# A Mammalian Genomic Signature Shaped by Single Nucleotide Variants Regulates Transcriptome Integrity and Diversity

**DOI:** 10.1101/2025.09.11.675687

**Authors:** Jian Yang, Samuel Ogunsola, Jason Wong, Aydan Wang, Roby Joehanes, Daniel Levy, Shalini Sharma, Chunyu Liu, Jiuyong Xie

## Abstract

**Background:** Many functional features of mammalian genomic sequences remain poorly defined, especially how sequence motifs and genetic variants within non-coding regions (NCRs) regulate transcriptome integrity and diversity. We have shown that G-tracts unusually positioned between the polypyrimidine tract and 3′ AG repress usage of the AG and are enriched at cryptic splice sites in cancer cells but their broader role across the extensive NCRs of mammalian genomes is unknown.

**Results:** Here, we identify a widely evolved genomic signature, G-tract-AG motifs consisting of guanine tracts closely upstream of AG dinucleotides, which is significantly associated with single-nucleotide variants (SNVs) identified in genome-wide association studies, particularly within NCRs. Approximately 9,000 such G-tracts within human genes are disrupted by variants of the *cis*-splicing quantitative trait loci in the Genotype-Tissue Expression project. Functionally, G-tracts repress splicing at the adjacent 3′ AG, primarily by stalling the second transesterification step. Disruption of G-tracts by SNVs relieves this repression, enabling splicing and generating novel transcript isoforms. These G-tract-disrupting SNVs are *in cis* across the majority of protein-coding genes and are among thousands of rare variants causing genetic diseases.

**Conclusions:** G-tract-AG signatures are widespread bipartite motifs with dual functions: G-tracts repress AG usage to safeguard transcriptome integrity, while SNV-induced disruption releases AGs for splicing to promote transcriptome diversity. Our findings provide mechanistic insights into the regulation of transcriptome integrity and diversity by a mammalian genomic signature, particularly for NCR SNVs associated with diverse traits and a new framework for their functional annotation.

## Background

Many sequence features of mammalian genomes, predominantly composed of non-coding regions (NCRs), remain largely unknown [1]. While cryptically spliced products from the NCR pose challenges to transcript integrity [2], novel mRNA isoforms driven by genetic variations could contribute to transcript diversity in evolution [3, 4]. It remains challenging, however, to identify the functional sequence motifs and single nucleotide variants (SNVs) in the NCR, particularly those reported in genome-wide association studies (GWAS) [5].

Several sequence motifs have been identified to prevent cryptic splicing of endogenous transcripts, such as pyrimidine-, UG- or CA-rich motifs [2, 6–10]. Lengthening uridine tracts also allow the emergence of new exons in primate evolution [4]. Distinct from these motifs, G-tracts are made of (G)_≥3_ purines. They are abundant in mammalian introns often as splicing enhancers [11–13], but not typically anticipated to be within the consensus sequence of the 3′ splice site (SS), (Y)_20_NY<u>AG</u> in humans (Y: C or T; N: A, C, G or T) [14]. When G-tracts do occur inside this region, they repress splicing as regulatory <u>e</u>lements between the polypyrimidine tract (<u>P</u>y) and 3′ <u>A</u>G dinucleotides (REPAG) [15–19]. REPAGs have been found within annotated 3′SS in eukaryotes from yeast to human and are enriched the most in mammals [18], likely contributing to the extensive use of alternative splicing [17, 20, 21].

Despite the abundance of G-tracts in mammalian introns [11], only about 2,000 REPAG-harbouring 3′SS have been identified in the human genome [17, 18]. Interestingly, in addition to annotated 3′SS, cryptic 3′SS in cancer tissues with splicing factor mutations also show REPAG enrichment [18]. This observation suggests that additional G-tract-AG signatures may lie deeper within introns or other non-coding regions, where they could serve additional roles.

Here, we searched the human genome and identified millions of these signatures, which are largely in the NCRs conserved in mammals. We show their potent ability to repress or de-repress splicing depending on the SNVs and highlight a distinct mechanism by which genetic variants in these regions could control splicing in their association with mammalian traits or diseases. These key sequences may also assist in the functional annotation of SNVs in NCRs [22–25].

## Results

### 1. Human G-tract-AG signatures: genome-wide presence, enrichment in GWAS-mapped regions and conservation among mammalian species

To identify genome-wide G-tract-AG signatures, we searched both strands of the human genome based on the criteria applied to the annotated 3’SS [18] (Fig. 1A). Initially, we identified and evaluated 111 millions of AG dinucleotides within splice site-like sequences (3’SSL) that had splicing signals greater than 0.1, the lower boundary of most constitutive exons (Additional file 1: Fig. S1), according to their MaxEnt scores [26]. Although numerous such 3’SSLs possess potential splicing signals, currently annotated 3’SS is only around 0.3 million [14]. This discrepancy suggests that the majority of these 3’SSLs remain either unused or actively repressed in cells. Consistent with this repression hypothesis, roughly 6 million of these 3′SSLs (about 5.4%) harbor the G-tract-AG signature - an enrichment of 2,630 times over that observed in annotated 3′SS [18]. This substantial enrichment underscores the potential functional importance of G-tract-AG signatures within these 3′SSL genomic regions.

**Figure 1.**
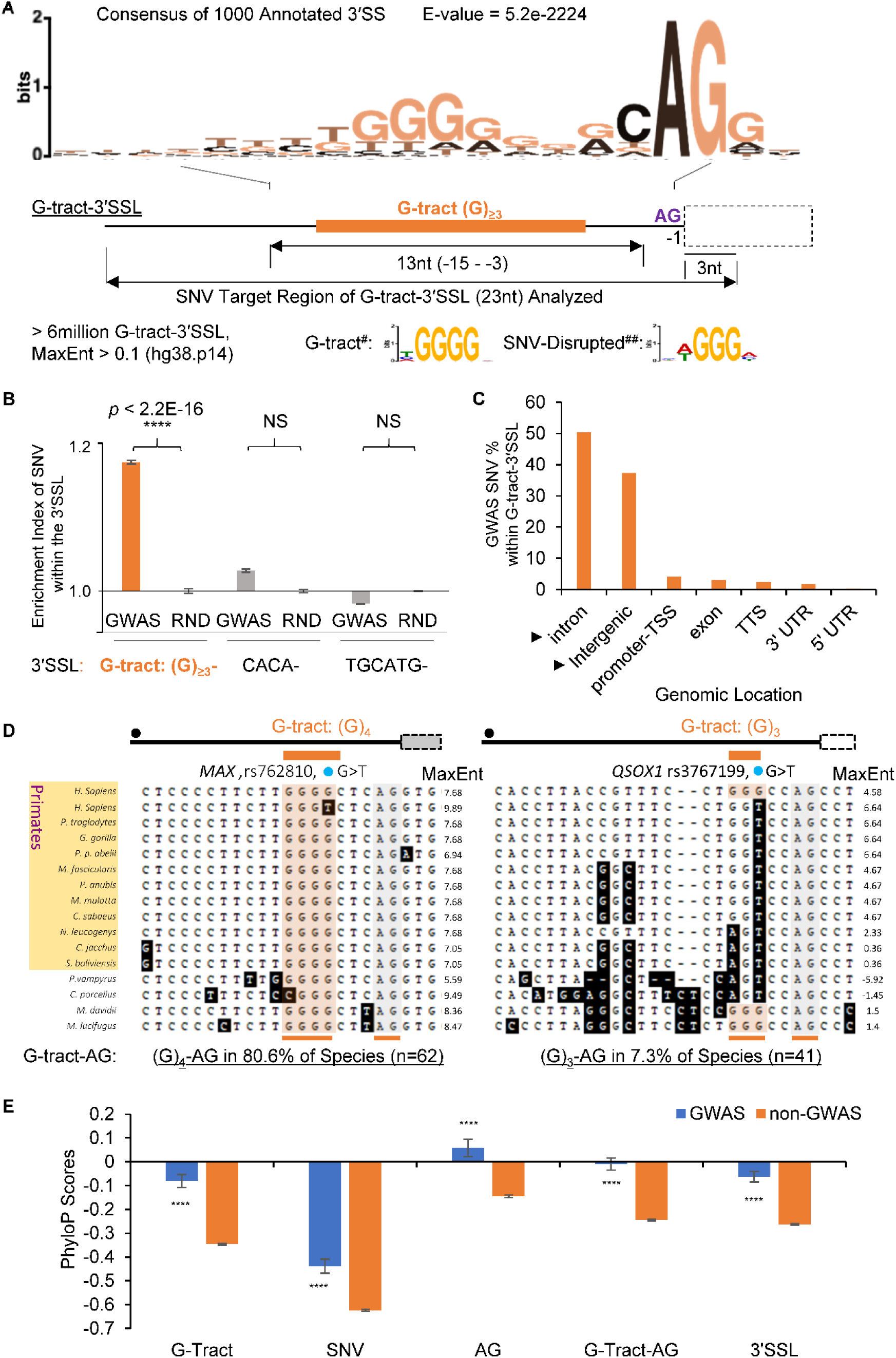
Distribution of G-tract-AG signatures in the human genome and GWAS-mapped regions, and in mammals. **A**. Diagram of the genome-wide search for G-tract-AG signatures. (Upper) A successful search within the Ensembl-annotated 3’SS with the criteria: the consensus motifs G-tract and AG of identified REPAG-3’SS (n = 1000). (Lower) Search criteria for the G-tract, AG positions/ranges within the 23nt 3’SS-like sequences in the genome, with the result summary and G-tract consensus motifs at the bottom. #: n = 700, E-value =1.9e-478; ##: n = 419, E-value =3.6e-122. **B**. Enrichment index of SNVs within G-tract-3’SSL among GWAS SNVs (input n = 128, 991, MAF ≥ 0.05, *p* < 5E-08). Random SNVs from the TOPMed (n = 1000 sets, each with 128, 991 SNVs (RND), matched by MAF bins and TSS distance, excluding GWAS SNVs), were used as background (Index = 1.0) for calculating the enrichment ratios of the abundance (GWAS/RND). *p*-value: by one-sample t-test. NS: non-significant. **C**. Genomic location profile of the GWAS SNVs within G-tract-3’SSL (n = 41,353), showing their mainly intron and intergenic locations. **D**. The G-tract-3’SSL of the *MAX* (Left) or *QSOX1* (Right) aligned among mammalian species, with human SNV variant sequences as the top two. 3’SSL MaxEnt scores are to the right of each sequence. Black dot: location of potential branch point. Dotted box: exon, and heavy black line: intron. Shaded box: exon already emerged in humans as a tissue-restricted alternative exon. **E**. PhyloP scores (470 mammals) of the motifs and SNVs from the GWAS database, at the human 3’SSL of G-tract-AG signatures. Scores are Means ± SEM, n = 3113, 3102, 3105, 3094, 3136 (GWAS); 104,705, 104,613, 104,486, 104,174, 105,758 (non-GWAS), for the G-tract, AG, SNV, G-Tract-AG and 3’SSL groups, respectively. ****: *p* < 3.5E-8.

To identify G-tract-AG signatures that are close to or disrupted by single nucleotide variants (SNVs), we first filtered their 3′SSL sequences (Fig. 1A, 23nt: 18nt + AG + 3nt, by the MaxEnt algorithm [26]) for the presence of any common SNVs from the TOPMed (Trans-Omics for Precision Medicine, <u>m</u>inor <u>a</u>llele frequency MAF ≥ 0.05) database. This analysis identified approximately 1.8 million unique SNVs, including 41,353 that overlap with the GWAS hits. These SNVs were significantly enriched in the GWAS loci (32.1%) compared to random SNVs in the human genome (30%, Fig. 1B, enrichment index, see also Methods). In contrast, there was no significant enrichment of SNVs in the CACA- or TGCATG-3′SSL in GWAS in parallel searches. Moreover, more than 95% of such GWAS SNVs were found in the NCR, mainly introns and intergenic regions (Fig. 1C). Further, within the genomic start-end boundaries of 3,202 GWAS genes, we identified 4,447 SNVs that directly disrupt the G-tracts [(G)_3-11_] in the sense strand (*in cis*, Fig. 1A, Additional file 2: Table S1a and Additional file 1: Fig. S2). Together, these data support a significant association of the G-tract-AG signatures with the GWAS SNVs.

Manual examination of a group of G-tract-AG signatures indicated that most of them were in mammals only (97%, n = 37), based on the MultiZ-alignment sequences of 100 vertebrates from the UCSC Genome Browser [27] (Additional file 1: Fig. S3). However, they exhibited higher divergence than those in the annotated 3′SS [15, 17]; for instance, about 25% of these signatures were found in primates only. In the two representative examples of different levels of conservation (Fig. 1D), the G-tract-AG signature in *MAX* is conserved in 80.6% of the species (n = 62), and the signature in *QSOX1* is conserved in 7.3% only (n = 41). The G-tract disruption occurred in other species as well, by A, C or T in guinea pig and primates, among other changes within the 3′SSLs (Additional file 1: Fig. S4). By MaxEnt scores, the G-tract disruption increased the splicing signal to or close to that of the human SNV-T variants in a few species (*MAX* in guinea pig and *QSOX1* in chimpanzee, gorilla and orangutan).

To systematically assess the conservation/divergence of the genomic signatures and G-tract-disrupting SNVs located between human gene start and end boundaries, we obtained their phyloP scores from the alignment of 470 mammalian genomes [27]. The mean phyloP scores for the G tract, G-tract-disrupting SNV, AG and G-tract-AG are −0.34, −0.61, −0.14 and −0.24, respectively (SEM = 0.004 each, n > 107K motifs, Additional file 1: Fig. S2B). The negative mean values indicate their accelerated evolution in mammals, particularly at SNV positions. However, within the GWAS subset, mean phyloP scores are significantly higher than those of the non-GWAS group (Fig. 1E), suggesting relative evolutionary constraint or reduced divergence at trait-associated sites. Based on phyloP scores and presence or absence of corresponding motifs in the MultiZ alignment [27], we estimate that about 63% of G-tract-AG signatures are primate-restricted (phyloP score ≤ 0), while the remaining signatures (phyloP score > 0) are conserved beyond primates in other mammals, with a very small fraction (< 0.1%) also in fish. Therefore, the G-tract-AG signatures and their G-tract-disruptive SNVs are predominantly mammalian features, exhibiting varying degrees of evolutionary divergence and conservation.

These data, together with the REPAG repression of splicing [15, 18], imply a much wider role for the G-tract-AG signature in the NCR of mammalian genomes, beyond its control of alternative splicing of annotated 3′SS [15–17].

### 2. G-tract repression and SNV-caused de-repression of splicing from the 3′ AG

To determine whether the G-tract-disruptive SNVs are globally associated with splicing, we examined156,380 such SNVs *in cis* within gene start-end boundaries in the human genome, and cross-referenced them with the *cis*-sQTLs (splicing quantitative trait loci, false discovery rate, FDR<0.05, n = 486,725 unique IDs) from the Genotype-Tissue Expression Project (GTEx) v10 [28]. Of these, approximately 9,000 SNVs were significant sQTLs (5.62%, Additional file 2: Table S1b), representing a 1.58-fold enrichment compared to randomly selected SNVs (n=1,000 sets of 156,380 SNVs, matched by their MAF and transcription start sites (TSS), *p* = 2.2E-16, one sample t-test). Of the 4447 GWAS SNVs, the sQTL proportion is even higher (14%). Thus, these G-tract-disruptive SNVs are significantly enriched for sQTLs, supporting their potential role in splicing and impact on the transcriptome.

To validate the potential effects of the G-tracts and SNVs on splicing, we chose the above two examples for their evolutionary conservation or divergence, increased MaxEnt scores upon disruption by SNVs and deep intron locations. We examined the effects in mini-gene splicing reporter assays [29] (Fig. 2 and Additional file 1: Fig. S5), which we have employed to consistently demonstrate the G-tract repression of splicing in ten REPAG reporters [15, 17, 19] (Additional file 3: Table S2). The *MAX* SNV rs762810-T is in the upstream 3′SS of exon 4 (E4), in a mammal-specific but much less conserved region than the E3 region (Fig. 2A). The E4 inclusion is at about 20% of the E3 by their RNA-Seq reads in blood cells, but invisible in the other 53 types of GTEx tissues including kidney and skin; Moreover, the intron 3 excision ratio is significantly higher in samples with the rs762810-T/T than those with the G/G genotype (Additional file 1: Fig. S6) [28]. The *MAX* G-tract repression and SNV de-repression effects were indeed validated in the splicing reporters (Fig. 2B): in human embryonic kidney (HEK293T) cells, the (G)_4_-carrying reporter skipped the E4. In contrast, the (G)_3_T-carrying reporter with a transversion of the last G to the rs762810-T resulted in 66% E4 usage (in molar ratio), and the (A)_4_ reporter with G to A transitions resulted in 98% E4 usage. Thus, it is the (G)_4_ that represses but the rs762810-T that de-represses splicing from the 3′ AG. The *QSOX1* rs3767199-T is in the first intron, 398nt downstream of exon 1 (Additional file 1: Fig. S5C). Like the rs762810-T, this variant also de-repressed the G-tract repression of splicing from its 3′ AG splice site, albeit to a smaller extent in the reporter assay.

**Figure 2.**
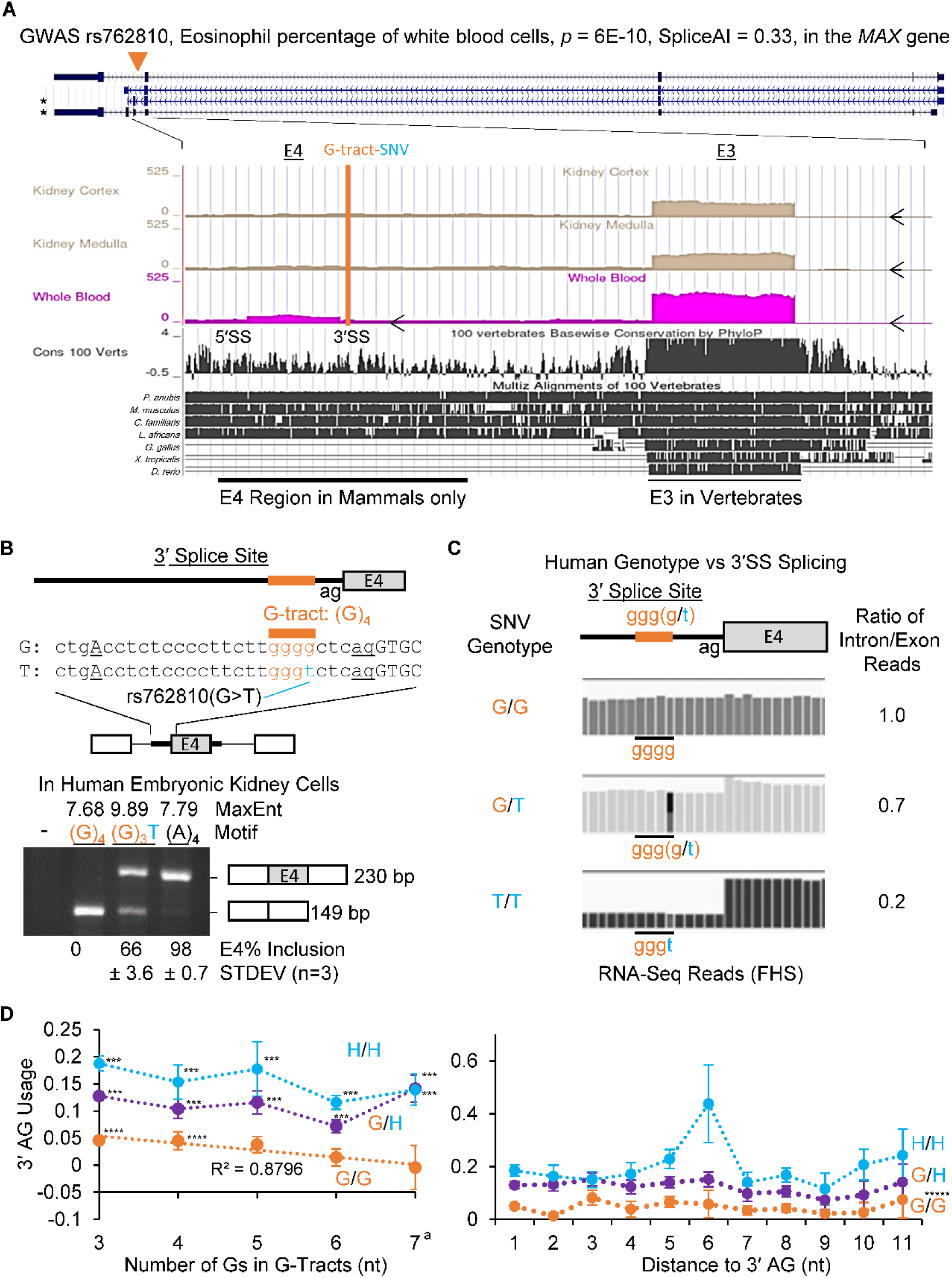
Functional validation of the NCR G-tract-AG signatures and SNVs: G-tract repression of splicing from the AG and disruption of the repression by SNV. **A**. Diagram of the SNV rs762810 in the genomic structure of the *MAX* gene transcripts, with the mammal-specific SNV region in the enlarged snapshots of GTEx RNA-Seq peaks and multiz alignment of 100 vertebrates from the UCSC Genome Browser. *: transcripts containing E4 (ENST00000557746.5 and NM_001407112.1). None of the other 53 GTEx tissues reaches the same level of E4 RNA-Seq peaks as in the blood. **B**. Splicing reporter assay of the G-tract-3’SS of the SNV allele in human embryonic kidney HEK293T cells. Intron/exon structure and sequence around the rs762810 loci with either the allele of G or T are above the agarose gels, which show the RT-PCR of spliced products. **C**. The SNV effect in genotype-RNA-Seq matched human samples of white blood cells from the FHS cohort, demonstrating the RNA-Seq reads at the G-tract-3’SS with different genotypes of the SNV alleles. **D**. Relationship between the 3′ AG usage (measured by the RNA-Seq reads, Mean ± SEM) and number of Gs in the G-tract (left panel, ^a^:7 or 8 Gs) or distance (nt) of the last G of (G)_3_ tracts to the 3′ AG (right panel) in 100 FHS participants of G/G genotypes. The corresponding 3′ AG usage among variant genotypes G/H or H/H (H: non-G nucleotides, A, C or T) is plotted in the same graphs for comparison. N = 171, 37, 19, 8, 8 (left panel), and 63, 25, 12, 13, 19, 7, 10, 15, 6, 5, 7 (right panel) participants. ***: *p* < 0.005, between G/G and variant genotype of the same number of Gs. ****: *p* < 0.002 between (G)_3_ or (G)_4_ and their respective longer G-tract G/G genotypes. *****: *p* < 3.5E-27 between G/G and non-G variant genotypes of the (G)_3_-tracts. Note: at 2nt or 6nt upstream (right panel), (G)_3_ or the H/H homozygote, respectively, exhibited stronger effects than that observed at their flanking positions (*p* < 0.05). However, these position effects need a larger sample size to validate.

To visualize the SNV genotype effect on splicing in humans, we examined the 3′SS intron/exon ratios of RNA-Seq reads, as described previously [30], in genotype- and RNA-Seq data-matched whole blood cell samples from participants in the Framingham Heart Study (FHS) (<u>50</u>). We found intron/exon junction-reads ratios of 1.0, 0.7, and 0.2 for individuals with G/G, G/T, or T/T genotypes of rs762810, respectively (Fig. 2C), indicating no splicing, partial splicing, or near-complete splicing of the 3′SS of the blood cell-specific exon. We further conducted association analyses with genotype or additive models between 3’ AG usage (measured as 1 - ratio of intron/exon reads within a 25 nt-window flanking the junction) and 3,302 SNVs located within G-tracts up to 13nt upstream of AG acceptor sites (Additional file 2: Table S1), using RNA-Seq data from 100 FHS participants. By the two models (Additional file 2: Table S1 and Additional files 4-5), we identified 241 pairs of SNV and G-tract-AG sites where the mean 3′ AG usage was increased in the non-G variant (H)-harboring genotype over the G/G homozygotes (Fig. 2D, *p* < 2.0E-18). The 3’ AG usage was not correlated with the 3’SS MaxEnt score (Pearson coefficient *r* = −0.05, *p* = 0.47), likely due to the inclusion of G-tract-AG as training decoy sites (∼20%) for the MaxEnt algorithm [26]. However, the mean value of the 3’ AG usage was correlated inversely with the G-tract lengths (3-8nt, *r* = −0.94, *p* < 0.02), where the additive effect by the H variants is evident in the (G)_3-6_ groups (Fig. 2D). In contrast, there was no such a correlation between the 3’ AG usage with the distance of the (G)_3_-tracts to the 3′ AG (*r* = −0.03, *p* = 0.92). Therefore, the human genotype and allele effects strongly support that a large proportion of the G-tracts repress and the G-tract-disrupting SNVs de-repress splicing for 3’ AG usage in a G-tract length-dependent way.

Taken together, these data support that the G-tract-AG signatures have dual functions in the NCR: (1) to repress splicing from the 3′ AG with the G-tract as a functional REPAG to protect transcript integrity and (2) to permit splicing for the emergence of novel exon (e.g., E4 in *MAX*) or transcript isoforms in mammals upon disruption of the G-tract by a SNV for transcript diversity.

### 3. Splicing repression at the two transesterification steps by the REPAG

For the mechanism of splicing repression by the G-tracts, we have shown previously that a REPAG element with its bound hnRNP (heterogeneous nuclear ribonucleoprotein) H1 represses splicing and reduces Py-binding by an early spliceosome factor U2AF65 [15, 17]. The hnRNP H1 binding is disrupted by mutating the G-tract [15], and by the *MAX* SNV-T variant in structure modelling (Additional file 1: Fig. S7). However, it is still unclear which of the two transesterification steps of splicing is repressed by such G-tracts. Our *in vitro* assay of splicing intermediates indicated that both steps were repressed but primarily at the 2^nd^ step.

In the *in vitro* splicing assay, we analyzed the kinetics of lariat formation in a pre-mRNA containing either the REPAG or its mutant as in our previous report (7) (Fig. 3A). Compared to the AdML vector, the presence of REPAG caused a significant delay by about 6 minutes in generating the lariat intermediate (lanes 1-3 vs. 4-6, and Fig. 3B). The delay was reversed upon mutating the REPAG sequence (lanes 7-9). Thus, REPAG delays the generation of the first step product, the lariat intermediate, which is consistent with the reduction of U2AF65 binding to the Py [15].

**Figure 3.**
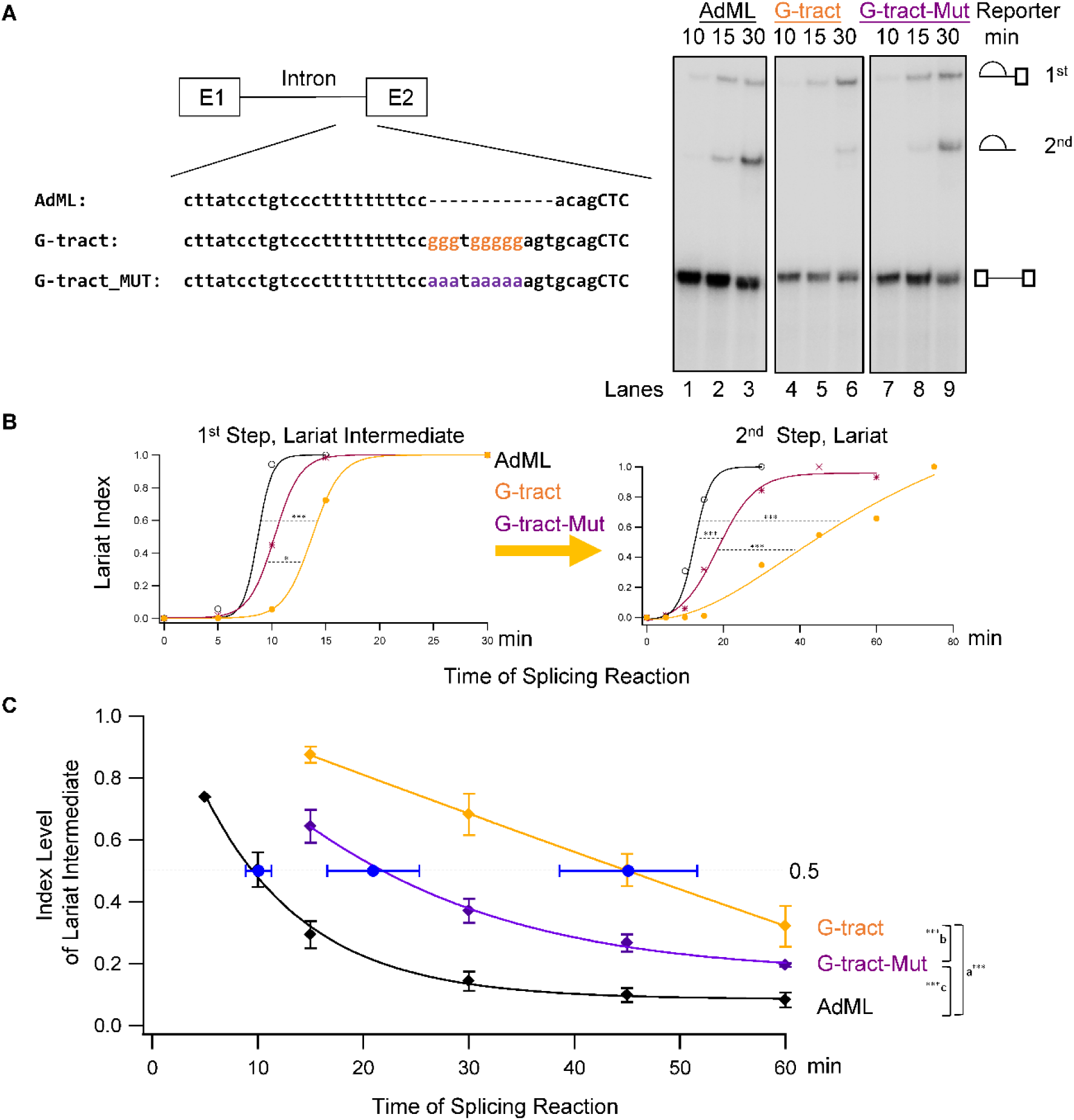
REPAG and SNV effects on the two transesterification steps of splicing. **A**. Left, diagram of the *in vitro* splicing reporters, AdML vector, REPAG-WT and REPAG-Mut and their 3’SS sequences. Right, examples of the *in vitro* splicing lariat products and pre-mRNA. N = 3-6 experiments of the three reporters. **B**. Time course of lariat product generation in the 1^st^ and 2^nd^ steps, after normalizing each product to its peak band intensities during the time course of the experiment. \*\*\**p* < 0.001, \**p* < 0.05, n = 8 pairs in total at two (10 and 15min) time points. **C**. Time course of the conversion of the 1^st^ step lariat intermediate to the 2^nd^ step lariat, after normalizing the lariat intermediate percentages of total lariats (1^st^ step lariat intermediate plus 2^nd^ step lariat) in each reaction to the highest percentage during the time course of the experiment. ^a^*p* < 9E-09, n = 12 pairs in total; ^b^*p* < 4E-06, ^c^*p* < 0.001, n = 9 pairs in total, at three (15, 30 and 45min) time points of 3-6 independent experiments. Blue dots and SEM (standard error of the mean) error bars: half conversion time (min). The best fitted curve (with minimal Chi-squared values, X^2^ < 0.007) was used to calculate the mean half conversion time of the 1^st^ step lariat intermediates in each experiment by their equations in Igor Pro (Wavemetrics).

Interestingly, the second step of splicing was also significantly delayed by REPAG but far more than would be expected from a simple delay of the first step (Fig. 3B). The half-life of lariat intermediates in conversion to the lariat was approximately 10, 21, and 45min for the AdML, REPAG mutant, and REPAG reporters, respectively (Mean, ±1.2, ±4.4, ±6.5, SEM, n = 6, 3 and 4, Fig. 3C), indicating a significant delay by the REPAG. Additionally, a less pronounced second-step repression was also observed for the REPAG mutant, likely due to the longer spacer between the Py and 3′ AG as reported [31].

Together, the results from the splicing intermediate assays demonstrate that the REPAG unusually represses both transesterification steps with a stronger effect on the 2^nd^ step of splicing.

### 4. Widespread impact of *in cis* disruption of the G-tracts of the genomic signatures by SNVs

The impact of the genomic G-tract-AG signature is likely widespread on mammalian genes. In the GWAS, the 3,202 genes that contain G-tract-disrupting SNVs *in cis* are strongly enriched for a variety of health-related traits including heart disease and cancer by DAVID functional clustering analysis (Fig. 4A, Additional file 2: Table S1a). About 77% of them are protein-coding genes. In the human genome, the 156,380 G-tract-disrupting SNVs are *in cis* within 23,479 unique genes, including about 71% of the 20,078 protein-coding genes in the hg38 (Fig. 4B, Additional file 2: Table S1b). These figures together with the sQTL enrichment and functional data on splicing (Figs. 2-3 and Additional file 1: Fig. S5) imply widespread impact by the G-tract-AG signatures and their SNVs on the transcriptomes and proteomes. For example, the *MAX* E4 usage contributes to two transcripts, resulting in truncated protein isoforms lacking the nuclear localization signal (NLS) or dimerization domain (Additional file 1: Fig. S8) [27, 32–35]. Several similarly truncated MAX isoforms showed cell growth-promoting instead of the tumor suppressor effect by the full length protein [33, 36–38]. The E4-promoting T-variant is associated with higher blood cell counts in GWAS [39], enriched among individuals with melanomas and associated with E4 ectopic inclusion in melanoma tumor cells (Additional file 1: Fig. S9), implying its association with cell growth as well. The usage of the *QSOX1* cryptic splice site caused by the rs3767199-T variant would lead to intron retention and nonsense-mediated mRNA decay reducing full-length protein levels (Additional file 1: Fig. S5A). Related to this anticipation, reduced protein level in *Qsox1* knockout mice caused lower pulse pressure, an effect also associated with the rs3767199-T variant in GWAS [40, 41]. Moreover, the sQTL of predominantly primate G-tract-AG signatures are highly enriched in cadherin family genes (*p* < 1E-10), which are important for cell-cell communications, while those also conserved in other mammals are enriched in genes encoding proteins with signal transduction domains (C2 or SH3, *p* < 1E-4). These distinct enrichment patterns suggest that the signatures may have contributed to the evolution of lineage-specific functional properties in primates and other mammals.

**Figure 4.**
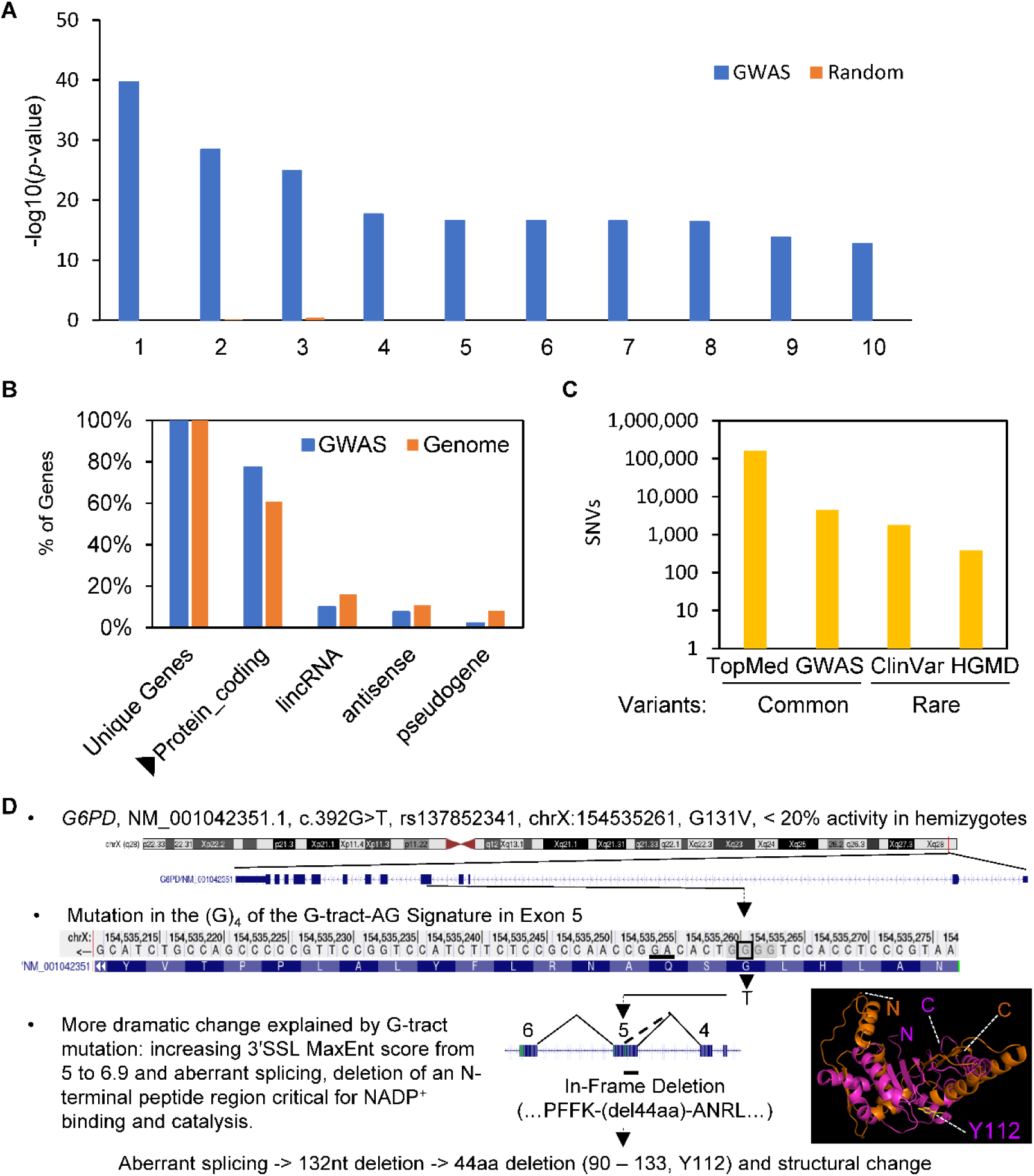
Disruption of the G-tracts of the signatures by human genetic variants *in cis* among GWAS-mapped genes, protein-coding genes and genetic disease genes. **A.** Significantly enriched GAD (Genetic Association Database) disease classes among the 3202 GWAS-mapped genes that harbor G-tract-AG signatures *in cis* with their G-tracts directly disrupted by GWAS SNVs, by DAVID functional clustering analysis. Random: mean of the − log10(*p*-value) of 3 sets of the same number of randomly picked human genes analyzed using the same parameters. 1. Chemdependency, 2. Metabolic, 3. Cardiovascular, 4. Pharmacogenomic, 5. Hematological, 6. Immune, 7. Renal, 8. Developmental, 9. Cancer, 10. Unknown. **B**. Percentages of the GWAS-mapped or genome-wide genes that harbor such signatures and SNVs. N = 3,202 and 23,479 genes for GWAS and the genome, respectively. **C**. Total number of such SNVs *in cis* within the host genes among the databases screened. **D**. A reasonable explanation provided by the G-tract-AG signature and its disruption to account for the unexplainable protein deficiency and genetic disease caused by a rare genetic variant, the SNV c.392G->T of the *G6PD* gene. Disruption of the G-tract motif by the variant causes partial exon skipping and protein truncation at the N-terminal. The AlphaFold3.0-predicted structure of the wild type or missense variant protein aligned well (RMSD = 0.925 or 0.885, respectively), but the truncated protein failed to do so (orange, RMSD = 14.500) with the corresponding N-terminal region (HQSD……FREDQI) of the electron microscopy structure (magenta, PDB #7UAG).

Beyond the genome-wide common SNVs, we also identified 2,145 rare genetic variants (MAF<0.01) that disrupt the G-tracts of the signatures in the human gene mutation database (HGMD) and ClinVar [42, 43] (Fig. 4B, Additional file 1: Fig. S10 and Additional file 6: Table S3). For instance, the c.392G>T variant of the glucose-6-phosphate dehydrogenase (*G6PD*) gene causes a neutral change of amino acids (G131V) but deficient enzyme activity in hemizygotes [44–46]. The discrepancy could be well explained by RNA level changes instead, by aberrant splicing due to T variant-disruption of the G-tract resulting in 44aa deletion at the N-terminal region (Fig. 4C). The variant enzyme will lose the Y112 for NADP^+^ binding and catalytic activity of the dehydrogenase [47].

## Discussion

Our finding provides not only a functional genomic signature to explain how transcriptome integrity and diversity are controlled in mammalian species by the signatures and their SNVs, but also a valuable tool for annotating similar sequence signatures and SNVs in the genomes. Identifying more of such signatures will help cover a much larger proportion of GWAS SNVs such as those in the CACA-3′SSL (Fig. 1B) or even coding regions (Fig. 1C, and 4D), or in other species (Fig. 1D). It will be particularly helpful for the identification of functional SNVs underestimated by existing prediction algorithms such as MaxEnt or SpliceAI [23]. For examples, the *MAX* and *QSOX1* SNVs were scored either below the confidence level 0.5 or not even scored by SpliceAI, as were 99% of the GWAS SNVs (Fig. 2A, Additional file 1: Fig. S11, and Additional file 7: Table S4). Further genome-wide functional annotation of the signatures will provide a global view of the balanced control between the maintenance of transcriptome integrity and the evolution of transcriptome diversity, as well as genetic differences among individuals or mammalian species. Moreover, detailed mechanistic studies on the motifs, splicing factors, and local structures will also provide further insights into the signature’s unusual effect on the 2^nd^ step of splicing, as found in other cases [48, 49]. Together, these studies on the signatures could help bridge the mammalian genome with transcriptome integrity and diversity but also advance personalized medicine through understanding and correcting cryptic splicing caused by SNVs [50].

## Conclusions

Taken together, our data define an evolved mammalian genomic signature (G-tract-AG) enriched for GWAS SNVs and GTEx sQTLs, and demonstrate that G-tract-disruptive SNVs act through a distinct molecular mechanism to control splicing. The G-tract, as a two-step tight splicing repressor, not only regulates alternative splicing of annotated 3′SS as previously reported [15], but also represses the latent AG splice sites from splicing in the introns of the genome. The 3′ AG becomes a splice site upon disruption of the G-tract by SNVs, leading to the generation of novel transcript isoforms. Therefore, it is likely that the widespread G-tract-AG bipartite motifs regulate mammalian transcriptome integrity and diversity depending on the SNV.

## Methods

### *In vitro* splicing assay

To the 3′ splice site of the adenovirus splicing reporter [51], we inserted the REPAG gggtgggggagtg upstream of the PRMT5 exon 3 and its mutant aaataaaaaagtg [15]. The *in vitro* splicing was carried out in HeLa nuclear extracts as reported [51]. The optimal time range for the time course measurements of lariat products was determined by pre-tests to avoid RNA degradation after prolonged incubation *in vitro*.

### Human genome search, G-tract-AG harbouring 3’SSL and phyloP score analysis

The GRCh38/hg38 reference genome was searched on both strands by identifying (G)_≥3_ (i.e., at least three consecutive guanines) within 1 to 13nt upstream of AG dinucleotides [18]. The 3’SS 23nt (18nt upstream plus 3nt downstream of AG) region was filtered for MaxEnt 3’SS strength measurement [26], and for the presence of TOPMed common SNVs (MAF ≥ 0.05) [52]. Briefly, to identify potential 3’SS AG sites, we developed a pipeline that first searched for genomic coordinates of AG dinucleotides on both strands across the human genome, then output the corresponding 23nt sequences covering the AG (5’- 18nt + AG + 3nt −3’, e.g. 5’-ttccaaacgaacttttgtaggga-3’). The likelihood of each sequence functioning as a valid 3’SS was quantified using MaxEntScan [26]. To map SNVs within the 23nt 3′SSL sequences, we used the TOPMed SNV annotation to retrieve allele and MAF information and integrated these with SNVs reported in the GWAS Catalog [5]. The TOPMed SNV annotation has been developed from deep sequencing on over 200,000 human genomes, with an average coverage of approximately 30-fold [52]. In this study, we selected SNVs with MAF ≥ 0.05 (approximately 3.5% of all TOPMed variants) from the TOPMed SNV annotation to identify variants within the 23nt region. To obtain MaxEnt scores for the 23nt sequences, we replaced the reference alleles with the alternative alleles and recalculated splice site strength using MaxEntScan.

To identify G-tract-AG-harbouring 3’SSL, we extracted 23nt sequences (Fig. 1A) containing (G)_≥3_ between the 6th and 18th nucleotides using our in-house pipeline. The consensus sequence motifs from these 23nt regions were then obtained using the online MEME motif analysis tool [53].

The phyloP scores of genomic motifs, SNVs and 3’SSL were obtained from the Hiller lab’s analysis of 470-mammal alignment through the UCSC Genome webserver (https://hgdownload.soe.ucsc.edu/goldenPath/hg38/phyloP470way/) [27]. The >107K scores of each group were averaged by their nucleotide lengths. The cutoff scores used to distinguish signatures present predominantly in primates (≤ 0) from those also in other mammals (> 0) was manually verified using the UCSC Genome Browser’s MultiZ alignment of 100 vertebrates.

### Enrichment analysis of G-tract-AG-harbouring 3′SSL among GWAS loci, GTEx sQTLs or health traits/diseases

Some SNVs in the GWAS Catalogue [5, 54] lack allele information or allele frequencies. Therefore, we annotated the GWAS SNVs using TopMed SNV data. We selected SNVs with MAF ≥ 0.05 and p-value ≤ 5E-08 from the GWAS Catalogue, resulting in 128,991 distinct GWAS SNVs. These SNVs were categorized into 20 groups according to five MAF bins (0.01, 0.05, 0.1, 0.2, 0.3, and 0.5) and four distance bins (0kb, 1kb, 10kb, 100kb, and 1Mb) from the transcription start site (TSS) of the closest genes, creating 20 matching groups [55]. Based on the same criteria, we generated 20 corresponding groups from all TOPMed SNVs. Specifically, TOPMed SNVs were categorized into five MAF bins (0.01, 0.05, 0.1, 0.2, 0.3, and 0.5) and four distance bins (0 kb, 1 kb, 10 kb, 100 kb, and 1 Mb) from the transcription start site (TSS) of the closest genes, yielding 20 matching groups parallel to those defined for the GWAS SNVs. From each group, we randomly selected the same number of SNPs as in the corresponding GWAS groups, yielding 128,991 SNPs in total. To minimize bias, we further generated 1000 random sets of 128,991 SNVs using the same procedure.

We calculated the proportion of GWAS SNVs located within G-tract-AG-harbouring 3′SSL (n = 29,284). We also calculated the proportion of the 128,991 randomly selected SNVs within these 3′SSL across 1000 sets. Fold-enrichment was calculated as the ratio of the proportion of GWAS SNVs within 3′SSL to the average proportion of matched SNVs within 3′SSLs across the 1000 sets. GTEx v10 sQTLs of 54 tissues were filtered to obtain 486,725 unique sQTLs, which was used to identify sQTLs located within the G-tract-AG-harbouring 3′SSL. One thousand random sets of SNVs matched by number, MAF and distance to TSS were generated using the same method described above. Enrichment ratio was calculated similarly as well.

Disease enrichment/cluster analysis was carried out using the DAVID functional clustering analysis at the NIH webserver (https://davidbioinformatics.nih.gov/) with the Genetic Association Database (GAD) [56, 57], which is a curated archive of human genetic association studies of complex diseases and disorders. GAD annotations include genes linked to reported disease associations, which allows enrichment analysis of gene–disease relationships. Genes from our list were tested against the GAD annotation set. Enrichment was assessed using DAVID’s modified Fisher’s exact test (EASE score). Three sets of control genes, 3202 per set, were randomly picked from 66,832 stable IDs of human genes for the same analyses.

### Study participants

This study used WGS and RNA-Seq data from selected samples of 1,544 Third Generation cohort participants (mean age 47±9 years, 52% women, 100% American White) [58], at the second examination cycle (2008–2011). Protocols for genetic material collection during regular health examinations were approved by the Institutional Review Board at Boston Medical Center. All participants provided written informed consent for genetic studies, and the research was conducted in accordance with relevant guidelines and regulations.

### RNA-Seq in the Framingham Heart Study

Peripheral whole blood samples were collected at the second examination cycle (2008-2011). RNA was isolated from peripheral whole blood using PAXgene™ tubes (PreAnalytiX, Hombrechtikon, Switzerland) to collect 2.5 mL samples from FHS participants [58]. Samples were incubated at room temperature for 4 hours for RNA stabilization and then stored at −80 °C. Prior to extraction, tubes were thawed at room temperature for 16 hours. Total RNA was extracted with a PAXgene Blood RNA Kit at the FHS Genetics Laboratory. Following centrifugation and washing, white blood cell pellets were lysed in guanidinium buffer. RNA quality was assessed by absorbance at 260 and 280 nm using a NanoDrop ND-1000, and integrity was evaluated with the Agilent Bioanalyzer 2100 microfluidic.

RNA sequencing was conducted at the University of Washington Northwest Genomics Center, an NHLBI TOPMed reference laboratory, following standard protocols. Data processing took place at the same facility. Base calls were generated on the NovaSeq 6000, and unaligned BAM files were converted to FASTQ format using Samtools bam2fq. Sequence quality was assessed with the FASTX-toolkit, and alignment to GRCh38 with GENCODE release 30 was performed using STAR. FastQC provided initial quality control metrics, and STAR created a genome index to generate BAM files at the gene and transcript levels [59]. The TOPMed RNA-Seq pipeline utilized RNA-SeQC for standard quality control metrics from aligned reads [60].

### Whole genome sequencing

DNA from peripheral whole blood from 7,215 FHS participants underwent whole genome sequencing (WGS) at the Broad Institute, a TOPMed contracting laboratory. Details on DNA extraction, processing, and variant identification followed standard procedures [52]. Consistent sequencing and data processing criteria were applied, aligning DNA sequences to human genome build GRCh38. The resulting BAM files were sent to TOPMed’s Informatics Research Center for re-alignment and standardized BAM file generation. This study utilized WGS data from Freeze 10a, which included approximately 208 million SNPs across TOPMed cohorts.

### Visualization of RNA-Seq reads corresponding to rs762810 (G>T) genotypes

We identified the same FHS participants with both RNA-Seq and WGS data. We then randomly selected three participants with G/G, G/T, and T/T genotypes of rs762810 (G > T, MAF = 0.25 for T). We applied Integrative Genomics Viewer (IGV) [61] to visualize the RNA-Seq reads count in response to the genotypes of rs762810, G/G, G/T, and T/T.

### Association analysis between SNVs and 3′ AG usage

We quantified 3′ AG usage of the 3′SSLs following previously described methods for intron/exon ratios of RNA-Seq reads within 25nt flanking the intron-exon junctions [62–64], in 100 randomly selected FHS participants. The 25nt downstream of the 3’ AG were considered as exon sequences here. Briefly, RNA-Seq reads were first aligned to the reference genome hg38 and reads mapped within 25nt flanking the intron-exon junctions overlapping the plus-strand G-tract-AG signatures were extracted using the featureCounts in Subread package [65]. The 3’ AG usage was quantified as: 1 - the ratio of (intron reads)/(exon reads), within 25nt flanking the junction. Additional quality control was applied to the intron-exon junction ratios. Specifically, we excluded AG sites with missing ratio values caused by zero-read counts on both sides of the AG site. We also removed sites with negative ratio values below −0.1. We focused on SNVs located within G-tracts up to 13nt upstream of AG sites (Additional file 2: Table S1). Two association models were applied to examine the relationship between SNV genotypes and 3’ AG usage in the 100 FHS participants. In the genotype model, genotypes were treated categorically (0/0, 0/1 or 1/1), using the homozygous reference (0/0) as the baseline when all three genotypes were present. If “0/0” was absent due to limited sample size, the genotype with the largest number of samples was used as the reference. SNVs with only one observed genotype were excluded. In the additive model, association was tested using alternative allele dosage (0, 1, or 2) as predictor for the ratios in the same 100 sample. Covariates included sex, blood cell counts, five genotype-based principal components (PCs) to account for population stratification, and 15 transcriptome-based PCs to adjust for unknown confounders affecting gene expression [66]. Given the limited sample size, statistical significance was assessed with a liberal threshold of *p* < 0.05. Violin plots were generated to show the associations between SNP genotypes and the ratios (S_files 1 and 2). To make sure the analysis method correctly reflects the splicing changes, we pre-tested it with the *MAX* E4 upstream 3’SS and obtained significant ratio changes consistent with the RNA-Seq and splicing reporter results in figure 2A-C.

### Structure modelling of protein-RNA complexes

We used AlphaFold 3.0 [67] to model the hnRNP H1 protein-RNA structure and changes by the SNV. We first confirmed the prediction accuracy by aligning the top-ranked AlphaFold models with the corresponding crystal or NMR structures of a group of RNA-binding proteins in complex with RNA oligos (≤ 30nt) from the protein data bank PDB [68, 69]. The root mean square deviation (RMSD) was 0.559 angstrom (± 0.049, mean ± SEM, n = 21 pairs of predicted and PDB structures) after alignment using PyMOL (Molecular Graphics System, Version 3.0 Schrödinger, LLC). This deviation is significantly less than the threshold of 3.0 (p < 0.0001, one sample two-tailed t-test) for structural differences [70]. We then modelled the protein-RNA complex structures between hnRNP H1 qRRM1 and qRRM2 and the REPAG-containing RNA oligos 5’-UGGGGCU-3’ and variant 5’-UGGGUCU-3’of the 3’SS of *MAX* intron 3.

### Mini-gene splicing reporter assay in HEK293T cells

G-tract, SNV or mutant reporters of *MAX* intron 3 or *QSOX1* intron 1 were made using the Gibson mutagenesis system [71], and verified by Sanger sequencing. Briefly, plasmids were transfected into HEK293T cells for overnight before RNA extraction for RT-PCR assay of spliced variants, as previously reported [72].

### Plasmid Construction

Minigene splicing reporters were constructed using the Gibson Assembly Master Mix (New England Biolabs [71]) following the manufacturer’s protocol. Linear PCR fragments were generated using primers with overlapping sequences corresponding to the ApaI/BglII linearized pDUP175 vector [29, 73]. The fragments were designed with the NEBuilder Assembly Tool.

For the pDUP-*MAX*-E4-WT plasmid, a genomic DNA fragment containing exon 4 (81bp) and partial flanking introns (27 bp upstream and 22 bp downstream) was PCR amplified from human genomic DNA. The primers used were *MAX*E4-WT-DUP175 (5′-CTTGGGTTTCTGATAGGGCCCTGACCTCTCCCCTTCTTGGGGCTCAGGTG-3′) and *MAX*_3′SS_DUP175_REV (5′-TCTCTGTCTCCACATGCCCAACCTGGAGTCTCTGGTAC-3′).

For the pDUP-STREX-*QSOX1*-3′SS-WT plasmid, the 3′ splice site of the STREX exon from the pDUPST1 vector was replaced by a 27-nt sequence from intron 1 of *QSOX1* using PCR with primers *QSOX1*_DupST1_3SS’_WT (5′-CTTGGGTTTCTGATAGGGCCCTGACACCTTACCGTTTCCTGGGCCAGCCAAGATGTCCATCTAC-3′) and DUP175_DupST1_3′SS_REV (5′-TCTCTGTCTCCACATGCCCAAGACATGGAACTGTGTGTG-3′). Mutant plasmids were created by amplifying the PCR fragments from the wild-type plasmid using complementary mutant-specific primers and cloned using the Gibson Assembly Master Mix. All plasmid constructs were verified through DNA sequencing.

### Cell Culture and Transfection

We verified the identity of the HEK293T cell line in the Xie laboratory using previously reported RNA-Seq data from our study [15]. This was supported in our dataset by significant enrichment of genes among the top 1,018 Harmonizome-HEK293T genes [74, 75], relative to GH_3_ cells [72] (*p* = 8.5E-7), and further by the expression, or lack of expression, of top 53 Harmonizome-HEK293T genes, in comparison with signature genes expressed in non-kidney tissues or cell lines used in the Xie laboratory [75] (*p* = 4.9E-15). The cells were also confirmed to be mycoplasma-free, as RNA-Seq analysis of triplicate samples each comprising at least 64 million paired-end reads [15], detected no mycoplasma-matching sequences [75]. Cells were cultured at 37°C with 5% CO₂. The cells were grown in Dulbecco’s Modified Eagle Medium supplemented with 10% fetal bovine serum and 1% penicillin-streptomycin-glutamine. For transfections, 1μg of plasmid DNA was used per well in a 6-well plate, and transfection was performed using Lipofectamine 3000 (Invitrogen), according to the manufacturer’s protocol.

### RT-PCR

Reverse transcription was carried out based on a previously described procedure [7]. Briefly, for reverse transcription, 1μg of total RNA (for *MAX*) or cytoplasmic RNA (for *QSOX1*) was added to a 10µl RT reaction and incubated at 45°C for 50min, followed by a 5-minute incubation at 90°C. For PCR amplification, 1μl of the RT product was used in a 24μl reaction. The PCR cycling was set for 30 cycles using primers DUP8a (5′-GACACCATGCATGGTGCACC-3′) and DUP10 (5′-CAAAGGACTCAAAGAACCTCTG-3′). PCR products were resolved on agarose gels, visualized under UV light, and photographed using a digital camera. Band intensities were quantified using ImageJ software [76]. Exon inclusion efficiency was calculated as the molar percentage of the included products by their intensity relative to the total intensity of both included and skipped products after normalization to the production lengths.

## Data Availability Statement

The whole genome sequencing (WGS) and RNA sequencing (RNA-Seq) data from the Framingham Heart Study (FHS) are available at the dbGaP database under accession code phs000007.v34.p15 (FHS) [https://www.ncbi.nlm.nih.gov/projects/gap/cgi-bin/study.cgi?study_id=phs000007.v34.p15] [77]. These data are available under restricted access to ensure confidentiality and protect participant privacy. Access can be obtained by submitting an ancillary study proposal and obtaining IRB approval. Timelines for the approval process range from 3-6 weeks. Researchers must first obtain an eRA Commons account, which is typically set up within 3-7 business days if the applicant’s institution is already registered with NIH (institutional registration may add an additional 2-4 weeks).

Our published raw RNA-Seq reads for the verification of HEK293T identity are freely available from the NCBI Sequence Read Archive with accession numbers SRR37830201, SRR37830202 and SRR37830203 [15]. Raw RNA-Seq reads for the control GH_3_ cell samples (X21-X23) are available under accession number SRR13670058 [72].

The IGV view of RNA-Seq peaks of 53 types of human tissues, from the GTEx Project [28], can be accessed through the GTEx portal IGV browser at https://www.gtexportal.org/home/ or the UCSC Genome Browser [27], at https://www.genome.ucsc.edu/. GTEx *cis*-sQTLs and summary statistics [78], are publicly available at the GTEx portal, https://www.gtexportal.org/home/downloads/adult-gtex/qtl, to download at https://storage.googleapis.com/adult-gtex/bulk-qtl/v10/susie-qtl/GTEx_v10_SuSiE_sQTL.tar [79].

All the other figure data including source code scripts and gel images are deposited in Figshare with accession #31743592 at https://doi.org/10.6084/m9.figshare.31743592 [75]. The source code is released under an open source license compliant with OSI (http://opensource.org/licenses) and also has a separate accession #31820986, at https://doi.org/10.6084/m9.figshare.31820986 [64].

## Competing Interests

The authors declare no competing interests.

## Funding

The work is supported by a discovery grant (RGPIN-2022-05023) by the Natural Sciences and Engineering Council of Canada (NSERC) to J.X., by NIH (GM127464 and P30CA023074) and Arizona Biomedical Research Centre (ABRC: RFGA2022-010-30) to S.S., and by NIH R01AA028263 and R01HL15569 to C.L. The Framingham Heart Study (FHS) was supported by NIH contracts N01-HC-25195, HHSN268201500001I, and 75N92019D00031. RNA-Seq was supported in part by the Division of Intramural Research (D. Levy, Principal Investigator) and an NIH Director’s Challenge Award (D. Levy, Principal Investigator).

## Ethical Approval

The Framingham Heart Study (FHS) was approved by the Institutional Review Board of Boston University Medical Campus (Protocol H-41461). All participants provided written informed consent for participation and for publication of research results. All procedures were conducted in accordance with applicable ethical guidelines and regulations and complied with the principles of the Declaration of Helsinki.

## Acknowledgements

We thank the donors for the Framingham Heart Study for sharing their genotype and RNA-Seq information, and Dr. Ren-Jang Lin for helpful comments.

## Author contributions

JX conceptualized the project with LC and SS. JY, SO and JW carried out the bioinformatics analyses, cell culture or *in vitro* splicing reporter assays, respectively. AW contributed to the REPAG MEME and AlphaFold3 structural analyses, RJ and DL contributed to the FHS RNA-Seq data. JX wrote the manuscript, with critical revisions by LC and SS. All authors have read and approved the manuscript.

## Declarations

Peer review information: William Fairbrother and Wenjing She were the primary editors of this article and managed its editorial process and peer review in collaboration with the rest of the editorial team. The peer-review history is available in the online version of this article.

## Additional Files

Additional file 1_Fig_S1-11.pdf

Additional file 2_Table_S1_G-tract-AG_SNV_in _GWAS_Genome.xlsx

Additional file 3_Table_S2_Functional REPAG_Validated_list.xlsx

Additional file 4_3’ Splice site usage_n374_pairs_additive_violin_plots.pdf

Additional file 5_3’ Splice site usage_n603_pairs_general_violin_plots.pdf

Additional file 6_Table_S3_ G-tract-AG_SNV_in ClinVar_HGMD.xlsx

Additional file 7_Table_S4_GWAS_SpliceAI.xlsx

## Notes

https://figshare.com/articles/dataset/_b_A_Mammalian_Genomic_Signature_Shaped_by_Single_Nucleotide_Variants_Regulating_Transcriptome_Integrity_and_Diversity_b_/31743592

### Competing Interest Statement

The authors have declared no competing interest.

### Summary of Updates

This version has been updated to reflect revisions made following peer review during journal submission.

