## Supplementary material for "A Mammalian Genomic Signature Shaped by Single Nucleotide Variants Regulates Transcriptome Integrity and Diversity": Additional file 1_Figures_S1-11.pdf

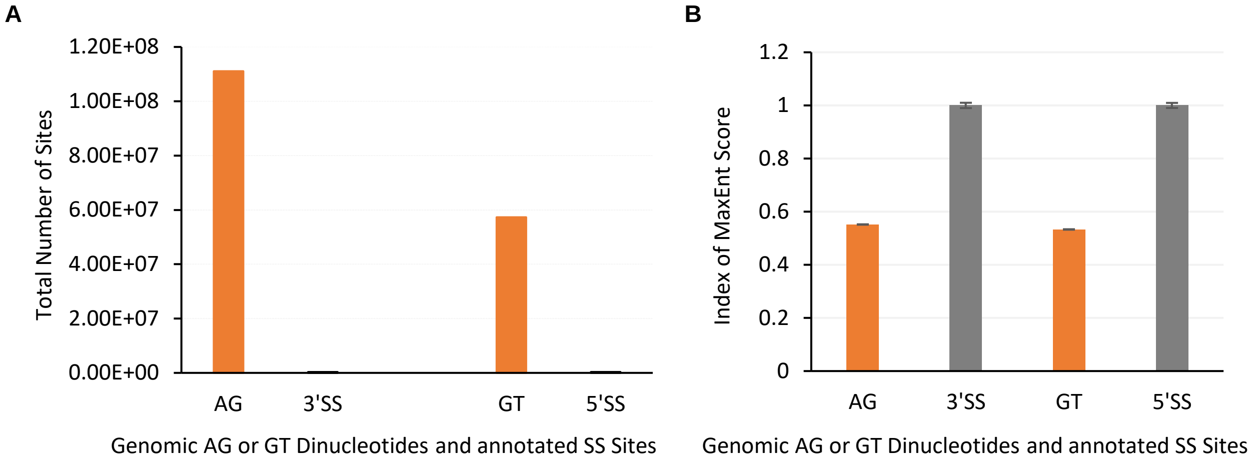


**Fig. S1**. **168 millions of AG/GT dinucleotides as potential cryptic splice sites in the human genome. A**. Total number of AG/GT dinucleotides within SS-like sequences with MaxEnt scores > 0.1 on both strands of the human genome hg38, with the relatively much smaller numbers of the annotated AG/GT 3′SS and 5′SS in the Ensembl for comparison. **B**. Index ratios of the MaxEnt scores (Mean ± SEM) of sequences harboring the dinucleotides on either strand of chromosomes, assuming the AG/GT as the SS dinucleotides, relative to a subset of annotated constitutive exon's SSs (n = 5,010 exons) in the Chr. 1 of the human genome hg38. The means of the 3′SS and 5′SS scores are 6.7 (± 0.07, SEM, minimal -33.57 – maximal 15.33, >0.1: 92.6%) and 6.96 (± 0.07, SEM, minimal -26.61 – maximal 11.81, >0.1: 94.0%), respectively. The scores of these AG/GT dinucleotide sites in the genome are half of the constitutive SS's scores on average, posing significant challenges with the potential for cryptic splicing under appropriate conditions (e.g. *trans*-acting factors, signaling or cell/tissue types, etc. spatial or temporal changes).

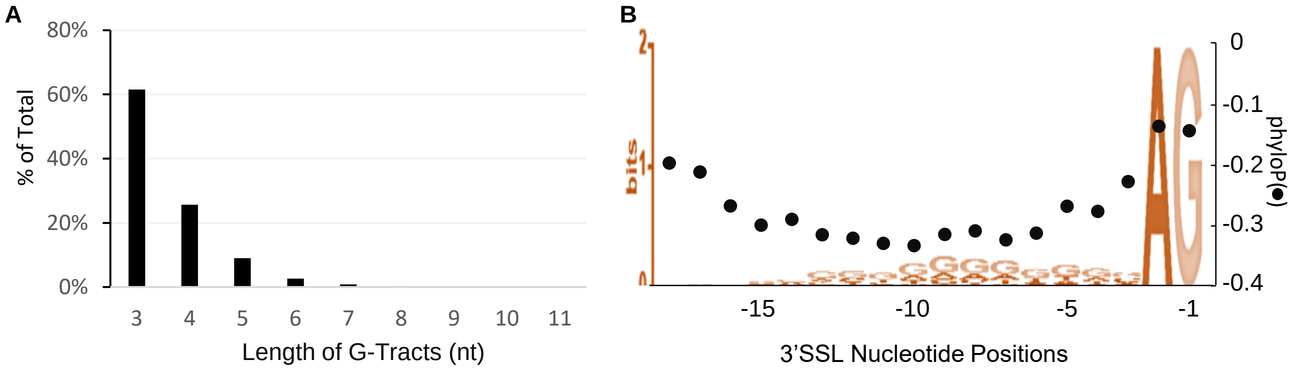


**Fig. S2**. **Length (nt), location and phyloP score of the G-tracts in the genomic signatures. A**. Length distribution of G-tracts. n = 7916 G-tracts, from 6114 GWAS ($p$ < 5E-08) SNV's G-tract-AG-3′SSL. **B**. MEME consensus motifs of G-tract-AG 3′SSL of 1998 sQTL within human genes, E value = 1.4E-442, showing the enrichment of Gs at the 3′SSL. Black dots indicate the phyloP scores from 470 mammalian species at each position of >107K sequences within human genes.

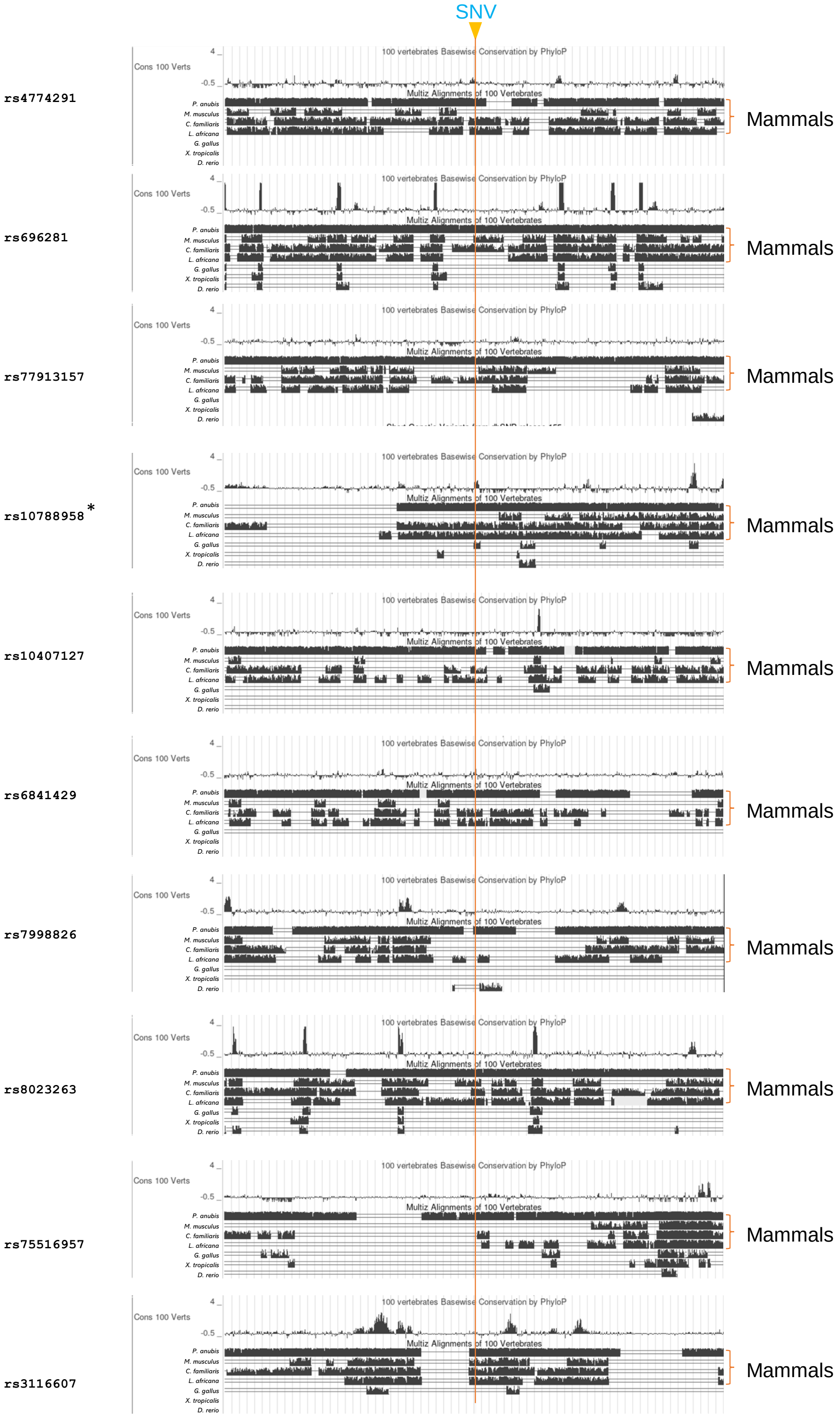


**Fig. S3. The human G-tract-AG signature is conserved in mammals only.** Shown are 10 examples of the manually examined 37 SNVs that disrupt the G-tracts of the signatures in the MultiZ alignment of 100 vertebrates. Screenshots, taken from the UCSC Genome Browser, indicate their conservation in mammals only, in contrast to their flanking exon regions that are also in the other vertebrates like birds or fish. *: The rs10788958 alignment in chicken does not contain the G-tract nor AG.

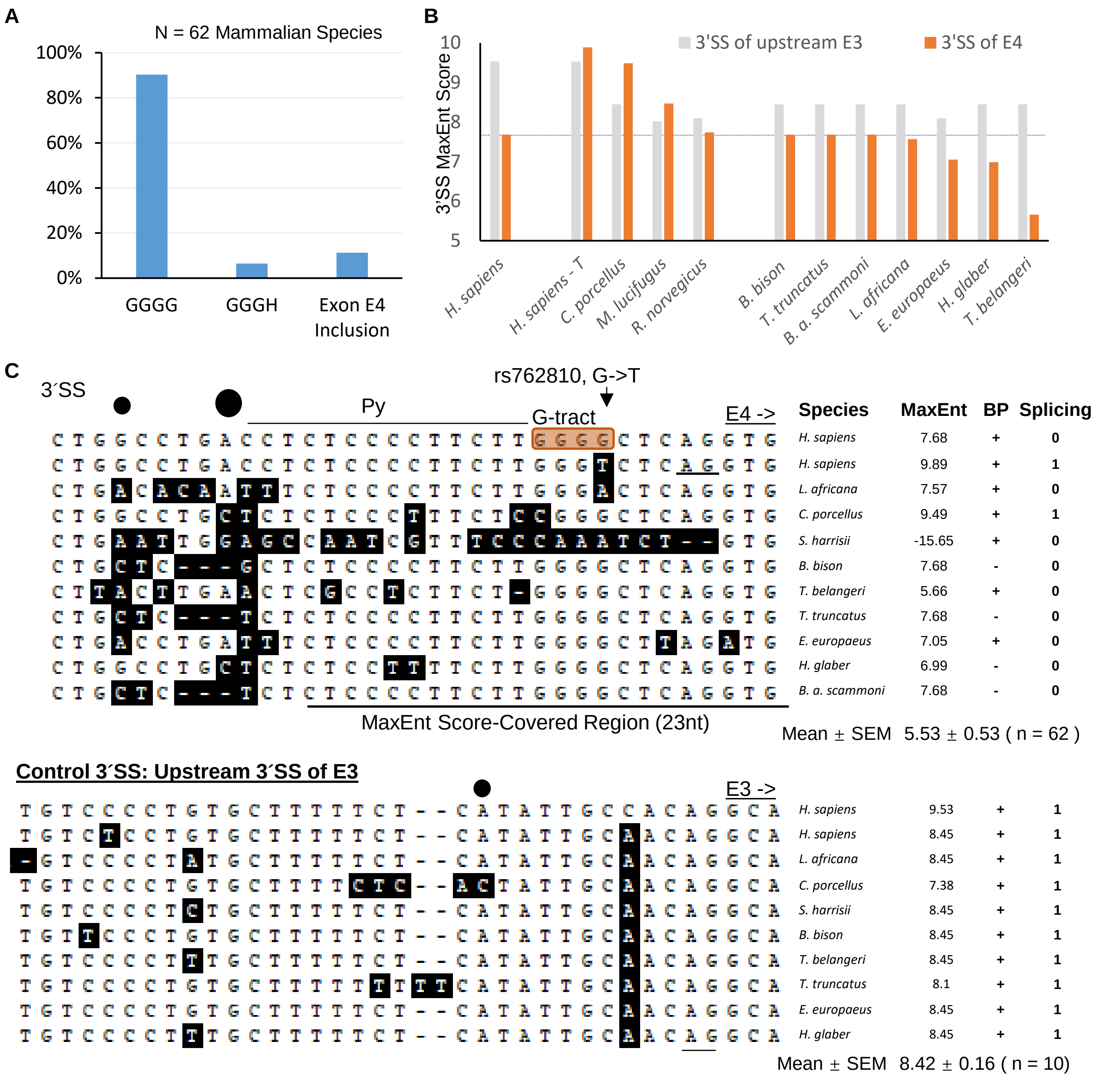


**Fig. S4. Divergent sequence and 3´SS strength changes of the G-tract-3´SS of the *MAX* gene in some mammalian species. A.** Percentages of species containing the G-tract, its disruption and predicted exon 4 (E4) inclusion among 6 of 62 mammalian species, based on: higher 3´ and 5´ SS-combined MaxEnt scores than the human reference sequence, presence of 3' AG and branch point motifs, shortened $(G)_4$- tract and intact ORF. H: A, C or T. **B.** In comparison with the human reference variant G (Left), 3´SS MaxEnt scores of E4 of representative mammalian species that are higher (Middle) or lower (Right), in comparison with those of the upstream E3. **C.** Additional examples of divergent 3´SS sequences of the *MAX* G-tract-3´SS motifs, besides SNV-disruption of the G-tract among mammals. Upper panel: G-tract-3´SS alignment. Lower panel: the less divergent upstream 3´SS of exon 3 of the same species for comparison. Nucleotides different from the human reference genome sequences are shaded in black. Black dots: positions of potential branch points in some species. Py: polypyrimidine tract. Splicing prediction: by MaxEnt scores, essential 3´SS motifs and/or transcript evidence. 1: positive or more likely; 0: negative or less likely. Note that here we use 3´SS since the evolved T-variant creates a splice site in humans (see functional validation later).

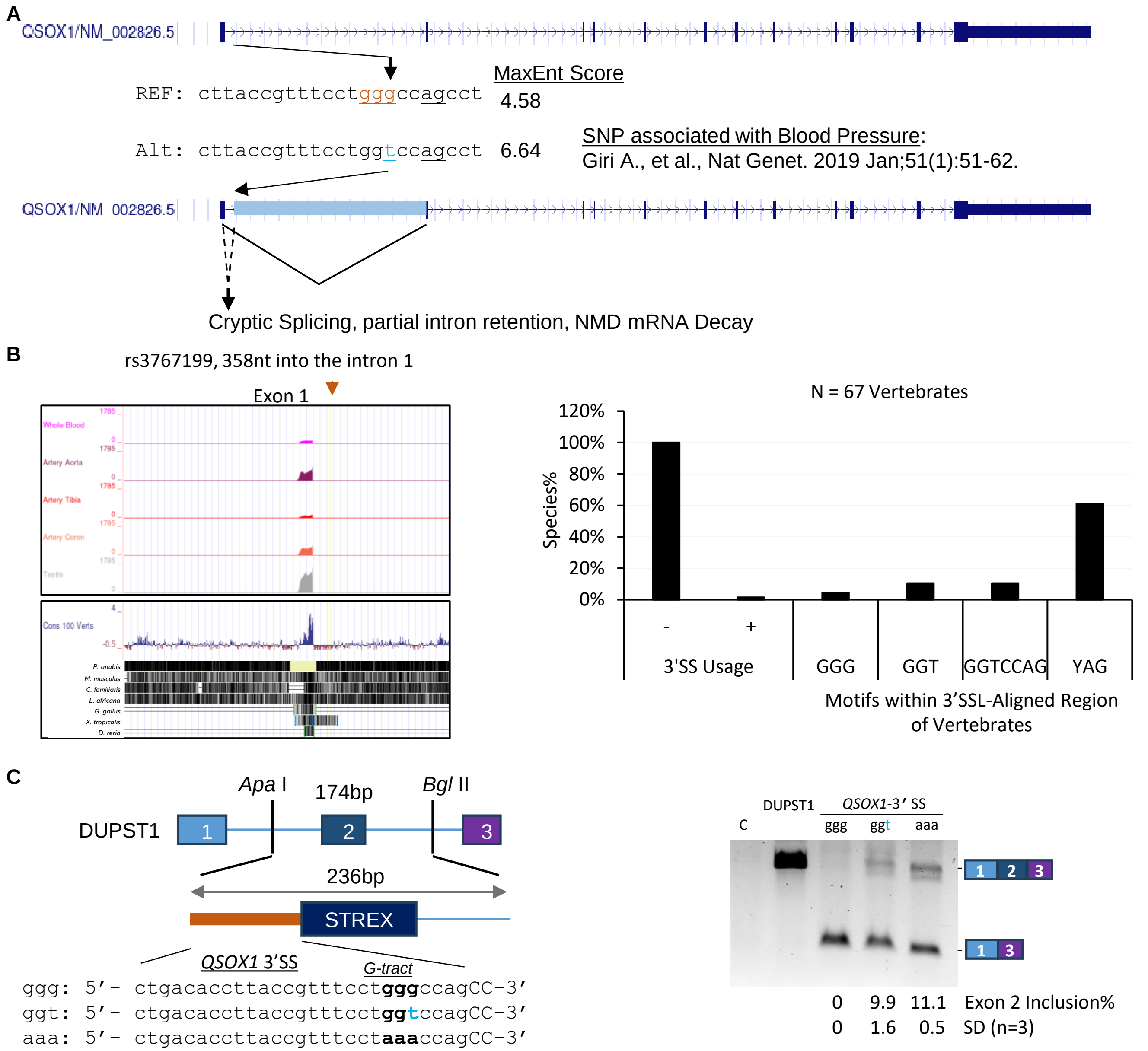


**Fig. S5**. Validation of the effect of SNV rs3767199-T on splicing through the intronic G-tract-AG signature of the *QSOX1* gene by minigene splicing reporter assay. **A**. Diagram of the SNV region in the intron 1 of the *QSOX1* gene, adapted from the UCSC genome browser. **B**. Left, repression of the SNV-3′ SSL from cryptic splicing in human tissues: snapshot of the SNV region in GTEx showing RNA-Seq peaks of the upstream exon 1 in several tissues but none downstream the SNV, nor in the rest of the 54 tissues, with the aligned 100 vertebrate conservation profile below, both from the UCSC genome browser. Note: the GGG-AG signature is not in *X. tropicalis* though it is also aligned locally. Right, percentages of 67 vertebrate species with potential 3′ SS usage or the 3′ SSL motifs. The GGG and GGT are in only a small percentages of the species and all are in mammals only. 3′ SS usage (only with human SNV) based on: presence of 3′ AG, shortened G-tract and higher MaxEnt 3′ SSL score than the human′s reference sequence. **C**. Left, diagram of the minigene splicing reporter, derived from the DUPST1 harboring the STREX exon (reference 29: Xie & Black, *Nature*′01). Right, agarose gel of the RT-PCR products of cytoplasmic RNA of the splicing reporters transiently expressed in HEK293T cells. C: negative control with untransfected-cellular RNA. The $(G)_3$ repression was also observed when in front of a 51nt beta-globin exon.

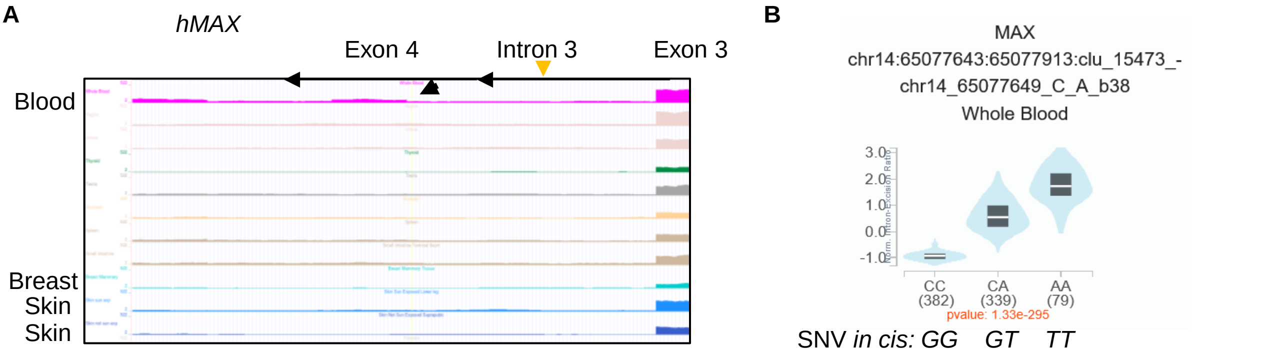


**Fig. S6**. SNV-association with *MAX* exon 4 usage in blood cells in GTEx. **A**. Snapshot of the IGV view of the RNA-Seq reads peaks of the rs762810-T allele-associated exon 4 in the intron 3 of *MAX* ENST00000557746.5, among a number of GTEx tissues in the UCSC Genome Browser. The arrowhead points to the location of the SNV and the upstream 3′SS of exon 4. Black arrows: direction of the *MAX* gene. **B**. GTEx result of the intron 3 excision ratios with different genotypes of the SNV rs762810.

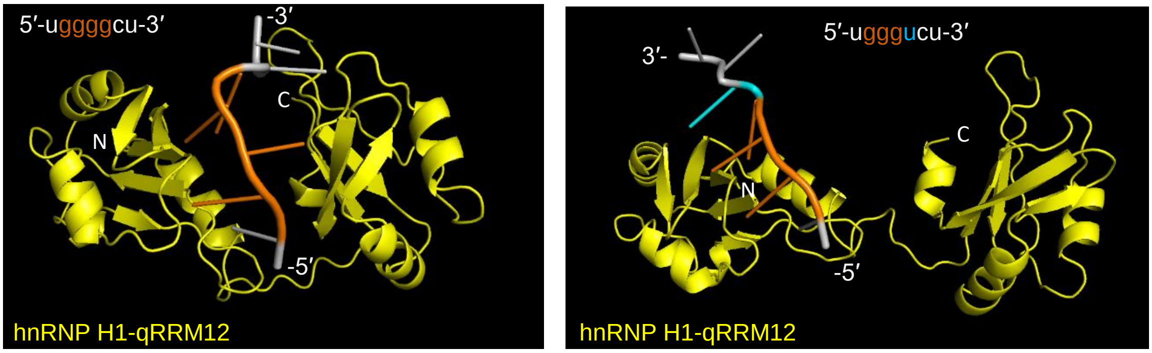


**Fig. S7**. AlphaFold 3.0-predicted structural conformation changes in the interaction between the G-tract-G or -U RNA variants and the trans-acting splicing repressor hnRNP H1 qRRM1 and 2 domains. Shown are the *MAX* G-tract 5′-UGGGGCU-3′ (Gs in oranges) or its rs762810-T variant 5′-UGGGUCU-3′(SNV U in cyan) in complex with the repressor. RMSD = 9.211 when aligned between the two complexes, indicating significant structural differences induced by the variants ($p < 0.0001$), in contrast to an average RMSD of 0.559 (±0.225, SEM) of 21 protein-RNA (≤ 30nt) complexes aligned with their experimental structures from the PDB in PyMol. The high-confidence RNA-protein complex show that the U-variant dramatically relaxed the G-tract RNA interaction with the trans-acting splicing repressor hnRNP H1. The complex switched from a compact key-lock pocket for the G-variant to a wide-open configuration for the U-variant, in a manner similar to the crystal structure changes for different G-tracts in reference 69 (Penumutchu et al., *J Am Chem Soc*, 2018), suggesting destabilization of the G-tract-repressor interaction by the U-variant.

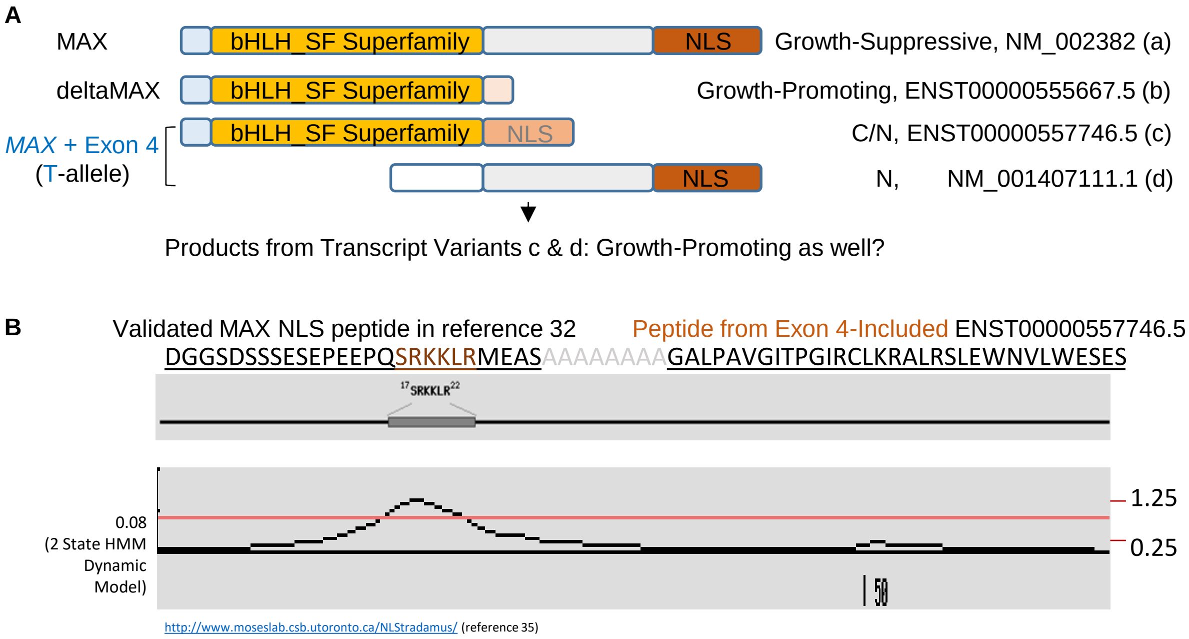


**Fig. S8**. Domain loss of the MAX protein isoforms generated from the transcripts with the SNV-T allele-promoted exon 4 usage. **A**. Truncation of the COOH or $NH_2$ terminals in the isoforms from the exon 4-included transcripts, in comparison with the representative full length or short (delta) isoforms with known effects on cell growth. **B**. Significant loss of the NLS signal strength from the transcript ENST00000557746.5 containing exon 4. Prediction based on reference 35 (Nguyen et al., *BMC Bioinformatics*, 2009) with the known NLS peptide and ENST00000557746.5-encoded NLS-candidate peptide input together with a poly(Alanine) linker.

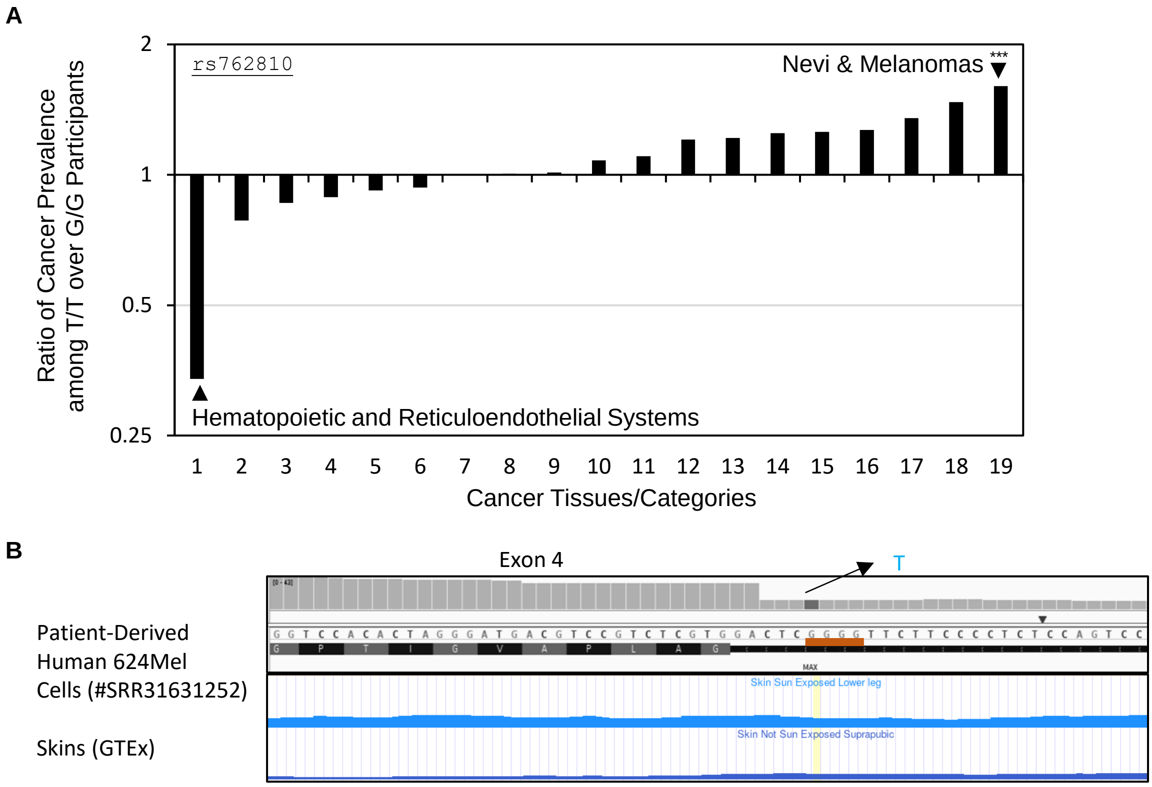


**Fig. S9. *MAX* variant genotype distribution among cancer patients of different tissues and exon 4 ectopic inclusion in melanoma cells with the T-variant. A.** Tissue-dependent ratio of cancer prevalence among the T/T over G/G genotypes of SNV rs762810 of the *MAX* gene. Shown is the log2 ratios of the cancer prevalence among different tissues or histological categories over that among the cohort after normalization to (divided by) the allele frequencies in the cohort. From left to right: 1. hematopoietic and reticuloendothelial systems, 2. trachea bronchus, and lung, 3. only one cancer, 4. colon, 5. lymph nodes, 6. rectum, rectosigmoid junction, anal canal and anus, not otherwise specified, 7. control: all FHS participants with genotype data for SNP rs762810, 8. healthy controls from FHS with genotype data for SNP rs762810, 9. prostate gland, 10. urinary bladder, 11. adenocarcinoma, 12. female breast (excludes skin of breast), 13. basal cell carcinoma, 14. corpus uteri or endometrium, 15. skin, excludes skin of labia majora, skin of vulva, skin of penis and skin of scrotum, 16. with multiple cancers, 17. squamous cell carcinoma, 18. duct carcinoma, 19. nevi & melanomas, in an FHS cohort with sex- and age-matched genotype data for the SNV rs762810 (***: $p$ < 4E-11, hypergeometric density test, n = 8,481 participants). **B**. Exon 4 ectopic inclusion (>50%) in patient melanoma-derived cells with the T variant only. Raw RNA-Seq sequence and BAM file derived from SRA #SRR31631252. Another sample with the T variant only and similar inclusion is #SRR31631251. The extra AG at the exon start is likely from the last two nts of exon 3 based on the gene sequence and RNA-Seq reads.

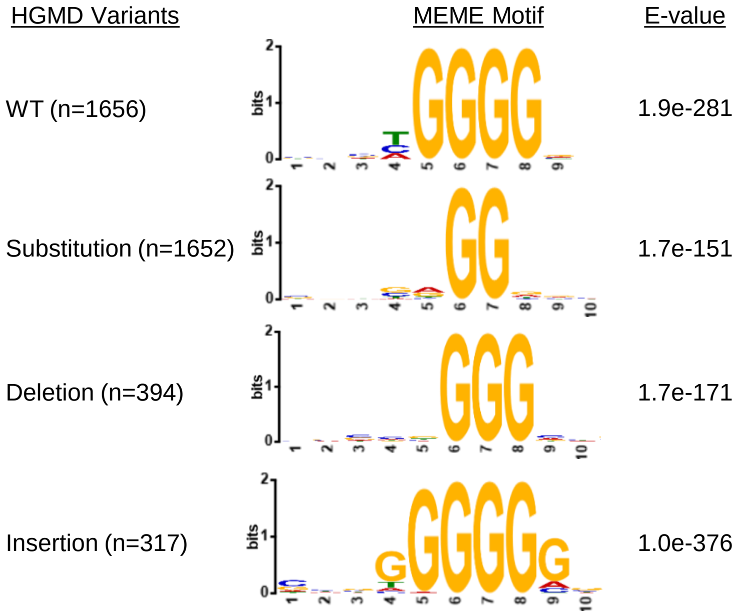


**Fig. S10.** A group of G-tract mutations/variants identified in human genetic diseases in the HGMD database. Shown are the changes of the consensus motif sequences by G-tract mutations in comparison with the wild type sequences, as analyzed using the MEME. The mutations are from reference 42 (Bacolla, et al., NAR'15).

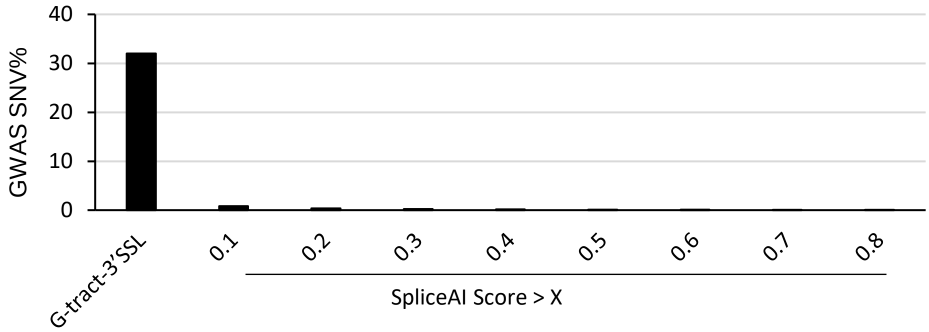


**Fig. S11.** Underrepresentation of the G-tract-AG signatures in current splicing-prediction software. Shown is the rarity of the G-tract-3′SSL SNVs among SpliceAI output results by comparison of the percentages of G-tract-3′SSL-overlapping SNVs with the percentages of the SpliceAI output among GWAS SNVs (n = 128,991 and 153,477 input GWAS SNVs in total, respectively). Of the 166 SpliceAI-scored SNVs above the confidence level of >0.5, only 7 are G-tract-disrupting SNVs (0.005% of GWAS). The G-tract-disrupting SNVs at 3′SSLs, 5.62% GTEx sQTLs reported here, are thus expected to add significantly to the list of splicing-regulating SNVs without the limitation of SNV distance to annotated splice sites by SpliceAI or other programs.
