## Supplementary material for "A Mammalian Genomic Signature Shaped by Single Nucleotide Variants Regulates Transcriptome Integrity and Diversity": Additional file 4_3' splice site usage_n374_pairs_additive_violin_plots.pdf

Violin plots of 3' splice site usage of 374 pairs of SNV and G-tract-AG-harboring 3’ splice site from the additive model, by analysis of whole genome sequencing and RNA-Seq - matched data among 100 FHS participants.

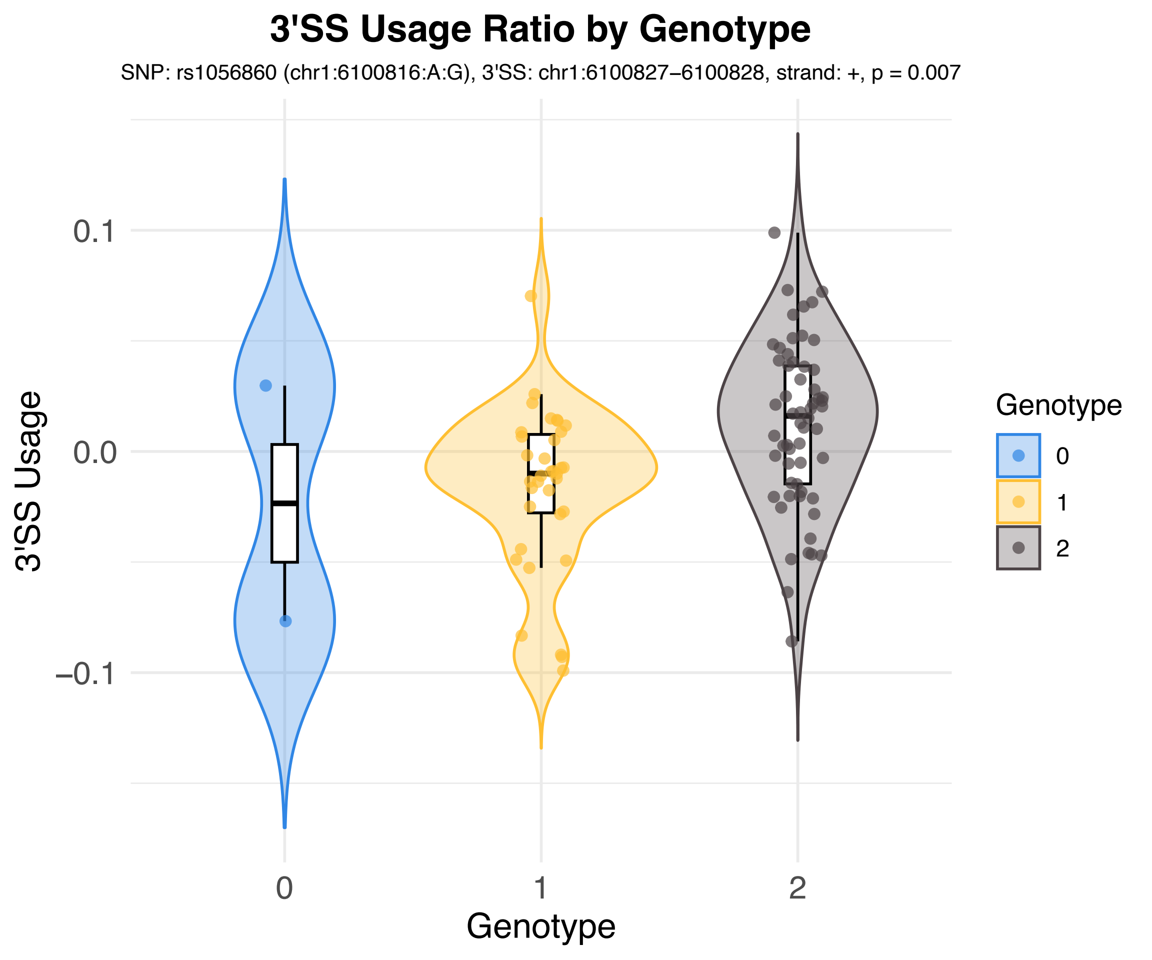
3'SS Usage Ratio by Genotype
SNP: rs1056860 (chr1:6100816:A:G), 3'SS: chr1:6100827–6100828, strand: +, p = 0.007
3'SS Usage
0.1
0.0
–0.1
0
1
2
Genotype
Genotype
0
1
2

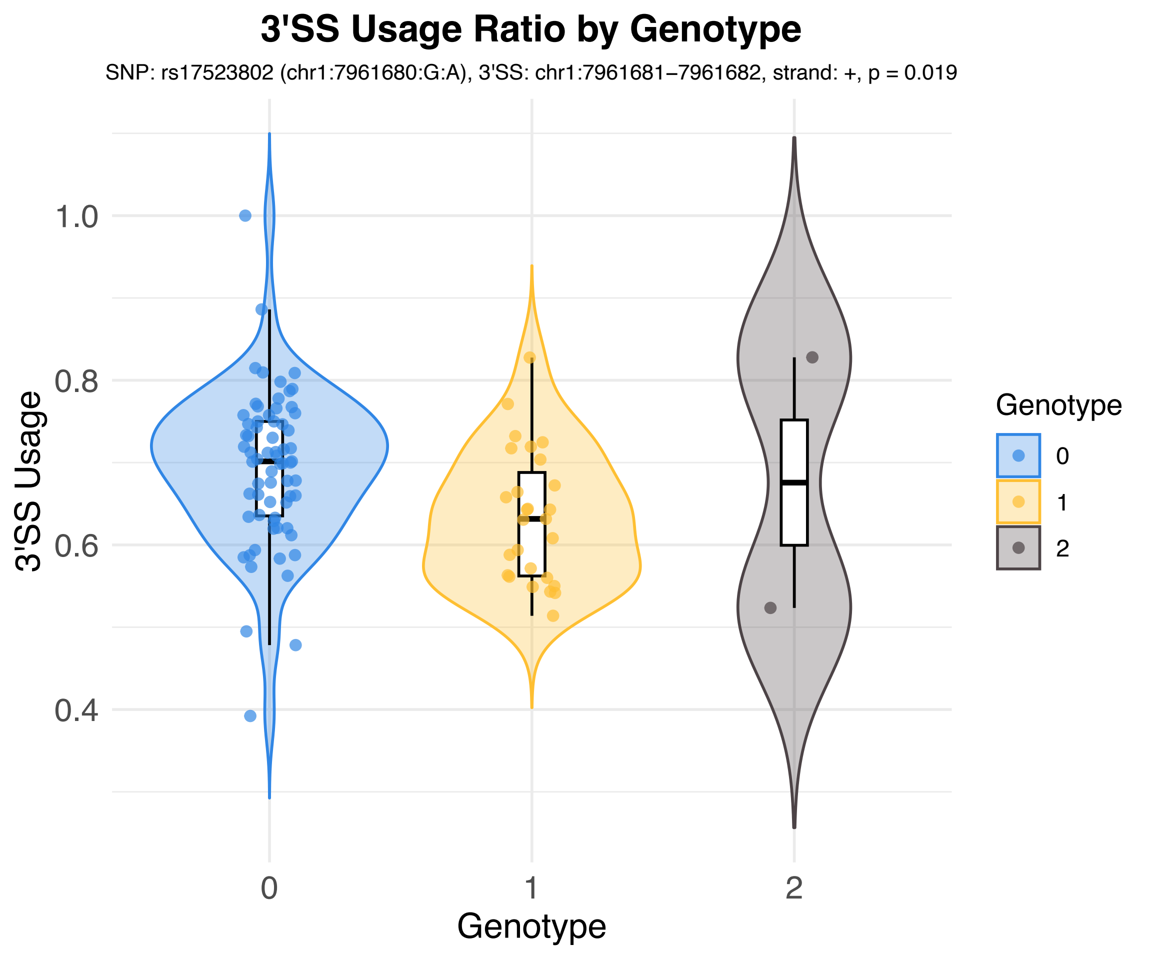
3'SS Usage Ratio by Genotype
SNP: rs17523802 (chr1:7961680:G:A), 3'SS: chr1:7961681−7961682, strand: +, p = 0.019
3'SS Usage
1.0
0.8
0.6
0.4
0
1
2
Genotype
Genotype
0
1
2

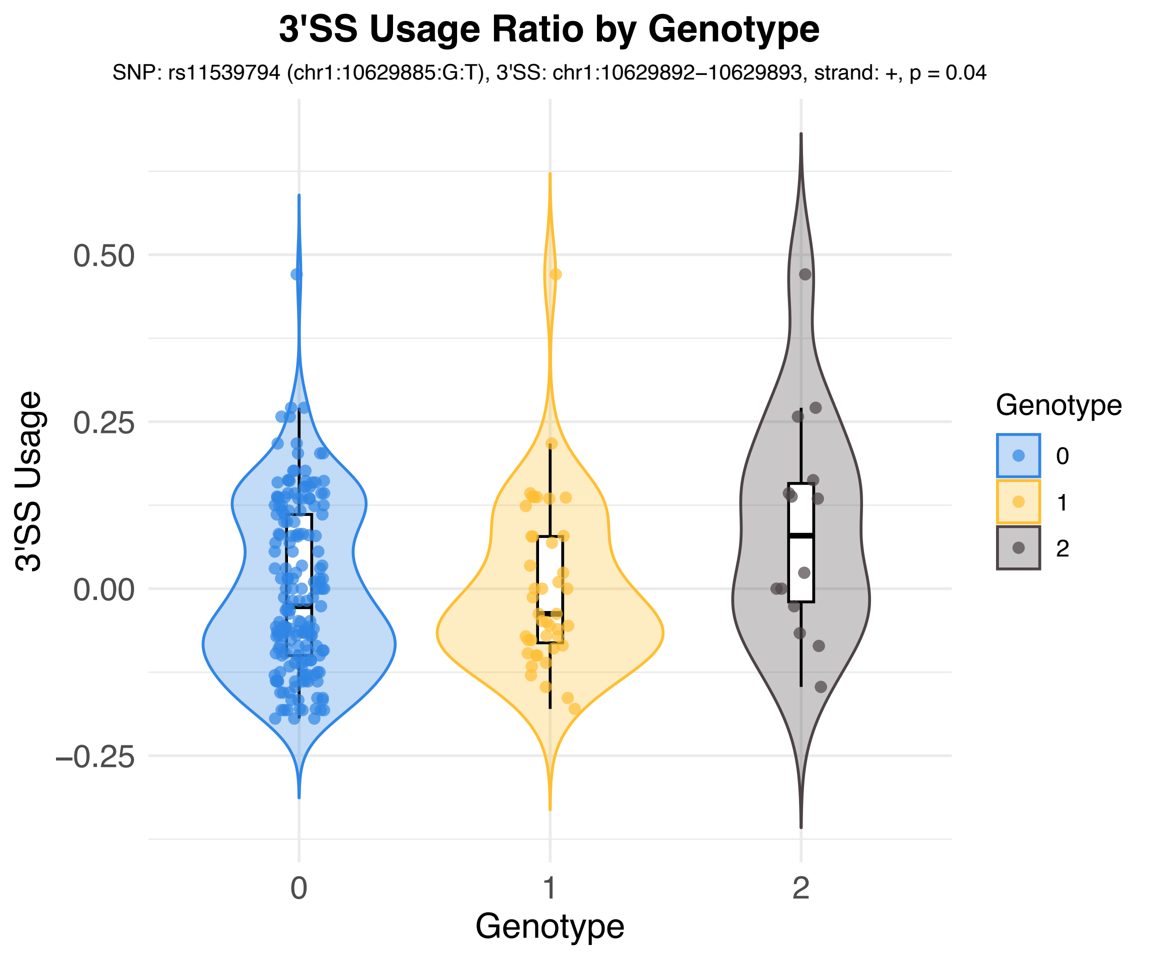
3'SS Usage Ratio by Genotype
SNP: rs11539794 (chr1:10629885:G:T), 3'SS: chr1:10629892−10629893, strand: +, p = 0.04
3'SS Usage
0.50
0.25
0.00
−0.25
0
1
2
Genotype
Genotype
0
1
2

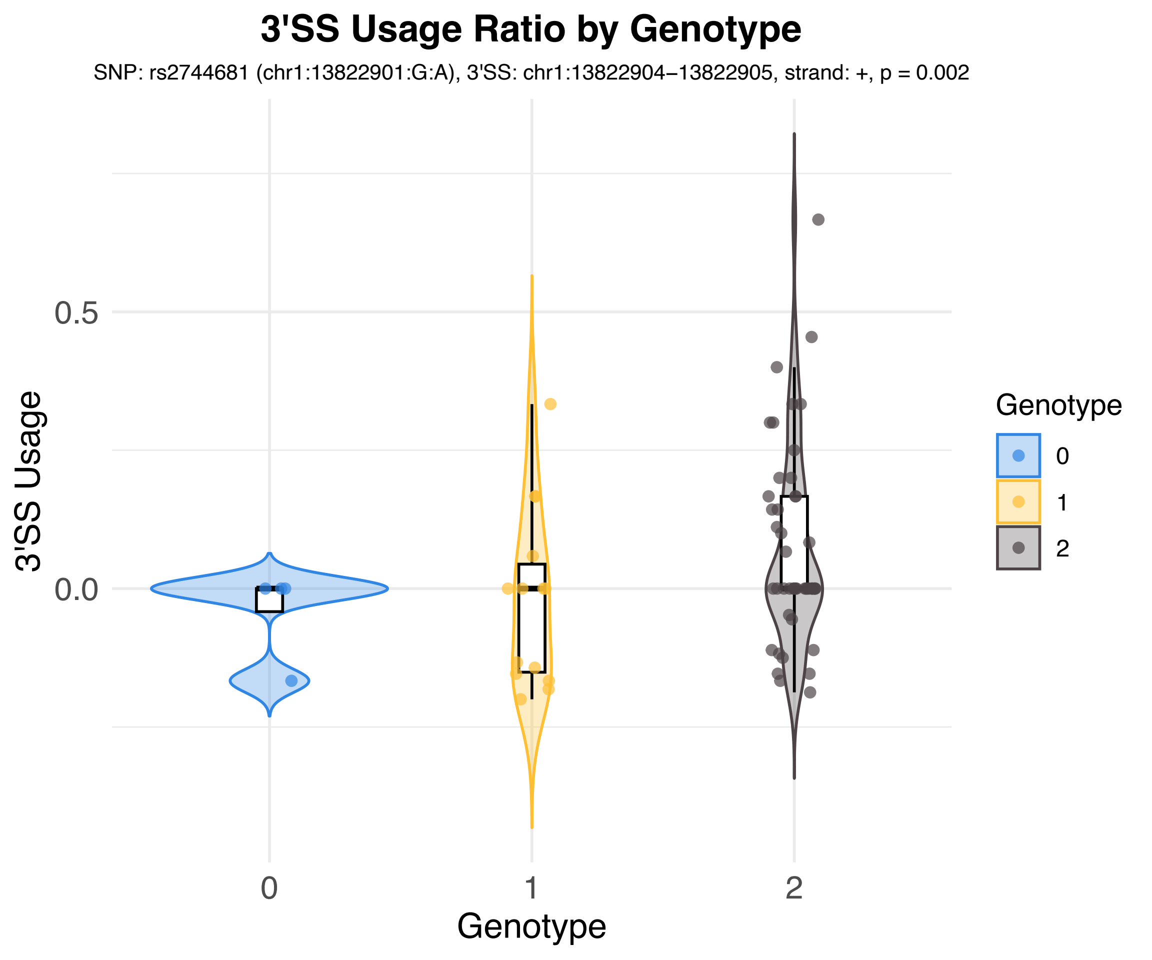
3'SS Usage Ratio by Genotype
SNP: rs2744681 (chr1:13822901:G:A), 3'SS: chr1:13822904−13822905, strand: +, p = 0.002
3'SS Usage
0.5
0.0
0
1
2
Genotype
Genotype
0
1
2

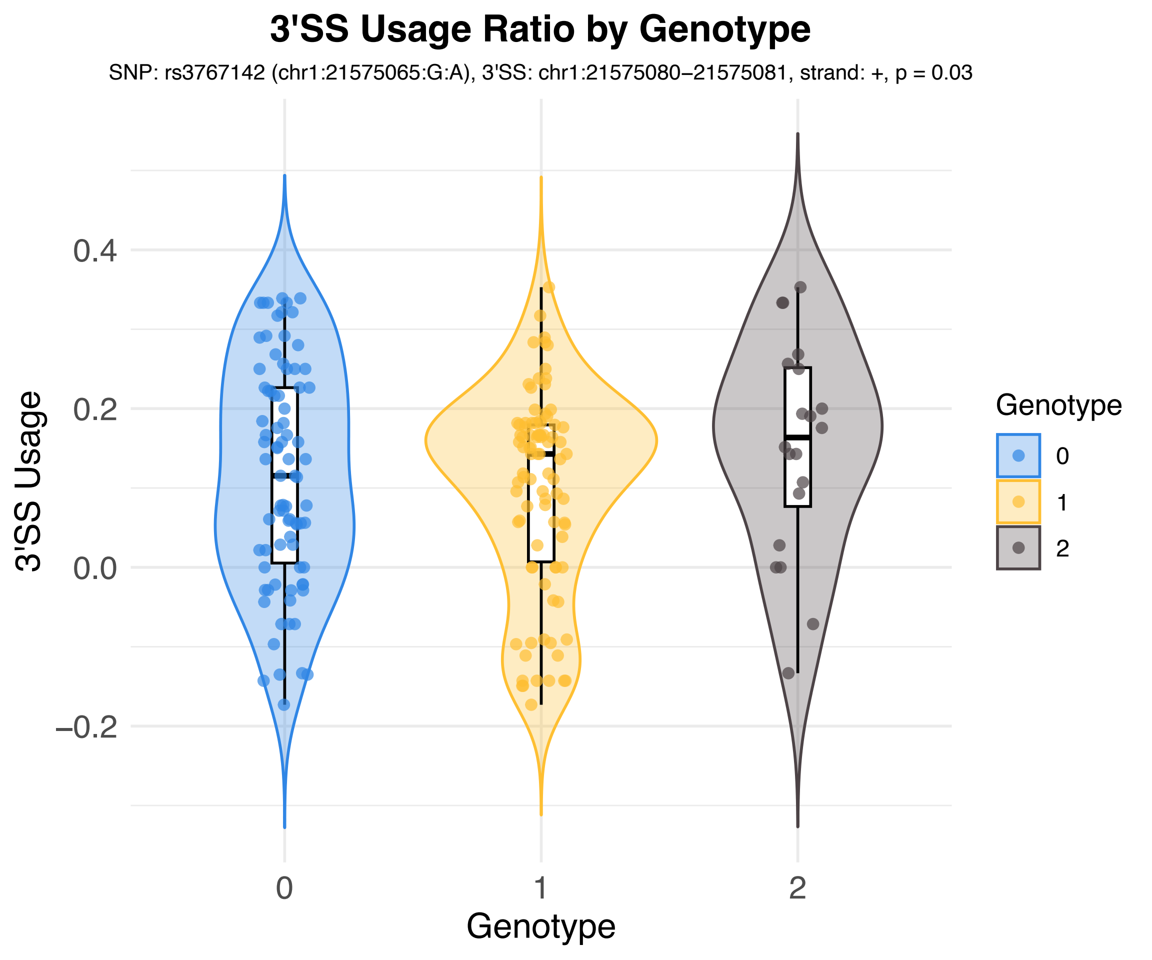
3'SS Usage Ratio by Genotype
SNP: rs3767142 (chr1:21575065:G:A), 3'SS: chr1:21575080−21575081, strand: +, p = 0.03
3'SS Usage
0.4
0.2
0.0
−0.2
0
1
2
Genotype
Genotype
0
1
2

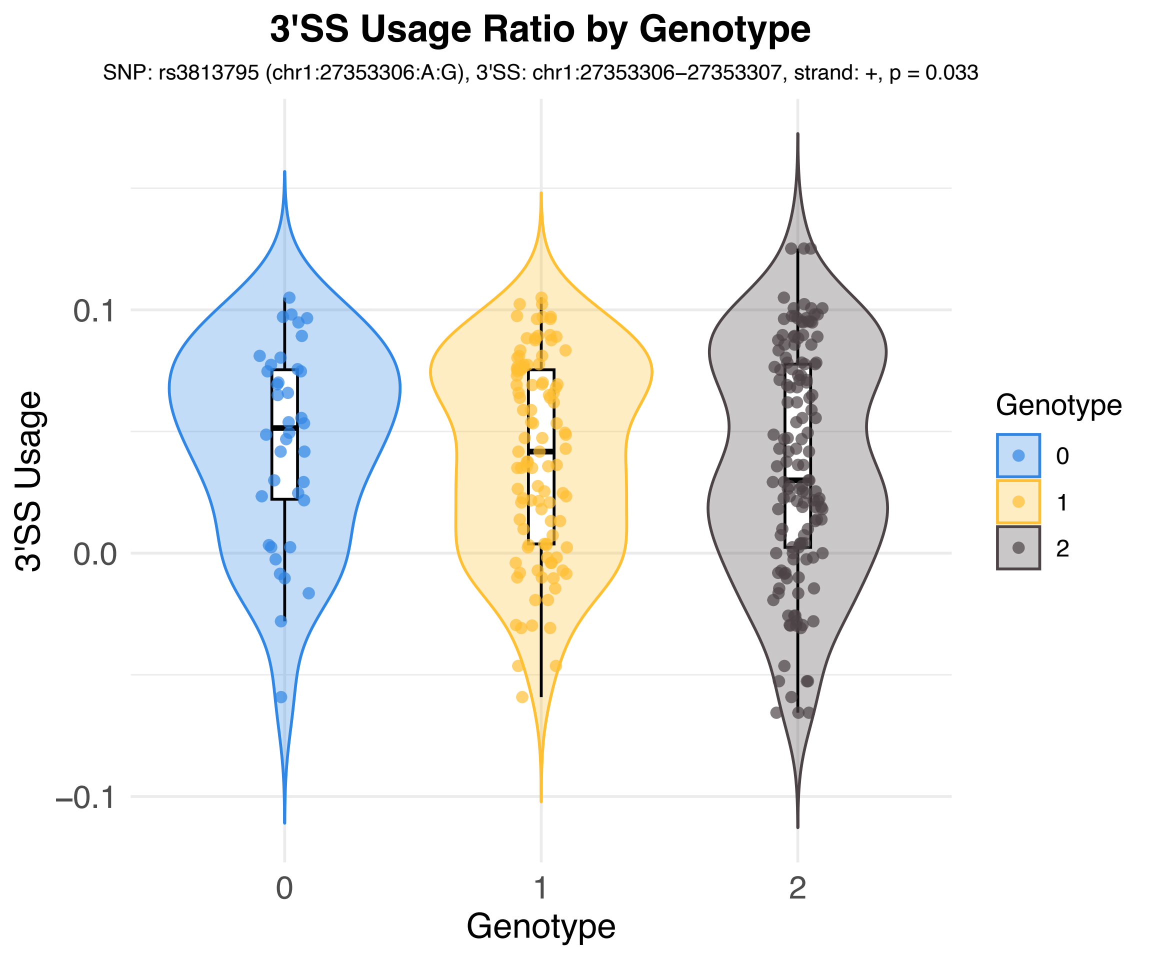
3'SS Usage Ratio by Genotype
SNP: rs3813795 (chr1:27353306:A:G), 3'SS: chr1:27353306–27353307, strand: +, p = 0.033
3'SS Usage
0.1
0.0
–0.1
0
1
2
Genotype
Genotype
0
1
2

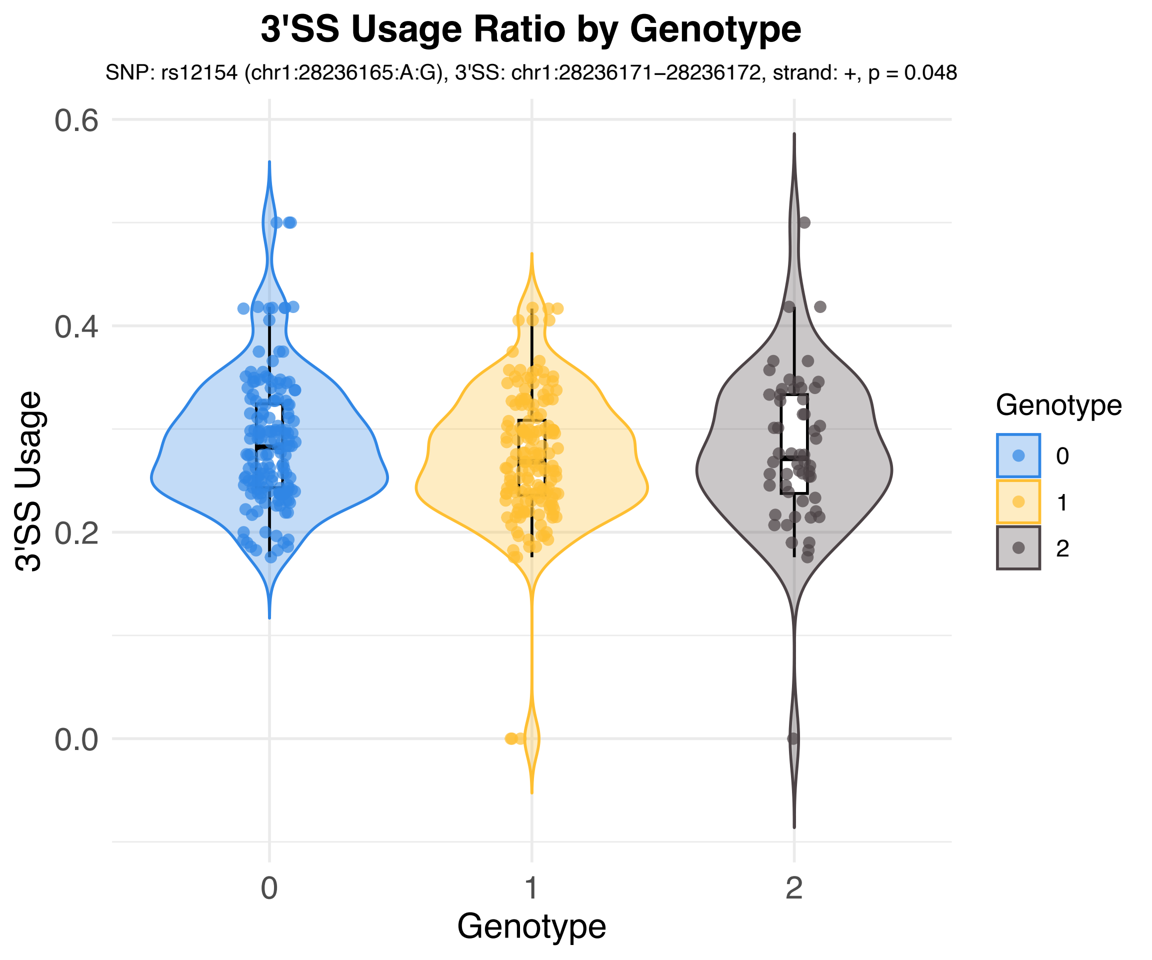
3'SS Usage Ratio by Genotype
SNP: rs12154 (chr1:28236165:A:G), 3'SS: chr1:28236171–28236172, strand: +, p = 0.048
3'SS Usage
0.6
0.4
0.2
0.0
0
1
2
Genotype
Genotype
0
1
2

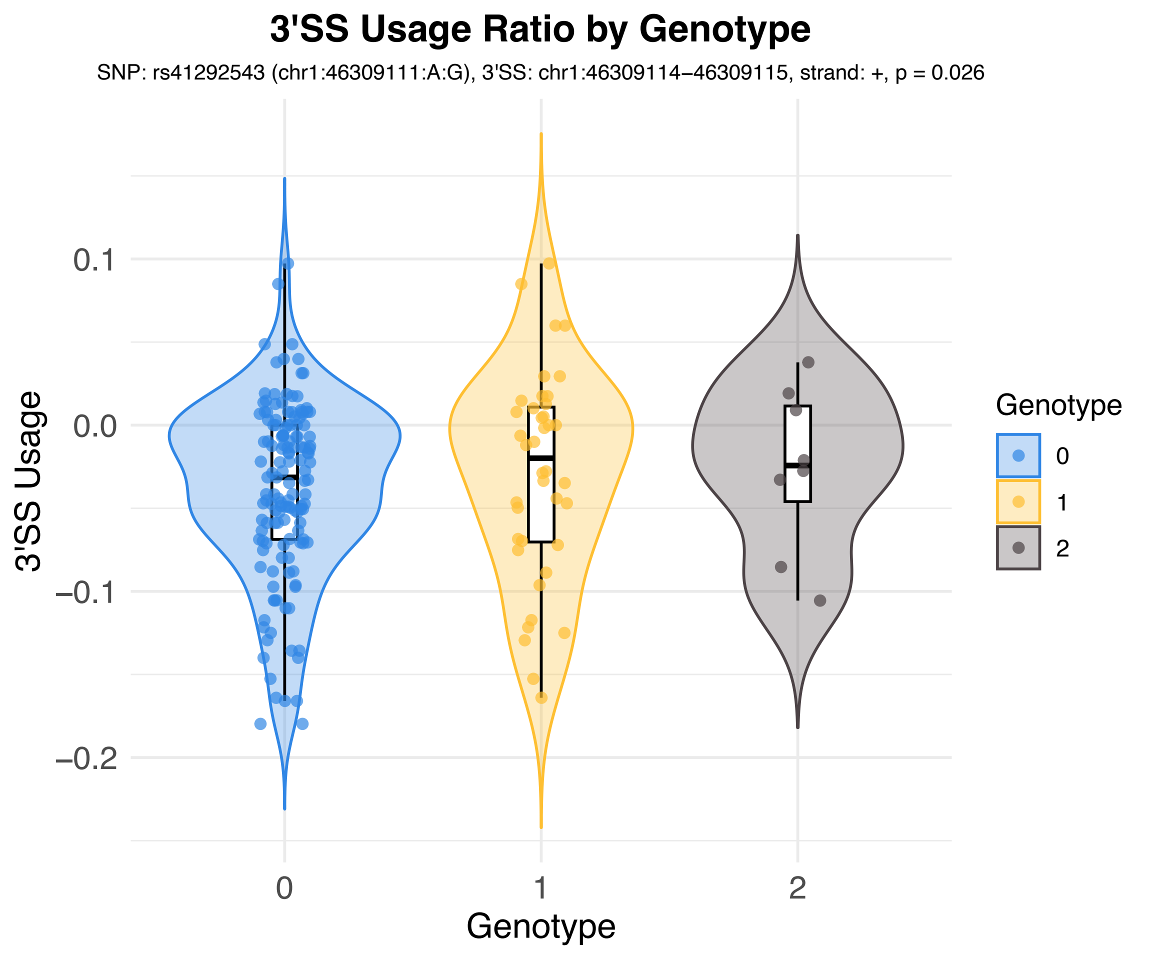
3'SS Usage Ratio by Genotype
SNP: rs41292543 (chr1:46309111:A:G), 3'SS: chr1:46309114–46309115, strand: +, p = 0.026
3'SS Usage
0.1
0.0
−0.1
−0.2
0
1
2
Genotype
Genotype
0
1
2

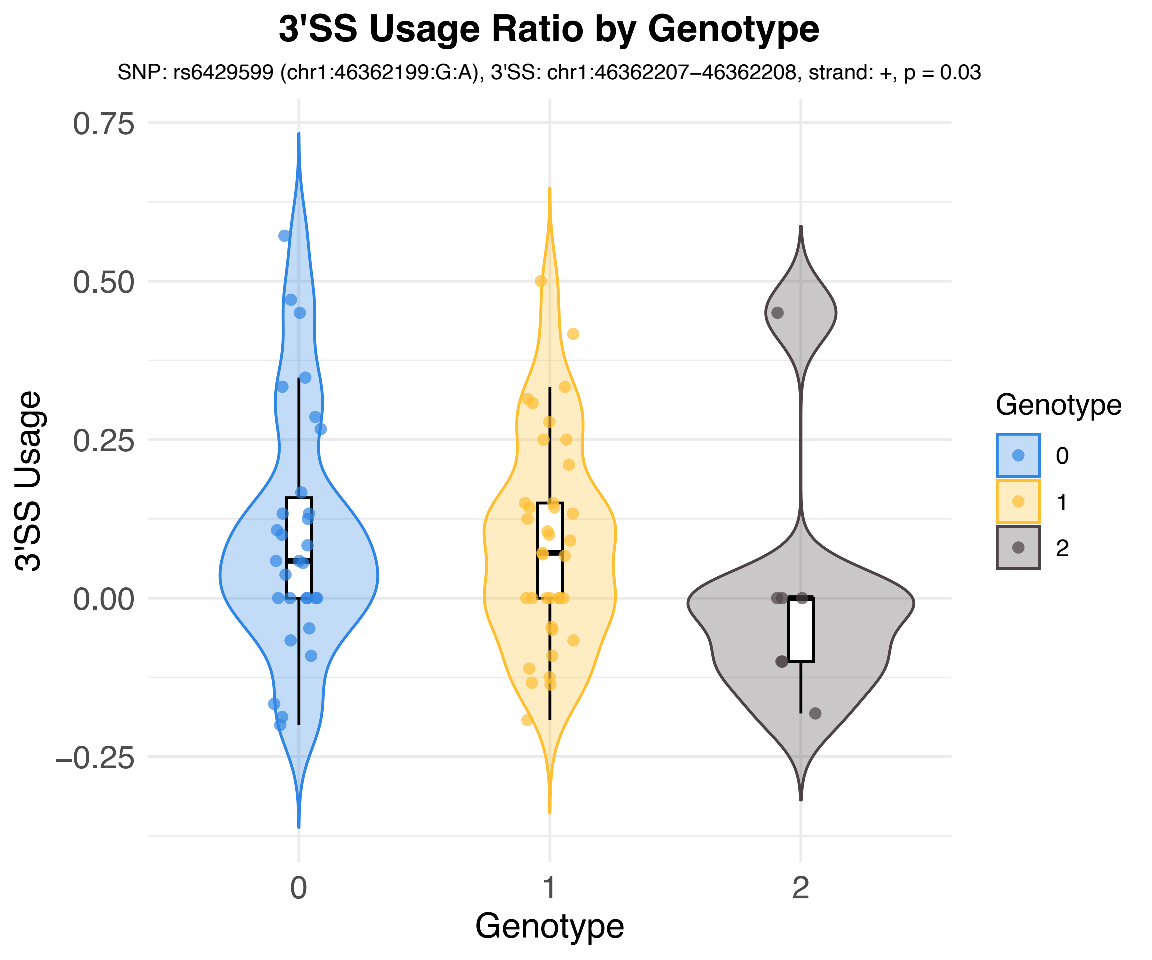
3'SS Usage Ratio by Genotype
SNP: rs6429599 (chr1:46362199:G:A), 3'SS: chr1:46362207−46362208, strand: +, p = 0.03
3'SS Usage
0.75
0.50
0.25
0.00
−0.25
0
1
2
Genotype
Genotype
0
1
2

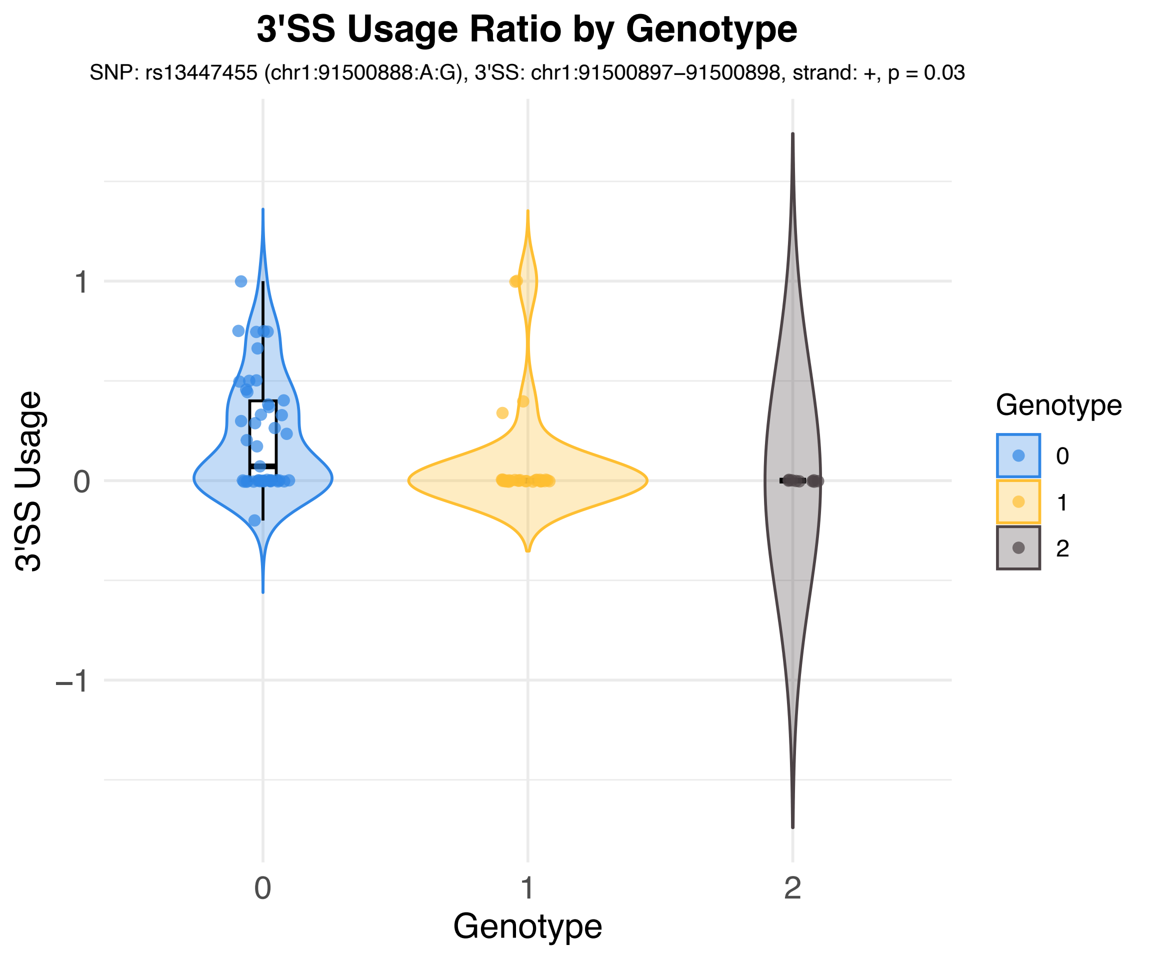
3'SS Usage Ratio by Genotype
SNP: rs13447455 (chr1:91500888:A:G), 3'SS: chr1:91500897–91500898, strand: +, p = 0.03
3'SS Usage
1
0
−1
0
1
2
Genotype
Genotype
0
1
2

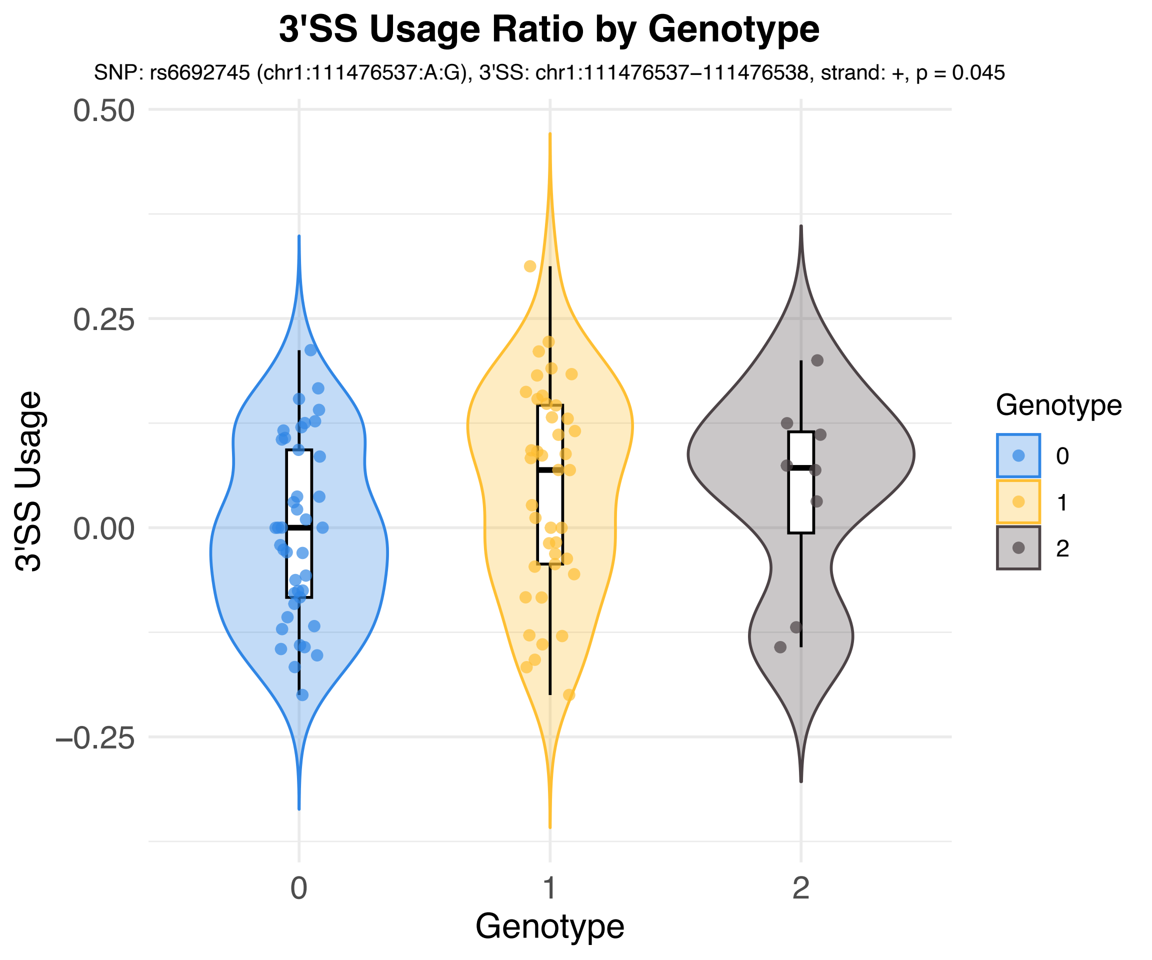
3'SS Usage Ratio by Genotype
SNP: rs6692745 (chr1:111476537:A:G), 3'SS: chr1:111476537−111476538, strand: +, p = 0.045
3'SS Usage
0.50
0.25
0.00
−0.25
0
1
2
Genotype
Genotype
0
1
2

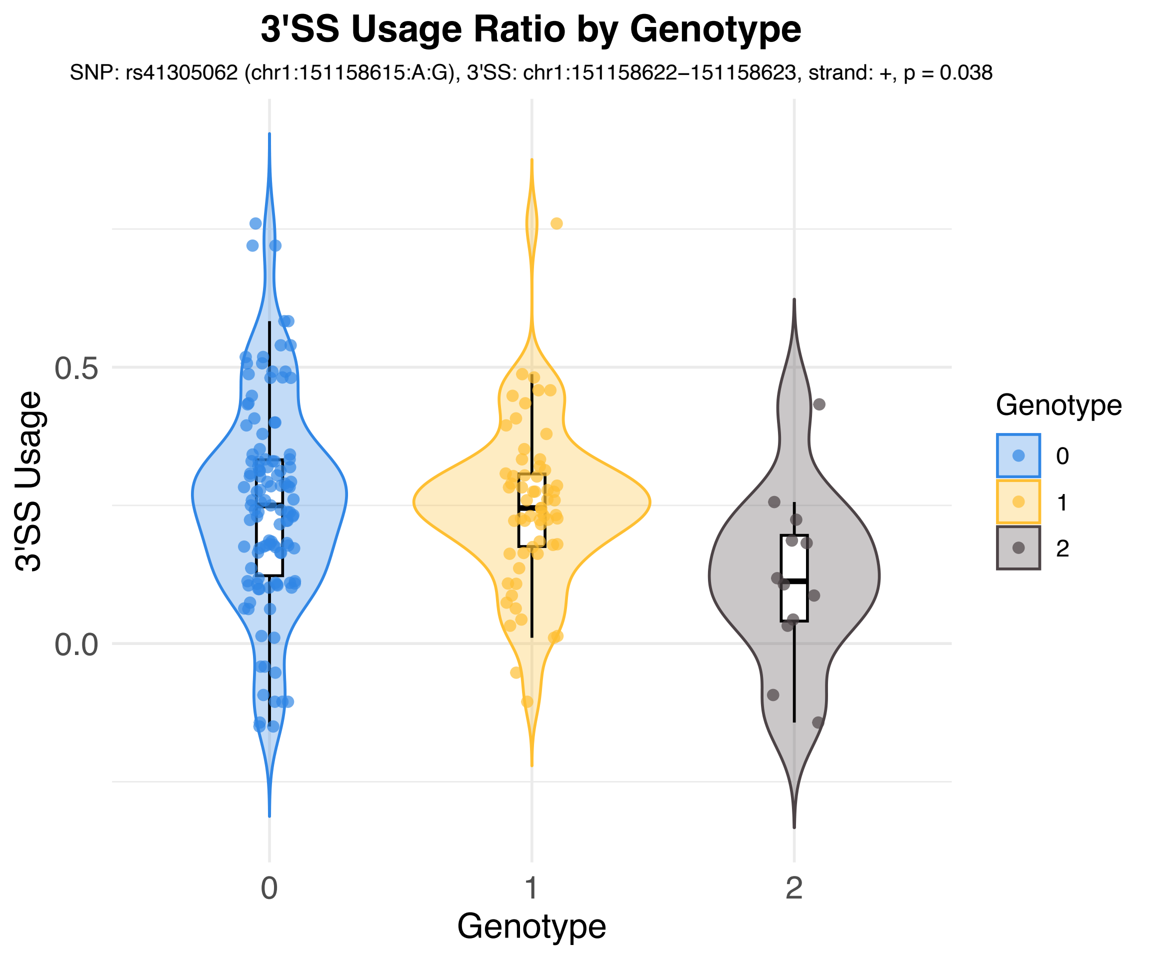
3'SS Usage Ratio by Genotype
SNP: rs41305062 (chr1:151158615:A:G), 3'SS: chr1:151158622–151158623, strand: +, p = 0.038
3'SS Usage
0.5
0.0
0
1
2
Genotype
Genotype
0
1
2

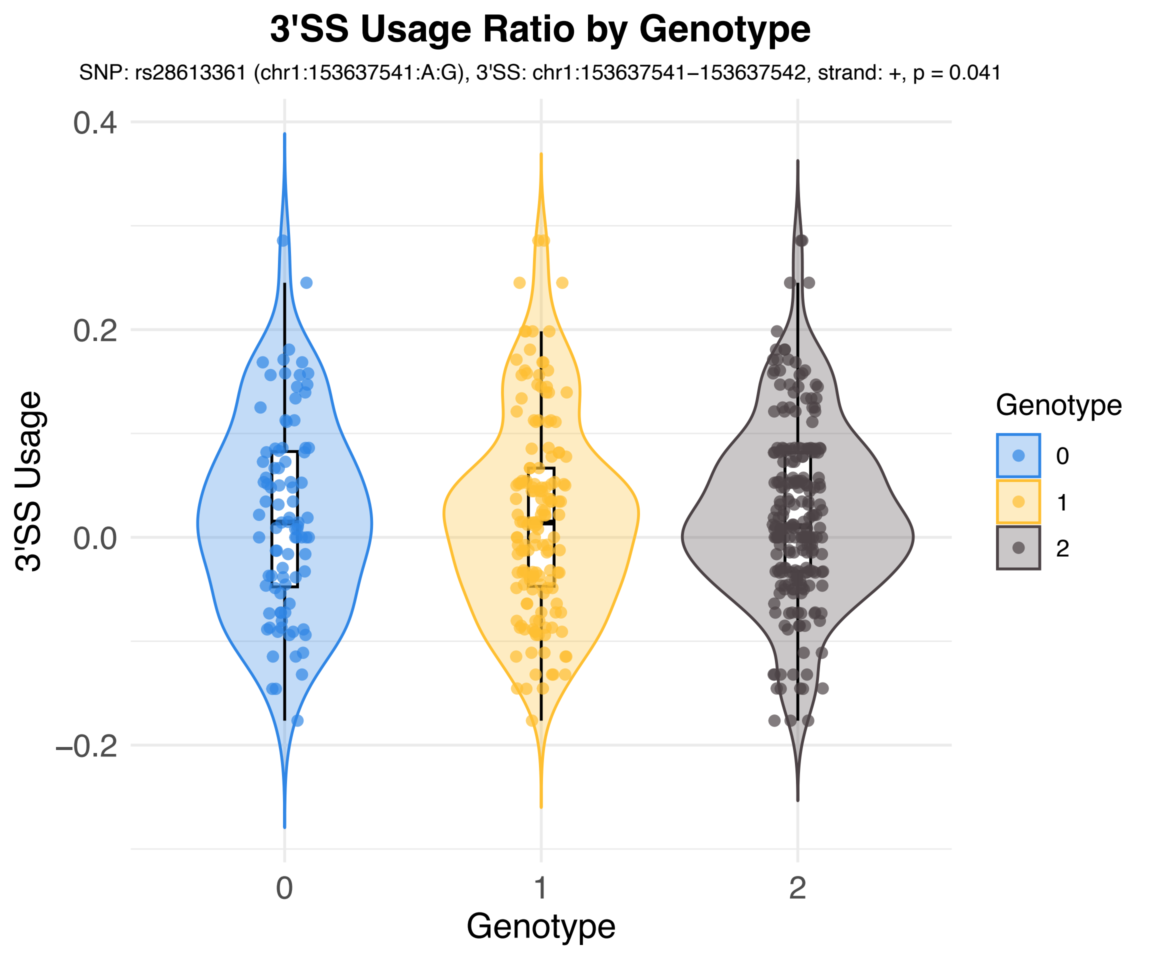
3'SS Usage Ratio by Genotype
SNP: rs28613361 (chr1:153637541:A:G), 3'SS: chr1:153637541−153637542, strand: +, p = 0.041
3'SS Usage
0.4
0.2
0.0
−0.2
0
1
2
Genotype
Genotype
0
1
2

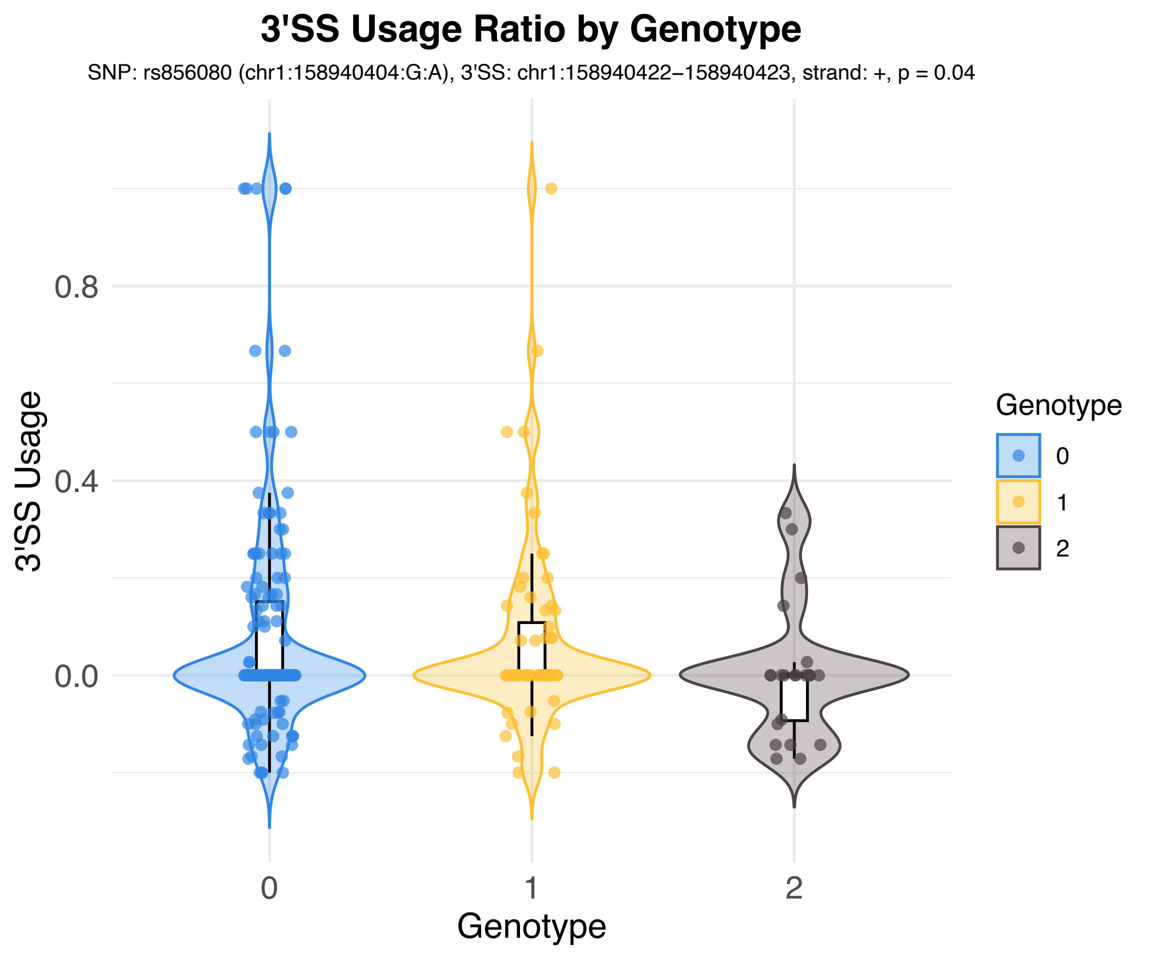
3'SS Usage Ratio by Genotype
SNP: rs856080 (chr1:158940404:G:A), 3'SS: chr1:158940422−158940423, strand: +, p = 0.04
3'SS Usage
0.8
0.4
0.0
0
1
2
Genotype
Genotype
0
1
2

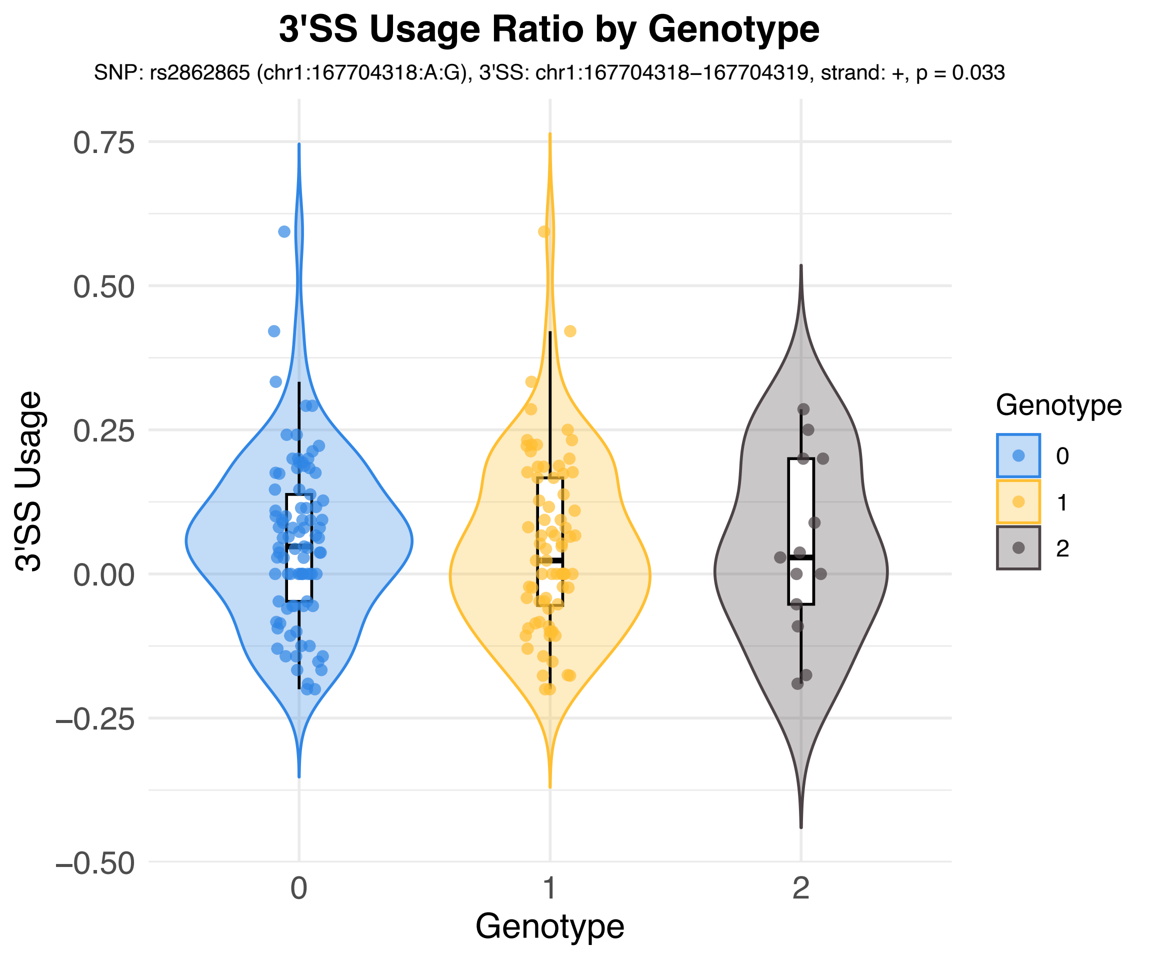
3'SS Usage Ratio by Genotype
SNP: rs2862865 (chr1:167704318:A:G), 3'SS: chr1:167704318−167704319, strand: +, p = 0.033
3'SS Usage
0.75
0.50
0.25
0.00
−0.25
−0.50
0
1
2
Genotype
Genotype
0
1
2

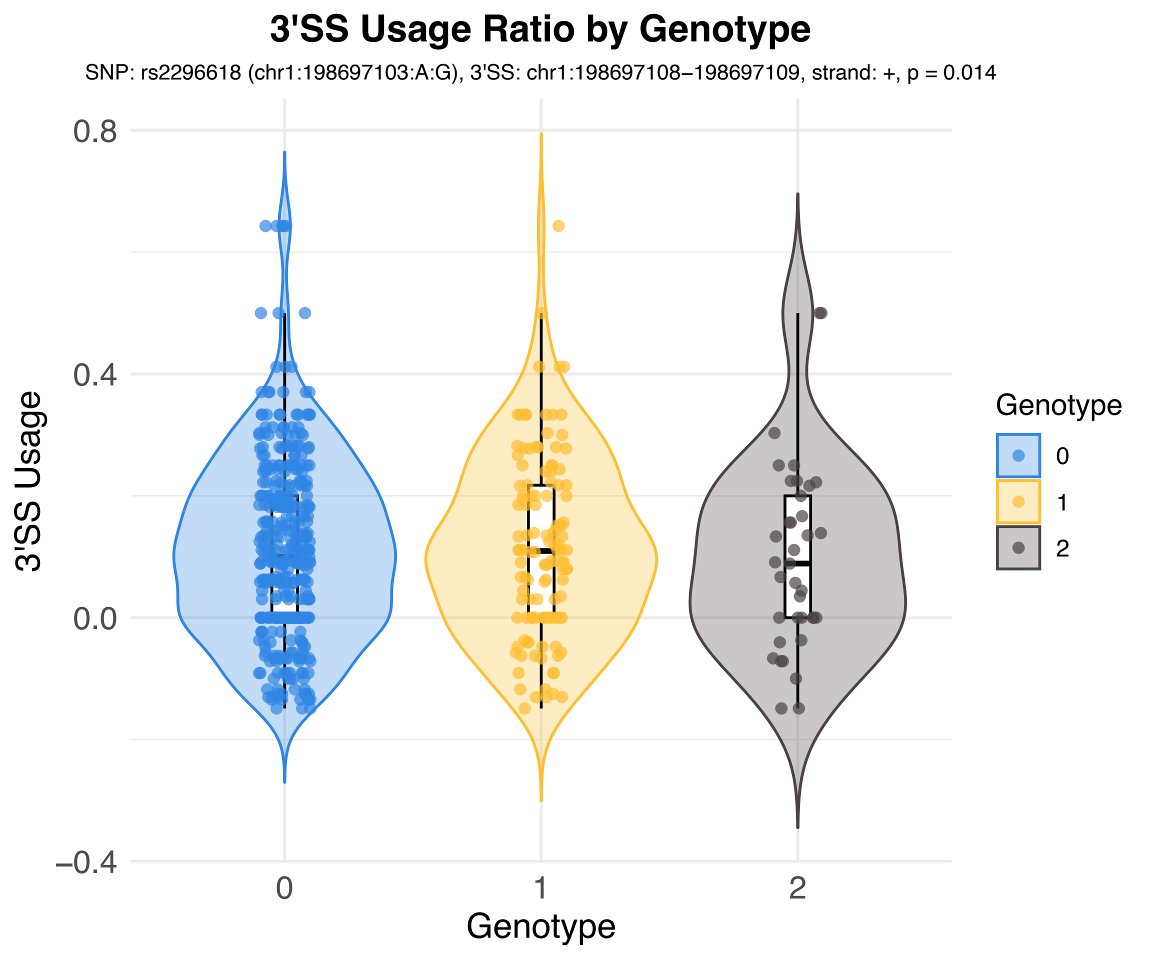

3'SS Usage Ratio by Genotype
SNP: rs2296618 (chr1:198697103:A:G), 3'SS: chr1:198697108−198697109, strand: +, p = 0.014
3'SS Usage
0.8
0.4
0.0
−0.4
0
1
2
Genotype
Genotype
0
1
2

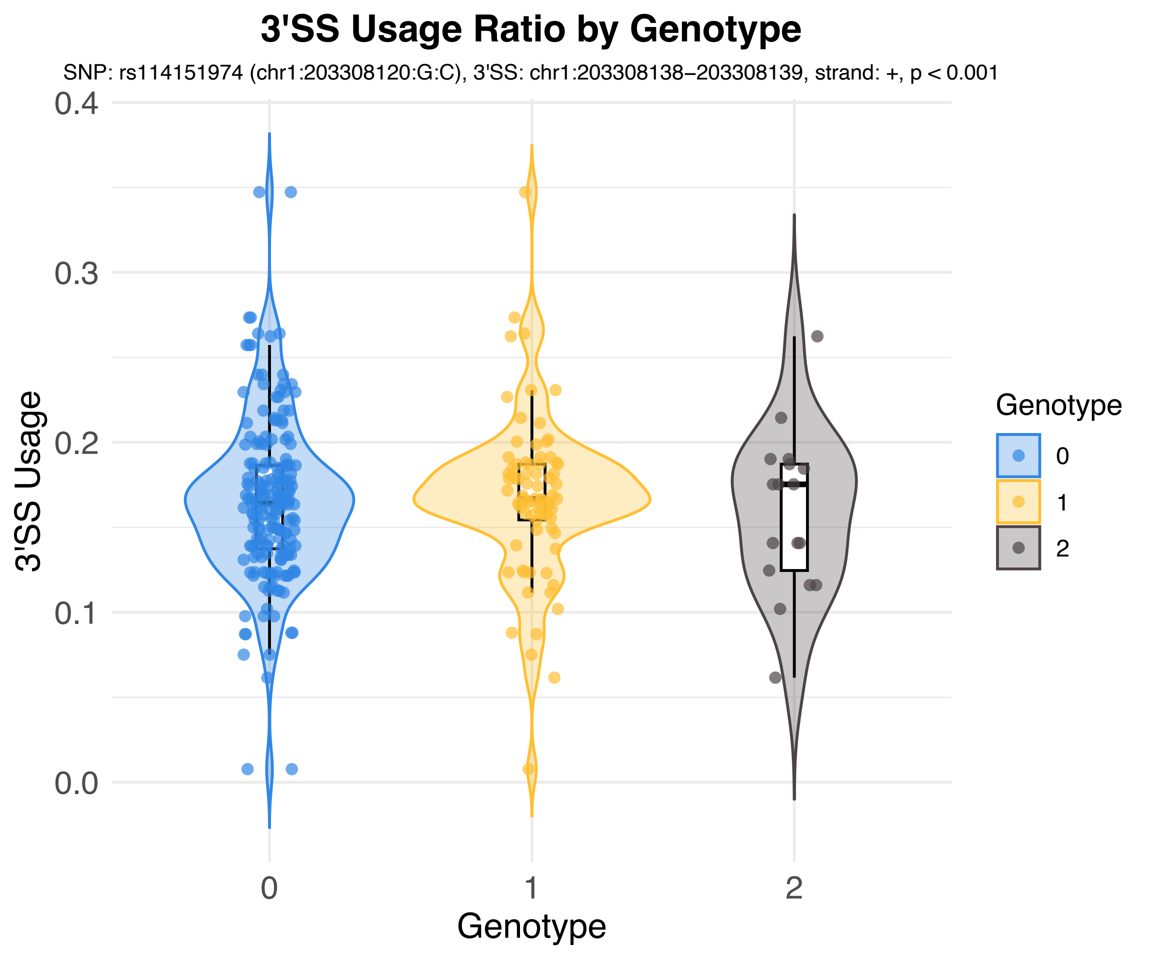
3'SS Usage Ratio by Genotype
SNP: rs114151974 (chr1:203308120:G:C), 3'SS: chr1:203308138–203308139, strand: +, p < 0.001
3'SS Usage
0.0
0.1
0.2
0.3
0.4
Genotype
0
1
2
Genotype
0
1
2

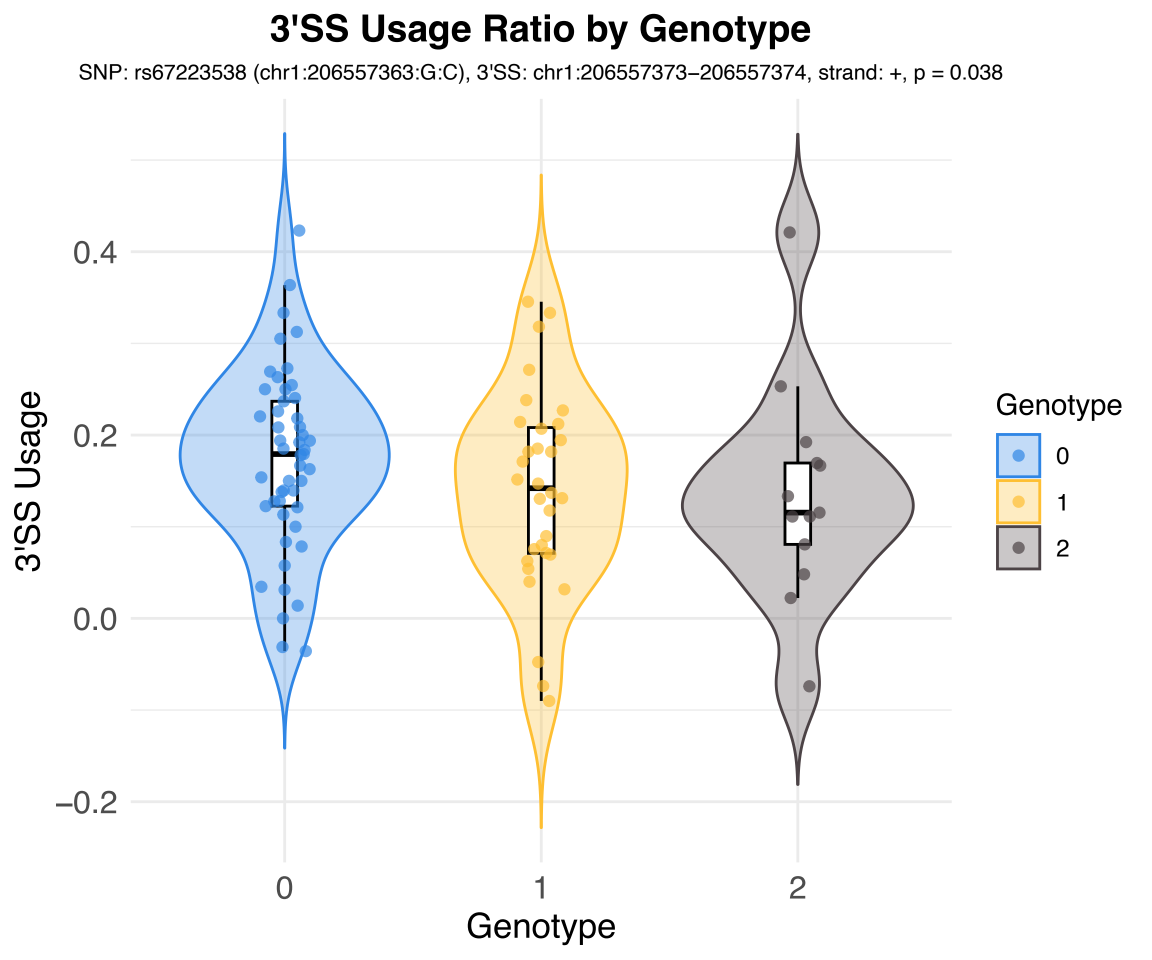

3'SS Usage Ratio by Genotype
SNP: rs67223538 (chr1:206557363:G:C), 3'SS: chr1:206557373–206557374, strand: +, p = 0.038
3'SS Usage
0.4
0.2
0.0
−0.2
0
1
2
Genotype
Genotype
0
1
2

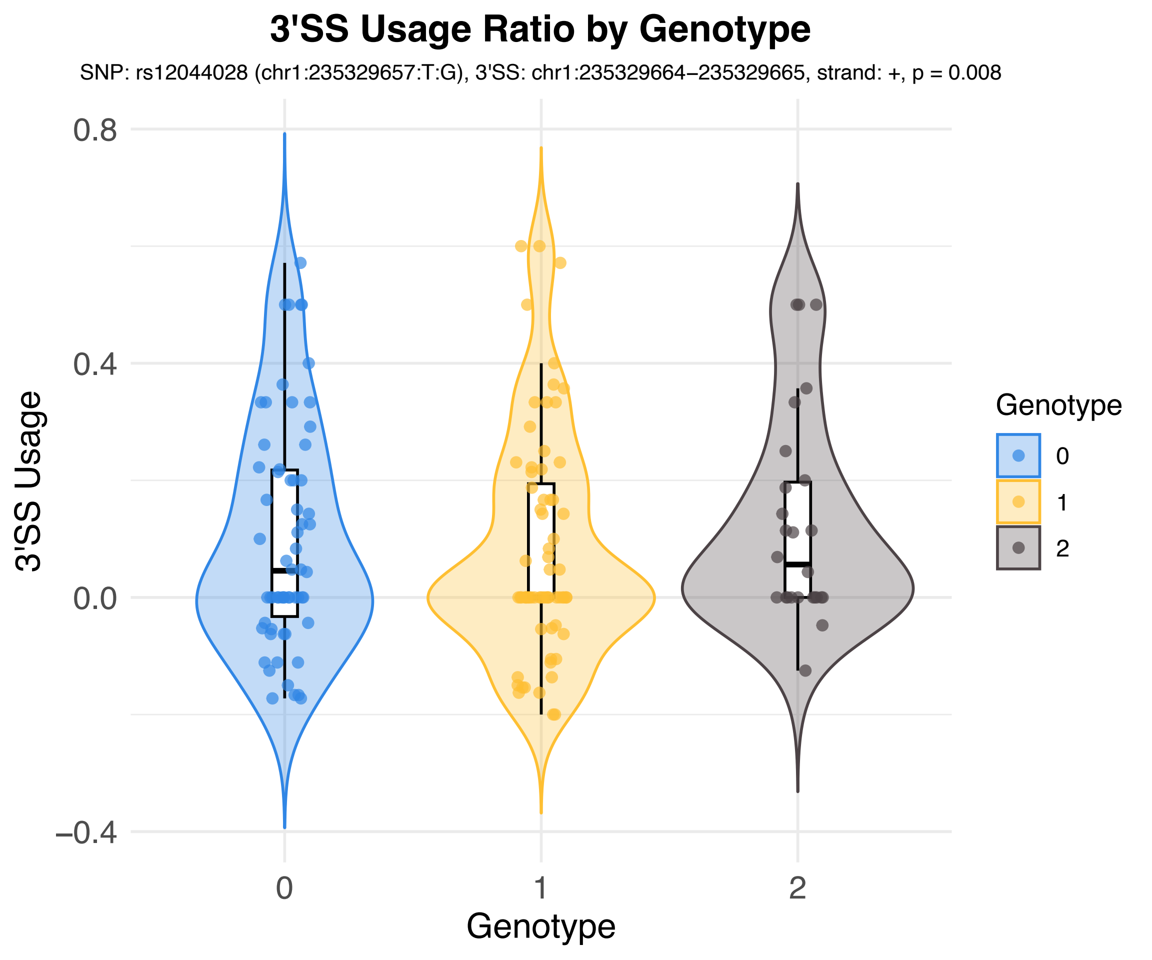
3'SS Usage Ratio by Genotype
SNP: rs12044028 (chr1:235329657:T:G), 3'SS: chr1:235329664–235329665, strand: +, p = 0.008
3'SS Usage
0.8
0.4
0.0
–0.4
0
1
2
Genotype
Genotype
0
1
2

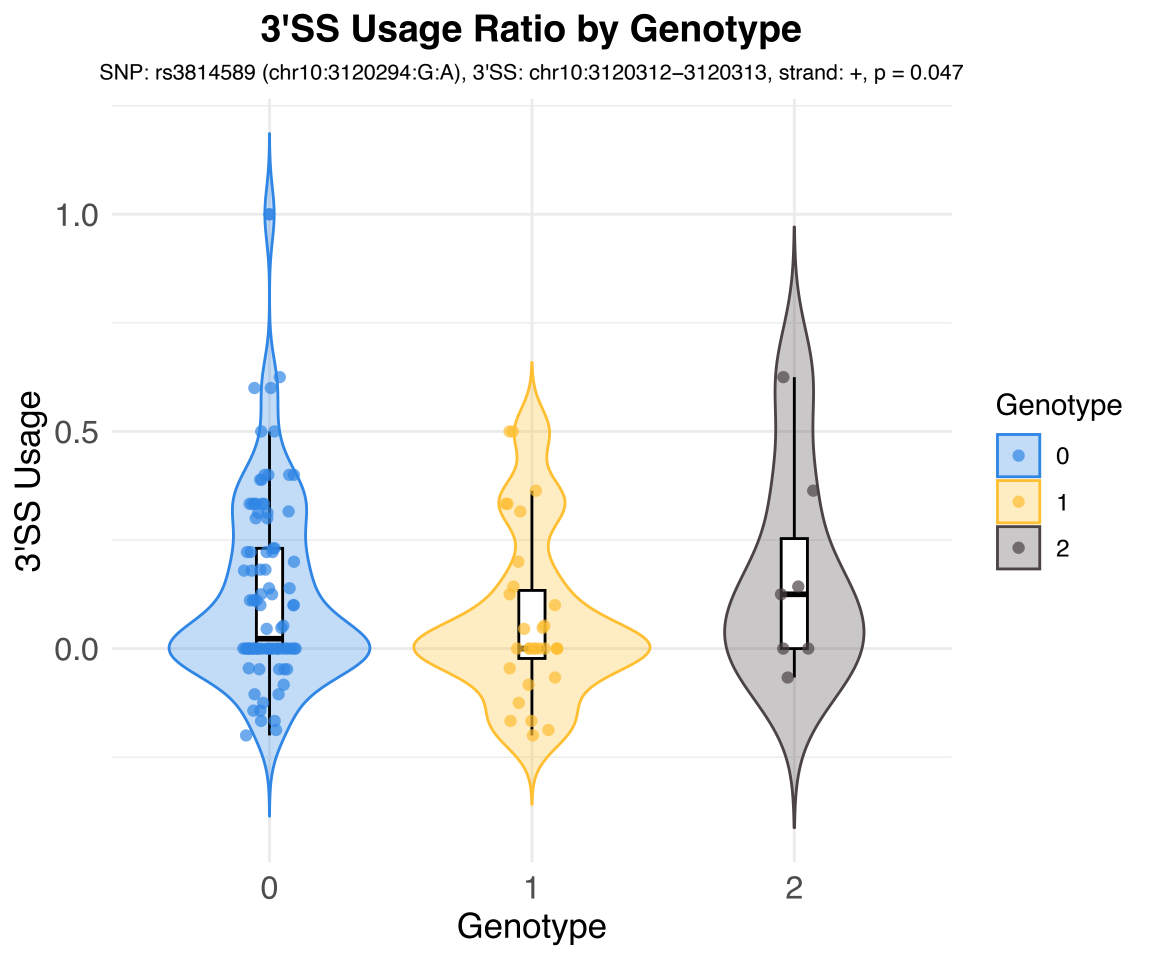
3'SS Usage Ratio by Genotype
SNP: rs3814589 (chr10:3120294:G:A), 3'SS: chr10:3120312–3120313, strand: +, p = 0.047
3'SS Usage
1.0
0.5
0.0
0
1
2
Genotype
Genotype
0
1
2

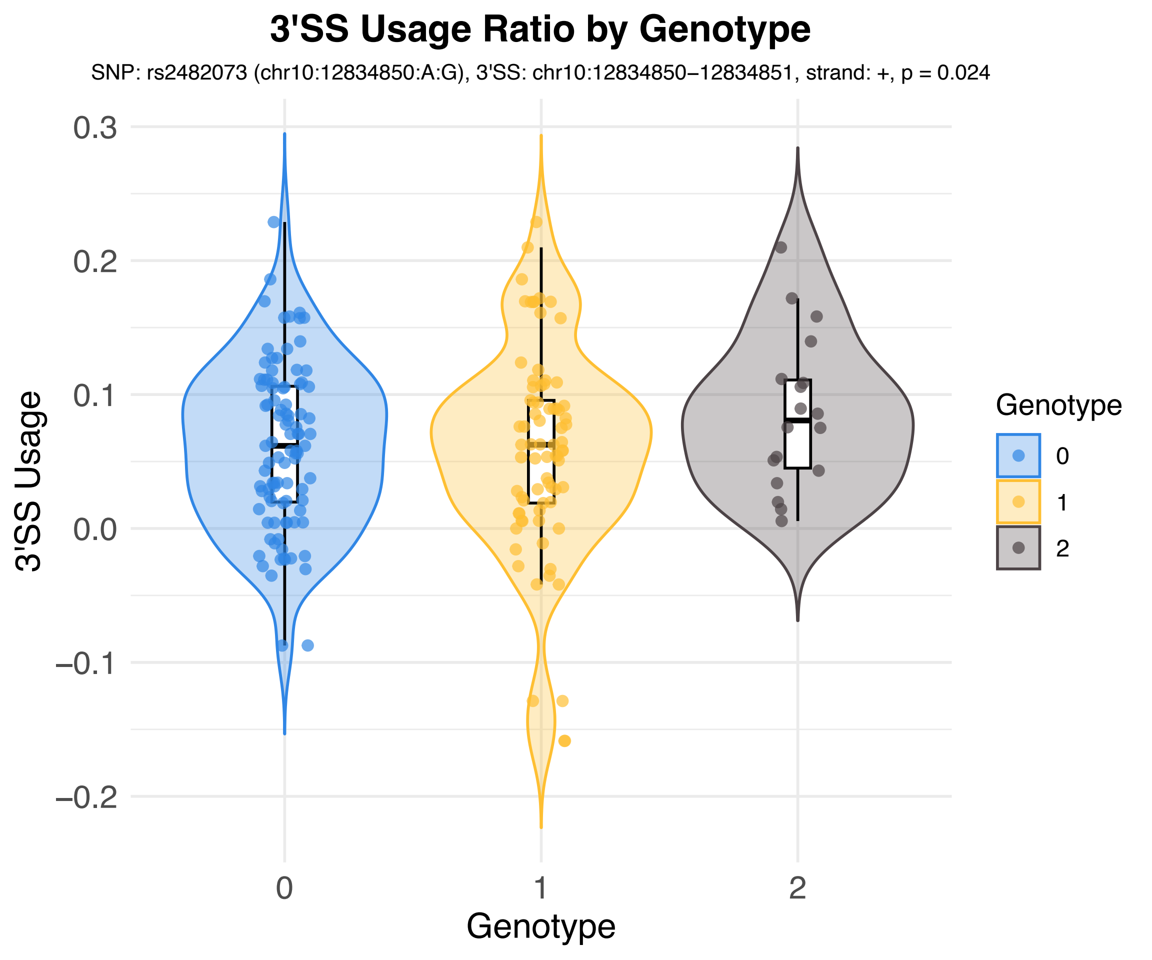

3'SS Usage Ratio by Genotype
SNP: rs2482073 (chr10:12834850:A:G), 3'SS: chr10:12834850−12834851, strand: +, p = 0.024
3'SS Usage
0.3
0.2
0.1
0.0
−0.1
−0.2
0
1
2
Genotype
Genotype
0
1
2

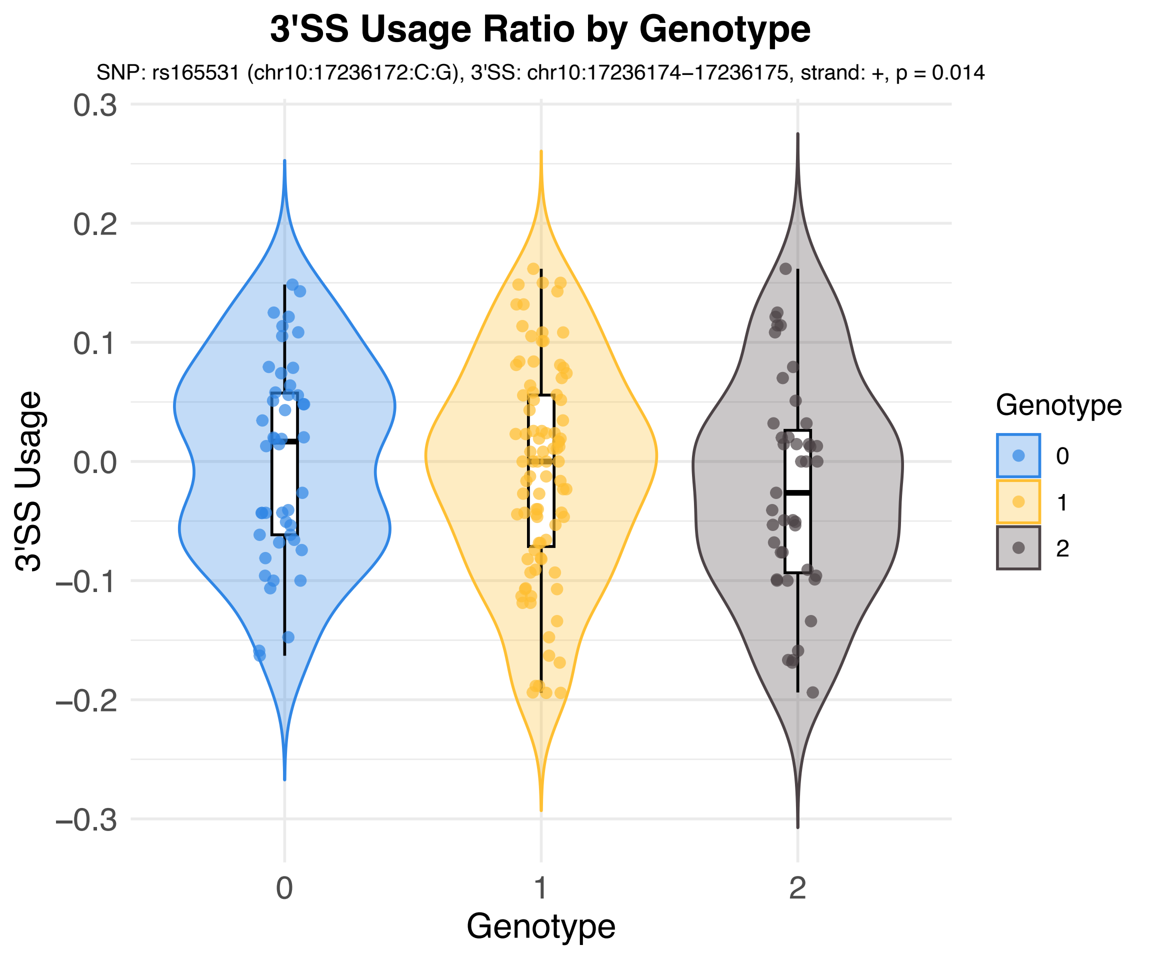
3'SS Usage Ratio by Genotype
SNP: rs165531 (chr10:17236172:C:G), 3'SS: chr10:17236174–17236175, strand: +, p = 0.014
3'SS Usage
0.3
0.2
0.1
0.0
−0.1
−0.2
−0.3
0
1
2
Genotype
Genotype
0
1
2

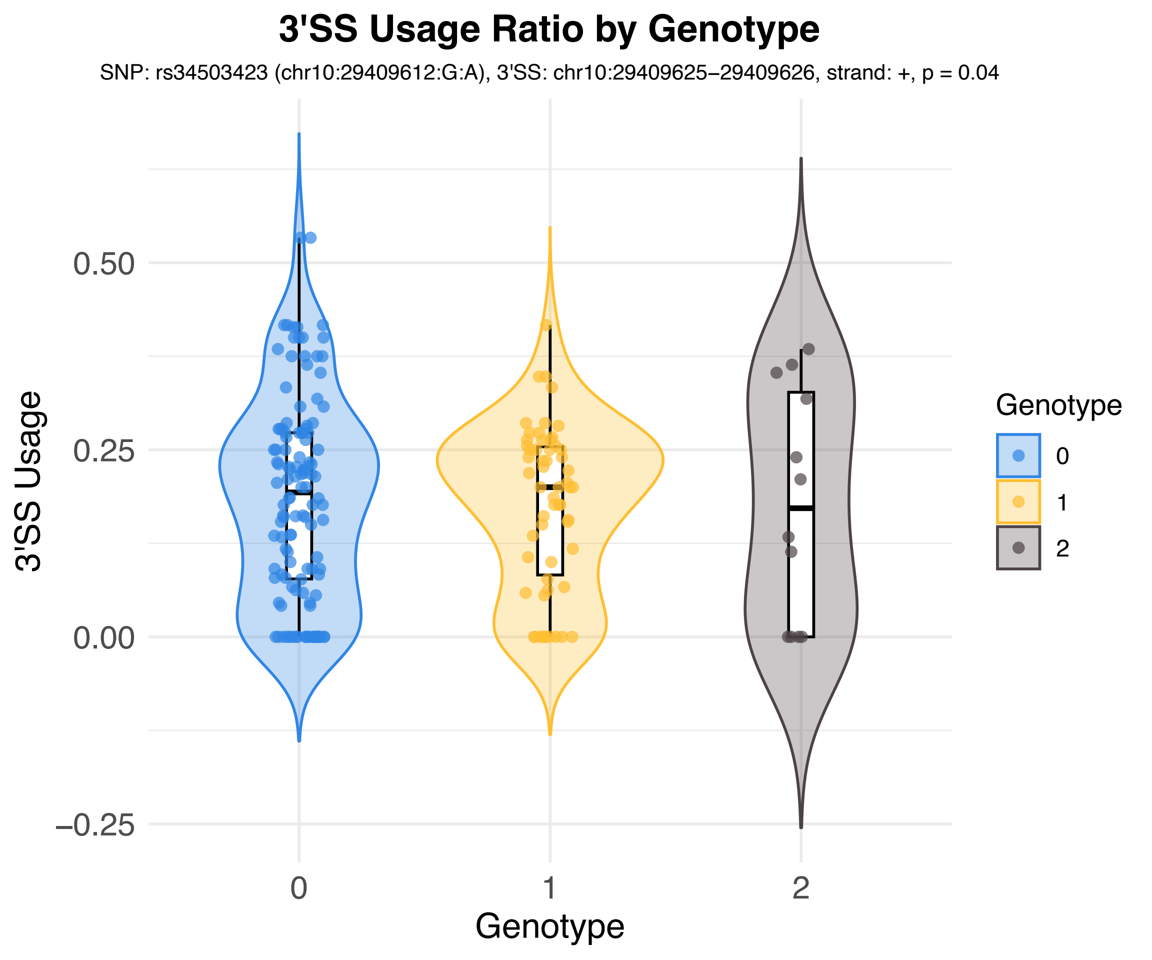
3'SS Usage Ratio by Genotype
SNP: rs34503423 (chr10:29409612:G:A), 3'SS: chr10:29409625–29409626, strand: +, p = 0.04
3'SS Usage
0.50
0.25
0.00
–0.25
0
1
2
Genotype
Genotype
0
1
2

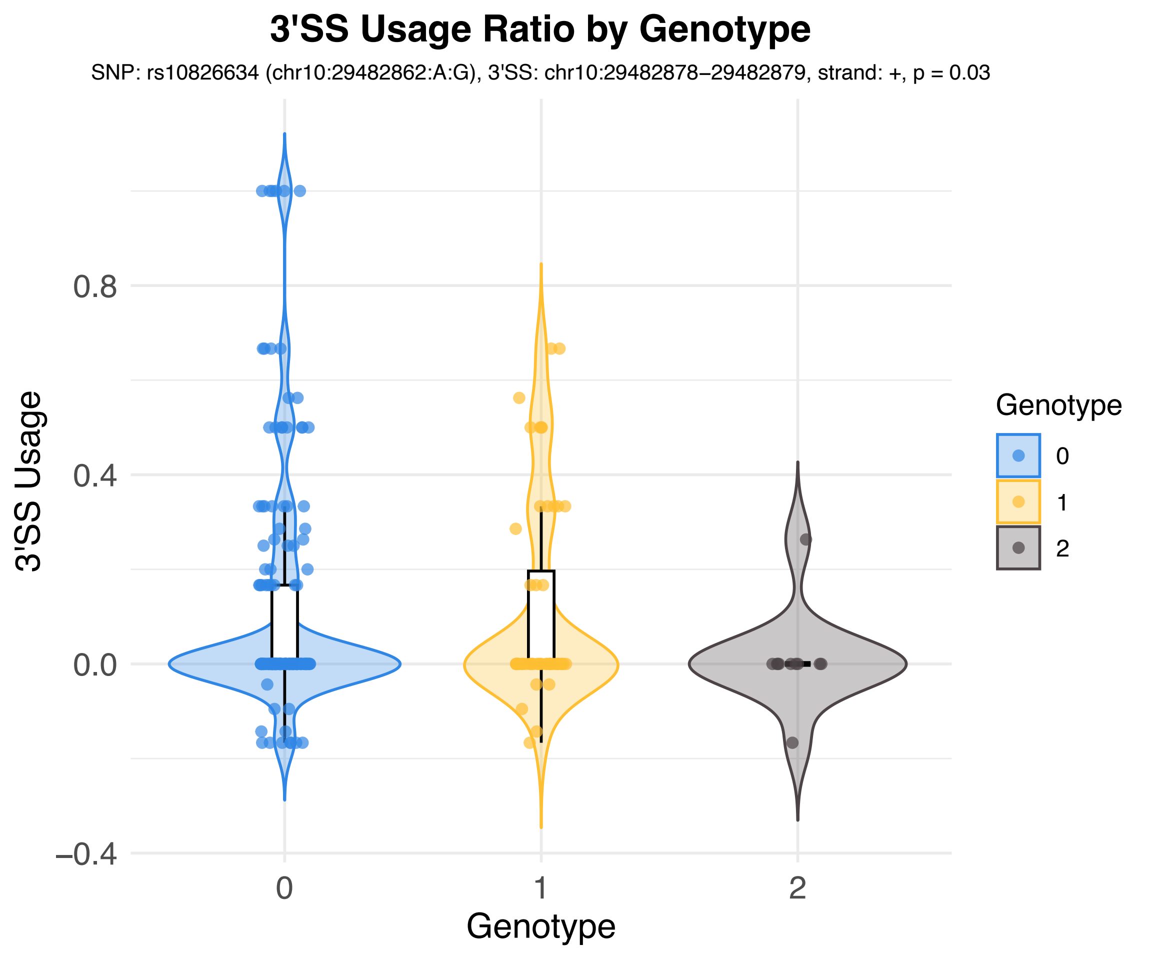
3'SS Usage Ratio by Genotype
SNP: rs10826634 (chr10:29482862:A:G), 3'SS: chr10:29482878–29482879, strand: +, p = 0.03
3'SS Usage
0.8
0.4
0.0
−0.4
0
1
2
Genotype
Genotype
0
1
2

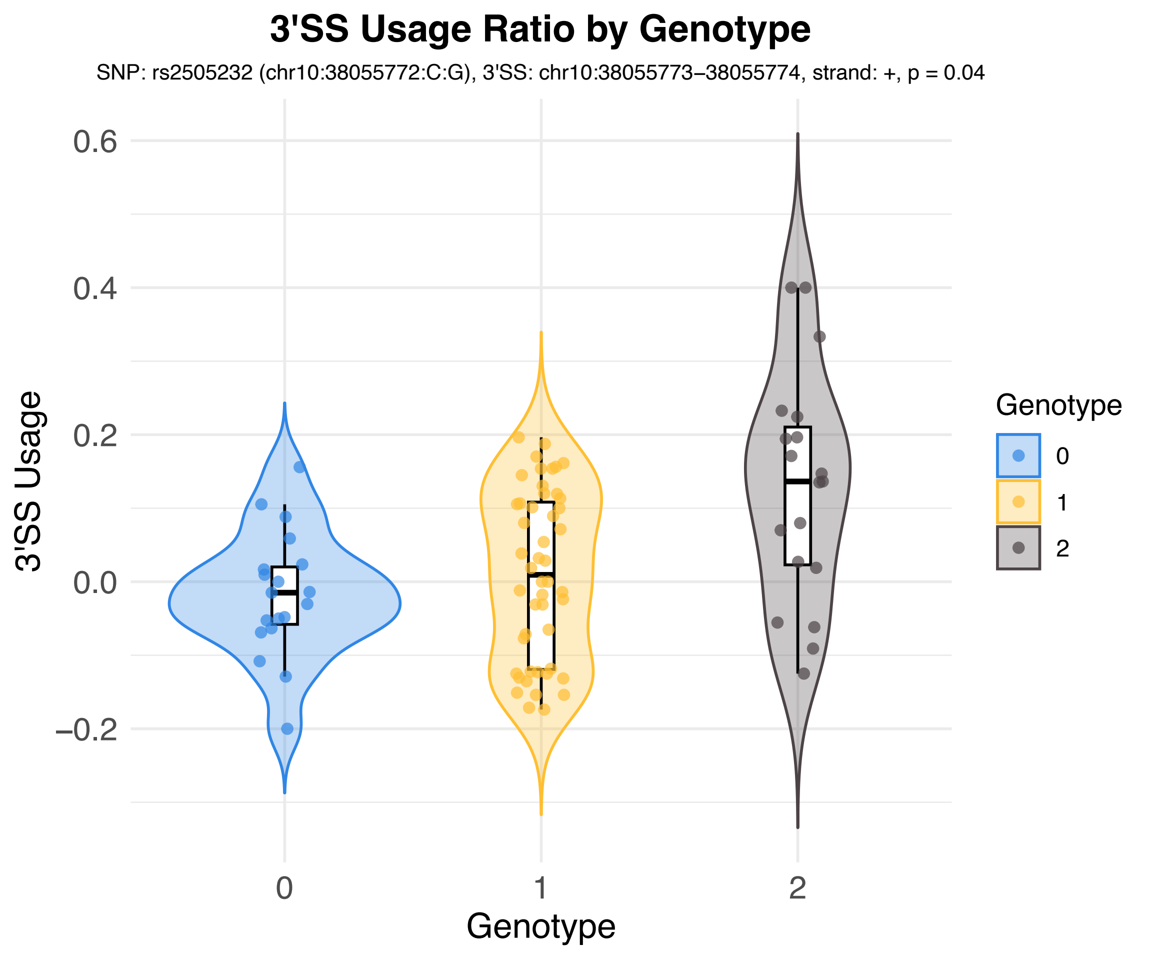
3'SS Usage Ratio by Genotype
SNP: rs2505232 (chr10:38055772:C:G), 3'SS: chr10:38055773–38055774, strand: +, p = 0.04
3'SS Usage
0.6
0.4
0.2
0.0
−0.2
0
1
2
Genotype
Genotype
0
1
2

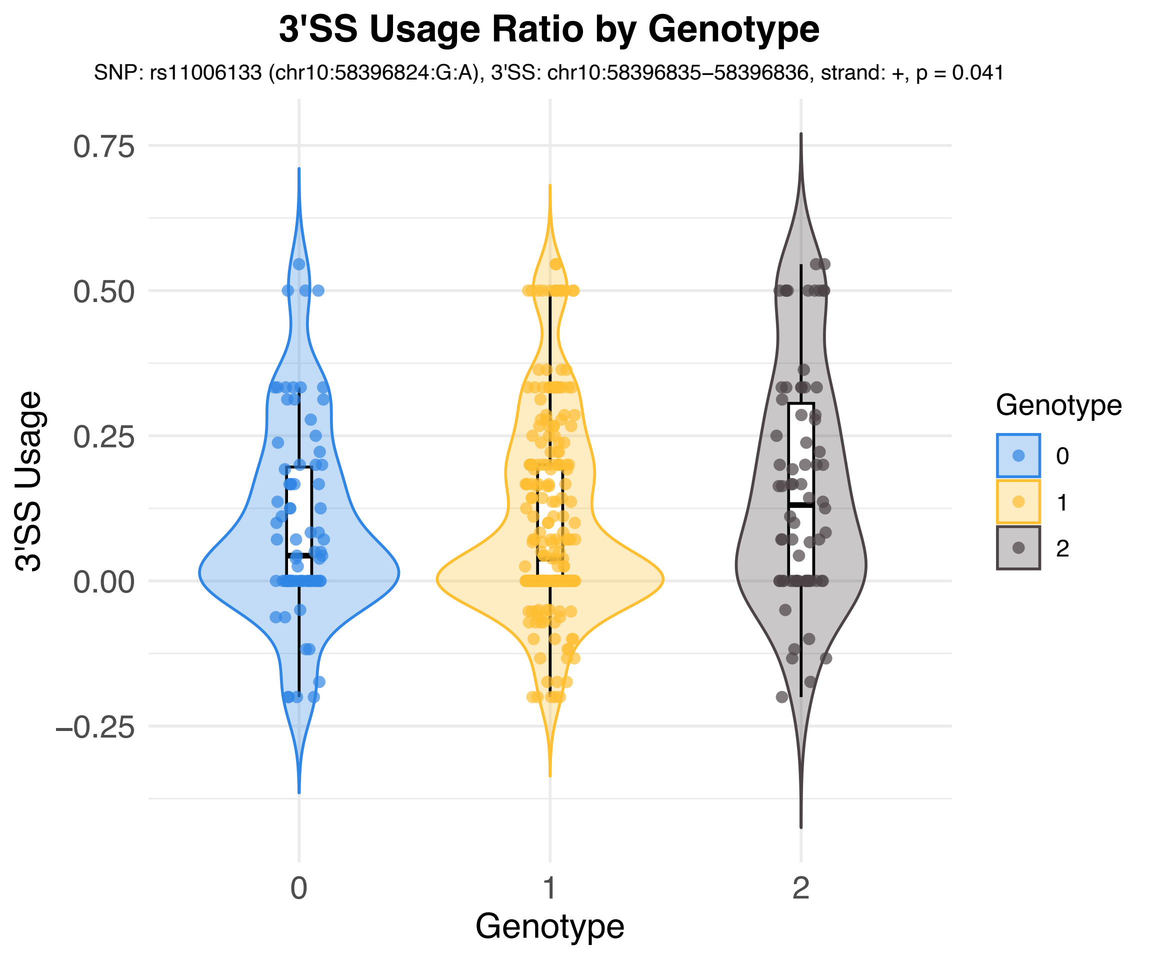
3'SS Usage Ratio by Genotype
SNP: rs11006133 (chr10:58396824:G:A), 3'SS: chr10:58396835–58396836, strand: +, p = 0.041
3'SS Usage
0.75
0.50
0.25
0.00
−0.25
0
1
2
Genotype
Genotype
0
1
2

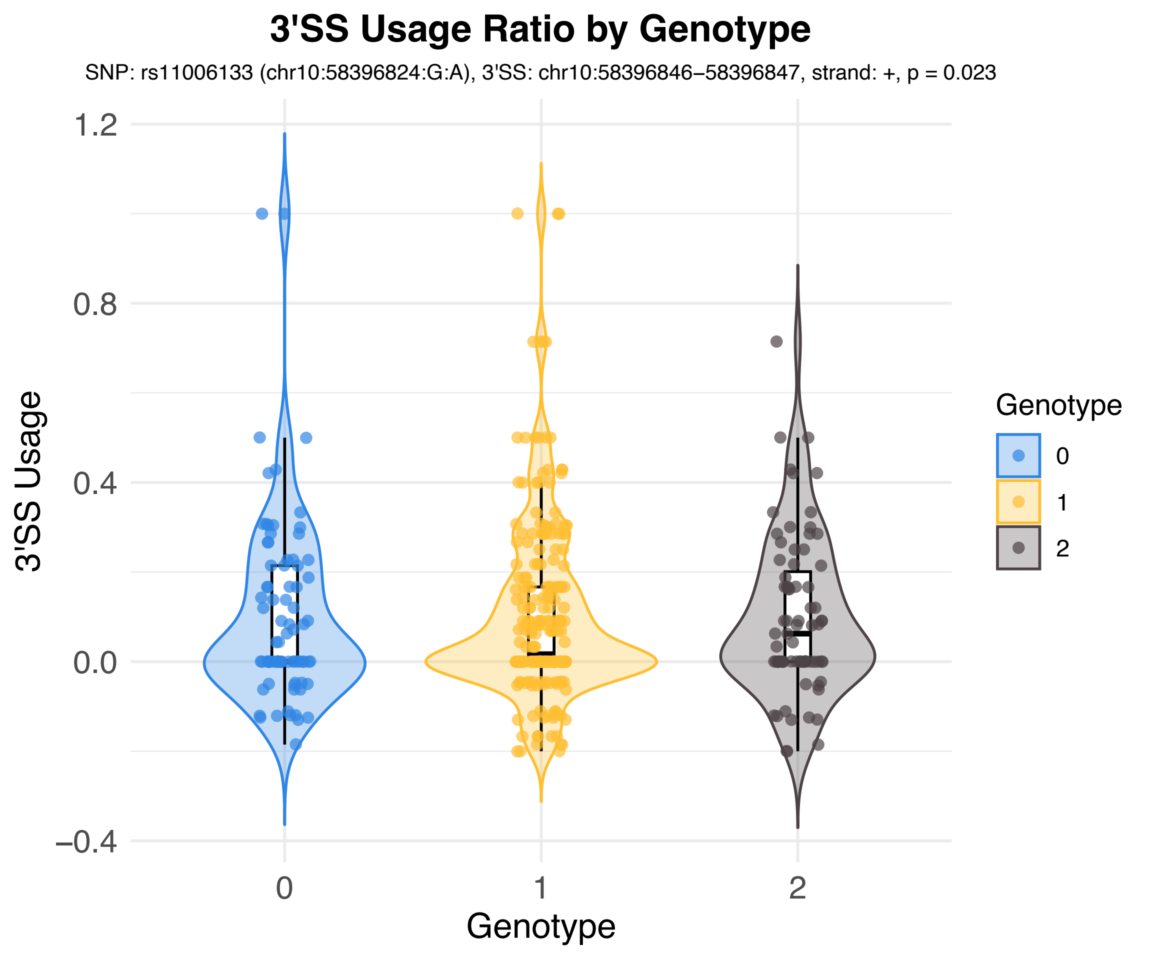
3'SS Usage Ratio by Genotype
SNP: rs11006133 (chr10:58396824:G:A), 3'SS: chr10:58396846–58396847, strand: +, p = 0.023
3'SS Usage
1.2
0.8
0.4
0.0
−0.4
0
1
2
Genotype
Genotype
0
1
2

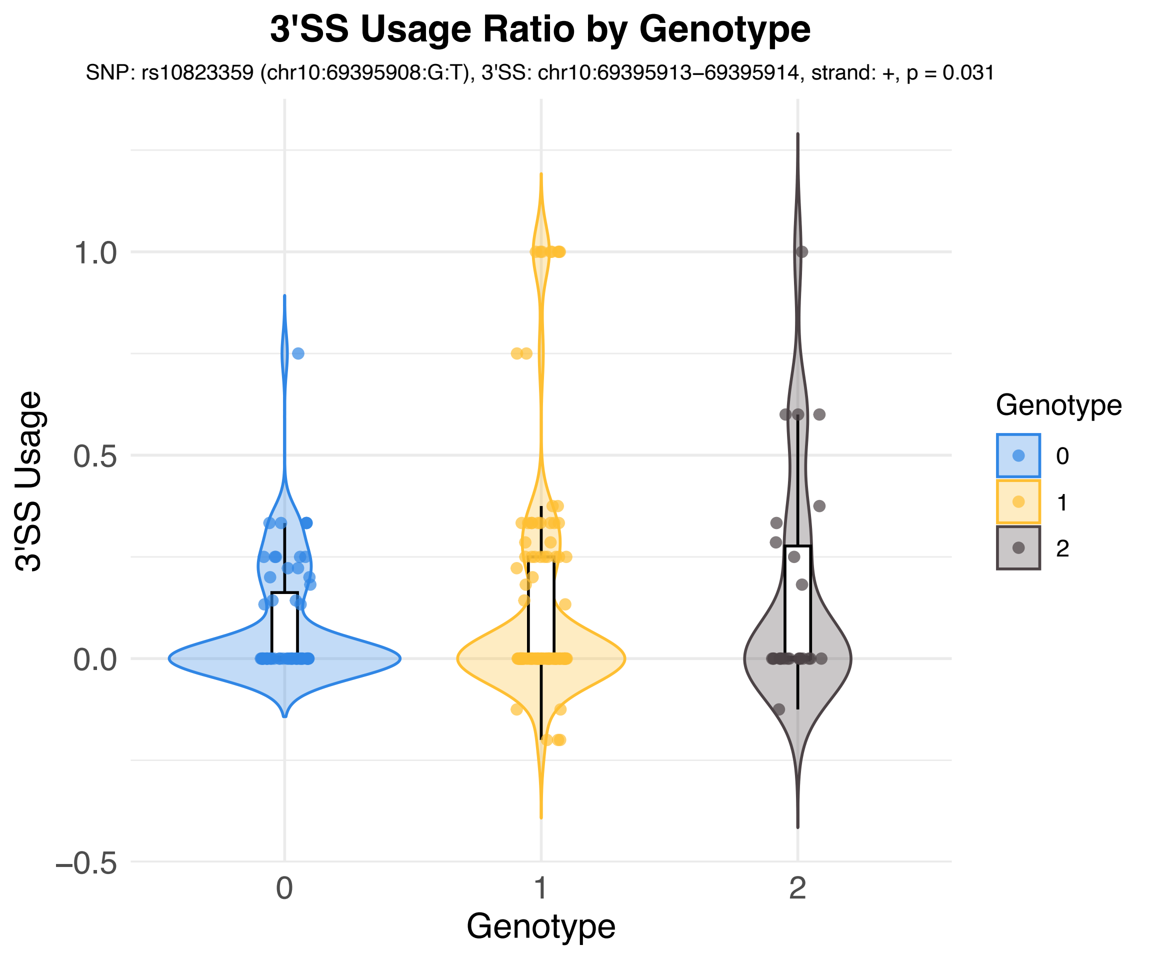

3'SS Usage Ratio by Genotype
SNP: rs10823359 (chr10:69395908:G:T), 3'SS: chr10:69395913–69395914, strand: +, p = 0.031
3'SS Usage
1.0
0.5
0.0
−0.5
0
1
2
Genotype
Genotype
0
1
2

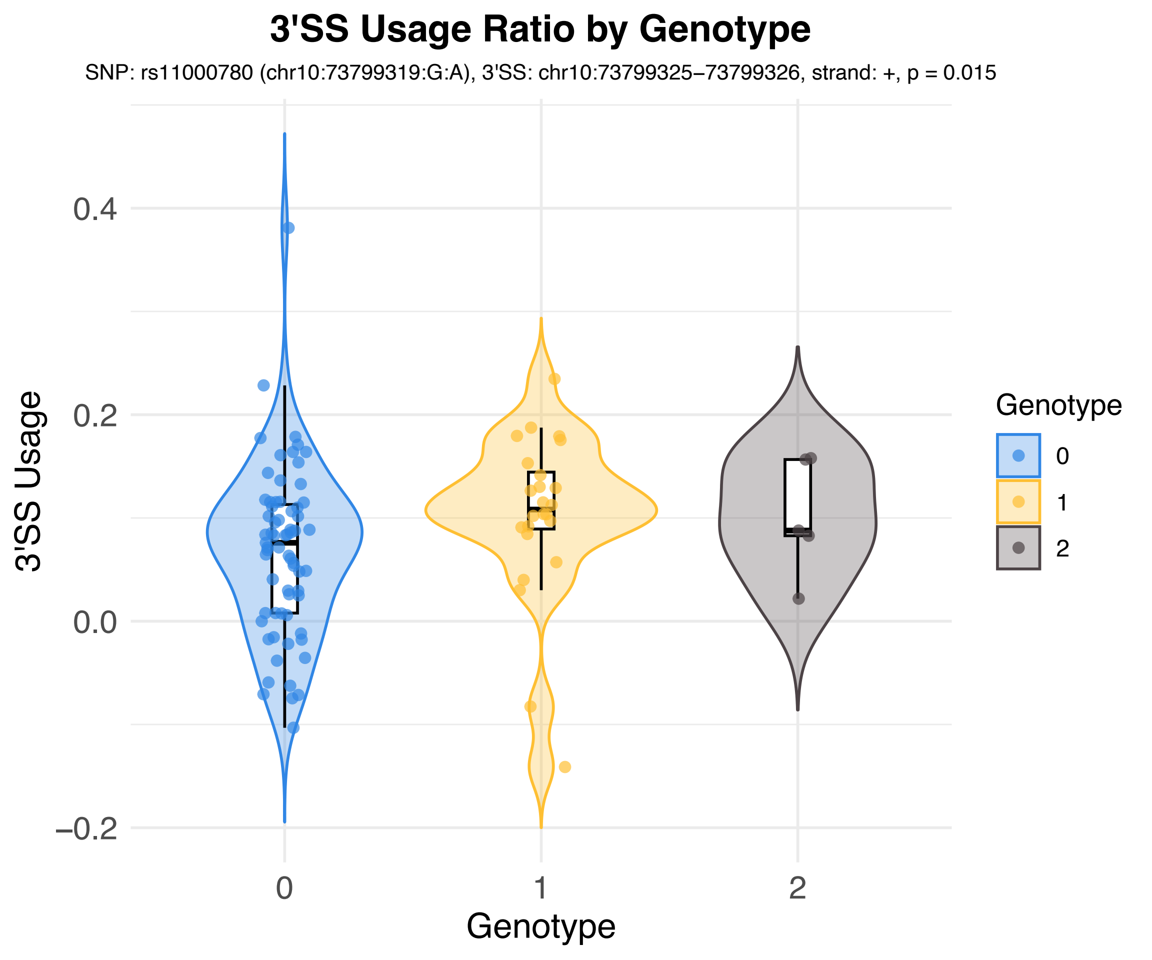
3'SS Usage Ratio by Genotype
SNP: rs11000780 (chr10:73799319:G:A), 3'SS: chr10:73799325–73799326, strand: +, p = 0.015
3'SS Usage
0.4
0.2
0.0
–0.2
0
1
2
Genotype
Genotype
0
1
2

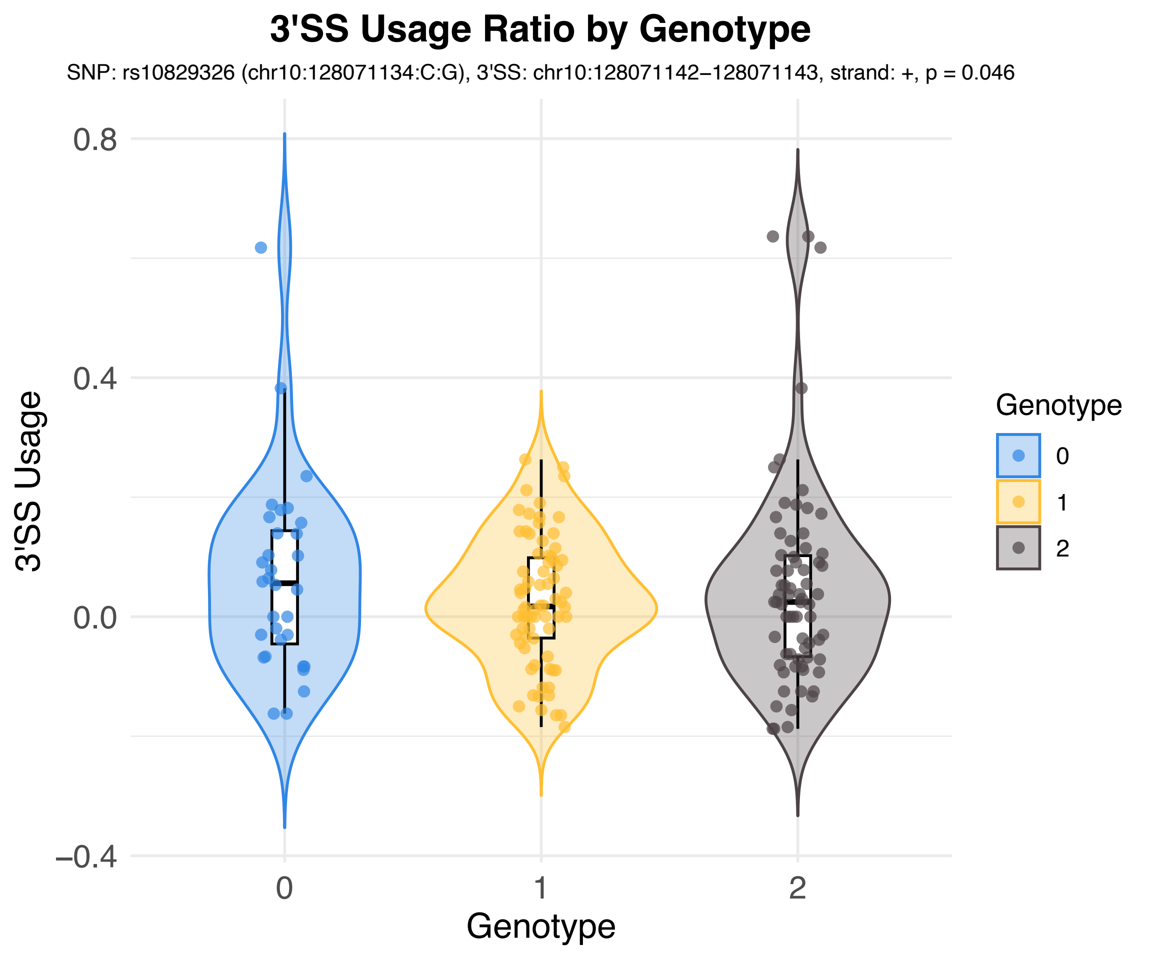
3'SS Usage Ratio by Genotype
SNP: rs10829326 (chr10:128071134:C:G), 3'SS: chr10:128071142−128071143, strand: +, p = 0.046
3'SS Usage
0.8
0.4
0.0
−0.4
0
1
2
Genotype
Genotype
0
1
2

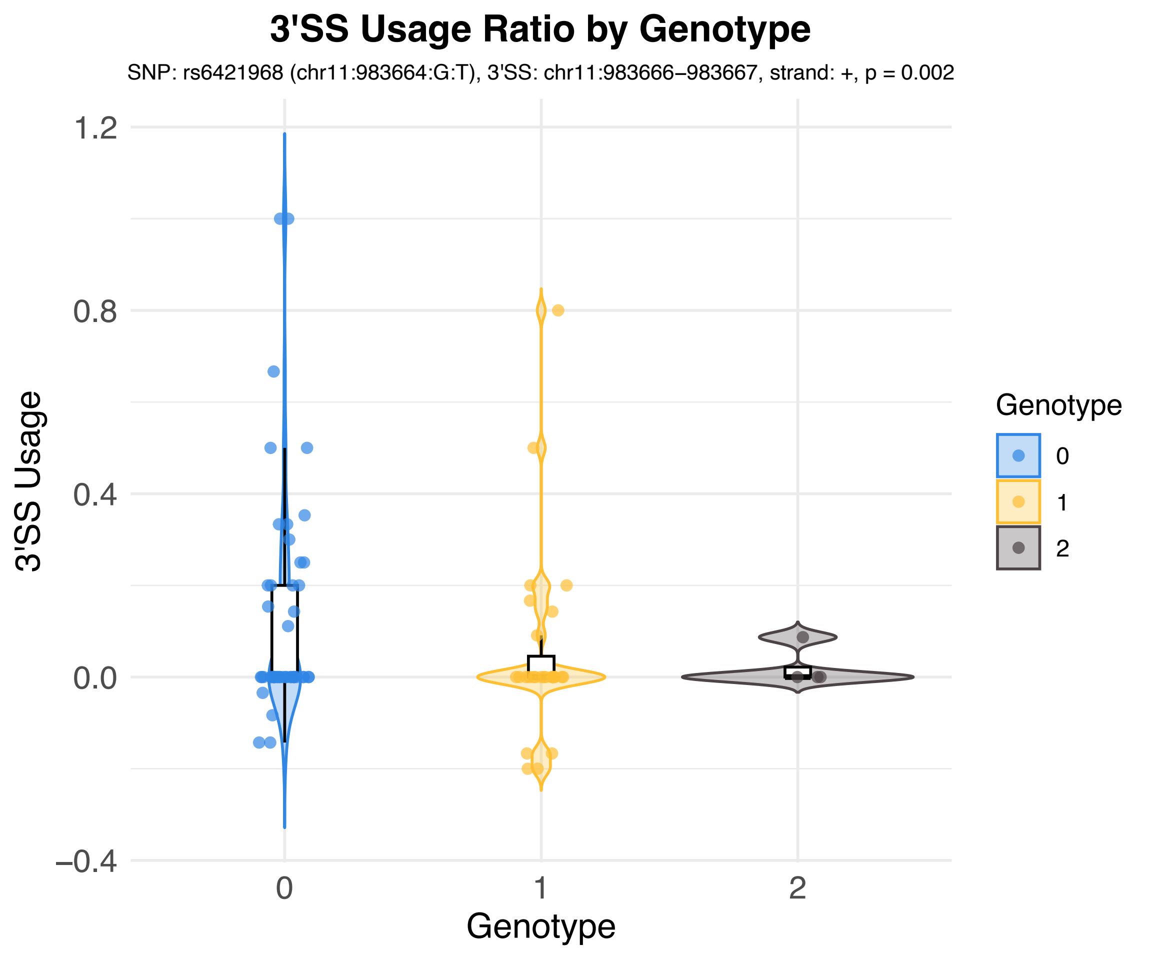
3'SS Usage Ratio by Genotype
SNP: rs6421968 (chr11:983664:G:T), 3'SS: chr11:983666−983667, strand: +, p = 0.002
3'SS Usage
1.2
0.8
0.4
0.0
−0.4
0
1
2
Genotype
Genotype
0
1
2

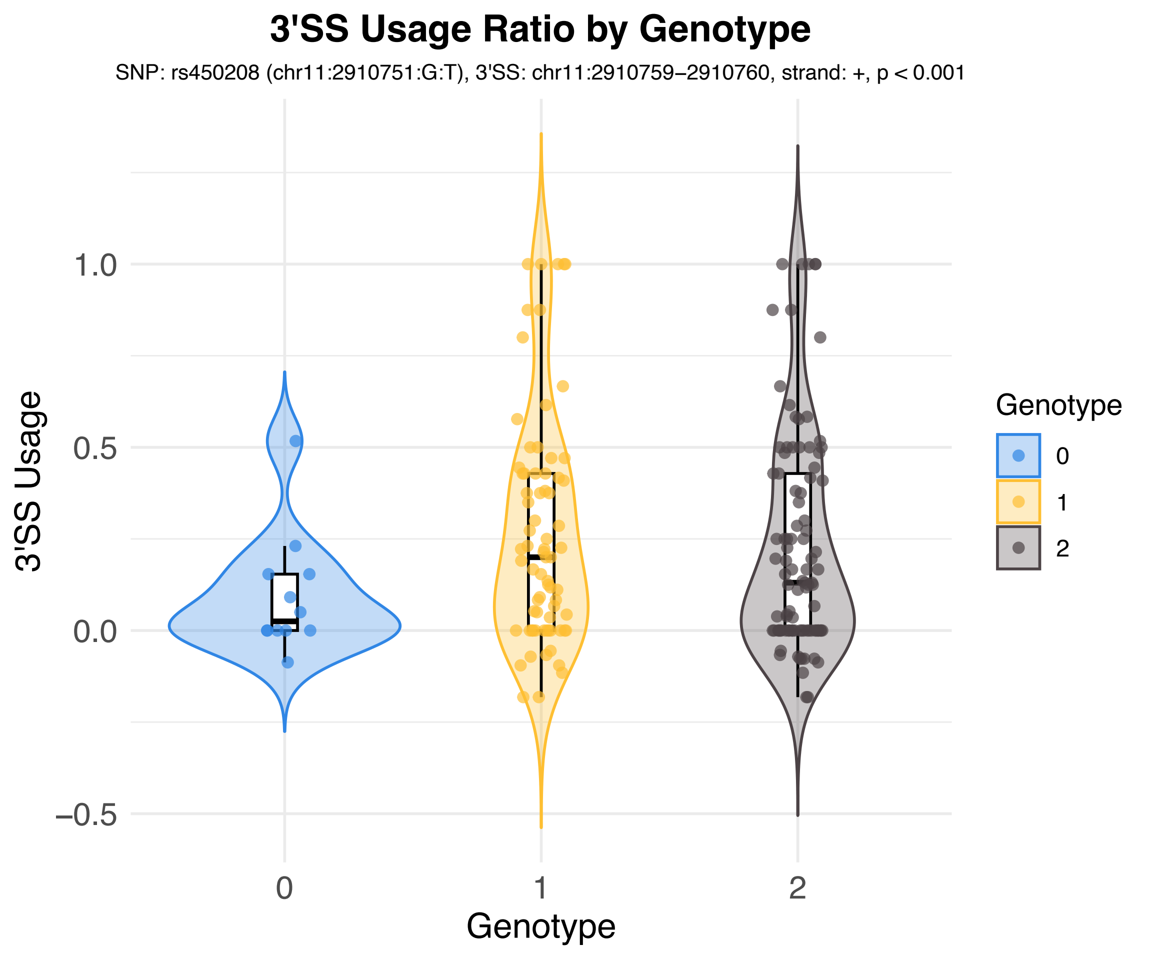
3'SS Usage Ratio by Genotype
SNP: rs450208 (chr11:2910751:G:T), 3'SS: chr11:2910759−2910760, strand: +, p < 0.001
3'SS Usage
1.0
0.5
0.0
−0.5
0
1
2
Genotype
Genotype
0
1
2

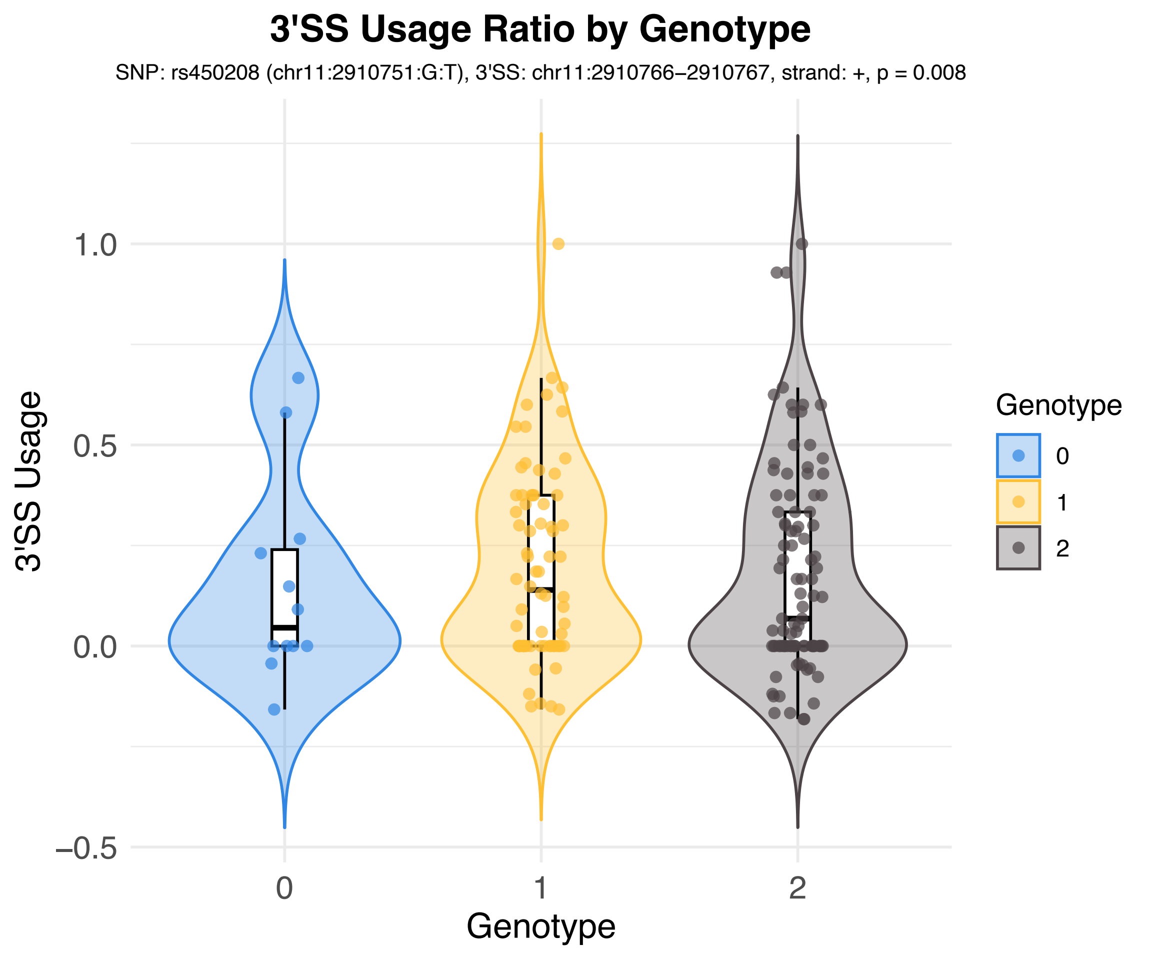
3'SS Usage Ratio by Genotype
SNP: rs450208 (chr11:2910751:G:T), 3'SS: chr11:2910766–2910767, strand: +, p = 0.008
3'SS Usage
1.0
0.5
0.0
−0.5
0
1
2
Genotype
Genotype
0
1
2

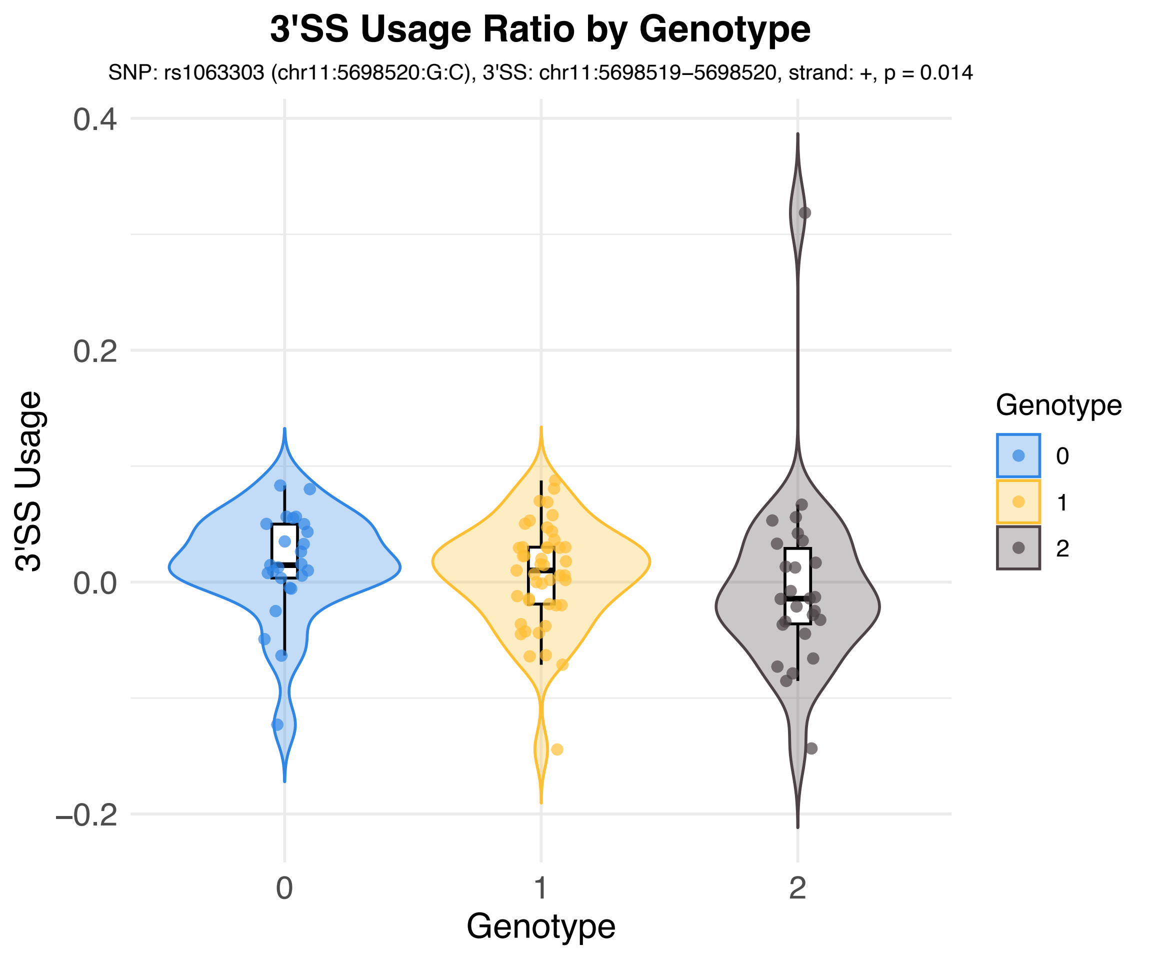
3'SS Usage Ratio by Genotype
SNP: rs1063303 (chr11:5698520:G:C), 3'SS: chr11:5698519−5698520, strand: +, p = 0.014
3'SS Usage
0.4
0.2
0.0
−0.2
0
1
2
Genotype
Genotype
0
1
2

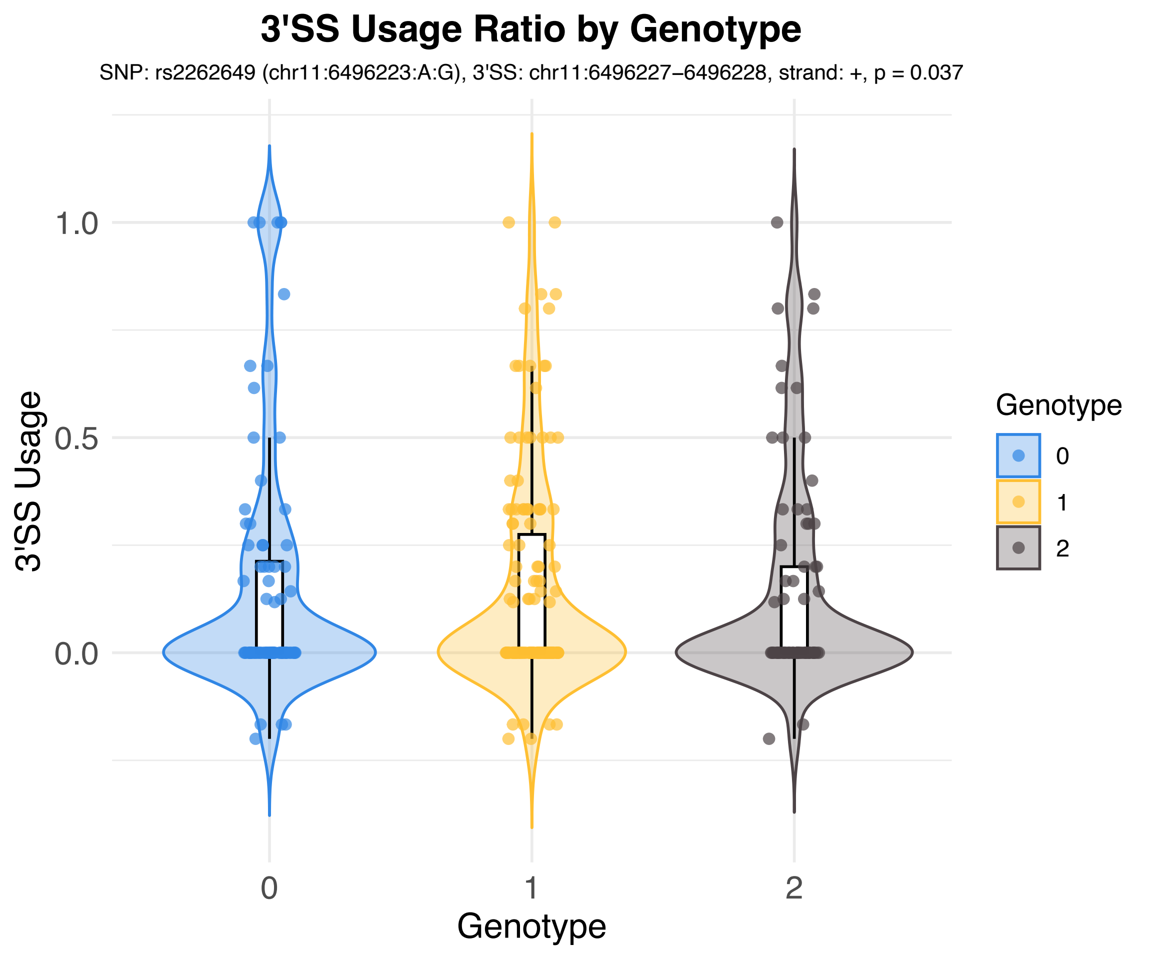
3'SS Usage Ratio by Genotype
SNP: rs2262649 (chr11:6496223:A:G), 3'SS: chr11:6496227–6496228, strand: +, p = 0.037
3'SS Usage
1.0
0.5
0.0
0
1
2
Genotype
Genotype
0
1
2

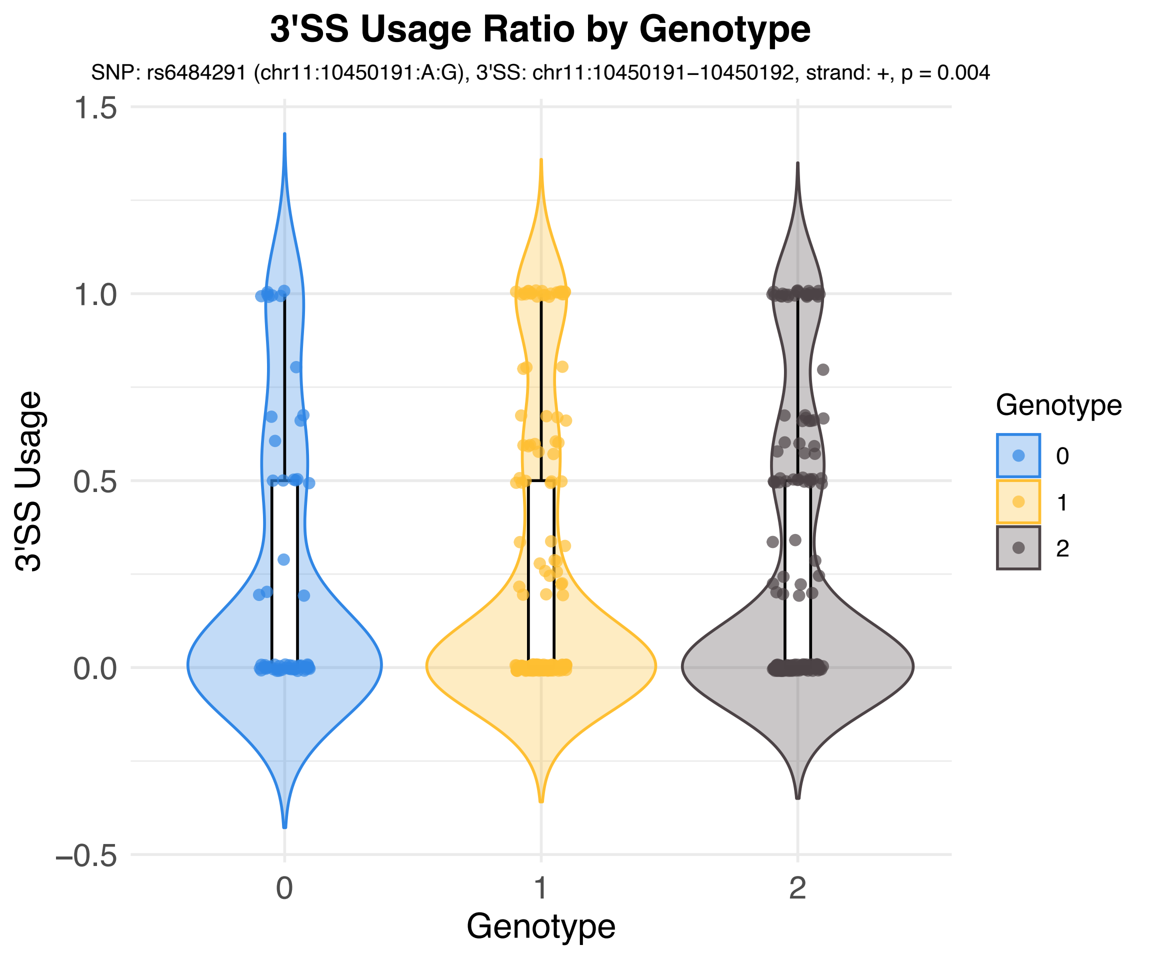
3'SS Usage Ratio by Genotype
SNP: rs6484291 (chr11:10450191:A:G), 3'SS: chr11:10450191−10450192, strand: +, p = 0.004
3'SS Usage
1.5
1.0
0.5
0.0
−0.5
0
1
2
Genotype
Genotype
0
1
2

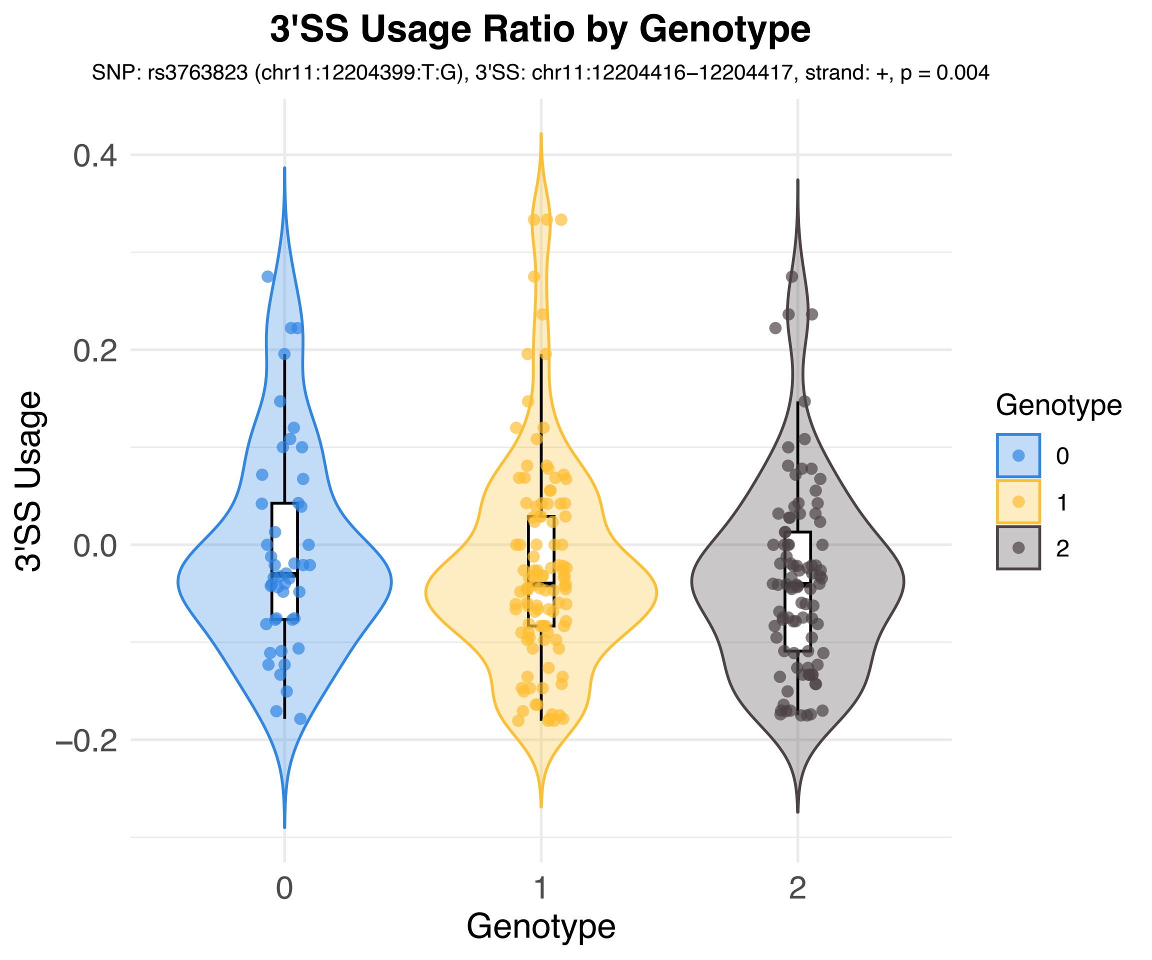

3'SS Usage Ratio by Genotype
SNP: rs3763823 (chr11:12204399:T:G), 3'SS: chr11:12204416−12204417, strand: +, p = 0.004
3'SS Usage
0.4
0.2
0.0
−0.2
0
1
2
Genotype
Genotype
0
1
2

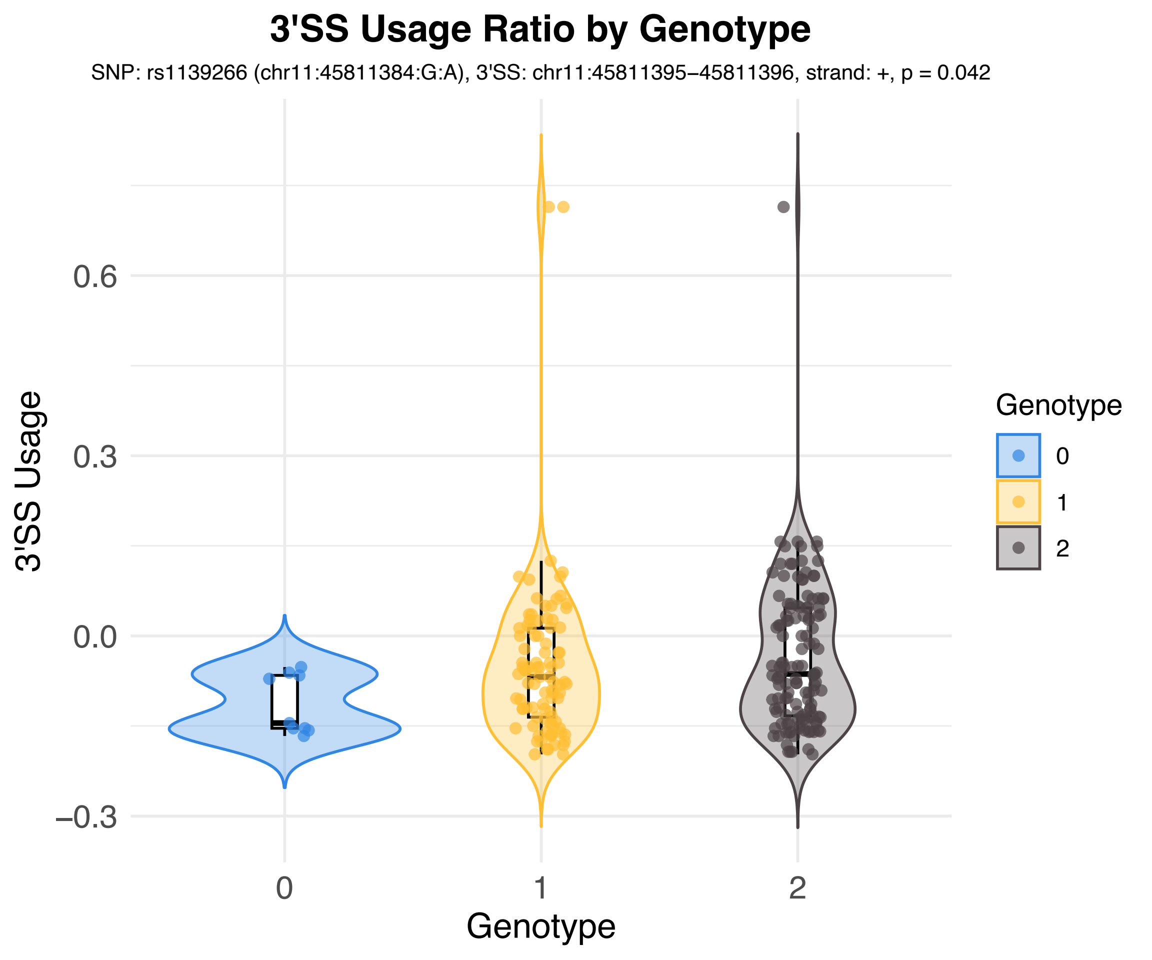
3'SS Usage Ratio by Genotype
SNP: rs1139266 (chr11:45811384:G:A), 3'SS: chr11:45811395–45811396, strand: +, p = 0.042
3'SS Usage
0.6
0.3
0.0
−0.3
0
1
2
Genotype
Genotype
0
1
2

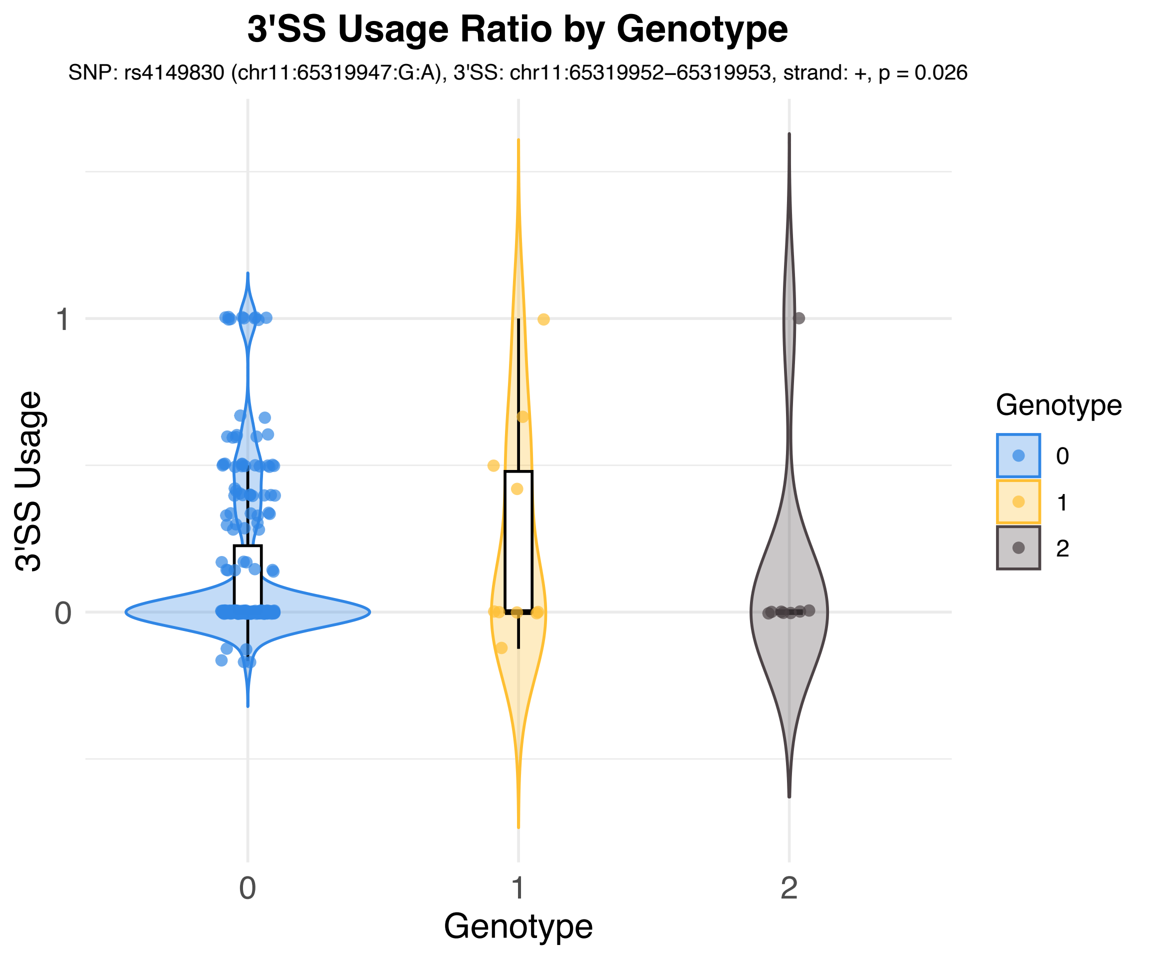
3'SS Usage Ratio by Genotype
SNP: rs4149830 (chr11:65319947:G:A), 3'SS: chr11:65319952–65319953, strand: +, p = 0.026
3'SS Usage
1
0
0
1
2
Genotype
Genotype
0
1
2

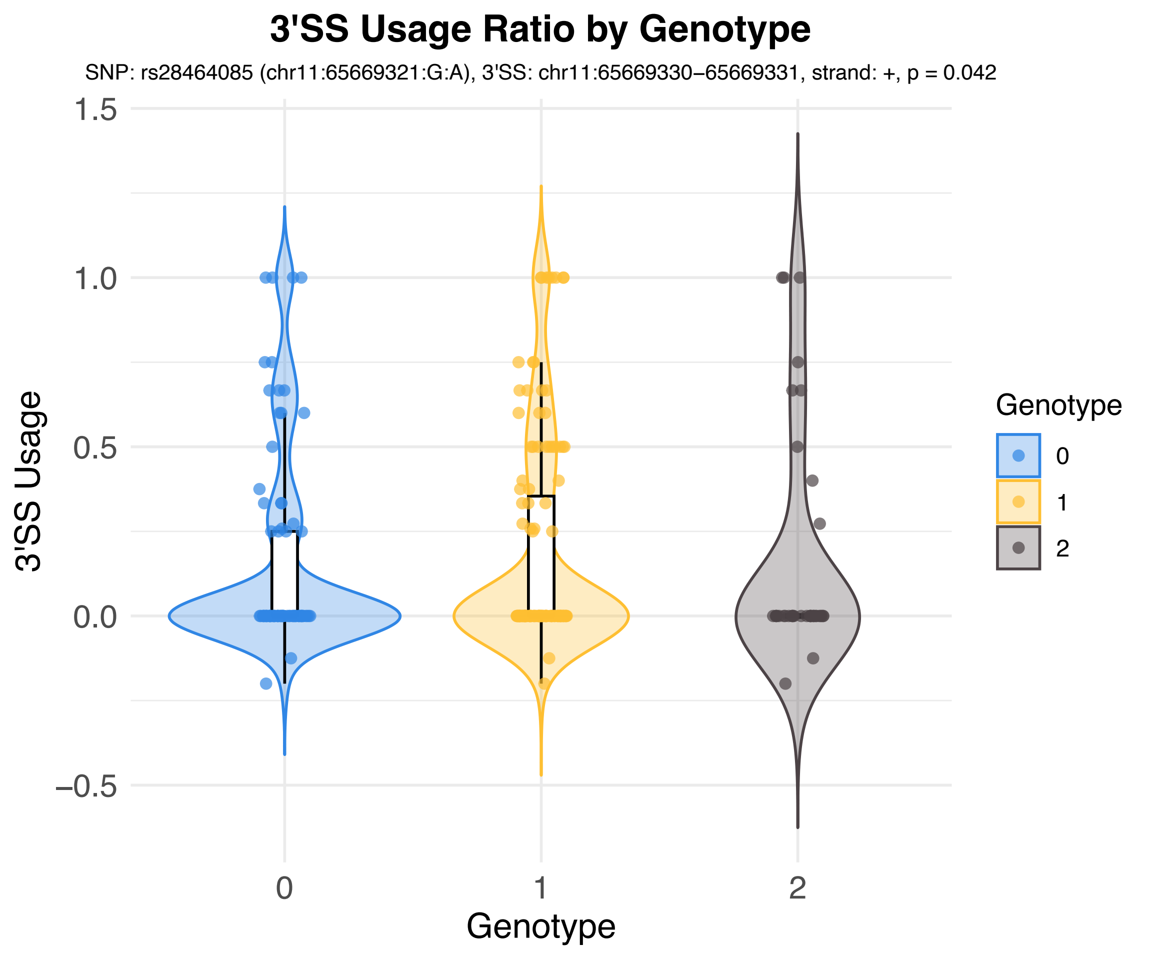
3'SS Usage Ratio by Genotype
SNP: rs28464085 (chr11:65669321:G:A), 3'SS: chr11:65669330−65669331, strand: +, p = 0.042
3'SS Usage
1.5
1.0
0.5
0.0
−0.5
0
1
2
Genotype
Genotype
0
1
2

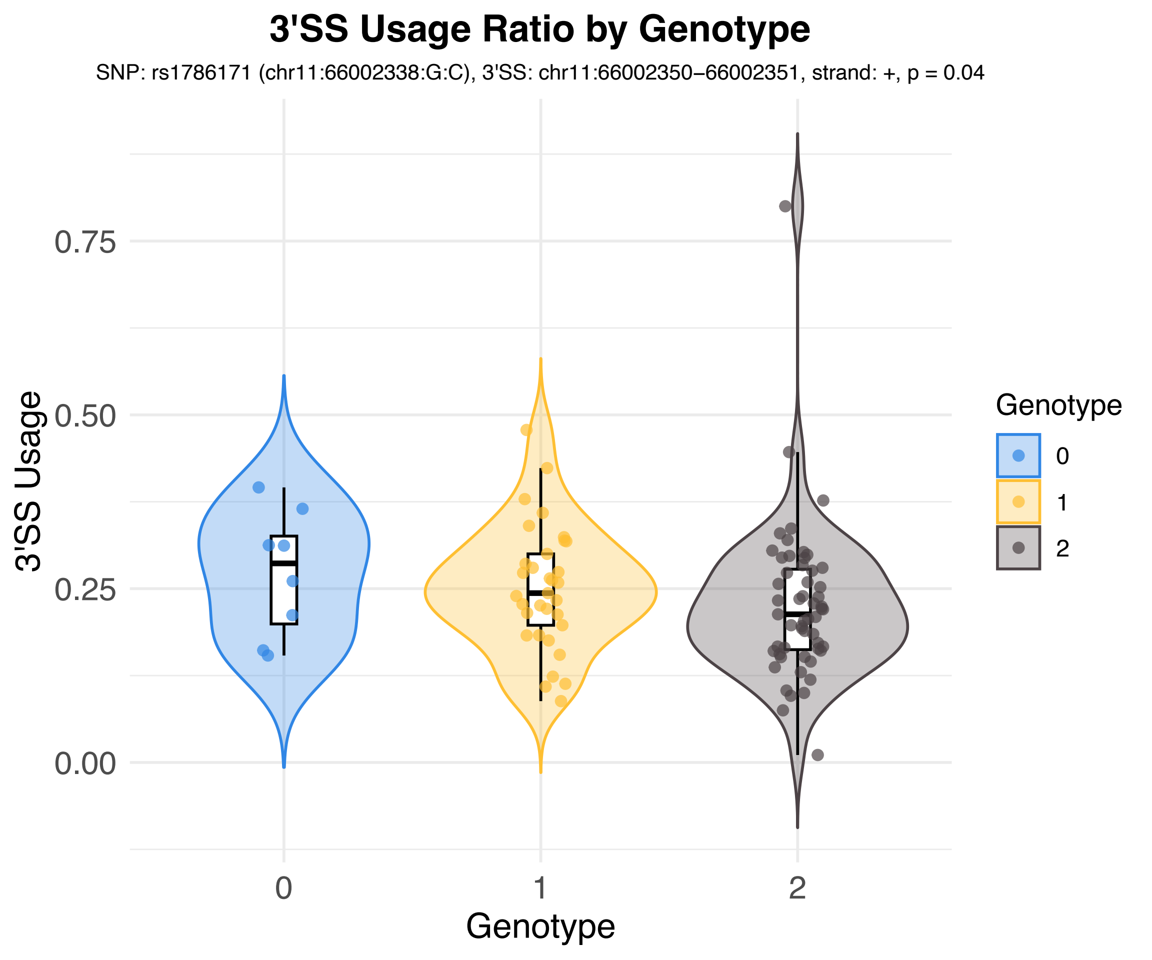
3'SS Usage Ratio by Genotype
SNP: rs1786171 (chr11:66002338:G:C), 3'SS: chr11:66002350−66002351, strand: +, p = 0.04
3'SS Usage
0.75
0.50
0.25
0.00
0
1
2
Genotype
Genotype
0
1
2

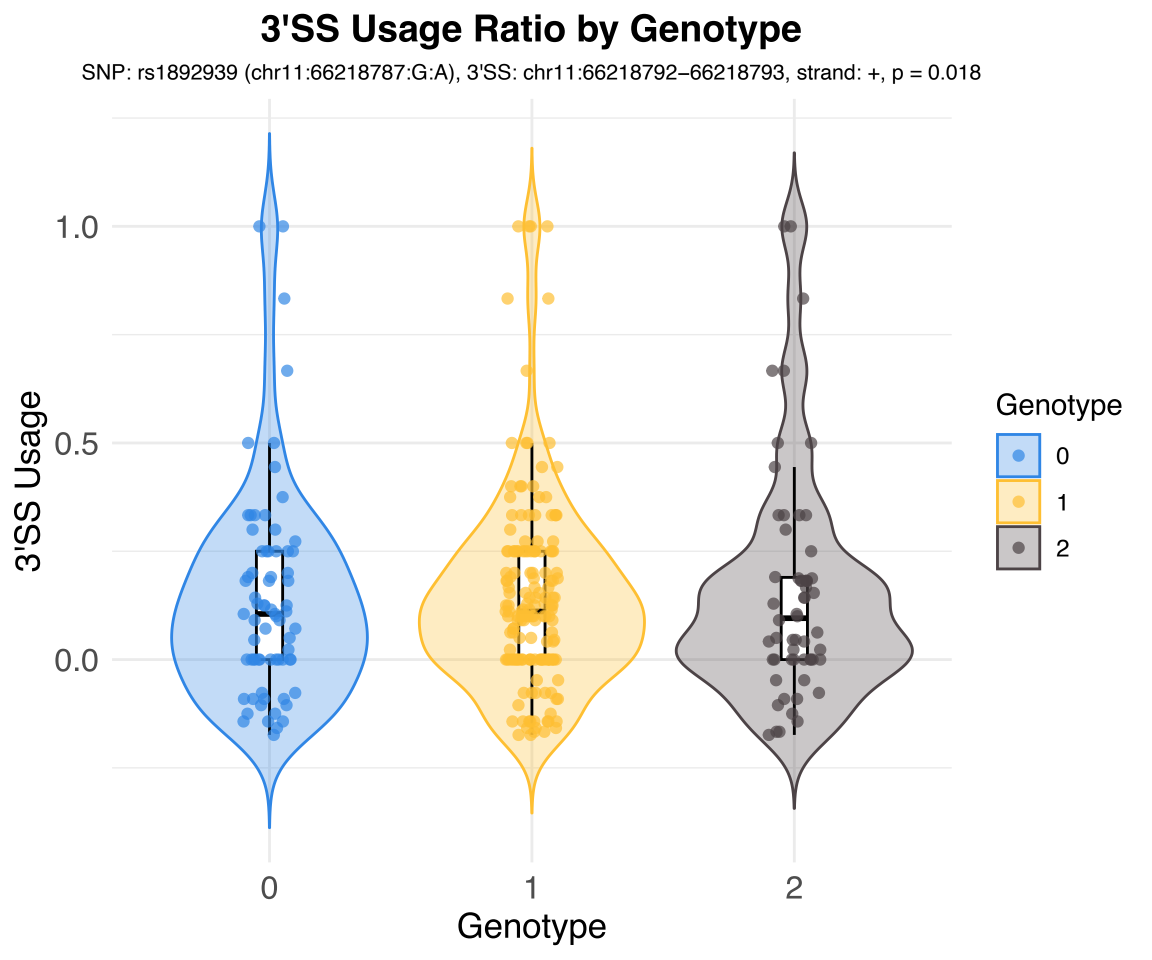
3'SS Usage Ratio by Genotype
SNP: rs1892939 (chr11:66218787:G:A), 3'SS: chr11:66218792–66218793, strand: +, p = 0.018
3'SS Usage
1.0
0.5
0.0
0
1
2
Genotype
Genotype
0
1
2

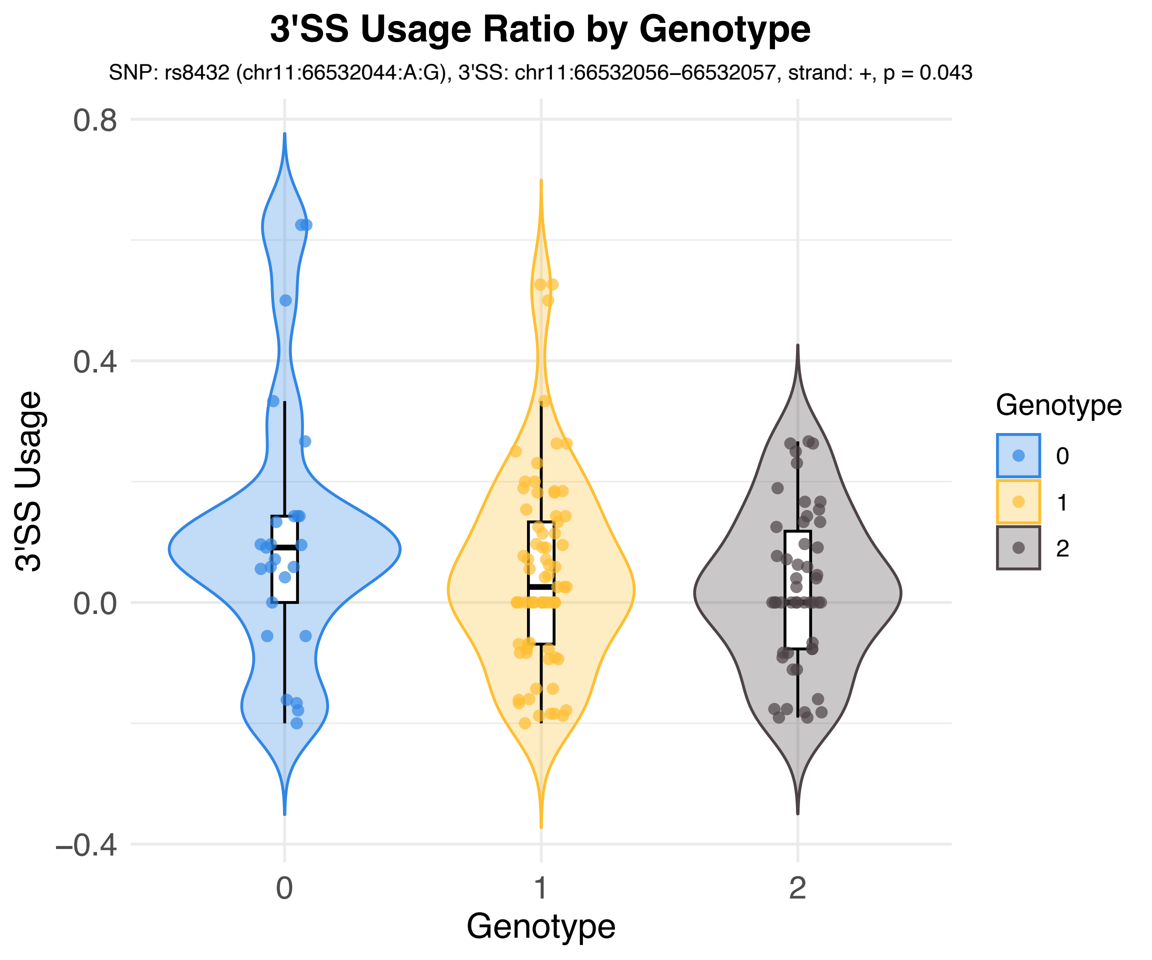
3'SS Usage Ratio by Genotype
SNP: rs8432 (chr11:66532044:A:G), 3'SS: chr11:66532056–66532057, strand: +, p = 0.043
3'SS Usage
0.8
0.4
0.0
−0.4
0
1
2
Genotype
Genotype
0
1
2

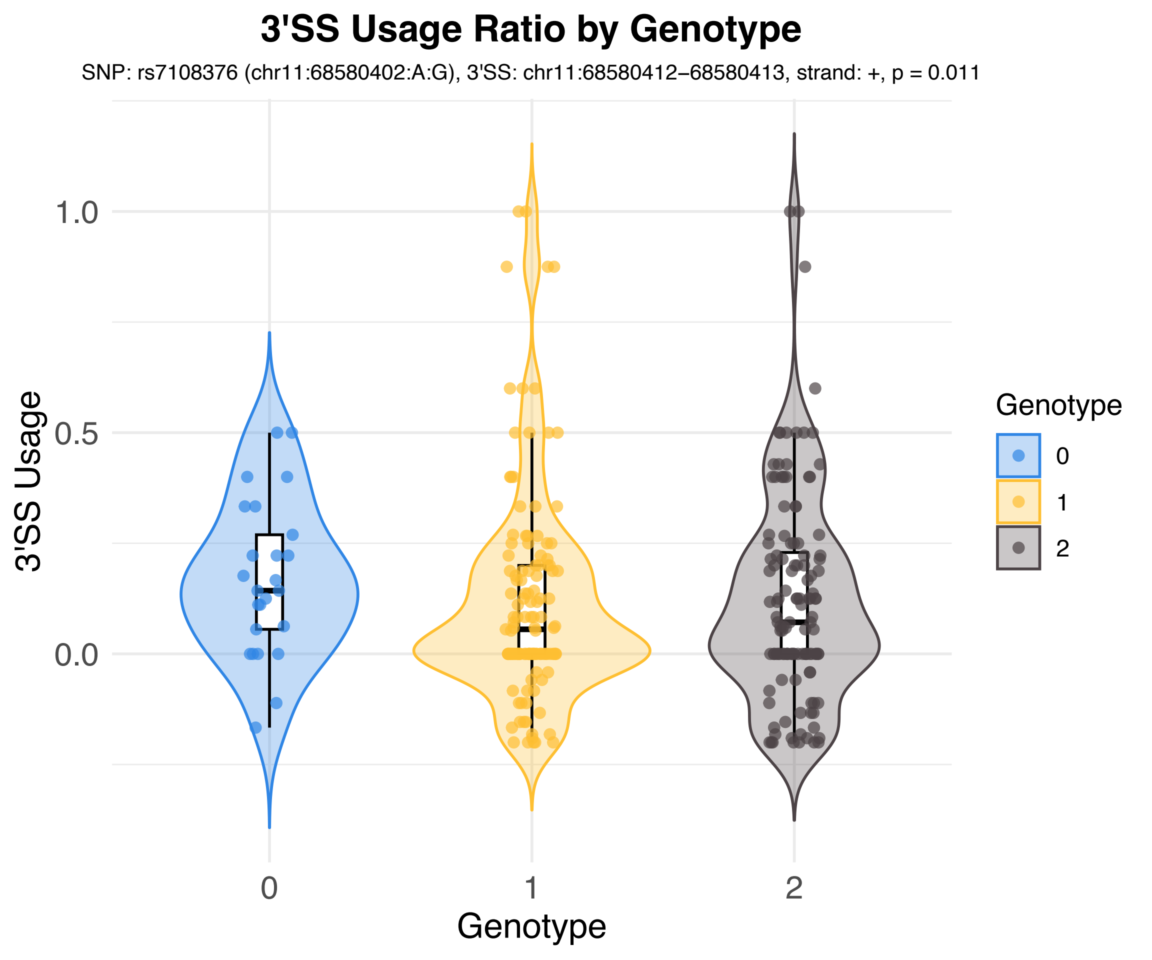

3'SS Usage Ratio by Genotype
SNP: rs7108376 (chr11:68580402:A:G), 3'SS: chr11:68580412–68580413, strand: +, p = 0.011
3'SS Usage
1.0
0.5
0.0
0
1
2
Genotype
Genotype
0
1
2

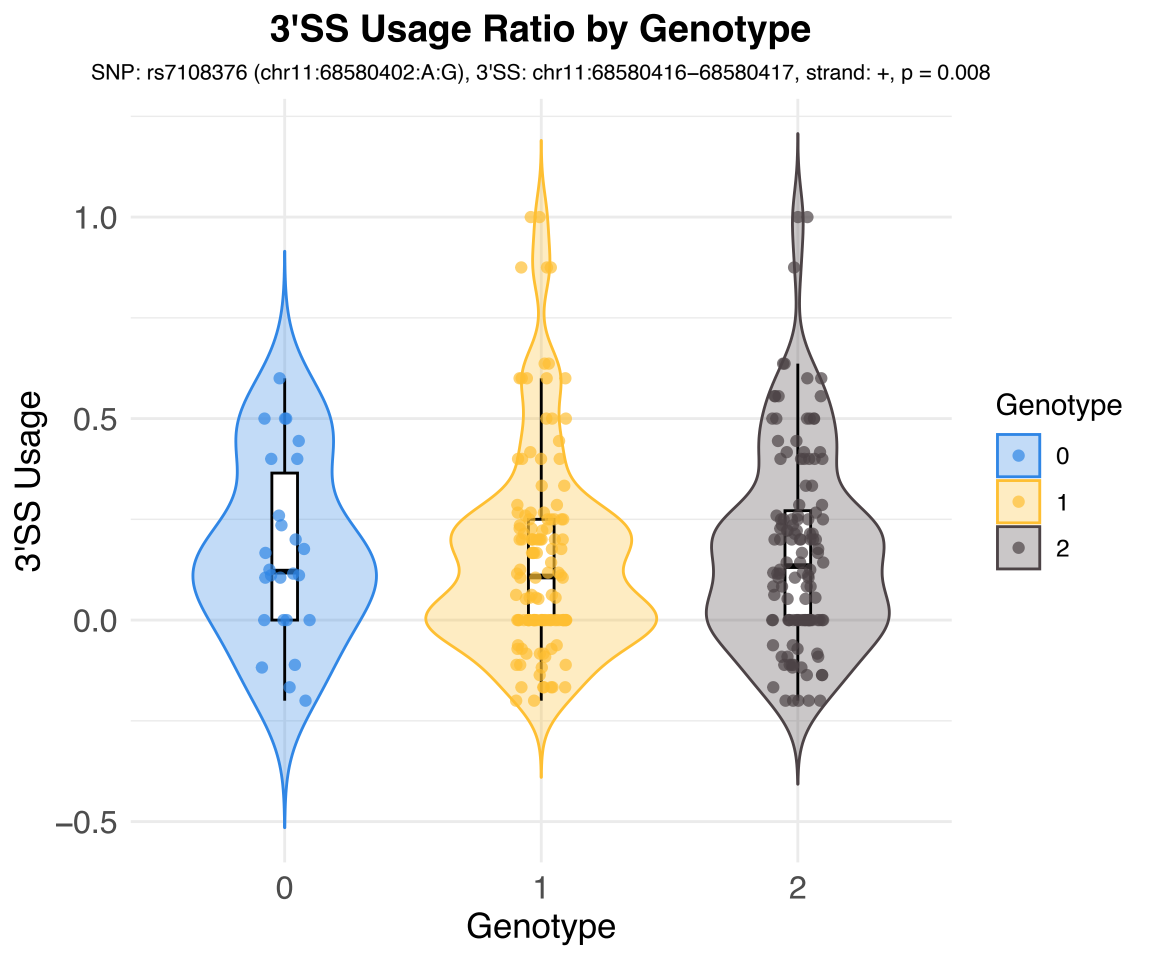

3'SS Usage Ratio by Genotype
SNP: rs7108376 (chr11:68580402:A:G), 3'SS: chr11:68580416–68580417, strand: +, p = 0.008
3'SS Usage
1.0
0.5
0.0
−0.5
0
1
2
Genotype
Genotype
0
1
2

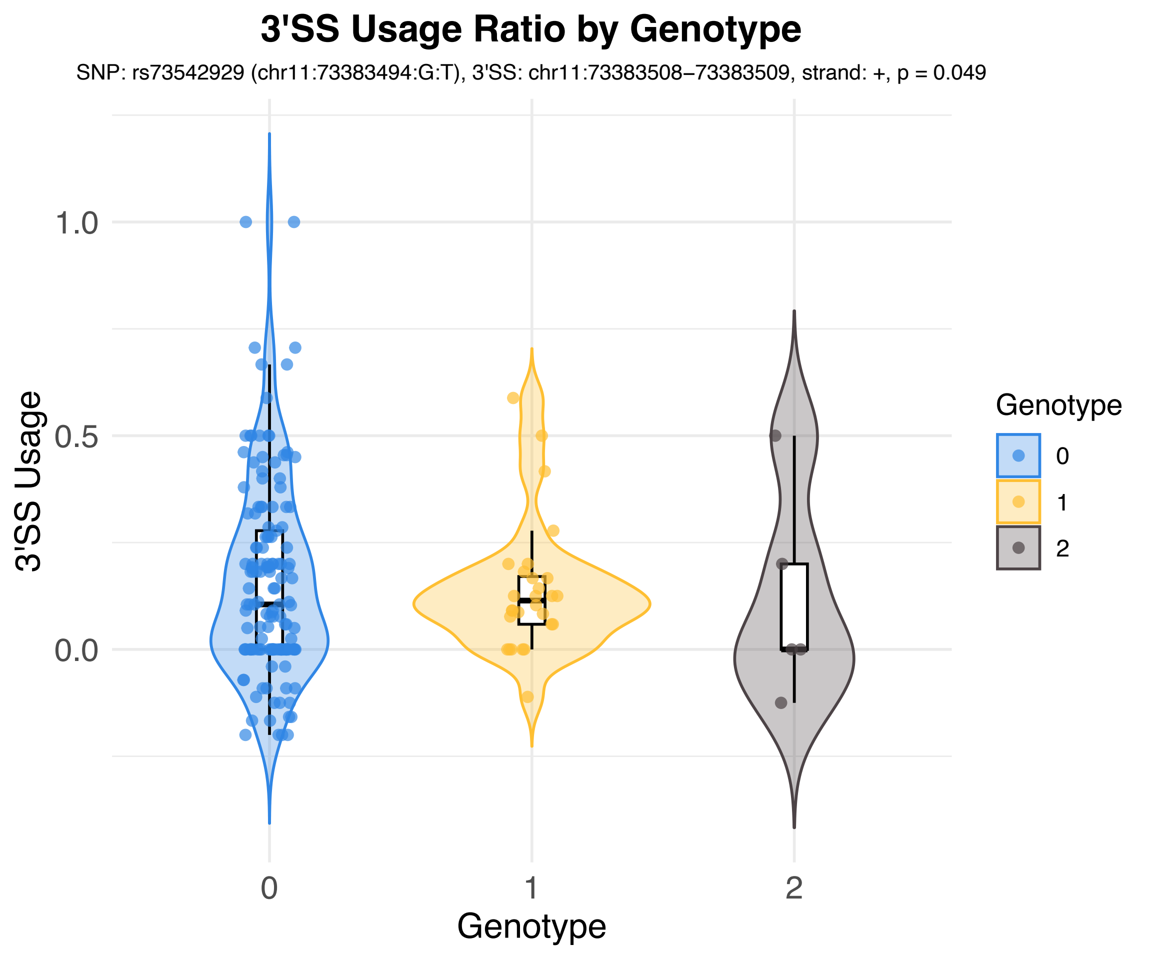
3'SS Usage Ratio by Genotype
SNP: rs73542929 (chr11:73383494:G:T), 3'SS: chr11:73383508–73383509, strand: +, p = 0.049
3'SS Usage
1.0
0.5
0.0
0
1
2
Genotype
Genotype
0
1
2

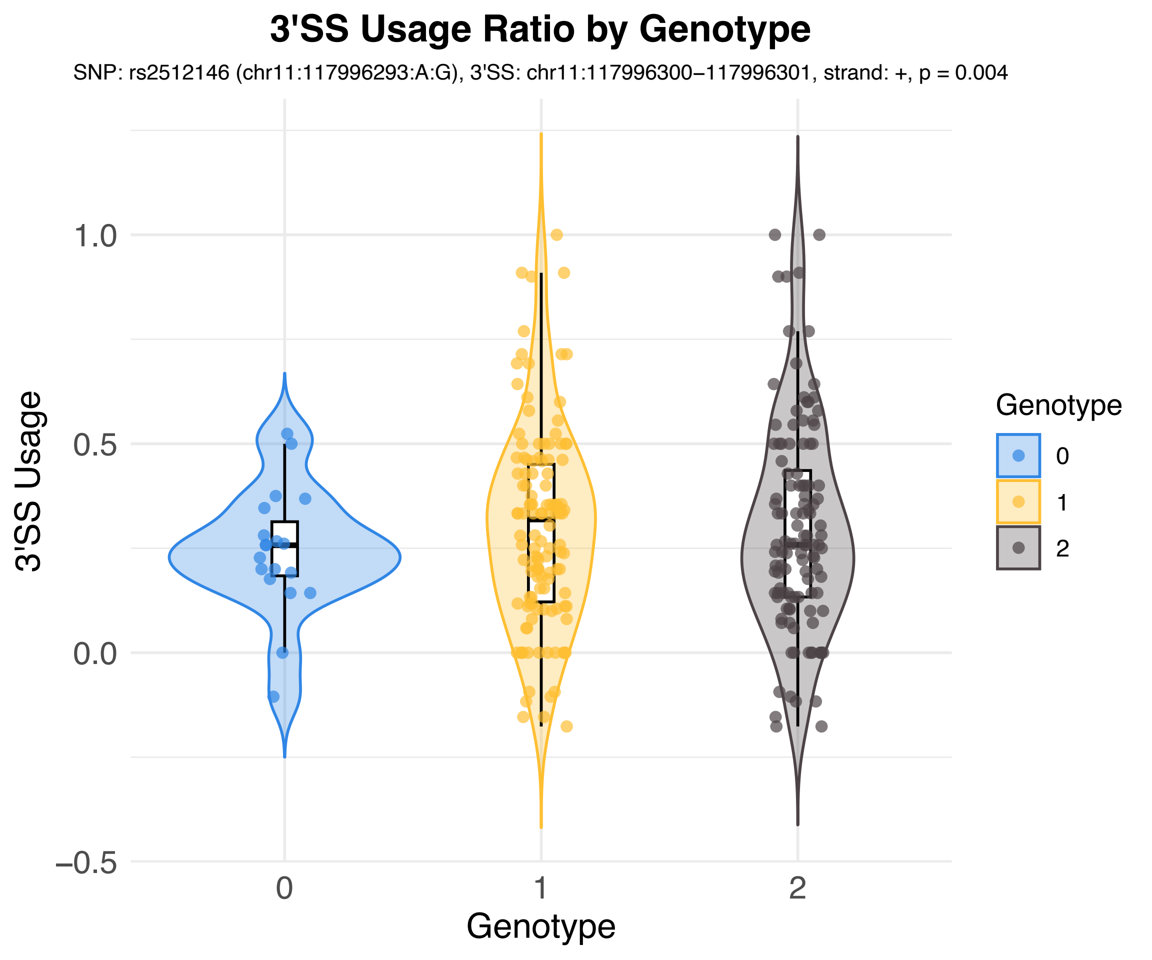

3'SS Usage Ratio by Genotype
SNP: rs2512146 (chr11:117996293:A:G), 3'SS: chr11:117996300−117996301, strand: +, p = 0.004
3'SS Usage
1.0
0.5
0.0
−0.5
0
1
2
Genotype
Genotype
0
1
2

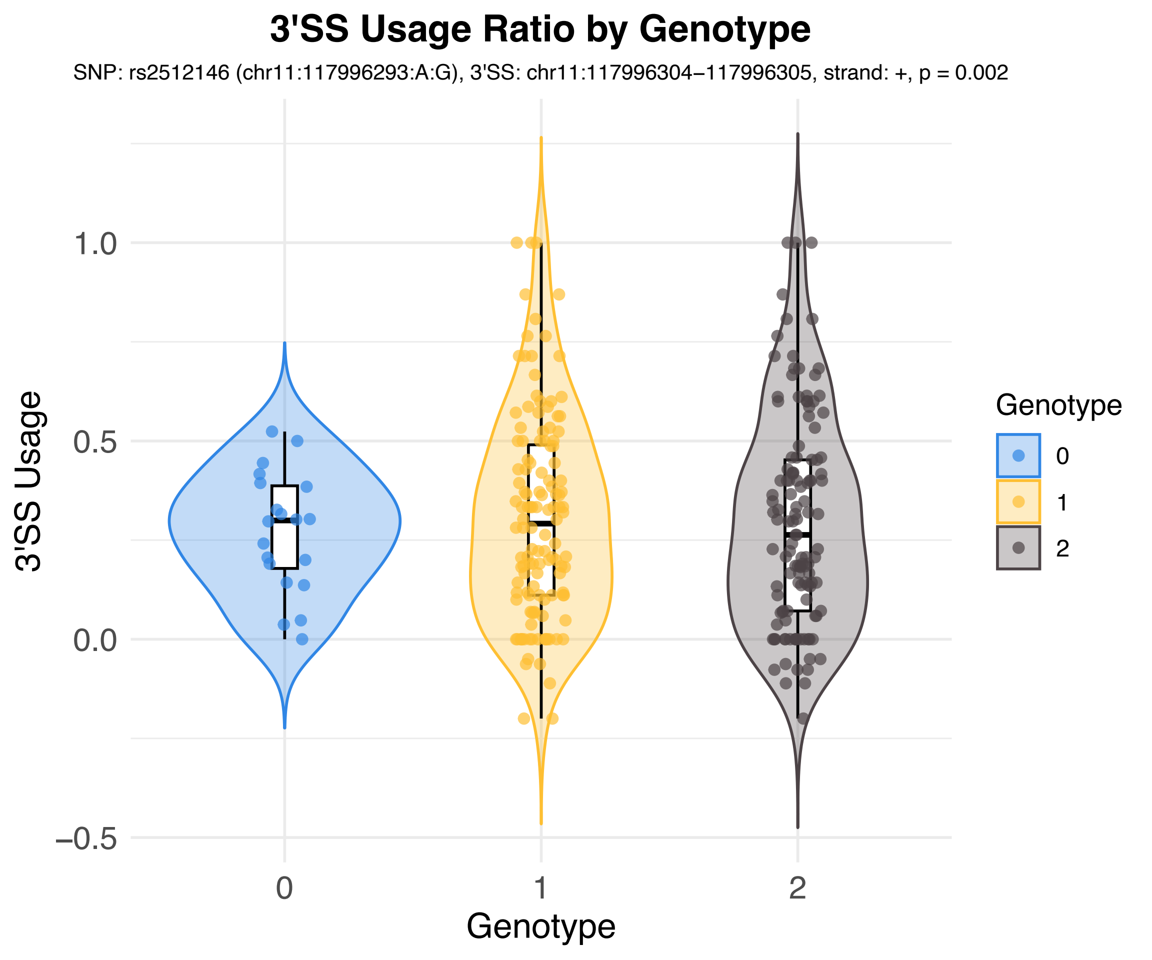
3'SS Usage Ratio by Genotype
SNP: rs2512146 (chr11:117996293:A:G), 3'SS: chr11:117996304–117996305, strand: +, p = 0.002
3'SS Usage
1.0
0.5
0.0
−0.5
0
1
2
Genotype
Genotype
0
1
2

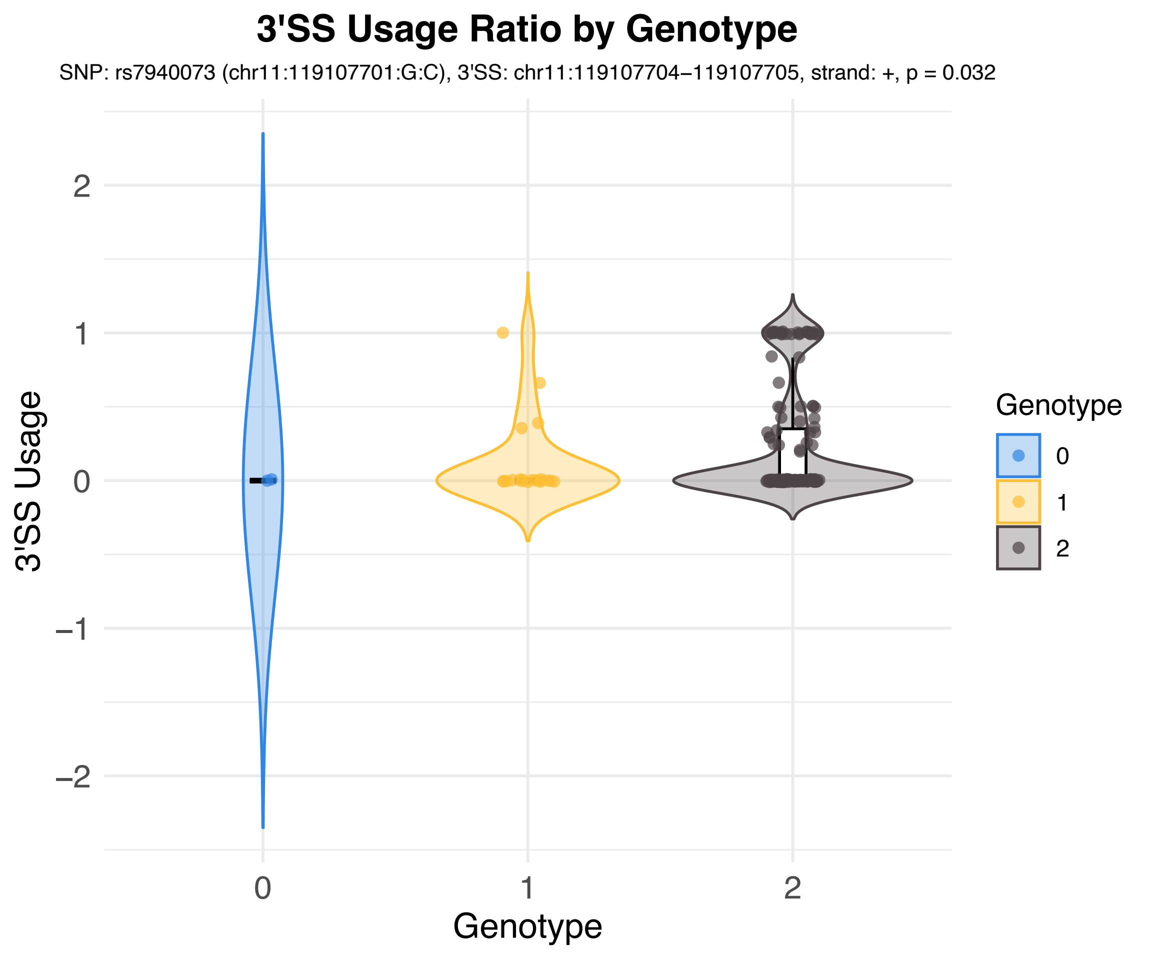
3'SS Usage Ratio by Genotype
SNP: rs7940073 (chr11:119107701:G:C), 3'SS: chr11:119107704–119107705, strand: +, p = 0.032
3'SS Usage
2
1
0
–1
–2
0
1
2
Genotype
Genotype
0
1
2

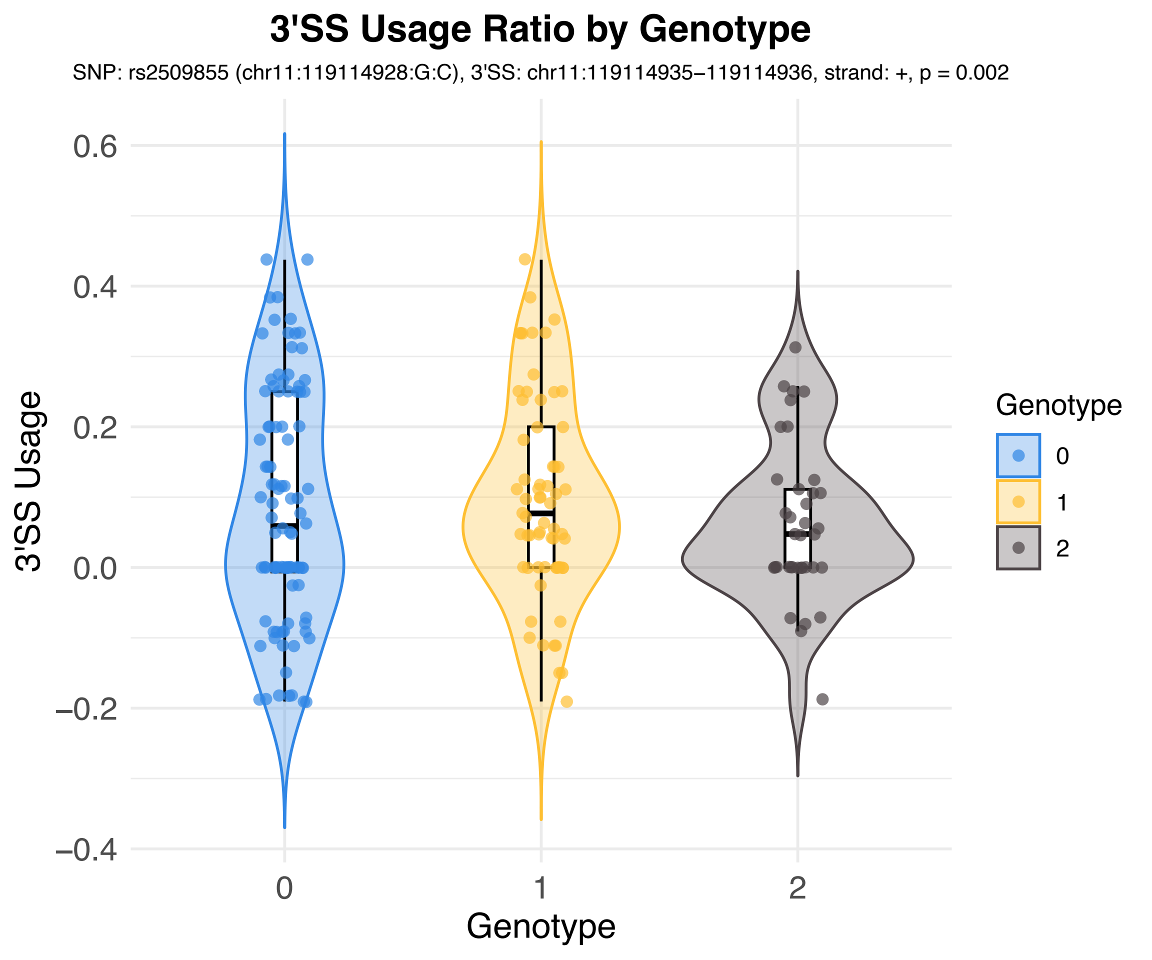
3'SS Usage Ratio by Genotype
SNP: rs2509855 (chr11:119114928:G:C), 3'SS: chr11:119114935−119114936, strand: +, p = 0.002
3'SS Usage
0.6
0.4
0.2
0.0
−0.2
−0.4
0
1
2
Genotype
Genotype
0
1
2

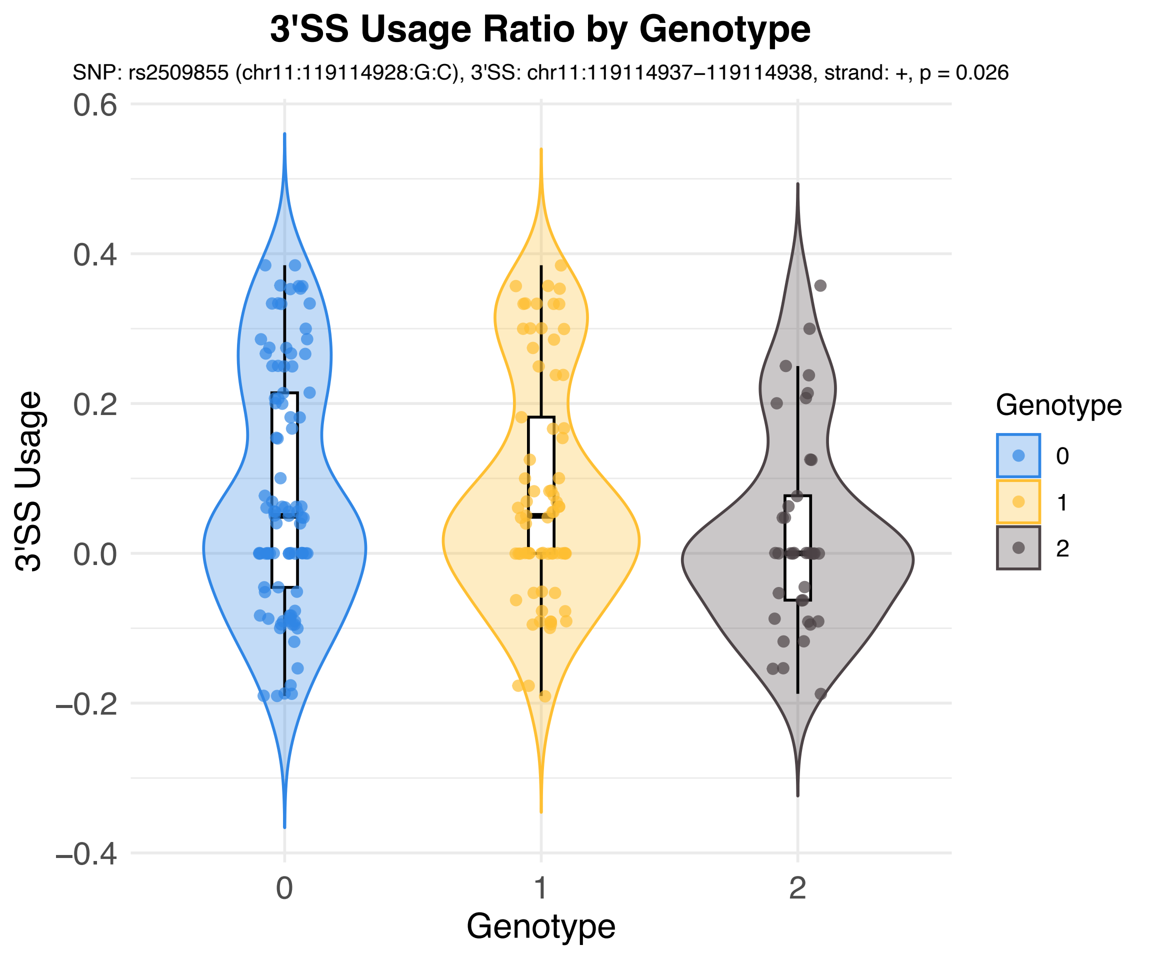
3'SS Usage Ratio by Genotype
SNP: rs2509855 (chr11:119114928:G:C), 3'SS: chr11:119114937−119114938, strand: +, p = 0.026
3'SS Usage
0.6
0.4
0.2
0.0
−0.2
−0.4
0
1
2
Genotype
Genotype
0
1
2

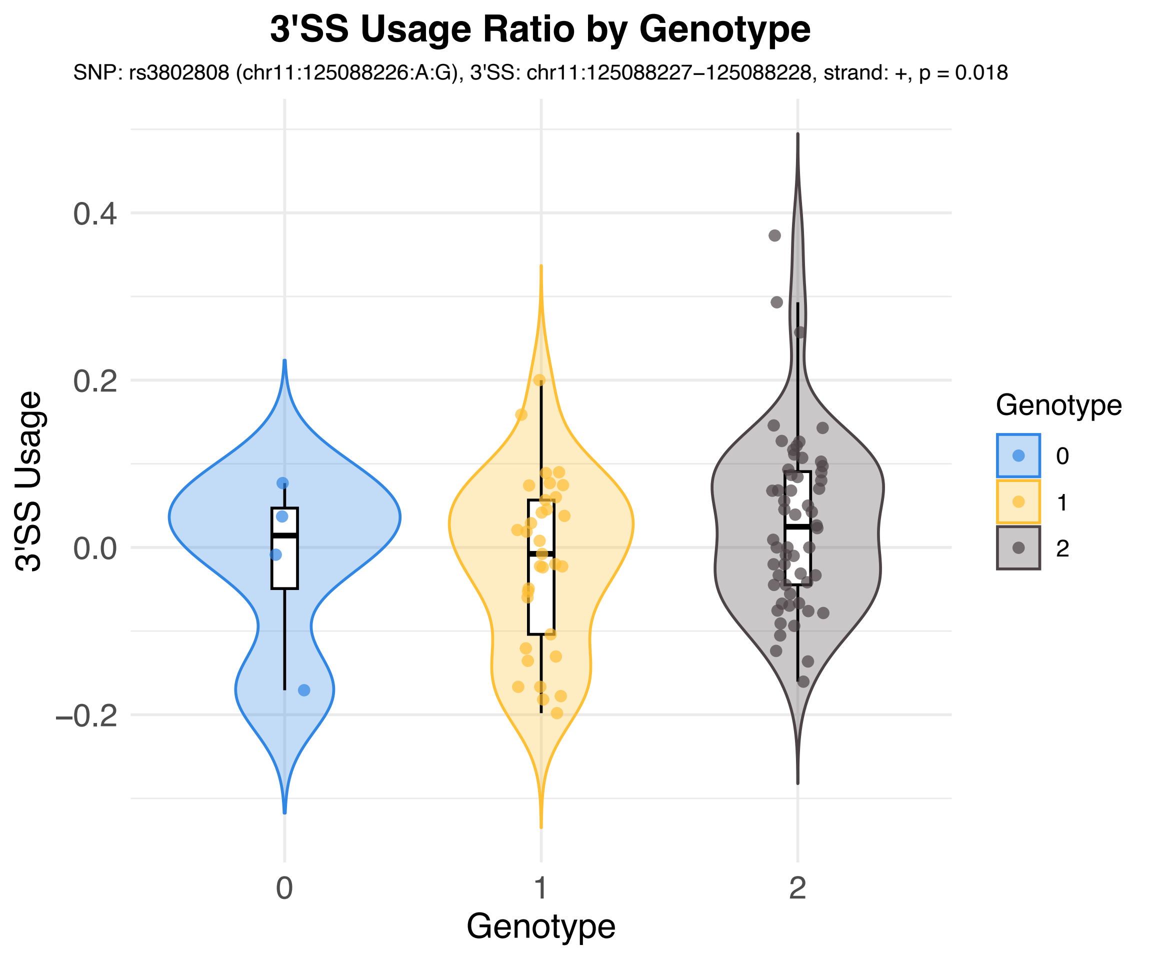
3'SS Usage Ratio by Genotype
SNP: rs3802808 (chr11:125088226:A:G), 3'SS: chr11:125088227−125088228, strand: +, p = 0.018
3'SS Usage
0.4
0.2
0.0
−0.2
0
1
2
Genotype
Genotype
0
1
2

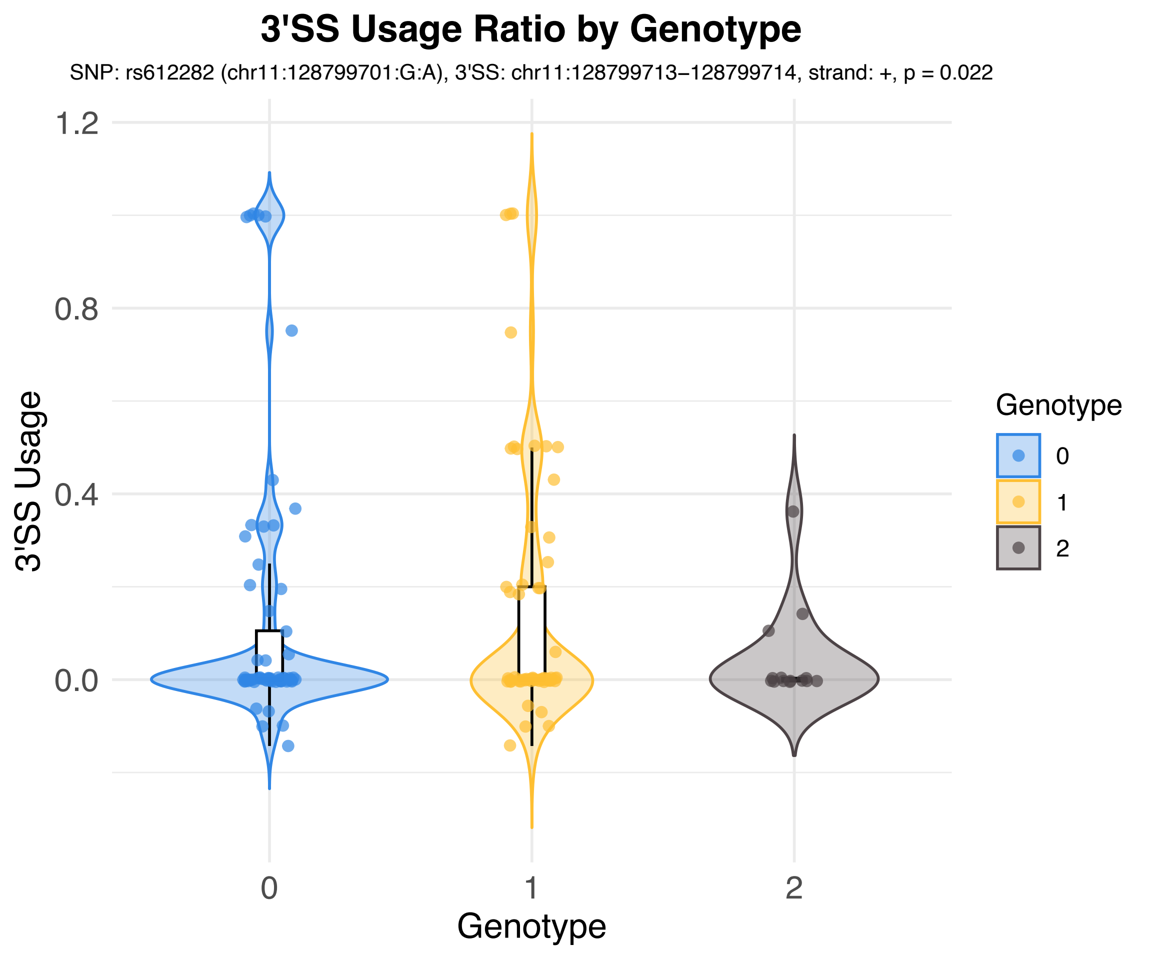

3'SS Usage Ratio by Genotype
SNP: rs612282 (chr11:128799701:G:A), 3'SS: chr11:128799713–128799714, strand: +, p = 0.022
3'SS Usage
1.2
0.8
0.4
0.0
0
1
2
Genotype
Genotype
0
1
2

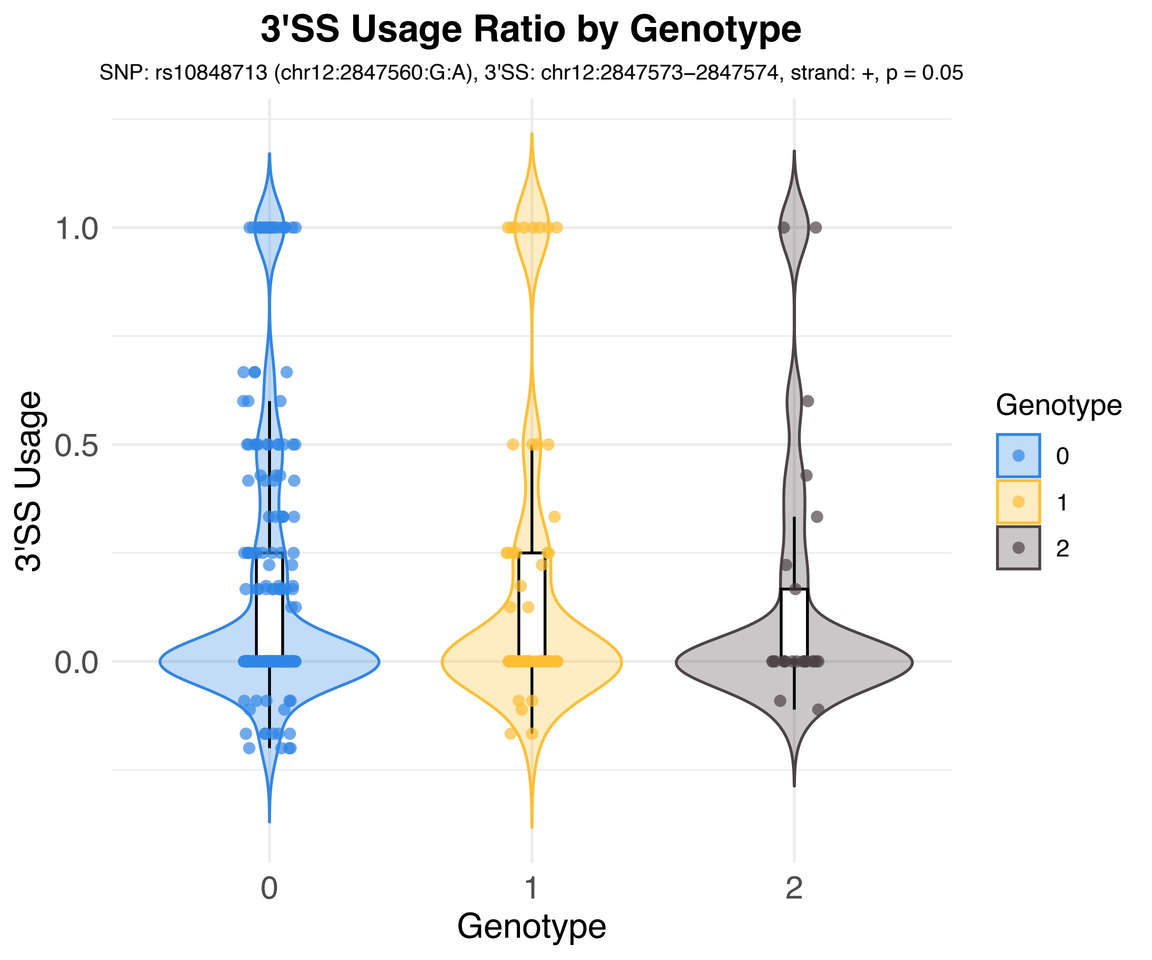

3'SS Usage Ratio by Genotype
SNP: rs10848713 (chr12:2847560:G:A), 3'SS: chr12:2847573–2847574, strand: +, p = 0.05
3'SS Usage
1.0
0.5
0.0
0
1
2
Genotype
Genotype
0
1
2

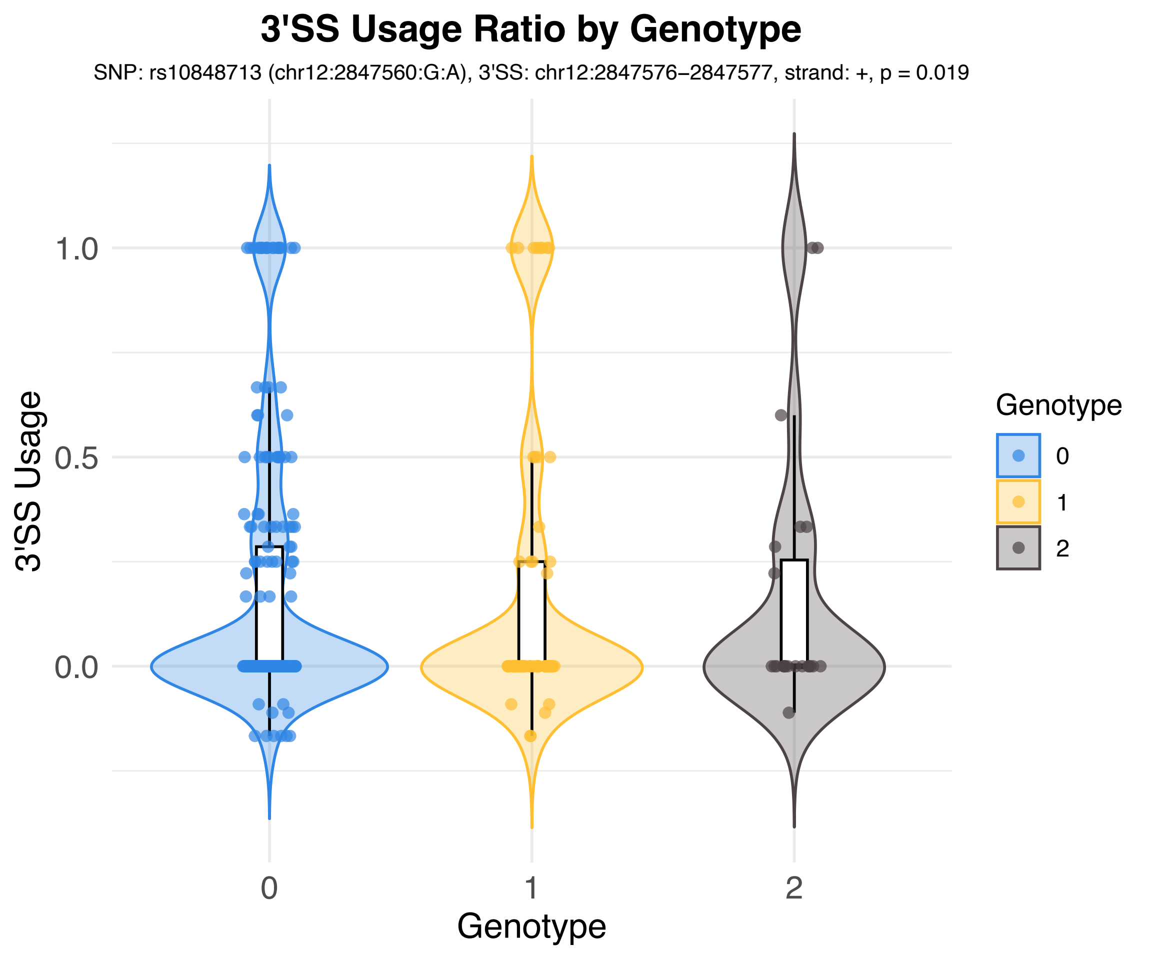
3'SS Usage Ratio by Genotype
SNP: rs10848713 (chr12:2847560:G:A), 3'SS: chr12:2847576–2847577, strand: +, p = 0.019
3'SS Usage
1.0
0.5
0.0
0
1
2
Genotype
Genotype
0
1
2

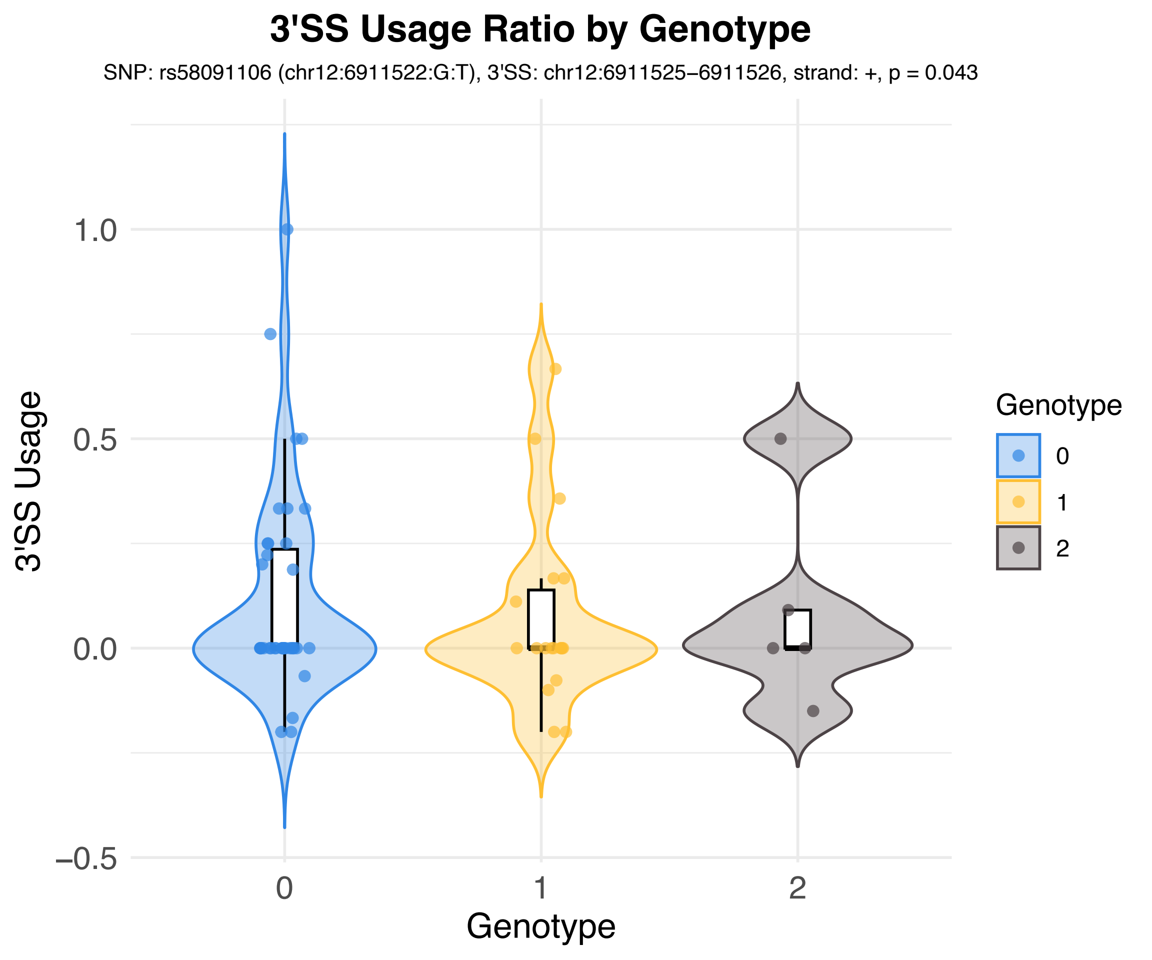
3'SS Usage Ratio by Genotype
SNP: rs58091106 (chr12:6911522:G:T), 3'SS: chr12:6911525–6911526, strand: +, p = 0.043
3'SS Usage
1.0
0.5
0.0
−0.5
0
1
2
Genotype
Genotype
0
1
2

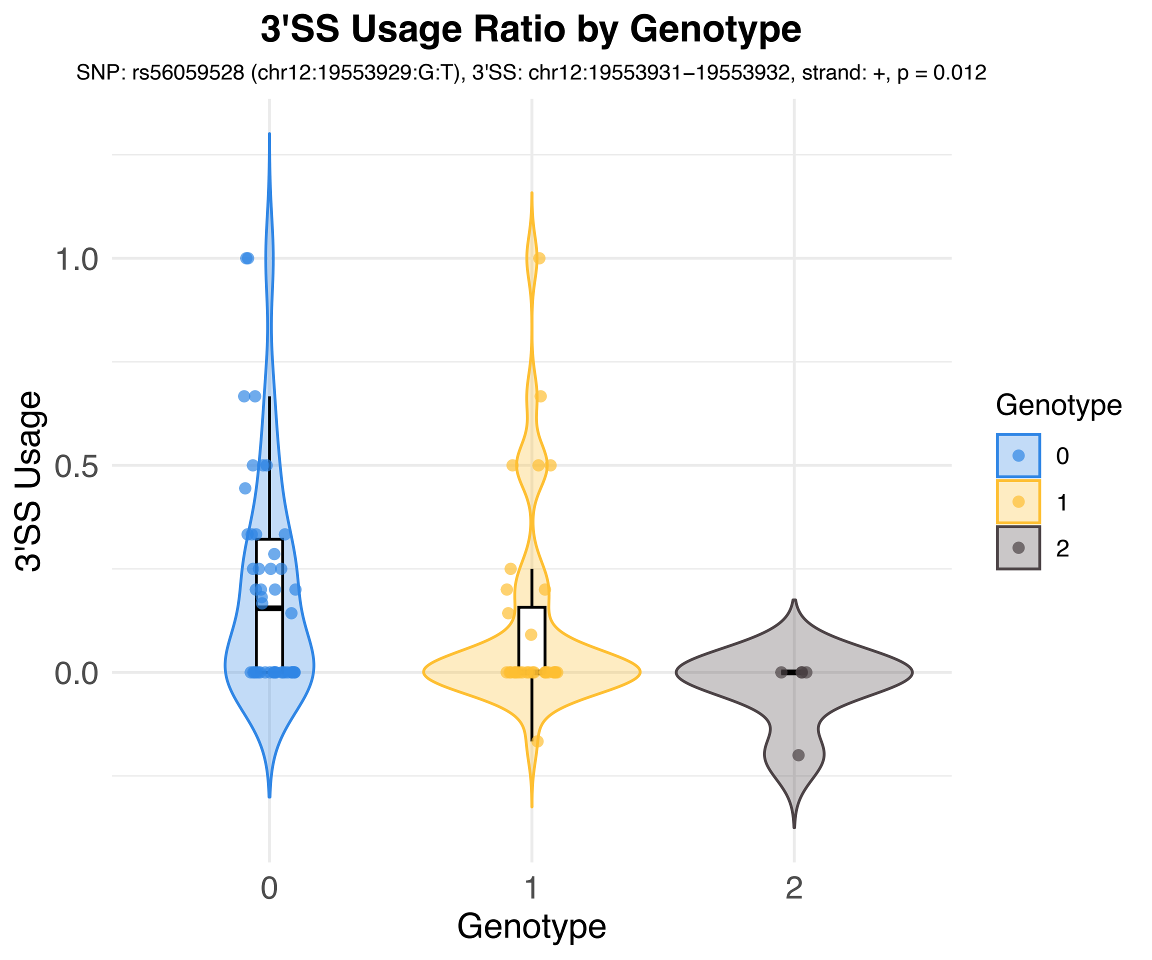
3'SS Usage Ratio by Genotype
SNP: rs56059528 (chr12:19553929:G:T), 3'SS: chr12:19553931−19553932, strand: +, p = 0.012
3'SS Usage
1.0
0.5
0.0
0
1
2
Genotype
Genotype
0
1
2

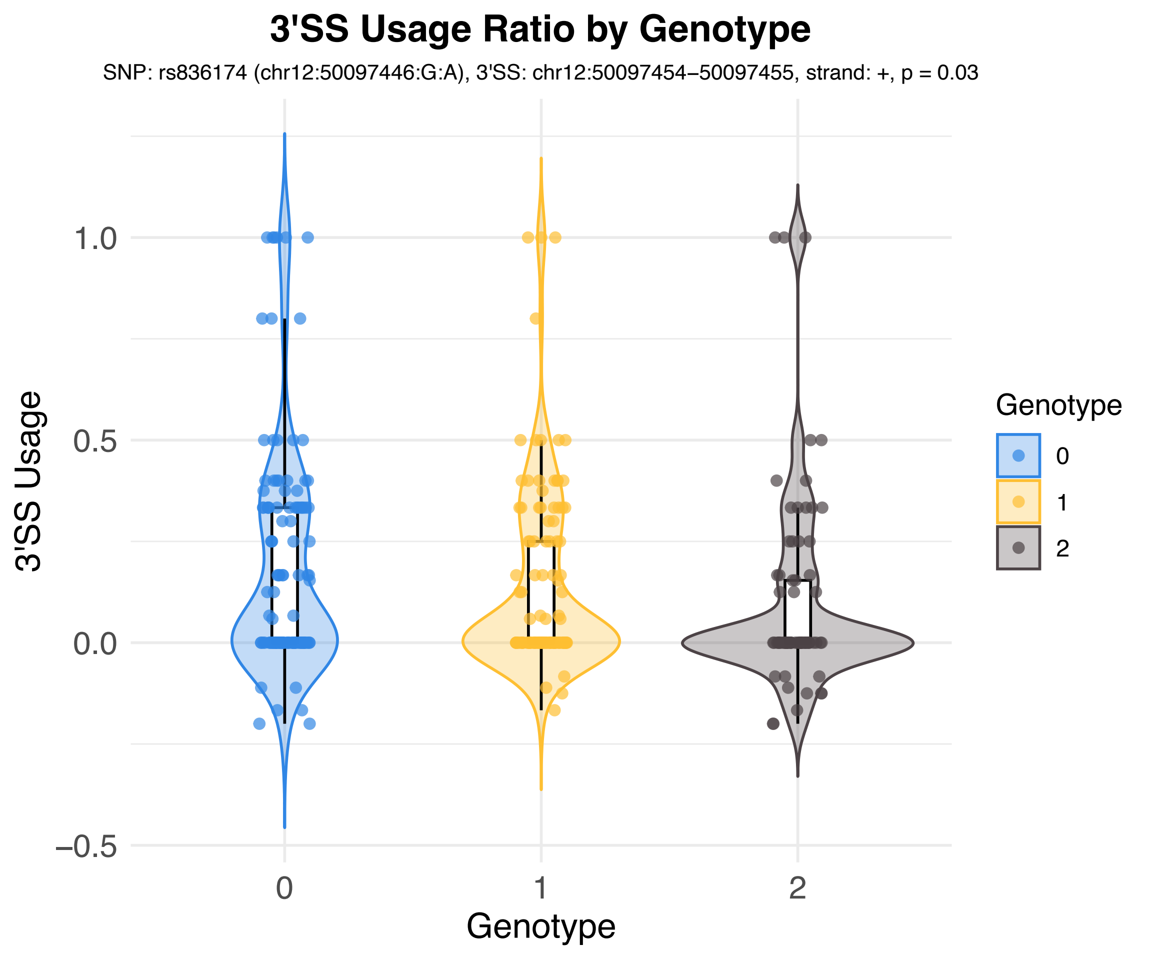

3'SS Usage Ratio by Genotype
SNP: rs836174 (chr12:50097446:G:A), 3'SS: chr12:50097454–50097455, strand: +, p = 0.03
3'SS Usage
1.0
0.5
0.0
−0.5
0
1
2
Genotype
Genotype
0
1
2

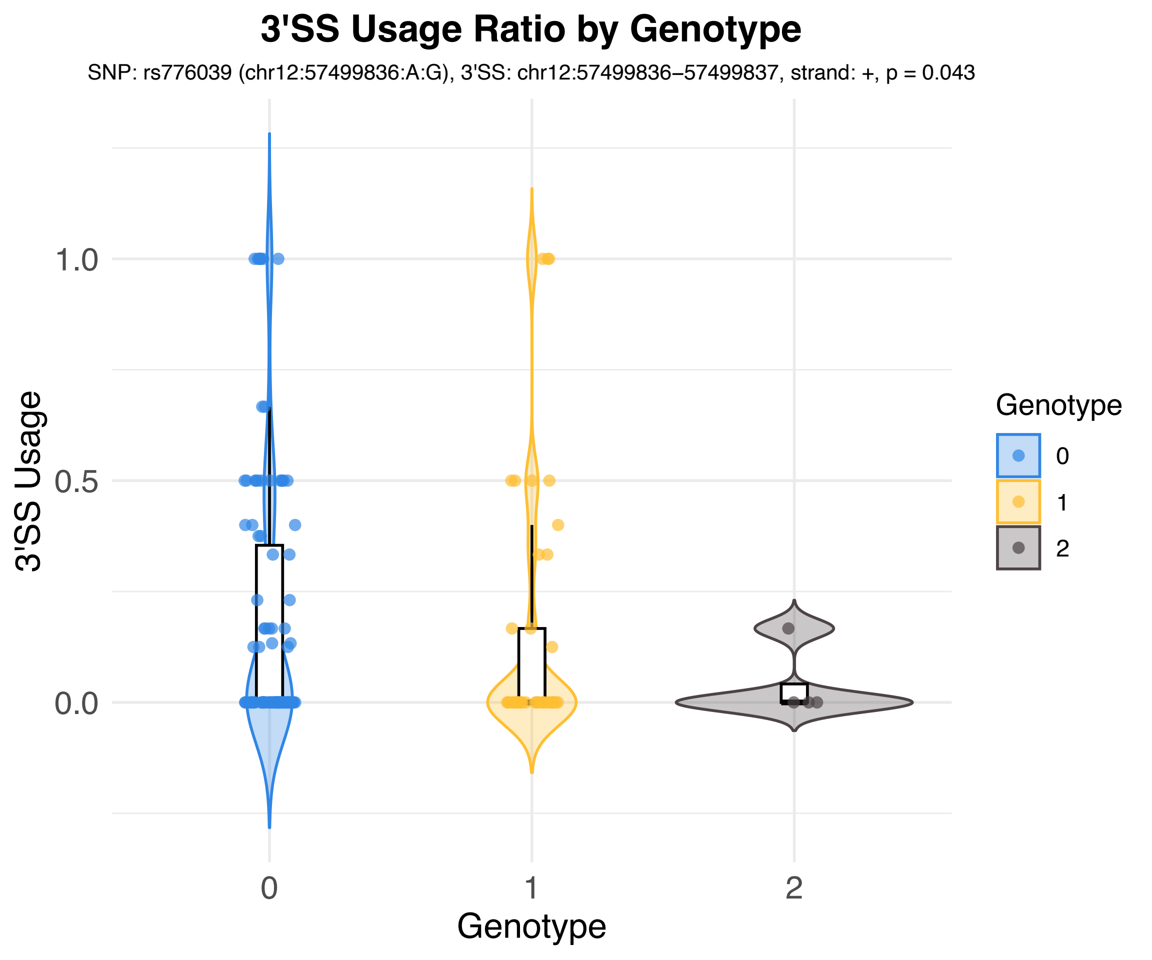
3'SS Usage Ratio by Genotype
SNP: rs776039 (chr12:57499836:A:G), 3'SS: chr12:57499836–57499837, strand: +, p = 0.043
3'SS Usage
1.0
0.5
0.0
0
1
2
Genotype
Genotype
0
1
2

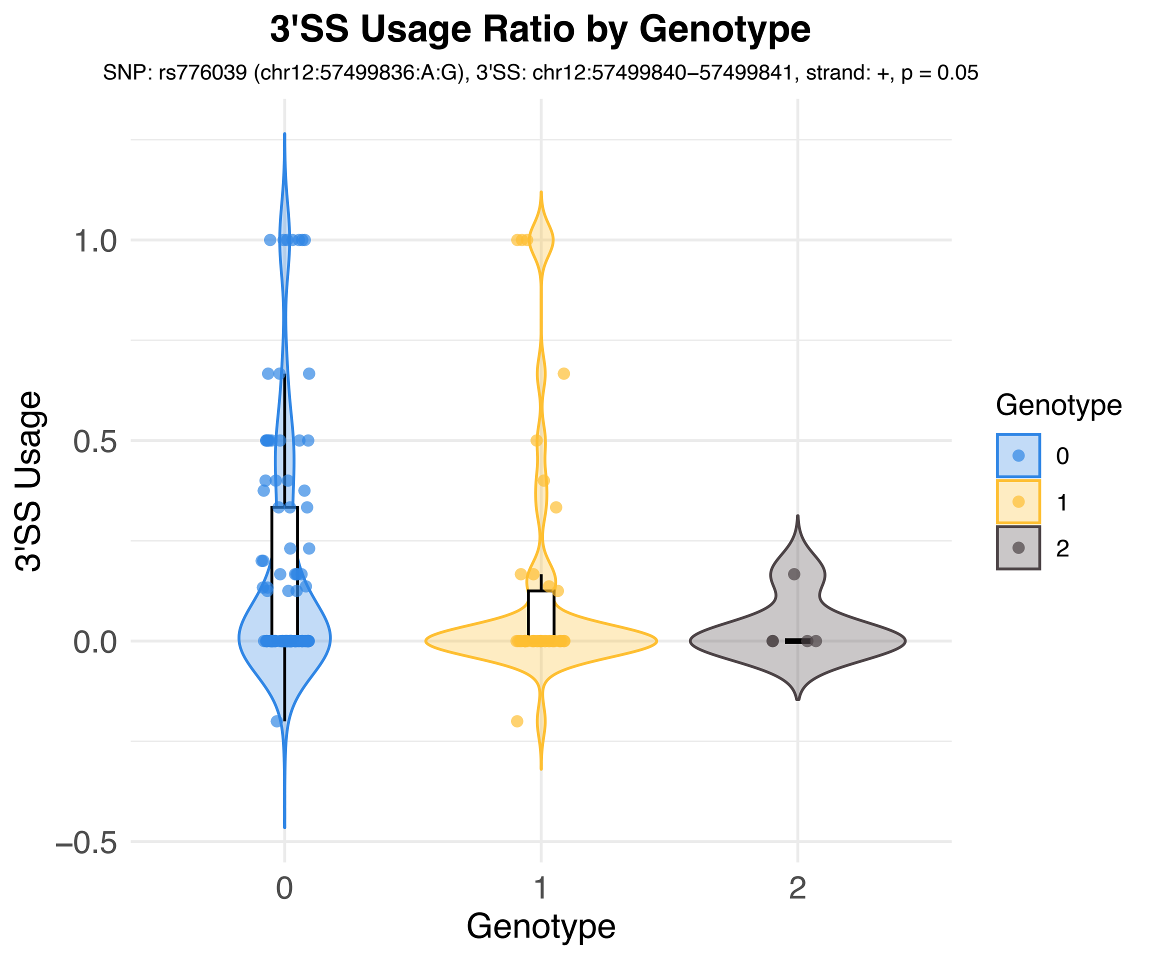
3'SS Usage Ratio by Genotype
SNP: rs776039 (chr12:57499836:A:G), 3'SS: chr12:57499840–57499841, strand: +, p = 0.05
3'SS Usage
1.0
0.5
0.0
−0.5
0
1
2
Genotype
Genotype
0
1
2

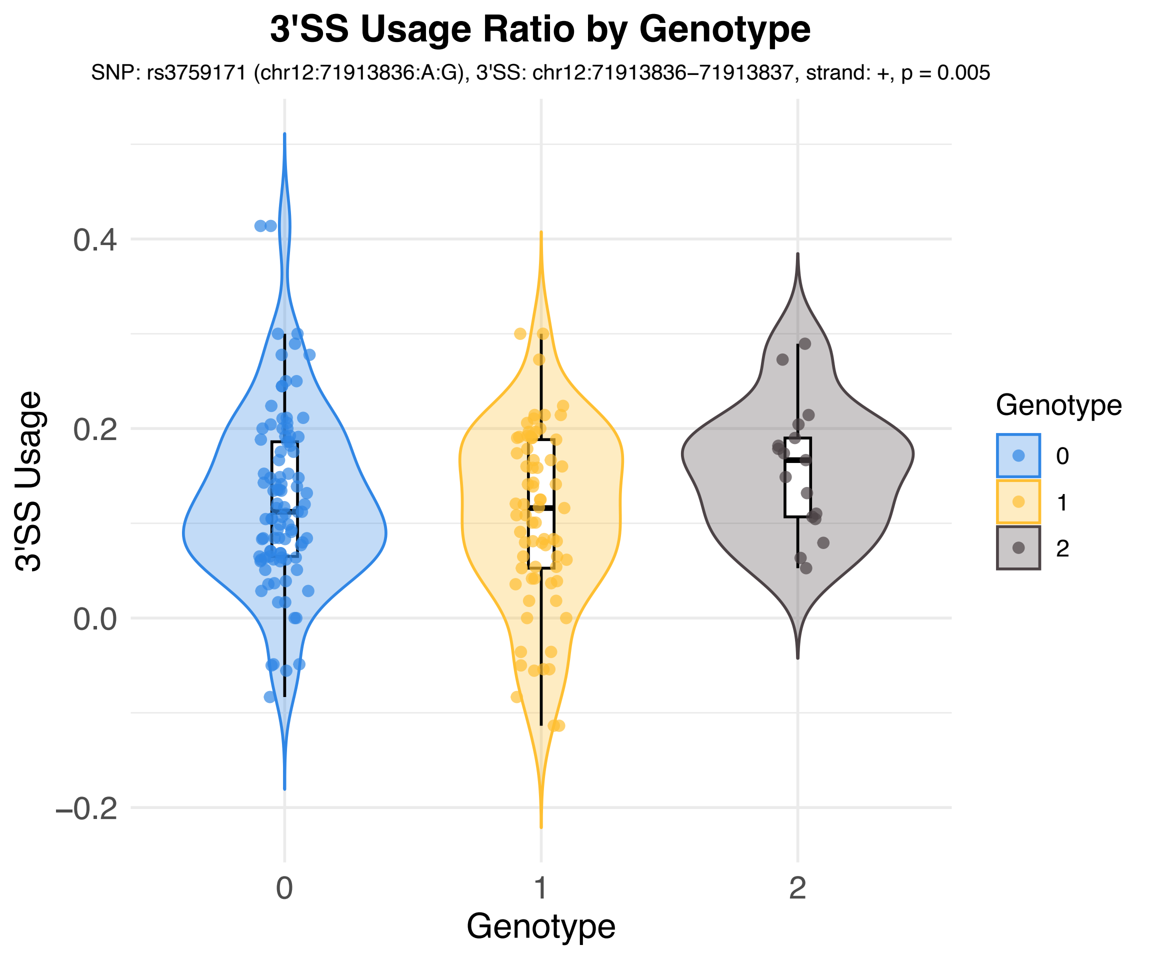
3'SS Usage Ratio by Genotype
SNP: rs3759171 (chr12:71913836:A:G), 3'SS: chr12:71913836–71913837, strand: +, p = 0.005
3'SS Usage
0.4
0.2
0.0
−0.2
0
1
2
Genotype
Genotype
0
1
2

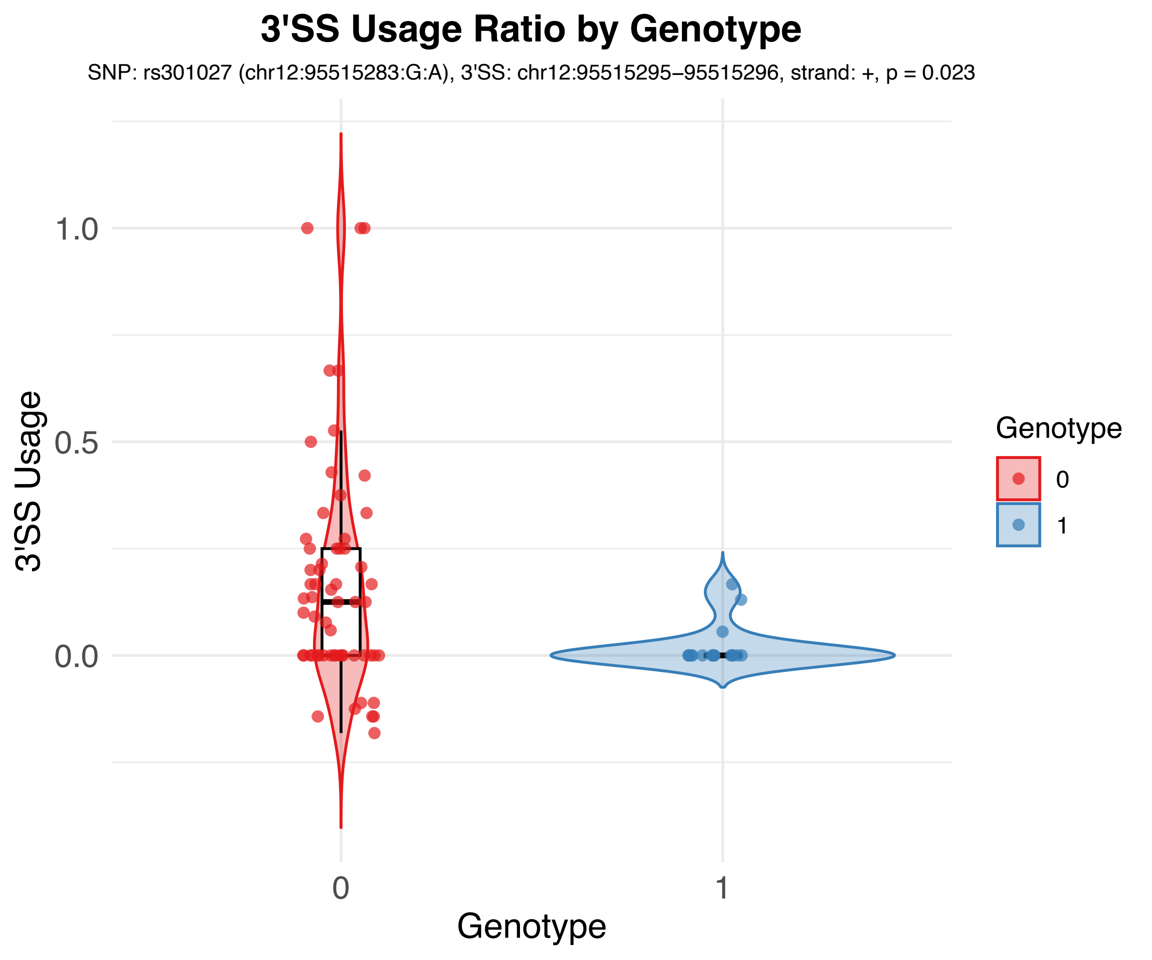

3'SS Usage Ratio by Genotype
SNP: rs301027 (chr12:95515283:G:A), 3'SS: chr12:95515295–95515296, strand: +, p = 0.023
3'SS Usage
1.0
0.5
0.0
0
1
Genotype
Genotype
0
1

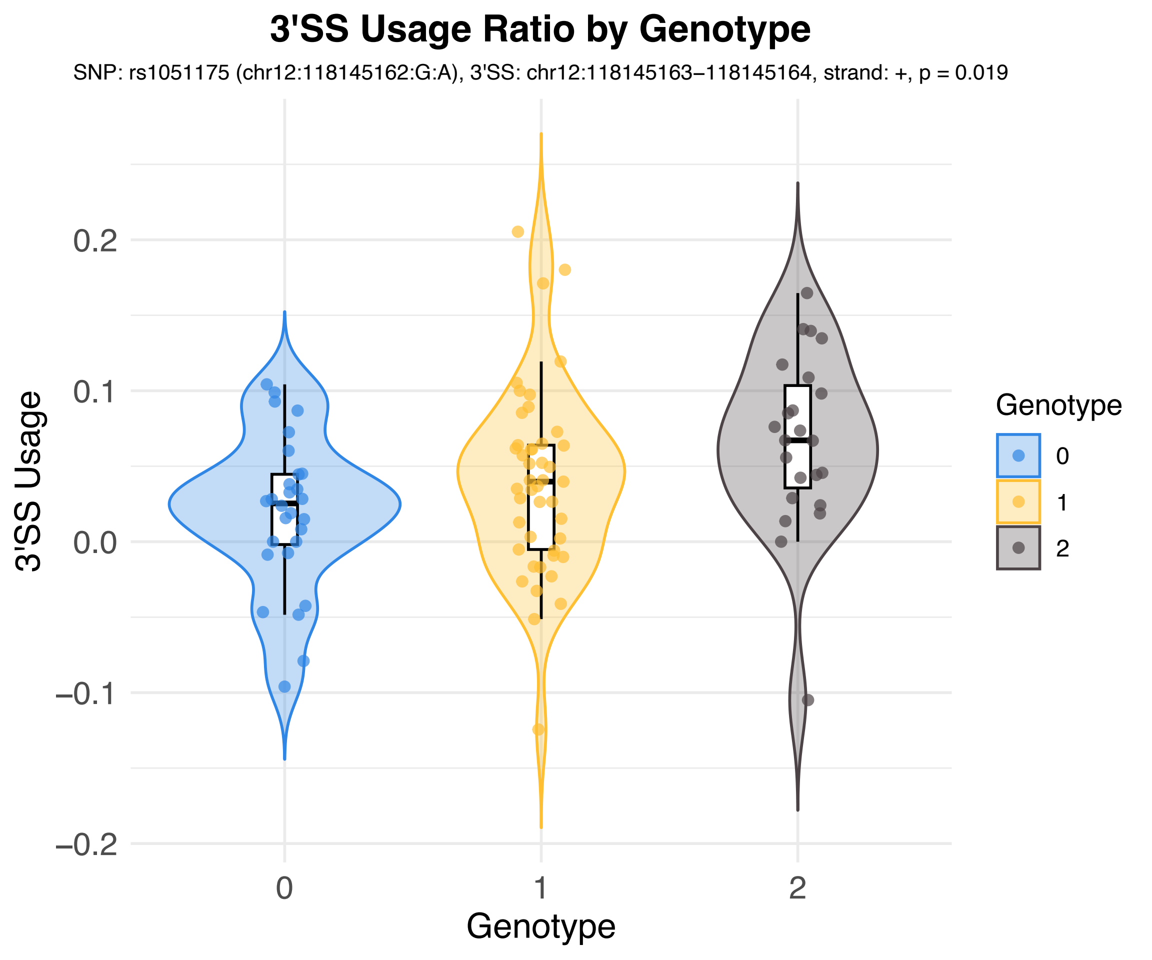

3'SS Usage Ratio by Genotype
SNP: rs1051175 (chr12:118145162:G:A), 3'SS: chr12:118145163–118145164, strand: +, p = 0.019
3'SS Usage
0.2
0.1
0.0
−0.1
−0.2
0
1
2
Genotype
Genotype
0
1
2

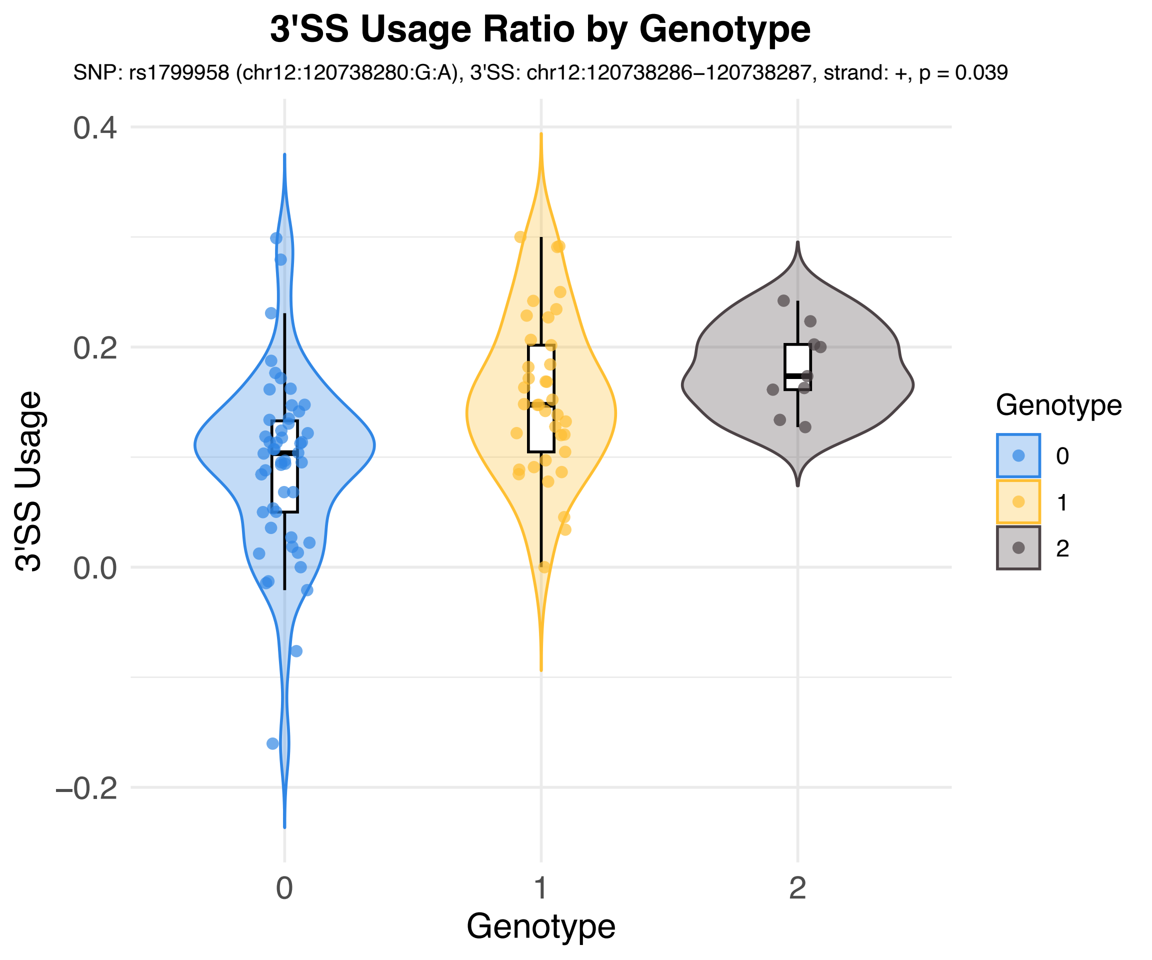
3'SS Usage Ratio by Genotype
SNP: rs1799958 (chr12:120738280:G:A), 3'SS: chr12:120738286–120738287, strand: +, p = 0.039
3'SS Usage
0.4
0.2
0.0
−0.2
Genotype
0
1
2
Genotype
0
1
2

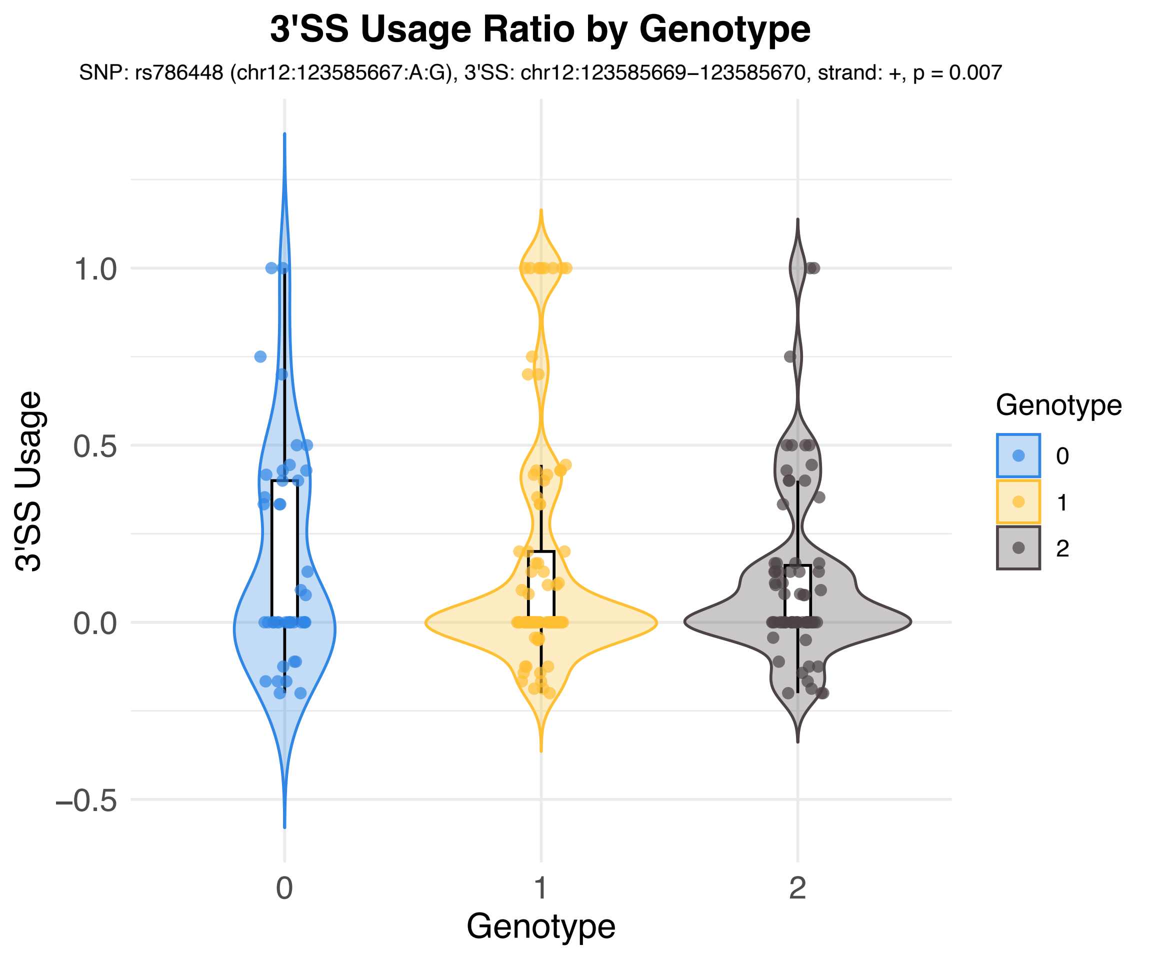
3'SS Usage Ratio by Genotype
SNP: rs786448 (chr12:123585667:A:G), 3'SS: chr12:123585669−123585670, strand: +, p = 0.007
3'SS Usage
1.0
0.5
0.0
−0.5
0
1
2
Genotype
Genotype
0
1
2

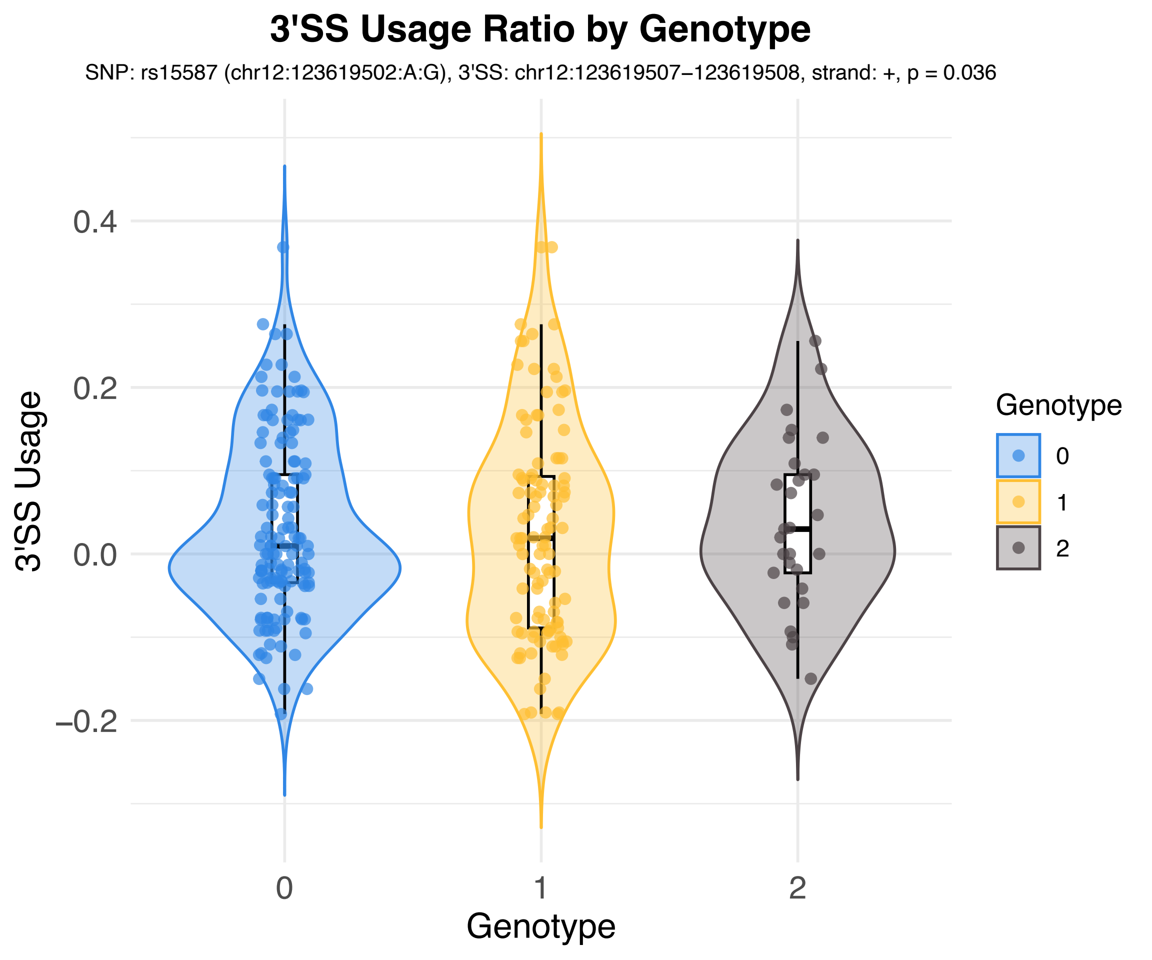
3'SS Usage Ratio by Genotype
SNP: rs15587 (chr12:123619502:A:G), 3'SS: chr12:123619507−123619508, strand: +, p = 0.036
3'SS Usage
0.4
0.2
0.0
−0.2
0
1
2
Genotype
Genotype
0
1
2

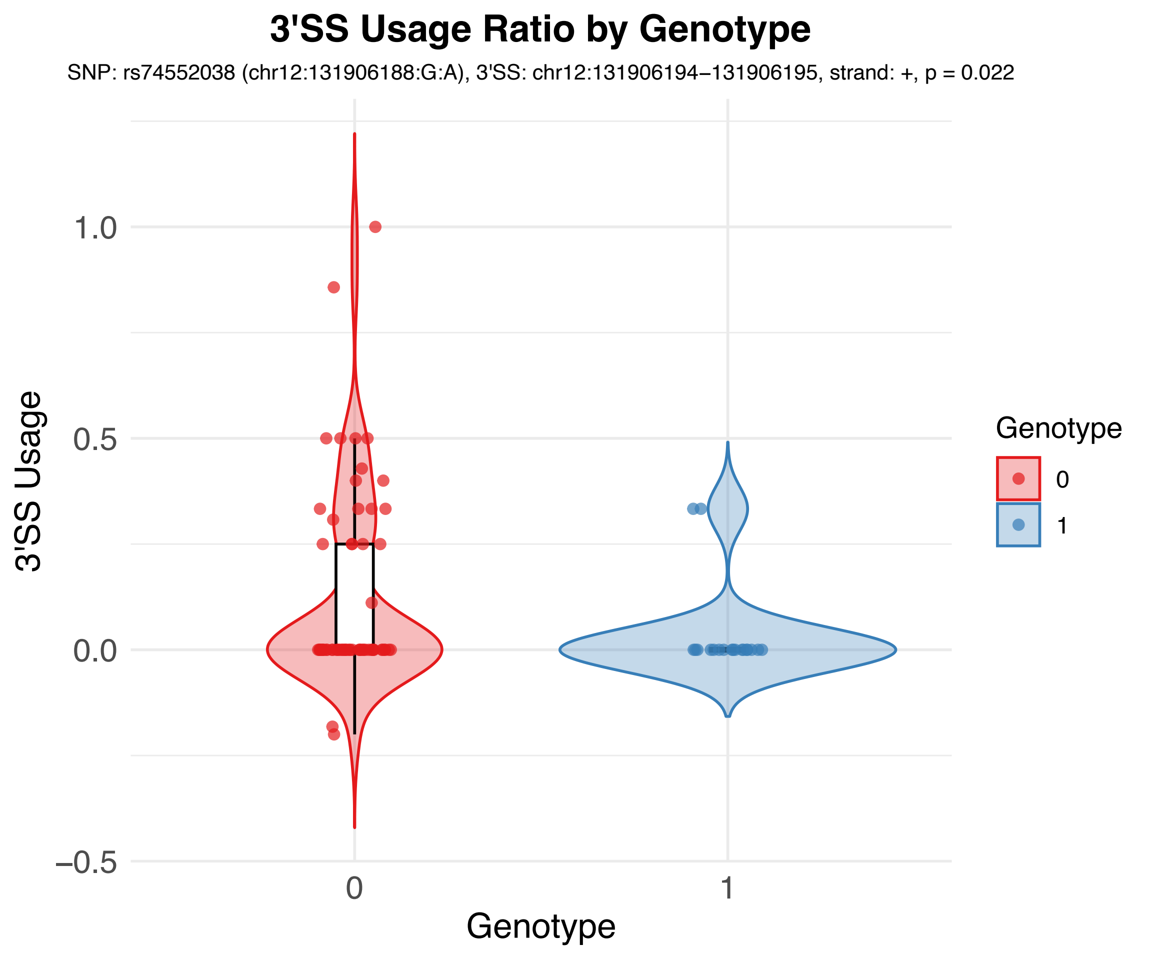

3'SS Usage Ratio by Genotype
SNP: rs74552038 (chr12:131906188:G:A), 3'SS: chr12:131906194−131906195, strand: +, p = 0.022
3'SS Usage
1.0
0.5
0.0
−0.5
0
1
Genotype
Genotype
0
1

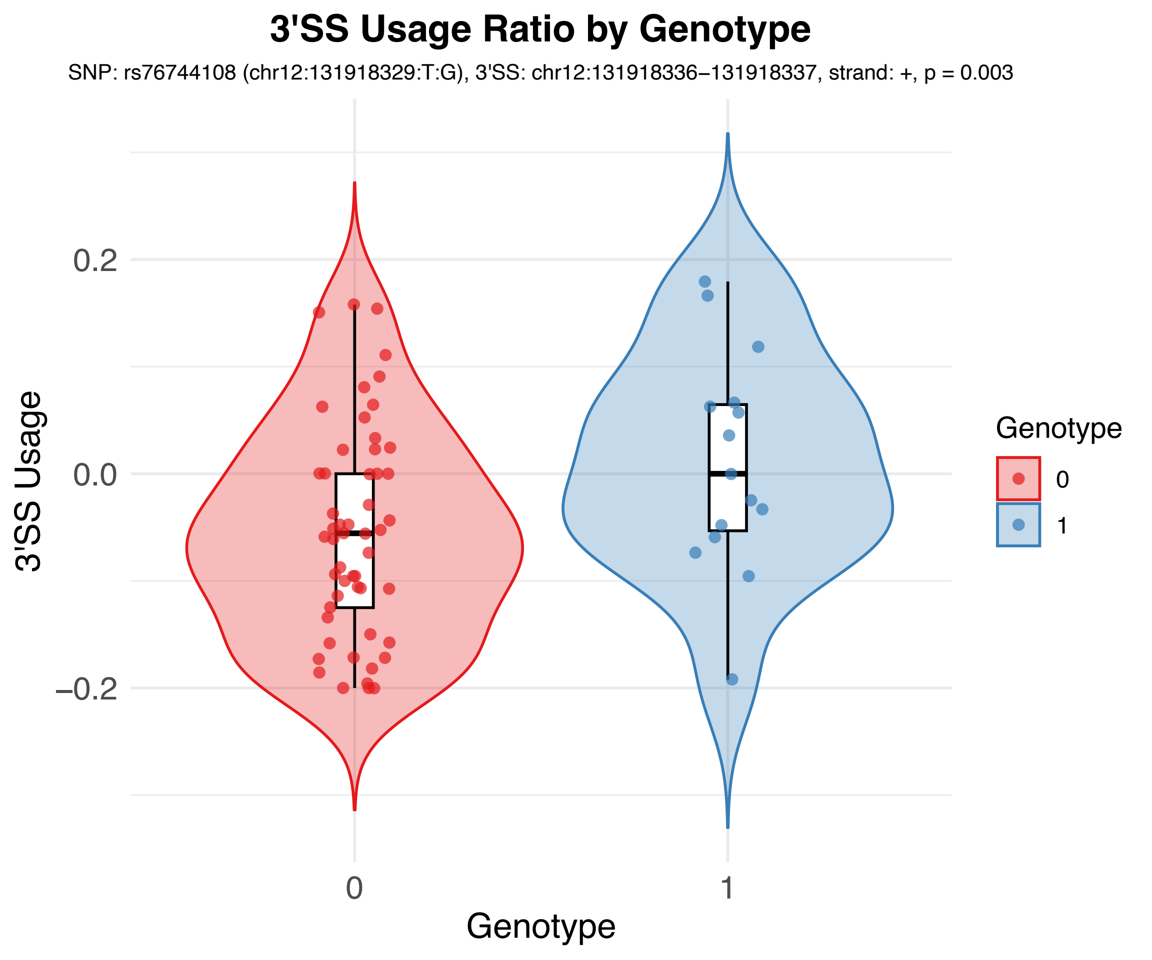
3'SS Usage Ratio by Genotype
SNP: rs76744108 (chr12:131918329:T:G), 3'SS: chr12:131918336−131918337, strand: +, p = 0.003
3'SS Usage
0.2
0.0
−0.2
0
1
Genotype
Genotype
0
1

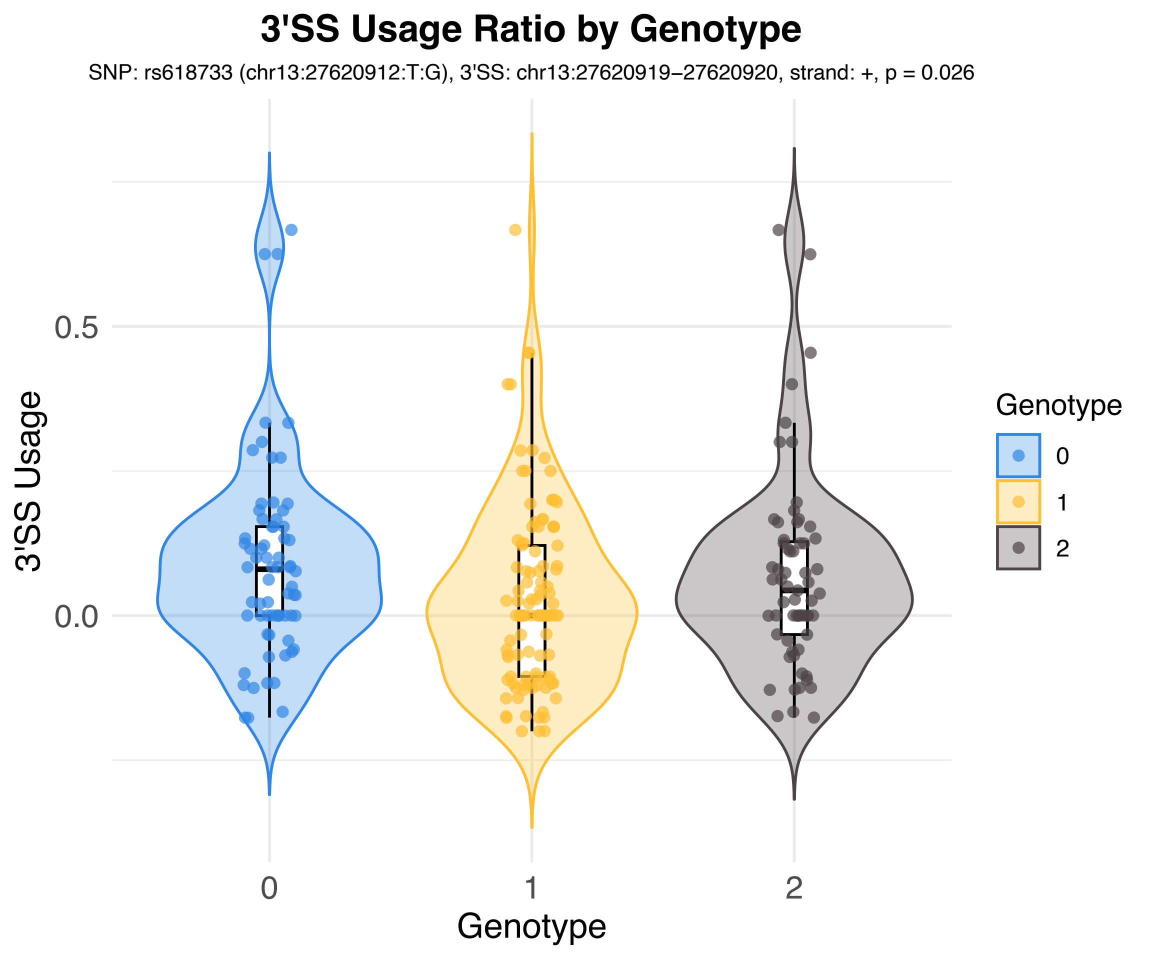

3'SS Usage Ratio by Genotype
SNP: rs618733 (chr13:27620912:T:G), 3'SS: chr13:27620919–27620920, strand: +, p = 0.026
3'SS Usage
0.5
0.0
0
1
2
Genotype
Genotype
0
1
2

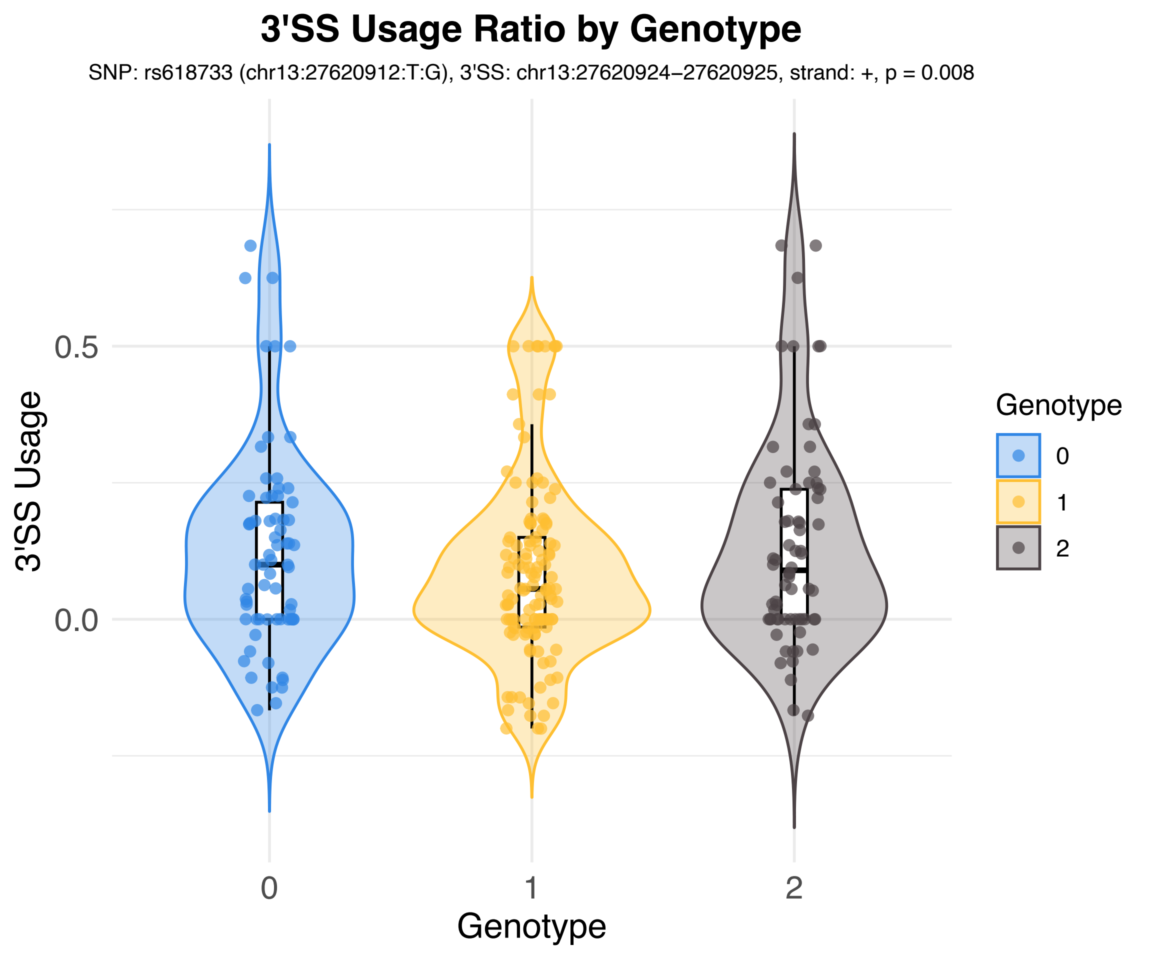

3'SS Usage Ratio by Genotype
SNP: rs618733 (chr13:27620912:T:G), 3'SS: chr13:27620924–27620925, strand: +, p = 0.008
3'SS Usage
0.5
0.0
0
1
2
Genotype
Genotype
0
1
2

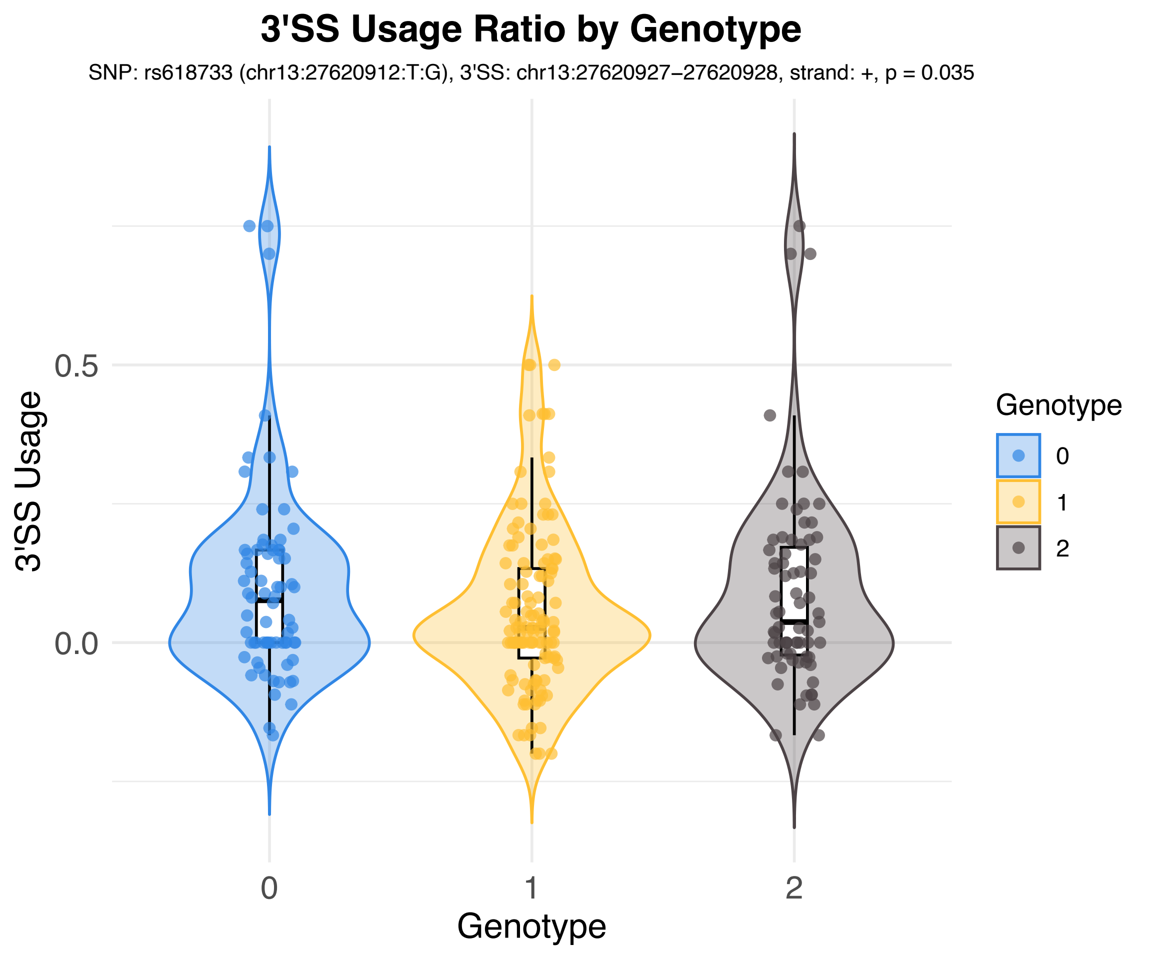

3'SS Usage Ratio by Genotype
SNP: rs618733 (chr13:27620912:T:G), 3'SS: chr13:27620927–27620928, strand: +, p = 0.035
3'SS Usage
0.5
0.0
0
1
2
Genotype
Genotype
0
1
2

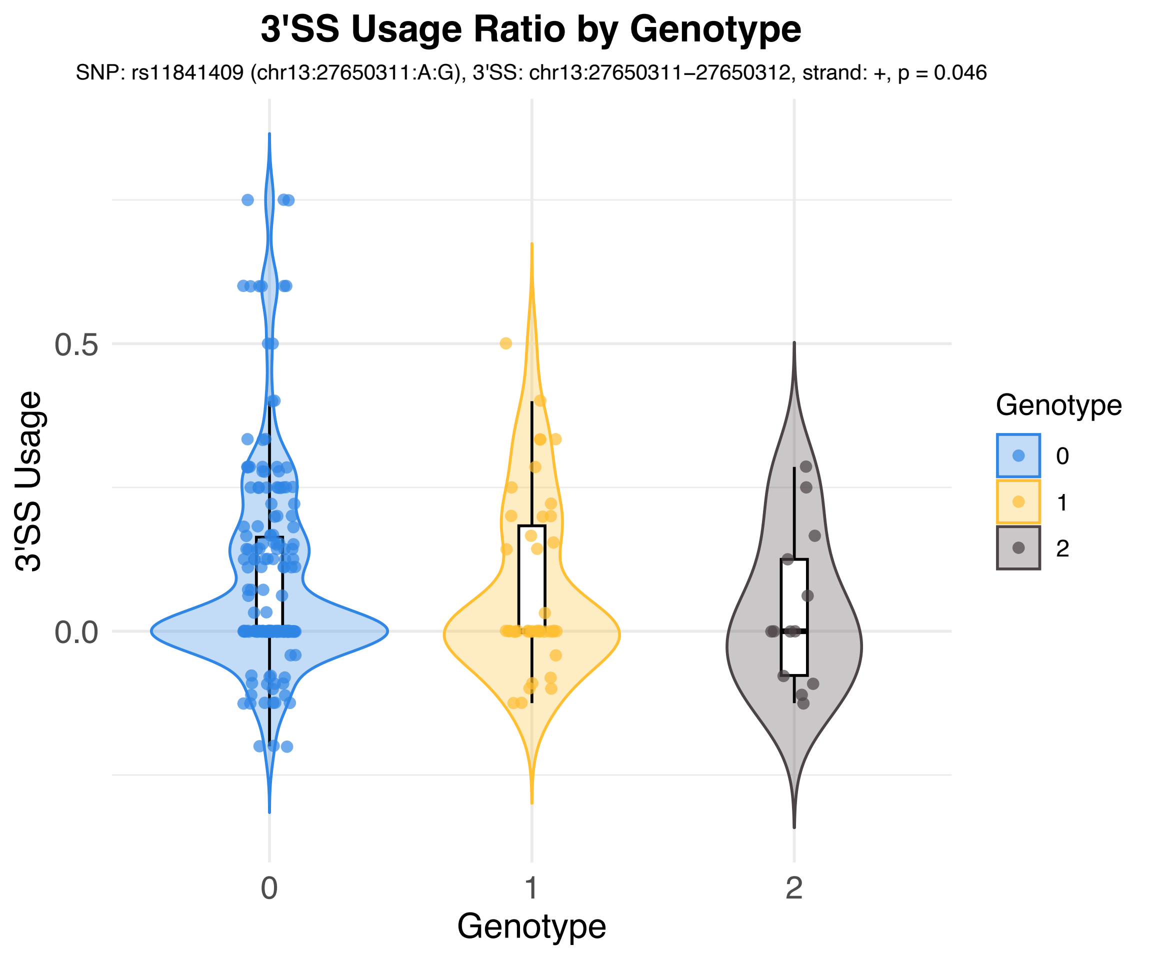
3'SS Usage Ratio by Genotype
SNP: rs11841409 (chr13:27650311:A:G), 3'SS: chr13:27650311–27650312, strand: +, p = 0.046
3'SS Usage
0.5
0.0
0
1
2
Genotype
Genotype
0
1
2

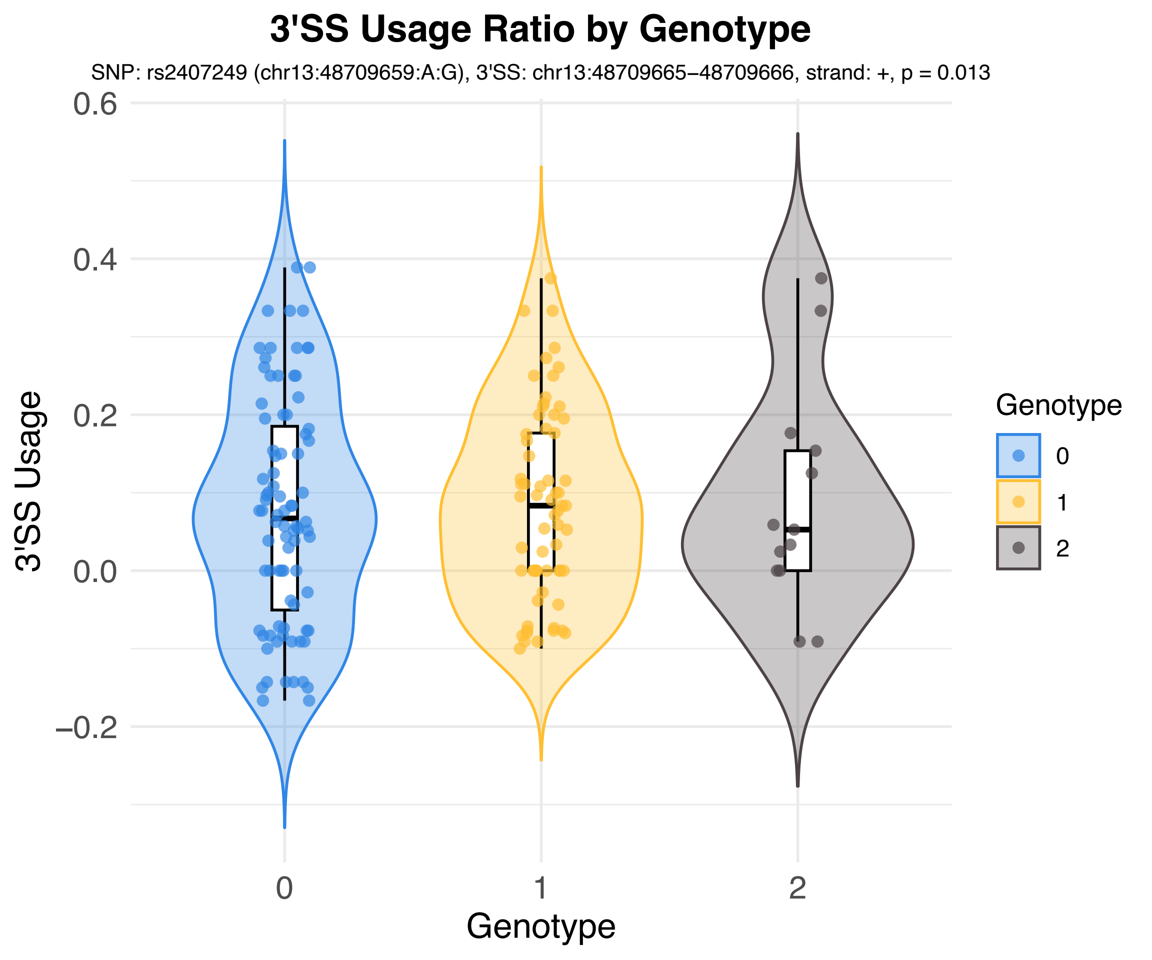
3'SS Usage Ratio by Genotype
SNP: rs2407249 (chr13:48709659:A:G), 3'SS: chr13:48709665–48709666, strand: +, p = 0.013
0.6
0.4
0.2
0.0
−0.2
3'SS Usage
0
1
2
Genotype
Genotype
0
1
2

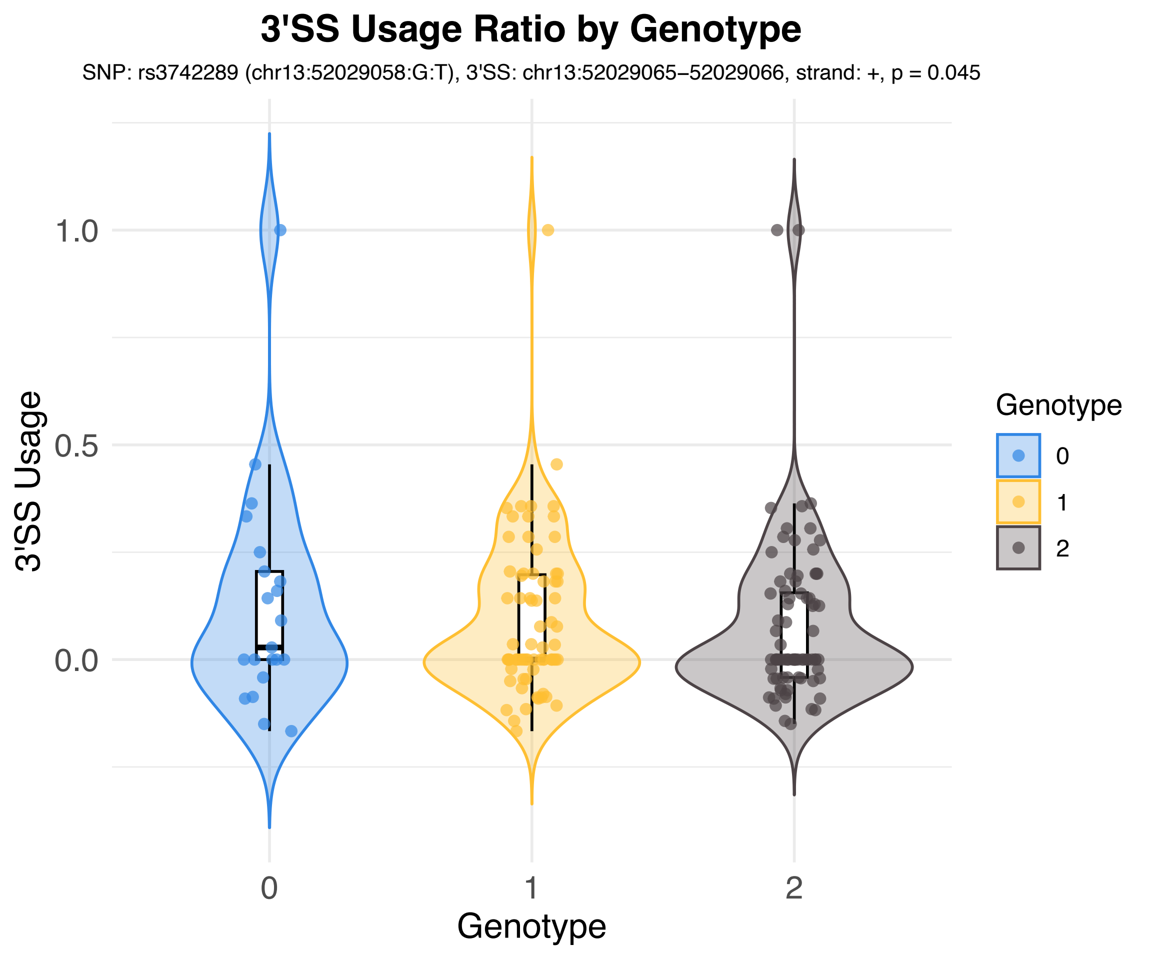

3'SS Usage Ratio by Genotype
SNP: rs3742289 (chr13:52029058:G:T), 3'SS: chr13:52029065–52029066, strand: +, p = 0.045
3'SS Usage
1.0
0.5
0.0
0
1
2
Genotype
Genotype
0
1
2

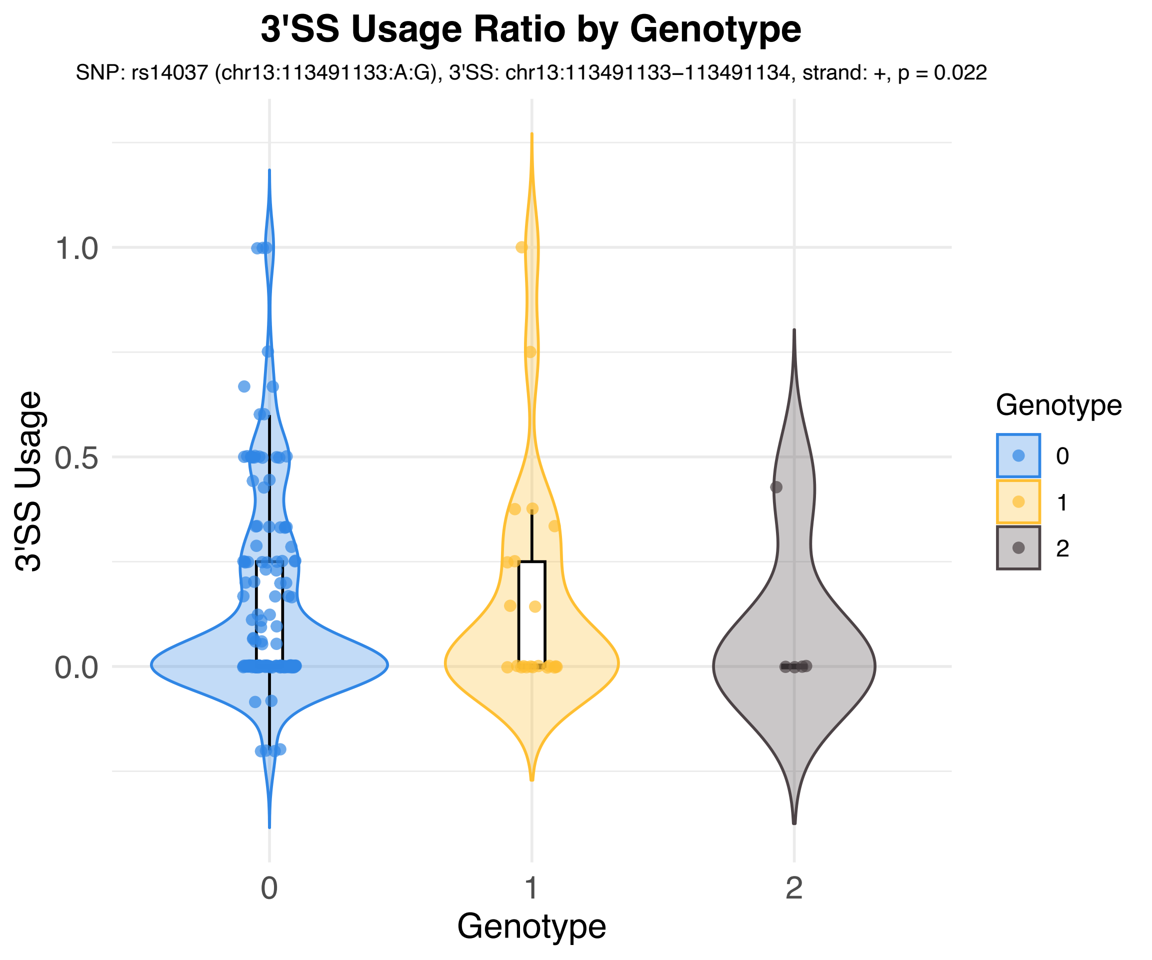
3'SS Usage Ratio by Genotype
SNP: rs14037 (chr13:113491133:A:G), 3'SS: chr13:113491133–113491134, strand: +, p = 0.022
3'SS Usage
1.0
0.5
0.0
0
1
2
Genotype
Genotype
0
1
2

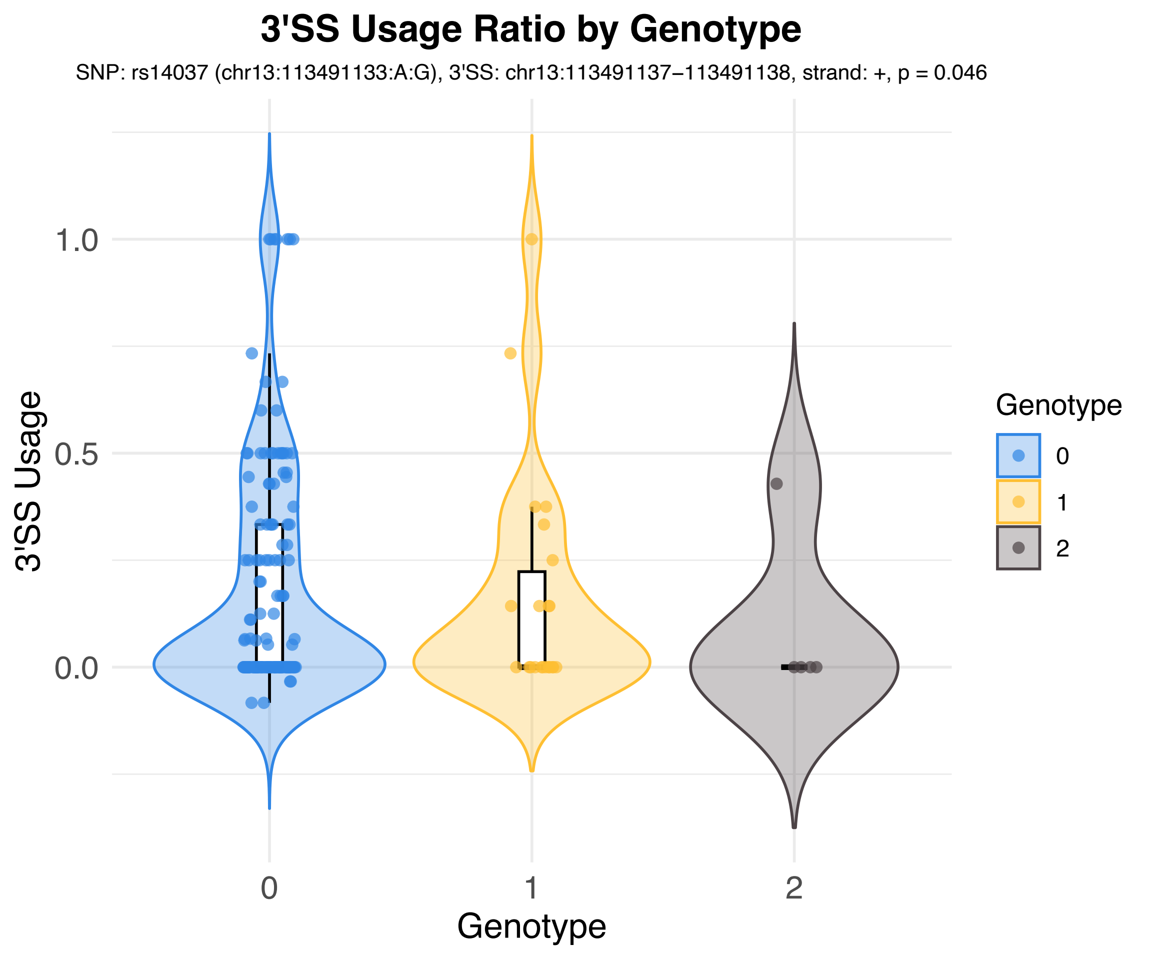
3'SS Usage Ratio by Genotype
SNP: rs14037 (chr13:113491133:A:G), 3'SS: chr13:113491137−113491138, strand: +, p = 0.046
3'SS Usage
1.0
0.5
0.0
0
1
2
Genotype
Genotype
0
1
2

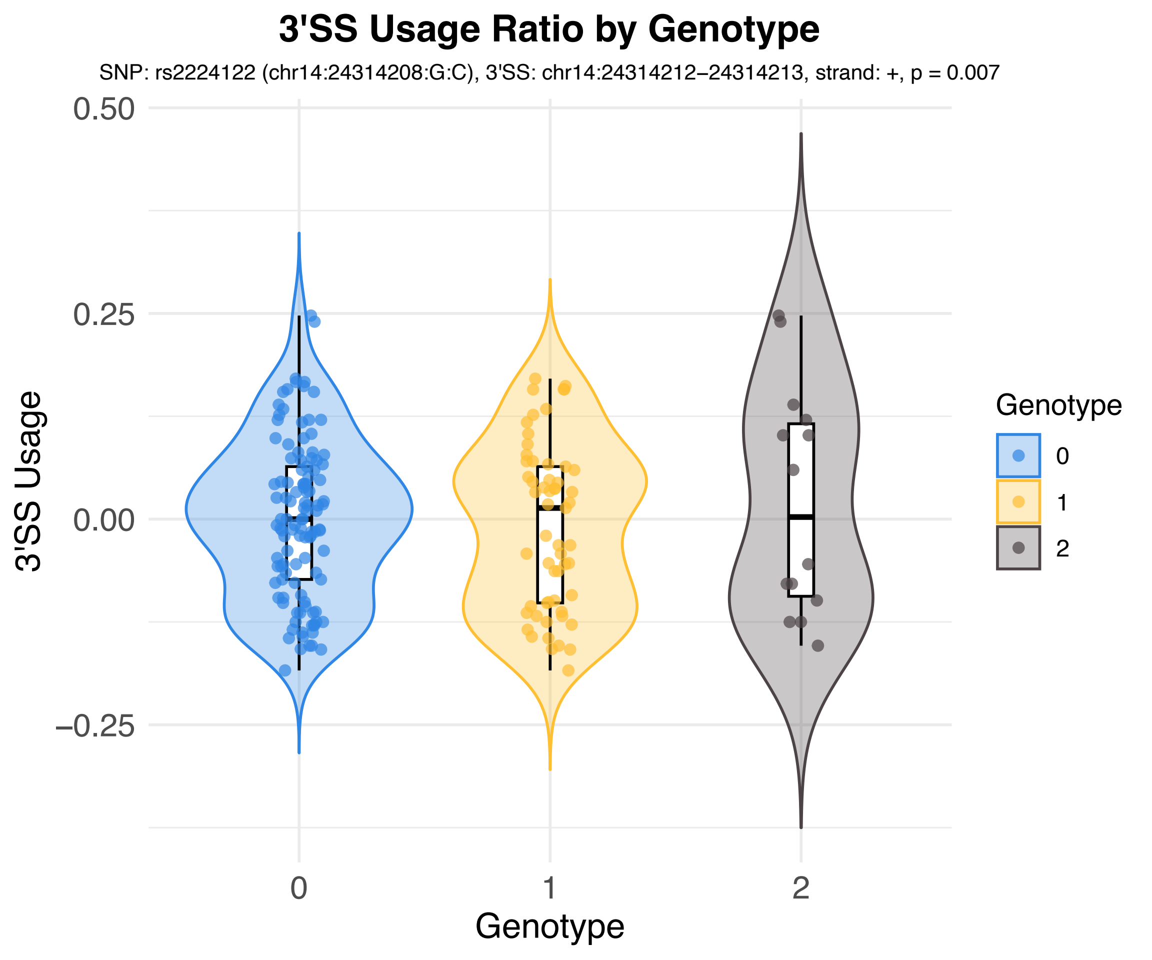

3'SS Usage Ratio by Genotype
SNP: rs2224122 (chr14:24314208:G:C), 3'SS: chr14:24314212−24314213, strand: +, p = 0.007
3'SS Usage
0.50
0.25
0.00
−0.25
0
1
2
Genotype
Genotype
0
1
2

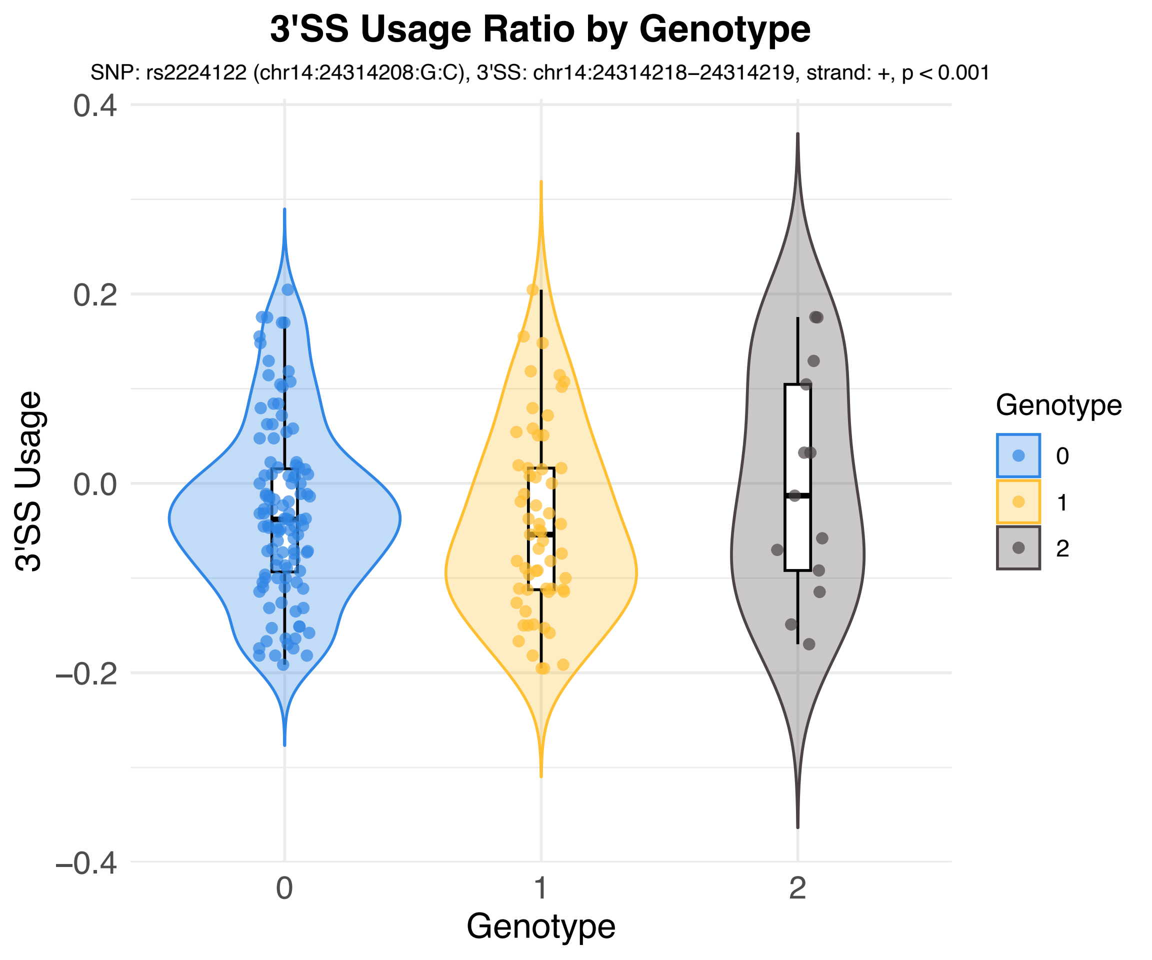
3'SS Usage Ratio by Genotype
SNP: rs2224122 (chr14:24314208:G:C), 3'SS: chr14:24314218–24314219, strand: +, p < 0.001
3'SS Usage
0.4
0.2
0.0
−0.2
−0.4
0
1
2
Genotype
Genotype
0
1
2

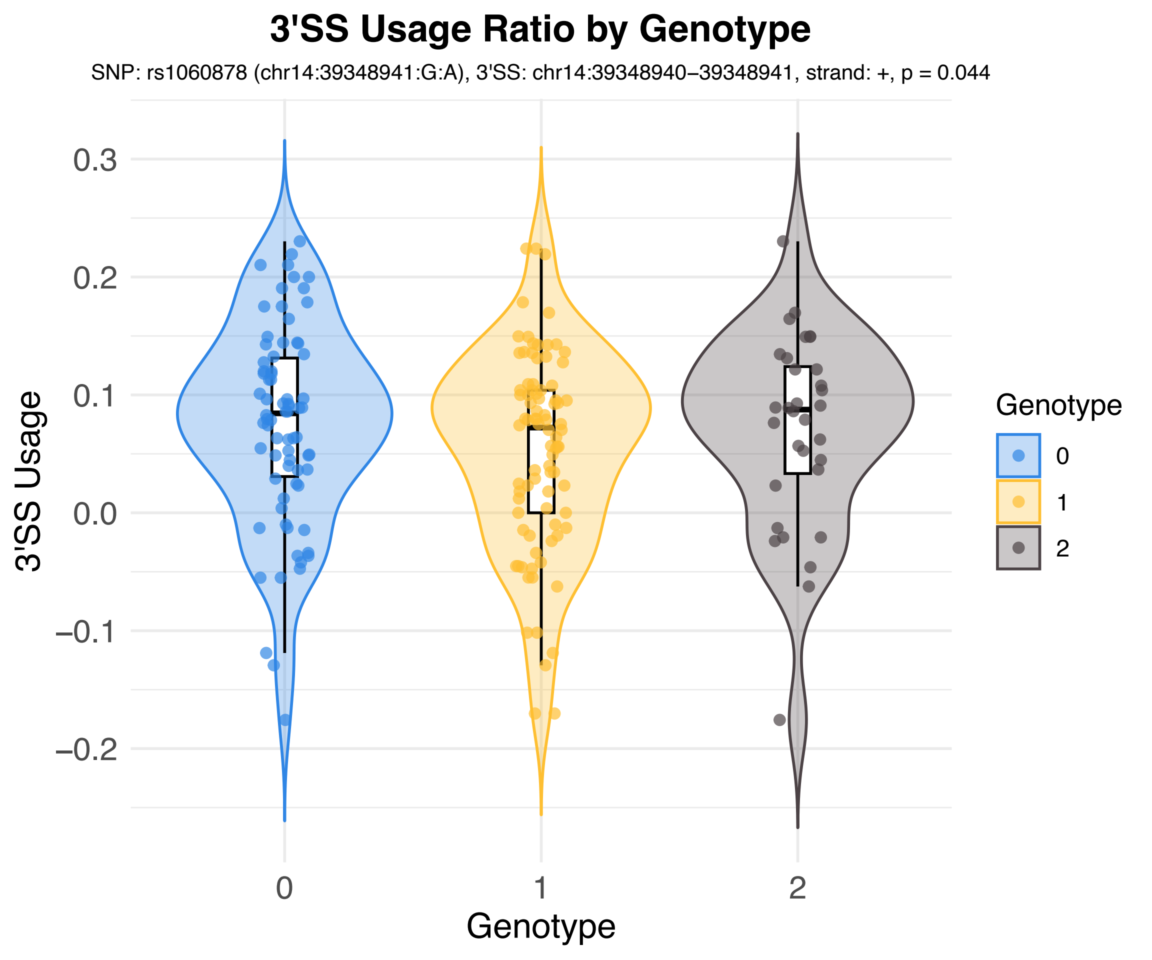
3'SS Usage Ratio by Genotype
SNP: rs1060878 (chr14:39348941:G:A), 3'SS: chr14:39348940–39348941, strand: +, p = 0.044
3'SS Usage
0.3
0.2
0.1
0.0
−0.1
−0.2
Genotype
0
1
2
Genotype
0
1
2

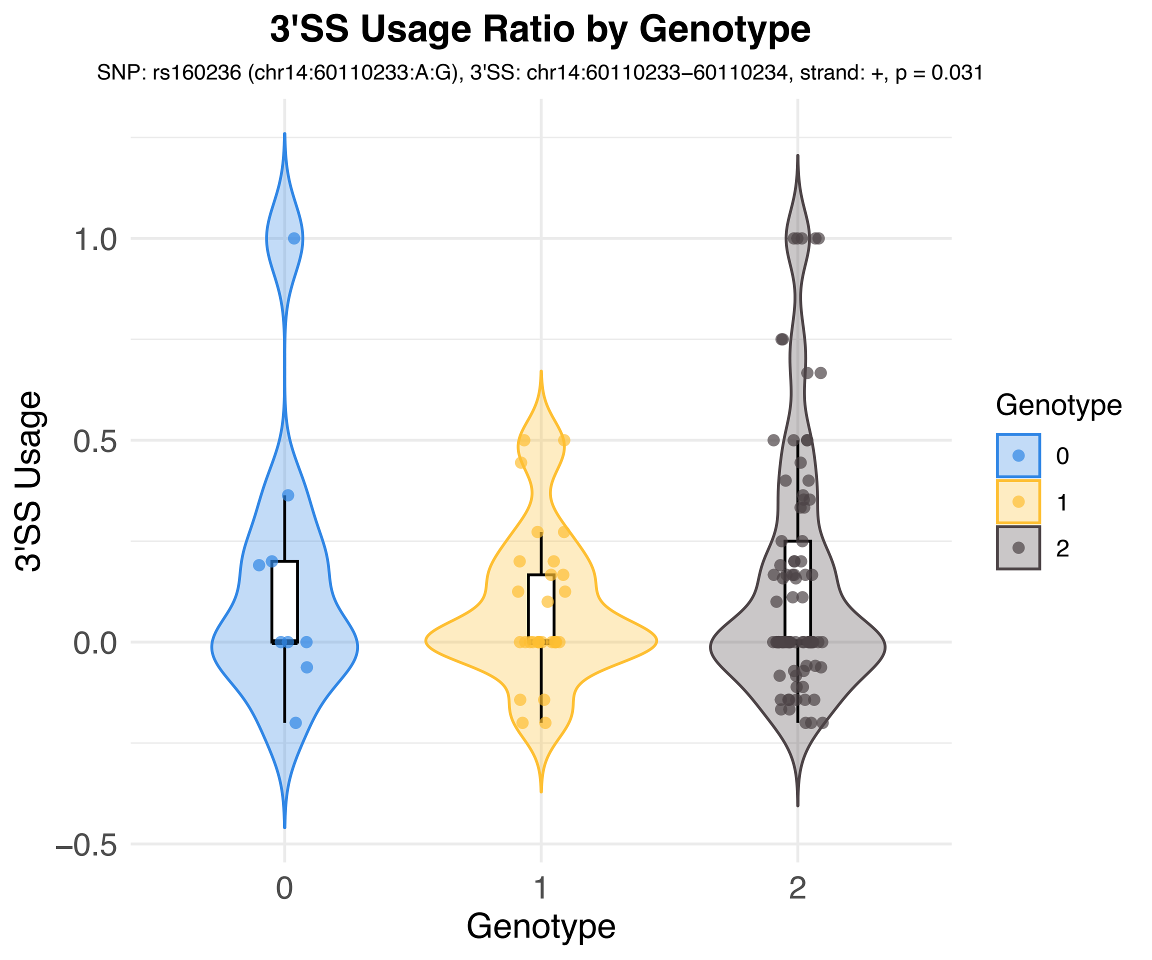
3'SS Usage Ratio by Genotype
SNP: rs160236 (chr14:60110233:A:G), 3'SS: chr14:60110233−60110234, strand: +, p = 0.031
3'SS Usage
1.0
0.5
0.0
−0.5
0
1
2
Genotype
Genotype
0
1
2

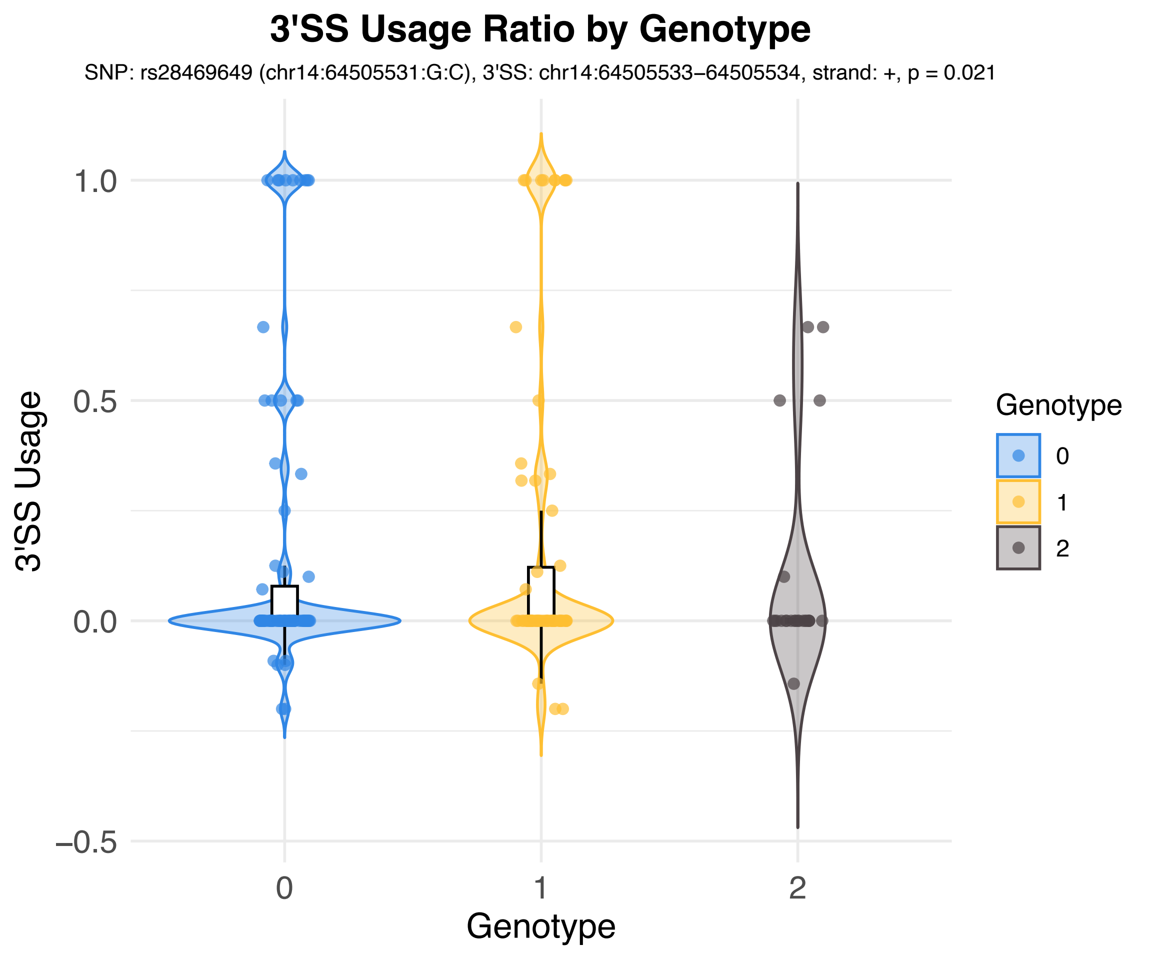

3'SS Usage Ratio by Genotype
SNP: rs28469649 (chr14:64505531:G:C), 3'SS: chr14:64505533–64505534, strand: +, p = 0.021
3'SS Usage
1.0
0.5
0.0
–0.5
0
1
2
Genotype
Genotype
0
1
2

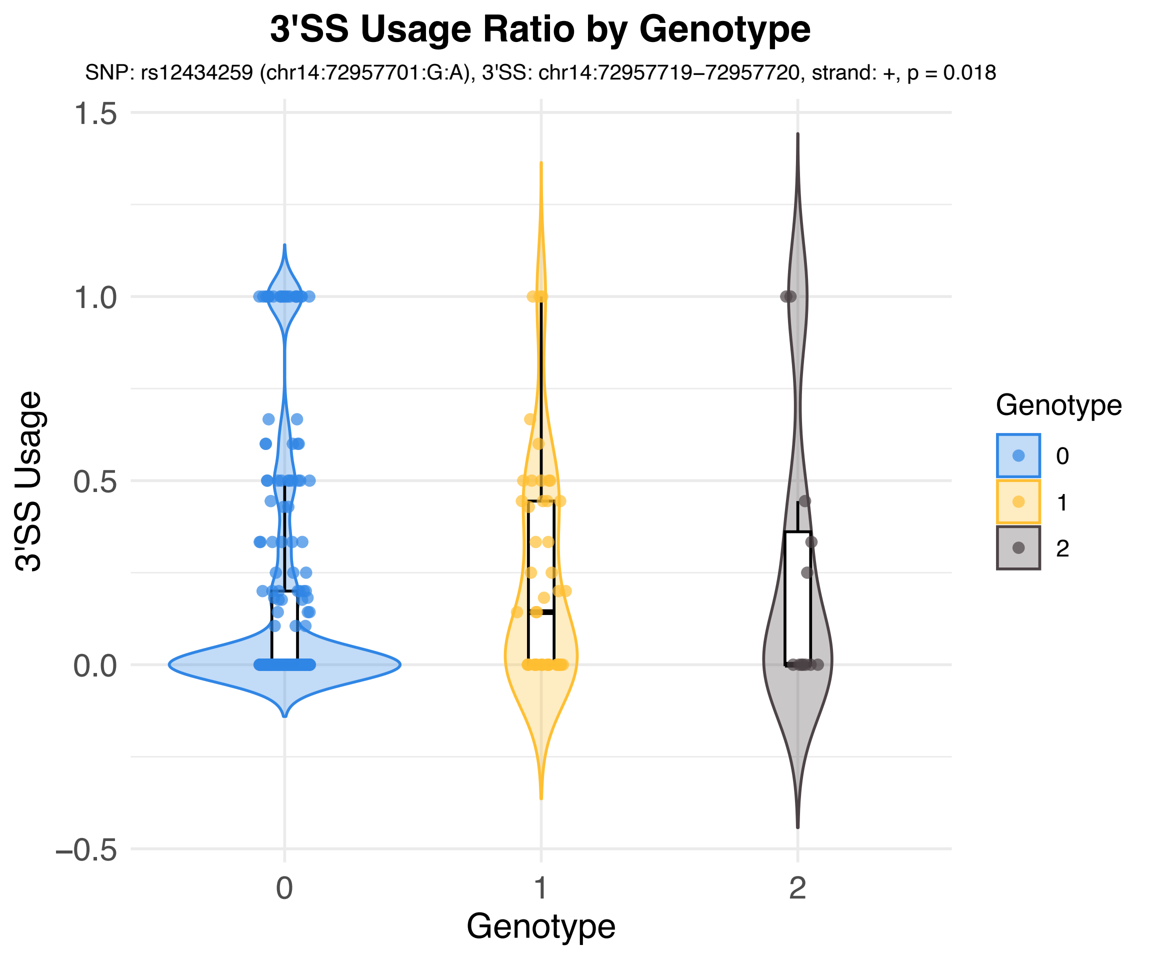

3'SS Usage Ratio by Genotype
SNP: rs12434259 (chr14:72957701:G:A), 3'SS: chr14:72957719–72957720, strand: +, p = 0.018
3'SS Usage
1.5
1.0
0.5
0.0
−0.5
0
1
2
Genotype
Genotype
0
1
2

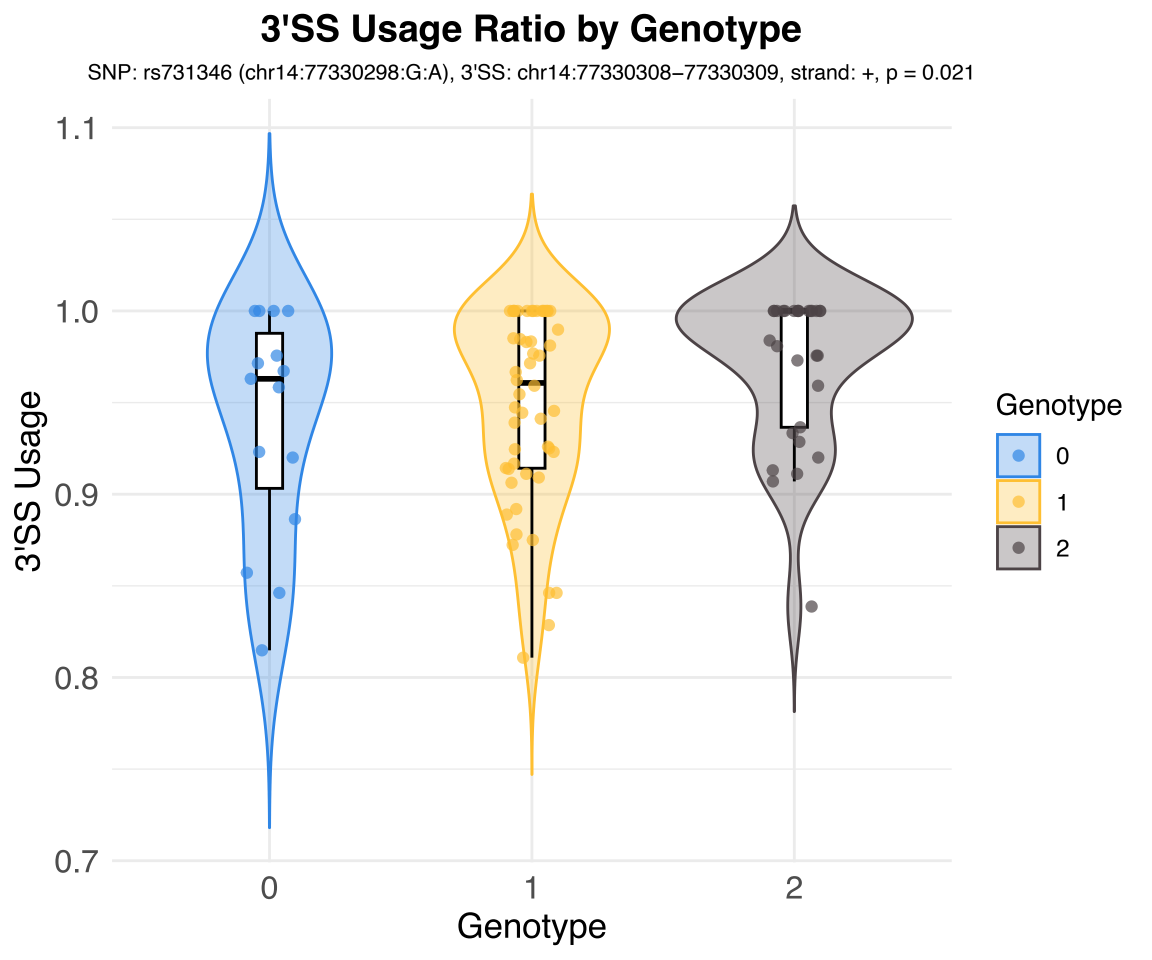

3'SS Usage Ratio by Genotype
SNP: rs731346 (chr14:77330298:G:A), 3'SS: chr14:77330308–77330309, strand: +, p = 0.021
3'SS Usage
1.1
1.0
0.9
0.8
0.7
0
1
2
Genotype
Genotype
0
1
2

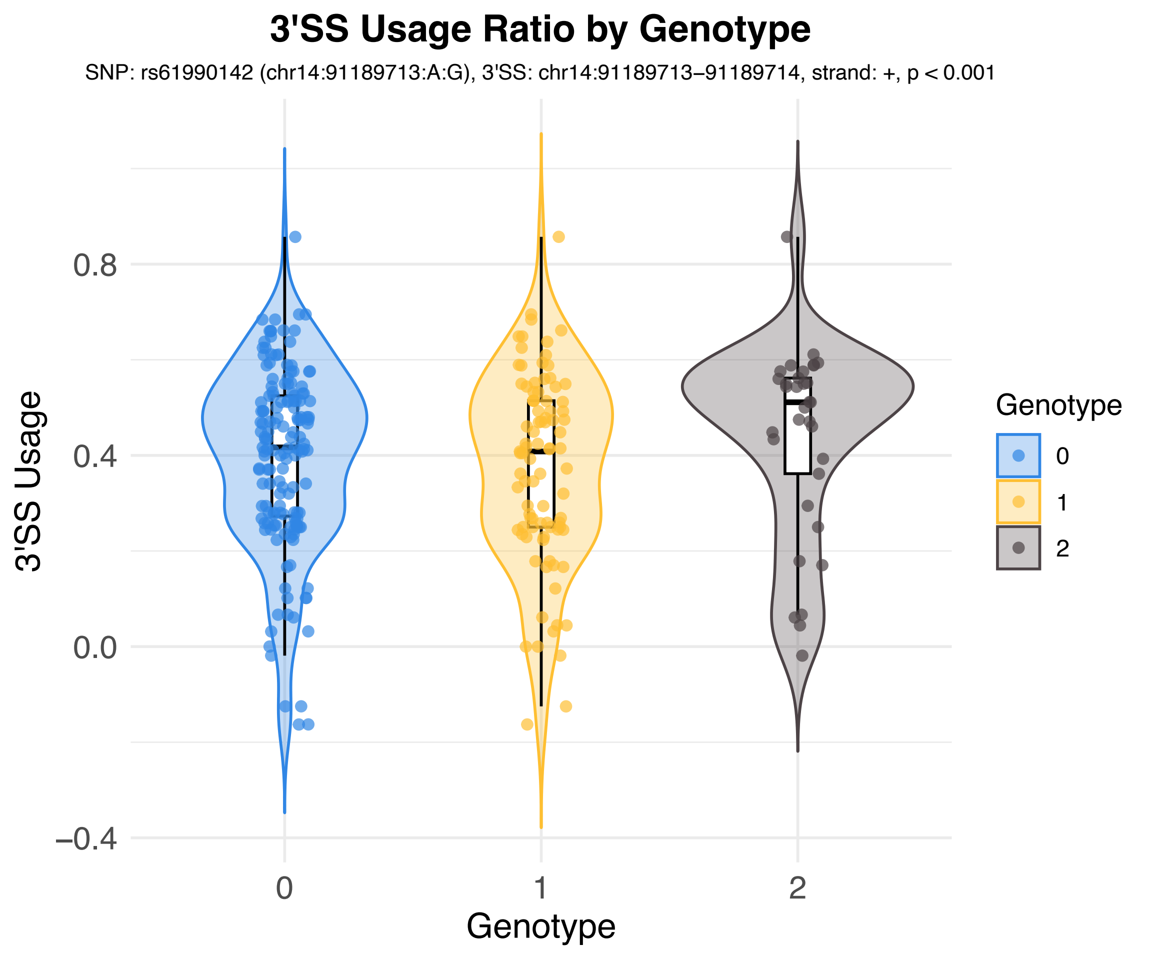
3'SS Usage Ratio by Genotype
SNP: rs61990142 (chr14:91189713:A:G), 3'SS: chr14:91189713–91189714, strand: +, p < 0.001
3'SS Usage
0.8
0.4
0.0
–0.4
0
1
2
Genotype
Genotype
0
1
2

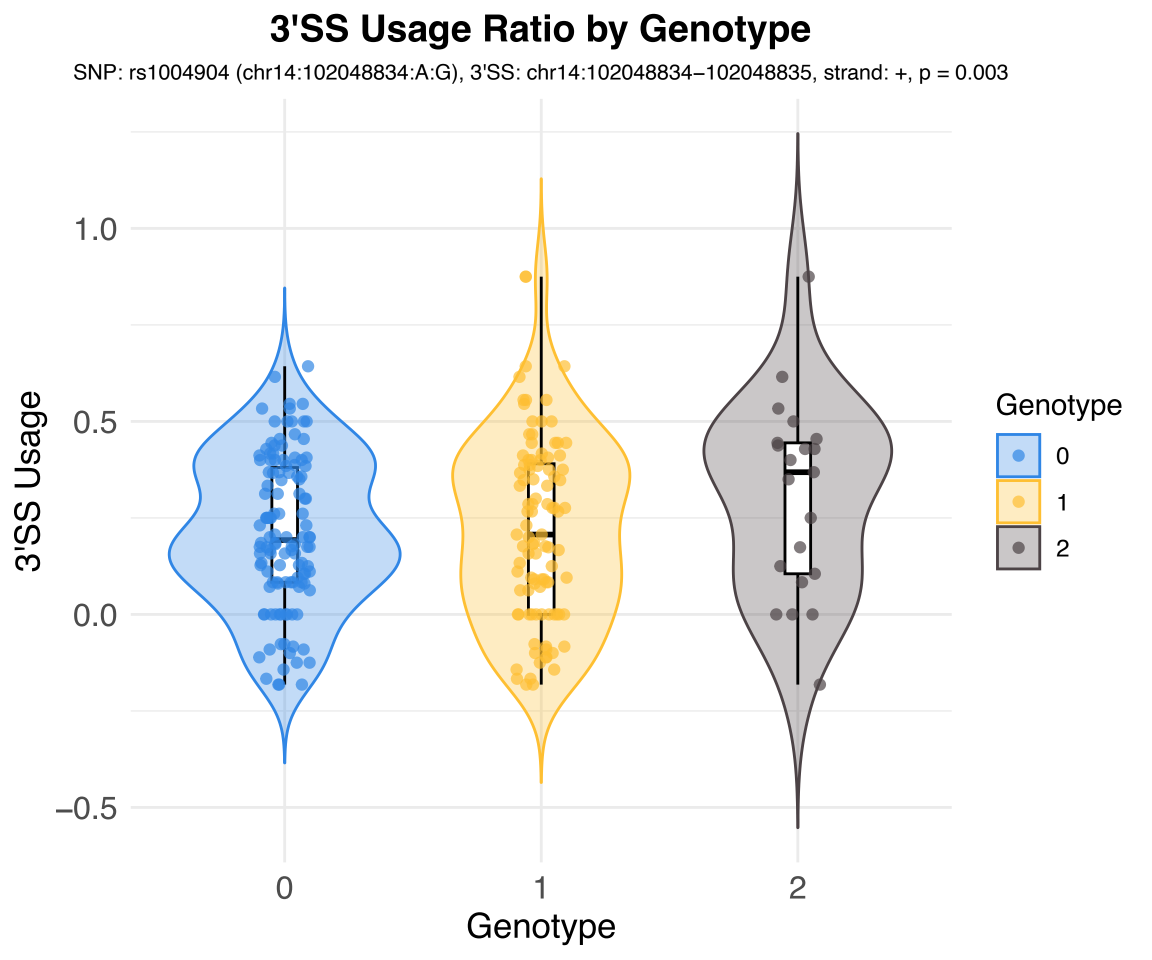
3'SS Usage Ratio by Genotype
SNP: rs1004904 (chr14:102048834:A:G), 3'SS: chr14:102048834−102048835, strand: +, p = 0.003
3'SS Usage
1.0
0.5
0.0
−0.5
0
1
2
Genotype
Genotype
0
1
2

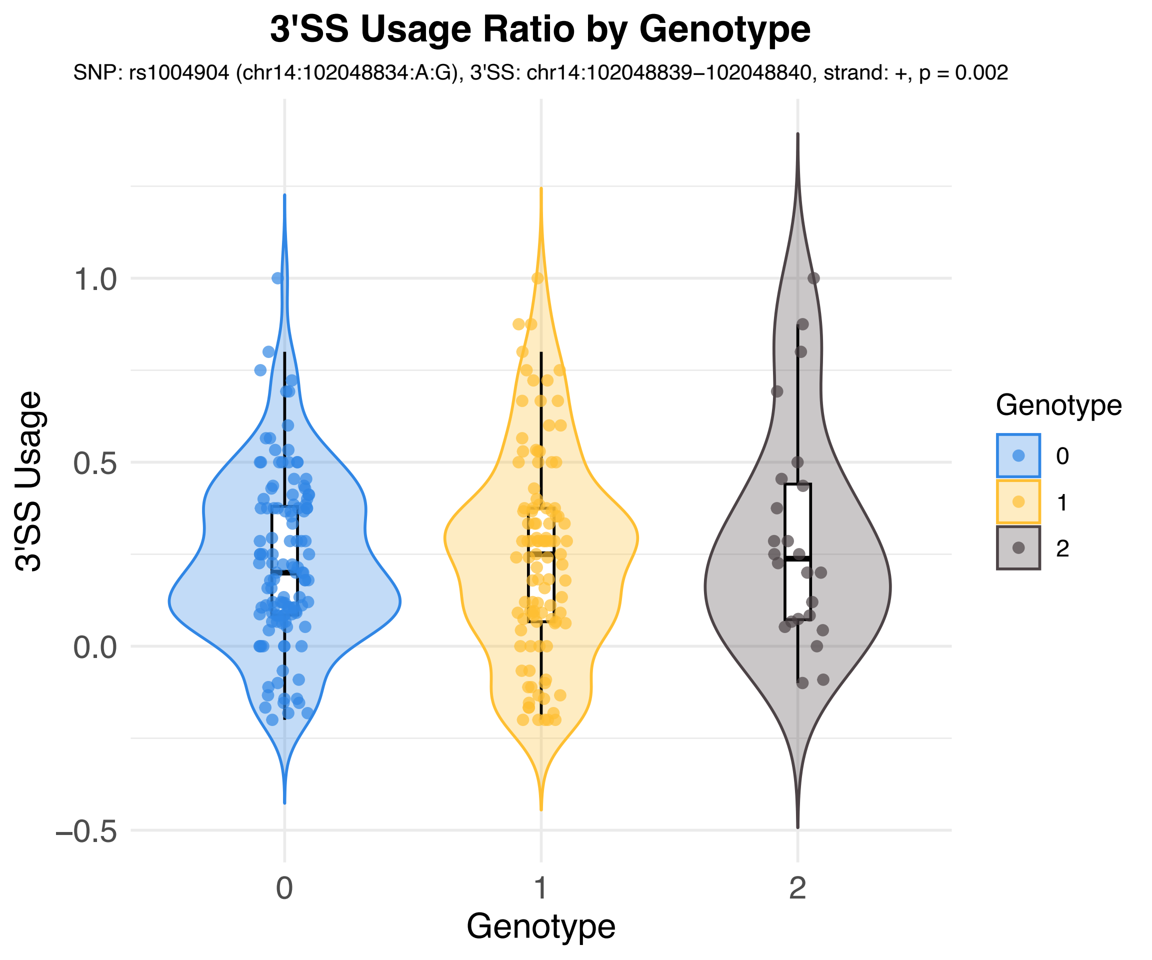
3'SS Usage Ratio by Genotype
SNP: rs1004904 (chr14:102048834:A:G), 3'SS: chr14:102048839−102048840, strand: +, p = 0.002
3'SS Usage
1.0
0.5
0.0
−0.5
0
1
2
Genotype
Genotype
0
1
2

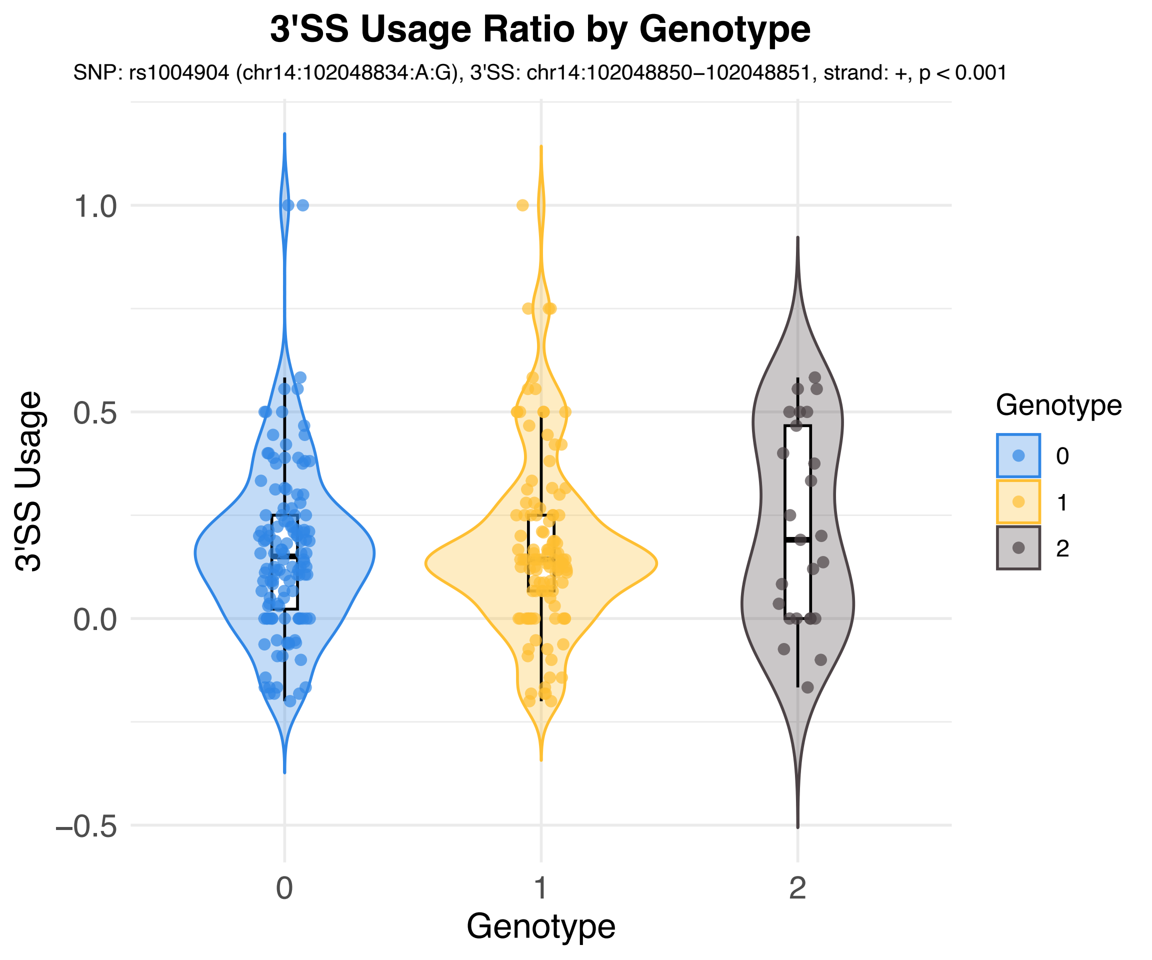
3'SS Usage Ratio by Genotype
SNP: rs1004904 (chr14:102048834:A:G), 3'SS: chr14:102048850−102048851, strand: +, p < 0.001
3'SS Usage
1.0
0.5
0.0
−0.5
0
1
2
Genotype
Genotype
0
1
2

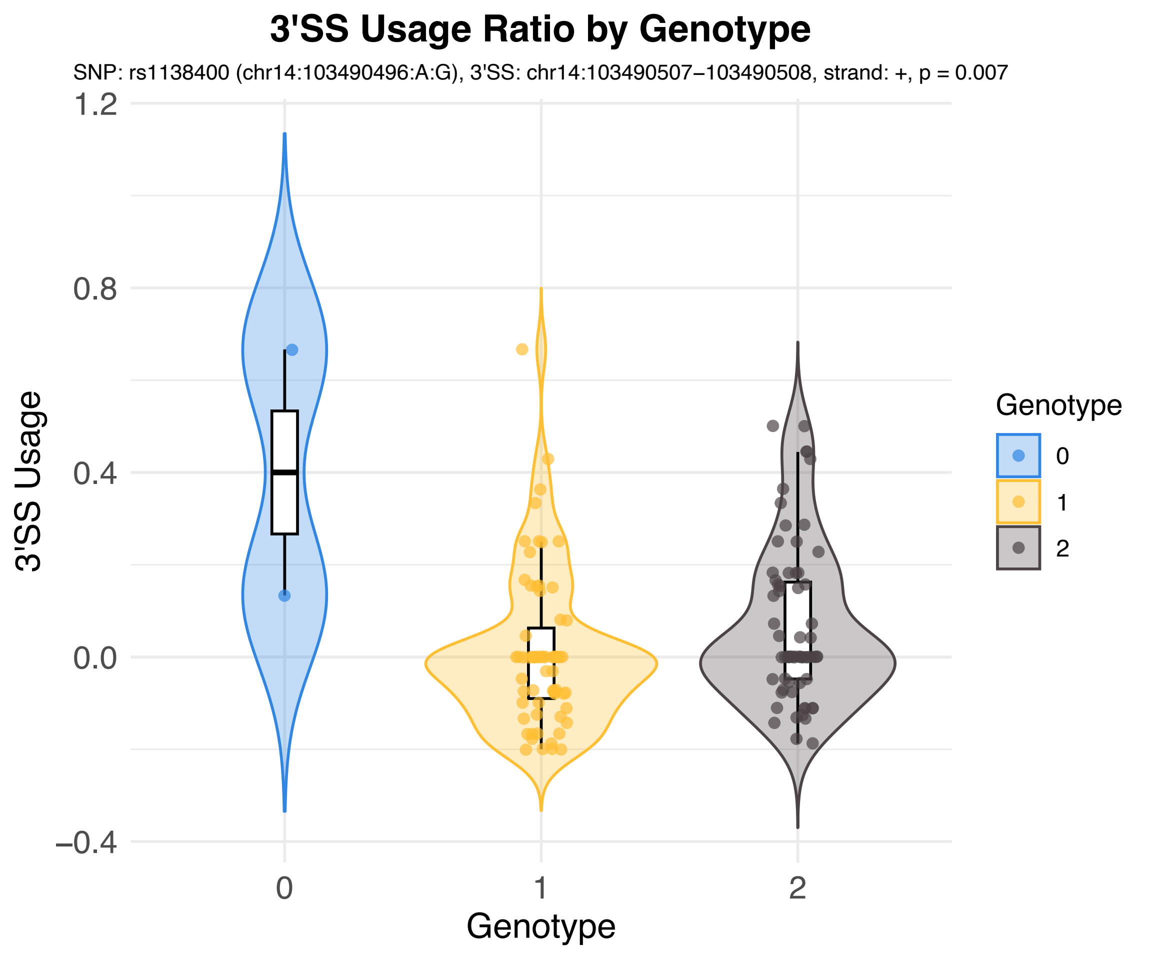

3'SS Usage Ratio by Genotype
SNP: rs1138400 (chr14:103490496:A:G), 3'SS: chr14:103490507−103490508, strand: +, p = 0.007
1.2
0.8
0.4
0.0
−0.4
3'SS Usage
0
1
2
Genotype
Genotype
0
1
2

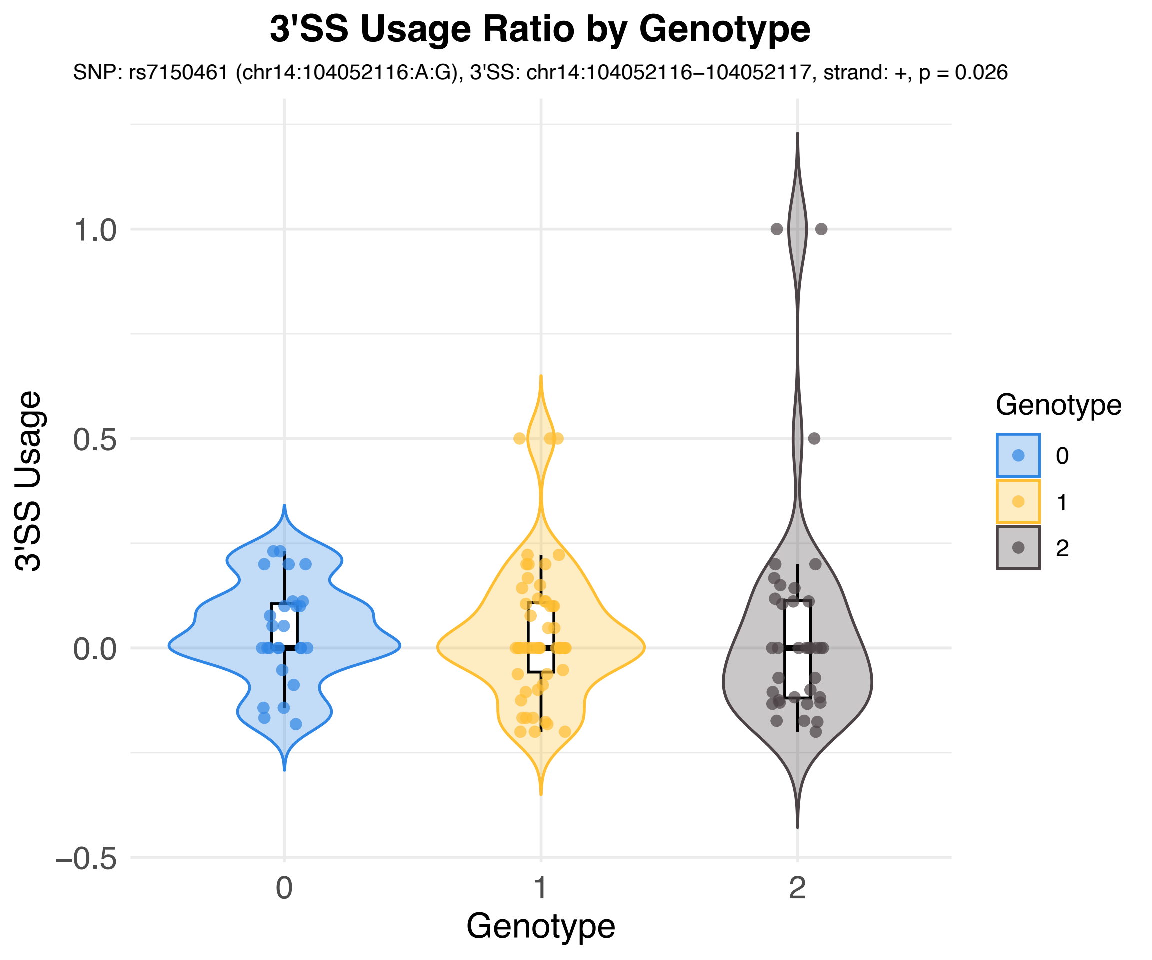
3'SS Usage Ratio by Genotype
SNP: rs7150461 (chr14:104052116:A:G), 3'SS: chr14:104052116–104052117, strand: +, p = 0.026
3'SS Usage
1.0
0.5
0.0
−0.5
0
1
2
Genotype
Genotype
0
1
2

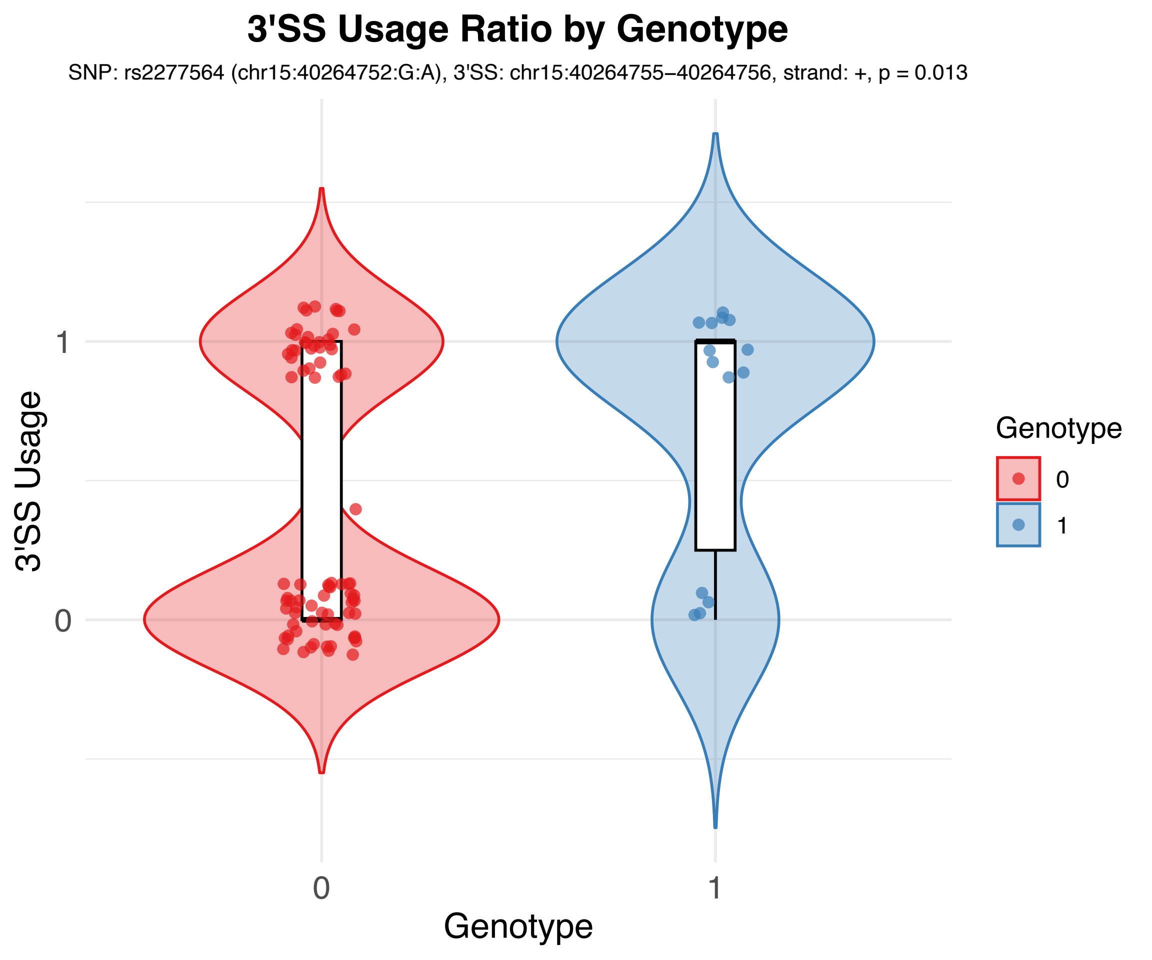
3'SS Usage Ratio by Genotype
SNP: rs2277564 (chr15:40264752:G:A), 3'SS: chr15:40264755–40264756, strand: +, p = 0.013
3'SS Usage
1
0
Genotype
0
1
Genotype
0
1

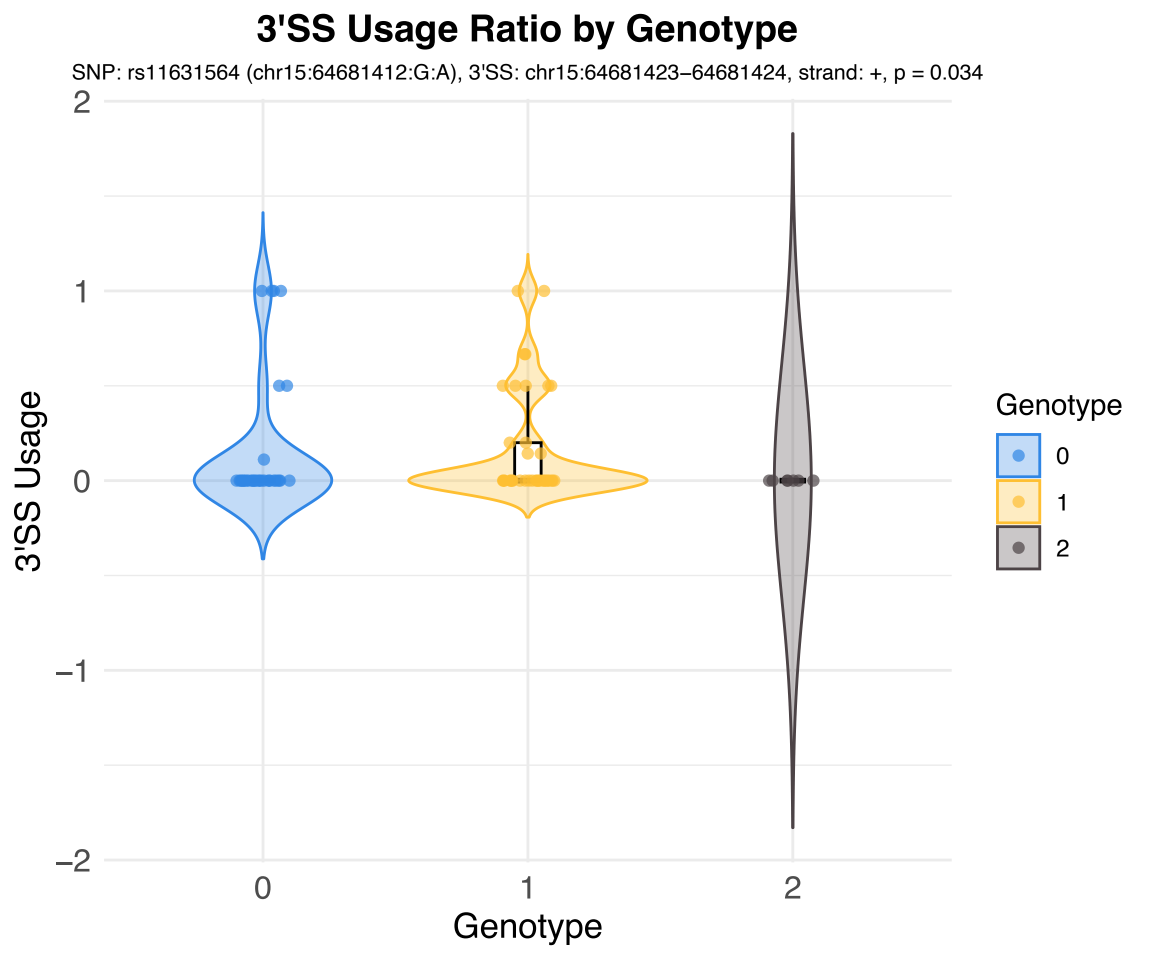
3'SS Usage Ratio by Genotype
SNP: rs11631564 (chr15:64681412:G:A), 3'SS: chr15:64681423–64681424, strand: +, p = 0.034
3'SS Usage
2
1
0
−1
−2
Genotype
0
1
2
Genotype
0
1
2

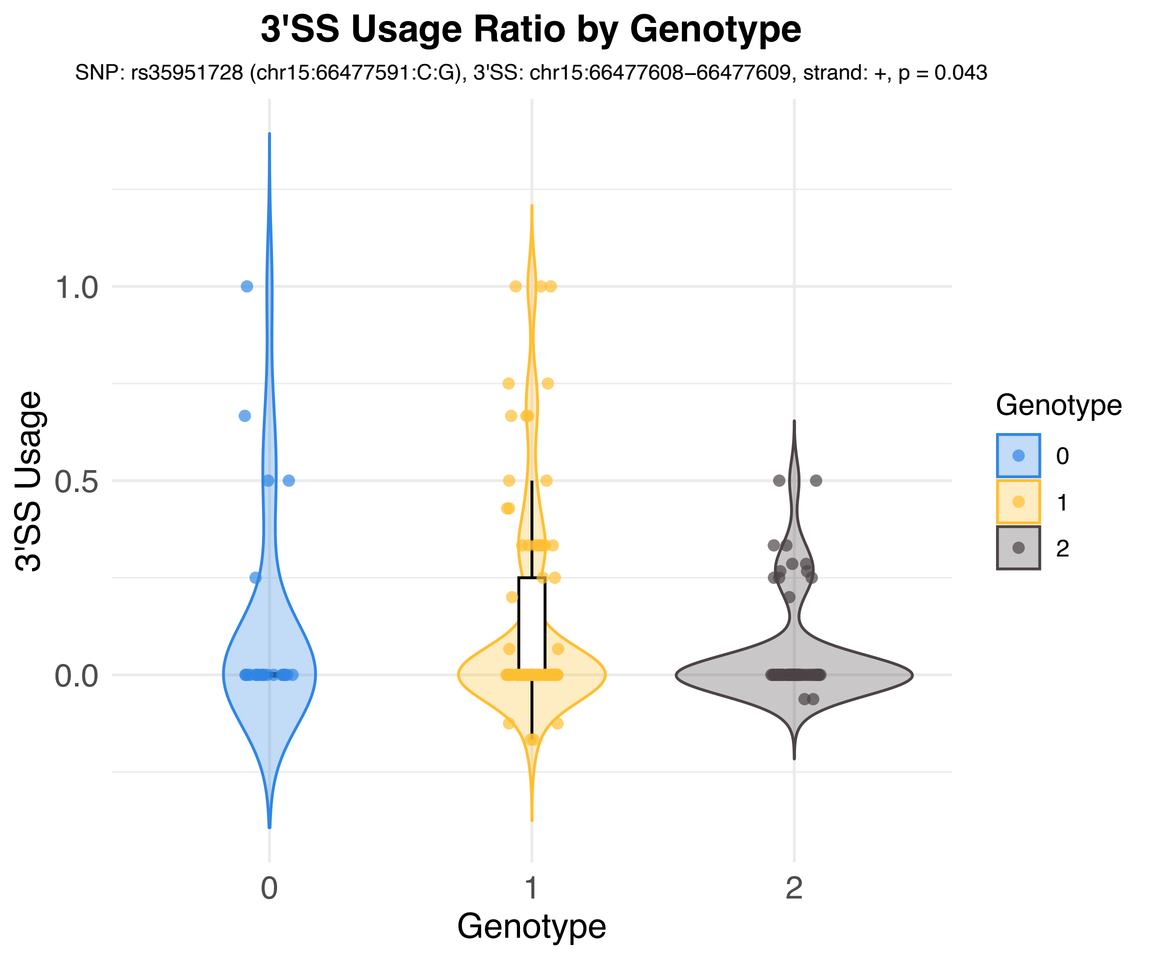

3'SS Usage Ratio by Genotype
SNP: rs35951728 (chr15:66477591:C:G), 3'SS: chr15:66477608–66477609, strand: +, p = 0.043
3'SS Usage
1.0
0.5
0.0
0
1
2
Genotype
Genotype
0
1
2

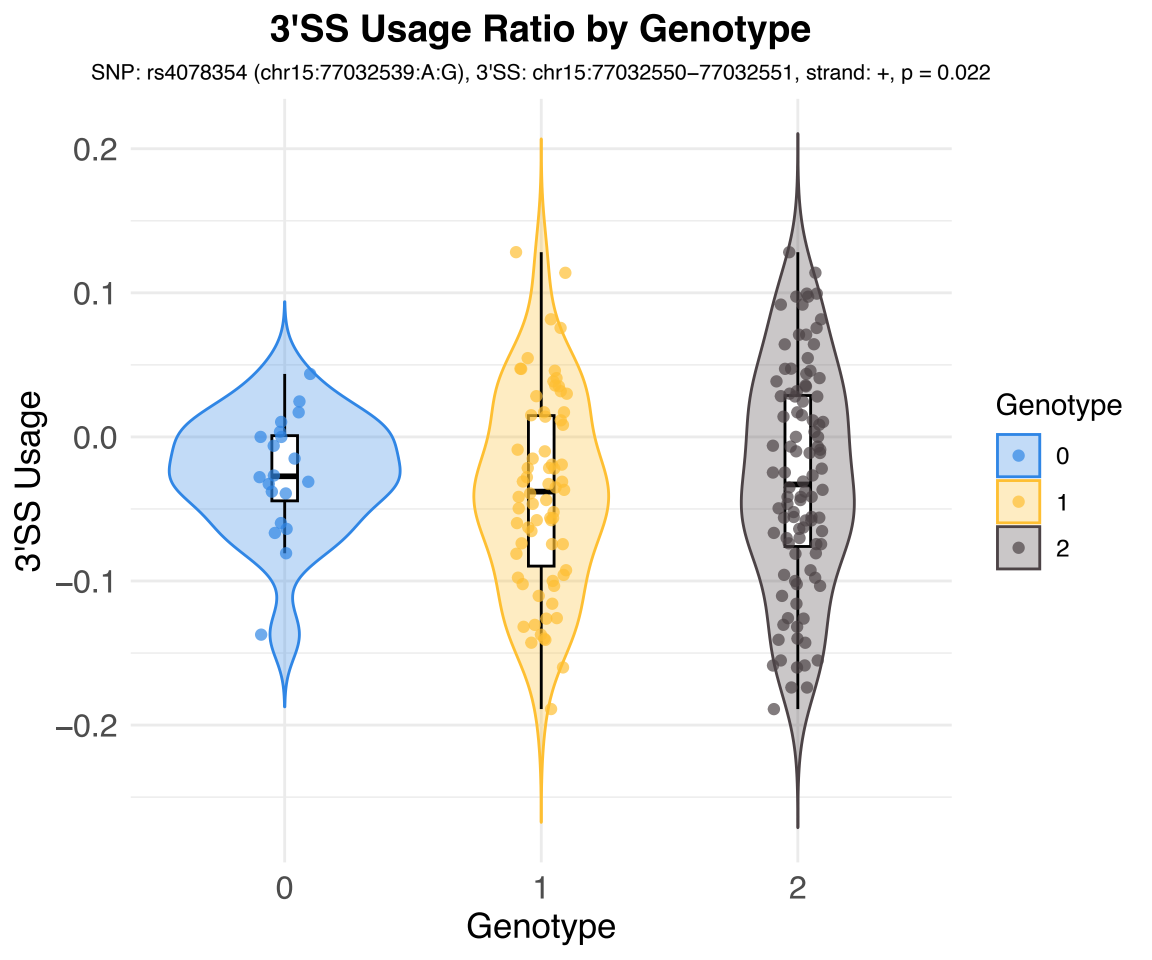
3'SS Usage Ratio by Genotype
SNP: rs4078354 (chr15:77032539:A:G), 3'SS: chr15:77032550–77032551, strand: +, p = 0.022
3'SS Usage
0.2
0.1
0.0
−0.1
−0.2
0
1
2
Genotype
Genotype
0
1
2

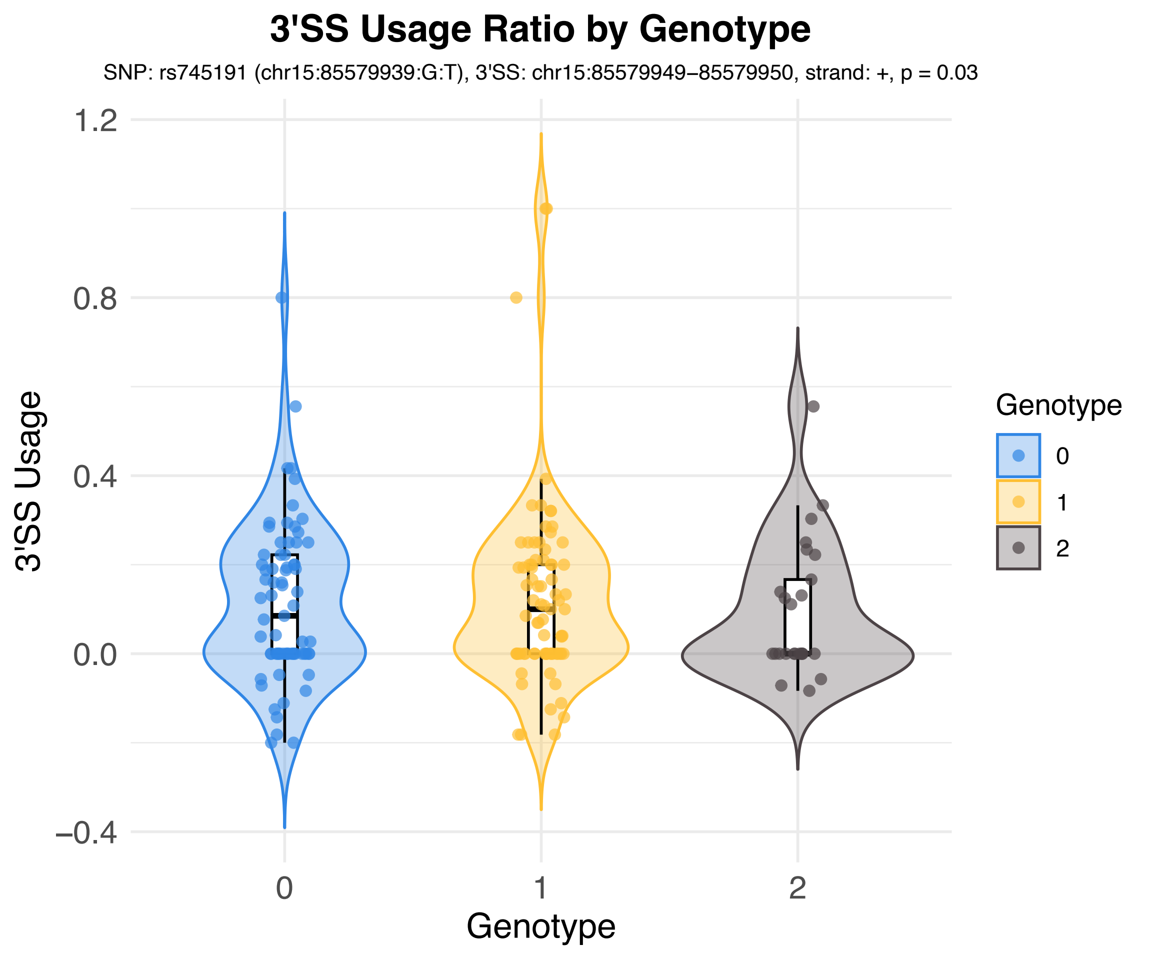
3'SS Usage Ratio by Genotype
SNP: rs745191 (chr15:85579939:G:T), 3'SS: chr15:85579949–85579950, strand: +, p = 0.03
3'SS Usage
1.2
0.8
0.4
0.0
–0.4
0
1
2
Genotype
Genotype
0
1
2

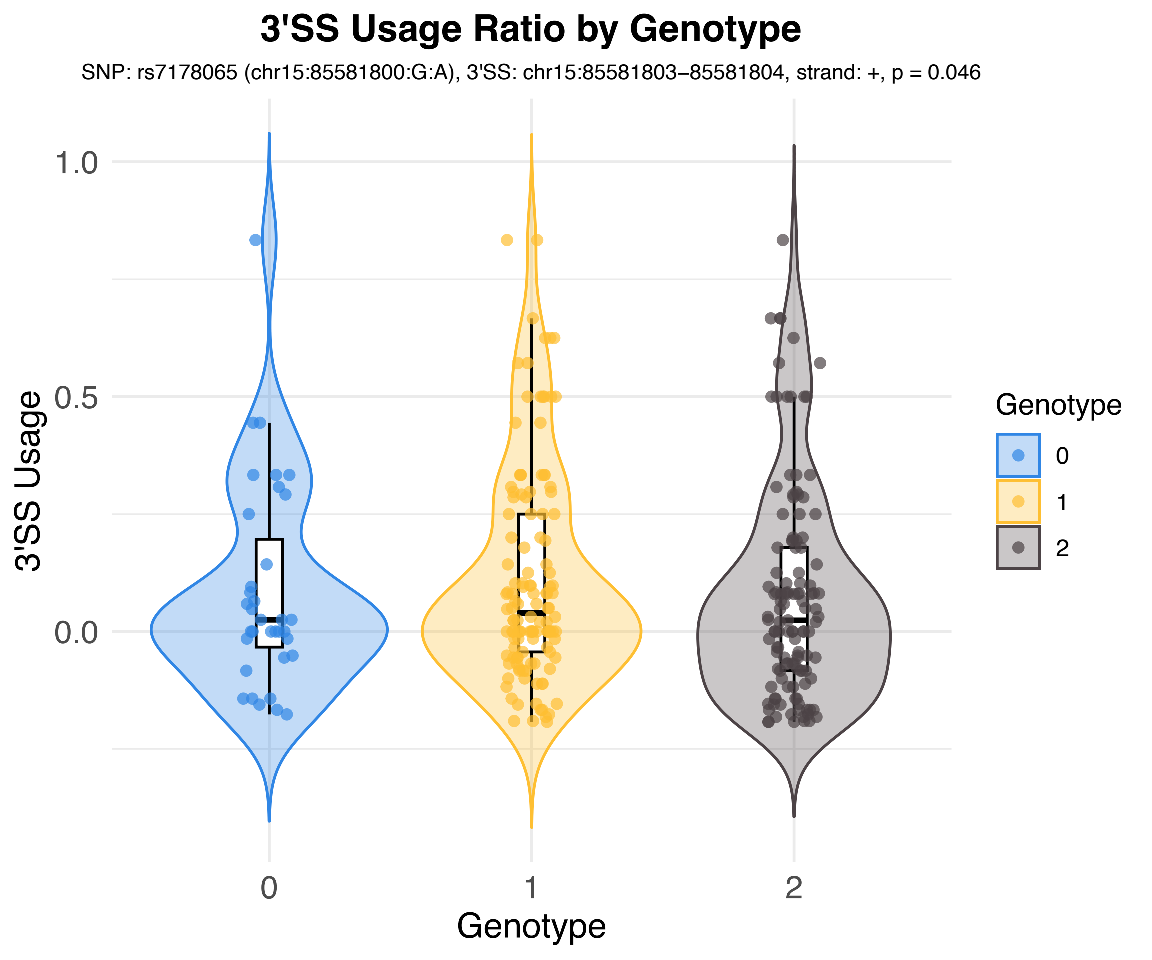

3'SS Usage Ratio by Genotype
SNP: rs7178065 (chr15:85581800:G:A), 3'SS: chr15:85581803–85581804, strand: +, p = 0.046
3'SS Usage
1.0
0.5
0.0
0
1
2
Genotype
Genotype
0
1
2

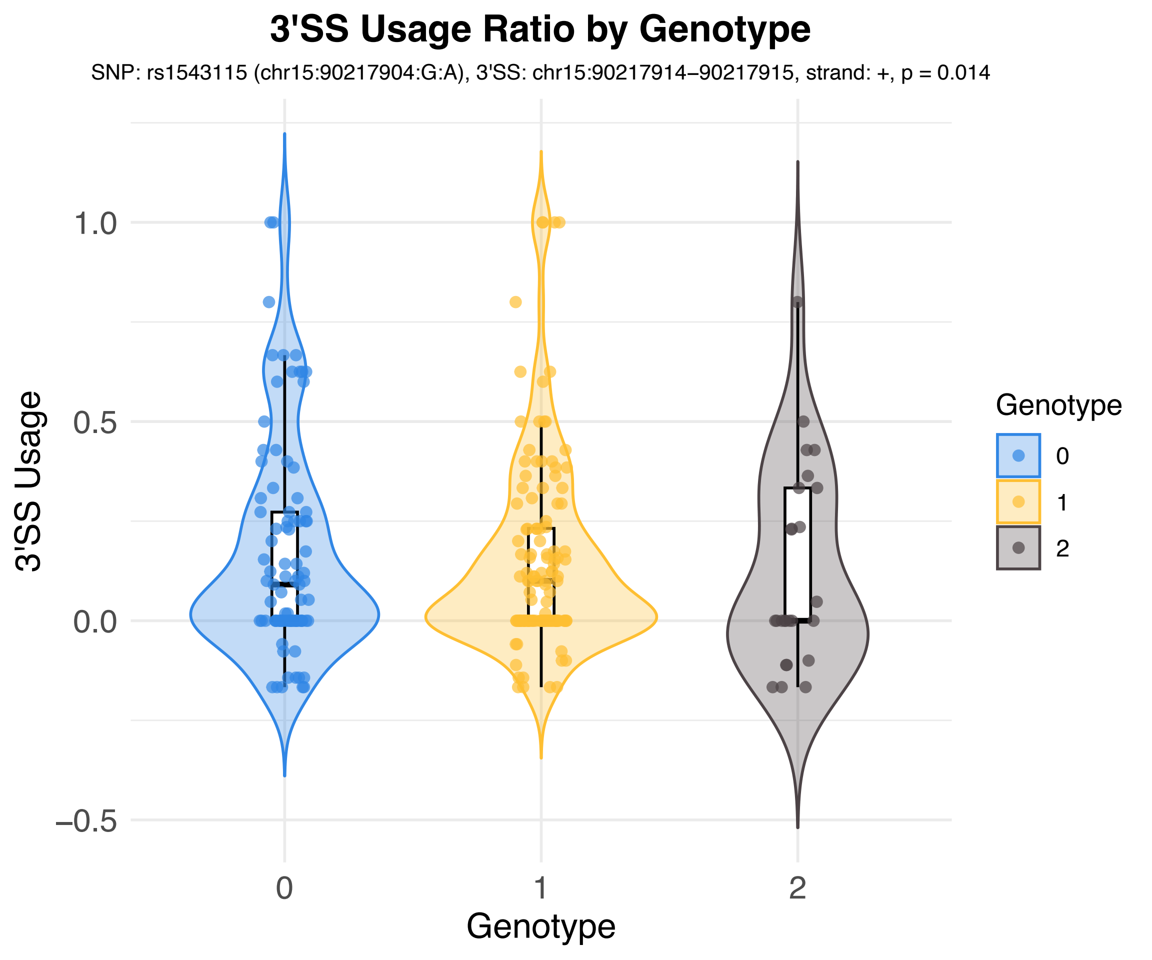
3'SS Usage Ratio by Genotype
SNP: rs1543115 (chr15:90217904:G:A), 3'SS: chr15:90217914–90217915, strand: +, p = 0.014
3'SS Usage
1.0
0.5
0.0
−0.5
0
1
2
Genotype
Genotype
0
1
2

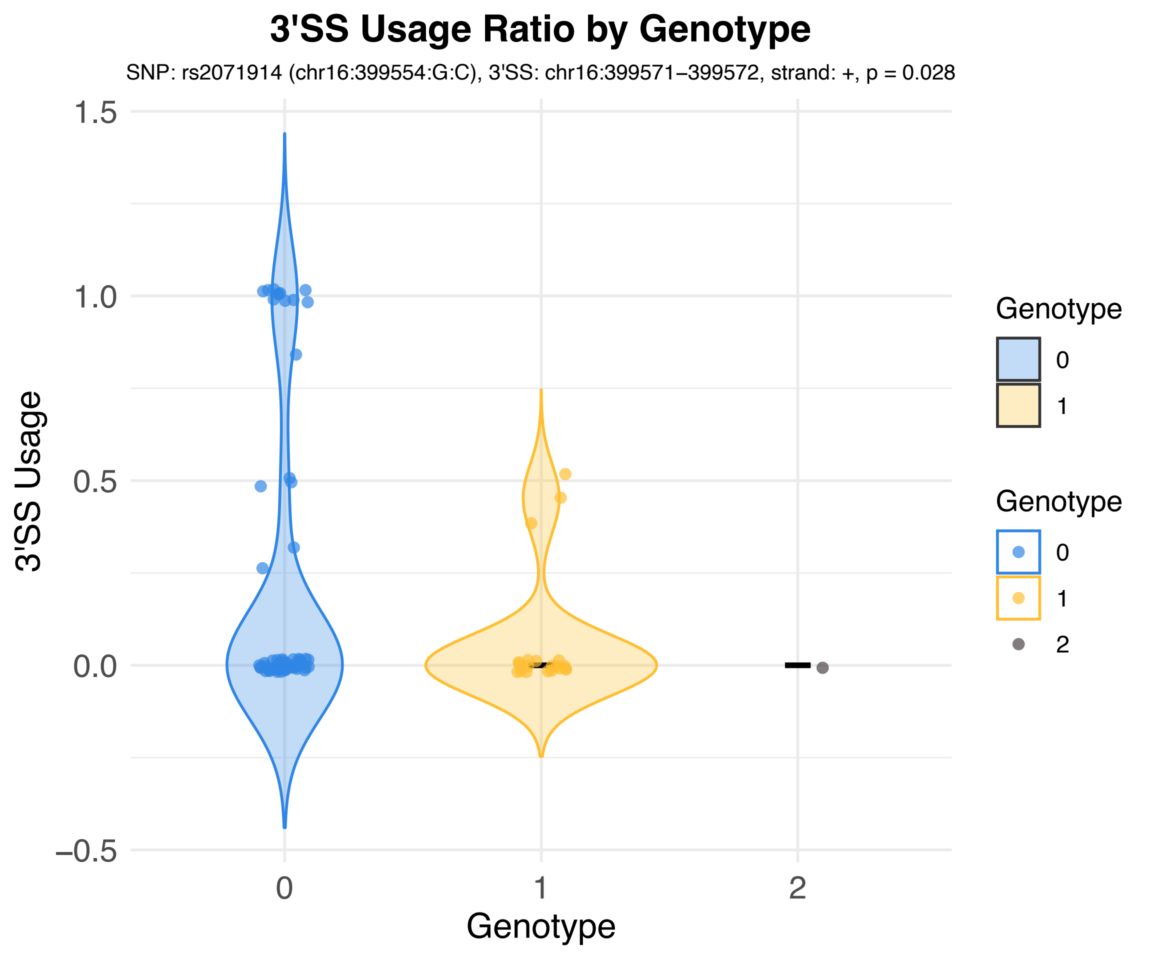
3'SS Usage Ratio by Genotype
SNP: rs2071914 (chr16:399554:G:C), 3'SS: chr16:399571–399572, strand: +, p = 0.028
3'SS Usage
1.5
1.0
0.5
0.0
−0.5
0
1
2
Genotype
Genotype
0
1
Genotype
0
1
2

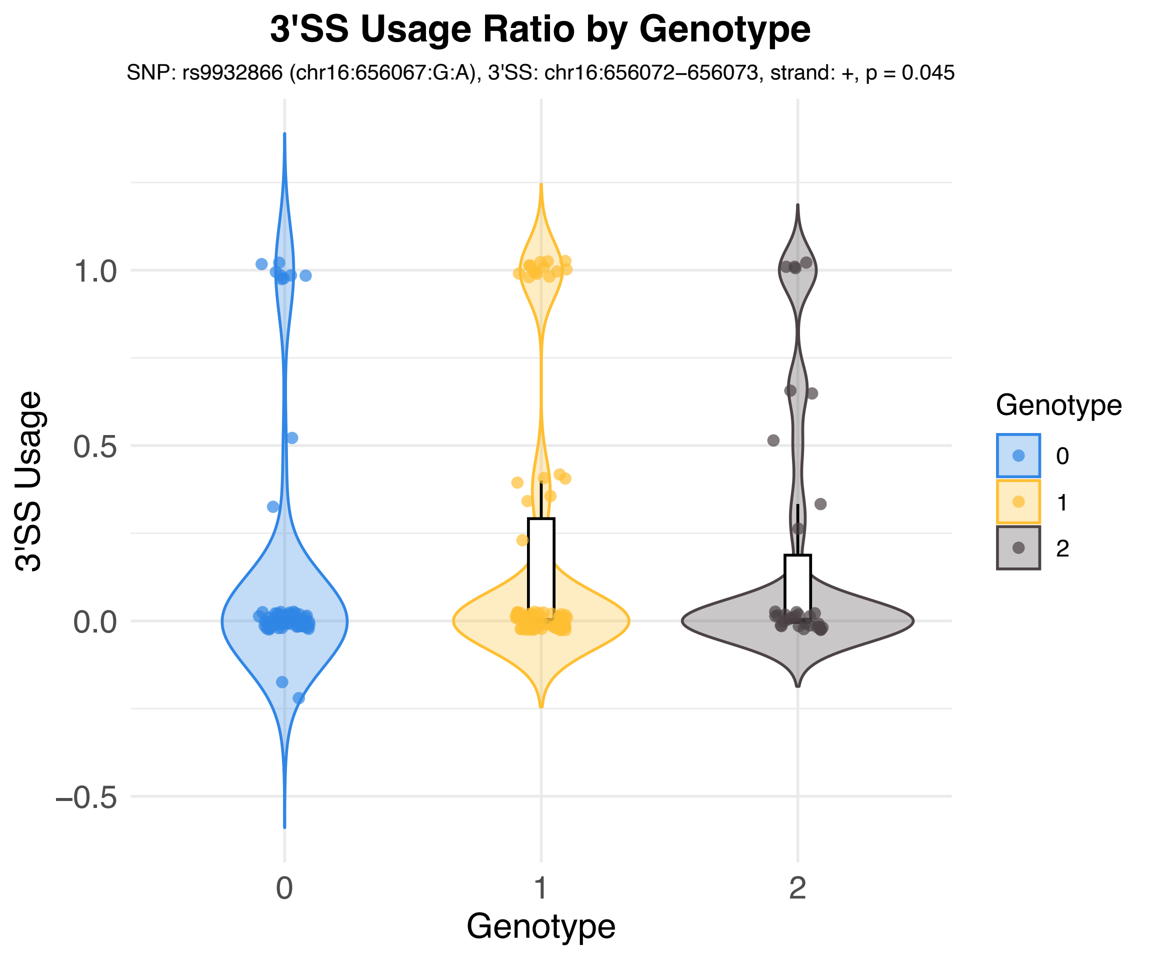

3'SS Usage Ratio by Genotype
SNP: rs9932866 (chr16:656067:G:A), 3'SS: chr16:656072–656073, strand: +, p = 0.045
3'SS Usage
1.0
0.5
0.0
−0.5
0
1
2
Genotype
Genotype
0
1
2

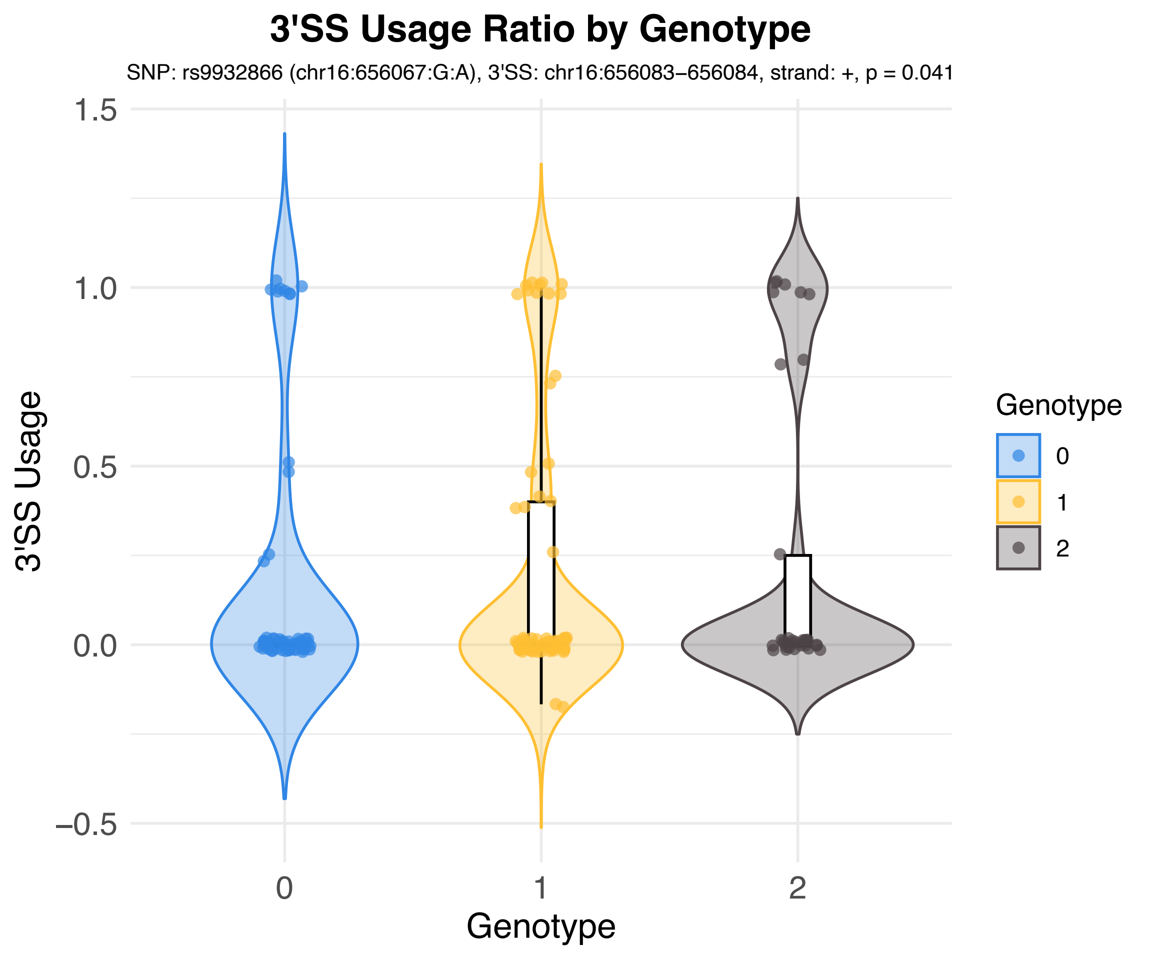
3'SS Usage Ratio by Genotype
SNP: rs9932866 (chr16:656067:G:A), 3'SS: chr16:656083–656084, strand: +, p = 0.041
3'SS Usage
1.5
1.0
0.5
0.0
–0.5
0
1
2
Genotype
Genotype
0
1
2

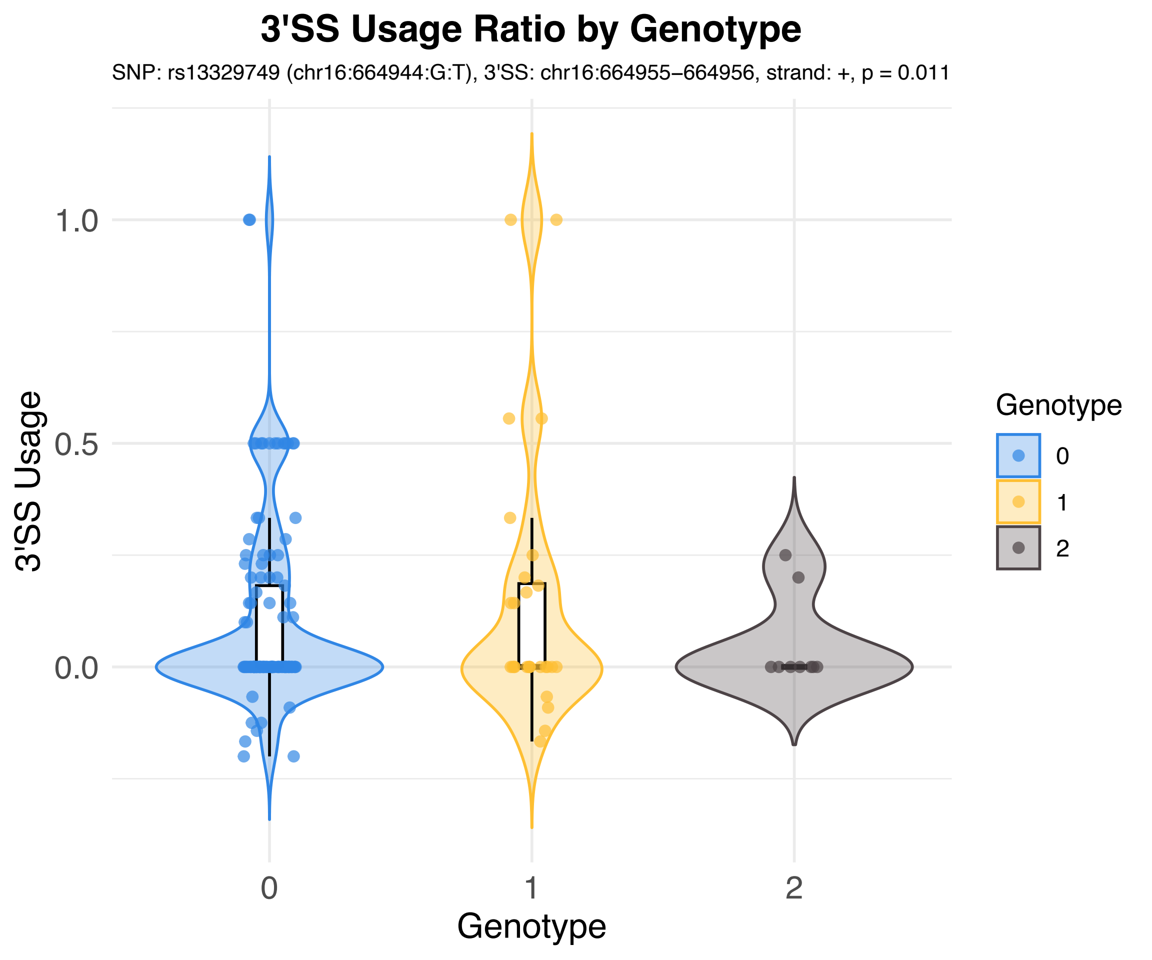
3'SS Usage Ratio by Genotype
SNP: rs13329749 (chr16:664944:G:T), 3'SS: chr16:664955–664956, strand: +, p = 0.011
3'SS Usage
1.0
0.5
0.0
0
1
2
Genotype
Genotype
0
1
2

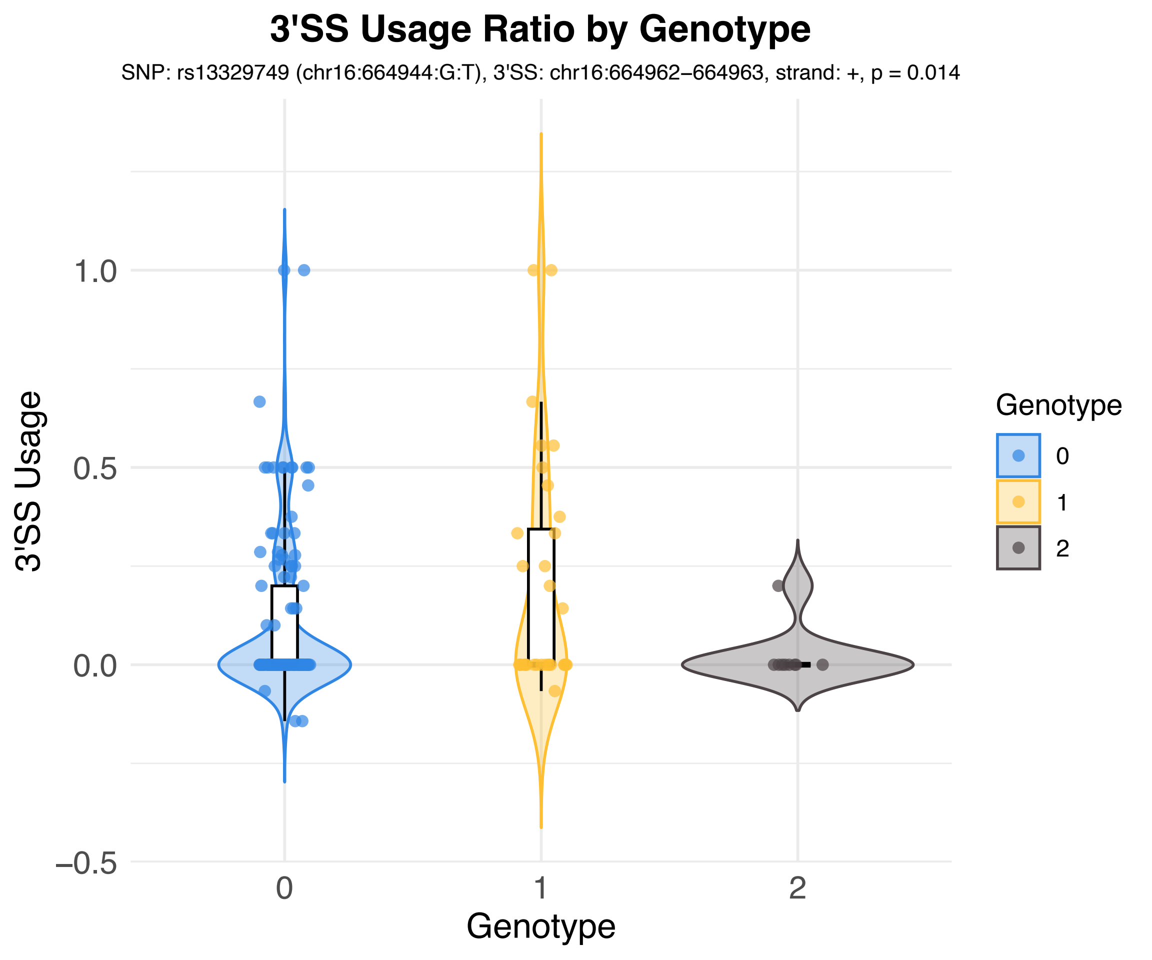
3'SS Usage Ratio by Genotype
SNP: rs13329749 (chr16:664944:G:T), 3'SS: chr16:664962−664963, strand: +, p = 0.014
3'SS Usage
1.0
0.5
0.0
−0.5
0
1
2
Genotype
Genotype
0
1
2

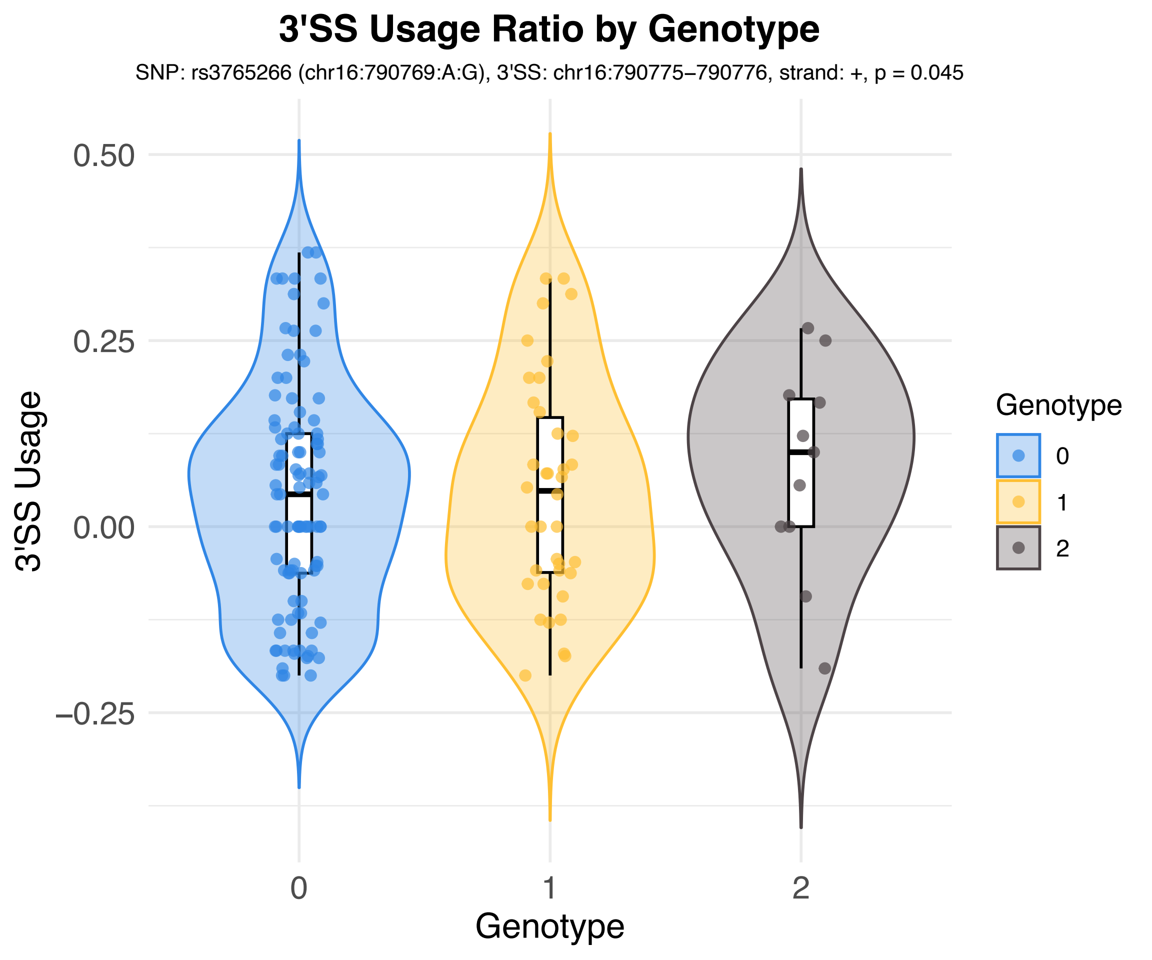
3'SS Usage Ratio by Genotype
SNP: rs3765266 (chr16:790769:A:G), 3'SS: chr16:790775−790776, strand: +, p = 0.045
3'SS Usage
0.50
0.25
0.00
−0.25
0
1
2
Genotype
Genotype
0
1
2

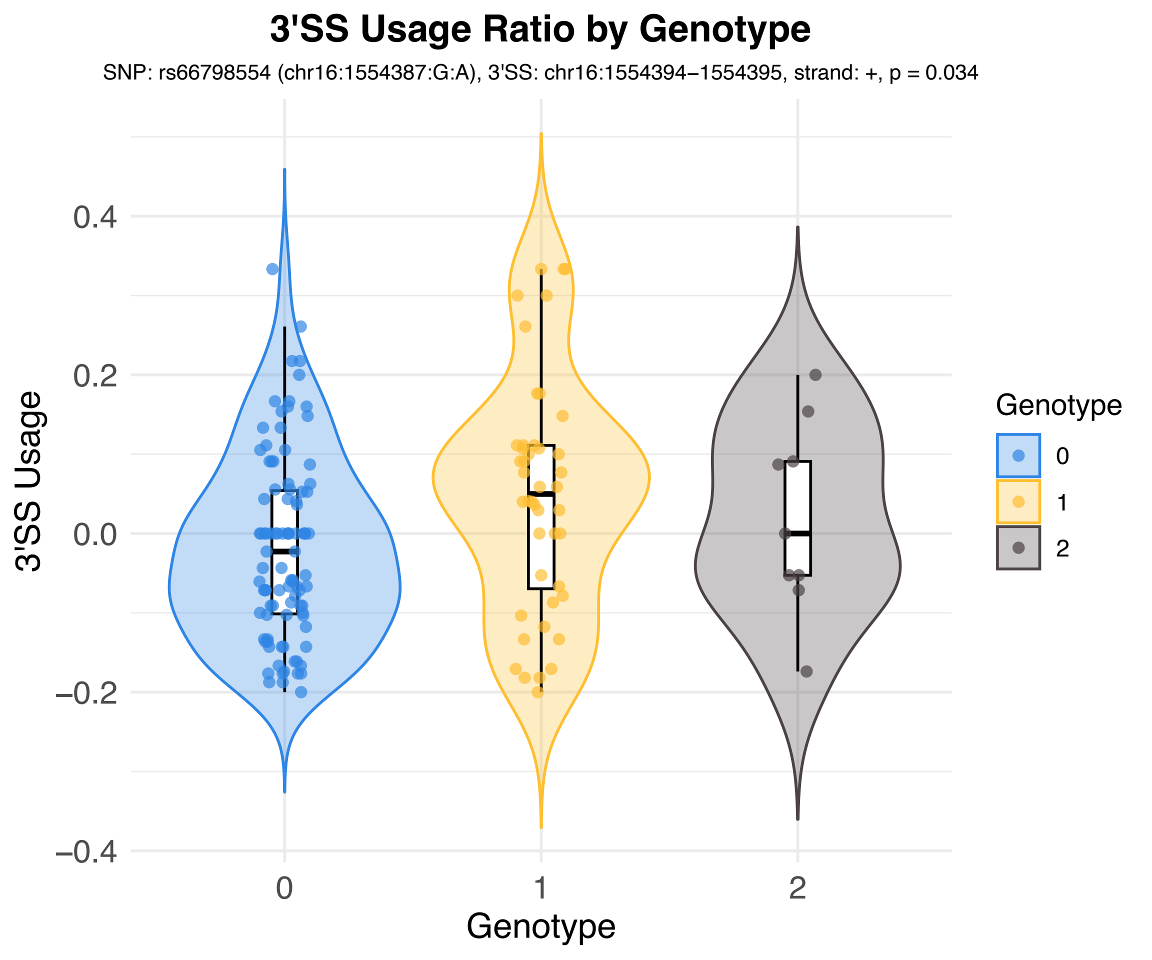

3'SS Usage Ratio by Genotype
SNP: rs66798554 (chr16:1554387:G:A), 3'SS: chr16:1554394–1554395, strand: +, p = 0.034
3'SS Usage
0.4
0.2
0.0
−0.2
−0.4
0
1
2
Genotype
Genotype
0
1
2

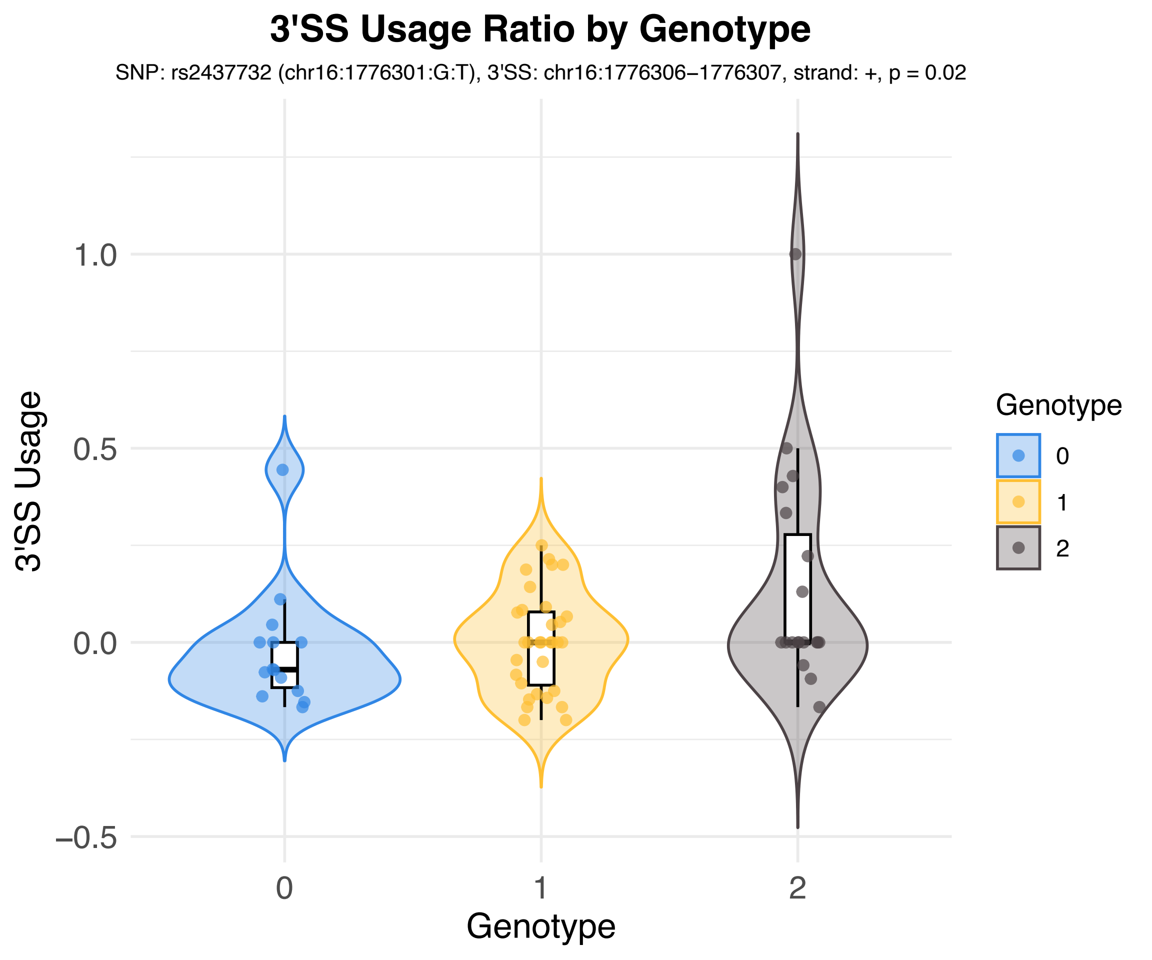
3'SS Usage Ratio by Genotype
SNP: rs2437732 (chr16:1776301:G:T), 3'SS: chr16:1776306−1776307, strand: +, p = 0.02
3'SS Usage
1.0
0.5
0.0
−0.5
0
1
2
Genotype
Genotype
0
1
2

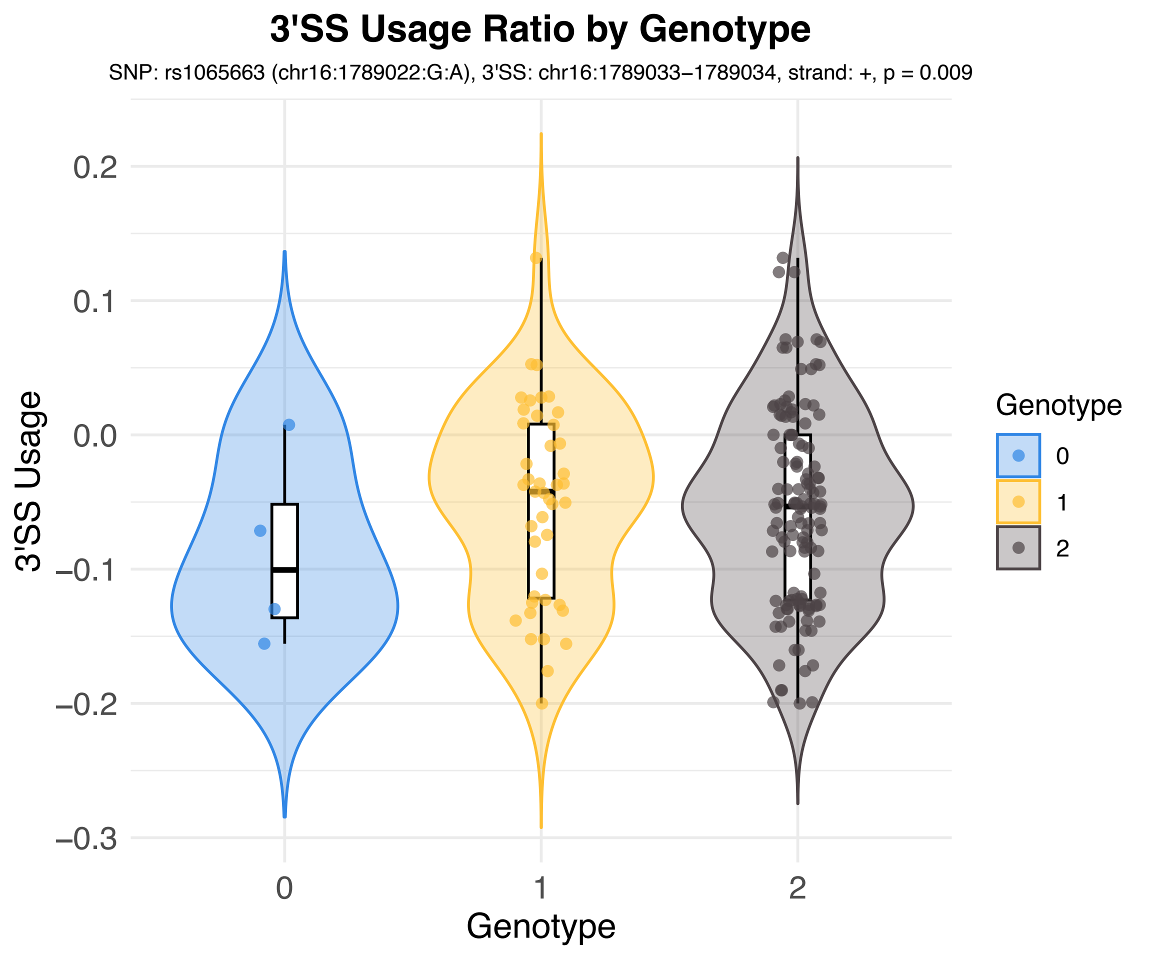
3'SS Usage Ratio by Genotype
SNP: rs1065663 (chr16:1789022:G:A), 3'SS: chr16:1789033−1789034, strand: +, p = 0.009
3'SS Usage
0.2
0.1
0.0
−0.1
−0.2
−0.3
0
1
2
Genotype
Genotype
0
1
2

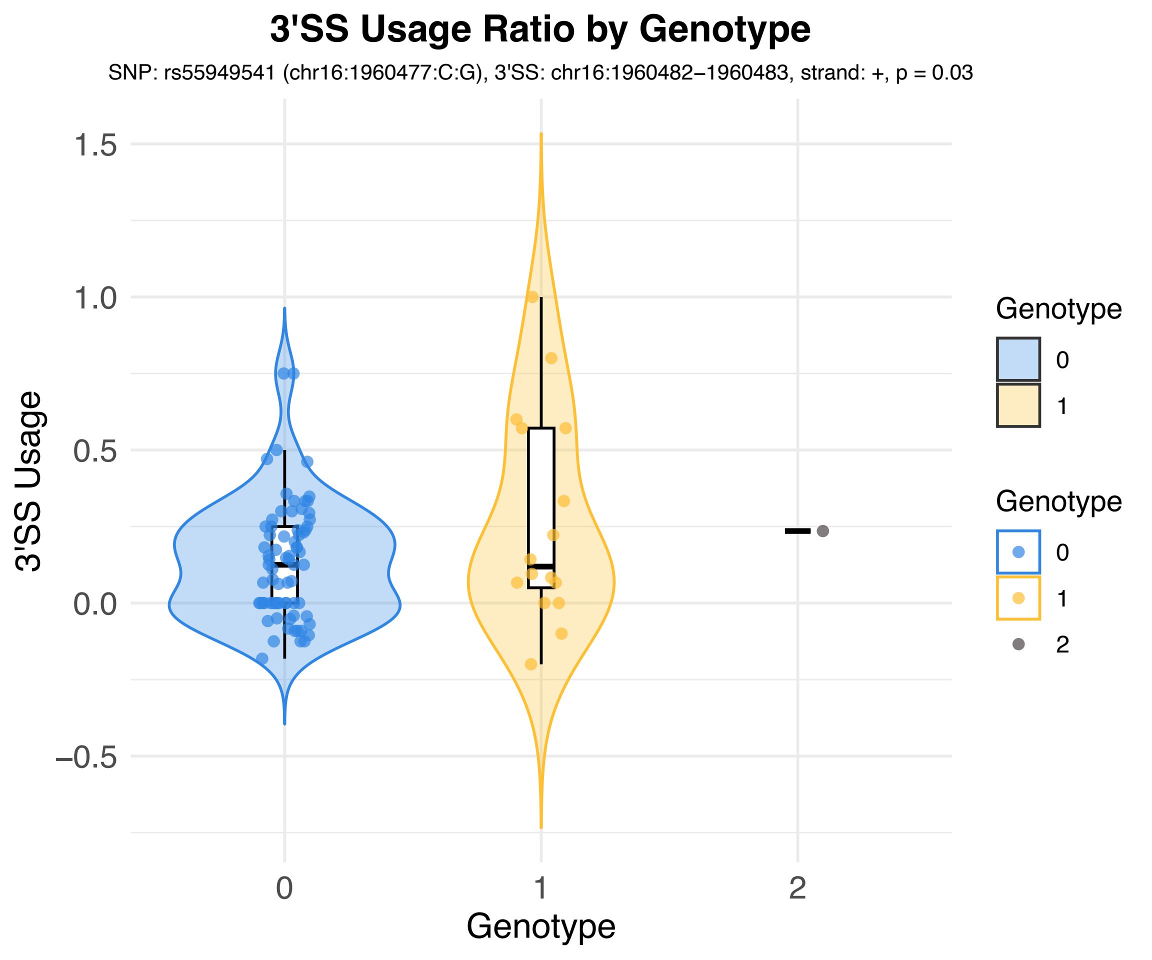
3'SS Usage Ratio by Genotype
SNP: rs55949541 (chr16:1960477:C:G), 3'SS: chr16:1960482−1960483, strand: +, p = 0.03
3'SS Usage
1.5
1.0
0.5
0.0
−0.5
0
1
2
Genotype
Genotype
0
1
Genotype
0
1
2

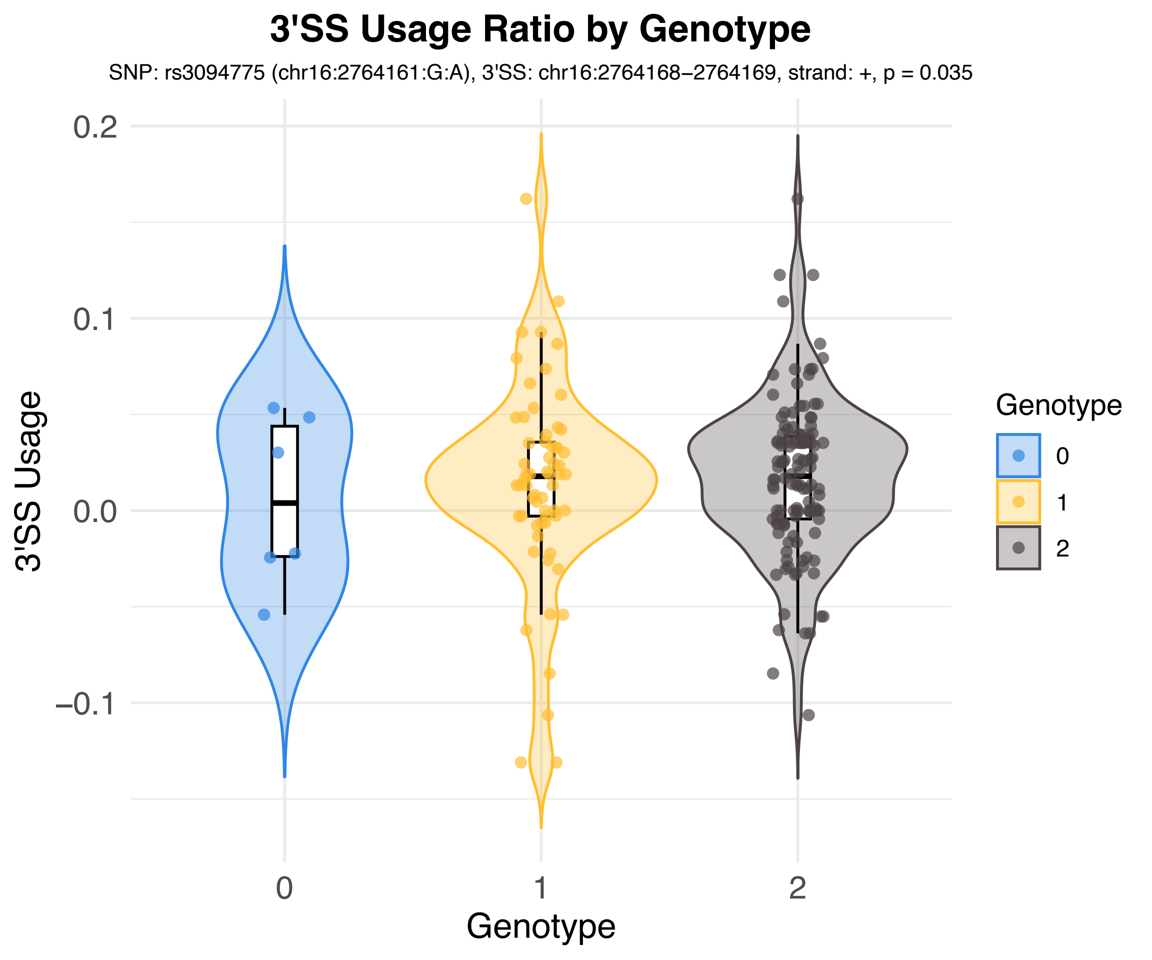
3'SS Usage Ratio by Genotype
SNP: rs3094775 (chr16:2764161:G:A), 3'SS: chr16:2764168–2764169, strand: +, p = 0.035
3'SS Usage
0.2
0.1
0.0
–0.1
0
1
2
Genotype
Genotype
0
1
2

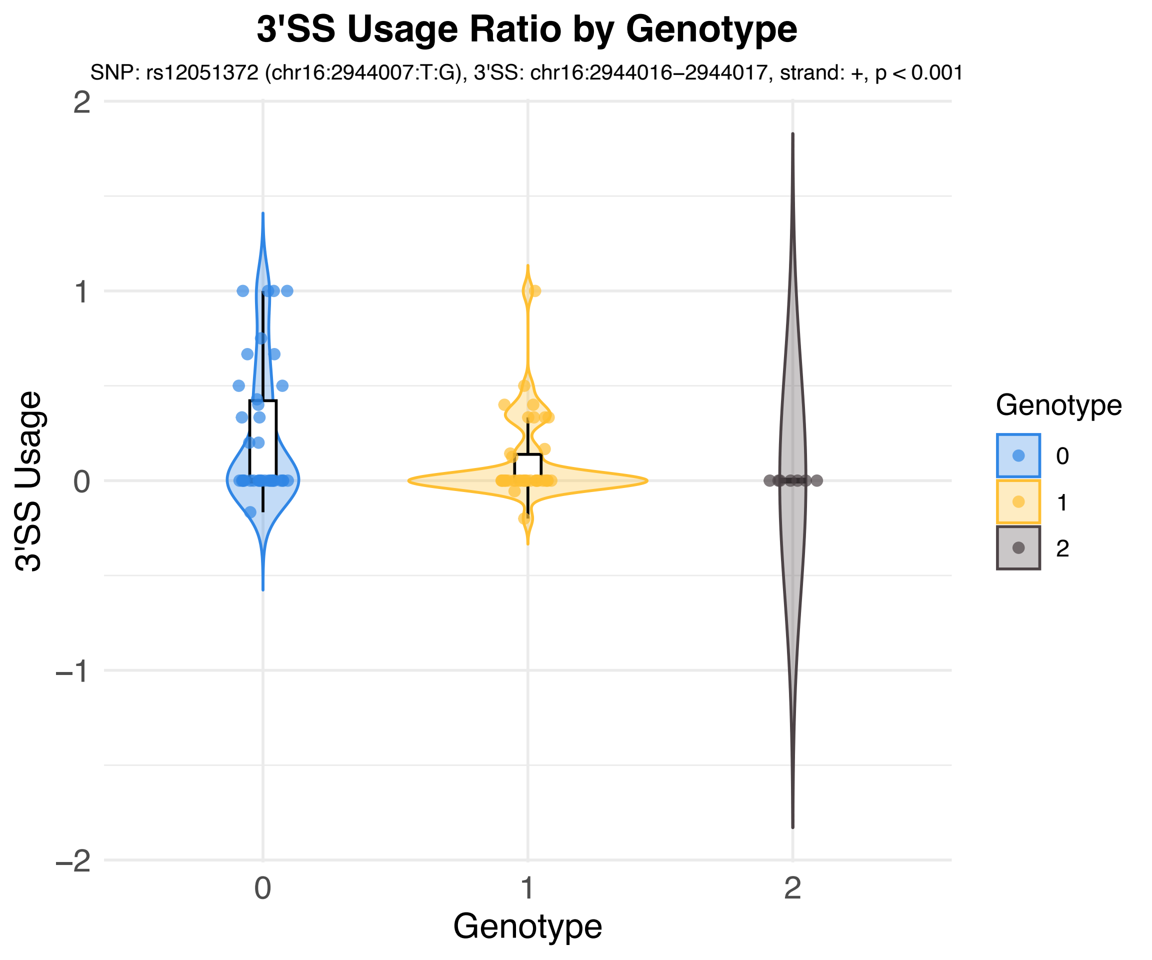
3'SS Usage Ratio by Genotype
SNP: rs12051372 (chr16:2944007:T:G), 3'SS: chr16:2944016–2944017, strand: +, p < 0.001
3'SS Usage
2
1
0
−1
−2
Genotype
0
1
2
Genotype
0
1
2

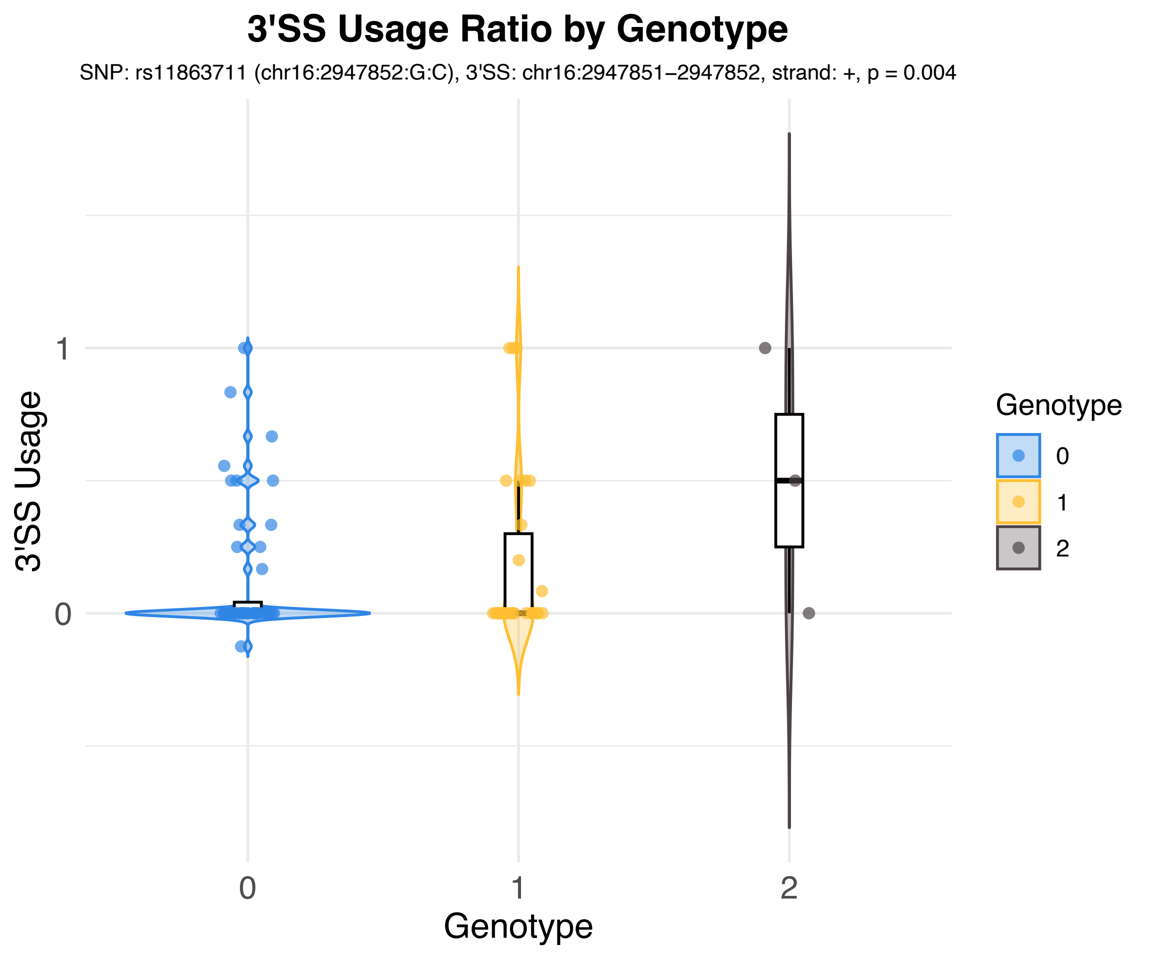
3'SS Usage Ratio by Genotype
SNP: rs11863711 (chr16:2947852:G:C), 3'SS: chr16:2947851−2947852, strand: +, p = 0.004
3'SS Usage
1
0
0
1
2
Genotype
Genotype
0
1
2

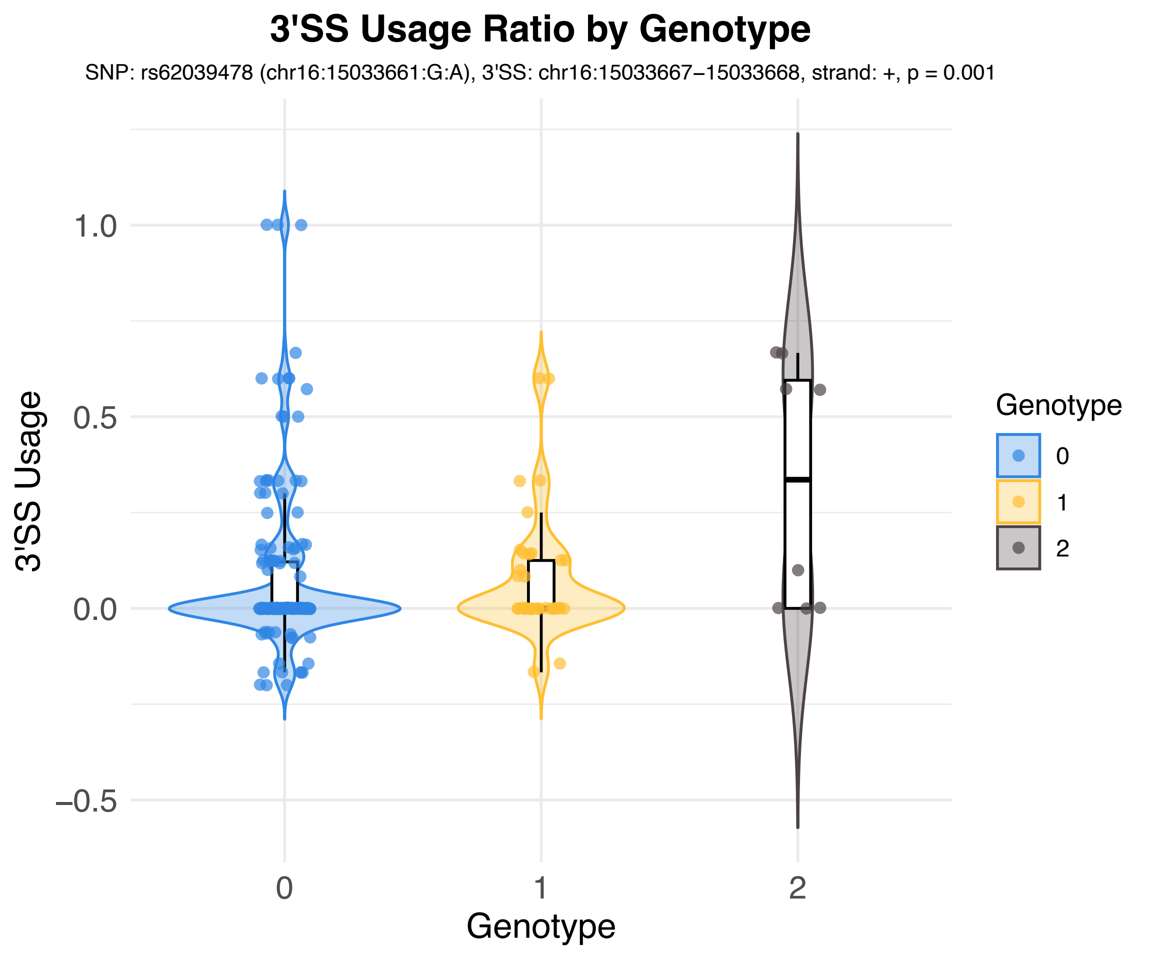

3'SS Usage Ratio by Genotype
SNP: rs62039478 (chr16:15033661:G:A), 3'SS: chr16:15033667−15033668, strand: +, p = 0.001
3'SS Usage
1.0
0.5
0.0
−0.5
0
1
2
Genotype
Genotype
0
1
2

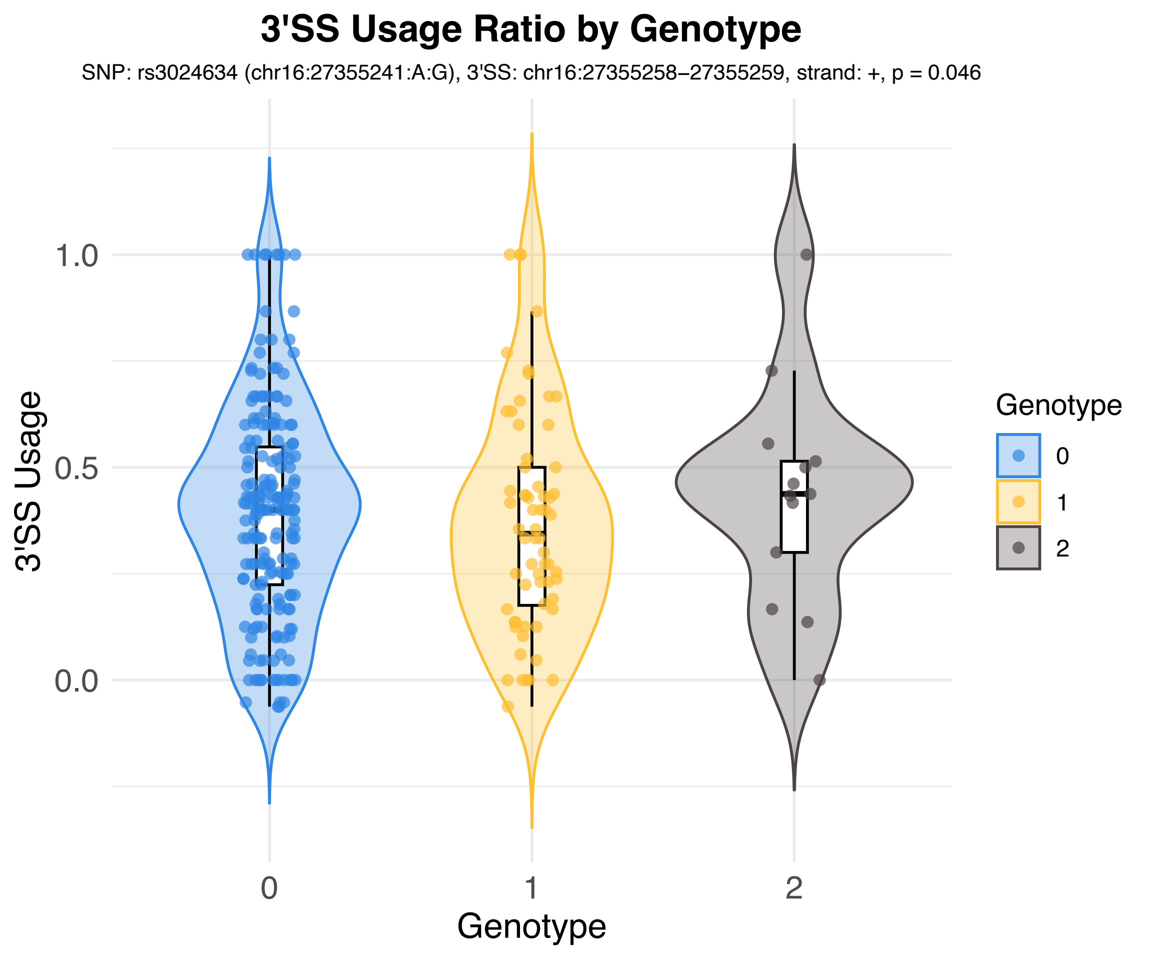

3'SS Usage Ratio by Genotype
SNP: rs3024634 (chr16:27355241:A:G), 3'SS: chr16:27355258–27355259, strand: +, p = 0.046
3'SS Usage
1.0
0.5
0.0
0
1
2
Genotype
Genotype
0
1
2

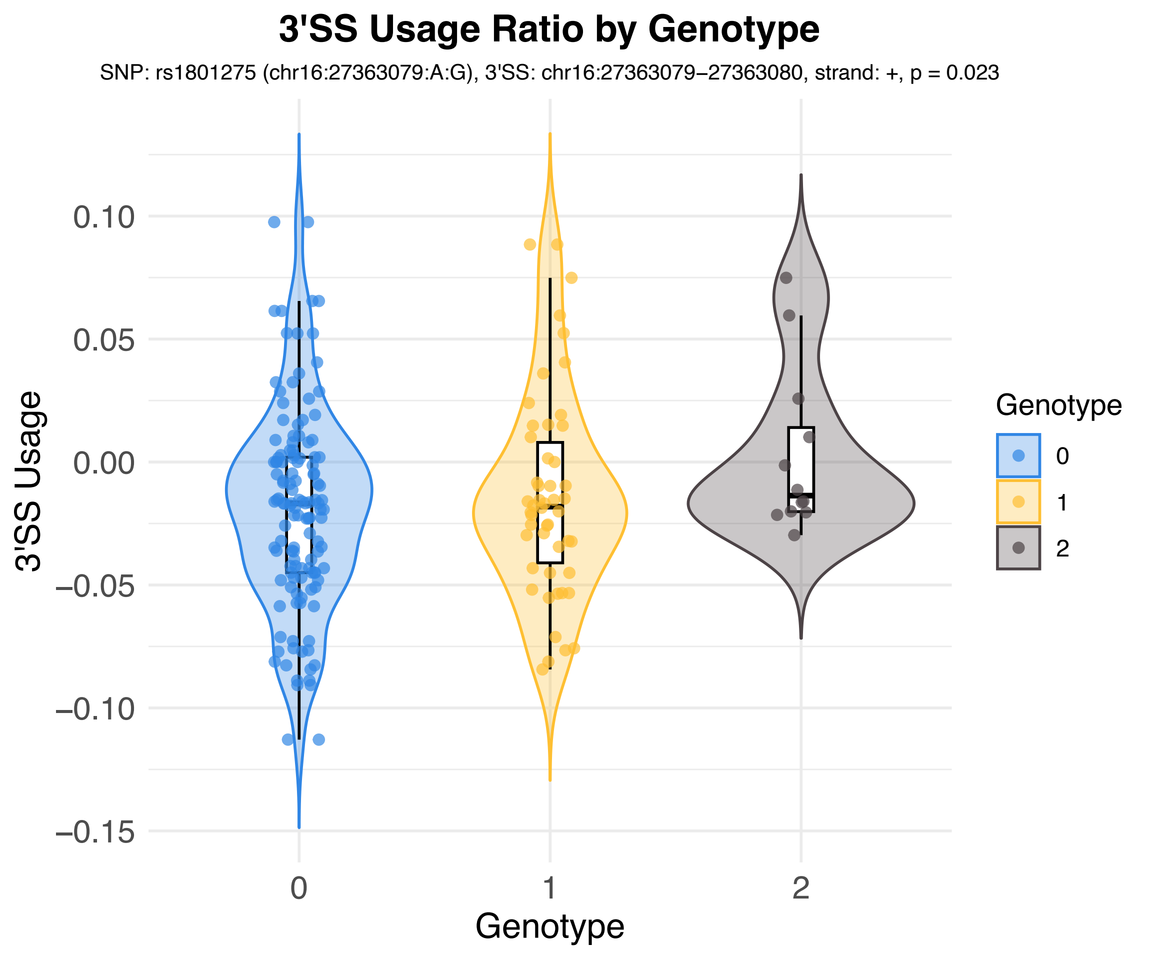
3'SS Usage Ratio by Genotype
SNP: rs1801275 (chr16:27363079:A:G), 3'SS: chr16:27363079−27363080, strand: +, p = 0.023
3'SS Usage
0.10
0.05
0.00
−0.05
−0.10
−0.15
0
1
2
Genotype
Genotype
0
1
2

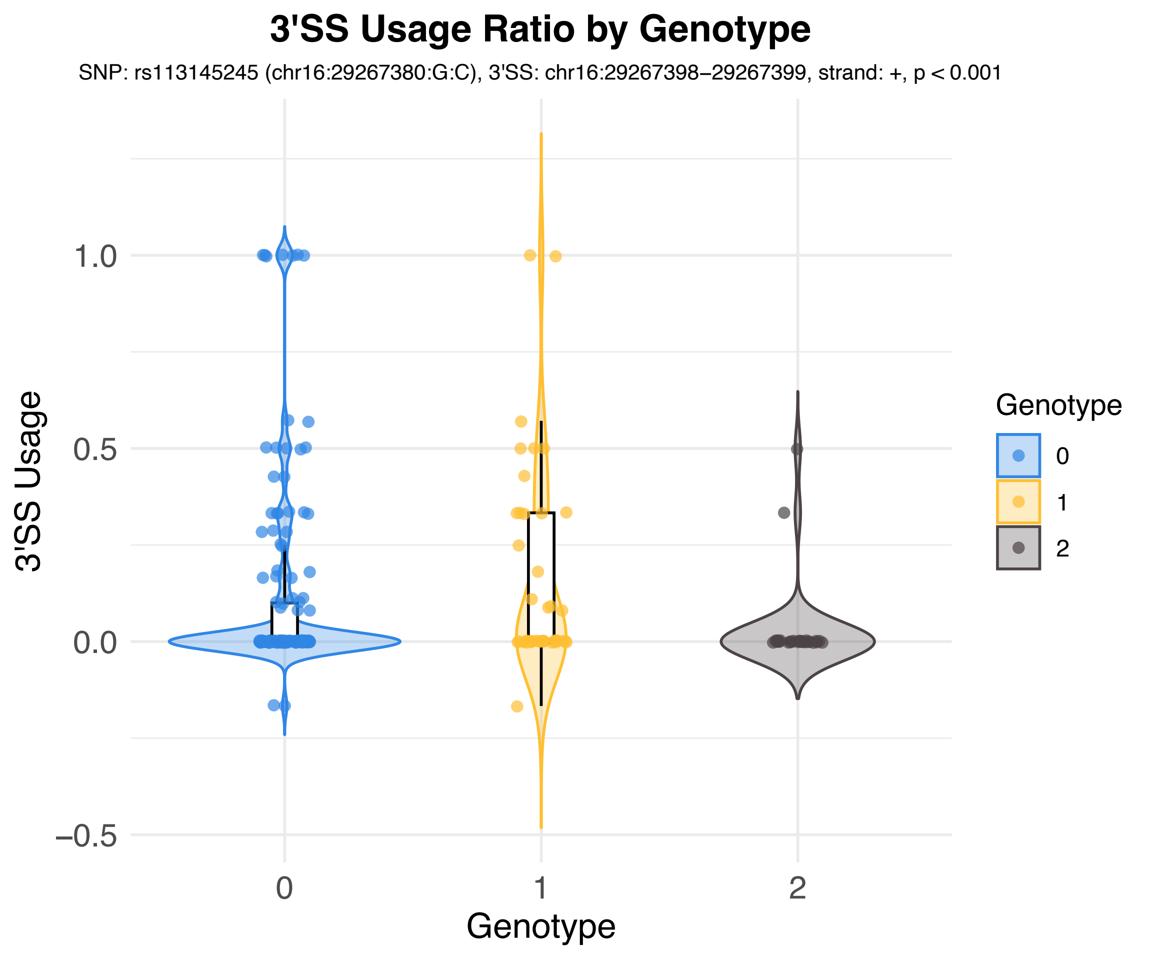
3'SS Usage Ratio by Genotype
SNP: rs113145245 (chr16:29267380:G:C), 3'SS: chr16:29267398–29267399, strand: +, p < 0.001
3'SS Usage
1.0
0.5
0.0
–0.5
0
1
2
Genotype
Genotype
0
1
2

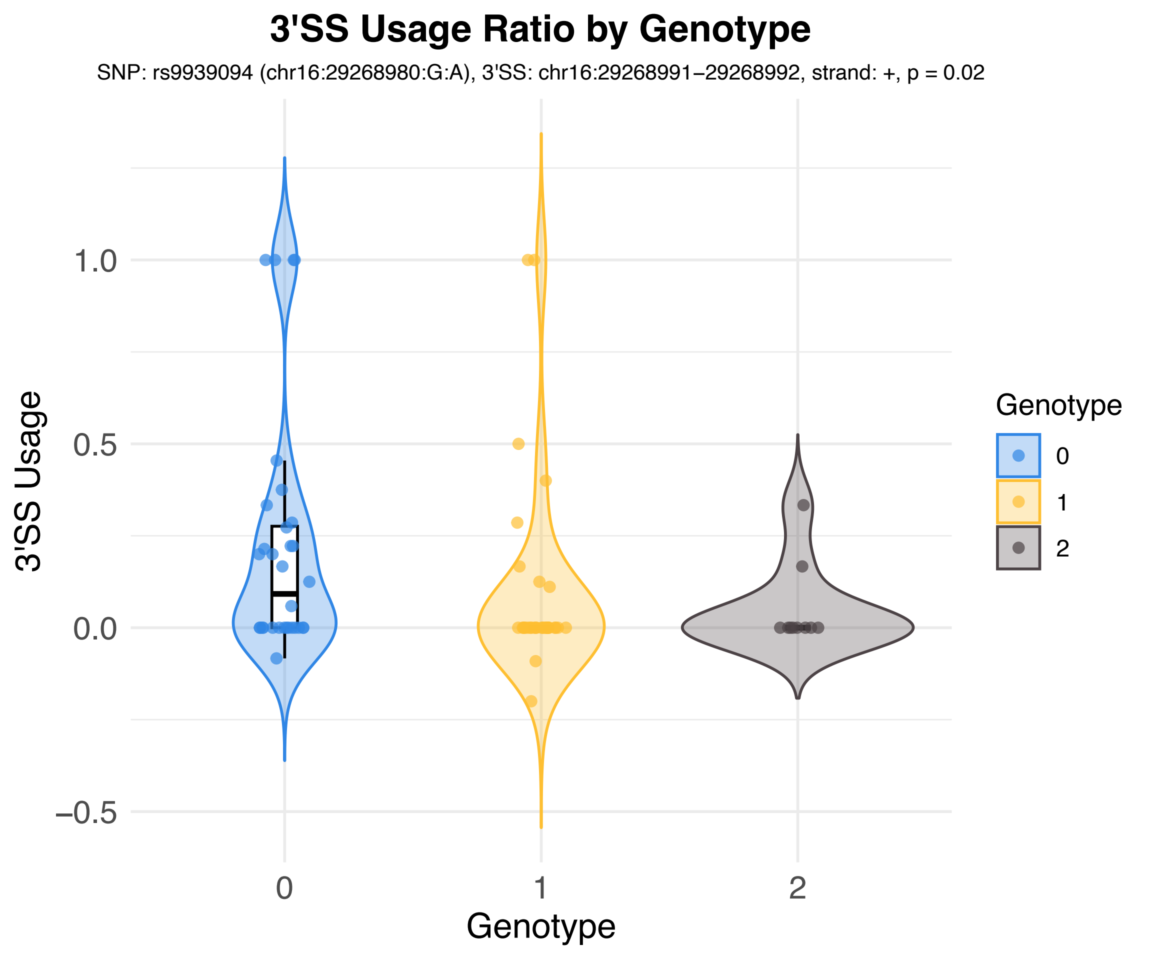
3'SS Usage Ratio by Genotype
SNP: rs9939094 (chr16:29268980:G:A), 3'SS: chr16:29268991–29268992, strand: +, p = 0.02
3'SS Usage
1.0
0.5
0.0
−0.5
0
1
2
Genotype
Genotype
0
1
2

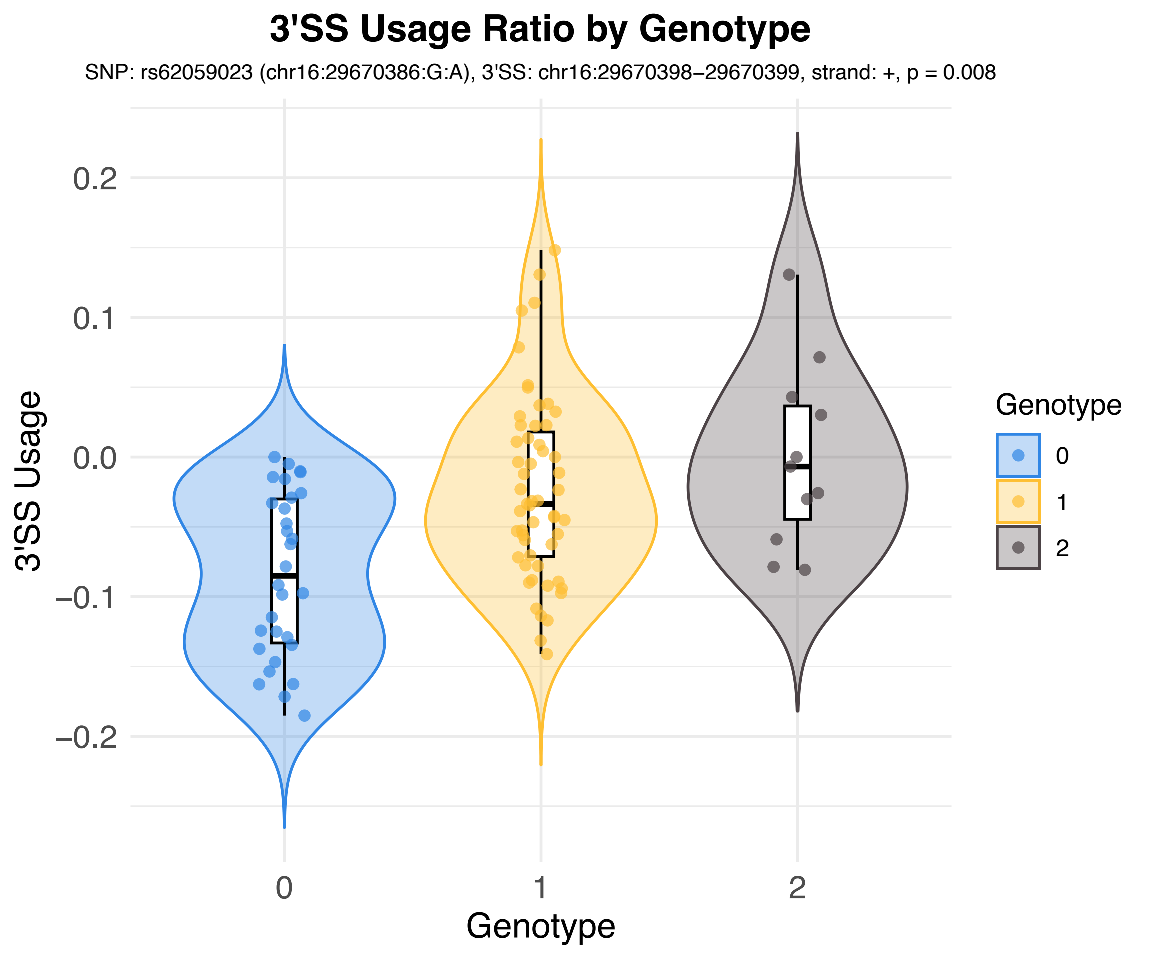
3'SS Usage Ratio by Genotype
SNP: rs62059023 (chr16:29670386:G:A), 3'SS: chr16:29670398–29670399, strand: +, p = 0.008
3'SS Usage
0.2
0.1
0.0
−0.1
−0.2
0
1
2
Genotype
Genotype
0
1
2

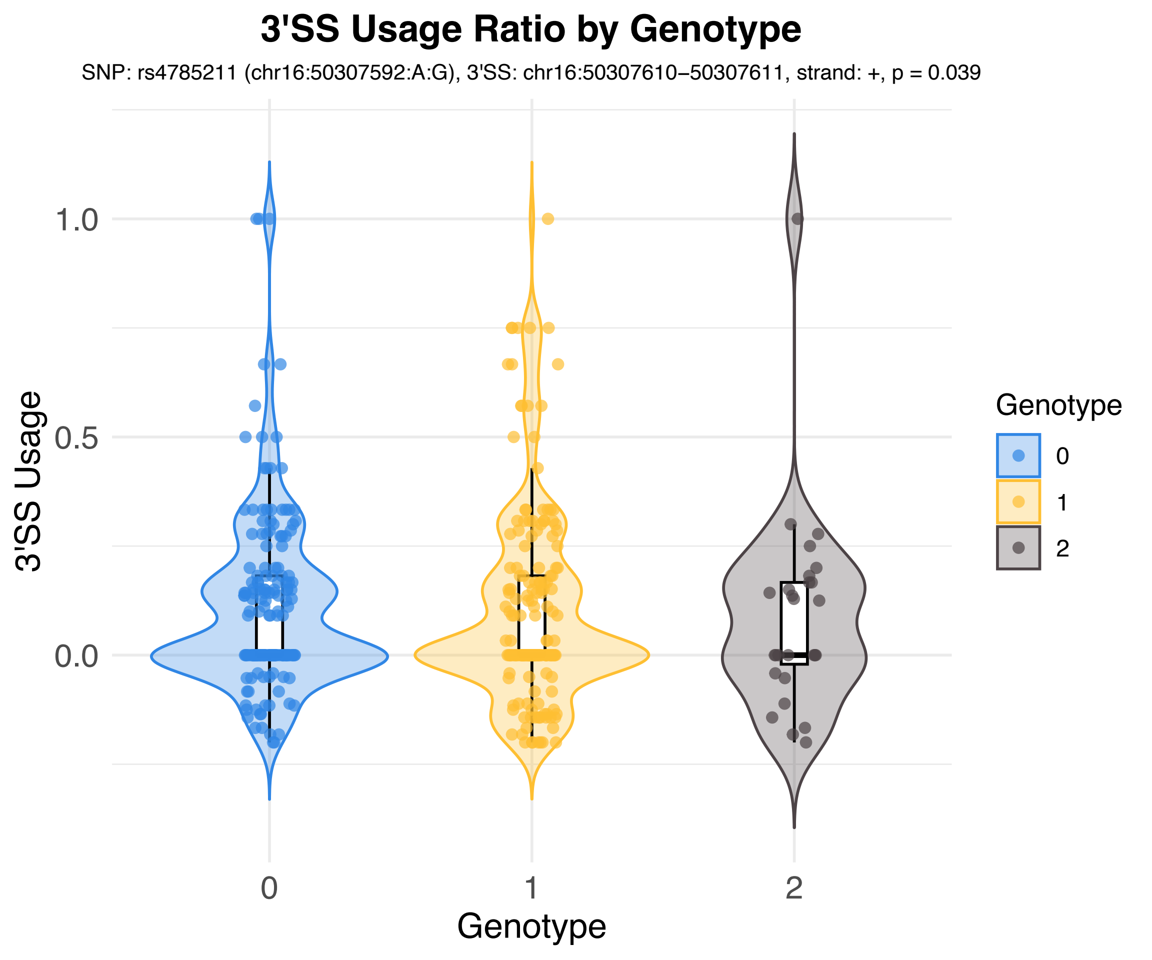

3'SS Usage Ratio by Genotype
SNP: rs4785211 (chr16:50307592:A:G), 3'SS: chr16:50307610−50307611, strand: +, p = 0.039
3'SS Usage
1.0
0.5
0.0
0
1
2
Genotype
Genotype
0
1
2

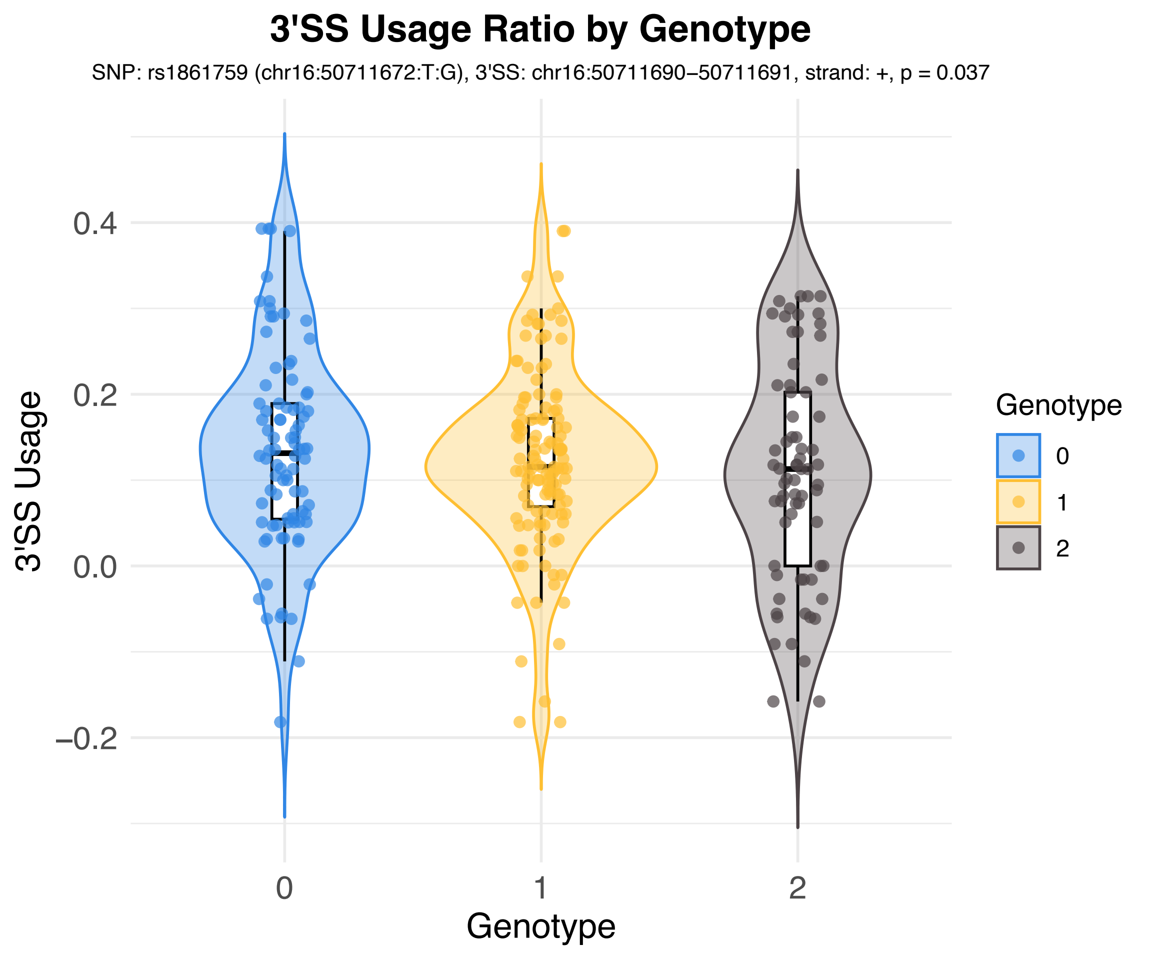
3'SS Usage Ratio by Genotype
SNP: rs1861759 (chr16:50711672:T:G), 3'SS: chr16:50711690–50711691, strand: +, p = 0.037
3'SS Usage
0.4
0.2
0.0
−0.2
0
1
2
Genotype
Genotype
0
1
2

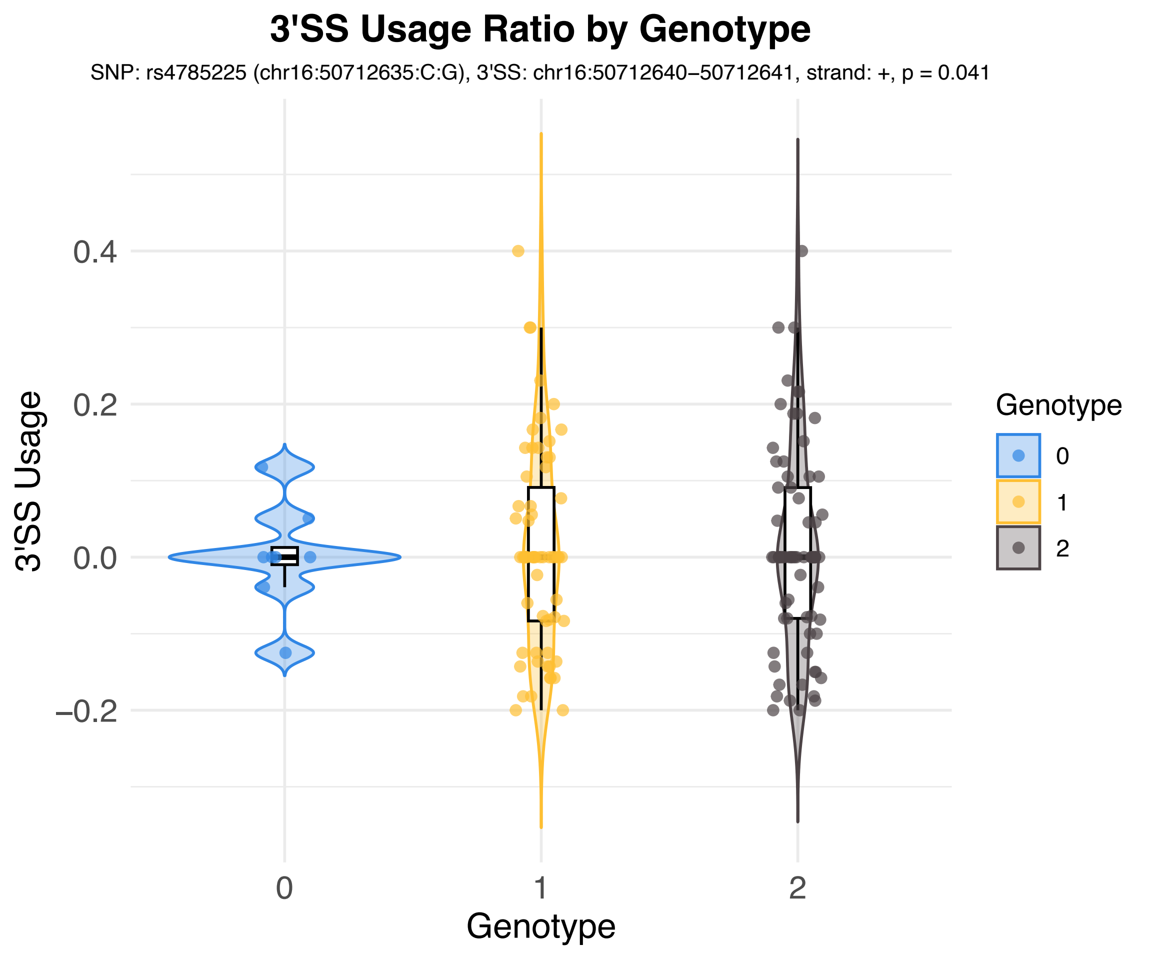
3'SS Usage Ratio by Genotype
SNP: rs4785225 (chr16:50712635:C:G), 3'SS: chr16:50712640–50712641, strand: +, p = 0.041
3'SS Usage
0.4
0.2
0.0
–0.2
0
1
2
Genotype
Genotype
0
1
2

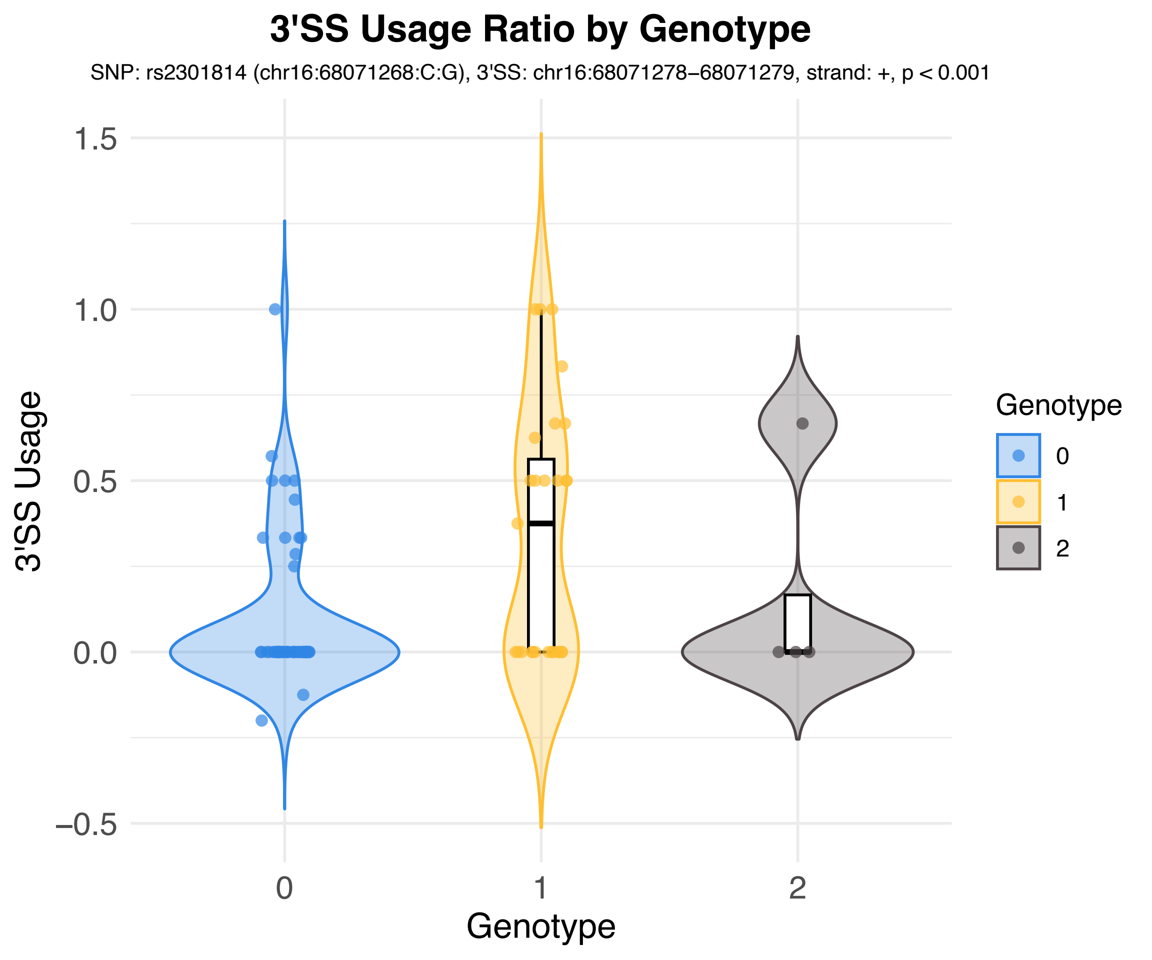

3'SS Usage Ratio by Genotype
SNP: rs2301814 (chr16:68071268:C:G), 3'SS: chr16:68071278–68071279, strand: +, p < 0.001
3'SS Usage
1.5
1.0
0.5
0.0
−0.5
0
1
2
Genotype
Genotype
0
1
2

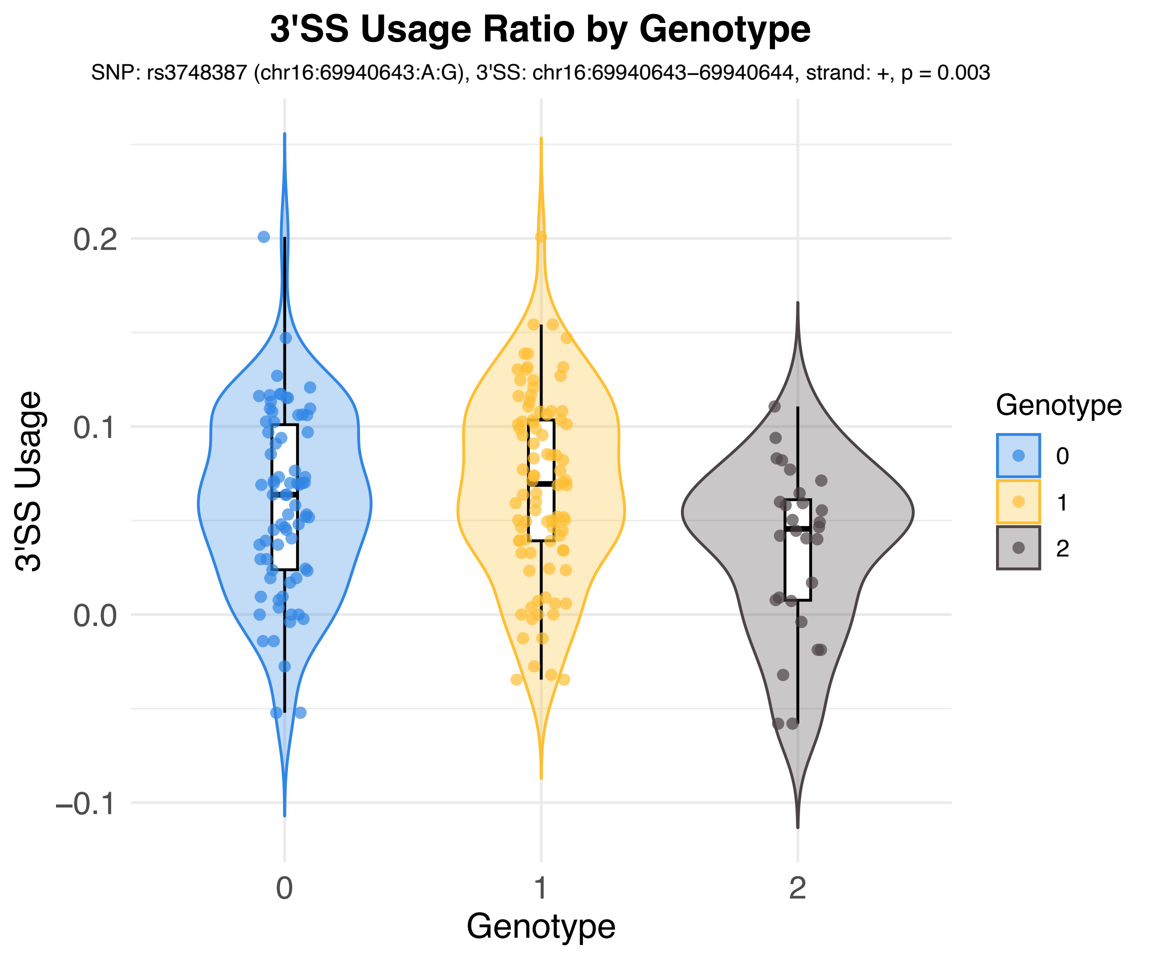
3'SS Usage Ratio by Genotype
SNP: rs3748387 (chr16:69940643:A:G), 3'SS: chr16:69940643–69940644, strand: +, p = 0.003
3'SS Usage
0.2
0.1
0.0
−0.1
0
1
2
Genotype
Genotype
0
1
2

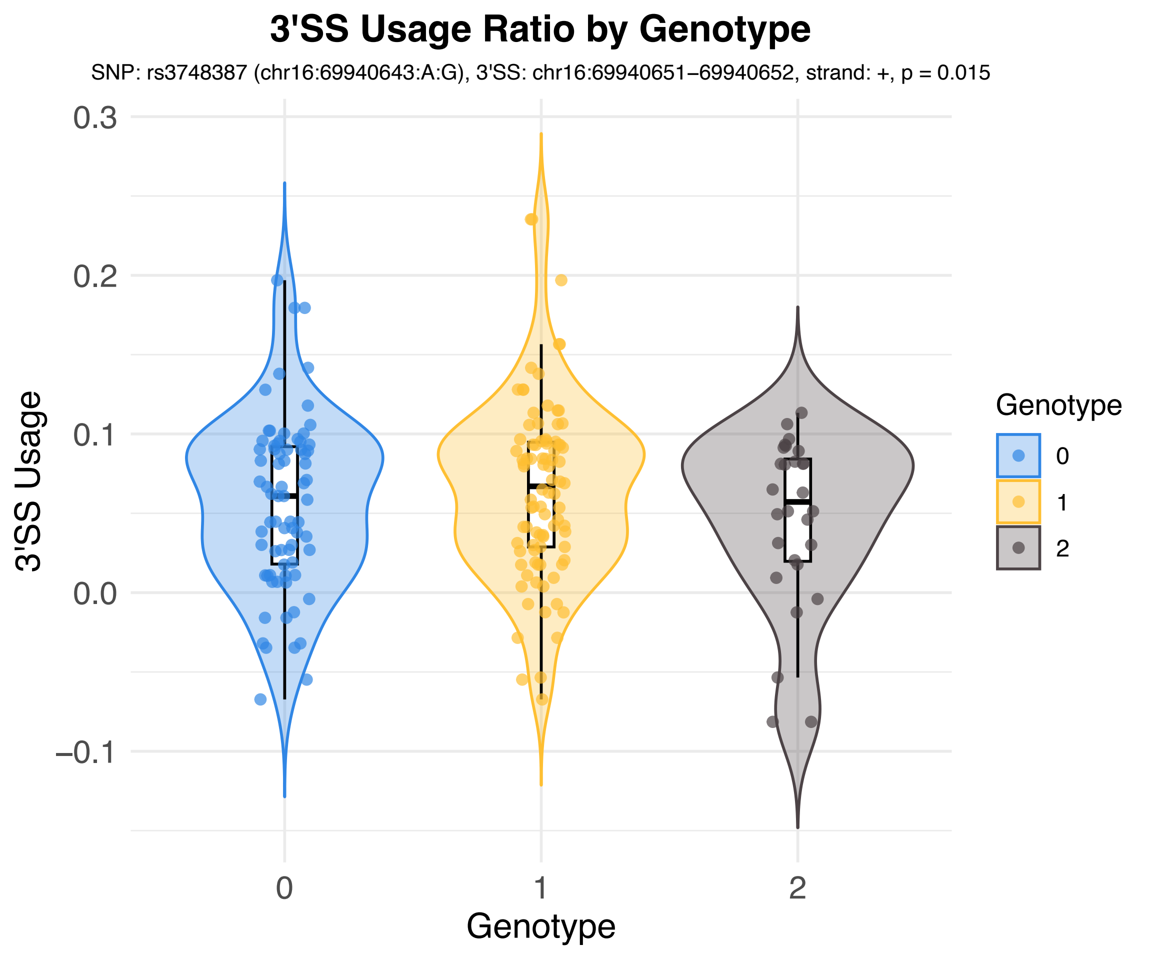

3'SS Usage Ratio by Genotype
SNP: rs3748387 (chr16:69940643:A:G), 3'SS: chr16:69940651–69940652, strand: +, p = 0.015
3'SS Usage
0.3
0.2
0.1
0.0
−0.1
0
1
2
Genotype
Genotype
0
1
2

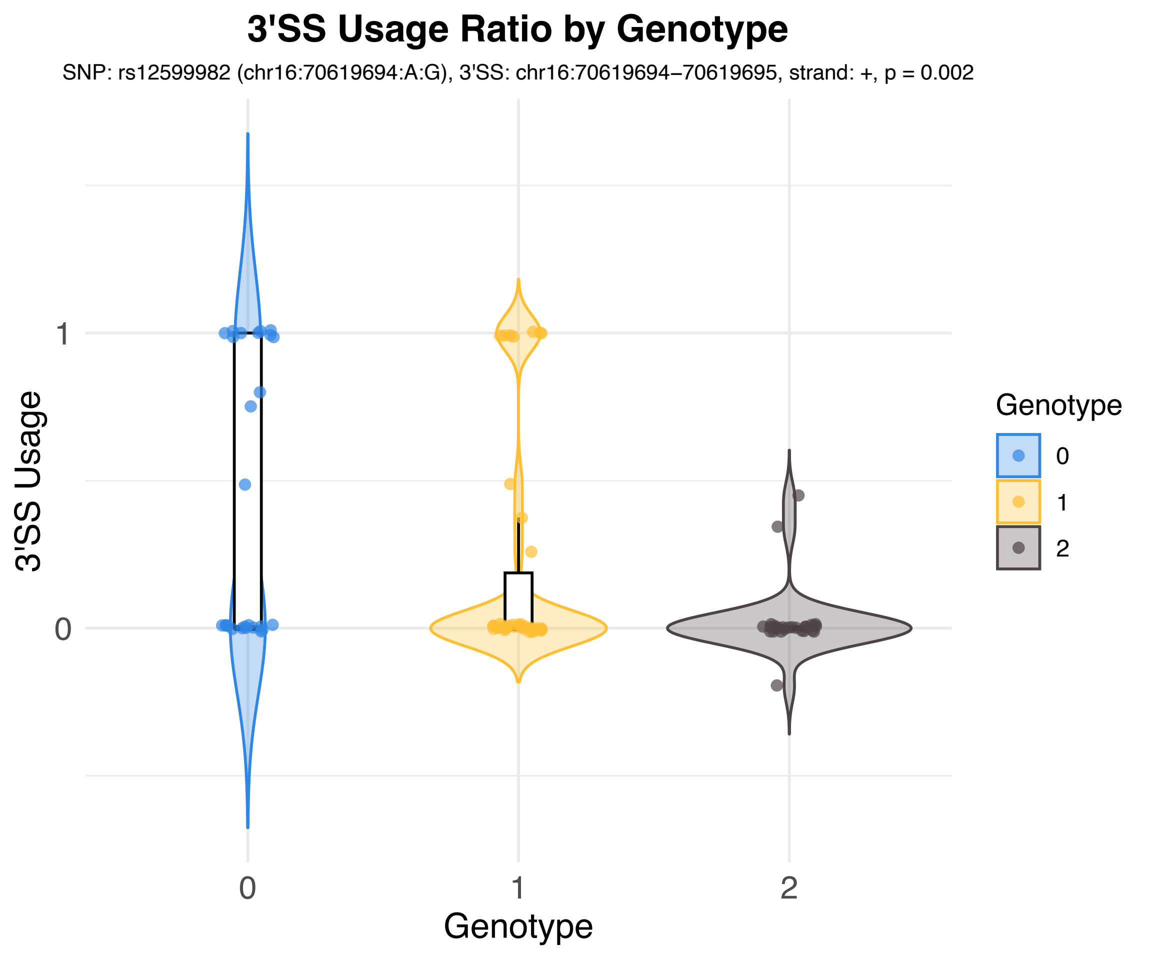
3'SS Usage Ratio by Genotype
SNP: rs12599982 (chr16:70619694:A:G), 3'SS: chr16:70619694–70619695, strand: +, p = 0.002
3'SS Usage
1
0
0
1
2
Genotype
Genotype
0
1
2

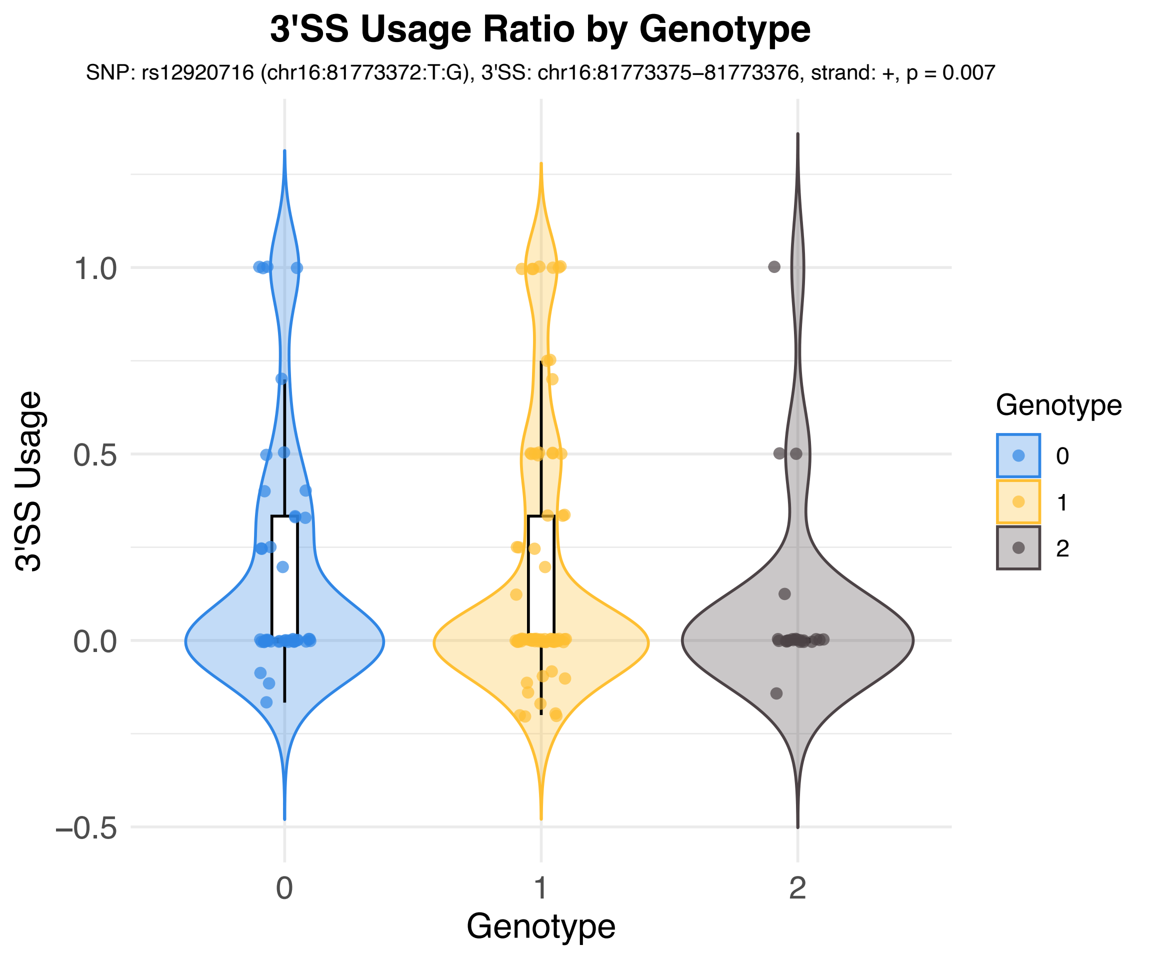
3'SS Usage Ratio by Genotype
SNP: rs12920716 (chr16:81773372:T:G), 3'SS: chr16:81773375–81773376, strand: +, p = 0.007
3'SS Usage
1.0
0.5
0.0
−0.5
0
1
2
Genotype
Genotype
0
1
2

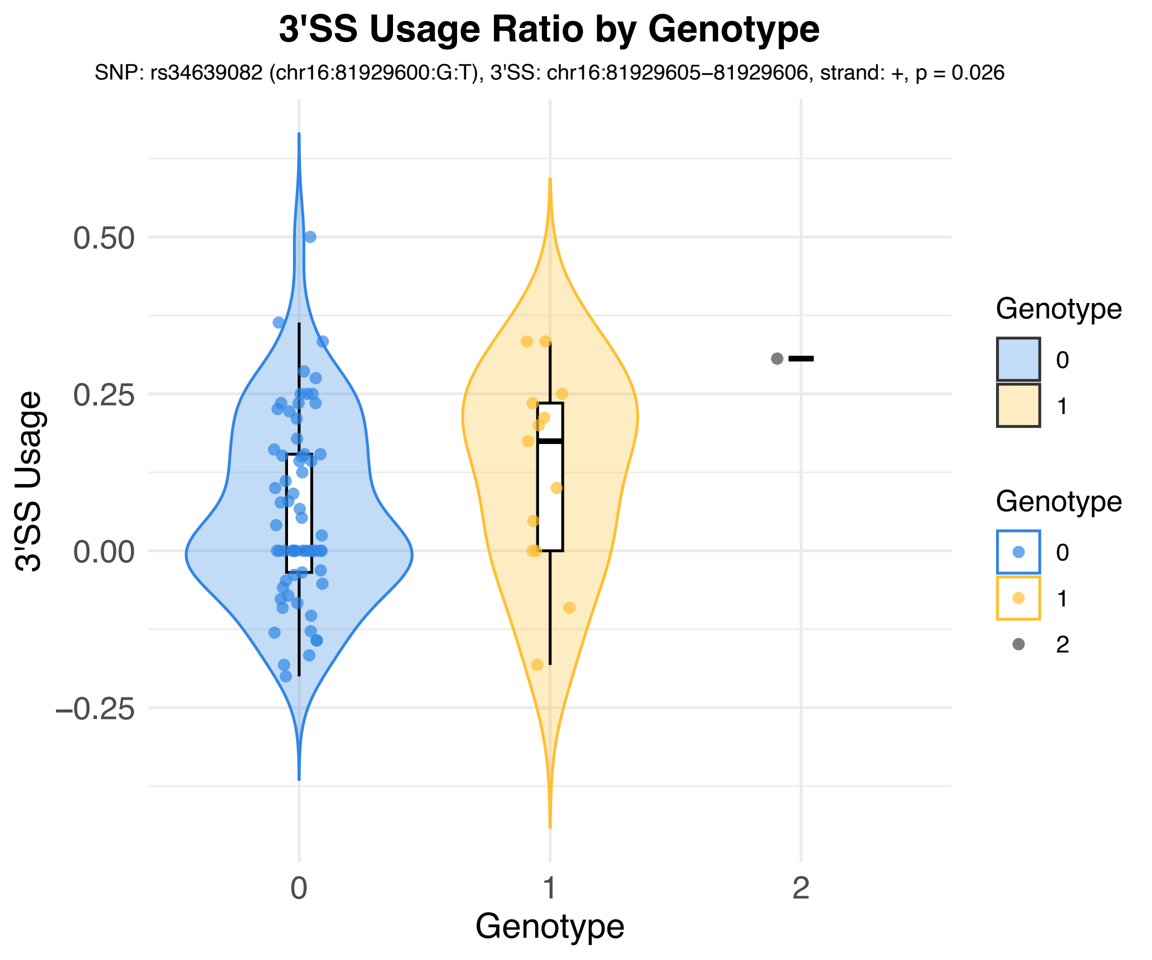

3'SS Usage Ratio by Genotype
SNP: rs34639082 (chr16:81929600:G:T), 3'SS: chr16:81929605–81929606, strand: +, p = 0.026
3'SS Usage
0.50
0.25
0.00
−0.25
0
1
2
Genotype
Genotype
0
1
Genotype
0
1
2

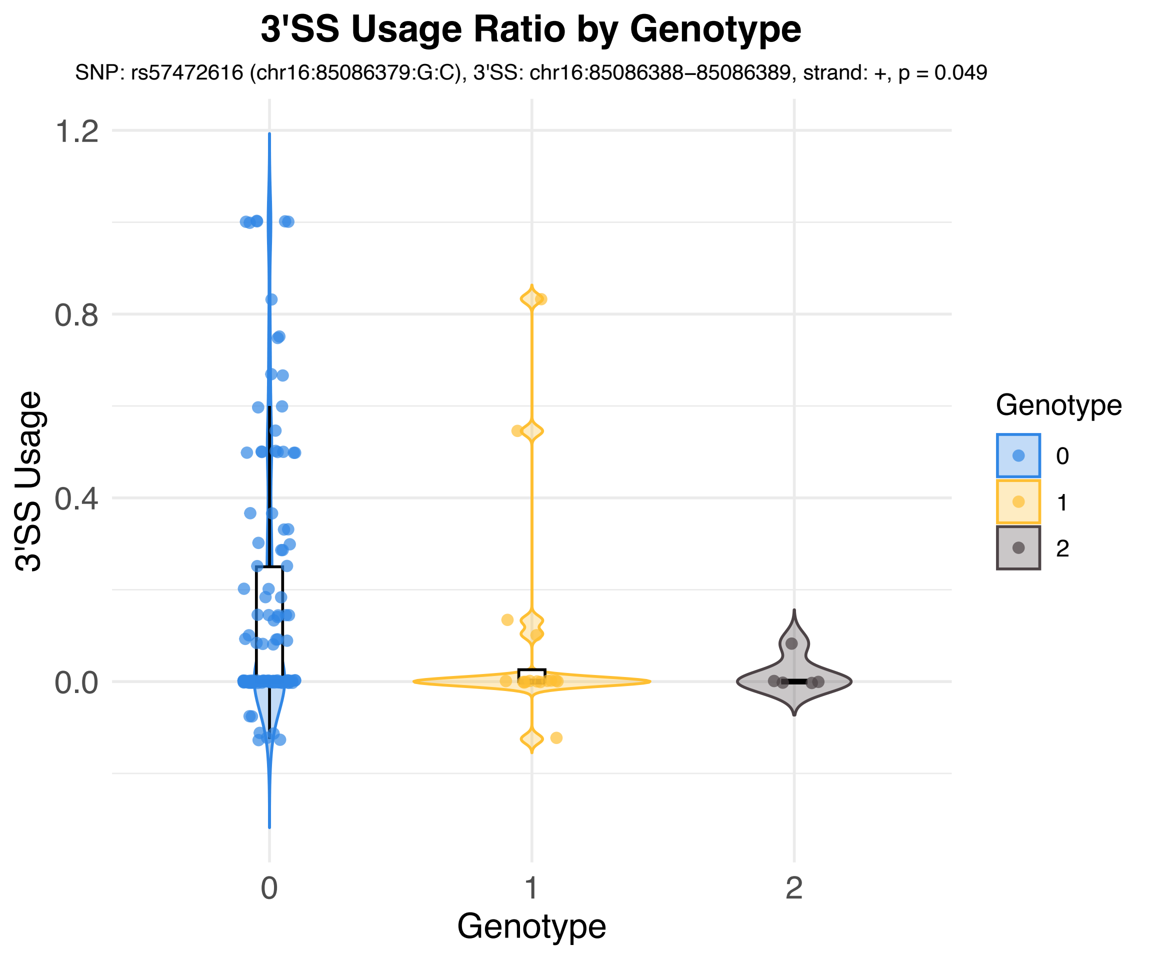
3'SS Usage Ratio by Genotype
SNP: rs57472616 (chr16:85086379:G:C), 3'SS: chr16:85086388–85086389, strand: +, p = 0.049
3'SS Usage
1.2
0.8
0.4
0.0
0
1
2
Genotype
Genotype
0
1
2

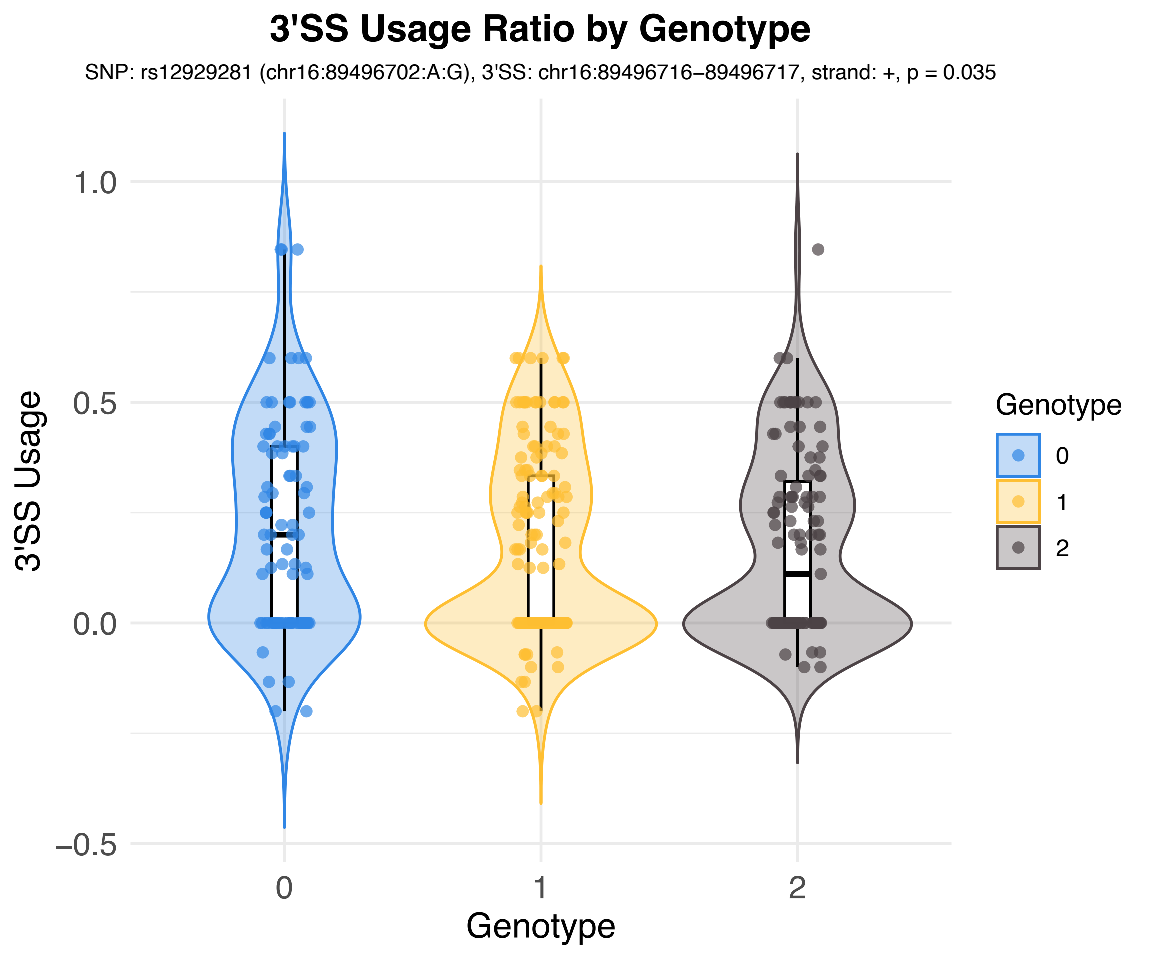
3'SS Usage Ratio by Genotype
SNP: rs12929281 (chr16:89496702:A:G), 3'SS: chr16:89496716–89496717, strand: +, p = 0.035
3'SS Usage
1.0
0.5
0.0
−0.5
0
1
2
Genotype
Genotype
0
1
2

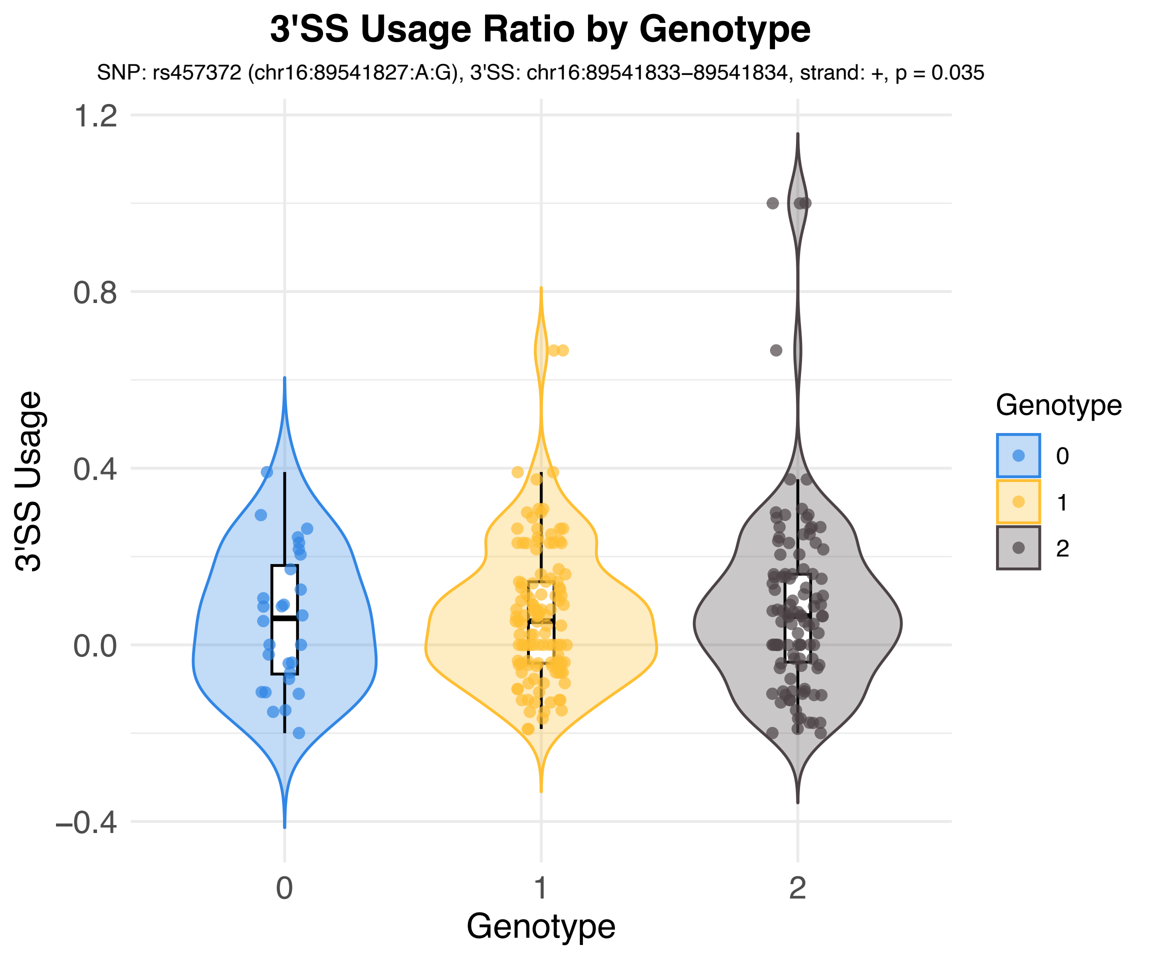

3'SS Usage Ratio by Genotype
SNP: rs457372 (chr16:89541827:A:G), 3'SS: chr16:89541833–89541834, strand: +, p = 0.035
3'SS Usage
1.2
0.8
0.4
0.0
−0.4
0
1
2
Genotype
Genotype
0
1
2

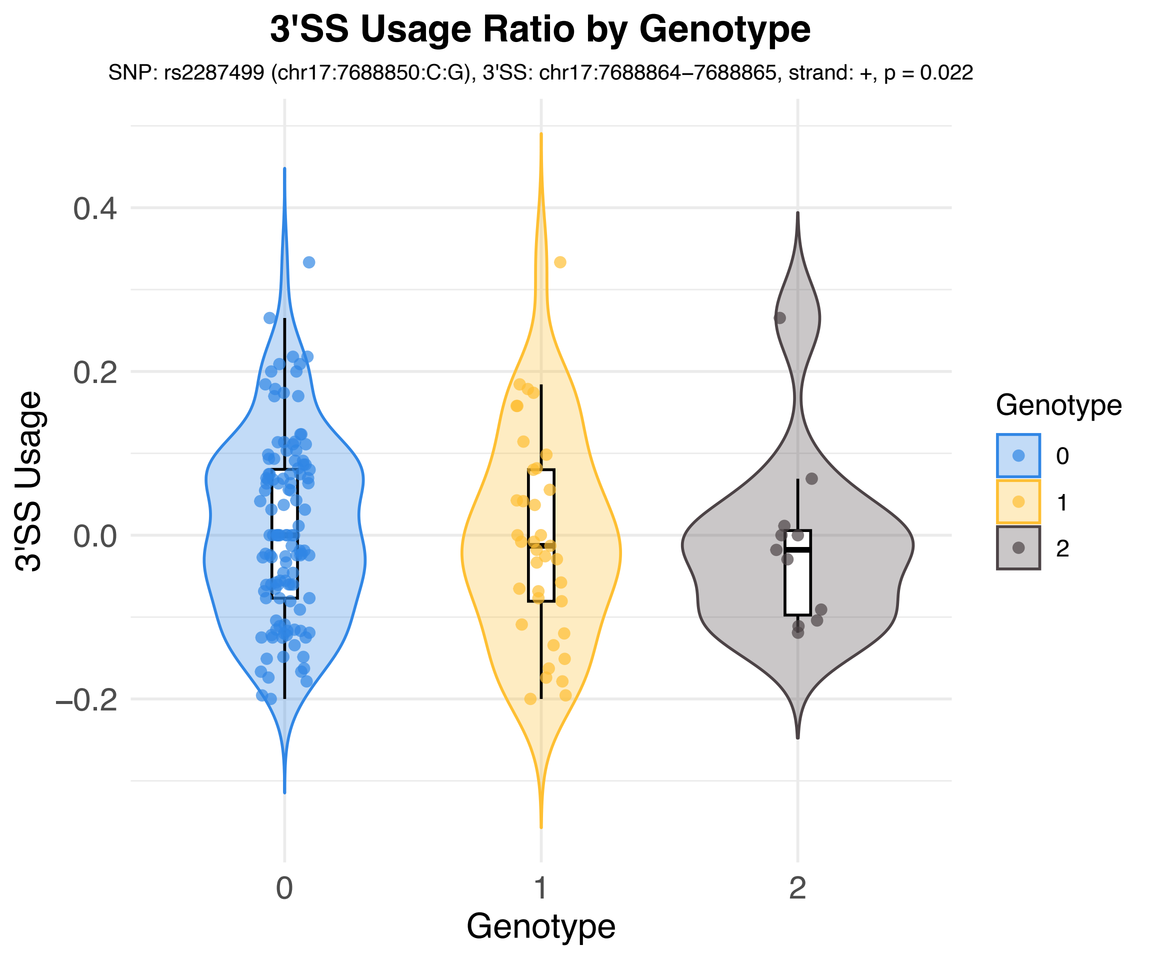
3'SS Usage Ratio by Genotype
SNP: rs2287499 (chr17:7688850:C:G), 3'SS: chr17:7688864–7688865, strand: +, p = 0.022
3'SS Usage
0.4
0.2
0.0
–0.2
0
1
2
Genotype
Genotype
0
1
2

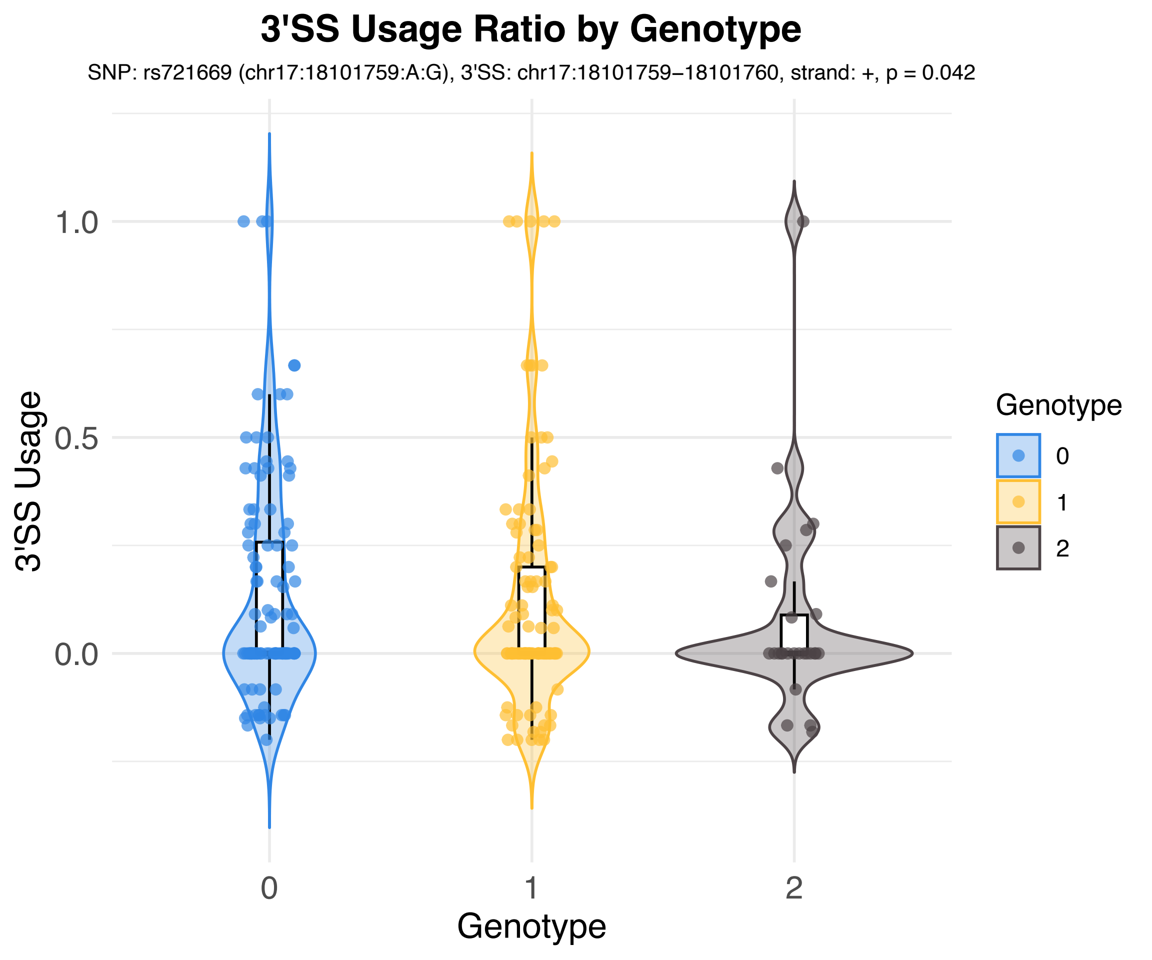

3'SS Usage Ratio by Genotype
SNP: rs721669 (chr17:18101759:A:G), 3'SS: chr17:18101759−18101760, strand: +, p = 0.042
3'SS Usage
1.0
0.5
0.0
0
1
2
Genotype
Genotype
0
1
2

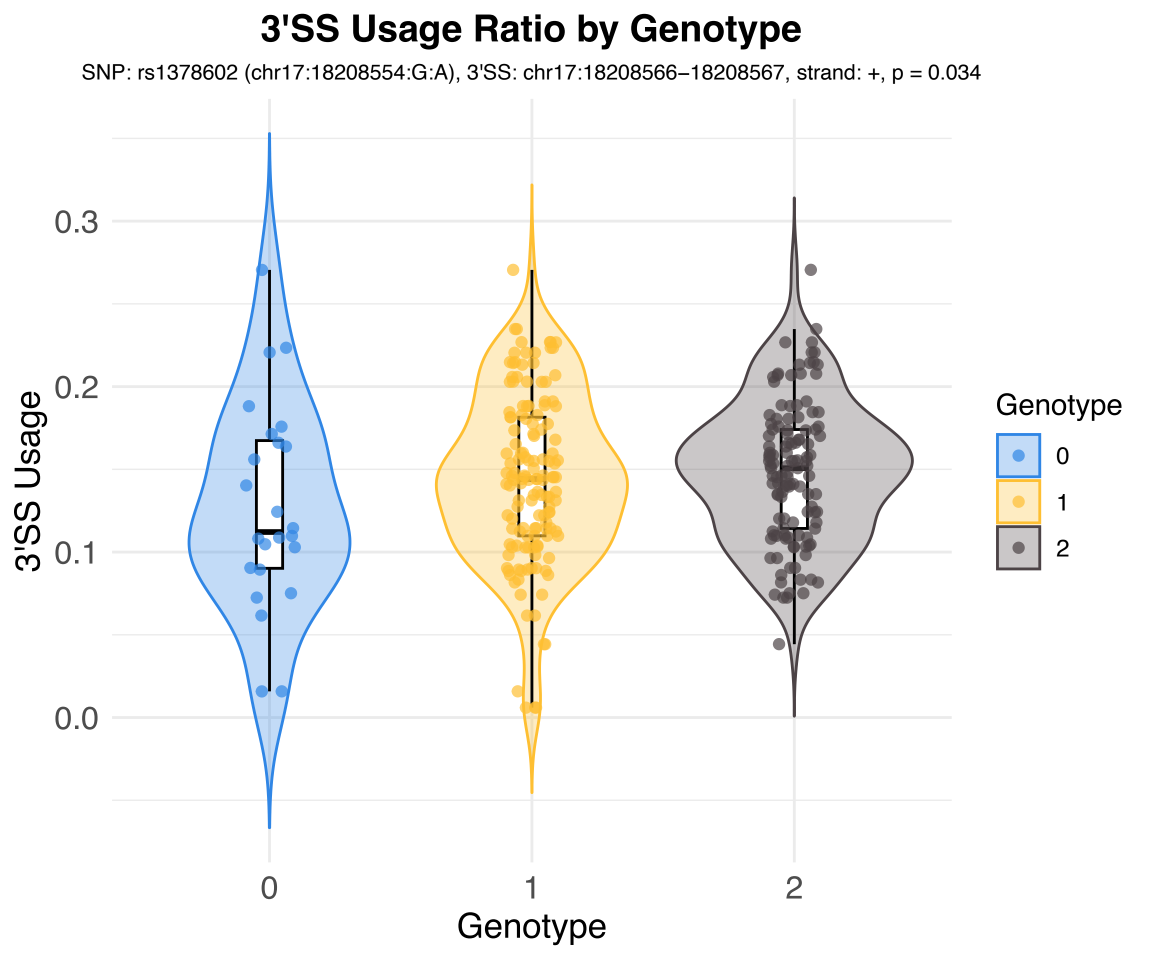
3'SS Usage Ratio by Genotype
SNP: rs1378602 (chr17:18208554:G:A), 3'SS: chr17:18208566−18208567, strand: +, p = 0.034
3'SS Usage
0.3
0.2
0.1
0.0
0
1
2
Genotype
Genotype
0
1
2

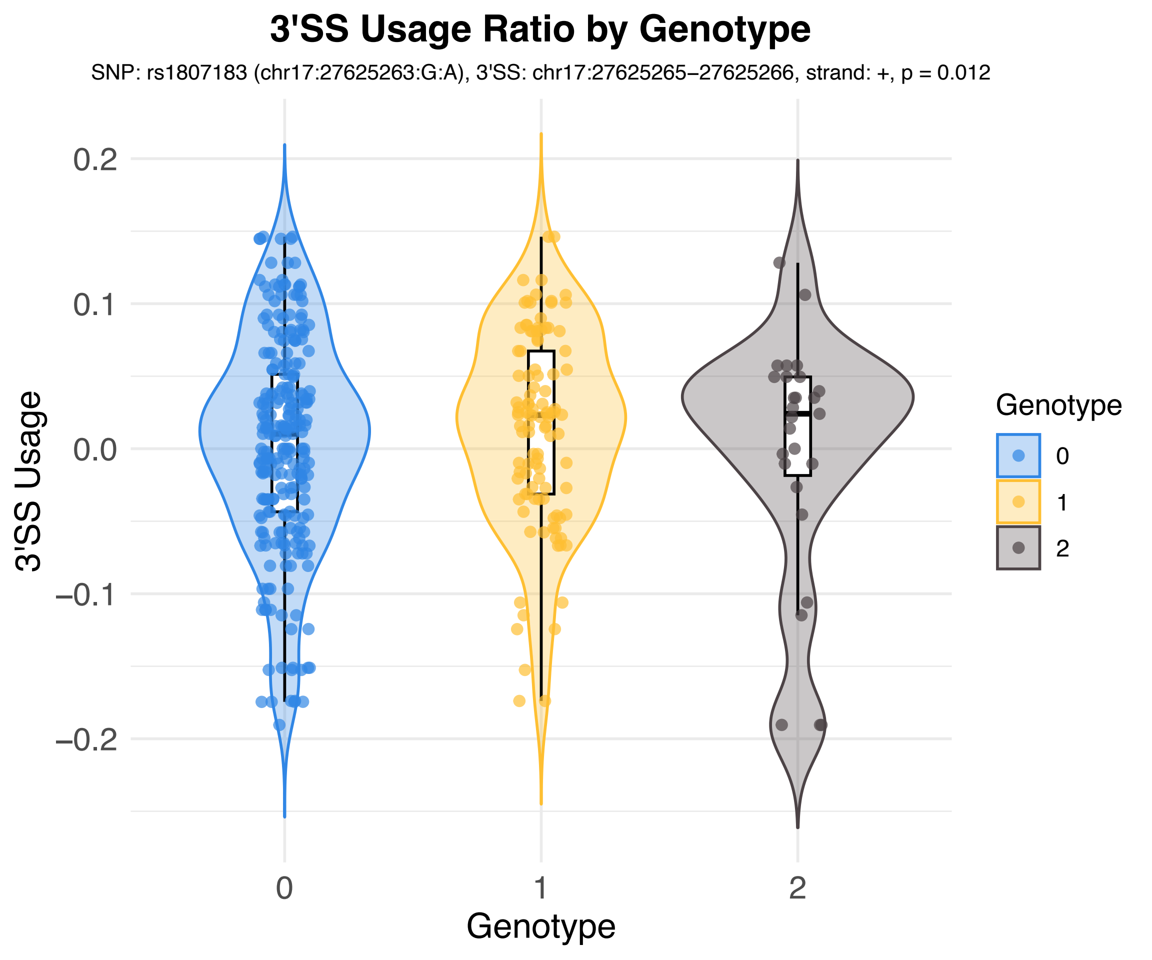
3'SS Usage Ratio by Genotype
SNP: rs1807183 (chr17:27625263:G:A), 3'SS: chr17:27625265–27625266, strand: +, p = 0.012
3'SS Usage
0.2
0.1
0.0
–0.1
–0.2
0
1
2
Genotype
Genotype
0
1
2

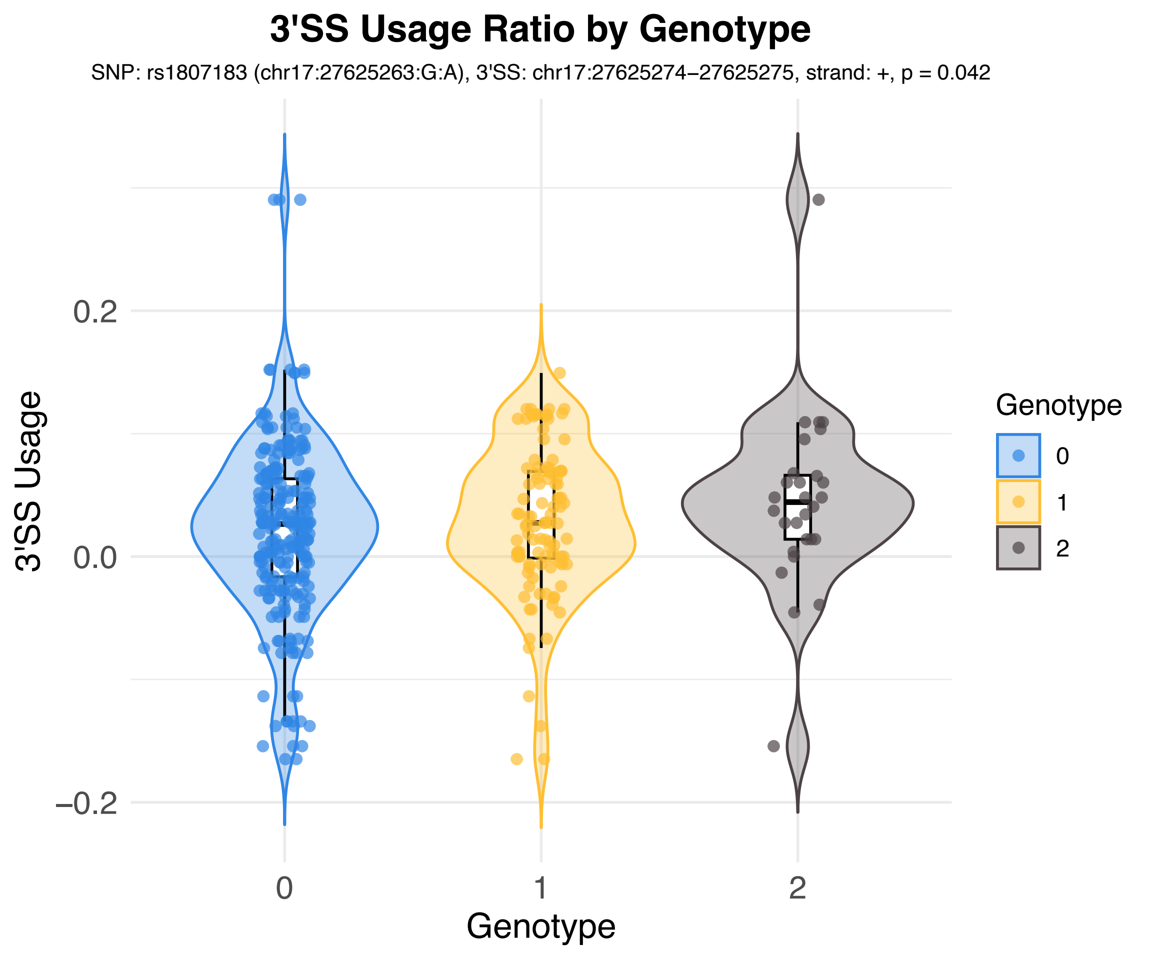
3'SS Usage Ratio by Genotype
SNP: rs1807183 (chr17:27625263:G:A), 3'SS: chr17:27625274–27625275, strand: +, p = 0.042
3'SS Usage
0.2
0.0
−0.2
0
1
2
Genotype
Genotype
0
1
2

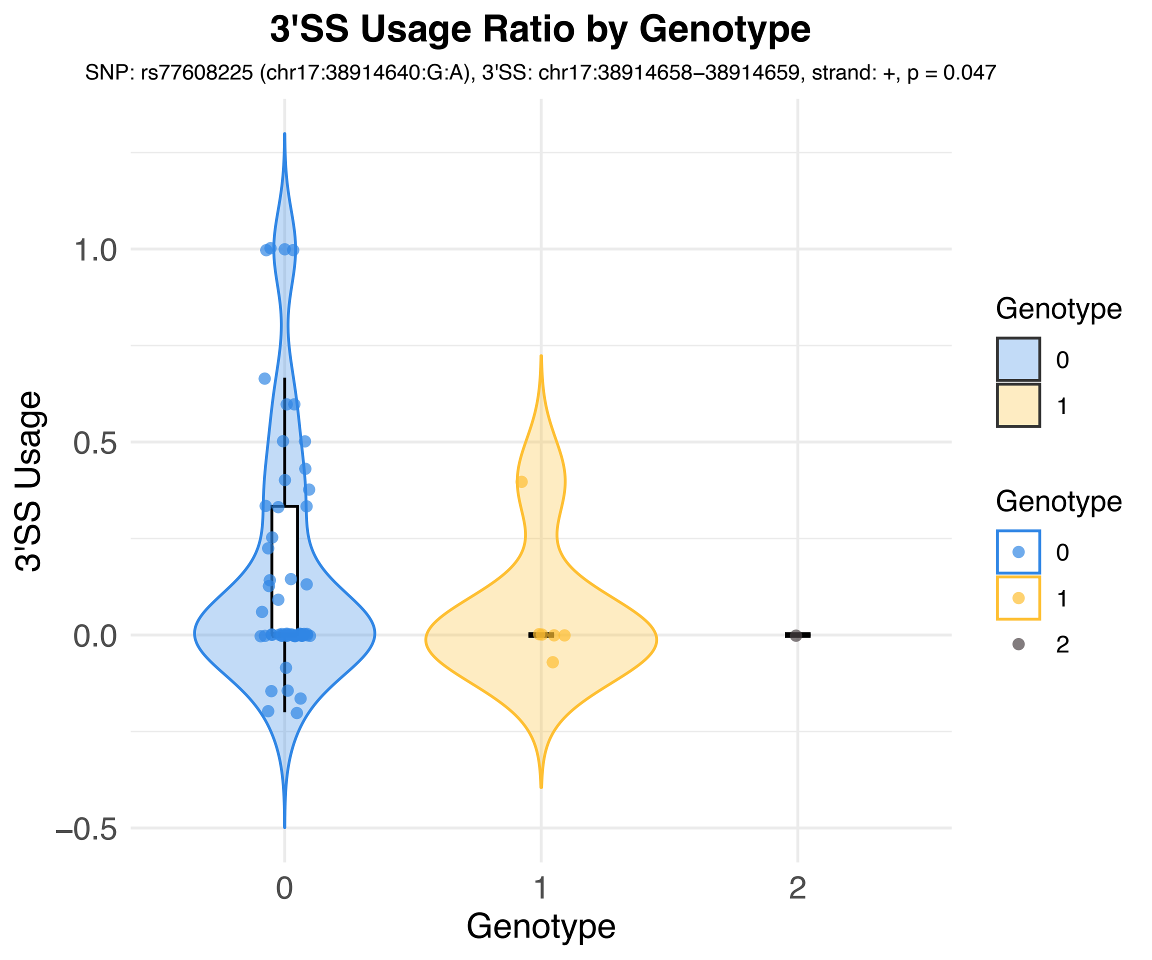
3'SS Usage Ratio by Genotype
SNP: rs77608225 (chr17:38914640:G:A), 3'SS: chr17:38914658–38914659, strand: +, p = 0.047
3'SS Usage
1.0
0.5
0.0
−0.5
0
1
2
Genotype
Genotype
0
1
Genotype
0
1
2

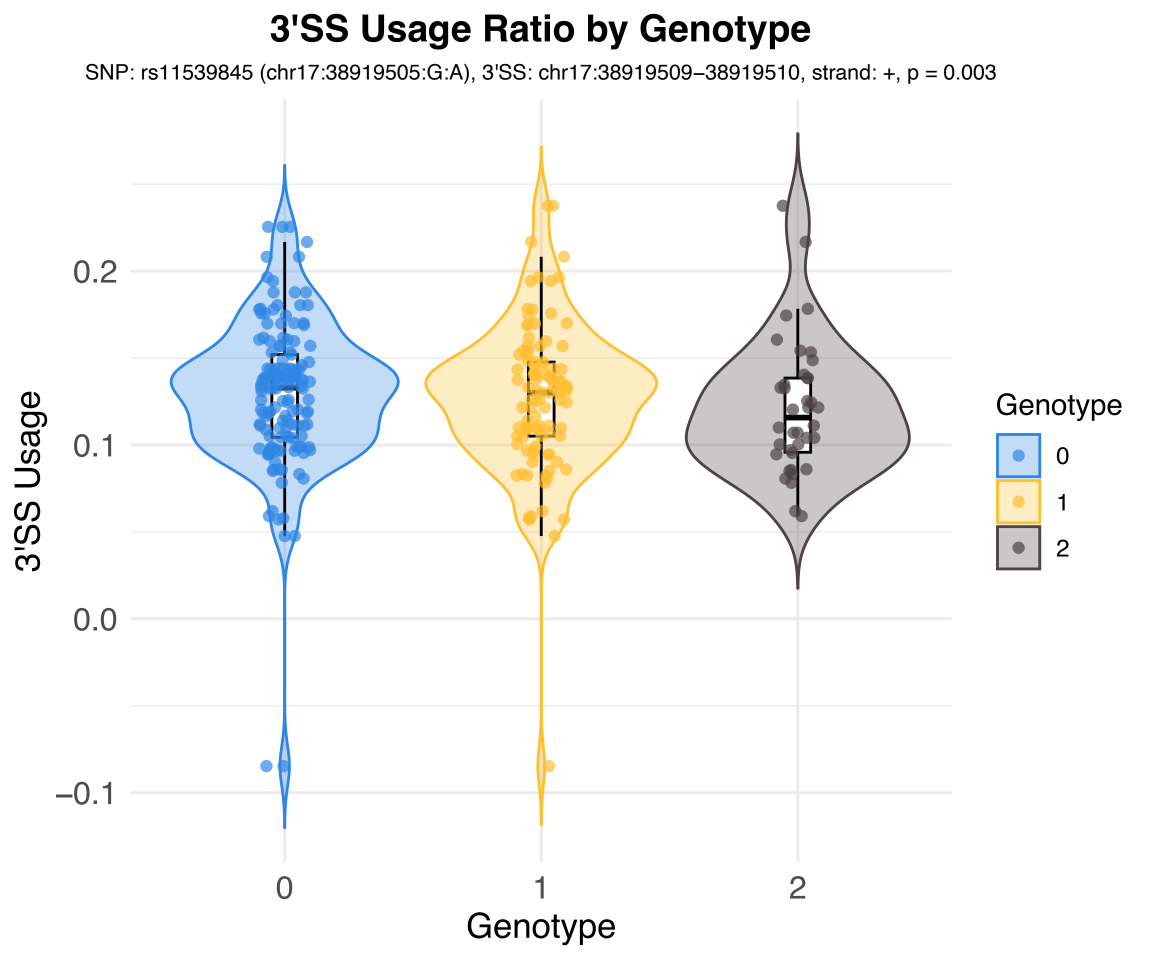
3'SS Usage Ratio by Genotype
SNP: rs11539845 (chr17:38919505:G:A), 3'SS: chr17:38919509–38919510, strand: +, p = 0.003
3'SS Usage
0.2
0.1
0.0
−0.1
0
1
2
Genotype
Genotype
0
1
2

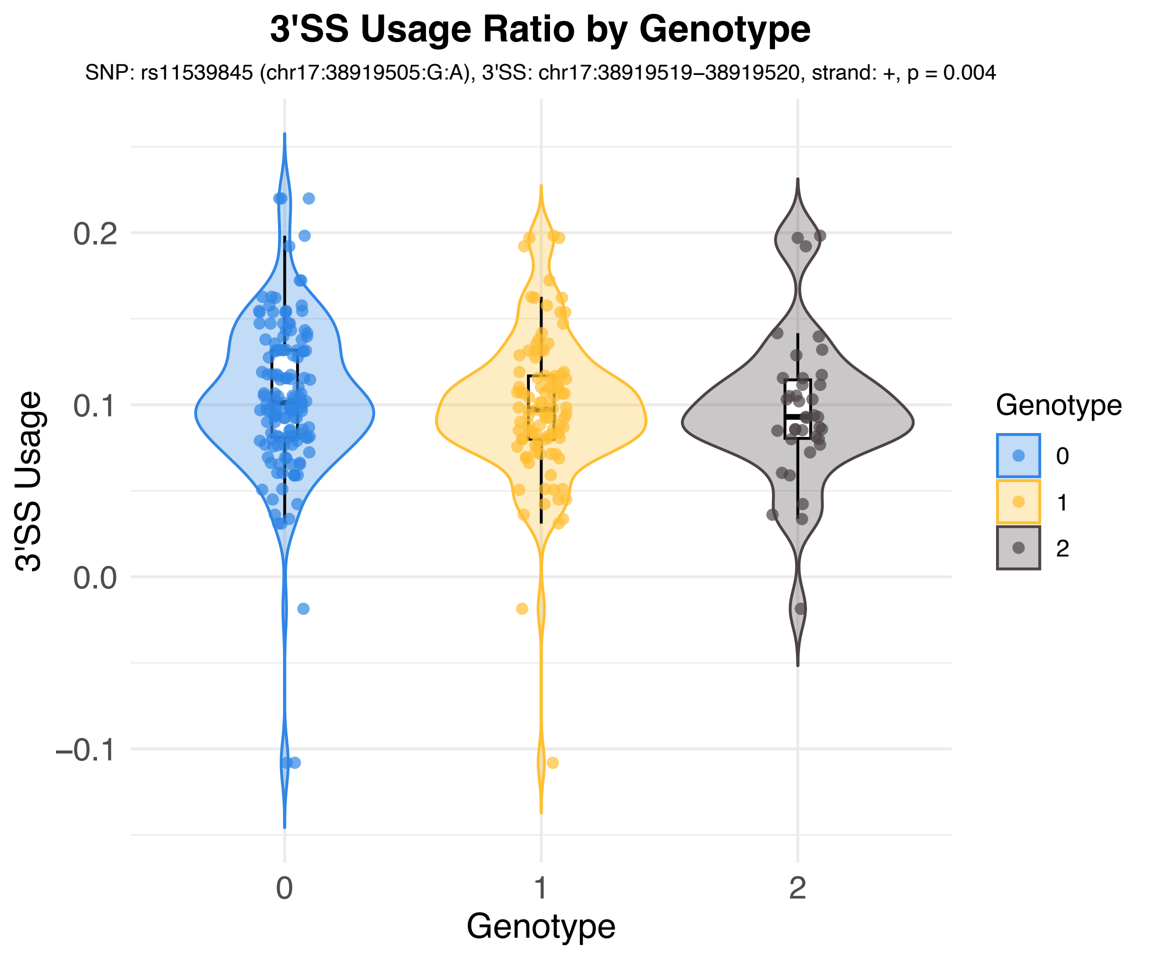

3'SS Usage Ratio by Genotype
SNP: rs11539845 (chr17:38919505:G:A), 3'SS: chr17:38919519–38919520, strand: +, p = 0.004
3'SS Usage
0.2
0.1
0.0
−0.1
0
1
2
Genotype
Genotype
0
1
2

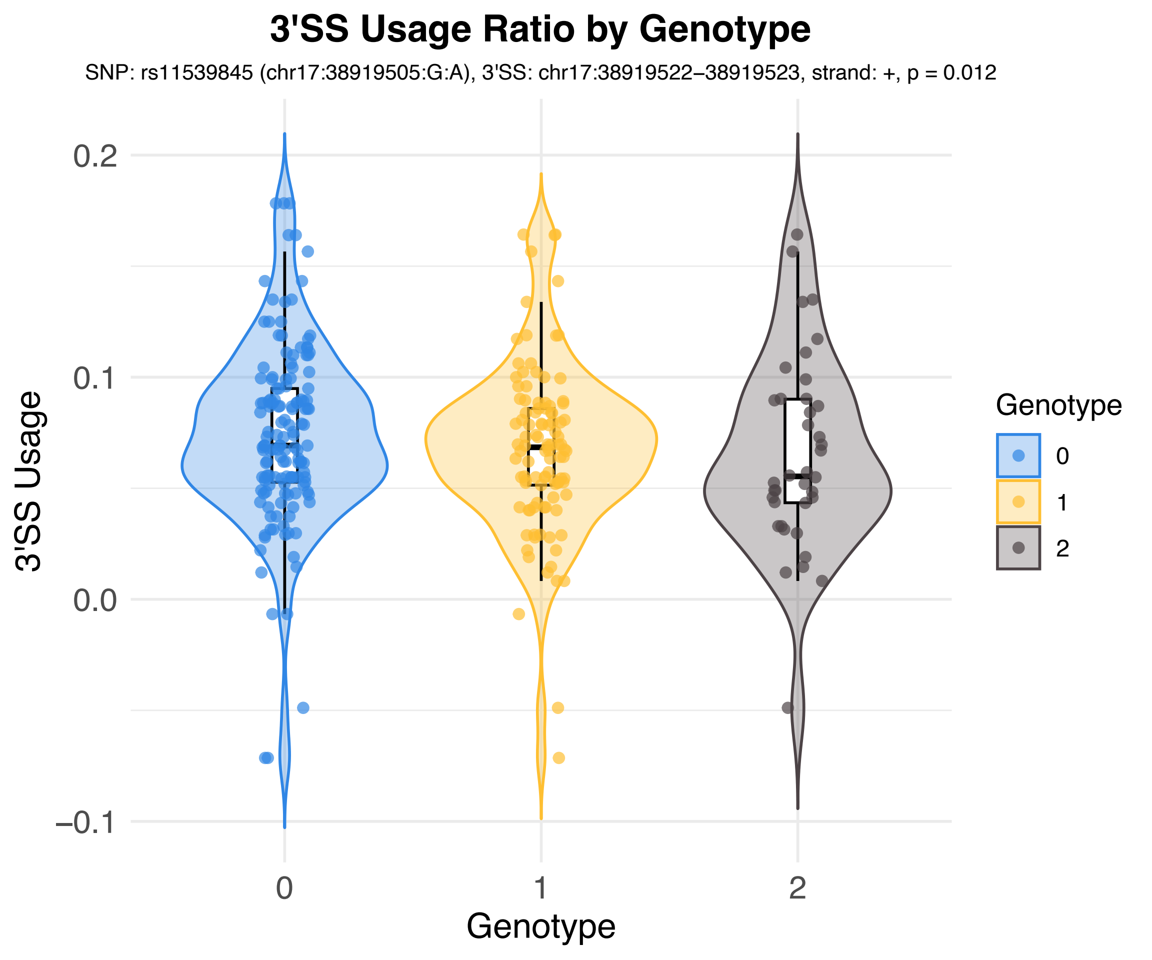
3'SS Usage Ratio by Genotype
SNP: rs11539845 (chr17:38919505:G:A), 3'SS: chr17:38919522–38919523, strand: +, p = 0.012
3'SS Usage
0.2
0.1
0.0
−0.1
0
1
2
Genotype
Genotype
0
1
2

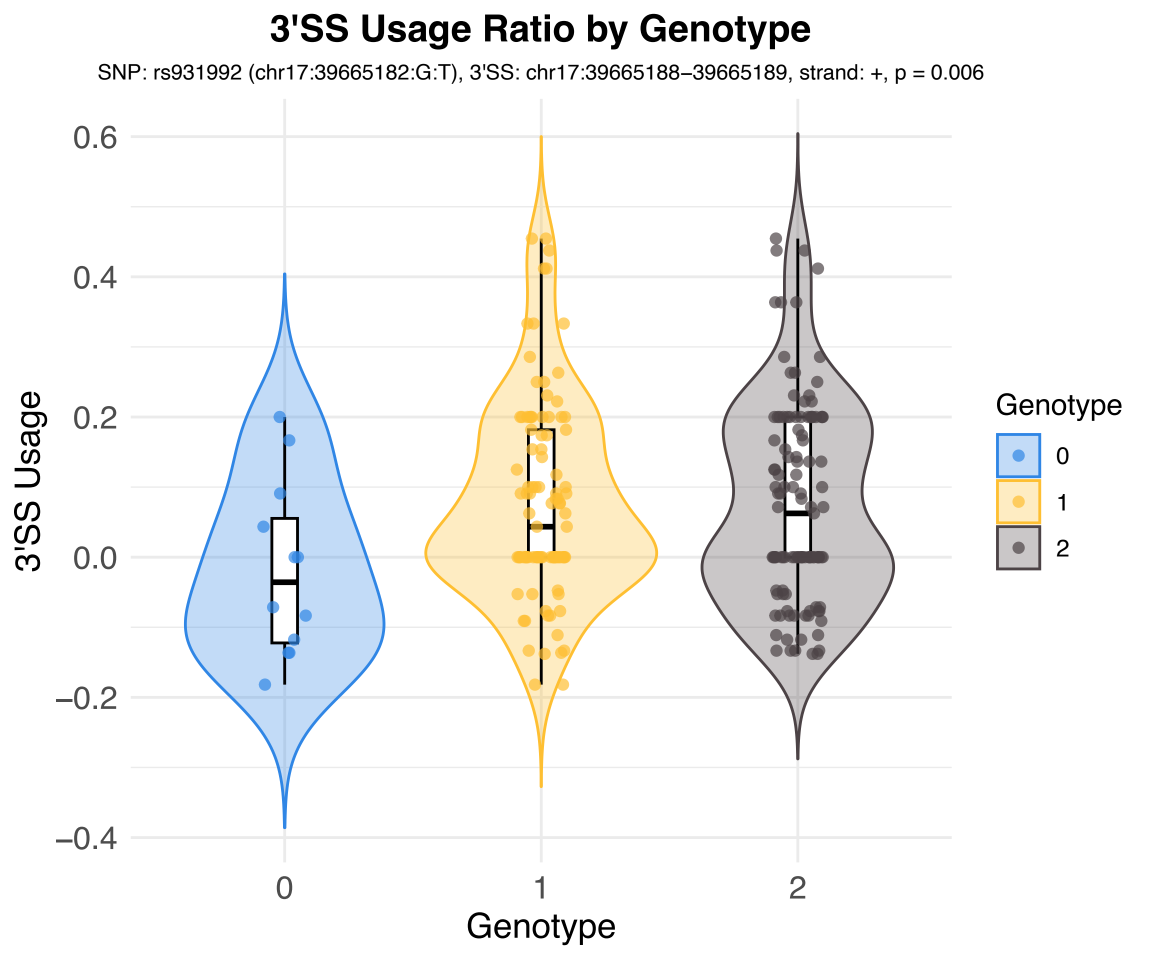
3'SS Usage Ratio by Genotype
SNP: rs931992 (chr17:39665182:G:T), 3'SS: chr17:39665188–39665189, strand: +, p = 0.006
3'SS Usage
0.6
0.4
0.2
0.0
−0.2
−0.4
0
1
2
Genotype
Genotype
0
1
2

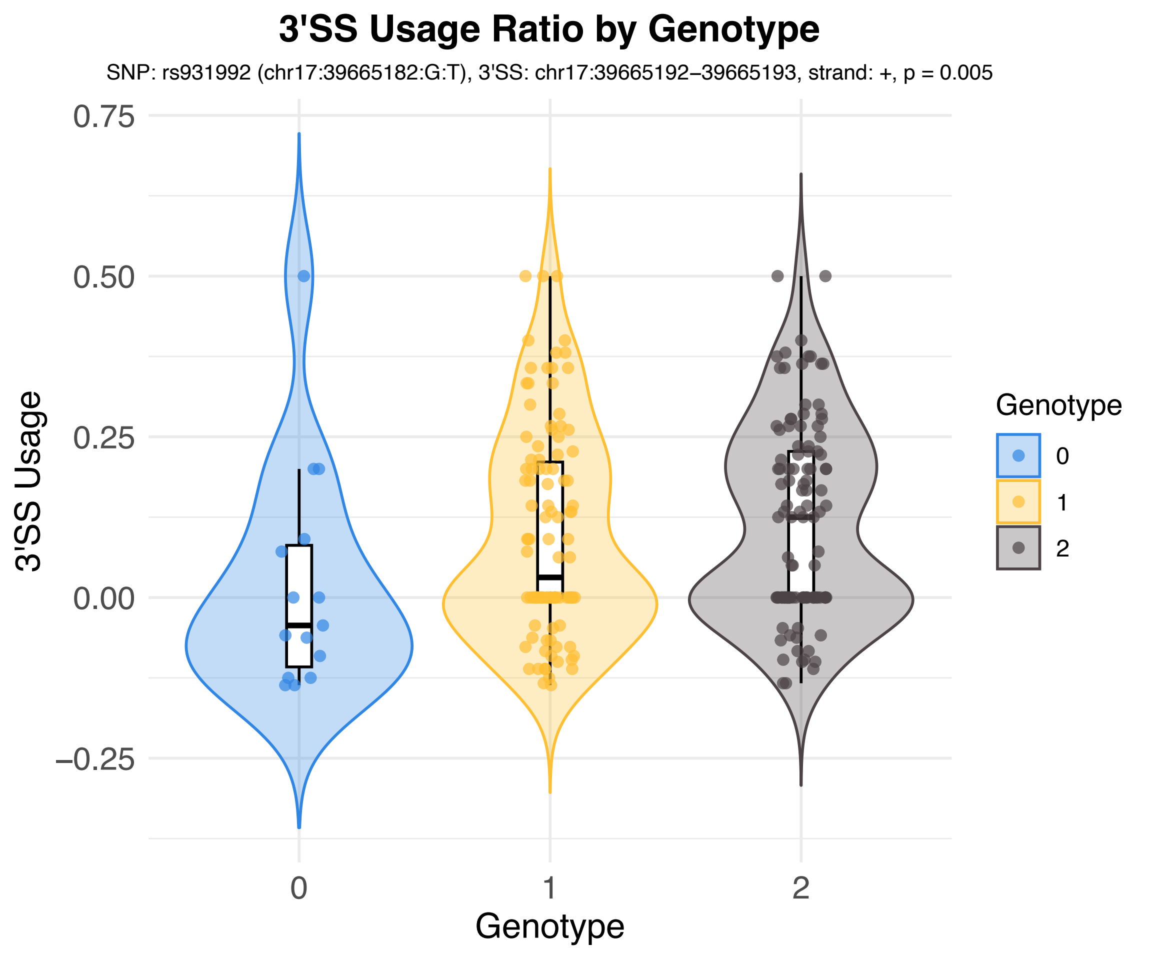

3'SS Usage Ratio by Genotype
SNP: rs931992 (chr17:39665182:G:T), 3'SS: chr17:39665192–39665193, strand: +, p = 0.005
0.75
0.50
0.25
0.00
−0.25
3'SS Usage
0
1
2
Genotype
Genotype
0
1
2

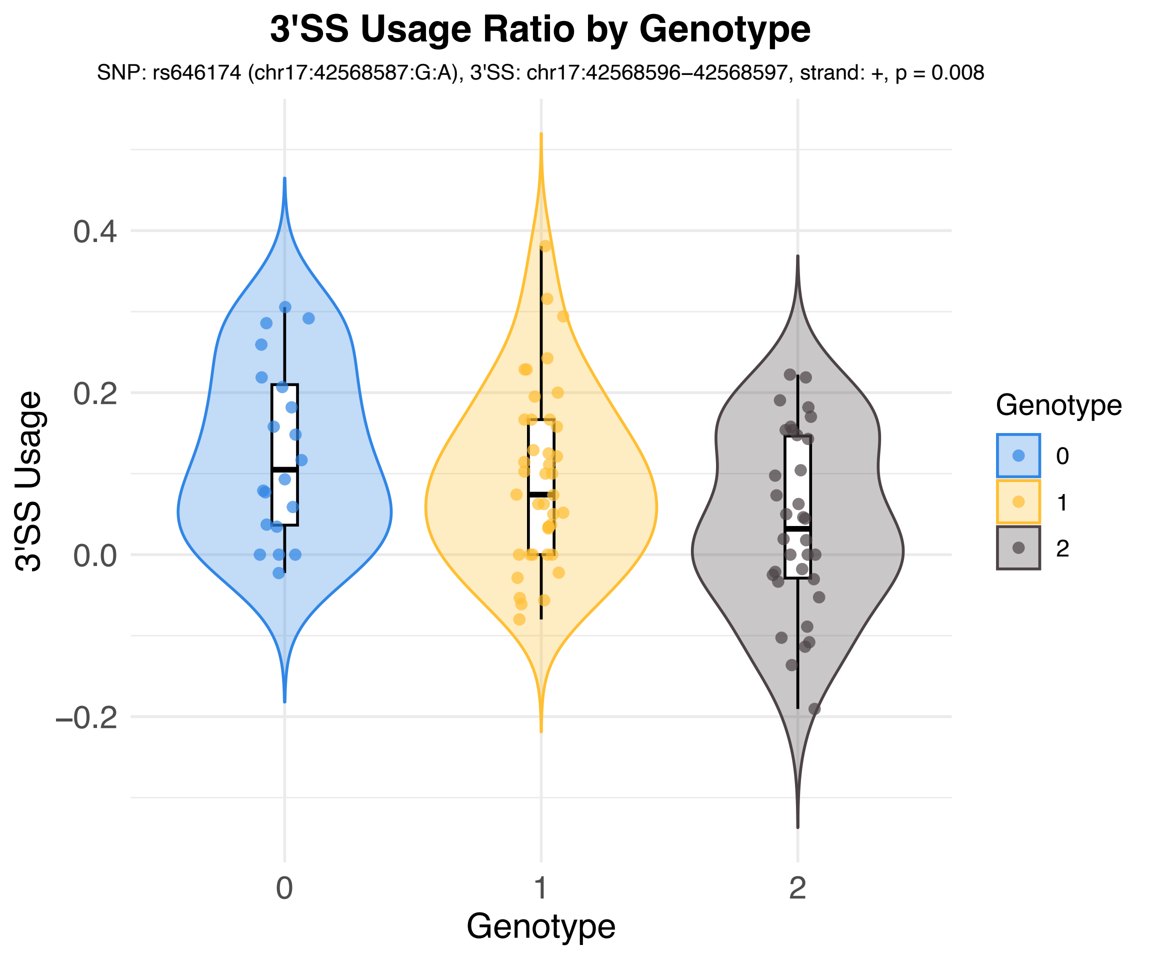
3'SS Usage Ratio by Genotype
SNP: rs646174 (chr17:42568587:G:A), 3'SS: chr17:42568596–42568597, strand: +, p = 0.008
3'SS Usage
0.4
0.2
0.0
–0.2
0
1
2
Genotype
Genotype
0
1
2

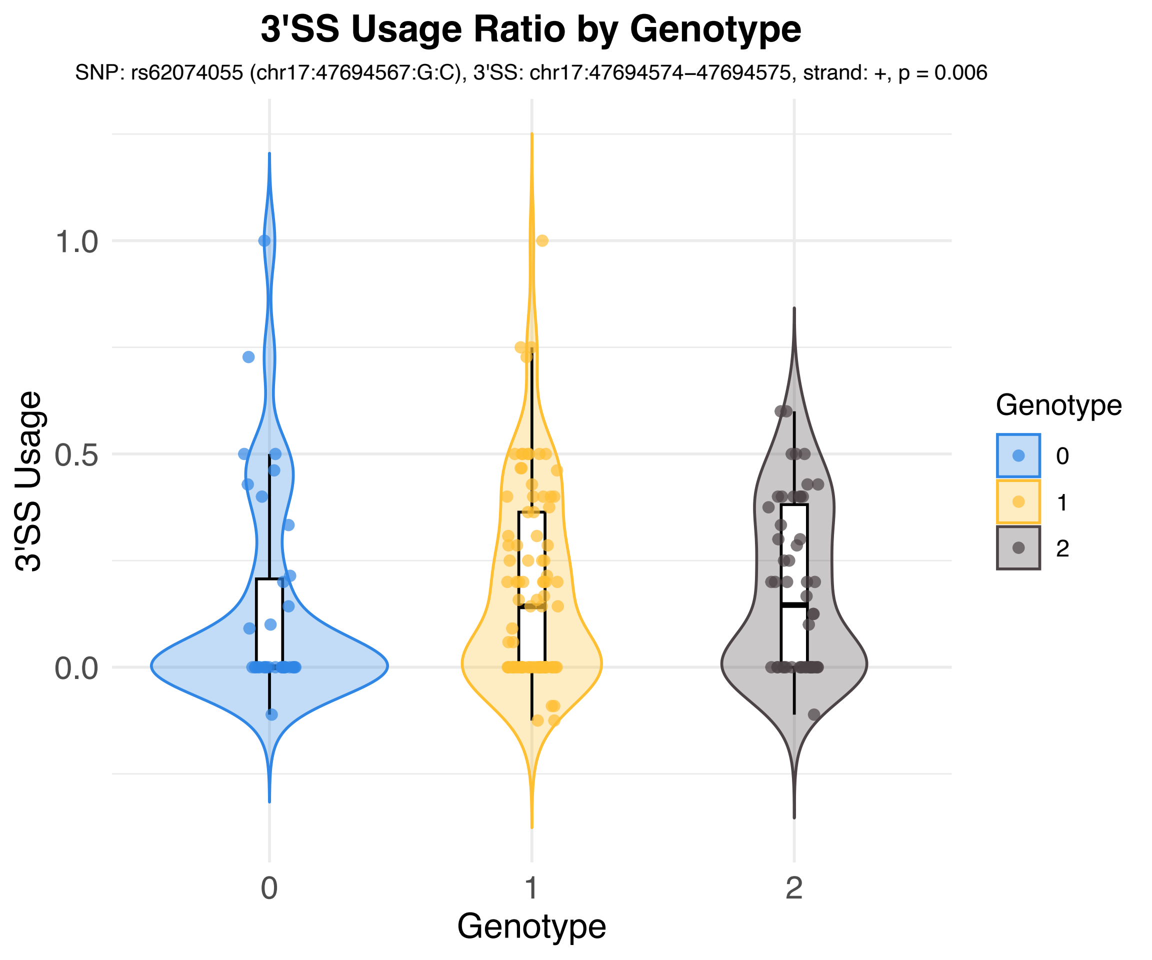
3'SS Usage Ratio by Genotype
SNP: rs62074055 (chr17:47694567:G:C), 3'SS: chr17:47694574–47694575, strand: +, p = 0.006
3'SS Usage
1.0
0.5
0.0
0
1
2
Genotype
Genotype
0
1
2

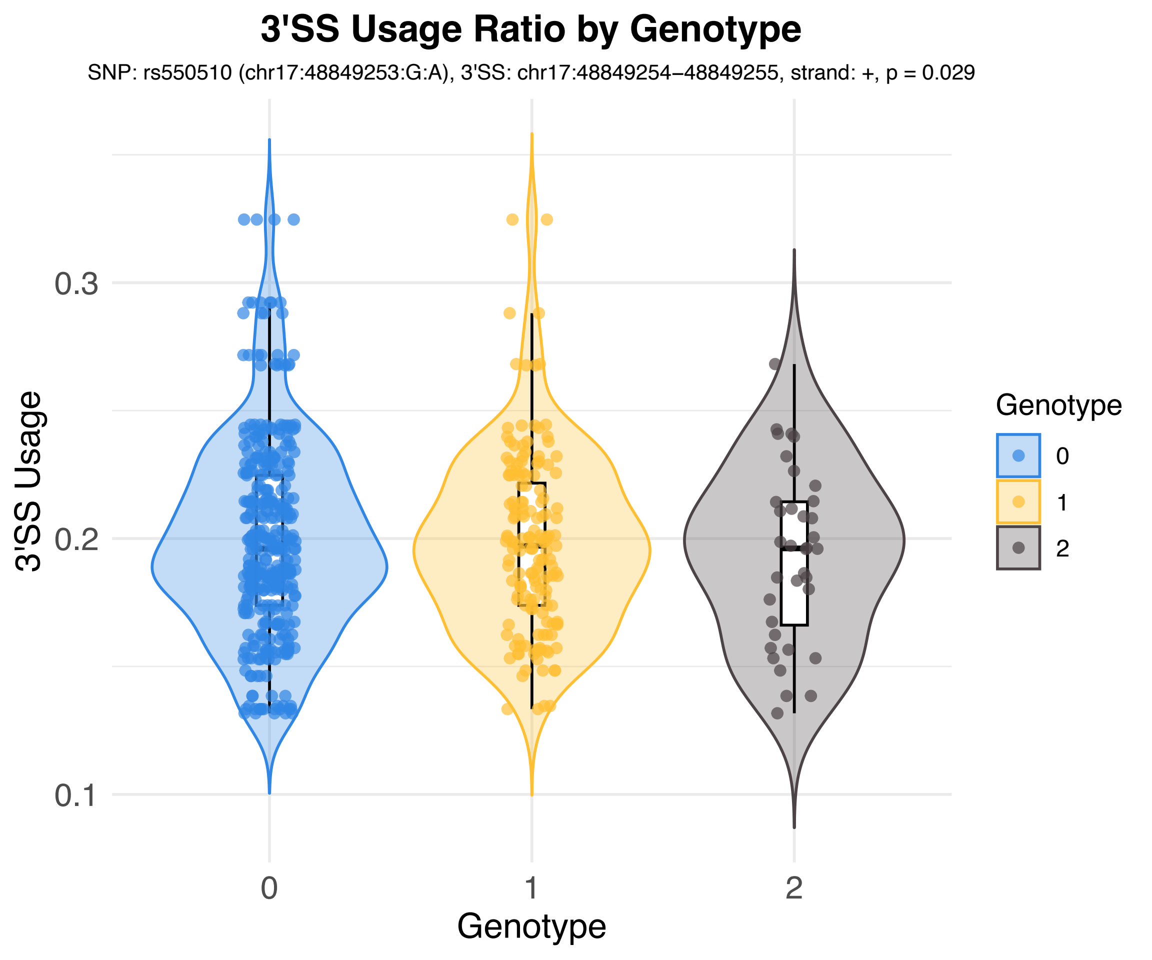

3'SS Usage Ratio by Genotype
SNP: rs550510 (chr17:48849253:G:A), 3'SS: chr17:48849254−48849255, strand: +, p = 0.029
3'SS Usage
0.1
0.2
0.3
0
1
2
Genotype
Genotype
0
1
2

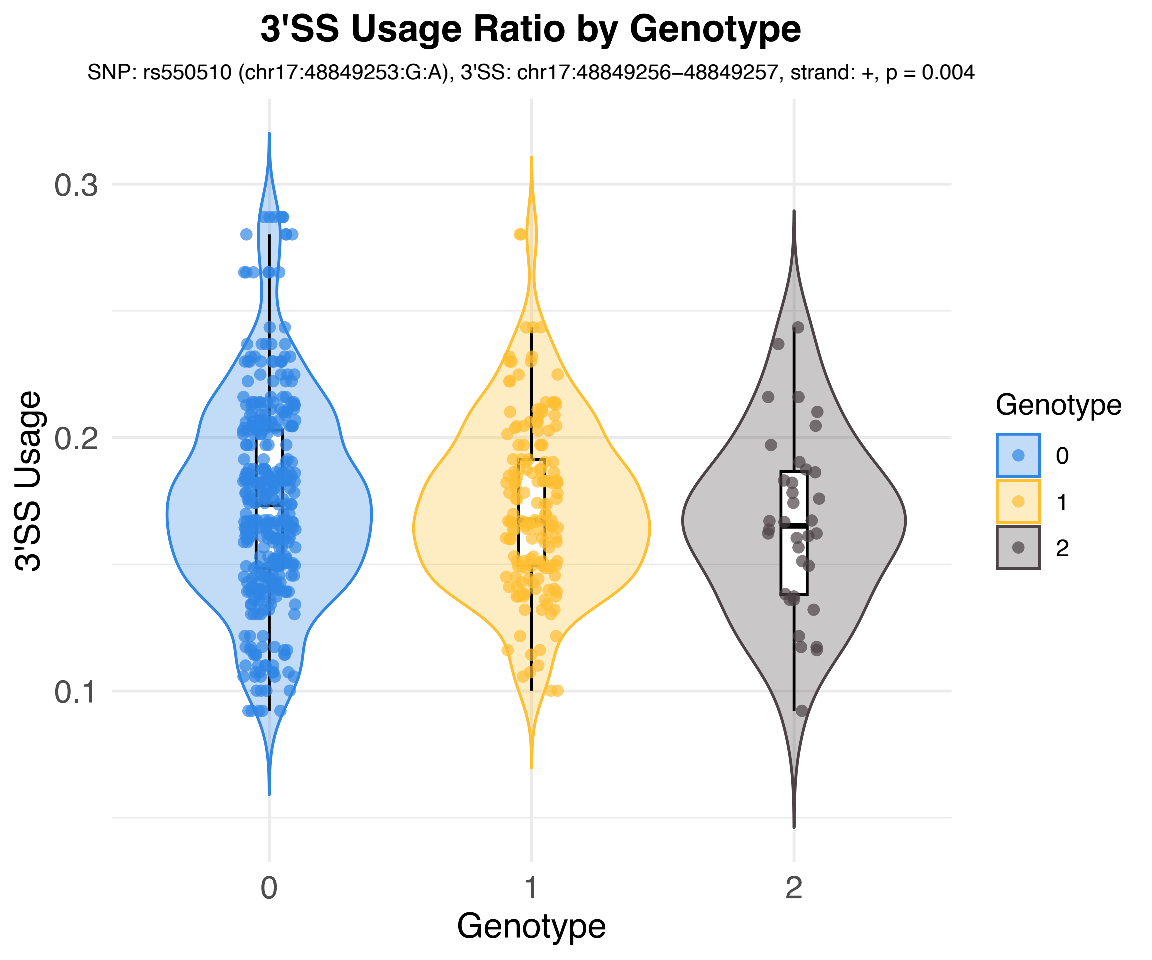

3'SS Usage Ratio by Genotype
SNP: rs550510 (chr17:48849253:G:A), 3'SS: chr17:48849256−48849257, strand: +, p = 0.004
3'SS Usage
0.3
0.2
0.1
0
1
2
Genotype
Genotype
0
1
2

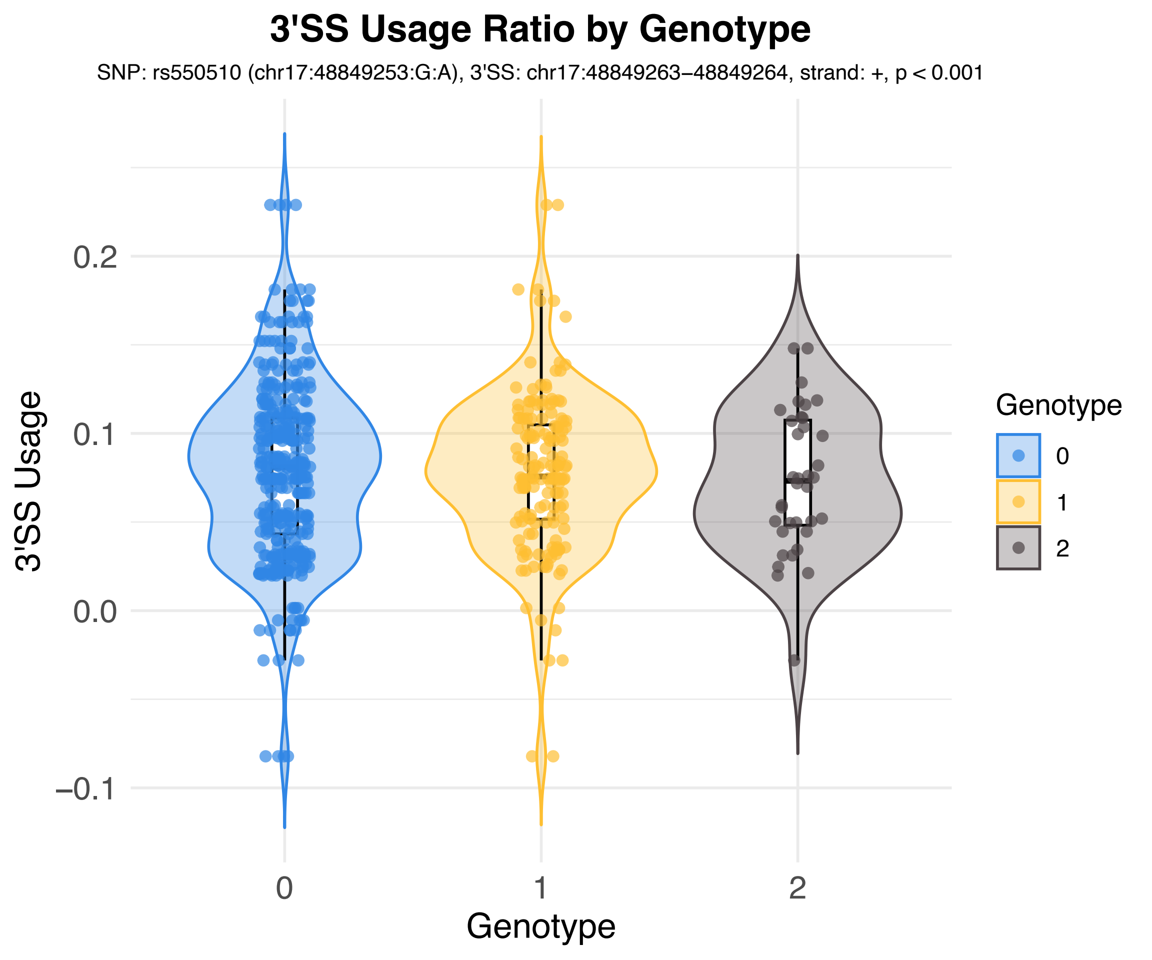
3'SS Usage Ratio by Genotype
SNP: rs550510 (chr17:48849253:G:A), 3'SS: chr17:48849263–48849264, strand: +, p < 0.001
3'SS Usage
0.2
0.1
0.0
−0.1
0
1
2
Genotype
Genotype
0
1
2

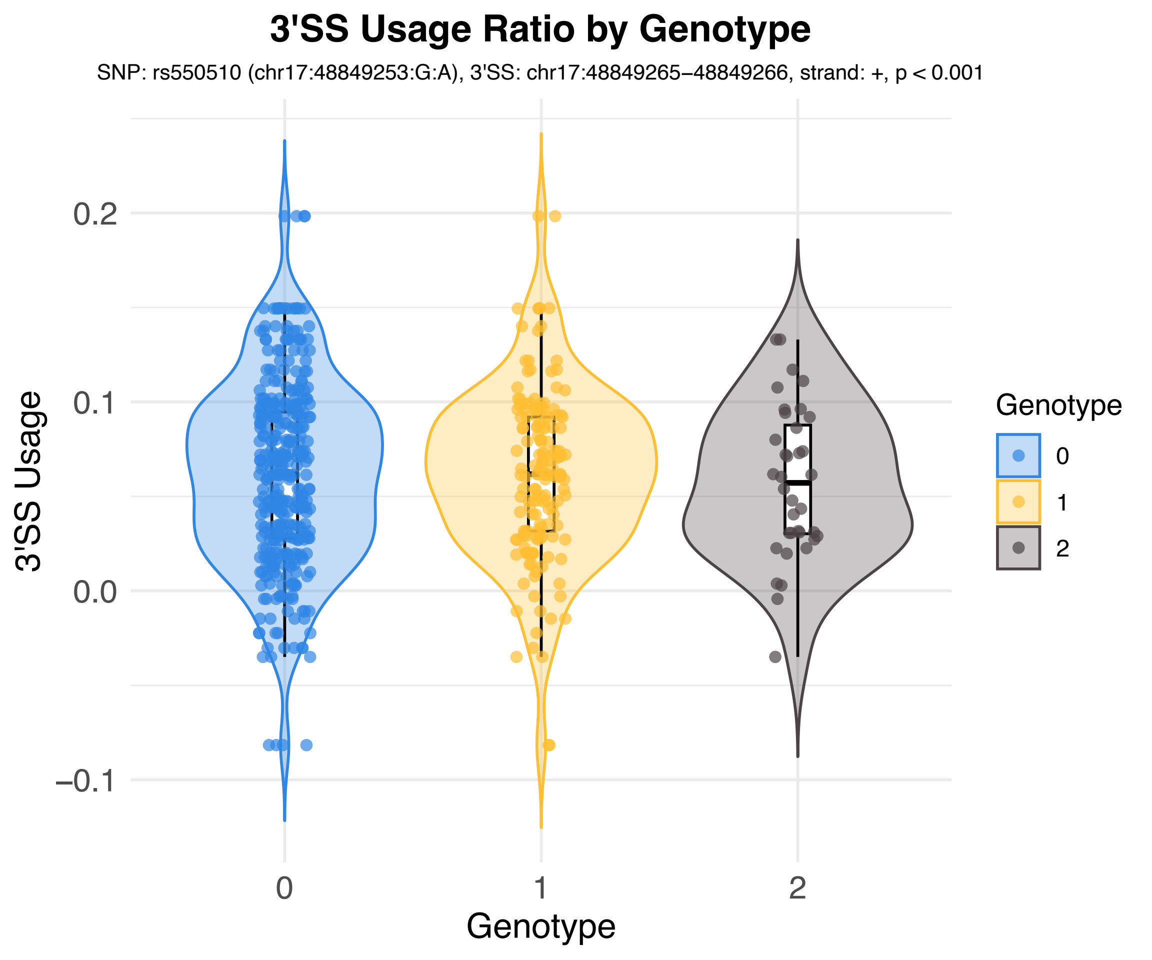
3'SS Usage Ratio by Genotype
SNP: rs550510 (chr17:48849253:G:A), 3'SS: chr17:48849265–48849266, strand: +, p < 0.001
3'SS Usage
0.2
0.1
0.0
−0.1
Genotype
0
1
2
Genotype
0
1
2

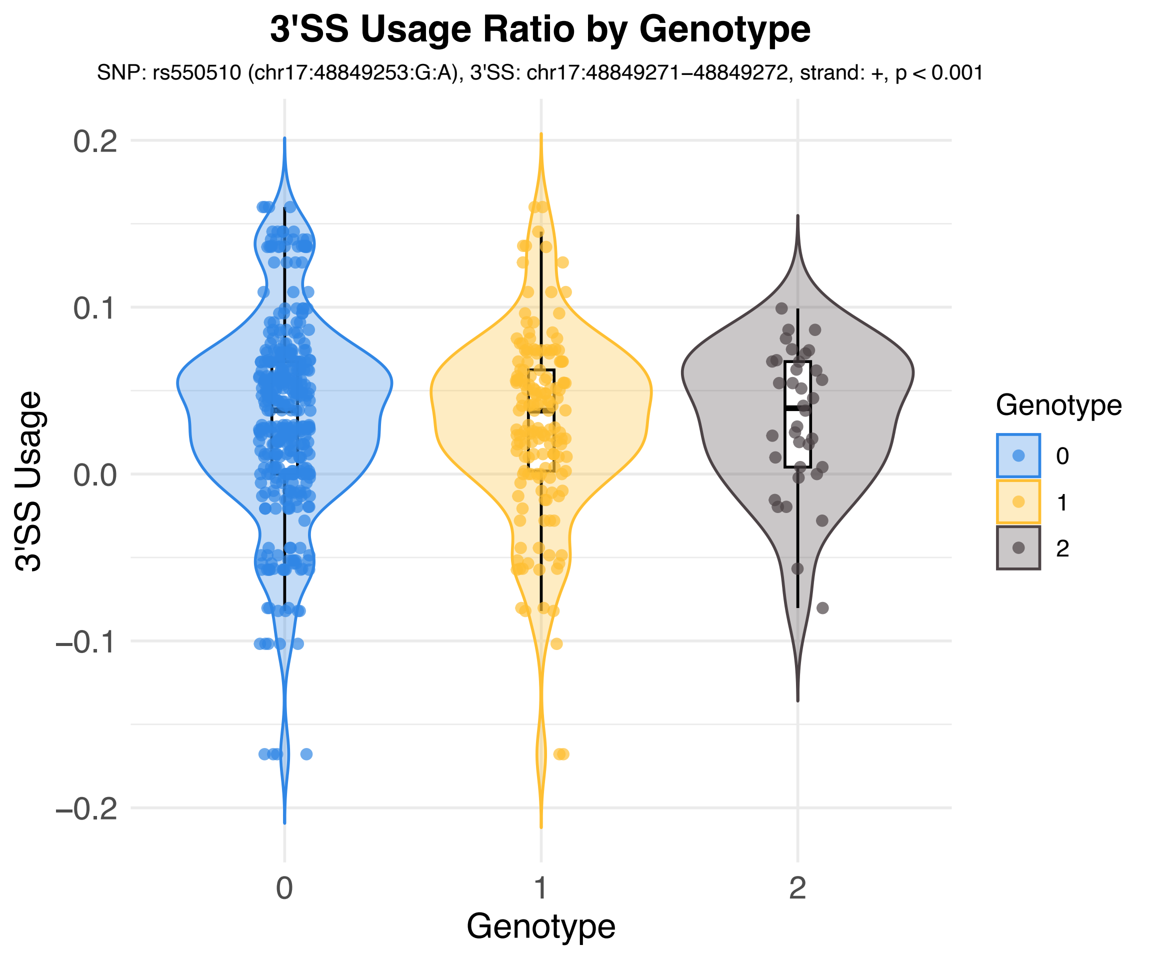
3'SS Usage Ratio by Genotype
SNP: rs550510 (chr17:48849253:G:A), 3'SS: chr17:48849271–48849272, strand: +, p < 0.001
3'SS Usage
0.2
0.1
0.0
–0.1
–0.2
0
1
2
Genotype
Genotype
0
1
2

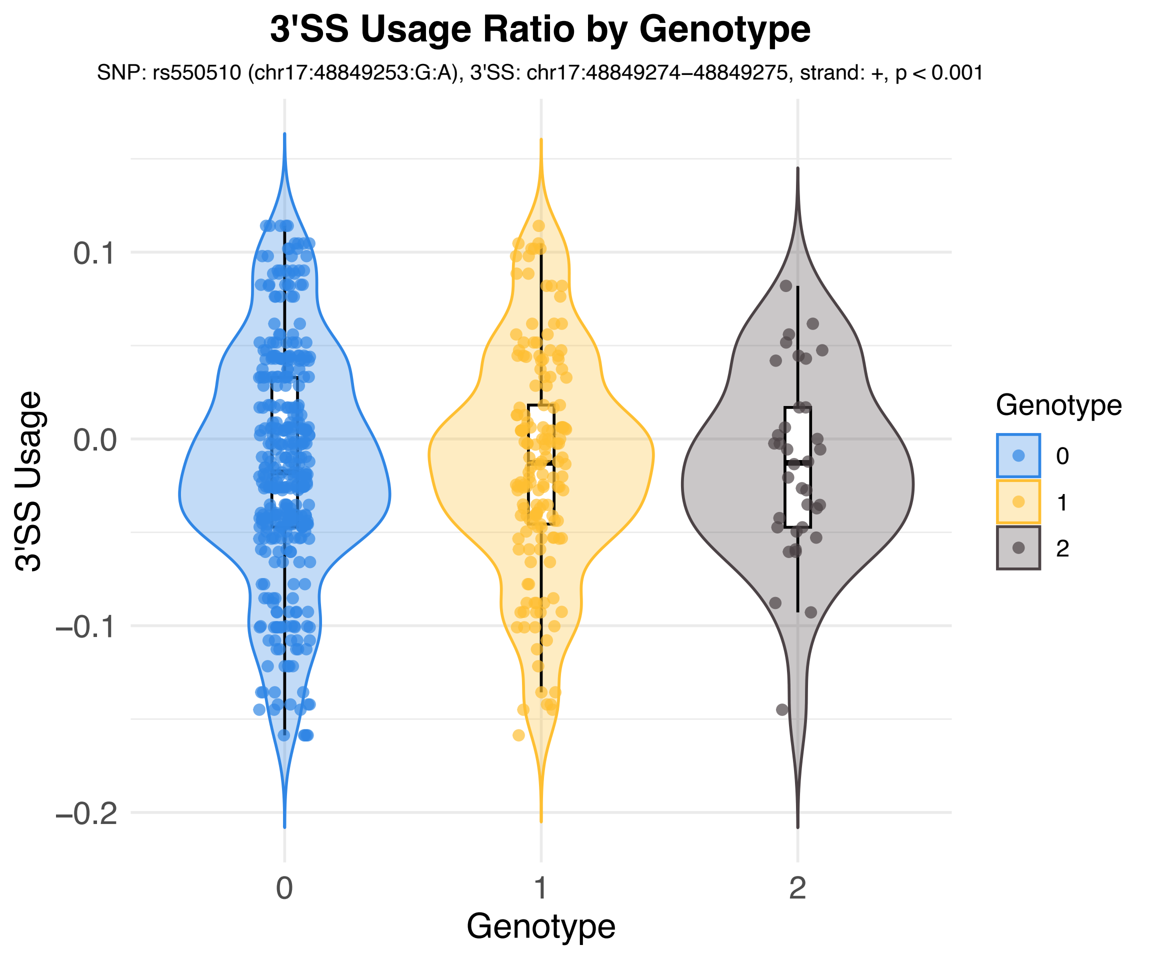
3'SS Usage Ratio by Genotype
SNP: rs550510 (chr17:48849253:G:A), 3'SS: chr17:48849274−48849275, strand: +, p < 0.001
3'SS Usage
0.1
0.0
−0.1
−0.2
0
1
2
Genotype
Genotype
0
1
2

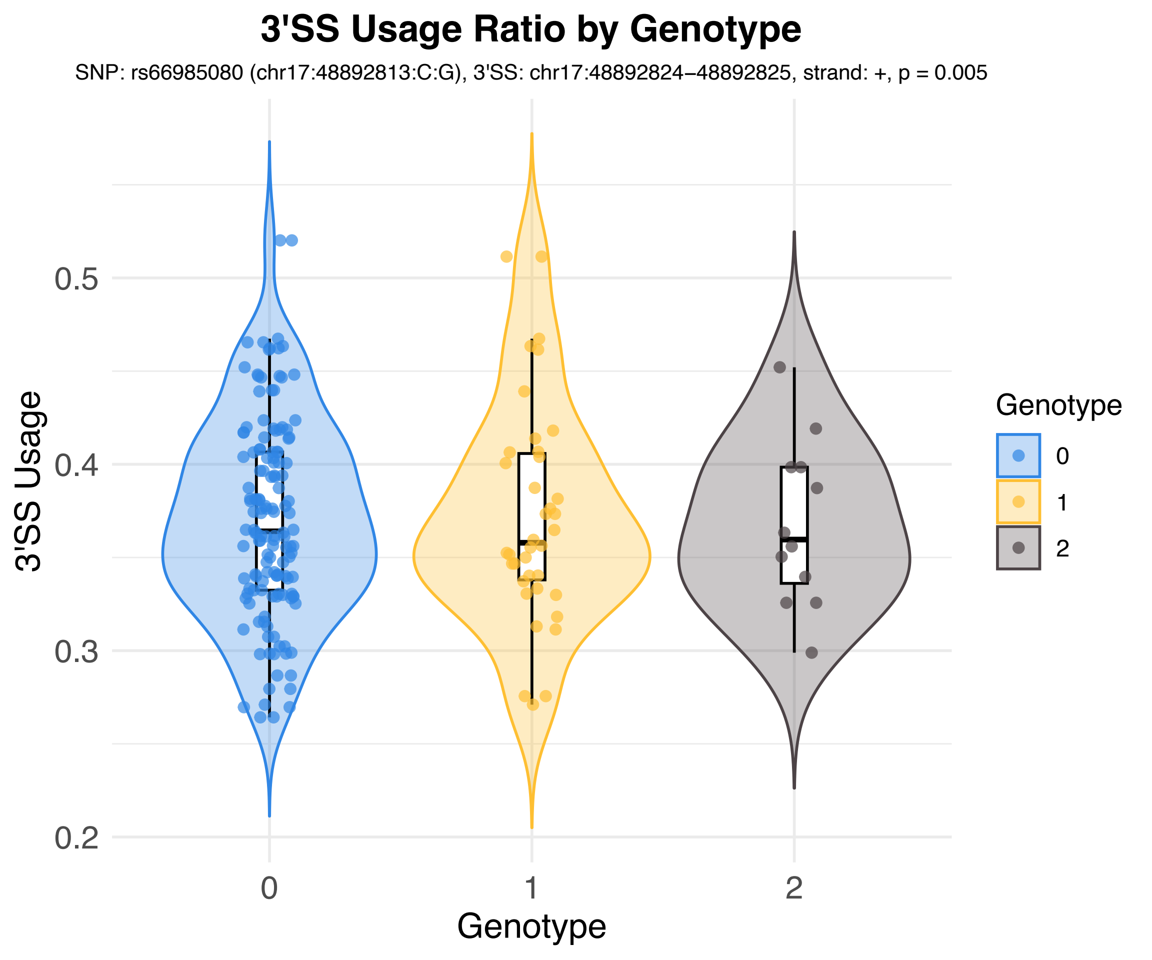
3'SS Usage Ratio by Genotype
SNP: rs66985080 (chr17:48892813:C:G), 3'SS: chr17:48892824–48892825, strand: +, p = 0.005
3'SS Usage
0.2
0.3
0.4
0.5
0
1
2
Genotype
Genotype
0
1
2

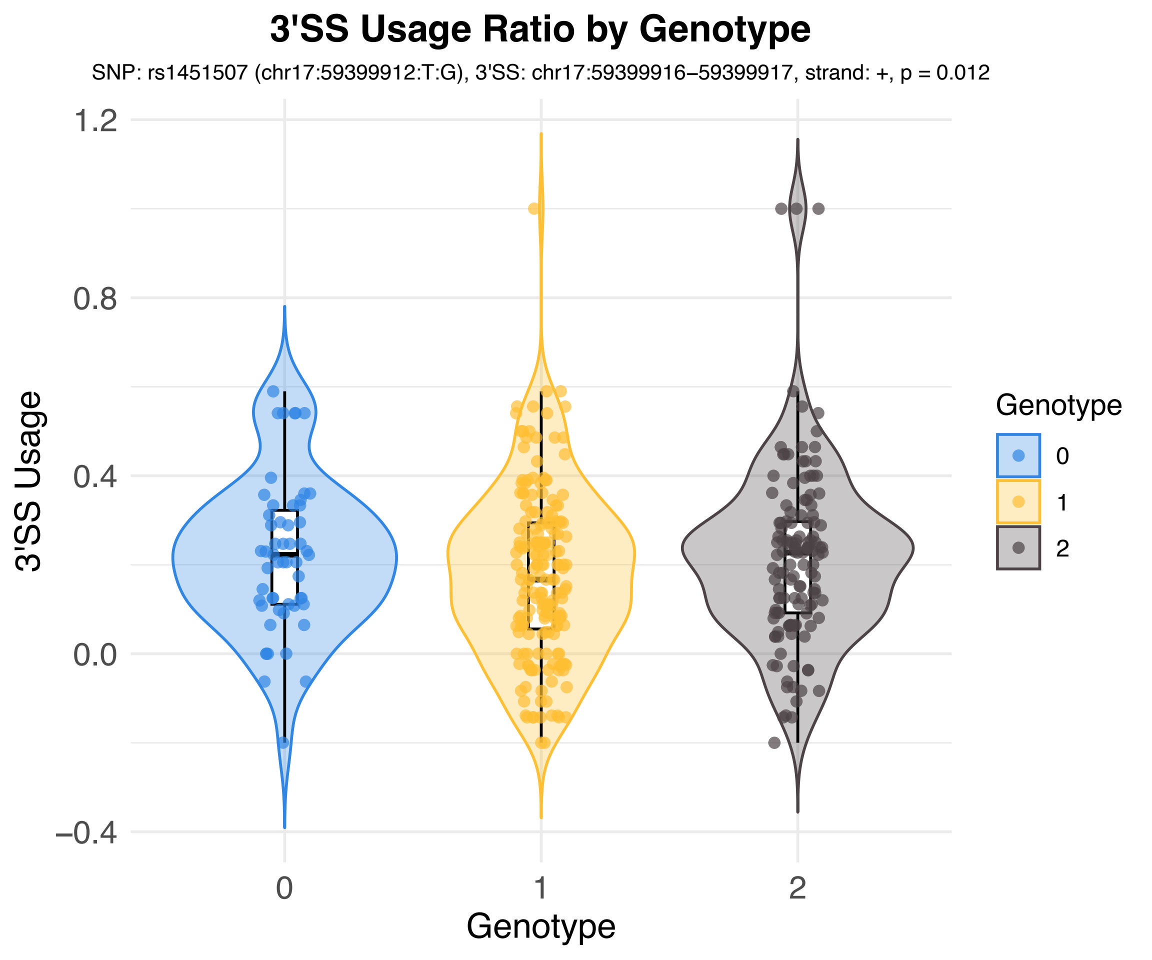

3'SS Usage Ratio by Genotype
SNP: rs1451507 (chr17:59399912:T:G), 3'SS: chr17:59399916–59399917, strand: +, p = 0.012
3'SS Usage
1.2
0.8
0.4
0.0
–0.4
0
1
2
Genotype
Genotype
0
1
2

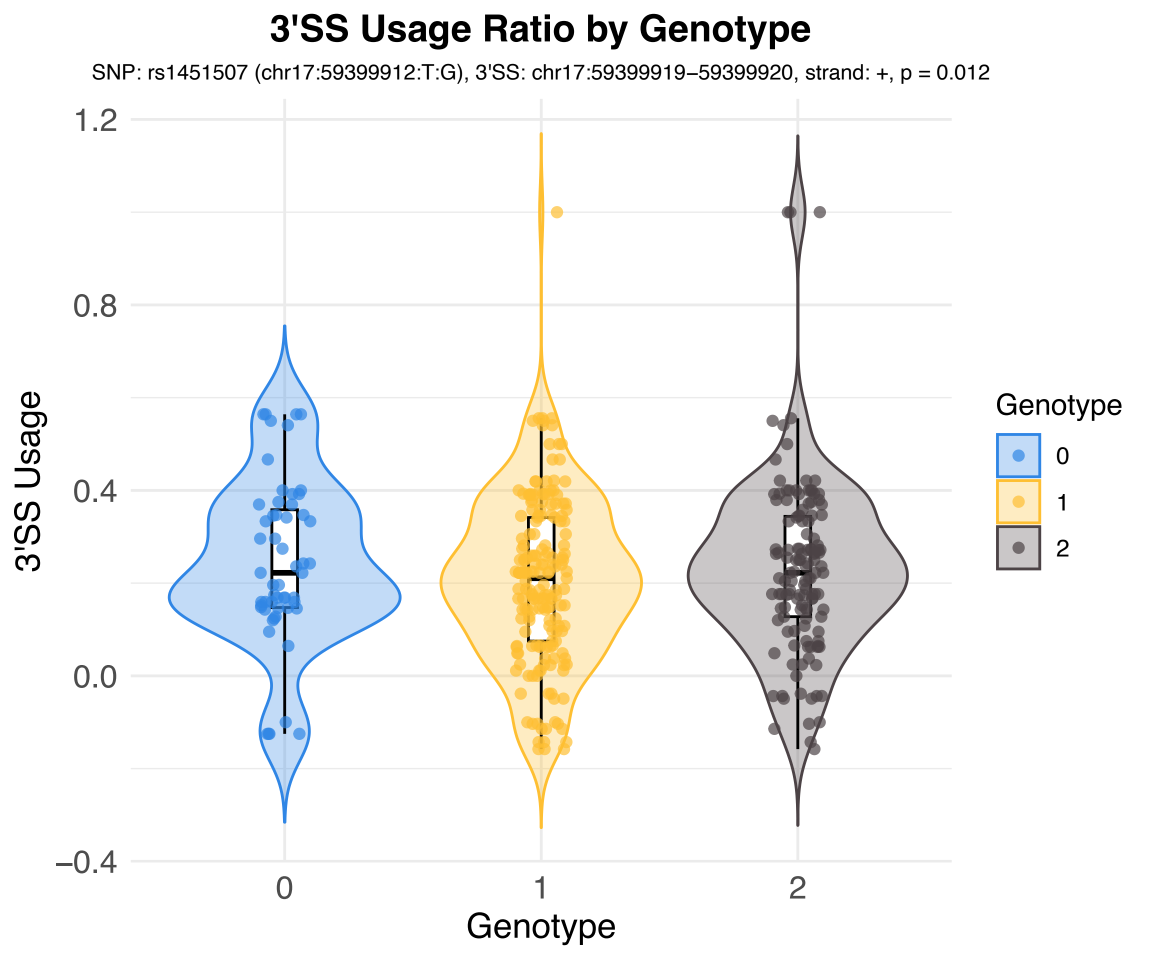
3'SS Usage Ratio by Genotype
SNP: rs1451507 (chr17:59399912:T:G), 3'SS: chr17:59399919–59399920, strand: +, p = 0.012
3'SS Usage
1.2
0.8
0.4
0.0
−0.4
0
1
2
Genotype
Genotype
0
1
2

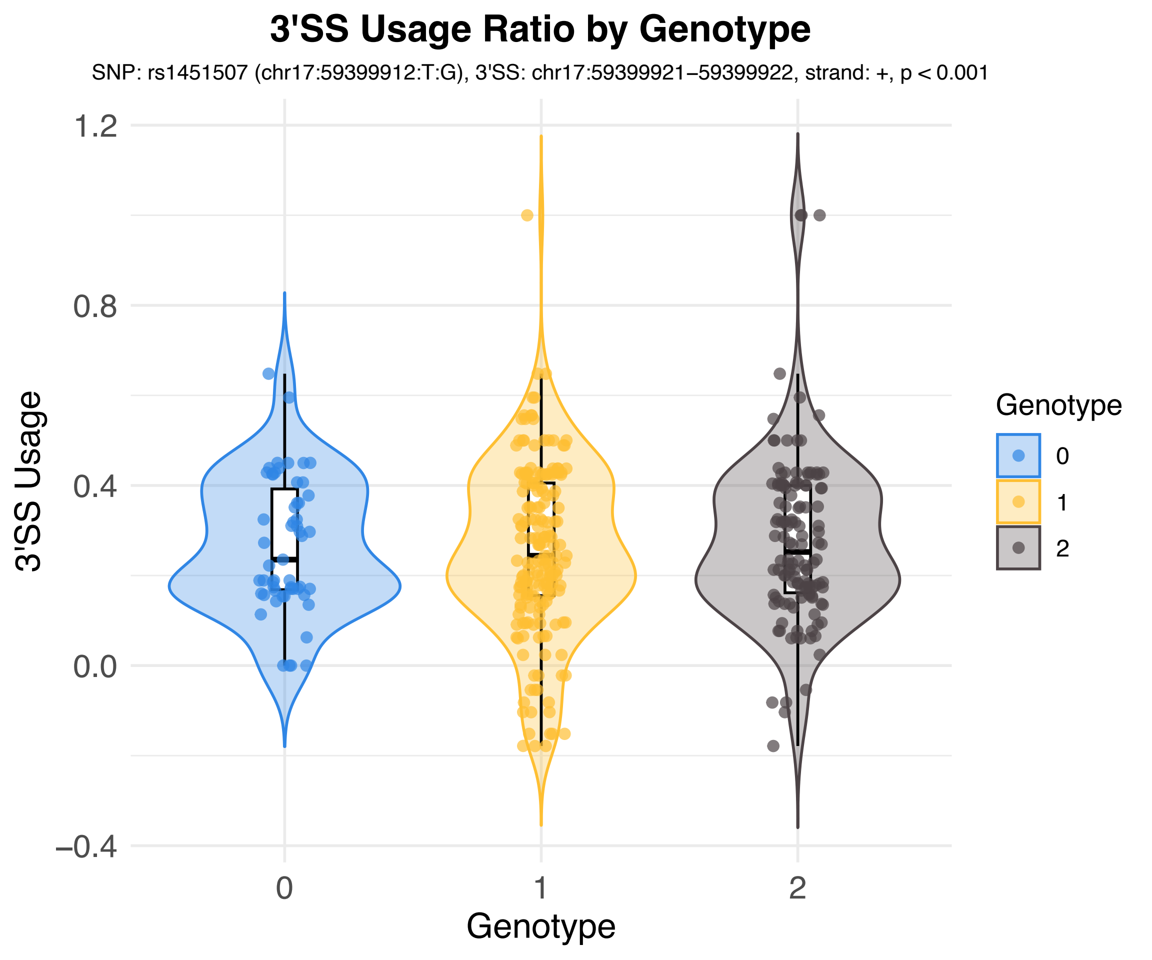

3'SS Usage Ratio by Genotype
SNP: rs1451507 (chr17:59399912:T:G), 3'SS: chr17:59399921–59399922, strand: +, p < 0.001
3'SS Usage
1.2
0.8
0.4
0.0
−0.4
0
1
2
Genotype
Genotype
0
1
2

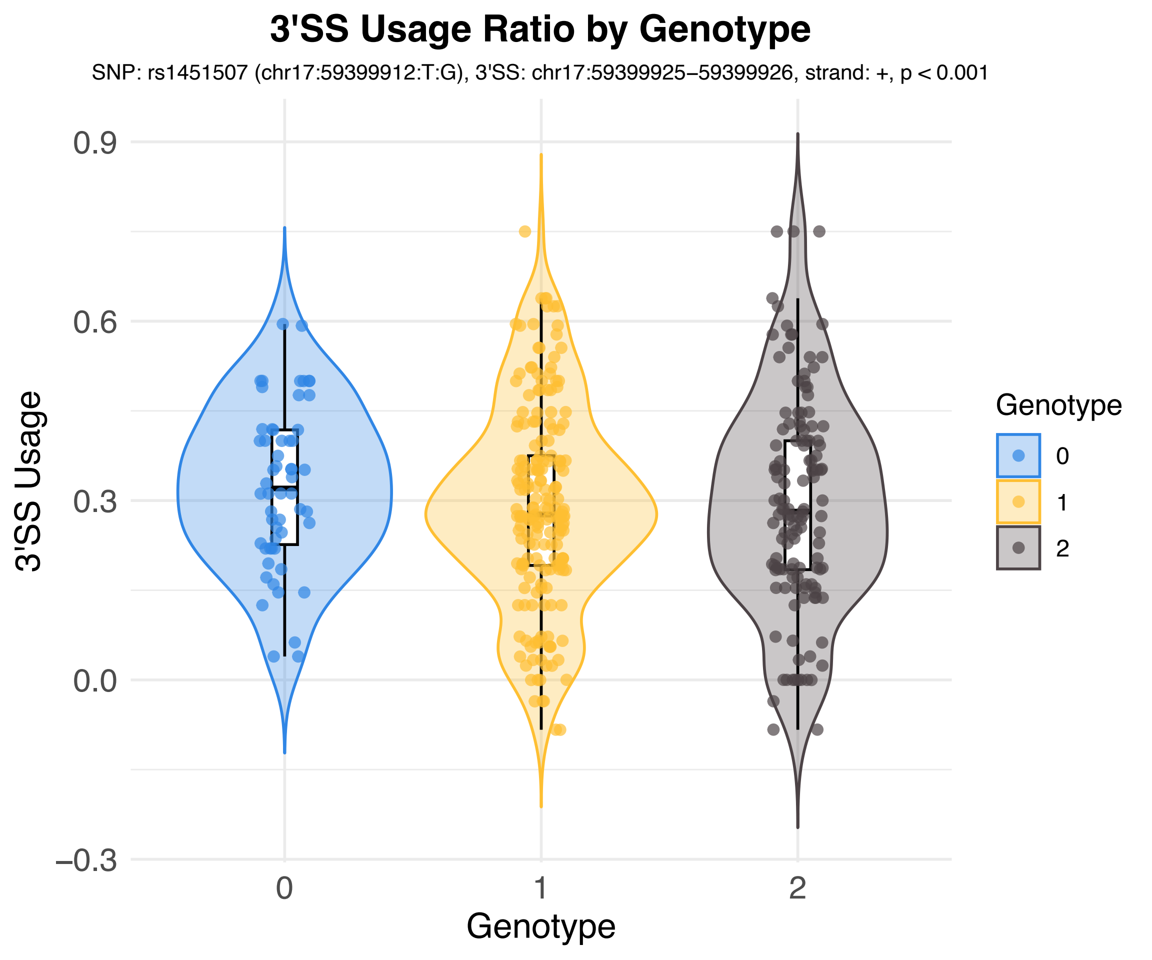
3'SS Usage Ratio by Genotype
SNP: rs1451507 (chr17:59399912:T:G), 3'SS: chr17:59399925–59399926, strand: +, p < 0.001
3'SS Usage
0.9
0.6
0.3
0.0
−0.3
0
1
2
Genotype
Genotype
0
1
2

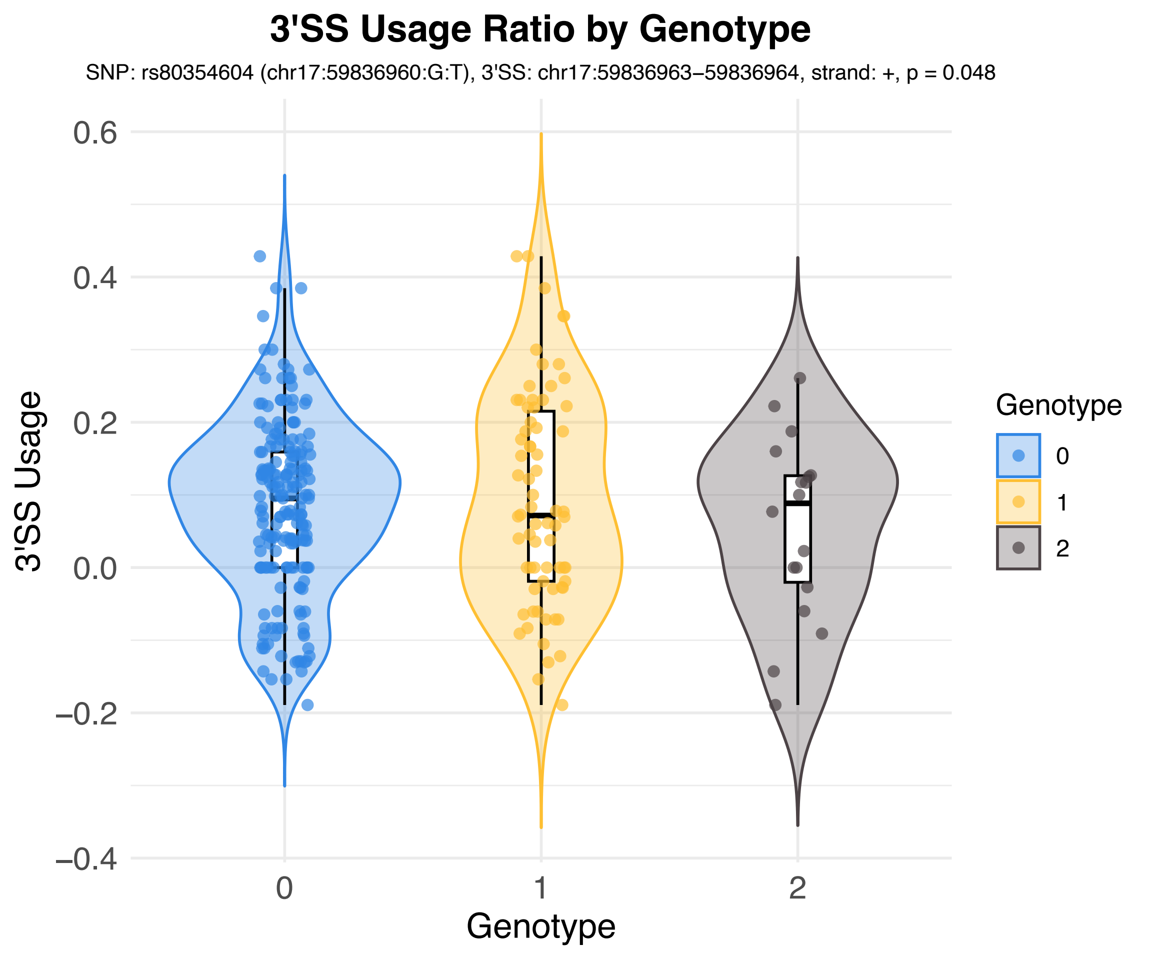

3'SS Usage Ratio by Genotype
SNP: rs80354604 (chr17:59836960:G:T), 3'SS: chr17:59836963–59836964, strand: +, p = 0.048
3'SS Usage
0.6
0.4
0.2
0.0
−0.2
−0.4
0
1
2
Genotype
Genotype
0
1
2

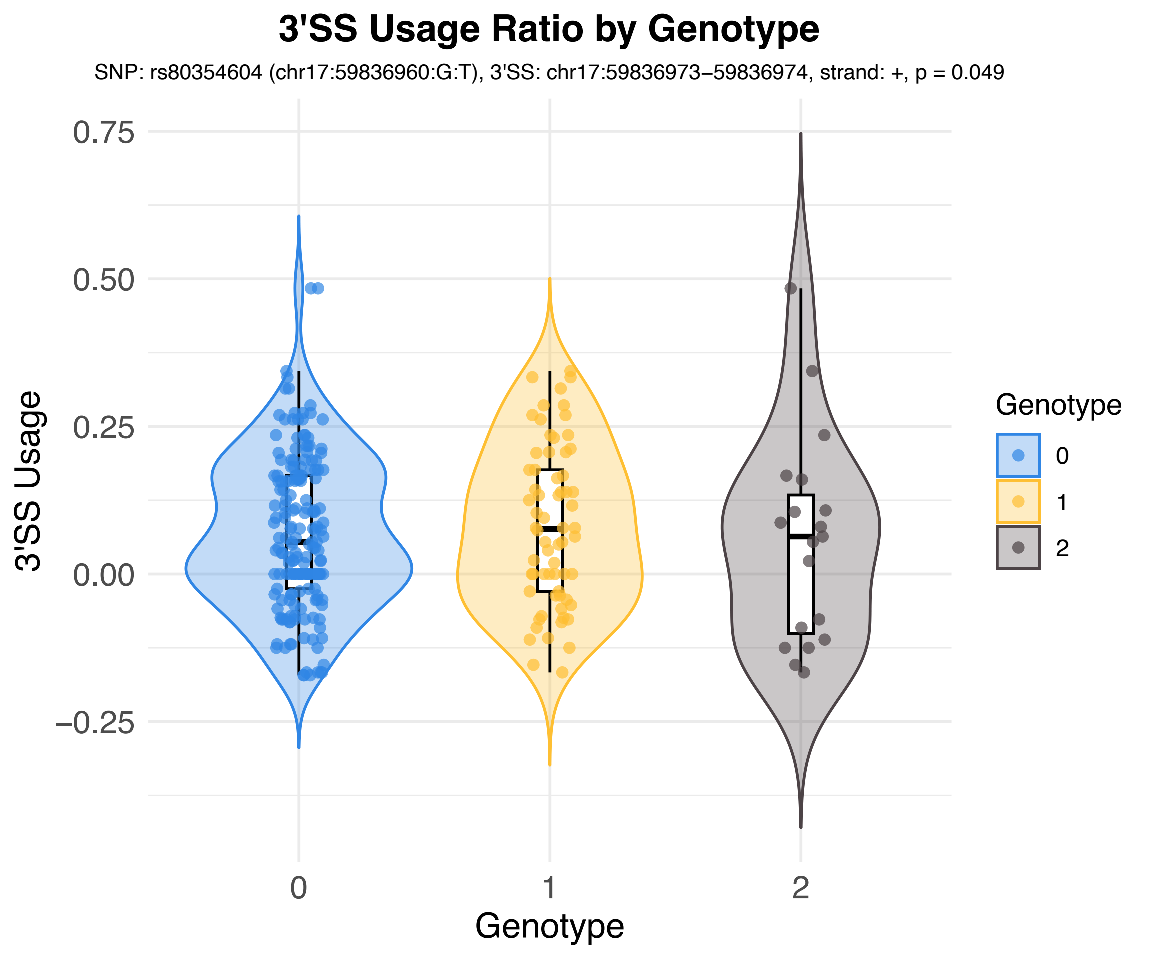

3'SS Usage Ratio by Genotype
SNP: rs80354604 (chr17:59836960:G:T), 3'SS: chr17:59836973–59836974, strand: +, p = 0.049
3'SS Usage
0.75
0.50
0.25
0.00
−0.25
0
1
2
Genotype
Genotype
0
1
2

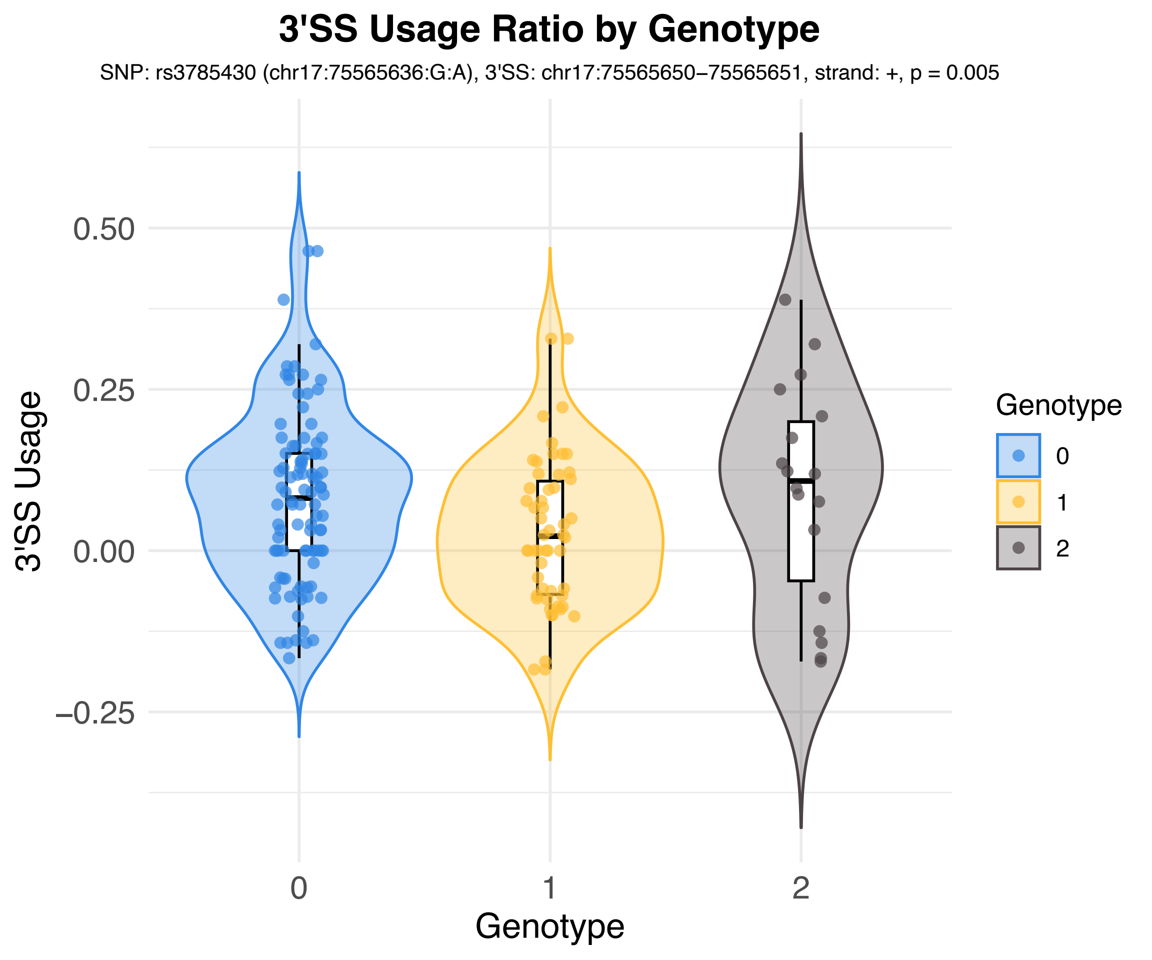
3'SS Usage Ratio by Genotype
SNP: rs3785430 (chr17:75565636:G:A), 3'SS: chr17:75565650–75565651, strand: +, p = 0.005
3'SS Usage
0.50
0.25
0.00
−0.25
0
1
2
Genotype
Genotype
0
1
2

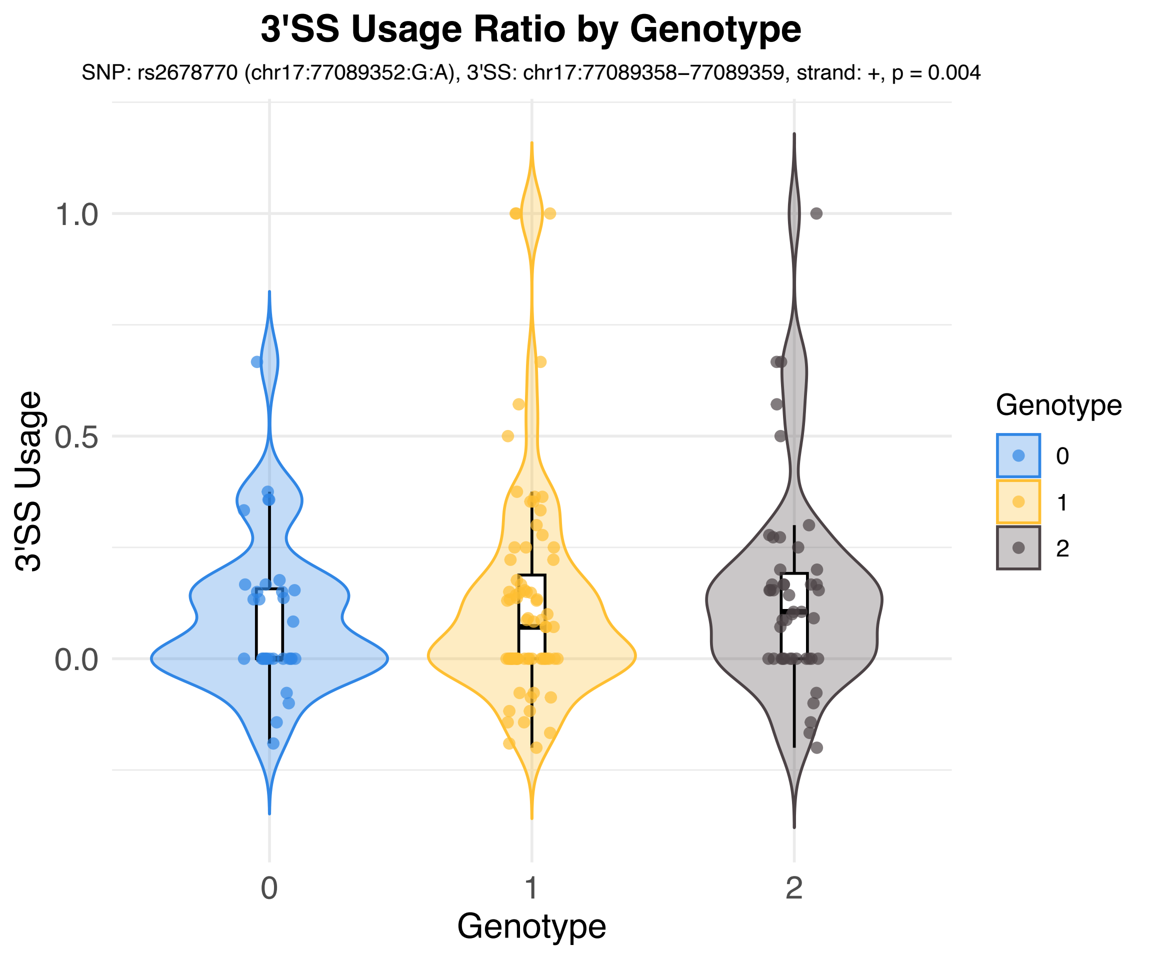
3'SS Usage Ratio by Genotype
SNP: rs2678770 (chr17:77089352:G:A), 3'SS: chr17:77089358−77089359, strand: +, p = 0.004
3'SS Usage
1.0
0.5
0.0
0
1
2
Genotype
Genotype
0
1
2

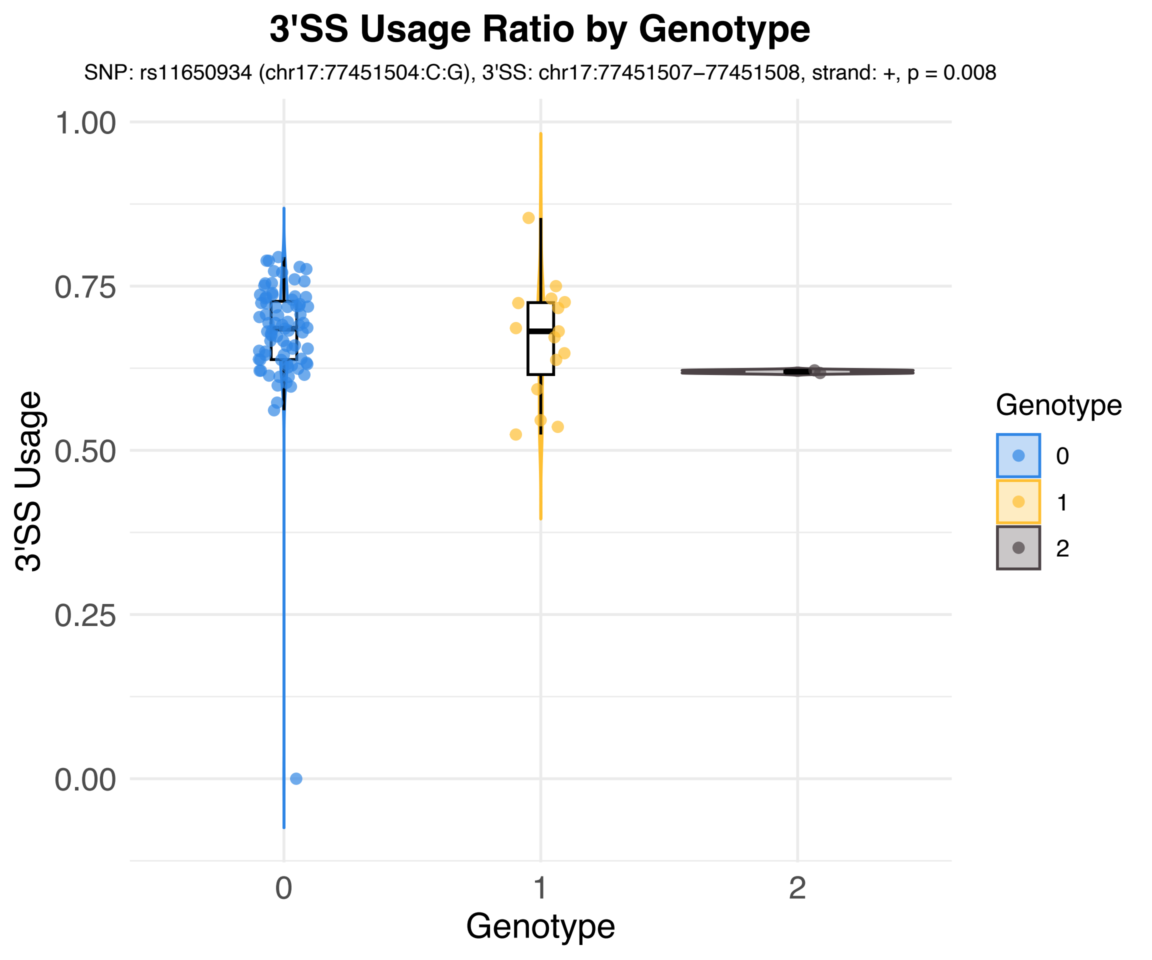
3'SS Usage Ratio by Genotype
SNP: rs11650934 (chr17:77451504:C:G), 3'SS: chr17:77451507−77451508, strand: +, p = 0.008
3'SS Usage
1.00
0.75
0.50
0.25
0.00
0
1
2
Genotype
Genotype
0
1
2

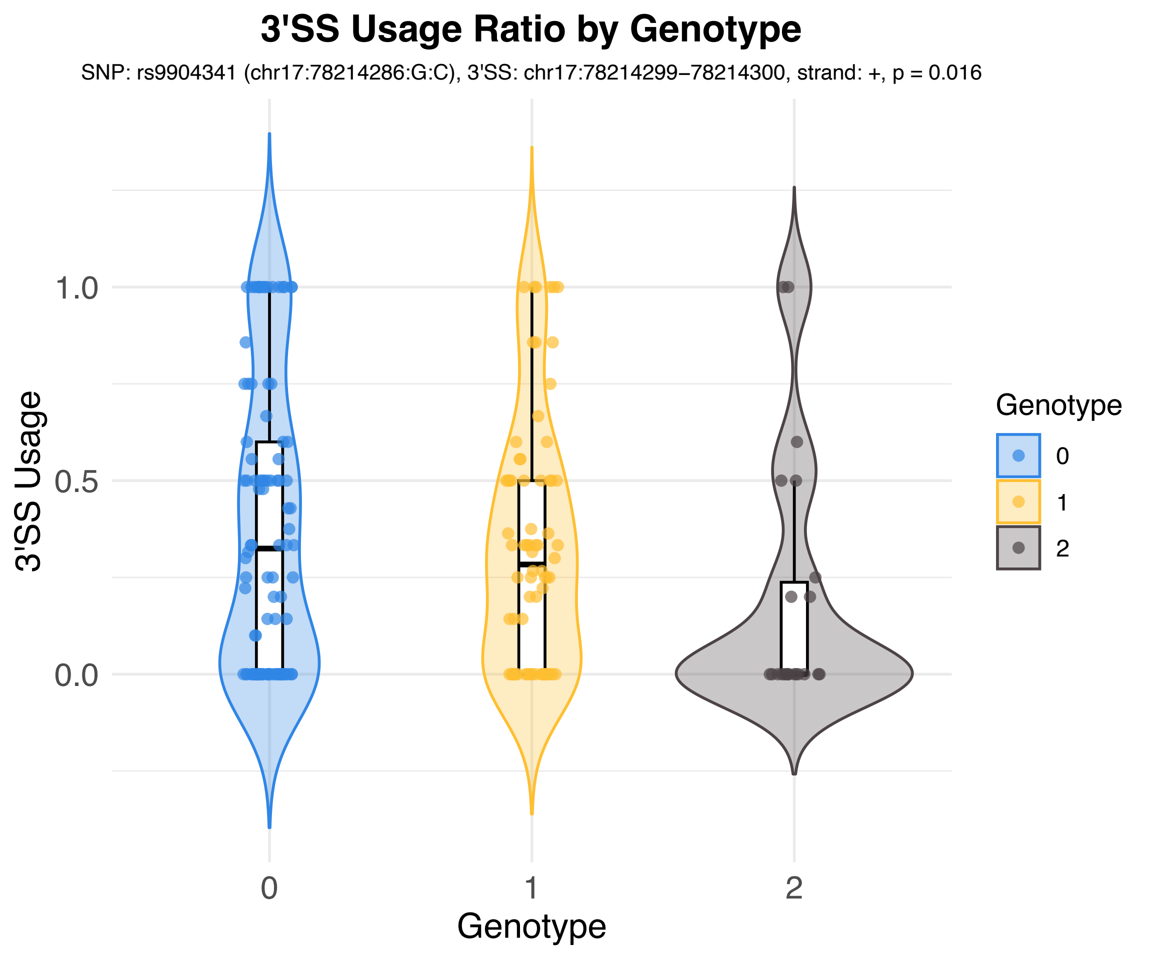
3'SS Usage Ratio by Genotype
SNP: rs9904341 (chr17:78214286:G:C), 3'SS: chr17:78214299−78214300, strand: +, p = 0.016
3'SS Usage
1.0
0.5
0.0
0
1
2
Genotype
Genotype
0
1
2

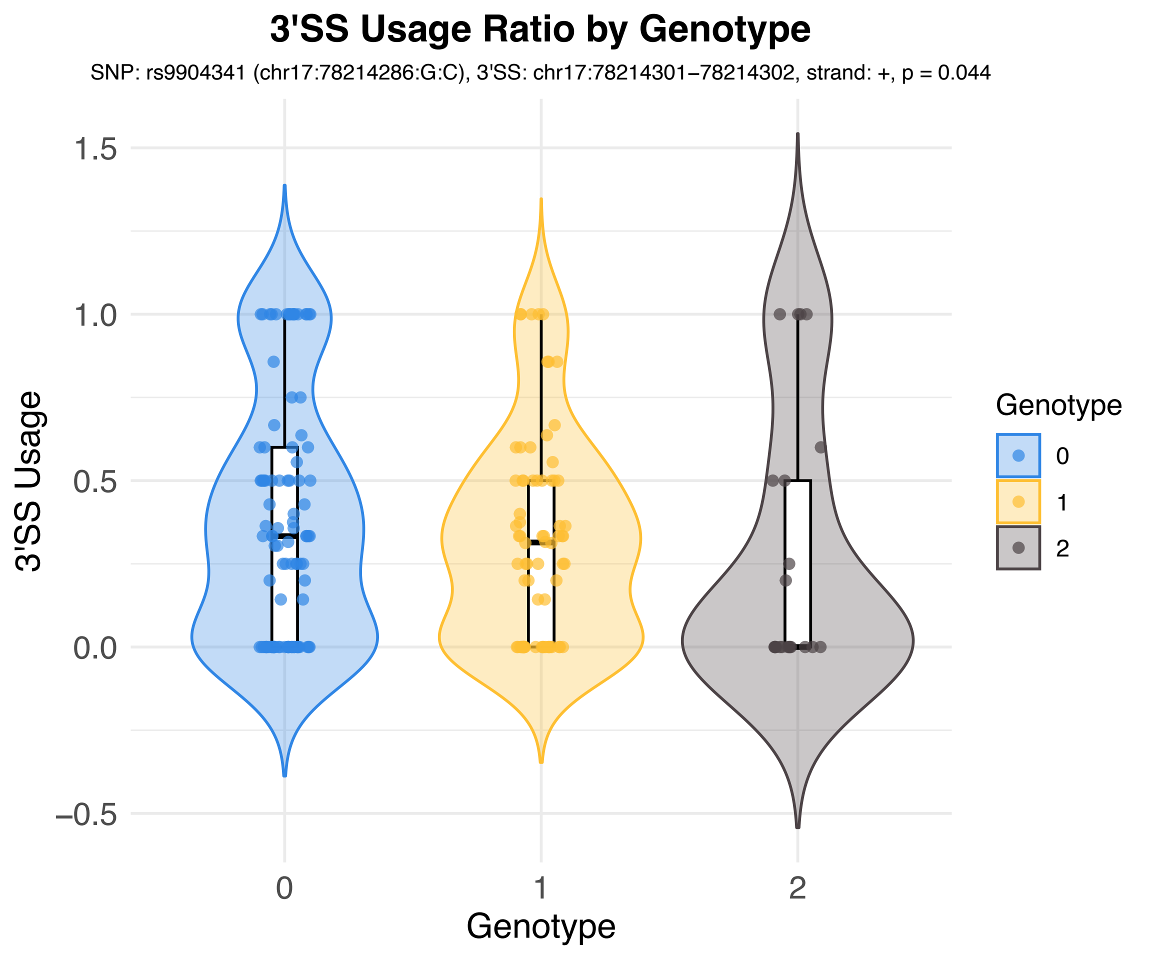
3'SS Usage Ratio by Genotype
SNP: rs9904341 (chr17:78214286:G:C), 3'SS: chr17:78214301–78214302, strand: +, p = 0.044
3'SS Usage
1.5
1.0
0.5
0.0
−0.5
0
1
2
Genotype
Genotype
0
1
2

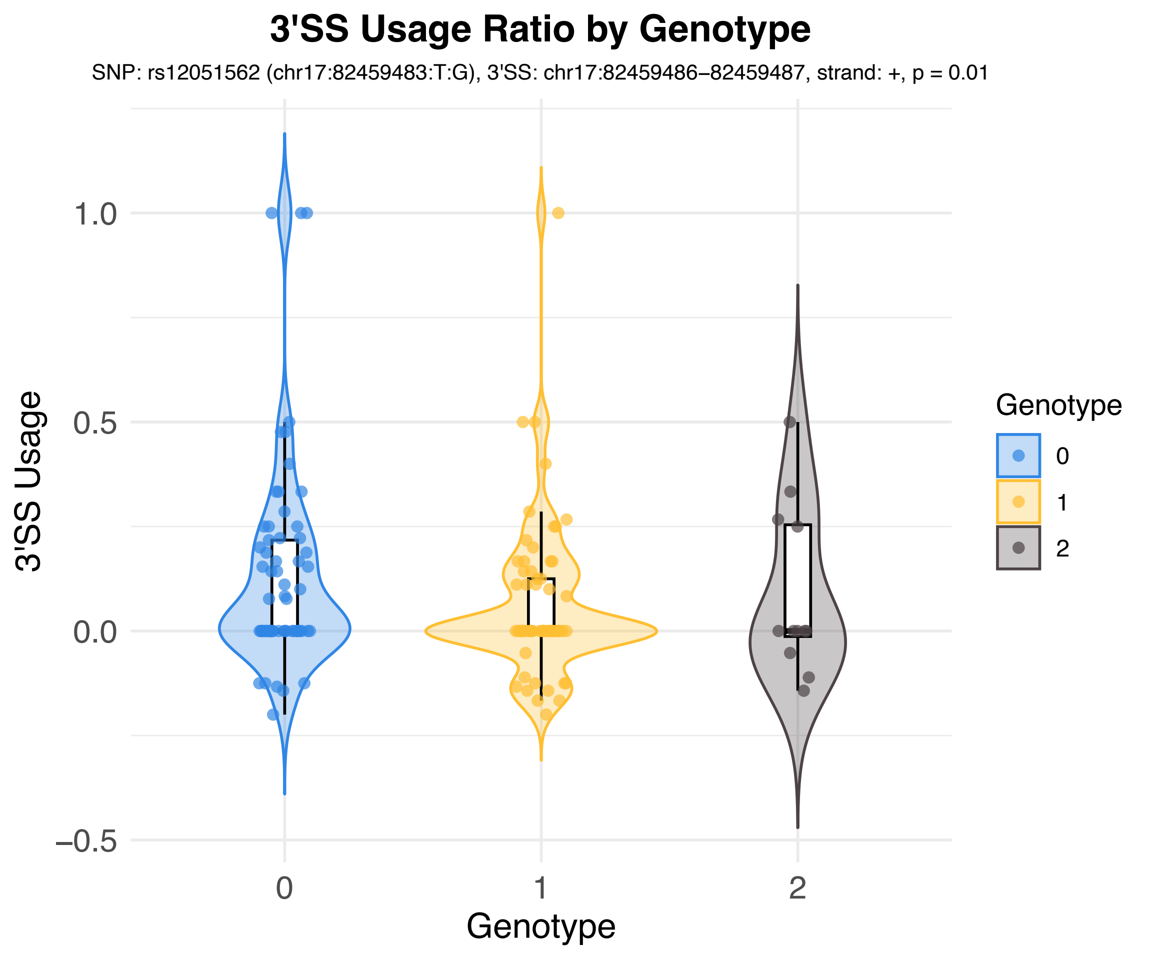
3'SS Usage Ratio by Genotype
SNP: rs12051562 (chr17:82459483:T:G), 3'SS: chr17:82459486–82459487, strand: +, p = 0.01
3'SS Usage
1.0
0.5
0.0
−0.5
0
1
2
Genotype
Genotype
0
1
2

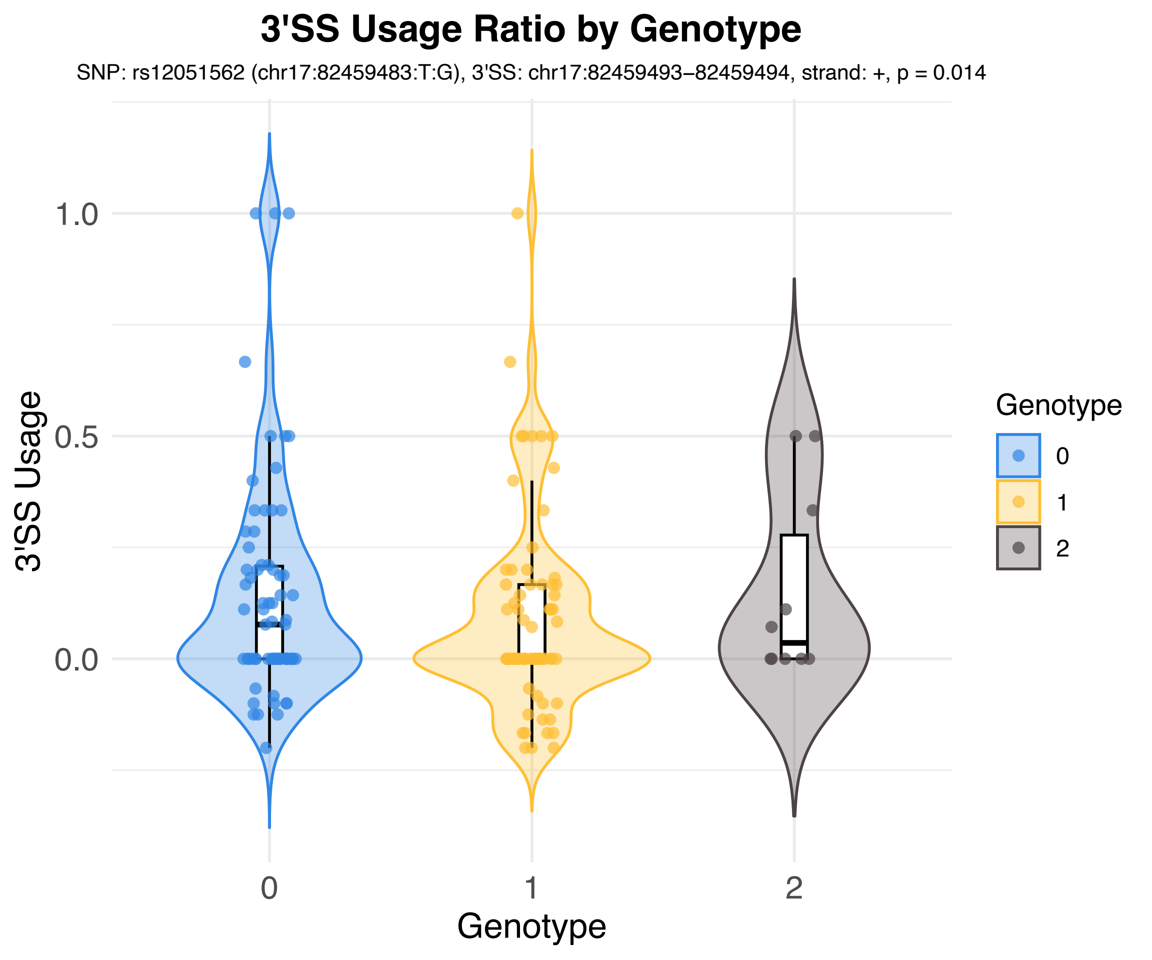

3'SS Usage Ratio by Genotype
SNP: rs12051562 (chr17:82459483:T:G), 3'SS: chr17:82459493–82459494, strand: +, p = 0.014
3'SS Usage
1.0
0.5
0.0
0
1
2
Genotype
Genotype
0
1
2

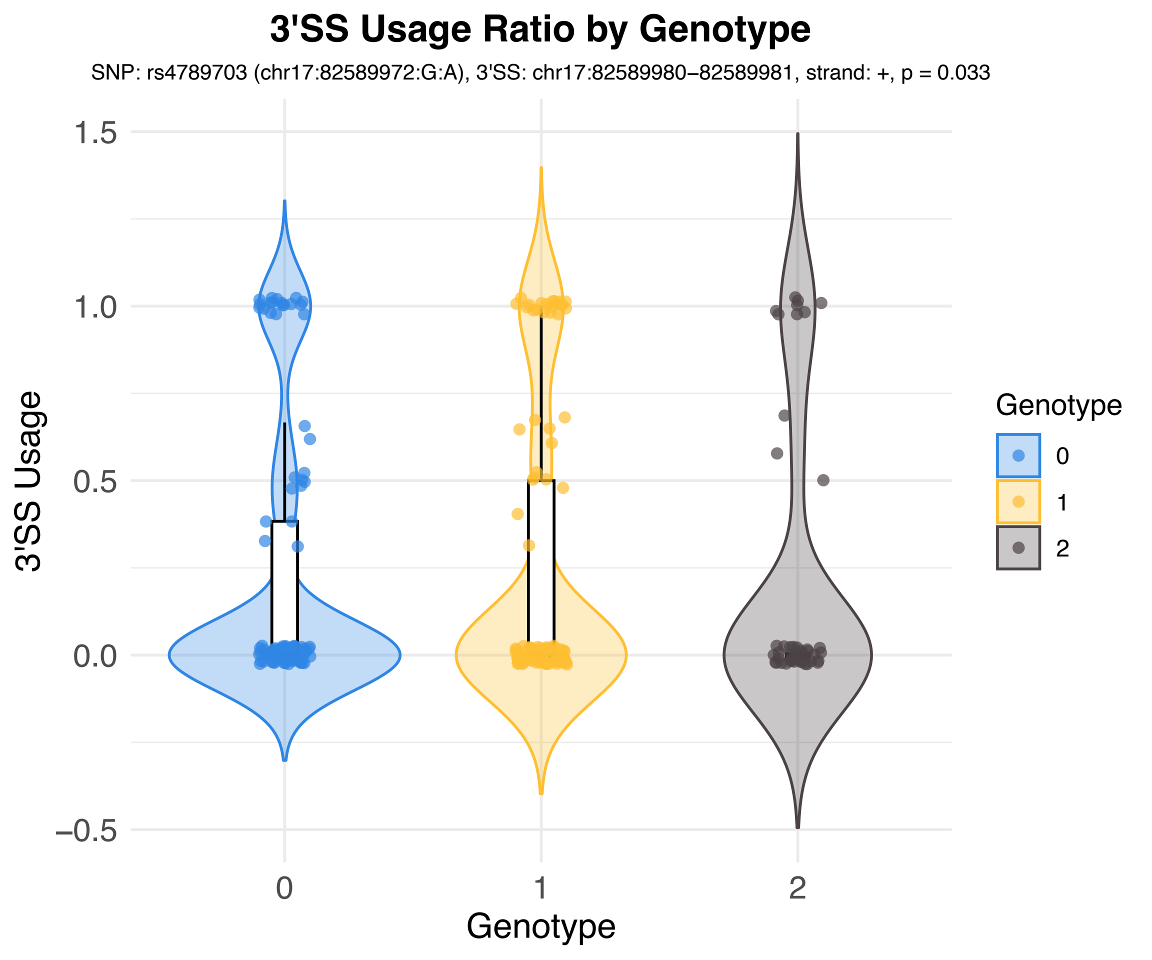
3'SS Usage Ratio by Genotype
SNP: rs4789703 (chr17:82589972:G:A), 3'SS: chr17:82589980–82589981, strand: +, p = 0.033
3'SS Usage
1.5
1.0
0.5
0.0
−0.5
0
1
2
Genotype
Genotype
0
1
2

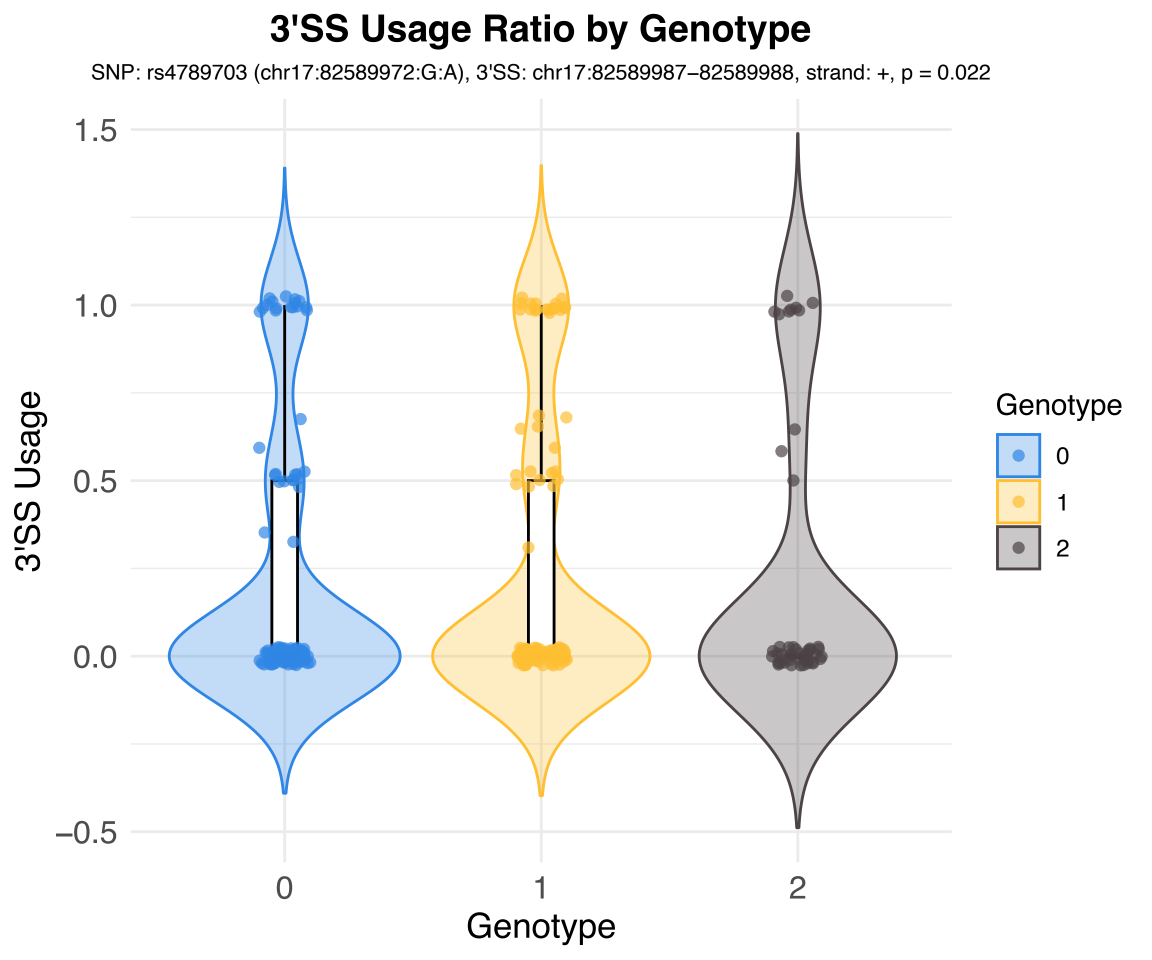

3'SS Usage Ratio by Genotype
SNP: rs4789703 (chr17:82589972:G:A), 3'SS: chr17:82589987–82589988, strand: +, p = 0.022
3'SS Usage
1.5
1.0
0.5
0.0
–0.5
0
1
2
Genotype
Genotype
0
1
2

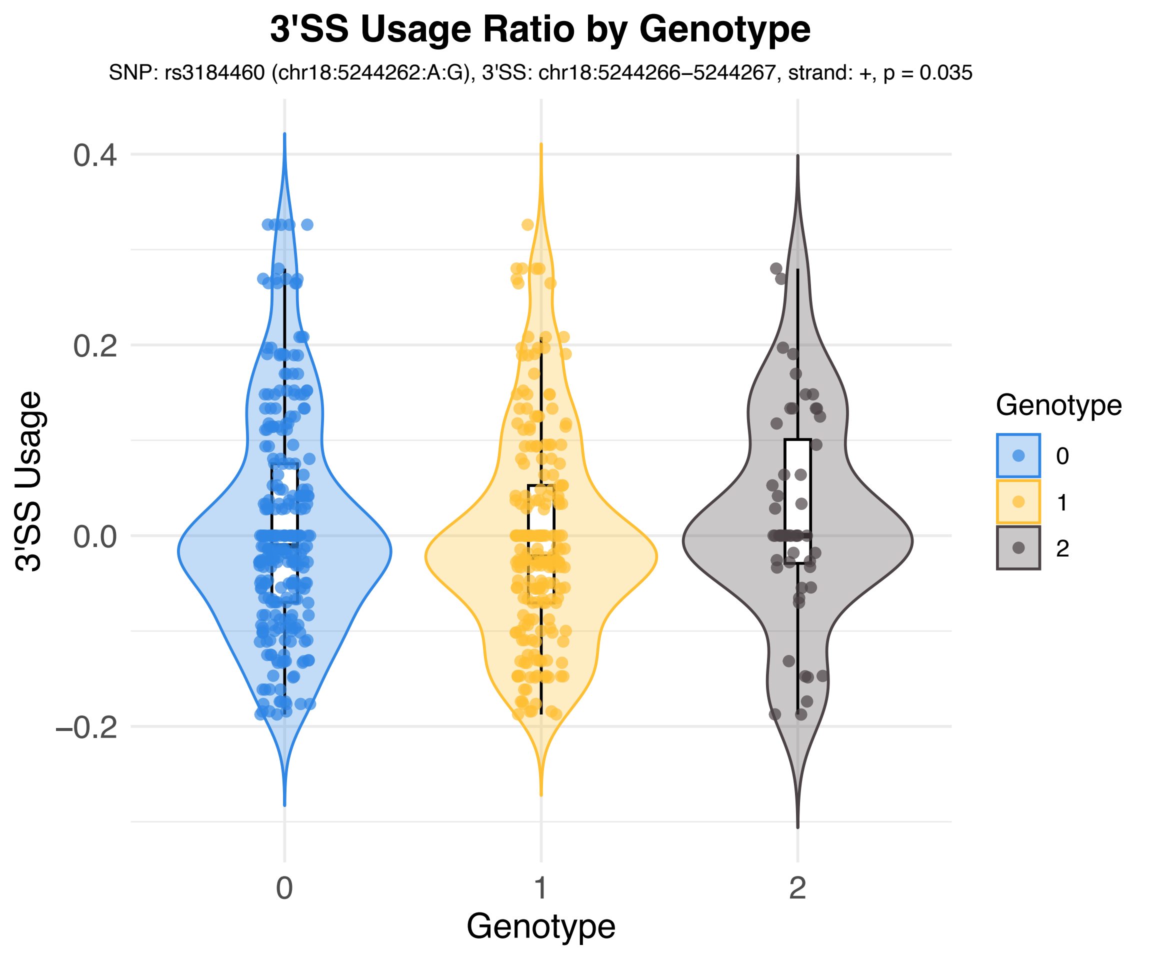
3'SS Usage Ratio by Genotype
SNP: rs3184460 (chr18:5244262:A:G), 3'SS: chr18:5244266–5244267, strand: +, p = 0.035
3'SS Usage
0.4
0.2
0.0
−0.2
0
1
2
Genotype
Genotype
0
1
2

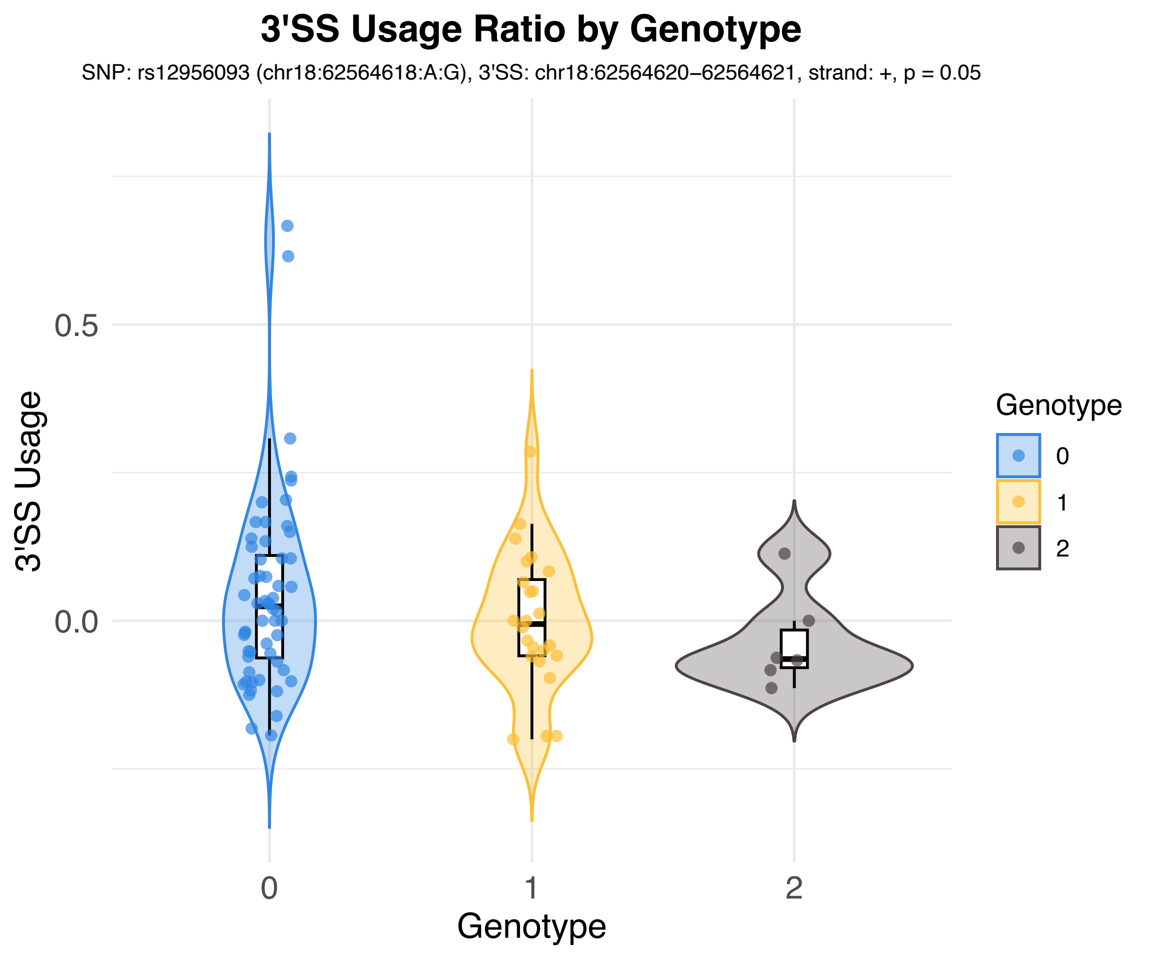
3'SS Usage Ratio by Genotype
SNP: rs12956093 (chr18:62564618:A:G), 3'SS: chr18:62564620−62564621, strand: +, p = 0.05
3'SS Usage
0.5
0.0
0
1
2
Genotype
Genotype
0
1
2

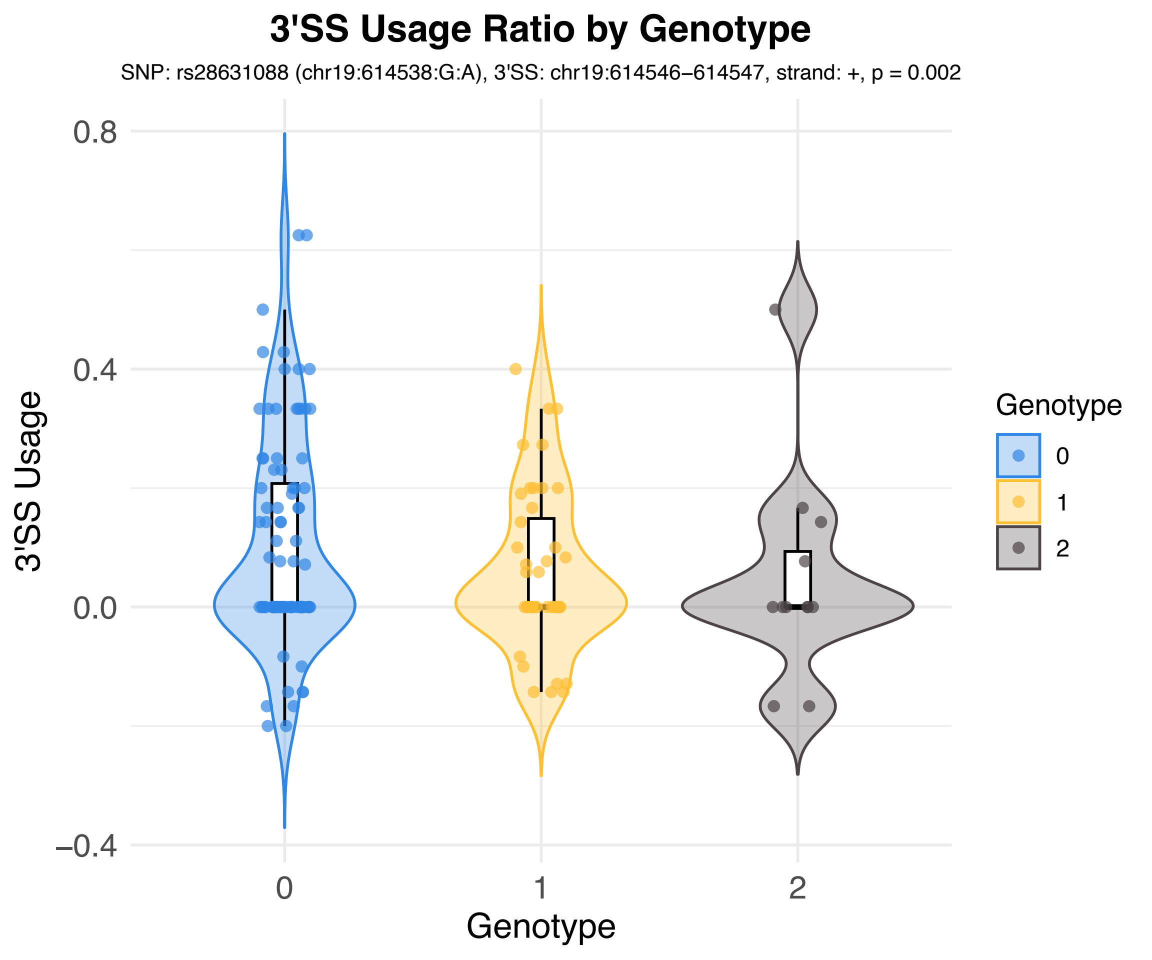

3'SS Usage Ratio by Genotype
SNP: rs28631088 (chr19:614538:G:A), 3'SS: chr19:614546–614547, strand: +, p = 0.002
3'SS Usage
0.8
0.4
0.0
−0.4
0
1
2
Genotype
Genotype
0
1
2

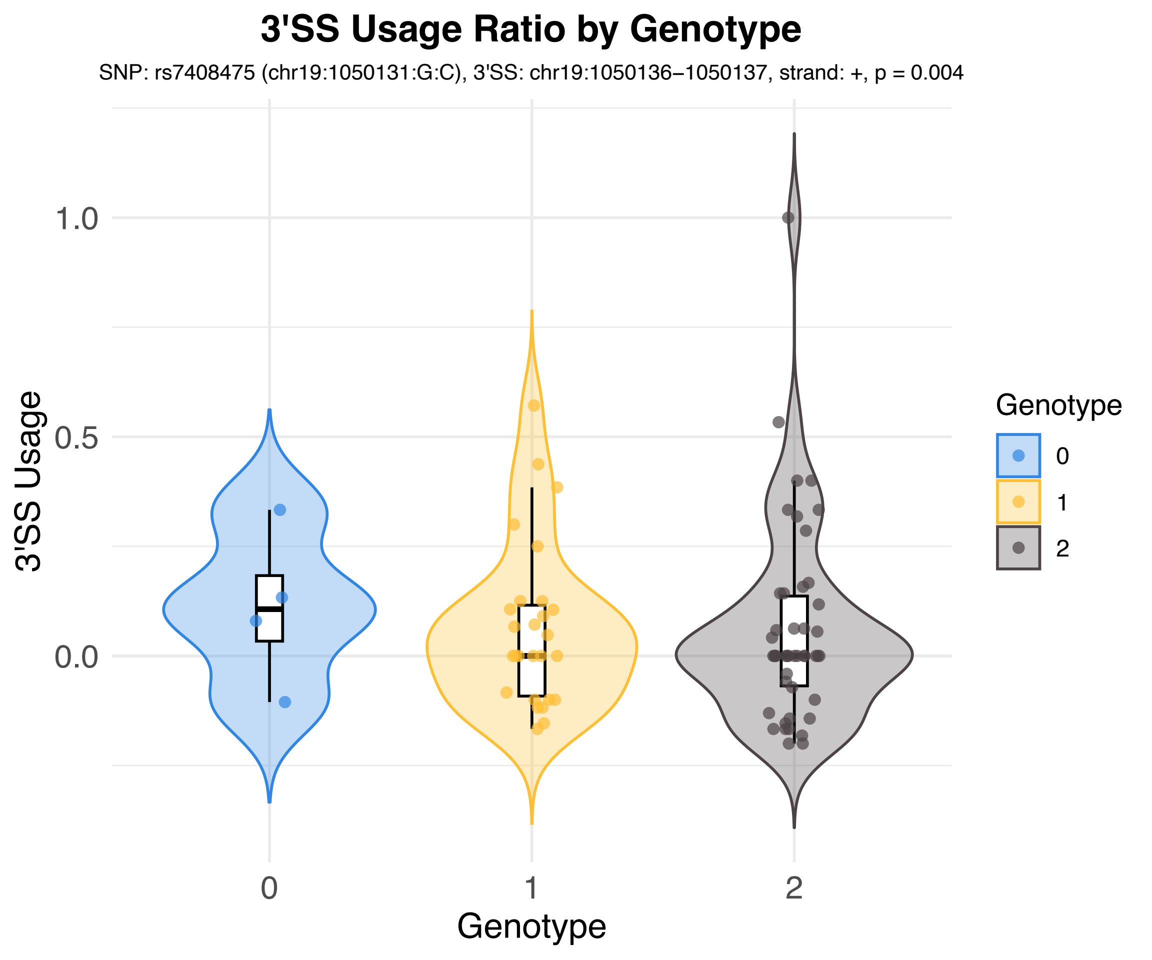
3'SS Usage Ratio by Genotype
SNP: rs7408475 (chr19:1050131:G:C), 3'SS: chr19:1050136–1050137, strand: +, p = 0.004
3'SS Usage
1.0
0.5
0.0
0
1
2
Genotype
Genotype
0
1
2

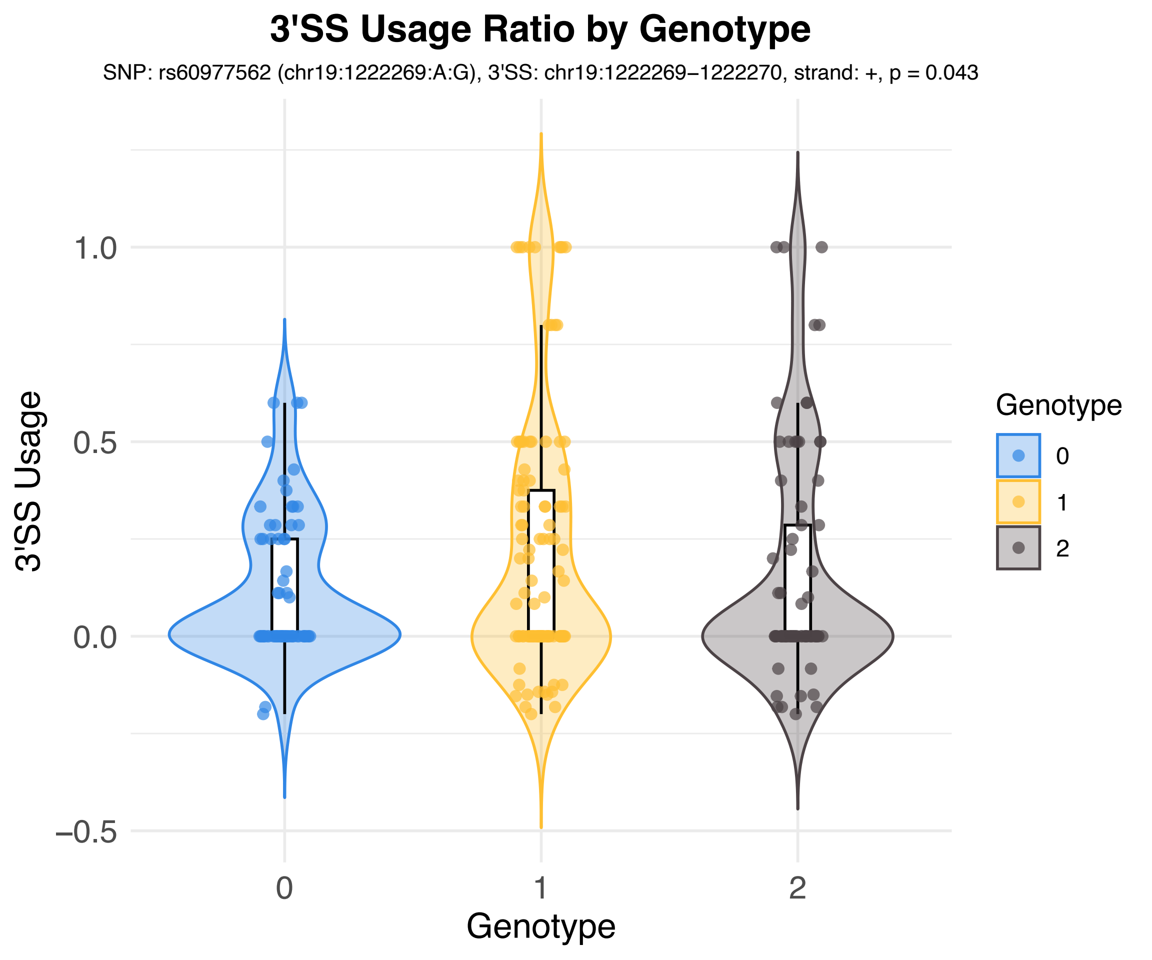
3'SS Usage Ratio by Genotype
SNP: rs60977562 (chr19:1222269:A:G), 3'SS: chr19:1222269−1222270, strand: +, p = 0.043
3'SS Usage
1.0
0.5
0.0
−0.5
0
1
2
Genotype
Genotype
0
1
2

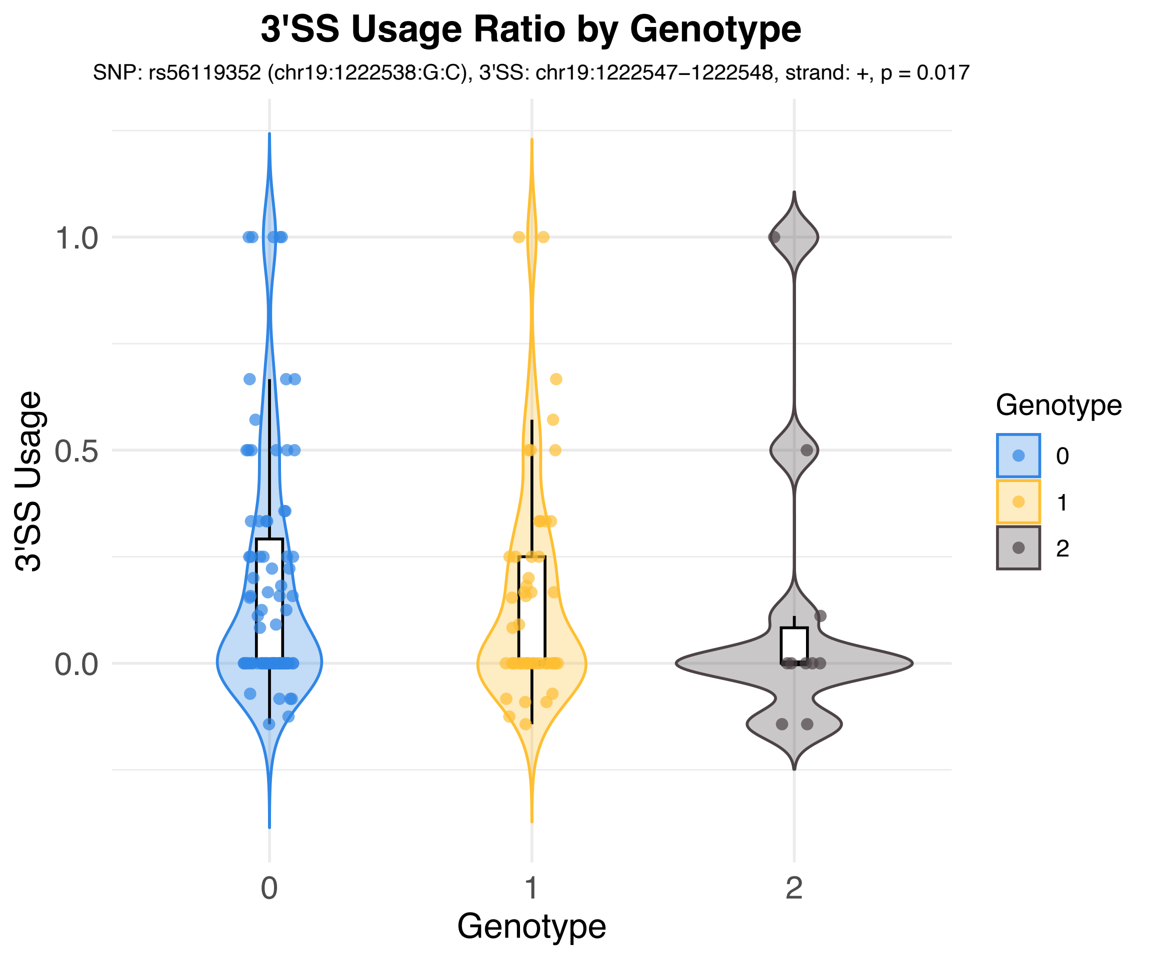
3'SS Usage Ratio by Genotype
SNP: rs56119352 (chr19:1222538:G:C), 3'SS: chr19:1222547−1222548, strand: +, p = 0.017
3'SS Usage
1.0
0.5
0.0
0
1
2
Genotype
Genotype
0
1
2

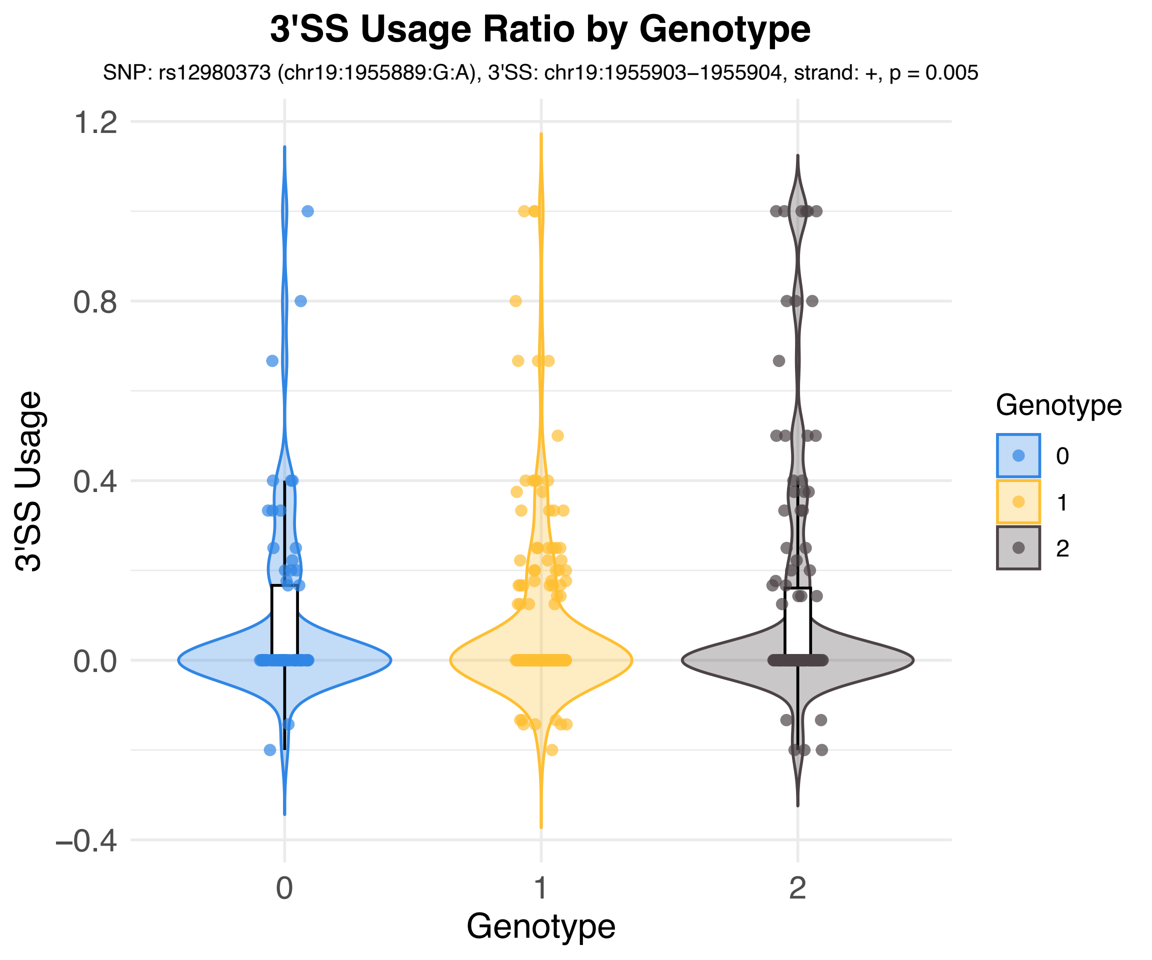
3'SS Usage Ratio by Genotype
SNP: rs12980373 (chr19:1955889:G:A), 3'SS: chr19:1955903−1955904, strand: +, p = 0.005
3'SS Usage
1.2
0.8
0.4
0.0
−0.4
0
1
2
Genotype
Genotype
0
1
2

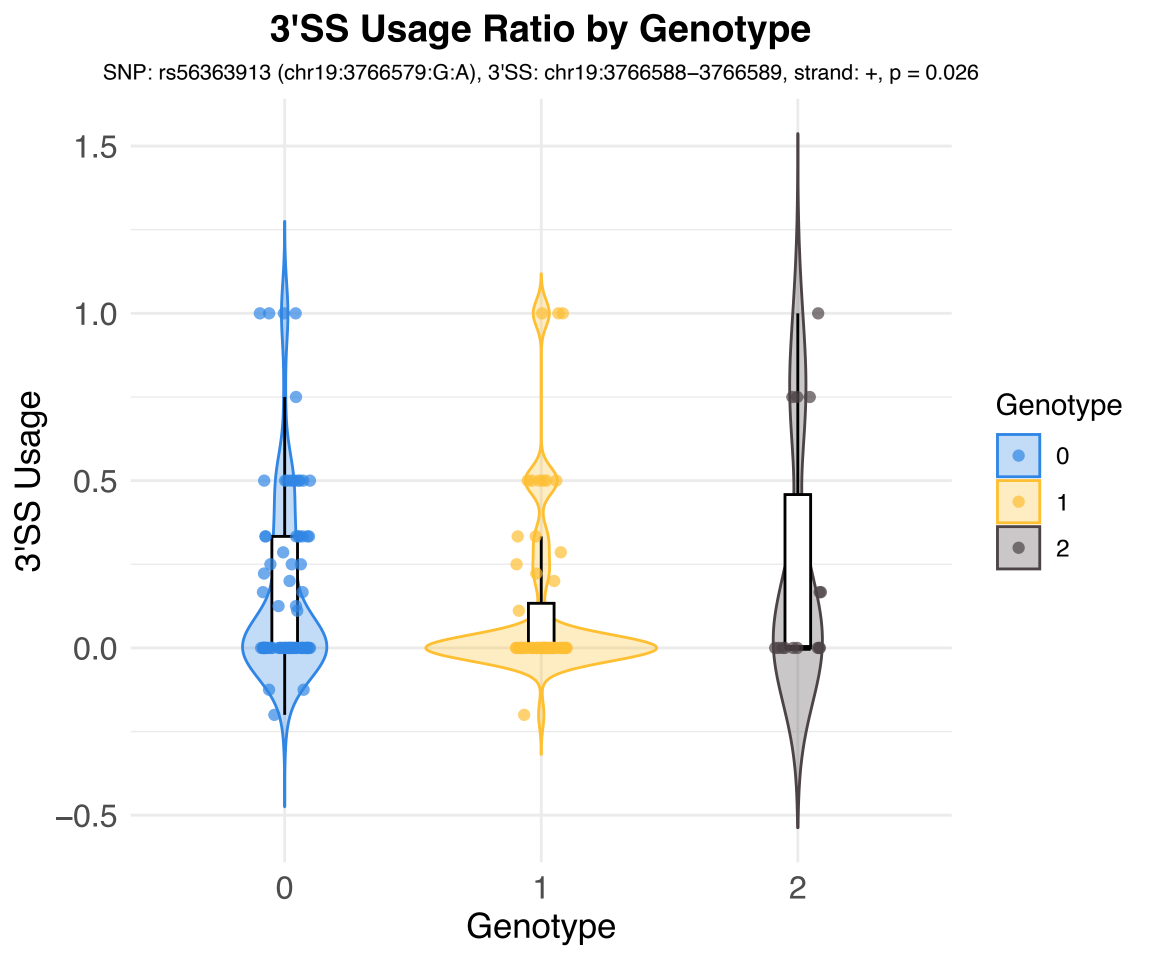

3'SS Usage Ratio by Genotype
SNP: rs56363913 (chr19:3766579:G:A), 3'SS: chr19:3766588–3766589, strand: +, p = 0.026
3'SS Usage
1.5
1.0
0.5
0.0
−0.5
0
1
2
Genotype
Genotype
0
1
2

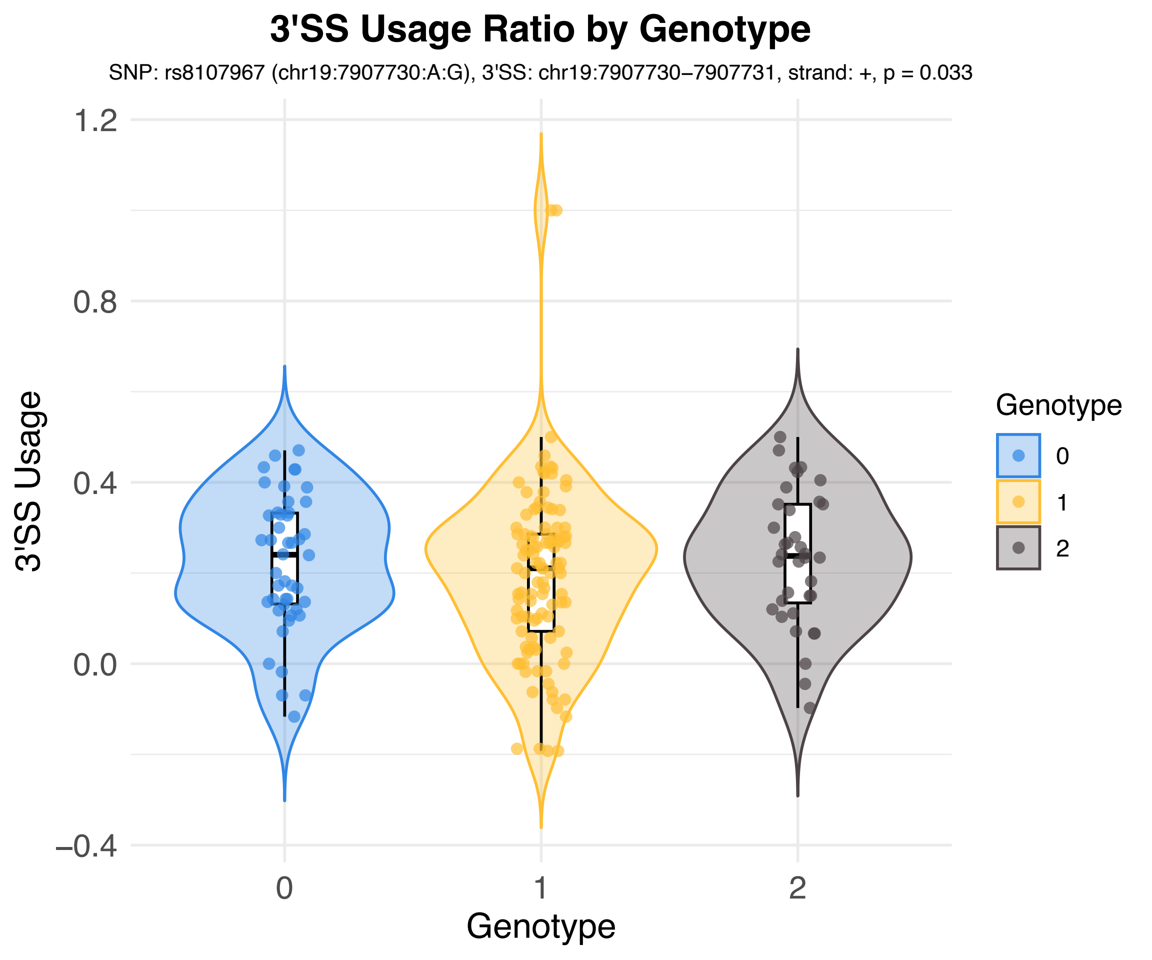

3'SS Usage Ratio by Genotype
SNP: rs8107967 (chr19:7907730:A:G), 3'SS: chr19:7907730–7907731, strand: +, p = 0.033
3'SS Usage
1.2
0.8
0.4
0.0
–0.4
0
1
2
Genotype
Genotype
0
1
2

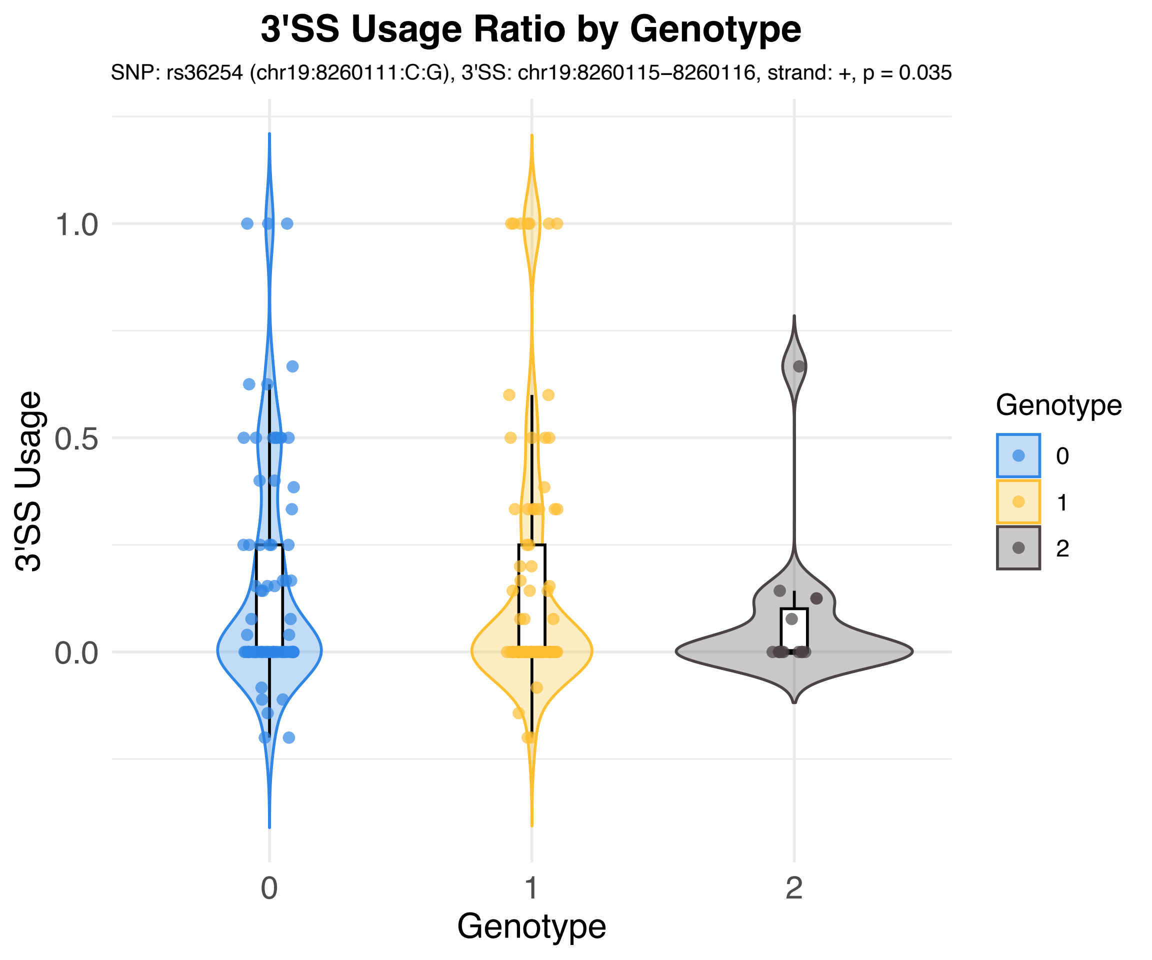
3'SS Usage Ratio by Genotype
SNP: rs36254 (chr19:8260111:C:G), 3'SS: chr19:8260115–8260116, strand: +, p = 0.035
3'SS Usage
1.0
0.5
0.0
0
1
2
Genotype
Genotype
0
1
2

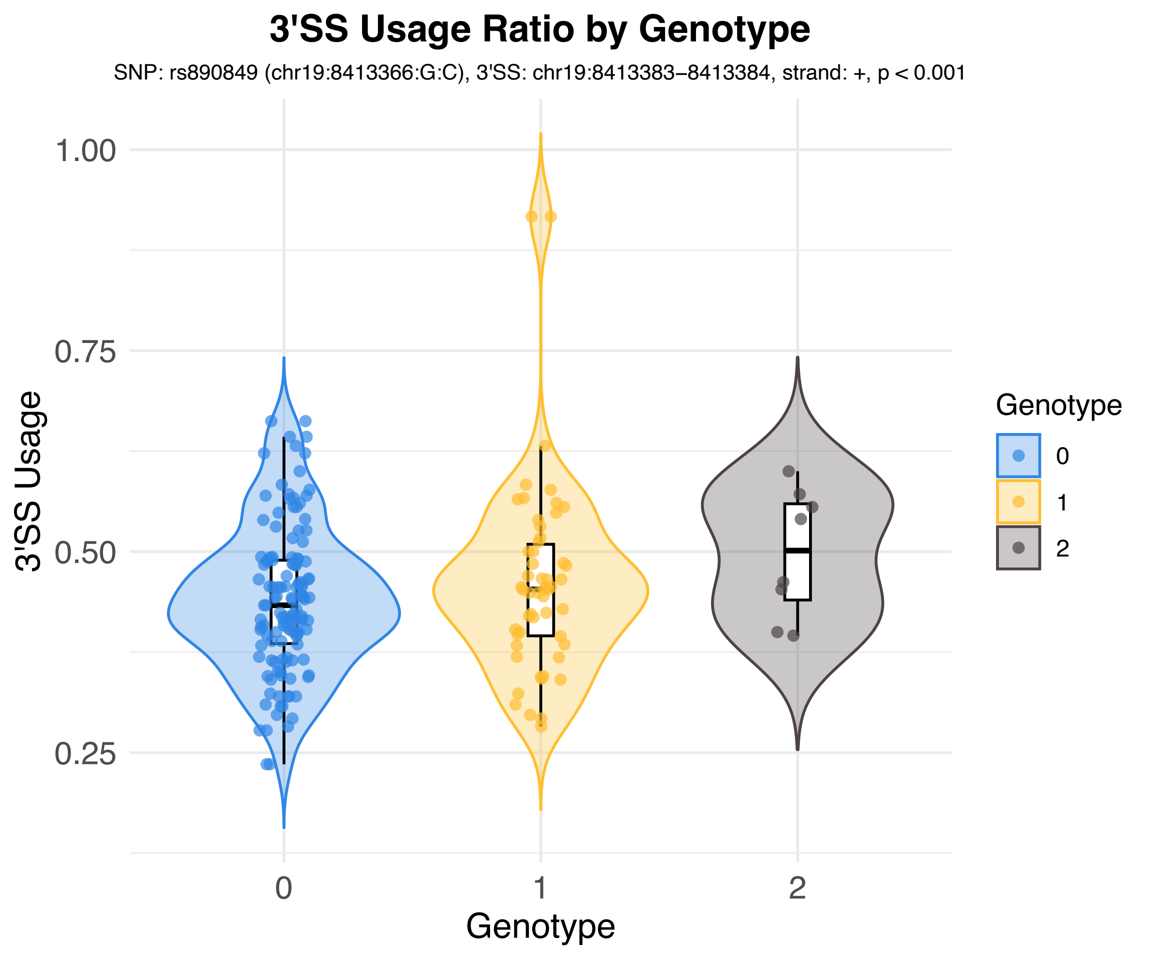
3'SS Usage Ratio by Genotype
SNP: rs890849 (chr19:8413366:G:C), 3'SS: chr19:8413383–8413384, strand: +, p < 0.001
3'SS Usage
1.00
0.75
0.50
0.25
0
1
2
Genotype
Genotype
0
1
2

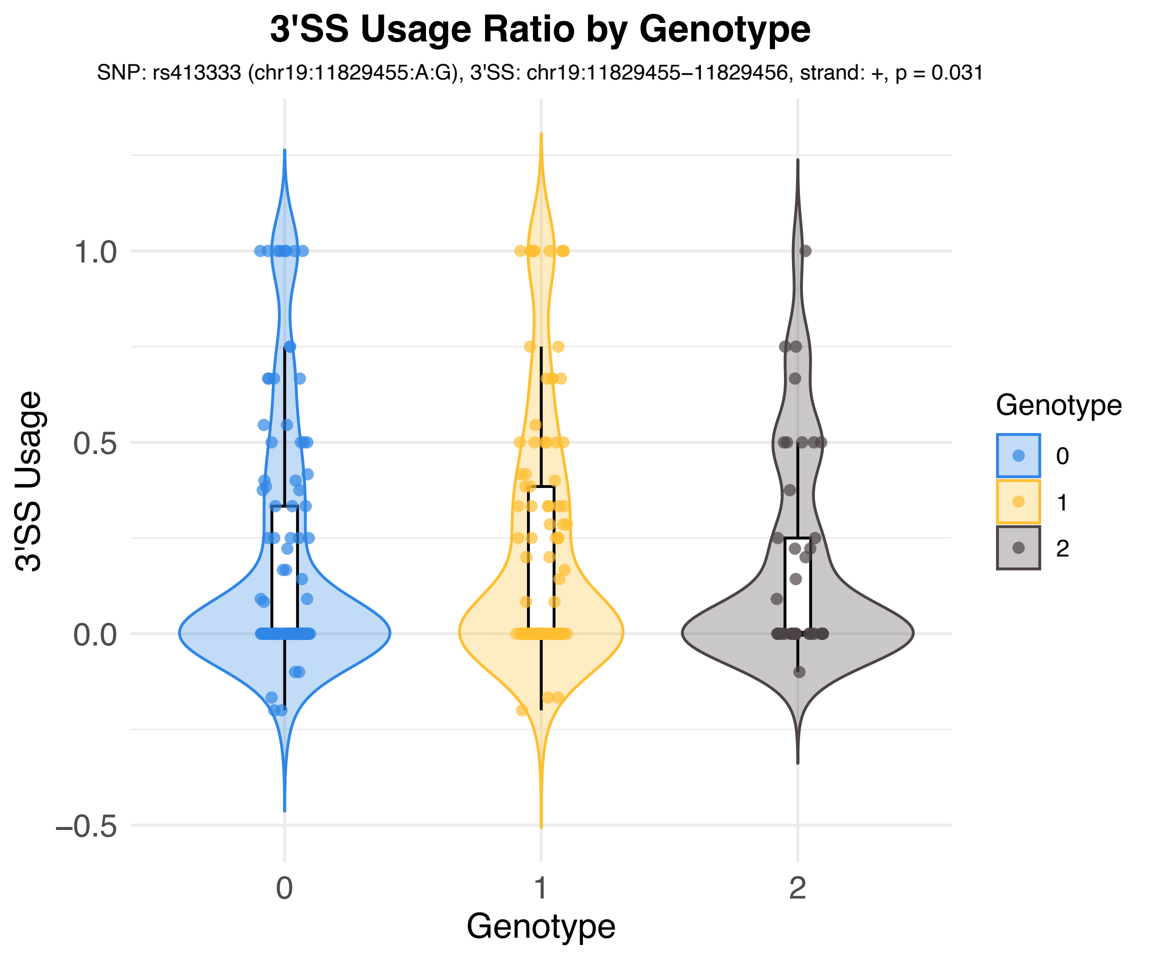
3'SS Usage Ratio by Genotype
SNP: rs413333 (chr19:11829455:A:G), 3'SS: chr19:11829455–11829456, strand: +, p = 0.031
3'SS Usage
1.0
0.5
0.0
–0.5
0
1
2
Genotype
Genotype
0
1
2

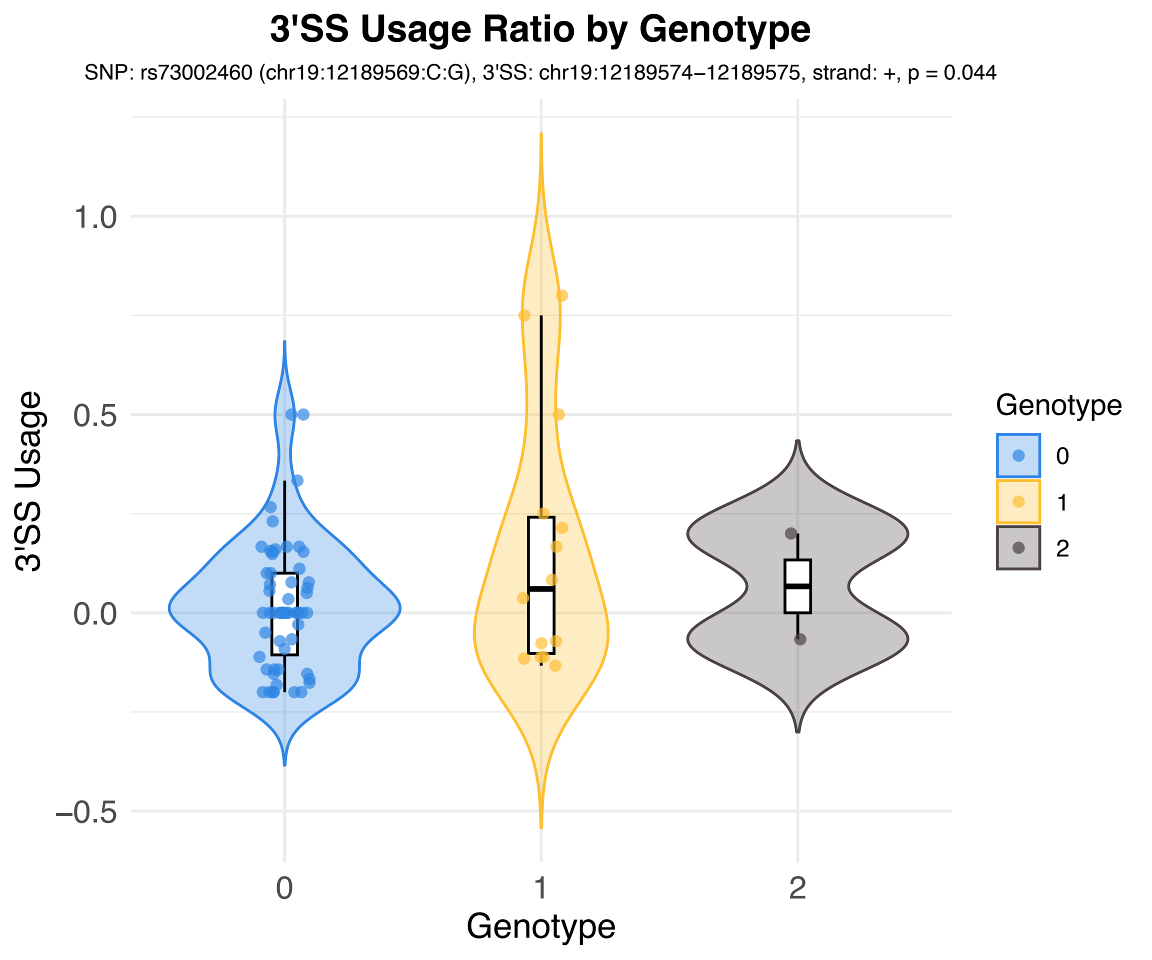
3'SS Usage Ratio by Genotype
SNP: rs73002460 (chr19:12189569:C:G), 3'SS: chr19:12189574–12189575, strand: +, p = 0.044
3'SS Usage
1.0
0.5
0.0
−0.5
0
1
2
Genotype
Genotype
0
1
2

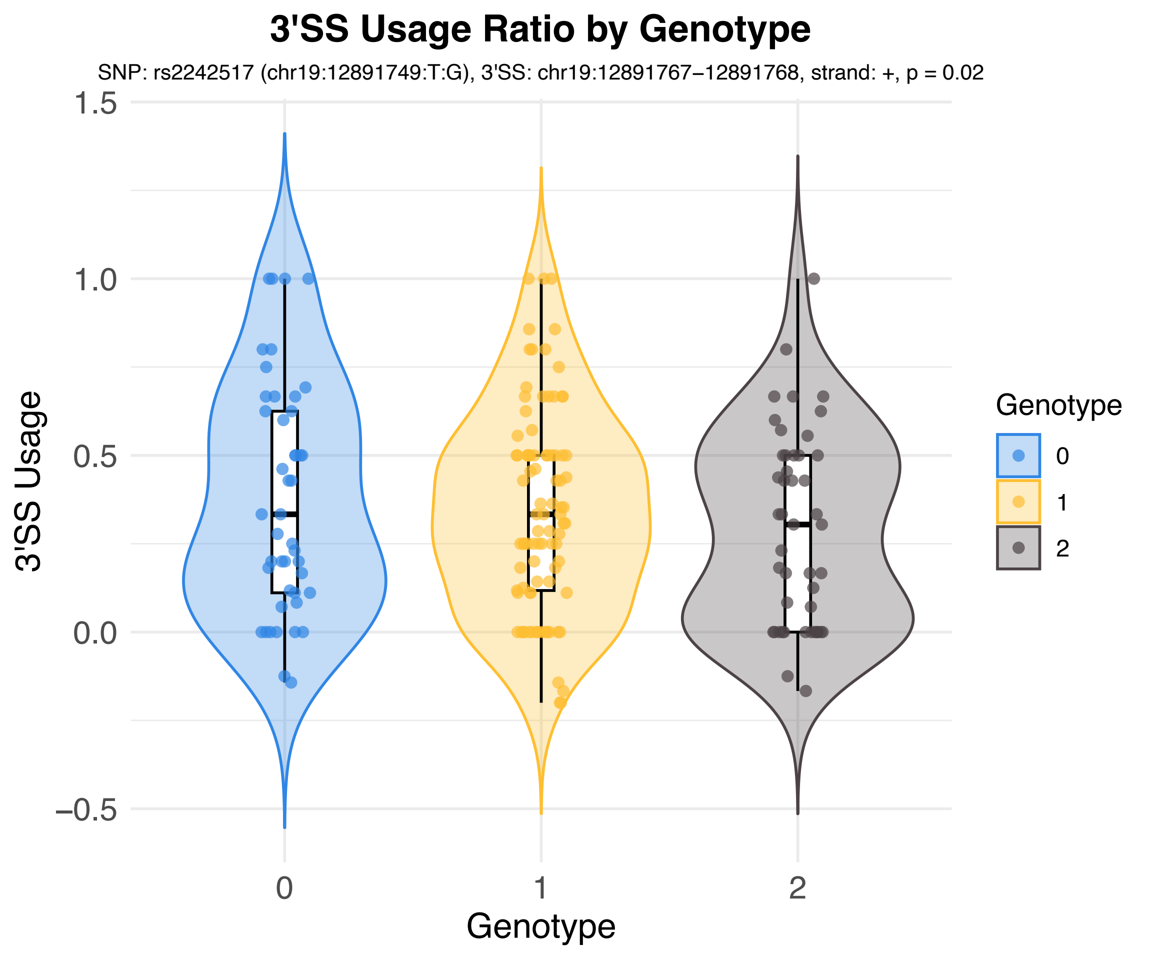

3'SS Usage Ratio by Genotype
SNP: rs2242517 (chr19:12891749:T:G), 3'SS: chr19:12891767−12891768, strand: +, p = 0.02
3'SS Usage
1.5
1.0
0.5
0.0
−0.5
0
1
2
Genotype
Genotype
0
1
2

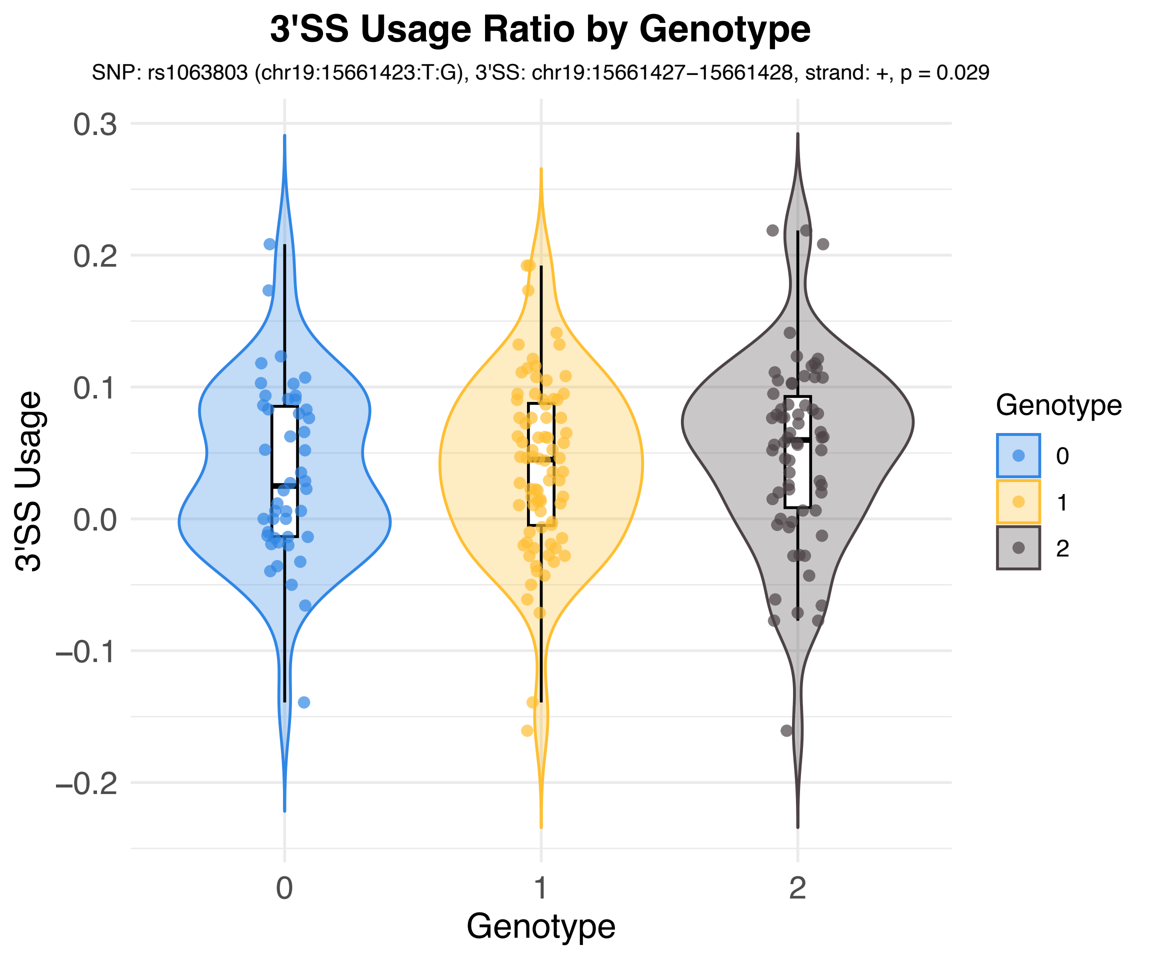
3'SS Usage Ratio by Genotype
SNP: rs1063803 (chr19:15661423:T:G), 3'SS: chr19:15661427−15661428, strand: +, p = 0.029
3'SS Usage
0.3
0.2
0.1
0.0
−0.1
−0.2
0
1
2
Genotype
Genotype
0
1
2

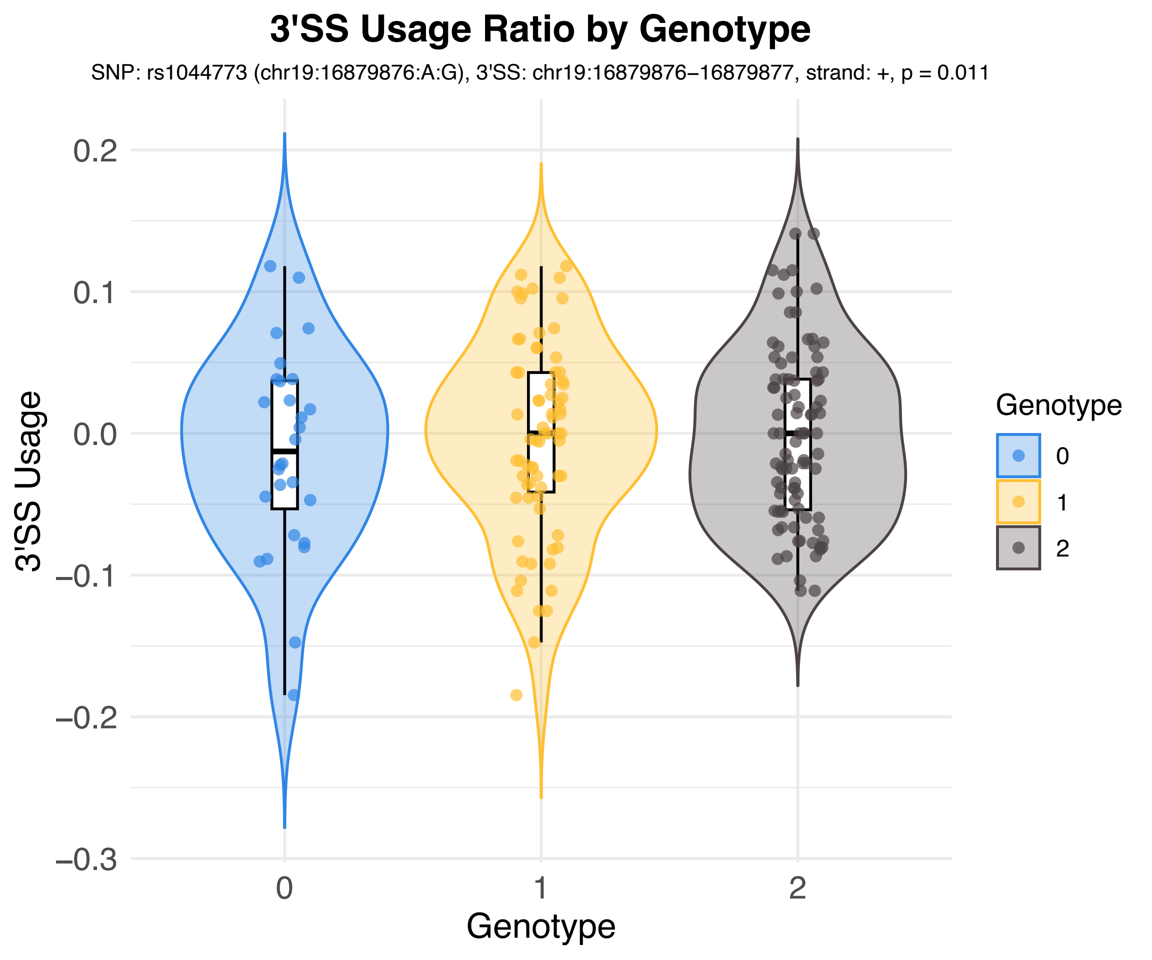
3'SS Usage Ratio by Genotype
SNP: rs1044773 (chr19:16879876:A:G), 3'SS: chr19:16879876–16879877, strand: +, p = 0.011
3'SS Usage
0.2
0.1
0.0
−0.1
−0.2
−0.3
0
1
2
Genotype
Genotype
0
1
2

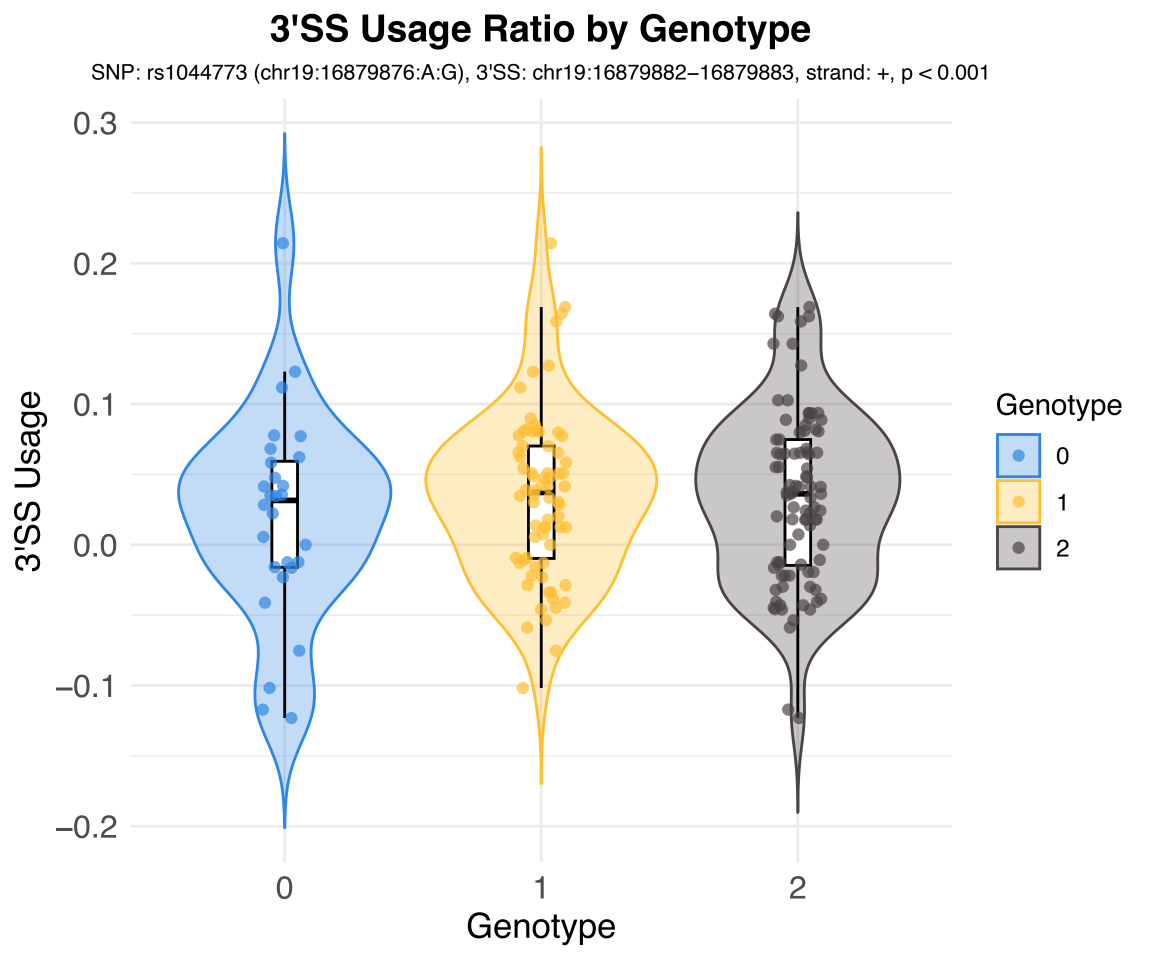

3'SS Usage Ratio by Genotype
SNP: rs1044773 (chr19:16879876:A:G), 3'SS: chr19:16879882–16879883, strand: +, p < 0.001
3'SS Usage
0.3
0.2
0.1
0.0
–0.1
–0.2
0
1
2
Genotype
Genotype
0
1
2

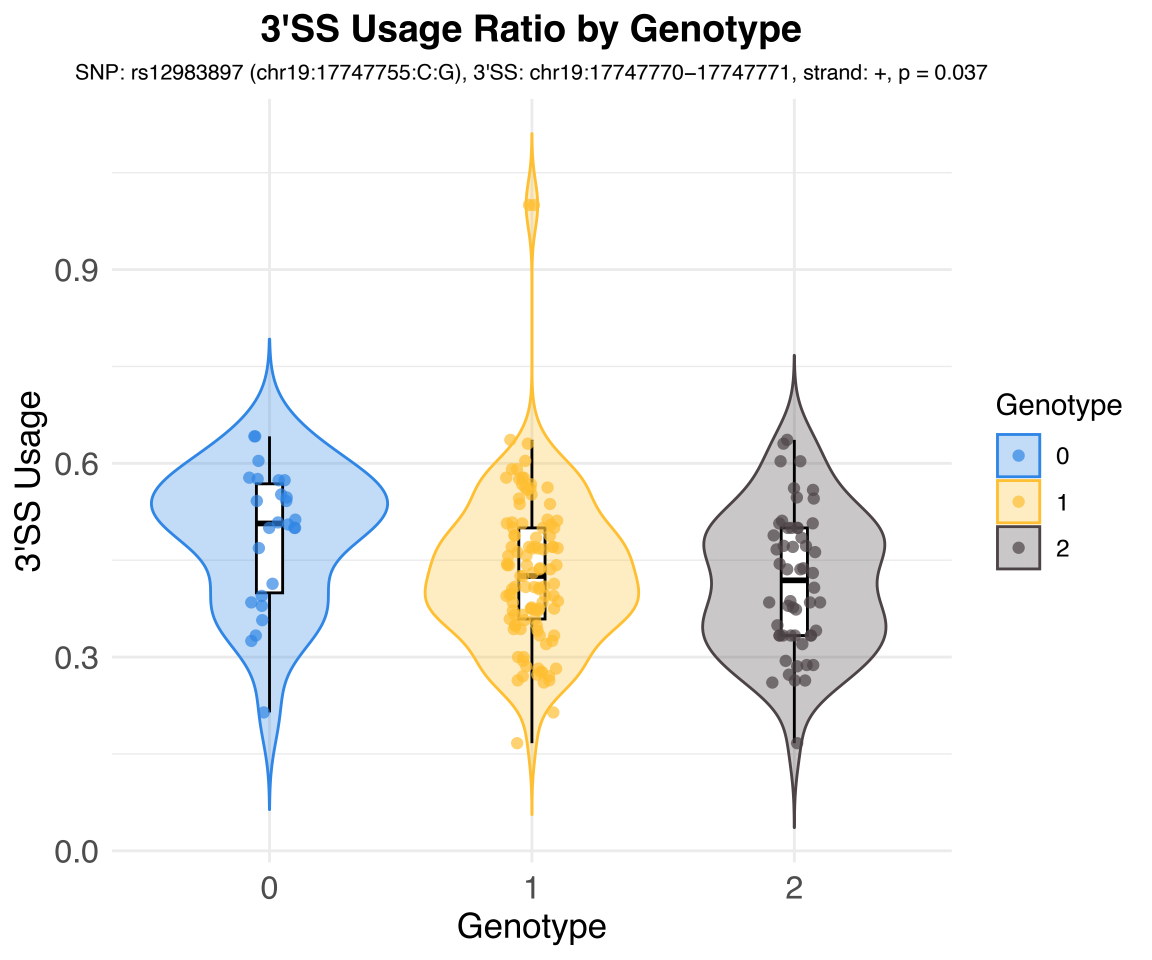
3'SS Usage Ratio by Genotype
SNP: rs12983897 (chr19:17747755:C:G), 3'SS: chr19:17747770−17747771, strand: +, p = 0.037
3'SS Usage
0.0
0.3
0.6
0.9
0
1
2
Genotype
Genotype
0
1
2

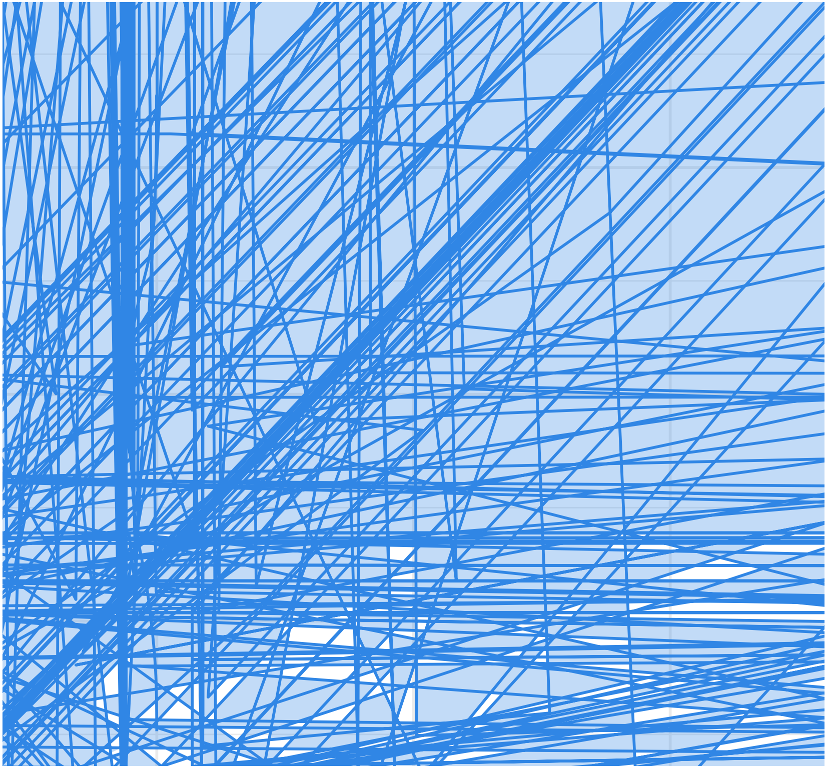

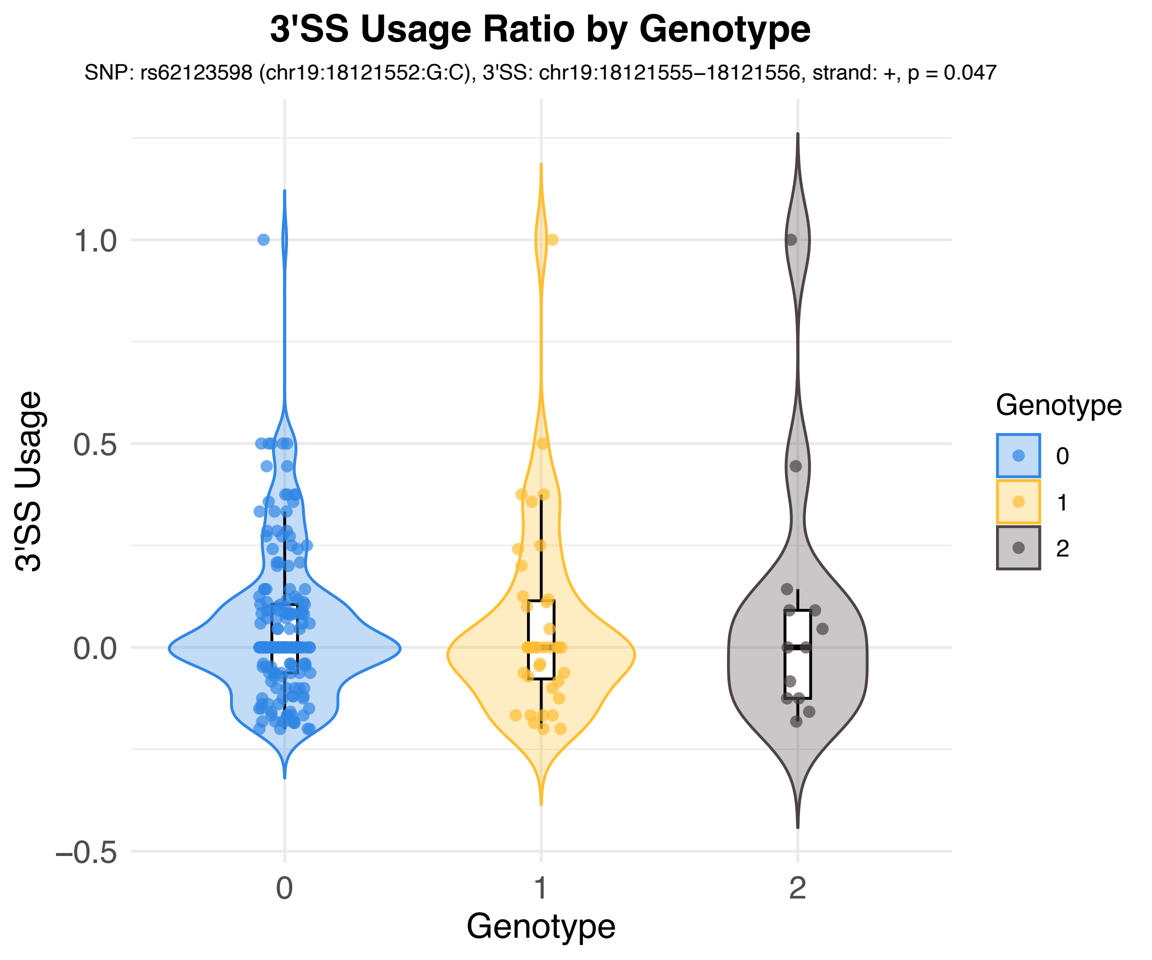
3'SS Usage Ratio by Genotype
SNP: rs62123598 (chr19:18121552:G:C), 3'SS: chr19:18121555−18121556, strand: +, p = 0.047
3'SS Usage
1.0
0.5
0.0
−0.5
0
1
2
Genotype
Genotype
0
1
2

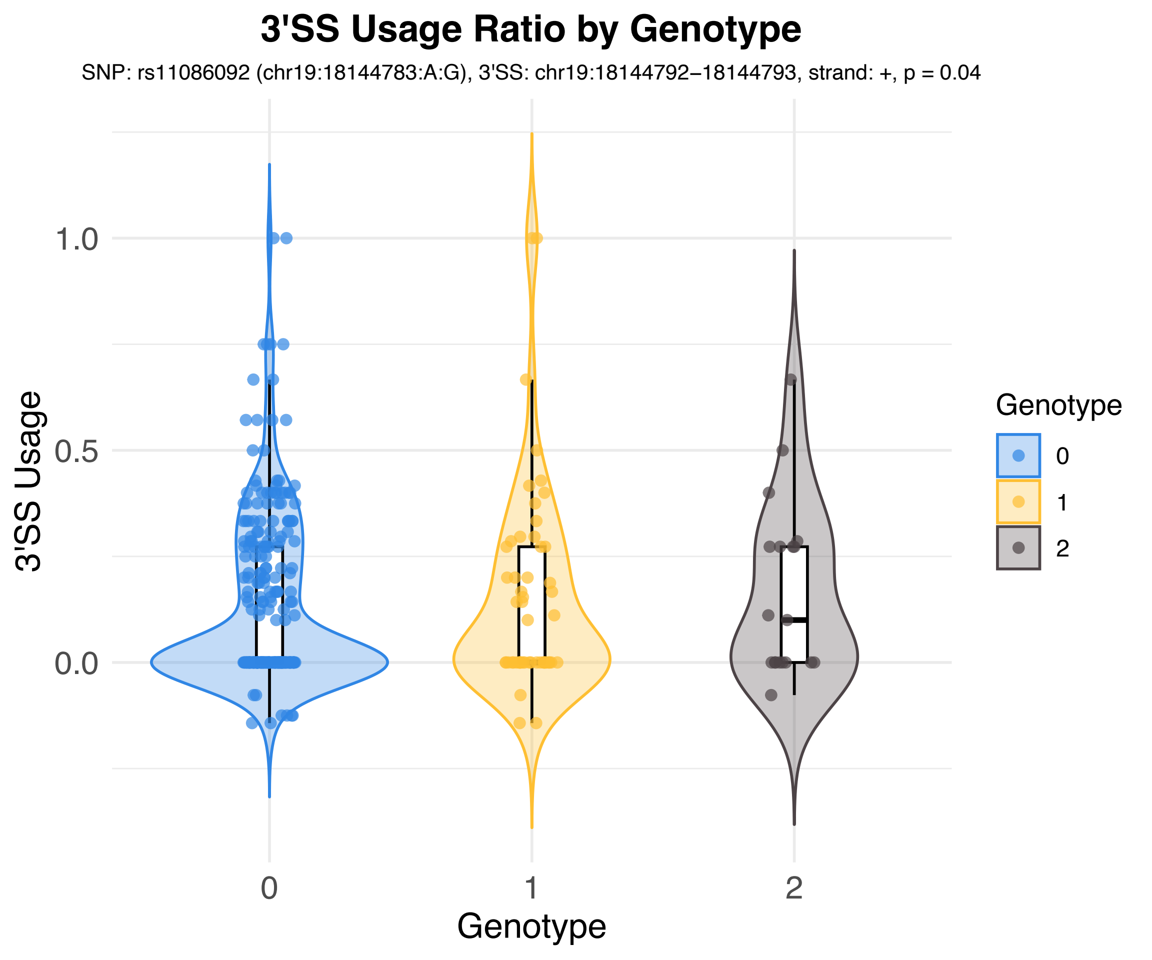

3'SS Usage Ratio by Genotype
SNP: rs11086092 (chr19:18144783:A:G), 3'SS: chr19:18144792–18144793, strand: +, p = 0.04
3'SS Usage
1.0
0.5
0.0
0
1
2
Genotype
Genotype
0
1
2

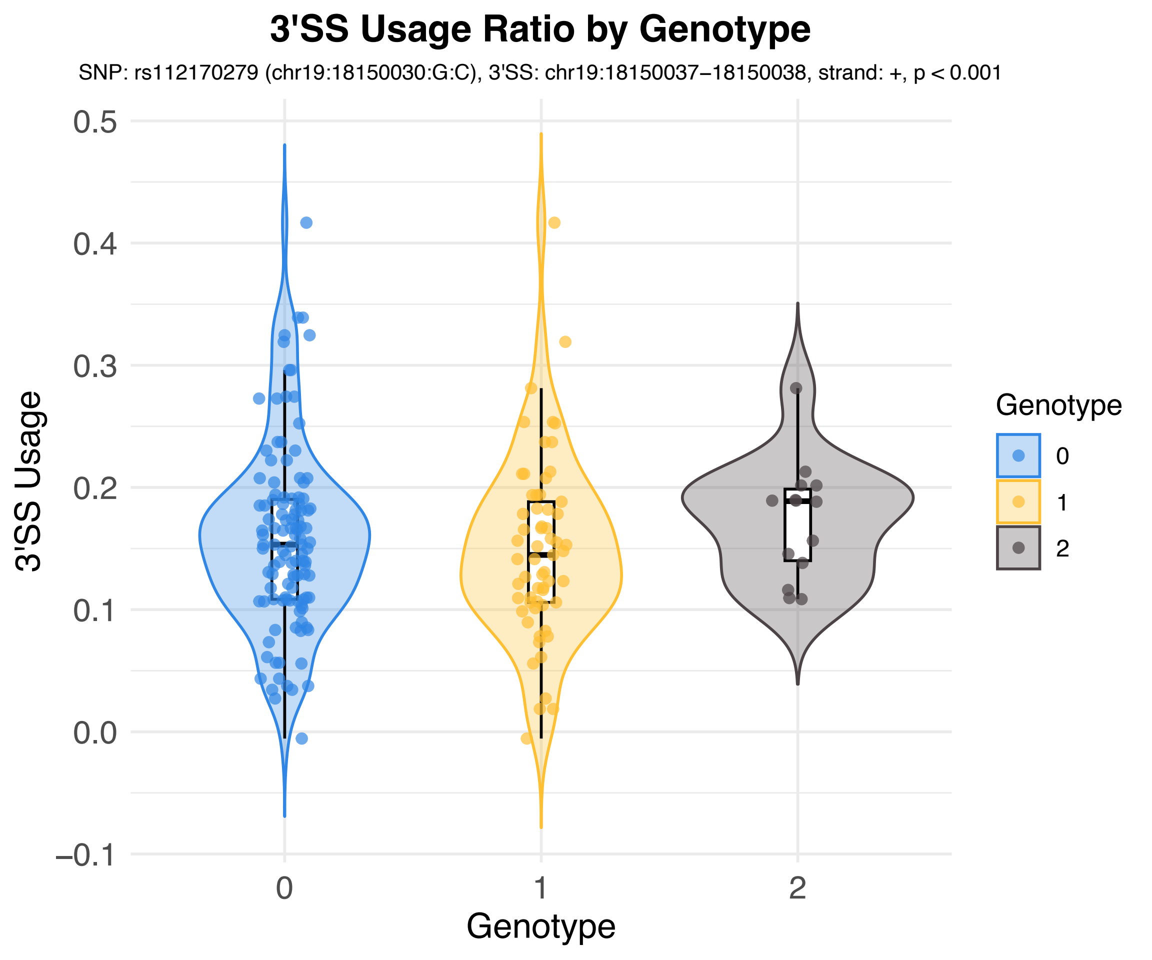

3'SS Usage Ratio by Genotype
SNP: rs112170279 (chr19:18150030:G:C), 3'SS: chr19:18150037−18150038, strand: +, p < 0.001
3'SS Usage
0.5
0.4
0.3
0.2
0.1
0.0
−0.1
0
1
2
Genotype
Genotype
0
1
2

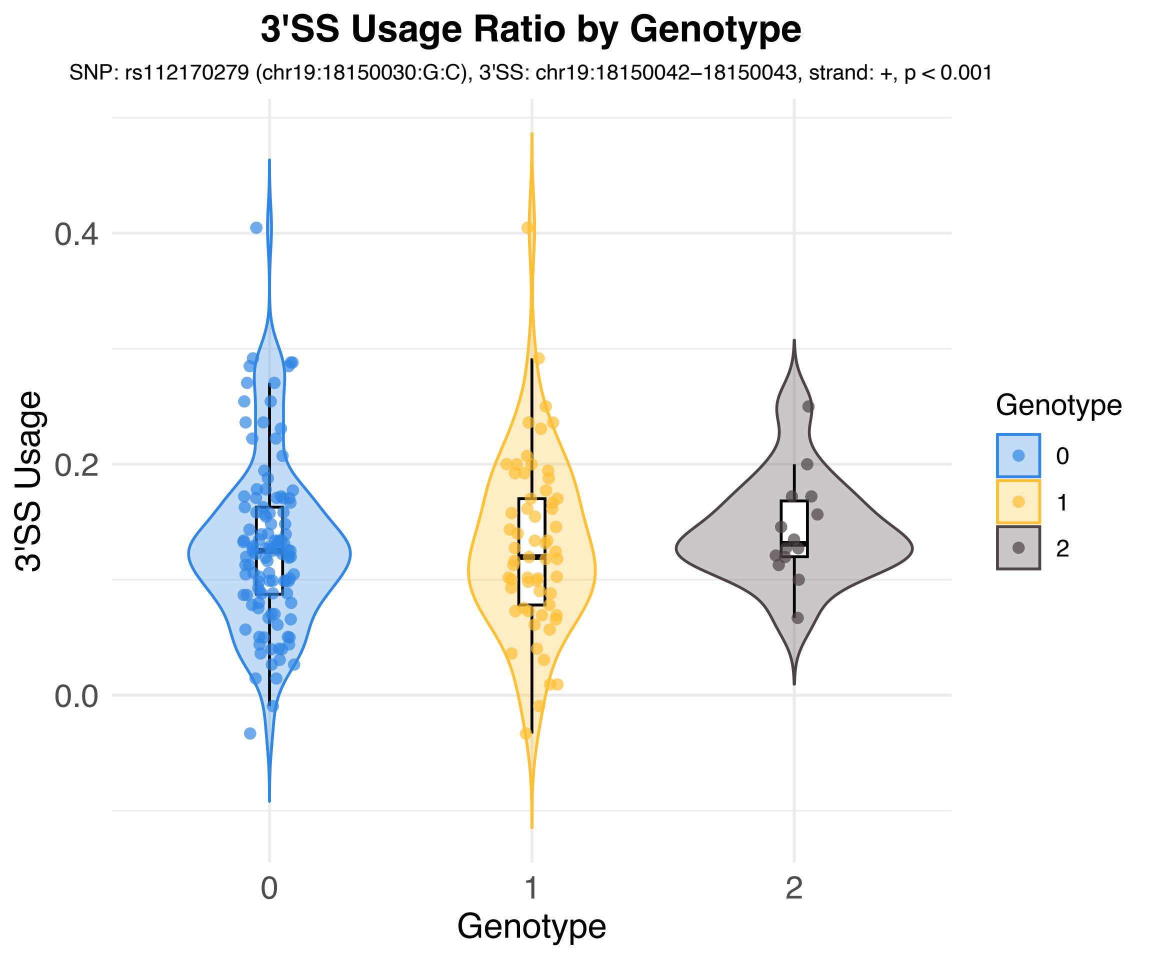
3'SS Usage Ratio by Genotype
SNP: rs112170279 (chr19:18150030:G:C), 3'SS: chr19:18150042–18150043, strand: +, p < 0.001
3'SS Usage
0.4
0.2
0.0
0
1
2
Genotype
Genotype
0
1
2

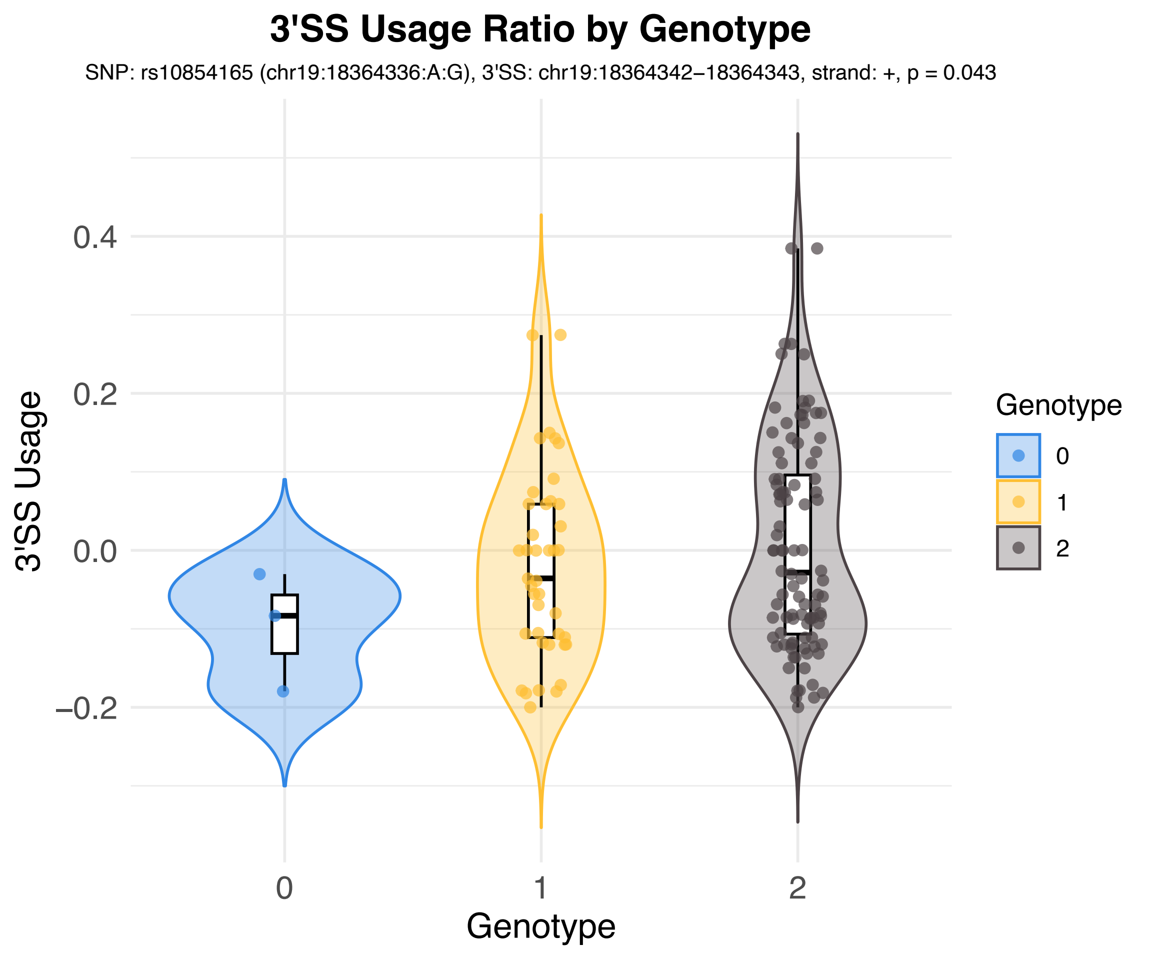
3'SS Usage Ratio by Genotype
SNP: rs10854165 (chr19:18364336:A:G), 3'SS: chr19:18364342–18364343, strand: +, p = 0.043
3'SS Usage
0.4
0.2
0.0
−0.2
0
1
2
Genotype
Genotype
0
1
2

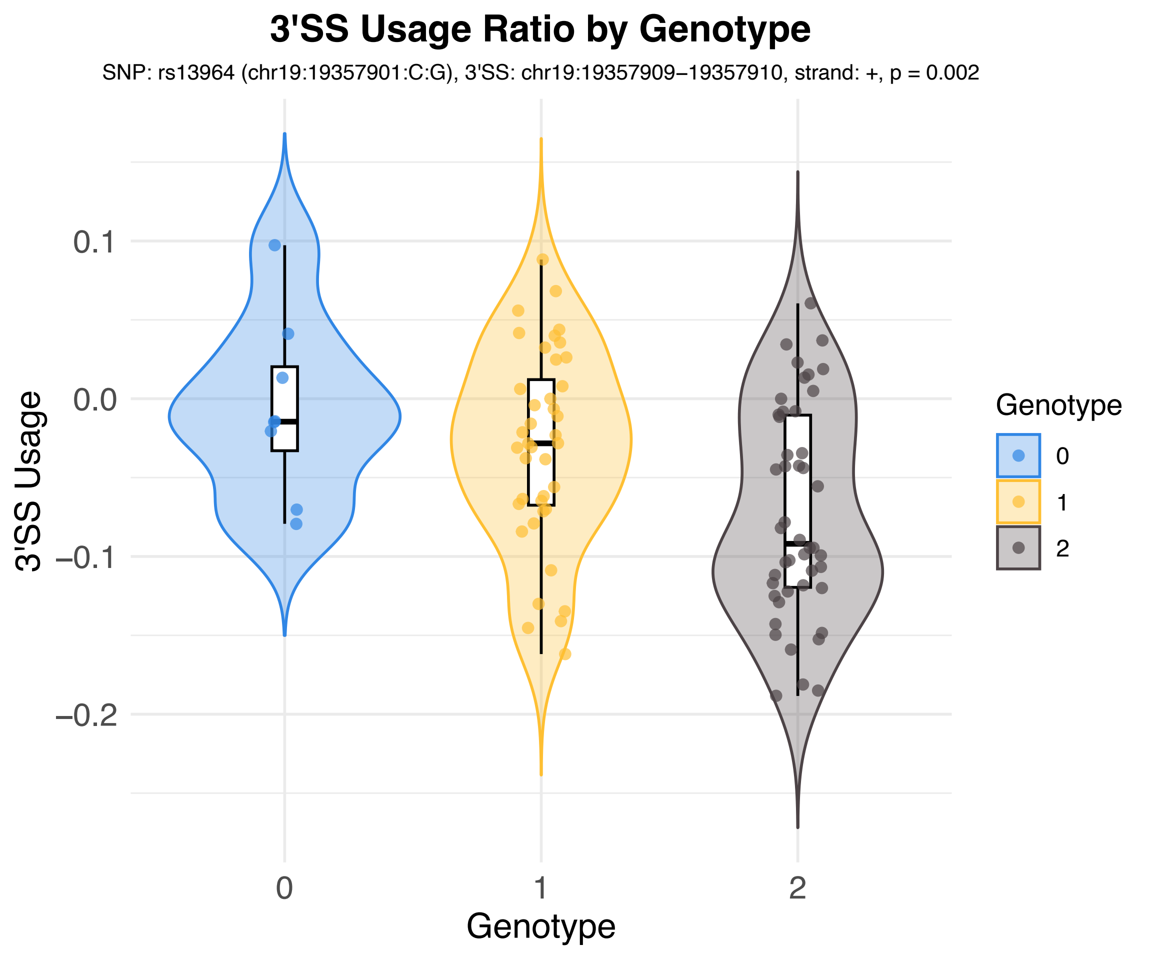

3'SS Usage Ratio by Genotype
SNP: rs13964 (chr19:19357901:C:G), 3'SS: chr19:19357909−19357910, strand: +, p = 0.002
3'SS Usage
0.1
0.0
−0.1
−0.2
0
1
2
Genotype
Genotype
0
1
2

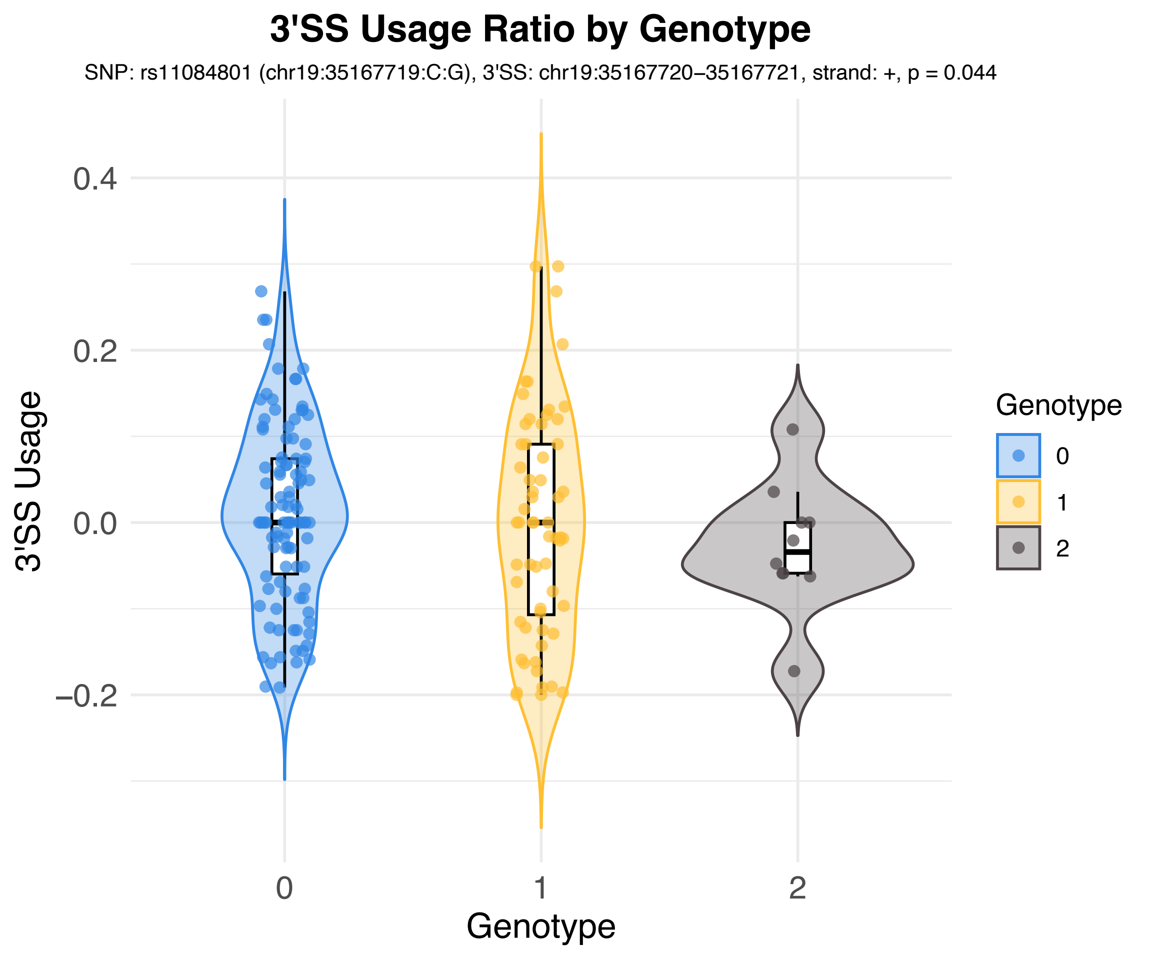

3'SS Usage Ratio by Genotype
SNP: rs11084801 (chr19:35167719:C:G), 3'SS: chr19:35167720–35167721, strand: +, p = 0.044
3'SS Usage
0.4
0.2
0.0
−0.2
0
1
2
Genotype
Genotype
0
1
2

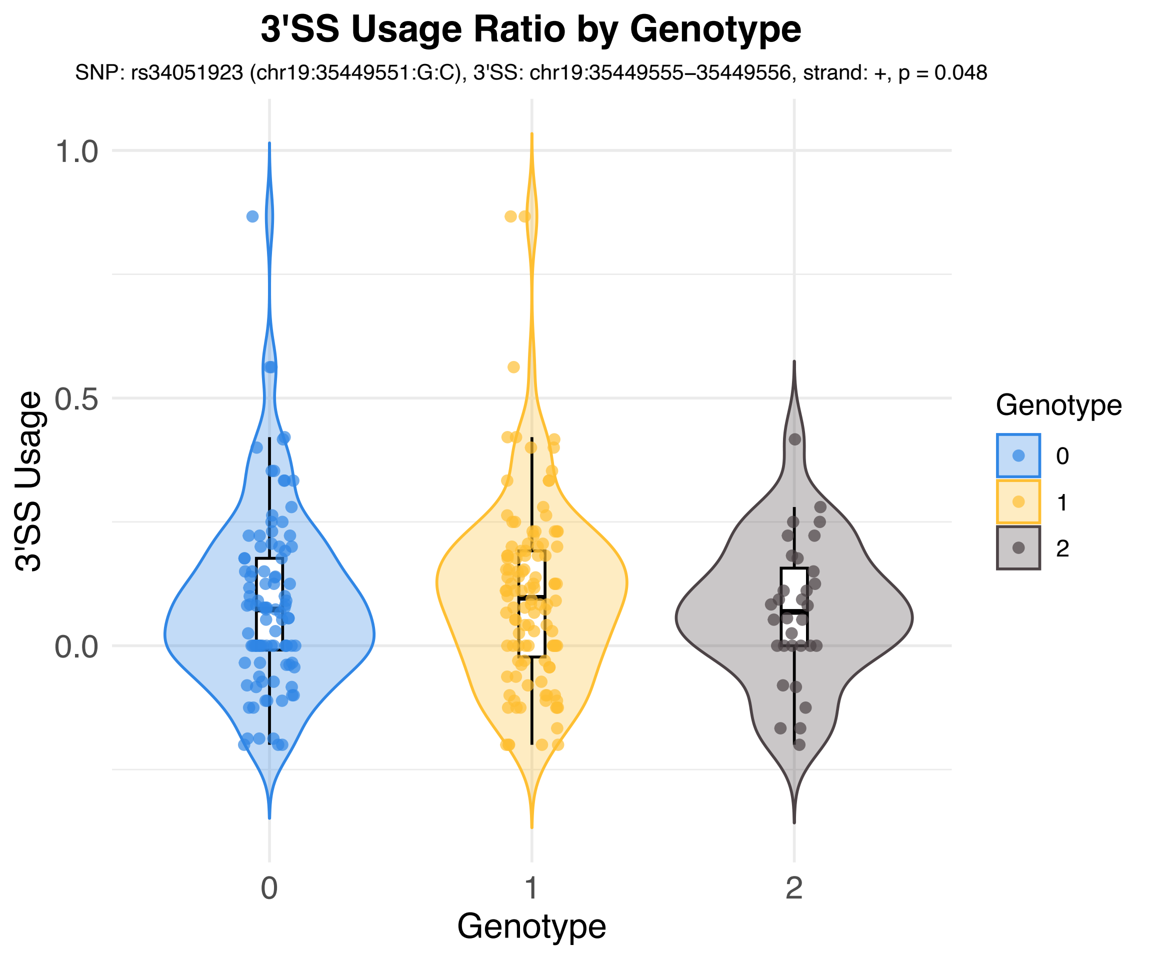
3'SS Usage Ratio by Genotype
SNP: rs34051923 (chr19:35449551:G:C), 3'SS: chr19:35449555–35449556, strand: +, p = 0.048
3'SS Usage
1.0
0.5
0.0
0
1
2
Genotype
Genotype
0
1
2

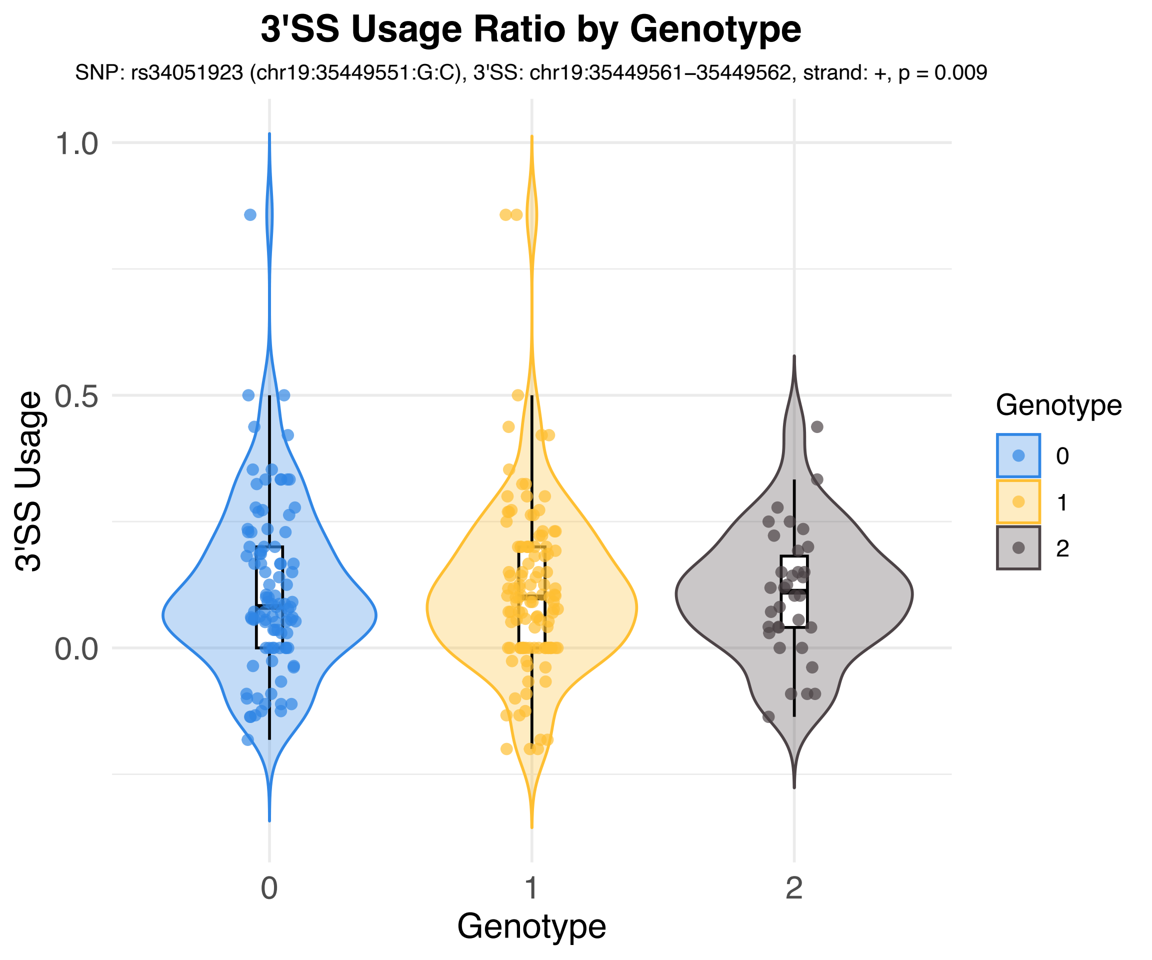

3'SS Usage Ratio by Genotype
SNP: rs34051923 (chr19:35449551:G:C), 3'SS: chr19:35449561–35449562, strand: +, p = 0.009
3'SS Usage
1.0
0.5
0.0
0
1
2
Genotype
Genotype
0
1
2

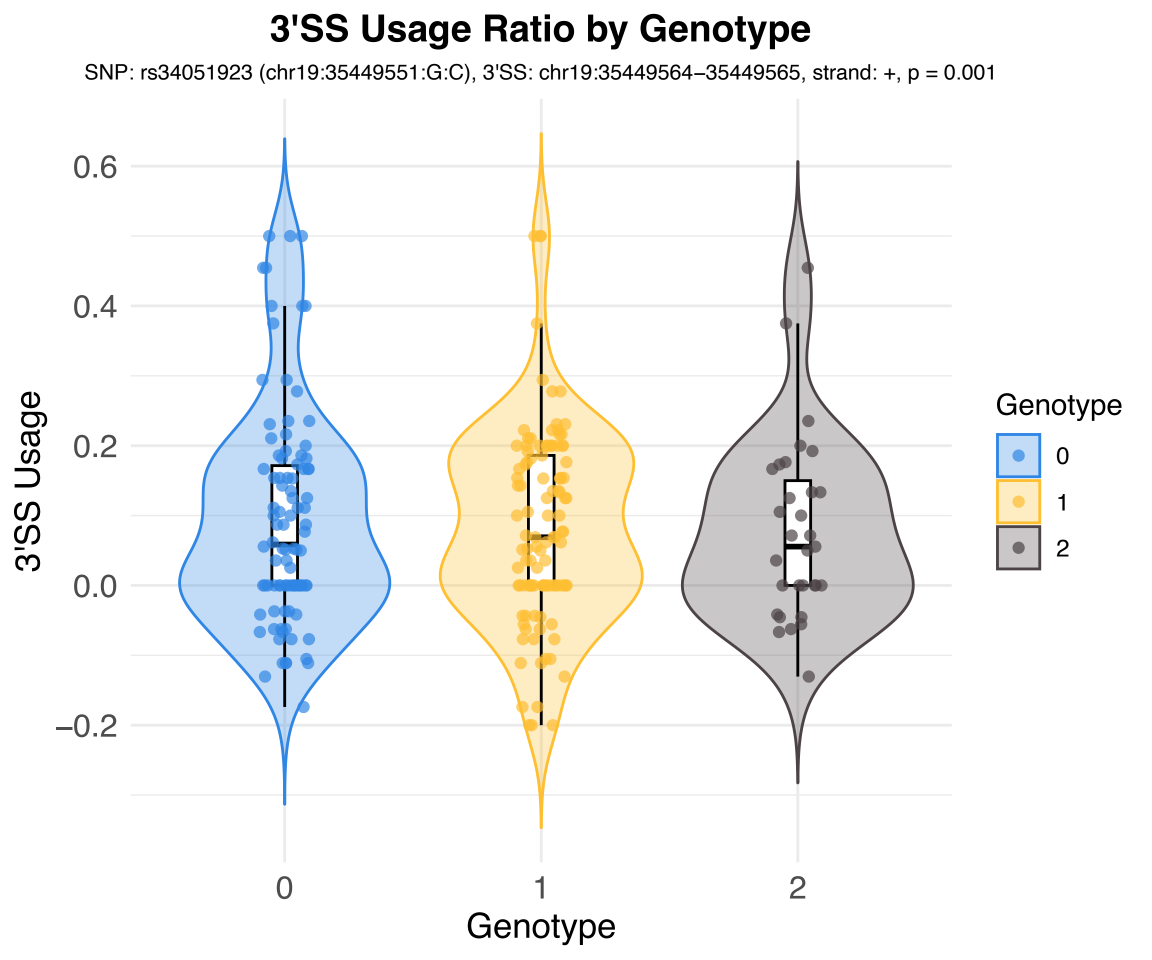

3'SS Usage Ratio by Genotype
SNP: rs34051923 (chr19:35449551:G:C), 3'SS: chr19:35449564–35449565, strand: +, p = 0.001
3'SS Usage
0.6
0.4
0.2
0.0
−0.2
0
1
2
Genotype
Genotype
0
1
2

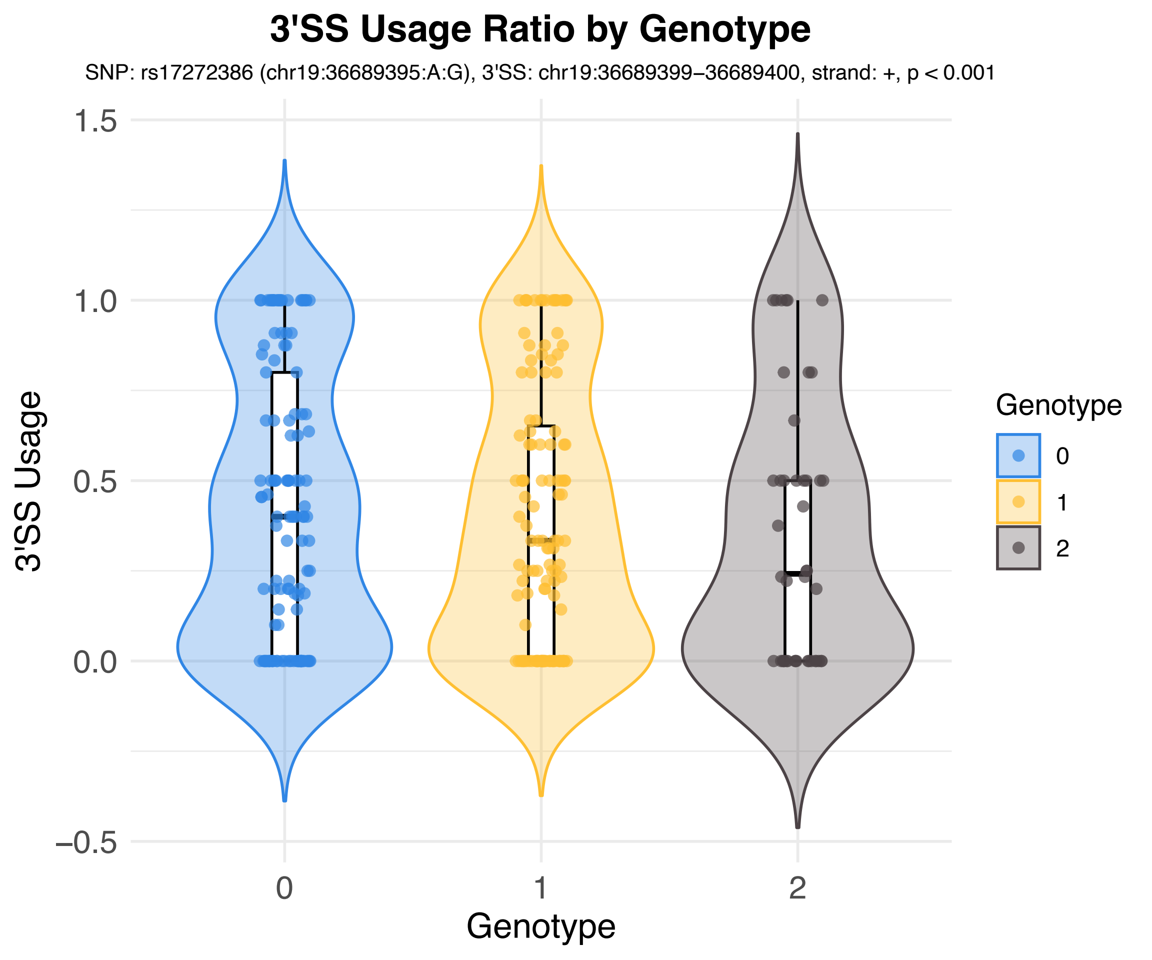

3'SS Usage Ratio by Genotype
SNP: rs17272386 (chr19:36689395:A:G), 3'SS: chr19:36689399–36689400, strand: +, p < 0.001
3'SS Usage
1.5
1.0
0.5
0.0
−0.5
0
1
2
Genotype
Genotype
0
1
2

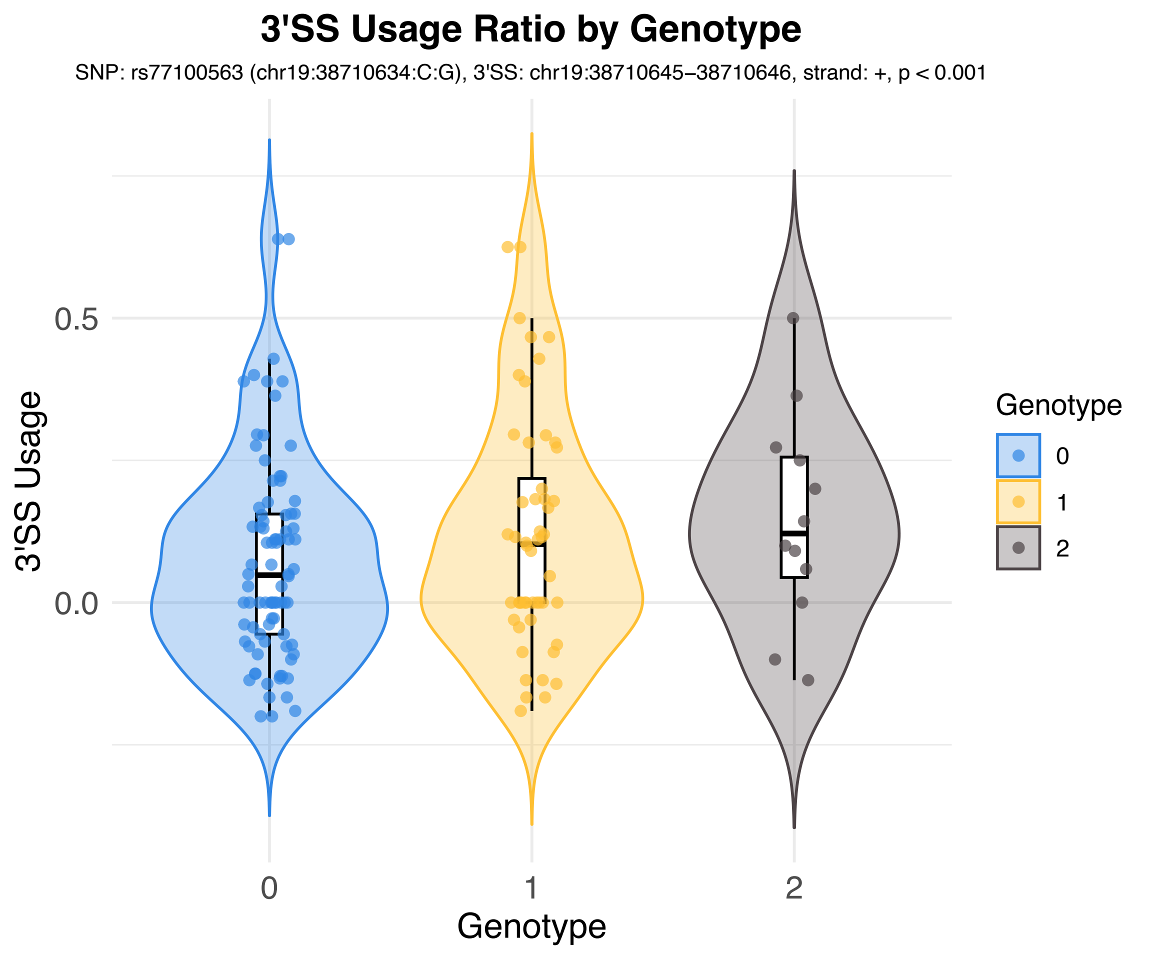
3'SS Usage Ratio by Genotype
SNP: rs77100563 (chr19:38710634:C:G), 3'SS: chr19:38710645–38710646, strand: +, p < 0.001
3'SS Usage
0.5
0.0
0
1
2
Genotype
Genotype
0
1
2

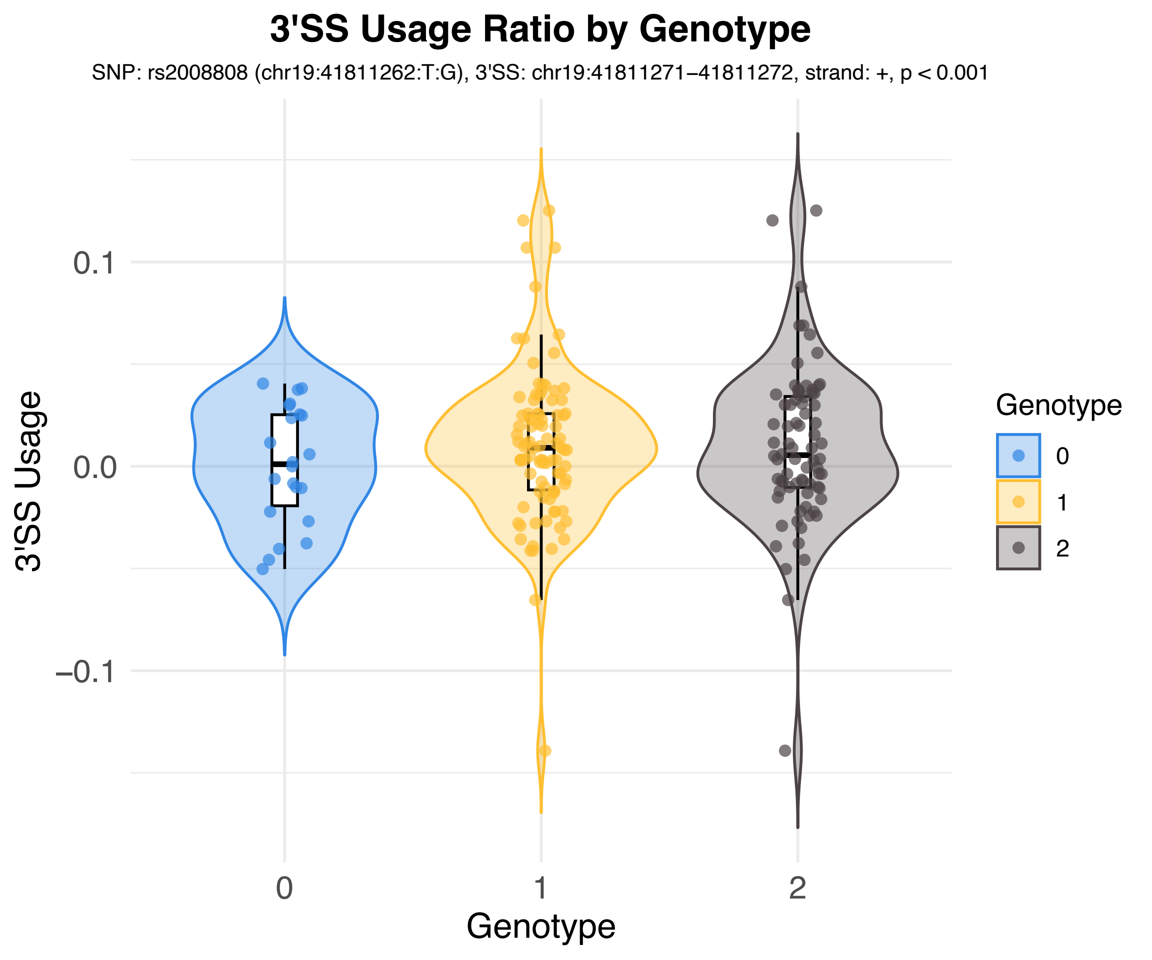

3'SS Usage Ratio by Genotype
SNP: rs2008808 (chr19:41811262:T:G), 3'SS: chr19:41811271–41811272, strand: +, p < 0.001
3'SS Usage
0.1
0.0
–0.1
0
1
2
Genotype
Genotype
0
1
2

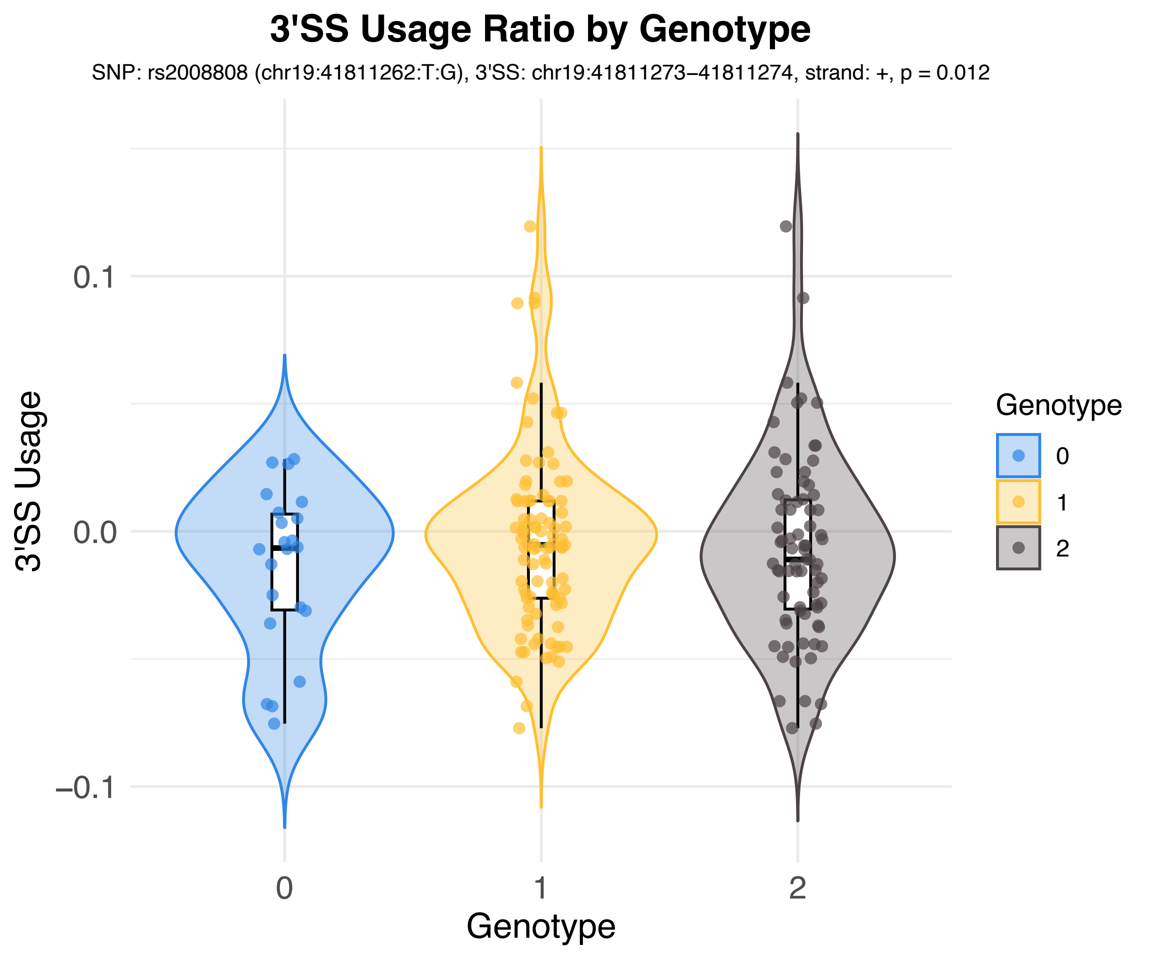
3'SS Usage Ratio by Genotype
SNP: rs2008808 (chr19:41811262:T:G), 3'SS: chr19:41811273–41811274, strand: +, p = 0.012
3'SS Usage
0.1
0.0
−0.1
0
1
2
Genotype
Genotype
0
1
2

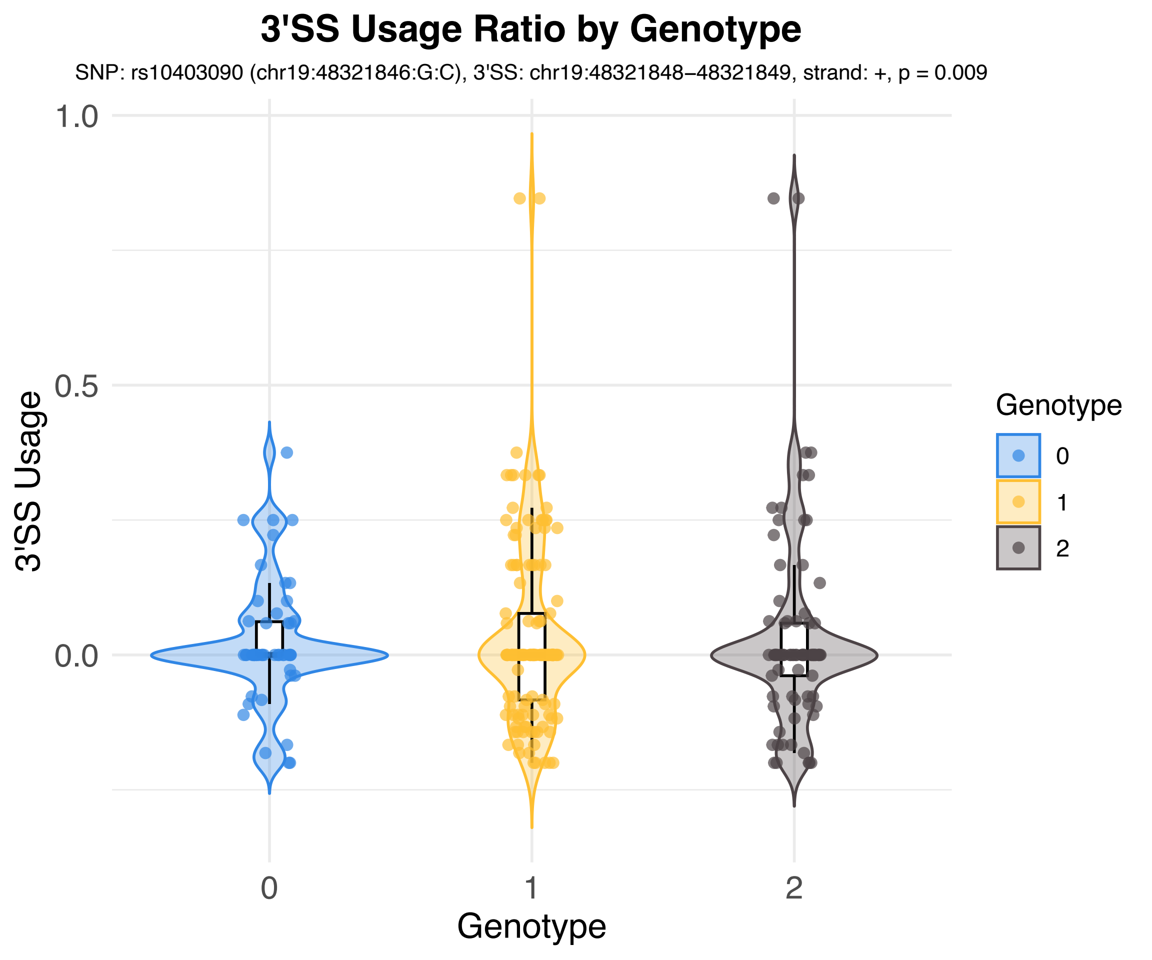

3'SS Usage Ratio by Genotype
SNP: rs10403090 (chr19:48321846:G:C), 3'SS: chr19:48321848–48321849, strand: +, p = 0.009
3'SS Usage
1.0
0.5
0.0
0
1
2
Genotype
Genotype
0
1
2

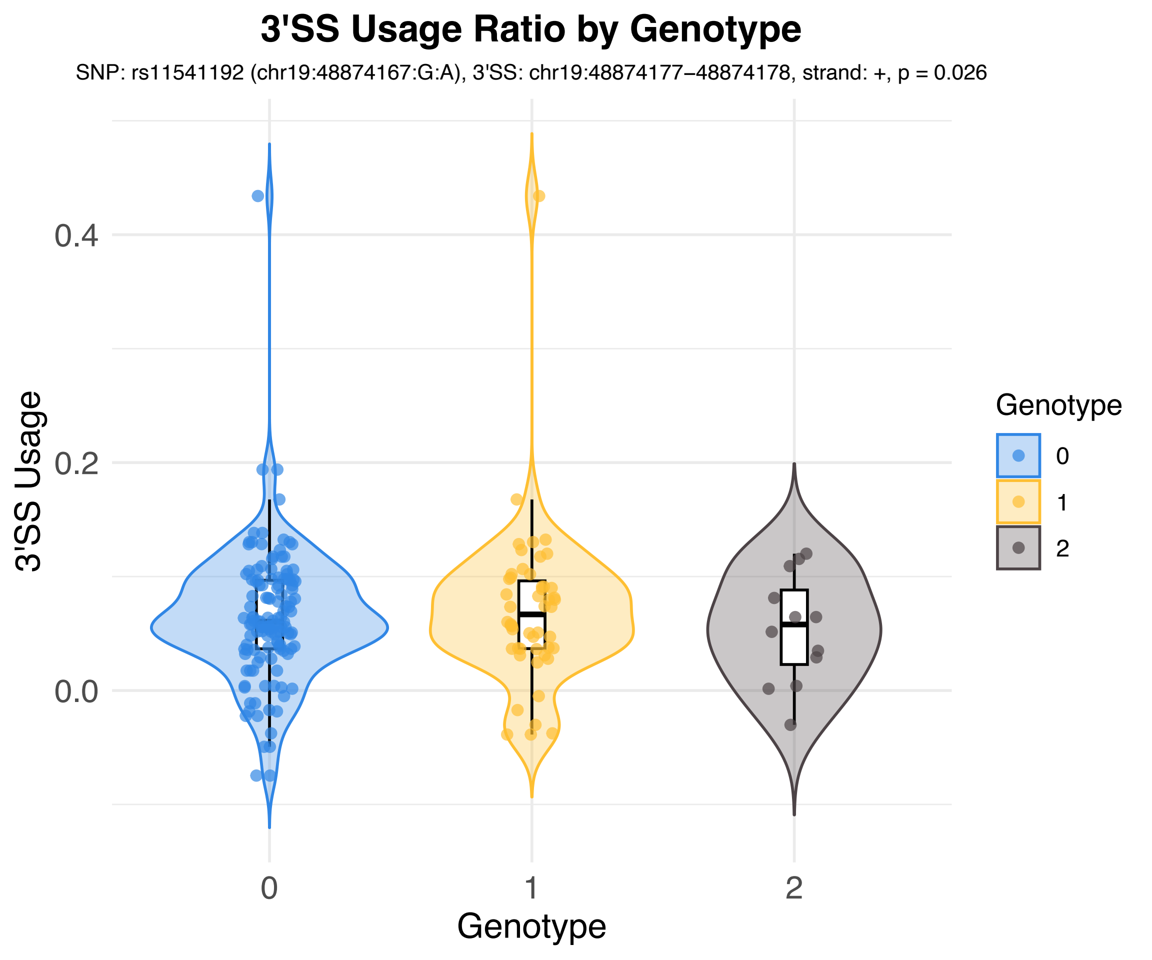

3'SS Usage Ratio by Genotype
SNP: rs11541192 (chr19:48874167:G:A), 3'SS: chr19:48874177−48874178, strand: +, p = 0.026
3'SS Usage
0.4
0.2
0.0
0
1
2
Genotype
Genotype
0
1
2

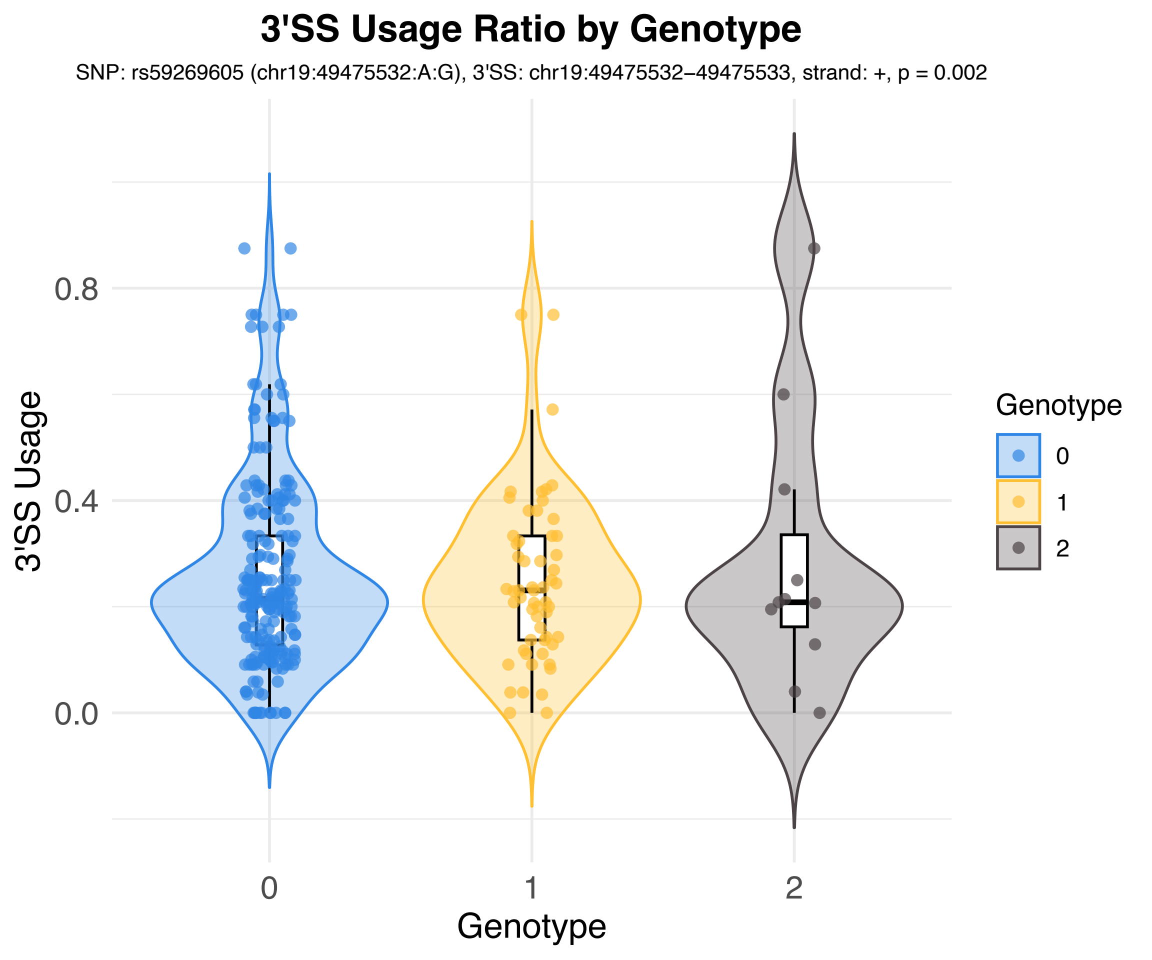
3'SS Usage Ratio by Genotype
SNP: rs59269605 (chr19:49475532:A:G), 3'SS: chr19:49475532–49475533, strand: +, p = 0.002
3'SS Usage
0.8
0.4
0.0
0
1
2
Genotype
Genotype
0
1
2

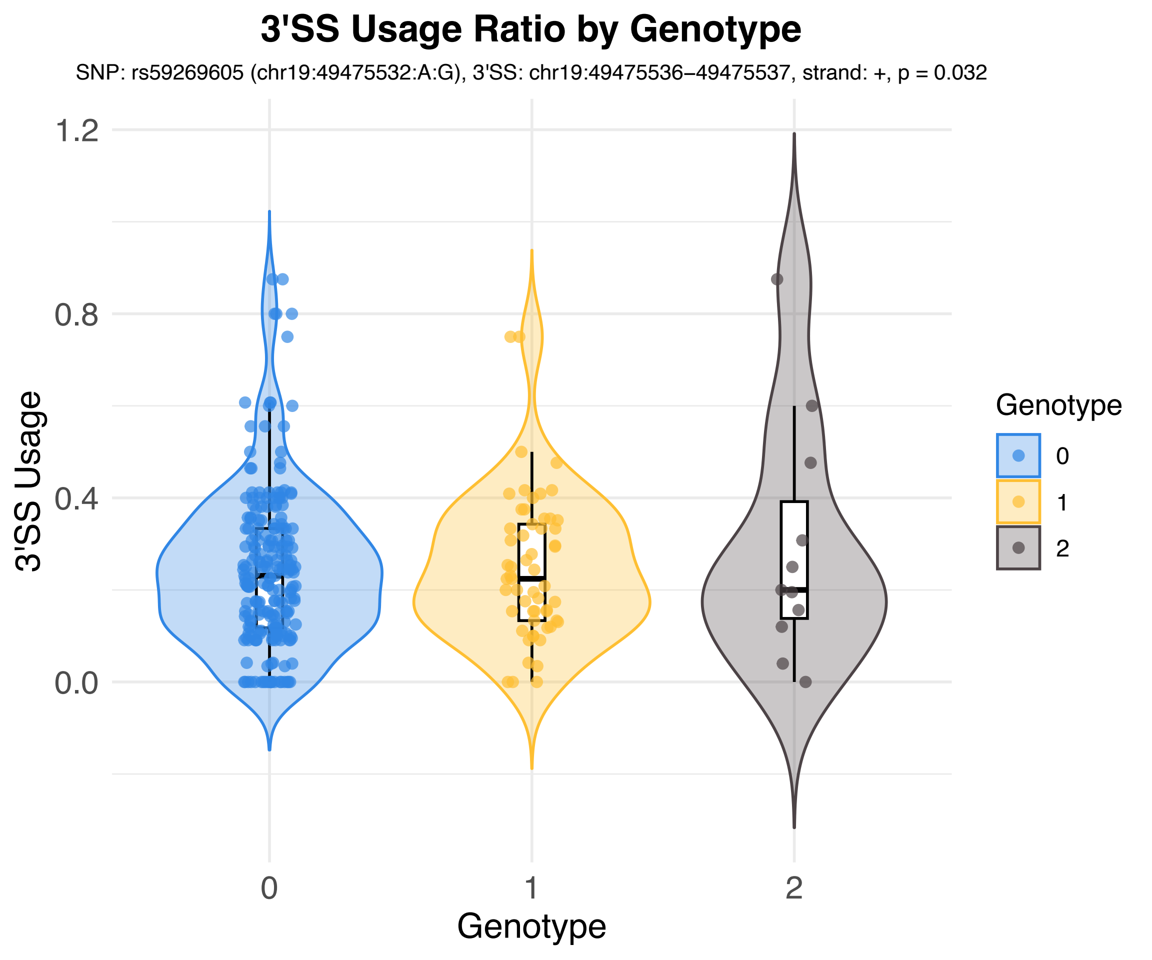
3'SS Usage Ratio by Genotype
SNP: rs59269605 (chr19:49475532:A:G), 3'SS: chr19:49475536–49475537, strand: +, p = 0.032
3'SS Usage
1.2
0.8
0.4
0.0
0
1
2
Genotype
Genotype
0
1
2

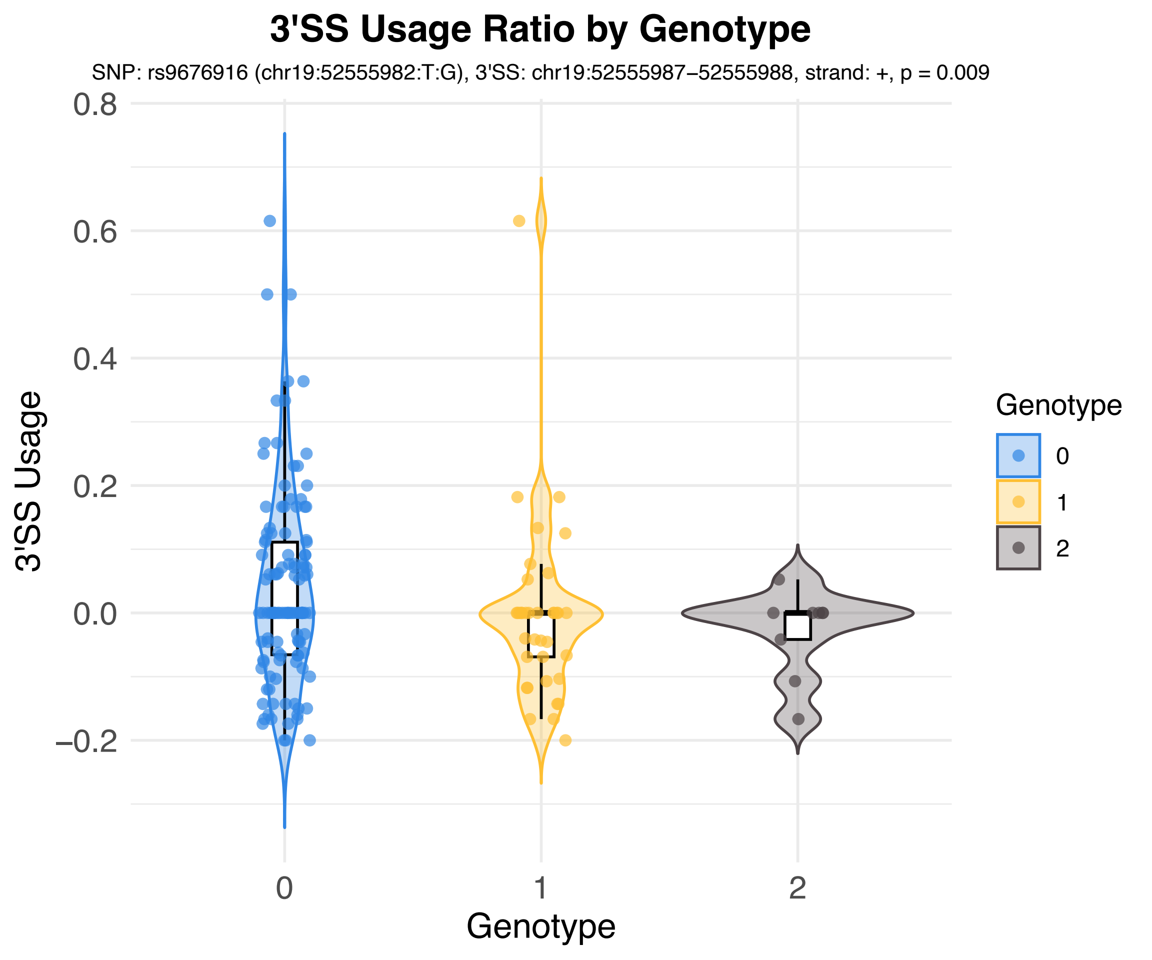

3'SS Usage Ratio by Genotype
SNP: rs9676916 (chr19:52555982:T:G), 3'SS: chr19:52555987–52555988, strand: +, p = 0.009
3'SS Usage
0.8
0.6
0.4
0.2
0.0
–0.2
0
1
2
Genotype
Genotype
0
1
2

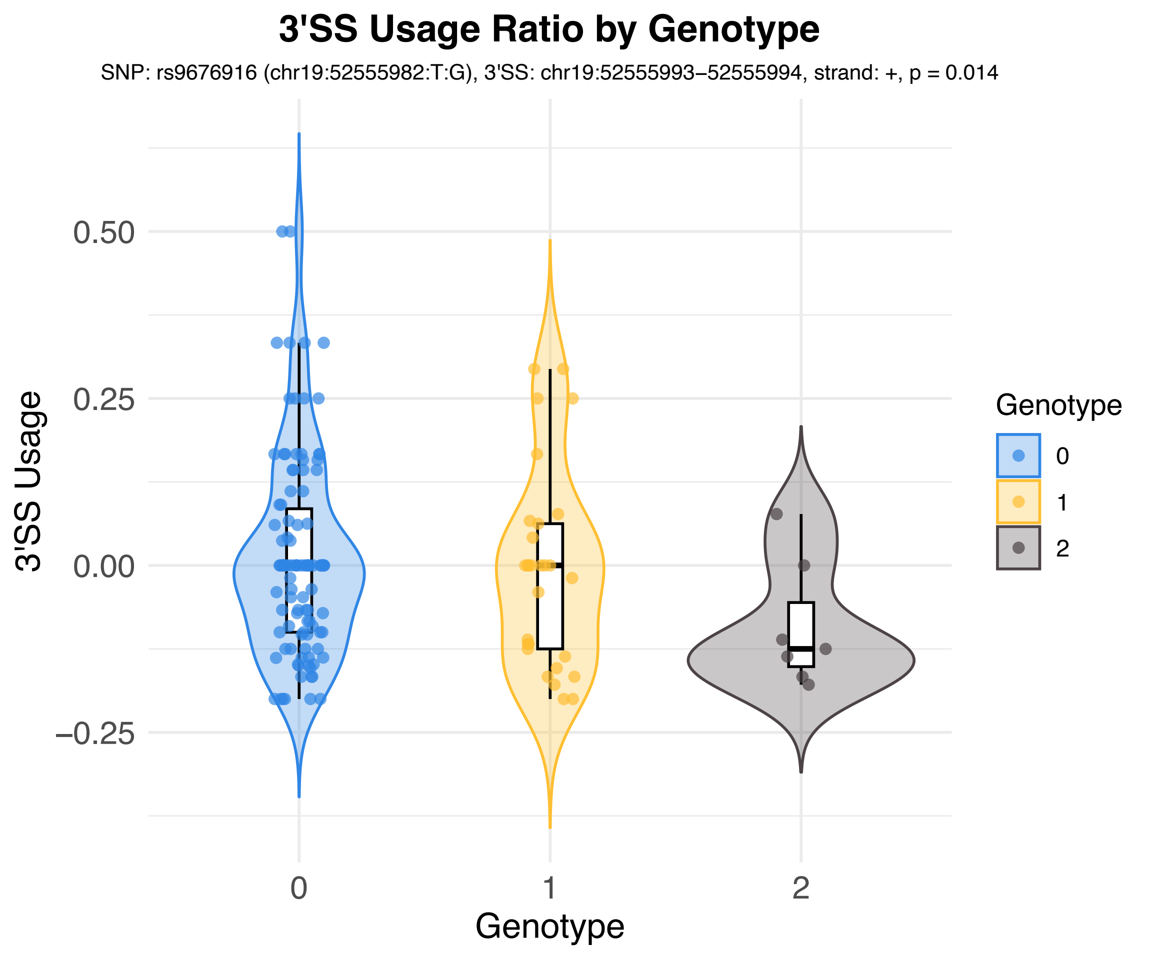
3'SS Usage Ratio by Genotype
SNP: rs9676916 (chr19:52555982:T:G), 3'SS: chr19:52555993–52555994, strand: +, p = 0.014
3'SS Usage
0.50
0.25
0.00
−0.25
0
1
2
Genotype
Genotype
0
1
2

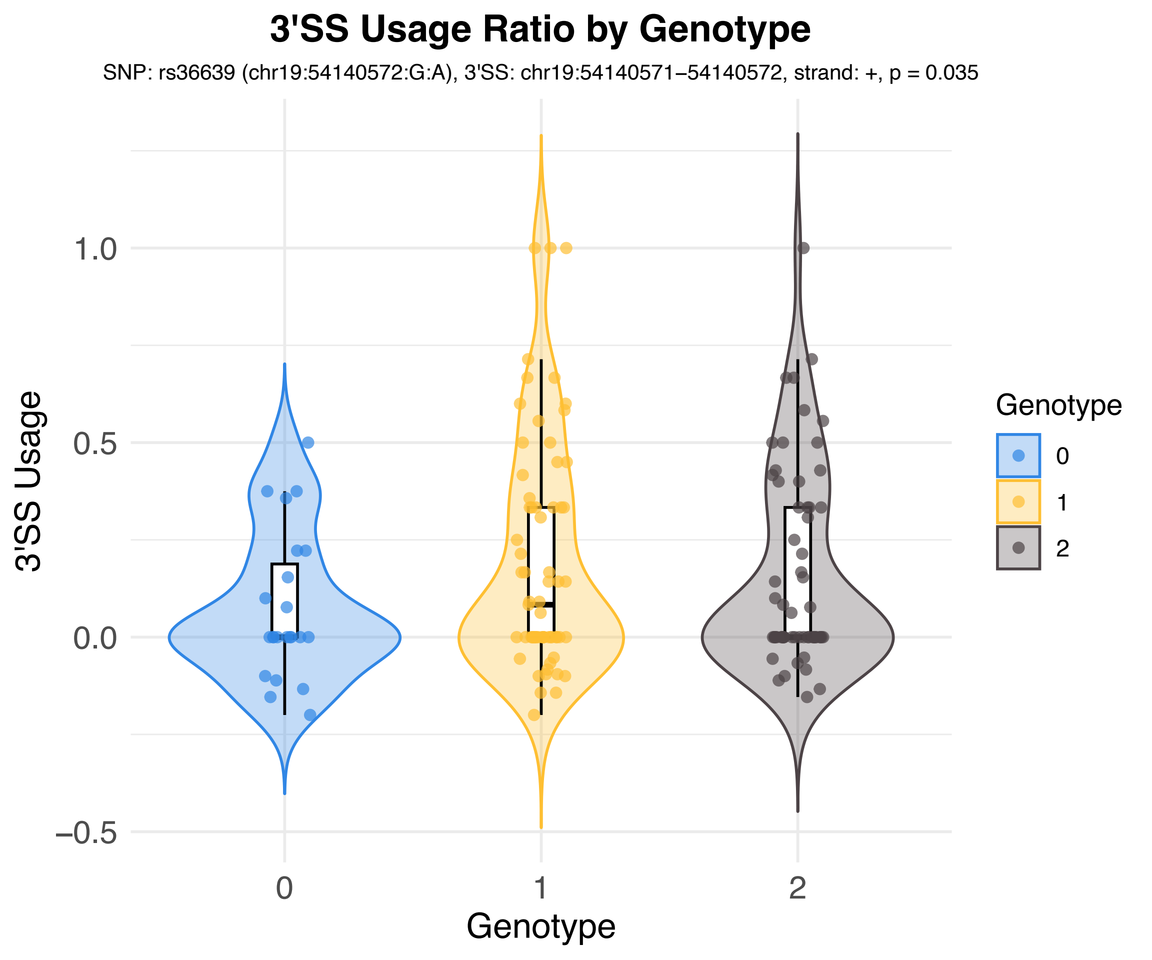
3'SS Usage Ratio by Genotype
SNP: rs36639 (chr19:54140572:G:A), 3'SS: chr19:54140571–54140572, strand: +, p = 0.035
3'SS Usage
1.0
0.5
0.0
−0.5
0
1
2
Genotype
Genotype
0
1
2

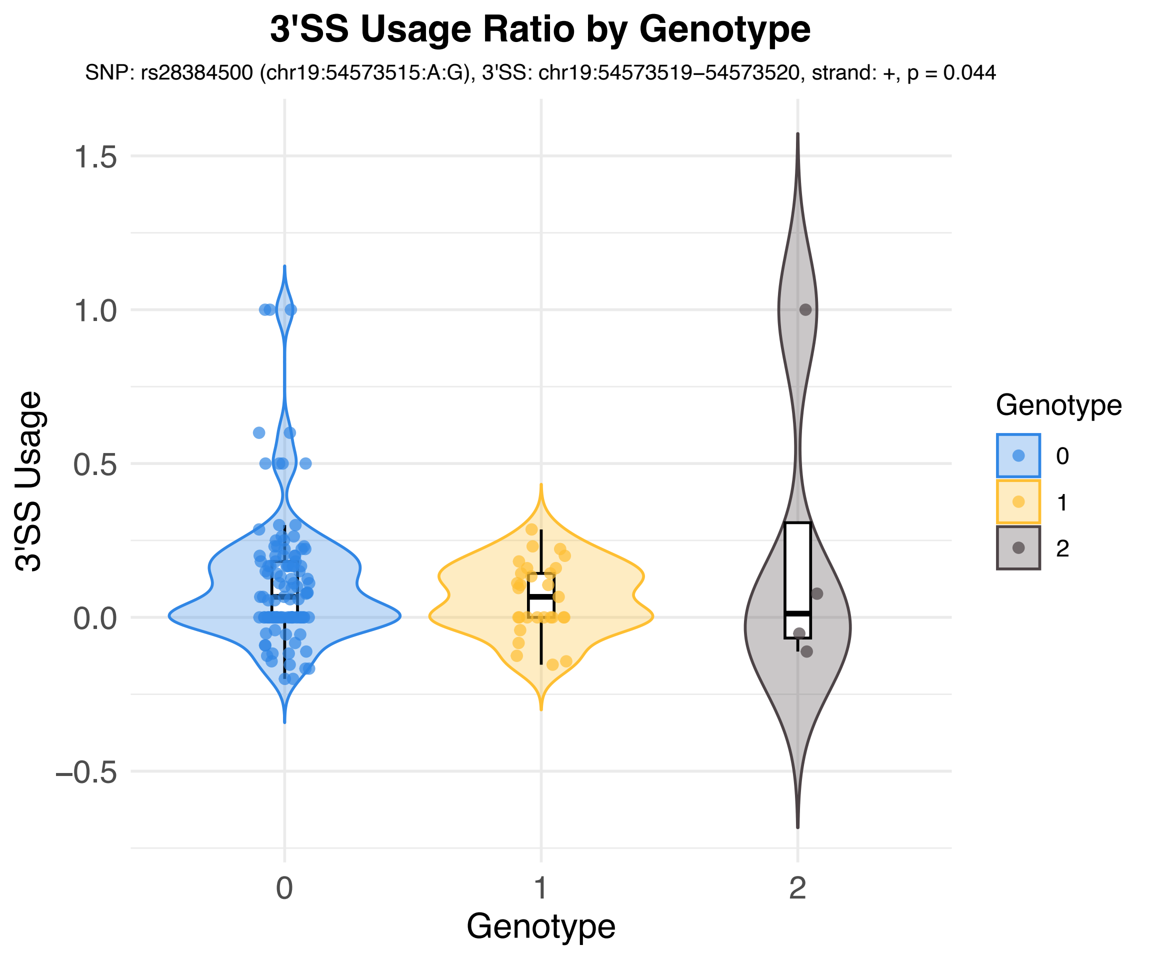
3'SS Usage Ratio by Genotype
SNP: rs28384500 (chr19:54573515:A:G), 3'SS: chr19:54573519–54573520, strand: +, p = 0.044
3'SS Usage
1.5
1.0
0.5
0.0
−0.5
0
1
2
Genotype
Genotype
0
1
2

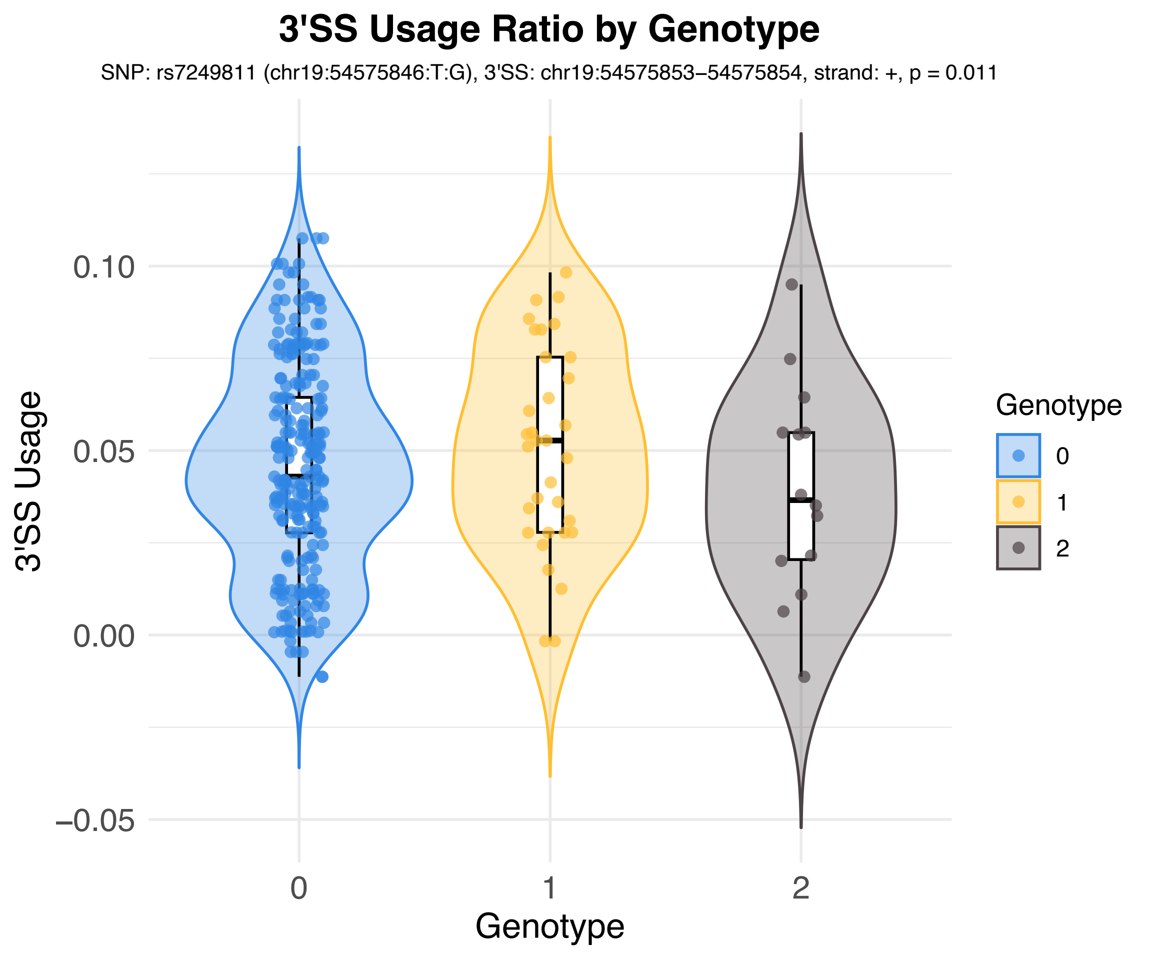
3'SS Usage Ratio by Genotype
SNP: rs7249811 (chr19:54575846:T:G), 3'SS: chr19:54575853–54575854, strand: +, p = 0.011
3'SS Usage
0.10
0.05
0.00
−0.05
0
1
2
Genotype
Genotype
0
1
2

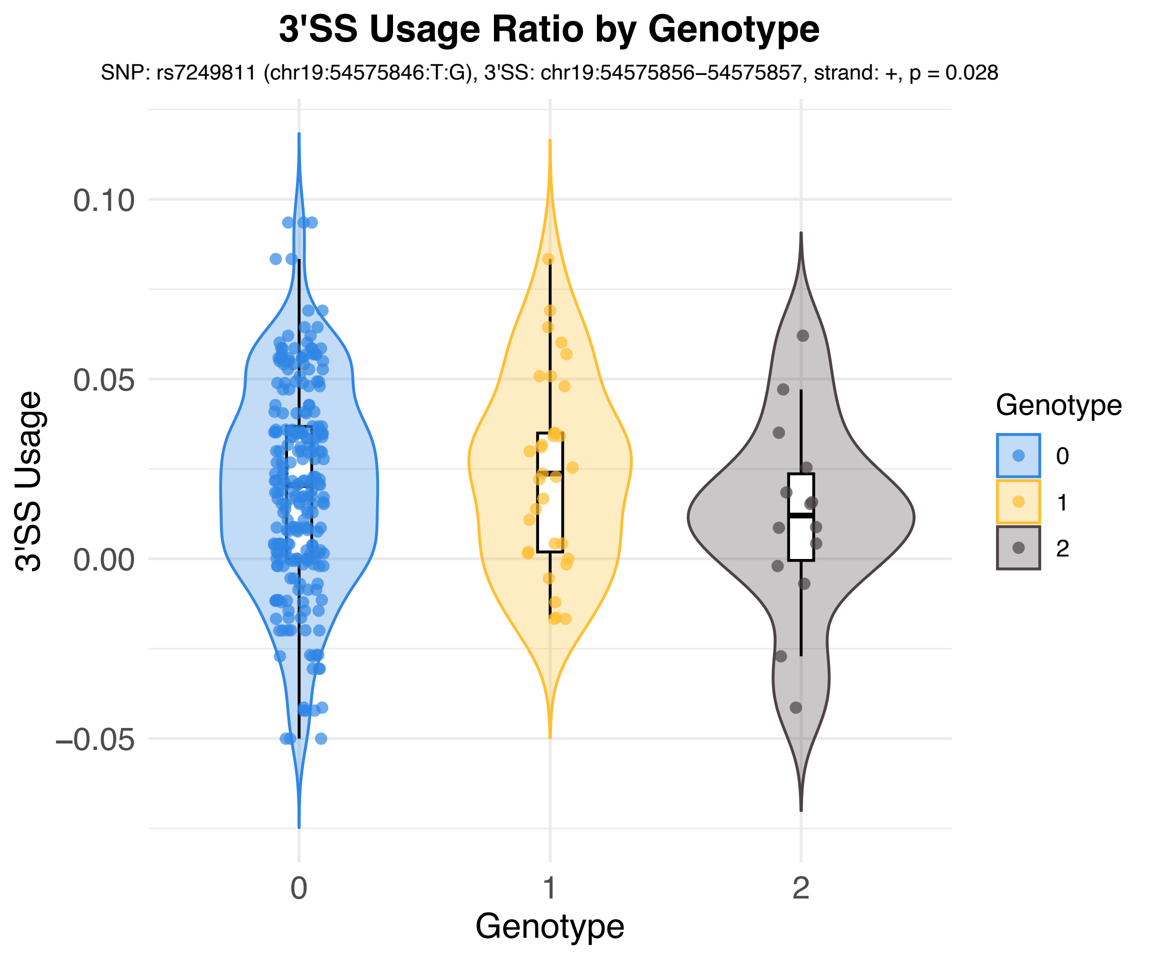
3'SS Usage Ratio by Genotype
SNP: rs7249811 (chr19:54575846:T:G), 3'SS: chr19:54575856–54575857, strand: +, p = 0.028
3'SS Usage
0.10
0.05
0.00
−0.05
0
1
2
Genotype
Genotype
0
1
2

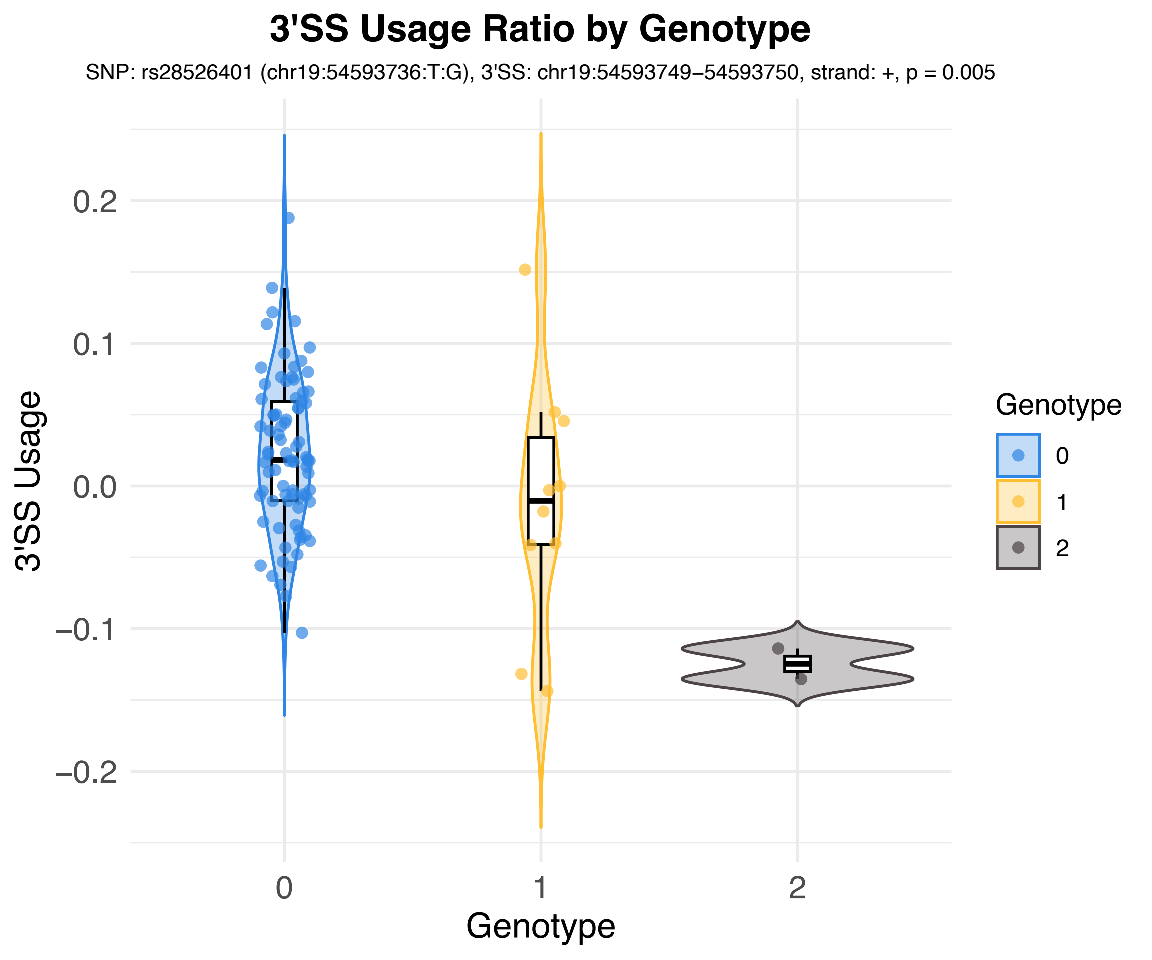

3'SS Usage Ratio by Genotype
SNP: rs28526401 (chr19:54593736:T:G), 3'SS: chr19:54593749−54593750, strand: +, p = 0.005
3'SS Usage
0.2
0.1
0.0
−0.1
−0.2
0
1
2
Genotype
Genotype
0
1
2

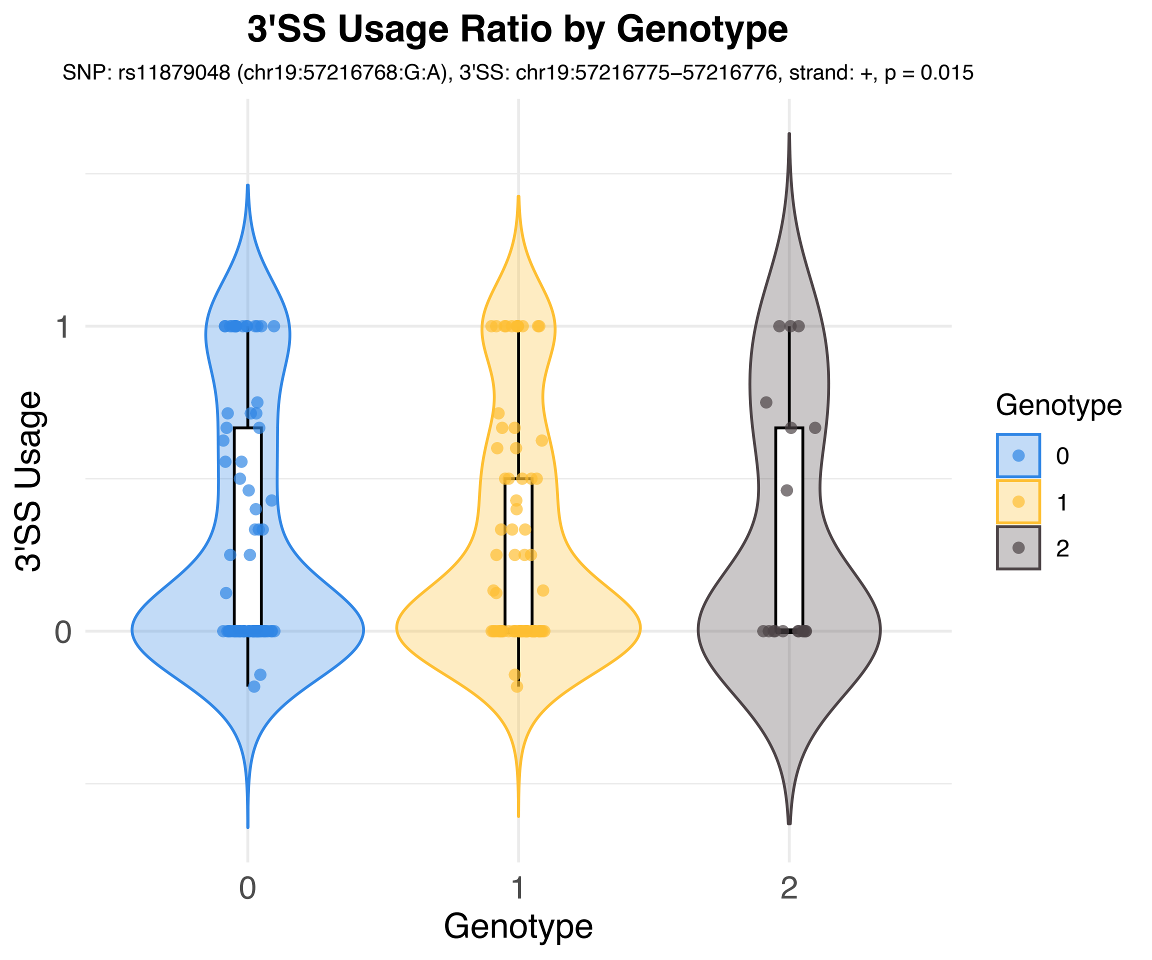

3'SS Usage Ratio by Genotype
SNP: rs11879048 (chr19:57216768:G:A), 3'SS: chr19:57216775–57216776, strand: +, p = 0.015
3'SS Usage
1
0
0
1
2
Genotype
Genotype
0
1
2

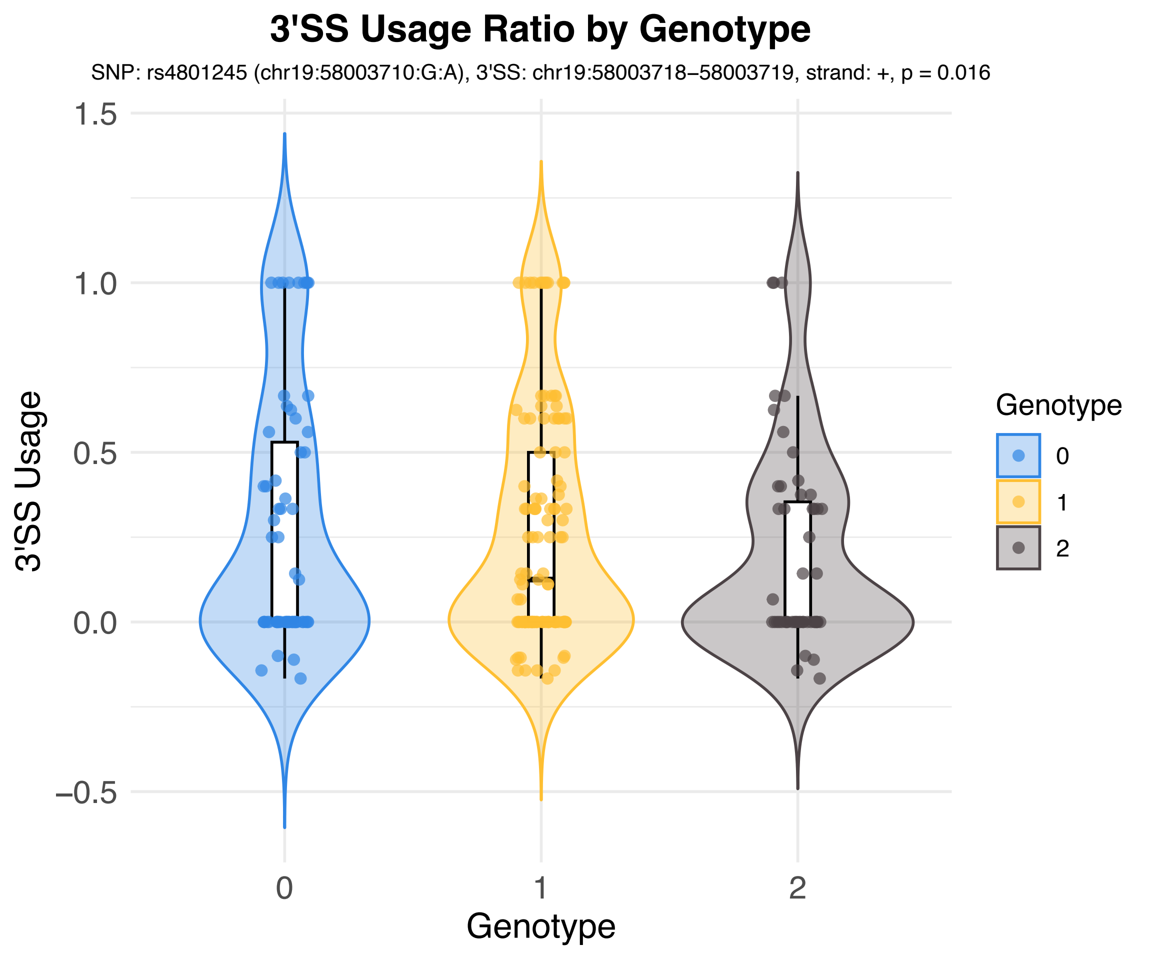
3'SS Usage Ratio by Genotype
SNP: rs4801245 (chr19:58003710:G:A), 3'SS: chr19:58003718–58003719, strand: +, p = 0.016
3'SS Usage
1.5
1.0
0.5
0.0
−0.5
0
1
2
Genotype
Genotype
0
1
2

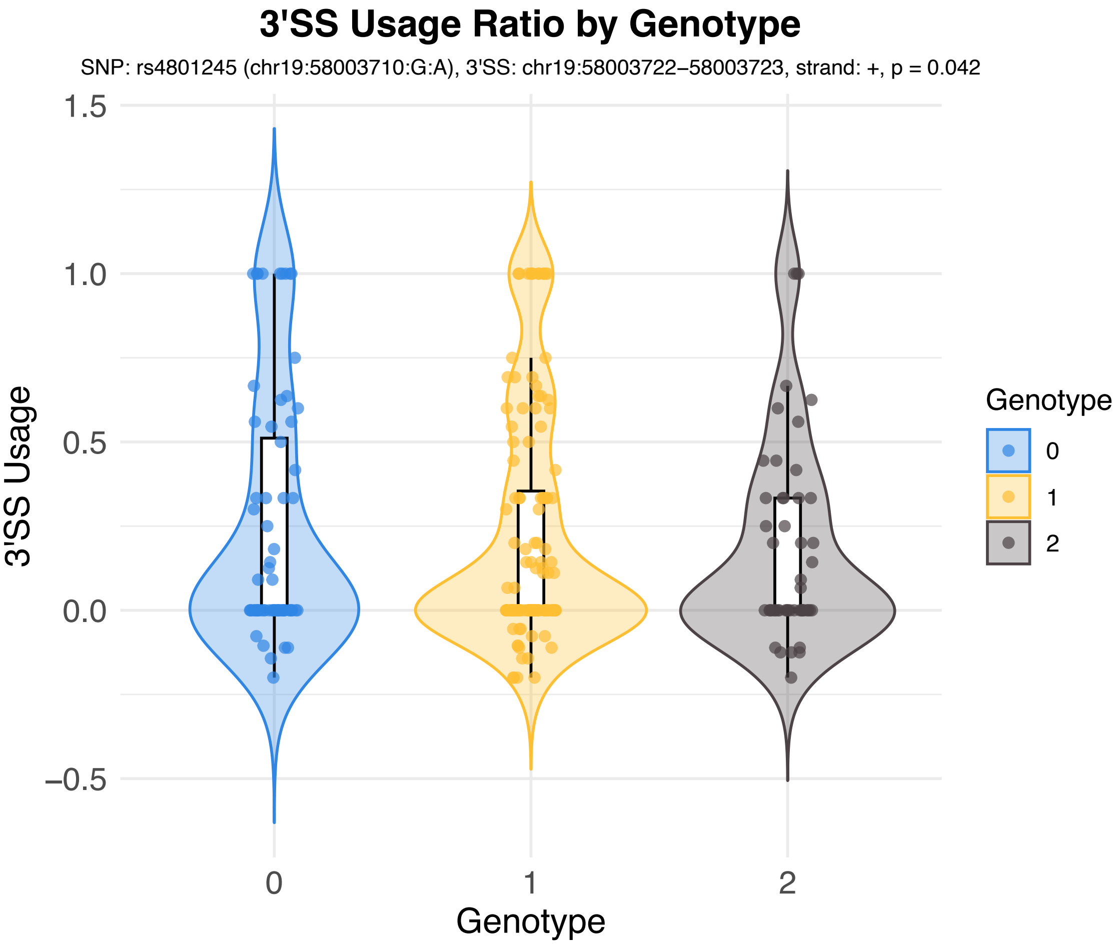

3'SS Usage Ratio by Genotype
SNP: rs4801245 (chr19:58003710:G:A), 3'SS: chr19:58003722–58003723, strand: +, p = 0.042
3'SS Usage
1.5
1.0
0.5
0.0
−0.5
0
1
2
Genotype
Genotype
0
1
2

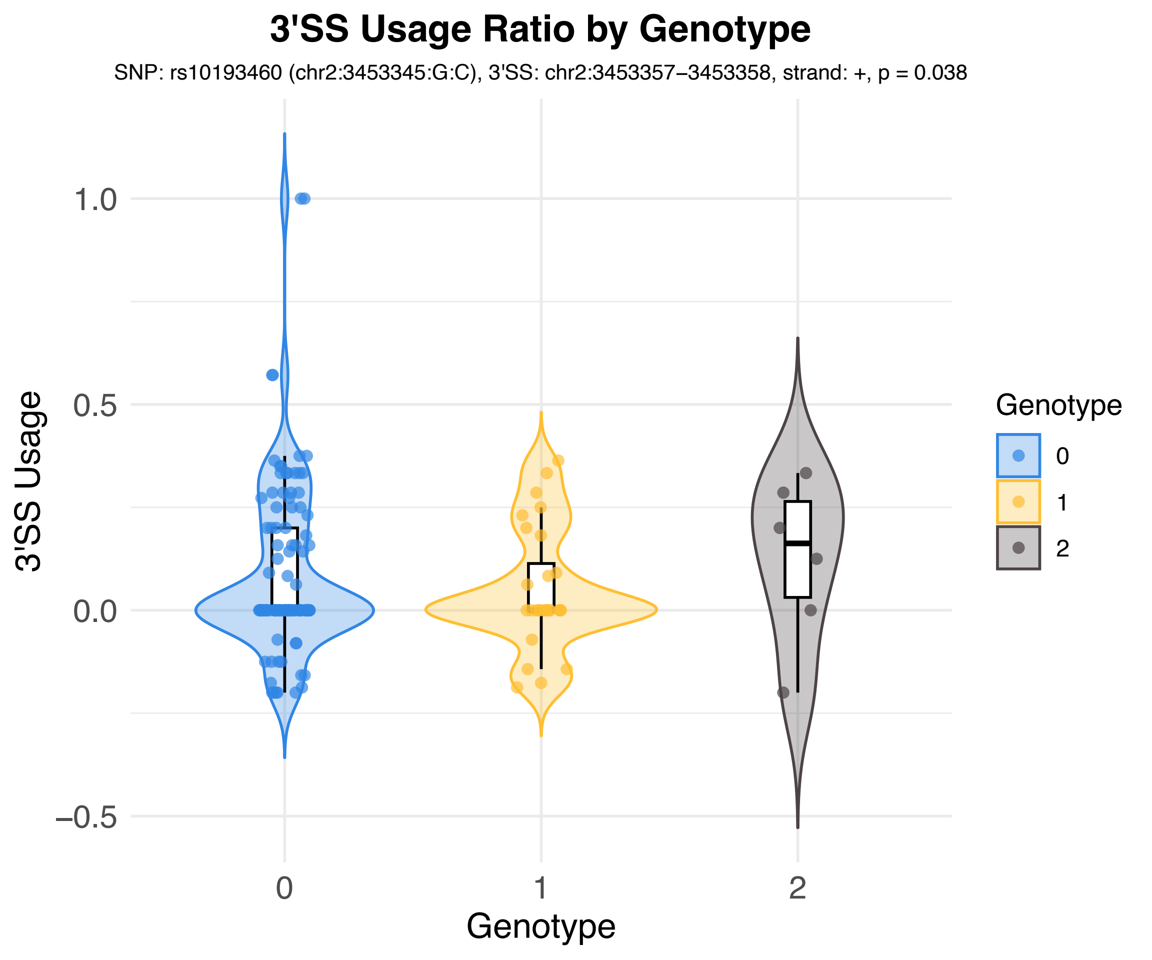

3'SS Usage Ratio by Genotype
SNP: rs10193460 (chr2:3453345:G:C), 3'SS: chr2:3453357–3453358, strand: +, p = 0.038
3'SS Usage
1.0
0.5
0.0
−0.5
0
1
2
Genotype
Genotype
0
1
2

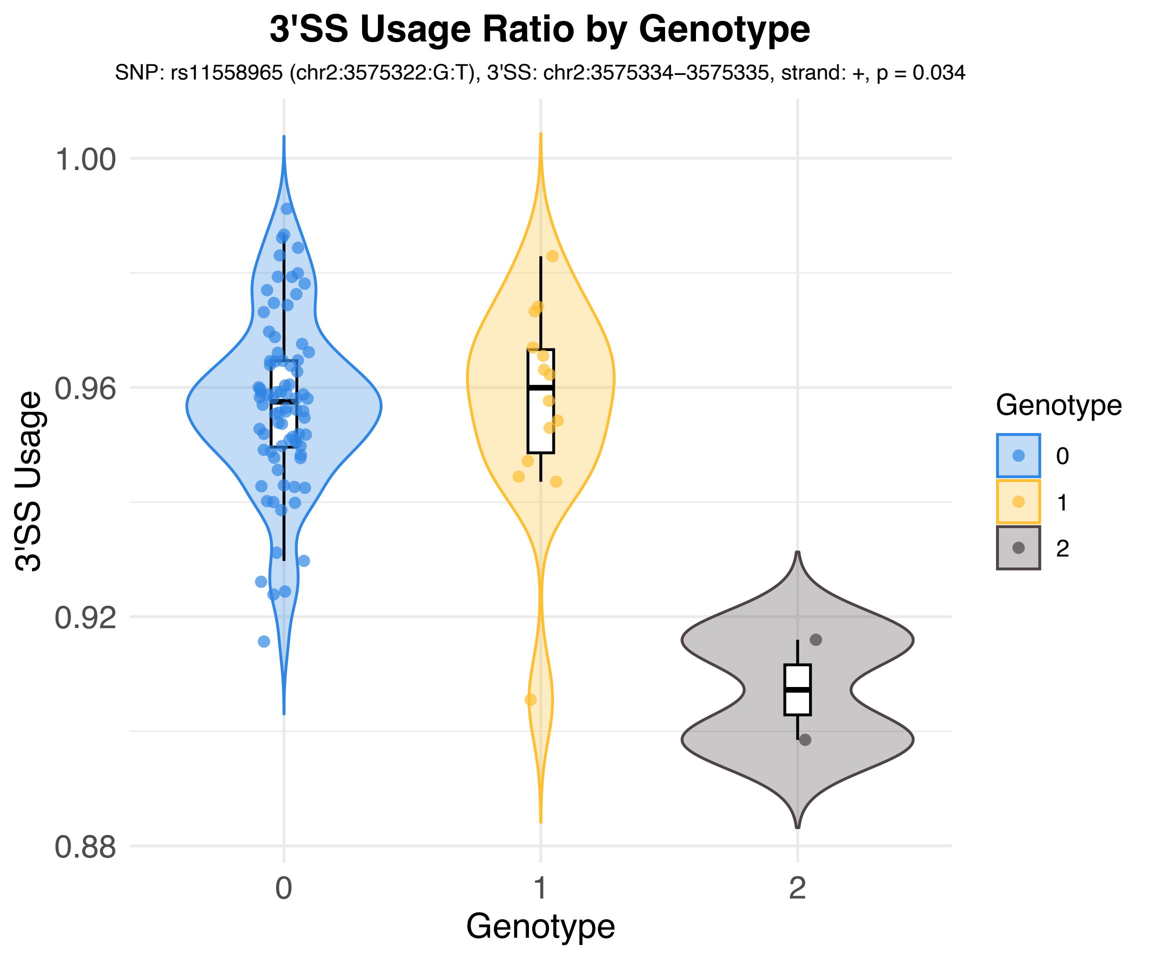
3'SS Usage Ratio by Genotype
SNP: rs11558965 (chr2:3575322:G:T), 3'SS: chr2:3575334–3575335, strand: +, p = 0.034
3'SS Usage
1.00
0.96
0.92
0.88
0
1
2
Genotype
Genotype
0
1
2

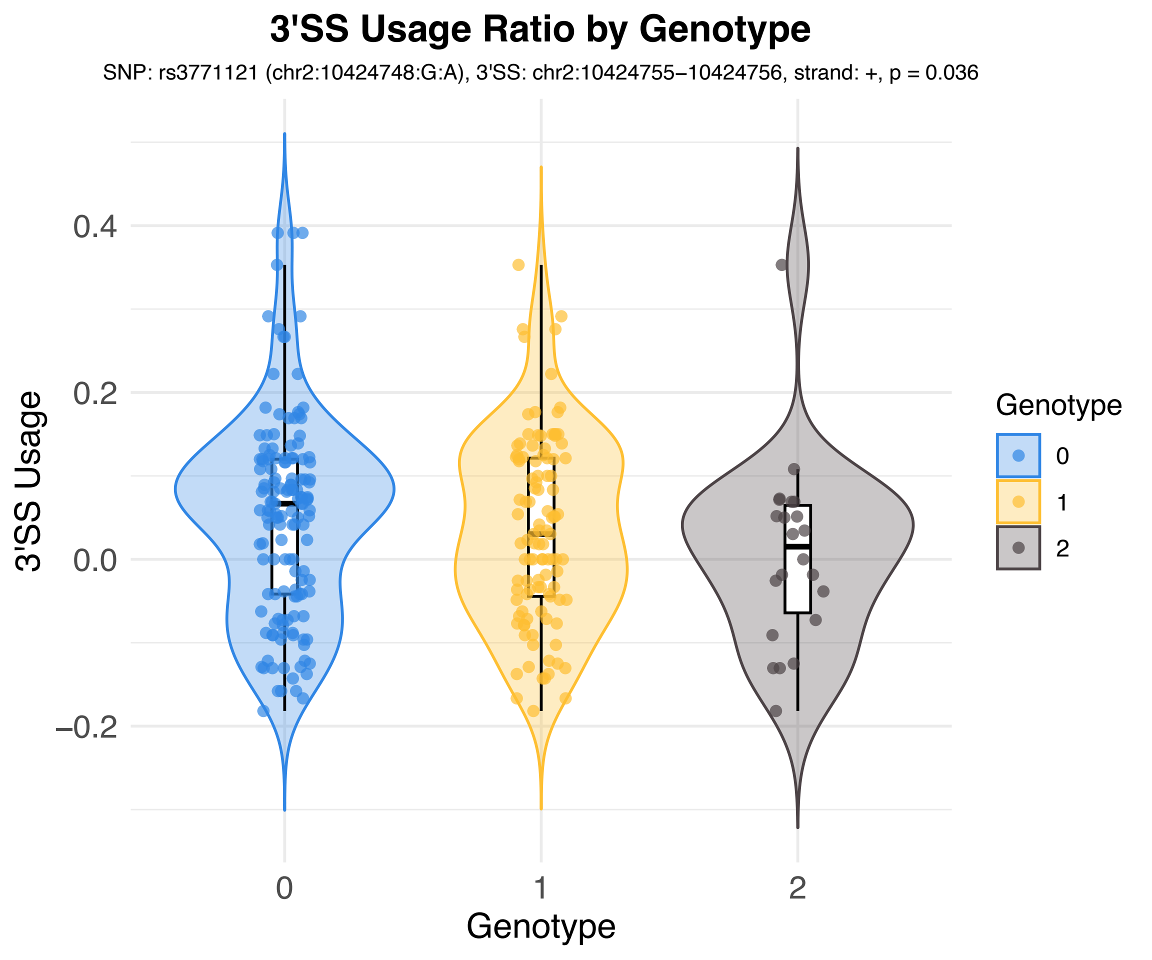

3'SS Usage Ratio by Genotype
SNP: rs3771121 (chr2:10424748:G:A), 3'SS: chr2:10424755−10424756, strand: +, p = 0.036
3'SS Usage
0.4
0.2
0.0
−0.2
0
1
2
Genotype
Genotype
0
1
2

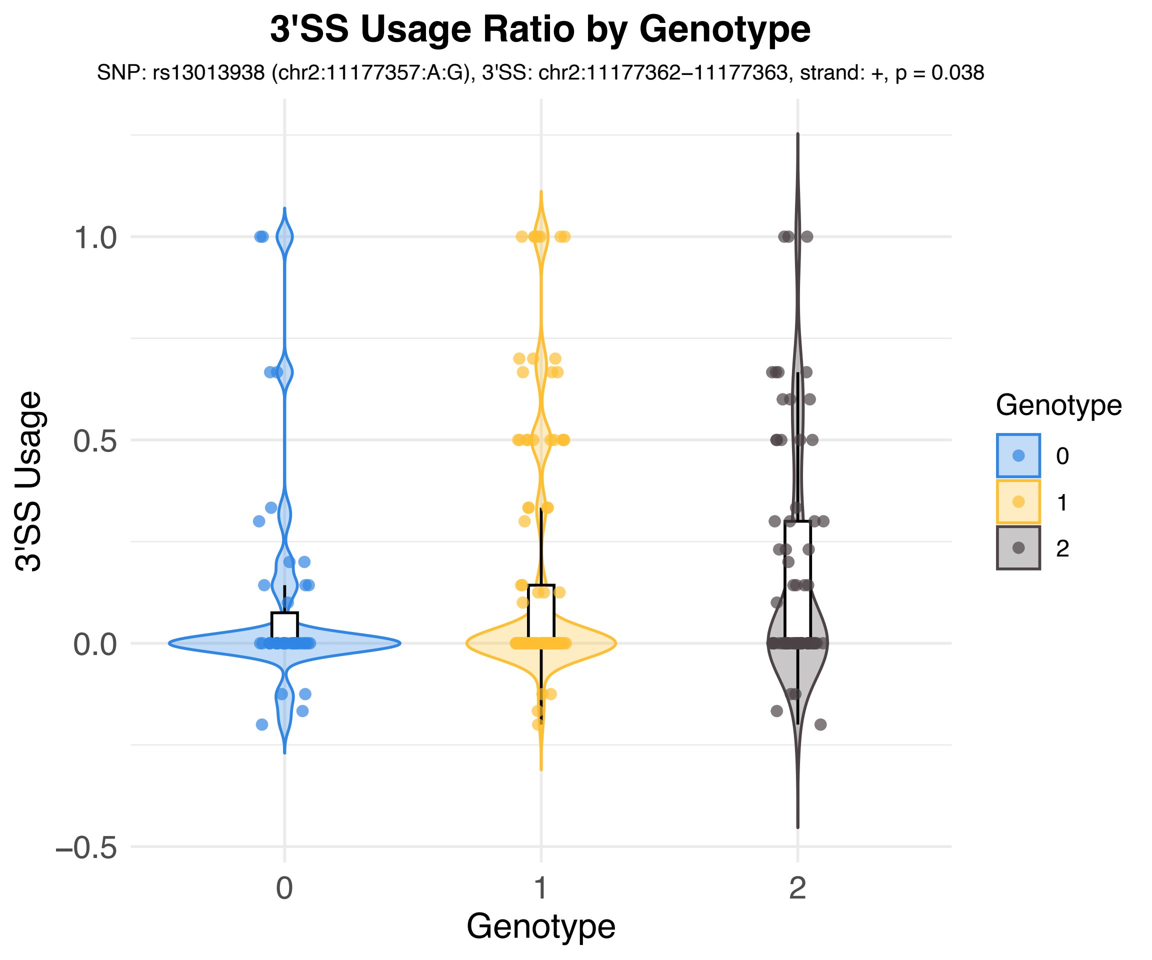
3'SS Usage Ratio by Genotype
SNP: rs13013938 (chr2:11177357:A:G), 3'SS: chr2:11177362−11177363, strand: +, p = 0.038
3'SS Usage
1.0
0.5
0.0
−0.5
0
1
2
Genotype
Genotype
0
1
2

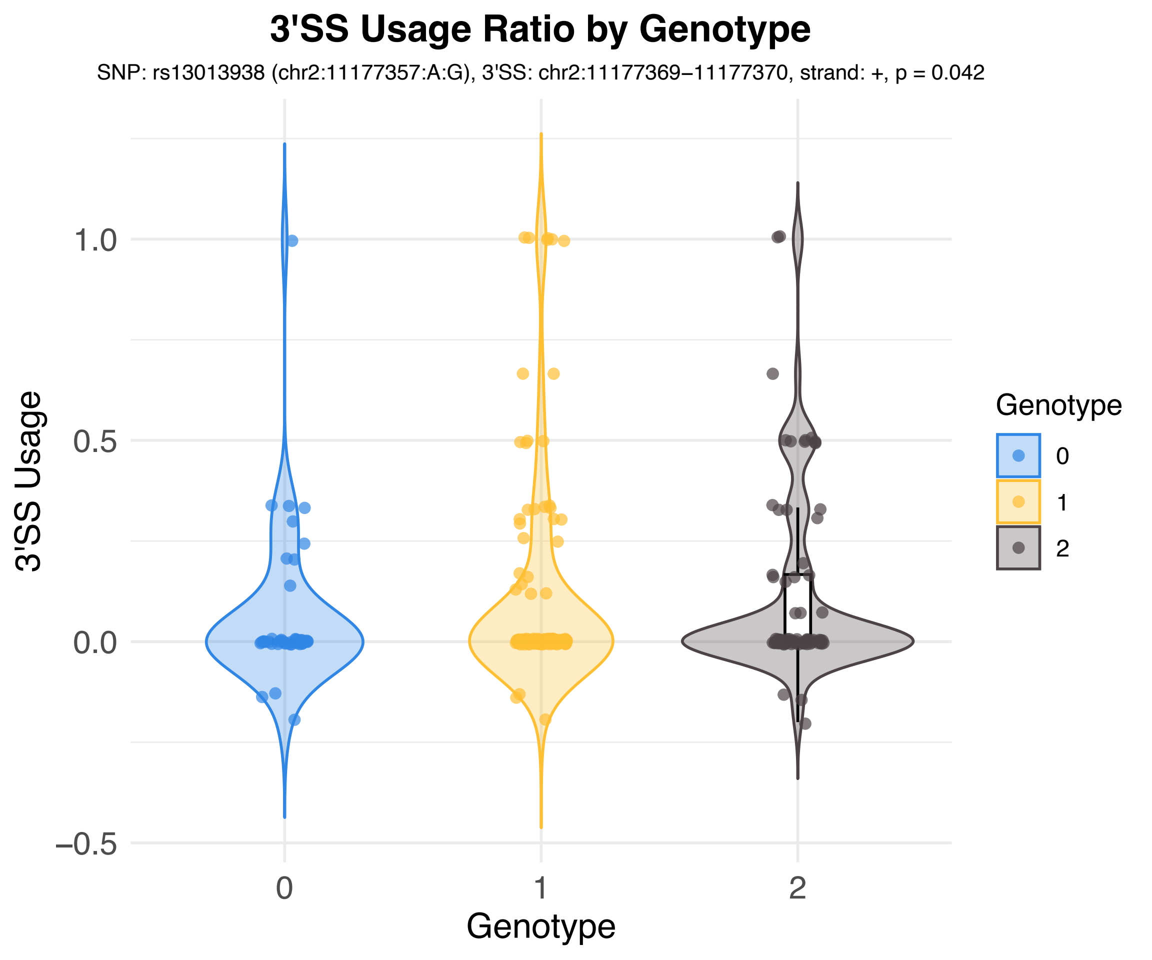
3'SS Usage Ratio by Genotype
SNP: rs13013938 (chr2:11177357:A:G), 3'SS: chr2:11177369−11177370, strand: +, p = 0.042
3'SS Usage
1.0
0.5
0.0
−0.5
0
1
2
Genotype
Genotype
0
1
2

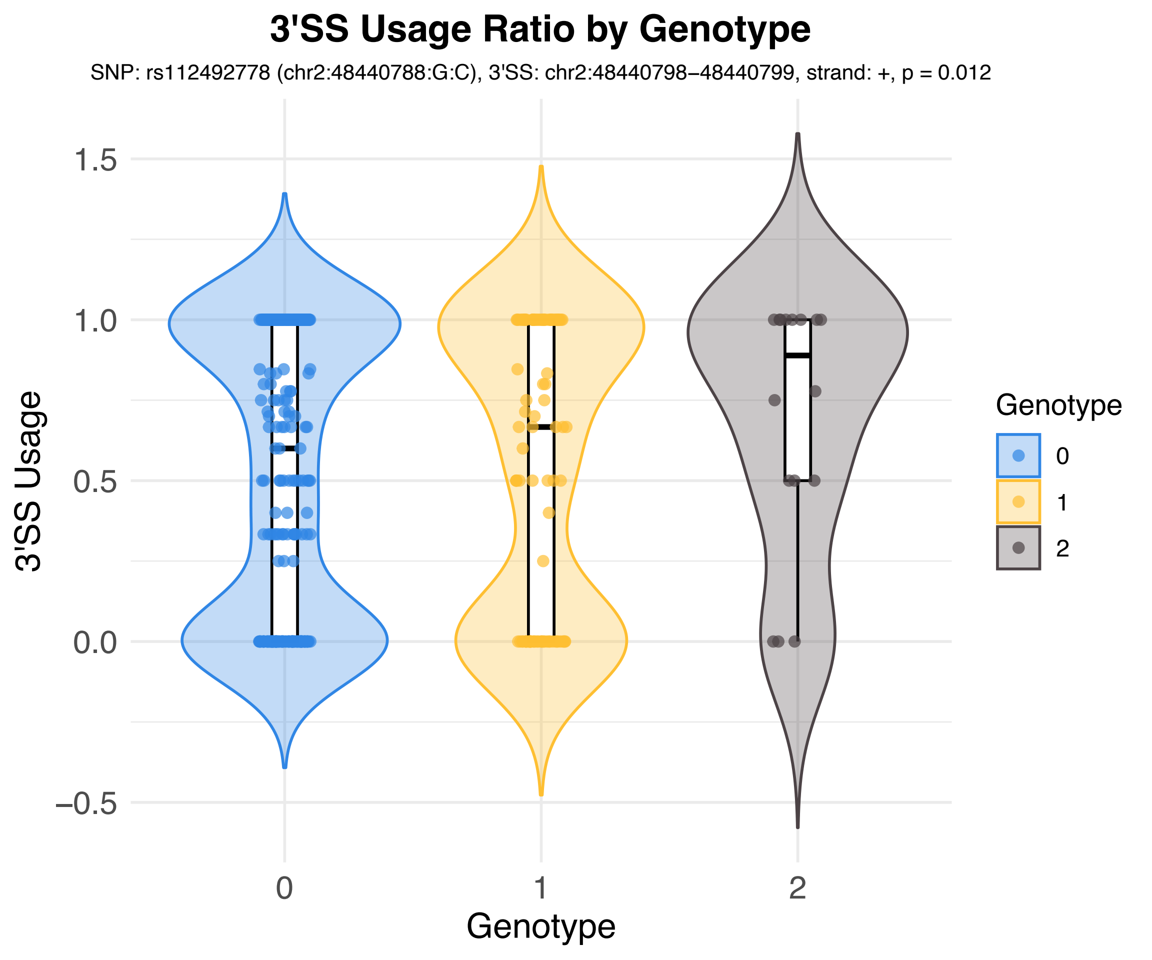

3'SS Usage Ratio by Genotype
SNP: rs112492778 (chr2:48440788:G:C), 3'SS: chr2:48440798−48440799, strand: +, p = 0.012
3'SS Usage
1.5
1.0
0.5
0.0
−0.5
0
1
2
Genotype
Genotype
0
1
2

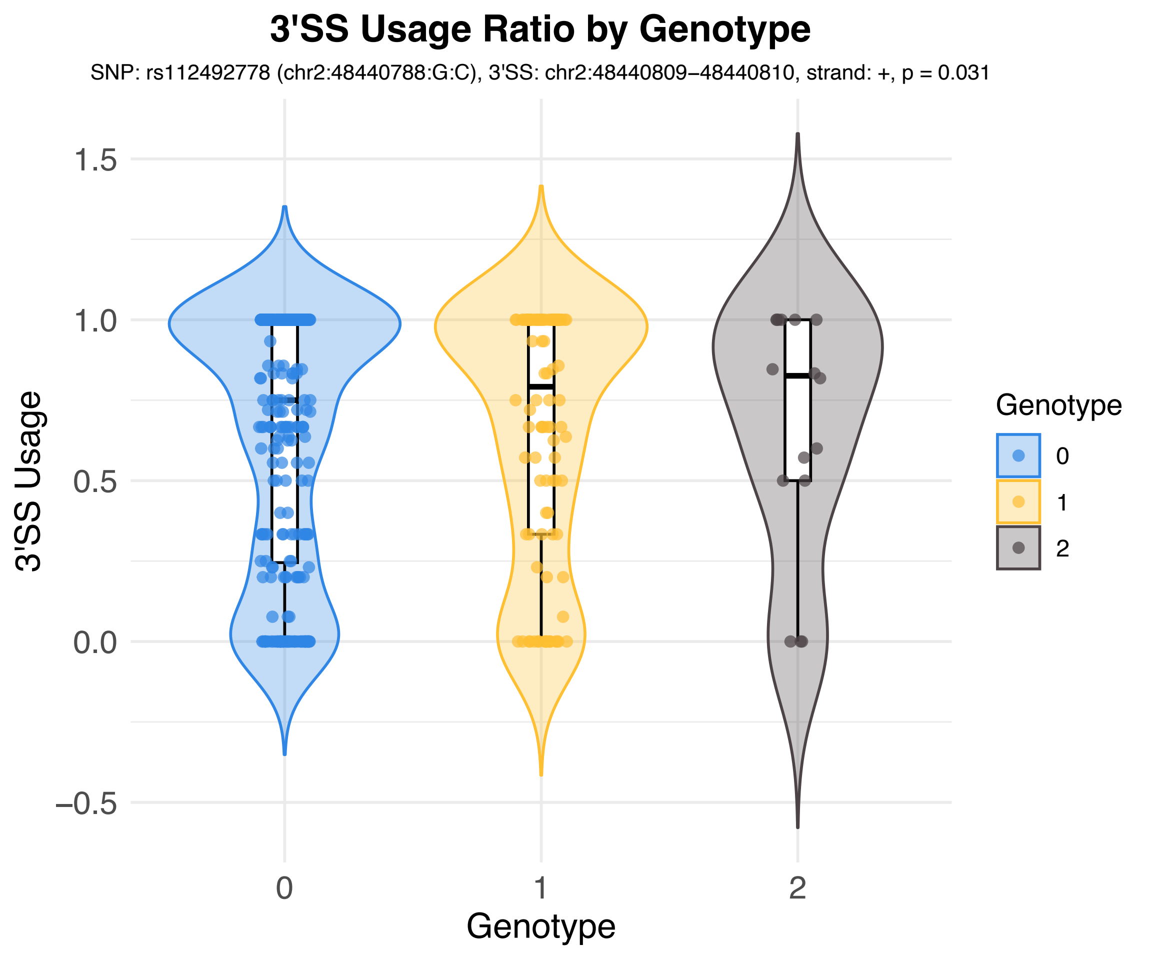

3'SS Usage Ratio by Genotype
SNP: rs112492778 (chr2:48440788:G:C), 3'SS: chr2:48440809−48440810, strand: +, p = 0.031
3'SS Usage
1.5
1.0
0.5
0.0
−0.5
0
1
2
Genotype
Genotype
0
1
2

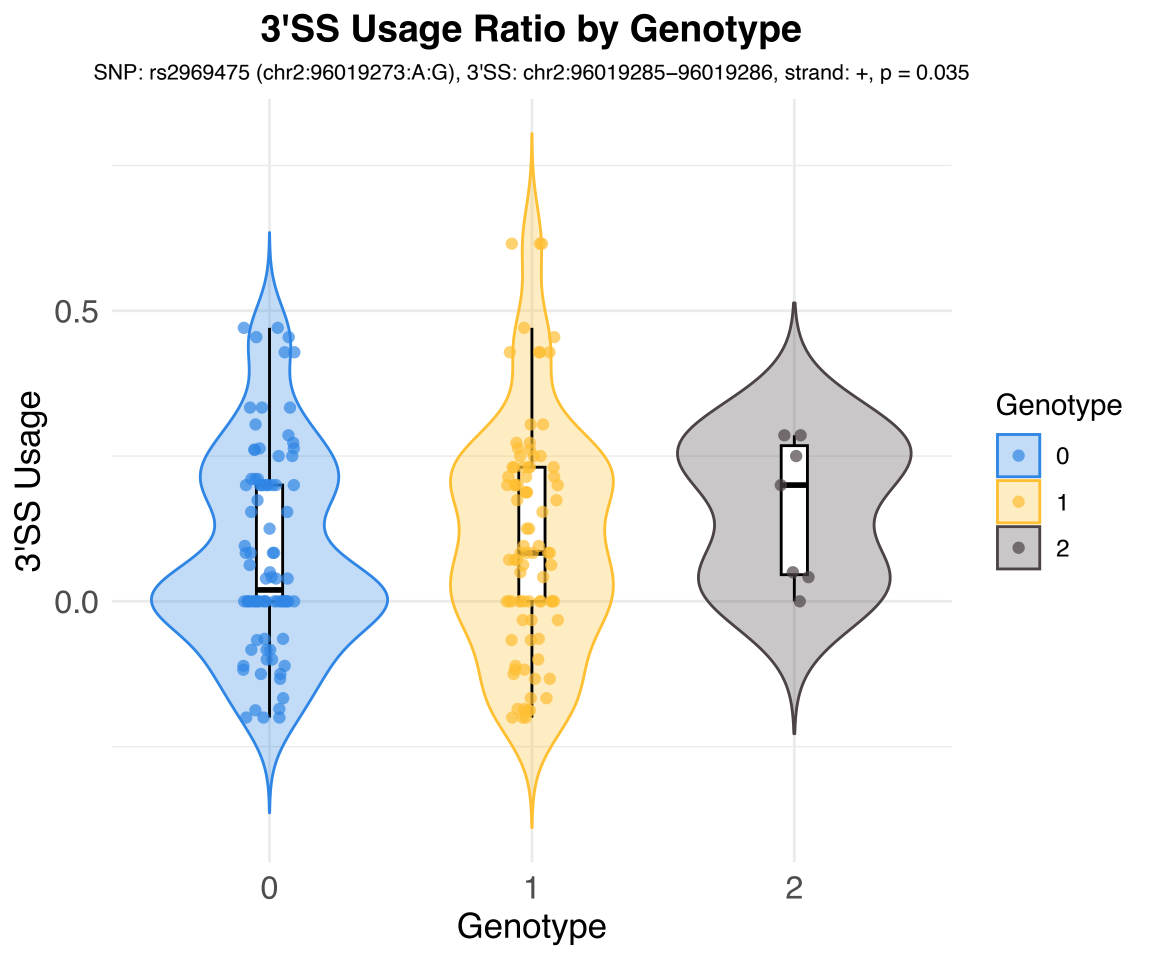

3'SS Usage Ratio by Genotype
SNP: rs2969475 (chr2:96019273:A:G), 3'SS: chr2:96019285–96019286, strand: +, p = 0.035
3'SS Usage
0.5
0.0
0
1
2
Genotype
Genotype
0
1
2

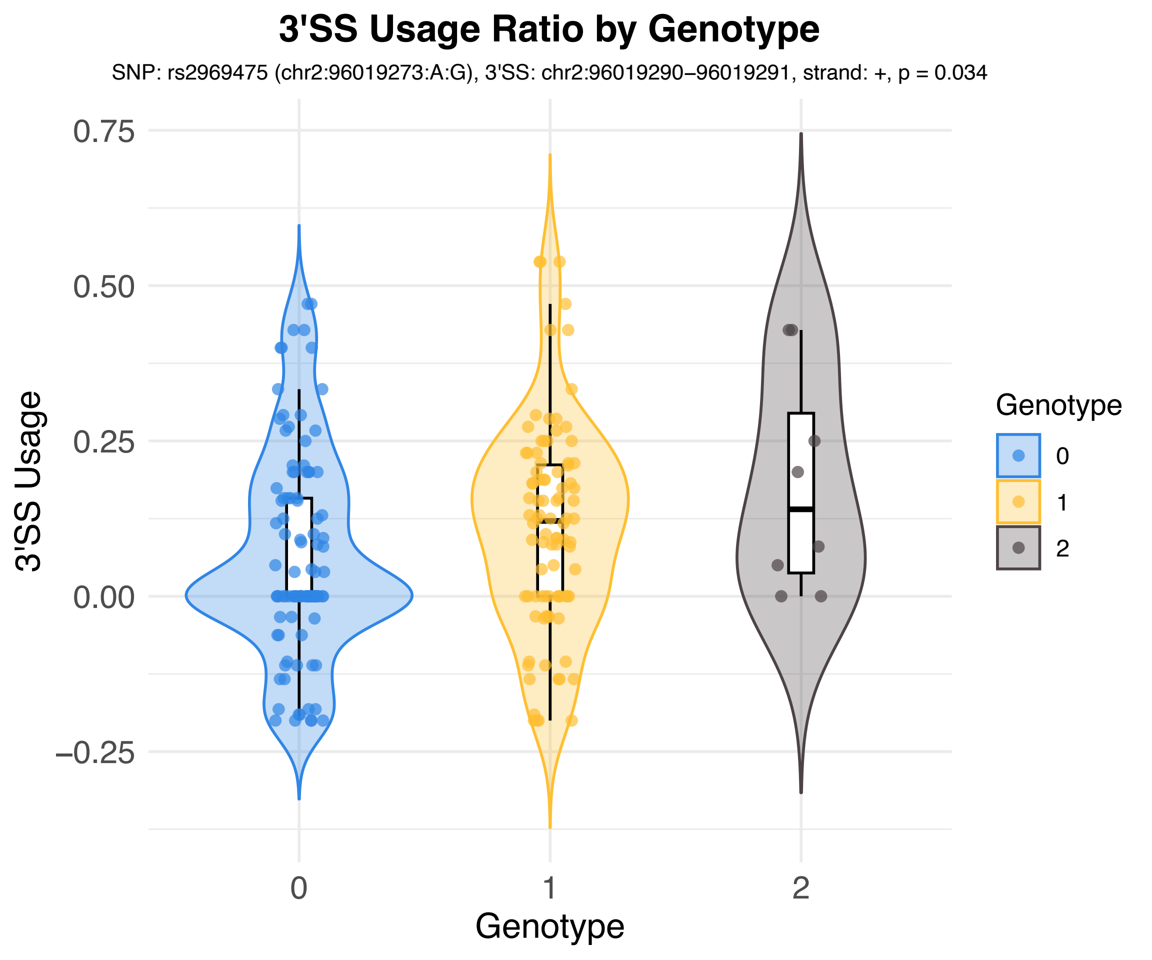
3'SS Usage Ratio by Genotype
SNP: rs2969475 (chr2:96019273:A:G), 3'SS: chr2:96019290–96019291, strand: +, p = 0.034
0.75
0.50
0.25
0.00
−0.25
3'SS Usage
0
1
2
Genotype
Genotype
0
1
2

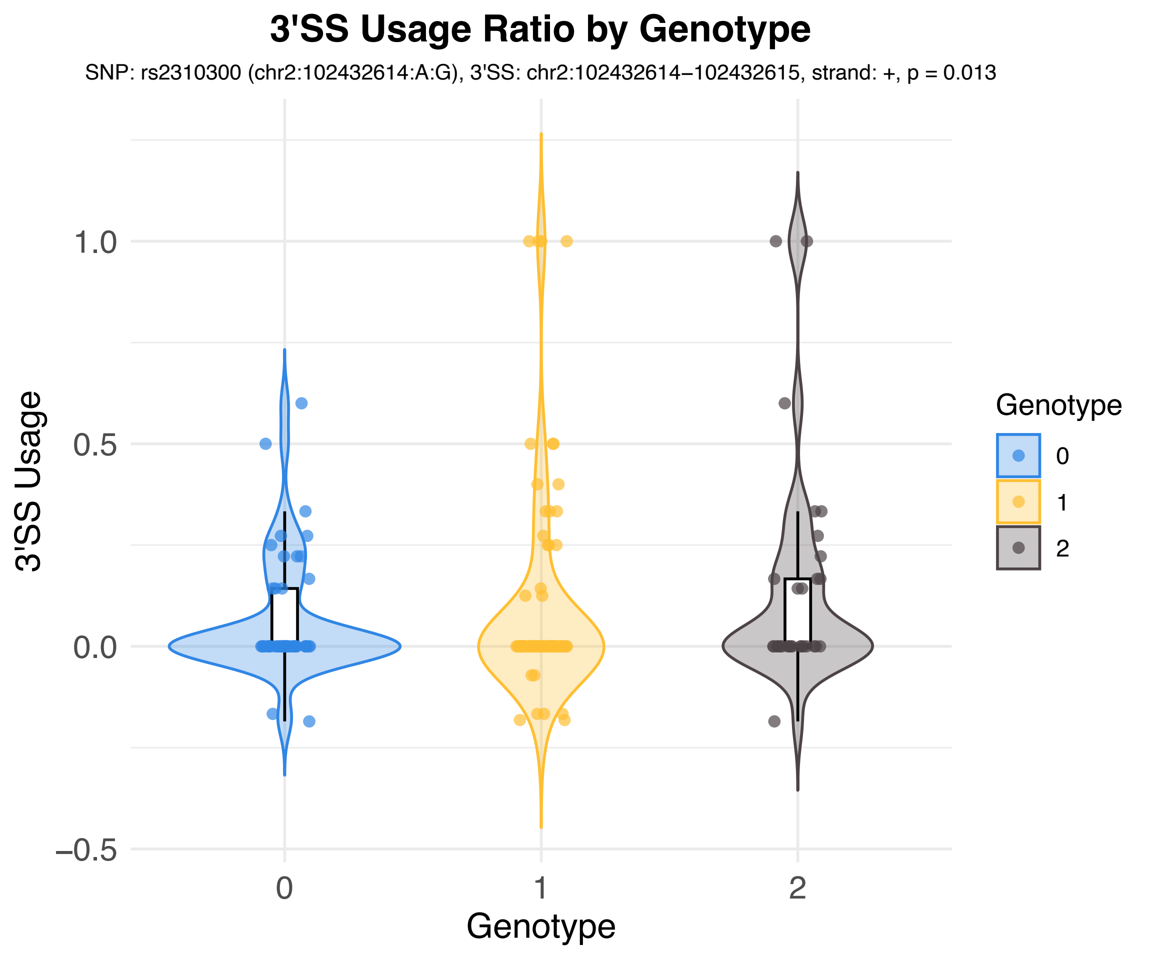

3'SS Usage Ratio by Genotype
SNP: rs2310300 (chr2:102432614:A:G), 3'SS: chr2:102432614–102432615, strand: +, p = 0.013
3'SS Usage
1.0
0.5
0.0
−0.5
0
1
2
Genotype
Genotype
0
1
2

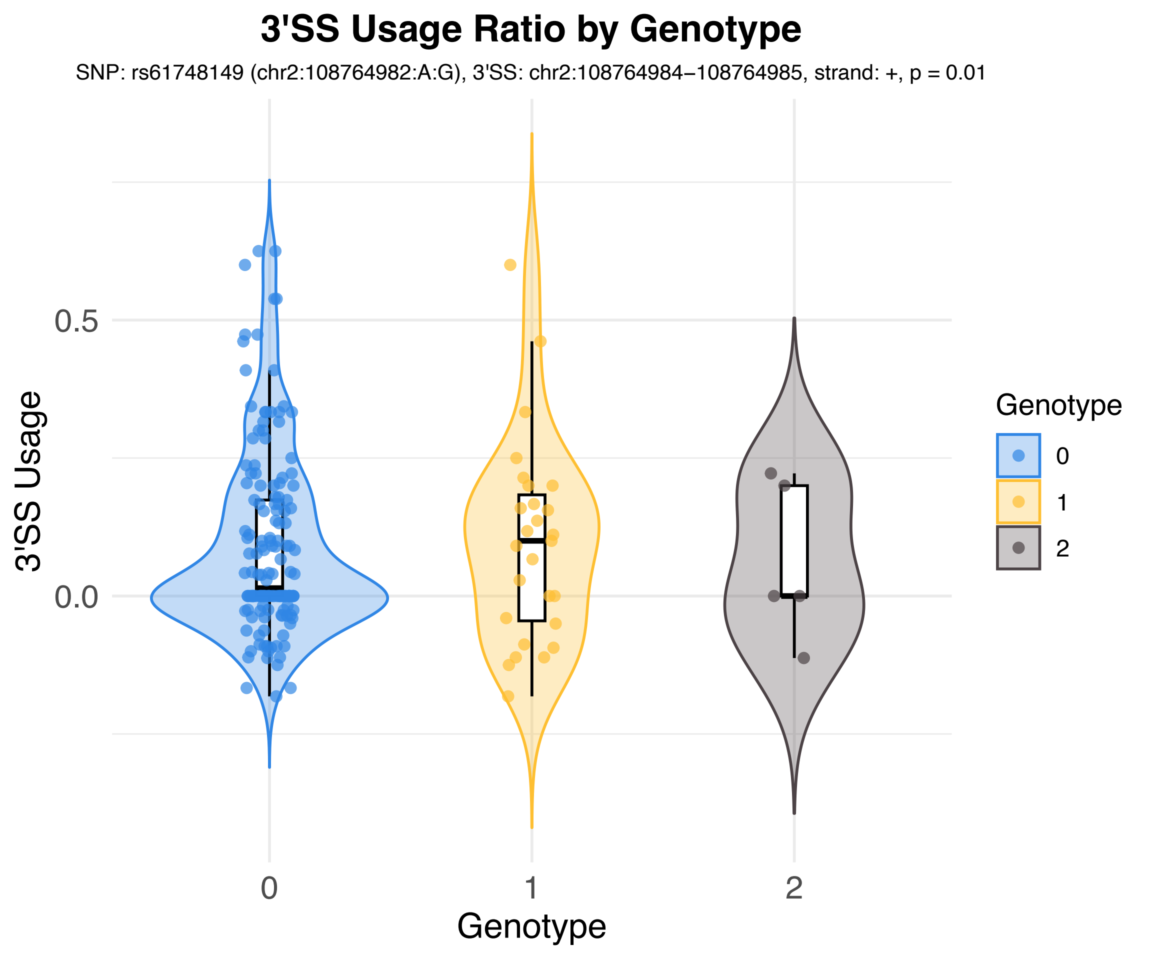
3'SS Usage Ratio by Genotype
SNP: rs61748149 (chr2:108764982:A:G), 3'SS: chr2:108764984−108764985, strand: +, p = 0.01
3'SS Usage
0.5
0.0
0
1
2
Genotype
Genotype
0
1
2

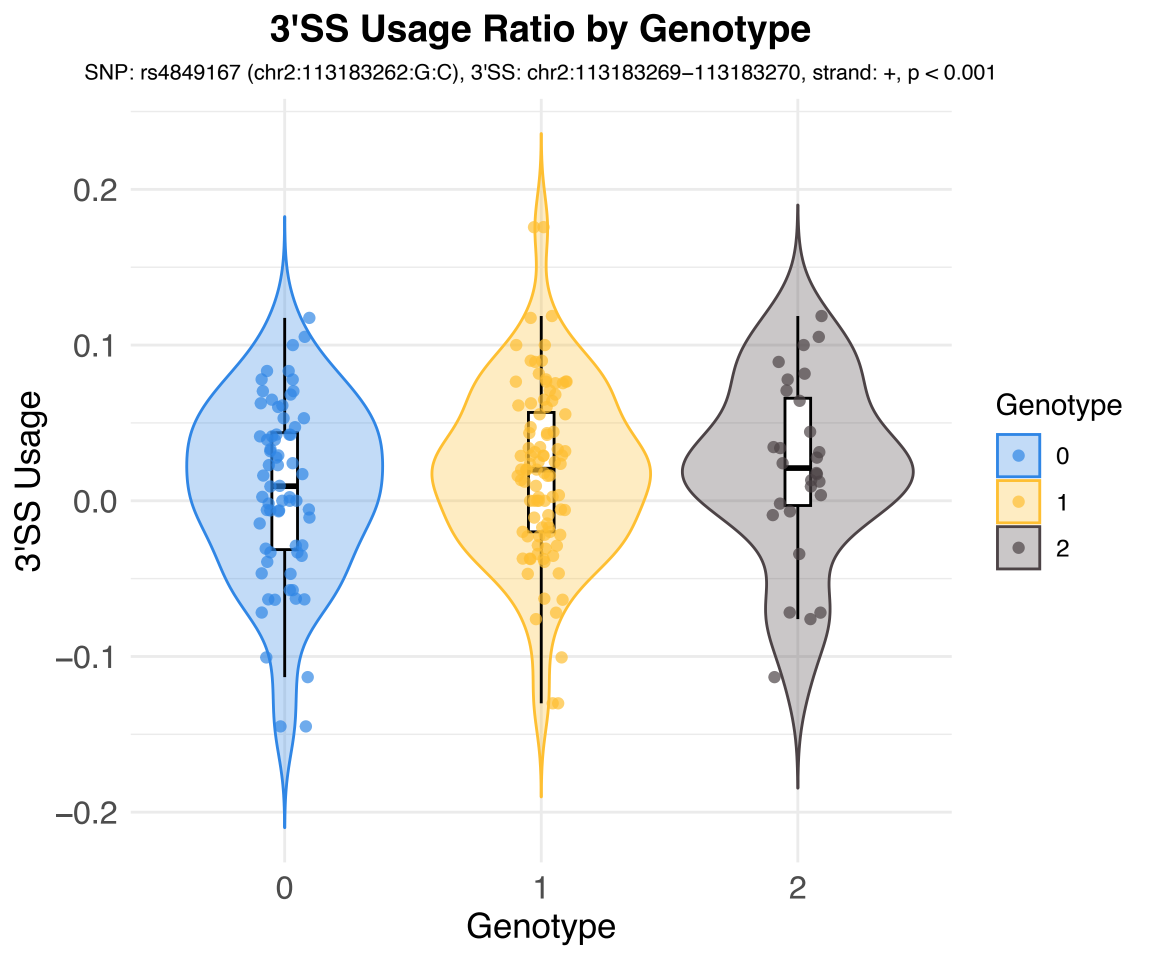
3'SS Usage Ratio by Genotype
SNP: rs4849167 (chr2:113183262:G:C), 3'SS: chr2:113183269−113183270, strand: +, p < 0.001
3'SS Usage
0.2
0.1
0.0
−0.1
−0.2
0
1
2
Genotype
Genotype
0
1
2

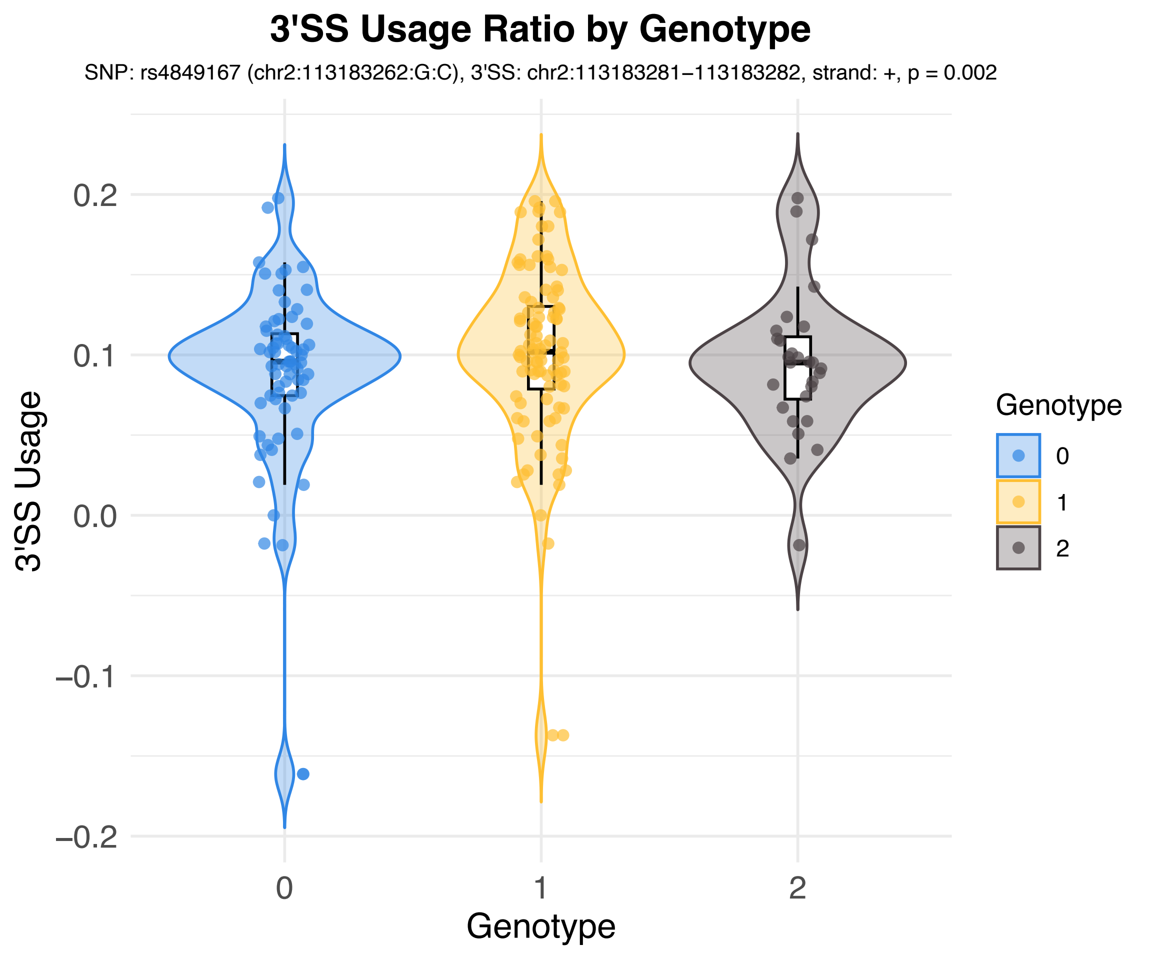

3'SS Usage Ratio by Genotype
SNP: rs4849167 (chr2:113183262:G:C), 3'SS: chr2:113183281−113183282, strand: +, p = 0.002
3'SS Usage
0.2
0.1
0.0
−0.1
−0.2
0
1
2
Genotype
Genotype
0
1
2

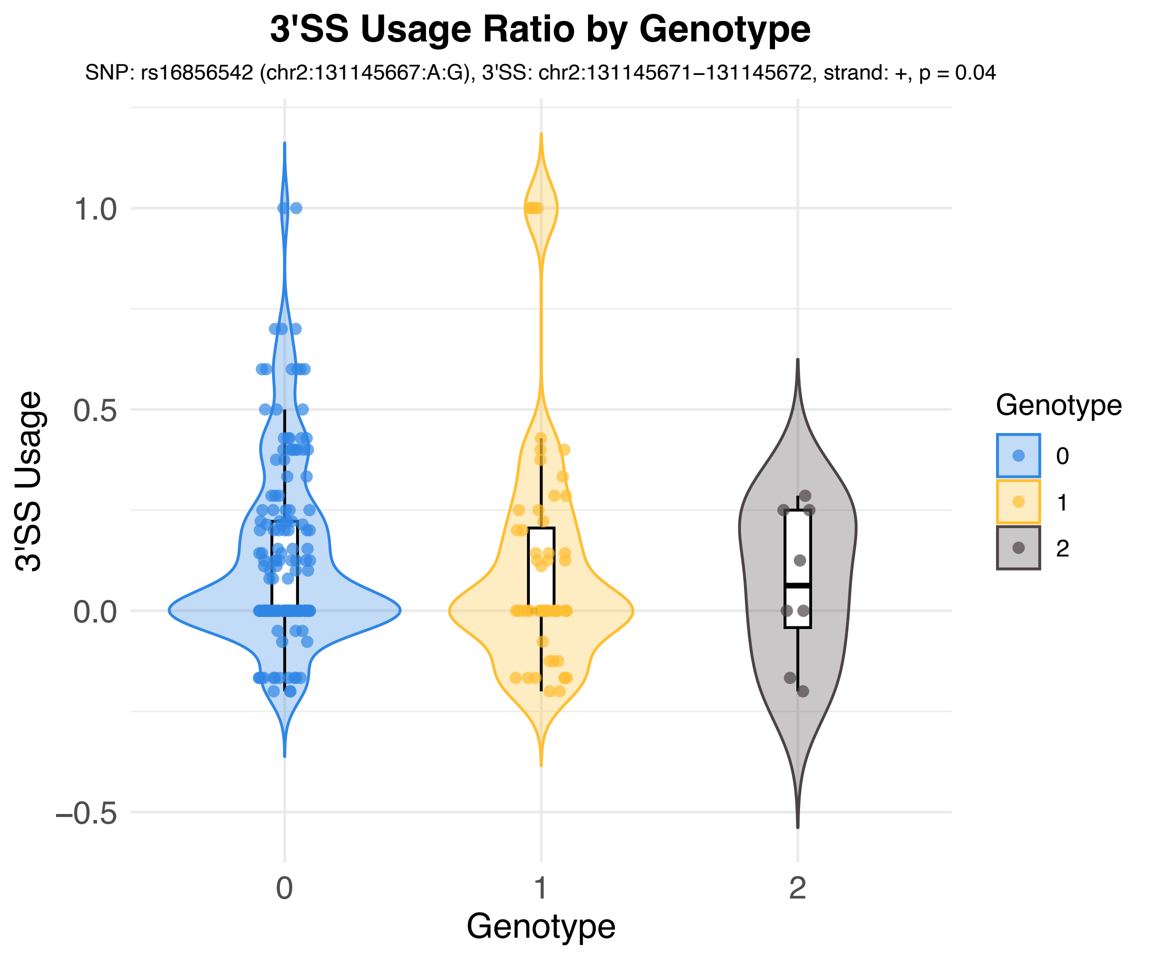
3'SS Usage Ratio by Genotype
SNP: rs16856542 (chr2:131145667:A:G), 3'SS: chr2:131145671−131145672, strand: +, p = 0.04
3'SS Usage
1.0
0.5
0.0
−0.5
0
1
2
Genotype
Genotype
0
1
2

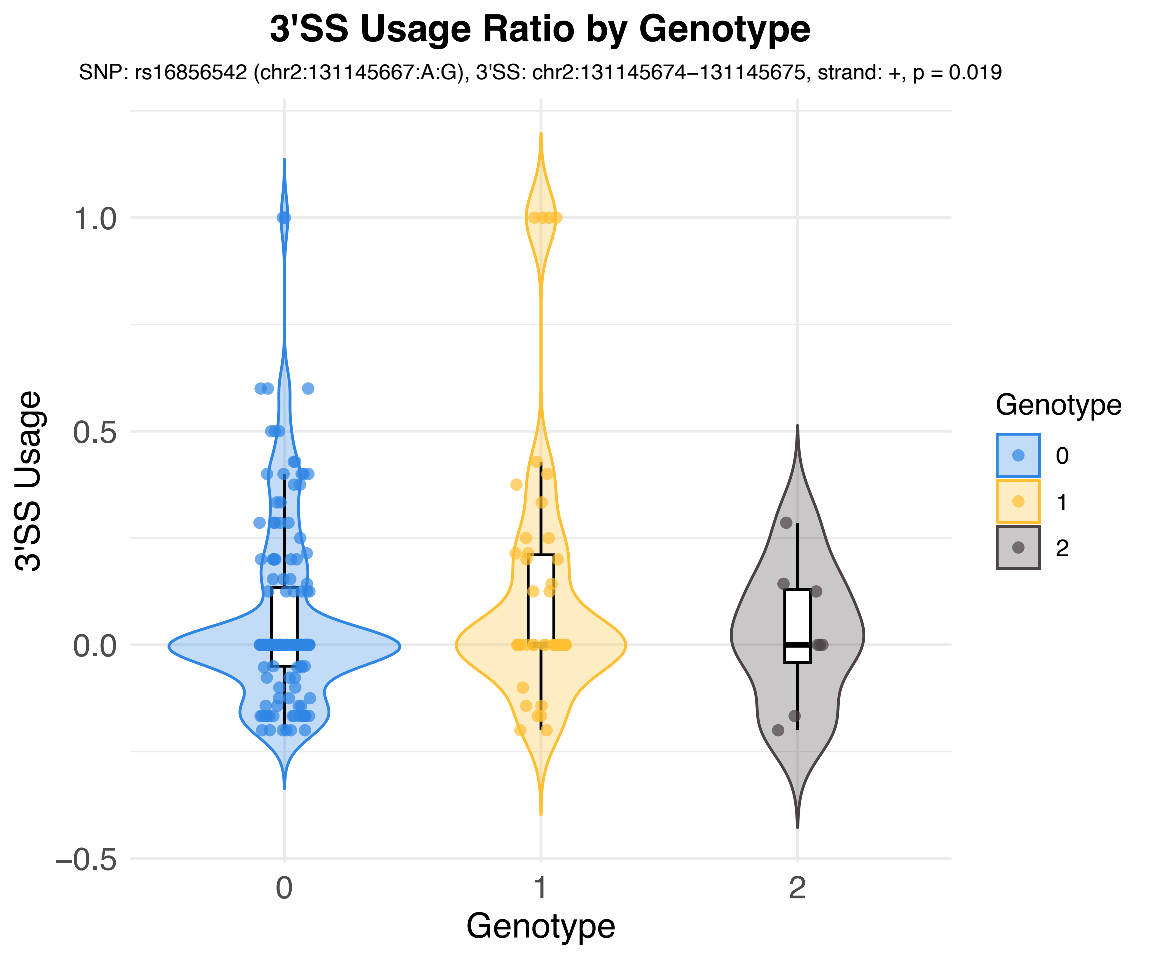
3'SS Usage Ratio by Genotype
SNP: rs16856542 (chr2:131145667:A:G), 3'SS: chr2:131145674–131145675, strand: +, p = 0.019
3'SS Usage
1.0
0.5
0.0
−0.5
0
1
2
Genotype
Genotype
0
1
2

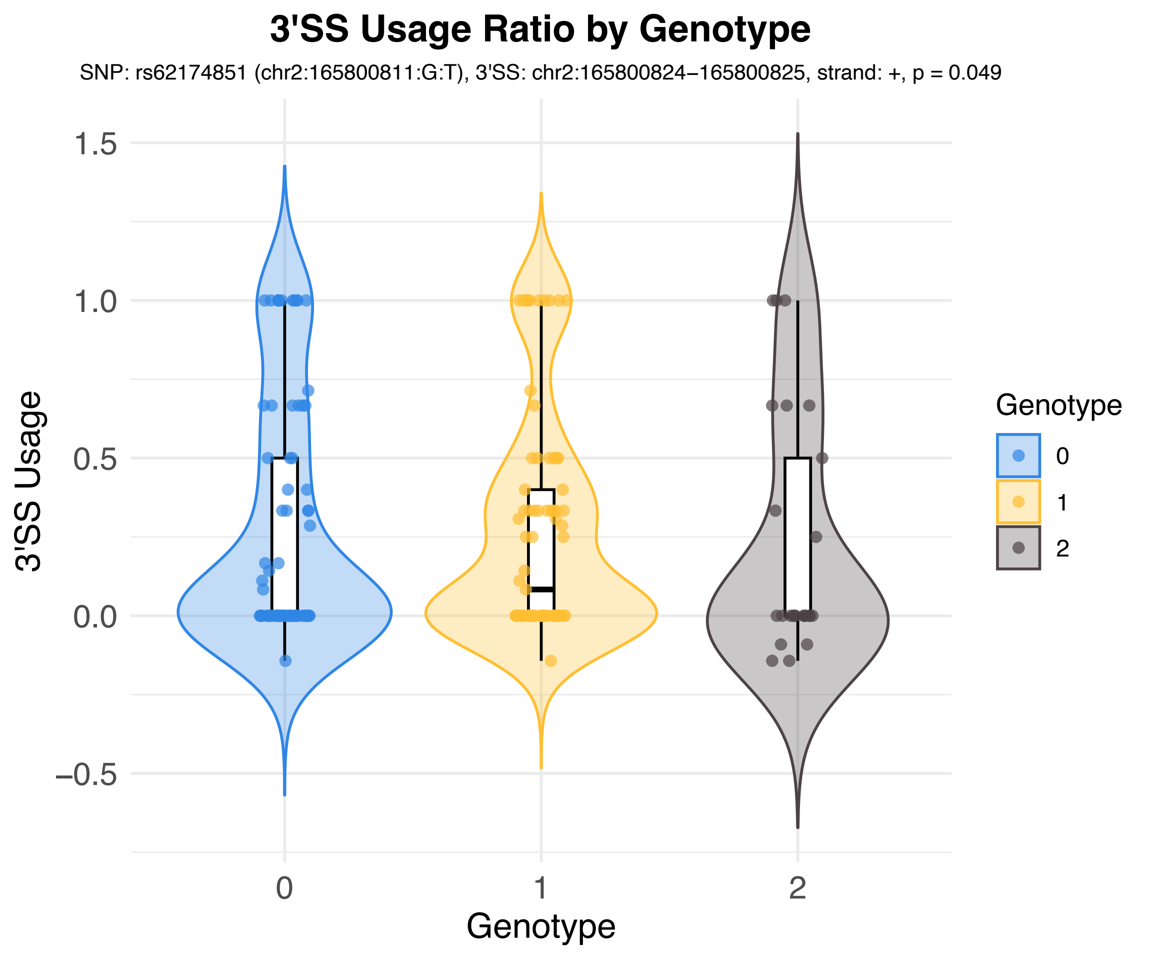

3'SS Usage Ratio by Genotype
SNP: rs62174851 (chr2:165800811:G:T), 3'SS: chr2:165800824−165800825, strand: +, p = 0.049
3'SS Usage
1.5
1.0
0.5
0.0
−0.5
0
1
2
Genotype
Genotype
0
1
2

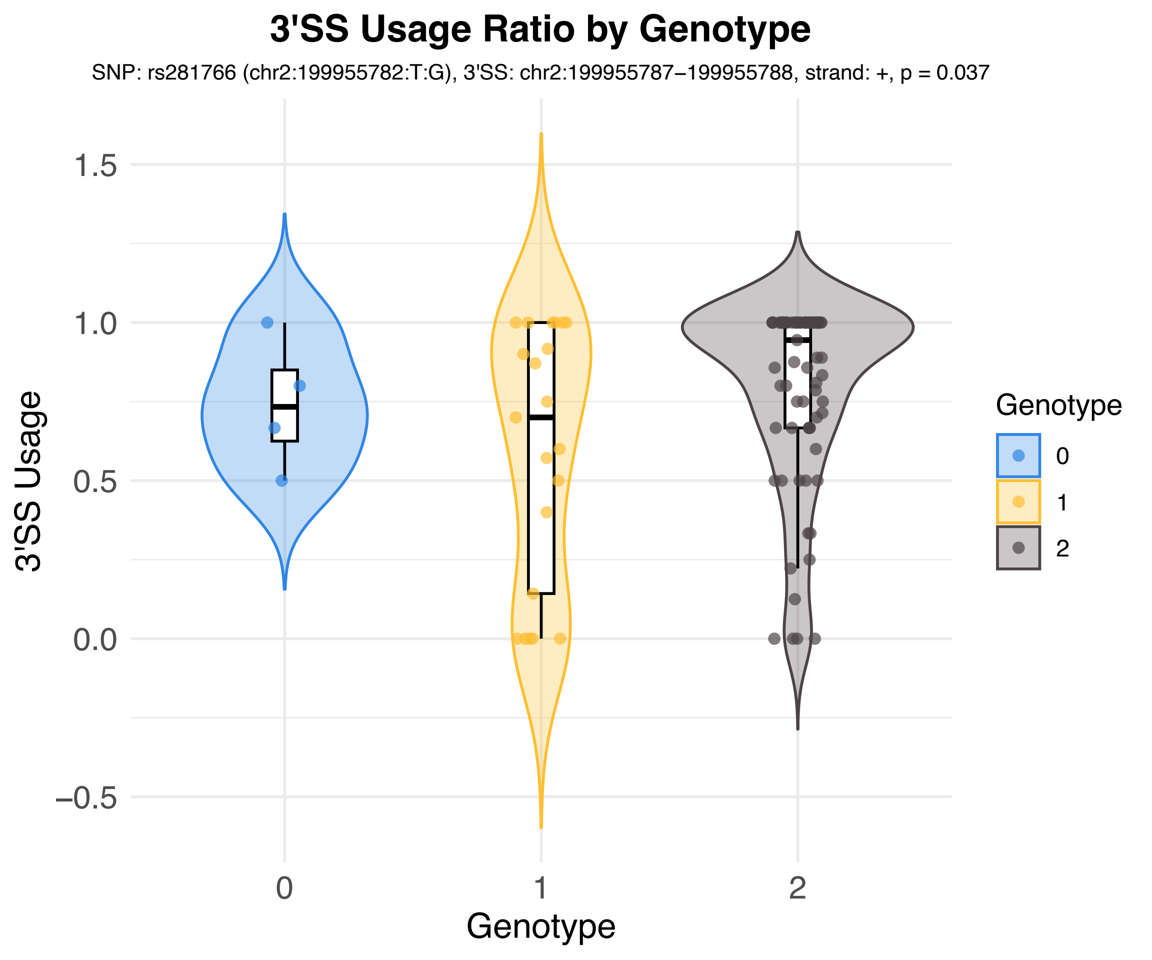

3'SS Usage Ratio by Genotype
SNP: rs281766 (chr2:199955782:T:G), 3'SS: chr2:199955787−199955788, strand: +, p = 0.037
3'SS Usage
1.5
1.0
0.5
0.0
−0.5
0
1
2
Genotype
Genotype
0
1
2

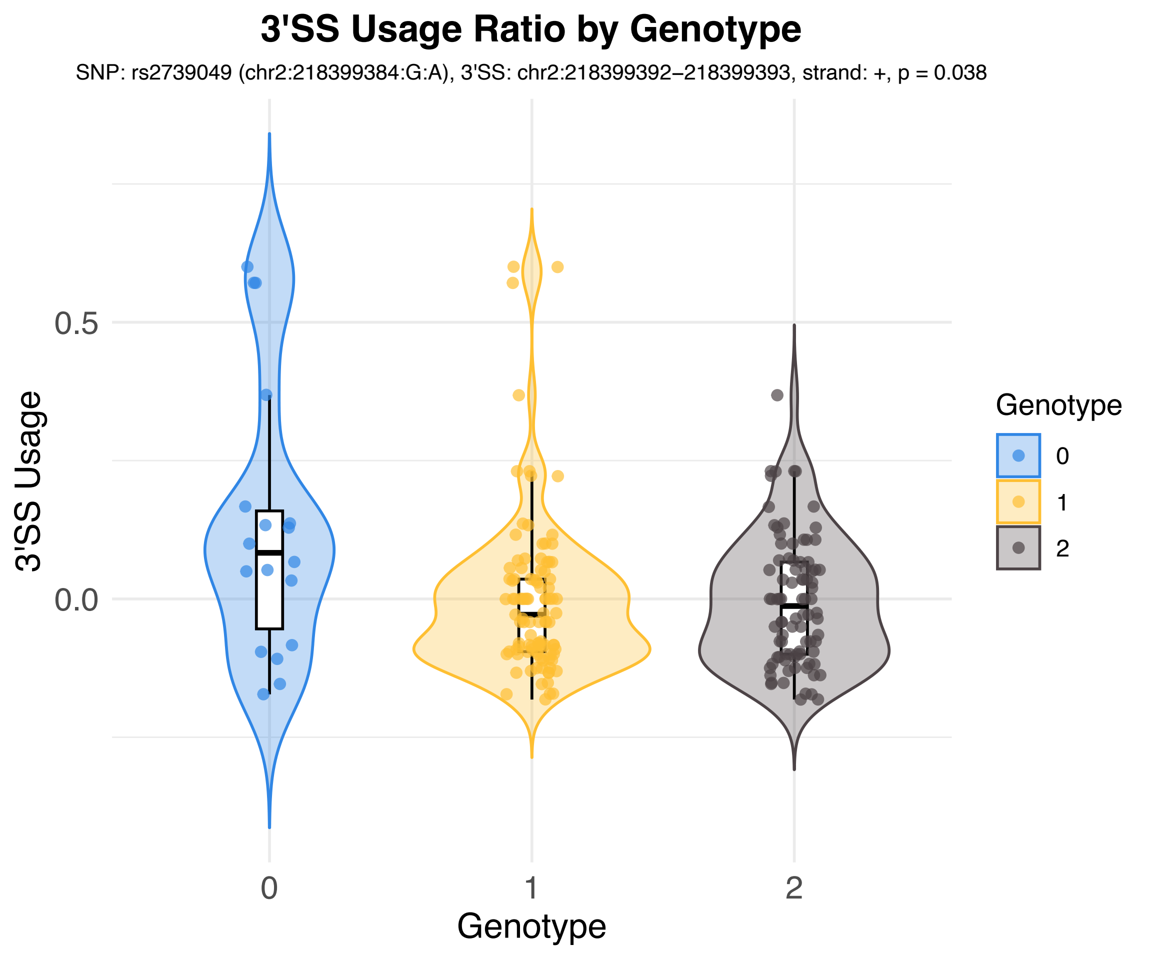

3'SS Usage Ratio by Genotype
SNP: rs2739049 (chr2:218399384:G:A), 3'SS: chr2:218399392–218399393, strand: +, p = 0.038
3'SS Usage
0.5
0.0
0
1
2
Genotype
Genotype
0
1
2

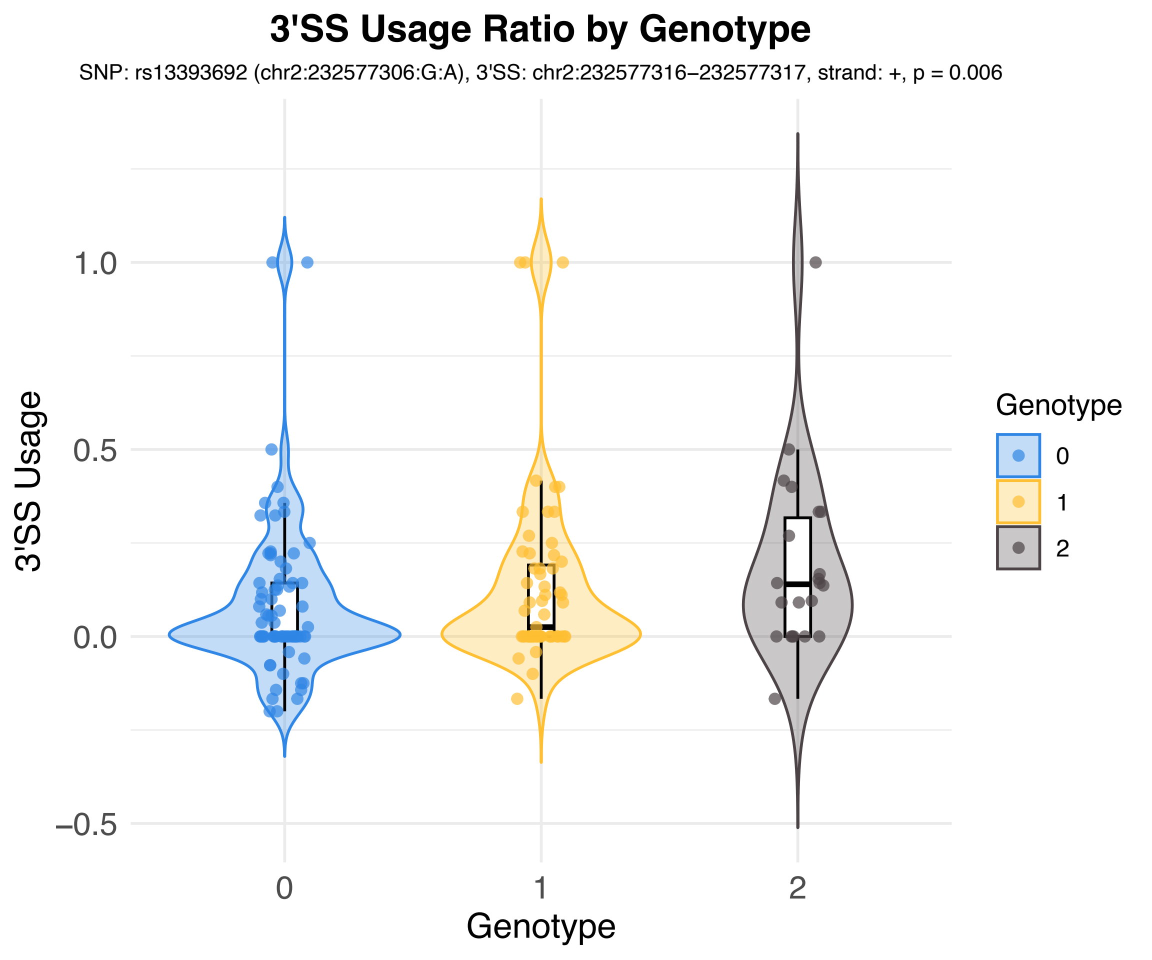
3'SS Usage Ratio by Genotype
SNP: rs13393692 (chr2:232577306:G:A), 3'SS: chr2:232577316–232577317, strand: +, p = 0.006
3'SS Usage
1.0
0.5
0.0
–0.5
0
1
2
Genotype
Genotype
0
1
2

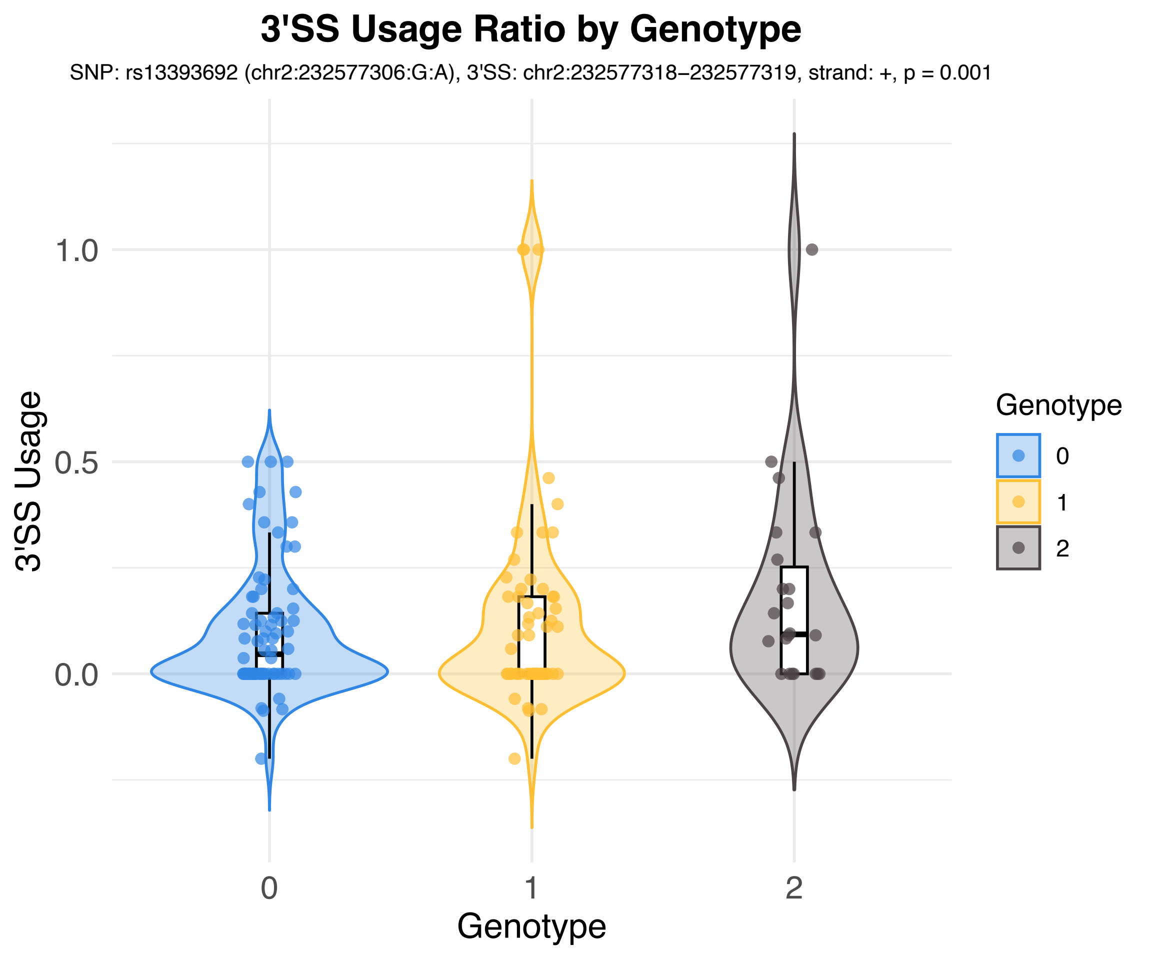
3'SS Usage Ratio by Genotype
SNP: rs13393692 (chr2:232577306:G:A), 3'SS: chr2:232577318–232577319, strand: +, p = 0.001
3'SS Usage
1.0
0.5
0.0
Genotype
0
1
2
Genotype
0
1
2

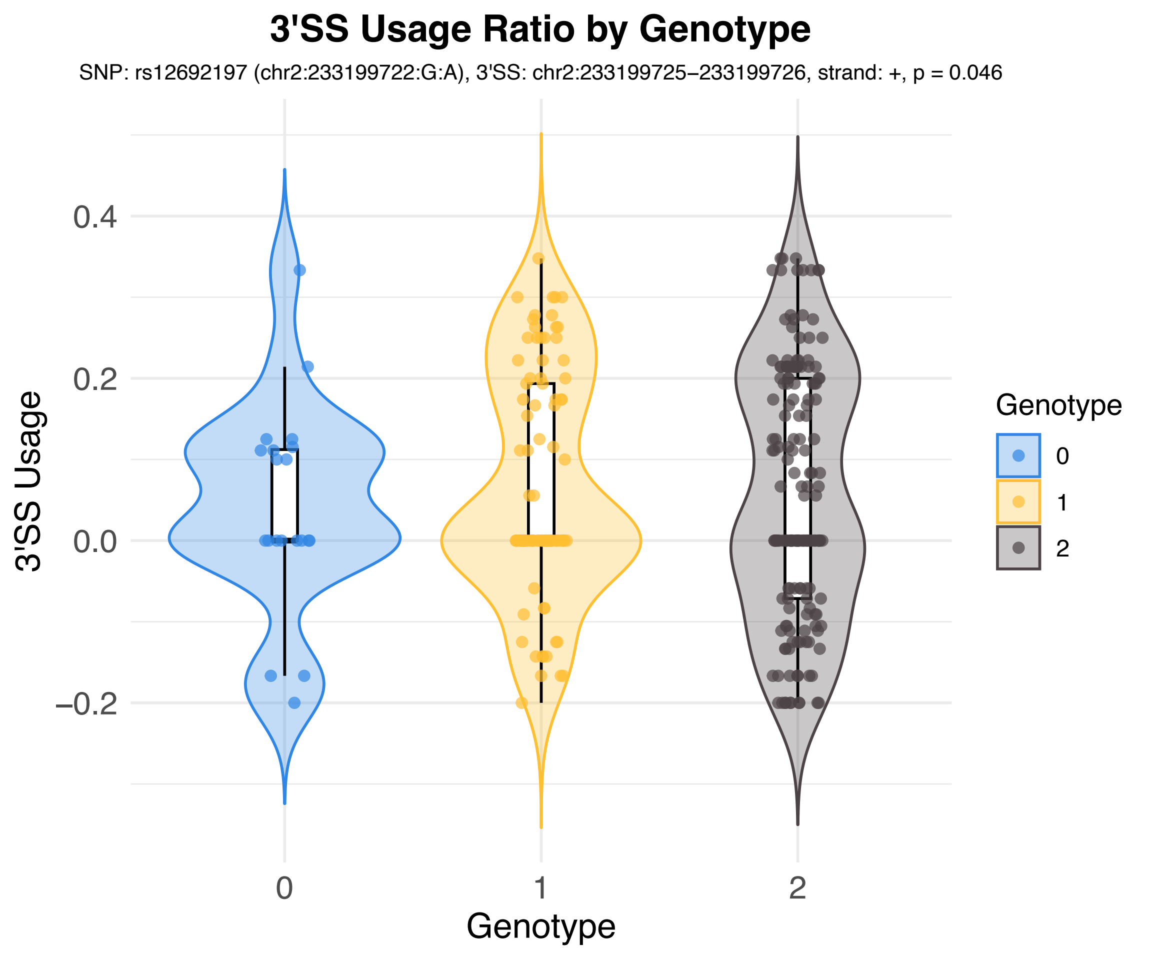
3'SS Usage Ratio by Genotype
SNP: rs12692197 (chr2:233199722:G:A), 3'SS: chr2:233199725–233199726, strand: +, p = 0.046
3'SS Usage
0.4
0.2
0.0
−0.2
Genotype
0
1
2
Genotype
0
1
2

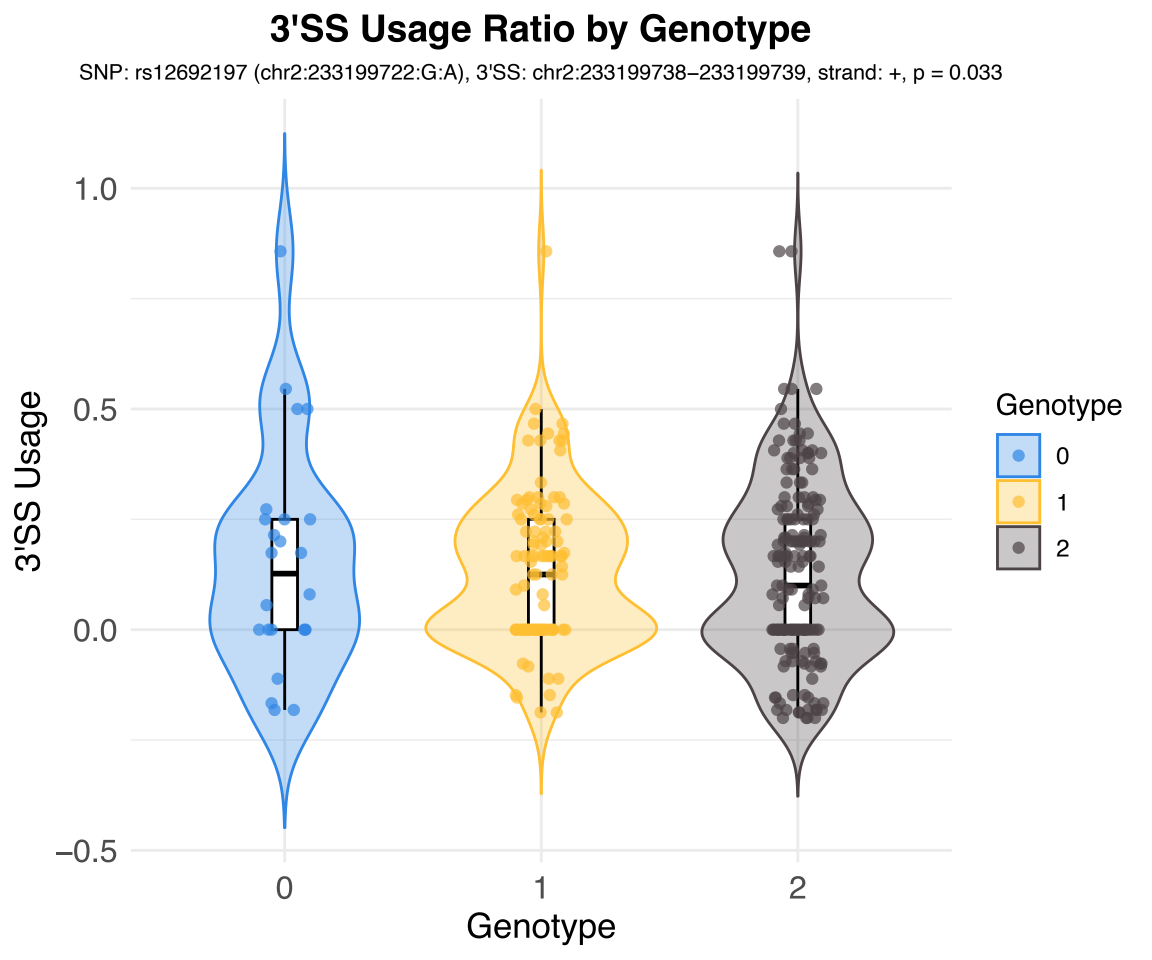
3'SS Usage Ratio by Genotype
SNP: rs12692197 (chr2:233199722:G:A), 3'SS: chr2:233199738–233199739, strand: +, p = 0.033
3'SS Usage
1.0
0.5
0.0
−0.5
0
1
2
Genotype
Genotype
0
1
2

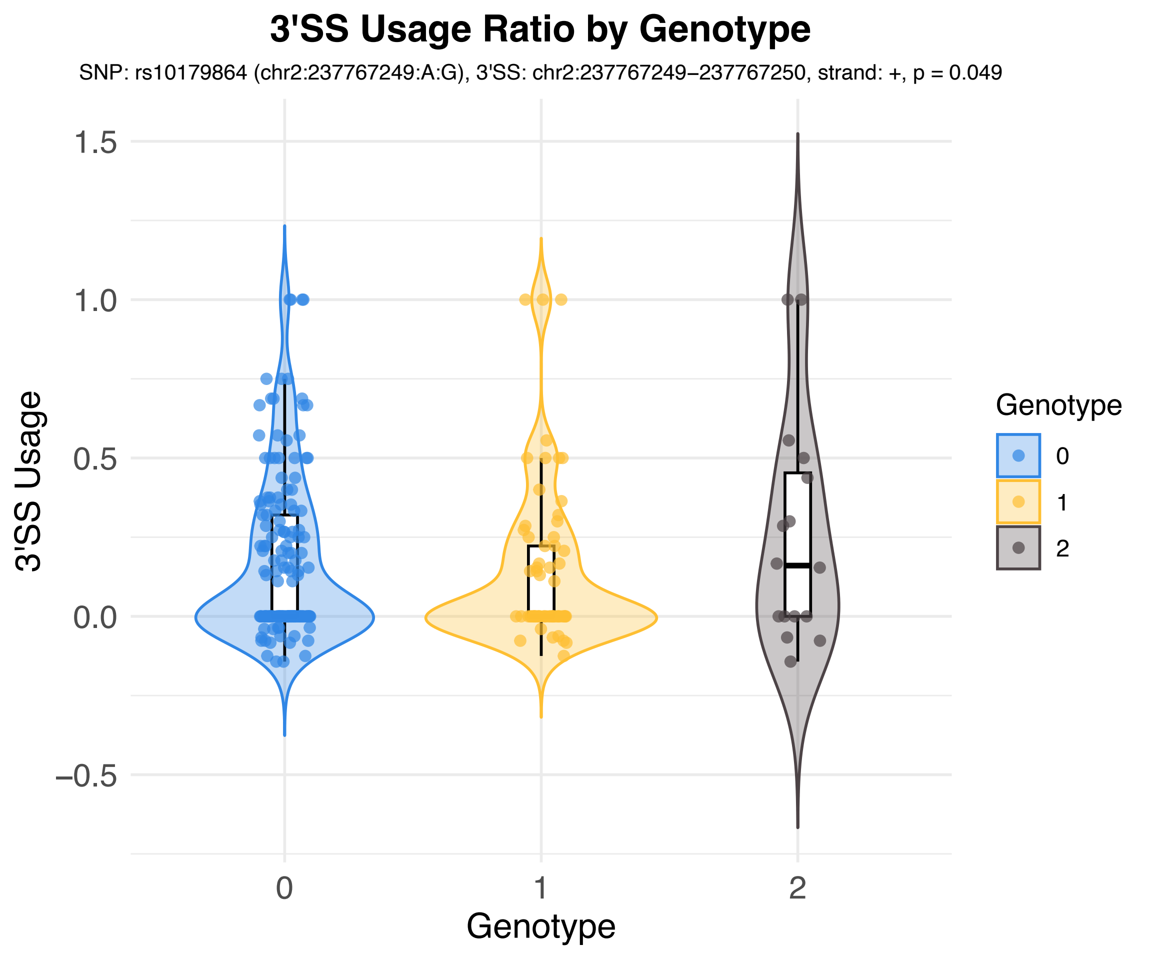

3'SS Usage Ratio by Genotype
SNP: rs10179864 (chr2:237767249:A:G), 3'SS: chr2:237767249–237767250, strand: +, p = 0.049
3'SS Usage
1.5
1.0
0.5
0.0
−0.5
0
1
2
Genotype
Genotype
0
1
2

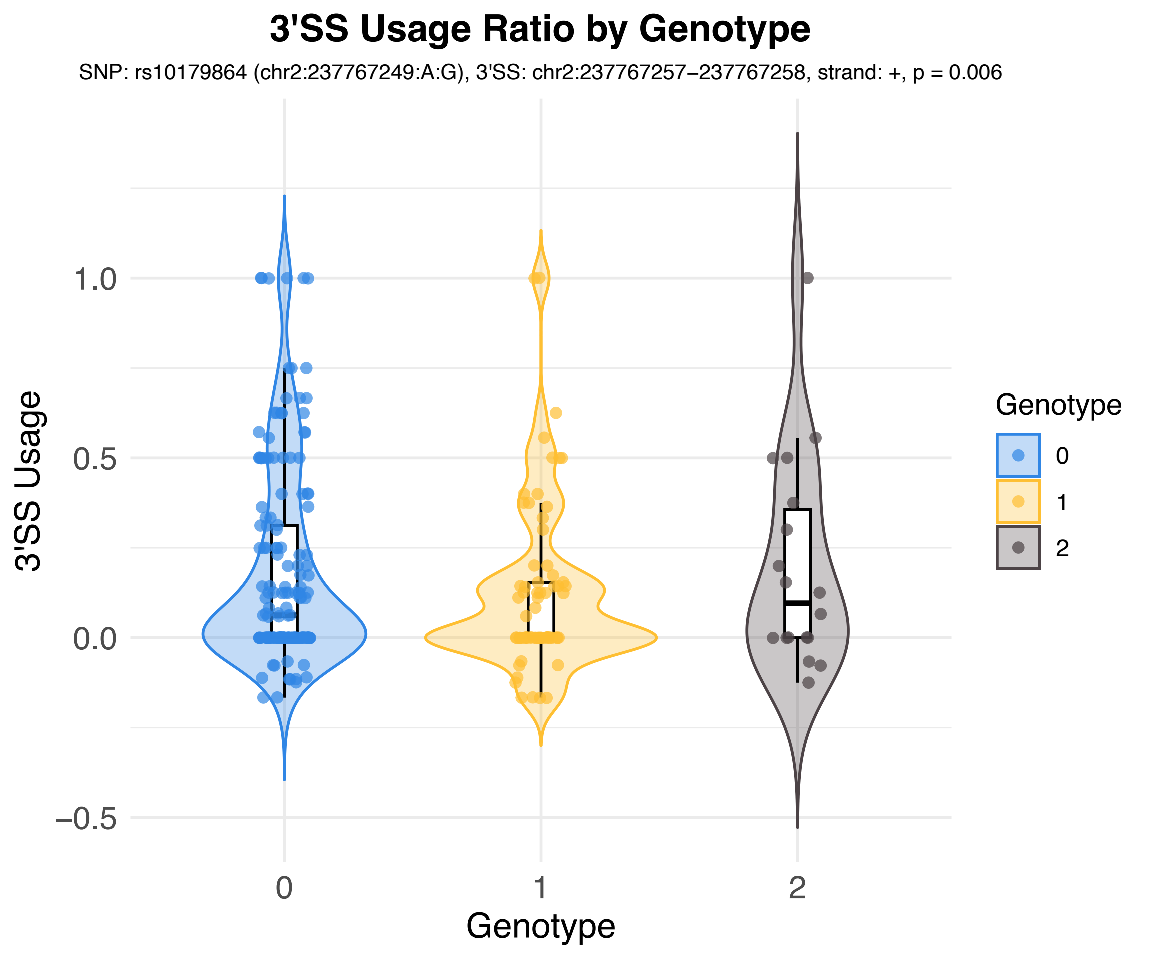
3'SS Usage Ratio by Genotype
SNP: rs10179864 (chr2:237767249:A:G), 3'SS: chr2:237767257–237767258, strand: +, p = 0.006
3'SS Usage
1.0
0.5
0.0
−0.5
0
1
2
Genotype
Genotype
0
1
2

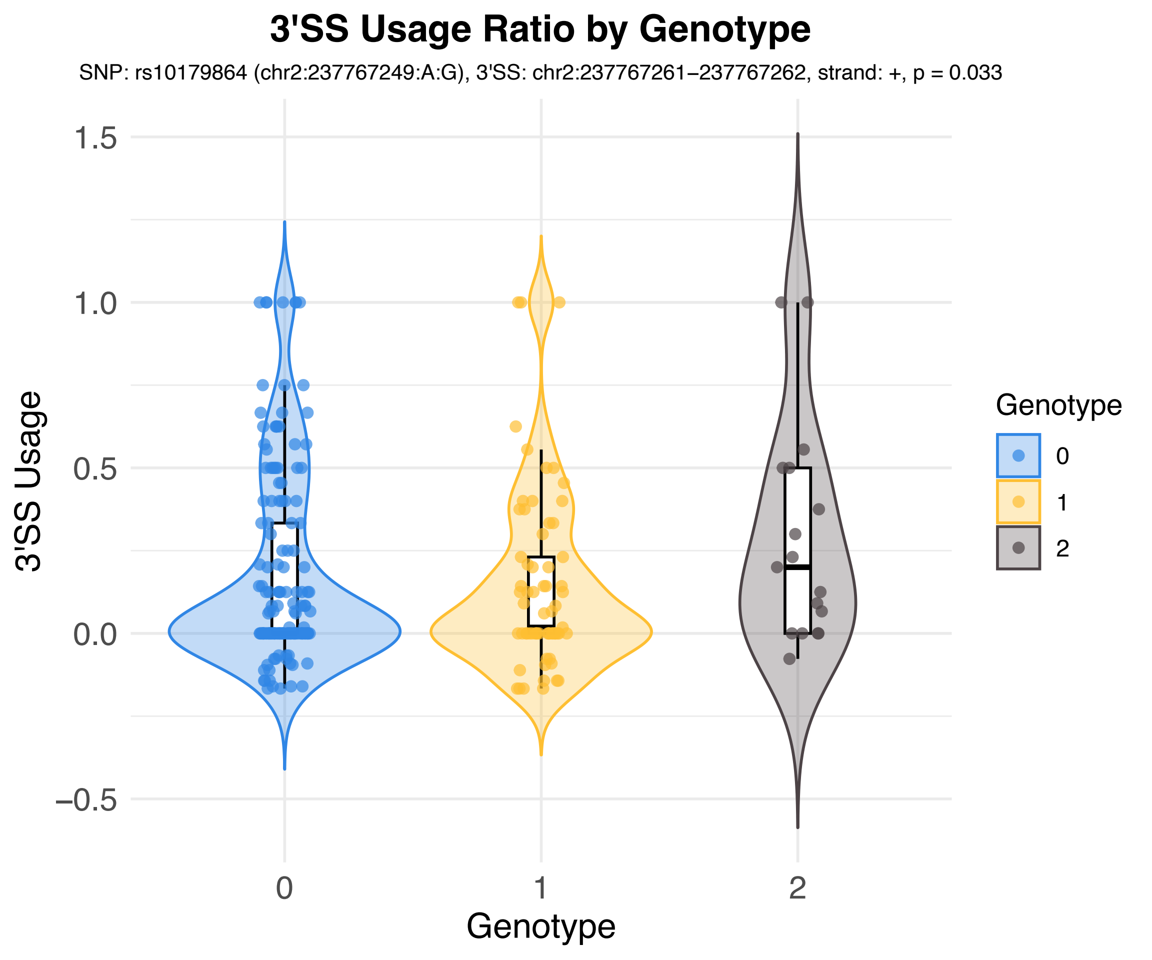

3'SS Usage Ratio by Genotype
SNP: rs10179864 (chr2:237767249:A:G), 3'SS: chr2:237767261–237767262, strand: +, p = 0.033
3'SS Usage
1.5
1.0
0.5
0.0
−0.5
0
1
2
Genotype
Genotype
0
1
2

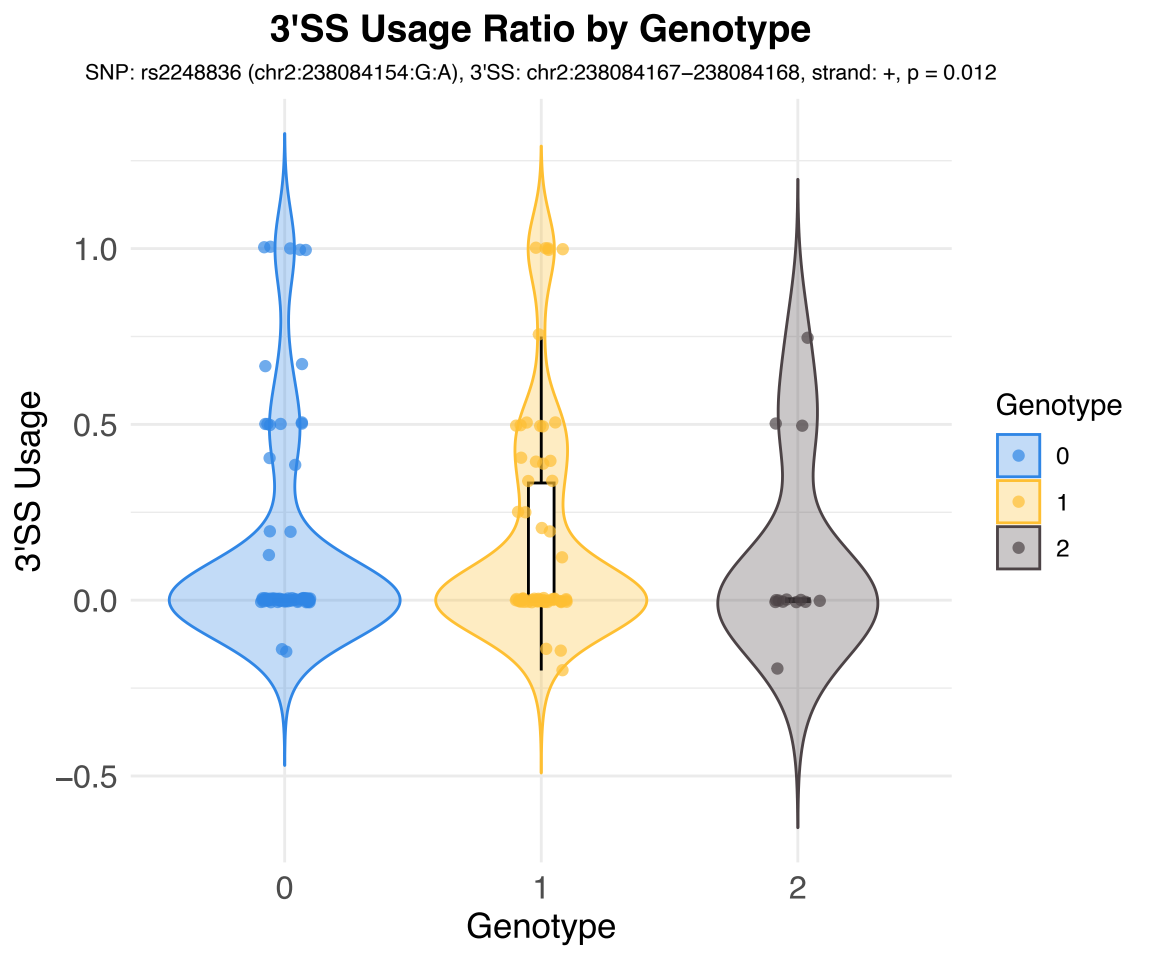
3'SS Usage Ratio by Genotype
SNP: rs2248836 (chr2:238084154:G:A), 3'SS: chr2:238084167−238084168, strand: +, p = 0.012
3'SS Usage
1.0
0.5
0.0
−0.5
0
1
2
Genotype
Genotype
0
1
2

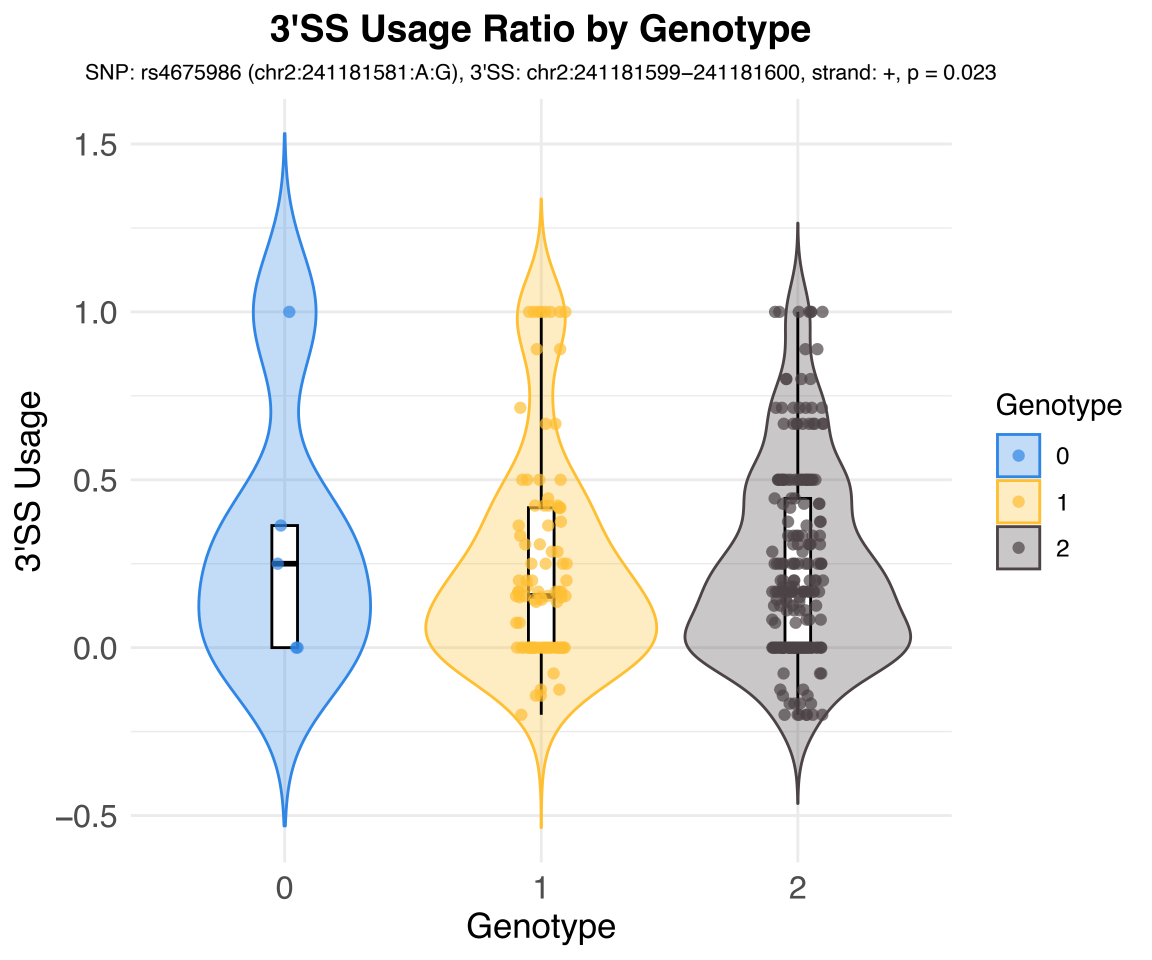
3'SS Usage Ratio by Genotype
SNP: rs4675986 (chr2:241181581:A:G), 3'SS: chr2:241181599−241181600, strand: +, p = 0.023
3'SS Usage
1.5
1.0
0.5
0.0
−0.5
0
1
2
Genotype
Genotype
0
1
2

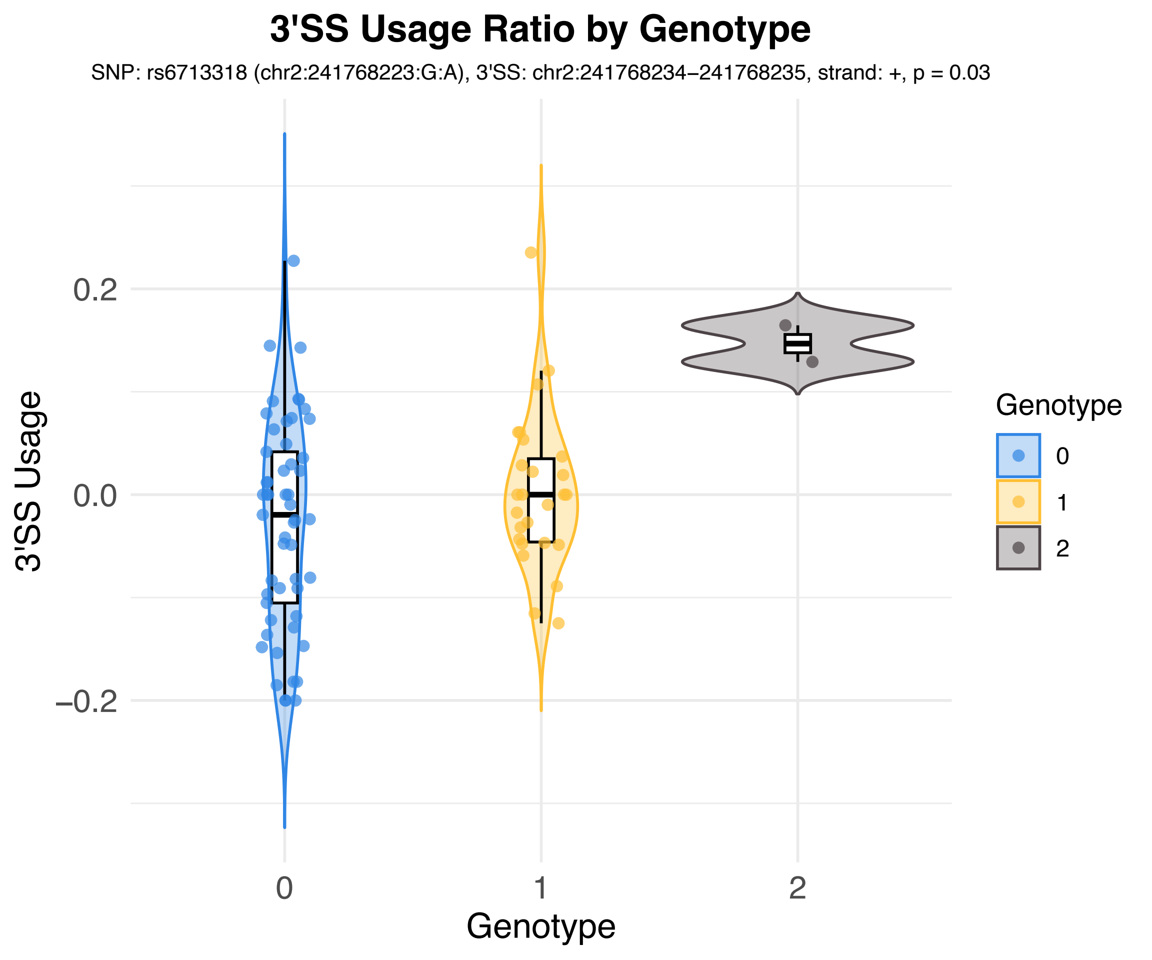
3'SS Usage Ratio by Genotype
SNP: rs6713318 (chr2:241768223:G:A), 3'SS: chr2:241768234–241768235, strand: +, p = 0.03
3'SS Usage
0.2
0.0
–0.2
0
1
2
Genotype
Genotype
0
1
2

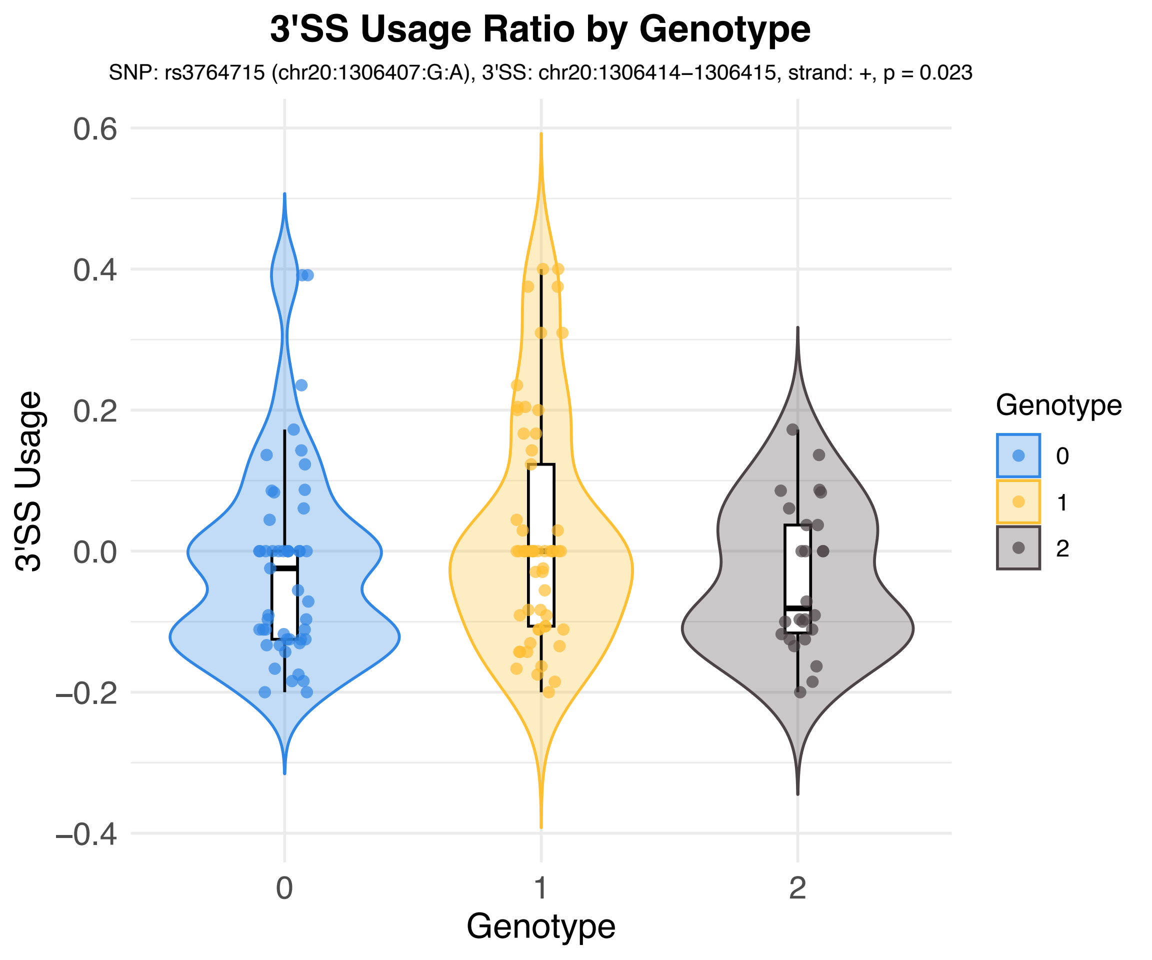

3'SS Usage Ratio by Genotype
SNP: rs3764715 (chr20:1306407:G:A), 3'SS: chr20:1306414–1306415, strand: +, p = 0.023
3'SS Usage
0.6
0.4
0.2
0.0
−0.2
−0.4
0
1
2
Genotype
Genotype
0
1
2

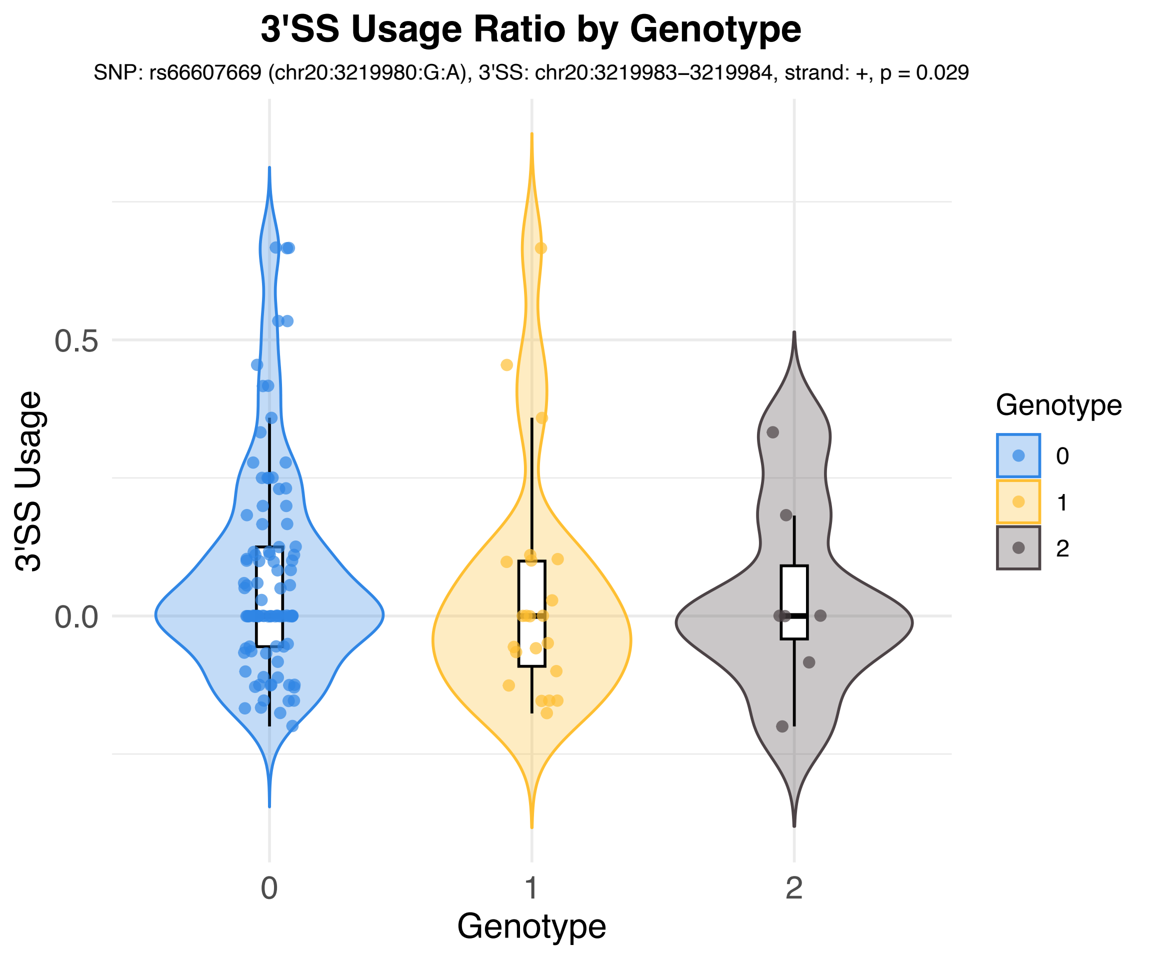
3'SS Usage Ratio by Genotype
SNP: rs66607669 (chr20:3219980:G:A), 3'SS: chr20:3219983–3219984, strand: +, p = 0.029
3'SS Usage
0.5
0.0
0
1
2
Genotype
Genotype
0
1
2

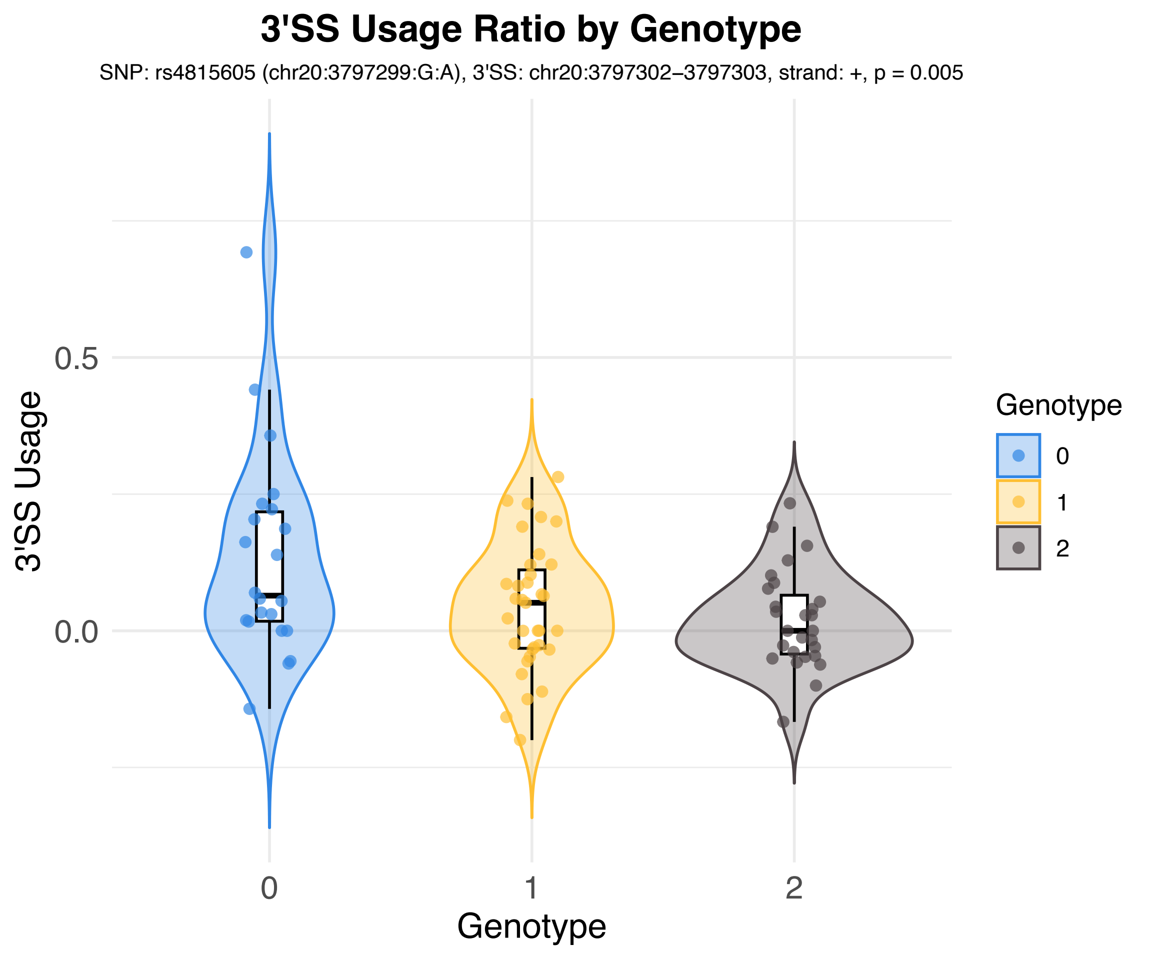
3'SS Usage Ratio by Genotype
SNP: rs4815605 (chr20:3797299:G:A), 3'SS: chr20:3797302–3797303, strand: +, p = 0.005
3'SS Usage
0.5
0.0
Genotype
0
1
2
Genotype
0
1
2

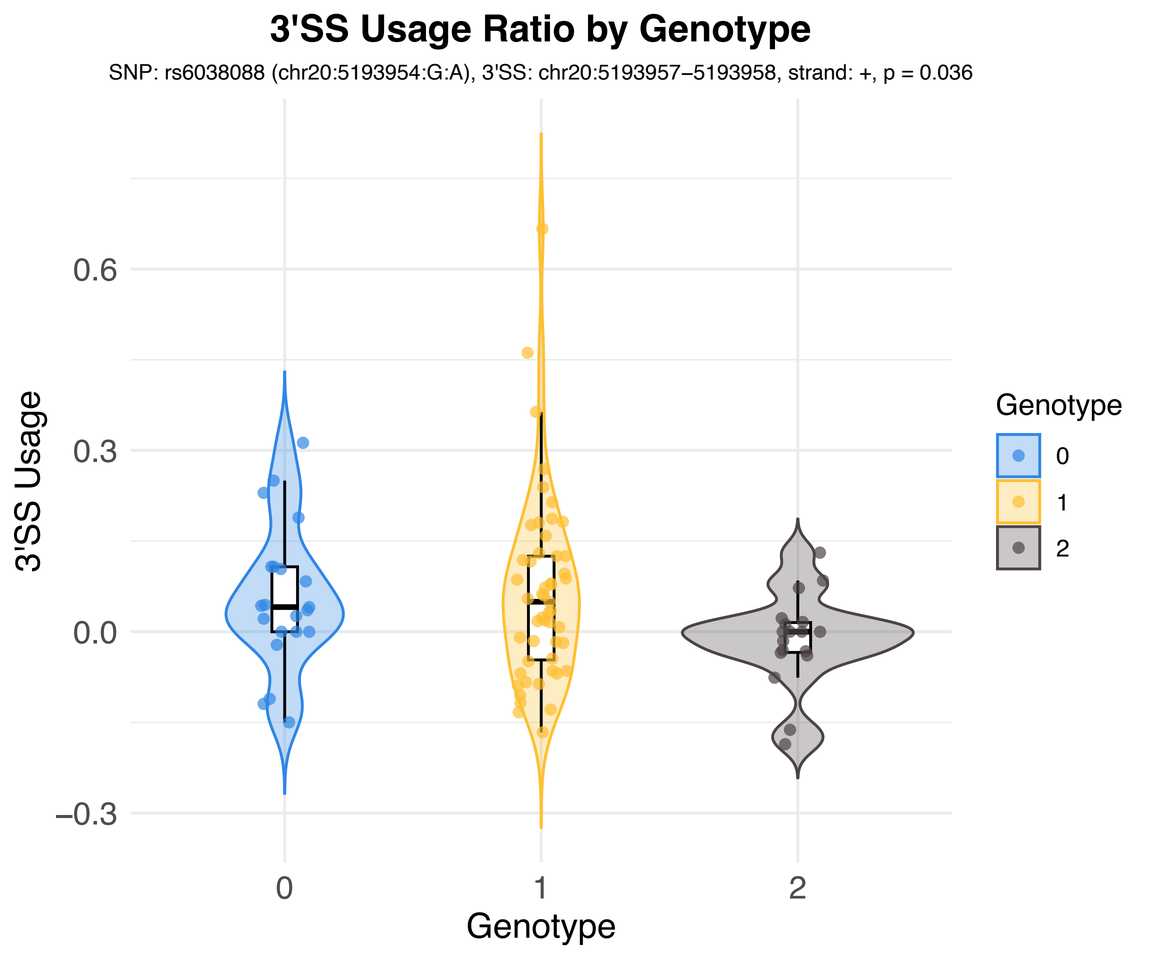

3'SS Usage Ratio by Genotype
SNP: rs6038088 (chr20:5193954:G:A), 3'SS: chr20:5193957−5193958, strand: +, p = 0.036
3'SS Usage
0.6
0.3
0.0
−0.3
0
1
2
Genotype
Genotype
0
1
2

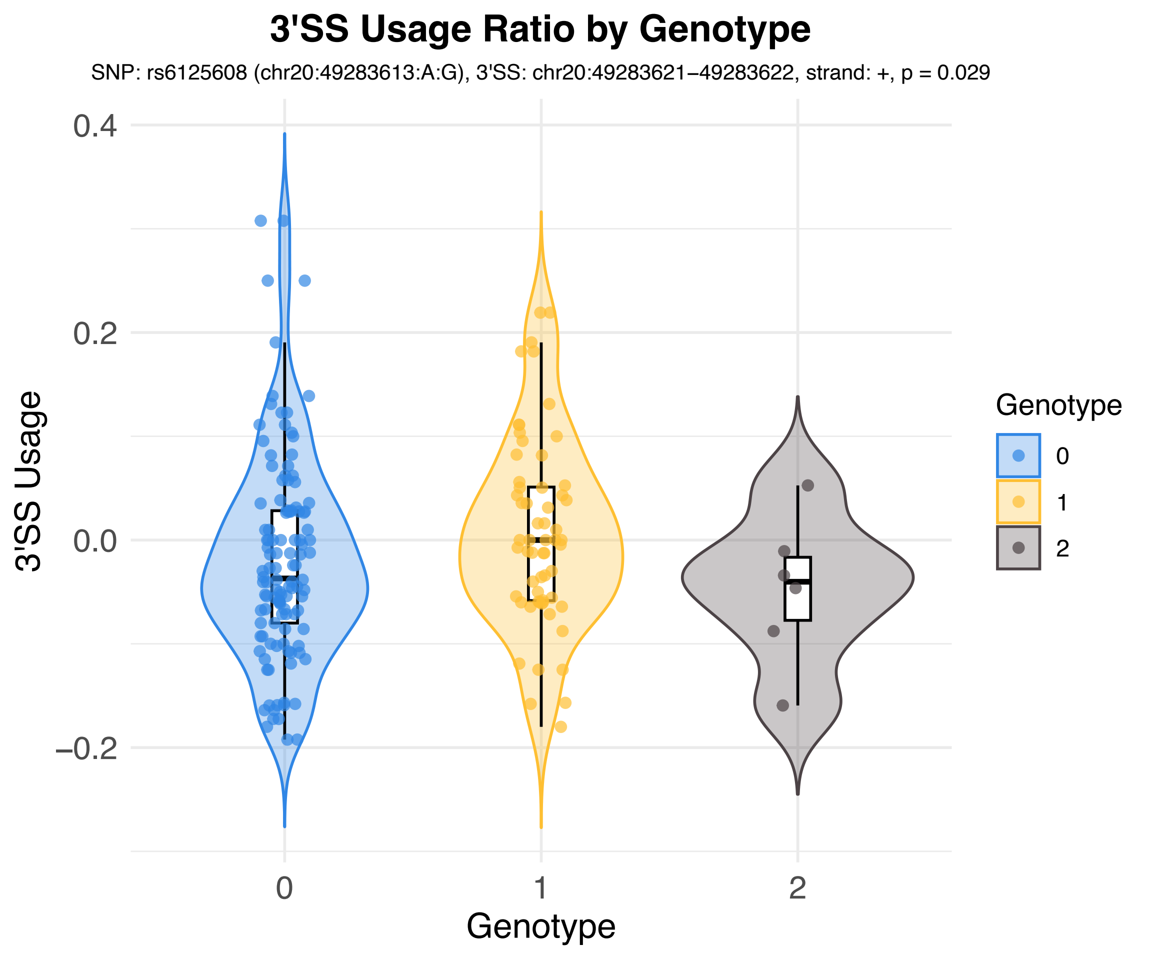
3'SS Usage Ratio by Genotype
SNP: rs6125608 (chr20:49283613:A:G), 3'SS: chr20:49283621–49283622, strand: +, p = 0.029
3'SS Usage
0.4
0.2
0.0
−0.2
0
1
2
Genotype
Genotype
0
1
2

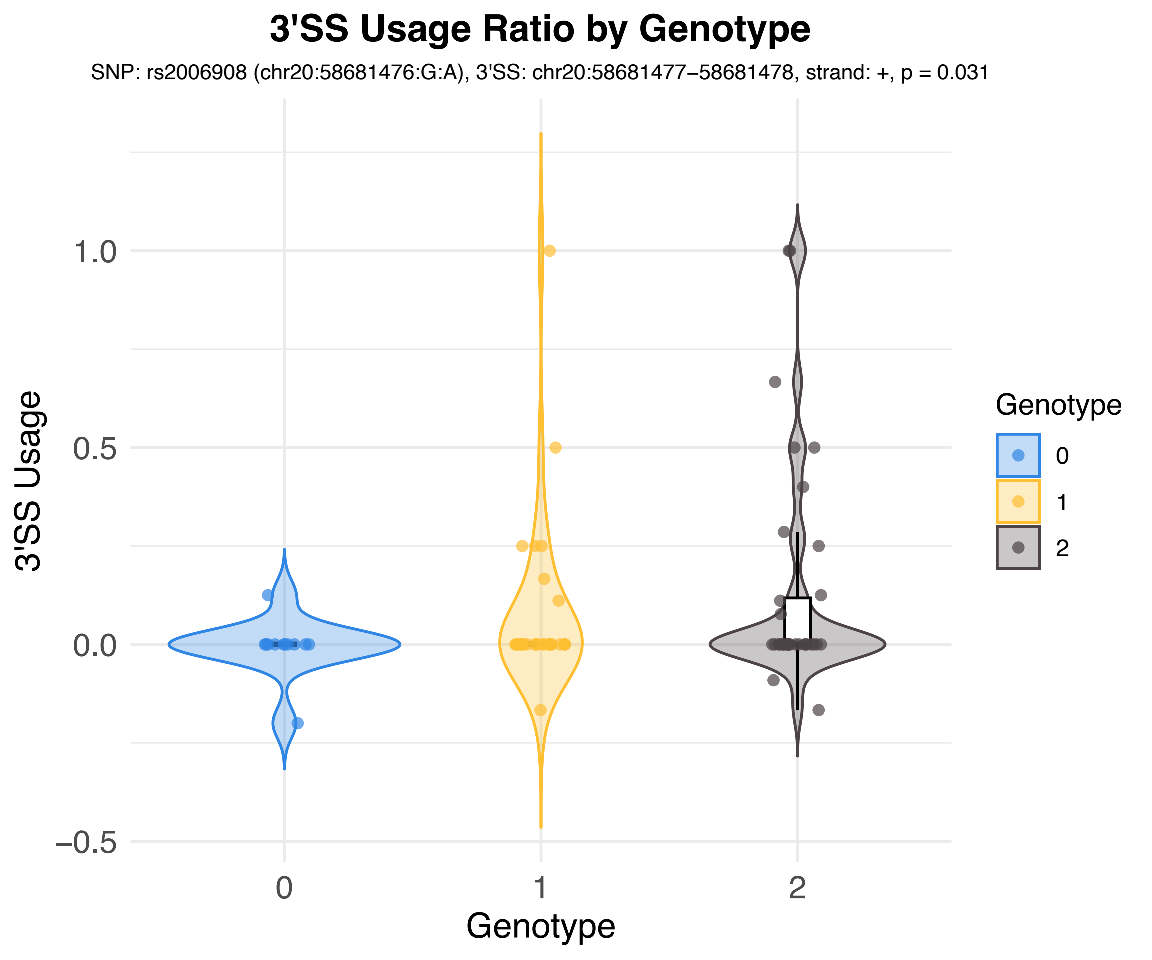
3'SS Usage Ratio by Genotype
SNP: rs2006908 (chr20:58681476:G:A), 3'SS: chr20:58681477–58681478, strand: +, p = 0.031
3'SS Usage
1.0
0.5
0.0
−0.5
0
1
2
Genotype
Genotype
0
1
2

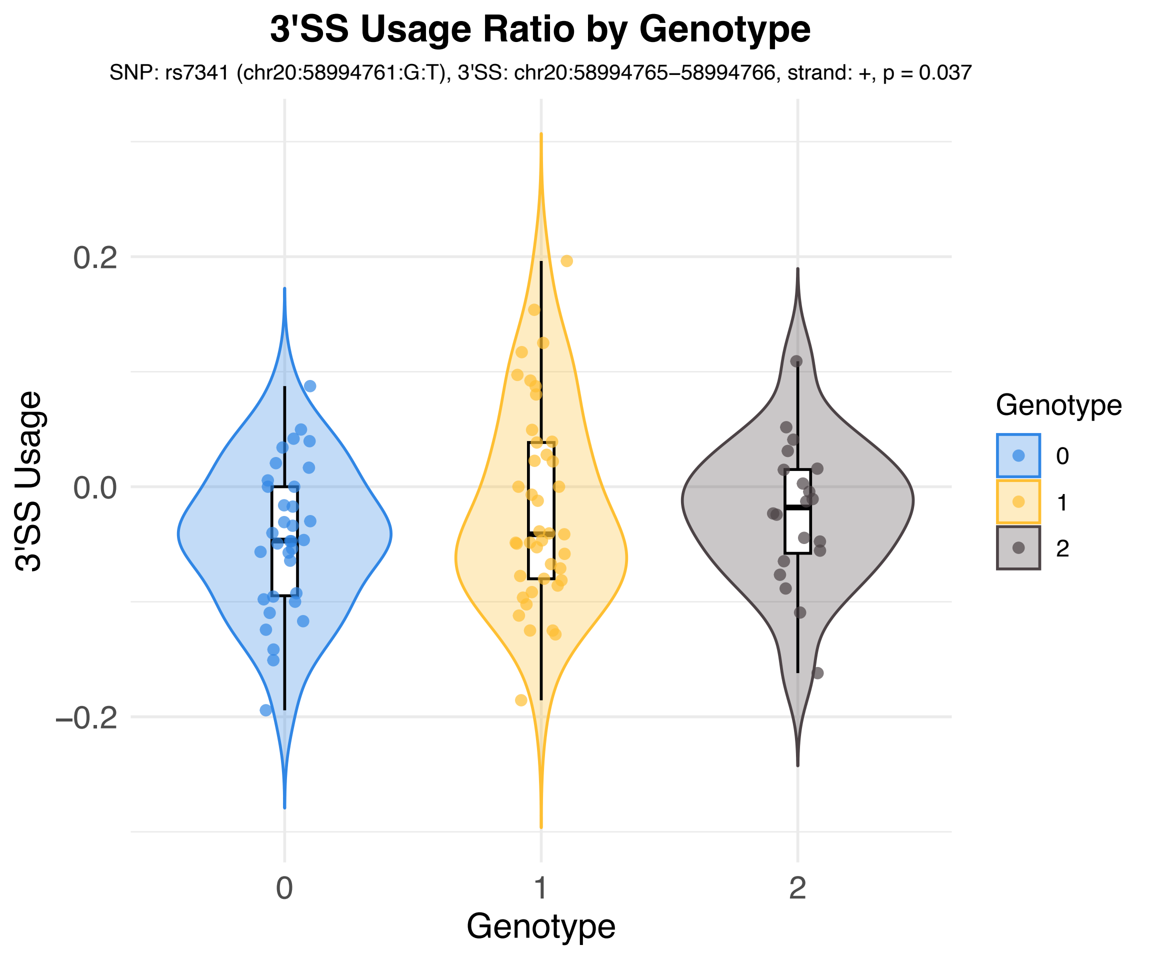
3'SS Usage Ratio by Genotype
SNP: rs7341 (chr20:58994761:G:T), 3'SS: chr20:58994765–58994766, strand: +, p = 0.037
3'SS Usage
0.2
0.0
−0.2
0
1
2
Genotype
Genotype
0
1
2

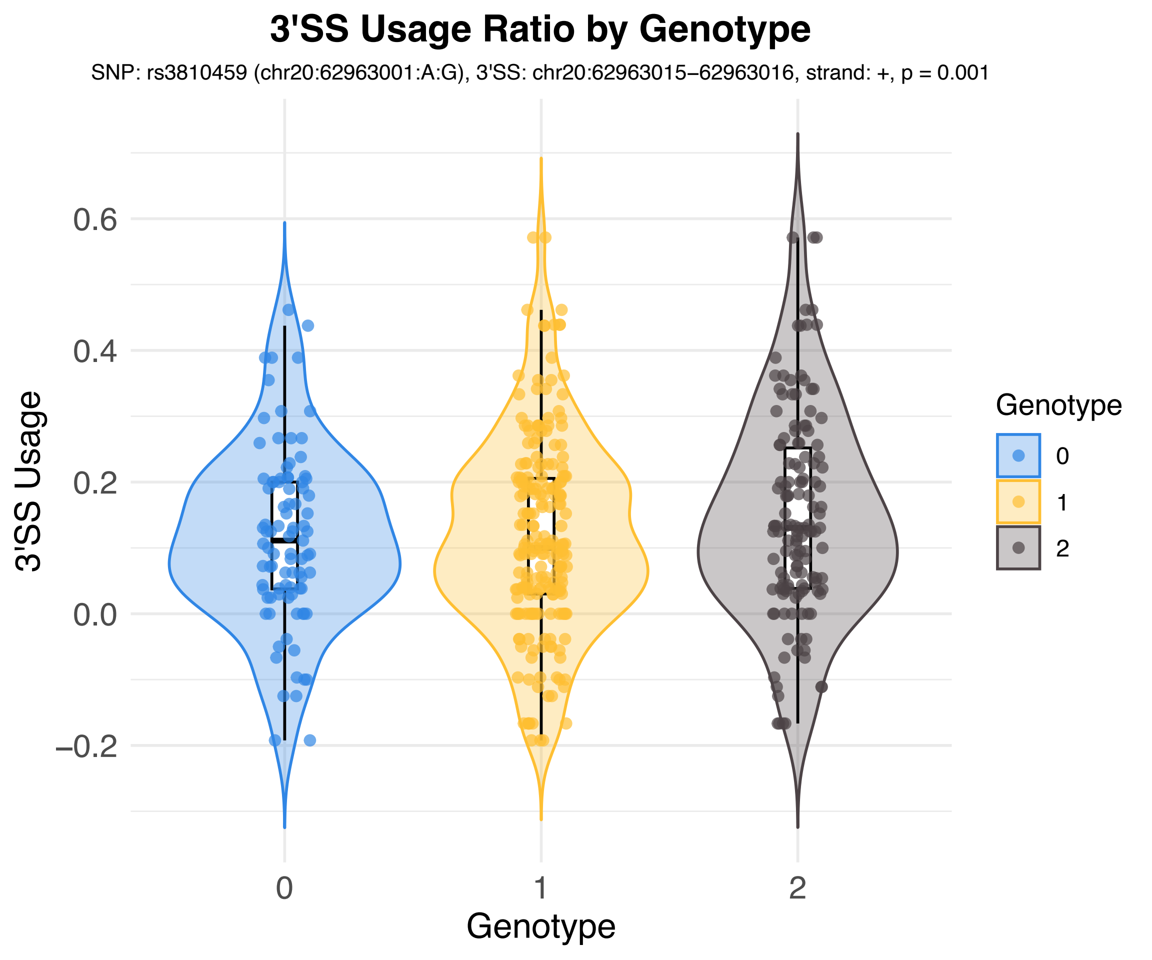
3'SS Usage Ratio by Genotype
SNP: rs3810459 (chr20:62963001:A:G), 3'SS: chr20:62963015–62963016, strand: +, p = 0.001
3'SS Usage
0.6
0.4
0.2
0.0
–0.2
0
1
2
Genotype
Genotype
0
1
2

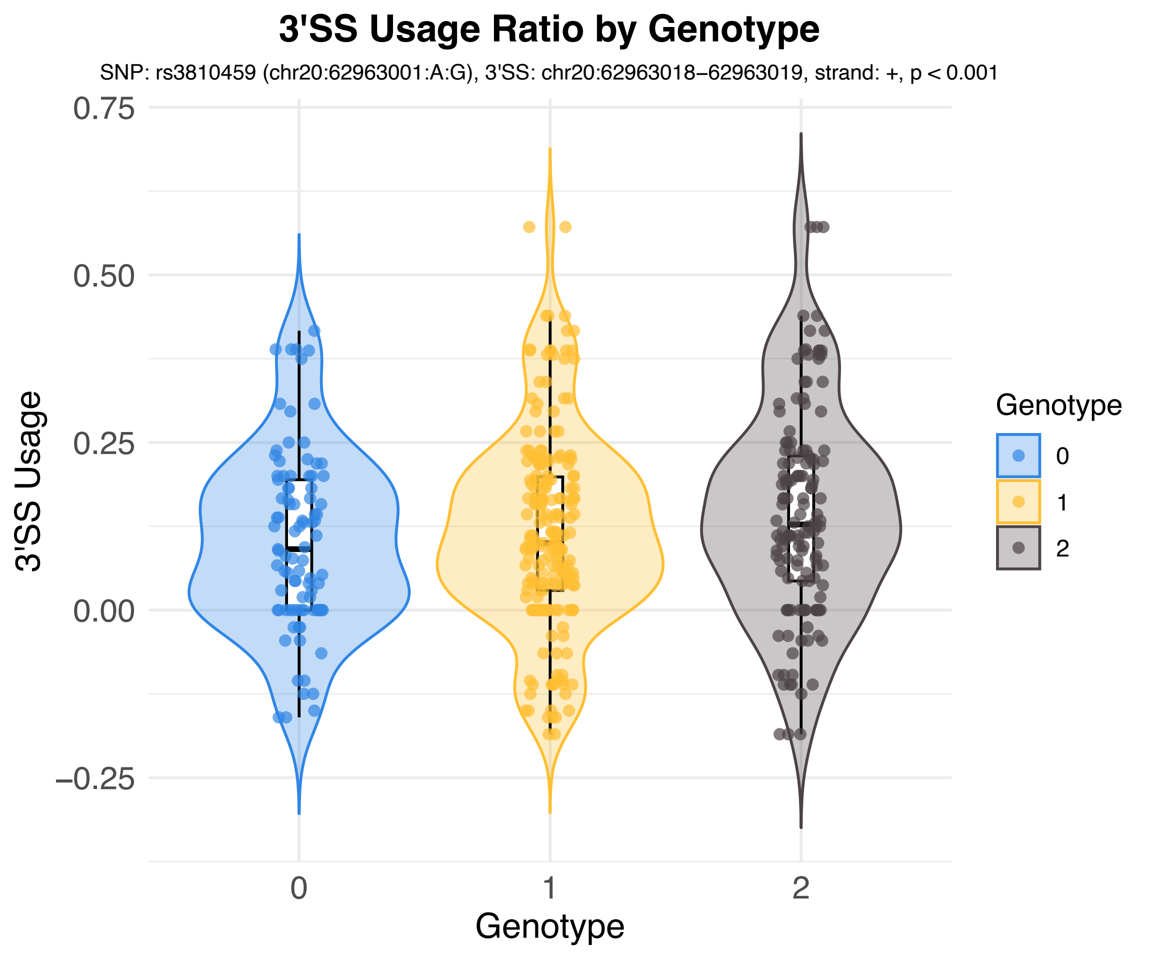
3'SS Usage Ratio by Genotype
SNP: rs3810459 (chr20:62963001:A:G), 3'SS: chr20:62963018–62963019, strand: +, p < 0.001
3'SS Usage
0.75
0.50
0.25
0.00
–0.25
0
1
2
Genotype
Genotype
0
1
2

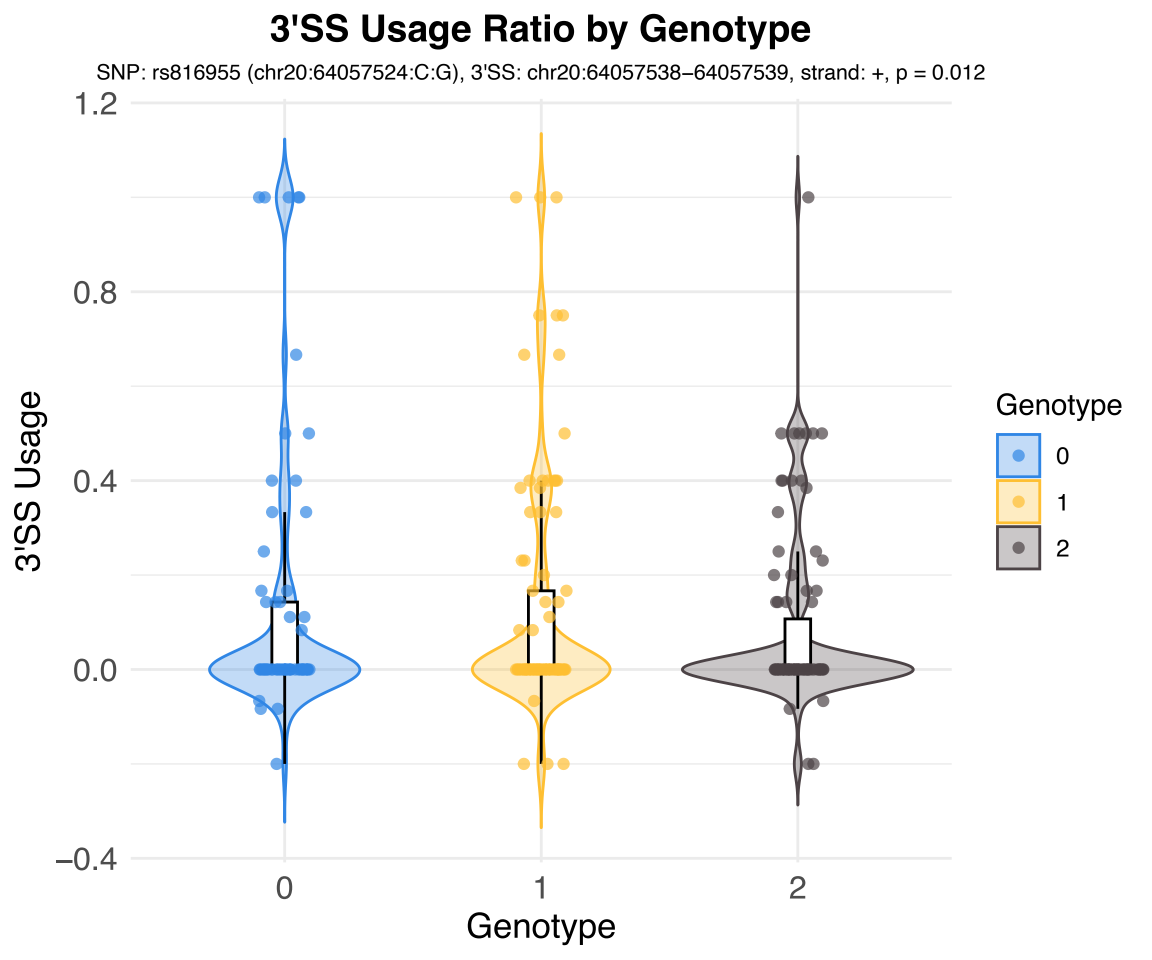

3'SS Usage Ratio by Genotype
SNP: rs816955 (chr20:64057524:C:G), 3'SS: chr20:64057538–64057539, strand: +, p = 0.012
3'SS Usage
1.2
0.8
0.4
0.0
−0.4
0
1
2
Genotype
Genotype
0
1
2

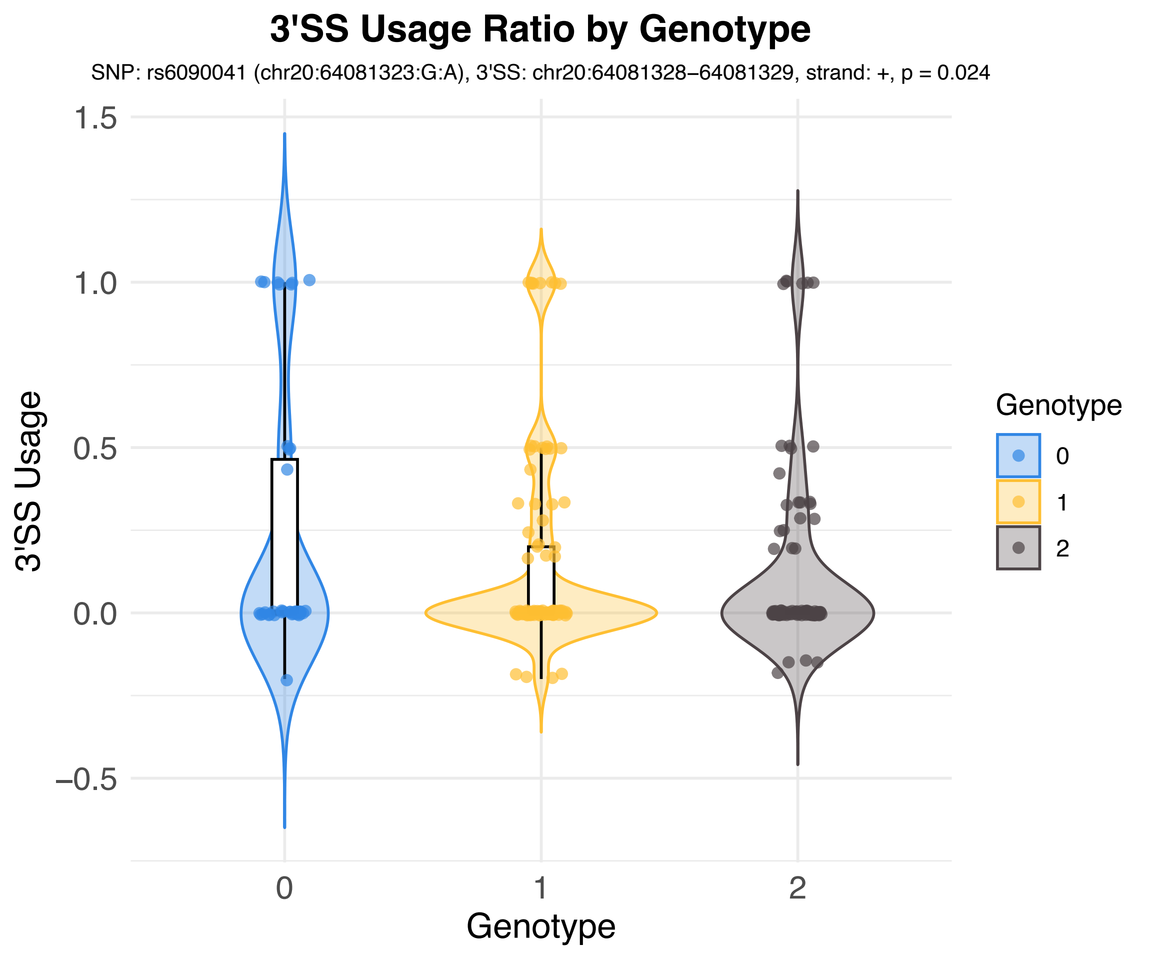
3'SS Usage Ratio by Genotype
SNP: rs6090041 (chr20:64081323:G:A), 3'SS: chr20:64081328–64081329, strand: +, p = 0.024
3'SS Usage
1.5
1.0
0.5
0.0
–0.5
0
1
2
Genotype
Genotype
0
1
2

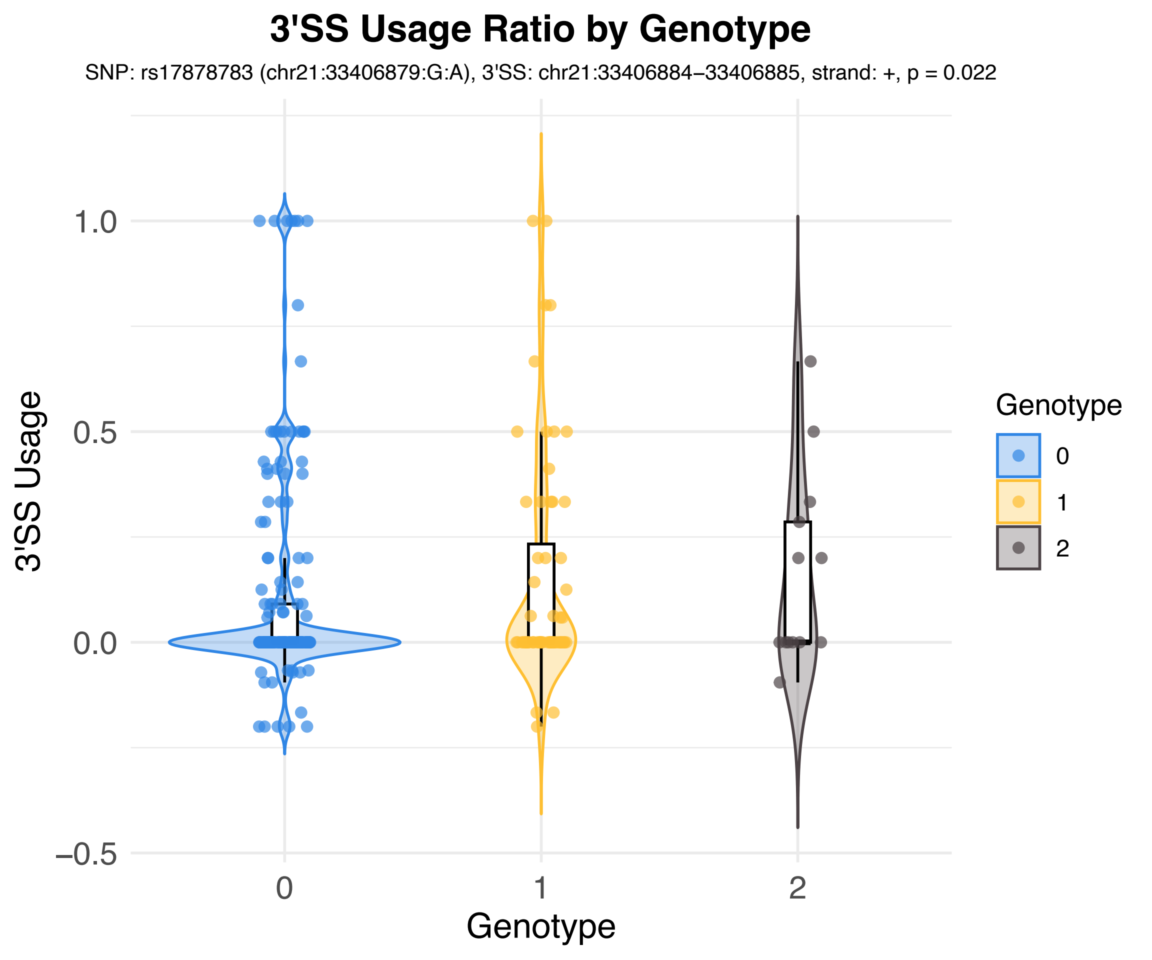
3'SS Usage Ratio by Genotype
SNP: rs17878783 (chr21:33406879:G:A), 3'SS: chr21:33406884–33406885, strand: +, p = 0.022
3'SS Usage
1.0
0.5
0.0
–0.5
0
1
2
Genotype
Genotype
0
1
2

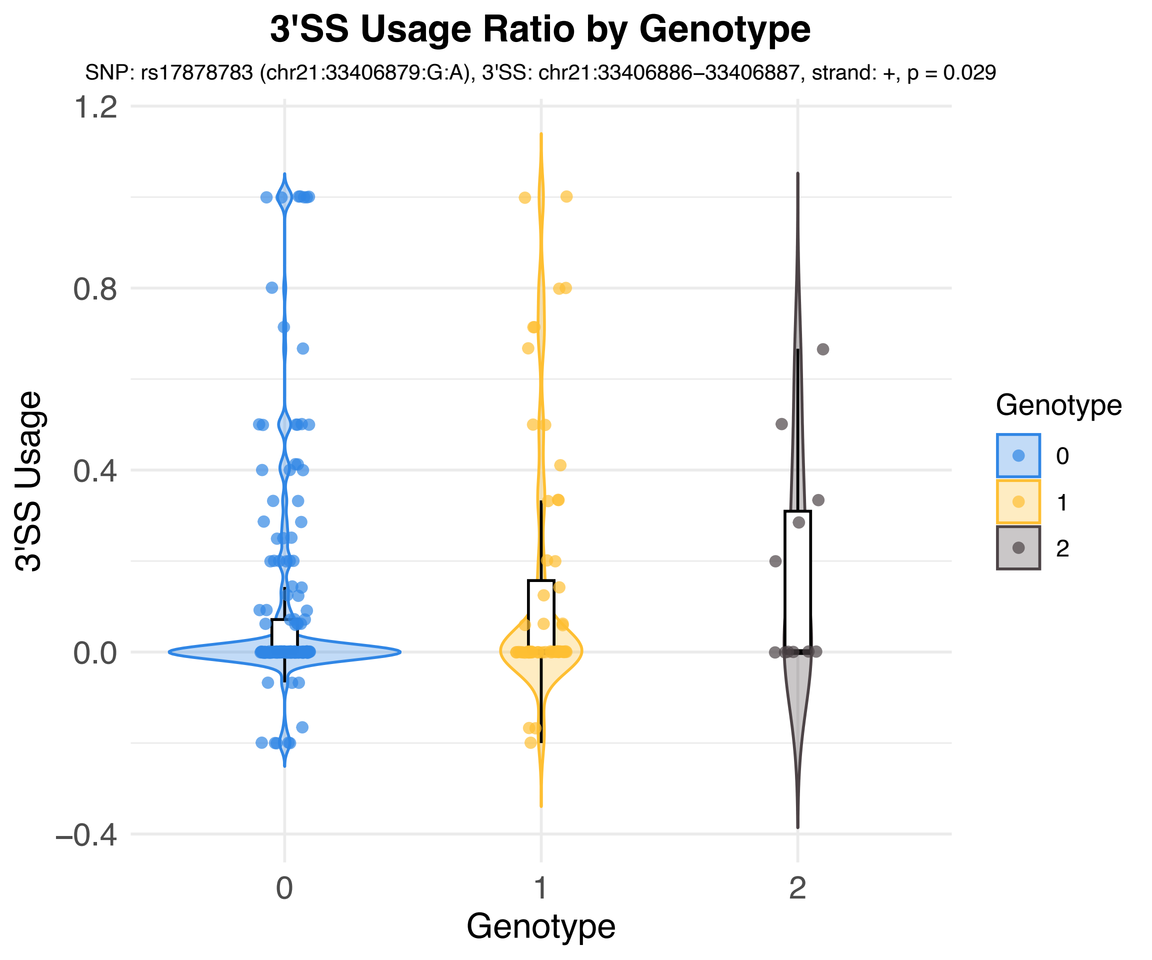
3'SS Usage Ratio by Genotype
SNP: rs17878783 (chr21:33406879:G:A), 3'SS: chr21:33406886–33406887, strand: +, p = 0.029
1.2
0.8
0.4
0.0
–0.4
3'SS Usage
0
1
2
Genotype
Genotype
0
1
2

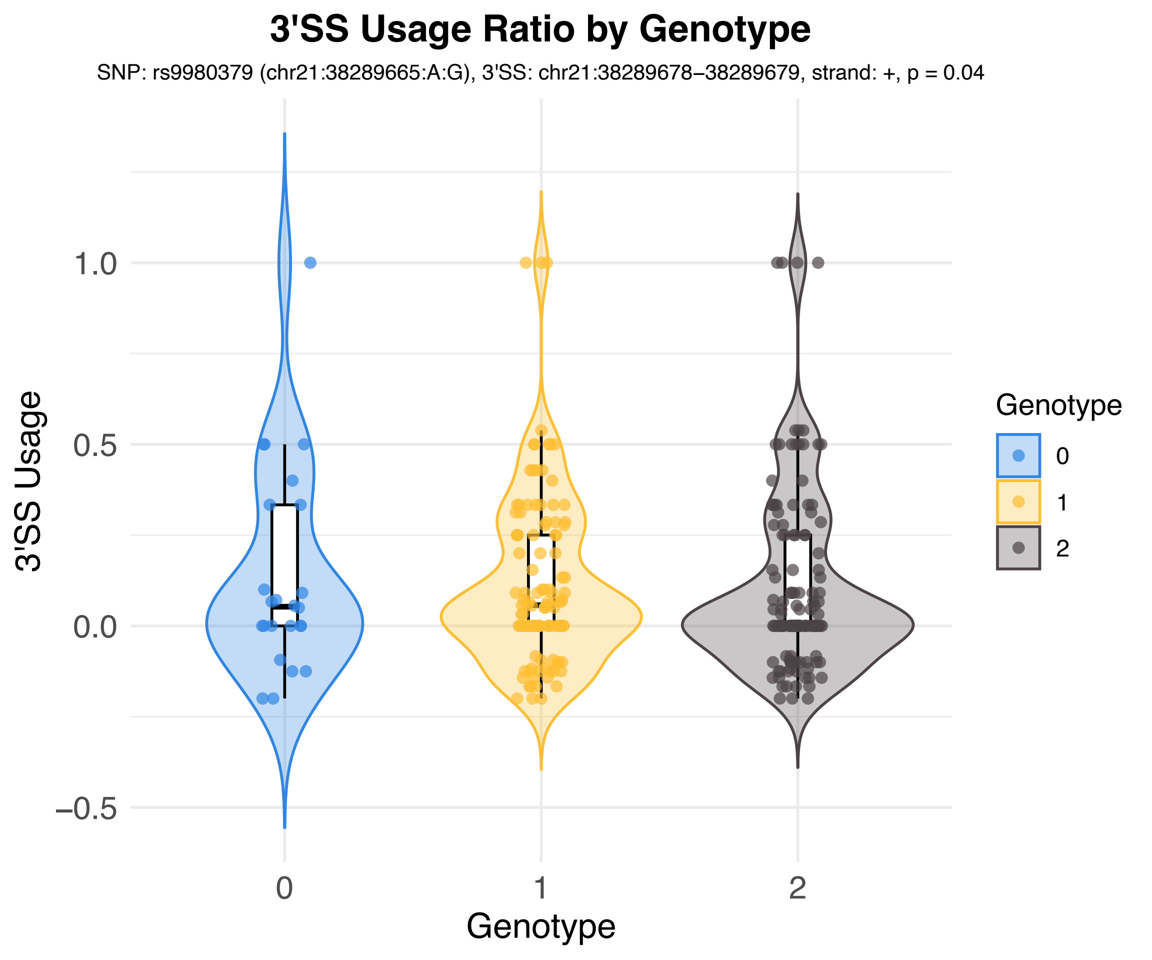

3'SS Usage Ratio by Genotype
SNP: rs9980379 (chr21:38289665:A:G), 3'SS: chr21:38289678–38289679, strand: +, p = 0.04
3'SS Usage
1.0
0.5
0.0
−0.5
0
1
2
Genotype
Genotype
0
1
2

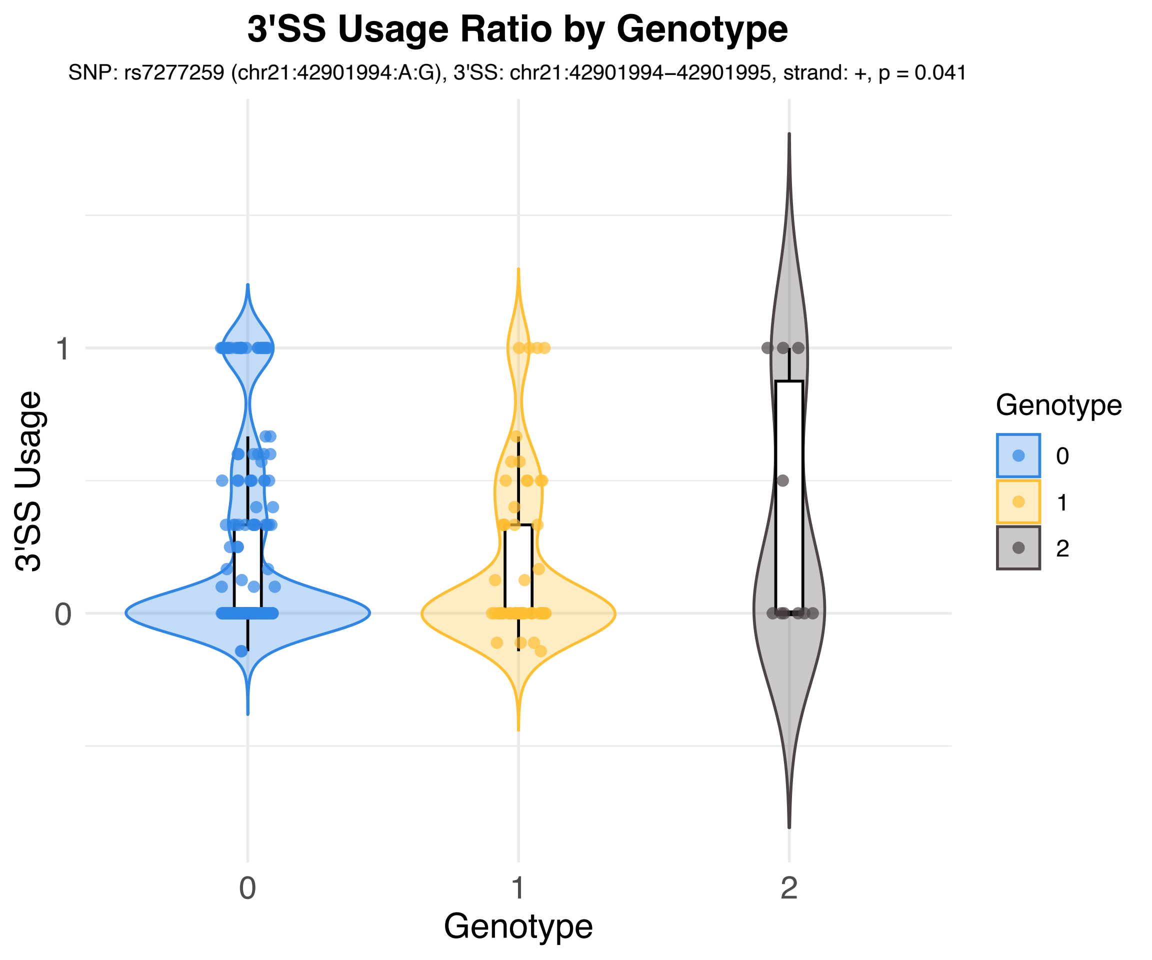
3'SS Usage Ratio by Genotype
SNP: rs7277259 (chr21:42901994:A:G), 3'SS: chr21:42901994–42901995, strand: +, p = 0.041
3'SS Usage
1
0
0
1
2
Genotype
Genotype
0
1
2

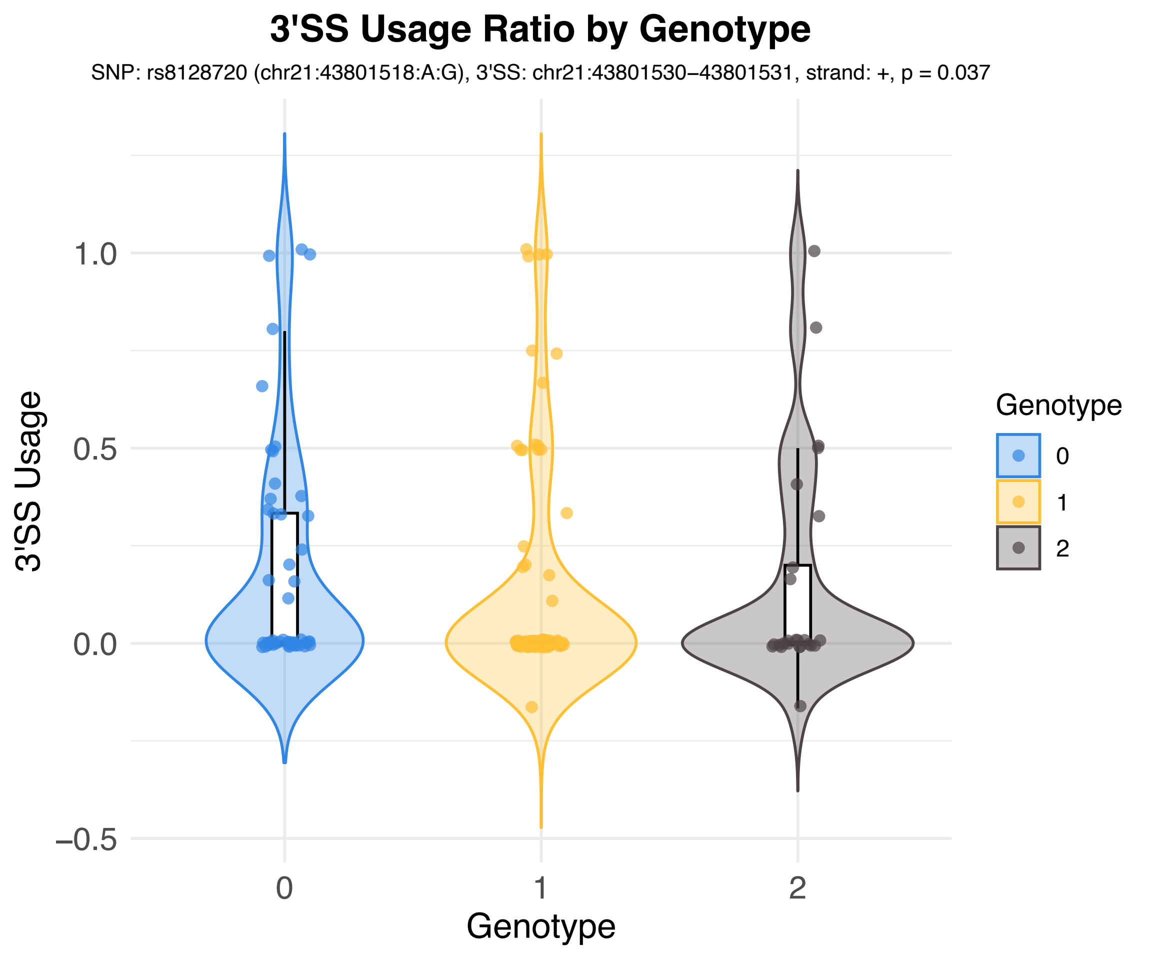
3'SS Usage Ratio by Genotype
SNP: rs8128720 (chr21:43801518:A:G), 3'SS: chr21:43801530−43801531, strand: +, p = 0.037
3'SS Usage
1.0
0.5
0.0
−0.5
0
1
2
Genotype
Genotype
0
1
2

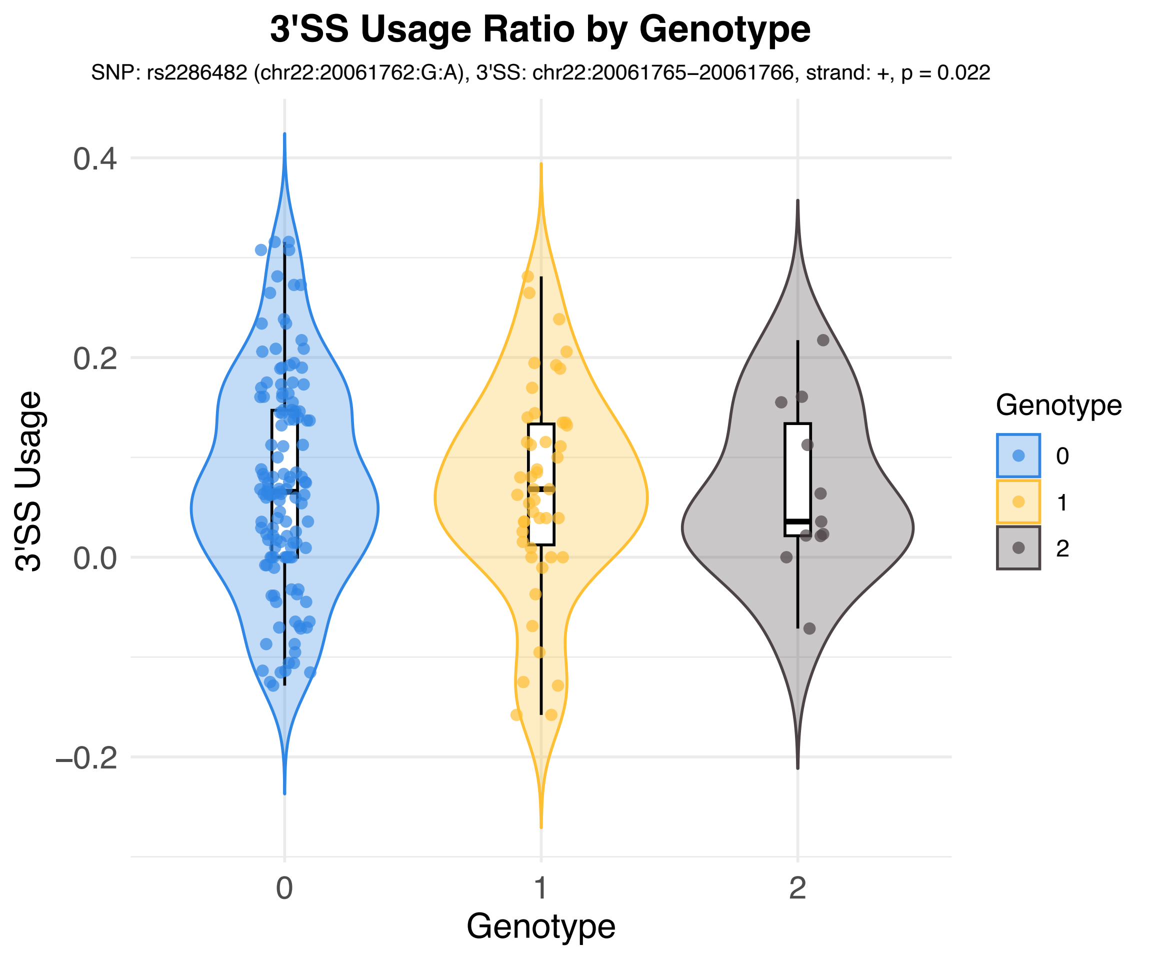
3'SS Usage Ratio by Genotype
SNP: rs2286482 (chr22:20061762:G:A), 3'SS: chr22:20061765–20061766, strand: +, p = 0.022
3'SS Usage
0.4
0.2
0.0
−0.2
0
1
2
Genotype
Genotype
0
1
2

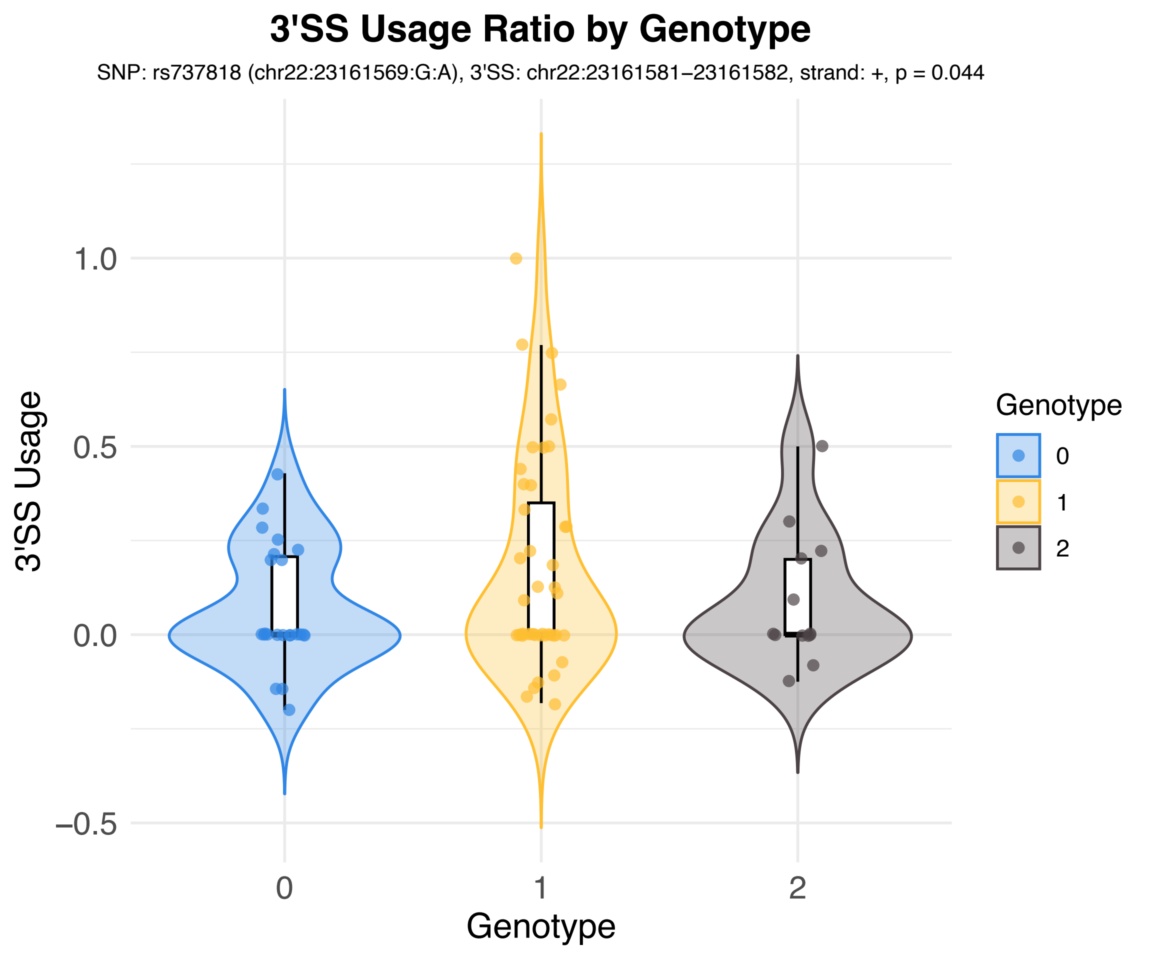
3'SS Usage Ratio by Genotype
SNP: rs737818 (chr22:23161569:G:A), 3'SS: chr22:23161581−23161582, strand: +, p = 0.044
3'SS Usage
1.0
0.5
0.0
−0.5
0
1
2
Genotype
Genotype
0
1
2

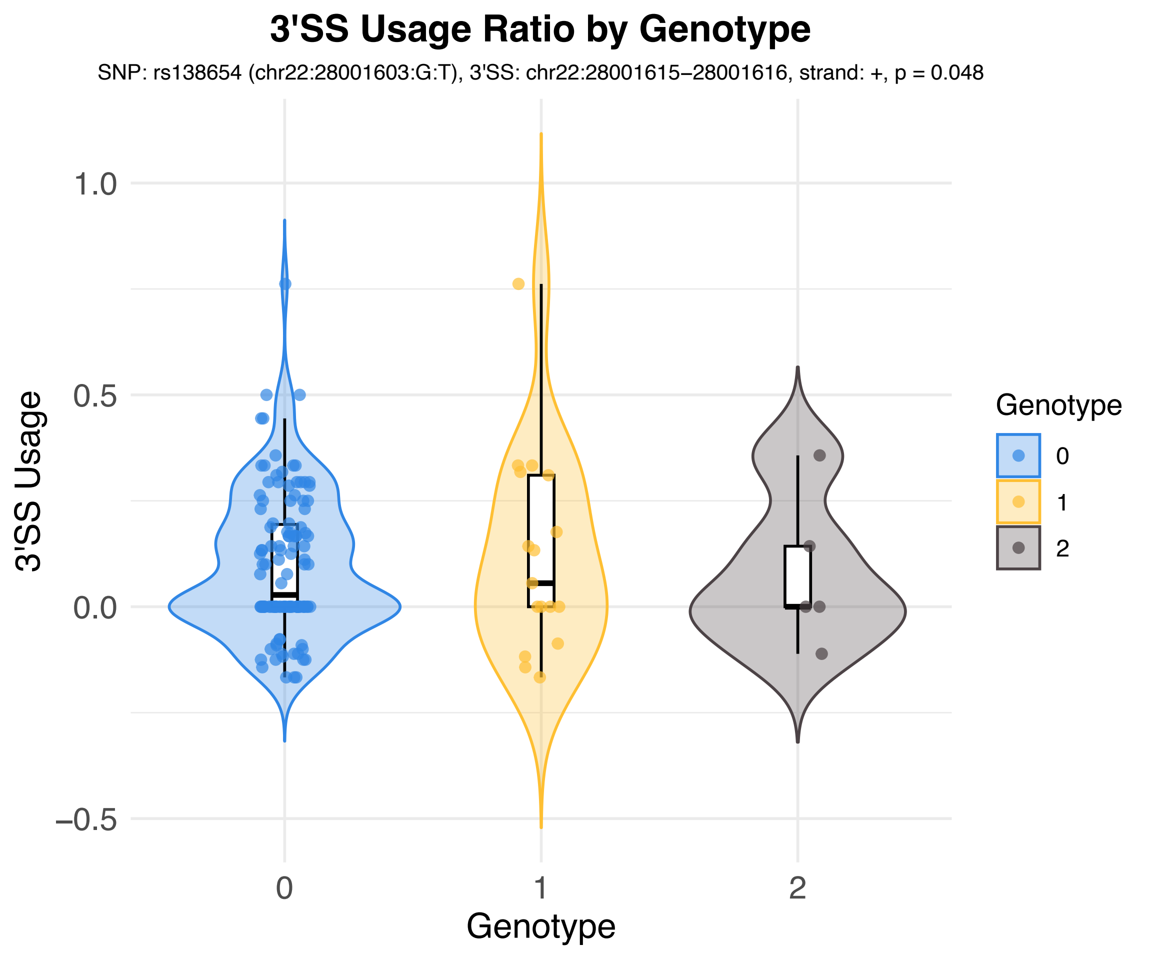
3'SS Usage Ratio by Genotype
SNP: rs138654 (chr22:28001603:G:T), 3'SS: chr22:28001615−28001616, strand: +, p = 0.048
3'SS Usage
1.0
0.5
0.0
−0.5
0
1
2
Genotype
Genotype
0
1
2

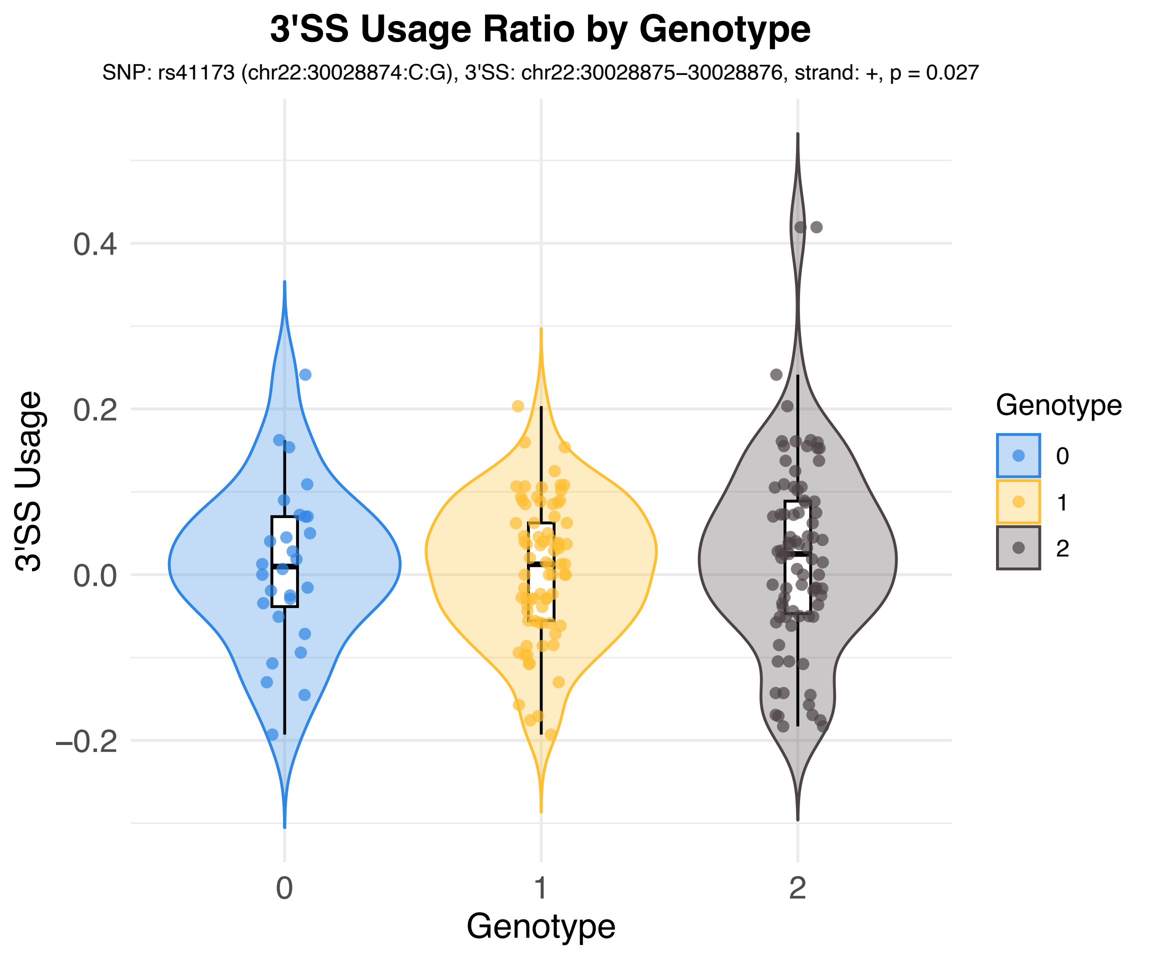

3'SS Usage Ratio by Genotype
SNP: rs41173 (chr22:30028874:C:G), 3'SS: chr22:30028875–30028876, strand: +, p = 0.027
3'SS Usage
0.4
0.2
0.0
−0.2
0
1
2
Genotype
Genotype
0
1
2

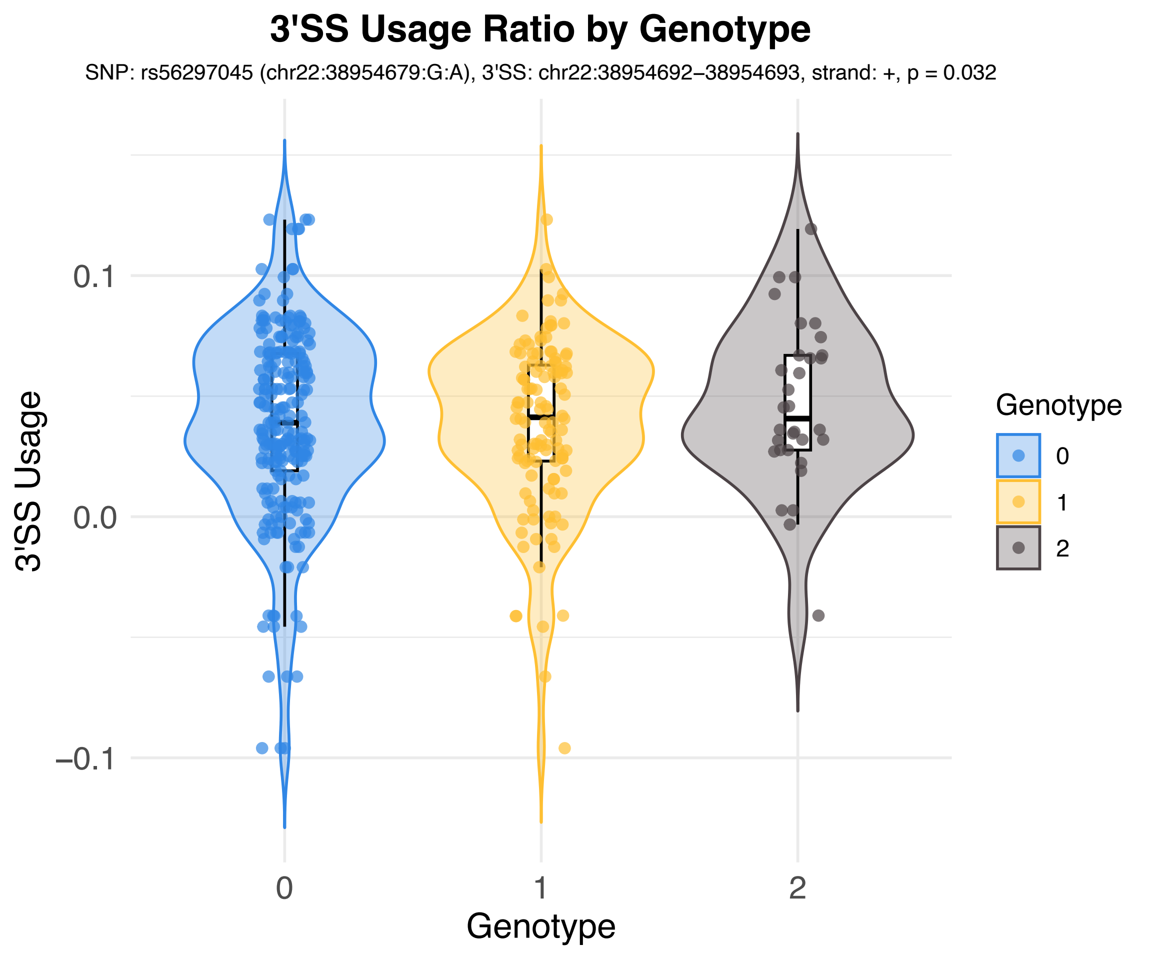
3'SS Usage Ratio by Genotype
SNP: rs56297045 (chr22:38954679:G:A), 3'SS: chr22:38954692–38954693, strand: +, p = 0.032
3'SS Usage
0.1
0.0
−0.1
0
1
2
Genotype
Genotype
0
1
2

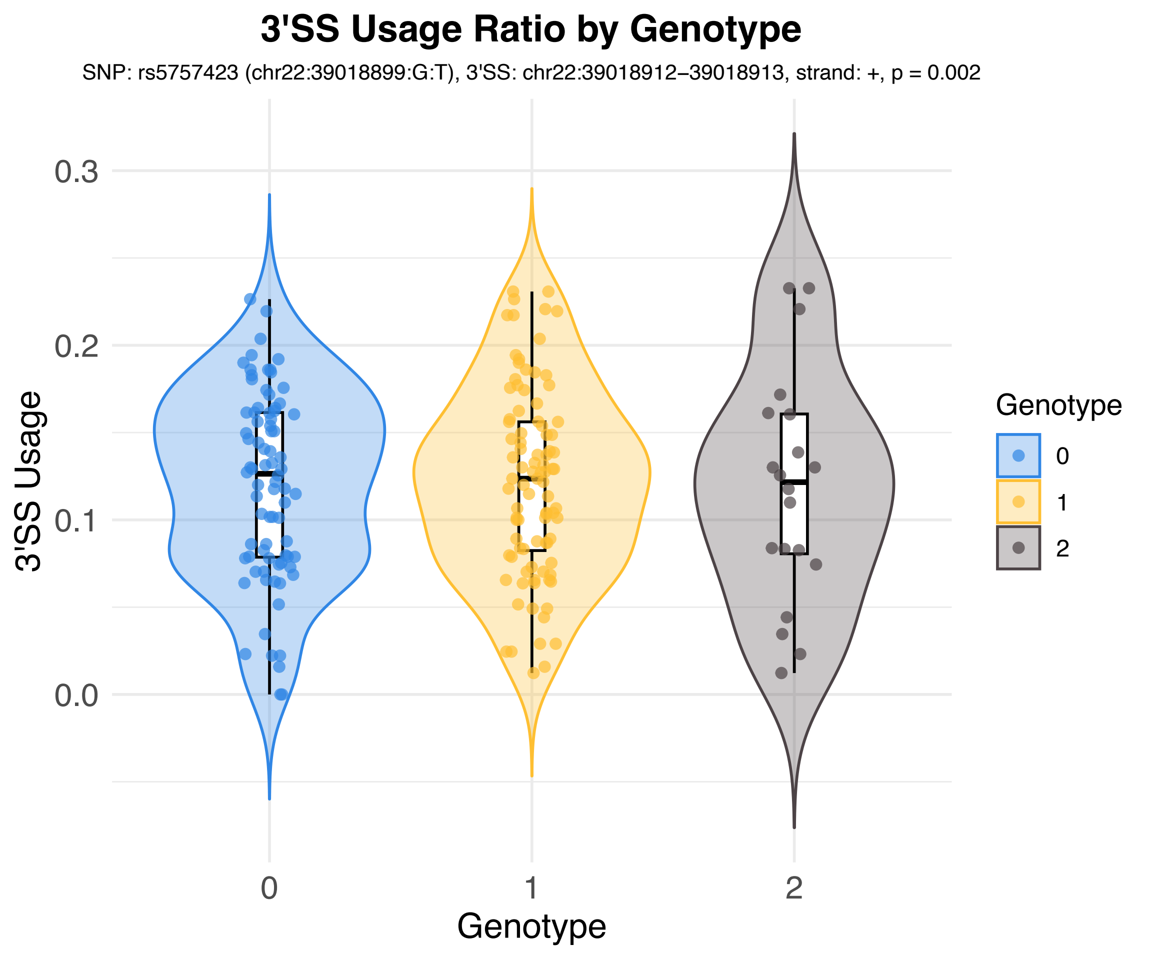
3'SS Usage Ratio by Genotype
SNP: rs5757423 (chr22:39018899:G:T), 3'SS: chr22:39018912–39018913, strand: +, p = 0.002
3'SS Usage
0.3
0.2
0.1
0.0
0
1
2
Genotype
Genotype
0
1
2

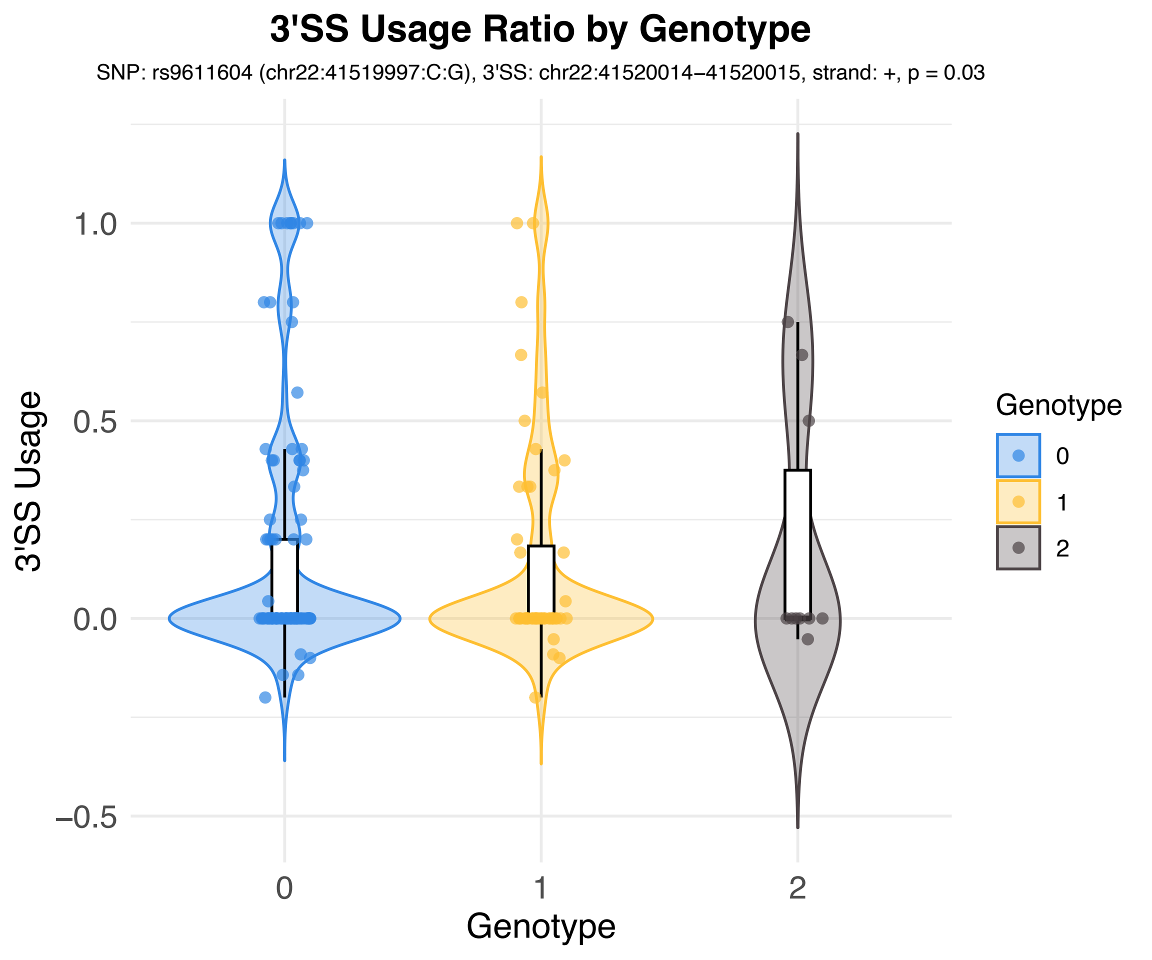
3'SS Usage Ratio by Genotype
SNP: rs9611604 (chr22:41519997:C:G), 3'SS: chr22:41520014–41520015, strand: +, p = 0.03
3'SS Usage
1.0
0.5
0.0
−0.5
0
1
2
Genotype
Genotype
0
1
2

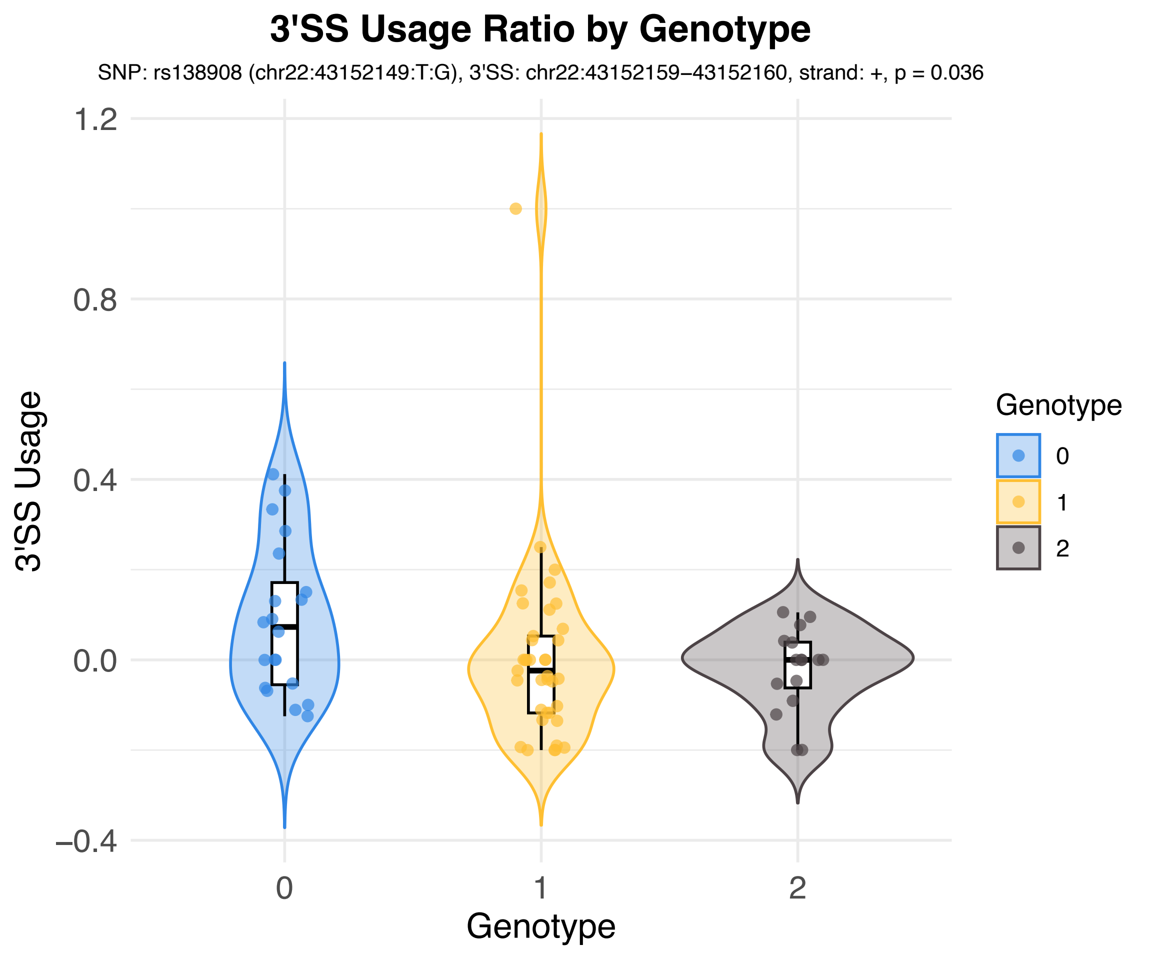
3'SS Usage Ratio by Genotype
SNP: rs138908 (chr22:43152149:T:G), 3'SS: chr22:43152159−43152160, strand: +, p = 0.036
3'SS Usage
1.2
0.8
0.4
0.0
−0.4
0
1
2
Genotype
Genotype
0
1
2

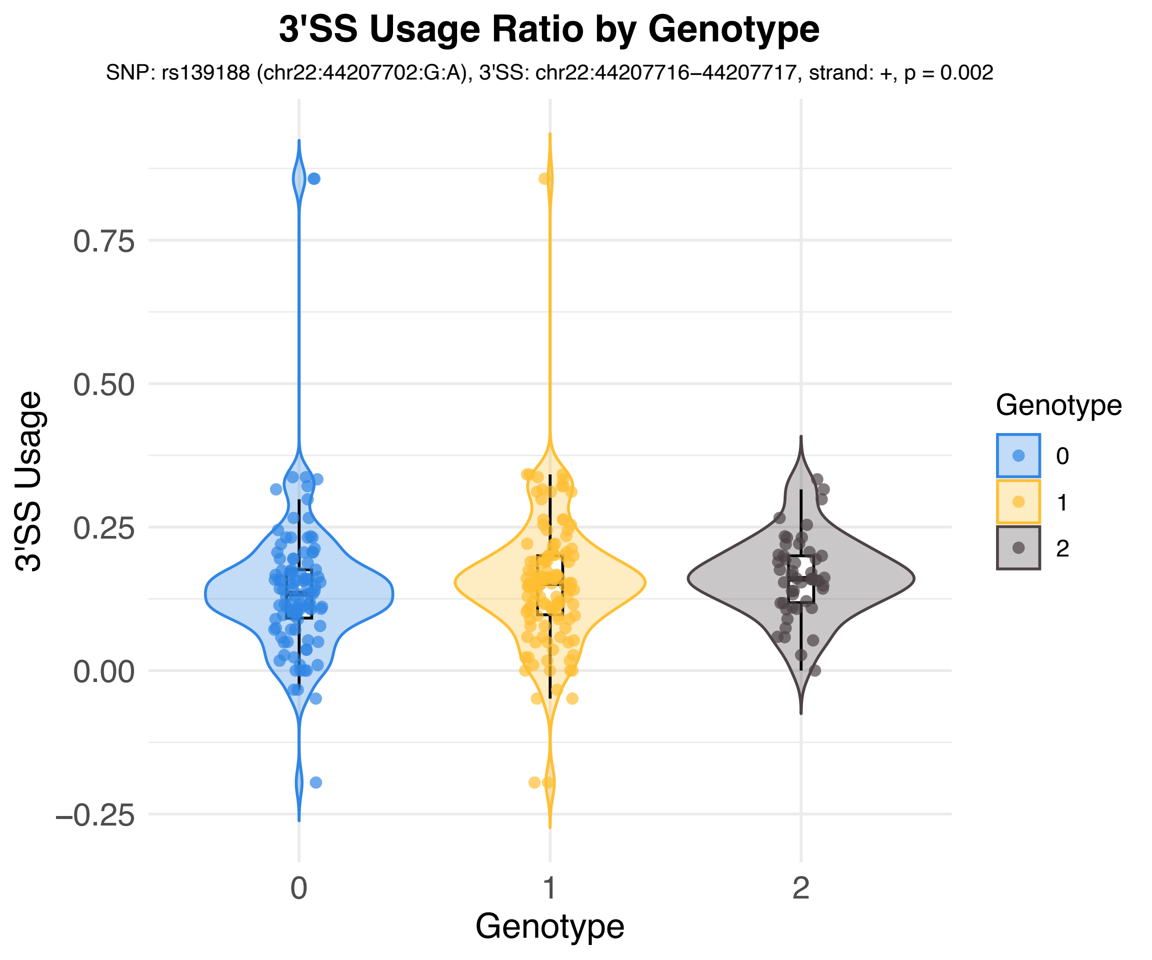
3'SS Usage Ratio by Genotype
SNP: rs139188 (chr22:44207702:G:A), 3'SS: chr22:44207716−44207717, strand: +, p = 0.002
3'SS Usage
0.75
0.50
0.25
0.00
−0.25
0
1
2
Genotype
Genotype
0
1
2

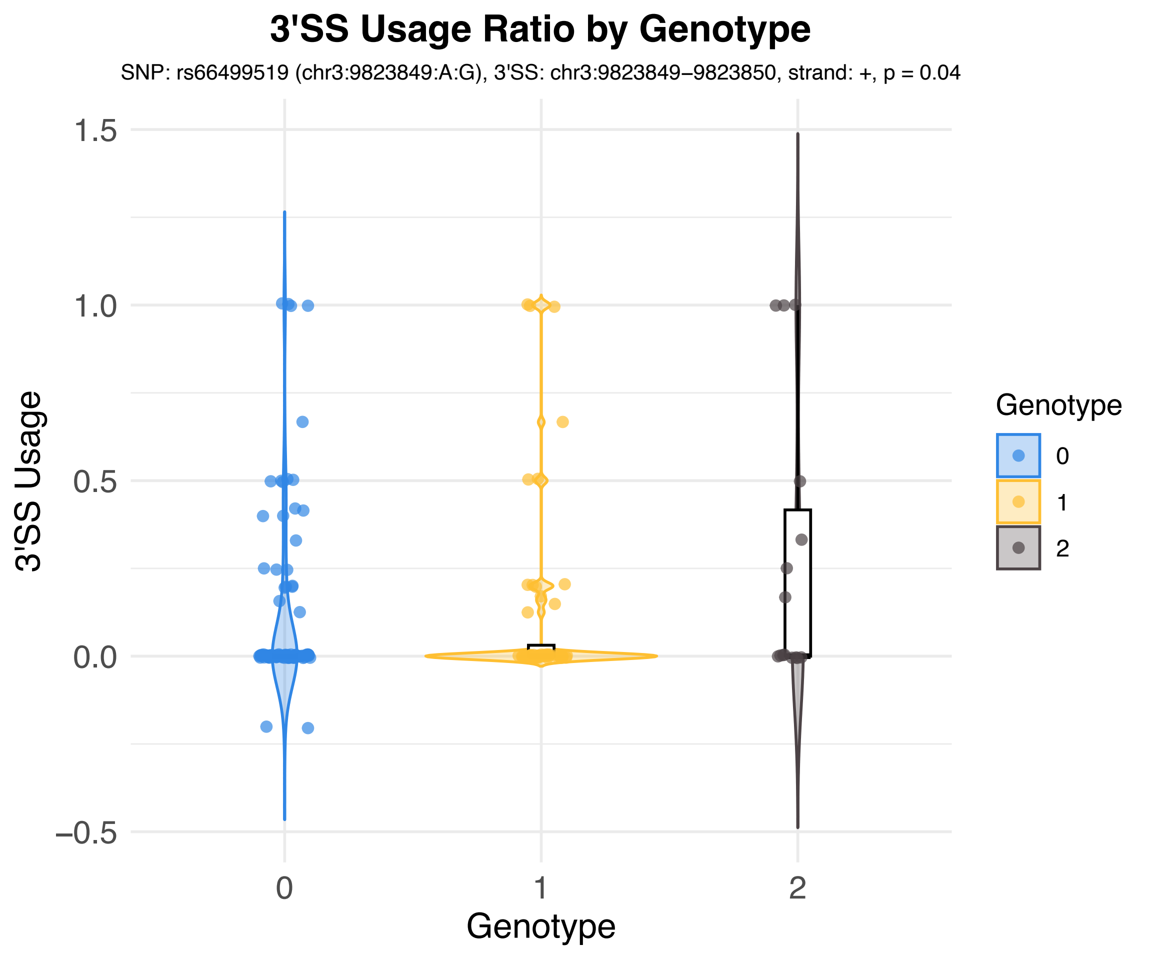

3'SS Usage Ratio by Genotype
SNP: rs66499519 (chr3:9823849:A:G), 3'SS: chr3:9823849−9823850, strand: +, p = 0.04
3'SS Usage
1.5
1.0
0.5
0.0
−0.5
0
1
2
Genotype
Genotype
0
1
2

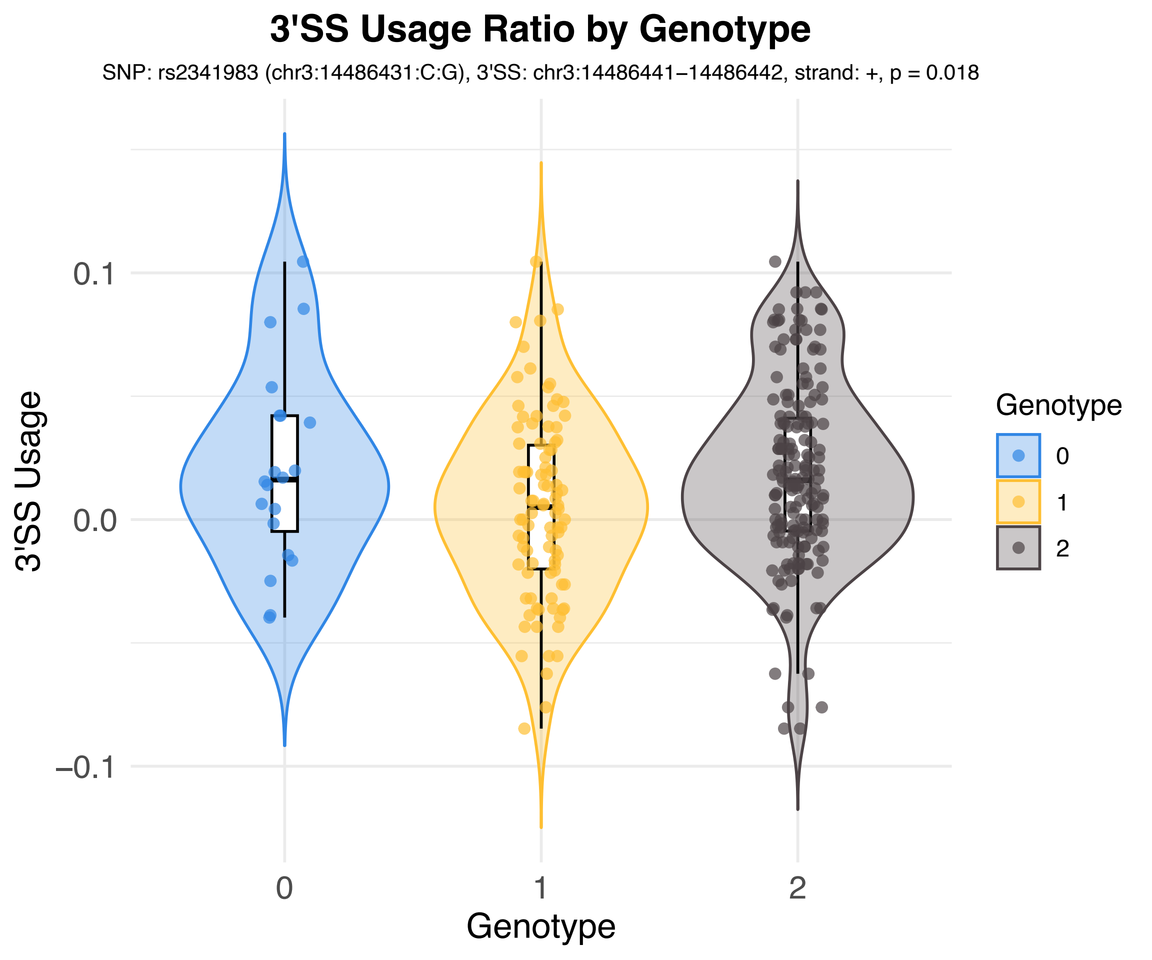
3'SS Usage Ratio by Genotype
SNP: rs2341983 (chr3:14486431:C:G), 3'SS: chr3:14486441−14486442, strand: +, p = 0.018
3'SS Usage
0.1
0.0
−0.1
0
1
2
Genotype
Genotype
0
1
2

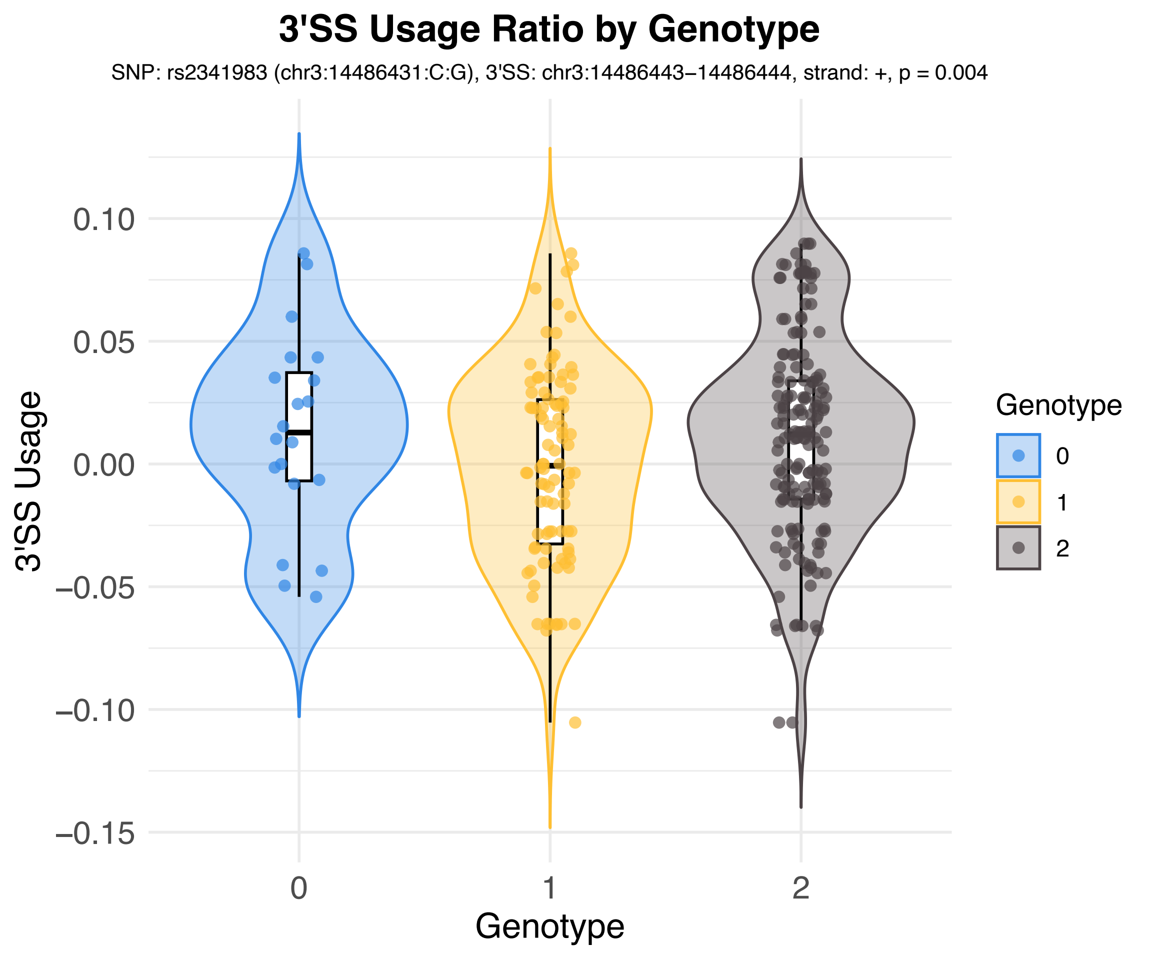
3'SS Usage Ratio by Genotype
SNP: rs2341983 (chr3:14486431:C:G), 3'SS: chr3:14486443−14486444, strand: +, p = 0.004
3'SS Usage
0.10
0.05
0.00
−0.05
−0.10
−0.15
0
1
2
Genotype
Genotype
0
1
2

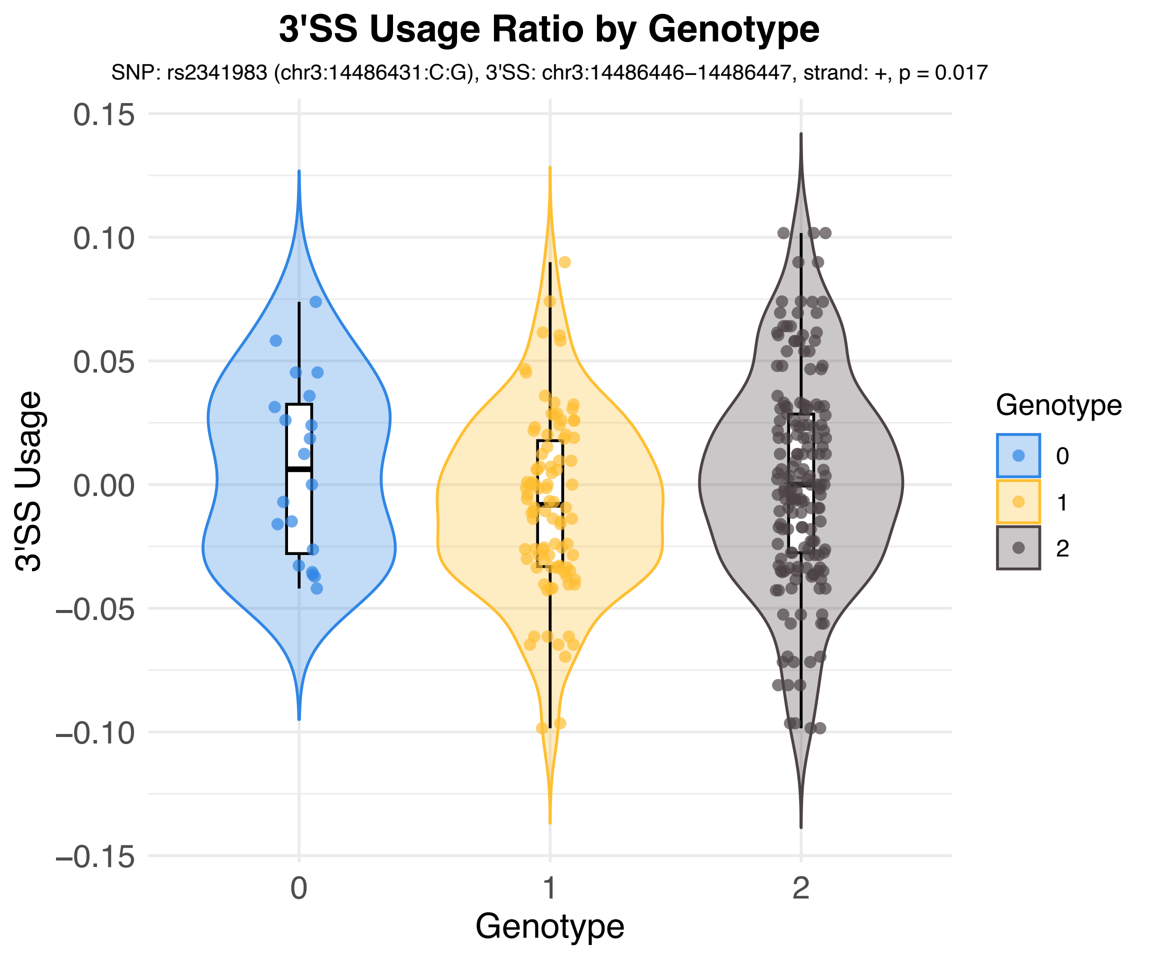

3'SS Usage Ratio by Genotype
SNP: rs2341983 (chr3:14486431:C:G), 3'SS: chr3:14486446−14486447, strand: +, p = 0.017
3'SS Usage
0.15
0.10
0.05
0.00
−0.05
−0.10
−0.15
0
1
2
Genotype
Genotype
0
1
2

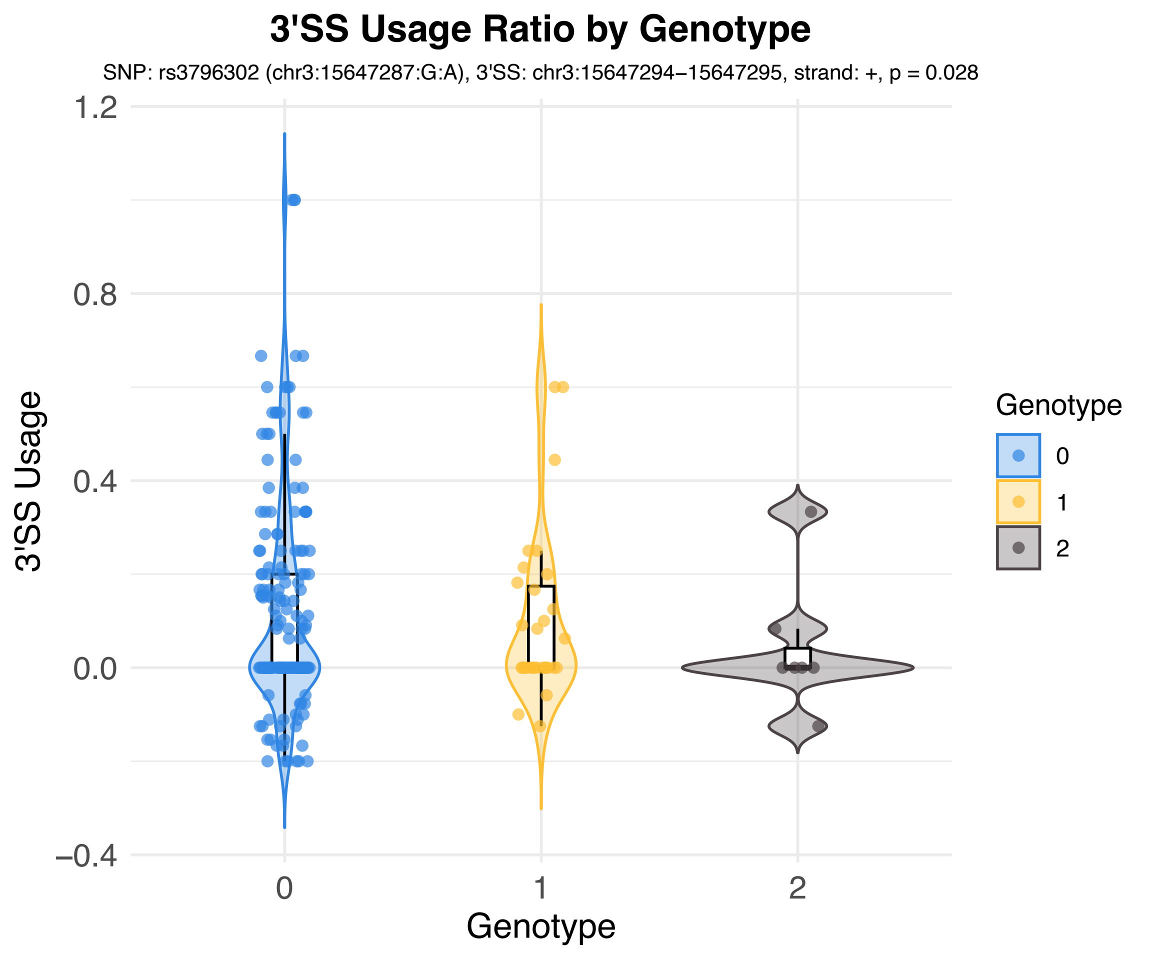
3'SS Usage Ratio by Genotype
SNP: rs3796302 (chr3:15647287:G:A), 3'SS: chr3:15647294−15647295, strand: +, p = 0.028
3'SS Usage
1.2
0.8
0.4
0.0
−0.4
0
1
2
Genotype
Genotype
0
1
2

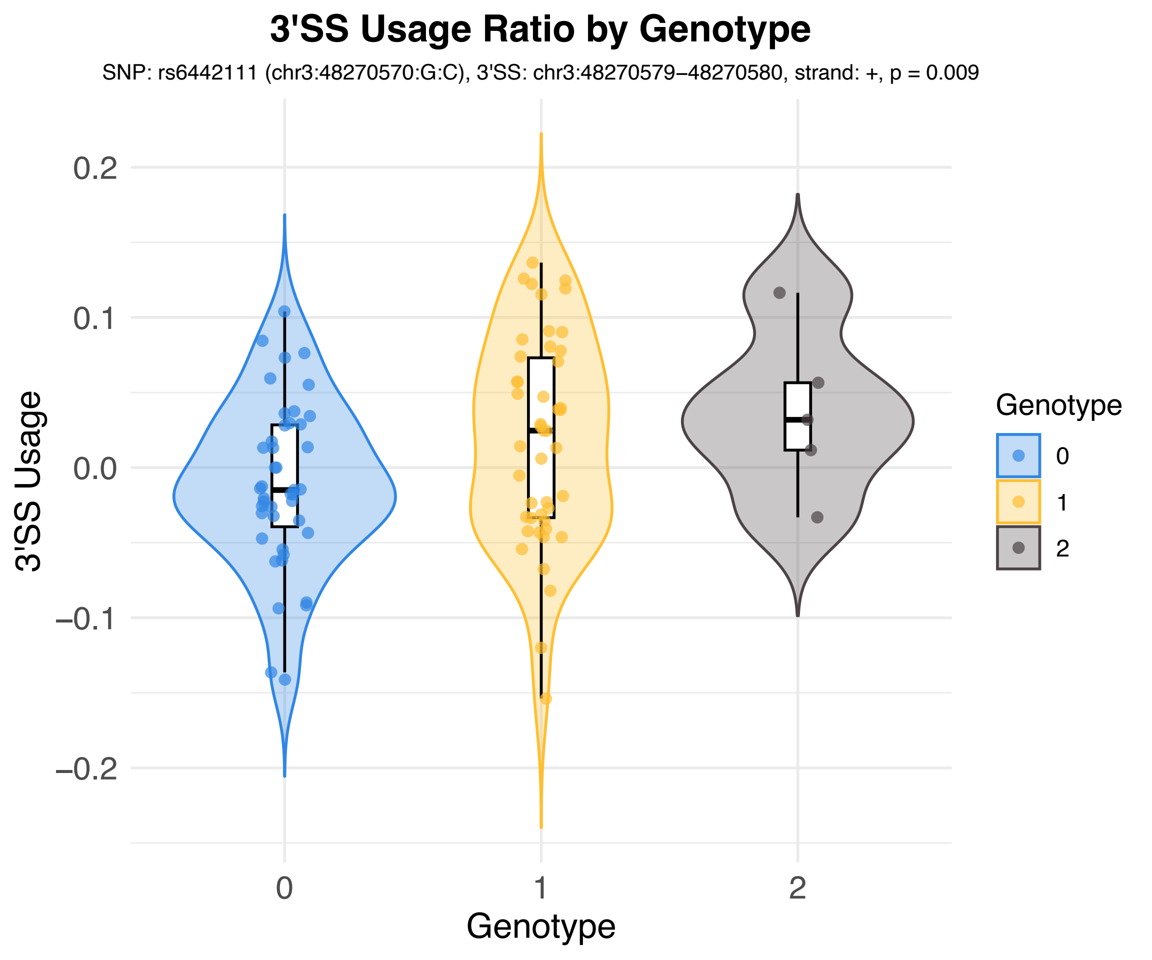
3'SS Usage Ratio by Genotype
SNP: rs6442111 (chr3:48270570:G:C), 3'SS: chr3:48270579–48270580, strand: +, p = 0.009
3'SS Usage
0.2
0.1
0.0
−0.1
−0.2
0
1
2
Genotype
Genotype
0
1
2

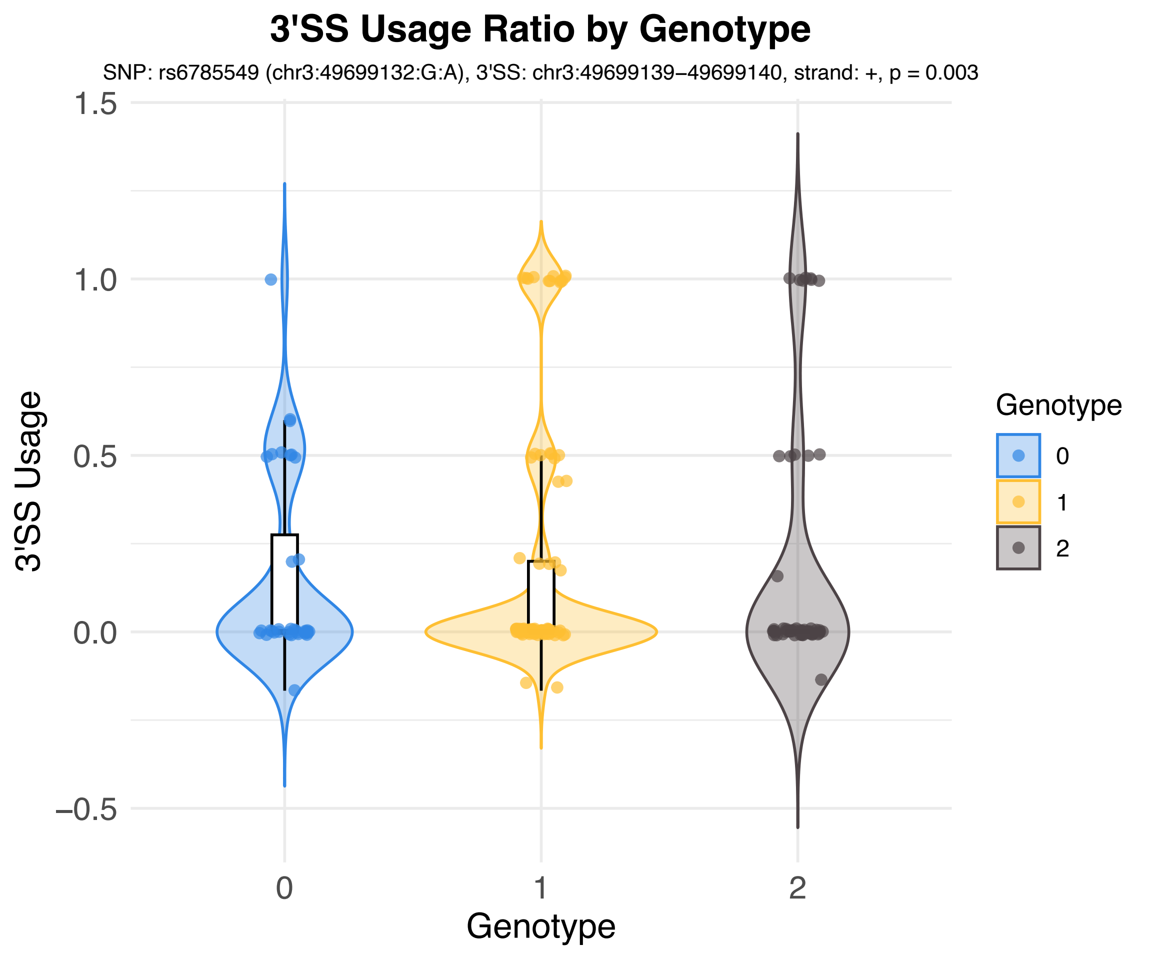
3'SS Usage Ratio by Genotype
SNP: rs6785549 (chr3:49699132:G:A), 3'SS: chr3:49699139−49699140, strand: +, p = 0.003
3'SS Usage
1.5
1.0
0.5
0.0
−0.5
0
1
2
Genotype
Genotype
0
1
2

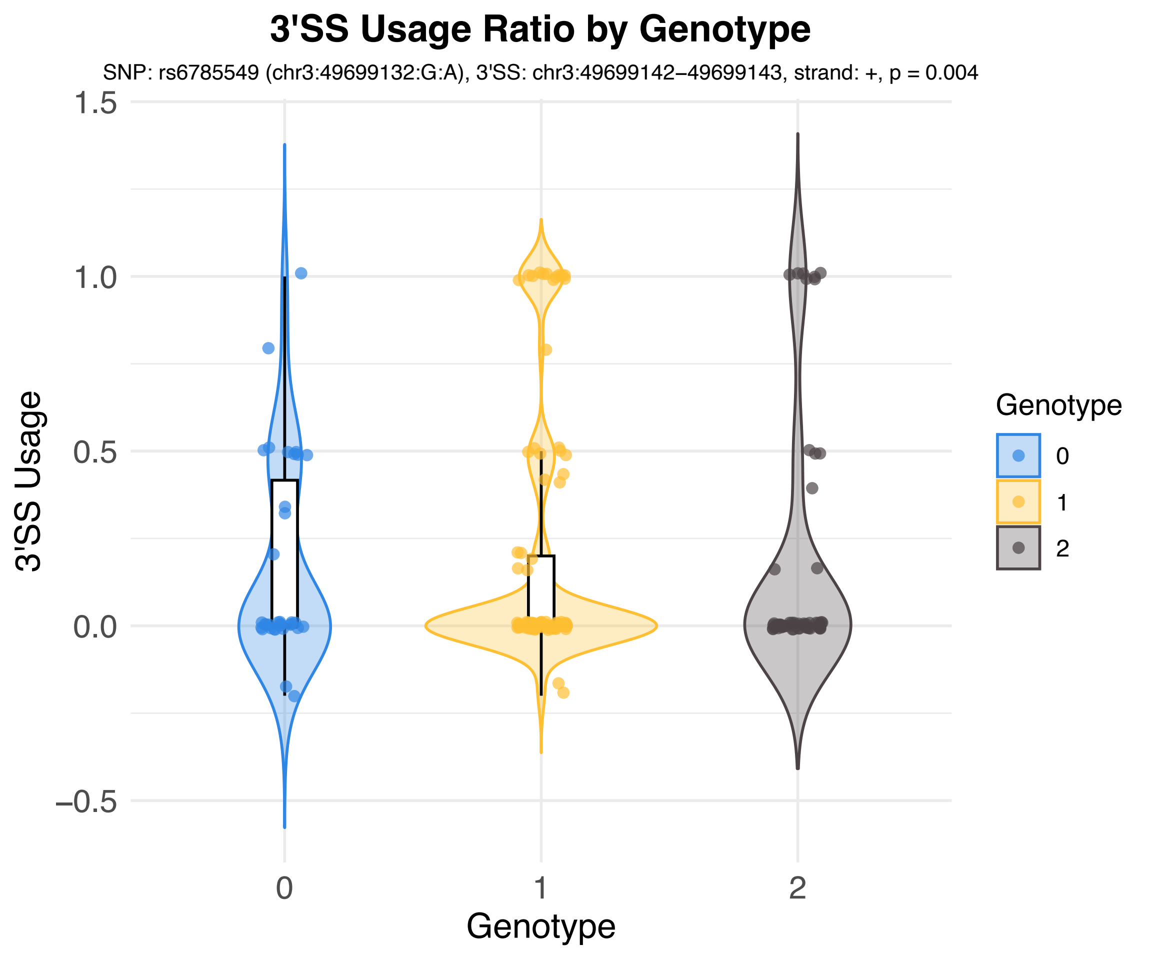
3'SS Usage Ratio by Genotype
SNP: rs6785549 (chr3:49699132:G:A), 3'SS: chr3:49699142−49699143, strand: +, p = 0.004
3'SS Usage
1.5
1.0
0.5
0.0
−0.5
0
1
2
Genotype
Genotype
0
1
2

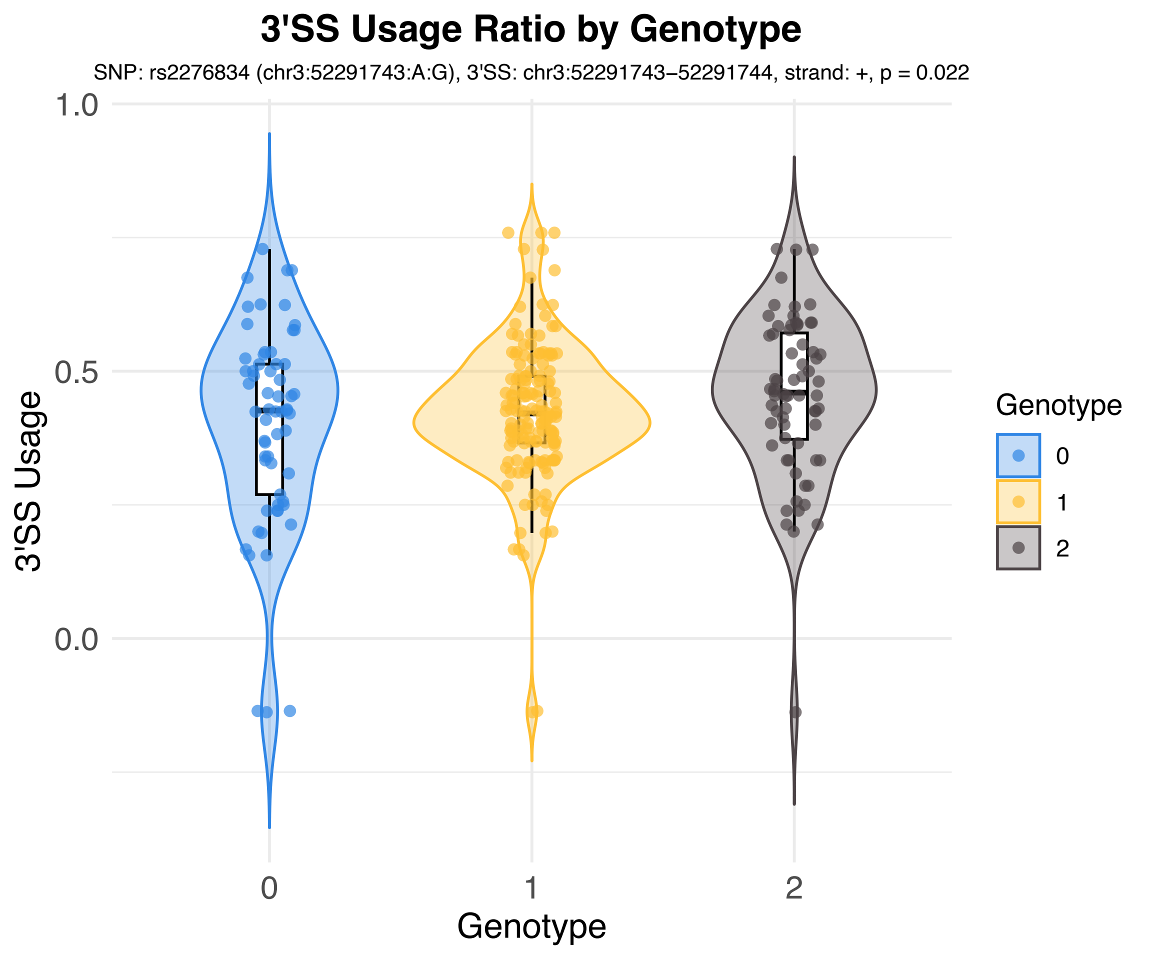

3'SS Usage Ratio by Genotype
SNP: rs2276834 (chr3:52291743:A:G), 3'SS: chr3:52291743−52291744, strand: +, p = 0.022
3'SS Usage
1.0
0.5
0.0
0
1
2
Genotype
Genotype
0
1
2

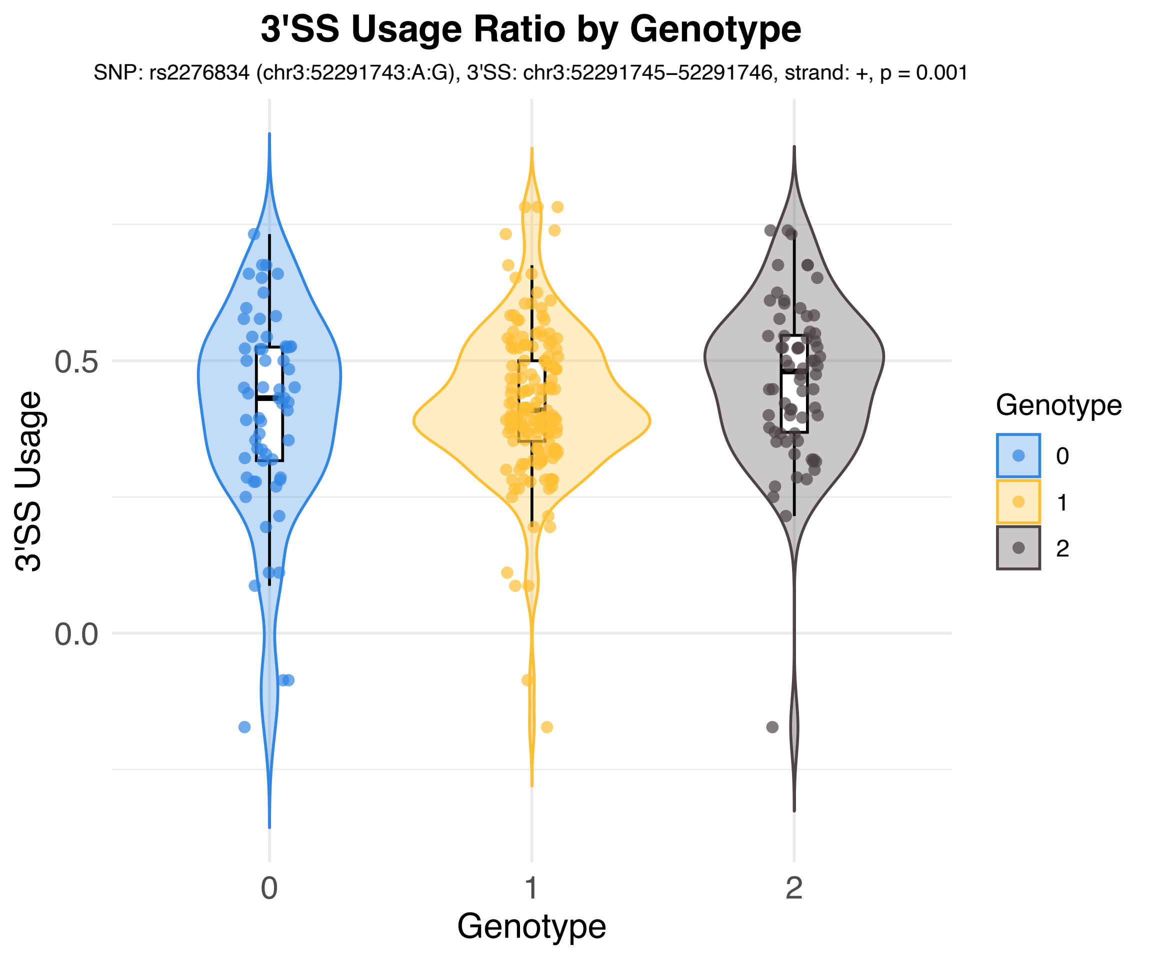
3'SS Usage Ratio by Genotype
SNP: rs2276834 (chr3:52291743:A:G), 3'SS: chr3:52291745−52291746, strand: +, p = 0.001
3'SS Usage
0.5
0.0
0
1
2
Genotype
Genotype
0
1
2

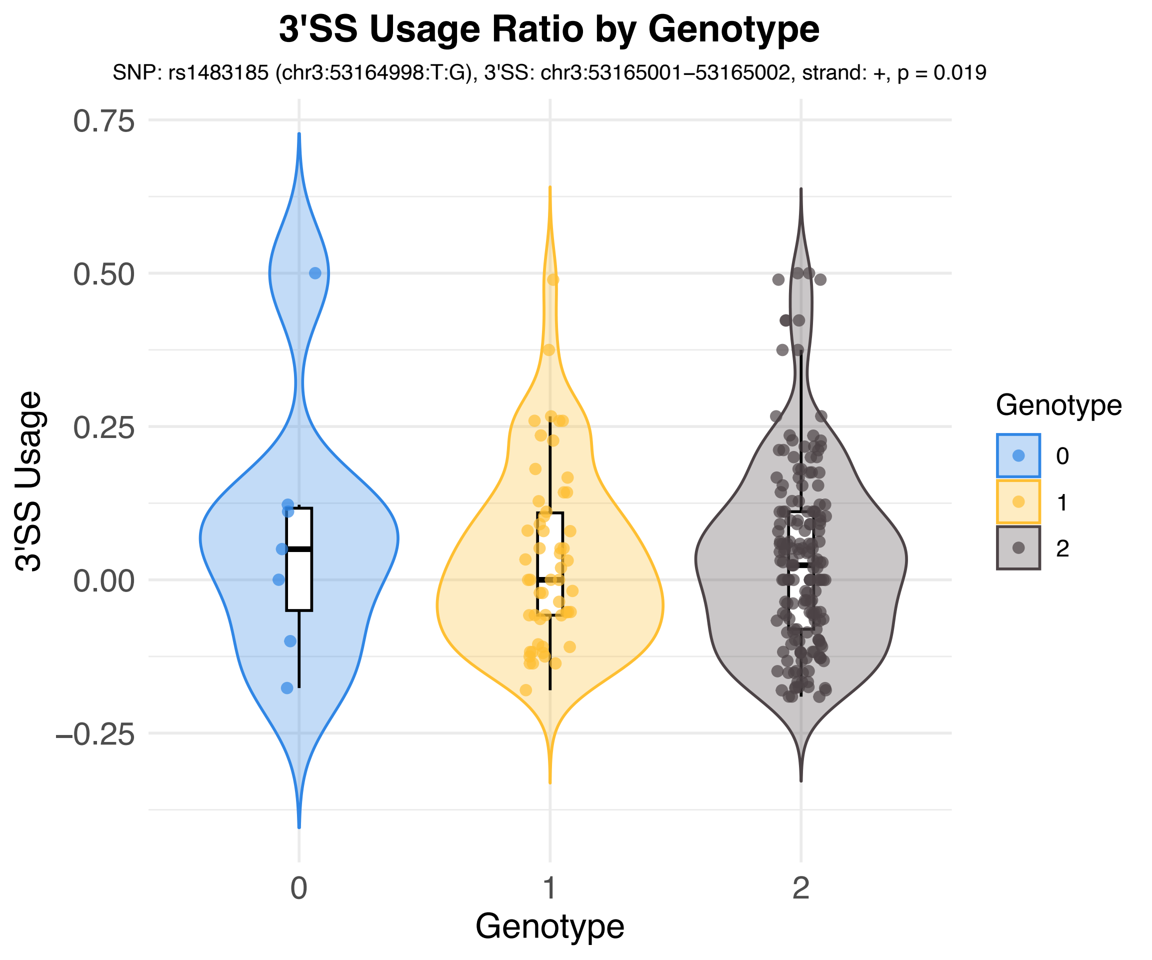

3'SS Usage Ratio by Genotype
SNP: rs1483185 (chr3:53164998:T:G), 3'SS: chr3:53165001–53165002, strand: +, p = 0.019
3'SS Usage
0.75
0.50
0.25
0.00
−0.25
0
1
2
Genotype
Genotype
0
1
2

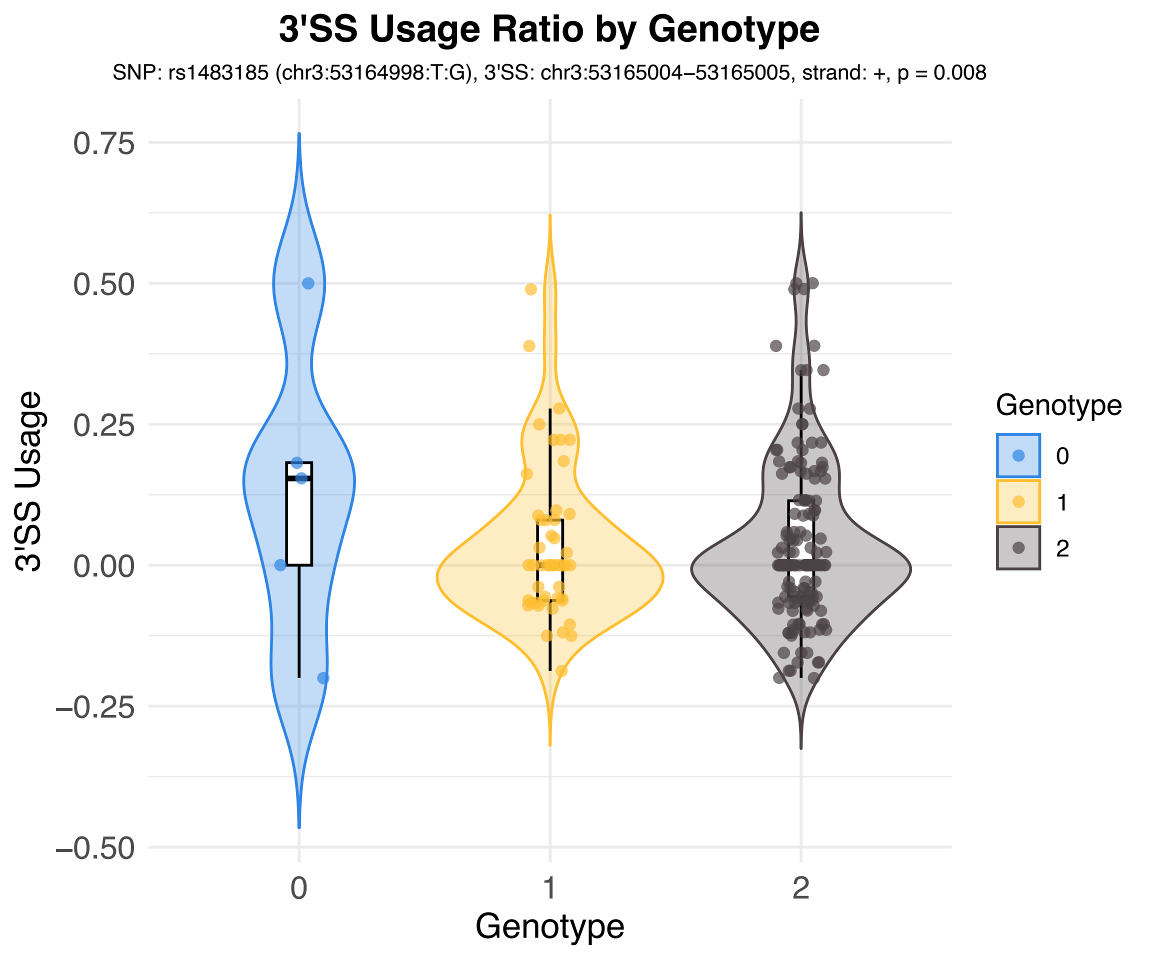

3'SS Usage Ratio by Genotype
SNP: rs1483185 (chr3:53164998:T:G), 3'SS: chr3:53165004–53165005, strand: +, p = 0.008
3'SS Usage
0.75
0.50
0.25
0.00
−0.25
−0.50
0
1
2
Genotype
Genotype
0
1
2

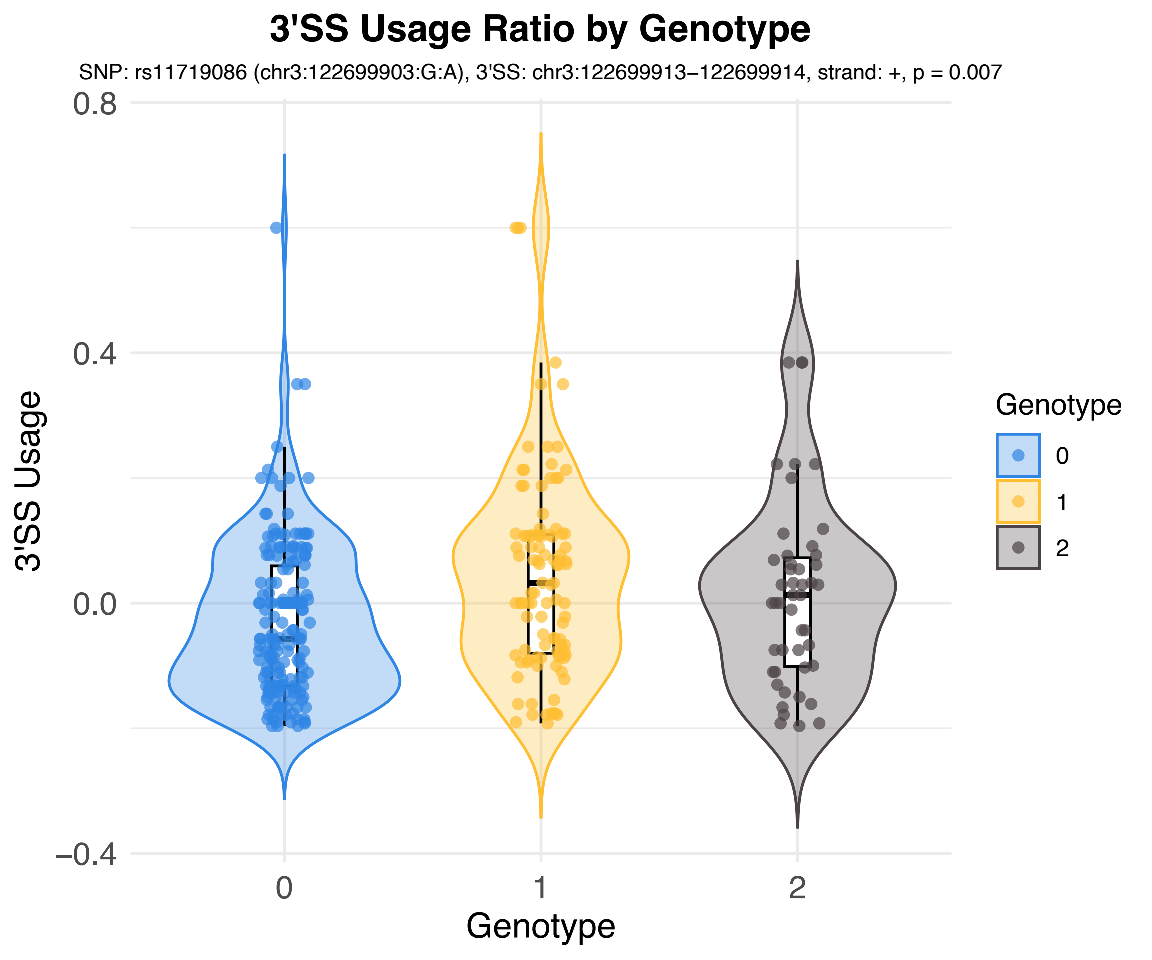
3'SS Usage Ratio by Genotype
SNP: rs11719086 (chr3:122699903:G:A), 3'SS: chr3:122699913−122699914, strand: +, p = 0.007
3'SS Usage
0.8
0.4
0.0
−0.4
0
1
2
Genotype
Genotype
0
1
2

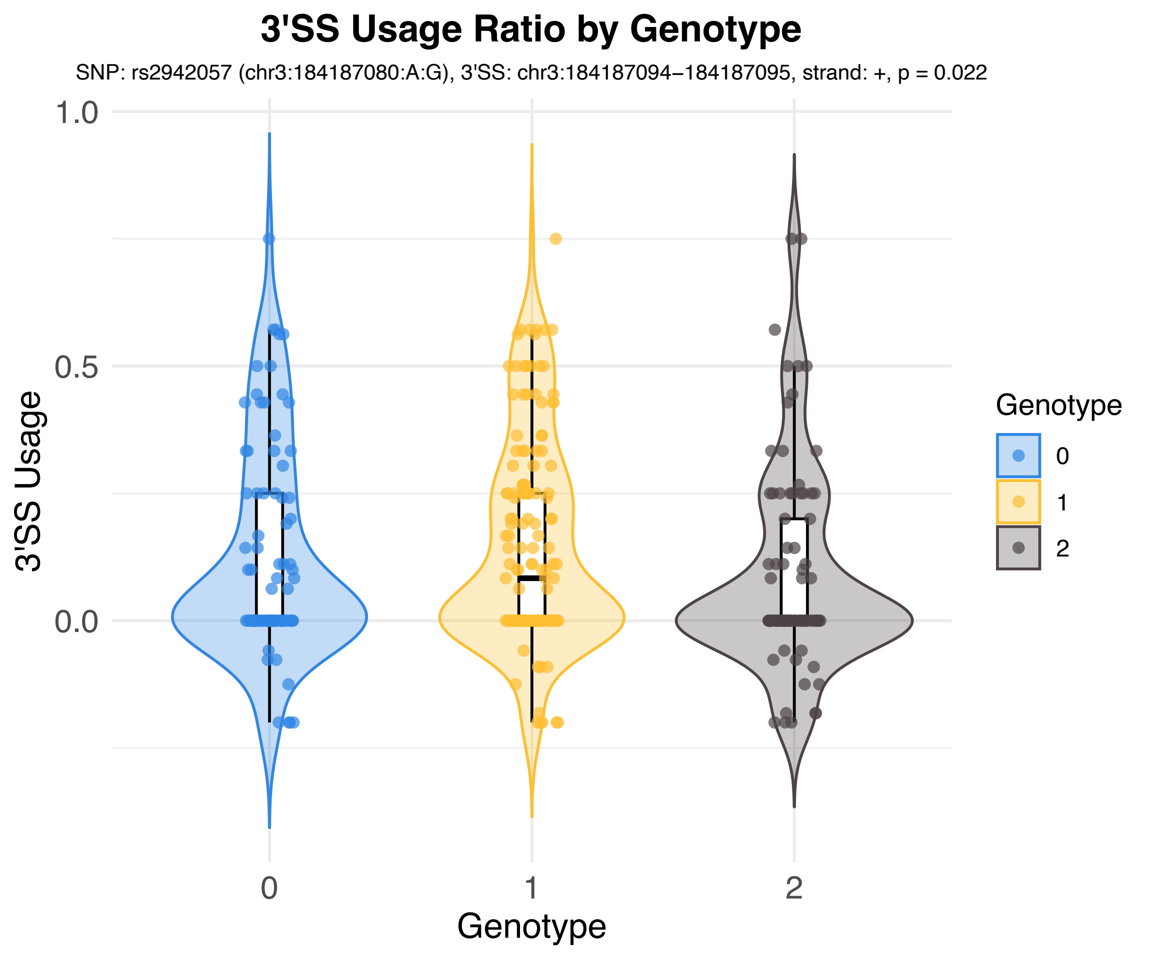
3'SS Usage Ratio by Genotype
SNP: rs2942057 (chr3:184187080:A:G), 3'SS: chr3:184187094−184187095, strand: +, p = 0.022
3'SS Usage
1.0
0.5
0.0
0
1
2
Genotype
Genotype
0
1
2

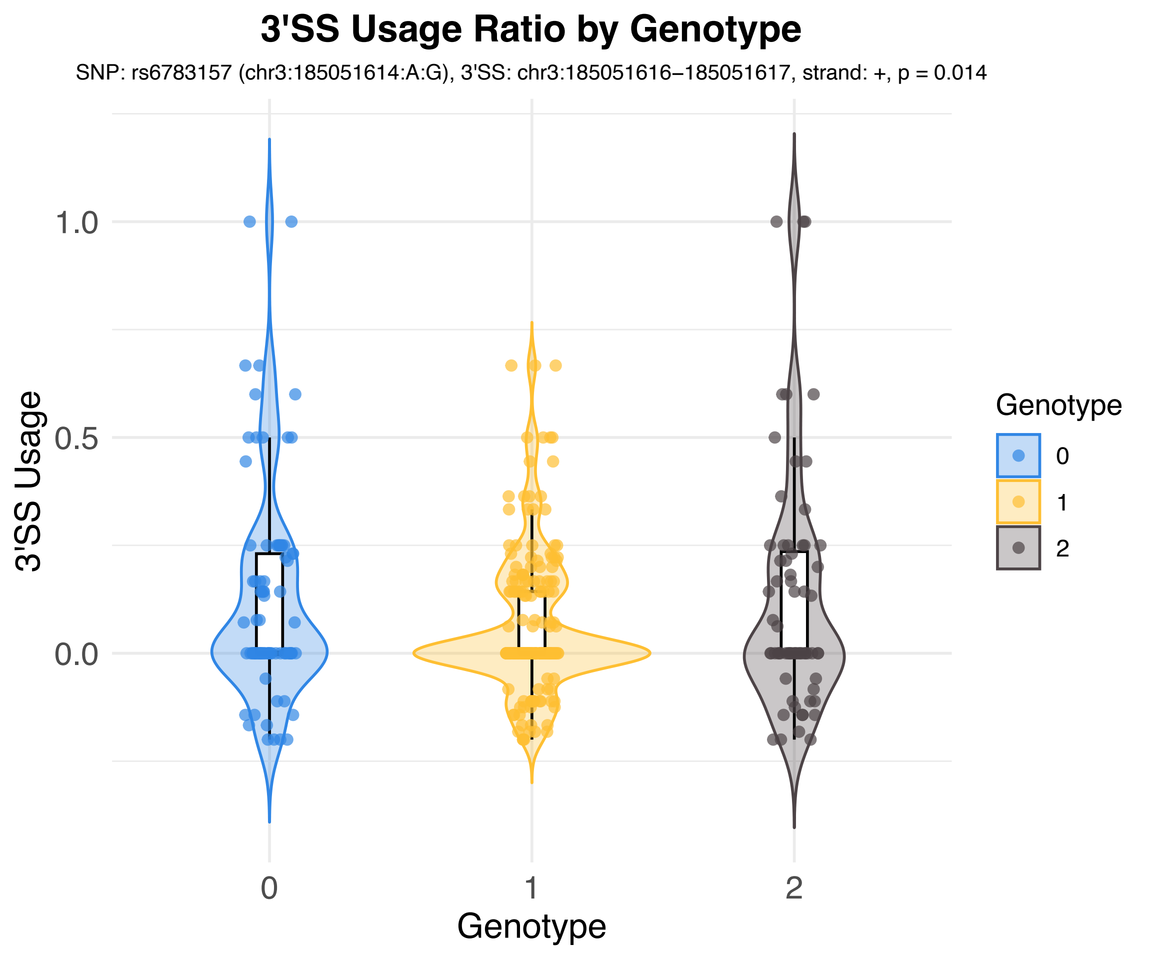
3'SS Usage Ratio by Genotype
SNP: rs6783157 (chr3:185051614:A:G), 3'SS: chr3:185051616–185051617, strand: +, p = 0.014
3'SS Usage
1.0
0.5
0.0
0
1
2
Genotype
Genotype
0
1
2

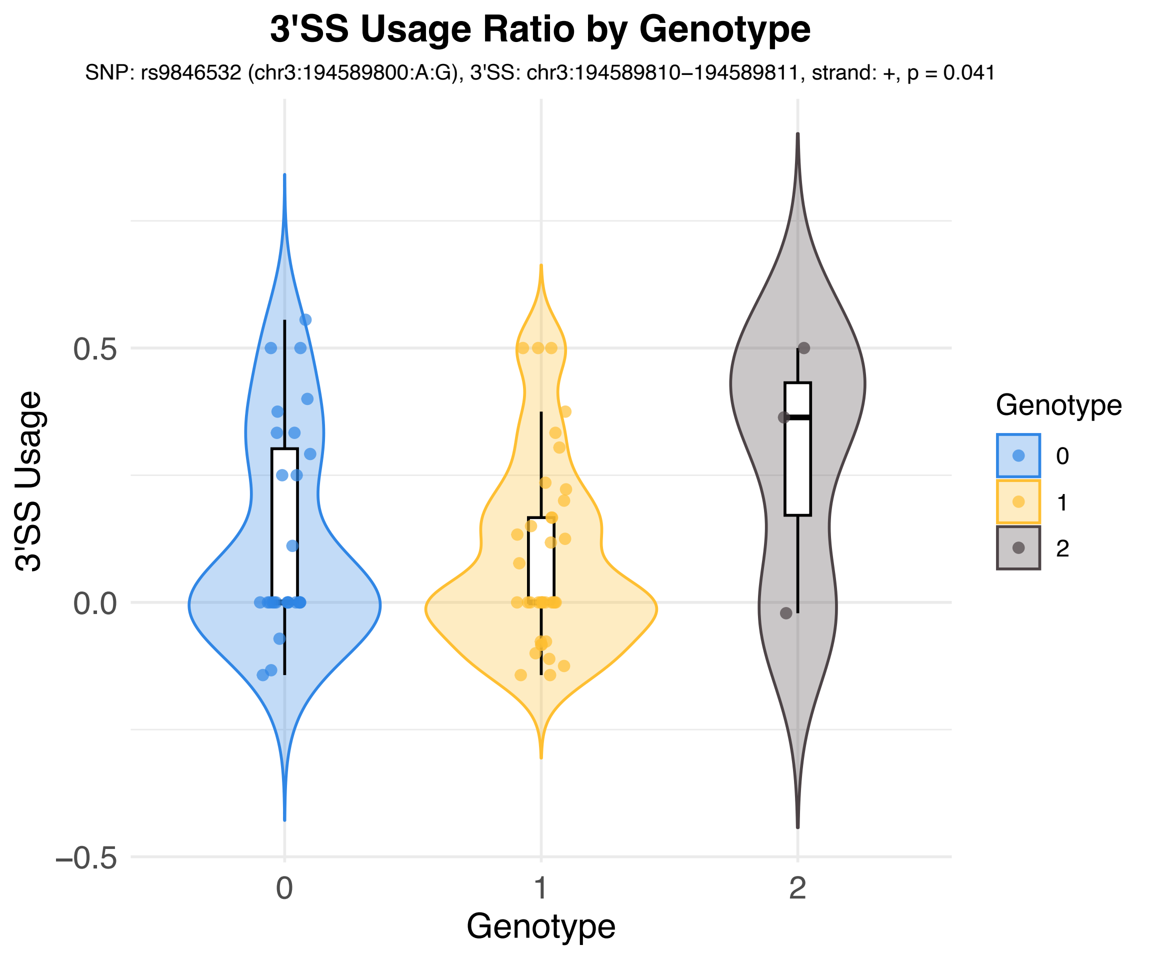
3'SS Usage Ratio by Genotype
SNP: rs9846532 (chr3:194589800:A:G), 3'SS: chr3:194589810−194589811, strand: +, p = 0.041
3'SS Usage
0.5
0.0
−0.5
0
1
2
Genotype
Genotype
0
1
2

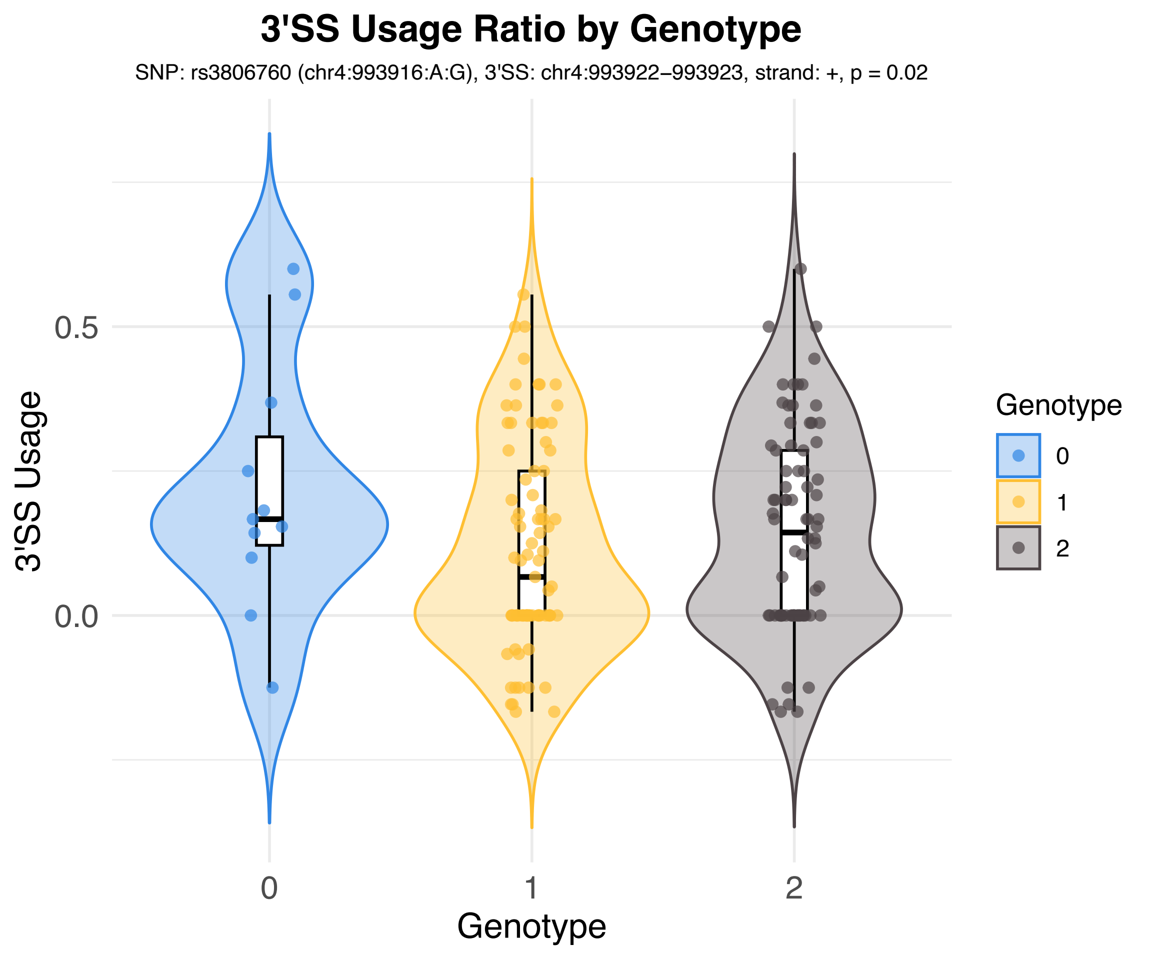
3'SS Usage Ratio by Genotype
SNP: rs3806760 (chr4:993916:A:G), 3'SS: chr4:993922−993923, strand: +, p = 0.02
3'SS Usage
0.5
0.0
0
1
2
Genotype
Genotype
0
1
2

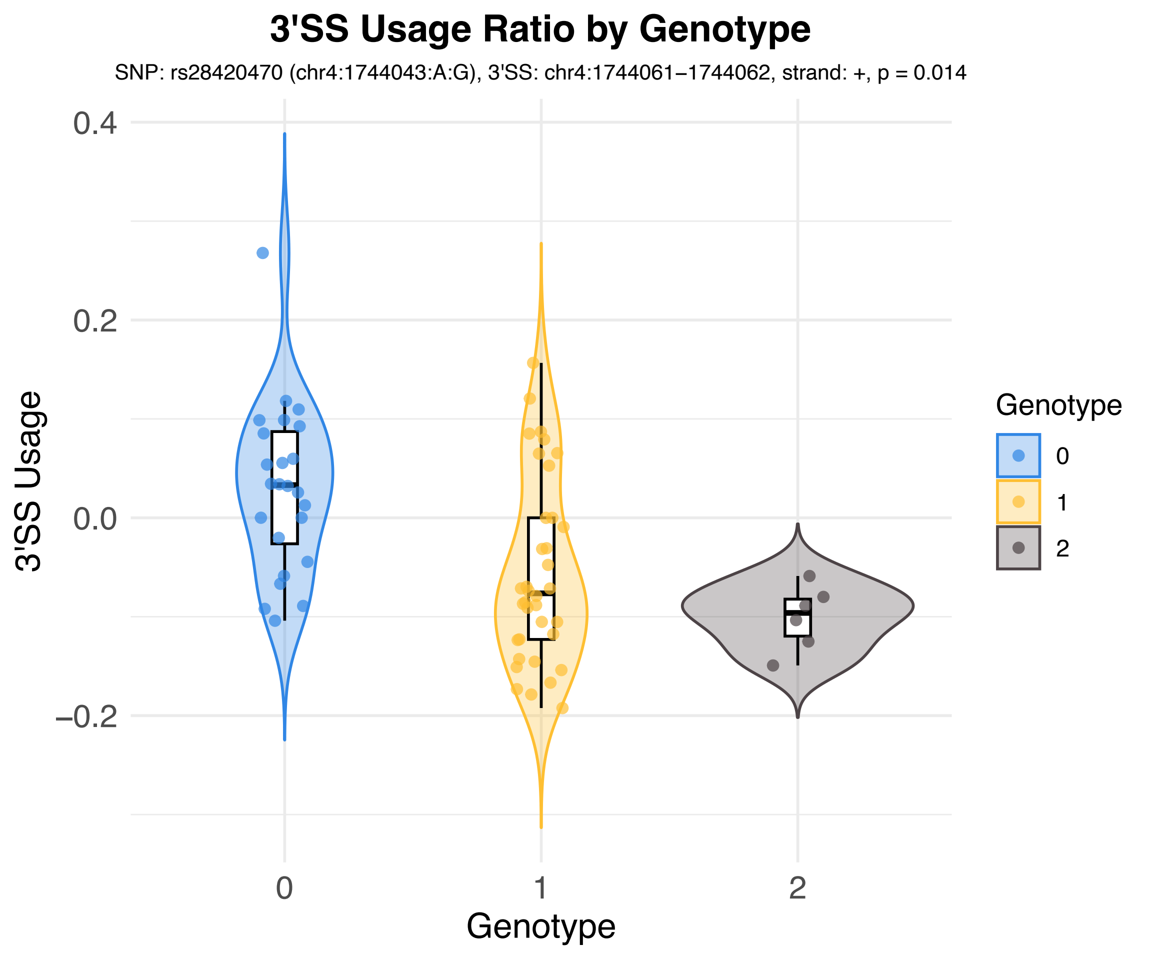

3'SS Usage Ratio by Genotype
SNP: rs28420470 (chr4:1744043:A:G), 3'SS: chr4:1744061–1744062, strand: +, p = 0.014
3'SS Usage
0.4
0.2
0.0
−0.2
Genotype
0
1
2
Genotype
0
1
2

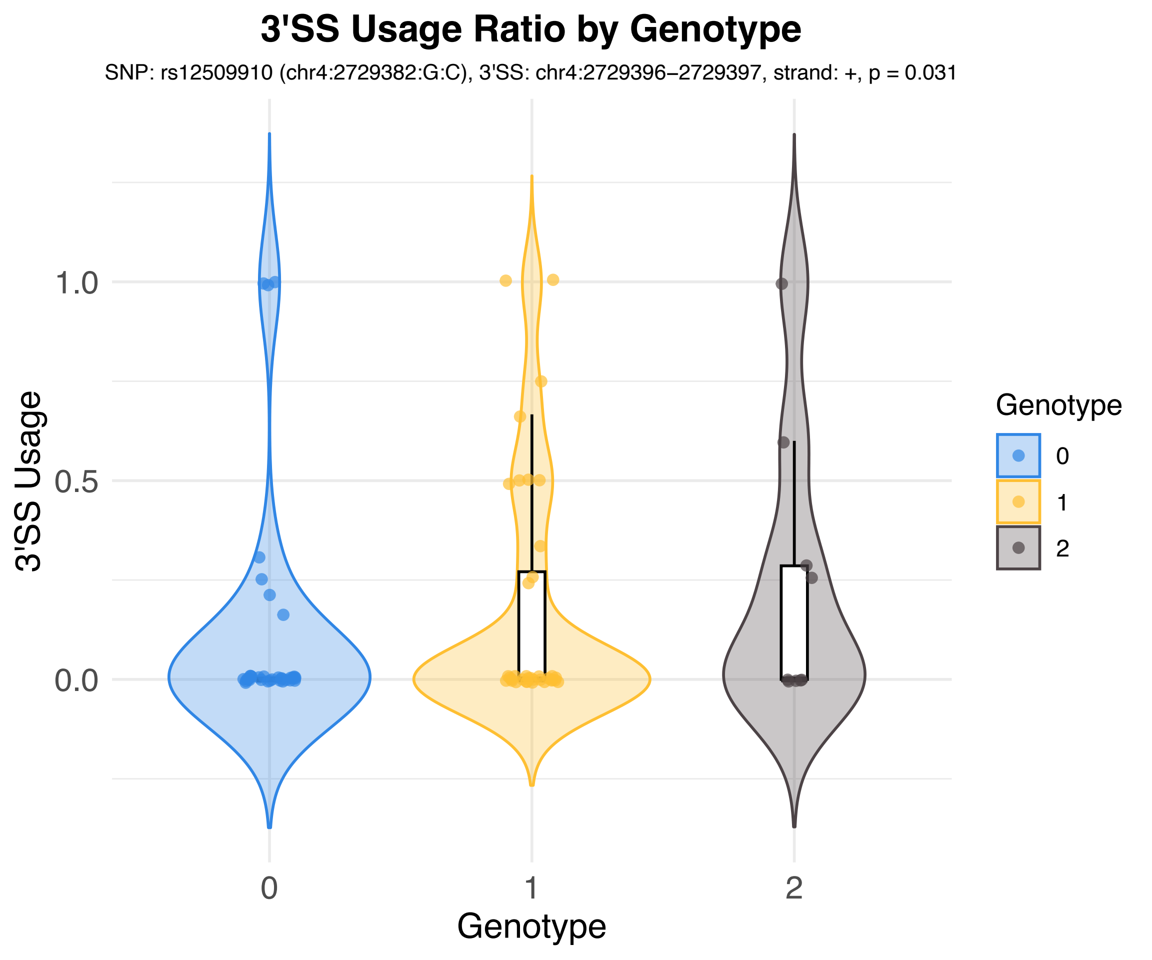
3'SS Usage Ratio by Genotype
SNP: rs12509910 (chr4:2729382:G:C), 3'SS: chr4:2729396–2729397, strand: +, p = 0.031
3'SS Usage
1.0
0.5
0.0
0
1
2
Genotype
Genotype
0
1
2

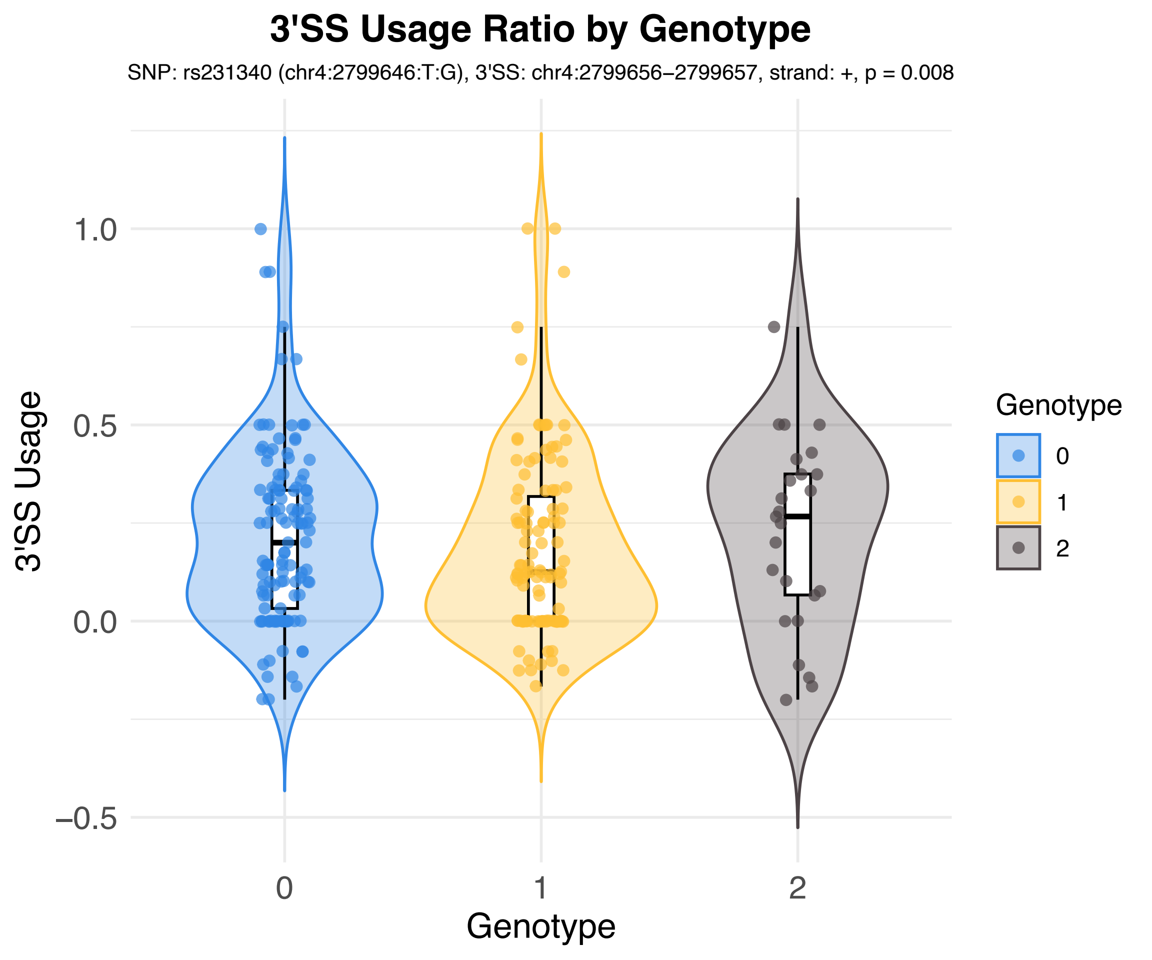

3'SS Usage Ratio by Genotype
SNP: rs231340 (chr4:2799646:T:G), 3'SS: chr4:2799656–2799657, strand: +, p = 0.008
3'SS Usage
1.0
0.5
0.0
−0.5
0
1
2
Genotype
Genotype
0
1
2

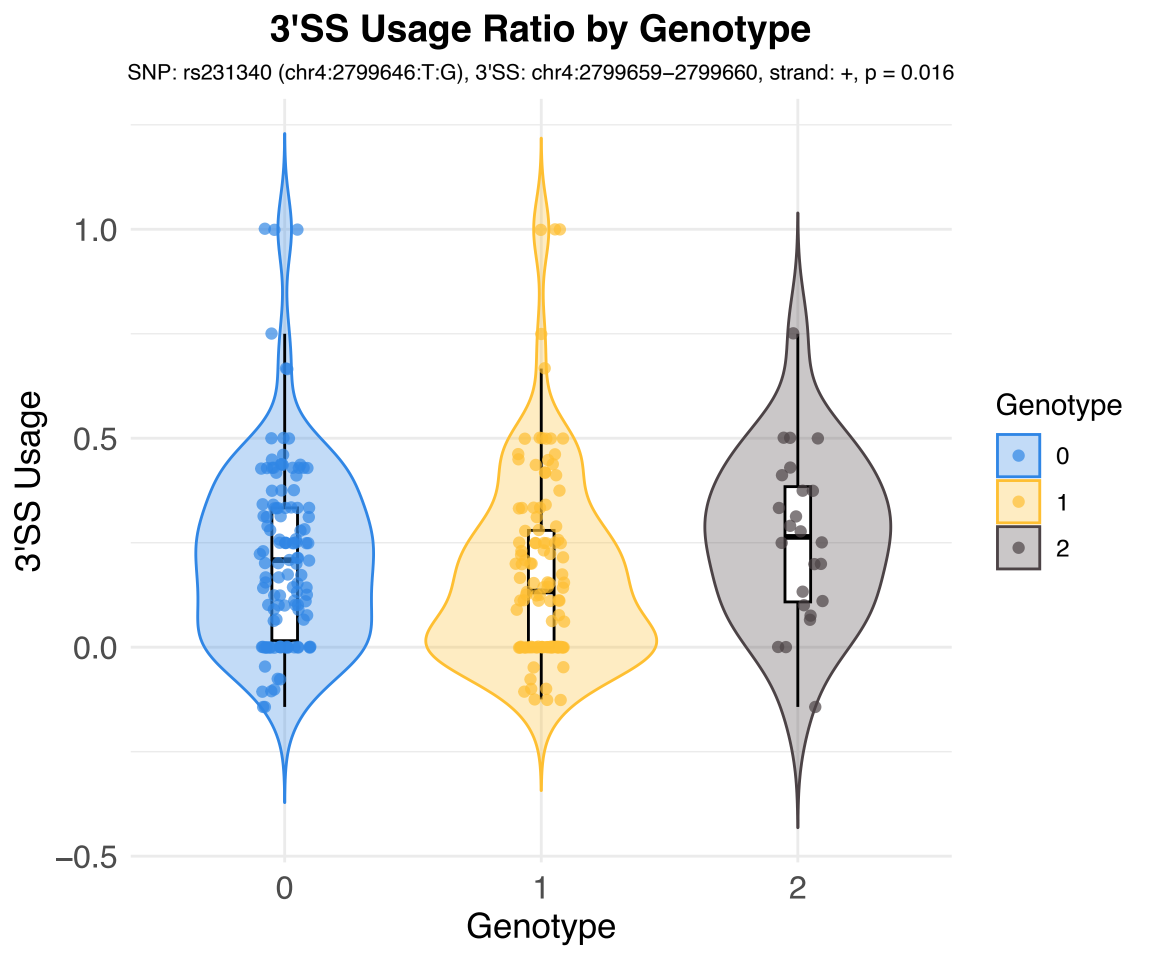

3'SS Usage Ratio by Genotype
SNP: rs231340 (chr4:2799646:T:G), 3'SS: chr4:2799659–2799660, strand: +, p = 0.016
3'SS Usage
1.0
0.5
0.0
−0.5
0
1
2
Genotype
Genotype
0
1
2

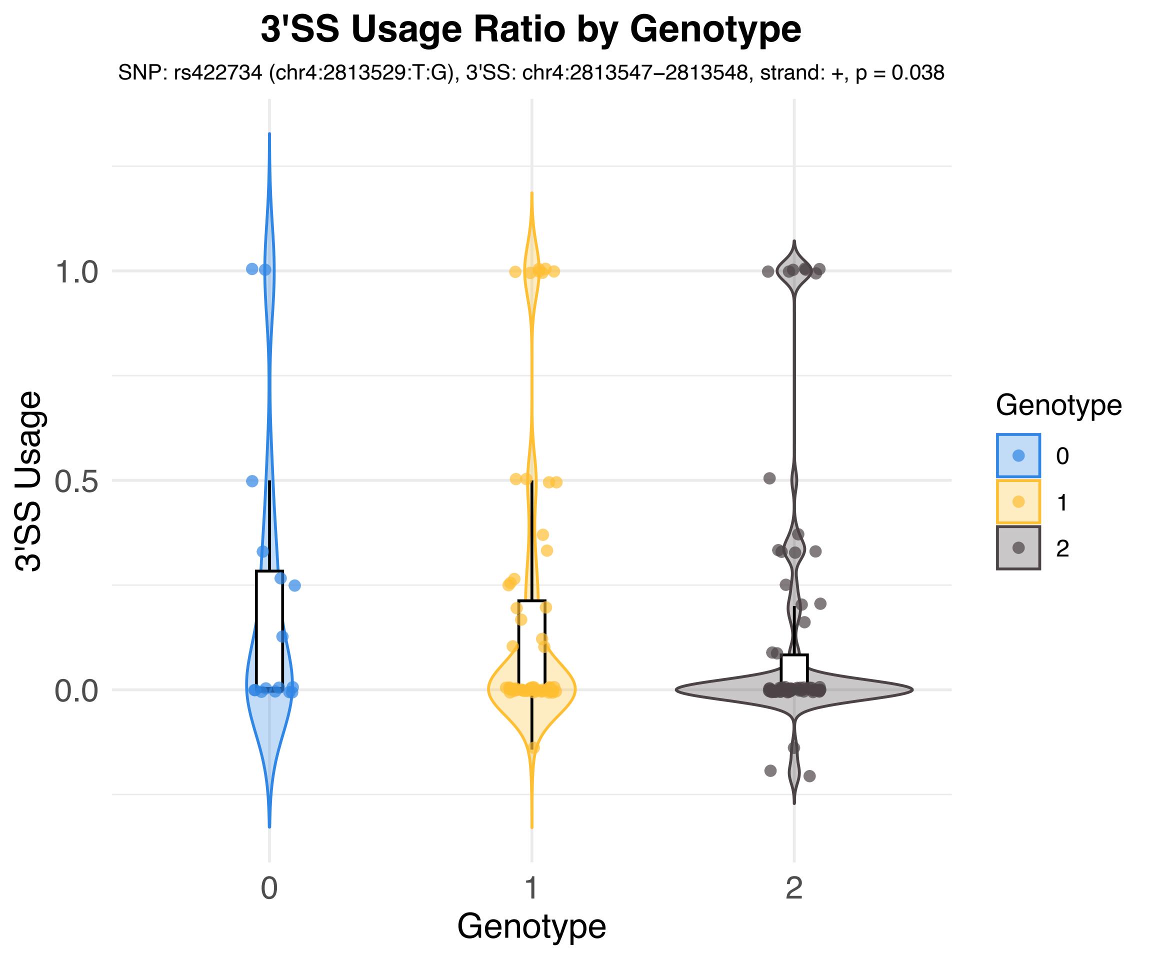

3'SS Usage Ratio by Genotype
SNP: rs422734 (chr4:2813529:T:G), 3'SS: chr4:2813547–2813548, strand: +, p = 0.038
3'SS Usage
1.0
0.5
0.0
0
1
2
Genotype
Genotype
0
1
2

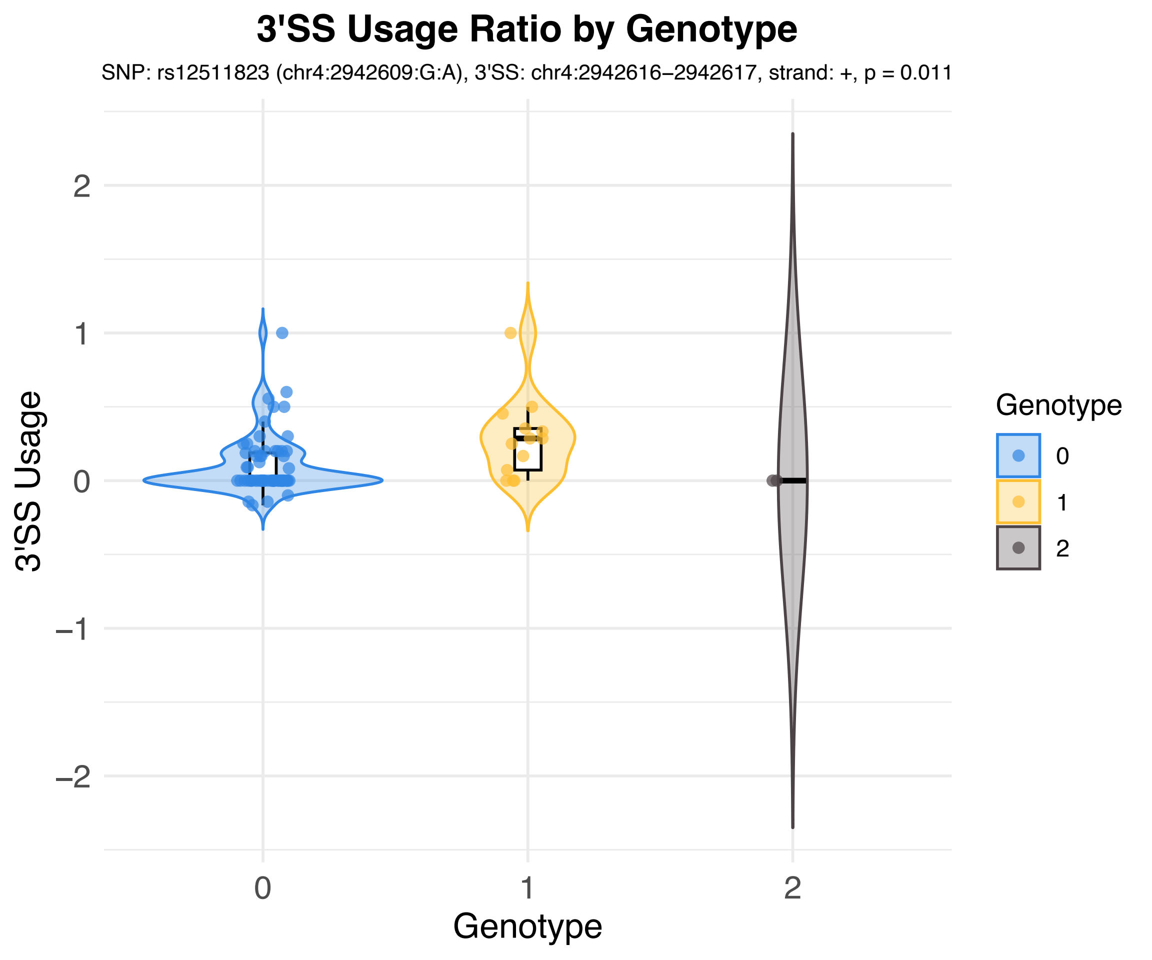
3'SS Usage Ratio by Genotype
SNP: rs12511823 (chr4:2942609:G:A), 3'SS: chr4:2942616−2942617, strand: +, p = 0.011
3'SS Usage
2
1
0
−1
−2
0
1
2
Genotype
Genotype
0
1
2

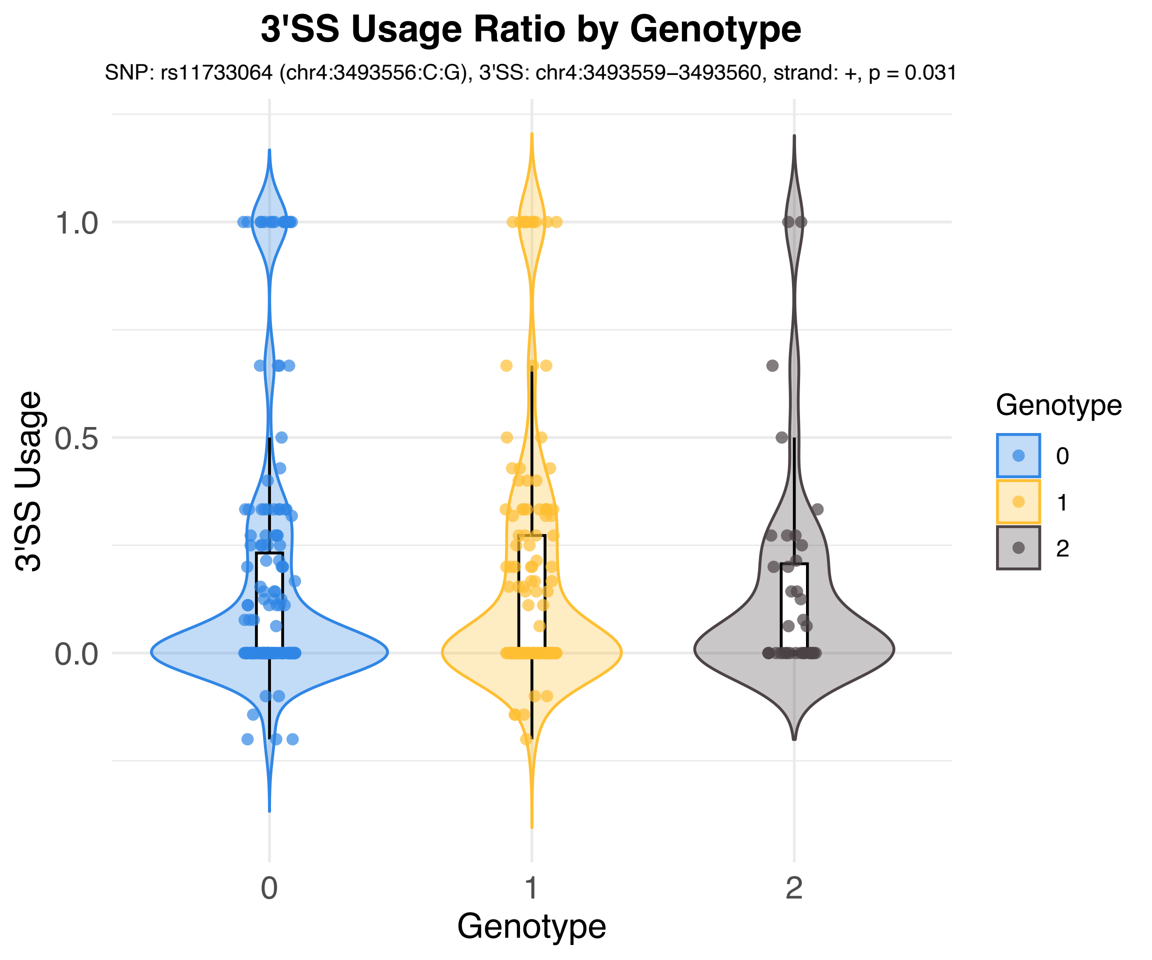
3'SS Usage Ratio by Genotype
SNP: rs11733064 (chr4:3493556:C:G), 3'SS: chr4:3493559–3493560, strand: +, p = 0.031
3'SS Usage
1.0
0.5
0.0
0
1
2
Genotype
Genotype
0
1
2

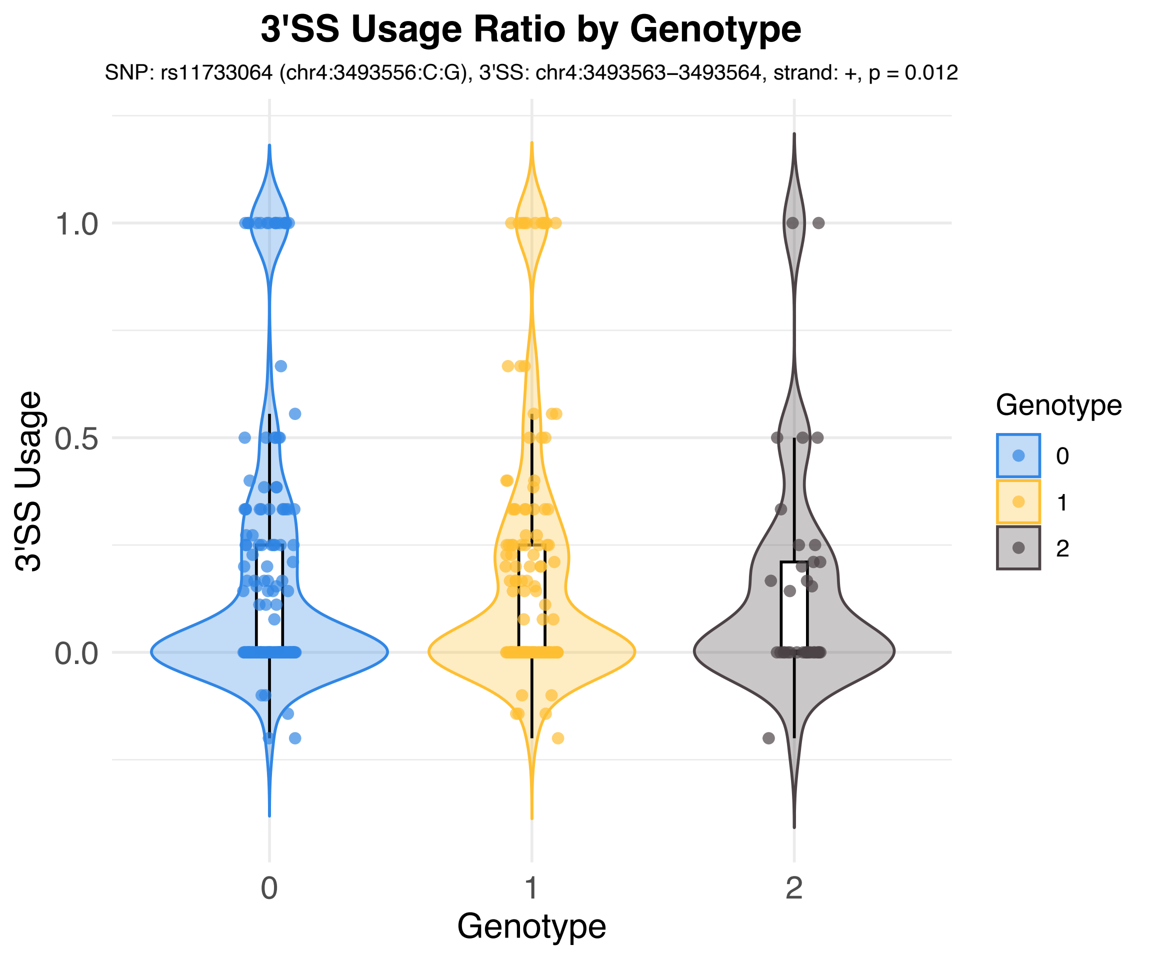

3'SS Usage Ratio by Genotype
SNP: rs11733064 (chr4:3493556:C:G), 3'SS: chr4:3493563–3493564, strand: +, p = 0.012
3'SS Usage
1.0
0.5
0.0
0
1
2
Genotype
Genotype
0
1
2

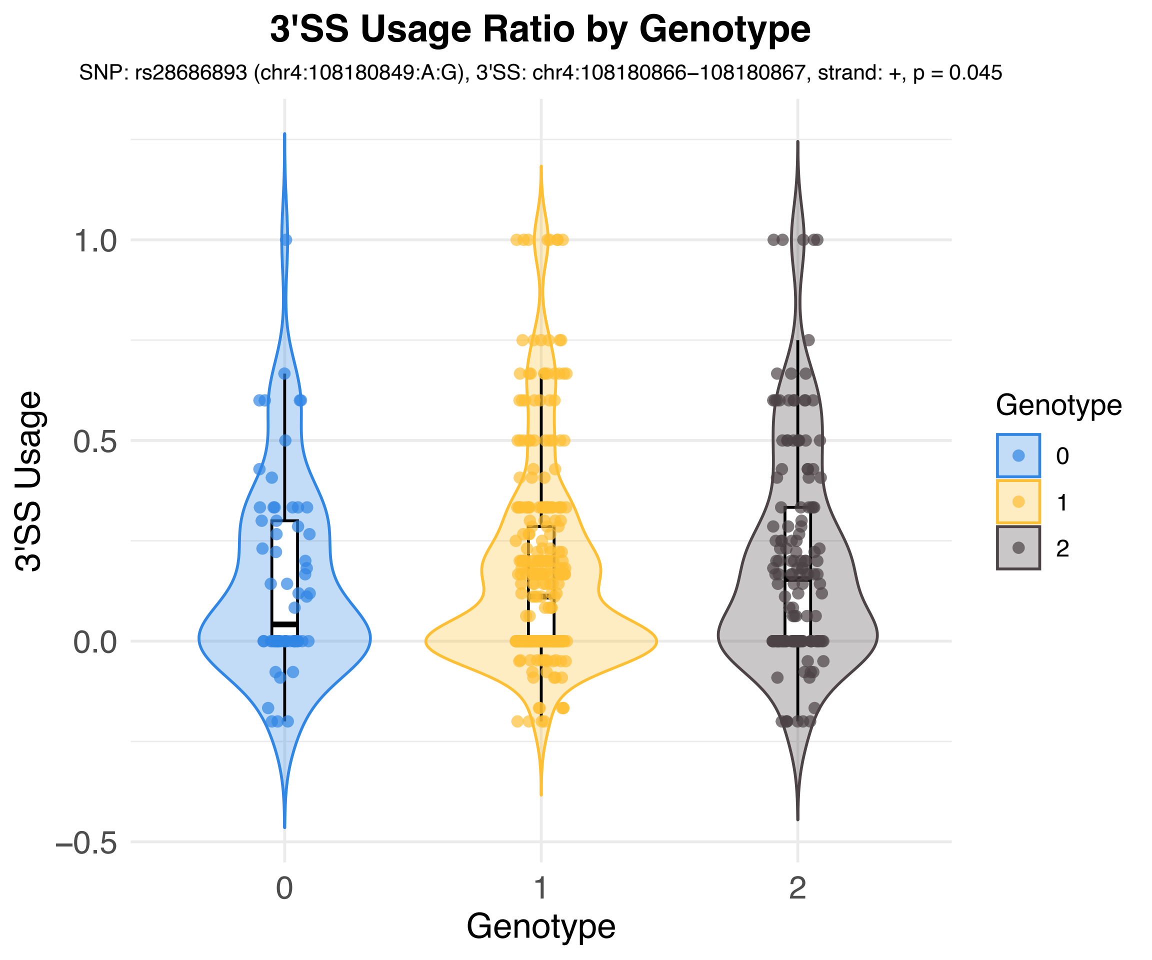
3'SS Usage Ratio by Genotype
SNP: rs28686893 (chr4:108180849:A:G), 3'SS: chr4:108180866–108180867, strand: +, p = 0.045
3'SS Usage
1.0
0.5
0.0
−0.5
0
1
2
Genotype
Genotype
0
1
2

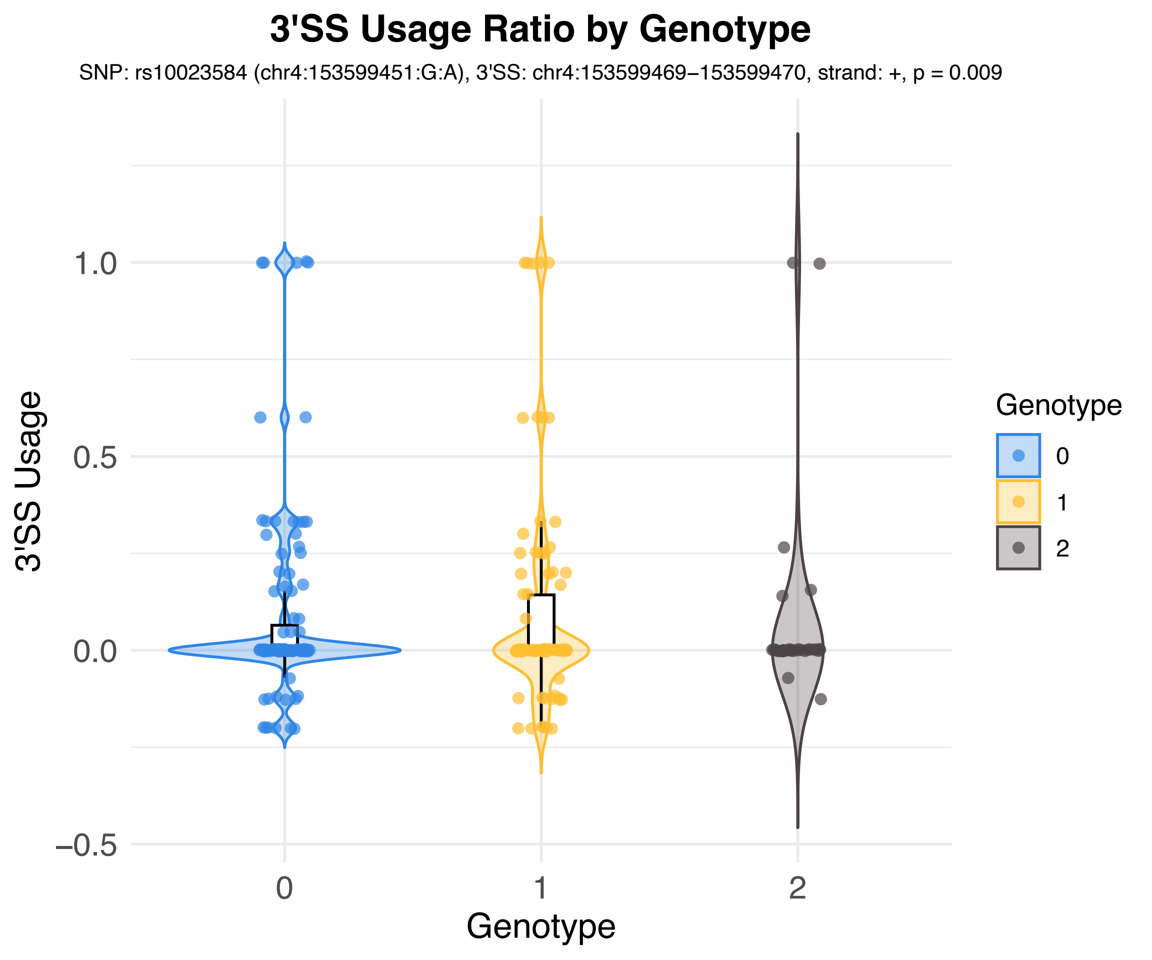

3'SS Usage Ratio by Genotype
SNP: rs10023584 (chr4:153599451:G:A), 3'SS: chr4:153599469−153599470, strand: +, p = 0.009
3'SS Usage
1.0
0.5
0.0
−0.5
0
1
2
Genotype
Genotype
0
1
2

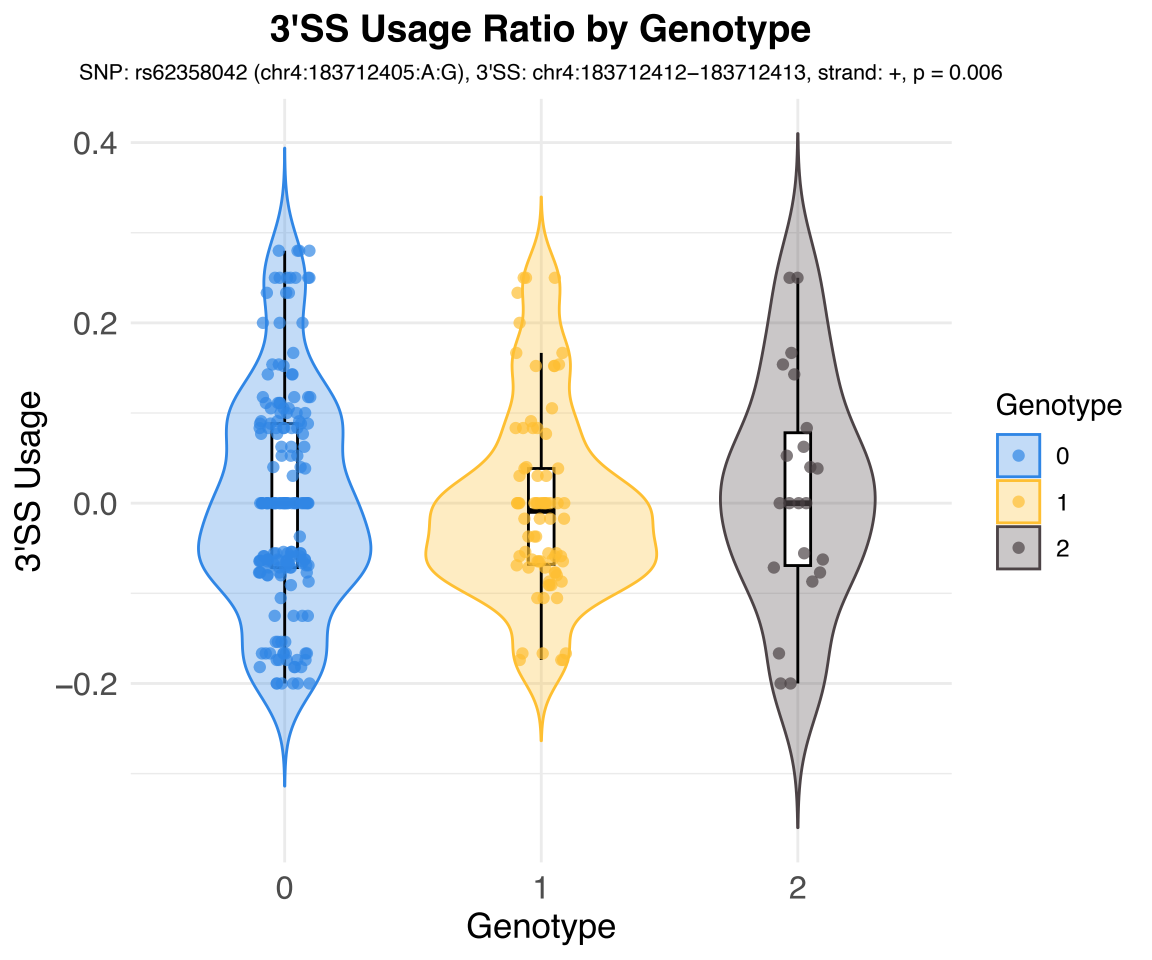

3'SS Usage Ratio by Genotype
SNP: rs62358042 (chr4:183712405:A:G), 3'SS: chr4:183712412–183712413, strand: +, p = 0.006
3'SS Usage
0.4
0.2
0.0
–0.2
0
1
2
Genotype
Genotype
0
1
2

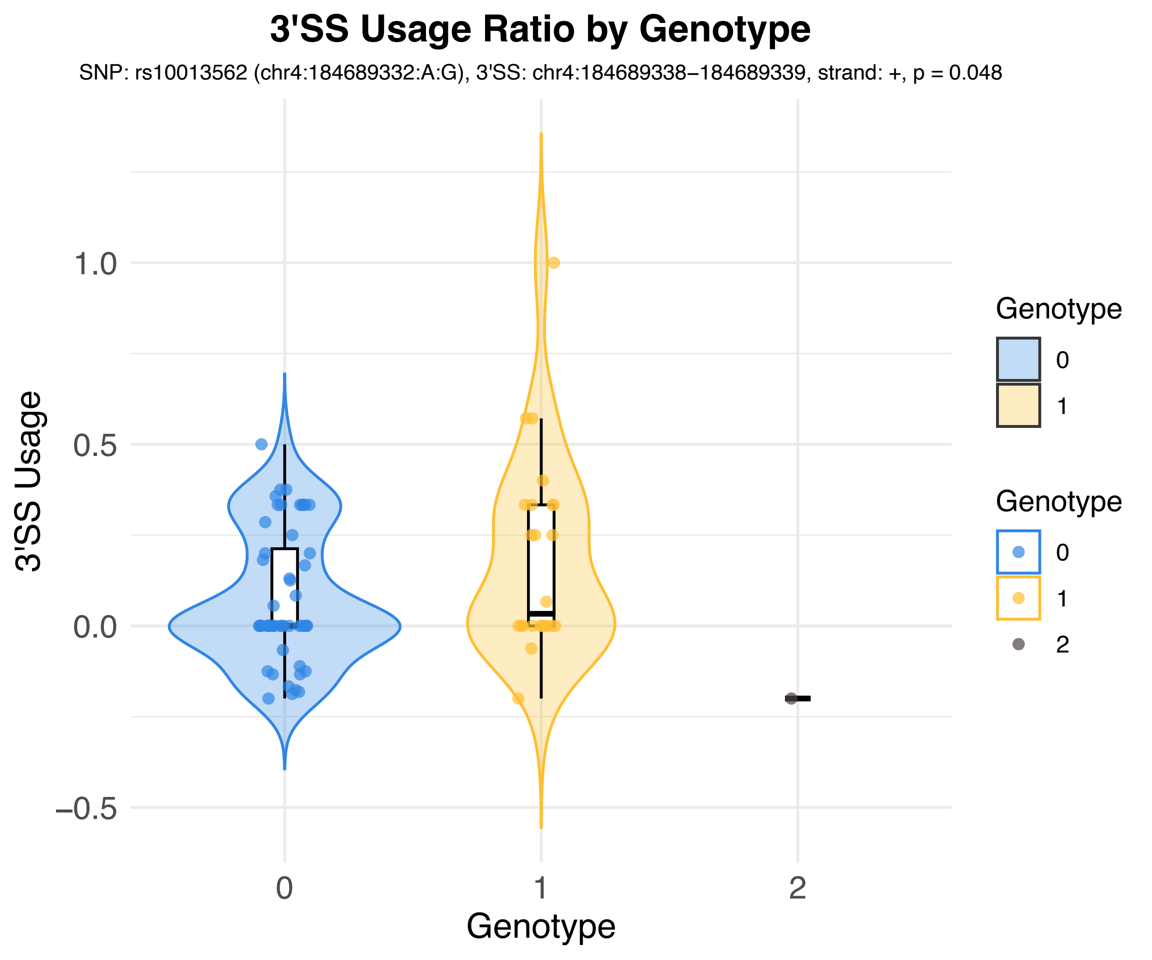
3'SS Usage Ratio by Genotype
SNP: rs10013562 (chr4:184689332:A:G), 3'SS: chr4:184689338−184689339, strand: +, p = 0.048
3'SS Usage
1.0
0.5
0.0
−0.5
0
1
2
Genotype
Genotype
0
1
Genotype
0
1
2

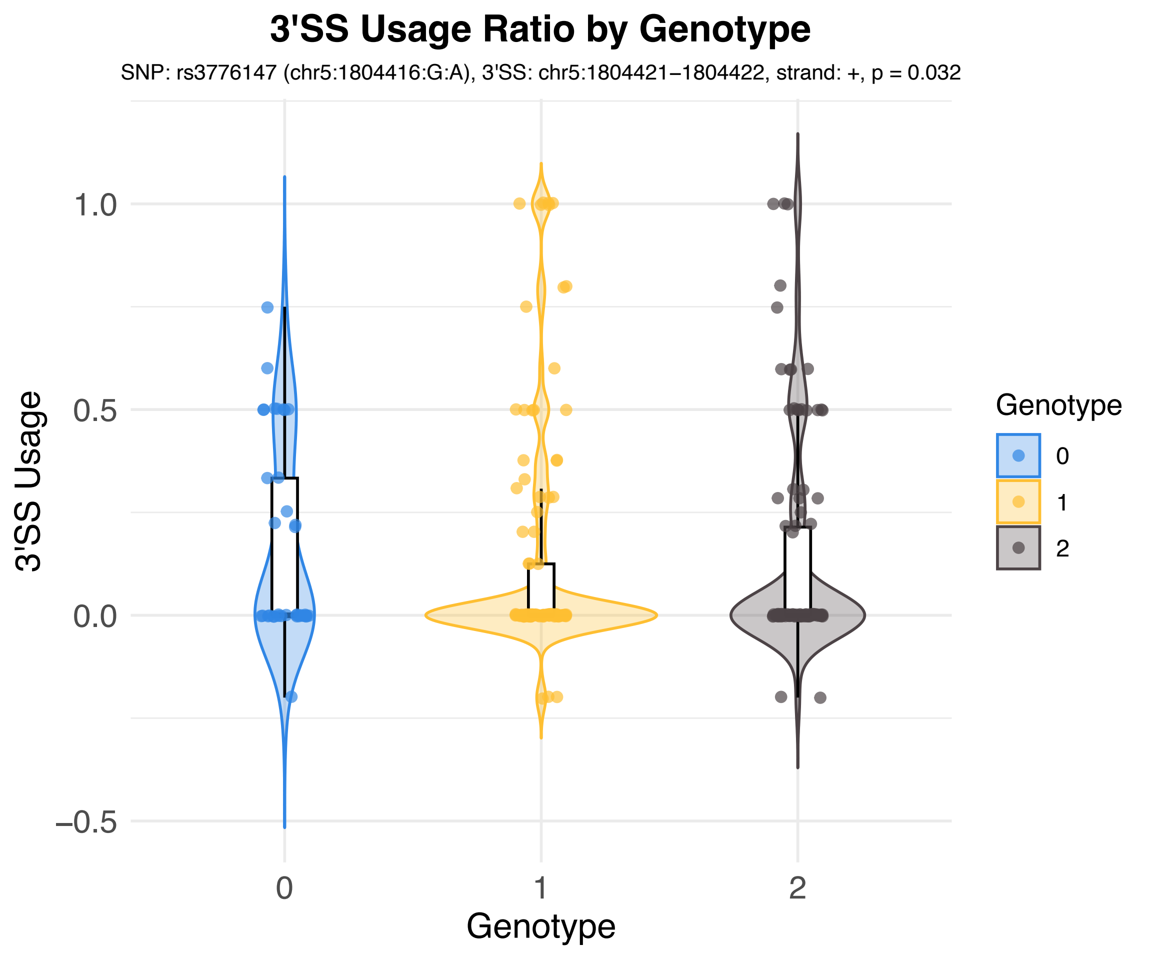
3'SS Usage Ratio by Genotype
SNP: rs3776147 (chr5:1804416:G:A), 3'SS: chr5:1804421−1804422, strand: +, p = 0.032
3'SS Usage
1.0
0.5
0.0
−0.5
0
1
2
Genotype
Genotype
0
1
2

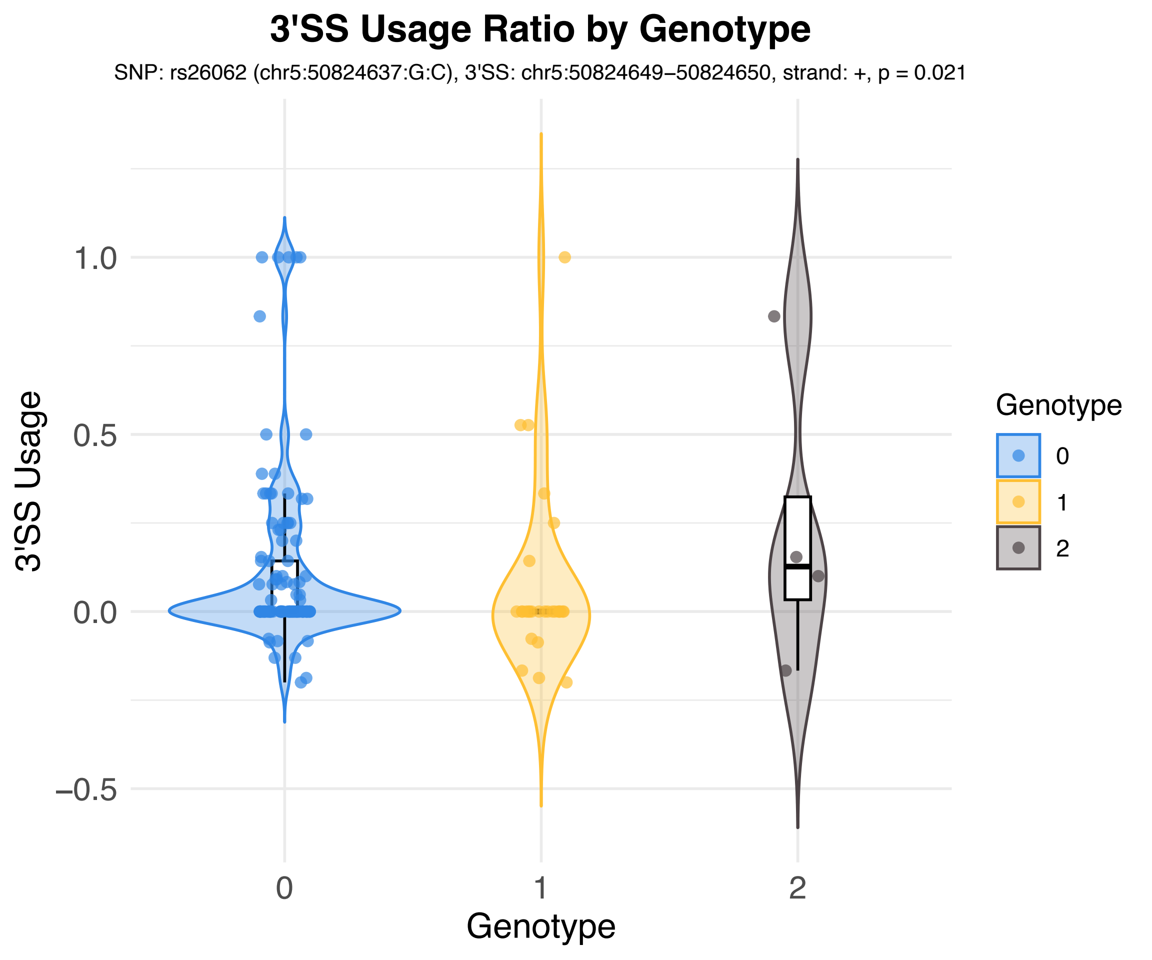
3'SS Usage Ratio by Genotype
SNP: rs26062 (chr5:50824637:G:C), 3'SS: chr5:50824649–50824650, strand: +, p = 0.021
3'SS Usage
1.0
0.5
0.0
−0.5
0
1
2
Genotype
Genotype
0
1
2

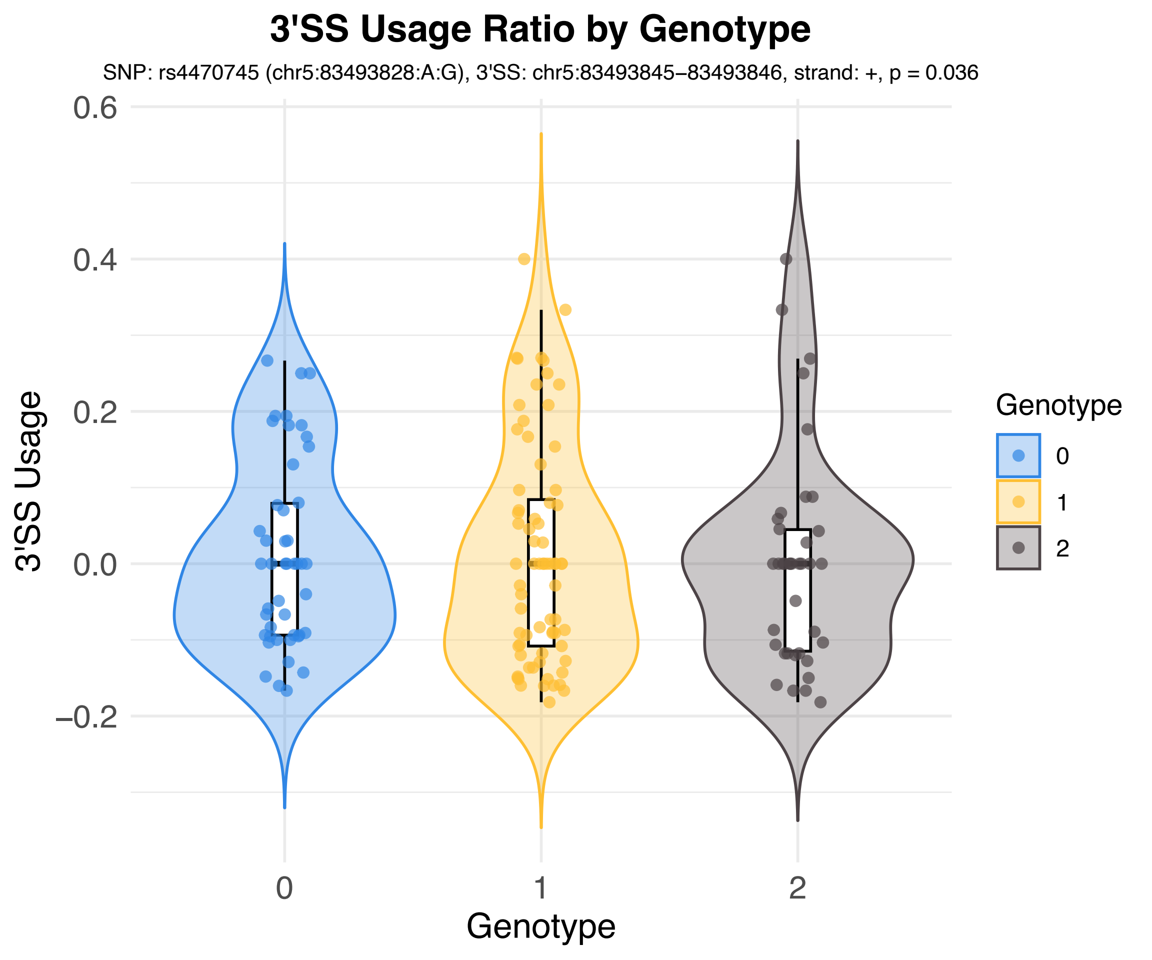
3'SS Usage Ratio by Genotype
SNP: rs4470745 (chr5:83493828:A:G), 3'SS: chr5:83493845–83493846, strand: +, p = 0.036
3'SS Usage
0.6
0.4
0.2
0.0
–0.2
0
1
2
Genotype
Genotype
0
1
2

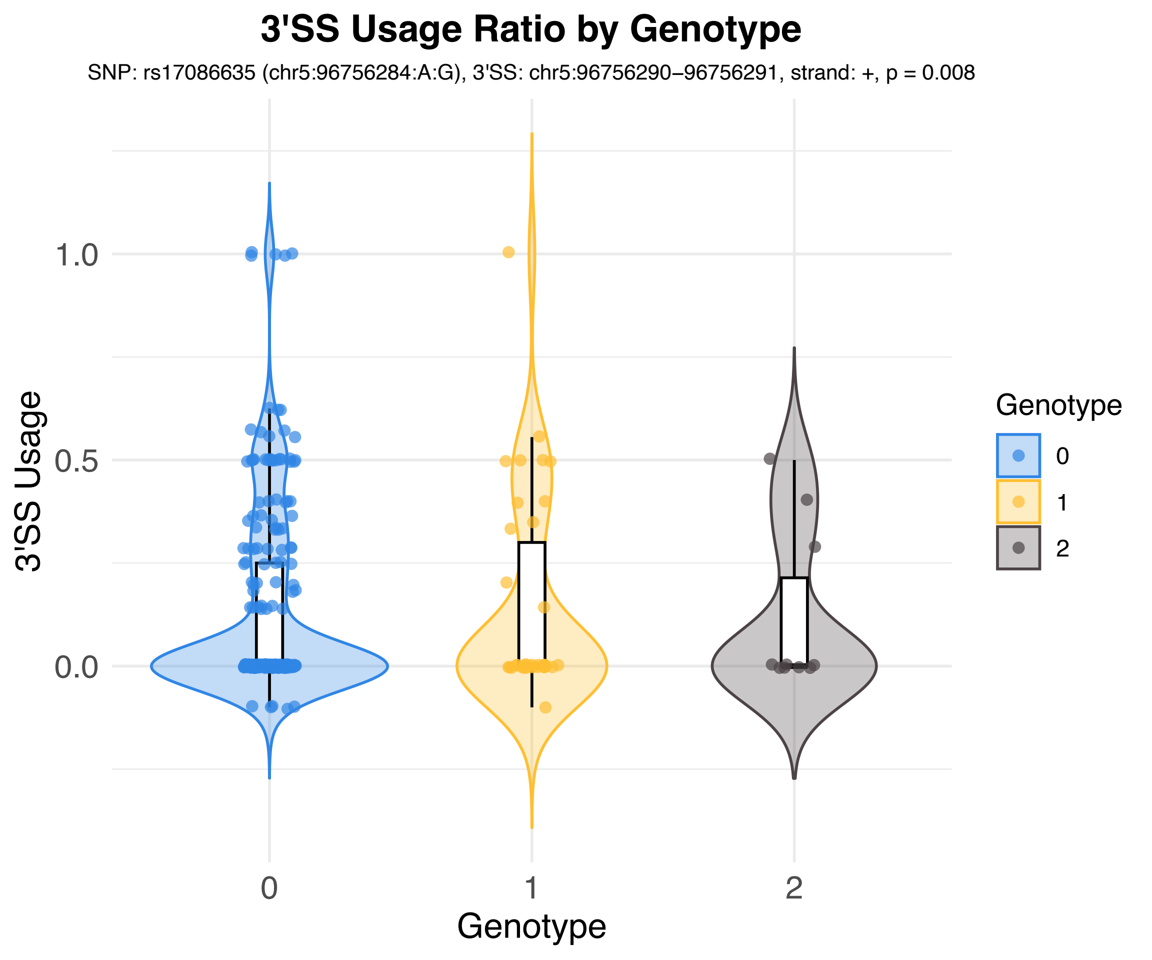
3'SS Usage Ratio by Genotype
SNP: rs17086635 (chr5:96756284:A:G), 3'SS: chr5:96756290−96756291, strand: +, p = 0.008
3'SS Usage
1.0
0.5
0.0
0
1
2
Genotype
Genotype
0
1
2

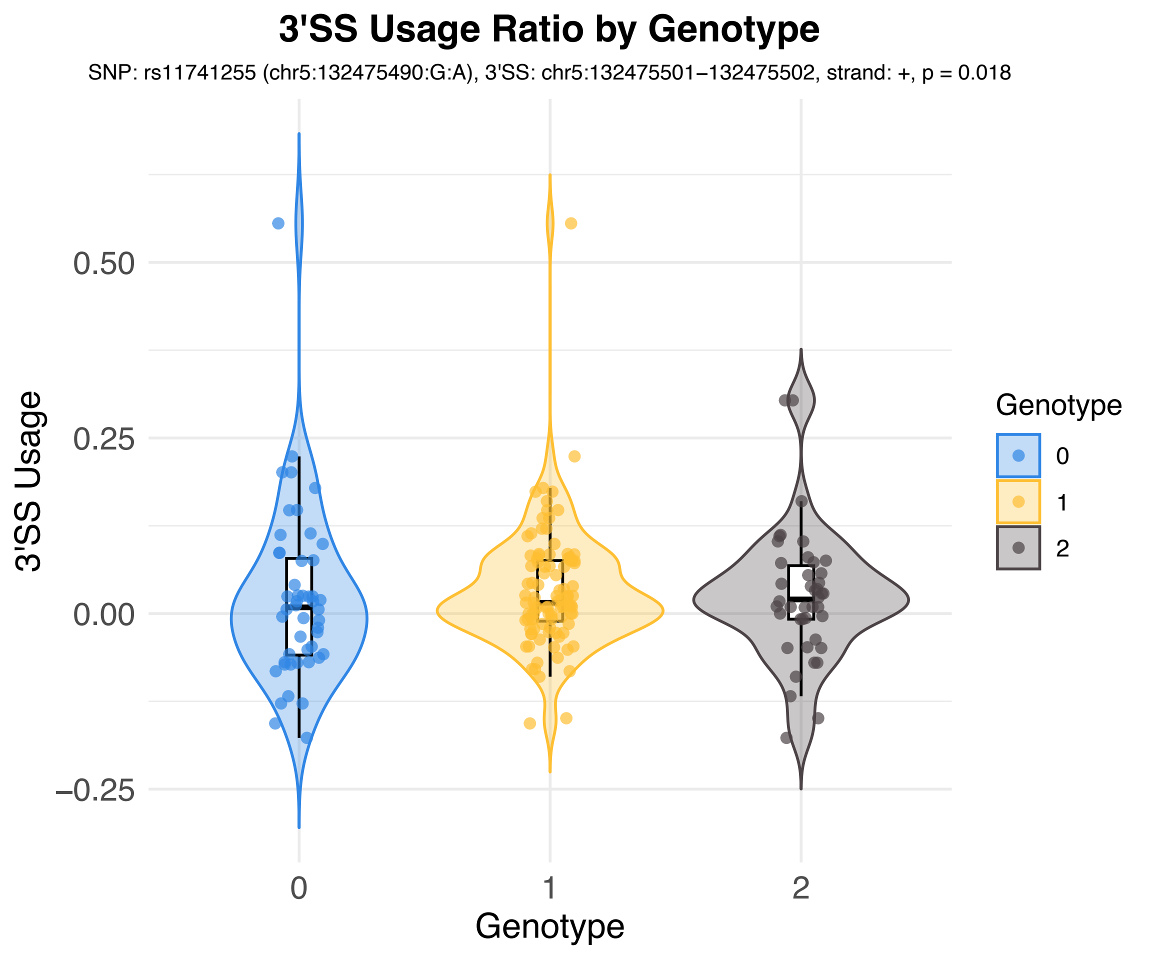
3'SS Usage Ratio by Genotype
SNP: rs11741255 (chr5:132475490:G:A), 3'SS: chr5:132475501−132475502, strand: +, p = 0.018
3'SS Usage
0.50
0.25
0.00
−0.25
0
1
2
Genotype
Genotype
0
1
2

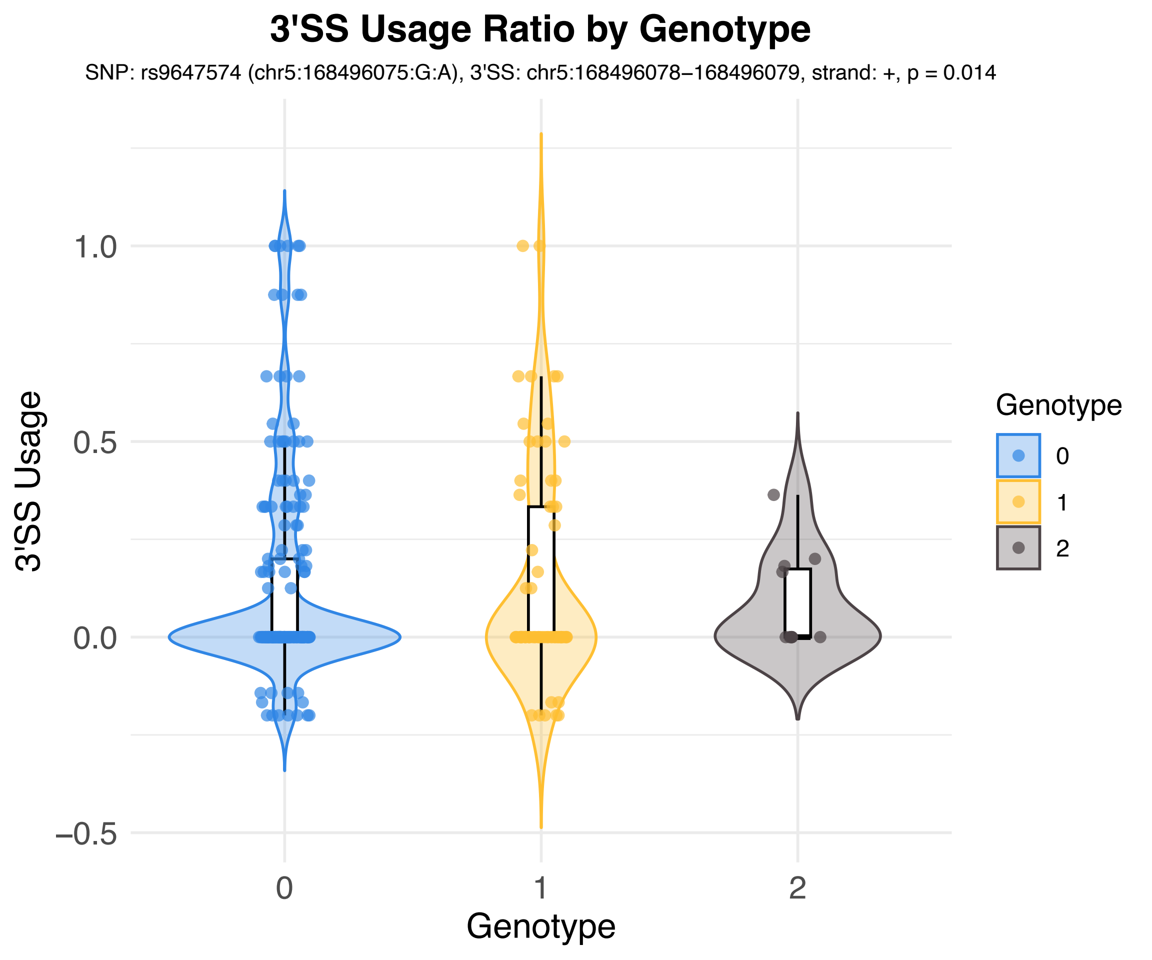
3'SS Usage Ratio by Genotype
SNP: rs9647574 (chr5:168496075:G:A), 3'SS: chr5:168496078–168496079, strand: +, p = 0.014
3'SS Usage
1.0
0.5
0.0
–0.5
0
1
2
Genotype
Genotype
0
1
2

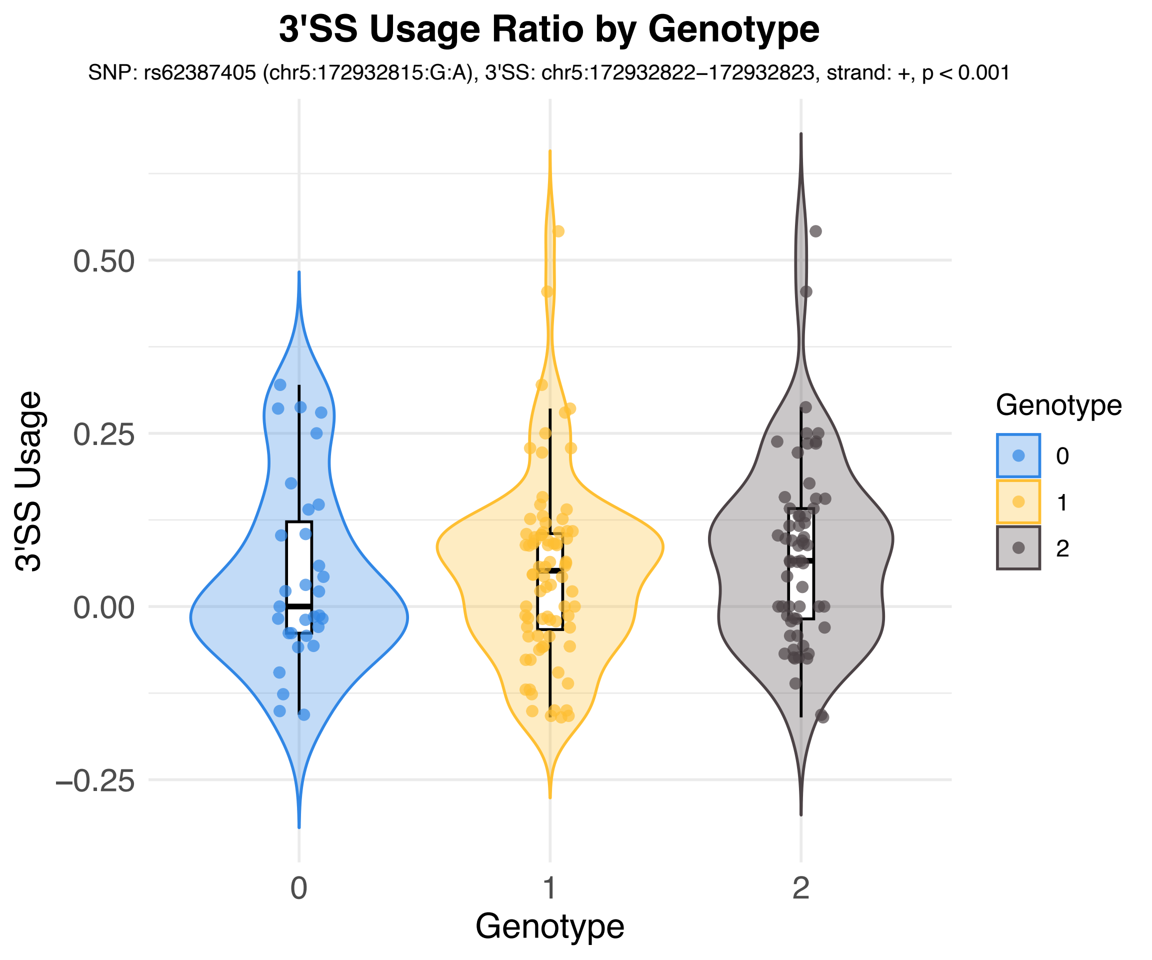
3'SS Usage Ratio by Genotype
SNP: rs62387405 (chr5:172932815:G:A), 3'SS: chr5:172932822−172932823, strand: +, p < 0.001
3'SS Usage
0.50
0.25
0.00
−0.25
0
1
2
Genotype
Genotype
0
1
2

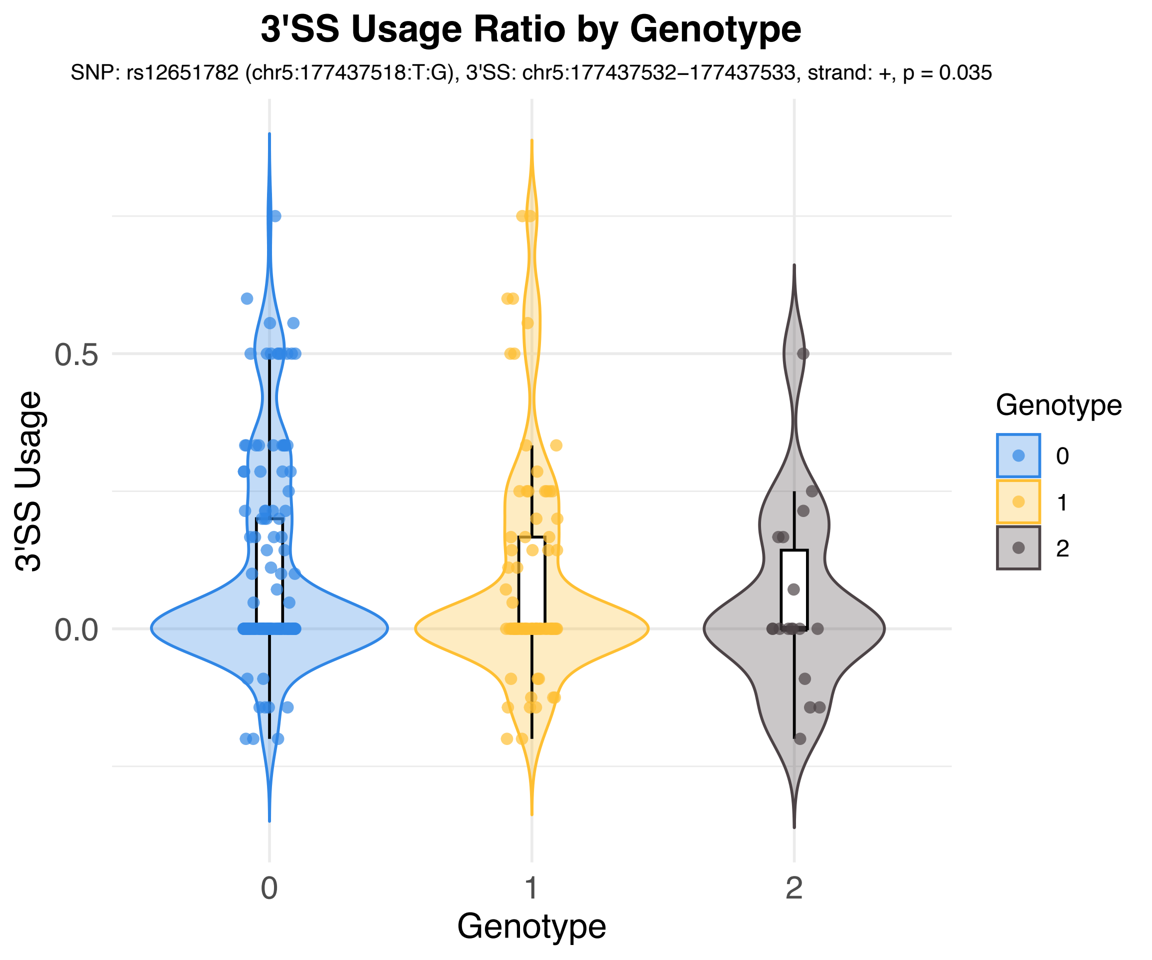
3'SS Usage Ratio by Genotype
SNP: rs12651782 (chr5:177437518:T:G), 3'SS: chr5:177437532−177437533, strand: +, p = 0.035
3'SS Usage
0.5
0.0
Genotype
0
1
2
Genotype
0
1
2

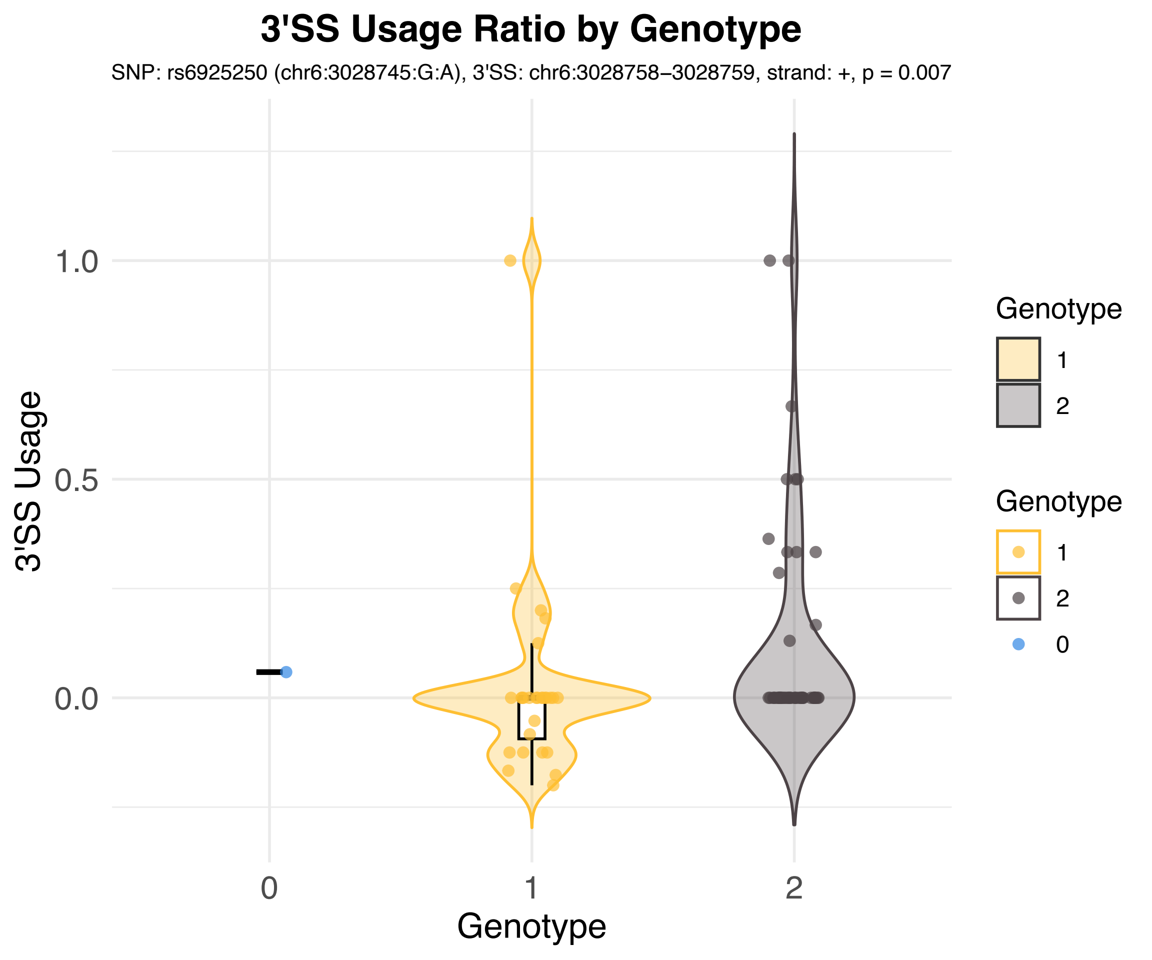
3'SS Usage Ratio by Genotype
SNP: rs6925250 (chr6:3028745:G:A), 3'SS: chr6:3028758–3028759, strand: +, p = 0.007
3'SS Usage
1.0
0.5
0.0
0
1
2
Genotype
Genotype
1
2
Genotype
1
2
0

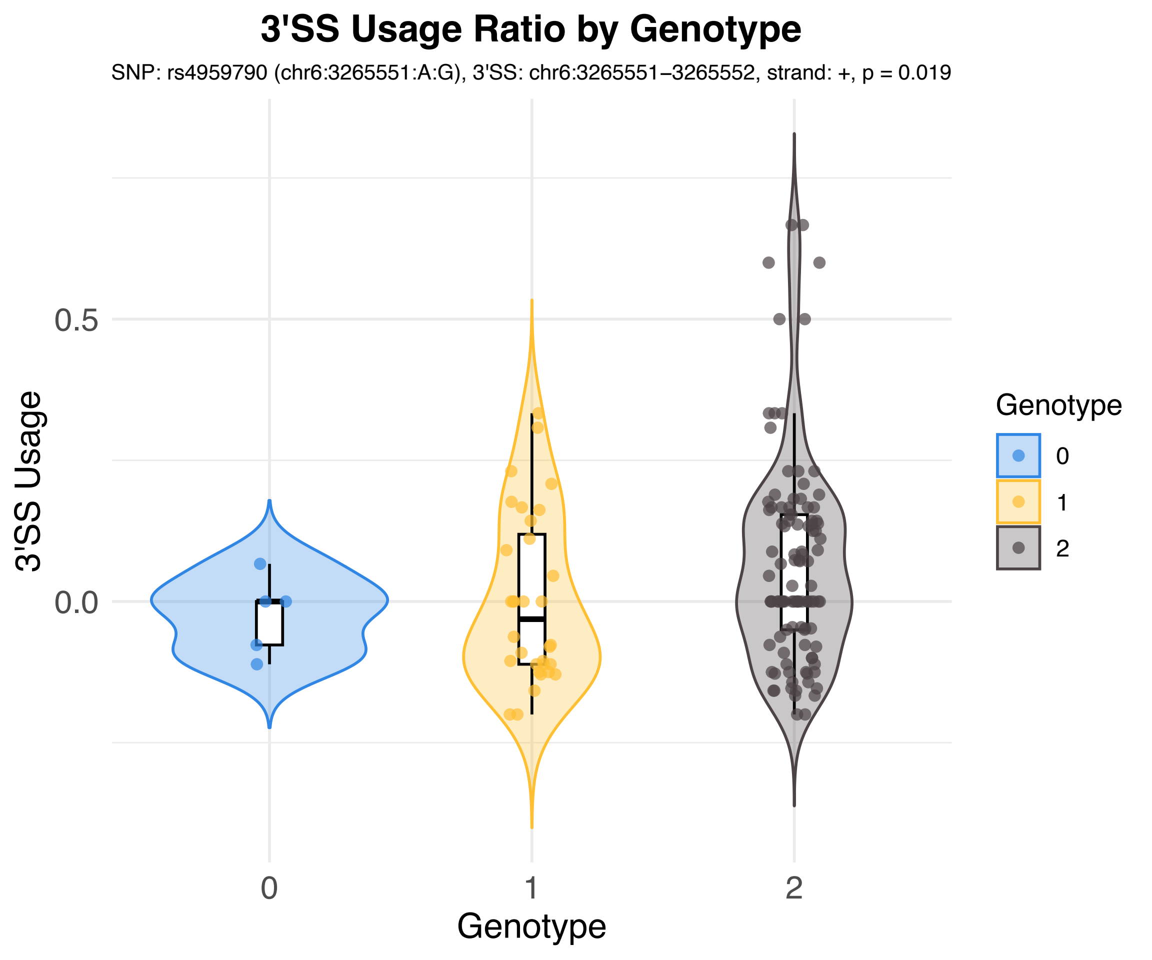

3'SS Usage Ratio by Genotype
SNP: rs4959790 (chr6:3265551:A:G), 3'SS: chr6:3265551–3265552, strand: +, p = 0.019
3'SS Usage
0.5
0.0
Genotype
0
1
2
Genotype
0
1
2

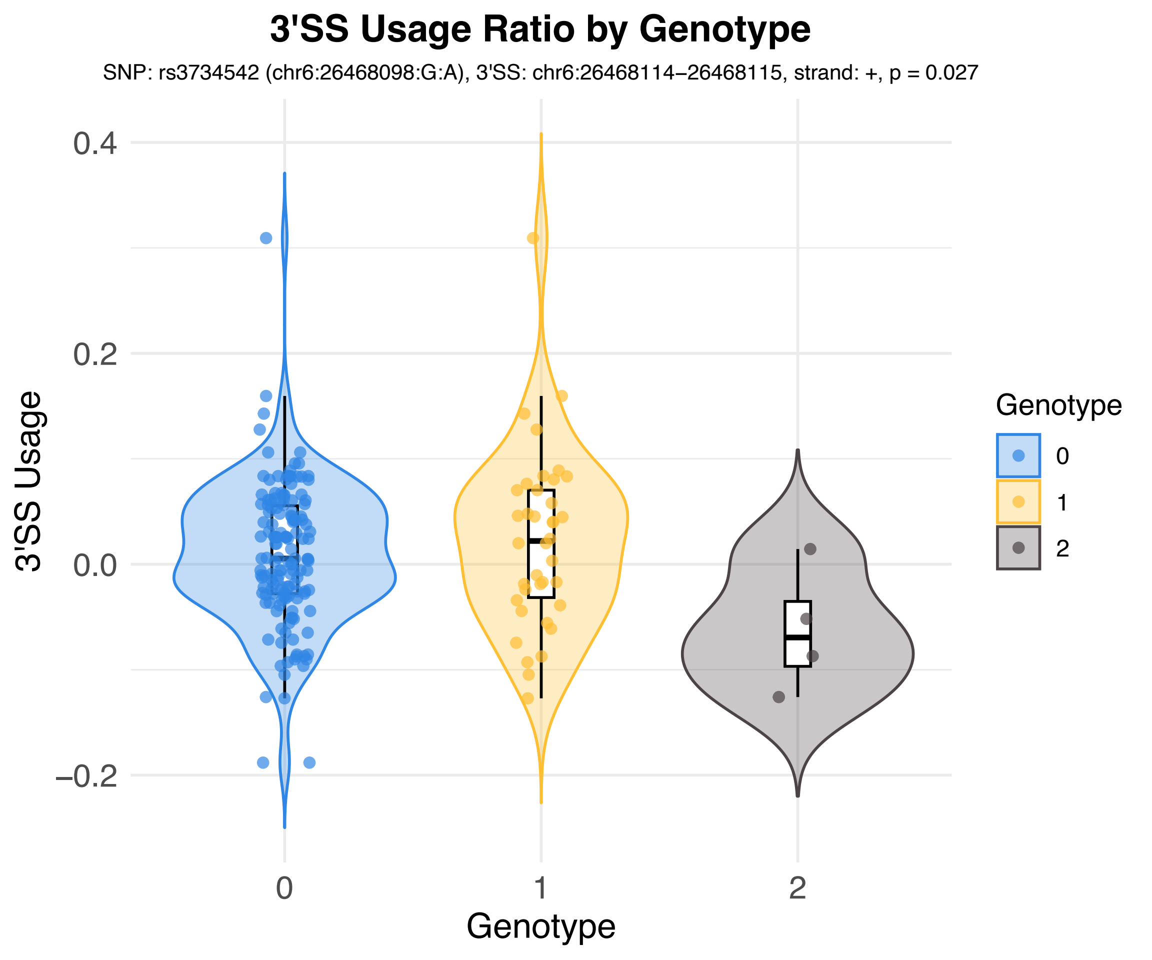
3'SS Usage Ratio by Genotype
SNP: rs3734542 (chr6:26468098:G:A), 3'SS: chr6:26468114−26468115, strand: +, p = 0.027
3'SS Usage
0.4
0.2
0.0
−0.2
0
1
2
Genotype
Genotype
0
1
2

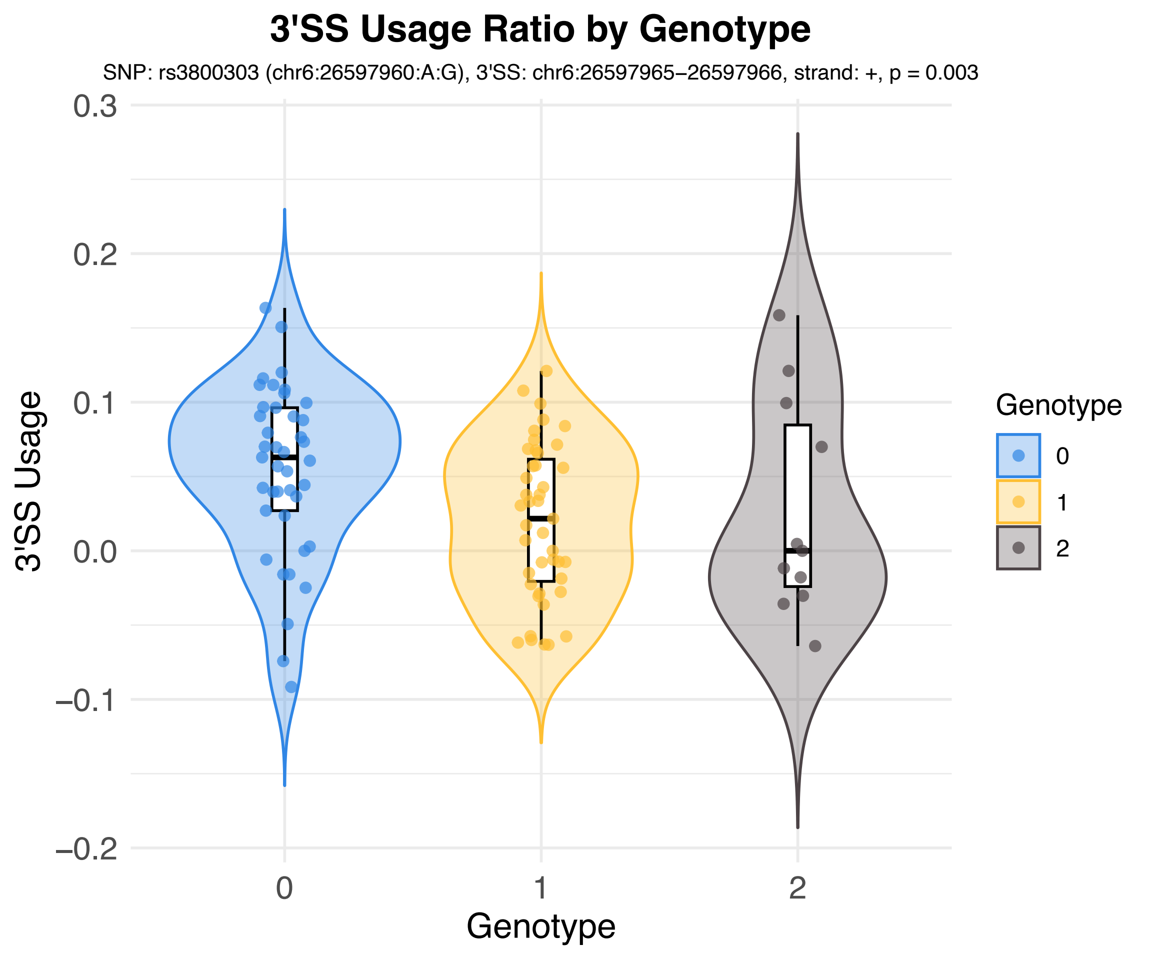

3'SS Usage Ratio by Genotype
SNP: rs3800303 (chr6:26597960:A:G), 3'SS: chr6:26597965−26597966, strand: +, p = 0.003
3'SS Usage
0.3
0.2
0.1
0.0
−0.1
−0.2
0
1
2
Genotype
Genotype
0
1
2

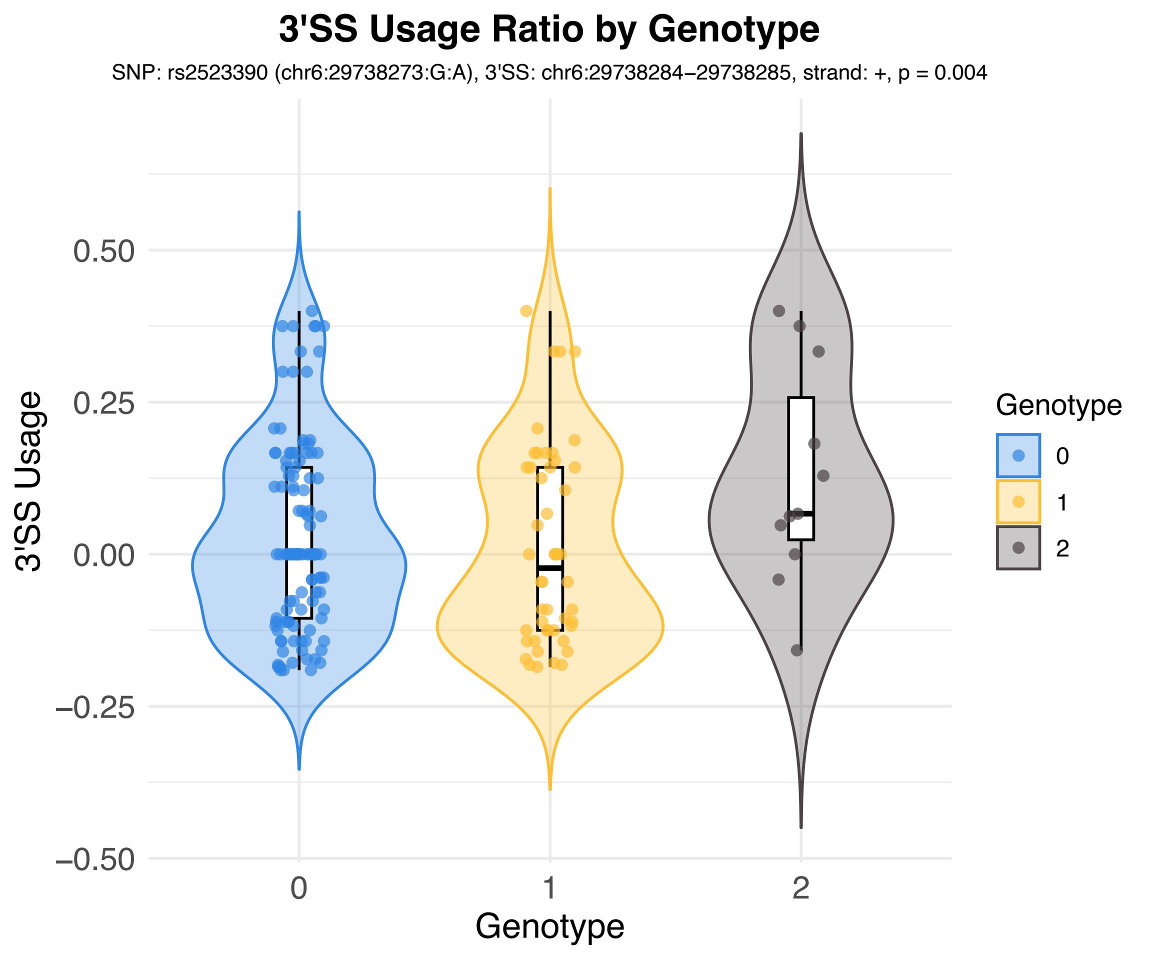
3'SS Usage Ratio by Genotype
SNP: rs2523390 (chr6:29738273:G:A), 3'SS: chr6:29738284−29738285, strand: +, p = 0.004
3'SS Usage
0.50
0.25
0.00
−0.25
−0.50
0
1
2
Genotype
Genotype
0
1
2

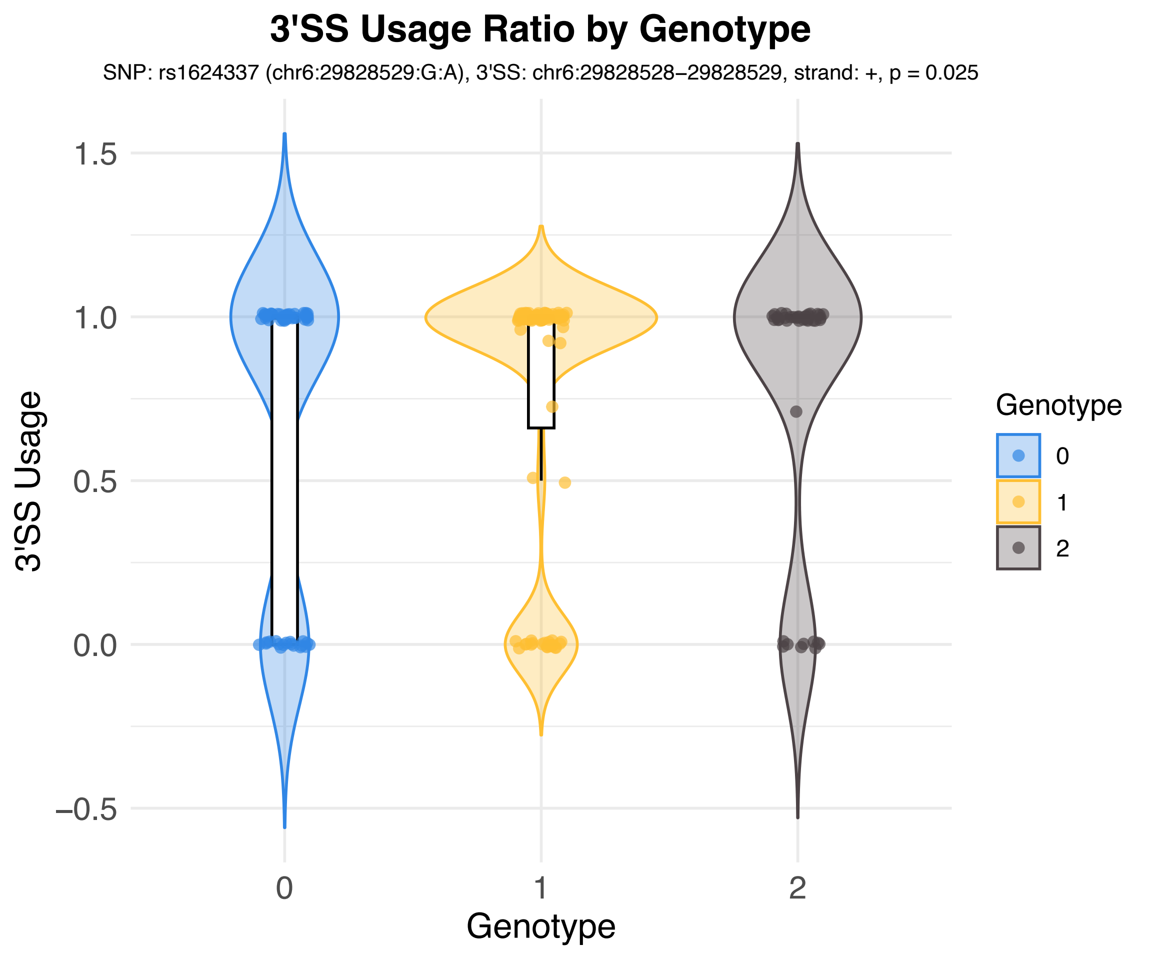

3'SS Usage Ratio by Genotype
SNP: rs1624337 (chr6:29828529:G:A), 3'SS: chr6:29828528−29828529, strand: +, p = 0.025
3'SS Usage
1.5
1.0
0.5
0.0
−0.5
0
1
2
Genotype
Genotype
0
1
2

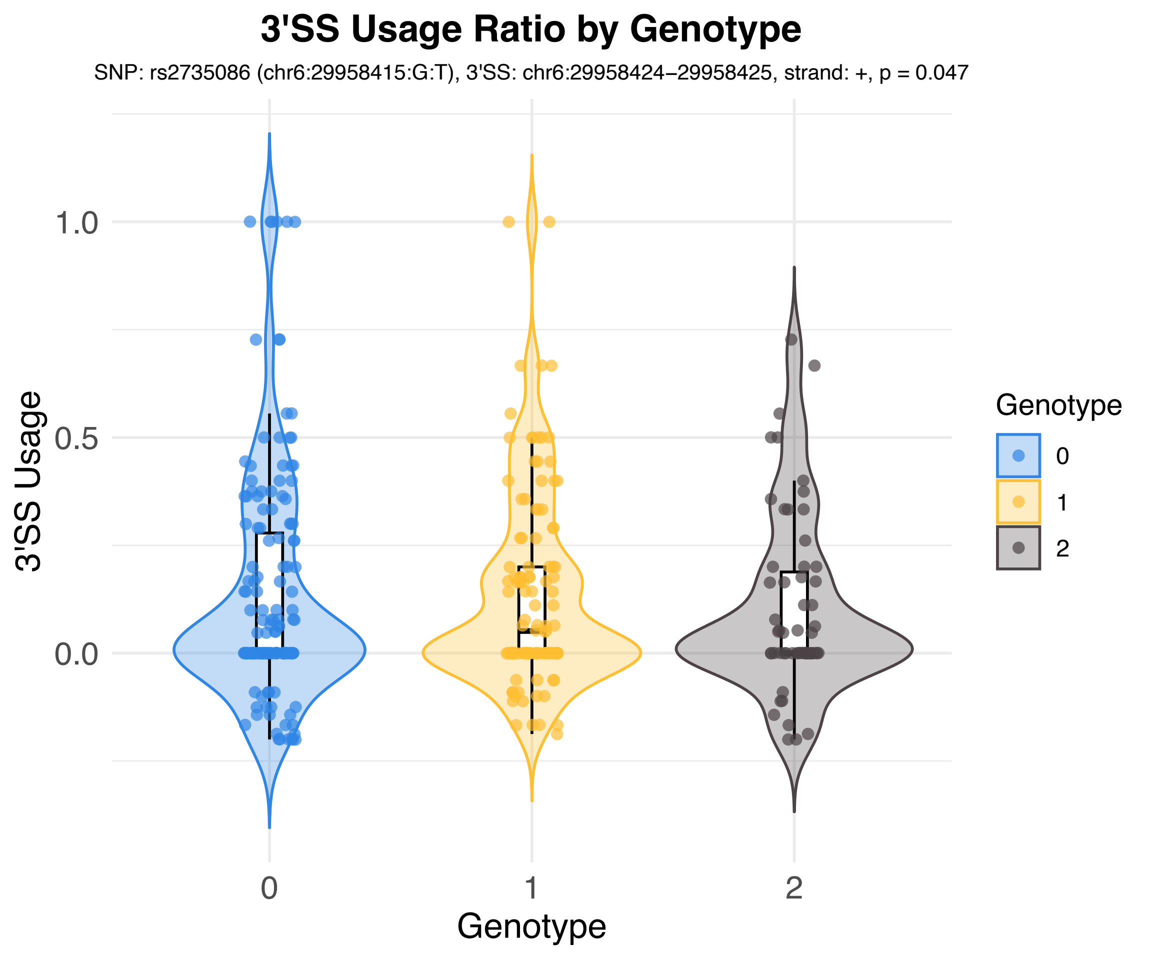

3'SS Usage Ratio by Genotype
SNP: rs2735086 (chr6:29958415:G:T), 3'SS: chr6:29958424−29958425, strand: +, p = 0.047
3'SS Usage
1.0
0.5
0.0
0
1
2
Genotype
Genotype
0
1
2

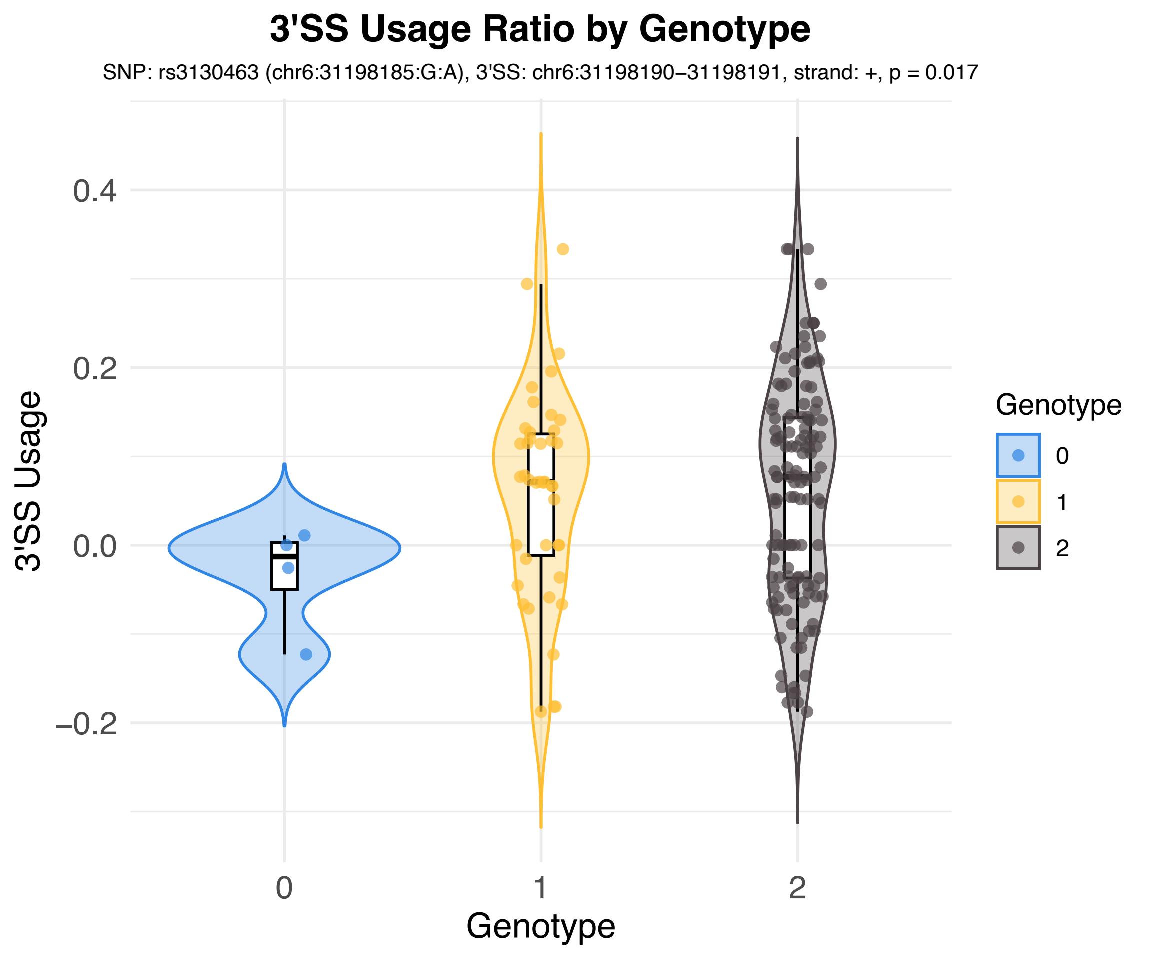
3'SS Usage Ratio by Genotype
SNP: rs3130463 (chr6:31198185:G:A), 3'SS: chr6:31198190–31198191, strand: +, p = 0.017
3'SS Usage
0.4
0.2
0.0
−0.2
0
1
2
Genotype
Genotype
0
1
2

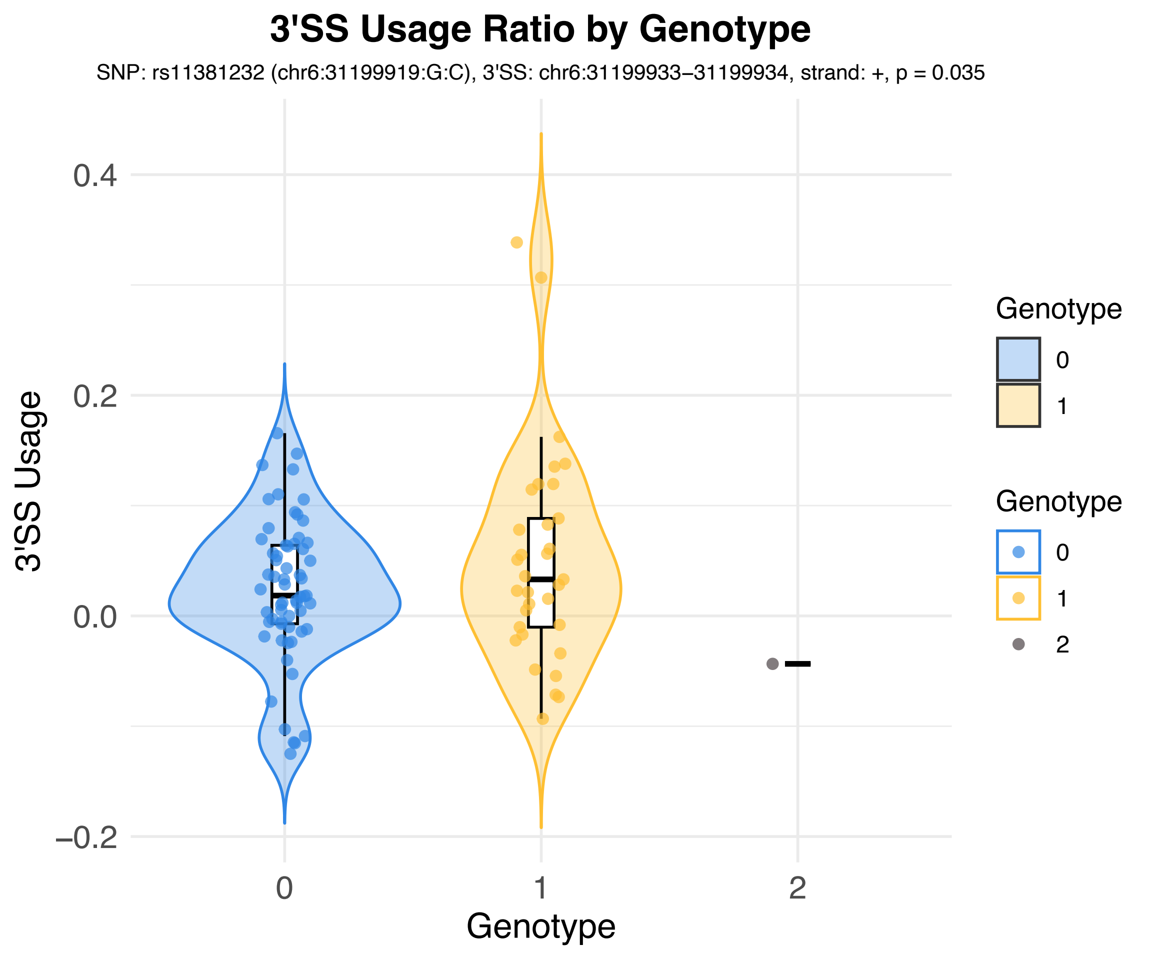

3'SS Usage Ratio by Genotype
SNP: rs11381232 (chr6:31199919:G:C), 3'SS: chr6:31199933–31199934, strand: +, p = 0.035
3'SS Usage
0.4
0.2
0.0
−0.2
0
1
2
Genotype
Genotype
0
1
Genotype
0
1
2

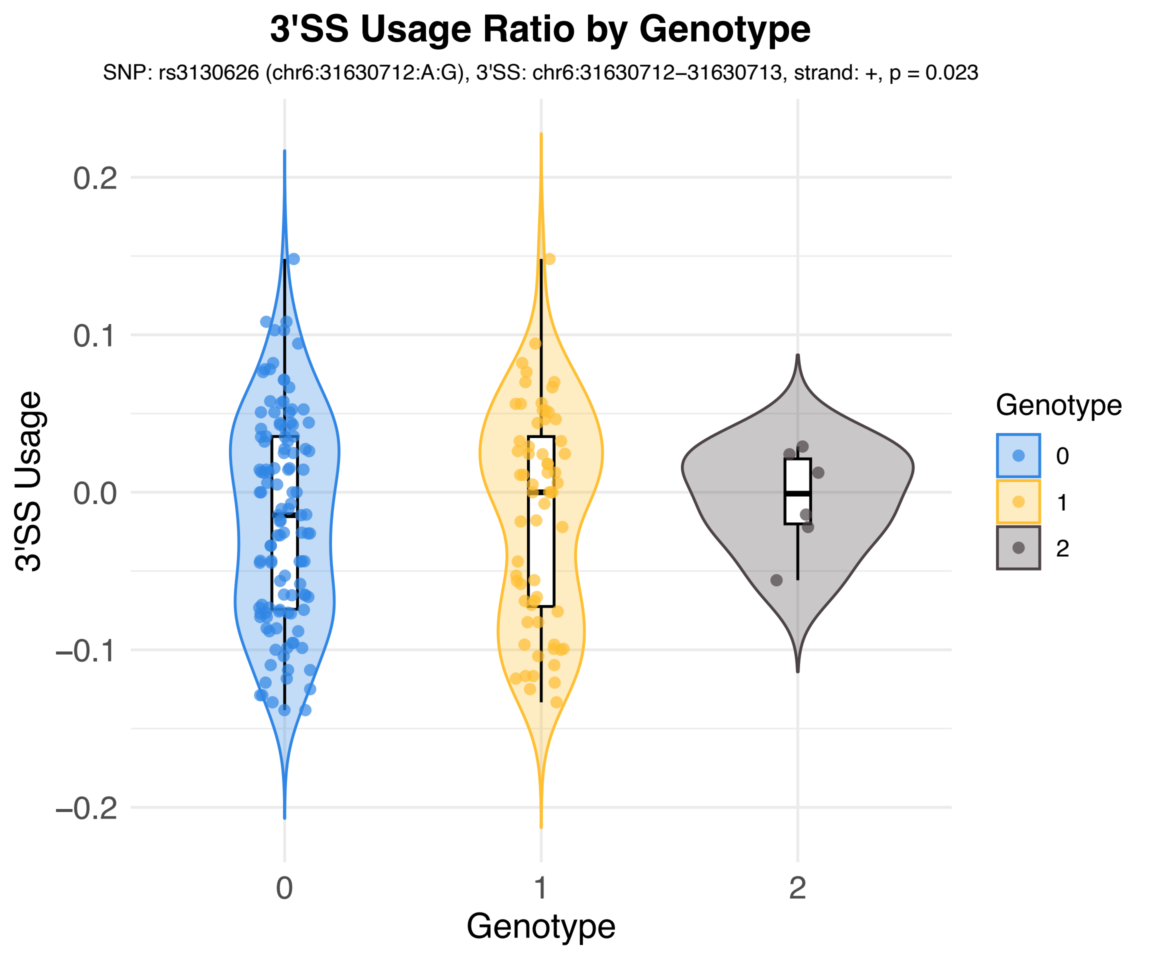
3'SS Usage Ratio by Genotype
SNP: rs3130626 (chr6:31630712:A:G), 3'SS: chr6:31630712–31630713, strand: +, p = 0.023
3'SS Usage
0.2
0.1
0.0
−0.1
−0.2
0
1
2
Genotype
Genotype
0
1
2

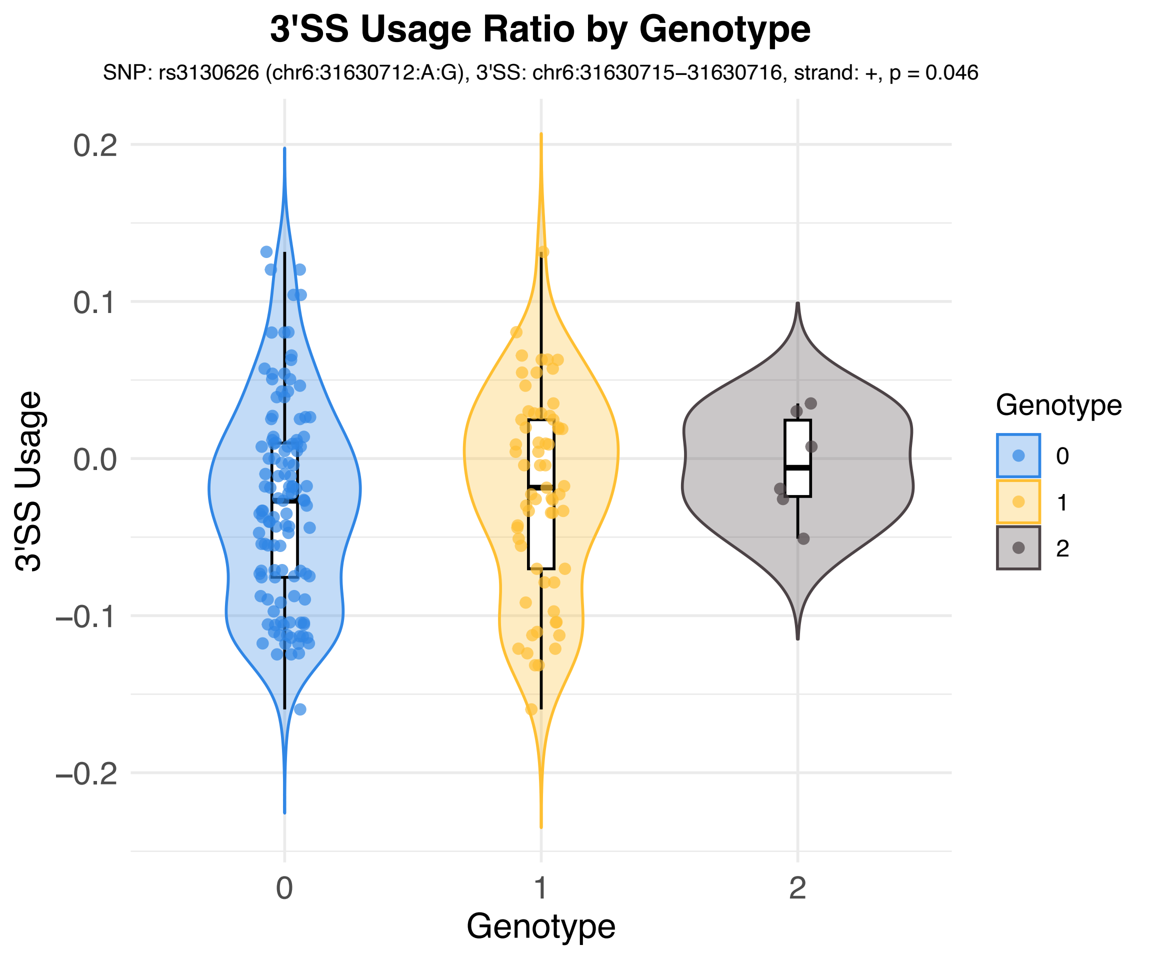
3'SS Usage Ratio by Genotype
SNP: rs3130626 (chr6:31630712:A:G), 3'SS: chr6:31630715–31630716, strand: +, p = 0.046
3'SS Usage
0.2
0.1
0.0
–0.1
–0.2
0
1
2
Genotype
Genotype
0
1
2

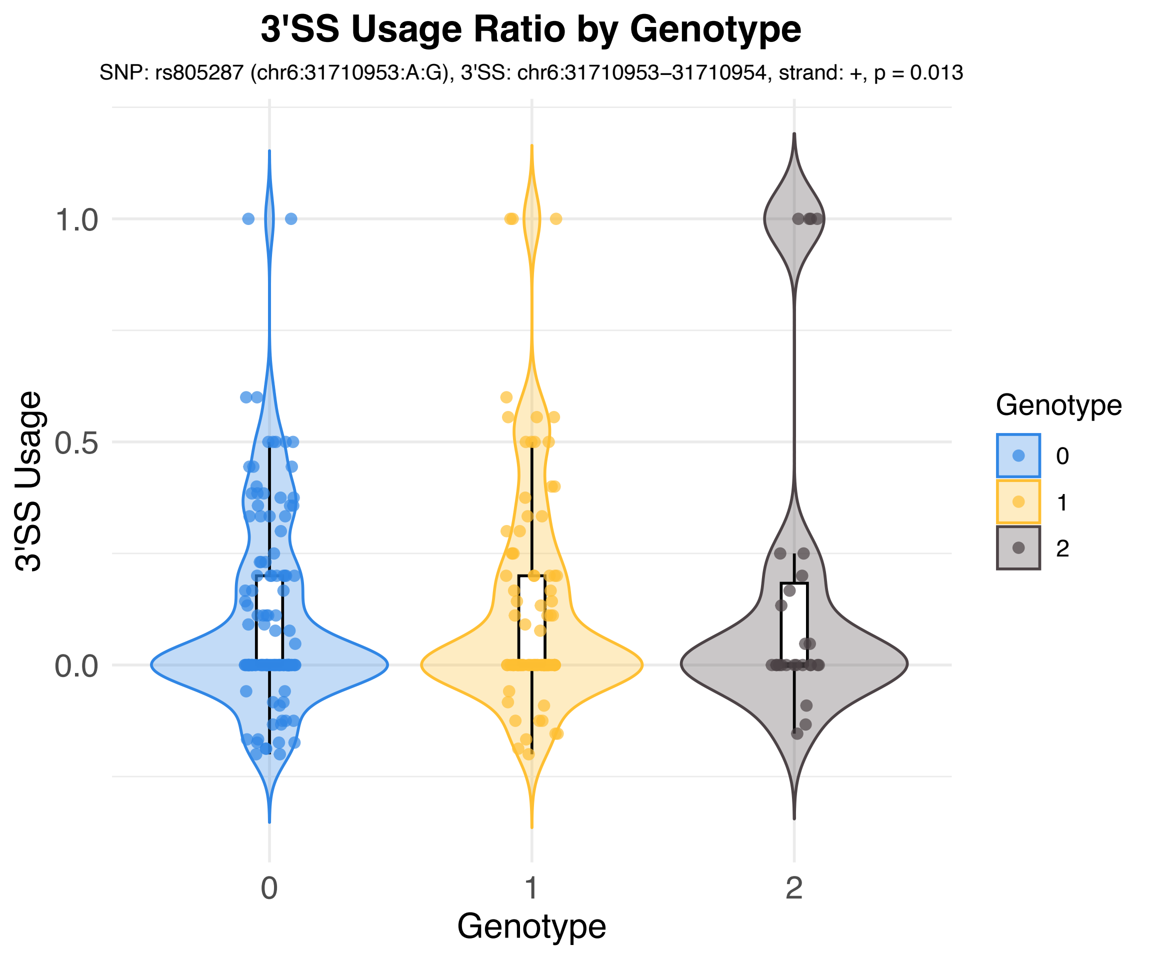
3'SS Usage Ratio by Genotype
SNP: rs805287 (chr6:31710953:A:G), 3'SS: chr6:31710953–31710954, strand: +, p = 0.013
3'SS Usage
1.0
0.5
0.0
0
1
2
Genotype
Genotype
0
1
2

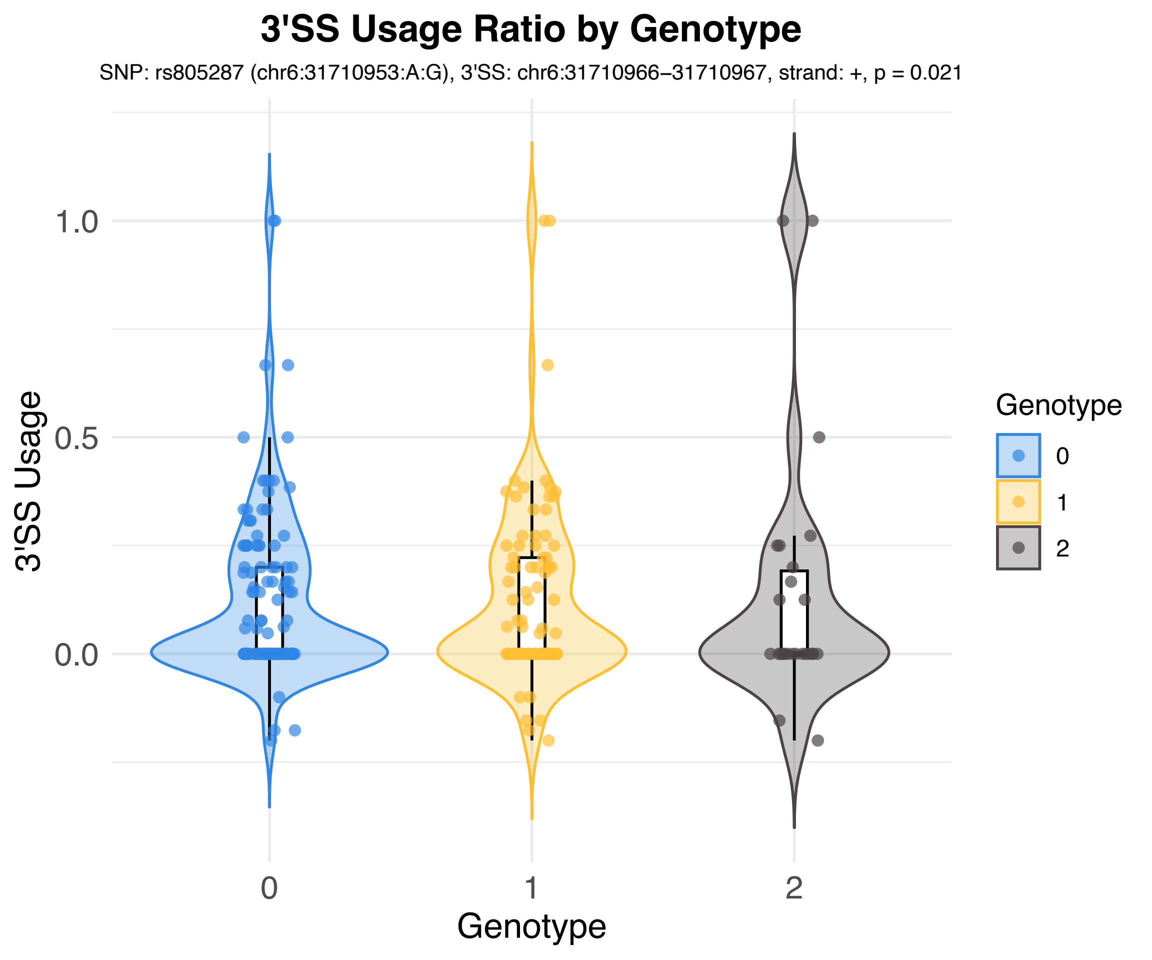
3'SS Usage Ratio by Genotype
SNP: rs805287 (chr6:31710953:A:G), 3'SS: chr6:31710966–31710967, strand: +, p = 0.021
3'SS Usage
1.0
0.5
0.0
0
1
2
Genotype
Genotype
0
1
2

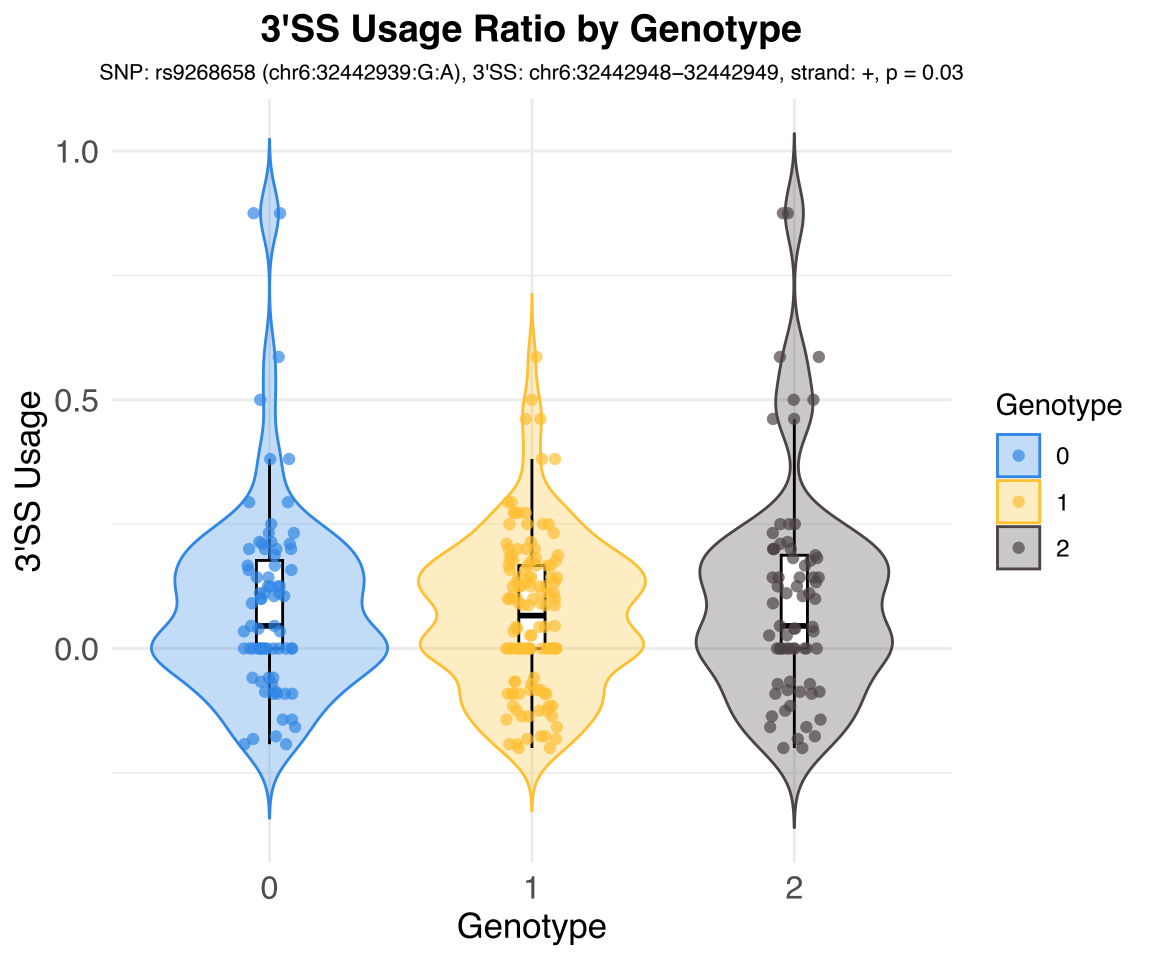

3'SS Usage Ratio by Genotype
SNP: rs9268658 (chr6:32442939:G:A), 3'SS: chr6:32442948–32442949, strand: +, p = 0.03
3'SS Usage
1.0
0.5
0.0
0
1
2
Genotype
Genotype
0
1
2

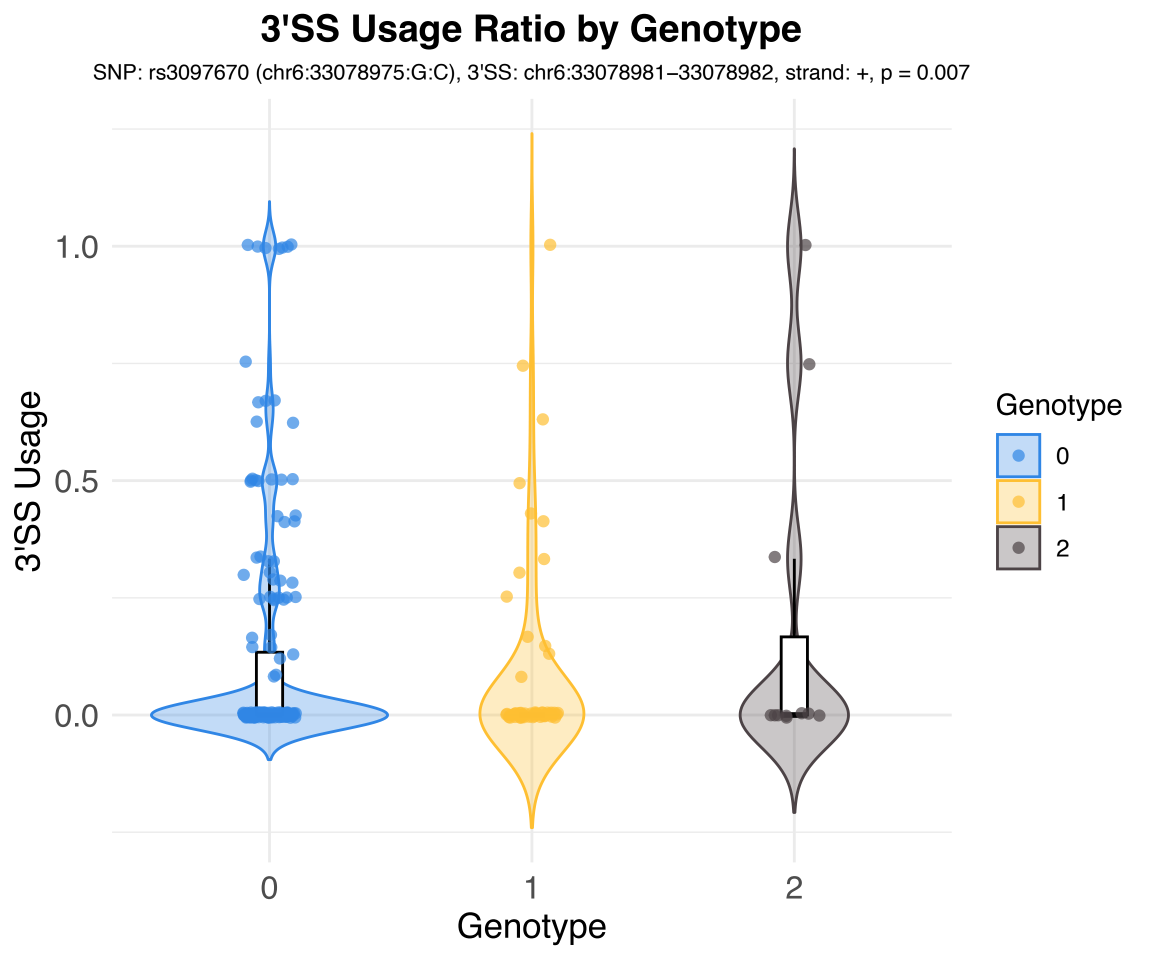
3'SS Usage Ratio by Genotype
SNP: rs3097670 (chr6:33078975:G:C), 3'SS: chr6:33078981–33078982, strand: +, p = 0.007
3'SS Usage
1.0
0.5
0.0
0
1
2
Genotype
Genotype
0
1
2

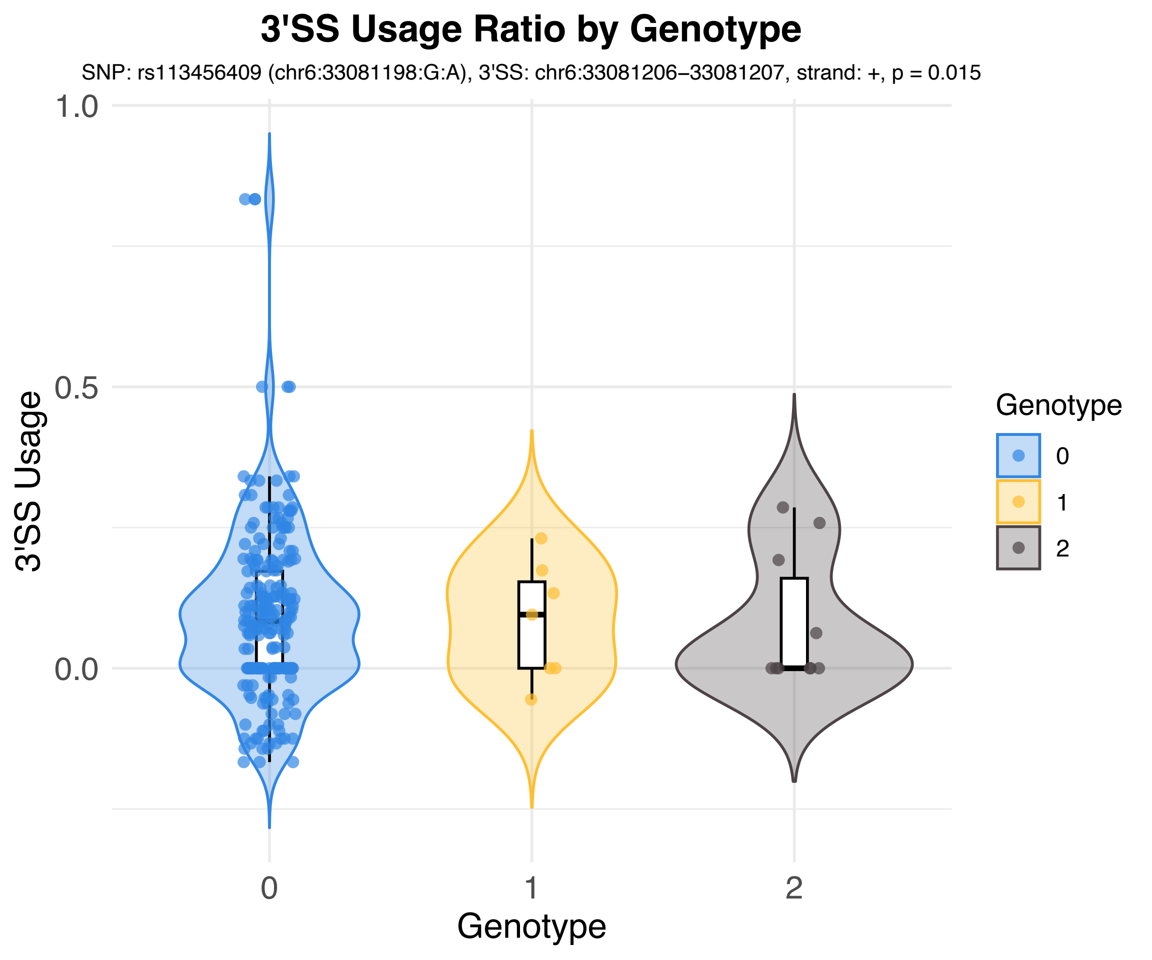
3'SS Usage Ratio by Genotype
SNP: rs113456409 (chr6:33081198:G:A), 3'SS: chr6:33081206–33081207, strand: +, p = 0.015
3'SS Usage
1.0
0.5
0.0
Genotype
0
1
2
Genotype
0
1
2

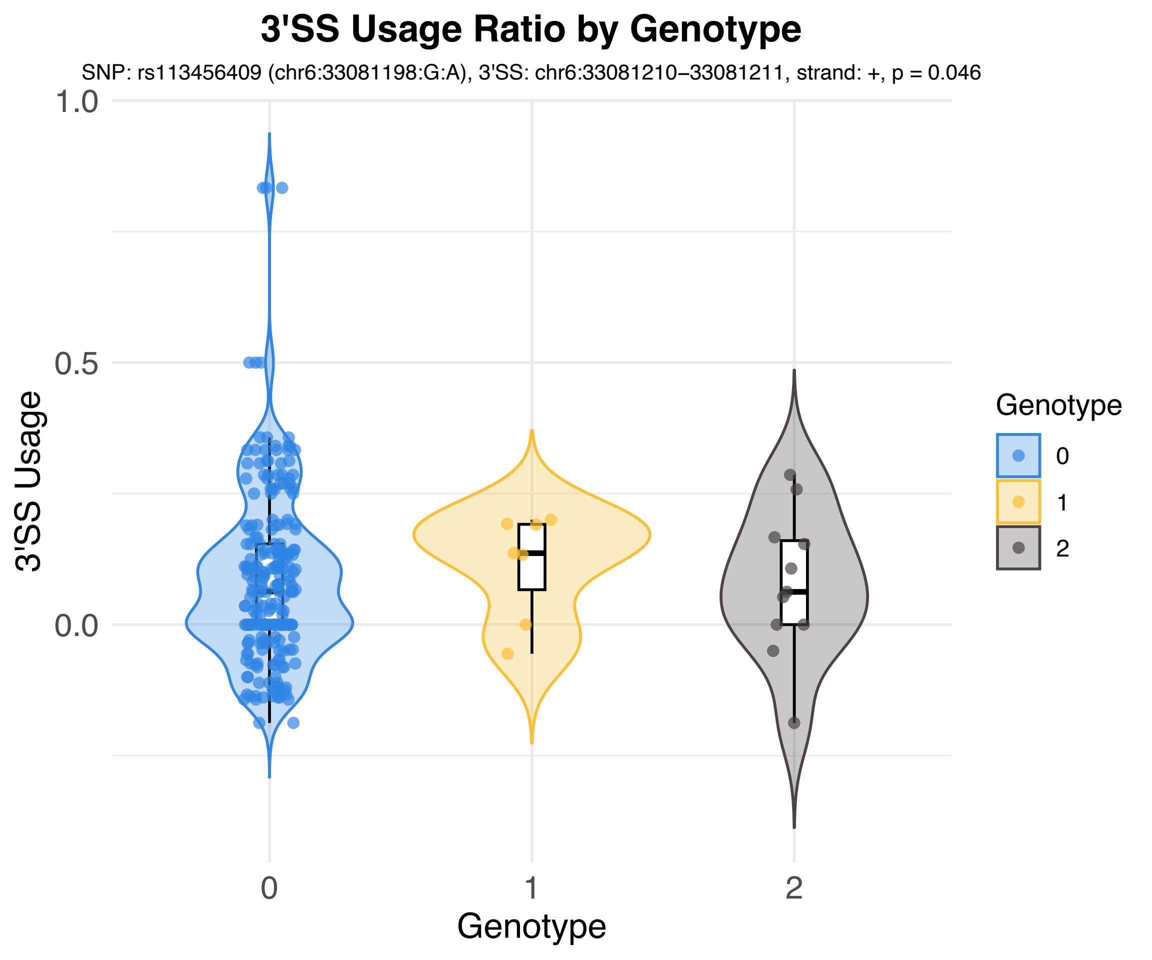
3'SS Usage Ratio by Genotype
SNP: rs113456409 (chr6:33081198:G:A), 3'SS: chr6:33081210–33081211, strand: +, p = 0.046
3'SS Usage
1.0
0.5
0.0
0
1
2
Genotype
Genotype
0
1
2

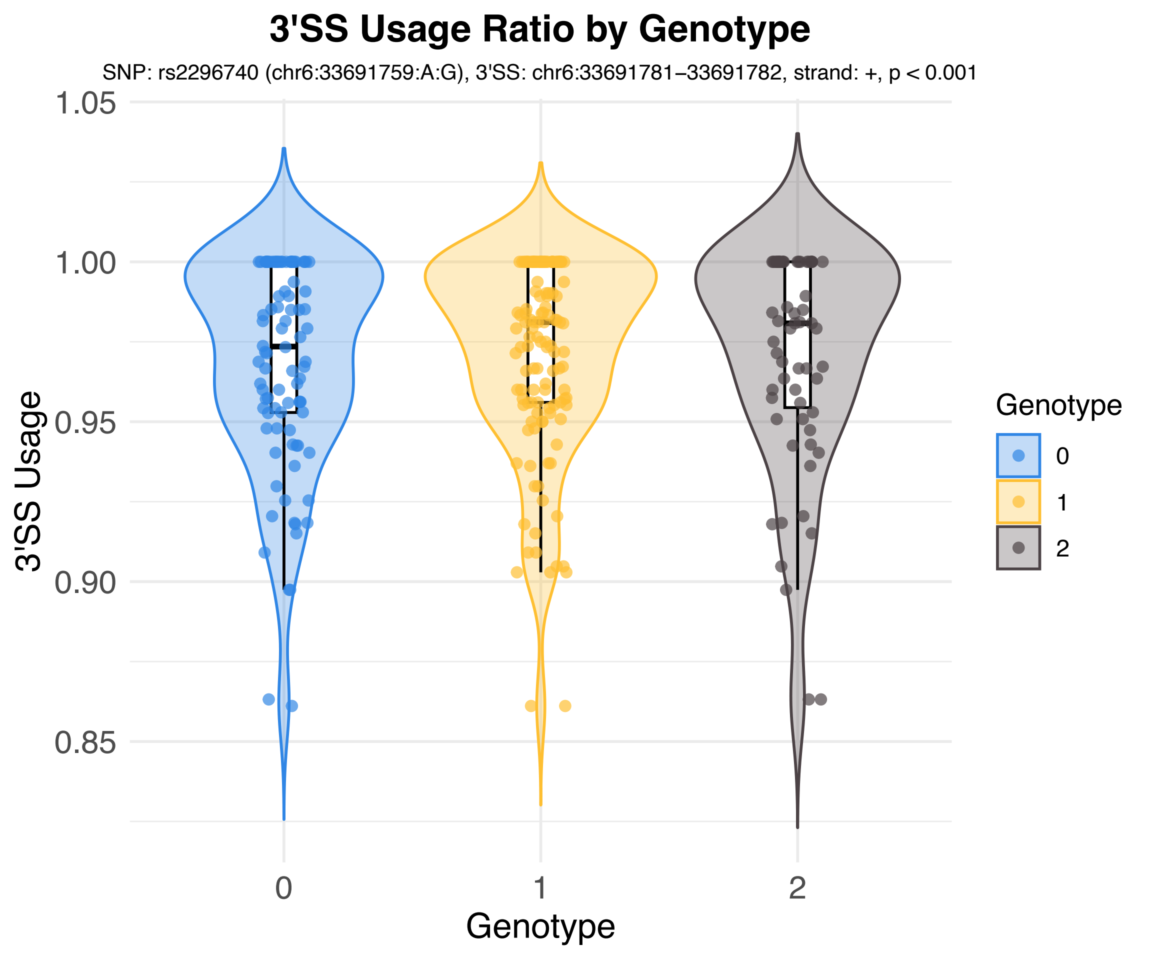

3'SS Usage Ratio by Genotype
SNP: rs2296740 (chr6:33691759:A:G), 3'SS: chr6:33691781–33691782, strand: +, p < 0.001
3'SS Usage
1.05
1.00
0.95
0.90
0.85
0
1
2
Genotype
Genotype
0
1
2

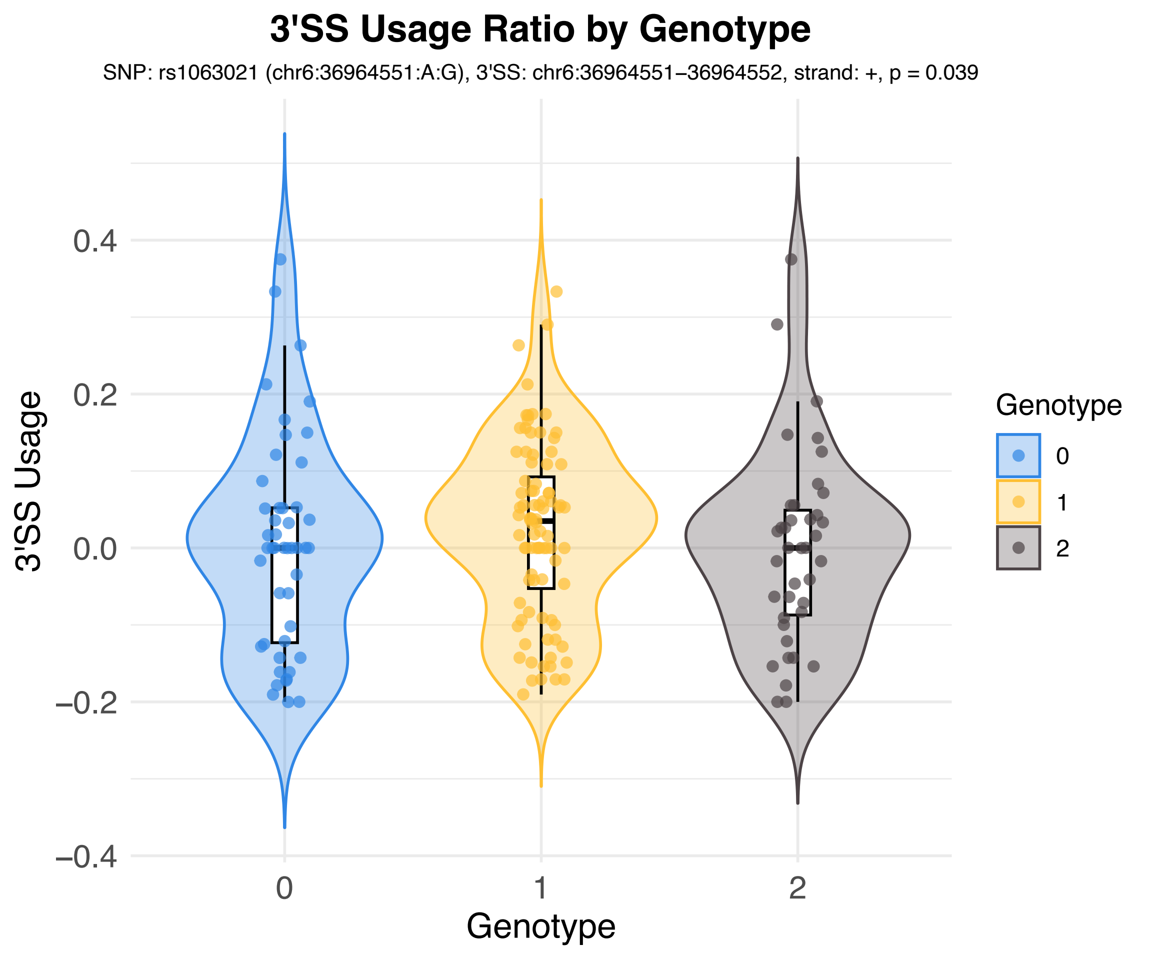
3'SS Usage Ratio by Genotype
SNP: rs1063021 (chr6:36964551:A:G), 3'SS: chr6:36964551–36964552, strand: +, p = 0.039
3'SS Usage
0.4
0.2
0.0
−0.2
−0.4
0
1
2
Genotype
Genotype
0
1
2

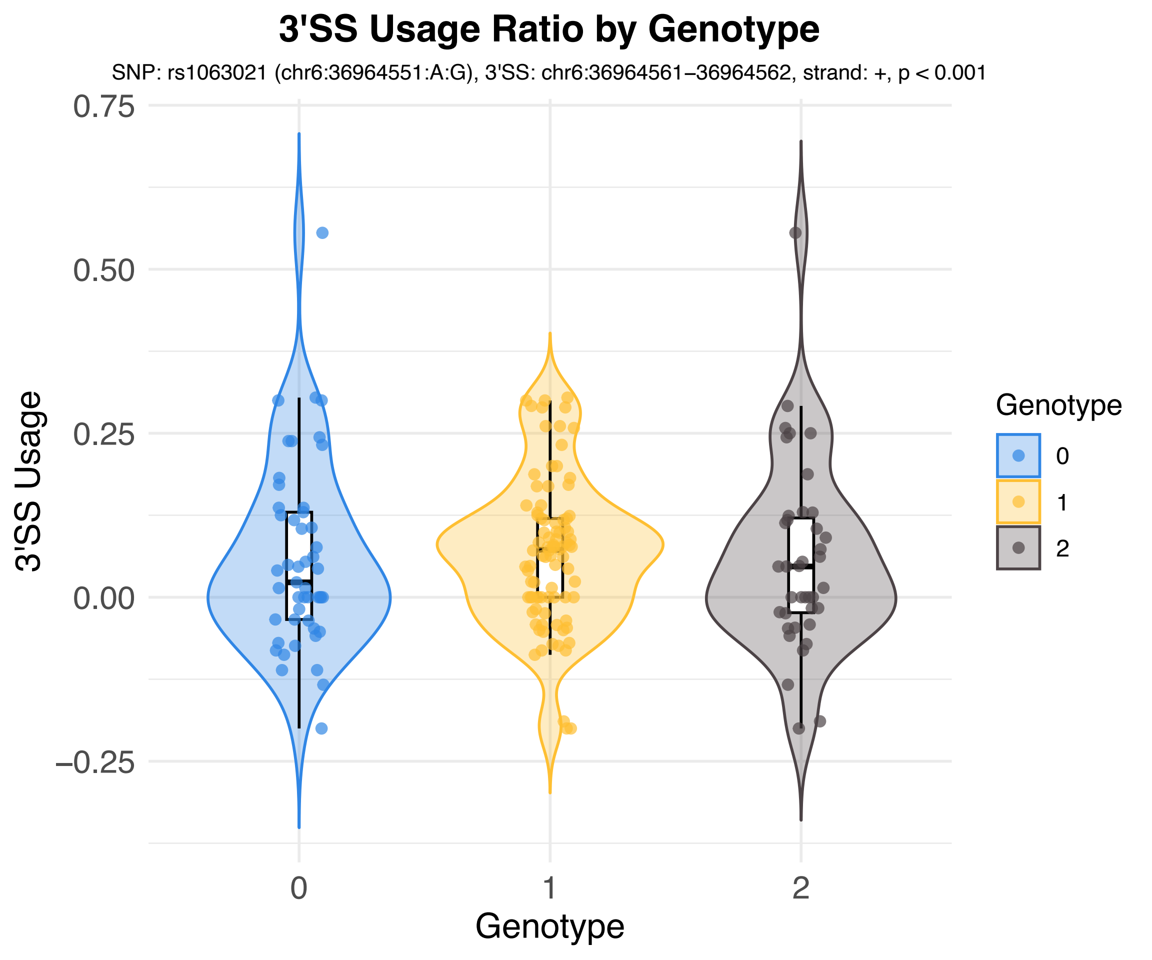

3'SS Usage Ratio by Genotype
SNP: rs1063021 (chr6:36964551:A:G), 3'SS: chr6:36964561–36964562, strand: +, p < 0.001
3'SS Usage
0.75
0.50
0.25
0.00
−0.25
0
1
2
Genotype
Genotype
0
1
2

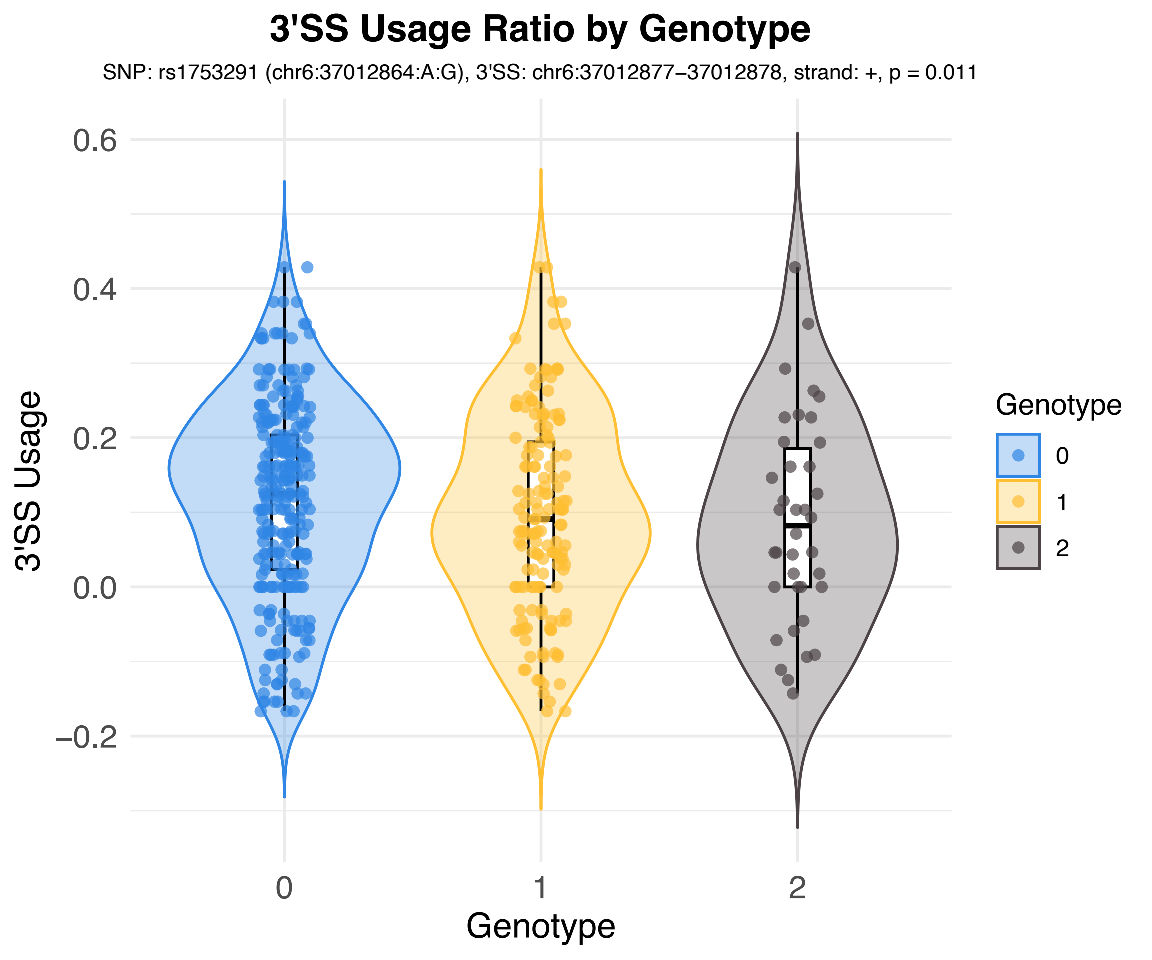
3'SS Usage Ratio by Genotype
SNP: rs1753291 (chr6:37012864:A:G), 3'SS: chr6:37012877–37012878, strand: +, p = 0.011
3'SS Usage
0.6
0.4
0.2
0.0
−0.2
0
1
2
Genotype
Genotype
0
1
2

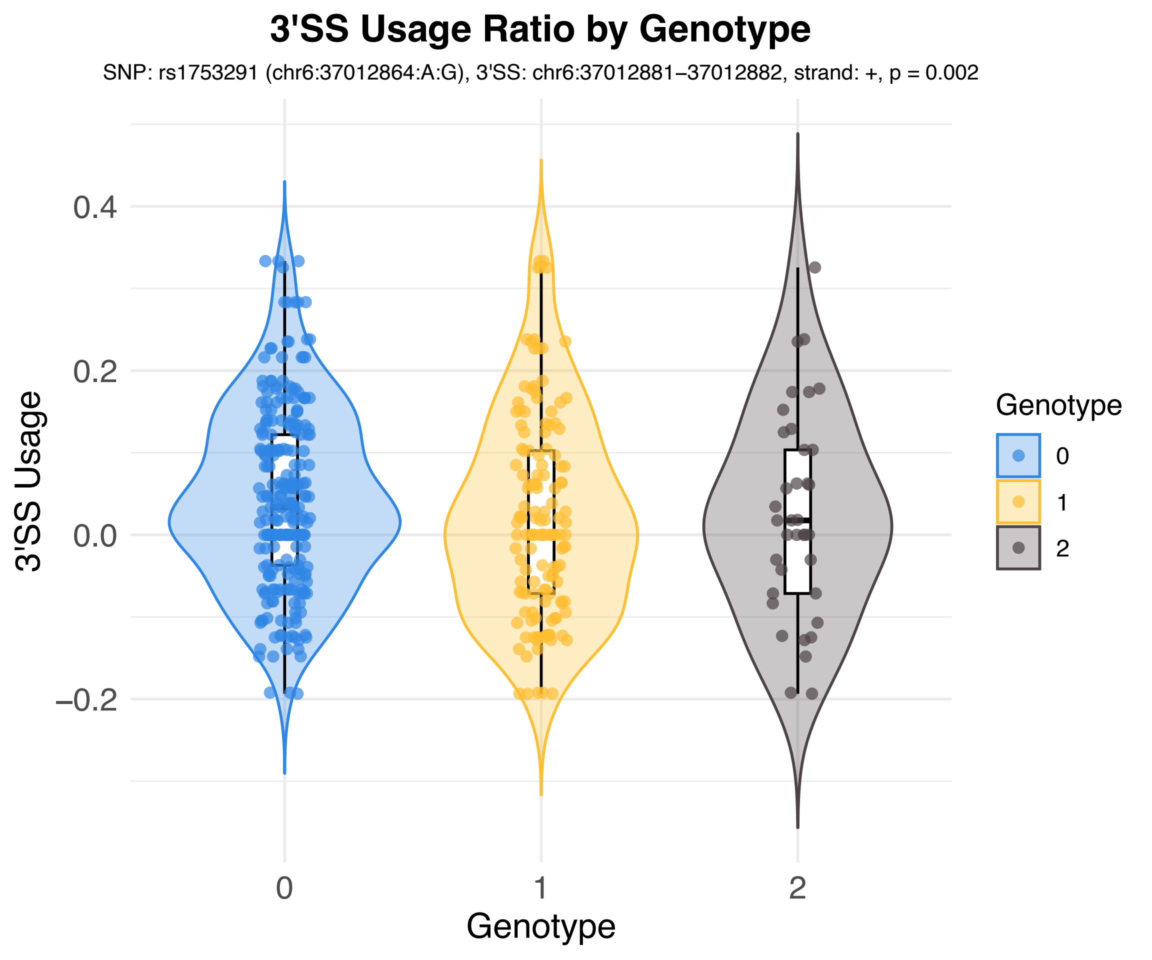
3'SS Usage Ratio by Genotype
SNP: rs1753291 (chr6:37012864:A:G), 3'SS: chr6:37012881–37012882, strand: +, p = 0.002
3'SS Usage
0.4
0.2
0.0
−0.2
0
1
2
Genotype
Genotype
0
1
2

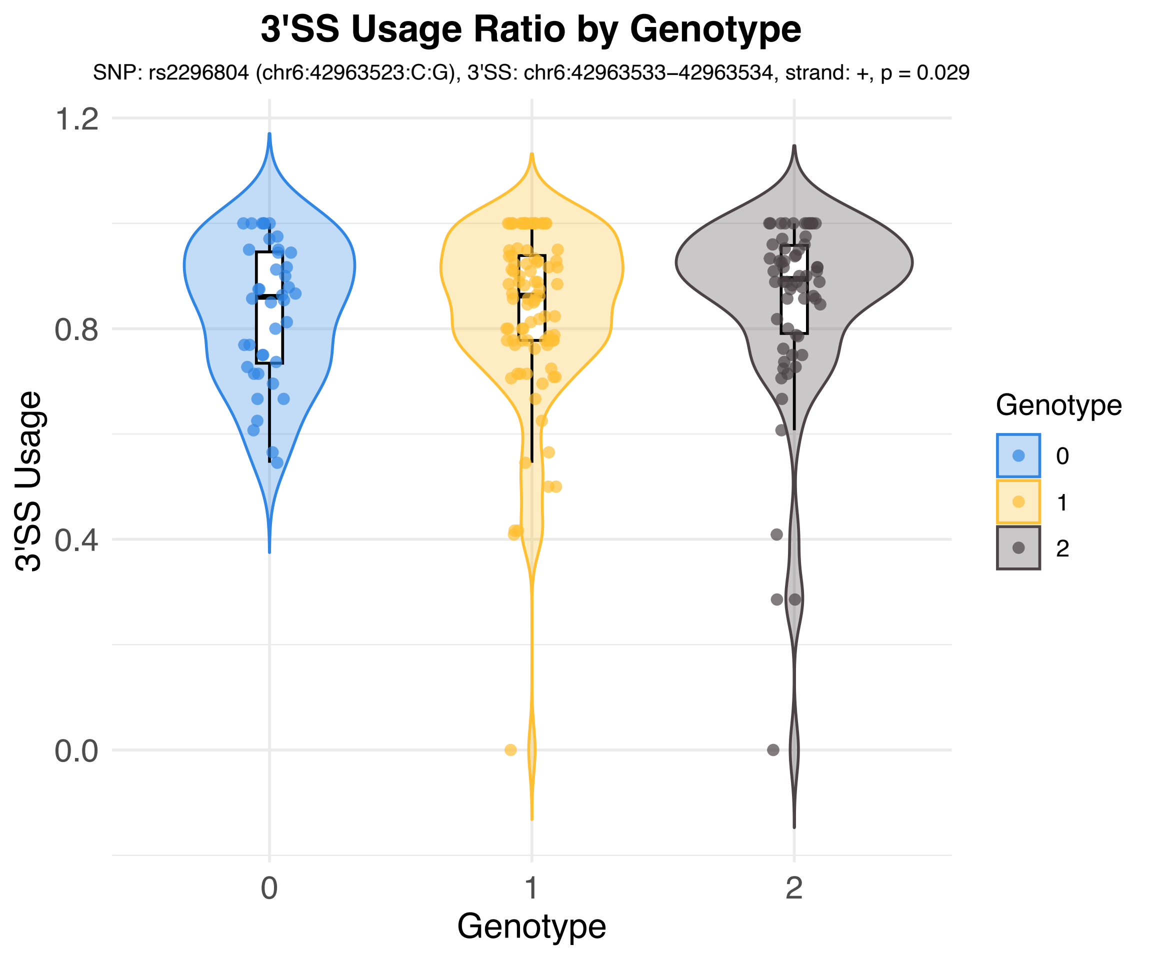

3'SS Usage Ratio by Genotype
SNP: rs2296804 (chr6:42963523:C:G), 3'SS: chr6:42963533–42963534, strand: +, p = 0.029
3'SS Usage
1.2
0.8
0.4
0.0
0
1
2
Genotype
Genotype
0
1
2

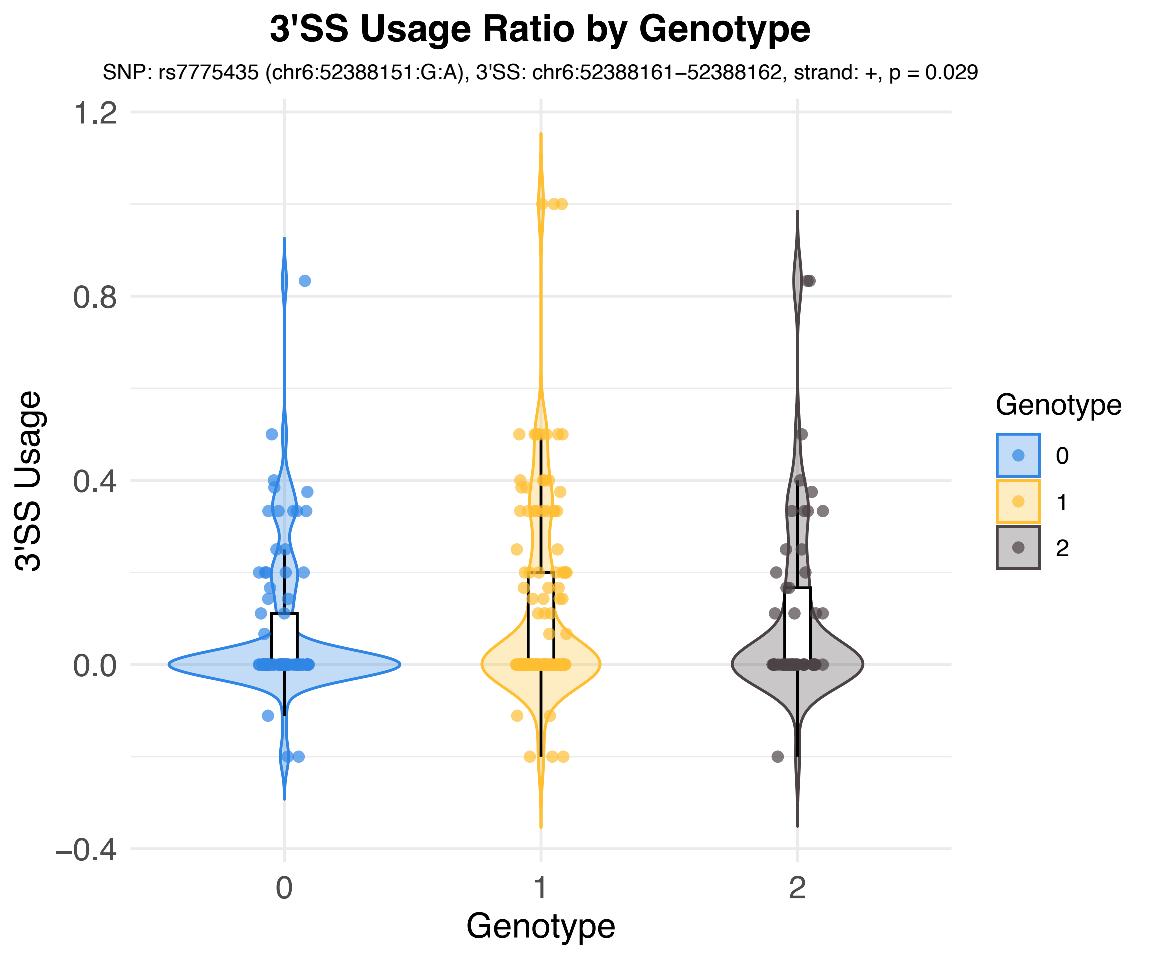

3'SS Usage Ratio by Genotype
SNP: rs7775435 (chr6:52388151:G:A), 3'SS: chr6:52388161–52388162, strand: +, p = 0.029
3'SS Usage
1.2
0.8
0.4
0.0
−0.4
0
1
2
Genotype
Genotype
0
1
2

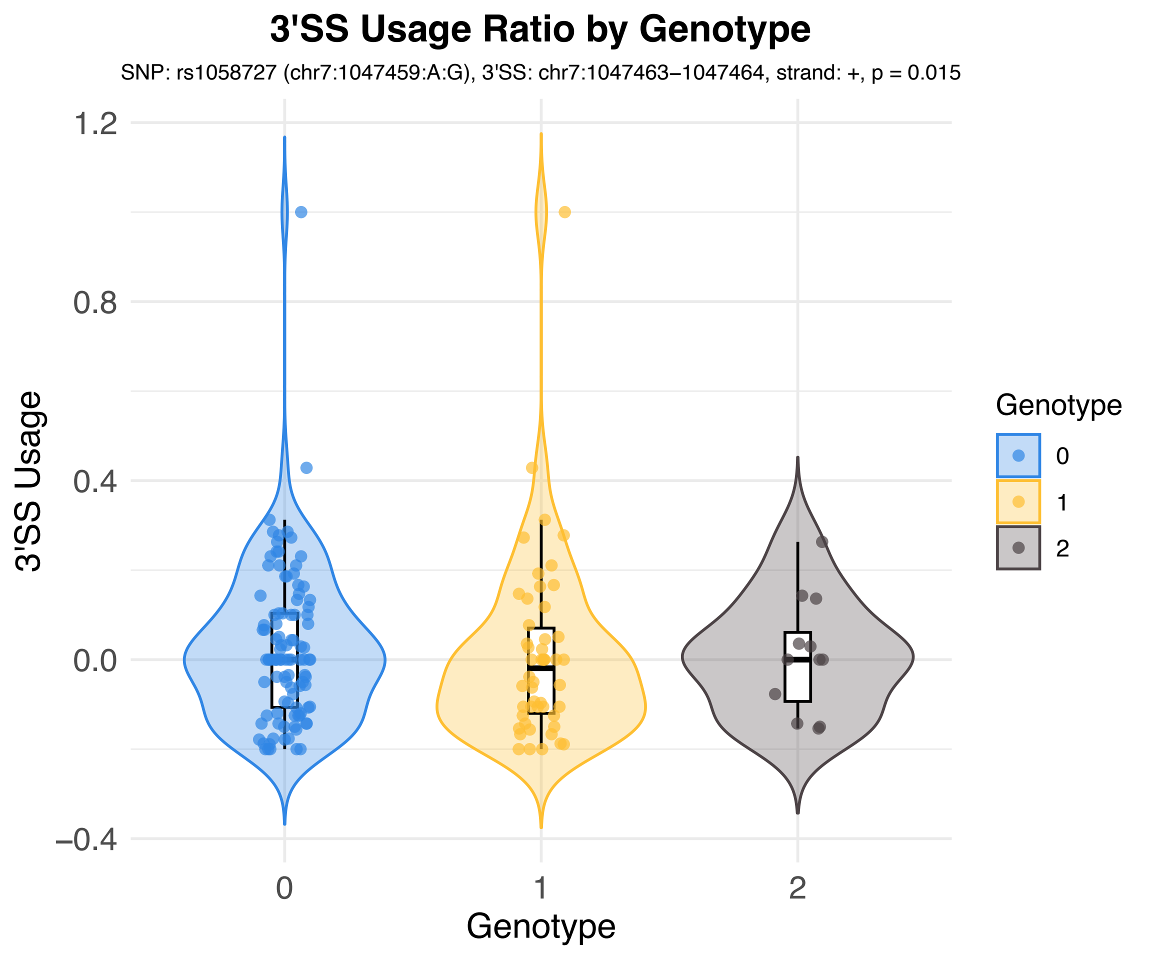
3'SS Usage Ratio by Genotype
SNP: rs1058727 (chr7:1047459:A:G), 3'SS: chr7:1047463–1047464, strand: +, p = 0.015
3'SS Usage
1.2
0.8
0.4
0.0
–0.4
0
1
2
Genotype
Genotype
0
1
2

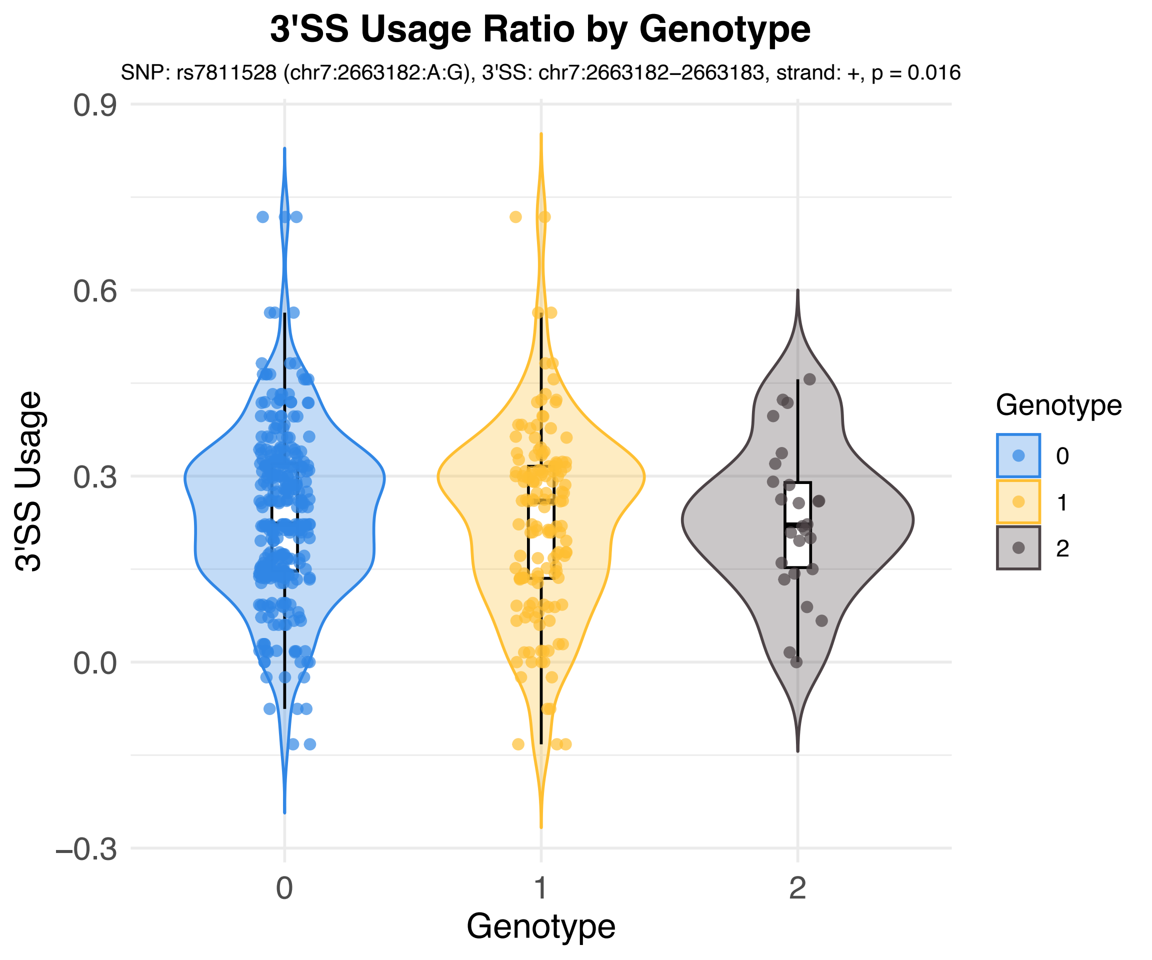

3'SS Usage Ratio by Genotype
SNP: rs7811528 (chr7:2663182:A:G), 3'SS: chr7:2663182–2663183, strand: +, p = 0.016
3'SS Usage
0.9
0.6
0.3
0.0
–0.3
0
1
2
Genotype
Genotype
0
1
2

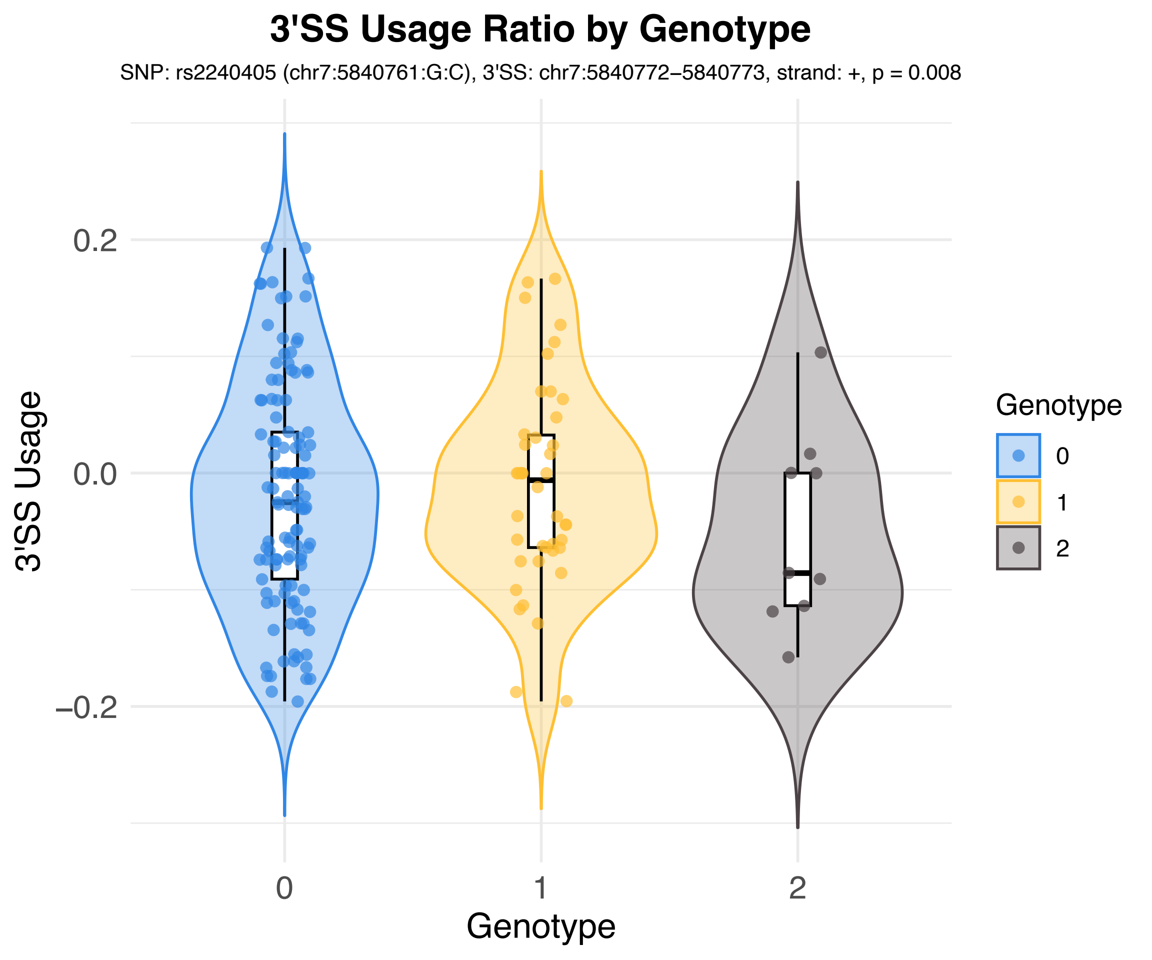
3'SS Usage Ratio by Genotype
SNP: rs2240405 (chr7:5840761:G:C), 3'SS: chr7:5840772–5840773, strand: +, p = 0.008
3'SS Usage
0.2
0.0
−0.2
0
1
2
Genotype
Genotype
0
1
2

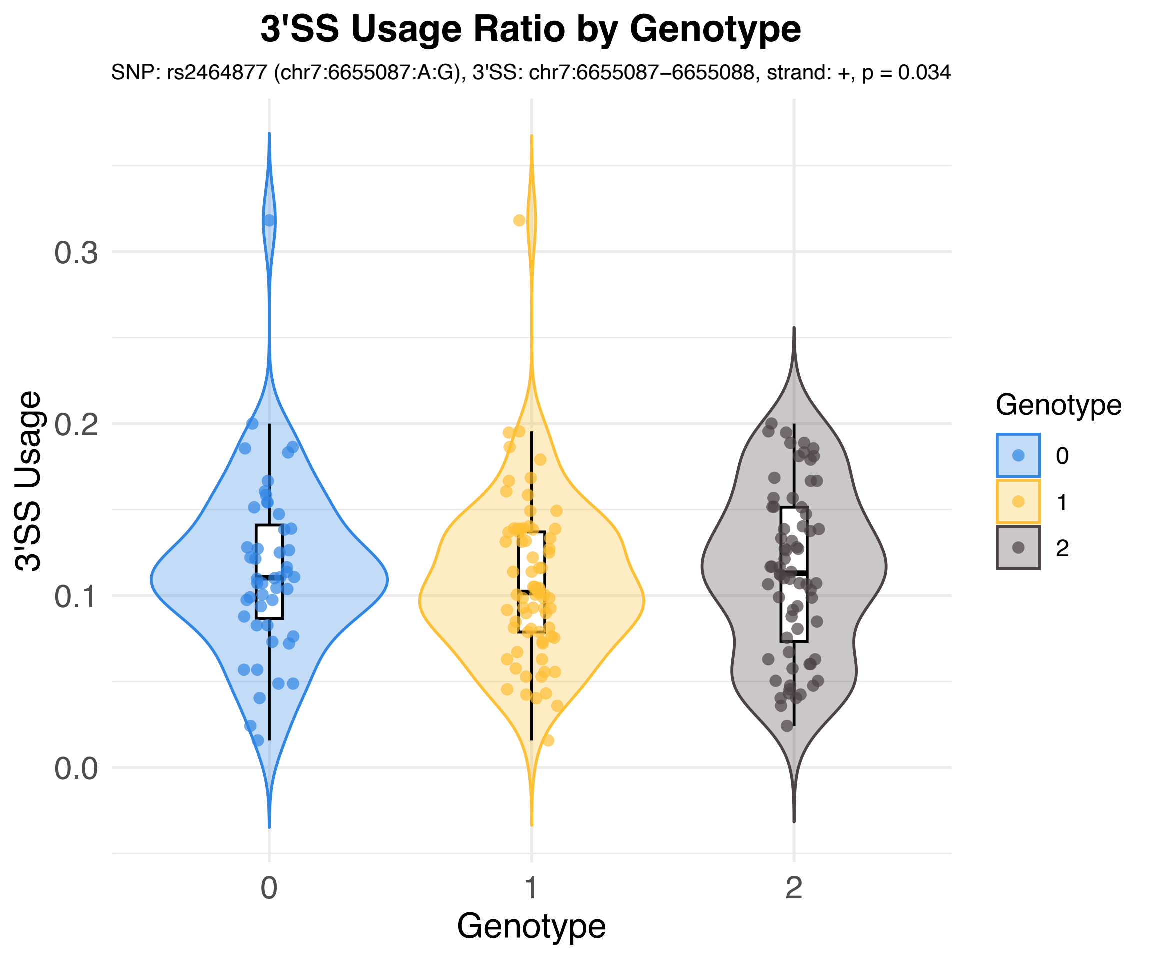

3'SS Usage Ratio by Genotype
SNP: rs2464877 (chr7:6655087:A:G), 3'SS: chr7:6655087−6655088, strand: +, p = 0.034
3'SS Usage
0.3
0.2
0.1
0.0
0
1
2
Genotype
Genotype
0
1
2

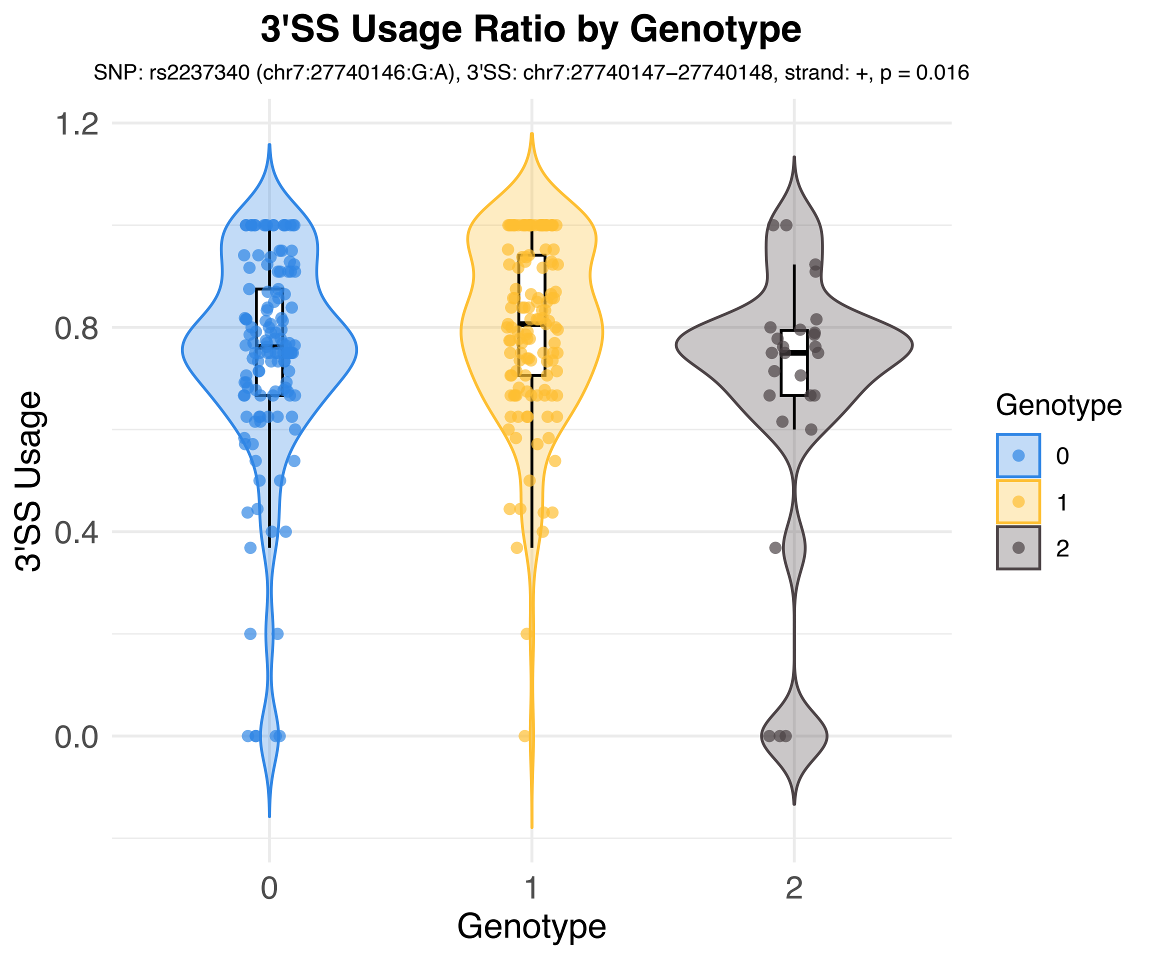
3'SS Usage Ratio by Genotype
SNP: rs2237340 (chr7:27740146:G:A), 3'SS: chr7:27740147–27740148, strand: +, p = 0.016
3'SS Usage
1.2
0.8
0.4
0.0
0
1
2
Genotype
Genotype
0
1
2

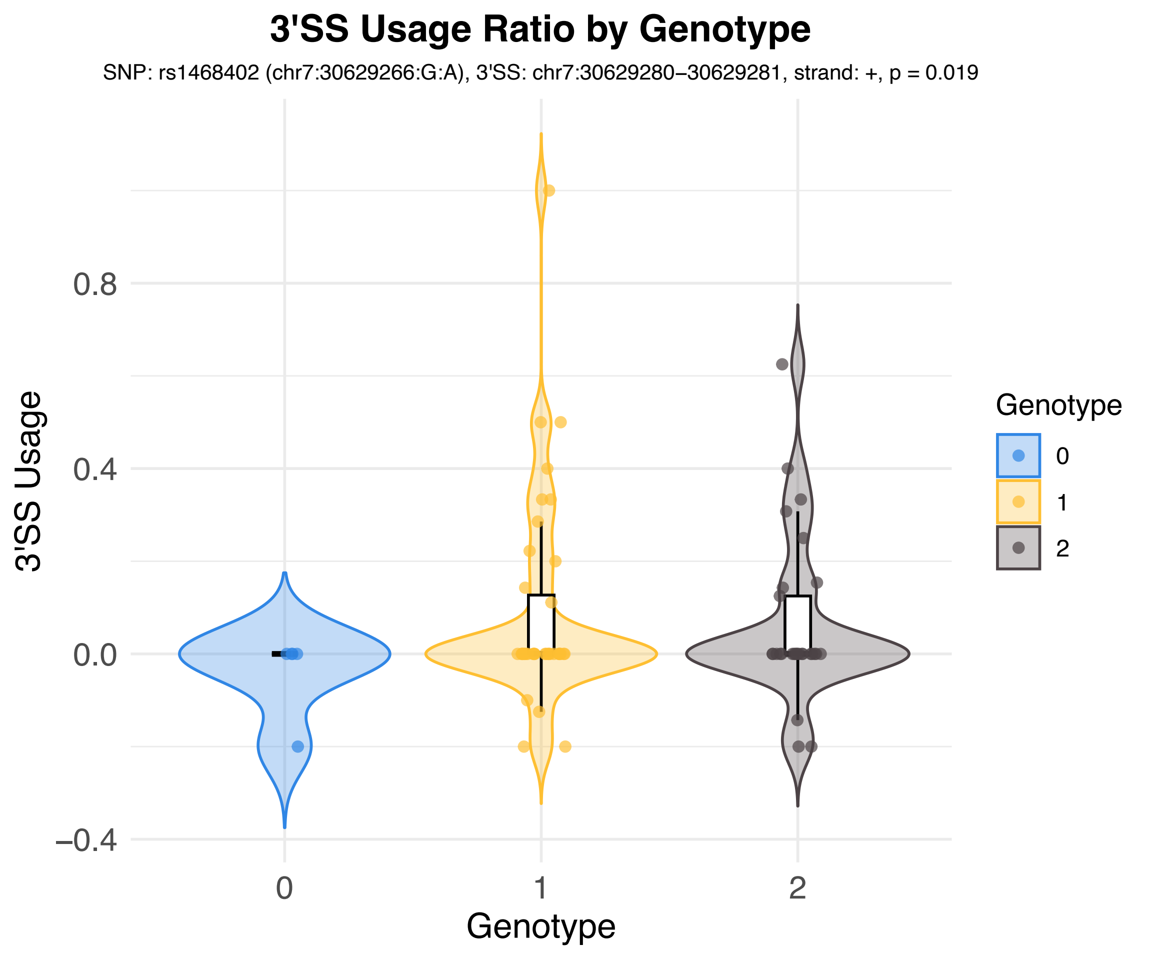

3'SS Usage Ratio by Genotype
SNP: rs1468402 (chr7:30629266:G:A), 3'SS: chr7:30629280–30629281, strand: +, p = 0.019
3'SS Usage
0.8
0.4
0.0
–0.4
0
1
2
Genotype
Genotype
0
1
2

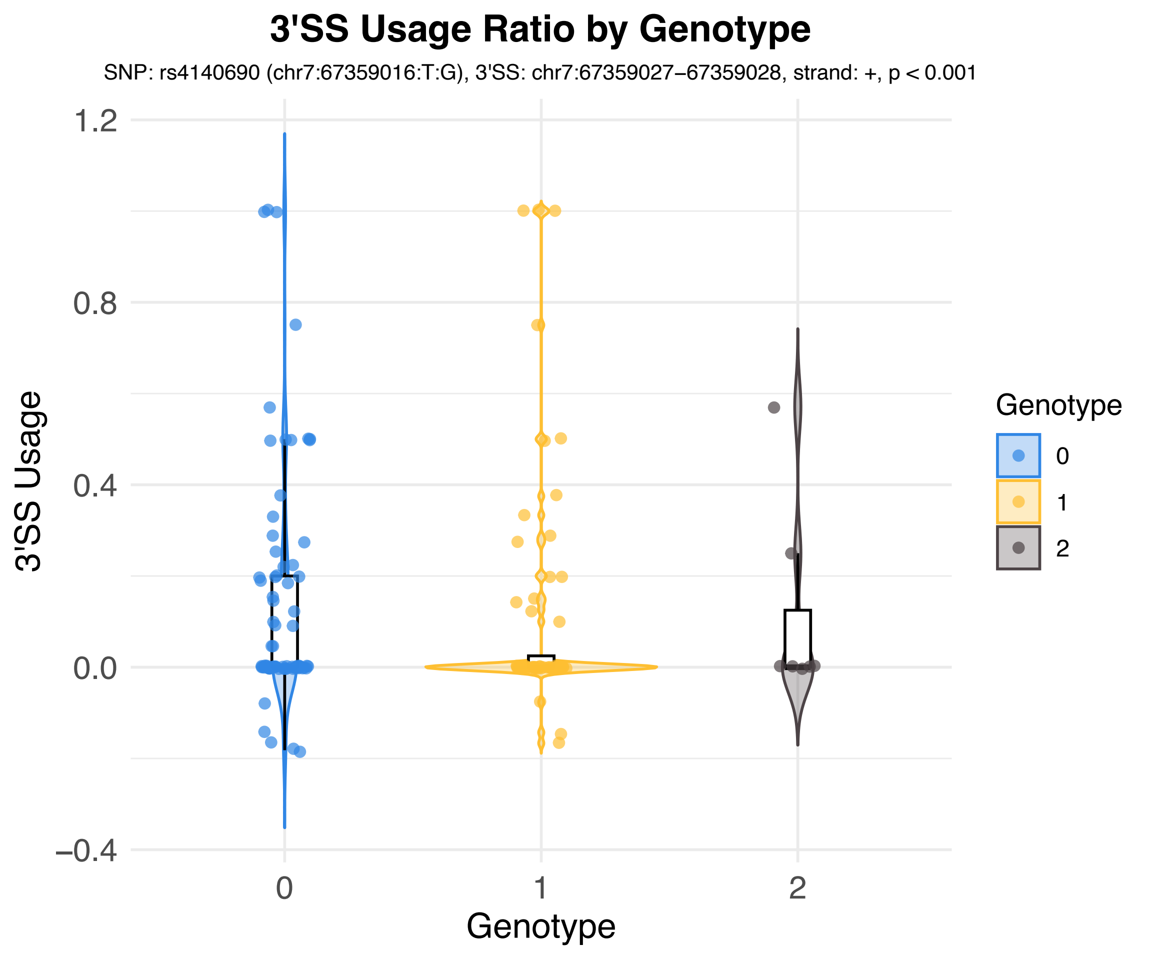
3'SS Usage Ratio by Genotype
SNP: rs4140690 (chr7:67359016:T:G), 3'SS: chr7:67359027−67359028, strand: +, p < 0.001
3'SS Usage
1.2
0.8
0.4
0.0
−0.4
0
1
2
Genotype
Genotype
0
1
2

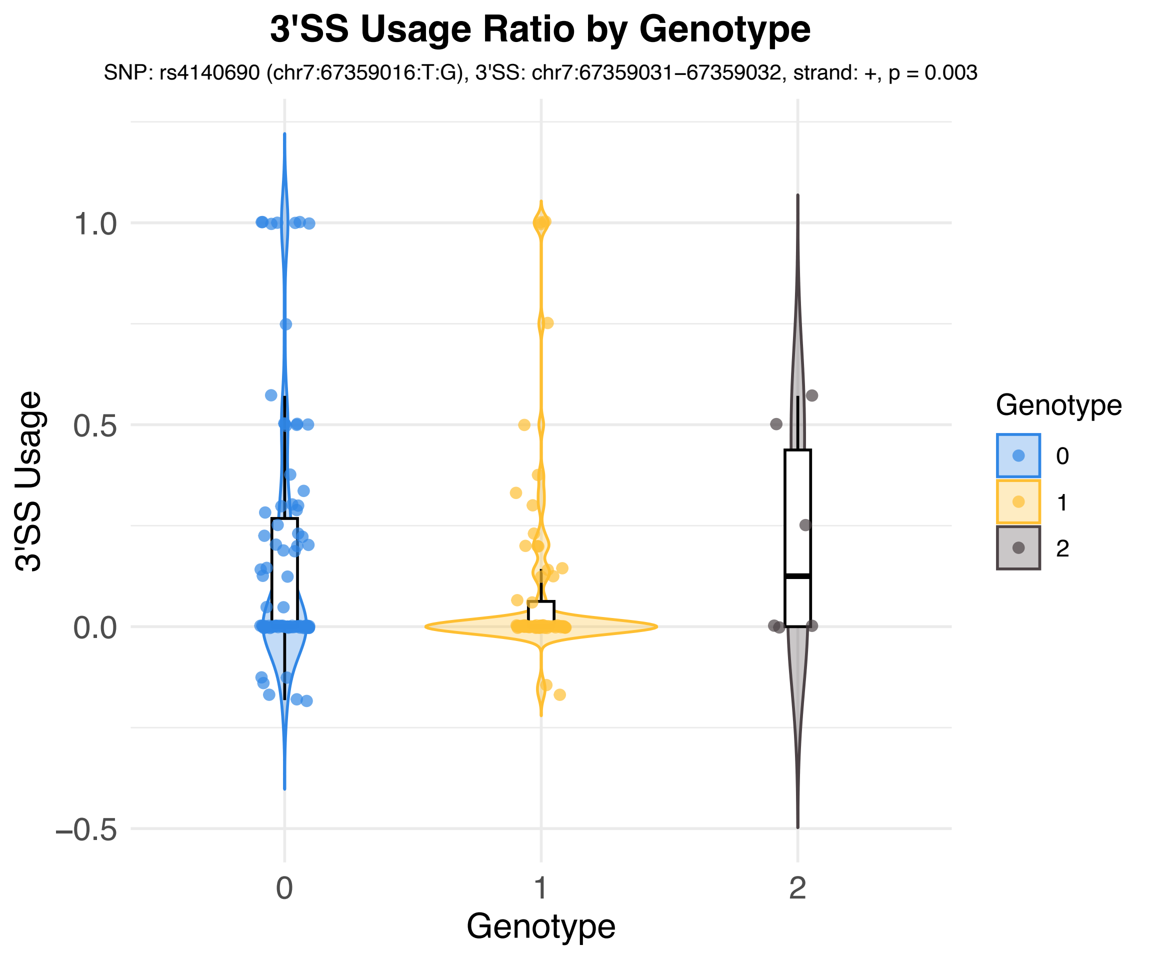

3'SS Usage Ratio by Genotype
SNP: rs4140690 (chr7:67359016:T:G), 3'SS: chr7:67359031–67359032, strand: +, p = 0.003
3'SS Usage
1.0
0.5
0.0
–0.5
0
1
2
Genotype
Genotype
0
1
2

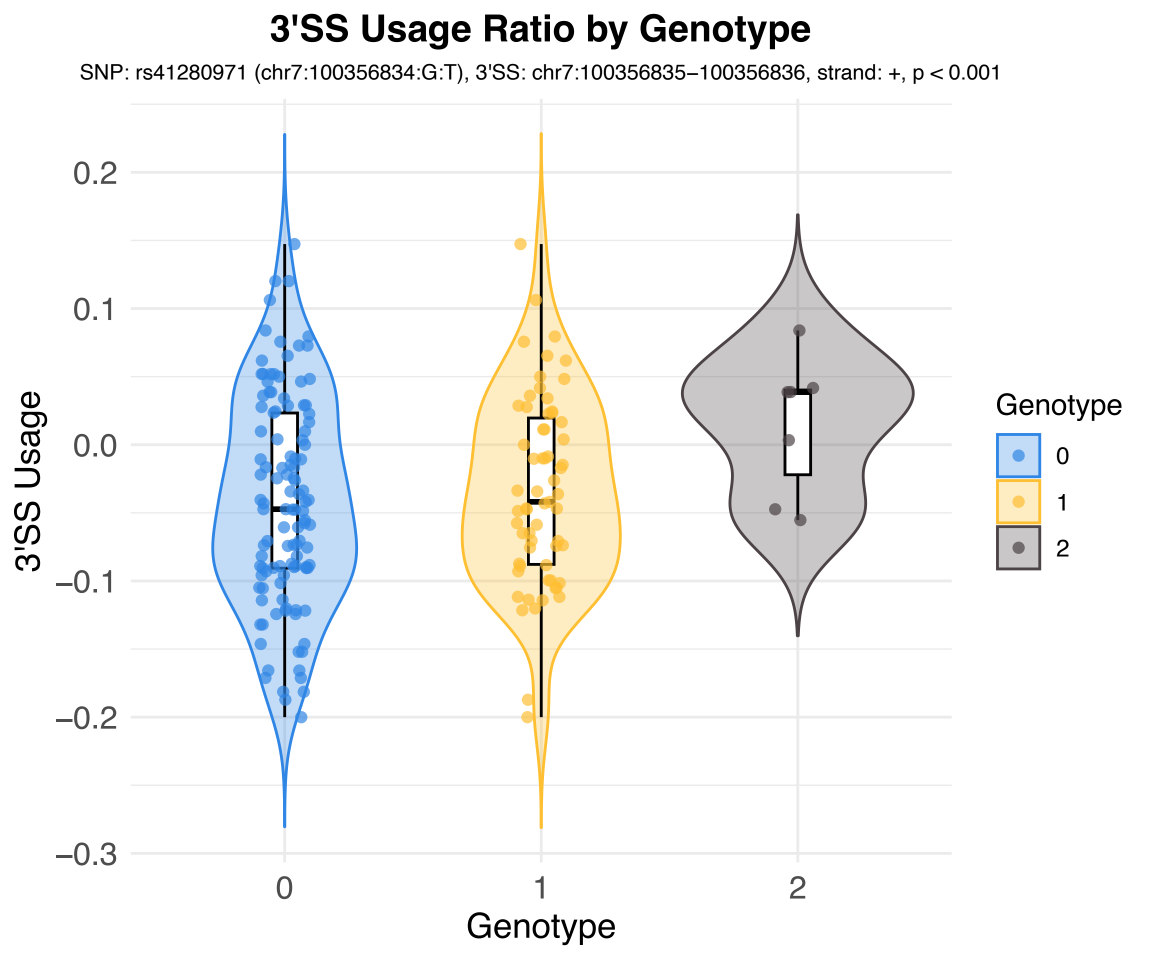
3'SS Usage Ratio by Genotype
SNP: rs41280971 (chr7:100356834:G:T), 3'SS: chr7:100356835–100356836, strand: +, p < 0.001
3'SS Usage
0.2
0.1
0.0
−0.1
−0.2
−0.3
0
1
2
Genotype
Genotype
0
1
2

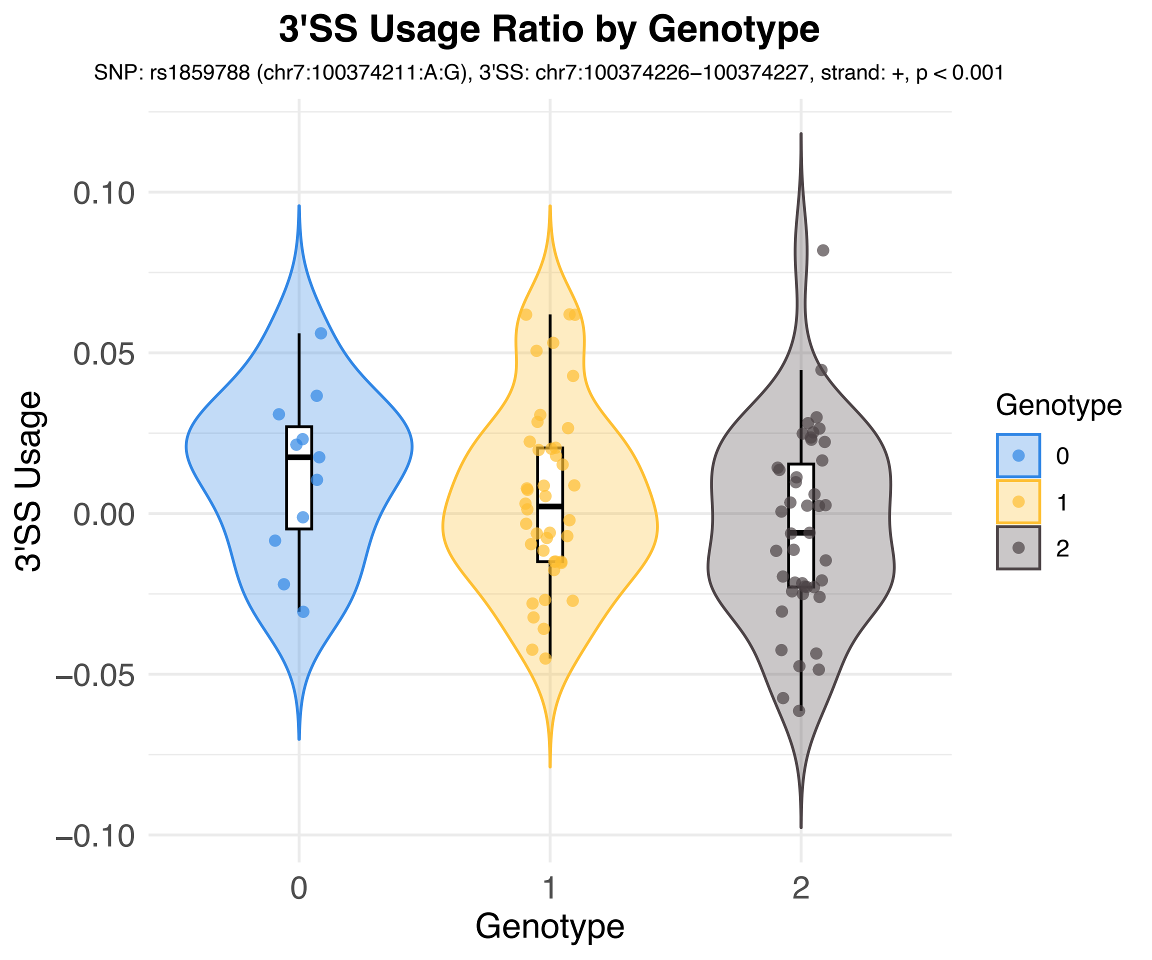
3'SS Usage Ratio by Genotype
SNP: rs1859788 (chr7:100374211:A:G), 3'SS: chr7:100374226–100374227, strand: +, p < 0.001
3'SS Usage
0.10
0.05
0.00
−0.05
−0.10
0
1
2
Genotype
Genotype
0
1
2

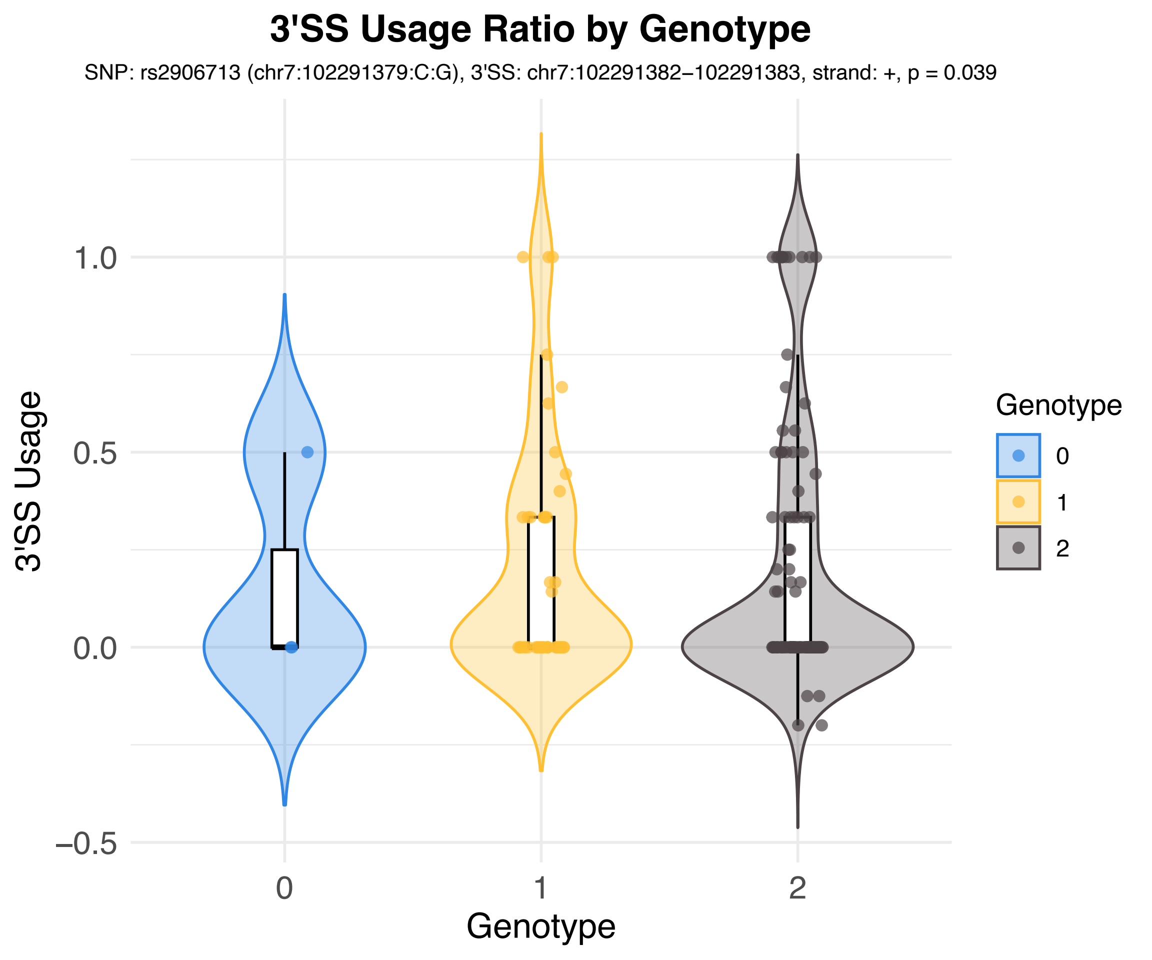
3'SS Usage Ratio by Genotype
SNP: rs2906713 (chr7:102291379:C:G), 3'SS: chr7:102291382−102291383, strand: +, p = 0.039
3'SS Usage
1.0
0.5
0.0
−0.5
0
1
2
Genotype
Genotype
0
1
2

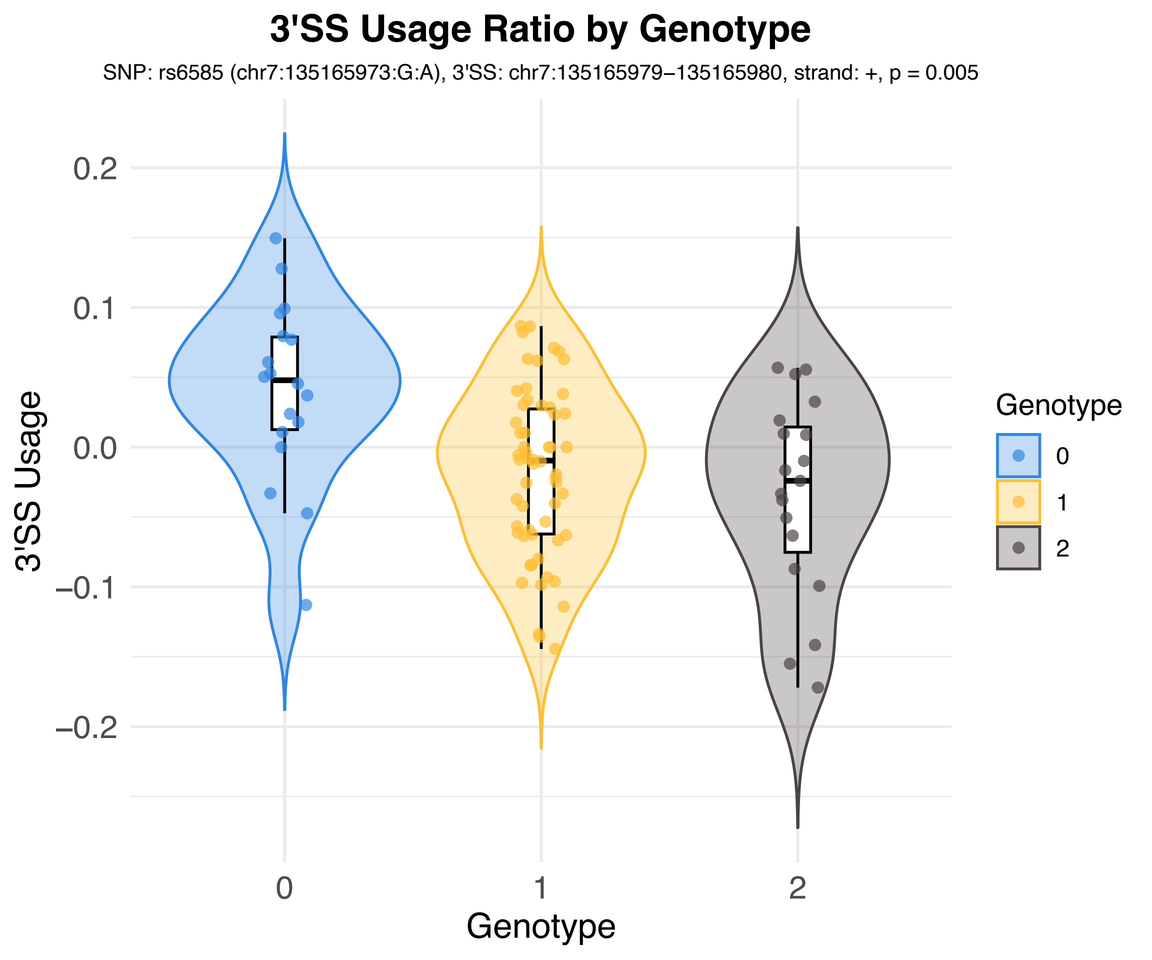
3'SS Usage Ratio by Genotype
SNP: rs6585 (chr7:135165973:G:A), 3'SS: chr7:135165979−135165980, strand: +, p = 0.005
3'SS Usage
0.2
0.1
0.0
−0.1
−0.2
0
1
2
Genotype
Genotype
0
1
2

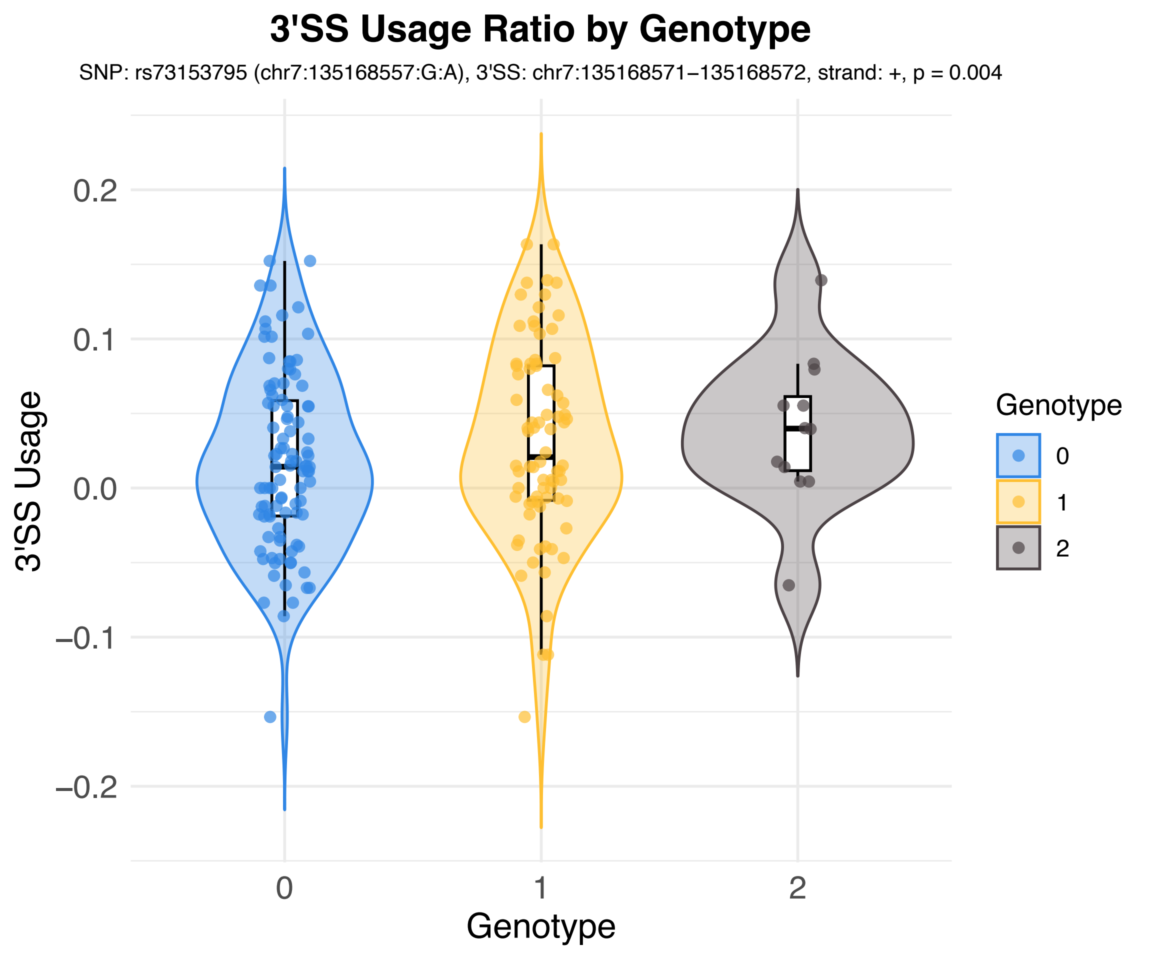
3'SS Usage Ratio by Genotype
SNP: rs73153795 (chr7:135168557:G:A), 3'SS: chr7:135168571–135168572, strand: +, p = 0.004
3'SS Usage
0.2
0.1
0.0
–0.1
–0.2
0
1
2
Genotype
Genotype
0
1
2

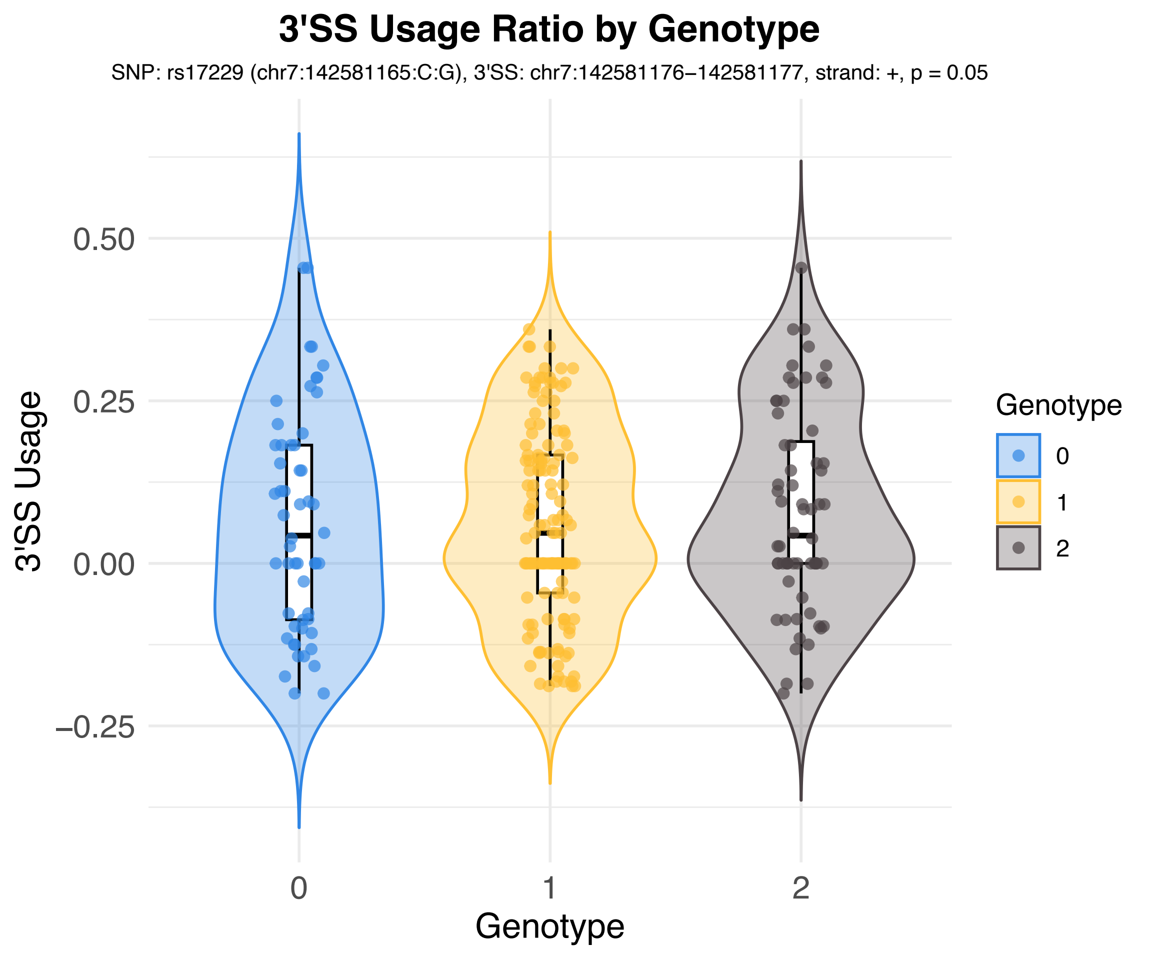

3'SS Usage Ratio by Genotype
SNP: rs17229 (chr7:142581165:C:G), 3'SS: chr7:142581176−142581177, strand: +, p = 0.05
3'SS Usage
0.50
0.25
0.00
−0.25
0
1
2
Genotype
Genotype
0
1
2

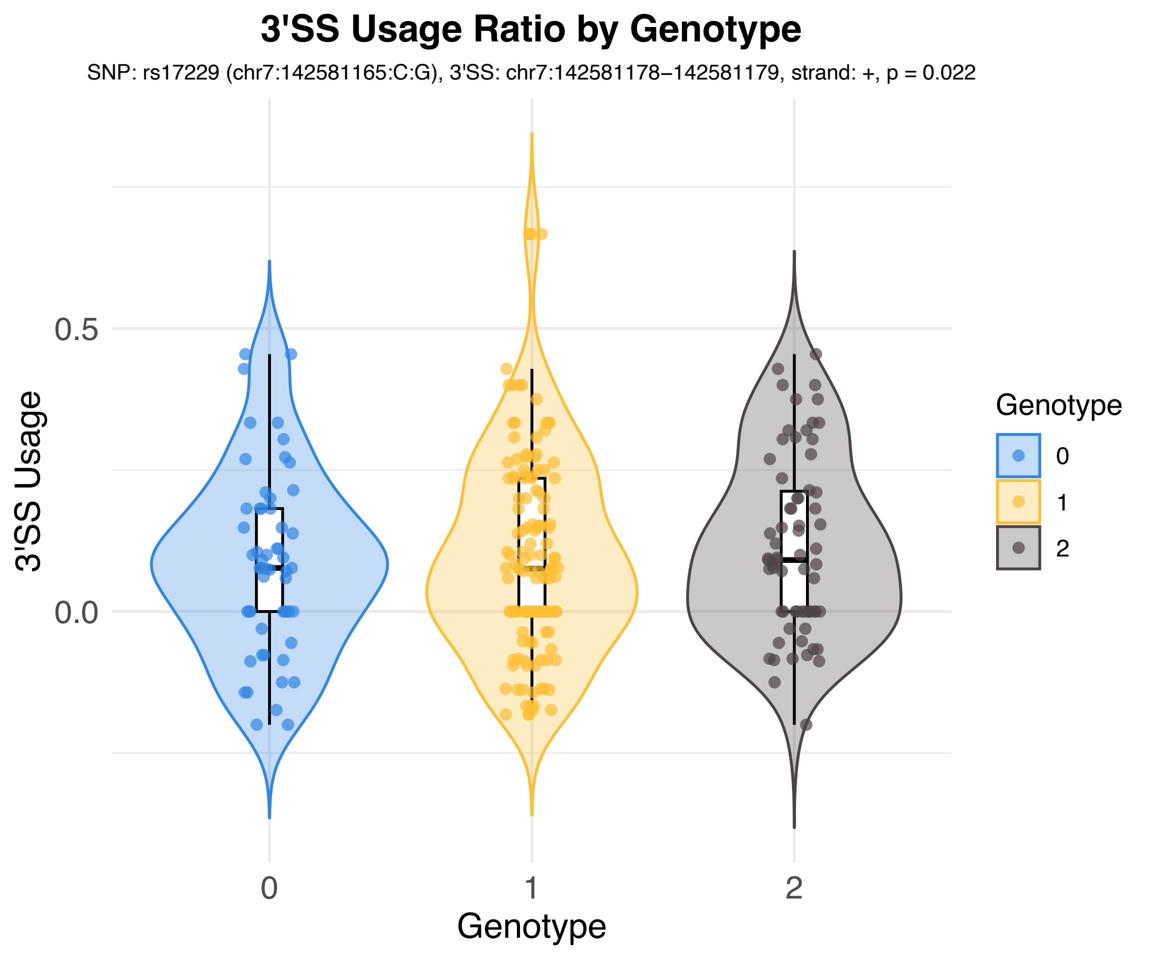

3'SS Usage Ratio by Genotype
SNP: rs17229 (chr7:142581165:C:G), 3'SS: chr7:142581178−142581179, strand: +, p = 0.022
3'SS Usage
0.5
0.0
0
1
2
Genotype
Genotype
0
1
2

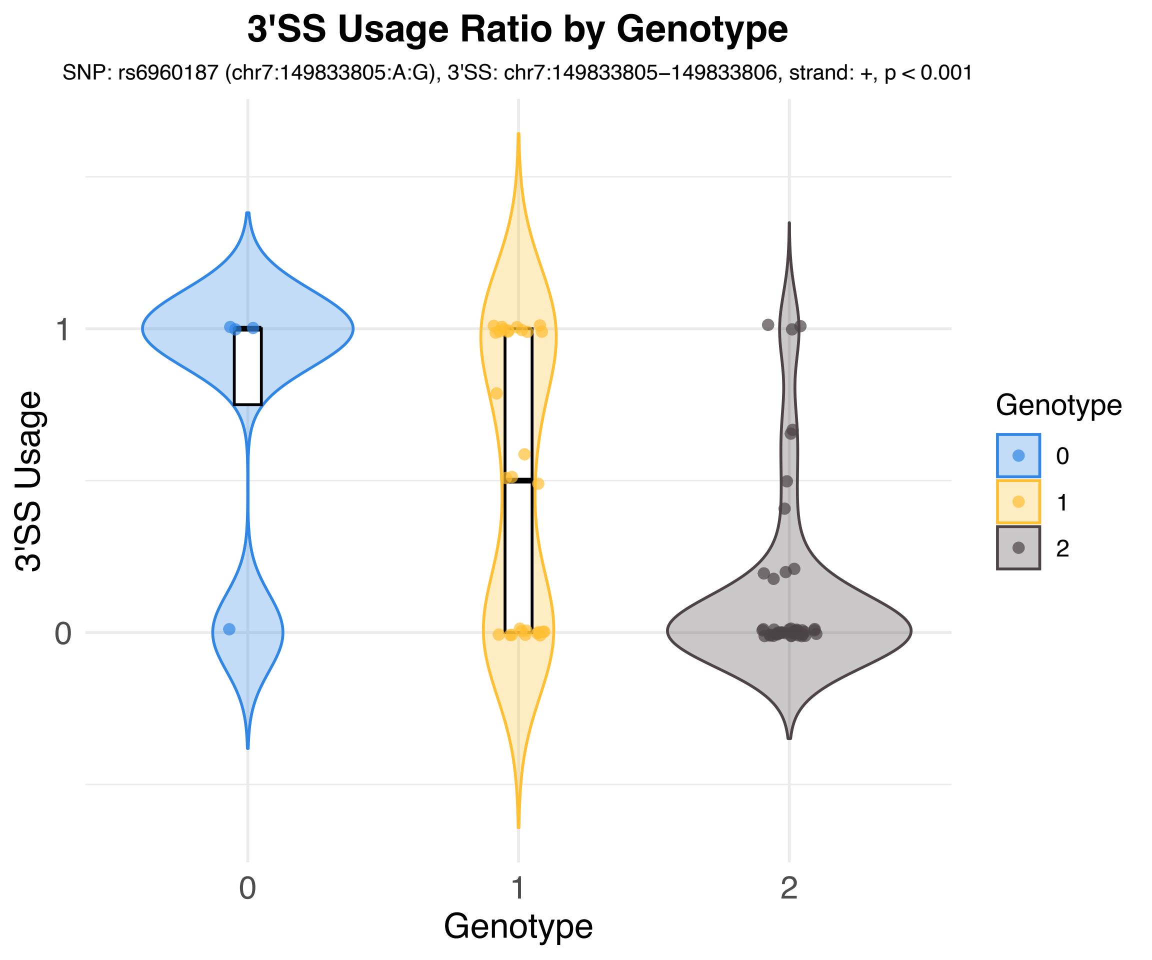
3'SS Usage Ratio by Genotype
SNP: rs6960187 (chr7:149833805:A:G), 3'SS: chr7:149833805−149833806, strand: +, p < 0.001
3'SS Usage
1
0
0
1
2
Genotype
Genotype
0
1
2

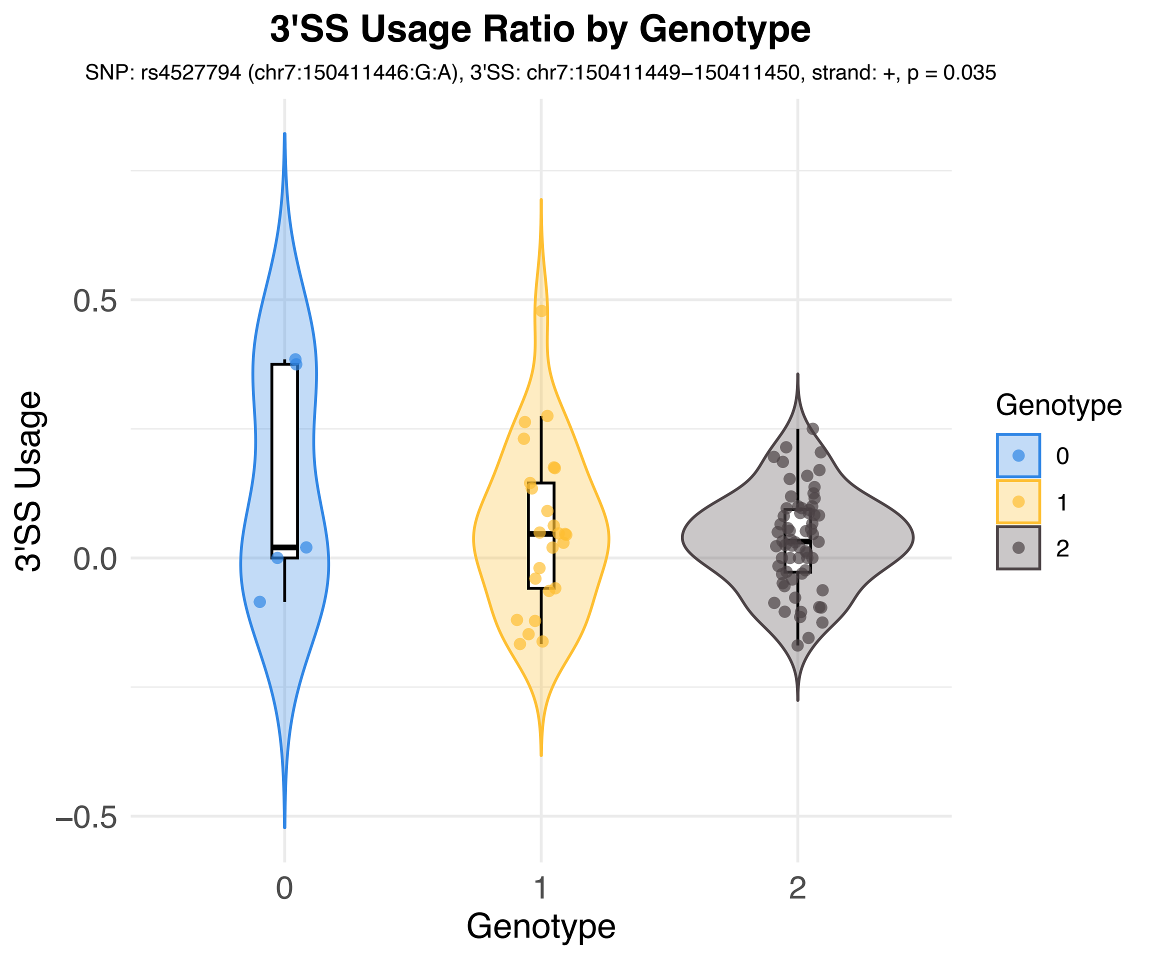

3'SS Usage Ratio by Genotype
SNP: rs4527794 (chr7:150411446:G:A), 3'SS: chr7:150411449−150411450, strand: +, p = 0.035
3'SS Usage
0.5
0.0
−0.5
0
1
2
Genotype
Genotype
0
1
2

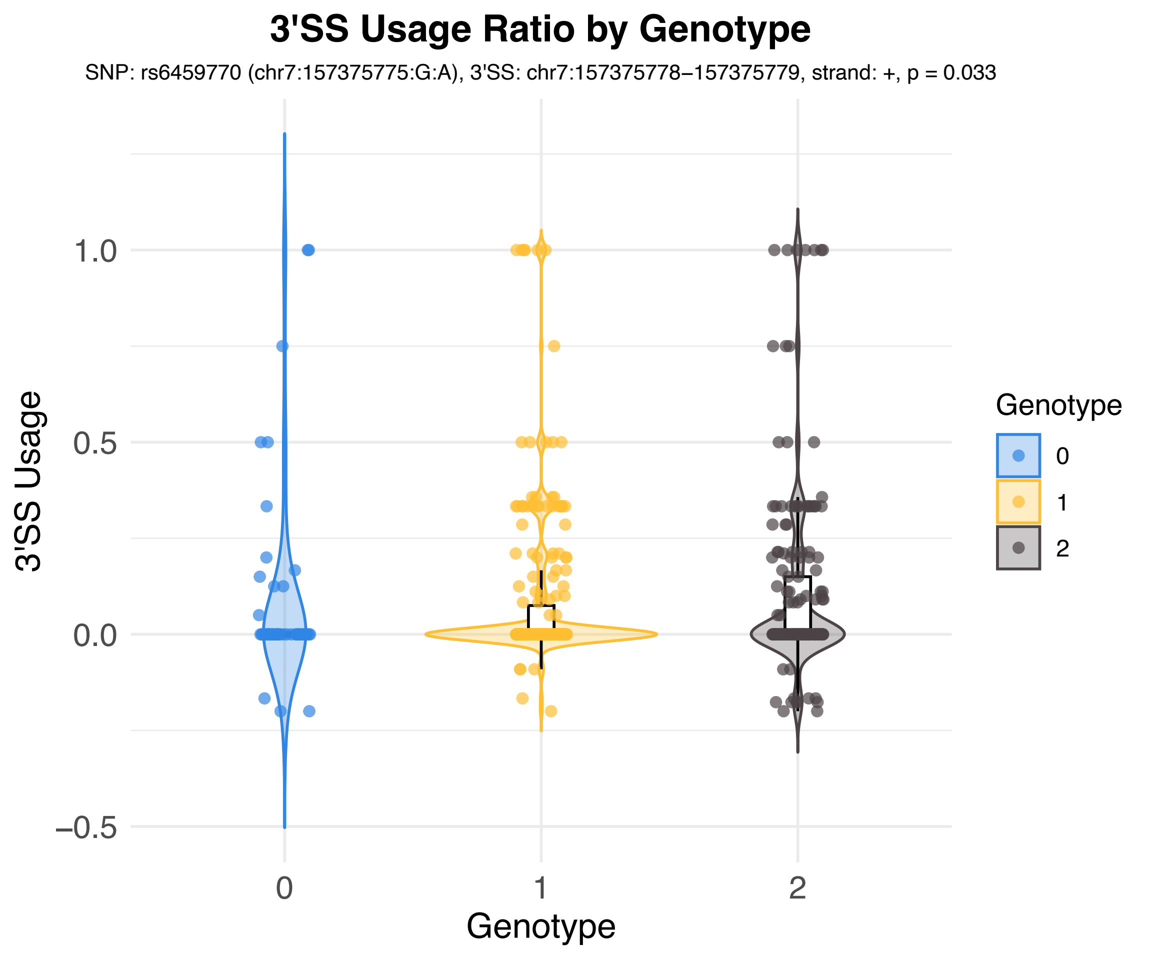
3'SS Usage Ratio by Genotype
SNP: rs6459770 (chr7:157375775:G:A), 3'SS: chr7:157375778–157375779, strand: +, p = 0.033
3'SS Usage
1.0
0.5
0.0
–0.5
0
1
2
Genotype
Genotype
0
1
2

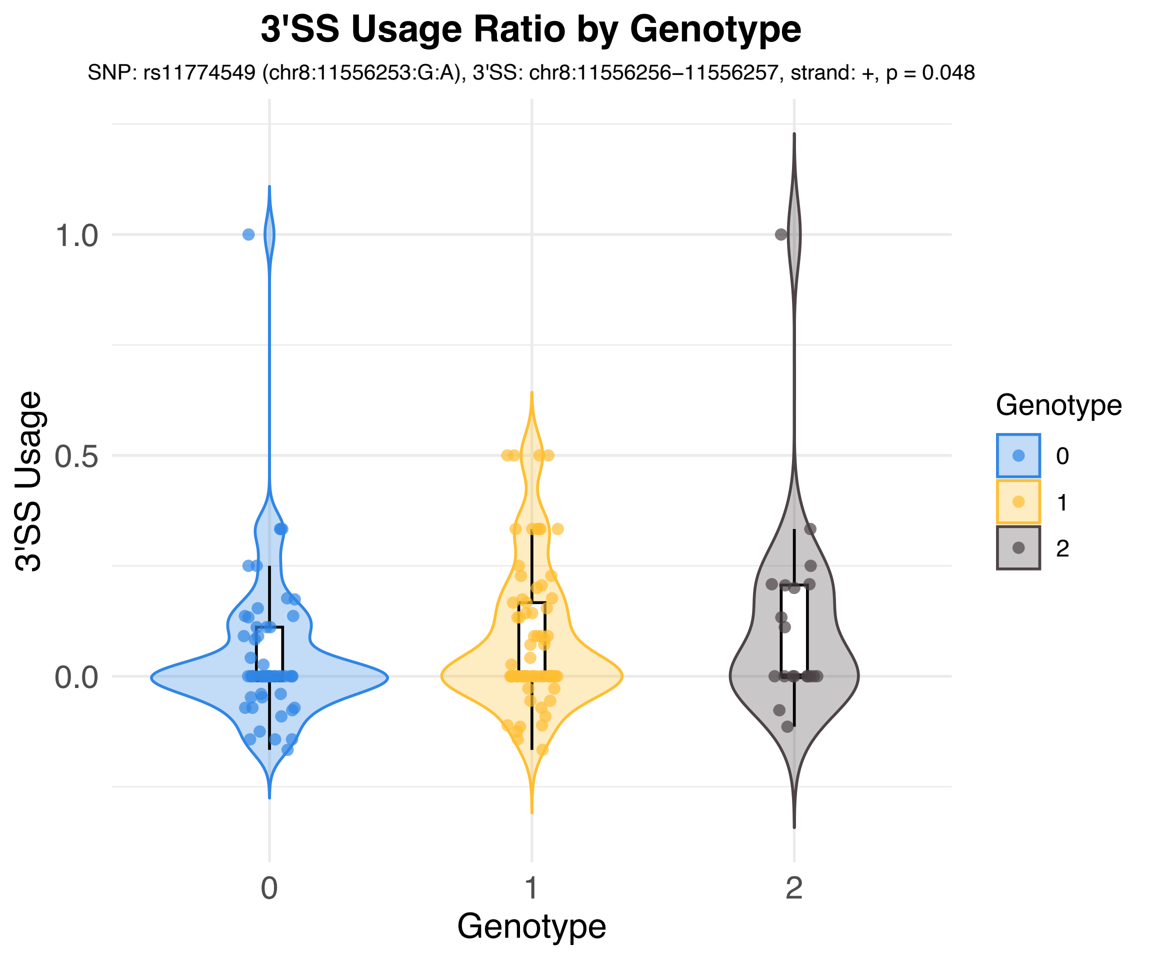
3'SS Usage Ratio by Genotype
SNP: rs11774549 (chr8:11556253:G:A), 3'SS: chr8:11556256−11556257, strand: +, p = 0.048
3'SS Usage
1.0
0.5
0.0
0
1
2
Genotype
Genotype
0
1
2

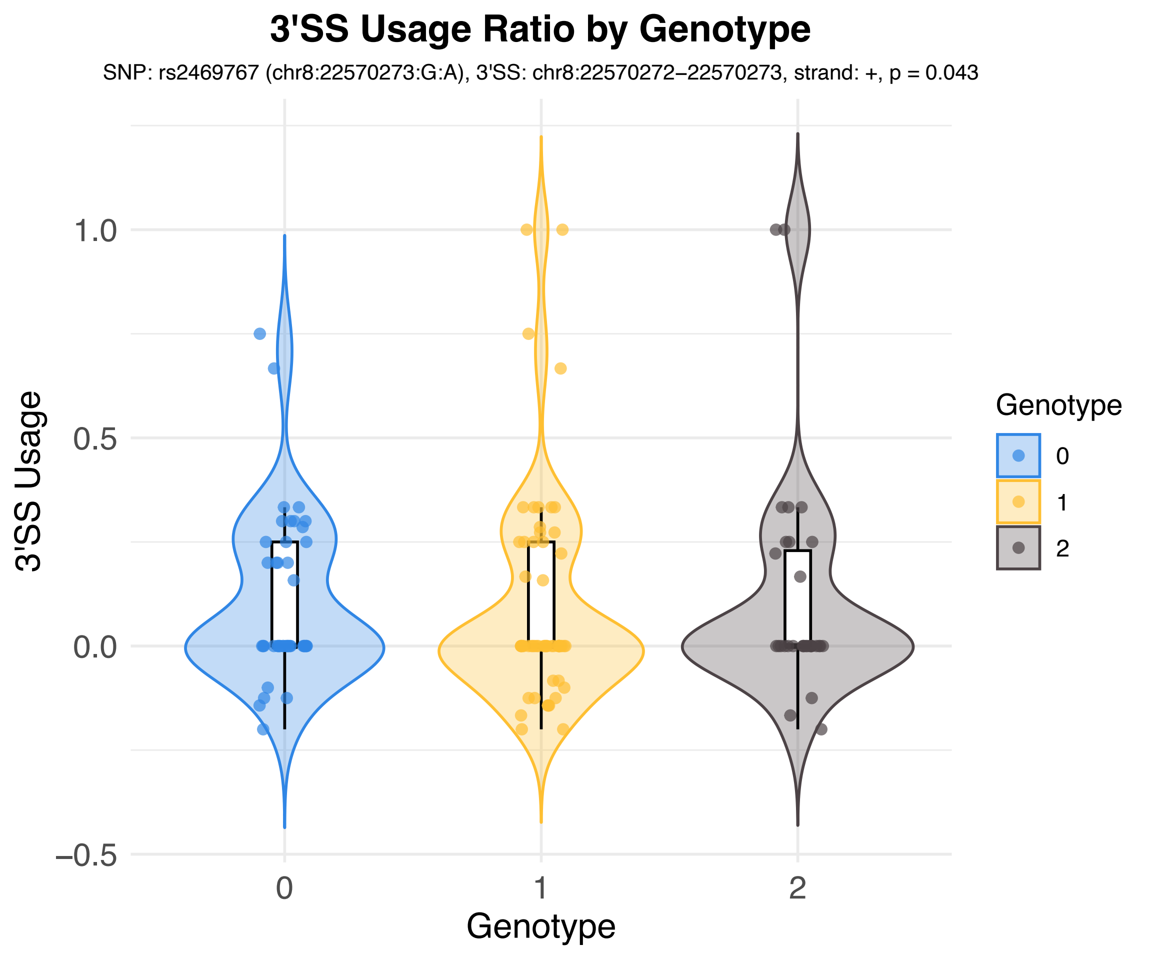
3'SS Usage Ratio by Genotype
SNP: rs2469767 (chr8:22570273:G:A), 3'SS: chr8:22570272−22570273, strand: +, p = 0.043
3'SS Usage
1.0
0.5
0.0
−0.5
0
1
2
Genotype
Genotype
0
1
2

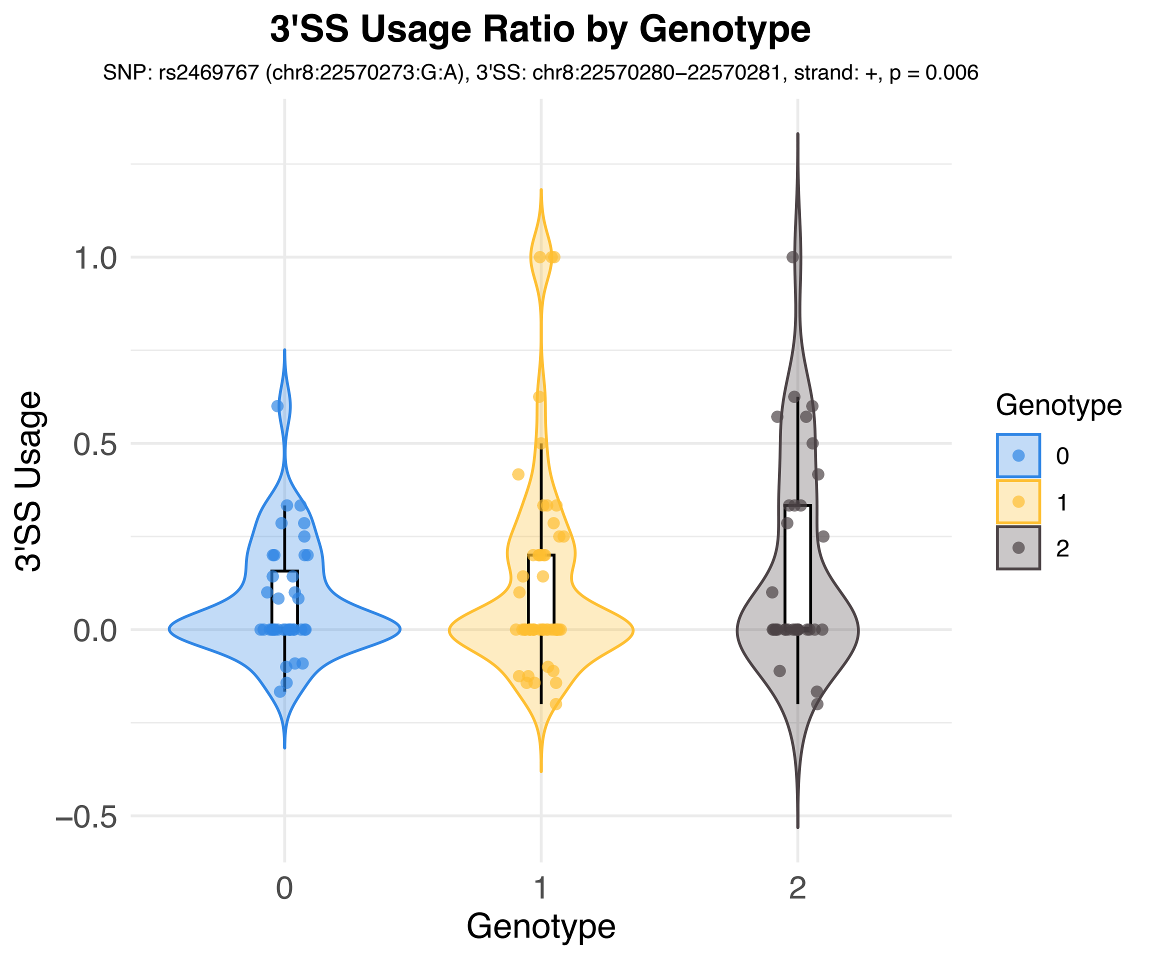
3'SS Usage Ratio by Genotype
SNP: rs2469767 (chr8:22570273:G:A), 3'SS: chr8:22570280−22570281, strand: +, p = 0.006
3'SS Usage
1.0
0.5
0.0
−0.5
0
1
2
Genotype
Genotype
0
1
2

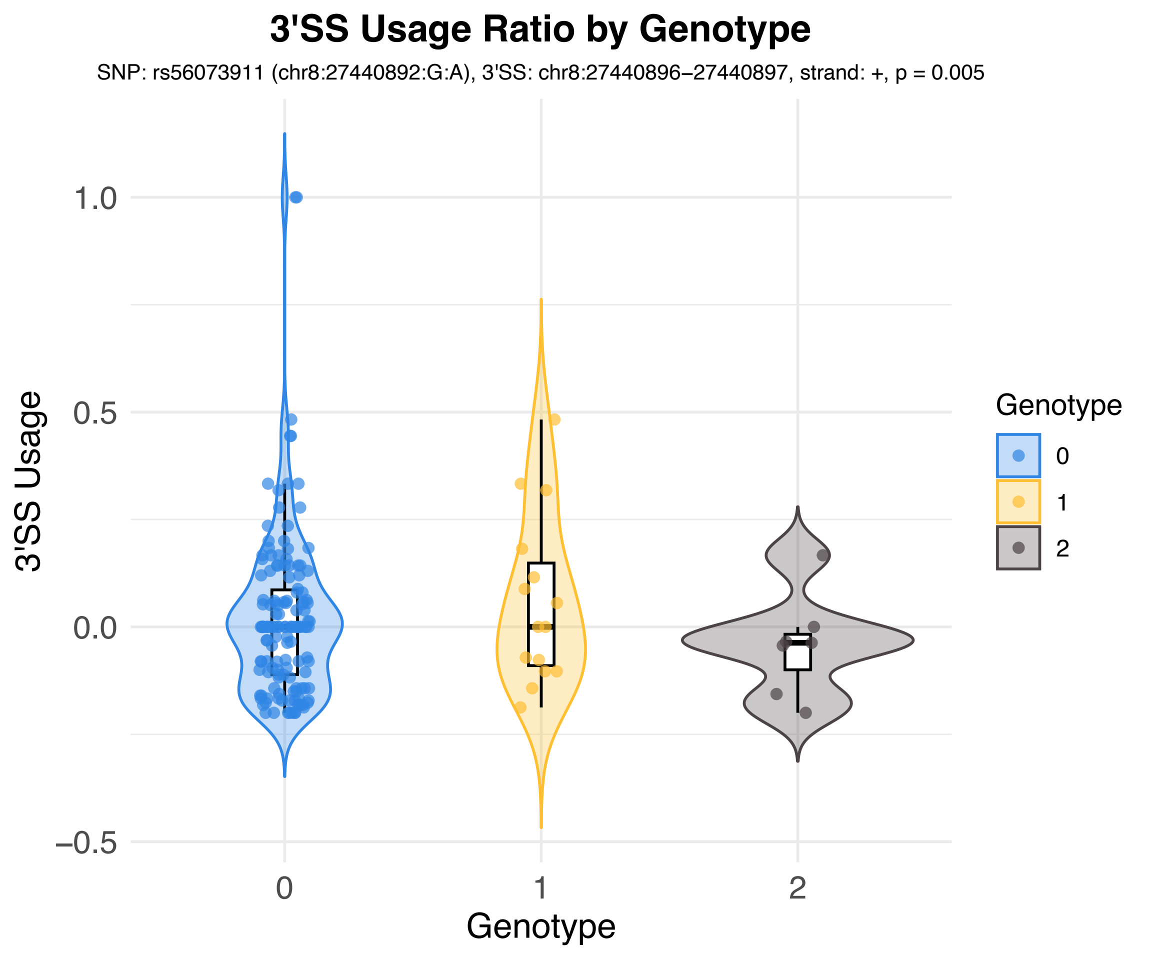
3'SS Usage Ratio by Genotype
SNP: rs56073911 (chr8:27440892:G:A), 3'SS: chr8:27440896–27440897, strand: +, p = 0.005
3'SS Usage
1.0
0.5
0.0
−0.5
0
1
2
Genotype
Genotype
0
1
2

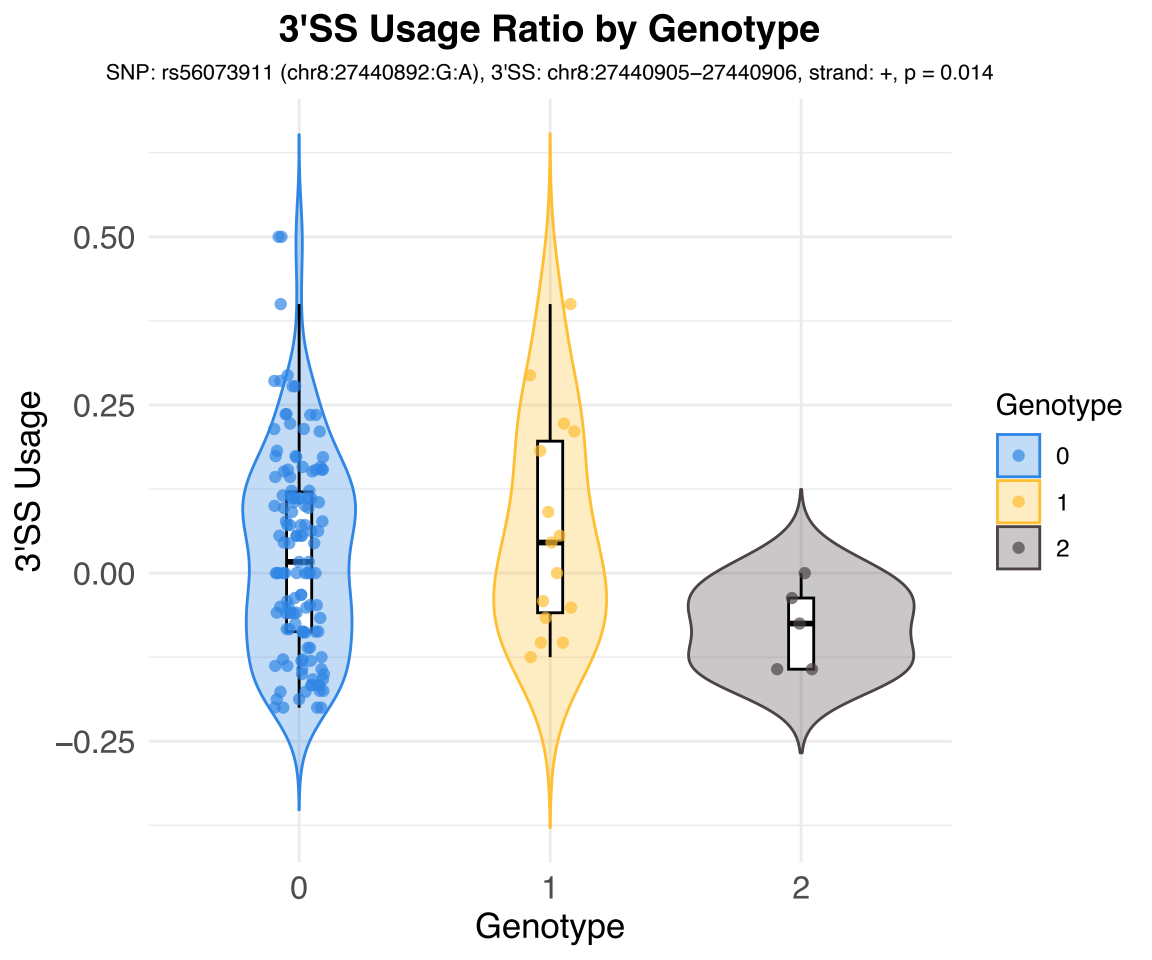
3'SS Usage Ratio by Genotype
SNP: rs56073911 (chr8:27440892:G:A), 3'SS: chr8:27440905−27440906, strand: +, p = 0.014
3'SS Usage
0.50
0.25
0.00
−0.25
0
1
2
Genotype
Genotype
0
1
2

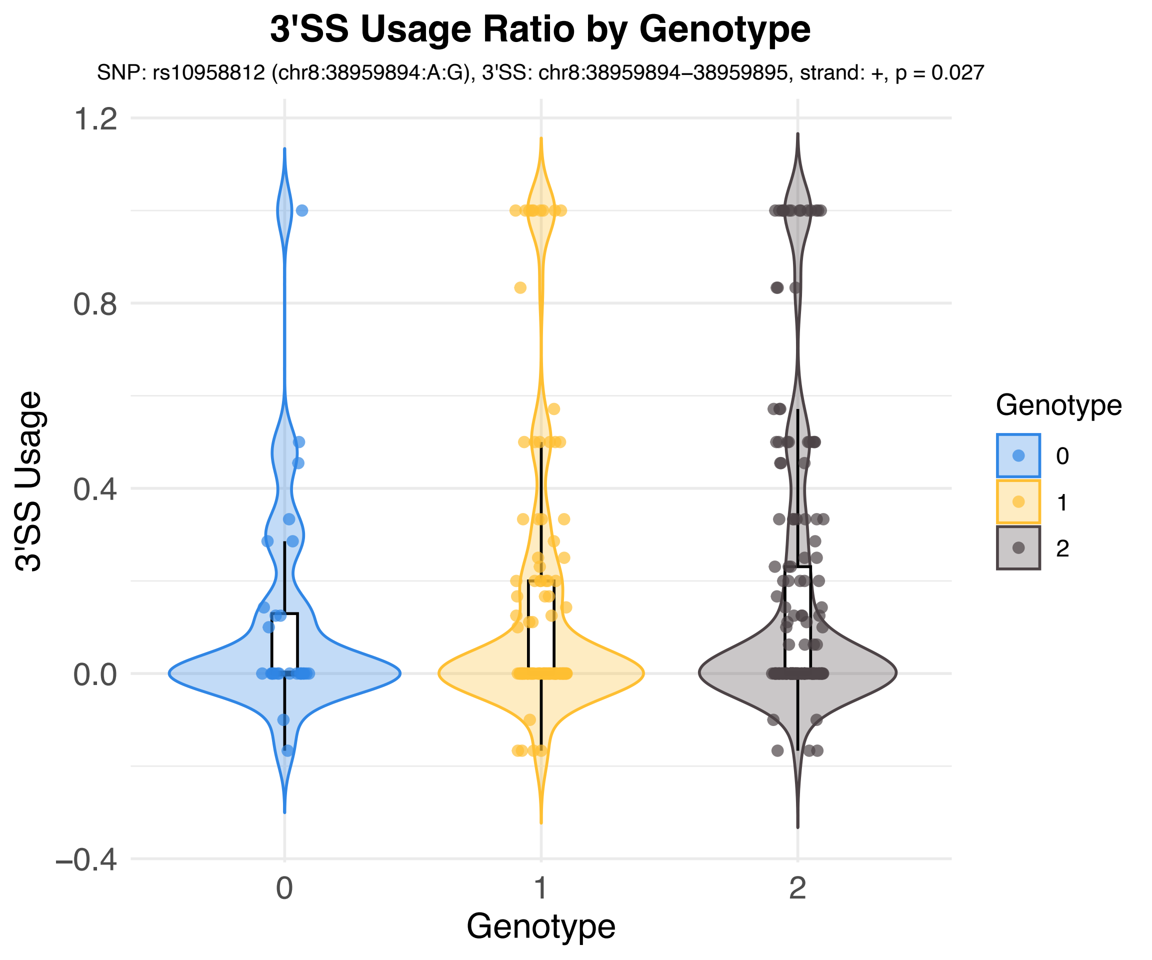
3'SS Usage Ratio by Genotype
SNP: rs10958812 (chr8:38959894:A:G), 3'SS: chr8:38959894–38959895, strand: +, p = 0.027
3'SS Usage
1.2
0.8
0.4
0.0
−0.4
0
1
2
Genotype
Genotype
0
1
2

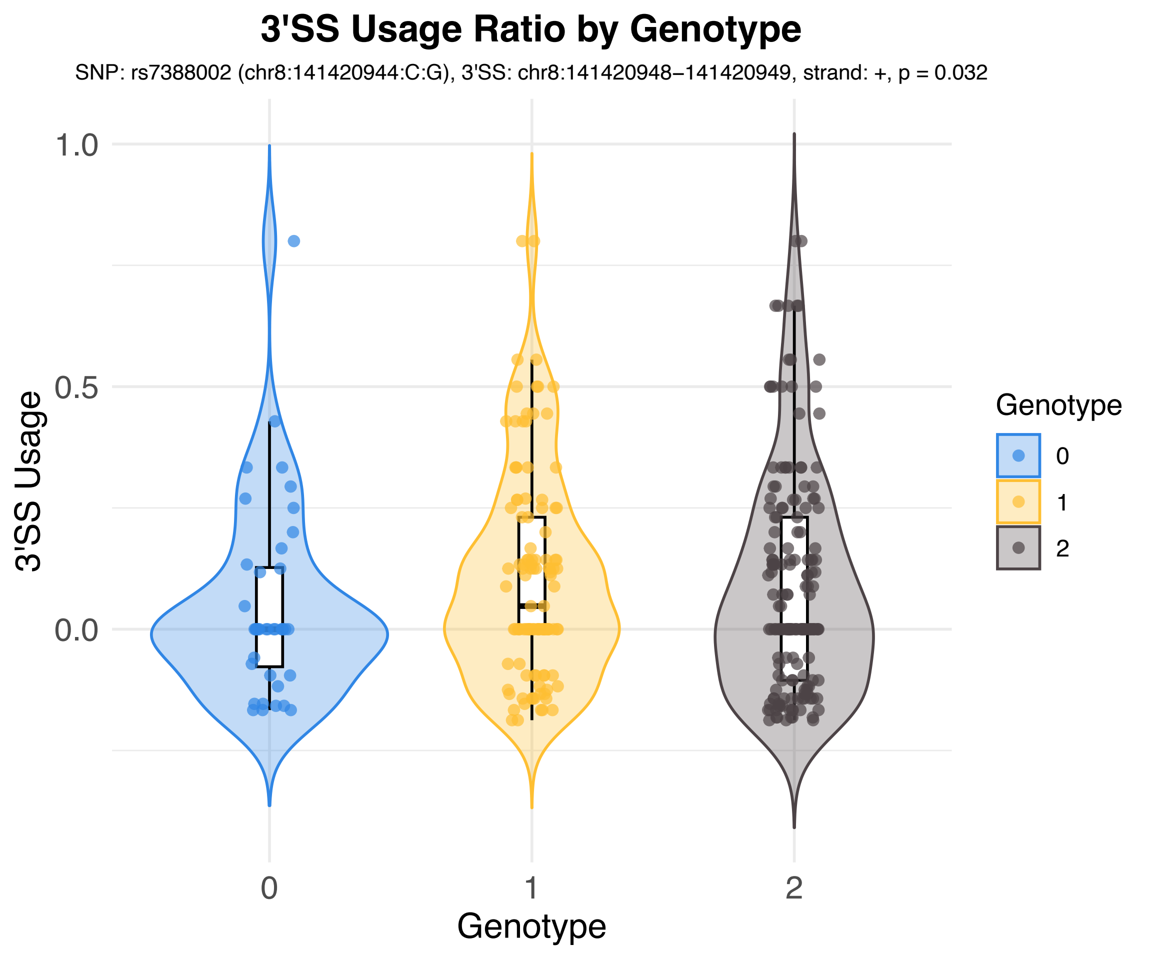
3'SS Usage Ratio by Genotype
SNP: rs7388002 (chr8:141420944:C:G), 3'SS: chr8:141420948−141420949, strand: +, p = 0.032
3'SS Usage
1.0
0.5
0.0
0
1
2
Genotype
Genotype
0
1
2

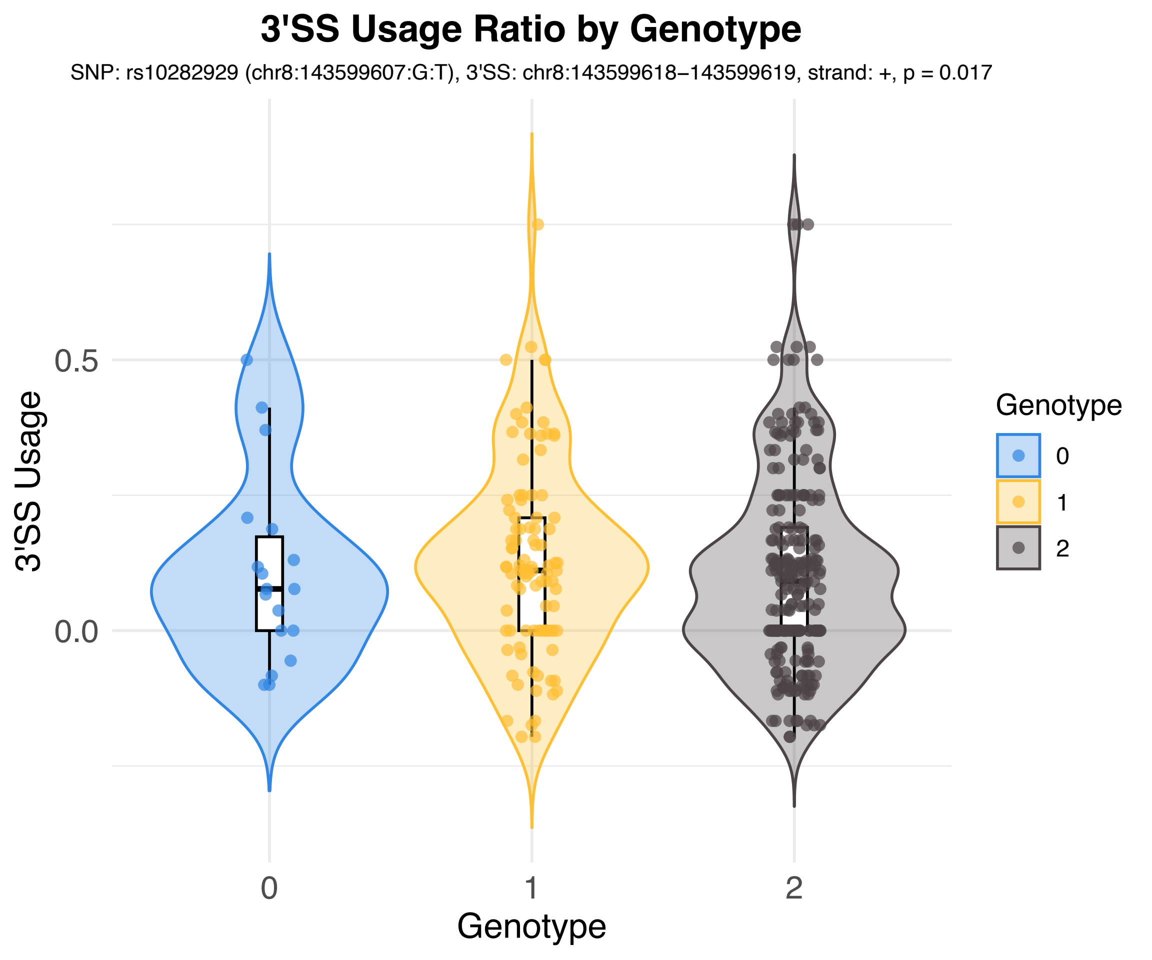
3'SS Usage Ratio by Genotype
SNP: rs10282929 (chr8:143599607:G:T), 3'SS: chr8:143599618−143599619, strand: +, p = 0.017
3'SS Usage
0.5
0.0
0
1
2
Genotype
Genotype
0
1
2

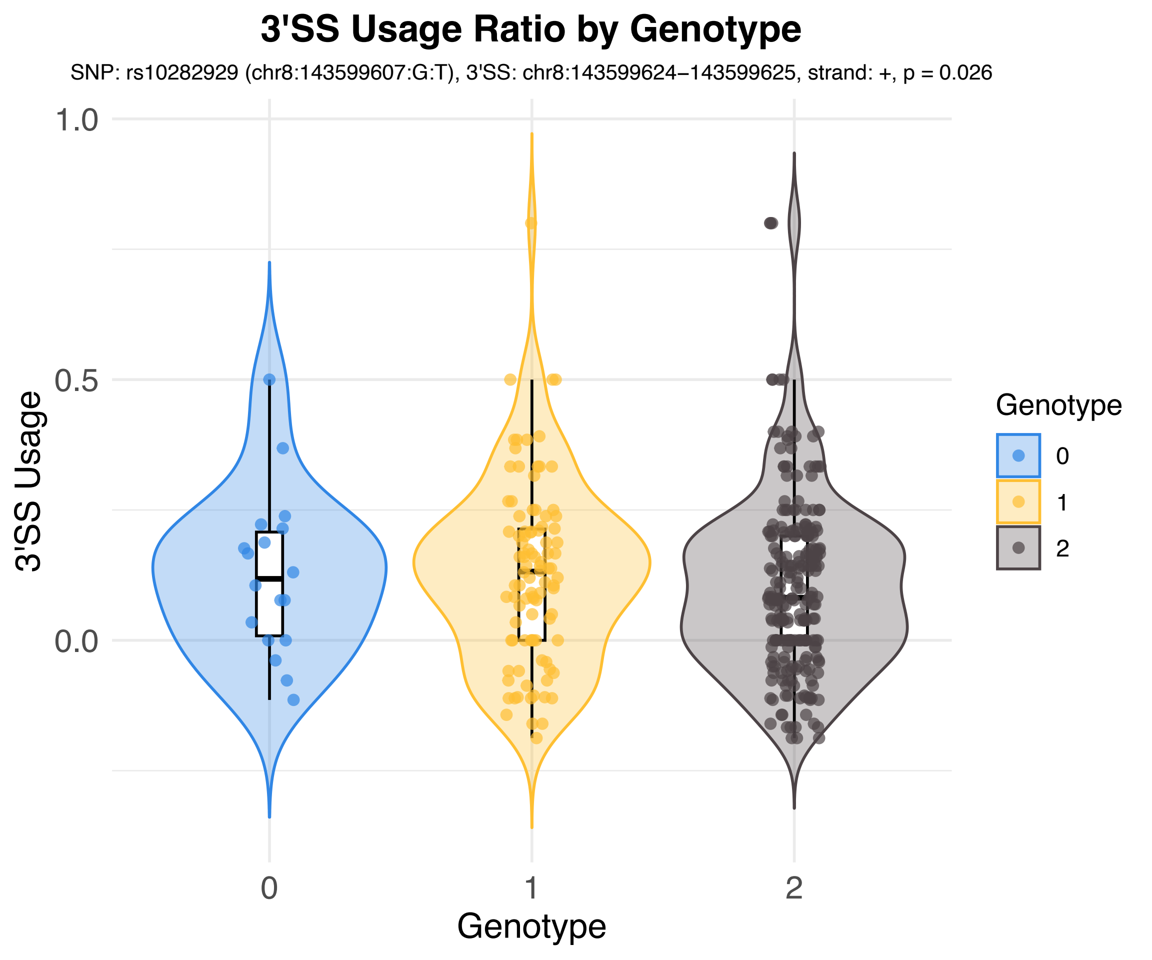
3'SS Usage Ratio by Genotype
SNP: rs10282929 (chr8:143599607:G:T), 3'SS: chr8:143599624−143599625, strand: +, p = 0.026
1.0
0.5
0.0
3'SS Usage
0
1
2
Genotype
Genotype
0
1
2

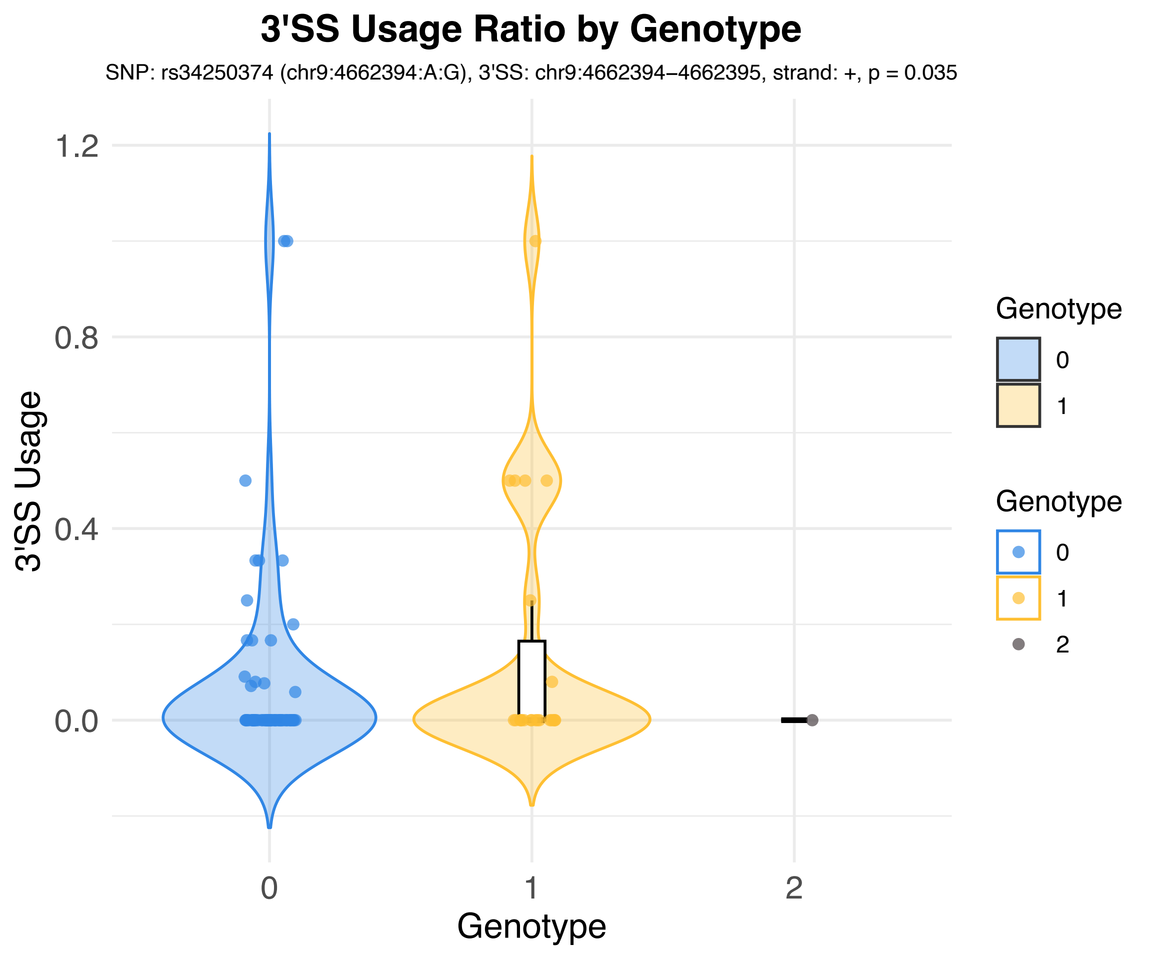
3'SS Usage Ratio by Genotype
SNP: rs34250374 (chr9:4662394:A:G), 3'SS: chr9:4662394−4662395, strand: +, p = 0.035
3'SS Usage
1.2
0.8
0.4
0.0
0
1
2
Genotype
Genotype
0
1
Genotype
0
1
2

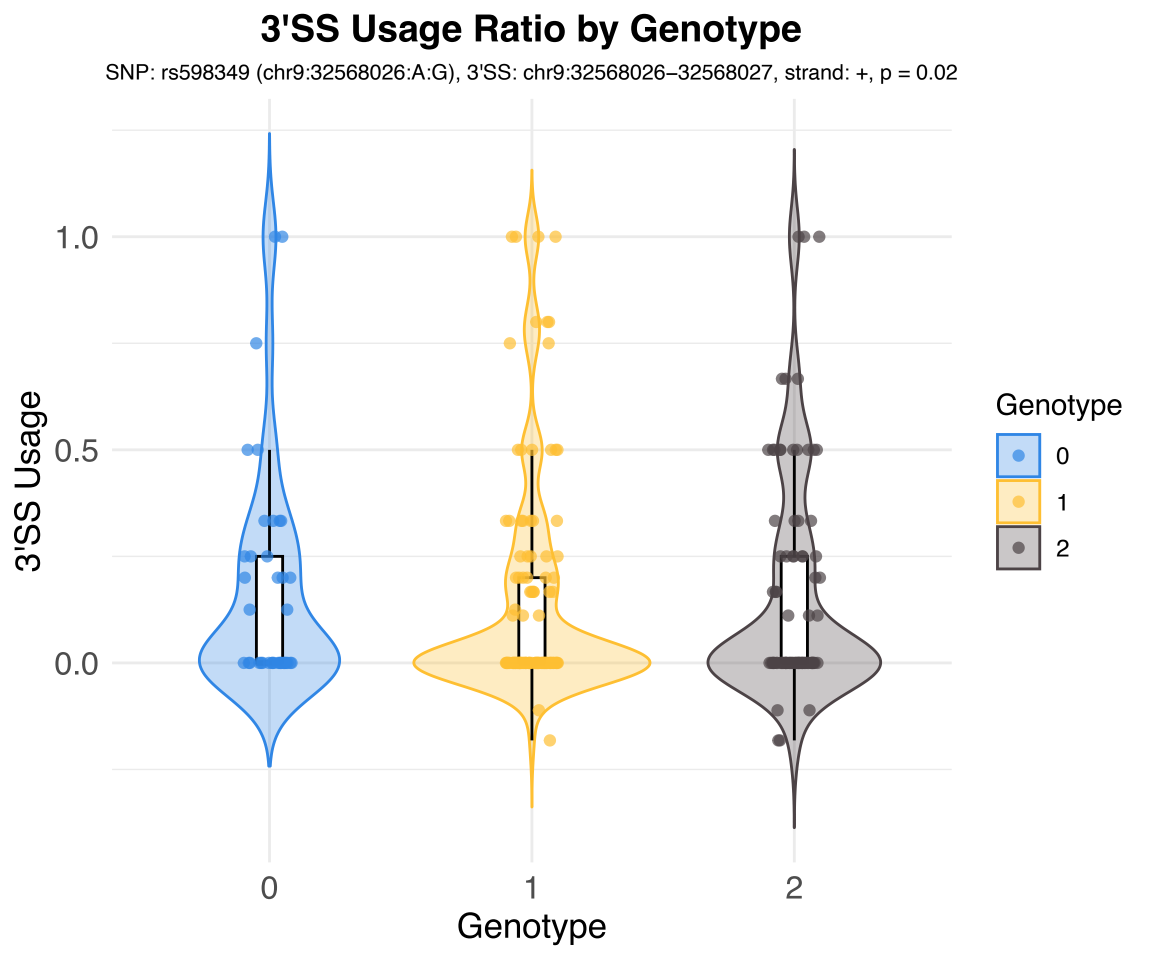
3'SS Usage Ratio by Genotype
SNP: rs598349 (chr9:32568026:A:G), 3'SS: chr9:32568026–32568027, strand: +, p = 0.02
3'SS Usage
1.0
0.5
0.0
0
1
2
Genotype
Genotype
0
1
2

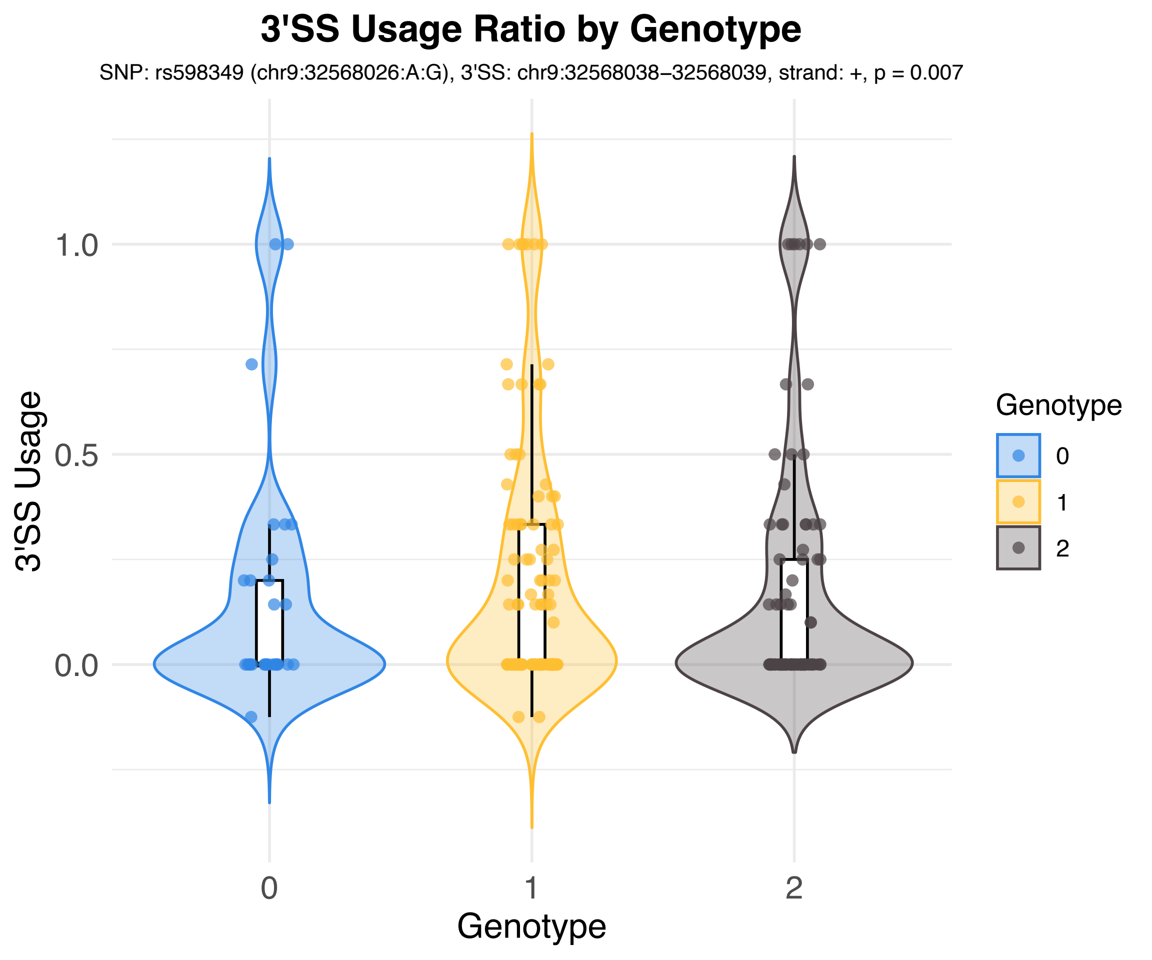

3'SS Usage Ratio by Genotype
SNP: rs598349 (chr9:32568026:A:G), 3'SS: chr9:32568038–32568039, strand: +, p = 0.007
3'SS Usage
1.0
0.5
0.0
0
1
2
Genotype
Genotype
0
1
2

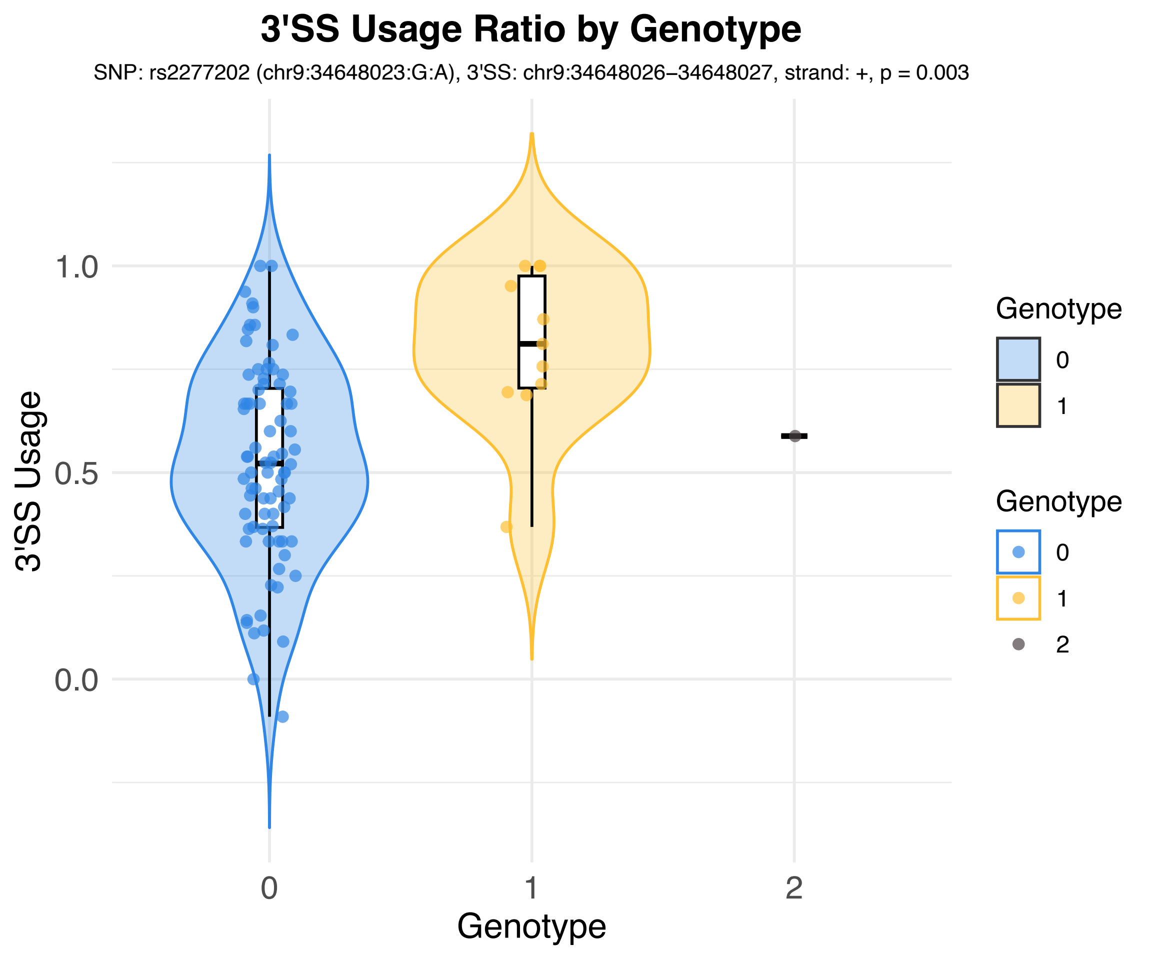
3'SS Usage Ratio by Genotype
SNP: rs2277202 (chr9:34648023:G:A), 3'SS: chr9:34648026–34648027, strand: +, p = 0.003
3'SS Usage
1.0
0.5
0.0
Genotype
0
1
2
Genotype
0
1
Genotype
0
1
2

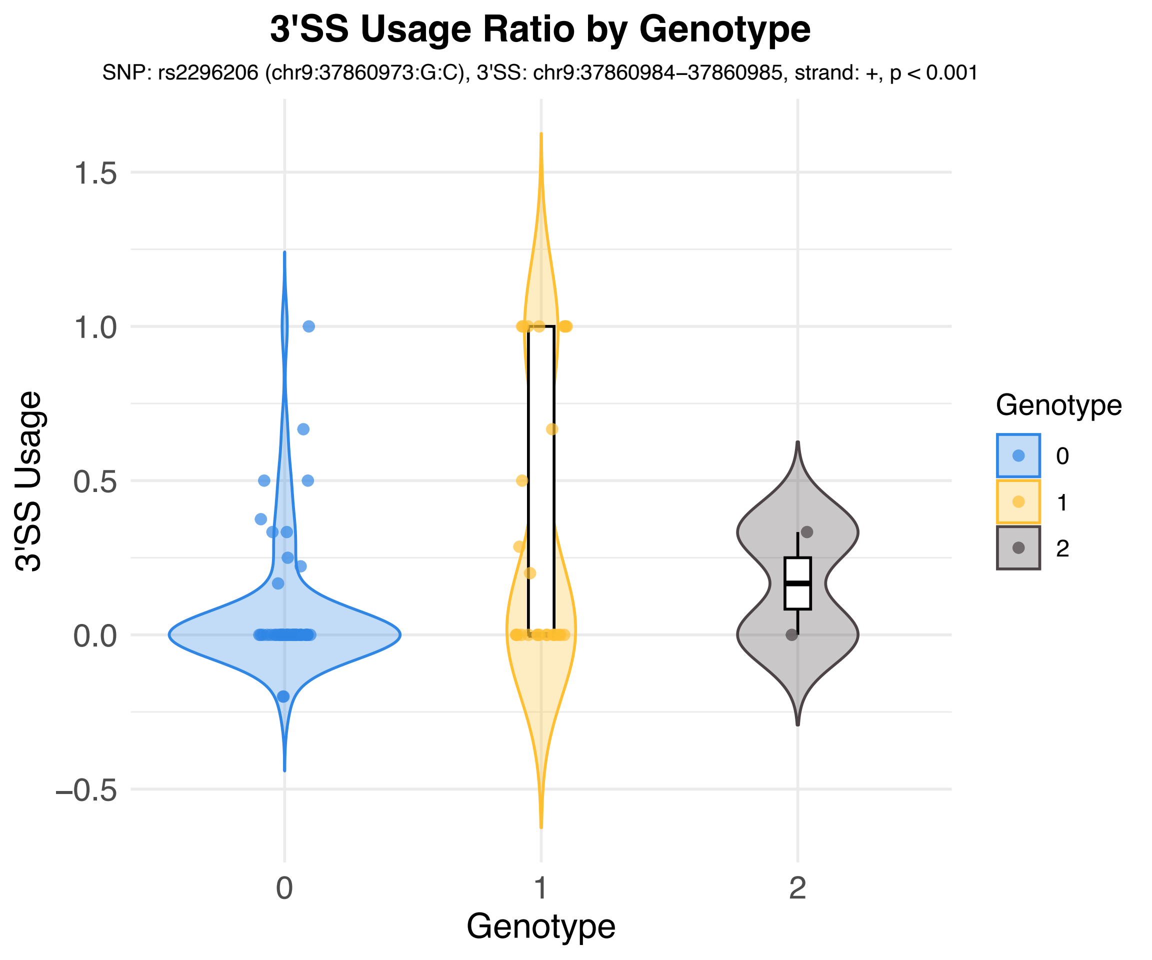
3'SS Usage Ratio by Genotype
SNP: rs2296206 (chr9:37860973:G:C), 3'SS: chr9:37860984–37860985, strand: +, p < 0.001
3'SS Usage
1.5
1.0
0.5
0.0
−0.5
0
1
2
Genotype
Genotype
0
1
2

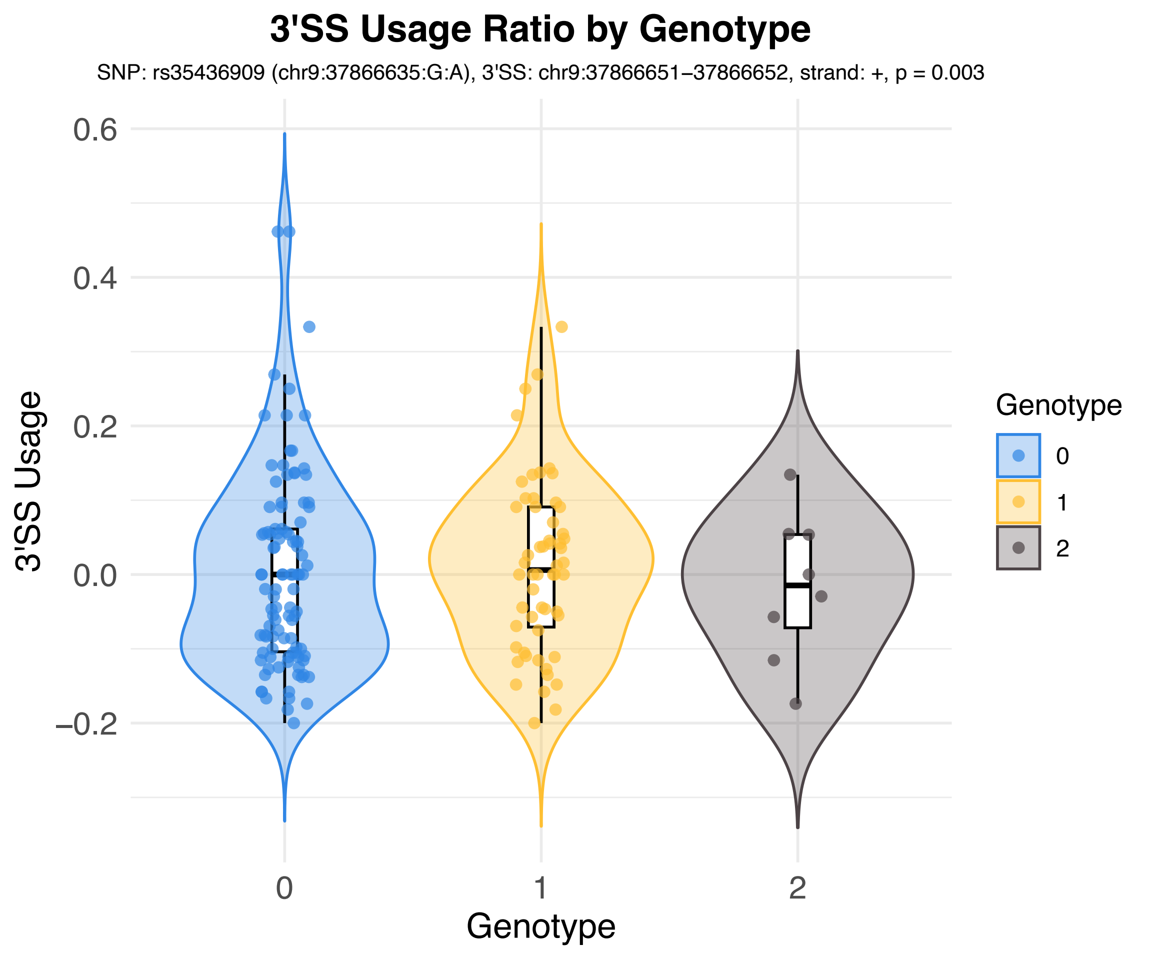
3'SS Usage Ratio by Genotype
SNP: rs35436909 (chr9:37866635:G:A), 3'SS: chr9:37866651–37866652, strand: +, p = 0.003
3'SS Usage
0.6
0.4
0.2
0.0
−0.2
0
1
2
Genotype
Genotype
0
1
2

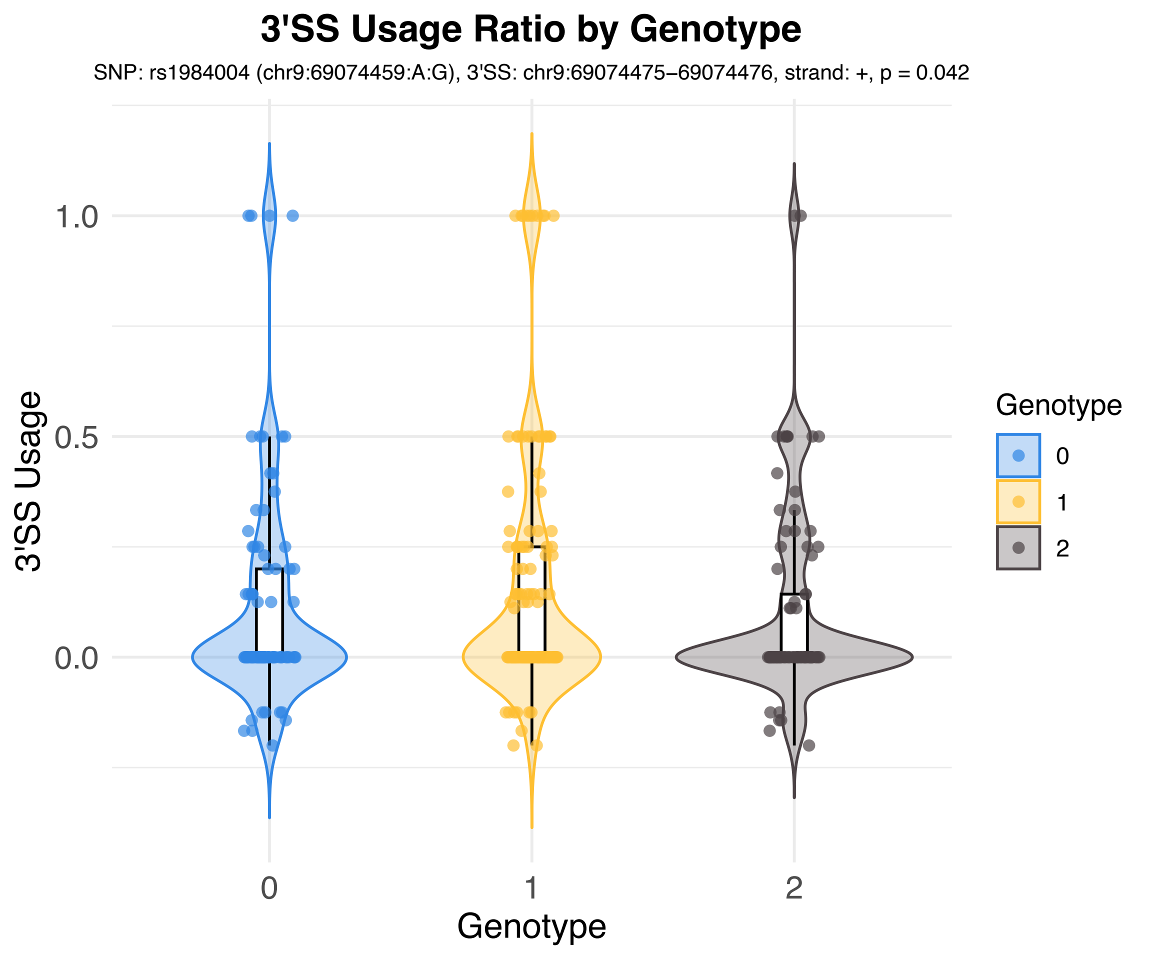
3'SS Usage Ratio by Genotype
SNP: rs1984004 (chr9:69074459:A:G), 3'SS: chr9:69074475–69074476, strand: +, p = 0.042
3'SS Usage
1.0
0.5
0.0
0
1
2
Genotype
Genotype
0
1
2

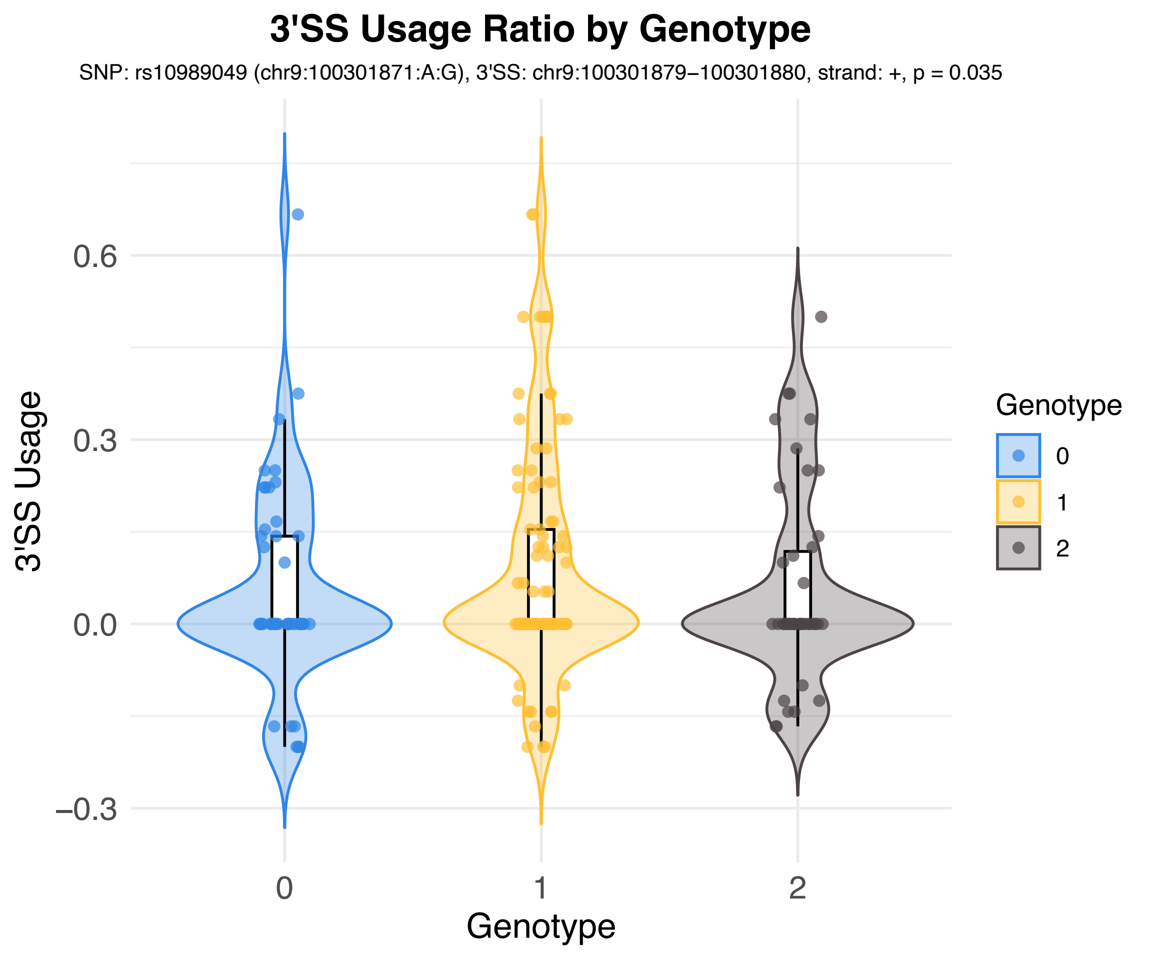

3'SS Usage Ratio by Genotype
SNP: rs10989049 (chr9:100301871:A:G), 3'SS: chr9:100301879−100301880, strand: +, p = 0.035
3'SS Usage
0.6
0.3
0.0
−0.3
0
1
2
Genotype
Genotype
0
1
2

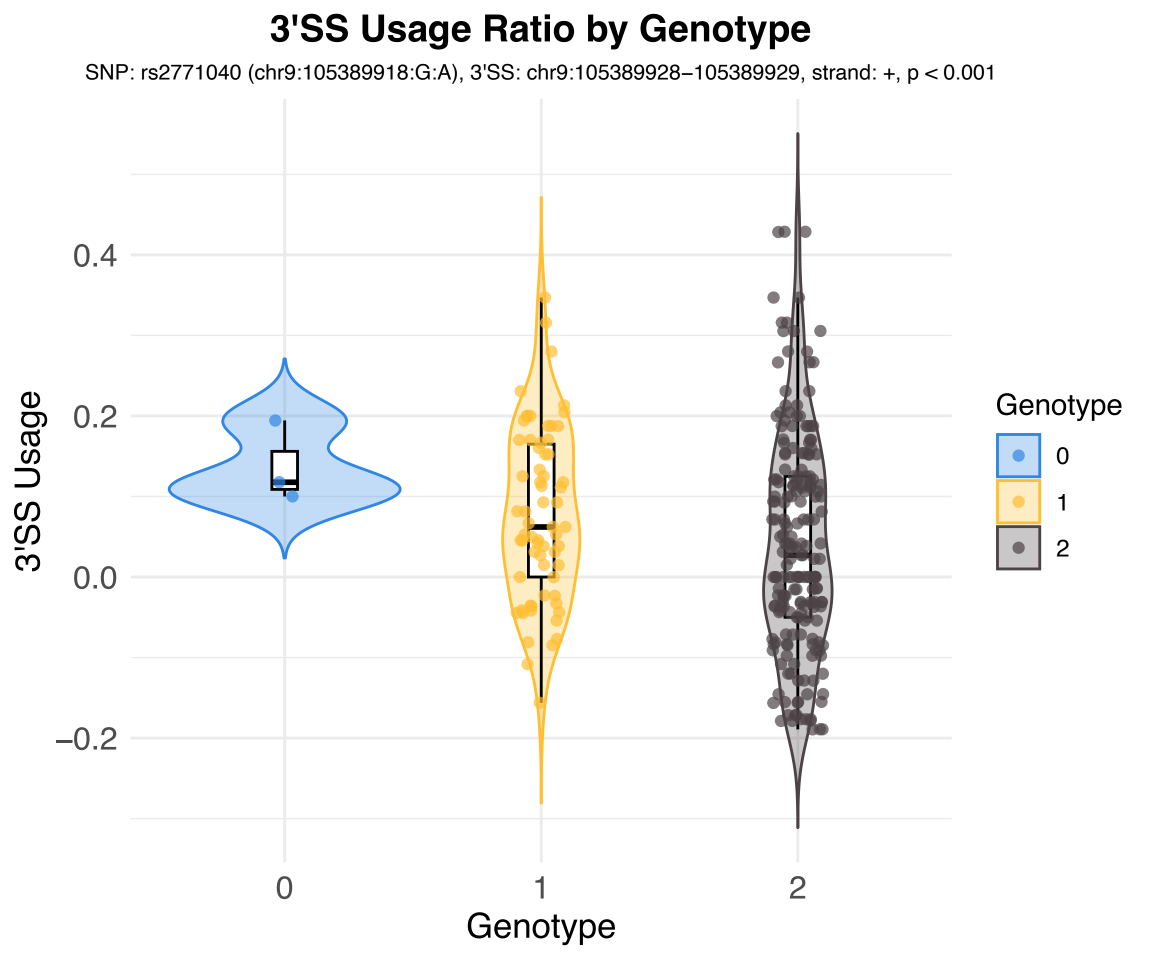

3'SS Usage Ratio by Genotype
SNP: rs2771040 (chr9:105389918:G:A), 3'SS: chr9:105389928−105389929, strand: +, p < 0.001
3'SS Usage
0.4
0.2
0.0
−0.2
0
1
2
Genotype
Genotype
0
1
2

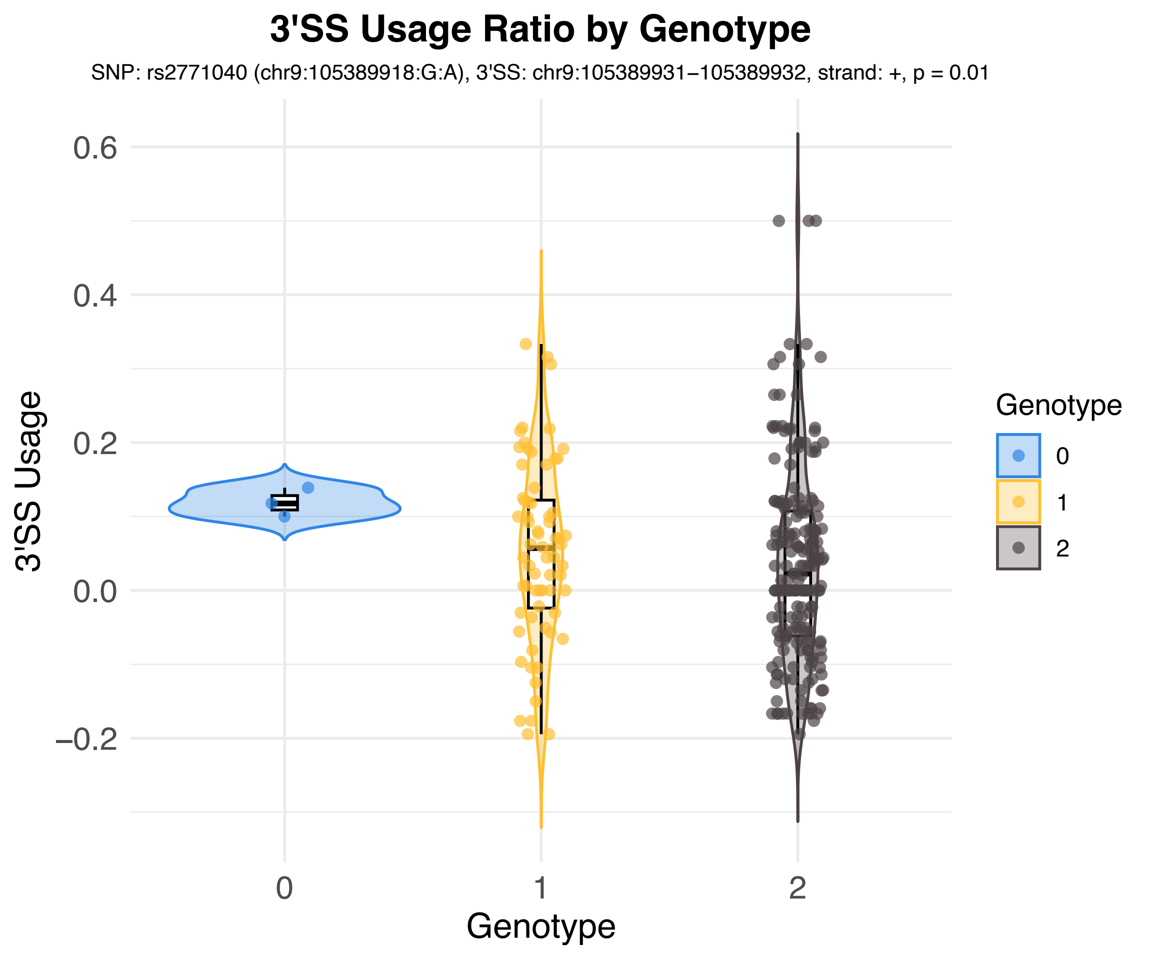
3'SS Usage Ratio by Genotype
SNP: rs2771040 (chr9:105389918:G:A), 3'SS: chr9:105389931−105389932, strand: +, p = 0.01
3'SS Usage
0.6
0.4
0.2
0.0
−0.2
0
1
2
Genotype
Genotype
0
1
2

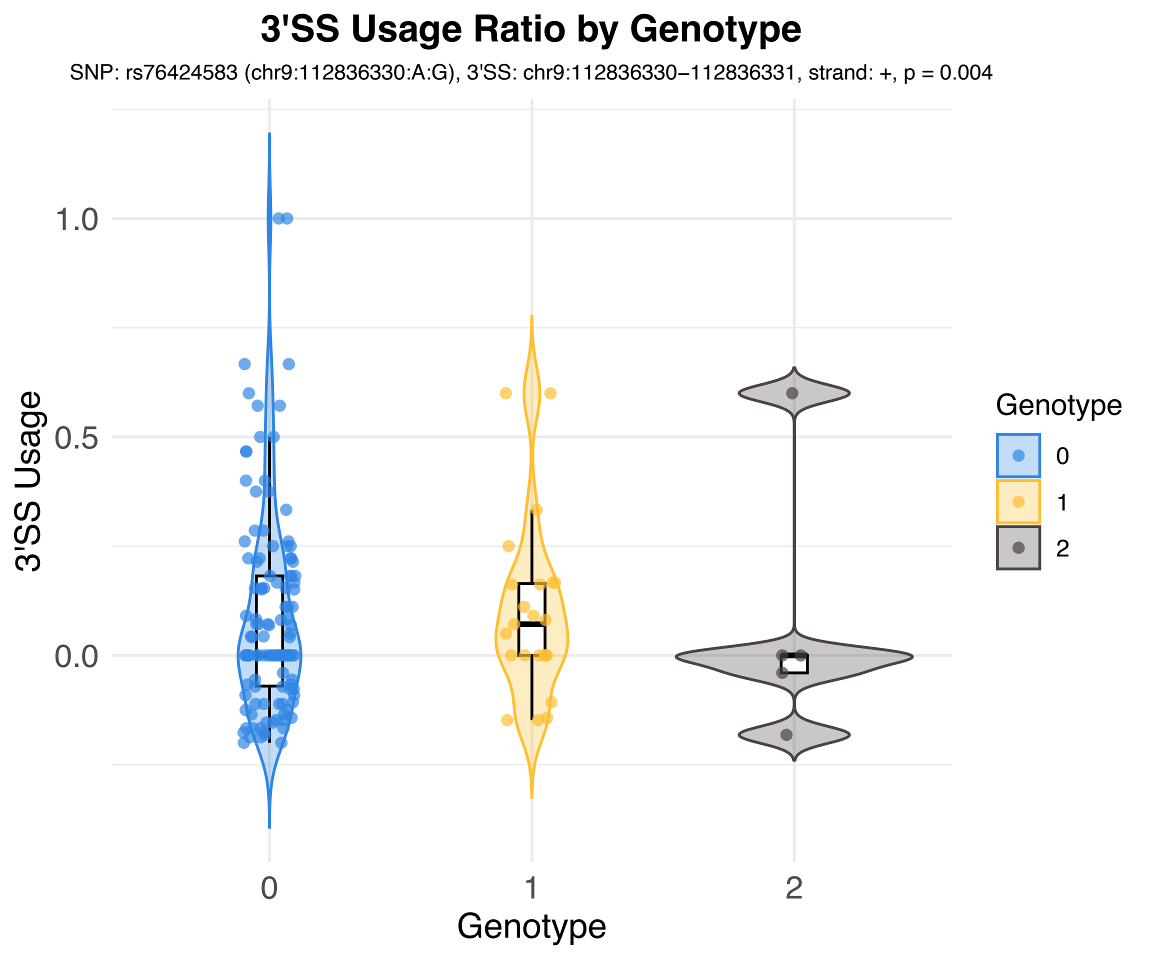

3'SS Usage Ratio by Genotype
SNP: rs76424583 (chr9:112836330:A:G), 3'SS: chr9:112836330−112836331, strand: +, p = 0.004
3'SS Usage
1.0
0.5
0.0
0
1
2
Genotype
Genotype
0
1
2

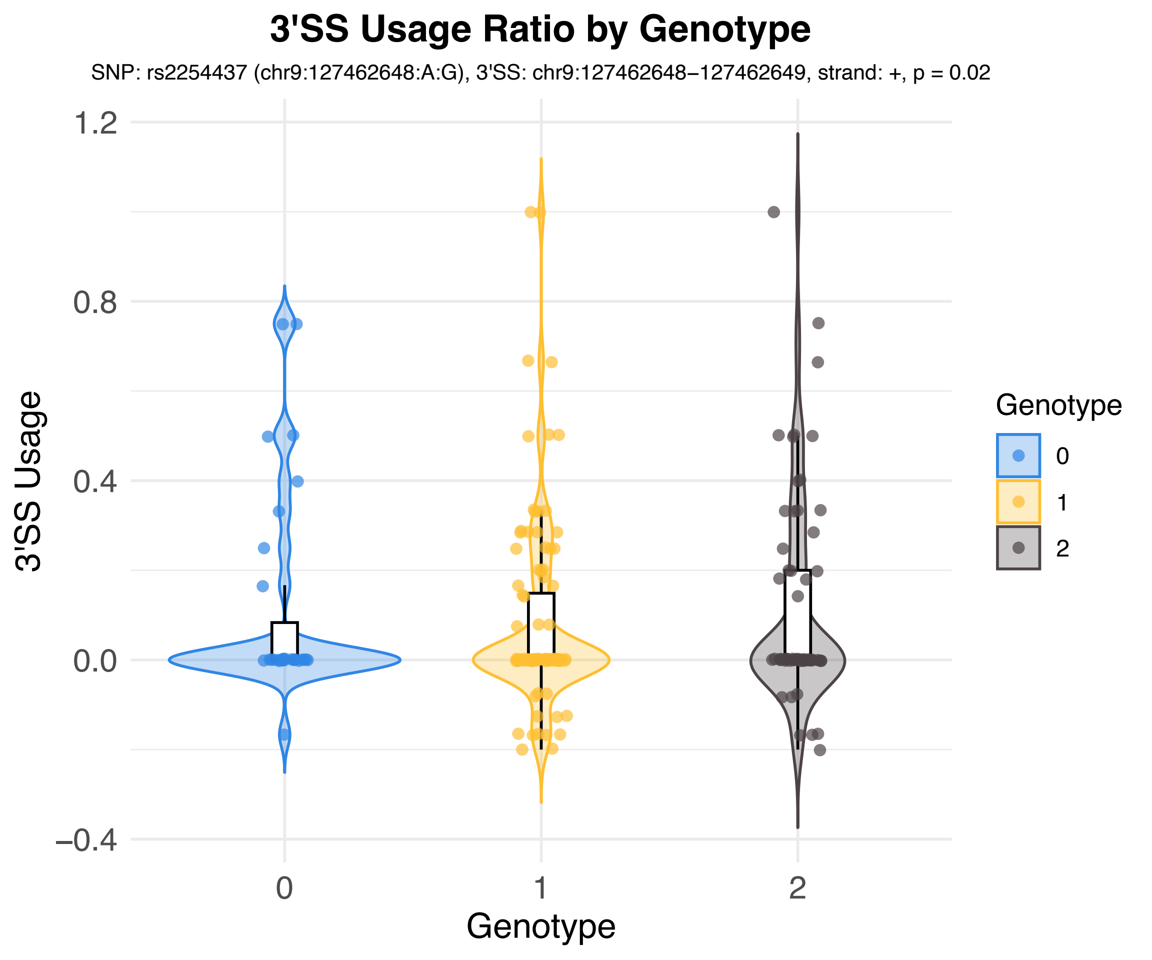
3'SS Usage Ratio by Genotype
SNP: rs2254437 (chr9:127462648:A:G), 3'SS: chr9:127462648−127462649, strand: +, p = 0.02
3'SS Usage
1.2
0.8
0.4
0.0
−0.4
0
1
2
Genotype
Genotype
0
1
2

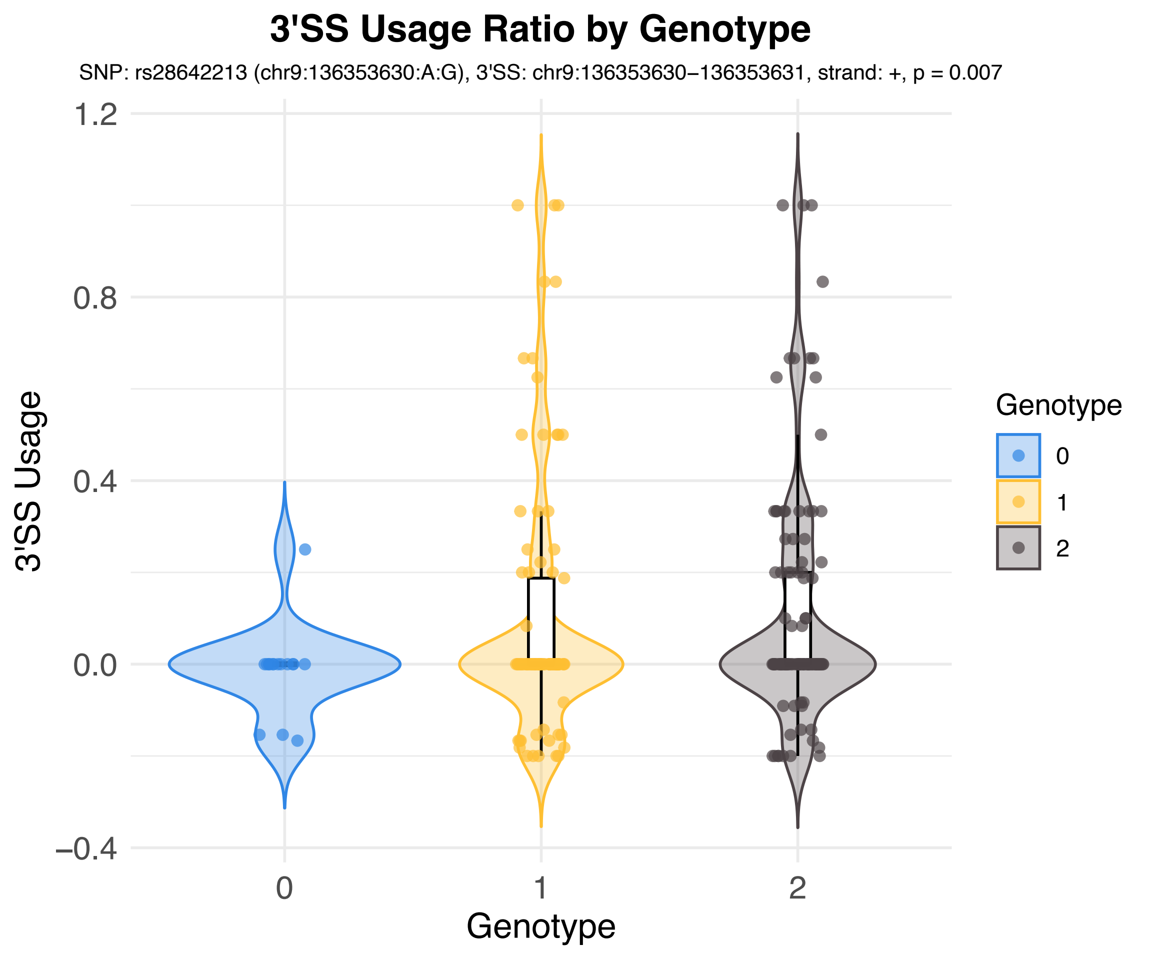

3'SS Usage Ratio by Genotype
SNP: rs28642213 (chr9:136353630:A:G), 3'SS: chr9:136353630−136353631, strand: +, p = 0.007
3'SS Usage
1.2
0.8
0.4
0.0
−0.4
0
1
2
Genotype
Genotype
0
1
2

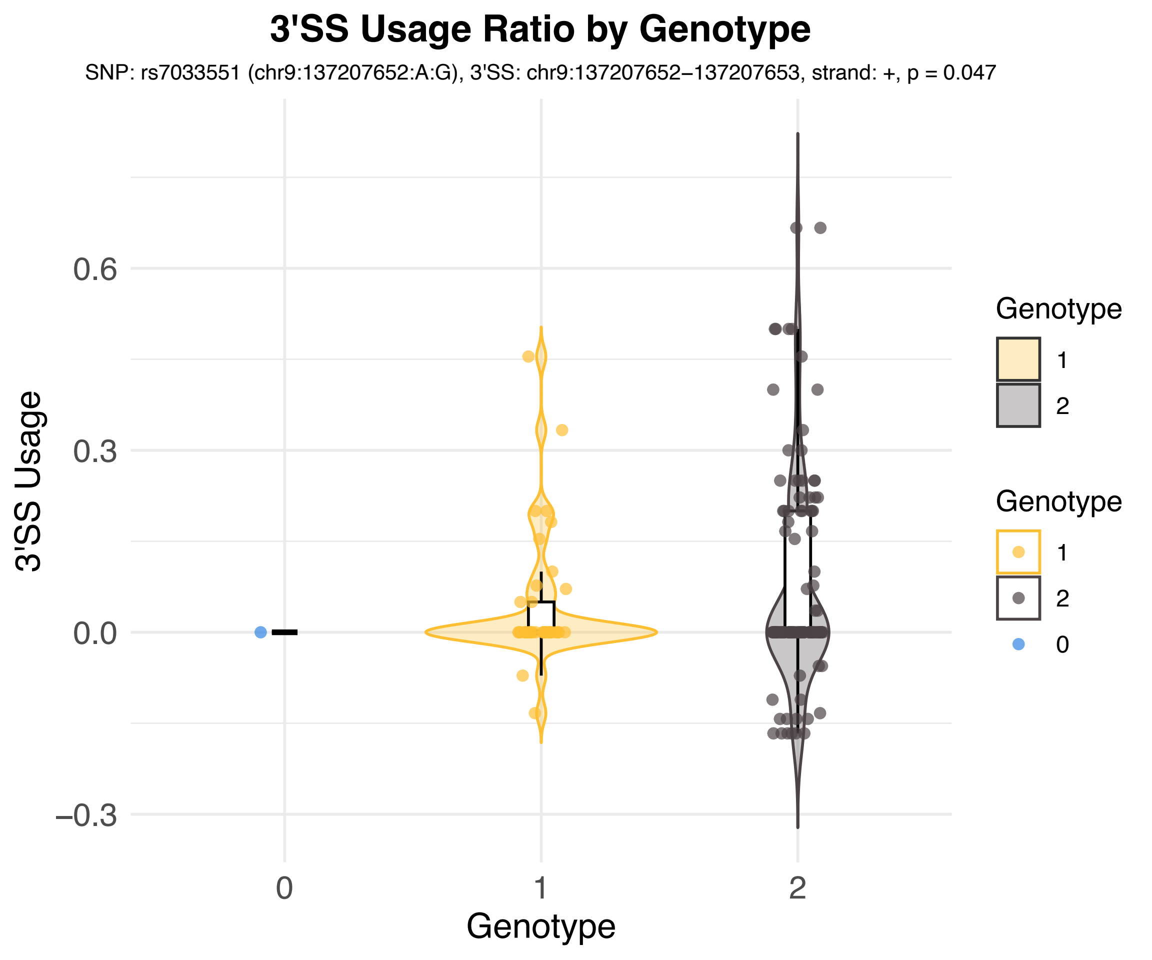

3'SS Usage Ratio by Genotype
SNP: rs7033551 (chr9:137207652:A:G), 3'SS: chr9:137207652−137207653, strand: +, p = 0.047
3'SS Usage
0.6
0.3
0.0
−0.3
0
1
2
Genotype
Genotype
1
2
Genotype
1
2
0

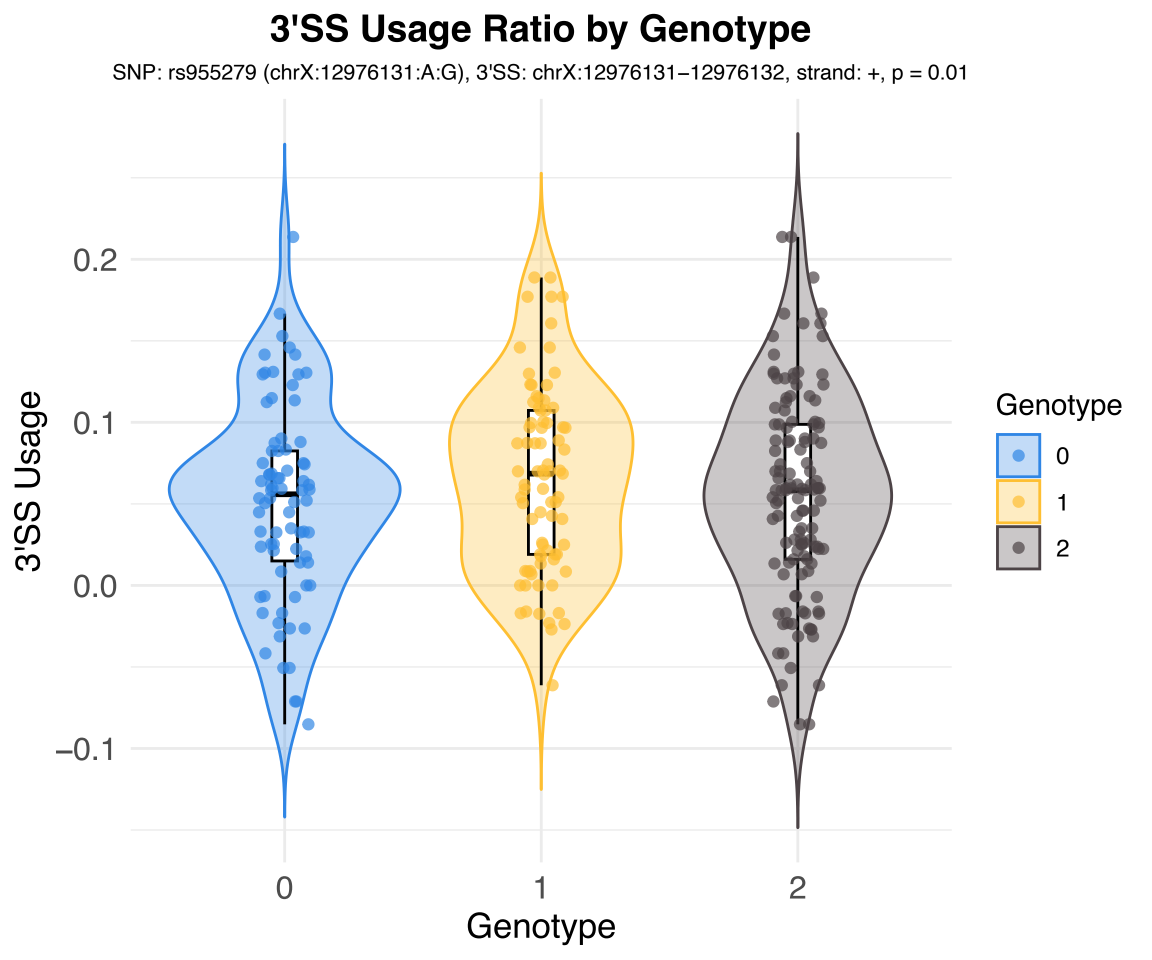
3'SS Usage Ratio by Genotype
SNP: rs955279 (chrX:12976131:A:G), 3'SS: chrX:12976131−12976132, strand: +, p = 0.01
3'SS Usage
0.2
0.1
0.0
−0.1
0
1
2
Genotype
Genotype
0
1
2

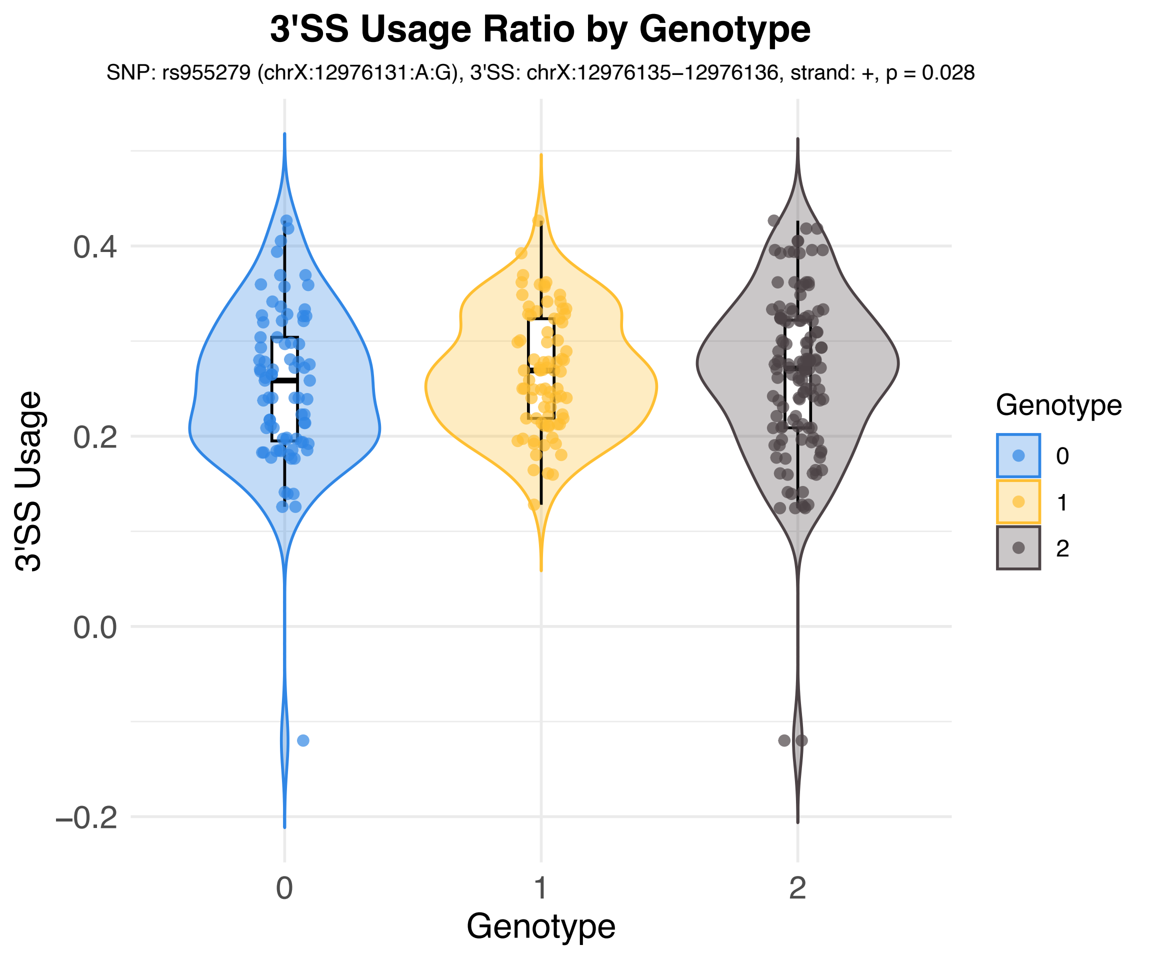
3'SS Usage Ratio by Genotype
SNP: rs955279 (chrX:12976131:A:G), 3'SS: chrX:12976135−12976136, strand: +, p = 0.028
3'SS Usage
0.4
0.2
0.0
−0.2
0
1
2
Genotype
Genotype
0
1
2

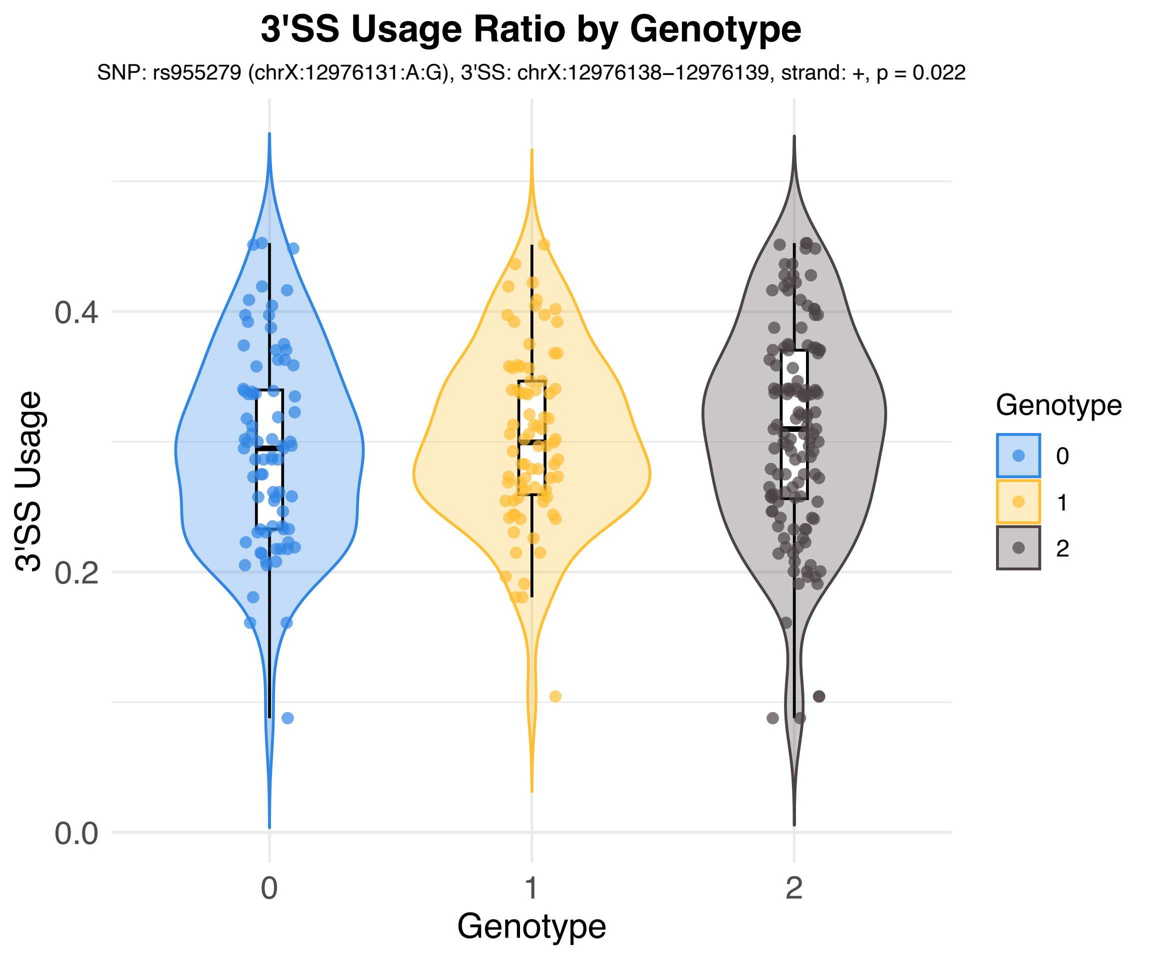

3'SS Usage Ratio by Genotype
SNP: rs955279 (chrX:12976131:A:G), 3'SS: chrX:12976138−12976139, strand: +, p = 0.022
3'SS Usage
0.0
0.2
0.4
0
1
2
Genotype
Genotype
0
1
2

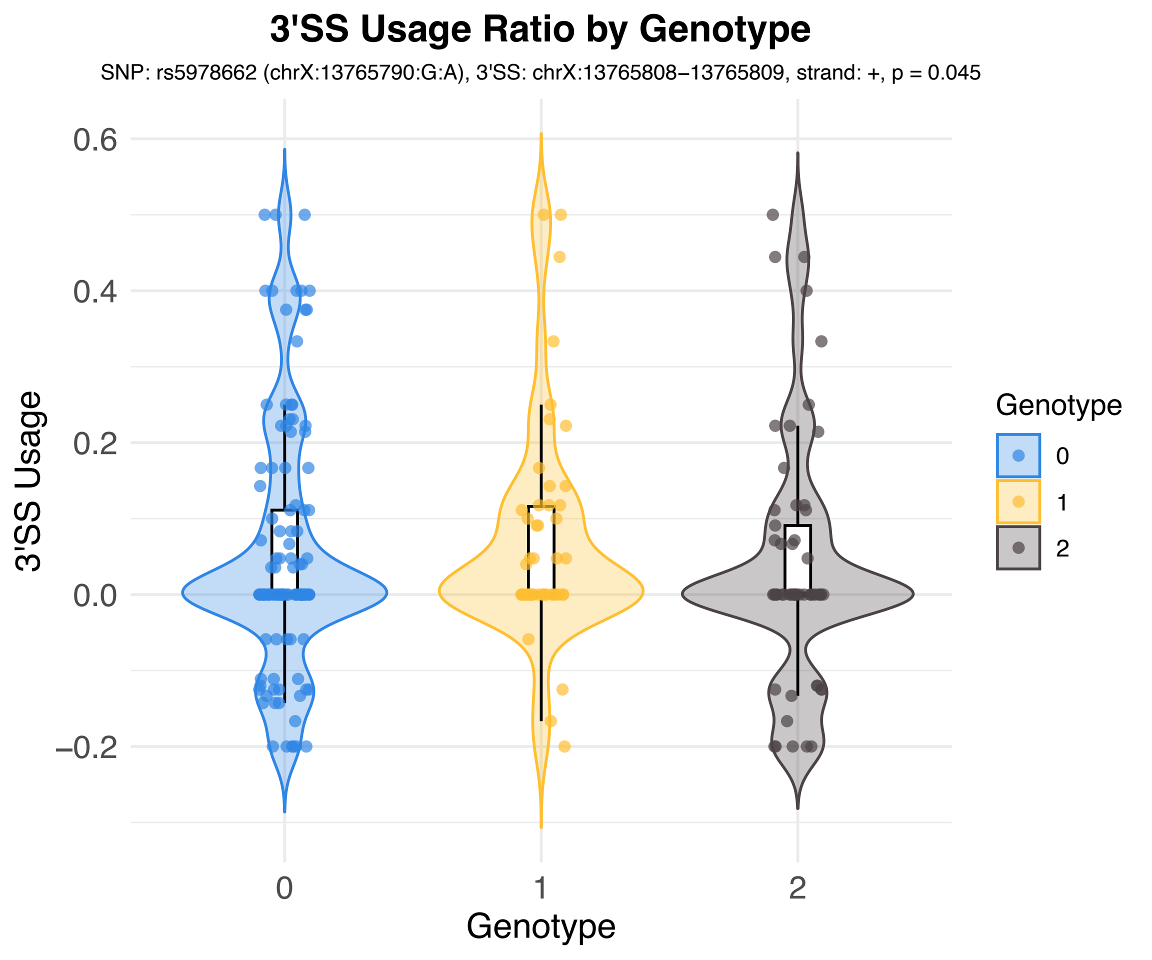
3'SS Usage Ratio by Genotype
SNP: rs5978662 (chrX:13765790:G:A), 3'SS: chrX:13765808–13765809, strand: +, p = 0.045
3'SS Usage
0.6
0.4
0.2
0.0
–0.2
0
1
2
Genotype
Genotype
0
1
2

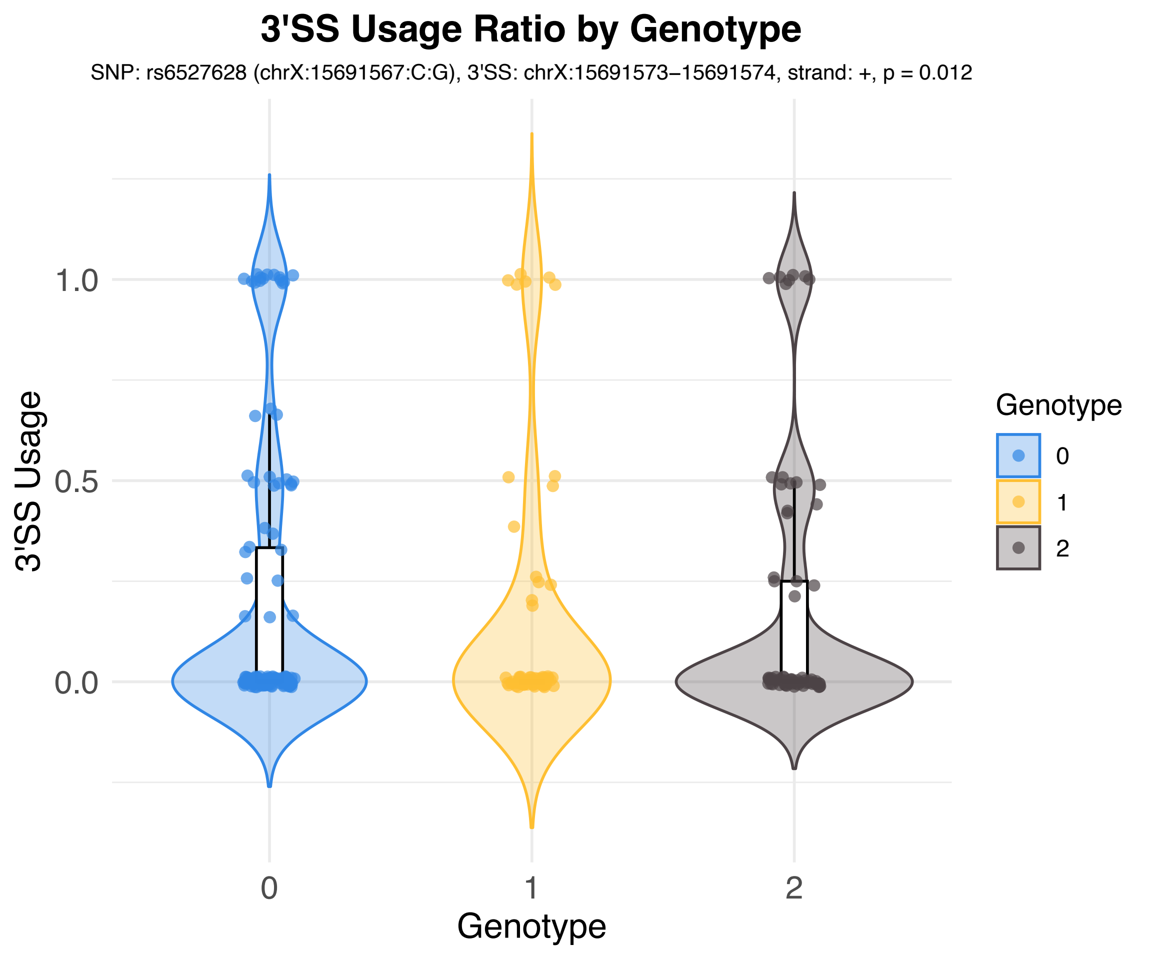
3'SS Usage Ratio by Genotype
SNP: rs6527628 (chrX:15691567:C:G), 3'SS: chrX:15691573−15691574, strand: +, p = 0.012
3'SS Usage
1.0
0.5
0.0
0
1
2
Genotype
Genotype
0
1
2

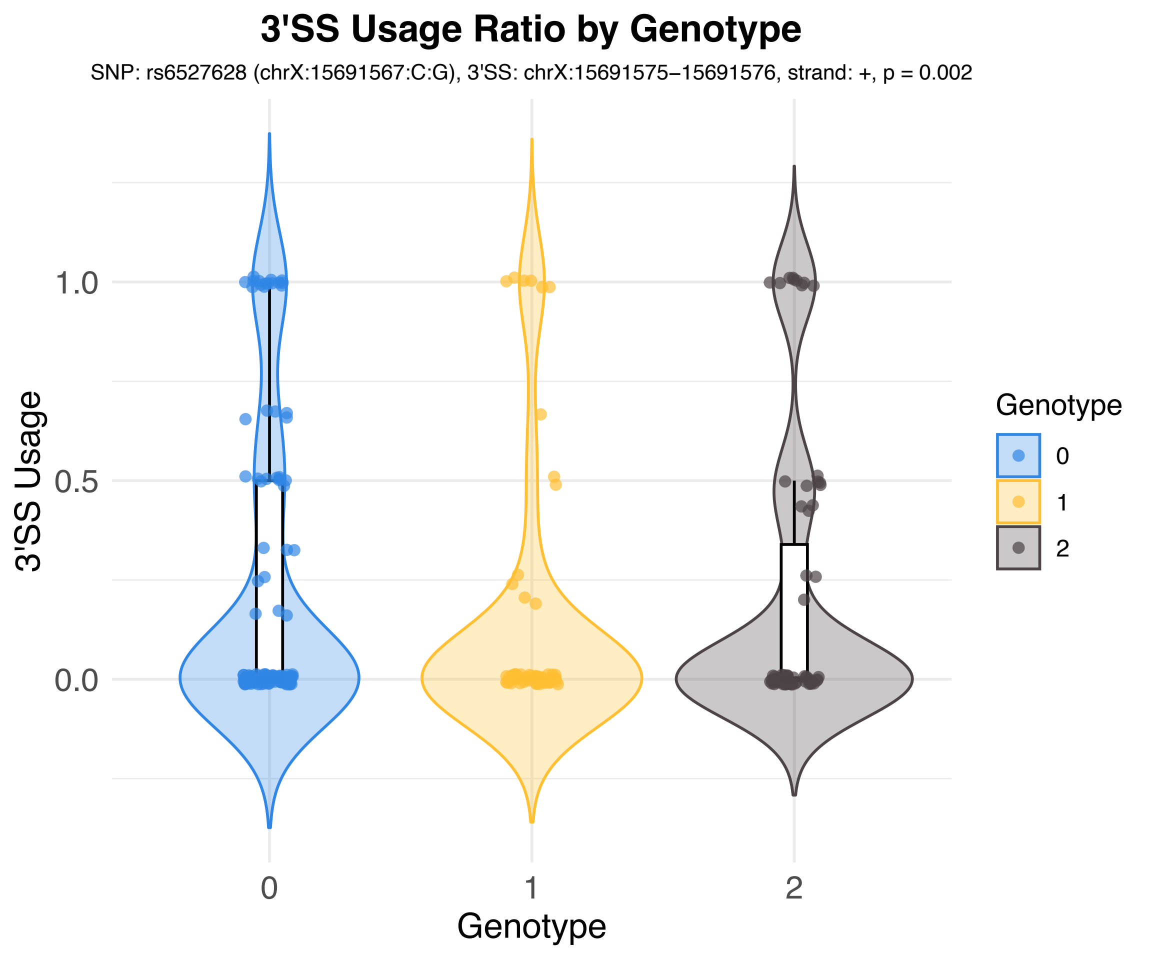

3'SS Usage Ratio by Genotype
SNP: rs6527628 (chrX:15691567:C:G), 3'SS: chrX:15691575−15691576, strand: +, p = 0.002
3'SS Usage
1.0
0.5
0.0
0
1
2
Genotype
Genotype
0
1
2

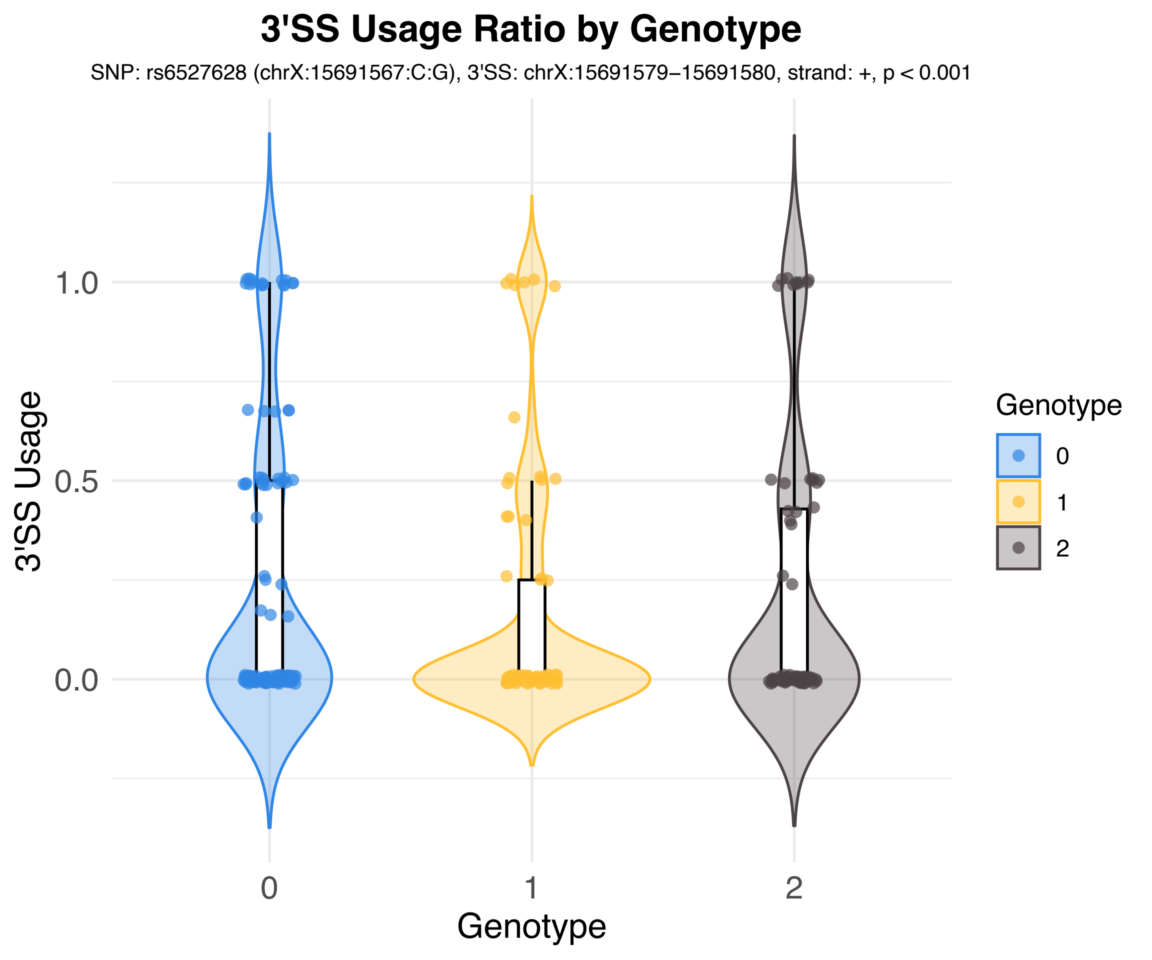

3'SS Usage Ratio by Genotype
SNP: rs6527628 (chrX:15691567:C:G), 3'SS: chrX:15691579−15691580, strand: +, p < 0.001
3'SS Usage
1.0
0.5
0.0
0
1
2
Genotype
Genotype
0
1
2

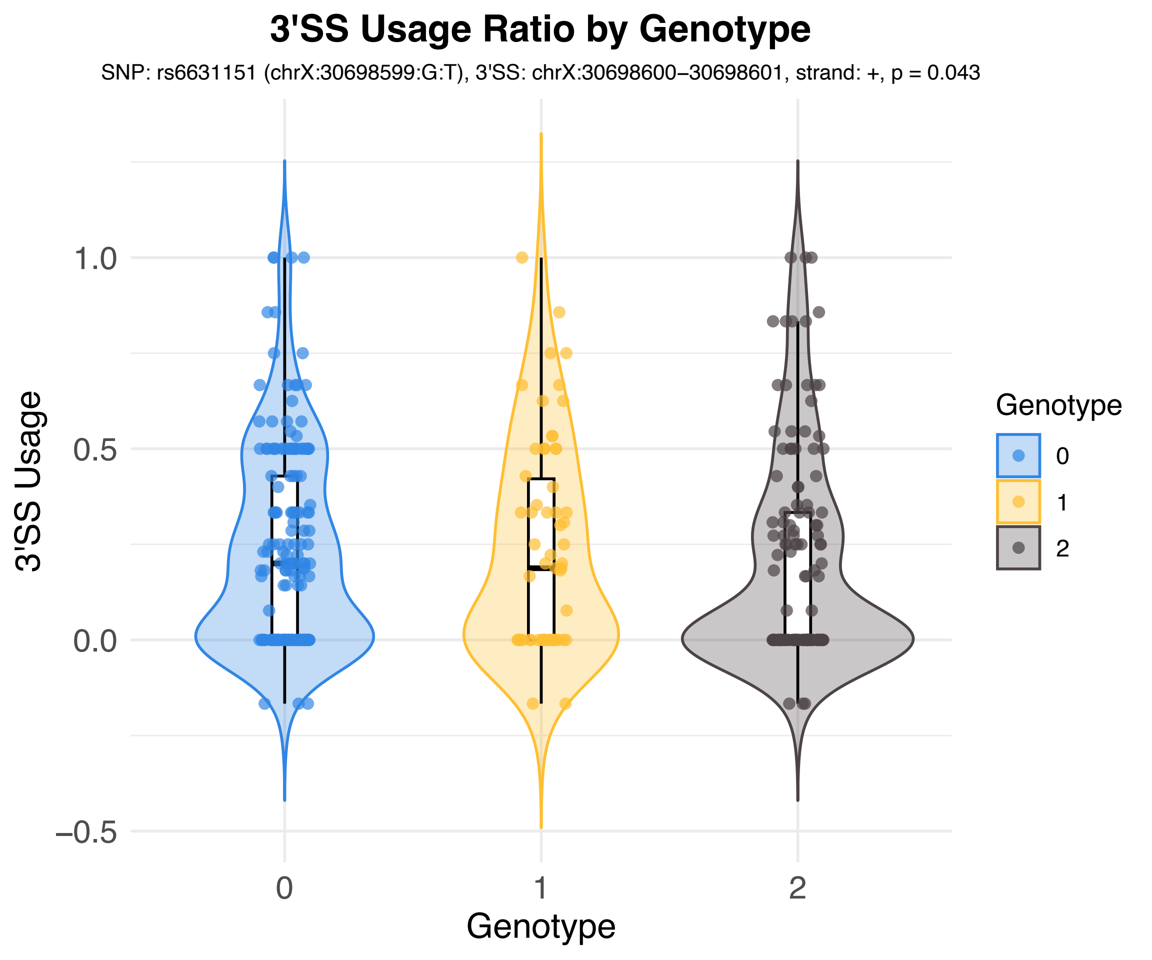

3'SS Usage Ratio by Genotype
SNP: rs6631151 (chrX:30698599:G:T), 3'SS: chrX:30698600–30698601, strand: +, p = 0.043
3'SS Usage
1.0
0.5
0.0
−0.5
0
1
2
Genotype
Genotype
0
1
2

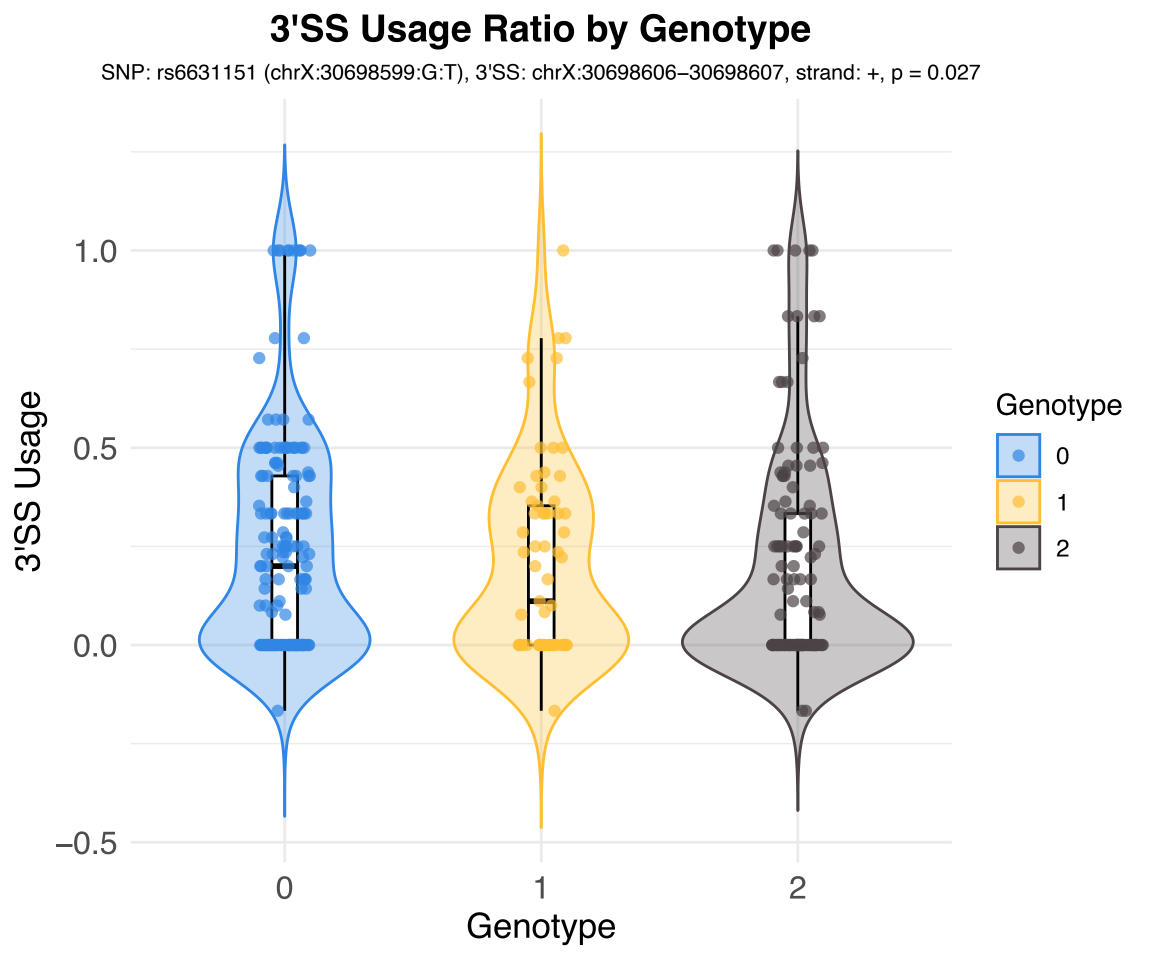
3'SS Usage Ratio by Genotype
SNP: rs6631151 (chrX:30698599:G:T), 3'SS: chrX:30698606–30698607, strand: +, p = 0.027
3'SS Usage
1.0
0.5
0.0
−0.5
0
1
2
Genotype
Genotype
0
1
2

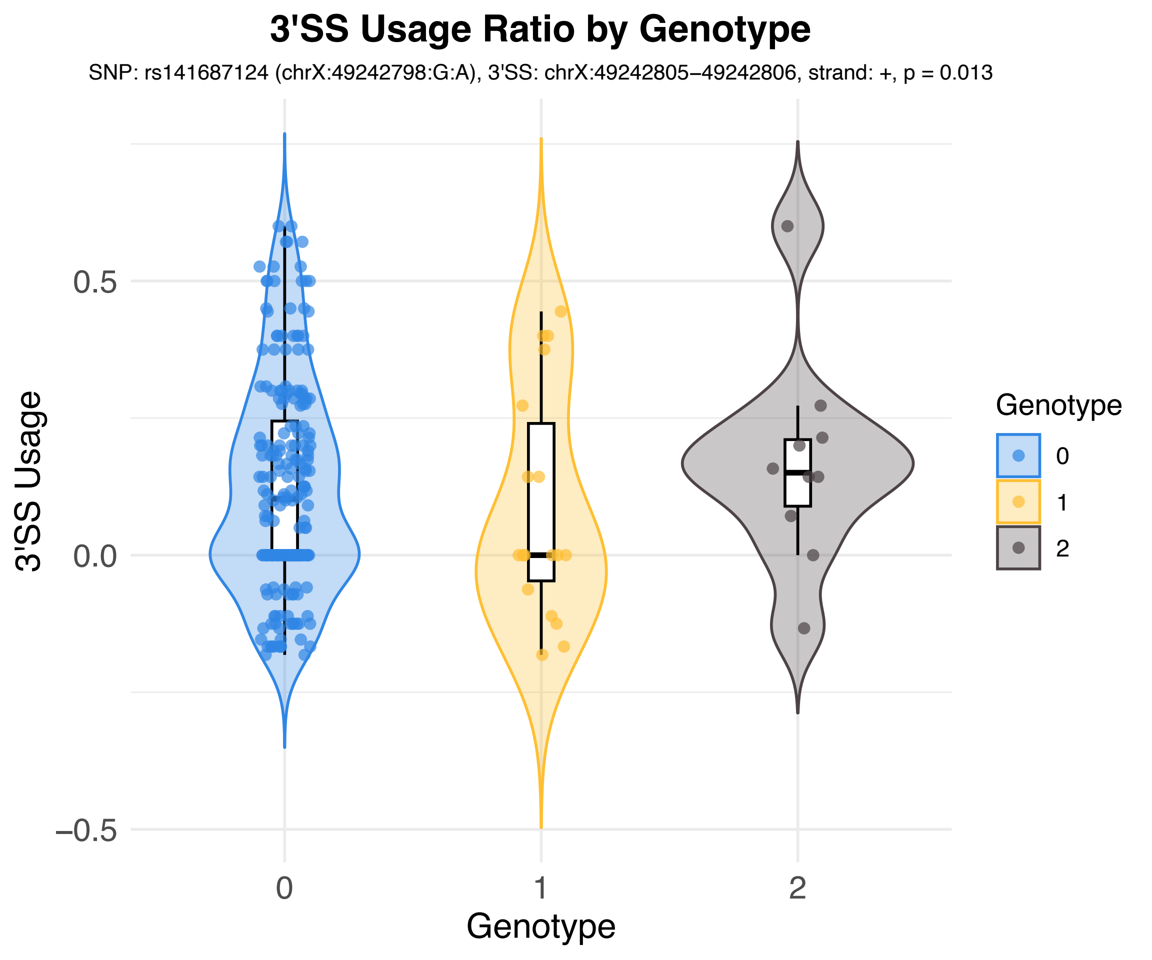
3'SS Usage Ratio by Genotype
SNP: rs141687124 (chrX:49242798:G:A), 3'SS: chrX:49242805−49242806, strand: +, p = 0.013
3'SS Usage
0.5
0.0
−0.5
0
1
2
Genotype
Genotype
0
1
2
