## Supplementary material for "A Mammalian Genomic Signature Shaped by Single Nucleotide Variants Regulates Transcriptome Integrity and Diversity": Additional file 5_3' splice site usage_n603_pairs_general_violin_plots.pdf

Violin plots of 3' splice site usage of 603 pairs of SNV and G-tract-AG-harboring 3’ splice site from the general genotype model, by analysis of whole genome sequencing and RNA-Seq - matched data among 100 FHS participants.

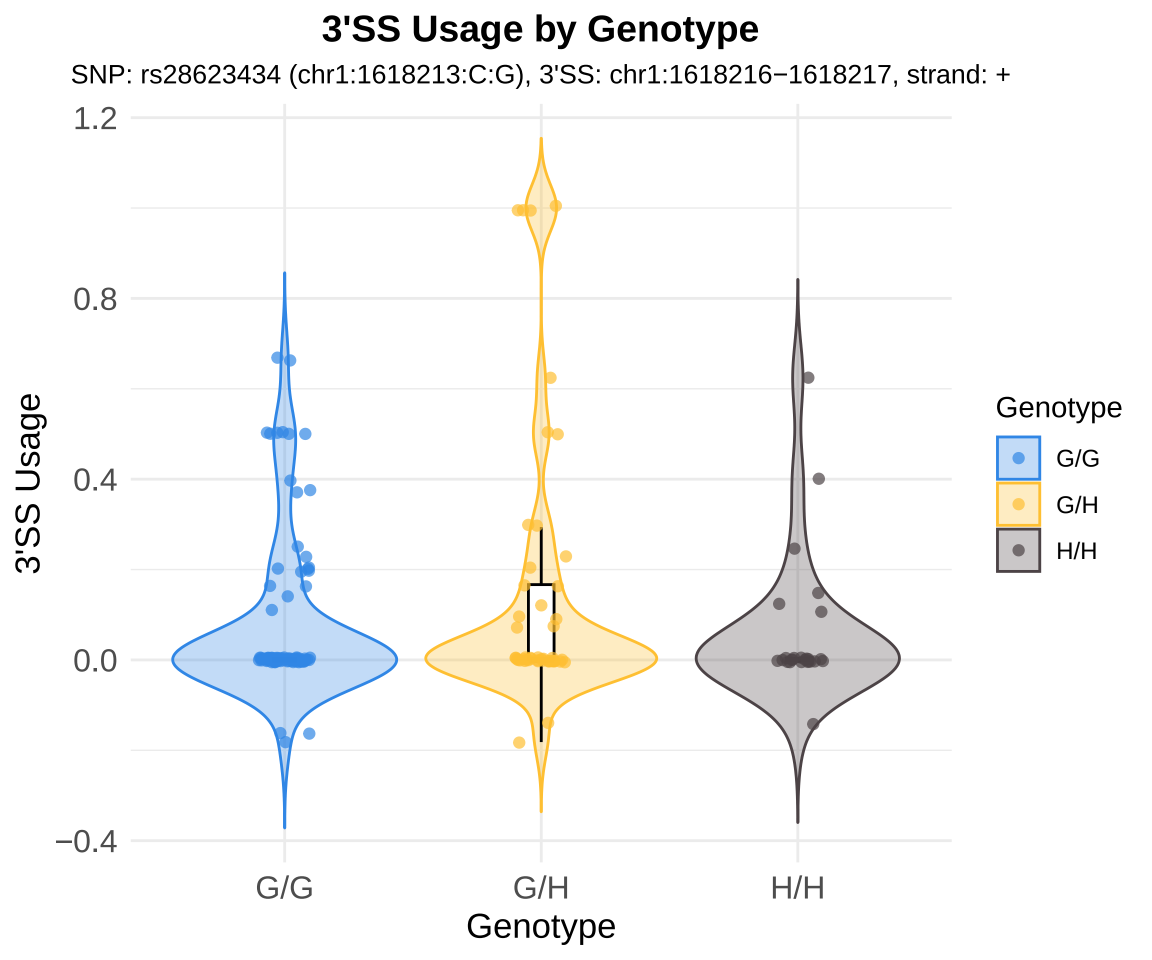
3'SS Usage by Genotype
SNP: rs28623434 (chr1:1618213:C:G), 3'SS: chr1:1618216−1618217, strand: +
3'SS Usage
1.2
0.8
0.4
0.0
−0.4
G/G
G/H
H/H
Genotype
Genotype
G/G
G/H
H/H

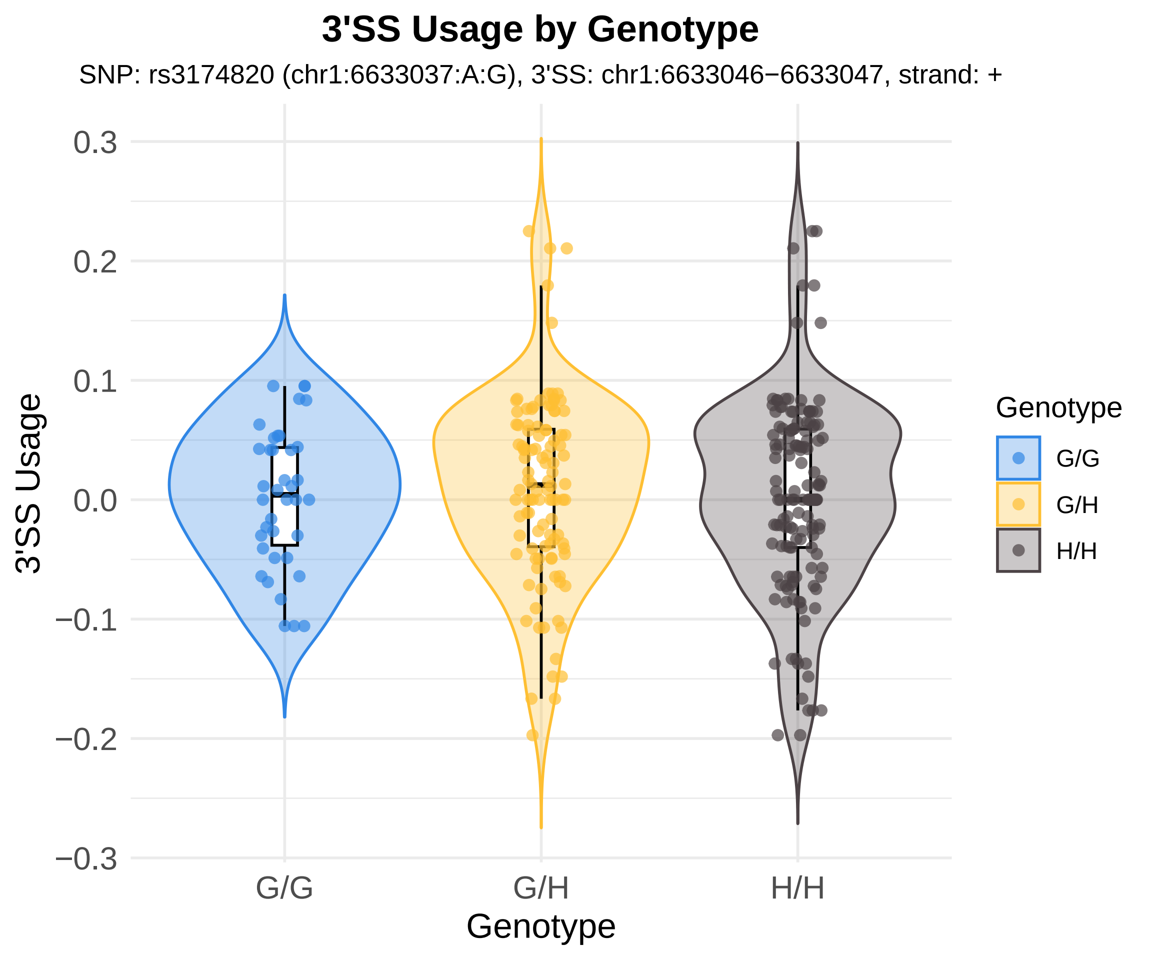
3'SS Usage by Genotype
SNP: rs3174820 (chr1:6633037:A:G), 3'SS: chr1:6633046−6633047, strand: +
3'SS Usage
0.3
0.2
0.1
0.0
−0.1
−0.2
−0.3
G/G
G/H
H/H
Genotype
Genotype
G/G
G/H
H/H

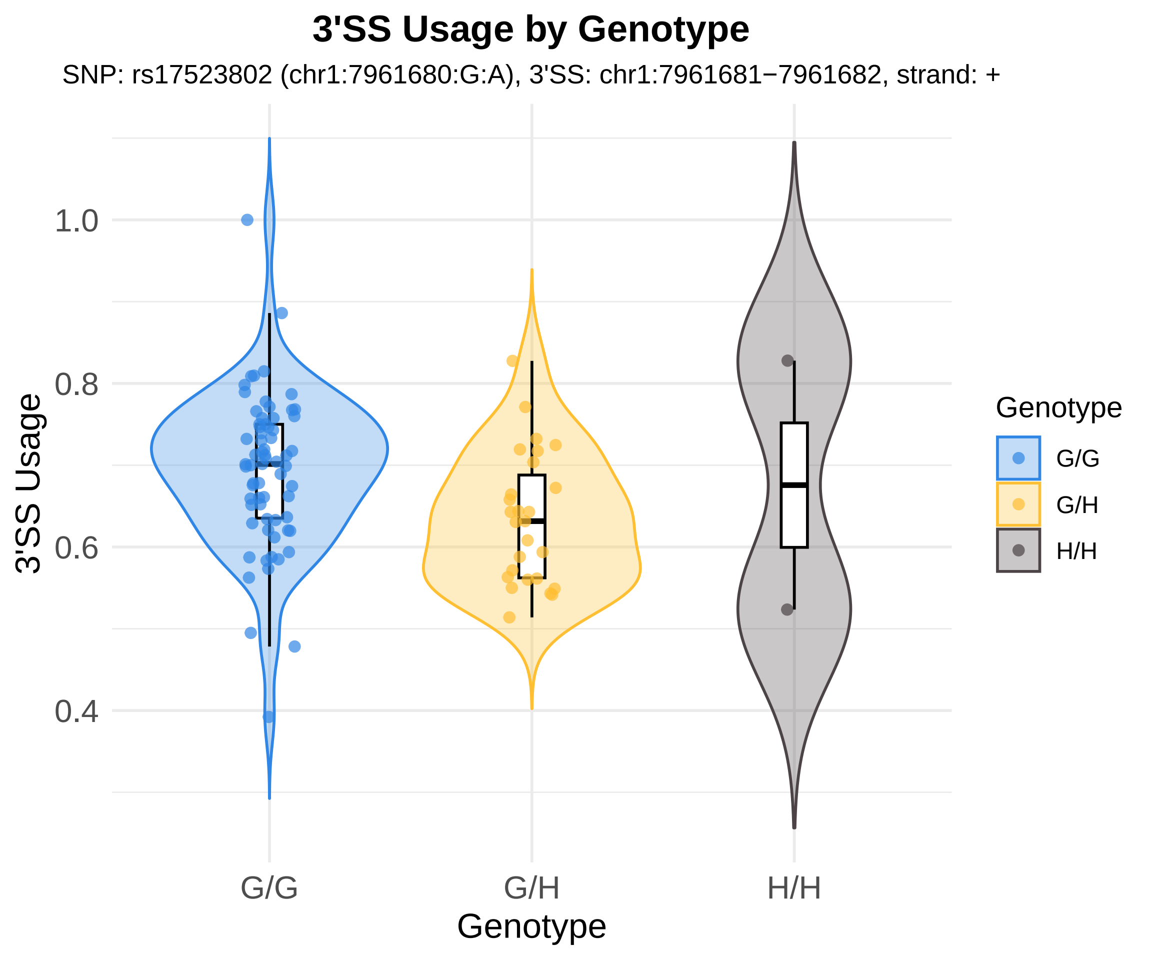
3'SS Usage by Genotype
SNP: rs17523802 (chr1:7961680:G:A), 3'SS: chr1:7961681−7961682, strand: +
3'SS Usage
1.0
0.8
0.6
0.4
G/G
G/H
H/H
Genotype
Genotype
G/G
G/H
H/H

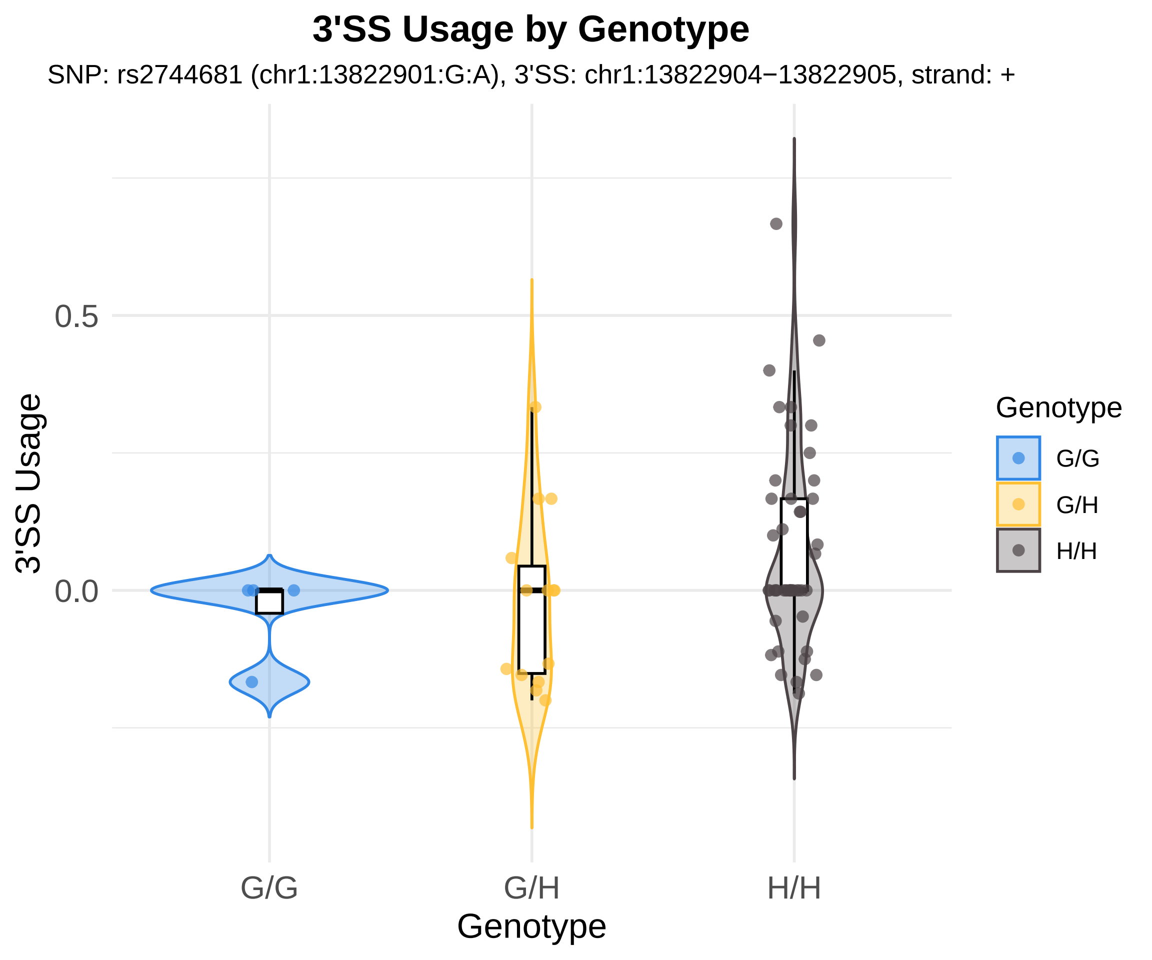

3'SS Usage by Genotype
SNP: rs2744681 (chr1:13822901:G:A), 3'SS: chr1:13822904−13822905, strand: +
3'SS Usage
0.5
0.0
G/G
G/H
H/H
Genotype
Genotype
G/G
G/H
H/H

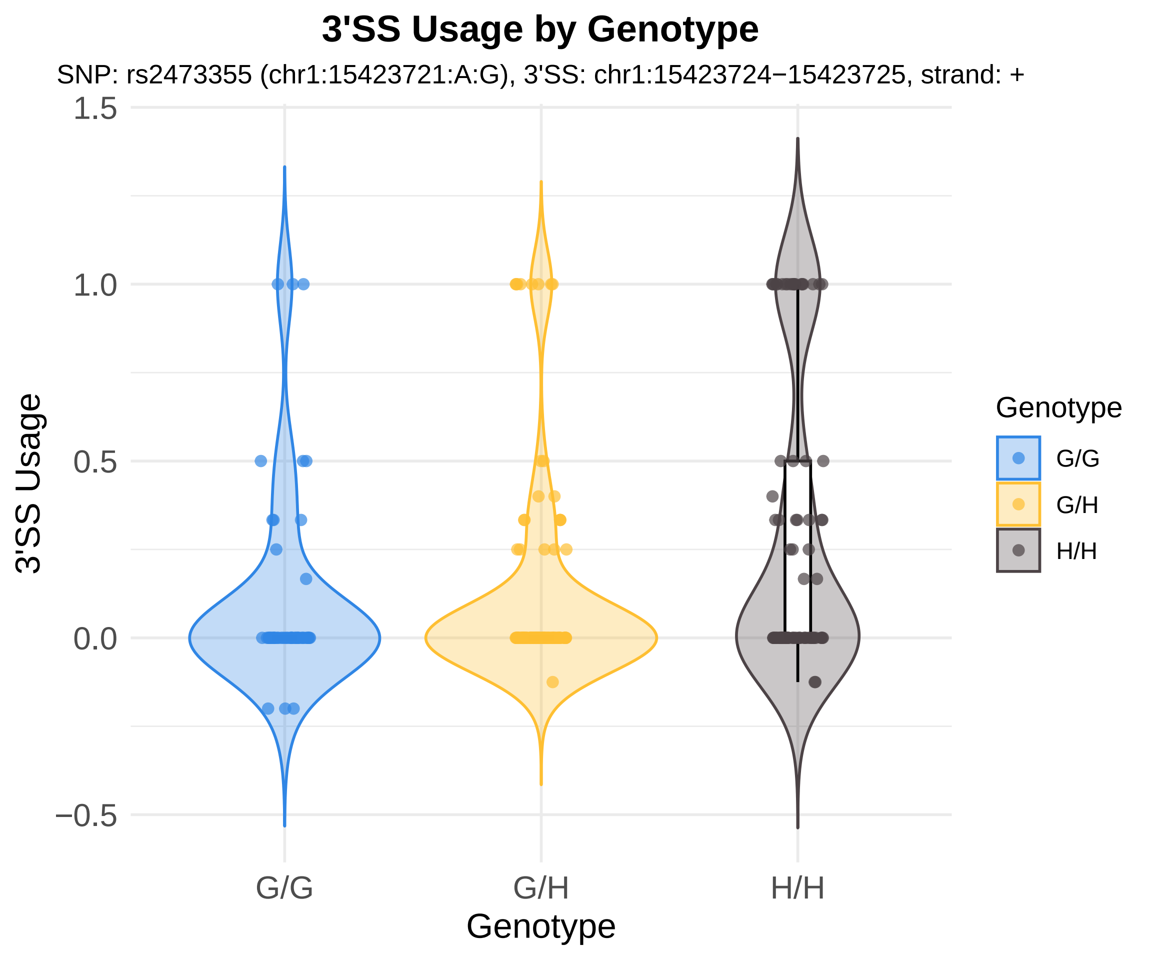

3'SS Usage by Genotype
SNP: rs2473355 (chr1:15423721:A:G), 3'SS: chr1:15423724−15423725, strand: +
3'SS Usage
1.5
1.0
0.5
0.0
−0.5
G/G
G/H
H/H
Genotype
Genotype
G/G
G/H
H/H

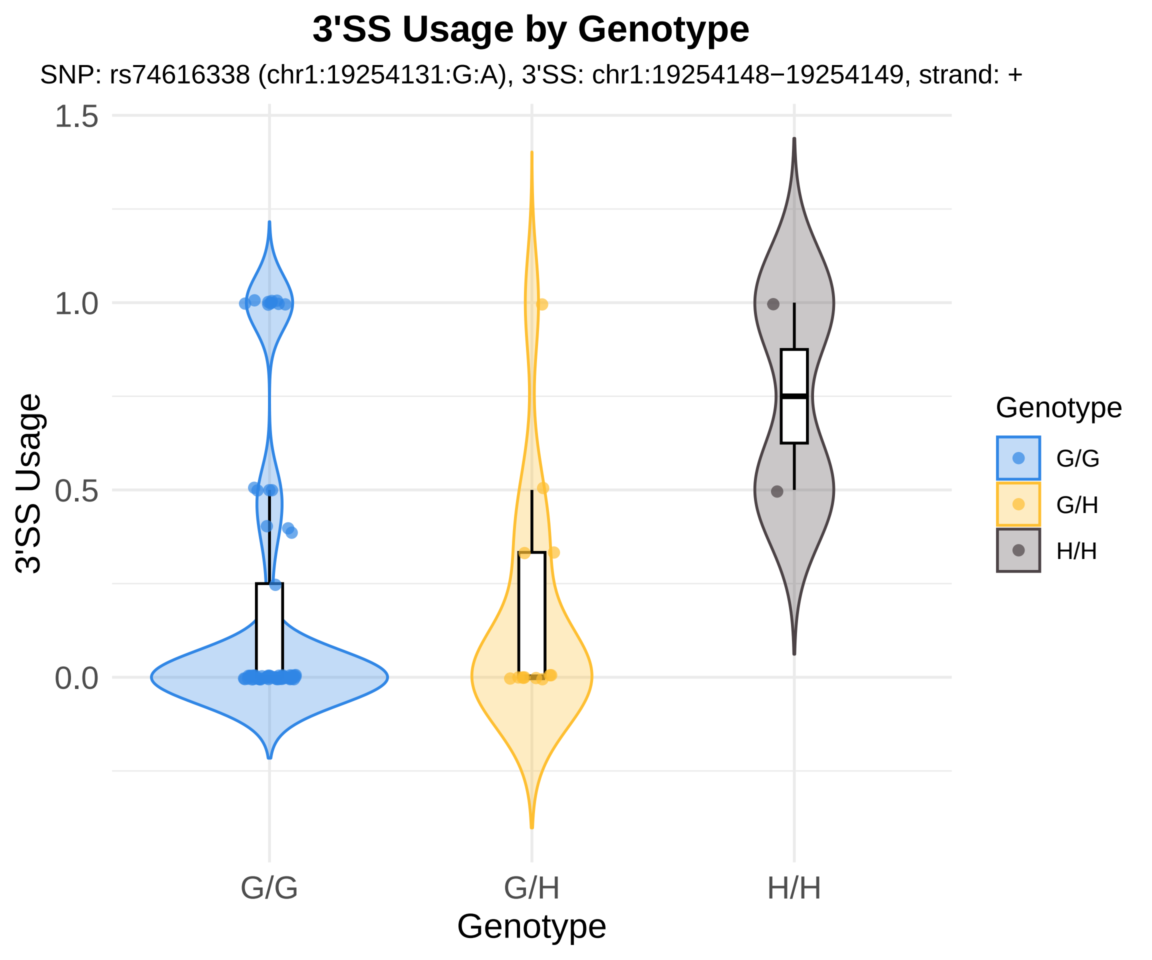
3'SS Usage by Genotype
SNP: rs74616338 (chr1:19254131:G:A), 3'SS: chr1:19254148−19254149, strand: +
3'SS Usage
1.5
1.0
0.5
0.0
G/G
G/H
H/H
Genotype
Genotype
G/G
G/H
H/H

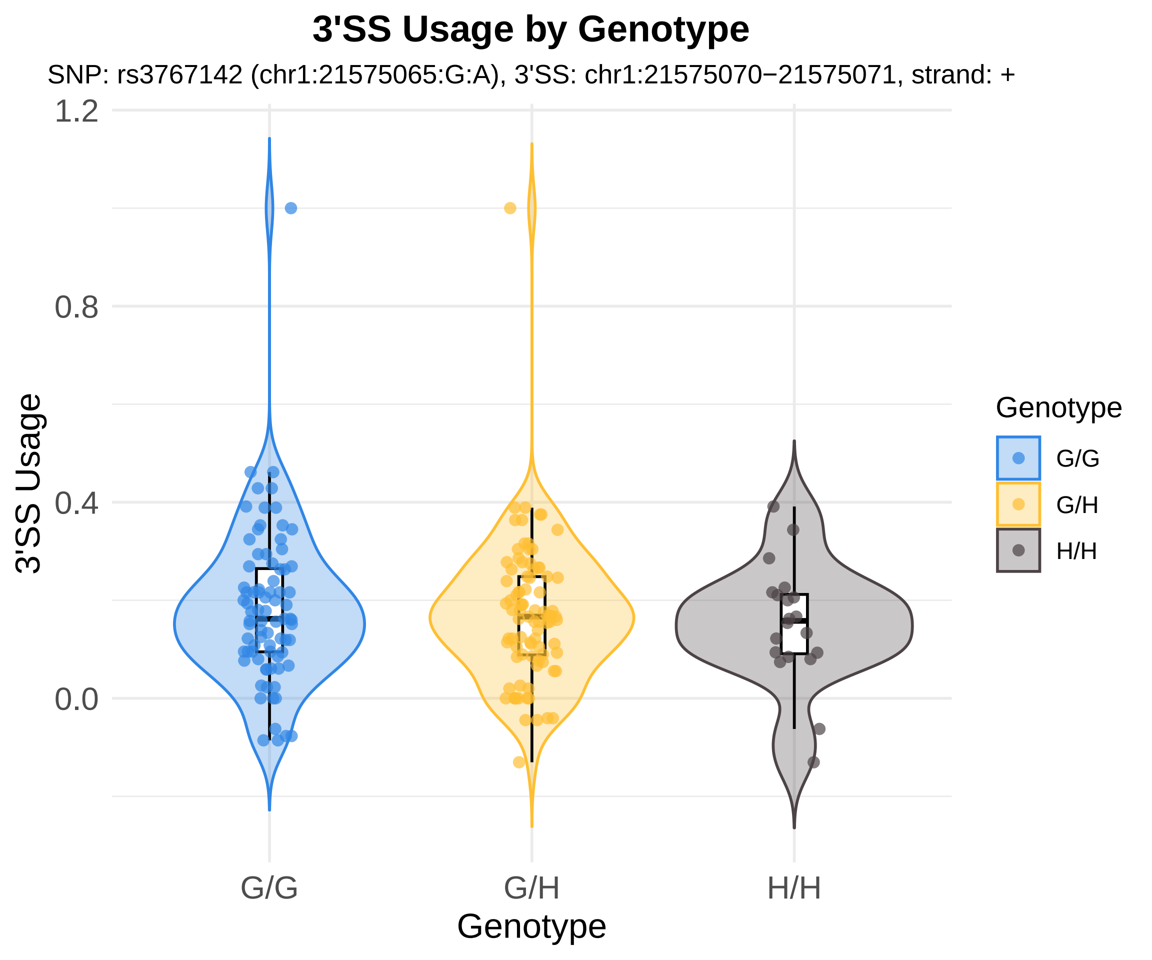
3'SS Usage by Genotype
SNP: rs3767142 (chr1:21575065:G:A), 3'SS: chr1:21575070−21575071, strand: +
3'SS Usage
1.2
0.8
0.4
0.0
G/G
G/H
H/H
Genotype
Genotype
G/G
G/H
H/H

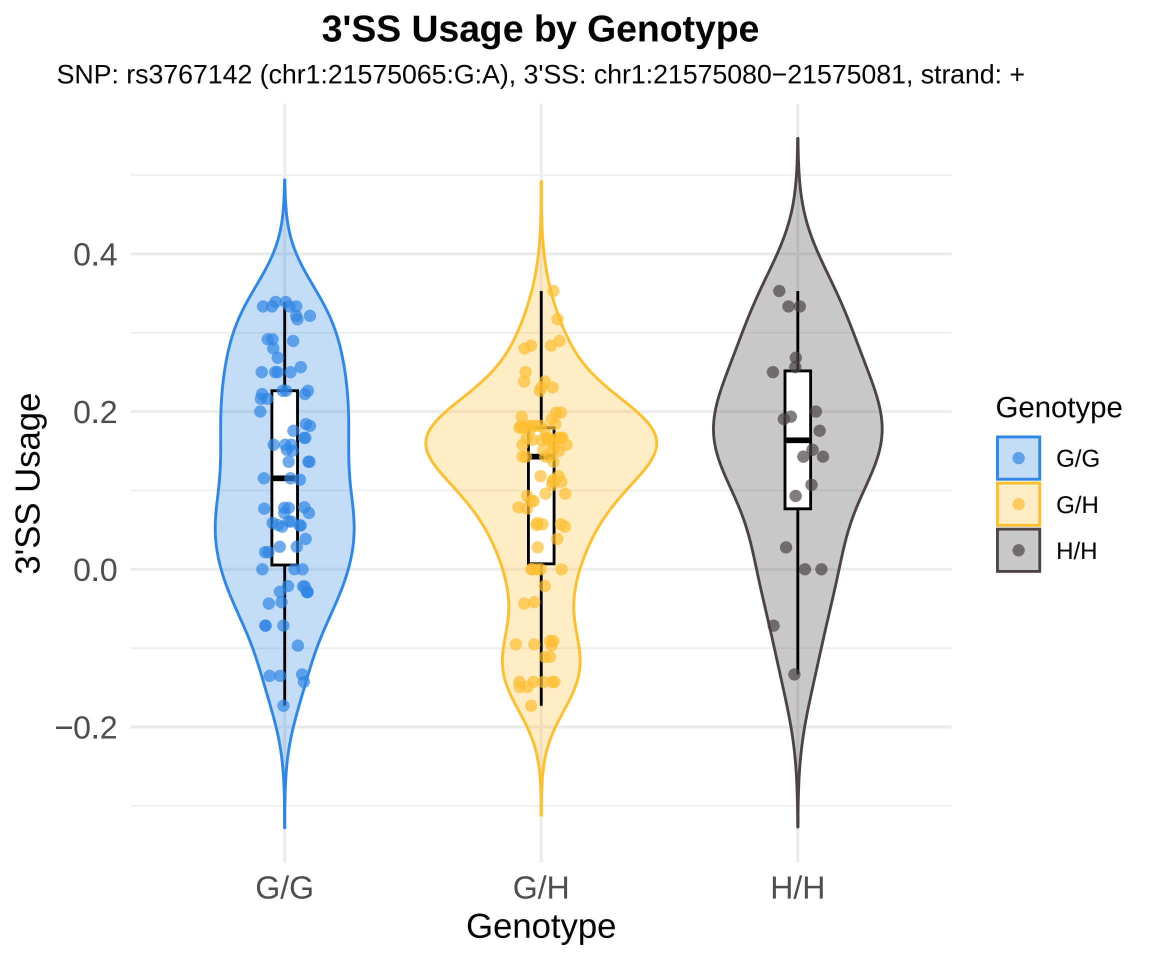

3'SS Usage by Genotype
SNP: rs3767142 (chr1:21575065:G:A), 3'SS: chr1:21575080−21575081, strand: +
3'SS Usage
0.4
0.2
0.0
−0.2
G/G
G/H
H/H
Genotype
Genotype
G/G
G/H
H/H

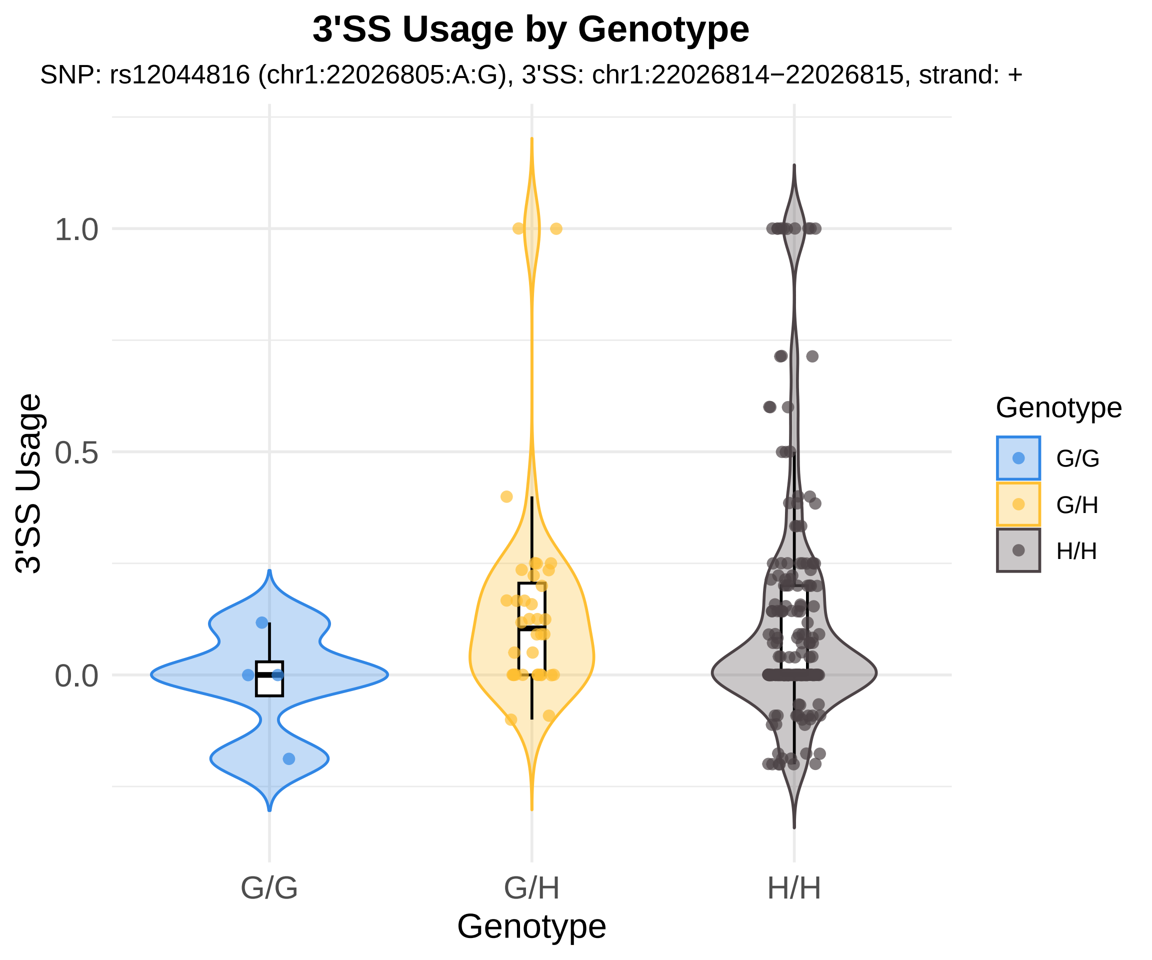
3'SS Usage by Genotype
SNP: rs12044816 (chr1:22026805:A:G), 3'SS: chr1:22026814−22026815, strand: +
3'SS Usage
1.0
0.5
0.0
G/G
G/H
H/H
Genotype
Genotype
G/G
G/H
H/H

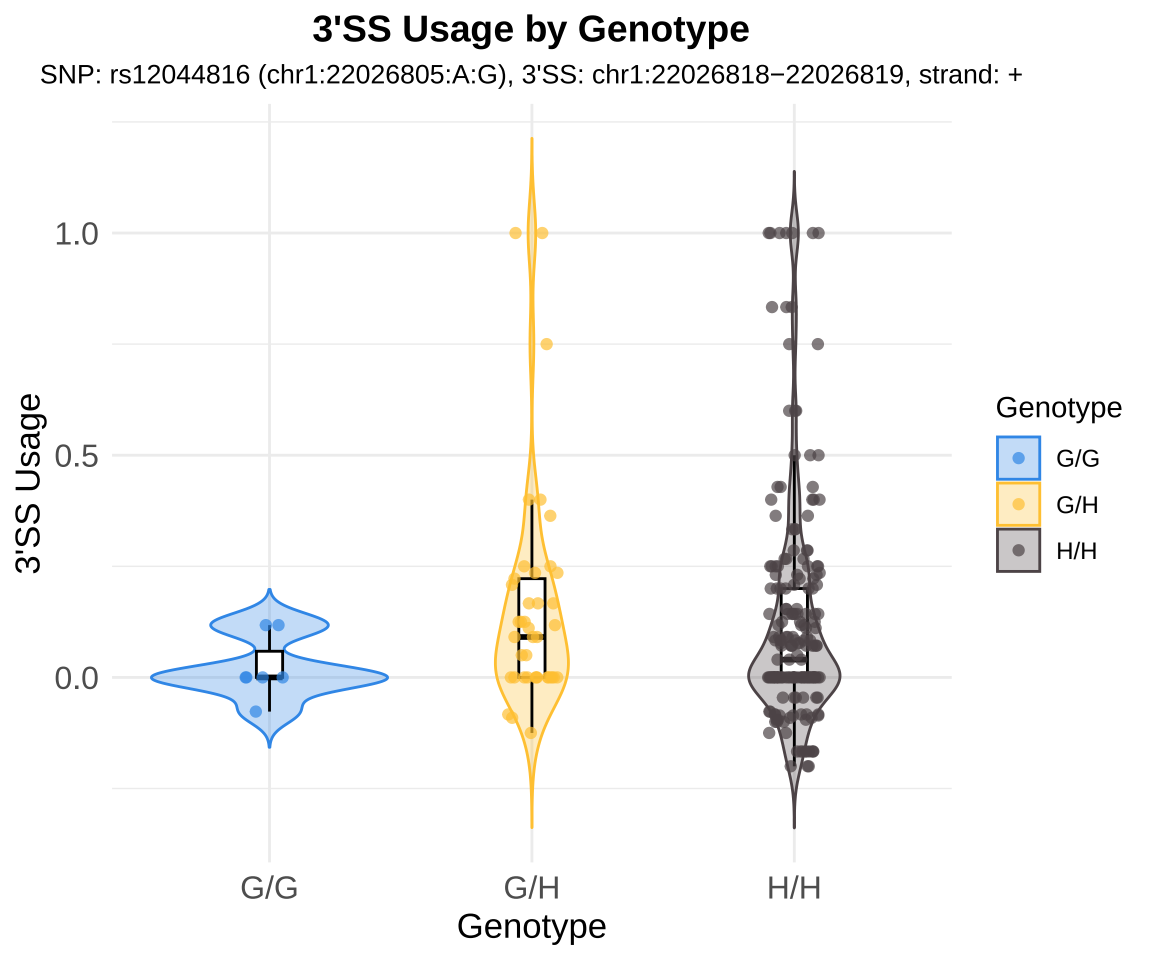
3'SS Usage by Genotype
SNP: rs12044816 (chr1:22026805:A:G), 3'SS: chr1:22026818−22026819, strand: +
3'SS Usage
1.0
0.5
0.0
G/G
G/H
H/H
Genotype
Genotype
G/G
G/H
H/H

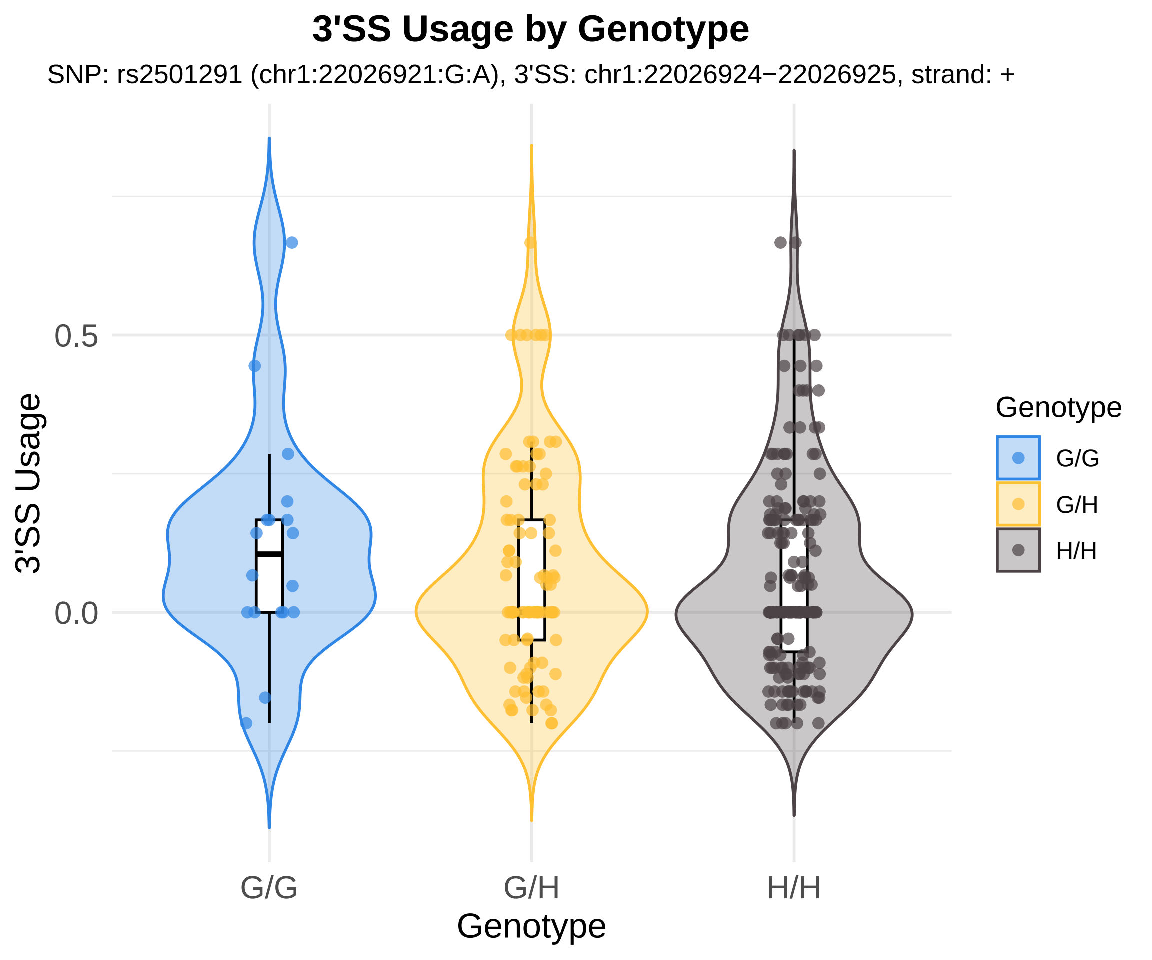
3'SS Usage by Genotype
SNP: rs2501291 (chr1:22026921:G:A), 3'SS: chr1:22026924−22026925, strand: +
3'SS Usage
0.5
0.0
G/G
G/H
H/H
Genotype
Genotype
G/G
G/H
H/H

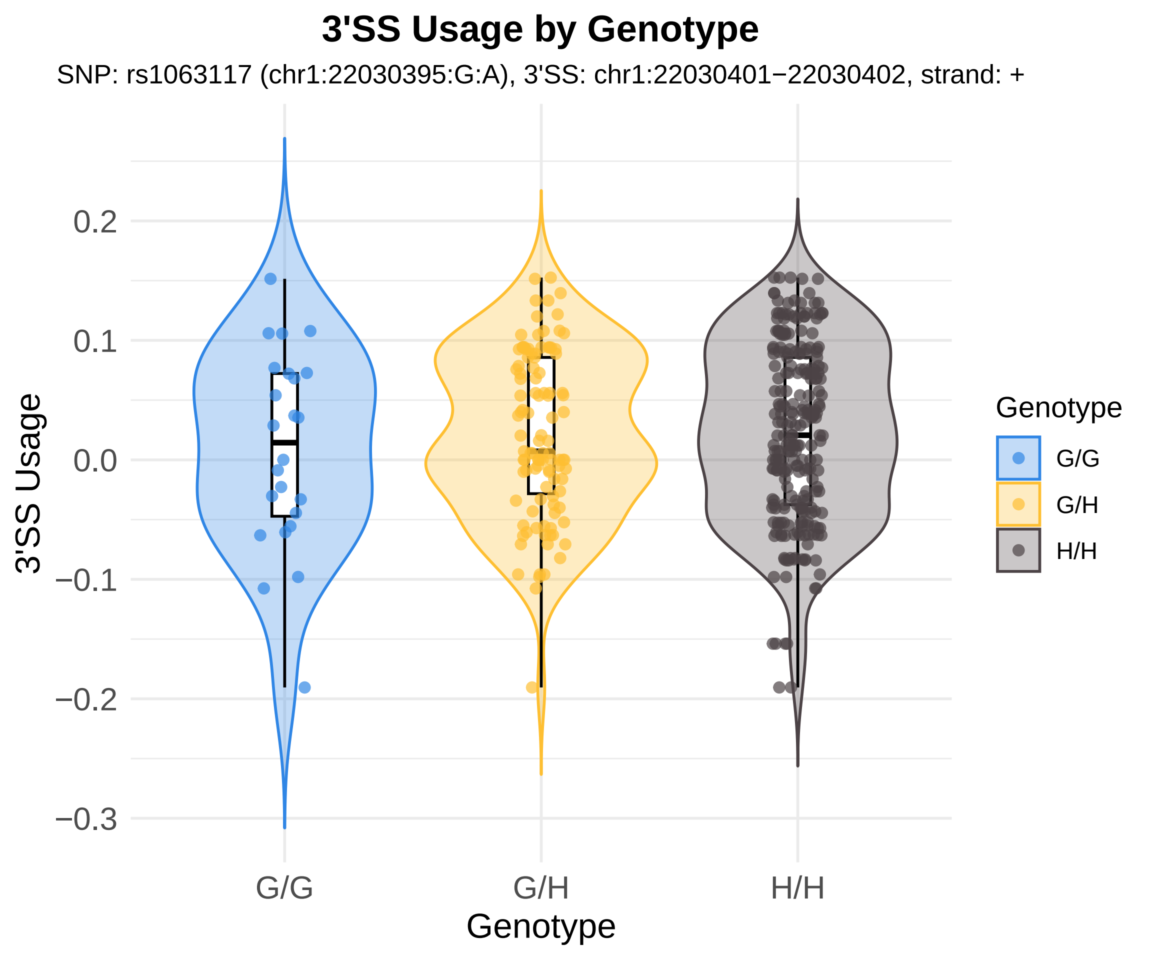
3'SS Usage by Genotype
SNP: rs1063117 (chr1:22030395:G:A), 3'SS: chr1:22030401−22030402, strand: +
0.2
0.1
0.0
−0.1
−0.2
−0.3
3'SS Usage
G/G
G/H
H/H
Genotype
Genotype
G/G
G/H
H/H

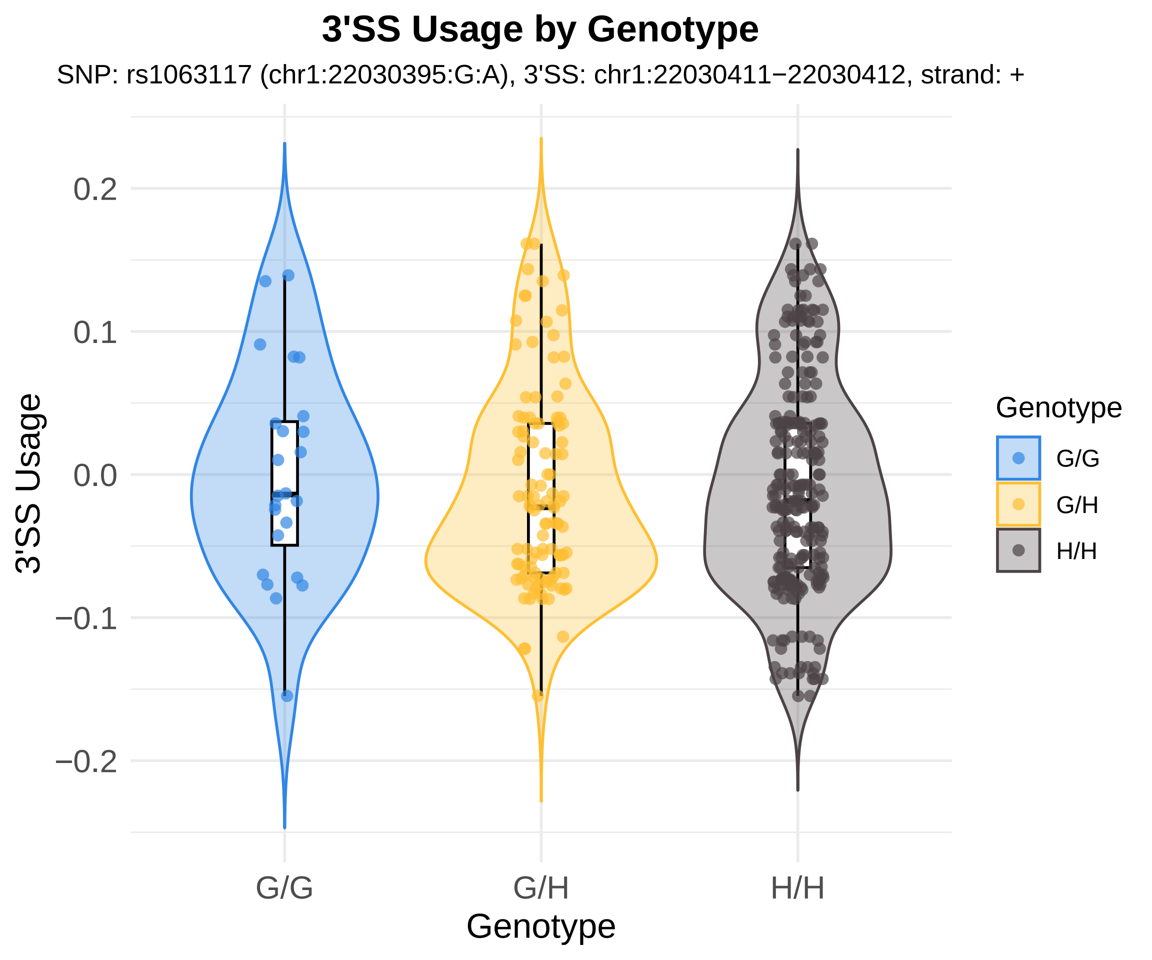
3'SS Usage by Genotype
SNP: rs1063117 (chr1:22030395:G:A), 3'SS: chr1:22030411−22030412, strand: +
3'SS Usage
0.2
0.1
0.0
−0.1
−0.2
G/G
G/H
H/H
Genotype
Genotype
G/G
G/H
H/H

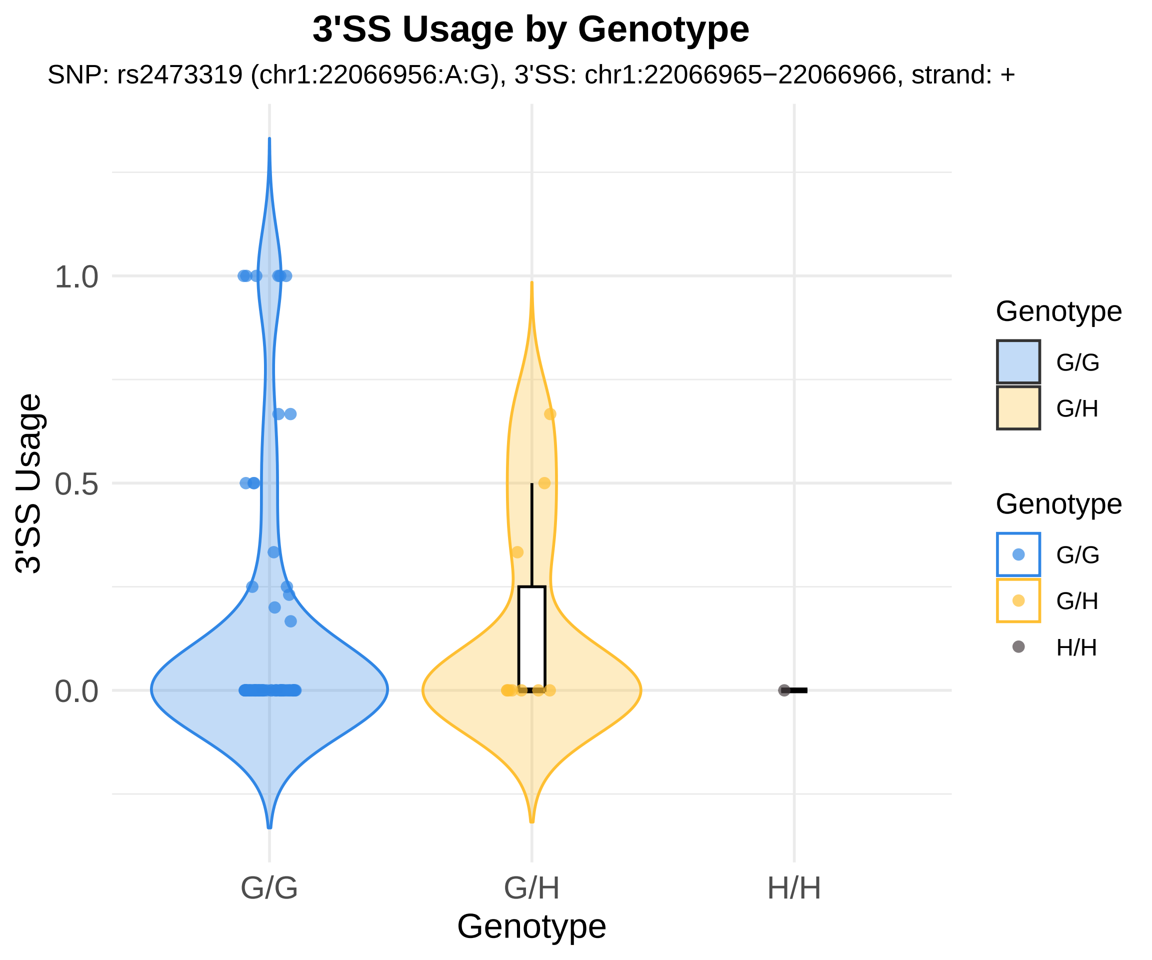

3'SS Usage by Genotype
SNP: rs2473319 (chr1:22066956:A:G), 3'SS: chr1:22066965−22066966, strand: +
3'SS Usage
1.0
0.5
0.0
G/G
G/H
H/H
Genotype
Genotype
G/G
G/H
Genotype
G/G
G/H
H/H

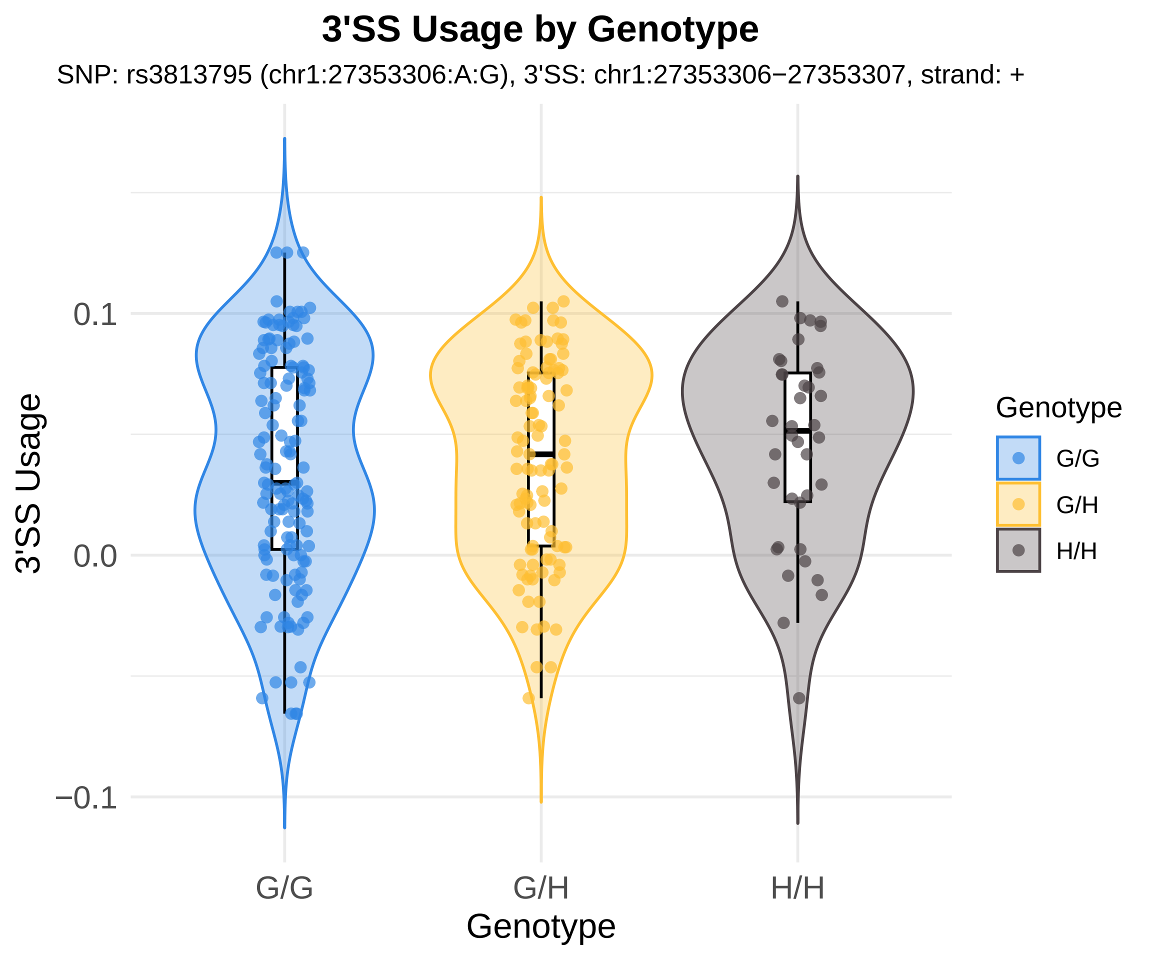
3'SS Usage by Genotype
SNP: rs3813795 (chr1:27353306:A:G), 3'SS: chr1:27353306−27353307, strand: +
3'SS Usage
0.1
0.0
−0.1
G/G
G/H
H/H
Genotype
Genotype
G/G
G/H
H/H

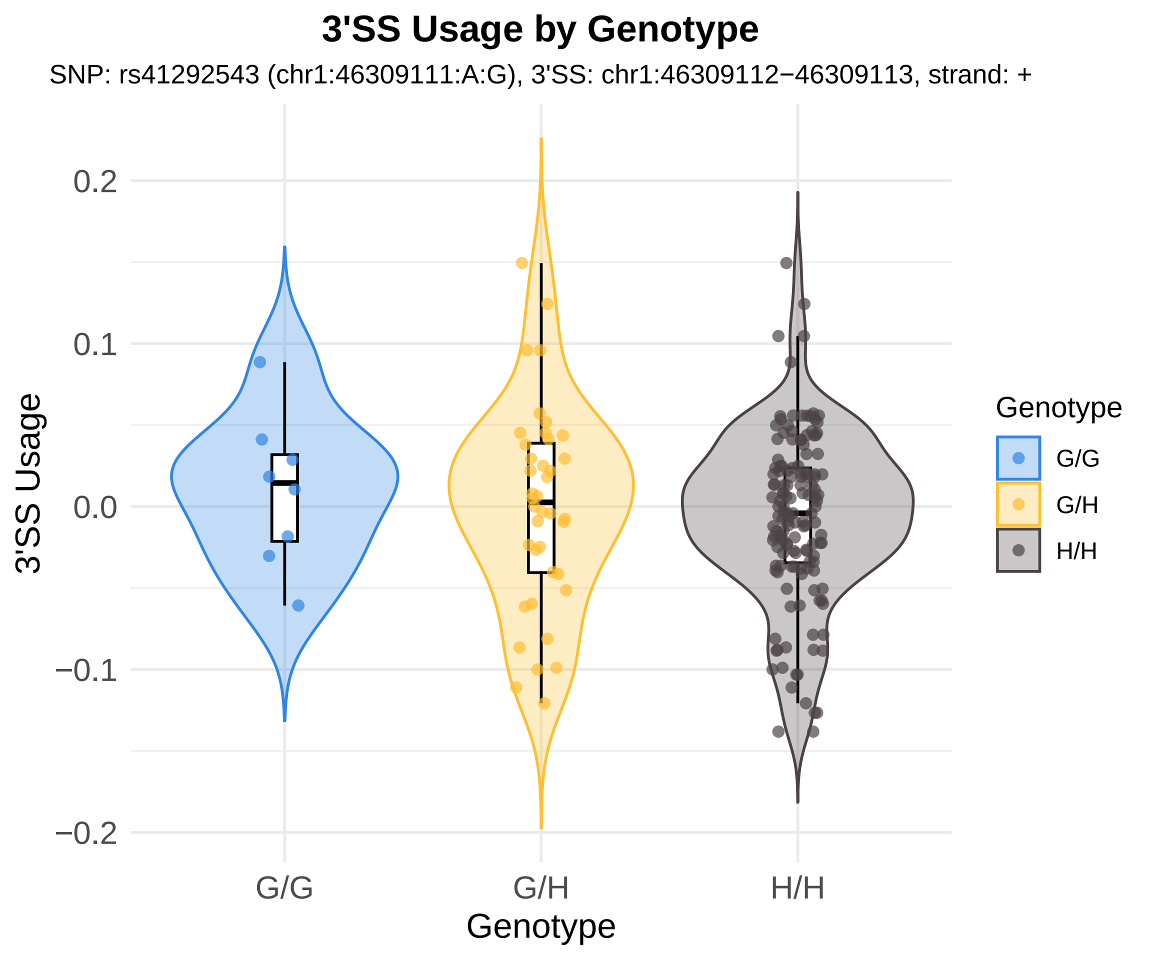

3'SS Usage by Genotype
SNP: rs41292543 (chr1:46309111:A:G), 3'SS: chr1:46309112−46309113, strand: +
3'SS Usage
0.2
0.1
0.0
−0.1
−0.2
G/G
G/H
H/H
Genotype
Genotype
G/G
G/H
H/H

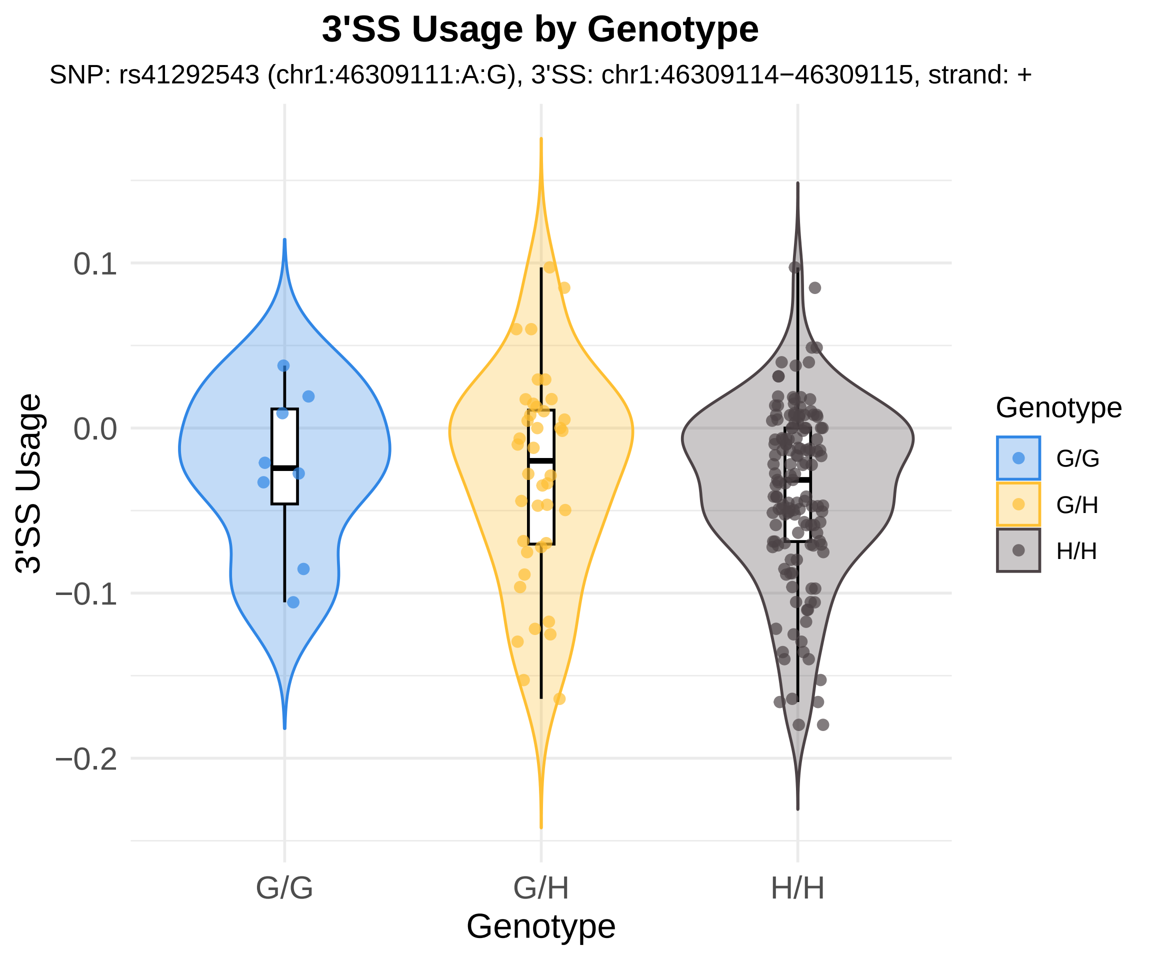

3'SS Usage by Genotype
SNP: rs41292543 (chr1:46309111:A:G), 3'SS: chr1:46309114−46309115, strand: +
3'SS Usage
0.1
0.0
−0.1
−0.2
G/G
G/H
H/H
Genotype
Genotype
G/G
G/H
H/H

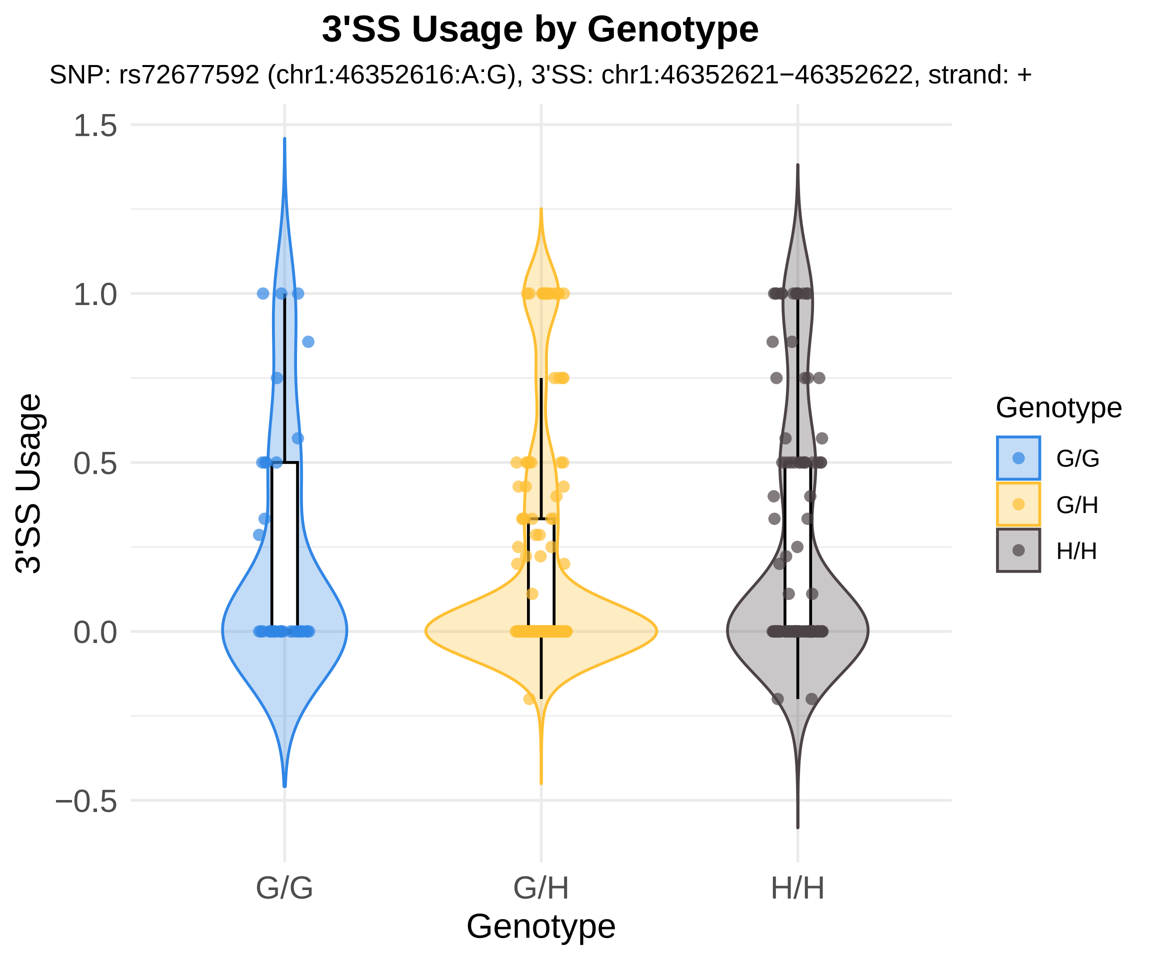
3'SS Usage by Genotype
SNP: rs72677592 (chr1:46352616:A:G), 3'SS: chr1:46352621−46352622, strand: +
1.5
1.0
0.5
0.0
−0.5
3'SS Usage
G/G
G/H
H/H
Genotype
Genotype
G/G
G/H
H/H

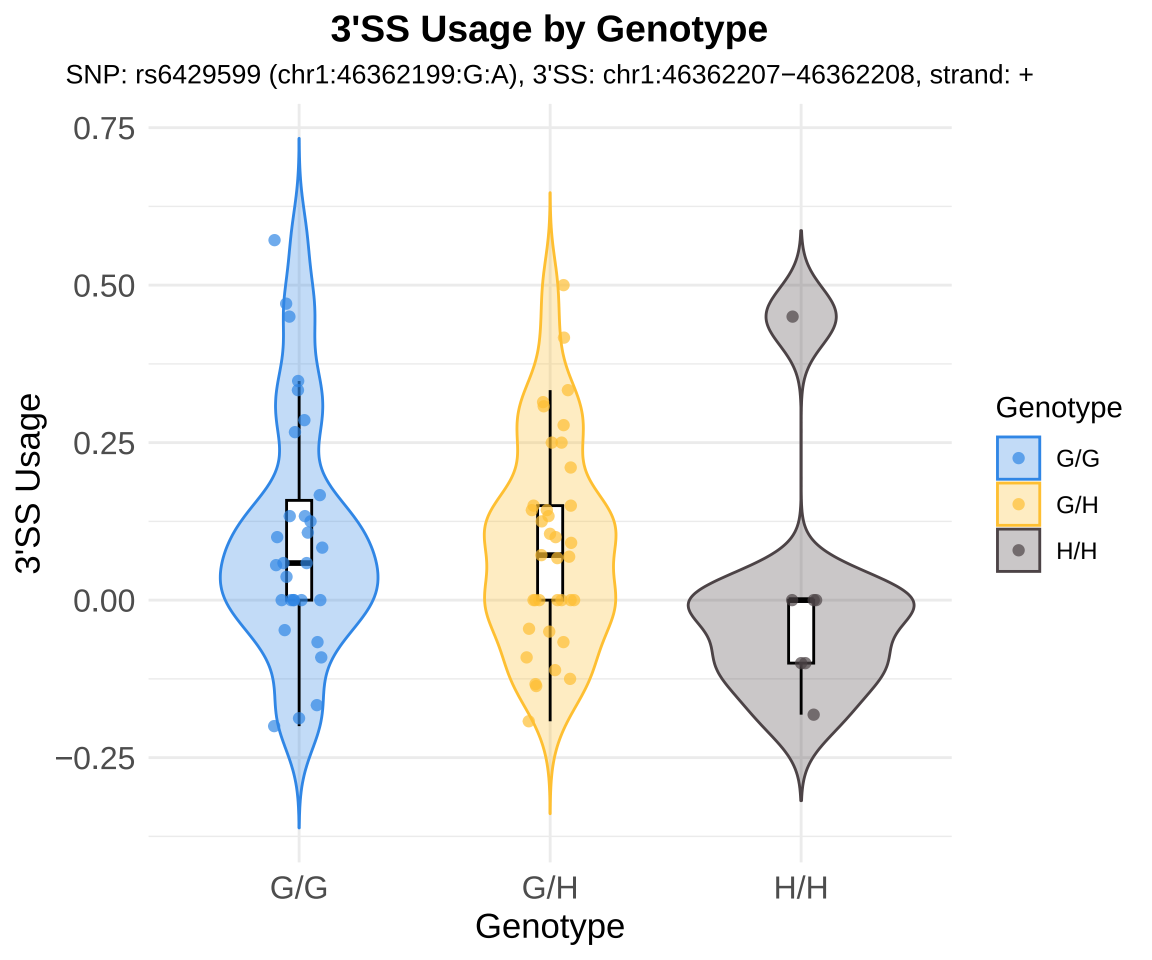

3'SS Usage by Genotype
SNP: rs6429599 (chr1:46362199:G:A), 3'SS: chr1:46362207−46362208, strand: +
3'SS Usage
0.75
0.50
0.25
0.00
−0.25
G/G
G/H
H/H
Genotype
Genotype
G/G
G/H
H/H

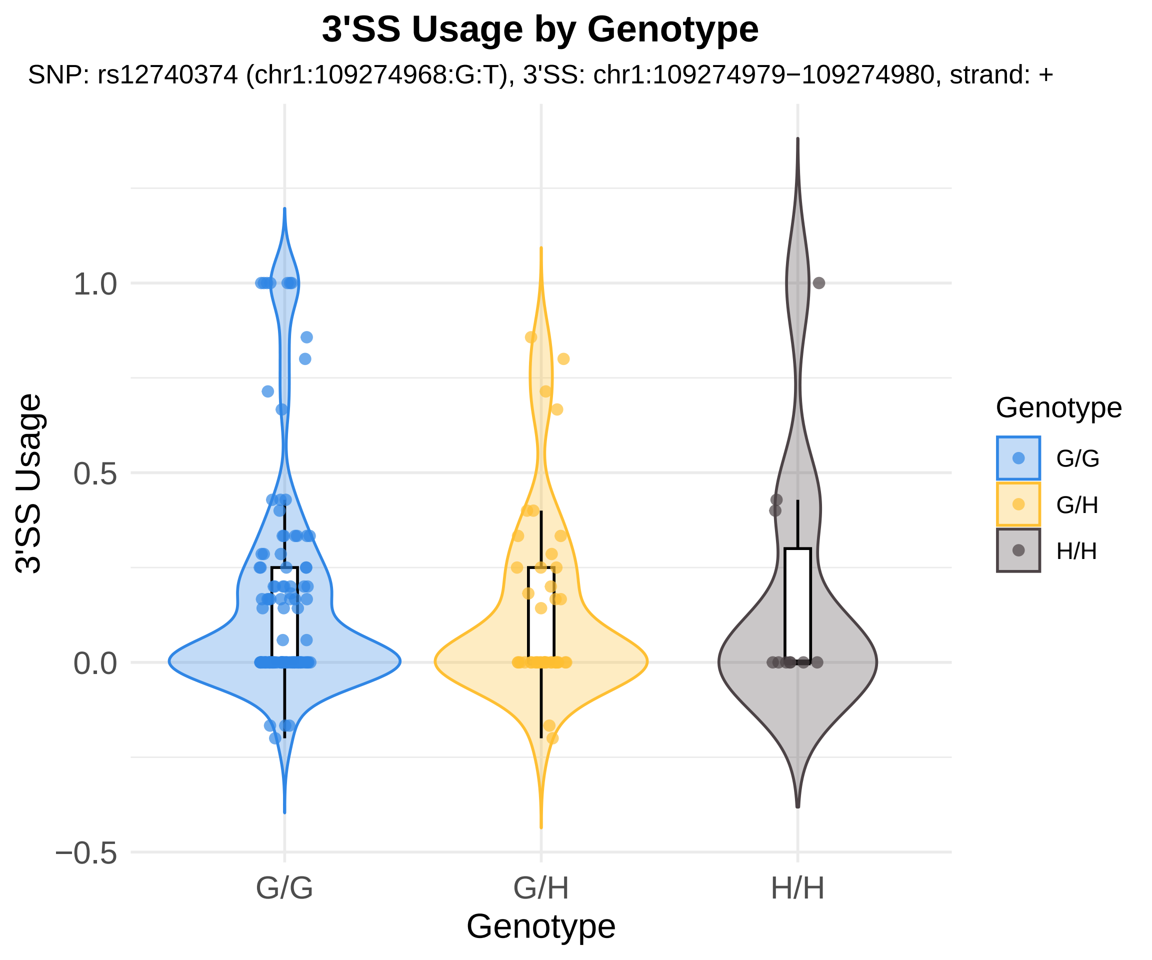
3'SS Usage by Genotype
SNP: rs12740374 (chr1:109274968:G:T), 3'SS: chr1:109274979−109274980, strand: +
3'SS Usage
1.0
0.5
0.0
−0.5
G/G
G/H
H/H
Genotype
Genotype
G/G
G/H
H/H

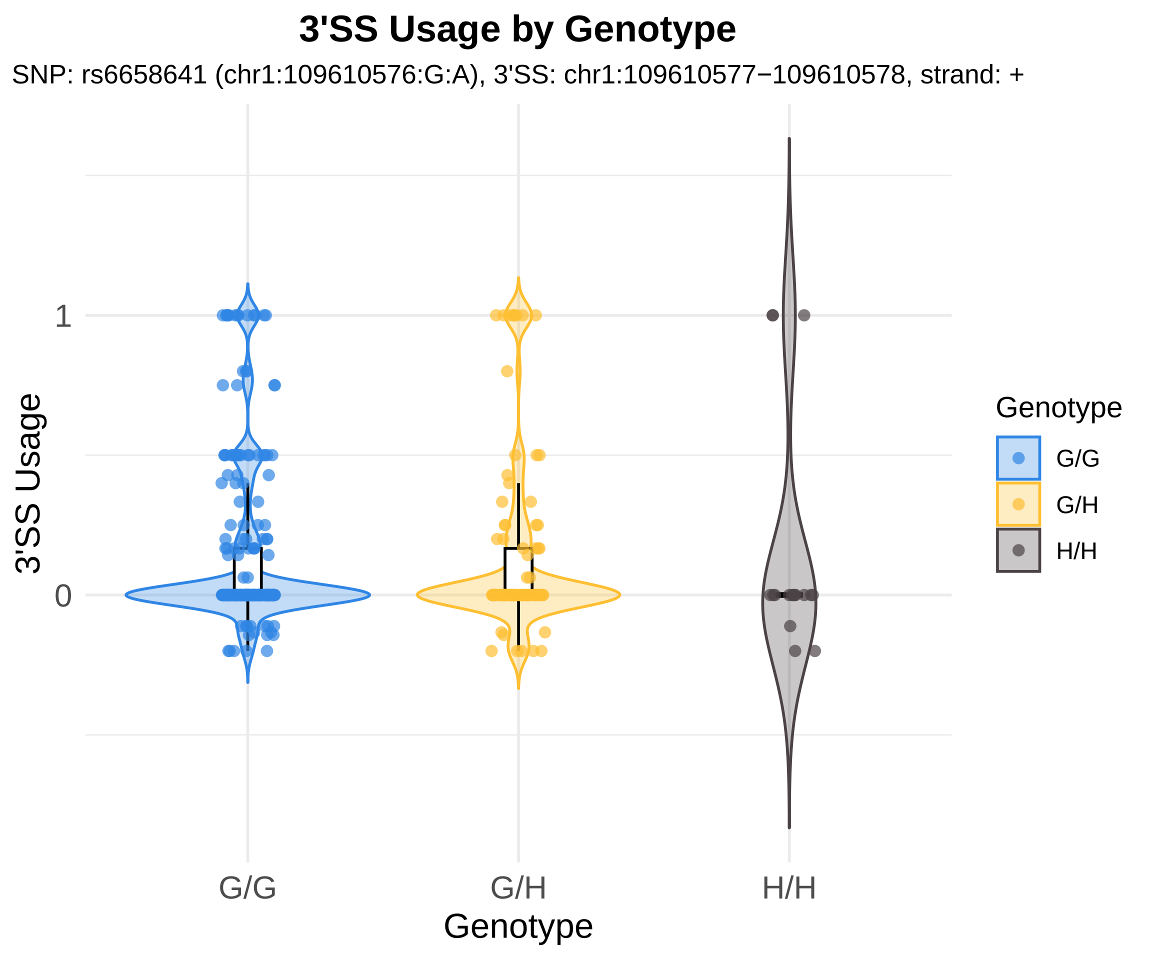
3'SS Usage by Genotype
SNP: rs6658641 (chr1:109610576:G:A), 3'SS: chr1:109610577−109610578, strand: +
3'SS Usage
1
0
G/G
G/H
H/H
Genotype
Genotype
G/G
G/H
H/H

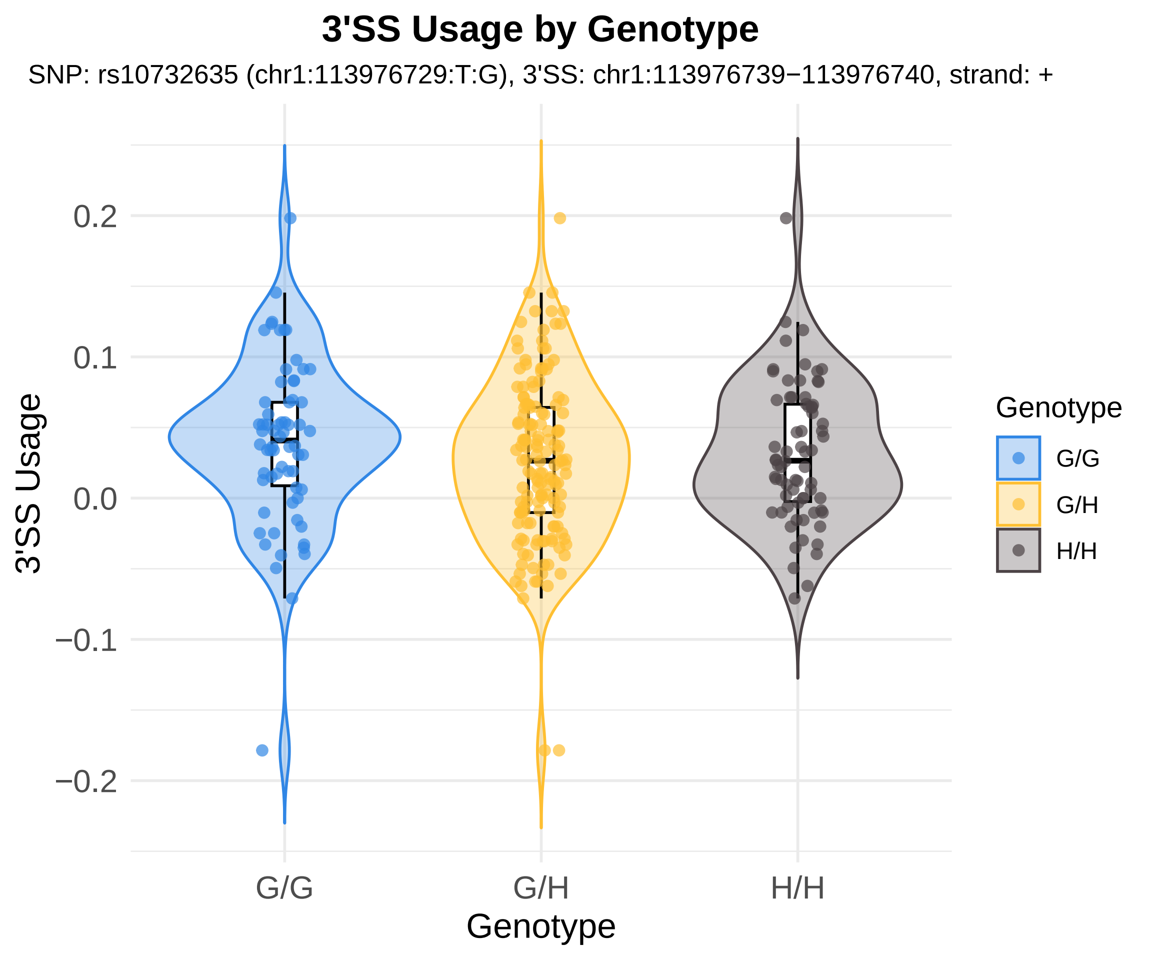
3'SS Usage by Genotype
SNP: rs10732635 (chr1:113976729:T:G), 3'SS: chr1:113976739−113976740, strand: +
3'SS Usage
0.2
0.1
0.0
−0.1
−0.2
G/G
G/H
H/H
Genotype
Genotype
G/G
G/H
H/H

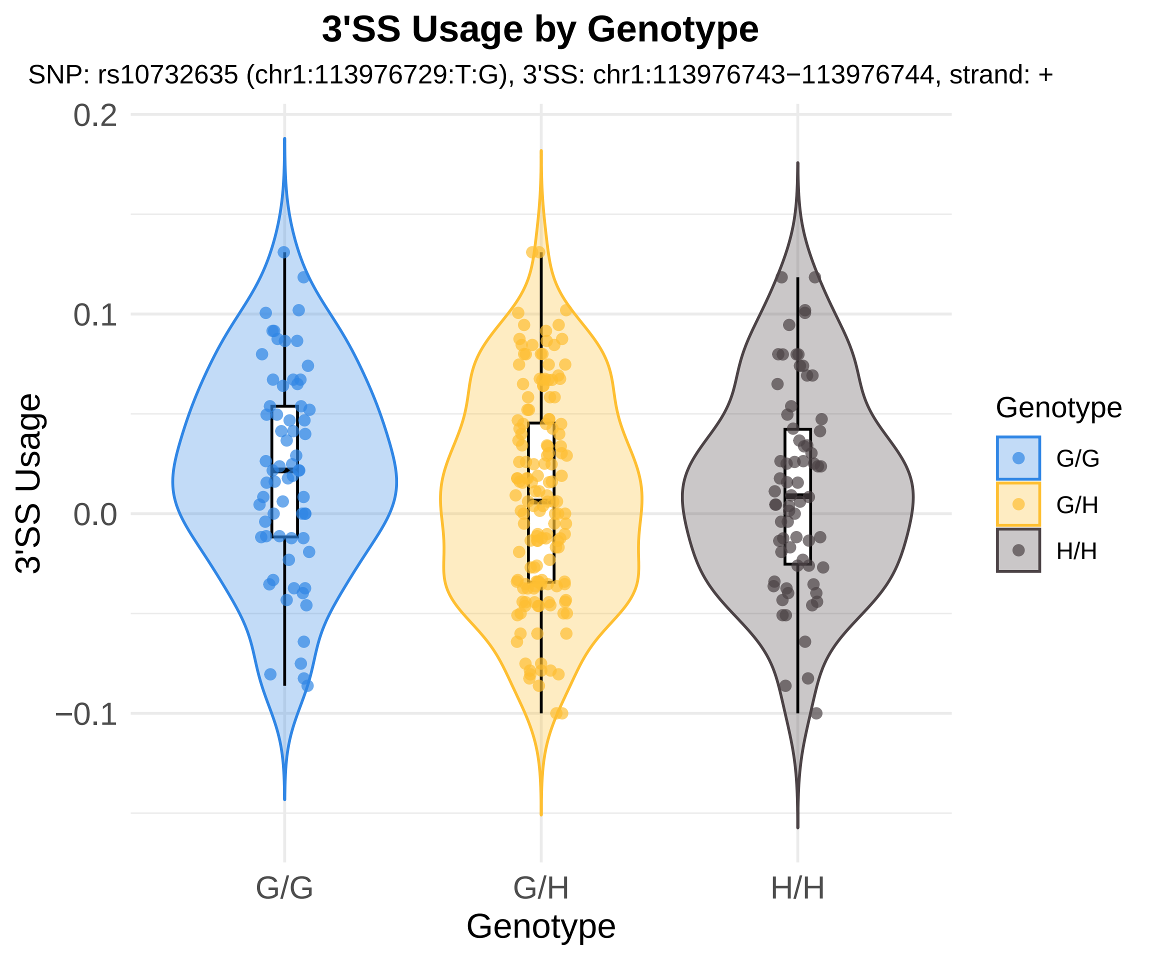
3'SS Usage by Genotype
SNP: rs10732635 (chr1:113976729:T:G), 3'SS: chr1:113976743−113976744, strand: +
0.2
0.1
0.0
−0.1
3'SS Usage
G/G
G/H
H/H
Genotype
Genotype
G/G
G/H
H/H

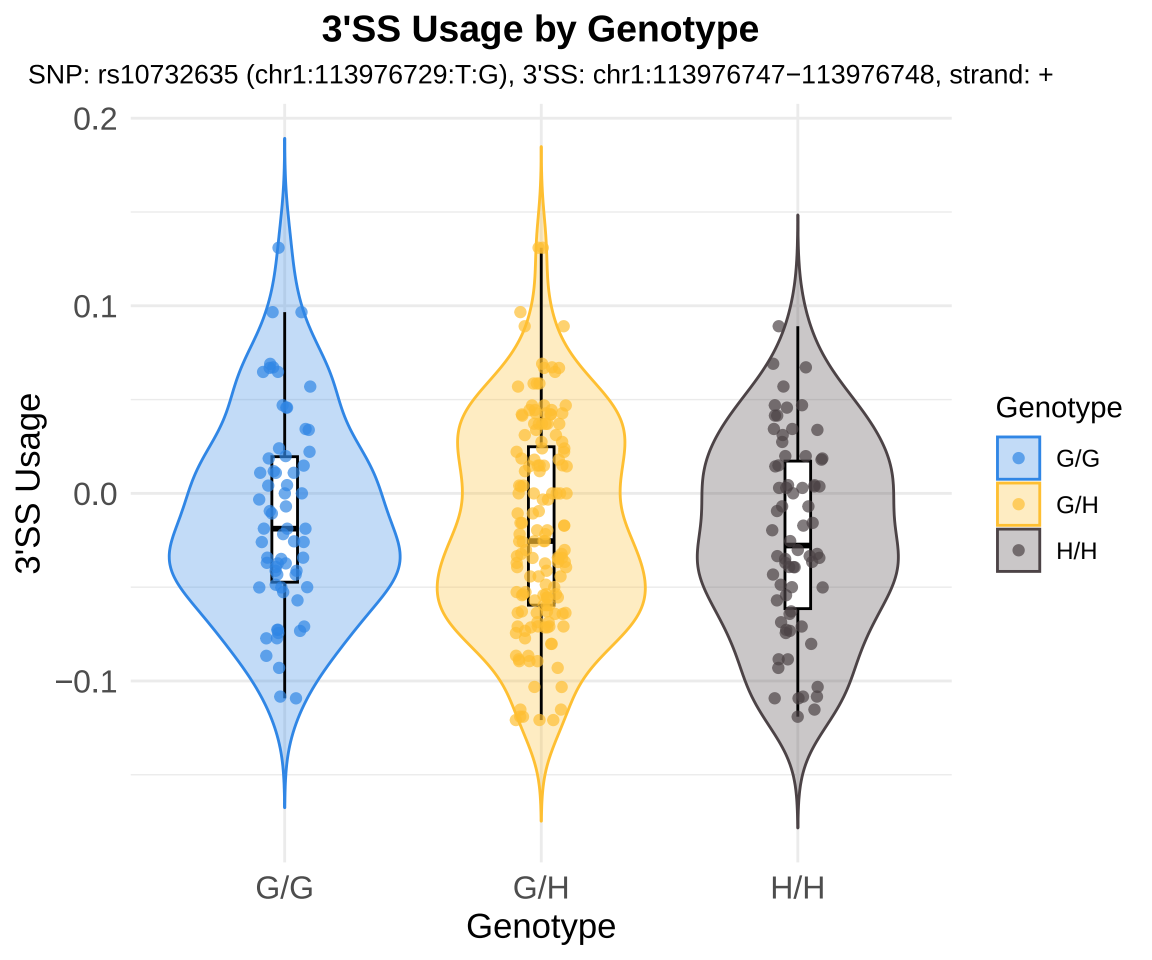

3'SS Usage by Genotype
SNP: rs10732635 (chr1:113976729:T:G), 3'SS: chr1:113976747−113976748, strand: +
0.2
0.1
0.0
−0.1
3'SS Usage
G/G
G/H
H/H
Genotype
Genotype
G/G
G/H
H/H

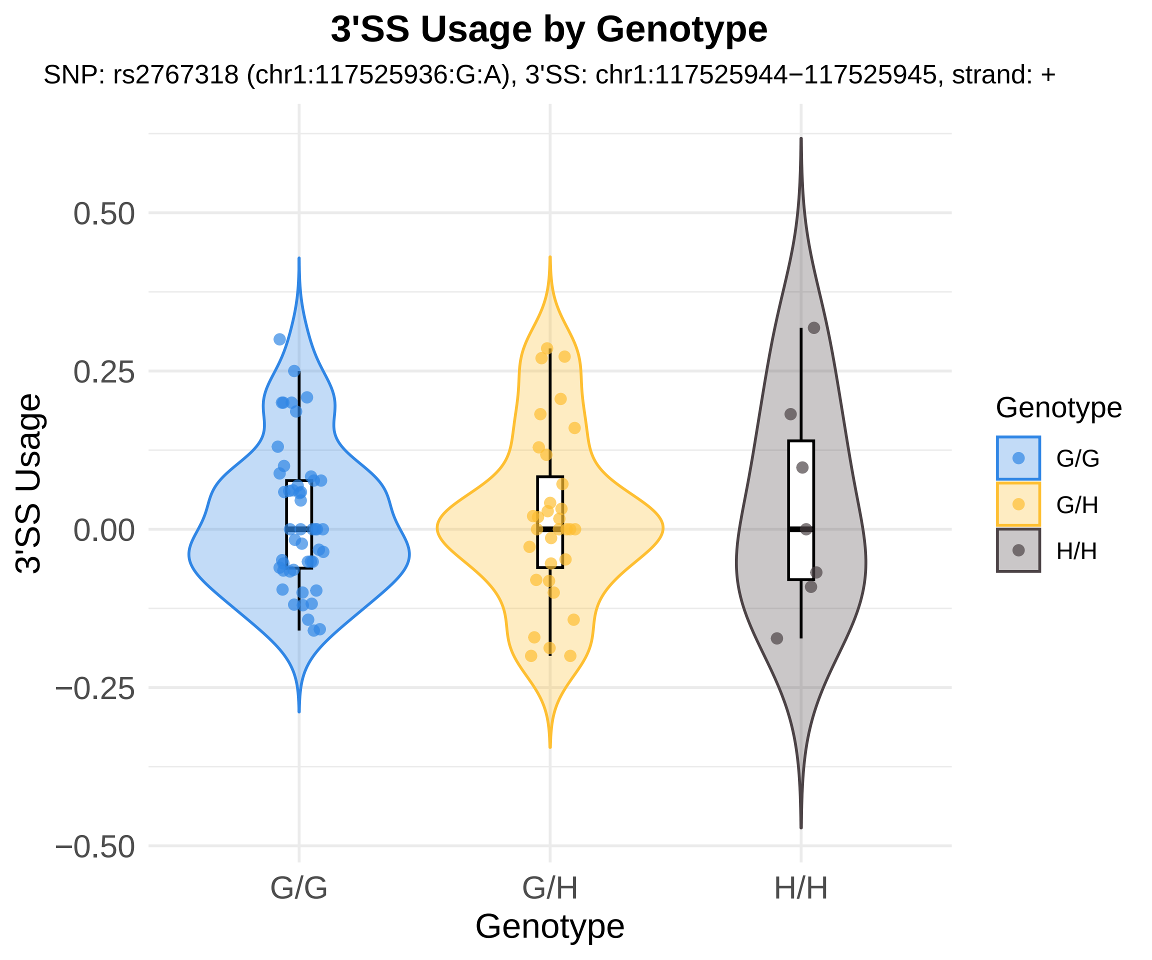
3'SS Usage by Genotype
SNP: rs2767318 (chr1:117525936:G:A), 3'SS: chr1:117525944−117525945, strand: +
3'SS Usage
0.50
0.25
0.00
−0.25
−0.50
G/G
G/H
H/H
Genotype
Genotype
G/G
G/H
H/H

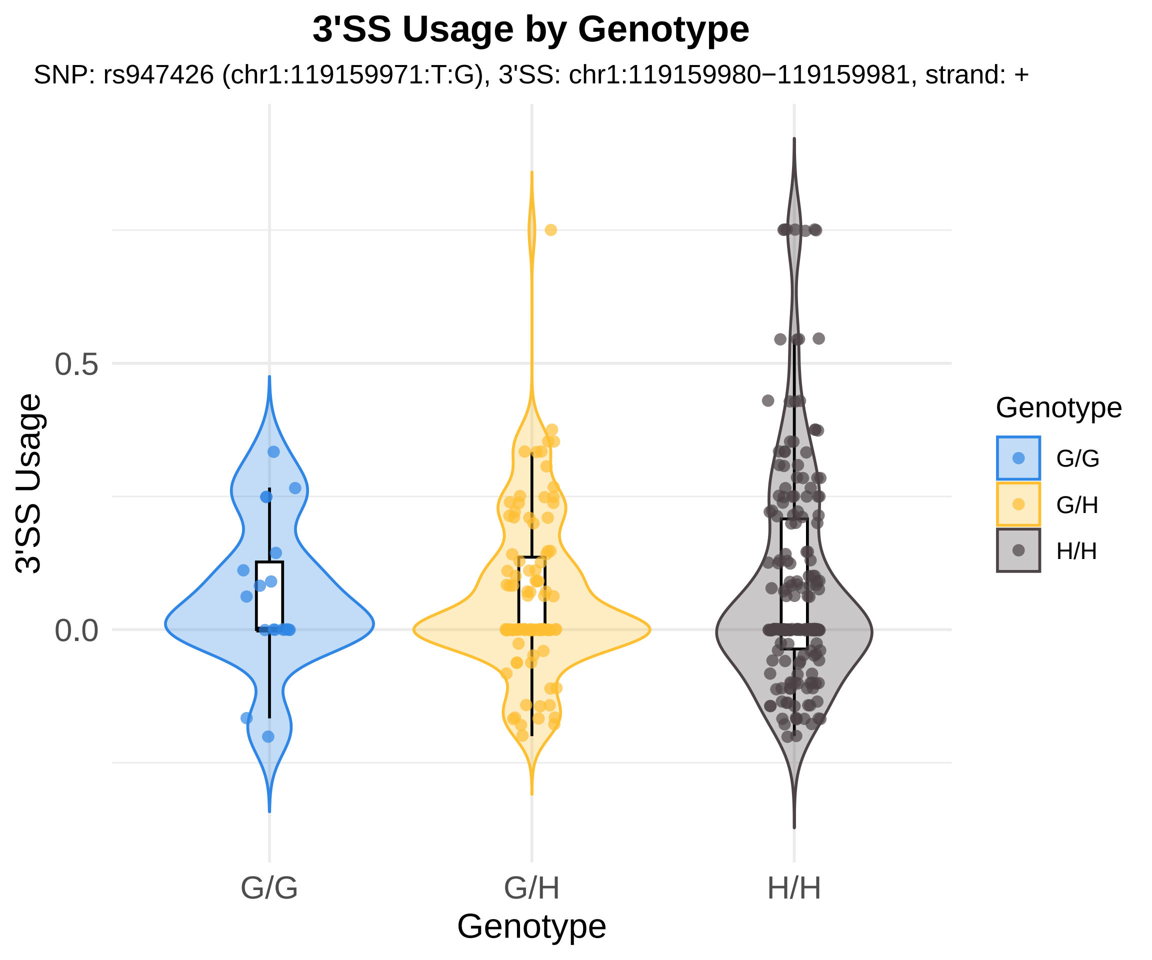
3'SS Usage by Genotype
SNP: rs947426 (chr1:119159971:T:G), 3'SS: chr1:119159980−119159981, strand: +
3'SS Usage
0.5
0.0
G/G
G/H
H/H
Genotype
Genotype
G/G
G/H
H/H

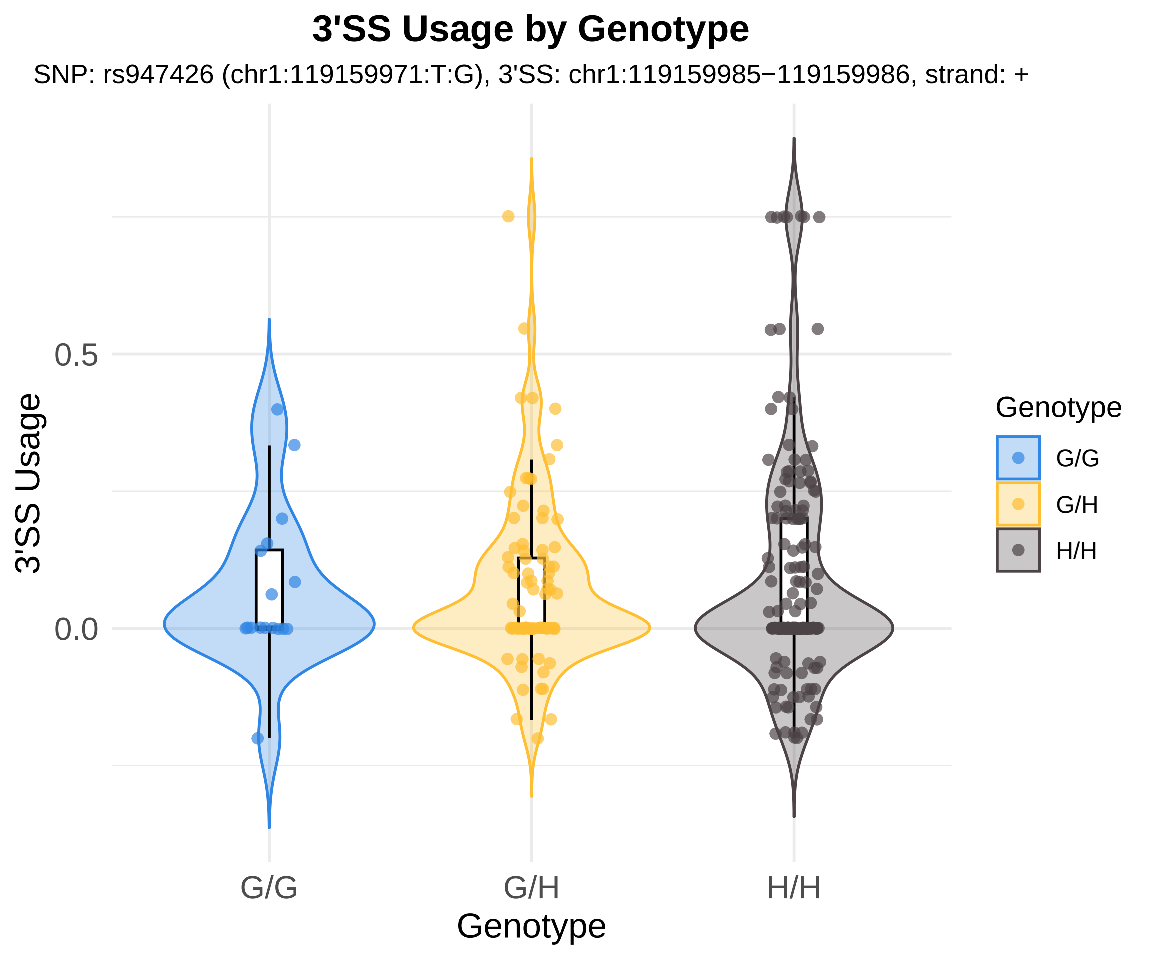

3'SS Usage by Genotype
SNP: rs947426 (chr1:119159971:T:G), 3'SS: chr1:119159985−119159986, strand: +
3'SS Usage
0.5
0.0
G/G
G/H
H/H
Genotype
Genotype
G/G
G/H
H/H

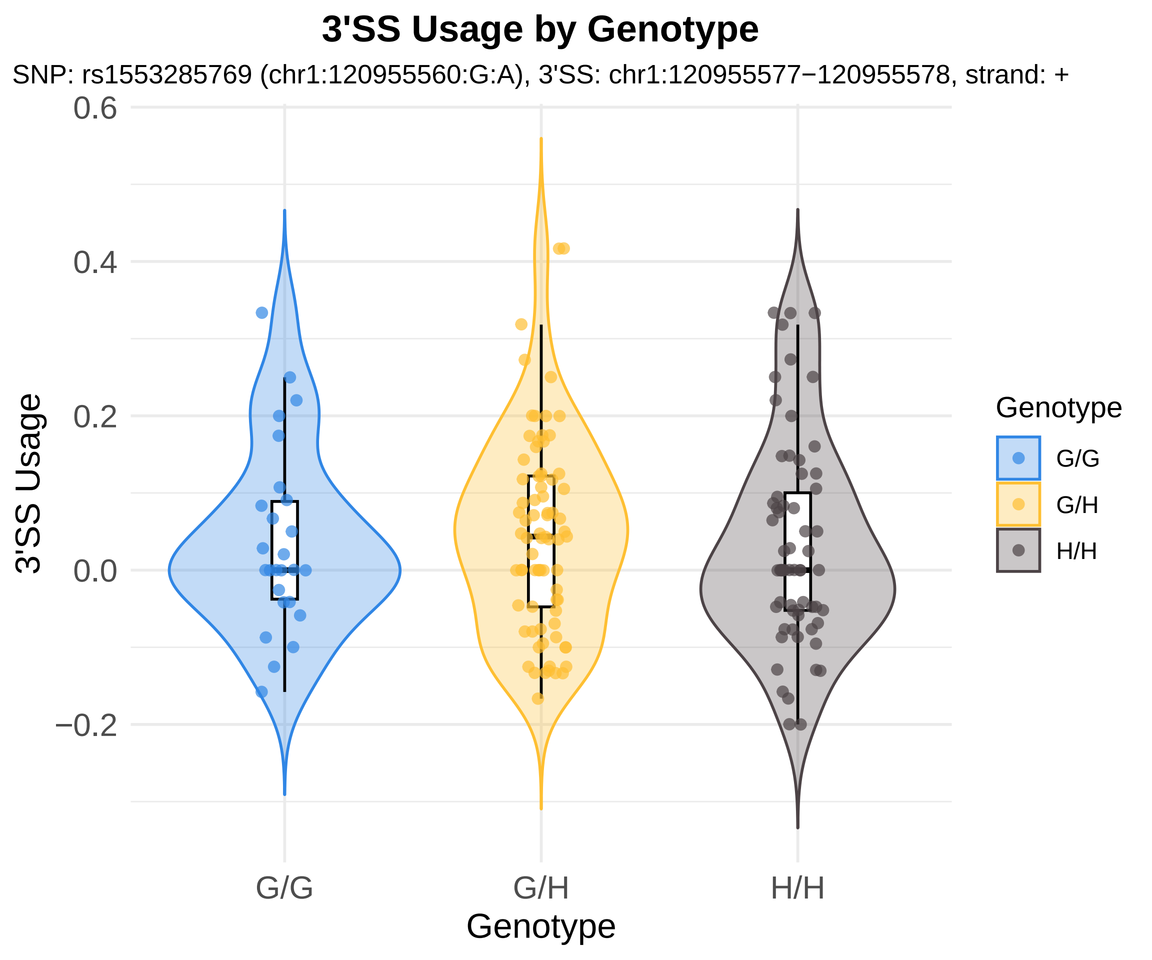
3'SS Usage by Genotype
SNP: rs1553285769 (chr1:120955560:G:A), 3'SS: chr1:120955577−120955578, strand: +
3'SS Usage
0.6
0.4
0.2
0.0
−0.2
G/G
G/H
H/H
Genotype
Genotype
G/G
G/H
H/H

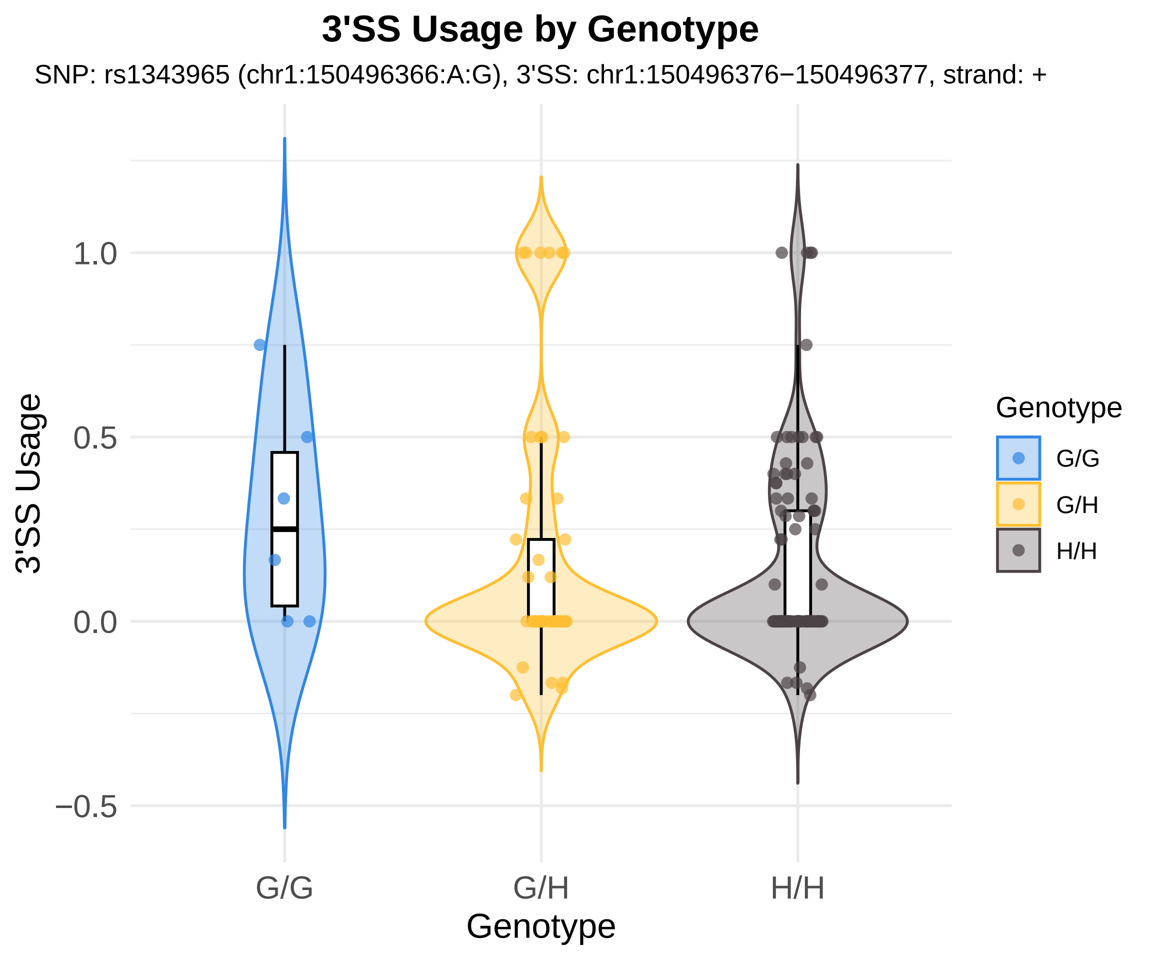
3'SS Usage by Genotype
SNP: rs1343965 (chr1:150496366:A:G), 3'SS: chr1:150496376−150496377, strand: +
3'SS Usage
1.0
0.5
0.0
−0.5
G/G
G/H
H/H
Genotype
Genotype
G/G
G/H
H/H

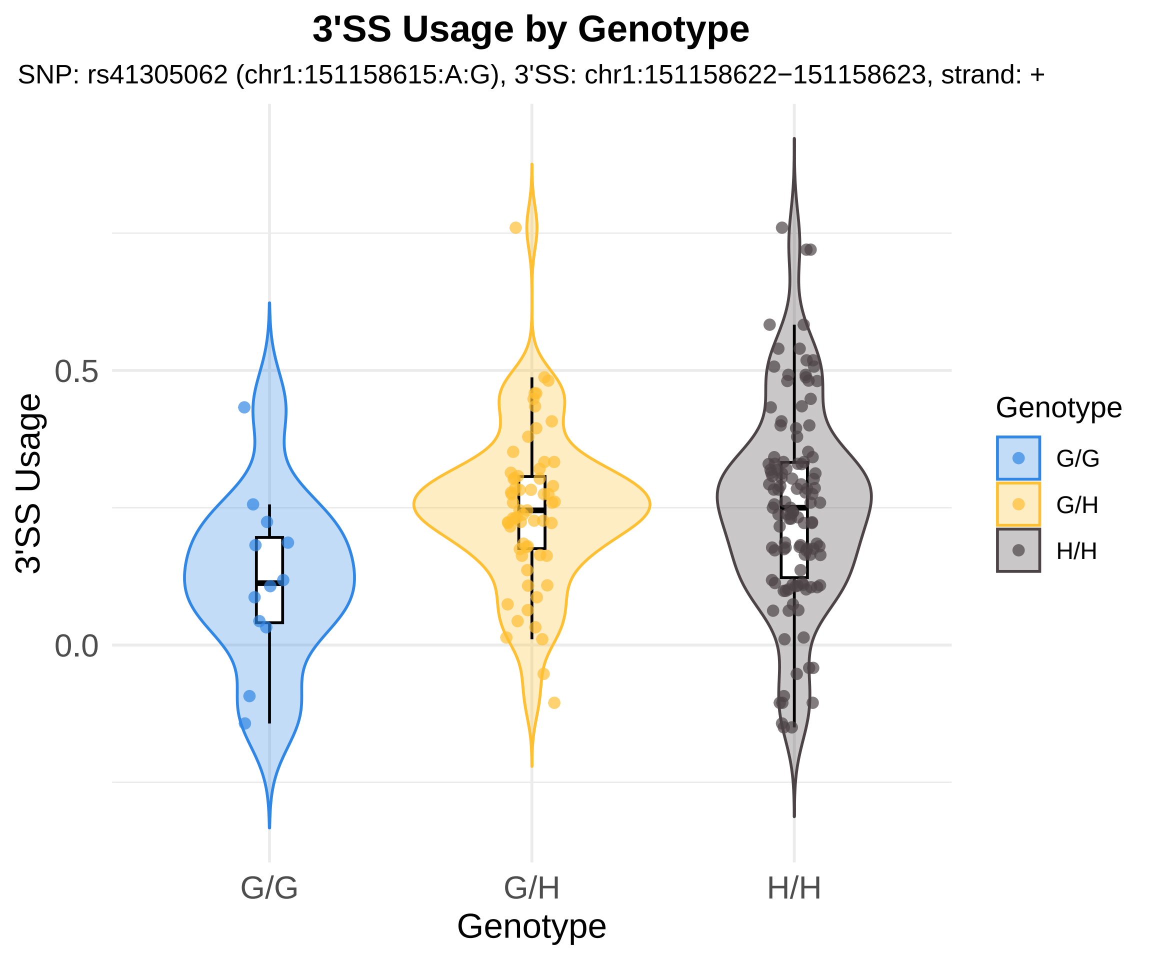
3'SS Usage by Genotype
SNP: rs41305062 (chr1:151158615:A:G), 3'SS: chr1:151158622−151158623, strand: +
3'SS Usage
0.5
0.0
G/G
G/H
H/H
Genotype
Genotype
G/G
G/H
H/H

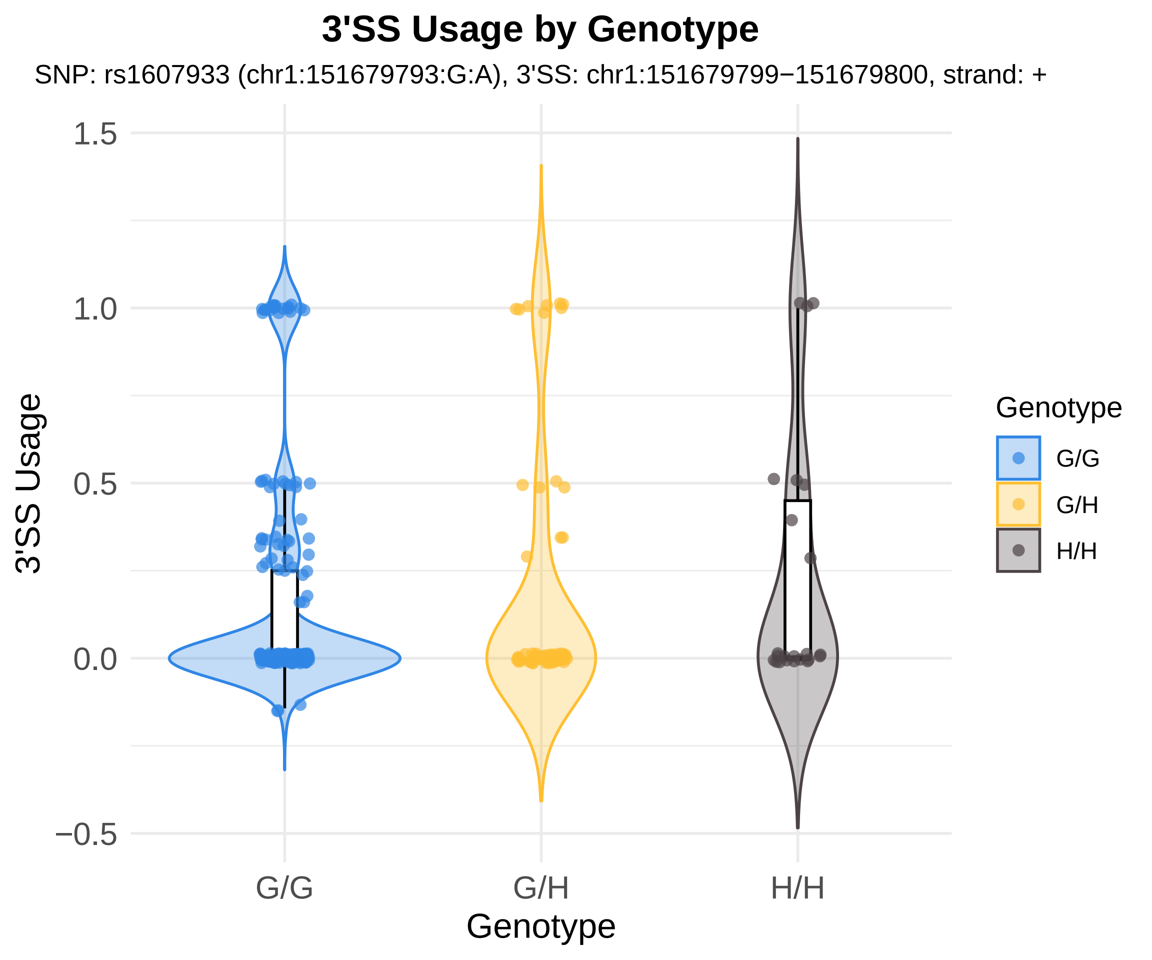
3'SS Usage by Genotype
SNP: rs1607933 (chr1:151679793:G:A), 3'SS: chr1:151679799−151679800, strand: +
1.5
1.0
0.5
0.0
−0.5
3'SS Usage
G/G
G/H
H/H
Genotype
Genotype
G/G
G/H
H/H

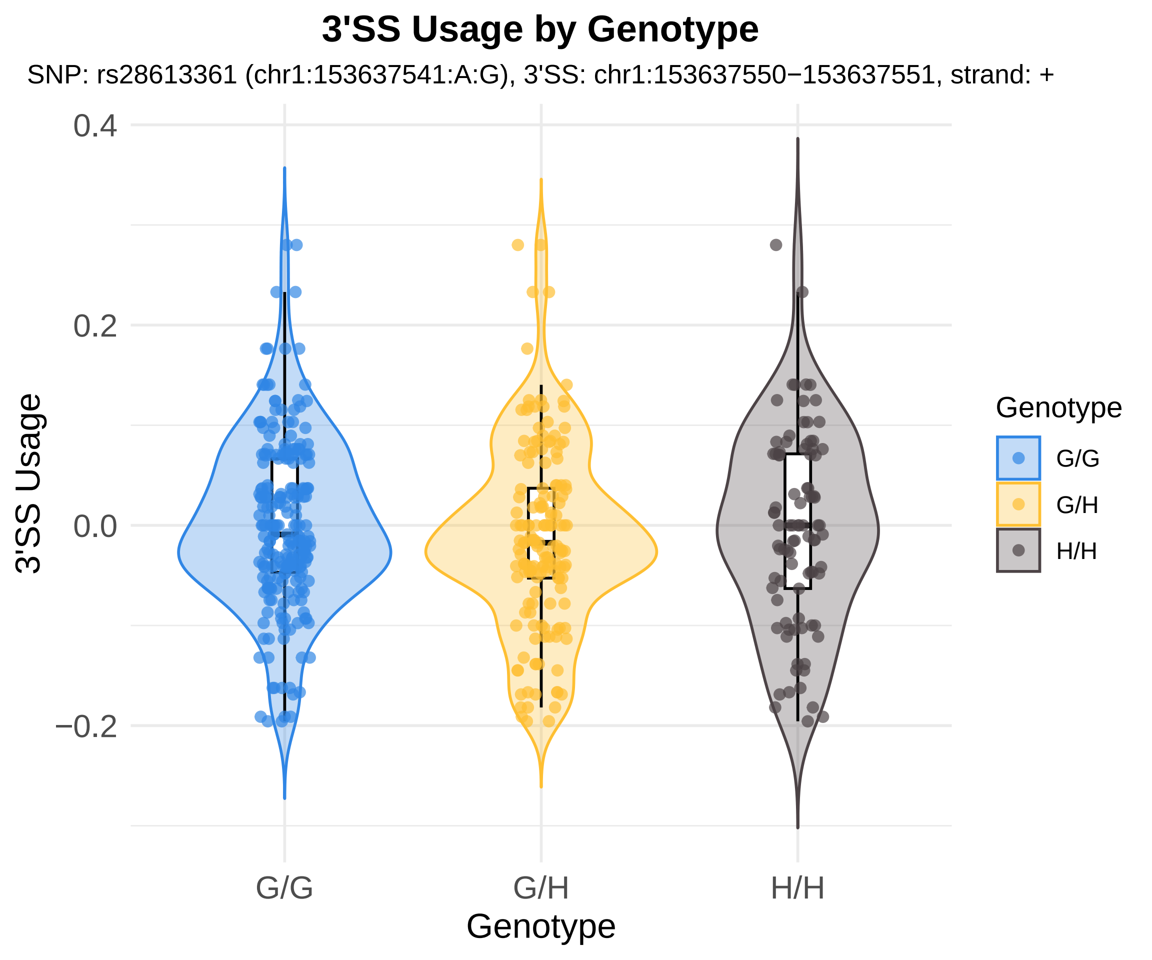

3'SS Usage by Genotype
SNP: rs28613361 (chr1:153637541:A:G), 3'SS: chr1:153637550−153637551, strand: +
3'SS Usage
0.4
0.2
0.0
−0.2
G/G
G/H
H/H
Genotype
Genotype
G/G
G/H
H/H

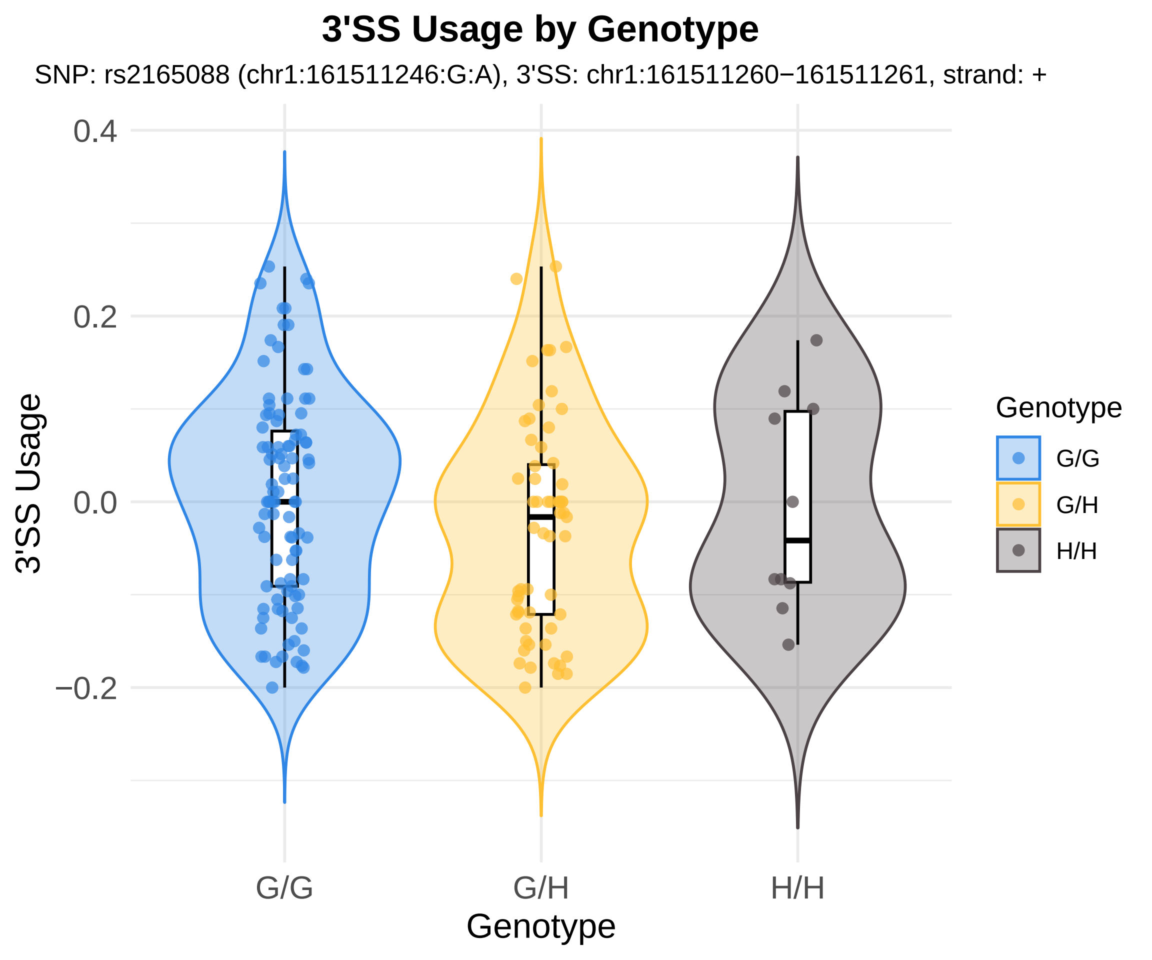

3'SS Usage by Genotype
SNP: rs2165088 (chr1:161511246:G:A), 3'SS: chr1:161511260−161511261, strand: +
3'SS Usage
0.4
0.2
0.0
−0.2
G/G
G/H
H/H
Genotype
Genotype
G/G
G/H
H/H

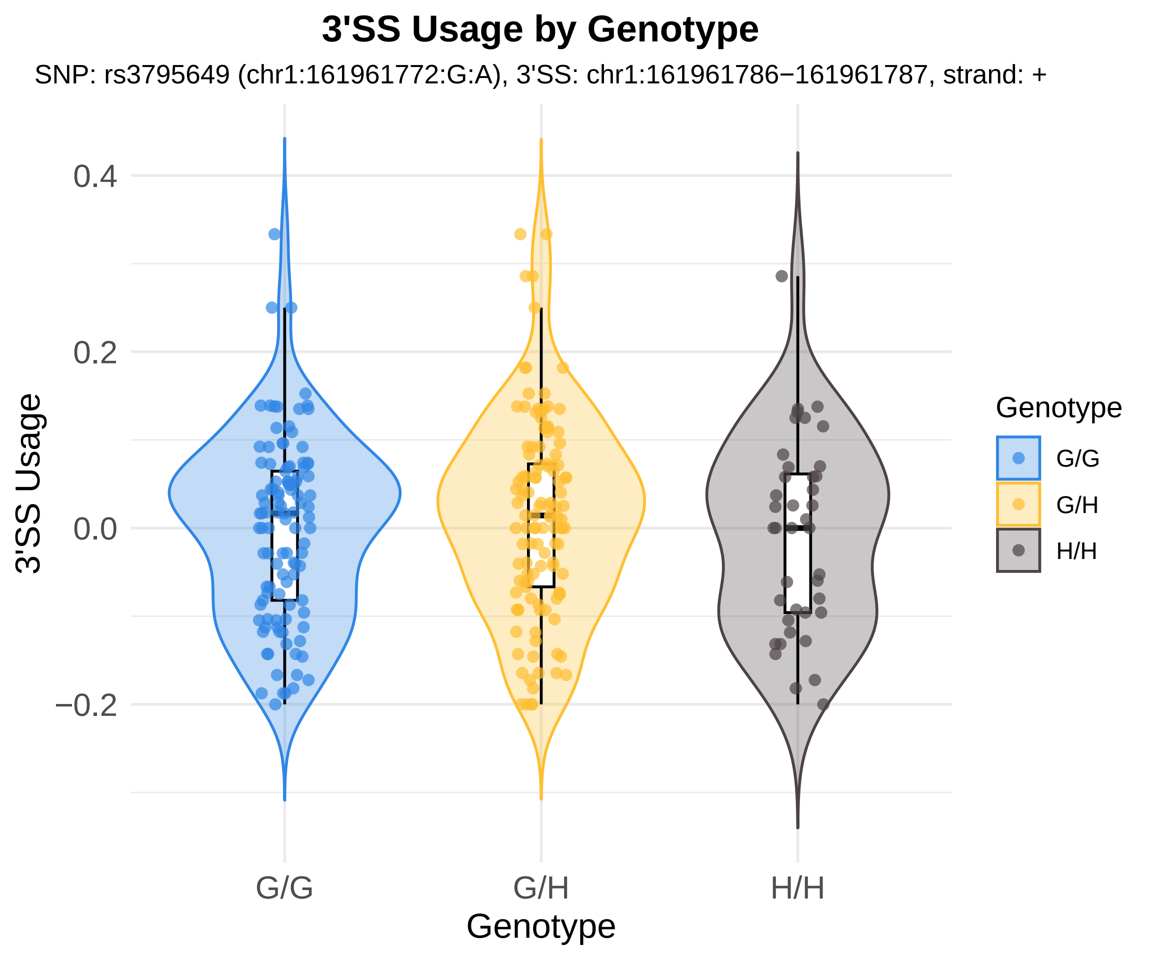

3'SS Usage by Genotype
SNP: rs3795649 (chr1:161961772:G:A), 3'SS: chr1:161961786−161961787, strand: +
3'SS Usage
0.4
0.2
0.0
−0.2
G/G
G/H
H/H
Genotype
Genotype
G/G
G/H
H/H

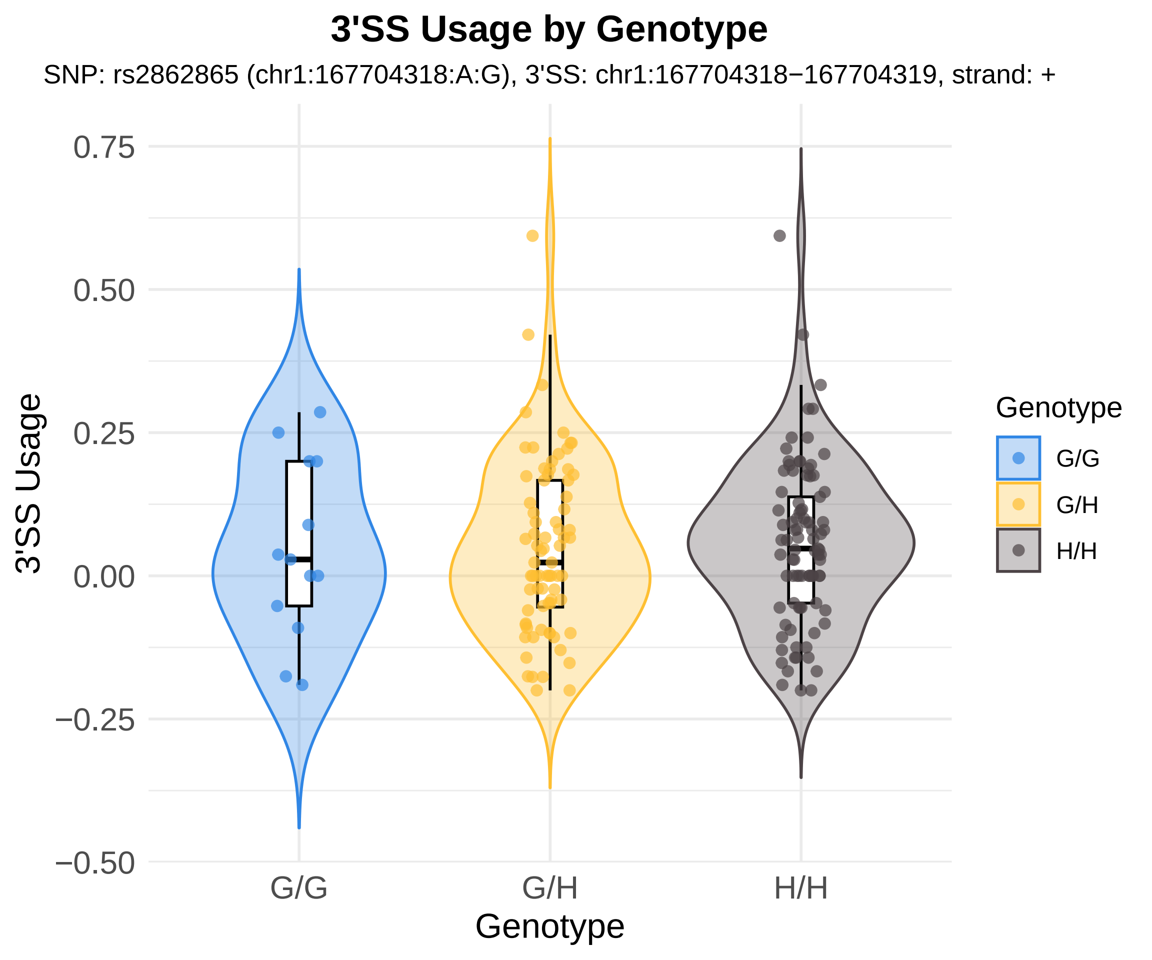

3'SS Usage by Genotype
SNP: rs2862865 (chr1:167704318:A:G), 3'SS: chr1:167704318−167704319, strand: +
3'SS Usage
0.75
0.50
0.25
0.00
−0.25
−0.50
G/G
G/H
H/H
Genotype
Genotype
G/G
G/H
H/H

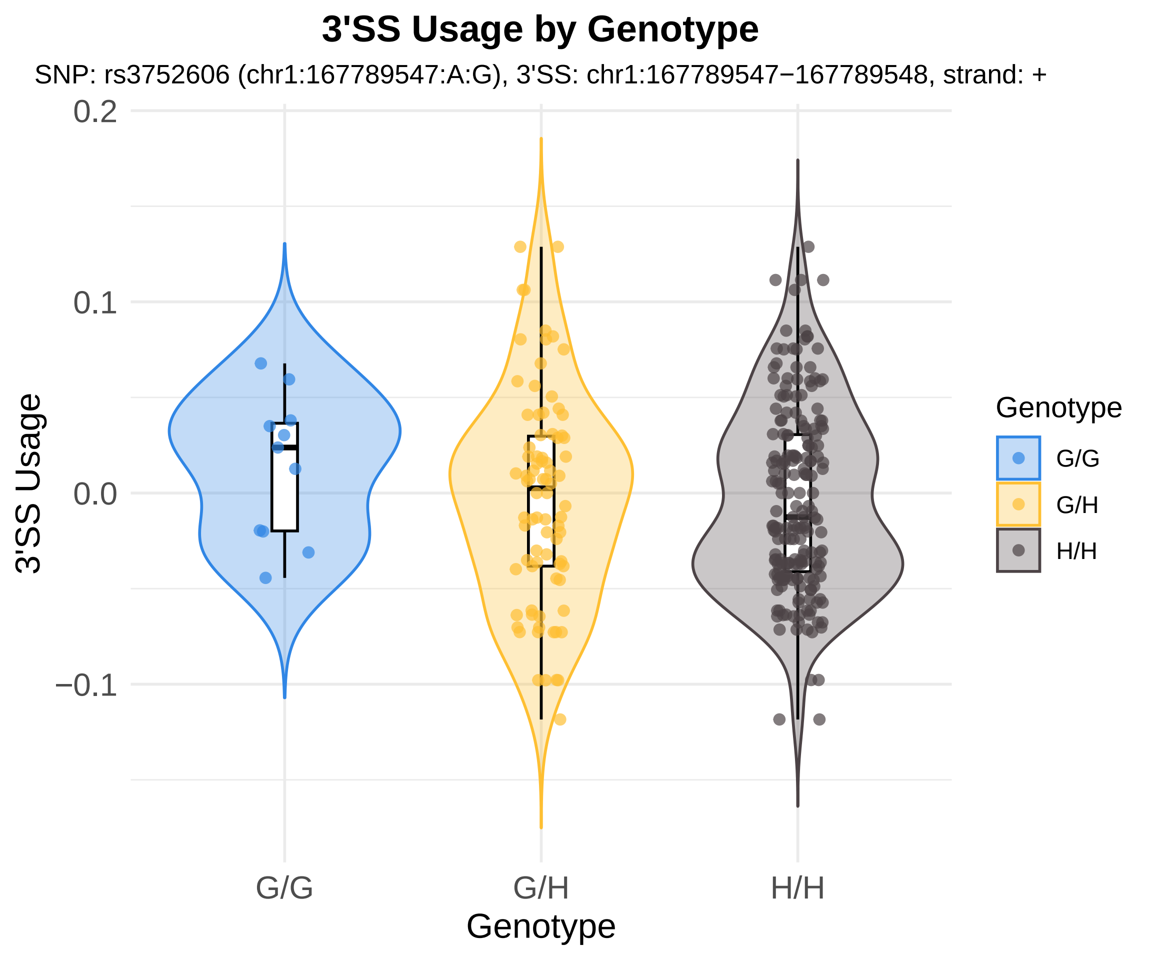

3'SS Usage by Genotype
SNP: rs3752606 (chr1:167789547:A:G), 3'SS: chr1:167789547−167789548, strand: +
0.2
0.1
0.0
−0.1
3'SS Usage
G/G
G/H
H/H
Genotype
Genotype
G/G
G/H
H/H

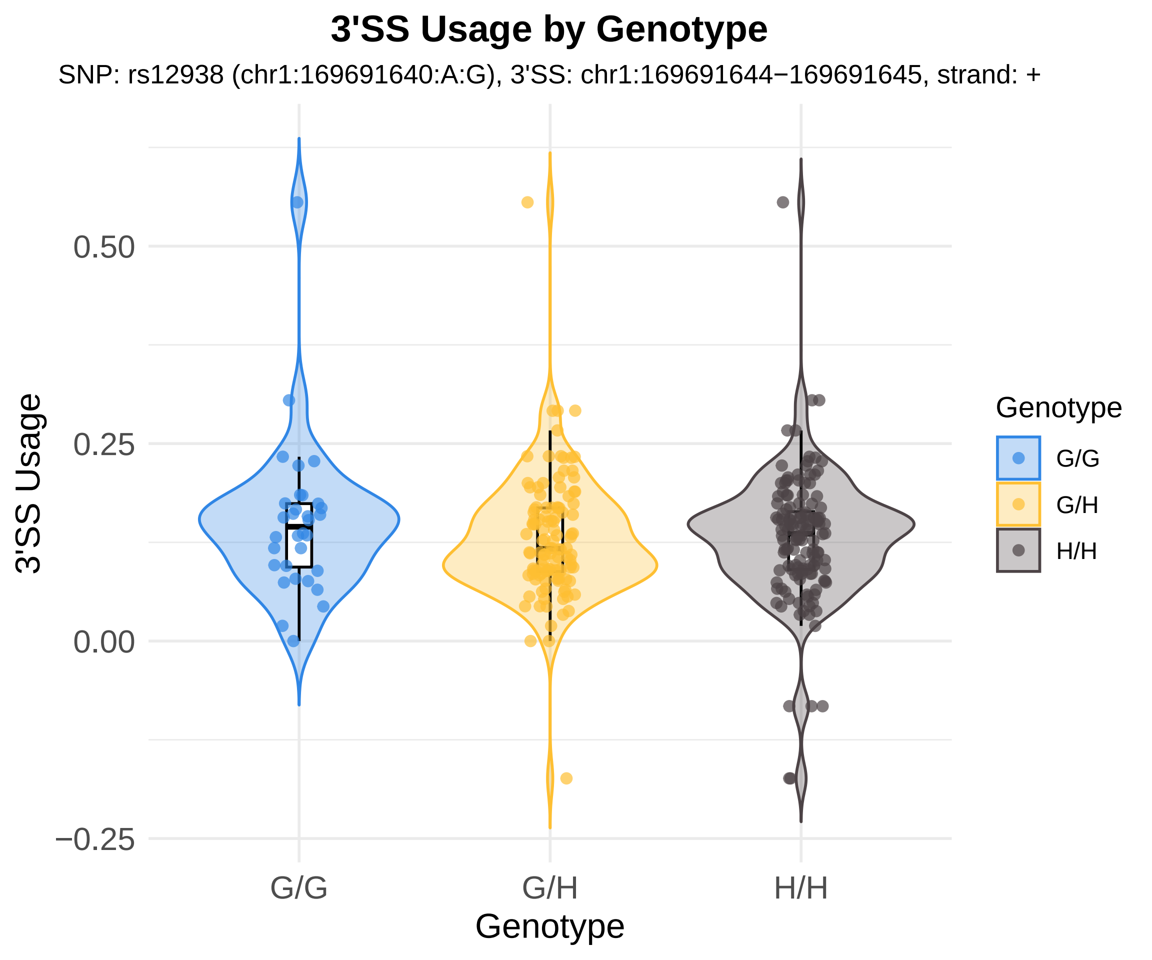
3'SS Usage by Genotype
SNP: rs12938 (chr1:169691640:A:G), 3'SS: chr1:169691644−169691645, strand: +
3'SS Usage
0.50
0.25
0.00
−0.25
G/G
G/H
H/H
Genotype
Genotype
G/G
G/H
H/H

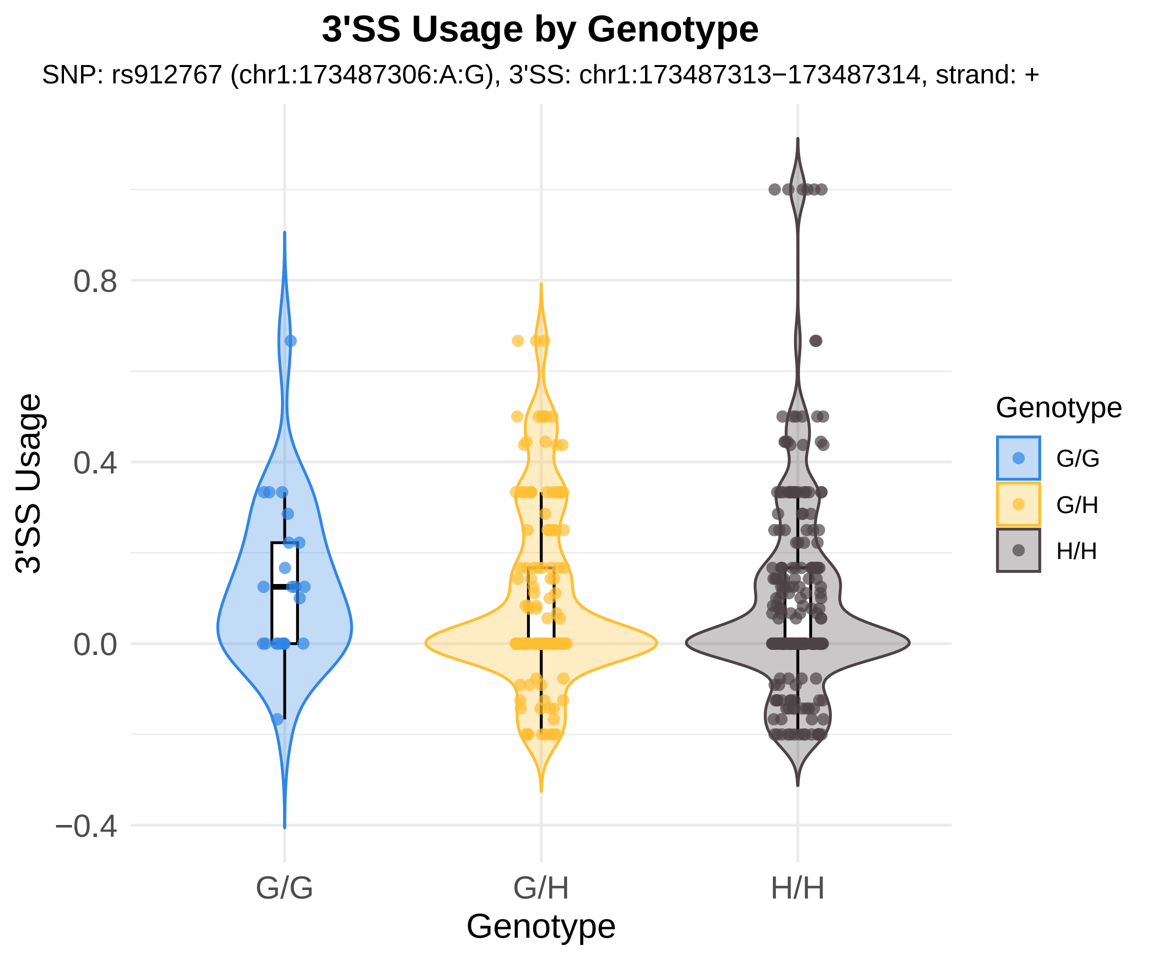
3'SS Usage by Genotype
SNP: rs912767 (chr1:173487306:A:G), 3'SS: chr1:173487313−173487314, strand: +
3'SS Usage
0.8
0.4
0.0
−0.4
G/G
G/H
H/H
Genotype
Genotype
G/G
G/H
H/H

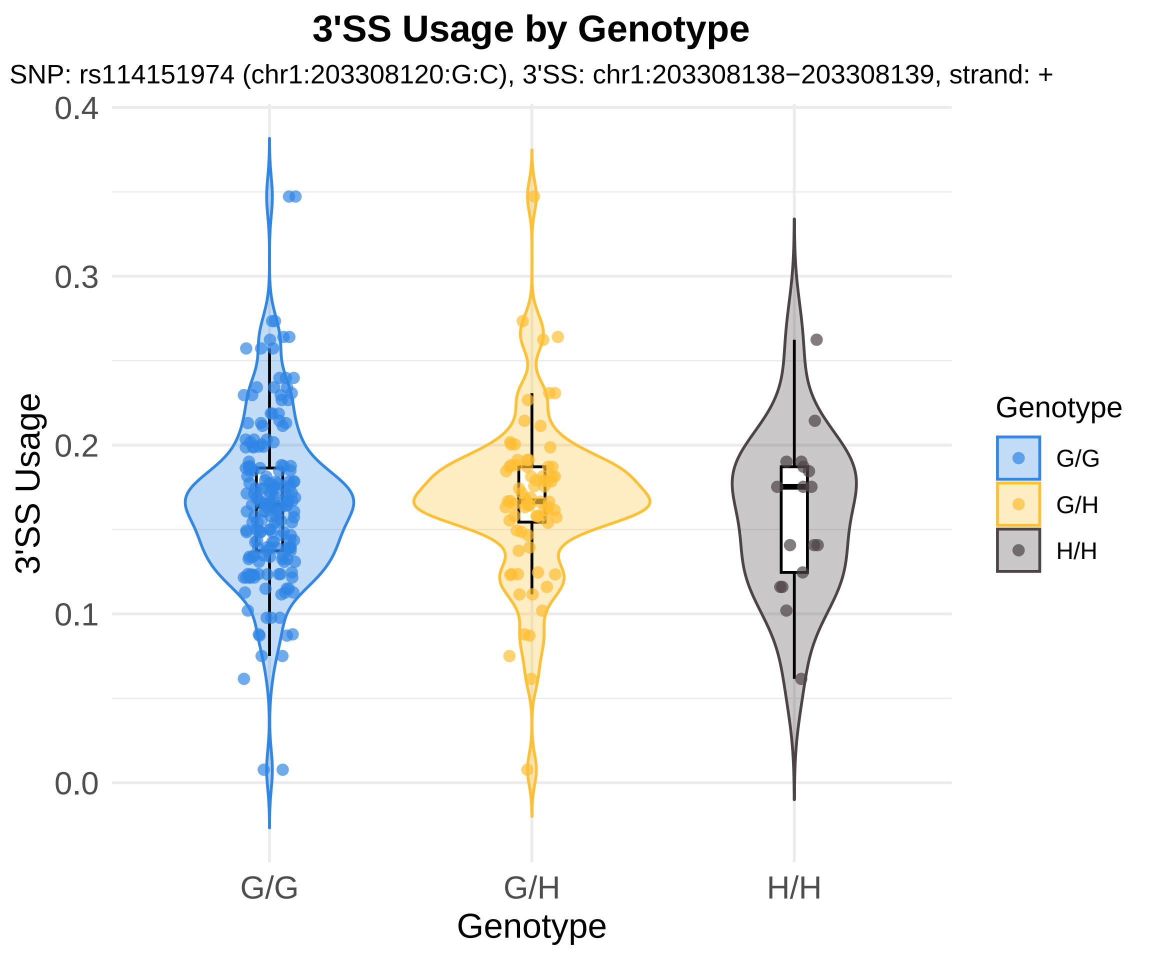

3'SS Usage by Genotype
SNP: rs114151974 (chr1:203308120:G:C), 3'SS: chr1:203308138−203308139, strand: +
0.4
0.3
0.2
0.1
0.0
3'SS Usage
G/G
G/H
H/H
Genotype
Genotype
G/G
G/H
H/H

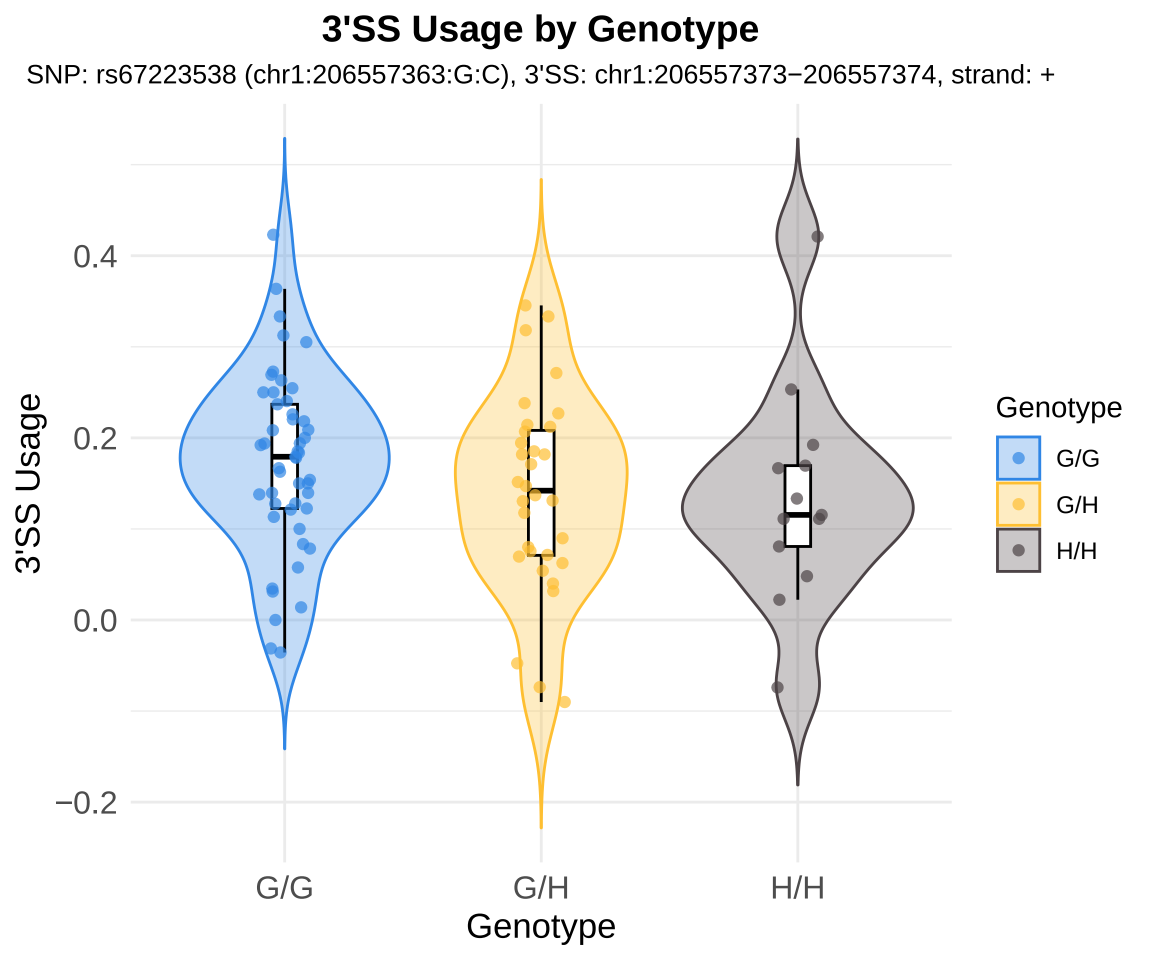

3'SS Usage by Genotype
SNP: rs67223538 (chr1:206557363:G:C), 3'SS: chr1:206557373−206557374, strand: +
3'SS Usage
0.4
0.2
0.0
−0.2
G/G
G/H
H/H
Genotype
Genotype
G/G
G/H
H/H

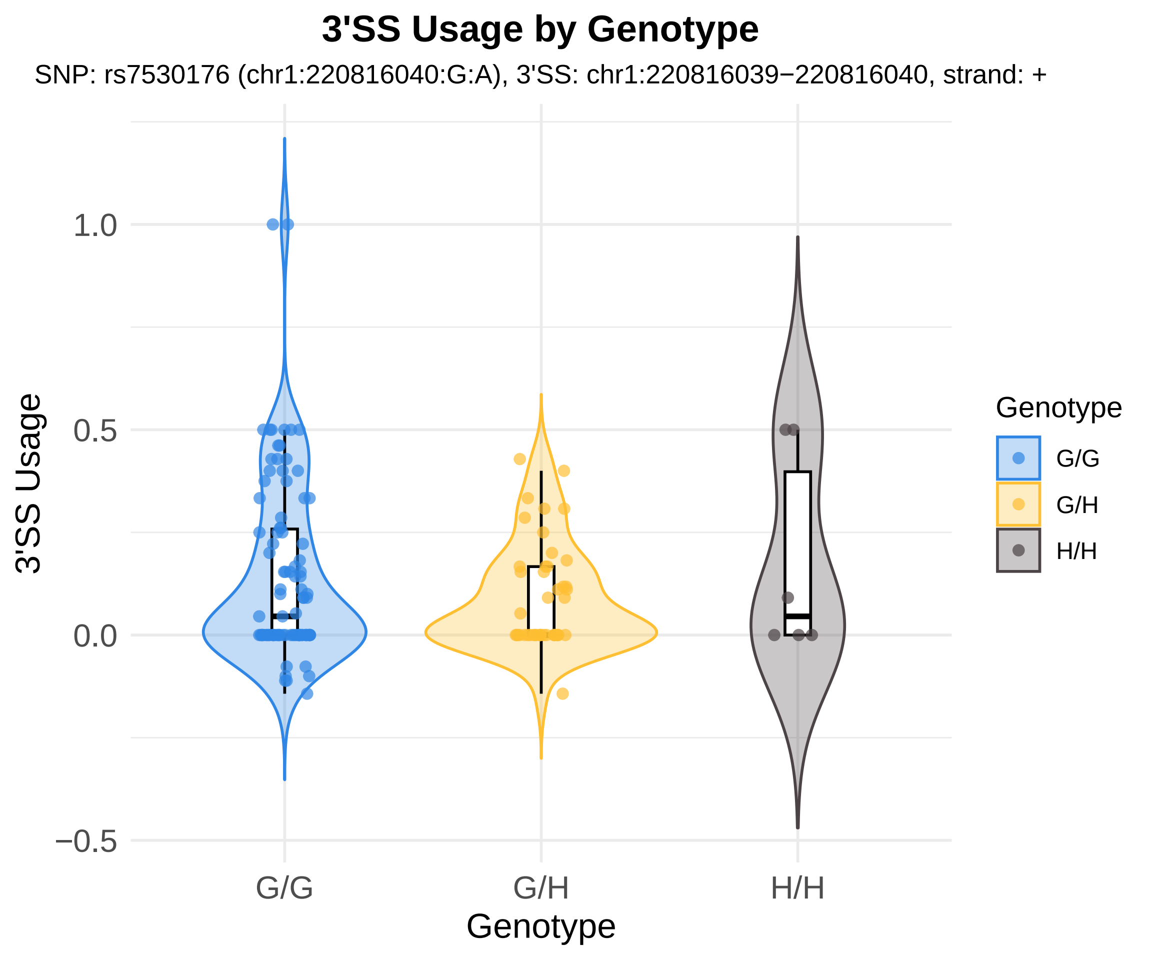

3'SS Usage by Genotype
SNP: rs7530176 (chr1:220816040:G:A), 3'SS: chr1:220816039−220816040, strand: +
3'SS Usage
1.0
0.5
0.0
−0.5
G/G
G/H
H/H
Genotype
Genotype
G/G
G/H
H/H

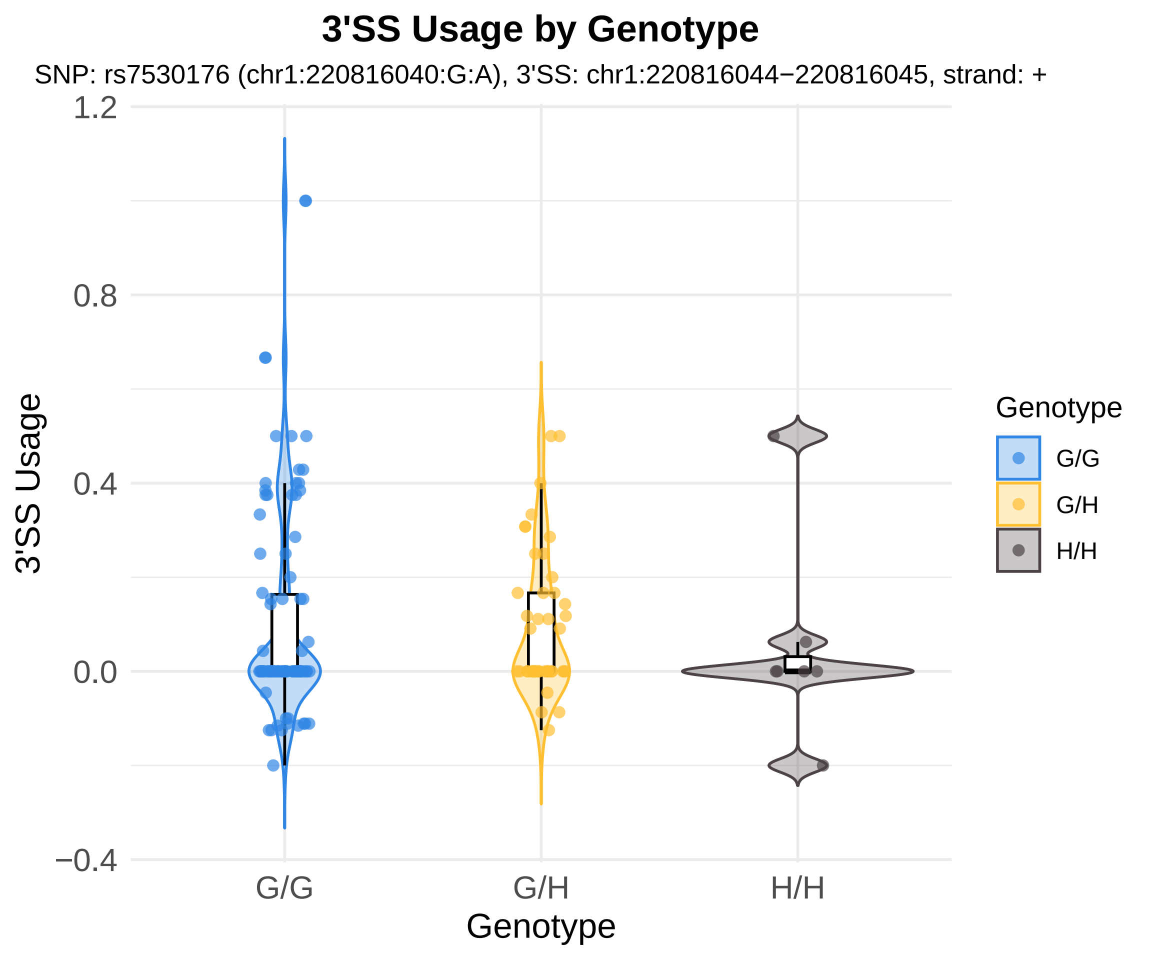
3'SS Usage by Genotype
SNP: rs7530176 (chr1:220816040:G:A), 3'SS: chr1:220816044−220816045, strand: +
3'SS Usage
1.2
0.8
0.4
0.0
−0.4
G/G
G/H
H/H
Genotype
Genotype
G/G
G/H
H/H

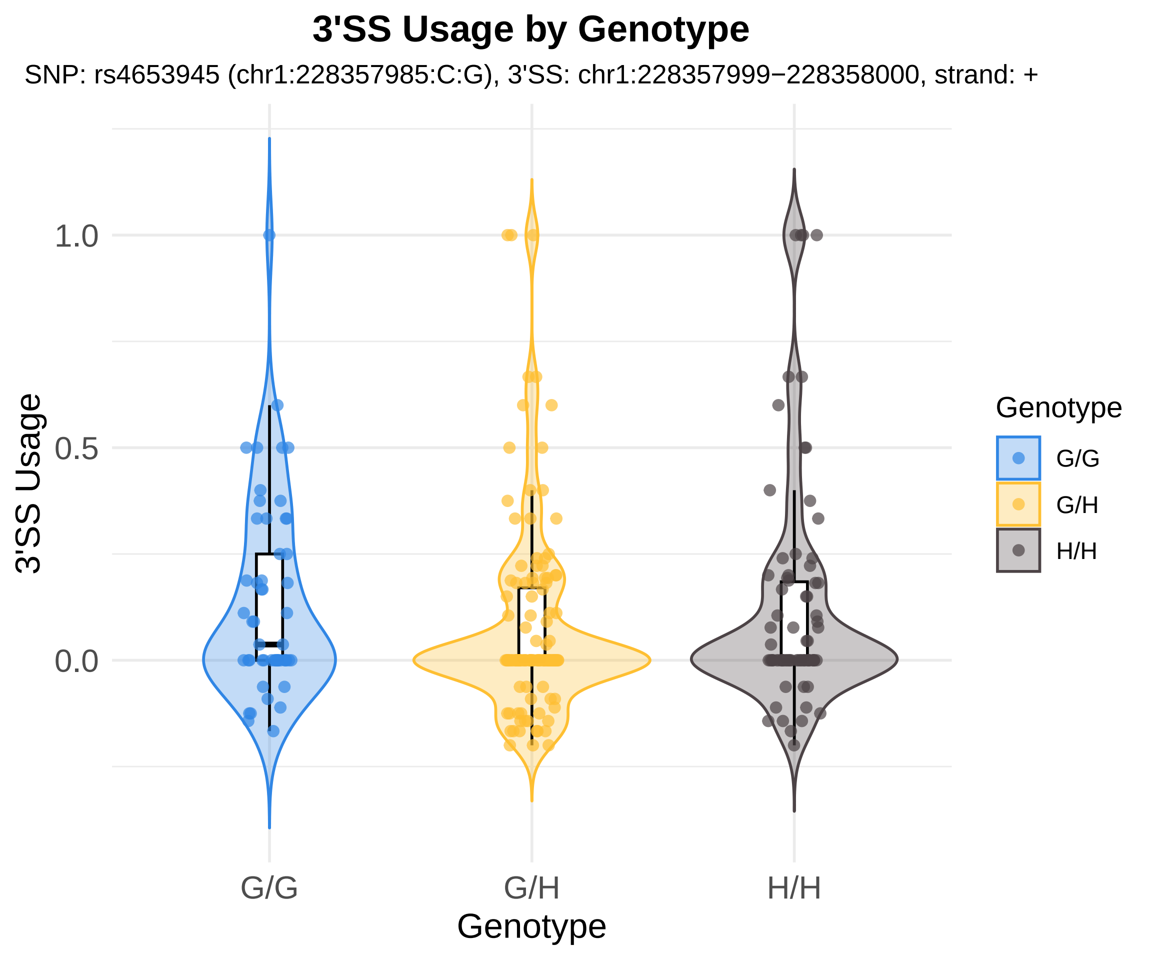

3'SS Usage by Genotype
SNP: rs4653945 (chr1:228357985:C:G), 3'SS: chr1:228357999−228358000, strand: +
3'SS Usage
1.0
0.5
0.0
G/G
G/H
H/H
Genotype
Genotype
G/G
G/H
H/H

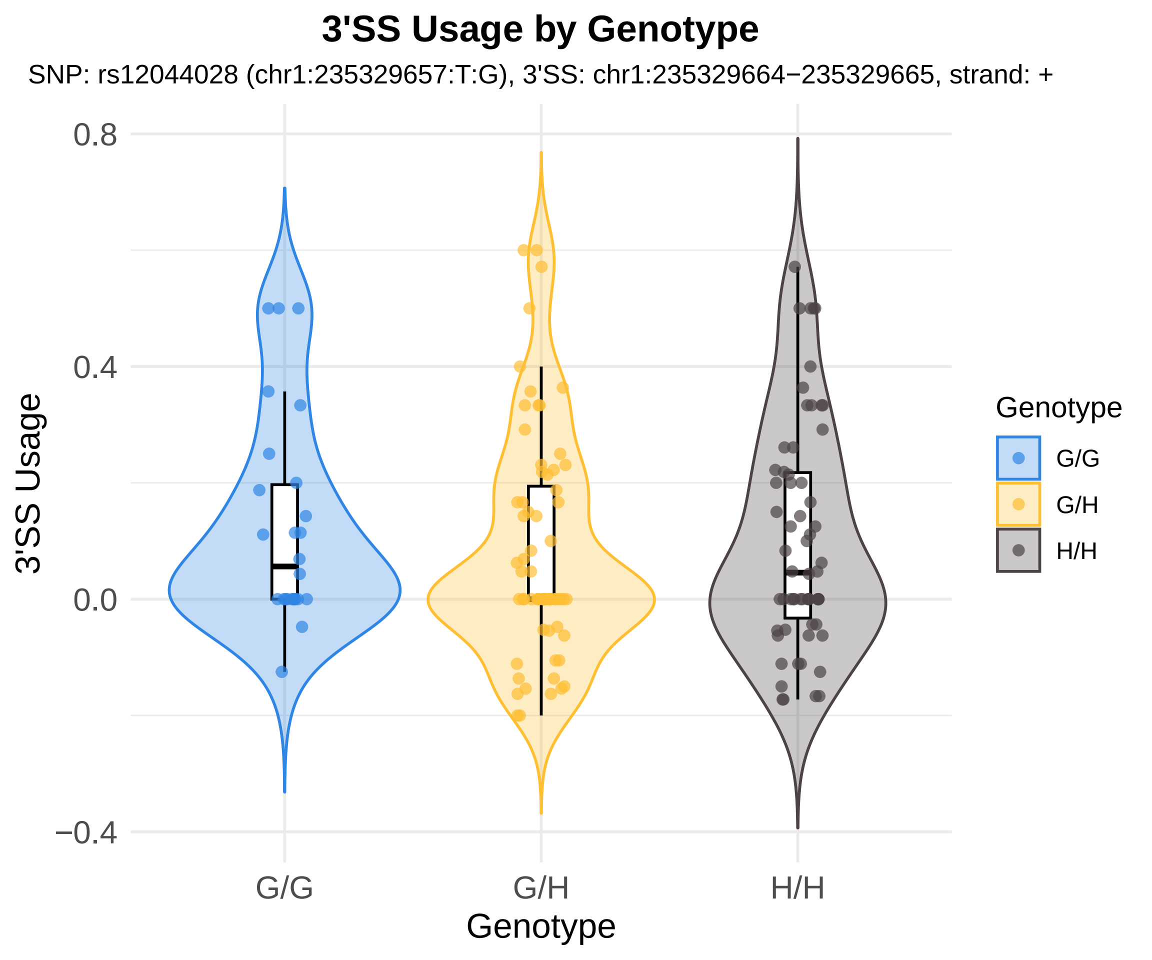

3'SS Usage by Genotype
SNP: rs12044028 (chr1:235329657:T:G), 3'SS: chr1:235329664−235329665, strand: +
0.8
0.4
0.0
−0.4
3'SS Usage
G/G
G/H
H/H
Genotype
Genotype
G/G
G/H
H/H

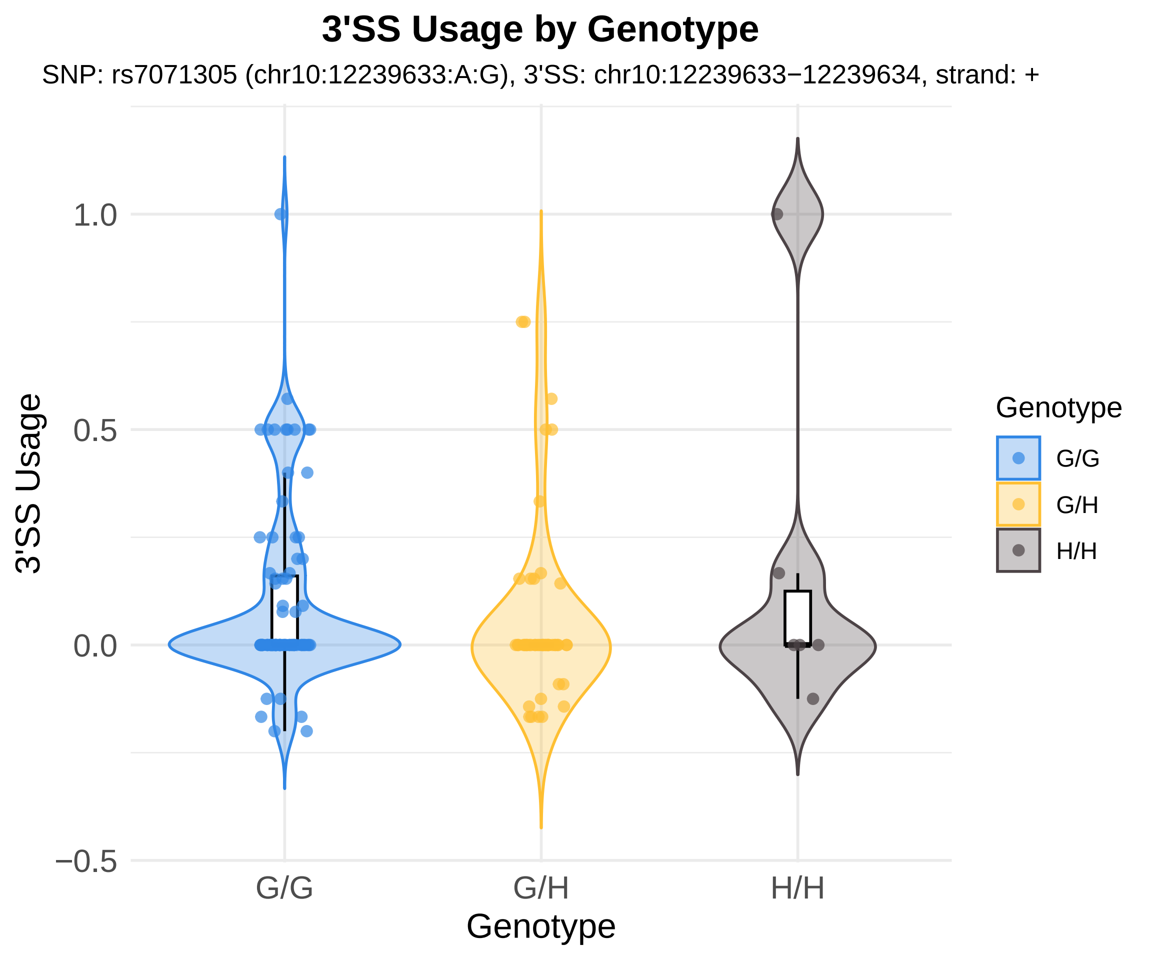

3'SS Usage by Genotype
SNP: rs7071305 (chr10:12239633:A:G), 3'SS: chr10:12239633−12239634, strand: +
3'SS Usage
1.0
0.5
0.0
−0.5
G/G
G/H
H/H
Genotype
Genotype
G/G
G/H
H/H

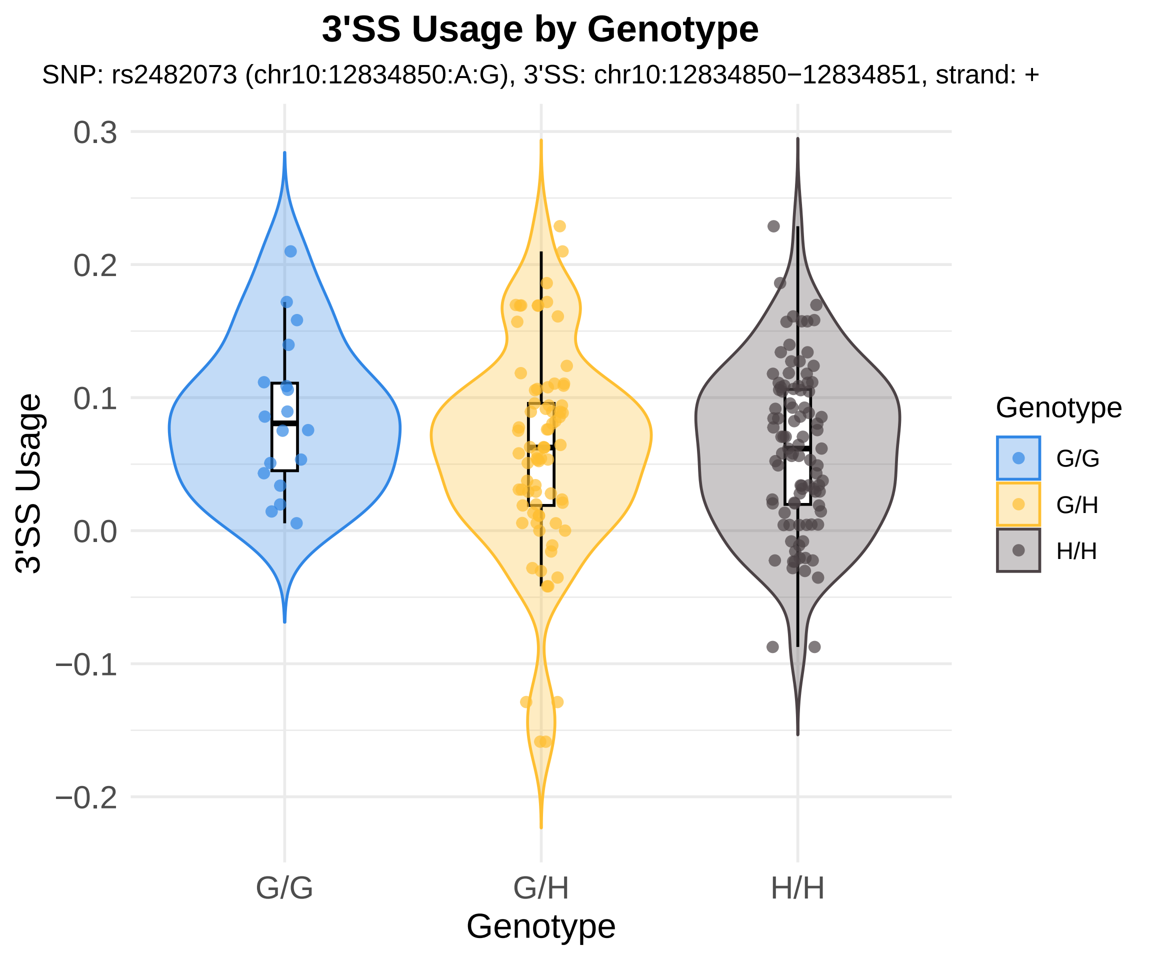
3'SS Usage by Genotype
SNP: rs2482073 (chr10:12834850:A:G), 3'SS: chr10:12834850−12834851, strand: +
0.3
0.2
0.1
0.0
−0.1
−0.2
3'SS Usage
G/G
G/H
H/H
Genotype
Genotype
G/G
G/H
H/H

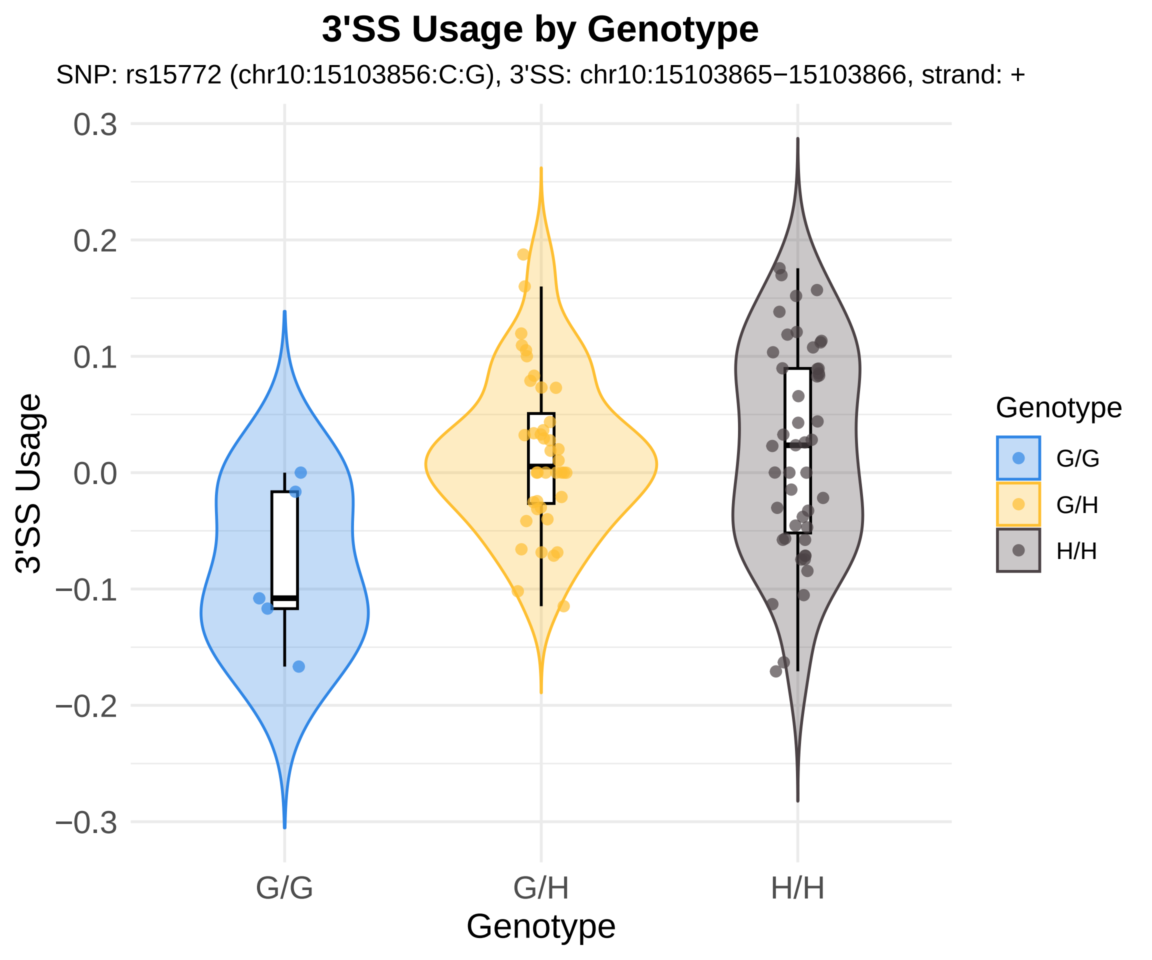

3'SS Usage by Genotype
SNP: rs15772 (chr10:15103856:C:G), 3'SS: chr10:15103865−15103866, strand: +
3'SS Usage
0.3
0.2
0.1
0.0
−0.1
−0.2
−0.3
G/G
G/H
H/H
Genotype
Genotype
G/G
G/H
H/H

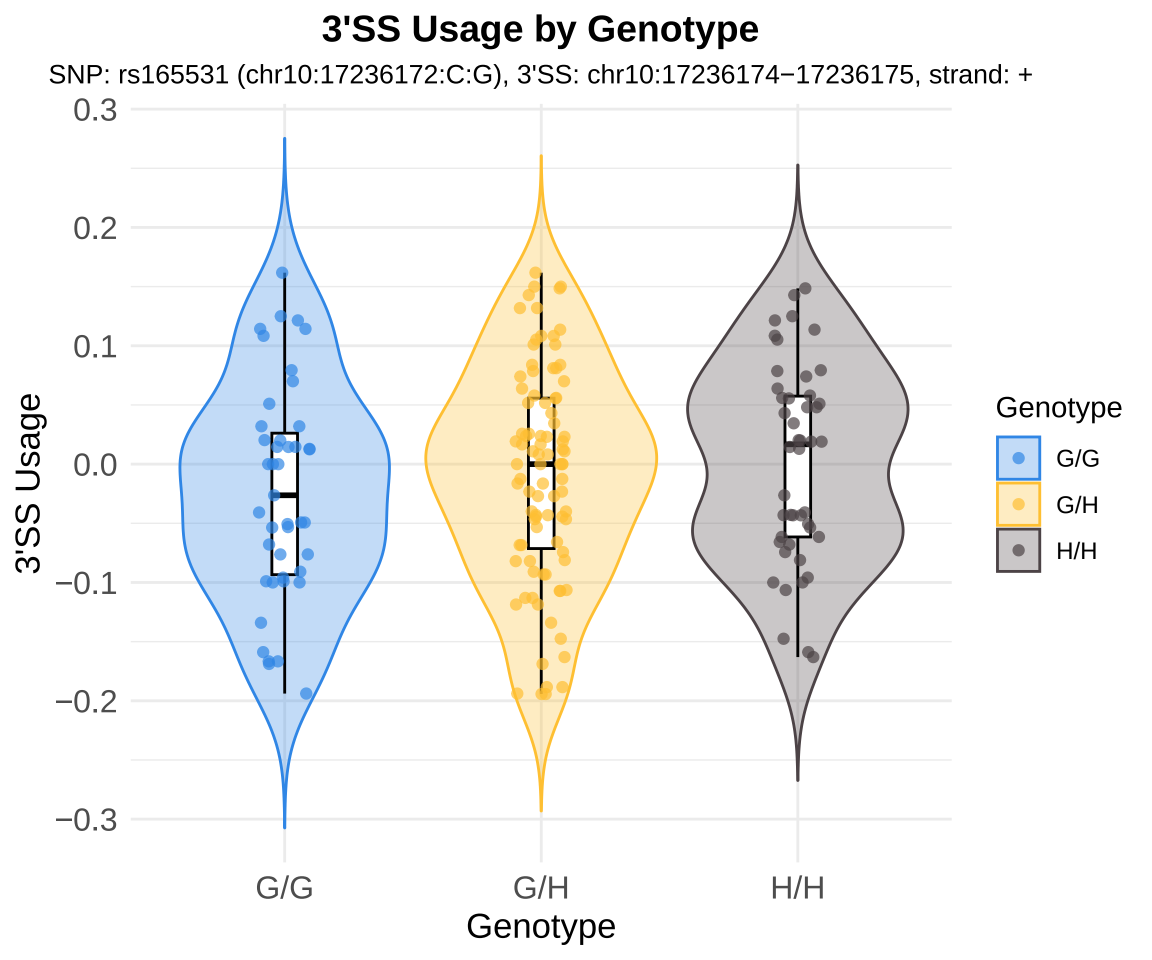
3'SS Usage by Genotype
SNP: rs165531 (chr10:17236172:C:G), 3'SS: chr10:17236174−17236175, strand: +
0.3
0.2
0.1
0.0
−0.1
−0.2
−0.3
3'SS Usage
G/G
G/H
H/H
Genotype
Genotype
G/G
G/H
H/H

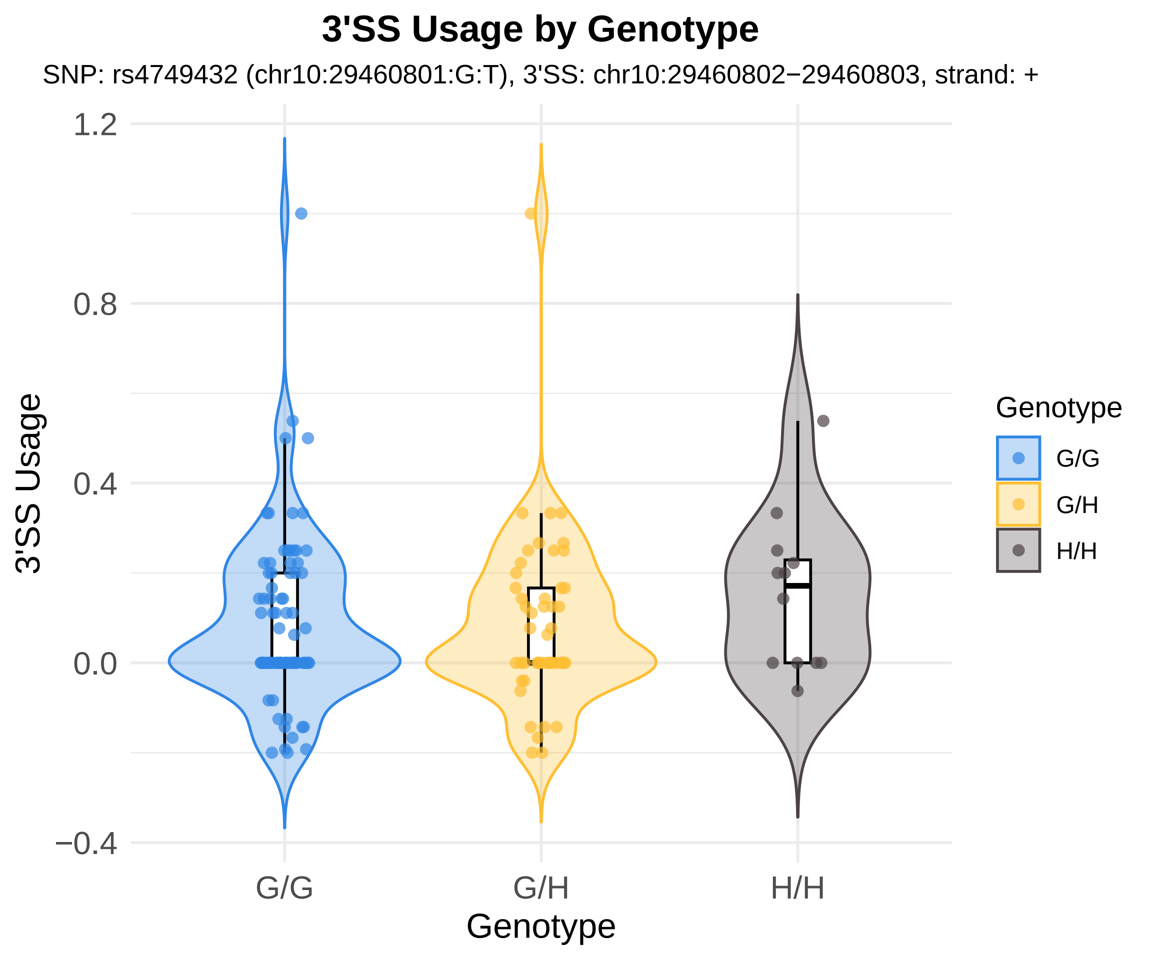
3'SS Usage by Genotype
SNP: rs4749432 (chr10:29460801:G:T), 3'SS: chr10:29460802−29460803, strand: +
3'SS Usage
1.2
0.8
0.4
0.0
−0.4
G/G
G/H
H/H
Genotype
Genotype
G/G
G/H
H/H

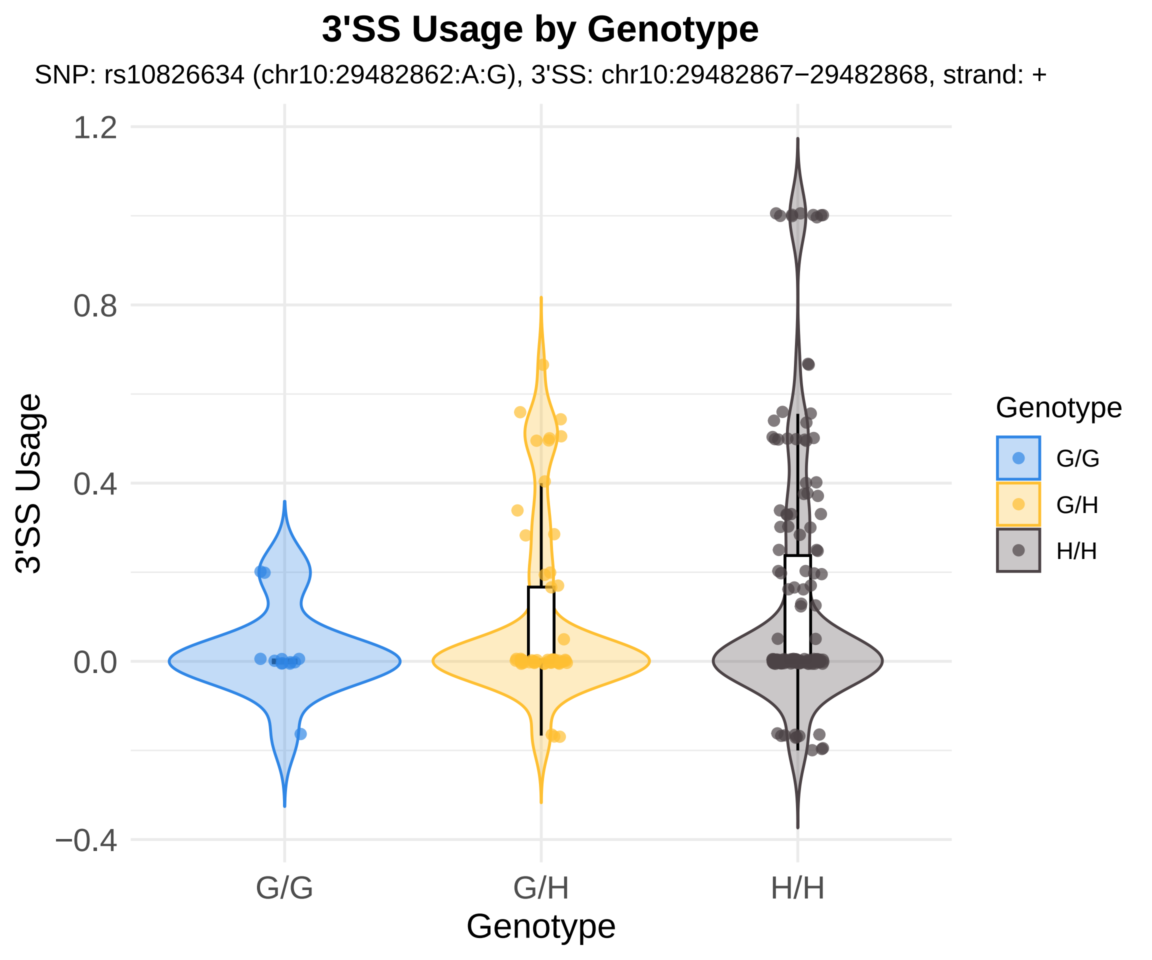
3'SS Usage by Genotype
SNP: rs10826634 (chr10:29482862:A:G), 3'SS: chr10:29482867−29482868, strand: +
1.2
0.8
0.4
0.0
−0.4
3'SS Usage
G/G
G/H
H/H
Genotype
Genotype
G/G
G/H
H/H

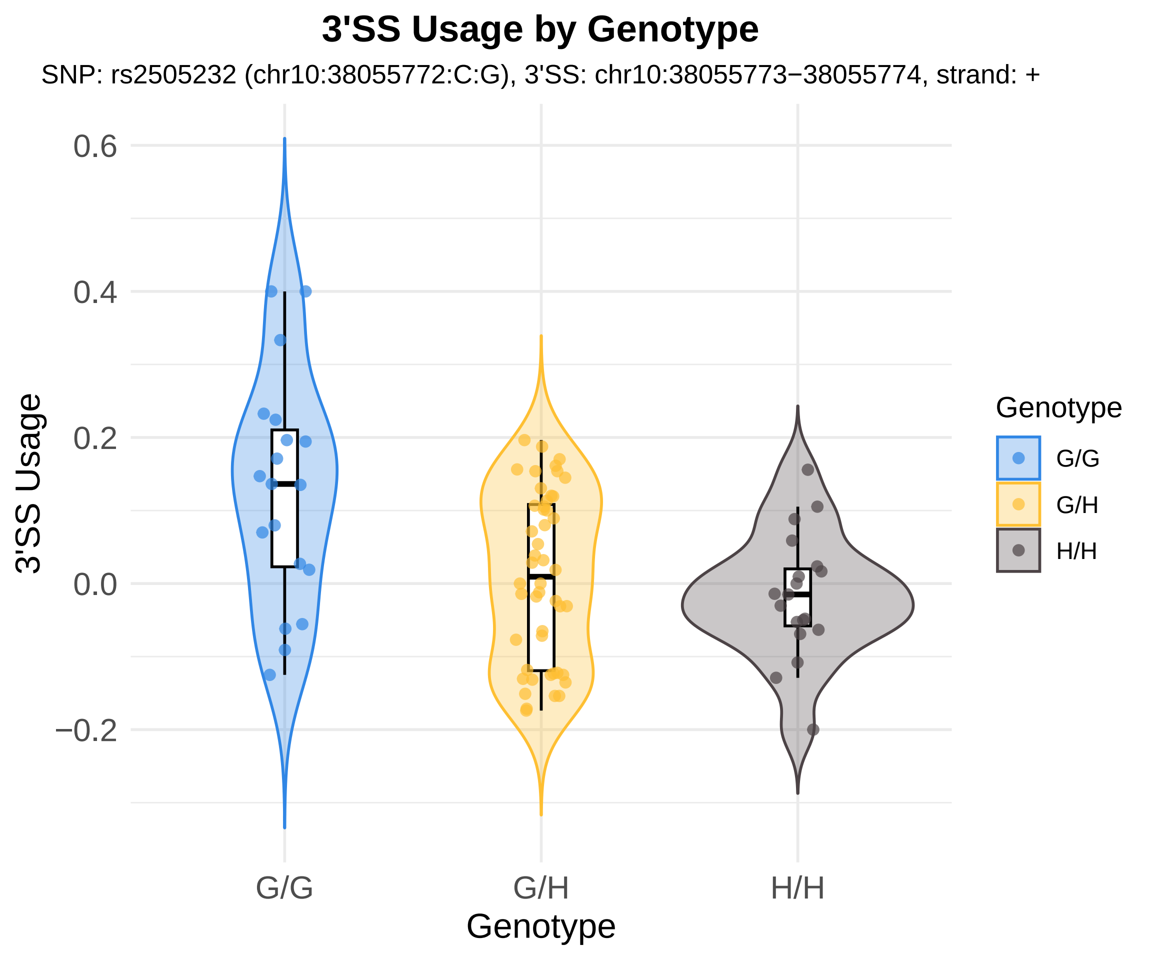

3'SS Usage by Genotype
SNP: rs2505232 (chr10:38055772:C:G), 3'SS: chr10:38055773−38055774, strand: +
3'SS Usage
0.6
0.4
0.2
0.0
−0.2
G/G
G/H
H/H
Genotype
Genotype
G/G
G/H
H/H

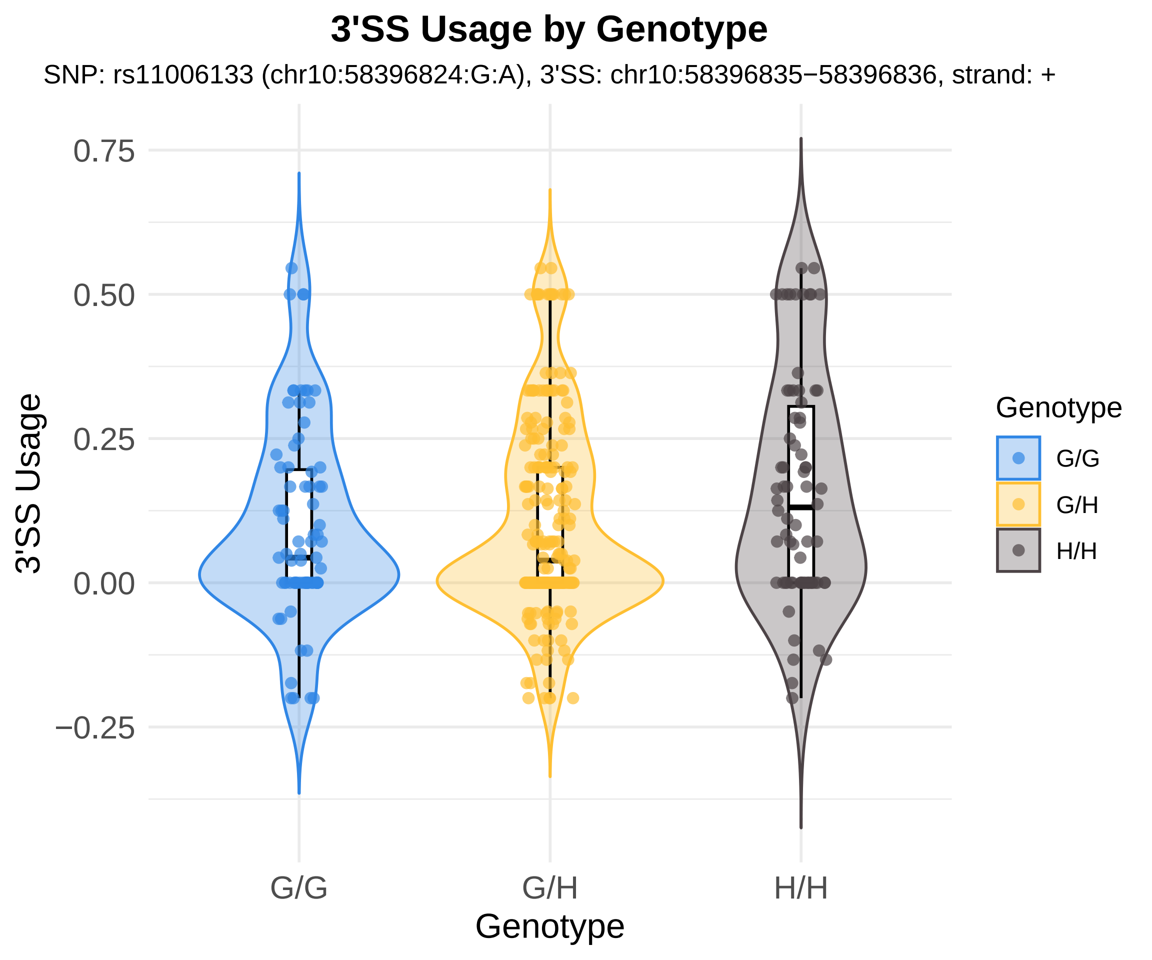
3'SS Usage by Genotype
SNP: rs11006133 (chr10:58396824:G:A), 3'SS: chr10:58396835−58396836, strand: +
3'SS Usage
0.75
0.50
0.25
0.00
−0.25
G/G
G/H
H/H
Genotype
Genotype
G/G
G/H
H/H

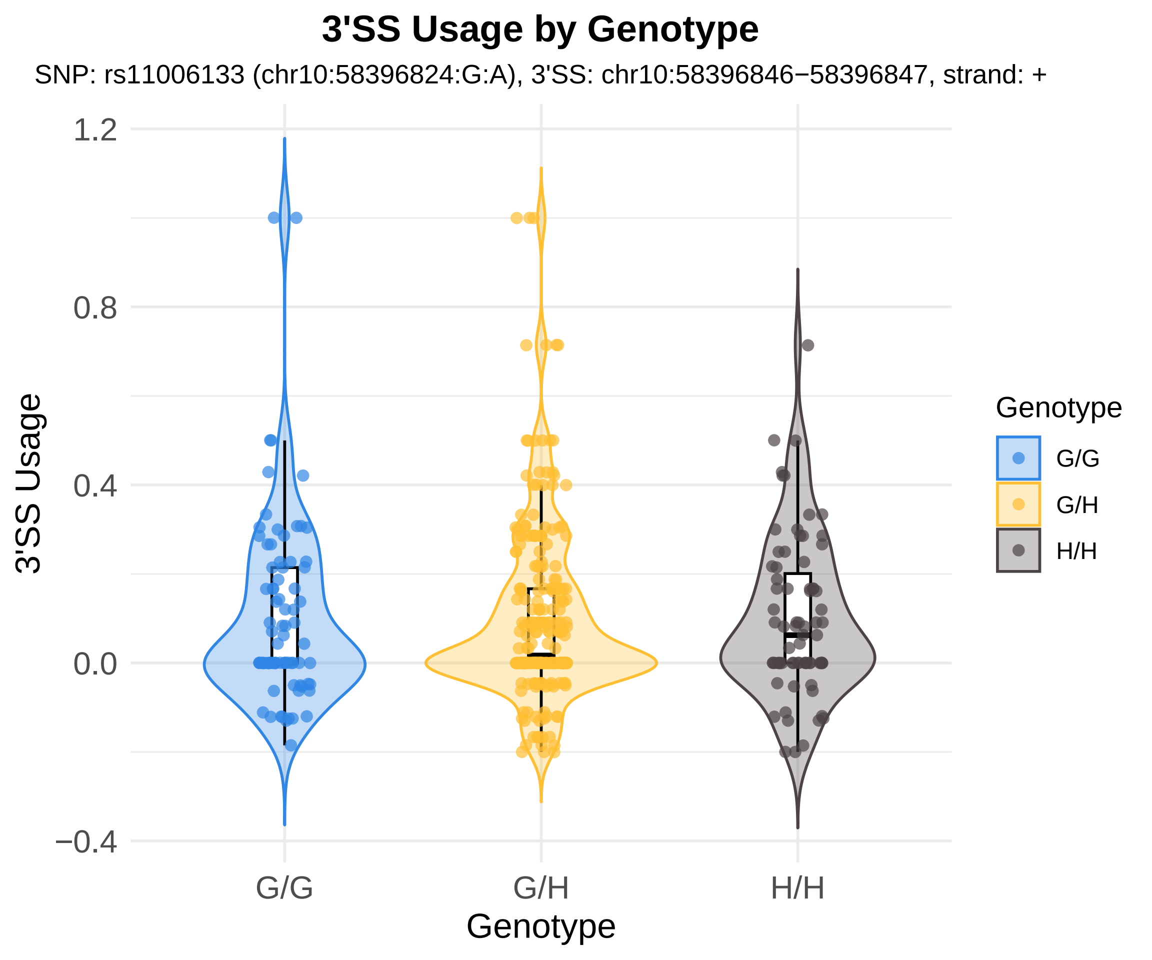
3'SS Usage by Genotype
SNP: rs11006133 (chr10:58396824:G:A), 3'SS: chr10:58396846−58396847, strand: +
3'SS Usage
1.2
0.8
0.4
0.0
−0.4
G/G
G/H
H/H
Genotype
Genotype
G/G
G/H
H/H

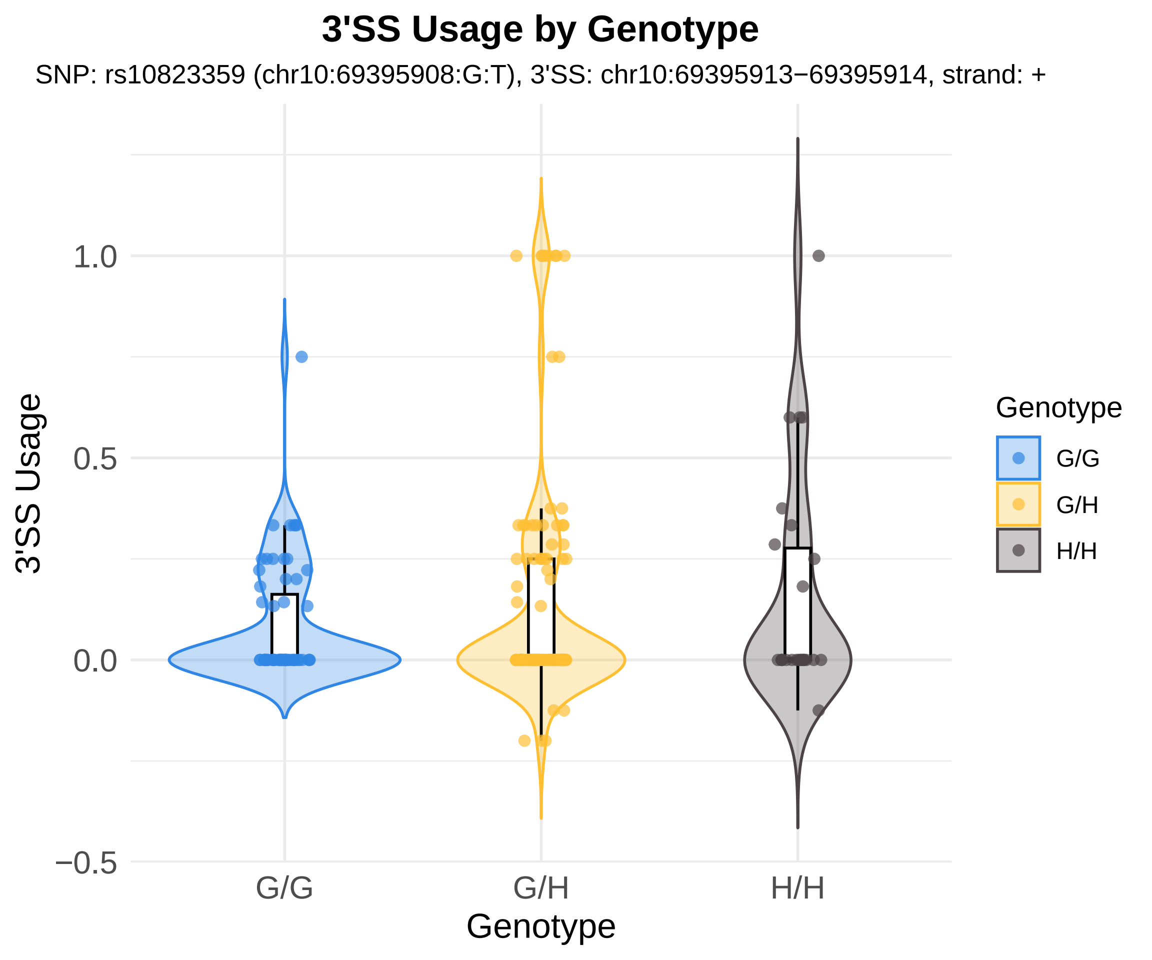
3'SS Usage by Genotype
SNP: rs10823359 (chr10:69395908:G:T), 3'SS: chr10:69395913−69395914, strand: +
3'SS Usage
1.0
0.5
0.0
−0.5
G/G
G/H
H/H
Genotype
Genotype
G/G
G/H
H/H

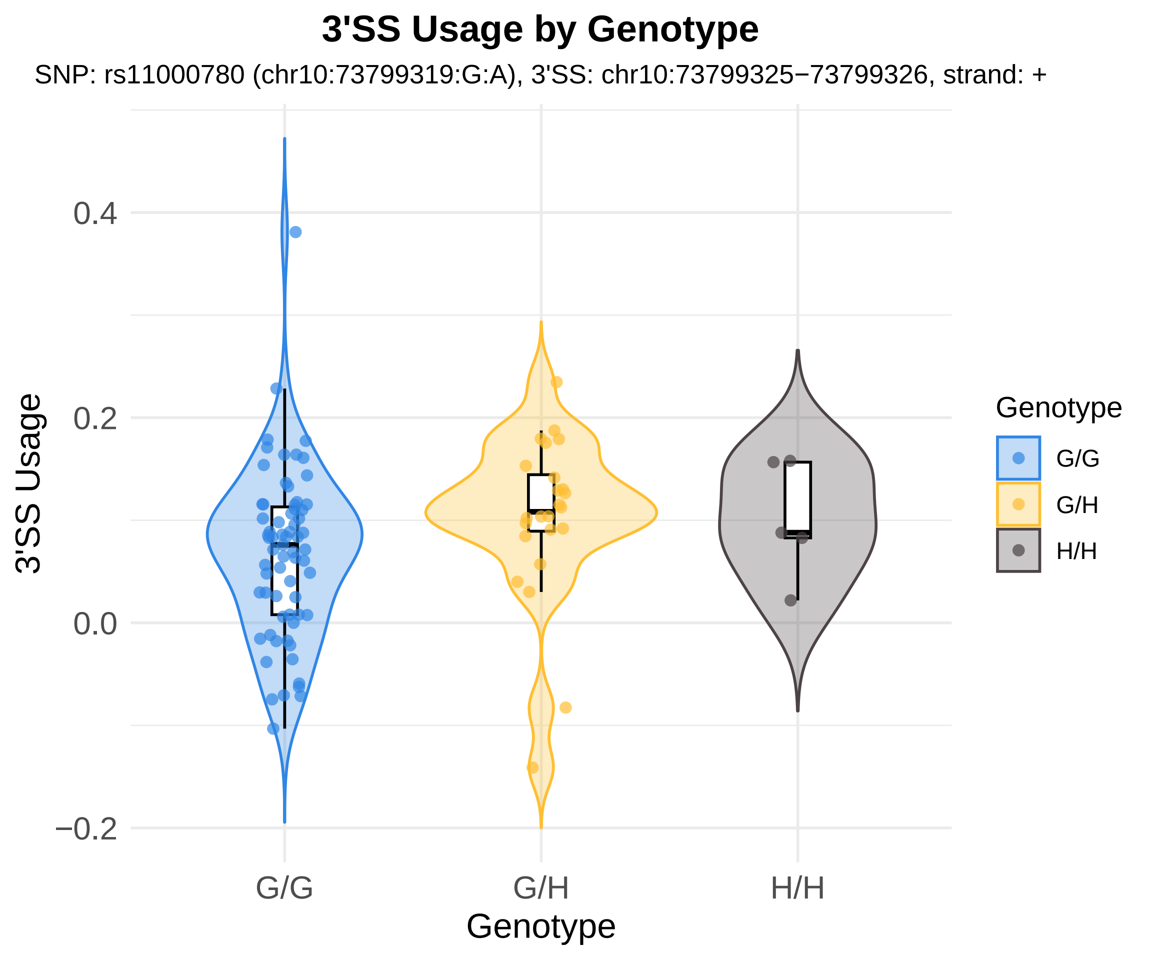
3'SS Usage by Genotype
SNP: rs11000780 (chr10:73799319:G:A), 3'SS: chr10:73799325−73799326, strand: +
3'SS Usage
0.4
0.2
0.0
−0.2
G/G
G/H
H/H
Genotype
Genotype
G/G
G/H
H/H

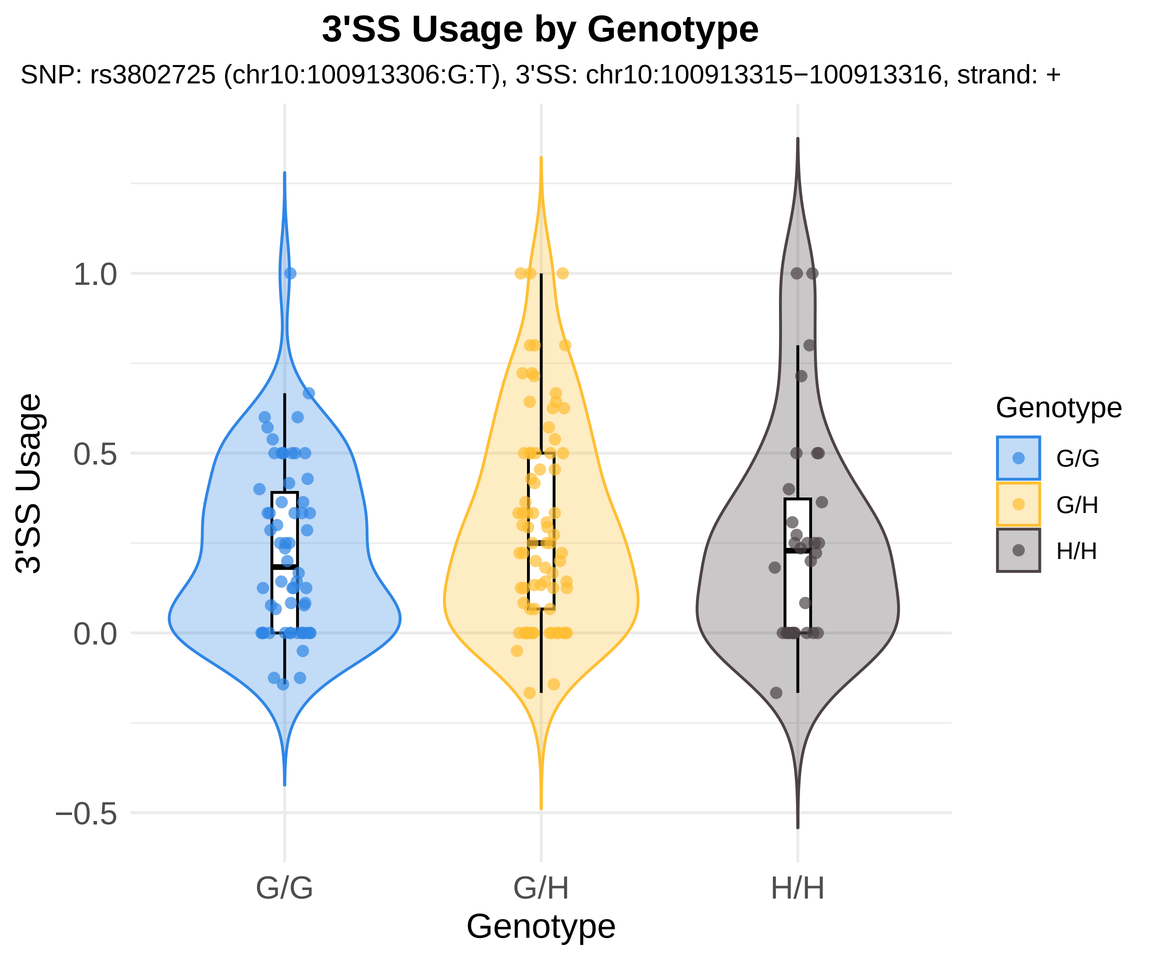

3'SS Usage by Genotype
SNP: rs3802725 (chr10:100913306:G:T), 3'SS: chr10:100913315−100913316, strand: +
3'SS Usage
1.0
0.5
0.0
−0.5
G/G
G/H
H/H
Genotype
Genotype
G/G
G/H
H/H

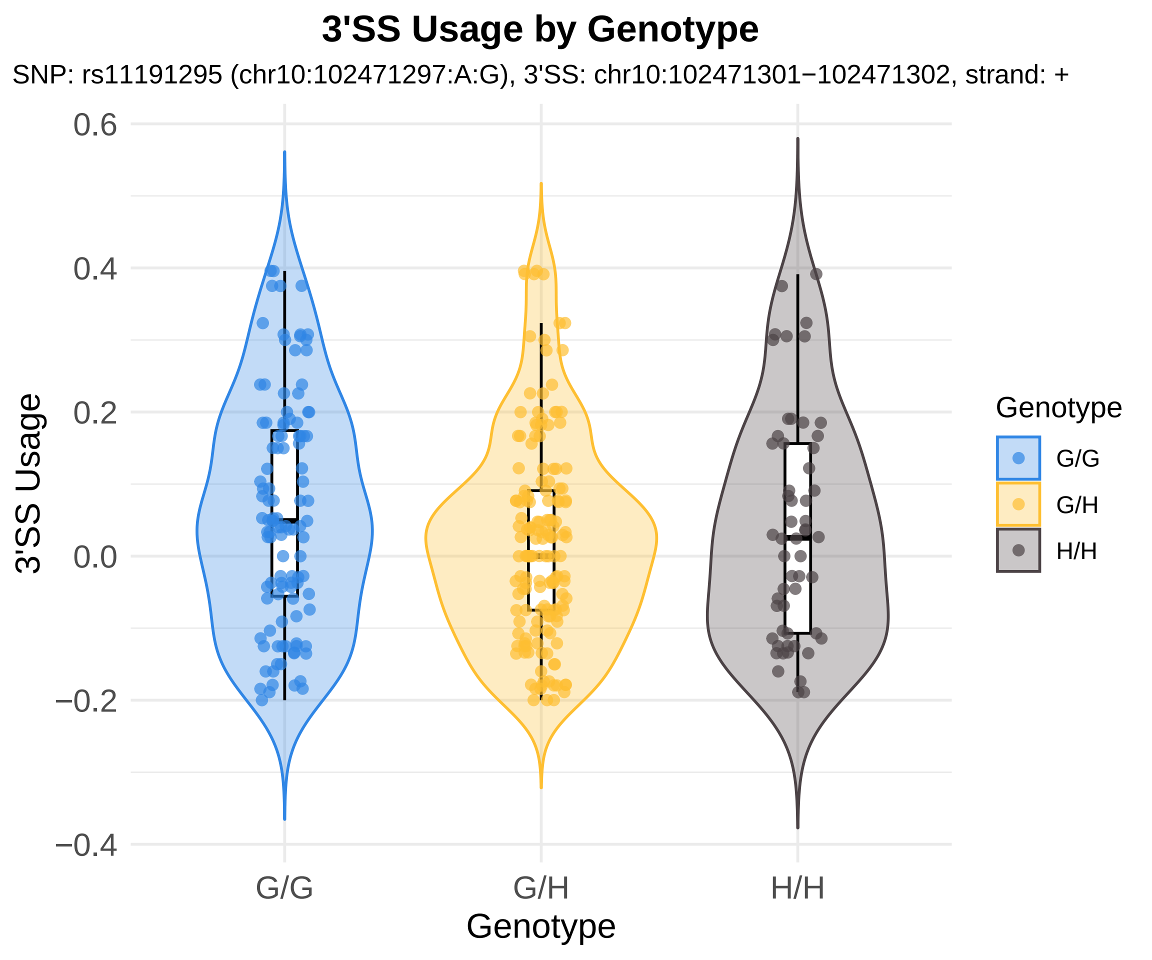
3'SS Usage by Genotype
SNP: rs11191295 (chr10:102471297:A:G), 3'SS: chr10:102471301−102471302, strand: +
3'SS Usage
0.6
0.4
0.2
0.0
−0.2
−0.4
G/G
G/H
H/H
Genotype
Genotype
G/G
G/H
H/H

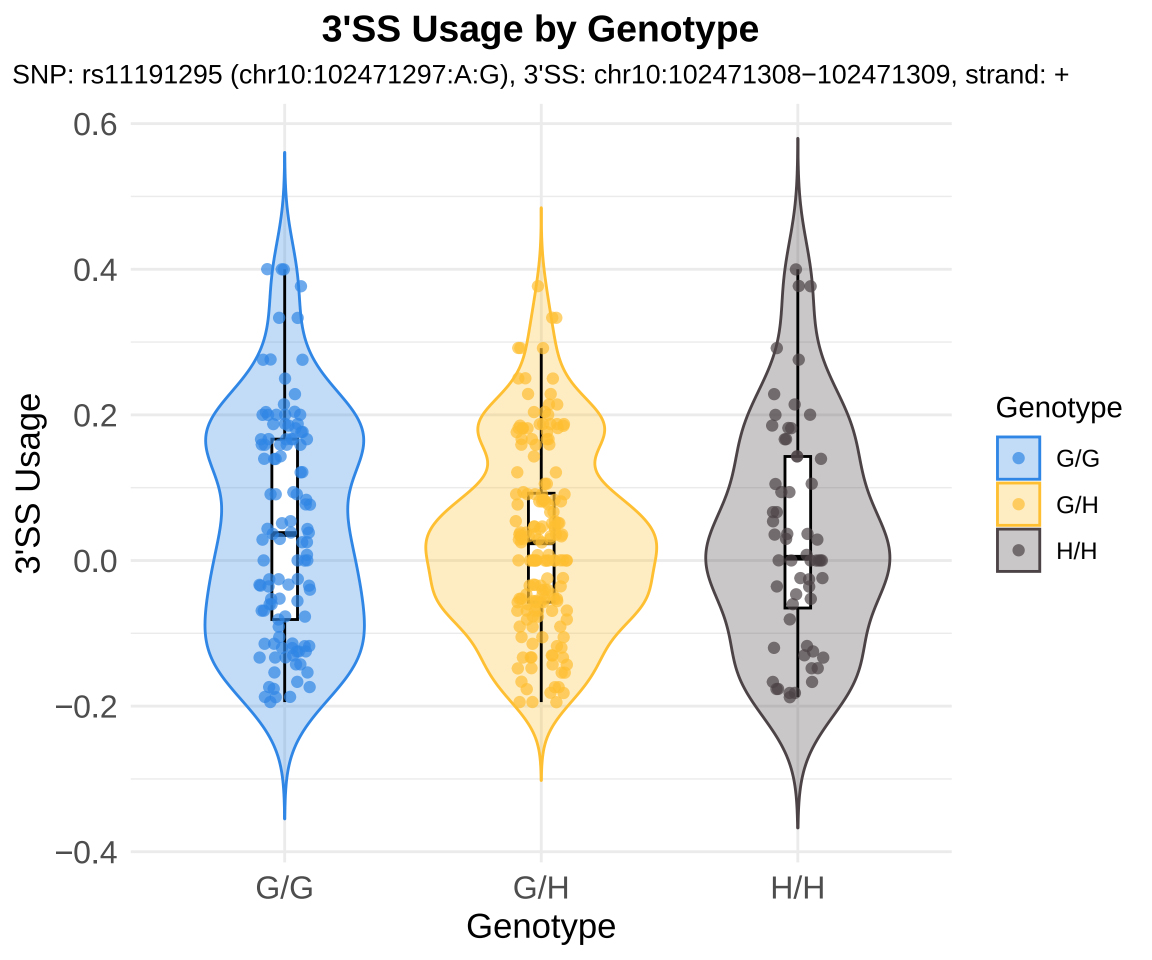
3'SS Usage by Genotype
SNP: rs11191295 (chr10:102471297:A:G), 3'SS: chr10:102471308−102471309, strand: +
3'SS Usage
0.6
0.4
0.2
0.0
−0.2
−0.4
G/G
G/H
H/H
Genotype
Genotype
G/G
G/H
H/H

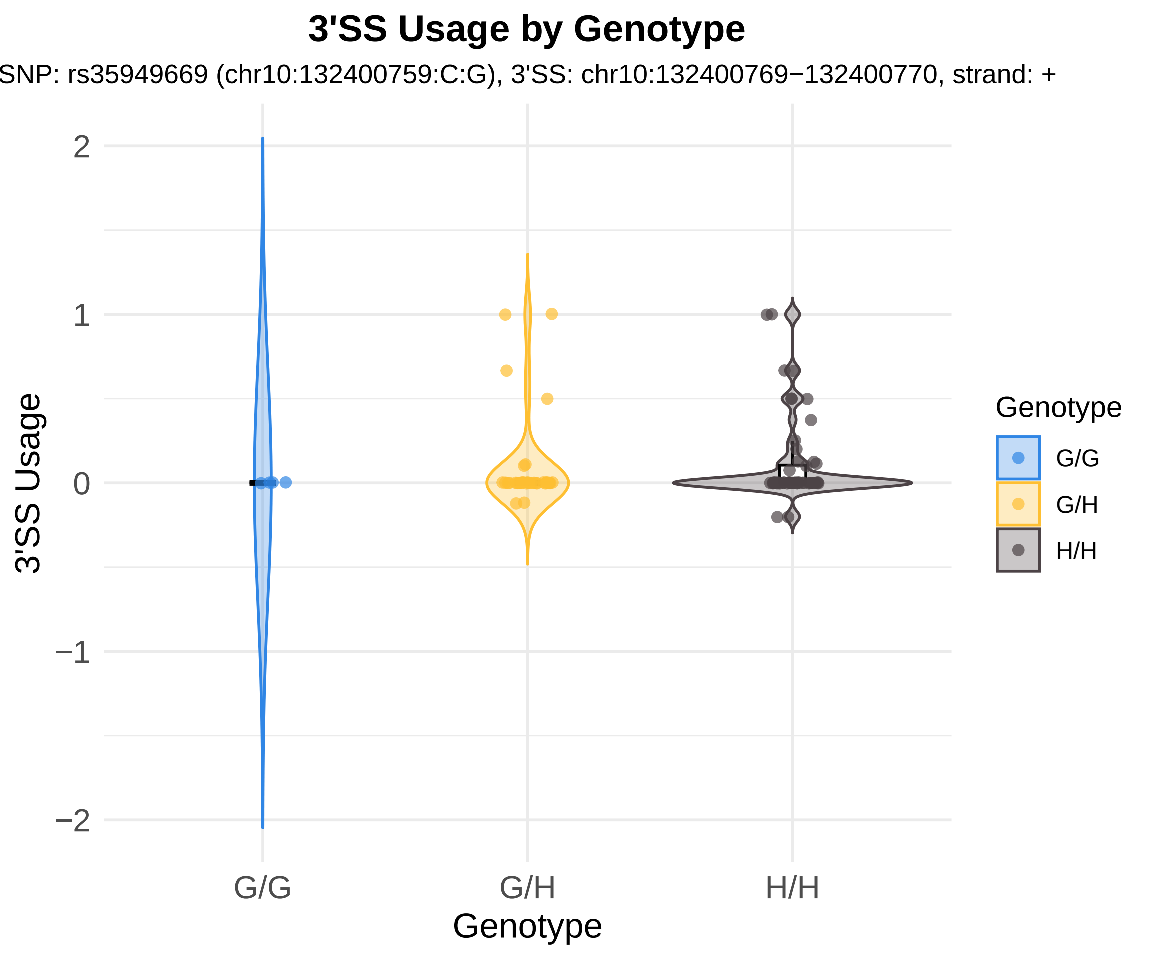
3'SS Usage by Genotype
SNP: rs35949669 (chr10:132400759:C:G), 3'SS: chr10:132400769−132400770, strand: +
3'SS Usage
2
1
0
−1
−2
G/G
G/H
H/H
Genotype
Genotype
G/G
G/H
H/H

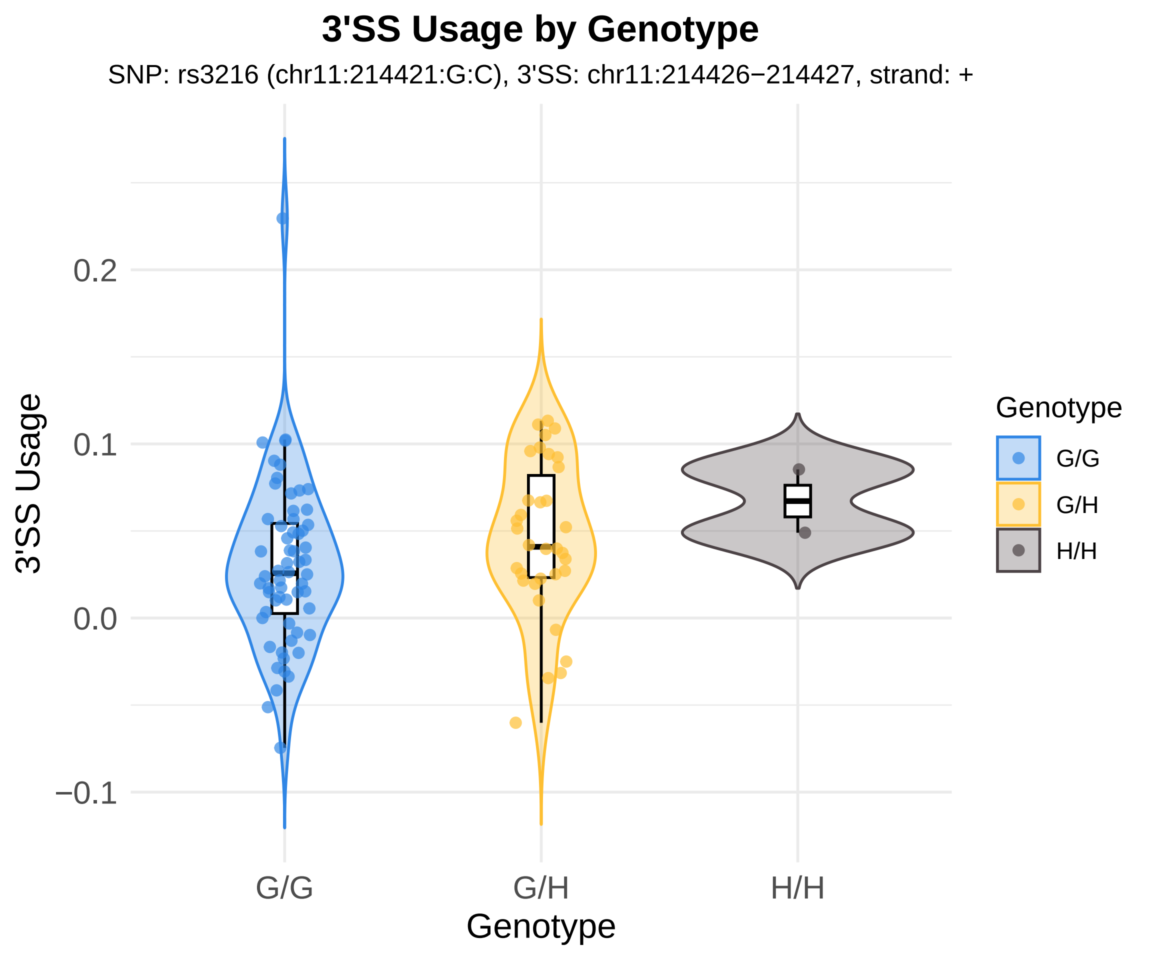
3'SS Usage by Genotype
SNP: rs3216 (chr11:214421:G:C), 3'SS: chr11:214426−214427, strand: +
3'SS Usage
0.2
0.1
0.0
−0.1
G/G
G/H
H/H
Genotype
Genotype
G/G
G/H
H/H

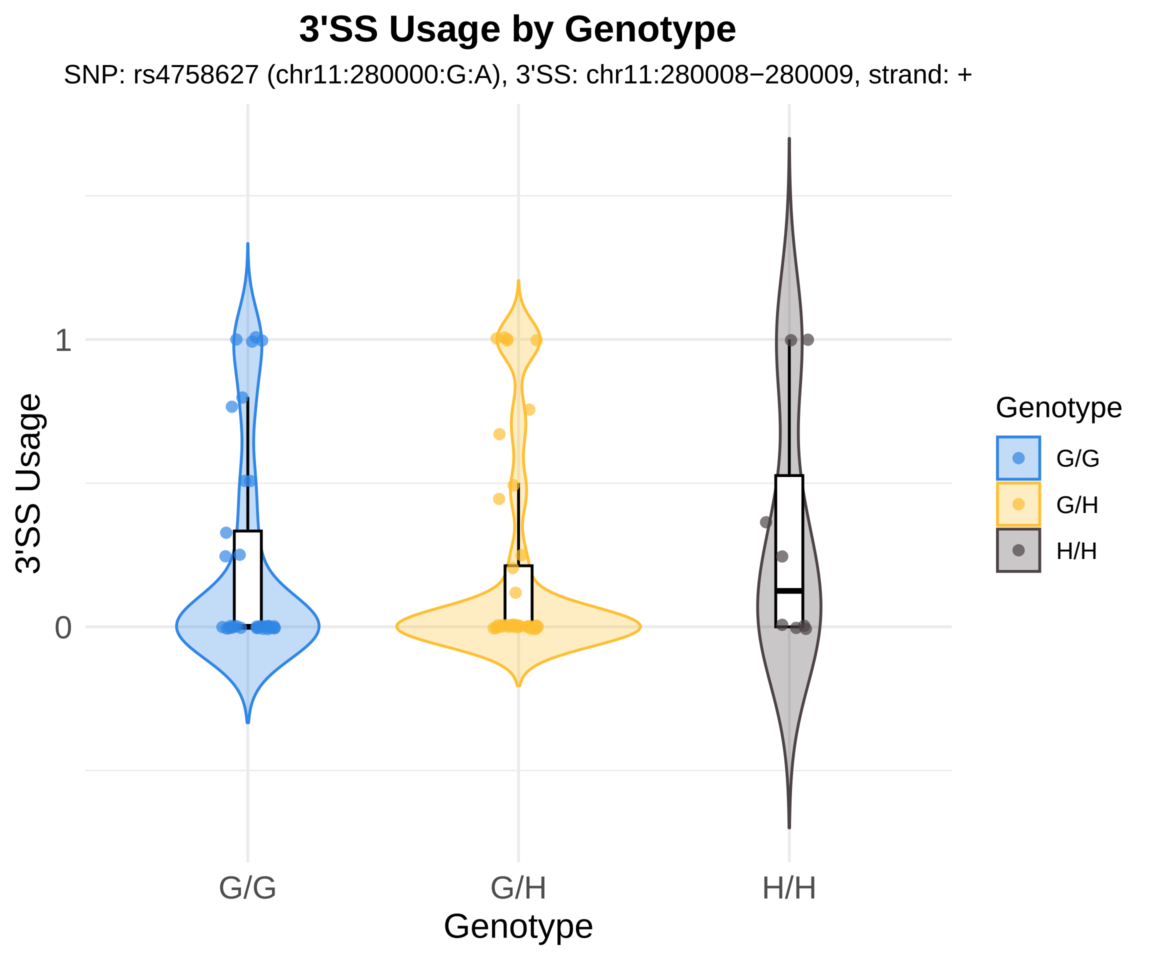
3'SS Usage by Genotype
SNP: rs4758627 (chr11:280000:G:A), 3'SS: chr11:280008−280009, strand: +
3'SS Usage
1
0
G/G
G/H
H/H
Genotype
Genotype
G/G
G/H
H/H

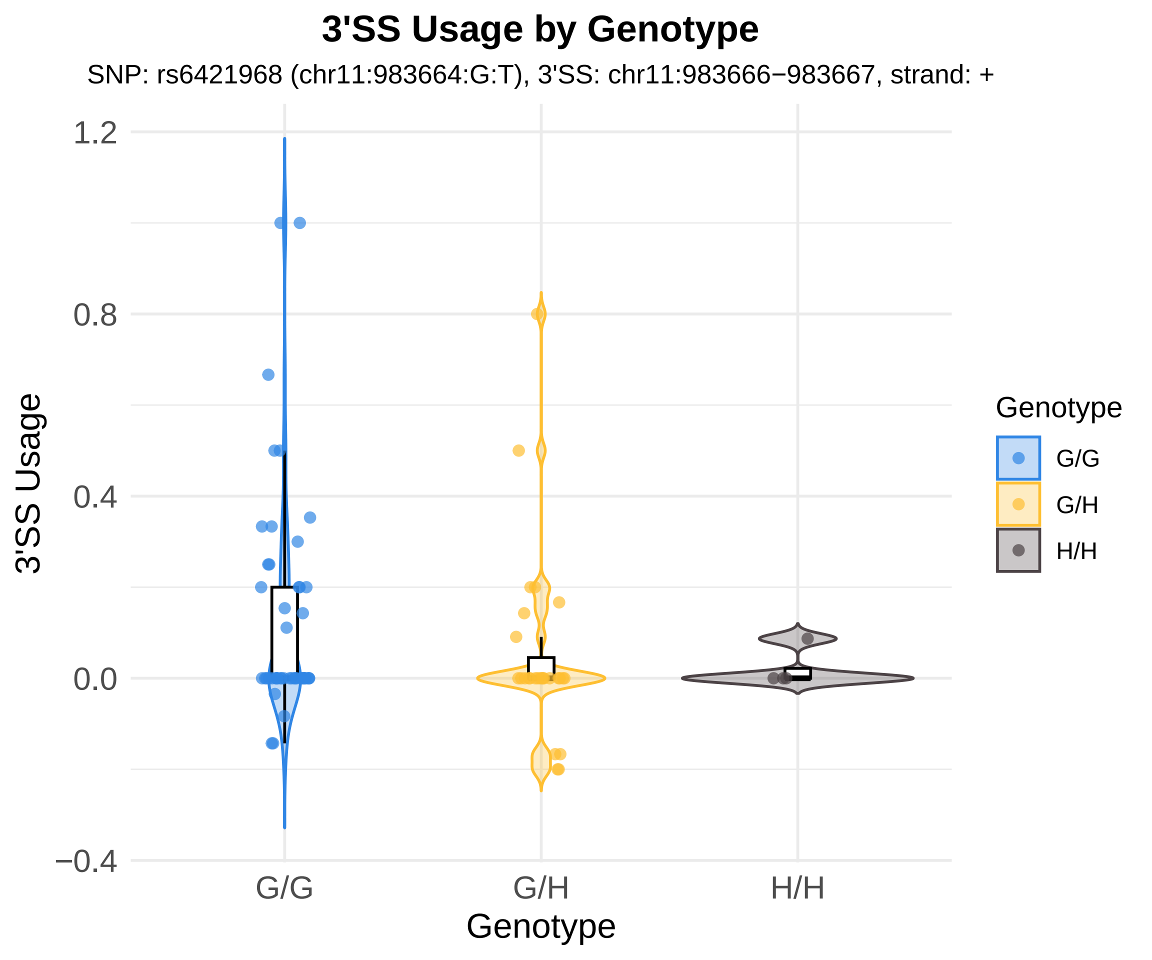
3'SS Usage by Genotype
SNP: rs6421968 (chr11:983664:G:T), 3'SS: chr11:983666−983667, strand: +
3'SS Usage
1.2
0.8
0.4
0.0
−0.4
G/G
G/H
H/H
Genotype
Genotype
G/G
G/H
H/H

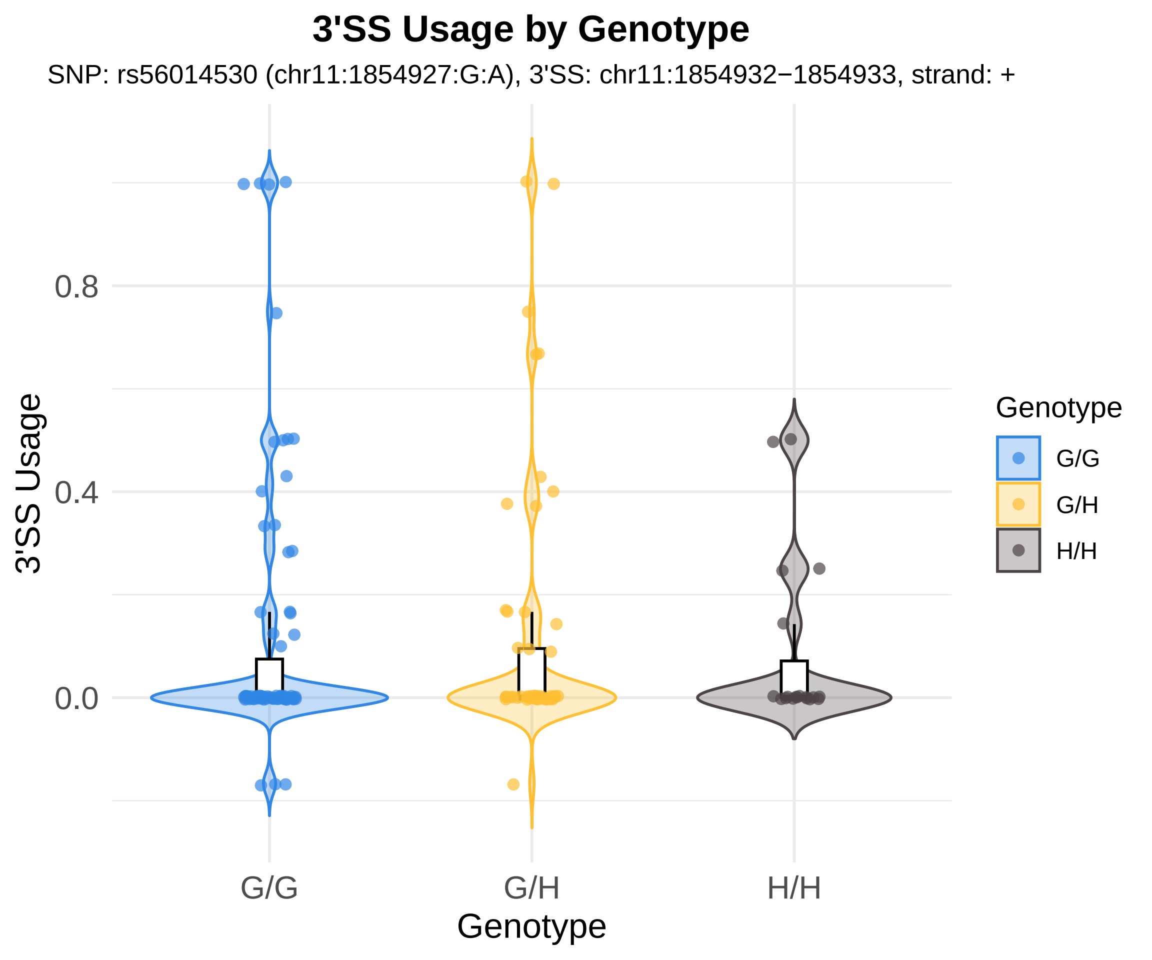
3'SS Usage by Genotype
SNP: rs56014530 (chr11:1854927:G:A), 3'SS: chr11:1854932−1854933, strand: +
3'SS Usage
0.8
0.4
0.0
G/G
G/H
H/H
Genotype
Genotype
G/G
G/H
H/H

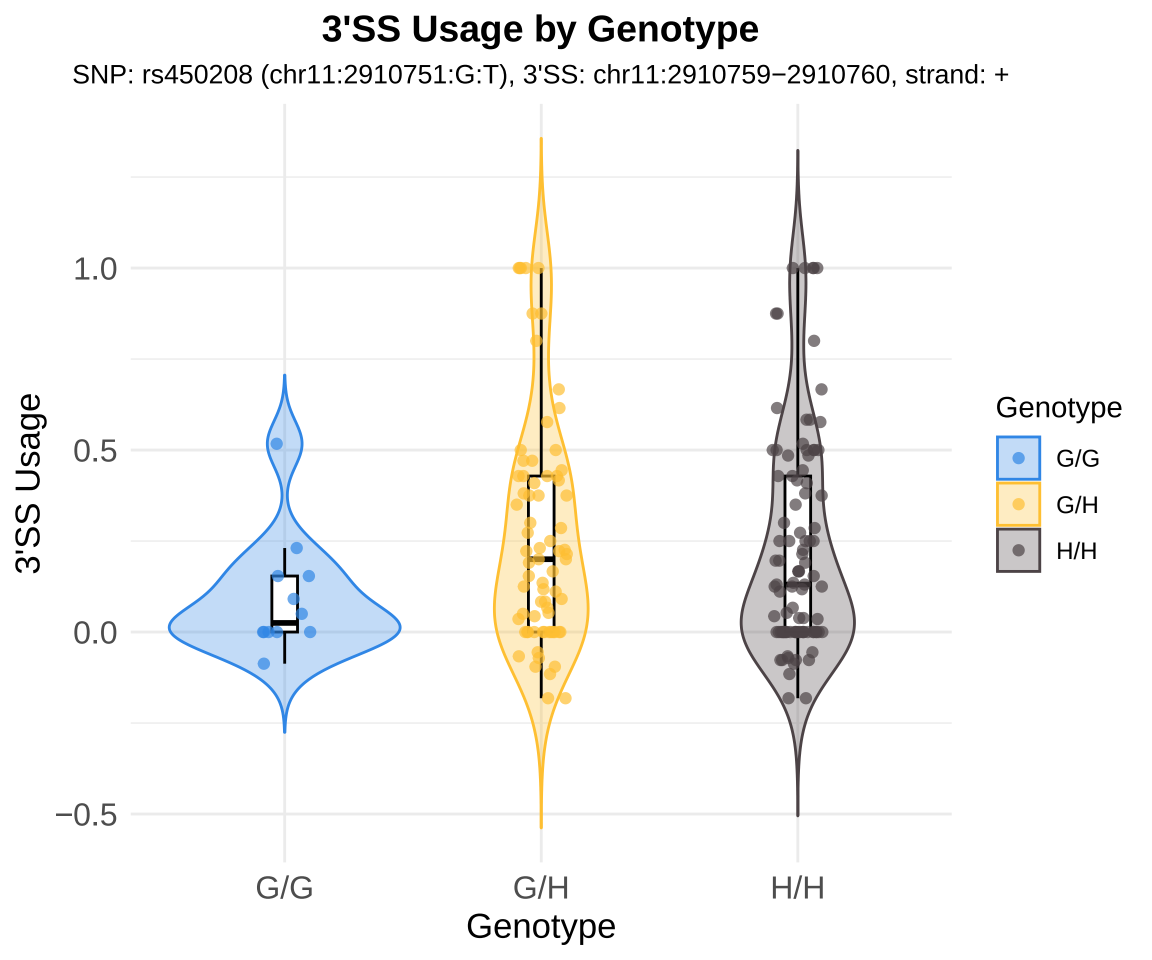
3'SS Usage by Genotype
SNP: rs450208 (chr11:2910751:G:T), 3'SS: chr11:2910759−2910760, strand: +
3'SS Usage
1.0
0.5
0.0
−0.5
G/G
G/H
H/H
Genotype
Genotype
G/G
G/H
H/H

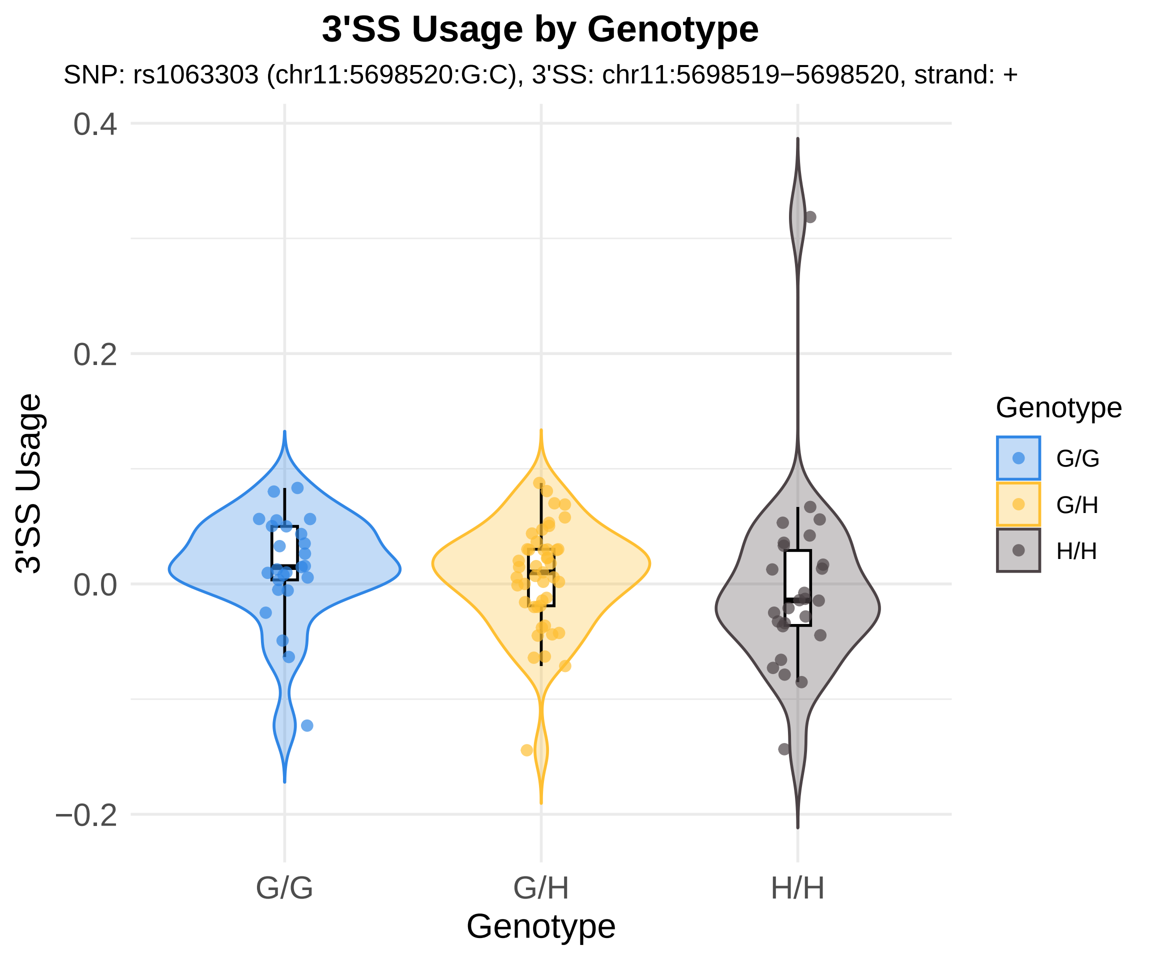
3'SS Usage by Genotype
SNP: rs1063303 (chr11:5698520:G:C), 3'SS: chr11:5698519−5698520, strand: +
3'SS Usage
0.4
0.2
0.0
−0.2
G/G
G/H
H/H
Genotype
Genotype
G/G
G/H
H/H

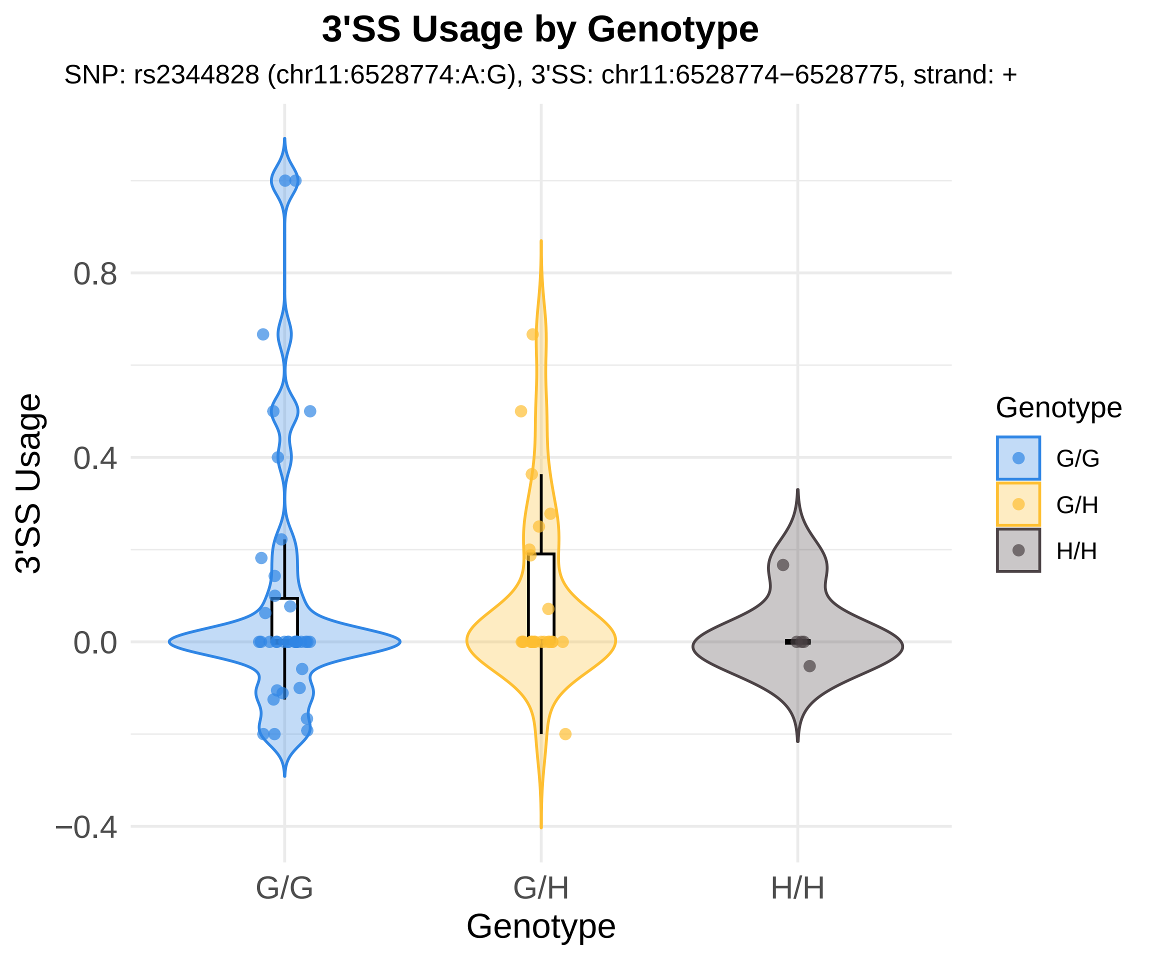
3'SS Usage by Genotype
SNP: rs2344828 (chr11:6528774:A:G), 3'SS: chr11:6528774−6528775, strand: +
3'SS Usage
0.8
0.4
0.0
−0.4
G/G
G/H
H/H
Genotype
Genotype
G/G
G/H
H/H

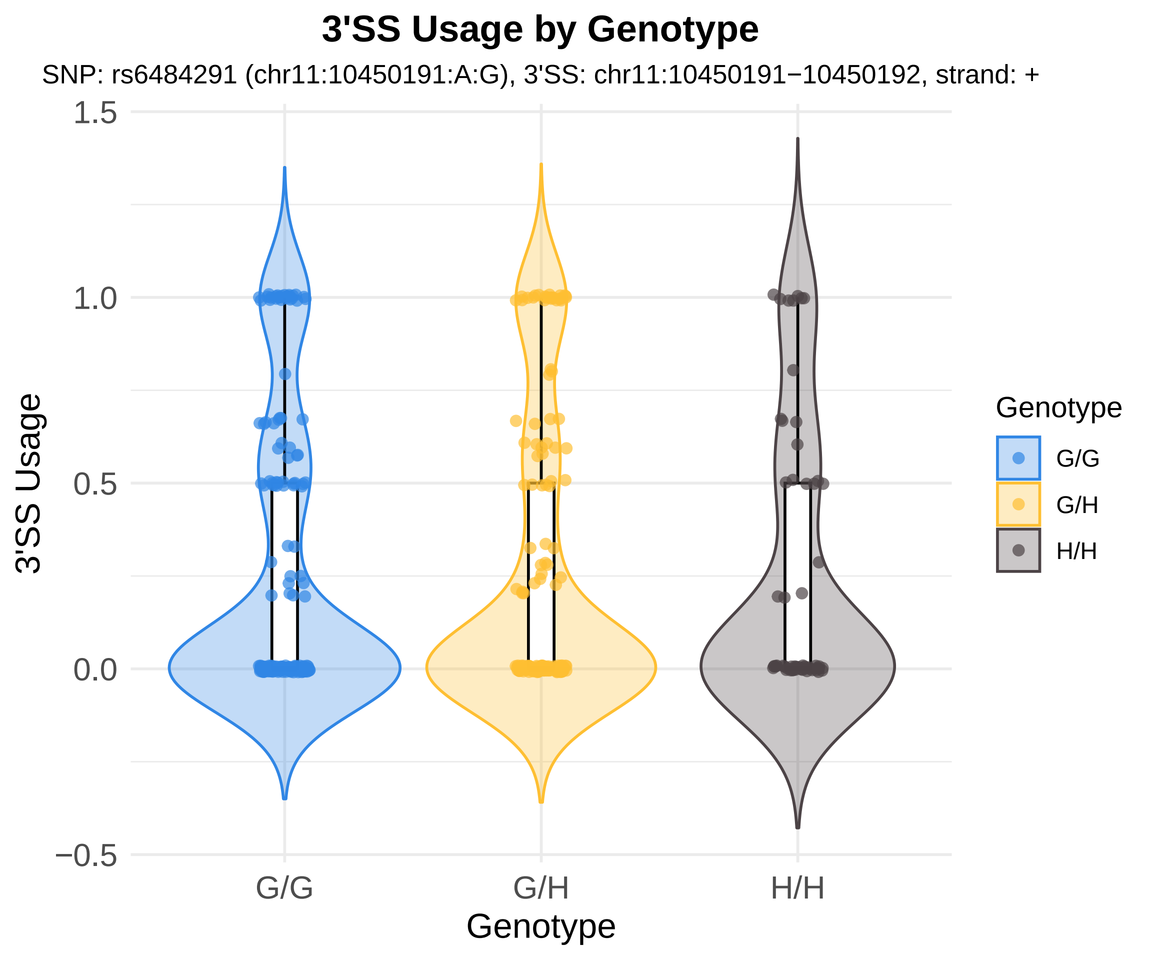

3'SS Usage by Genotype
SNP: rs6484291 (chr11:10450191:A:G), 3'SS: chr11:10450191−10450192, strand: +
3'SS Usage
1.5
1.0
0.5
0.0
−0.5
G/G
G/H
H/H
Genotype
Genotype
G/G
G/H
H/H

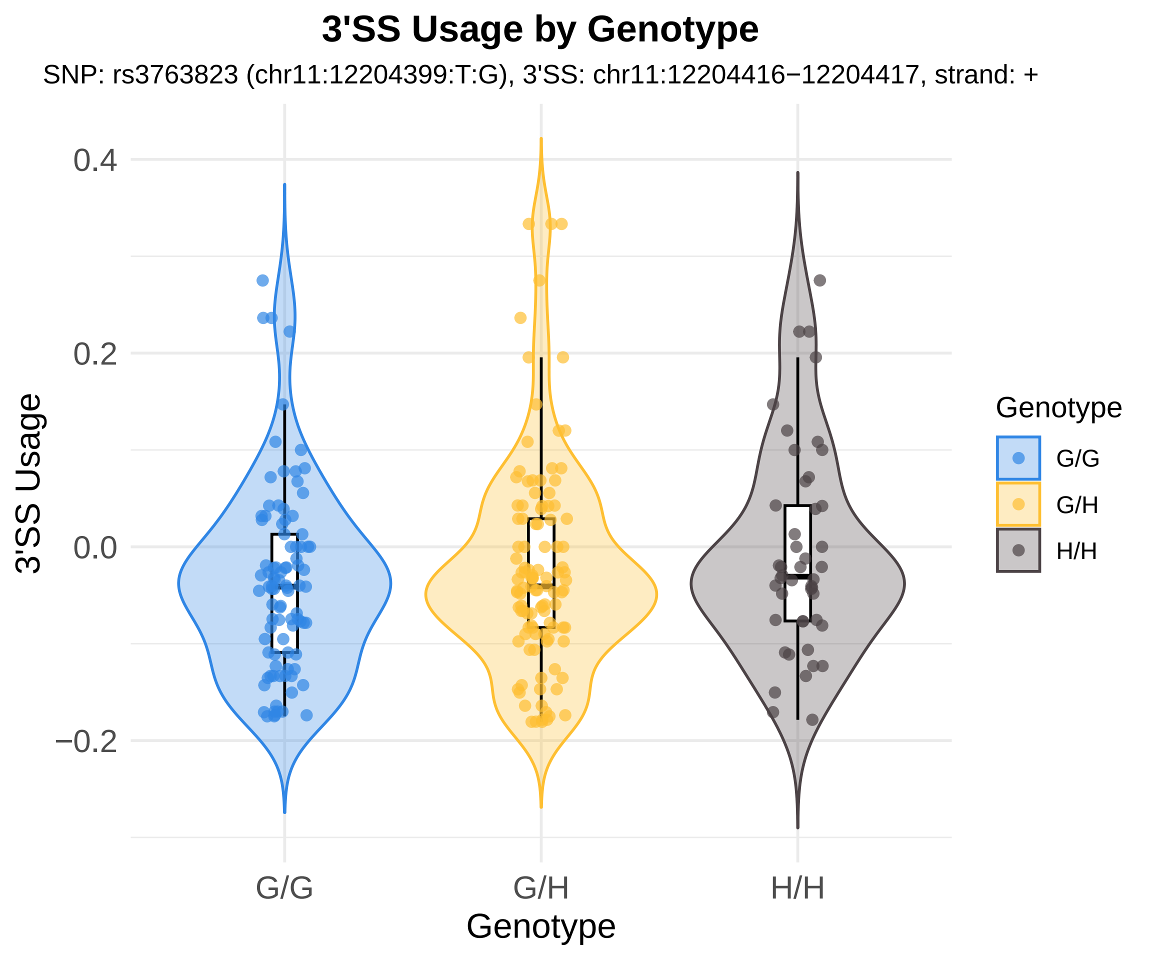
3'SS Usage by Genotype
SNP: rs3763823 (chr11:12204399:T:G), 3'SS: chr11:12204416−12204417, strand: +
3'SS Usage
0.4
0.2
0.0
−0.2
G/G
G/H
H/H
Genotype
Genotype
G/G
G/H
H/H

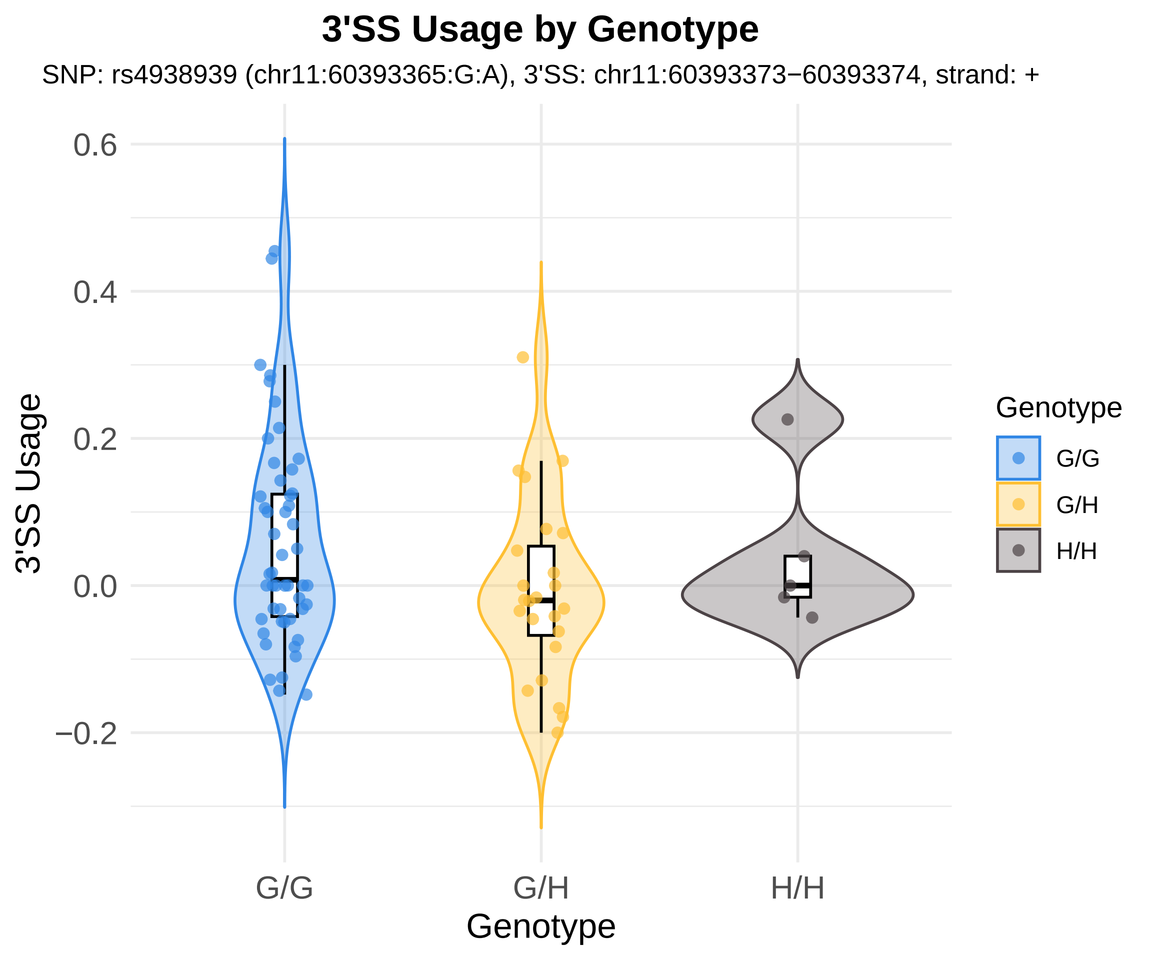
3'SS Usage by Genotype
SNP: rs4938939 (chr11:60393365:G:A), 3'SS: chr11:60393373−60393374, strand: +
3'SS Usage
0.6
0.4
0.2
0.0
−0.2
G/G
G/H
H/H
Genotype
Genotype
G/G
G/H
H/H

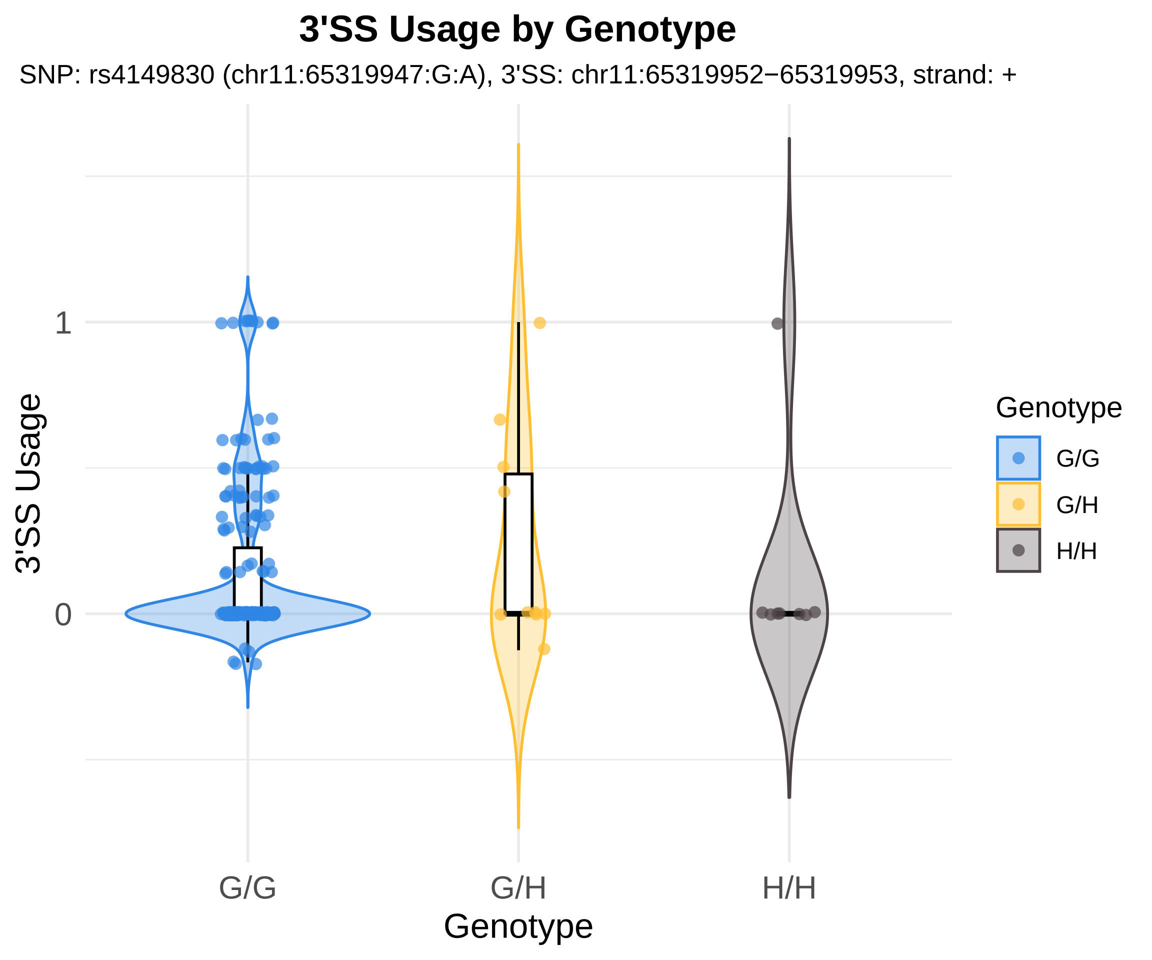
3'SS Usage by Genotype
SNP: rs4149830 (chr11:65319947:G:A), 3'SS: chr11:65319952−65319953, strand: +
3'SS Usage
1
0
G/G
G/H
H/H
Genotype
Genotype
G/G
G/H
H/H

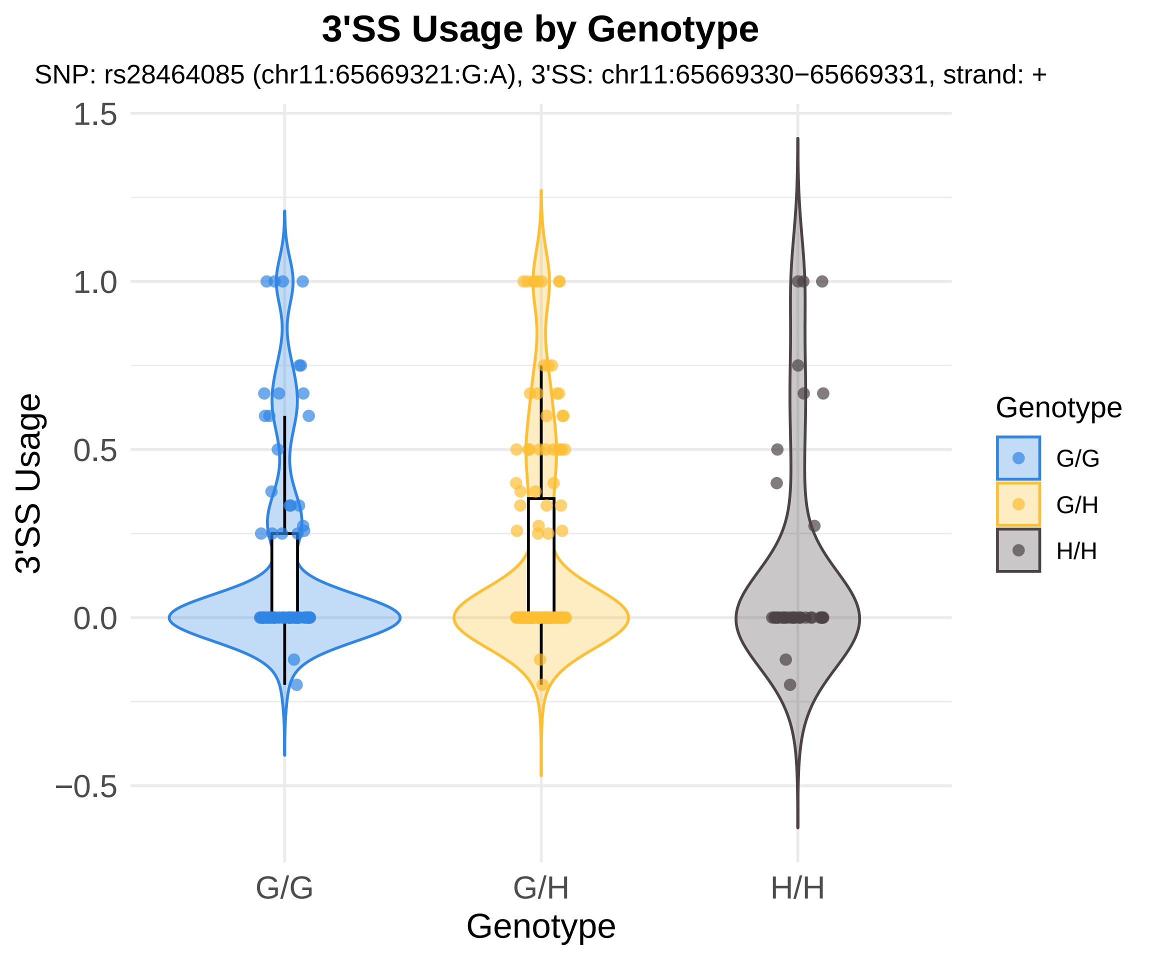

3'SS Usage by Genotype
SNP: rs28464085 (chr11:65669321:G:A), 3'SS: chr11:65669330−65669331, strand: +
1.5
1.0
0.5
0.0
−0.5
3'SS Usage
G/G
G/H
H/H
Genotype
Genotype
G/G
G/H
H/H

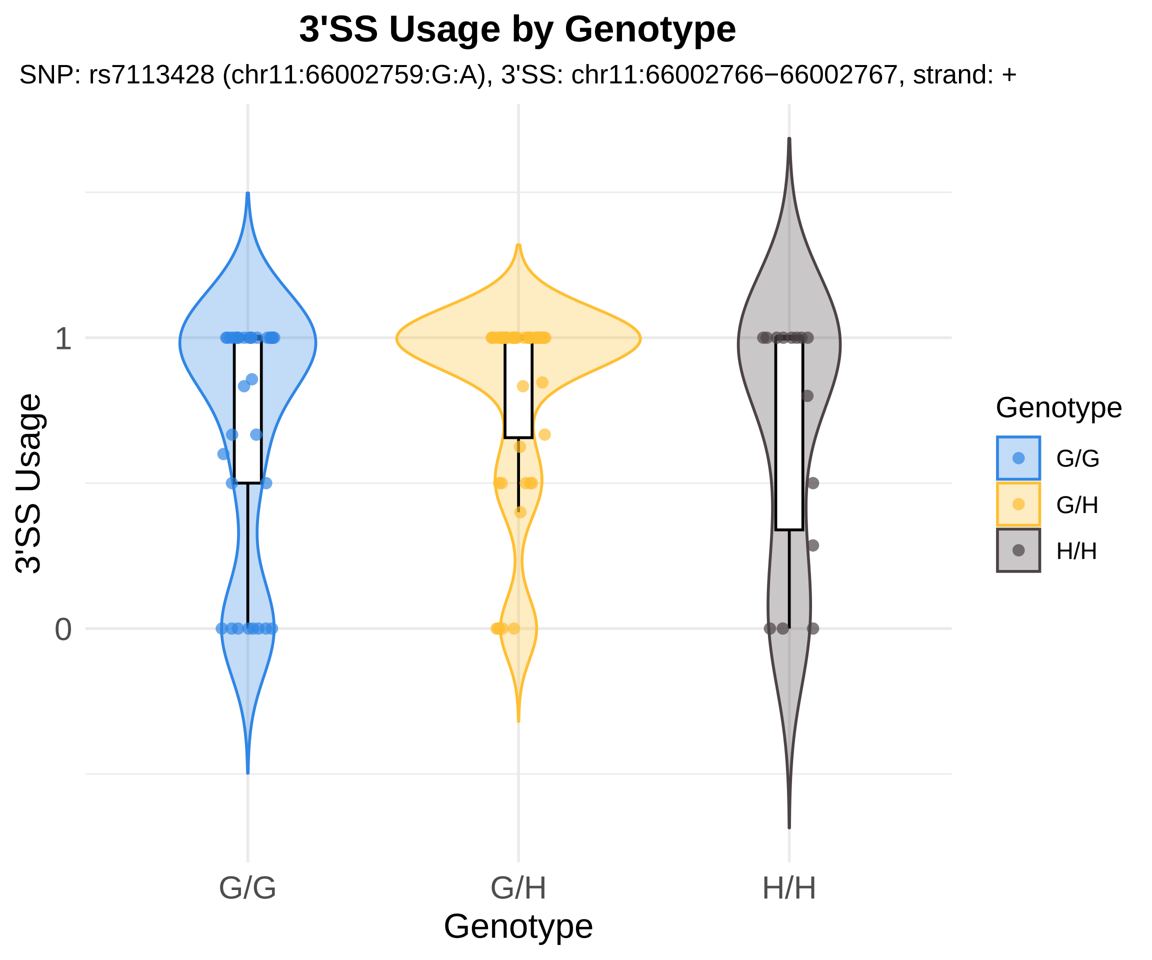
3'SS Usage by Genotype
SNP: rs7113428 (chr11:66002759:G:A), 3'SS: chr11:66002766−66002767, strand: +
3'SS Usage
1
0
G/G
G/H
H/H
Genotype
Genotype
G/G
G/H
H/H

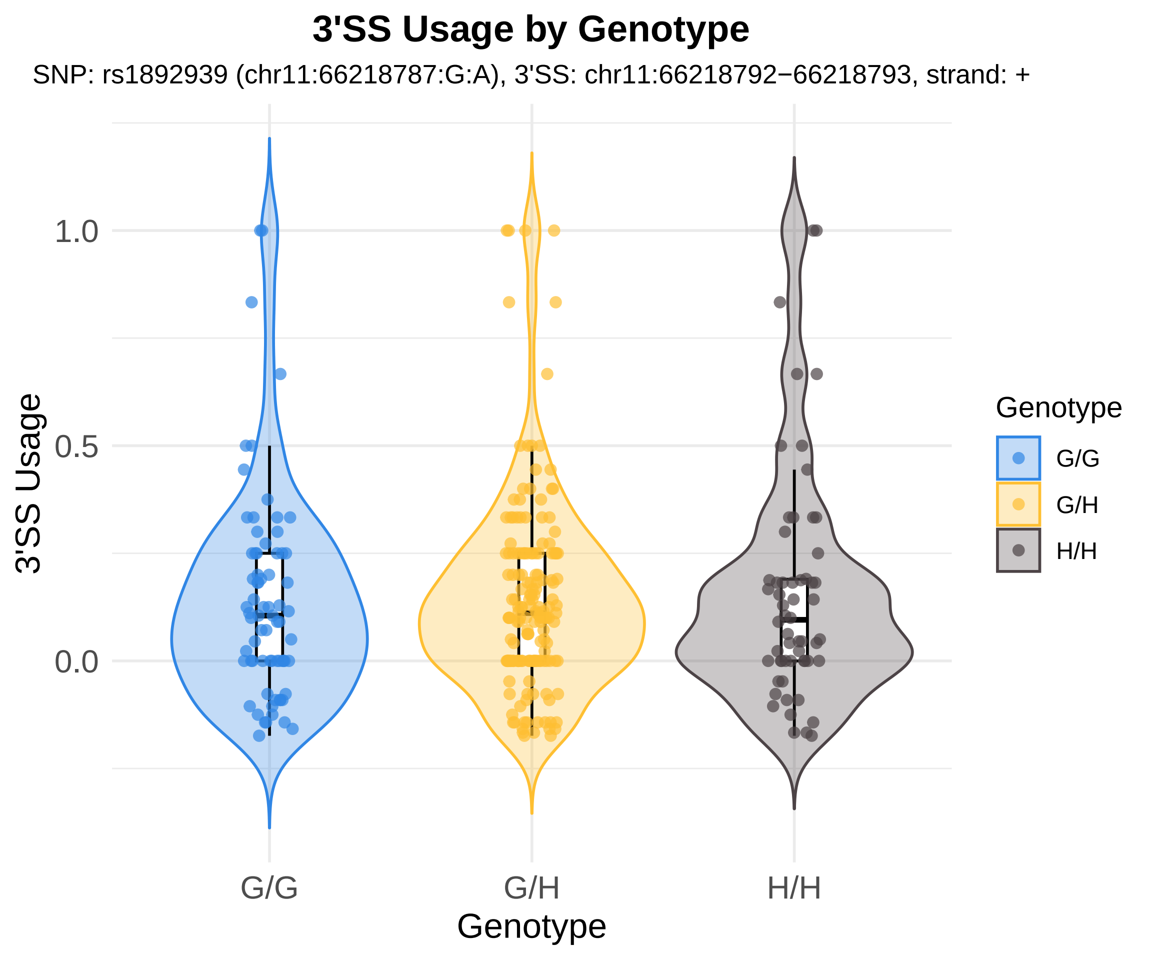
3'SS Usage by Genotype
SNP: rs1892939 (chr11:66218787:G:A), 3'SS: chr11:66218792−66218793, strand: +
3'SS Usage
1.0
0.5
0.0
G/G
G/H
H/H
Genotype
Genotype
G/G
G/H
H/H

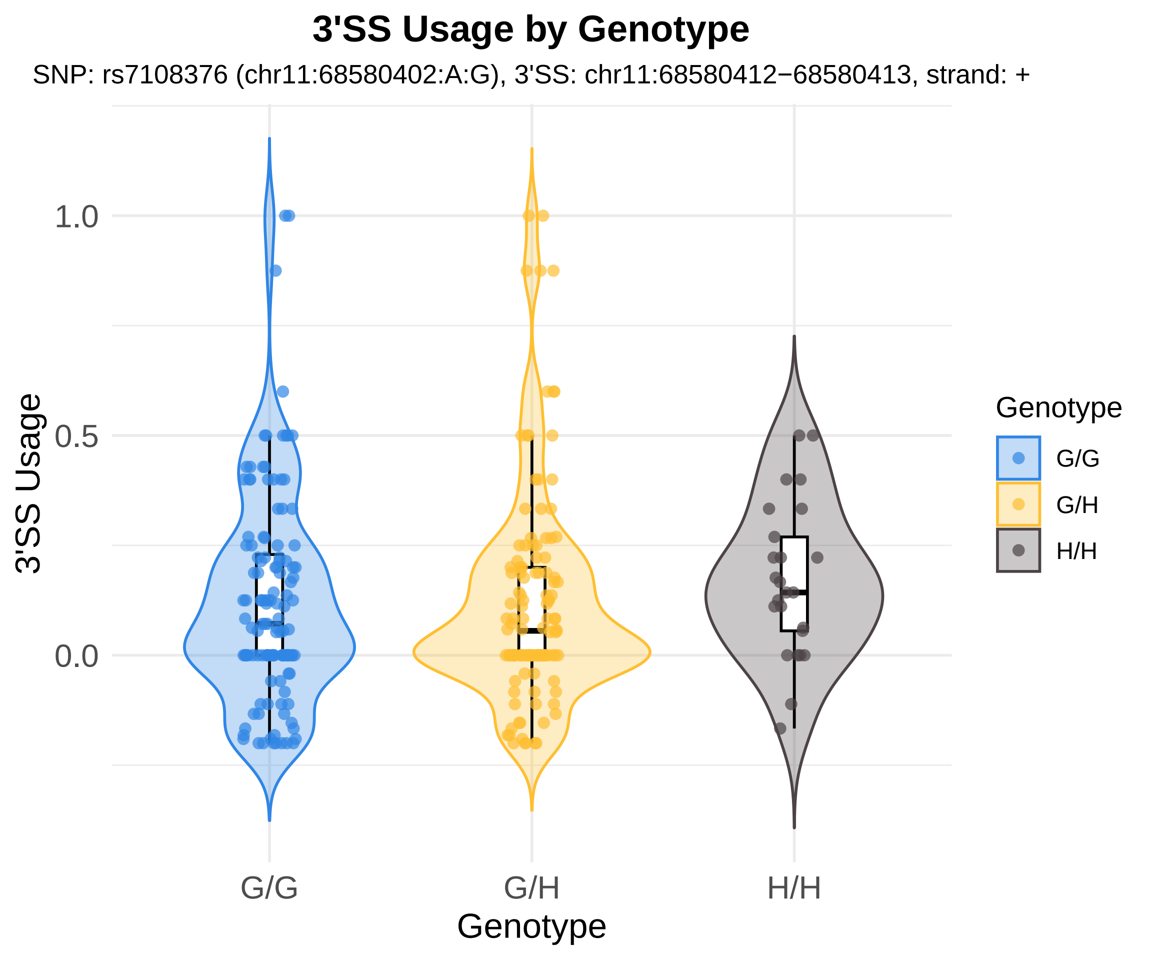
3'SS Usage by Genotype
SNP: rs7108376 (chr11:68580402:A:G), 3'SS: chr11:68580412−68580413, strand: +
1.0
0.5
0.0
3'SS Usage
G/G
G/H
H/H
Genotype
Genotype
G/G
G/H
H/H

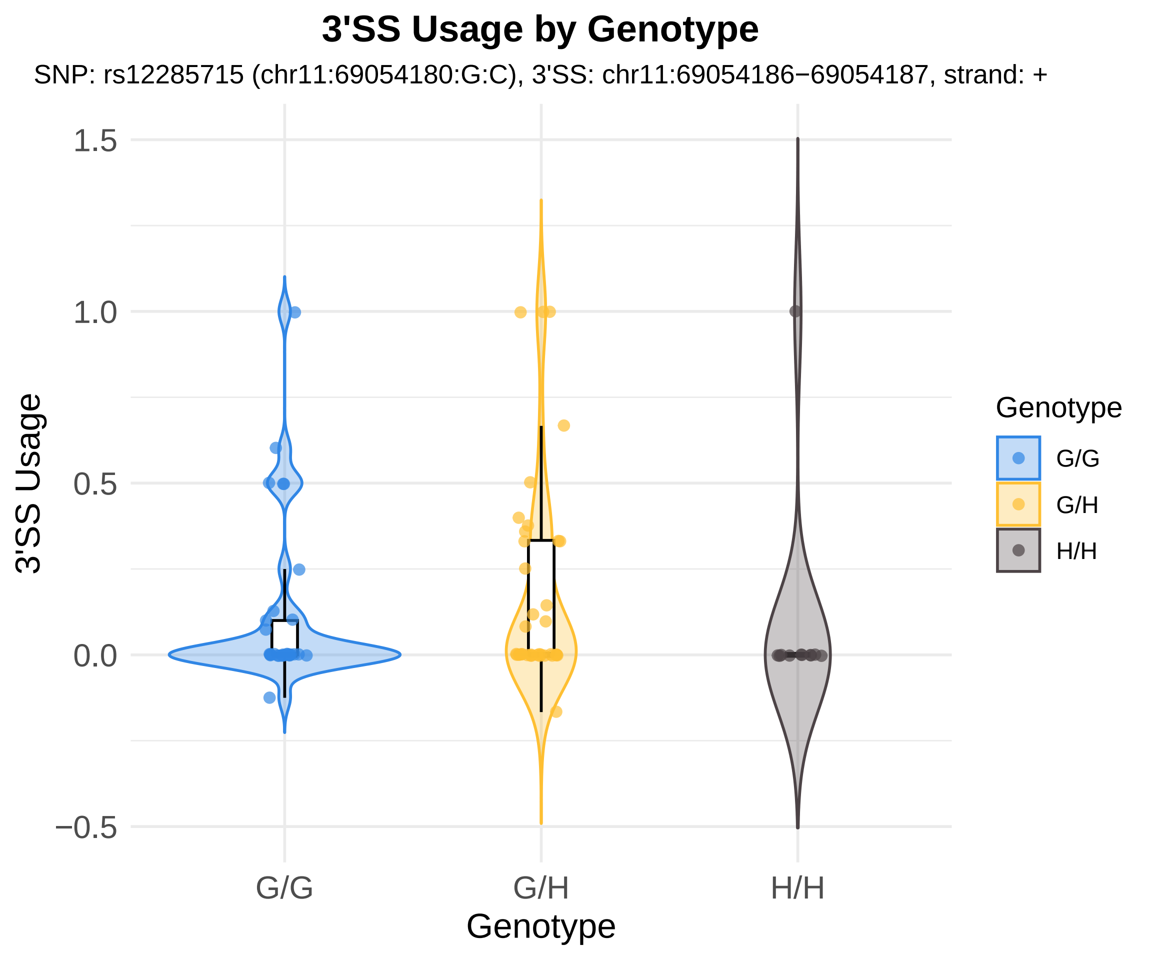
3'SS Usage by Genotype
SNP: rs12285715 (chr11:69054180:G:C), 3'SS: chr11:69054186−69054187, strand: +
3'SS Usage
1.5
1.0
0.5
0.0
−0.5
G/G
G/H
H/H
Genotype
Genotype
G/G
G/H
H/H

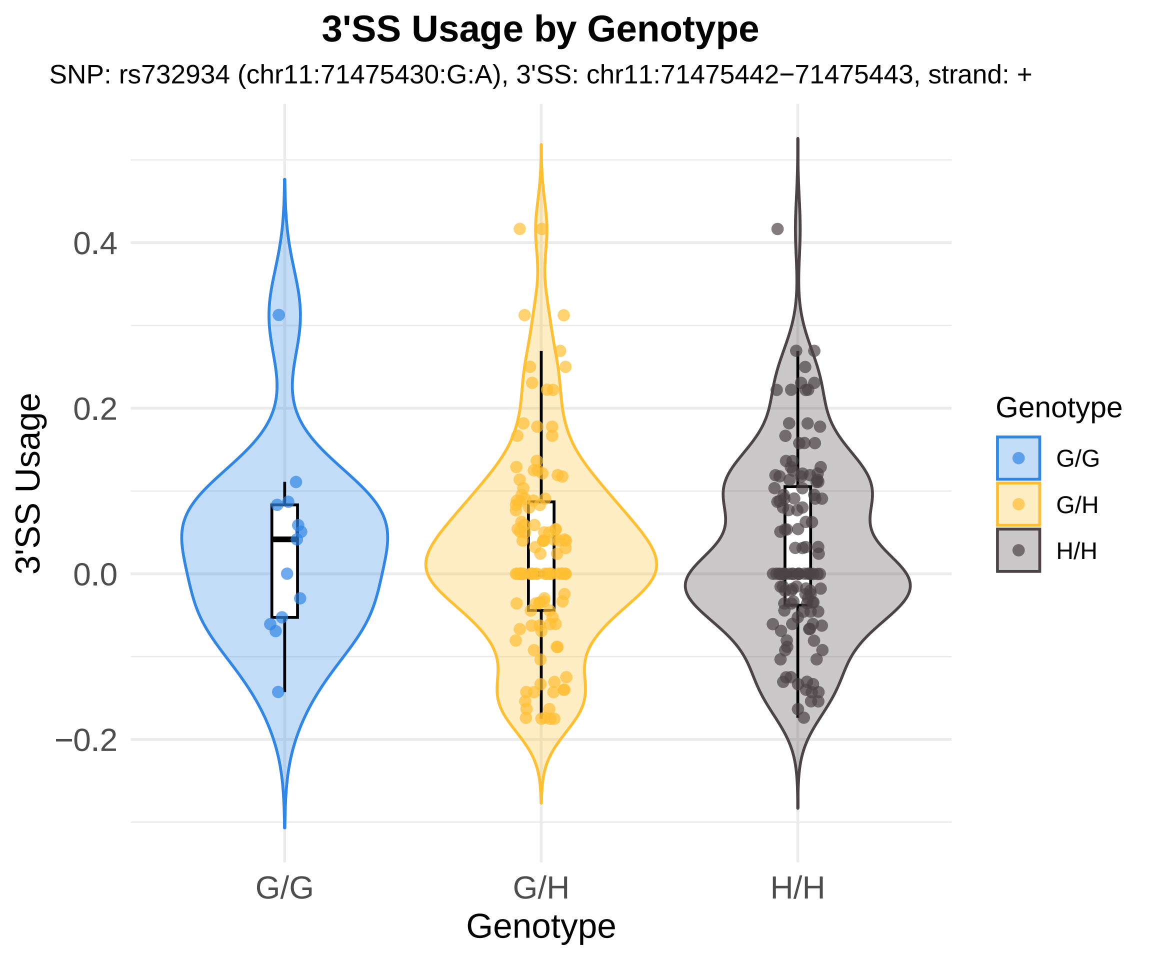
3'SS Usage by Genotype
SNP: rs732934 (chr11:71475430:G:A), 3'SS: chr11:71475442−71475443, strand: +
3'SS Usage
0.4
0.2
0.0
−0.2
G/G
G/H
H/H
Genotype
Genotype
G/G
G/H
H/H

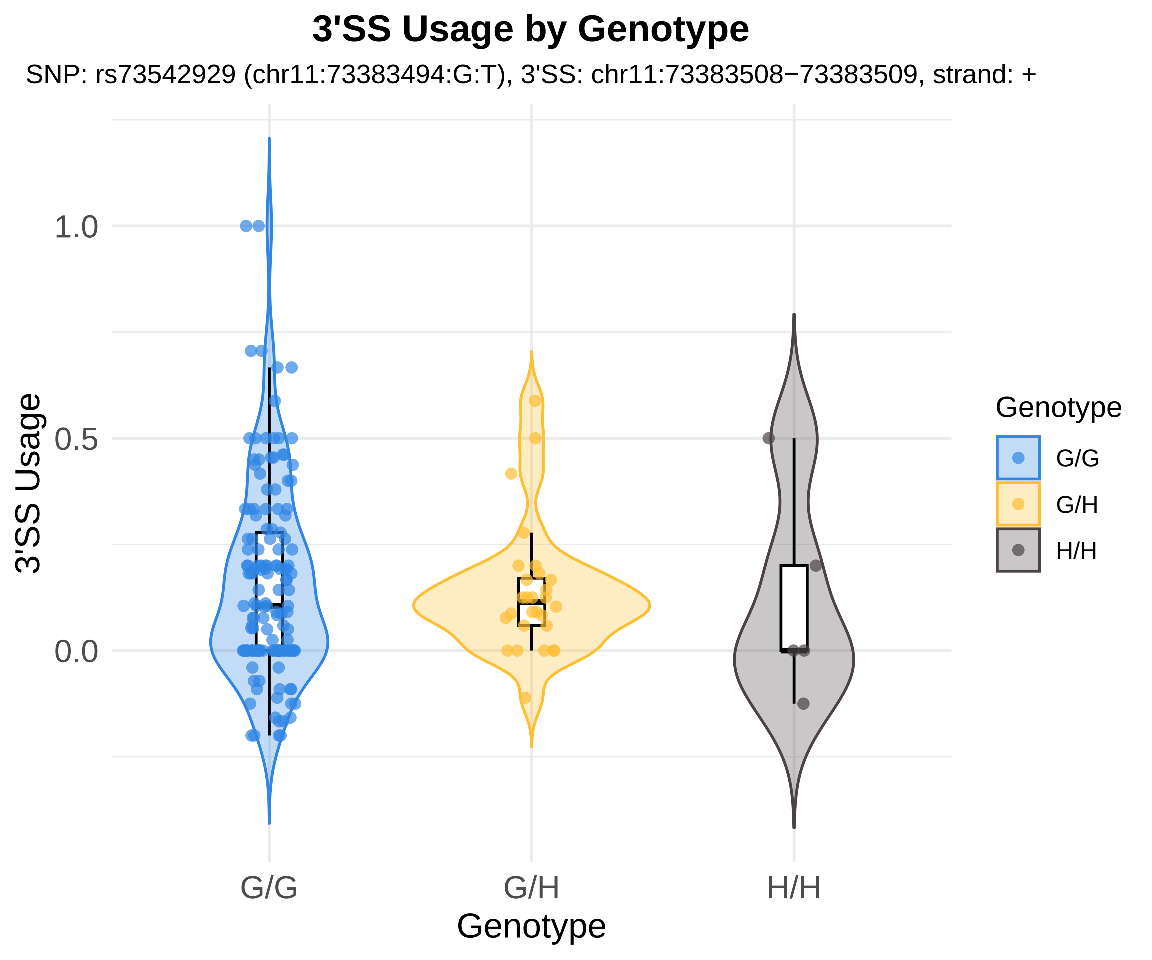

3'SS Usage by Genotype
SNP: rs73542929 (chr11:73383494:G:T), 3'SS: chr11:73383508−73383509, strand: +
3'SS Usage
1.0
0.5
0.0
G/G
G/H
H/H
Genotype
Genotype
G/G
G/H
H/H

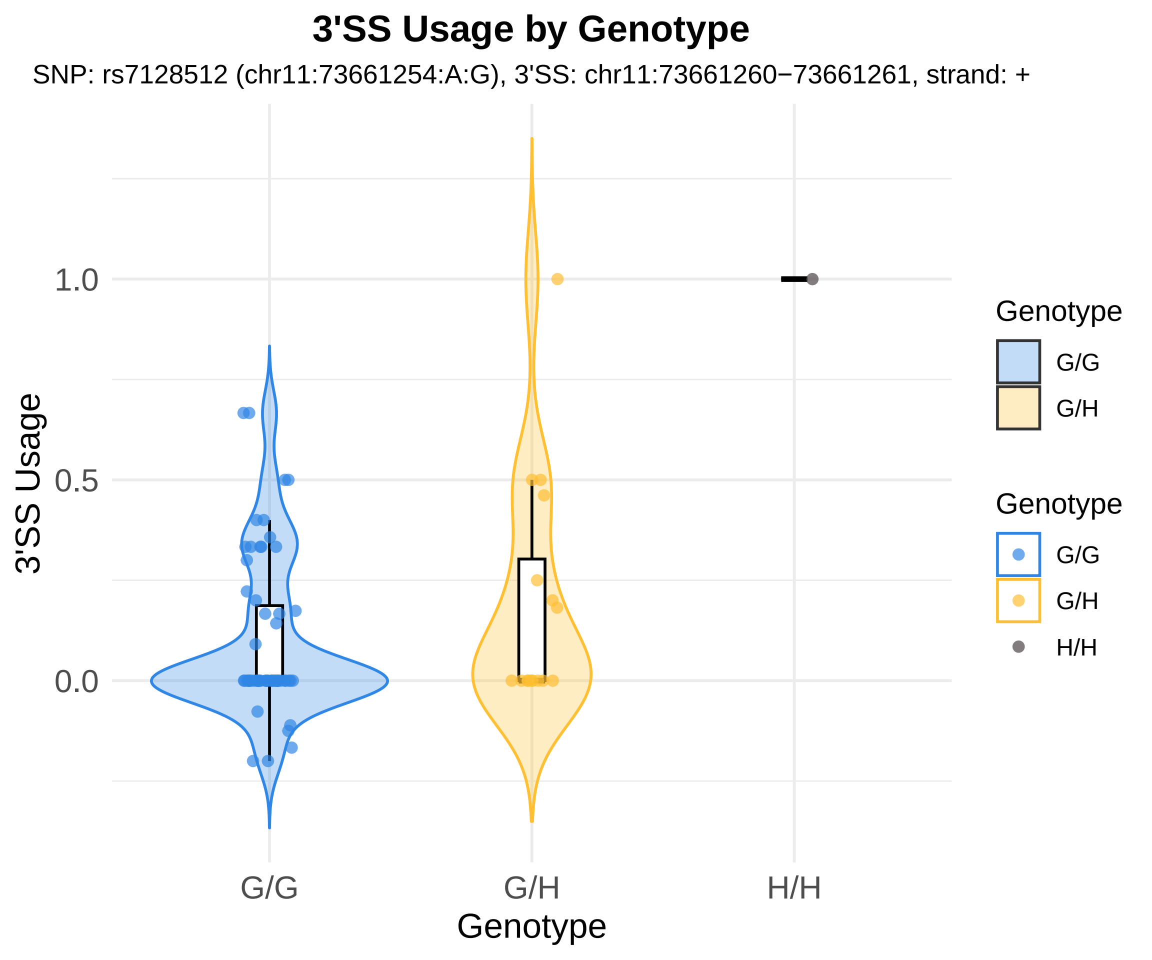
3'SS Usage by Genotype
SNP: rs7128512 (chr11:73661254:A:G), 3'SS: chr11:73661260−73661261, strand: +
3'SS Usage
1.0
0.5
0.0
G/G
G/H
H/H
Genotype
Genotype
G/G
G/H
Genotype
G/G
G/H
H/H

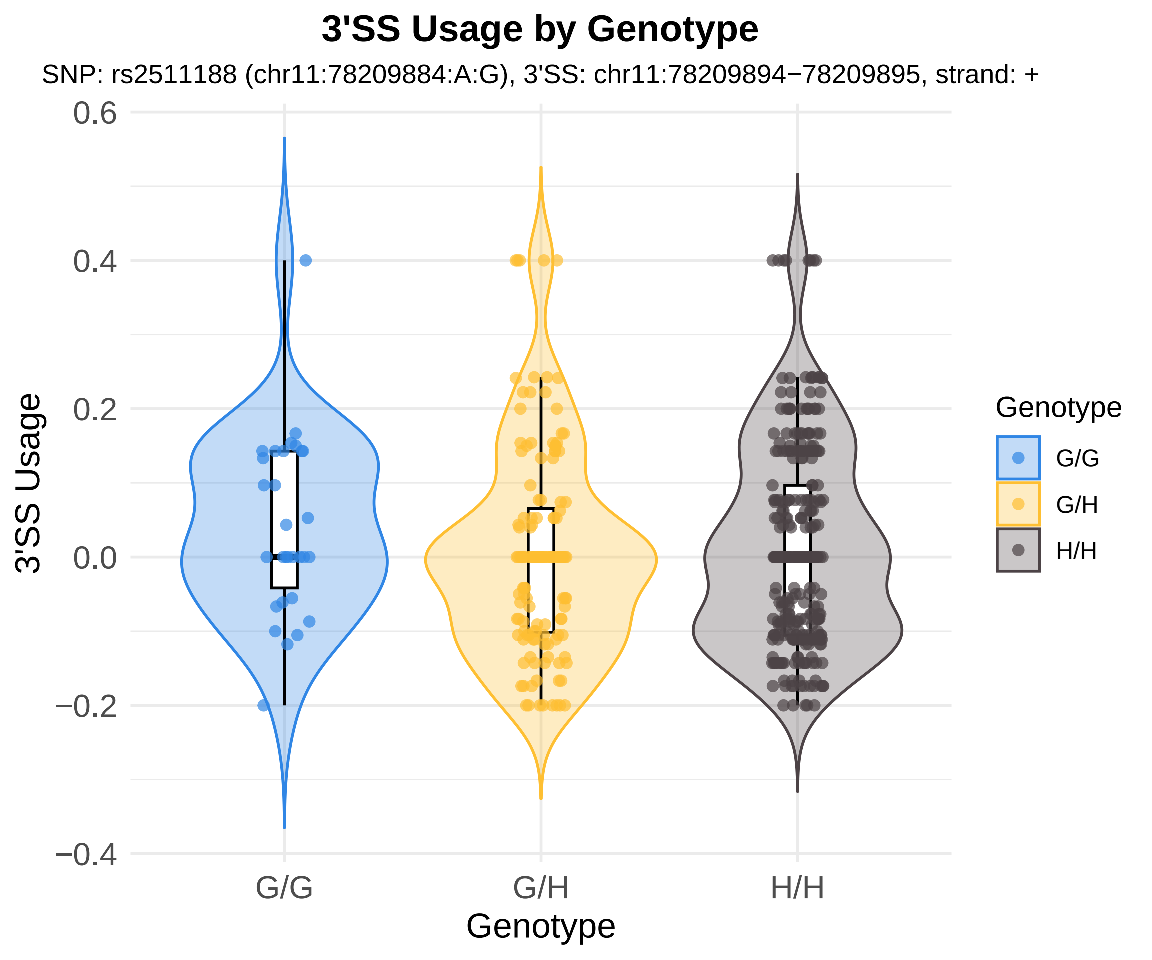

3'SS Usage by Genotype
SNP: rs2511188 (chr11:78209884:A:G), 3'SS: chr11:78209894−78209895, strand: +
0.6
0.4
0.2
0.0
−0.2
−0.4
3'SS Usage
G/G
G/H
H/H
Genotype
Genotype
G/G
G/H
H/H

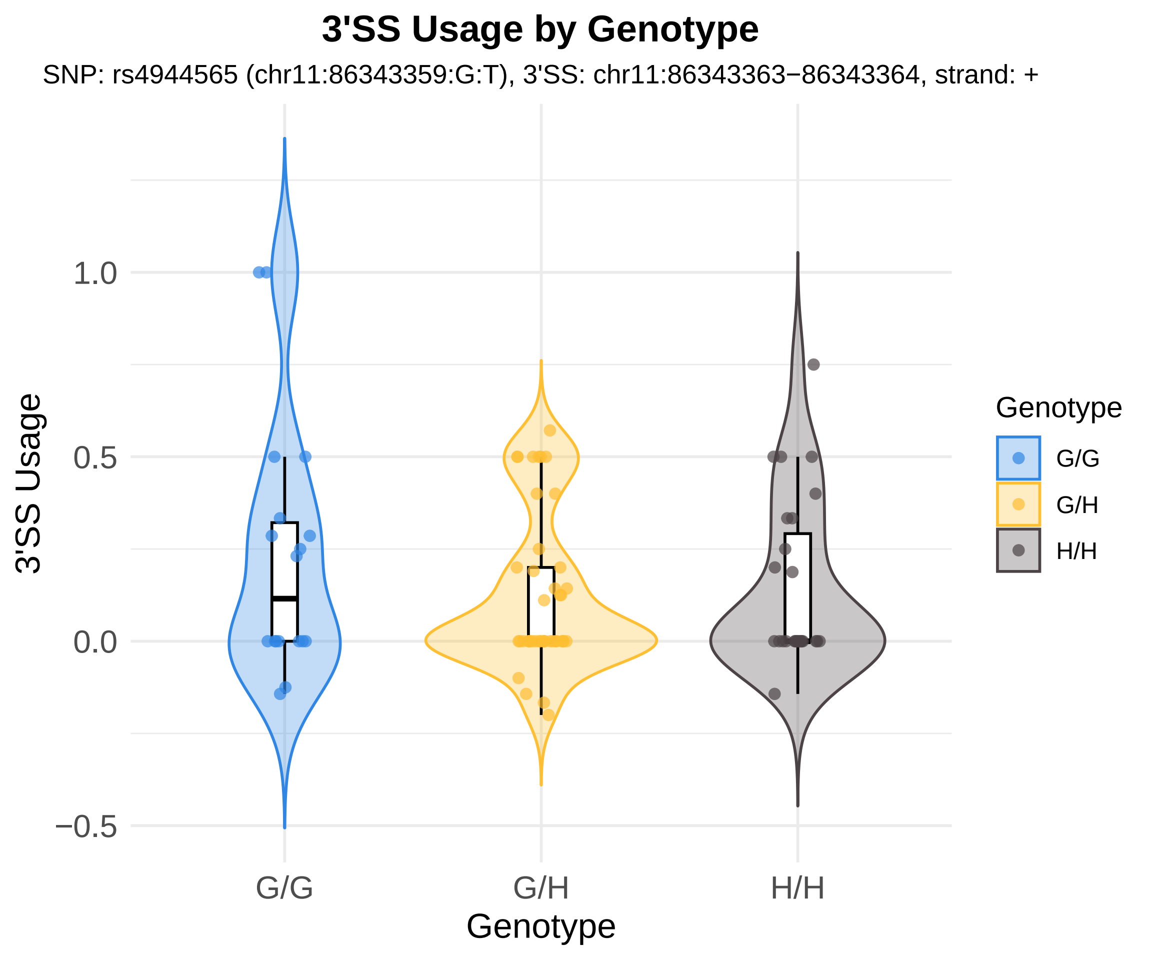
3'SS Usage by Genotype
SNP: rs4944565 (chr11:86343359:G:T), 3'SS: chr11:86343363−86343364, strand: +
3'SS Usage
1.0
0.5
0.0
−0.5
G/G
G/H
H/H
Genotype
Genotype
G/G
G/H
H/H

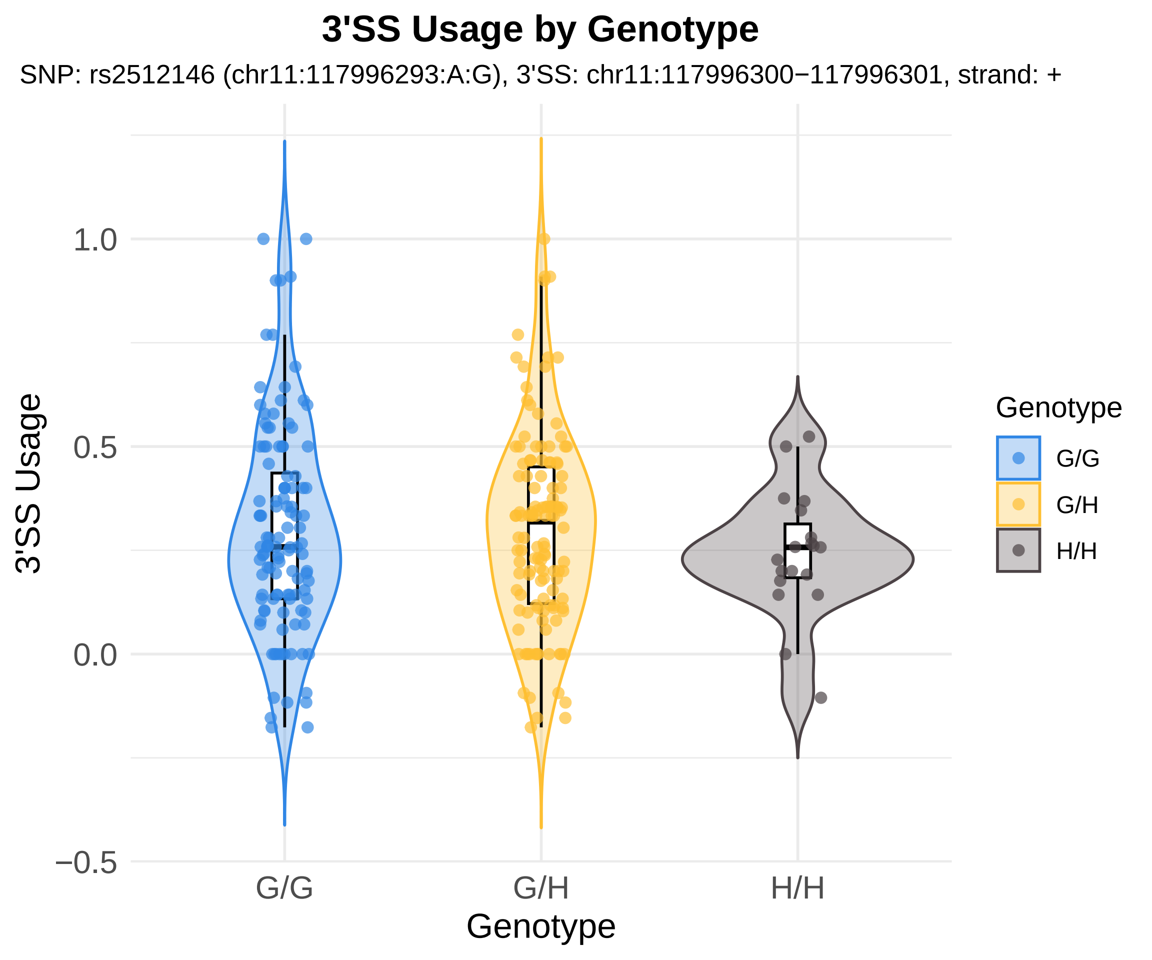
3'SS Usage by Genotype
SNP: rs2512146 (chr11:117996293:A:G), 3'SS: chr11:117996300−117996301, strand: +
3'SS Usage
1.0
0.5
0.0
−0.5
G/G
G/H
H/H
Genotype
Genotype
G/G
G/H
H/H

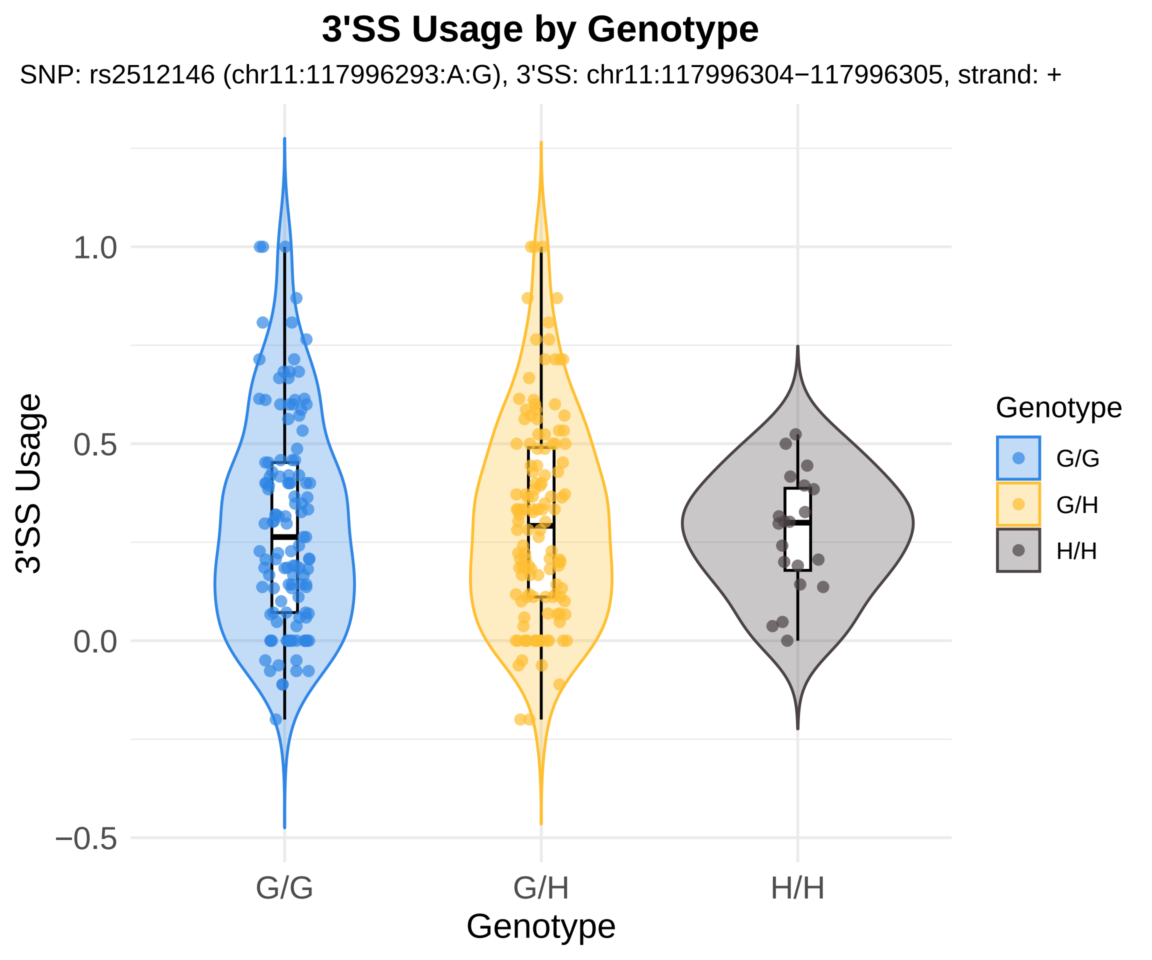
3'SS Usage by Genotype
SNP: rs2512146 (chr11:117996293:A:G), 3'SS: chr11:117996304−117996305, strand: +
3'SS Usage
1.0
0.5
0.0
−0.5
G/G
G/H
H/H
Genotype
Genotype
G/G
G/H
H/H

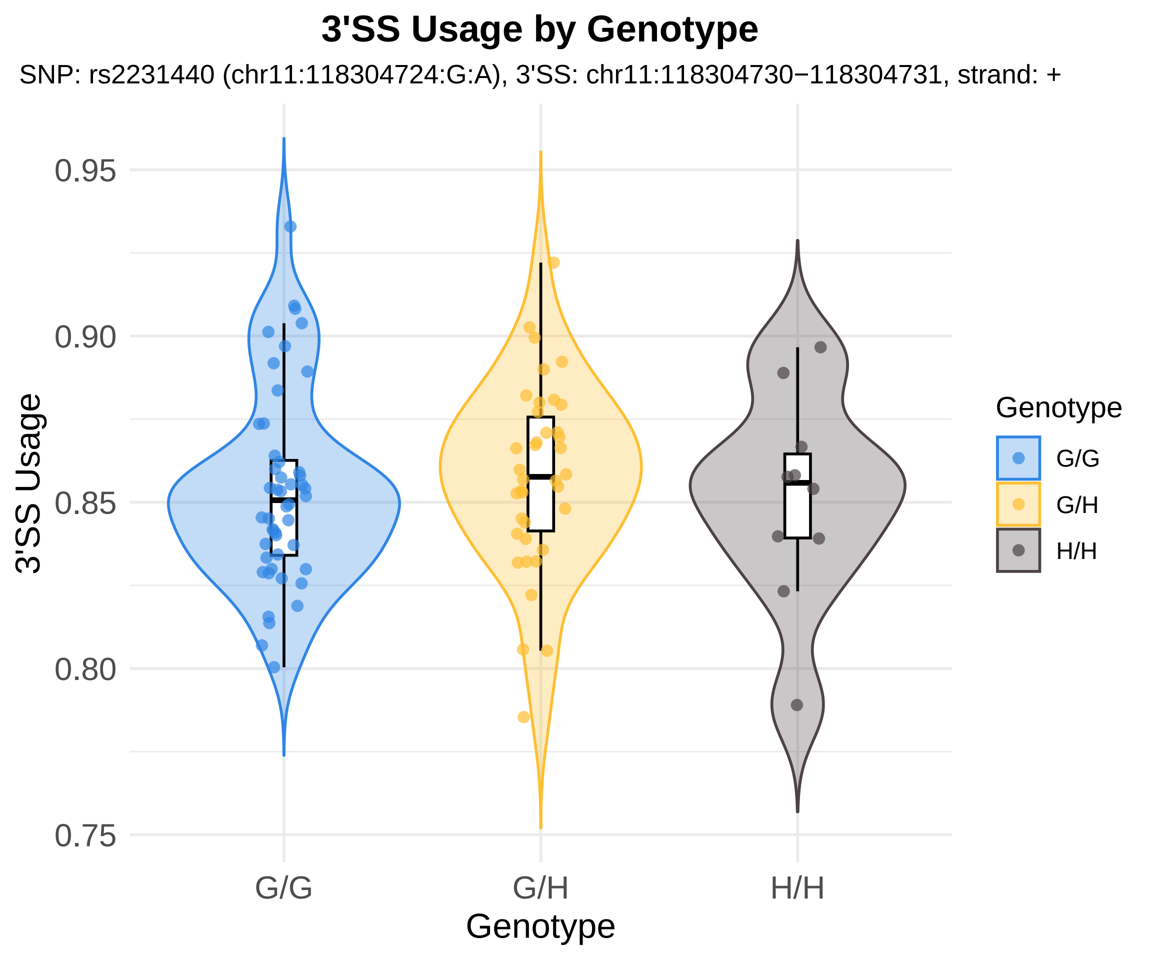
3'SS Usage by Genotype
SNP: rs2231440 (chr11:118304724:G:A), 3'SS: chr11:118304730−118304731, strand: +
3'SS Usage
0.95
0.90
0.85
0.80
0.75
G/G
G/H
H/H
Genotype
Genotype
G/G
G/H
H/H

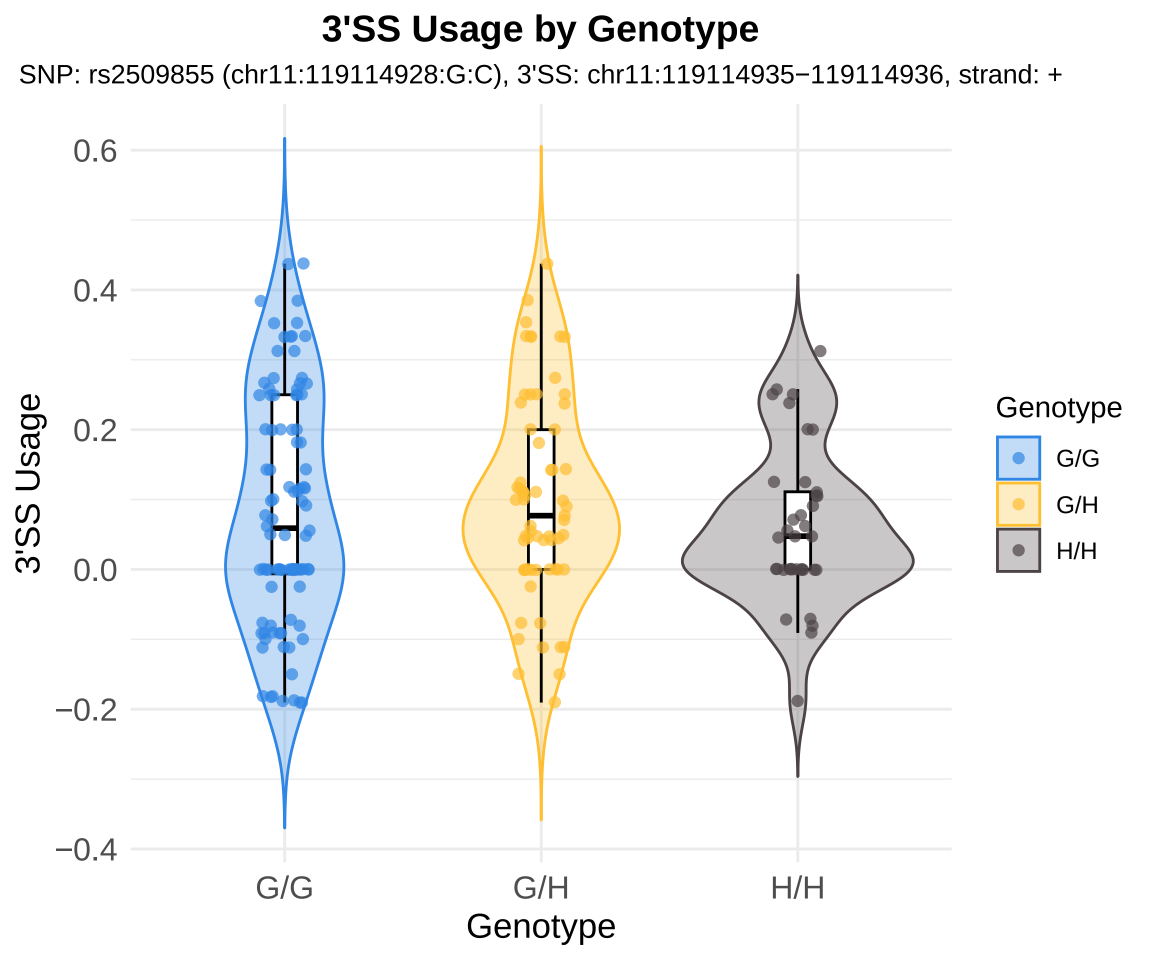
3'SS Usage by Genotype
SNP: rs2509855 (chr11:119114928:G:C), 3'SS: chr11:119114935−119114936, strand: +
0.6
0.4
0.2
0.0
−0.2
−0.4
3'SS Usage
G/G
G/H
H/H
Genotype
Genotype
G/G
G/H
H/H

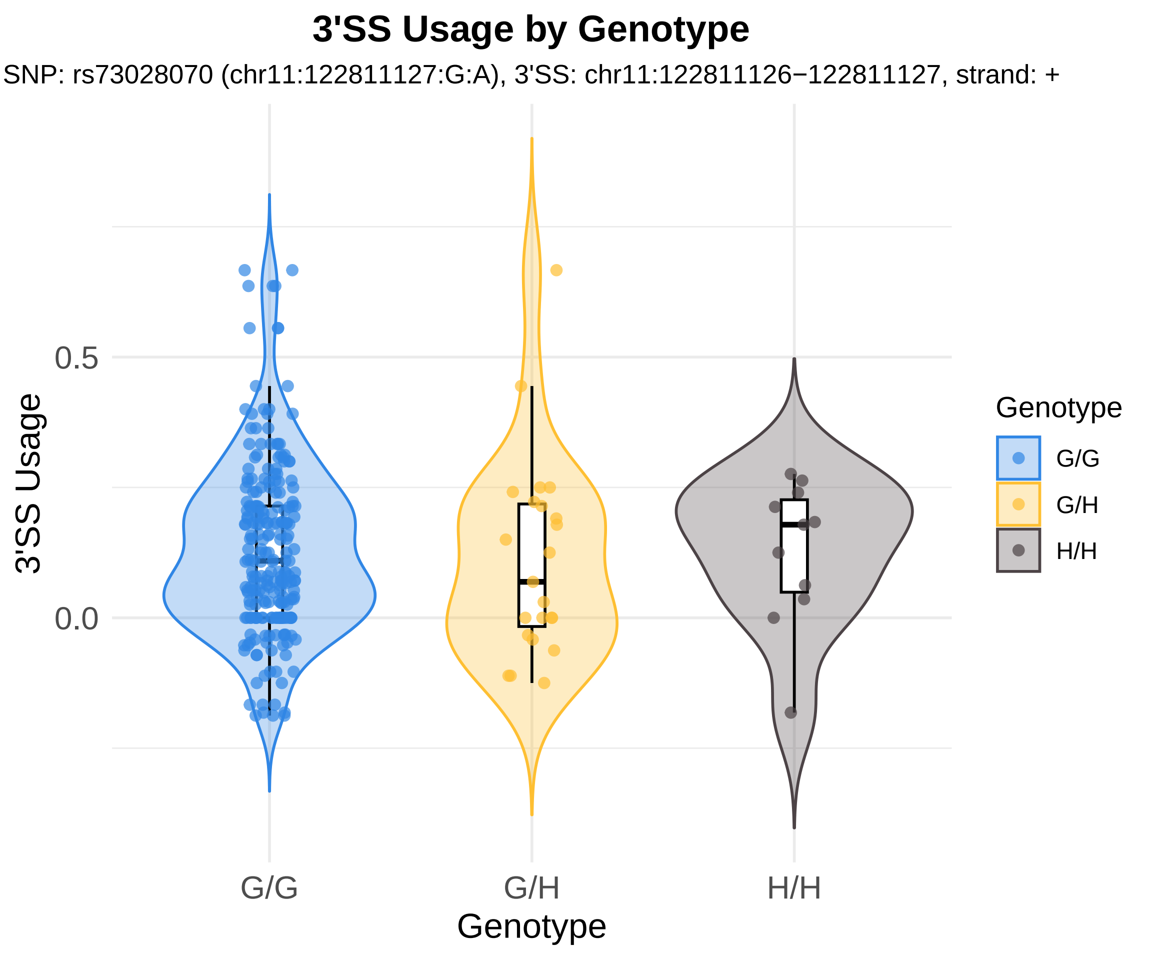

3'SS Usage by Genotype
SNP: rs73028070 (chr11:122811127:G:A), 3'SS: chr11:122811126−122811127, strand: +
3'SS Usage
0.5
0.0
G/G
G/H
H/H
Genotype
Genotype
G/G
G/H
H/H

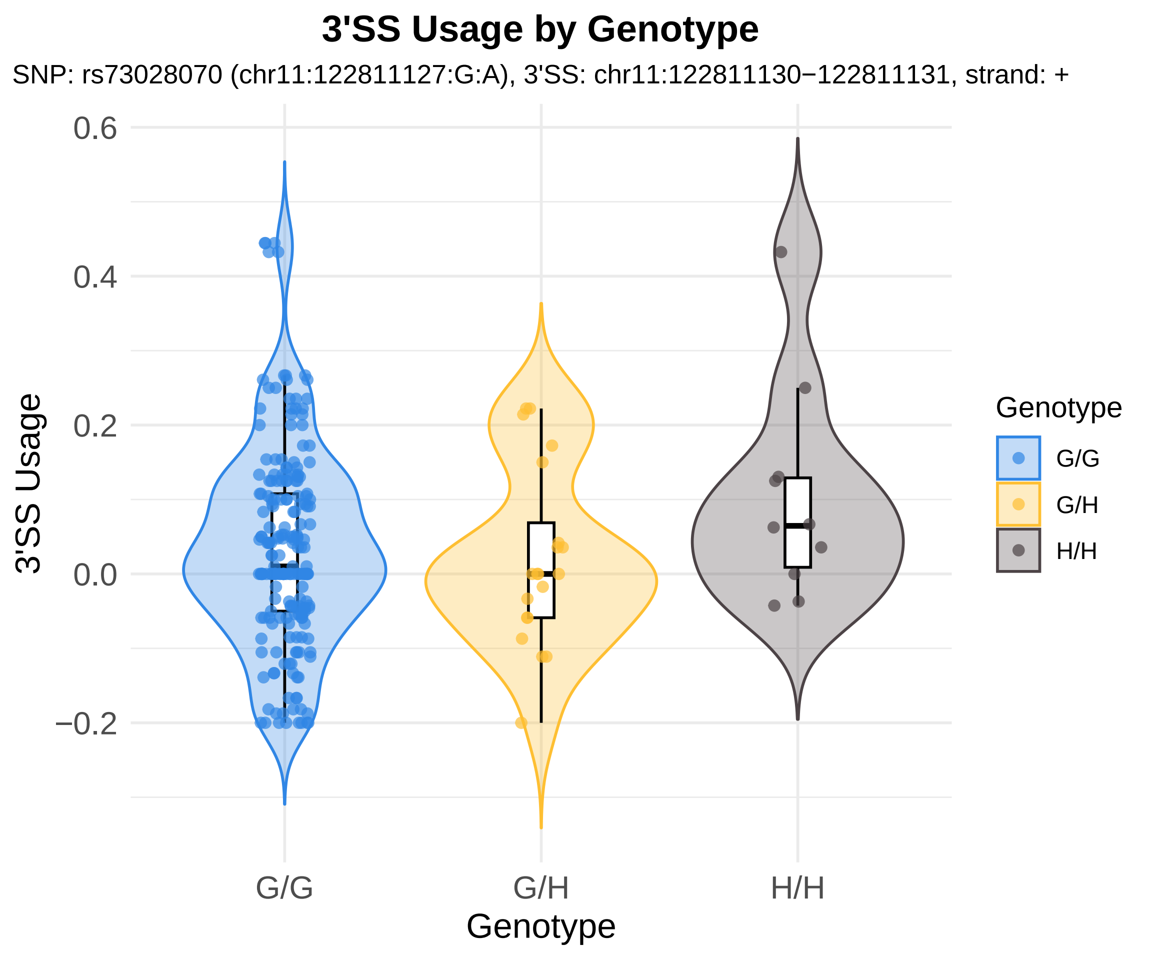
3'SS Usage by Genotype
SNP: rs73028070 (chr11:122811127:G:A), 3'SS: chr11:122811130−122811131, strand: +
3'SS Usage
0.6
0.4
0.2
0.0
−0.2
G/G
G/H
H/H
Genotype
Genotype
G/G
G/H
H/H

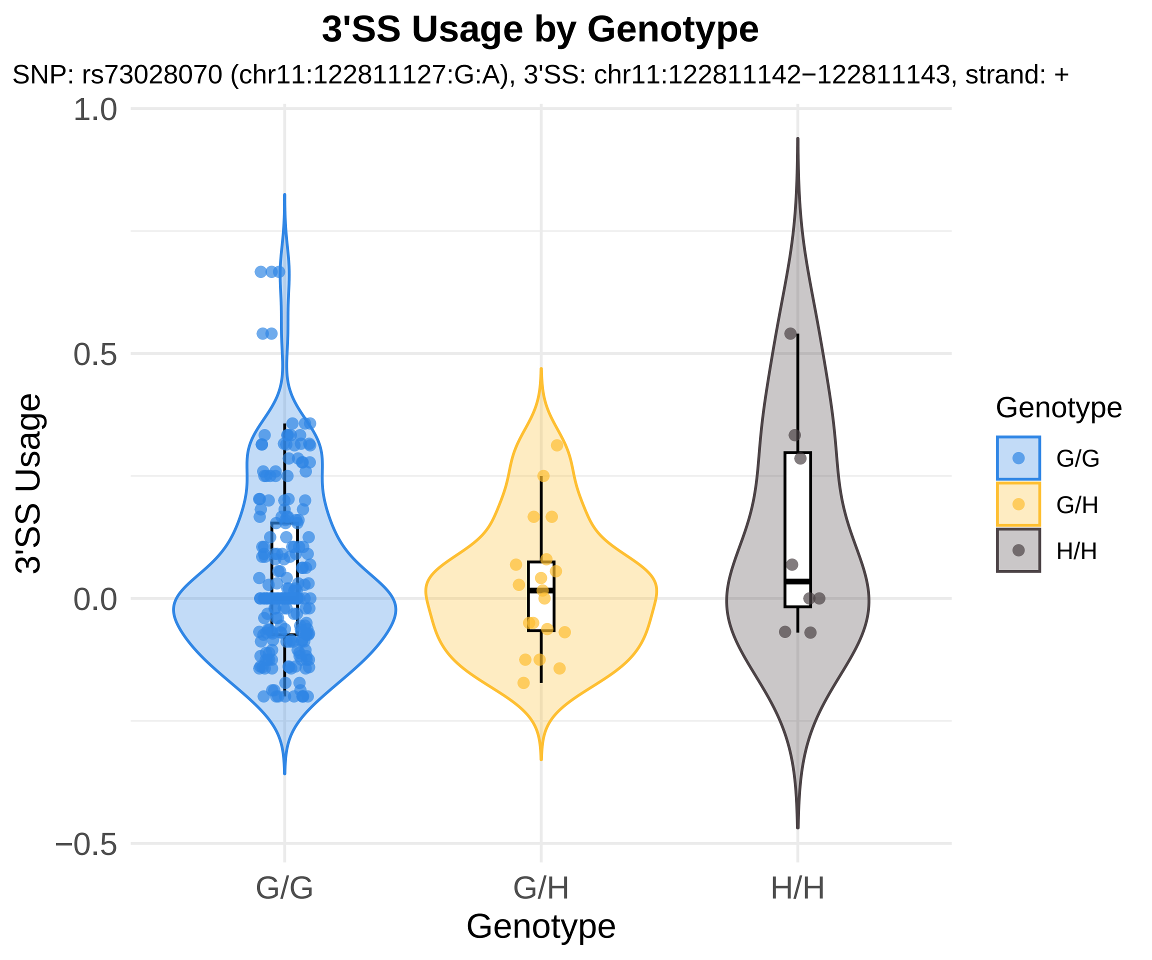
3'SS Usage by Genotype
SNP: rs73028070 (chr11:122811127:G:A), 3'SS: chr11:122811142−122811143, strand: +
3'SS Usage
1.0
0.5
0.0
−0.5
G/G
G/H
H/H
Genotype
Genotype
G/G
G/H
H/H

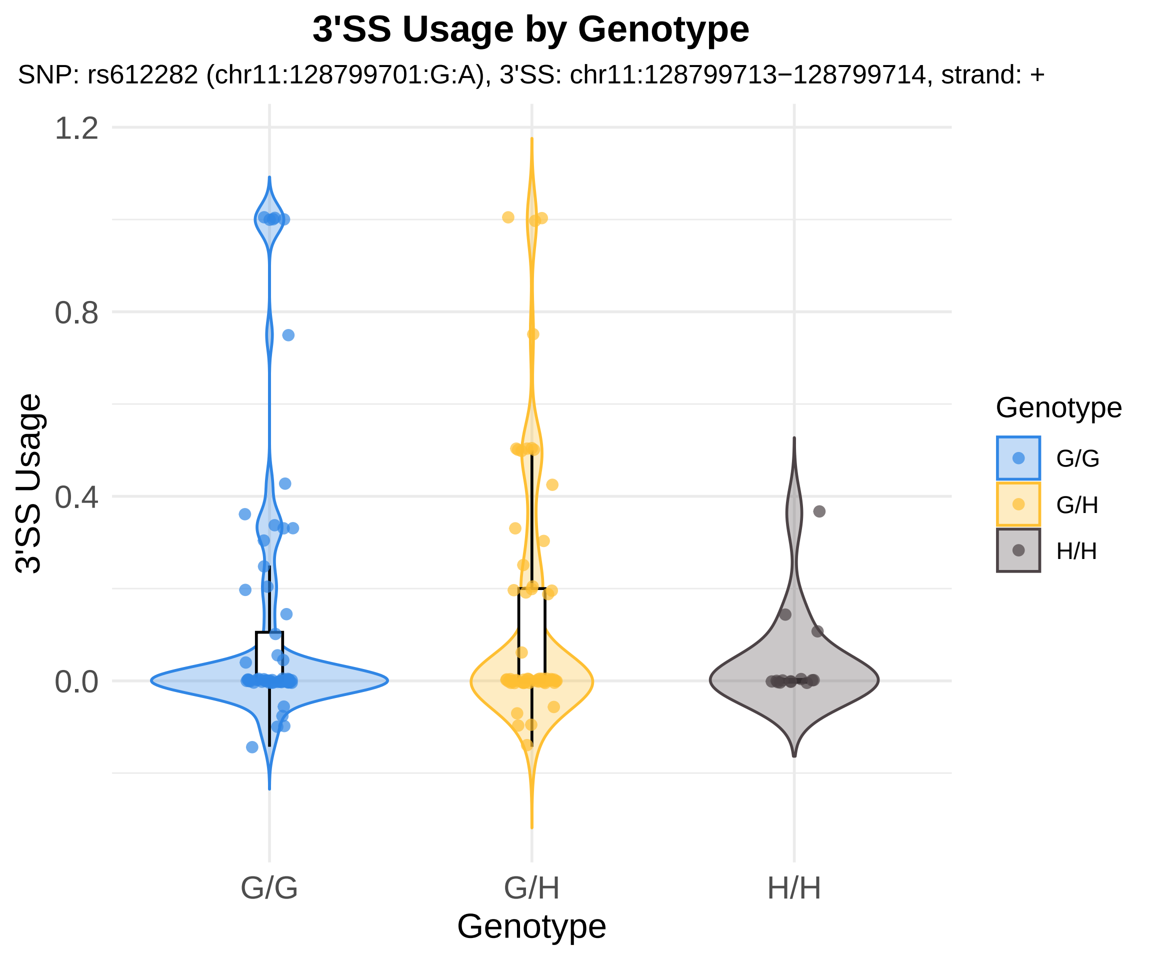
3'SS Usage by Genotype
SNP: rs612282 (chr11:128799701:G:A), 3'SS: chr11:128799713−128799714, strand: +
1.2
0.8
0.4
0.0
3'SS Usage
G/G
G/H
H/H
Genotype
Genotype
G/G
G/H
H/H

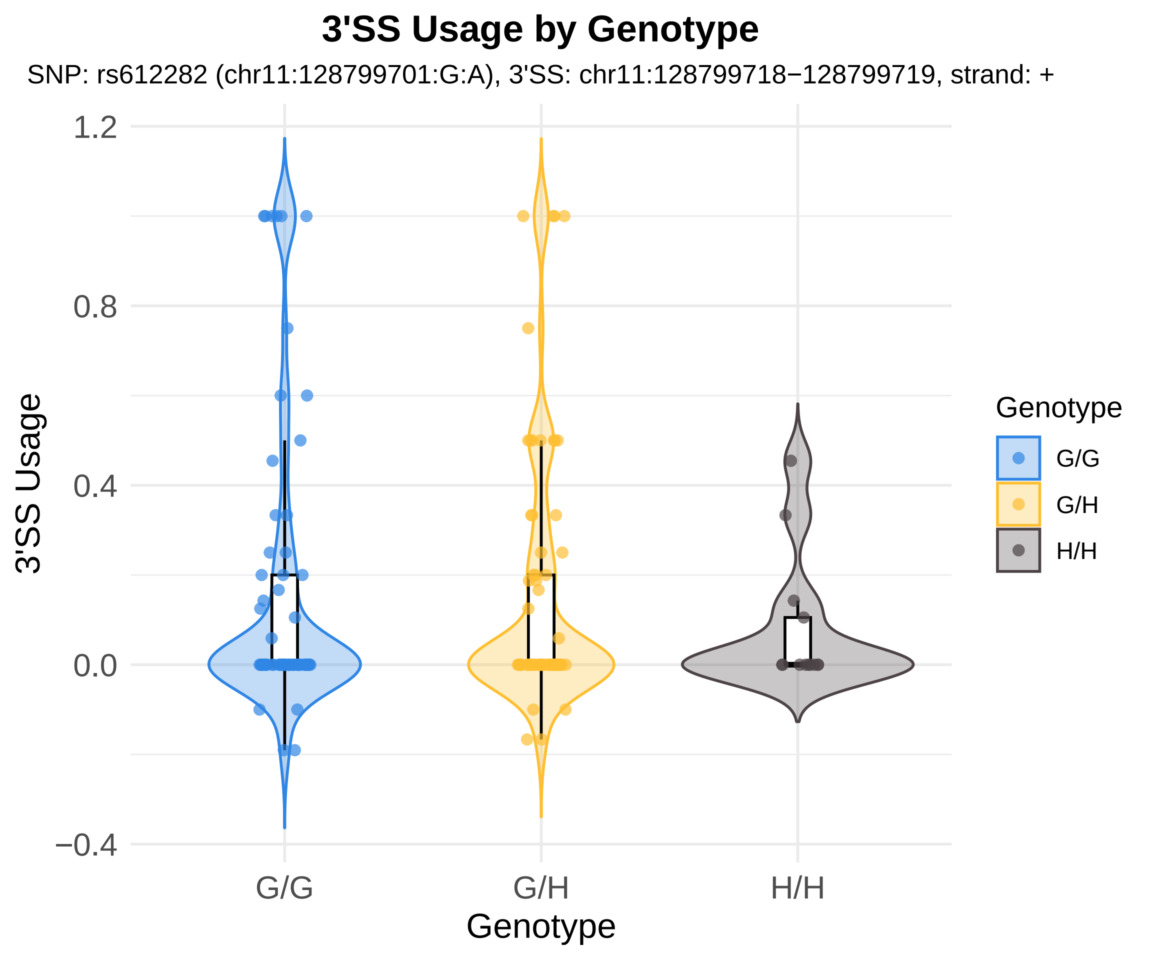
3'SS Usage by Genotype
SNP: rs612282 (chr11:128799701:G:A), 3'SS: chr11:128799718−128799719, strand: +
3'SS Usage
1.2
0.8
0.4
0.0
−0.4
G/G
G/H
H/H
Genotype
Genotype
G/G
G/H
H/H

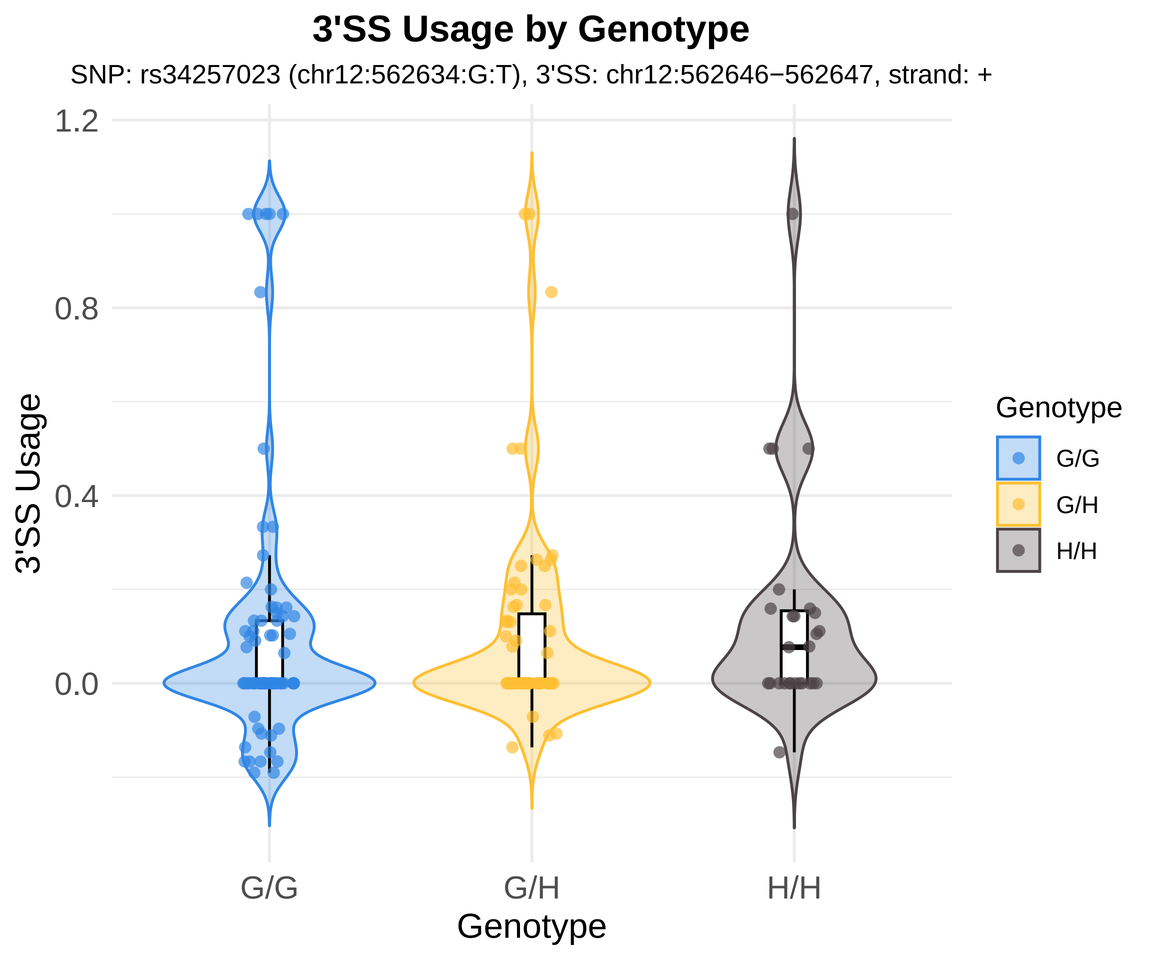
3'SS Usage by Genotype
SNP: rs34257023 (chr12:562634:G:T), 3'SS: chr12:562646−562647, strand: +
1.2
0.8
0.4
0.0
3'SS Usage
G/G
G/H
H/H
Genotype
Genotype
G/G
G/H
H/H

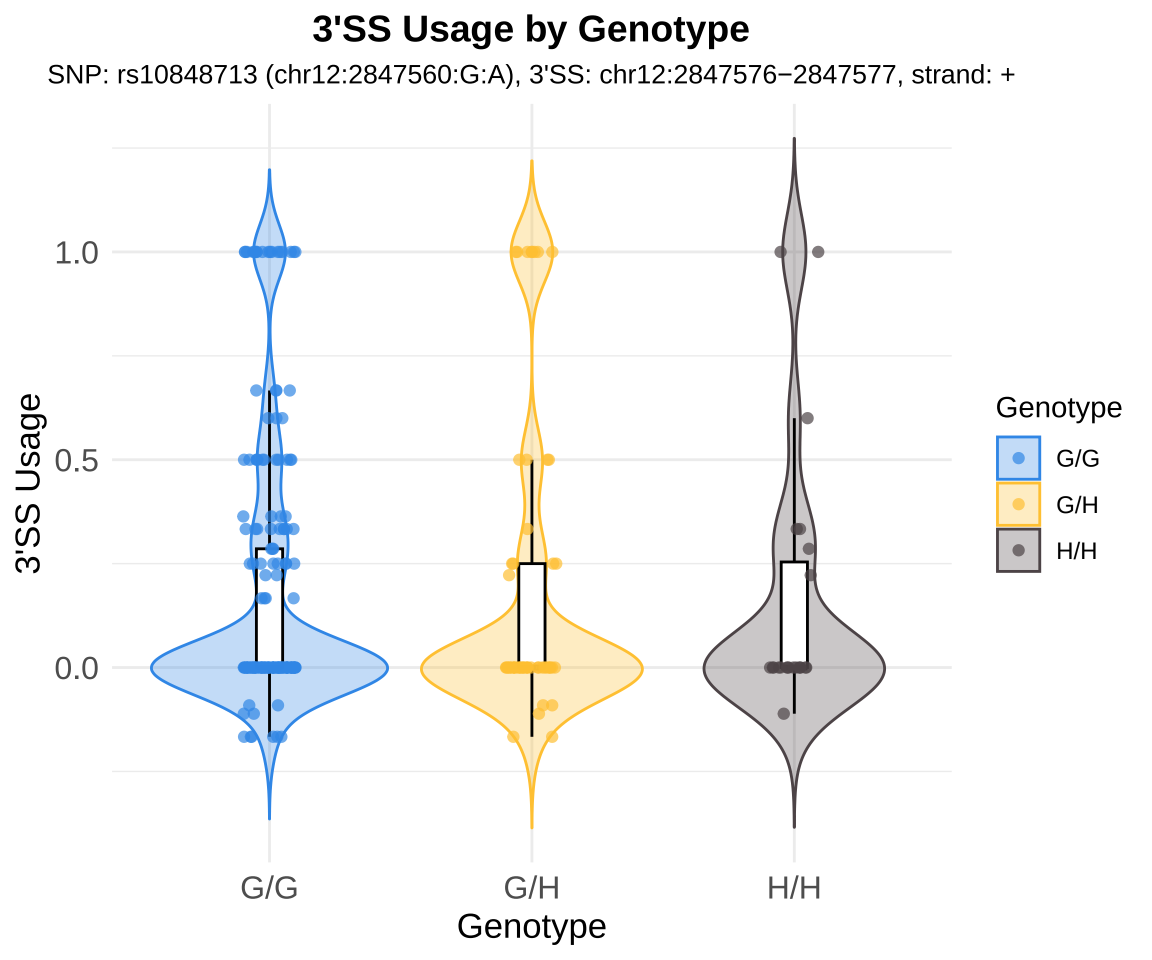

3'SS Usage by Genotype
SNP: rs10848713 (chr12:2847560:G:A), 3'SS: chr12:2847576−2847577, strand: +
3'SS Usage
1.0
0.5
0.0
G/G
G/H
H/H
Genotype
Genotype
G/G
G/H
H/H

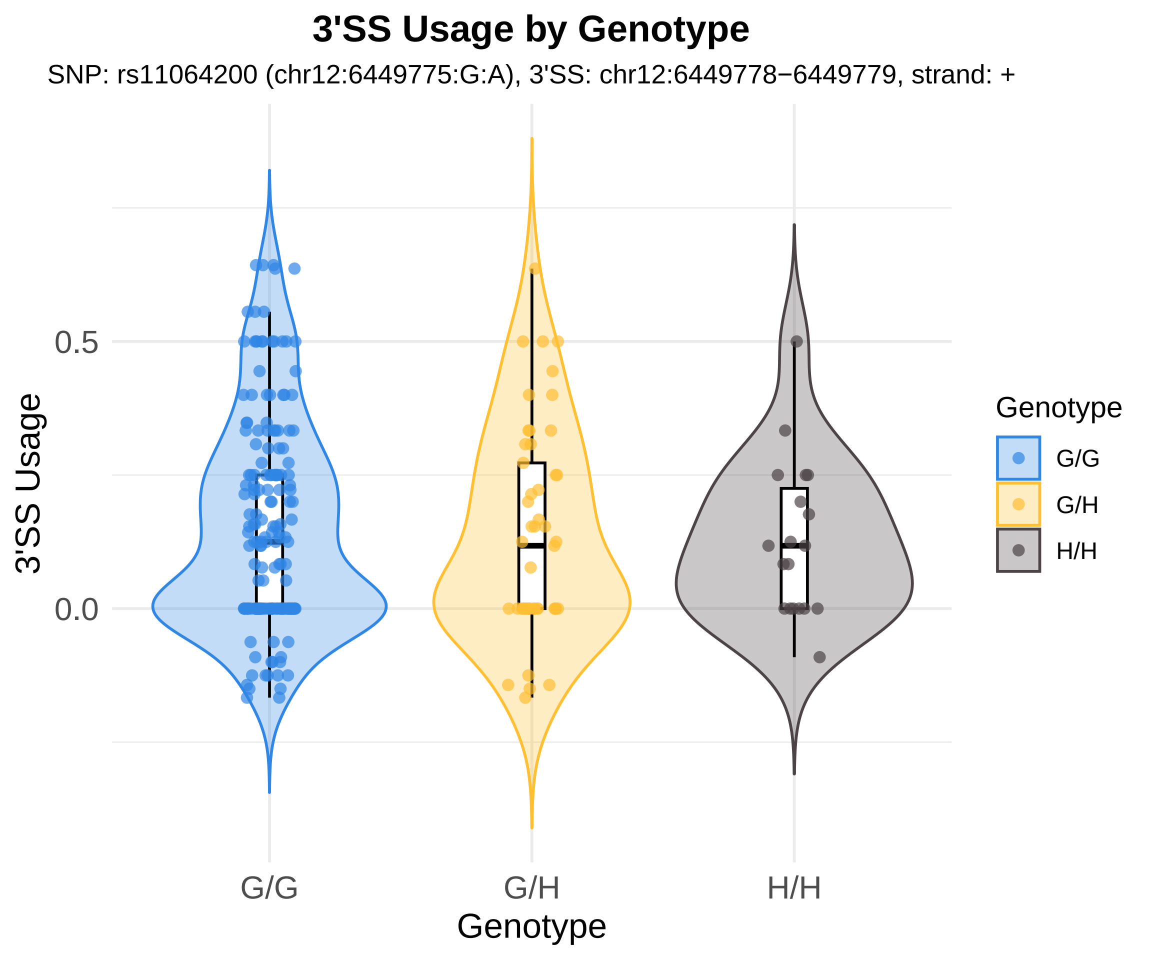
3'SS Usage by Genotype
SNP: rs11064200 (chr12:6449775:G:A), 3'SS: chr12:6449778−6449779, strand: +
3'SS Usage
0.5
0.0
G/G
G/H
H/H
Genotype
Genotype
G/G
G/H
H/H

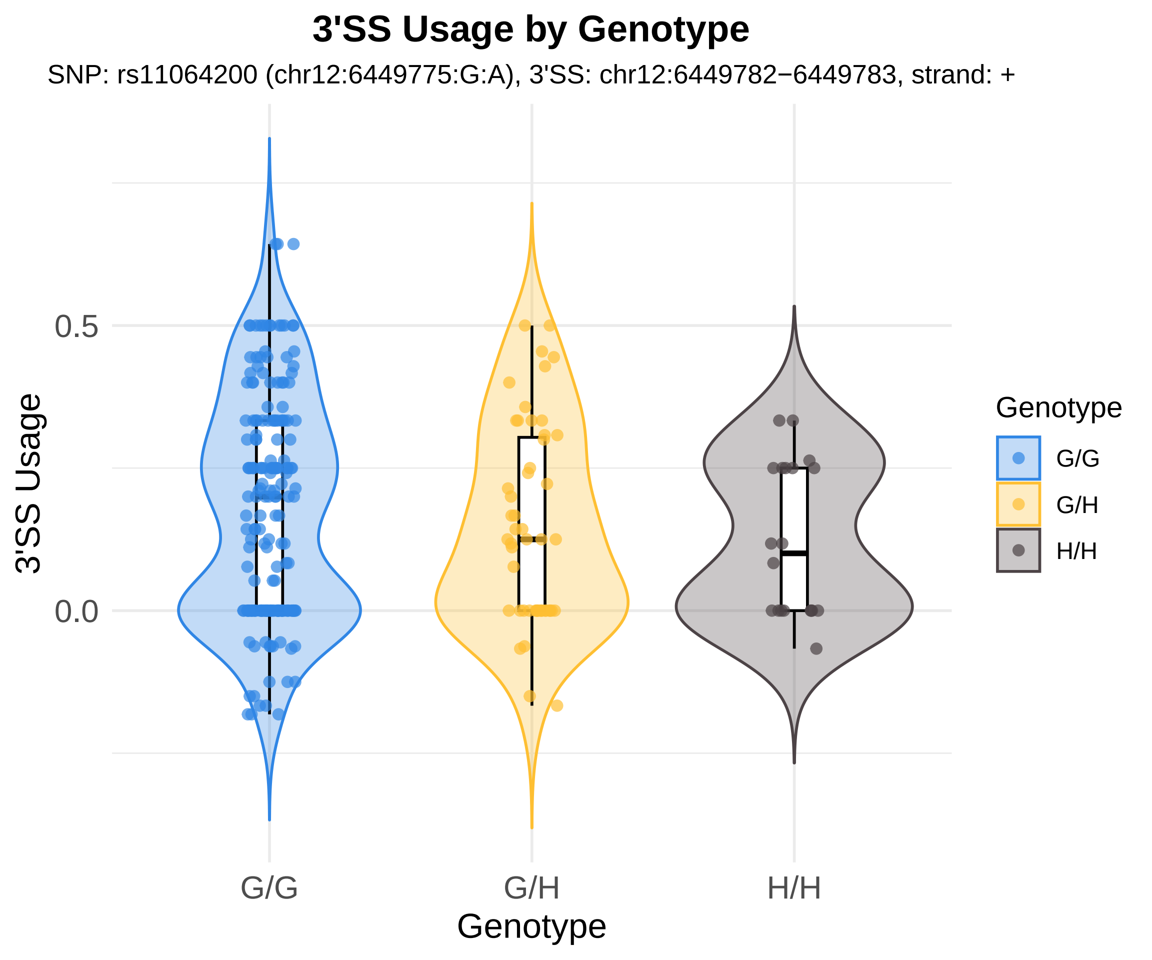

3'SS Usage by Genotype
SNP: rs11064200 (chr12:6449775:G:A), 3'SS: chr12:6449782−6449783, strand: +
3'SS Usage
0.5
0.0
G/G
G/H
H/H
Genotype
Genotype
G/G
G/H
H/H

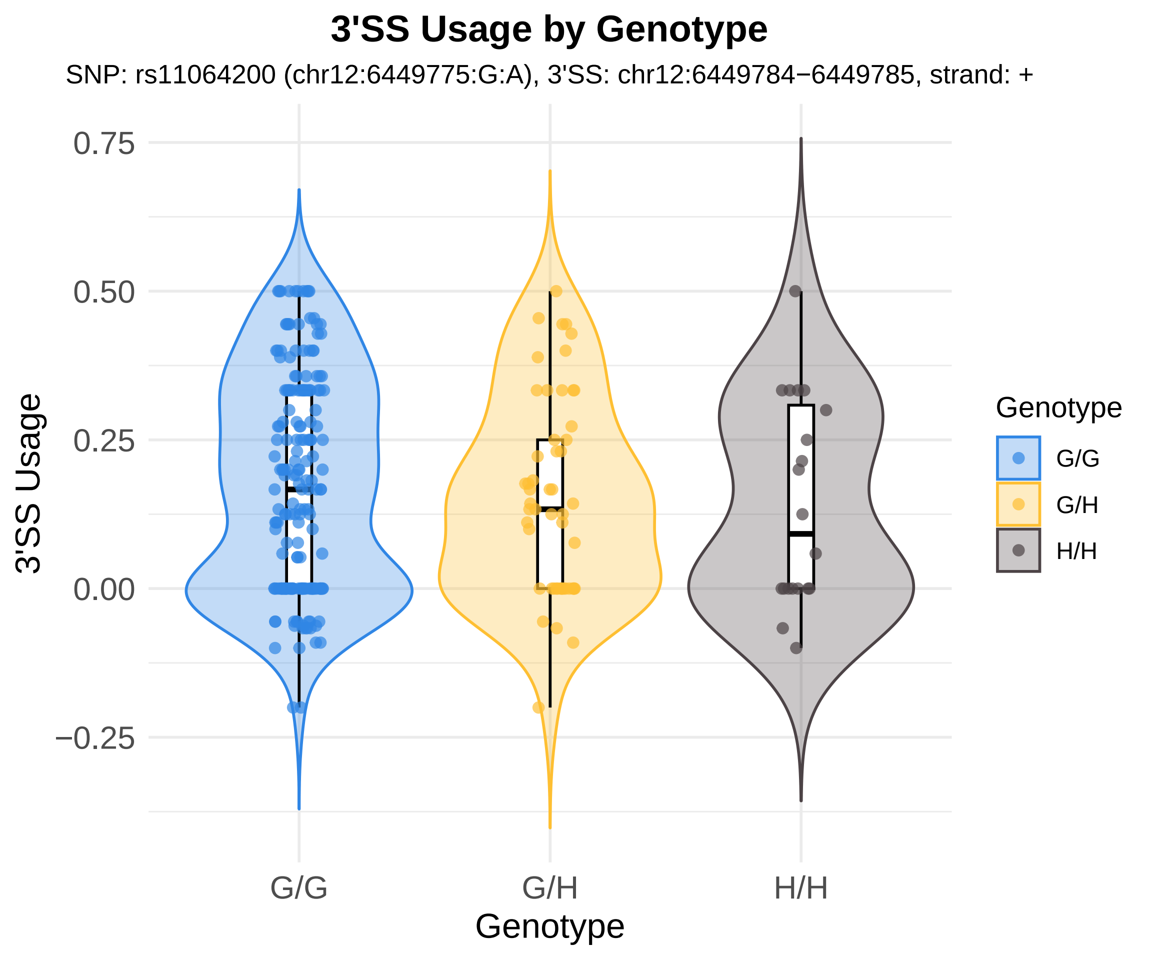
3'SS Usage by Genotype
SNP: rs11064200 (chr12:6449775:G:A), 3'SS: chr12:6449784−6449785, strand: +
3'SS Usage
0.75
0.50
0.25
0.00
−0.25
G/G
G/H
H/H
Genotype
Genotype
G/G
G/H
H/H

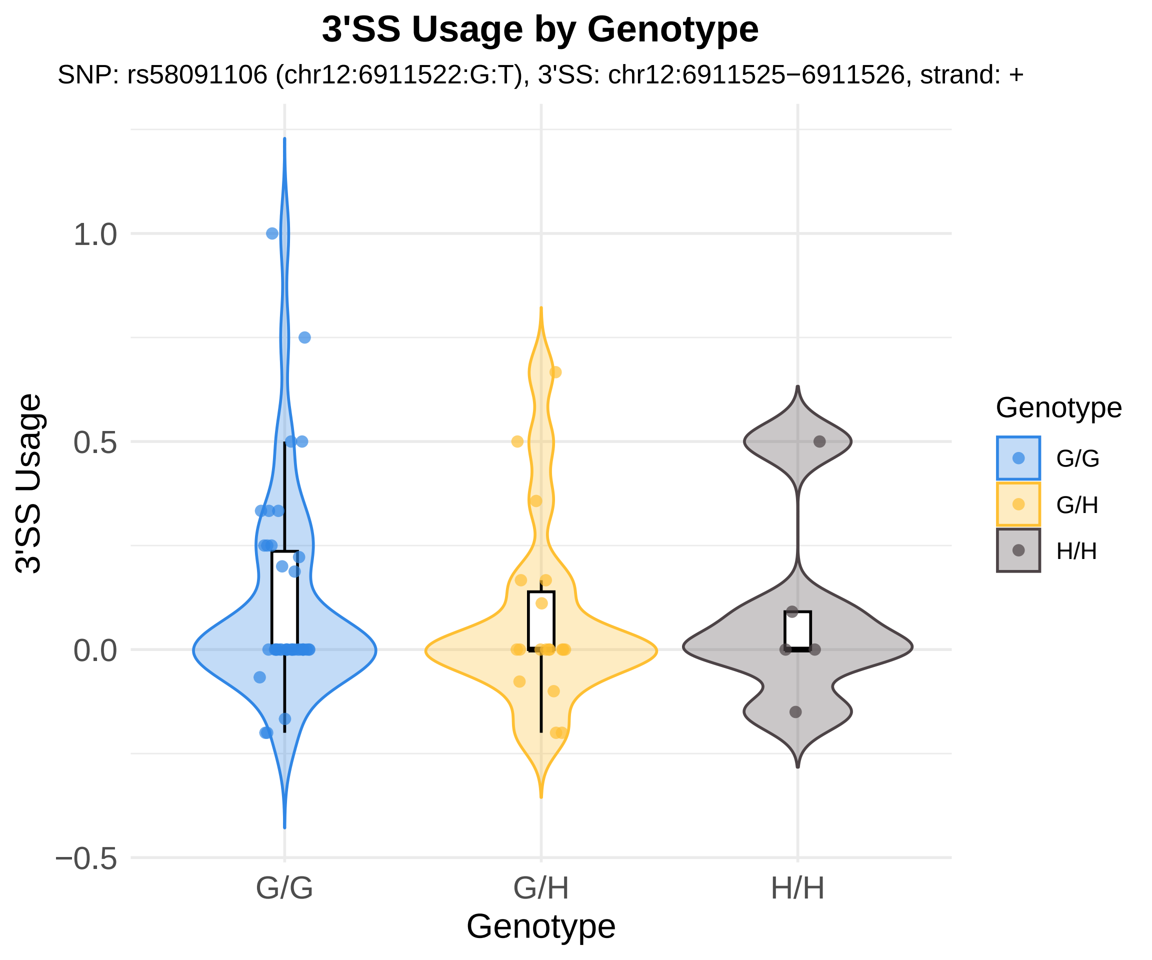
3'SS Usage by Genotype
SNP: rs58091106 (chr12:6911522:G:T), 3'SS: chr12:6911525−6911526, strand: +
3'SS Usage
1.0
0.5
0.0
−0.5
G/G
G/H
H/H
Genotype
Genotype
G/G
G/H
H/H

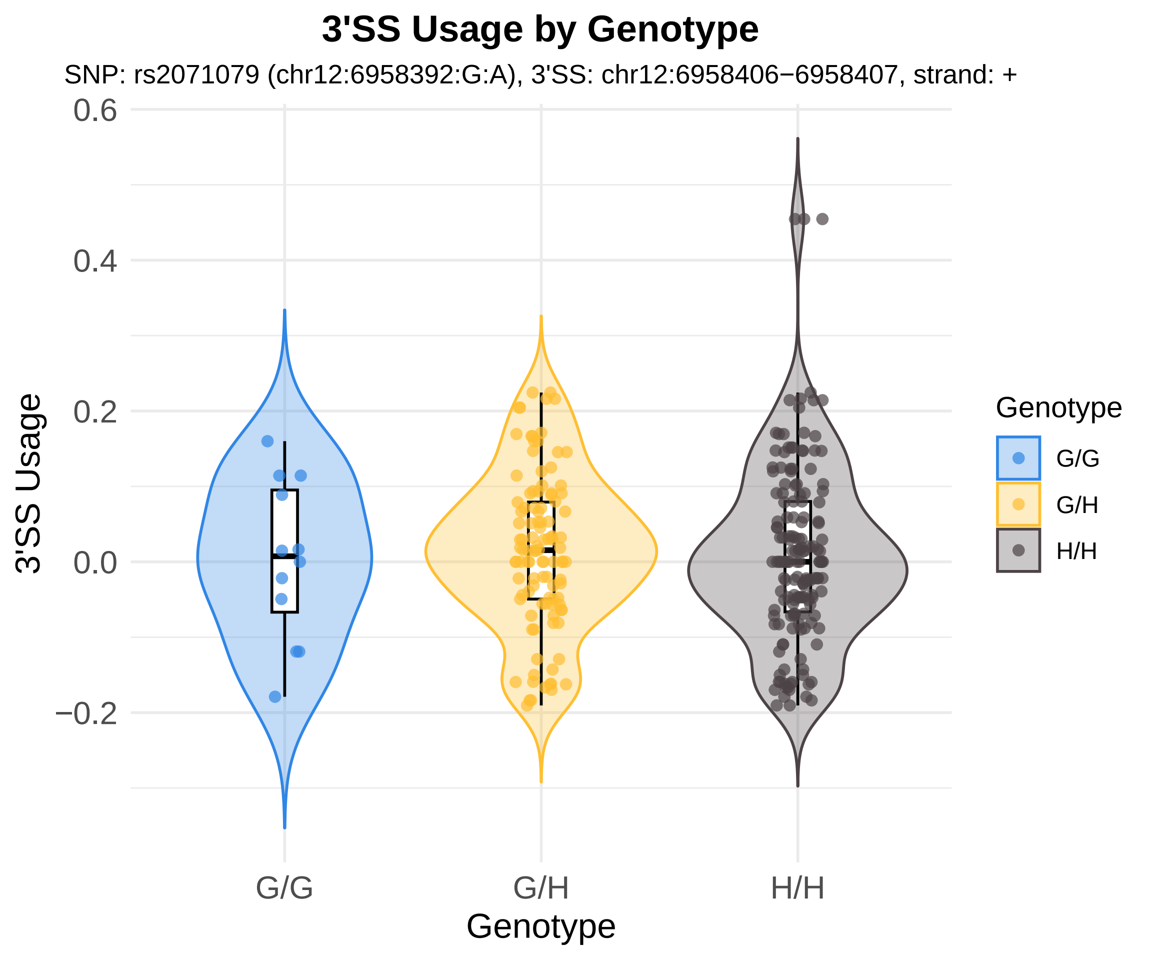

3'SS Usage by Genotype
SNP: rs2071079 (chr12:6958392:G:A), 3'SS: chr12:6958406−6958407, strand: +
0.6
0.4
0.2
0.0
−0.2
3'SS Usage
G/G
G/H
H/H
Genotype
Genotype
G/G
G/H
H/H

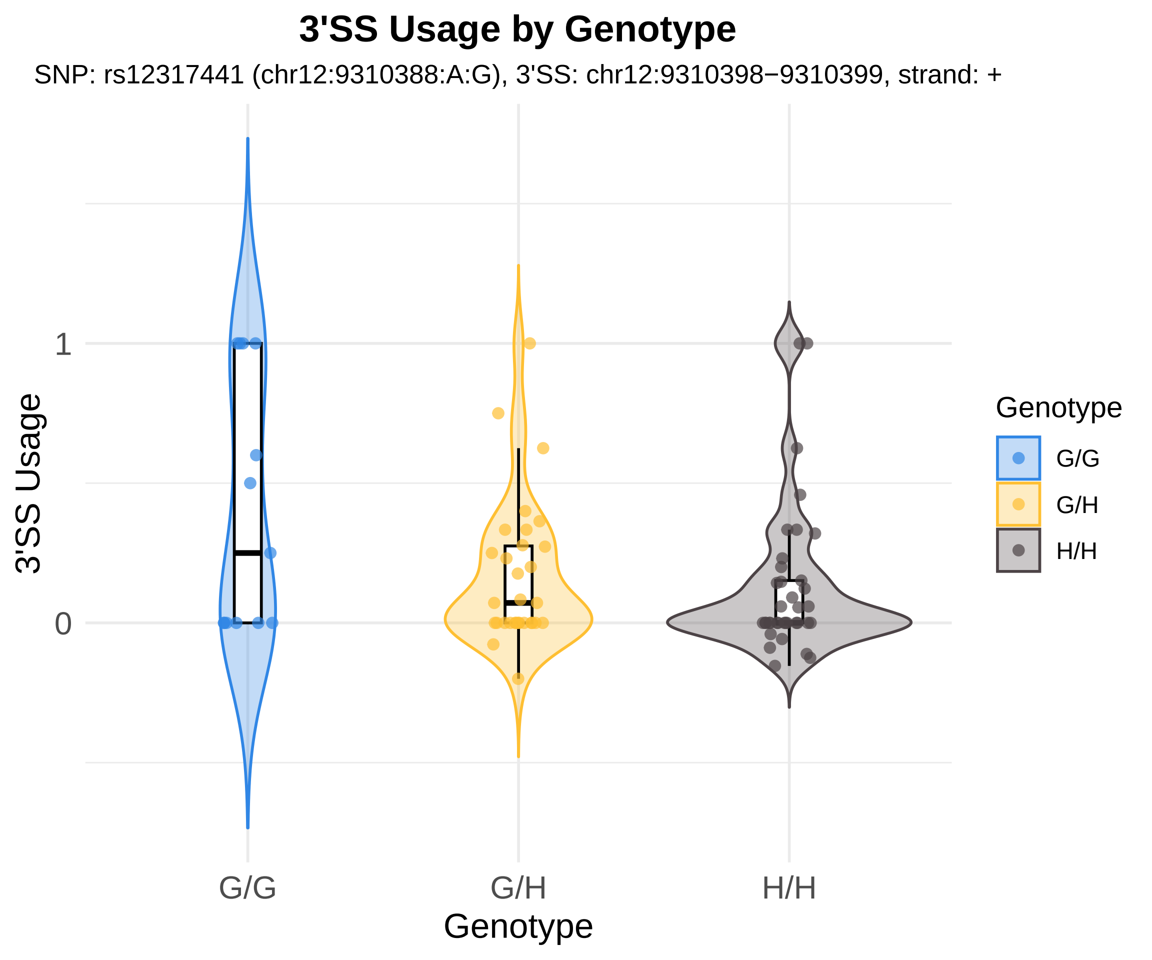
3'SS Usage by Genotype
SNP: rs12317441 (chr12:9310388:A:G), 3'SS: chr12:9310398−9310399, strand: +
3'SS Usage
1
0
G/G
G/H
H/H
Genotype
Genotype
G/G
G/H
H/H

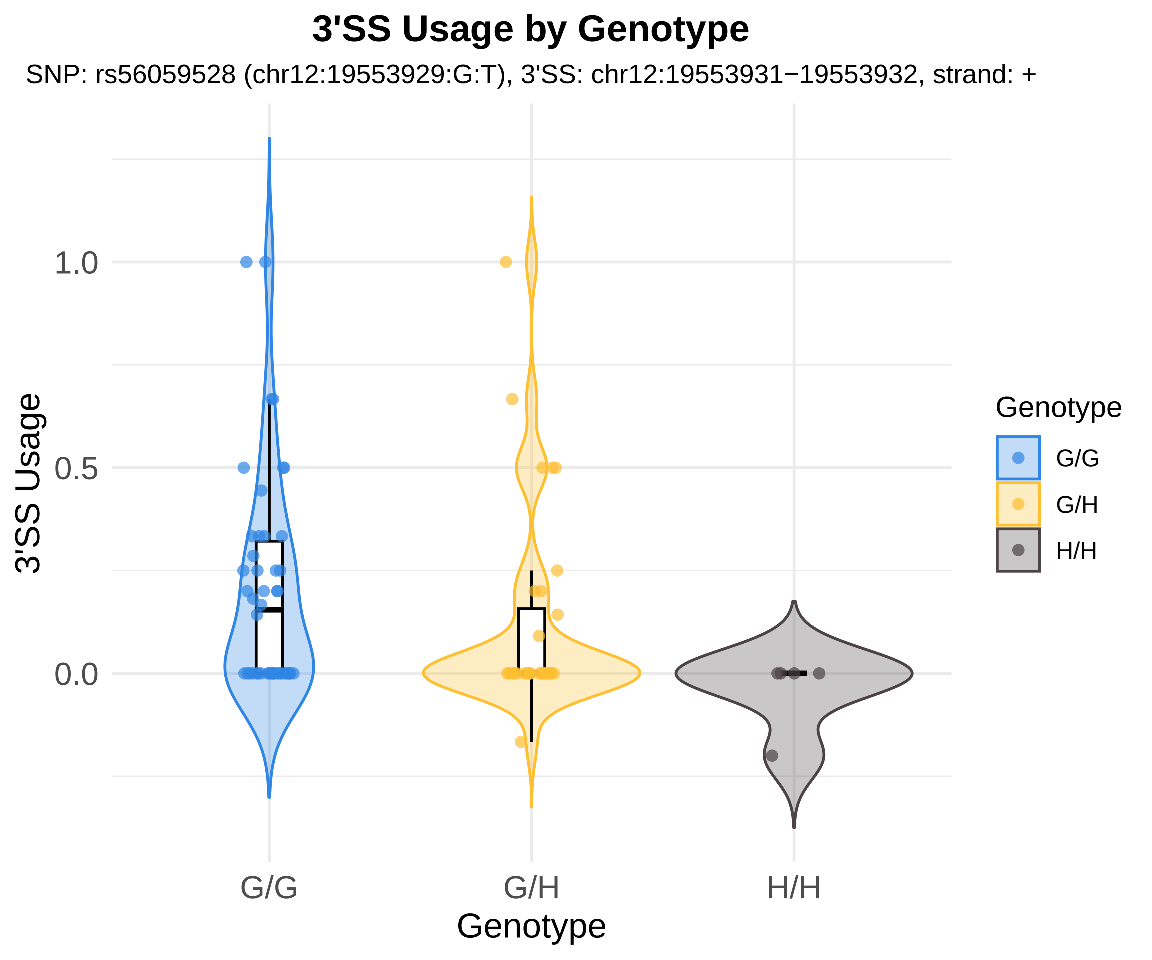
3'SS Usage by Genotype
SNP: rs56059528 (chr12:19553929:G:T), 3'SS: chr12:19553931−19553932, strand: +
3'SS Usage
1.0
0.5
0.0
G/G
G/H
H/H
Genotype
Genotype
G/G
G/H
H/H

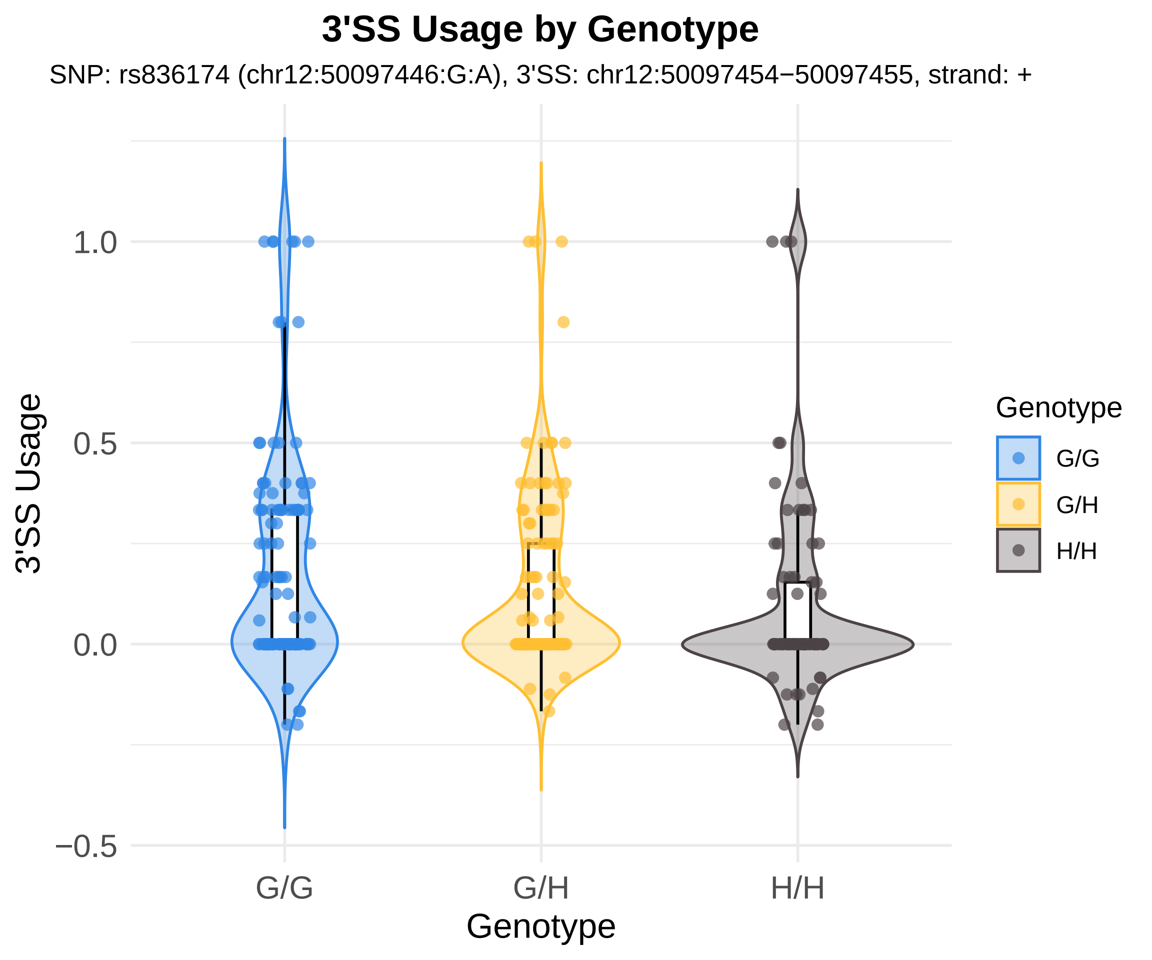

3'SS Usage by Genotype
SNP: rs836174 (chr12:50097446:G:A), 3'SS: chr12:50097454−50097455, strand: +
3'SS Usage
1.0
0.5
0.0
−0.5
G/G
G/H
H/H
Genotype
Genotype
G/G
G/H
H/H

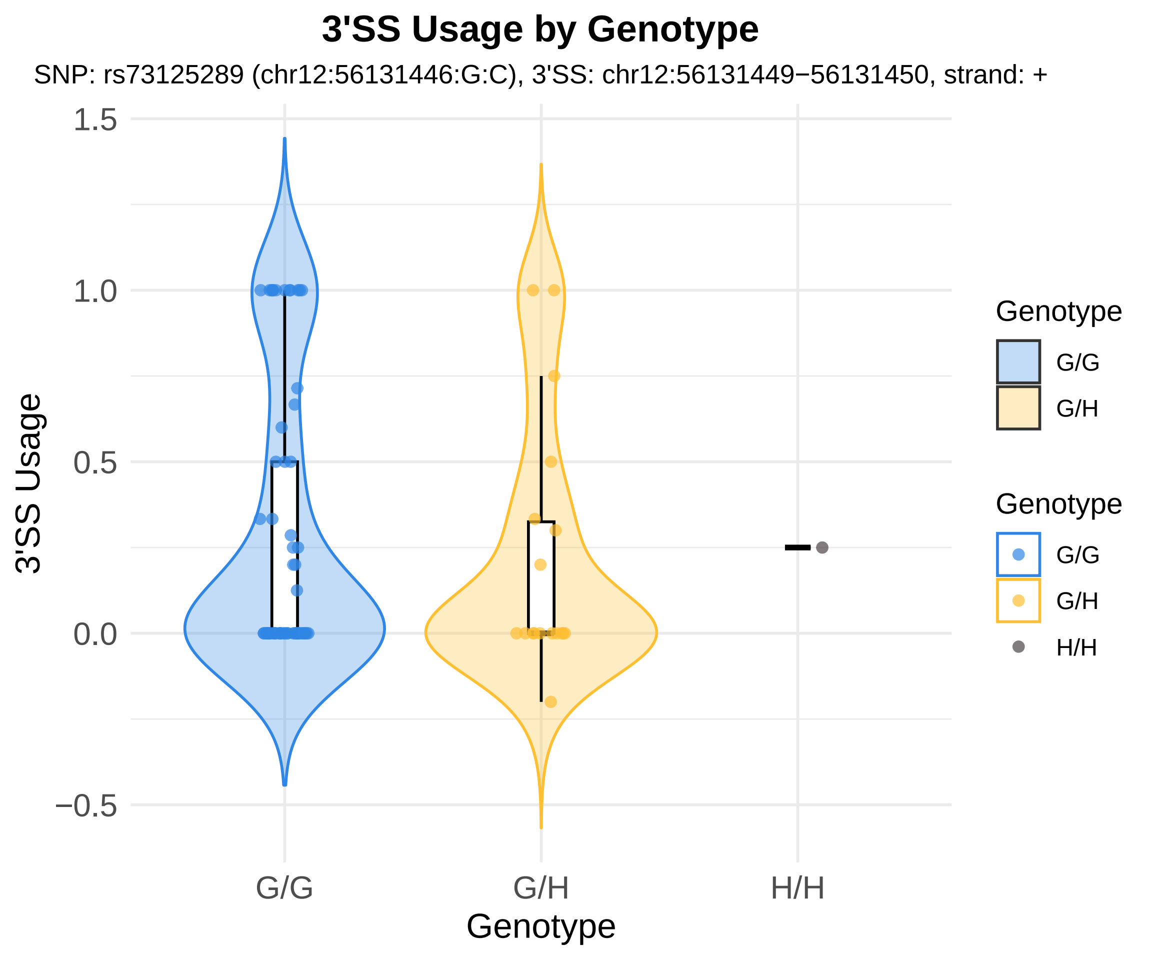
3'SS Usage by Genotype
SNP: rs73125289 (chr12:56131446:G:C), 3'SS: chr12:56131449−56131450, strand: +
1.5
1.0
0.5
0.0
−0.5
3'SS Usage
G/G
G/H
H/H
Genotype
Genotype
G/G
G/H
Genotype
G/G
G/H
H/H

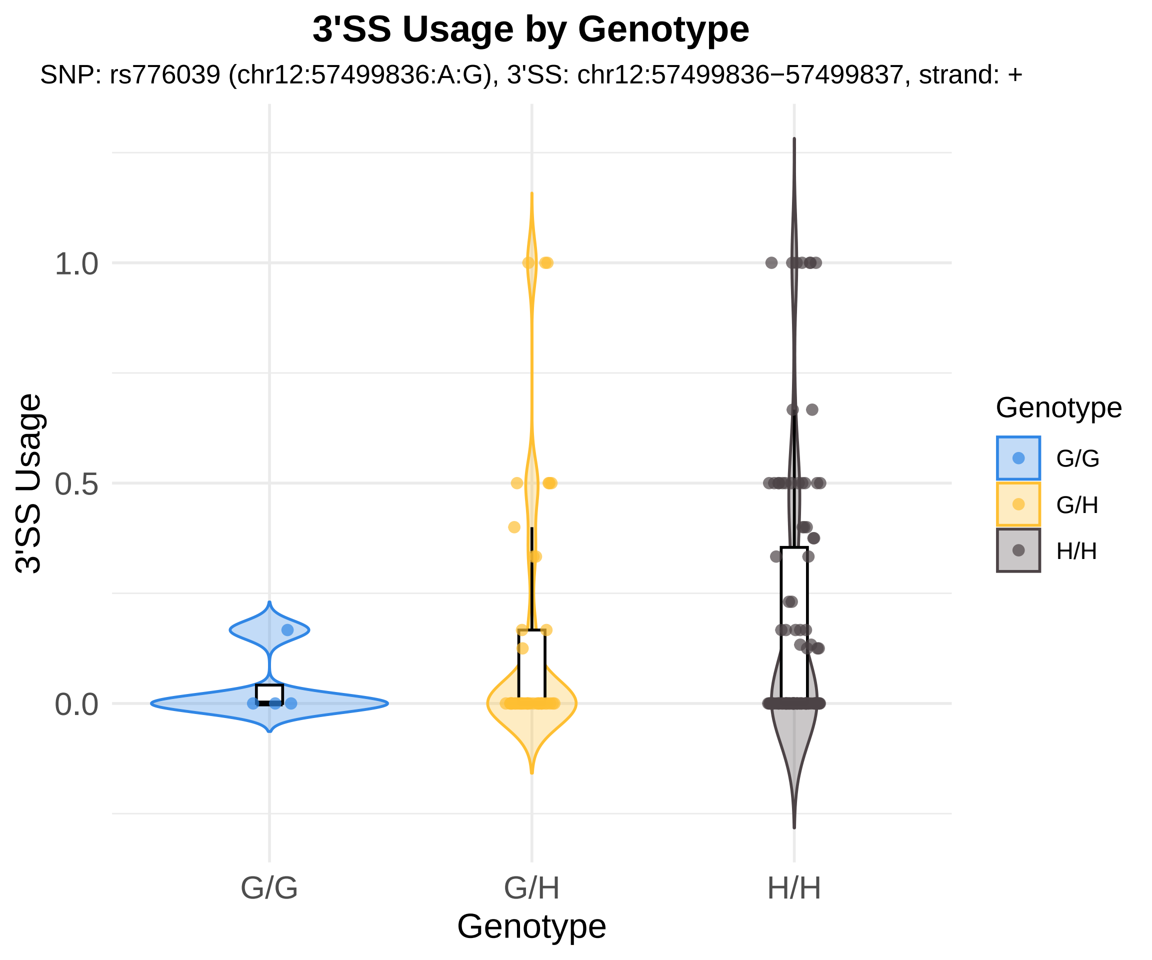
3'SS Usage by Genotype
SNP: rs776039 (chr12:57499836:A:G), 3'SS: chr12:57499836−57499837, strand: +
3'SS Usage
1.0
0.5
0.0
G/G
G/H
H/H
Genotype
Genotype
G/G
G/H
H/H

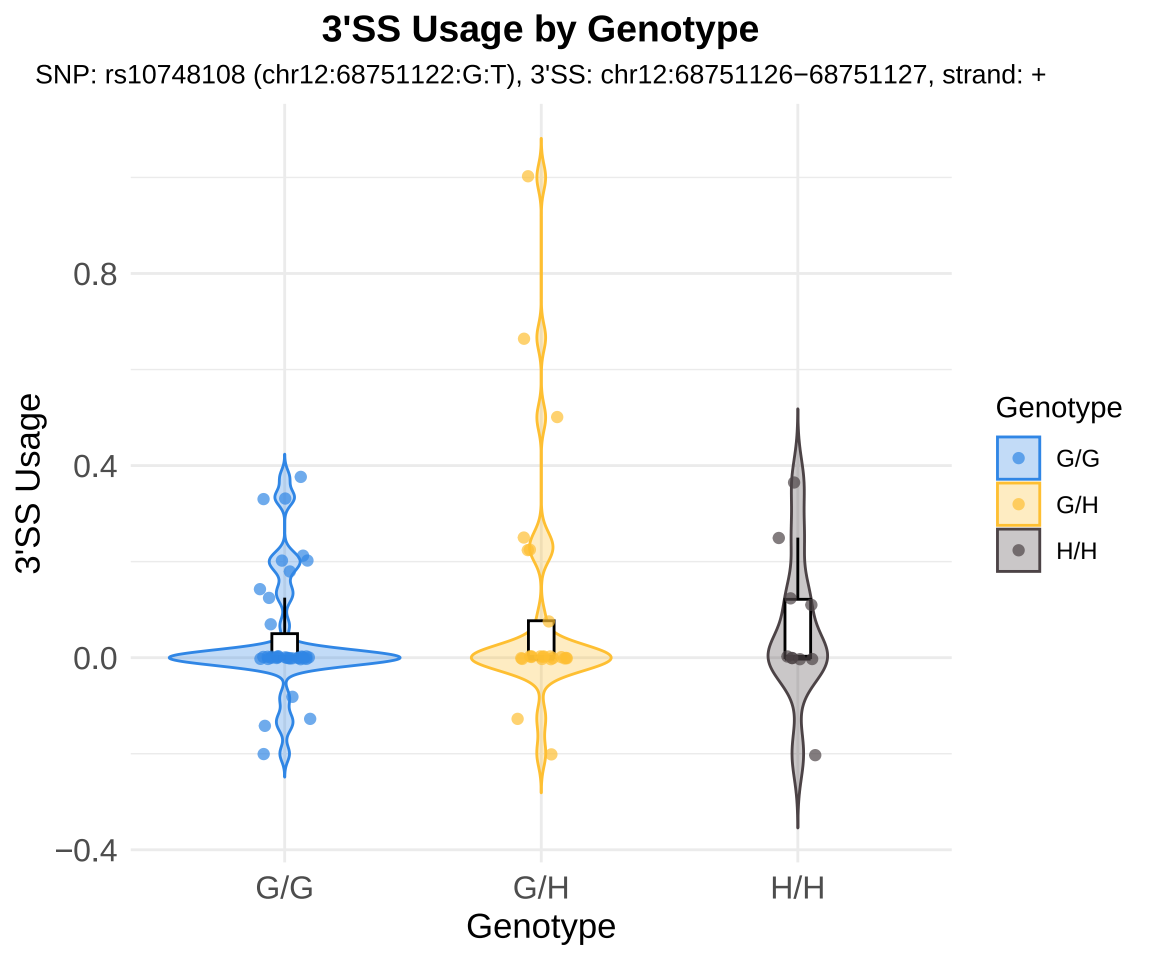
3'SS Usage by Genotype
SNP: rs10748108 (chr12:68751122:G:T), 3'SS: chr12:68751126−68751127, strand: +
3'SS Usage
0.8
0.4
0.0
−0.4
G/G
G/H
H/H
Genotype
Genotype
G/G
G/H
H/H

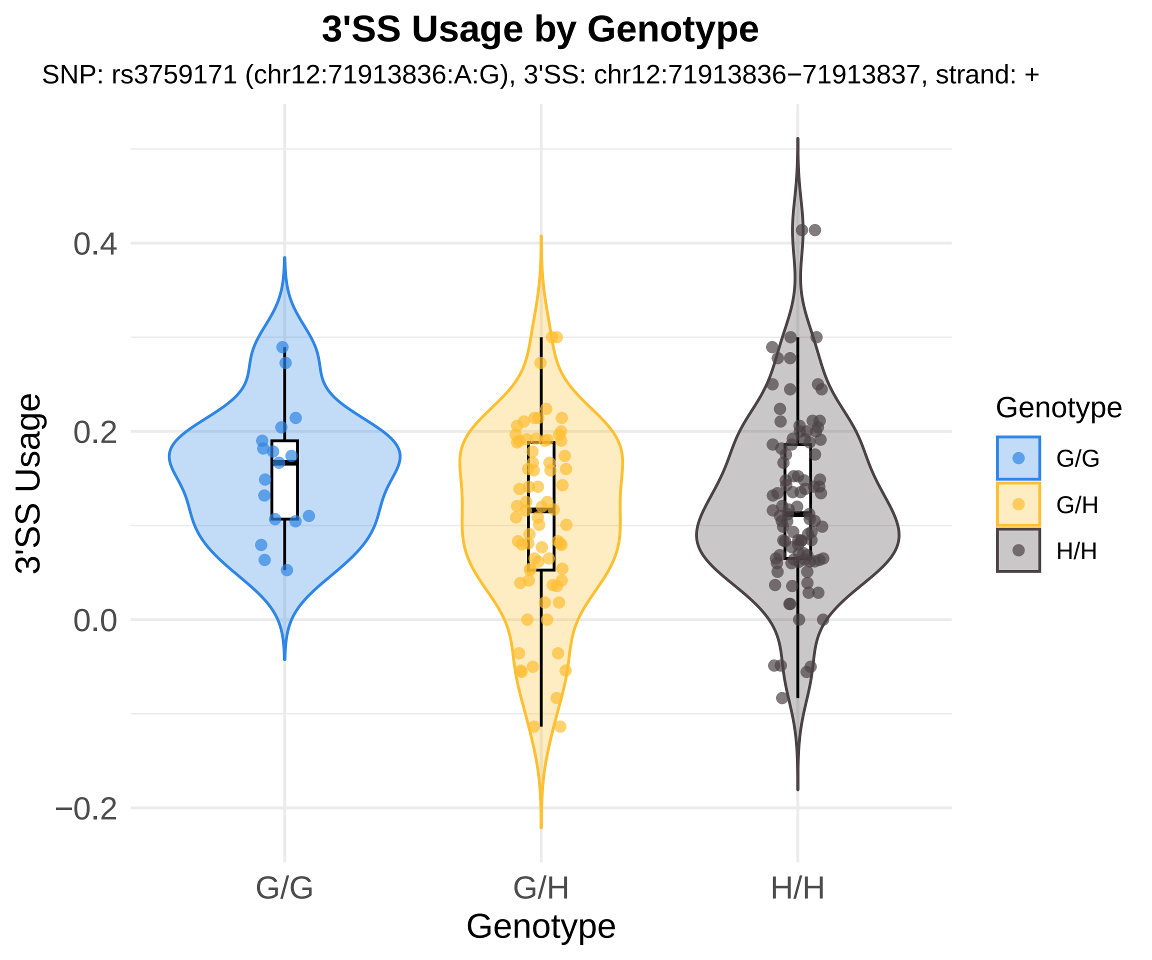
3'SS Usage by Genotype
SNP: rs3759171 (chr12:71913836:A:G), 3'SS: chr12:71913836−71913837, strand: +
3'SS Usage
0.4
0.2
0.0
−0.2
G/G
G/H
H/H
Genotype
Genotype
G/G
G/H
H/H

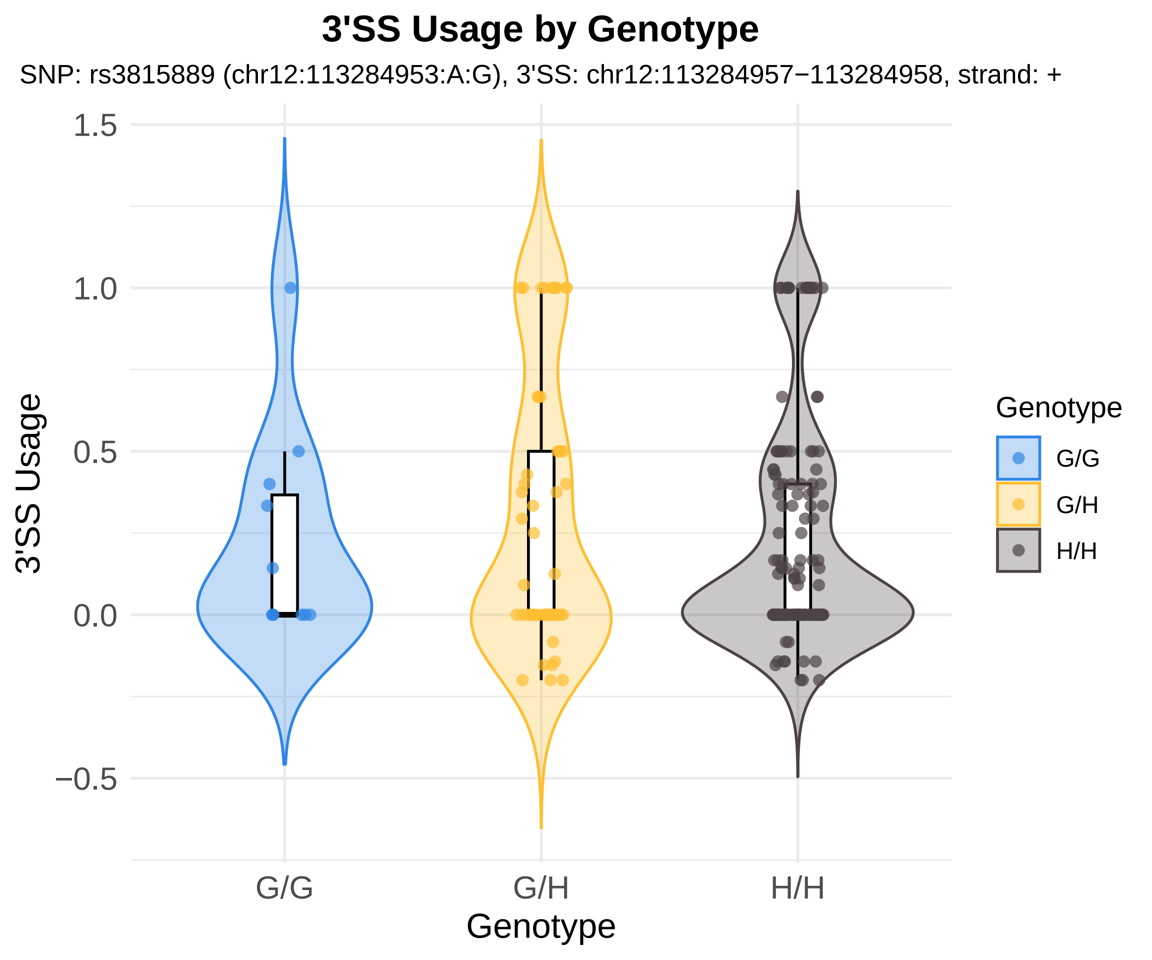

3'SS Usage by Genotype
SNP: rs3815889 (chr12:113284953:A:G), 3'SS: chr12:113284957−113284958, strand: +
1.5
1.0
0.5
0.0
−0.5
3'SS Usage
G/G
G/H
H/H
Genotype
Genotype
G/G
G/H
H/H

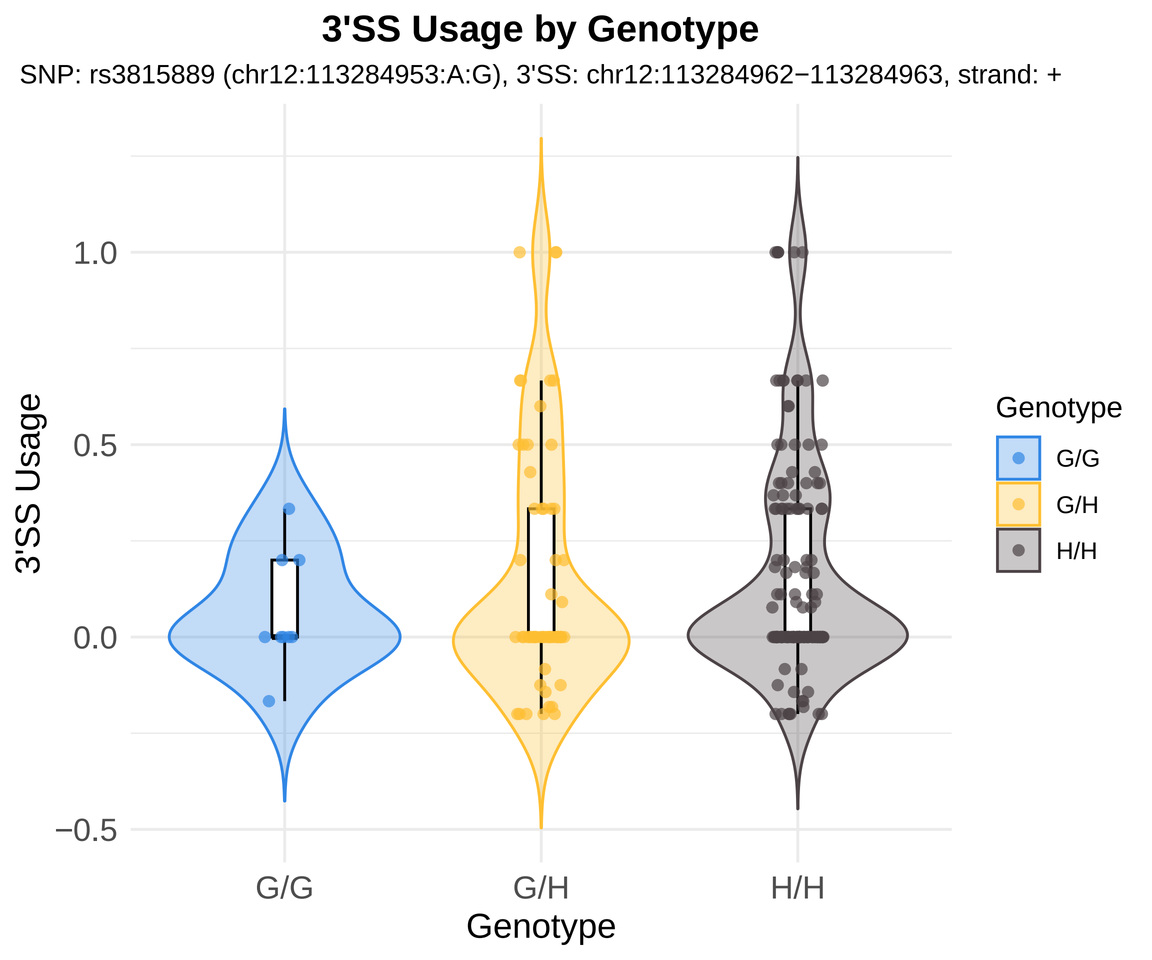

3'SS Usage by Genotype
SNP: rs3815889 (chr12:113284953:A:G), 3'SS: chr12:113284962−113284963, strand: +
3'SS Usage
1.0
0.5
0.0
−0.5
G/G
G/H
H/H
Genotype
Genotype
G/G
G/H
H/H

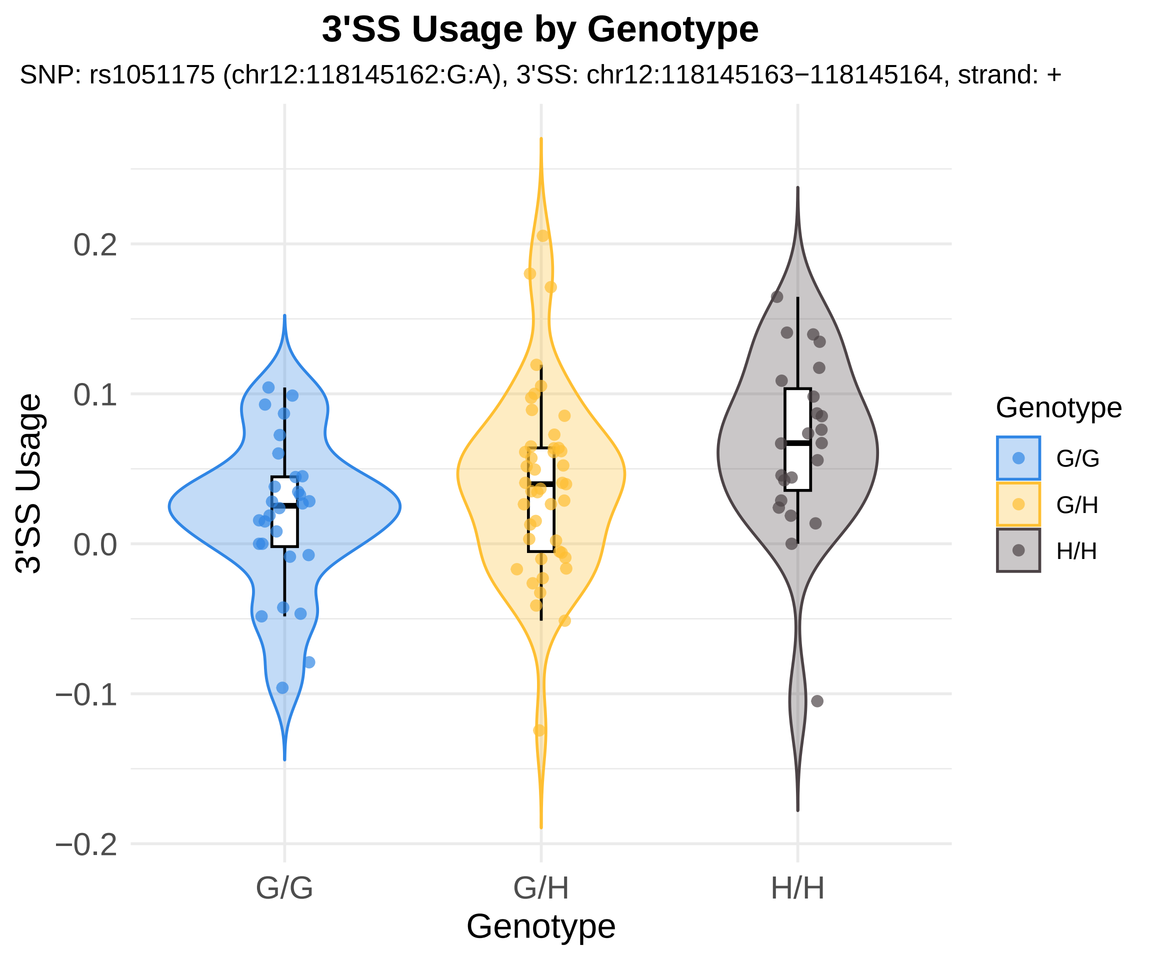
3'SS Usage by Genotype
SNP: rs1051175 (chr12:118145162:G:A), 3'SS: chr12:118145163−118145164, strand: +
3'SS Usage
0.2
0.1
0.0
−0.1
−0.2
G/G
G/H
H/H
Genotype
Genotype
G/G
G/H
H/H

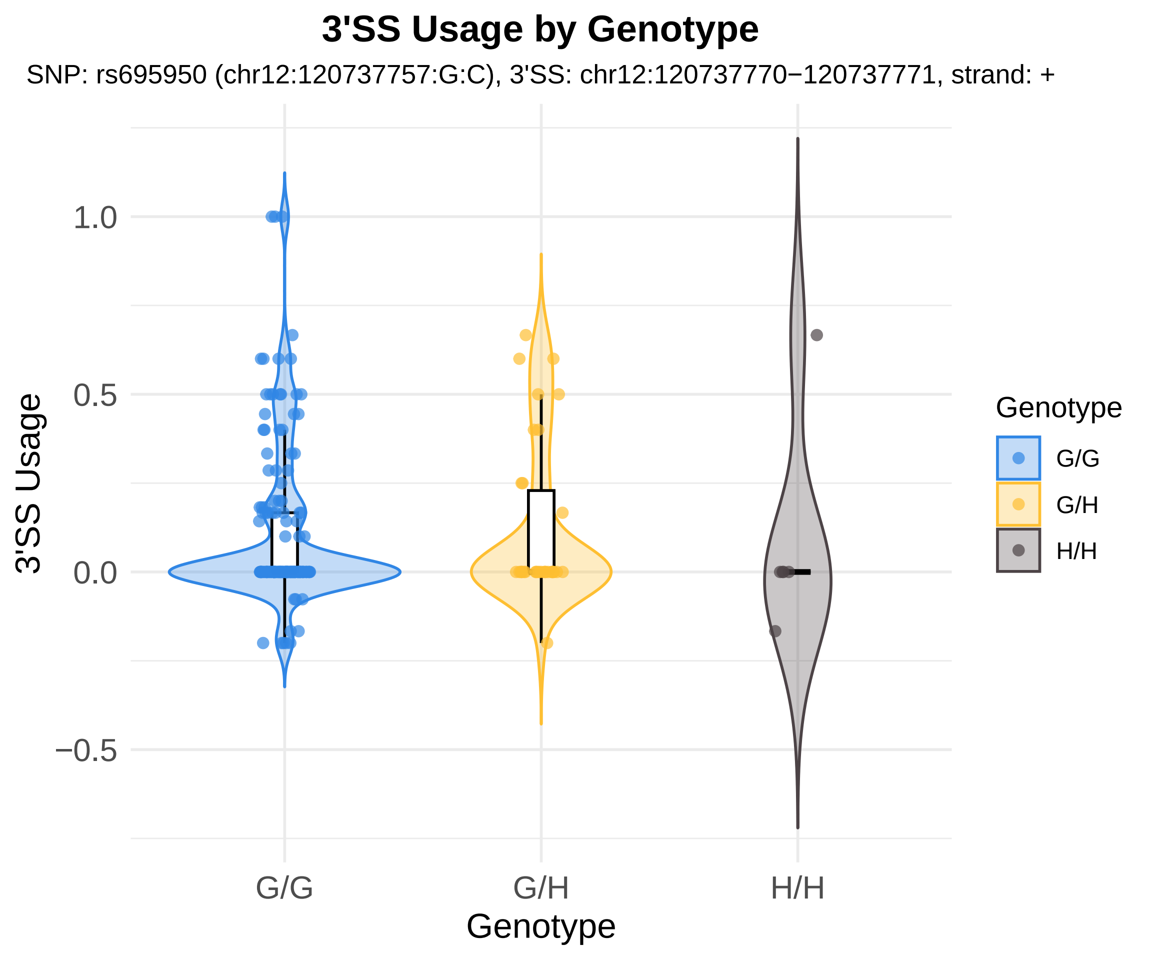
3'SS Usage by Genotype
SNP: rs695950 (chr12:120737757:G:C), 3'SS: chr12:120737770−120737771, strand: +
3'SS Usage
1.0
0.5
0.0
−0.5
G/G
G/H
H/H
Genotype
Genotype
G/G
G/H
H/H

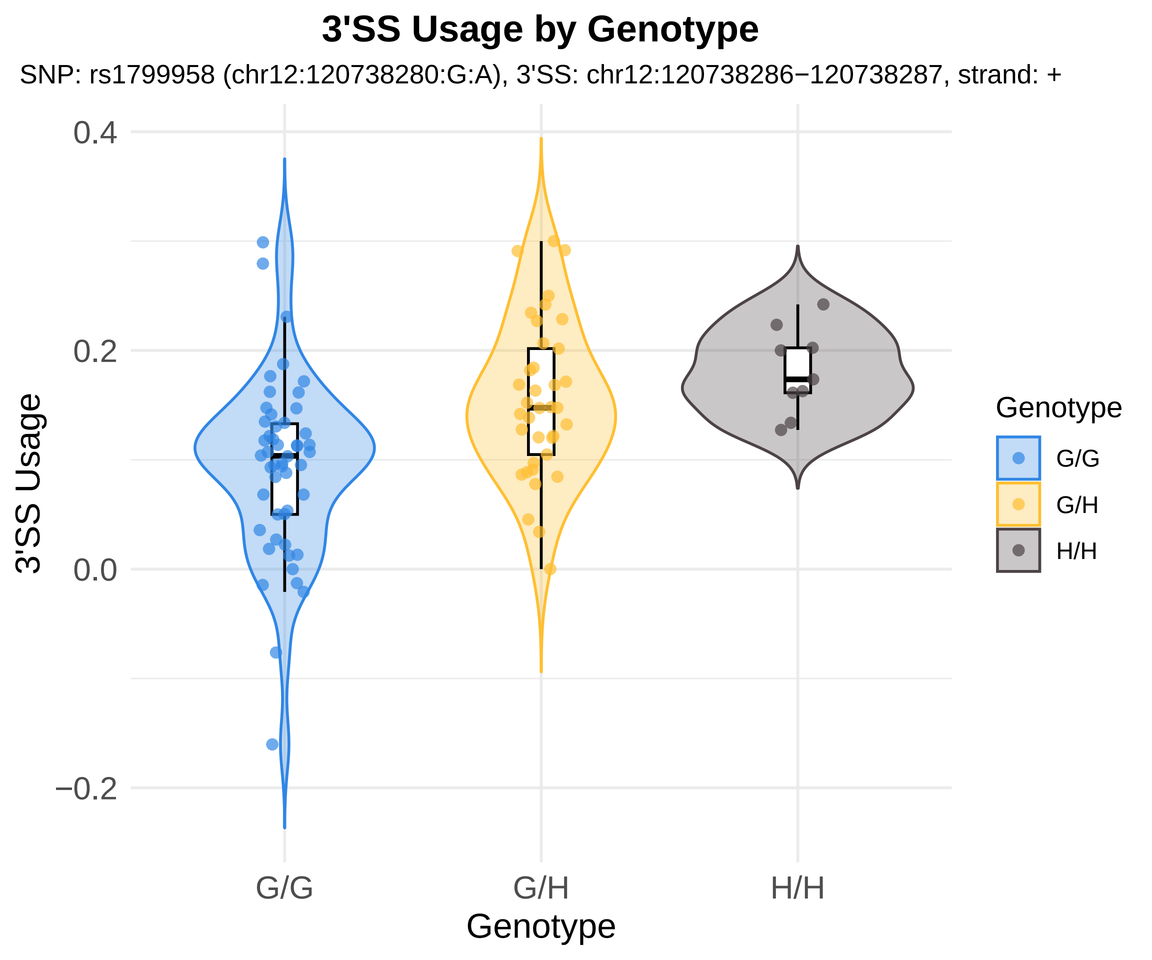
3'SS Usage by Genotype
SNP: rs1799958 (chr12:120738280:G:A), 3'SS: chr12:120738286−120738287, strand: +
3'SS Usage
0.4
0.2
0.0
−0.2
G/G
G/H
H/H
Genotype
Genotype
G/G
G/H
H/H

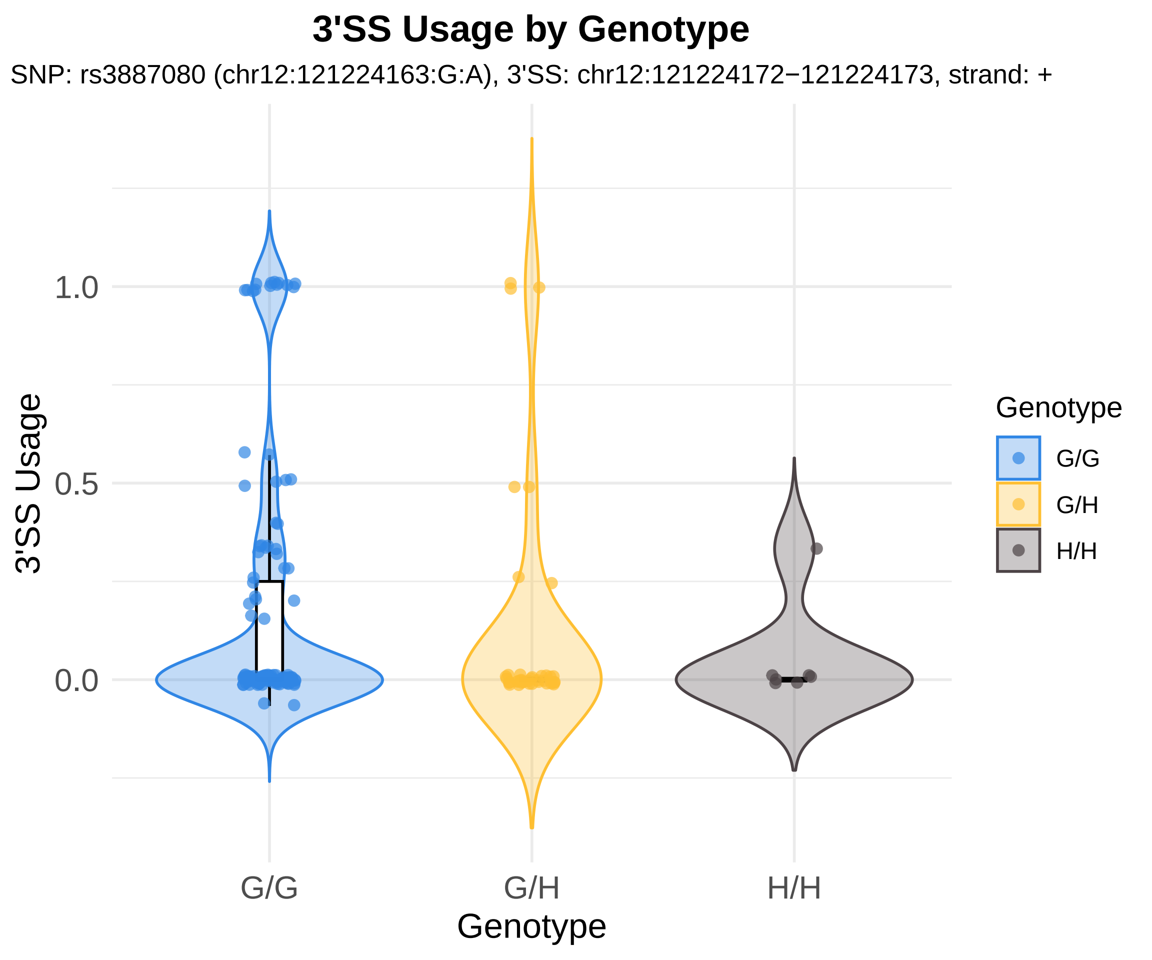
3'SS Usage by Genotype
SNP: rs3887080 (chr12:121224163:G:A), 3'SS: chr12:121224172−121224173, strand: +
3'SS Usage
1.0
0.5
0.0
G/G
G/H
H/H
Genotype
Genotype
G/G
G/H
H/H

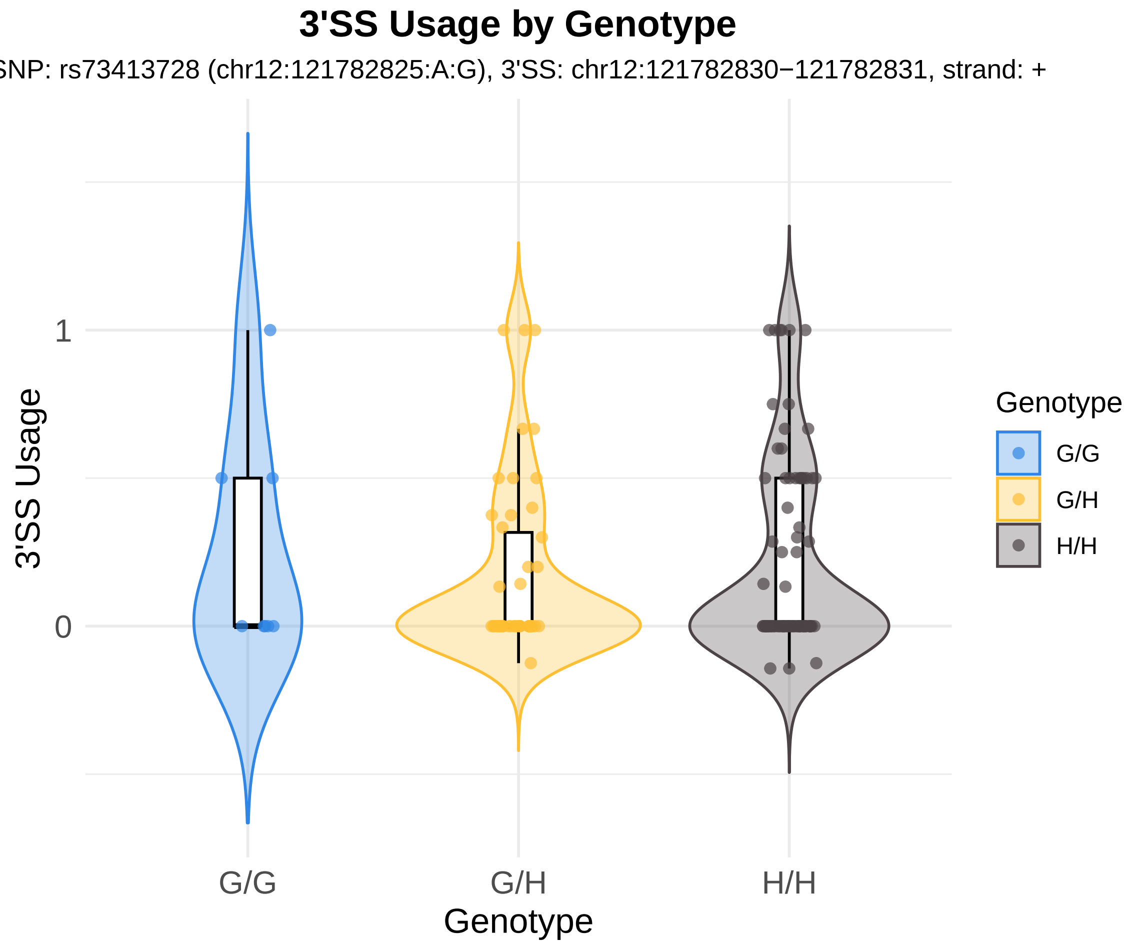
3'SS Usage by Genotype
SNP: rs73413728 (chr12:121782825:A:G), 3'SS: chr12:121782830−121782831, strand: +
3'SS Usage
1
0
G/G
G/H
H/H
Genotype
Genotype
G/G
G/H
H/H

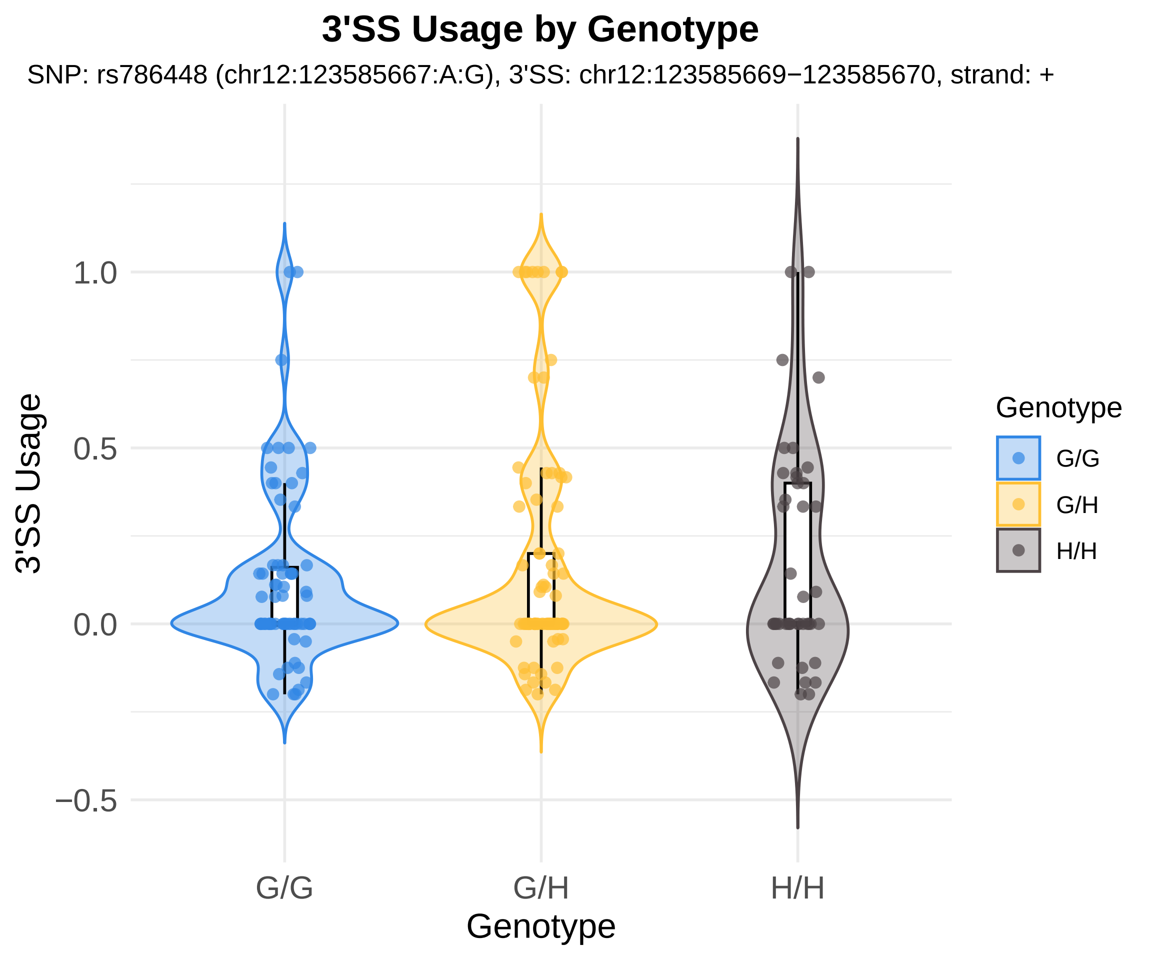

3'SS Usage by Genotype
SNP: rs786448 (chr12:123585667:A:G), 3'SS: chr12:123585669−123585670, strand: +
3'SS Usage
1.0
0.5
0.0
−0.5
G/G
G/H
H/H
Genotype
Genotype
G/G
G/H
H/H

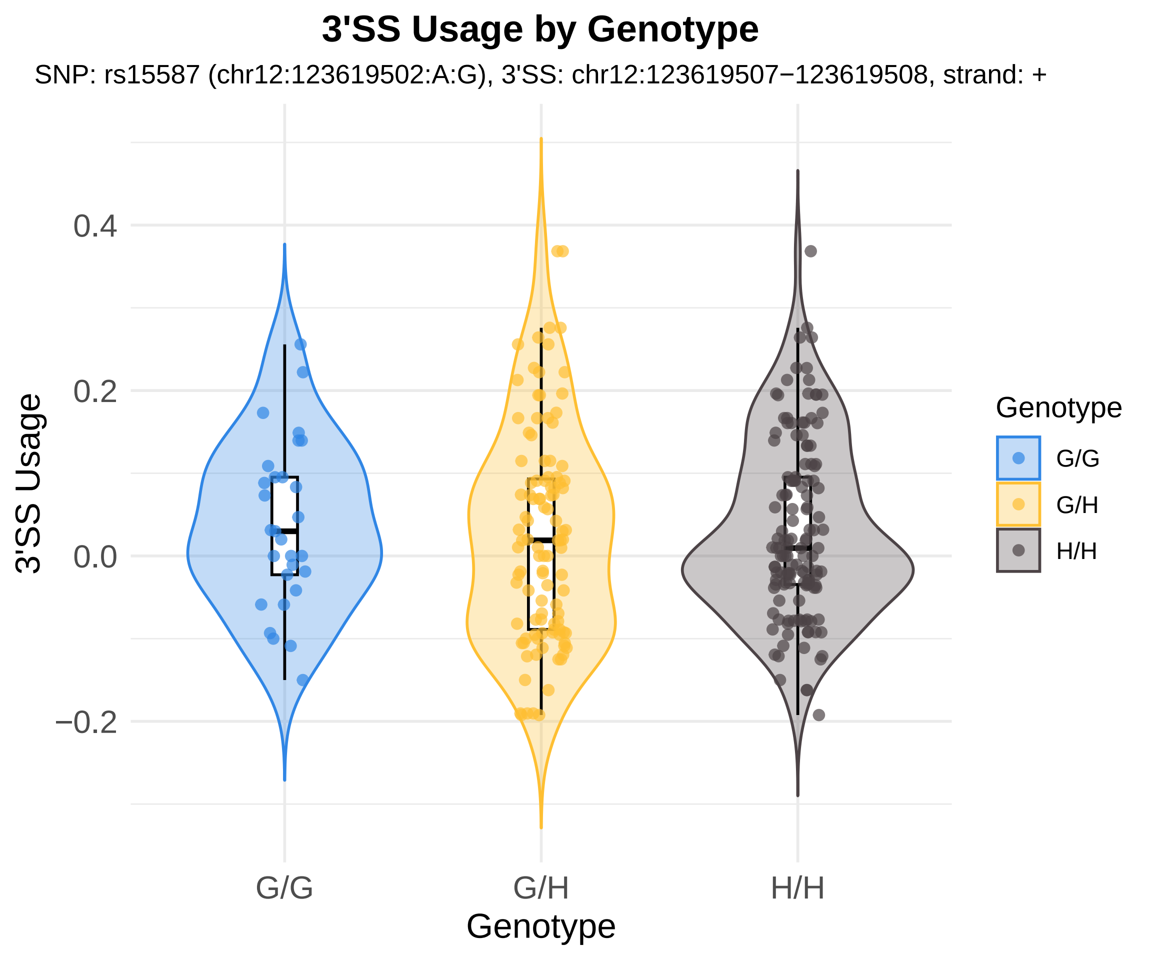

3'SS Usage by Genotype
SNP: rs15587 (chr12:123619502:A:G), 3'SS: chr12:123619507−123619508, strand: +
3'SS Usage
0.4
0.2
0.0
−0.2
G/G
G/H
H/H
Genotype
Genotype
G/G
G/H
H/H

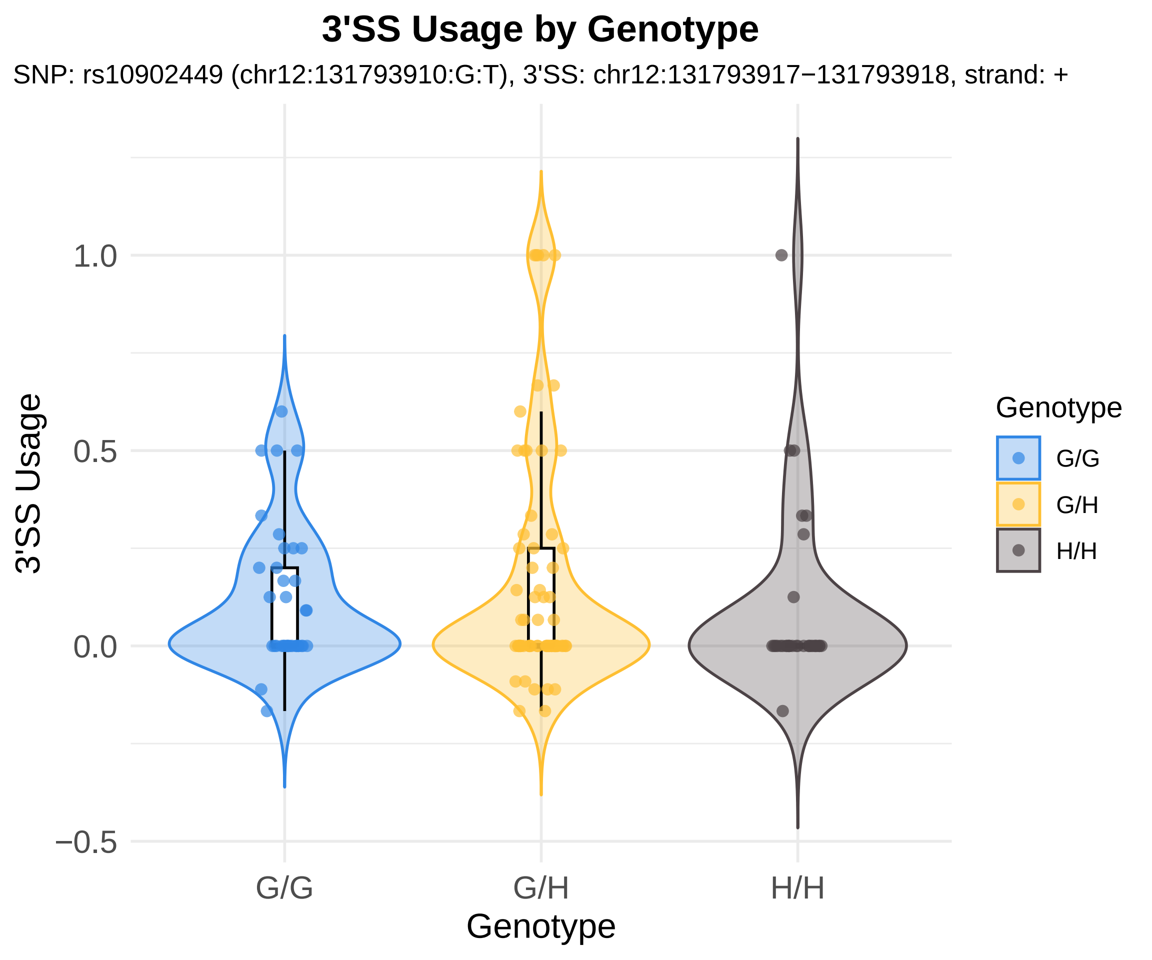

3'SS Usage by Genotype
SNP: rs10902449 (chr12:131793910:G:T), 3'SS: chr12:131793917−131793918, strand: +
3'SS Usage
1.0
0.5
0.0
−0.5
G/G
G/H
H/H
Genotype
Genotype
G/G
G/H
H/H

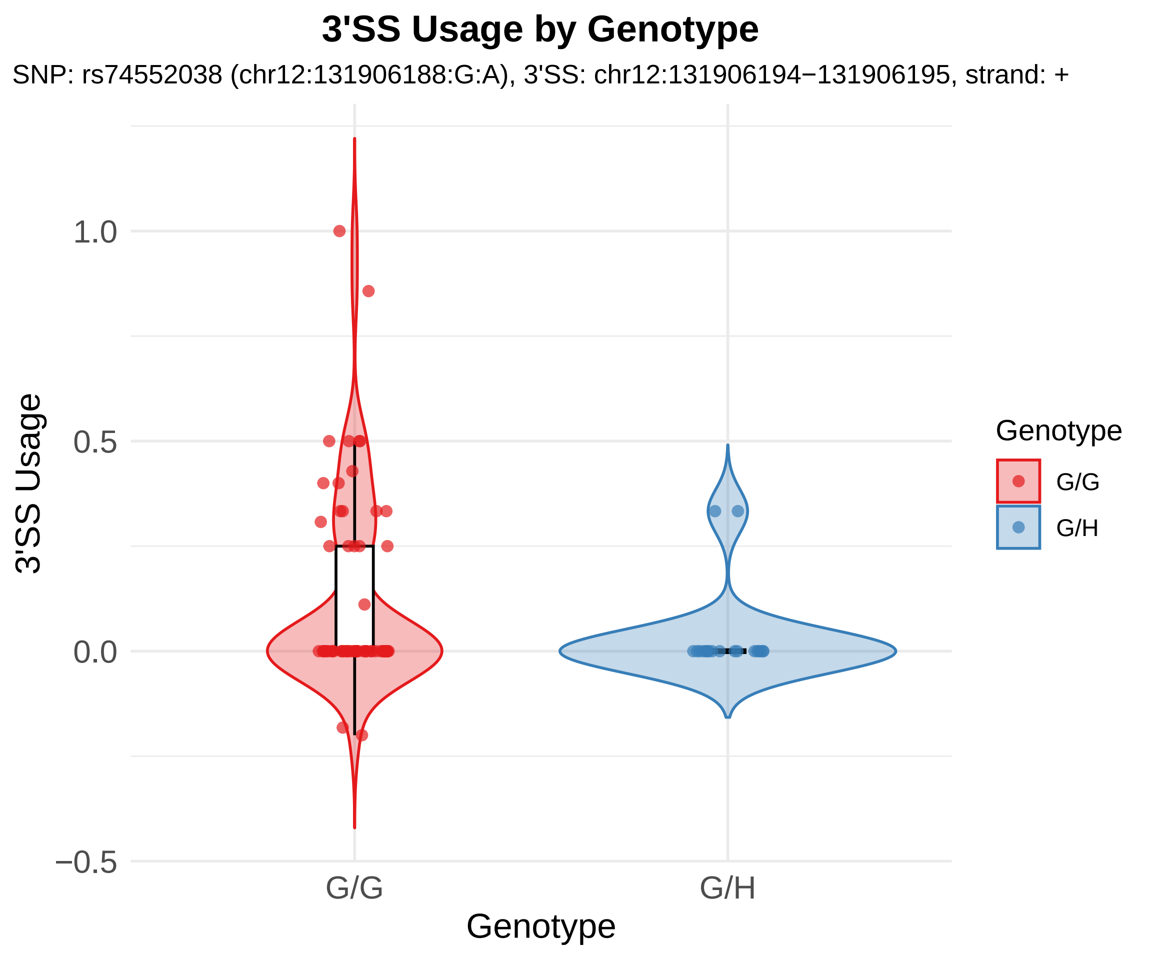
3'SS Usage by Genotype
SNP: rs74552038 (chr12:131906188:G:A), 3'SS: chr12:131906194−131906195, strand: +
3'SS Usage
1.0
0.5
0.0
−0.5
G/G
G/H
Genotype
Genotype
G/G
G/H

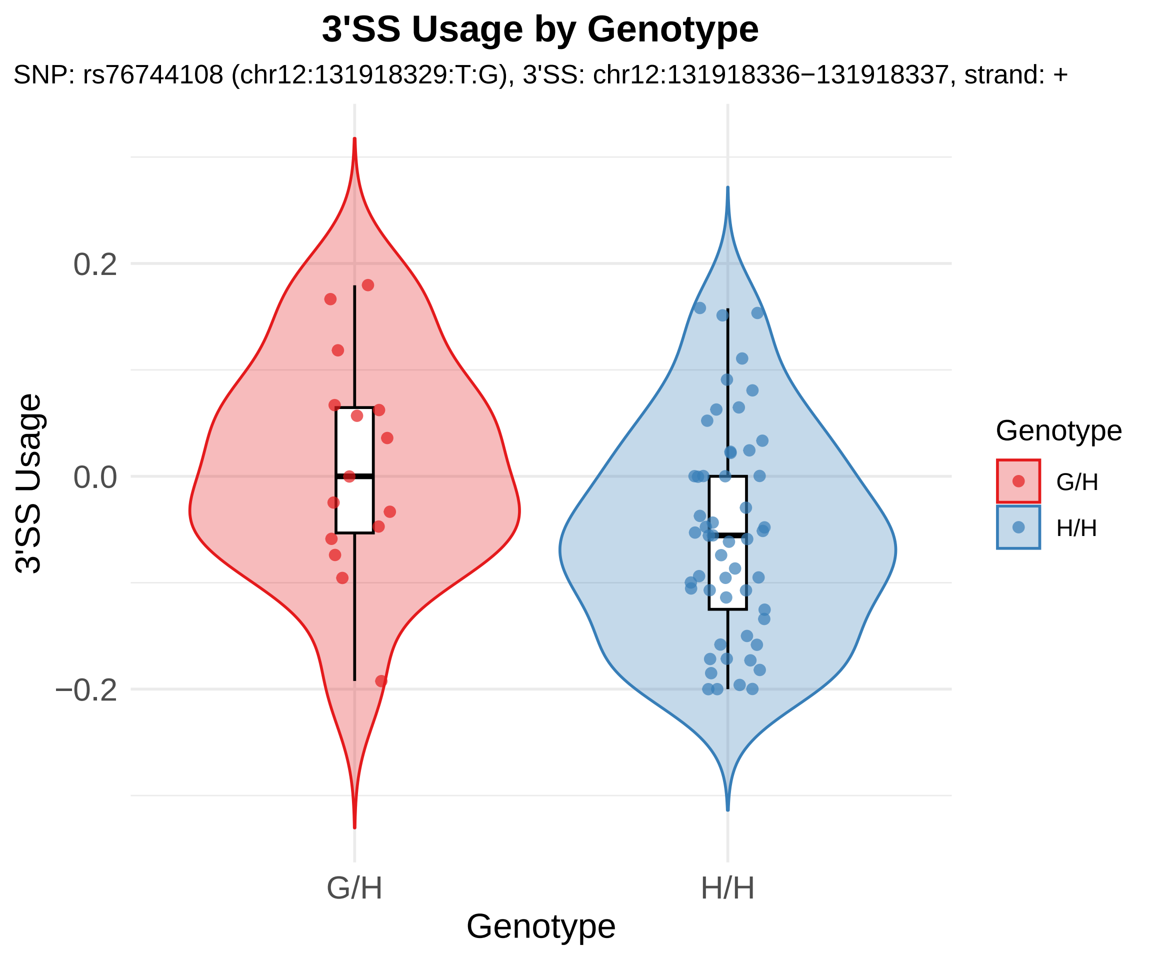
3'SS Usage by Genotype
SNP: rs76744108 (chr12:131918329:T:G), 3'SS: chr12:131918336−131918337, strand: +
3'SS Usage
0.2
0.0
−0.2
G/H
H/H
Genotype
Genotype
G/H
H/H

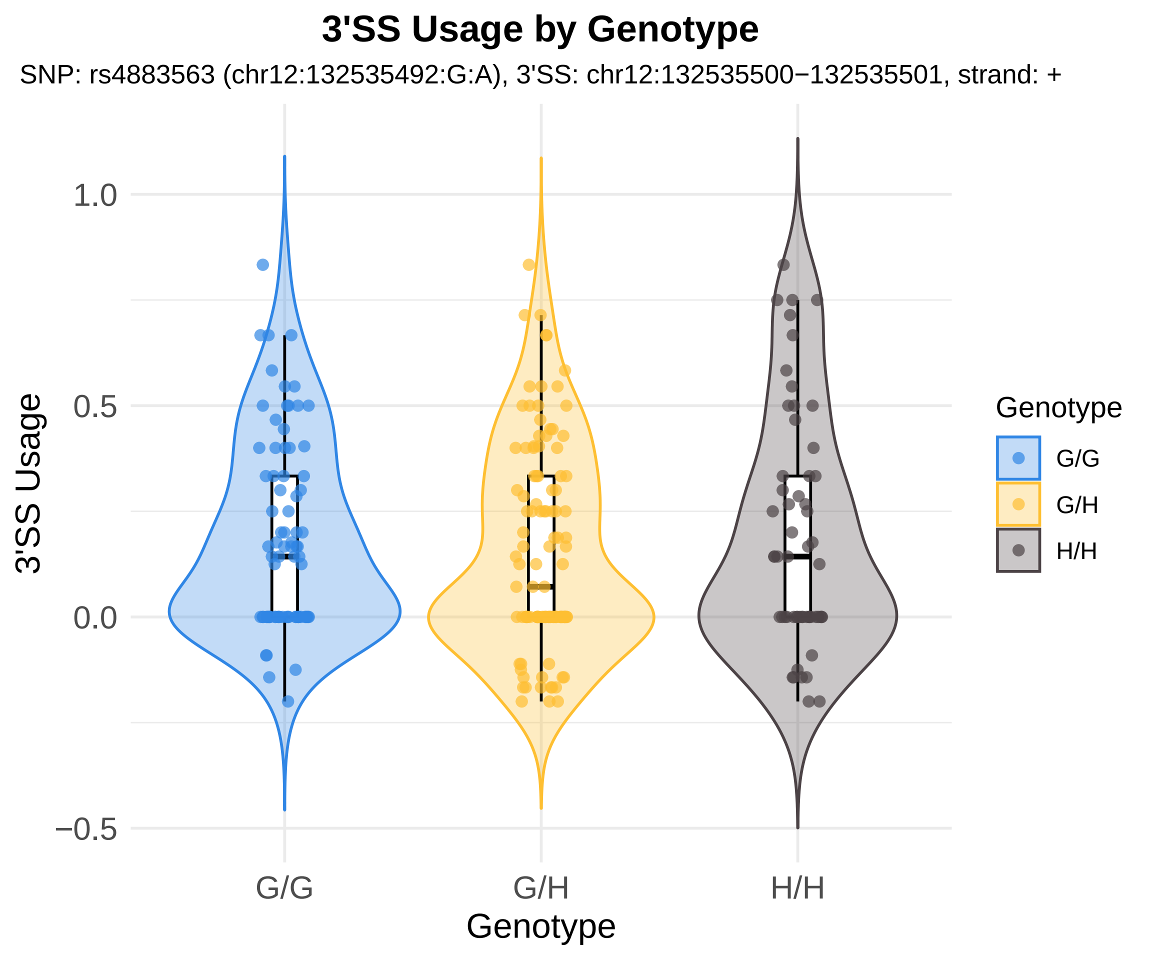
3'SS Usage by Genotype
SNP: rs4883563 (chr12:132535492:G:A), 3'SS: chr12:132535500−132535501, strand: +
3'SS Usage
1.0
0.5
0.0
−0.5
G/G
G/H
H/H
Genotype
Genotype
G/G
G/H
H/H

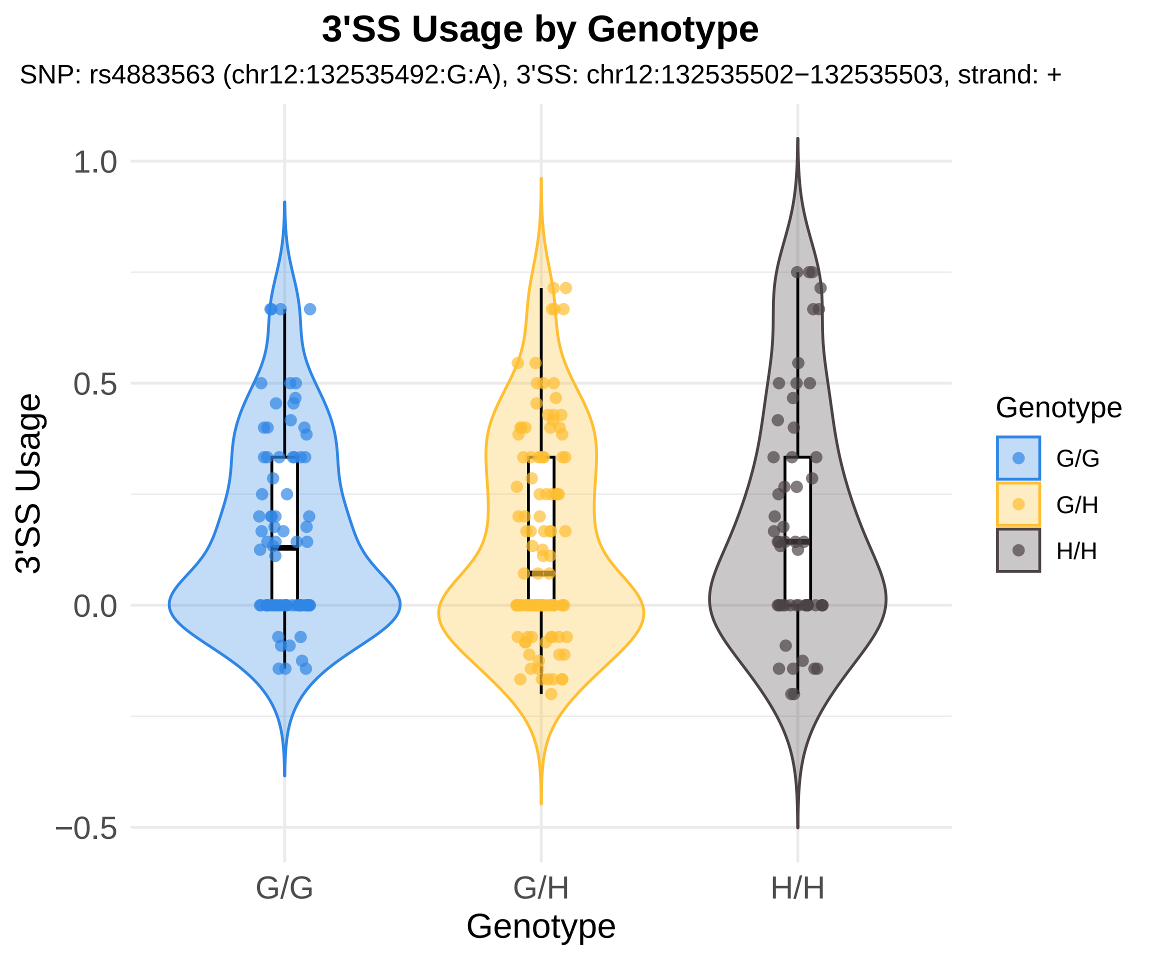
3'SS Usage by Genotype
SNP: rs4883563 (chr12:132535492:G:A), 3'SS: chr12:132535502−132535503, strand: +
1.0
0.5
0.0
−0.5
3'SS Usage
G/G
G/H
H/H
Genotype
Genotype
G/G
G/H
H/H

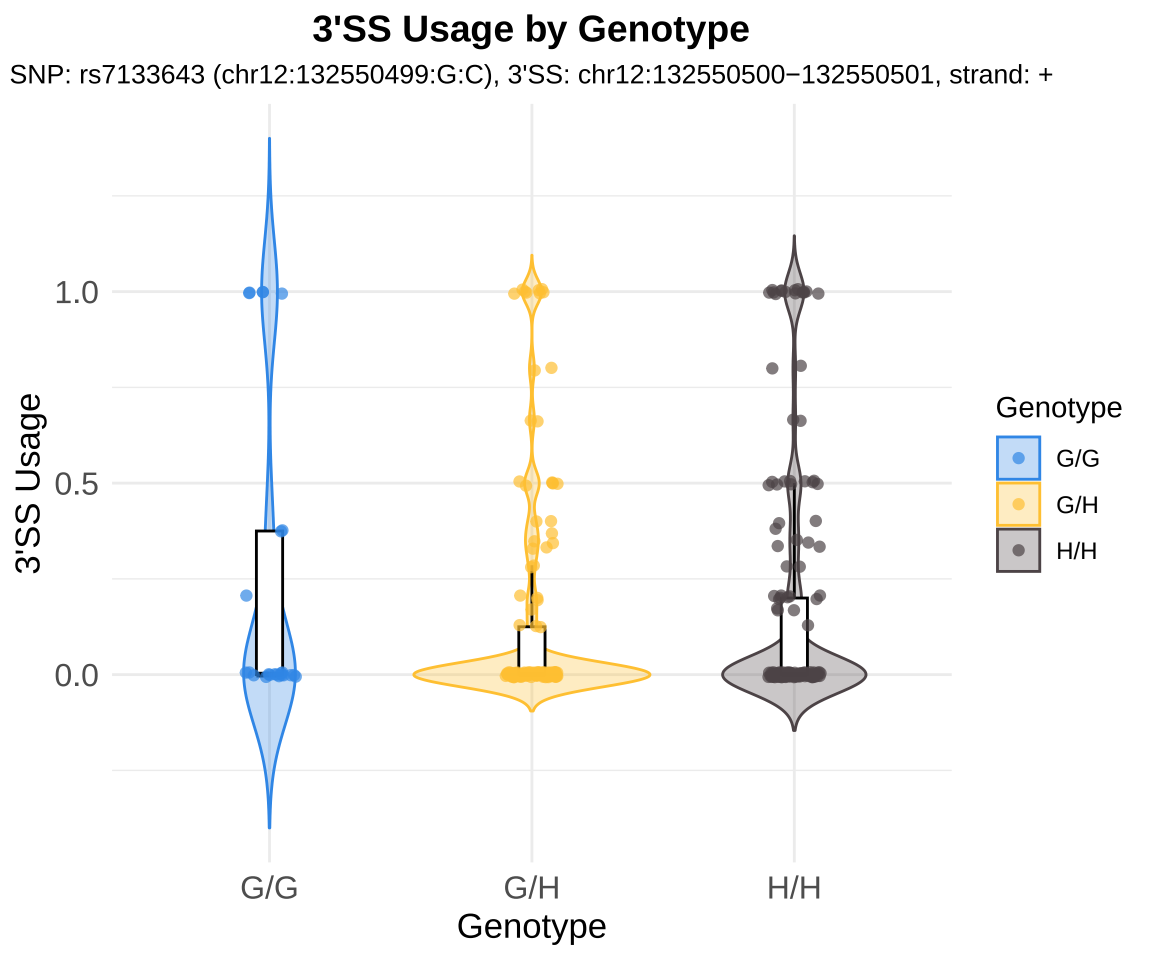

3'SS Usage by Genotype
SNP: rs7133643 (chr12:132550499:G:C), 3'SS: chr12:132550500−132550501, strand: +
3'SS Usage
1.0
0.5
0.0
G/G
G/H
H/H
Genotype
Genotype
G/G
G/H
H/H

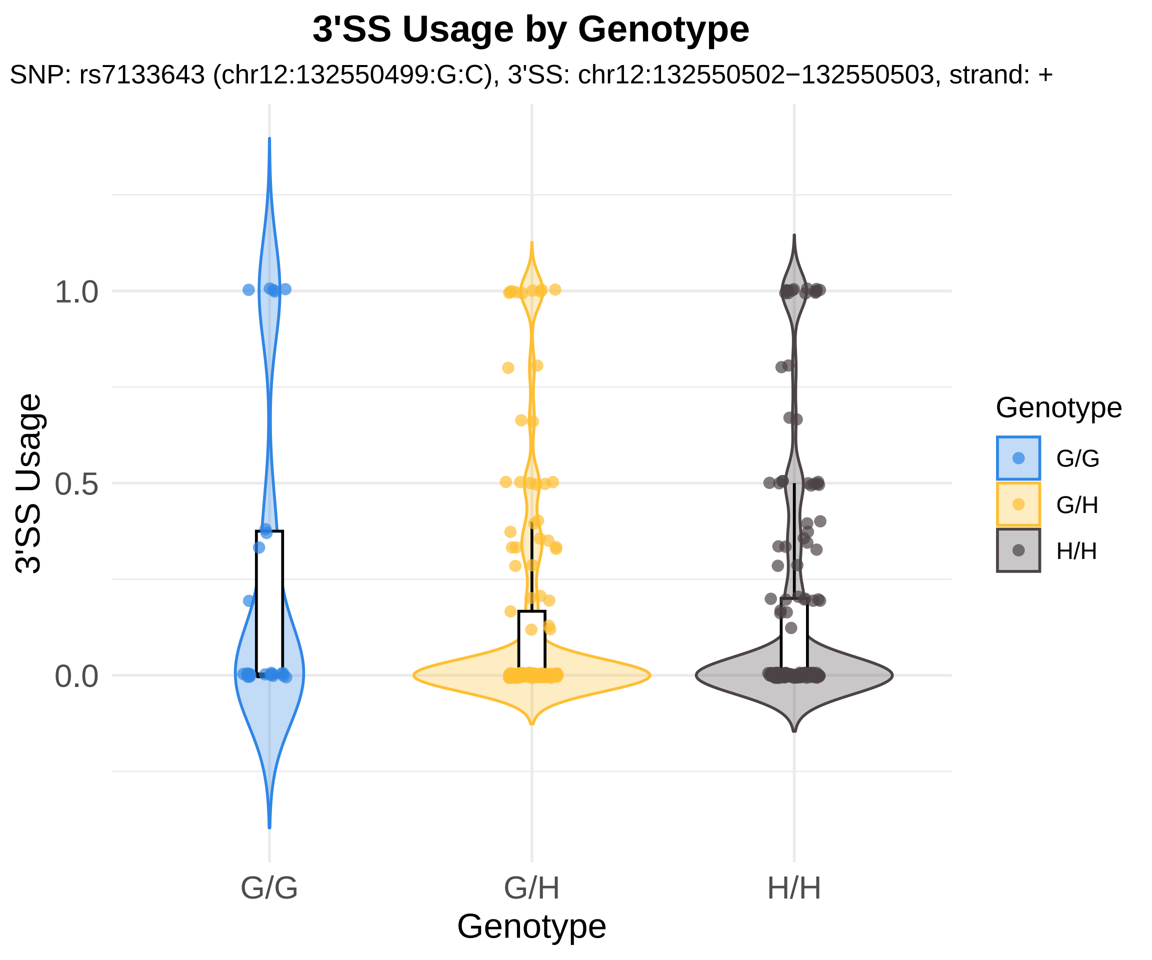
3'SS Usage by Genotype
SNP: rs7133643 (chr12:132550499:G:C), 3'SS: chr12:132550502−132550503, strand: +
3'SS Usage
1.0
0.5
0.0
G/G
G/H
H/H
Genotype
Genotype
G/G
G/H
H/H

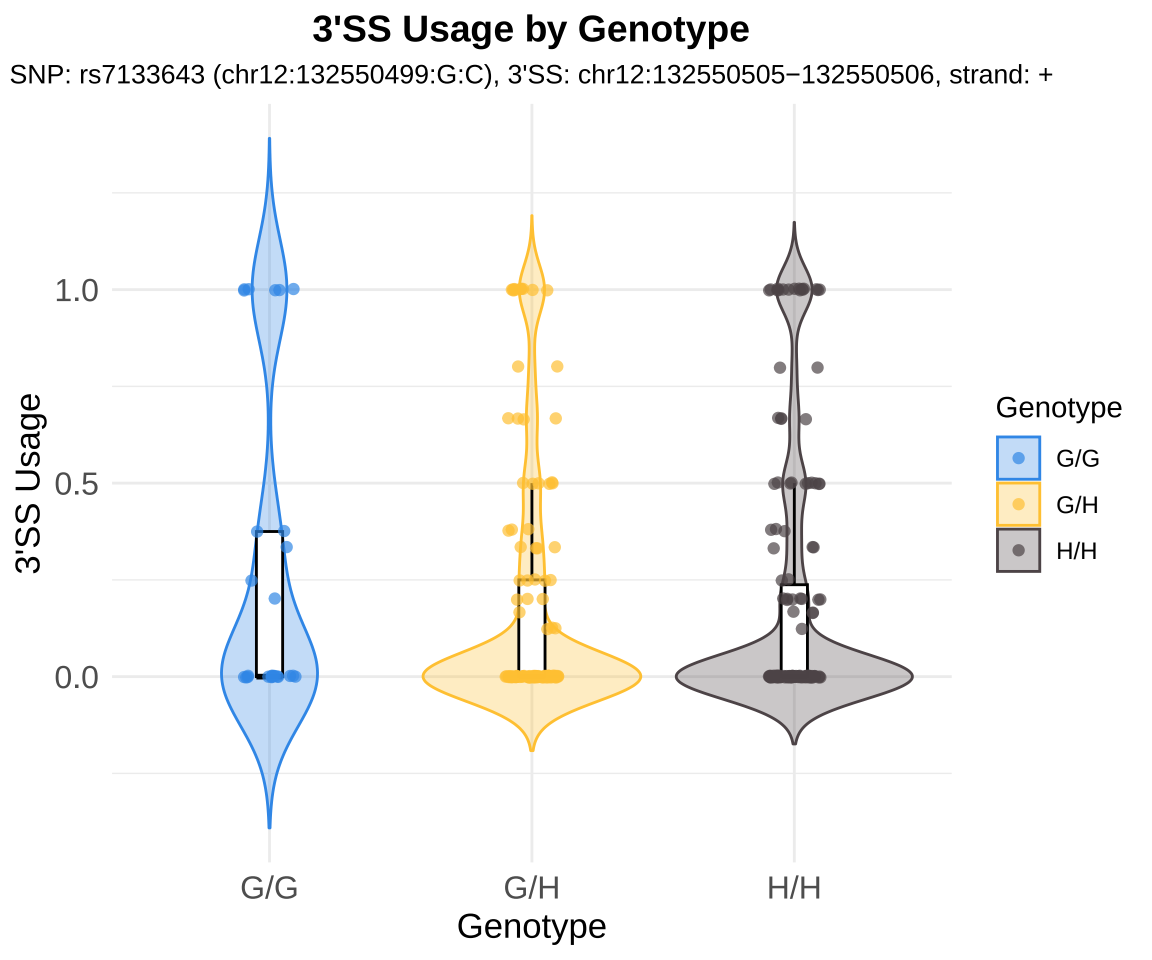
3'SS Usage by Genotype
SNP: rs7133643 (chr12:132550499:G:C), 3'SS: chr12:132550505−132550506, strand: +
3'SS Usage
1.0
0.5
0.0
G/G
G/H
H/H
Genotype
Genotype
G/G
G/H
H/H

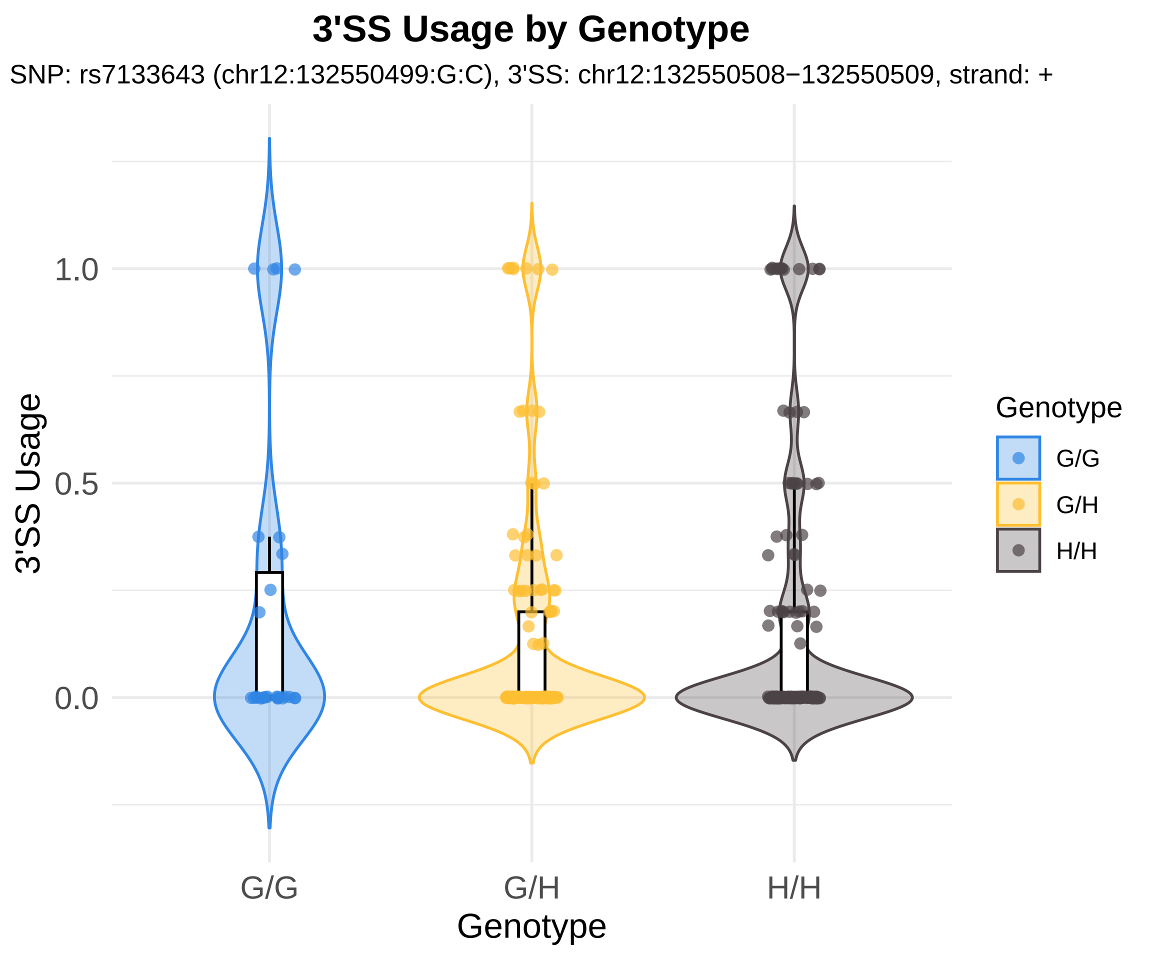
3'SS Usage by Genotype
SNP: rs7133643 (chr12:132550499:G:C), 3'SS: chr12:132550508−132550509, strand: +
3'SS Usage
1.0
0.5
0.0
G/G
G/H
H/H
Genotype
Genotype
G/G
G/H
H/H

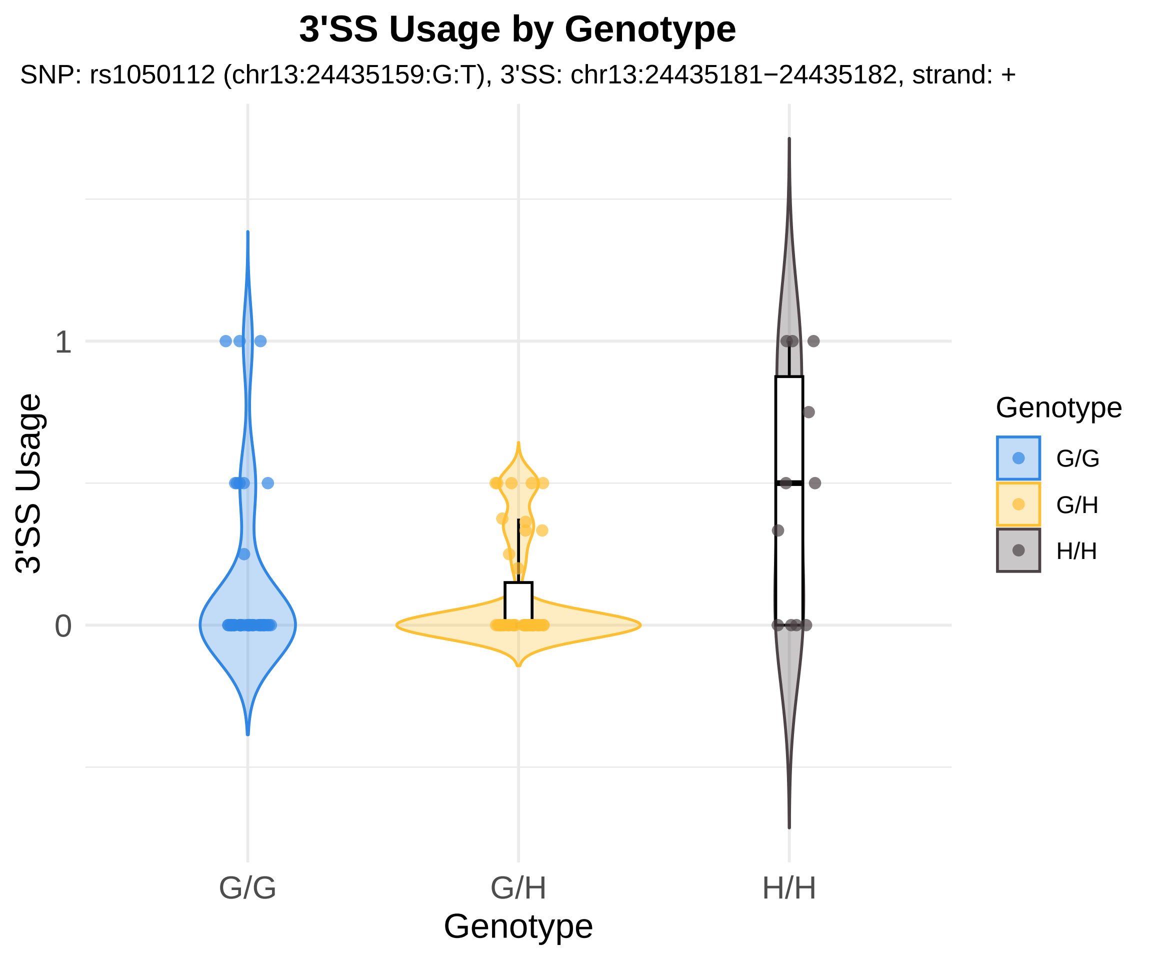
3'SS Usage by Genotype
SNP: rs1050112 (chr13:24435159:G:T), 3'SS: chr13:24435181−24435182, strand: +
3'SS Usage
1
0
G/G
G/H
H/H
Genotype
Genotype
G/G
G/H
H/H

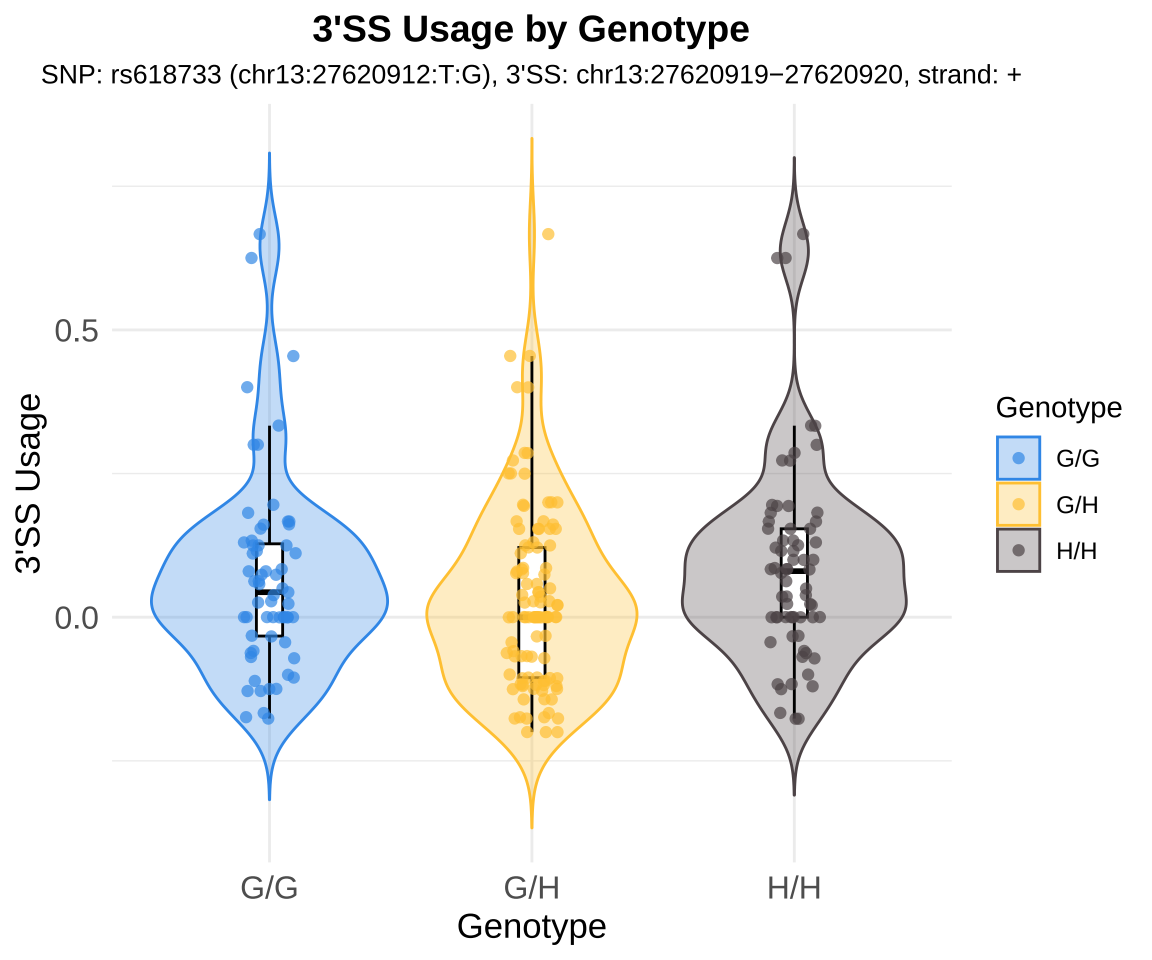

3'SS Usage by Genotype
SNP: rs618733 (chr13:27620912:T:G), 3'SS: chr13:27620919−27620920, strand: +
3'SS Usage
0.5
0.0
G/G
G/H
H/H
Genotype
Genotype
G/G
G/H
H/H

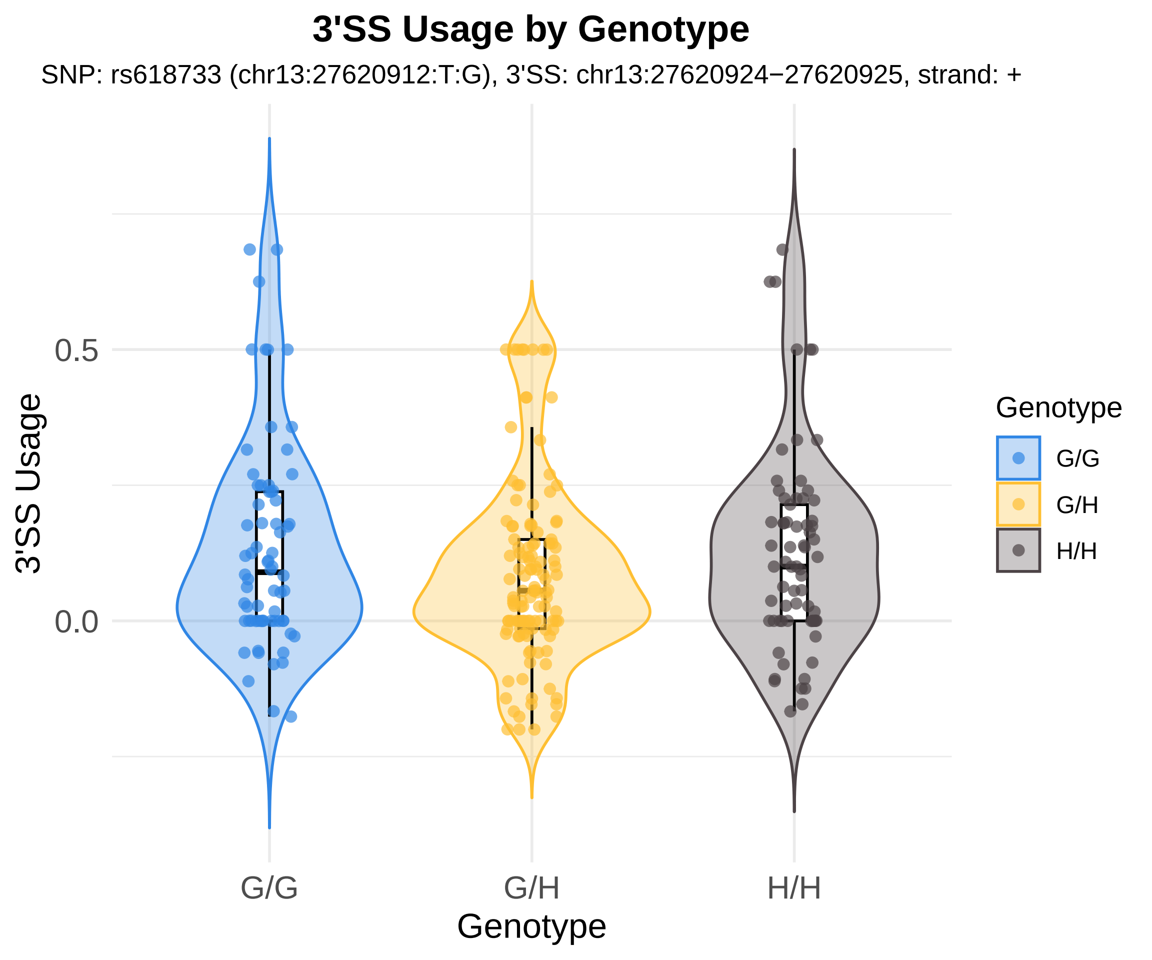
3'SS Usage by Genotype
SNP: rs618733 (chr13:27620912:T:G), 3'SS: chr13:27620924−27620925, strand: +
3'SS Usage
0.5
0.0
G/G
G/H
H/H
Genotype
Genotype
G/G
G/H
H/H

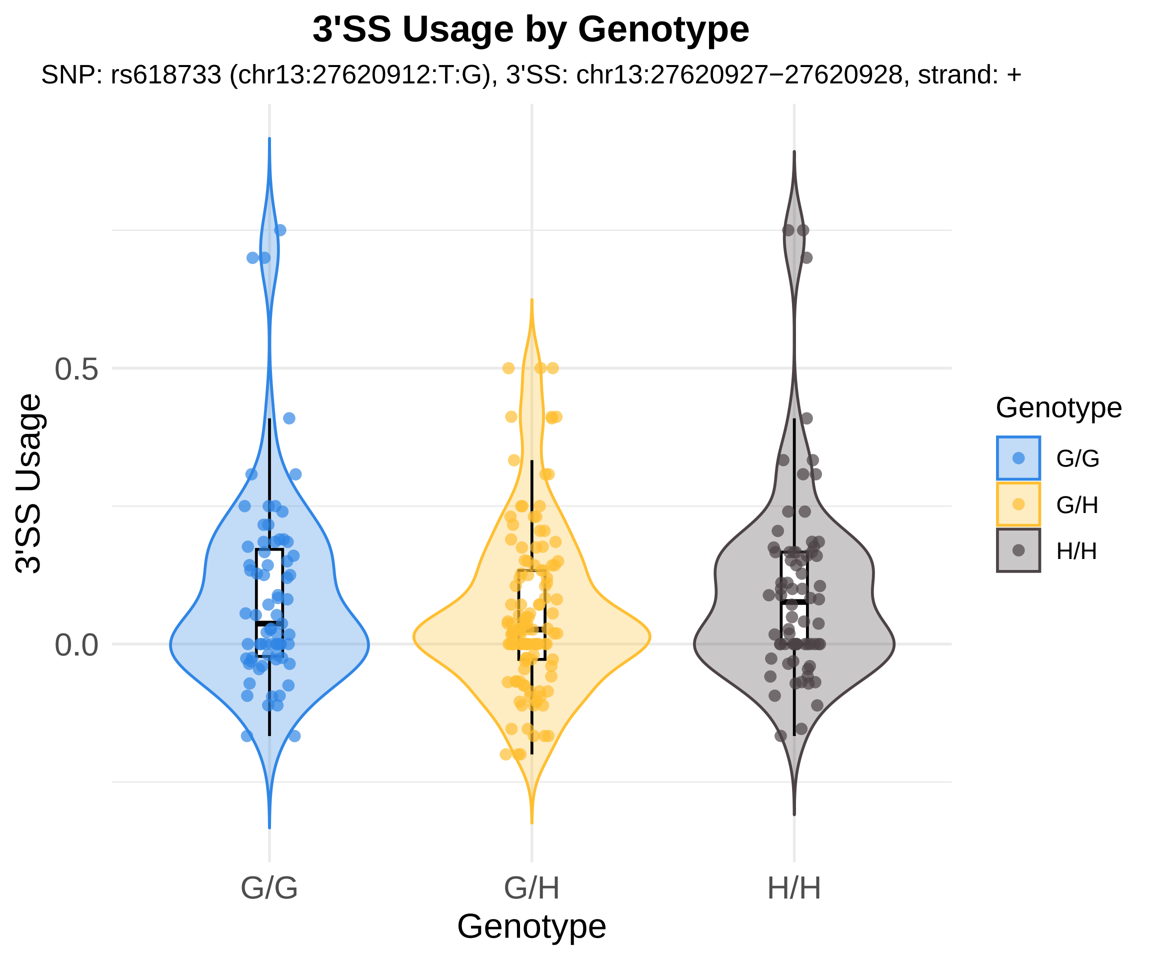
3'SS Usage by Genotype
SNP: rs618733 (chr13:27620912:T:G), 3'SS: chr13:27620927−27620928, strand: +
3'SS Usage
0.5
0.0
G/G
G/H
H/H
Genotype
Genotype
G/G
G/H
H/H

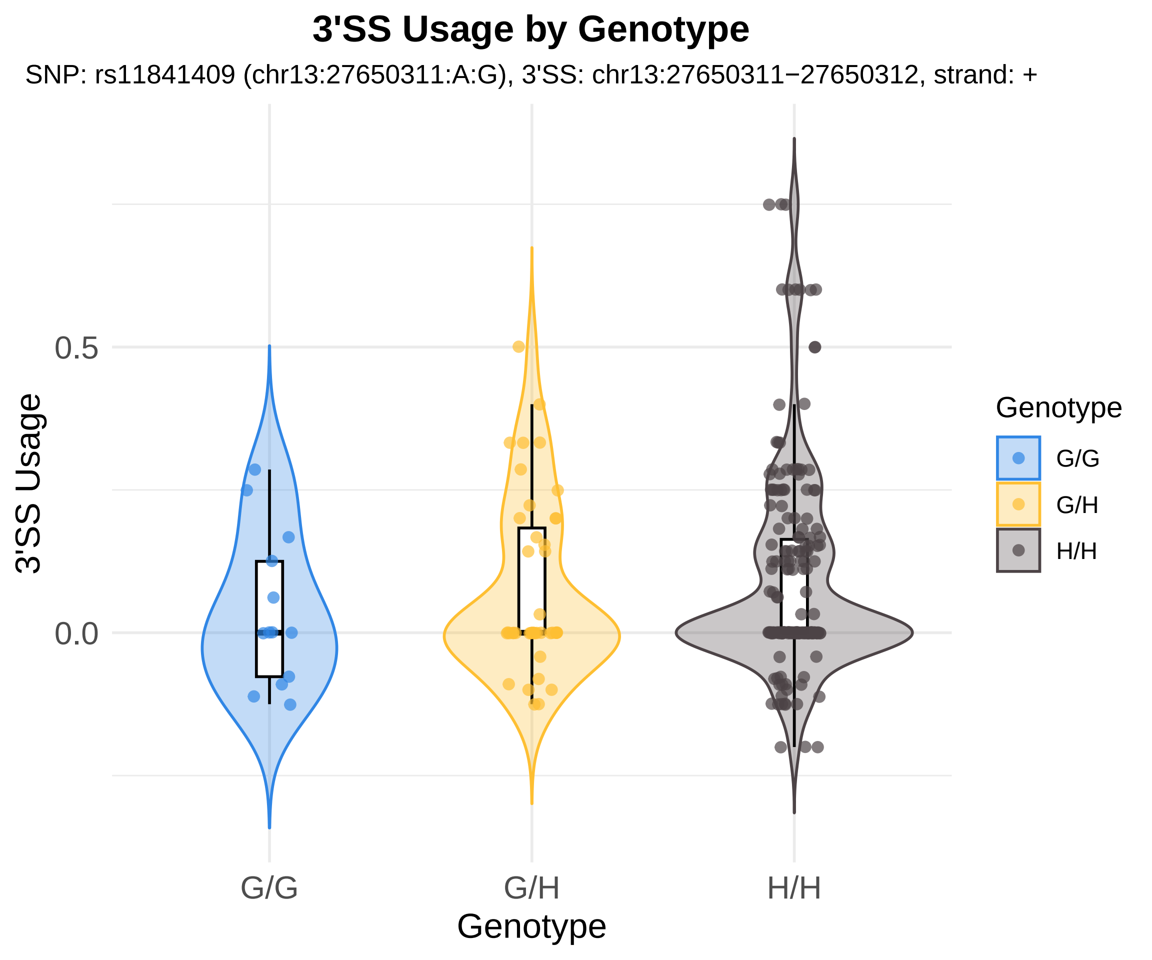
3'SS Usage by Genotype
SNP: rs11841409 (chr13:27650311:A:G), 3'SS: chr13:27650311−27650312, strand: +
3'SS Usage
0.5
0.0
G/G
G/H
H/H
Genotype
Genotype
G/G
G/H
H/H

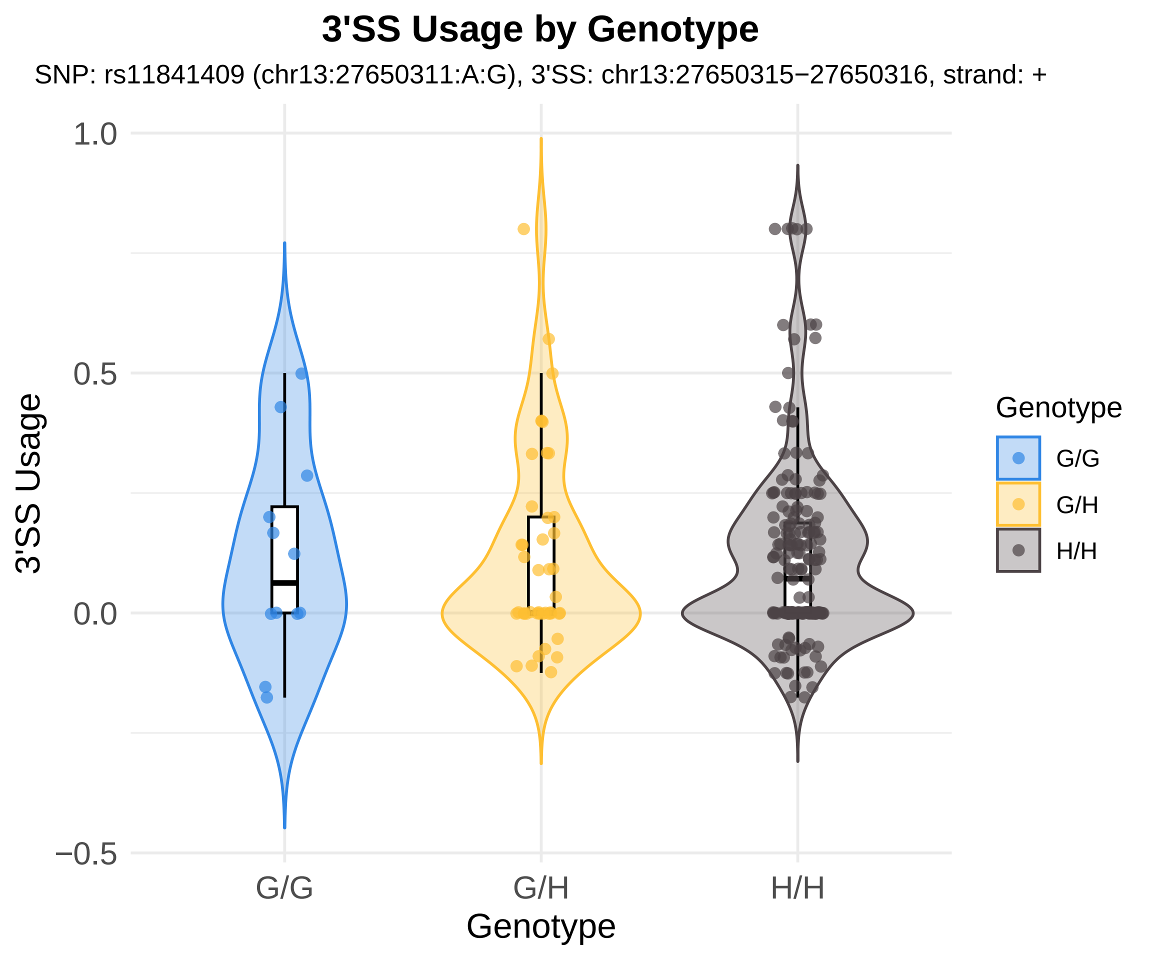

3'SS Usage by Genotype
SNP: rs11841409 (chr13:27650311:A:G), 3'SS: chr13:27650315−27650316, strand: +
1.0
0.5
0.0
−0.5
3'SS Usage
G/G
G/H
H/H
Genotype
Genotype
G/G
G/H
H/H

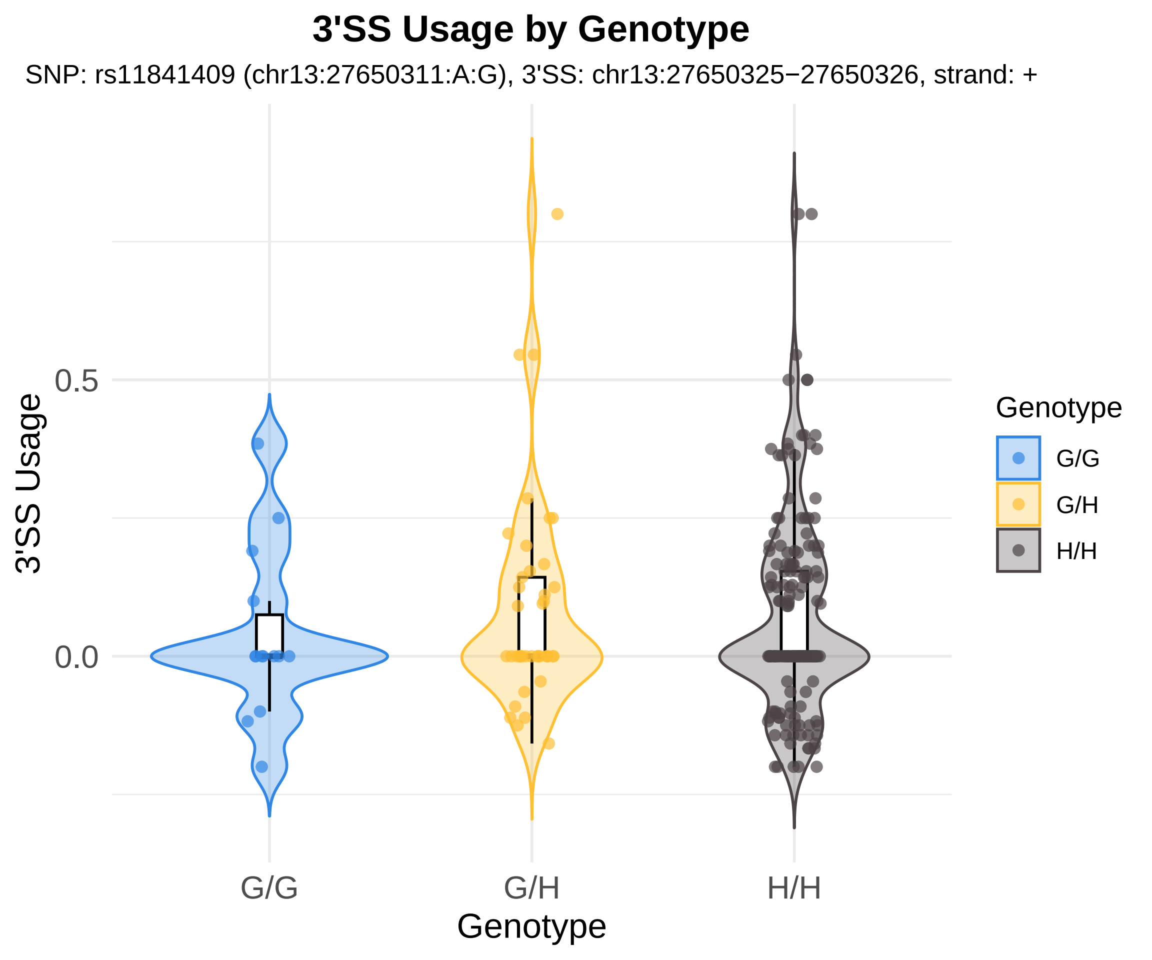
3'SS Usage by Genotype
SNP: rs11841409 (chr13:27650311:A:G), 3'SS: chr13:27650325−27650326, strand: +
3'SS Usage
0.5
0.0
G/G
G/H
H/H
Genotype
Genotype
G/G
G/H
H/H

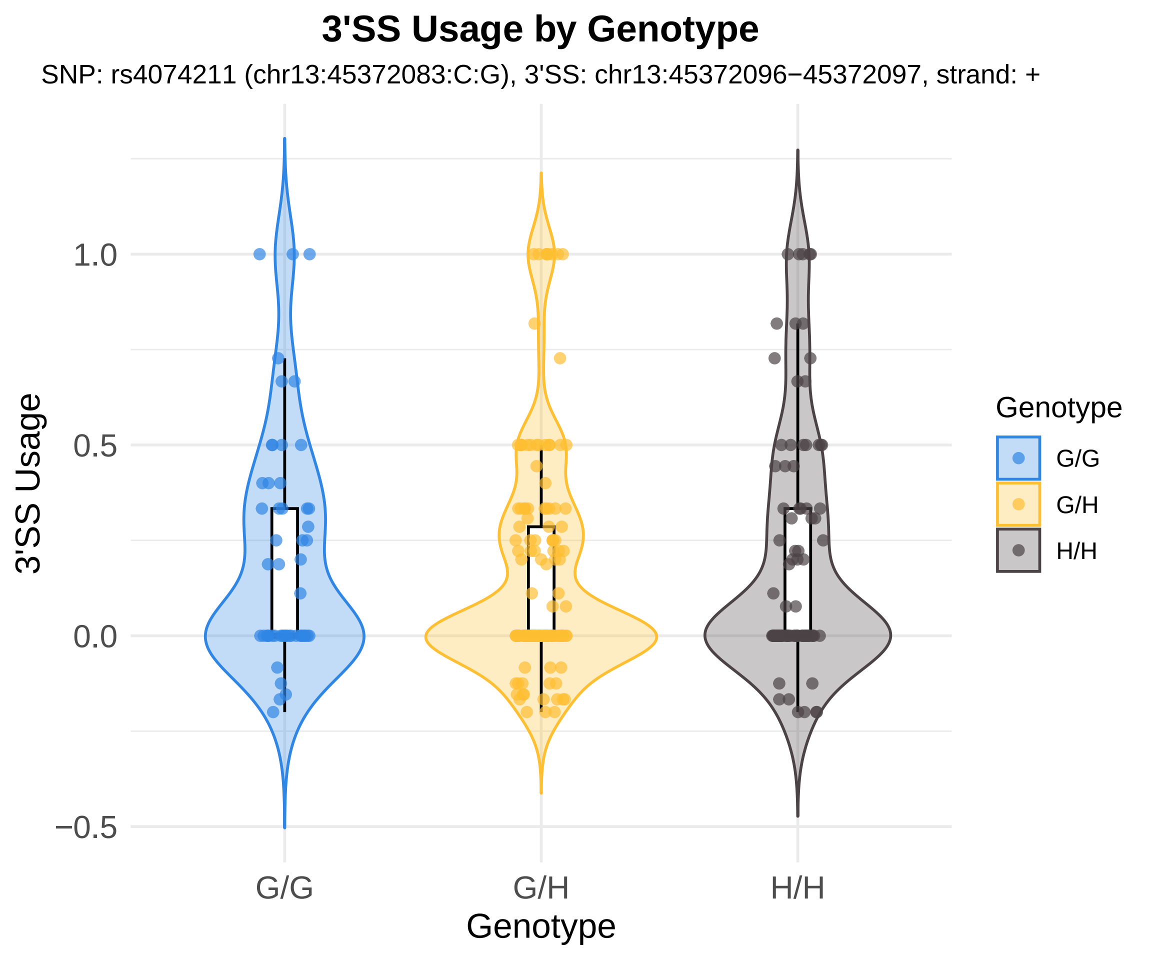

3'SS Usage by Genotype
SNP: rs4074211 (chr13:45372083:C:G), 3'SS: chr13:45372096−45372097, strand: +
3'SS Usage
1.0
0.5
0.0
−0.5
G/G
G/H
H/H
Genotype
Genotype
G/G
G/H
H/H

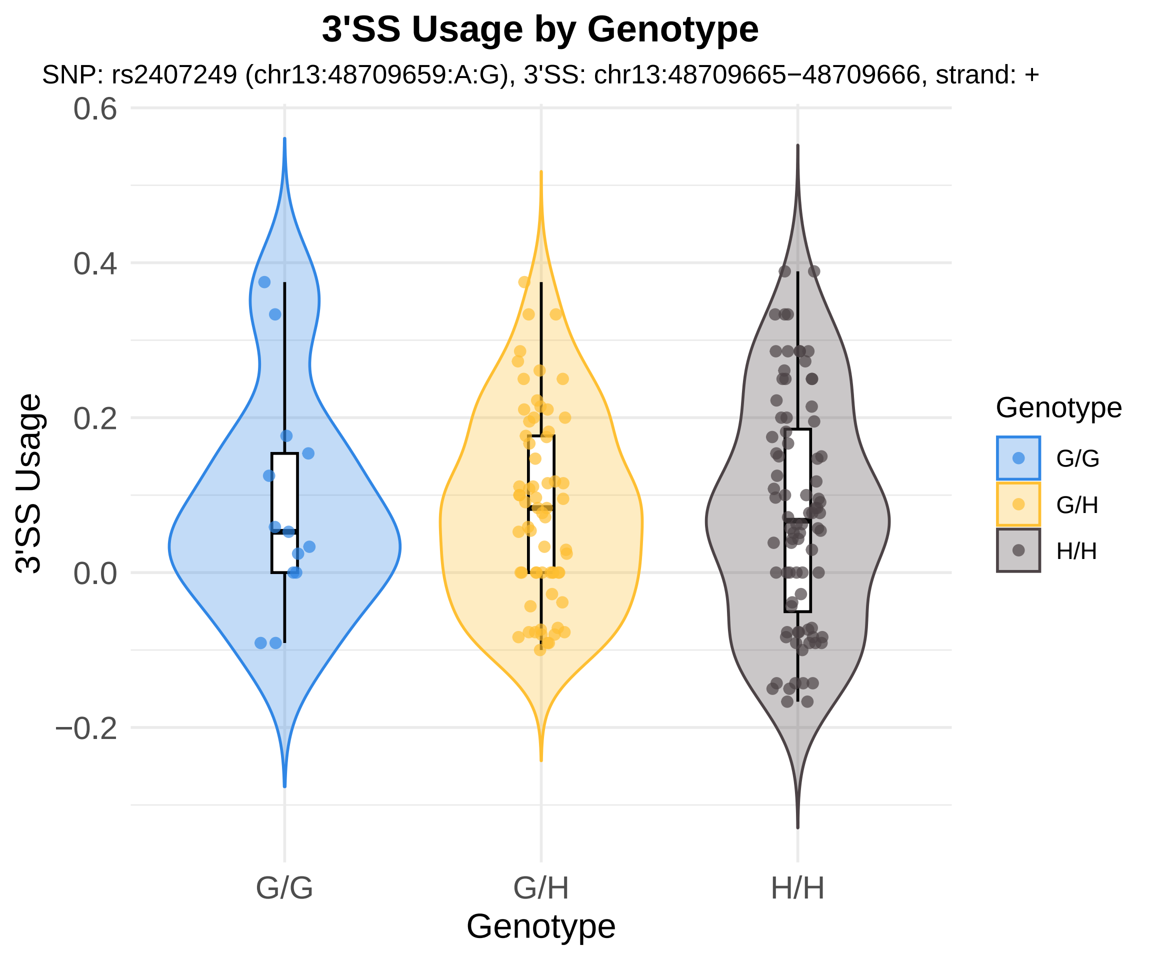

3'SS Usage by Genotype
SNP: rs2407249 (chr13:48709659:A:G), 3'SS: chr13:48709665−48709666, strand: +
0.6
0.4
0.2
0.0
−0.2
3'SS Usage
G/G
G/H
H/H
Genotype
Genotype
G/G
G/H
H/H

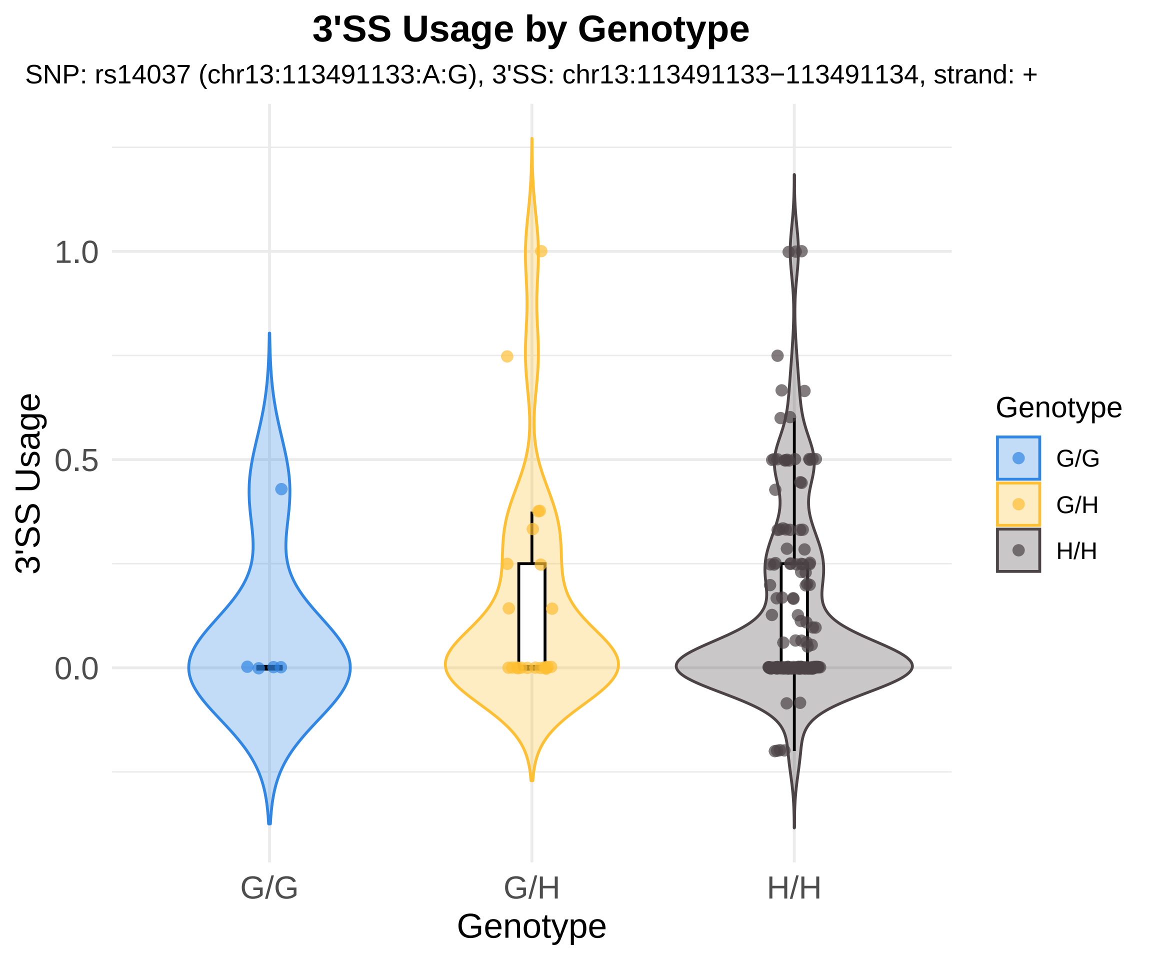
3'SS Usage by Genotype
SNP: rs14037 (chr13:113491133:A:G), 3'SS: chr13:113491133−113491134, strand: +
3'SS Usage
1.0
0.5
0.0
G/G
G/H
H/H
Genotype
Genotype
G/G
G/H
H/H

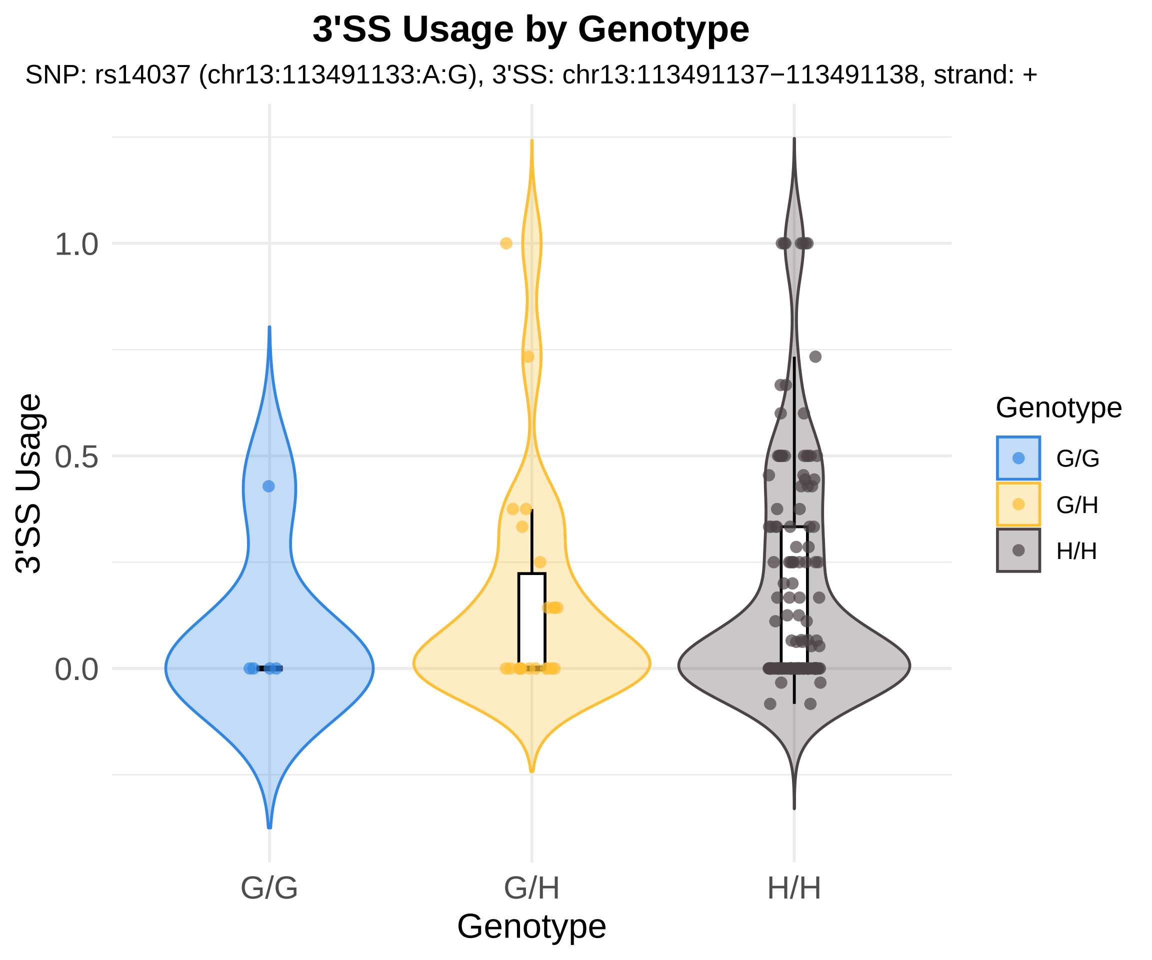

3'SS Usage by Genotype
SNP: rs14037 (chr13:113491133:A:G), 3'SS: chr13:113491137−113491138, strand: +
3'SS Usage
1.0
0.5
0.0
G/G
G/H
H/H
Genotype
Genotype
G/G
G/H
H/H

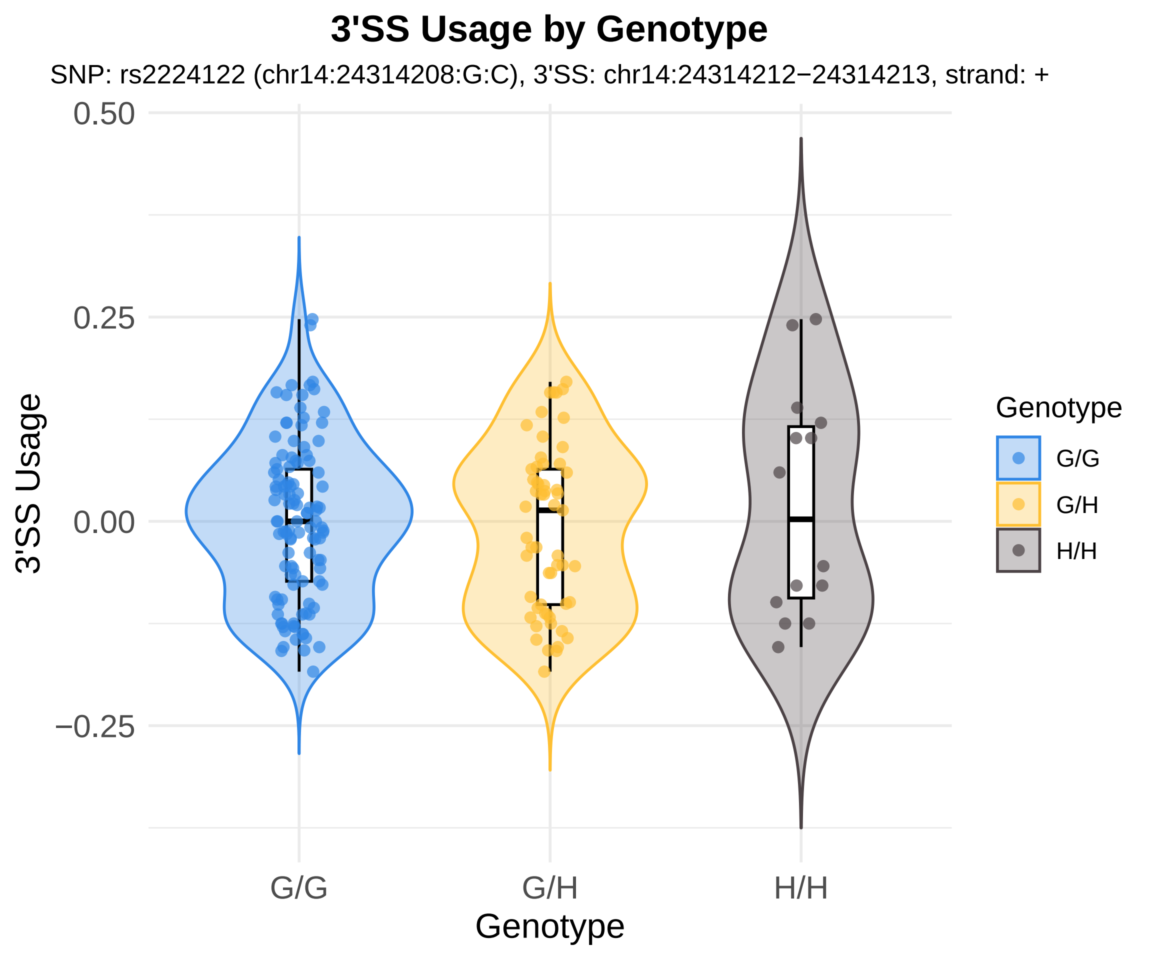

3'SS Usage by Genotype
SNP: rs2224122 (chr14:24314208:G:C), 3'SS: chr14:24314212−24314213, strand: +
0.50
0.25
0.00
−0.25
3'SS Usage
G/G
G/H
H/H
Genotype
Genotype
G/G
G/H
H/H

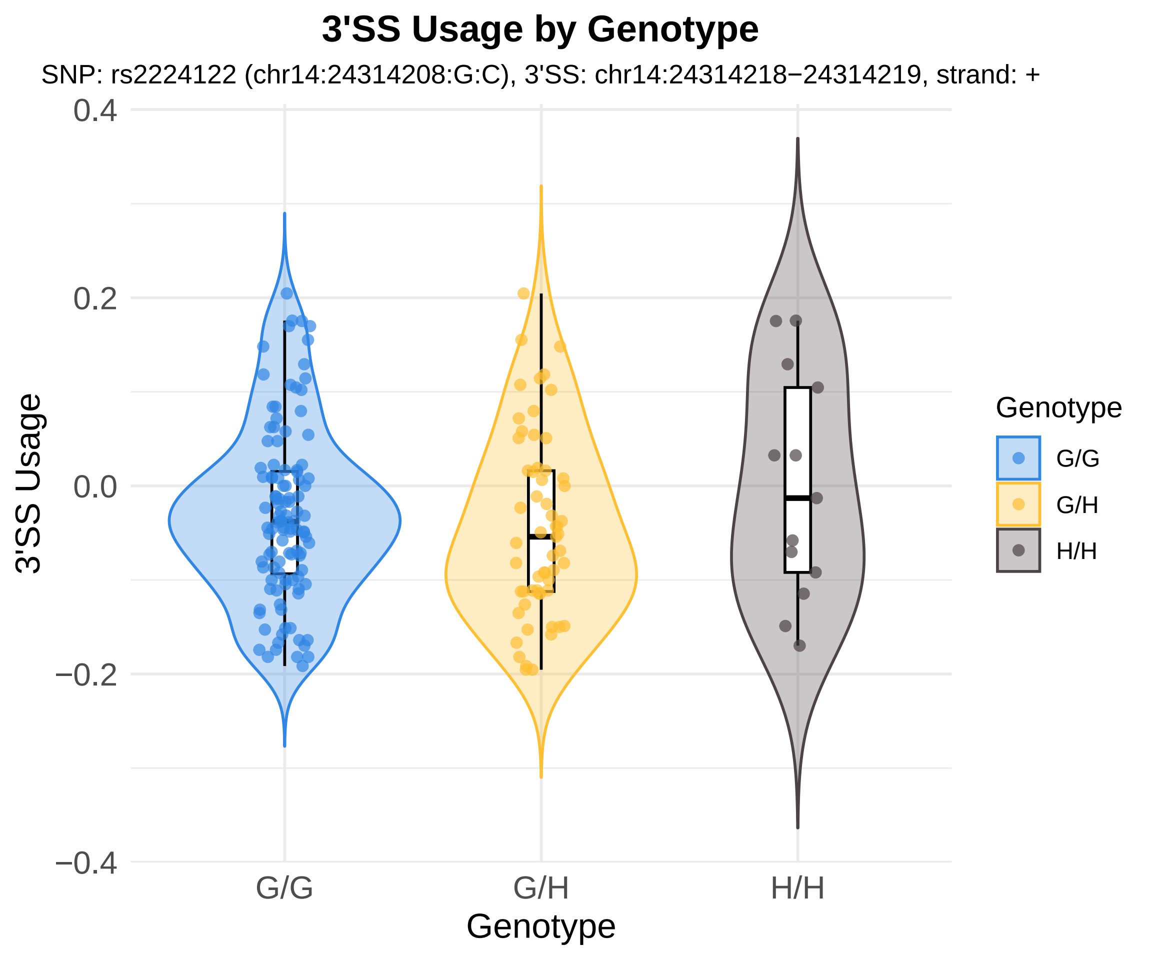
3'SS Usage by Genotype
SNP: rs2224122 (chr14:24314208:G:C), 3'SS: chr14:24314218−24314219, strand: +
3'SS Usage
0.4
0.2
0.0
−0.2
−0.4
G/G
G/H
H/H
Genotype
Genotype
G/G
G/H
H/H

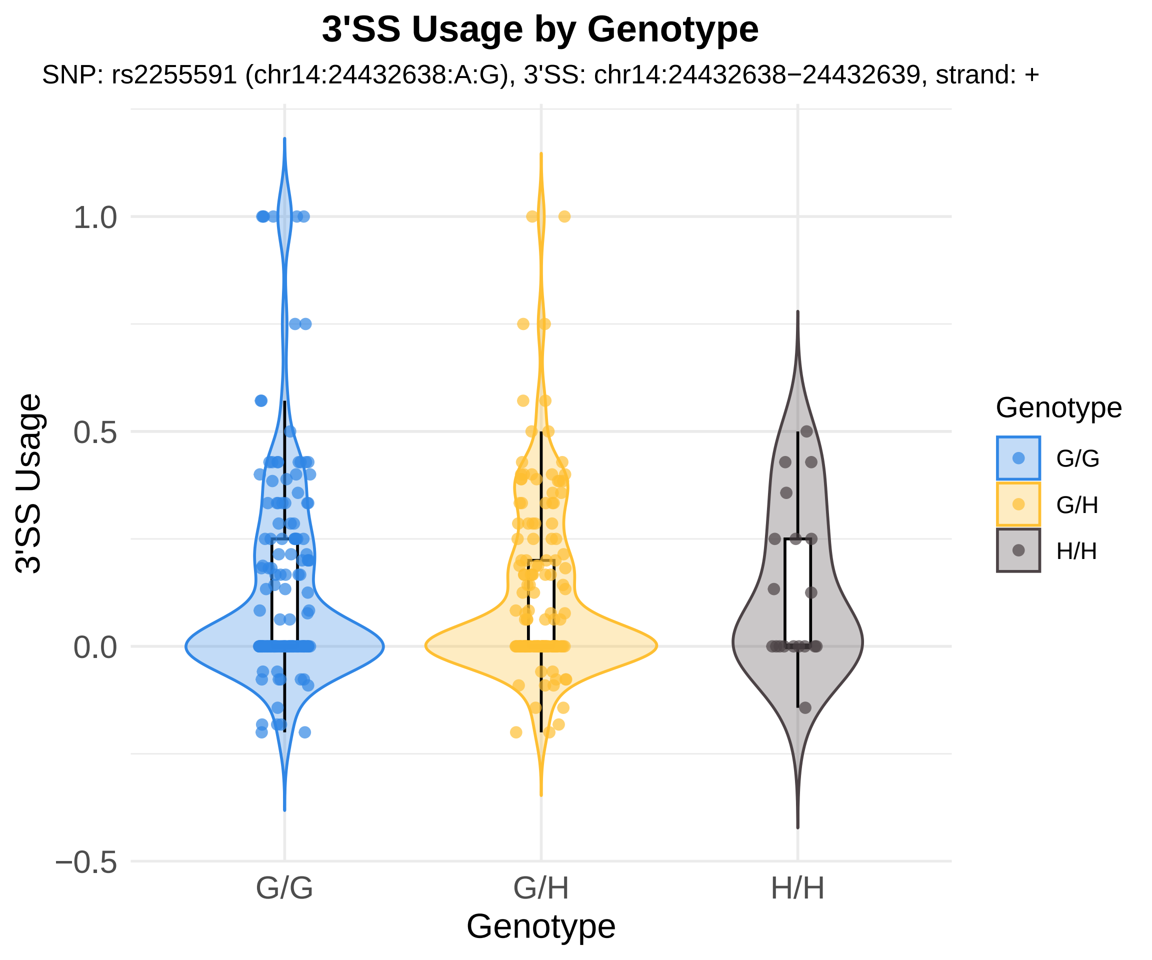
3'SS Usage by Genotype
SNP: rs2255591 (chr14:24432638:A:G), 3'SS: chr14:24432638−24432639, strand: +
3'SS Usage
1.0
0.5
0.0
−0.5
G/G
G/H
H/H
Genotype
Genotype
G/G
G/H
H/H

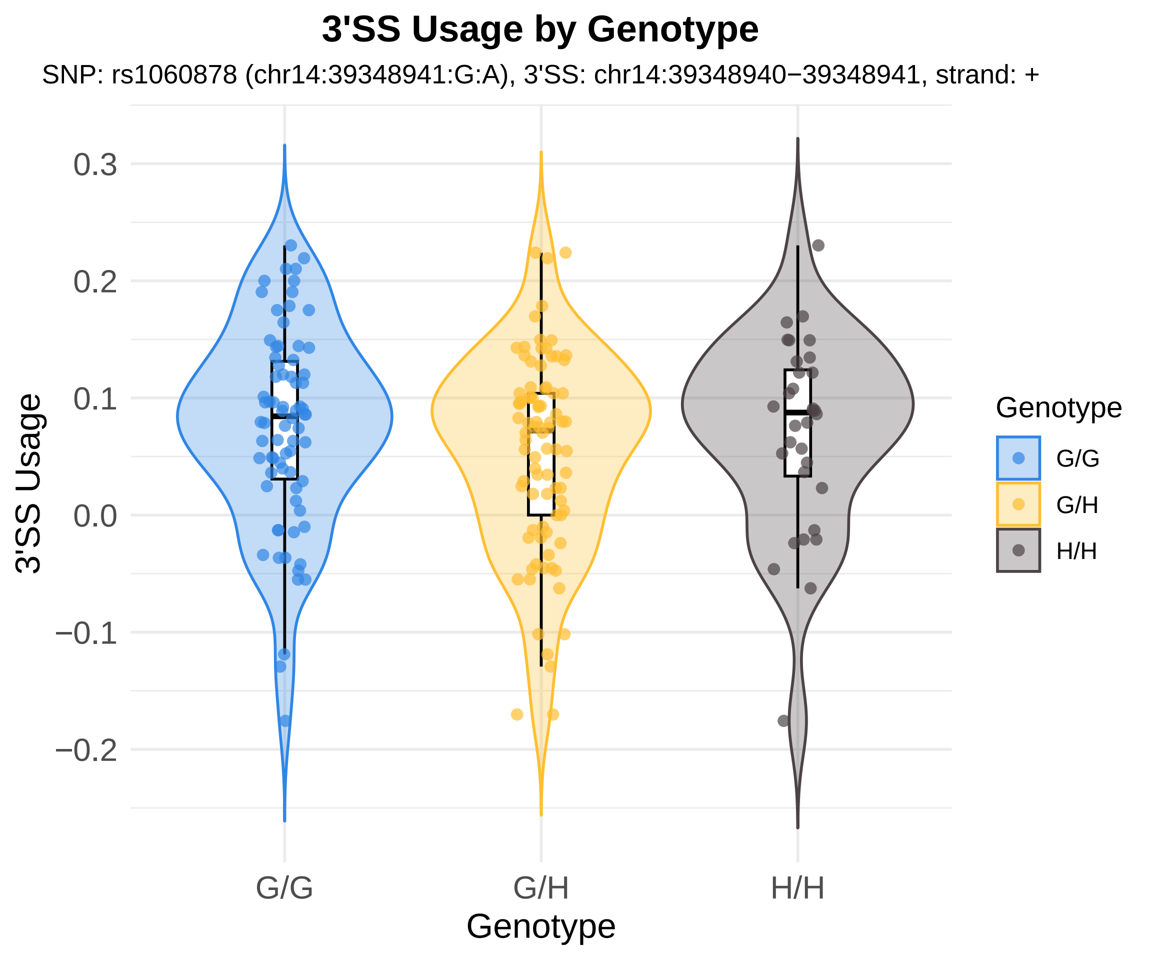
3'SS Usage by Genotype
SNP: rs1060878 (chr14:39348941:G:A), 3'SS: chr14:39348940−39348941, strand: +
3'SS Usage
0.3
0.2
0.1
0.0
−0.1
−0.2
G/G
G/H
H/H
Genotype
Genotype
G/G
G/H
H/H

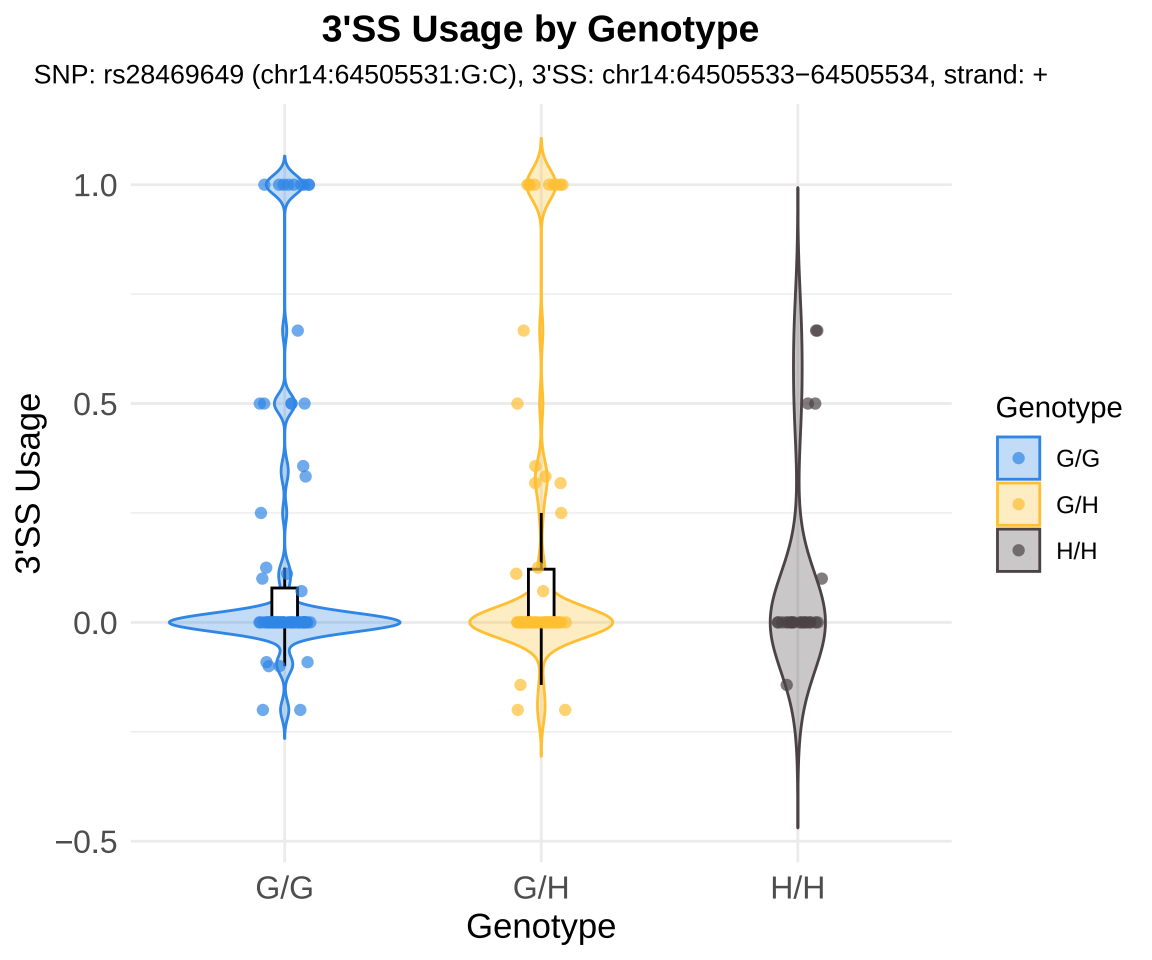

3'SS Usage by Genotype
SNP: rs28469649 (chr14:64505531:G:C), 3'SS: chr14:64505533−64505534, strand: +
1.0
0.5
0.0
−0.5
3'SS Usage
G/G
G/H
H/H
Genotype
Genotype
G/G
G/H
H/H

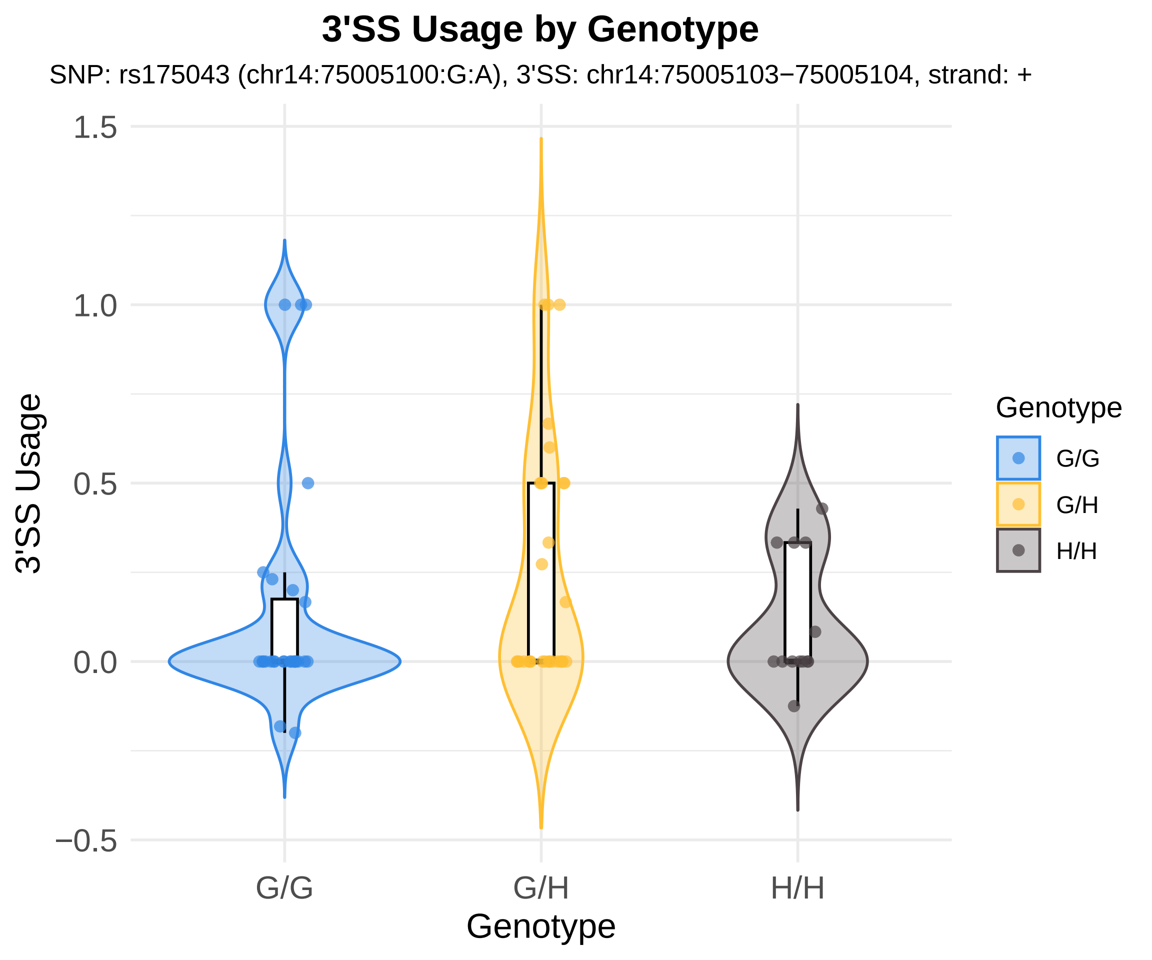
3'SS Usage by Genotype
SNP: rs175043 (chr14:75005100:G:A), 3'SS: chr14:75005103−75005104, strand: +
3'SS Usage
1.5
1.0
0.5
0.0
−0.5
G/G
G/H
H/H
Genotype
Genotype
G/G
G/H
H/H

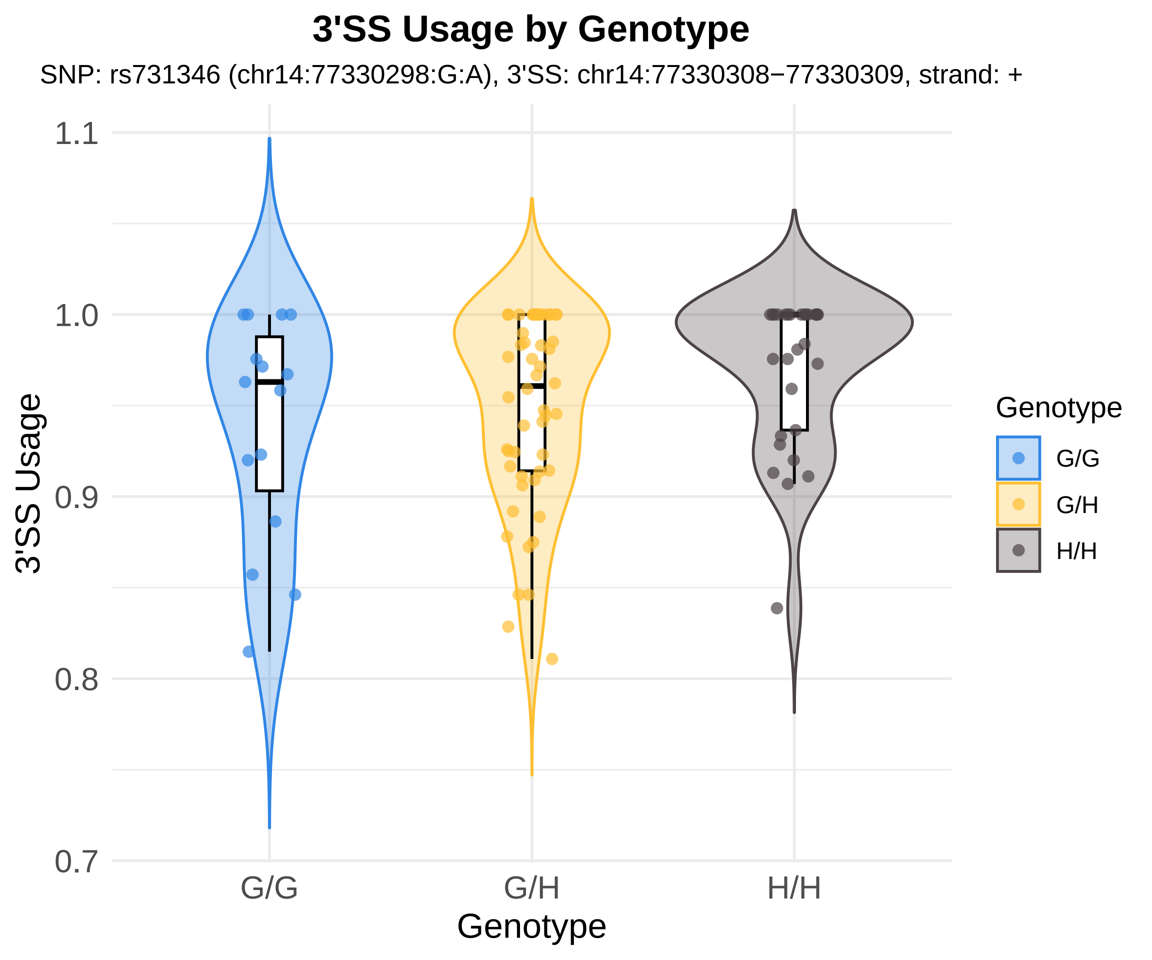
3'SS Usage by Genotype
SNP: rs731346 (chr14:77330298:G:A), 3'SS: chr14:77330308−77330309, strand: +
3'SS Usage
1.1
1.0
0.9
0.8
0.7
G/G
G/H
H/H
Genotype
Genotype
G/G
G/H
H/H

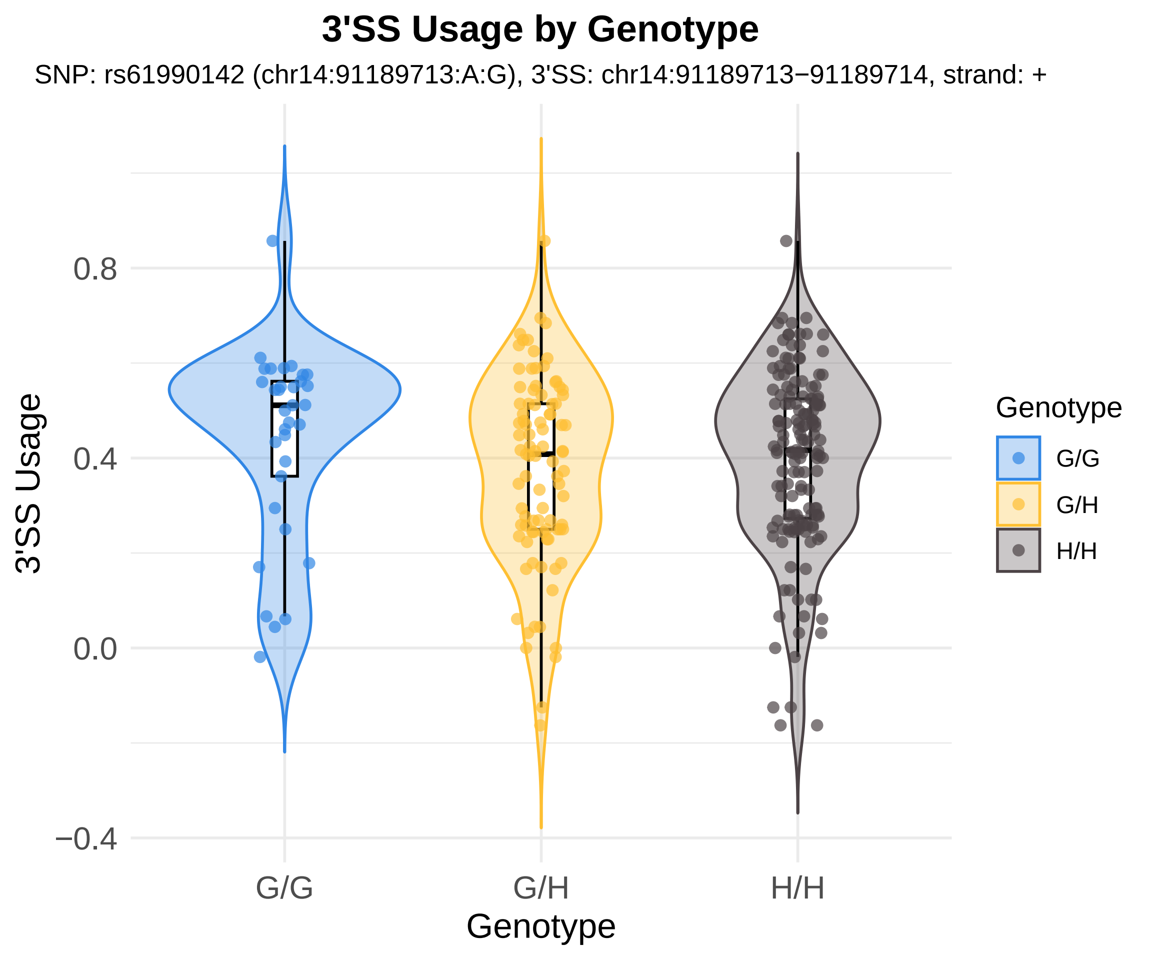

3'SS Usage by Genotype
SNP: rs61990142 (chr14:91189713:A:G), 3'SS: chr14:91189713−91189714, strand: +
3'SS Usage
0.8
0.4
0.0
−0.4
G/G
G/H
H/H
Genotype
Genotype
G/G
G/H
H/H

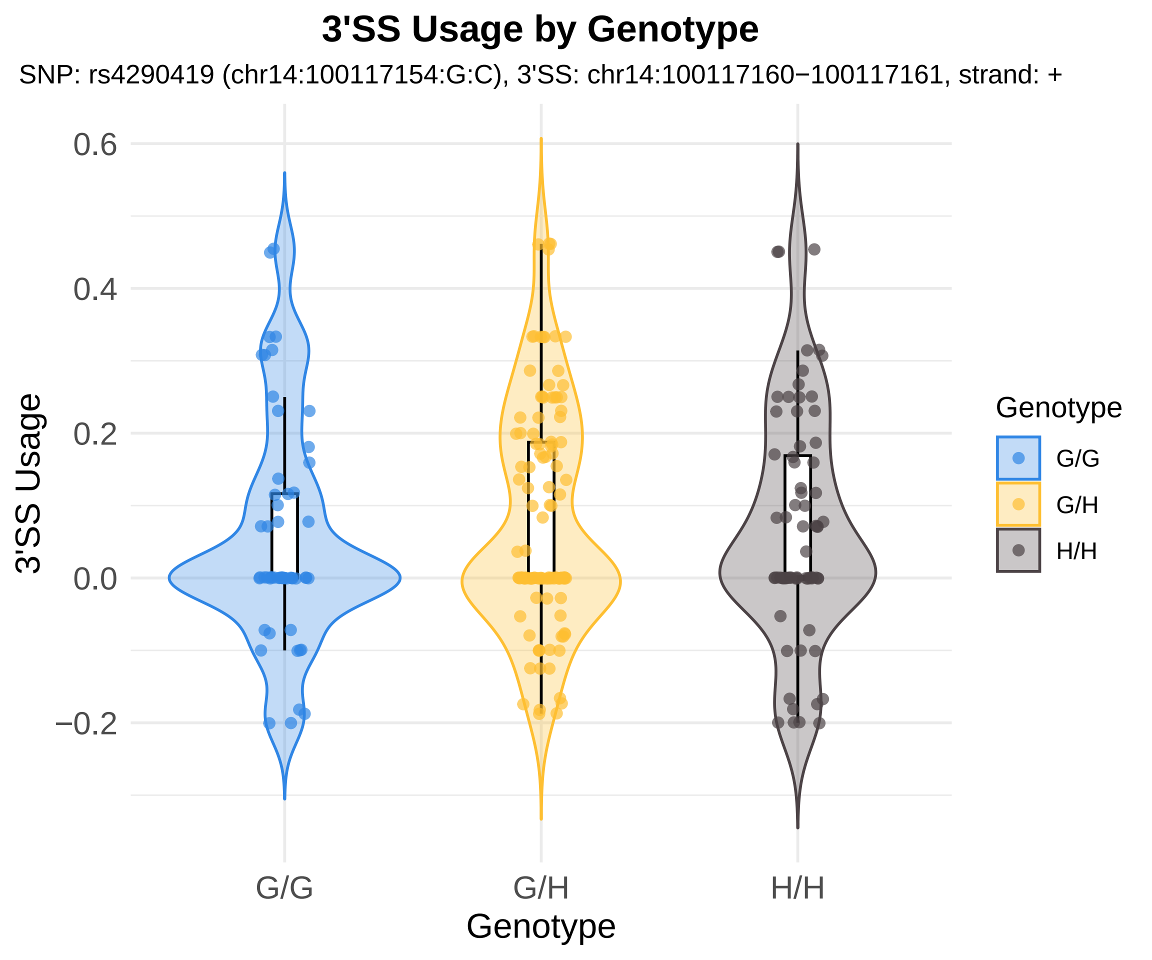

3'SS Usage by Genotype
SNP: rs4290419 (chr14:100117154:G:C), 3'SS: chr14:100117160−100117161, strand: +
0.6
0.4
0.2
0.0
−0.2
3'SS Usage
G/G
G/H
H/H
Genotype
Genotype
G/G
G/H
H/H

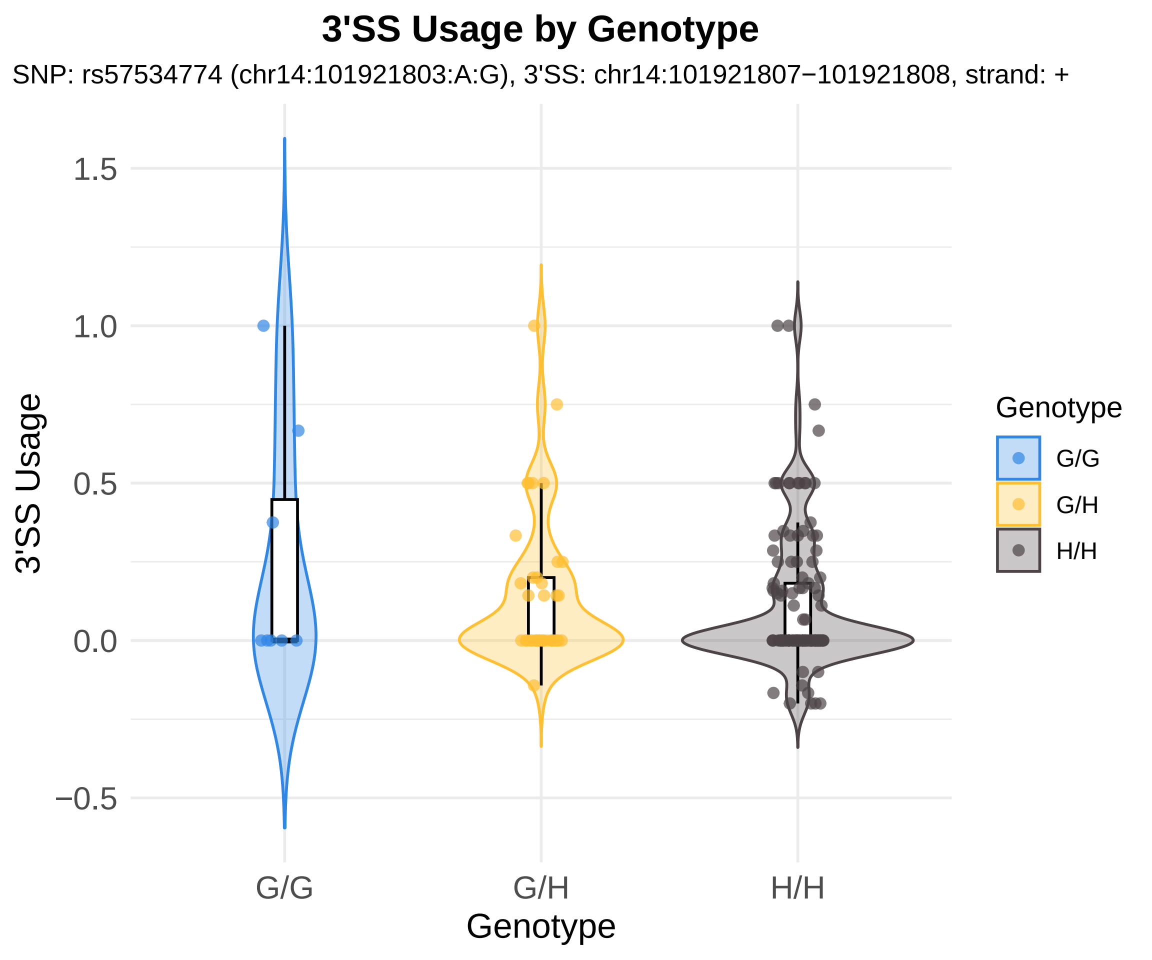

3'SS Usage by Genotype
SNP: rs57534774 (chr14:101921803:A:G), 3'SS: chr14:101921807−101921808, strand: +
3'SS Usage
1.5
1.0
0.5
0.0
−0.5
G/G
G/H
H/H
Genotype
Genotype
G/G
G/H
H/H

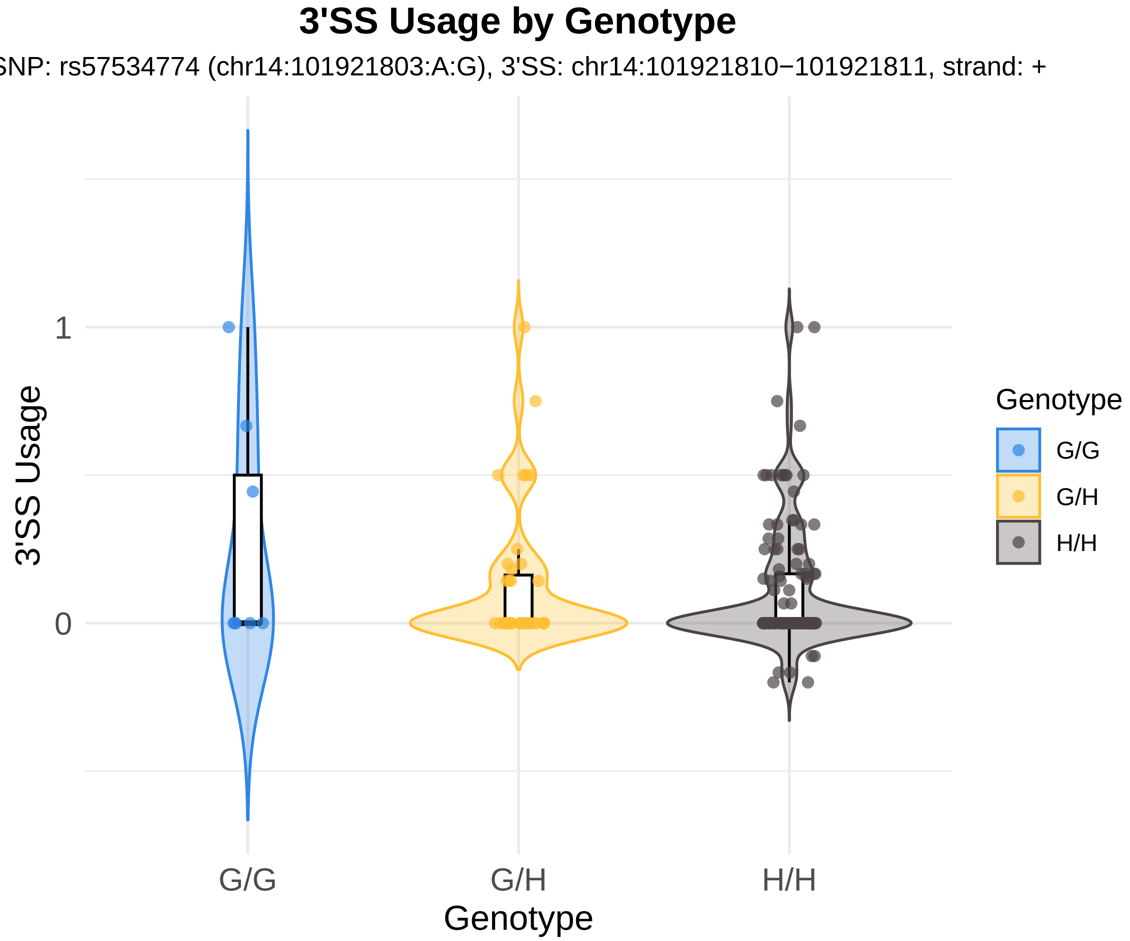

3'SS Usage by Genotype
SNP: rs57534774 (chr14:101921803:A:G), 3'SS: chr14:101921810−101921811, strand: +
3'SS Usage
1
0
G/G
G/H
H/H
Genotype
Genotype
G/G
G/H
H/H

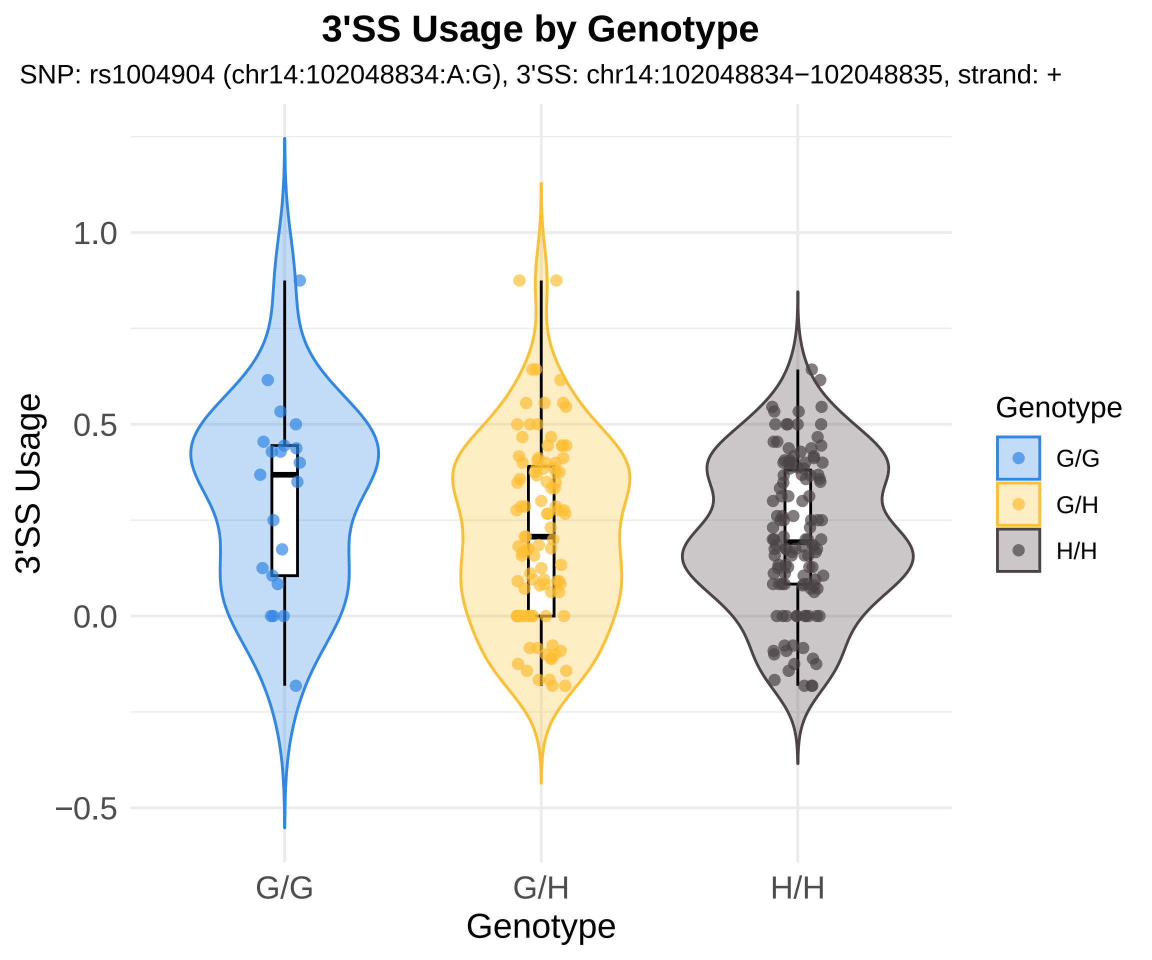
3'SS Usage by Genotype
SNP: rs1004904 (chr14:102048834:A:G), 3'SS: chr14:102048834−102048835, strand: +
3'SS Usage
1.0
0.5
0.0
−0.5
G/G
G/H
H/H
Genotype
Genotype
G/G
G/H
H/H

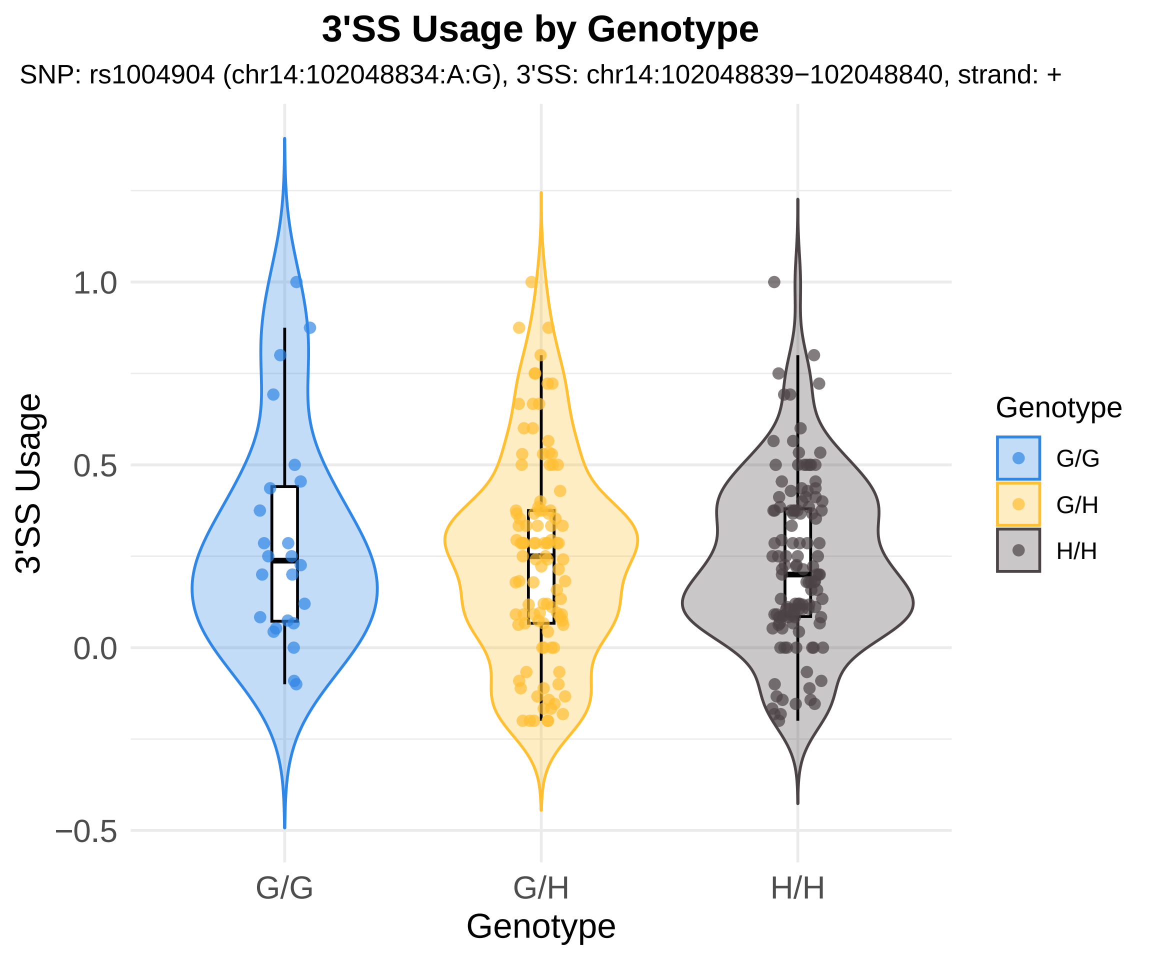

3'SS Usage by Genotype
SNP: rs1004904 (chr14:102048834:A:G), 3'SS: chr14:102048839−102048840, strand: +
3'SS Usage
1.0
0.5
0.0
−0.5
G/G
G/H
H/H
Genotype
Genotype
G/G
G/H
H/H

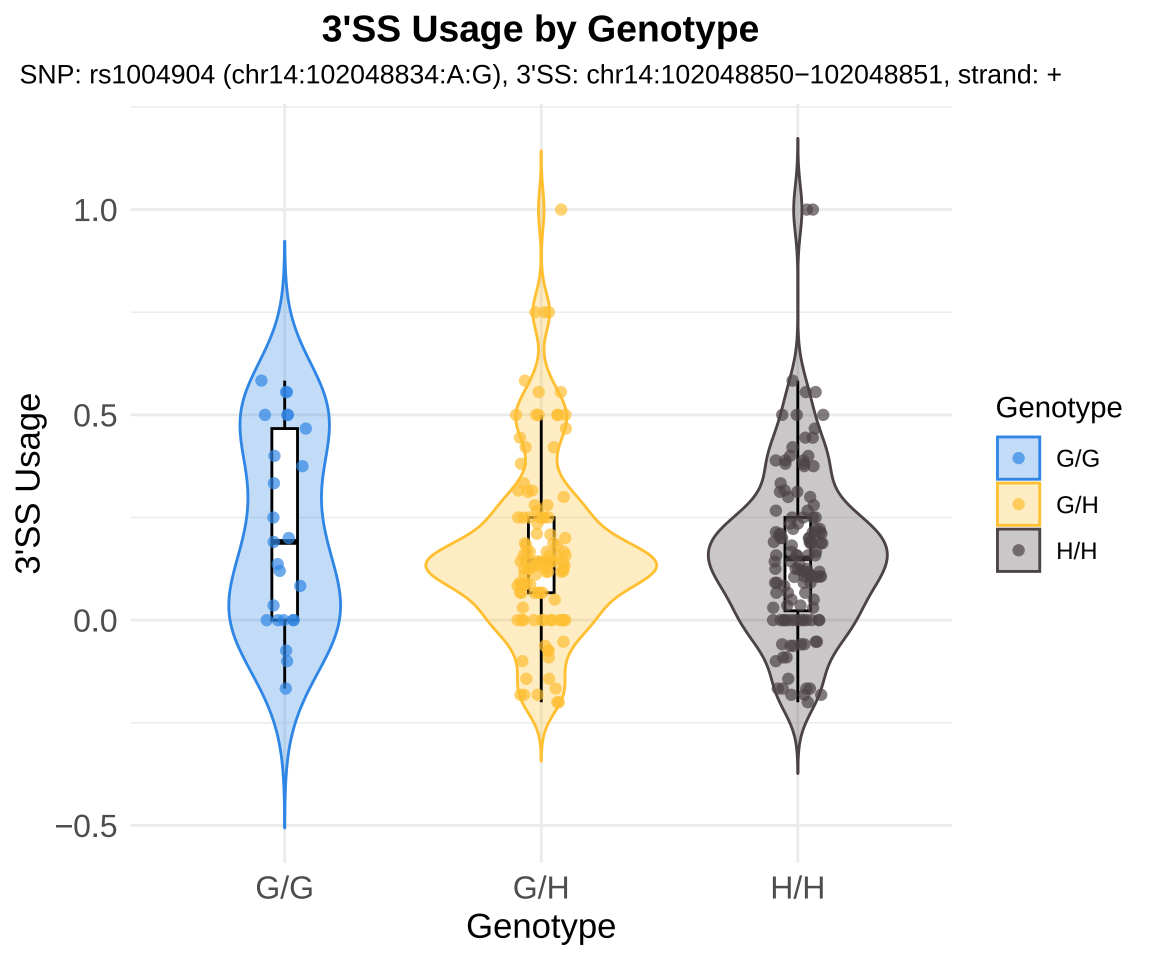
3'SS Usage by Genotype
SNP: rs1004904 (chr14:102048834:A:G), 3'SS: chr14:102048850−102048851, strand: +
3'SS Usage
1.0
0.5
0.0
−0.5
G/G
G/H
H/H
Genotype
Genotype
G/G
G/H
H/H

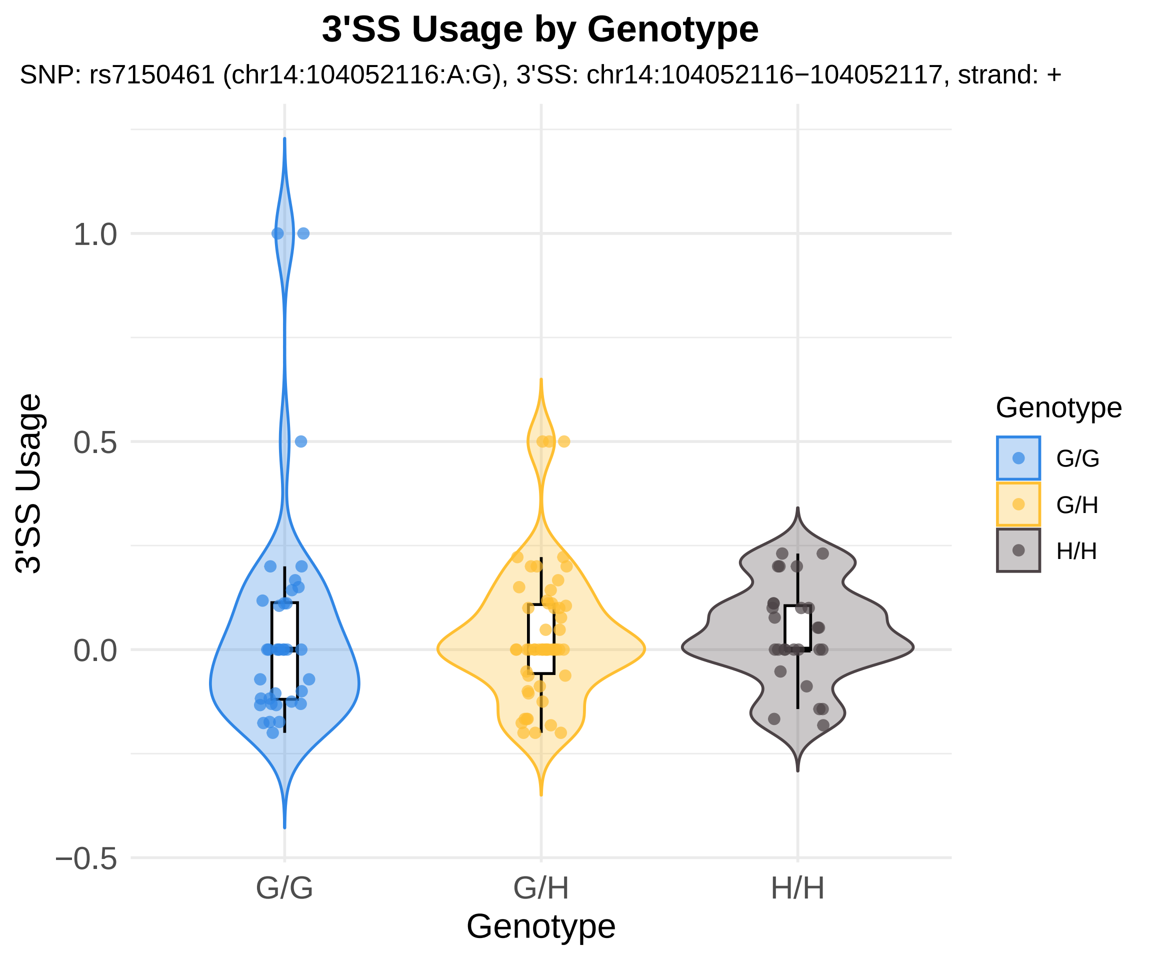
3'SS Usage by Genotype
SNP: rs7150461 (chr14:104052116:A:G), 3'SS: chr14:104052116−104052117, strand: +
3'SS Usage
1.0
0.5
0.0
−0.5
G/G
G/H
H/H
Genotype
Genotype
G/G
G/H
H/H

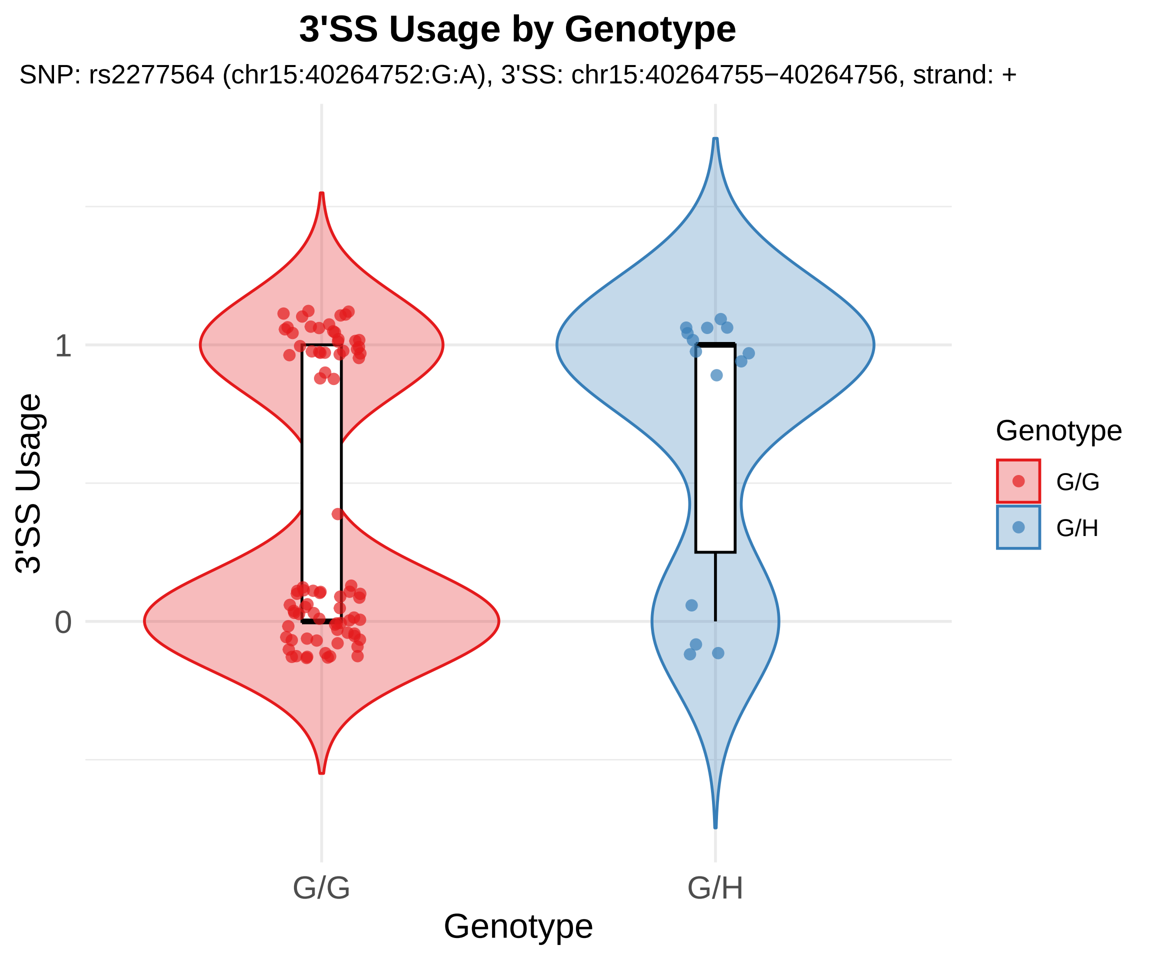

3'SS Usage by Genotype
SNP: rs2277564 (chr15:40264752:G:A), 3'SS: chr15:40264755−40264756, strand: +
3'SS Usage
1
0
G/G
G/H
Genotype
Genotype
G/G
G/H

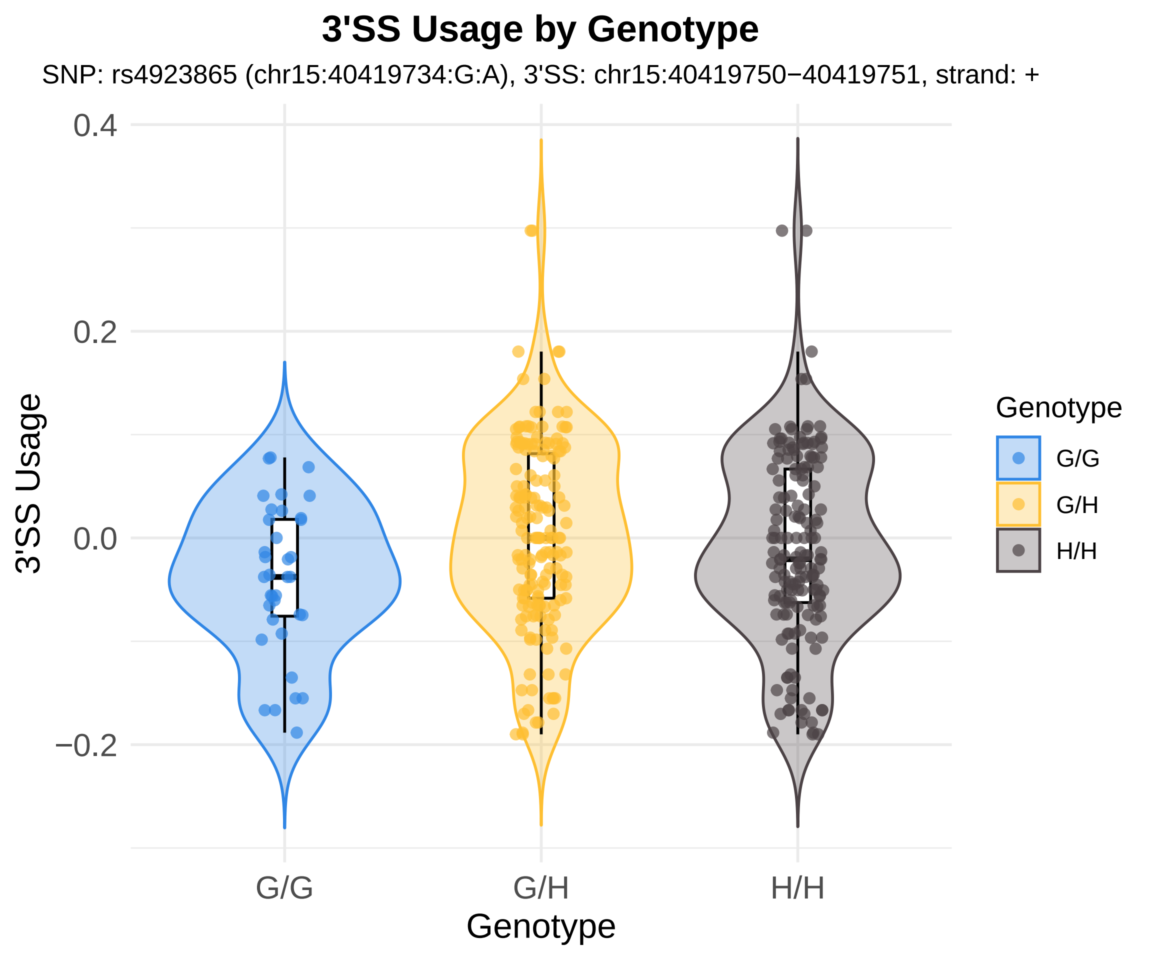
3'SS Usage by Genotype
SNP: rs4923865 (chr15:40419734:G:A), 3'SS: chr15:40419750−40419751, strand: +
0.4
0.2
0.0
−0.2
3'SS Usage
G/G
G/H
H/H
Genotype
Genotype
G/G
G/H
H/H

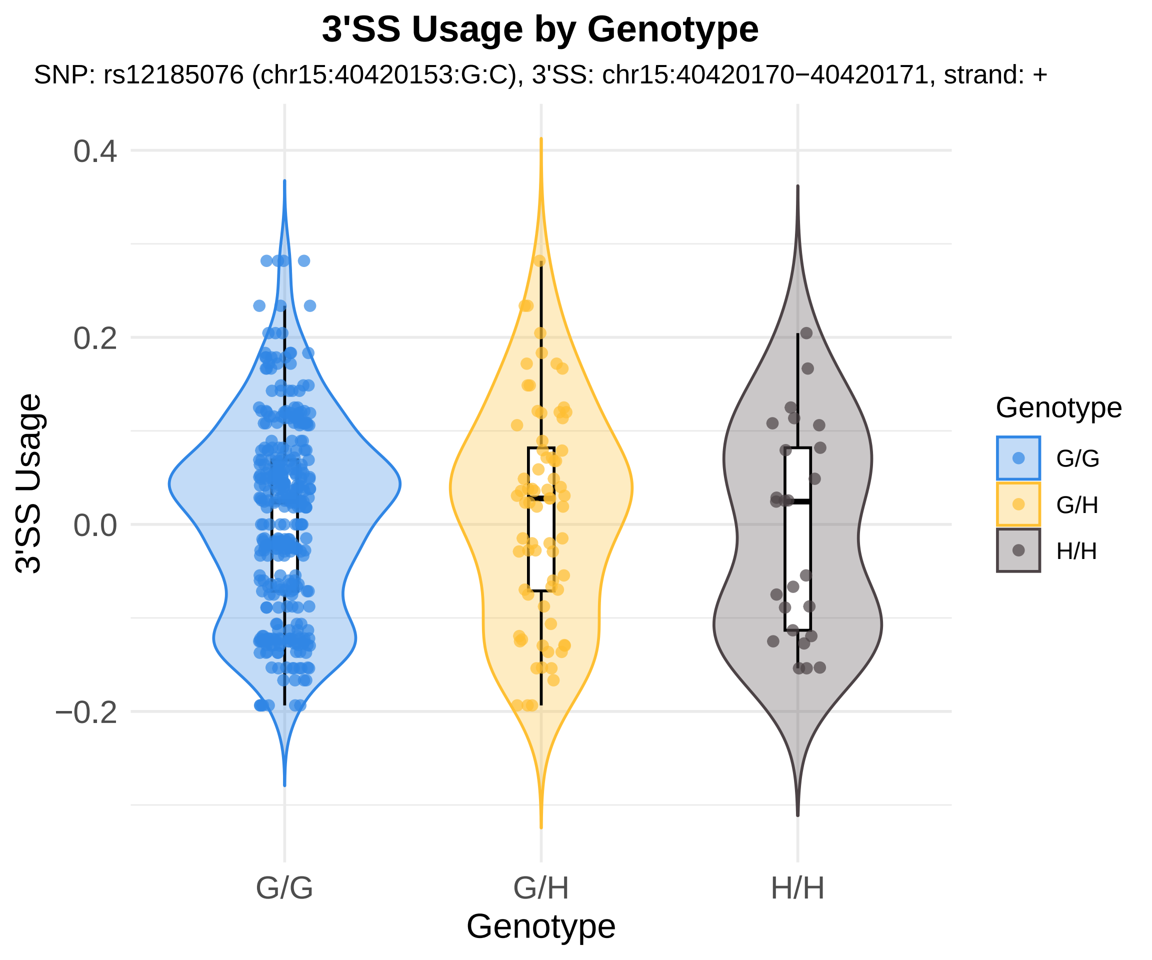
3'SS Usage by Genotype
SNP: rs12185076 (chr15:40420153:G:C), 3'SS: chr15:40420170−40420171, strand: +
3'SS Usage
0.4
0.2
0.0
−0.2
G/G
G/H
H/H
Genotype
Genotype
G/G
G/H
H/H

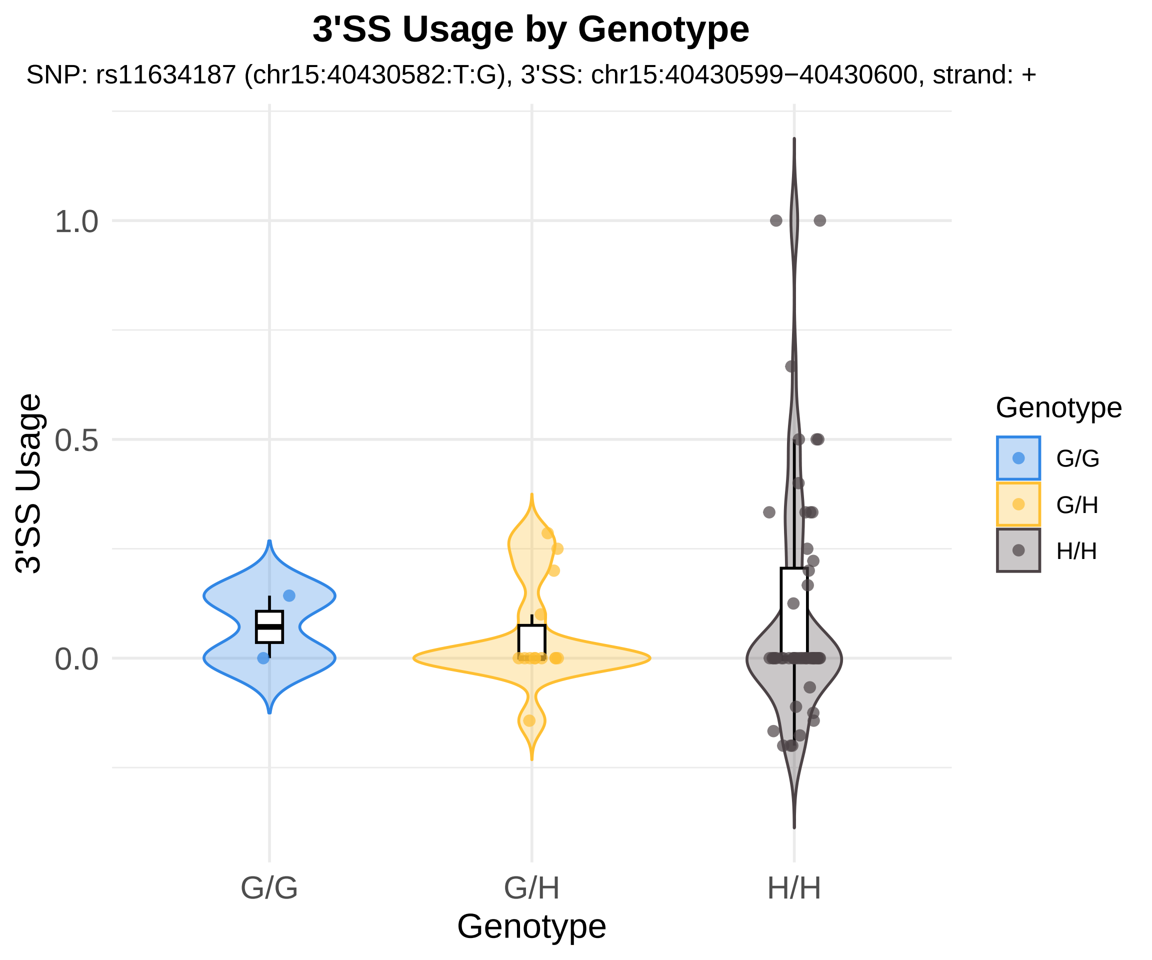

3'SS Usage by Genotype
SNP: rs11634187 (chr15:40430582:T:G), 3'SS: chr15:40430599−40430600, strand: +
3'SS Usage
1.0
0.5
0.0
G/G
G/H
H/H
Genotype
Genotype
G/G
G/H
H/H

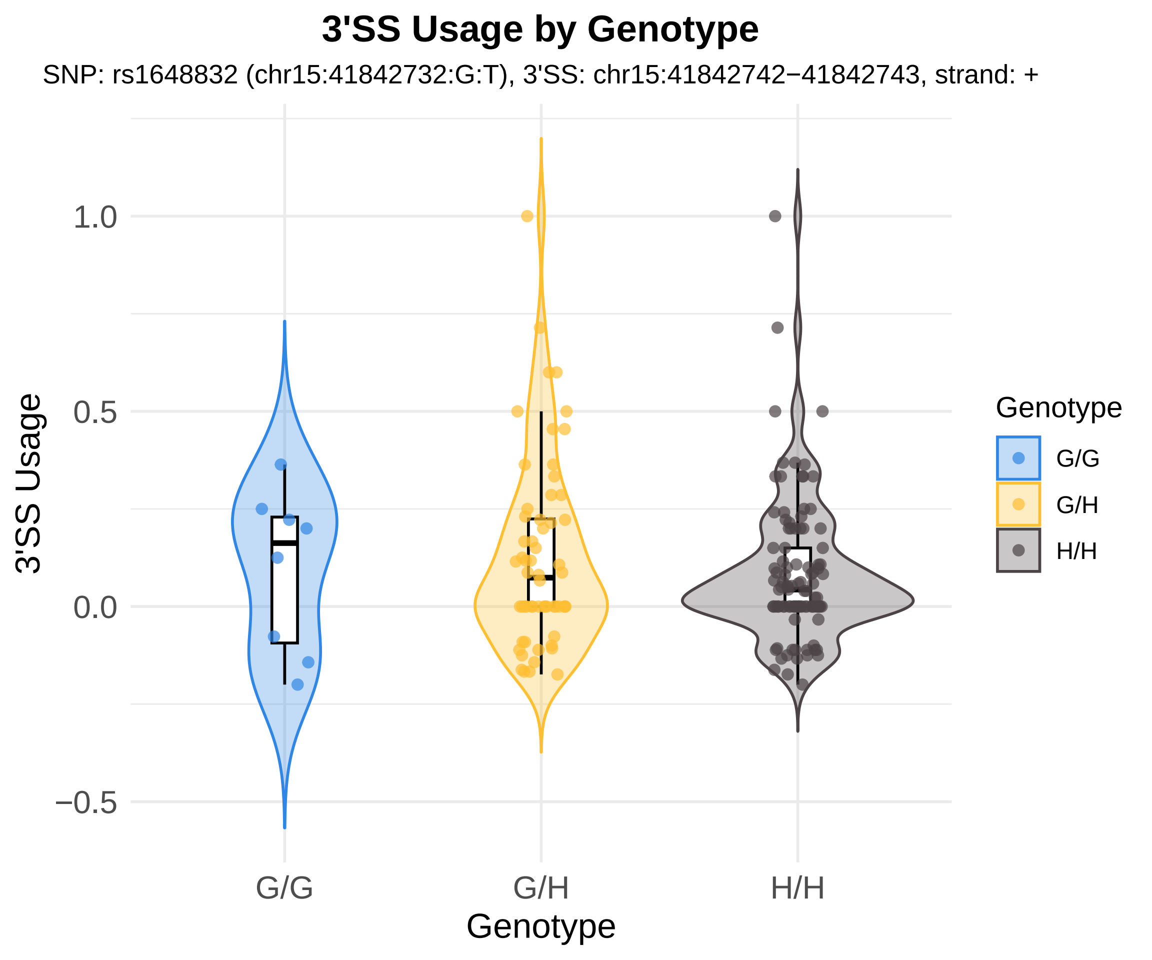
3'SS Usage by Genotype
SNP: rs1648832 (chr15:41842732:G:T), 3'SS: chr15:41842742−41842743, strand: +
3'SS Usage
1.0
0.5
0.0
−0.5
G/G
G/H
H/H
Genotype
Genotype
G/G
G/H
H/H

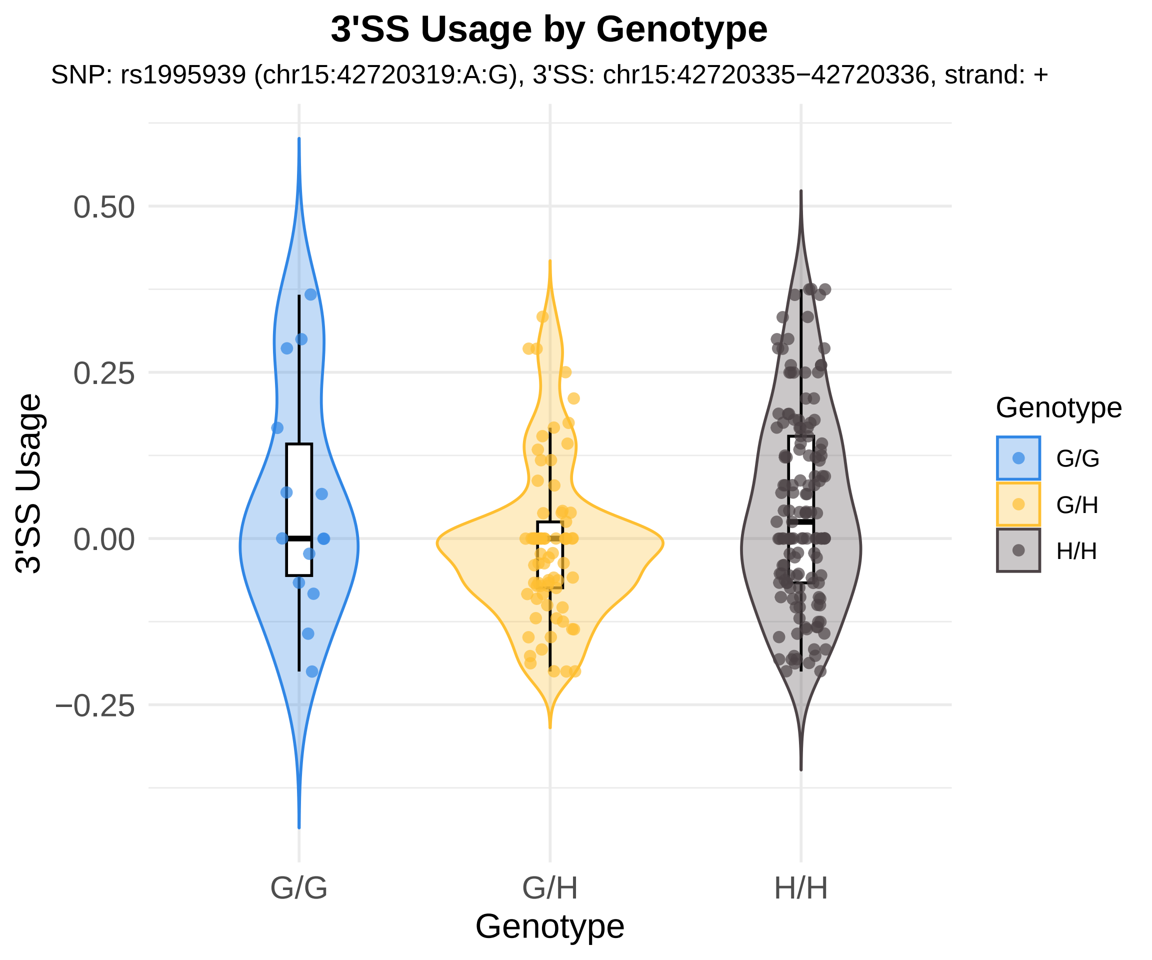
3'SS Usage by Genotype
SNP: rs1995939 (chr15:42720319:A:G), 3'SS: chr15:42720335−42720336, strand: +
0.50
0.25
0.00
−0.25
3'SS Usage
G/G
G/H
H/H
Genotype
Genotype
G/G
G/H
H/H

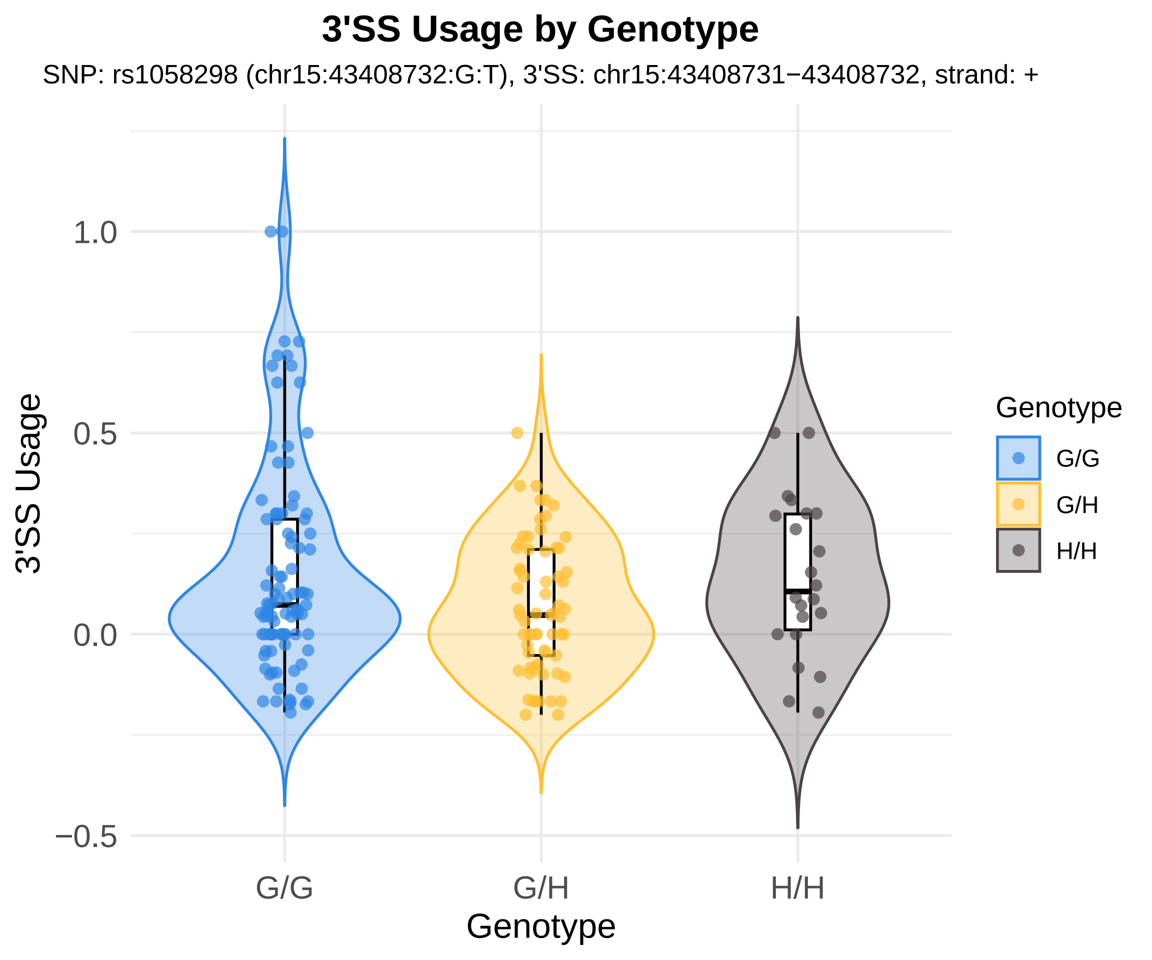

3'SS Usage by Genotype
SNP: rs1058298 (chr15:43408732:G:T), 3'SS: chr15:43408731−43408732, strand: +
3'SS Usage
1.0
0.5
0.0
−0.5
G/G
G/H
H/H
Genotype
Genotype
G/G
G/H
H/H

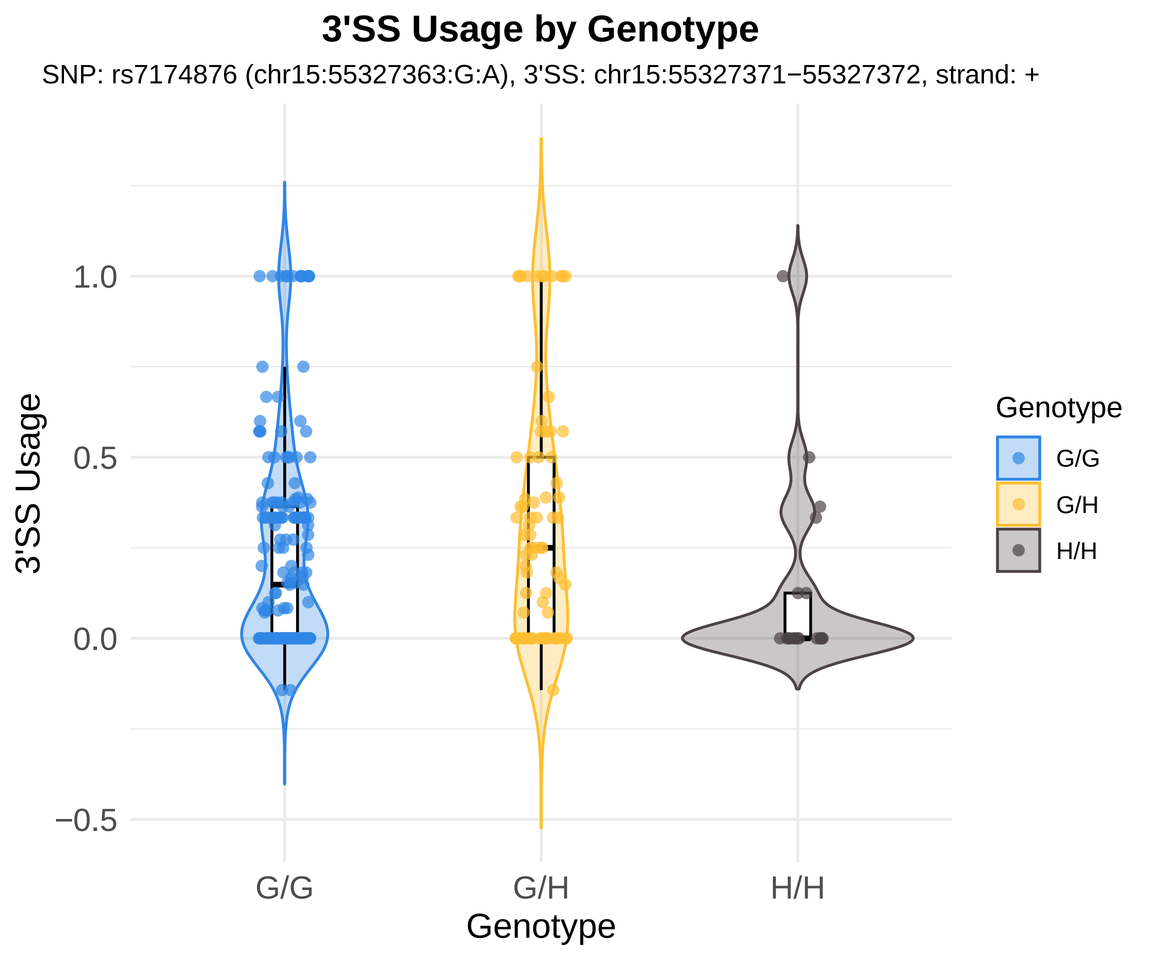

3'SS Usage by Genotype
SNP: rs7174876 (chr15:55327363:G:A), 3'SS: chr15:55327371−55327372, strand: +
3'SS Usage
1.0
0.5
0.0
−0.5
G/G
G/H
H/H
Genotype
Genotype
G/G
G/H
H/H

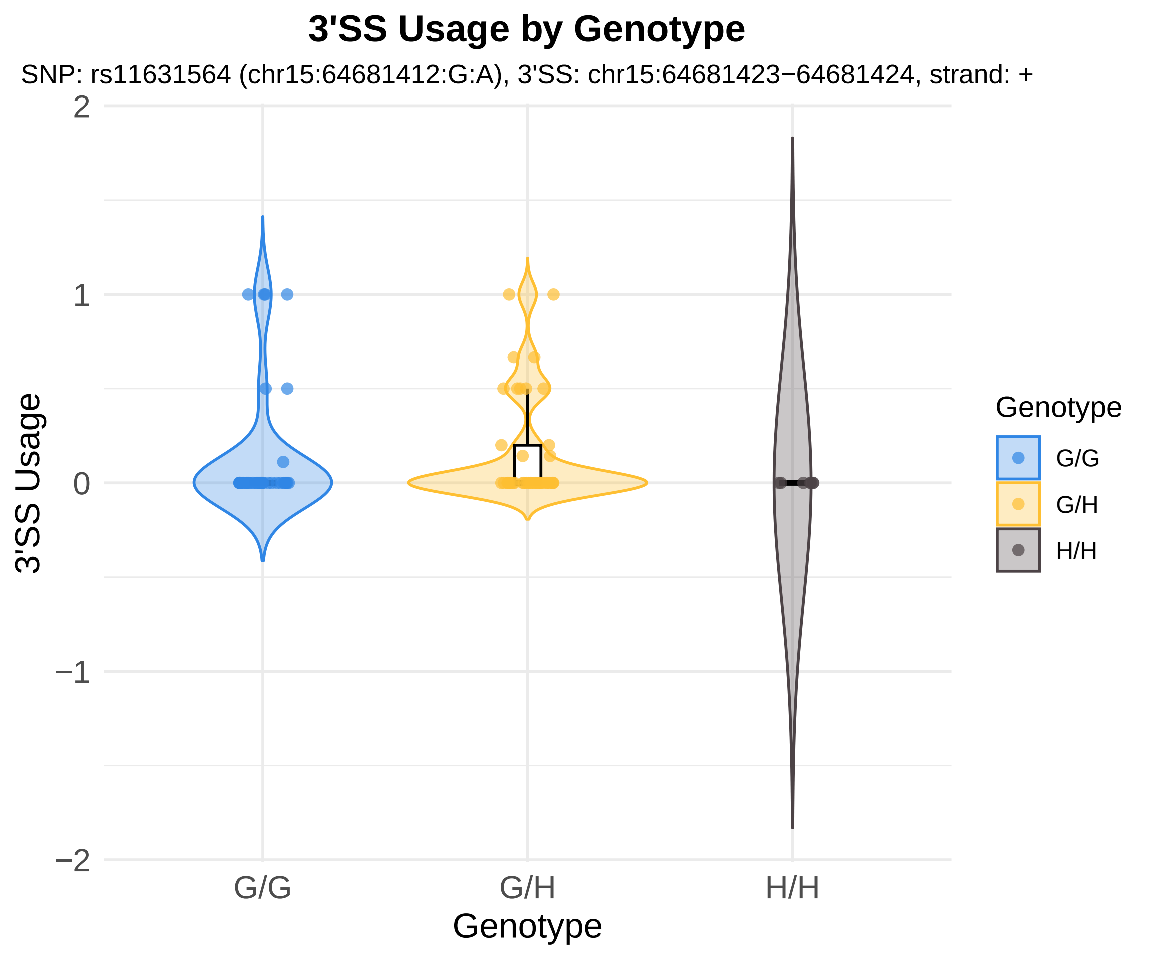

3'SS Usage by Genotype
SNP: rs11631564 (chr15:64681412:G:A), 3'SS: chr15:64681423−64681424, strand: +
2
1
0
−1
−2
3'SS Usage
G/G
G/H
H/H
Genotype
Genotype
G/G
G/H
H/H

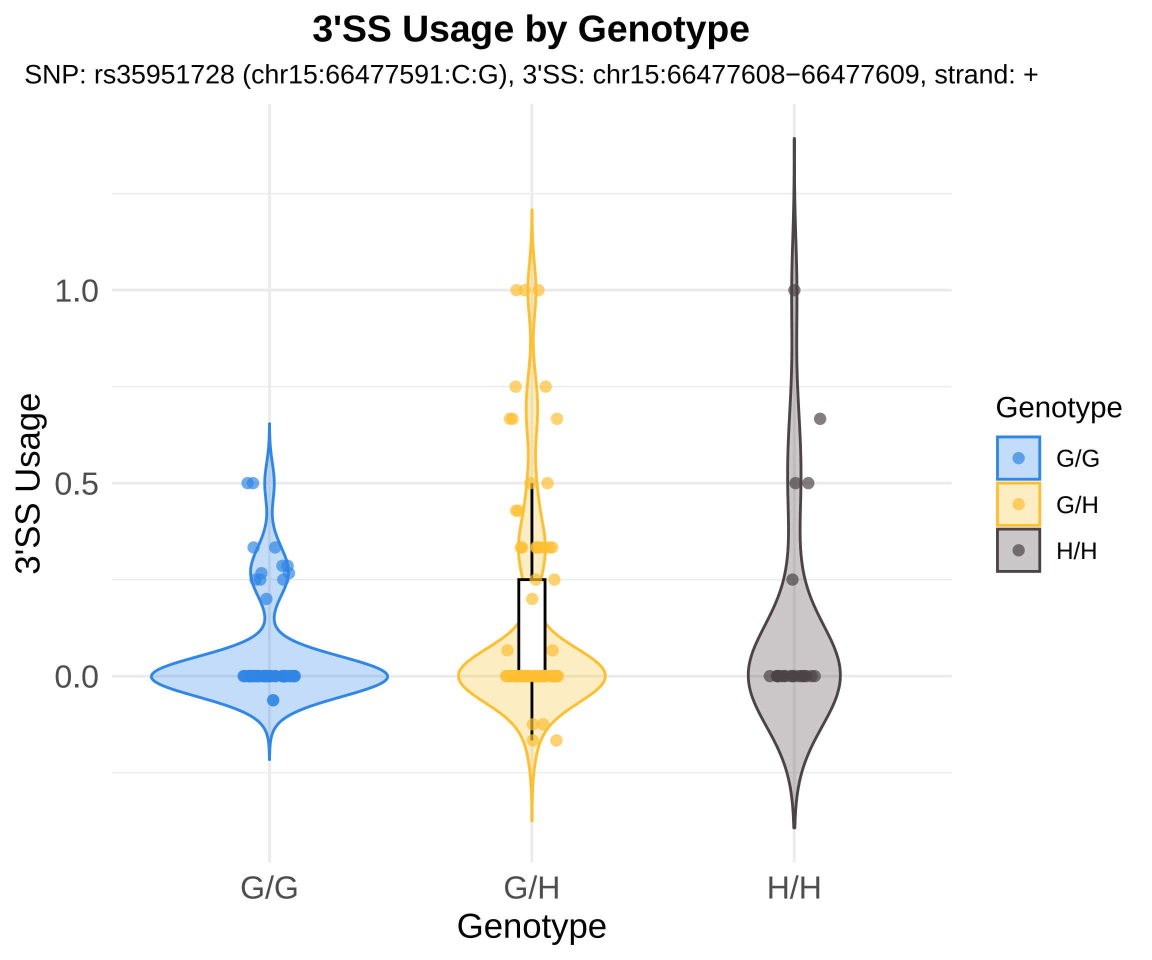

3'SS Usage by Genotype
SNP: rs35951728 (chr15:66477591:C:G), 3'SS: chr15:66477608−66477609, strand: +
3'SS Usage
1.0
0.5
0.0
G/G
G/H
H/H
Genotype
Genotype
G/G
G/H
H/H

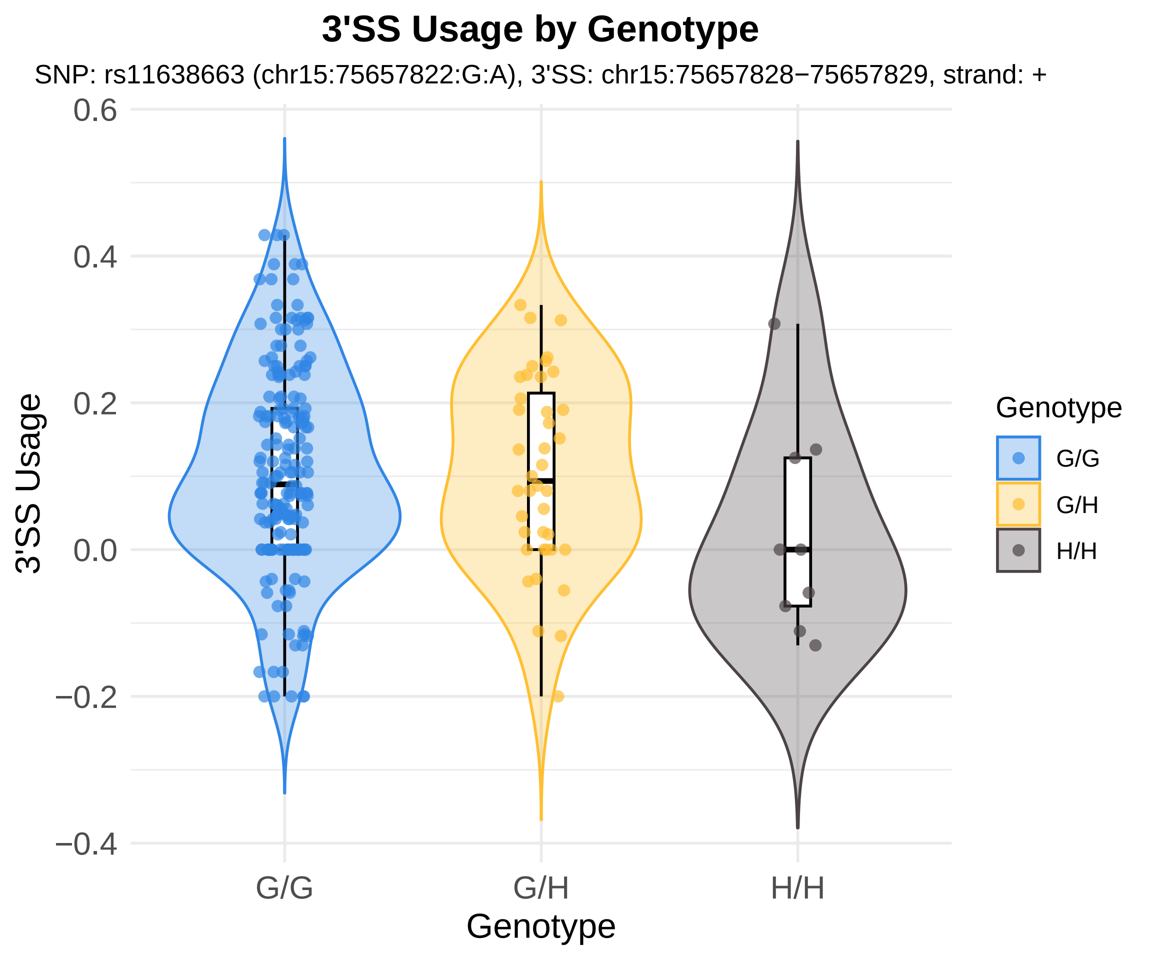
3'SS Usage by Genotype
SNP: rs11638663 (chr15:75657822:G:A), 3'SS: chr15:75657828−75657829, strand: +
3'SS Usage
0.6
0.4
0.2
0.0
−0.2
−0.4
G/G
G/H
H/H
Genotype
Genotype
G/G
G/H
H/H

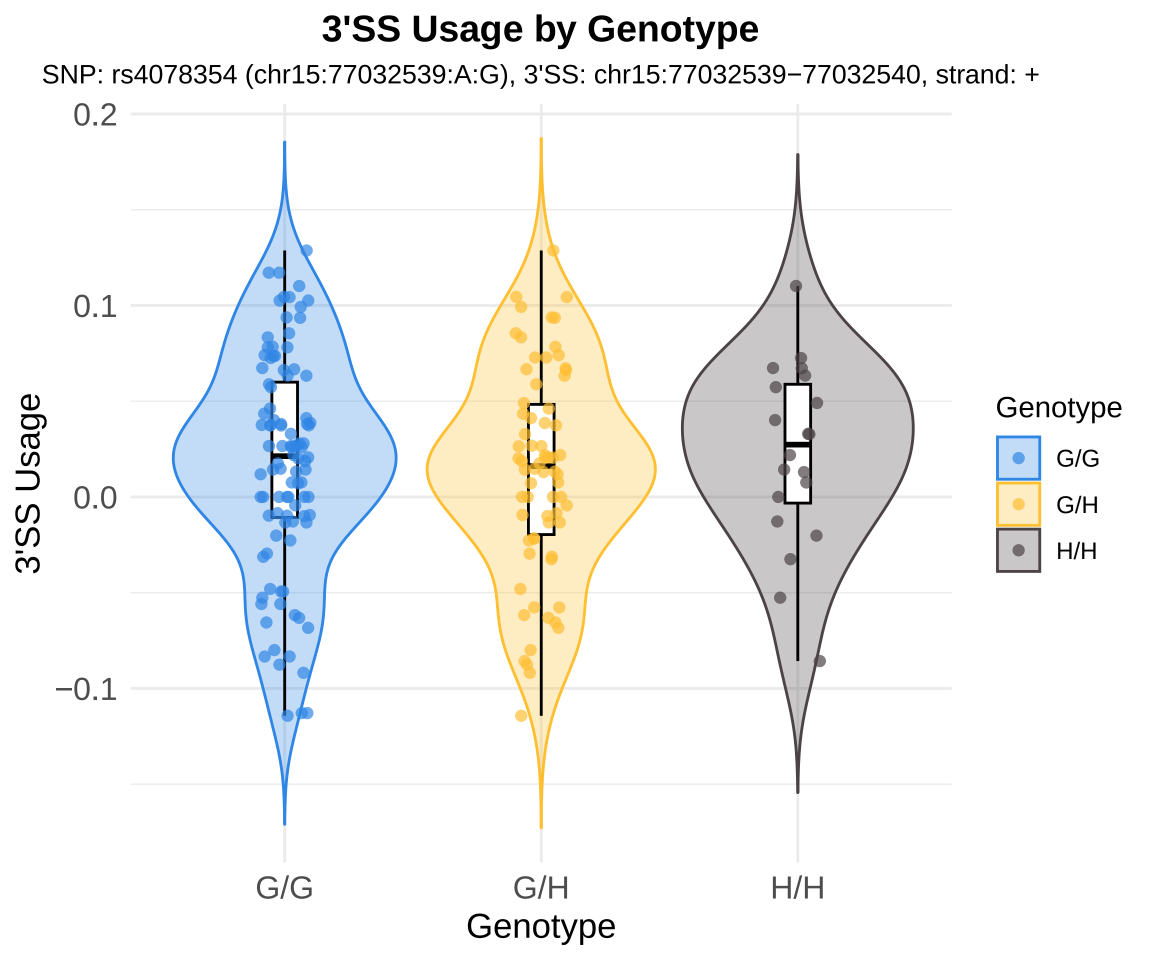

3'SS Usage by Genotype
SNP: rs4078354 (chr15:77032539:A:G), 3'SS: chr15:77032539−77032540, strand: +
3'SS Usage
0.2
0.1
0.0
−0.1
G/G
G/H
H/H
Genotype
Genotype
G/G
G/H
H/H

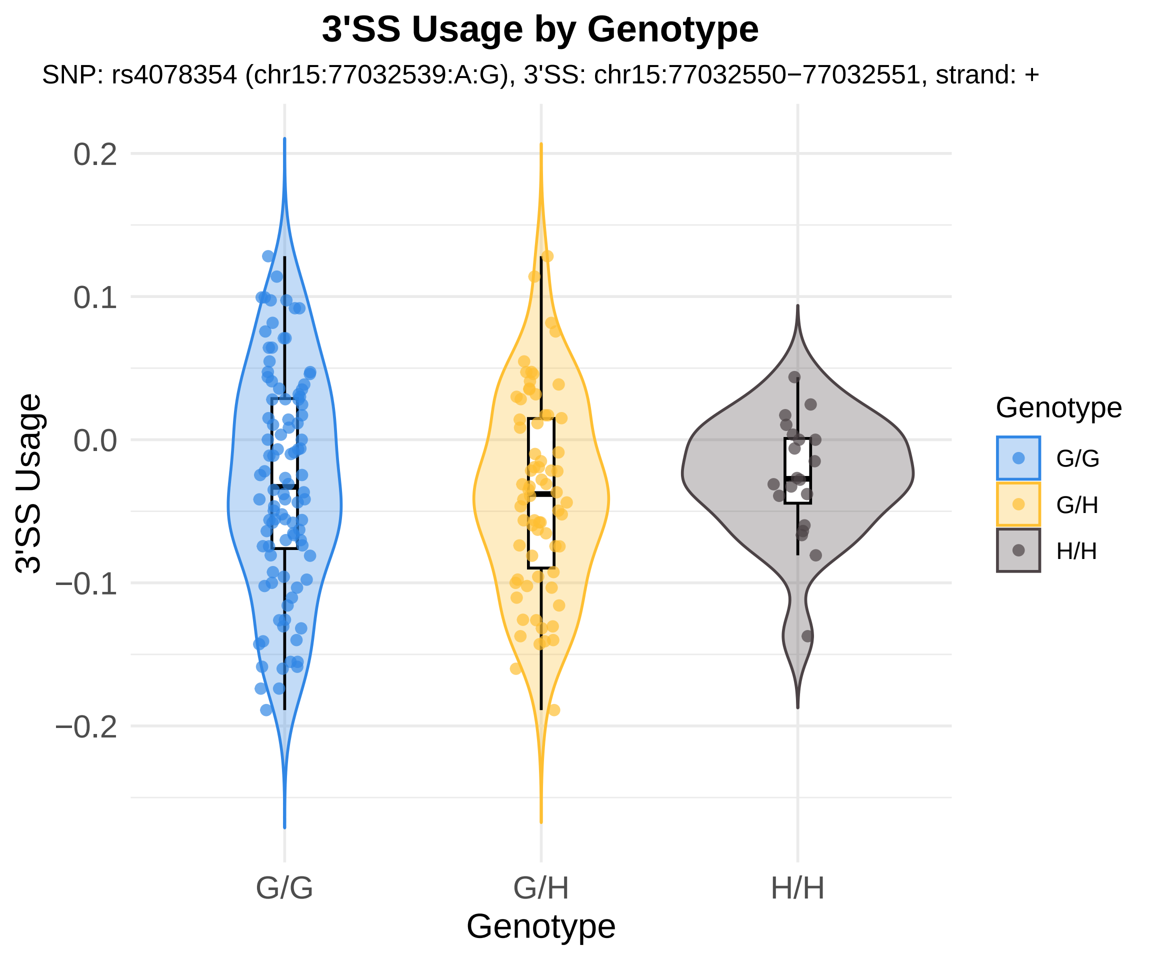
3'SS Usage by Genotype
SNP: rs4078354 (chr15:77032539:A:G), 3'SS: chr15:77032550−77032551, strand: +
0.2
0.1
0.0
−0.1
−0.2
3'SS Usage
G/G
G/H
H/H
Genotype
Genotype
G/G
G/H
H/H

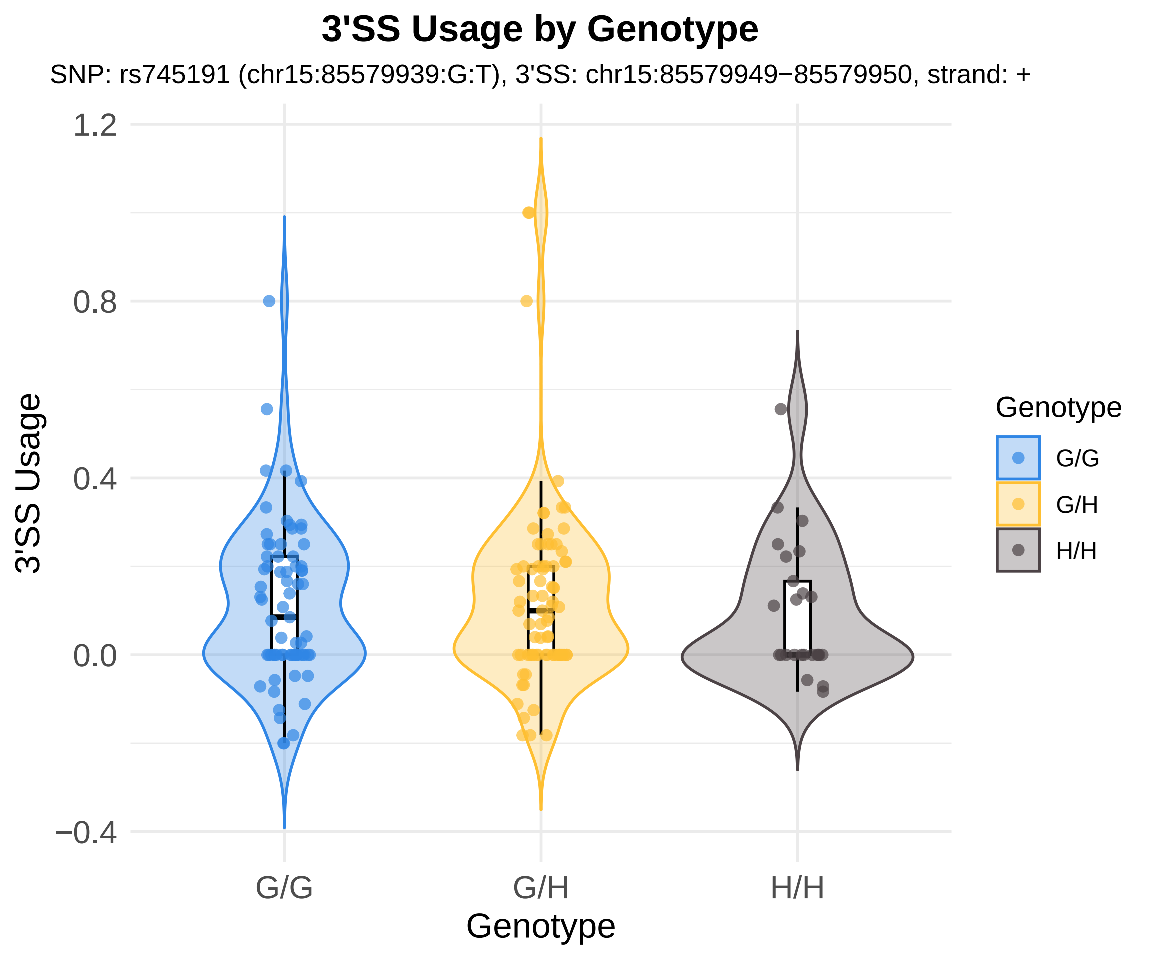
3'SS Usage by Genotype
SNP: rs745191 (chr15:85579939:G:T), 3'SS: chr15:85579949−85579950, strand: +
3'SS Usage
1.2
0.8
0.4
0.0
−0.4
G/G
G/H
H/H
Genotype
Genotype
G/G
G/H
H/H

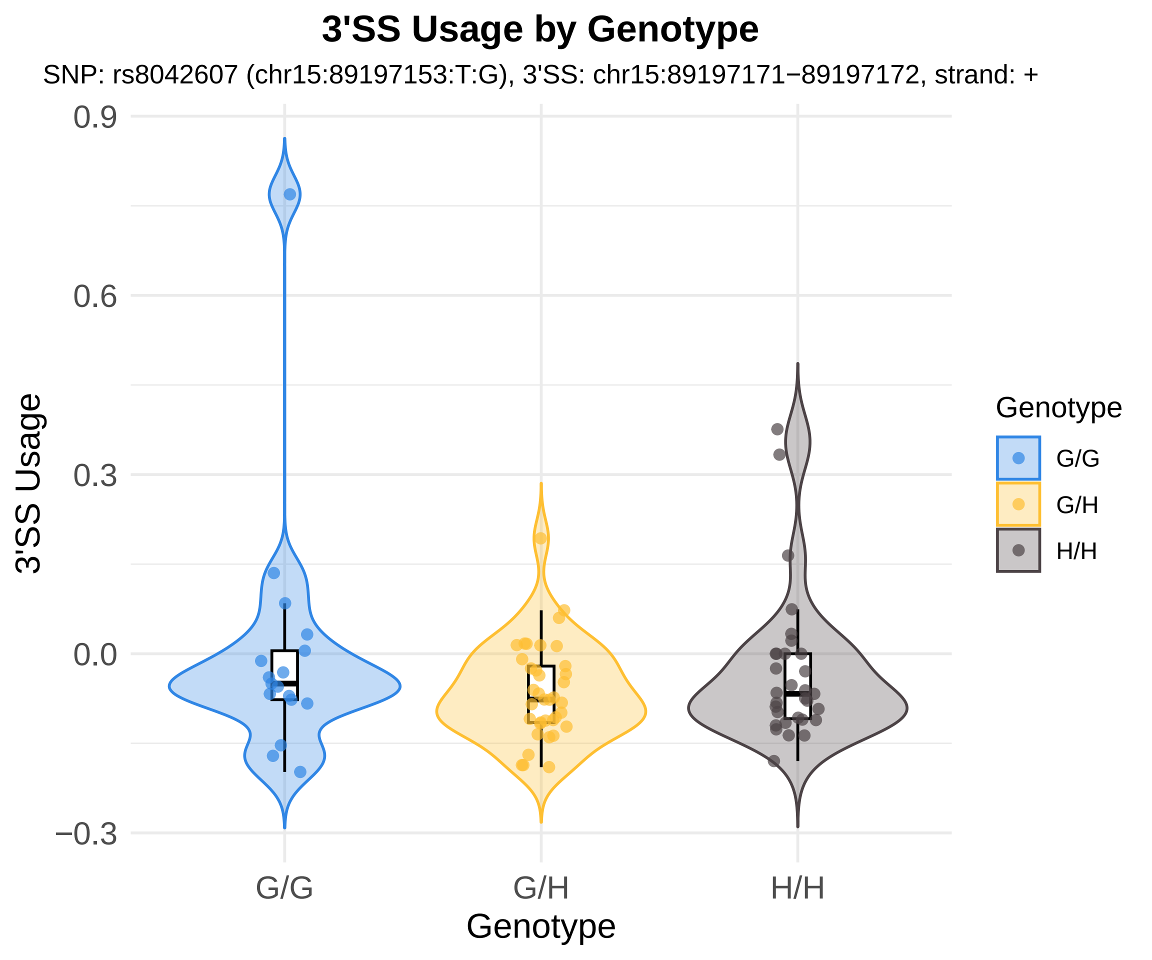
3'SS Usage by Genotype
SNP: rs8042607 (chr15:89197153:T:G), 3'SS: chr15:89197171−89197172, strand: +
0.9
0.6
0.3
0.0
−0.3
3'SS Usage
G/G
G/H
H/H
Genotype
Genotype
G/G
G/H
H/H

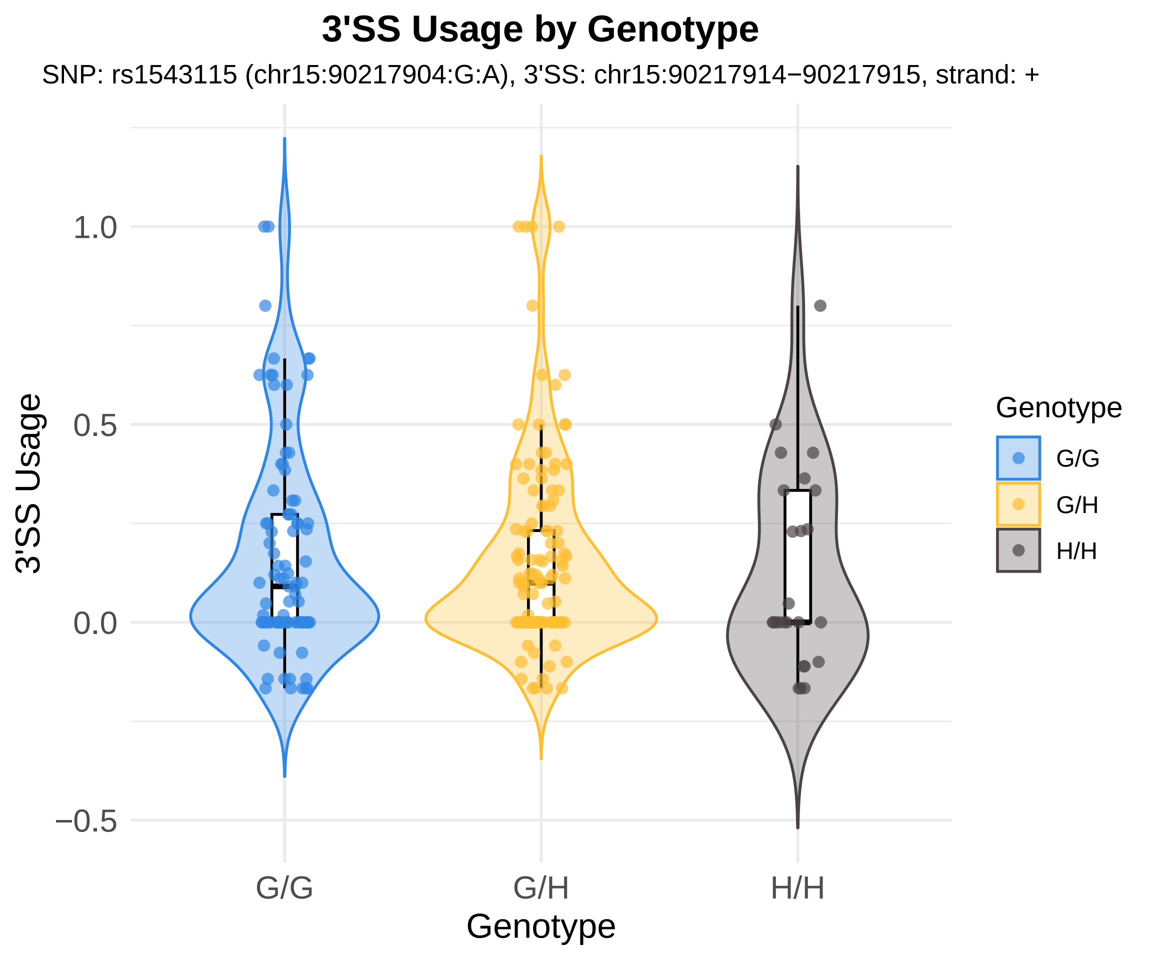
3'SS Usage by Genotype
SNP: rs1543115 (chr15:90217904:G:A), 3'SS: chr15:90217914−90217915, strand: +
3'SS Usage
1.0
0.5
0.0
−0.5
G/G
G/H
H/H
Genotype
Genotype
G/G
G/H
H/H

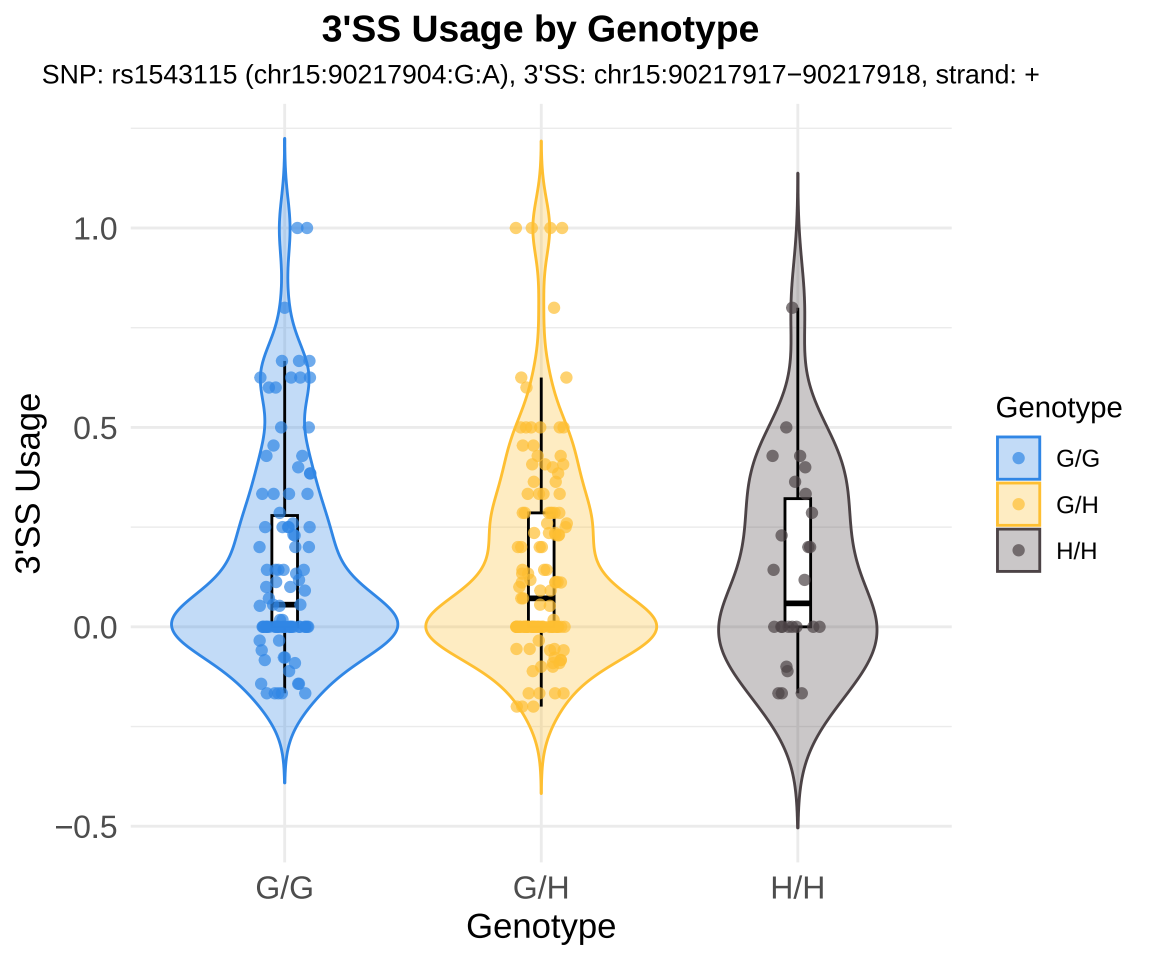
3'SS Usage by Genotype
SNP: rs1543115 (chr15:90217904:G:A), 3'SS: chr15:90217917−90217918, strand: +
3'SS Usage
1.0
0.5
0.0
−0.5
G/G
G/H
H/H
Genotype
Genotype
G/G
G/H
H/H

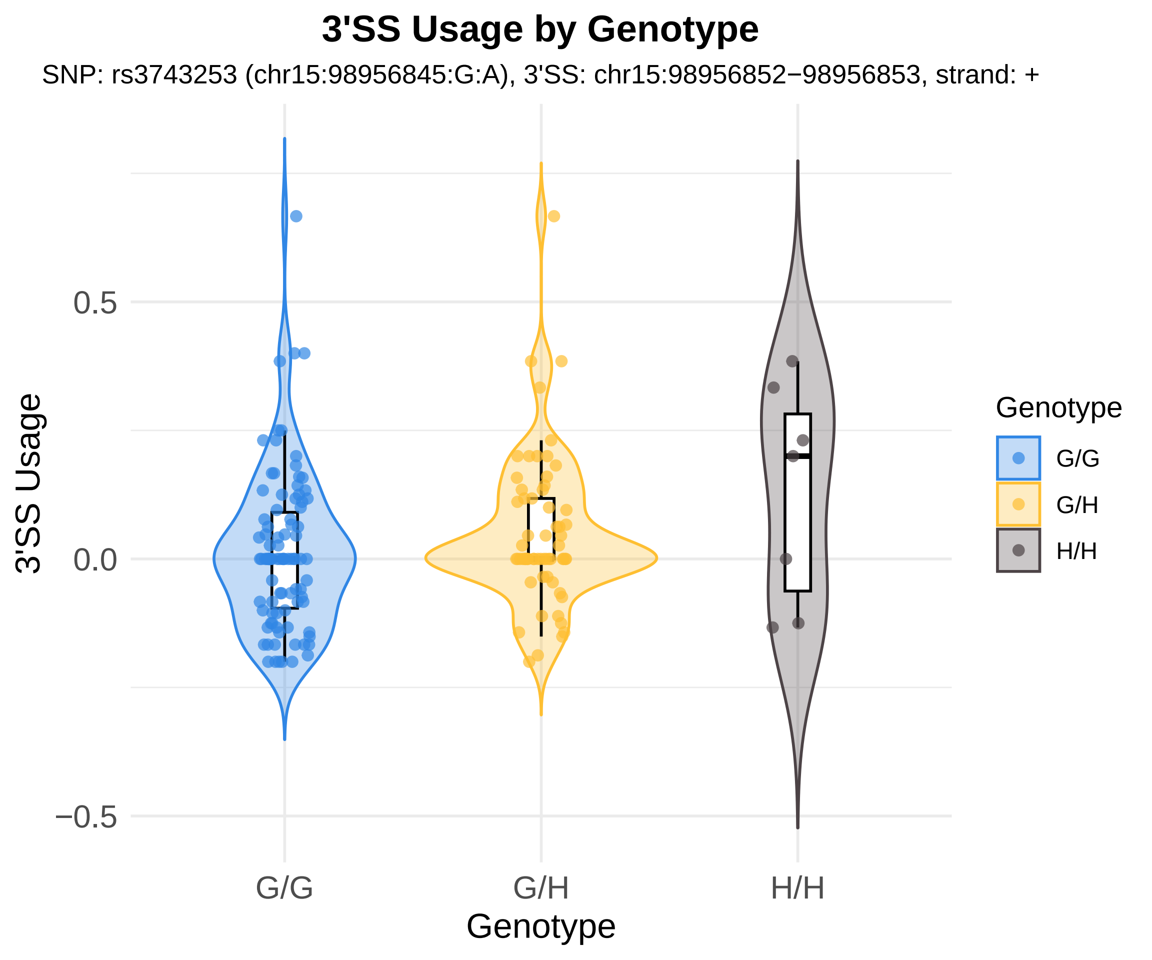
3'SS Usage by Genotype
SNP: rs3743253 (chr15:98956845:G:A), 3'SS: chr15:98956852−98956853, strand: +
3'SS Usage
0.5
0.0
−0.5
G/G
G/H
H/H
Genotype
Genotype
G/G
G/H
H/H

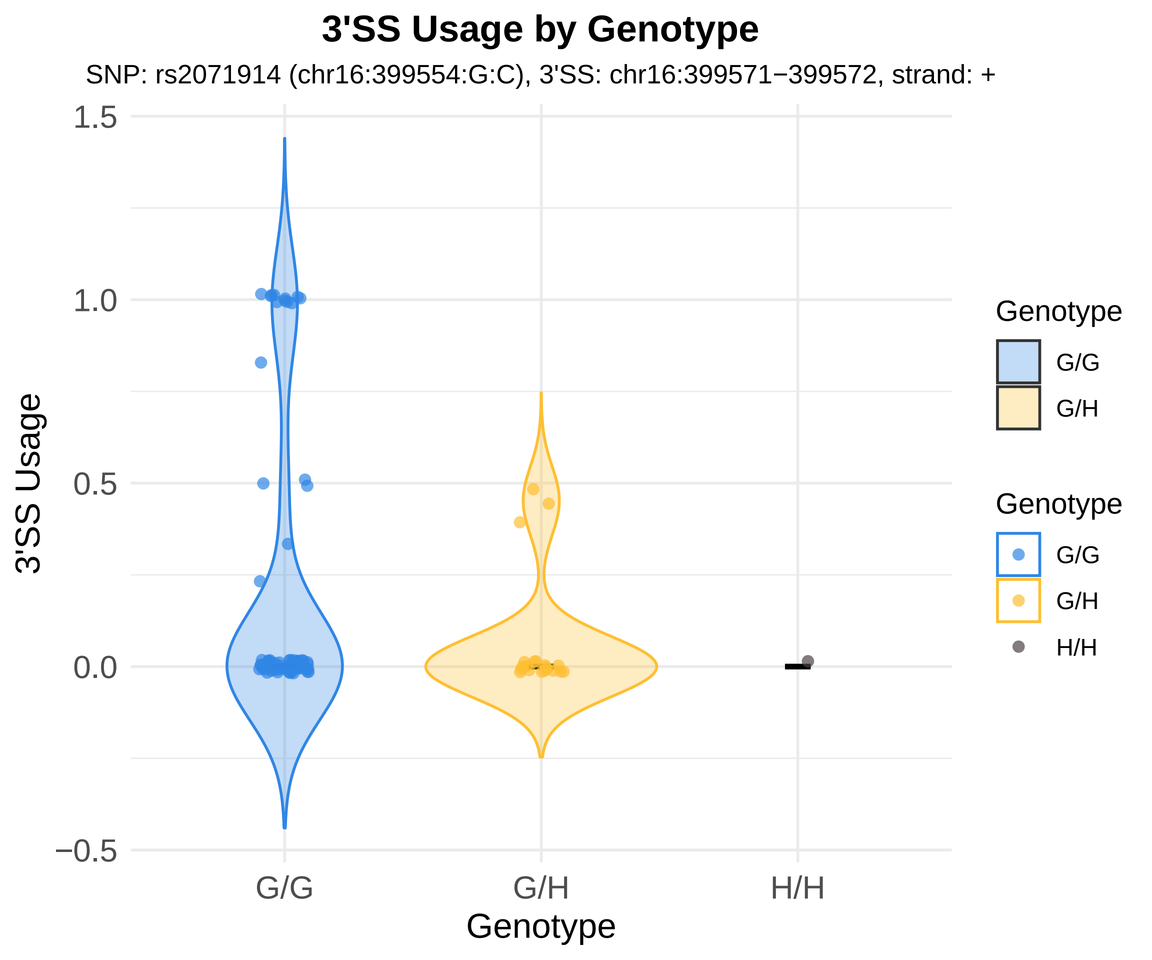

3'SS Usage by Genotype
SNP: rs2071914 (chr16:399554:G:C), 3'SS: chr16:399571−399572, strand: +
1.5
1.0
0.5
0.0
−0.5
3'SS Usage
G/G
G/H
H/H
Genotype
Genotype
G/G
G/H
Genotype
G/G
G/H
H/H

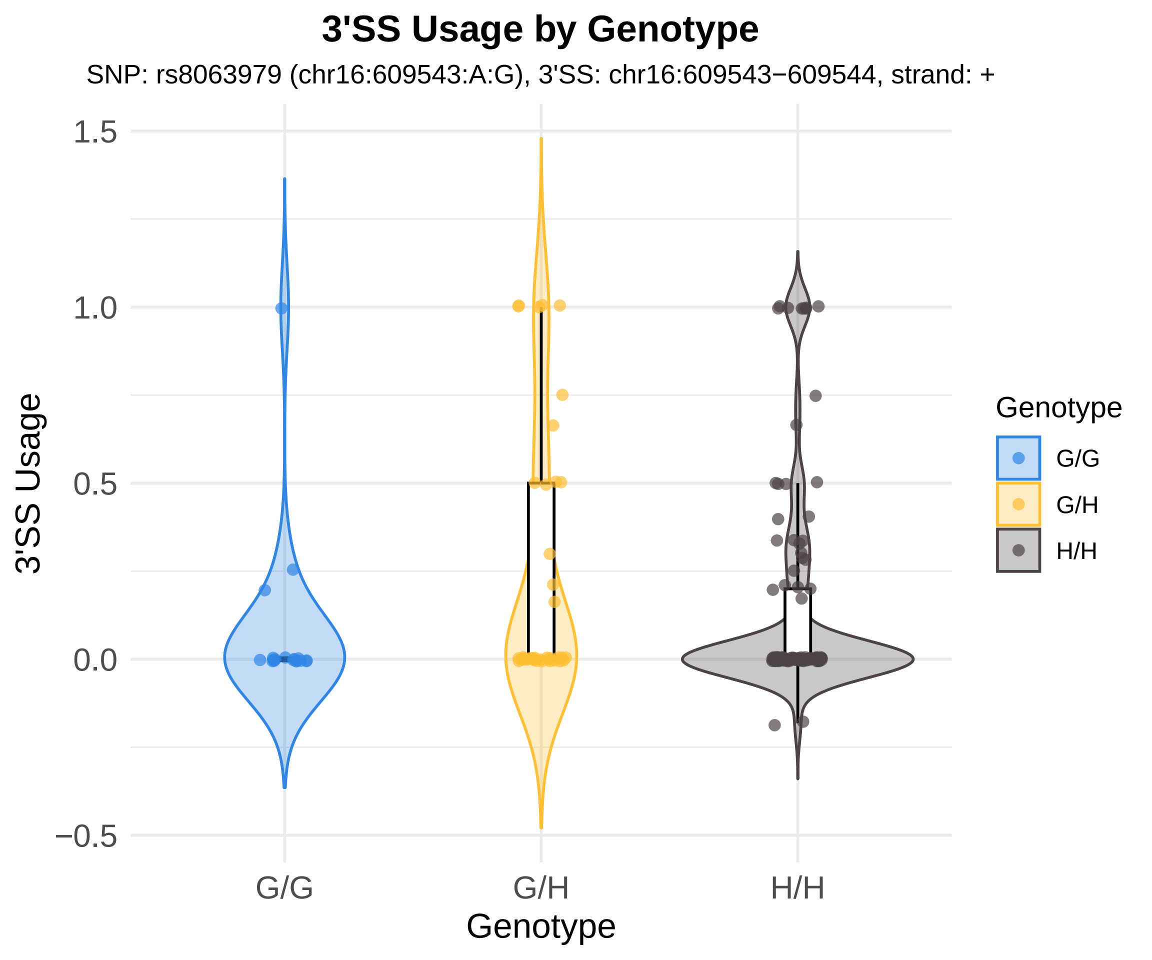
3'SS Usage by Genotype
SNP: rs8063979 (chr16:609543:A:G), 3'SS: chr16:609543−609544, strand: +
3'SS Usage
1.5
1.0
0.5
0.0
−0.5
G/G
G/H
H/H
Genotype
Genotype
G/G
G/H
H/H

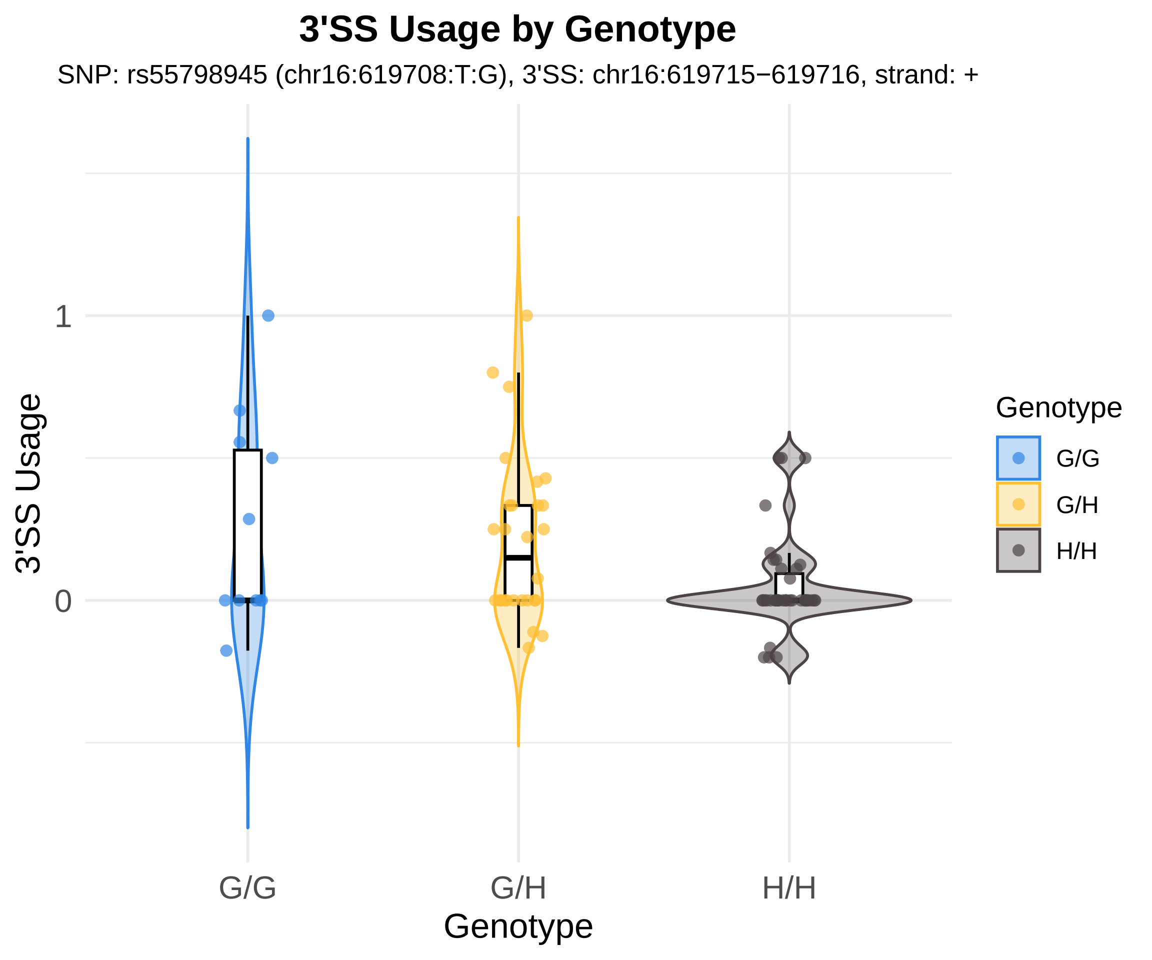
3'SS Usage by Genotype
SNP: rs55798945 (chr16:619708:T:G), 3'SS: chr16:619715−619716, strand: +
3'SS Usage
1
0
G/G
G/H
H/H
Genotype
Genotype
G/G
G/H
H/H

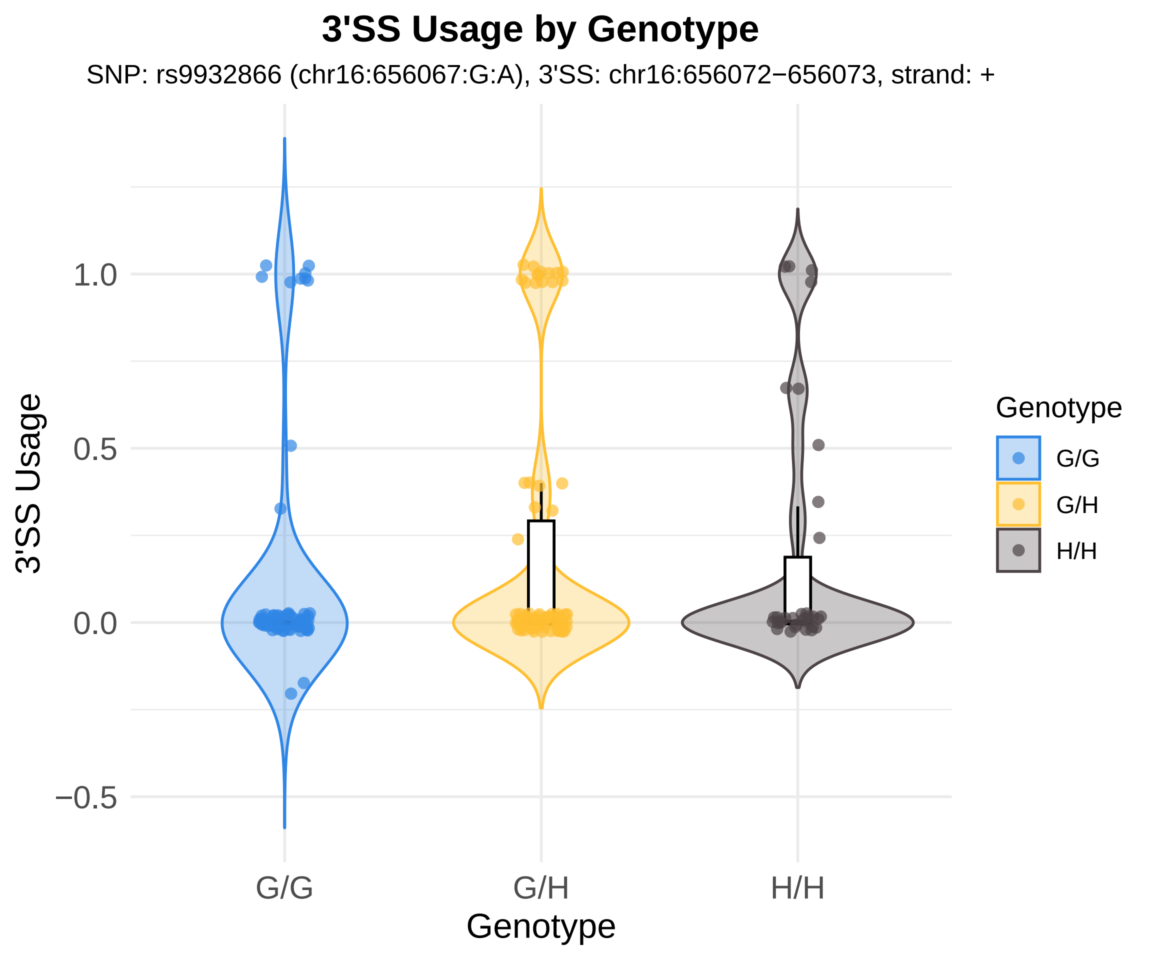
3'SS Usage by Genotype
SNP: rs9932866 (chr16:656067:G:A), 3'SS: chr16:656072−656073, strand: +
3'SS Usage
1.0
0.5
0.0
−0.5
G/G
G/H
H/H
Genotype
Genotype
G/G
G/H
H/H

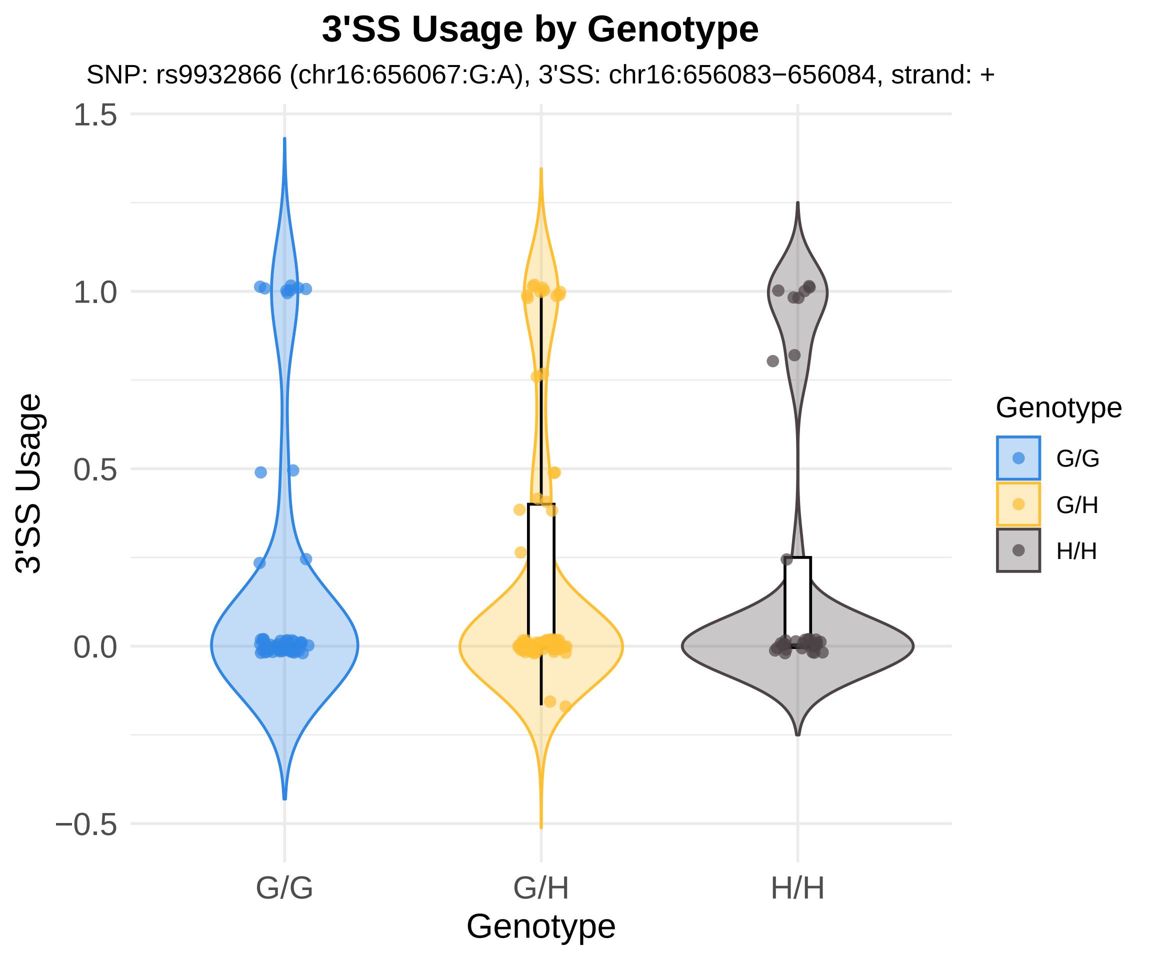
3'SS Usage by Genotype
SNP: rs9932866 (chr16:656067:G:A), 3'SS: chr16:656083−656084, strand: +
1.5
1.0
0.5
0.0
−0.5
3'SS Usage
G/G
G/H
H/H
Genotype
Genotype
G/G
G/H
H/H

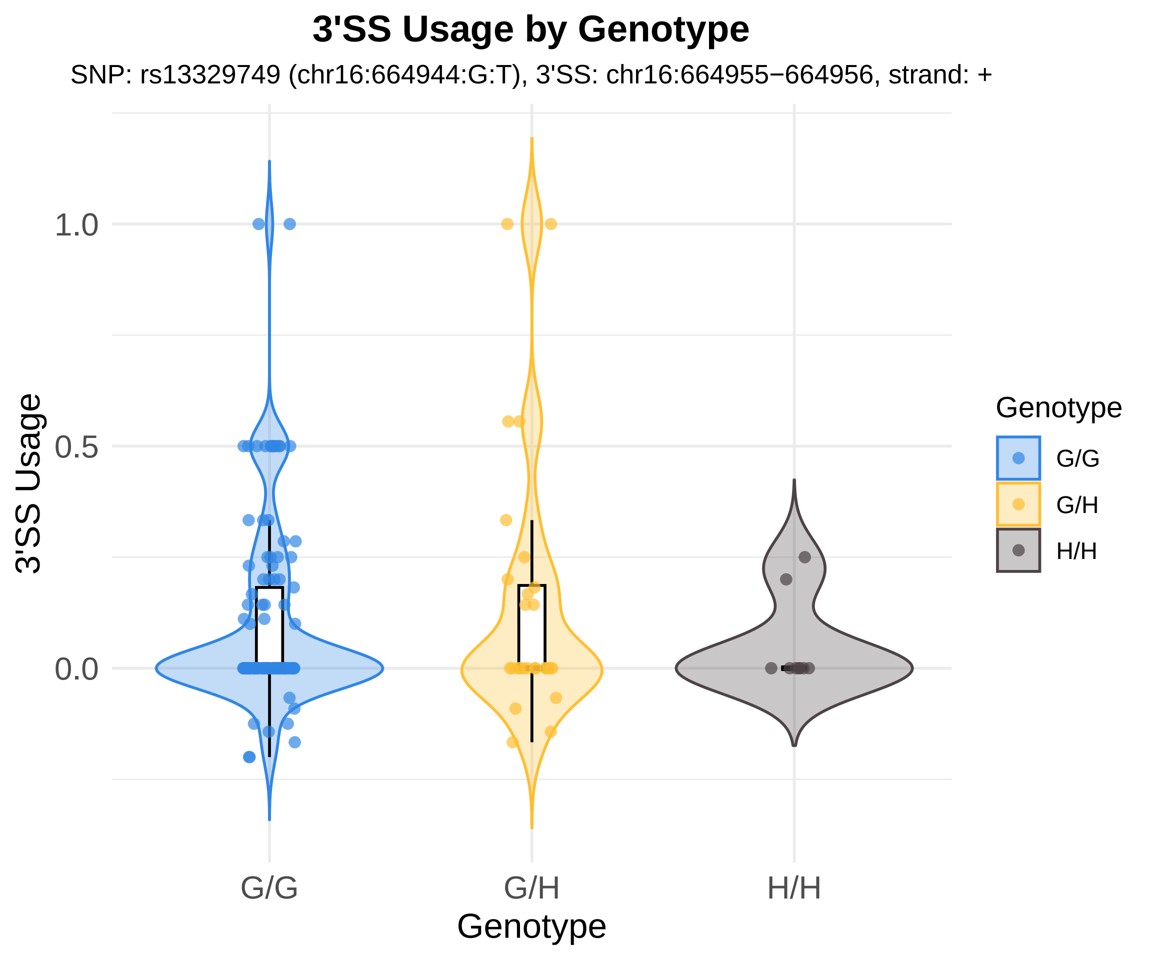

3'SS Usage by Genotype
SNP: rs13329749 (chr16:664944:G:T), 3'SS: chr16:664955−664956, strand: +
3'SS Usage
1.0
0.5
0.0
G/G
G/H
H/H
Genotype
Genotype
G/G
G/H
H/H

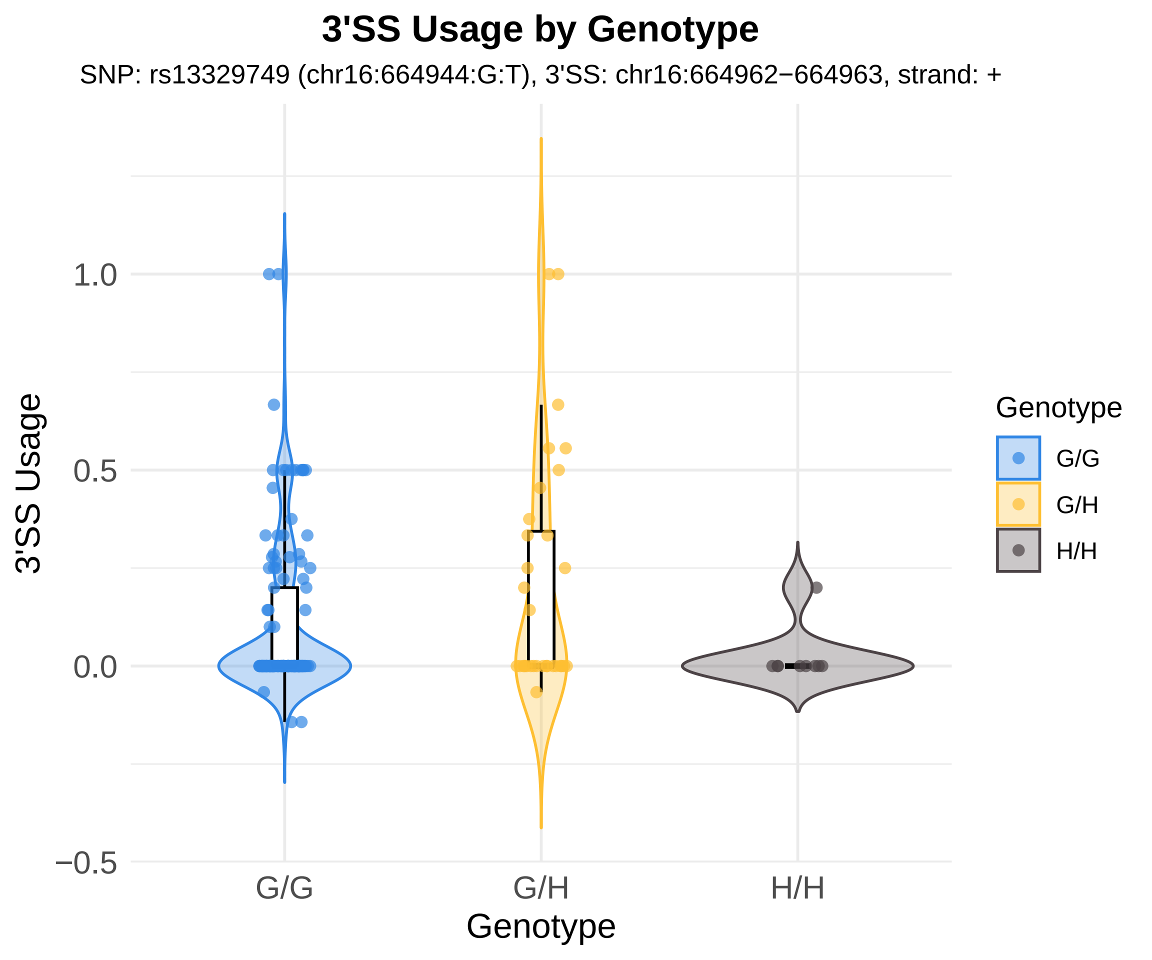

3'SS Usage by Genotype
SNP: rs13329749 (chr16:664944:G:T), 3'SS: chr16:664962−664963, strand: +
3'SS Usage
1.0
0.5
0.0
−0.5
G/G
G/H
H/H
Genotype
Genotype
G/G
G/H
H/H

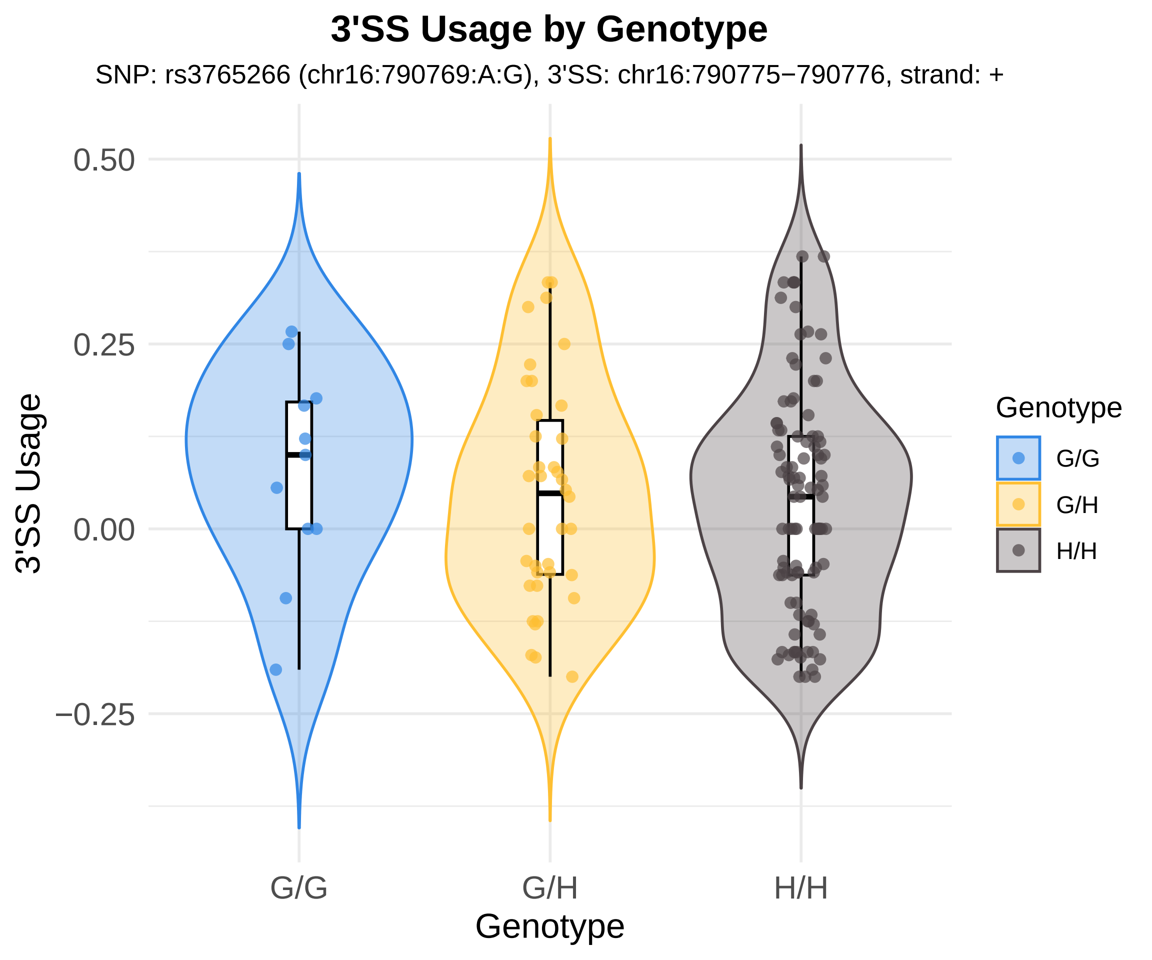
3'SS Usage by Genotype
SNP: rs3765266 (chr16:790769:A:G), 3'SS: chr16:790775−790776, strand: +
3'SS Usage
0.50
0.25
0.00
−0.25
G/G
G/H
H/H
Genotype
Genotype
G/G
G/H
H/H

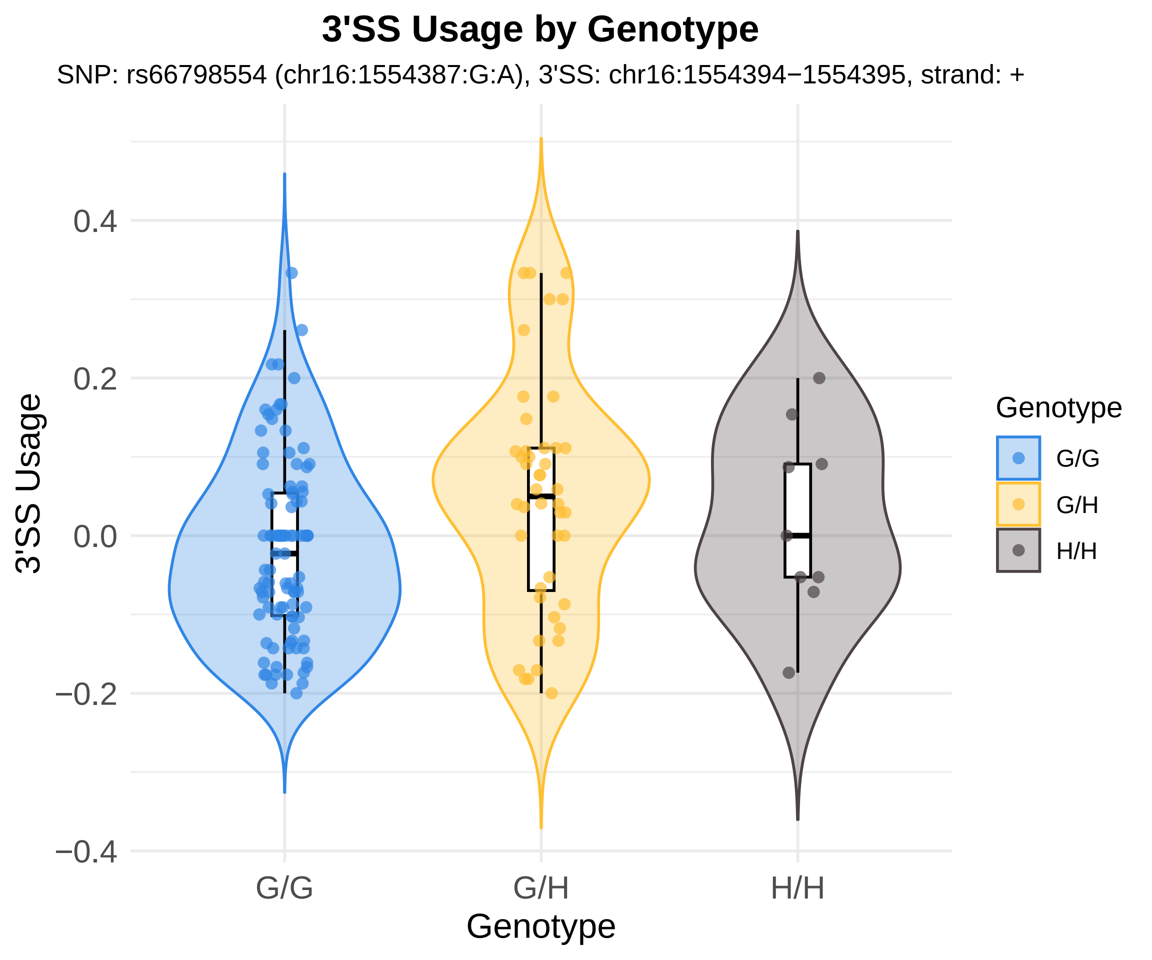
3'SS Usage by Genotype
SNP: rs66798554 (chr16:1554387:G:A), 3'SS: chr16:1554394−1554395, strand: +
3'SS Usage
0.4
0.2
0.0
−0.2
−0.4
G/G
G/H
H/H
Genotype
Genotype
G/G
G/H
H/H

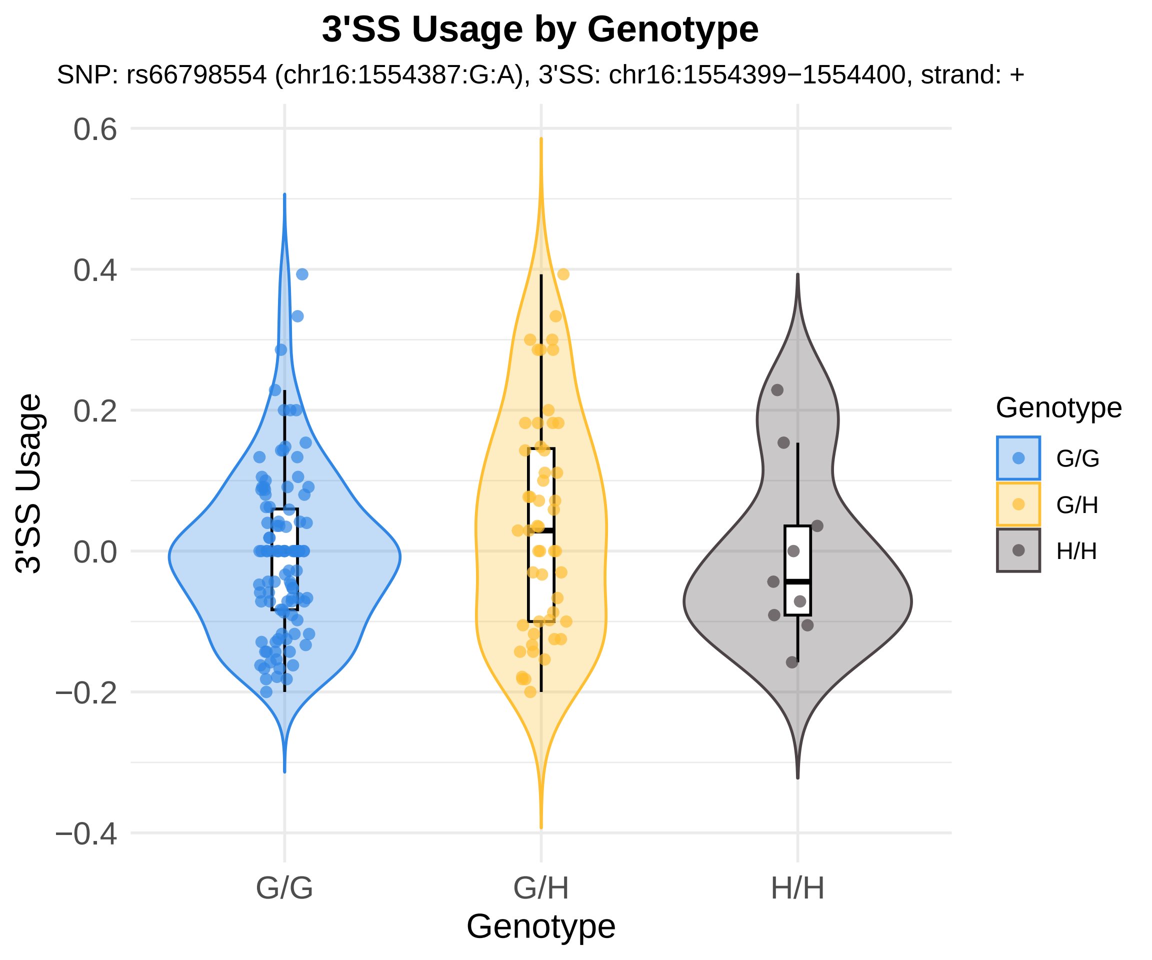
3'SS Usage by Genotype
SNP: rs66798554 (chr16:1554387:G:A), 3'SS: chr16:1554399−1554400, strand: +
3'SS Usage
0.6
0.4
0.2
0.0
−0.2
−0.4
G/G
G/H
H/H
Genotype
Genotype
G/G
G/H
H/H

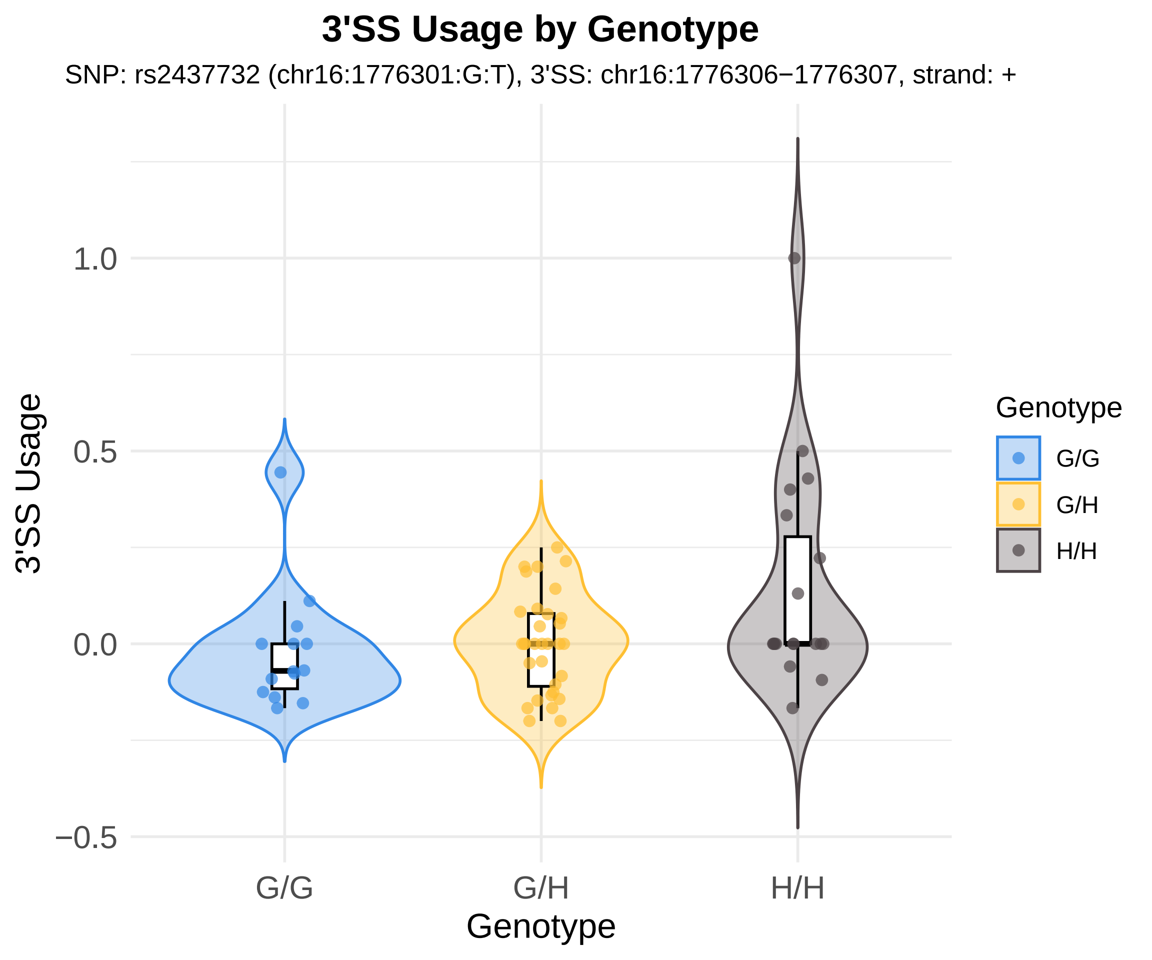
3'SS Usage by Genotype
SNP: rs2437732 (chr16:1776301:G:T), 3'SS: chr16:1776306−1776307, strand: +
3'SS Usage
1.0
0.5
0.0
−0.5
G/G
G/H
H/H
Genotype
Genotype
G/G
G/H
H/H

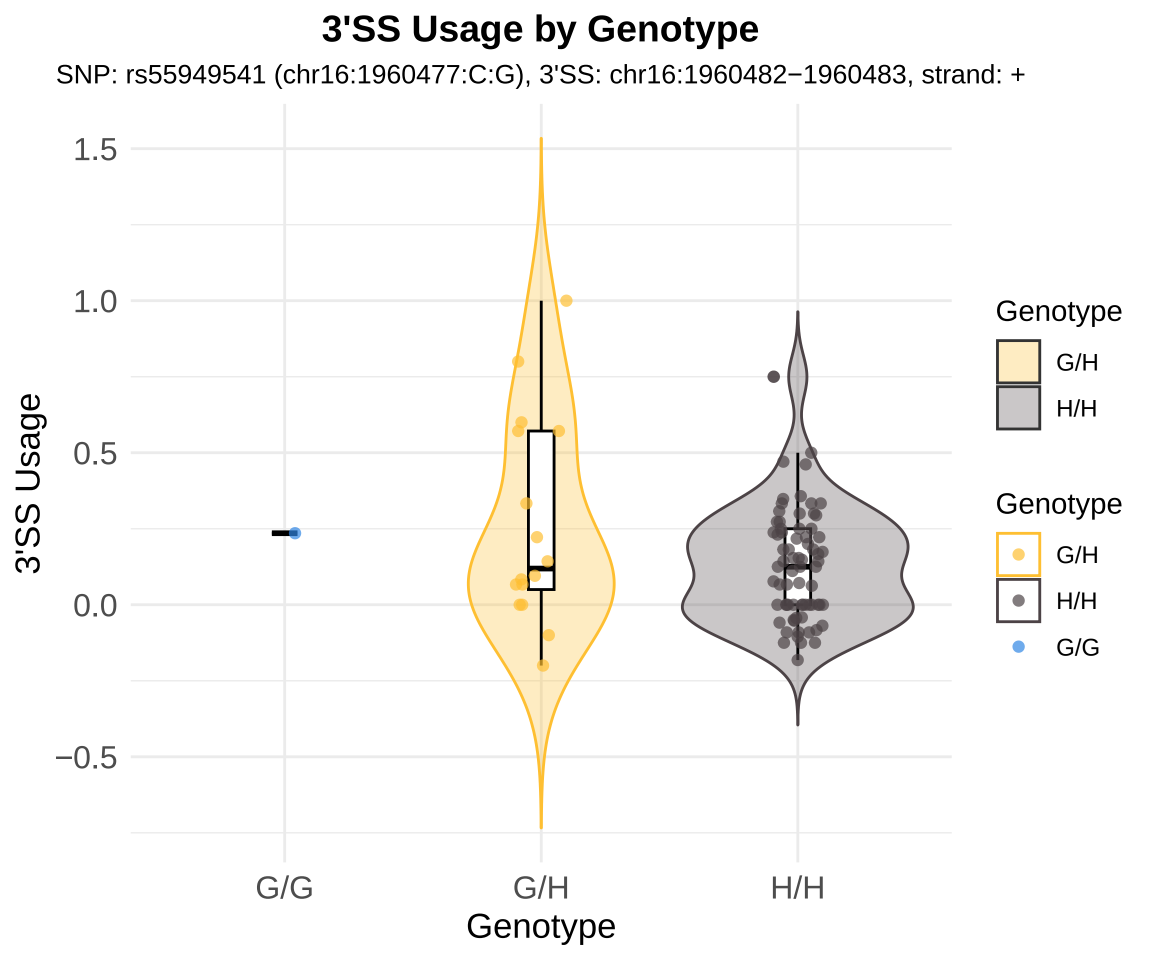
3'SS Usage by Genotype
SNP: rs55949541 (chr16:1960477:C:G), 3'SS: chr16:1960482−1960483, strand: +
3'SS Usage
1.5
1.0
0.5
0.0
−0.5
G/G
G/H
H/H
Genotype
Genotype
G/H
H/H
Genotype
G/H
H/H
G/G

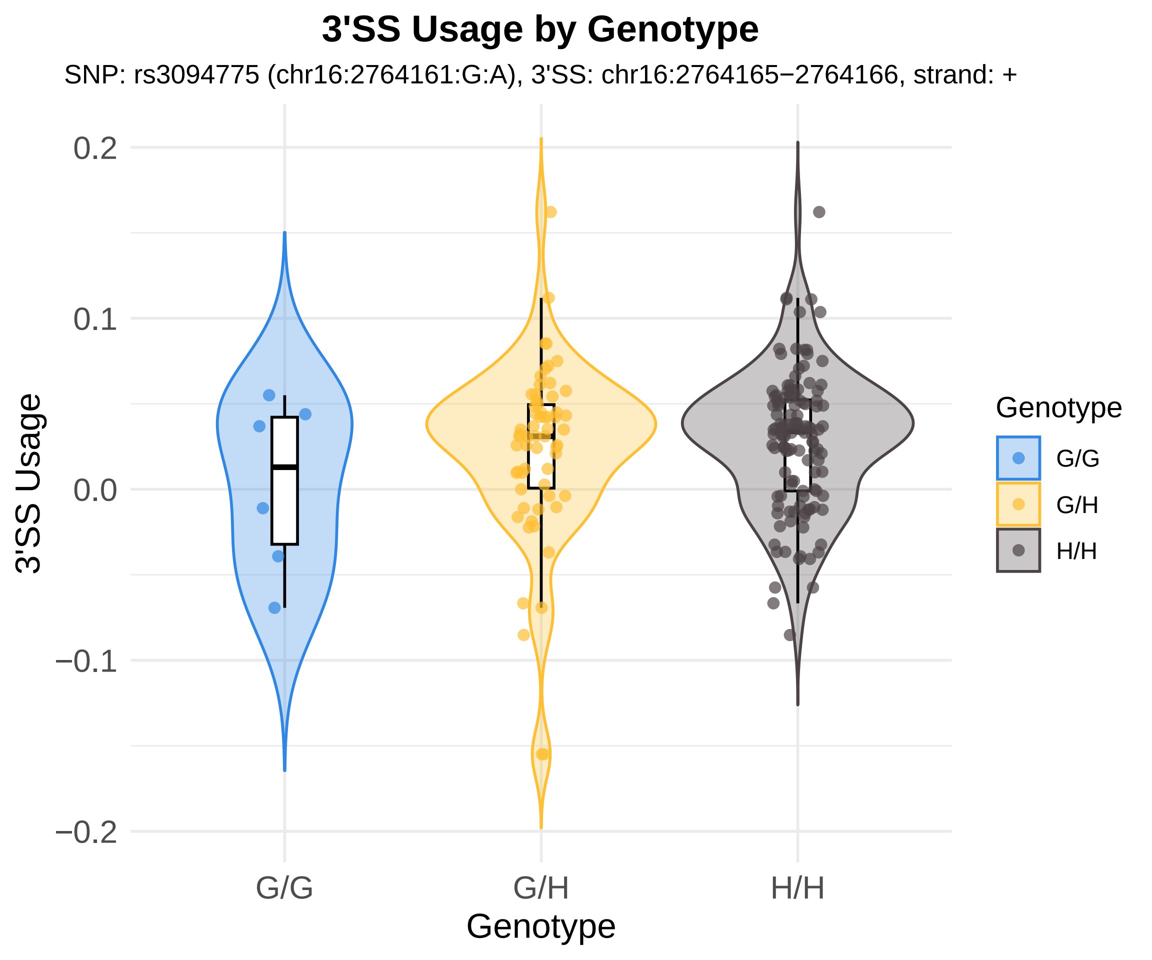

3'SS Usage by Genotype
SNP: rs3094775 (chr16:2764161:G:A), 3'SS: chr16:2764165−2764166, strand: +
3'SS Usage
0.2
0.1
0.0
−0.1
−0.2
G/G
G/H
H/H
Genotype
Genotype
G/G
G/H
H/H

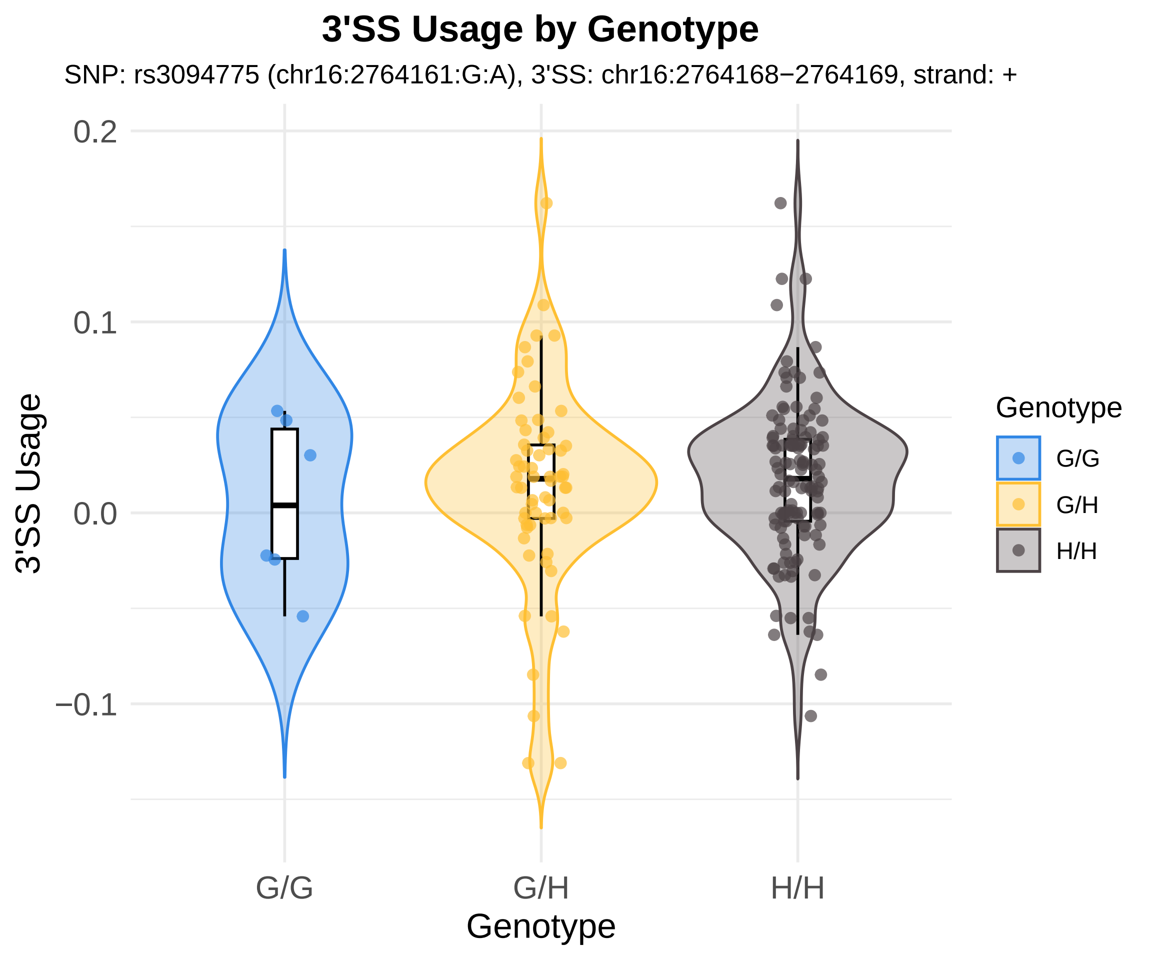
3'SS Usage by Genotype
SNP: rs3094775 (chr16:2764161:G:A), 3'SS: chr16:2764168−2764169, strand: +
0.2
0.1
0.0
−0.1
3'SS Usage
G/G
G/H
H/H
Genotype
Genotype
G/G
G/H
H/H

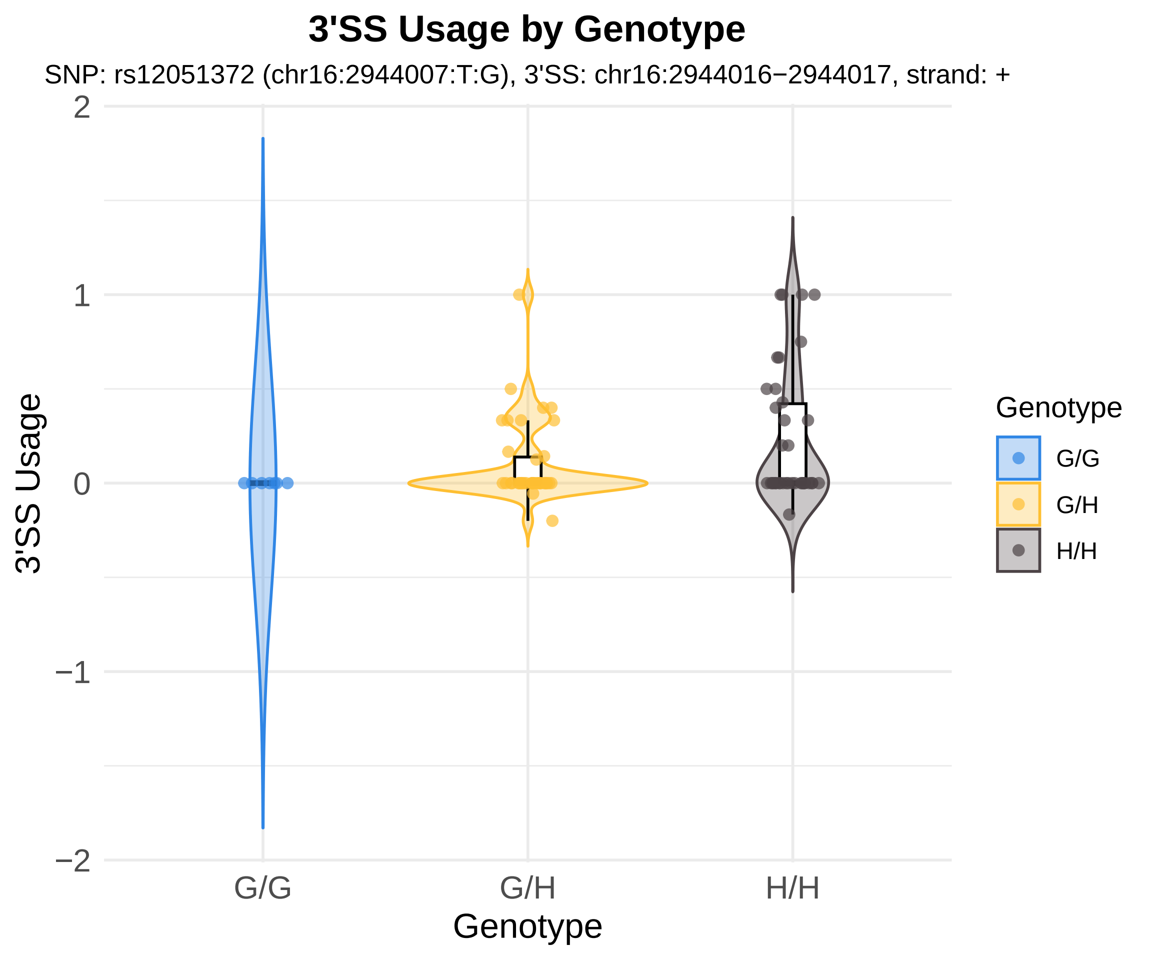
3'SS Usage by Genotype
SNP: rs12051372 (chr16:2944007:T:G), 3'SS: chr16:2944016−2944017, strand: +
3'SS Usage
2
1
0
−1
−2
G/G
G/H
H/H
Genotype
Genotype
G/G
G/H
H/H

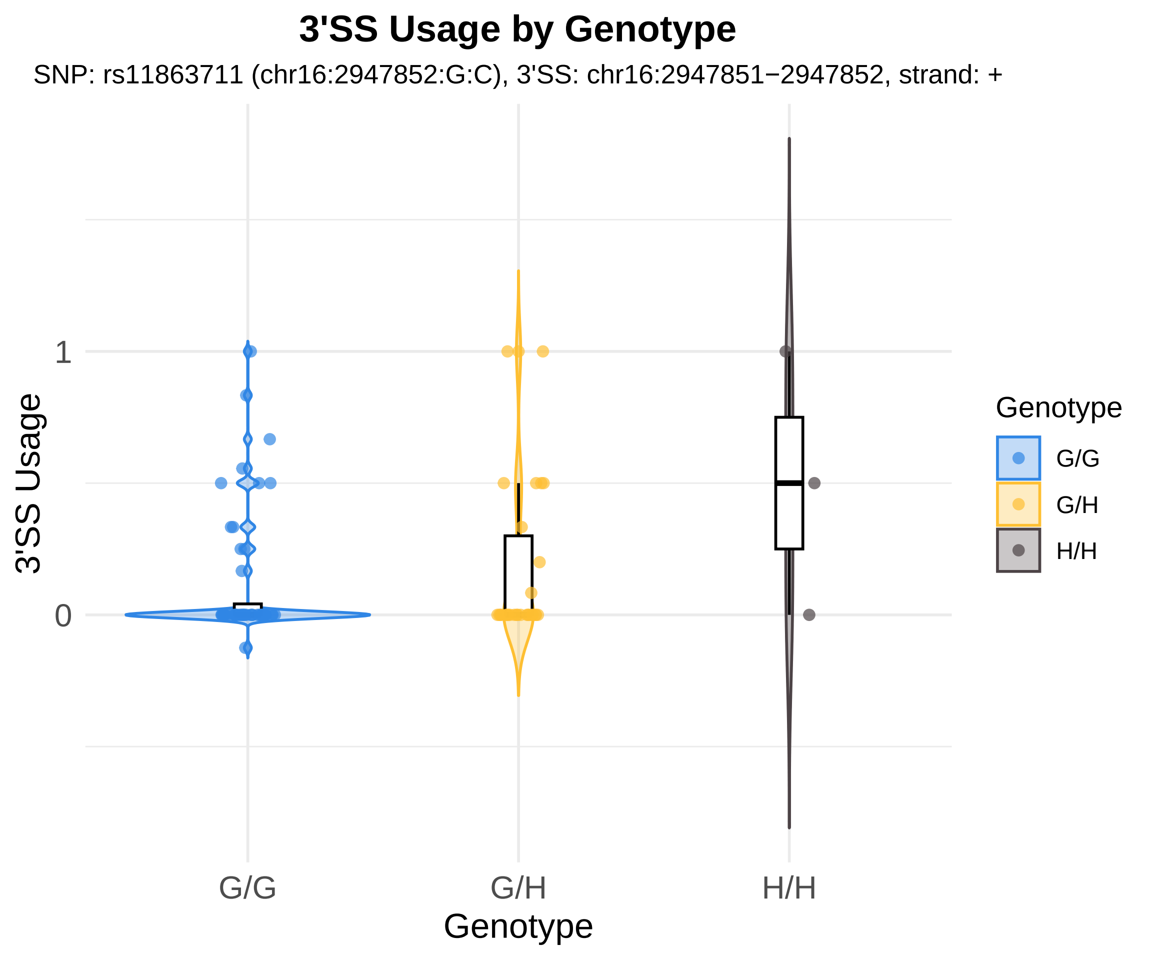

3'SS Usage by Genotype
SNP: rs11863711 (chr16:2947852:G:C), 3'SS: chr16:2947851−2947852, strand: +
3'SS Usage
1
0
G/G
G/H
H/H
Genotype
Genotype
G/G
G/H
H/H

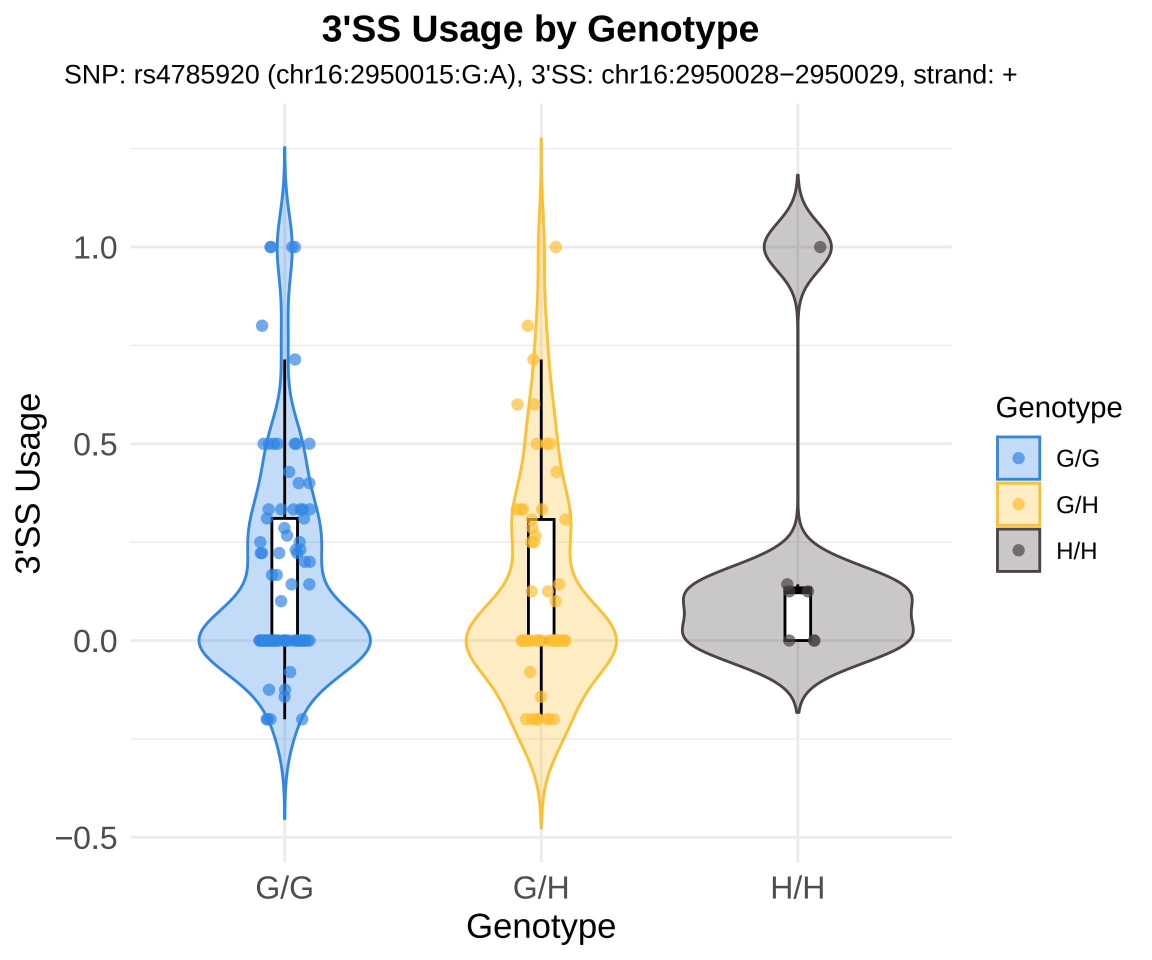

3'SS Usage by Genotype
SNP: rs4785920 (chr16:2950015:G:A), 3'SS: chr16:2950028−2950029, strand: +
3'SS Usage
1.0
0.5
0.0
−0.5
G/G
G/H
H/H
Genotype
Genotype
G/G
G/H
H/H

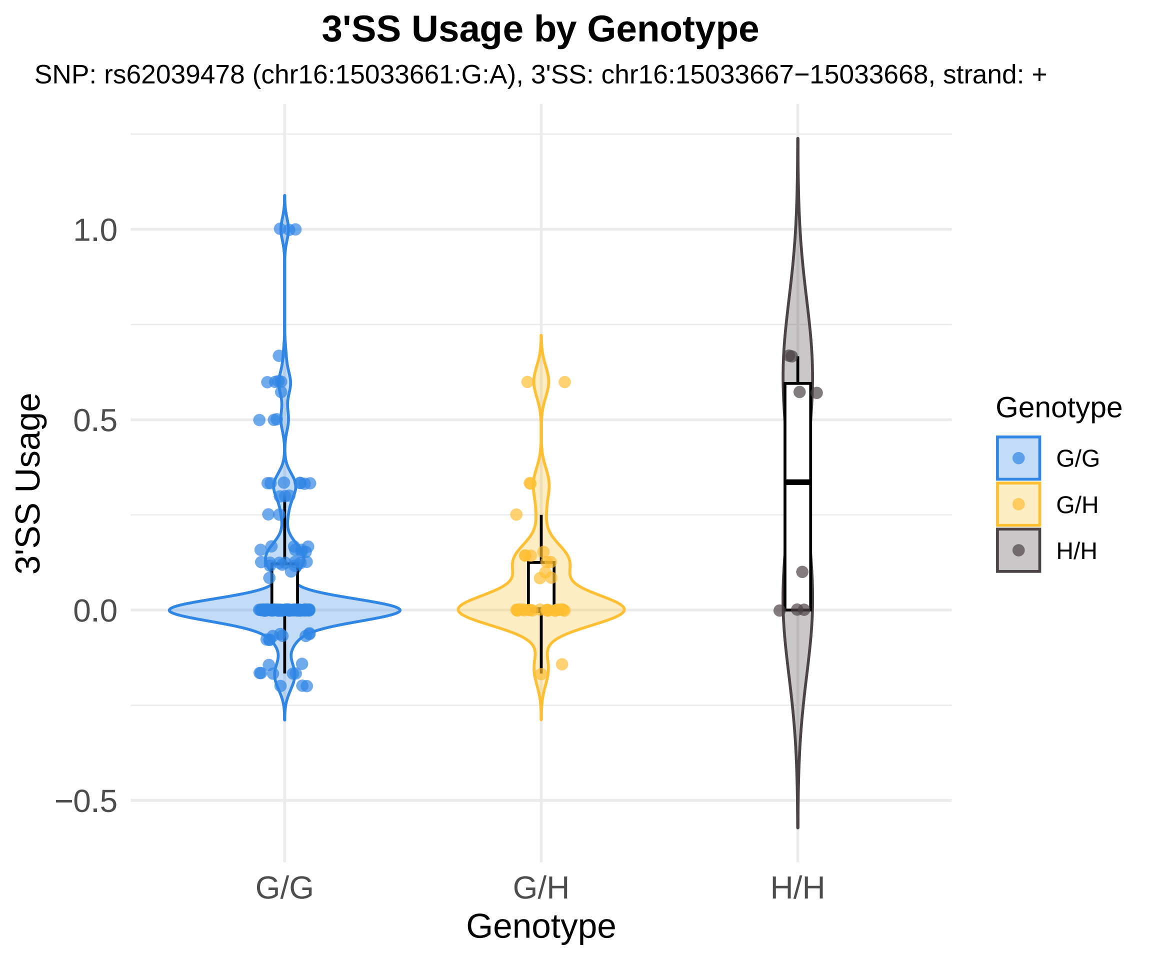
3'SS Usage by Genotype
SNP: rs62039478 (chr16:15033661:G:A), 3'SS: chr16:15033667−15033668, strand: +
3'SS Usage
1.0
0.5
0.0
−0.5
G/G
G/H
H/H
Genotype
Genotype
G/G
G/H
H/H

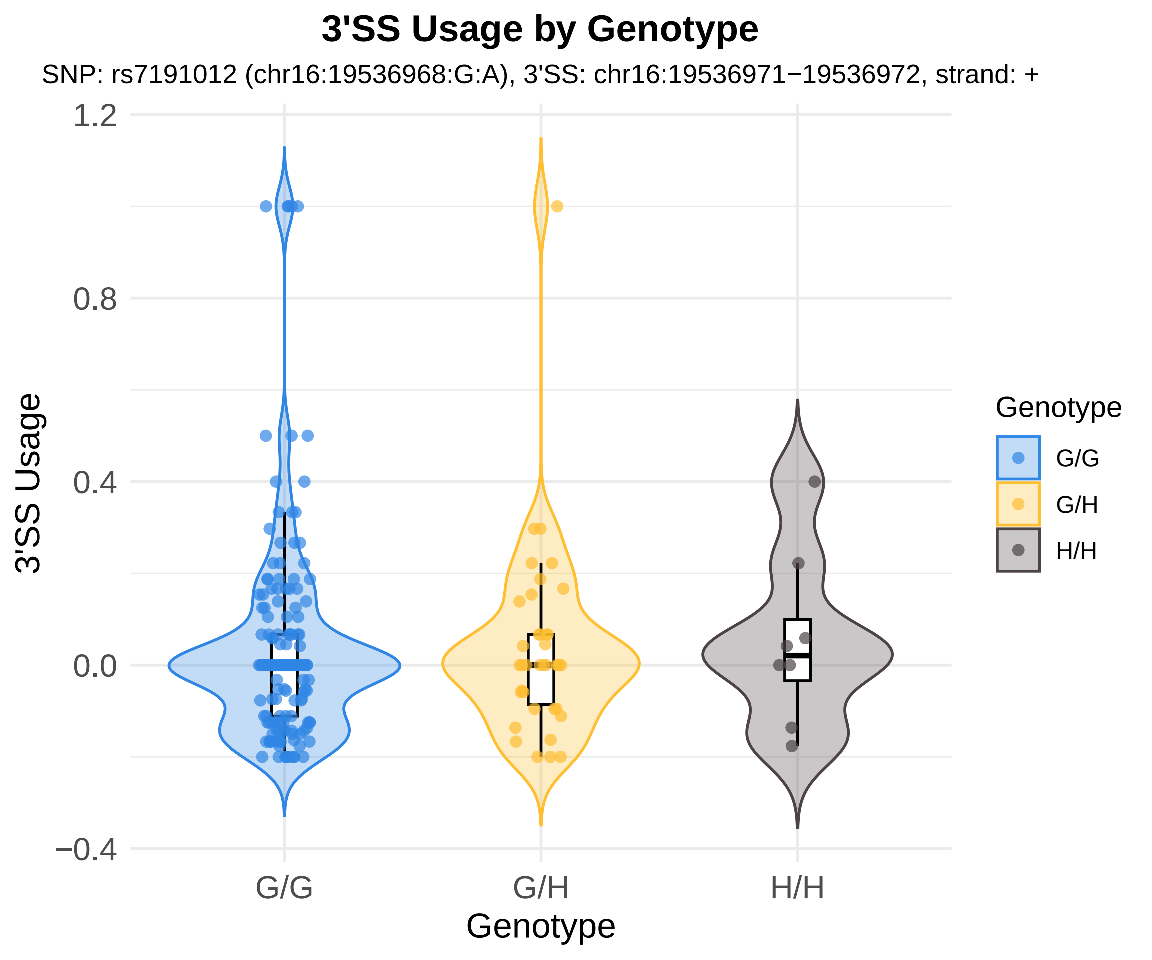

3'SS Usage by Genotype
SNP: rs7191012 (chr16:19536968:G:A), 3'SS: chr16:19536971−19536972, strand: +
3'SS Usage
1.2
0.8
0.4
0.0
−0.4
G/G
G/H
H/H
Genotype
Genotype
G/G
G/H
H/H

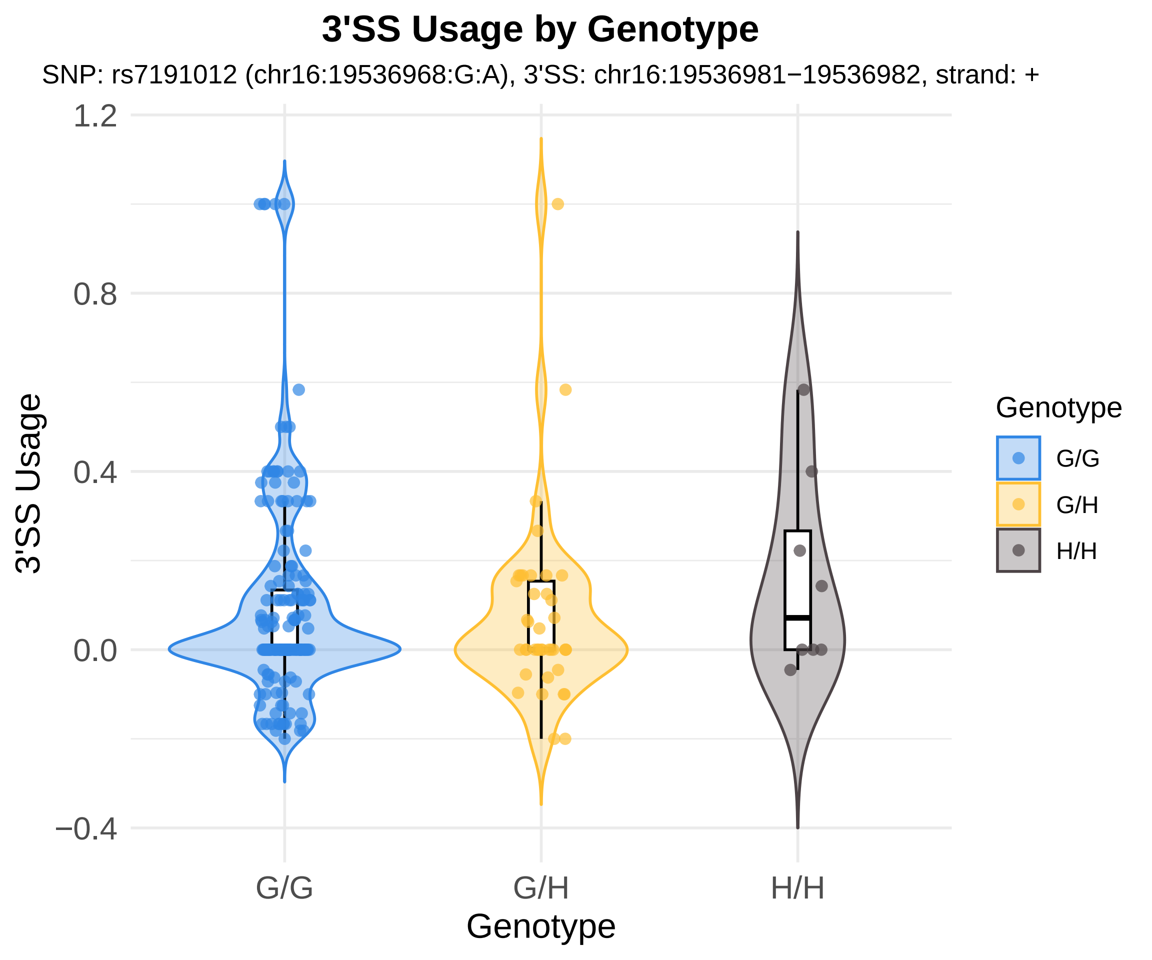

3'SS Usage by Genotype
SNP: rs7191012 (chr16:19536968:G:A), 3'SS: chr16:19536981−19536982, strand: +
3'SS Usage
1.2
0.8
0.4
0.0
−0.4
G/G
G/H
H/H
Genotype
Genotype
G/G
G/H
H/H

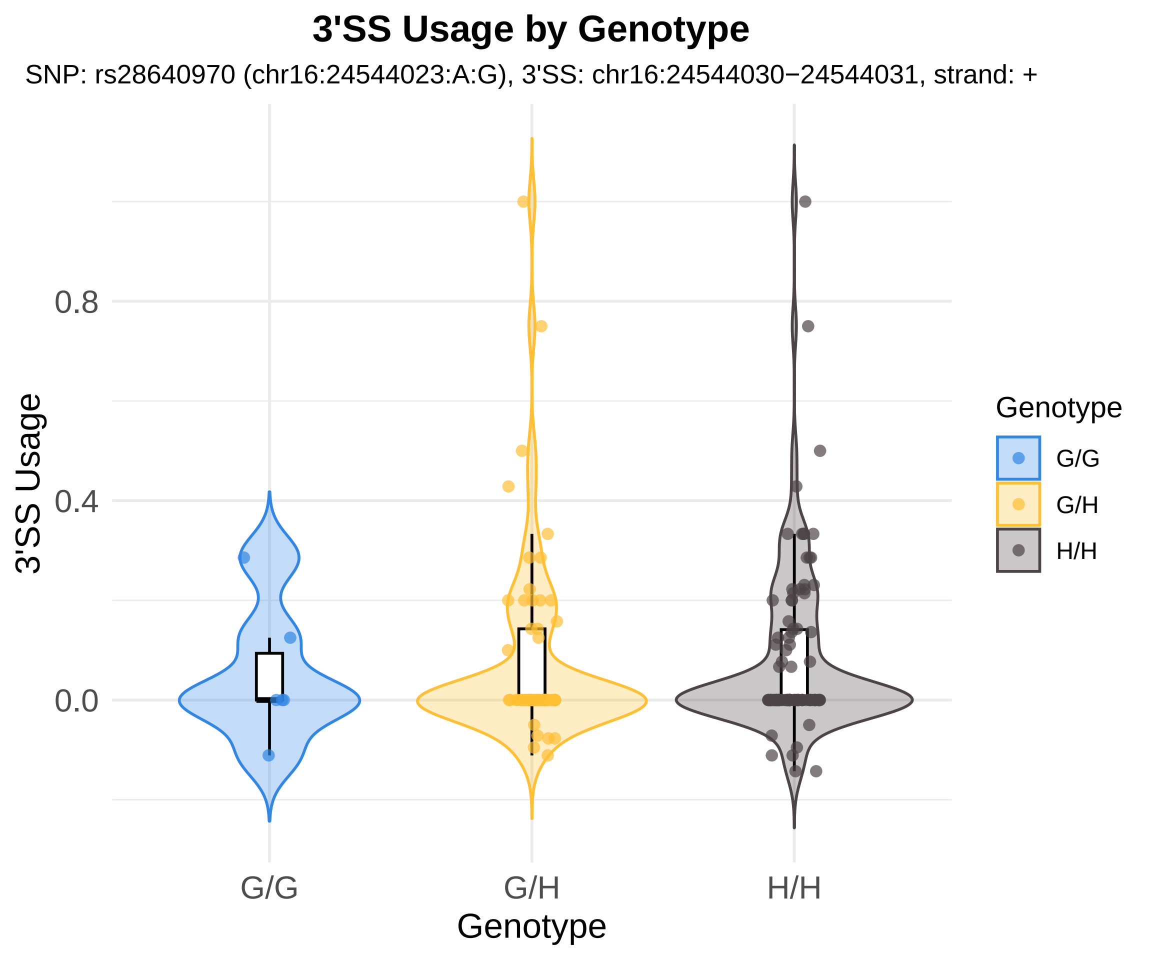
3'SS Usage by Genotype
SNP: rs28640970 (chr16:24544023:A:G), 3'SS: chr16:24544030−24544031, strand: +
3'SS Usage
0.8
0.4
0.0
G/G
G/H
H/H
Genotype
Genotype
G/G
G/H
H/H

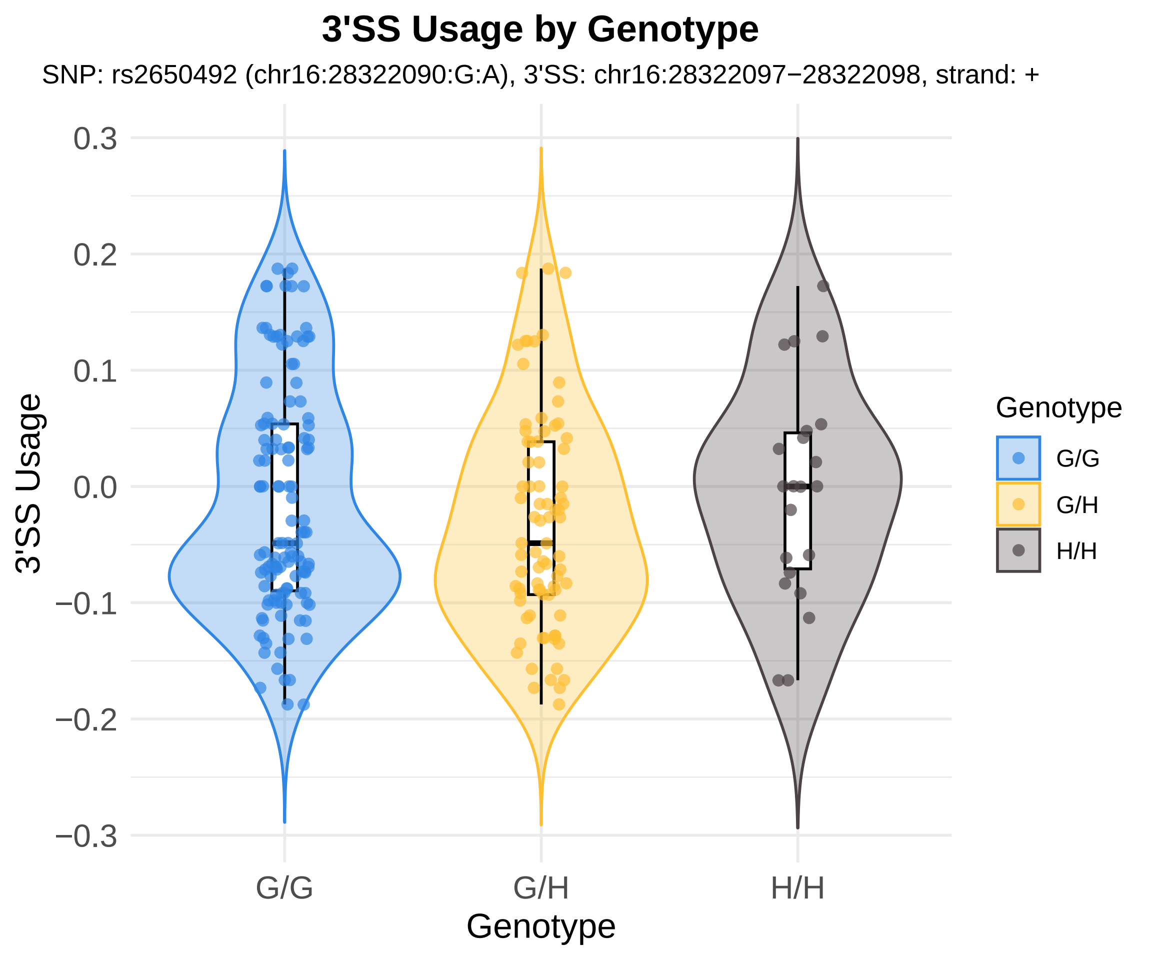
3'SS Usage by Genotype
SNP: rs2650492 (chr16:28322090:G:A), 3'SS: chr16:28322097−28322098, strand: +
3'SS Usage
0.3
0.2
0.1
0.0
−0.1
−0.2
−0.3
G/G
G/H
H/H
Genotype
Genotype
G/G
G/H
H/H

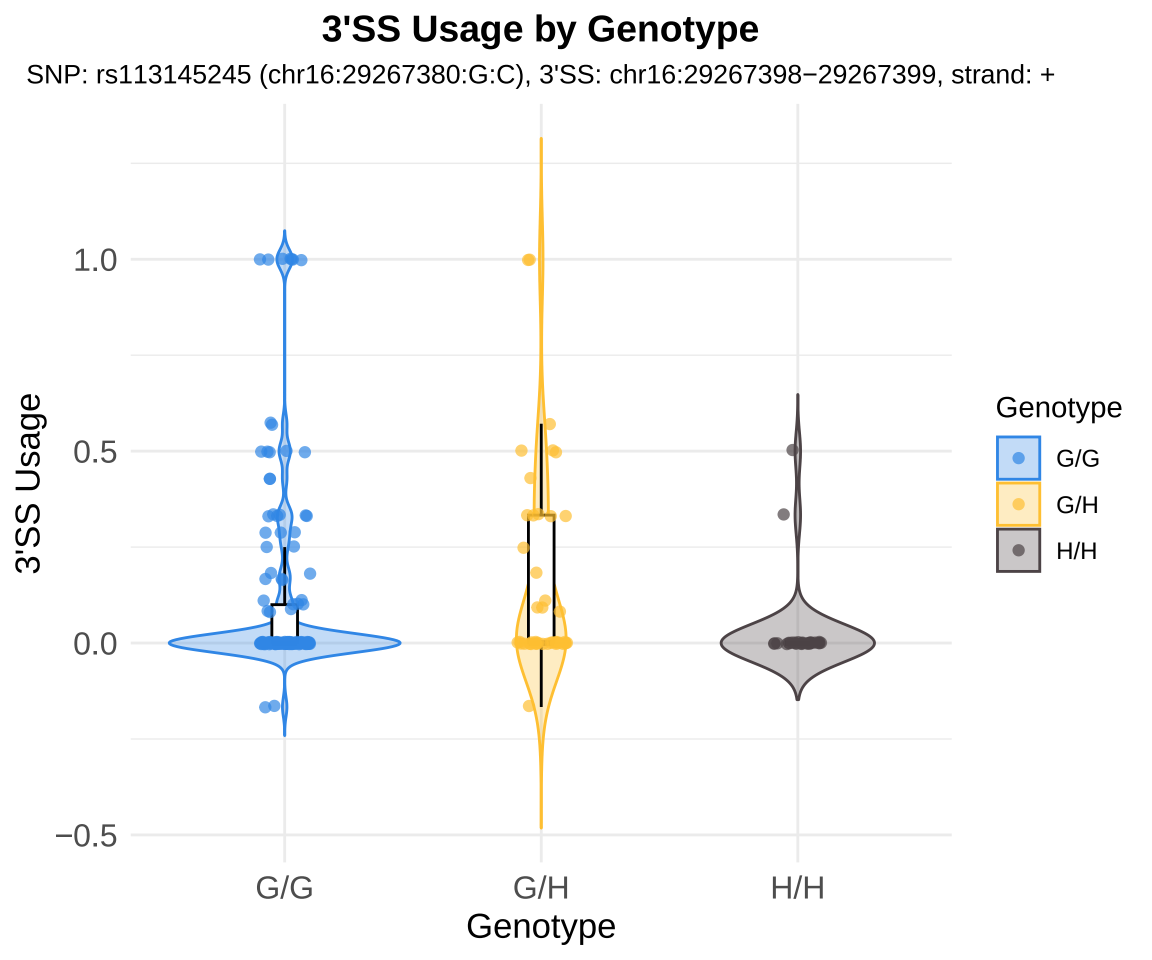
3'SS Usage by Genotype
SNP: rs113145245 (chr16:29267380:G:C), 3'SS: chr16:29267398−29267399, strand: +
3'SS Usage
1.0
0.5
0.0
−0.5
G/G
G/H
H/H
Genotype
Genotype
G/G
G/H
H/H

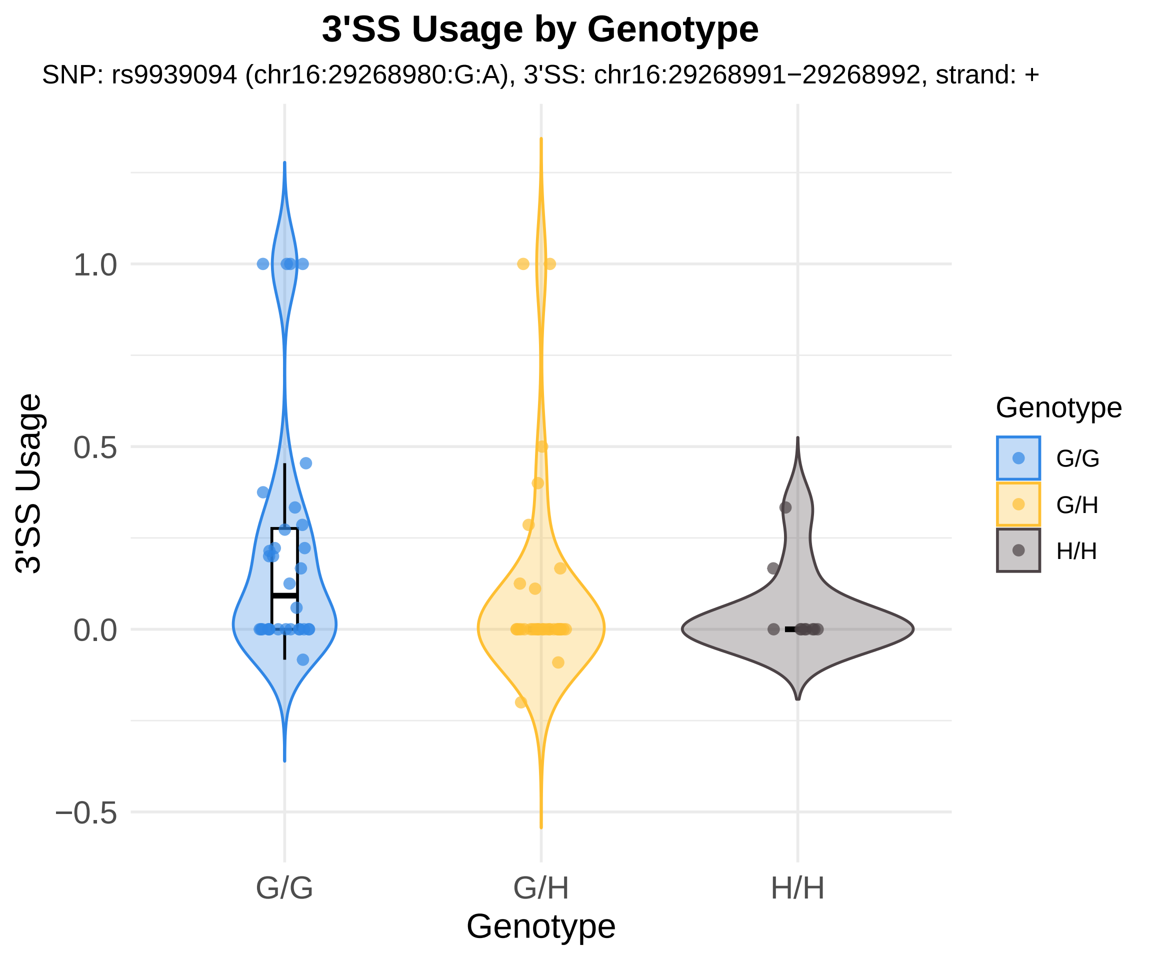

3'SS Usage by Genotype
SNP: rs9939094 (chr16:29268980:G:A), 3'SS: chr16:29268991−29268992, strand: +
3'SS Usage
1.0
0.5
0.0
−0.5
G/G
G/H
H/H
Genotype
Genotype
G/G
G/H
H/H

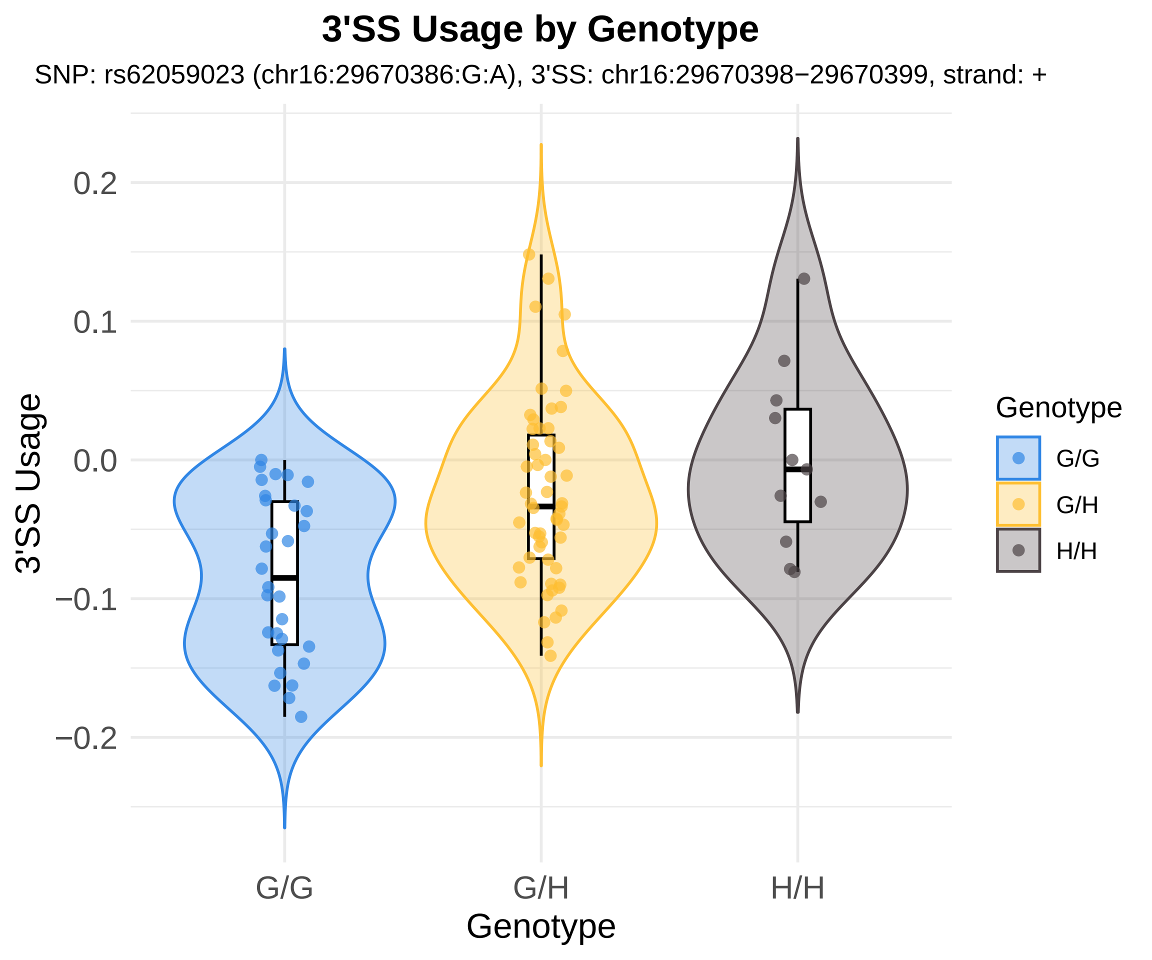
3'SS Usage by Genotype
SNP: rs62059023 (chr16:29670386:G:A), 3'SS: chr16:29670398−29670399, strand: +
3'SS Usage
0.2
0.1
0.0
−0.1
−0.2
G/G
G/H
H/H
Genotype
Genotype
G/G
G/H
H/H

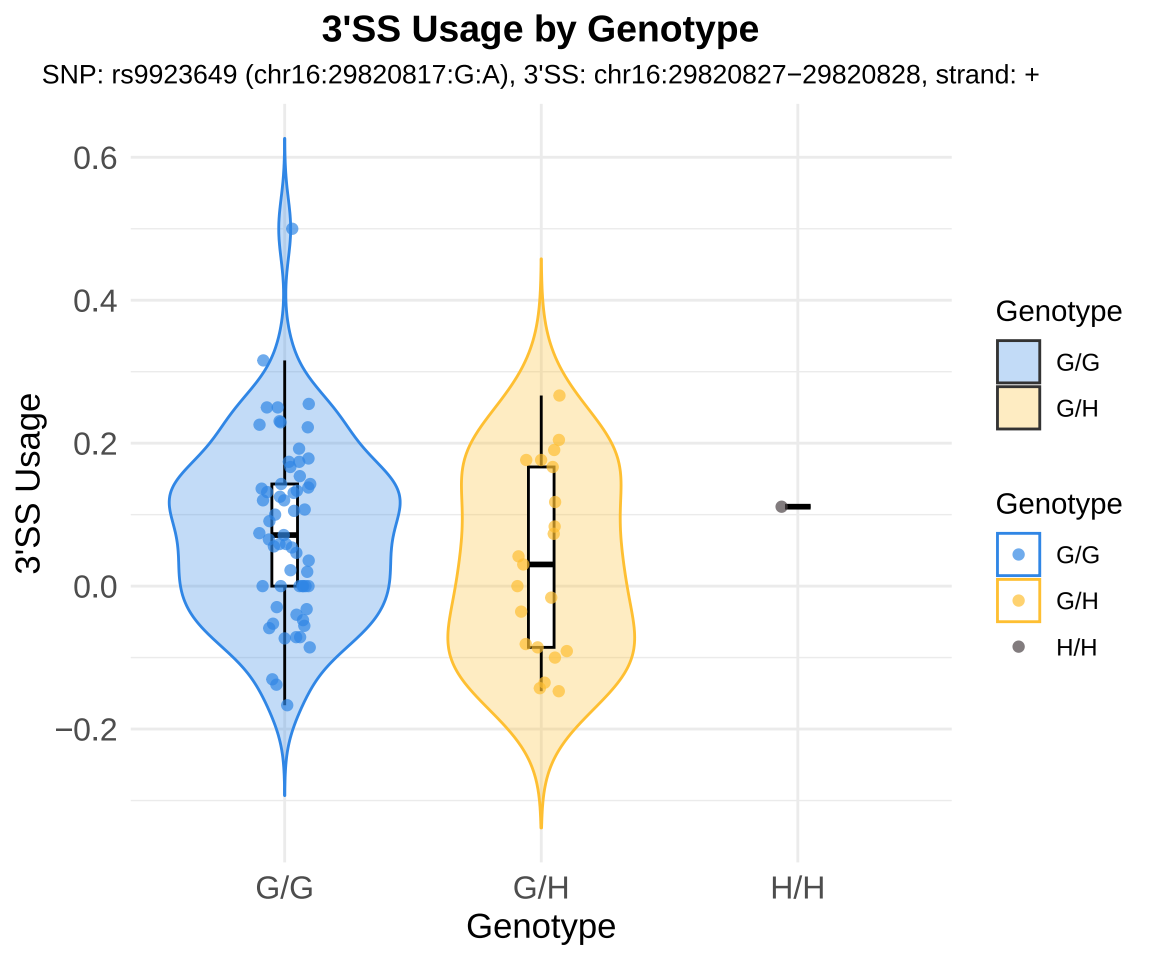

3'SS Usage by Genotype
SNP: rs9923649 (chr16:29820817:G:A), 3'SS: chr16:29820827−29820828, strand: +
3'SS Usage
0.6
0.4
0.2
0.0
−0.2
G/G
G/H
H/H
Genotype
Genotype
G/G
G/H
Genotype
G/G
G/H
H/H

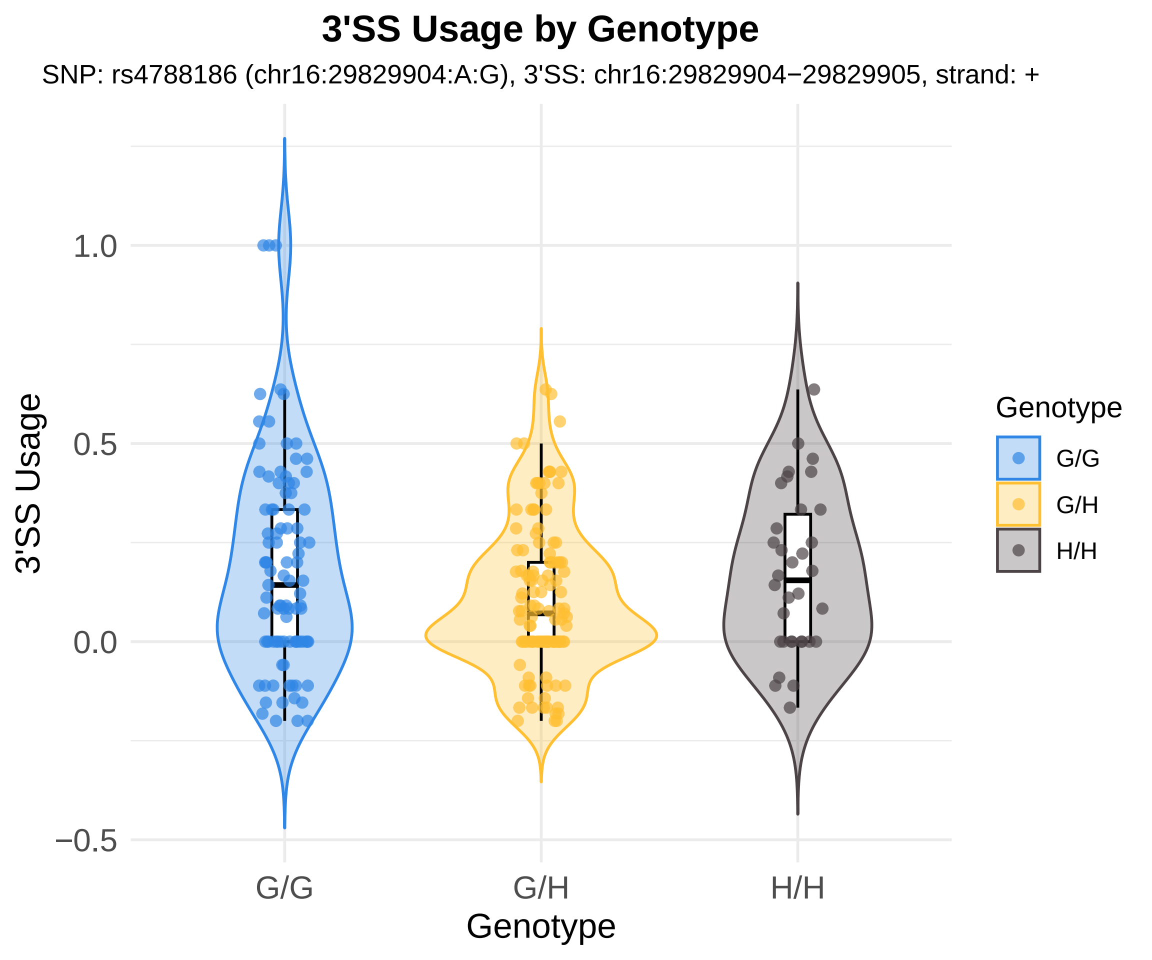
3'SS Usage by Genotype
SNP: rs4788186 (chr16:29829904:A:G), 3'SS: chr16:29829904−29829905, strand: +
3'SS Usage
1.0
0.5
0.0
−0.5
G/G
G/H
H/H
Genotype
Genotype
G/G
G/H
H/H

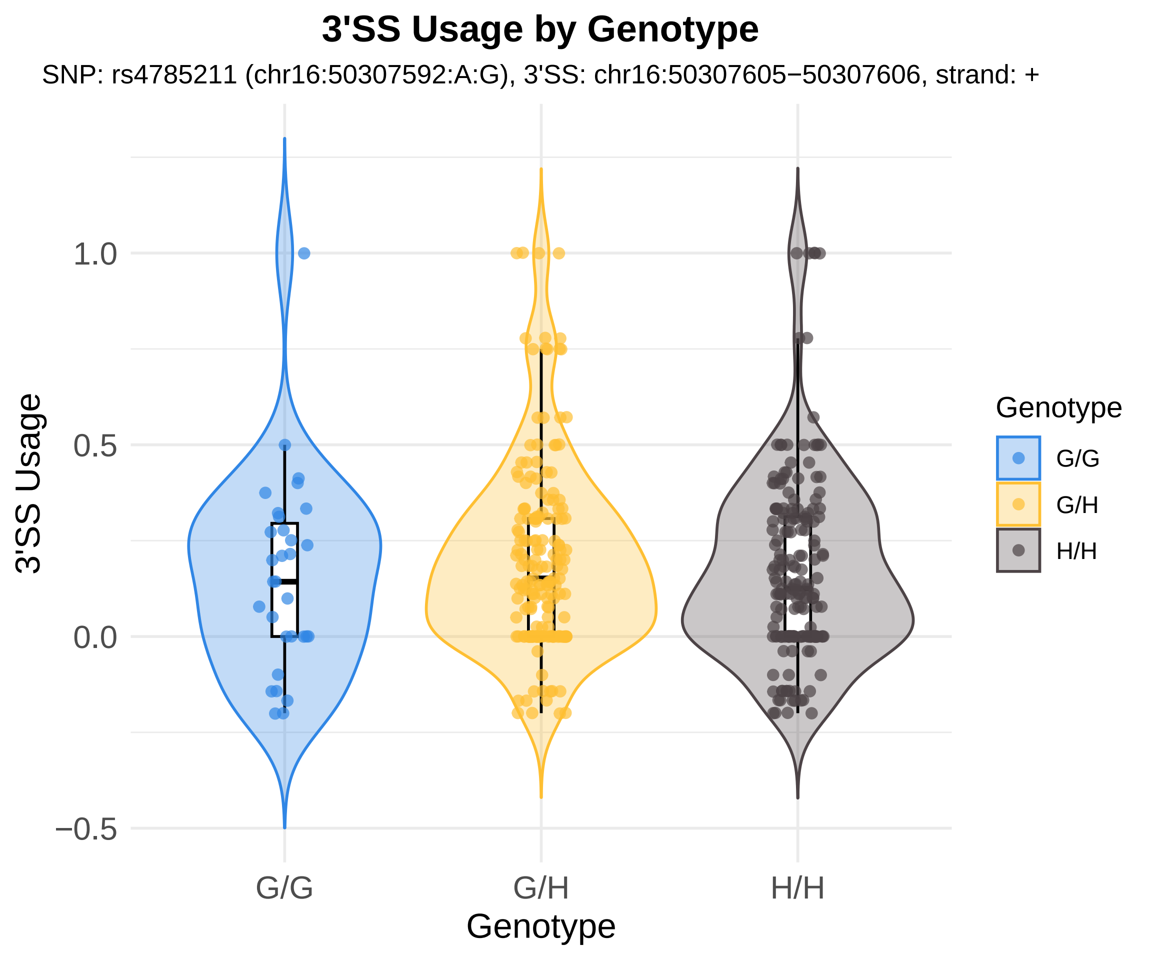
3'SS Usage by Genotype
SNP: rs4785211 (chr16:50307592:A:G), 3'SS: chr16:50307605−50307606, strand: +
3'SS Usage
1.0
0.5
0.0
−0.5
G/G
G/H
H/H
Genotype
Genotype
G/G
G/H
H/H

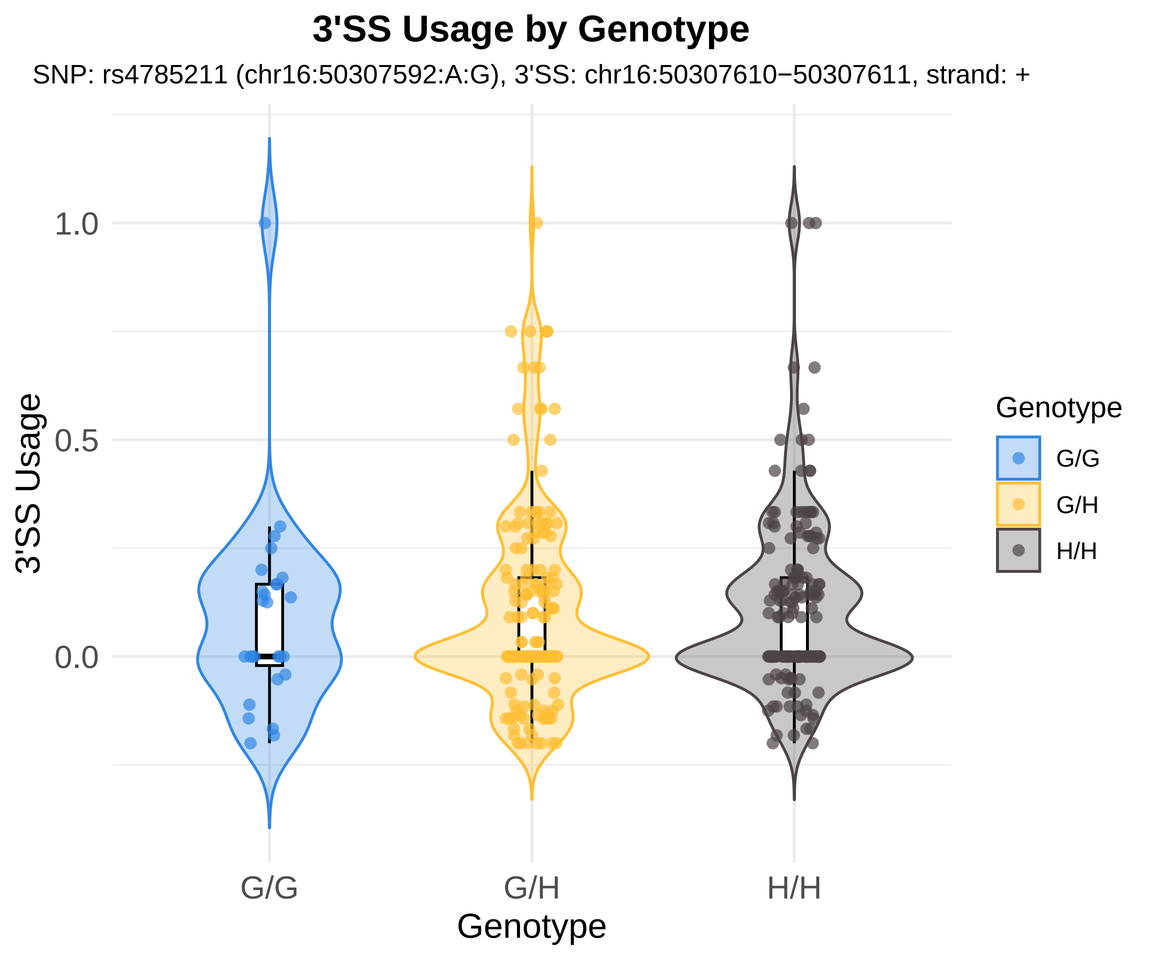
3'SS Usage by Genotype
SNP: rs4785211 (chr16:50307592:A:G), 3'SS: chr16:50307610−50307611, strand: +
3'SS Usage
1.0
0.5
0.0
G/G
G/H
H/H
Genotype
Genotype
G/G
G/H
H/H

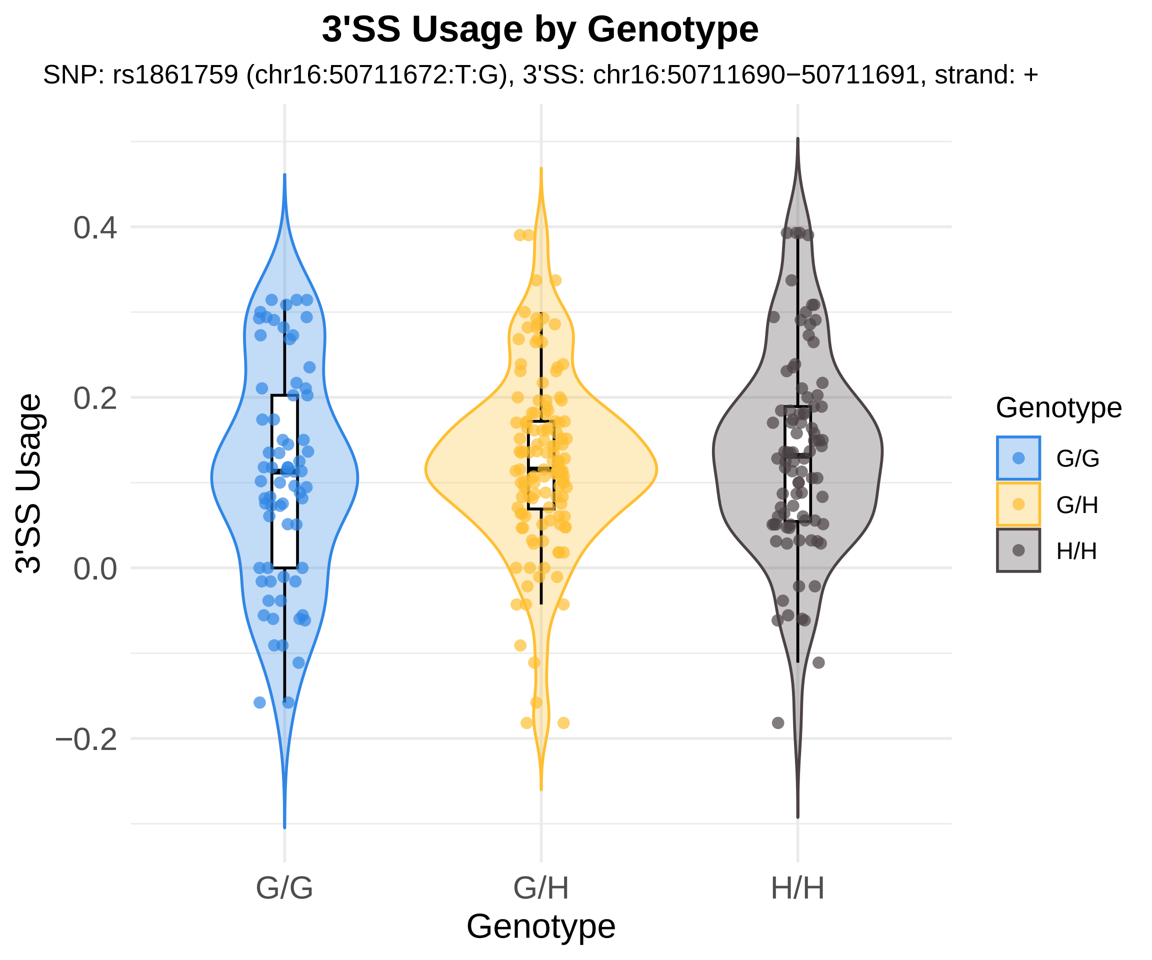
3'SS Usage by Genotype
SNP: rs1861759 (chr16:50711672:T:G), 3'SS: chr16:50711690−50711691, strand: +
3'SS Usage
0.4
0.2
0.0
−0.2
G/G
G/H
H/H
Genotype
Genotype
G/G
G/H
H/H

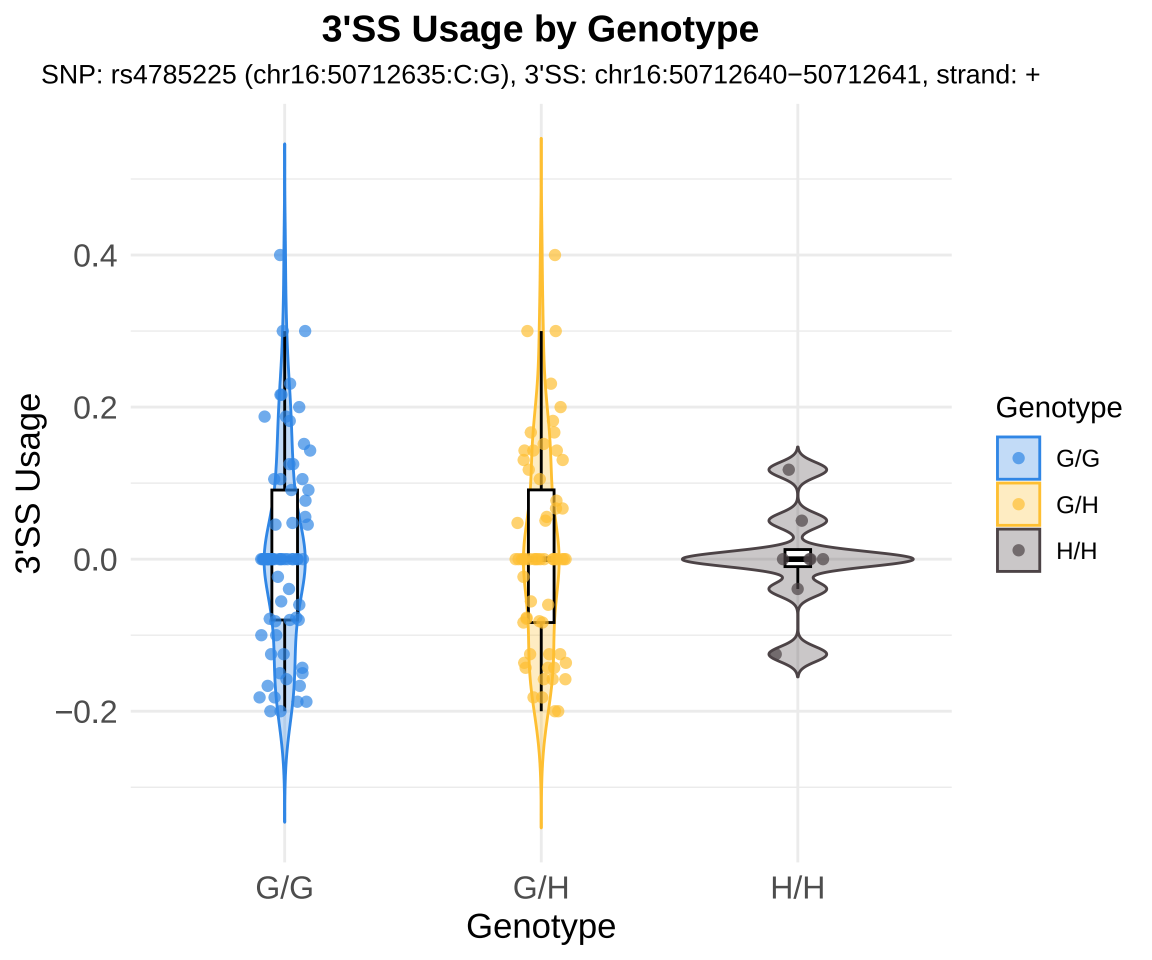
3'SS Usage by Genotype
SNP: rs4785225 (chr16:50712635:C:G), 3'SS: chr16:50712640−50712641, strand: +
3'SS Usage
0.4
0.2
0.0
−0.2
G/G
G/H
H/H
Genotype
Genotype
G/G
G/H
H/H

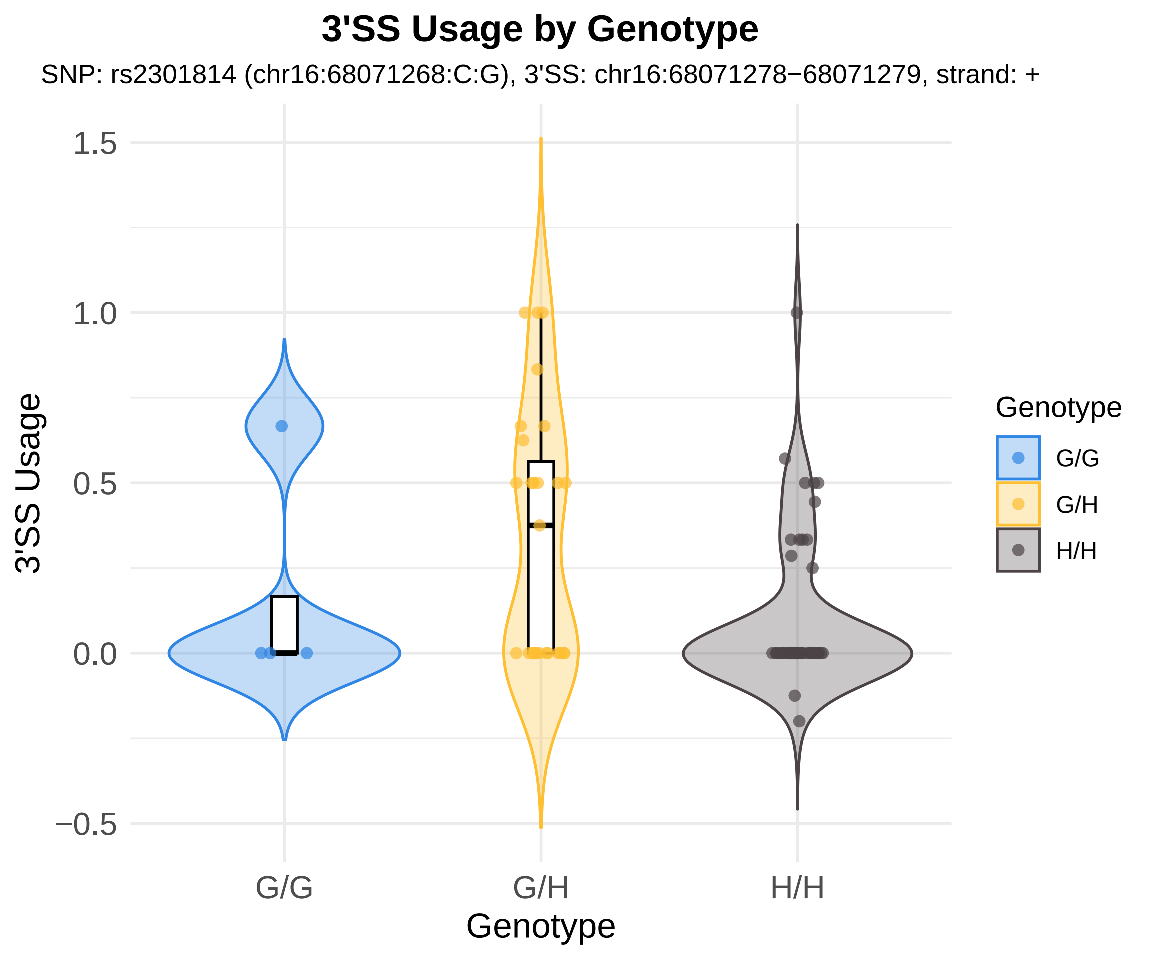
3'SS Usage by Genotype
SNP: rs2301814 (chr16:68071268:C:G), 3'SS: chr16:68071278−68071279, strand: +
3'SS Usage
1.5
1.0
0.5
0.0
−0.5
G/G
G/H
H/H
Genotype
Genotype
G/G
G/H
H/H

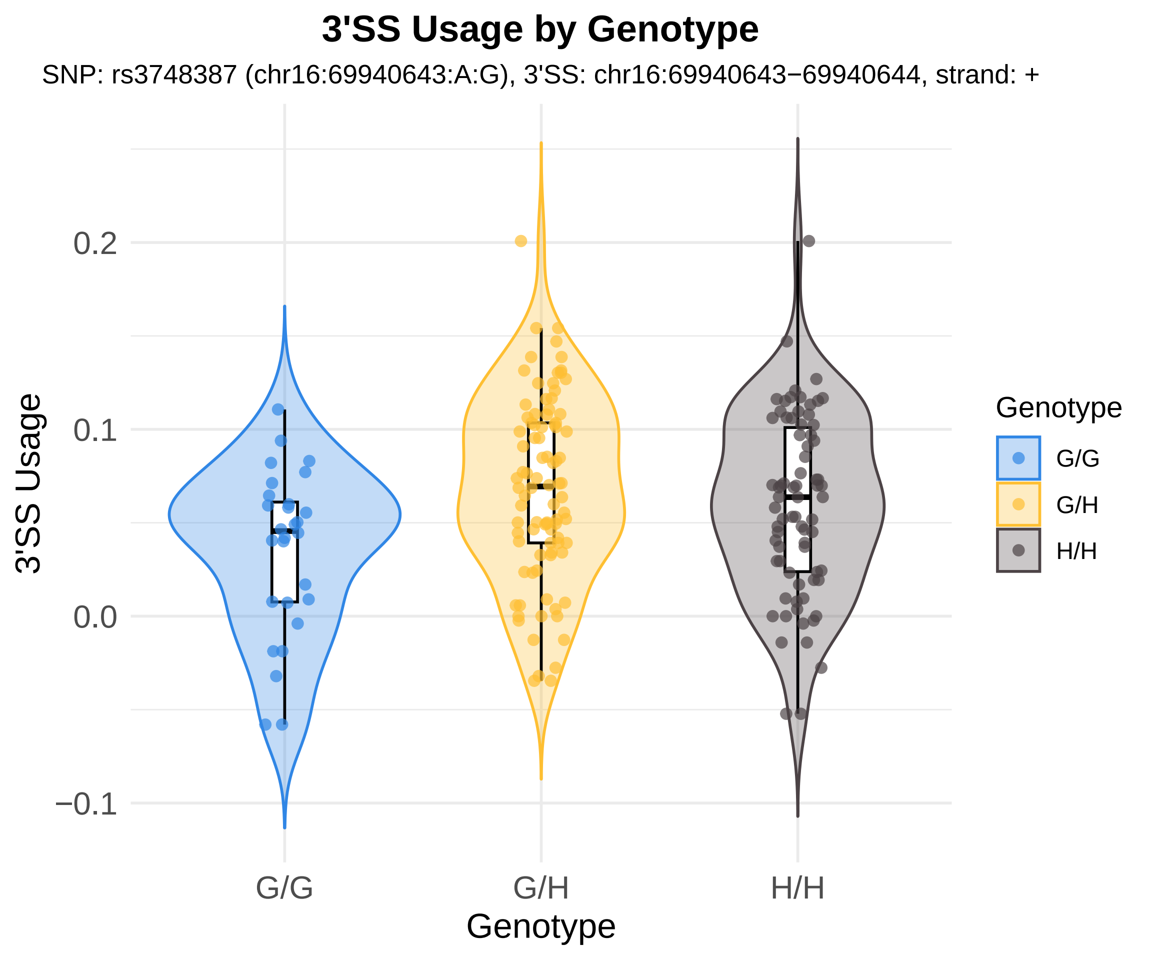

3'SS Usage by Genotype
SNP: rs3748387 (chr16:69940643:A:G), 3'SS: chr16:69940643−69940644, strand: +
3'SS Usage
0.2
0.1
0.0
−0.1
G/G
G/H
H/H
Genotype
Genotype
G/G
G/H
H/H

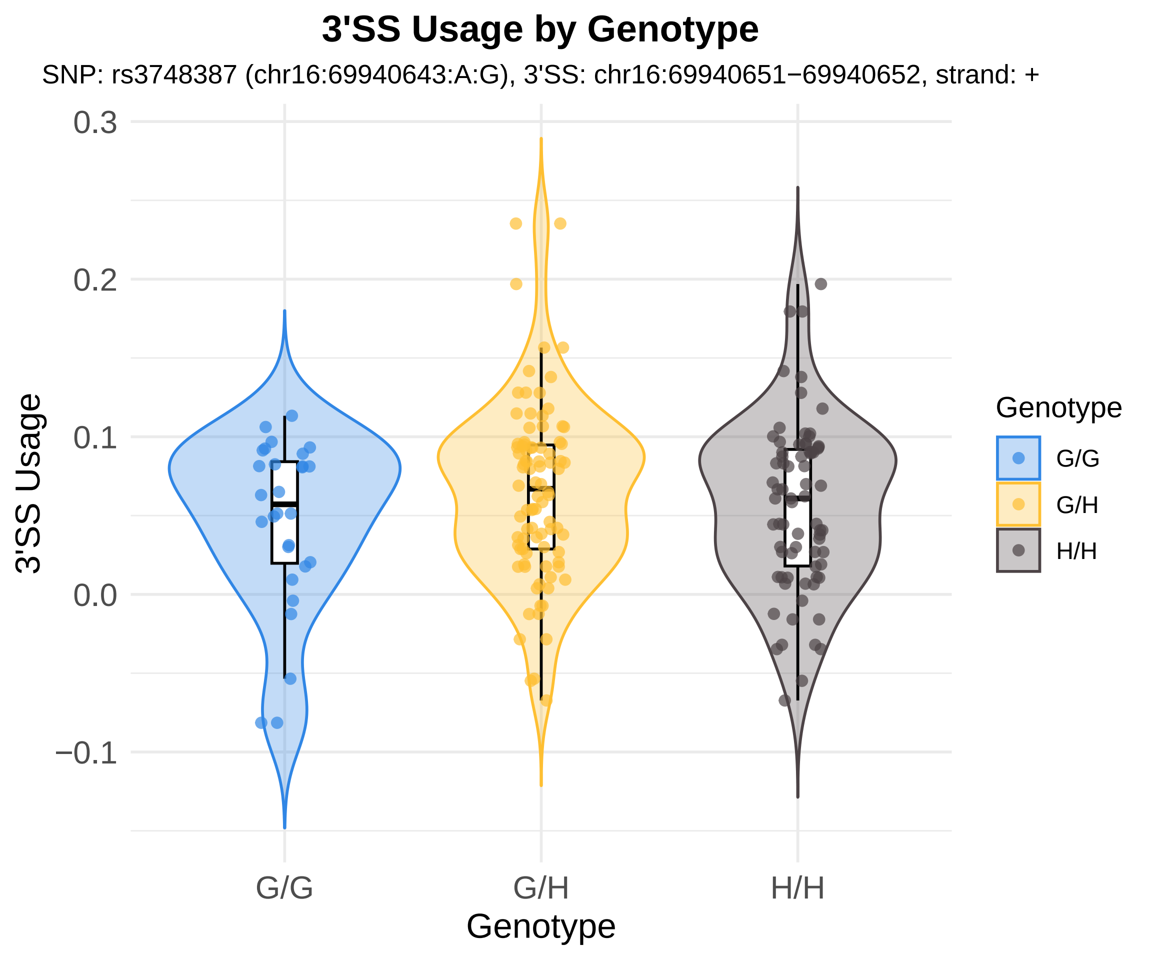
3'SS Usage by Genotype
SNP: rs3748387 (chr16:69940643:A:G), 3'SS: chr16:69940651−69940652, strand: +
3'SS Usage
0.3
0.2
0.1
0.0
−0.1
G/G
G/H
H/H
Genotype
Genotype
G/G
G/H
H/H

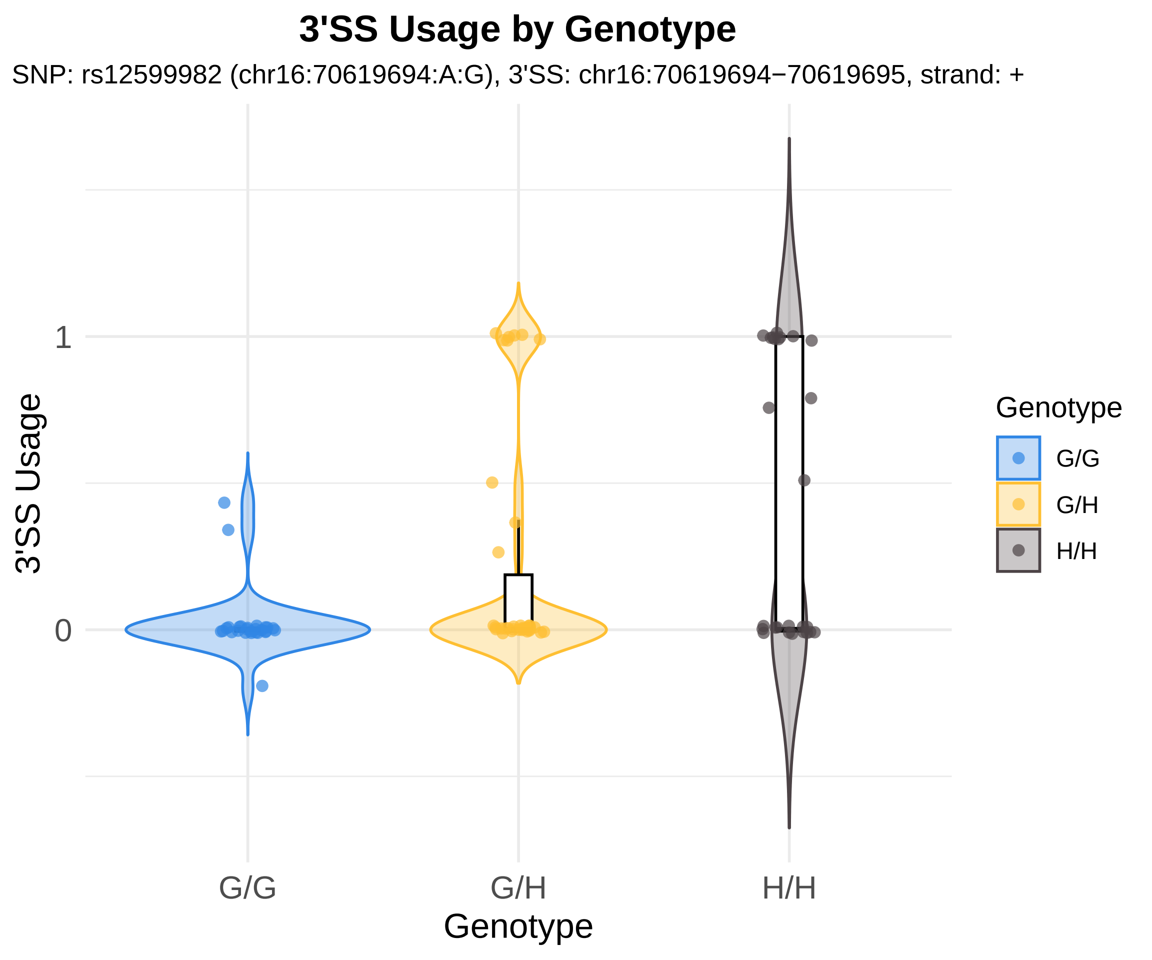

3'SS Usage by Genotype
SNP: rs12599982 (chr16:70619694:A:G), 3'SS: chr16:70619694−70619695, strand: +
3'SS Usage
1
0
G/G
G/H
H/H
Genotype
Genotype
G/G
G/H
H/H

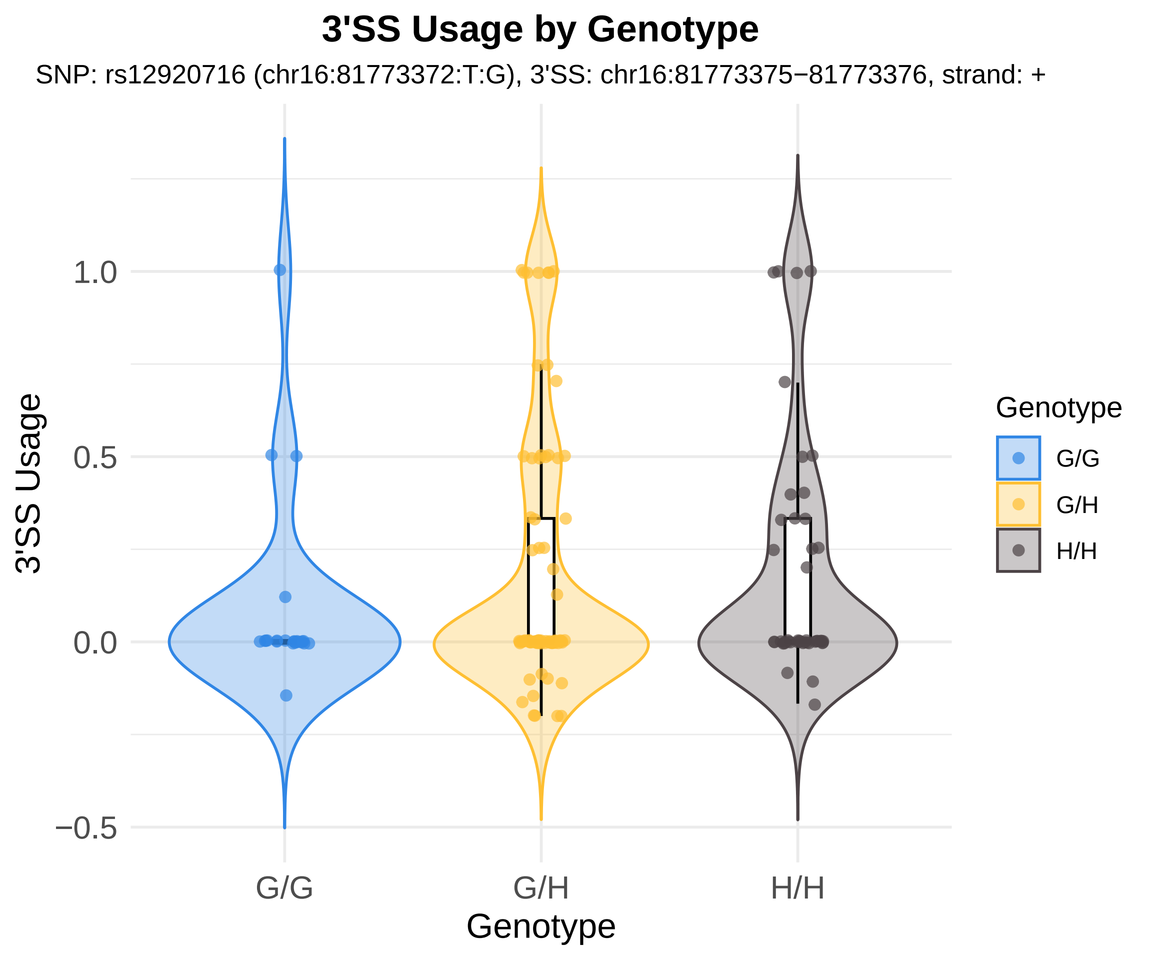
3'SS Usage by Genotype
SNP: rs12920716 (chr16:81773372:T:G), 3'SS: chr16:81773375−81773376, strand: +
3'SS Usage
1.0
0.5
0.0
−0.5
G/G
G/H
H/H
Genotype
Genotype
G/G
G/H
H/H

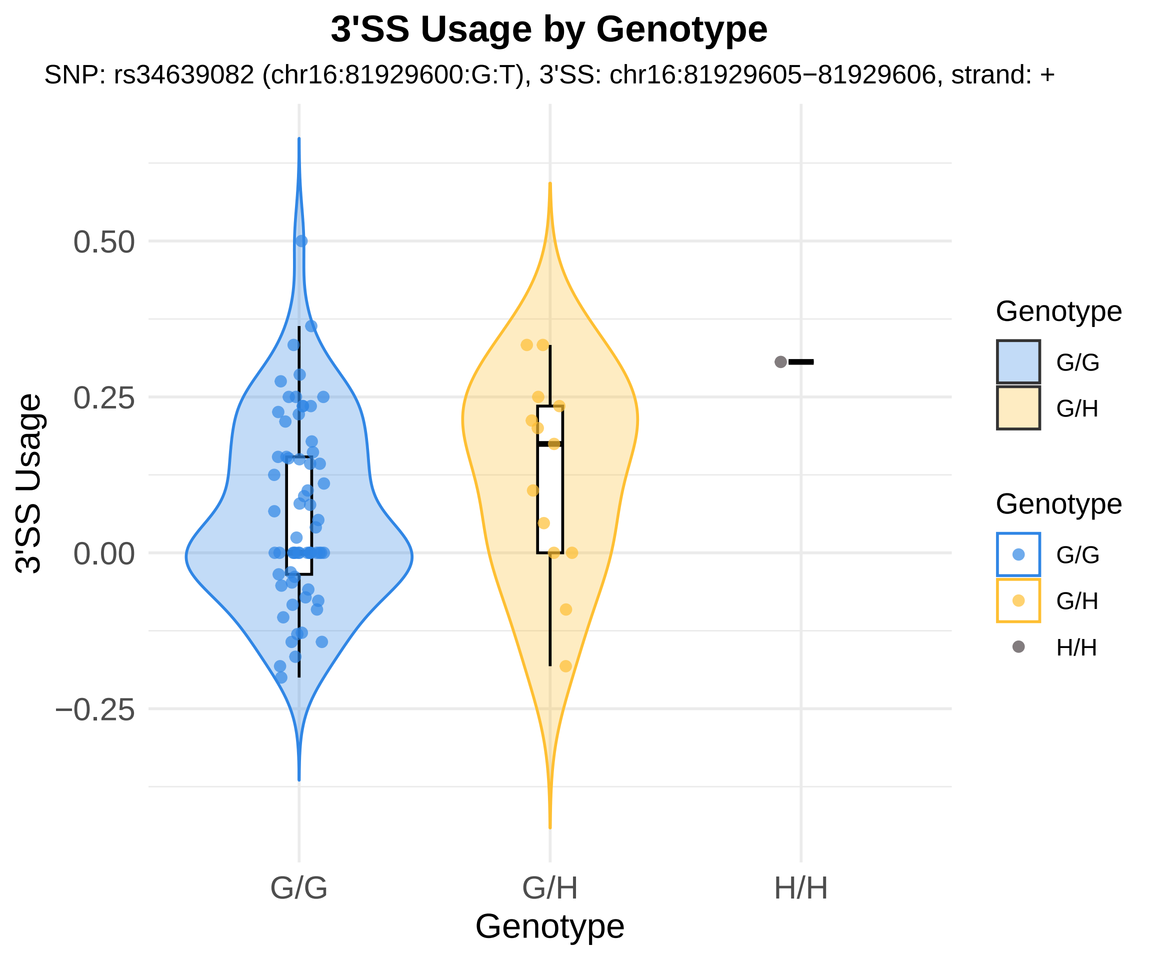
3'SS Usage by Genotype
SNP: rs34639082 (chr16:81929600:G:T), 3'SS: chr16:81929605−81929606, strand: +
3'SS Usage
0.50
0.25
0.00
−0.25
G/G
G/H
H/H
Genotype
Genotype
G/G
G/H
Genotype
G/G
G/H
H/H

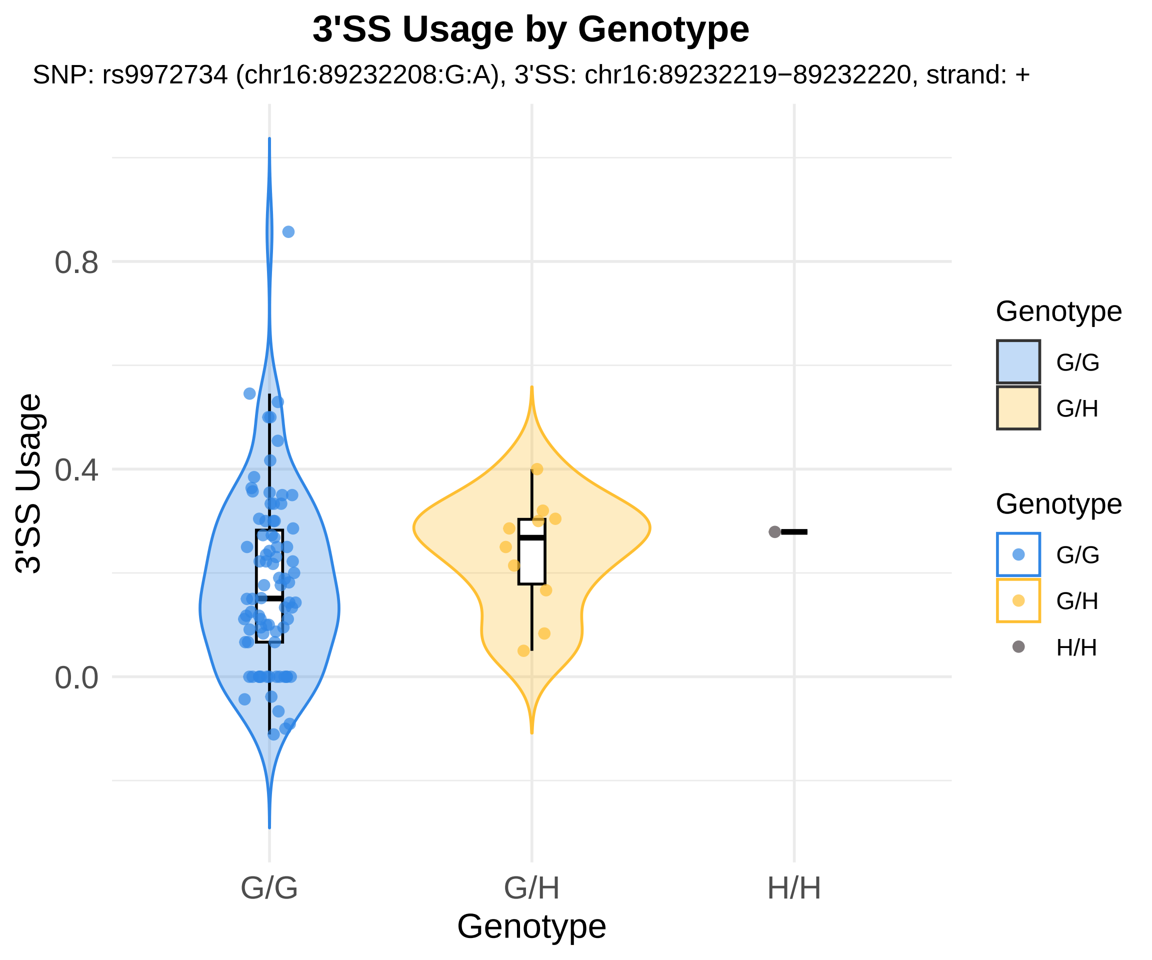

3'SS Usage by Genotype
SNP: rs9972734 (chr16:89232208:G:A), 3'SS: chr16:89232219−89232220, strand: +
3'SS Usage
0.8
0.4
0.0
G/G
G/H
H/H
Genotype
Genotype
G/G
G/H
Genotype
G/G
G/H
H/H

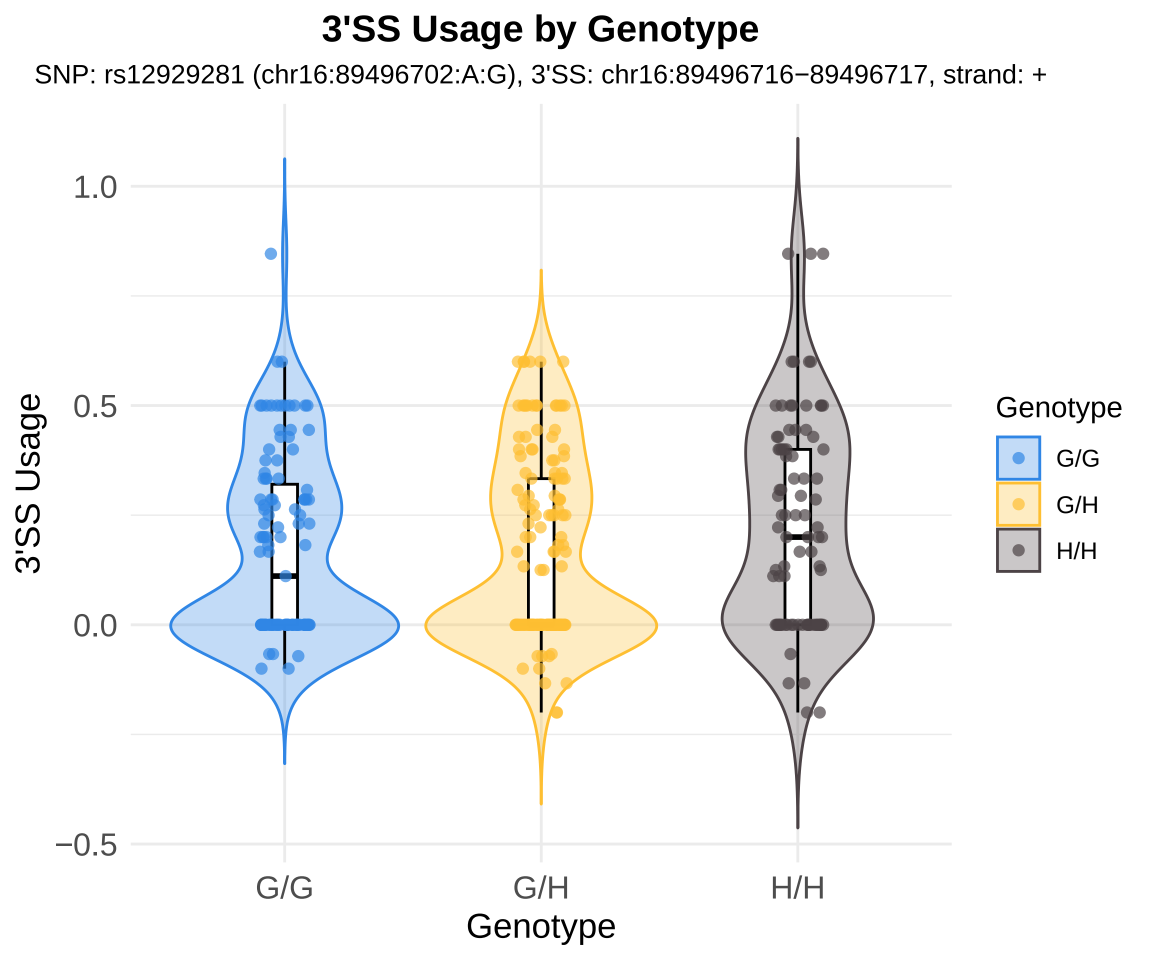

3'SS Usage by Genotype
SNP: rs12929281 (chr16:89496702:A:G), 3'SS: chr16:89496716−89496717, strand: +
3'SS Usage
1.0
0.5
0.0
−0.5
G/G
G/H
H/H
Genotype
Genotype
G/G
G/H
H/H

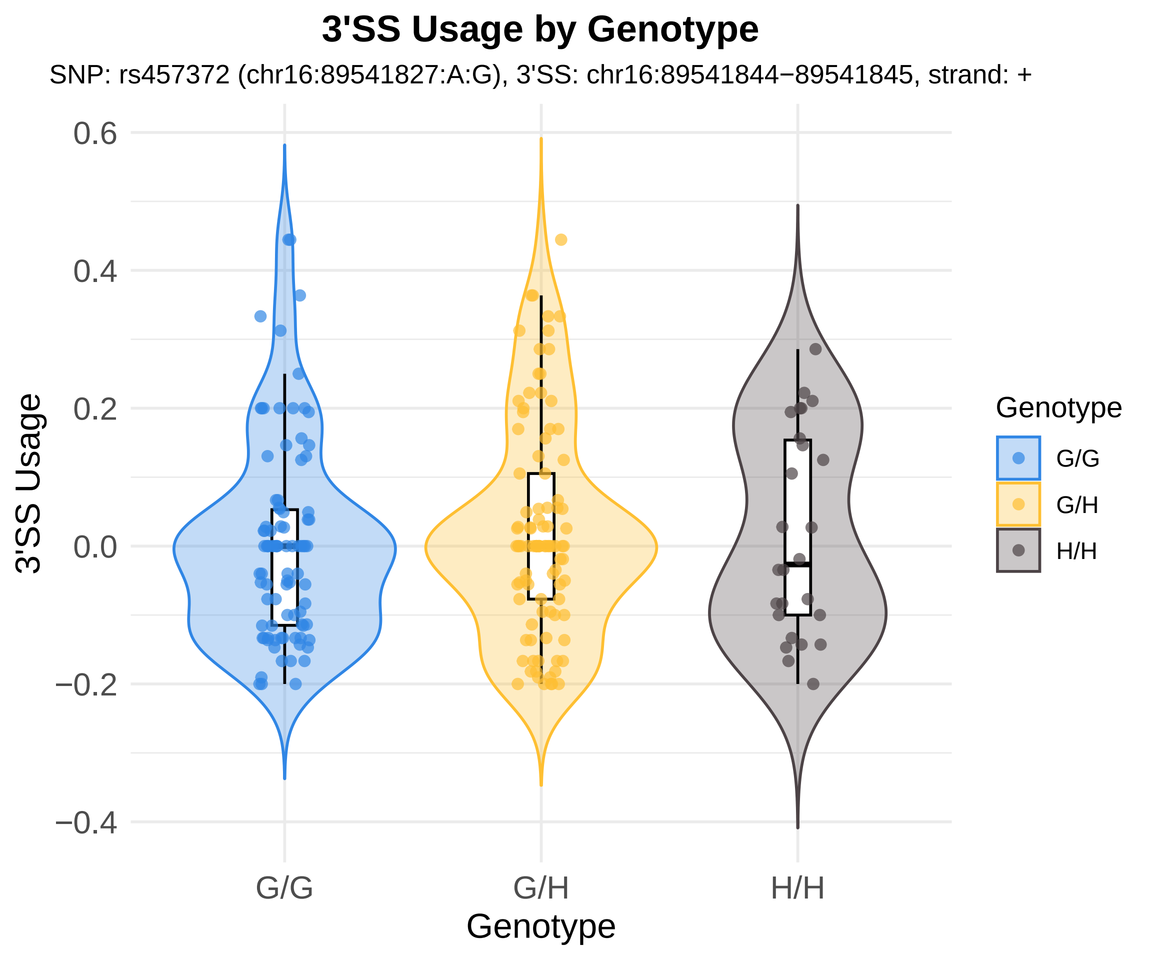
3'SS Usage by Genotype
SNP: rs457372 (chr16:89541827:A:G), 3'SS: chr16:89541844−89541845, strand: +
3'SS Usage
0.6
0.4
0.2
0.0
−0.2
−0.4
G/G
G/H
H/H
Genotype
Genotype
G/G
G/H
H/H

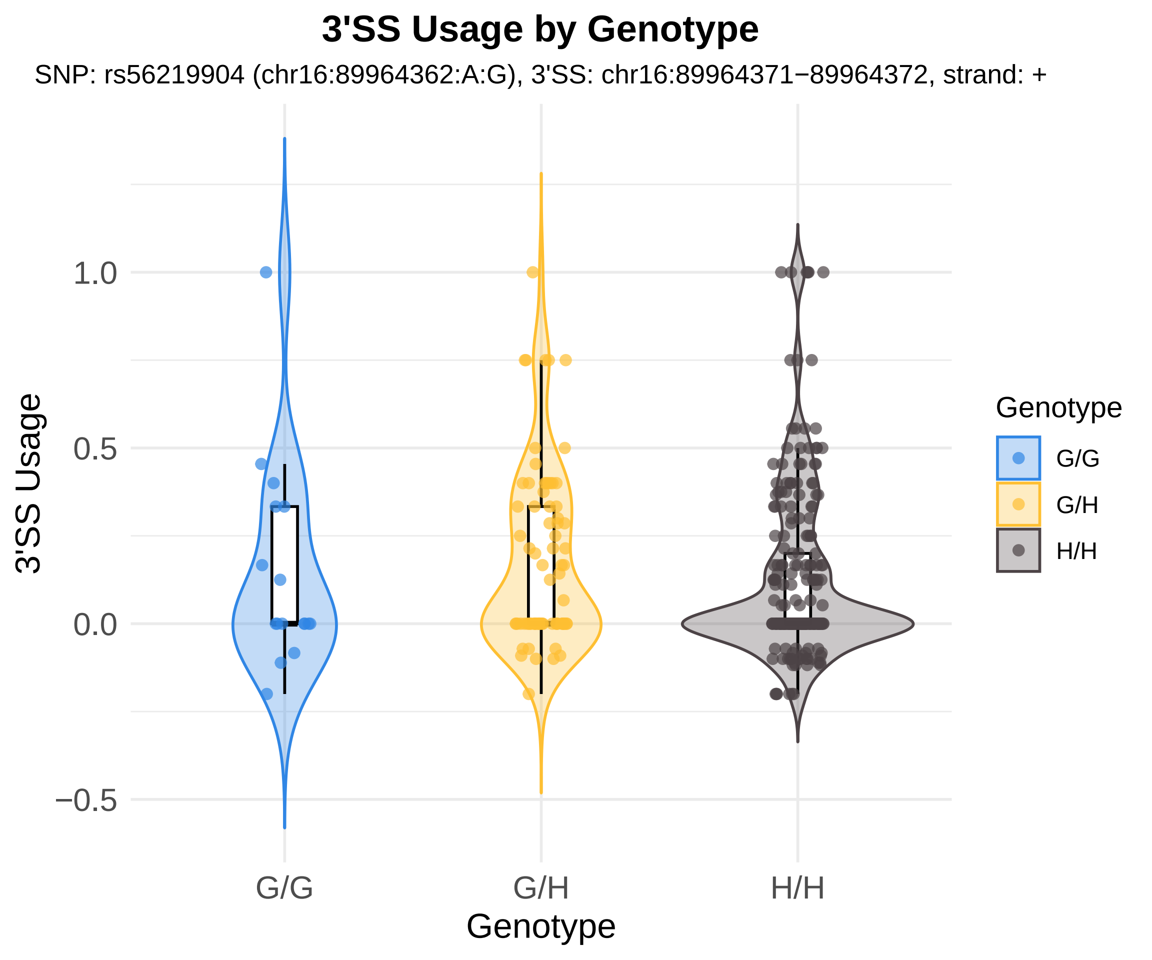
3'SS Usage by Genotype
SNP: rs56219904 (chr16:89964362:A:G), 3'SS: chr16:89964371−89964372, strand: +
3'SS Usage
1.0
0.5
0.0
−0.5
G/G
G/H
H/H
Genotype
Genotype
G/G
G/H
H/H

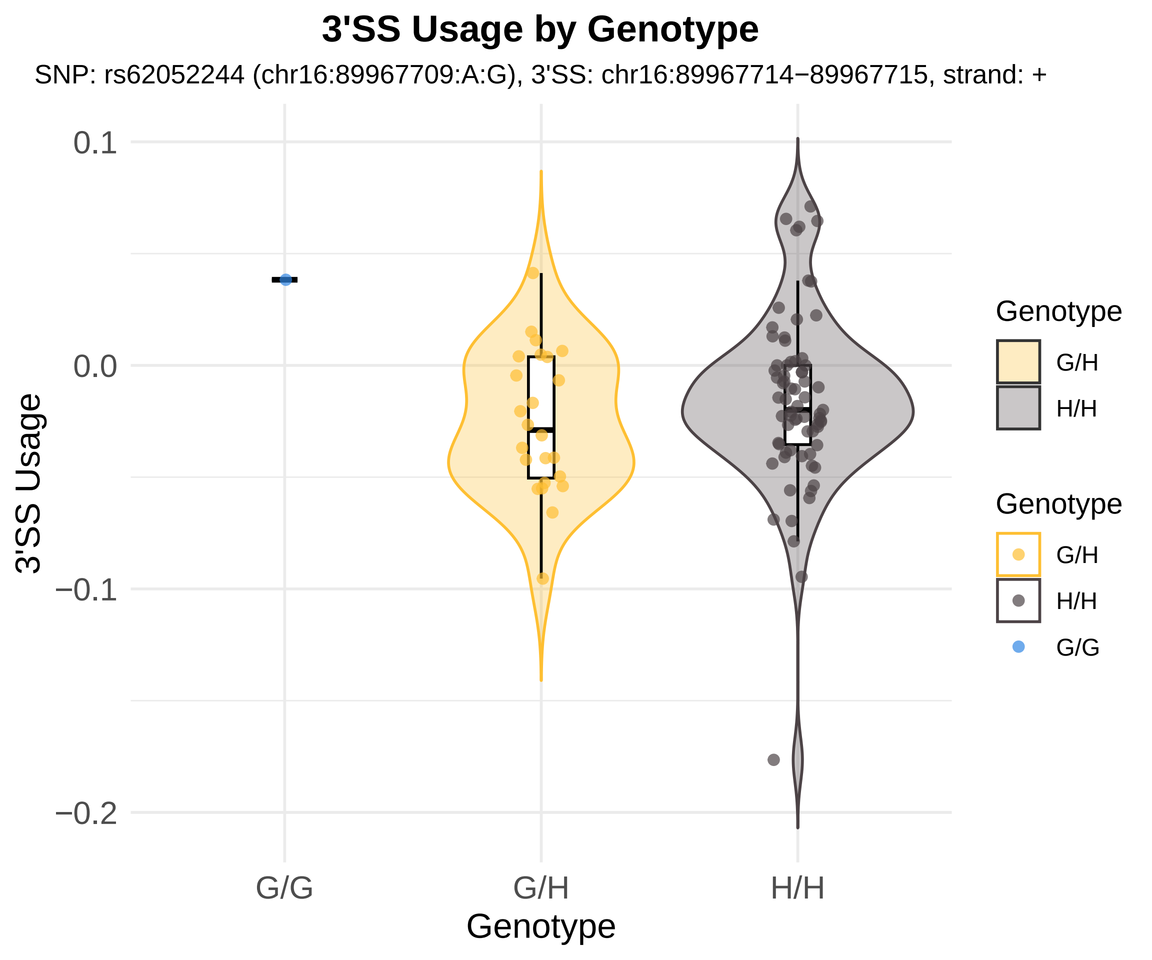

3'SS Usage by Genotype
SNP: rs62052244 (chr16:89967709:A:G), 3'SS: chr16:89967714−89967715, strand: +
0.1
0.0
−0.1
−0.2
3'SS Usage
G/G
G/H
H/H
Genotype
Genotype
G/H
H/H
Genotype
G/H
H/H
G/G

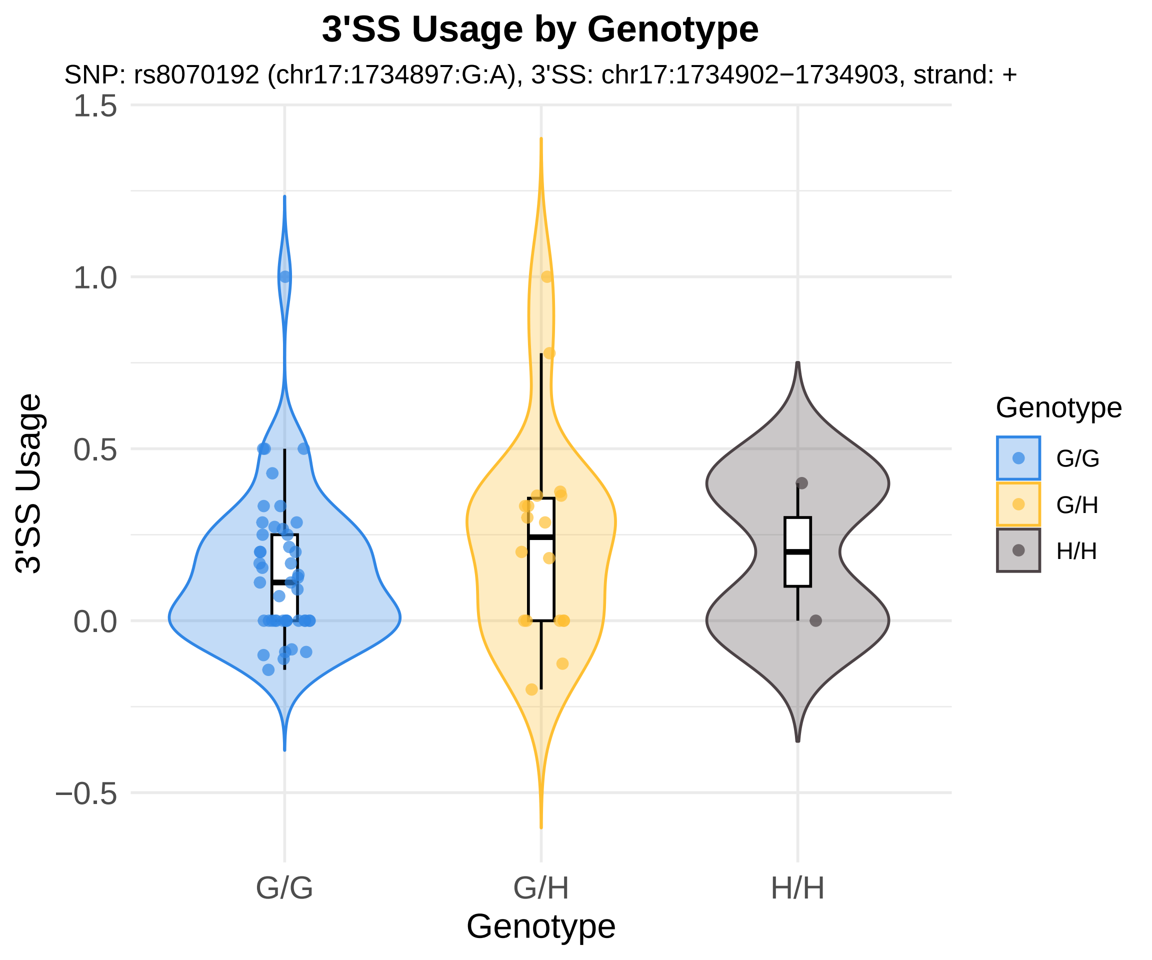
3'SS Usage by Genotype
SNP: rs8070192 (chr17:1734897:G:A), 3'SS: chr17:1734902−1734903, strand: +
3'SS Usage
1.5
1.0
0.5
0.0
−0.5
G/G
G/H
H/H
Genotype
Genotype
G/G
G/H
H/H

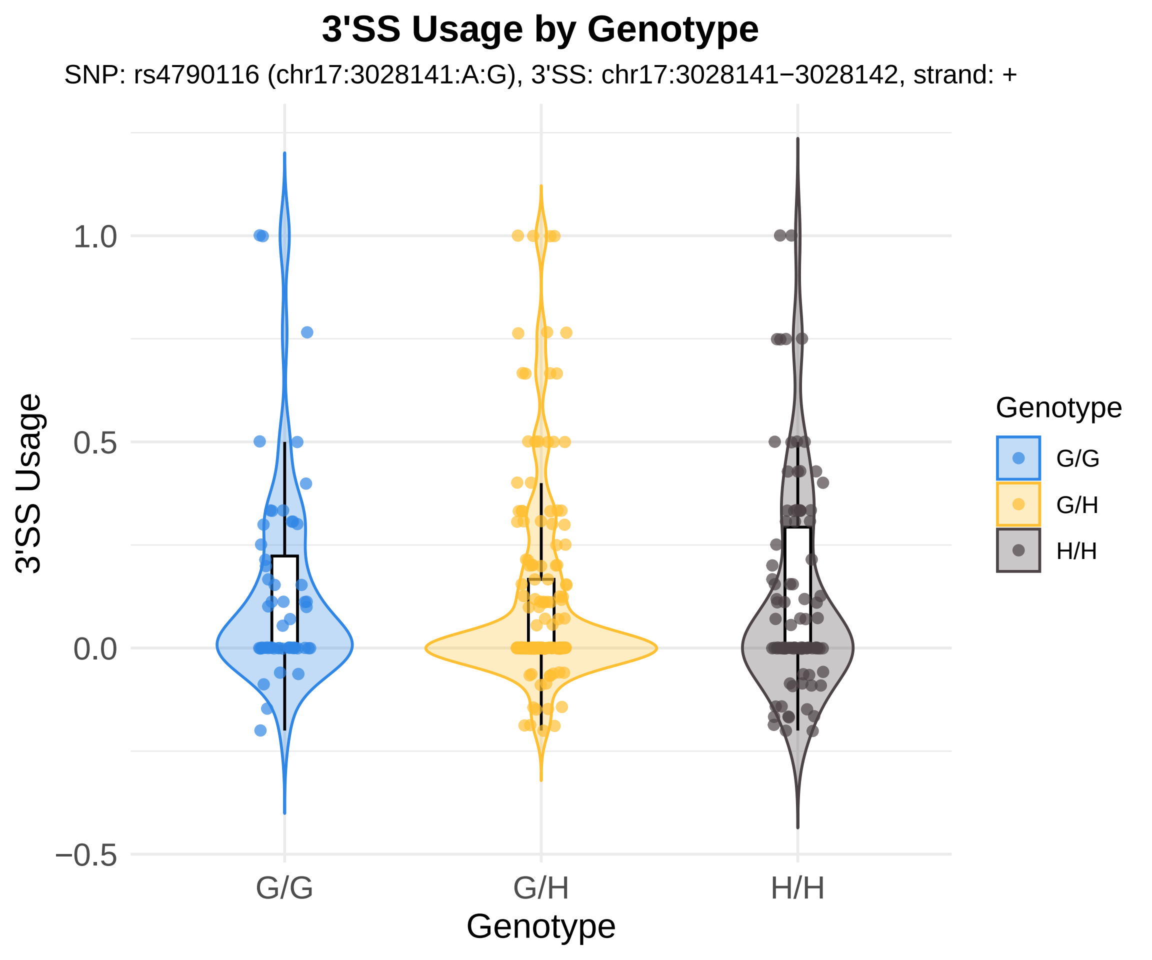
3'SS Usage by Genotype
SNP: rs4790116 (chr17:3028141:A:G), 3'SS: chr17:3028141−3028142, strand: +
3'SS Usage
1.0
0.5
0.0
−0.5
G/G
G/H
H/H
Genotype
Genotype
G/G
G/H
H/H

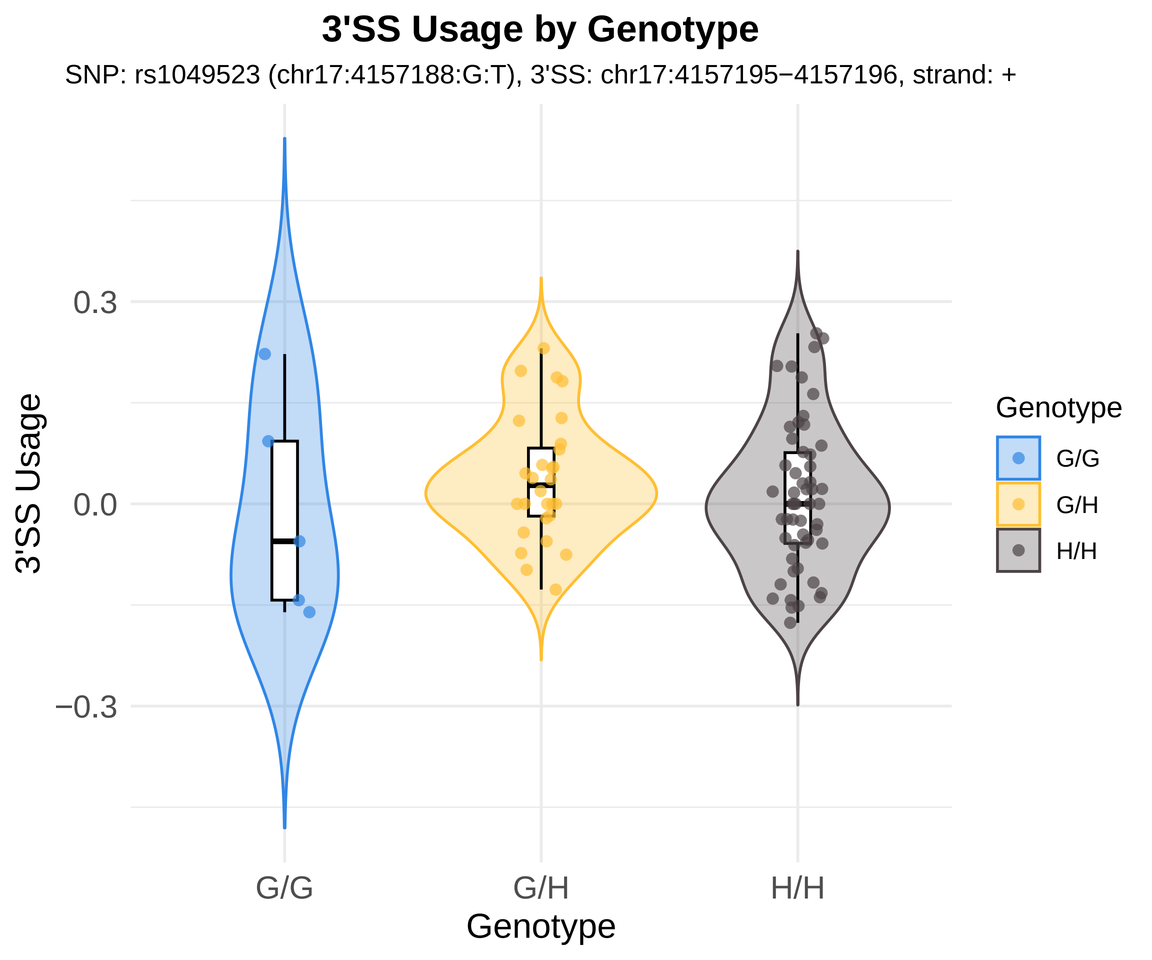
3'SS Usage by Genotype
SNP: rs1049523 (chr17:4157188:G:T), 3'SS: chr17:4157195−4157196, strand: +
3'SS Usage
0.3
0.0
−0.3
G/G
G/H
H/H
Genotype
Genotype
G/G
G/H
H/H

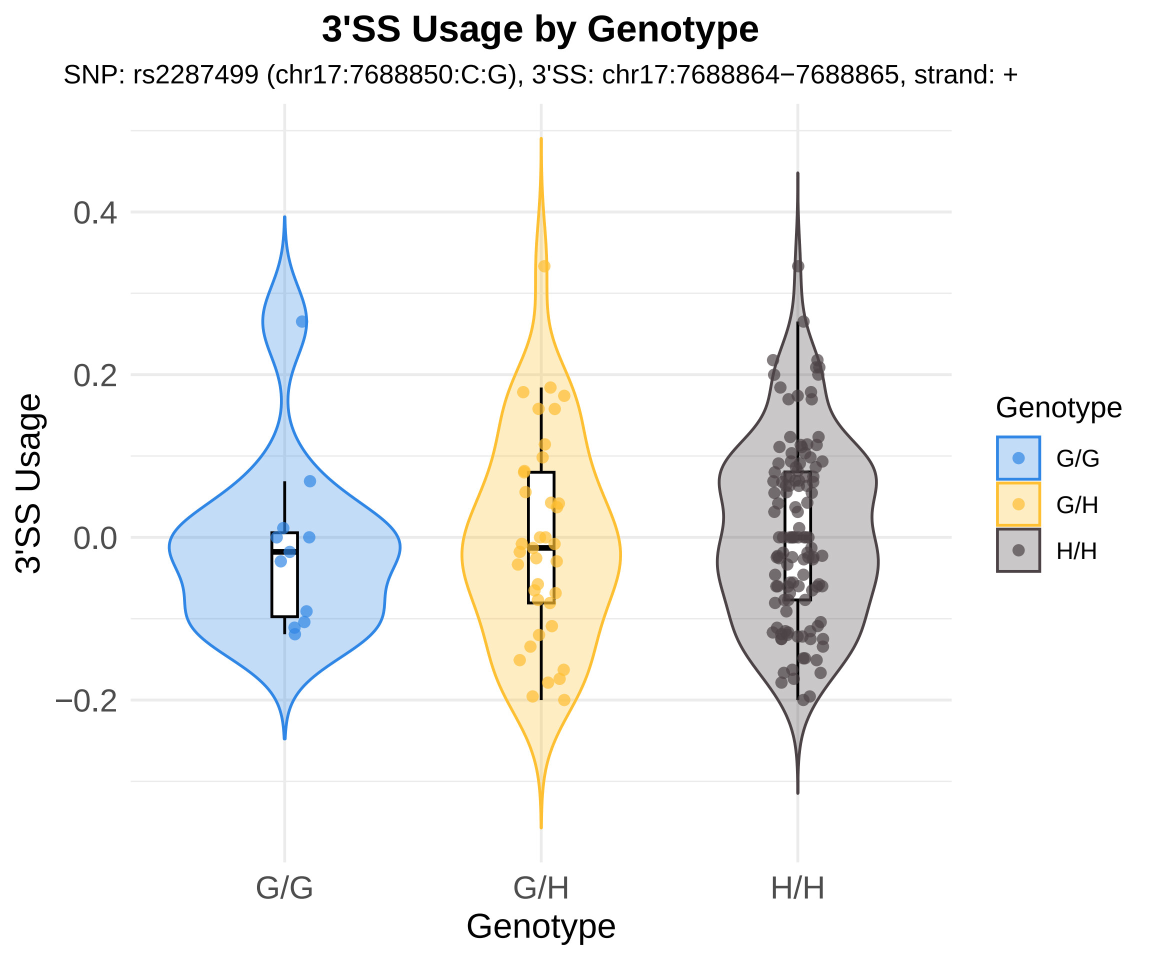

3'SS Usage by Genotype
SNP: rs2287499 (chr17:7688850:C:G), 3'SS: chr17:7688864−7688865, strand: +
3'SS Usage
0.4
0.2
0.0
−0.2
G/G
G/H
H/H
Genotype
Genotype
G/G
G/H
H/H

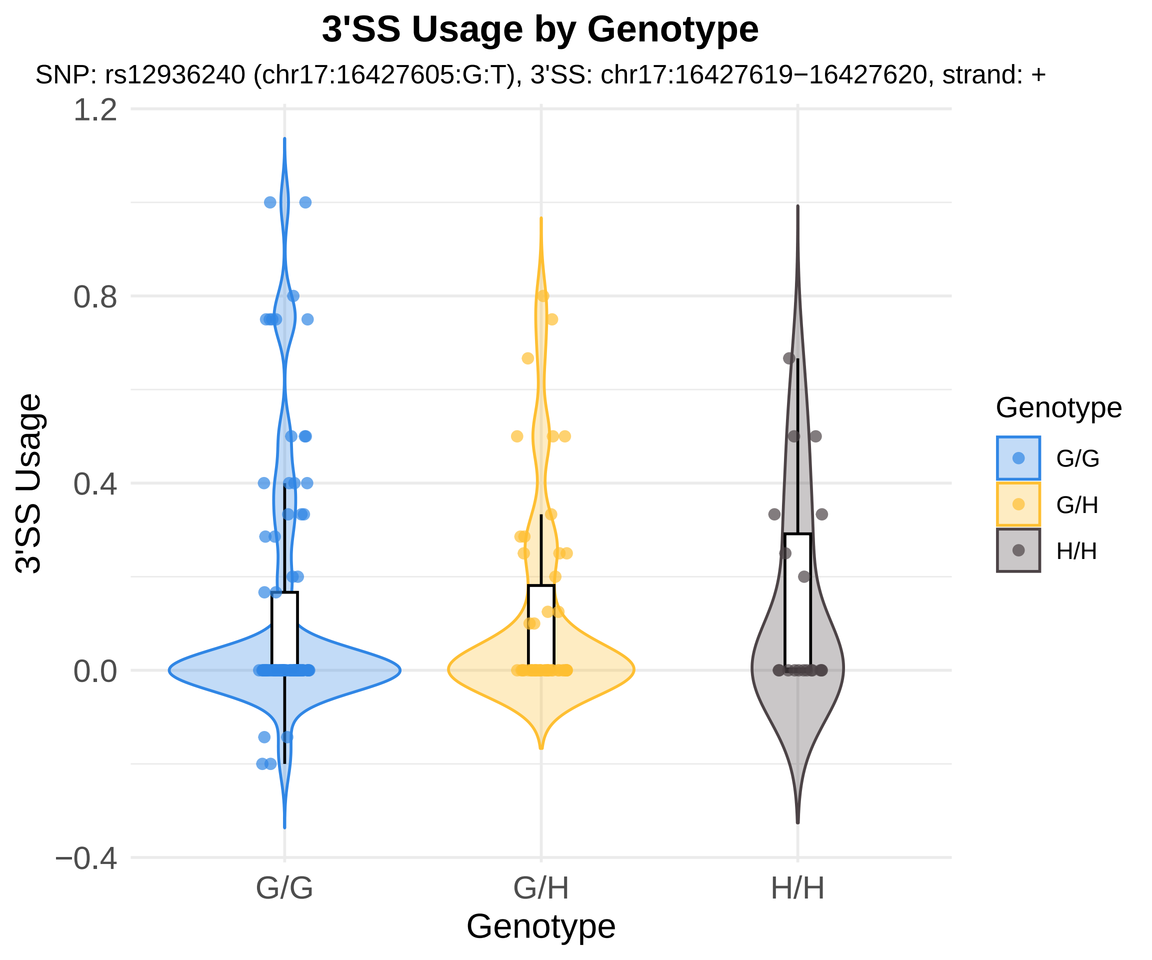

3'SS Usage by Genotype
SNP: rs12936240 (chr17:16427605:G:T), 3'SS: chr17:16427619−16427620, strand: +
1.2
0.8
0.4
0.0
−0.4
3'SS Usage
G/G
G/H
H/H
Genotype
Genotype
G/G
G/H
H/H

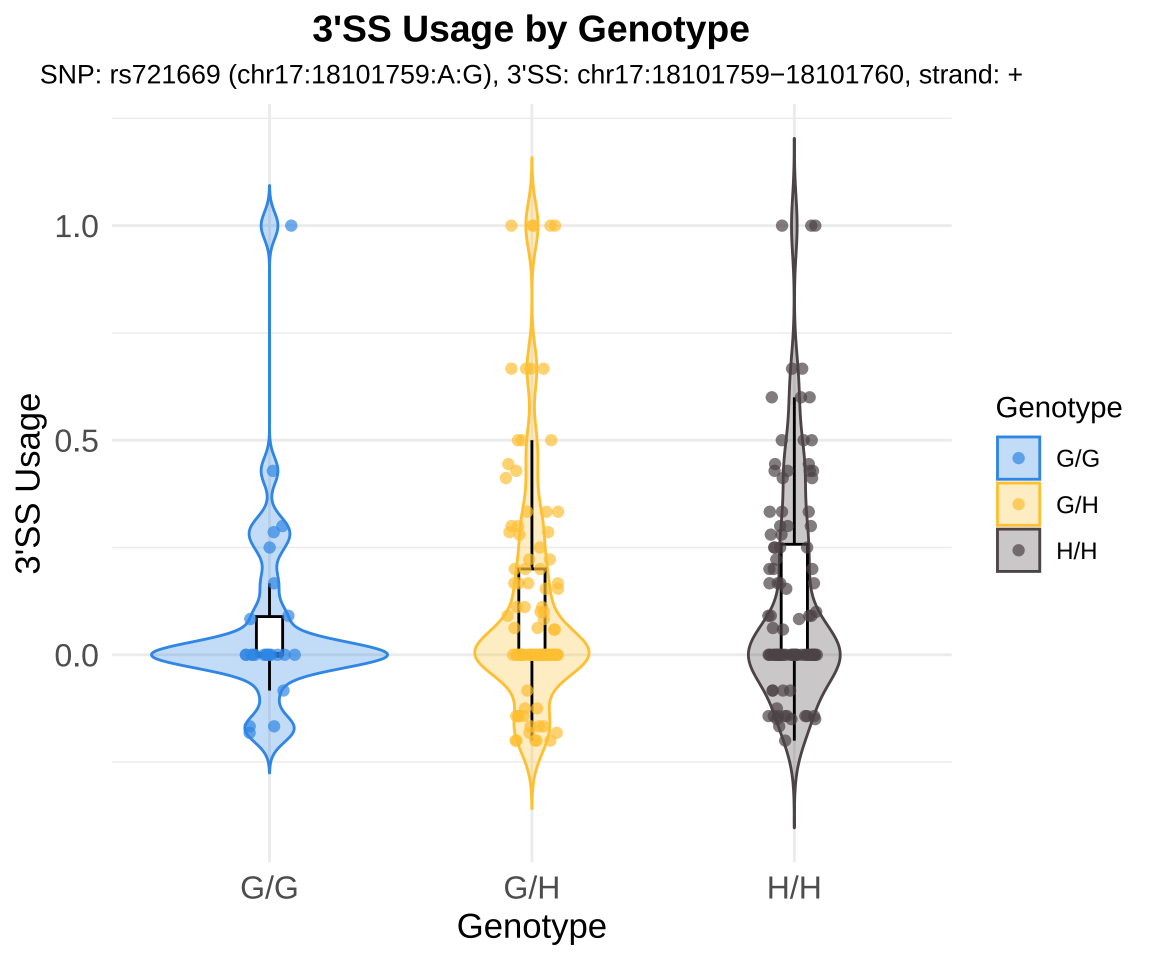

3'SS Usage by Genotype
SNP: rs721669 (chr17:18101759:A:G), 3'SS: chr17:18101759−18101760, strand: +
3'SS Usage
1.0
0.5
0.0
G/G
G/H
H/H
Genotype
Genotype
G/G
G/H
H/H

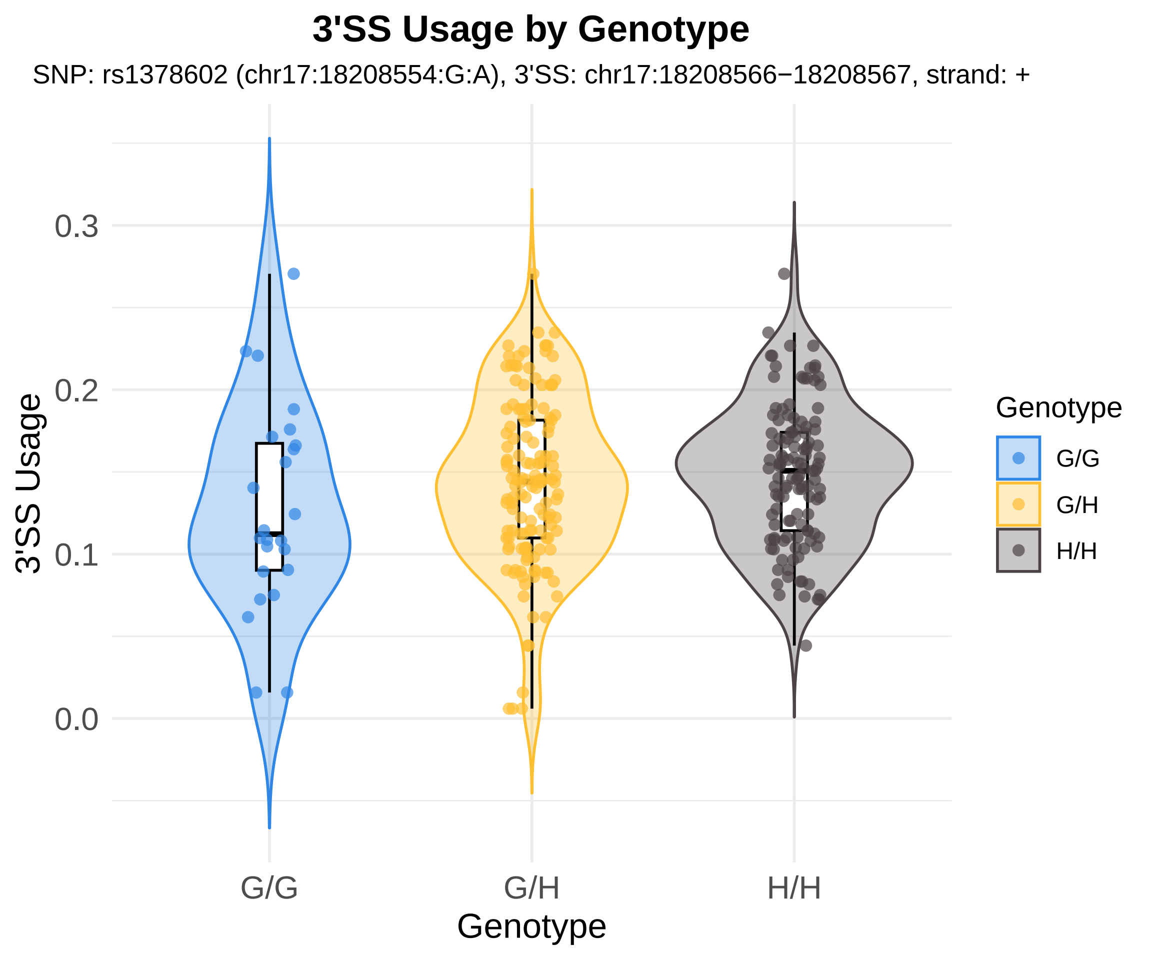

3'SS Usage by Genotype
SNP: rs1378602 (chr17:18208554:G:A), 3'SS: chr17:18208566−18208567, strand: +
3'SS Usage
0.0
0.1
0.2
0.3
G/G
G/H
H/H
Genotype
Genotype
G/G
G/H
H/H

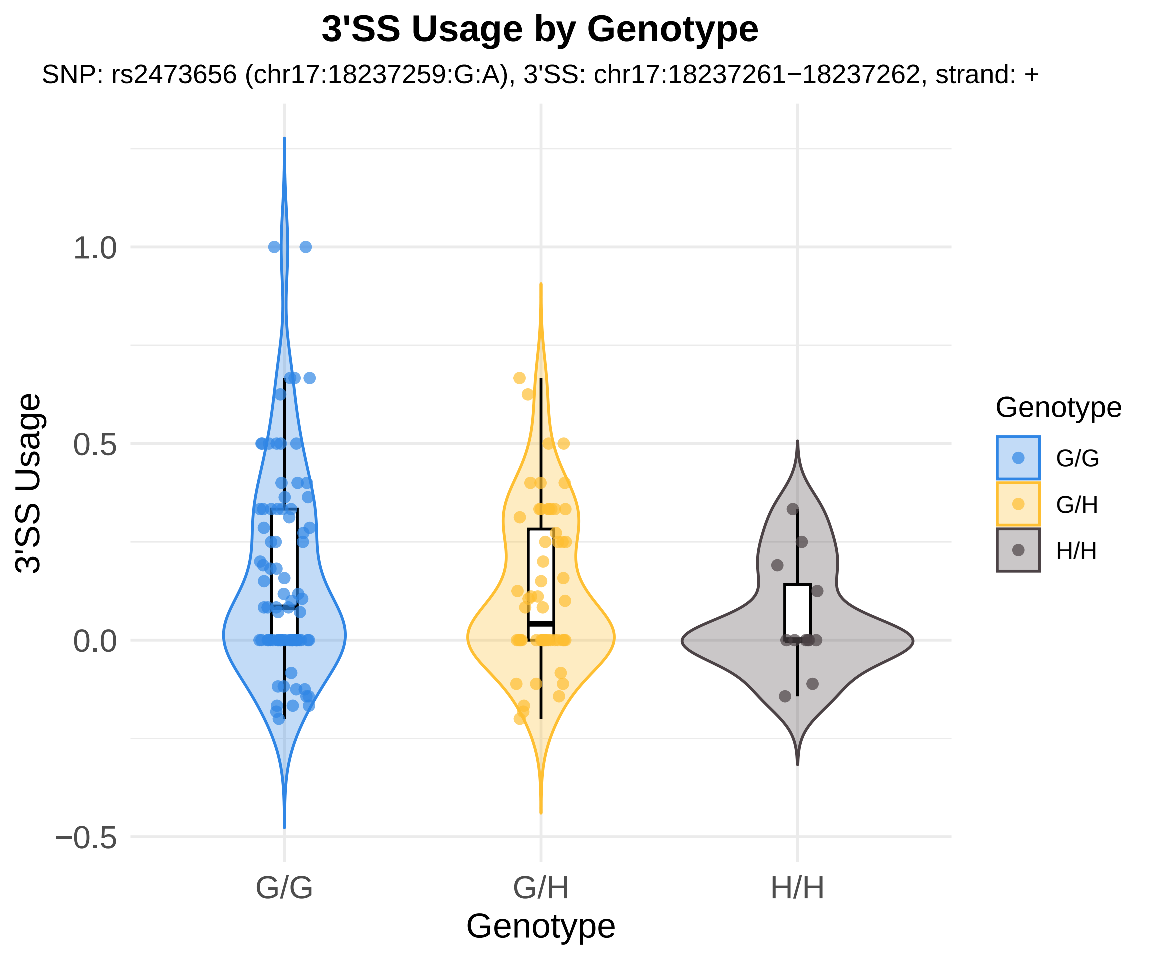
3'SS Usage by Genotype
SNP: rs2473656 (chr17:18237259:G:A), 3'SS: chr17:18237261−18237262, strand: +
3'SS Usage
1.0
0.5
0.0
−0.5
G/G
G/H
H/H
Genotype
Genotype
G/G
G/H
H/H

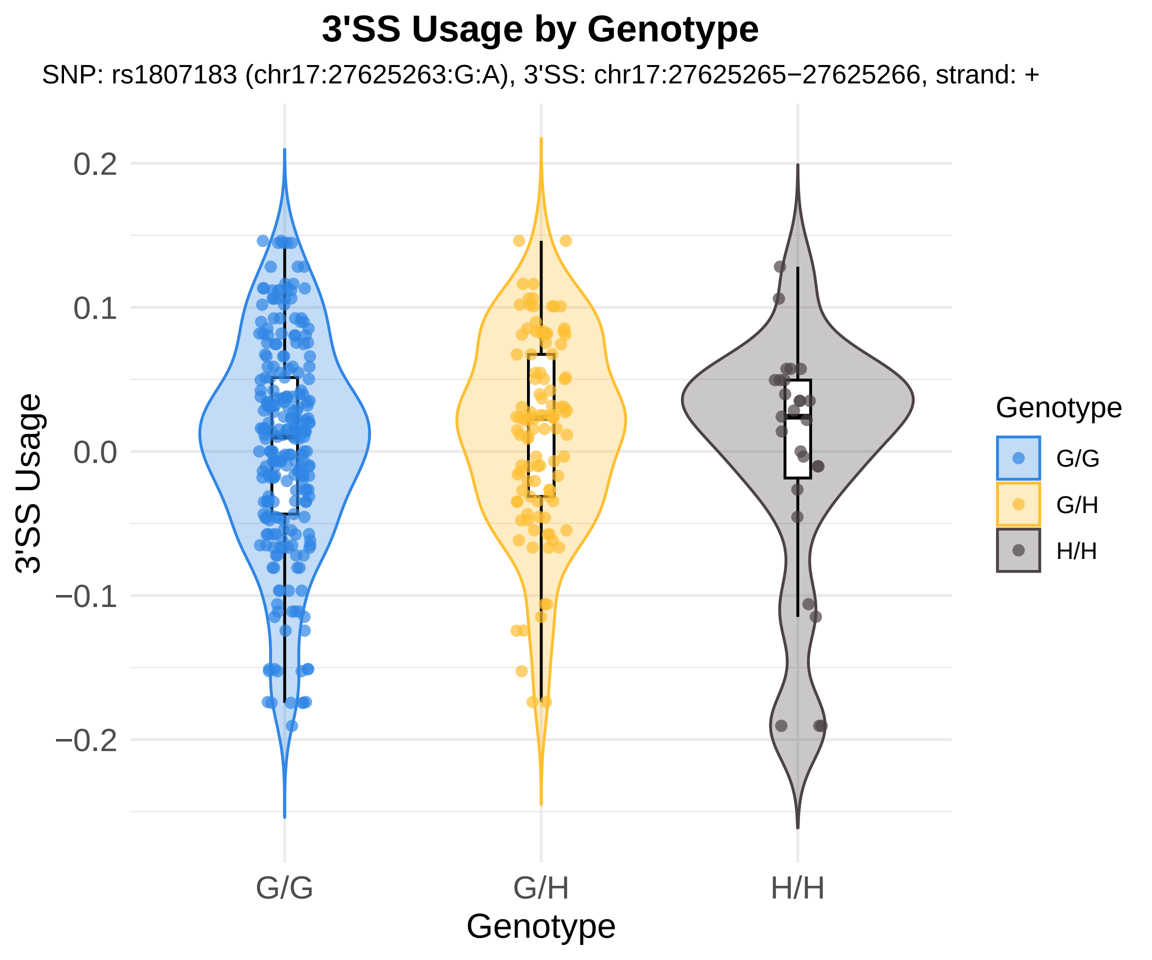
3'SS Usage by Genotype
SNP: rs1807183 (chr17:27625263:G:A), 3'SS: chr17:27625265−27625266, strand: +
3'SS Usage
0.2
0.1
0.0
−0.1
−0.2
G/G
G/H
H/H
Genotype
Genotype
G/G
G/H
H/H

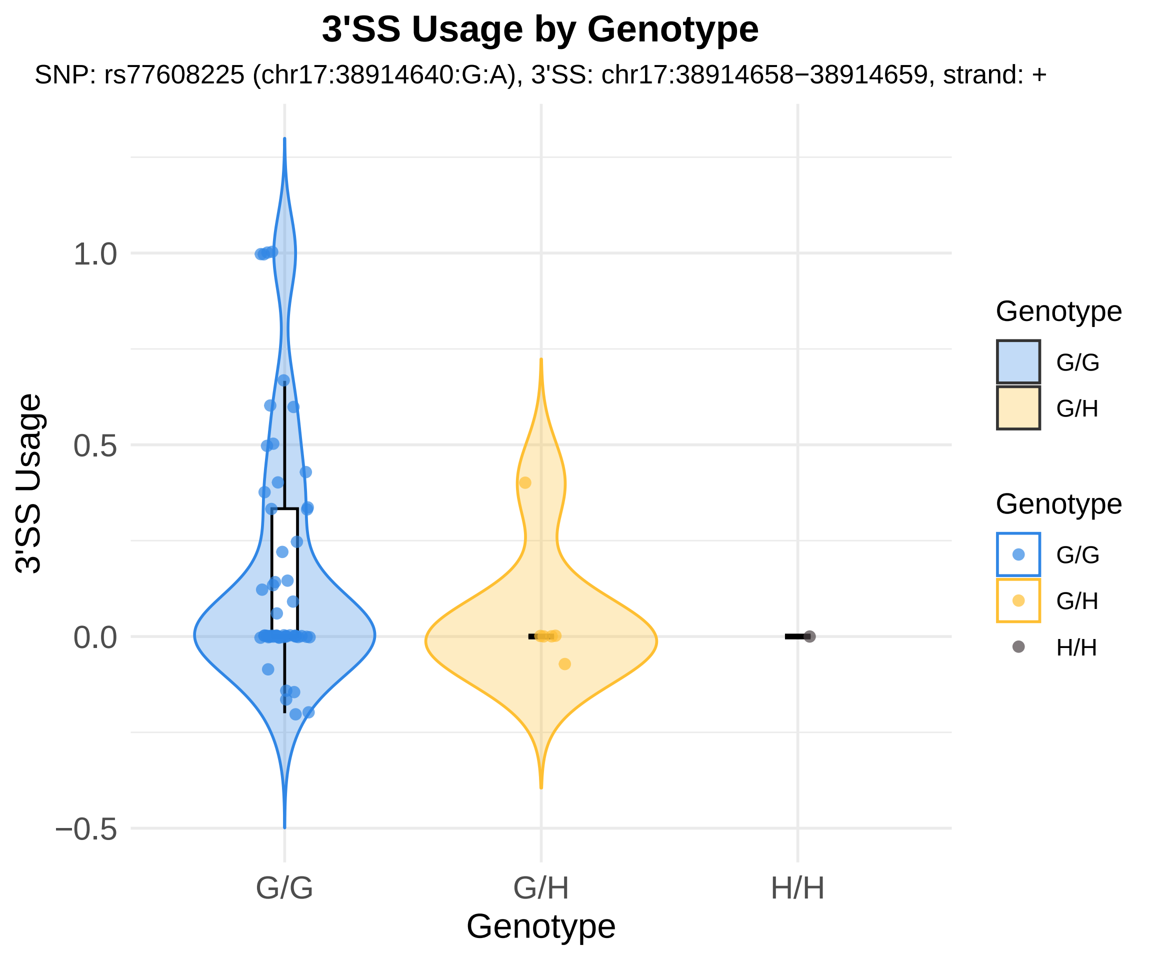
3'SS Usage by Genotype
SNP: rs77608225 (chr17:38914640:G:A), 3'SS: chr17:38914658−38914659, strand: +
3'SS Usage
1.0
0.5
0.0
−0.5
G/G
G/H
H/H
Genotype
Genotype
G/G
G/H
Genotype
G/G
G/H
H/H

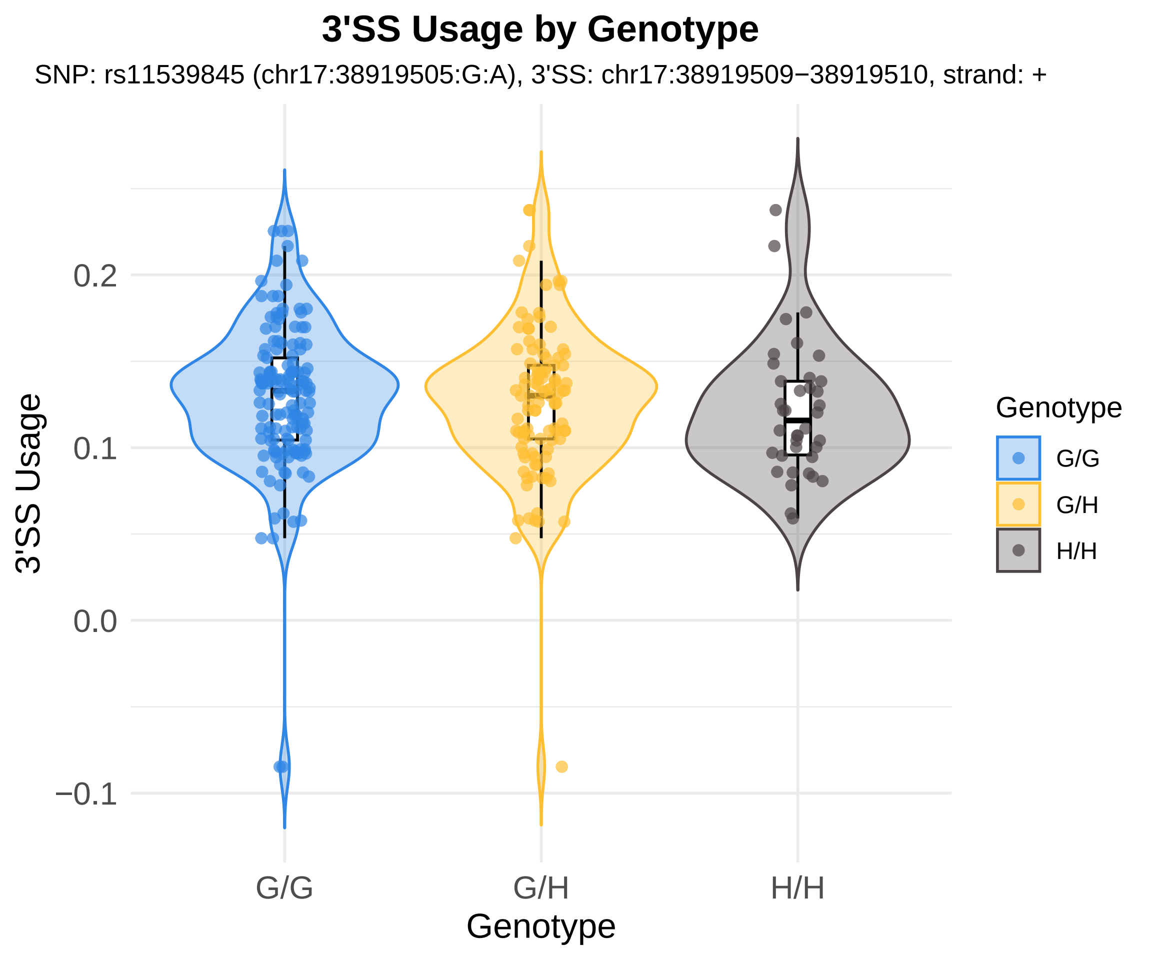
3'SS Usage by Genotype
SNP: rs11539845 (chr17:38919505:G:A), 3'SS: chr17:38919509−38919510, strand: +
3'SS Usage
0.2
0.1
0.0
−0.1
G/G
G/H
H/H
Genotype
Genotype
G/G
G/H
H/H

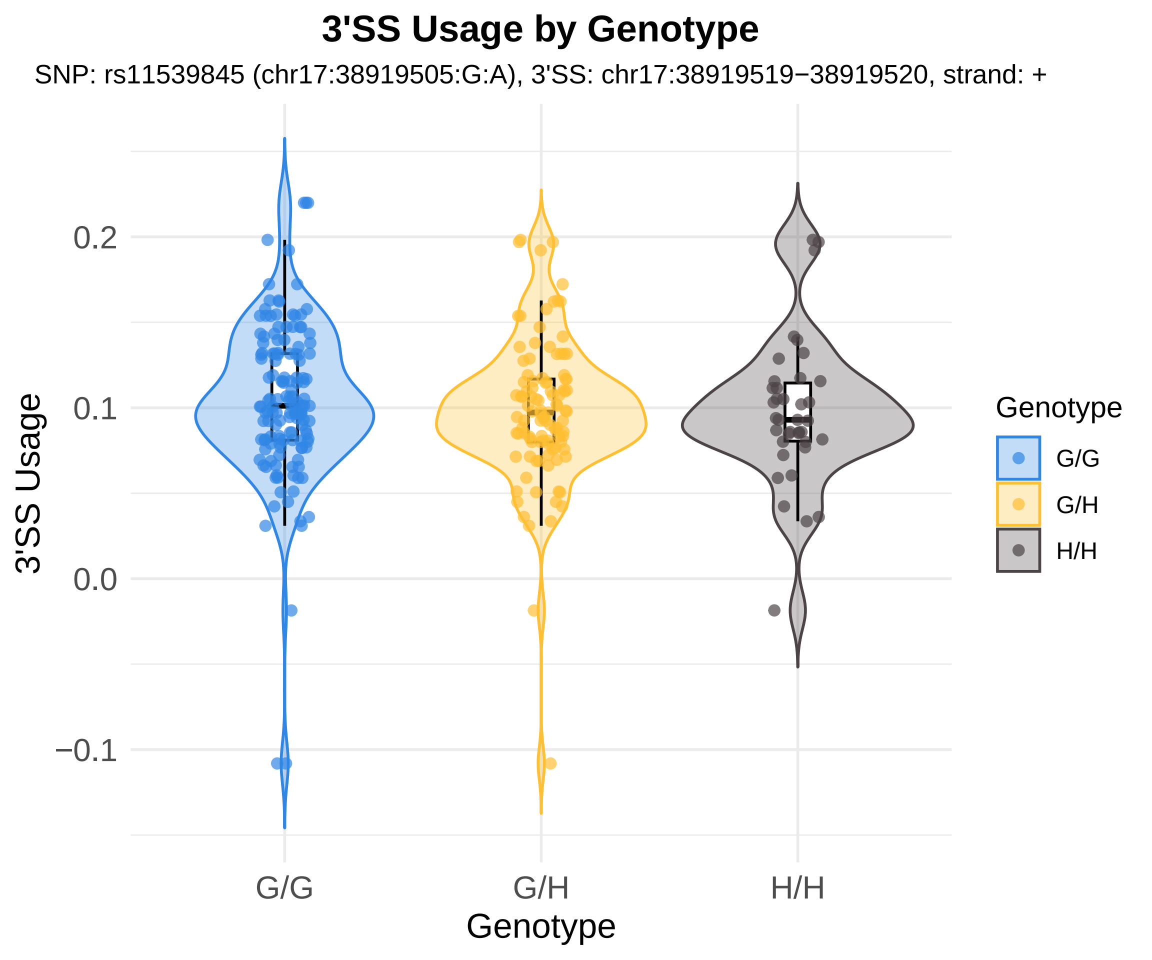
3'SS Usage by Genotype
SNP: rs11539845 (chr17:38919505:G:A), 3'SS: chr17:38919519−38919520, strand: +
3'SS Usage
0.2
0.1
0.0
−0.1
G/G
G/H
H/H
Genotype
Genotype
G/G
G/H
H/H

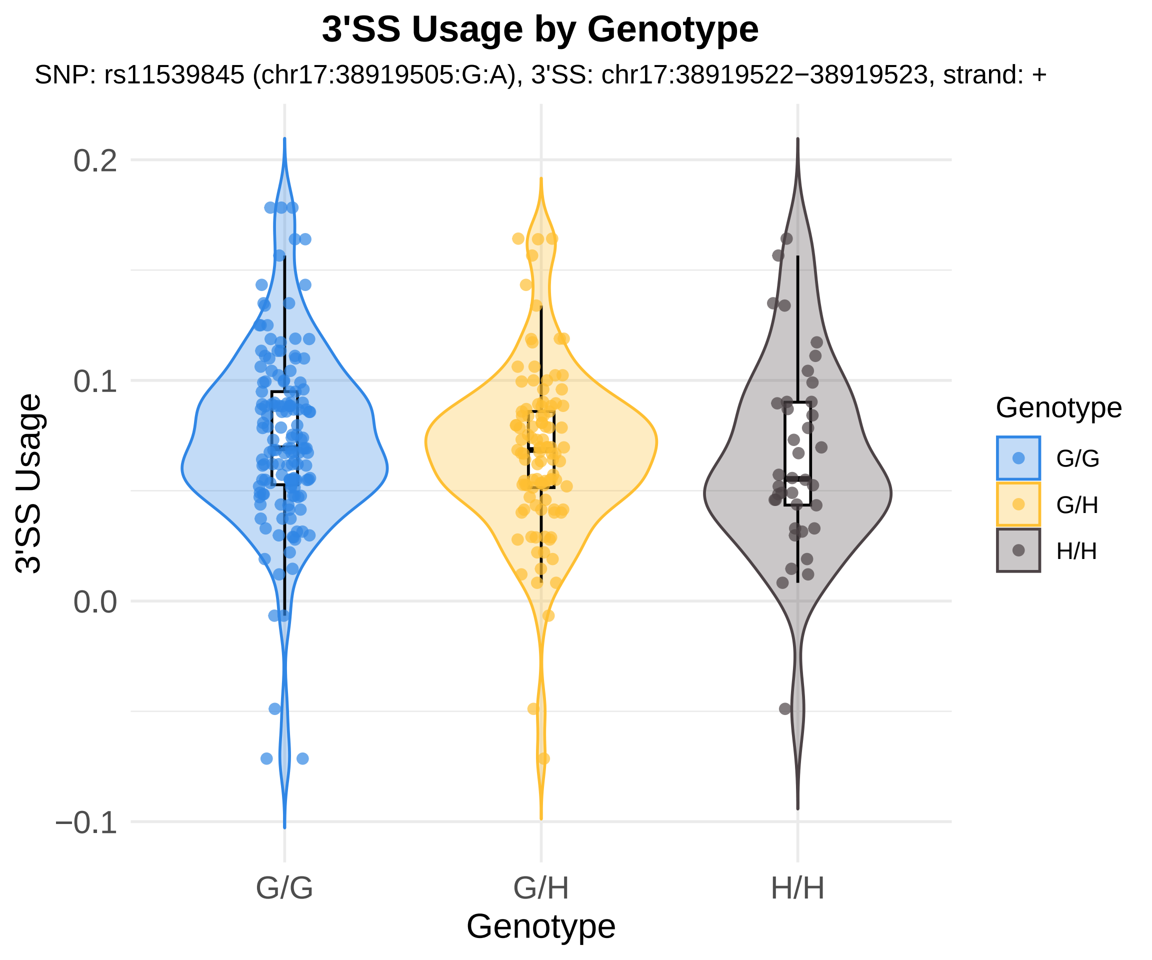

3'SS Usage by Genotype
SNP: rs11539845 (chr17:38919505:G:A), 3'SS: chr17:38919522−38919523, strand: +
3'SS Usage
0.2
0.1
0.0
−0.1
G/G
G/H
H/H
Genotype
Genotype
G/G
G/H
H/H

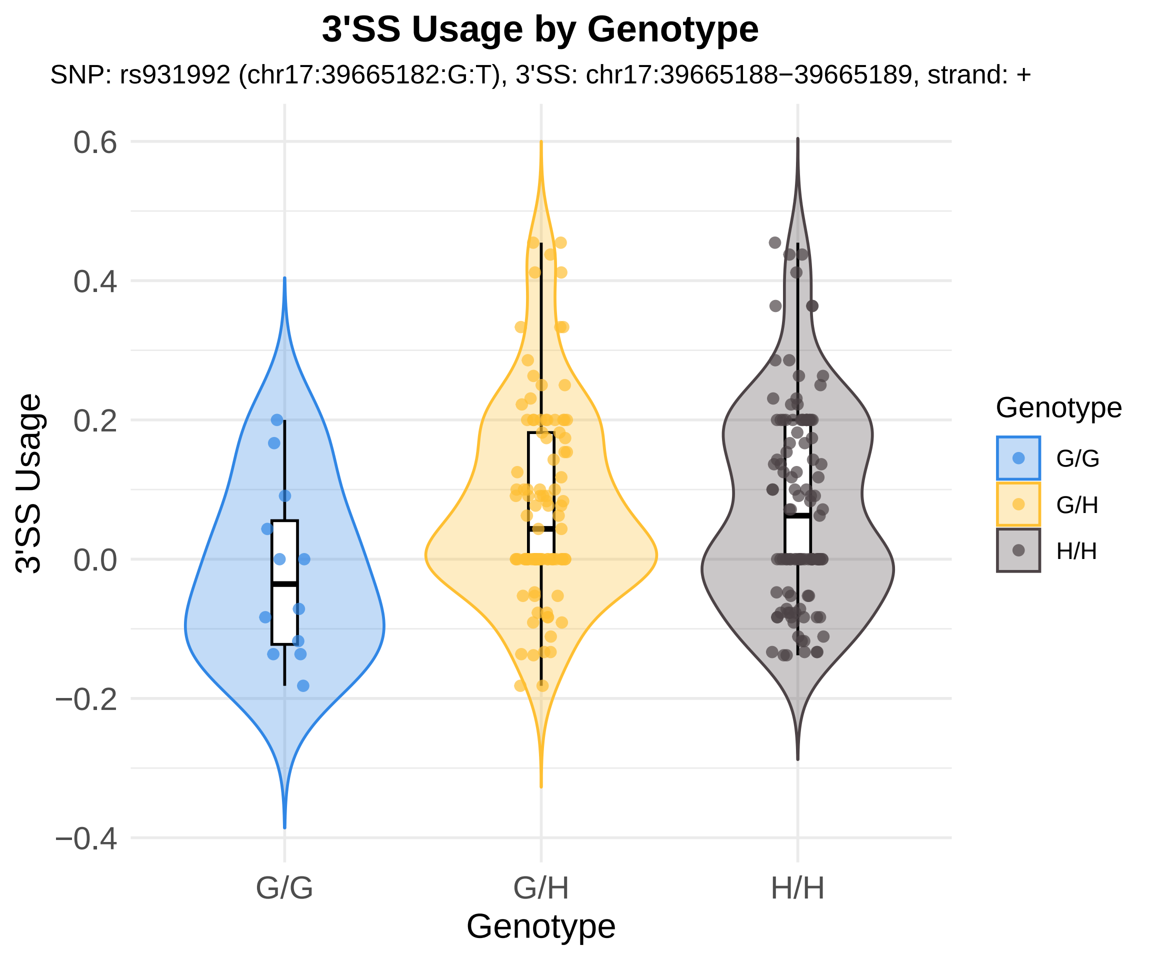
3'SS Usage by Genotype
SNP: rs931992 (chr17:39665182:G:T), 3'SS: chr17:39665188−39665189, strand: +
3'SS Usage
0.6
0.4
0.2
0.0
−0.2
−0.4
G/G
G/H
H/H
Genotype
Genotype
G/G
G/H
H/H

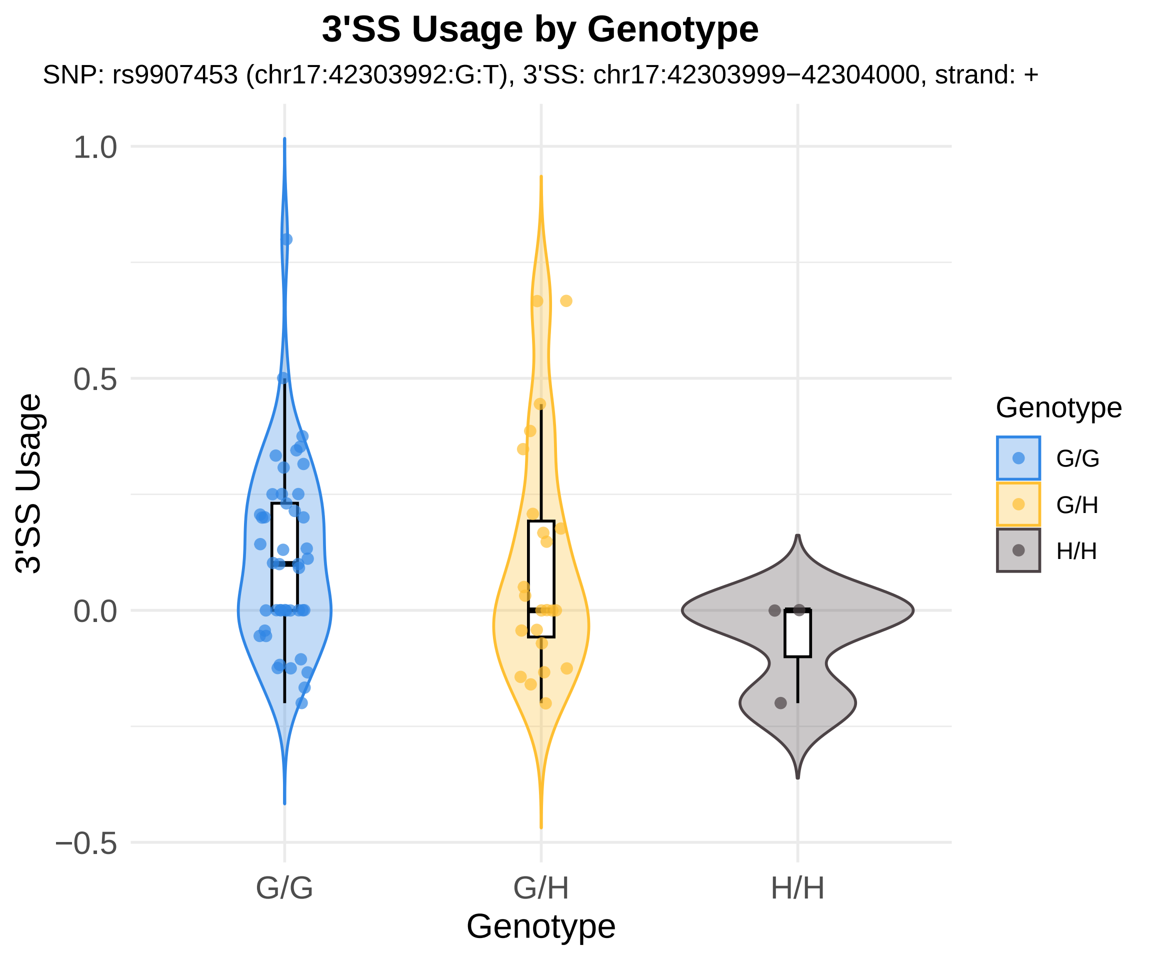
3'SS Usage by Genotype
SNP: rs9907453 (chr17:42303992:G:T), 3'SS: chr17:42303999−42304000, strand: +
3'SS Usage
1.0
0.5
0.0
−0.5
G/G
G/H
H/H
Genotype
Genotype
G/G
G/H
H/H

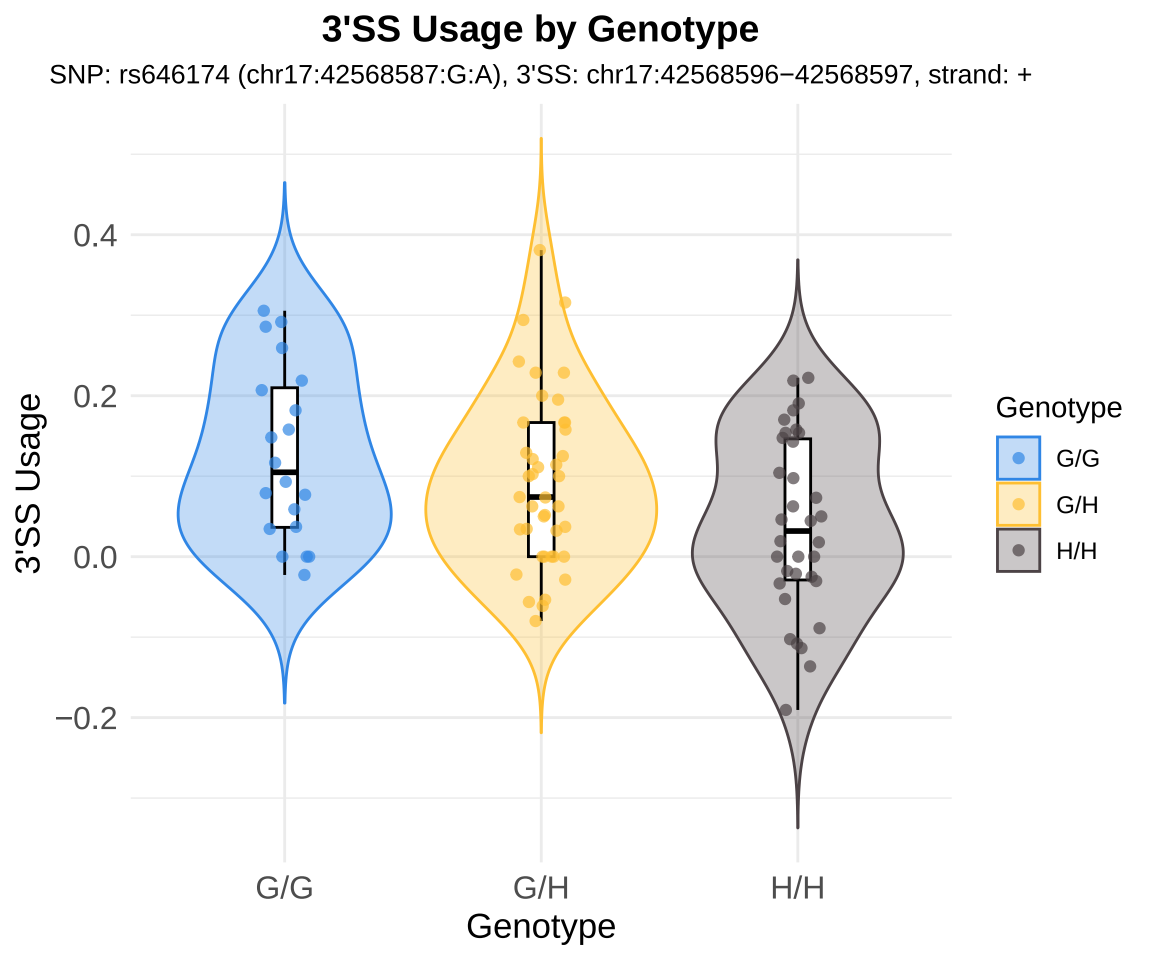

3'SS Usage by Genotype
SNP: rs646174 (chr17:42568587:G:A), 3'SS: chr17:42568596−42568597, strand: +
3'SS Usage
0.4
0.2
0.0
−0.2
G/G
G/H
H/H
Genotype
Genotype
G/G
G/H
H/H

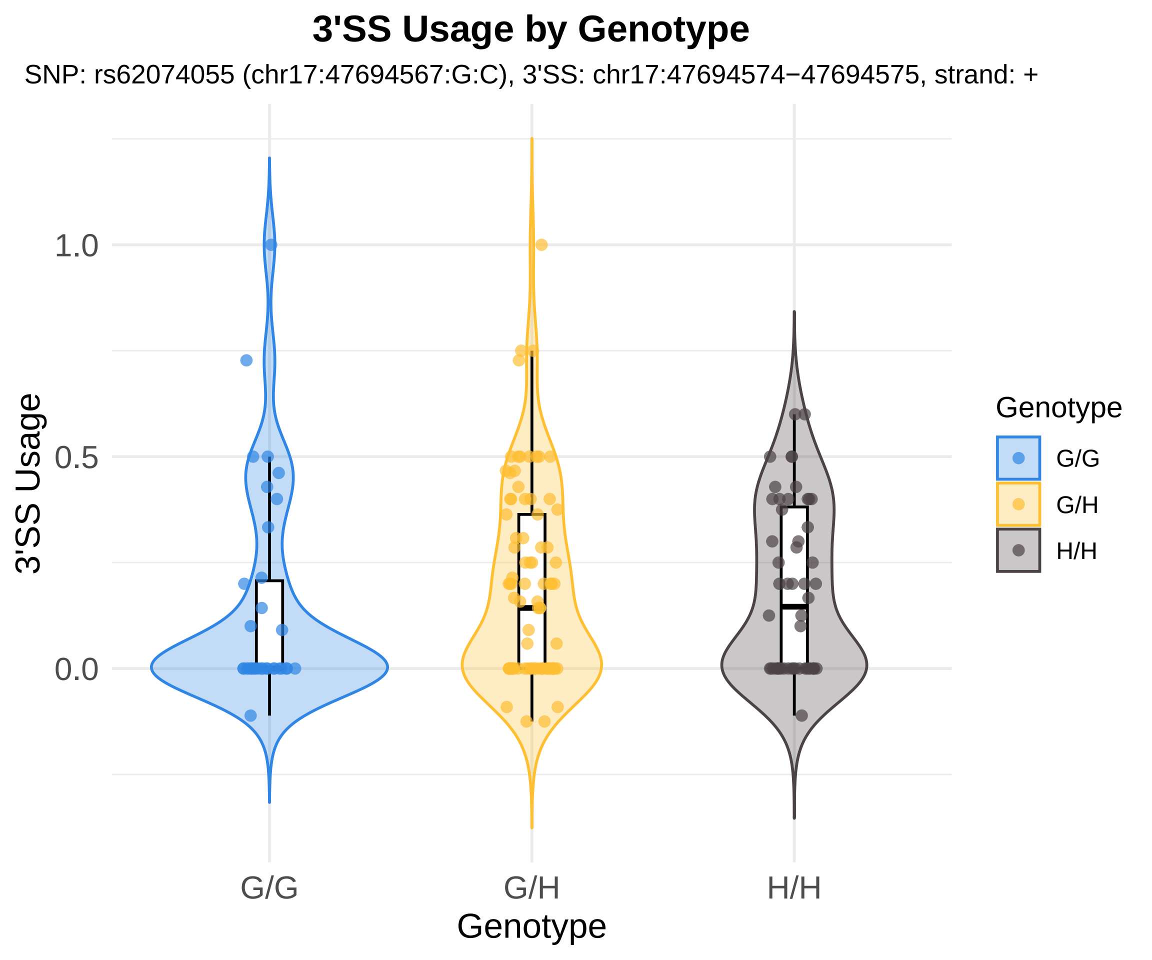
3'SS Usage by Genotype
SNP: rs62074055 (chr17:47694567:G:C), 3'SS: chr17:47694574−47694575, strand: +
3'SS Usage
1.0
0.5
0.0
G/G
G/H
H/H
Genotype
Genotype
G/G
G/H
H/H

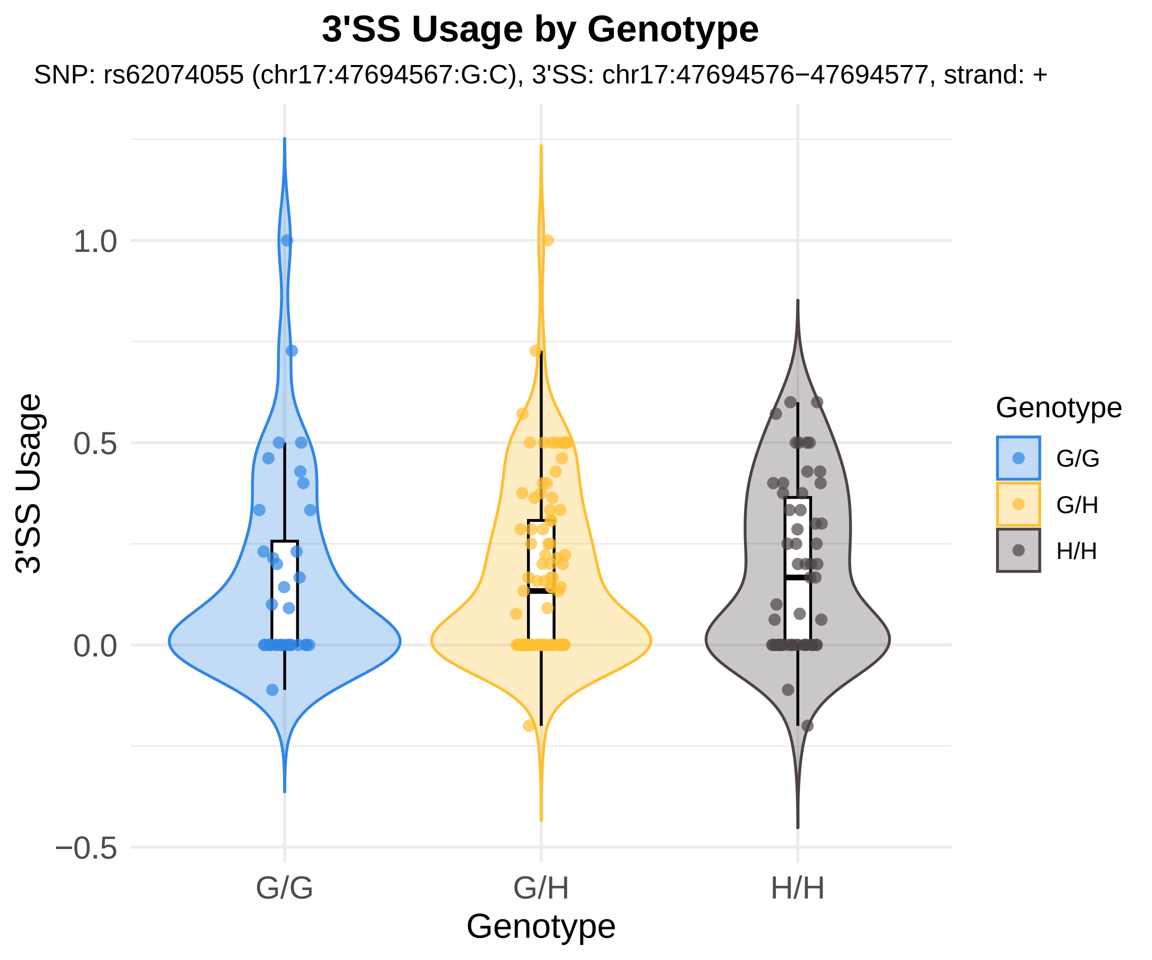
3'SS Usage by Genotype
SNP: rs62074055 (chr17:47694567:G:C), 3'SS: chr17:47694576−47694577, strand: +
3'SS Usage
1.0
0.5
0.0
−0.5
G/G
G/H
H/H
Genotype
Genotype
G/G
G/H
H/H

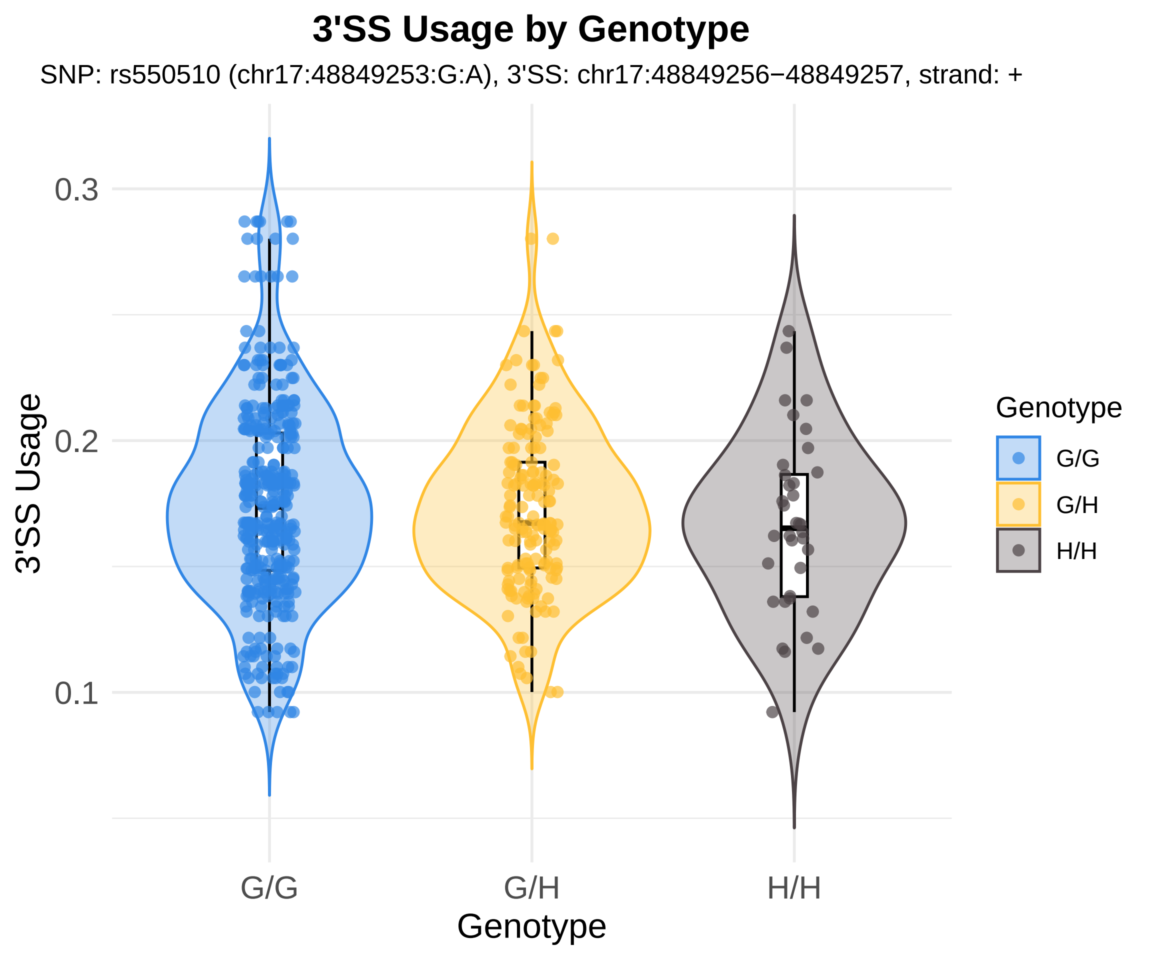

3'SS Usage by Genotype
SNP: rs550510 (chr17:48849253:G:A), 3'SS: chr17:48849256−48849257, strand: +
3'SS Usage
0.3
0.2
0.1
G/G
G/H
H/H
Genotype
Genotype
G/G
G/H
H/H

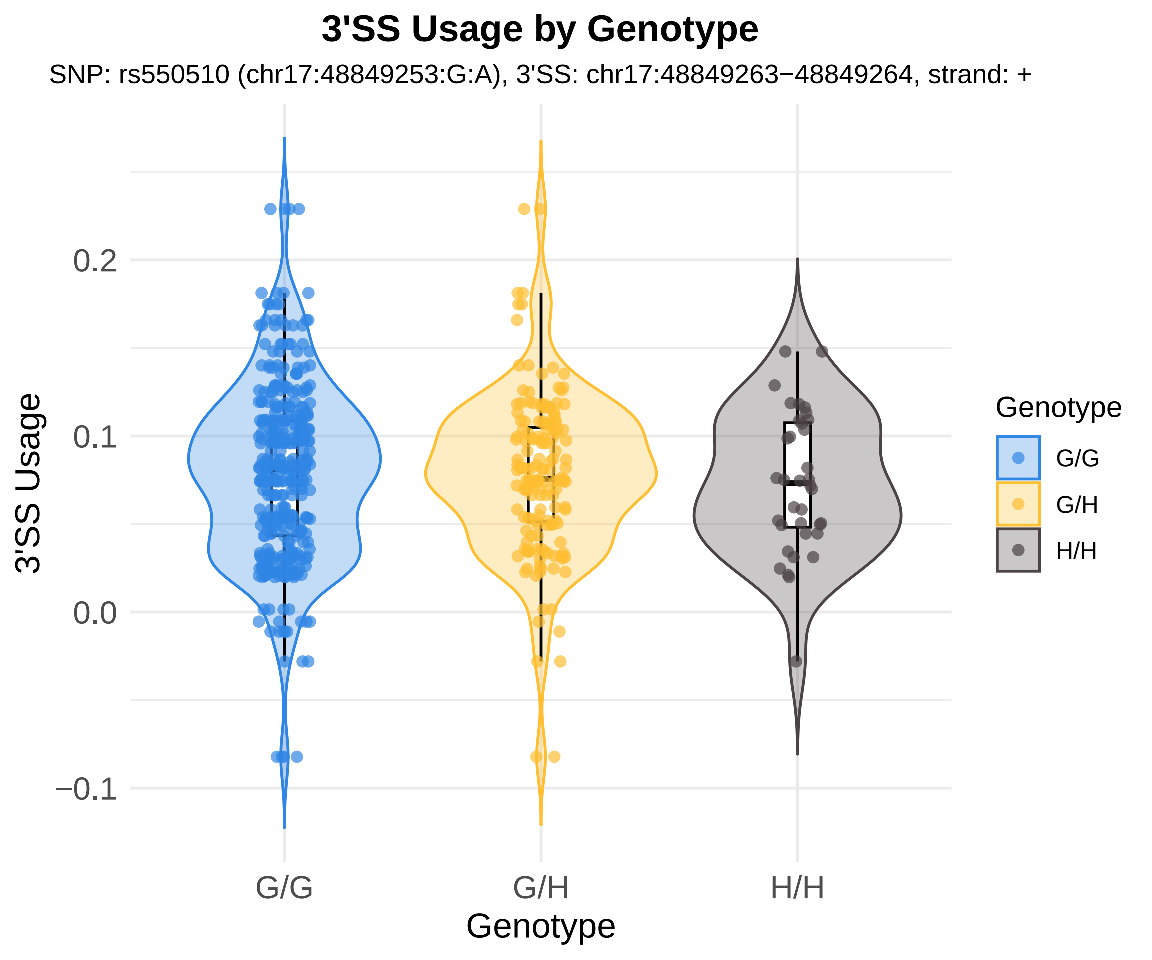
3'SS Usage by Genotype
SNP: rs550510 (chr17:48849253:G:A), 3'SS: chr17:48849263−48849264, strand: +
3'SS Usage
0.2
0.1
0.0
−0.1
G/G
G/H
H/H
Genotype
Genotype
G/G
G/H
H/H

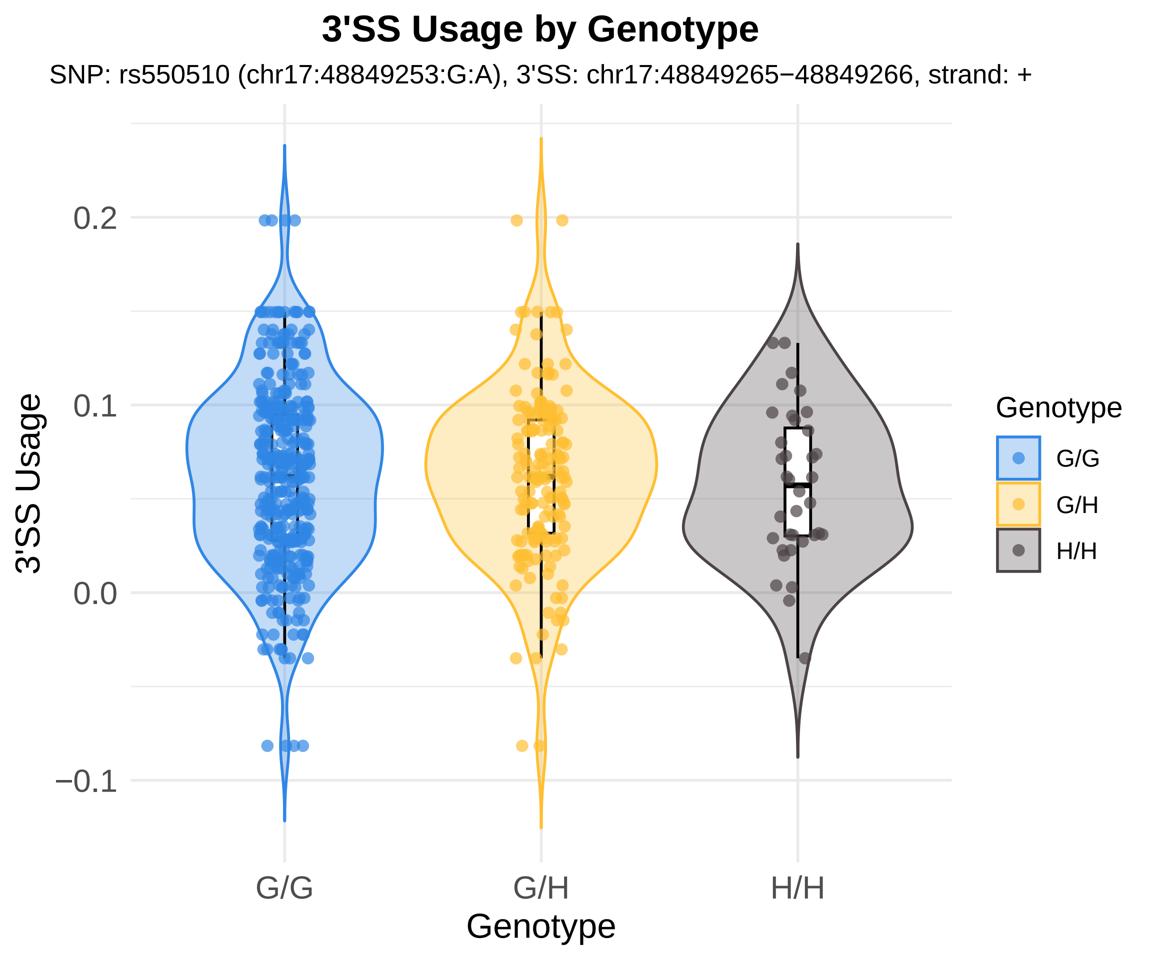

3'SS Usage by Genotype
SNP: rs550510 (chr17:48849253:G:A), 3'SS: chr17:48849265−48849266, strand: +
3'SS Usage
0.2
0.1
0.0
−0.1
G/G
G/H
H/H
Genotype
Genotype
G/G
G/H
H/H

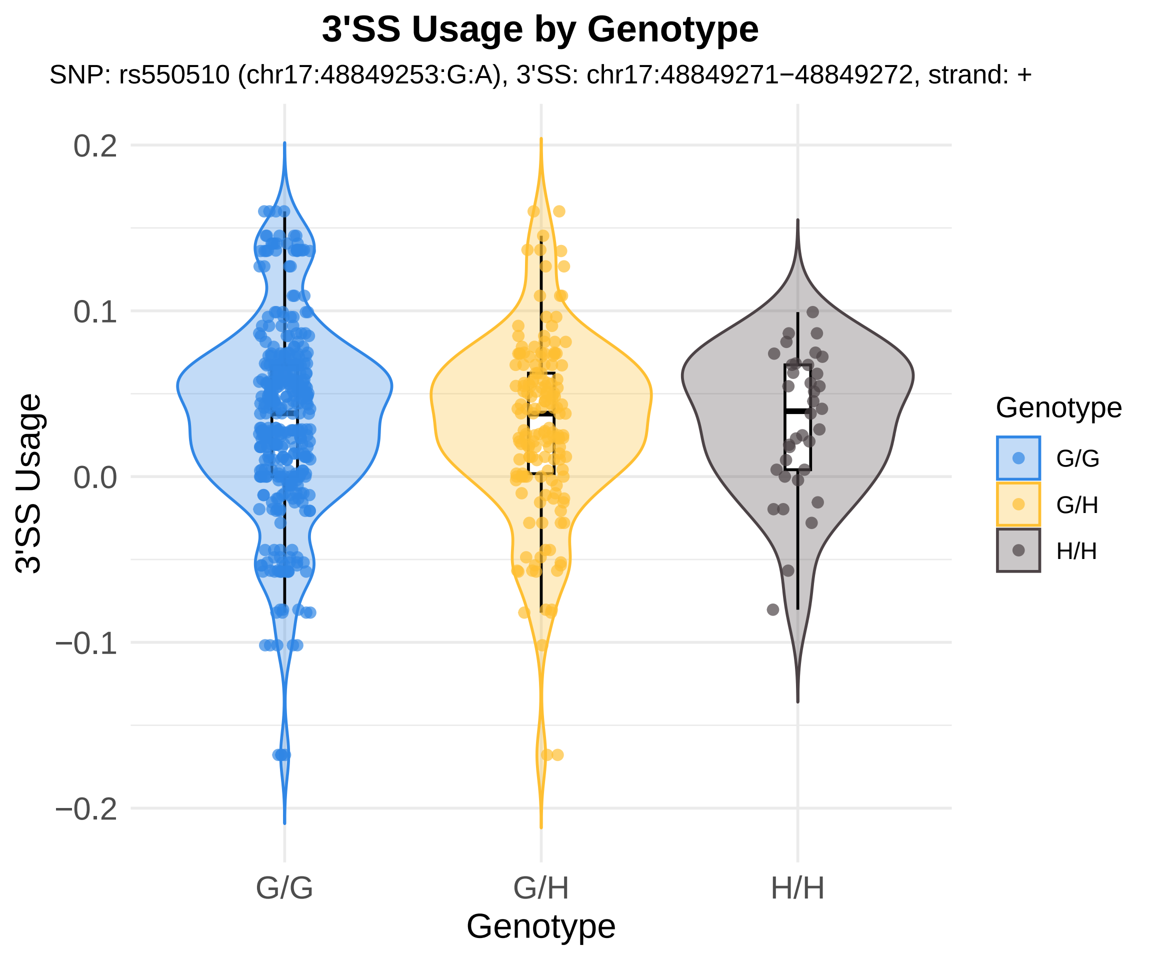
3'SS Usage by Genotype
SNP: rs550510 (chr17:48849253:G:A), 3'SS: chr17:48849271−48849272, strand: +
3'SS Usage
0.2
0.1
0.0
−0.1
−0.2
G/G
G/H
H/H
Genotype
Genotype
G/G
G/H
H/H

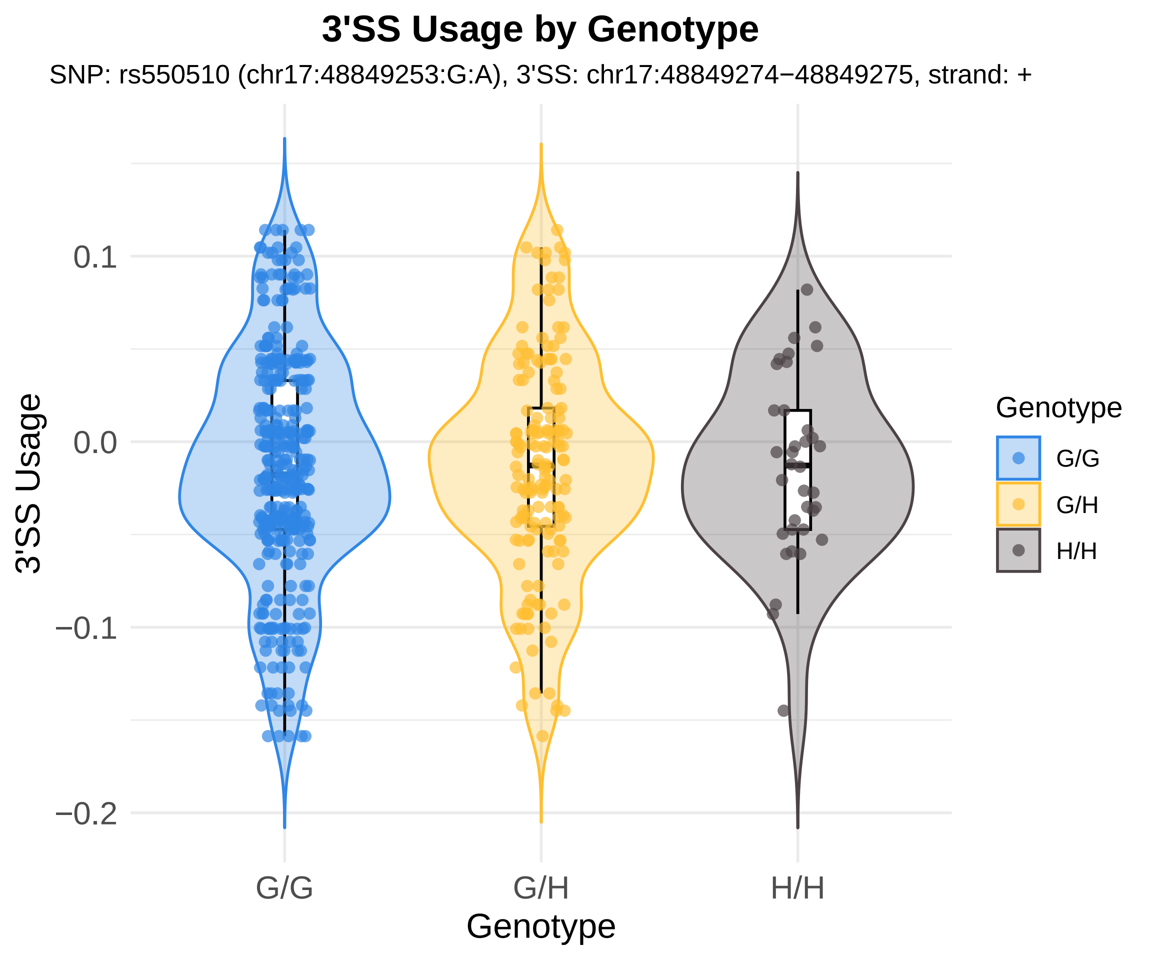
3'SS Usage by Genotype
SNP: rs550510 (chr17:48849253:G:A), 3'SS: chr17:48849274−48849275, strand: +
3'SS Usage
0.1
0.0
−0.1
−0.2
G/G
G/H
H/H
Genotype
Genotype
G/G
G/H
H/H

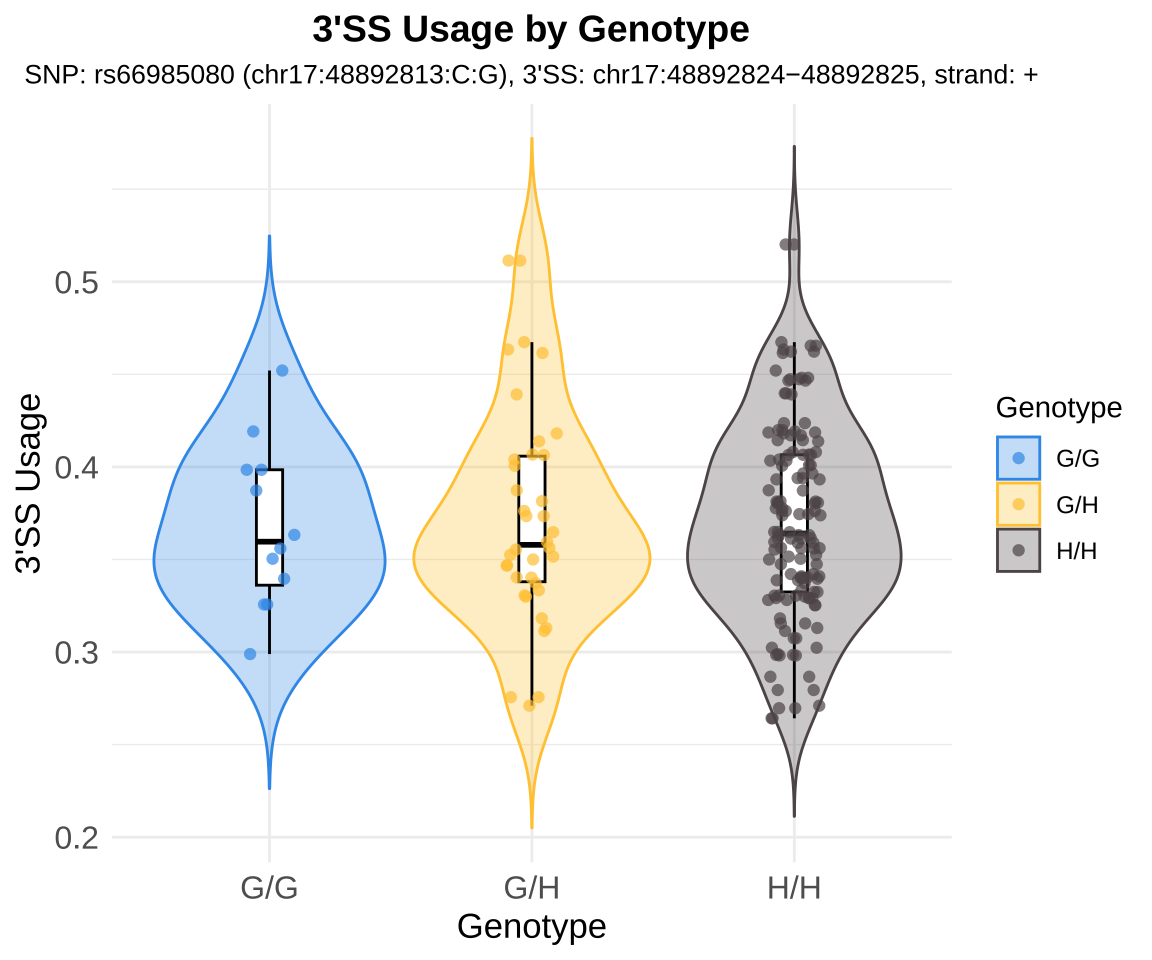

3'SS Usage by Genotype
SNP: rs66985080 (chr17:48892813:C:G), 3'SS: chr17:48892824−48892825, strand: +
3'SS Usage
0.5
0.4
0.3
0.2
G/G
G/H
H/H
Genotype
Genotype
G/G
G/H
H/H

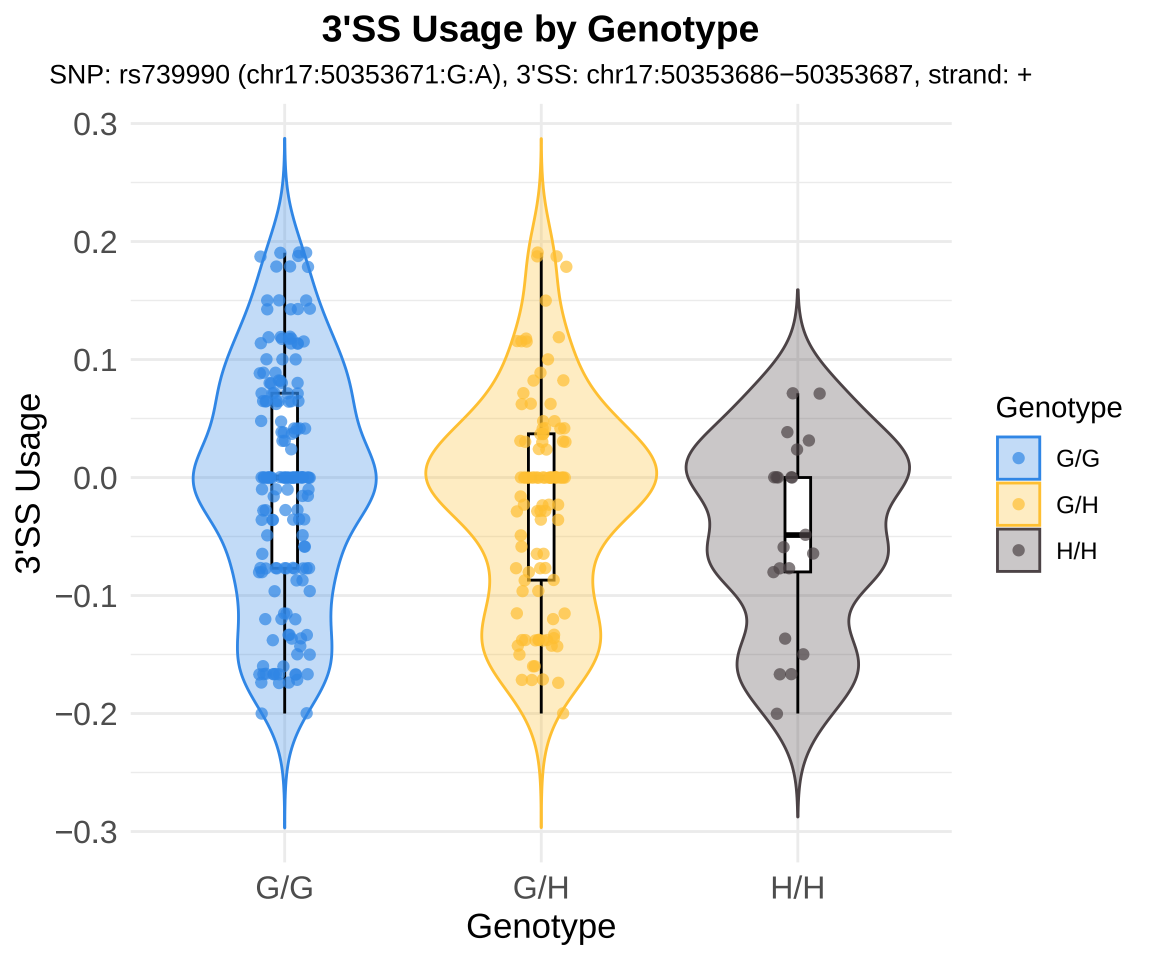
3'SS Usage by Genotype
SNP: rs739990 (chr17:50353671:G:A), 3'SS: chr17:50353686−50353687, strand: +
0.3
0.2
0.1
0.0
−0.1
−0.2
−0.3
3'SS Usage
G/G
G/H
H/H
Genotype
Genotype
G/G
G/H
H/H

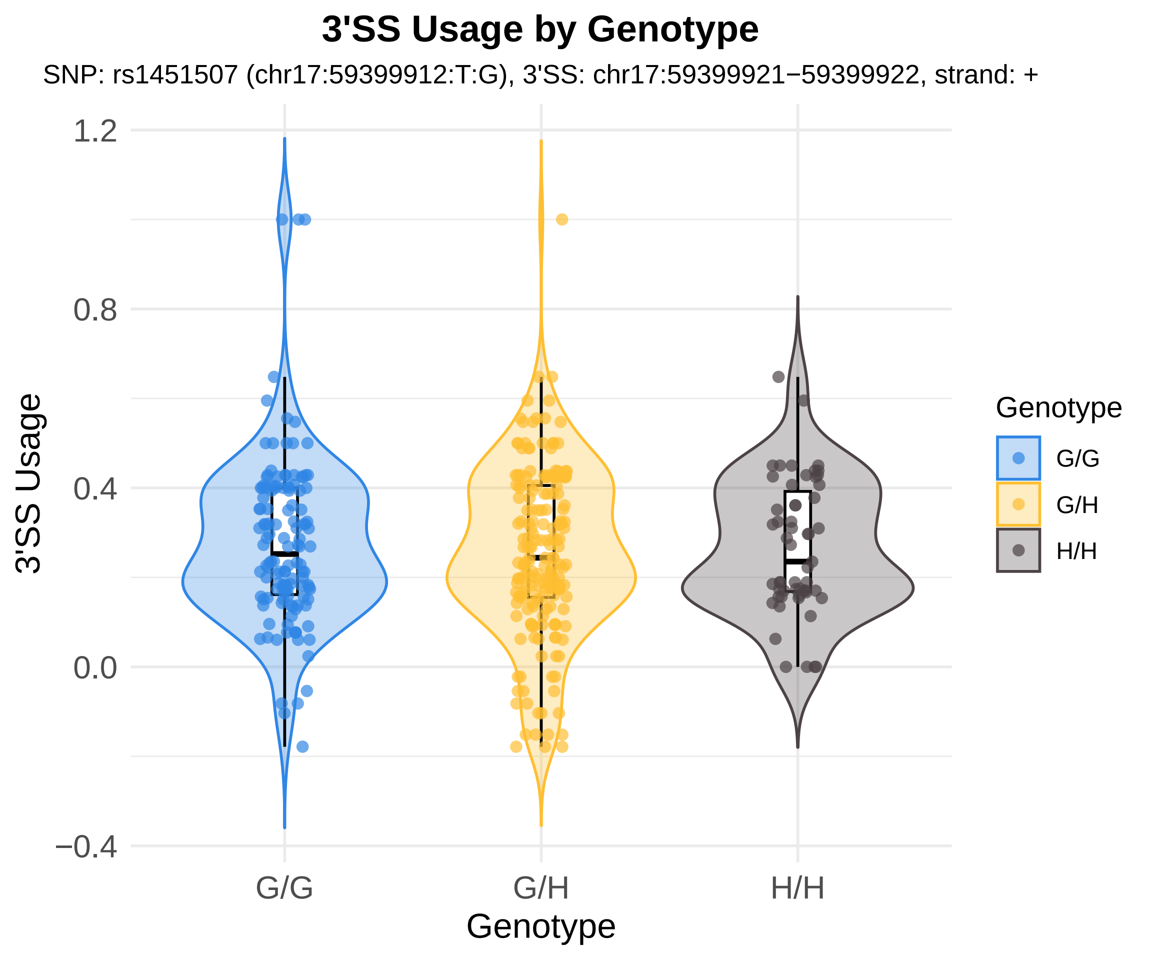
3'SS Usage by Genotype
SNP: rs1451507 (chr17:59399912:T:G), 3'SS: chr17:59399921−59399922, strand: +
1.2
0.8
0.4
0.0
−0.4
3'SS Usage
G/G
G/H
H/H
Genotype
Genotype
G/G
G/H
H/H

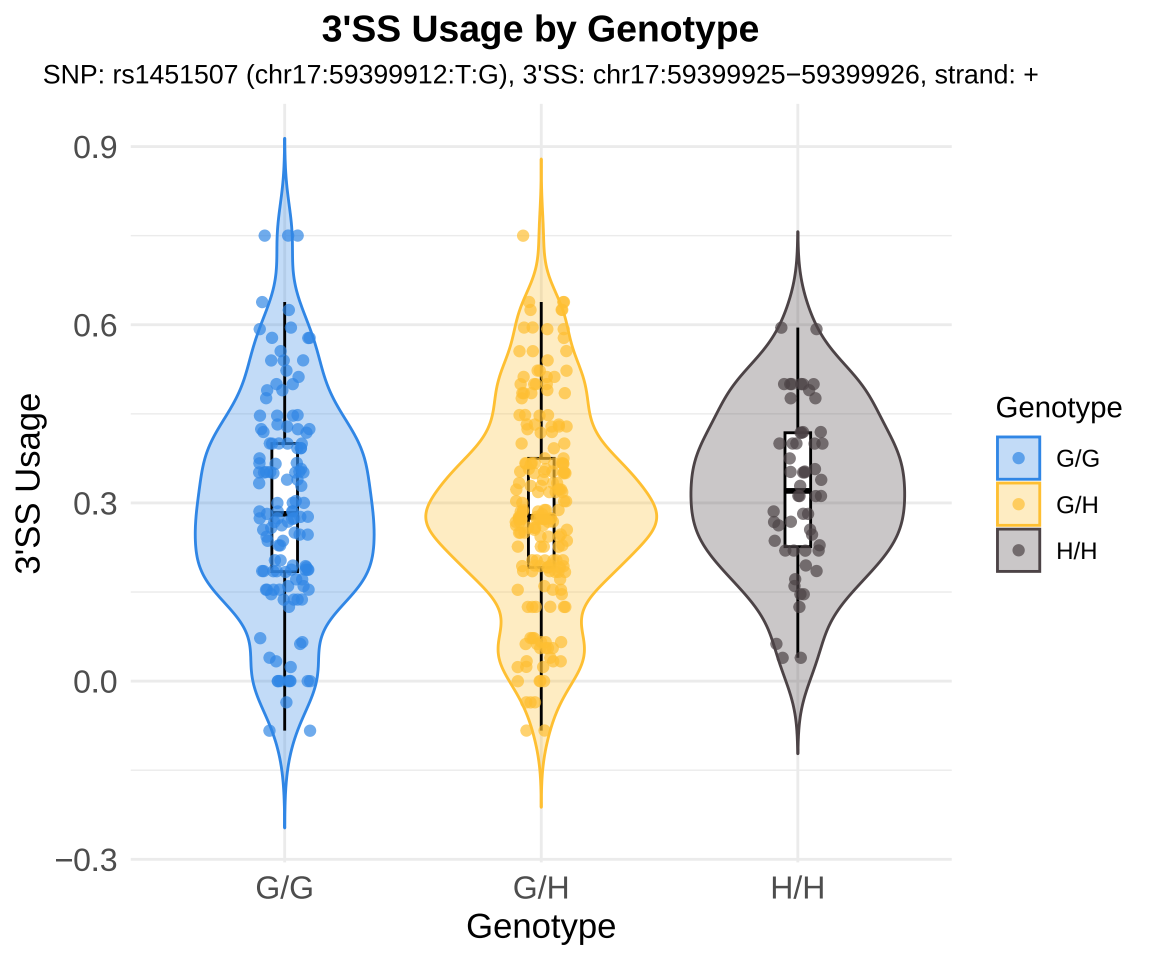
3'SS Usage by Genotype
SNP: rs1451507 (chr17:59399912:T:G), 3'SS: chr17:59399925−59399926, strand: +
3'SS Usage
0.9
0.6
0.3
0.0
−0.3
G/G
G/H
H/H
Genotype
Genotype
G/G
G/H
H/H

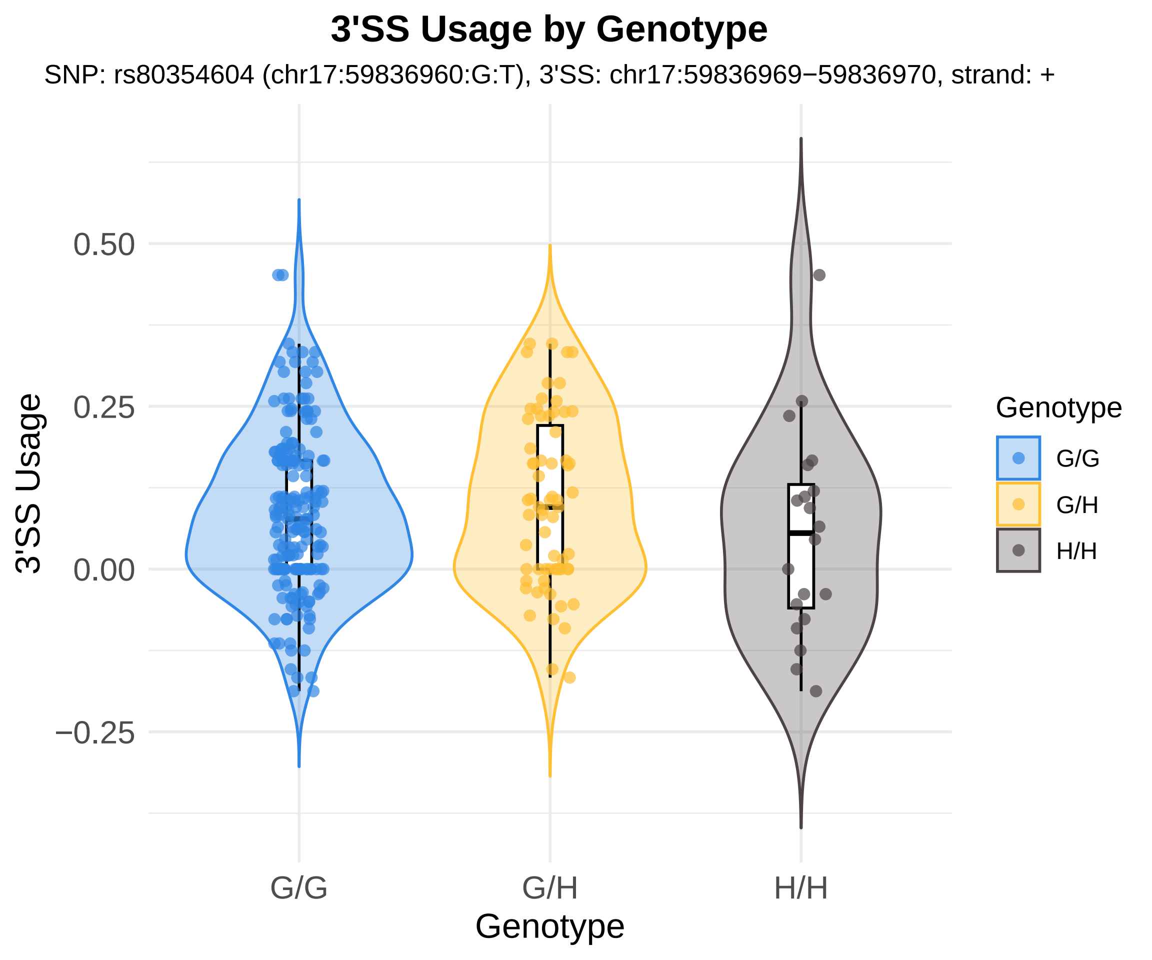
3'SS Usage by Genotype
SNP: rs80354604 (chr17:59836960:G:T), 3'SS: chr17:59836969−59836970, strand: +
3'SS Usage
0.50
0.25
0.00
−0.25
G/G
G/H
H/H
Genotype
Genotype
G/G
G/H
H/H

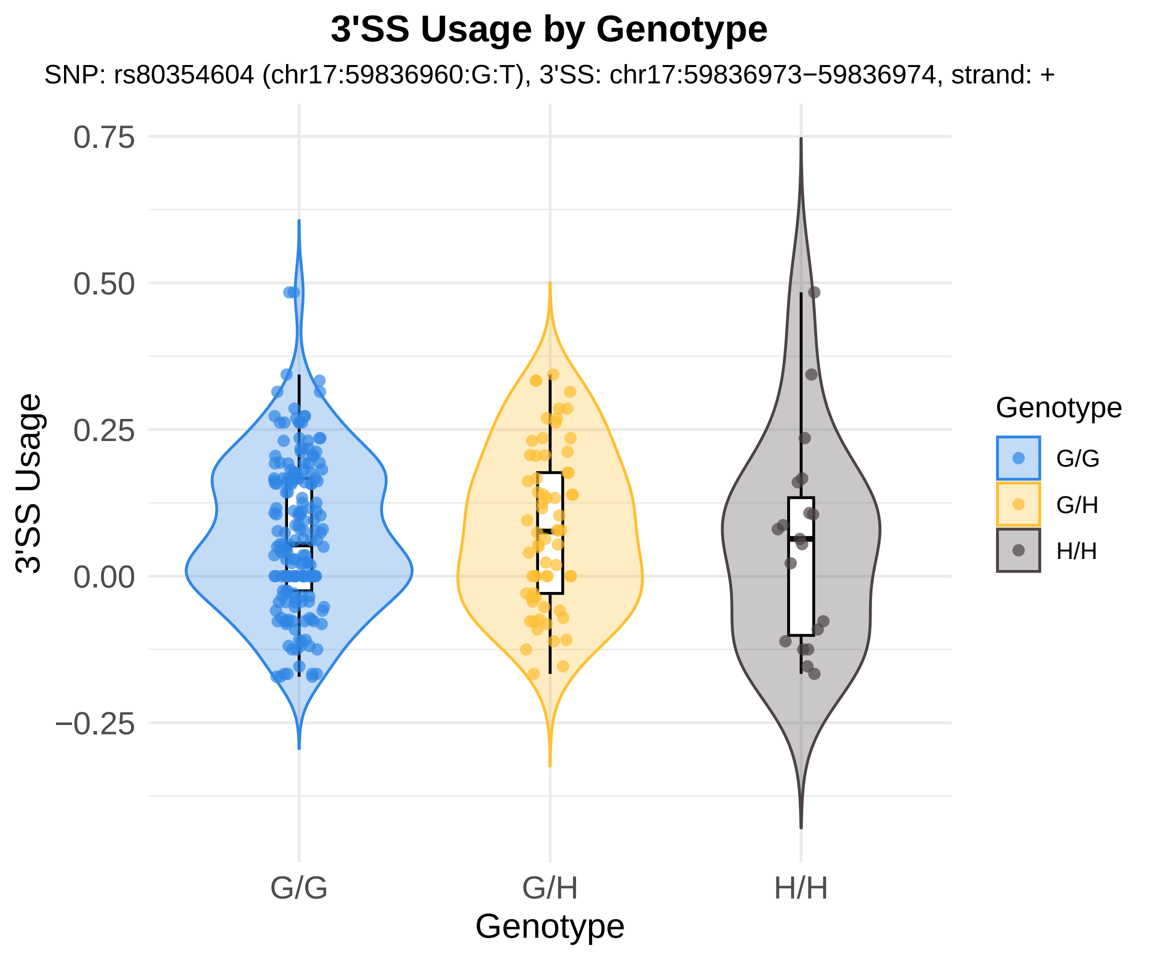
3'SS Usage by Genotype
SNP: rs80354604 (chr17:59836960:G:T), 3'SS: chr17:59836973−59836974, strand: +
3'SS Usage
0.75
0.50
0.25
0.00
−0.25
G/G
G/H
H/H
Genotype
Genotype
G/G
G/H
H/H

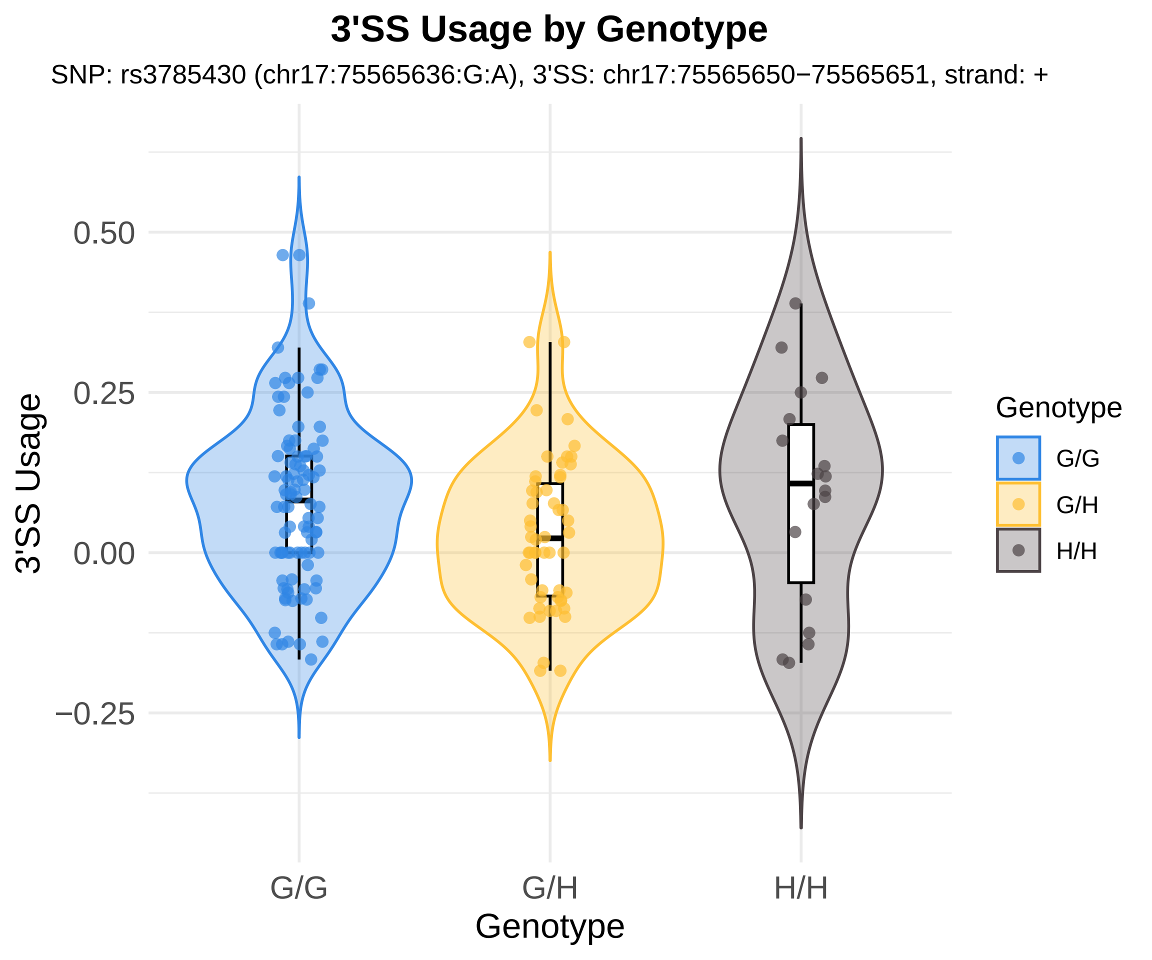
3'SS Usage by Genotype
SNP: rs3785430 (chr17:75565636:G:A), 3'SS: chr17:75565650−75565651, strand: +
3'SS Usage
0.50
0.25
0.00
−0.25
G/G
G/H
H/H
Genotype
Genotype
G/G
G/H
H/H

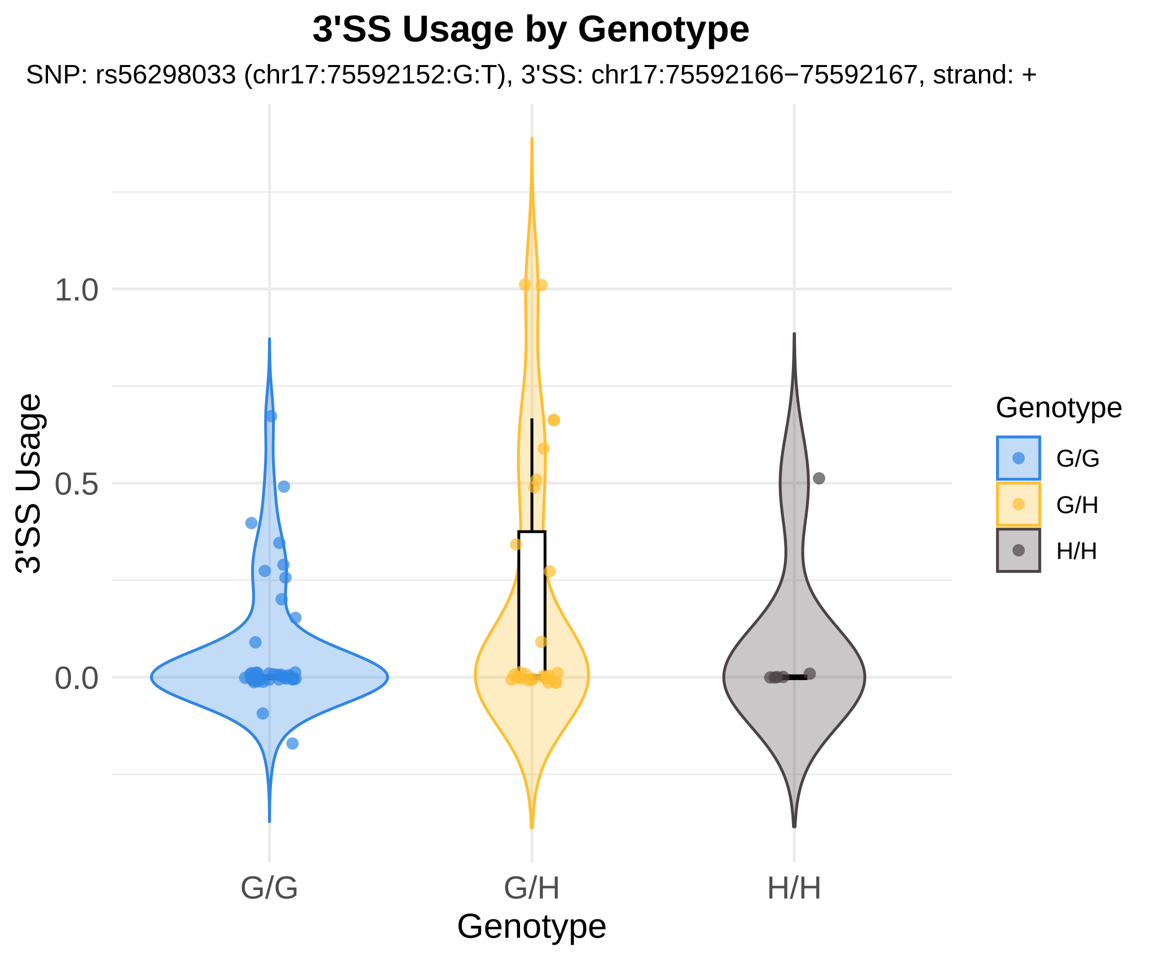
3'SS Usage by Genotype
SNP: rs56298033 (chr17:75592152:G:T), 3'SS: chr17:75592166−75592167, strand: +
3'SS Usage
1.0
0.5
0.0
G/G
G/H
H/H
Genotype
Genotype
G/G
G/H
H/H

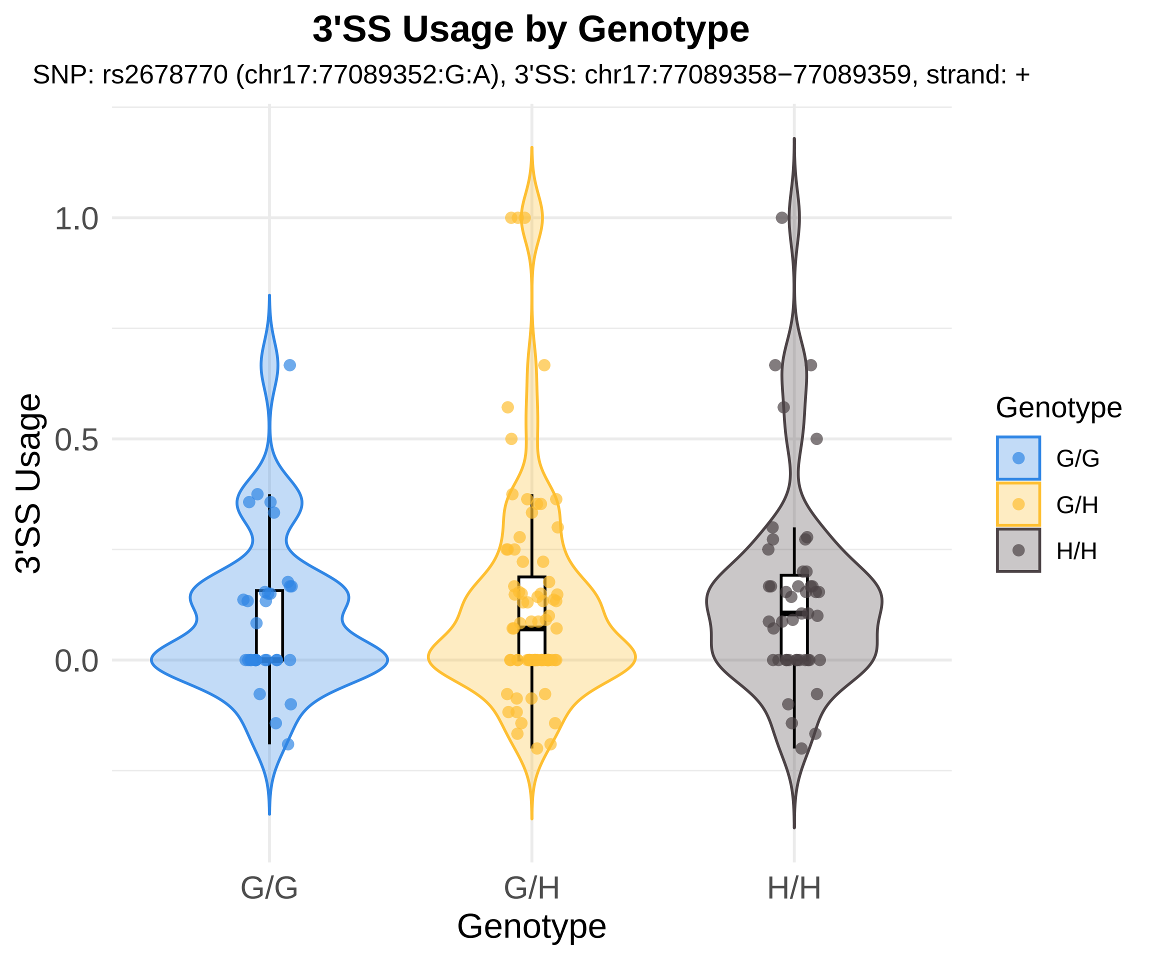

3'SS Usage by Genotype
SNP: rs2678770 (chr17:77089352:G:A), 3'SS: chr17:77089358−77089359, strand: +
1.0
0.5
0.0
3'SS Usage
G/G
G/H
H/H
Genotype
Genotype
G/G
G/H
H/H

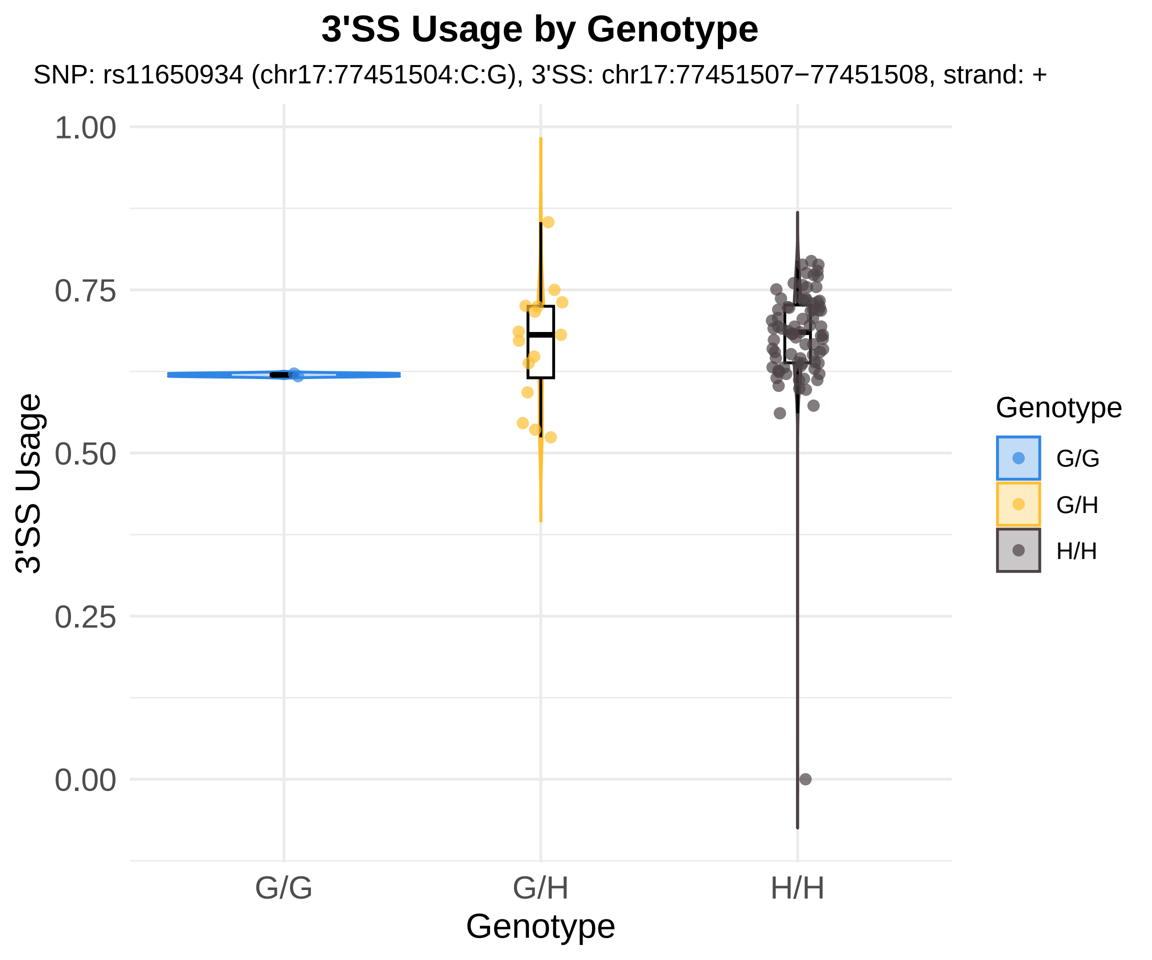
3'SS Usage by Genotype
SNP: rs11650934 (chr17:77451504:C:G), 3'SS: chr17:77451507−77451508, strand: +
1.00
0.75
0.50
0.25
0.00
3'SS Usage
G/G
G/H
H/H
Genotype
Genotype
G/G
G/H
H/H

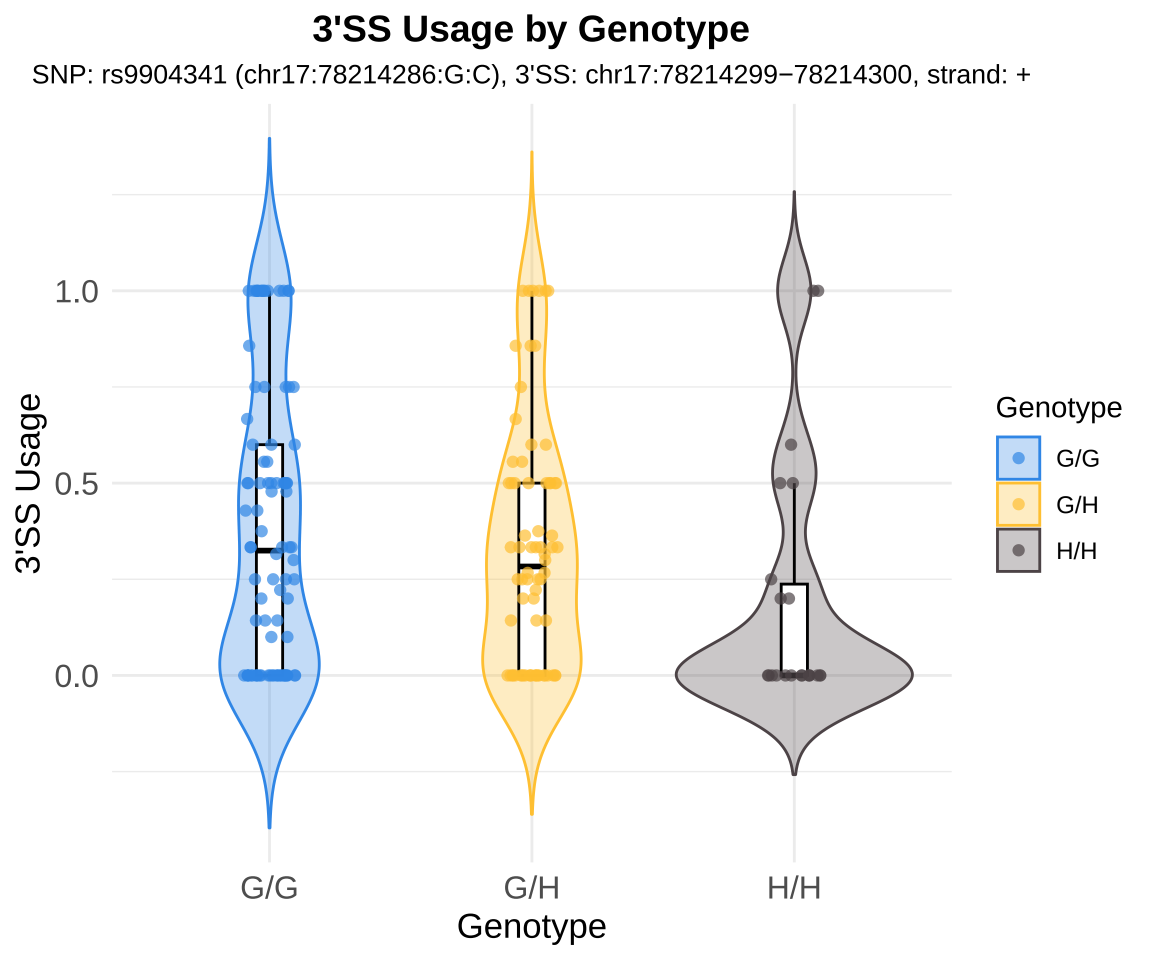
3'SS Usage by Genotype
SNP: rs9904341 (chr17:78214286:G:C), 3'SS: chr17:78214299−78214300, strand: +
3'SS Usage
1.0
0.5
0.0
G/G
G/H
H/H
Genotype
Genotype
G/G
G/H
H/H

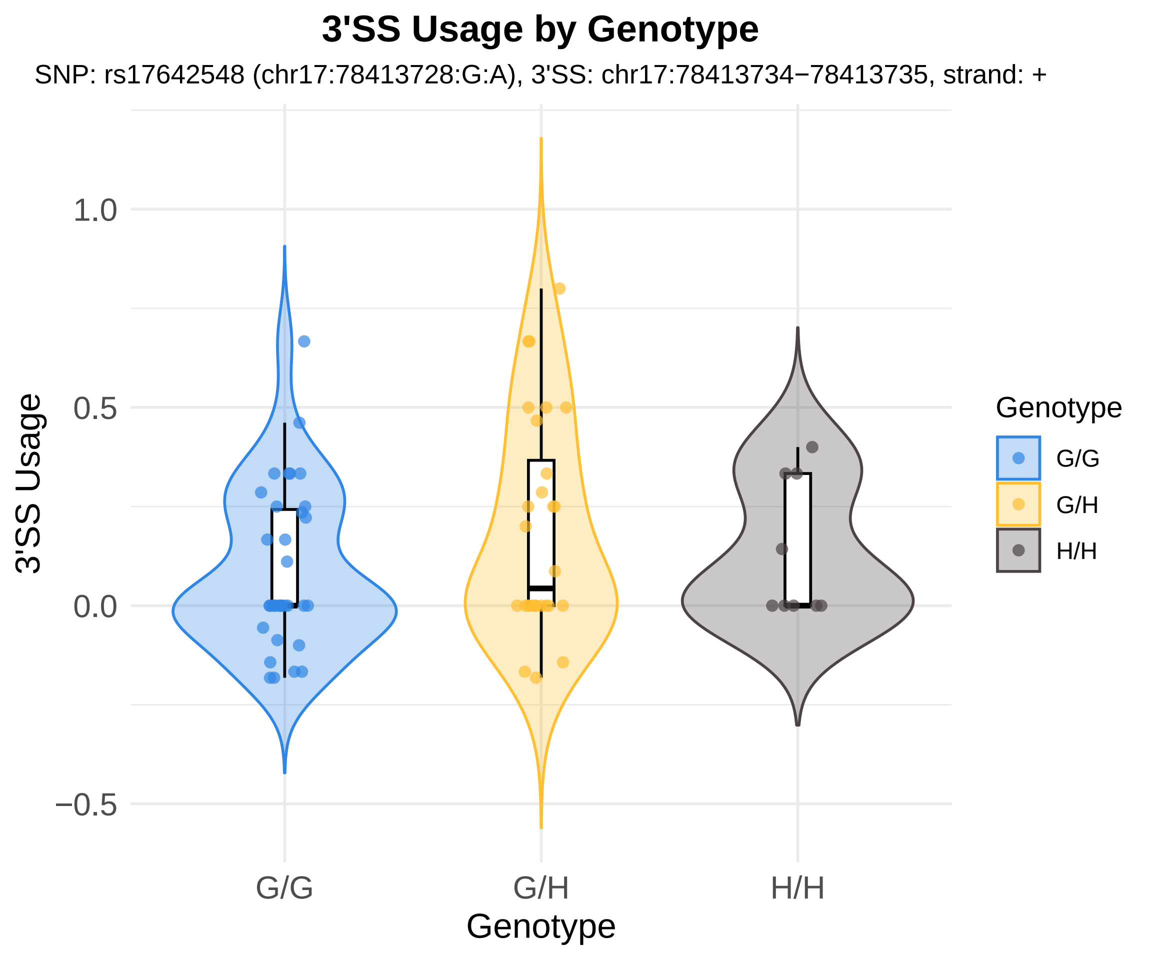
3'SS Usage by Genotype
SNP: rs17642548 (chr17:78413728:G:A), 3'SS: chr17:78413734−78413735, strand: +
3'SS Usage
1.0
0.5
0.0
−0.5
G/G
G/H
H/H
Genotype
Genotype
G/G
G/H
H/H

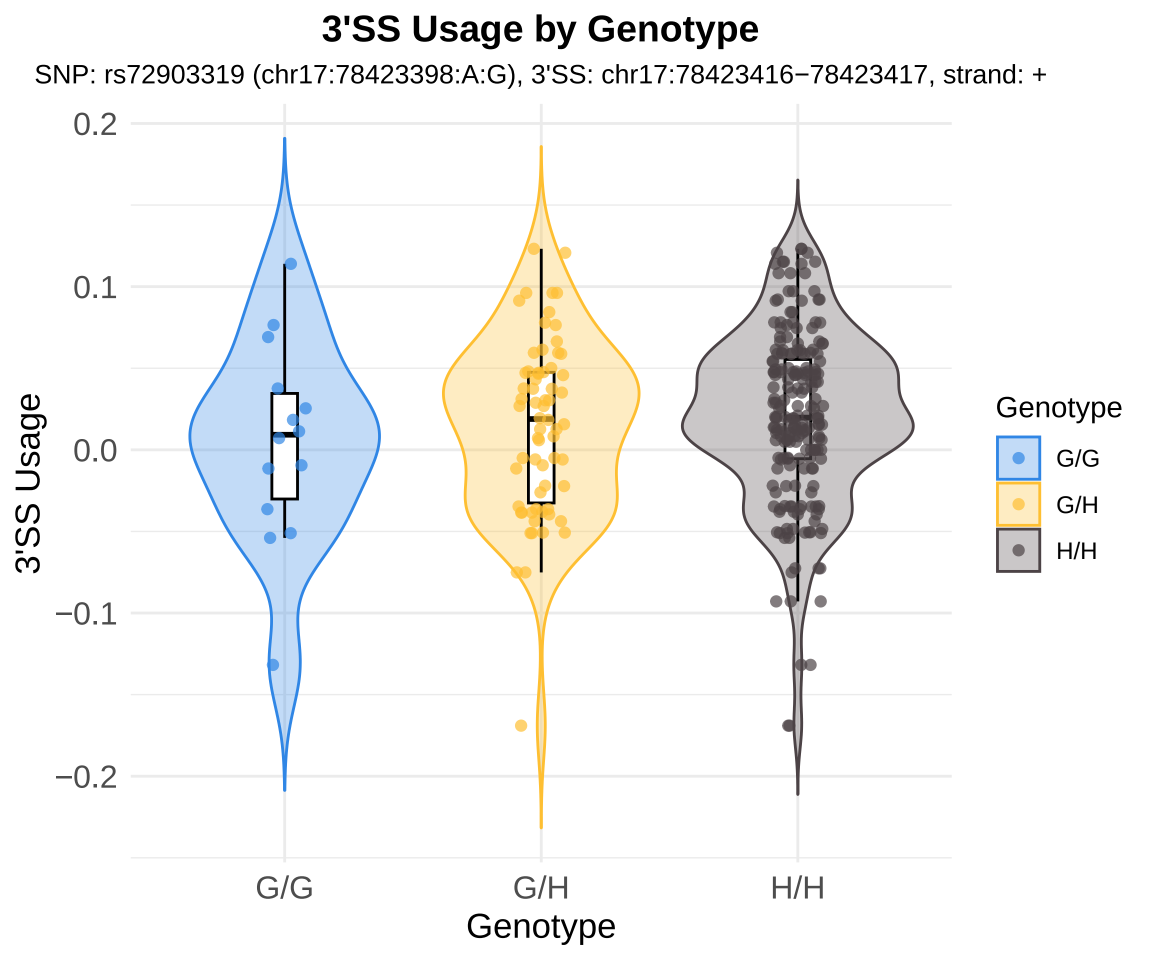

3'SS Usage by Genotype
SNP: rs72903319 (chr17:78423398:A:G), 3'SS: chr17:78423416−78423417, strand: +
3'SS Usage
0.2
0.1
0.0
−0.1
−0.2
G/G
G/H
H/H
Genotype
Genotype
G/G
G/H
H/H

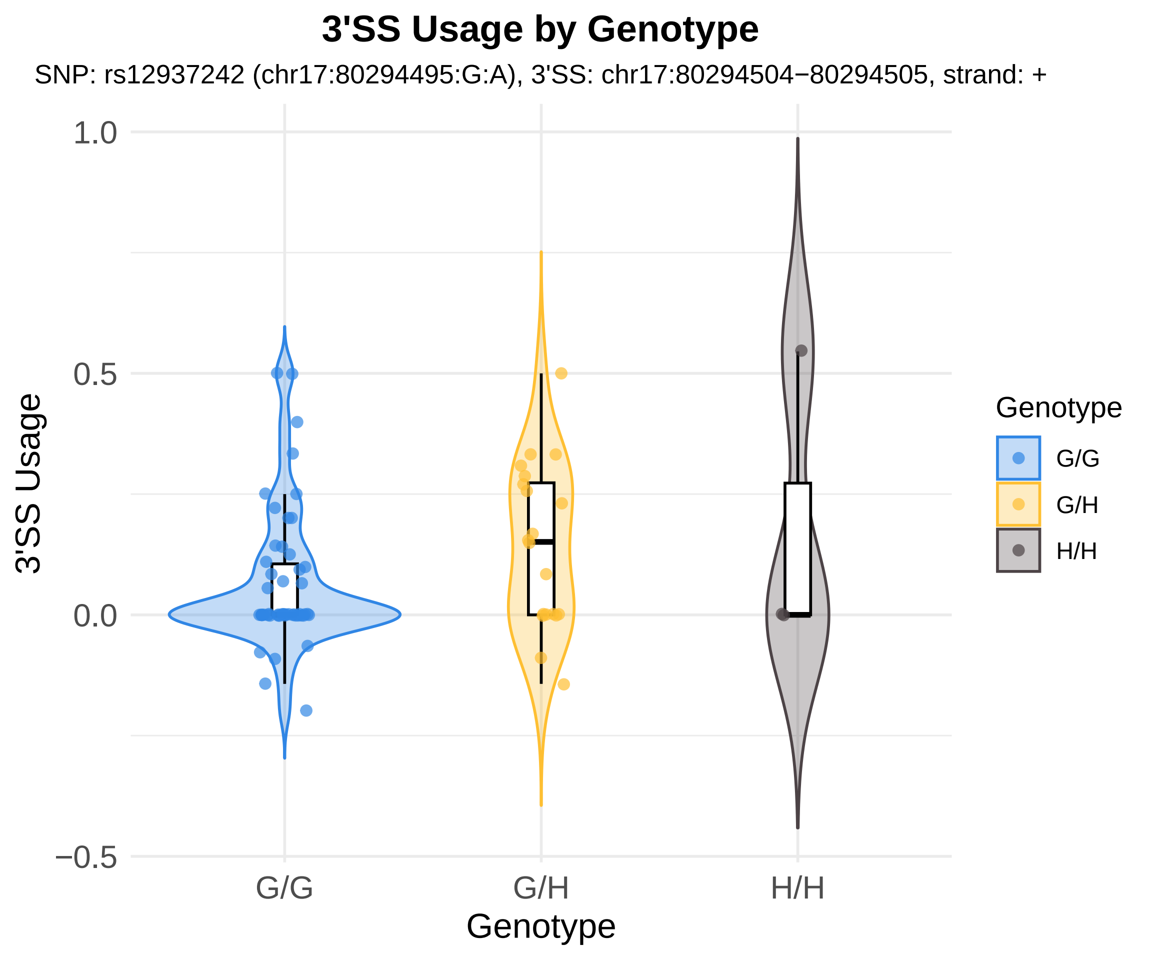

3'SS Usage by Genotype
SNP: rs12937242 (chr17:80294495:G:A), 3'SS: chr17:80294504−80294505, strand: +
1.0
0.5
0.0
−0.5
3'SS Usage
G/G
G/H
H/H
Genotype
Genotype
G/G
G/H
H/H

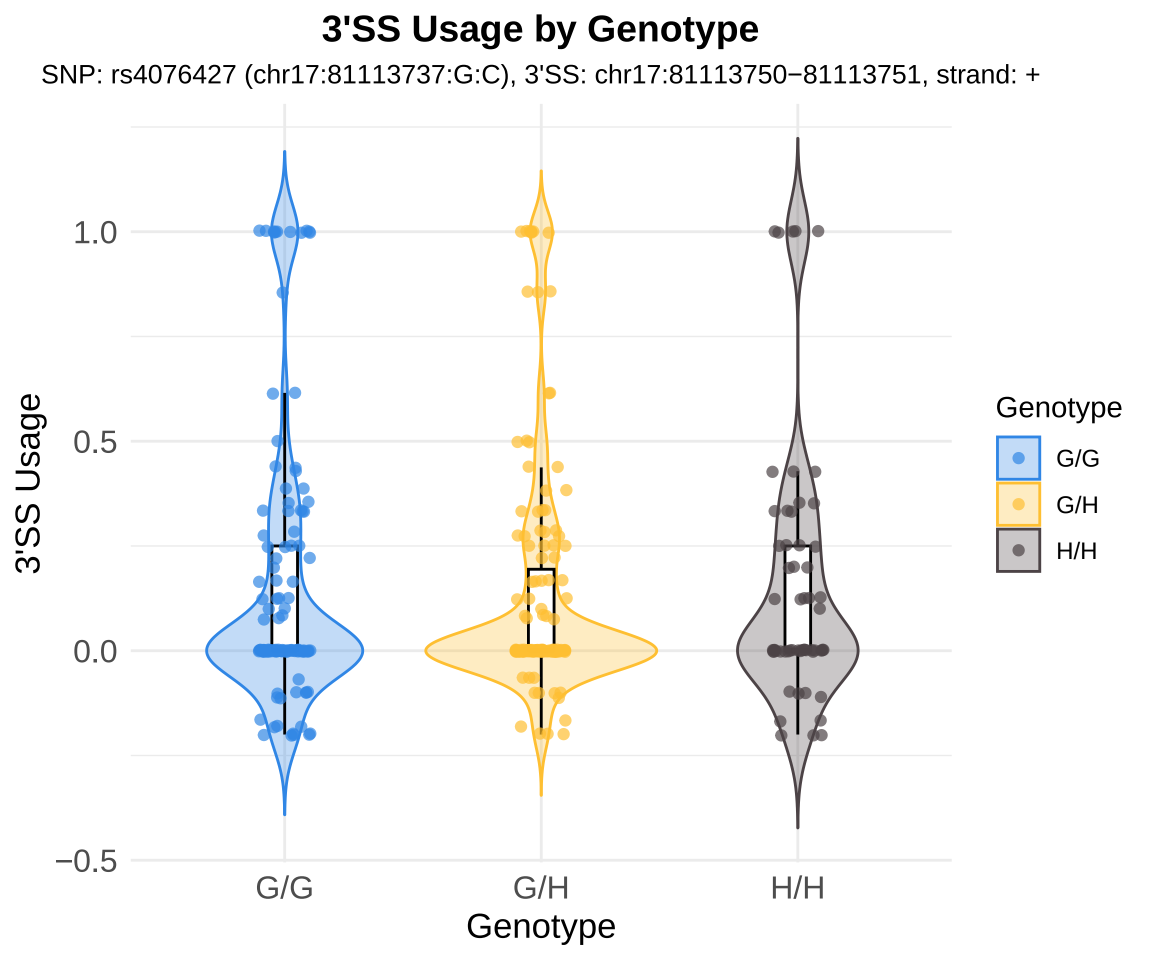
3'SS Usage by Genotype
SNP: rs4076427 (chr17:81113737:G:C), 3'SS: chr17:81113750−81113751, strand: +
3'SS Usage
1.0
0.5
0.0
−0.5
G/G
G/H
H/H
Genotype
Genotype
G/G
G/H
H/H

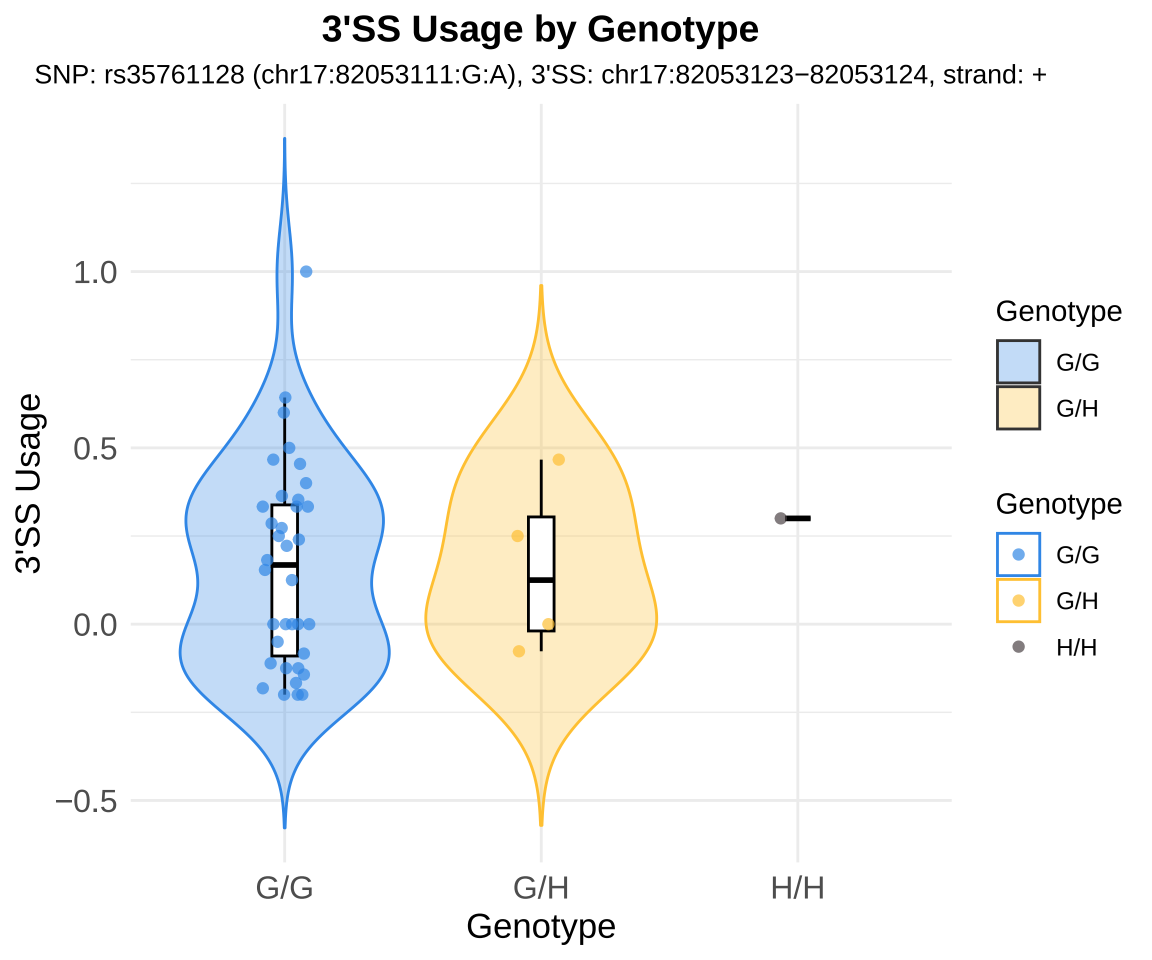
3'SS Usage by Genotype
SNP: rs35761128 (chr17:82053111:G:A), 3'SS: chr17:82053123−82053124, strand: +
3'SS Usage
1.0
0.5
0.0
−0.5
G/G
G/H
H/H
Genotype
Genotype
G/G
G/H
Genotype
G/G
G/H
H/H

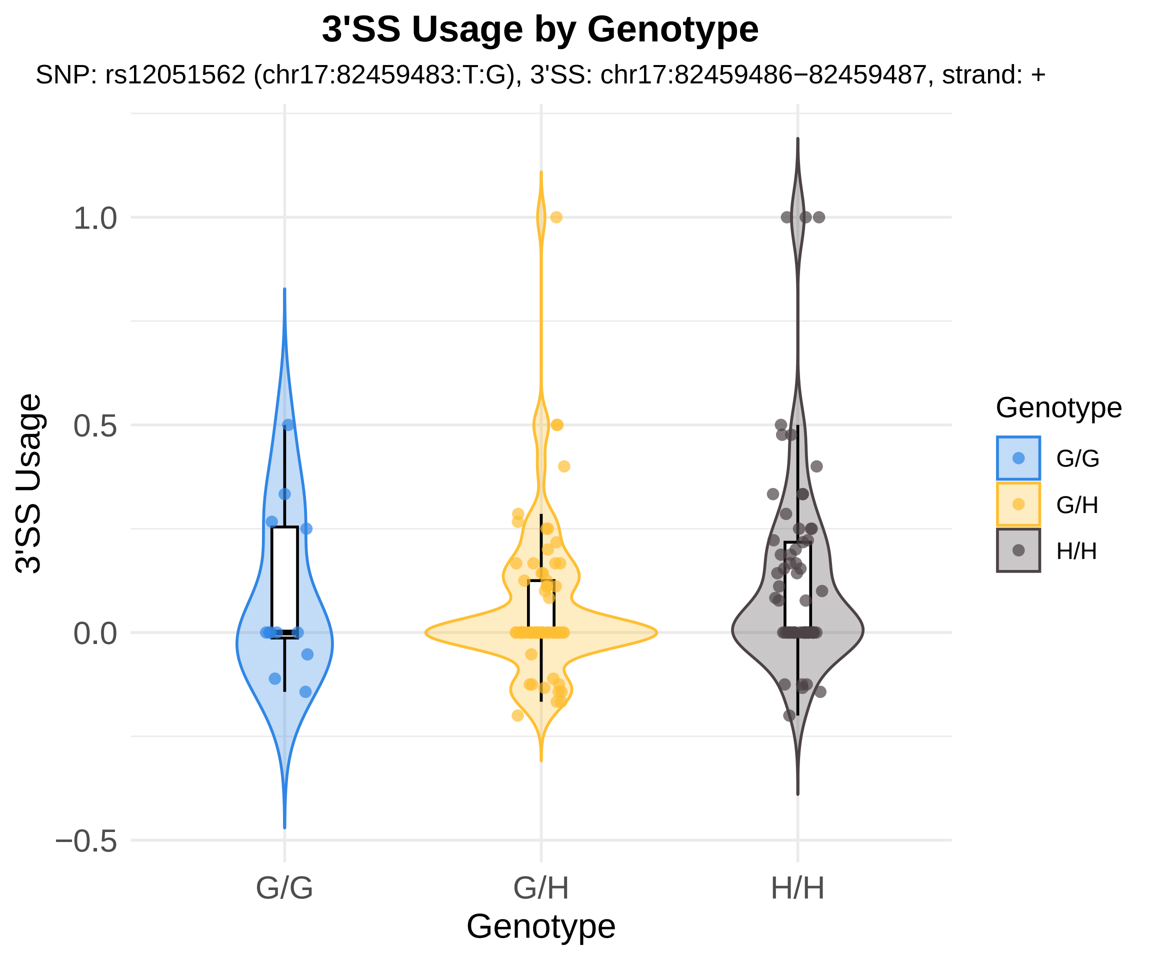

3'SS Usage by Genotype
SNP: rs12051562 (chr17:82459483:T:G), 3'SS: chr17:82459486−82459487, strand: +
3'SS Usage
1.0
0.5
0.0
−0.5
G/G
G/H
H/H
Genotype
Genotype
G/G
G/H
H/H

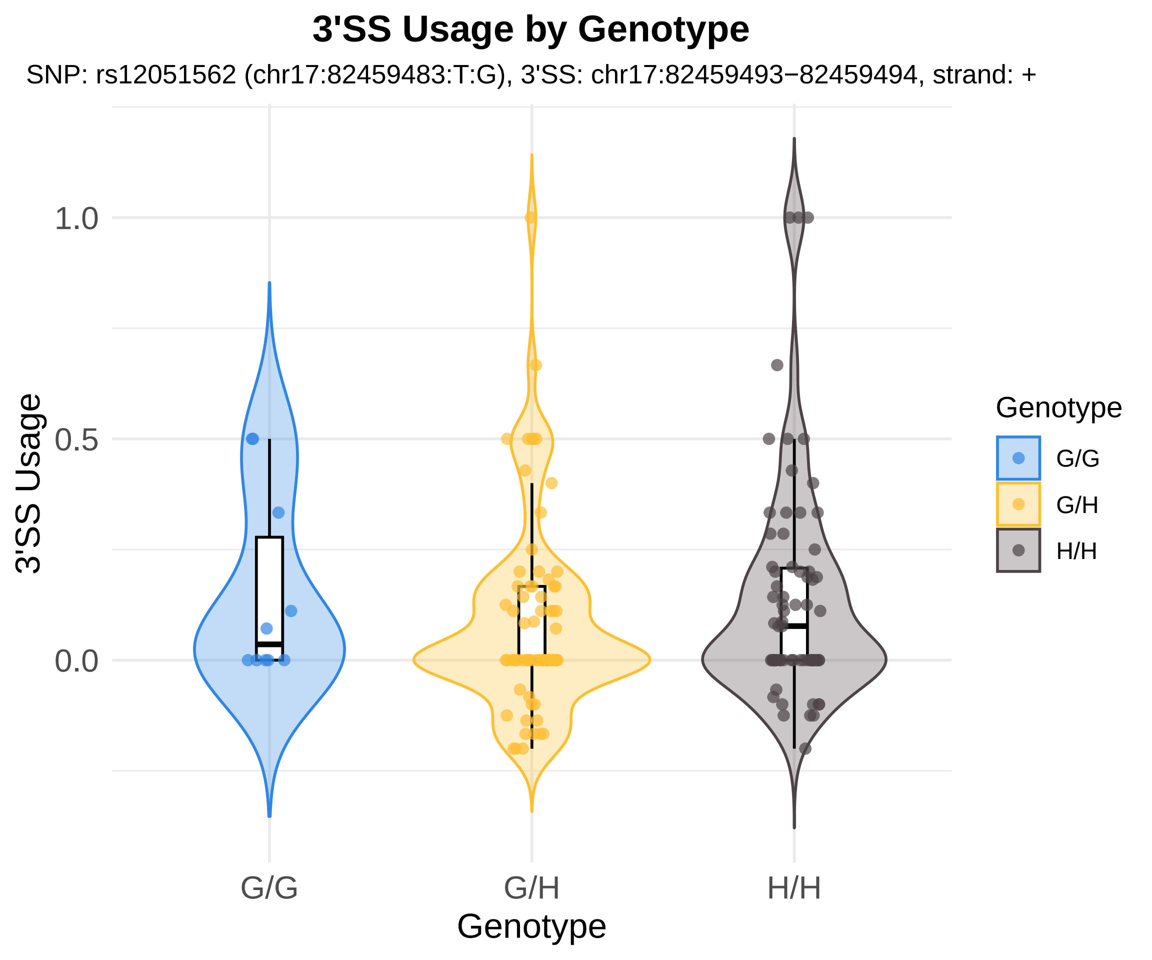
3'SS Usage by Genotype
SNP: rs12051562 (chr17:82459483:T:G), 3'SS: chr17:82459493−82459494, strand: +
3'SS Usage
1.0
0.5
0.0
G/G
G/H
H/H
Genotype
Genotype
G/G
G/H
H/H

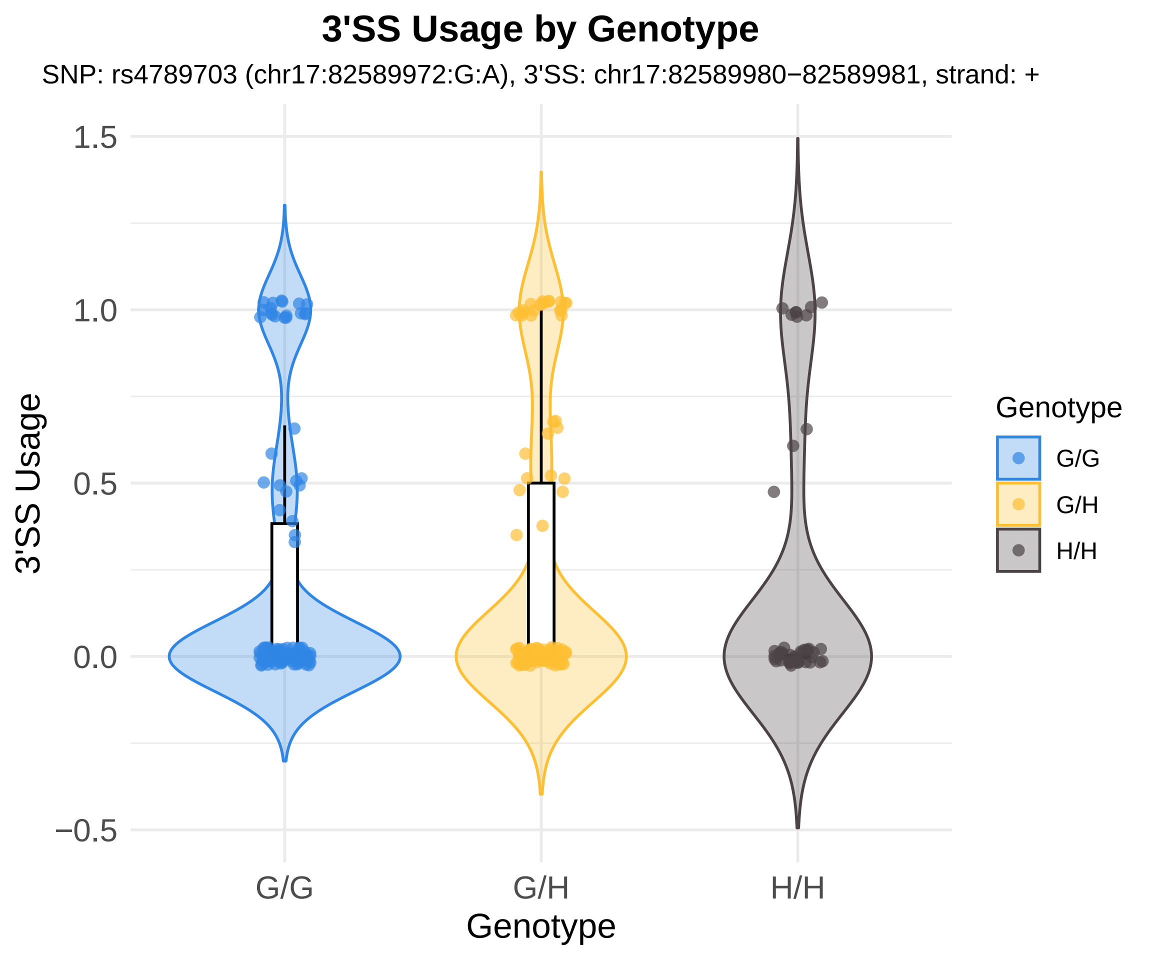

3'SS Usage by Genotype
SNP: rs4789703 (chr17:82589972:G:A), 3'SS: chr17:82589980−82589981, strand: +
3'SS Usage
1.5
1.0
0.5
0.0
−0.5
G/G
G/H
H/H
Genotype
Genotype
G/G
G/H
H/H

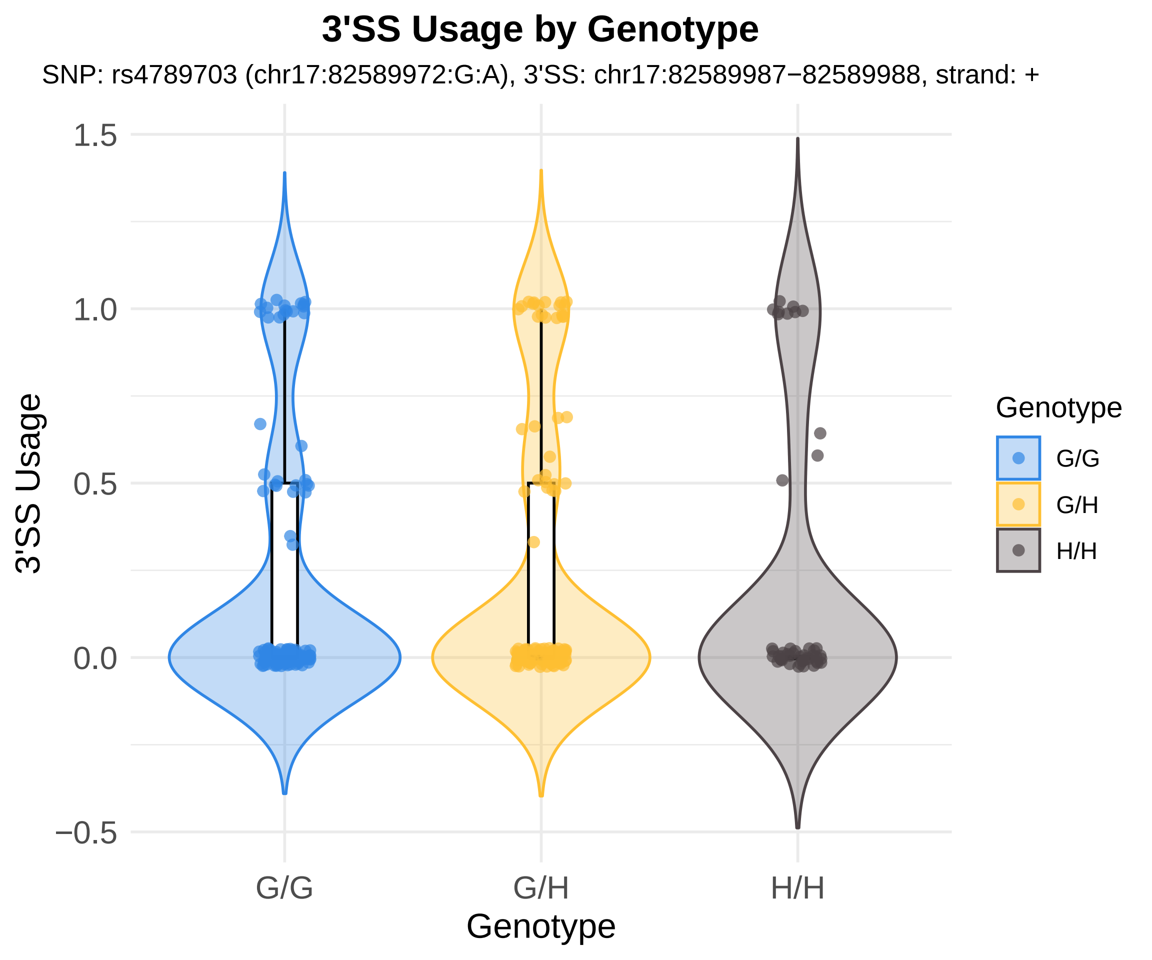
3'SS Usage by Genotype
SNP: rs4789703 (chr17:82589972:G:A), 3'SS: chr17:82589987−82589988, strand: +
3'SS Usage
1.5
1.0
0.5
0.0
−0.5
G/G
G/H
H/H
Genotype
Genotype
G/G
G/H
H/H

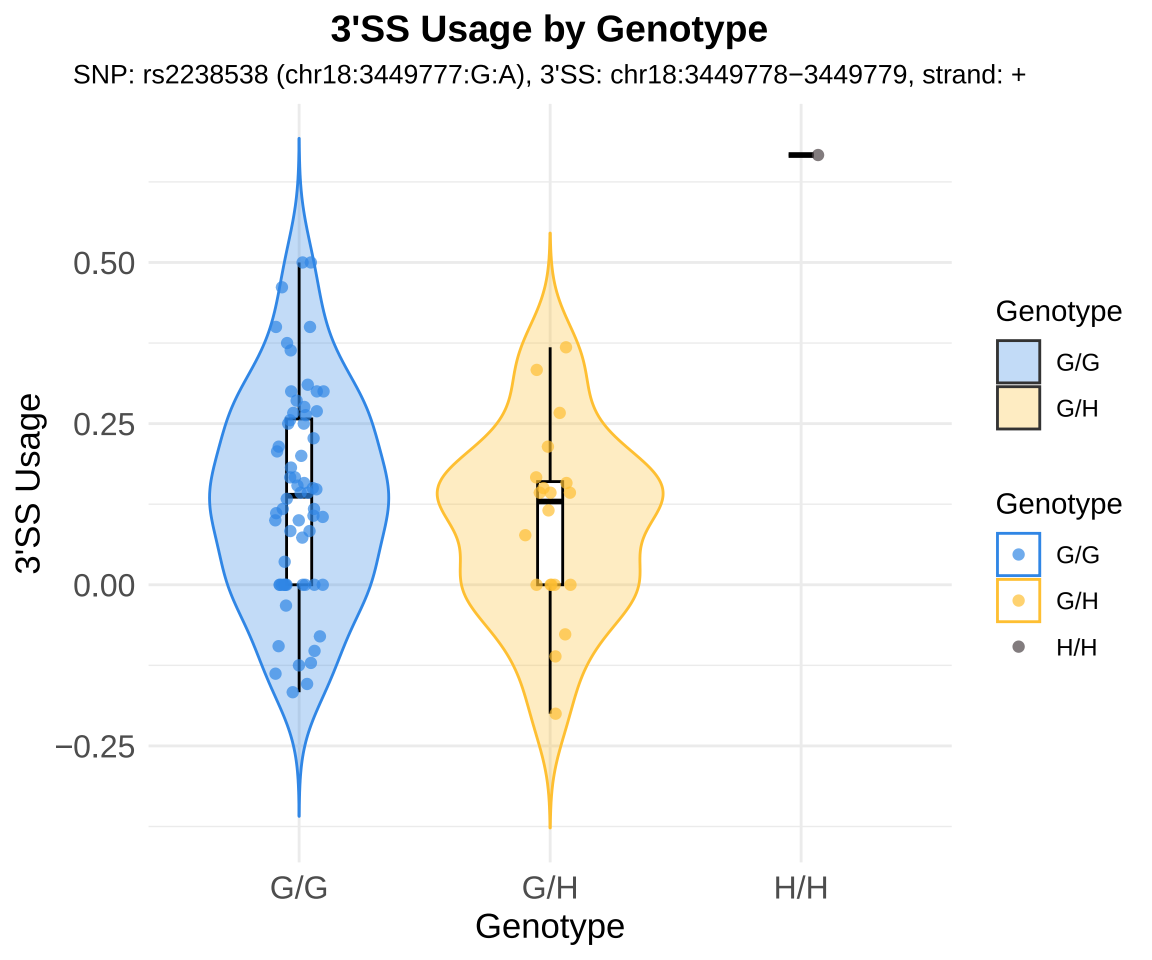
3'SS Usage by Genotype
SNP: rs2238538 (chr18:3449777:G:A), 3'SS: chr18:3449778−3449779, strand: +
3'SS Usage
0.50
0.25
0.00
−0.25
G/G
G/H
H/H
Genotype
Genotype
G/G
G/H
Genotype
G/G
G/H
H/H

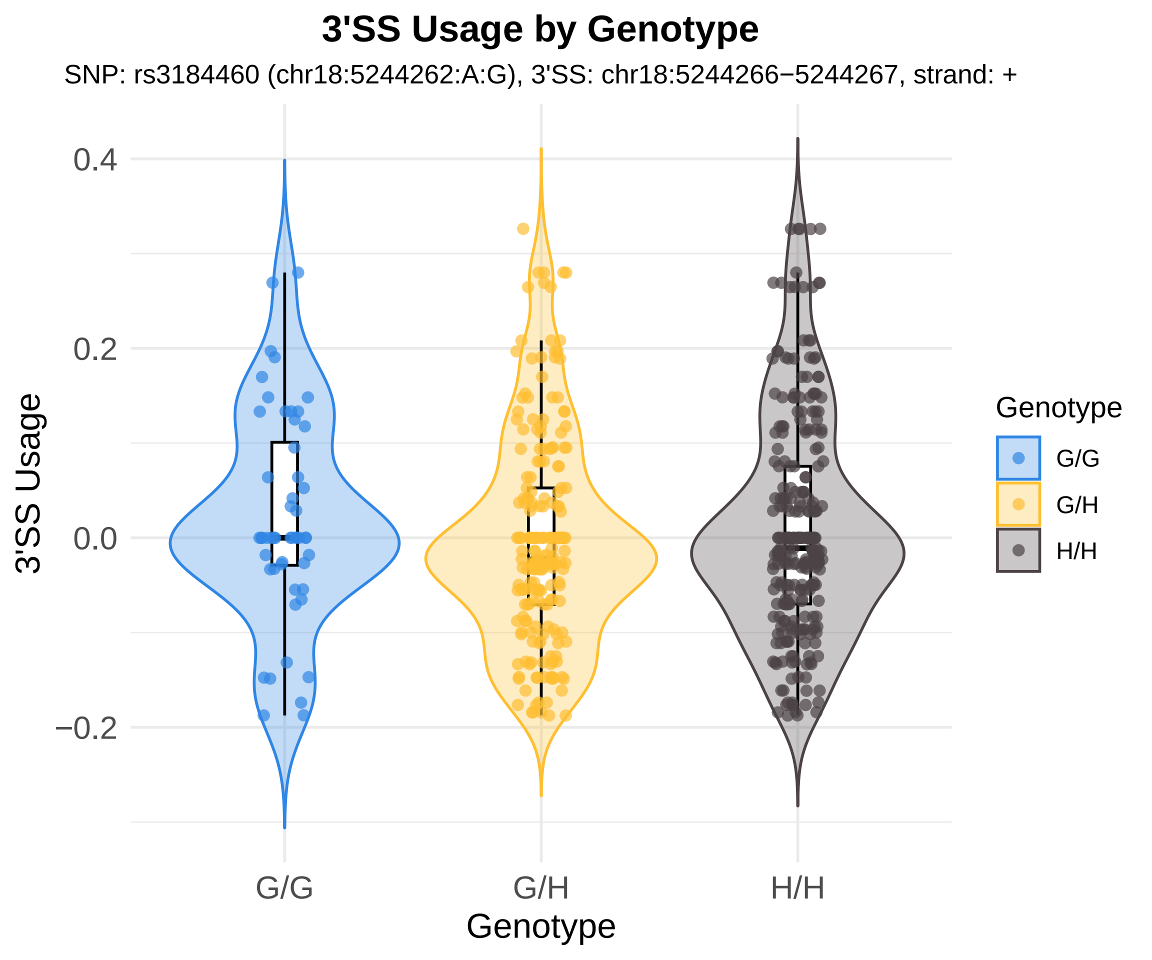

3'SS Usage by Genotype
SNP: rs3184460 (chr18:5244262:A:G), 3'SS: chr18:5244266−5244267, strand: +
0.4
0.2
0.0
−0.2
3'SS Usage
G/G
G/H
H/H
Genotype
Genotype
G/G
G/H
H/H

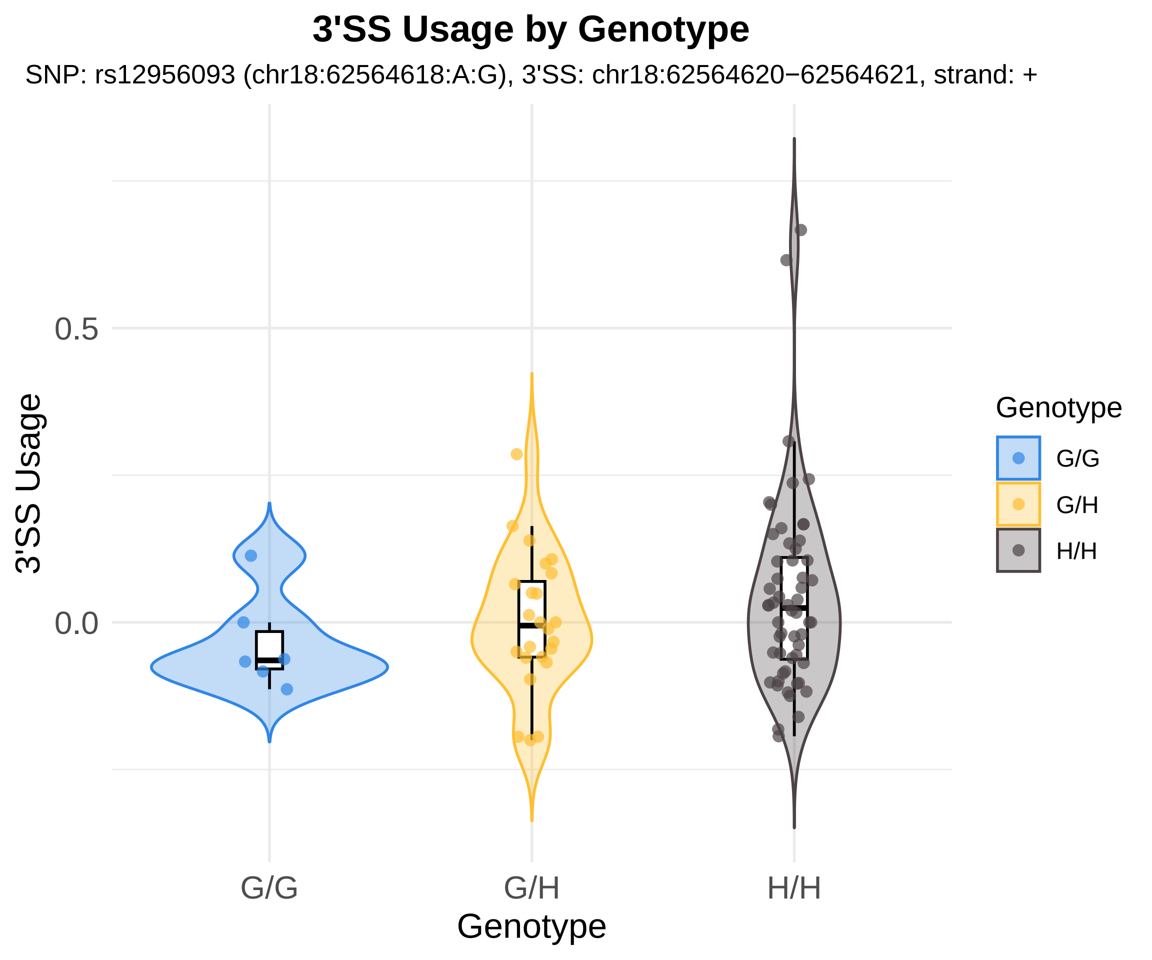

3'SS Usage by Genotype
SNP: rs12956093 (chr18:62564618:A:G), 3'SS: chr18:62564620−62564621, strand: +
3'SS Usage
0.5
0.0
G/G
G/H
H/H
Genotype
Genotype
G/G
G/H
H/H

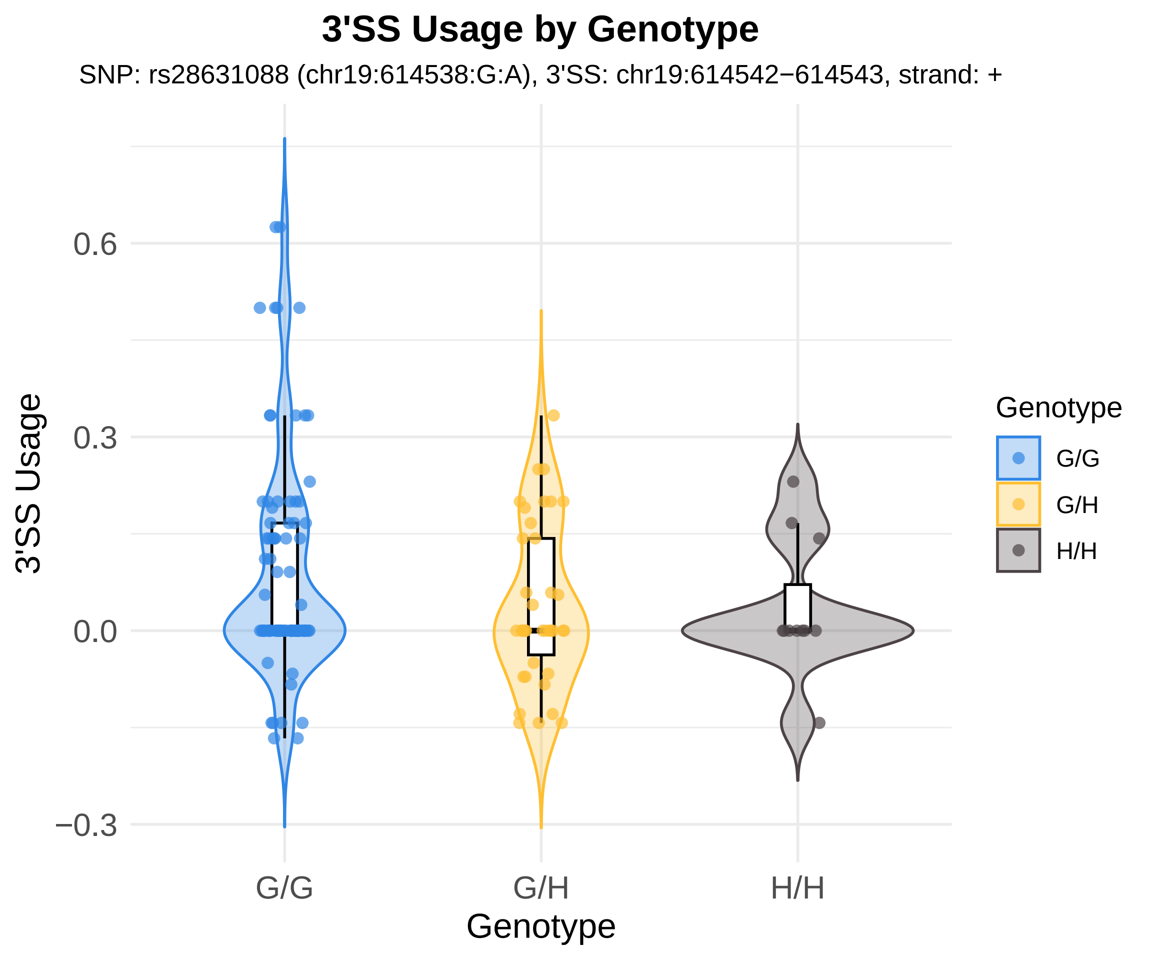
3'SS Usage by Genotype
SNP: rs28631088 (chr19:614538:G:A), 3'SS: chr19:614542−614543, strand: +
3'SS Usage
0.6
0.3
0.0
−0.3
G/G
G/H
H/H
Genotype
Genotype
G/G
G/H
H/H

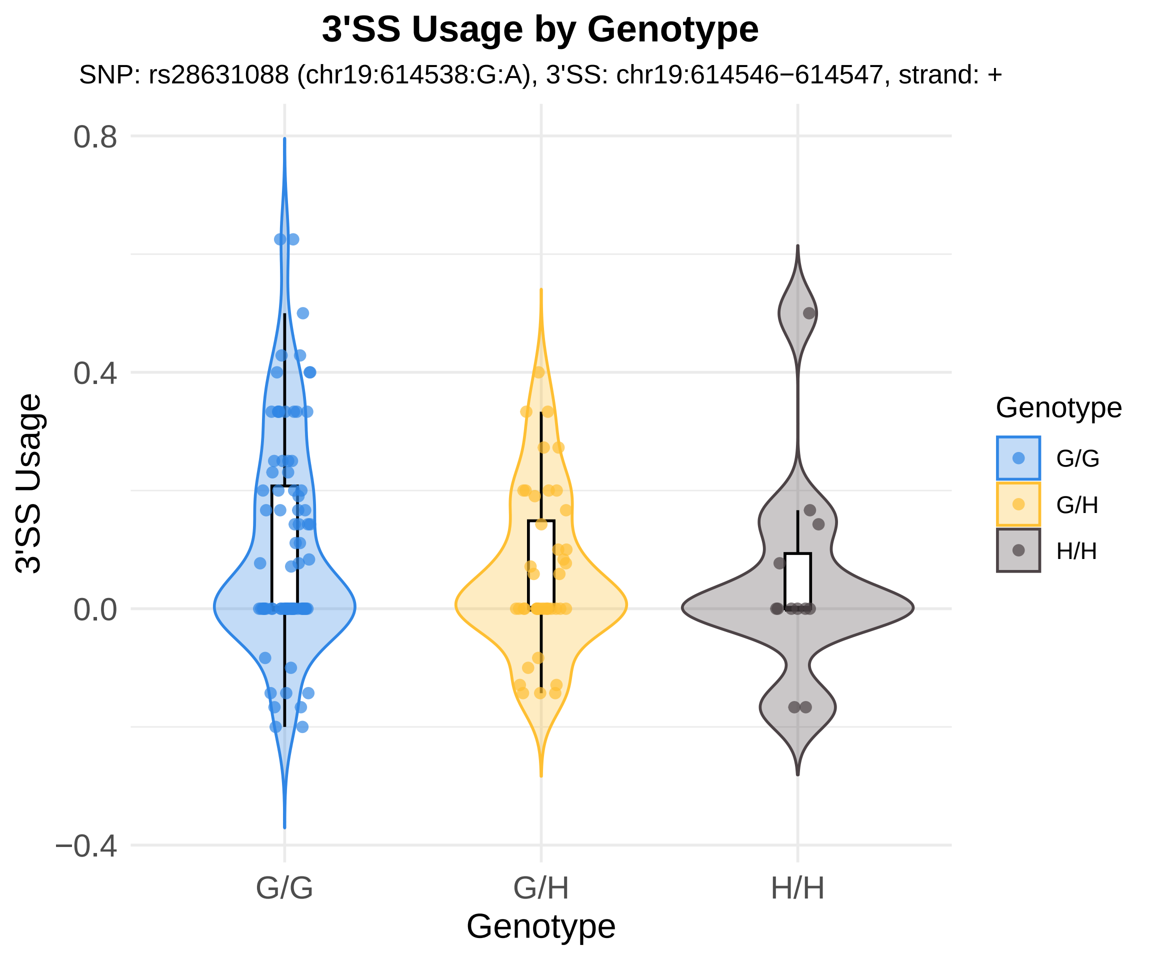
3'SS Usage by Genotype
SNP: rs28631088 (chr19:614538:G:A), 3'SS: chr19:614546−614547, strand: +
0.8
0.4
0.0
−0.4
3'SS Usage
G/G
G/H
H/H
Genotype
Genotype
G/G
G/H
H/H

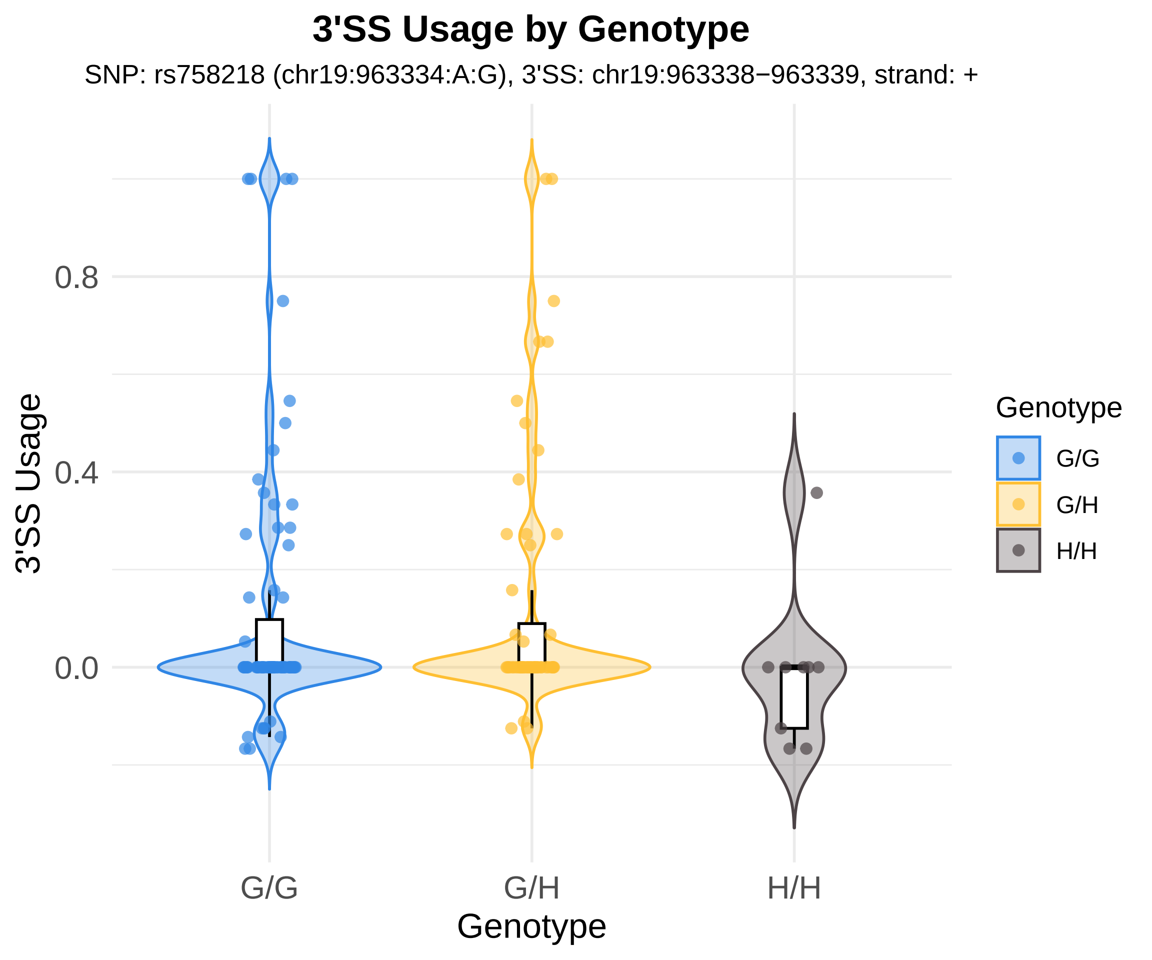
3'SS Usage by Genotype
SNP: rs758218 (chr19:963334:A:G), 3'SS: chr19:963338−963339, strand: +
3'SS Usage
0.8
0.4
0.0
G/G
G/H
H/H
Genotype
Genotype
G/G
G/H
H/H

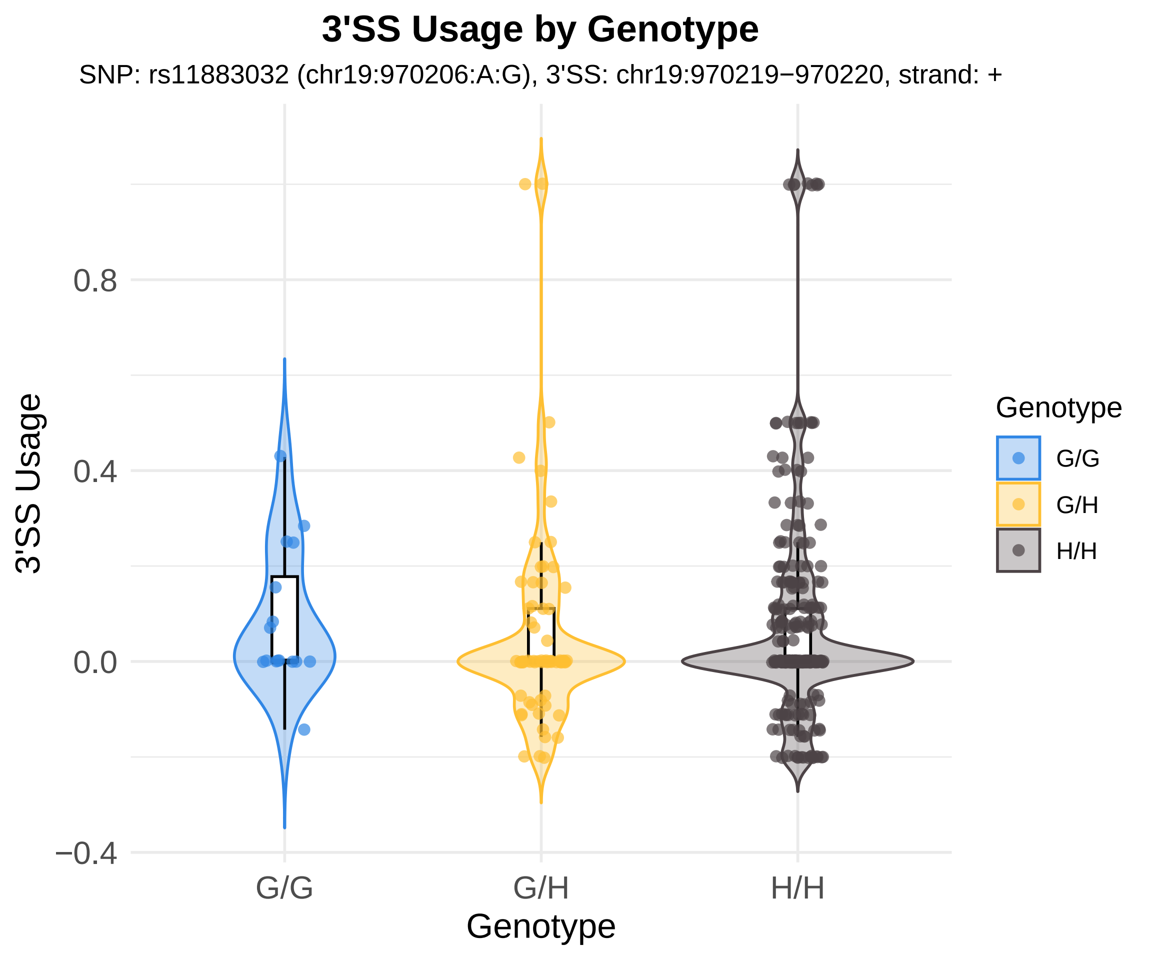
3'SS Usage by Genotype
SNP: rs11883032 (chr19:970206:A:G), 3'SS: chr19:970219−970220, strand: +
3'SS Usage
0.8
0.4
0.0
−0.4
G/G
G/H
H/H
Genotype
Genotype
G/G
G/H
H/H

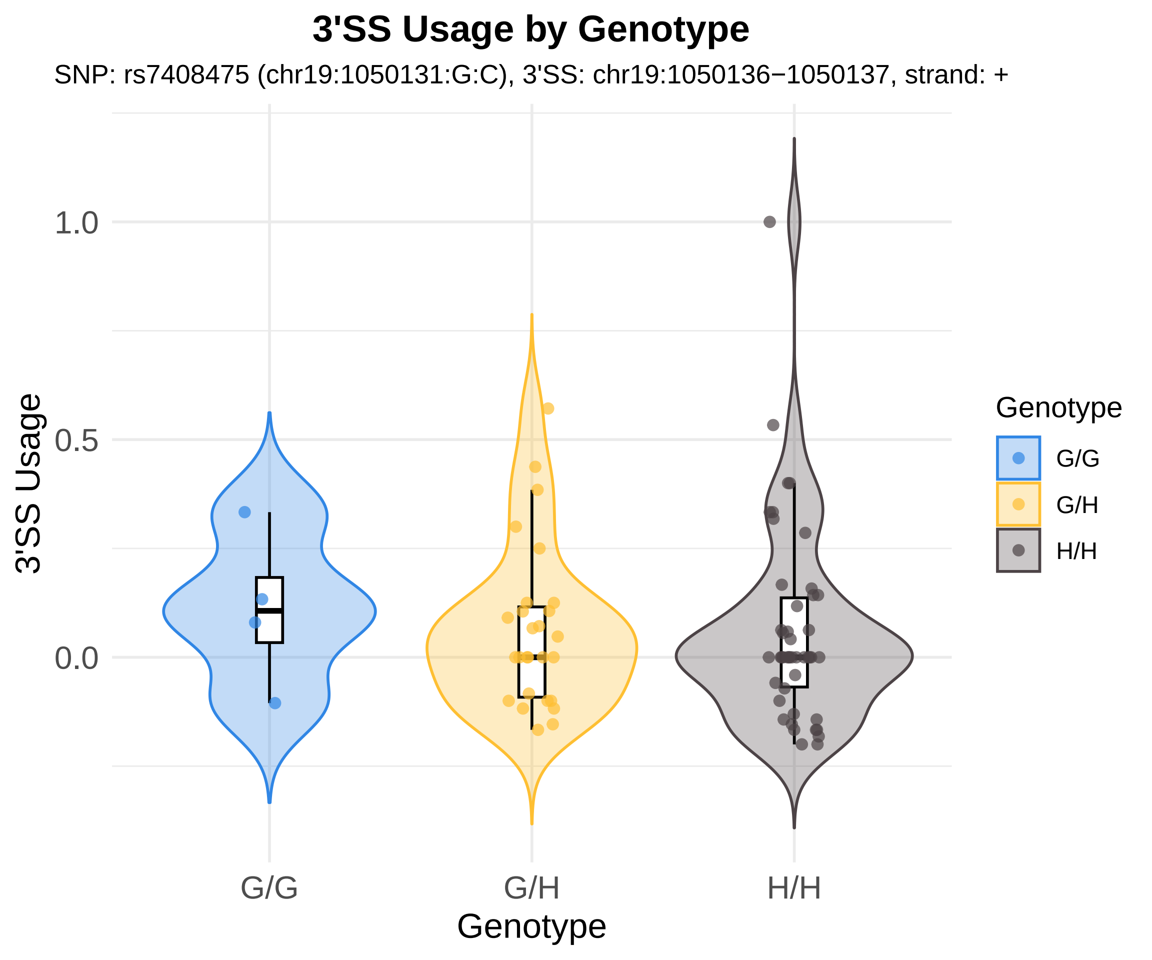
3'SS Usage by Genotype
SNP: rs7408475 (chr19:1050131:G:C), 3'SS: chr19:1050136−1050137, strand: +
3'SS Usage
1.0
0.5
0.0
G/G
G/H
H/H
Genotype
Genotype
G/G
G/H
H/H

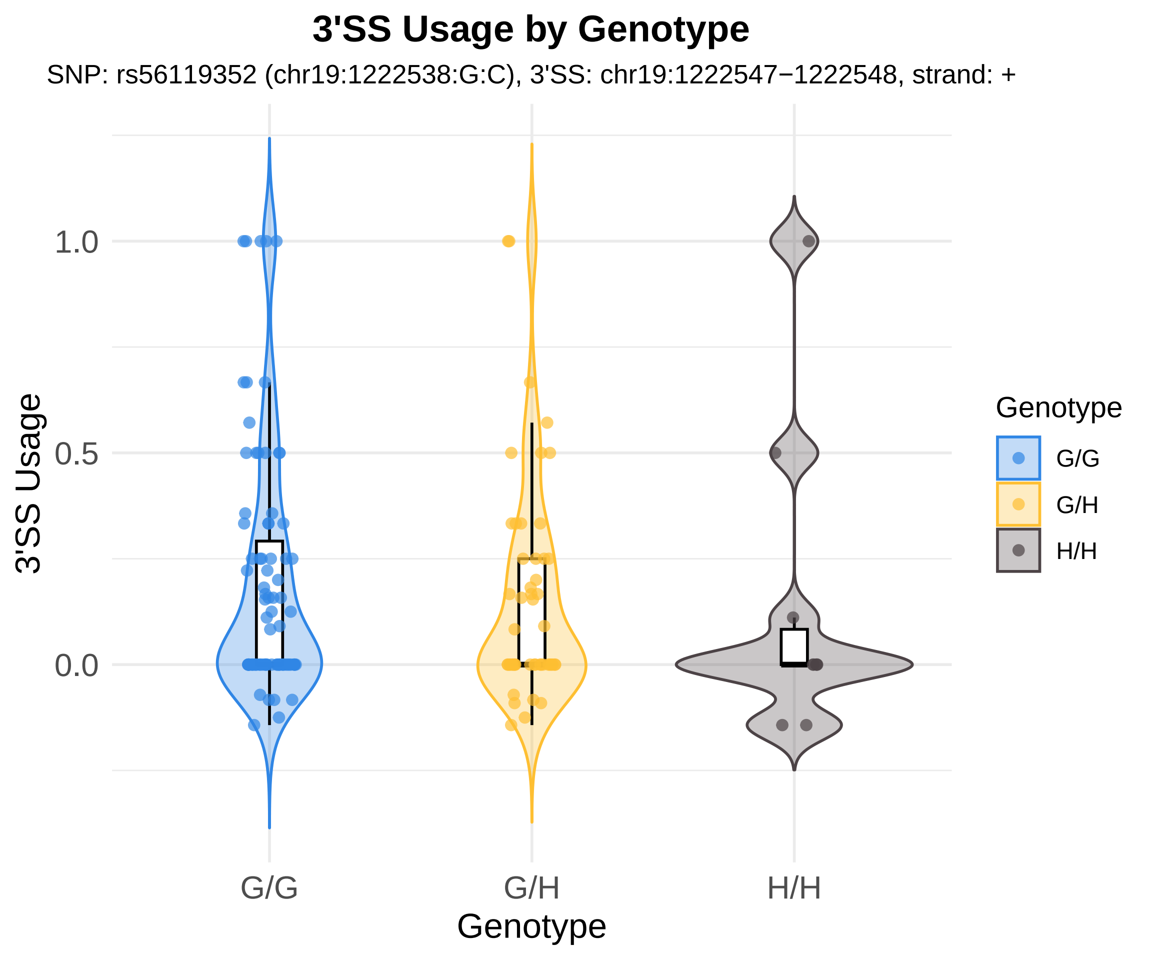

3'SS Usage by Genotype
SNP: rs56119352 (chr19:1222538:G:C), 3'SS: chr19:1222547−1222548, strand: +
3'SS Usage
1.0
0.5
0.0
G/G
G/H
H/H
Genotype
Genotype
G/G
G/H
H/H

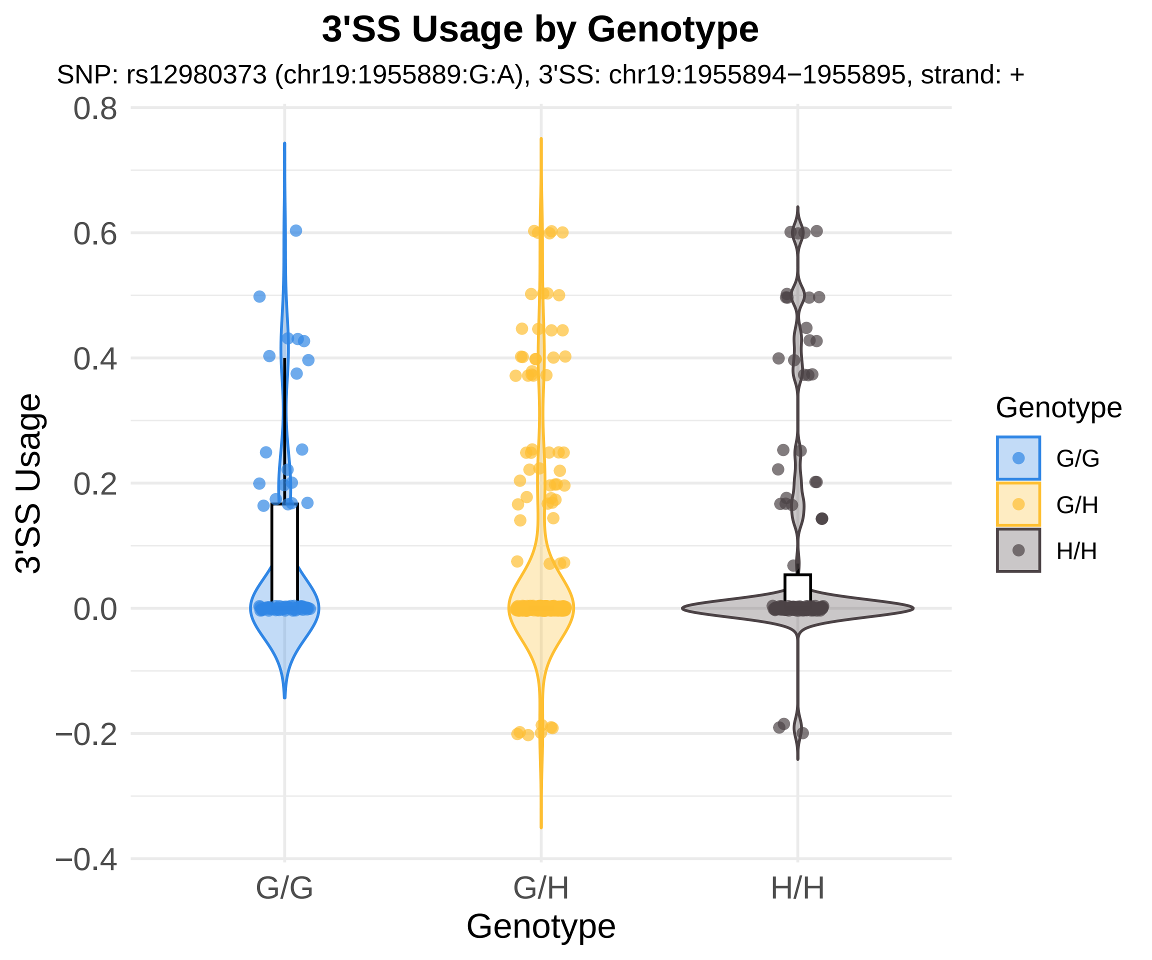
3'SS Usage by Genotype
SNP: rs12980373 (chr19:1955889:G:A), 3'SS: chr19:1955894−1955895, strand: +
3'SS Usage
0.8
0.6
0.4
0.2
0.0
−0.2
−0.4
G/G
G/H
H/H
Genotype
Genotype
G/G
G/H
H/H

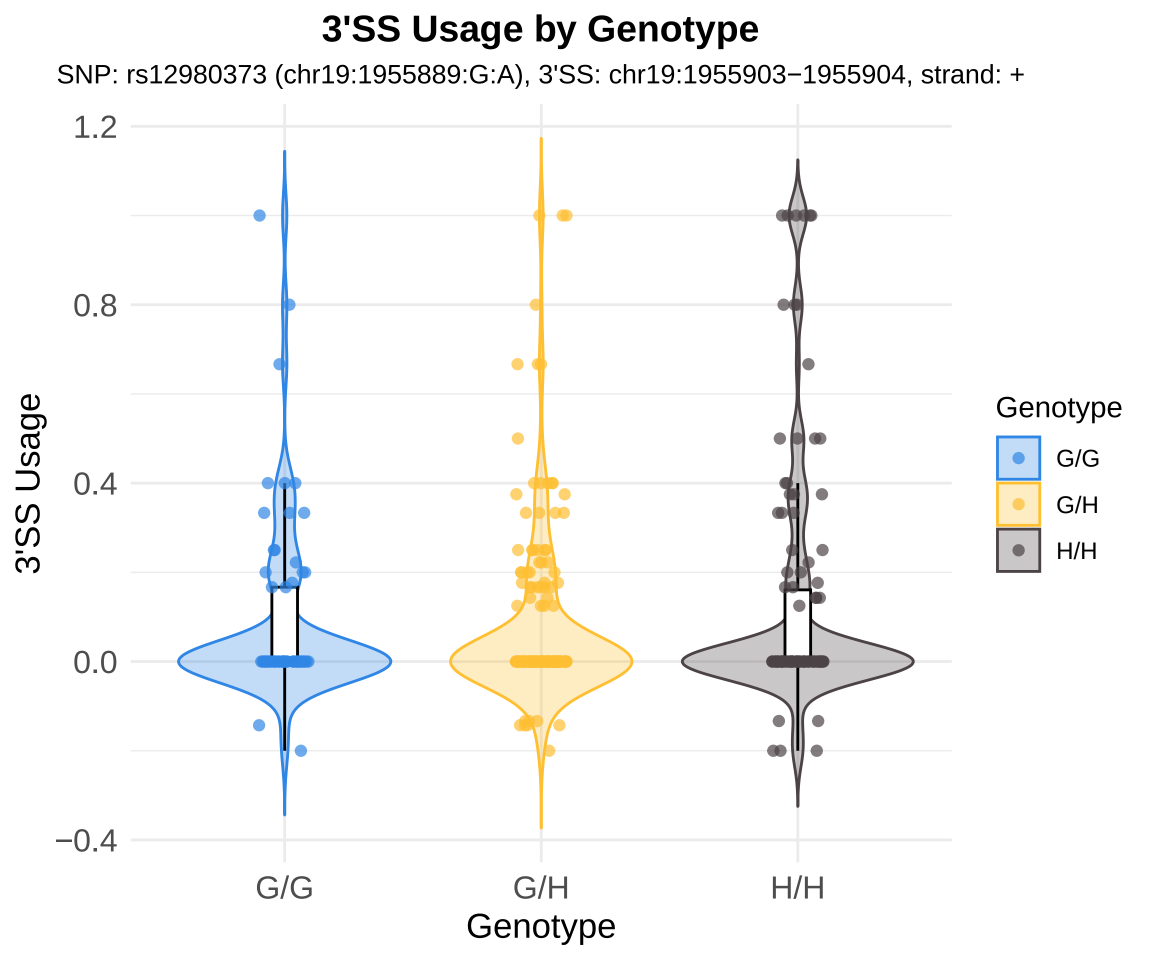
3'SS Usage by Genotype
SNP: rs12980373 (chr19:1955889:G:A), 3'SS: chr19:1955903−1955904, strand: +
3'SS Usage
1.2
0.8
0.4
0.0
−0.4
G/G
G/H
H/H
Genotype
Genotype
G/G
G/H
H/H

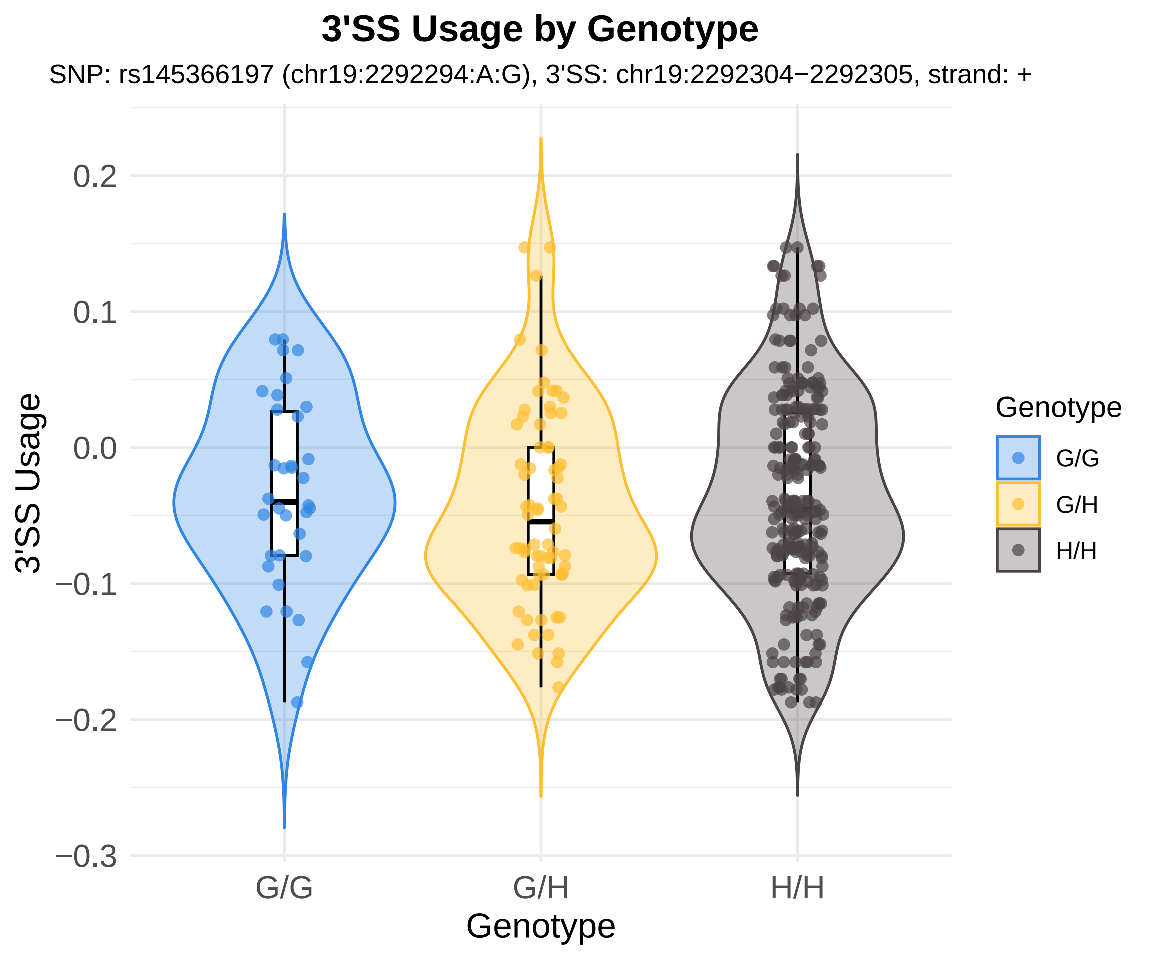
3'SS Usage by Genotype
SNP: rs145366197 (chr19:2292294:A:G), 3'SS: chr19:2292304−2292305, strand: +
3'SS Usage
0.2
0.1
0.0
−0.1
−0.2
−0.3
G/G
G/H
H/H
Genotype
Genotype
G/G
G/H
H/H

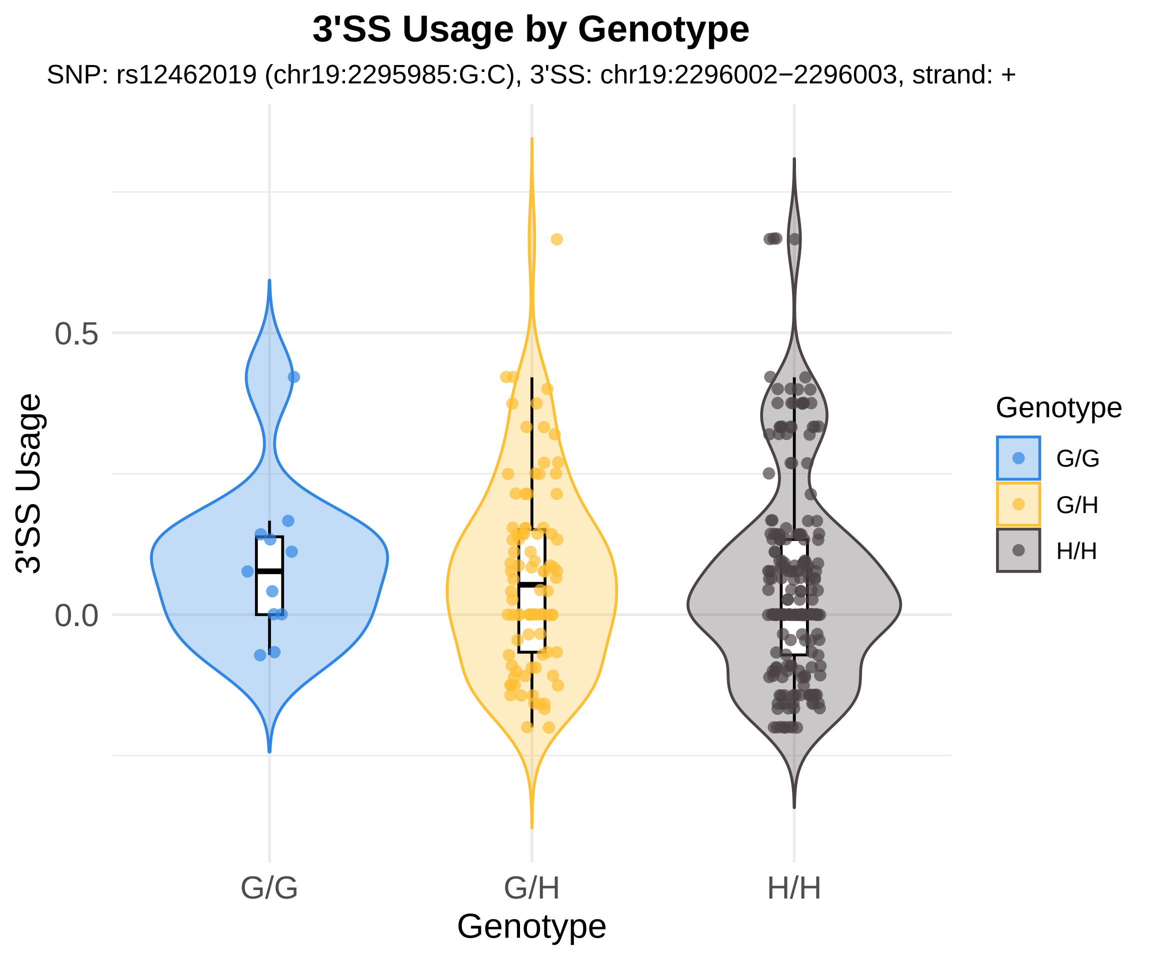

3'SS Usage by Genotype
SNP: rs12462019 (chr19:2295985:G:C), 3'SS: chr19:2296002−2296003, strand: +
3'SS Usage
0.5
0.0
G/G
G/H
H/H
Genotype
Genotype
G/G
G/H
H/H

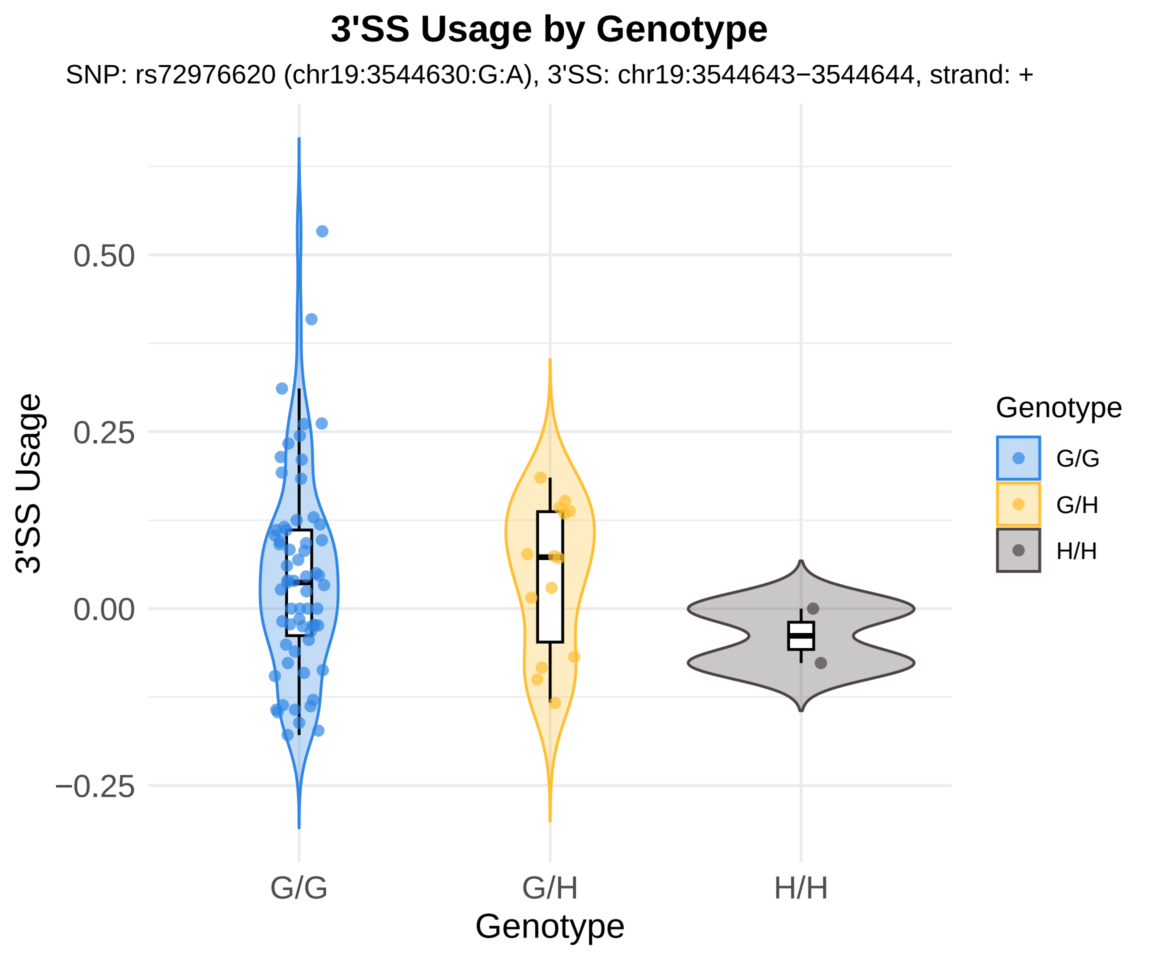
3'SS Usage by Genotype
SNP: rs72976620 (chr19:3544630:G:A), 3'SS: chr19:3544643−3544644, strand: +
3'SS Usage
0.50
0.25
0.00
−0.25
G/G
G/H
H/H
Genotype
Genotype
G/G
G/H
H/H

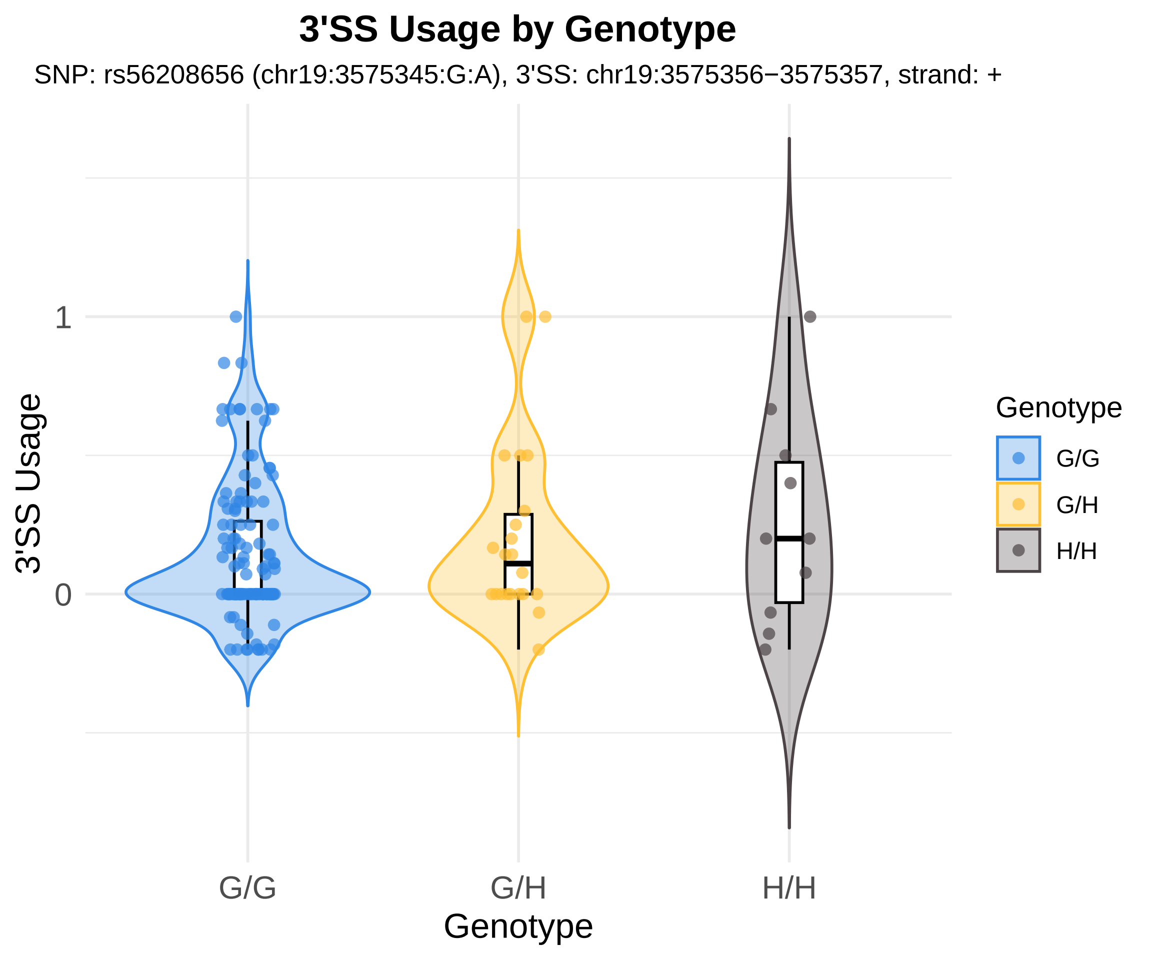

3'SS Usage by Genotype
SNP: rs56208656 (chr19:3575345:G:A), 3'SS: chr19:3575356−3575357, strand: +
3'SS Usage
1
0
G/G
G/H
H/H
Genotype
Genotype
G/G
G/H
H/H

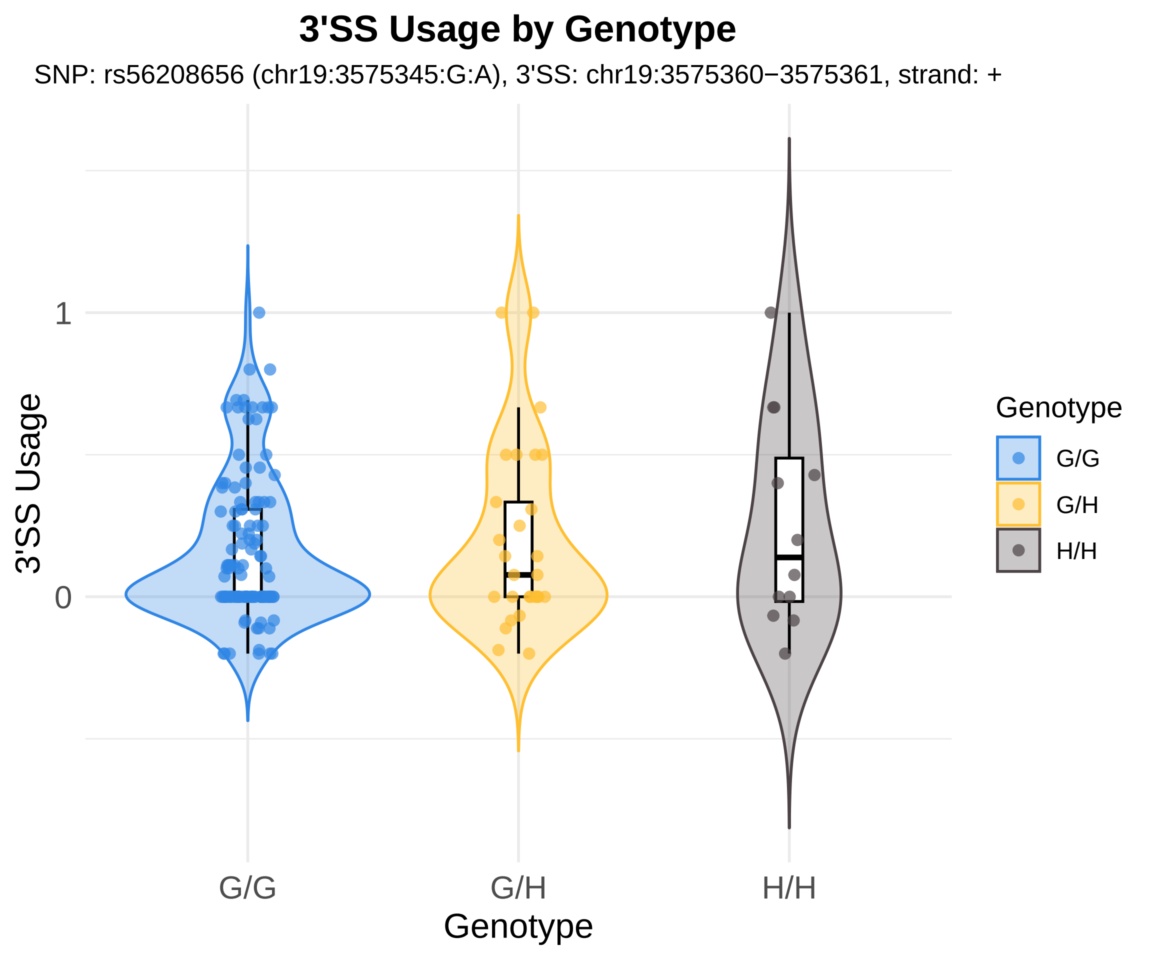
3'SS Usage by Genotype
SNP: rs56208656 (chr19:3575345:G:A), 3'SS: chr19:3575360−3575361, strand: +
3'SS Usage
1
0
G/G
G/H
H/H
Genotype
Genotype
G/G
G/H
H/H

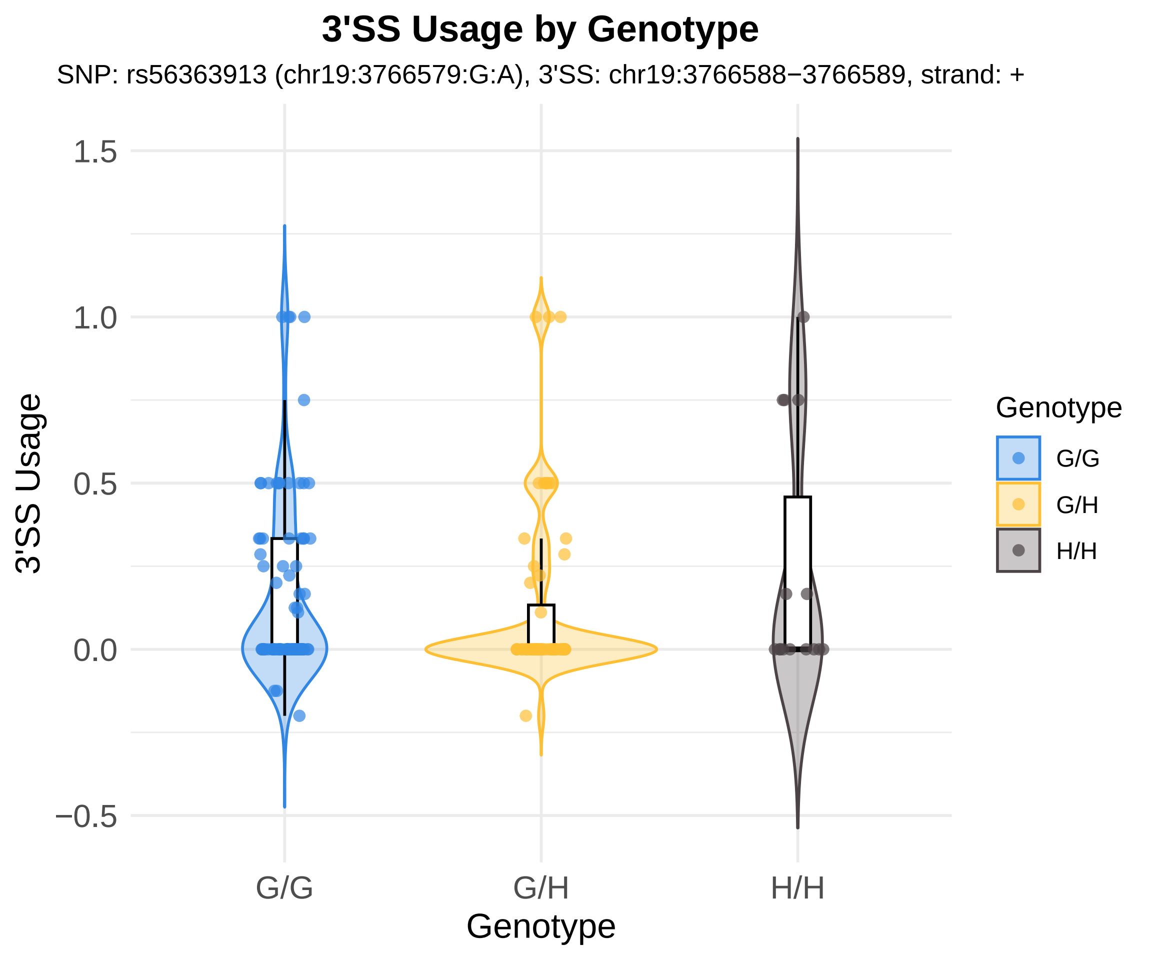
3'SS Usage by Genotype
SNP: rs56363913 (chr19:3766579:G:A), 3'SS: chr19:3766588−3766589, strand: +
3'SS Usage
1.5
1.0
0.5
0.0
−0.5
G/G
G/H
H/H
Genotype
Genotype
G/G
G/H
H/H

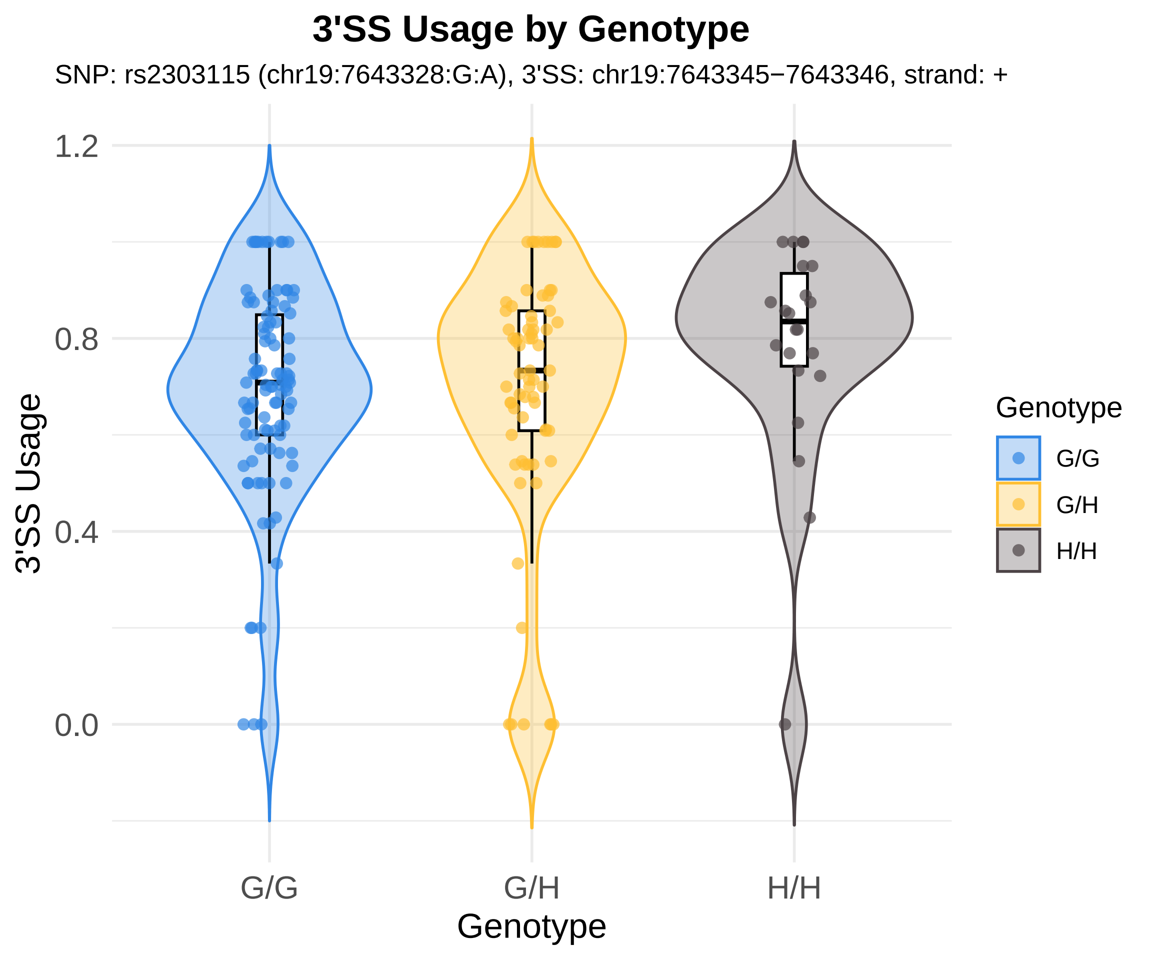
3'SS Usage by Genotype
SNP: rs2303115 (chr19:7643328:G:A), 3'SS: chr19:7643345−7643346, strand: +
3'SS Usage
1.2
0.8
0.4
0.0
G/G
G/H
H/H
Genotype
Genotype
G/G
G/H
H/H

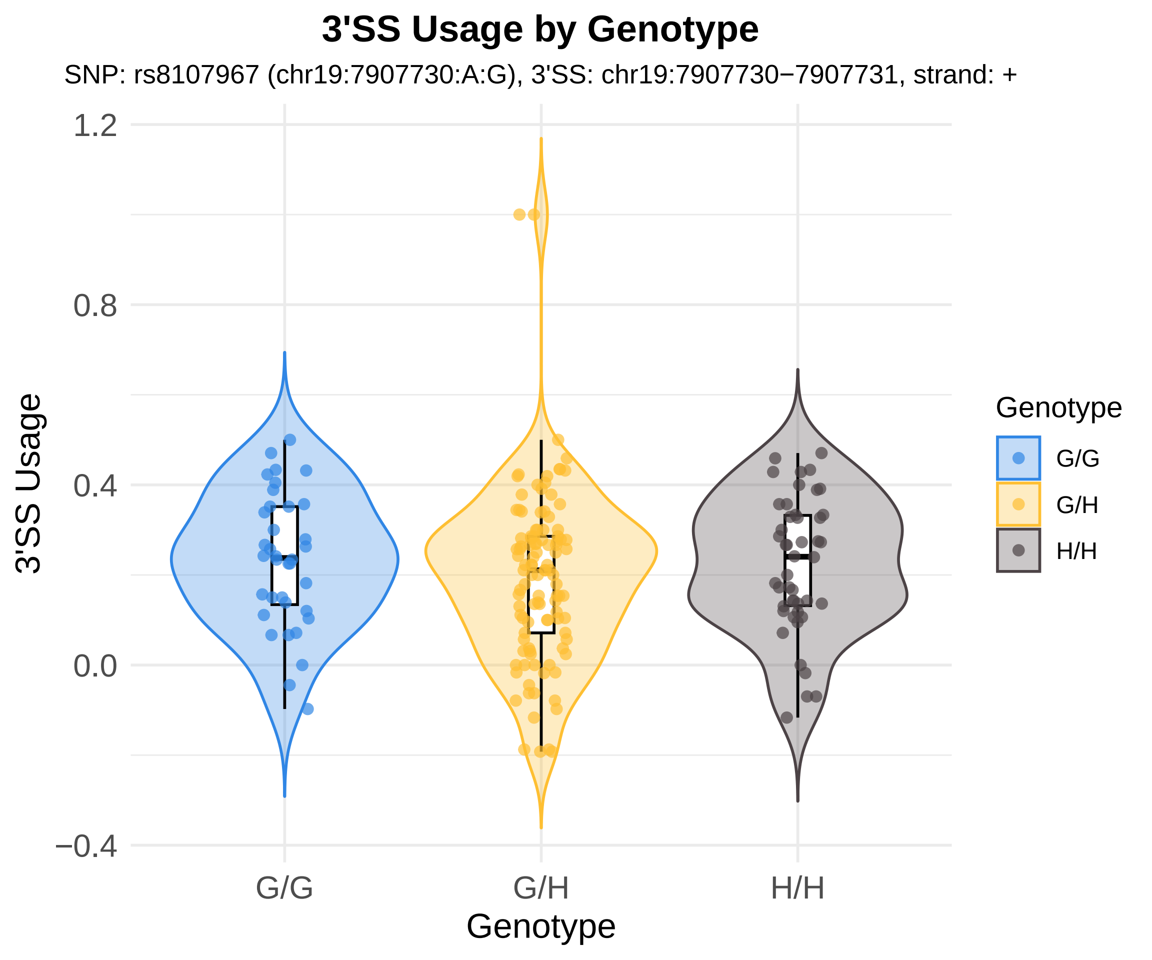

3'SS Usage by Genotype
SNP: rs8107967 (chr19:7907730:A:G), 3'SS: chr19:7907730−7907731, strand: +
3'SS Usage
1.2
0.8
0.4
0.0
−0.4
G/G
G/H
H/H
Genotype
Genotype
G/G
G/H
H/H

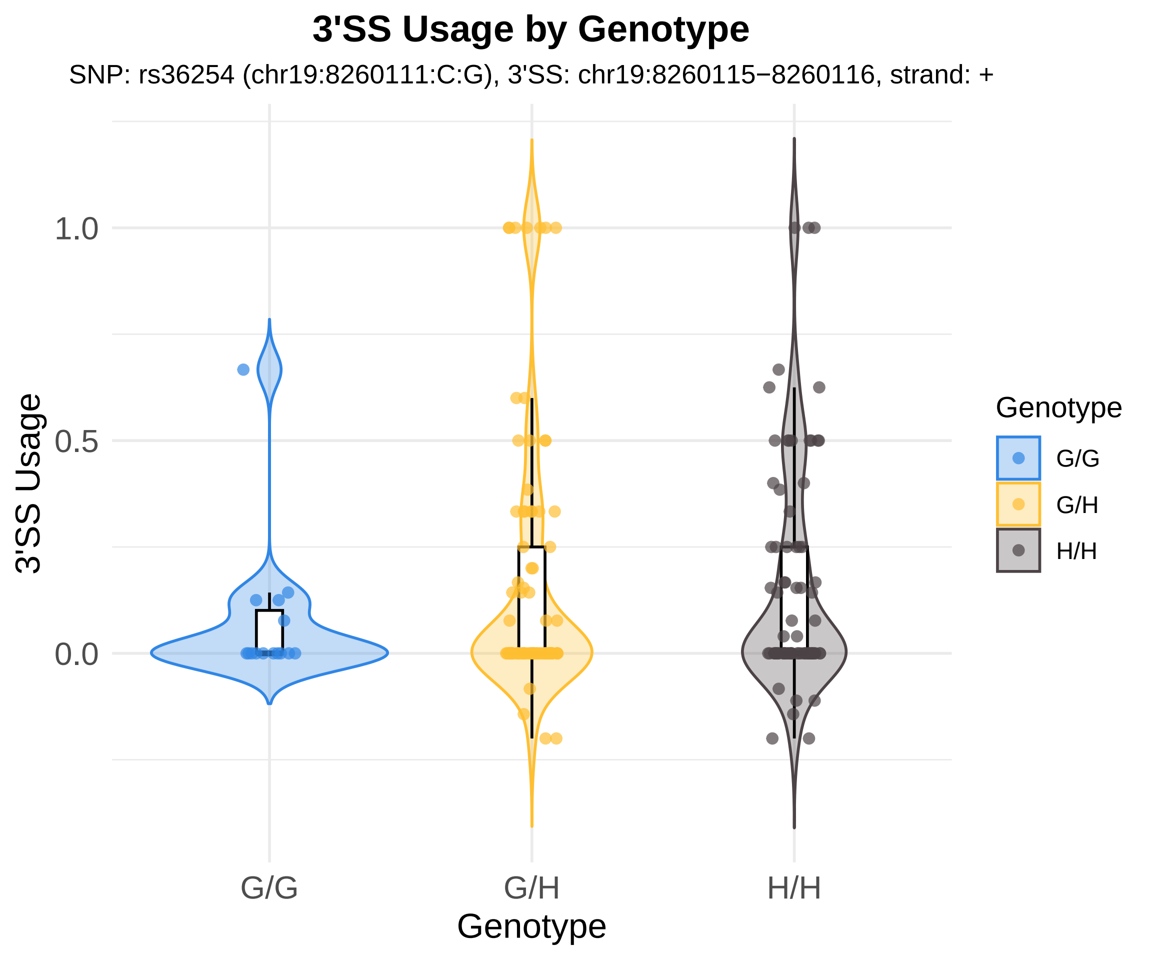

3'SS Usage by Genotype
SNP: rs36254 (chr19:8260111:C:G), 3'SS: chr19:8260115−8260116, strand: +
3'SS Usage
1.0
0.5
0.0
G/G
G/H
H/H
Genotype
Genotype
G/G
G/H
H/H

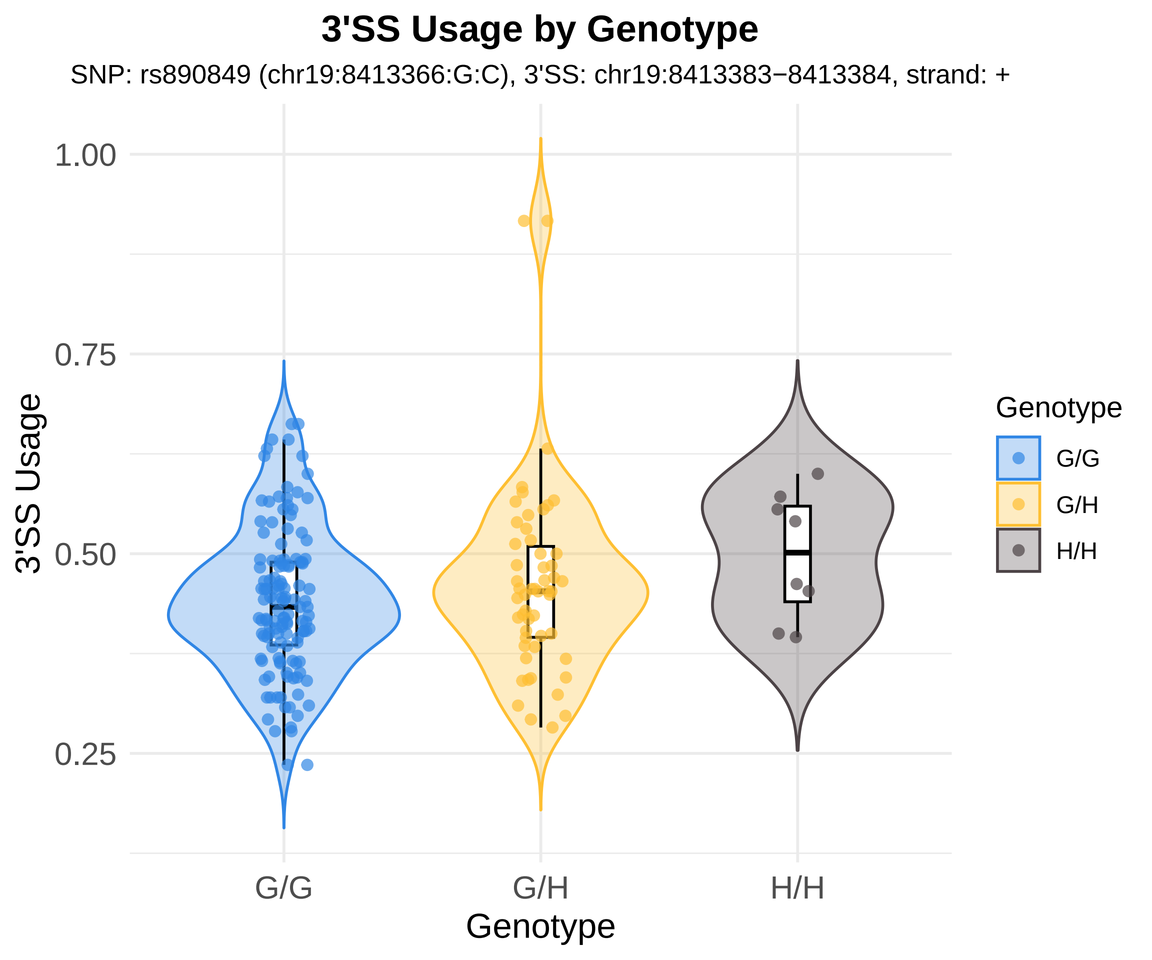

3'SS Usage by Genotype
SNP: rs890849 (chr19:8413366:G:C), 3'SS: chr19:8413383−8413384, strand: +
3'SS Usage
1.00
0.75
0.50
0.25
G/G
G/H
H/H
Genotype
Genotype
G/G
G/H
H/H

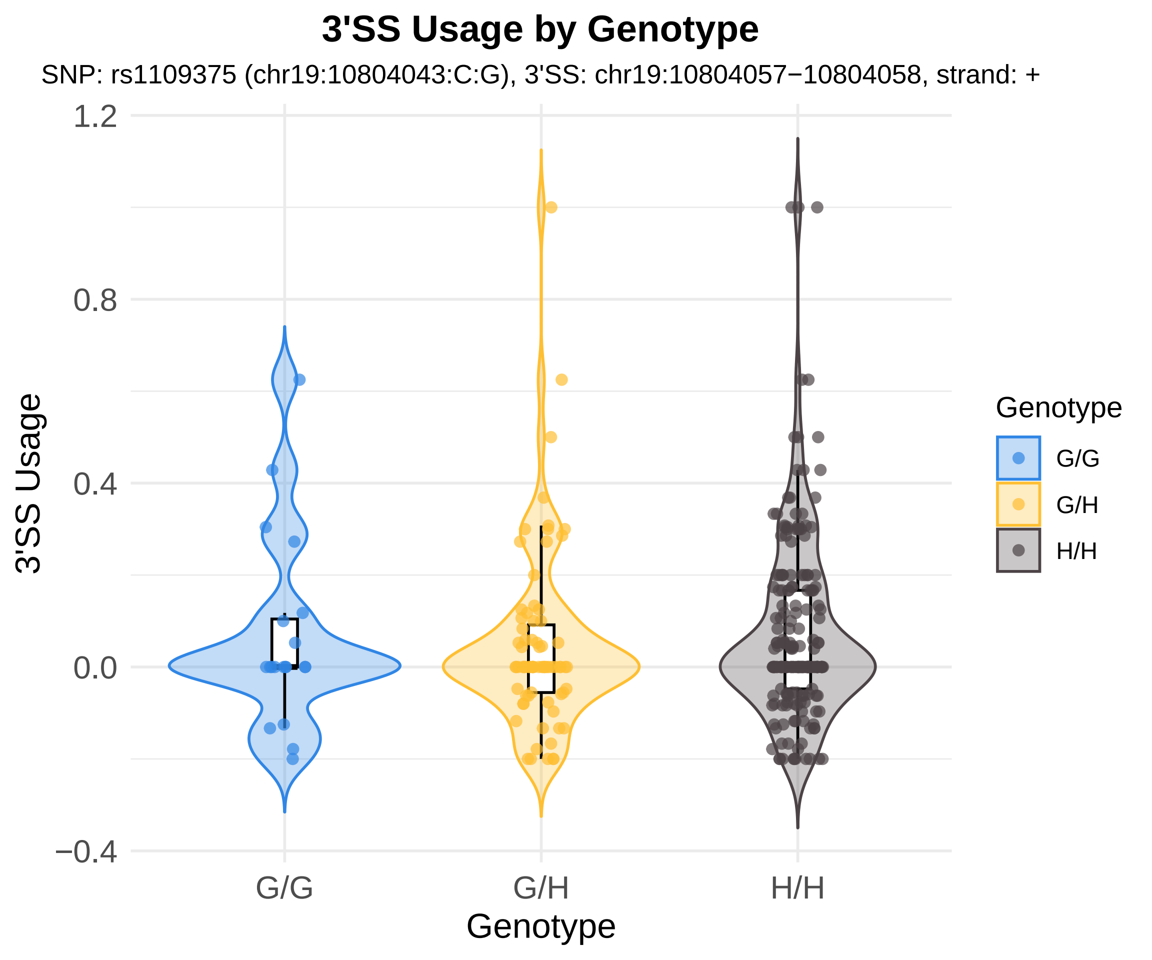

3'SS Usage by Genotype
SNP: rs1109375 (chr19:10804043:C:G), 3'SS: chr19:10804057−10804058, strand: +
1.2
0.8
0.4
0.0
−0.4
3'SS Usage
G/G
G/H
H/H
Genotype
Genotype
G/G
G/H
H/H

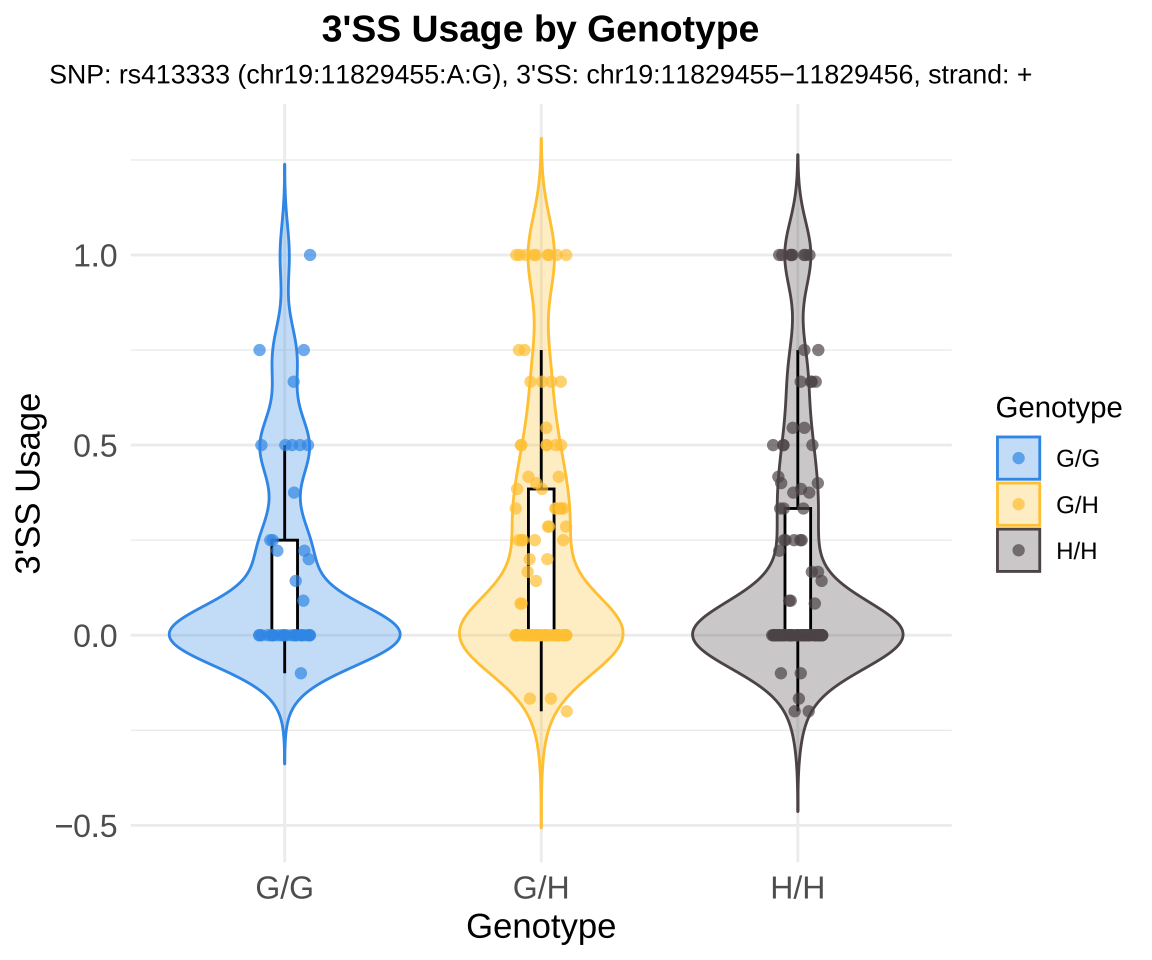
3'SS Usage by Genotype
SNP: rs413333 (chr19:11829455:A:G), 3'SS: chr19:11829455−11829456, strand: +
3'SS Usage
1.0
0.5
0.0
−0.5
G/G
G/H
H/H
Genotype
Genotype
G/G
G/H
H/H

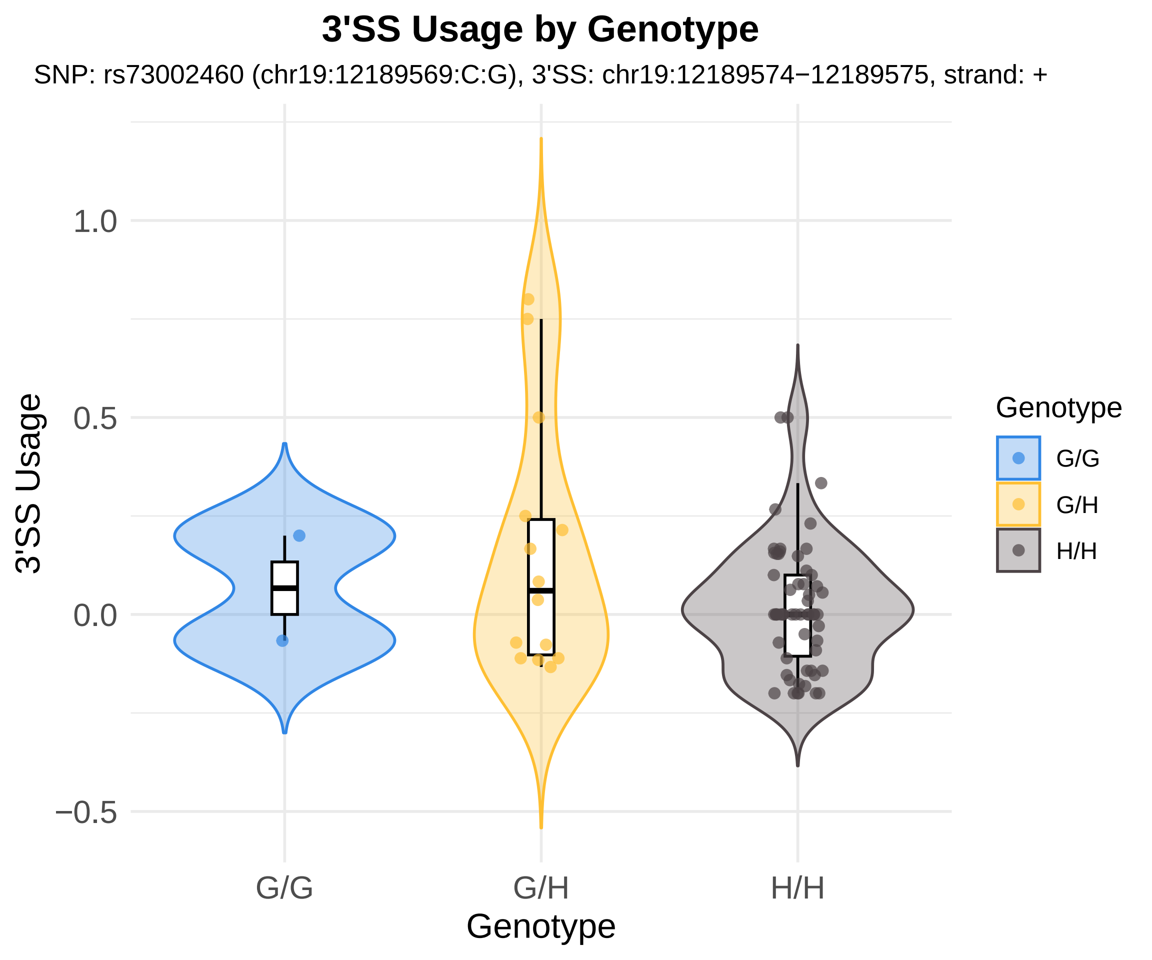

3'SS Usage by Genotype
SNP: rs73002460 (chr19:12189569:C:G), 3'SS: chr19:12189574−12189575, strand: +
3'SS Usage
1.0
0.5
0.0
−0.5
G/G
G/H
H/H
Genotype
Genotype
G/G
G/H
H/H

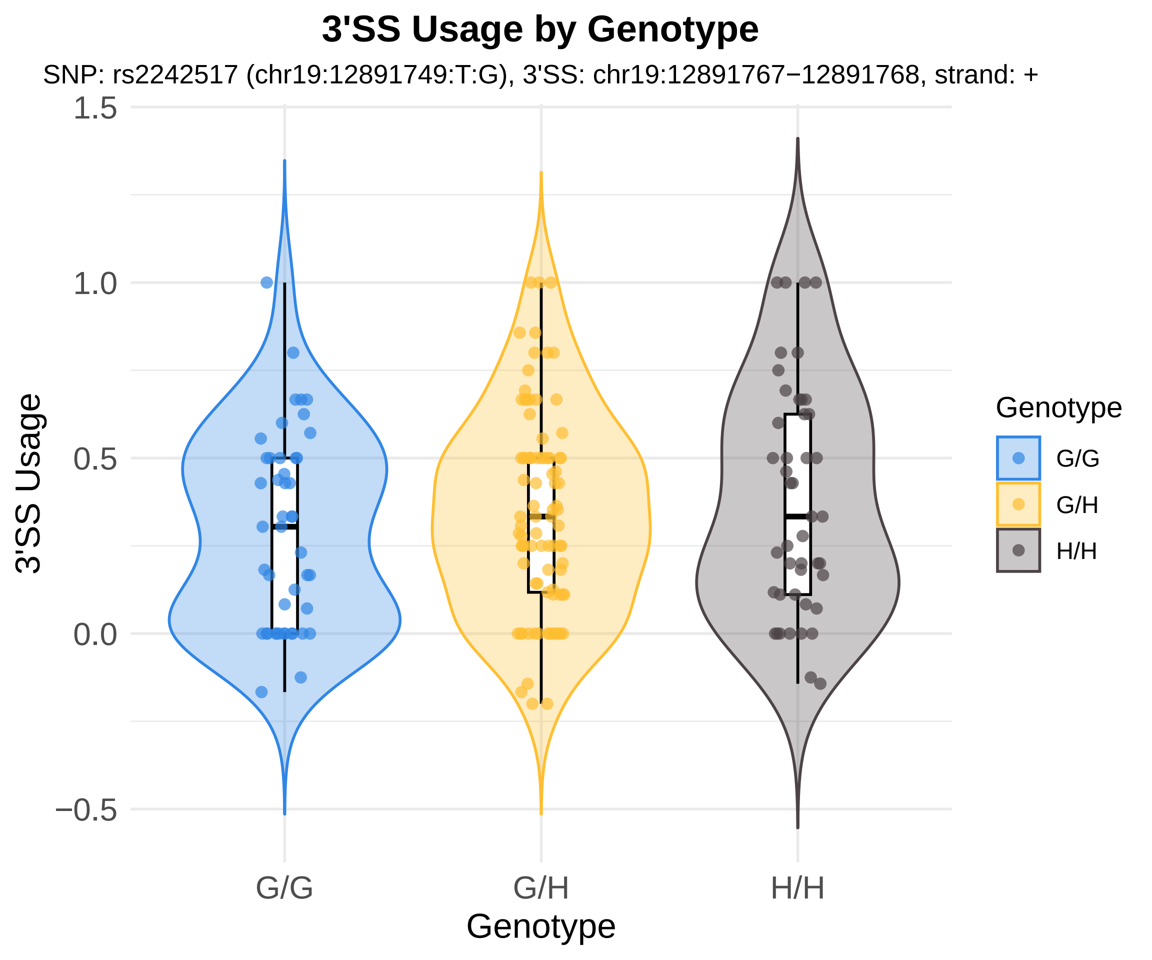

3'SS Usage by Genotype
SNP: rs2242517 (chr19:12891749:T:G), 3'SS: chr19:12891767−12891768, strand: +
1.5
1.0
0.5
0.0
−0.5
3'SS Usage
G/G
G/H
H/H
Genotype
Genotype
G/G
G/H
H/H

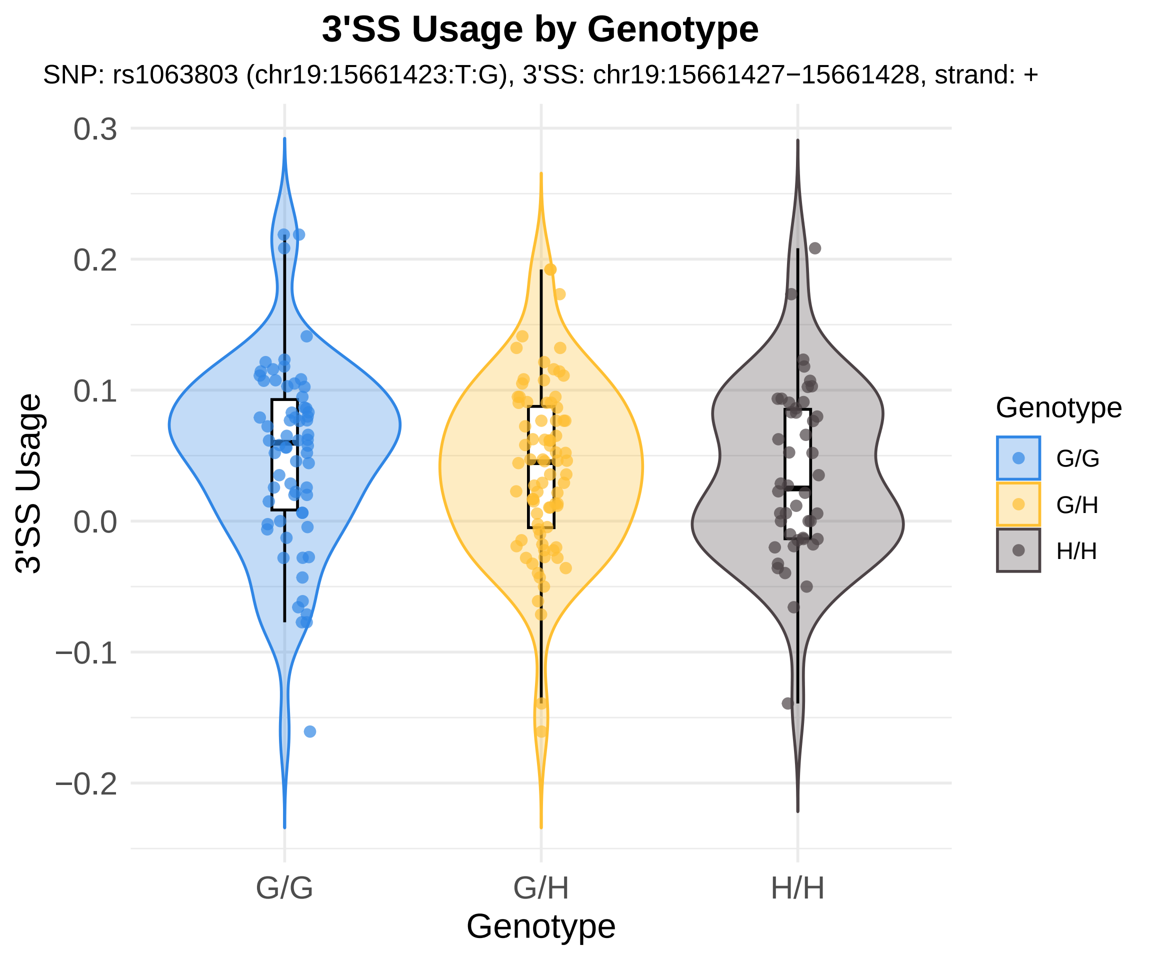
3'SS Usage by Genotype
SNP: rs1063803 (chr19:15661423:T:G), 3'SS: chr19:15661427−15661428, strand: +
3'SS Usage
0.3
0.2
0.1
0.0
−0.1
−0.2
G/G
G/H
H/H
Genotype
Genotype
G/G
G/H
H/H

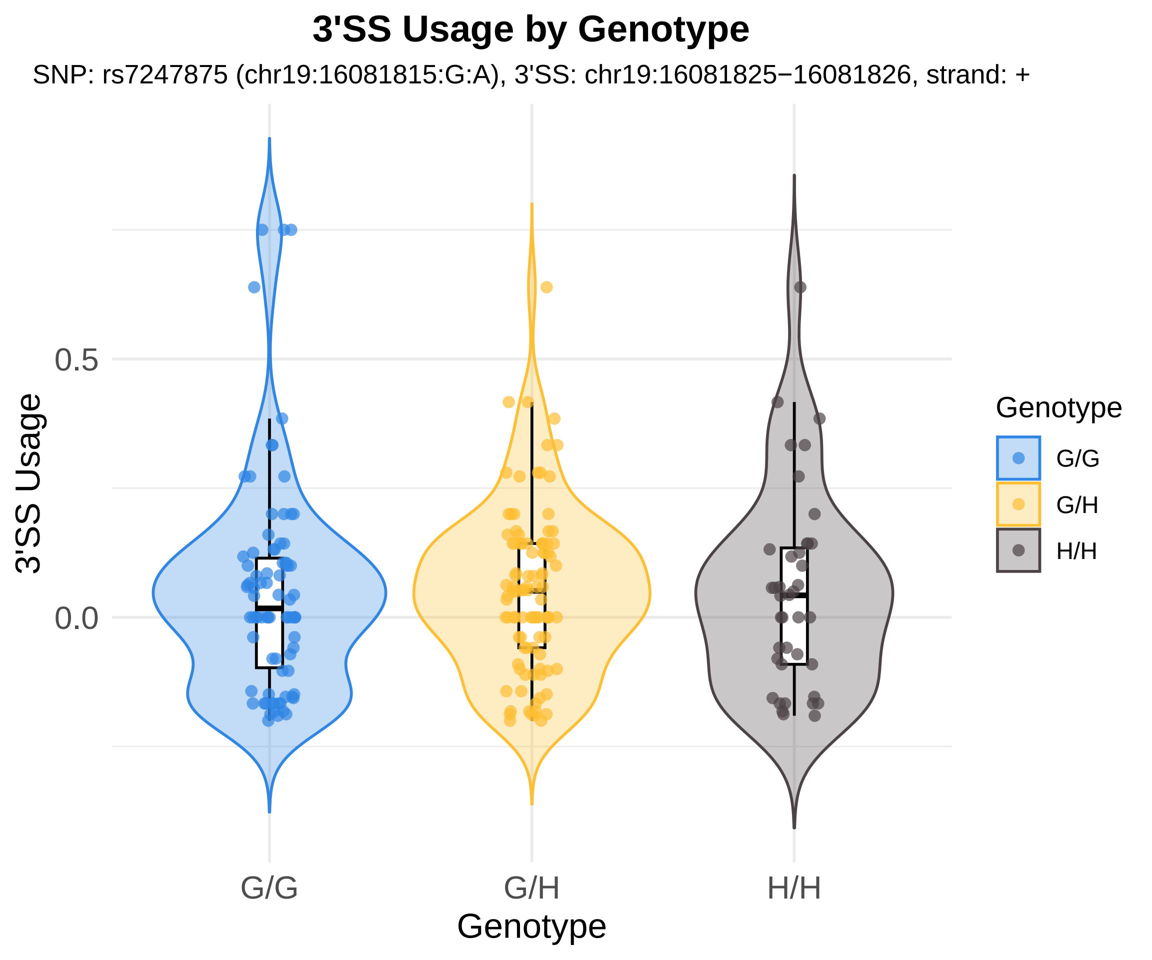

3'SS Usage by Genotype
SNP: rs7247875 (chr19:16081815:G:A), 3'SS: chr19:16081825−16081826, strand: +
3'SS Usage
0.5
0.0
G/G
G/H
H/H
Genotype
Genotype
G/G
G/H
H/H

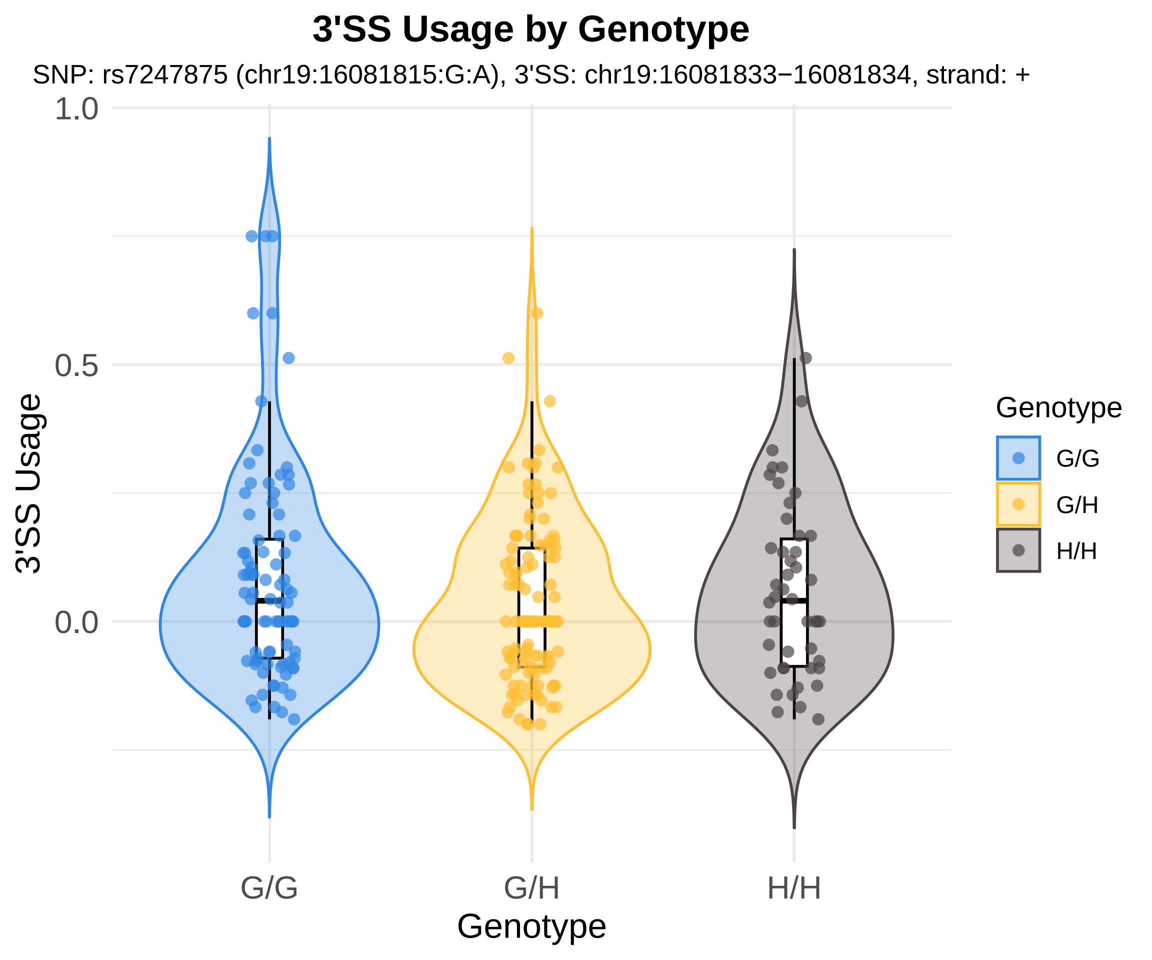
3'SS Usage by Genotype
SNP: rs7247875 (chr19:16081815:G:A), 3'SS: chr19:16081833−16081834, strand: +
1.0
0.5
0.0
3'SS Usage
G/G
G/H
H/H
Genotype
Genotype
G/G
G/H
H/H

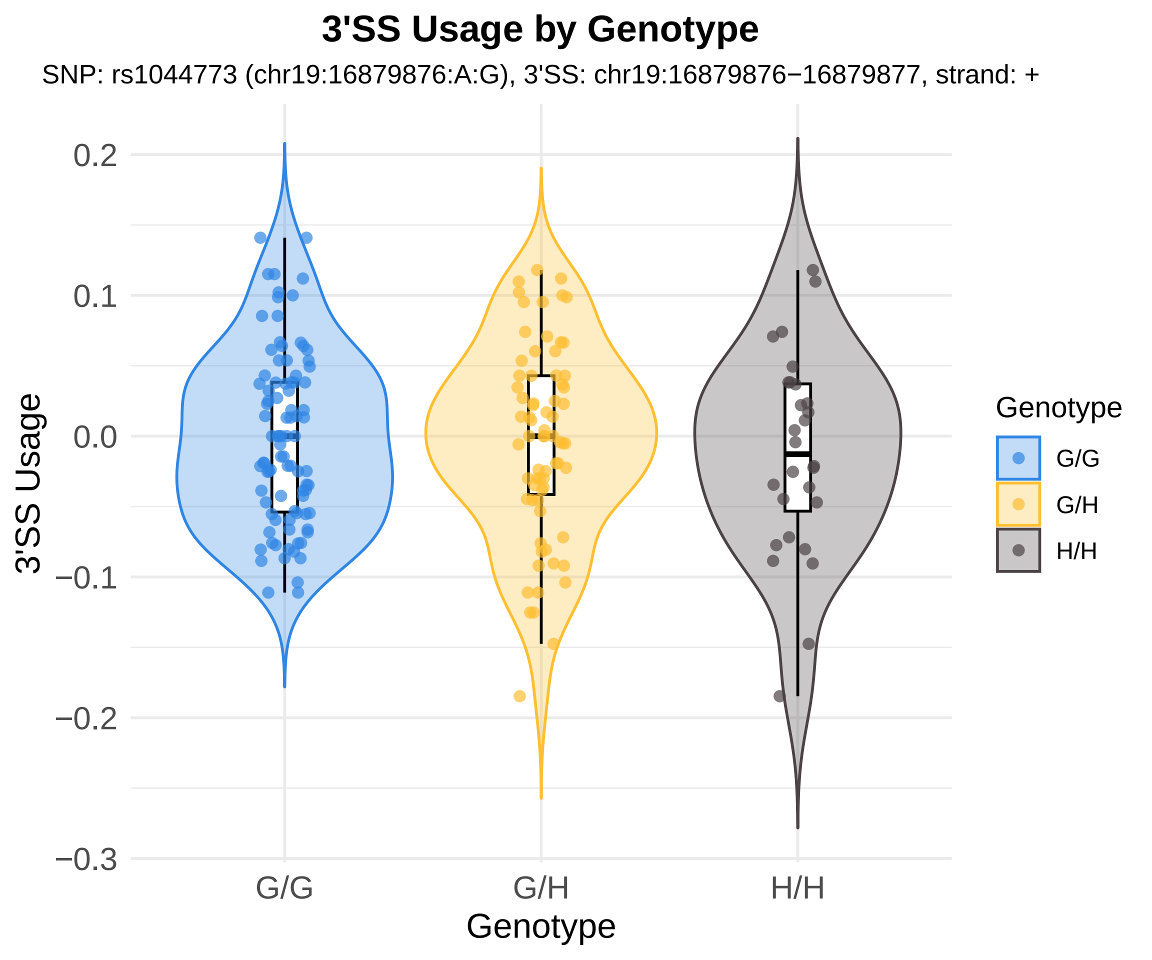
3'SS Usage by Genotype
SNP: rs1044773 (chr19:16879876:A:G), 3'SS: chr19:16879876−16879877, strand: +
0.2
0.1
0.0
−0.1
−0.2
−0.3
3'SS Usage
G/G
G/H
H/H
Genotype
Genotype
G/G
G/H
H/H

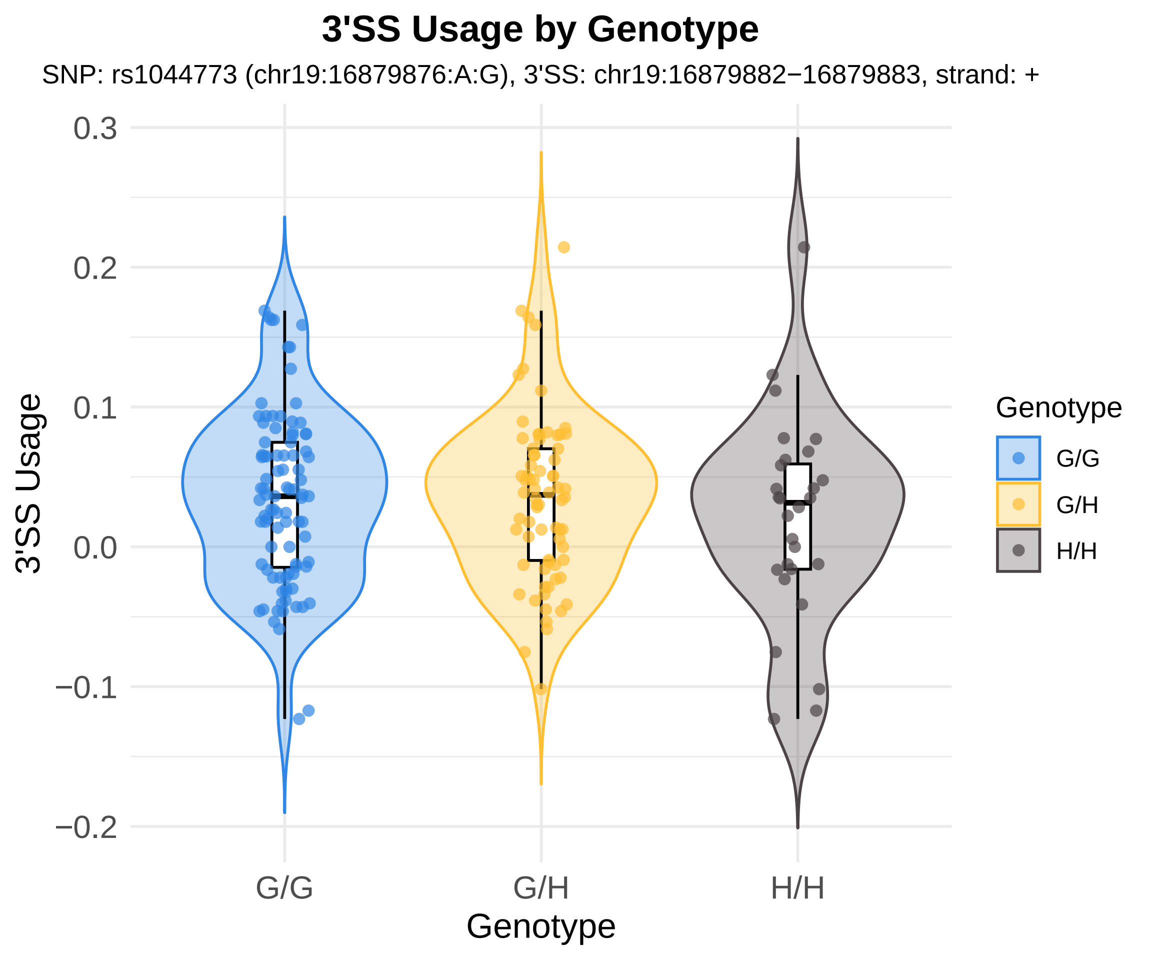
3'SS Usage by Genotype
SNP: rs1044773 (chr19:16879876:A:G), 3'SS: chr19:16879882−16879883, strand: +
3'SS Usage
0.3
0.2
0.1
0.0
−0.1
−0.2
G/G
G/H
H/H
Genotype
Genotype
G/G
G/H
H/H

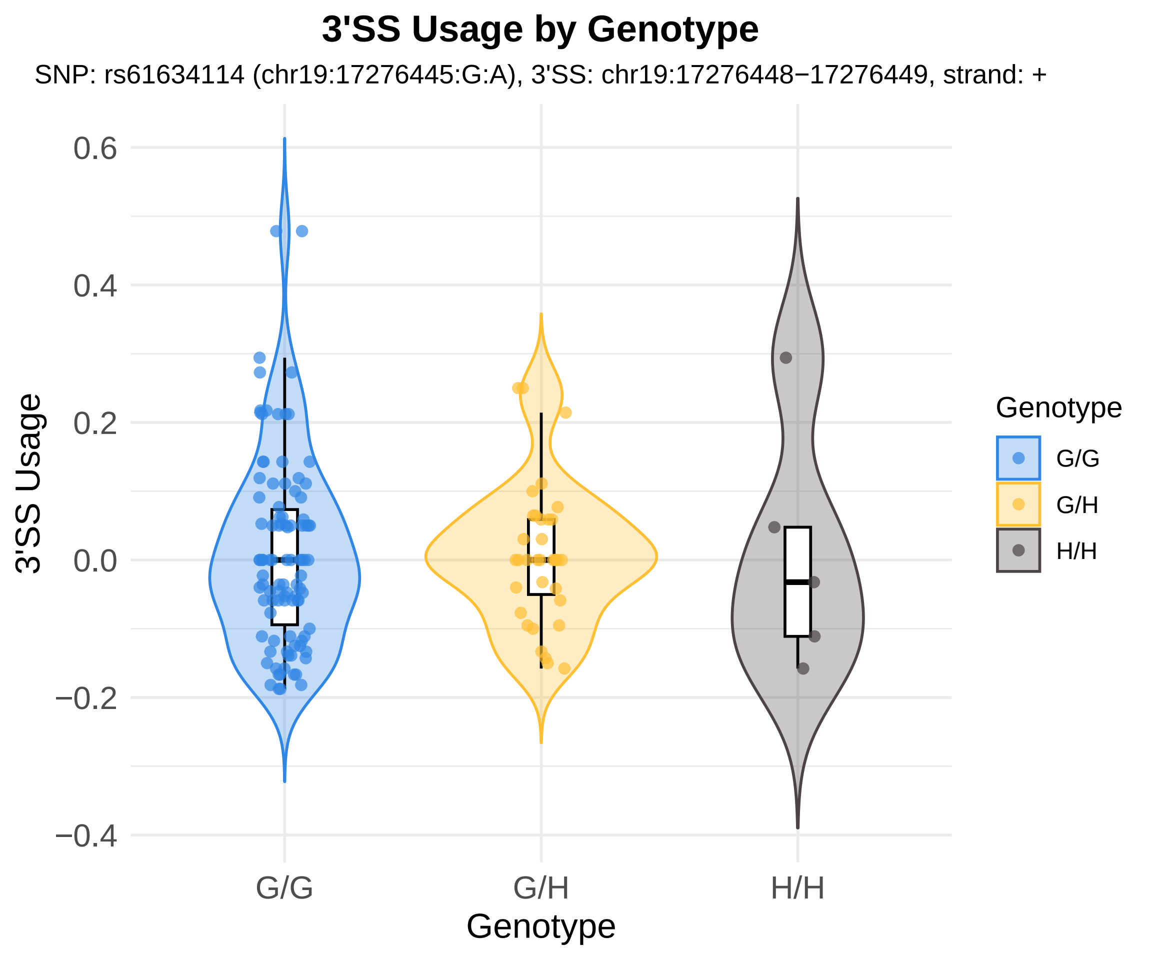
3'SS Usage by Genotype
SNP: rs61634114 (chr19:17276445:G:A), 3'SS: chr19:17276448−17276449, strand: +
3'SS Usage
0.6
0.4
0.2
0.0
−0.2
−0.4
G/G
G/H
H/H
Genotype
Genotype
G/G
G/H
H/H

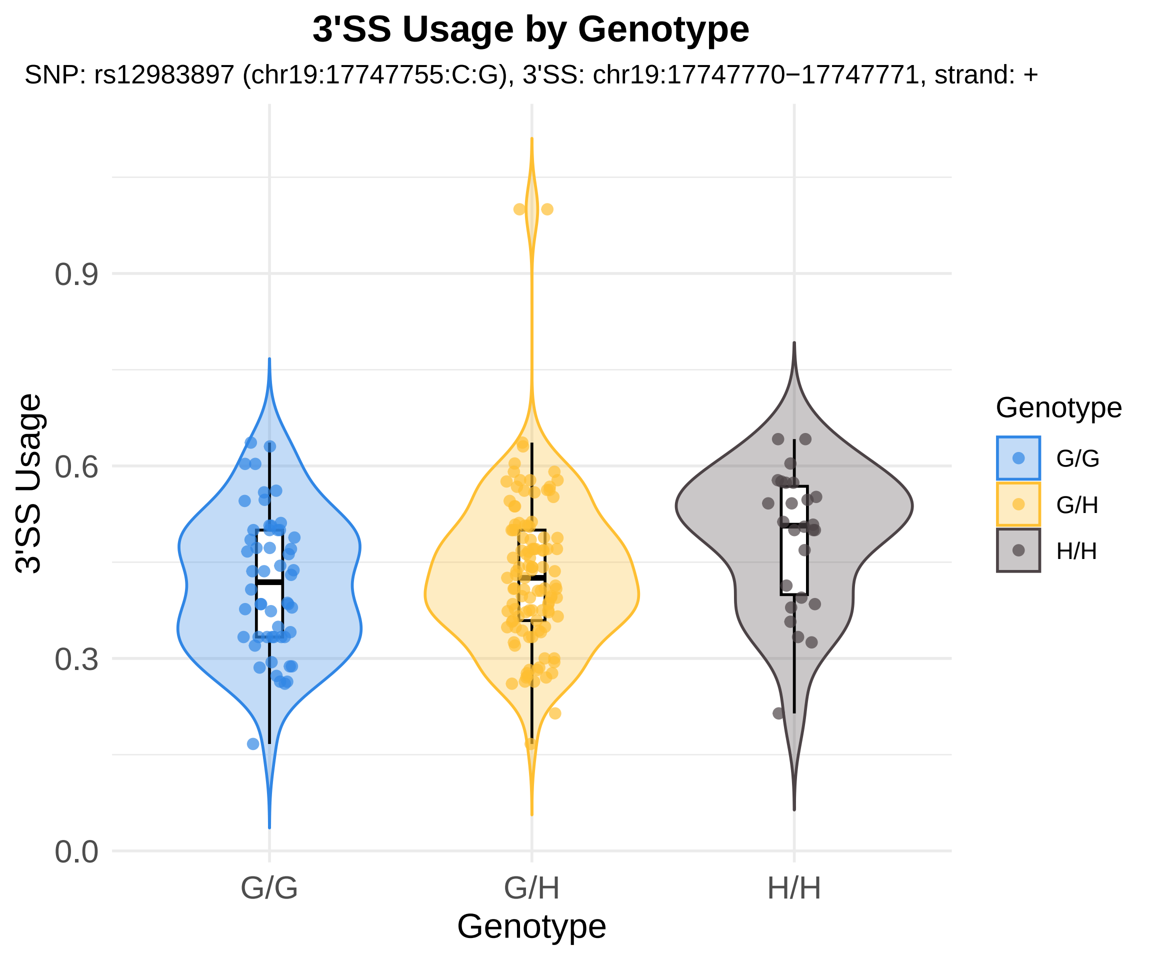
3'SS Usage by Genotype
SNP: rs12983897 (chr19:17747755:C:G), 3'SS: chr19:17747770−17747771, strand: +
3'SS Usage
0.0
0.3
0.6
0.9
G/G
G/H
H/H
Genotype
Genotype
G/G
G/H
H/H

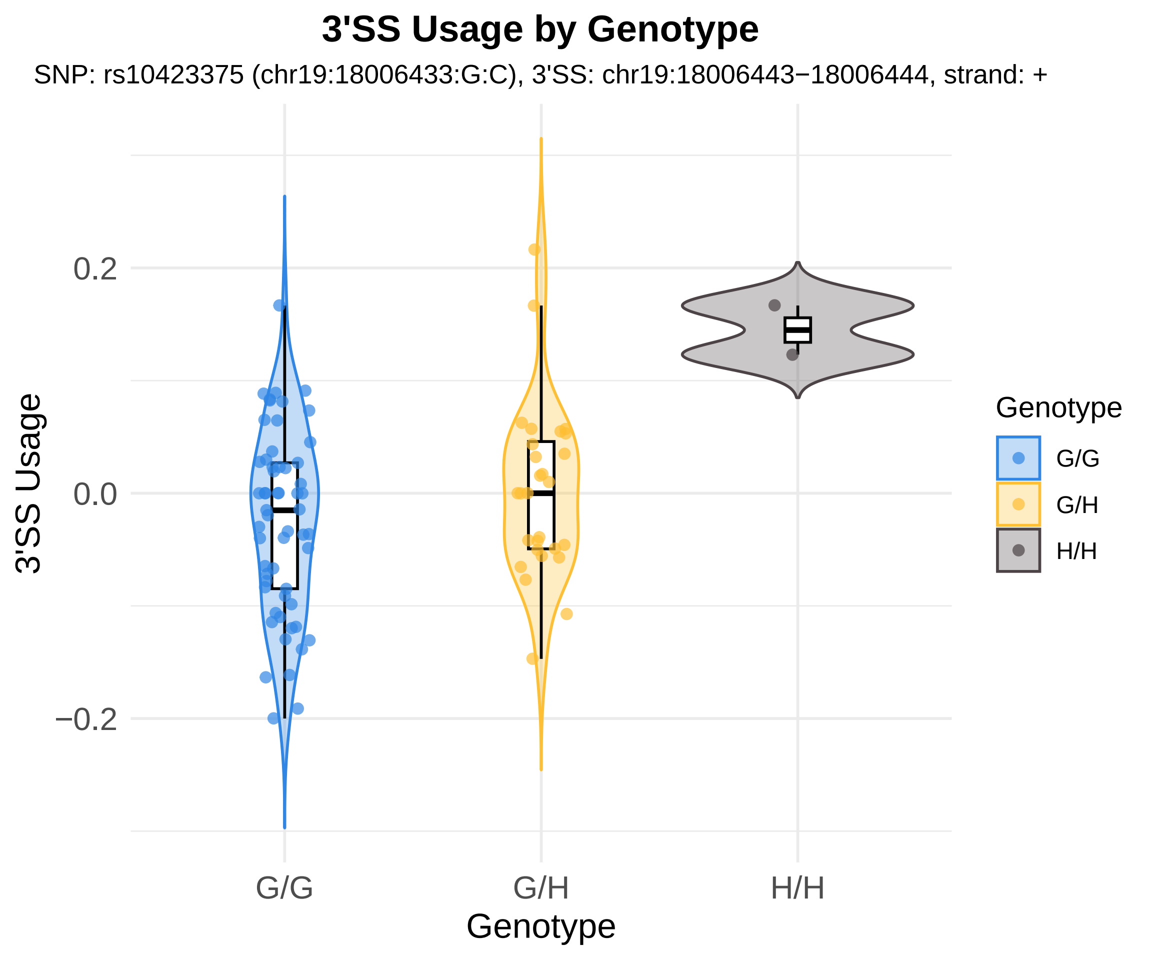
3'SS Usage by Genotype
SNP: rs10423375 (chr19:18006433:G:C), 3'SS: chr19:18006443−18006444, strand: +
3'SS Usage
0.2
0.0
−0.2
G/G
G/H
H/H
Genotype
Genotype
G/G
G/H
H/H

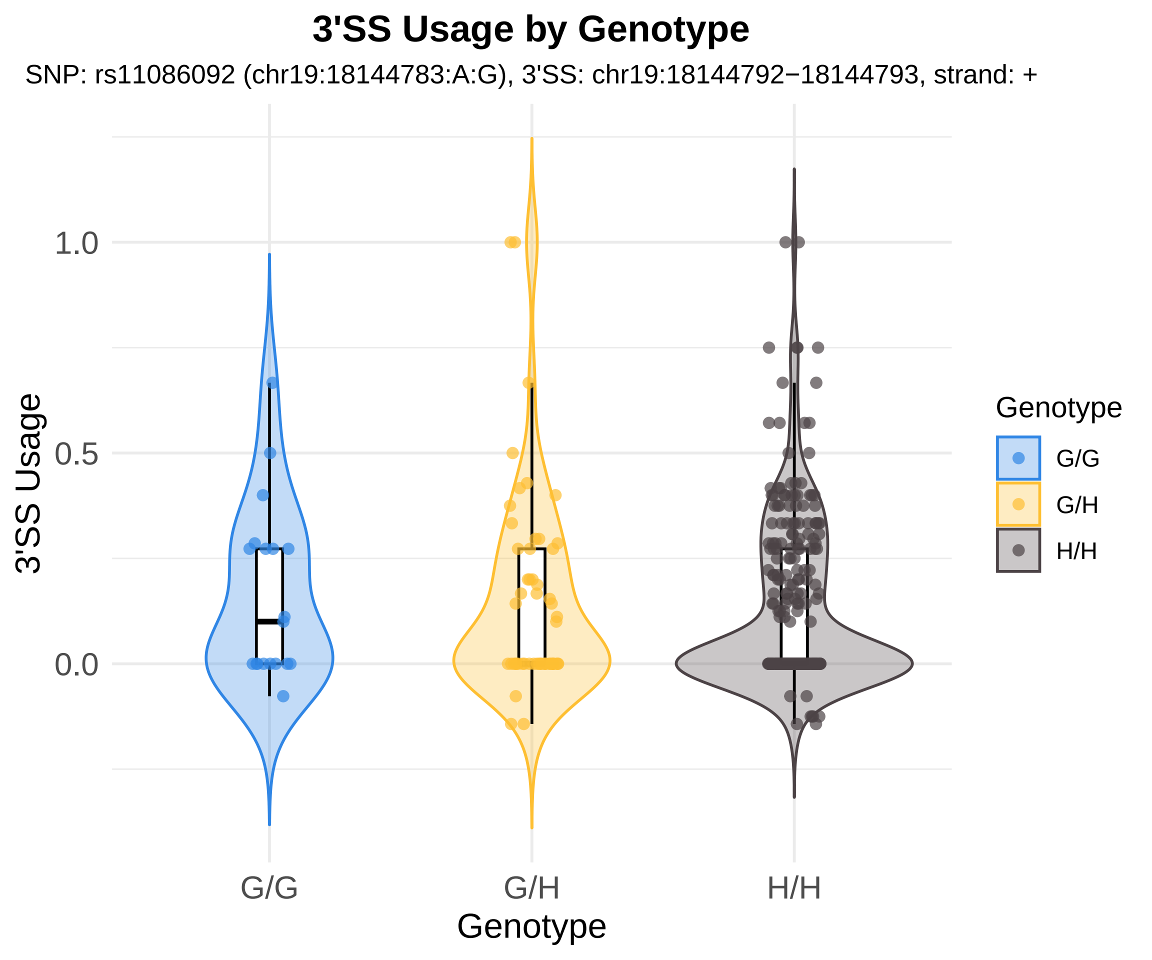
3'SS Usage by Genotype
SNP: rs11086092 (chr19:18144783:A:G), 3'SS: chr19:18144792−18144793, strand: +
1.0
0.5
0.0
3'SS Usage
G/G
G/H
H/H
Genotype
Genotype
G/G
G/H
H/H

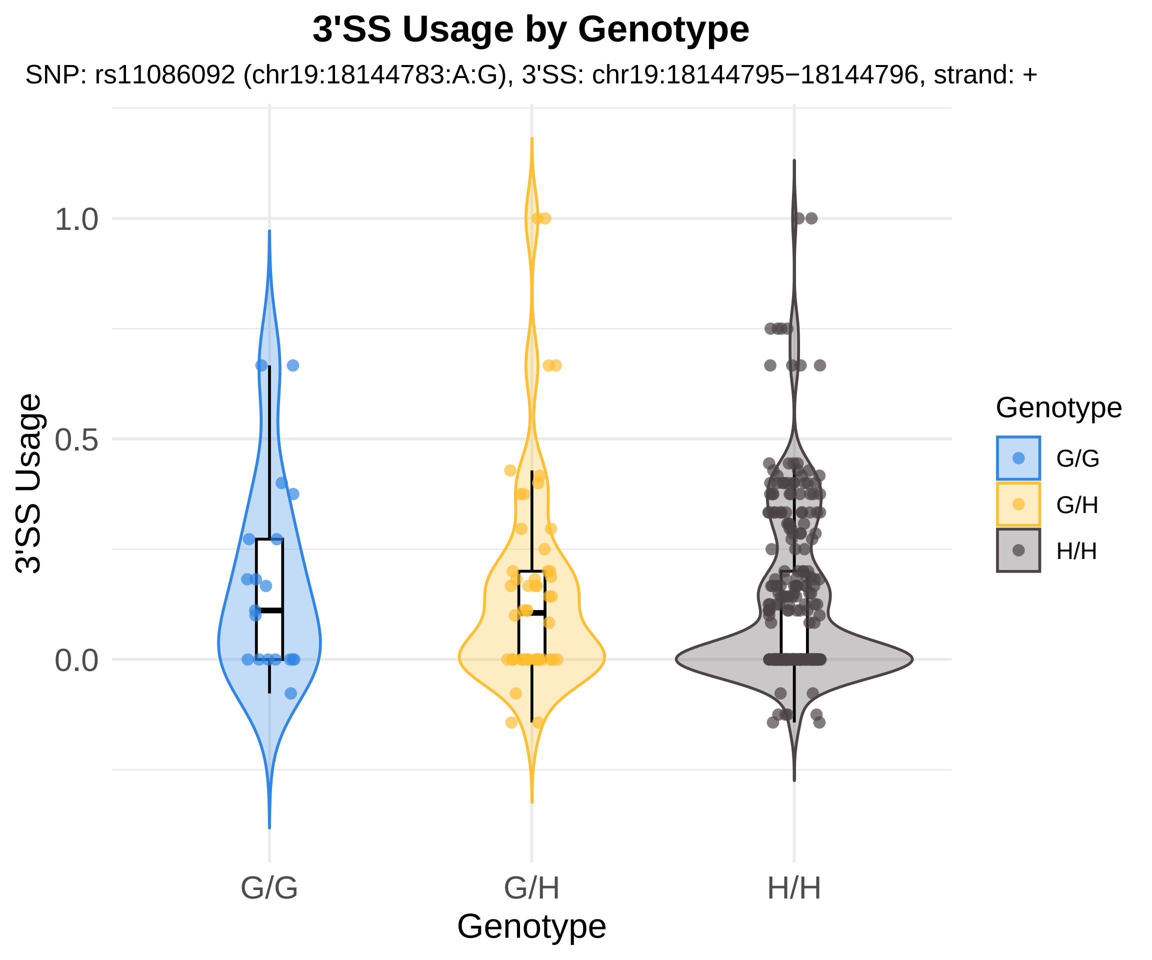

3'SS Usage by Genotype
SNP: rs11086092 (chr19:18144783:A:G), 3'SS: chr19:18144795−18144796, strand: +
3'SS Usage
1.0
0.5
0.0
G/G
G/H
H/H
Genotype
Genotype
G/G
G/H
H/H

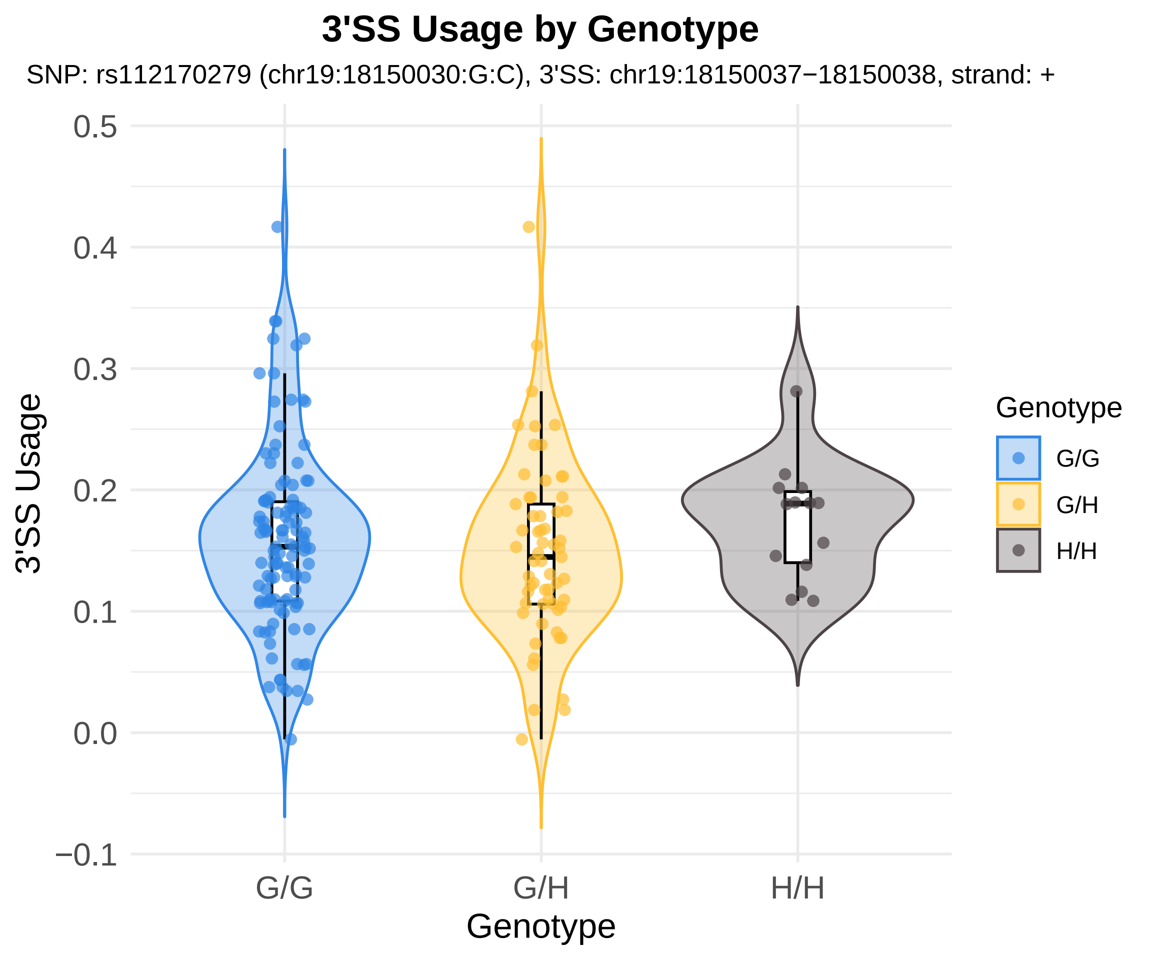
3'SS Usage by Genotype
SNP: rs112170279 (chr19:18150030:G:C), 3'SS: chr19:18150037−18150038, strand: +
3'SS Usage
0.5
0.4
0.3
0.2
0.1
0.0
−0.1
G/G
G/H
H/H
Genotype
Genotype
G/G
G/H
H/H

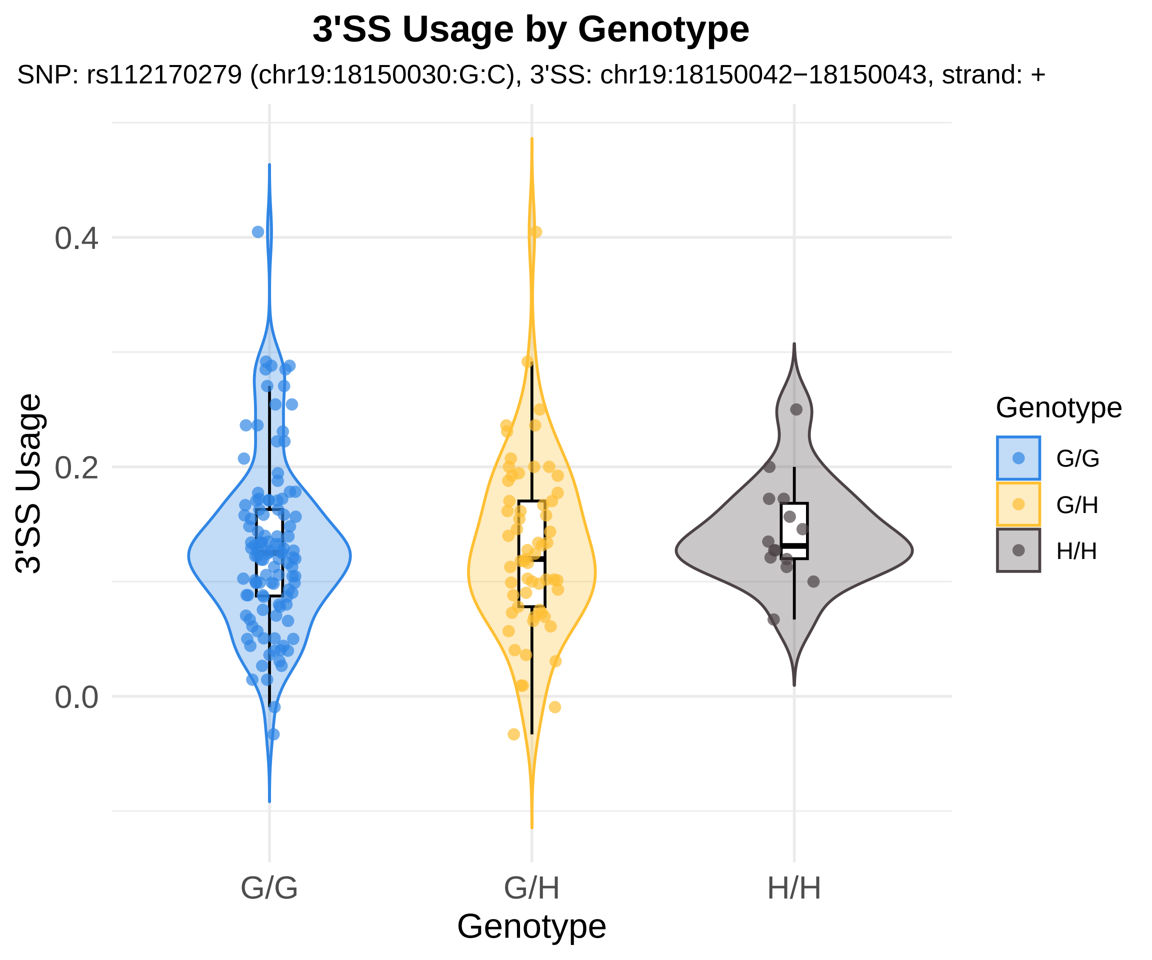
3'SS Usage by Genotype
SNP: rs112170279 (chr19:18150030:G:C), 3'SS: chr19:18150042−18150043, strand: +
3'SS Usage
0.4
0.2
0.0
G/G
G/H
H/H
Genotype
Genotype
G/G
G/H
H/H

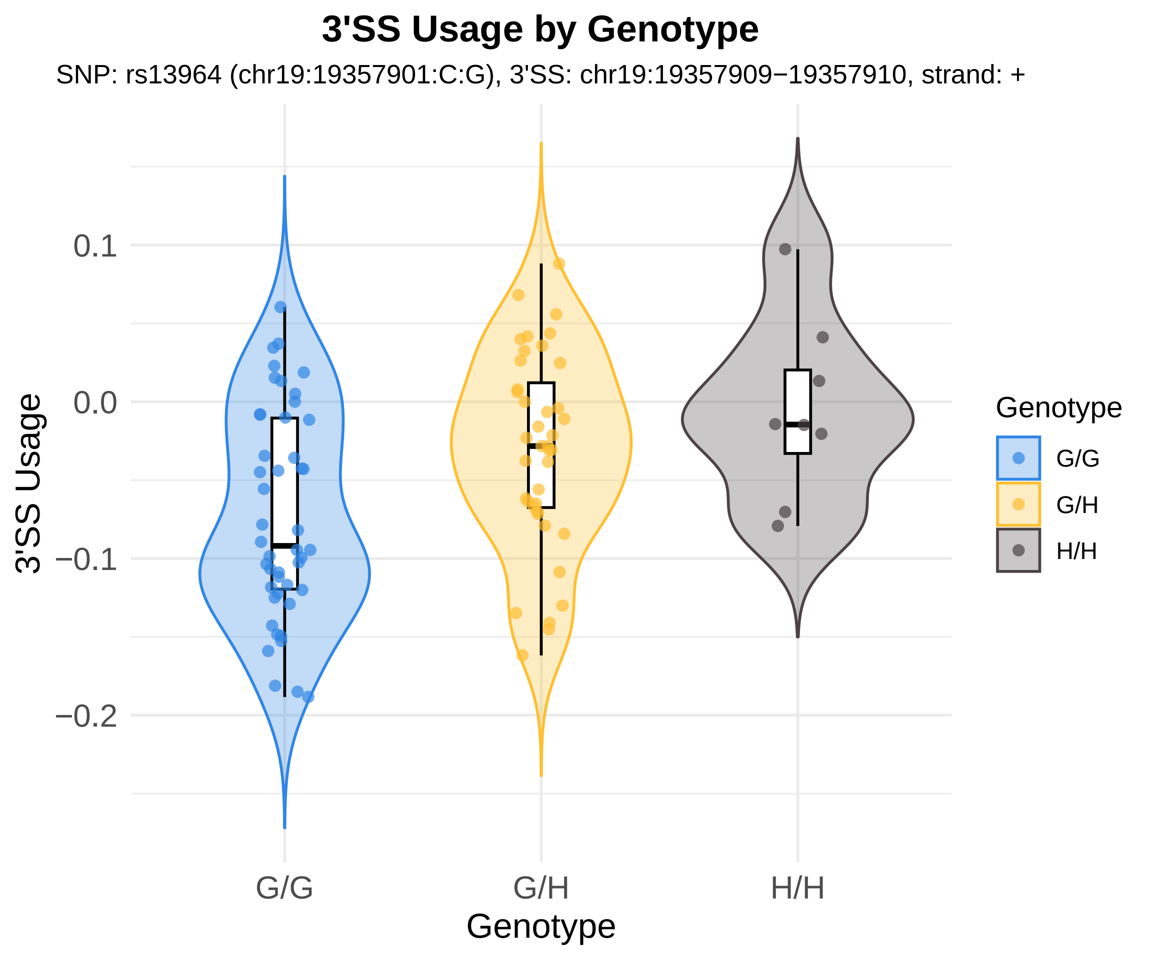
3'SS Usage by Genotype
SNP: rs13964 (chr19:19357901:C:G), 3'SS: chr19:19357909−19357910, strand: +
3'SS Usage
0.1
0.0
−0.1
−0.2
G/G
G/H
H/H
Genotype
Genotype
G/G
G/H
H/H

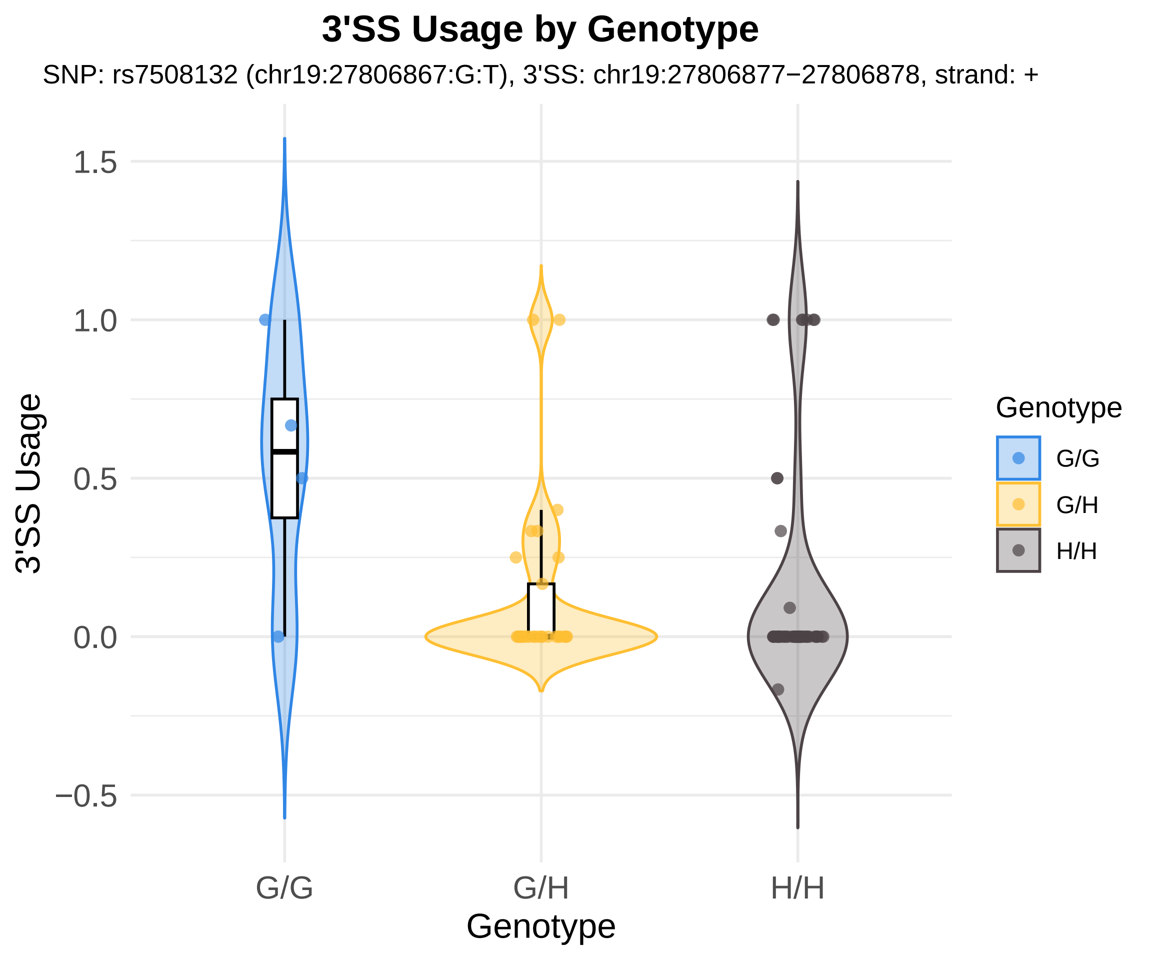

3'SS Usage by Genotype
SNP: rs7508132 (chr19:27806867:G:T), 3'SS: chr19:27806877−27806878, strand: +
3'SS Usage
1.5
1.0
0.5
0.0
−0.5
G/G
G/H
H/H
Genotype
Genotype
G/G
G/H
H/H

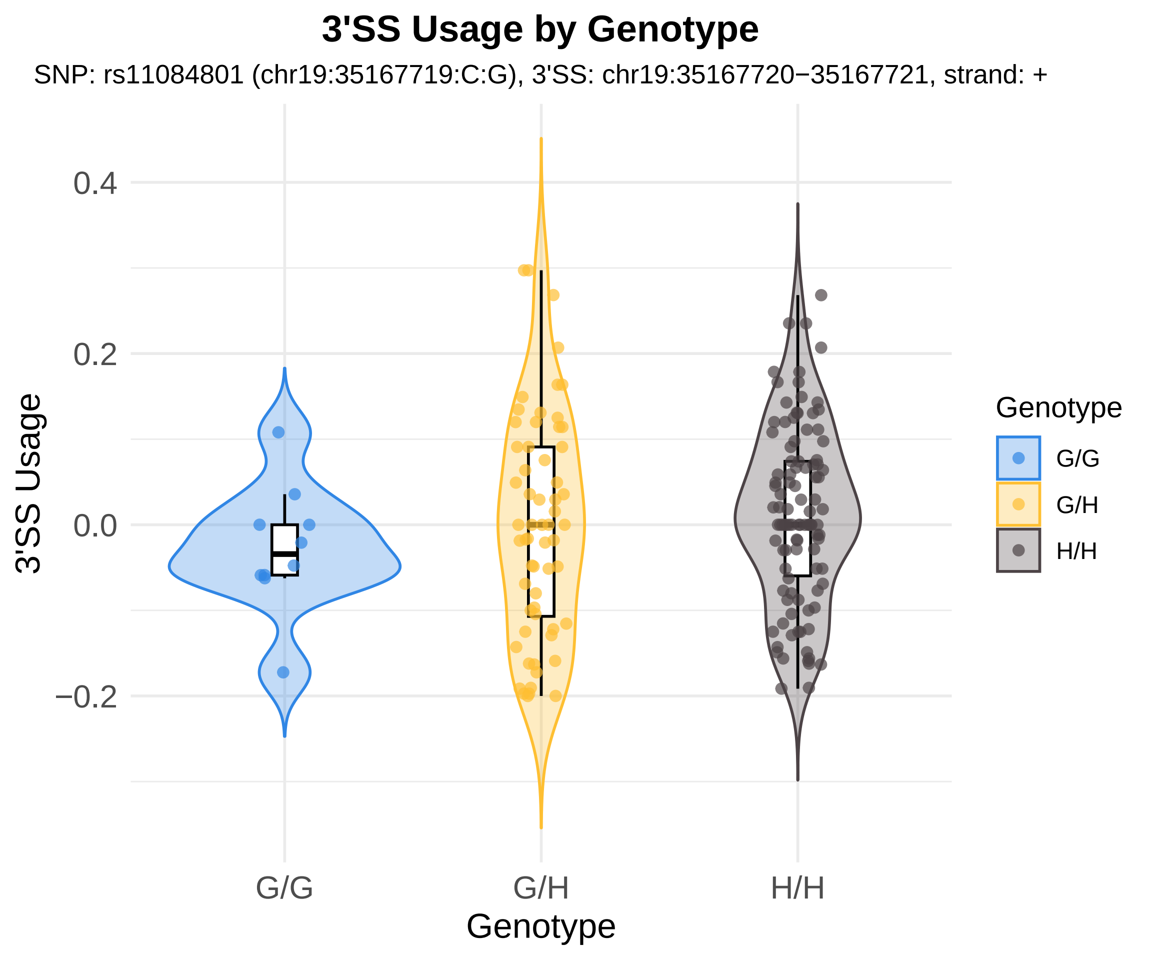
3'SS Usage by Genotype
SNP: rs11084801 (chr19:35167719:C:G), 3'SS: chr19:35167720−35167721, strand: +
3'SS Usage
0.4
0.2
0.0
−0.2
G/G
G/H
H/H
Genotype
Genotype
G/G
G/H
H/H

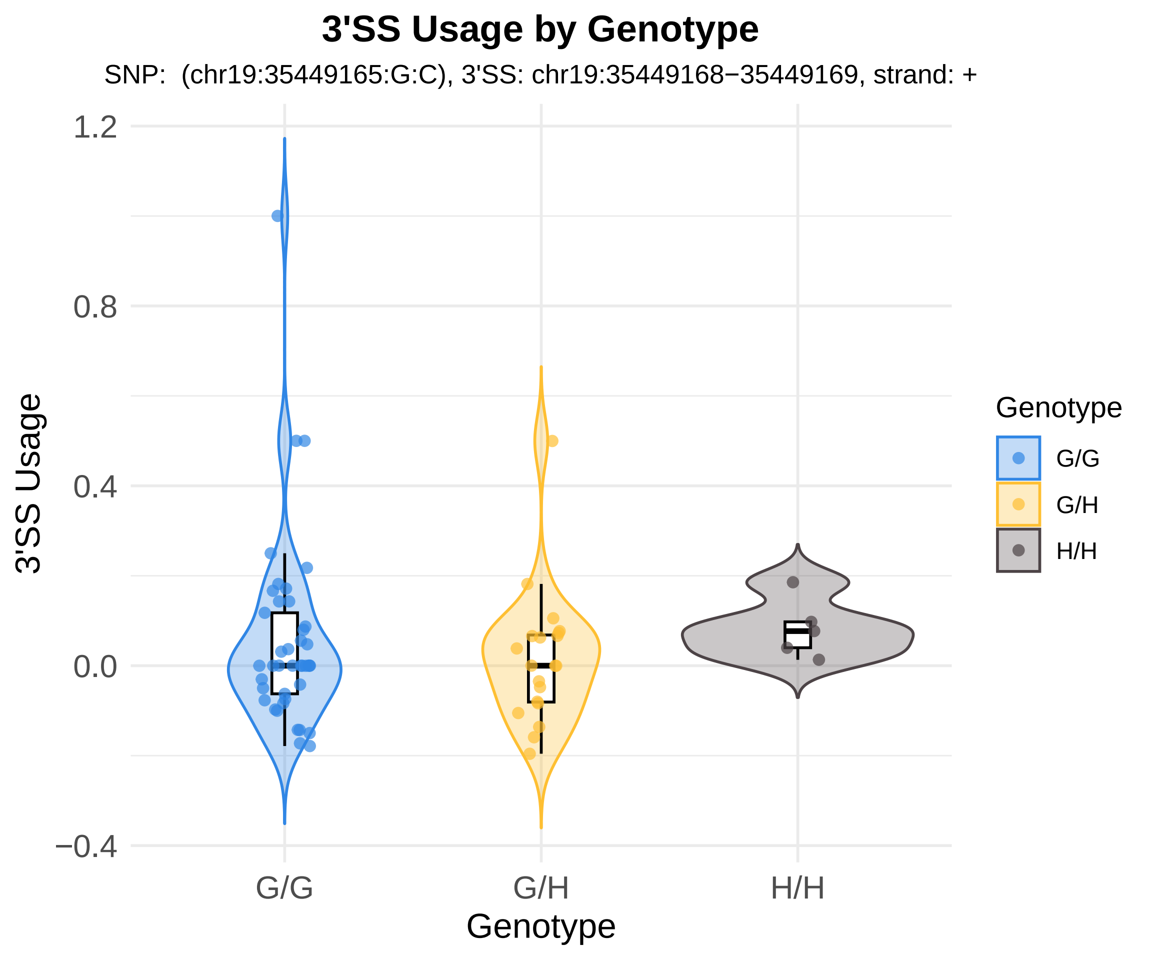
3'SS Usage by Genotype
SNP: (chr19:35449165:G:C), 3'SS: chr19:35449168−35449169, strand: +
3'SS Usage
1.2
0.8
0.4
0.0
−0.4
G/G
G/H
H/H
Genotype
Genotype
G/G
G/H
H/H

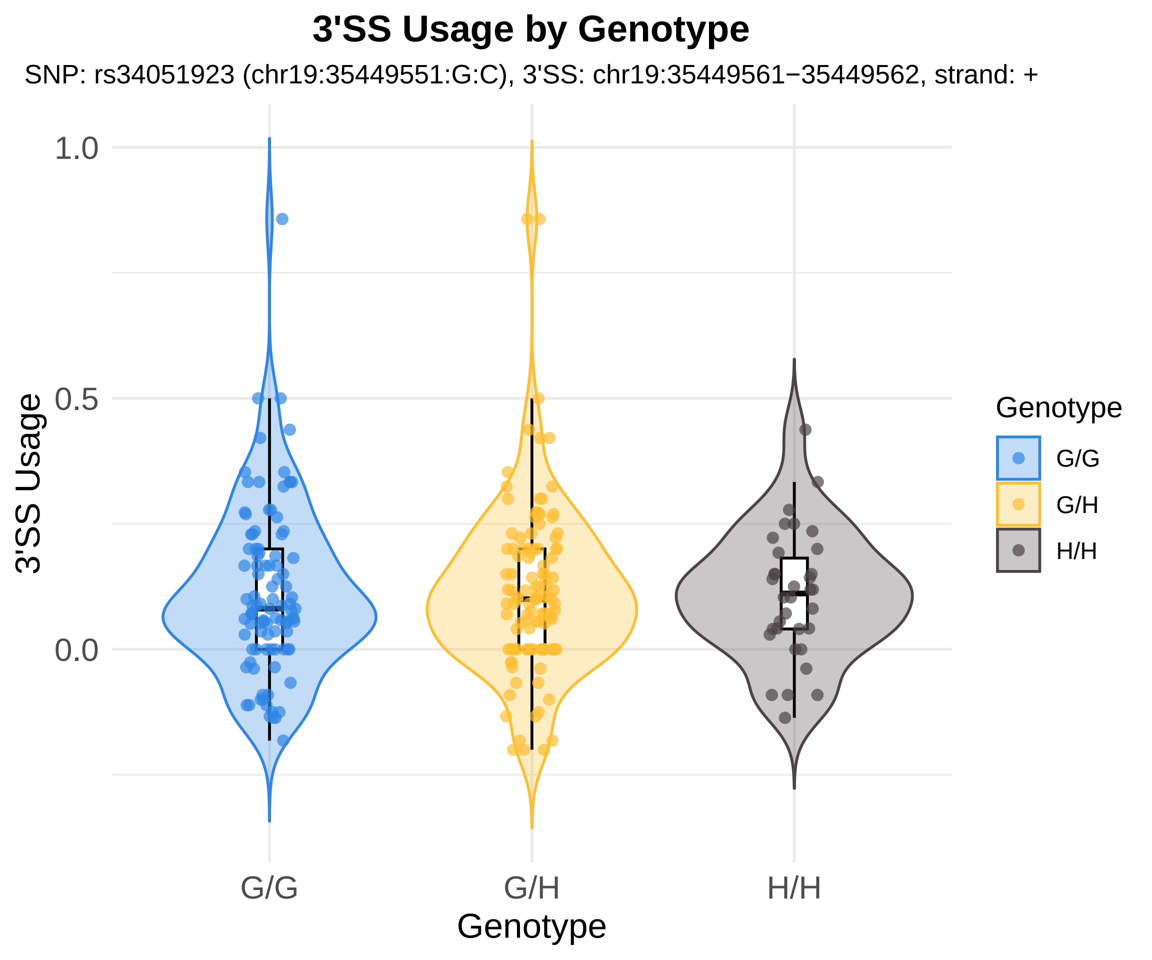

3'SS Usage by Genotype
SNP: rs34051923 (chr19:35449551:G:C), 3'SS: chr19:35449561−35449562, strand: +
1.0
0.5
0.0
3'SS Usage
G/G
G/H
H/H
Genotype
Genotype
G/G
G/H
H/H

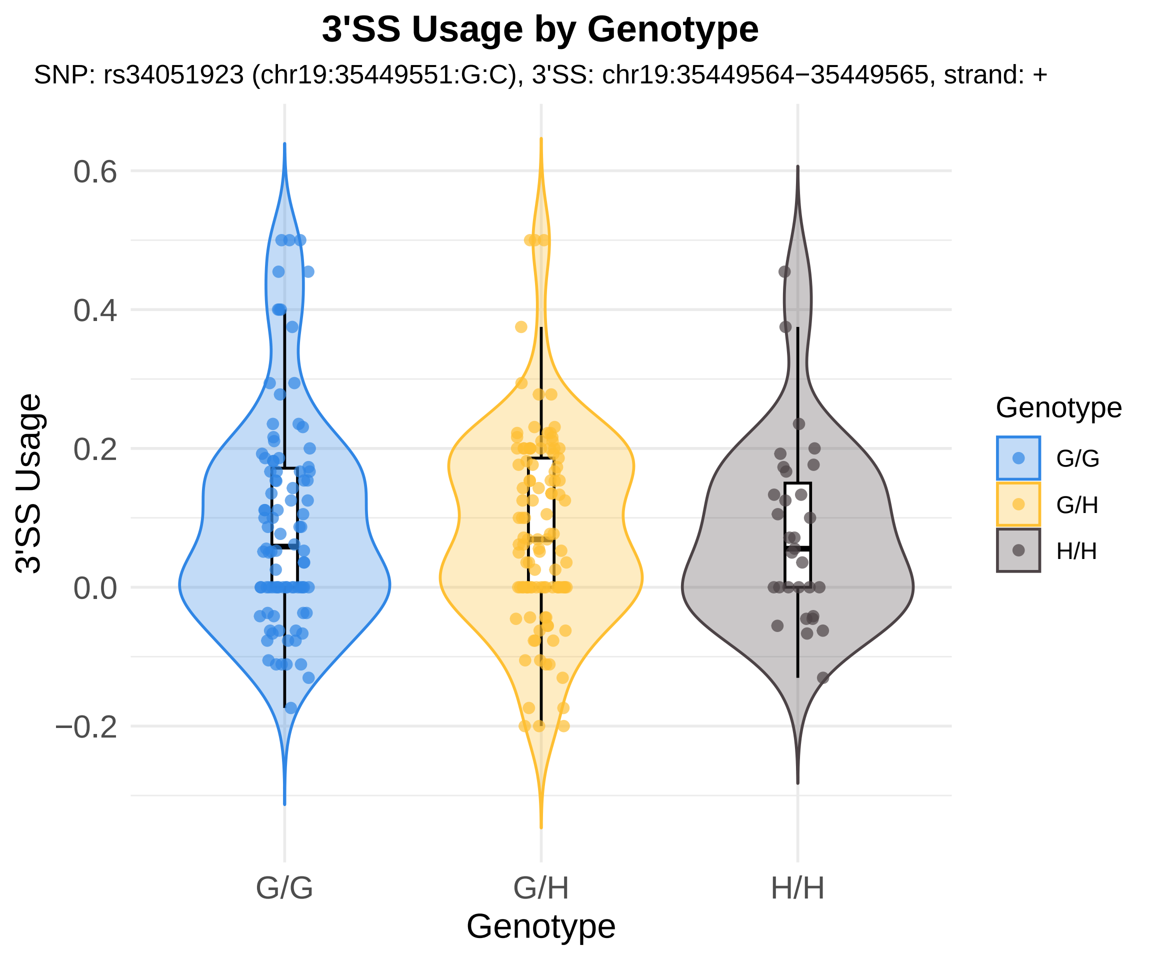

3'SS Usage by Genotype
SNP: rs34051923 (chr19:35449551:G:C), 3'SS: chr19:35449564−35449565, strand: +
3'SS Usage
0.6
0.4
0.2
0.0
−0.2
G/G
G/H
H/H
Genotype
Genotype
G/G
G/H
H/H

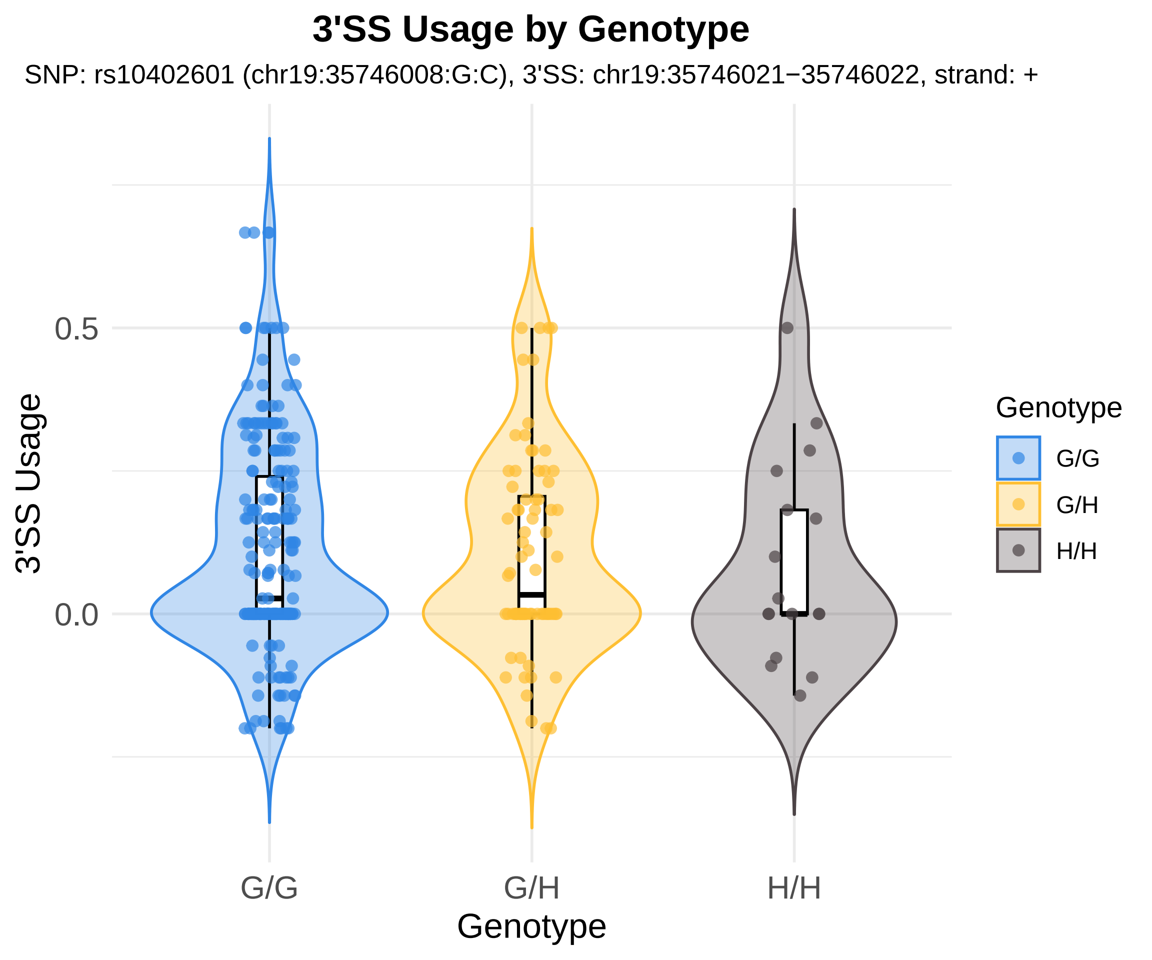

3'SS Usage by Genotype
SNP: rs10402601 (chr19:35746008:G:C), 3'SS: chr19:35746021−35746022, strand: +
3'SS Usage
0.5
0.0
G/G
G/H
H/H
Genotype
Genotype
G/G
G/H
H/H

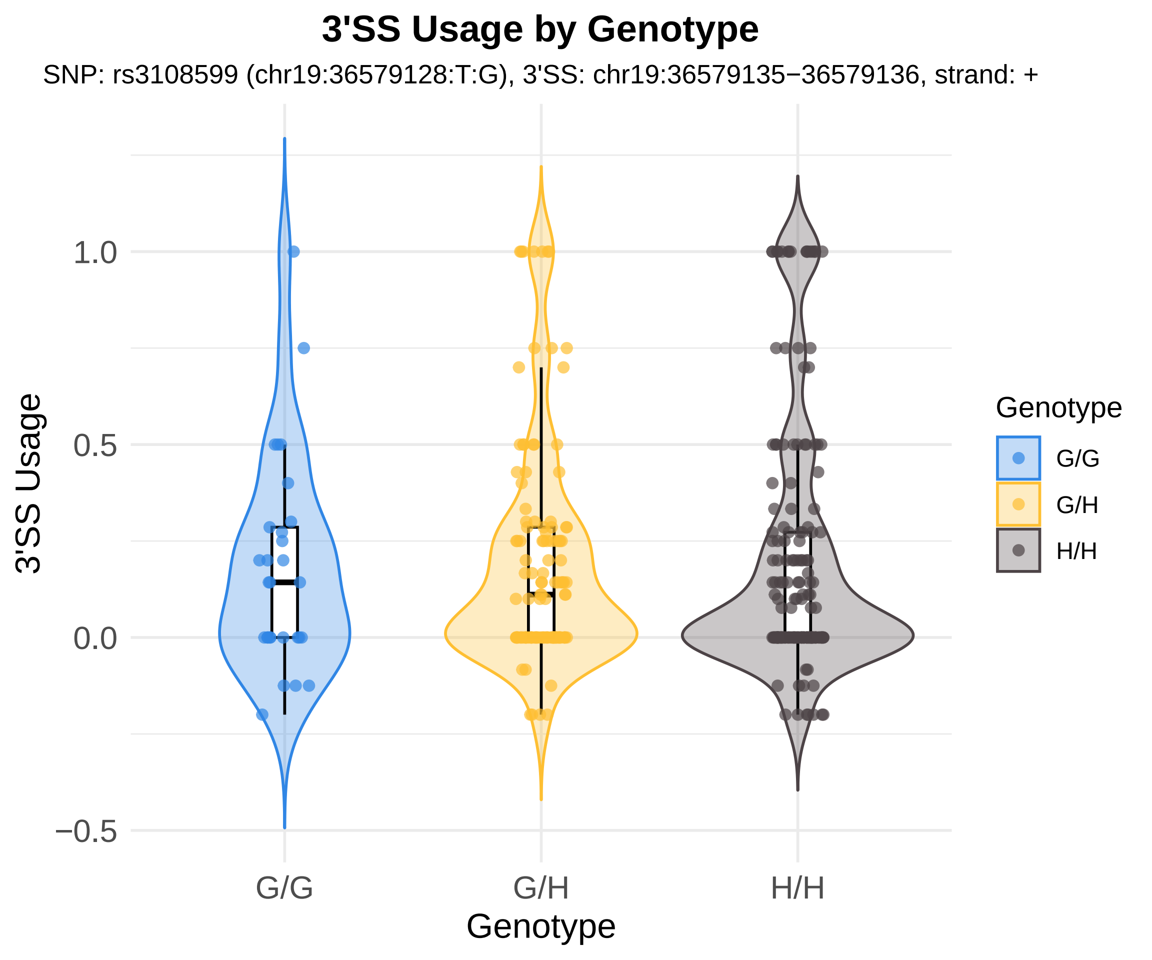
3'SS Usage by Genotype
SNP: rs3108599 (chr19:36579128:T:G), 3'SS: chr19:36579135−36579136, strand: +
3'SS Usage
1.0
0.5
0.0
−0.5
G/G
G/H
H/H
Genotype
Genotype
G/G
G/H
H/H

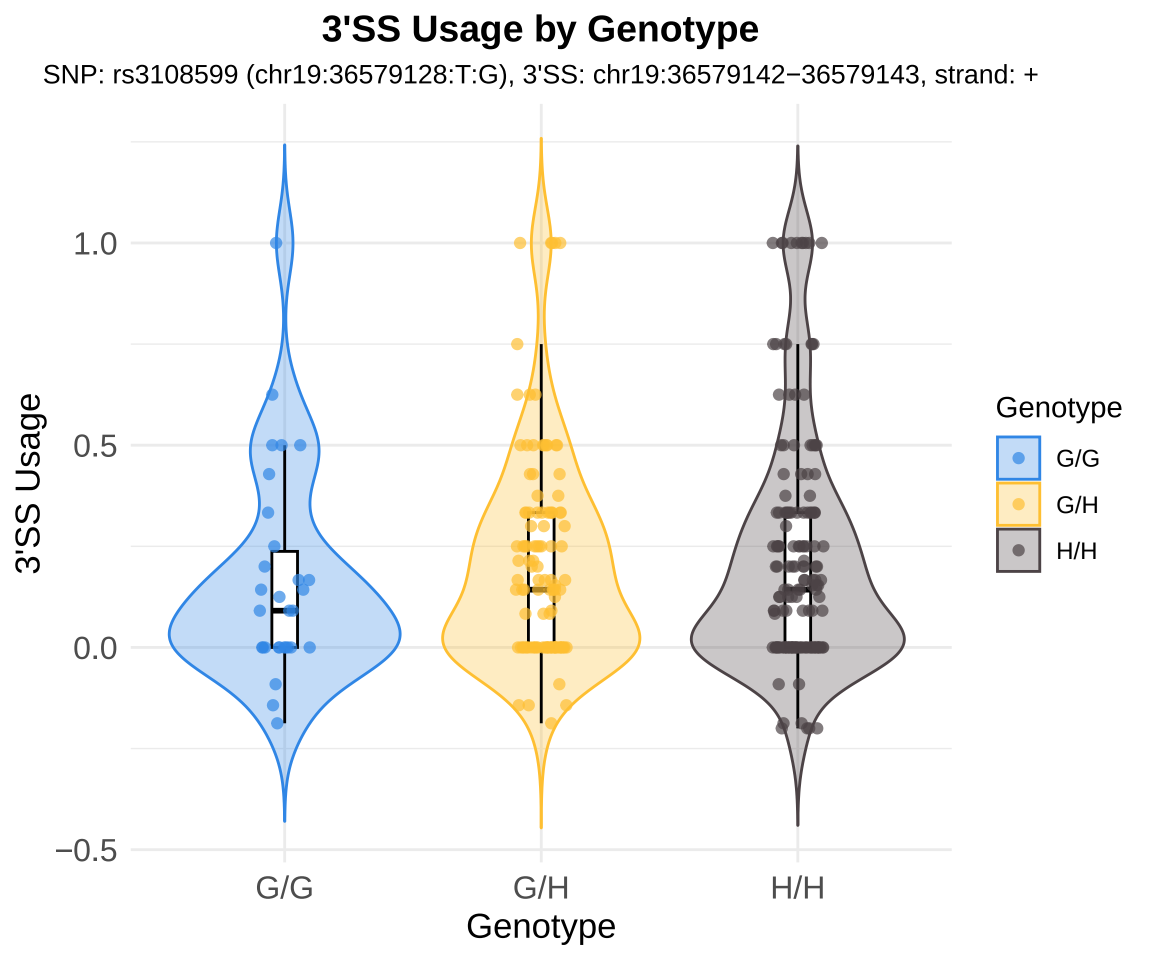
3'SS Usage by Genotype
SNP: rs3108599 (chr19:36579128:T:G), 3'SS: chr19:36579142−36579143, strand: +
3'SS Usage
1.0
0.5
0.0
−0.5
G/G
G/H
H/H
Genotype
Genotype
G/G
G/H
H/H

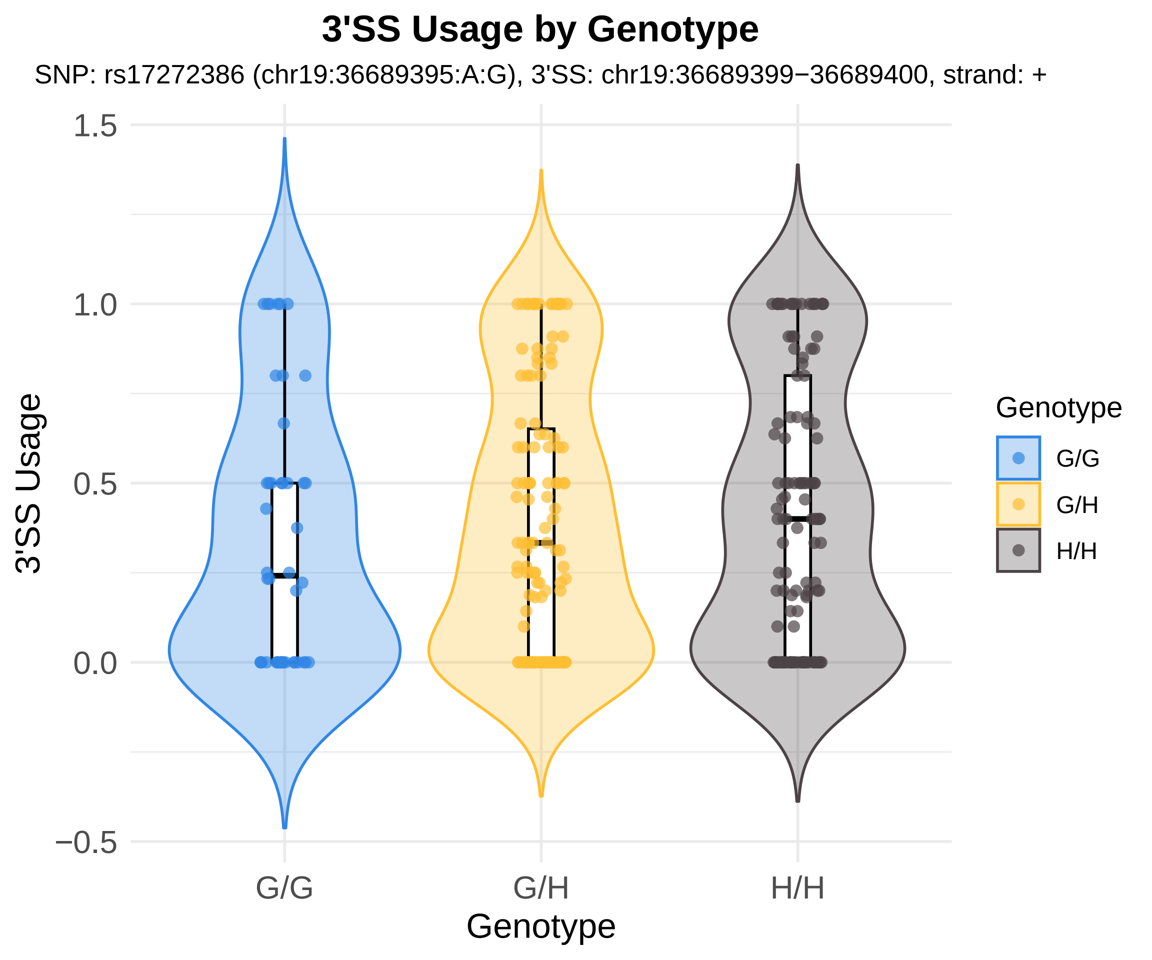

3'SS Usage by Genotype
SNP: rs17272386 (chr19:36689395:A:G), 3'SS: chr19:36689399−36689400, strand: +
1.5
1.0
0.5
0.0
−0.5
3'SS Usage
G/G
G/H
H/H
Genotype
Genotype
G/G
G/H
H/H

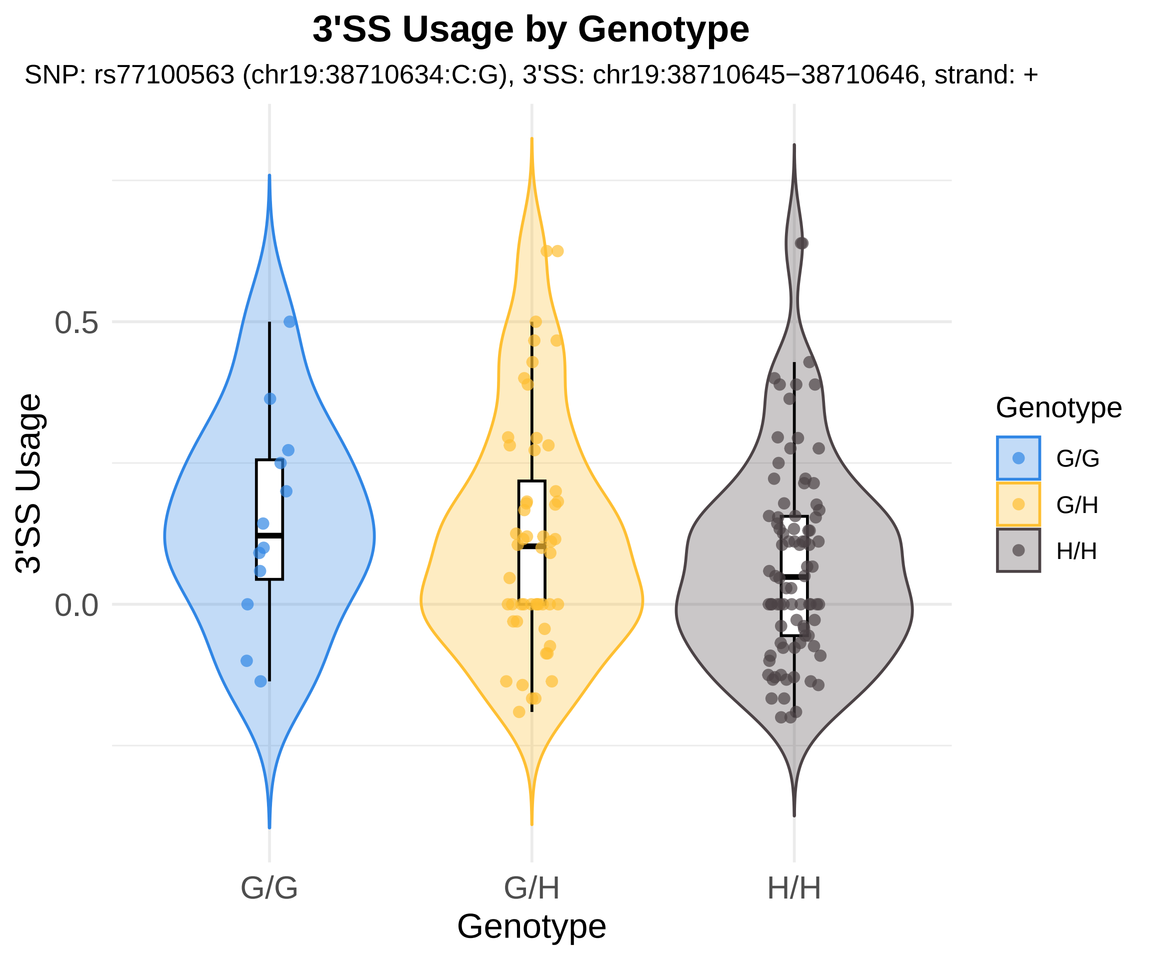
3'SS Usage by Genotype
SNP: rs77100563 (chr19:38710634:C:G), 3'SS: chr19:38710645−38710646, strand: +
3'SS Usage
0.5
0.0
G/G
G/H
H/H
Genotype
Genotype
G/G
G/H
H/H

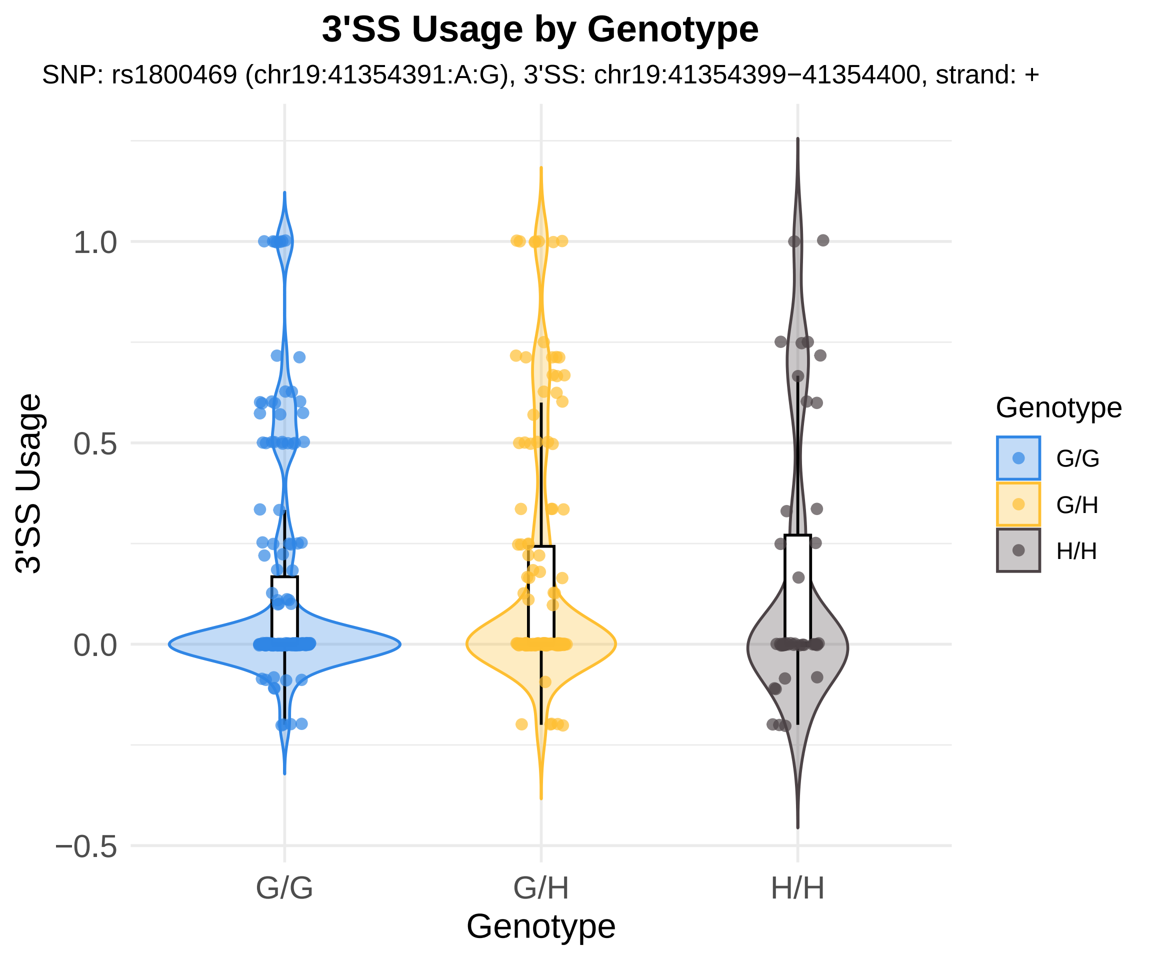
3'SS Usage by Genotype
SNP: rs1800469 (chr19:41354391:A:G), 3'SS: chr19:41354399−41354400, strand: +
3'SS Usage
1.0
0.5
0.0
−0.5
G/G
G/H
H/H
Genotype
Genotype
G/G
G/H
H/H

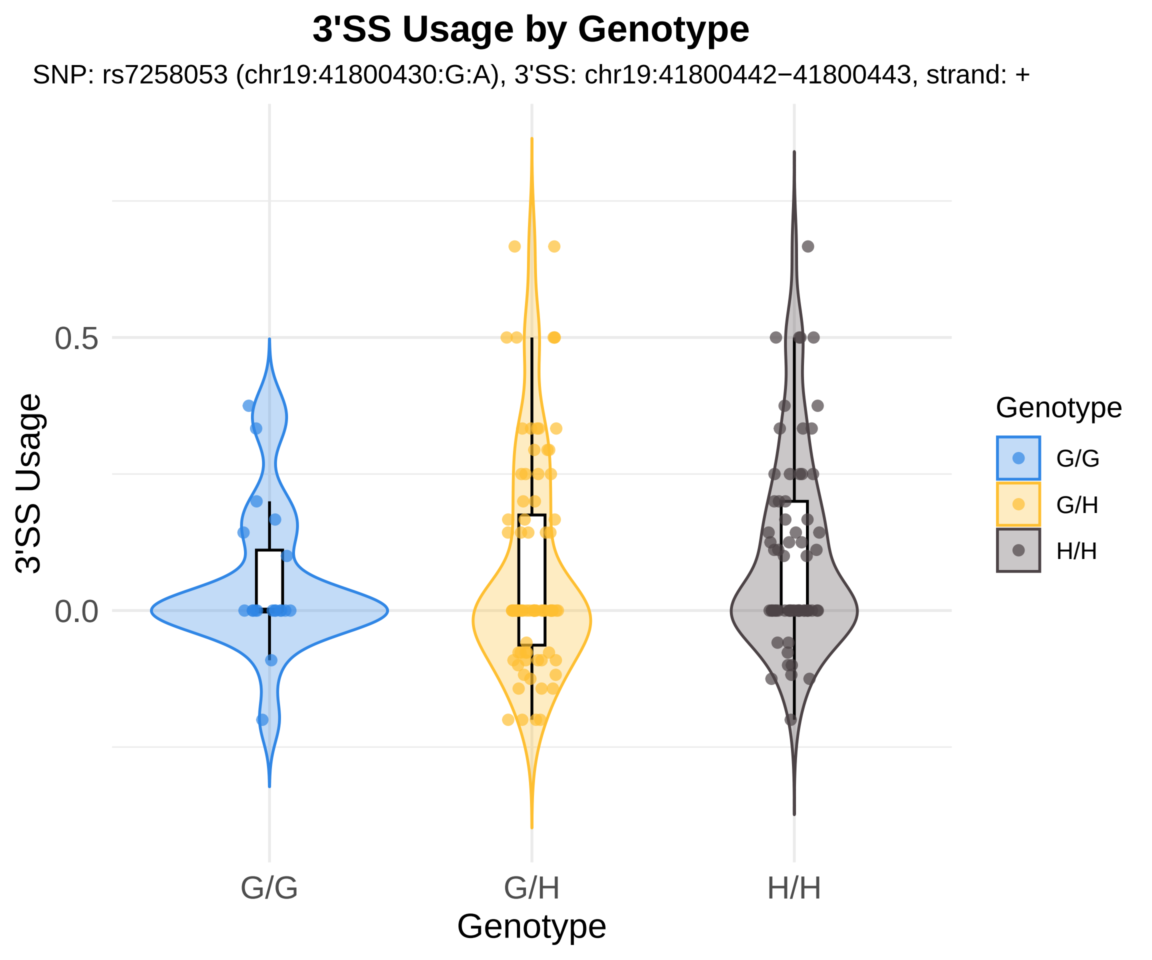

3'SS Usage by Genotype
SNP: rs7258053 (chr19:41800430:G:A), 3'SS: chr19:41800442−41800443, strand: +
3'SS Usage
0.5
0.0
G/G
G/H
H/H
Genotype
Genotype
G/G
G/H
H/H

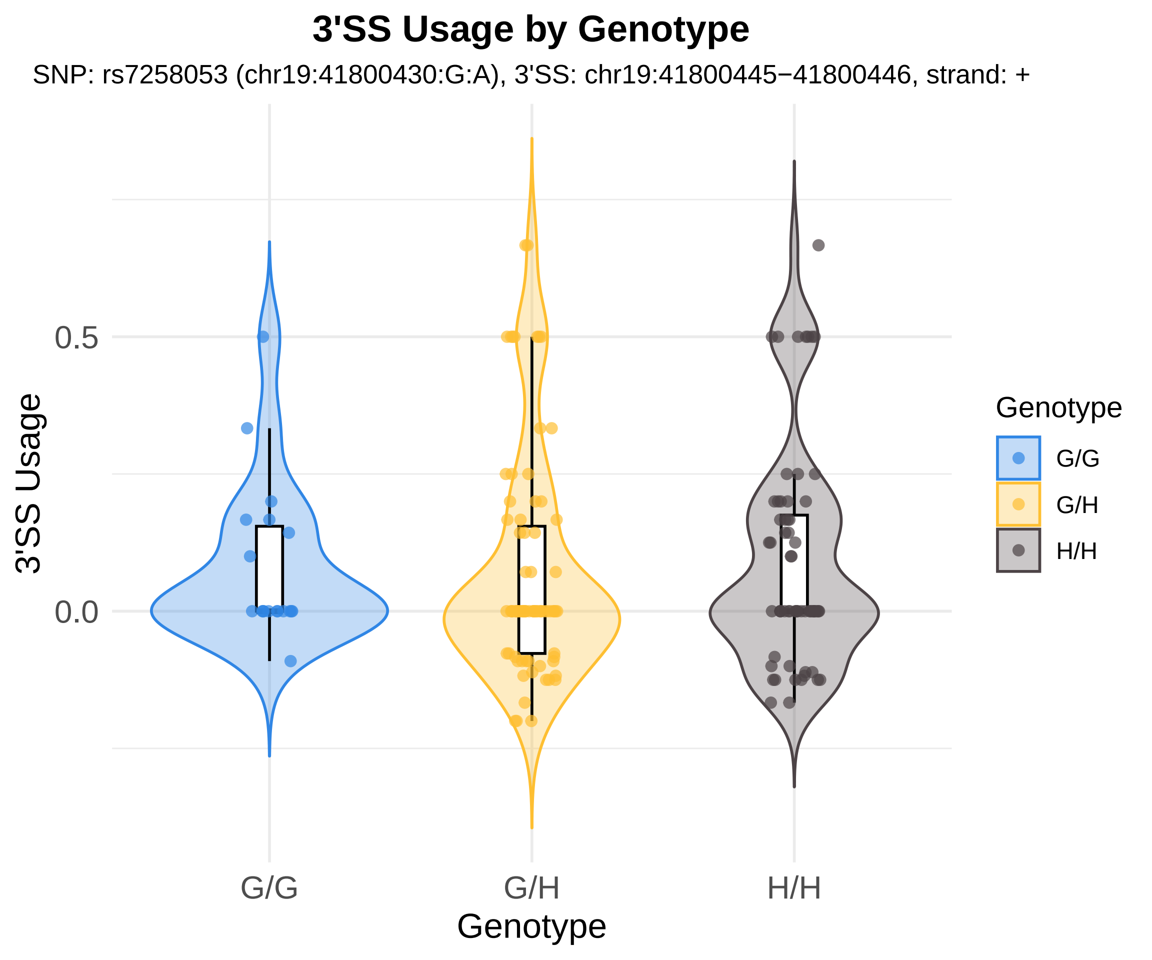

3'SS Usage by Genotype
SNP: rs7258053 (chr19:41800430:G:A), 3'SS: chr19:41800445−41800446, strand: +
3'SS Usage
0.5
0.0
G/G
G/H
H/H
Genotype
Genotype
G/G
G/H
H/H

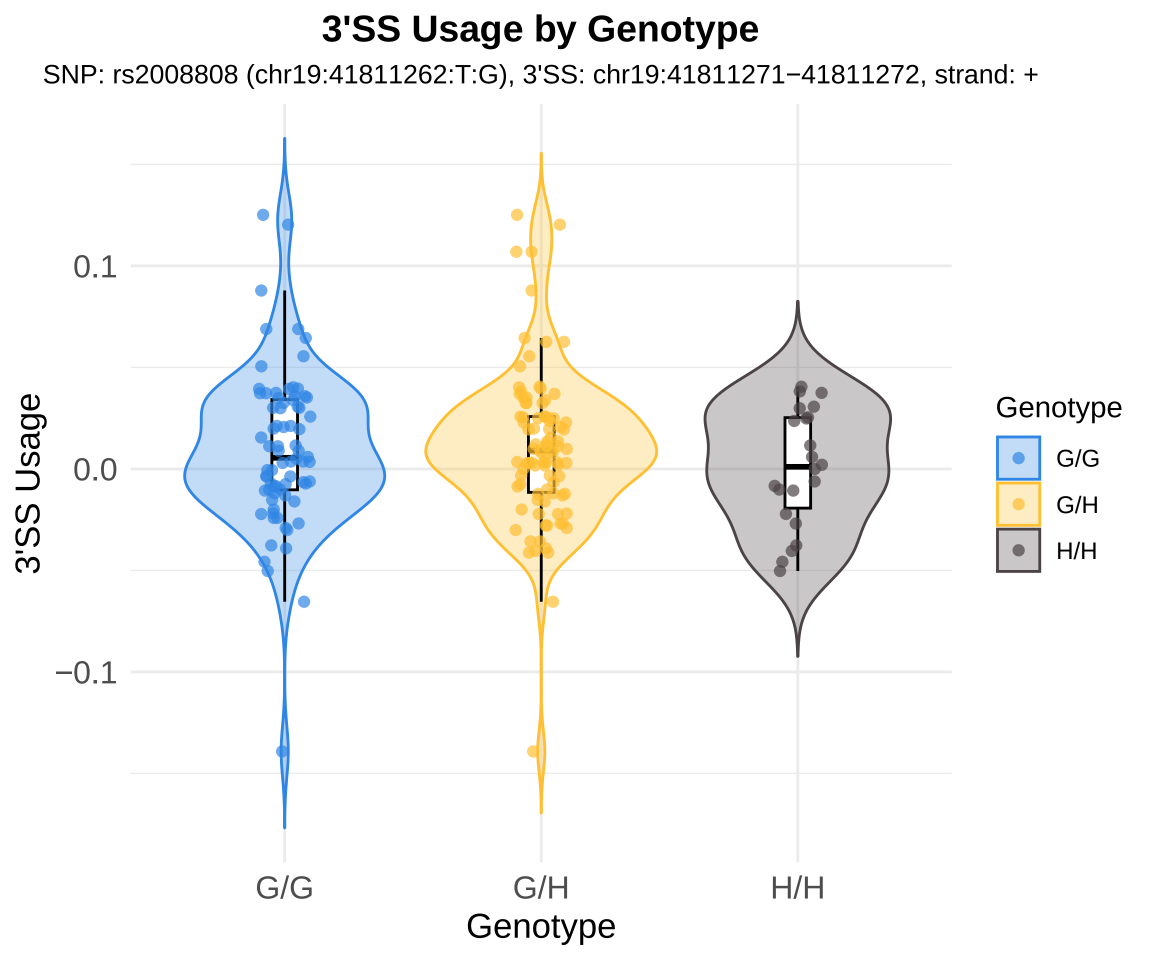
3'SS Usage by Genotype
SNP: rs2008808 (chr19:41811262:T:G), 3'SS: chr19:41811271−41811272, strand: +
3'SS Usage
0.1
0.0
−0.1
G/G
G/H
H/H
Genotype
Genotype
G/G
G/H
H/H

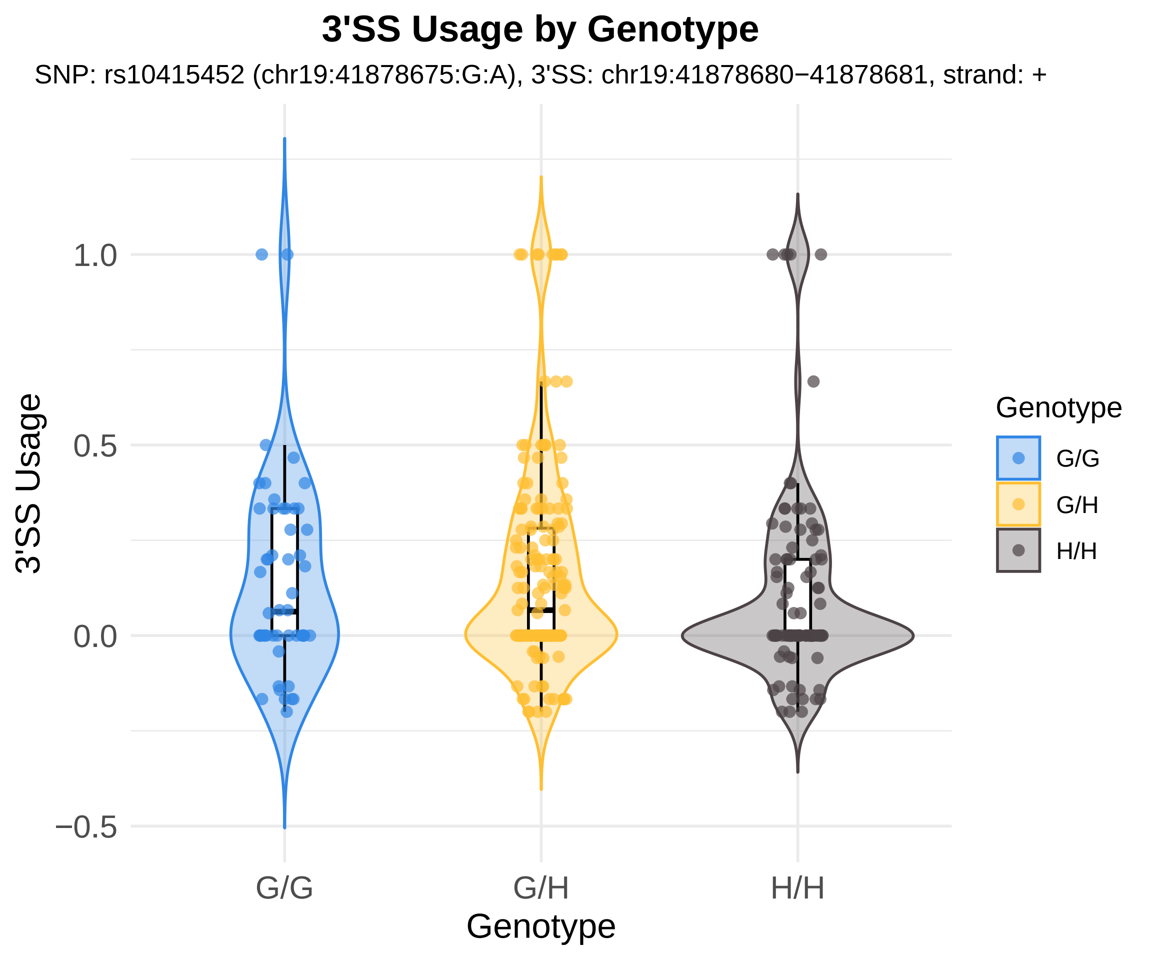

3'SS Usage by Genotype
SNP: rs10415452 (chr19:41878675:G:A), 3'SS: chr19:41878680−41878681, strand: +
3'SS Usage
1.0
0.5
0.0
−0.5
G/G
G/H
H/H
Genotype
Genotype
G/G
G/H
H/H

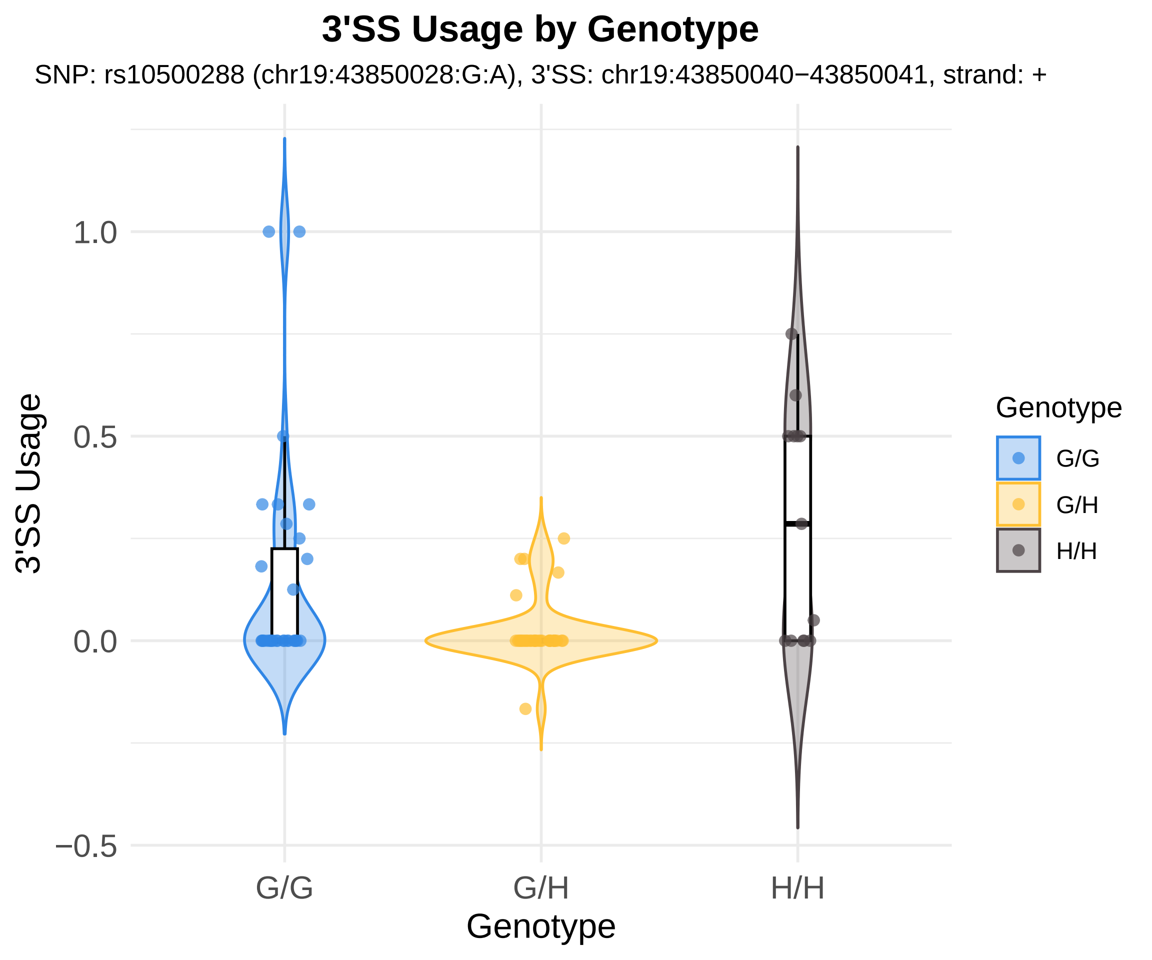
3'SS Usage by Genotype
SNP: rs10500288 (chr19:43850028:G:A), 3'SS: chr19:43850040−43850041, strand: +
3'SS Usage
1.0
0.5
0.0
−0.5
G/G
G/H
H/H
Genotype
Genotype
G/G
G/H
H/H

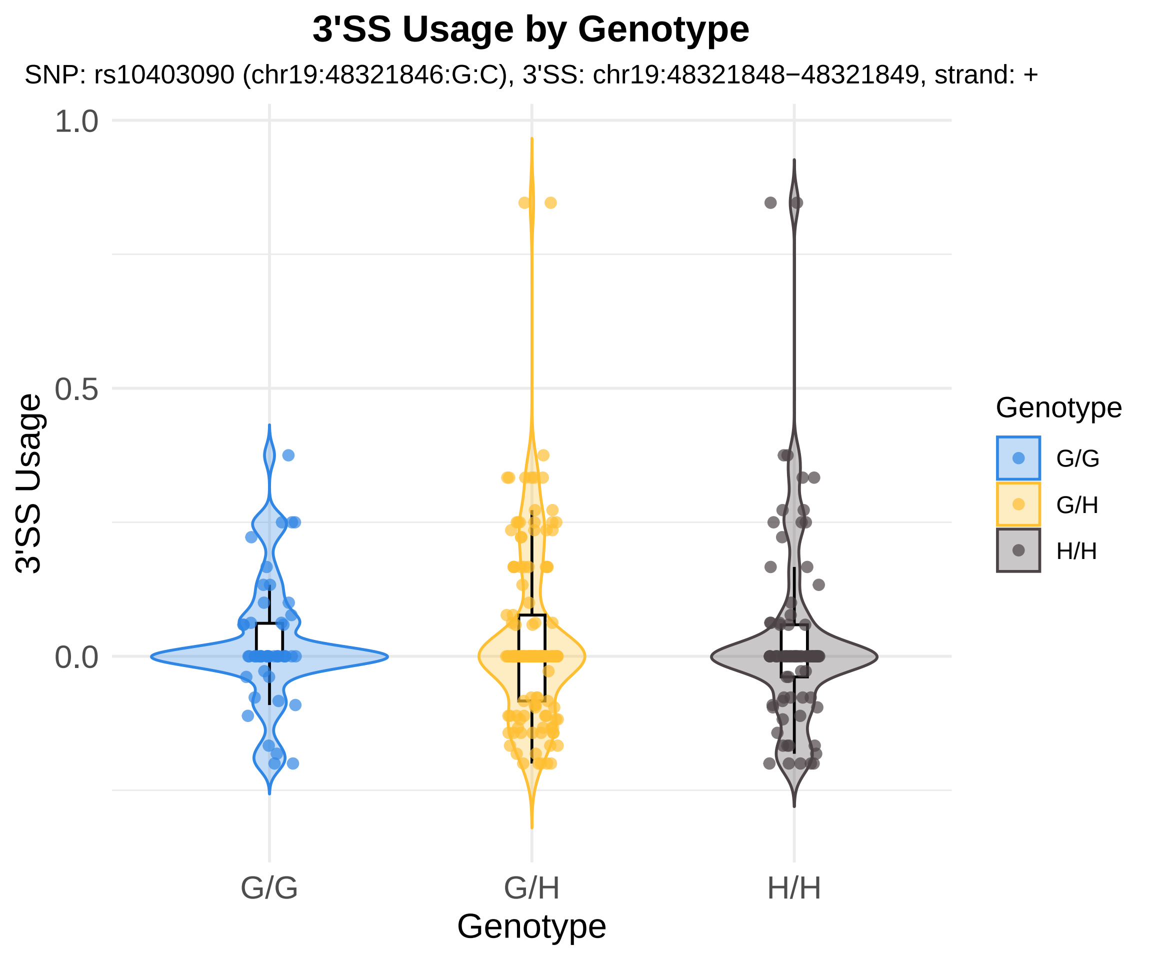

3'SS Usage by Genotype
SNP: rs10403090 (chr19:48321846:G:C), 3'SS: chr19:48321848−48321849, strand: +
1.0
0.5
0.0
3'SS Usage
G/G
G/H
H/H
Genotype
Genotype
G/G
G/H
H/H

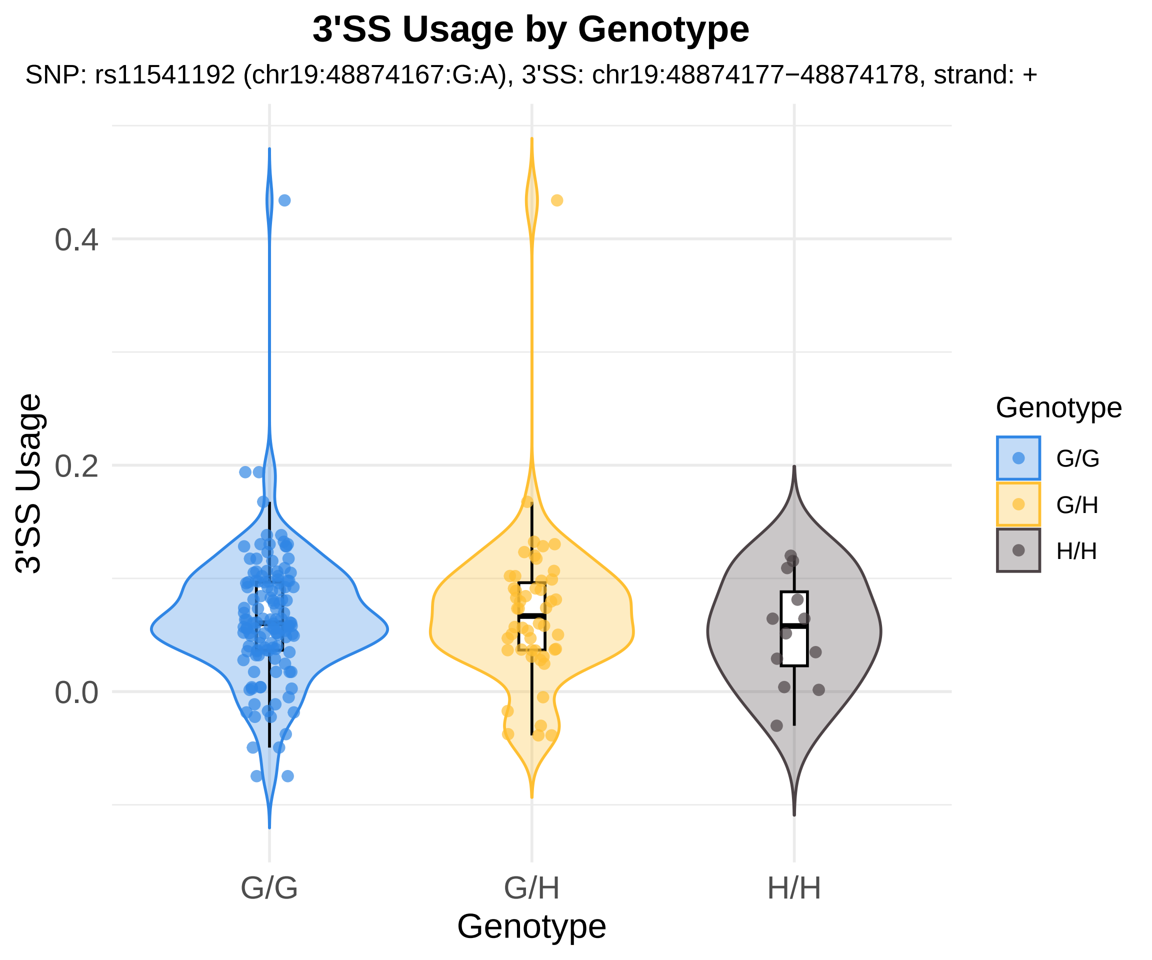

3'SS Usage by Genotype
SNP: rs11541192 (chr19:48874167:G:A), 3'SS: chr19:48874177−48874178, strand: +
3'SS Usage
0.4
0.2
0.0
G/G
G/H
H/H
Genotype
Genotype
G/G
G/H
H/H

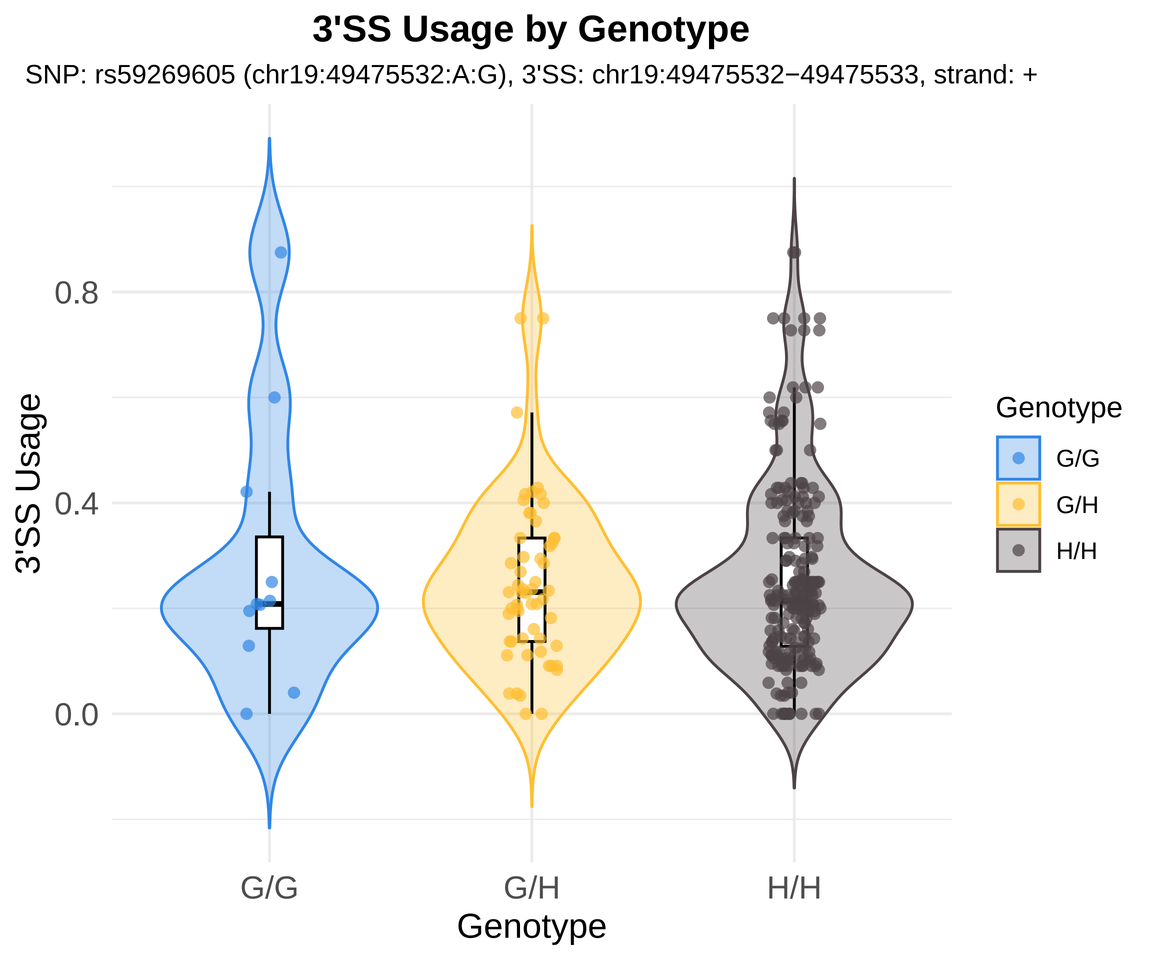
3'SS Usage by Genotype
SNP: rs59269605 (chr19:49475532:A:G), 3'SS: chr19:49475532−49475533, strand: +
3'SS Usage
0.8
0.4
0.0
G/G
G/H
H/H
Genotype
Genotype
G/G
G/H
H/H

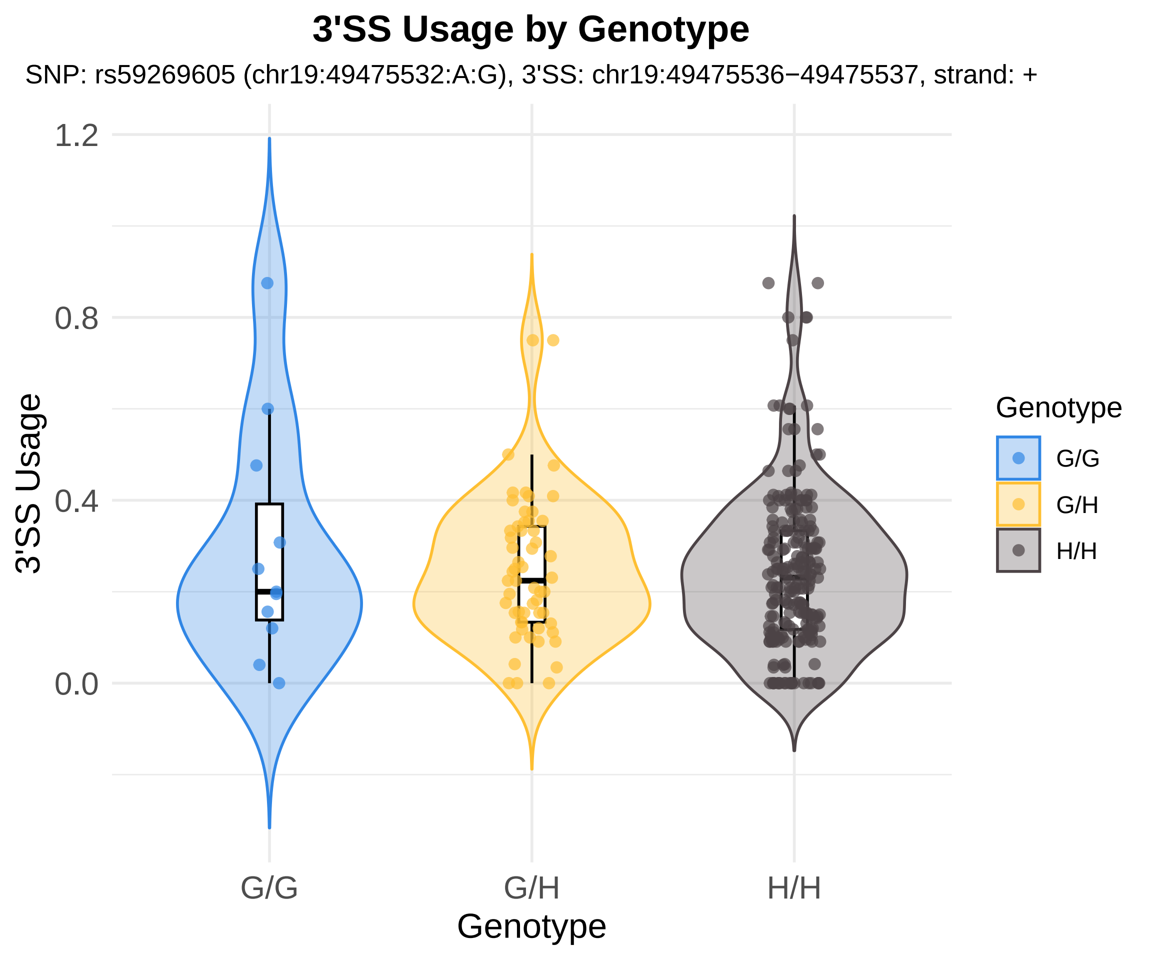

3'SS Usage by Genotype
SNP: rs59269605 (chr19:49475532:A:G), 3'SS: chr19:49475536−49475537, strand: +
1.2
0.8
0.4
0.0
3'SS Usage
G/G
G/H
H/H
Genotype
Genotype
G/G
G/H
H/H

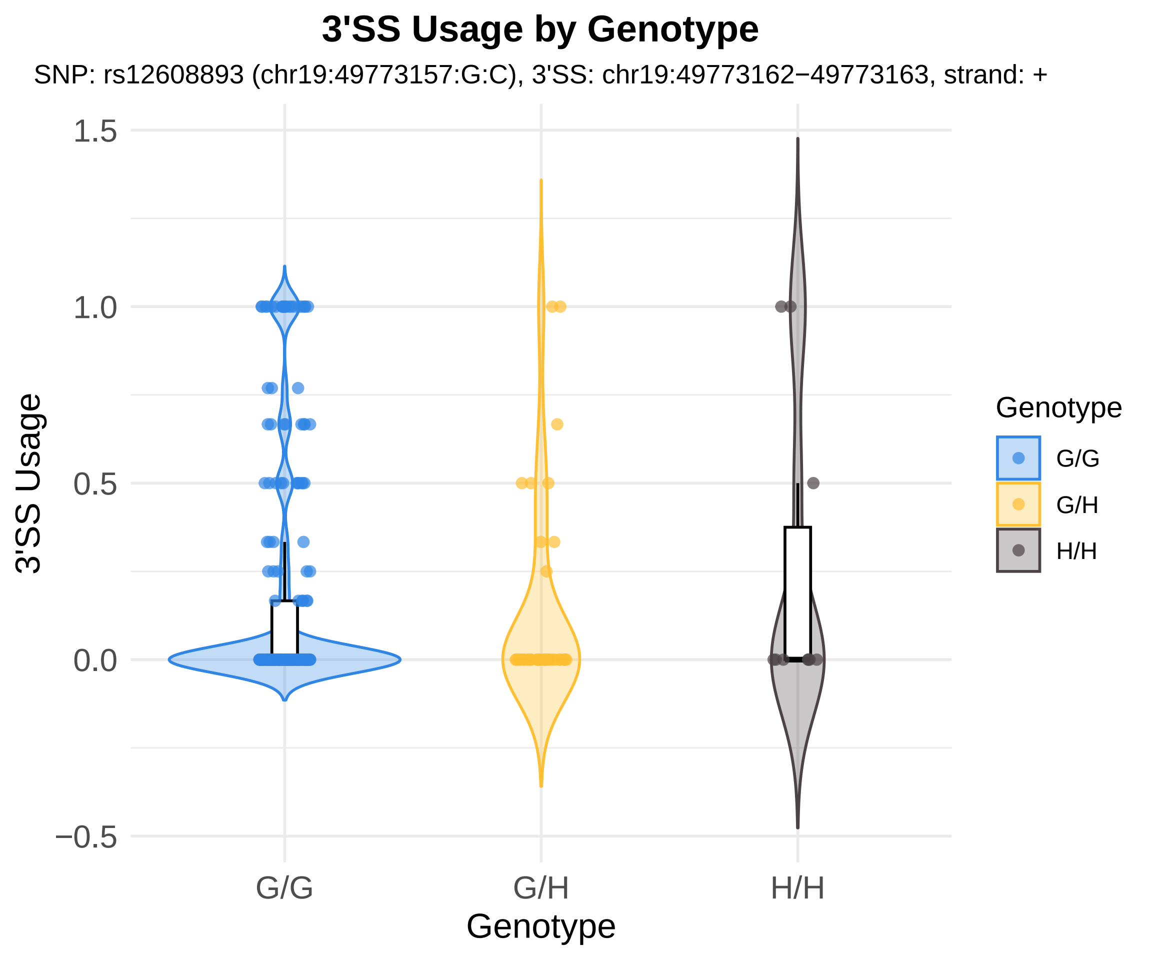
3'SS Usage by Genotype
SNP: rs12608893 (chr19:49773157:G:C), 3'SS: chr19:49773162−49773163, strand: +
3'SS Usage
1.5
1.0
0.5
0.0
−0.5
G/G
G/H
H/H
Genotype
Genotype
G/G
G/H
H/H

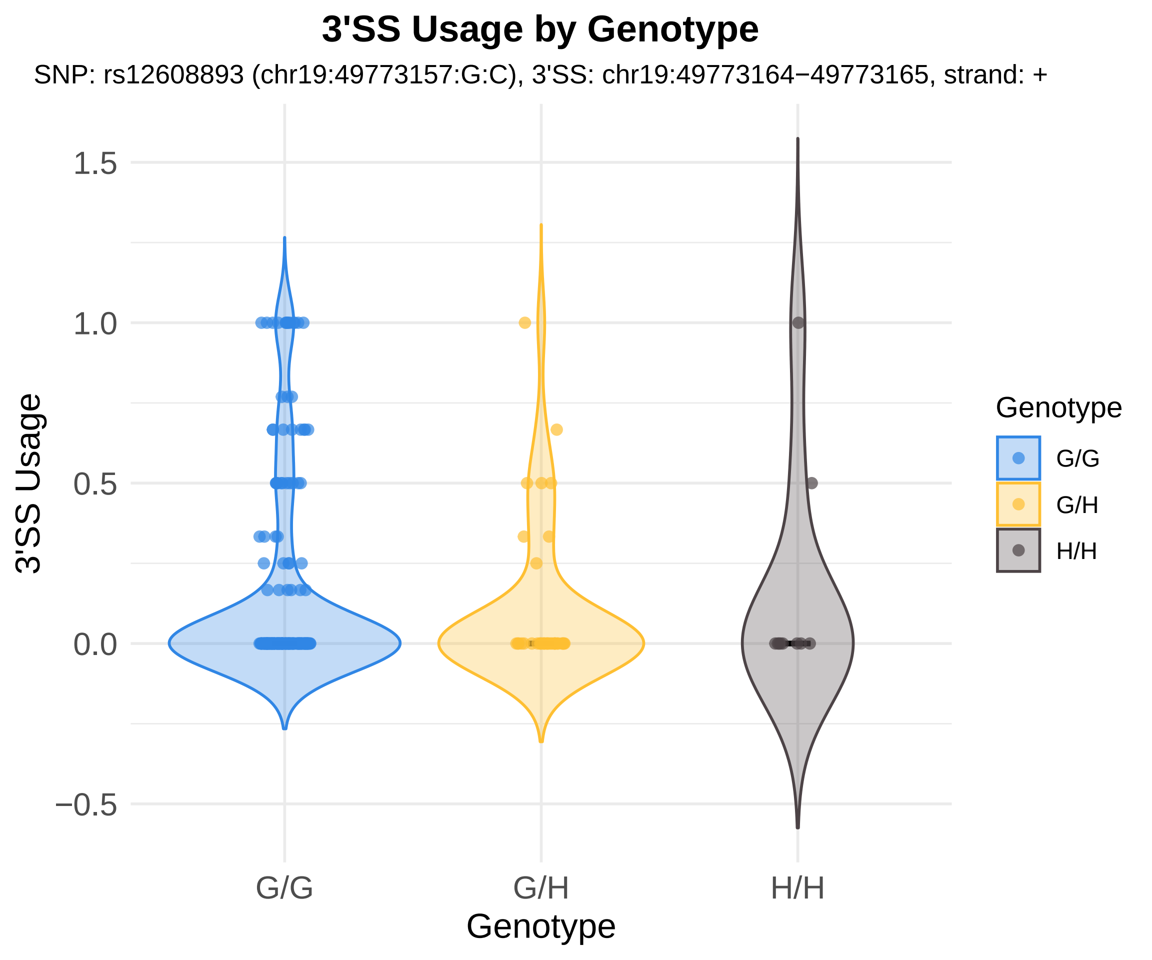

3'SS Usage by Genotype
SNP: rs12608893 (chr19:49773157:G:C), 3'SS: chr19:49773164−49773165, strand: +
3'SS Usage
1.5
1.0
0.5
0.0
−0.5
G/G
G/H
H/H
Genotype
Genotype
G/G
G/H
H/H

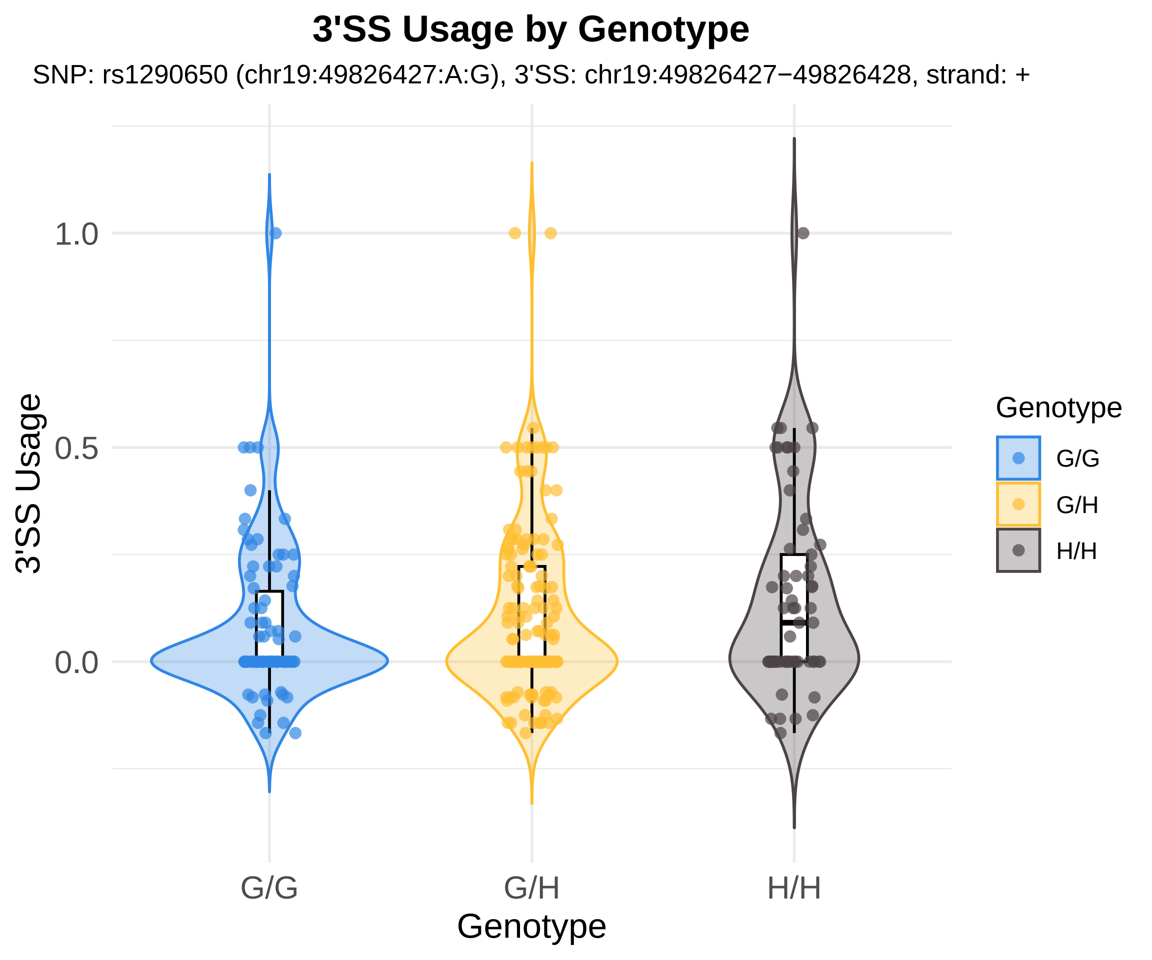
3'SS Usage by Genotype
SNP: rs1290650 (chr19:49826427:A:G), 3'SS: chr19:49826427−49826428, strand: +
3'SS Usage
1.0
0.5
0.0
G/G
G/H
H/H
Genotype
Genotype
G/G
G/H
H/H

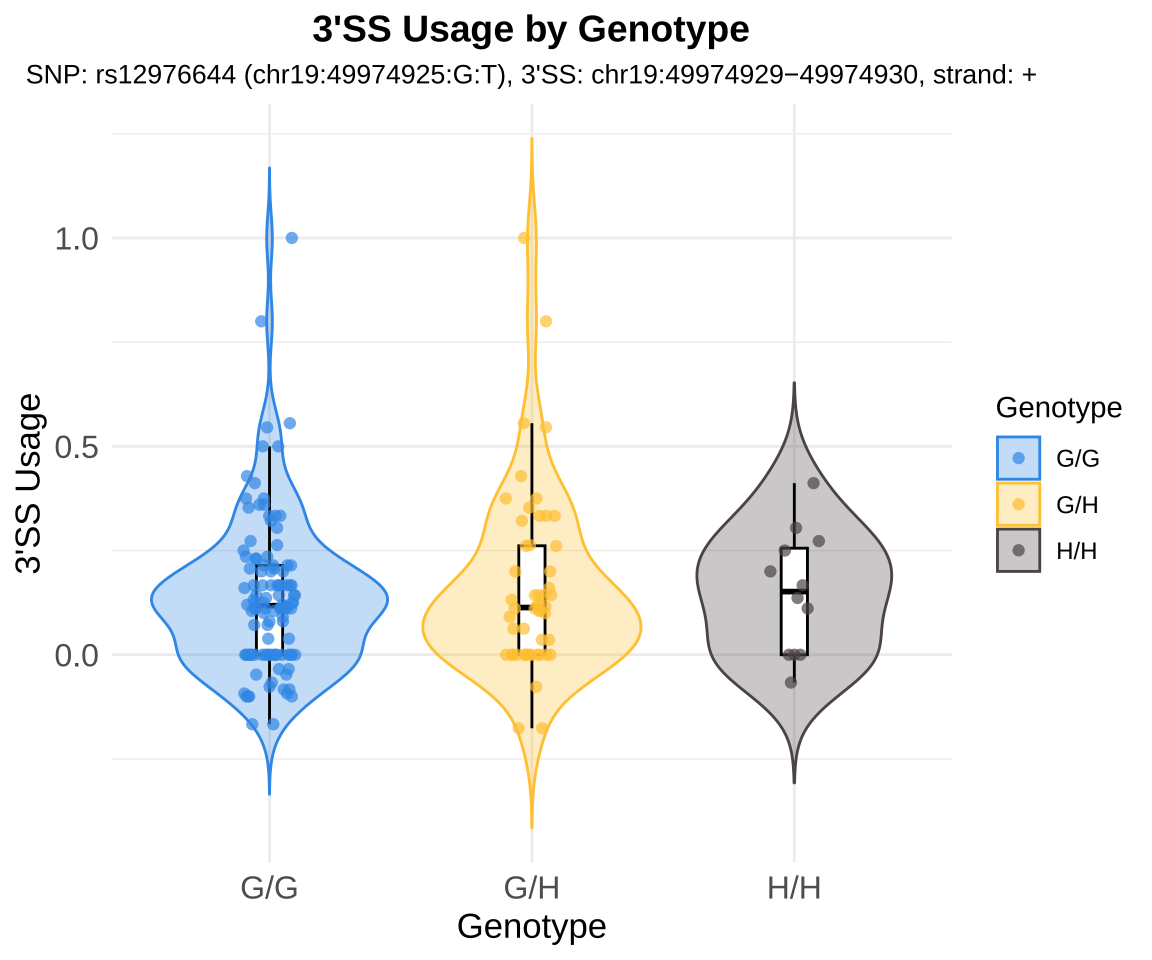

3'SS Usage by Genotype
SNP: rs12976644 (chr19:49974925:G:T), 3'SS: chr19:49974929−49974930, strand: +
3'SS Usage
1.0
0.5
0.0
G/G
G/H
H/H
Genotype
Genotype
G/G
G/H
H/H

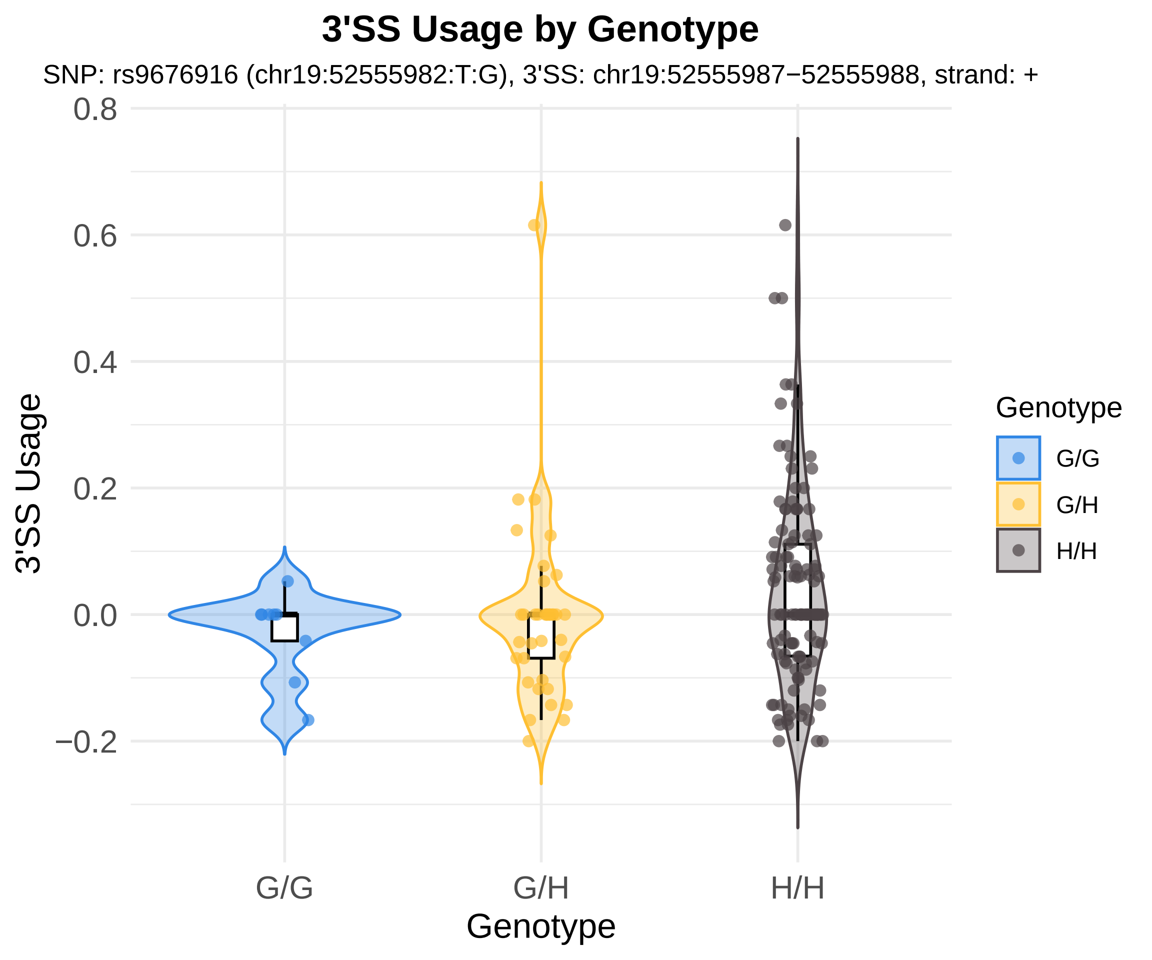
3'SS Usage by Genotype
SNP: rs9676916 (chr19:52555982:T:G), 3'SS: chr19:52555987−52555988, strand: +
0.8
0.6
0.4
0.2
0.0
−0.2
3'SS Usage
G/G
G/H
H/H
Genotype
Genotype
G/G
G/H
H/H

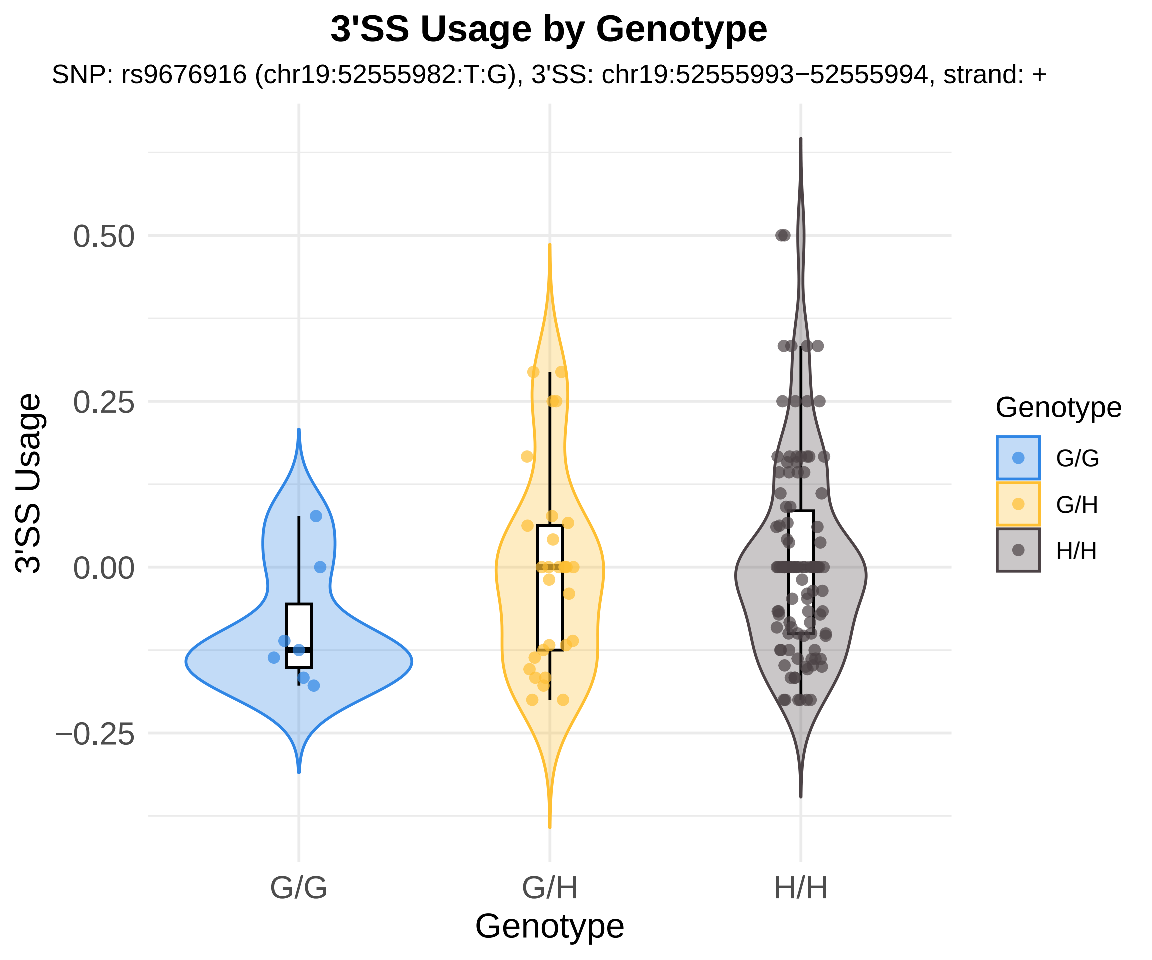
3'SS Usage by Genotype
SNP: rs9676916 (chr19:52555982:T:G), 3'SS: chr19:52555993−52555994, strand: +
3'SS Usage
0.50
0.25
0.00
−0.25
G/G
G/H
H/H
Genotype
Genotype
G/G
G/H
H/H

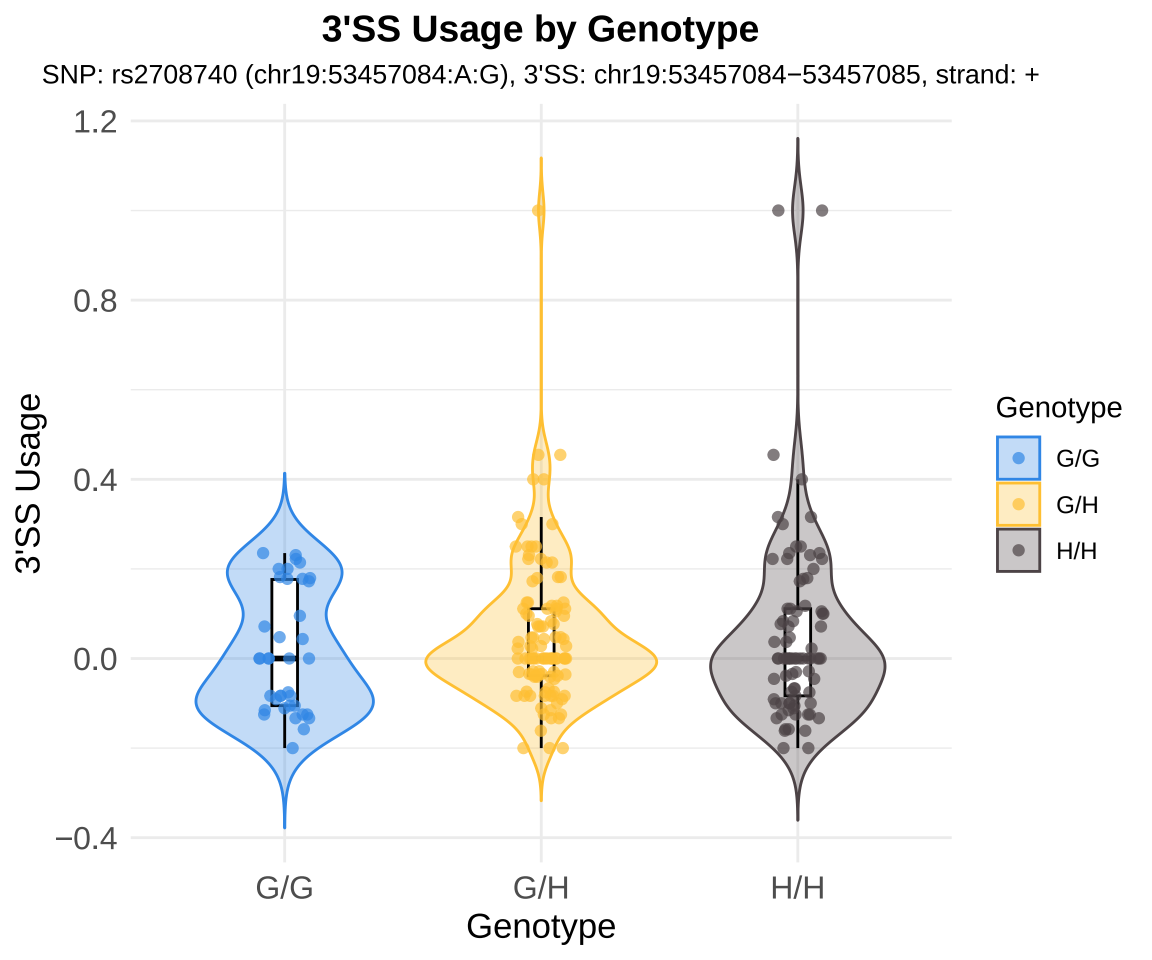
3'SS Usage by Genotype
SNP: rs2708740 (chr19:53457084:A:G), 3'SS: chr19:53457084−53457085, strand: +
1.2
0.8
0.4
0.0
−0.4
3'SS Usage
G/G
G/H
H/H
Genotype
Genotype
G/G
G/H
H/H

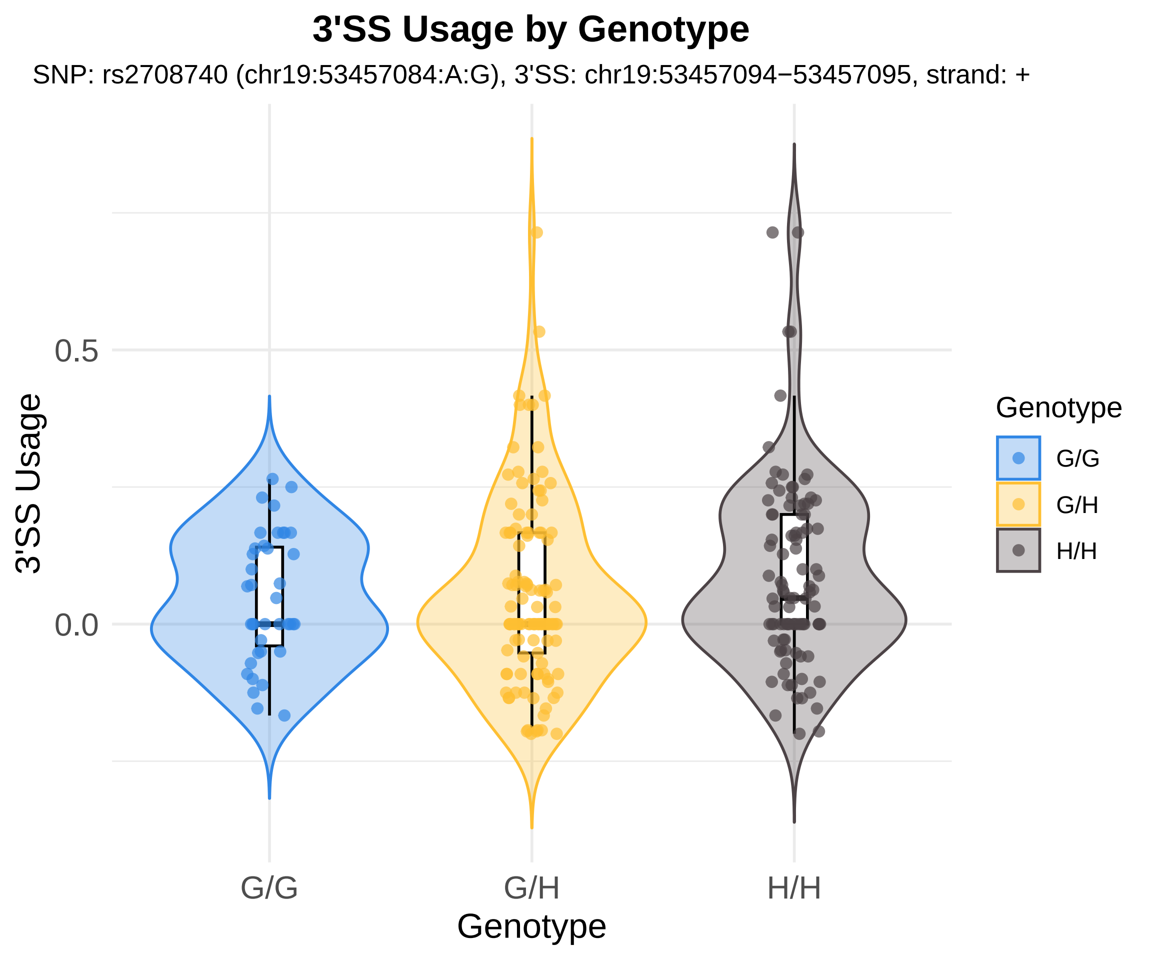
3'SS Usage by Genotype
SNP: rs2708740 (chr19:53457084:A:G), 3'SS: chr19:53457094−53457095, strand: +
3'SS Usage
0.5
0.0
G/G
G/H
H/H
Genotype
Genotype
G/G
G/H
H/H

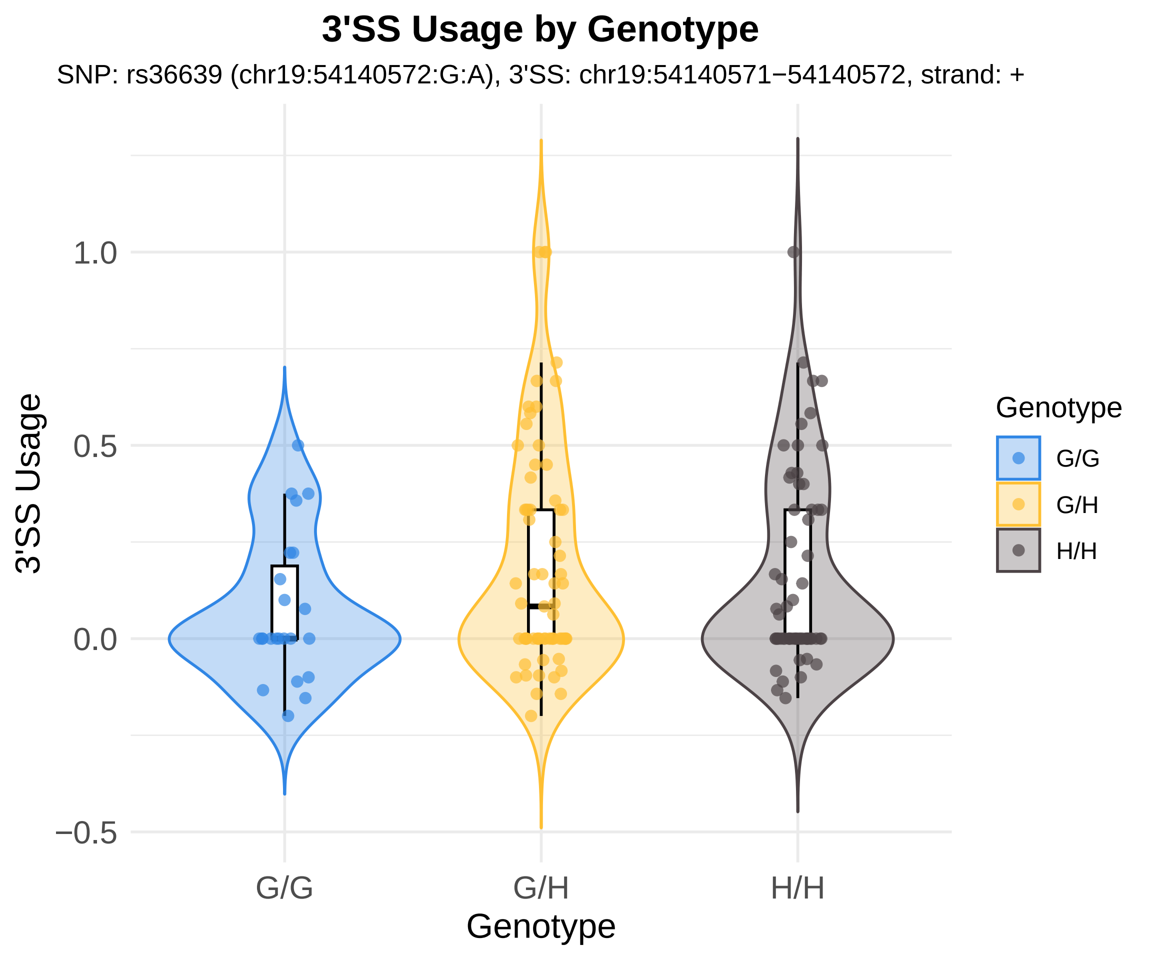
3'SS Usage by Genotype
SNP: rs36639 (chr19:54140572:G:A), 3'SS: chr19:54140571−54140572, strand: +
3'SS Usage
1.0
0.5
0.0
−0.5
G/G
G/H
H/H
Genotype
Genotype
G/G
G/H
H/H

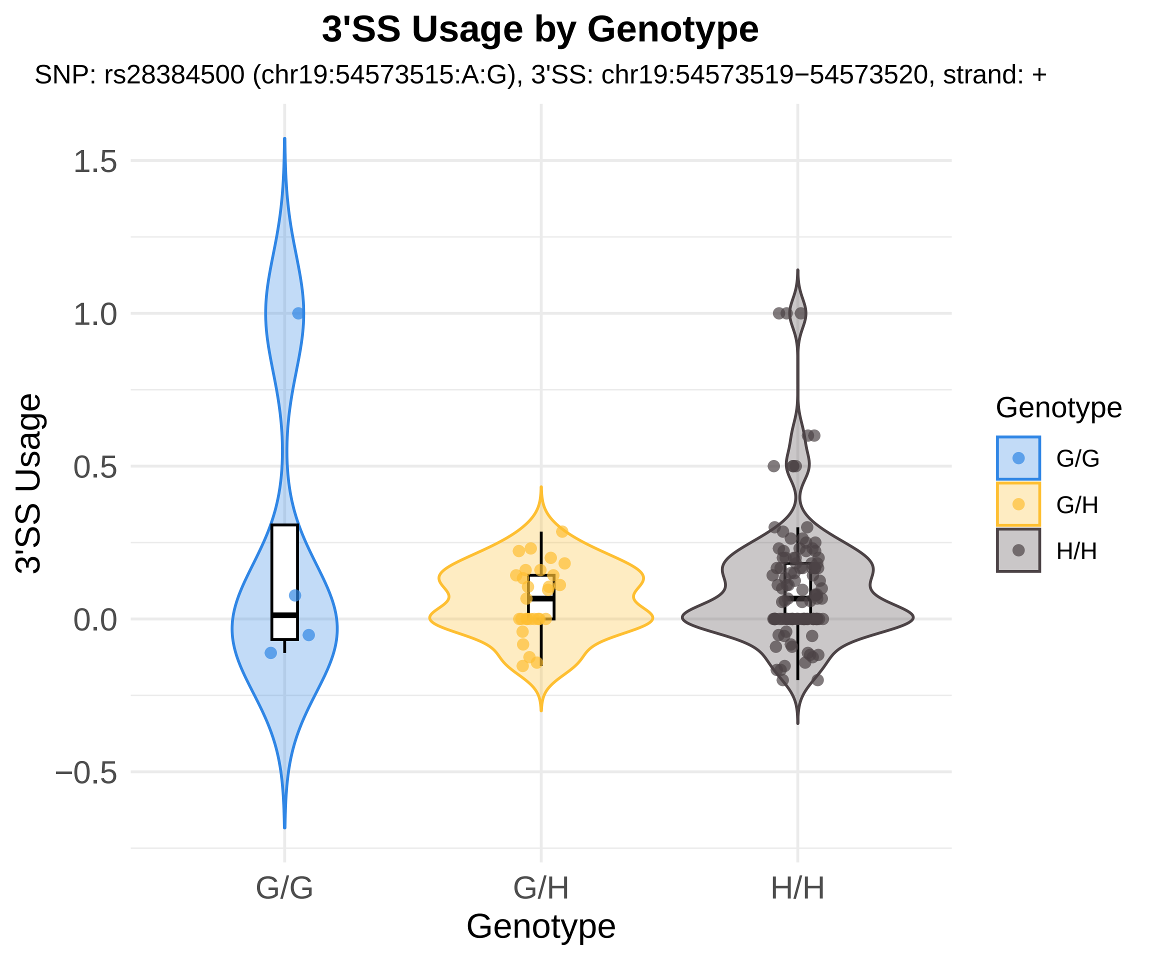
3'SS Usage by Genotype
SNP: rs28384500 (chr19:54573515:A:G), 3'SS: chr19:54573519−54573520, strand: +
3'SS Usage
1.5
1.0
0.5
0.0
−0.5
G/G
G/H
H/H
Genotype
Genotype
G/G
G/H
H/H

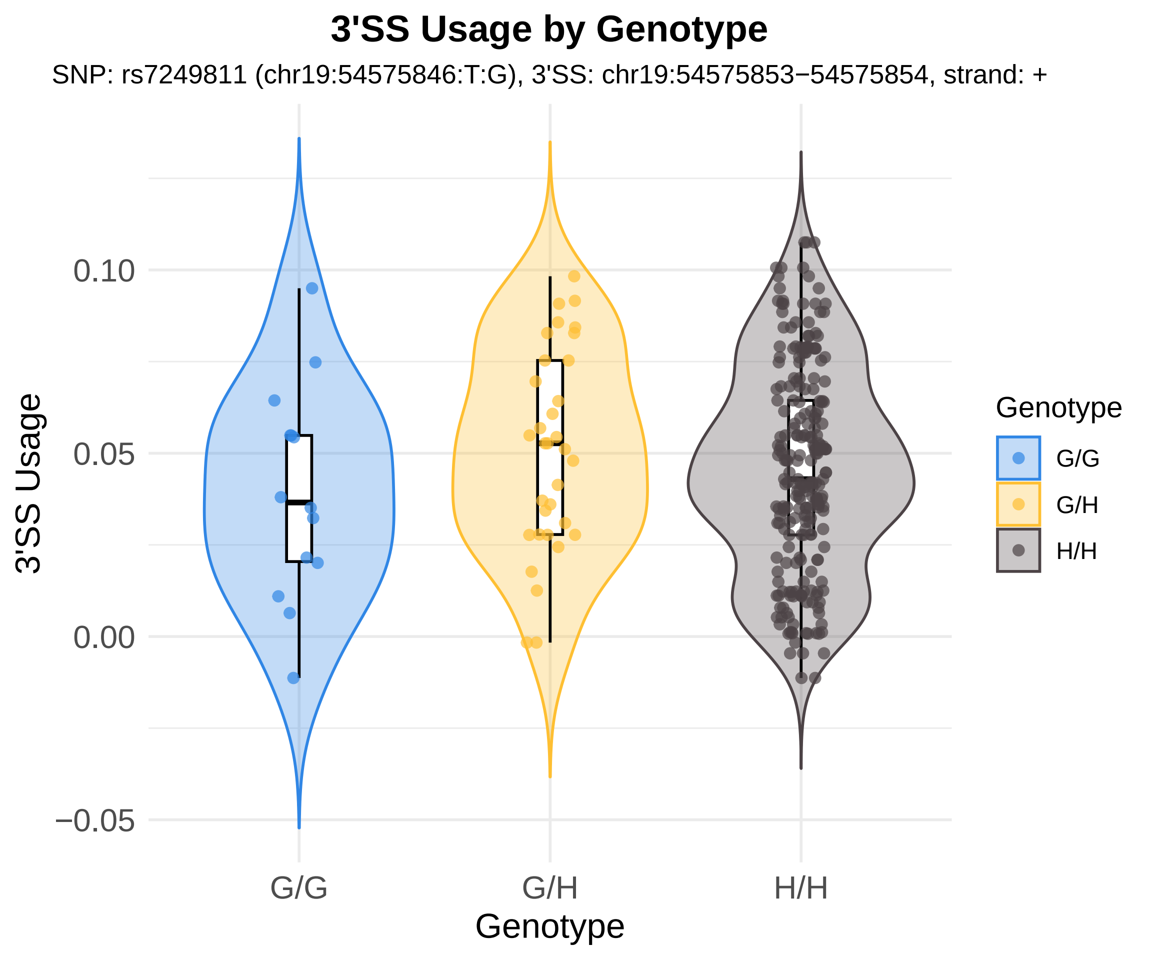
3'SS Usage by Genotype
SNP: rs7249811 (chr19:54575846:T:G), 3'SS: chr19:54575853−54575854, strand: +
3'SS Usage
0.10
0.05
0.00
−0.05
G/G
G/H
H/H
Genotype
Genotype
G/G
G/H
H/H

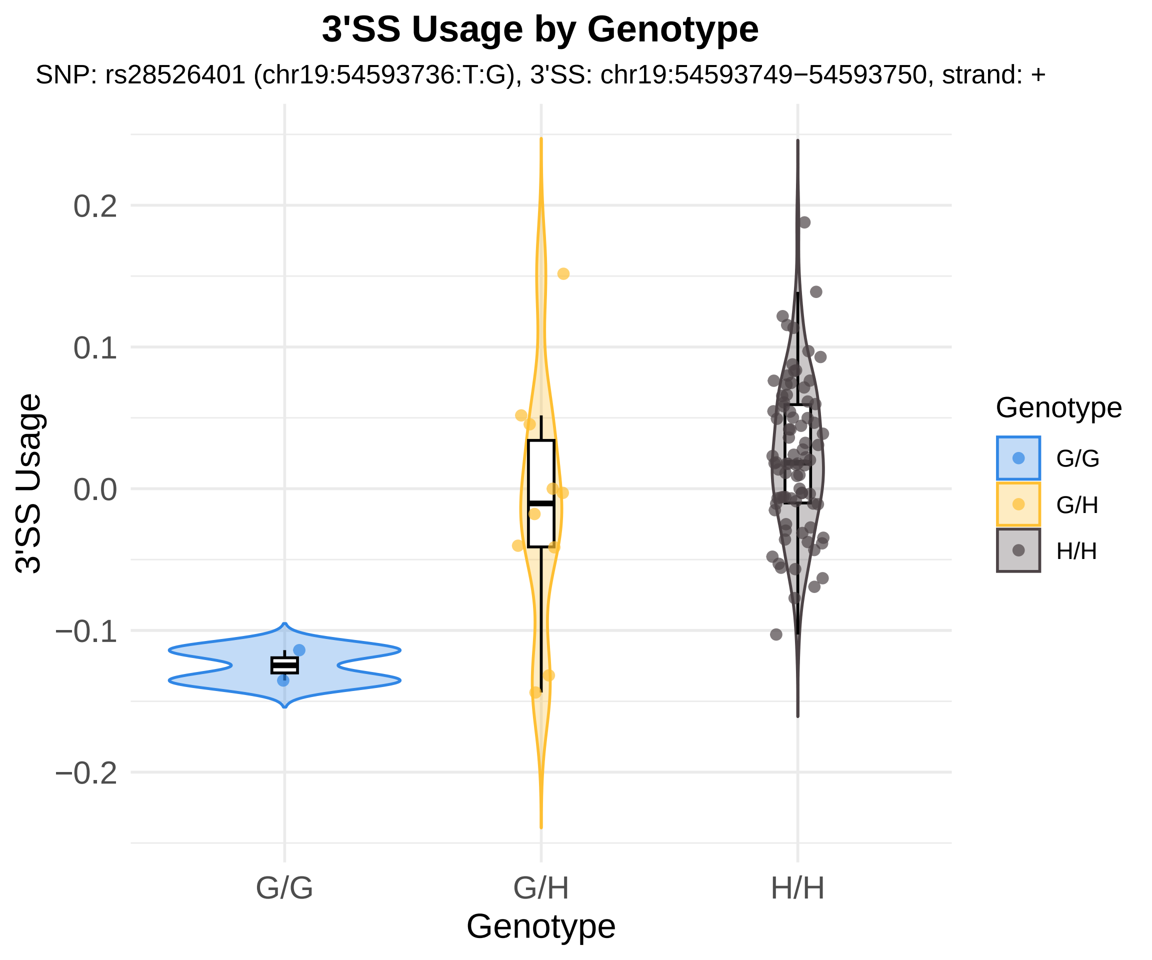
3'SS Usage by Genotype
SNP: rs28526401 (chr19:54593736:T:G), 3'SS: chr19:54593749−54593750, strand: +
3'SS Usage
0.2
0.1
0.0
−0.1
−0.2
G/G
G/H
H/H
Genotype
Genotype
G/G
G/H
H/H

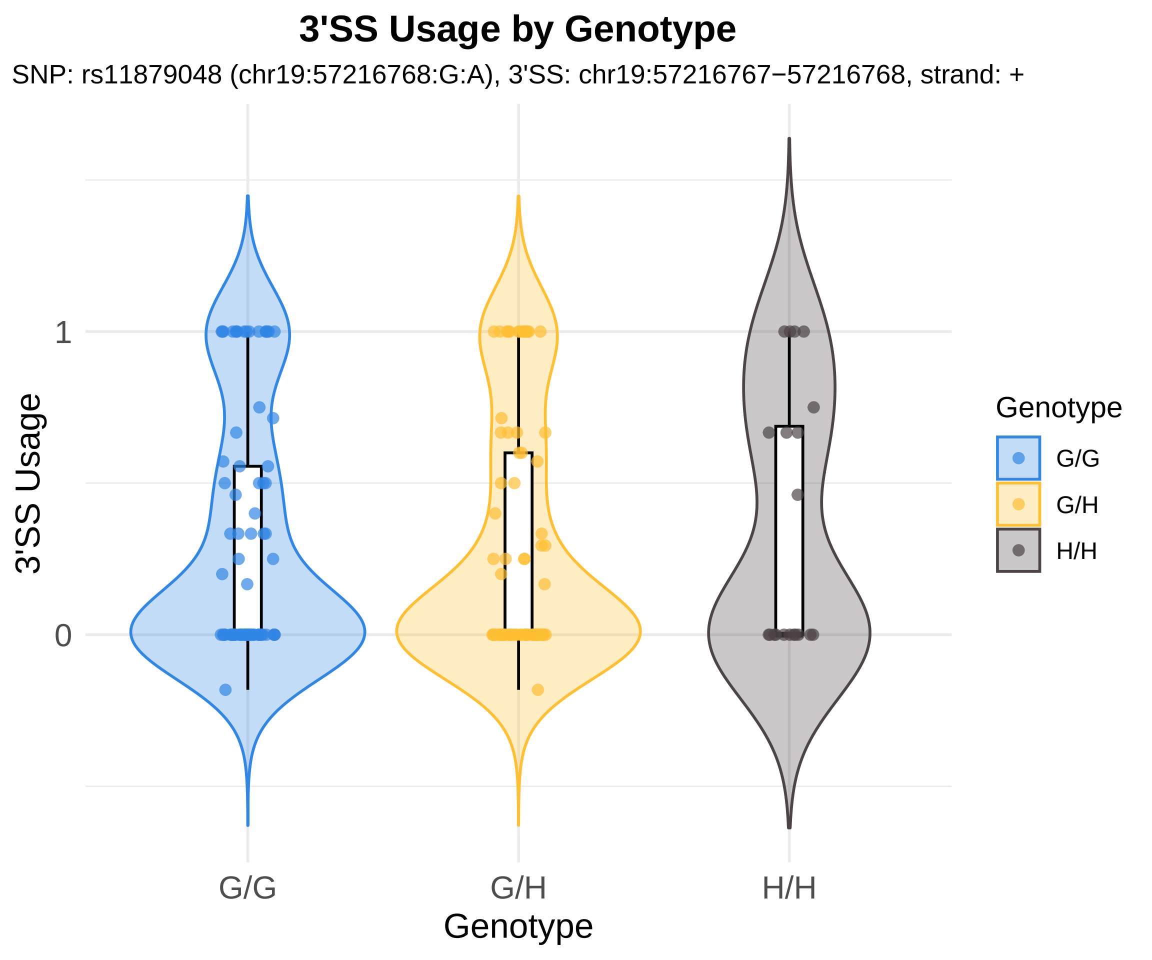

3'SS Usage by Genotype
SNP: rs11879048 (chr19:57216768:G:A), 3'SS: chr19:57216767−57216768, strand: +
3'SS Usage
1
0
G/G
G/H
H/H
Genotype
Genotype
G/G
G/H
H/H

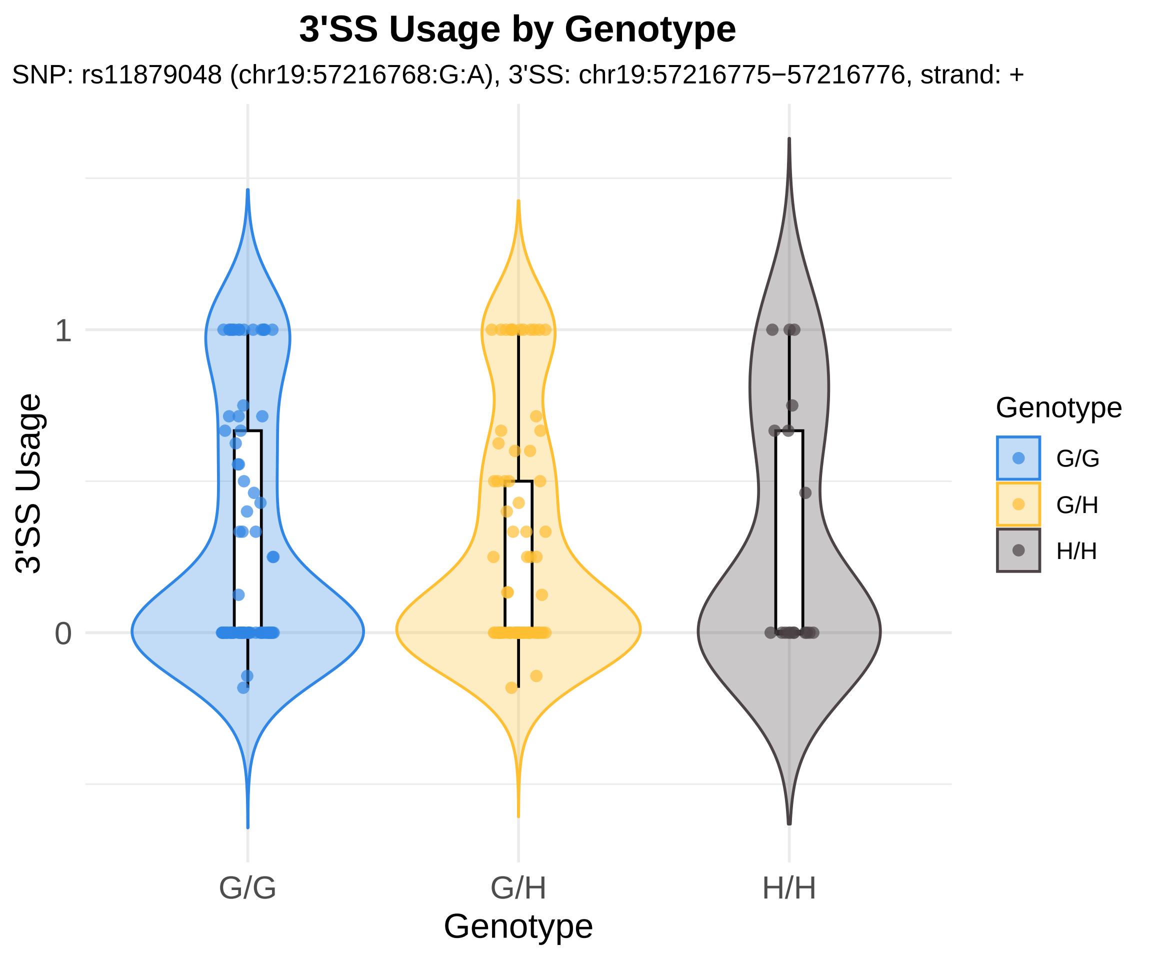
3'SS Usage by Genotype
SNP: rs11879048 (chr19:57216768:G:A), 3'SS: chr19:57216775−57216776, strand: +
3'SS Usage
1
0
G/G
G/H
H/H
Genotype
Genotype
G/G
G/H
H/H

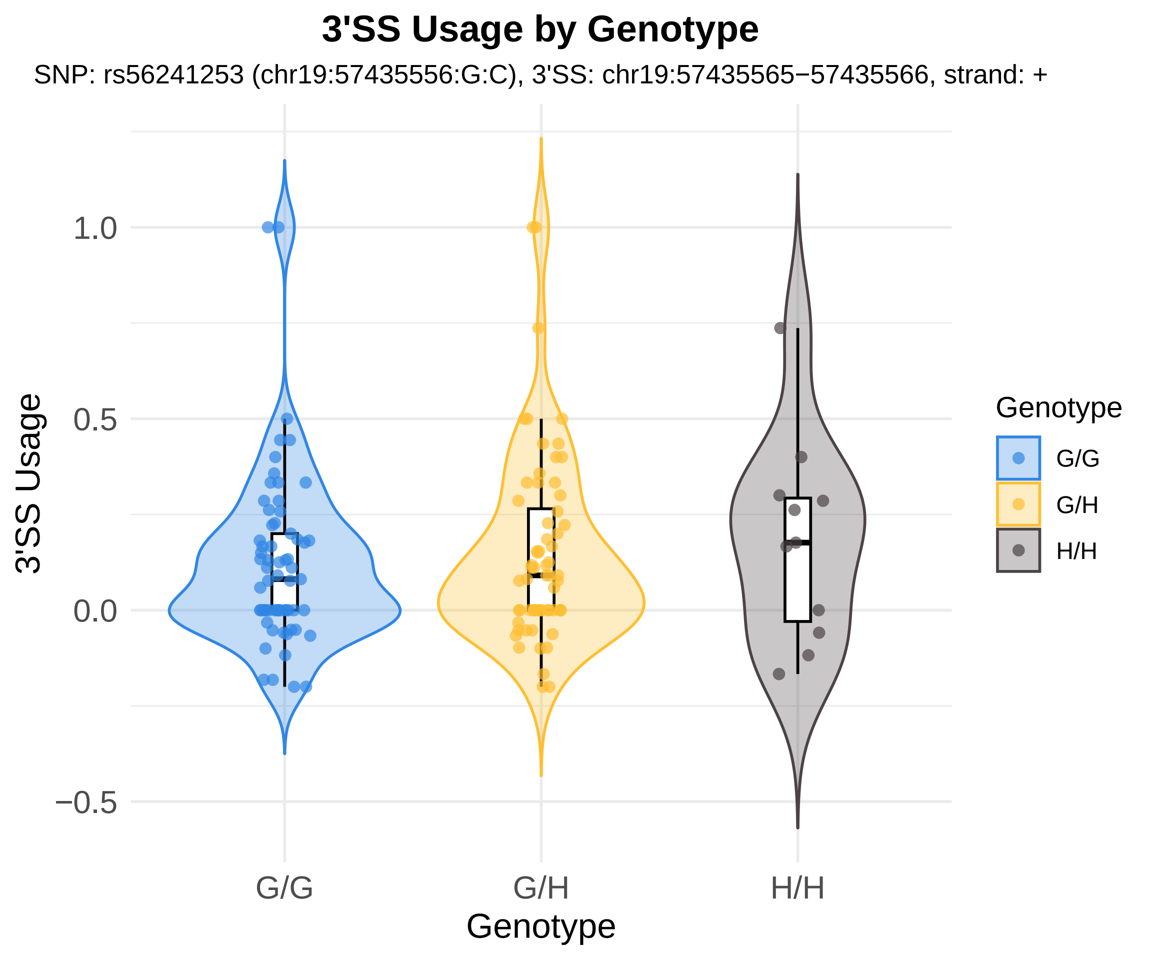

3'SS Usage by Genotype
SNP: rs56241253 (chr19:57435556:G:C), 3'SS: chr19:57435565−57435566, strand: +
3'SS Usage
1.0
0.5
0.0
−0.5
G/G
G/H
H/H
Genotype
Genotype
G/G
G/H
H/H

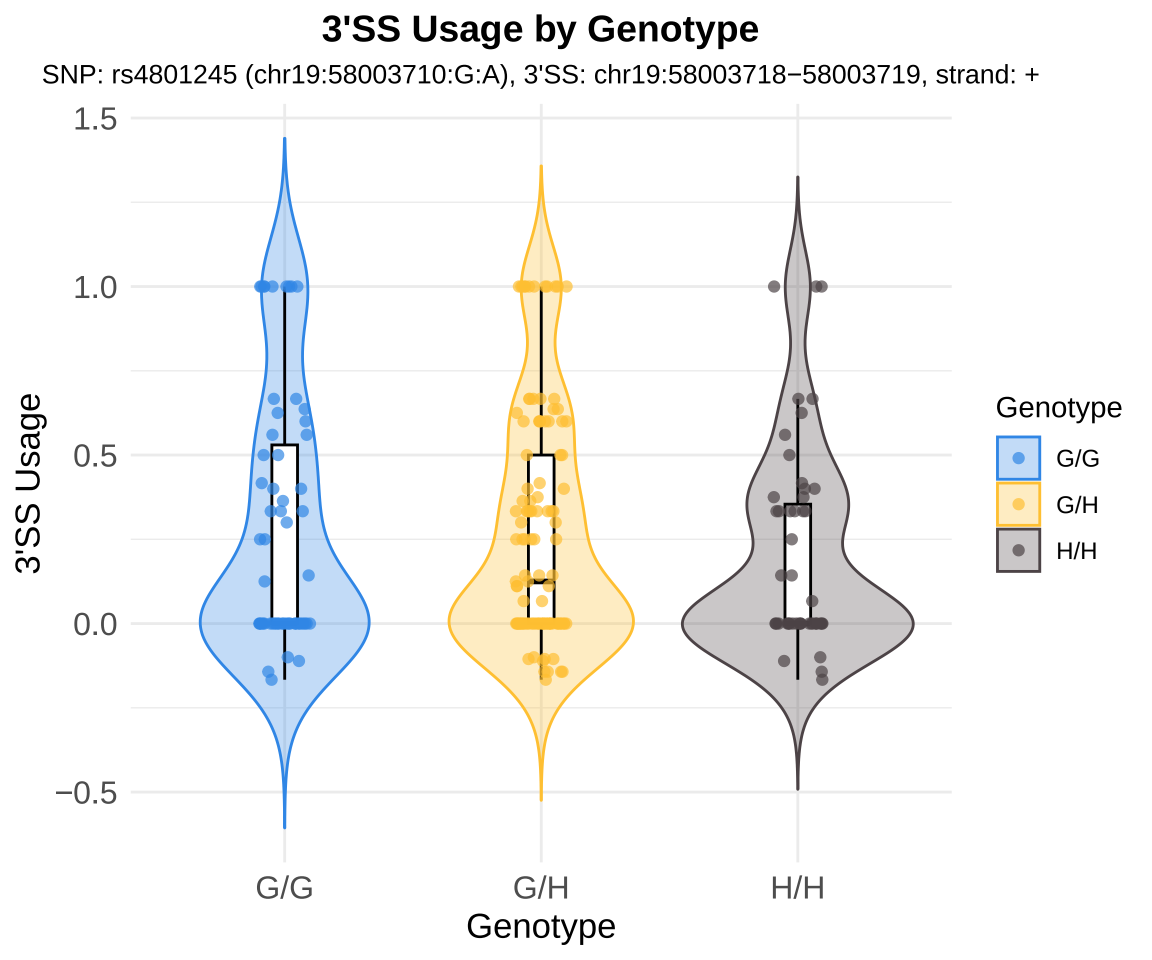
3'SS Usage by Genotype
SNP: rs4801245 (chr19:58003710:G:A), 3'SS: chr19:58003718−58003719, strand: +
3'SS Usage
1.5
1.0
0.5
0.0
−0.5
G/G
G/H
H/H
Genotype
Genotype
G/G
G/H
H/H

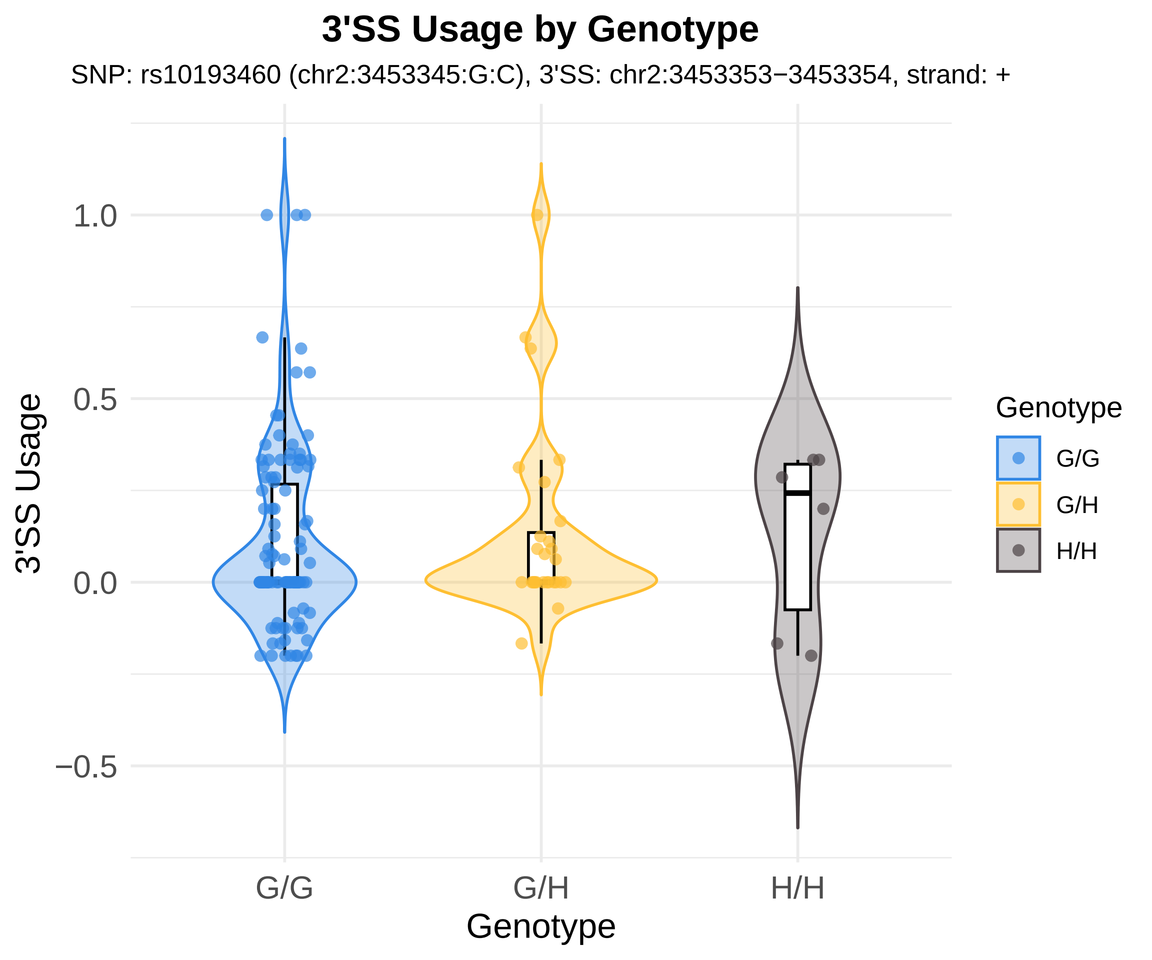
3'SS Usage by Genotype
SNP: rs10193460 (chr2:3453345:G:C), 3'SS: chr2:3453353−3453354, strand: +
1.0
0.5
0.0
−0.5
3'SS Usage
G/G
G/H
H/H
Genotype
Genotype
G/G
G/H
H/H

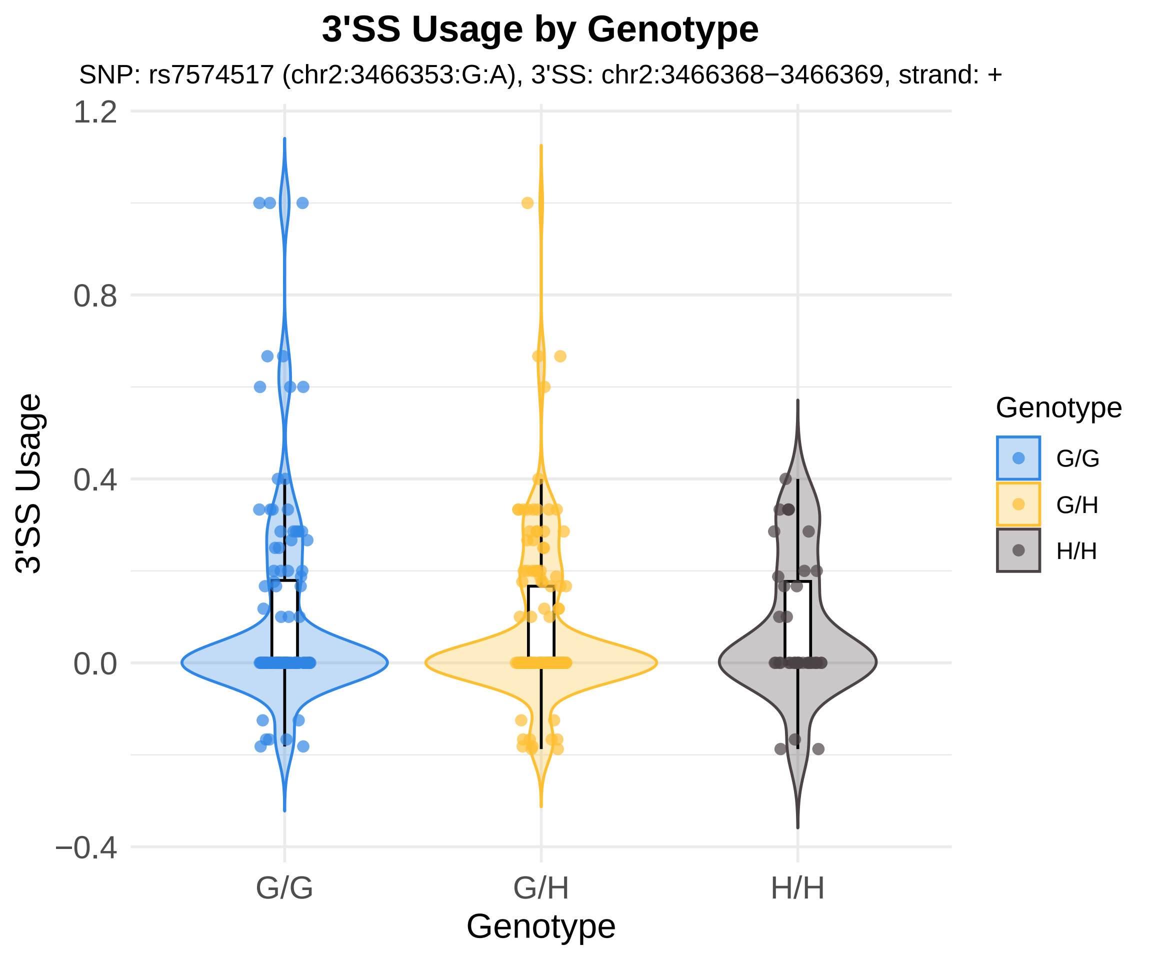
3'SS Usage by Genotype
SNP: rs7574517 (chr2:3466353:G:A), 3'SS: chr2:3466368−3466369, strand: +
3'SS Usage
1.2
0.8
0.4
0.0
−0.4
G/G
G/H
H/H
Genotype
Genotype
G/G
G/H
H/H

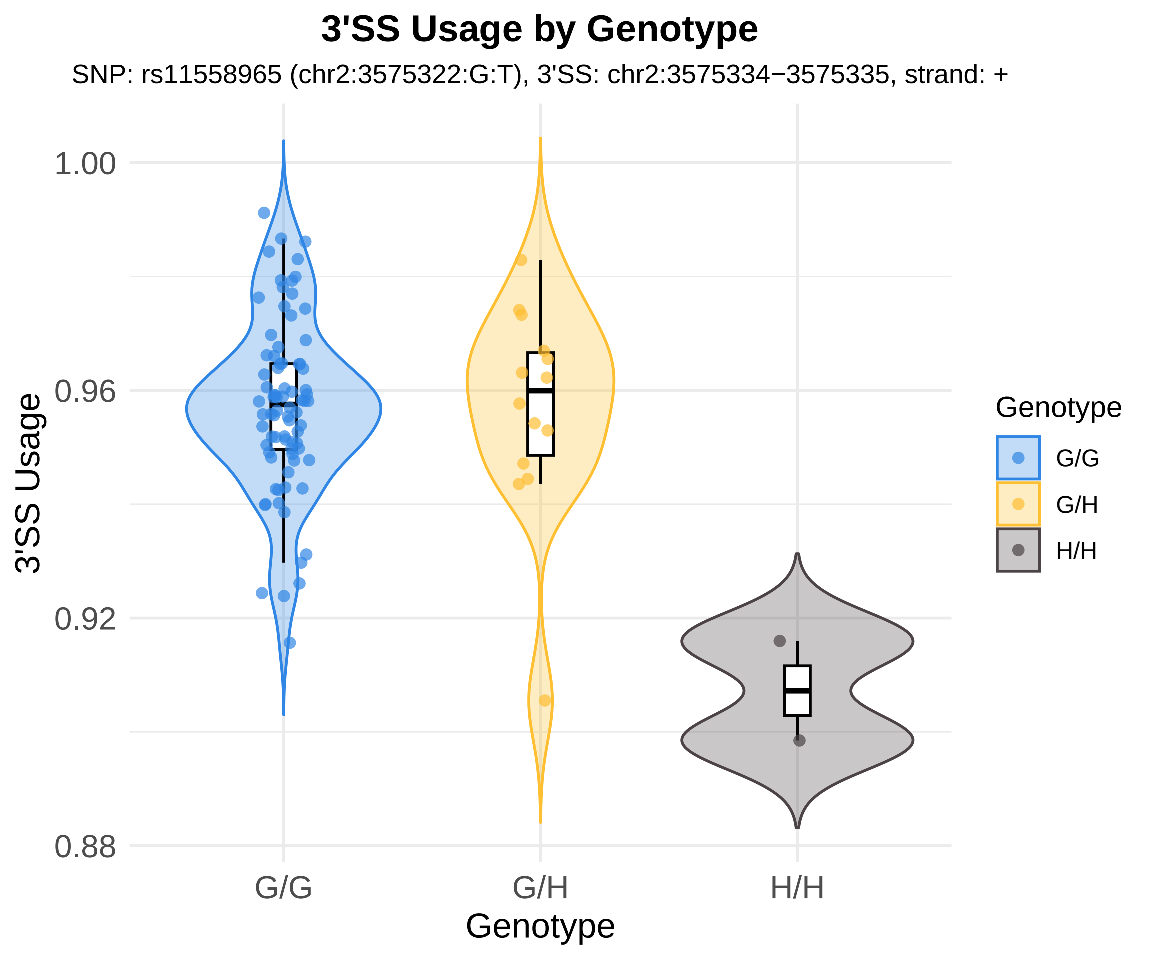
3'SS Usage by Genotype
SNP: rs11558965 (chr2:3575322:G:T), 3'SS: chr2:3575334−3575335, strand: +
3'SS Usage
1.00
0.96
0.92
0.88
G/G
G/H
H/H
Genotype
Genotype
G/G
G/H
H/H

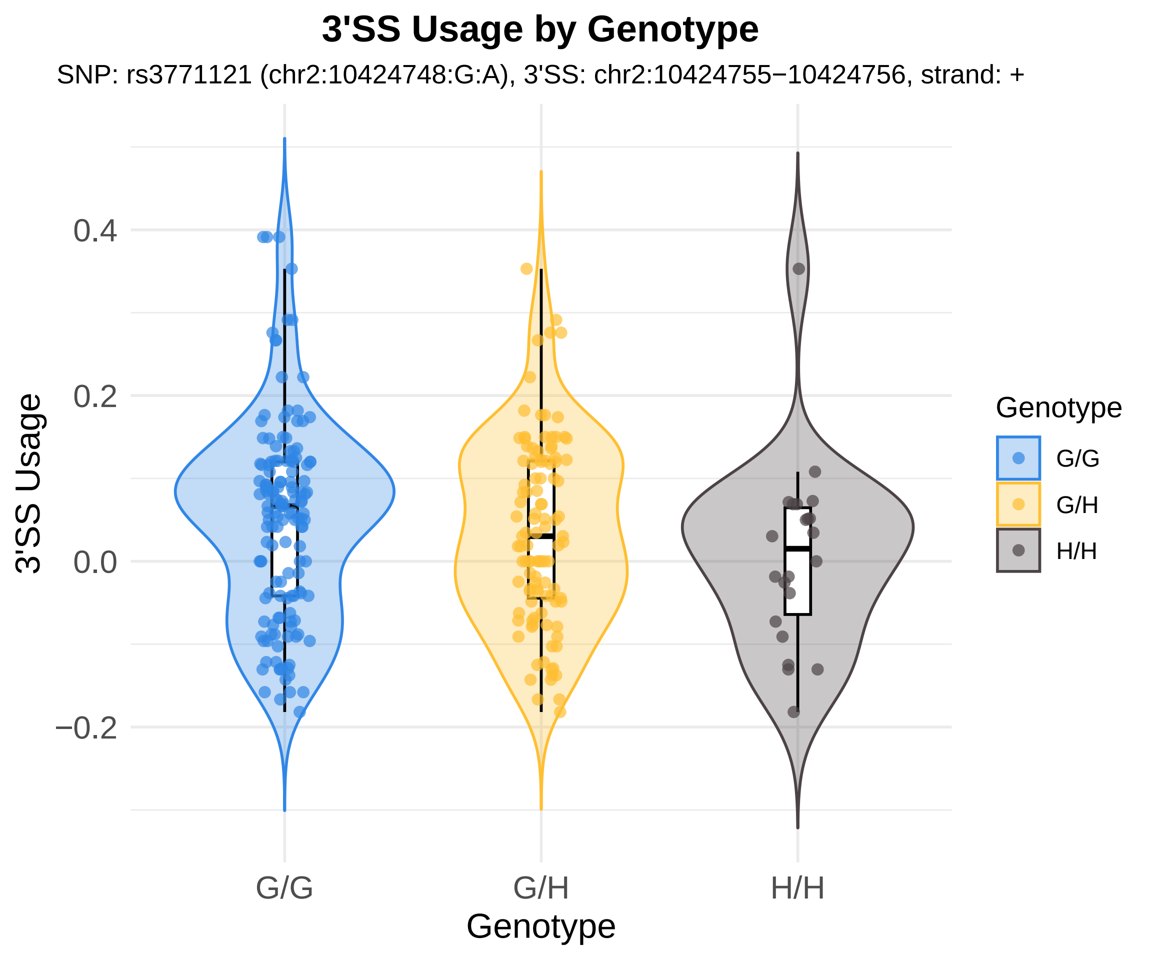

3'SS Usage by Genotype
SNP: rs3771121 (chr2:10424748:G:A), 3'SS: chr2:10424755−10424756, strand: +
3'SS Usage
0.4
0.2
0.0
−0.2
G/G
G/H
H/H
Genotype
Genotype
G/G
G/H
H/H

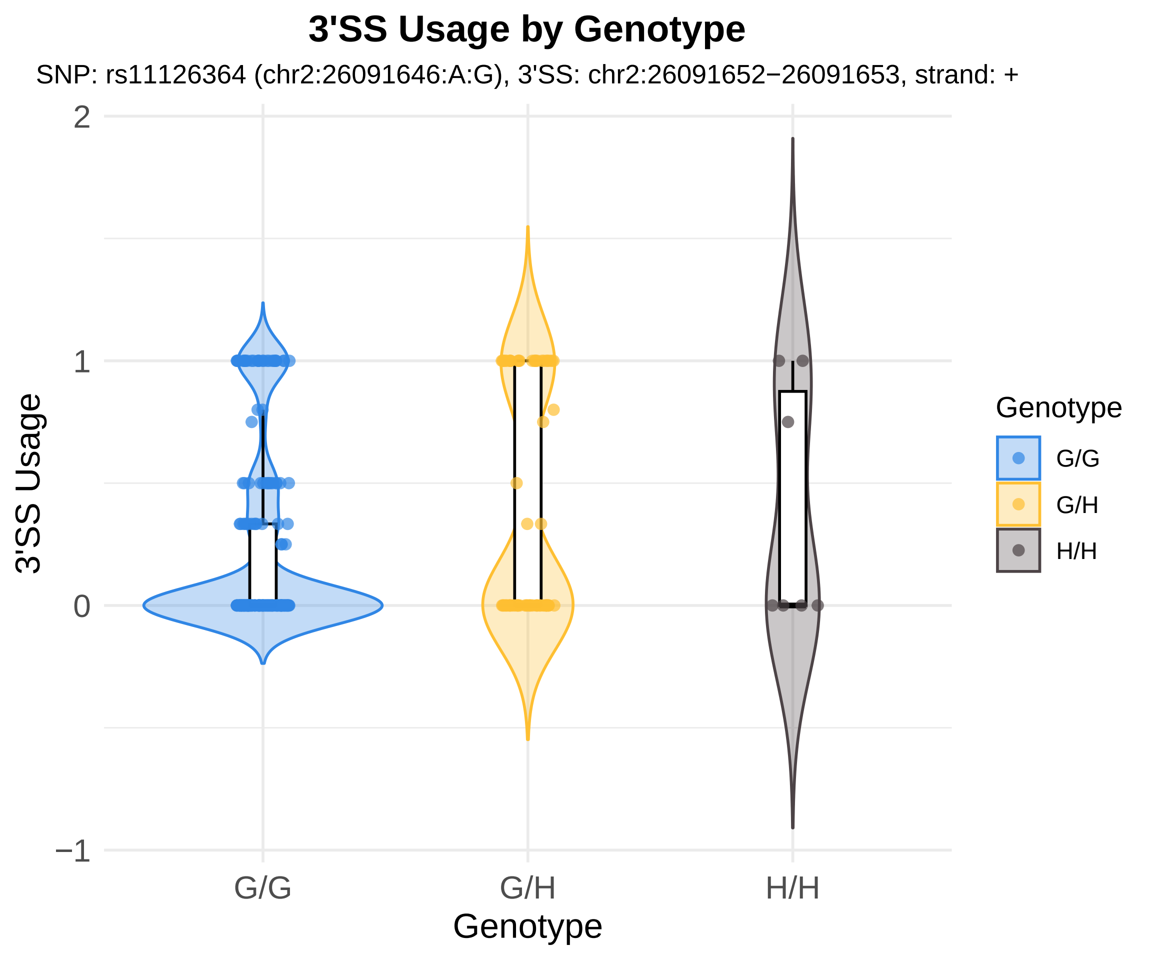
3'SS Usage by Genotype
SNP: rs11126364 (chr2:26091646:A:G), 3'SS: chr2:26091652−26091653, strand: +
3'SS Usage
2
1
0
−1
G/G
G/H
H/H
Genotype
Genotype
G/G
G/H
H/H

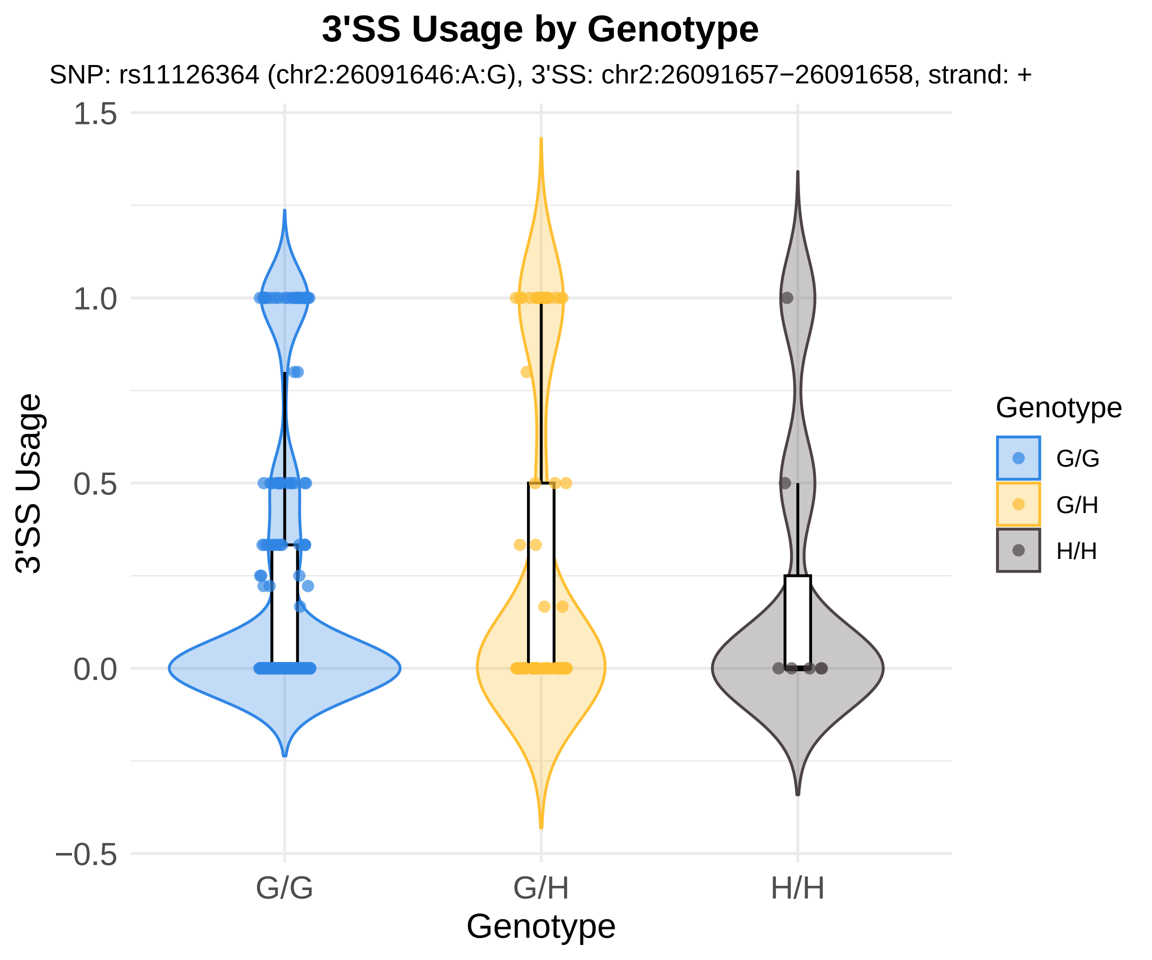

3'SS Usage by Genotype
SNP: rs11126364 (chr2:26091646:A:G), 3'SS: chr2:26091657−26091658, strand: +
3'SS Usage
1.5
1.0
0.5
0.0
−0.5
G/G
G/H
H/H
Genotype
Genotype
G/G
G/H
H/H

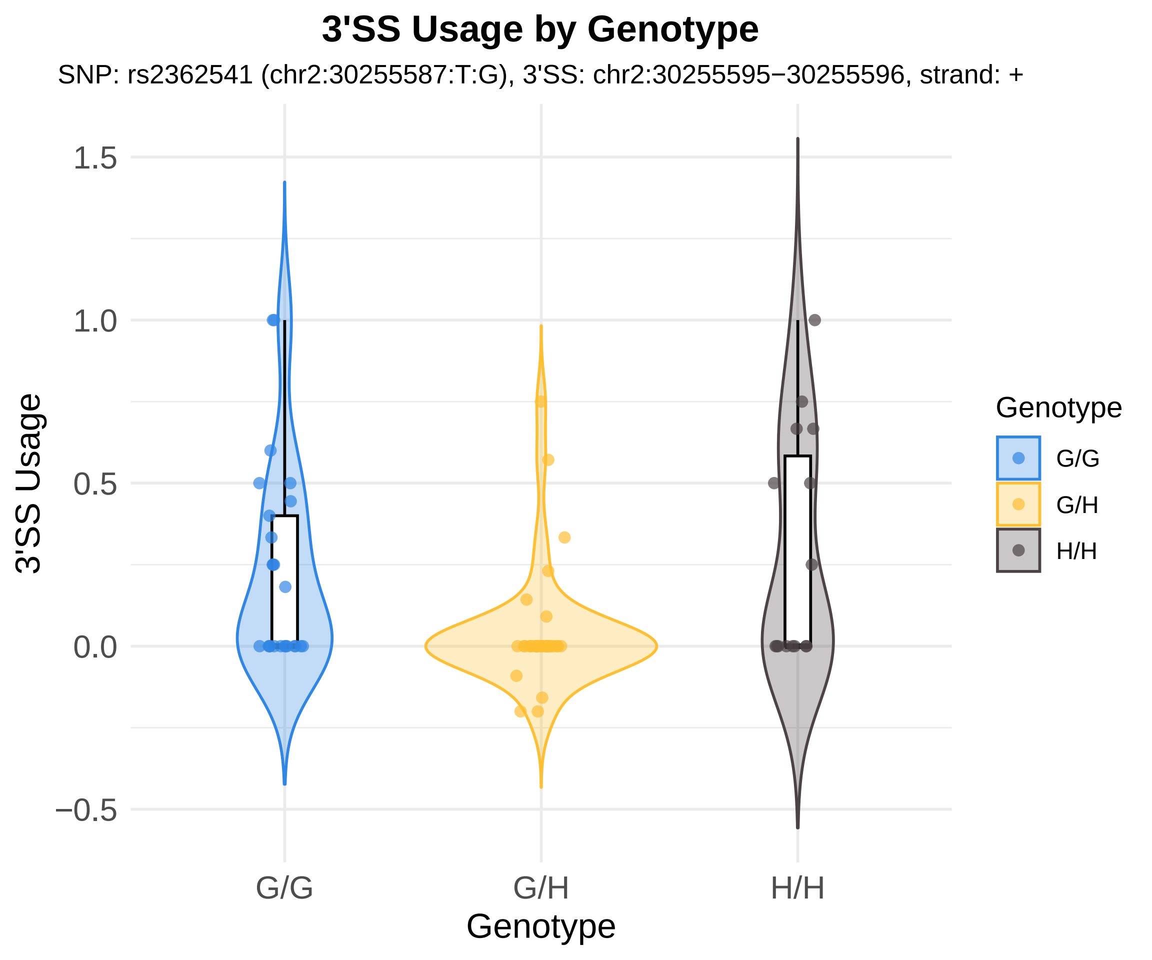
3'SS Usage by Genotype
SNP: rs2362541 (chr2:30255587:T:G), 3'SS: chr2:30255595−30255596, strand: +
3'SS Usage
1.5
1.0
0.5
0.0
−0.5
G/G
G/H
H/H
Genotype
Genotype
G/G
G/H
H/H

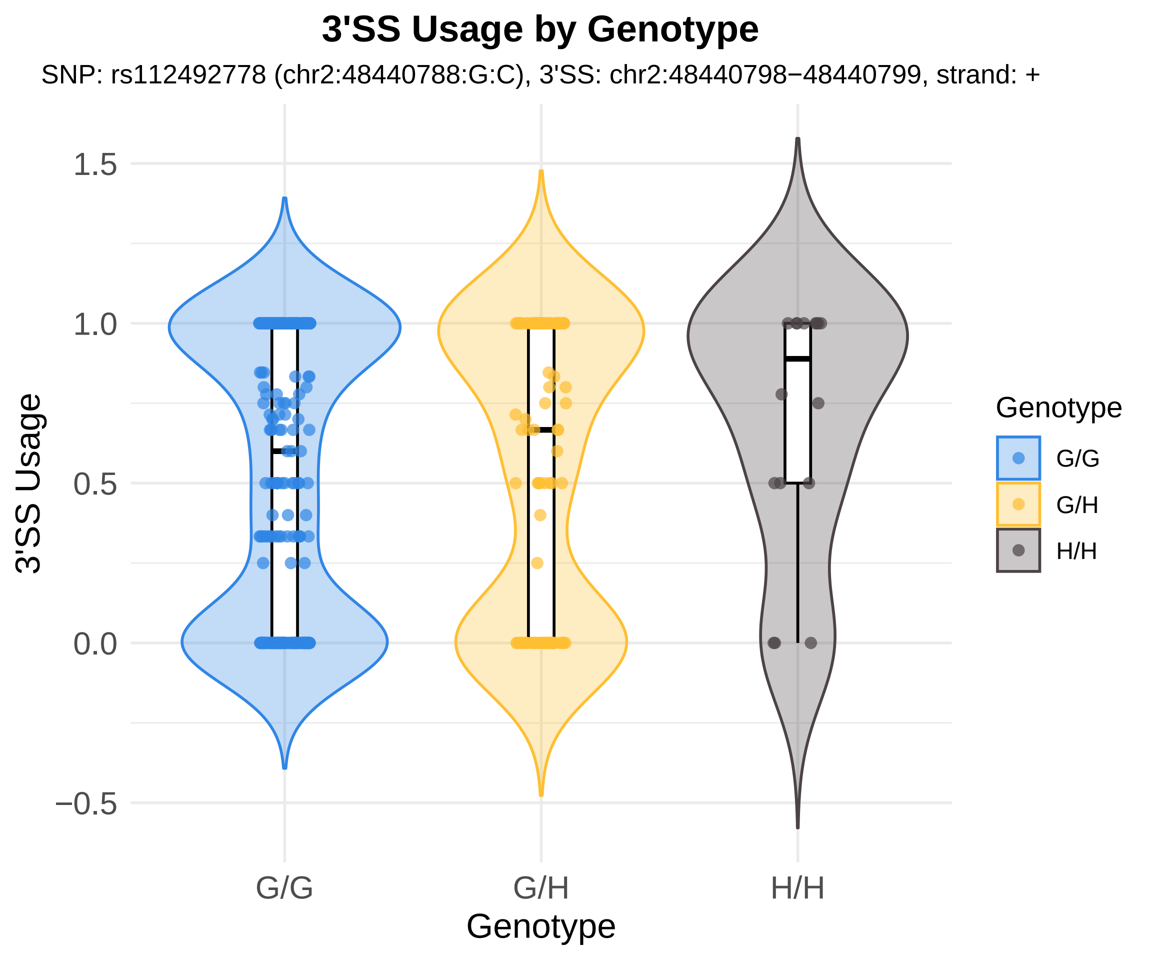

3'SS Usage by Genotype
SNP: rs112492778 (chr2:48440788:G:C), 3'SS: chr2:48440798−48440799, strand: +
3'SS Usage
1.5
1.0
0.5
0.0
−0.5
G/G
G/H
H/H
Genotype
Genotype
G/G
G/H
H/H

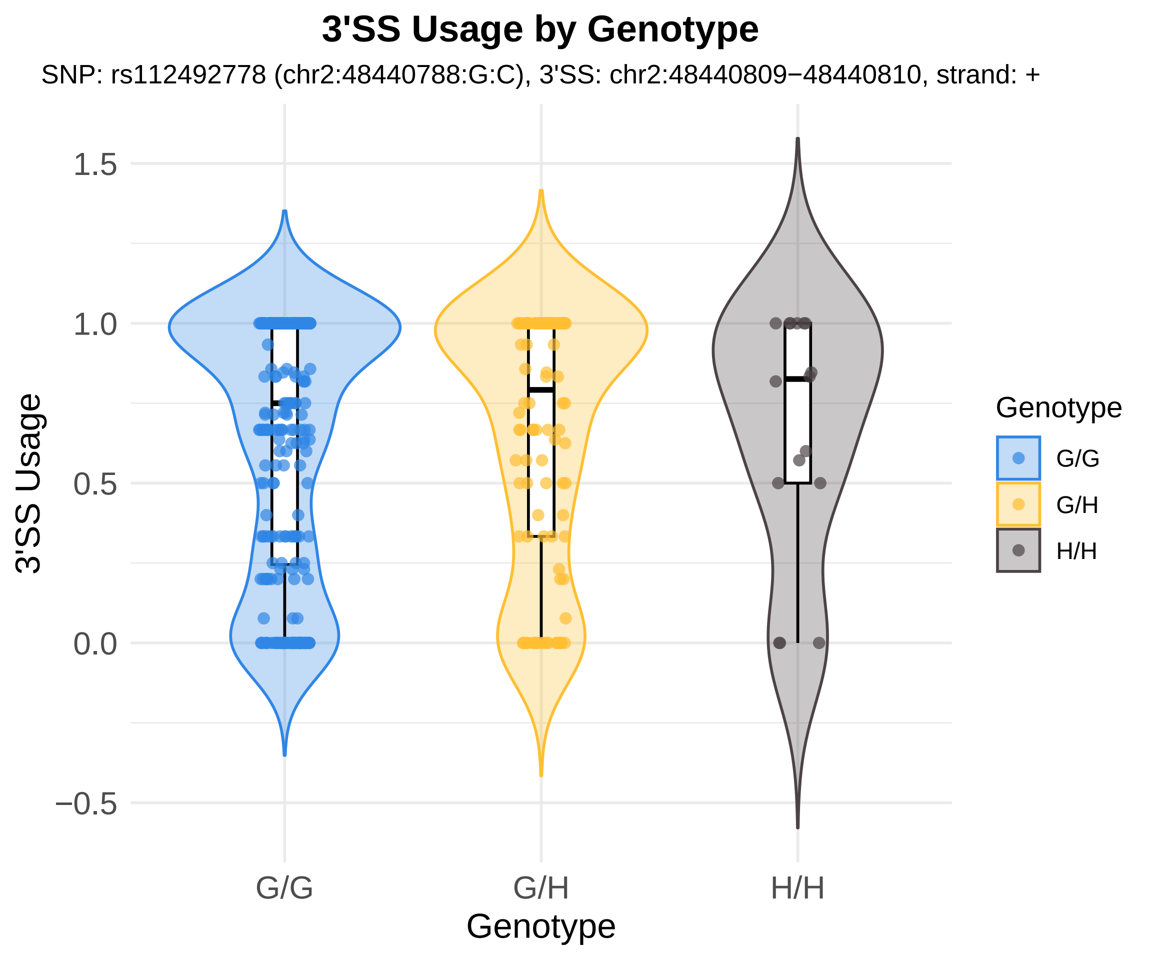

3'SS Usage by Genotype
SNP: rs112492778 (chr2:48440788:G:C), 3'SS: chr2:48440809−48440810, strand: +
1.5
1.0
0.5
0.0
−0.5
3'SS Usage
G/G
G/H
H/H
Genotype
Genotype
G/G
G/H
H/H

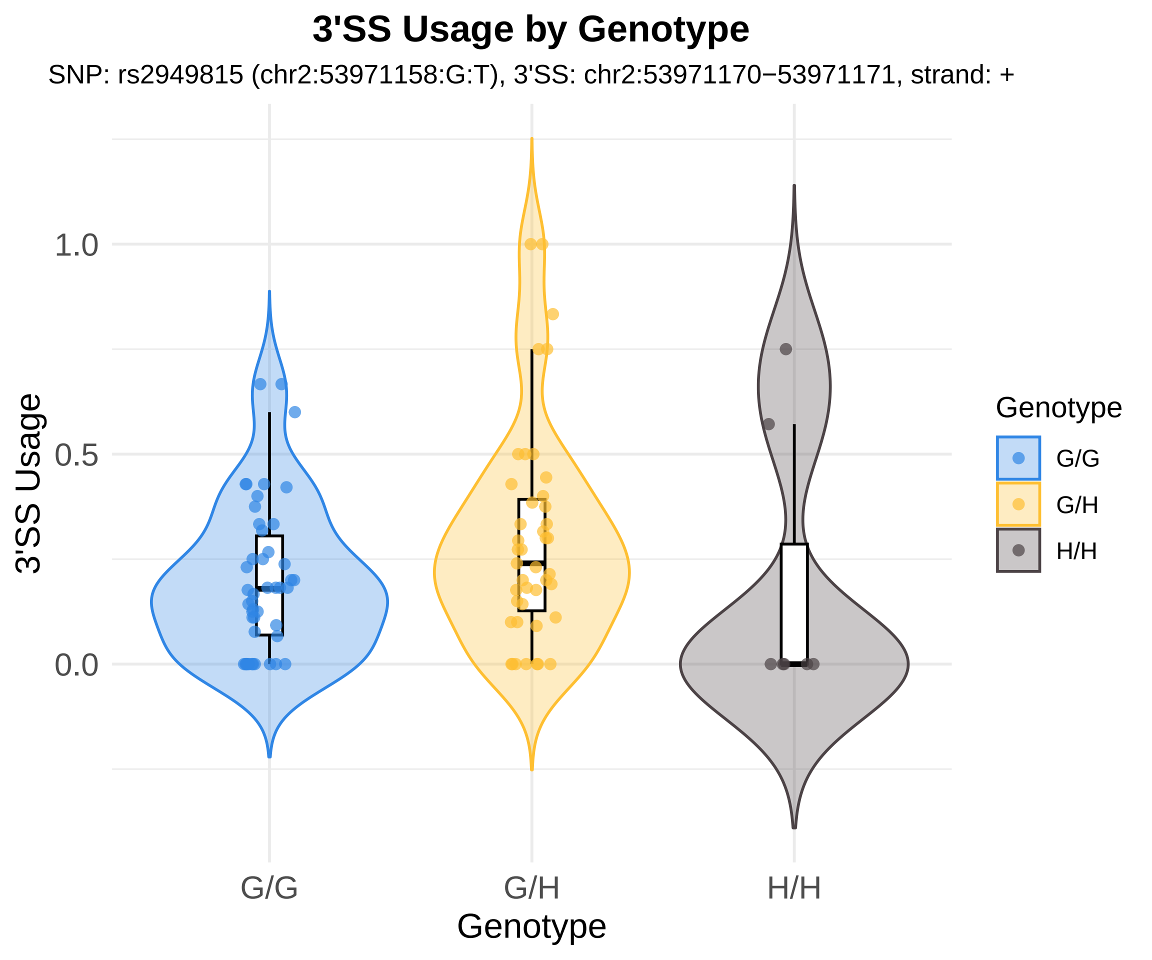

3'SS Usage by Genotype
SNP: rs2949815 (chr2:53971158:G:T), 3'SS: chr2:53971170−53971171, strand: +
3'SS Usage
1.0
0.5
0.0
G/G
G/H
H/H
Genotype
Genotype
G/G
G/H
H/H

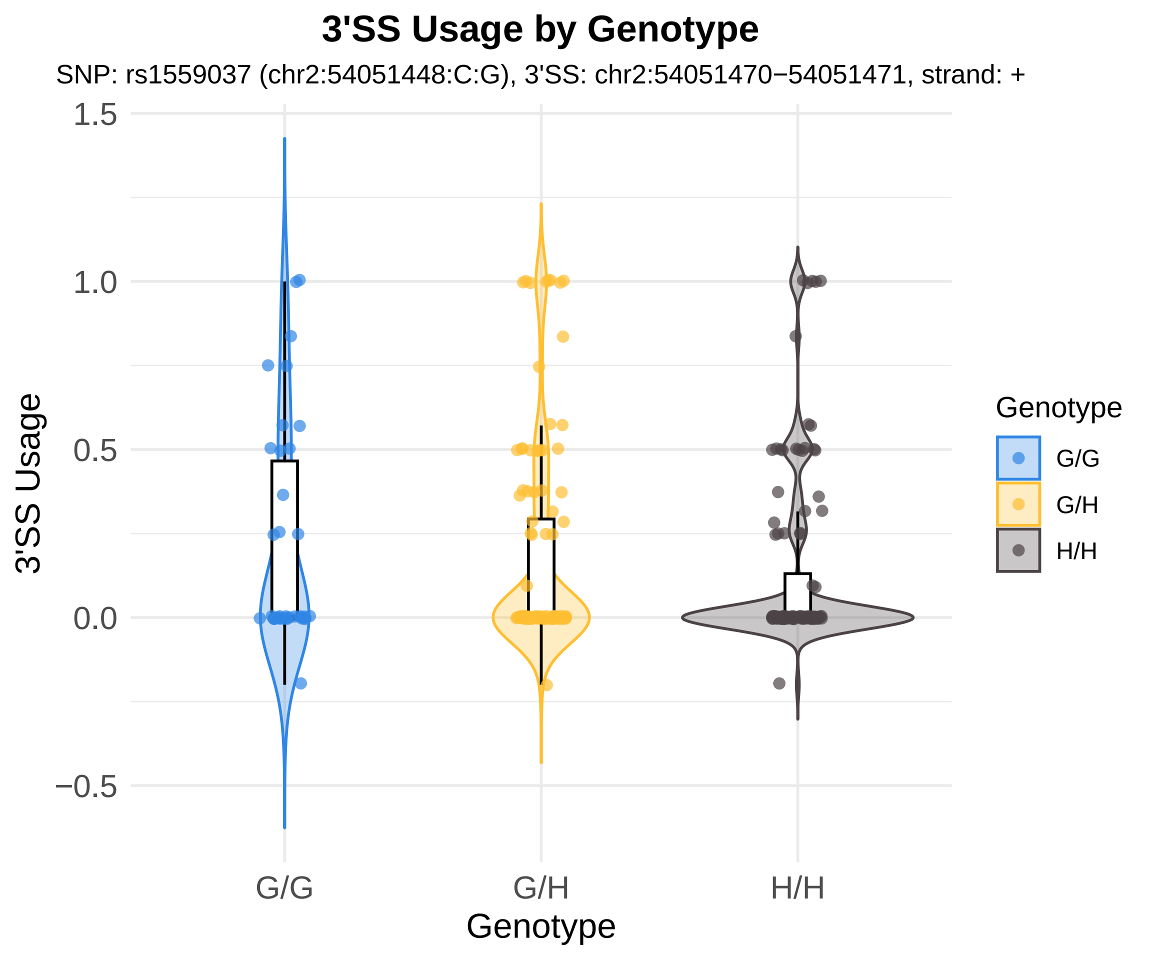
3'SS Usage by Genotype
SNP: rs1559037 (chr2:54051448:C:G), 3'SS: chr2:54051470−54051471, strand: +
1.5
1.0
0.5
0.0
−0.5
3'SS Usage
G/G
G/H
H/H
Genotype
Genotype
G/G
G/H
H/H

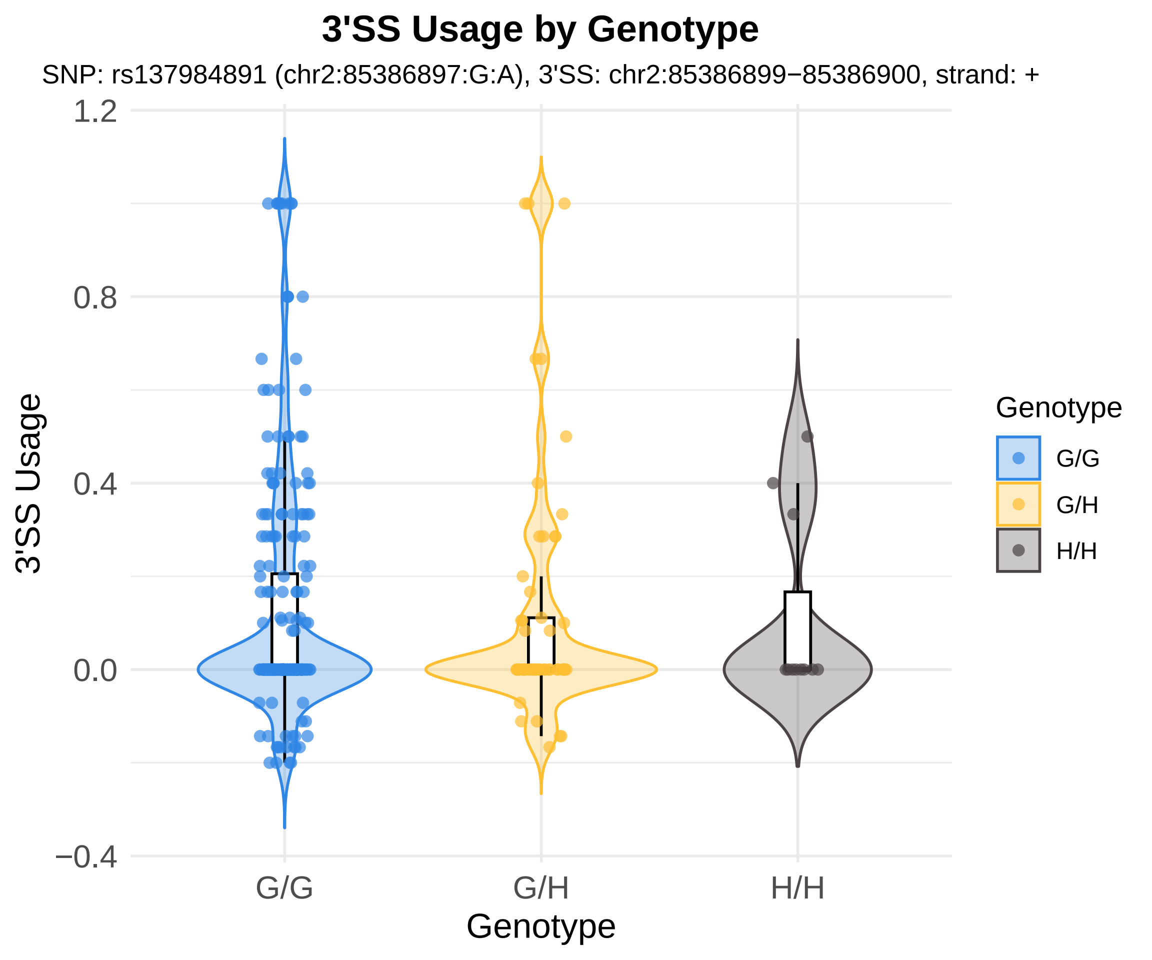
3'SS Usage by Genotype
SNP: rs137984891 (chr2:85386897:G:A), 3'SS: chr2:85386899−85386900, strand: +
3'SS Usage
1.2
0.8
0.4
0.0
−0.4
G/G
G/H
H/H
Genotype
Genotype
G/G
G/H
H/H

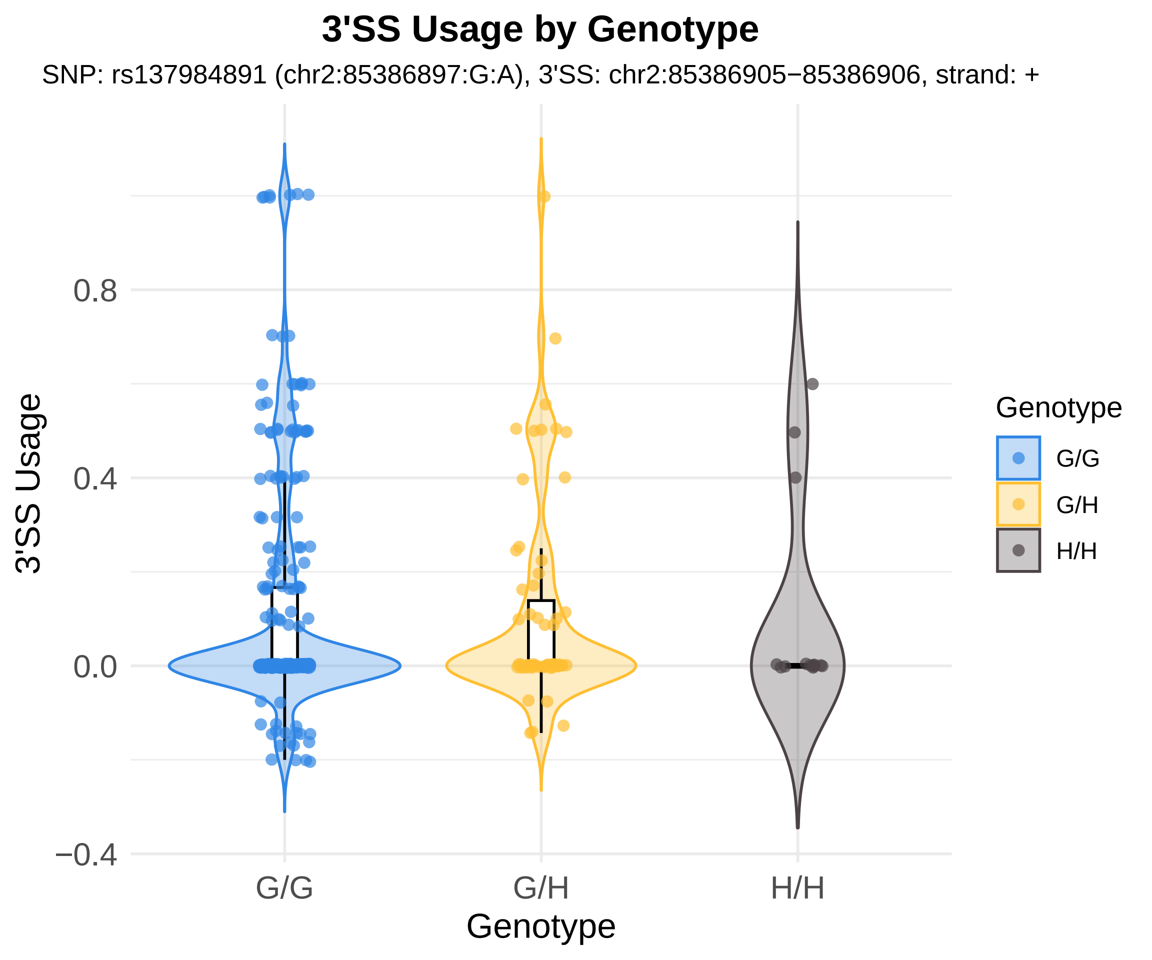
3'SS Usage by Genotype
SNP: rs137984891 (chr2:85386897:G:A), 3'SS: chr2:85386905−85386906, strand: +
3'SS Usage
0.8
0.4
0.0
−0.4
G/G
G/H
H/H
Genotype
Genotype
G/G
G/H
H/H

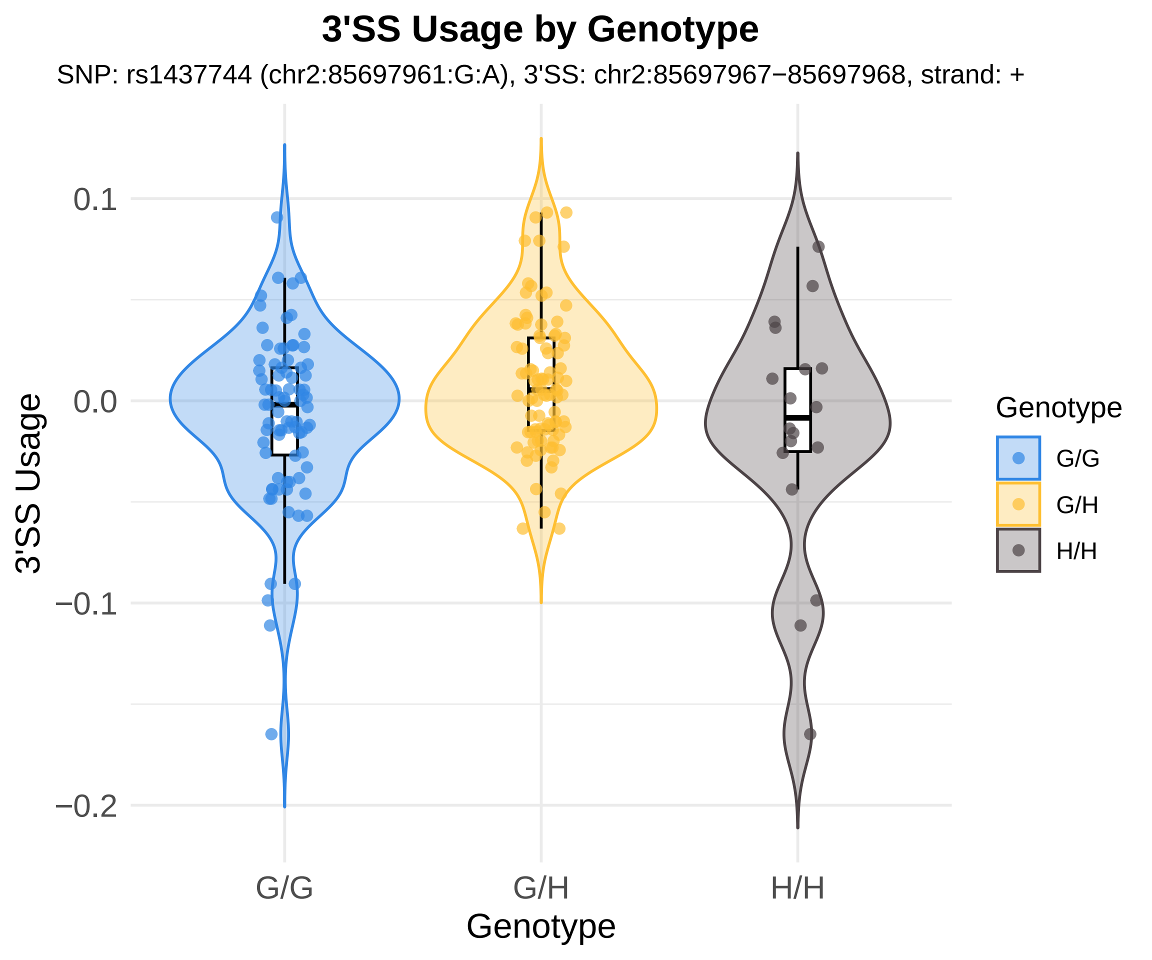
3'SS Usage by Genotype
SNP: rs1437744 (chr2:85697961:G:A), 3'SS: chr2:85697967−85697968, strand: +
0.1
0.0
−0.1
−0.2
3'SS Usage
G/G
G/H
H/H
Genotype
Genotype
G/G
G/H
H/H

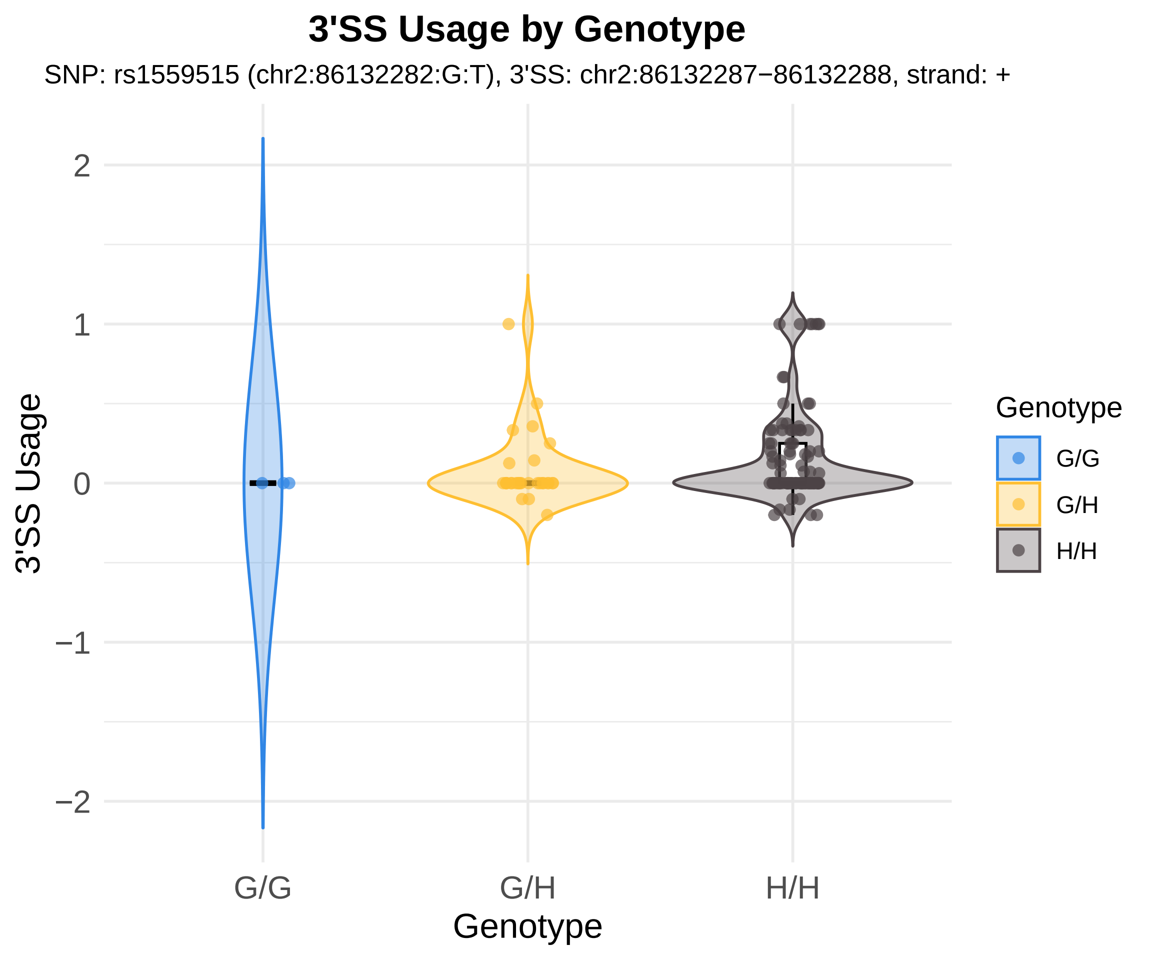

3'SS Usage by Genotype
SNP: rs1559515 (chr2:86132282:G:T), 3'SS: chr2:86132287−86132288, strand: +
3'SS Usage
2
1
0
−1
−2
G/G
G/H
H/H
Genotype
Genotype
G/G
G/H
H/H

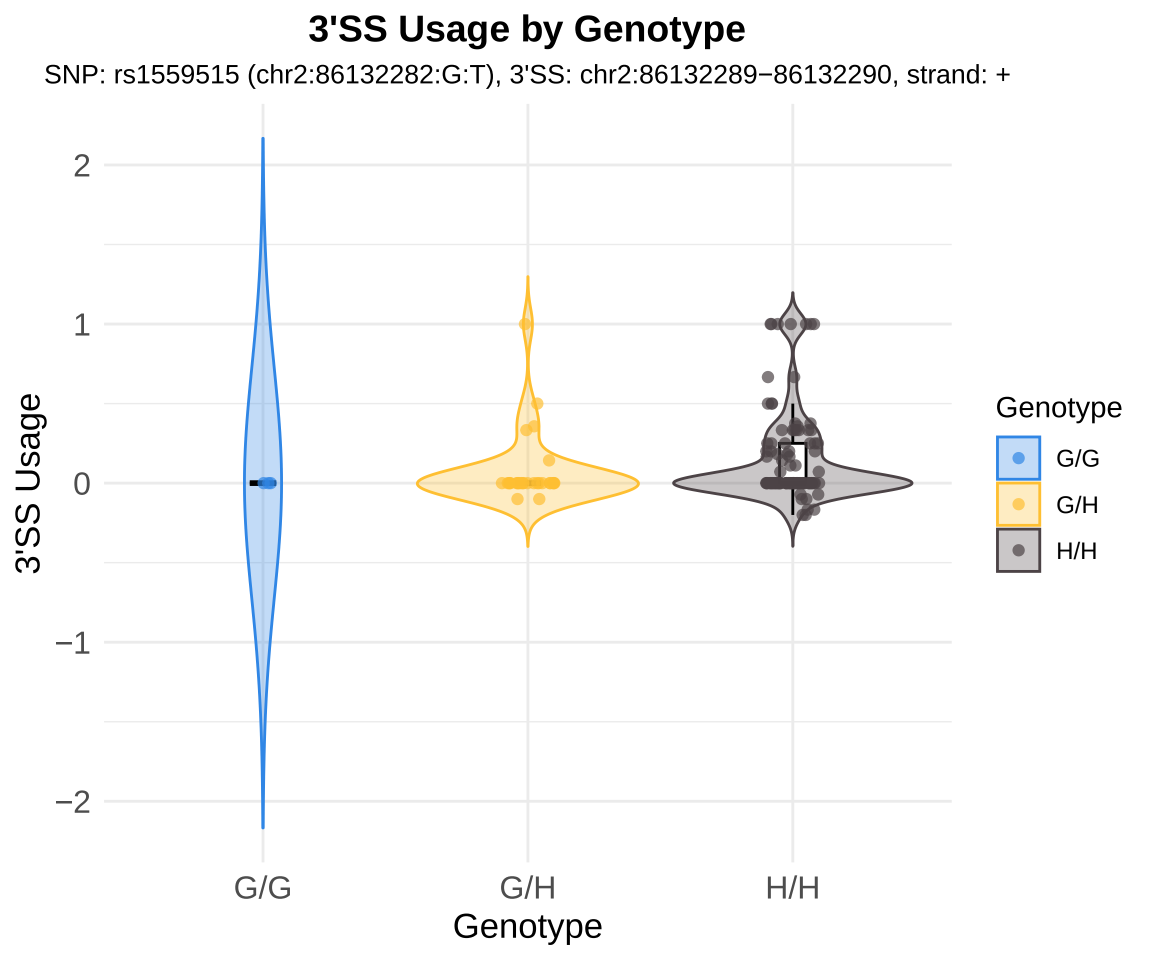
3'SS Usage by Genotype
SNP: rs1559515 (chr2:86132282:G:T), 3'SS: chr2:86132289−86132290, strand: +
3'SS Usage
2
1
0
−1
−2
G/G
G/H
H/H
Genotype
Genotype
G/G
G/H
H/H

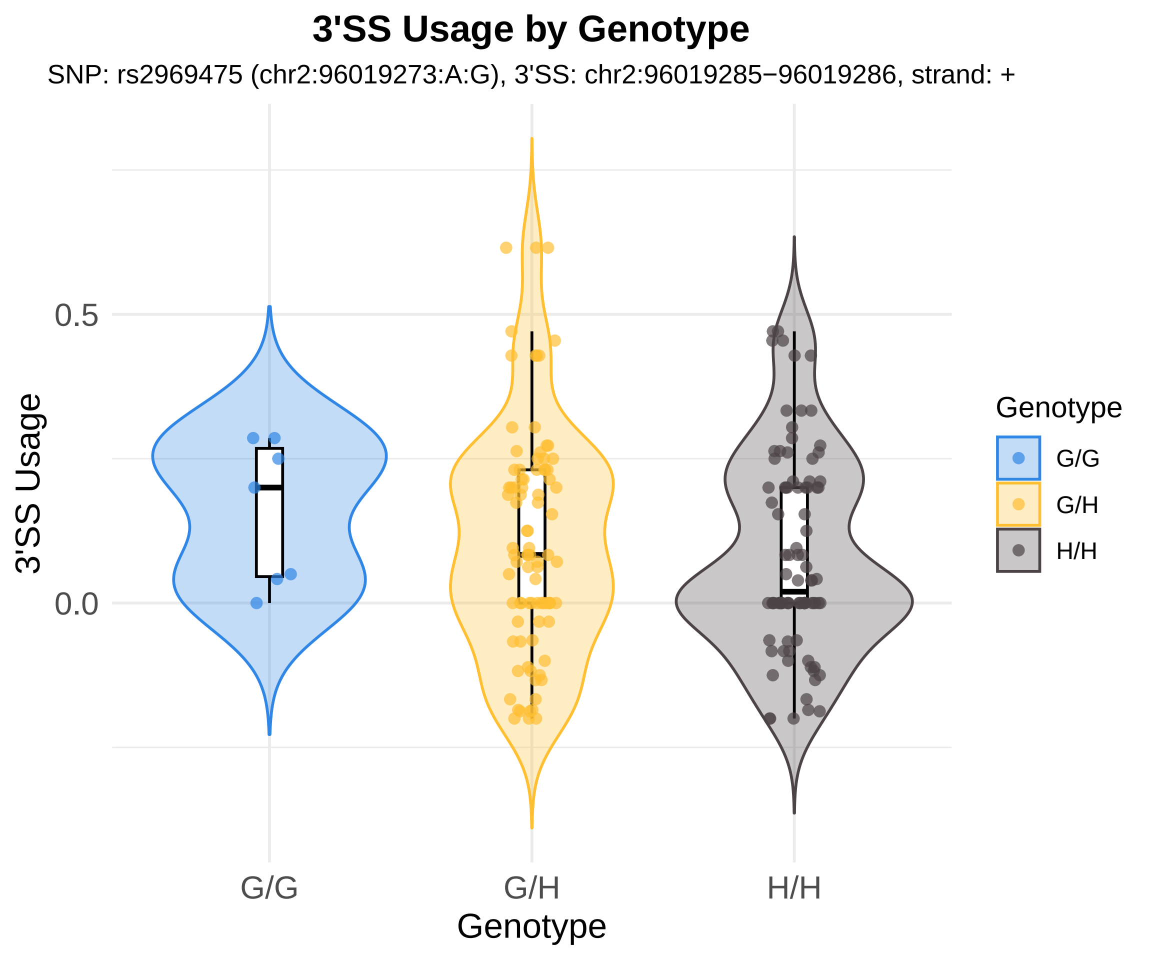
3'SS Usage by Genotype
SNP: rs2969475 (chr2:96019273:A:G), 3'SS: chr2:96019285−96019286, strand: +
3'SS Usage
0.5
0.0
G/G
G/H
H/H
Genotype
Genotype
G/G
G/H
H/H

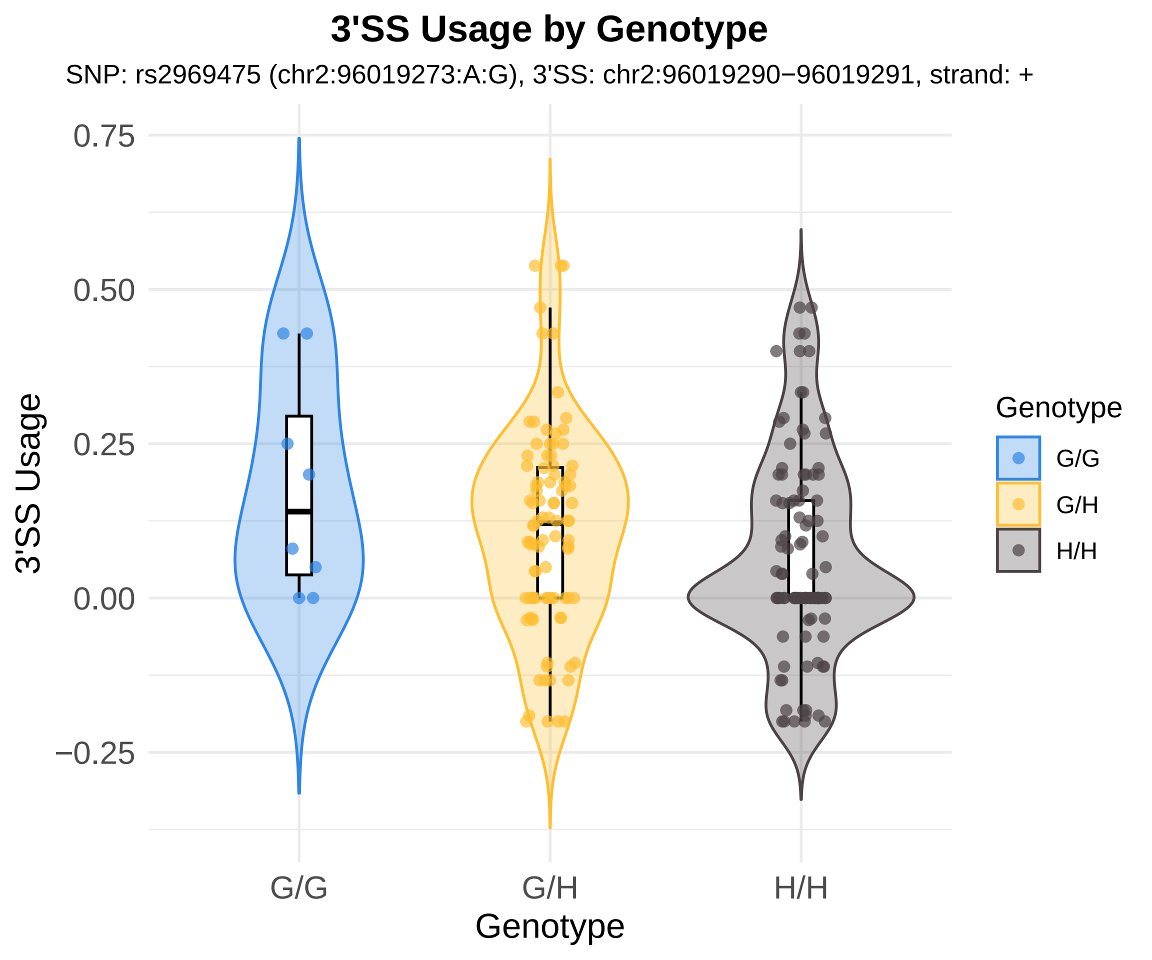

3'SS Usage by Genotype
SNP: rs2969475 (chr2:96019273:A:G), 3'SS: chr2:96019290−96019291, strand: +
3'SS Usage
0.75
0.50
0.25
0.00
−0.25
G/G
G/H
H/H
Genotype
Genotype
G/G
G/H
H/H

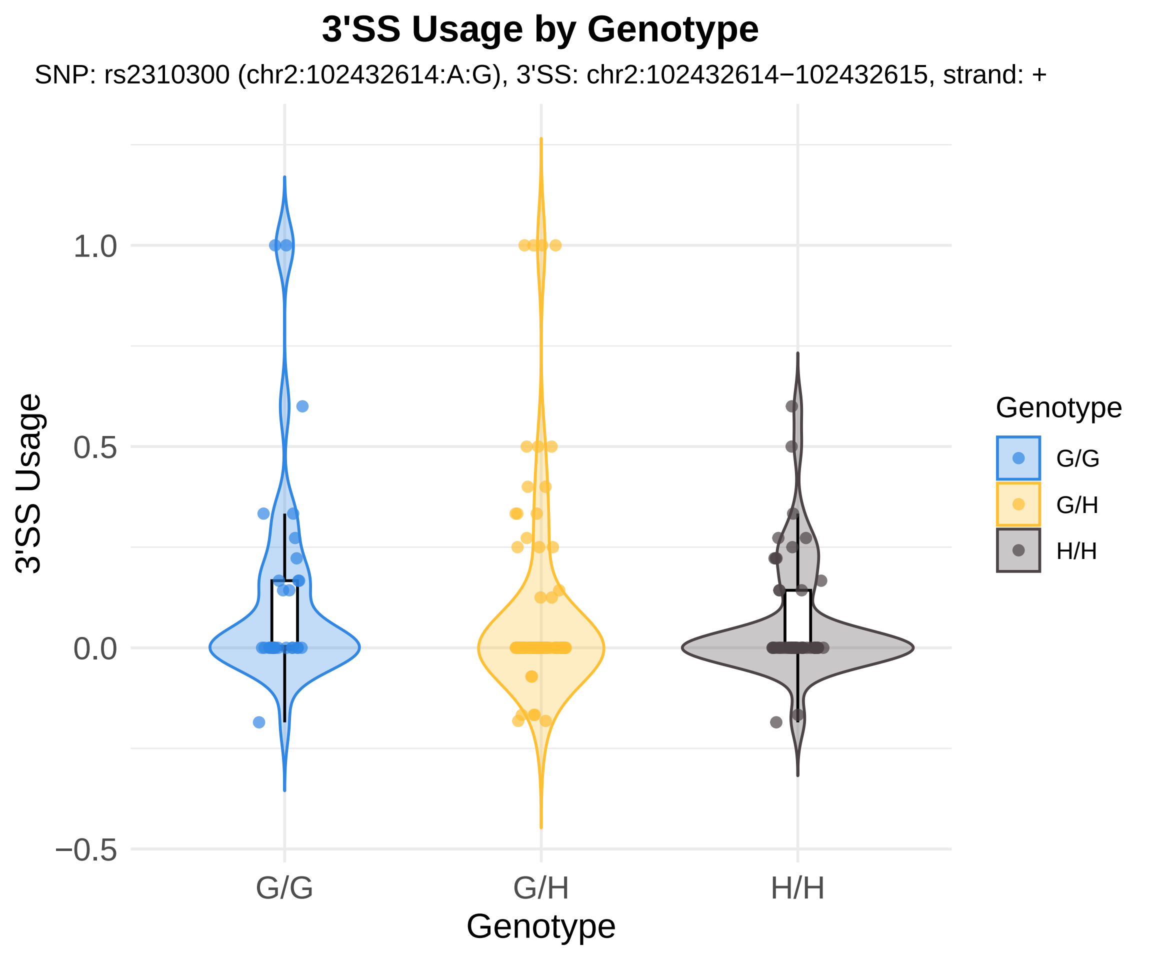
3'SS Usage by Genotype
SNP: rs2310300 (chr2:102432614:A:G), 3'SS: chr2:102432614−102432615, strand: +
3'SS Usage
1.0
0.5
0.0
−0.5
G/G
G/H
H/H
Genotype
Genotype
G/G
G/H
H/H

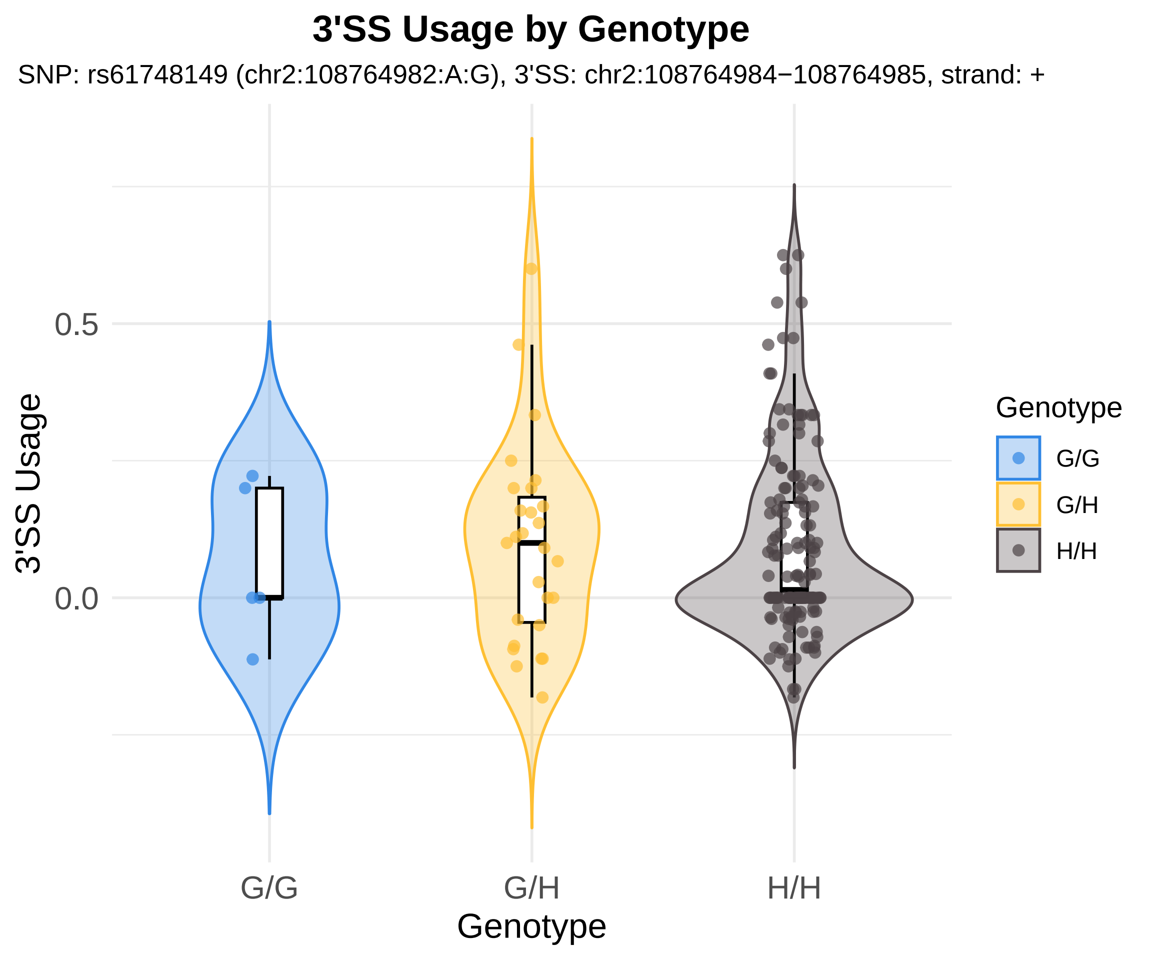

3'SS Usage by Genotype
SNP: rs61748149 (chr2:108764982:A:G), 3'SS: chr2:108764984−108764985, strand: +
3'SS Usage
0.5
0.0
G/G
G/H
H/H
Genotype
Genotype
G/G
G/H
H/H

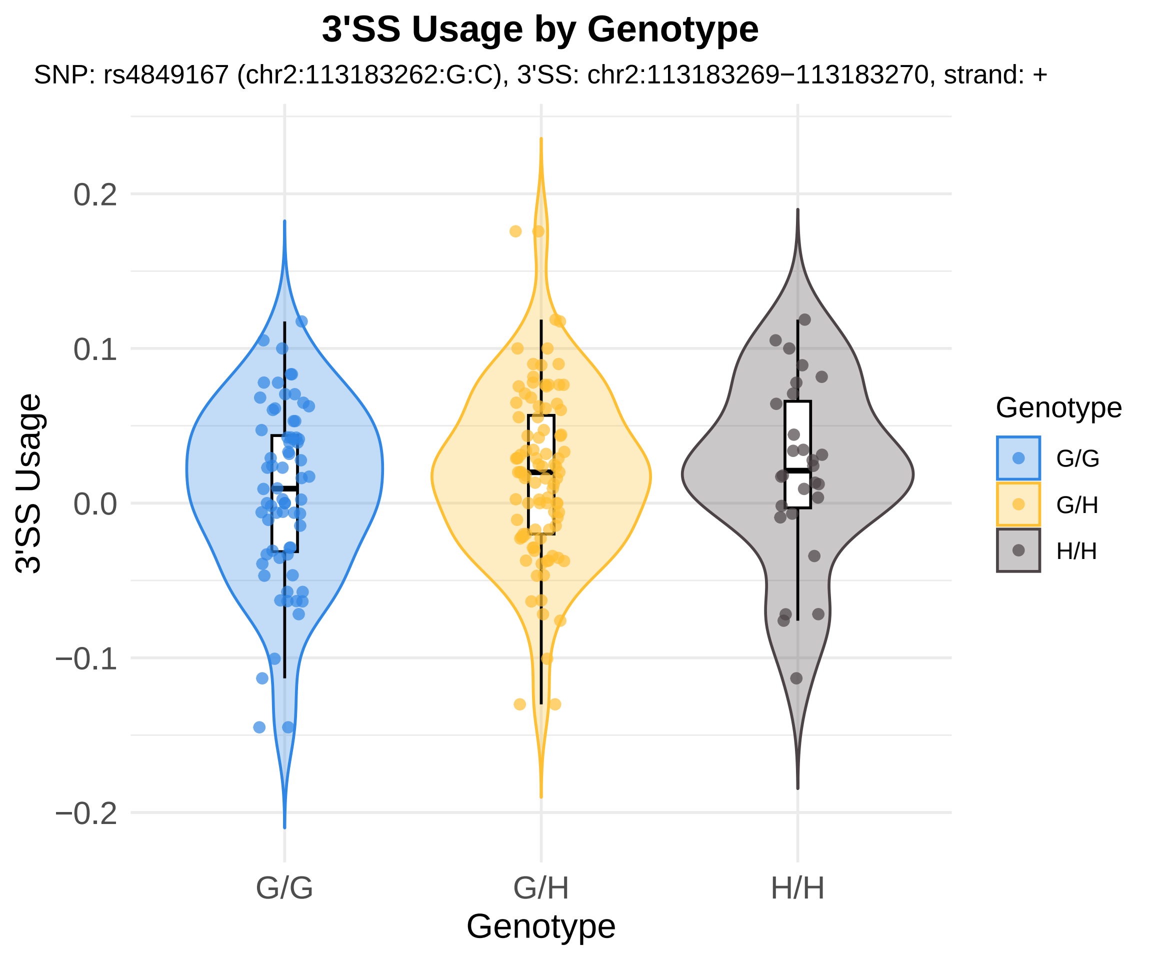

3'SS Usage by Genotype
SNP: rs4849167 (chr2:113183262:G:C), 3'SS: chr2:113183269−113183270, strand: +
3'SS Usage
0.2
0.1
0.0
−0.1
−0.2
G/G
G/H
H/H
Genotype
Genotype
G/G
G/H
H/H

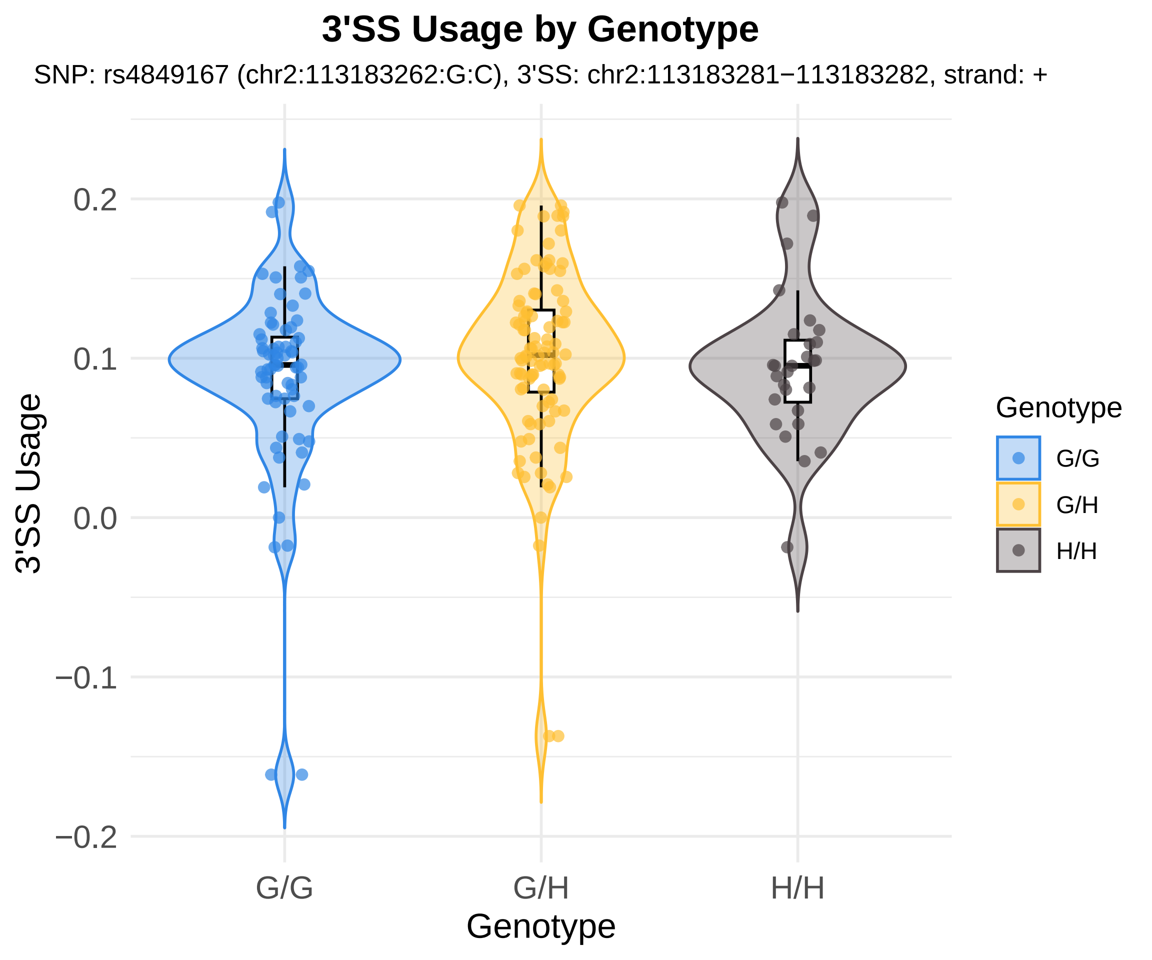
3'SS Usage by Genotype
SNP: rs4849167 (chr2:113183262:G:C), 3'SS: chr2:113183281−113183282, strand: +
3'SS Usage
0.2
0.1
0.0
−0.1
−0.2
G/G
G/H
H/H
Genotype
Genotype
G/G
G/H
H/H

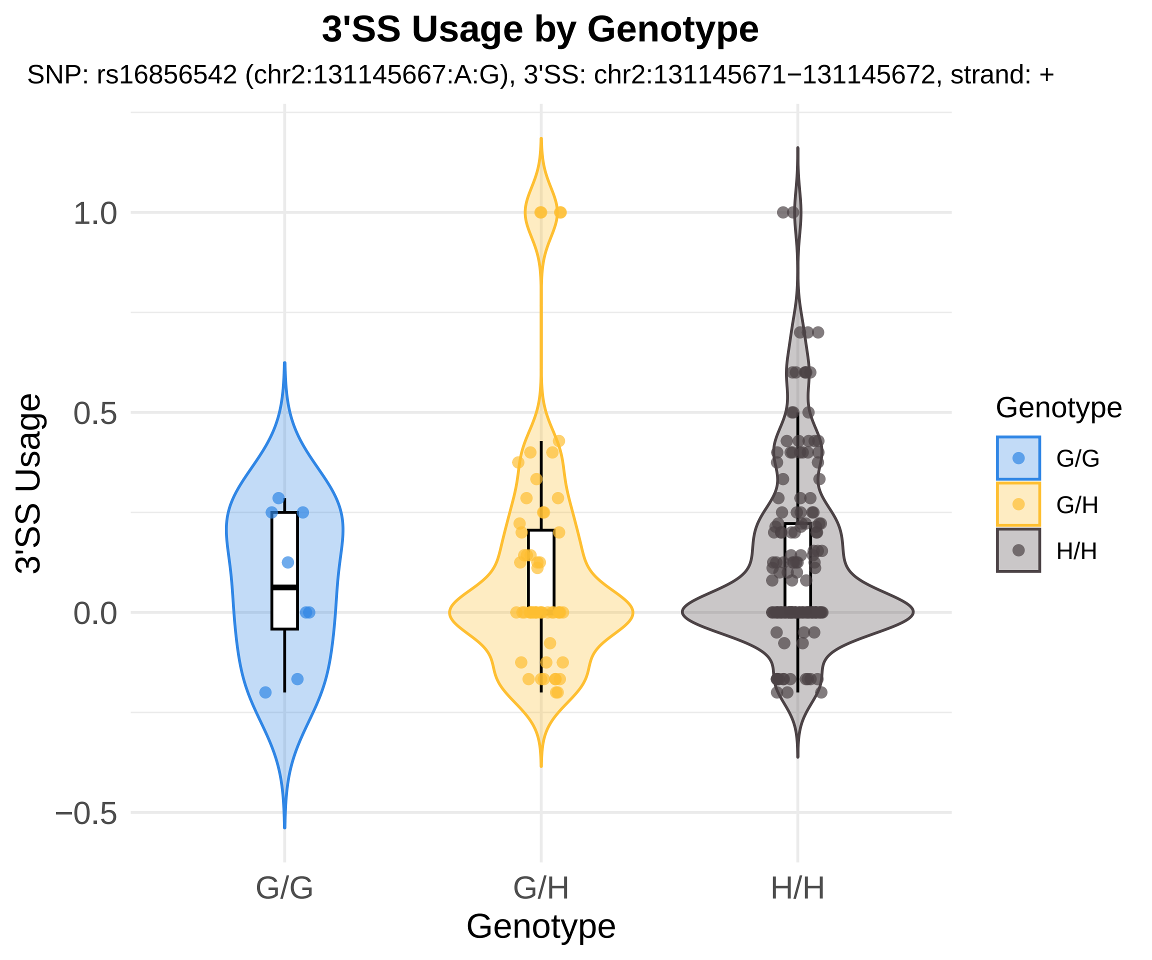
3'SS Usage by Genotype
SNP: rs16856542 (chr2:131145667:A:G), 3'SS: chr2:131145671−131145672, strand: +
3'SS Usage
1.0
0.5
0.0
−0.5
G/G
G/H
H/H
Genotype
Genotype
G/G
G/H
H/H

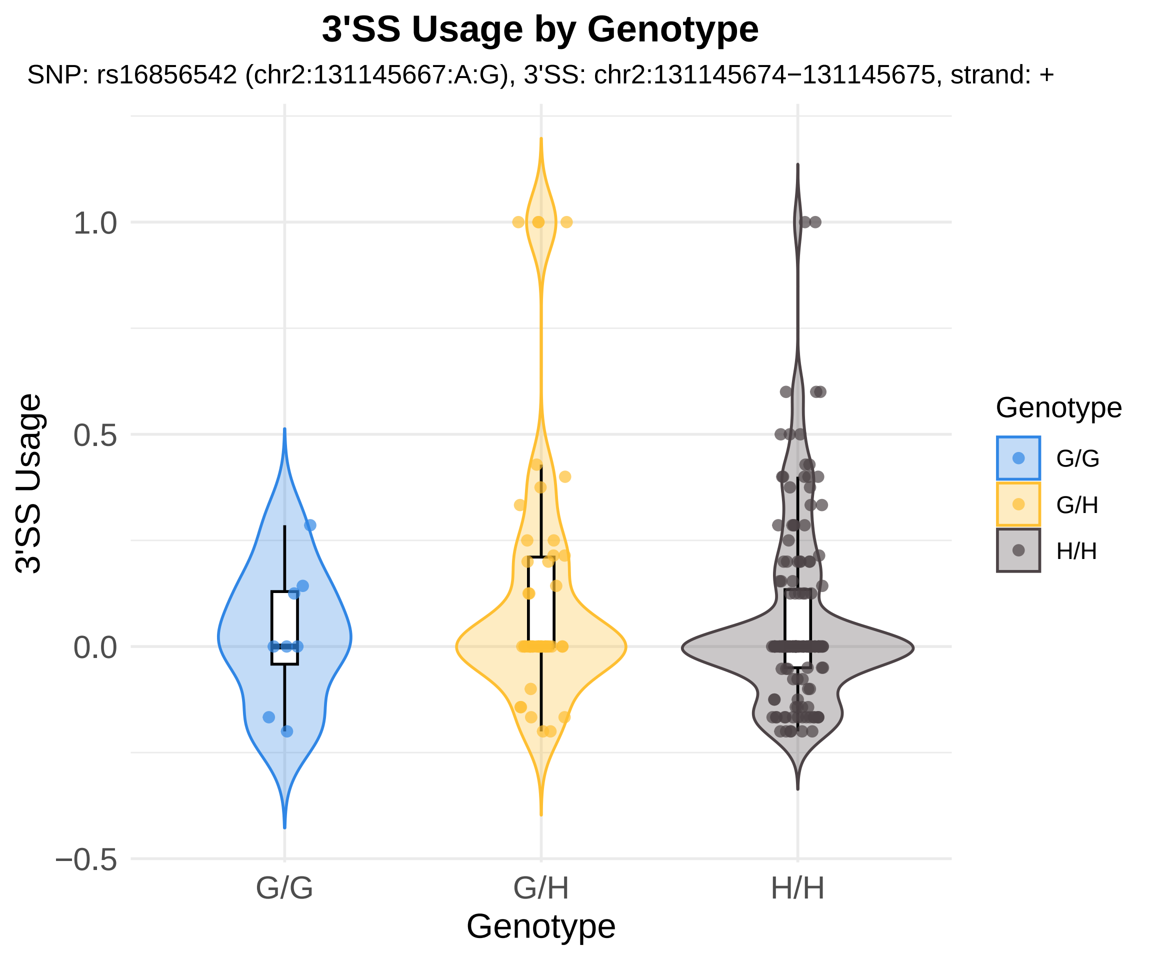

3'SS Usage by Genotype
SNP: rs16856542 (chr2:131145667:A:G), 3'SS: chr2:131145674−131145675, strand: +
3'SS Usage
1.0
0.5
0.0
−0.5
G/G
G/H
H/H
Genotype
Genotype
G/G
G/H
H/H

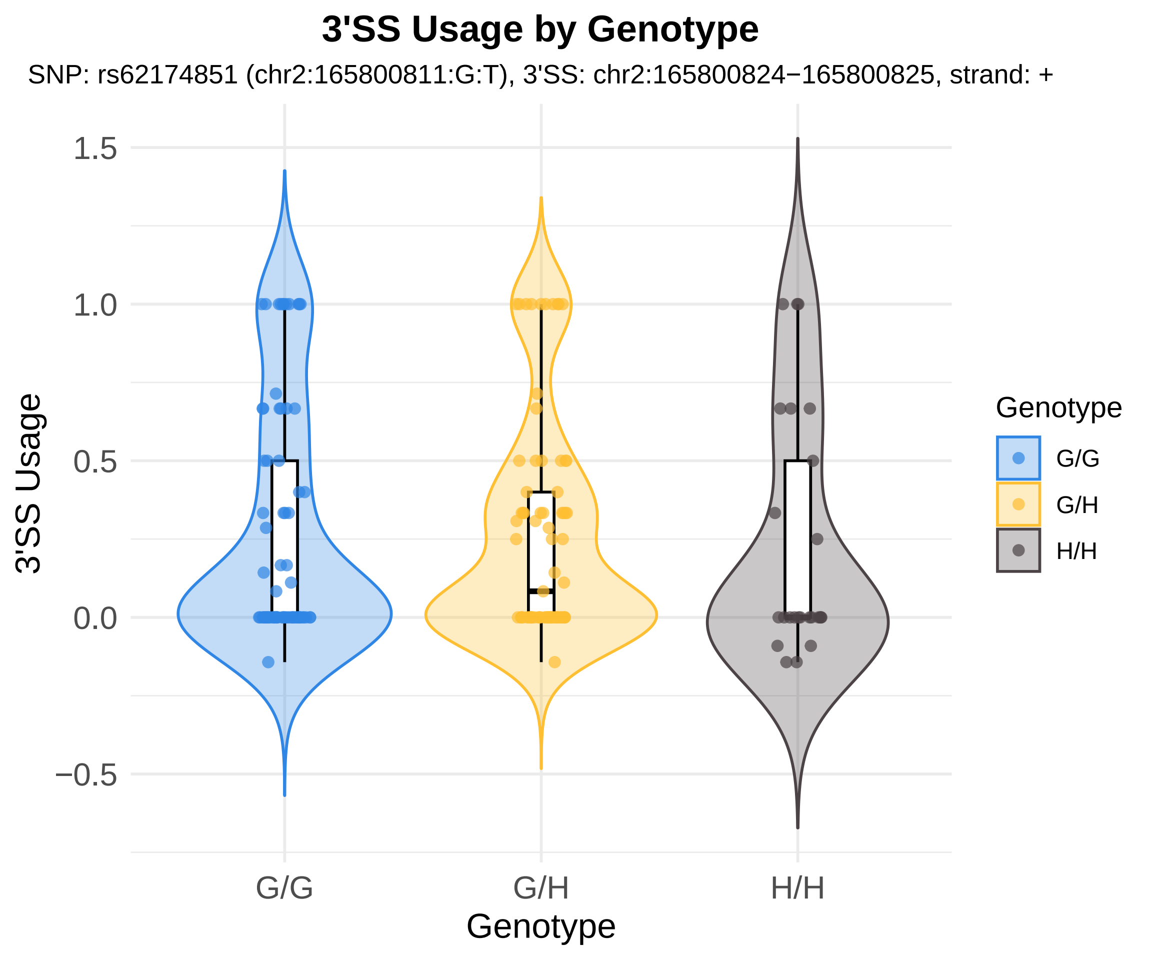
3'SS Usage by Genotype
SNP: rs62174851 (chr2:165800811:G:T), 3'SS: chr2:165800824−165800825, strand: +
3'SS Usage
1.5
1.0
0.5
0.0
−0.5
G/G
G/H
H/H
Genotype
Genotype
G/G
G/H
H/H

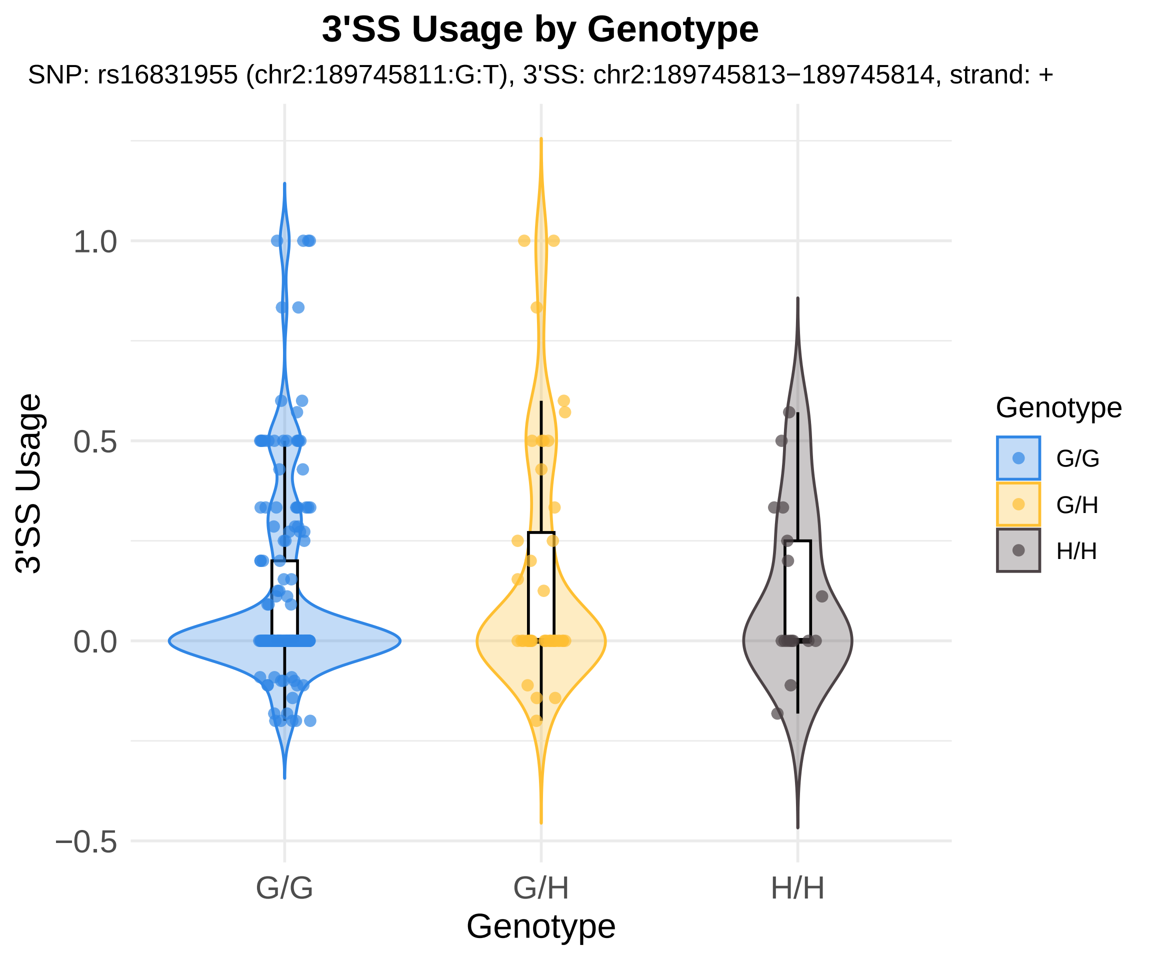
3'SS Usage by Genotype
SNP: rs16831955 (chr2:189745811:G:T), 3'SS: chr2:189745813−189745814, strand: +
3'SS Usage
1.0
0.5
0.0
−0.5
G/G
G/H
H/H
Genotype
Genotype
G/G
G/H
H/H

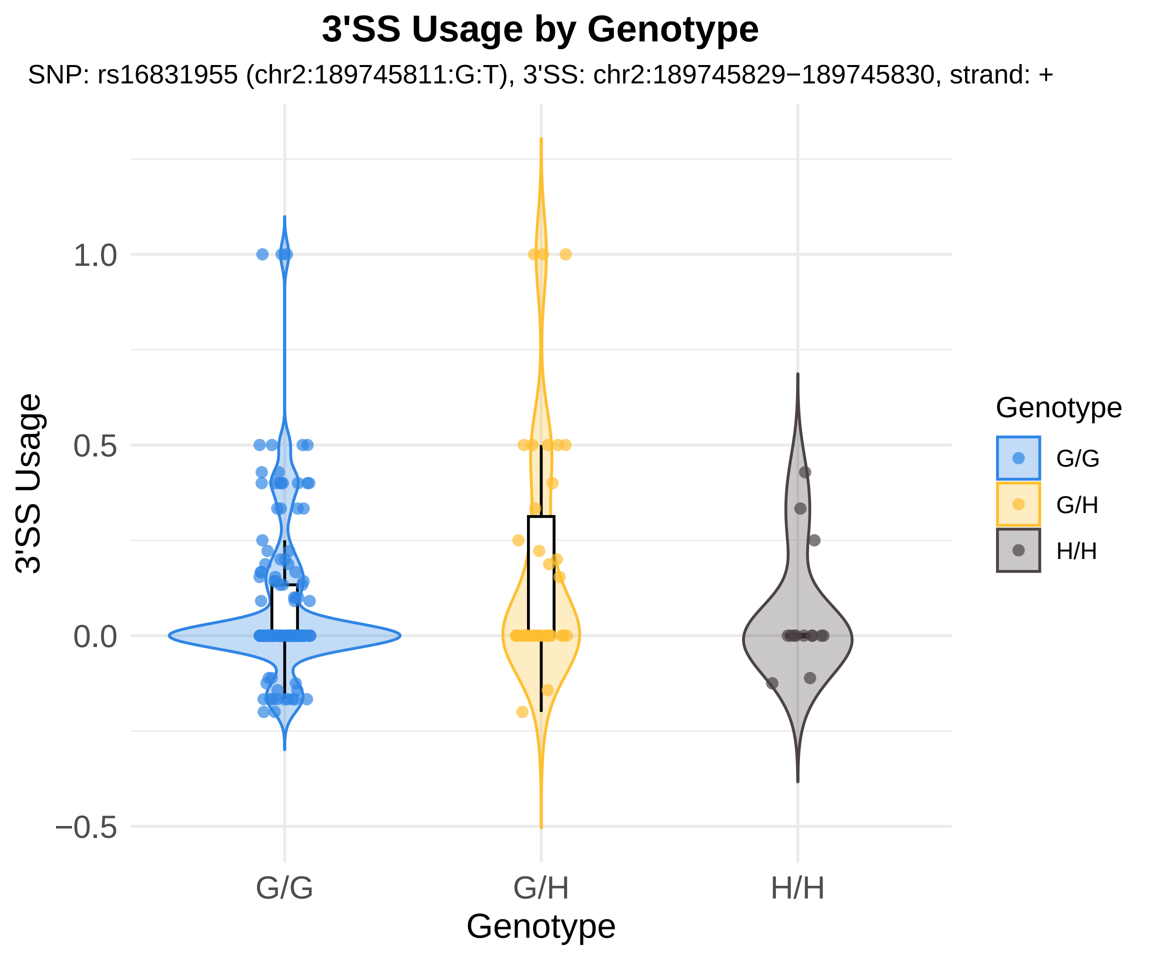

3'SS Usage by Genotype
SNP: rs16831955 (chr2:189745811:G:T), 3'SS: chr2:189745829−189745830, strand: +
3'SS Usage
1.0
0.5
0.0
−0.5
G/G
G/H
H/H
Genotype
Genotype
G/G
G/H
H/H

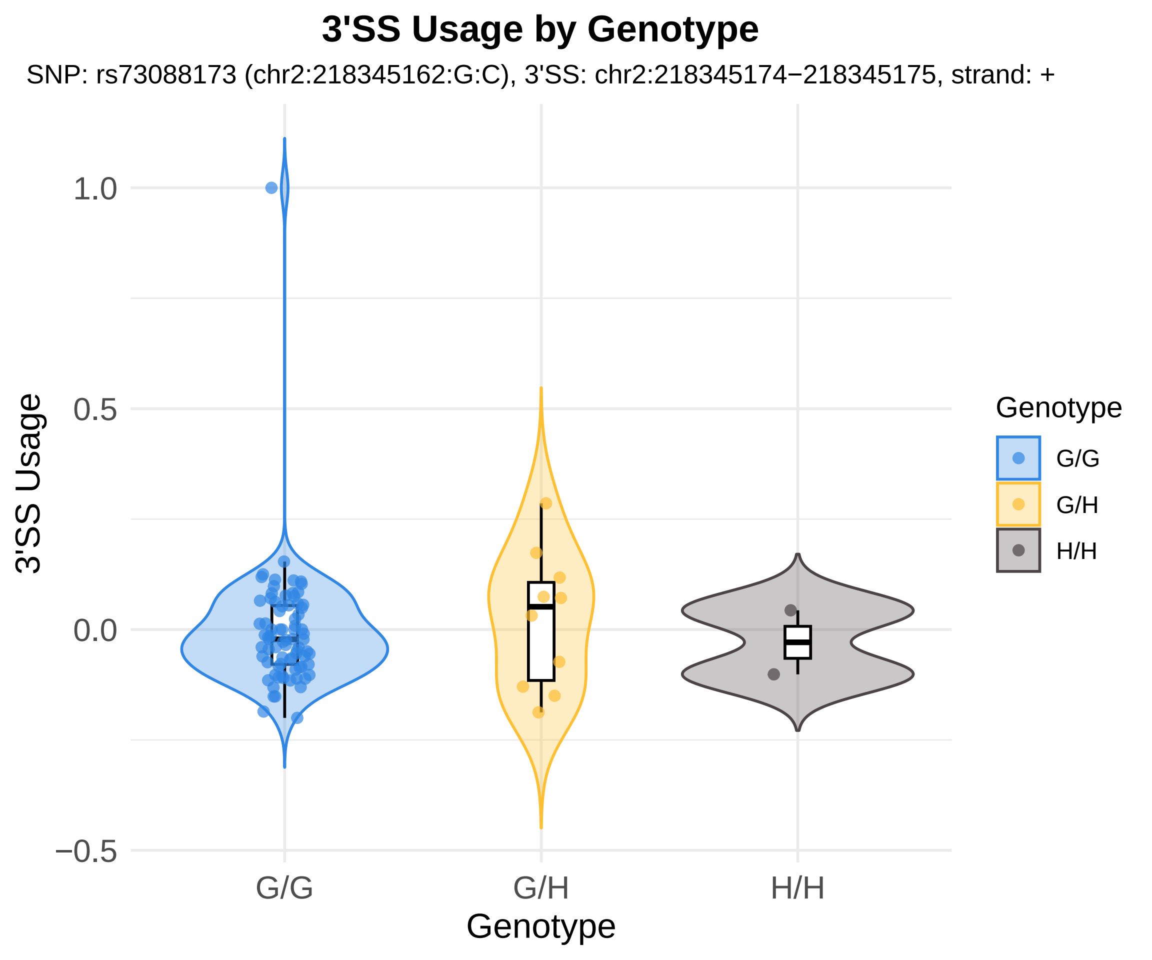

3'SS Usage by Genotype
SNP: rs73088173 (chr2:218345162:G:C), 3'SS: chr2:218345174−218345175, strand: +
3'SS Usage
1.0
0.5
0.0
−0.5
G/G
G/H
H/H
Genotype
Genotype
G/G
G/H
H/H

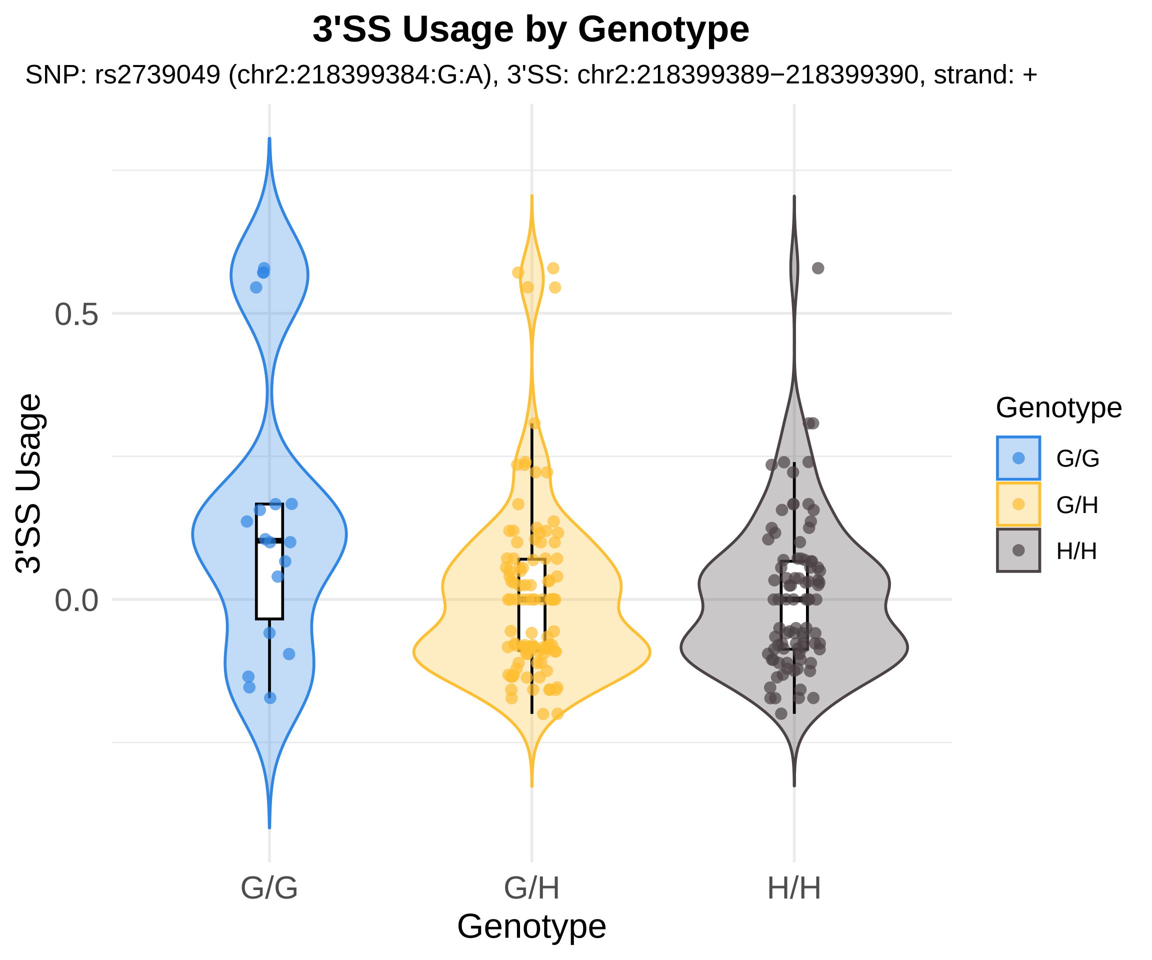
3'SS Usage by Genotype
SNP: rs2739049 (chr2:218399384:G:A), 3'SS: chr2:218399389−218399390, strand: +
3'SS Usage
0.5
0.0
G/G
G/H
H/H
Genotype
Genotype
G/G
G/H
H/H

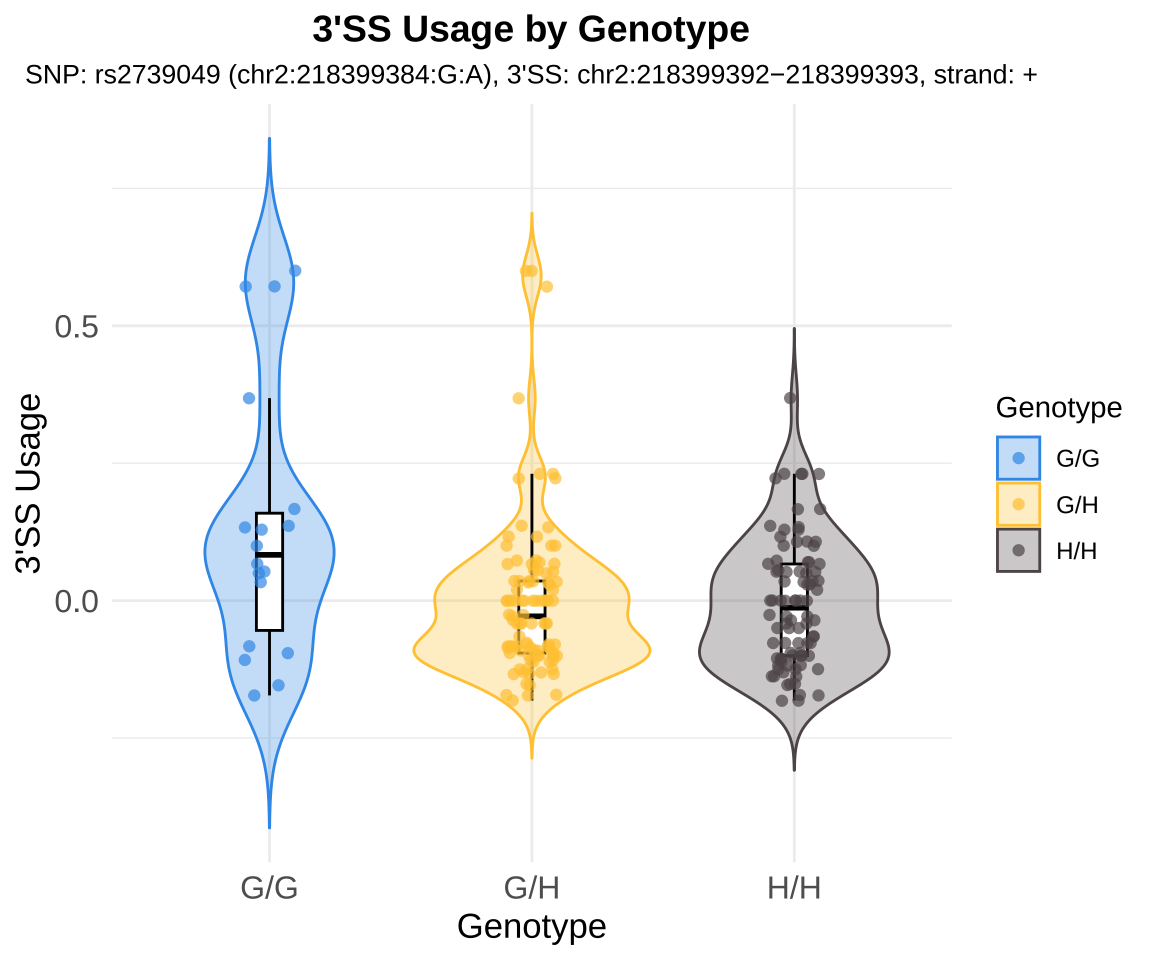

3'SS Usage by Genotype
SNP: rs2739049 (chr2:218399384:G:A), 3'SS: chr2:218399392−218399393, strand: +
3'SS Usage
0.5
0.0
G/G
G/H
H/H
Genotype
Genotype
G/G
G/H
H/H

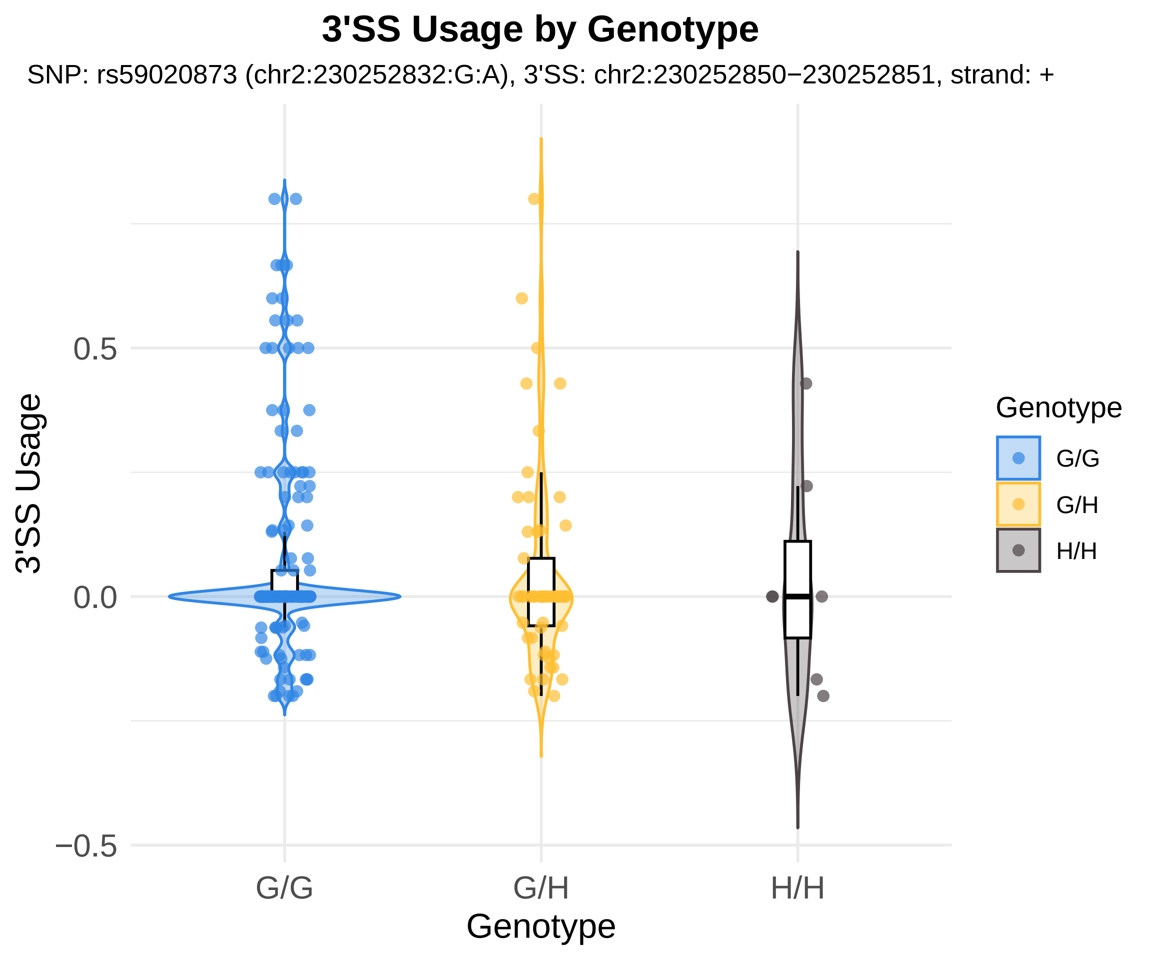
3'SS Usage by Genotype
SNP: rs59020873 (chr2:230252832:G:A), 3'SS: chr2:230252850−230252851, strand: +
3'SS Usage
0.5
0.0
−0.5
G/G
G/H
H/H
Genotype
Genotype
G/G
G/H
H/H

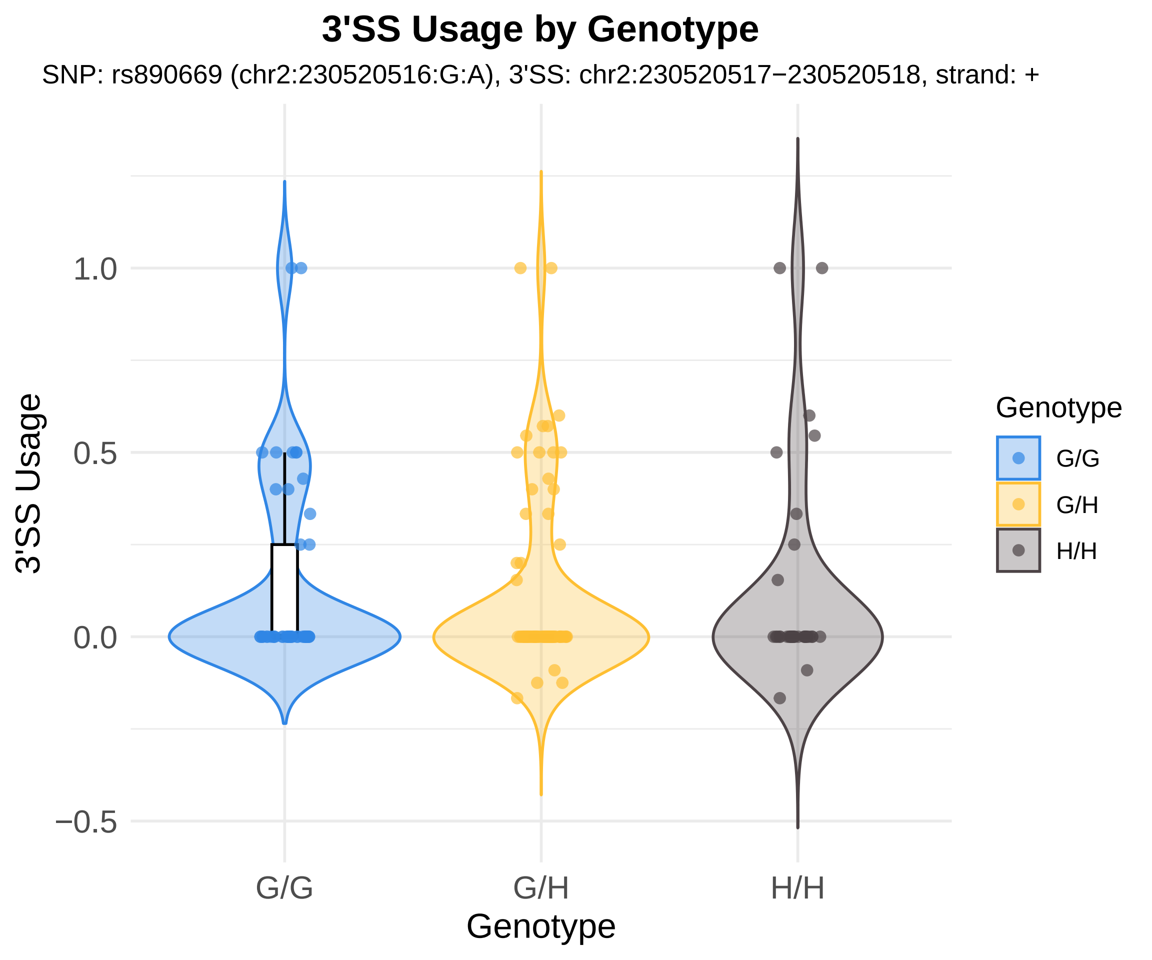

3'SS Usage by Genotype
SNP: rs890669 (chr2:230520516:G:A), 3'SS: chr2:230520517−230520518, strand: +
3'SS Usage
1.0
0.5
0.0
−0.5
G/G
G/H
H/H
Genotype
Genotype
G/G
G/H
H/H

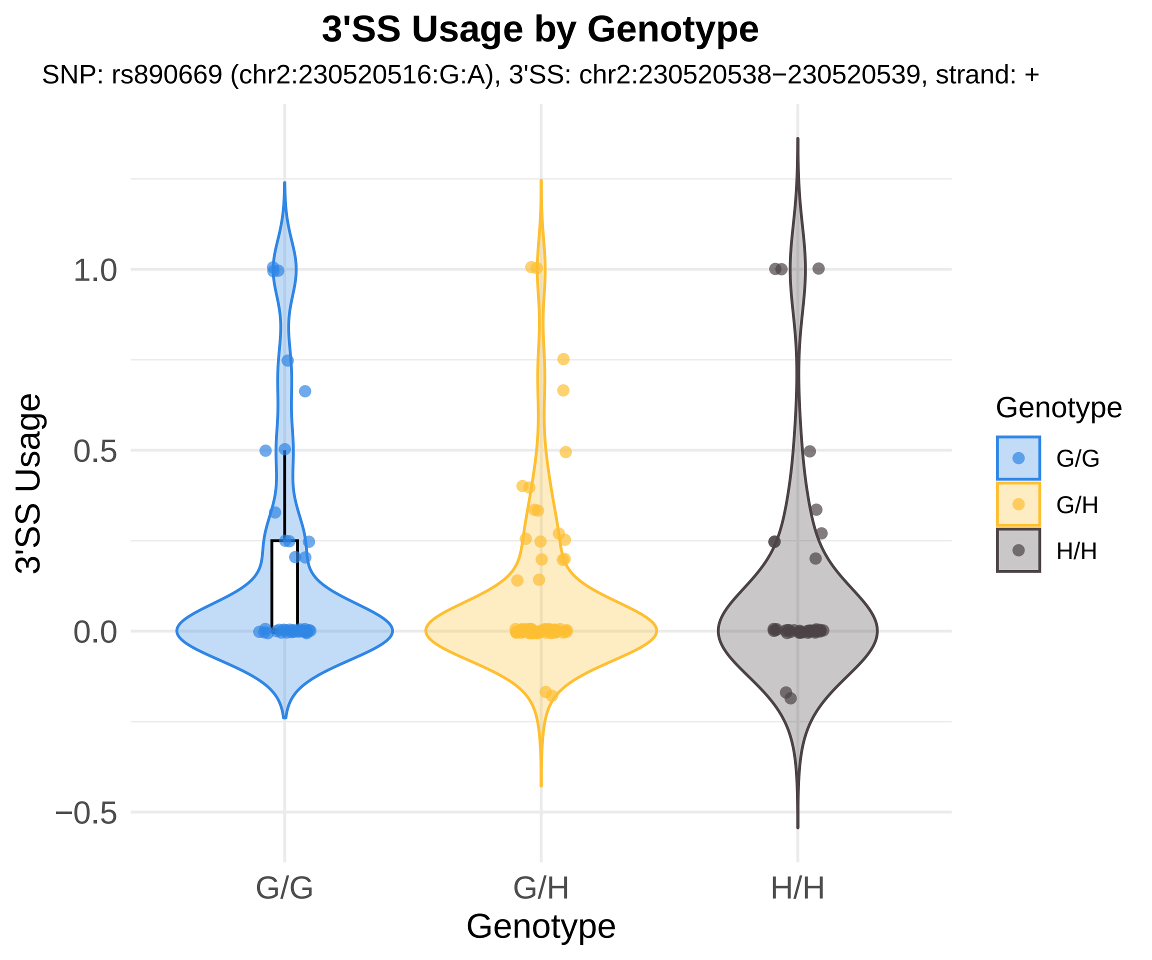
3'SS Usage by Genotype
SNP: rs890669 (chr2:230520516:G:A), 3'SS: chr2:230520538−230520539, strand: +
3'SS Usage
1.0
0.5
0.0
−0.5
G/G
G/H
H/H
Genotype
Genotype
G/G
G/H
H/H

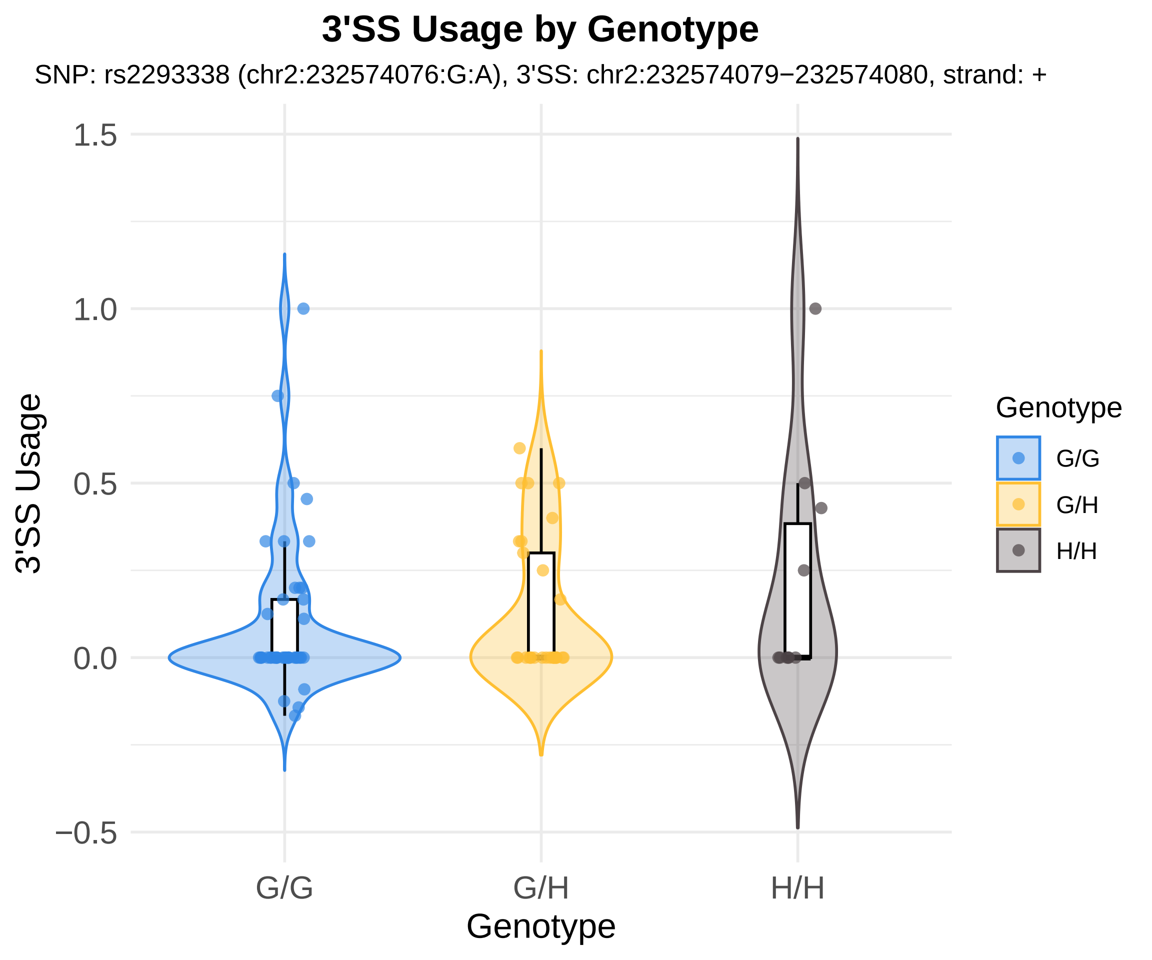
3'SS Usage by Genotype
SNP: rs2293338 (chr2:232574076:G:A), 3'SS: chr2:232574079−232574080, strand: +
1.5
1.0
0.5
0.0
−0.5
3'SS Usage
G/G
G/H
H/H
Genotype
Genotype
G/G
G/H
H/H

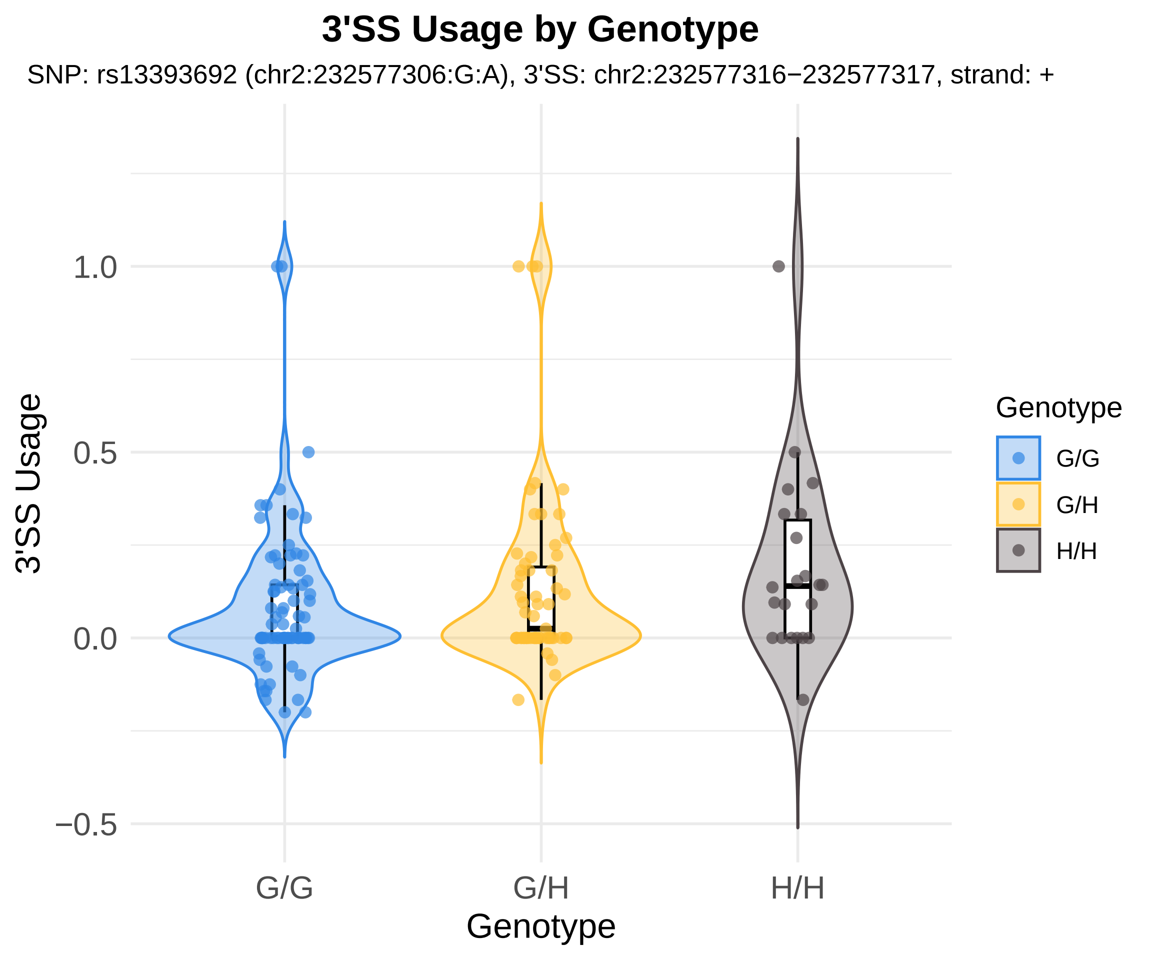
3'SS Usage by Genotype
SNP: rs13393692 (chr2:232577306:G:A), 3'SS: chr2:232577316−232577317, strand: +
3'SS Usage
1.0
0.5
0.0
−0.5
G/G
G/H
H/H
Genotype
Genotype
G/G
G/H
H/H

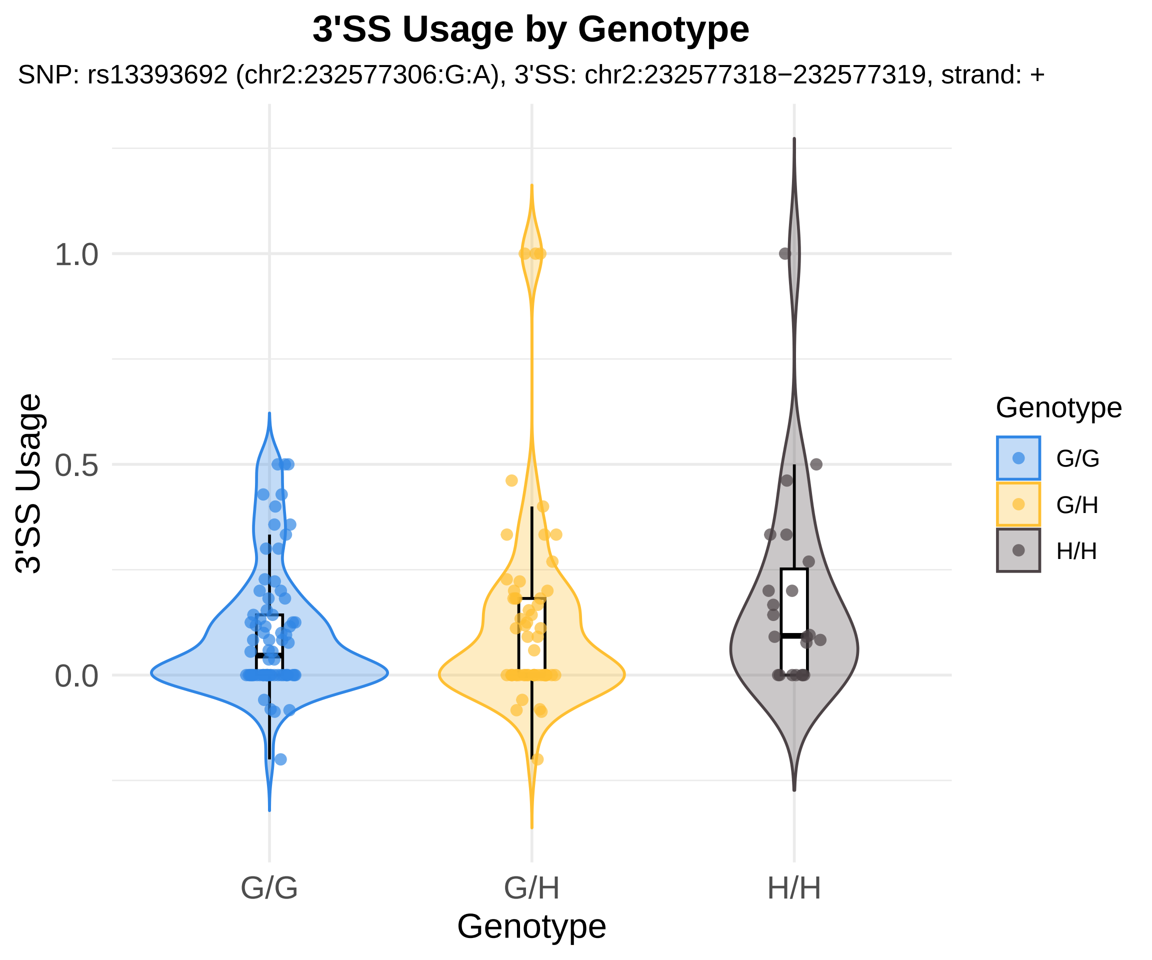

3'SS Usage by Genotype
SNP: rs13393692 (chr2:232577306:G:A), 3'SS: chr2:232577318−232577319, strand: +
3'SS Usage
1.0
0.5
0.0
G/G
G/H
H/H
Genotype
Genotype
G/G
G/H
H/H

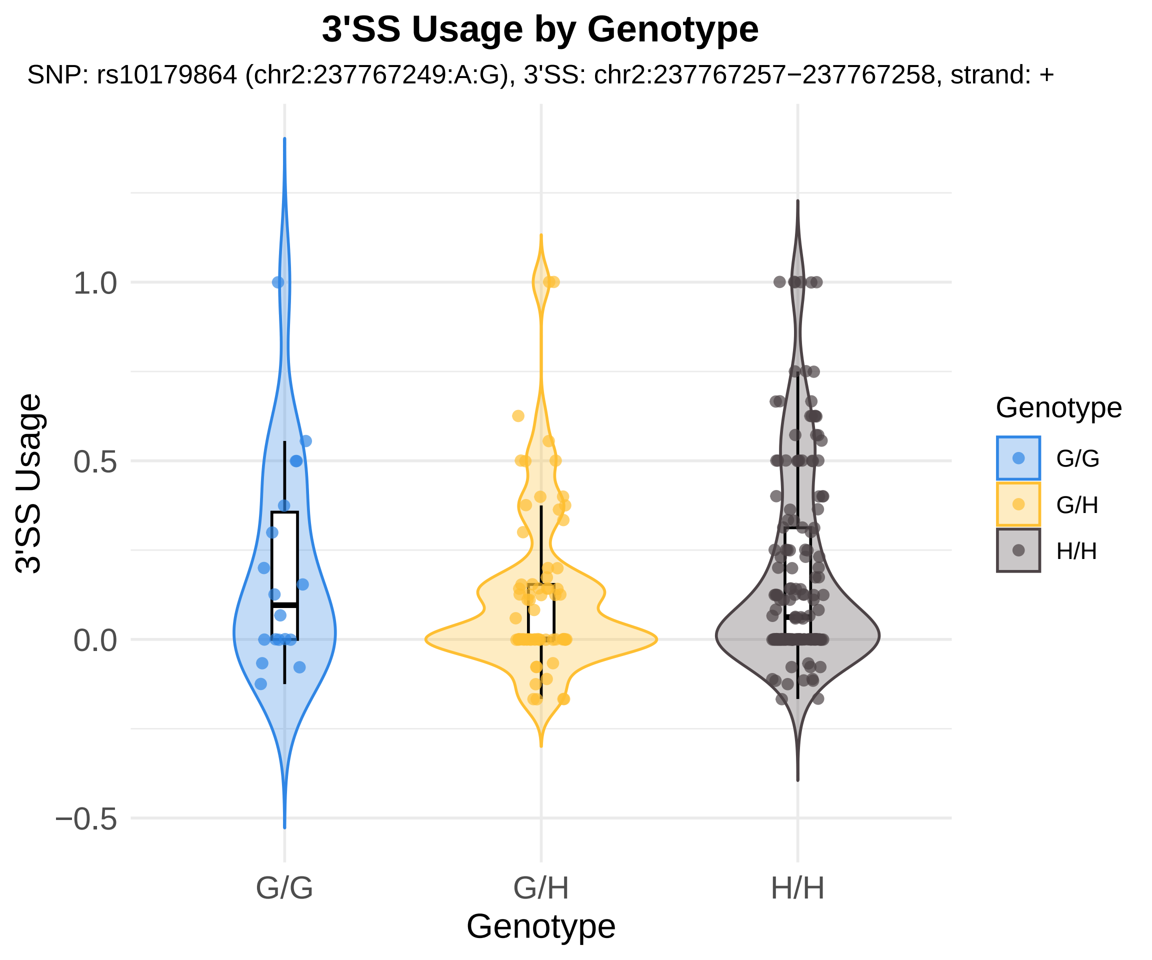
3'SS Usage by Genotype
SNP: rs10179864 (chr2:237767249:A:G), 3'SS: chr2:237767257−237767258, strand: +
3'SS Usage
1.0
0.5
0.0
−0.5
G/G
G/H
H/H
Genotype
Genotype
G/G
G/H
H/H

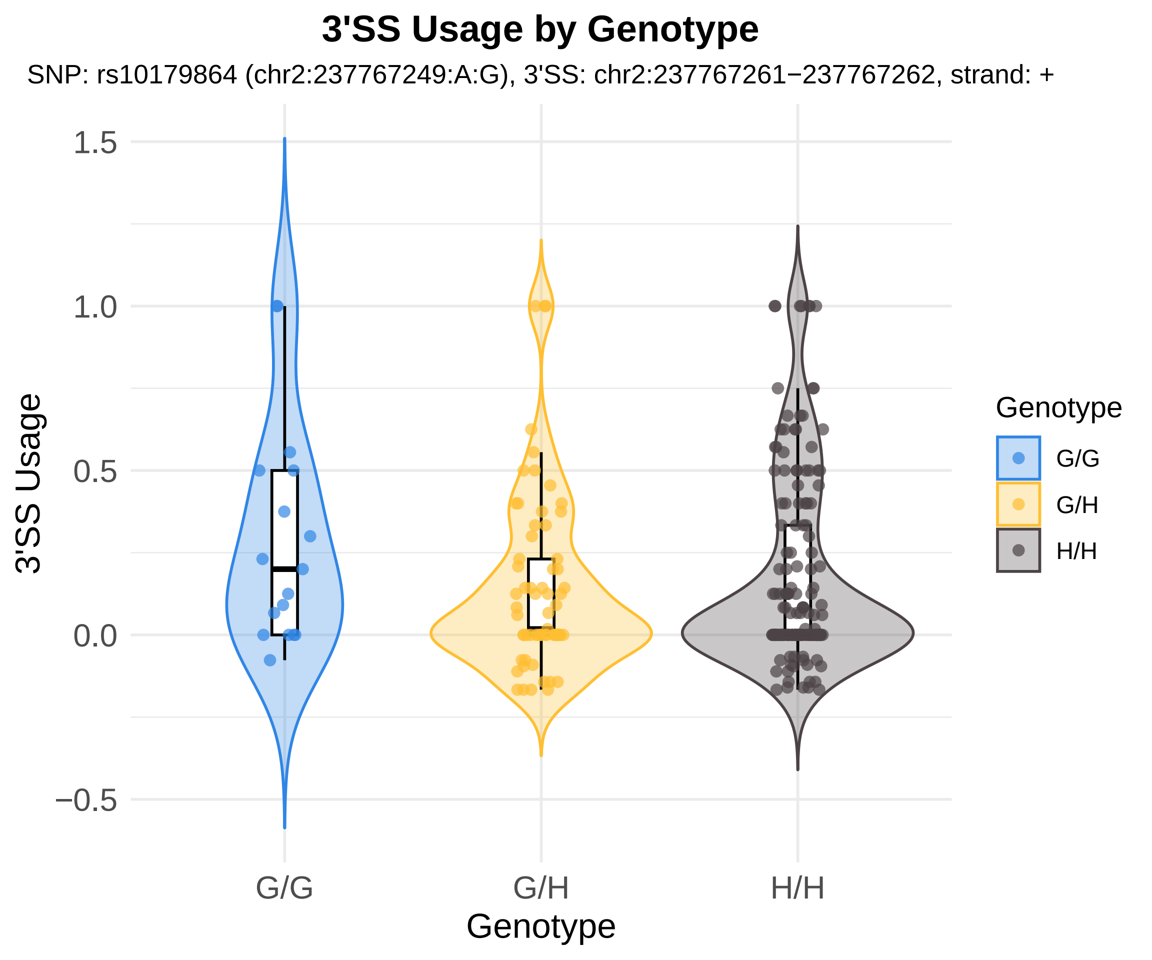
3'SS Usage by Genotype
SNP: rs10179864 (chr2:237767249:A:G), 3'SS: chr2:237767261−237767262, strand: +
3'SS Usage
1.5
1.0
0.5
0.0
−0.5
G/G
G/H
H/H
Genotype
Genotype
G/G
G/H
H/H

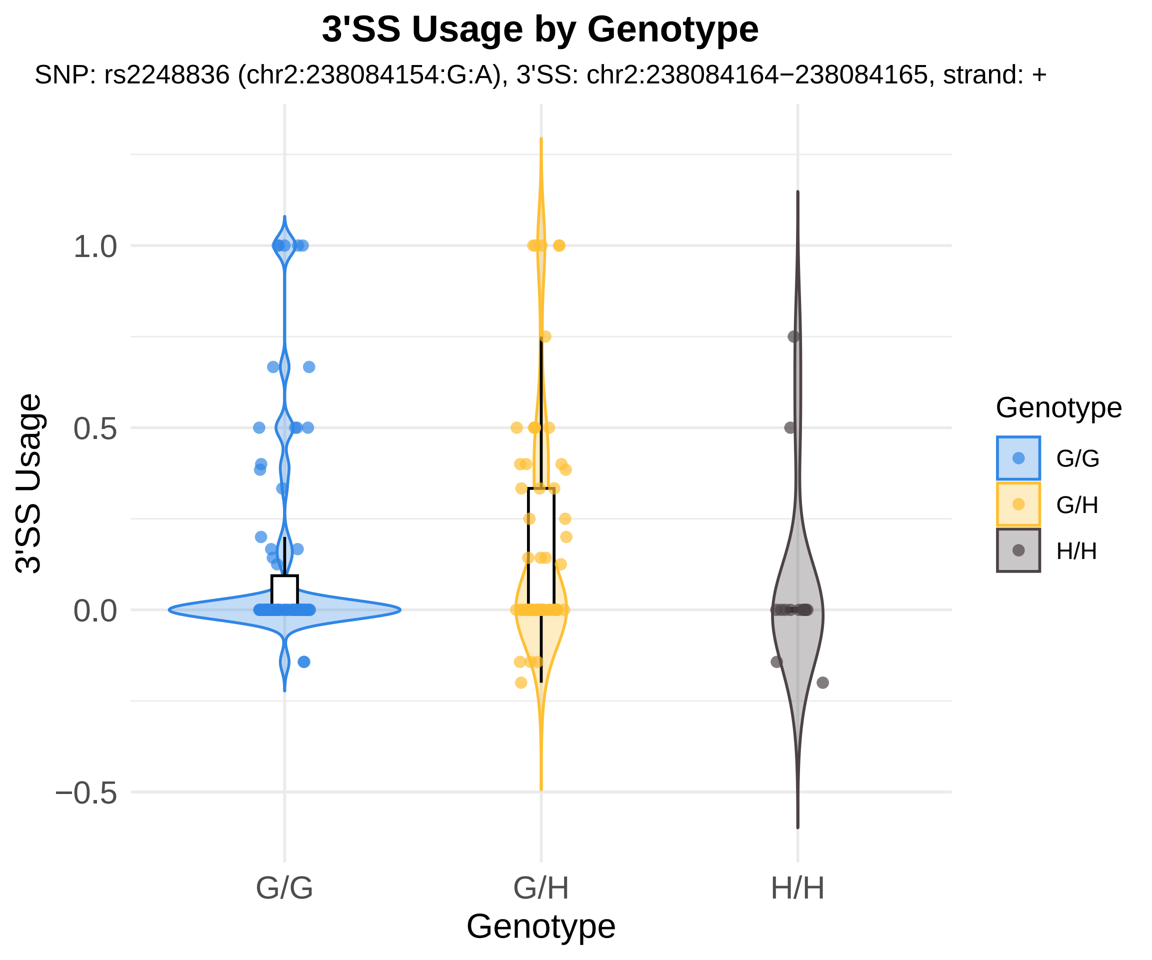
3'SS Usage by Genotype
SNP: rs2248836 (chr2:238084154:G:A), 3'SS: chr2:238084164−238084165, strand: +
3'SS Usage
1.0
0.5
0.0
−0.5
G/G
G/H
H/H
Genotype
Genotype
G/G
G/H
H/H

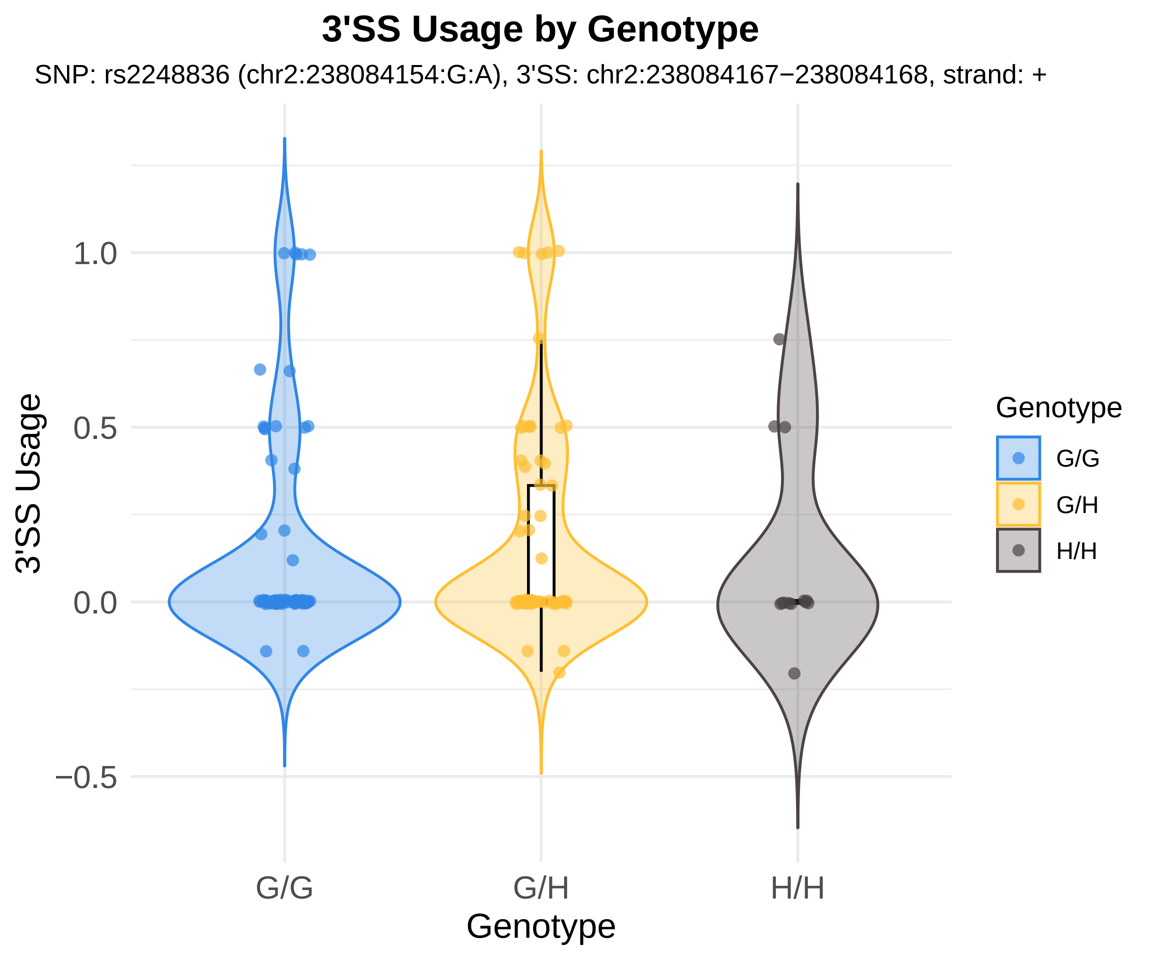

3'SS Usage by Genotype
SNP: rs2248836 (chr2:238084154:G:A), 3'SS: chr2:238084167−238084168, strand: +
3'SS Usage
1.0
0.5
0.0
−0.5
G/G
G/H
H/H
Genotype
Genotype
G/G
G/H
H/H

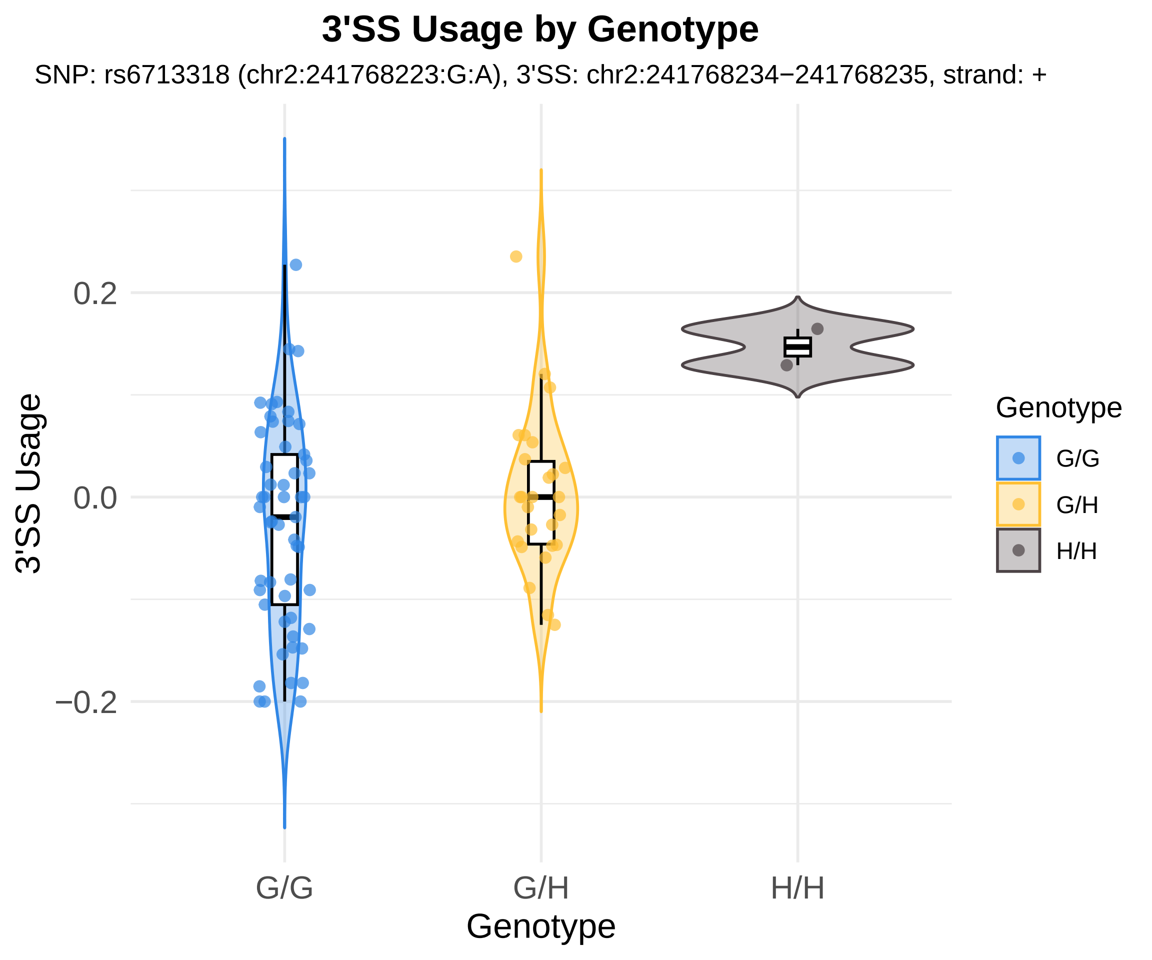
3'SS Usage by Genotype
SNP: rs6713318 (chr2:241768223:G:A), 3'SS: chr2:241768234−241768235, strand: +
3'SS Usage
0.2
0.0
−0.2
G/G
G/H
H/H
Genotype
Genotype
G/G
G/H
H/H

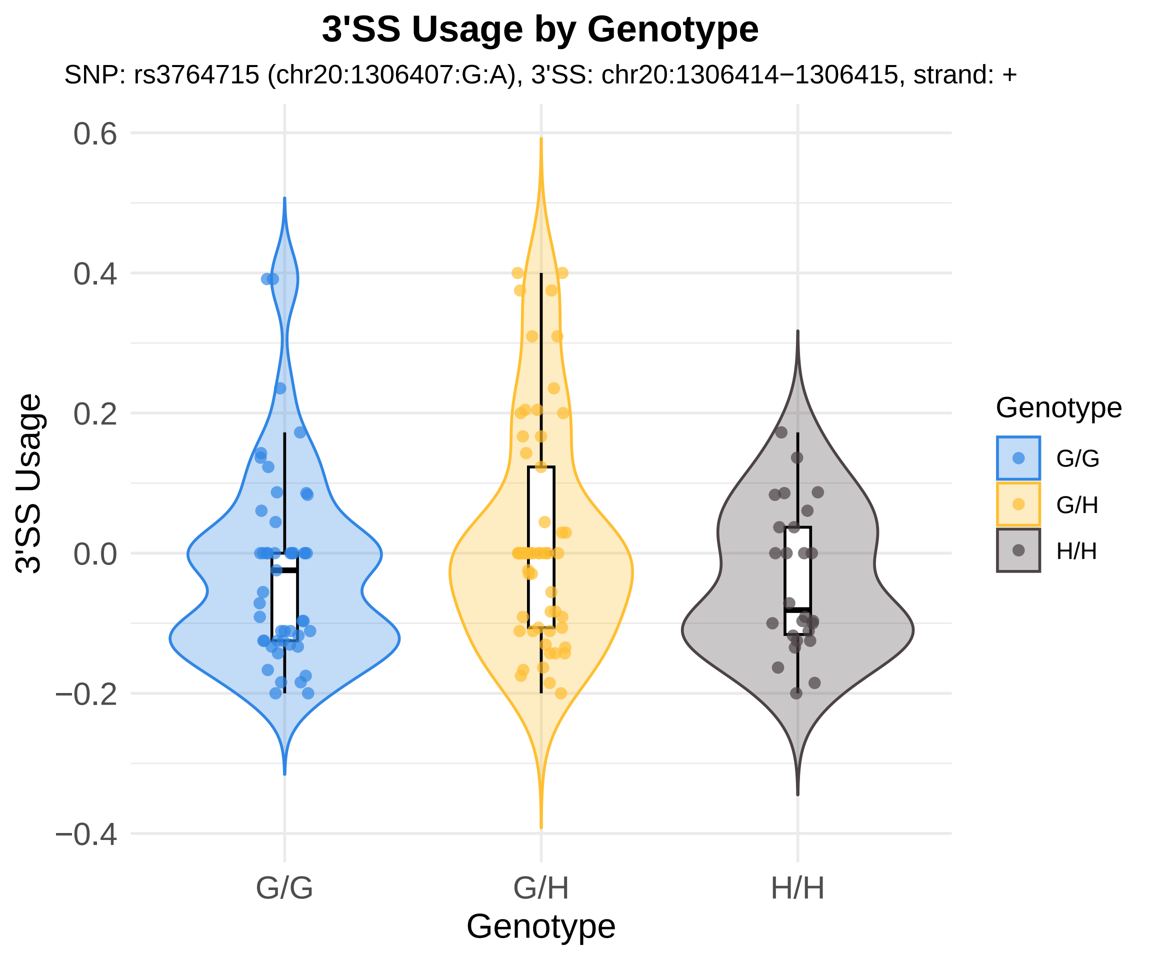
3'SS Usage by Genotype
SNP: rs3764715 (chr20:1306407:G:A), 3'SS: chr20:1306414−1306415, strand: +
3'SS Usage
0.6
0.4
0.2
0.0
−0.2
−0.4
G/G
G/H
H/H
Genotype
Genotype
G/G
G/H
H/H

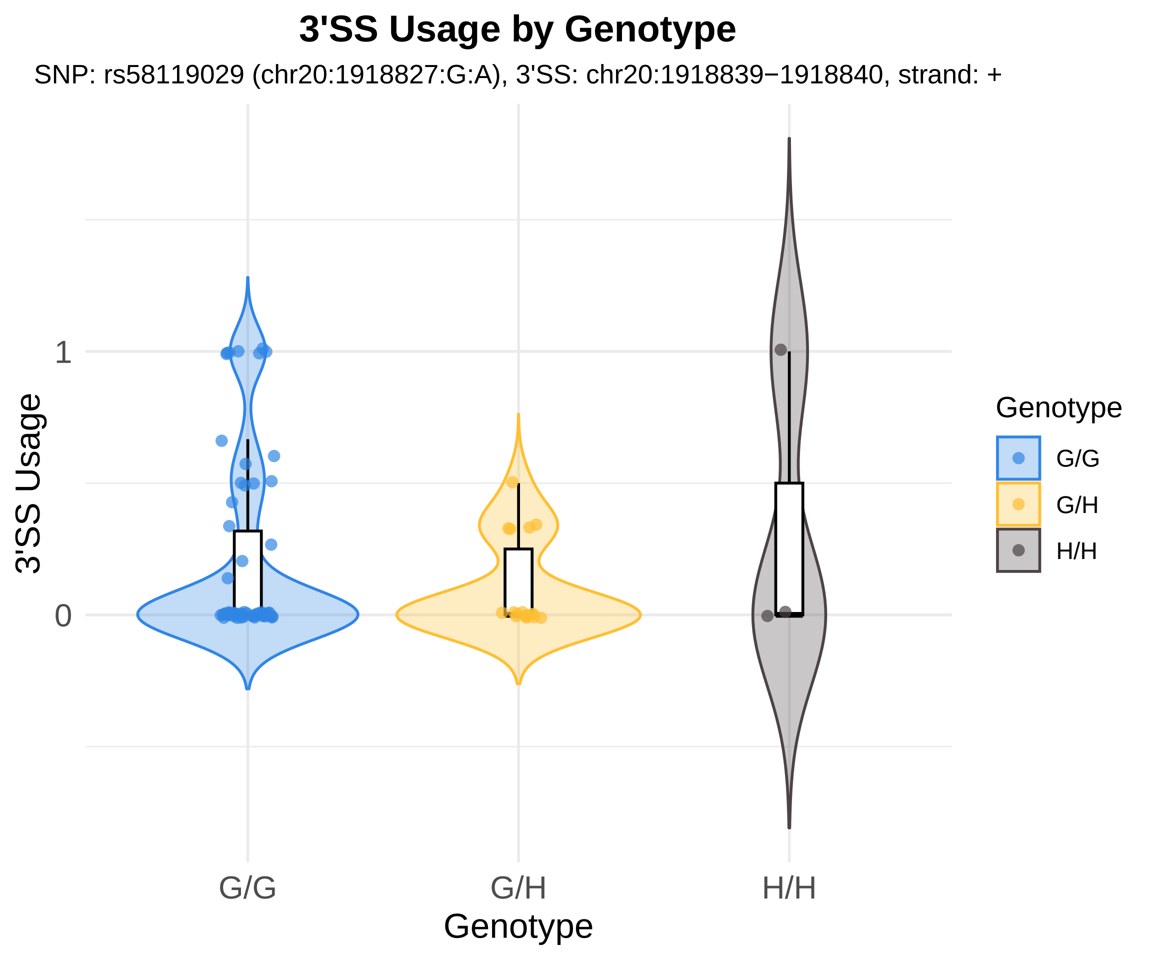
3'SS Usage by Genotype
SNP: rs58119029 (chr20:1918827:G:A), 3'SS: chr20:1918839−1918840, strand: +
3'SS Usage
1
0
G/G
G/H
H/H
Genotype
Genotype
G/G
G/H
H/H

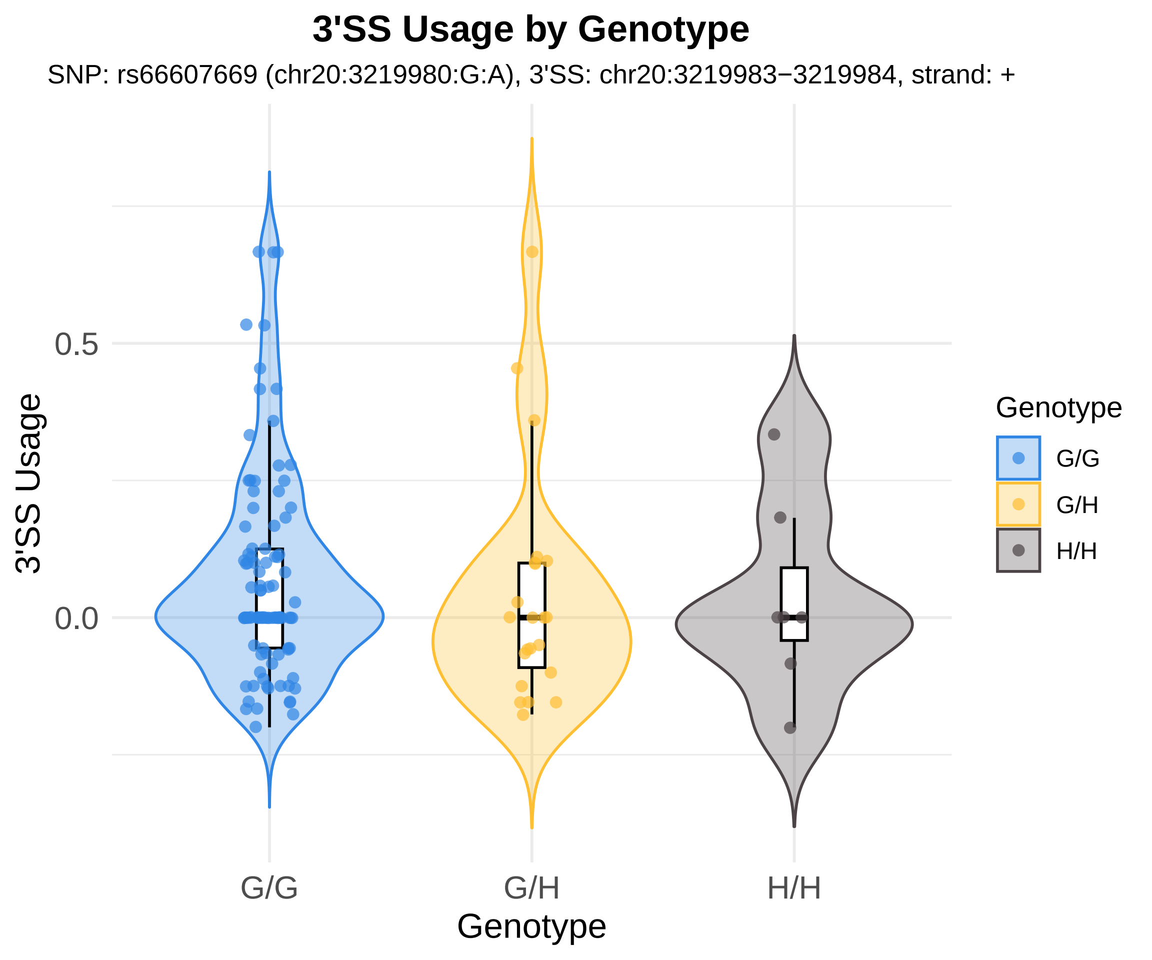
3'SS Usage by Genotype
SNP: rs66607669 (chr20:3219980:G:A), 3'SS: chr20:3219983−3219984, strand: +
3'SS Usage
0.5
0.0
G/G
G/H
H/H
Genotype
Genotype
G/G
G/H
H/H

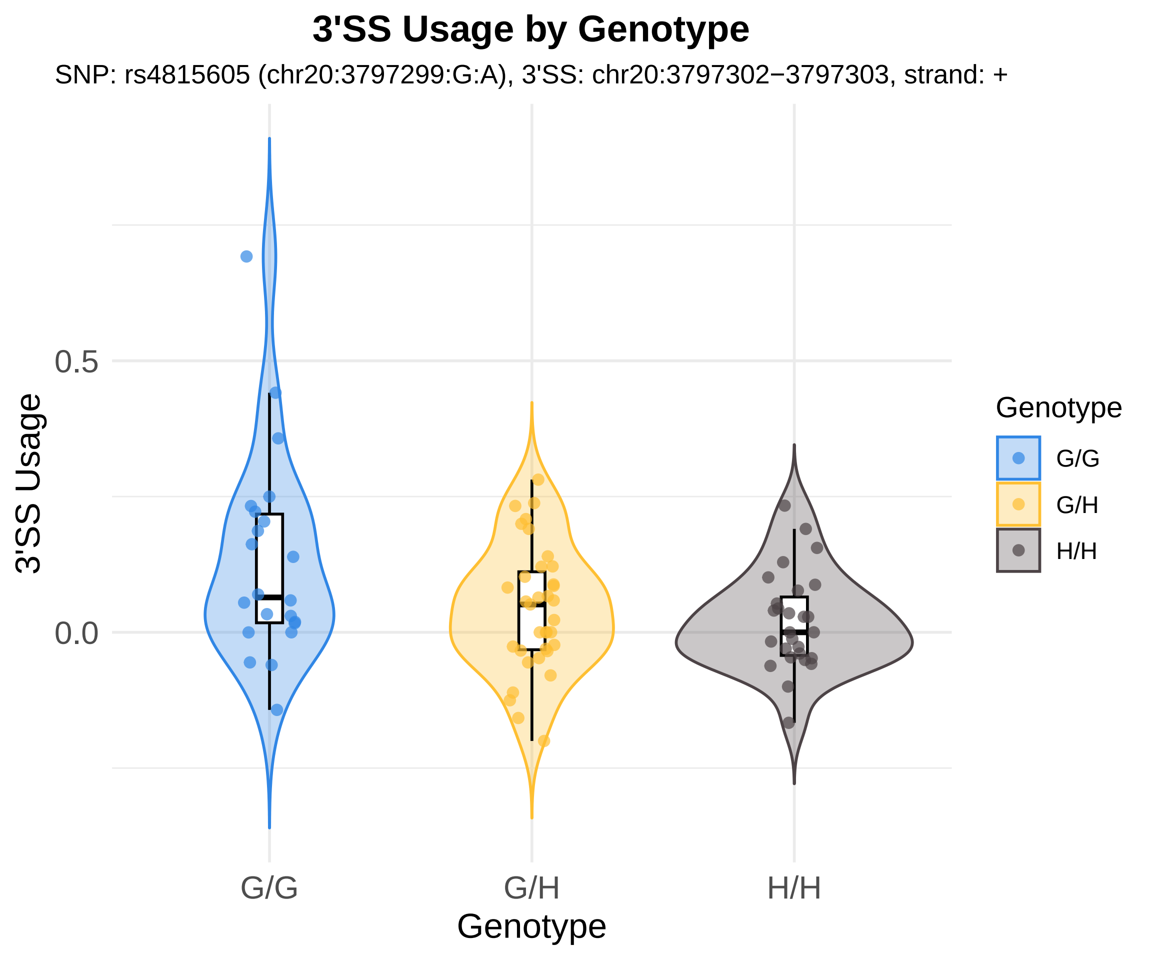
3'SS Usage by Genotype
SNP: rs4815605 (chr20:3797299:G:A), 3'SS: chr20:3797302−3797303, strand: +
3'SS Usage
0.5
0.0
G/G
G/H
H/H
Genotype
Genotype
G/G
G/H
H/H

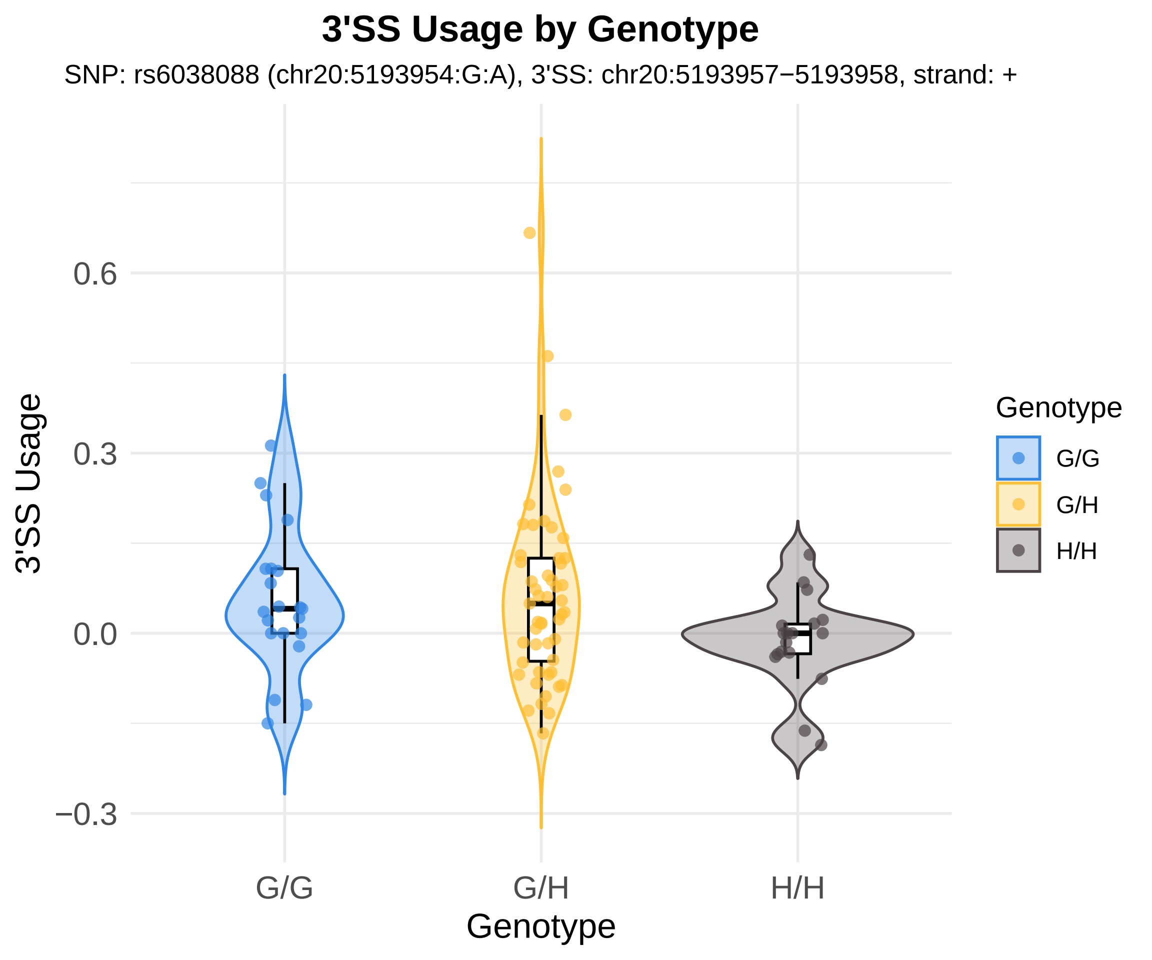
3'SS Usage by Genotype
SNP: rs6038088 (chr20:5193954:G:A), 3'SS: chr20:5193957−5193958, strand: +
3'SS Usage
0.6
0.3
0.0
−0.3
G/G
G/H
H/H
Genotype
Genotype
G/G
G/H
H/H

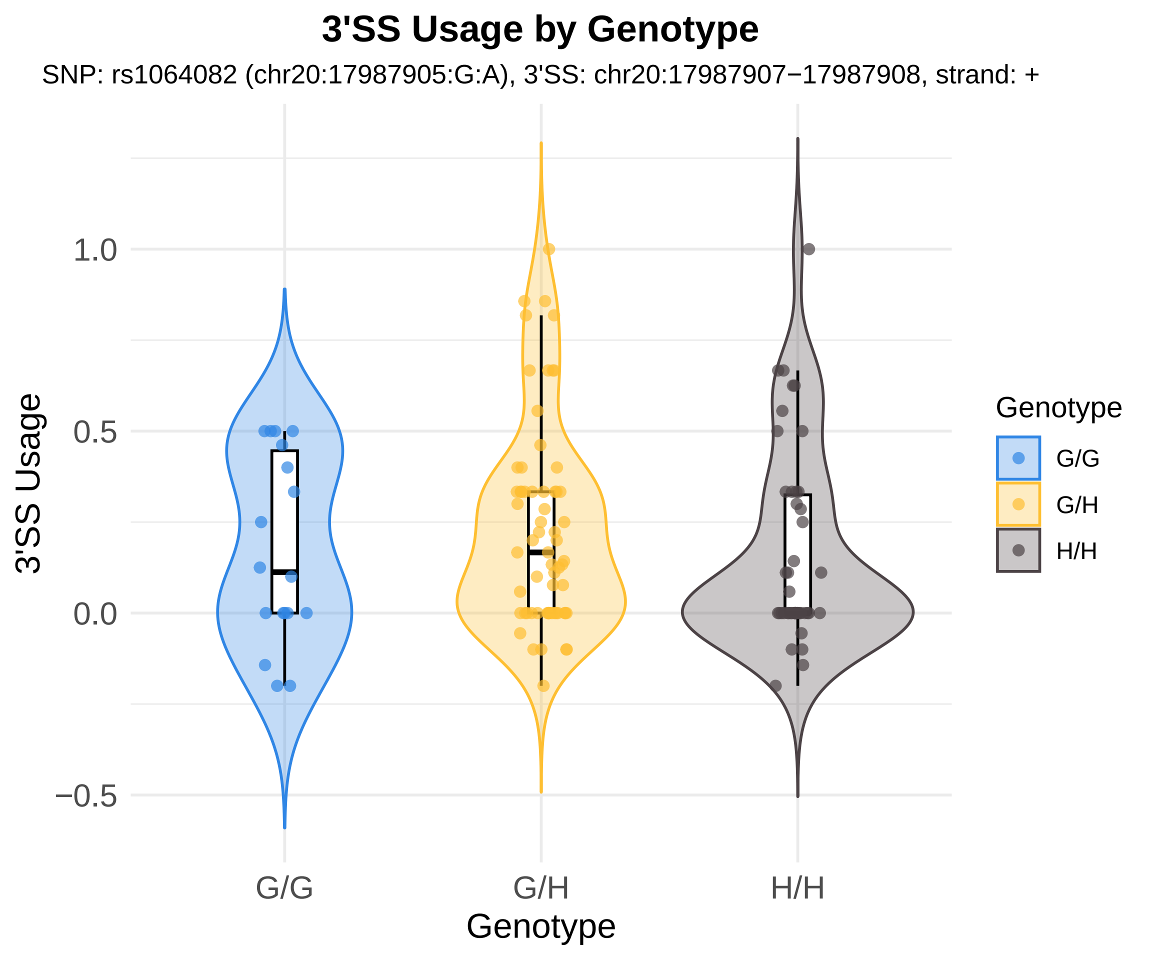
3'SS Usage by Genotype
SNP: rs1064082 (chr20:17987905:G:A), 3'SS: chr20:17987907−17987908, strand: +
3'SS Usage
1.0
0.5
0.0
−0.5
G/G
G/H
H/H
Genotype
Genotype
G/G
G/H
H/H

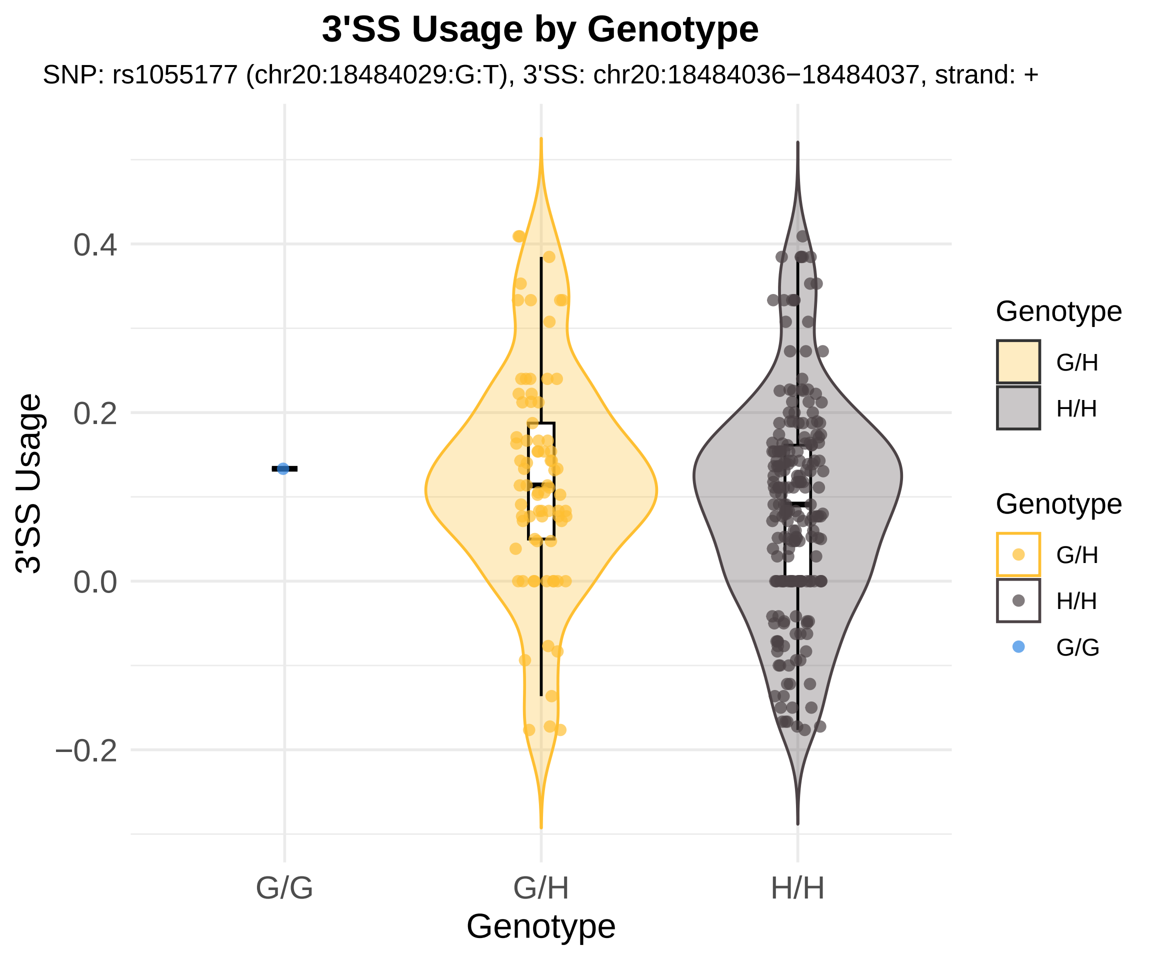

3'SS Usage by Genotype
SNP: rs1055177 (chr20:18484029:G:T), 3'SS: chr20:18484036−18484037, strand: +
3'SS Usage
0.4
0.2
0.0
−0.2
G/G
G/H
H/H
Genotype
Genotype
G/H
H/H
Genotype
G/H
H/H
G/G

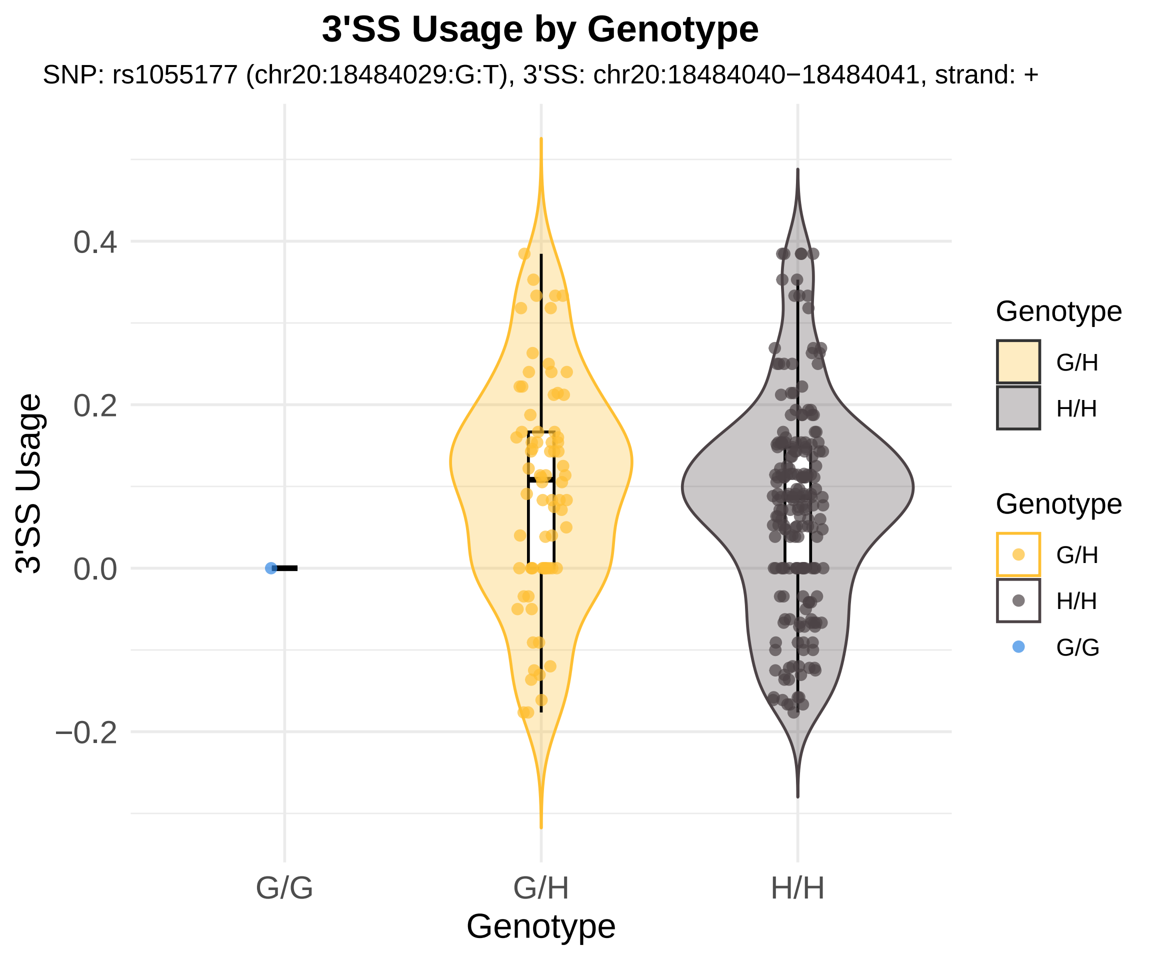

3'SS Usage by Genotype
SNP: rs1055177 (chr20:18484029:G:T), 3'SS: chr20:18484040−18484041, strand: +
3'SS Usage
0.4
0.2
0.0
−0.2
G/G
G/H
H/H
Genotype
Genotype
G/H
H/H
Genotype
G/H
H/H
G/G

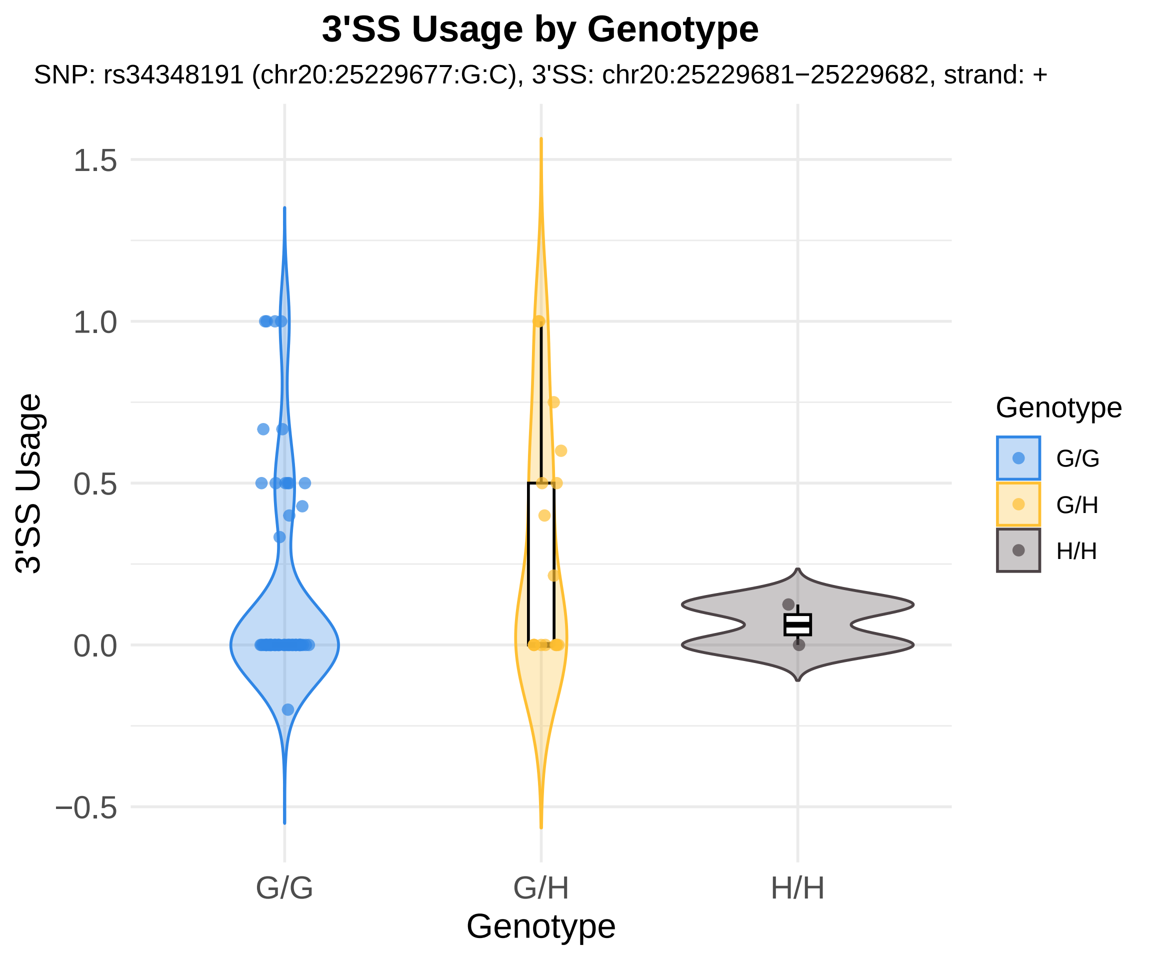
3'SS Usage by Genotype
SNP: rs34348191 (chr20:25229677:G:C), 3'SS: chr20:25229681−25229682, strand: +
3'SS Usage
1.5
1.0
0.5
0.0
−0.5
G/G
G/H
H/H
Genotype
Genotype
G/G
G/H
H/H

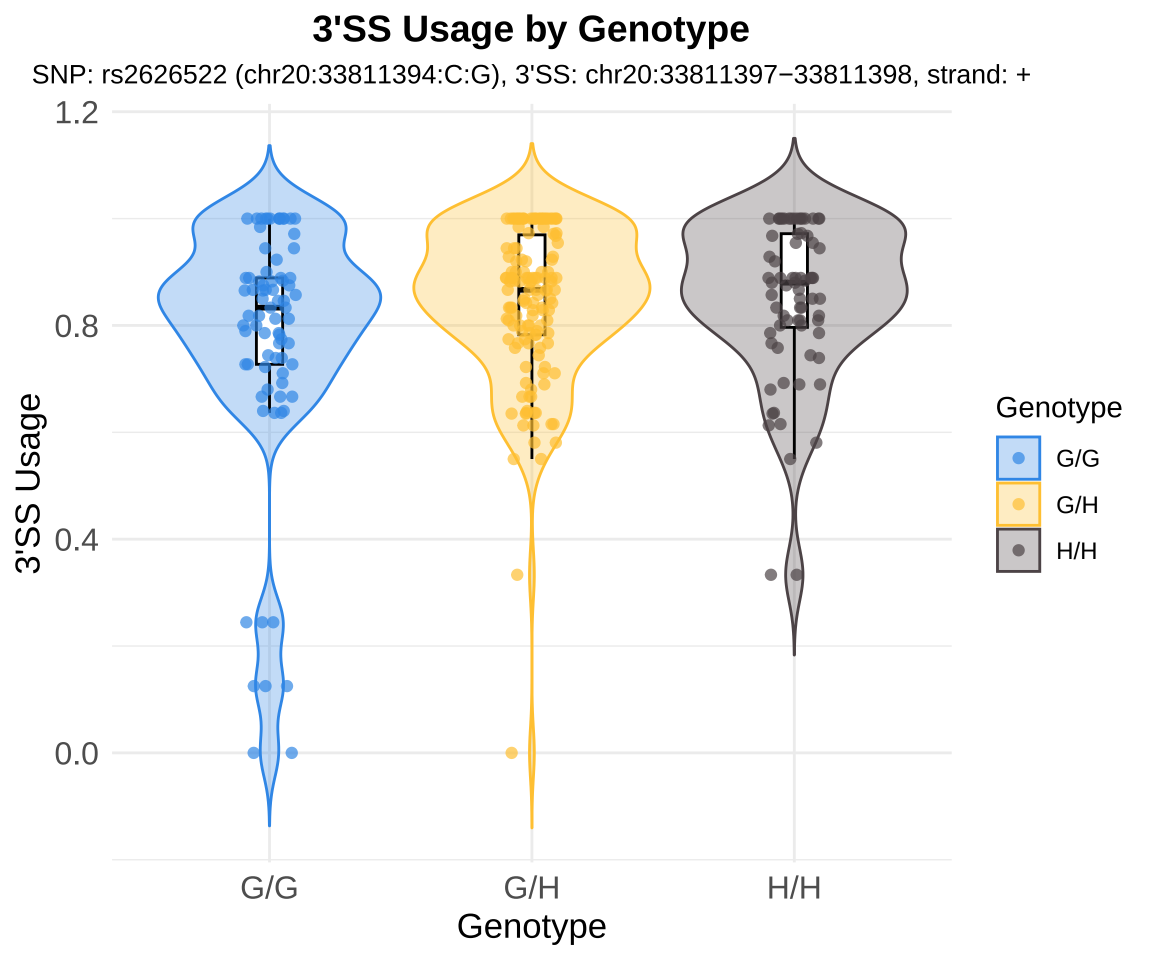
3'SS Usage by Genotype
SNP: rs2626522 (chr20:33811394:C:G), 3'SS: chr20:33811397−33811398, strand: +
1.2
0.8
0.4
0.0
3'SS Usage
G/G
G/H
H/H
Genotype
Genotype
G/G
G/H
H/H

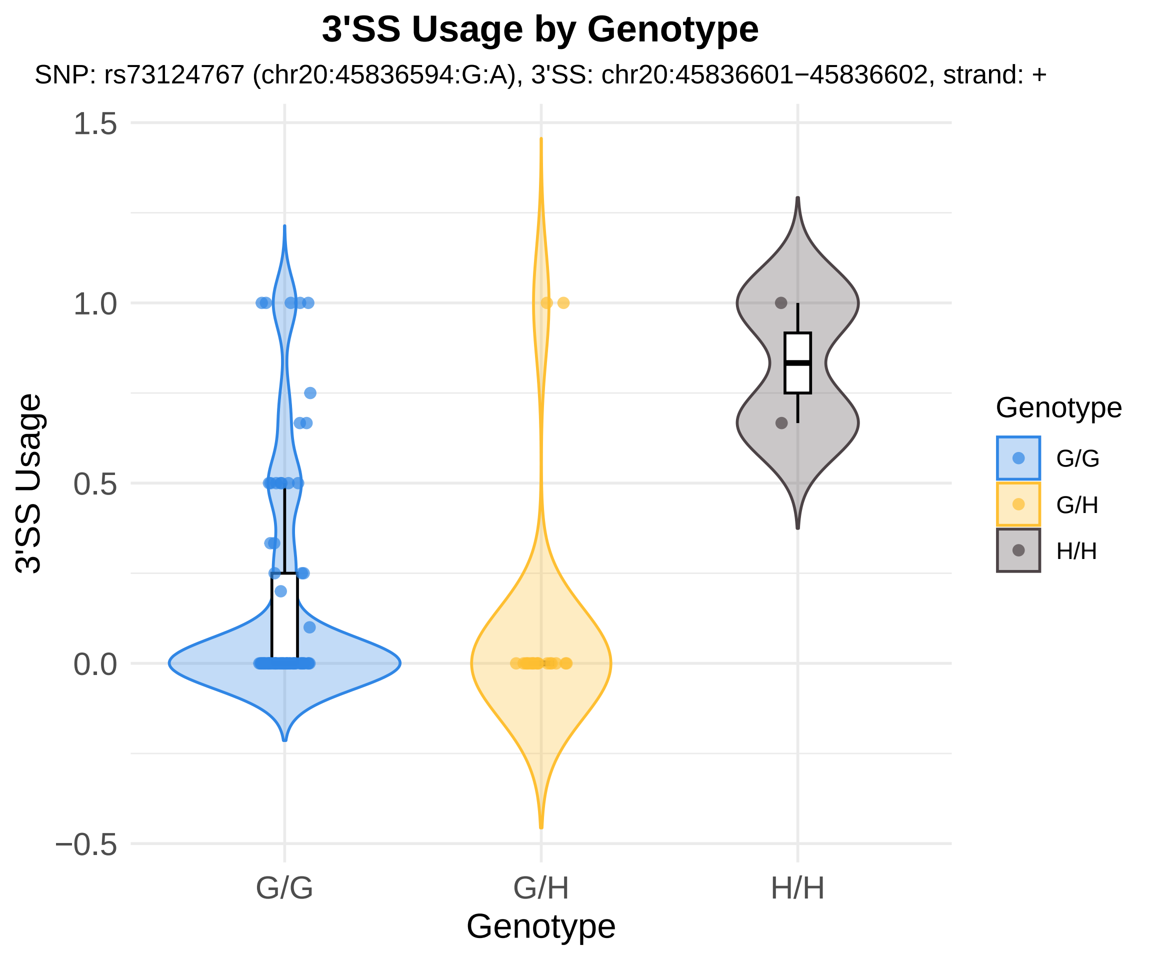

3'SS Usage by Genotype
SNP: rs73124767 (chr20:45836594:G:A), 3'SS: chr20:45836601−45836602, strand: +
1.5
1.0
0.5
0.0
−0.5
3'SS Usage
G/G
G/H
H/H
Genotype
Genotype
G/G
G/H
H/H

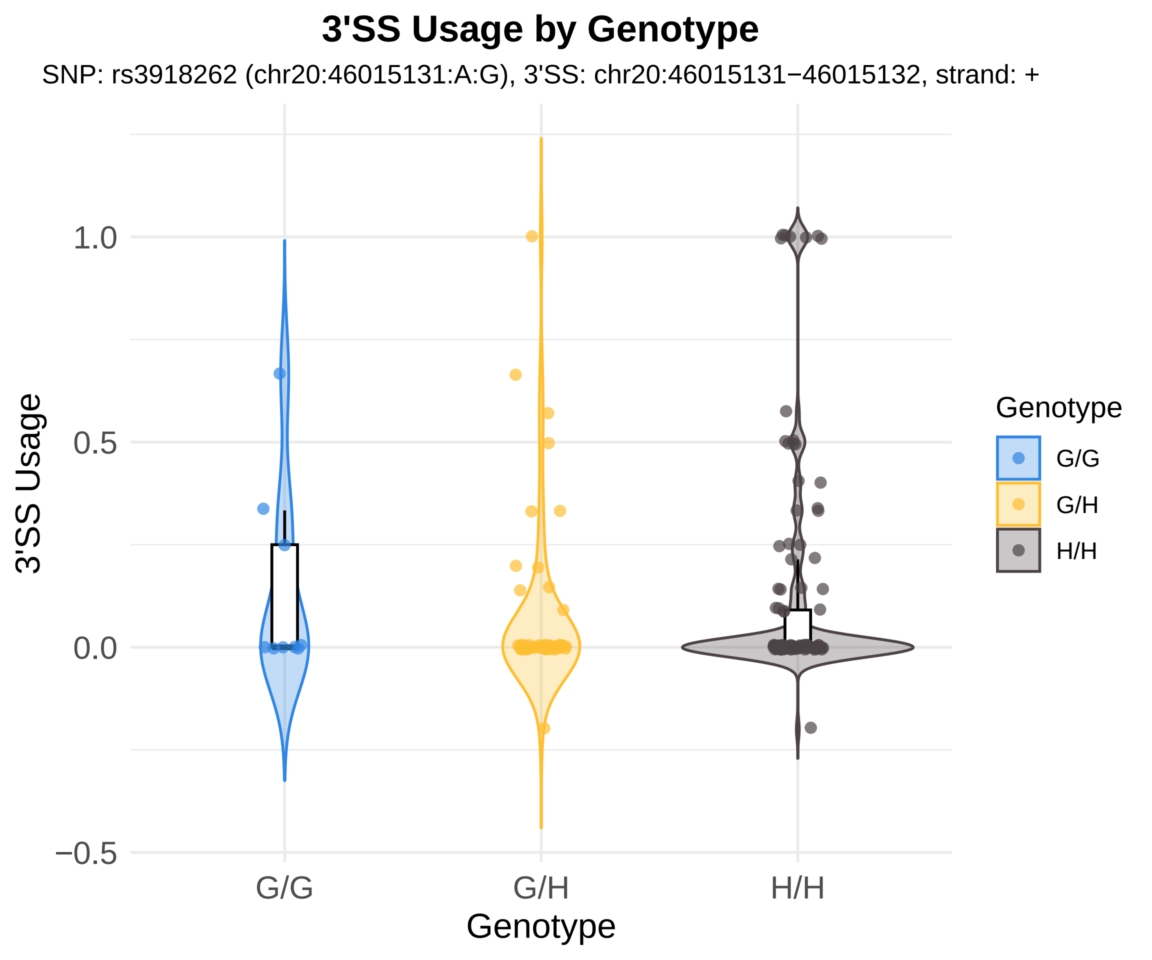
3'SS Usage by Genotype
SNP: rs3918262 (chr20:46015131:A:G), 3'SS: chr20:46015131−46015132, strand: +
3'SS Usage
1.0
0.5
0.0
−0.5
G/G
G/H
H/H
Genotype
Genotype
G/G
G/H
H/H

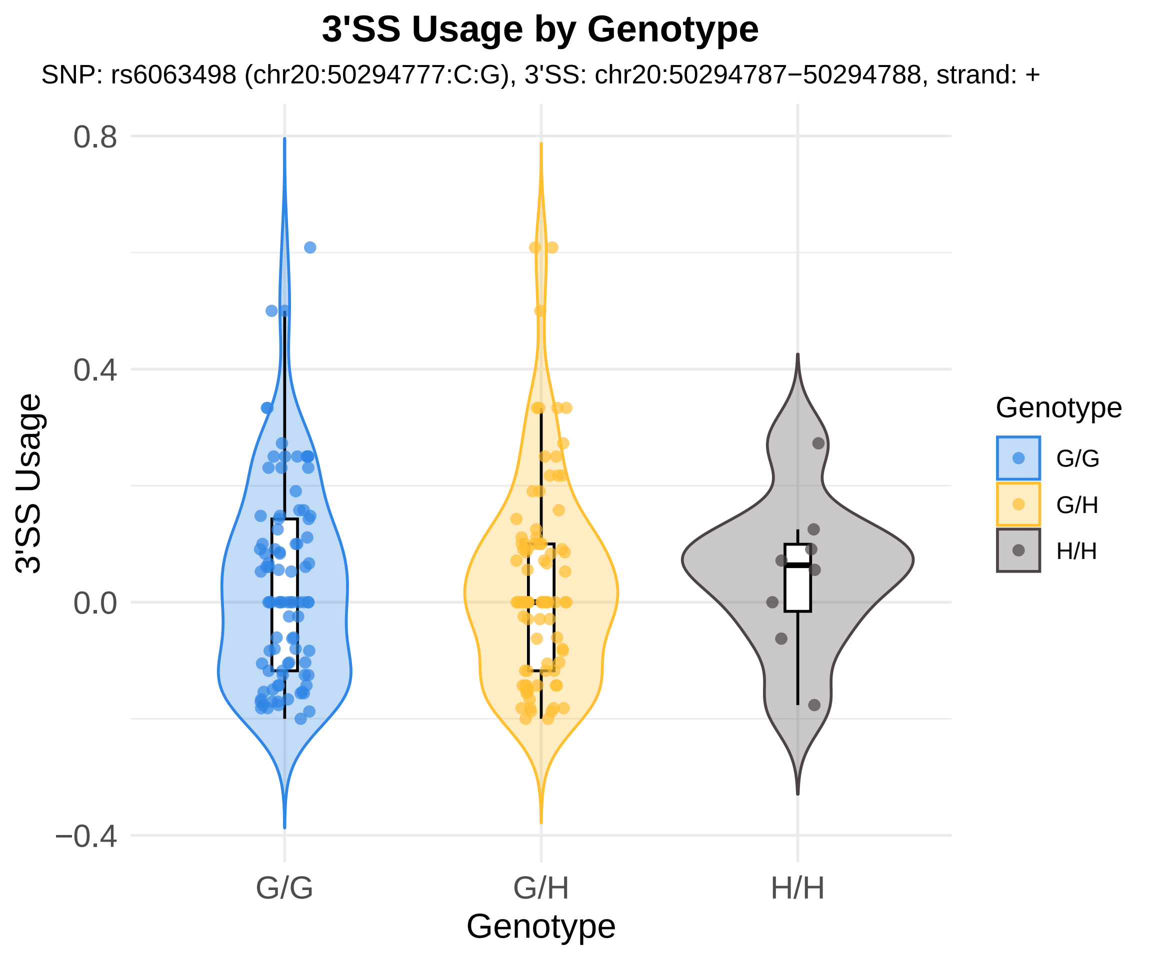
3'SS Usage by Genotype
SNP: rs6063498 (chr20:50294777:C:G), 3'SS: chr20:50294787−50294788, strand: +
0.8
0.4
0.0
−0.4
3'SS Usage
G/G
G/H
H/H
Genotype
Genotype
G/G
G/H
H/H

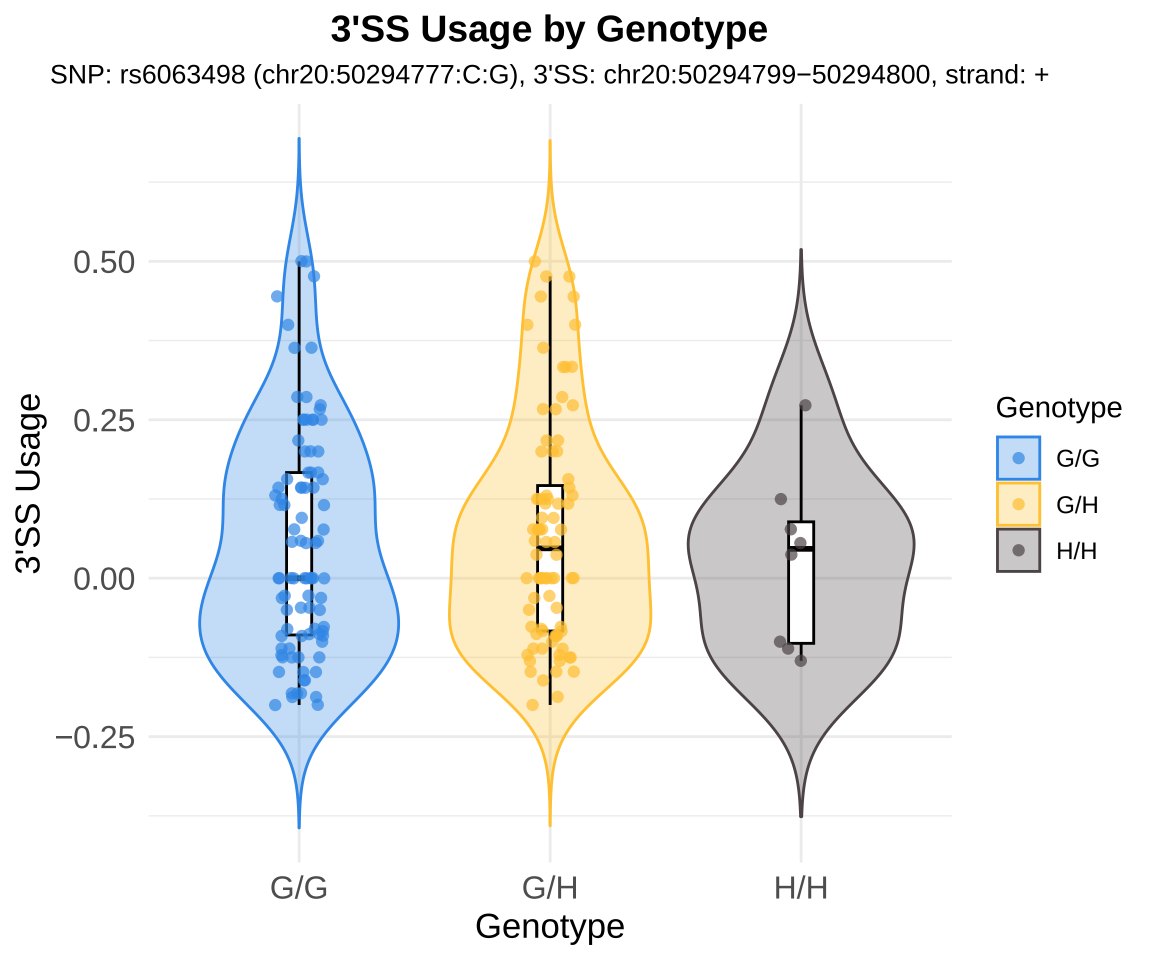

3'SS Usage by Genotype
SNP: rs6063498 (chr20:50294777:C:G), 3'SS: chr20:50294799−50294800, strand: +
3'SS Usage
0.50
0.25
0.00
−0.25
G/G
G/H
H/H
Genotype
Genotype
G/G
G/H
H/H

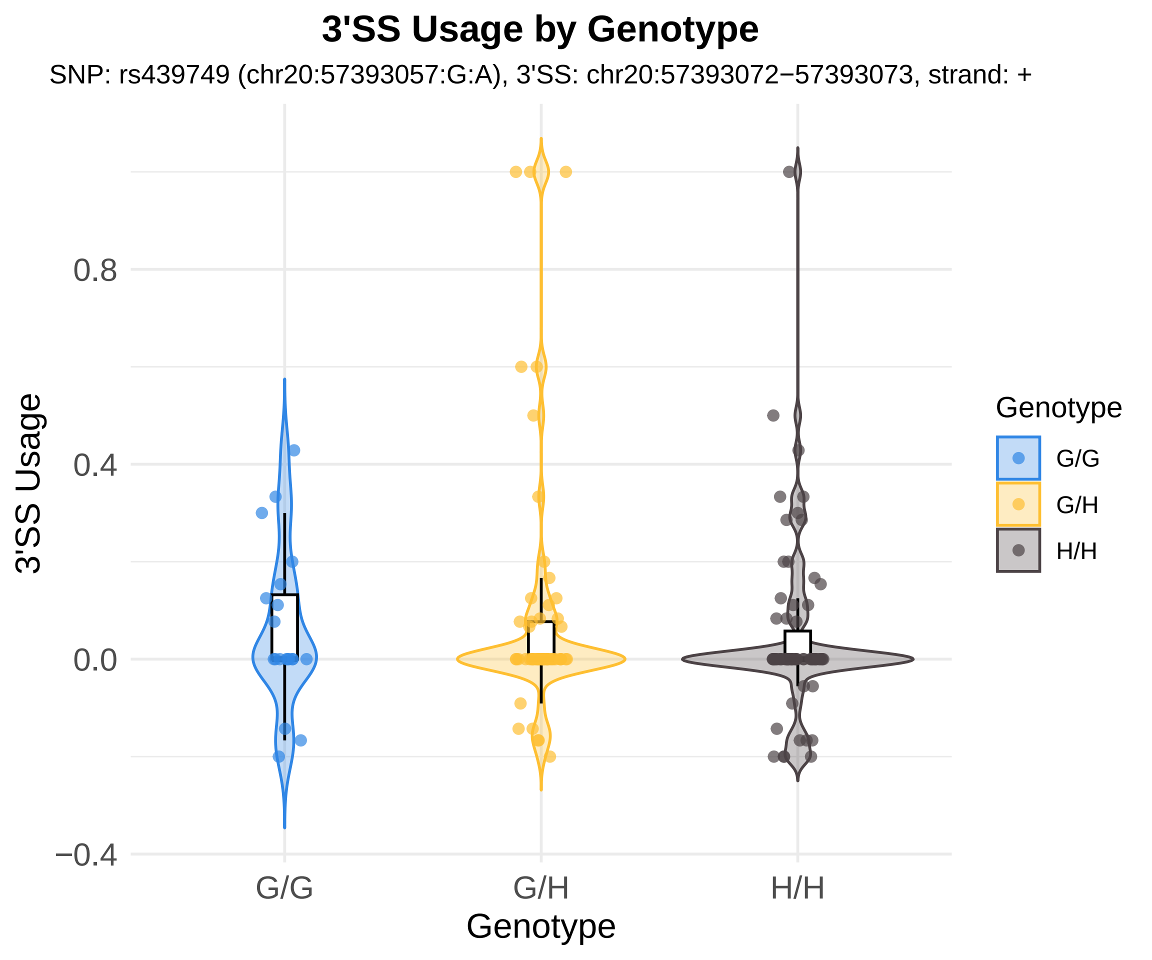

3'SS Usage by Genotype
SNP: rs439749 (chr20:57393057:G:A), 3'SS: chr20:57393072−57393073, strand: +
3'SS Usage
0.8
0.4
0.0
−0.4
G/G
G/H
H/H
Genotype
Genotype
G/G
G/H
H/H

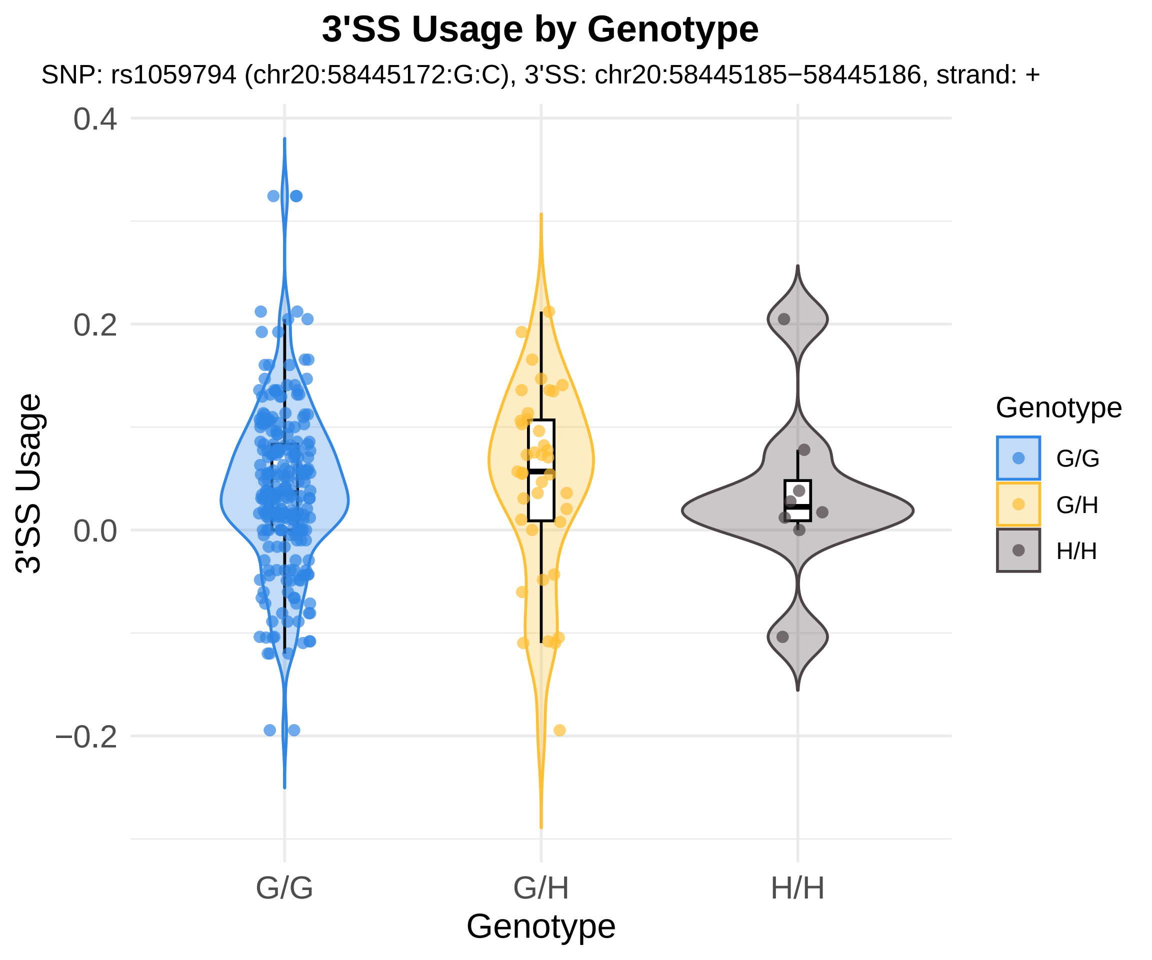
3'SS Usage by Genotype
SNP: rs1059794 (chr20:58445172:G:C), 3'SS: chr20:58445185−58445186, strand: +
0.4
0.2
0.0
−0.2
3'SS Usage
G/G
G/H
H/H
Genotype
Genotype
G/G
G/H
H/H

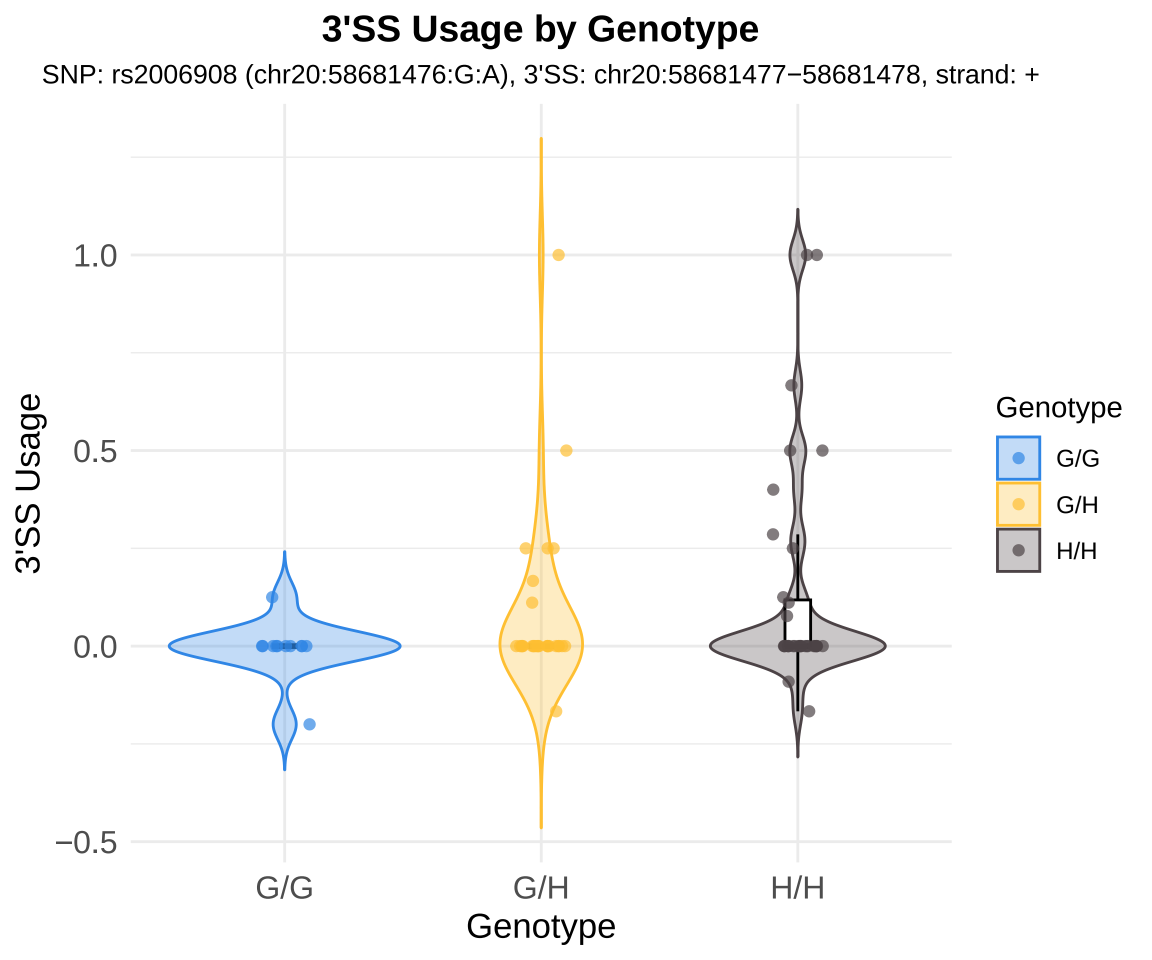

3'SS Usage by Genotype
SNP: rs2006908 (chr20:58681476:G:A), 3'SS: chr20:58681477−58681478, strand: +
3'SS Usage
1.0
0.5
0.0
−0.5
G/G
G/H
H/H
Genotype
Genotype
G/G
G/H
H/H

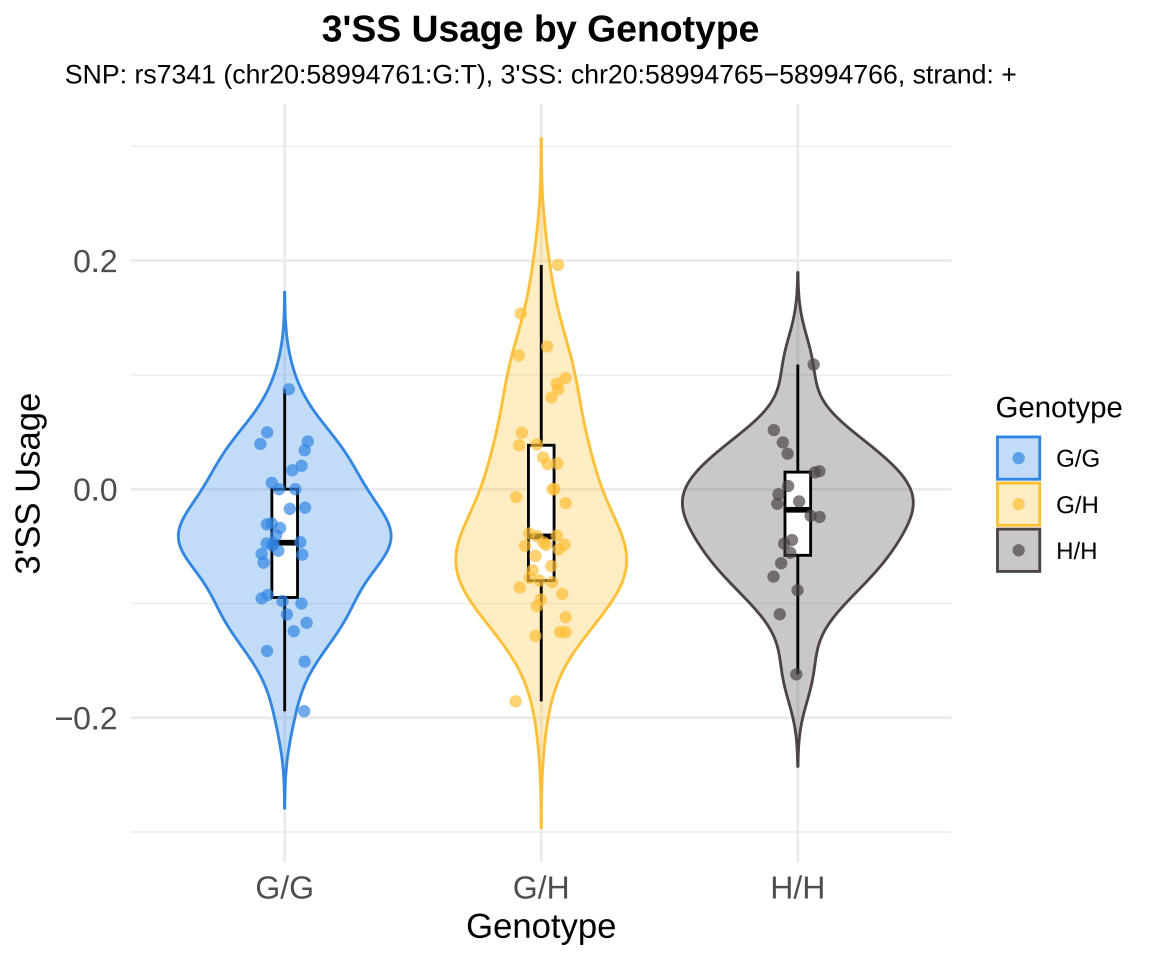
3'SS Usage by Genotype
SNP: rs7341 (chr20:58994761:G:T), 3'SS: chr20:58994765−58994766, strand: +
3'SS Usage
0.2
0.0
−0.2
G/G
G/H
H/H
Genotype
Genotype
G/G
G/H
H/H

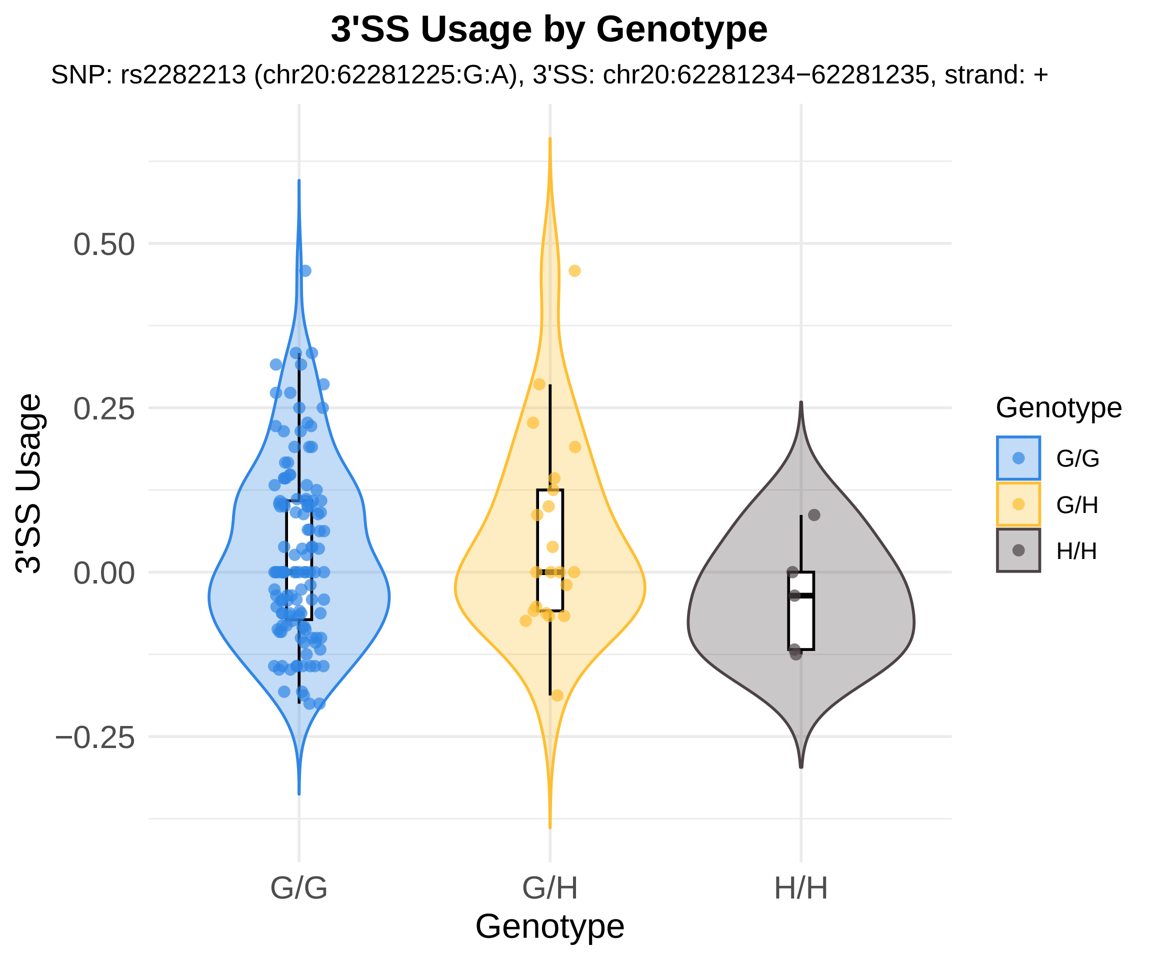

3'SS Usage by Genotype
SNP: rs2282213 (chr20:62281225:G:A), 3'SS: chr20:62281234−62281235, strand: +
3'SS Usage
0.50
0.25
0.00
−0.25
G/G
G/H
H/H
Genotype
Genotype
G/G
G/H
H/H

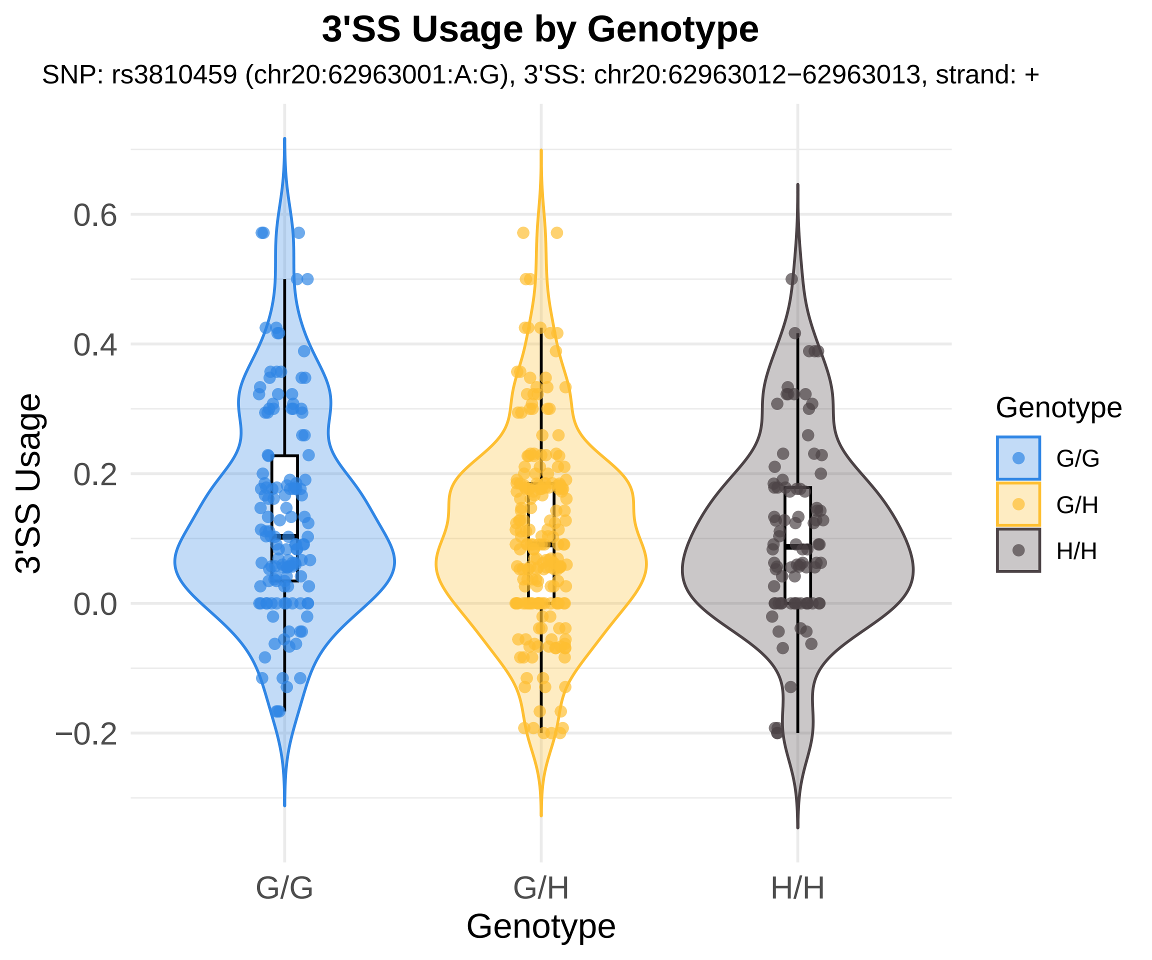

3'SS Usage by Genotype
SNP: rs3810459 (chr20:62963001:A:G), 3'SS: chr20:62963012−62963013, strand: +
3'SS Usage
0.6
0.4
0.2
0.0
−0.2
G/G
G/H
H/H
Genotype
Genotype
G/G
G/H
H/H

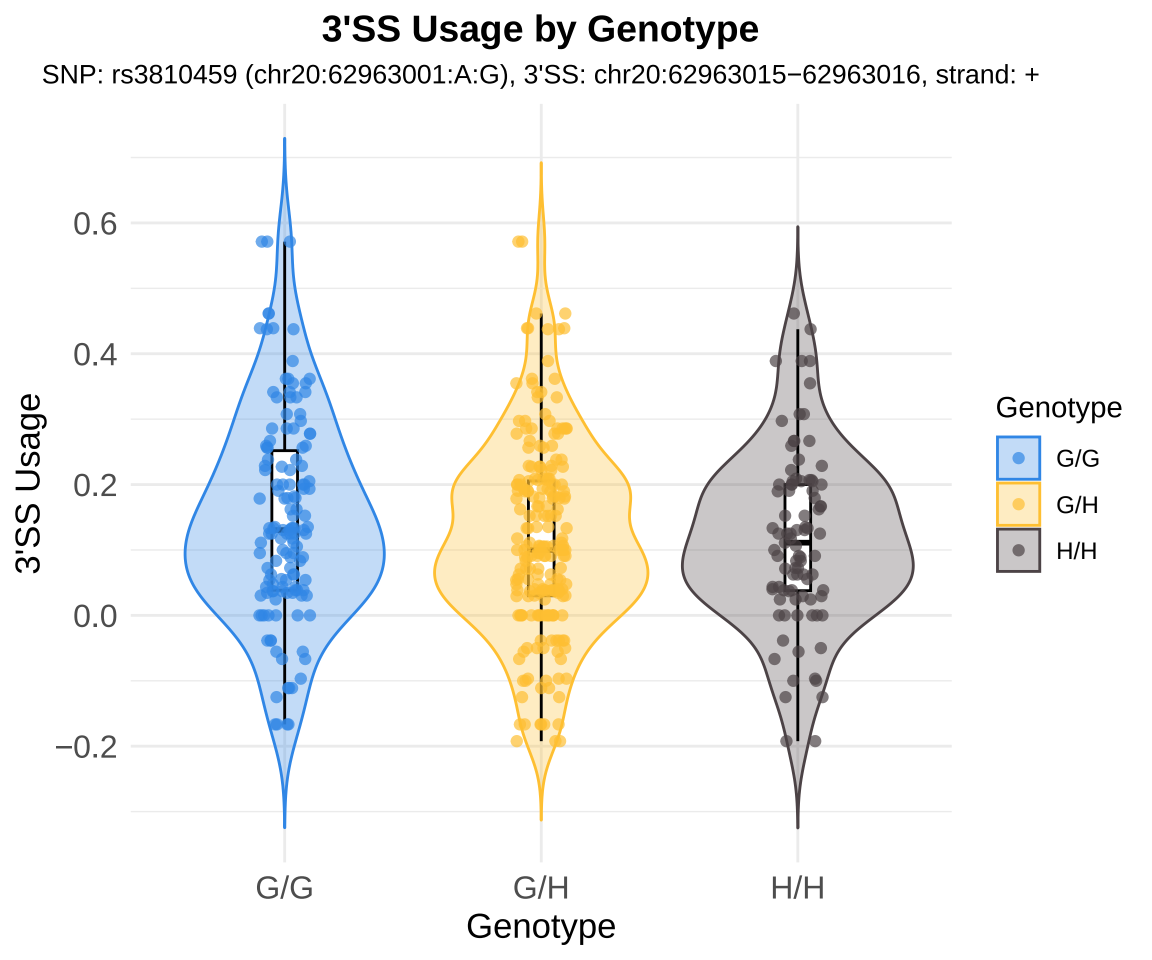
3'SS Usage by Genotype
SNP: rs3810459 (chr20:62963001:A:G), 3'SS: chr20:62963015−62963016, strand: +
3'SS Usage
0.6
0.4
0.2
0.0
−0.2
G/G
G/H
H/H
Genotype
Genotype
G/G
G/H
H/H

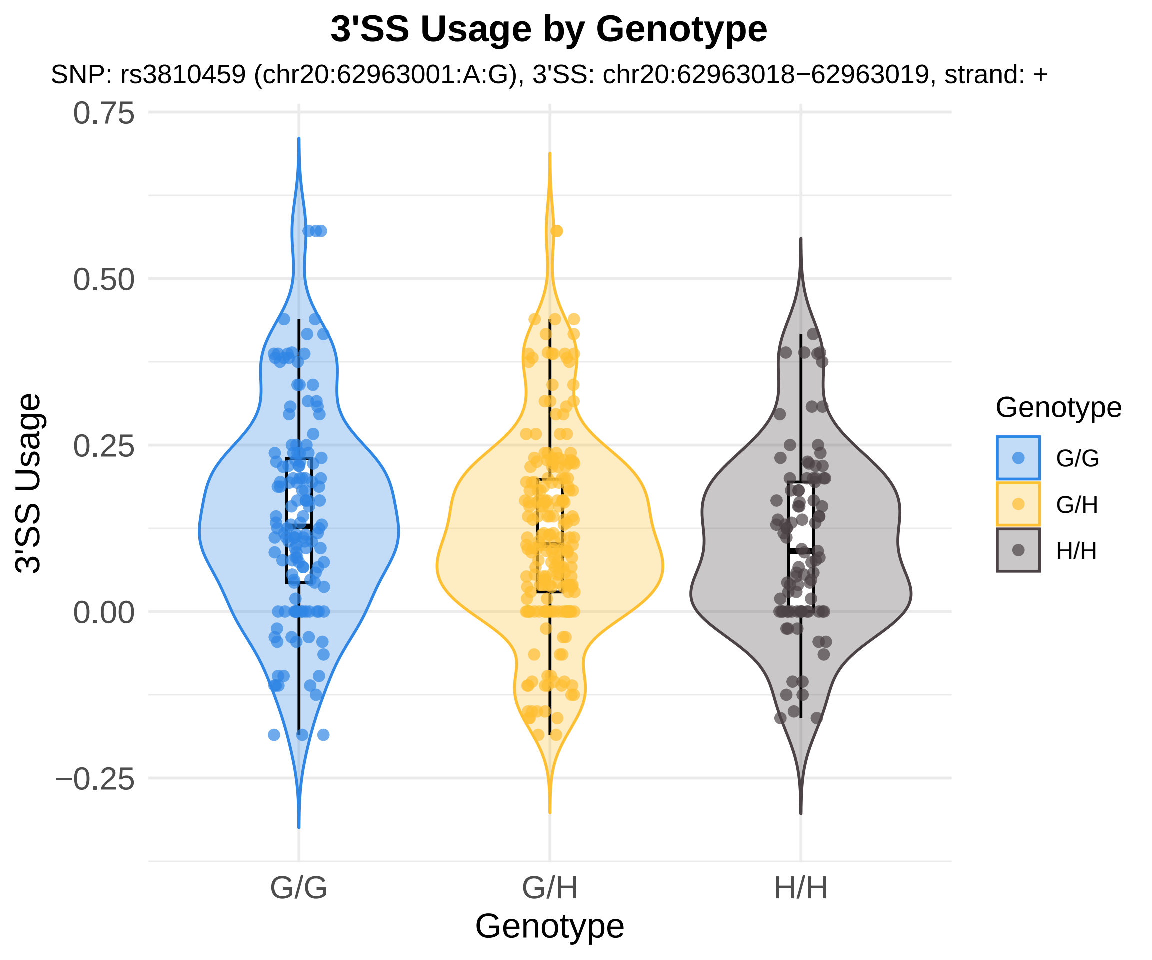
3'SS Usage by Genotype
SNP: rs3810459 (chr20:62963001:A:G), 3'SS: chr20:62963018−62963019, strand: +
0.75
0.50
0.25
0.00
−0.25
3'SS Usage
G/G
G/H
H/H
Genotype
Genotype
G/G
G/H
H/H

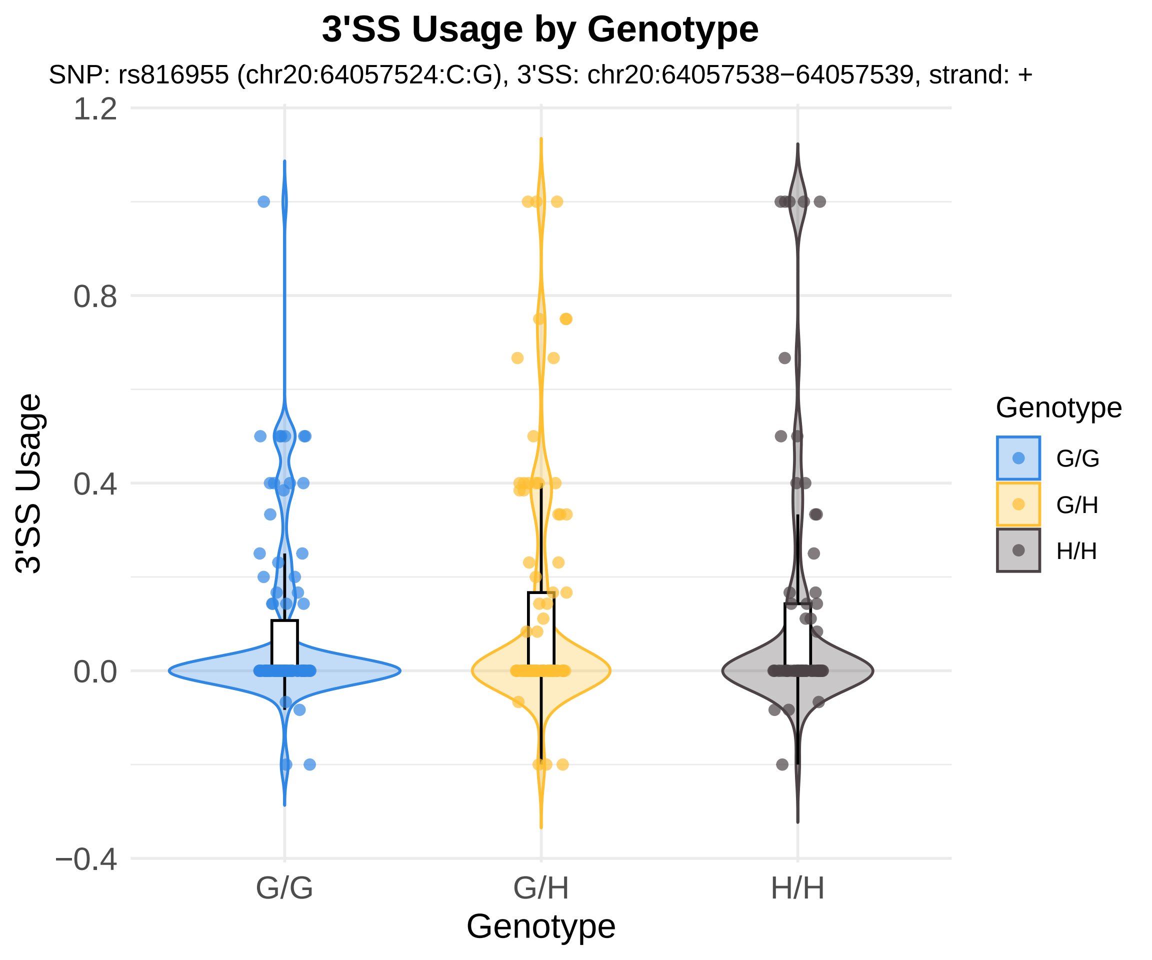
3'SS Usage by Genotype
SNP: rs816955 (chr20:64057524:C:G), 3'SS: chr20:64057538−64057539, strand: +
1.2
0.8
0.4
0.0
−0.4
3'SS Usage
G/G
G/H
H/H
Genotype
Genotype
G/G
G/H
H/H

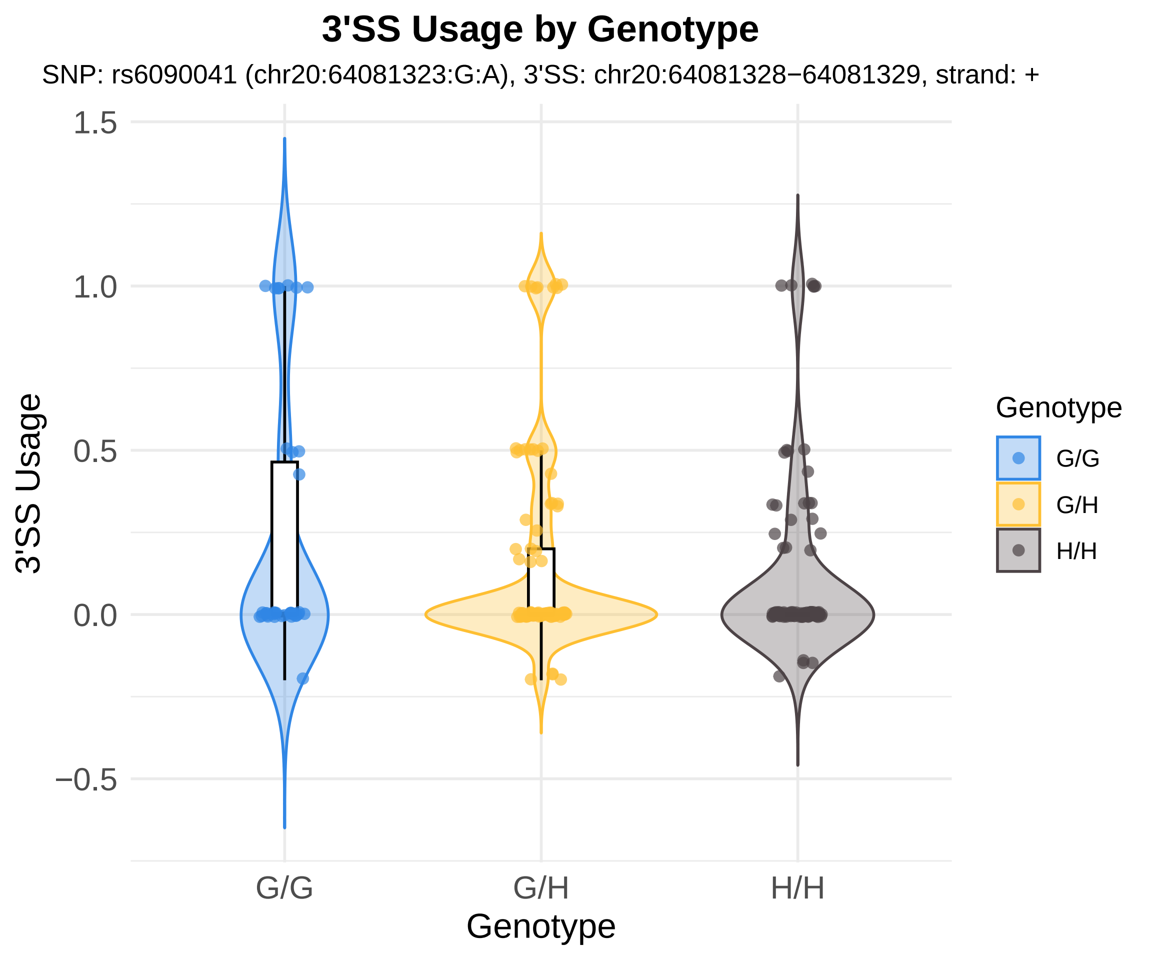
3'SS Usage by Genotype
SNP: rs6090041 (chr20:64081323:G:A), 3'SS: chr20:64081328−64081329, strand: +
1.5
1.0
0.5
0.0
−0.5
3'SS Usage
G/G
G/H
H/H
Genotype
Genotype
G/G
G/H
H/H

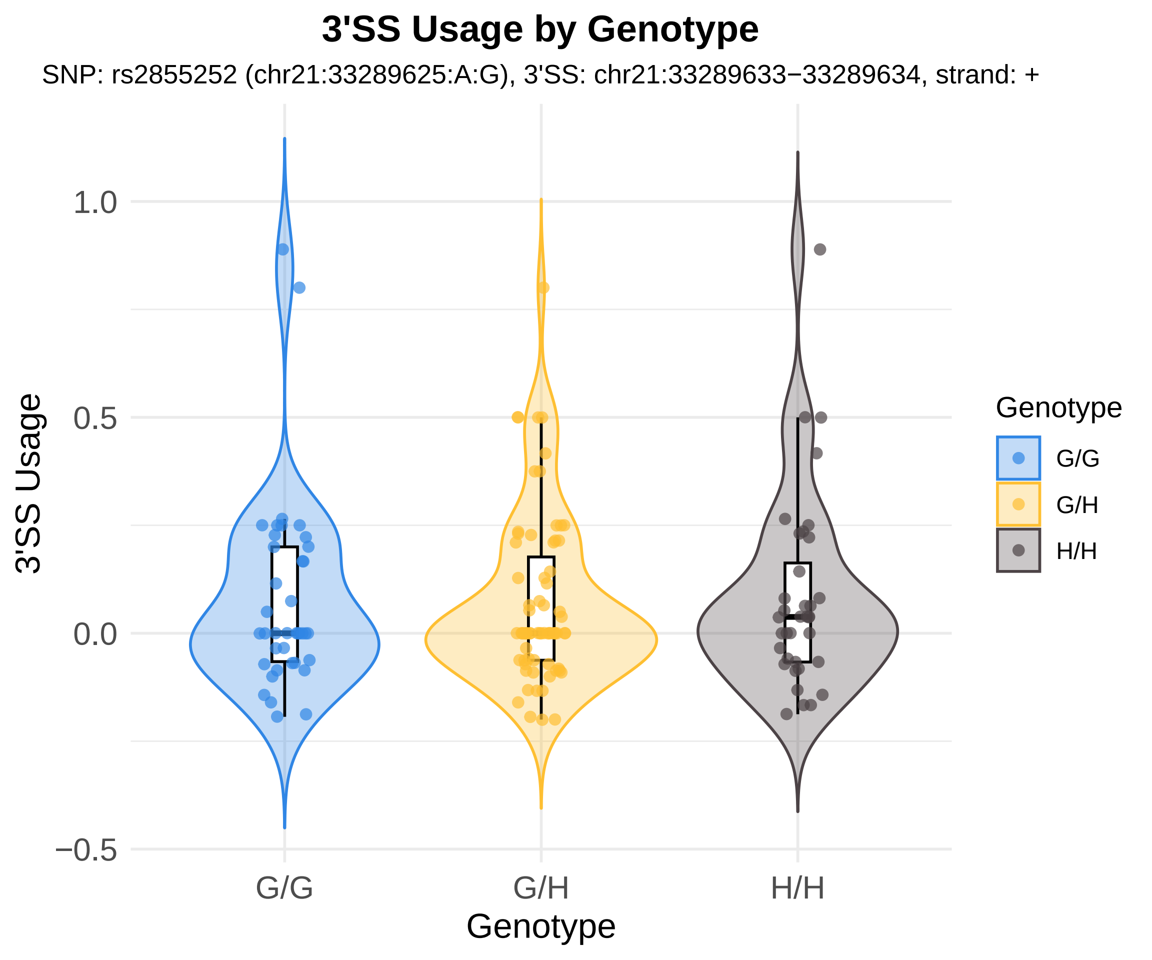
3'SS Usage by Genotype
SNP: rs2855252 (chr21:33289625:A:G), 3'SS: chr21:33289633−33289634, strand: +
3'SS Usage
1.0
0.5
0.0
−0.5
G/G
G/H
H/H
Genotype
Genotype
G/G
G/H
H/H

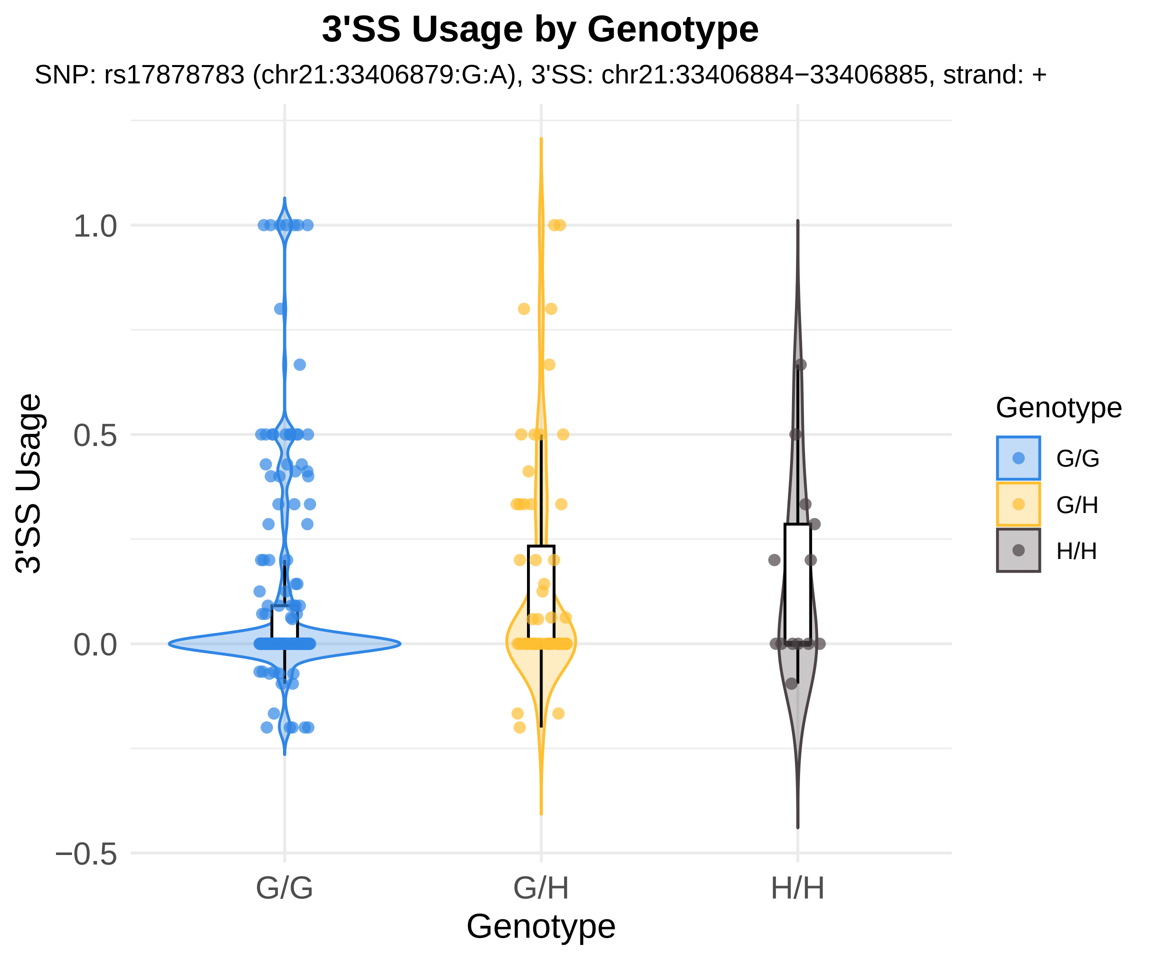

3'SS Usage by Genotype
SNP: rs17878783 (chr21:33406879:G:A), 3'SS: chr21:33406884−33406885, strand: +
3'SS Usage
1.0
0.5
0.0
−0.5
G/G
G/H
H/H
Genotype
Genotype
G/G
G/H
H/H

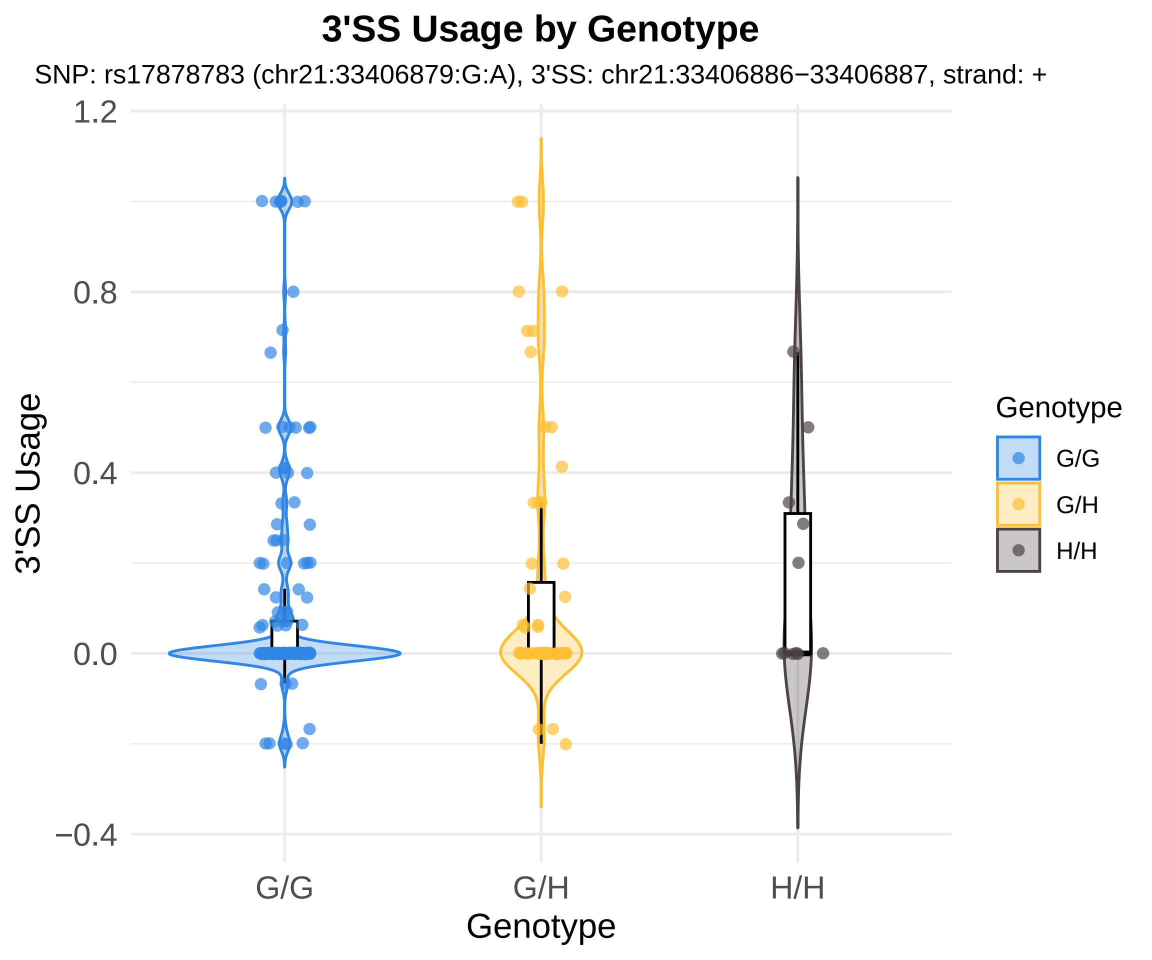
3'SS Usage by Genotype
SNP: rs17878783 (chr21:33406879:G:A), 3'SS: chr21:33406886−33406887, strand: +
1.2
0.8
0.4
0.0
−0.4
3'SS Usage
G/G
G/H
H/H
Genotype
Genotype
G/G
G/H
H/H

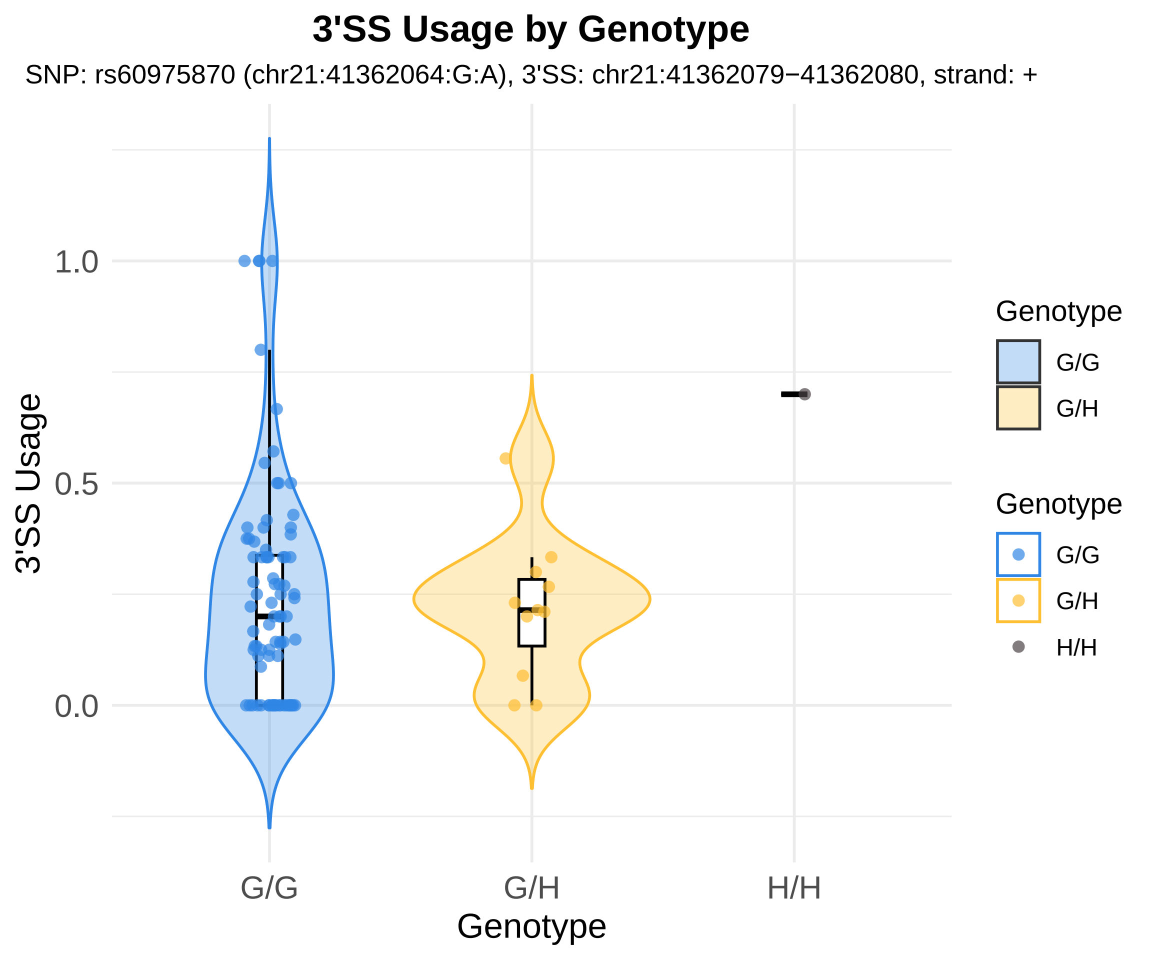
3'SS Usage by Genotype
SNP: rs60975870 (chr21:41362064:G:A), 3'SS: chr21:41362079−41362080, strand: +
3'SS Usage
1.0
0.5
0.0
G/G
G/H
H/H
Genotype
Genotype
G/G
G/H
Genotype
G/G
G/H
H/H

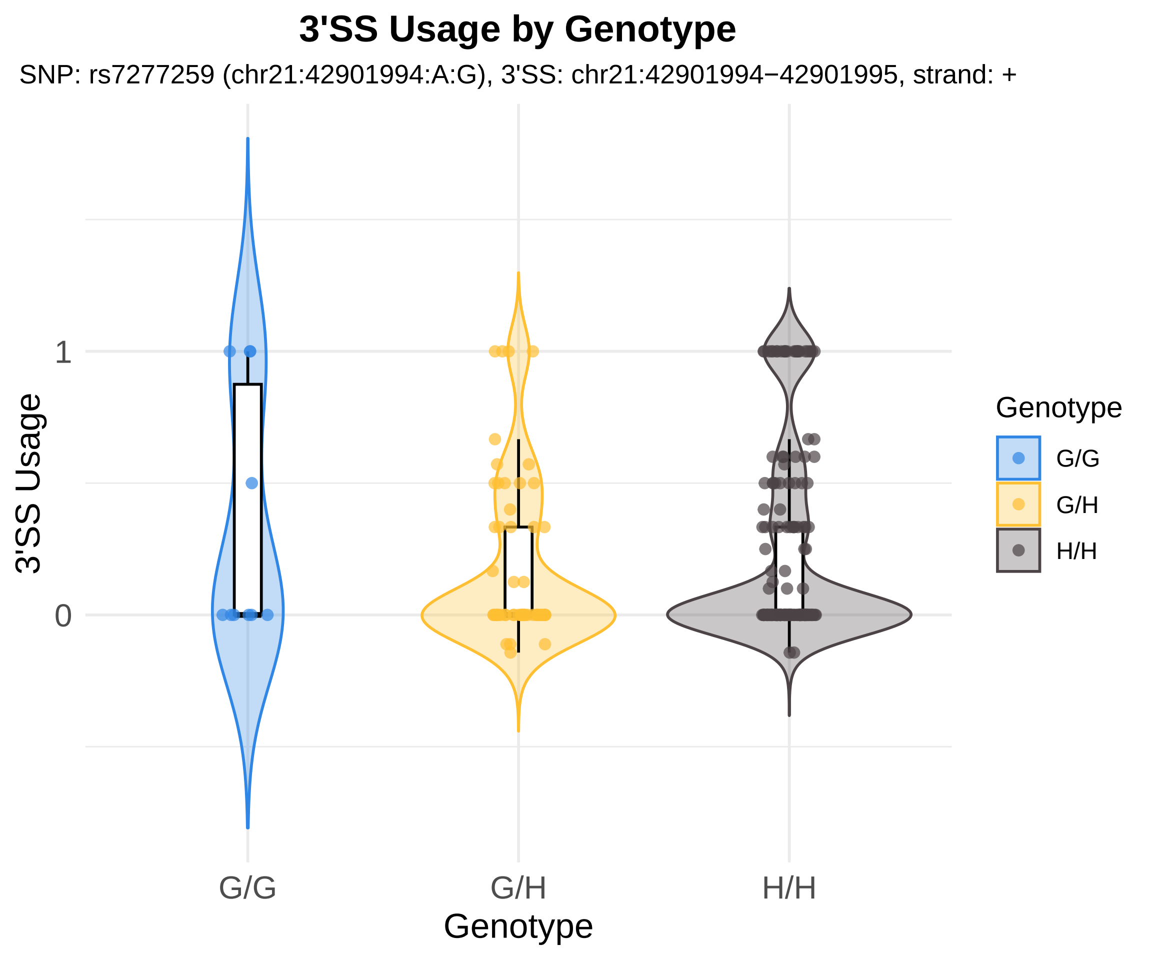

3'SS Usage by Genotype
SNP: rs7277259 (chr21:42901994:A:G), 3'SS: chr21:42901994−42901995, strand: +
3'SS Usage
1
0
G/G
G/H
H/H
Genotype
Genotype
G/G
G/H
H/H

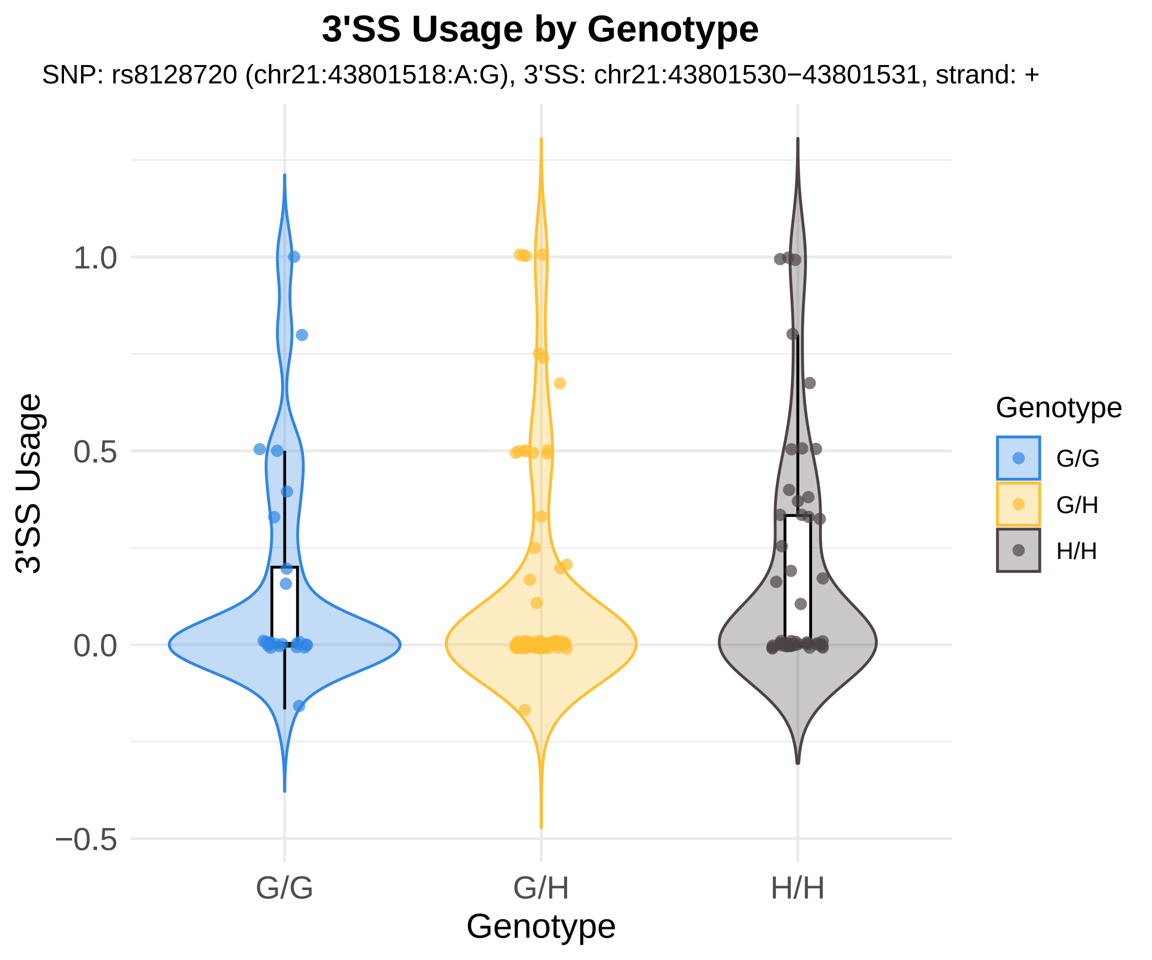
3'SS Usage by Genotype
SNP: rs8128720 (chr21:43801518:A:G), 3'SS: chr21:43801530−43801531, strand: +
3'SS Usage
1.0
0.5
0.0
−0.5
G/G
G/H
H/H
Genotype
Genotype
G/G
G/H
H/H

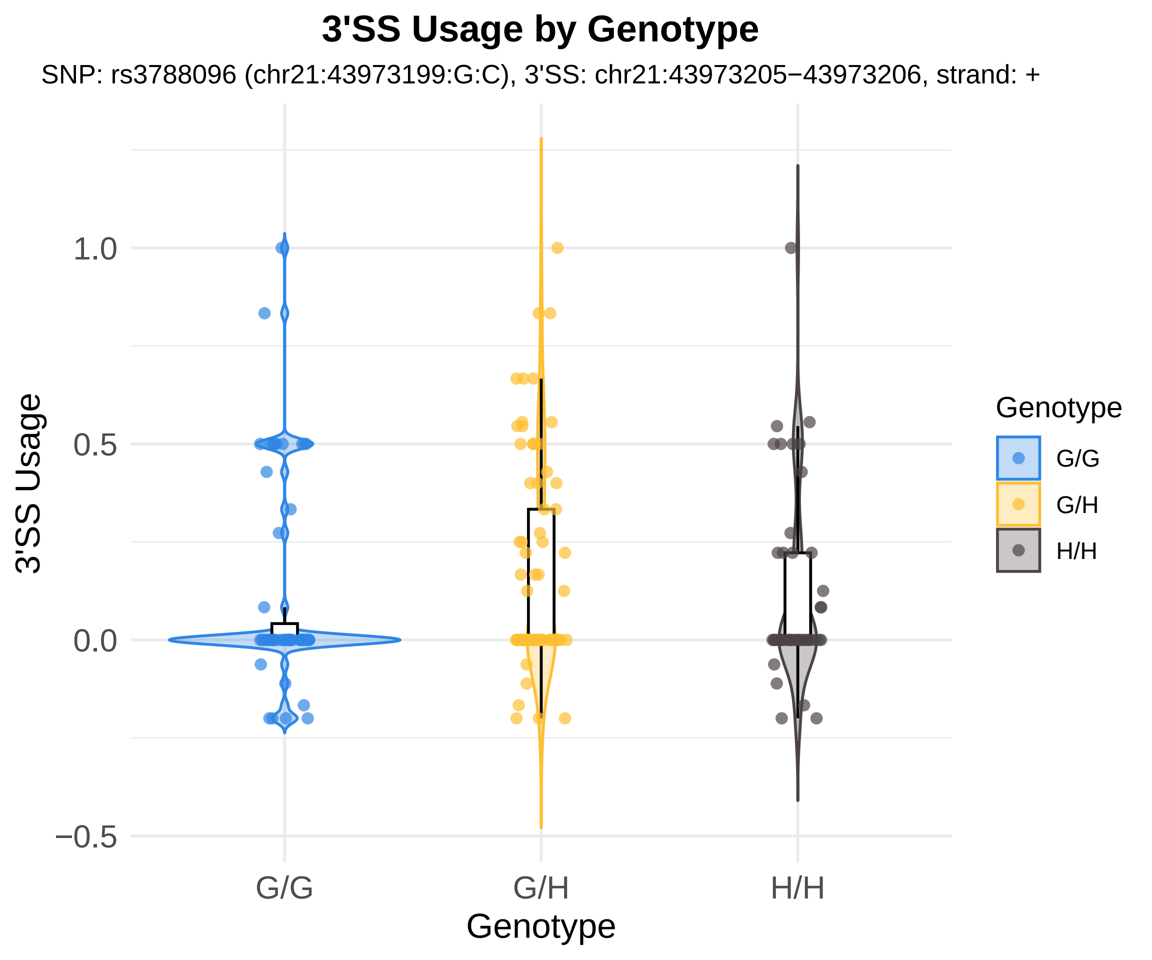

3'SS Usage by Genotype
SNP: rs3788096 (chr21:43973199:G:C), 3'SS: chr21:43973205−43973206, strand: +
3'SS Usage
1.0
0.5
0.0
−0.5
G/G
G/H
H/H
Genotype
Genotype
G/G
G/H
H/H

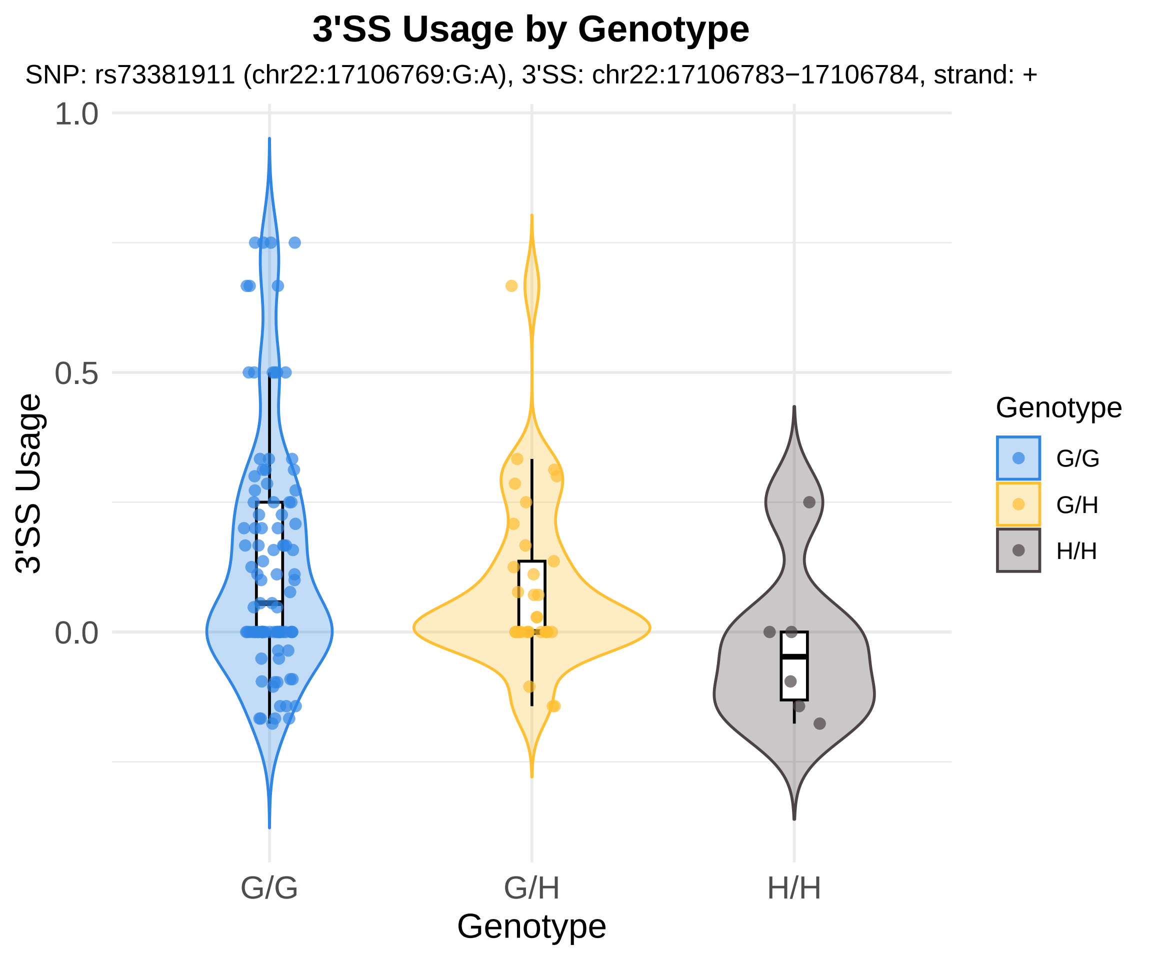
3'SS Usage by Genotype
SNP: rs73381911 (chr22:17106769:G:A), 3'SS: chr22:17106783−17106784, strand: +
3'SS Usage
1.0
0.5
0.0
G/G
G/H
H/H
Genotype
Genotype
G/G
G/H
H/H

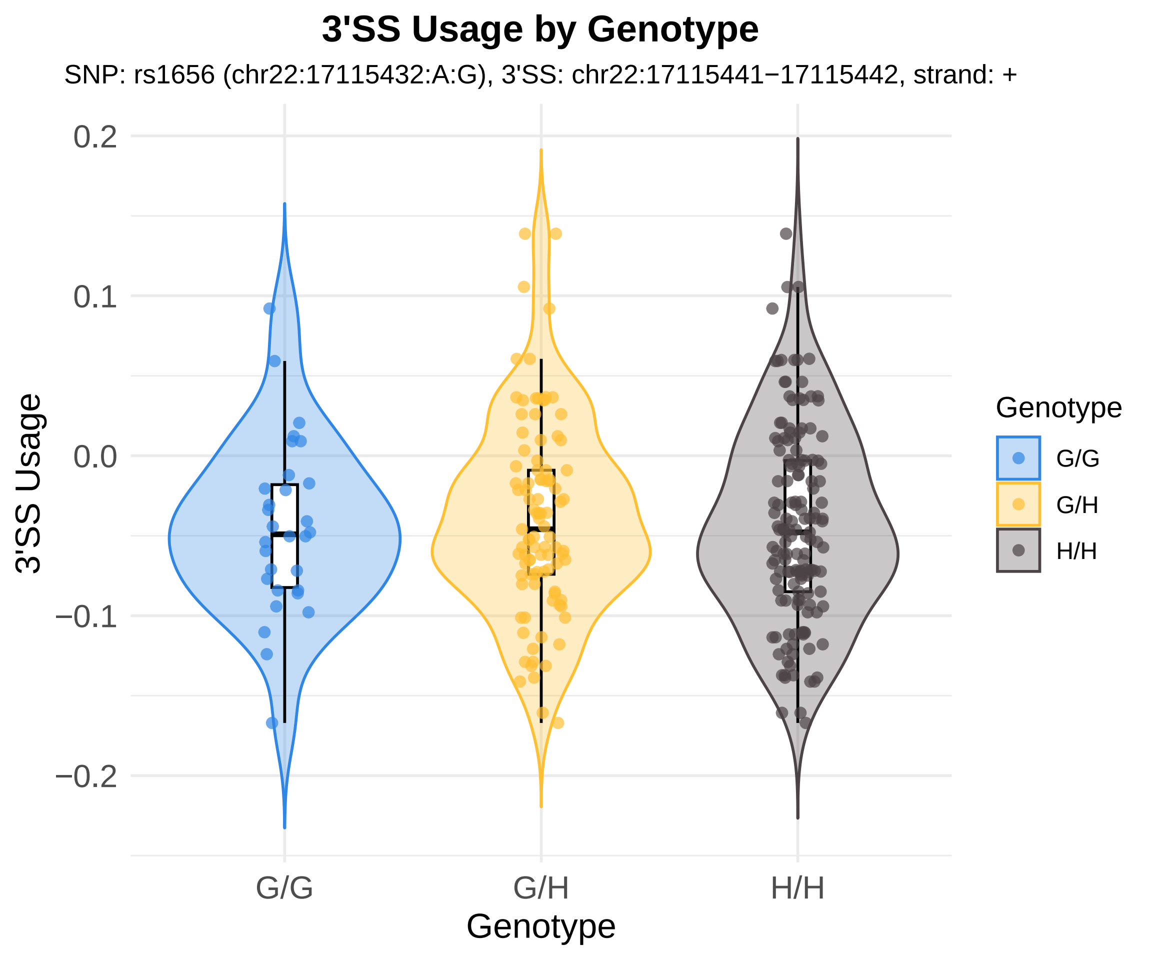
3'SS Usage by Genotype
SNP: rs1656 (chr22:17115432:A:G), 3'SS: chr22:17115441−17115442, strand: +
0.2
0.1
0.0
−0.1
−0.2
3'SS Usage
G/G
G/H
H/H
Genotype
Genotype
G/G
G/H
H/H

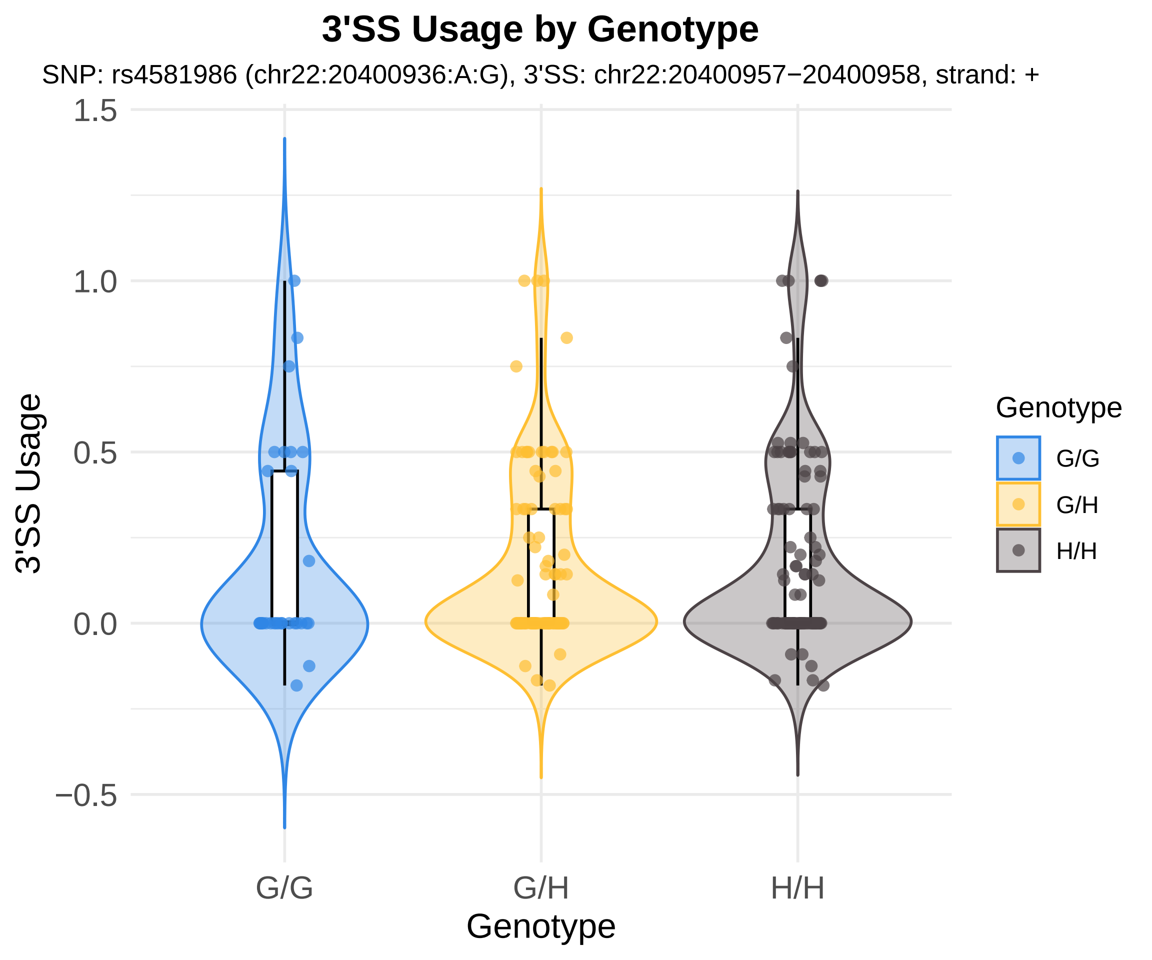

3'SS Usage by Genotype
SNP: rs4581986 (chr22:20400936:A:G), 3'SS: chr22:20400957−20400958, strand: +
1.5
1.0
0.5
0.0
−0.5
3'SS Usage
G/G
G/H
H/H
Genotype
Genotype
G/G
G/H
H/H

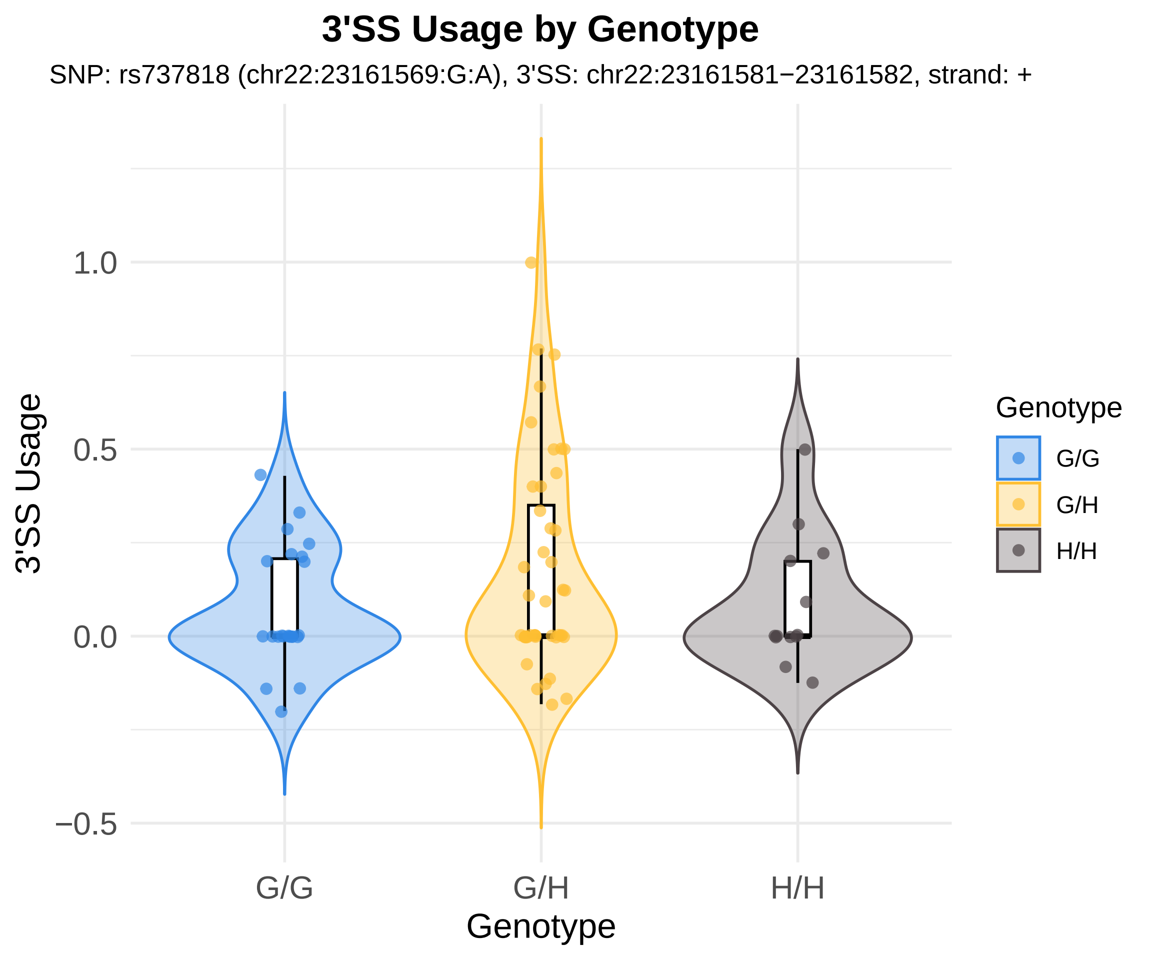

3'SS Usage by Genotype
SNP: rs737818 (chr22:23161569:G:A), 3'SS: chr22:23161581−23161582, strand: +
3'SS Usage
1.0
0.5
0.0
−0.5
G/G
G/H
H/H
Genotype
Genotype
G/G
G/H
H/H

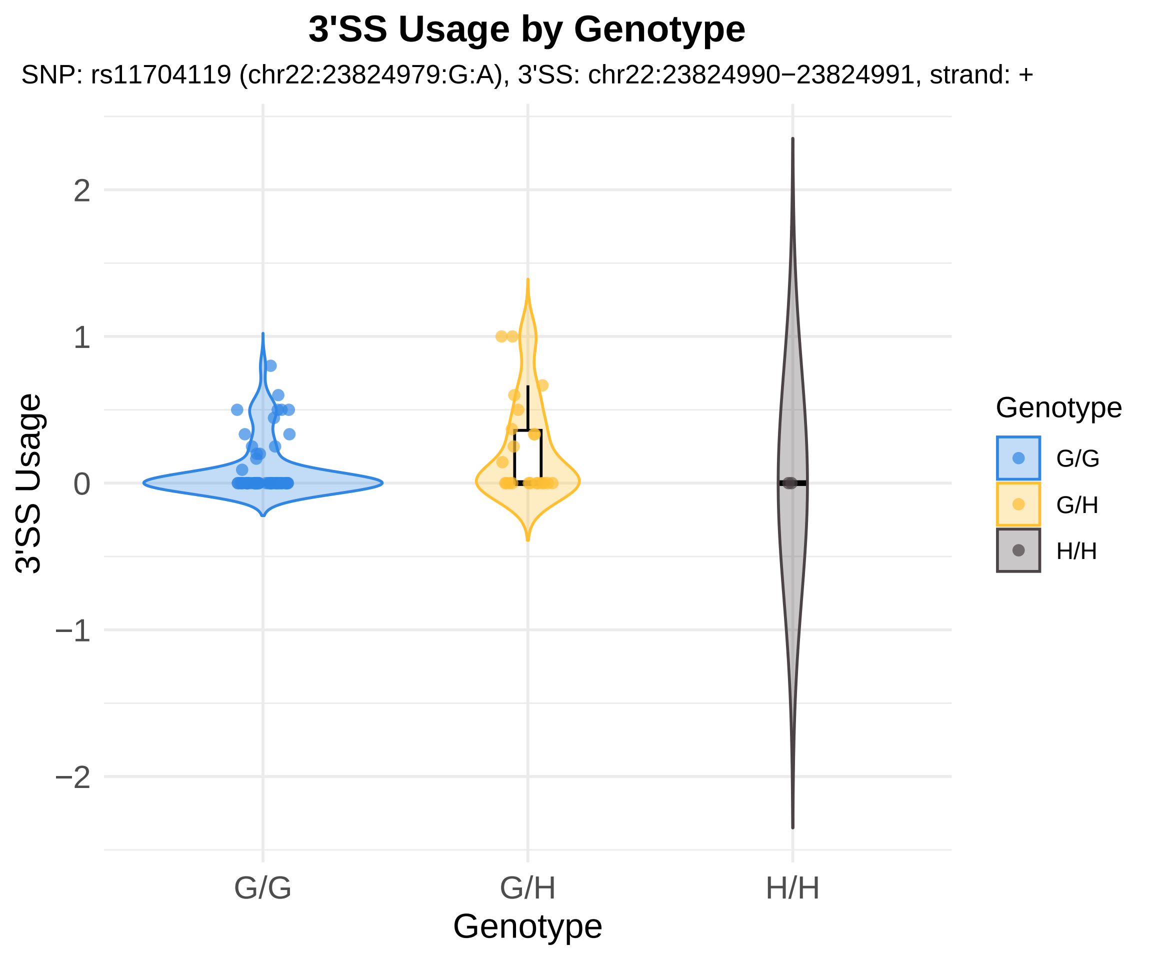
3'SS Usage by Genotype
SNP: rs11704119 (chr22:23824979:G:A), 3'SS: chr22:23824990−23824991, strand: +
3'SS Usage
2
1
0
−1
−2
G/G
G/H
H/H
Genotype
Genotype
G/G
G/H
H/H

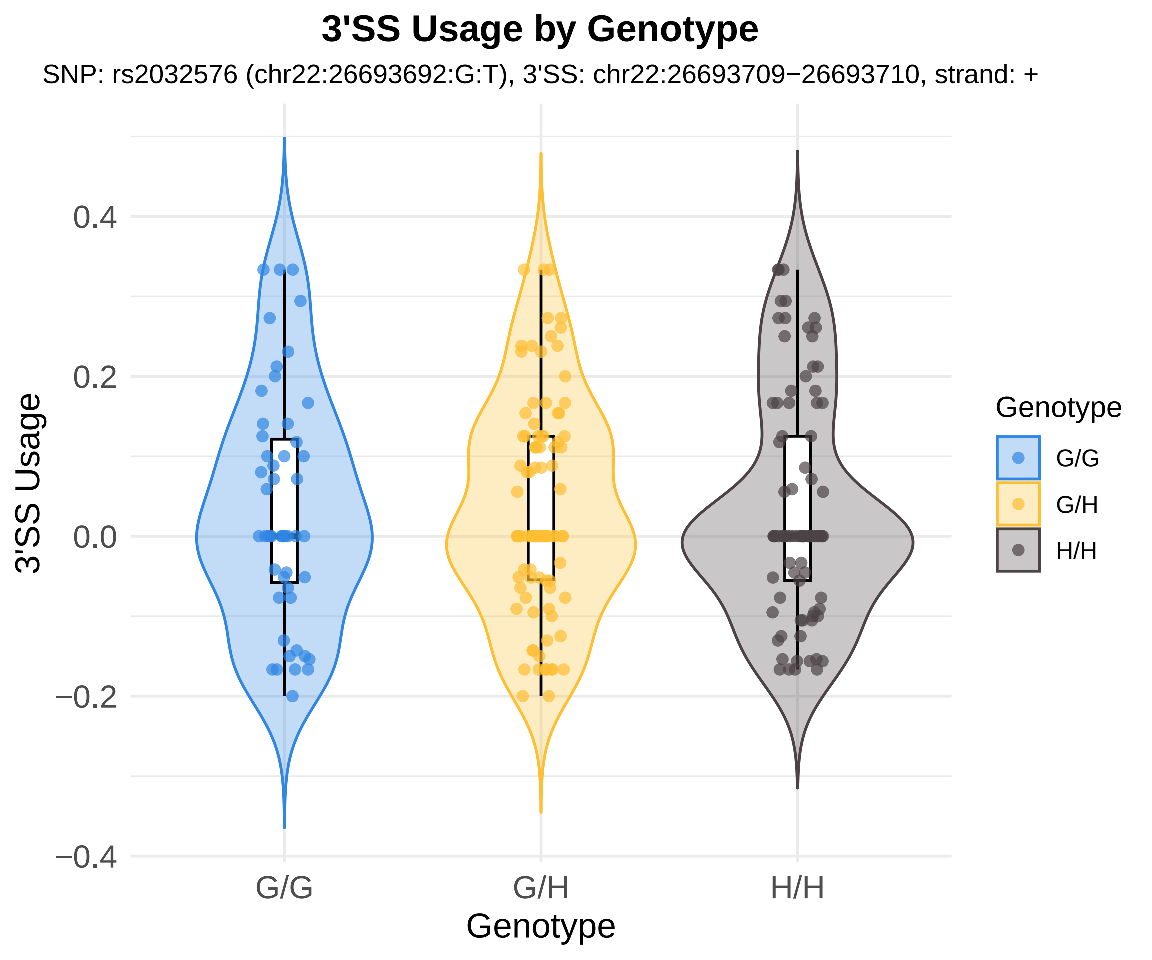
3'SS Usage by Genotype
SNP: rs2032576 (chr22:26693692:G:T), 3'SS: chr22:26693709−26693710, strand: +
3'SS Usage
0.4
0.2
0.0
−0.2
−0.4
G/G
G/H
H/H
Genotype
Genotype
G/G
G/H
H/H

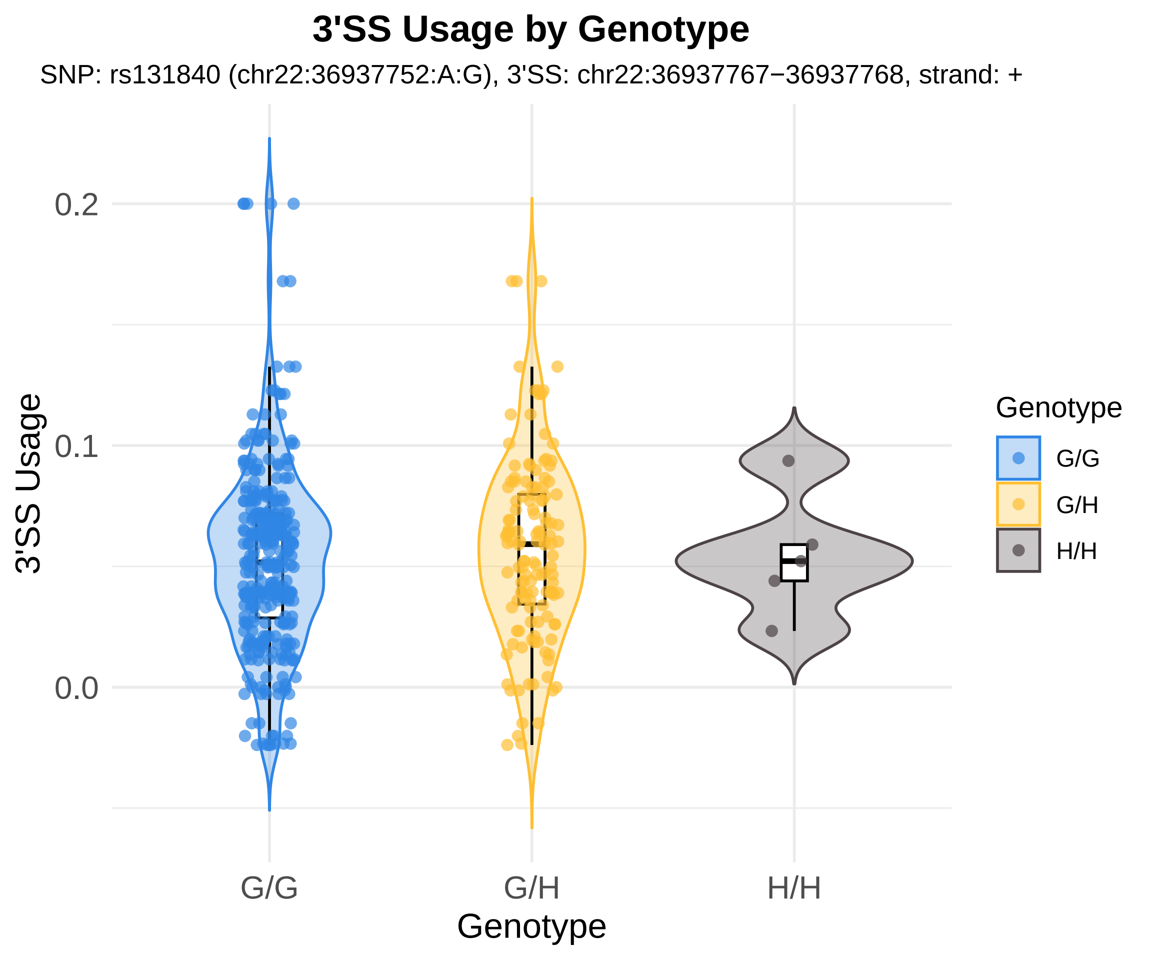

3'SS Usage by Genotype
SNP: rs131840 (chr22:36937752:A:G), 3'SS: chr22:36937767−36937768, strand: +
3'SS Usage
0.2
0.1
0.0
G/G
G/H
H/H
Genotype
Genotype
G/G
G/H
H/H

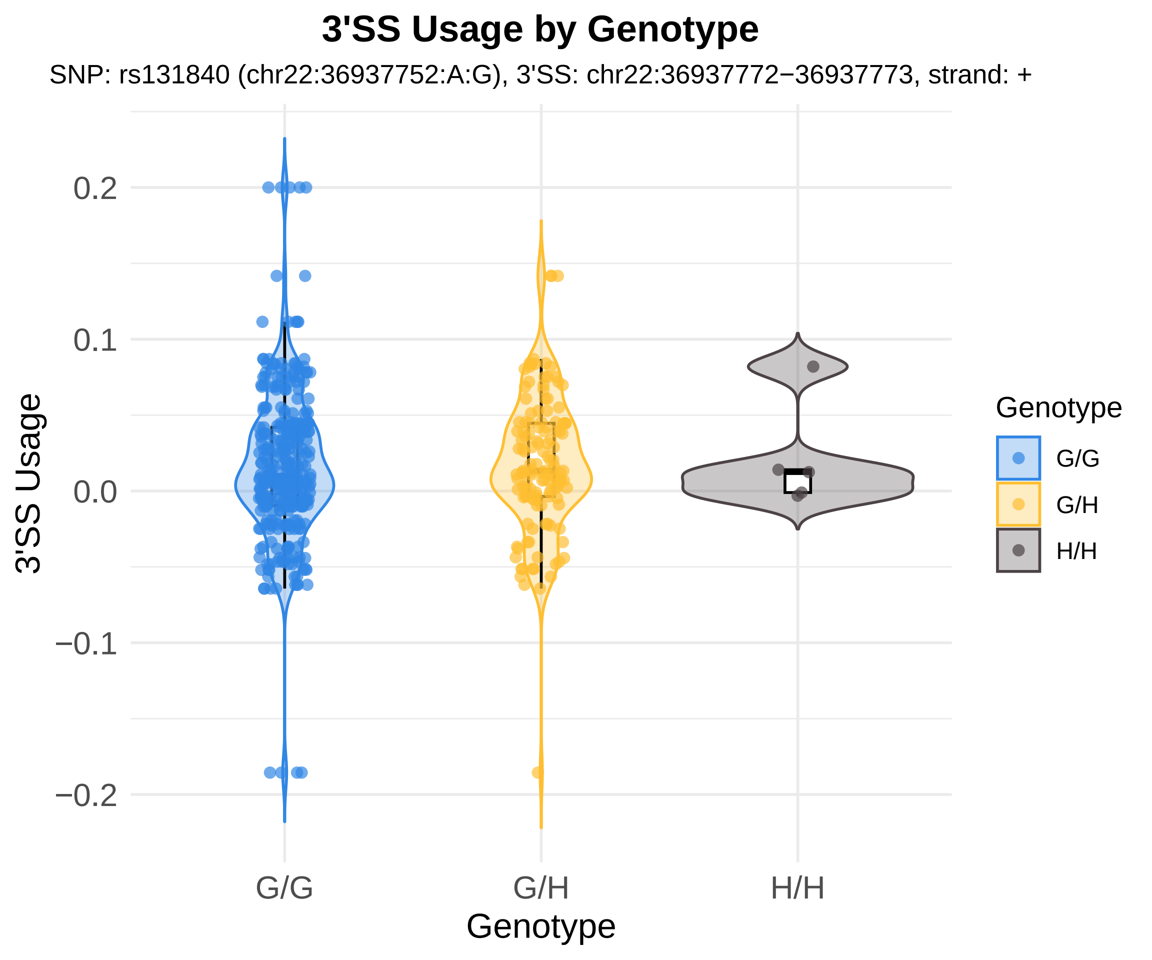

3'SS Usage by Genotype
SNP: rs131840 (chr22:36937752:A:G), 3'SS: chr22:36937772−36937773, strand: +
3'SS Usage
0.2
0.1
0.0
−0.1
−0.2
G/G
G/H
H/H
Genotype
Genotype
G/G
G/H
H/H

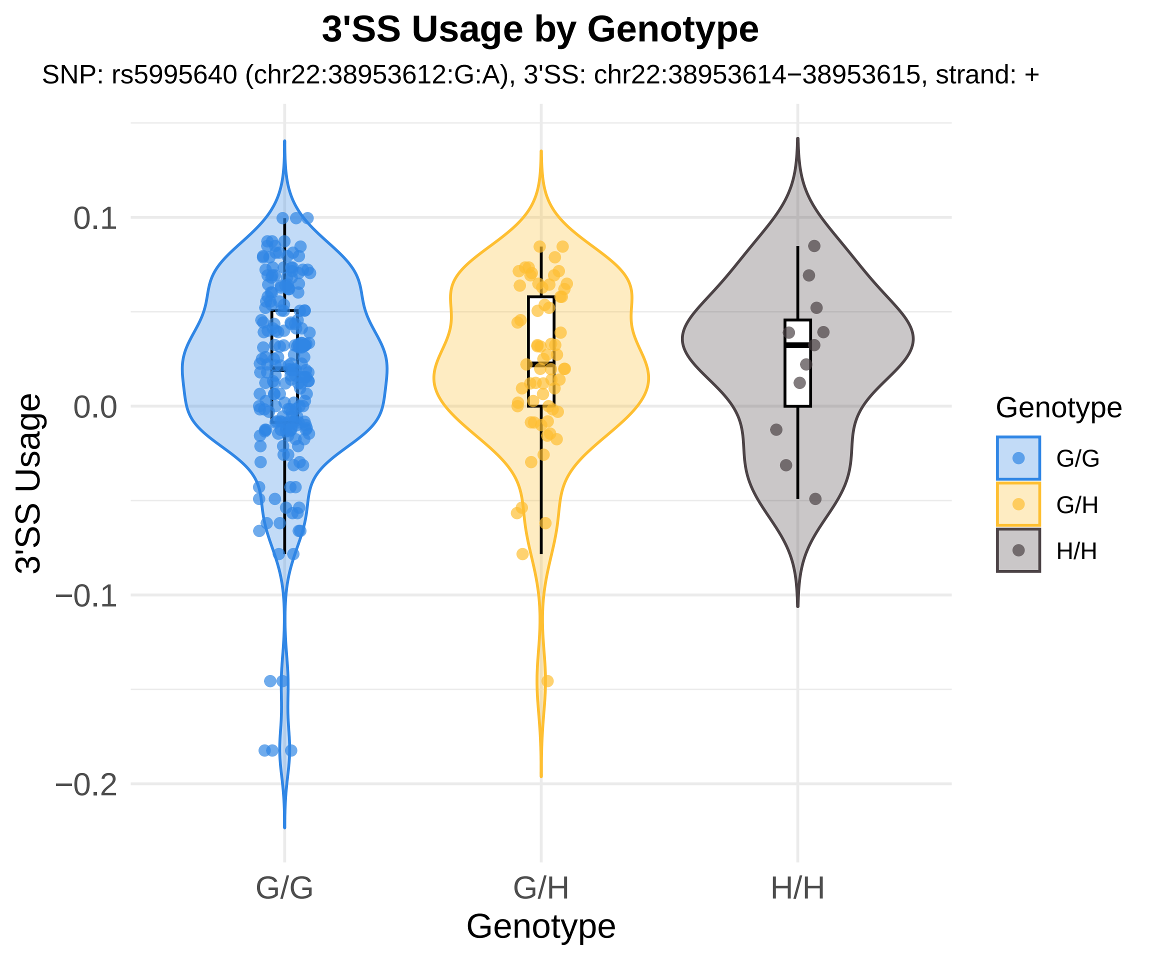

3'SS Usage by Genotype
SNP: rs5995640 (chr22:38953612:G:A), 3'SS: chr22:38953614−38953615, strand: +
3'SS Usage
0.1
0.0
−0.1
−0.2
G/G
G/H
H/H
Genotype
Genotype
G/G
G/H
H/H

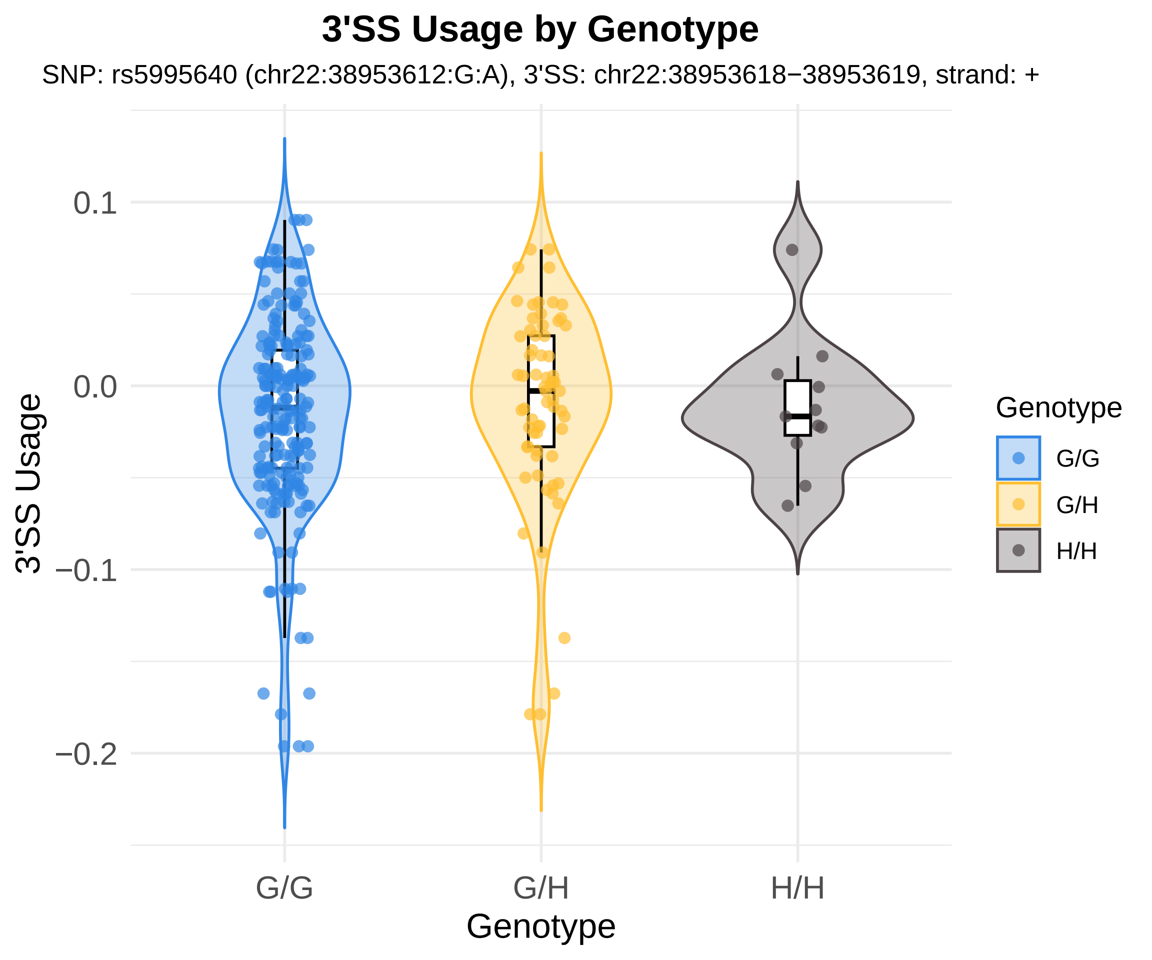
3'SS Usage by Genotype
SNP: rs5995640 (chr22:38953612:G:A), 3'SS: chr22:38953618−38953619, strand: +
3'SS Usage
0.1
0.0
−0.1
−0.2
G/G
G/H
H/H
Genotype
Genotype
G/G
G/H
H/H

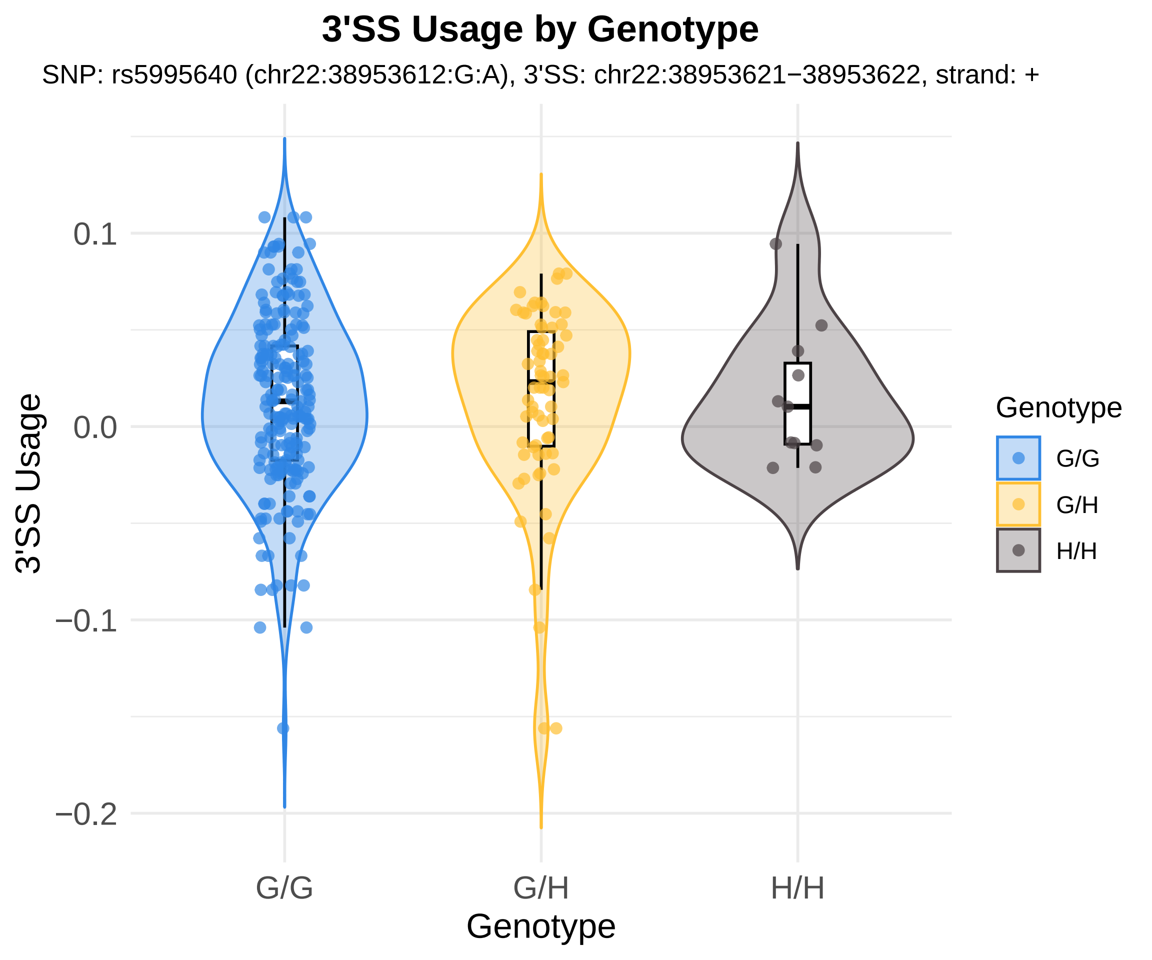
3'SS Usage by Genotype
SNP: rs5995640 (chr22:38953612:G:A), 3'SS: chr22:38953621−38953622, strand: +
3'SS Usage
0.1
0.0
−0.1
−0.2
G/G
G/H
H/H
Genotype
Genotype
G/G
G/H
H/H

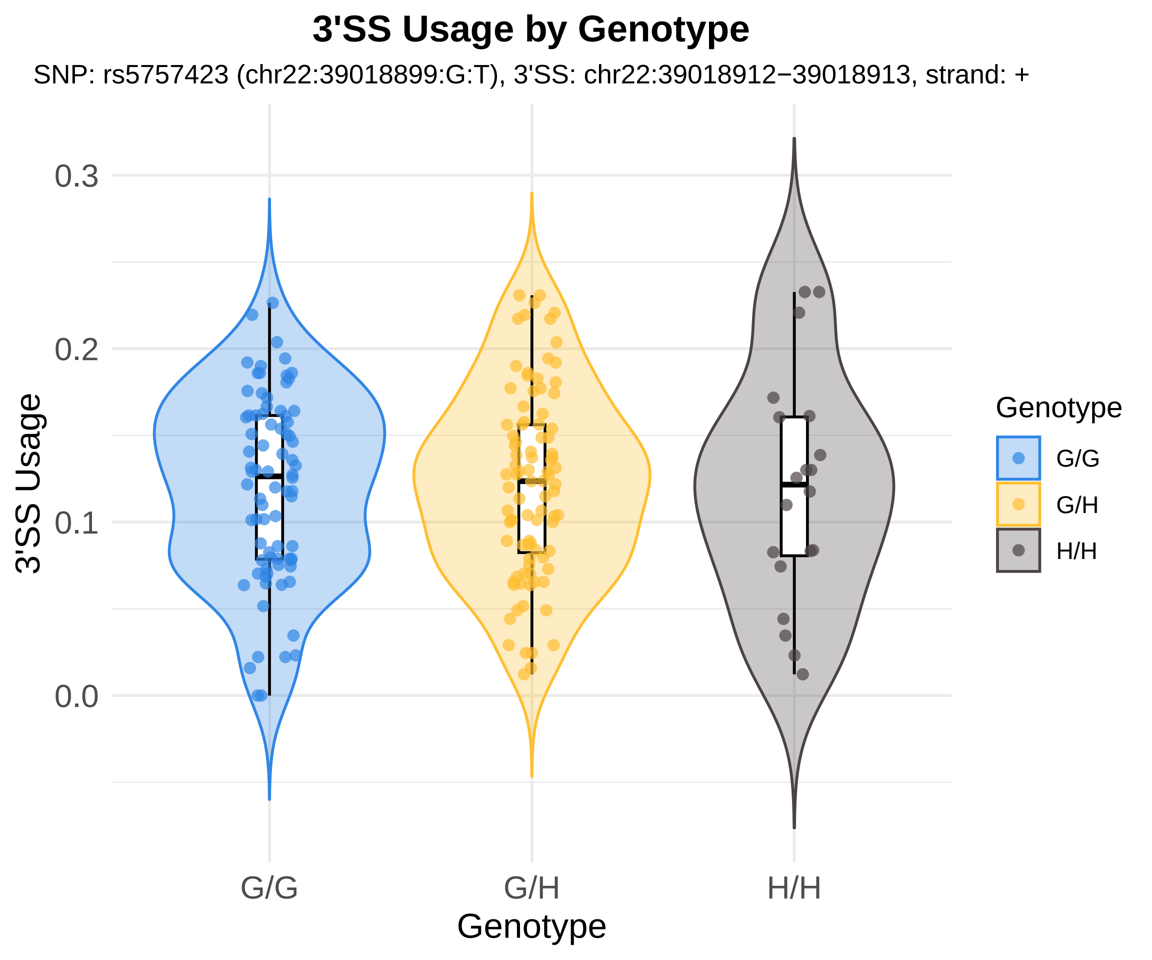
3'SS Usage by Genotype
SNP: rs5757423 (chr22:39018899:G:T), 3'SS: chr22:39018912−39018913, strand: +
0.3
0.2
0.1
0.0
3'SS Usage
G/G
G/H
H/H
Genotype
Genotype
G/G
G/H
H/H

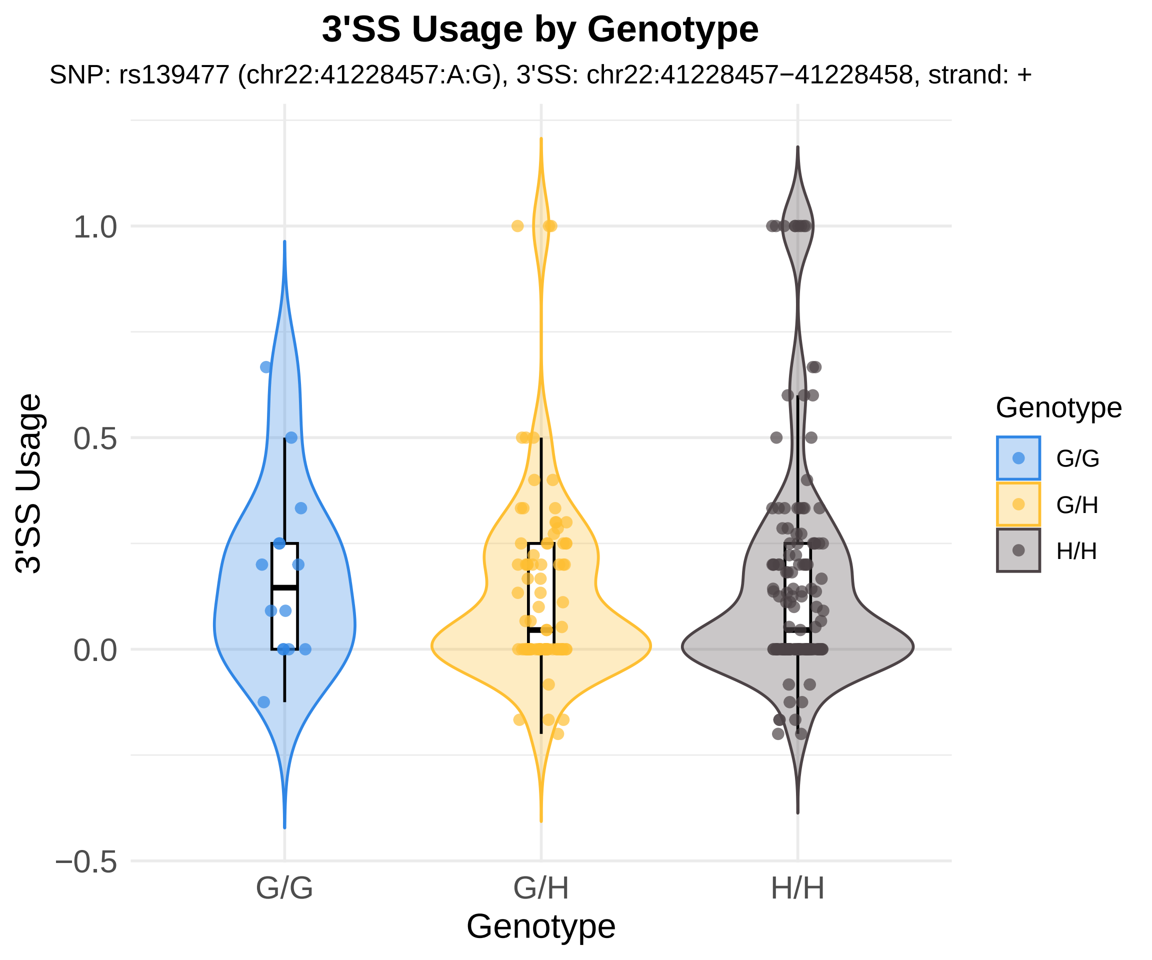

3'SS Usage by Genotype
SNP: rs139477 (chr22:41228457:A:G), 3'SS: chr22:41228457−41228458, strand: +
3'SS Usage
1.0
0.5
0.0
−0.5
G/G
G/H
H/H
Genotype
Genotype
G/G
G/H
H/H

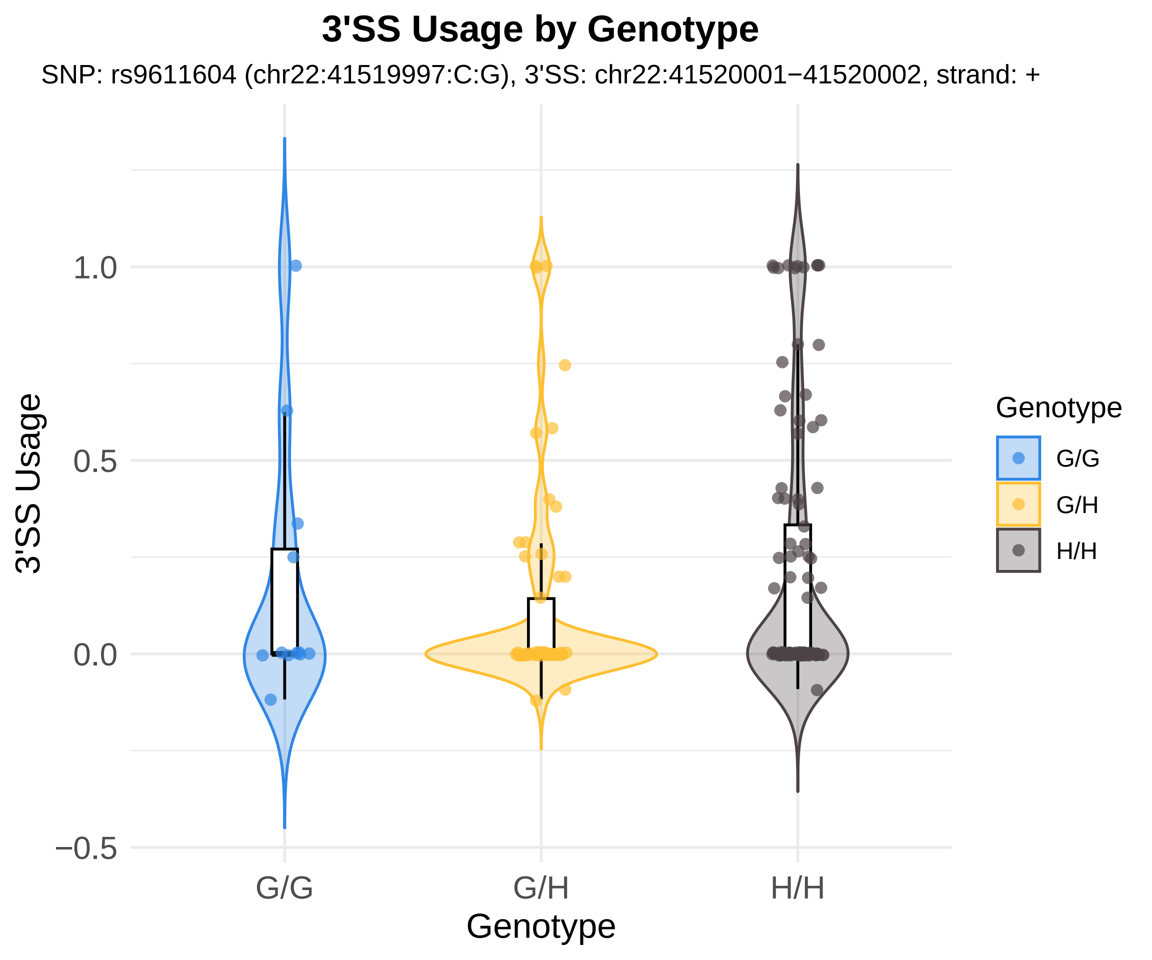
3'SS Usage by Genotype
SNP: rs9611604 (chr22:41519997:C:G), 3'SS: chr22:41520001−41520002, strand: +
3'SS Usage
1.0
0.5
0.0
−0.5
G/G
G/H
H/H
Genotype
Genotype
G/G
G/H
H/H

### 3'SS Usage by Genotype

SNP: rs9611604 (chr22:41519997:C:G), 3'SS: chr22:41520014−41520015, strand: +

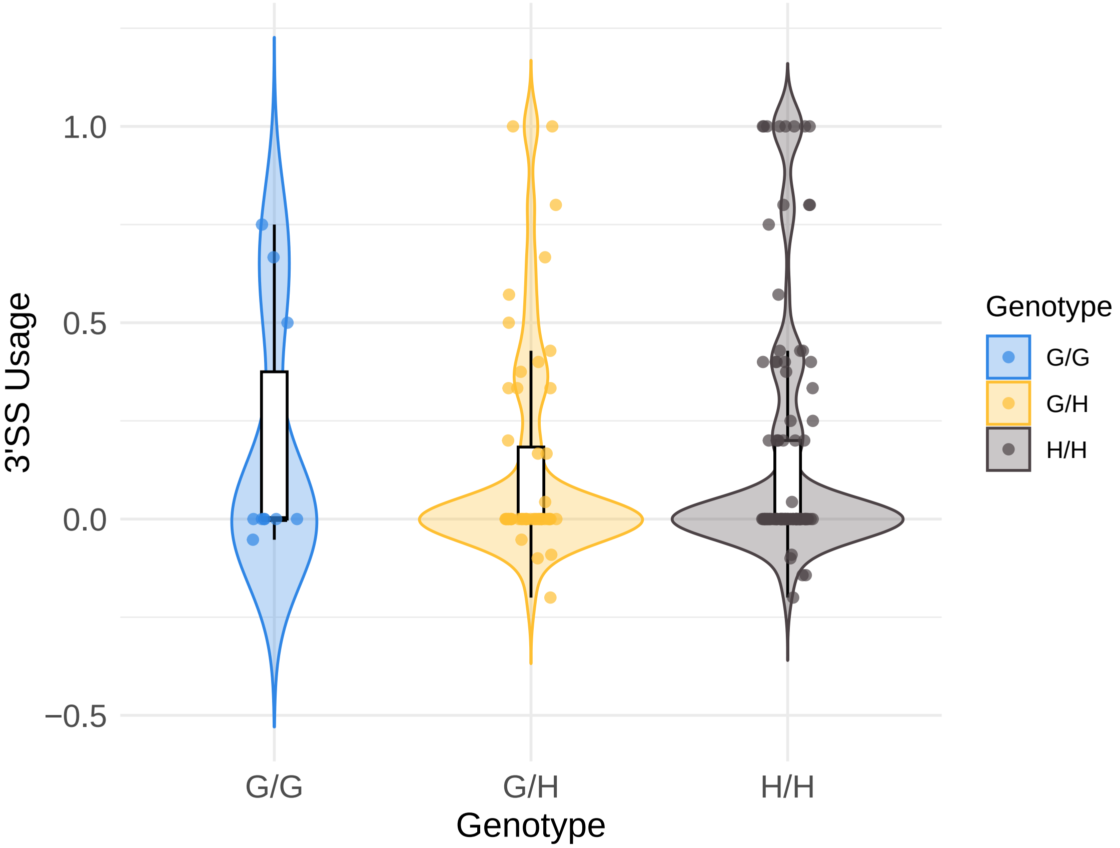

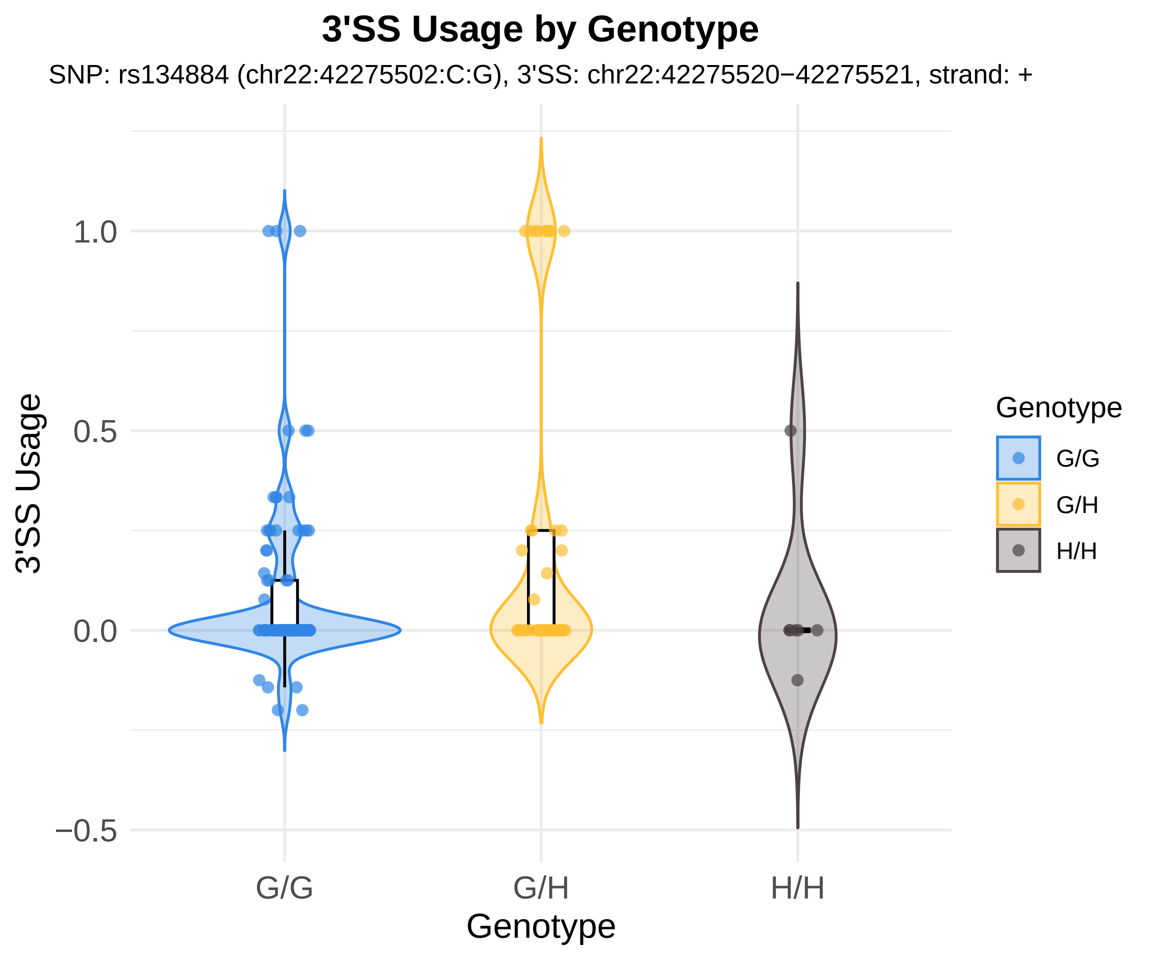
3'SS Usage by Genotype
SNP: rs134884 (chr22:42275502:C:G), 3'SS: chr22:42275520−42275521, strand: +
3'SS Usage
1.0
0.5
0.0
−0.5
G/G
G/H
H/H
Genotype
Genotype
G/G
G/H
H/H

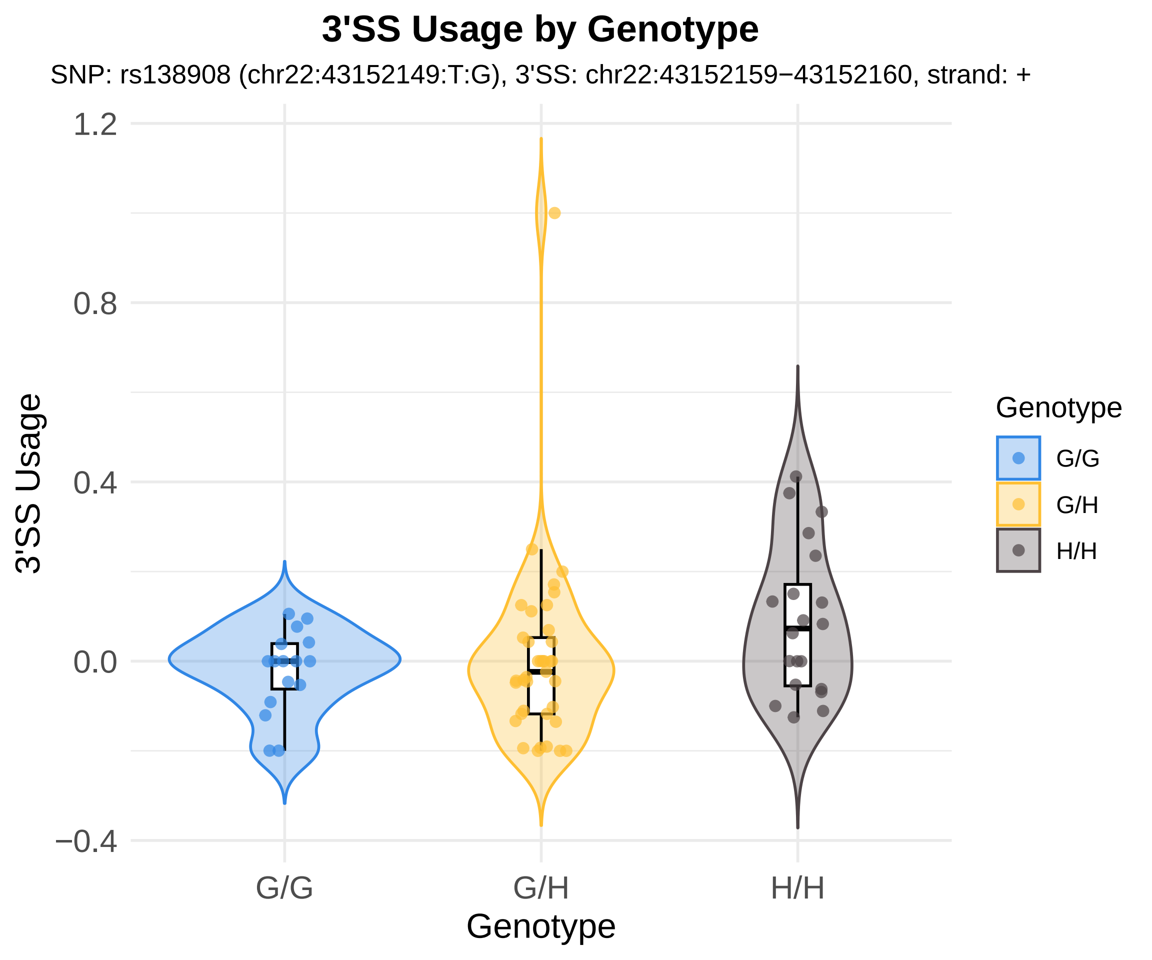
3'SS Usage by Genotype
SNP: rs138908 (chr22:43152149:T:G), 3'SS: chr22:43152159−43152160, strand: +
1.2
0.8
0.4
0.0
−0.4
3'SS Usage
G/G
G/H
H/H
Genotype
Genotype
G/G
G/H
H/H

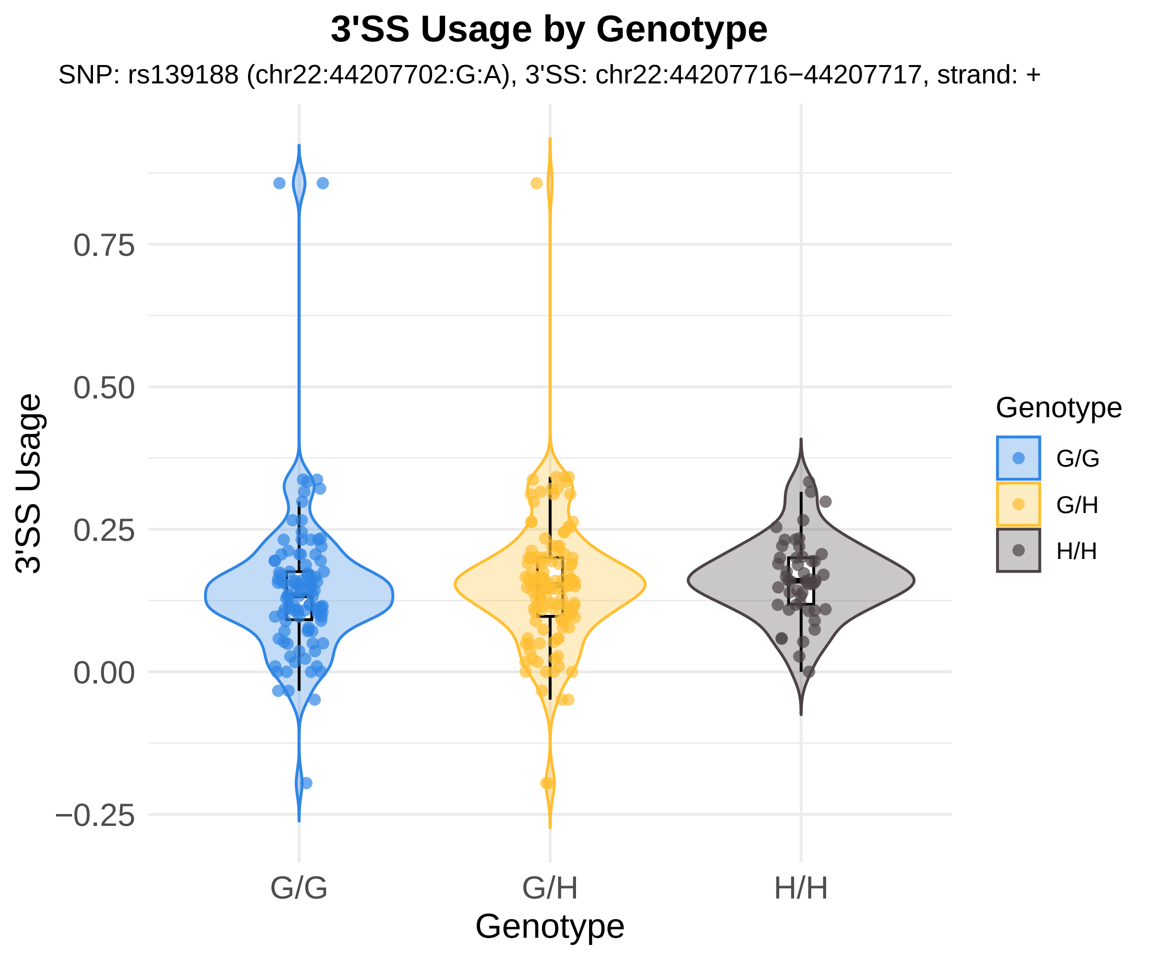

3'SS Usage by Genotype
SNP: rs139188 (chr22:44207702:G:A), 3'SS: chr22:44207716−44207717, strand: +
0.75
0.50
0.25
0.00
−0.25
3'SS Usage
G/G
G/H
H/H
Genotype
Genotype
G/G
G/H
H/H

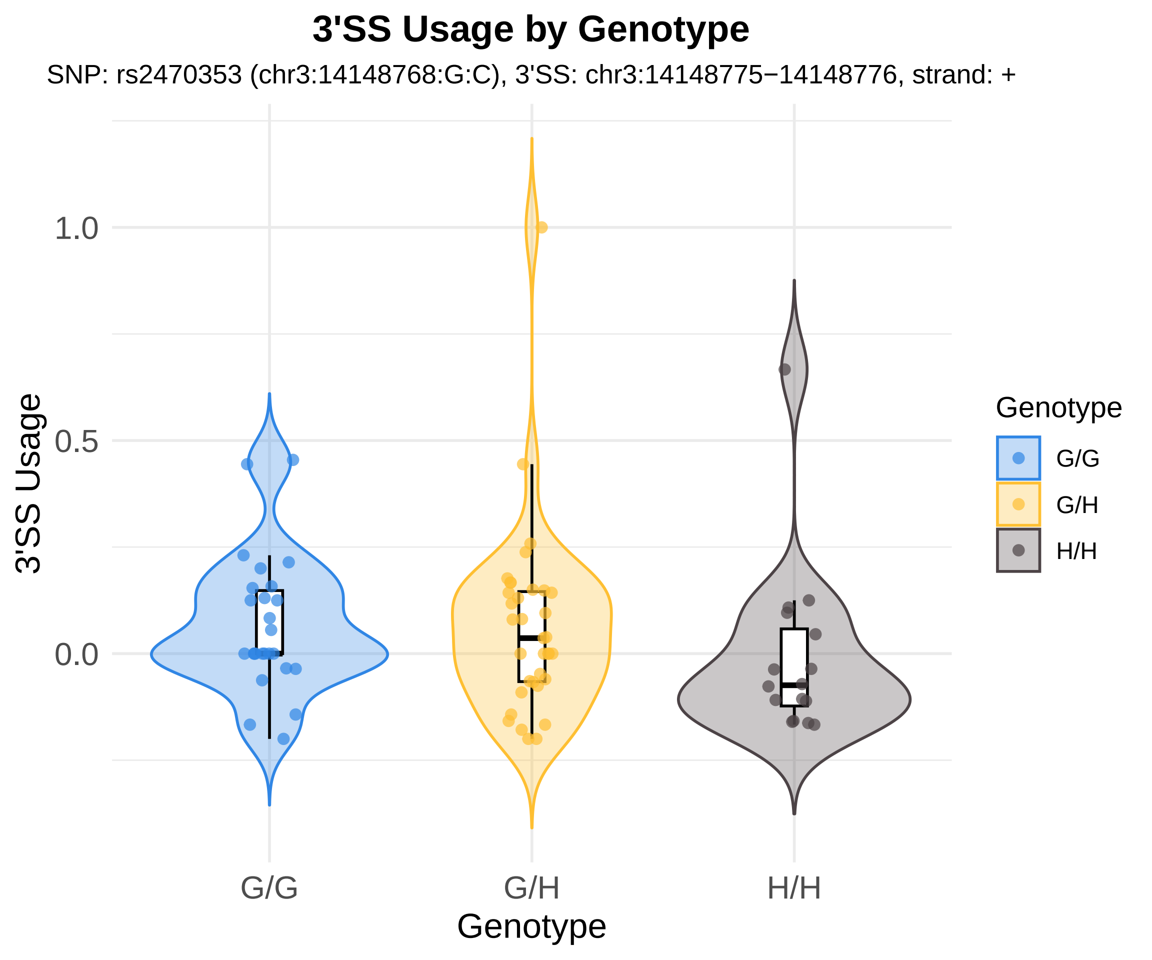

3'SS Usage by Genotype
SNP: rs2470353 (chr3:14148768:G:C), 3'SS: chr3:14148775−14148776, strand: +
3'SS Usage
1.0
0.5
0.0
G/G
G/H
H/H
Genotype
Genotype
G/G
G/H
H/H

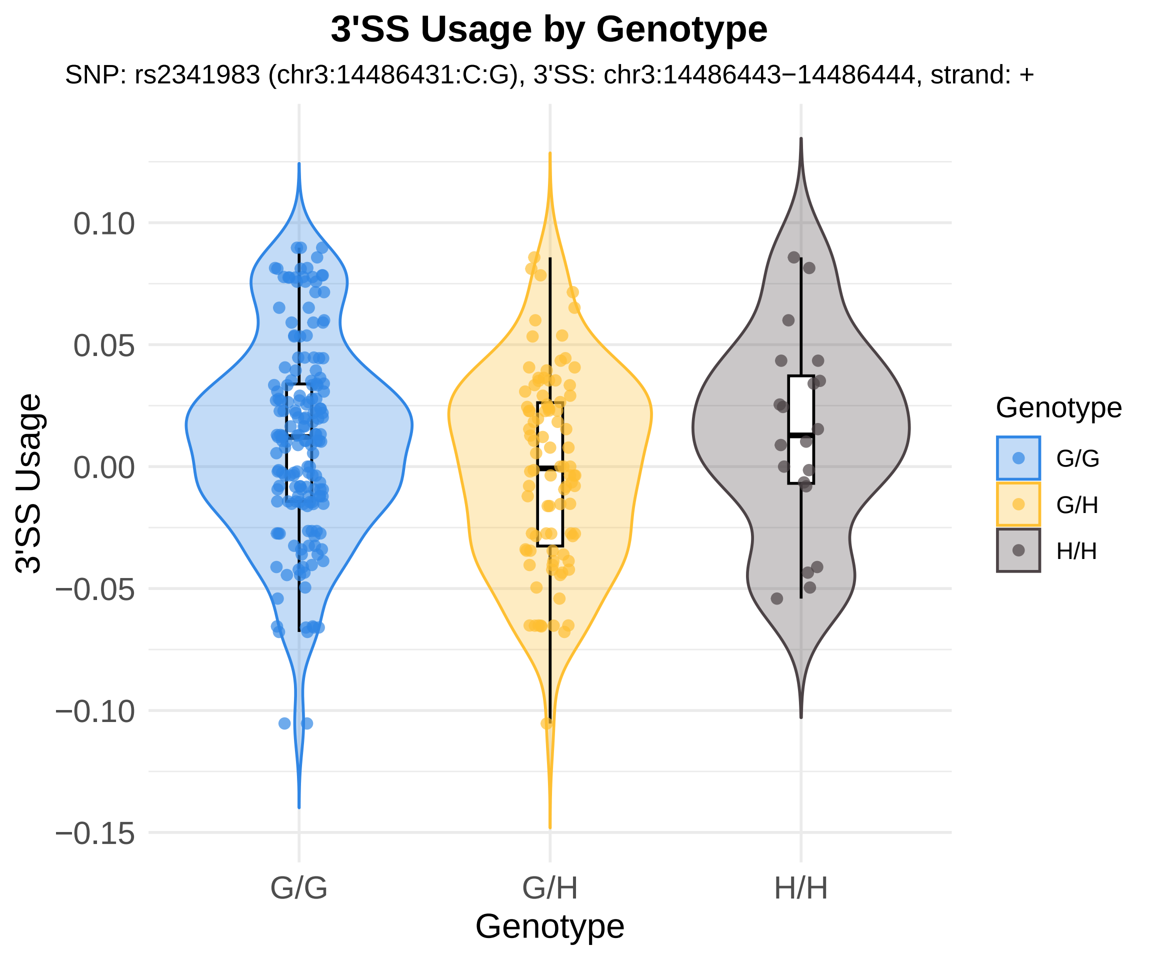
3'SS Usage by Genotype
SNP: rs2341983 (chr3:14486431:C:G), 3'SS: chr3:14486443−14486444, strand: +
3'SS Usage
0.10
0.05
0.00
−0.05
−0.10
−0.15
G/G
G/H
H/H
Genotype
Genotype
G/G
G/H
H/H

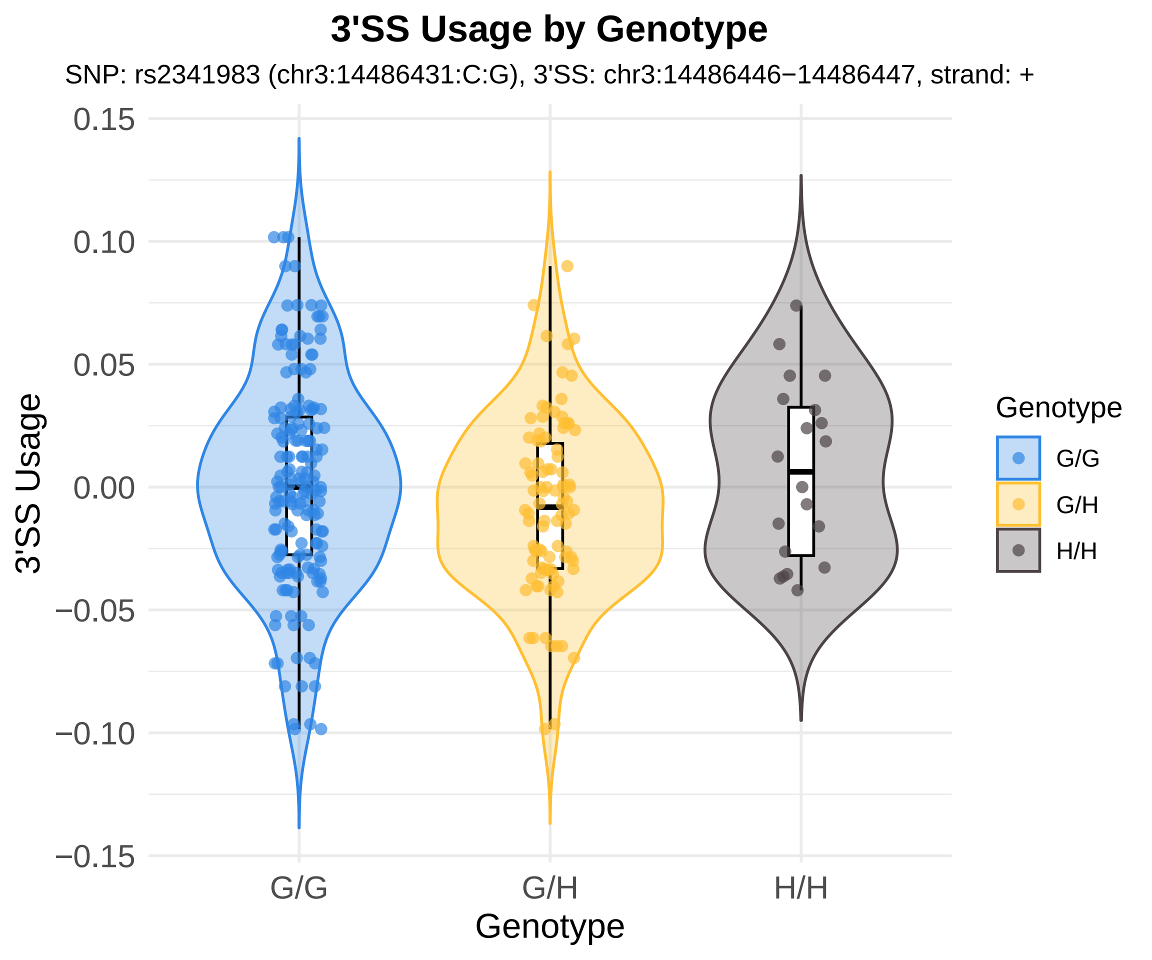
3'SS Usage by Genotype
SNP: rs2341983 (chr3:14486431:C:G), 3'SS: chr3:14486446−14486447, strand: +
3'SS Usage
0.15
0.10
0.05
0.00
−0.05
−0.10
−0.15
G/G
G/H
H/H
Genotype
Genotype
G/G
G/H
H/H

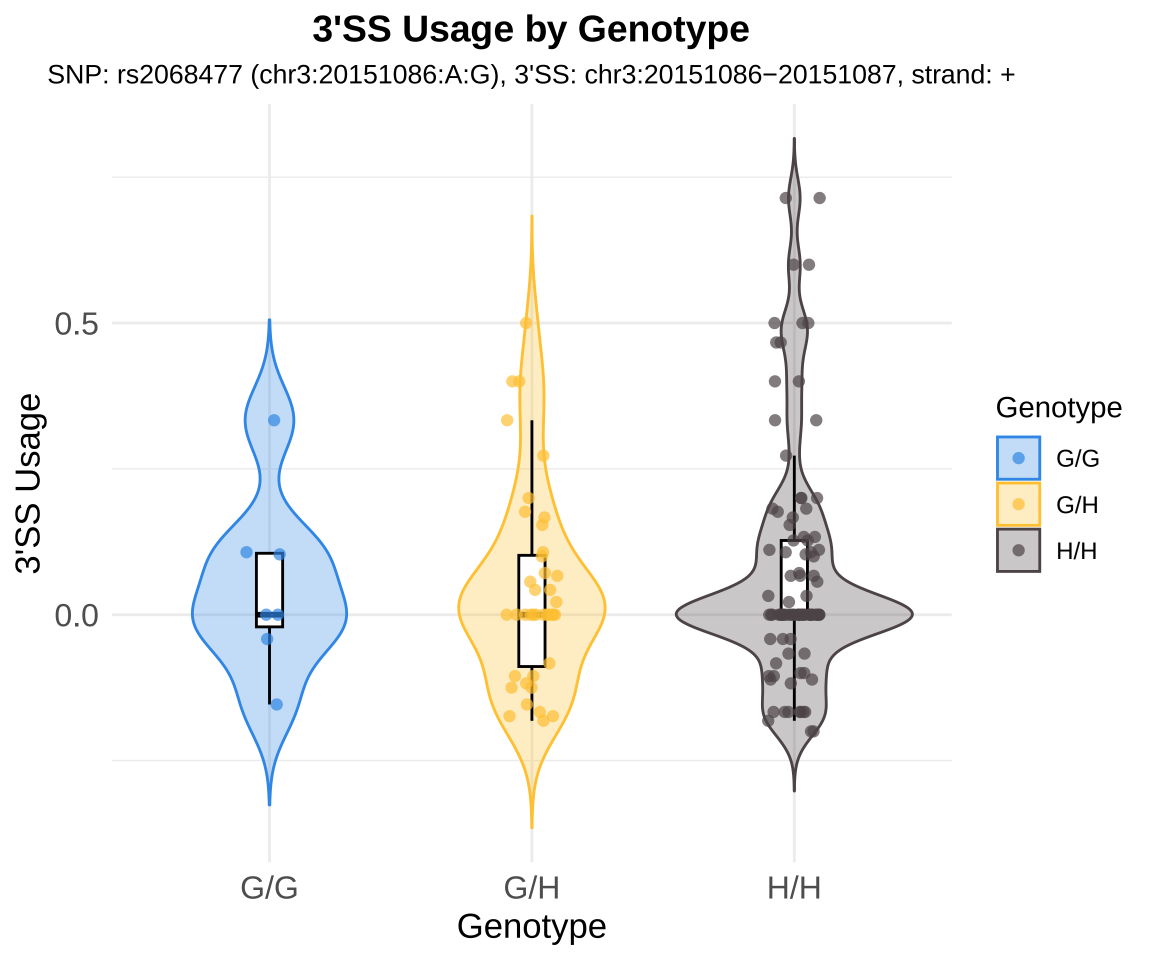
3'SS Usage by Genotype
SNP: rs2068477 (chr3:20151086:A:G), 3'SS: chr3:20151086−20151087, strand: +
3'SS Usage
0.5
0.0
G/G
G/H
H/H
Genotype
Genotype
G/G
G/H
H/H

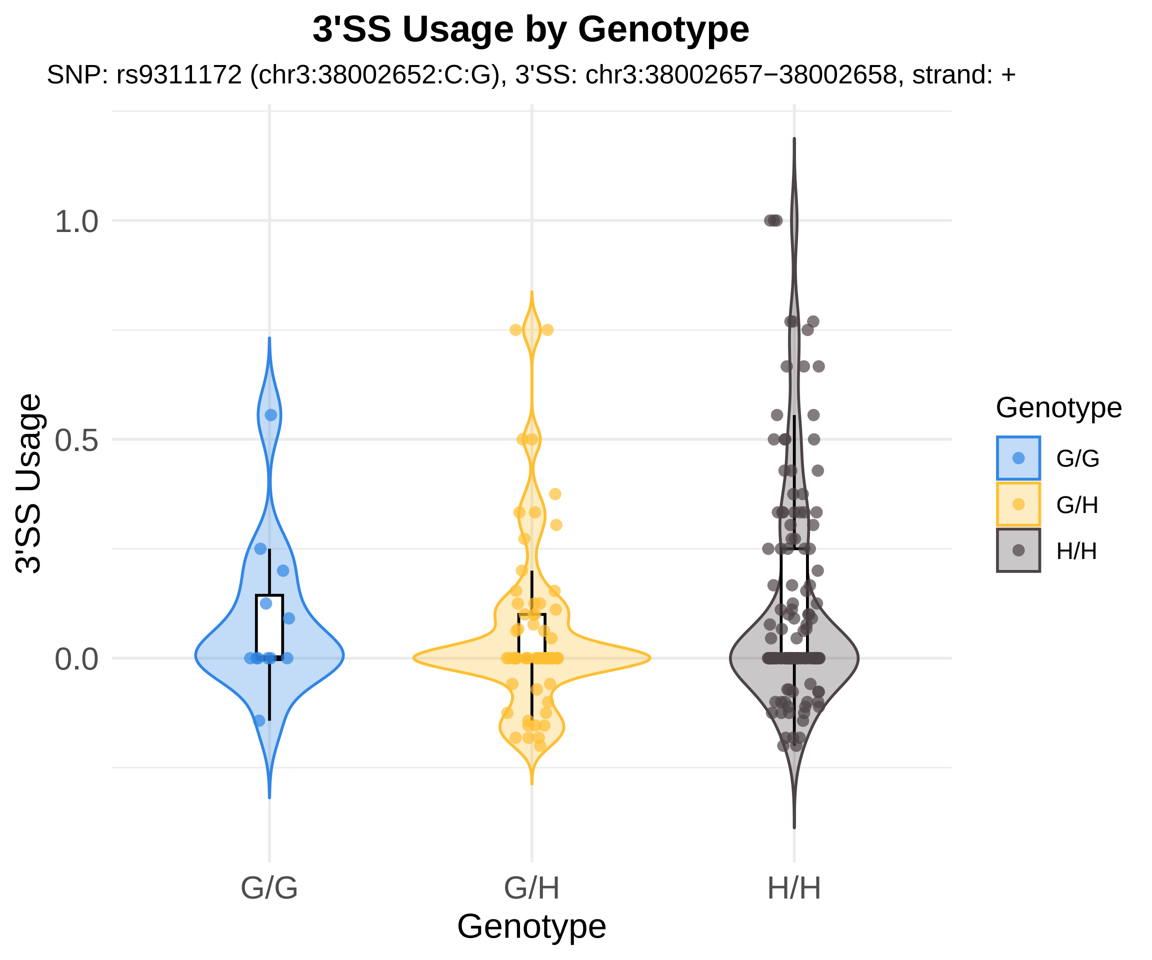
3'SS Usage by Genotype
SNP: rs9311172 (chr3:38002652:C:G), 3'SS: chr3:38002657−38002658, strand: +
3'SS Usage
1.0
0.5
0.0
G/G
G/H
H/H
Genotype
Genotype
G/G
G/H
H/H

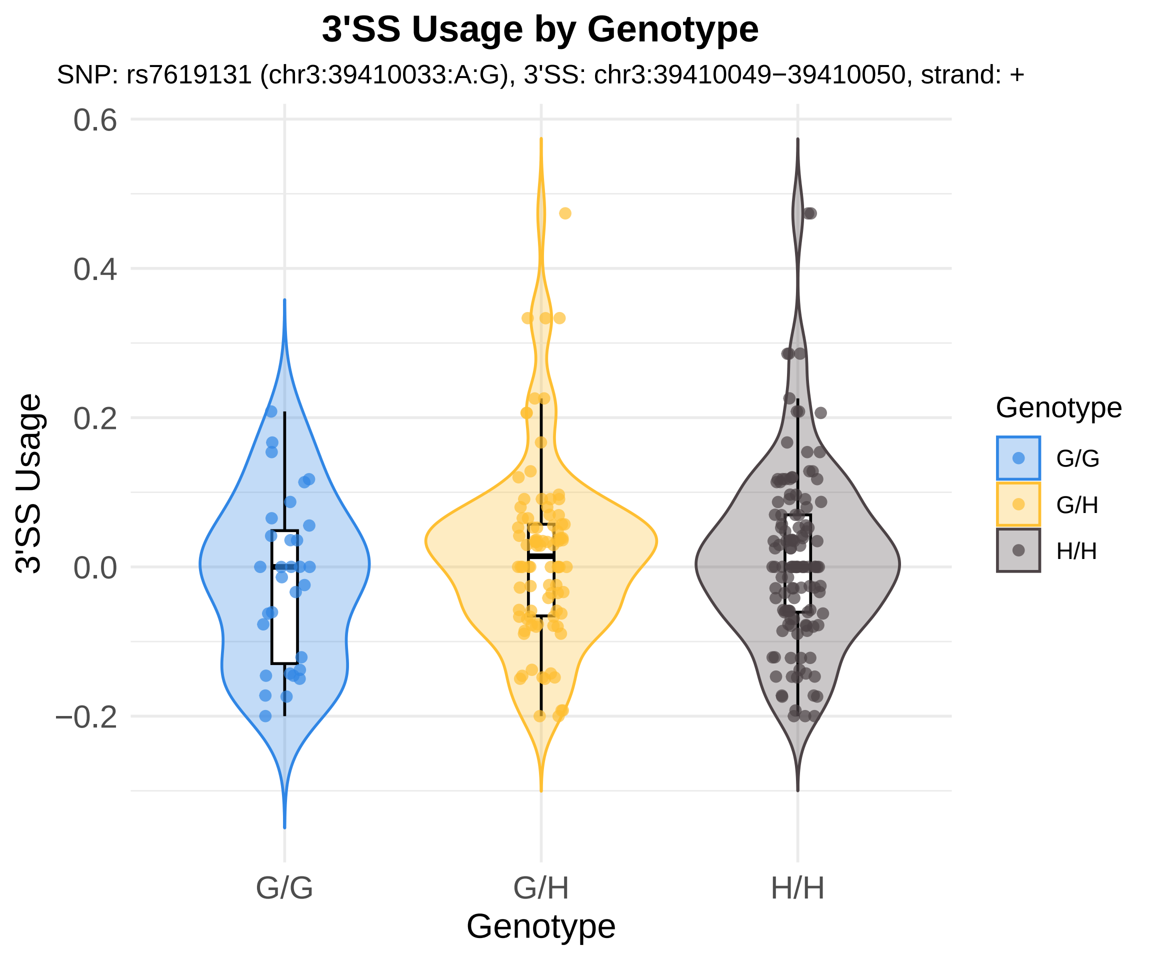
3'SS Usage by Genotype
SNP: rs7619131 (chr3:39410033:A:G), 3'SS: chr3:39410049−39410050, strand: +
3'SS Usage
0.6
0.4
0.2
0.0
−0.2
G/G
G/H
H/H
Genotype
Genotype
G/G
G/H
H/H

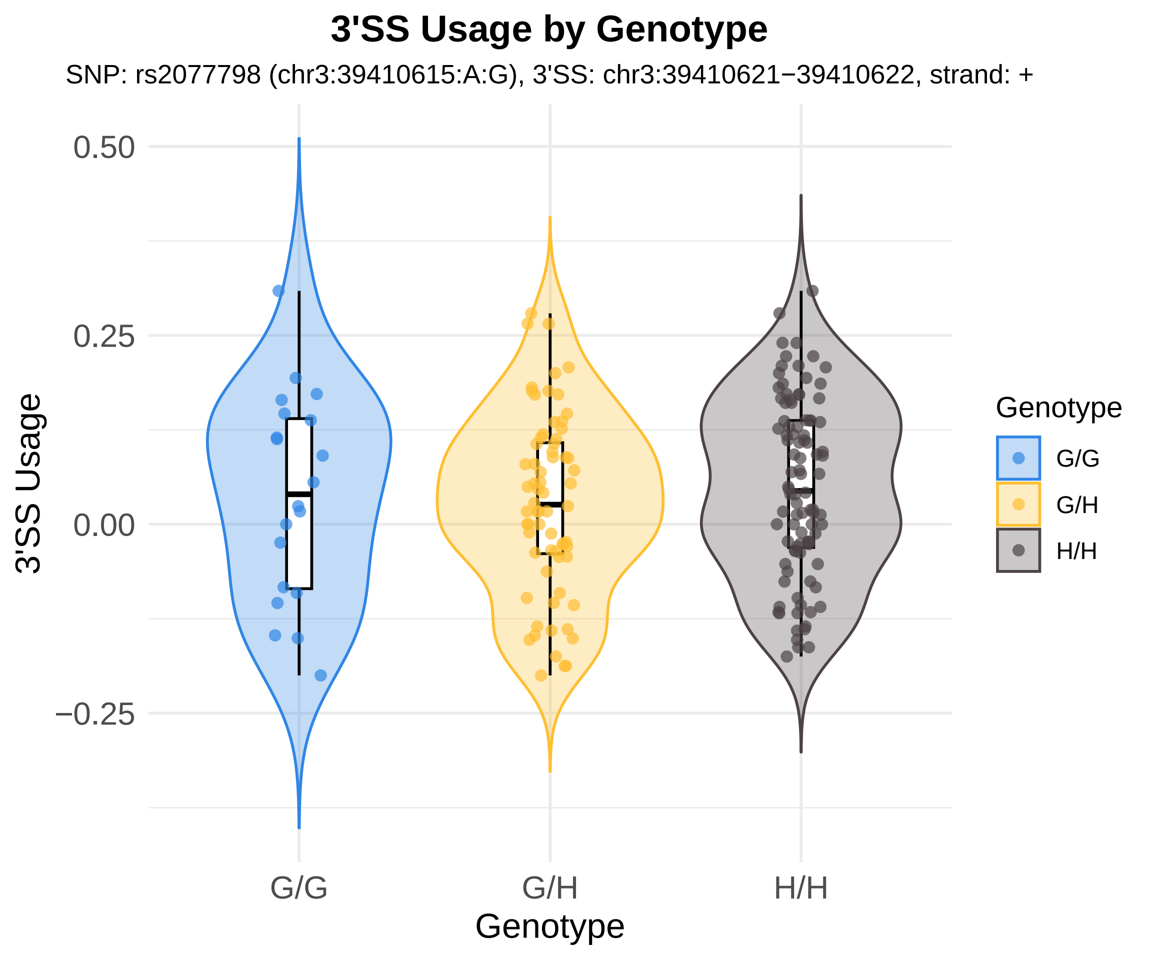
3'SS Usage by Genotype
SNP: rs2077798 (chr3:39410615:A:G), 3'SS: chr3:39410621−39410622, strand: +
0.50
0.25
0.00
−0.25
3'SS Usage
G/G
G/H
H/H
Genotype
Genotype
G/G
G/H
H/H

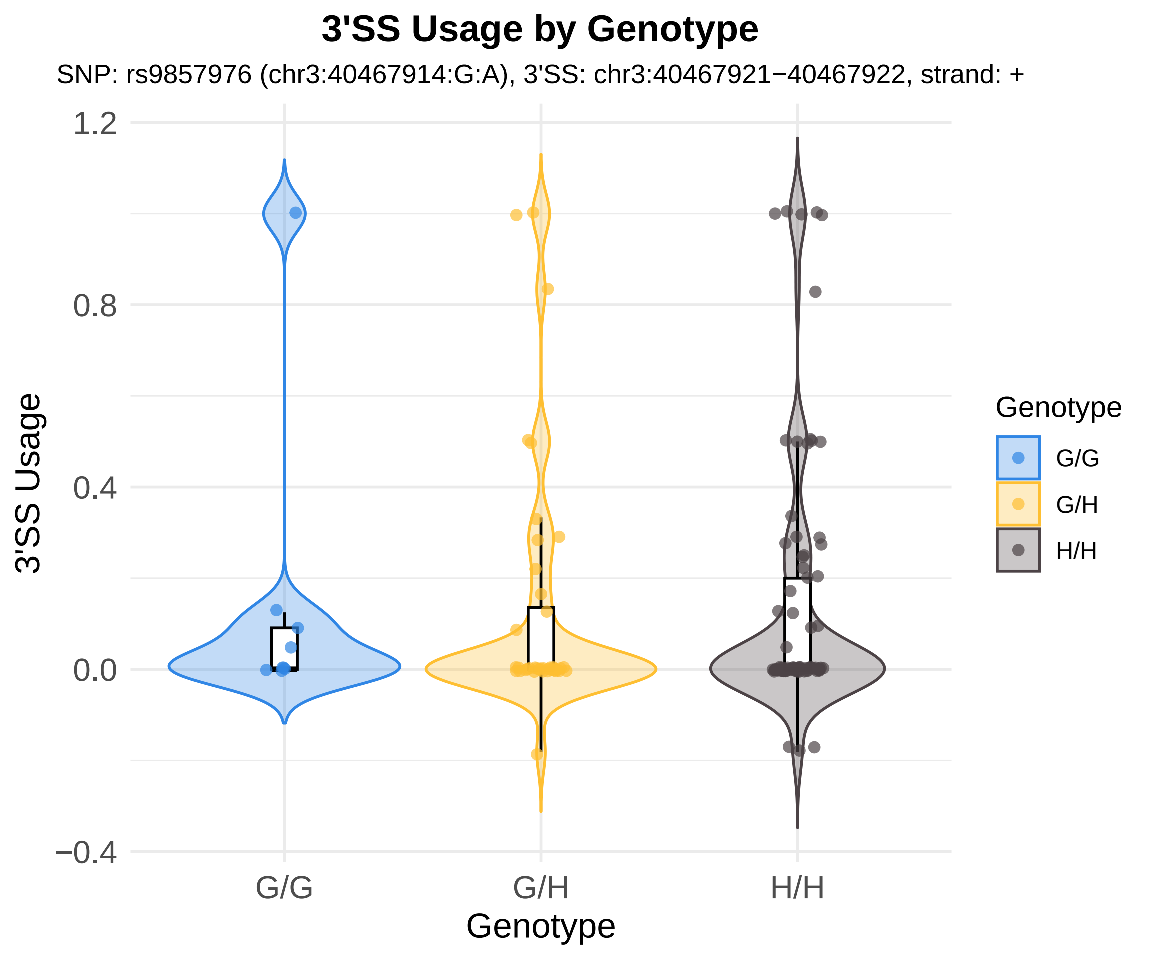
3'SS Usage by Genotype
SNP: rs9857976 (chr3:40467914:G:A), 3'SS: chr3:40467921−40467922, strand: +
3'SS Usage
1.2
0.8
0.4
0.0
−0.4
G/G
G/H
H/H
Genotype
Genotype
G/G
G/H
H/H

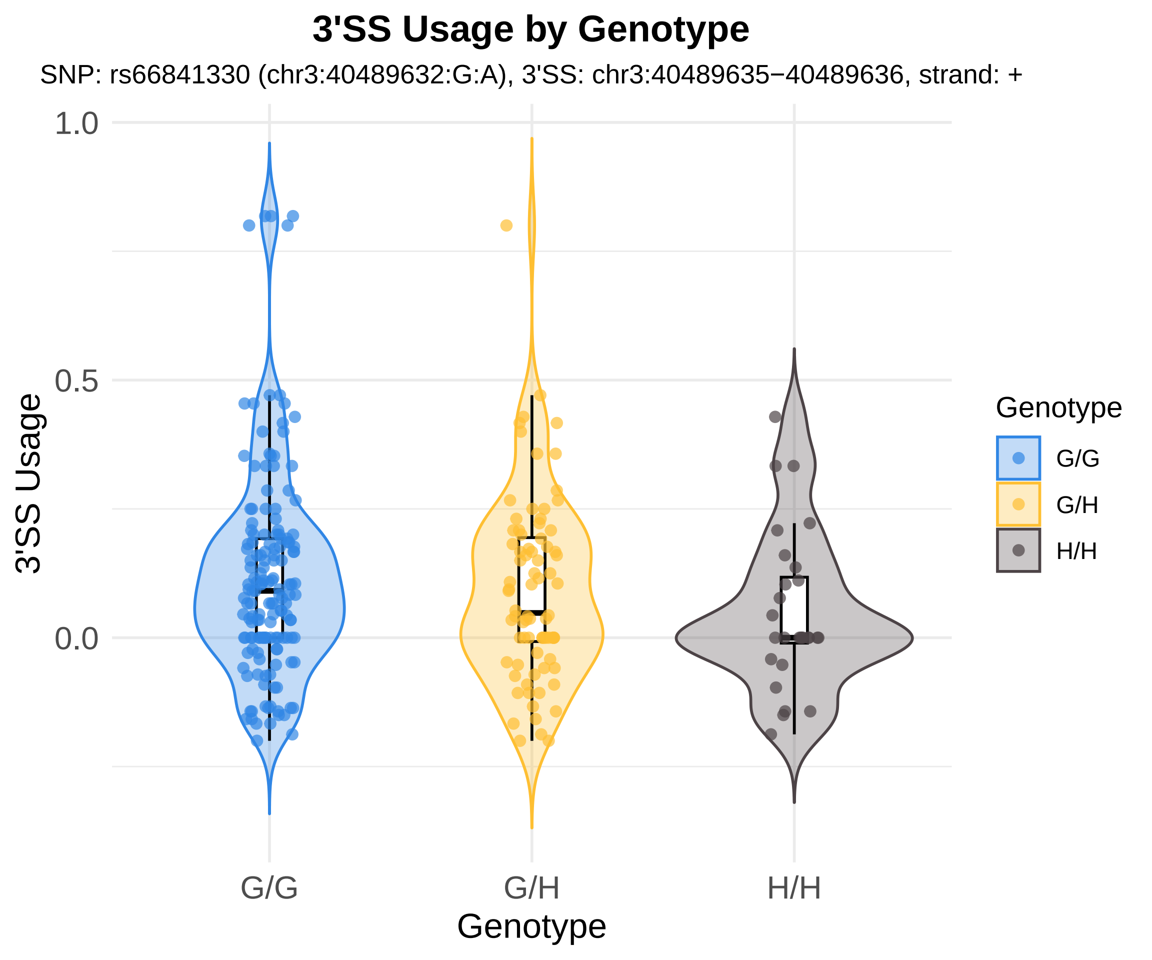

3'SS Usage by Genotype
SNP: rs66841330 (chr3:40489632:G:A), 3'SS: chr3:40489635−40489636, strand: +
1.0
0.5
0.0
3'SS Usage
G/G
G/H
H/H
Genotype
Genotype
G/G
G/H
H/H

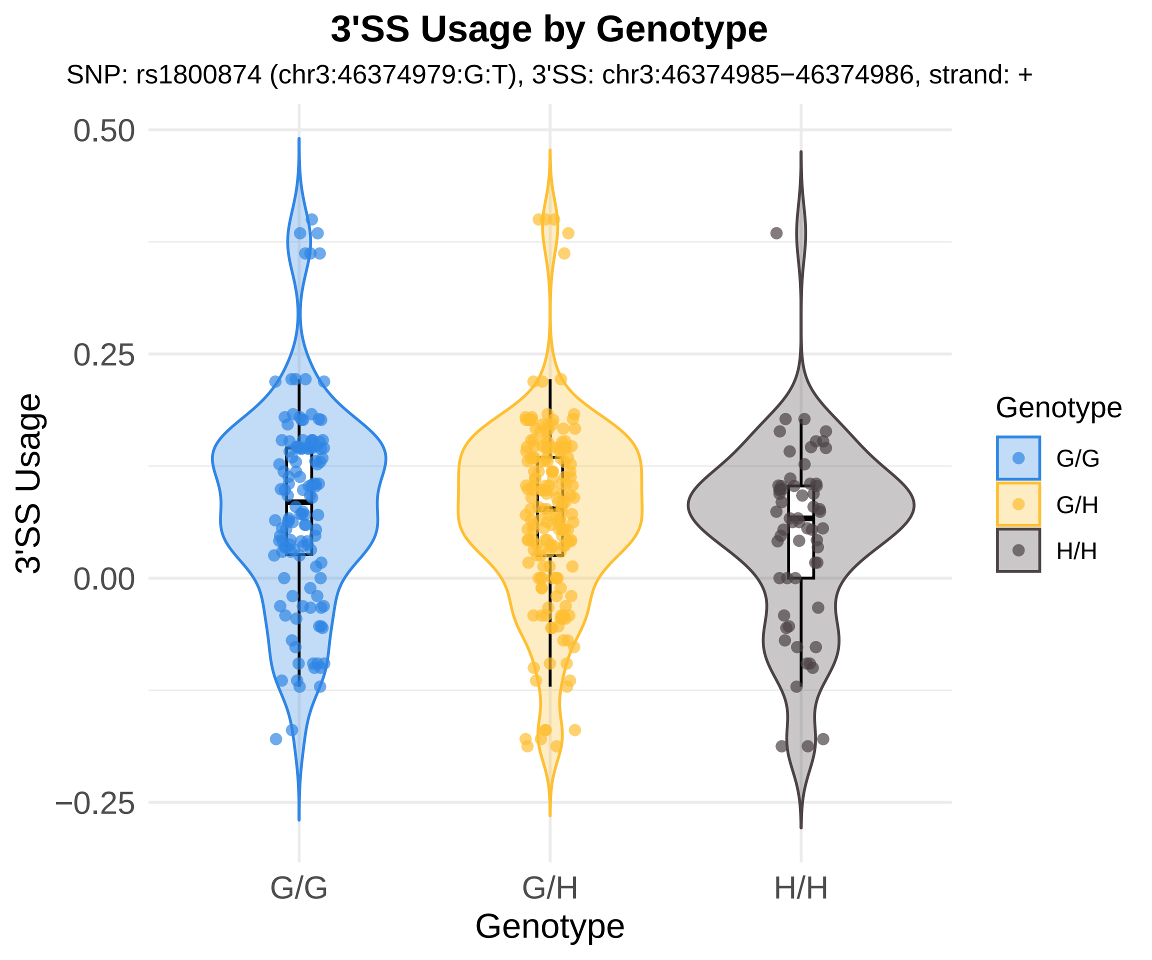
3'SS Usage by Genotype
SNP: rs1800874 (chr3:46374979:G:T), 3'SS: chr3:46374985−46374986, strand: +
3'SS Usage
0.50
0.25
0.00
−0.25
G/G
G/H
H/H
Genotype
Genotype
G/G
G/H
H/H

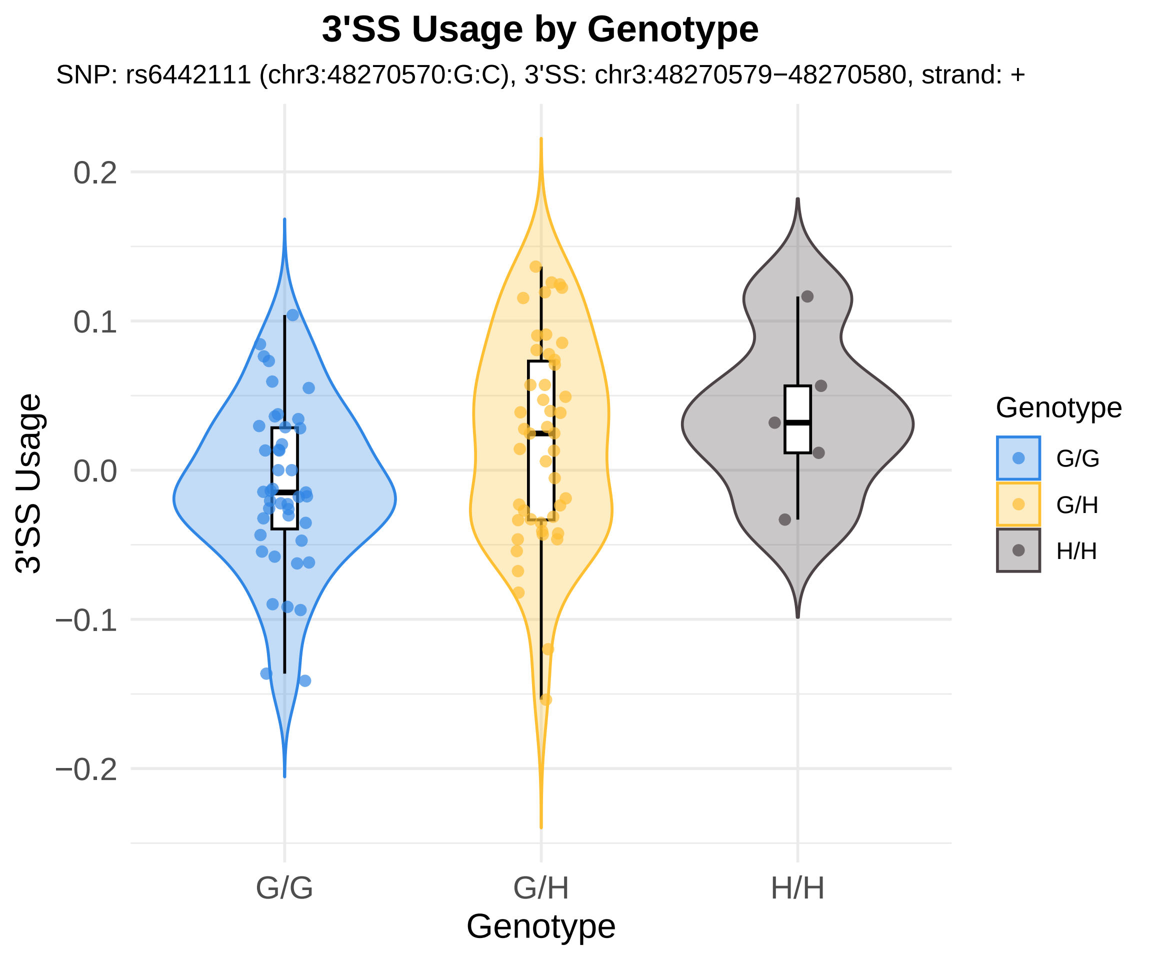
3'SS Usage by Genotype
SNP: rs6442111 (chr3:48270570:G:C), 3'SS: chr3:48270579−48270580, strand: +
3'SS Usage
0.2
0.1
0.0
−0.1
−0.2
G/G
G/H
H/H
Genotype
Genotype
G/G
G/H
H/H

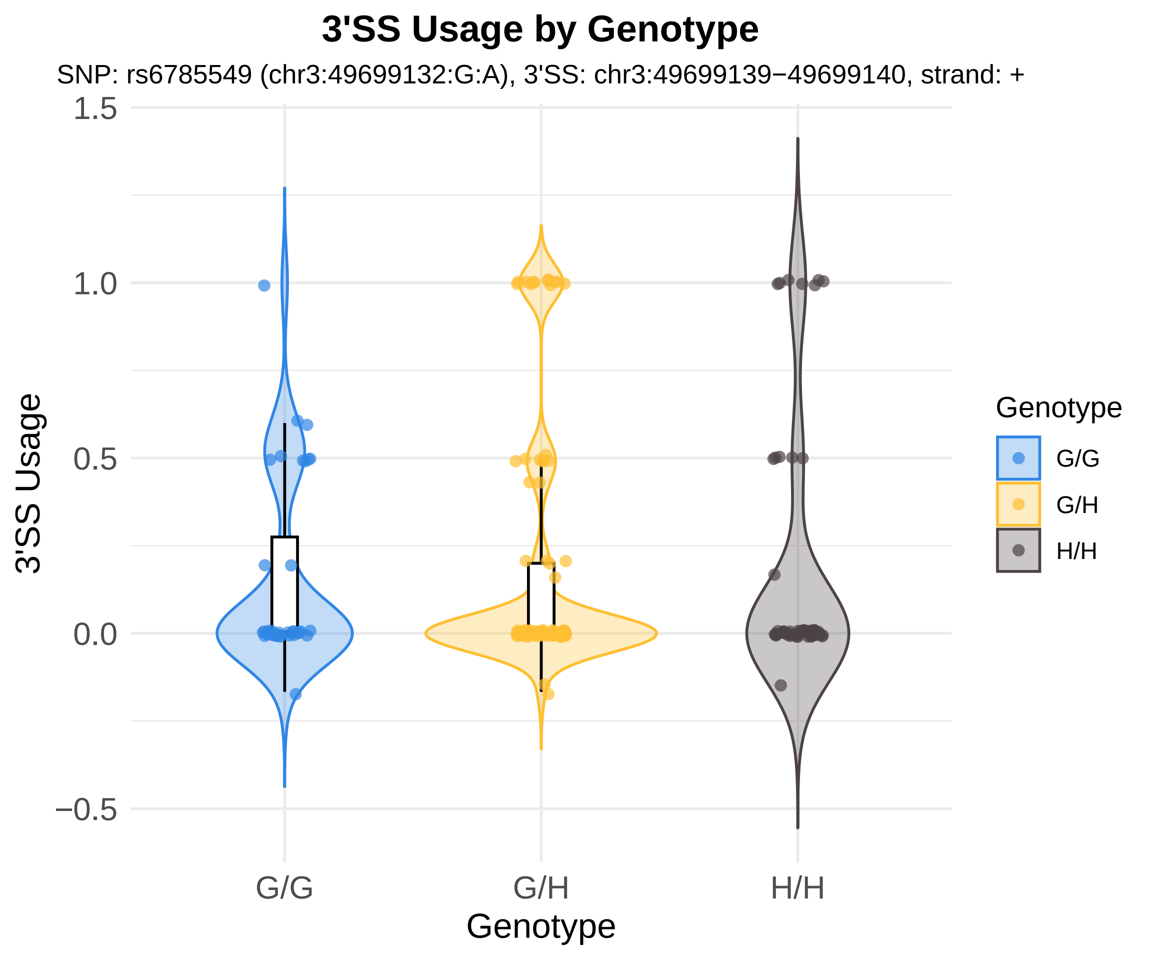
3'SS Usage by Genotype
SNP: rs6785549 (chr3:49699132:G:A), 3'SS: chr3:49699139−49699140, strand: +
1.5
1.0
0.5
0.0
−0.5
3'SS Usage
G/G
G/H
H/H
Genotype
Genotype
G/G
G/H
H/H

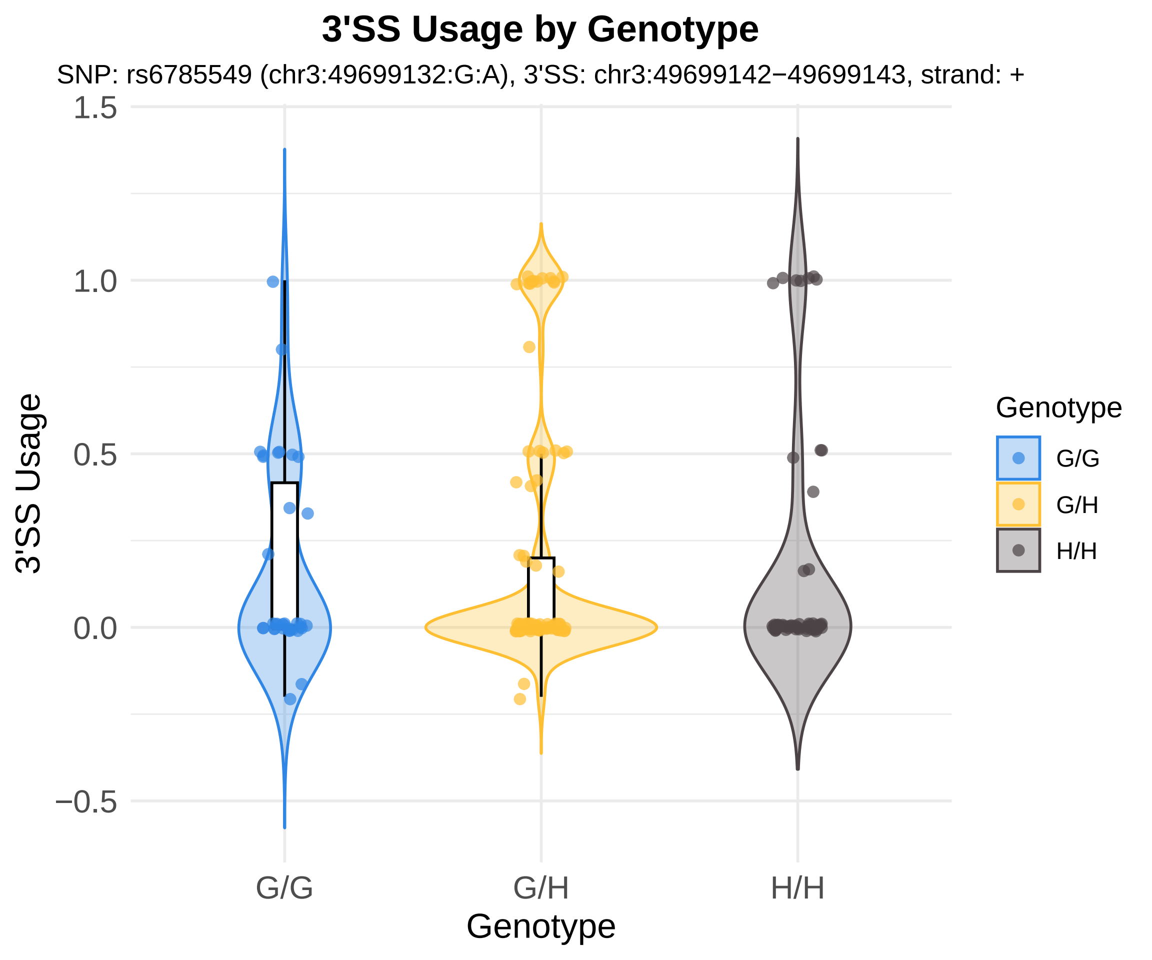

3'SS Usage by Genotype
SNP: rs6785549 (chr3:49699132:G:A), 3'SS: chr3:49699142−49699143, strand: +
3'SS Usage
1.5
1.0
0.5
0.0
−0.5
G/G
G/H
H/H
Genotype
Genotype
G/G
G/H
H/H

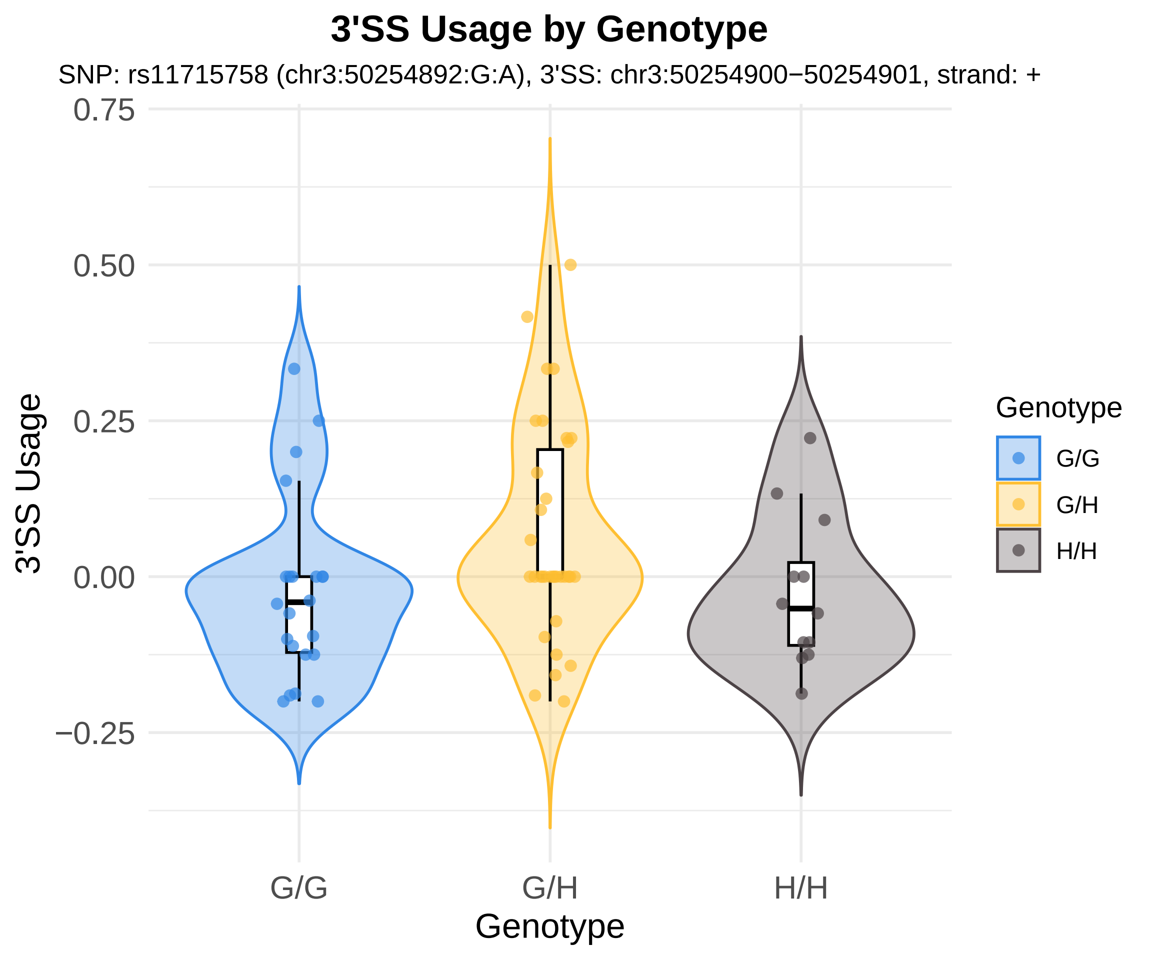

3'SS Usage by Genotype
SNP: rs11715758 (chr3:50254892:G:A), 3'SS: chr3:50254900−50254901, strand: +
0.75
0.50
0.25
0.00
−0.25
3'SS Usage
G/G
G/H
H/H
Genotype
Genotype
G/G
G/H
H/H

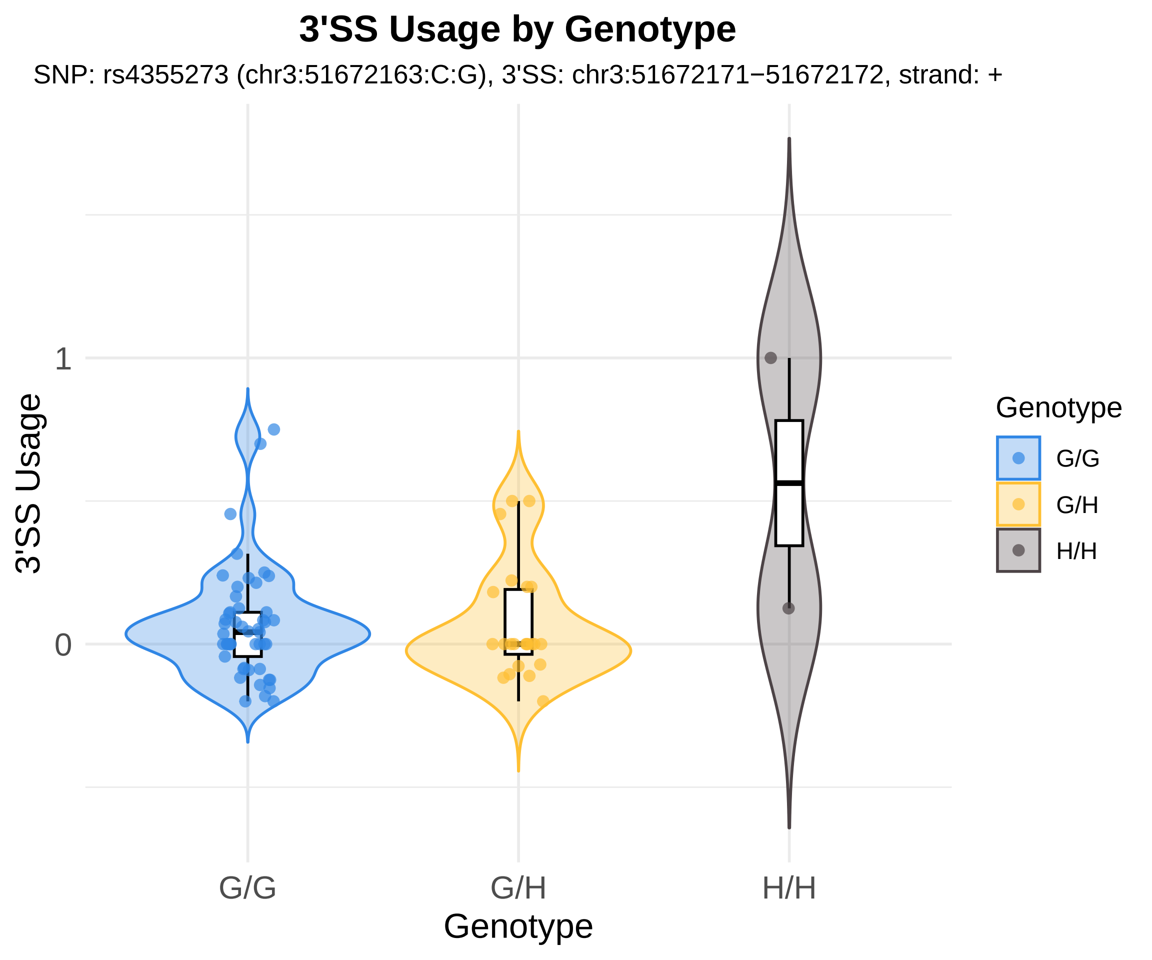

3'SS Usage by Genotype
SNP: rs4355273 (chr3:51672163:C:G), 3'SS: chr3:51672171−51672172, strand: +
3'SS Usage
1
0
G/G
G/H
H/H
Genotype
Genotype
G/G
G/H
H/H

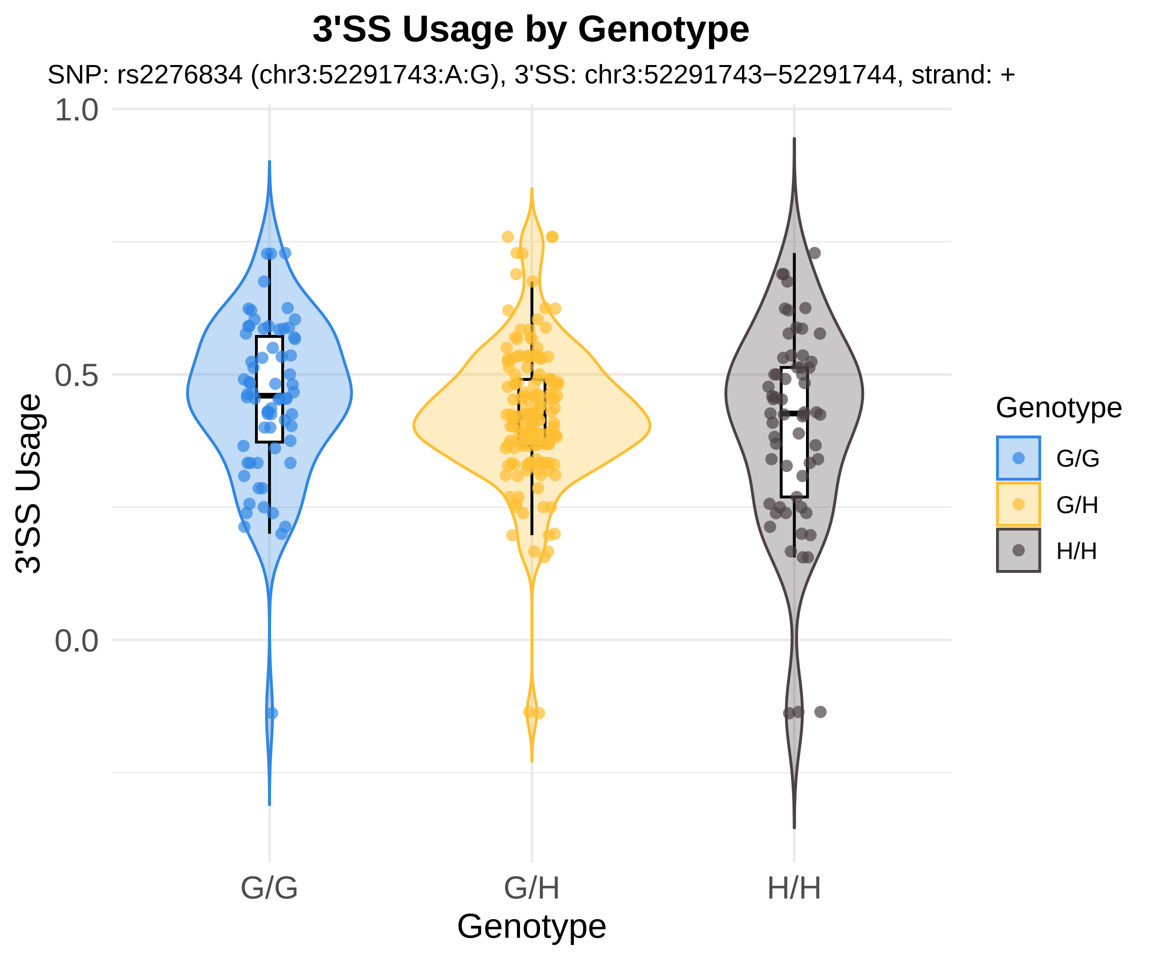
3'SS Usage by Genotype
SNP: rs2276834 (chr3:52291743:A:G), 3'SS: chr3:52291743−52291744, strand: +
1.0
0.5
0.0
3'SS Usage
G/G
G/H
H/H
Genotype
Genotype
G/G
G/H
H/H

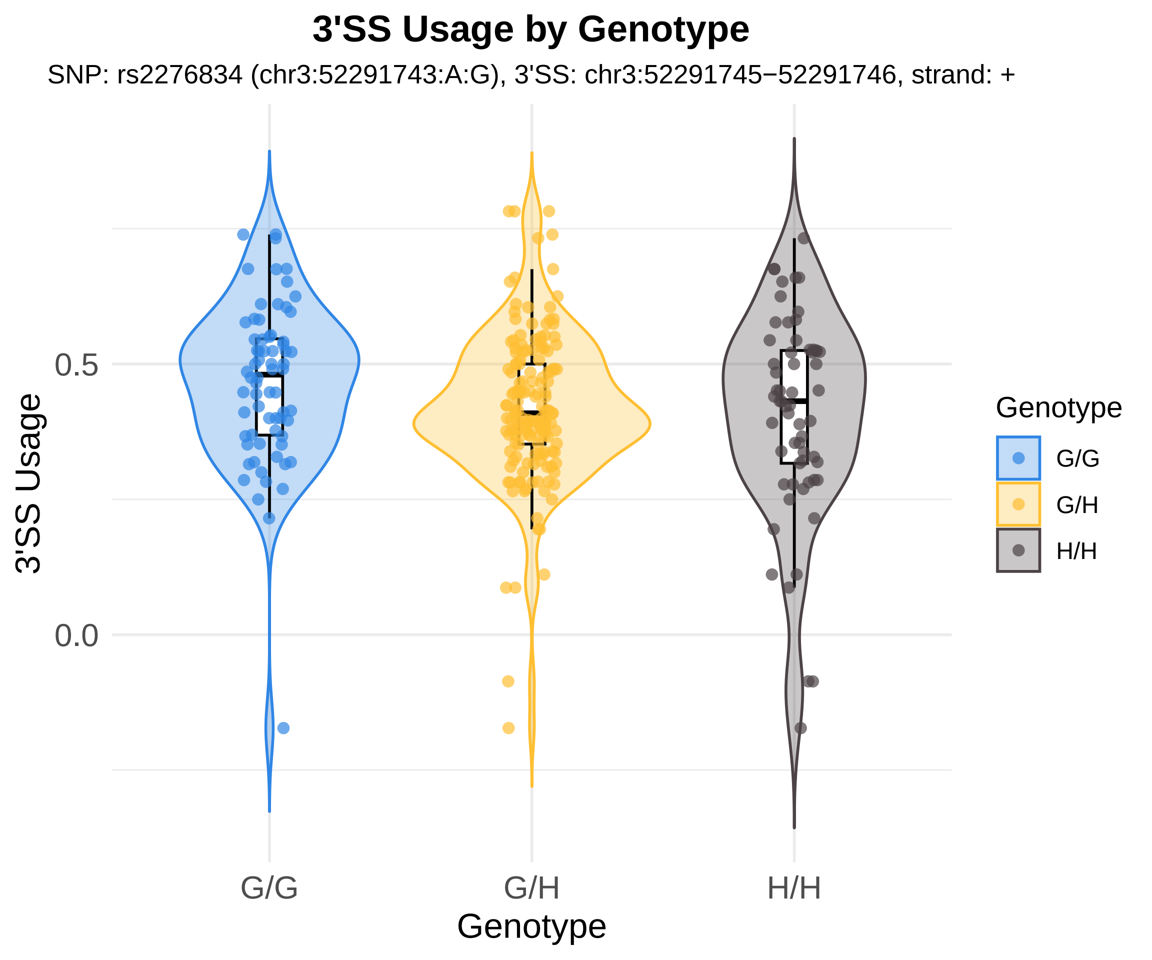
3'SS Usage by Genotype
SNP: rs2276834 (chr3:52291743:A:G), 3'SS: chr3:52291745−52291746, strand: +
3'SS Usage
0.5
0.0
G/G
G/H
H/H
Genotype
Genotype
G/G
G/H
H/H

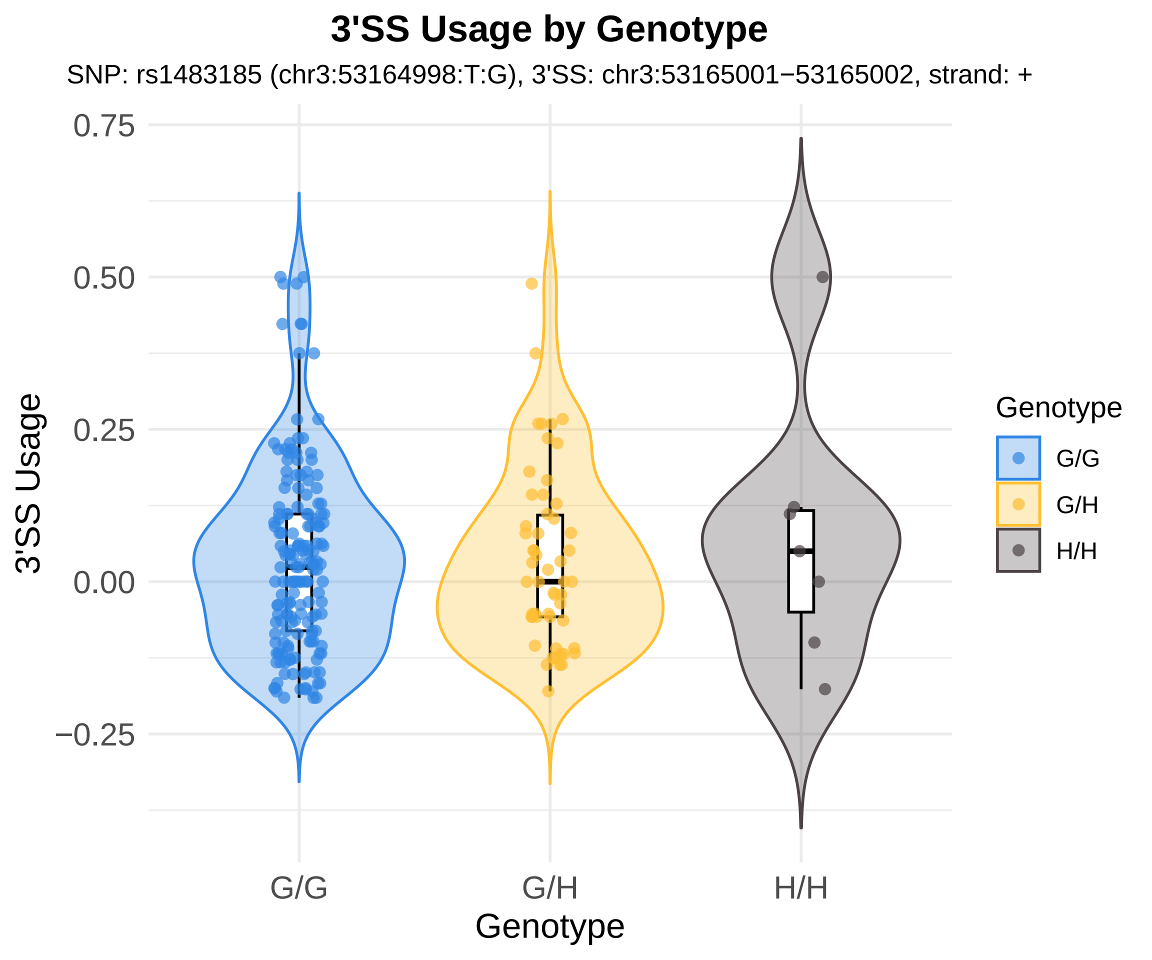

3'SS Usage by Genotype
SNP: rs1483185 (chr3:53164998:T:G), 3'SS: chr3:53165001−53165002, strand: +
0.75
0.50
0.25
0.00
−0.25
3'SS Usage
G/G
G/H
H/H
Genotype
Genotype
G/G
G/H
H/H

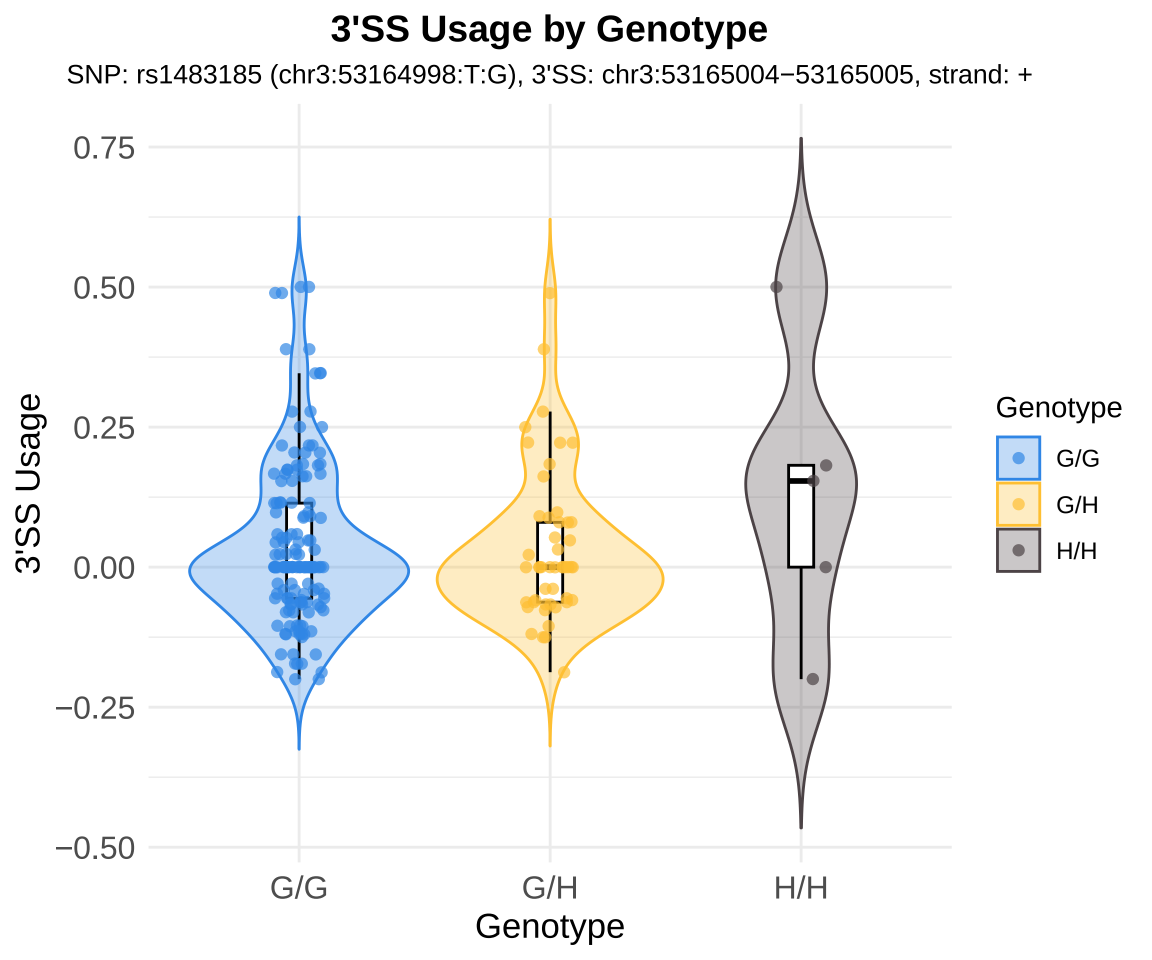
3'SS Usage by Genotype
SNP: rs1483185 (chr3:53164998:T:G), 3'SS: chr3:53165004−53165005, strand: +
3'SS Usage
0.75
0.50
0.25
0.00
−0.25
−0.50
G/G
G/H
H/H
Genotype
Genotype
G/G
G/H
H/H

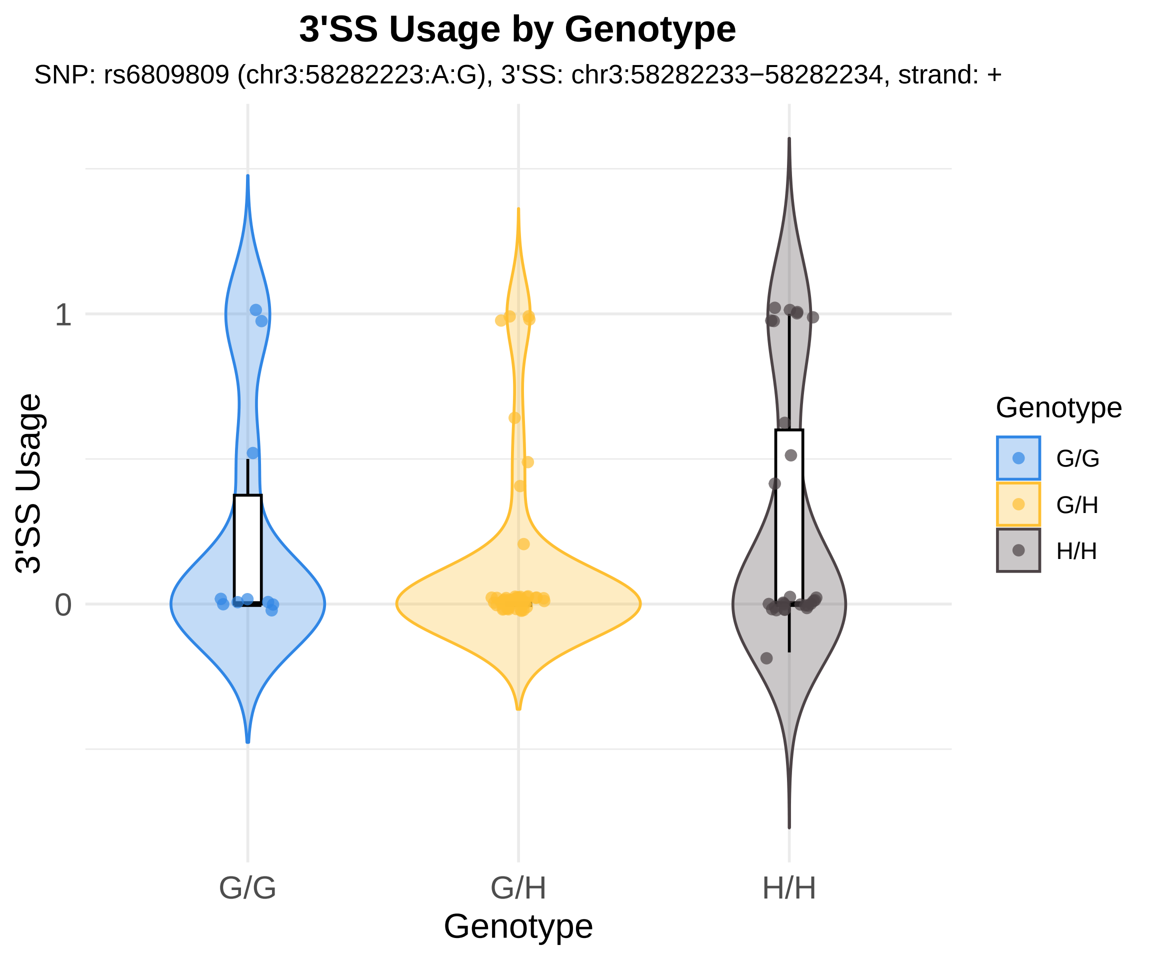

3'SS Usage by Genotype
SNP: rs6809809 (chr3:58282223:A:G), 3'SS: chr3:58282233−58282234, strand: +
3'SS Usage
1
0
G/G
G/H
H/H
Genotype
Genotype
G/G
G/H
H/H

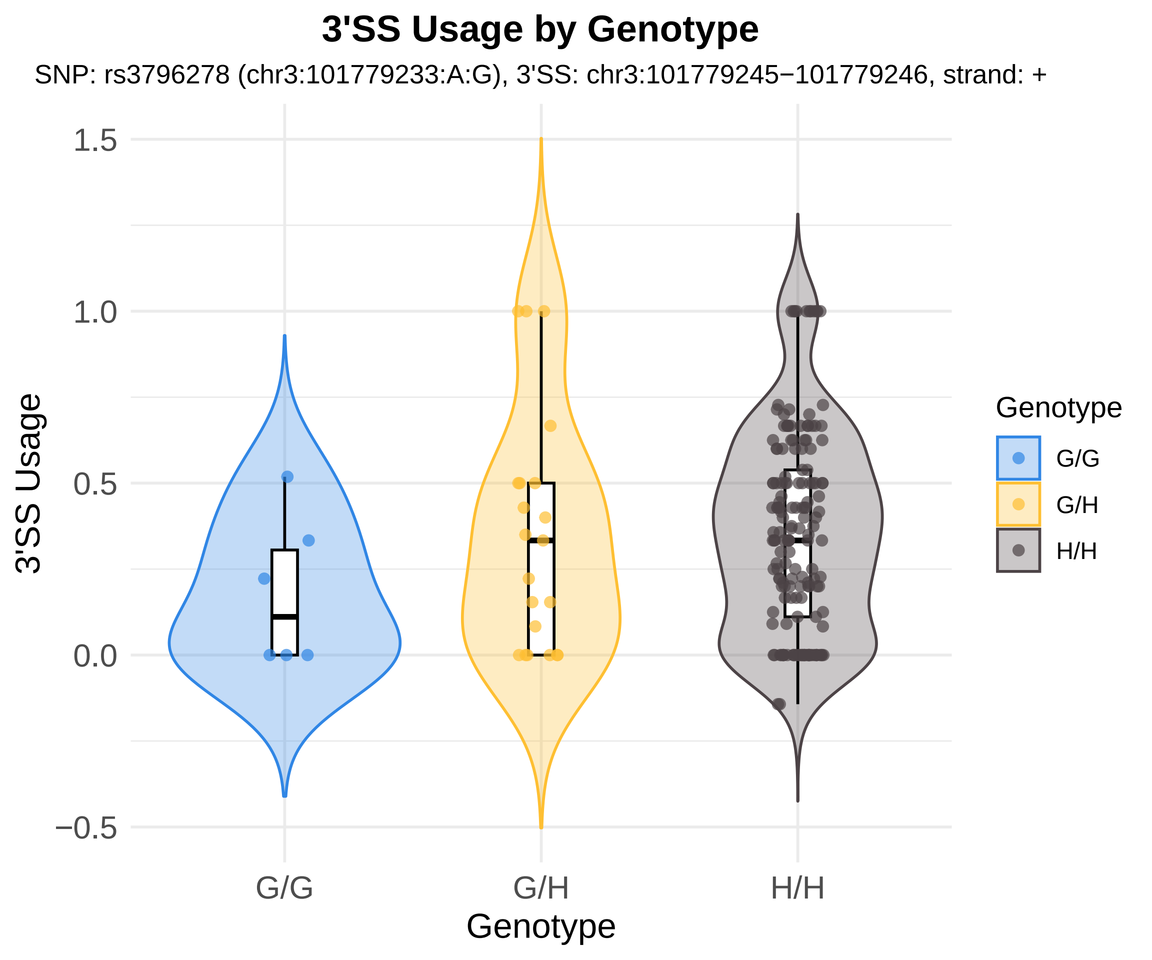

3'SS Usage by Genotype
SNP: rs3796278 (chr3:101779233:A:G), 3'SS: chr3:101779245−101779246, strand: +
3'SS Usage
1.5
1.0
0.5
0.0
−0.5
G/G
G/H
H/H
Genotype
Genotype
G/G
G/H
H/H

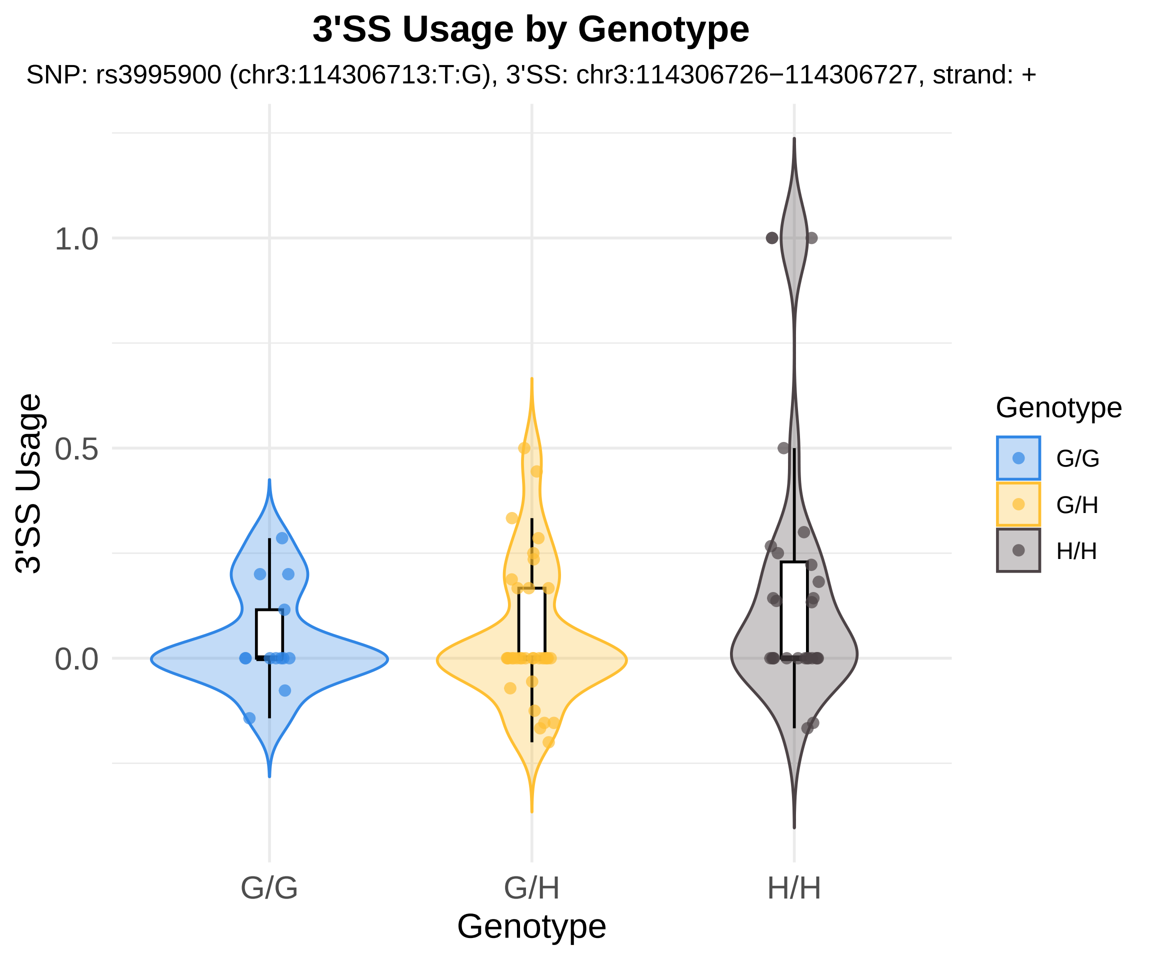
3'SS Usage by Genotype
SNP: rs3995900 (chr3:114306713:T:G), 3'SS: chr3:114306726−114306727, strand: +
3'SS Usage
1.0
0.5
0.0
G/G
G/H
H/H
Genotype
Genotype
G/G
G/H
H/H

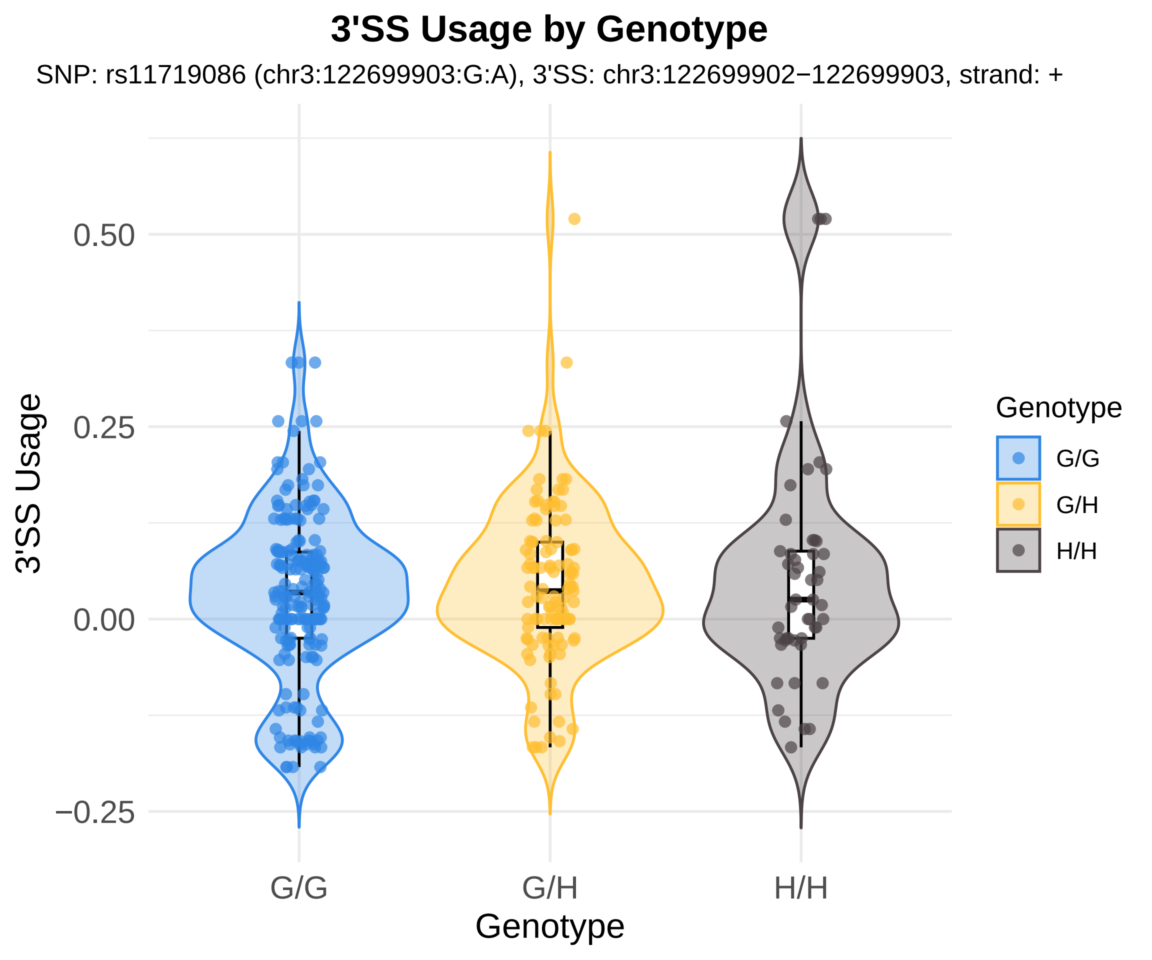
3'SS Usage by Genotype
SNP: rs11719086 (chr3:122699903:G:A), 3'SS: chr3:122699902−122699903, strand: +
3'SS Usage
0.50
0.25
0.00
−0.25
G/G
G/H
H/H
Genotype
Genotype
G/G
G/H
H/H

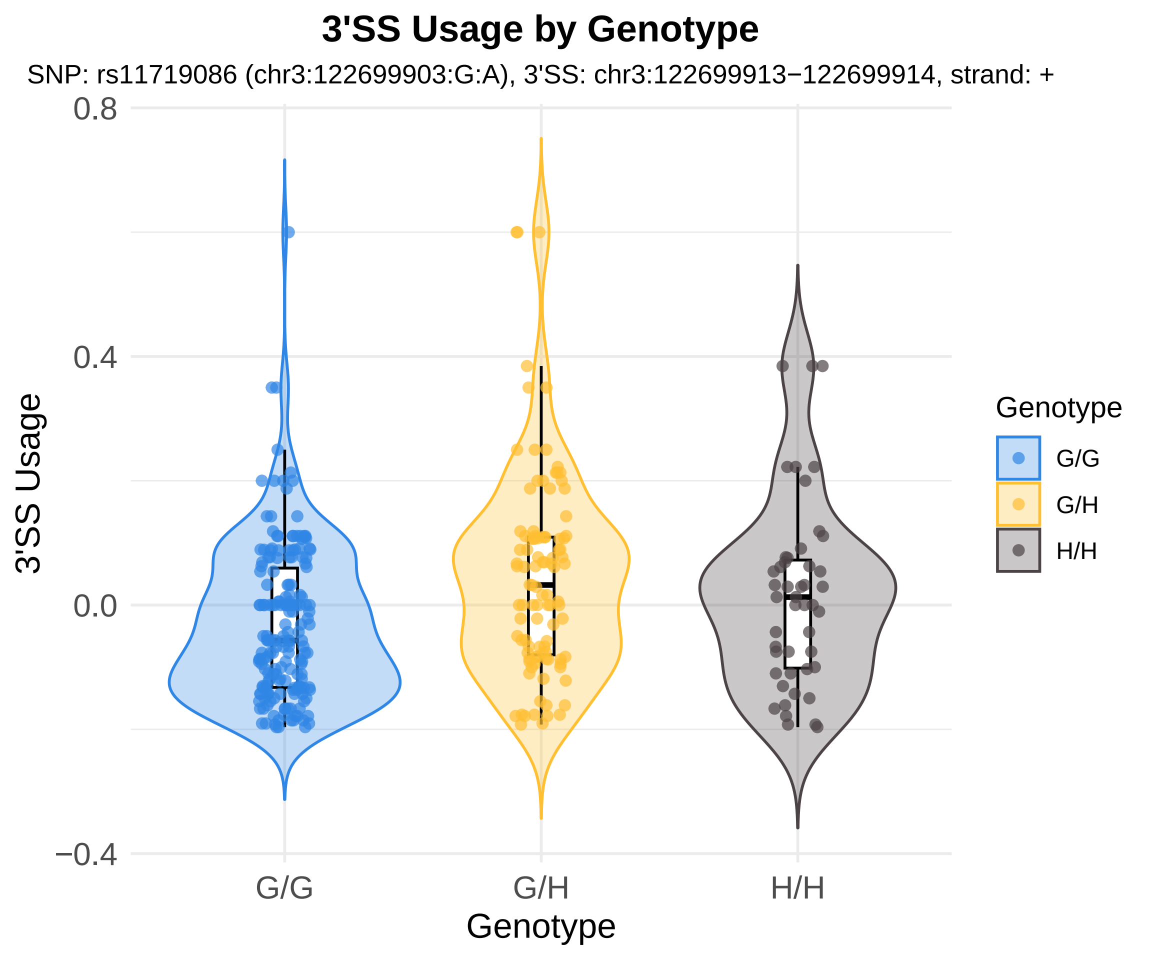

3'SS Usage by Genotype
SNP: rs11719086 (chr3:122699903:G:A), 3'SS: chr3:122699913−122699914, strand: +
0.8
0.4
0.0
−0.4
3'SS Usage
G/G
G/H
H/H
Genotype
Genotype
G/G
G/H
H/H

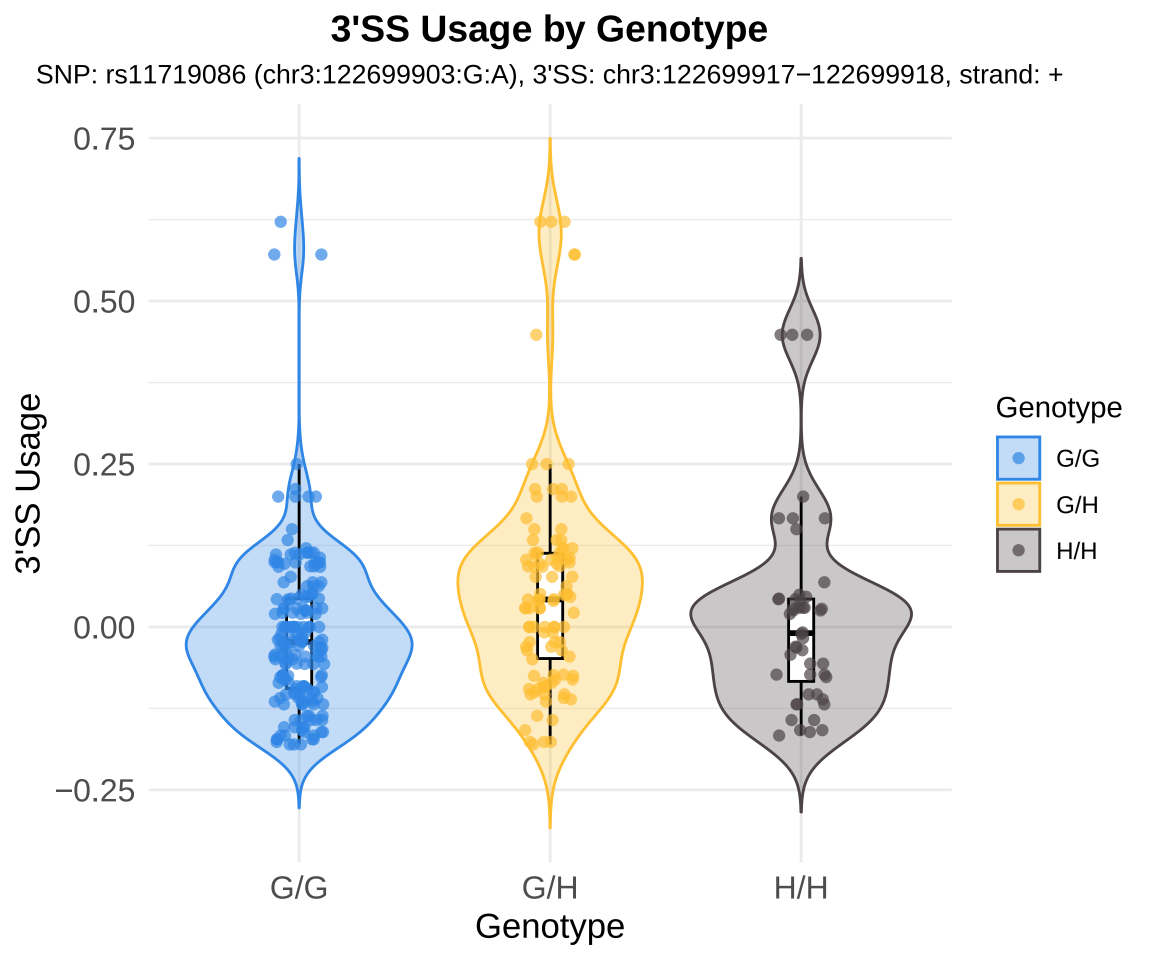
3'SS Usage by Genotype
SNP: rs11719086 (chr3:122699903:G:A), 3'SS: chr3:122699917−122699918, strand: +
3'SS Usage
0.75
0.50
0.25
0.00
−0.25
G/G
G/H
H/H
Genotype
Genotype
G/G
G/H
H/H

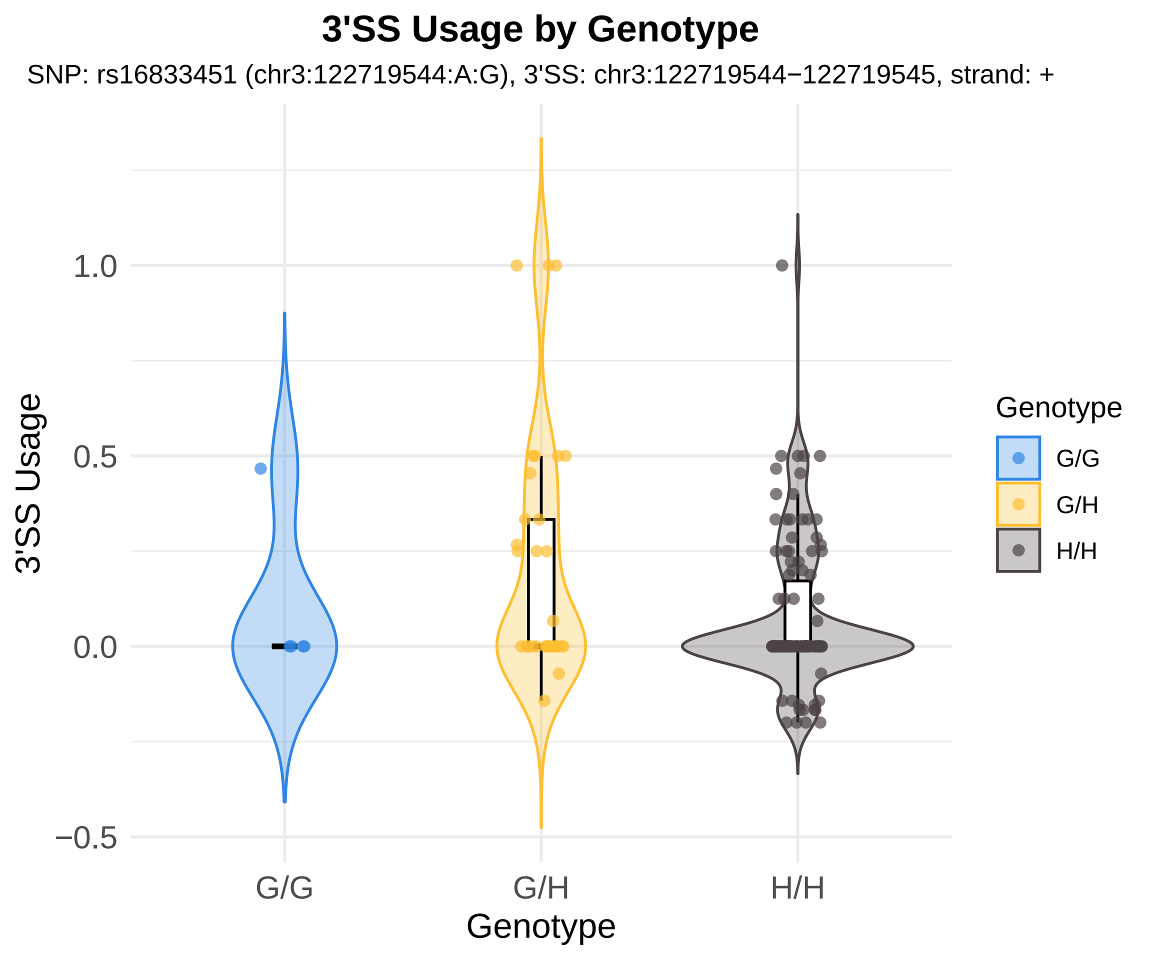

3'SS Usage by Genotype
SNP: rs16833451 (chr3:122719544:A:G), 3'SS: chr3:122719544−122719545, strand: +
3'SS Usage
1.0
0.5
0.0
−0.5
G/G
G/H
H/H
Genotype
Genotype
G/G
G/H
H/H

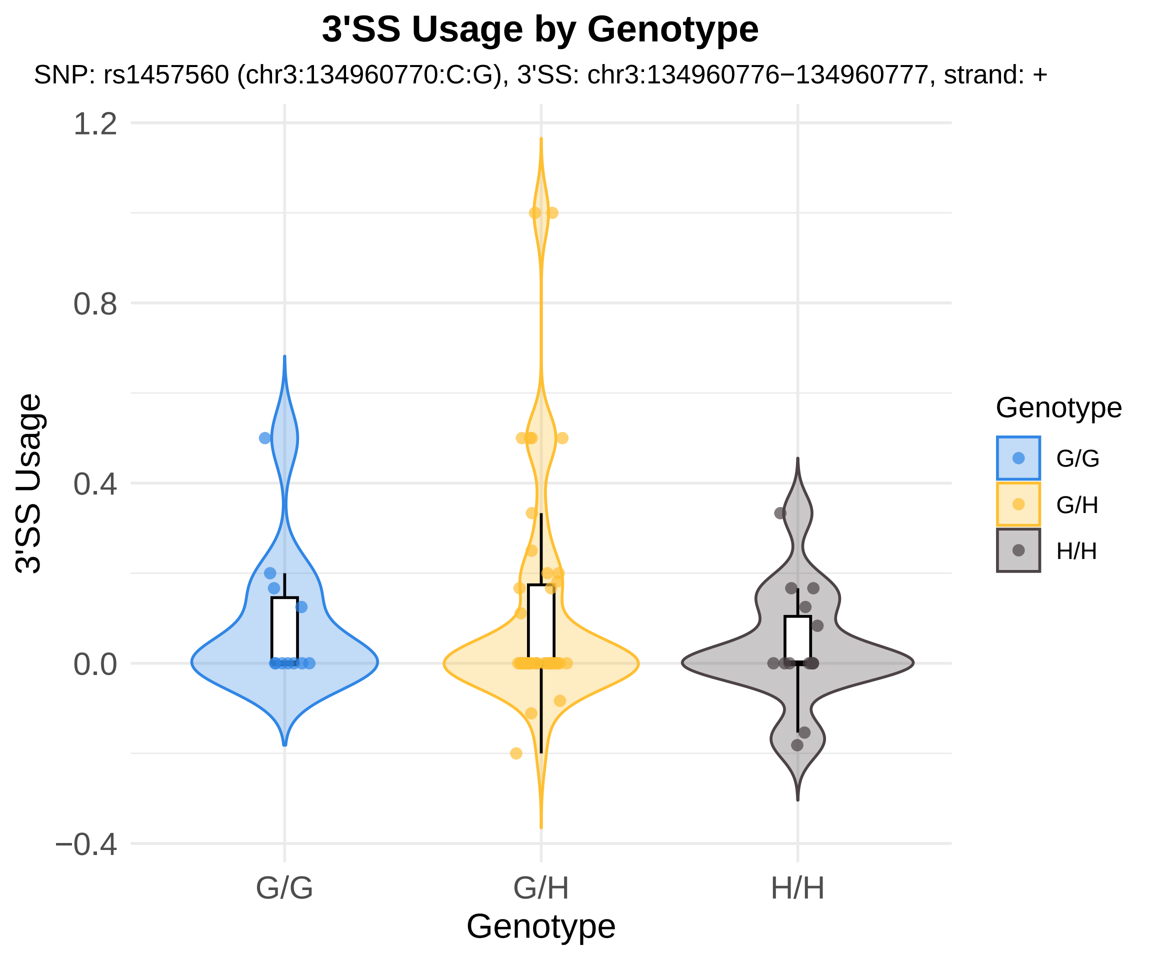

3'SS Usage by Genotype
SNP: rs1457560 (chr3:134960770:C:G), 3'SS: chr3:134960776−134960777, strand: +
3'SS Usage
1.2
0.8
0.4
0.0
−0.4
G/G
G/H
H/H
Genotype
Genotype
G/G
G/H
H/H

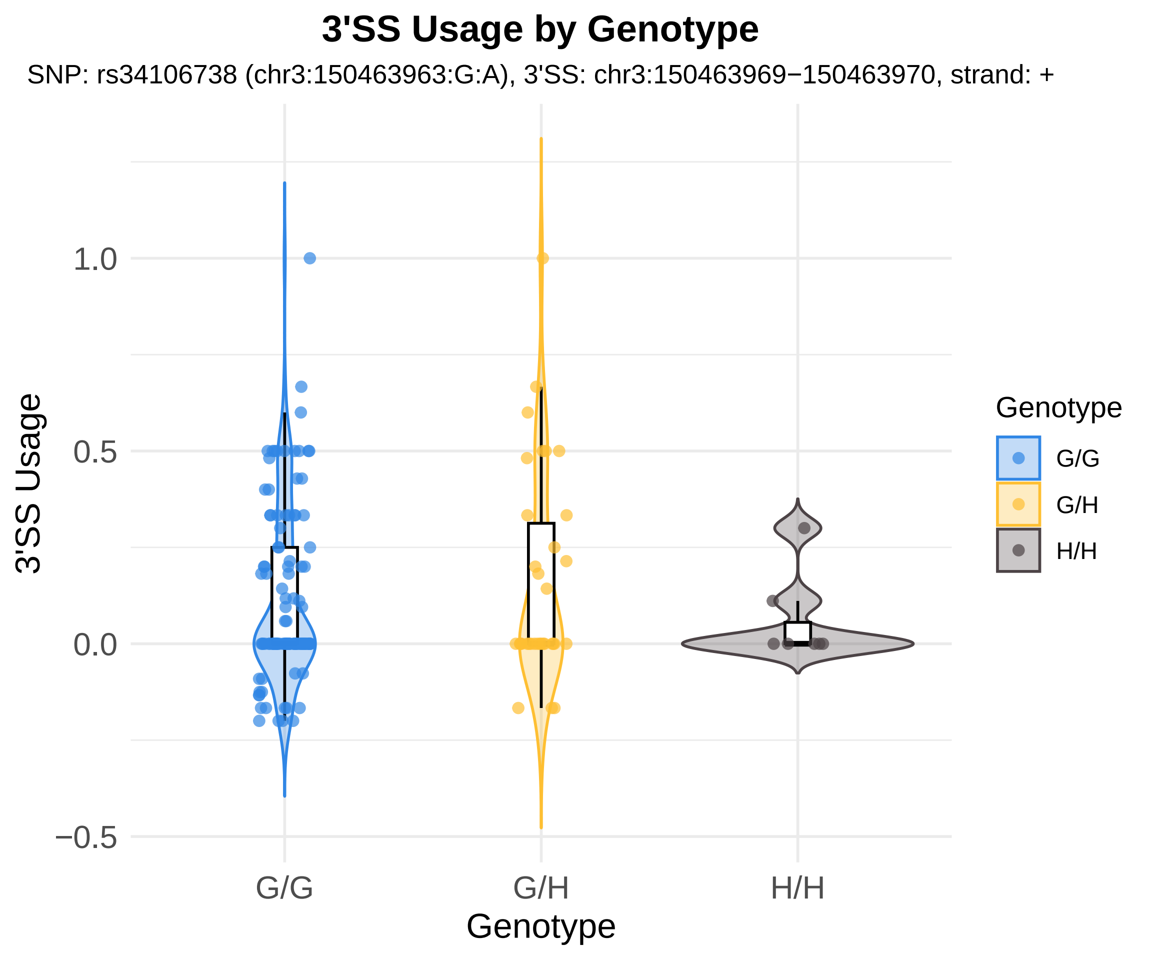

3'SS Usage by Genotype
SNP: rs34106738 (chr3:150463963:G:A), 3'SS: chr3:150463969−150463970, strand: +
3'SS Usage
1.0
0.5
0.0
−0.5
G/G
G/H
H/H
Genotype
Genotype
G/G
G/H
H/H

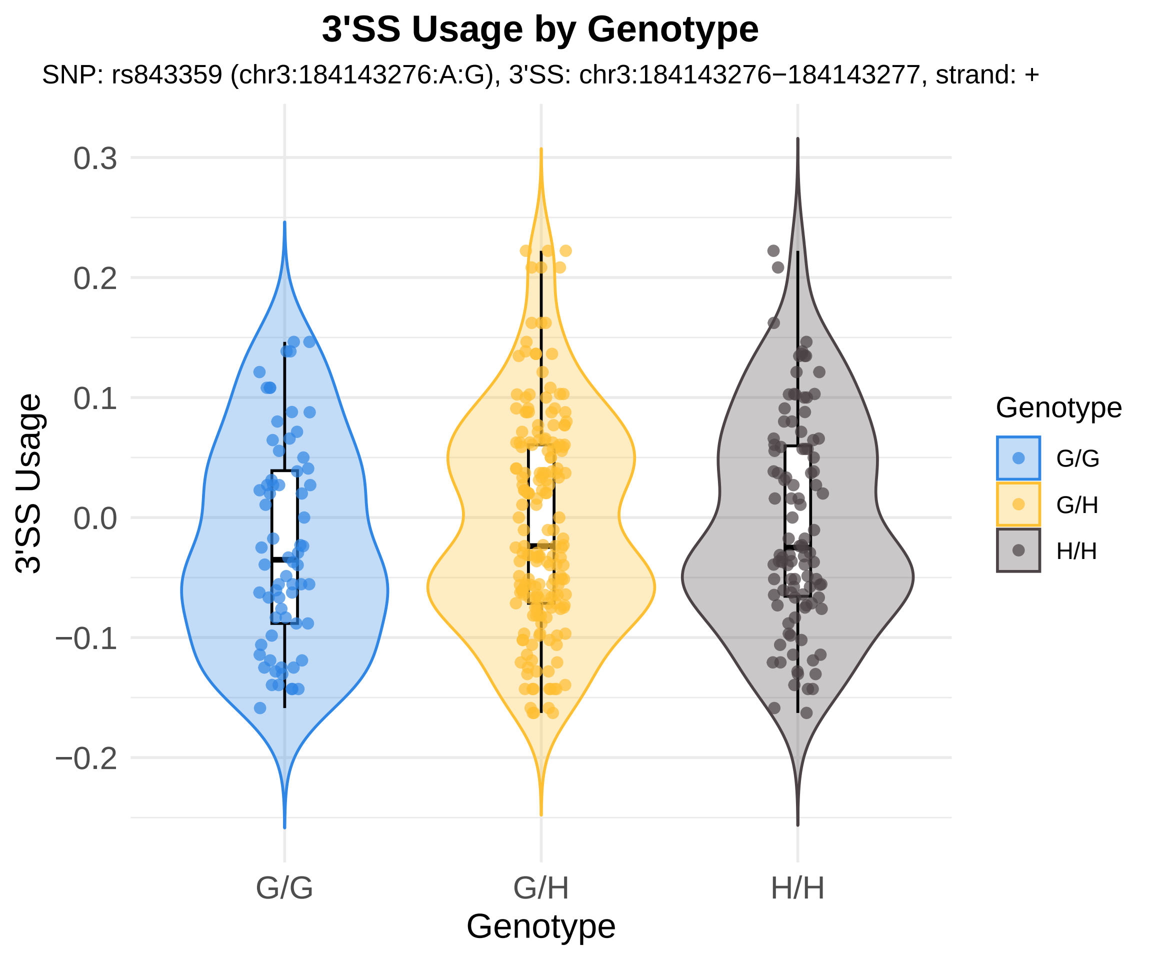

3'SS Usage by Genotype
SNP: rs843359 (chr3:184143276:A:G), 3'SS: chr3:184143276−184143277, strand: +
3'SS Usage
0.3
0.2
0.1
0.0
−0.1
−0.2
G/G
G/H
H/H
Genotype
Genotype
G/G
G/H
H/H

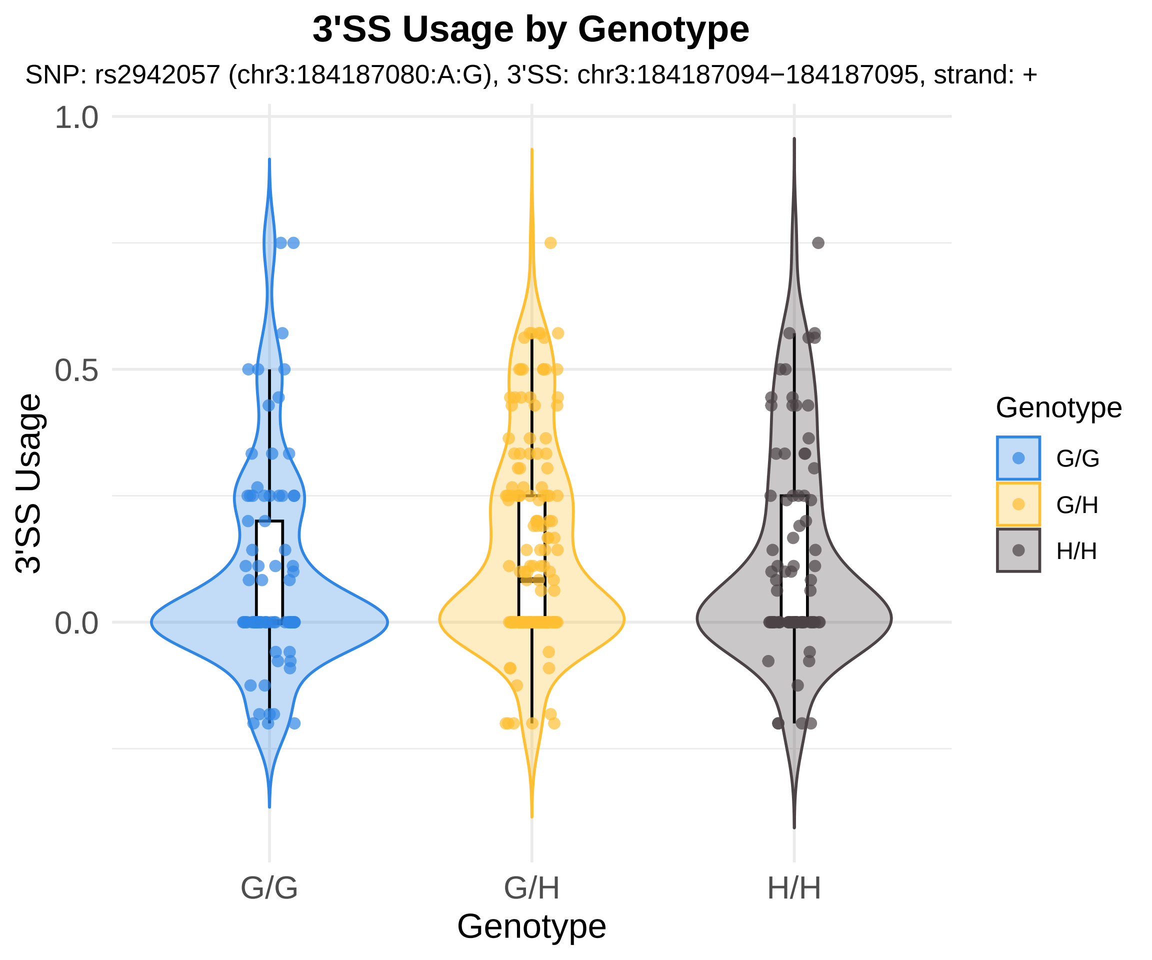
3'SS Usage by Genotype
SNP: rs2942057 (chr3:184187080:A:G), 3'SS: chr3:184187094−184187095, strand: +
1.0
0.5
0.0
3'SS Usage
G/G
G/H
H/H
Genotype
Genotype
G/G
G/H
H/H

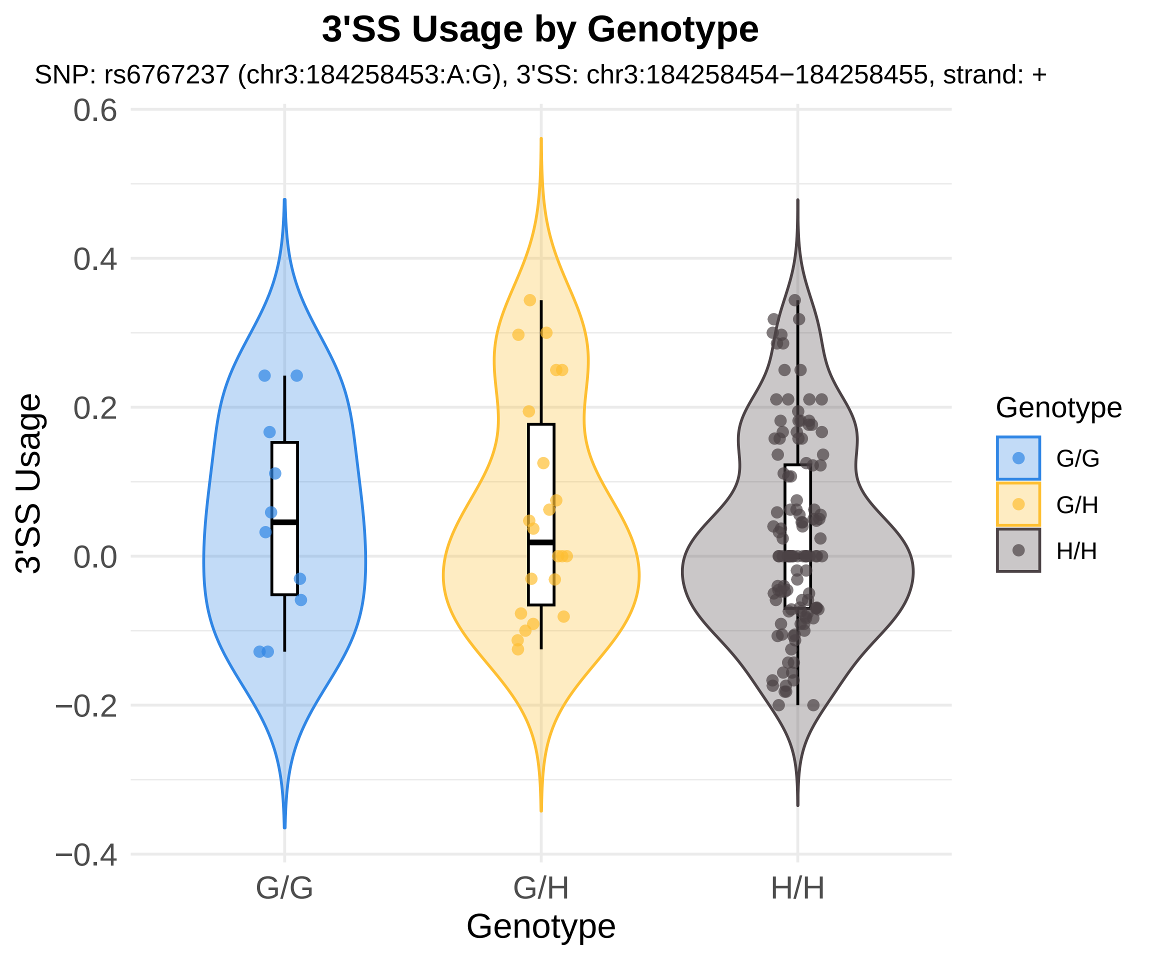

3'SS Usage by Genotype
SNP: rs6767237 (chr3:184258453:A:G), 3'SS: chr3:184258454−184258455, strand: +
3'SS Usage
0.6
0.4
0.2
0.0
−0.2
−0.4
G/G
G/H
H/H
Genotype
Genotype
G/G
G/H
H/H

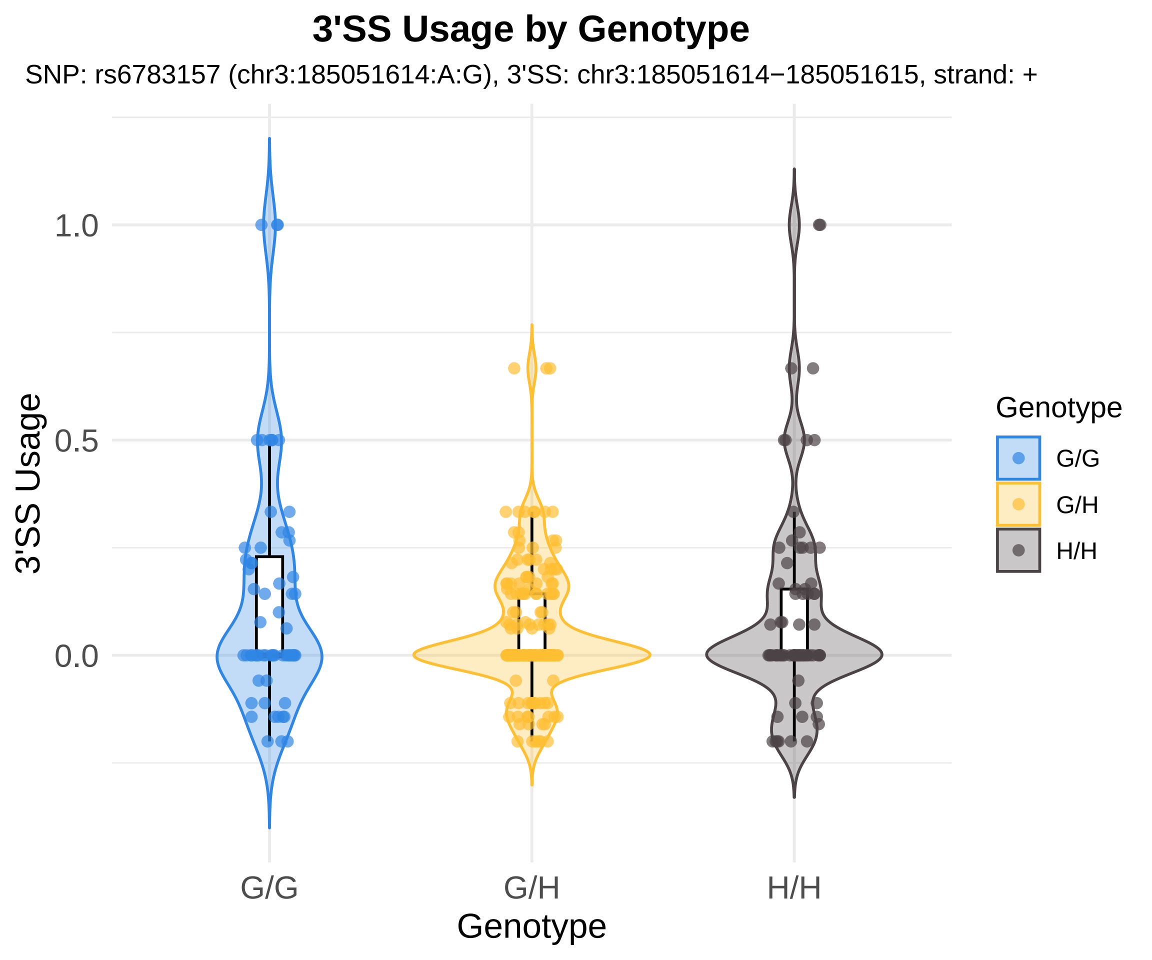
3'SS Usage by Genotype
SNP: rs6783157 (chr3:185051614:A:G), 3'SS: chr3:185051614−185051615, strand: +
1.0
0.5
0.0
3'SS Usage
G/G
G/H
H/H
Genotype
Genotype
G/G
G/H
H/H

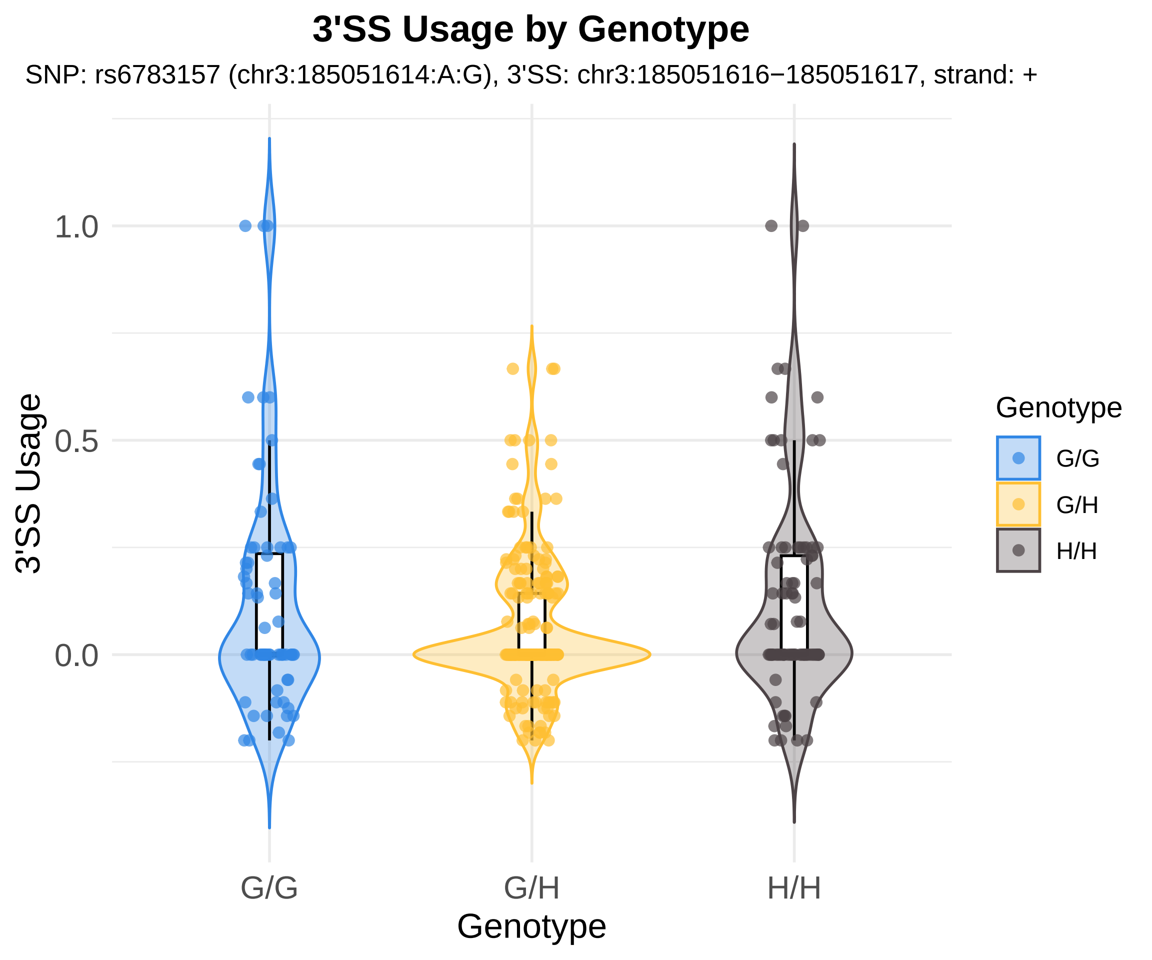
3'SS Usage by Genotype
SNP: rs6783157 (chr3:185051614:A:G), 3'SS: chr3:185051616−185051617, strand: +
3'SS Usage
1.0
0.5
0.0
G/G
G/H
H/H
Genotype
Genotype
G/G
G/H
H/H

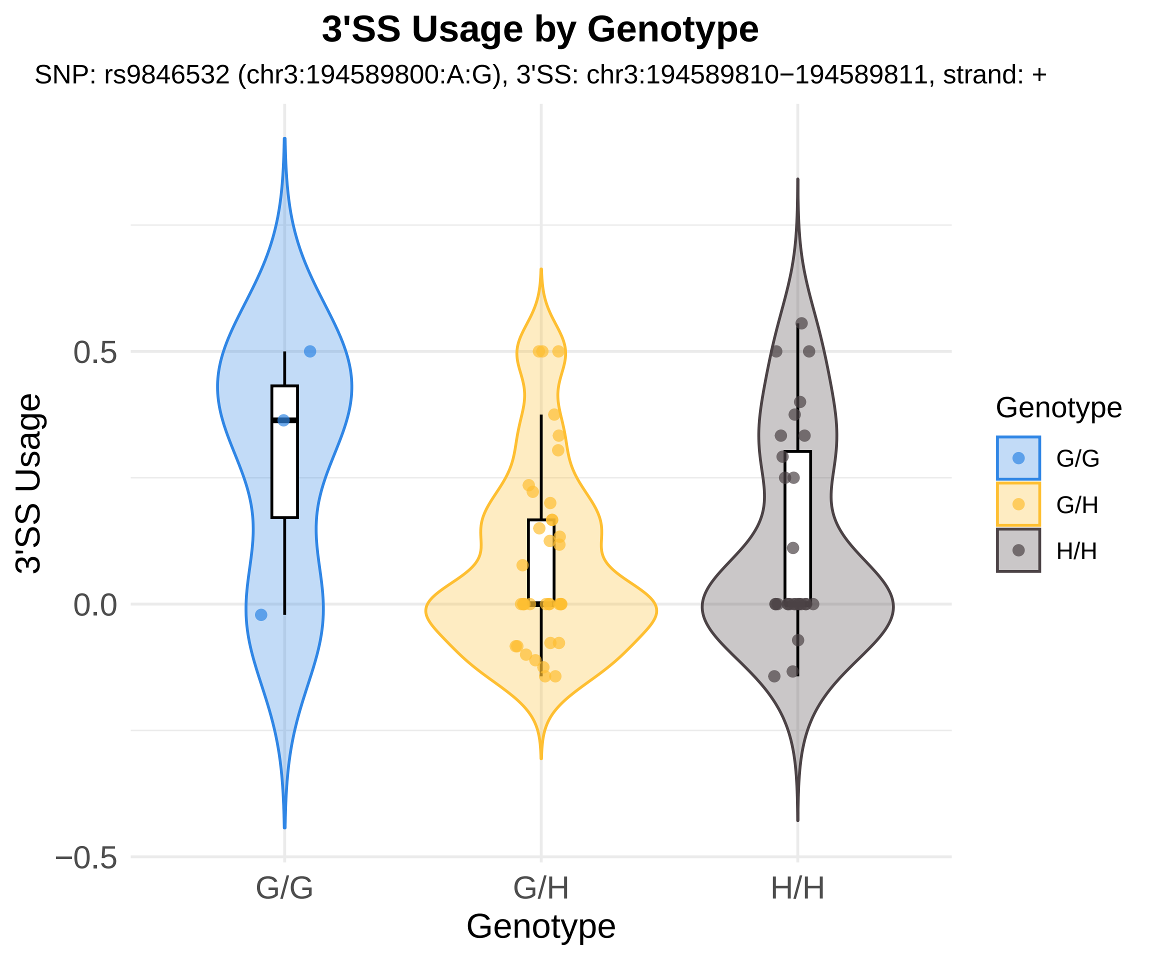

3'SS Usage by Genotype
SNP: rs9846532 (chr3:194589800:A:G), 3'SS: chr3:194589810−194589811, strand: +
3'SS Usage
0.5
0.0
−0.5
G/G
G/H
H/H
Genotype
Genotype
G/G
G/H
H/H

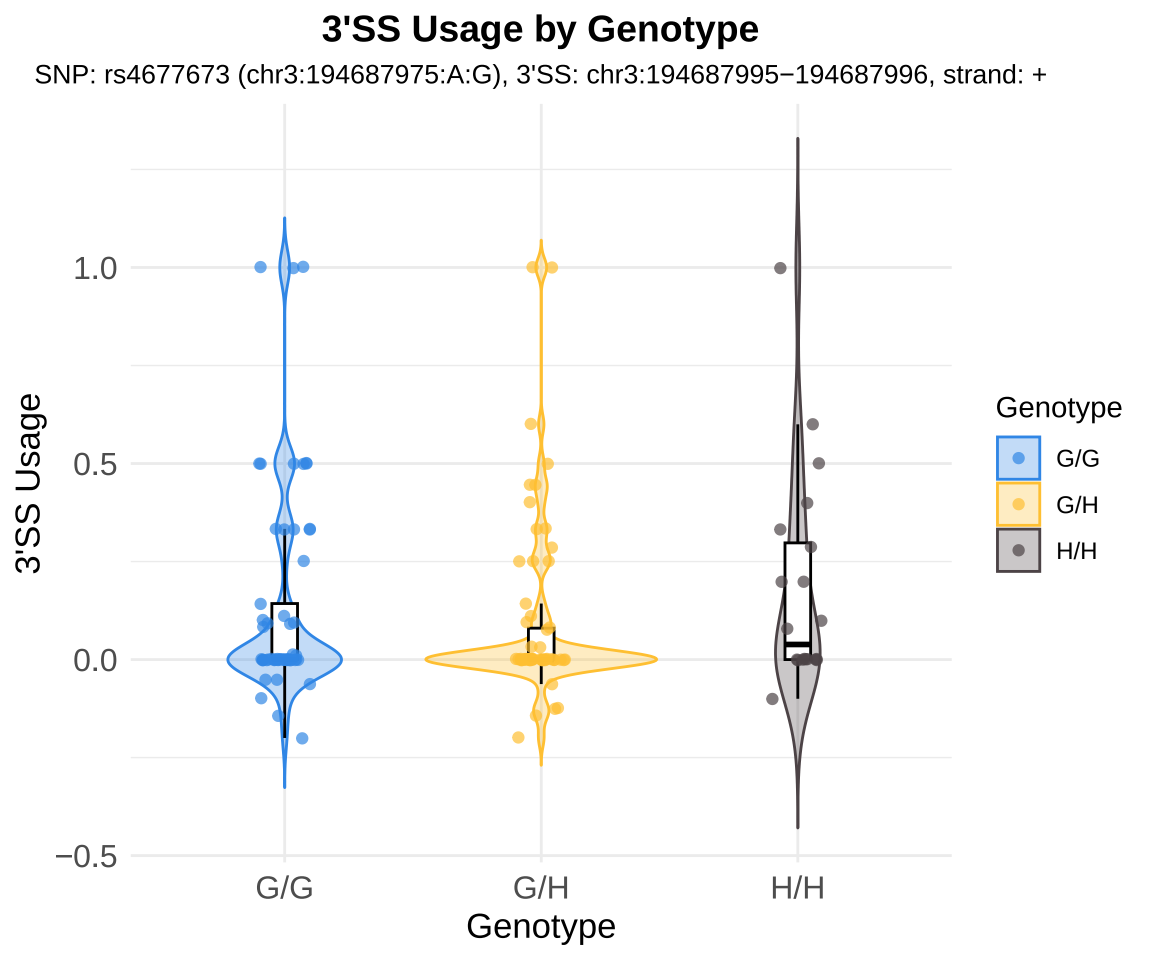
3'SS Usage by Genotype
SNP: rs4677673 (chr3:194687975:A:G), 3'SS: chr3:194687995−194687996, strand: +
3'SS Usage
1.0
0.5
0.0
−0.5
G/G
G/H
H/H
Genotype
Genotype
G/G
G/H
H/H

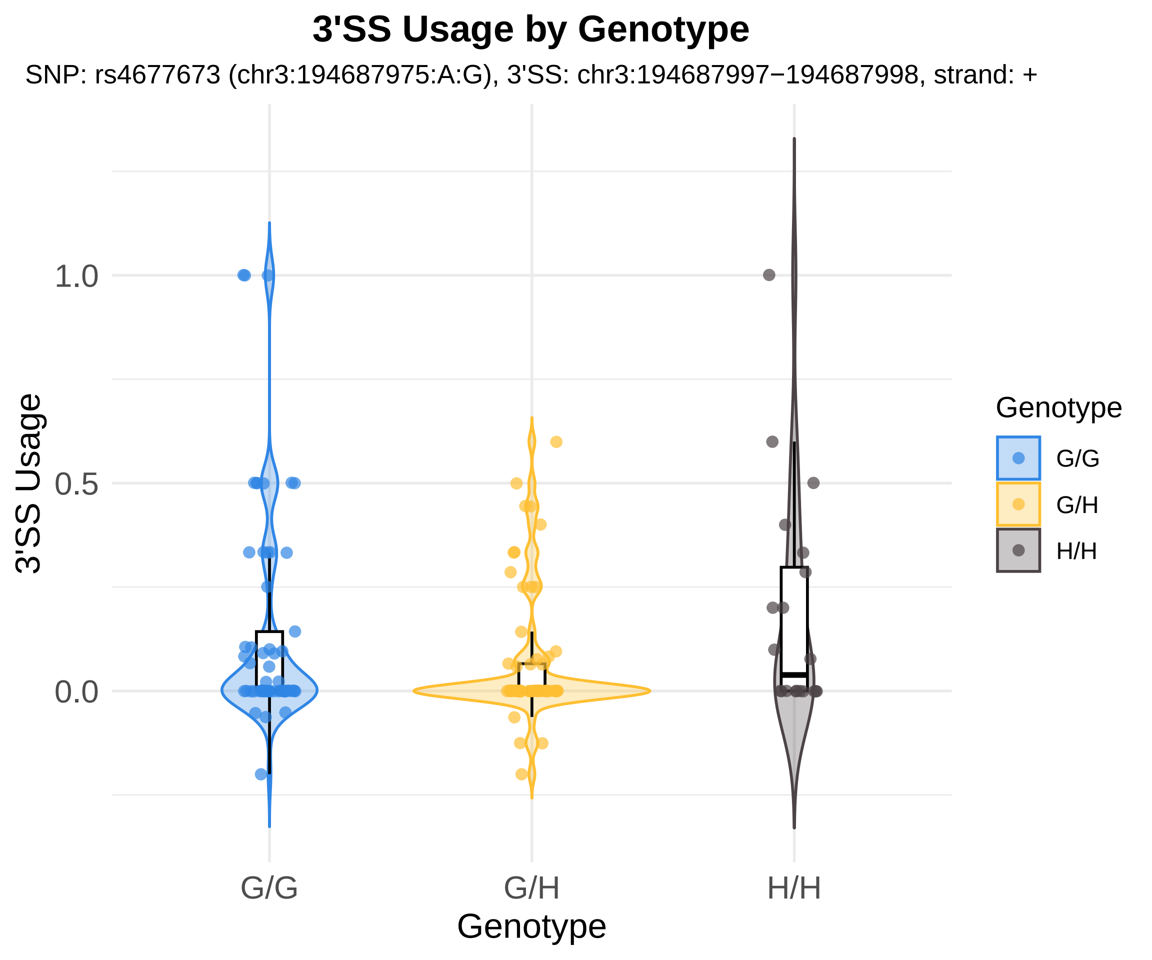

3'SS Usage by Genotype
SNP: rs4677673 (chr3:194687975:A:G), 3'SS: chr3:194687997−194687998, strand: +
3'SS Usage
1.0
0.5
0.0
G/G
G/H
H/H
Genotype
Genotype
G/G
G/H
H/H

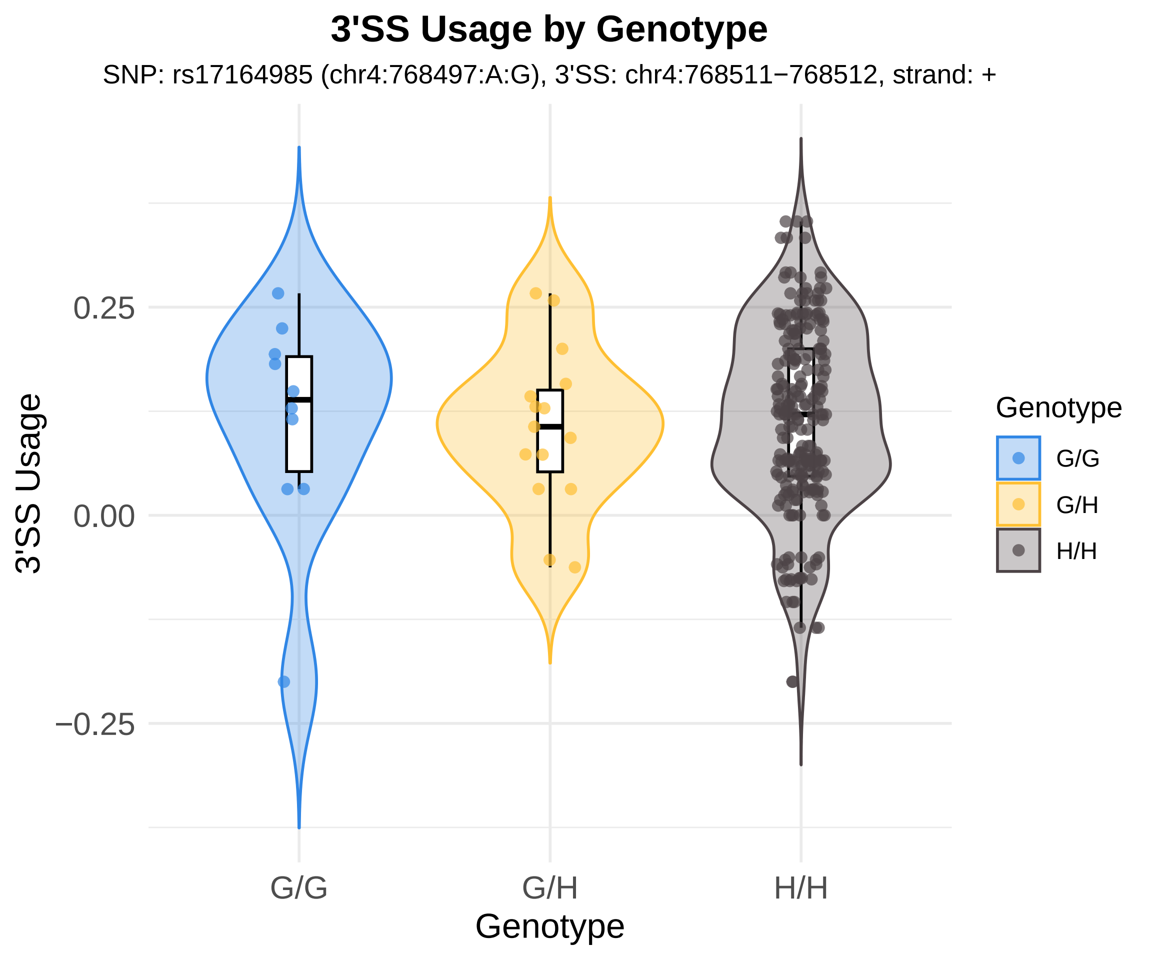
3'SS Usage by Genotype
SNP: rs17164985 (chr4:768497:A:G), 3'SS: chr4:768511−768512, strand: +
3'SS Usage
0.25
0.00
−0.25
G/G
G/H
H/H
Genotype
Genotype
G/G
G/H
H/H

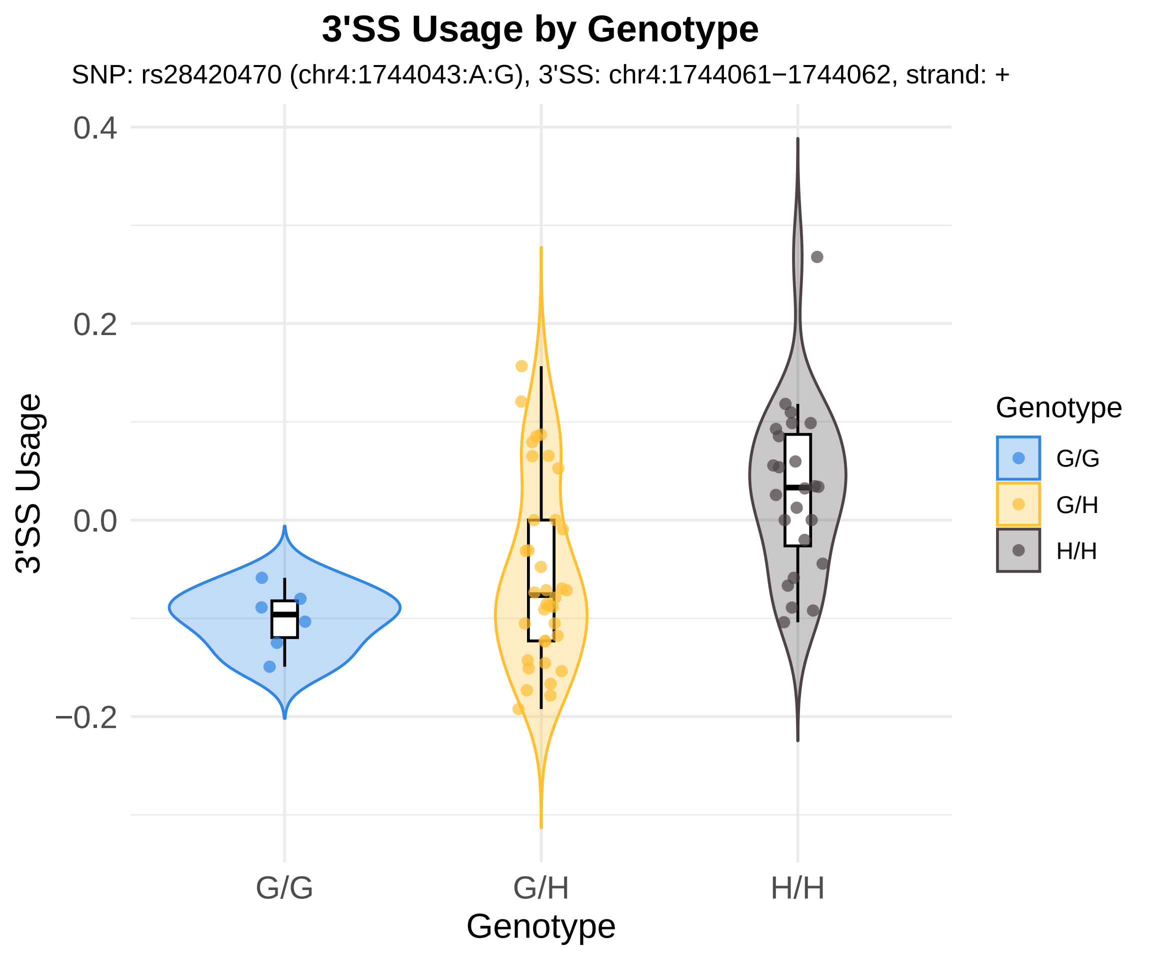

3'SS Usage by Genotype
SNP: rs28420470 (chr4:1744043:A:G), 3'SS: chr4:1744061−1744062, strand: +
0.4
0.2
0.0
−0.2
3'SS Usage
G/G
G/H
H/H
Genotype
Genotype
G/G
G/H
H/H

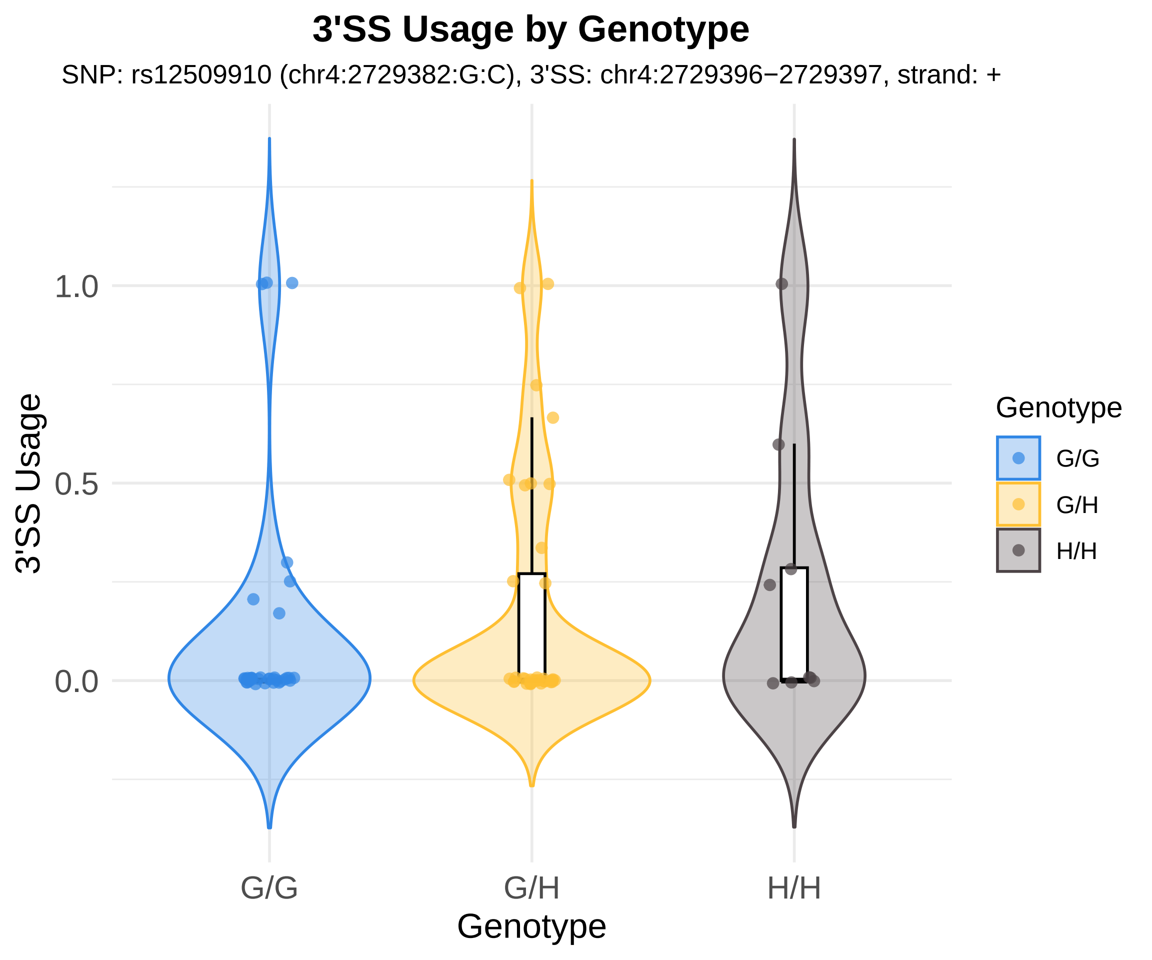
3'SS Usage by Genotype
SNP: rs12509910 (chr4:2729382:G:C), 3'SS: chr4:2729396−2729397, strand: +
3'SS Usage
1.0
0.5
0.0
G/G
G/H
H/H
Genotype
Genotype
G/G
G/H
H/H

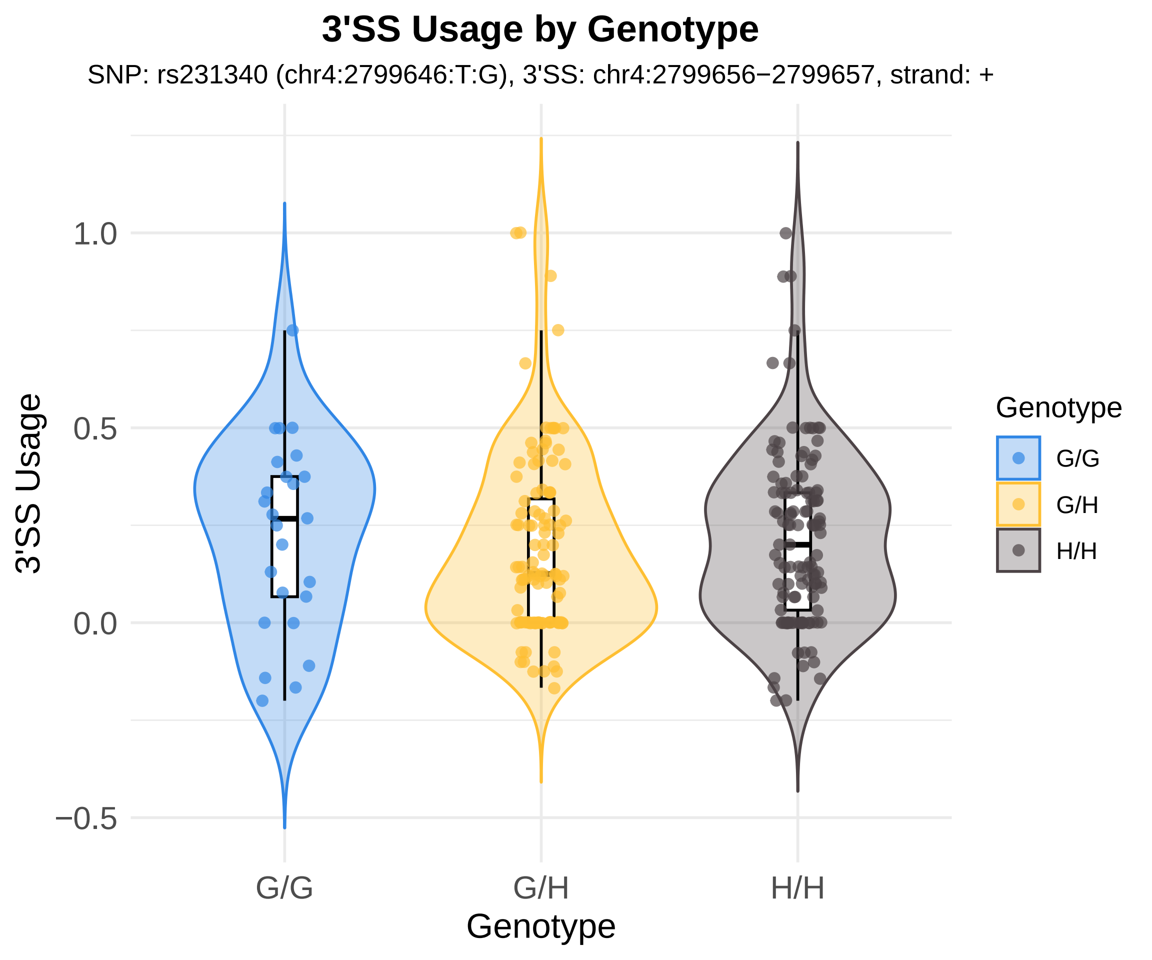

3'SS Usage by Genotype
SNP: rs231340 (chr4:2799646:T:G), 3'SS: chr4:2799656−2799657, strand: +
3'SS Usage
1.0
0.5
0.0
−0.5
G/G
G/H
H/H
Genotype
Genotype
G/G
G/H
H/H

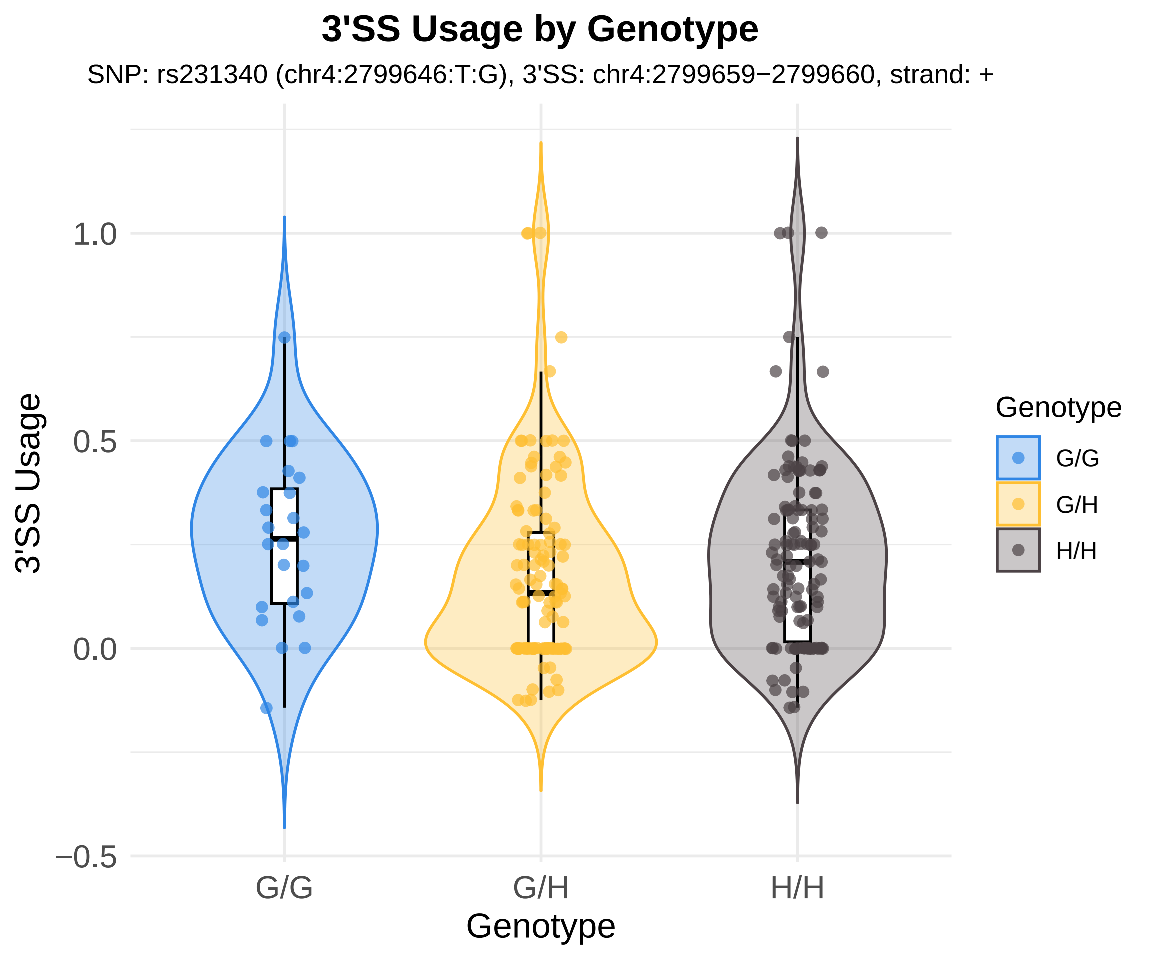
3'SS Usage by Genotype
SNP: rs231340 (chr4:2799646:T:G), 3'SS: chr4:2799659−2799660, strand: +
3'SS Usage
1.0
0.5
0.0
−0.5
G/G
G/H
H/H
Genotype
Genotype
G/G
G/H
H/H

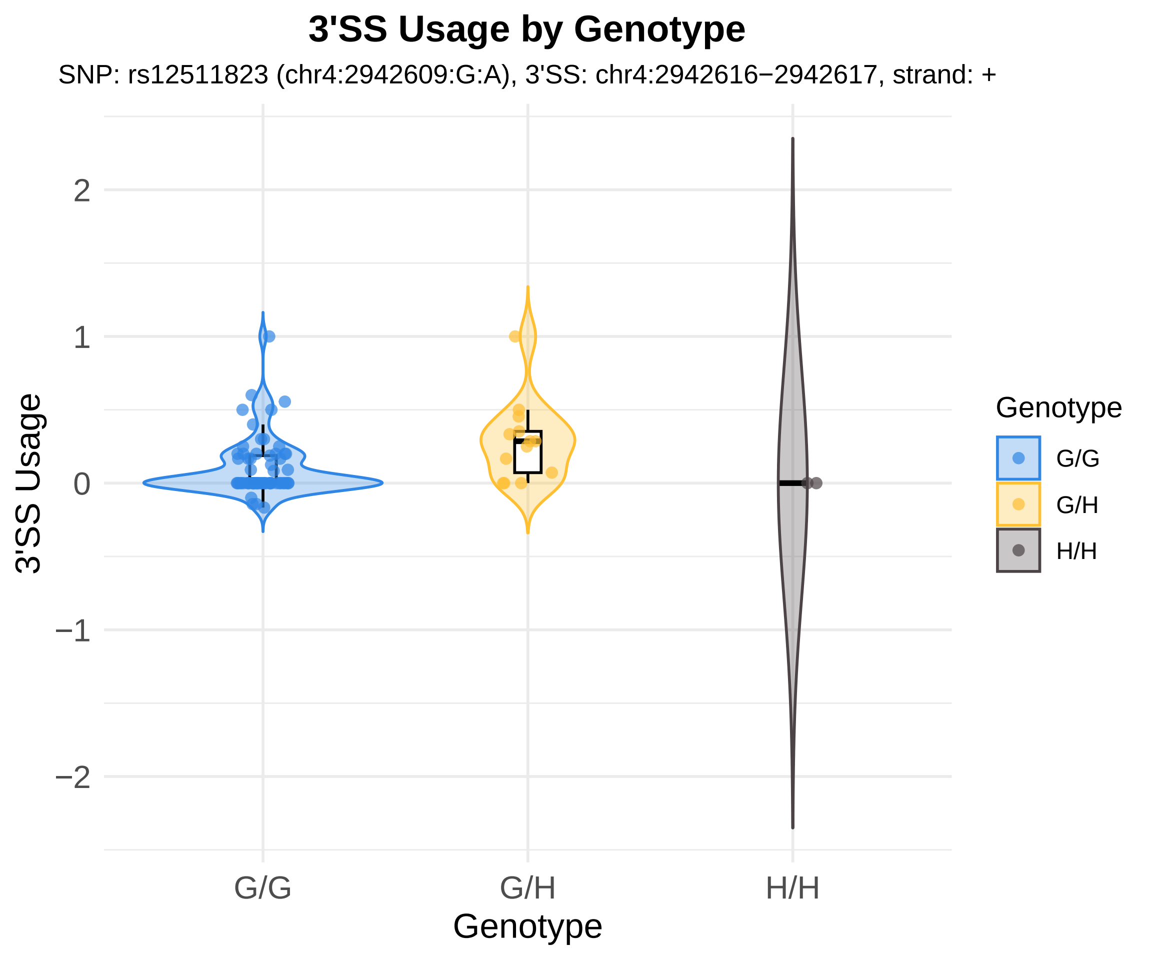
3'SS Usage by Genotype
SNP: rs12511823 (chr4:2942609:G:A), 3'SS: chr4:2942616−2942617, strand: +
3'SS Usage
2
1
0
−1
−2
G/G
G/H
H/H
Genotype
Genotype
G/G
G/H
H/H

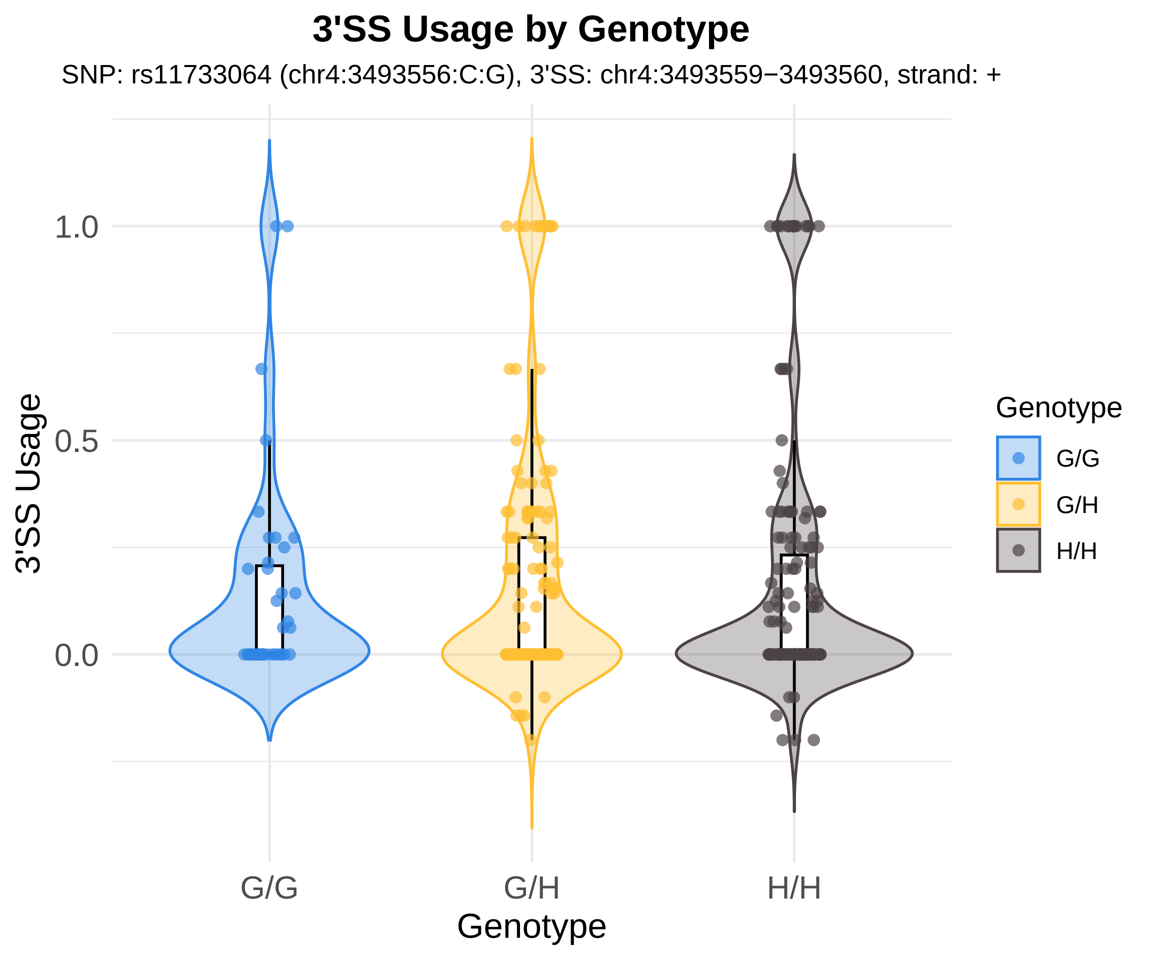
3'SS Usage by Genotype
SNP: rs11733064 (chr4:3493556:C:G), 3'SS: chr4:3493559−3493560, strand: +
3'SS Usage
1.0
0.5
0.0
G/G
G/H
H/H
Genotype
Genotype
G/G
G/H
H/H

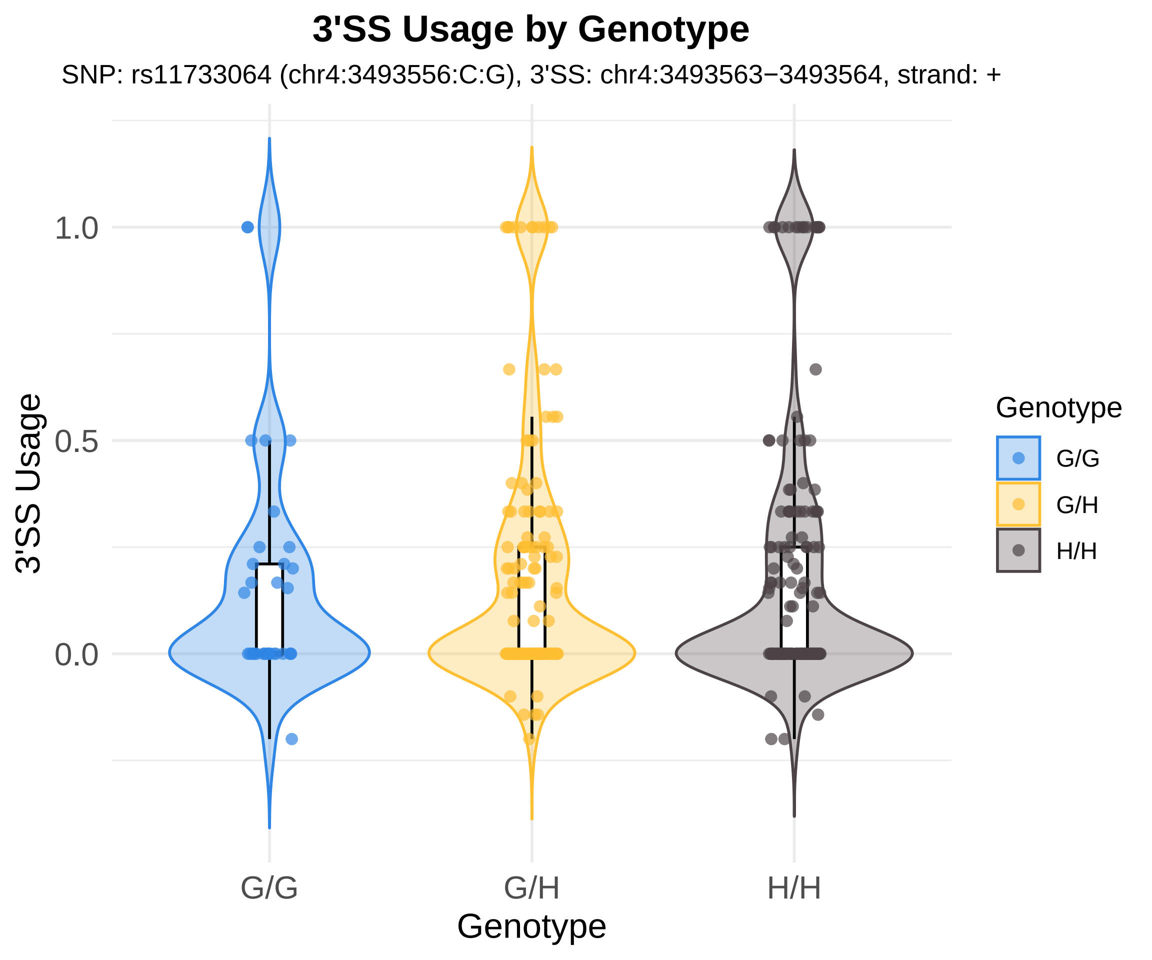
3'SS Usage by Genotype
SNP: rs11733064 (chr4:3493556:C:G), 3'SS: chr4:3493563−3493564, strand: +
3'SS Usage
1.0
0.5
0.0
G/G
G/H
H/H
Genotype
Genotype
G/G
G/H
H/H

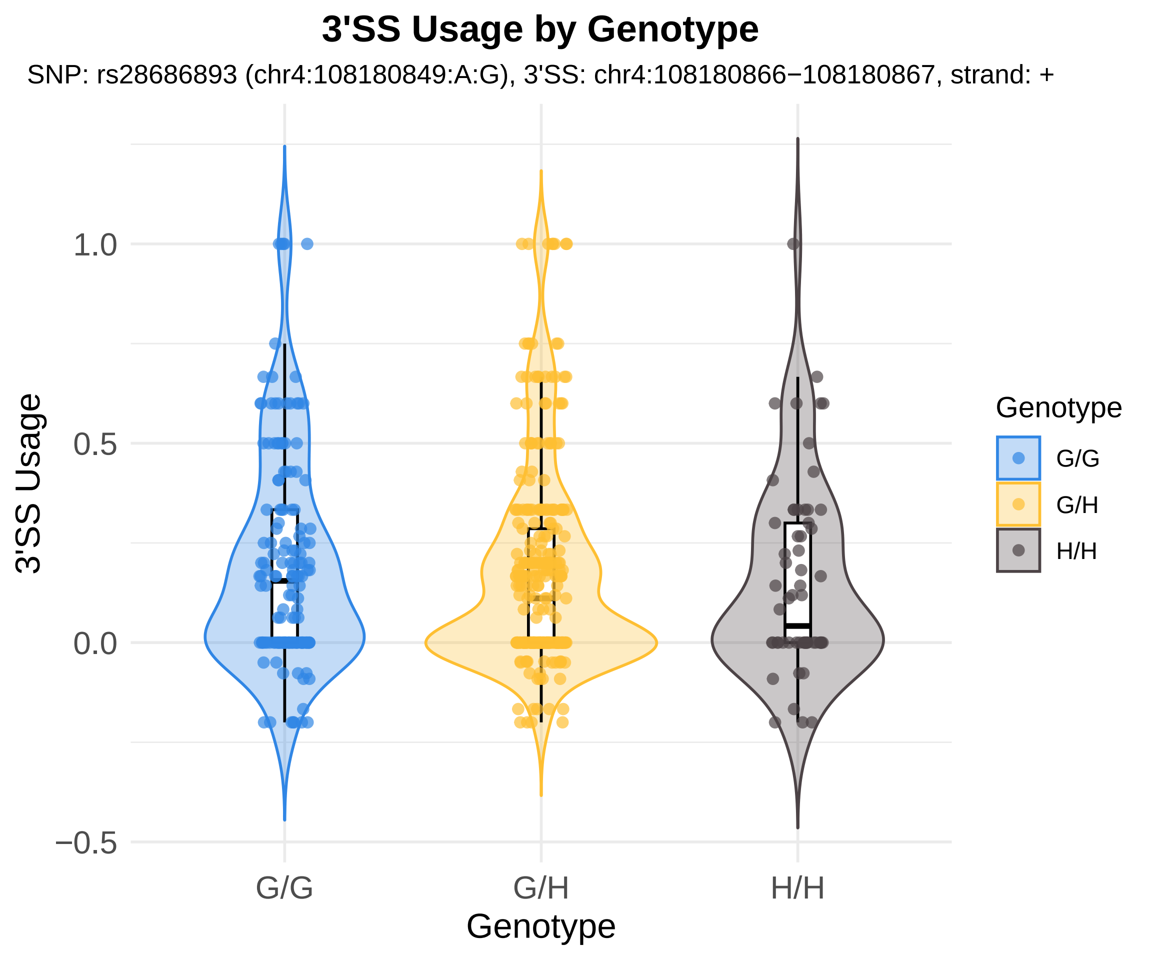
3'SS Usage by Genotype
SNP: rs28686893 (chr4:108180849:A:G), 3'SS: chr4:108180866−108180867, strand: +
3'SS Usage
1.0
0.5
0.0
−0.5
G/G
G/H
H/H
Genotype
Genotype
G/G
G/H
H/H

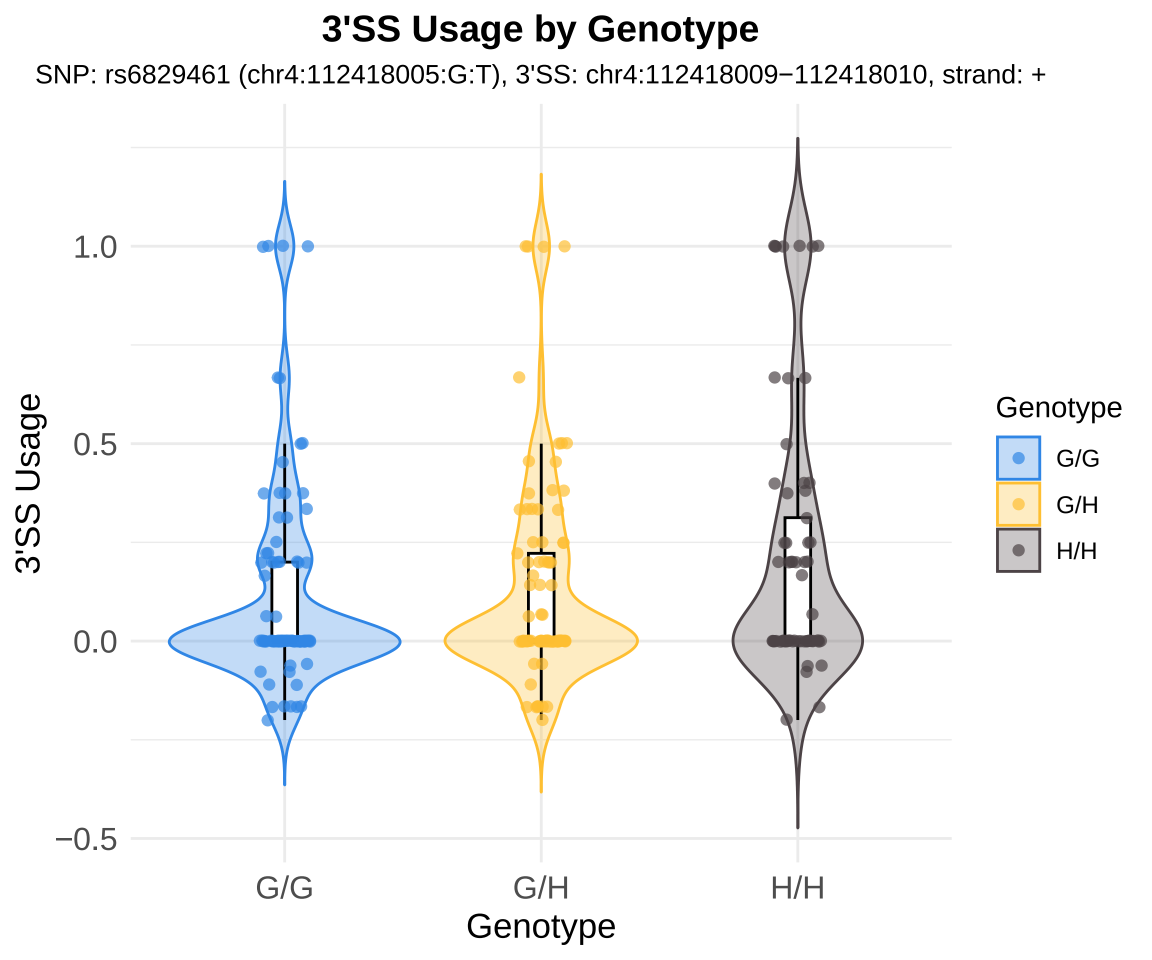
3'SS Usage by Genotype
SNP: rs6829461 (chr4:112418005:G:T), 3'SS: chr4:112418009−112418010, strand: +
3'SS Usage
1.0
0.5
0.0
−0.5
G/G
G/H
H/H
Genotype
Genotype
G/G
G/H
H/H

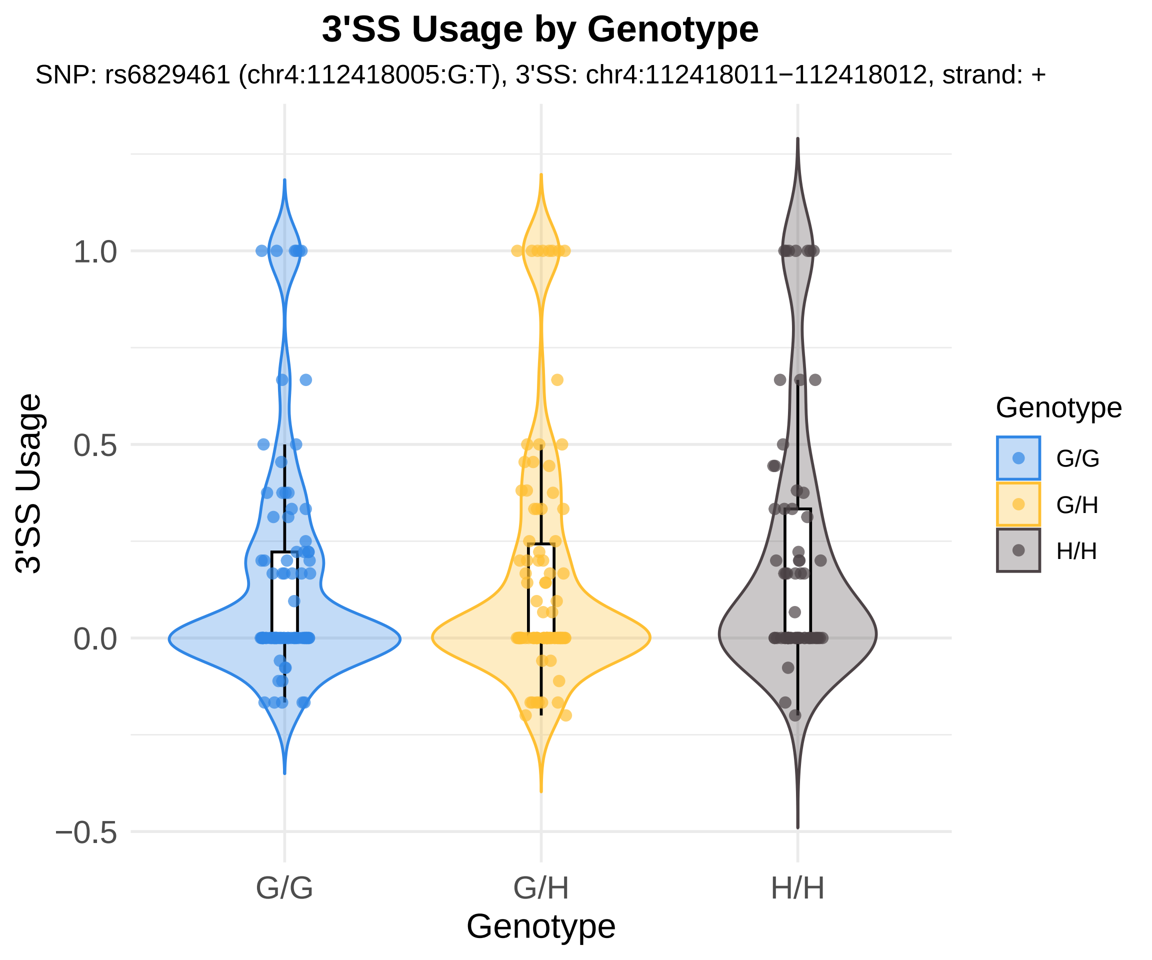

3'SS Usage by Genotype
SNP: rs6829461 (chr4:112418005:G:T), 3'SS: chr4:112418011−112418012, strand: +
3'SS Usage
1.0
0.5
0.0
−0.5
G/G
G/H
H/H
Genotype
Genotype
G/G
G/H
H/H

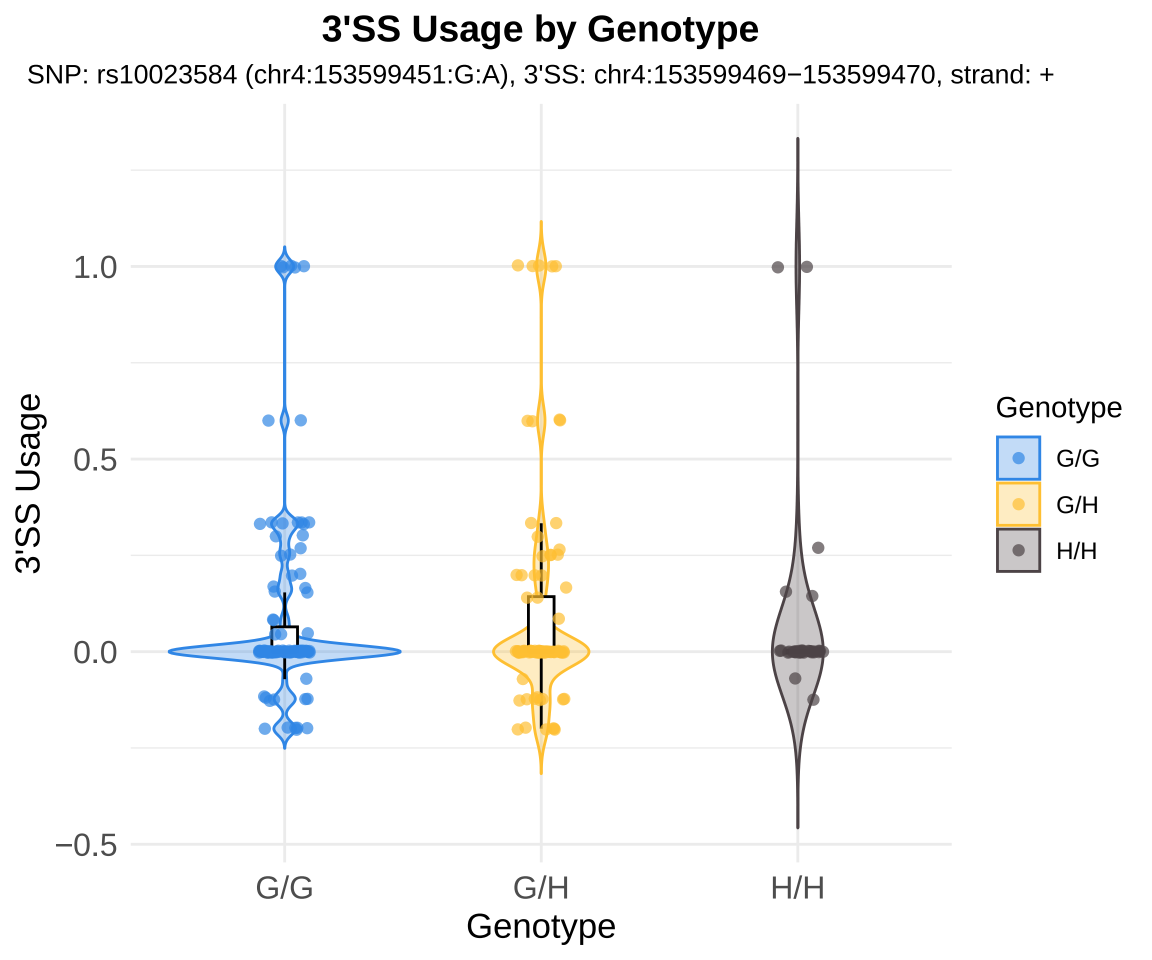
3'SS Usage by Genotype
SNP: rs10023584 (chr4:153599451:G:A), 3'SS: chr4:153599469−153599470, strand: +
3'SS Usage
1.0
0.5
0.0
−0.5
G/G
G/H
H/H
Genotype
Genotype
G/G
G/H
H/H

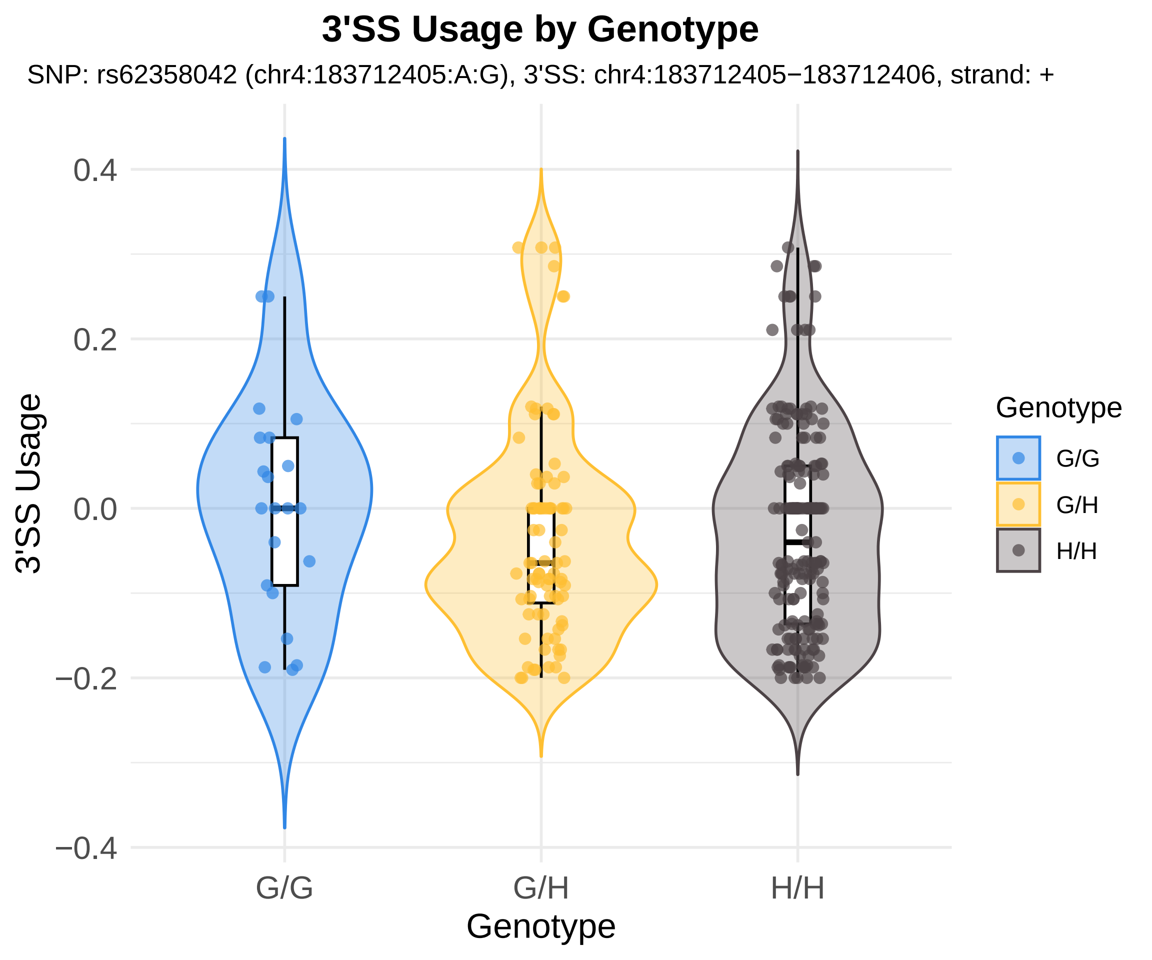
3'SS Usage by Genotype
SNP: rs62358042 (chr4:183712405:A:G), 3'SS: chr4:183712405−183712406, strand: +
0.4
0.2
0.0
−0.2
−0.4
3'SS Usage
G/G
G/H
H/H
Genotype
Genotype
G/G
G/H
H/H

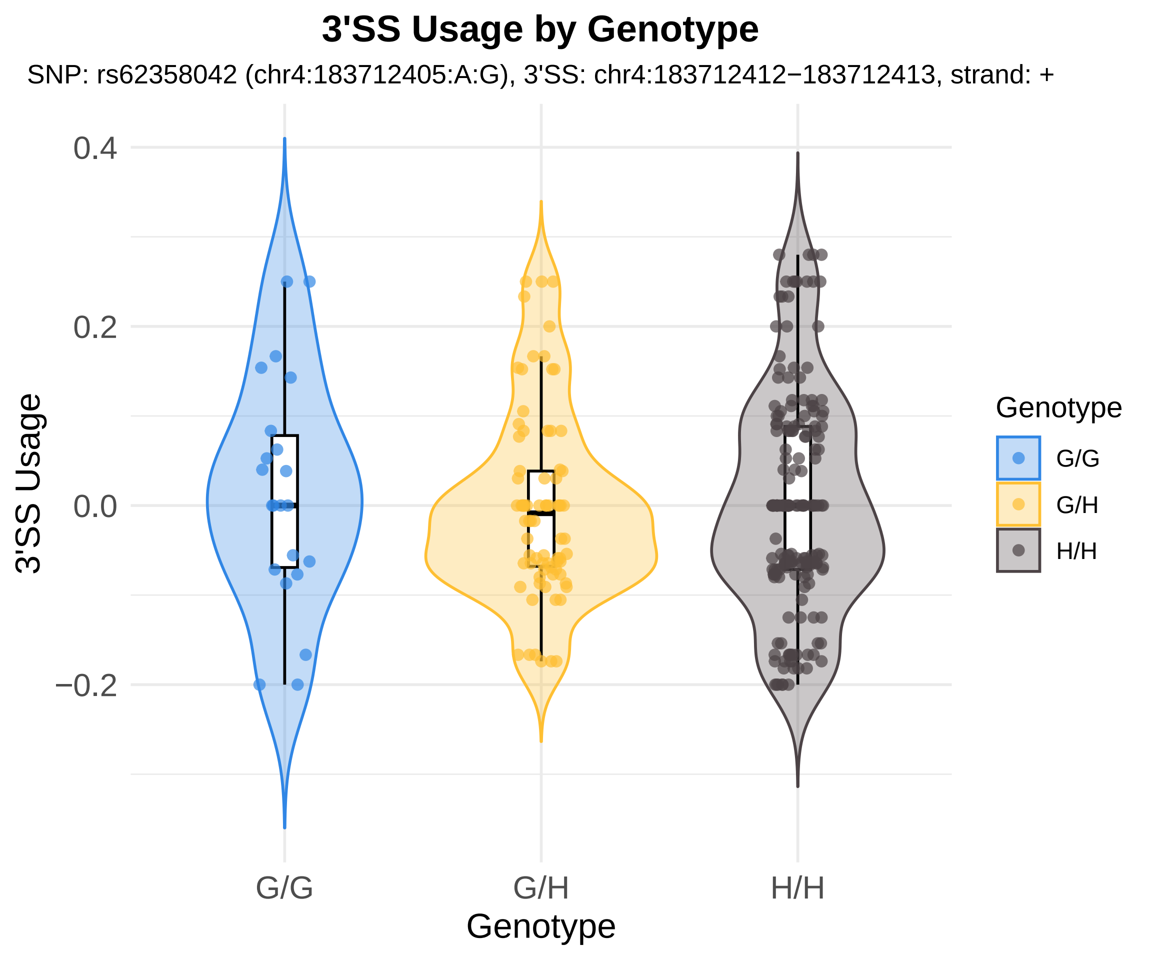

3'SS Usage by Genotype
SNP: rs62358042 (chr4:183712405:A:G), 3'SS: chr4:183712412−183712413, strand: +
0.4
0.2
0.0
−0.2
3'SS Usage
G/G
G/H
H/H
Genotype
Genotype
G/G
G/H
H/H

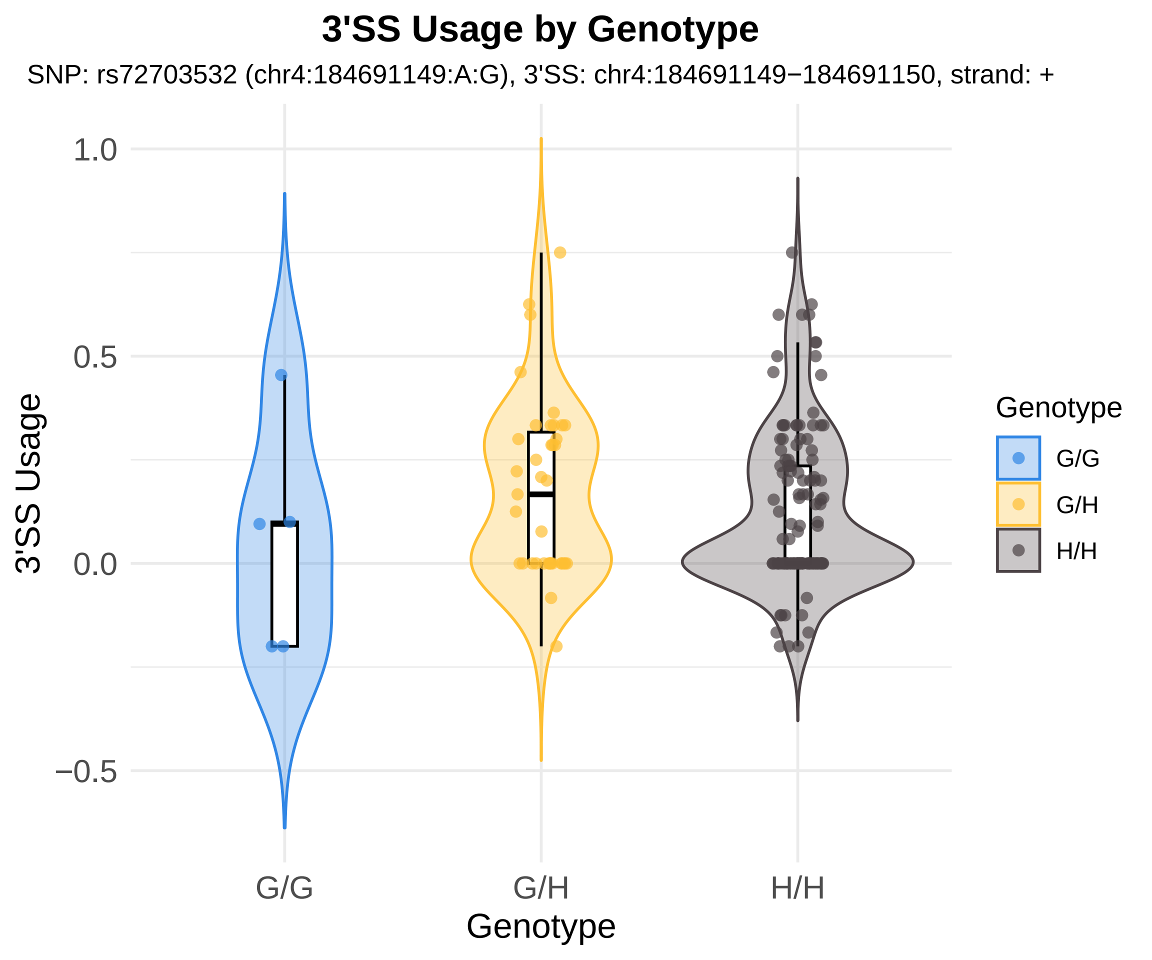
3'SS Usage by Genotype
SNP: rs72703532 (chr4:184691149:A:G), 3'SS: chr4:184691149−184691150, strand: +
1.0
0.5
0.0
−0.5
3'SS Usage
G/G
G/H
H/H
Genotype
Genotype
G/G
G/H
H/H

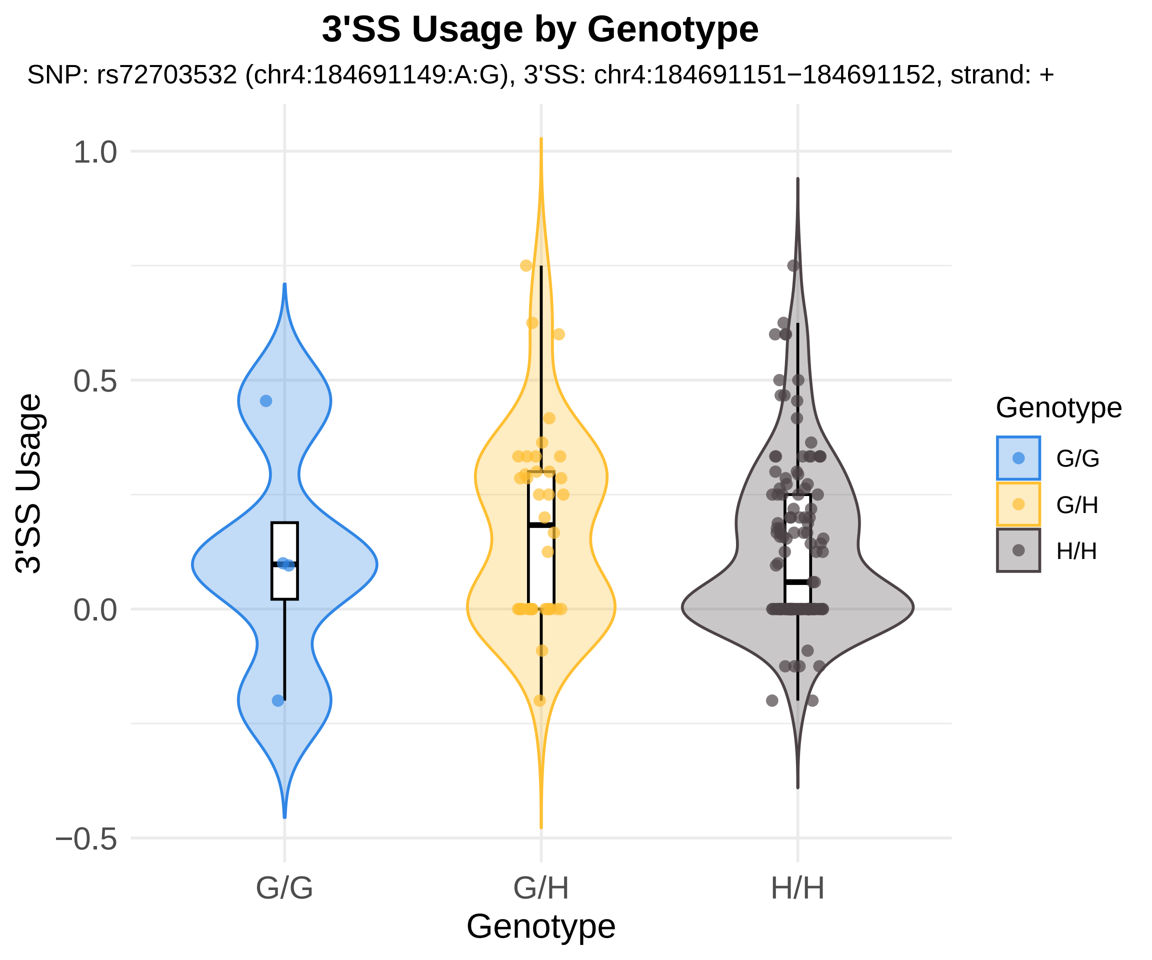

3'SS Usage by Genotype
SNP: rs72703532 (chr4:184691149:A:G), 3'SS: chr4:184691151−184691152, strand: +
3'SS Usage
1.0
0.5
0.0
−0.5
G/G
G/H
H/H
Genotype
Genotype
G/G
G/H
H/H

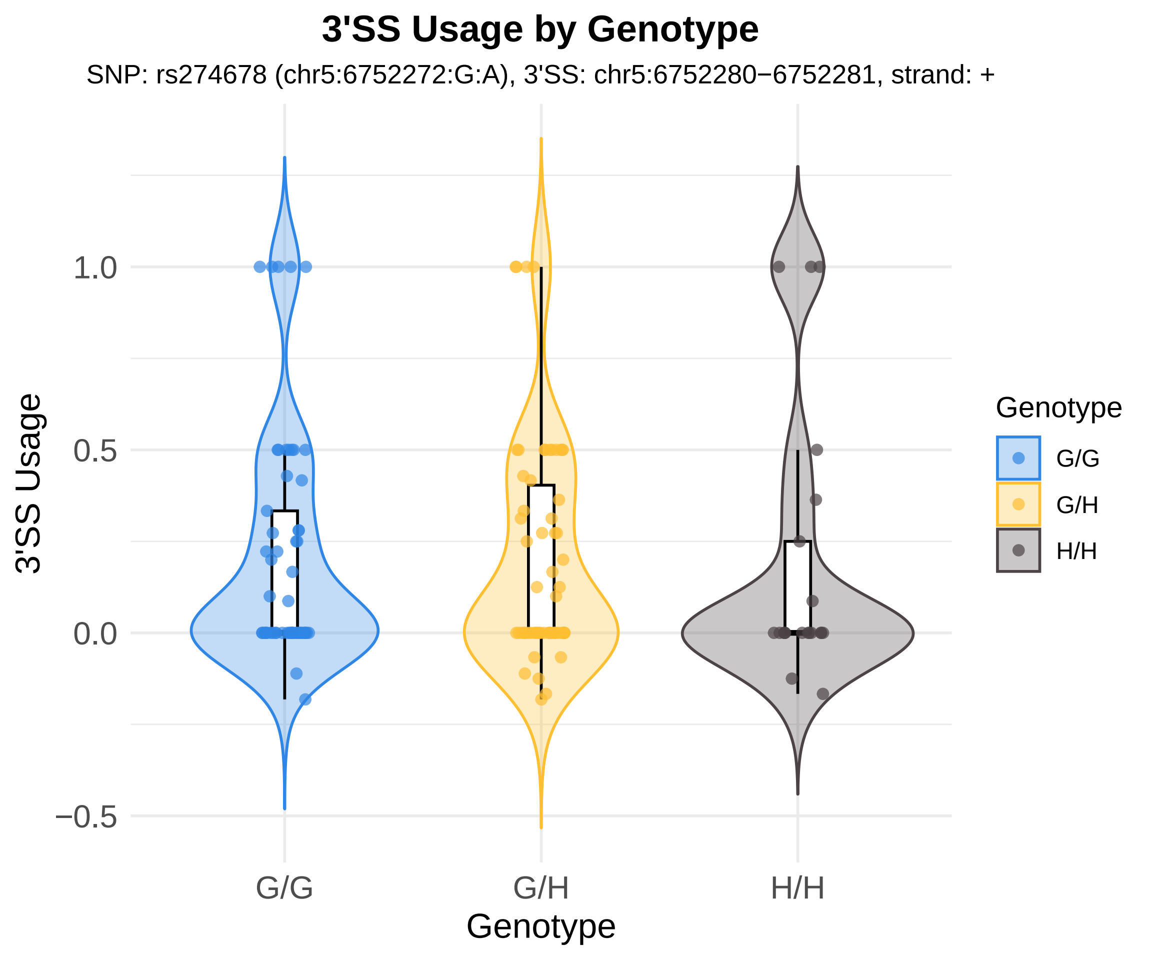
3'SS Usage by Genotype
SNP: rs274678 (chr5:6752272:G:A), 3'SS: chr5:6752280−6752281, strand: +
3'SS Usage
1.0
0.5
0.0
−0.5
G/G
G/H
H/H
Genotype
Genotype
G/G
G/H
H/H

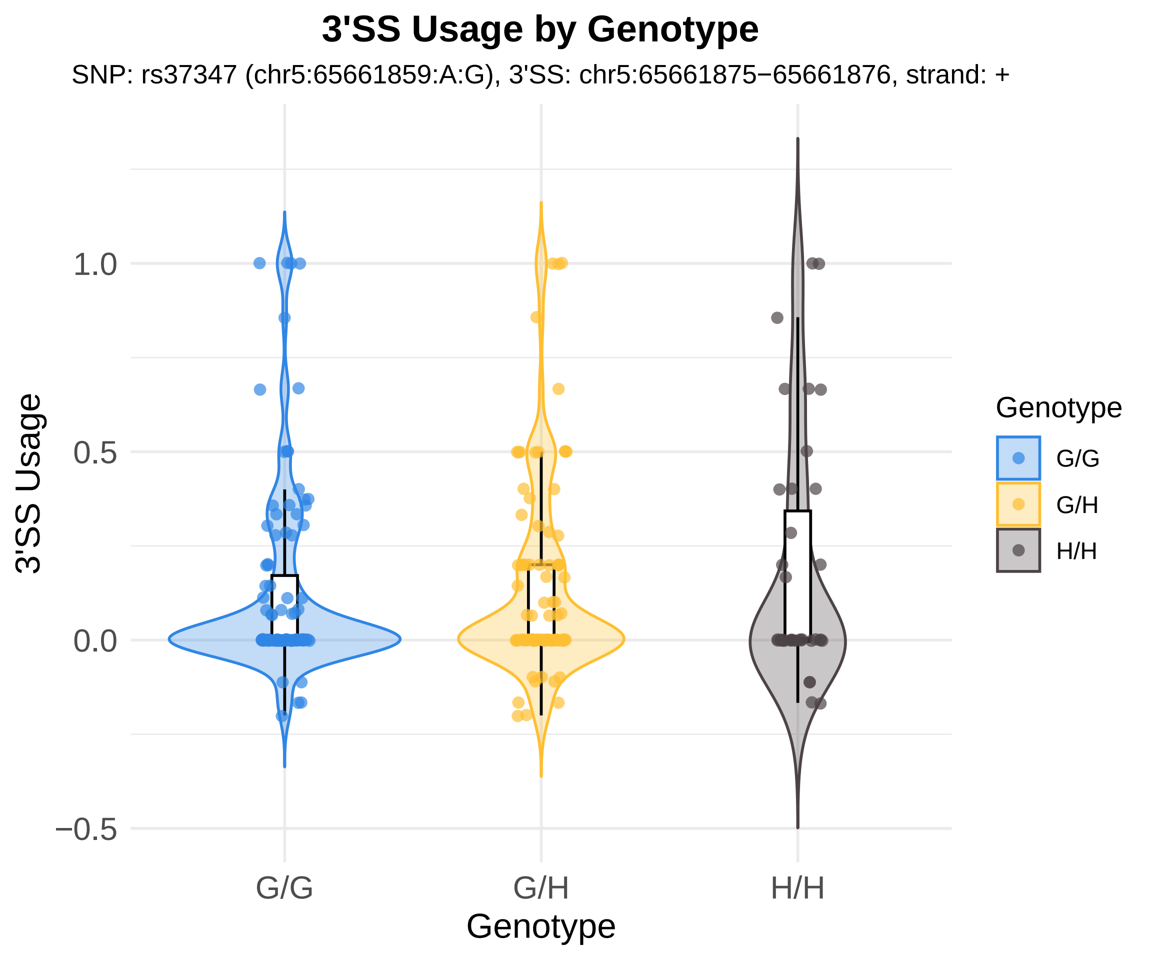
3'SS Usage by Genotype
SNP: rs37347 (chr5:65661859:A:G), 3'SS: chr5:65661875−65661876, strand: +
3'SS Usage
1.0
0.5
0.0
−0.5
G/G
G/H
H/H
Genotype
Genotype
G/G
G/H
H/H

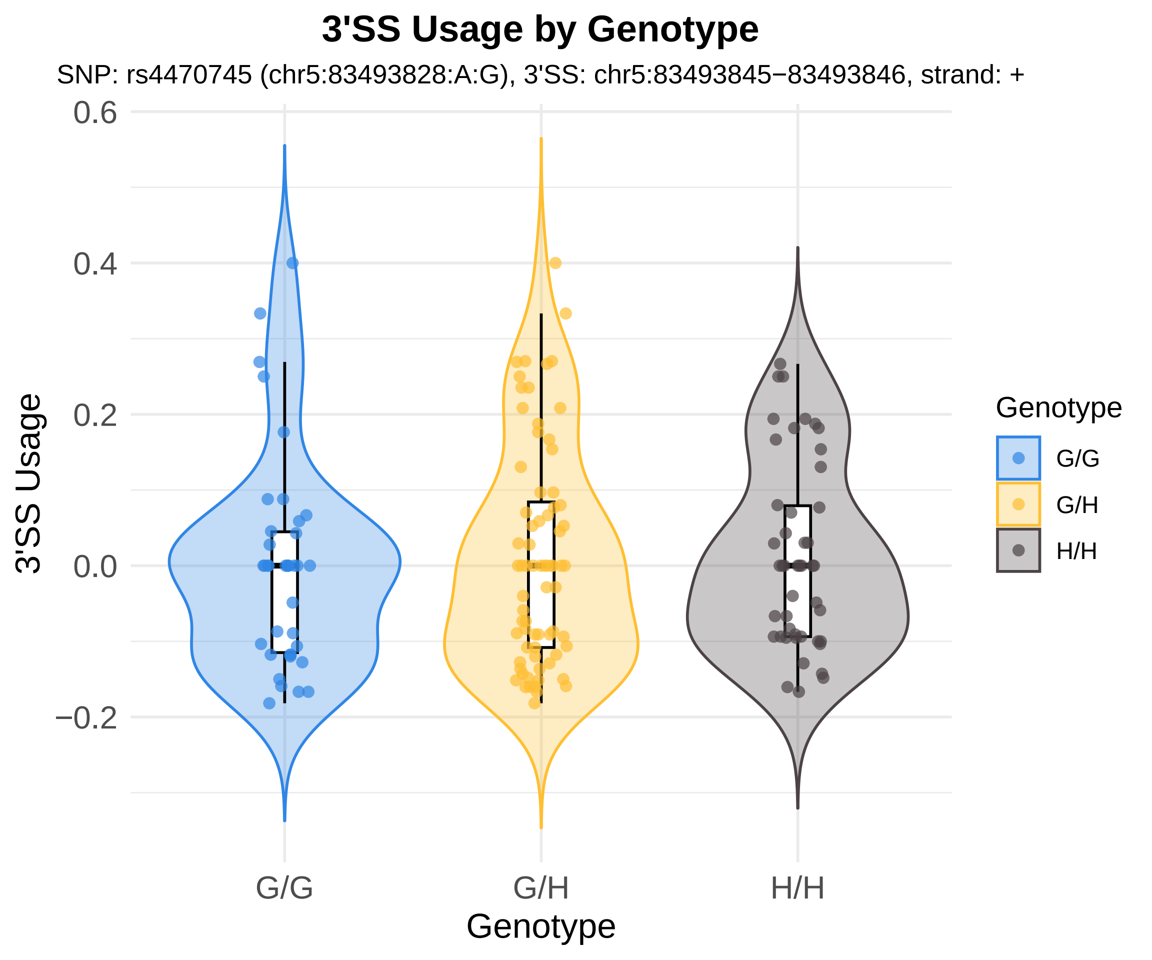
3'SS Usage by Genotype
SNP: rs4470745 (chr5:83493828:A:G), 3'SS: chr5:83493845−83493846, strand: +
0.6
0.4
0.2
0.0
−0.2
3'SS Usage
G/G
G/H
H/H
Genotype
Genotype
G/G
G/H
H/H

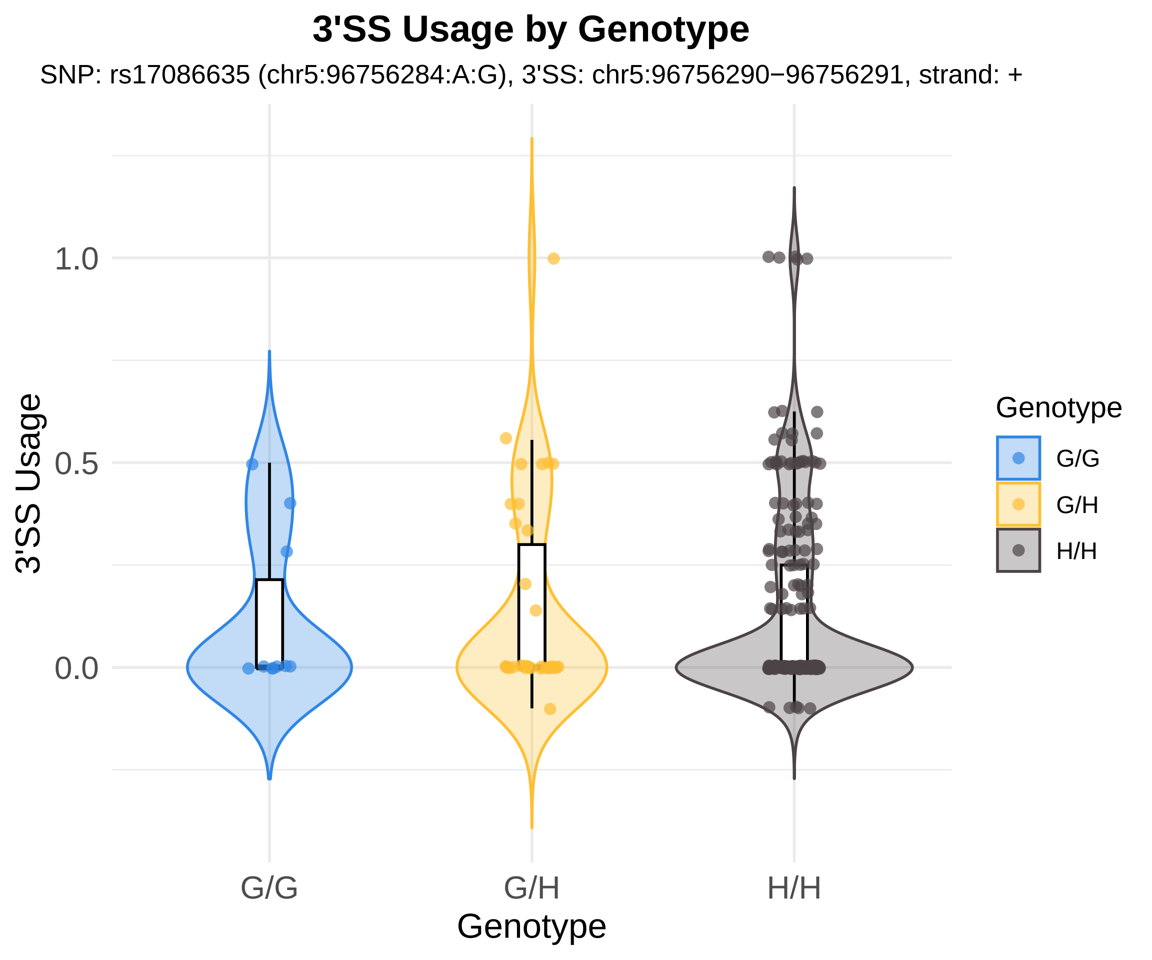

3'SS Usage by Genotype
SNP: rs17086635 (chr5:96756284:A:G), 3'SS: chr5:96756290−96756291, strand: +
3'SS Usage
1.0
0.5
0.0
G/G
G/H
H/H
Genotype
Genotype
G/G
G/H
H/H

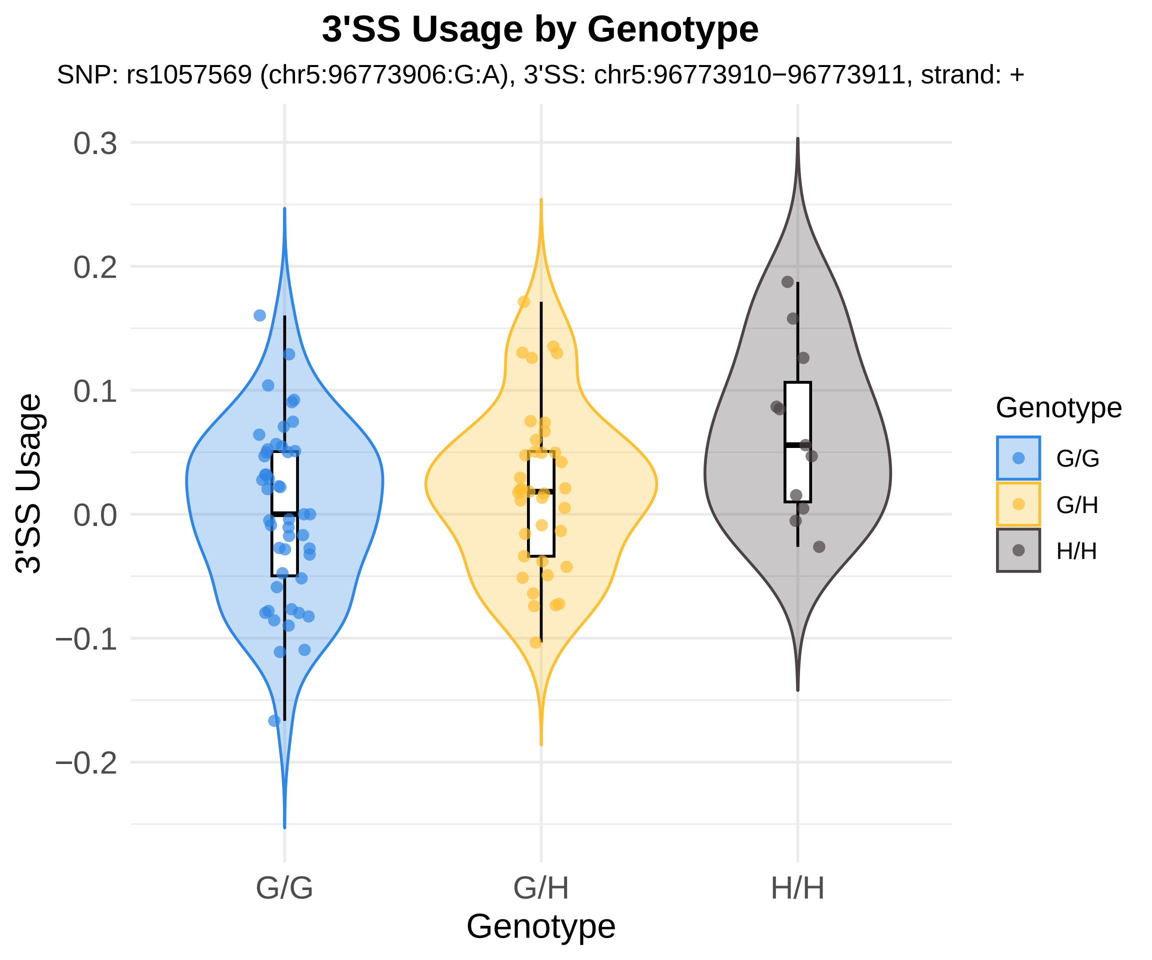
3'SS Usage by Genotype
SNP: rs1057569 (chr5:96773906:G:A), 3'SS: chr5:96773910−96773911, strand: +
3'SS Usage
0.3
0.2
0.1
0.0
−0.1
−0.2
G/G
G/H
H/H
Genotype
Genotype
G/G
G/H
H/H

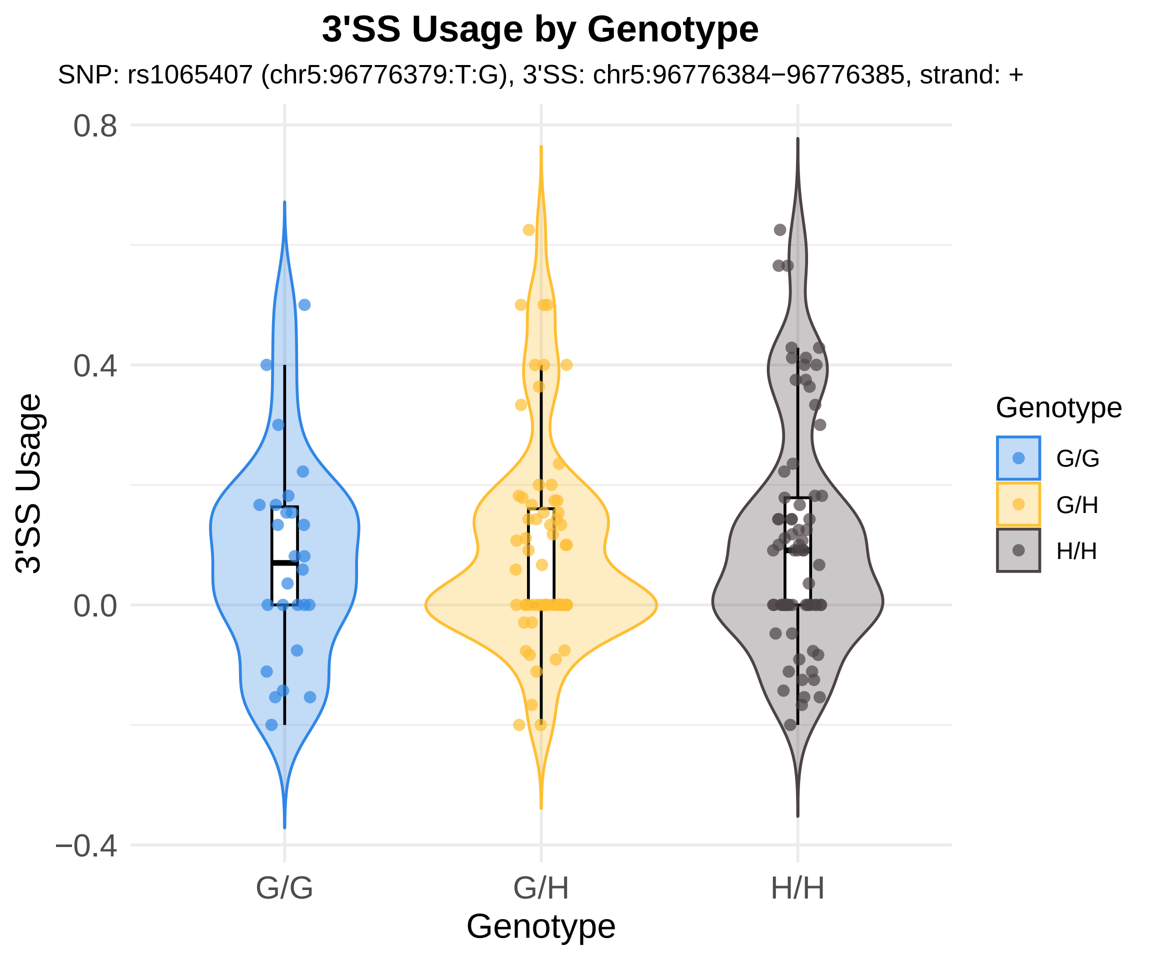

3'SS Usage by Genotype
SNP: rs1065407 (chr5:96776379:T:G), 3'SS: chr5:96776384−96776385, strand: +
0.8
0.4
0.0
−0.4
3'SS Usage
G/G
G/H
H/H
Genotype
Genotype
G/G
G/H
H/H

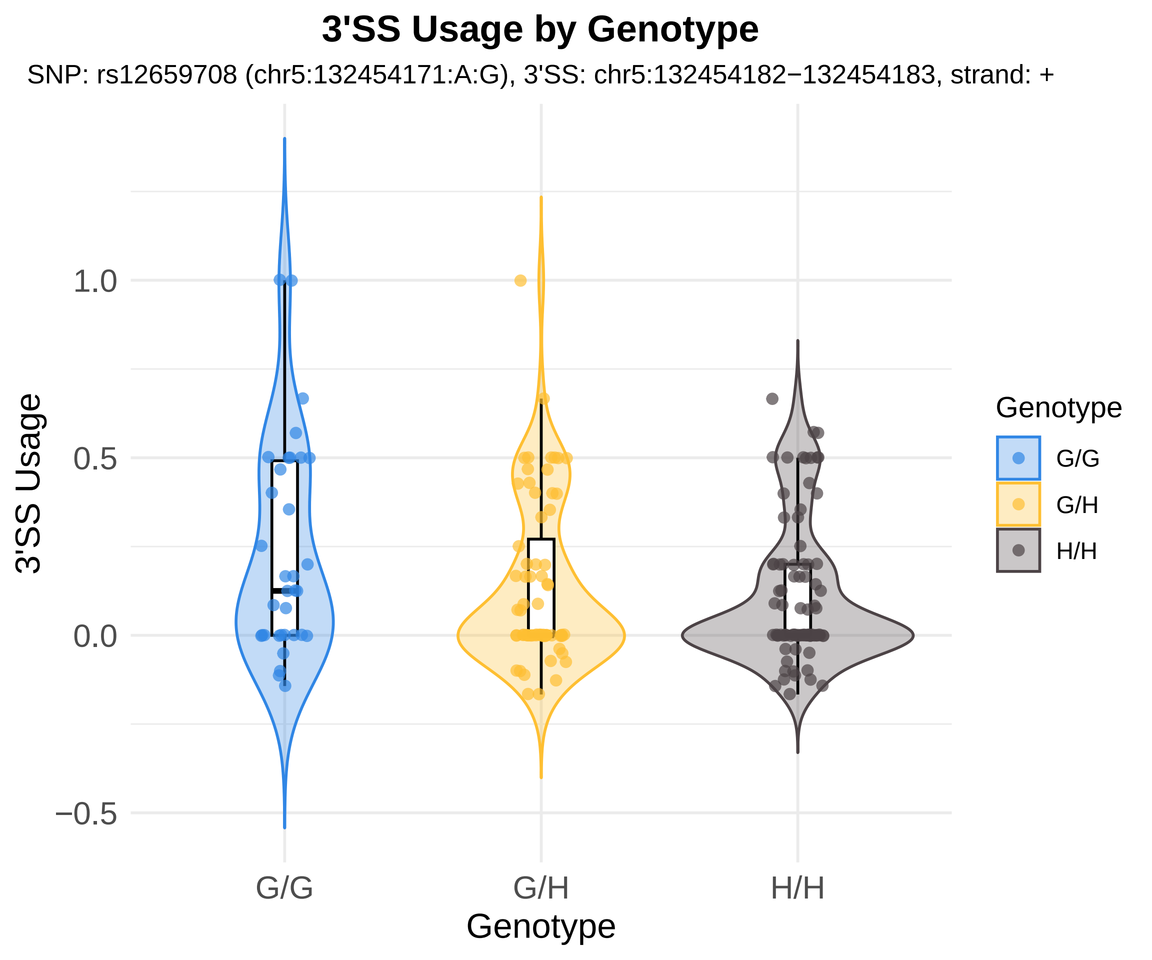

3'SS Usage by Genotype
SNP: rs12659708 (chr5:132454171:A:G), 3'SS: chr5:132454182−132454183, strand: +
3'SS Usage
1.0
0.5
0.0
−0.5
G/G
G/H
H/H
Genotype
Genotype
G/G
G/H
H/H

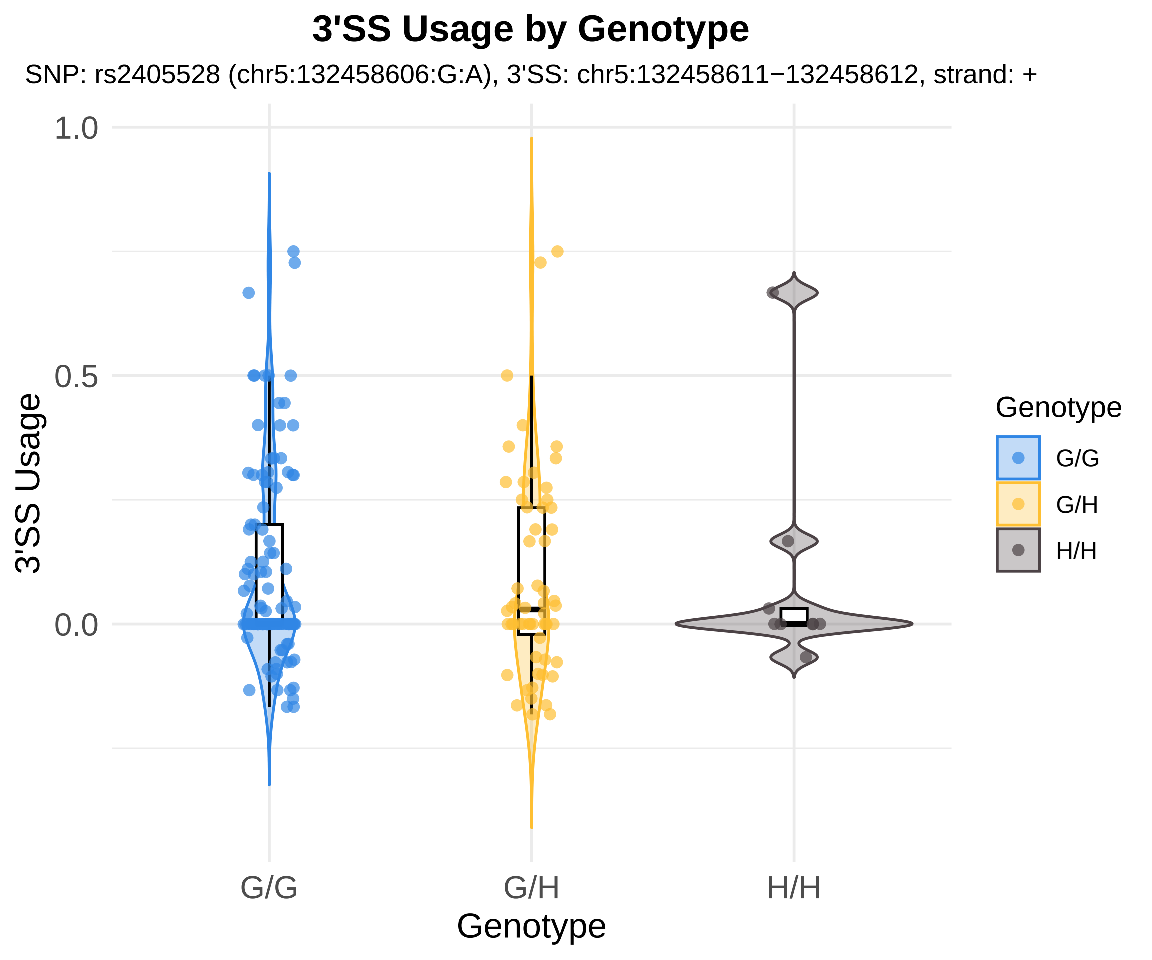
3'SS Usage by Genotype
SNP: rs2405528 (chr5:132458606:G:A), 3'SS: chr5:132458611−132458612, strand: +
3'SS Usage
1.0
0.5
0.0
G/G
G/H
H/H
Genotype
Genotype
G/G
G/H
H/H

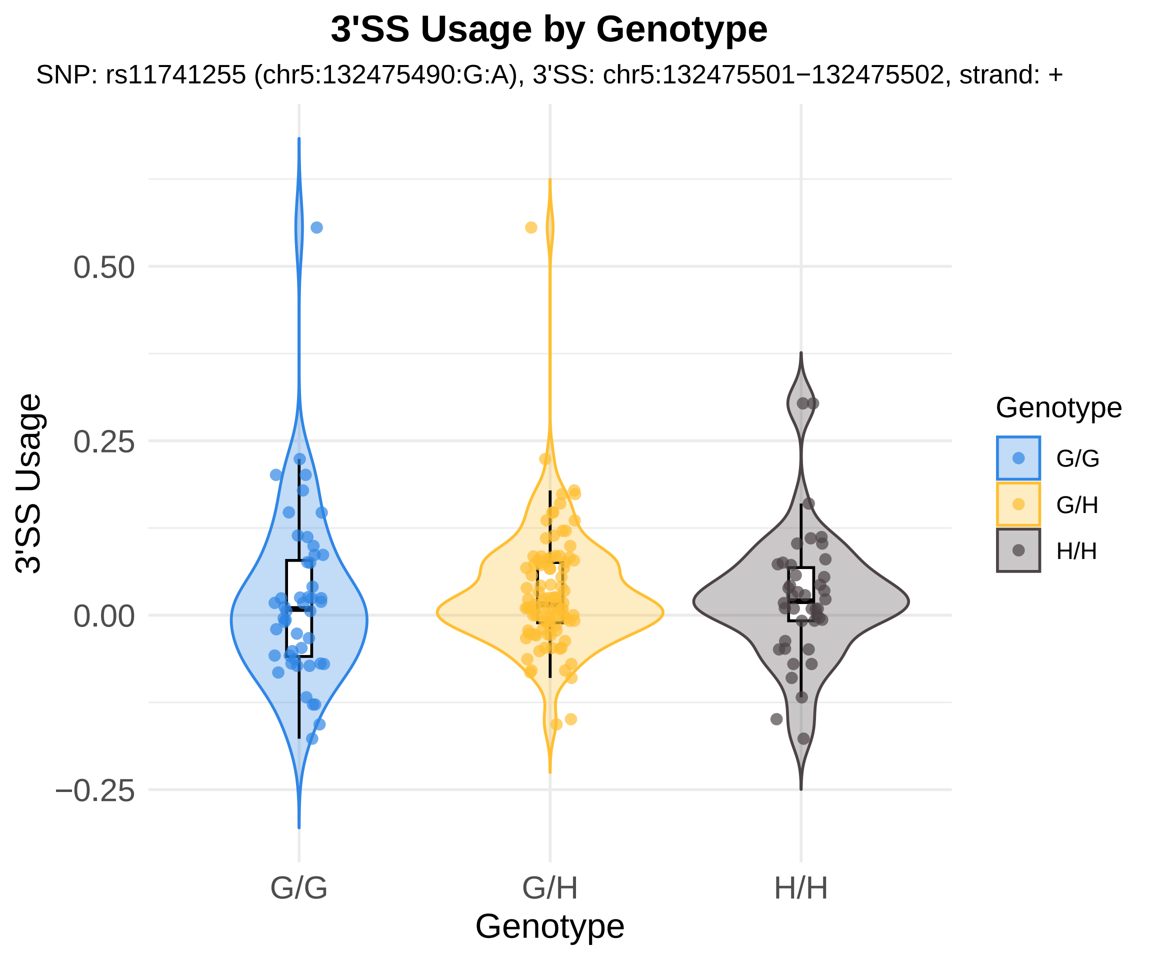

3'SS Usage by Genotype
SNP: rs11741255 (chr5:132475490:G:A), 3'SS: chr5:132475501−132475502, strand: +
3'SS Usage
0.50
0.25
0.00
−0.25
G/G
G/H
H/H
Genotype
Genotype
G/G
G/H
H/H

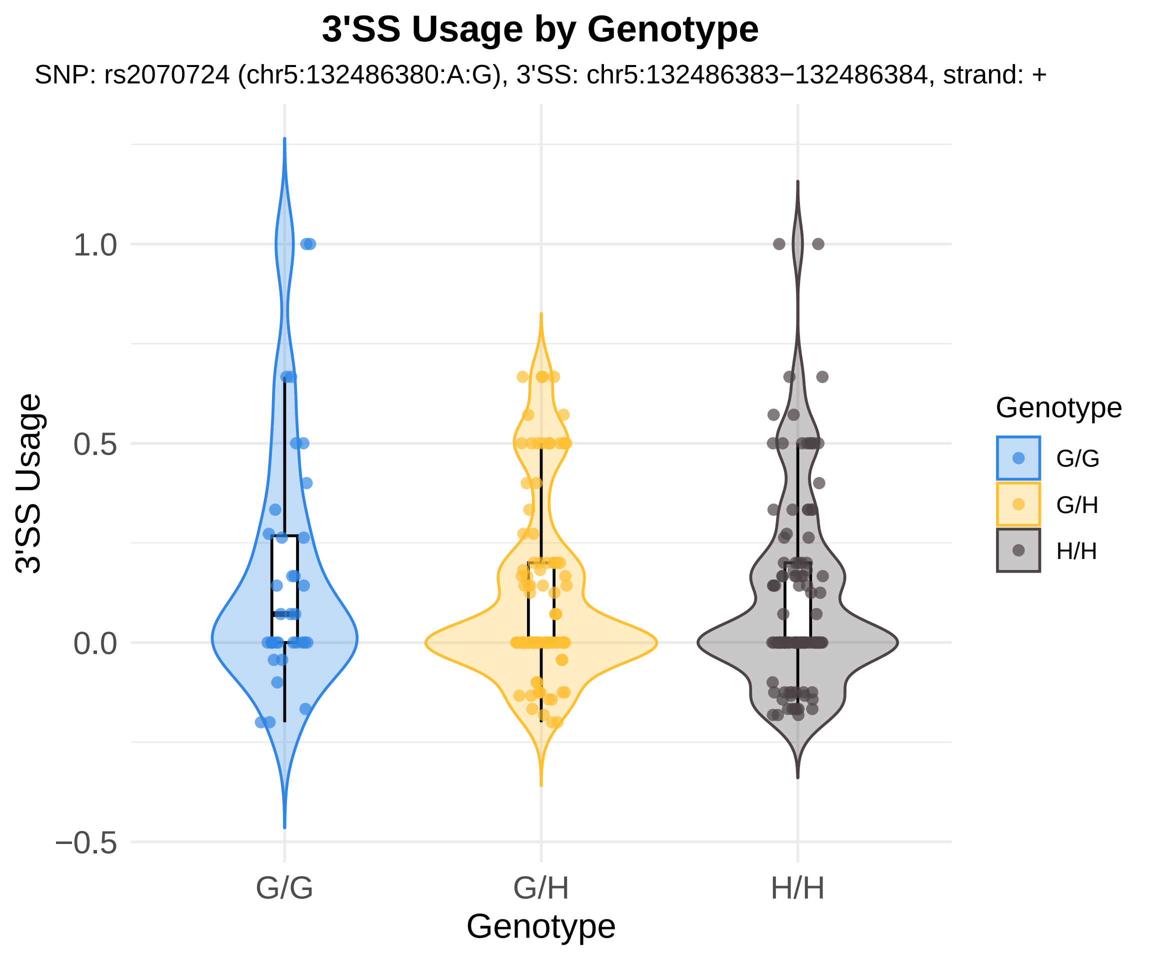

3'SS Usage by Genotype
SNP: rs2070724 (chr5:132486380:A:G), 3'SS: chr5:132486383−132486384, strand: +
3'SS Usage
1.0
0.5
0.0
−0.5
G/G
G/H
H/H
Genotype
Genotype
G/G
G/H
H/H

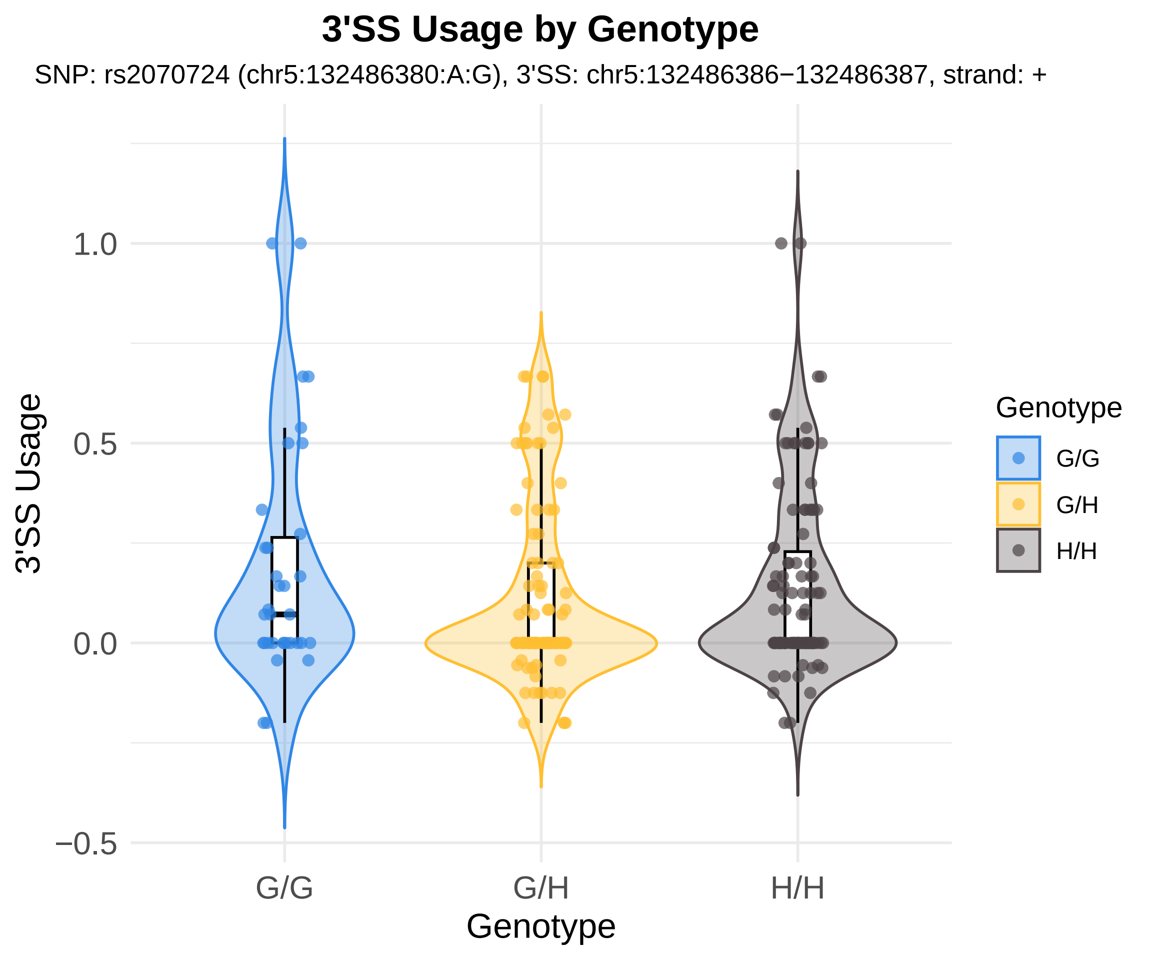

3'SS Usage by Genotype
SNP: rs2070724 (chr5:132486380:A:G), 3'SS: chr5:132486386−132486387, strand: +
3'SS Usage
1.0
0.5
0.0
−0.5
G/G
G/H
H/H
Genotype
Genotype
G/G
G/H
H/H

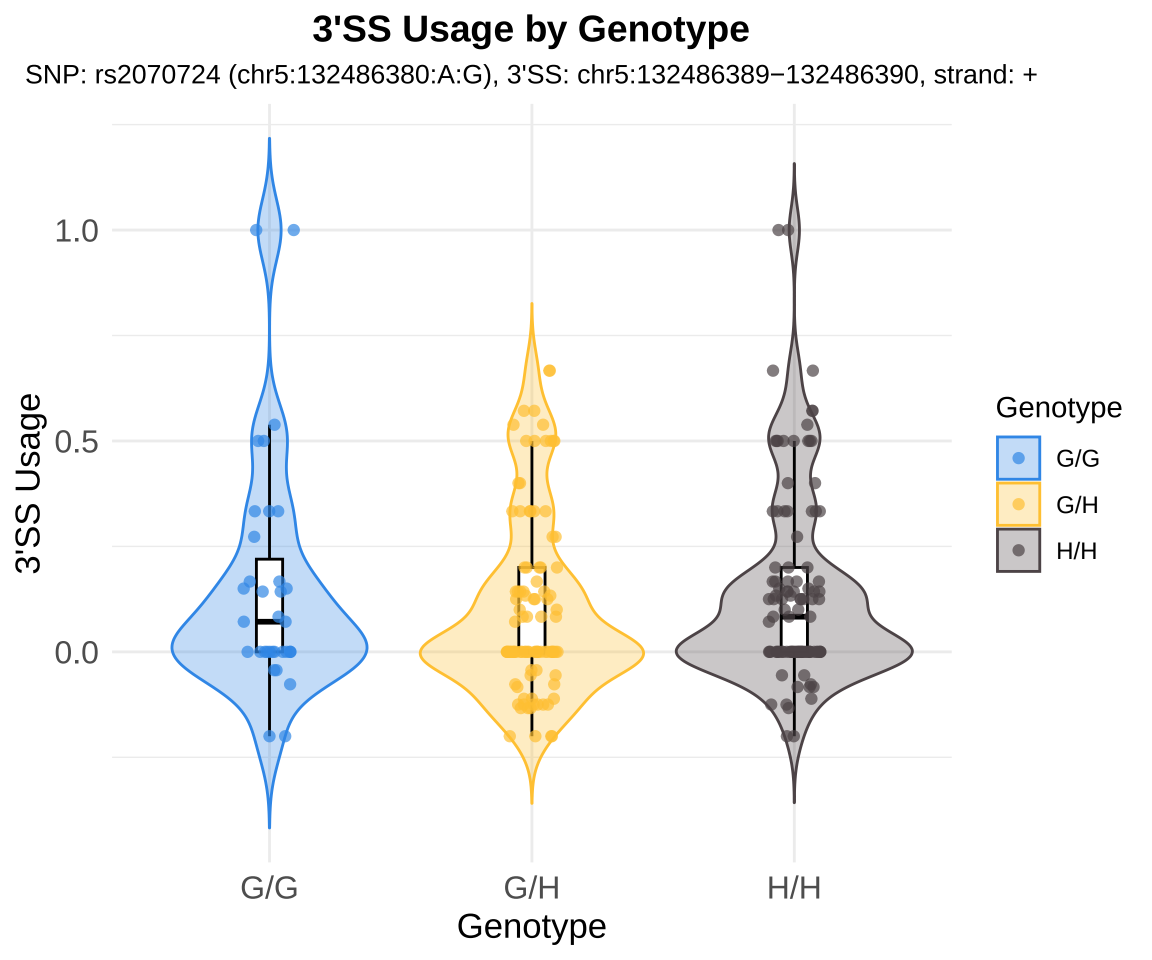

3'SS Usage by Genotype
SNP: rs2070724 (chr5:132486380:A:G), 3'SS: chr5:132486389−132486390, strand: +
3'SS Usage
1.0
0.5
0.0
G/G
G/H
H/H
Genotype
Genotype
G/G
G/H
H/H

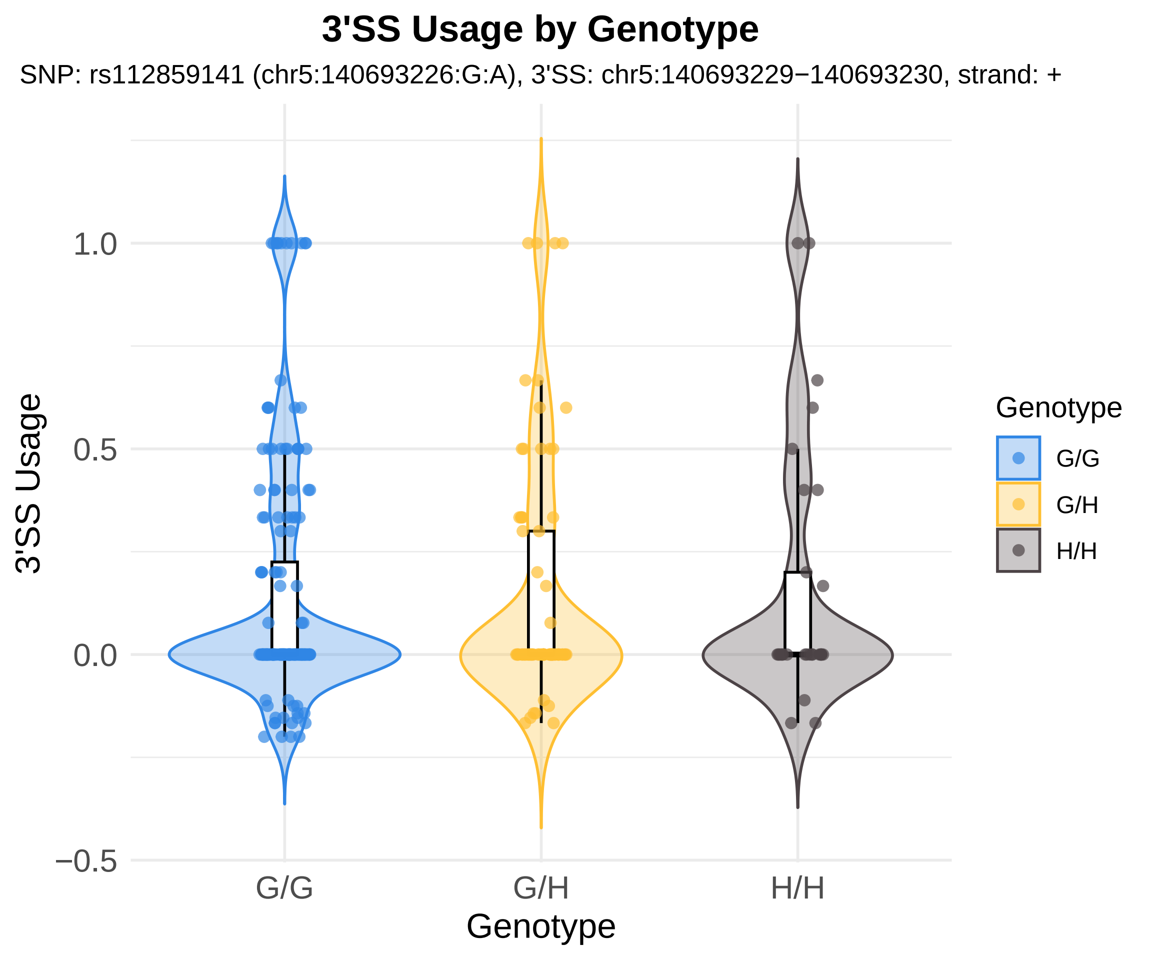
3'SS Usage by Genotype
SNP: rs112859141 (chr5:140693226:G:A), 3'SS: chr5:140693229−140693230, strand: +
3'SS Usage
1.0
0.5
0.0
−0.5
G/G
G/H
H/H
Genotype
Genotype
G/G
G/H
H/H

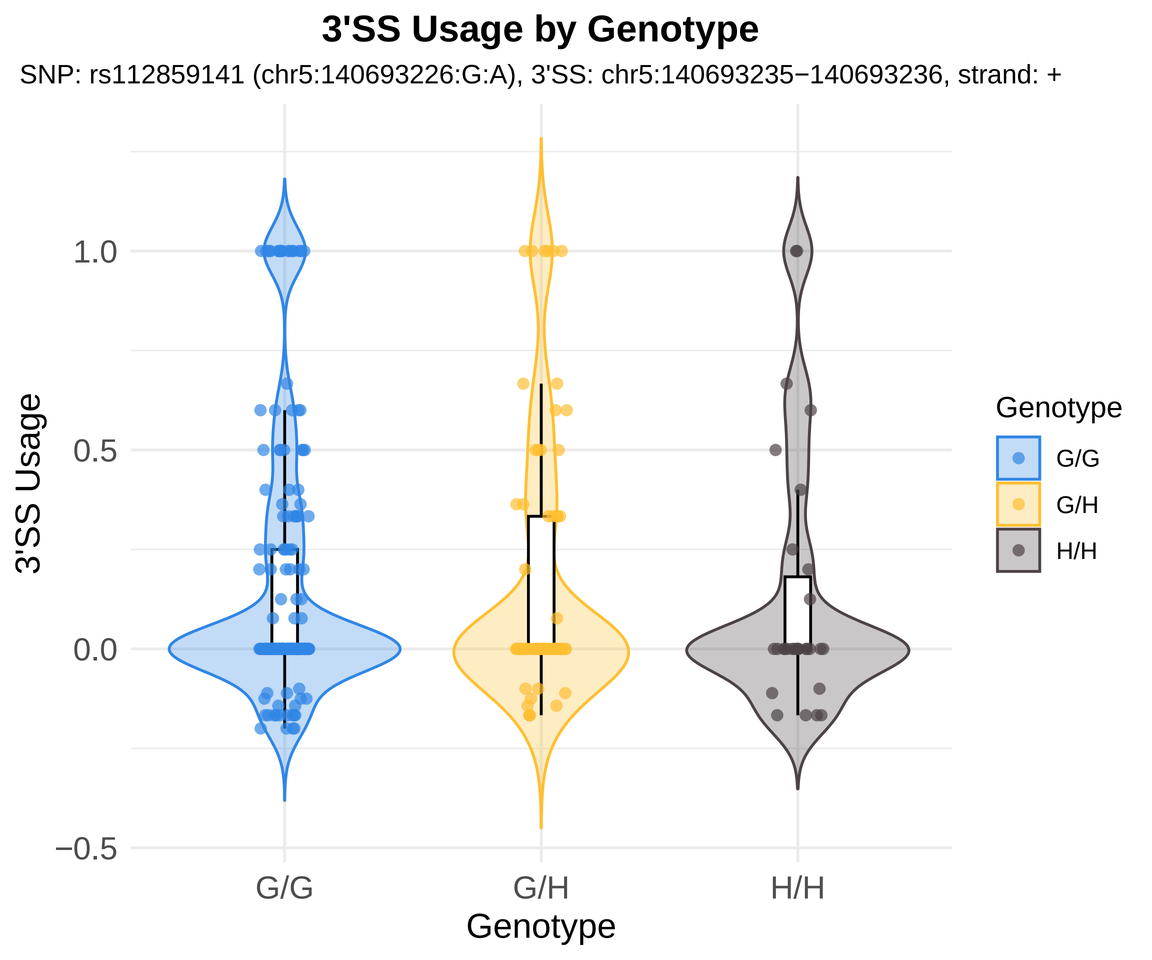

3'SS Usage by Genotype
SNP: rs112859141 (chr5:140693226:G:A), 3'SS: chr5:140693235−140693236, strand: +
3'SS Usage
1.0
0.5
0.0
−0.5
G/G
G/H
H/H
Genotype
Genotype
G/G
G/H
H/H

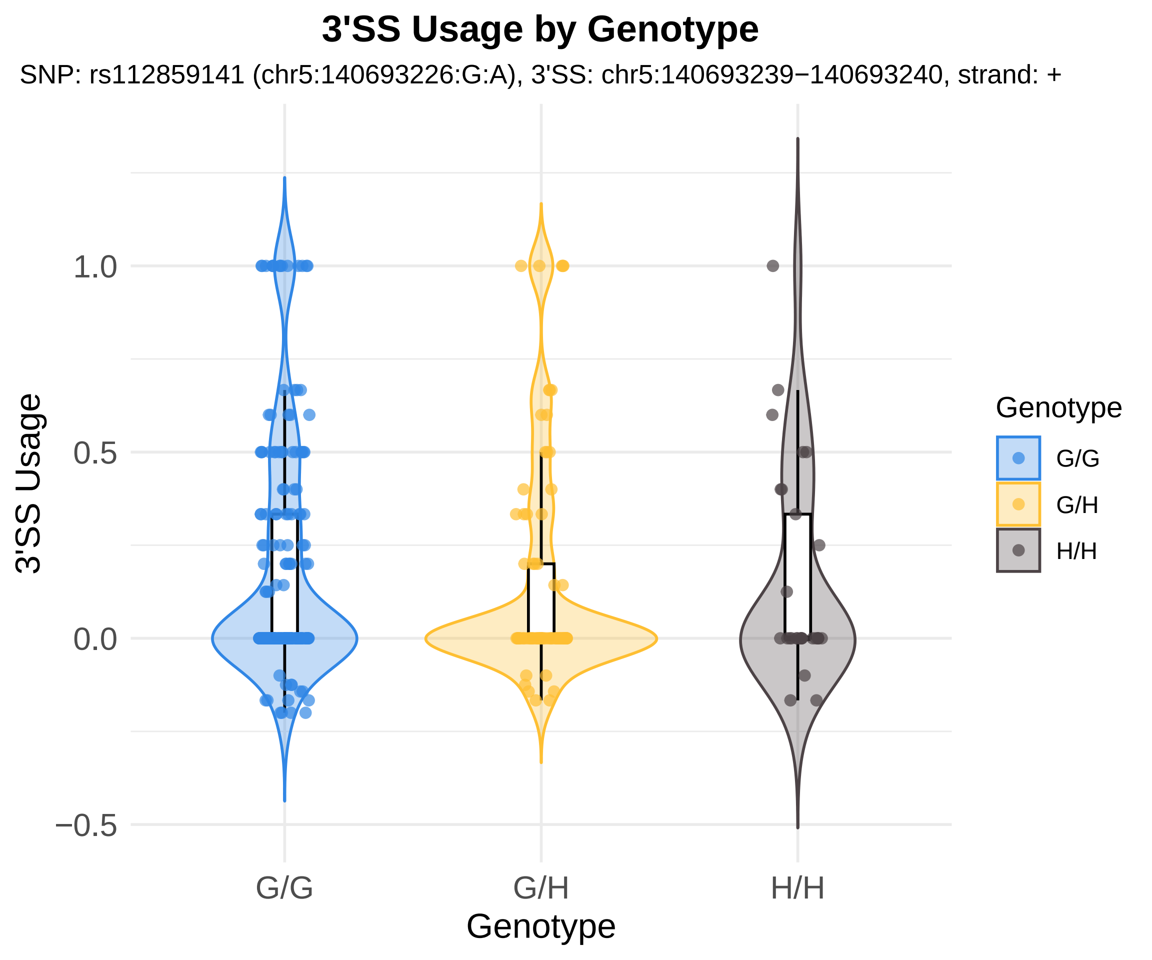
3'SS Usage by Genotype
SNP: rs112859141 (chr5:140693226:G:A), 3'SS: chr5:140693239−140693240, strand: +
3'SS Usage
1.0
0.5
0.0
−0.5
G/G
G/H
H/H
Genotype
Genotype
G/G
G/H
H/H

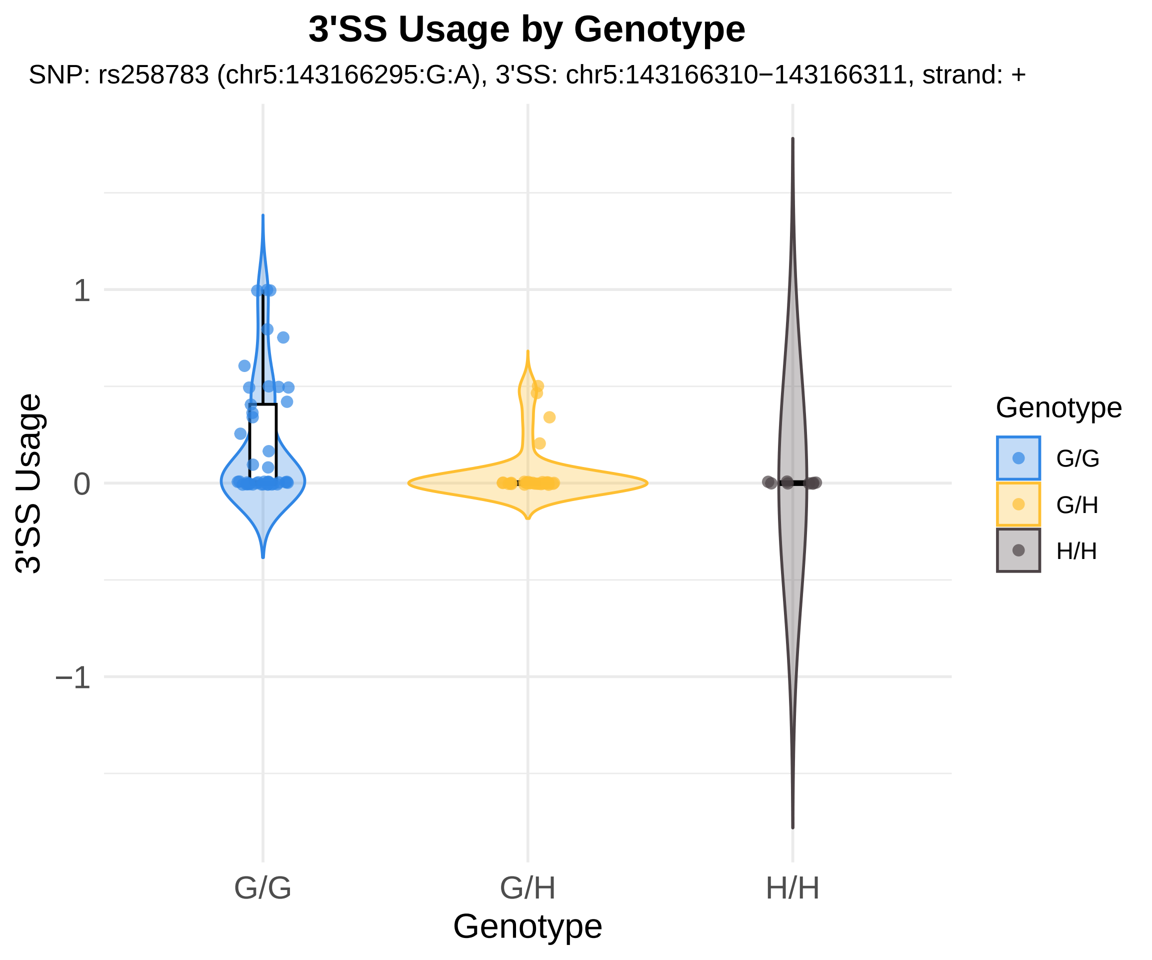

3'SS Usage by Genotype
SNP: rs258783 (chr5:143166295:G:A), 3'SS: chr5:143166310−143166311, strand: +
3'SS Usage
1
0
−1
G/G
G/H
H/H
Genotype
Genotype
G/G
G/H
H/H

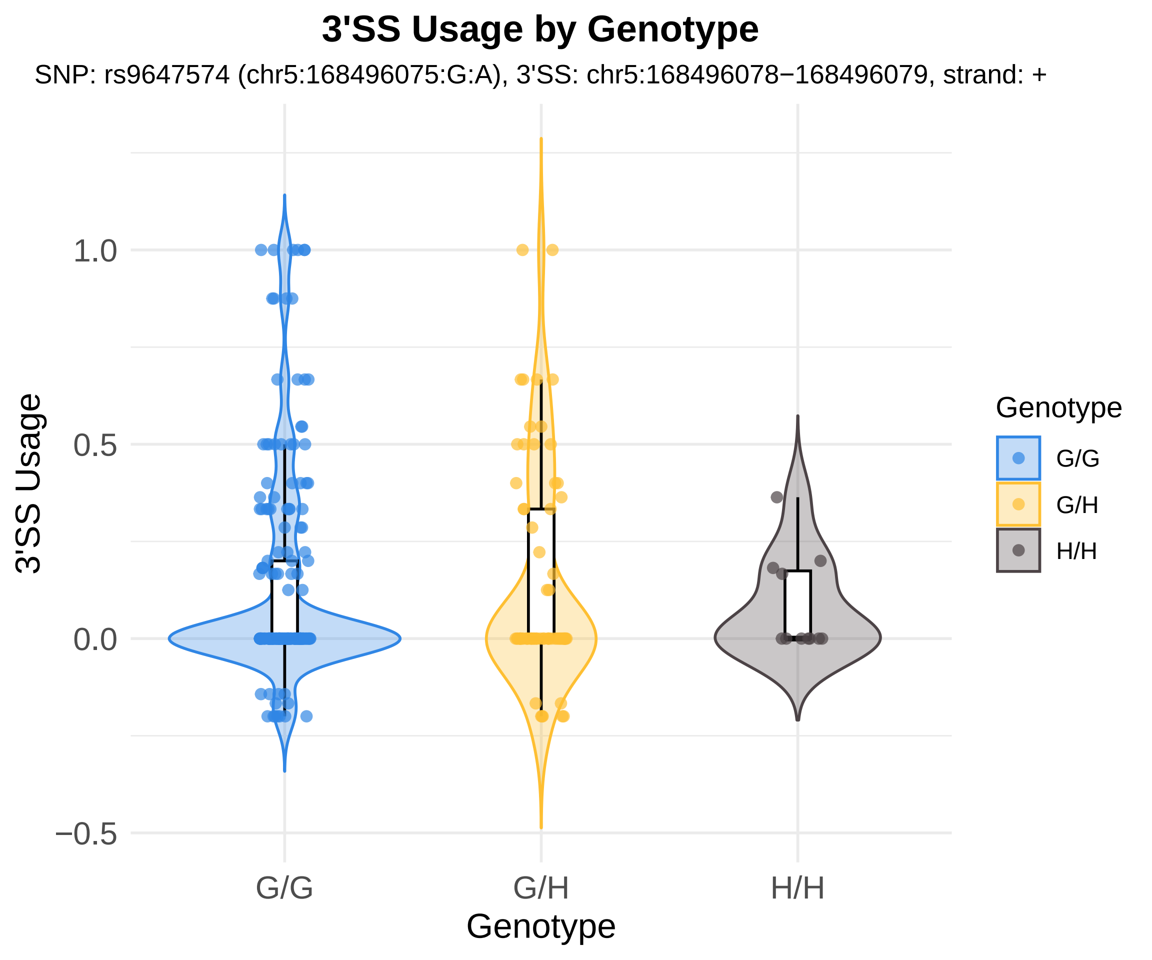

3'SS Usage by Genotype
SNP: rs9647574 (chr5:168496075:G:A), 3'SS: chr5:168496078−168496079, strand: +
3'SS Usage
1.0
0.5
0.0
−0.5
G/G
G/H
H/H
Genotype
Genotype
G/G
G/H
H/H

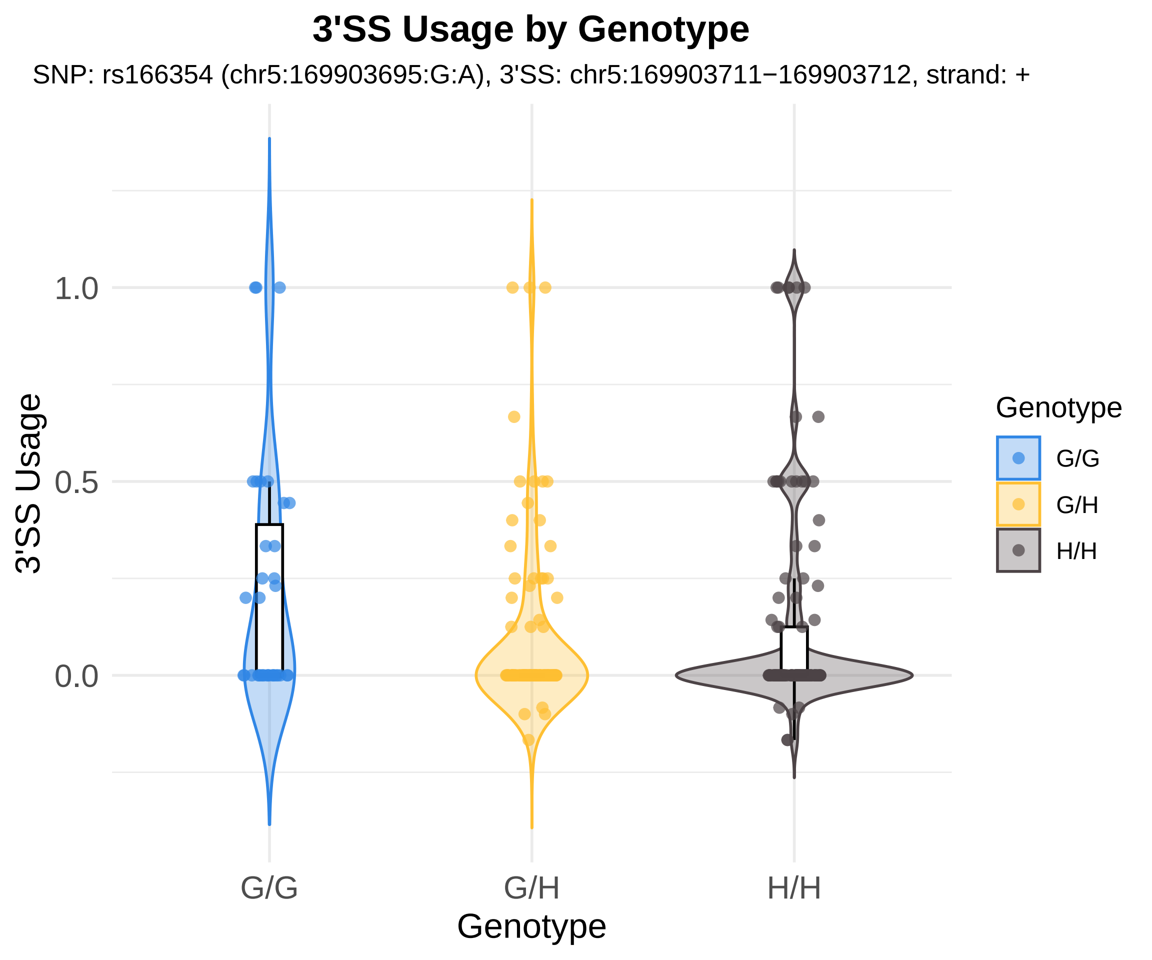

3'SS Usage by Genotype
SNP: rs166354 (chr5:169903695:G:A), 3'SS: chr5:169903711−169903712, strand: +
3'SS Usage
1.0
0.5
0.0
G/G
G/H
H/H
Genotype
Genotype
G/G
G/H
H/H

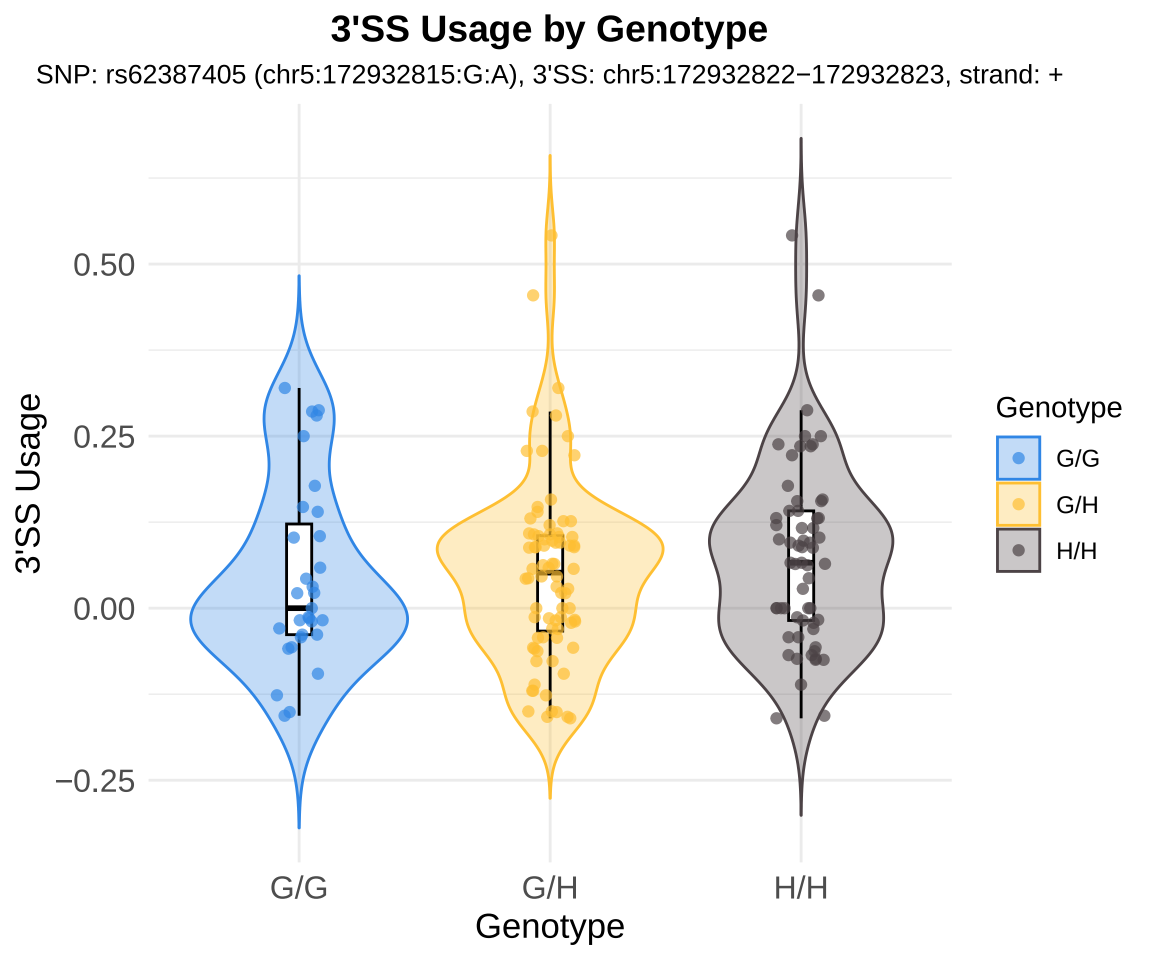

3'SS Usage by Genotype
SNP: rs62387405 (chr5:172932815:G:A), 3'SS: chr5:172932822−172932823, strand: +
3'SS Usage
0.50
0.25
0.00
−0.25
G/G
G/H
H/H
Genotype
Genotype
G/G
G/H
H/H

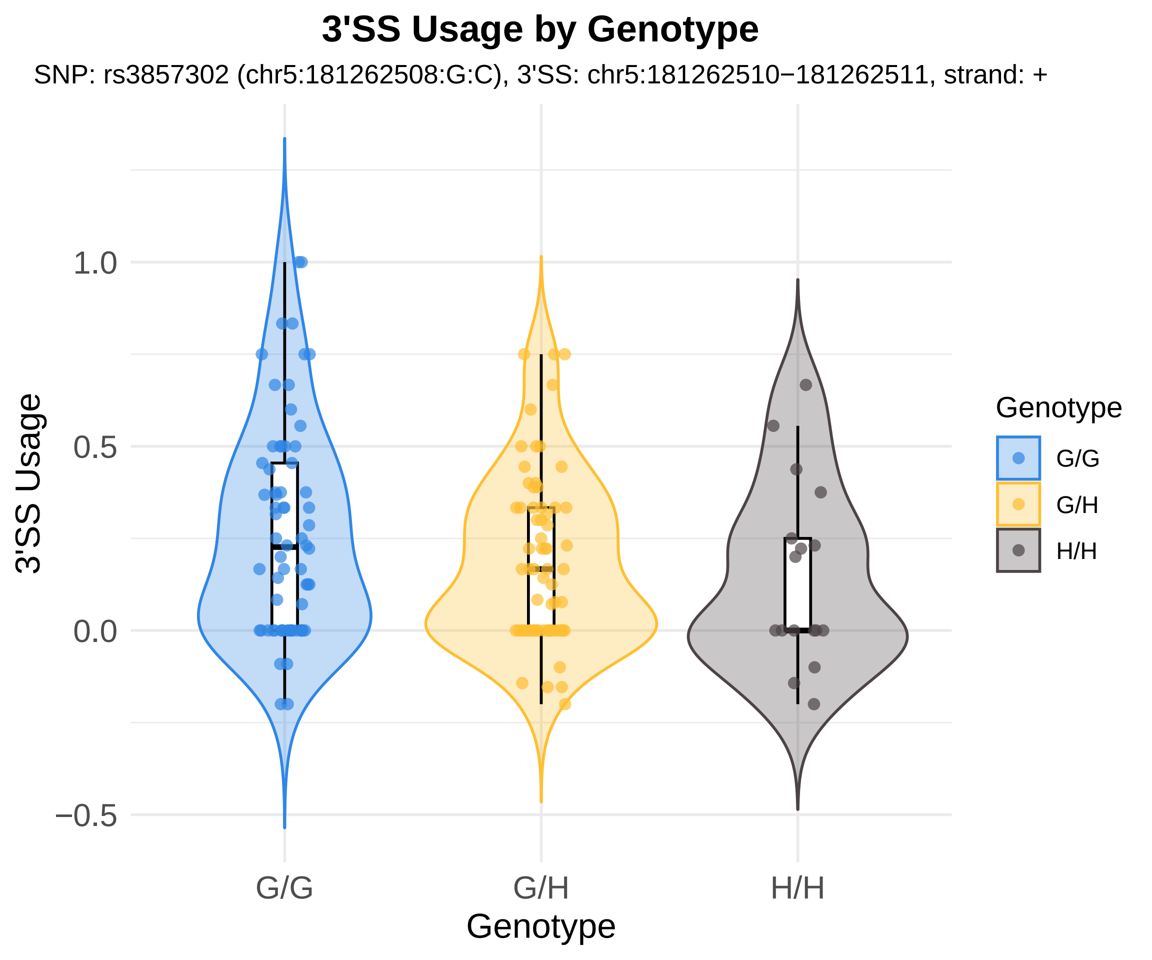
3'SS Usage by Genotype
SNP: rs3857302 (chr5:181262508:G:C), 3'SS: chr5:181262510−181262511, strand: +
3'SS Usage
1.0
0.5
0.0
−0.5
G/G
G/H
H/H
Genotype
Genotype
G/G
G/H
H/H

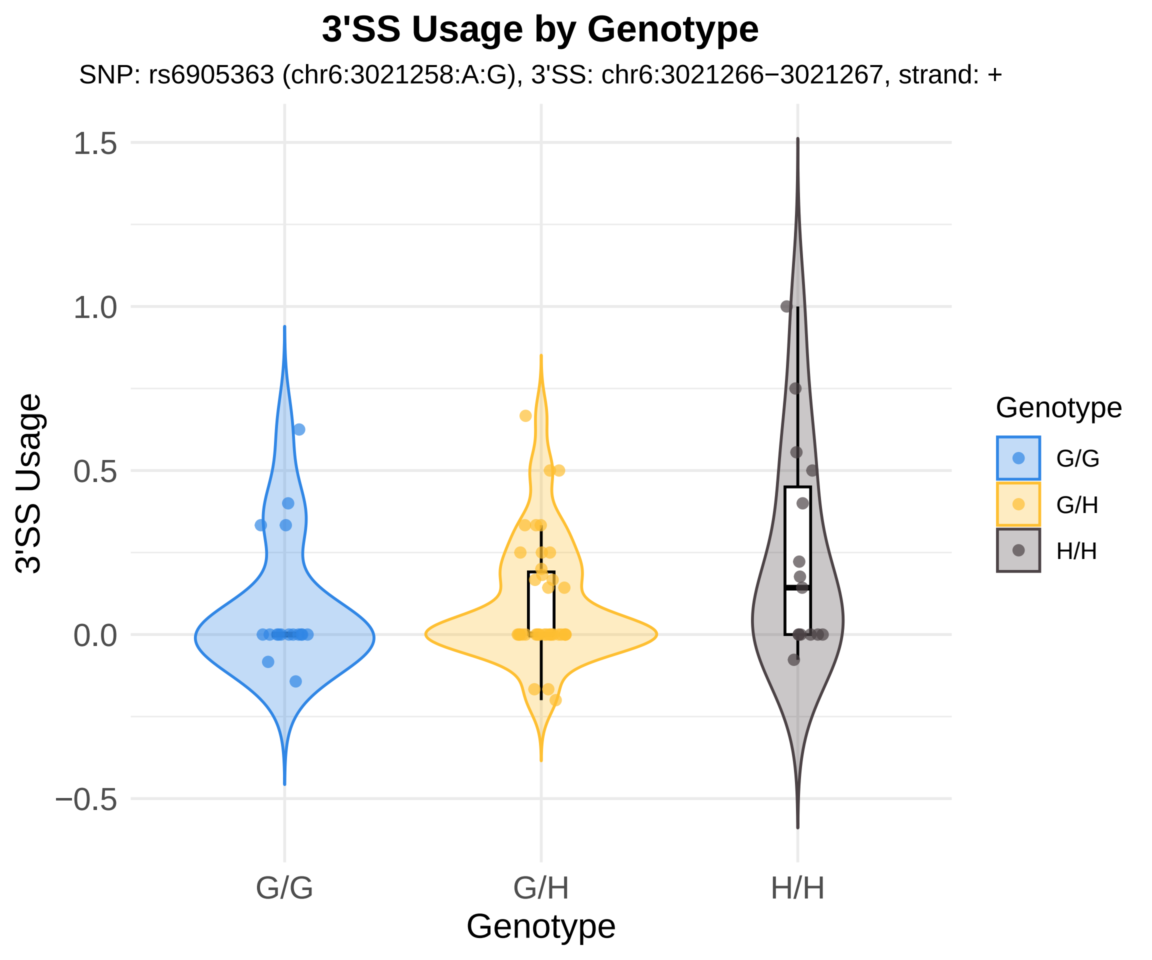
3'SS Usage by Genotype
SNP: rs6905363 (chr6:3021258:A:G), 3'SS: chr6:3021266−3021267, strand: +
3'SS Usage
1.5
1.0
0.5
0.0
−0.5
G/G
G/H
H/H
Genotype
Genotype
G/G
G/H
H/H

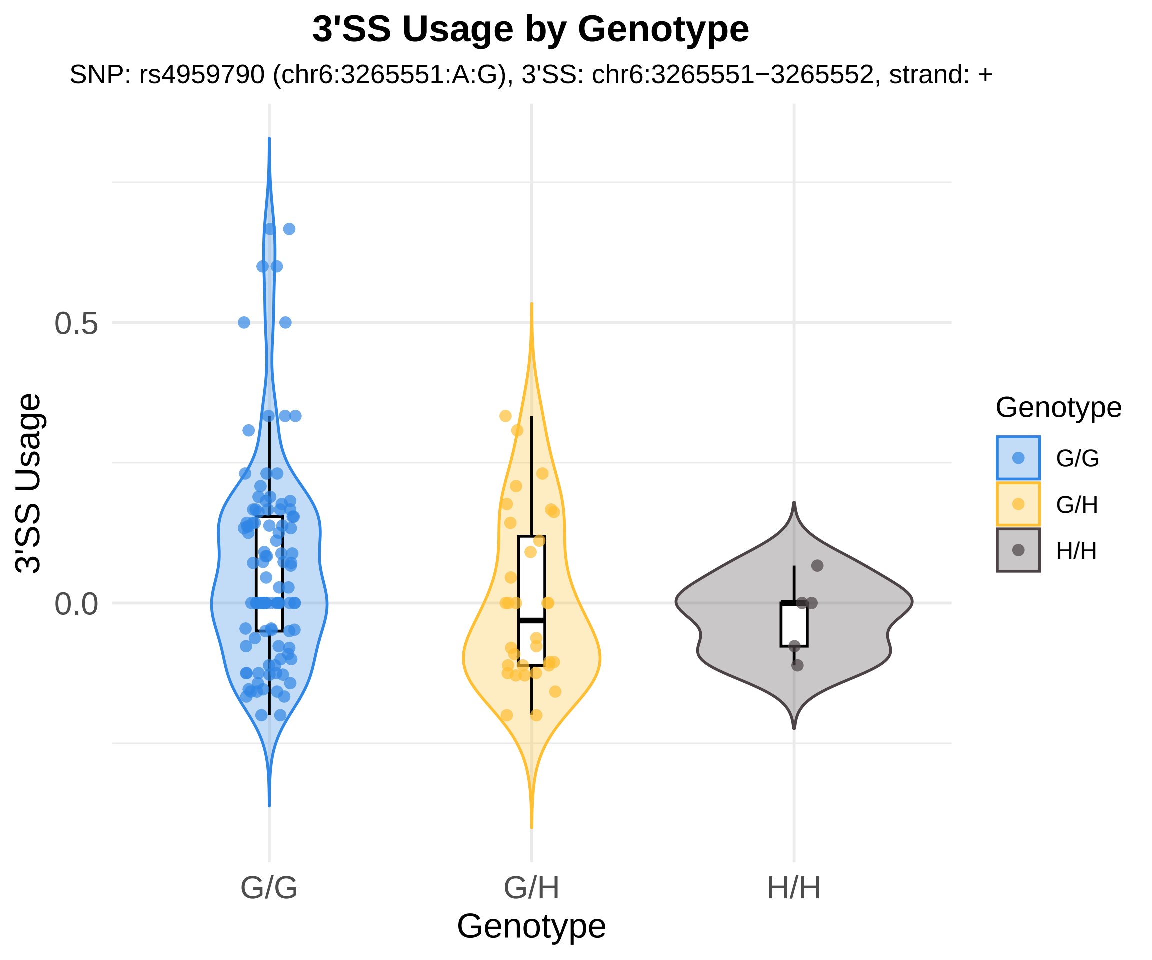
3'SS Usage by Genotype
SNP: rs4959790 (chr6:3265551:A:G), 3'SS: chr6:3265551−3265552, strand: +
3'SS Usage
0.5
0.0
G/G
G/H
H/H
Genotype
Genotype
G/G
G/H
H/H

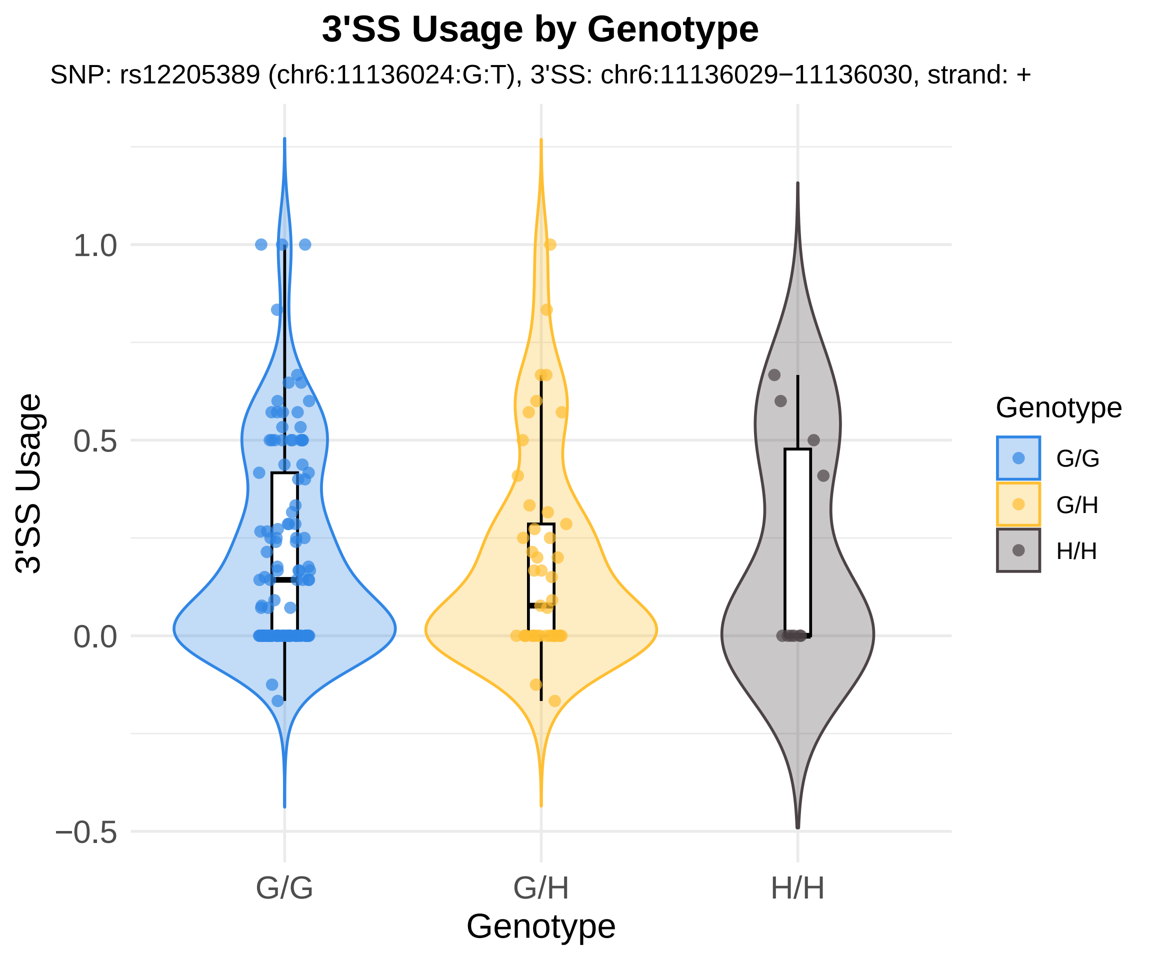
3'SS Usage by Genotype
SNP: rs12205389 (chr6:11136024:G:T), 3'SS: chr6:11136029−11136030, strand: +
3'SS Usage
1.0
0.5
0.0
−0.5
G/G
G/H
H/H
Genotype
Genotype
G/G
G/H
H/H

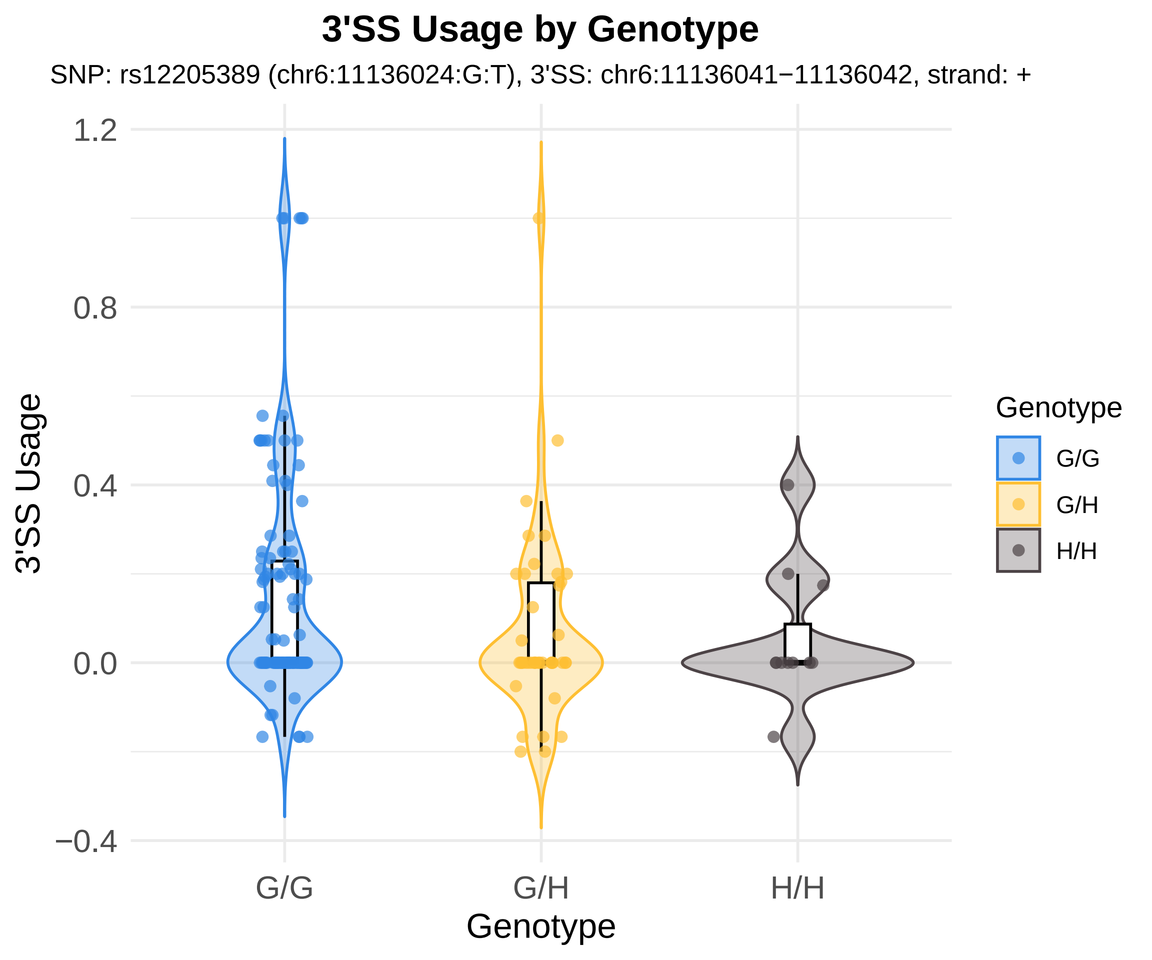

3'SS Usage by Genotype
SNP: rs12205389 (chr6:11136024:G:T), 3'SS: chr6:11136041−11136042, strand: +
1.2
0.8
0.4
0.0
−0.4
3'SS Usage
G/G
G/H
H/H
Genotype
Genotype
G/G
G/H
H/H

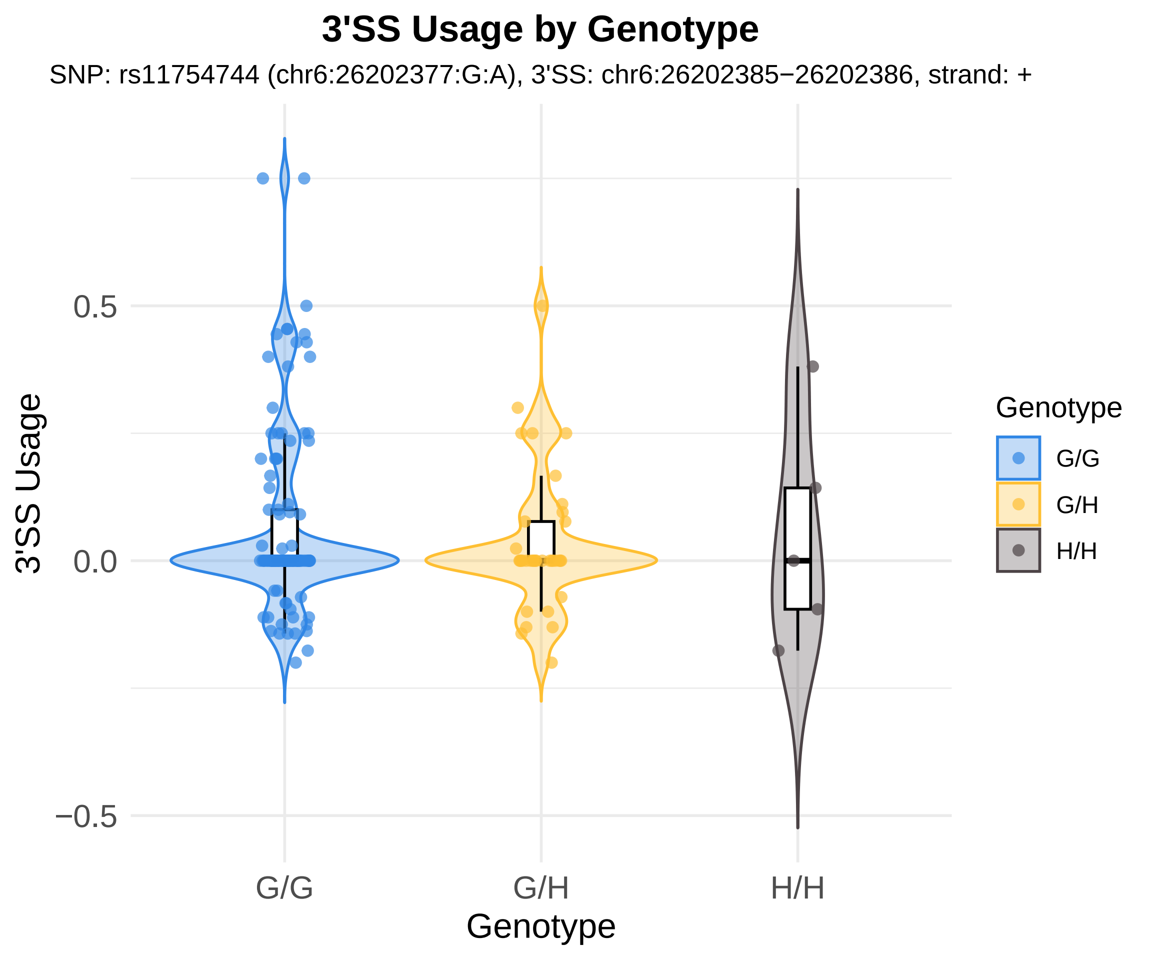
3'SS Usage by Genotype
SNP: rs11754744 (chr6:26202377:G:A), 3'SS: chr6:26202385−26202386, strand: +
3'SS Usage
0.5
0.0
−0.5
G/G
G/H
H/H
Genotype
Genotype
G/G
G/H
H/H

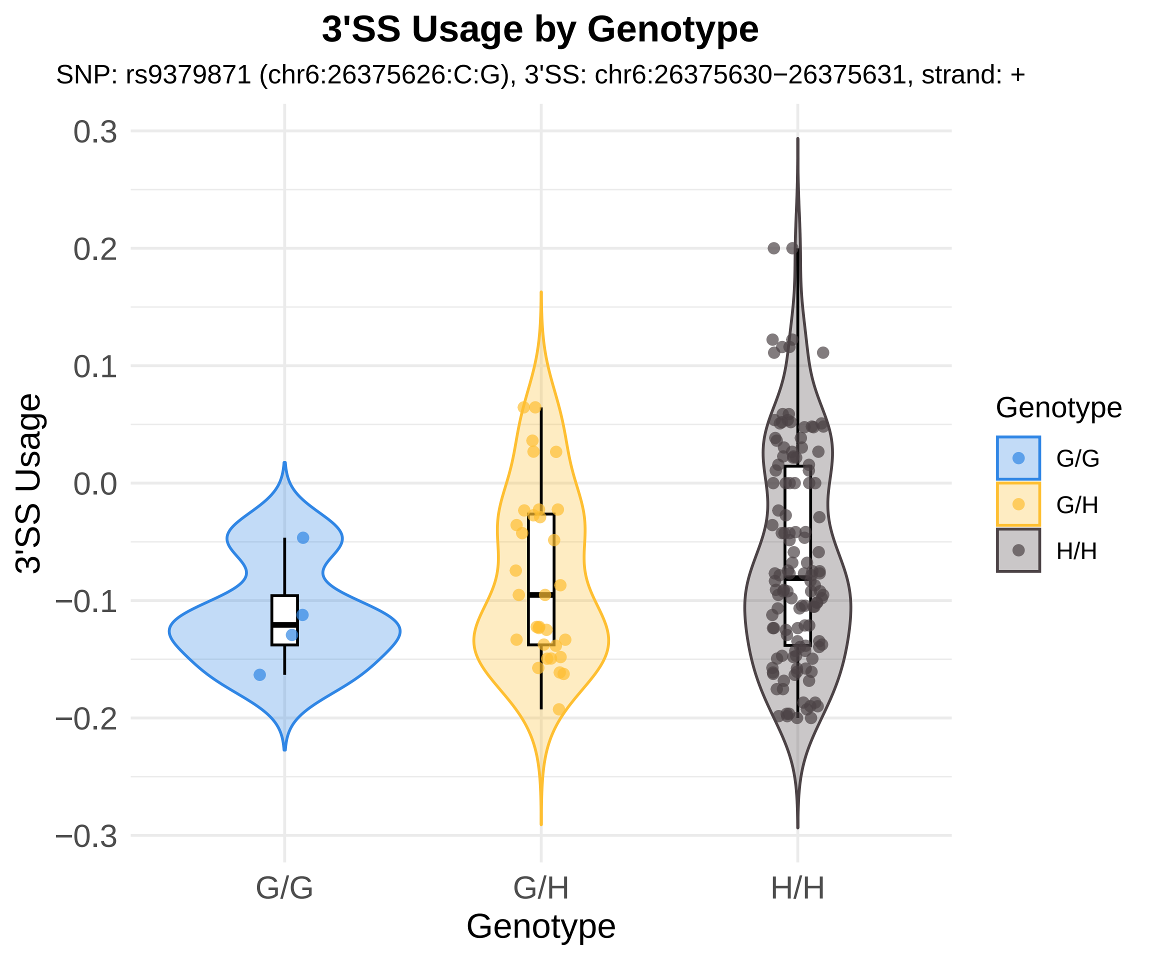
3'SS Usage by Genotype
SNP: rs9379871 (chr6:26375626:C:G), 3'SS: chr6:26375630−26375631, strand: +
3'SS Usage
0.3
0.2
0.1
0.0
−0.1
−0.2
−0.3
G/G
G/H
H/H
Genotype
Genotype
G/G
G/H
H/H

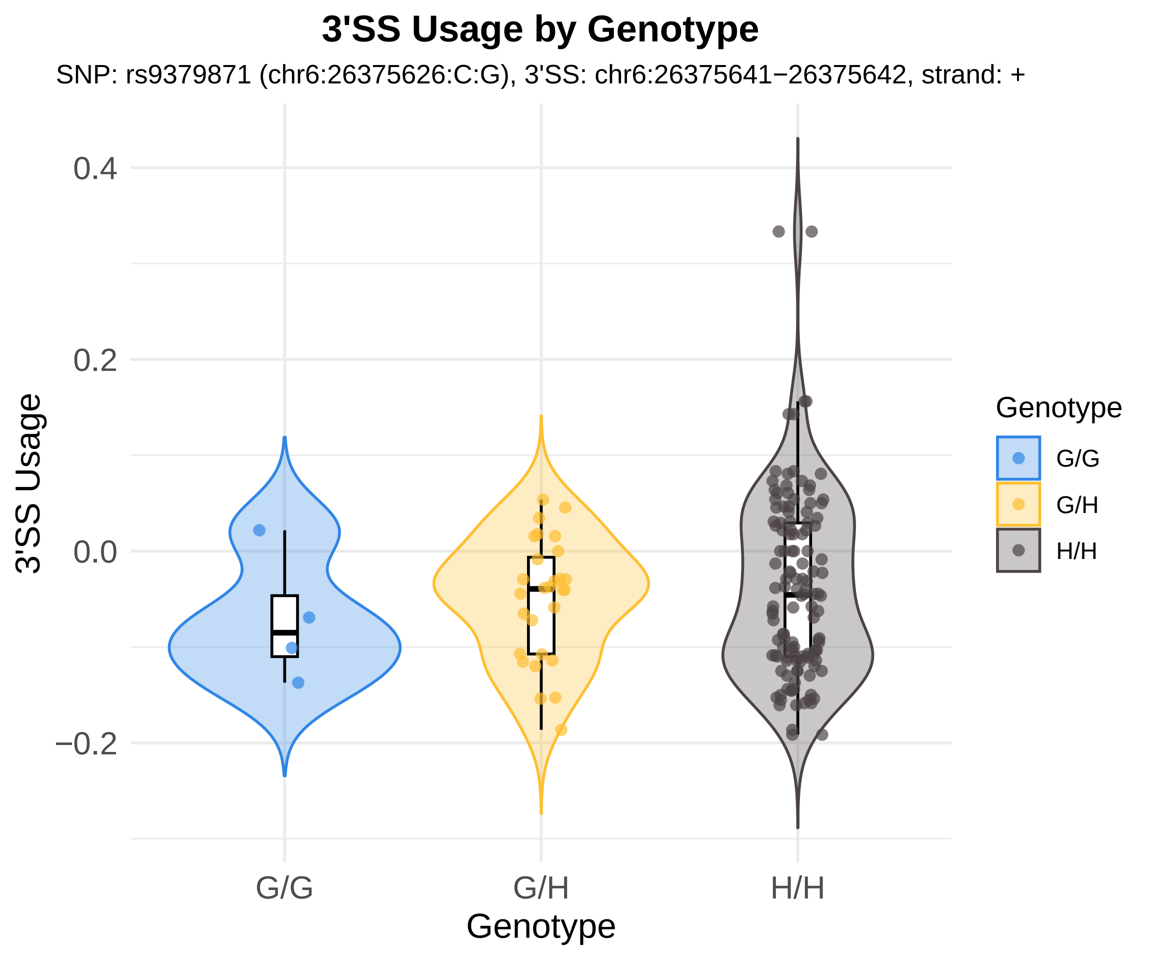

3'SS Usage by Genotype
SNP: rs9379871 (chr6:26375626:C:G), 3'SS: chr6:26375641−26375642, strand: +
3'SS Usage
0.4
0.2
0.0
−0.2
G/G
G/H
H/H
Genotype
Genotype
G/G
G/H
H/H

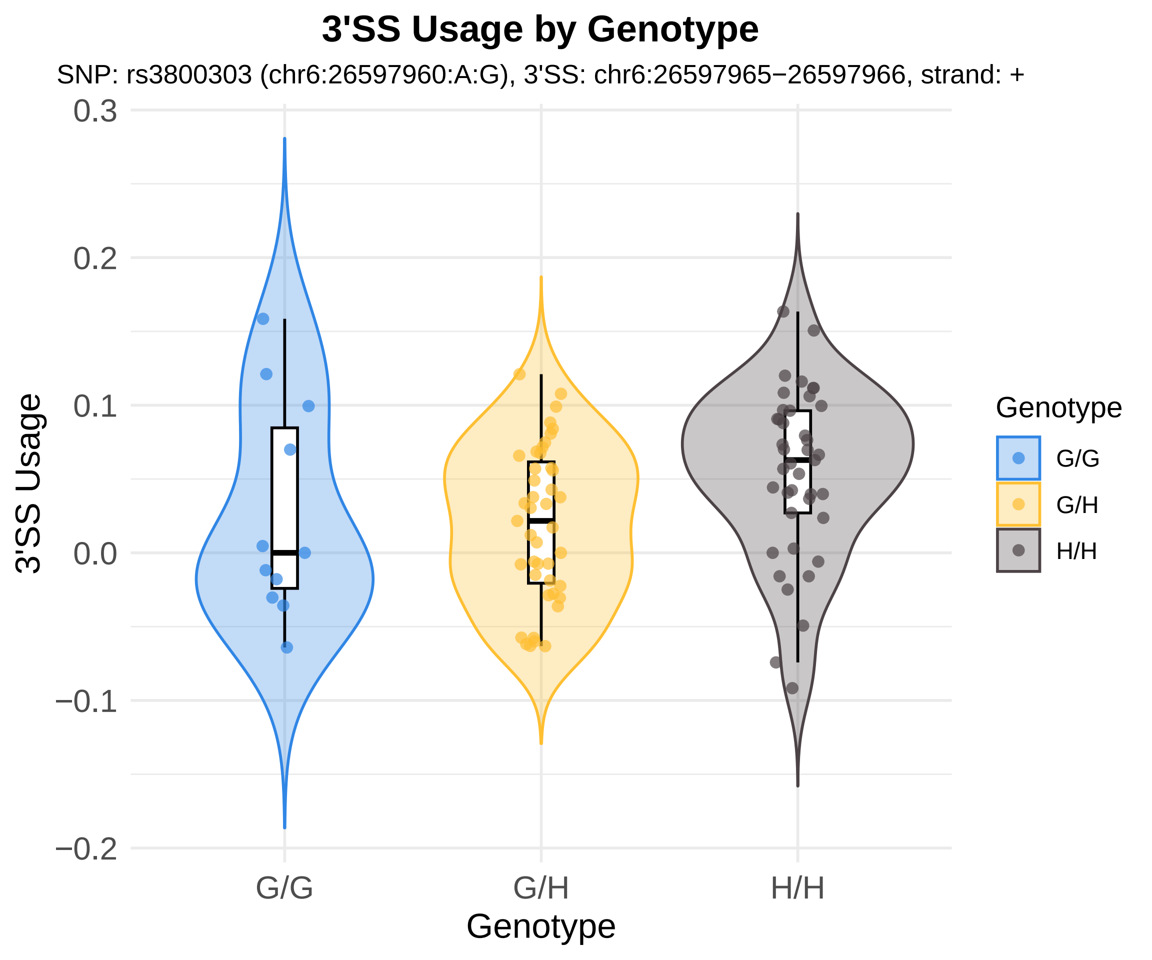

3'SS Usage by Genotype
SNP: rs3800303 (chr6:26597960:A:G), 3'SS: chr6:26597965−26597966, strand: +
3'SS Usage
0.3
0.2
0.1
0.0
−0.1
−0.2
G/G
G/H
H/H
Genotype
Genotype
G/G
G/H
H/H

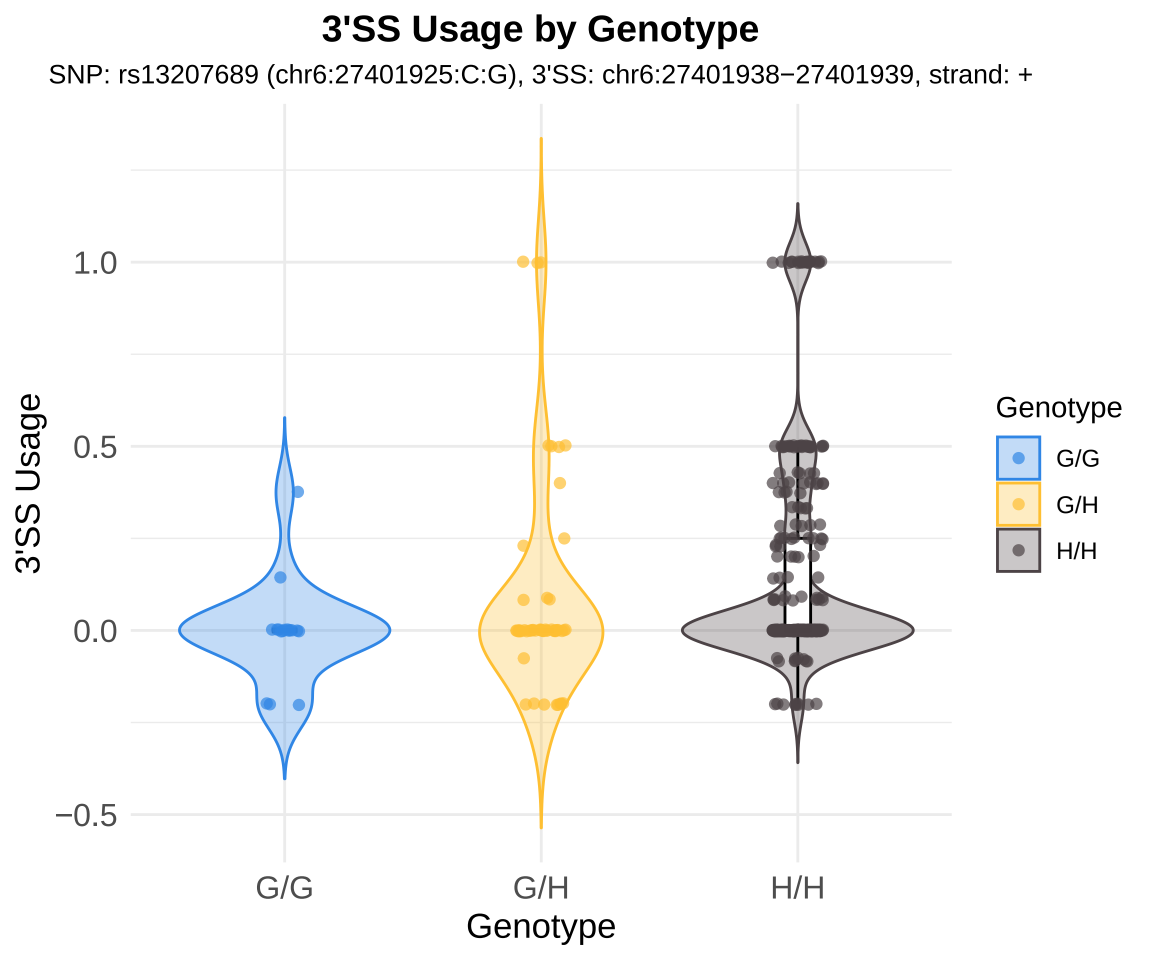
3'SS Usage by Genotype
SNP: rs13207689 (chr6:27401925:C:G), 3'SS: chr6:27401938−27401939, strand: +
3'SS Usage
1.0
0.5
0.0
−0.5
G/G
G/H
H/H
Genotype
Genotype
G/G
G/H
H/H

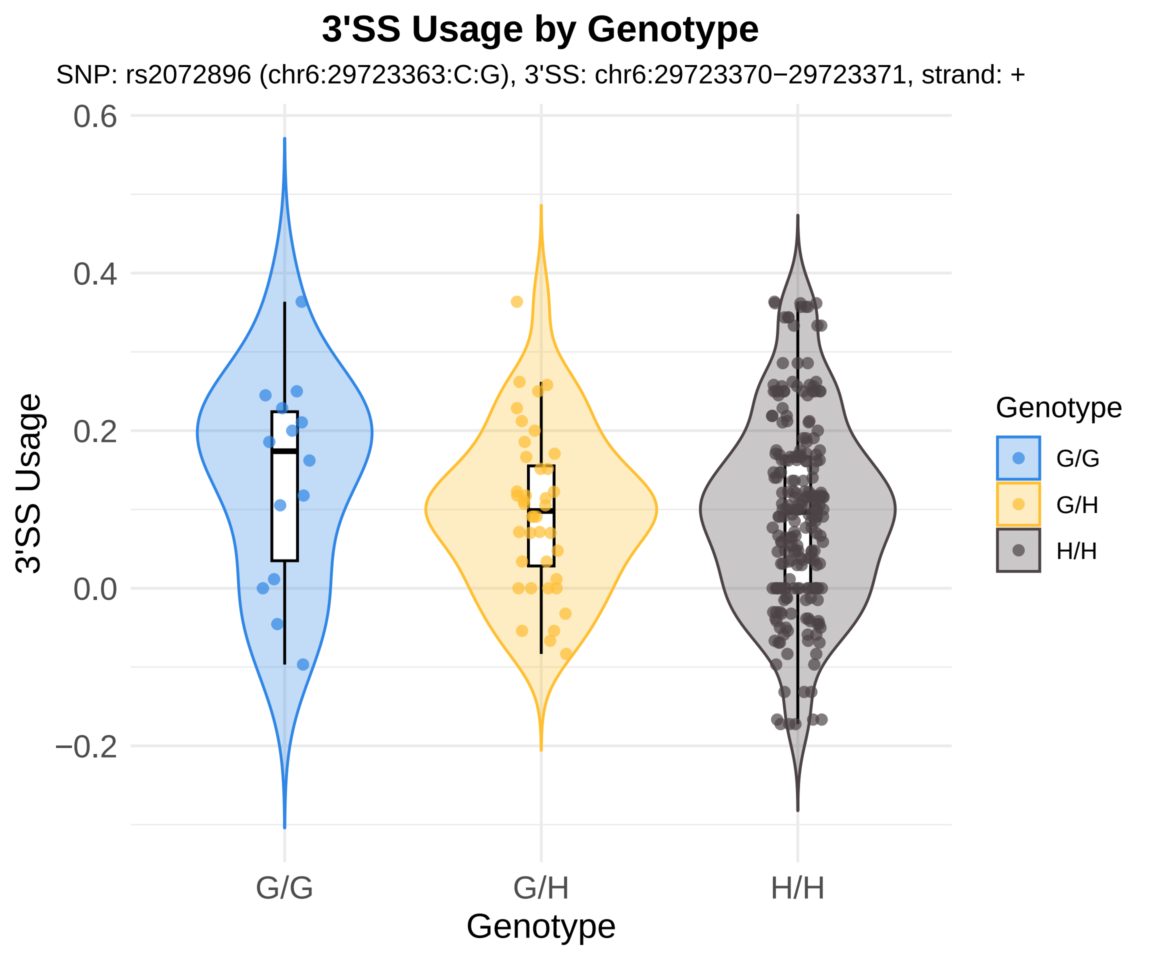
3'SS Usage by Genotype
SNP: rs2072896 (chr6:29723363:C:G), 3'SS: chr6:29723370−29723371, strand: +
3'SS Usage
0.6
0.4
0.2
0.0
−0.2
G/G
G/H
H/H
Genotype
Genotype
G/G
G/H
H/H

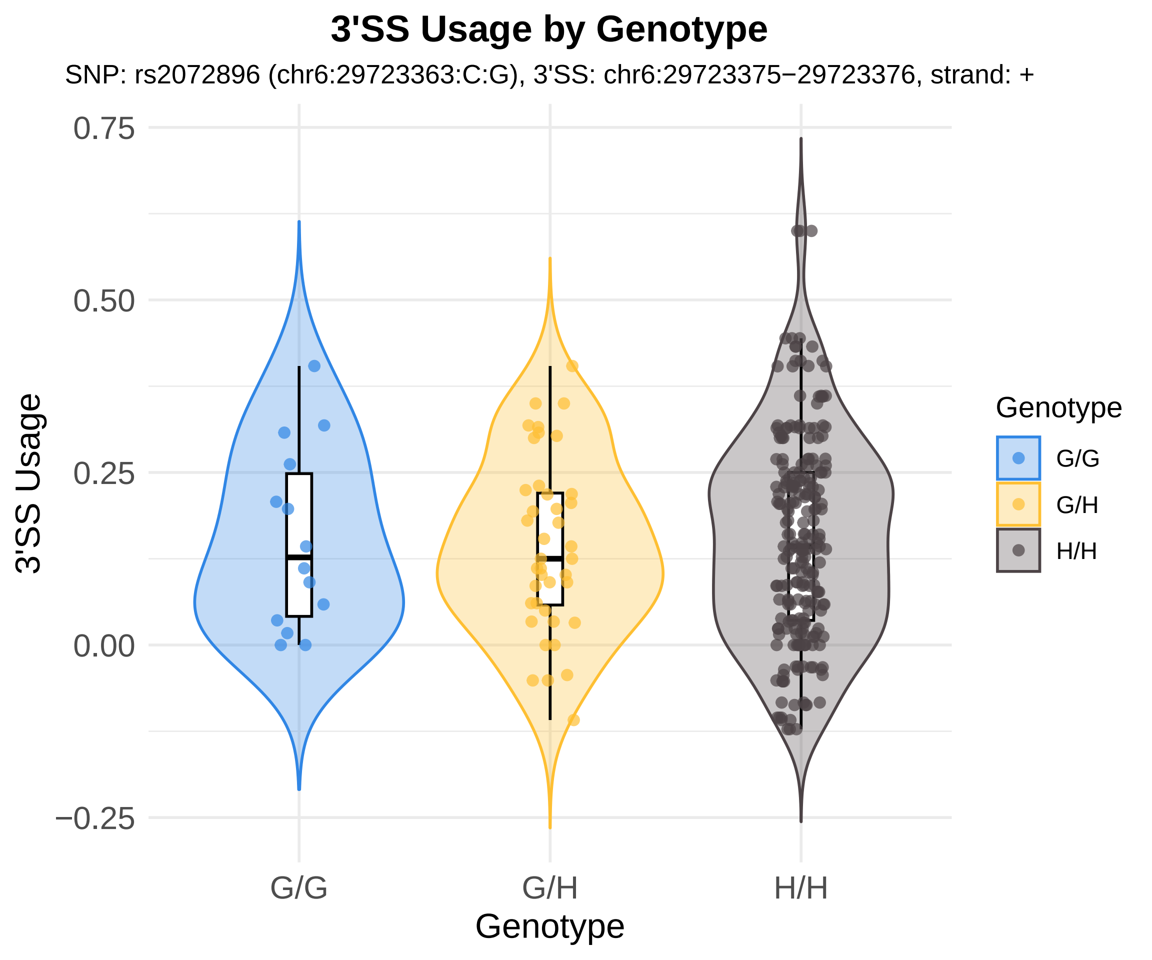
3'SS Usage by Genotype
SNP: rs2072896 (chr6:29723363:C:G), 3'SS: chr6:29723375−29723376, strand: +
3'SS Usage
0.75
0.50
0.25
0.00
−0.25
G/G
G/H
H/H
Genotype
Genotype
G/G
G/H
H/H

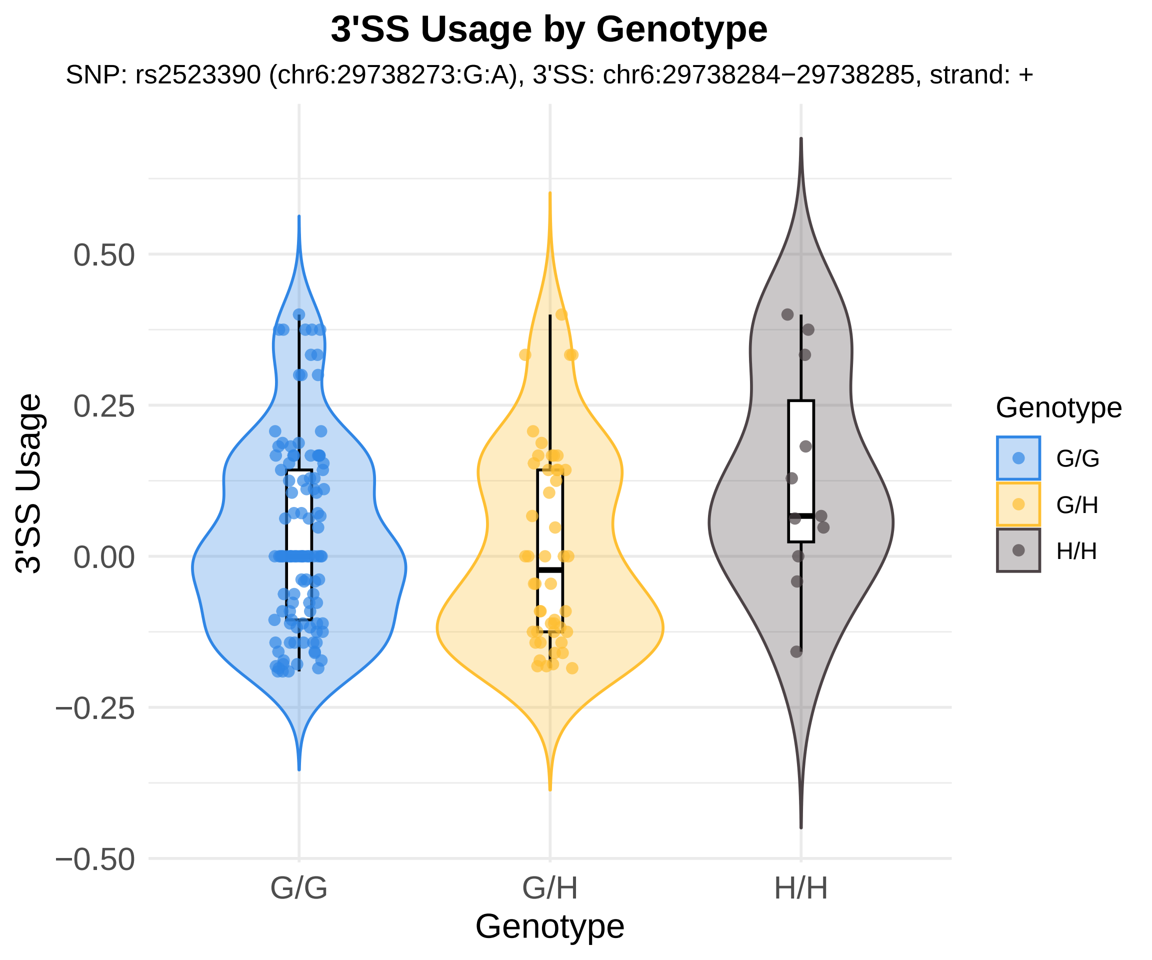
3'SS Usage by Genotype
SNP: rs2523390 (chr6:29738273:G:A), 3'SS: chr6:29738284−29738285, strand: +
3'SS Usage
0.50
0.25
0.00
−0.25
−0.50
G/G
G/H
H/H
Genotype
Genotype
G/G
G/H
H/H

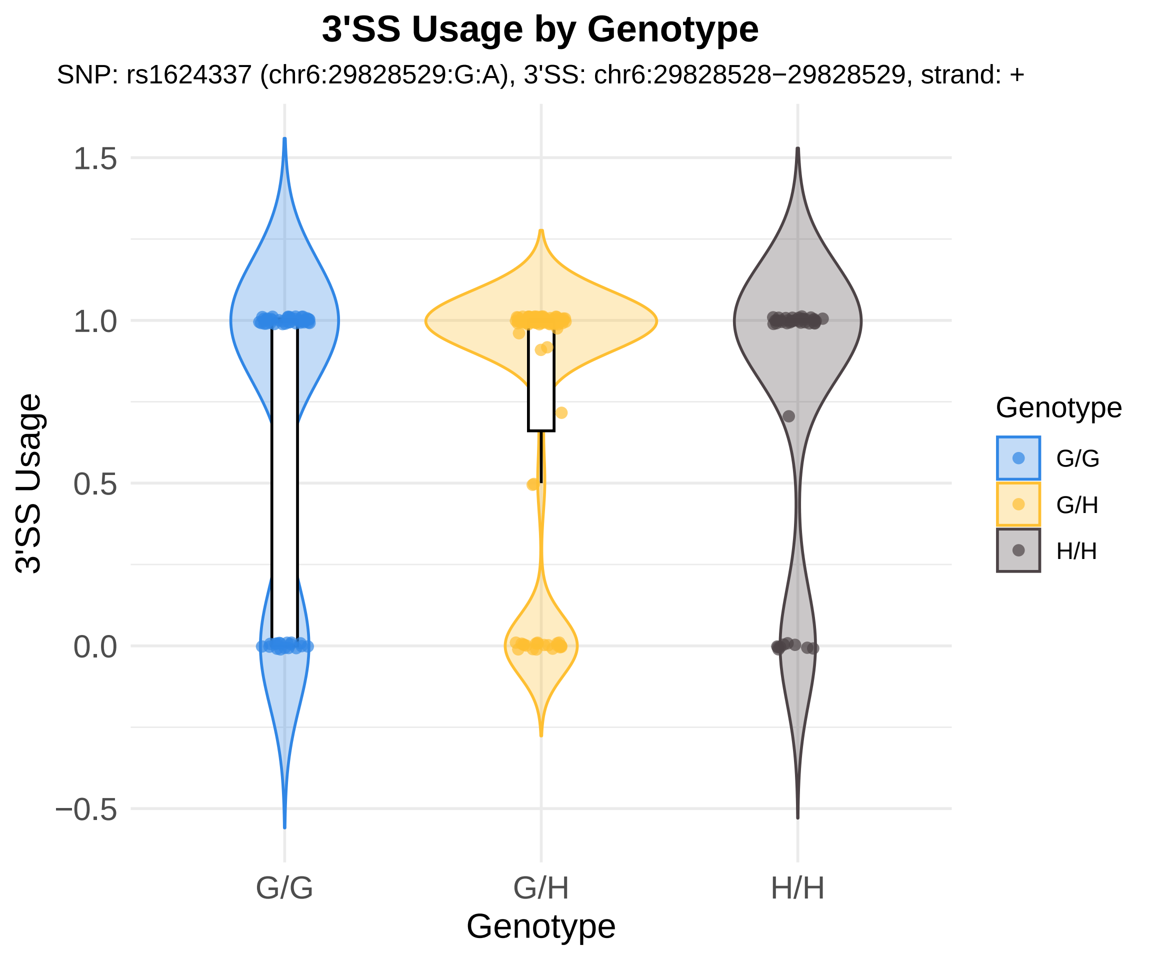
3'SS Usage by Genotype
SNP: rs1624337 (chr6:29828529:G:A), 3'SS: chr6:29828528−29828529, strand: +
3'SS Usage
1.5
1.0
0.5
0.0
−0.5
G/G
G/H
H/H
Genotype
Genotype
G/G
G/H
H/H

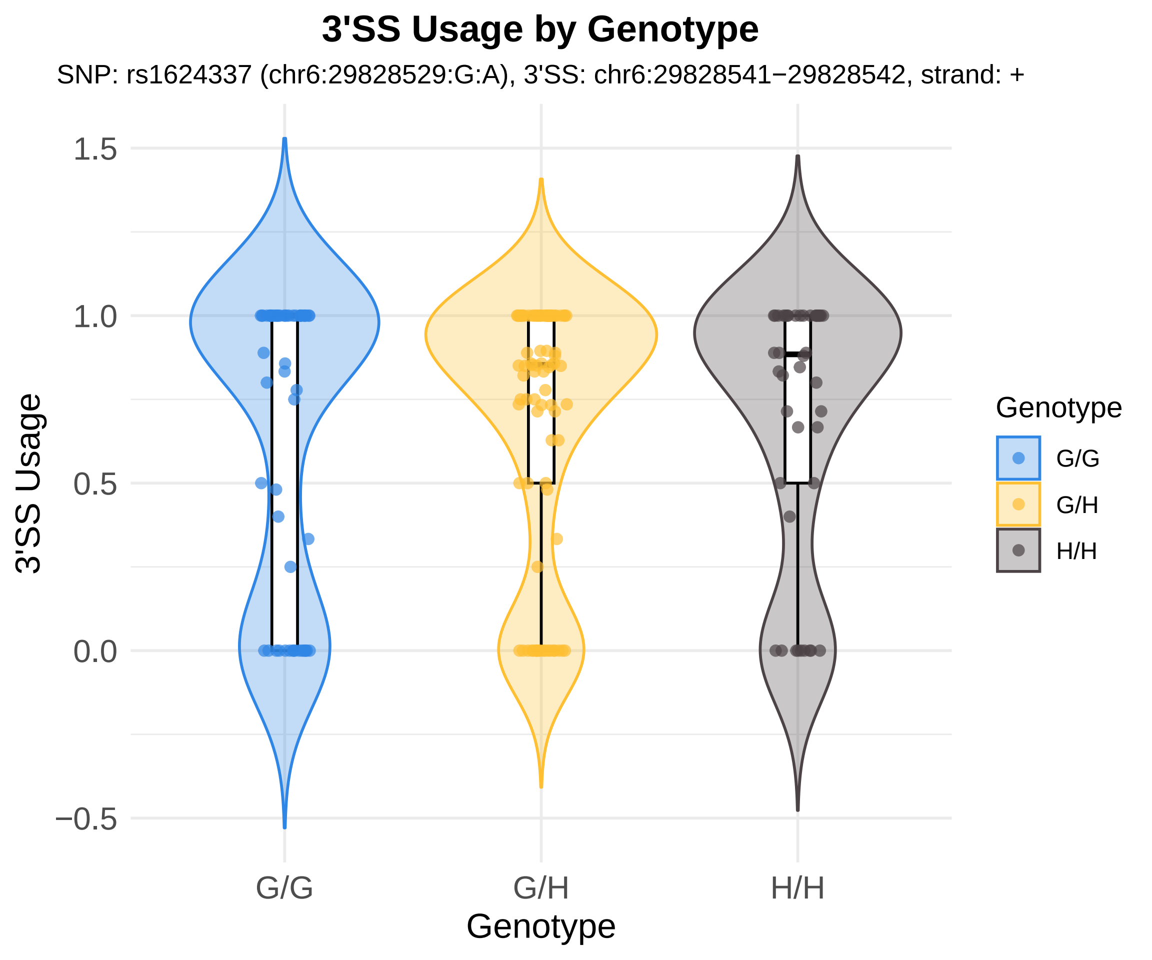

3'SS Usage by Genotype
SNP: rs1624337 (chr6:29828529:G:A), 3'SS: chr6:29828541−29828542, strand: +
3'SS Usage
1.5
1.0
0.5
0.0
−0.5
G/G
G/H
H/H
Genotype
Genotype
G/G
G/H
H/H

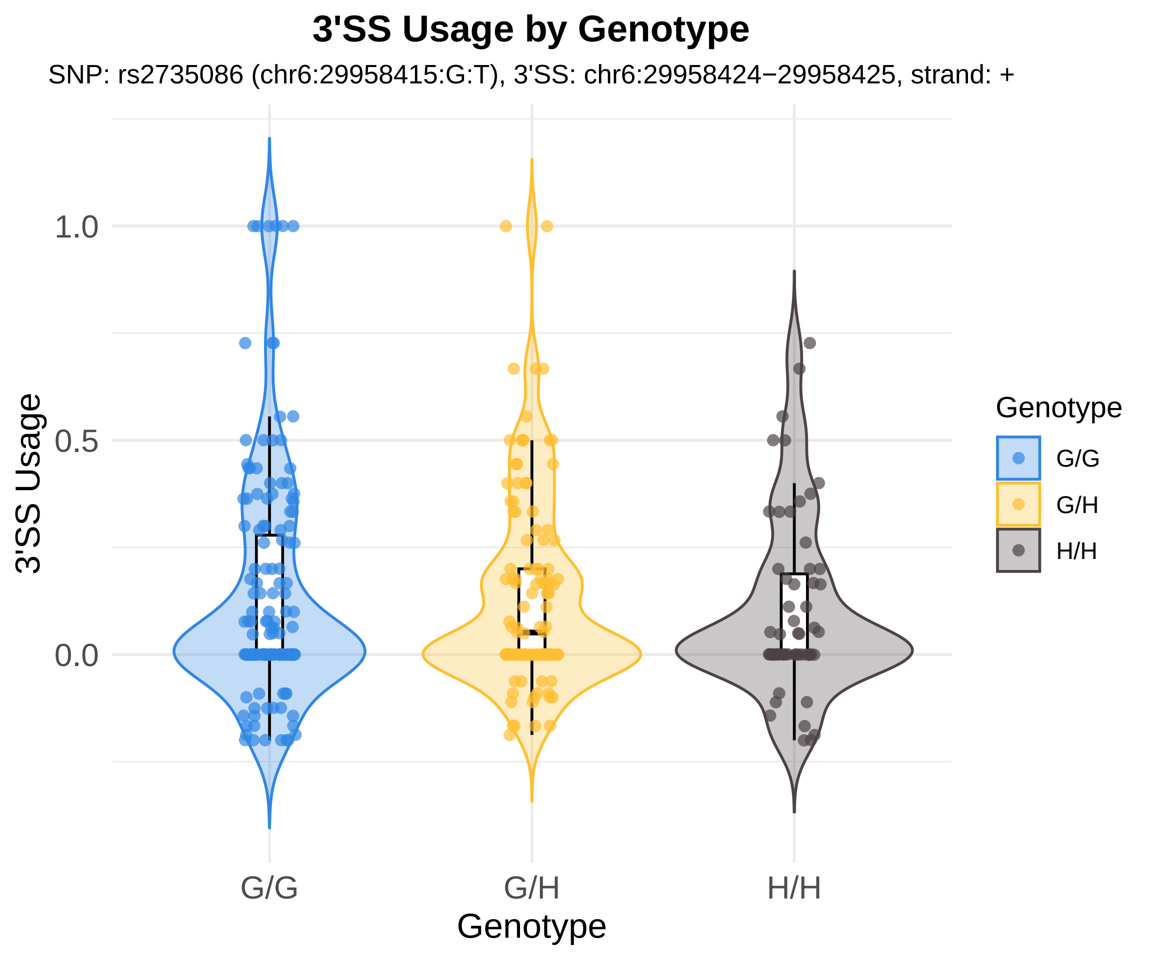
3'SS Usage by Genotype
SNP: rs2735086 (chr6:29958415:G:T), 3'SS: chr6:29958424−29958425, strand: +
3'SS Usage
1.0
0.5
0.0
G/G
G/H
H/H
Genotype
Genotype
G/G
G/H
H/H

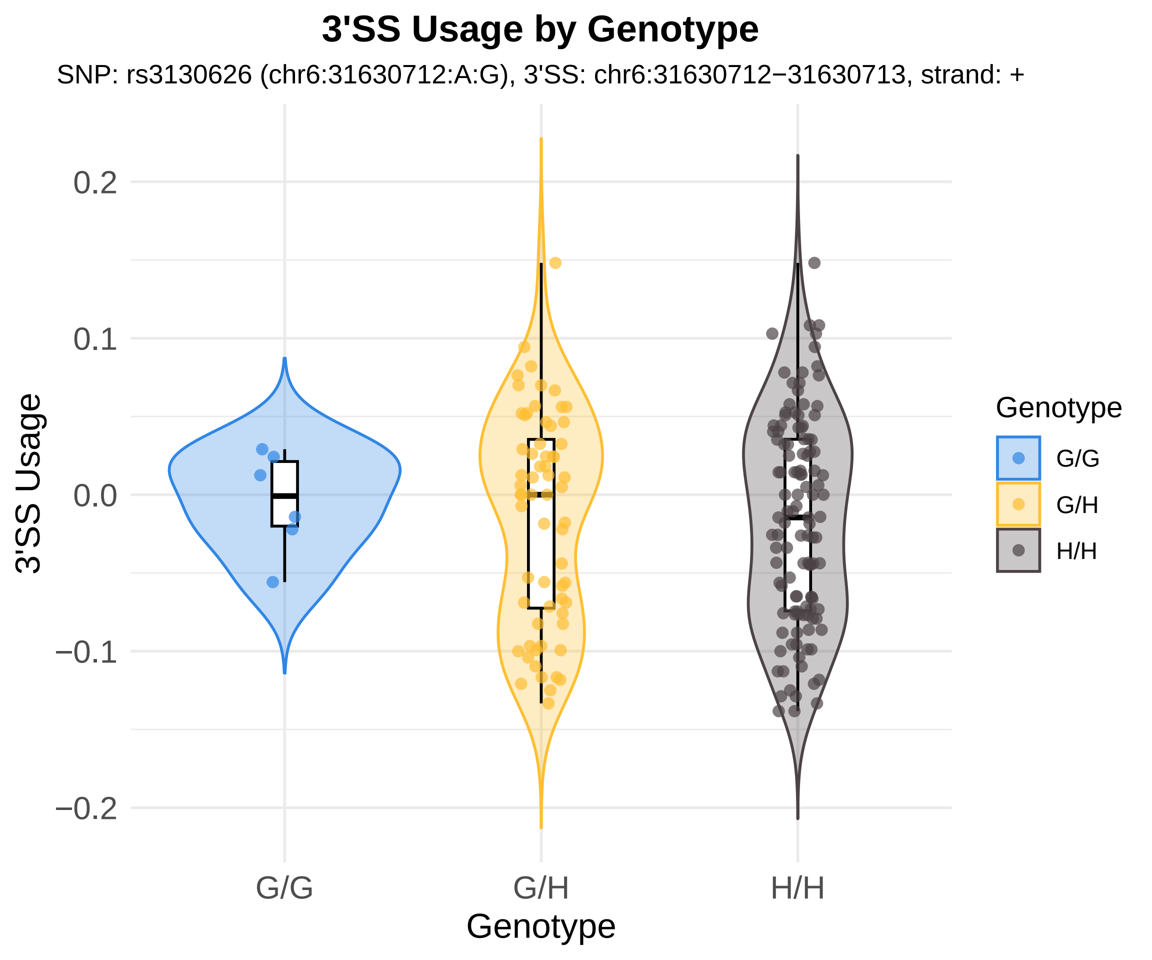

3'SS Usage by Genotype
SNP: rs3130626 (chr6:31630712:A:G), 3'SS: chr6:31630712−31630713, strand: +
3'SS Usage
0.2
0.1
0.0
−0.1
−0.2
G/G
G/H
H/H
Genotype
Genotype
G/G
G/H
H/H

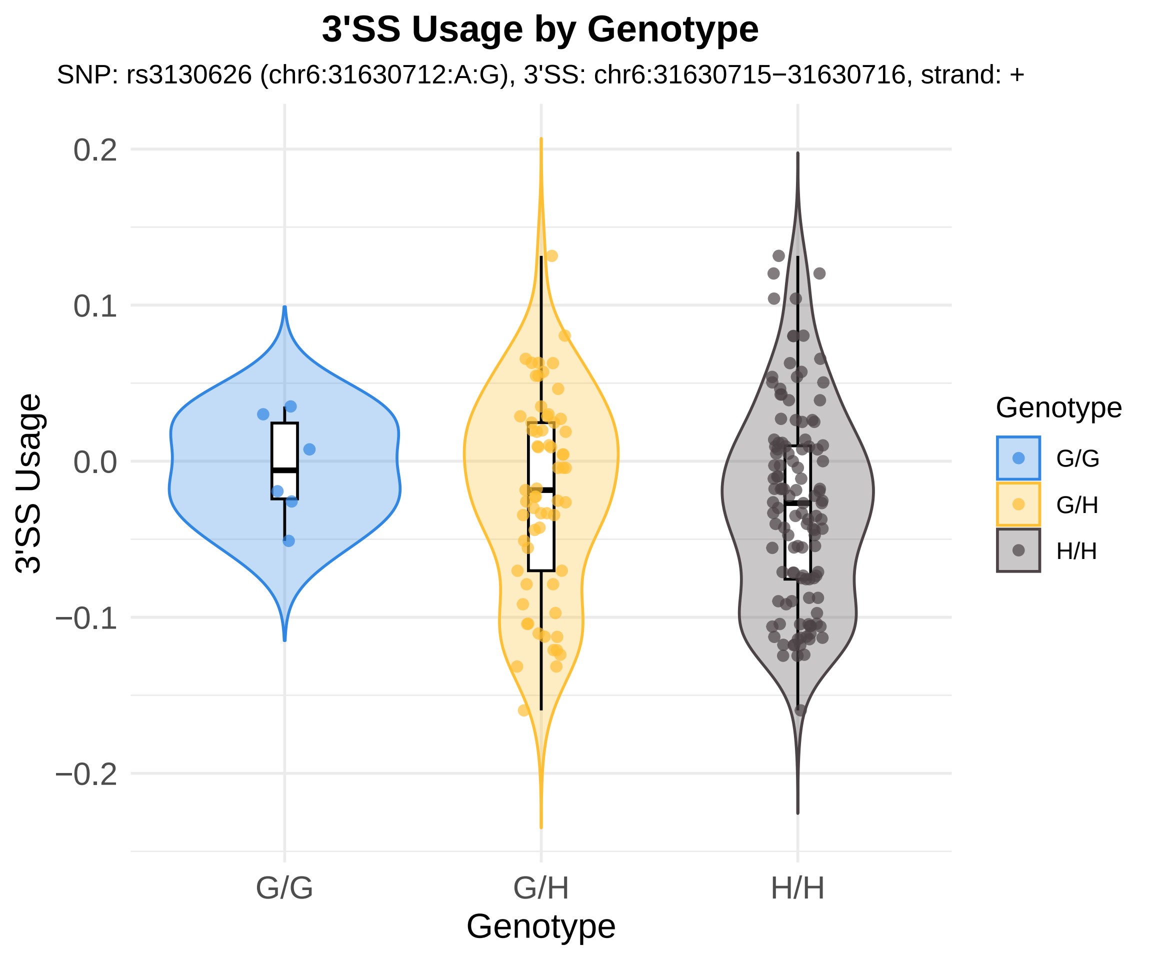
3'SS Usage by Genotype
SNP: rs3130626 (chr6:31630712:A:G), 3'SS: chr6:31630715−31630716, strand: +
0.2
0.1
0.0
−0.1
−0.2
3'SS Usage
G/G
G/H
H/H
Genotype
Genotype
G/G
G/H
H/H

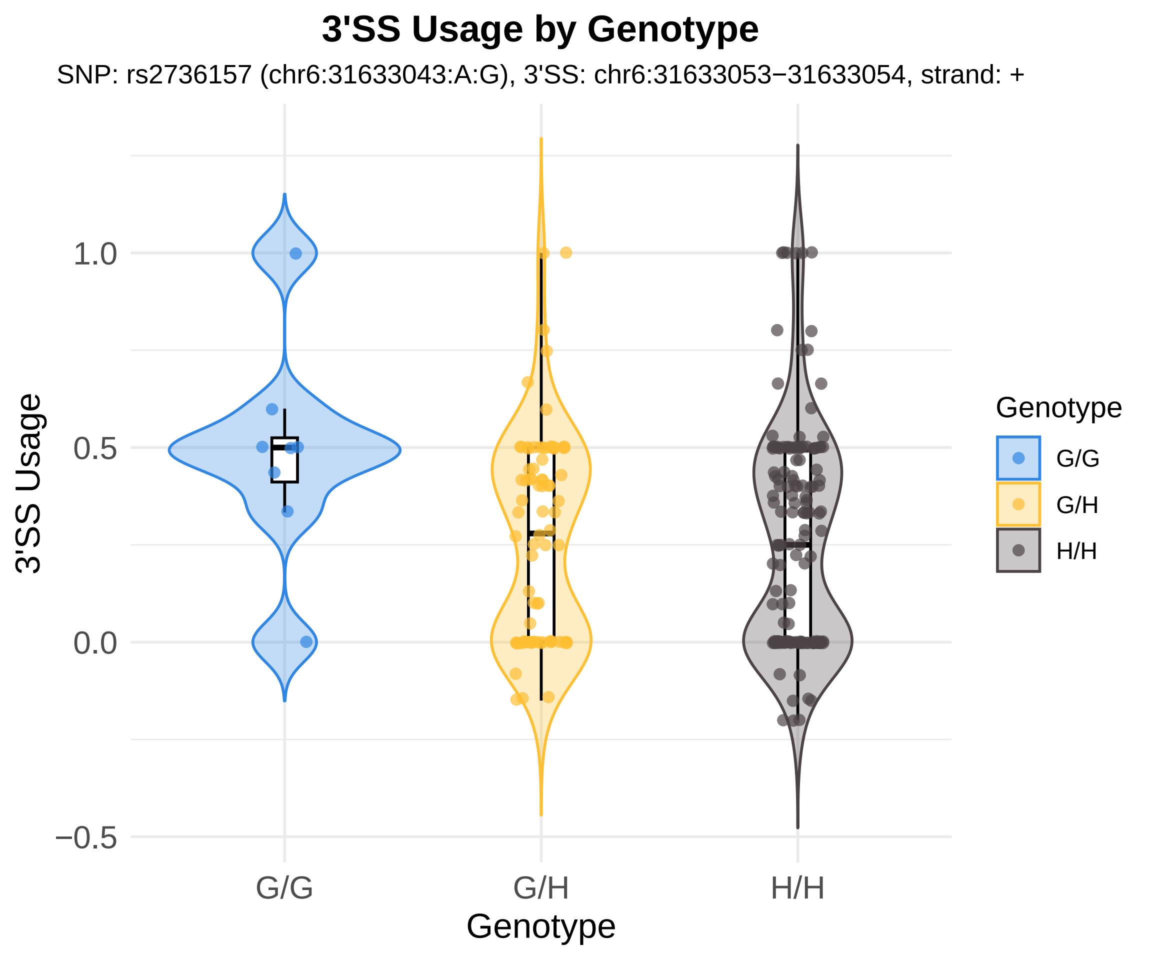

3'SS Usage by Genotype
SNP: rs2736157 (chr6:31633043:A:G), 3'SS: chr6:31633053−31633054, strand: +
3'SS Usage
1.0
0.5
0.0
−0.5
G/G
G/H
H/H
Genotype
Genotype
G/G
G/H
H/H

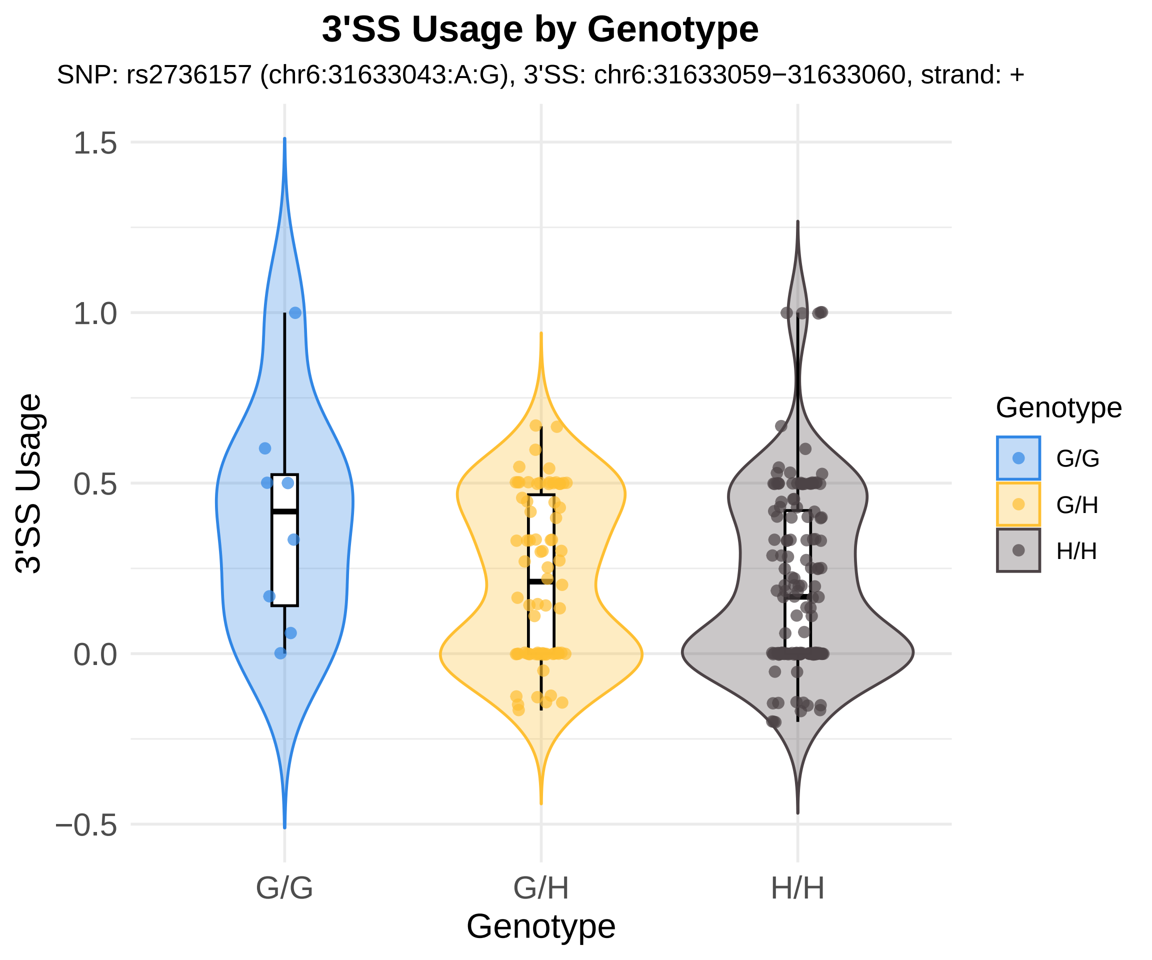

3'SS Usage by Genotype
SNP: rs2736157 (chr6:31633043:A:G), 3'SS: chr6:31633059−31633060, strand: +
3'SS Usage
1.5
1.0
0.5
0.0
−0.5
G/G
G/H
H/H
Genotype
Genotype
G/G
G/H
H/H

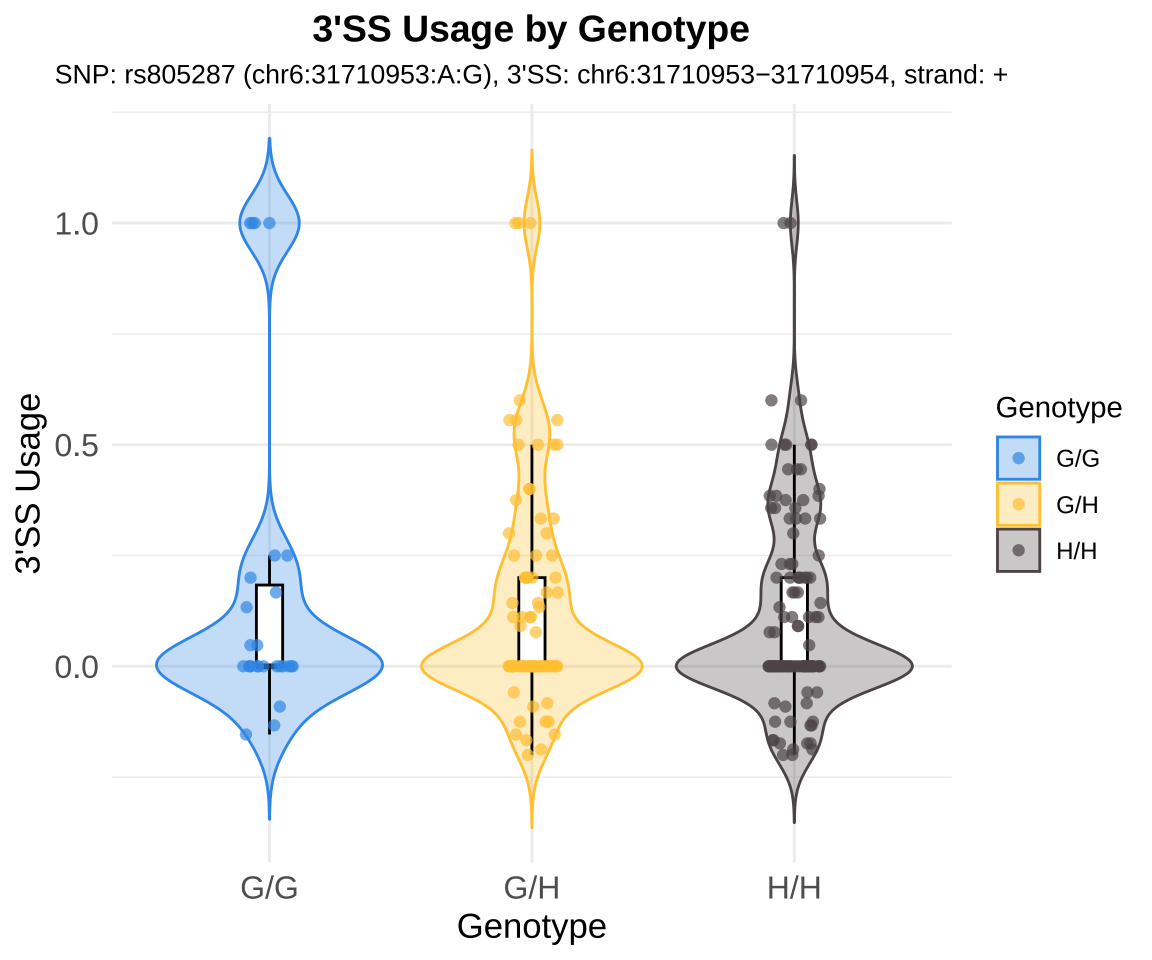
3'SS Usage by Genotype
SNP: rs805287 (chr6:31710953:A:G), 3'SS: chr6:31710953−31710954, strand: +
3'SS Usage
1.0
0.5
0.0
G/G
G/H
H/H
Genotype
Genotype
G/G
G/H
H/H

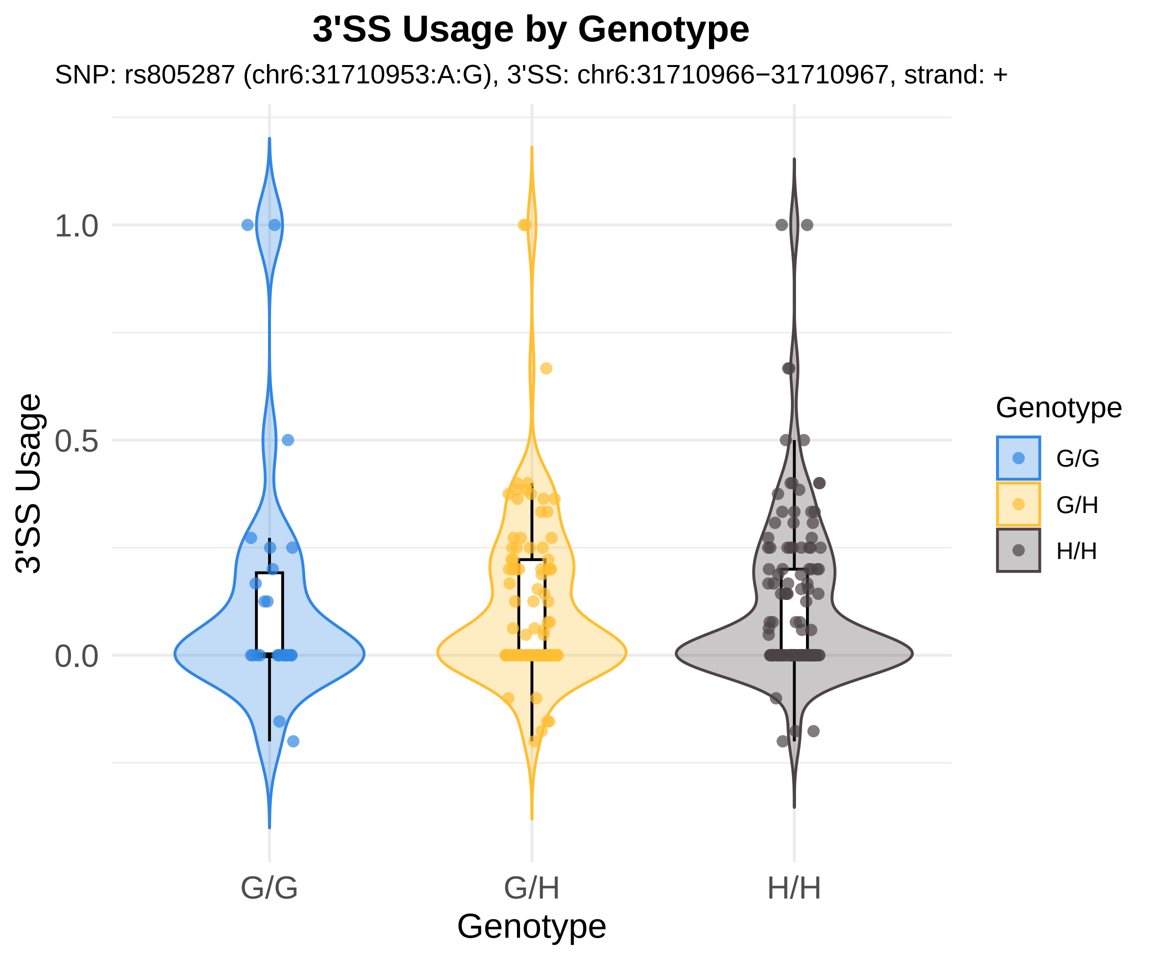
3'SS Usage by Genotype
SNP: rs805287 (chr6:31710953:A:G), 3'SS: chr6:31710966−31710967, strand: +
3'SS Usage
1.0
0.5
0.0
G/G
G/H
H/H
Genotype
Genotype
G/G
G/H
H/H

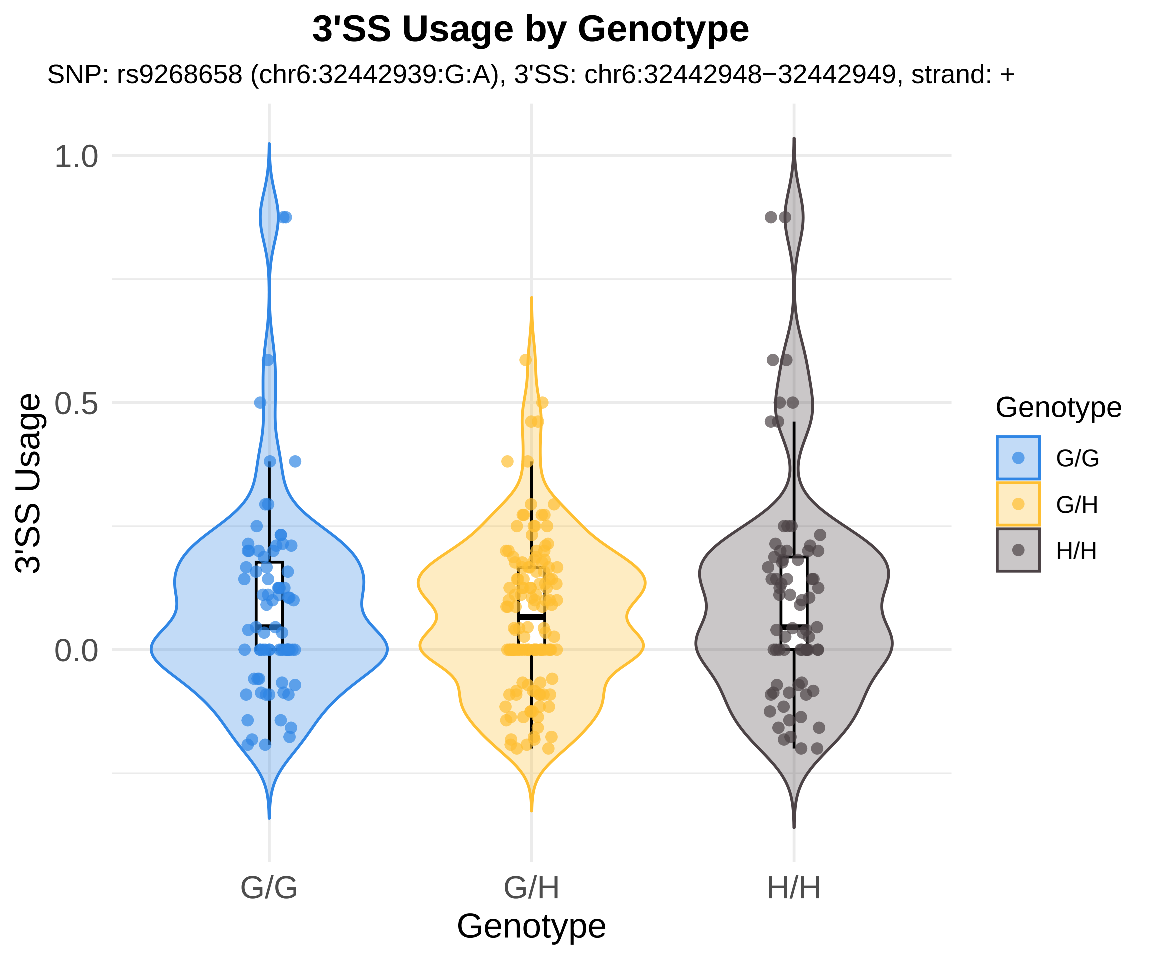
3'SS Usage by Genotype
SNP: rs9268658 (chr6:32442939:G:A), 3'SS: chr6:32442948−32442949, strand: +
3'SS Usage
1.0
0.5
0.0
G/G
G/H
H/H
Genotype
Genotype
G/G
G/H
H/H

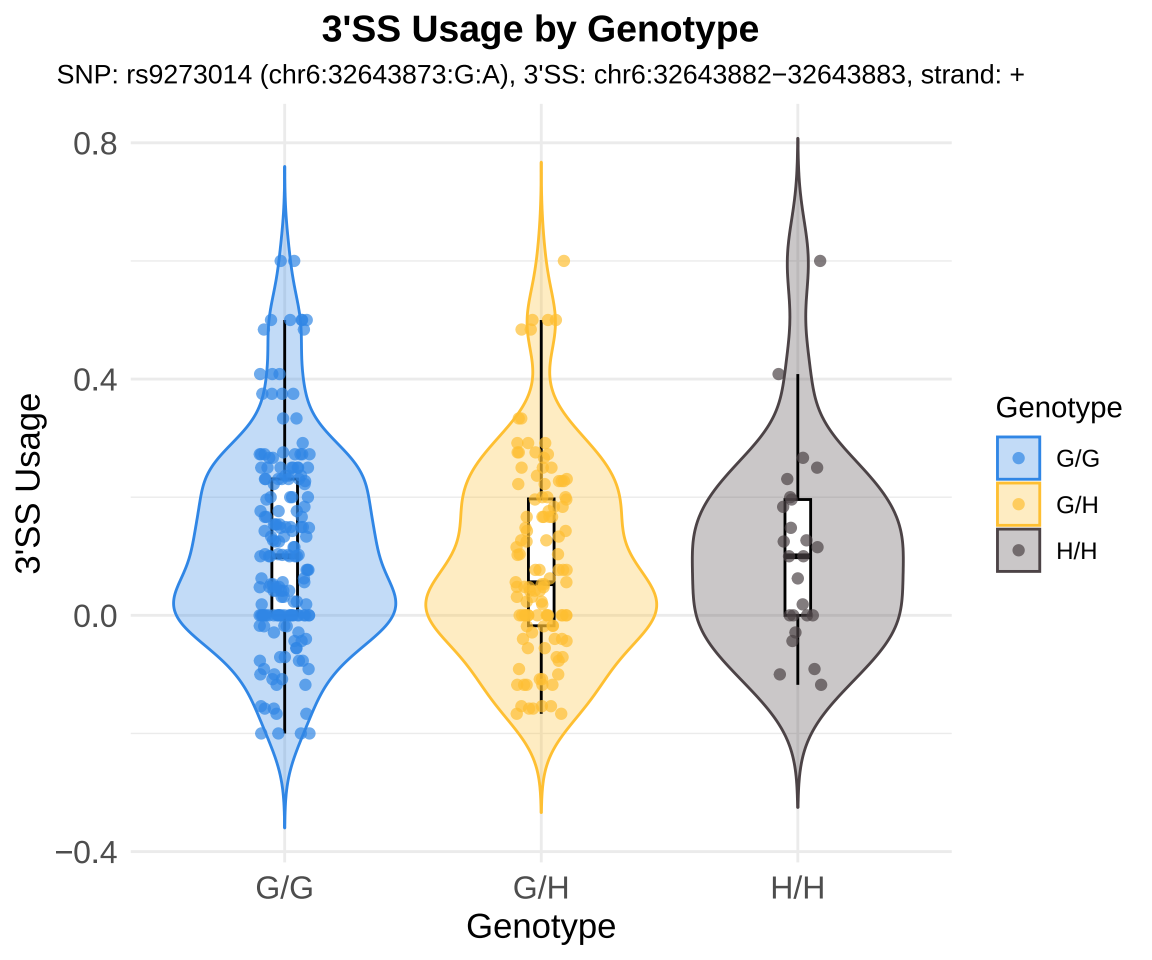
3'SS Usage by Genotype
SNP: rs9273014 (chr6:32643873:G:A), 3'SS: chr6:32643882−32643883, strand: +
3'SS Usage
0.8
0.4
0.0
−0.4
G/G
G/H
H/H
Genotype
Genotype
G/G
G/H
H/H

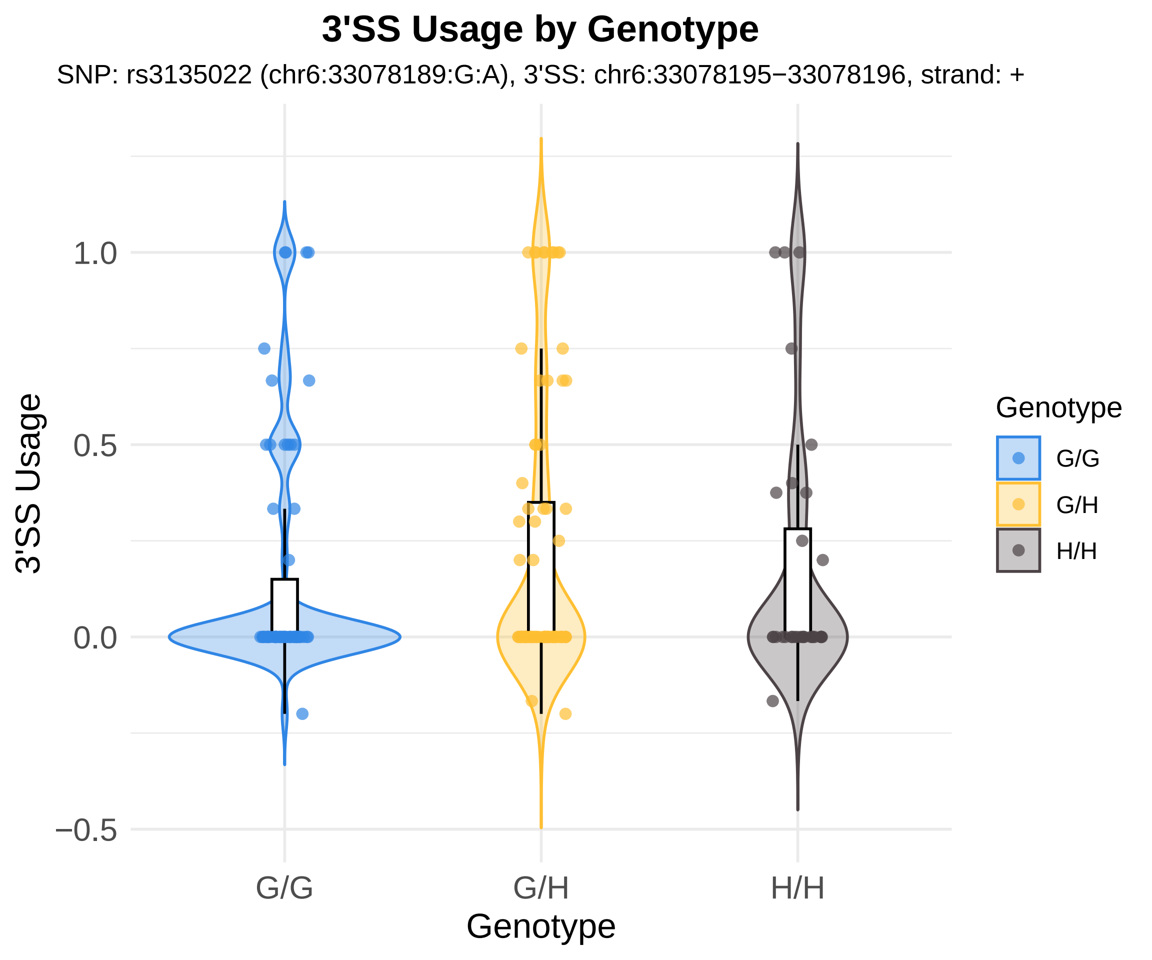

3'SS Usage by Genotype
SNP: rs3135022 (chr6:33078189:G:A), 3'SS: chr6:33078195−33078196, strand: +
3'SS Usage
1.0
0.5
0.0
−0.5
G/G
G/H
H/H
Genotype
Genotype
G/G
G/H
H/H

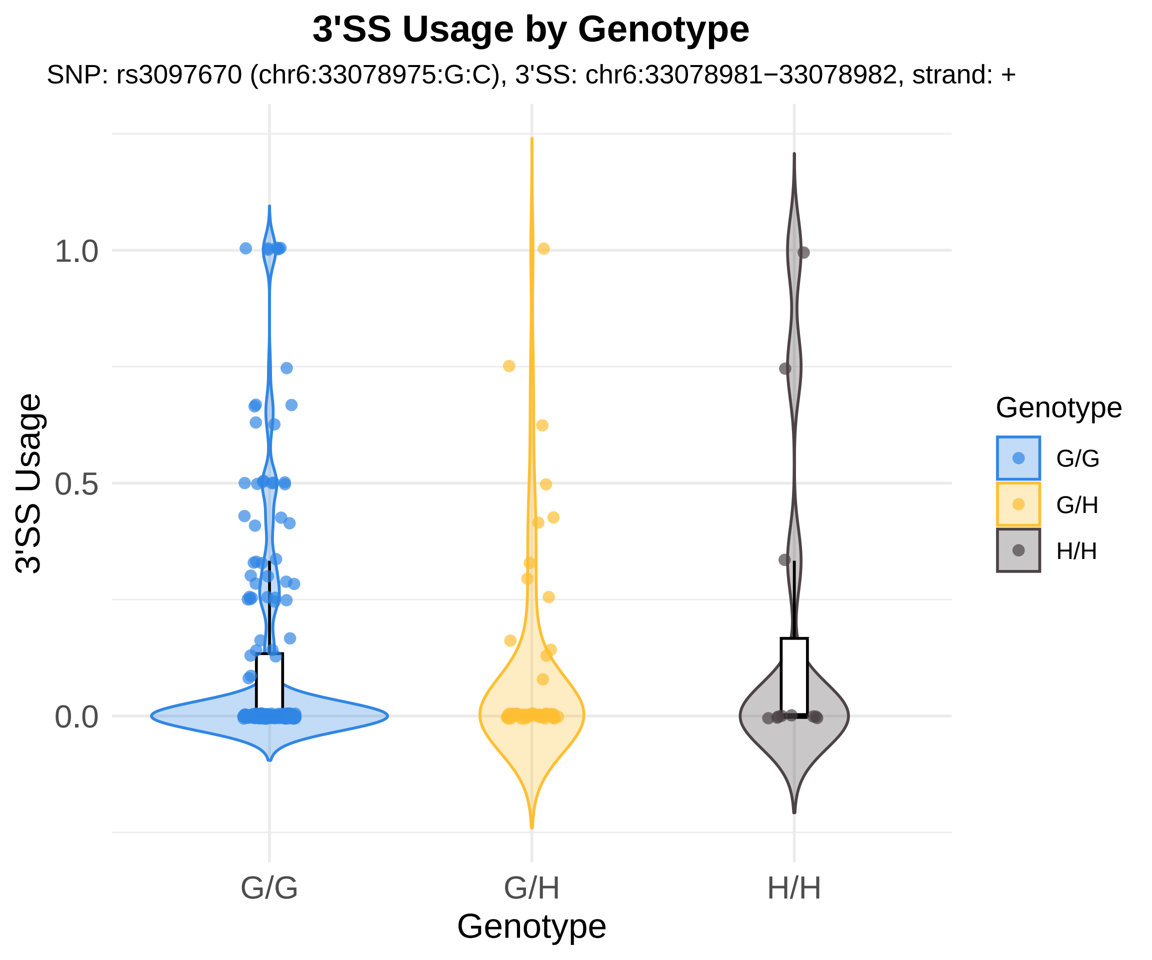
3'SS Usage by Genotype
SNP: rs3097670 (chr6:33078975:G:C), 3'SS: chr6:33078981−33078982, strand: +
3'SS Usage
1.0
0.5
0.0
G/G
G/H
H/H
Genotype
Genotype
G/G
G/H
H/H

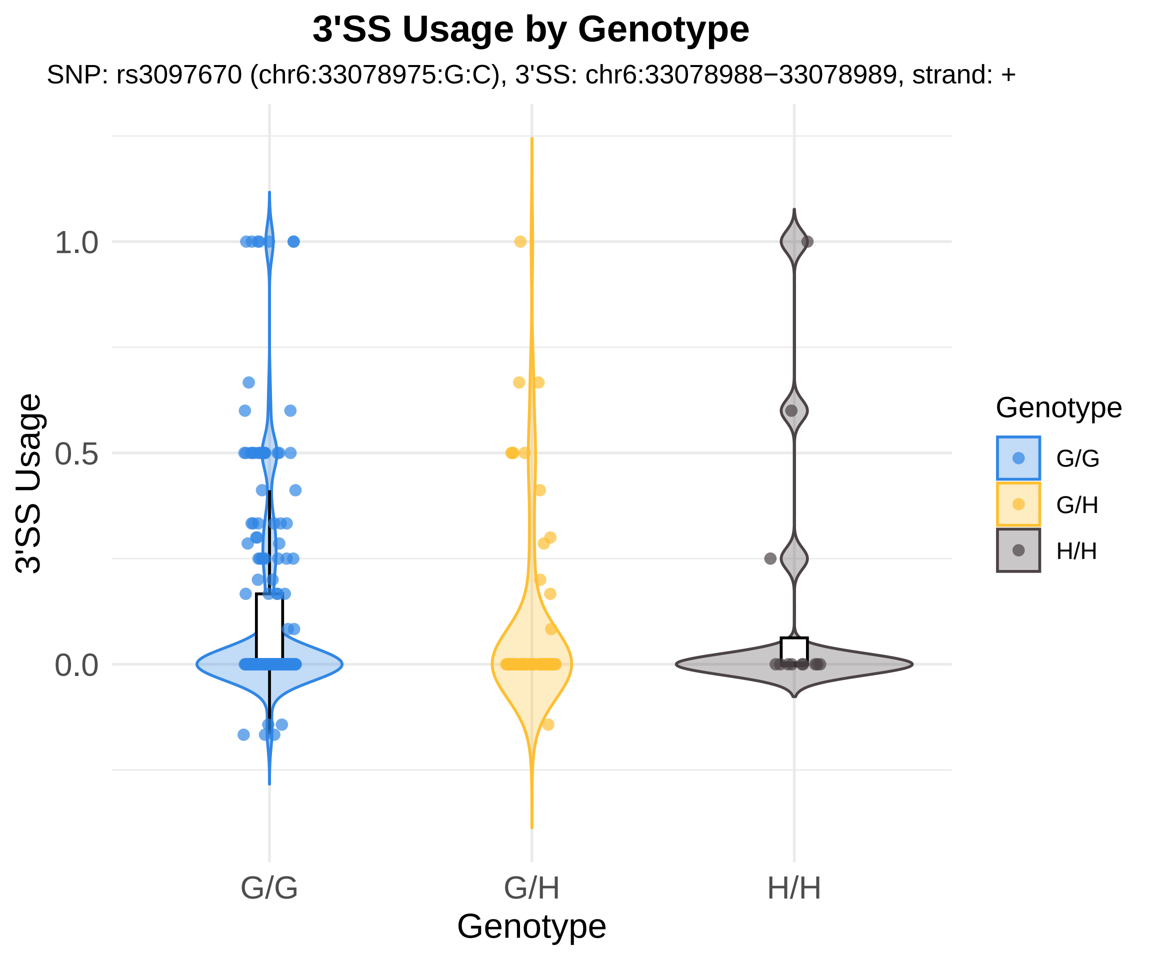
3'SS Usage by Genotype
SNP: rs3097670 (chr6:33078975:G:C), 3'SS: chr6:33078988−33078989, strand: +
3'SS Usage
1.0
0.5
0.0
G/G
G/H
H/H
Genotype
Genotype
G/G
G/H
H/H

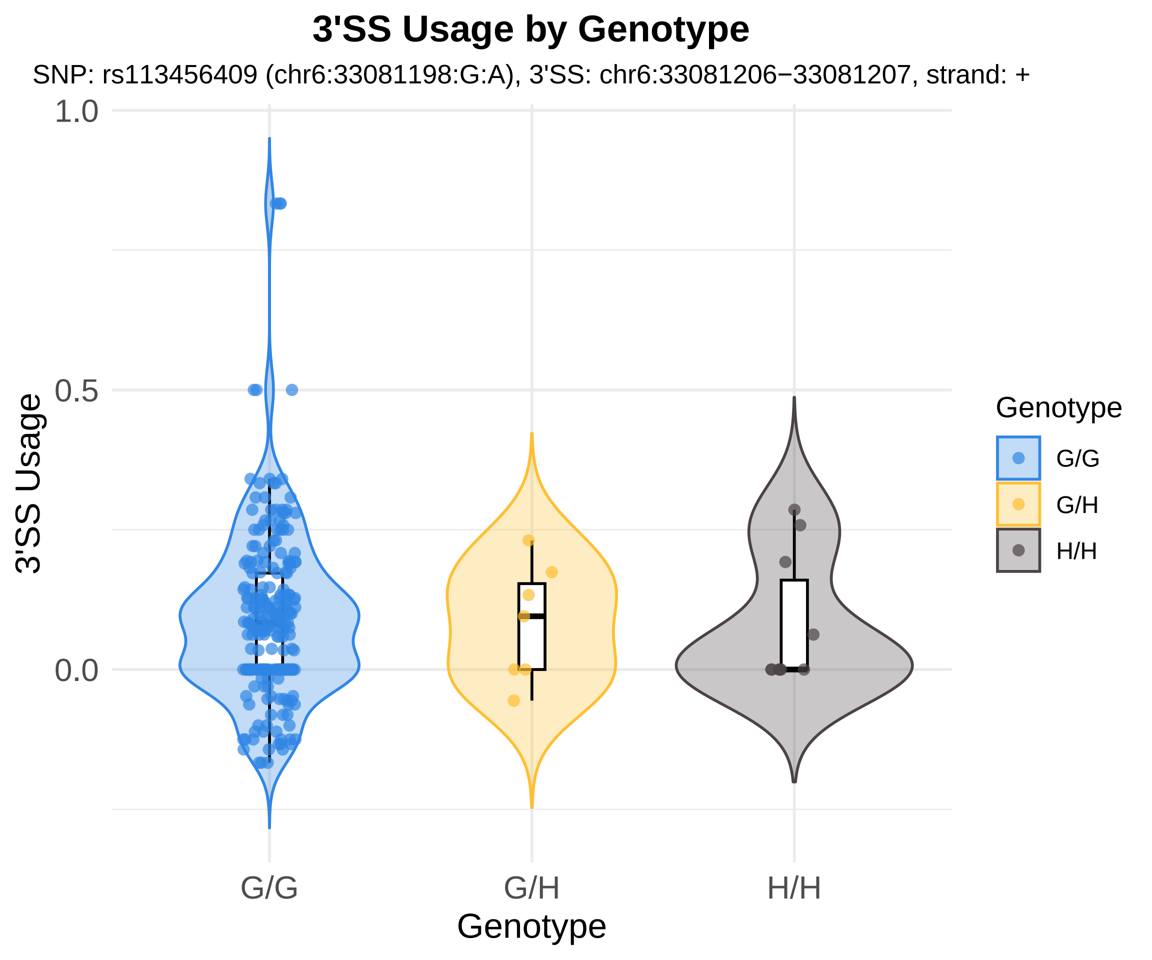
3'SS Usage by Genotype
SNP: rs113456409 (chr6:33081198:G:A), 3'SS: chr6:33081206−33081207, strand: +
1.0
0.5
0.0
3'SS Usage
G/G
G/H
H/H
Genotype
Genotype
G/G
G/H
H/H

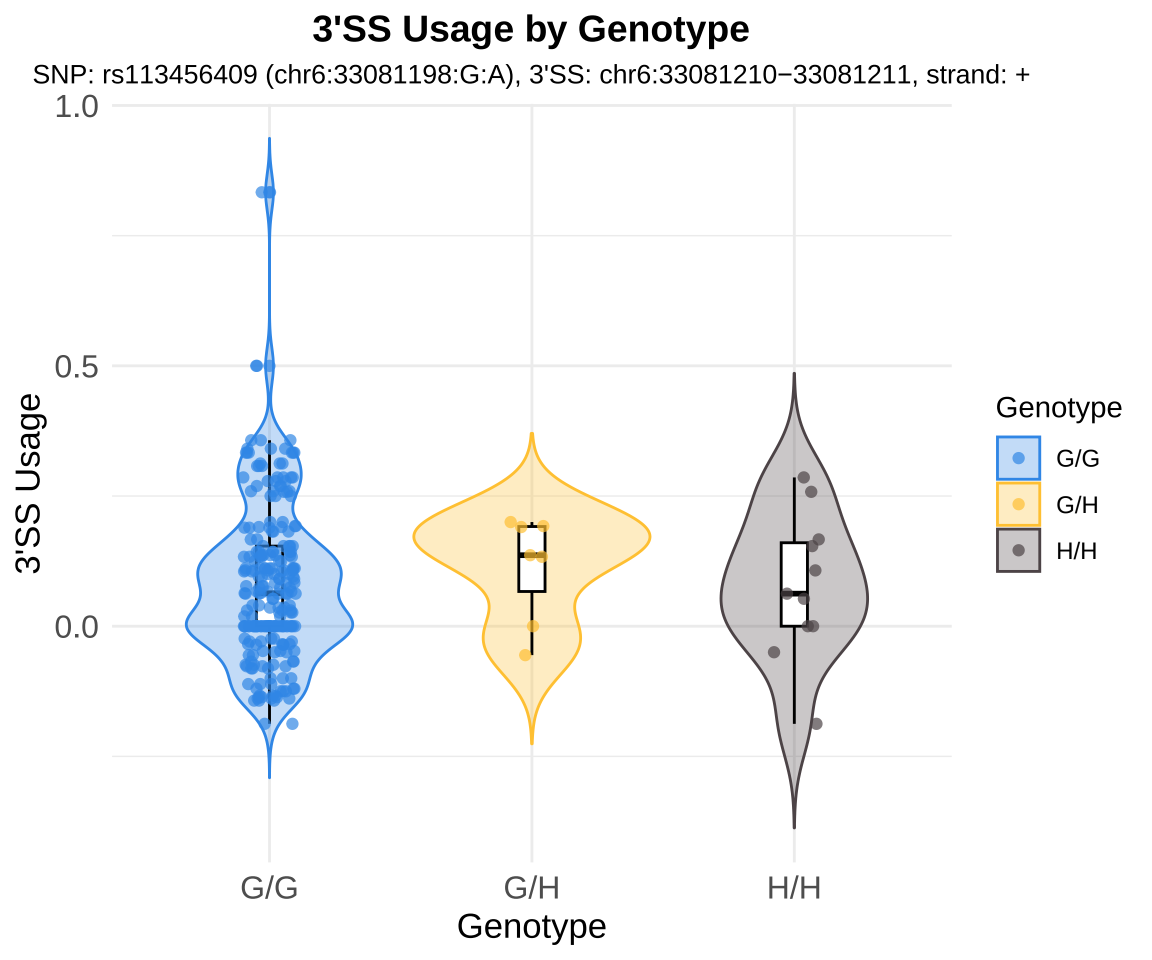
3'SS Usage by Genotype
SNP: rs113456409 (chr6:33081198:G:A), 3'SS: chr6:33081210−33081211, strand: +
1.0
0.5
0.0
3'SS Usage
G/G
G/H
H/H
Genotype
Genotype
G/G
G/H
H/H

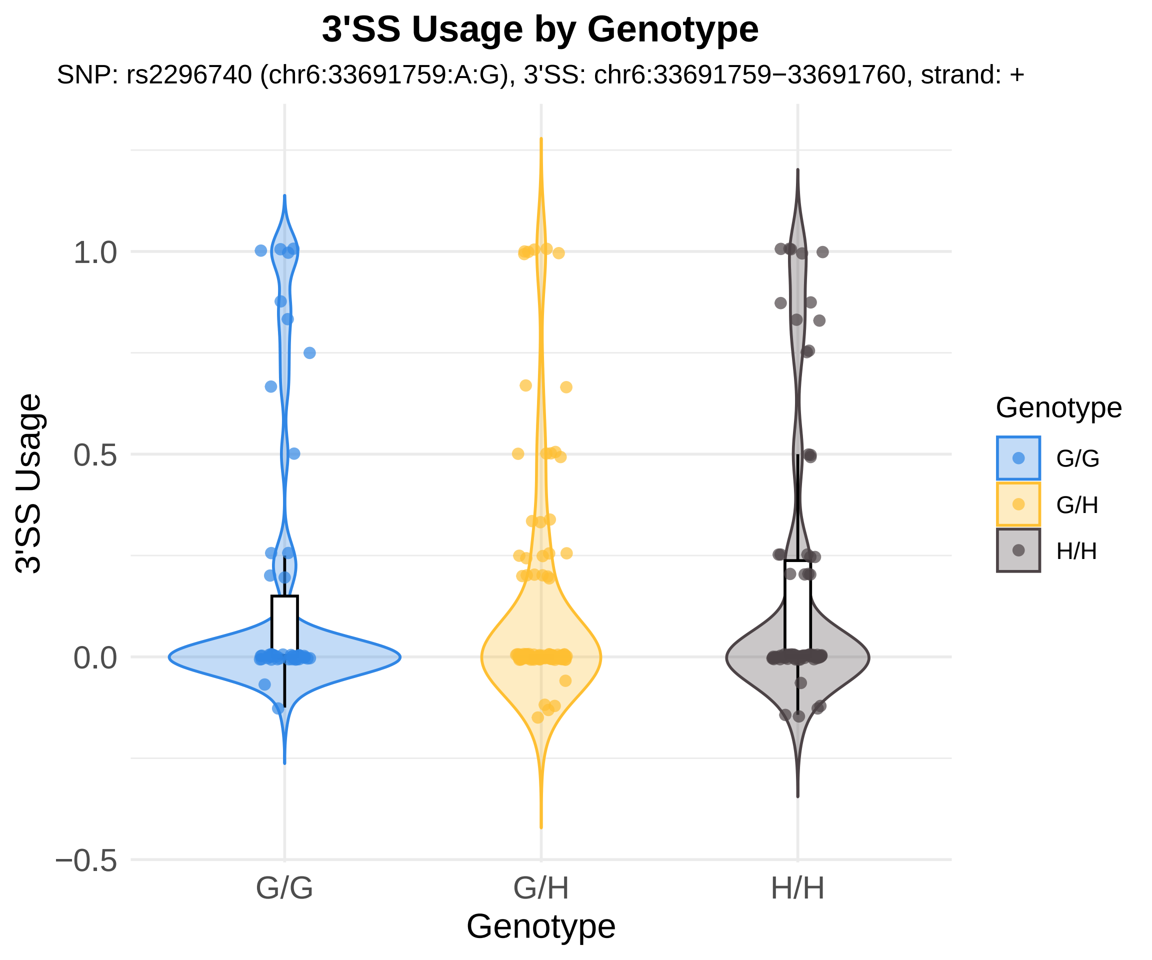
3'SS Usage by Genotype
SNP: rs2296740 (chr6:33691759:A:G), 3'SS: chr6:33691759−33691760, strand: +
3'SS Usage
1.0
0.5
0.0
−0.5
G/G
G/H
H/H
Genotype
Genotype
G/G
G/H
H/H

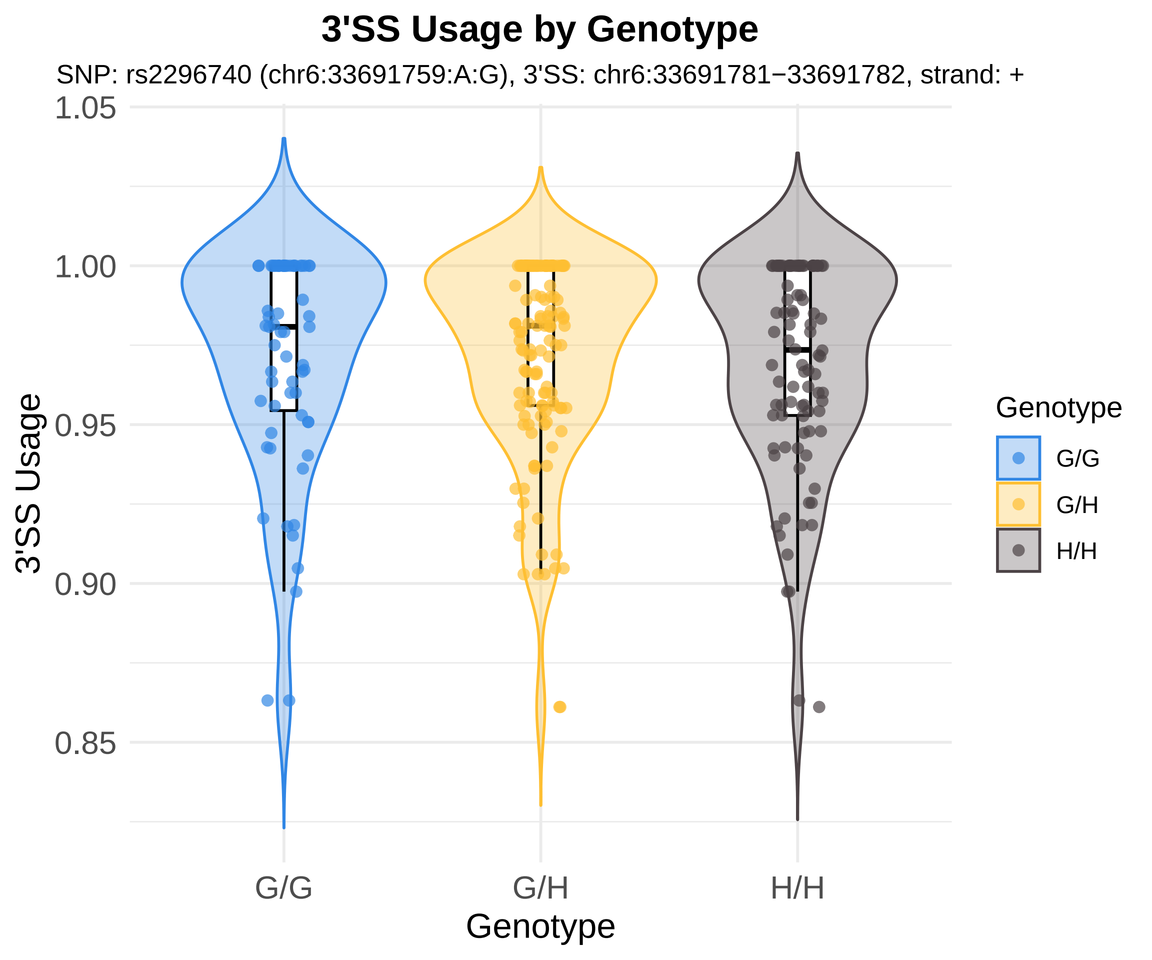
3'SS Usage by Genotype
SNP: rs2296740 (chr6:33691759:A:G), 3'SS: chr6:33691781−33691782, strand: +
3'SS Usage
1.05
1.00
0.95
0.90
0.85
G/G
G/H
H/H
Genotype
Genotype
G/G
G/H
H/H

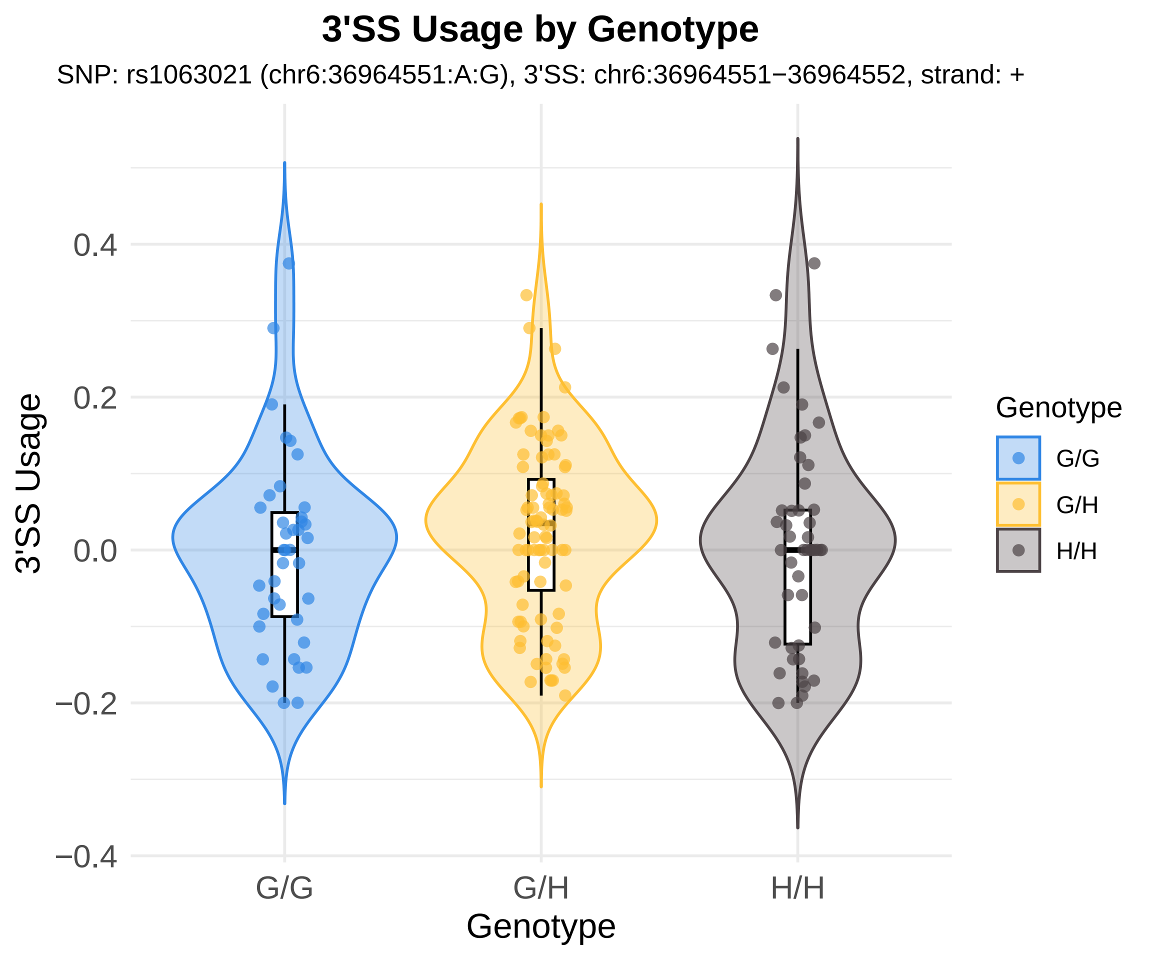

3'SS Usage by Genotype
SNP: rs1063021 (chr6:36964551:A:G), 3'SS: chr6:36964551−36964552, strand: +
3'SS Usage
0.4
0.2
0.0
−0.2
−0.4
G/G
G/H
H/H
Genotype
Genotype
G/G
G/H
H/H

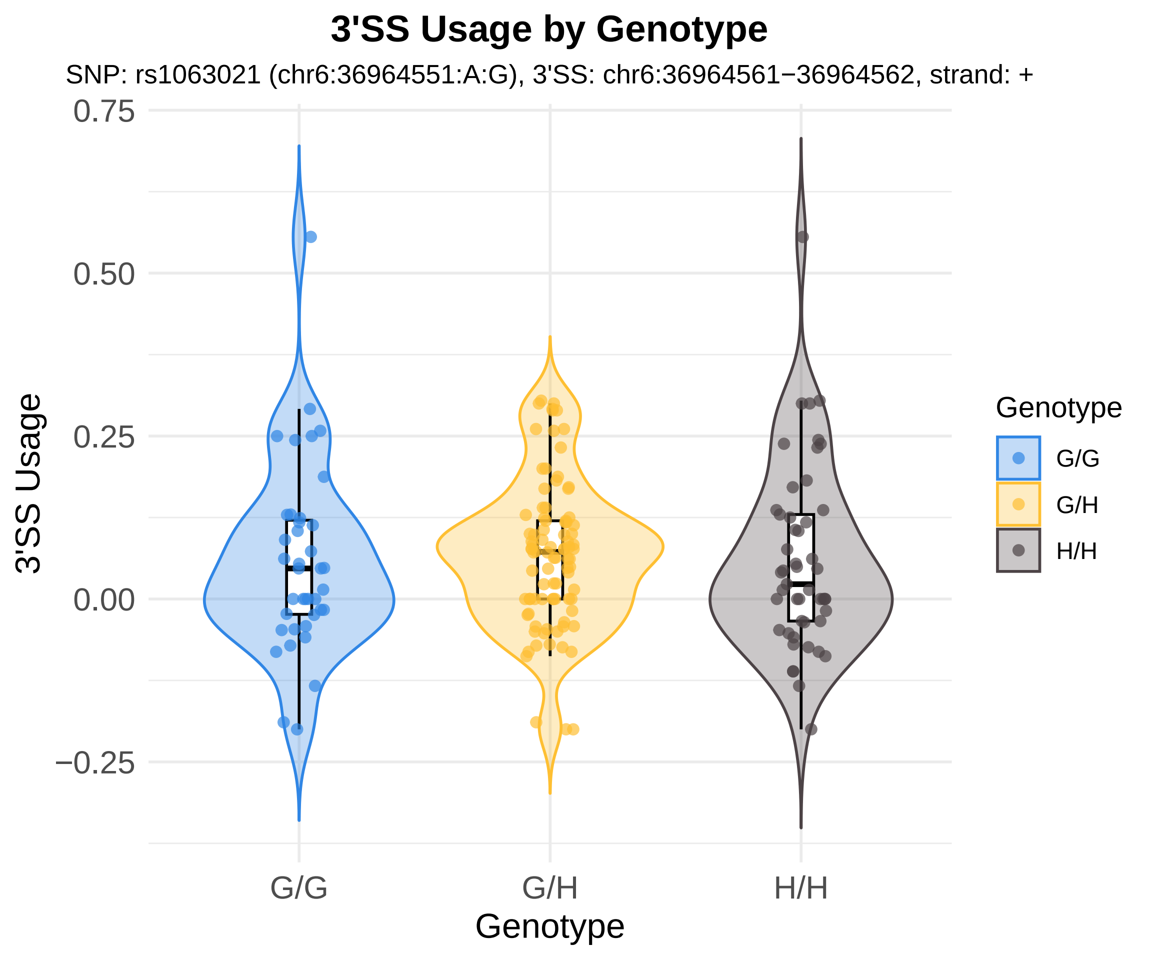
3'SS Usage by Genotype
SNP: rs1063021 (chr6:36964551:A:G), 3'SS: chr6:36964561−36964562, strand: +
0.75
0.50
0.25
0.00
−0.25
3'SS Usage
G/G
G/H
H/H
Genotype
Genotype
G/G
G/H
H/H

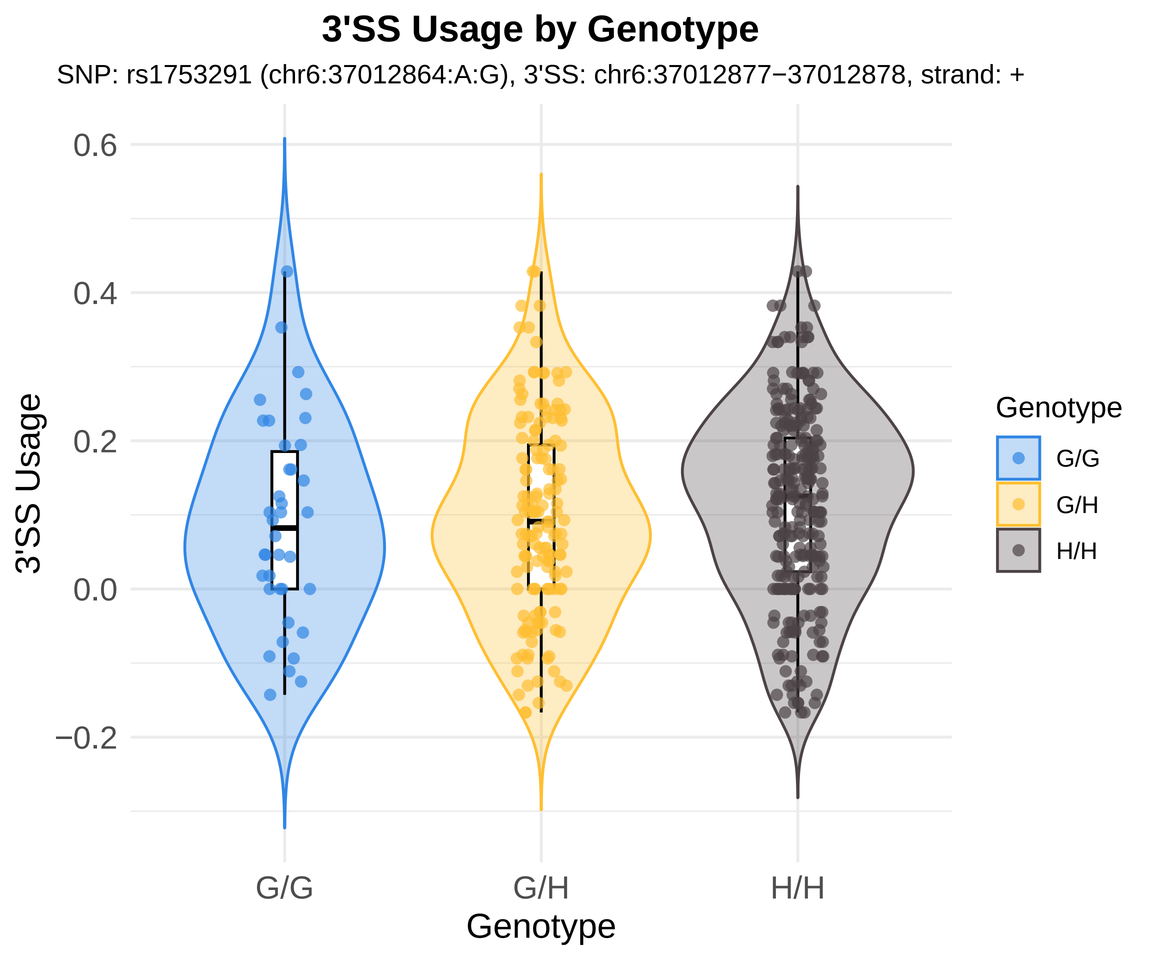
3'SS Usage by Genotype
SNP: rs1753291 (chr6:37012864:A:G), 3'SS: chr6:37012877−37012878, strand: +
3'SS Usage
0.6
0.4
0.2
0.0
−0.2
G/G
G/H
H/H
Genotype
Genotype
G/G
G/H
H/H

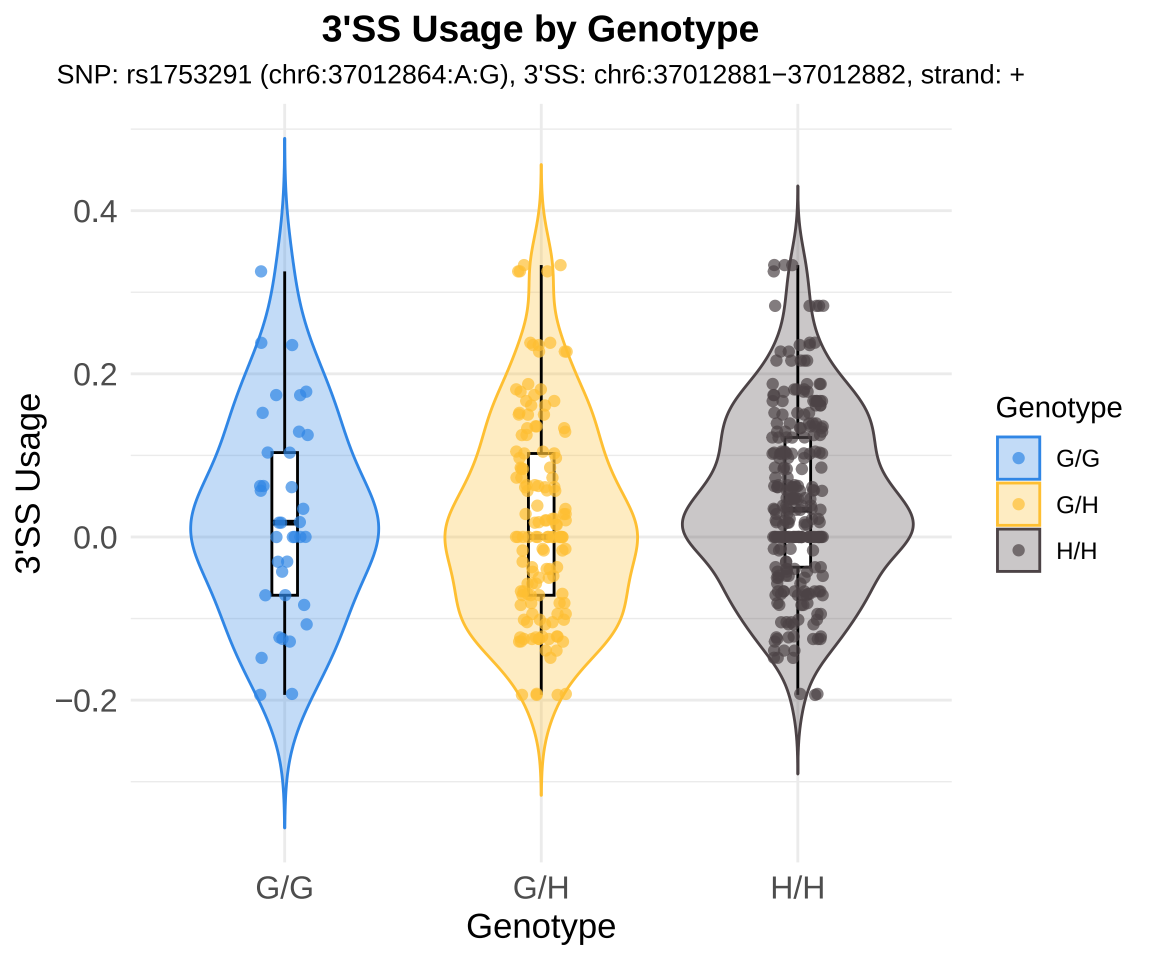
3'SS Usage by Genotype
SNP: rs1753291 (chr6:37012864:A:G), 3'SS: chr6:37012881−37012882, strand: +
3'SS Usage
0.4
0.2
0.0
−0.2
G/G
G/H
H/H
Genotype
Genotype
G/G
G/H
H/H

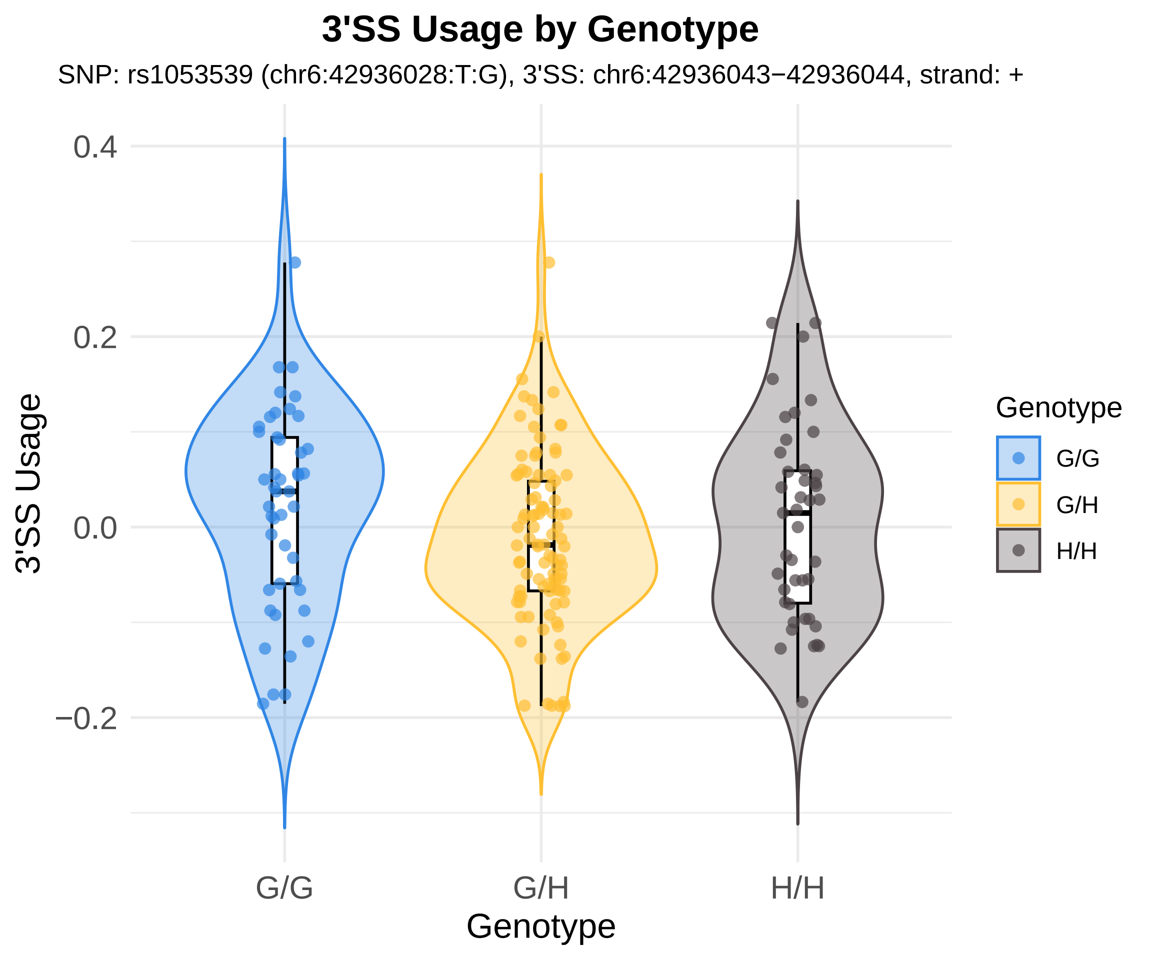

3'SS Usage by Genotype
SNP: rs1053539 (chr6:42936028:T:G), 3'SS: chr6:42936043−42936044, strand: +
3'SS Usage
0.4
0.2
0.0
−0.2
G/G
G/H
H/H
Genotype
Genotype
G/G
G/H
H/H

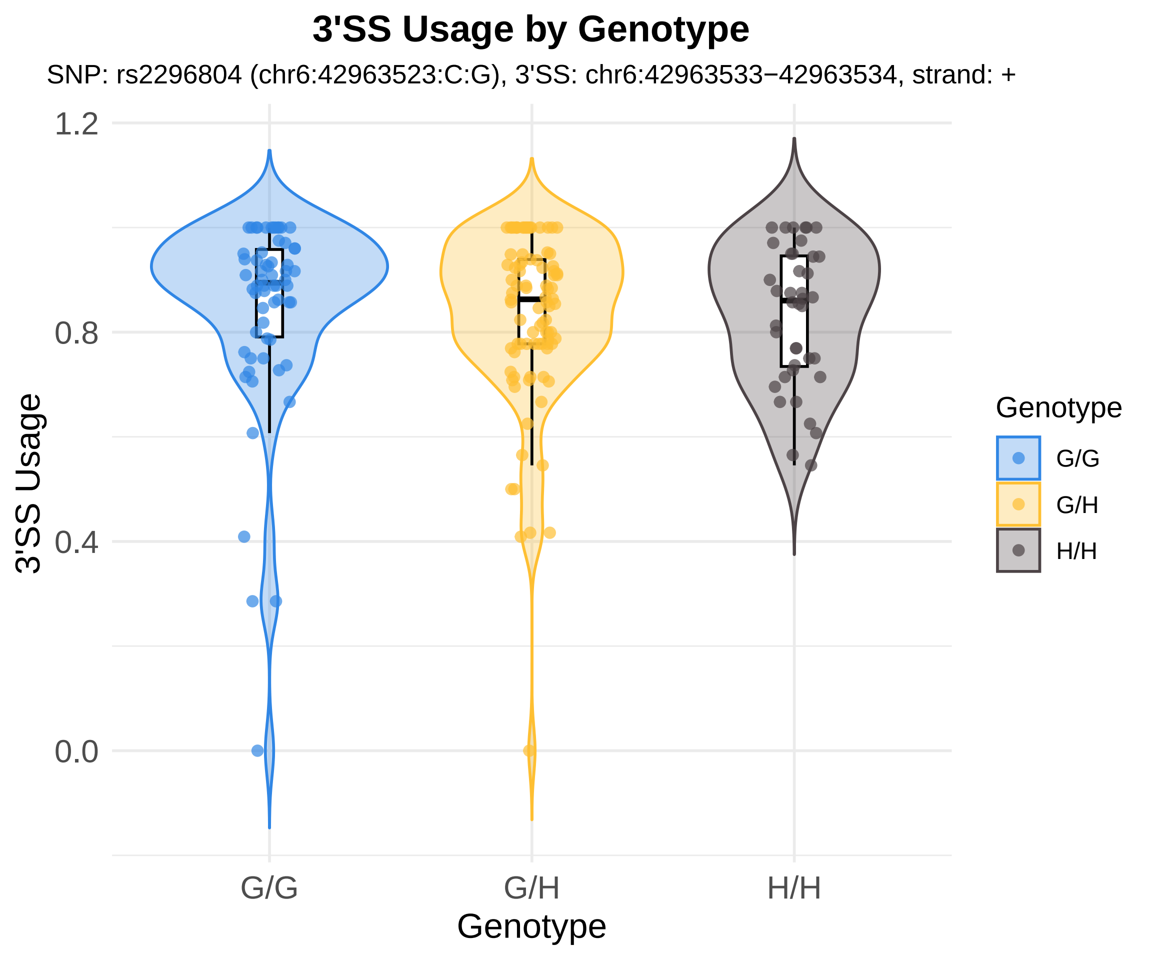
3'SS Usage by Genotype
SNP: rs2296804 (chr6:42963523:C:G), 3'SS: chr6:42963533−42963534, strand: +
1.2
0.8
0.4
0.0
3'SS Usage
G/G
G/H
H/H
Genotype
Genotype
G/G
G/H
H/H

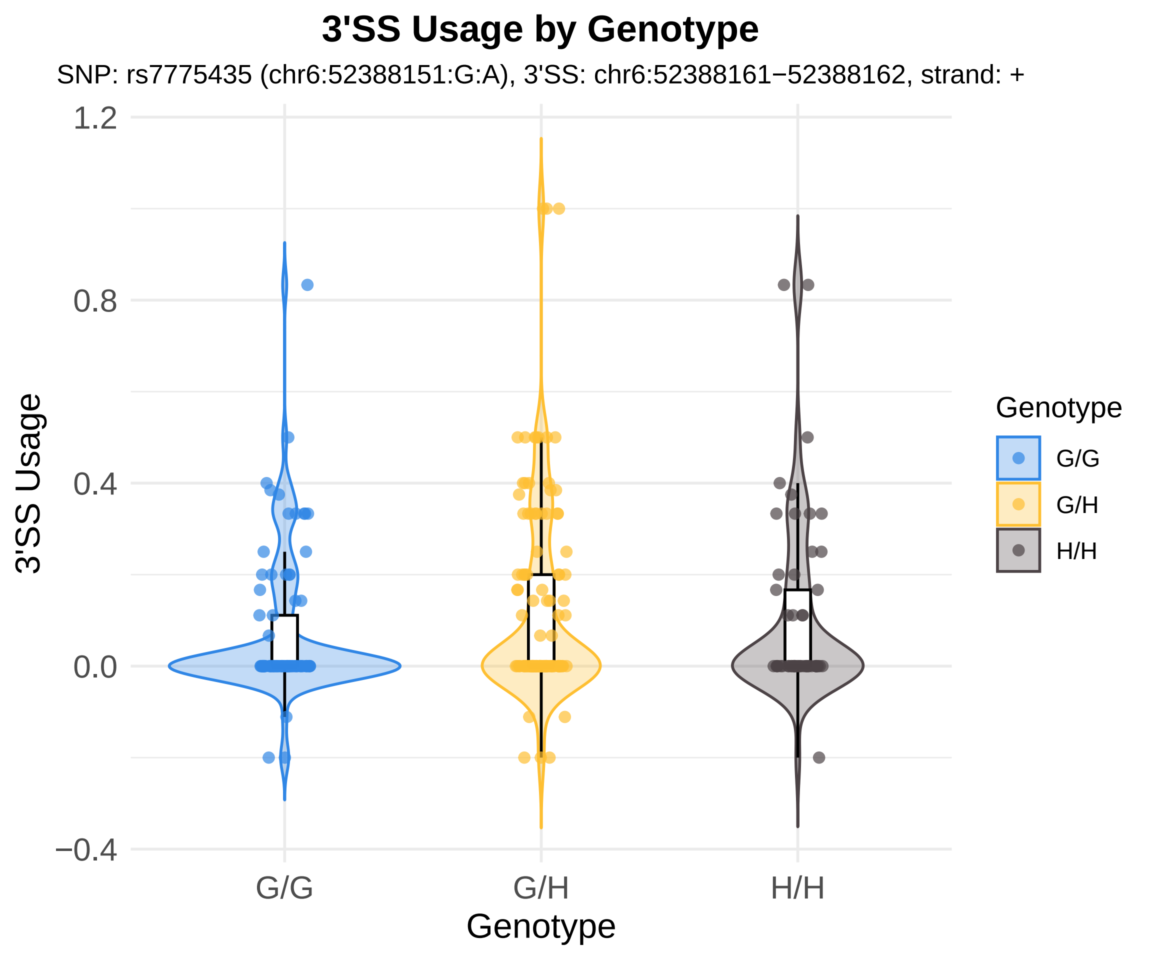

3'SS Usage by Genotype
SNP: rs7775435 (chr6:52388151:G:A), 3'SS: chr6:52388161−52388162, strand: +
3'SS Usage
1.2
0.8
0.4
0.0
−0.4
G/G
G/H
H/H
Genotype
Genotype
G/G
G/H
H/H

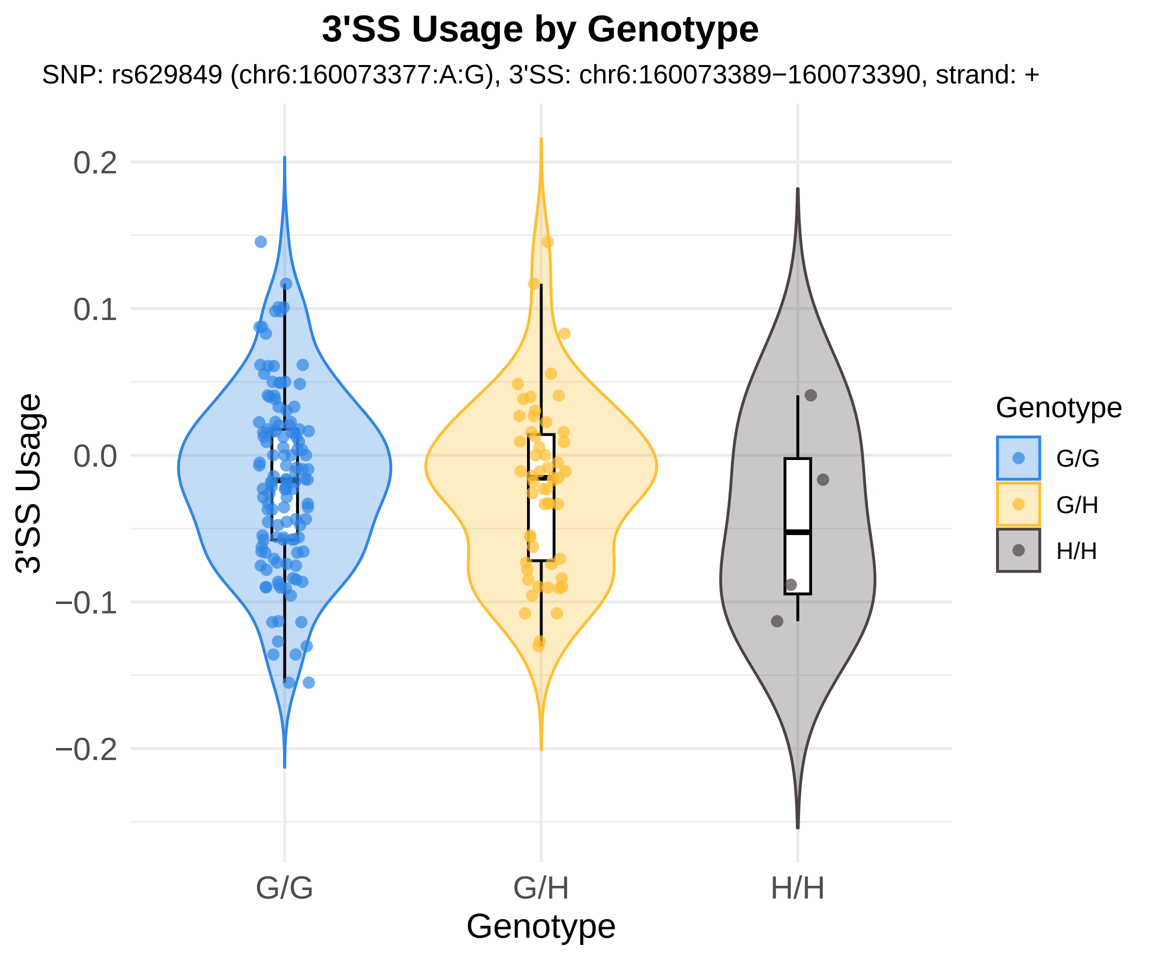
3'SS Usage by Genotype
SNP: rs629849 (chr6:160073377:A:G), 3'SS: chr6:160073389−160073390, strand: +
0.2
0.1
0.0
−0.1
−0.2
3'SS Usage
G/G
G/H
H/H
Genotype
Genotype
G/G
G/H
H/H

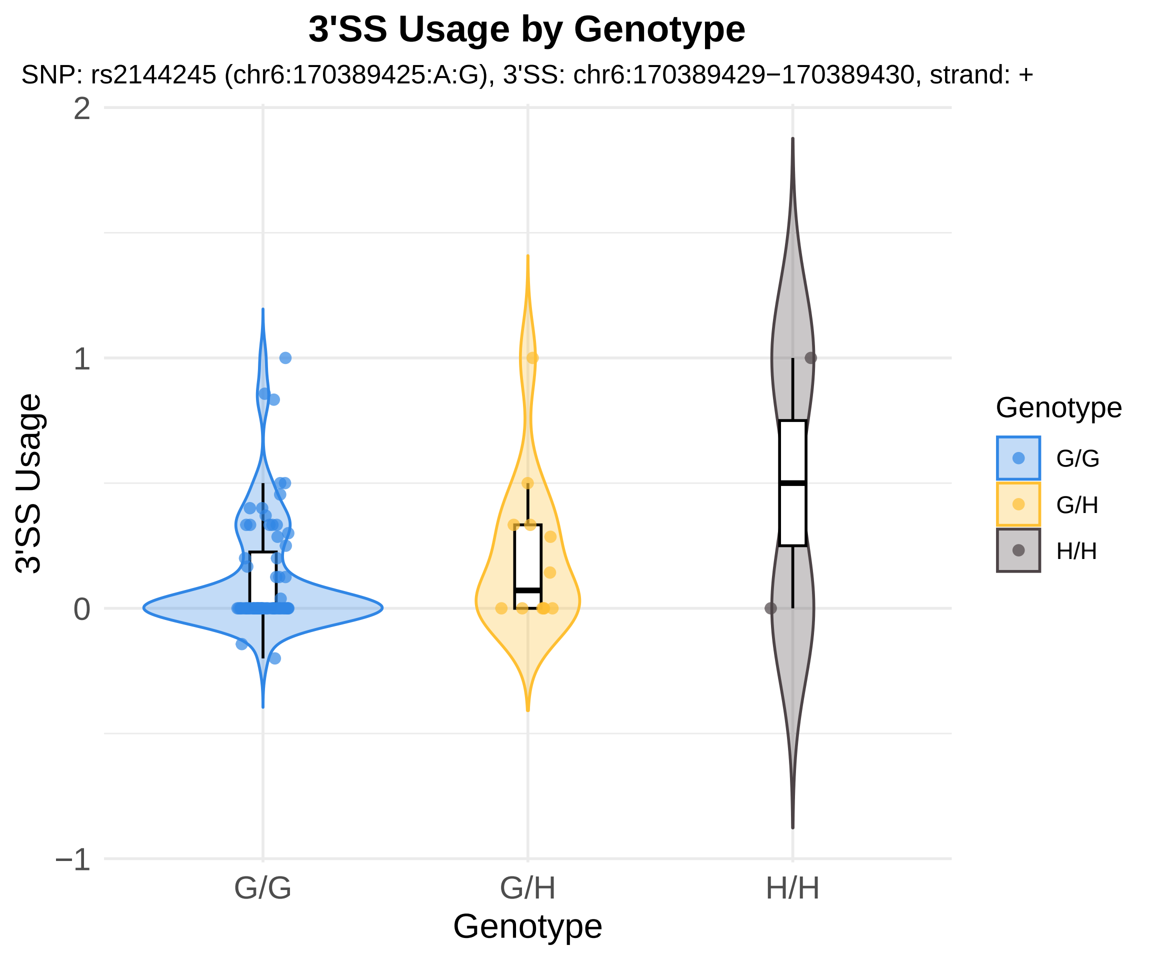
3'SS Usage by Genotype
SNP: rs2144245 (chr6:170389425:A:G), 3'SS: chr6:170389429−170389430, strand: +
3'SS Usage
2
1
0
−1
G/G
G/H
H/H
Genotype
Genotype
G/G
G/H
H/H

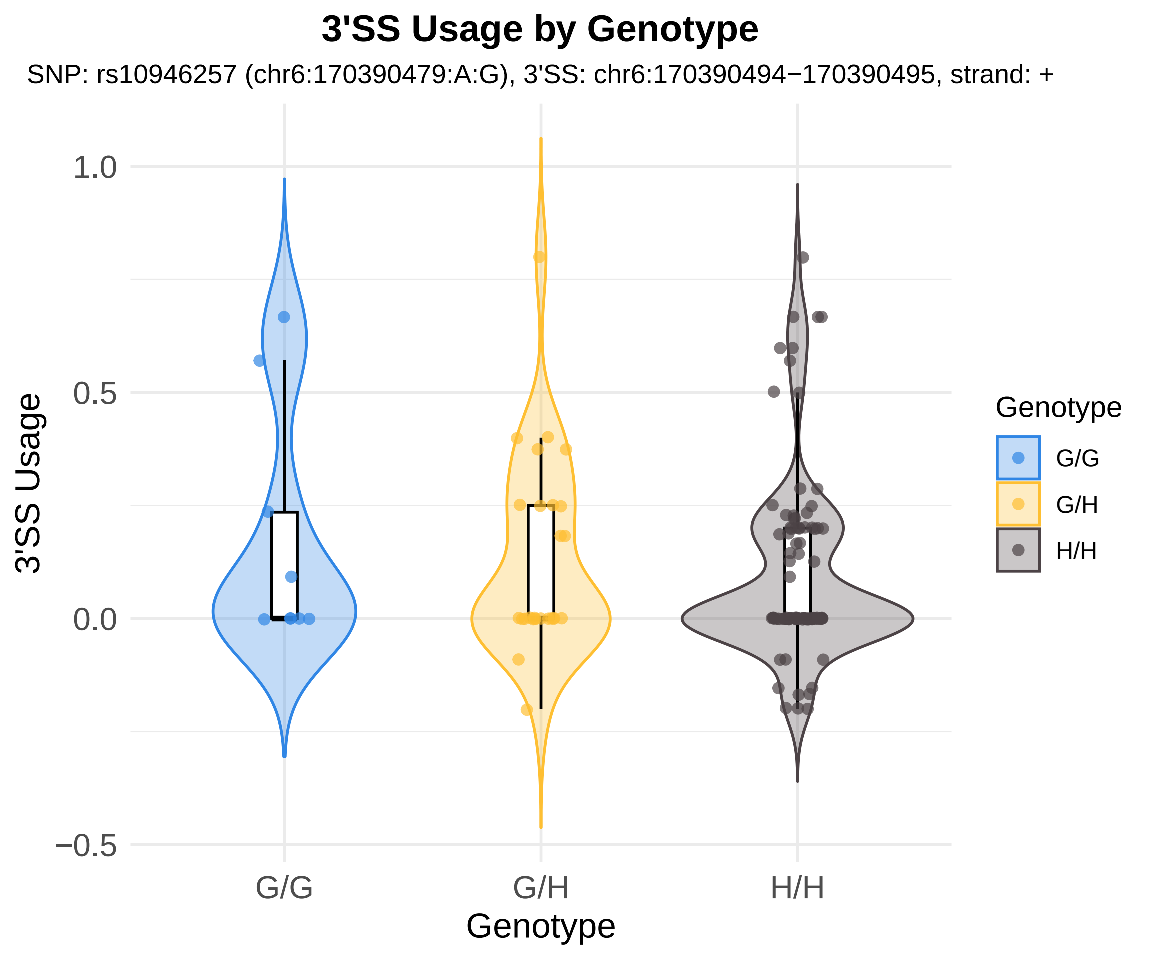

3'SS Usage by Genotype
SNP: rs10946257 (chr6:170390479:A:G), 3'SS: chr6:170390494−170390495, strand: +
3'SS Usage
1.0
0.5
0.0
−0.5
G/G
G/H
H/H
Genotype
Genotype
G/G
G/H
H/H

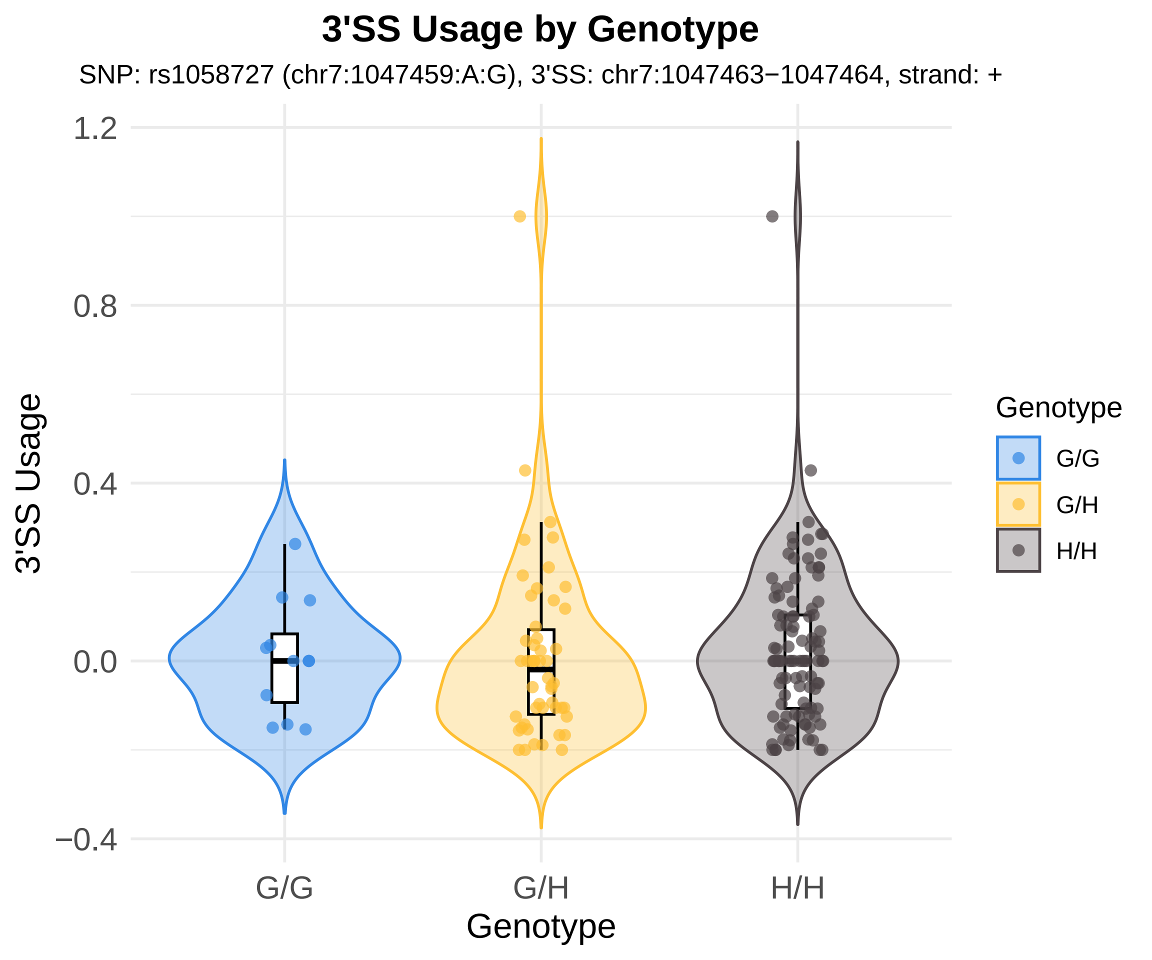
3'SS Usage by Genotype
SNP: rs1058727 (chr7:1047459:A:G), 3'SS: chr7:1047463−1047464, strand: +
3'SS Usage
1.2
0.8
0.4
0.0
−0.4
G/G
G/H
H/H
Genotype
Genotype
G/G
G/H
H/H

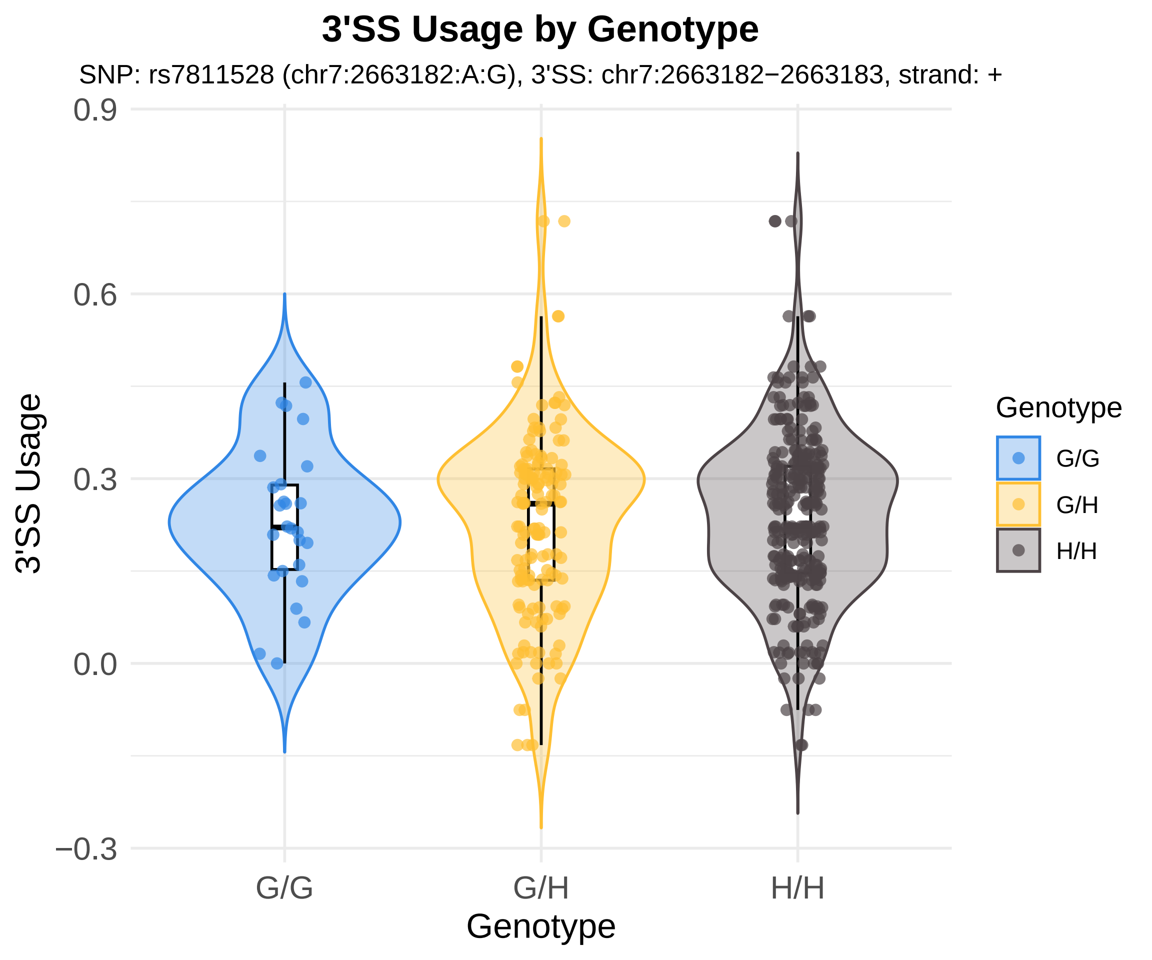
3'SS Usage by Genotype
SNP: rs7811528 (chr7:2663182:A:G), 3'SS: chr7:2663182−2663183, strand: +
3'SS Usage
0.9
0.6
0.3
0.0
−0.3
G/G
G/H
H/H
Genotype
Genotype
G/G
G/H
H/H

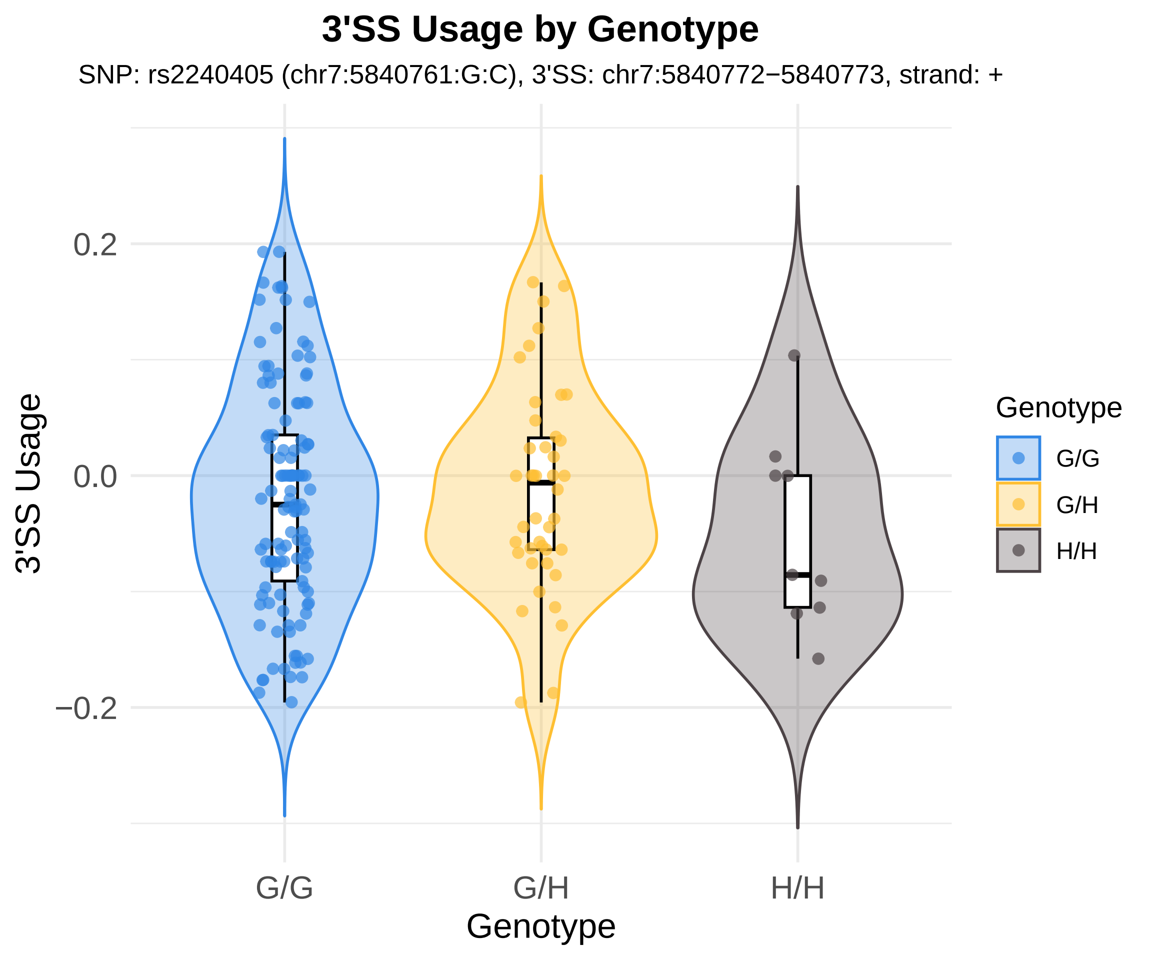
3'SS Usage by Genotype
SNP: rs2240405 (chr7:5840761:G:C), 3'SS: chr7:5840772−5840773, strand: +
3'SS Usage
0.2
0.0
−0.2
G/G
G/H
H/H
Genotype
Genotype
G/G
G/H
H/H

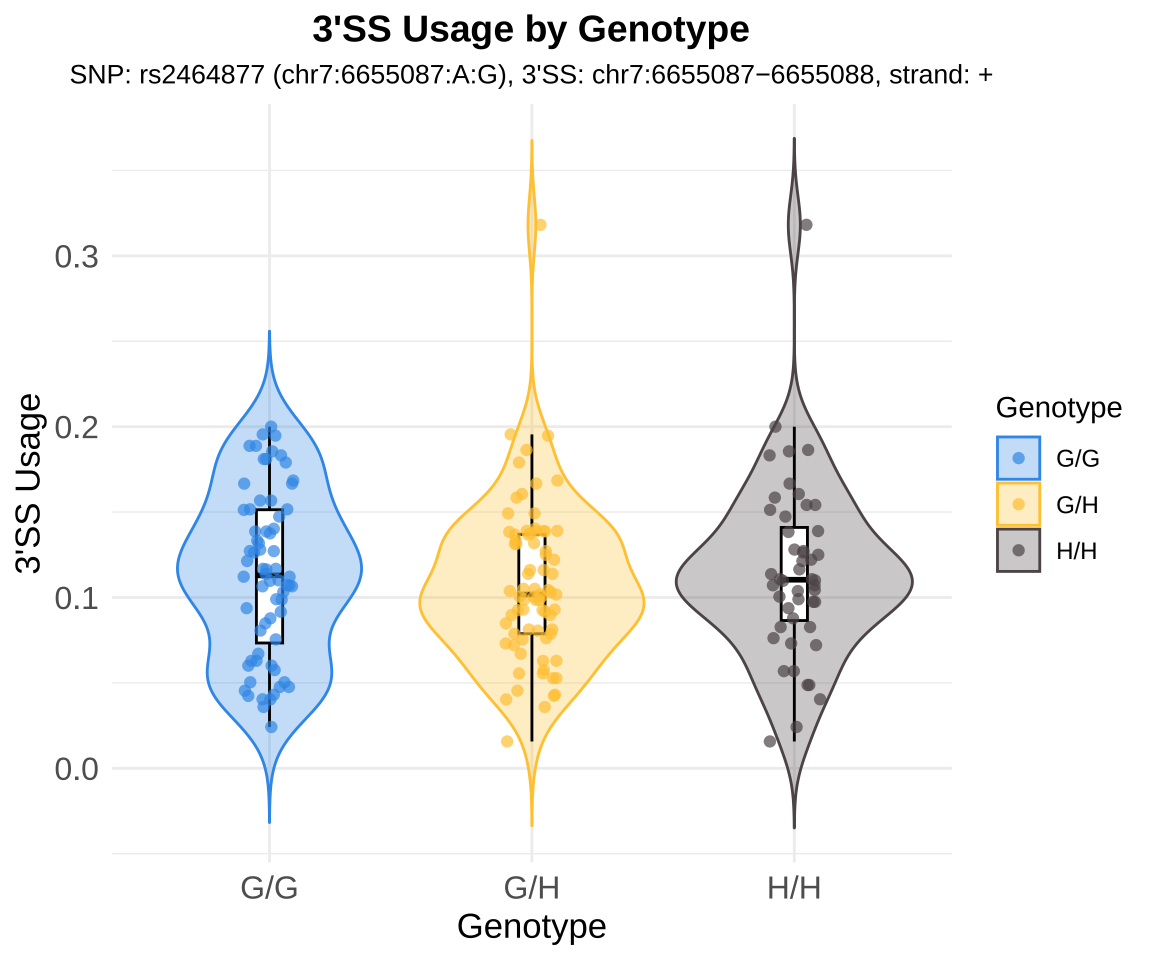

3'SS Usage by Genotype
SNP: rs2464877 (chr7:6655087:A:G), 3'SS: chr7:6655087−6655088, strand: +
3'SS Usage
0.0
0.1
0.2
0.3
G/G
G/H
H/H
Genotype
Genotype
G/G
G/H
H/H

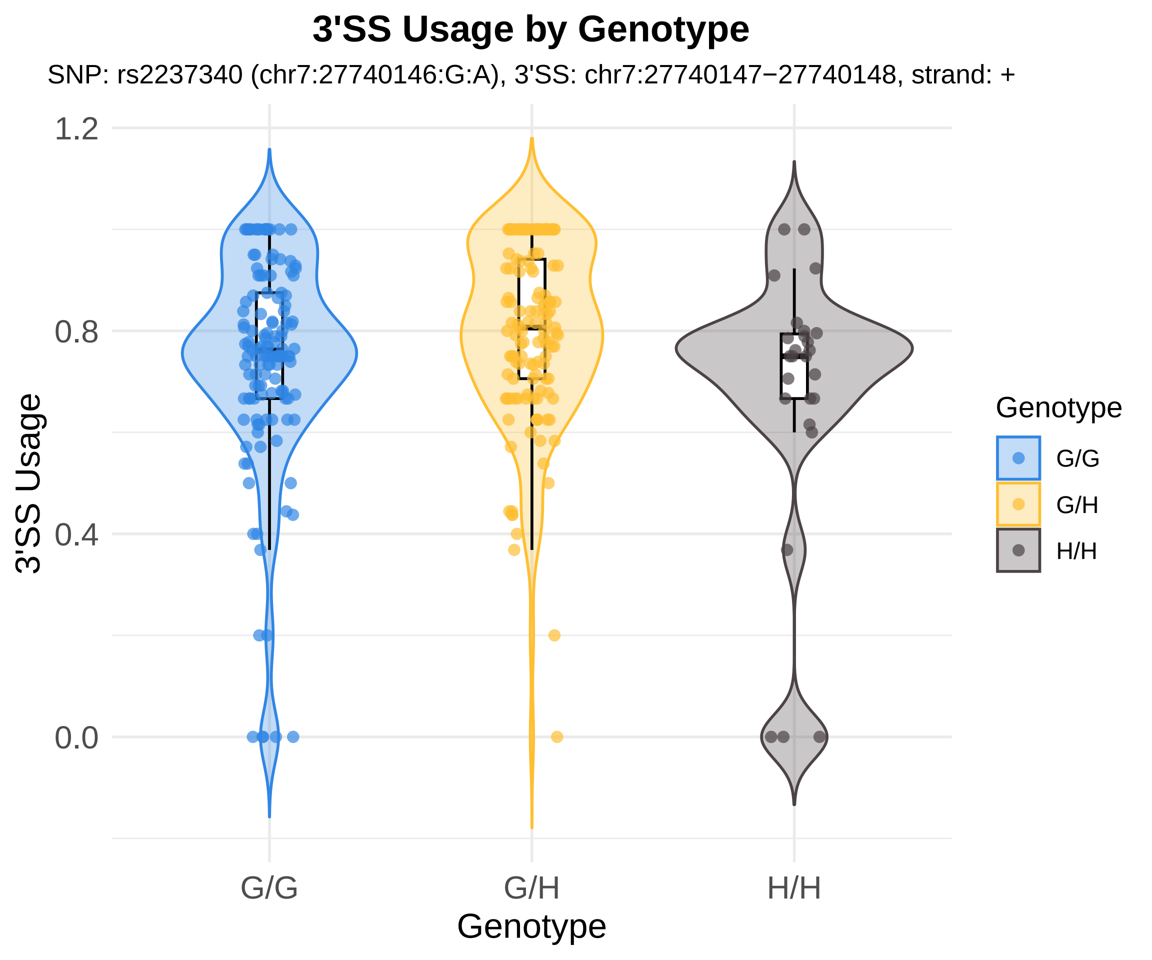

3'SS Usage by Genotype
SNP: rs2237340 (chr7:27740146:G:A), 3'SS: chr7:27740147−27740148, strand: +
3'SS Usage
1.2
0.8
0.4
0.0
G/G
G/H
H/H
Genotype
Genotype
G/G
G/H
H/H

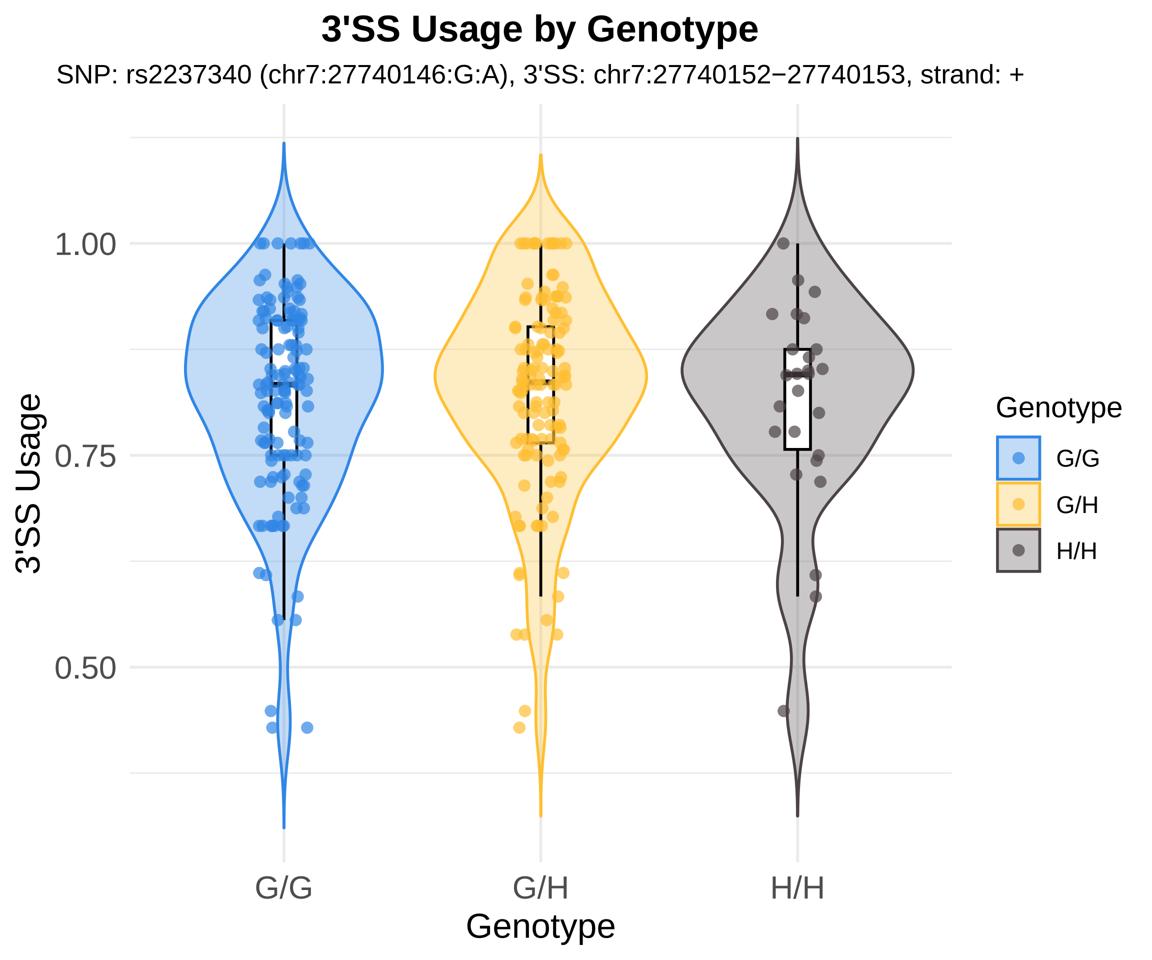
3'SS Usage by Genotype
SNP: rs2237340 (chr7:27740146:G:A), 3'SS: chr7:27740152−27740153, strand: +
3'SS Usage
1.00
0.75
0.50
G/G
G/H
H/H
Genotype
Genotype
G/G
G/H
H/H

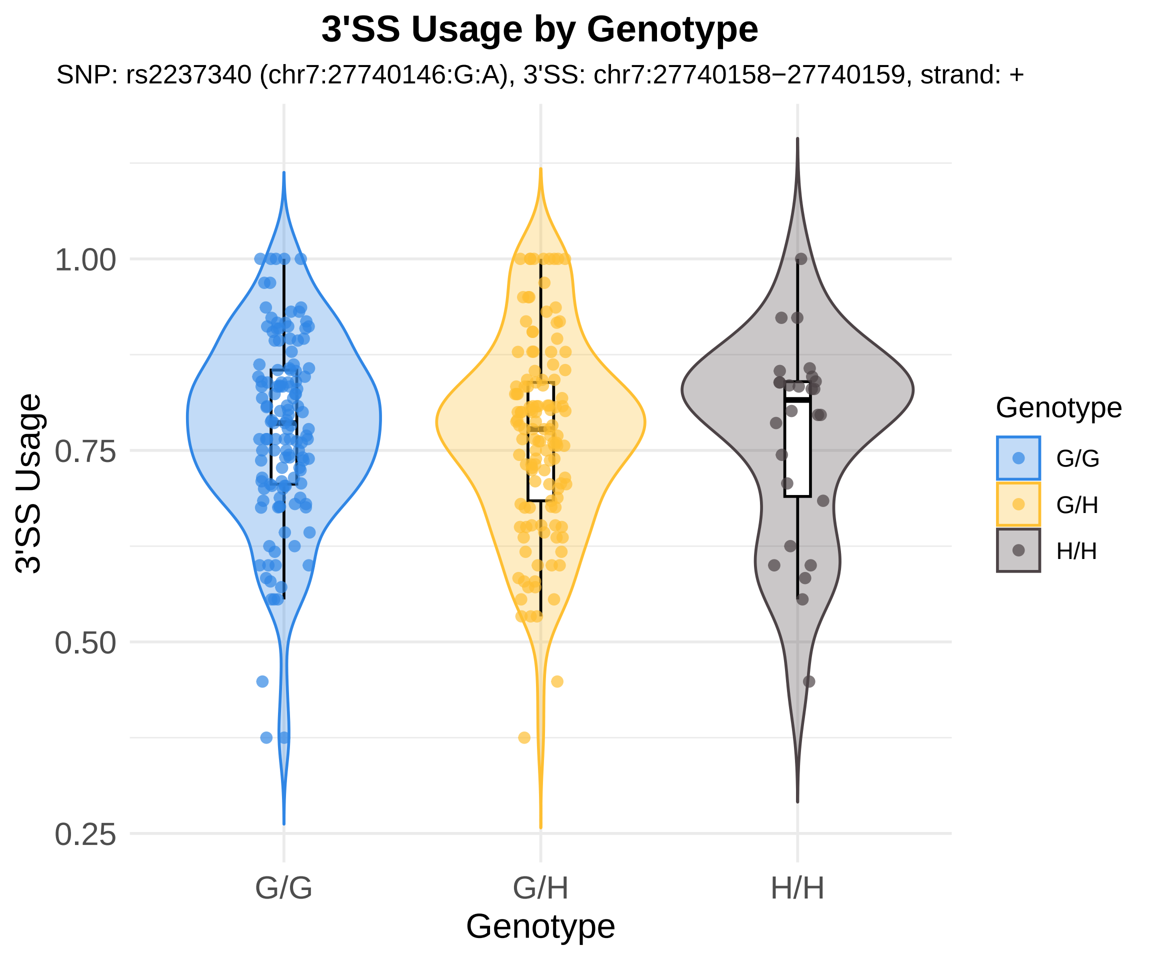

3'SS Usage by Genotype
SNP: rs2237340 (chr7:27740146:G:A), 3'SS: chr7:27740158−27740159, strand: +
3'SS Usage
1.00
0.75
0.50
0.25
G/G
G/H
H/H
Genotype
Genotype
G/G
G/H
H/H

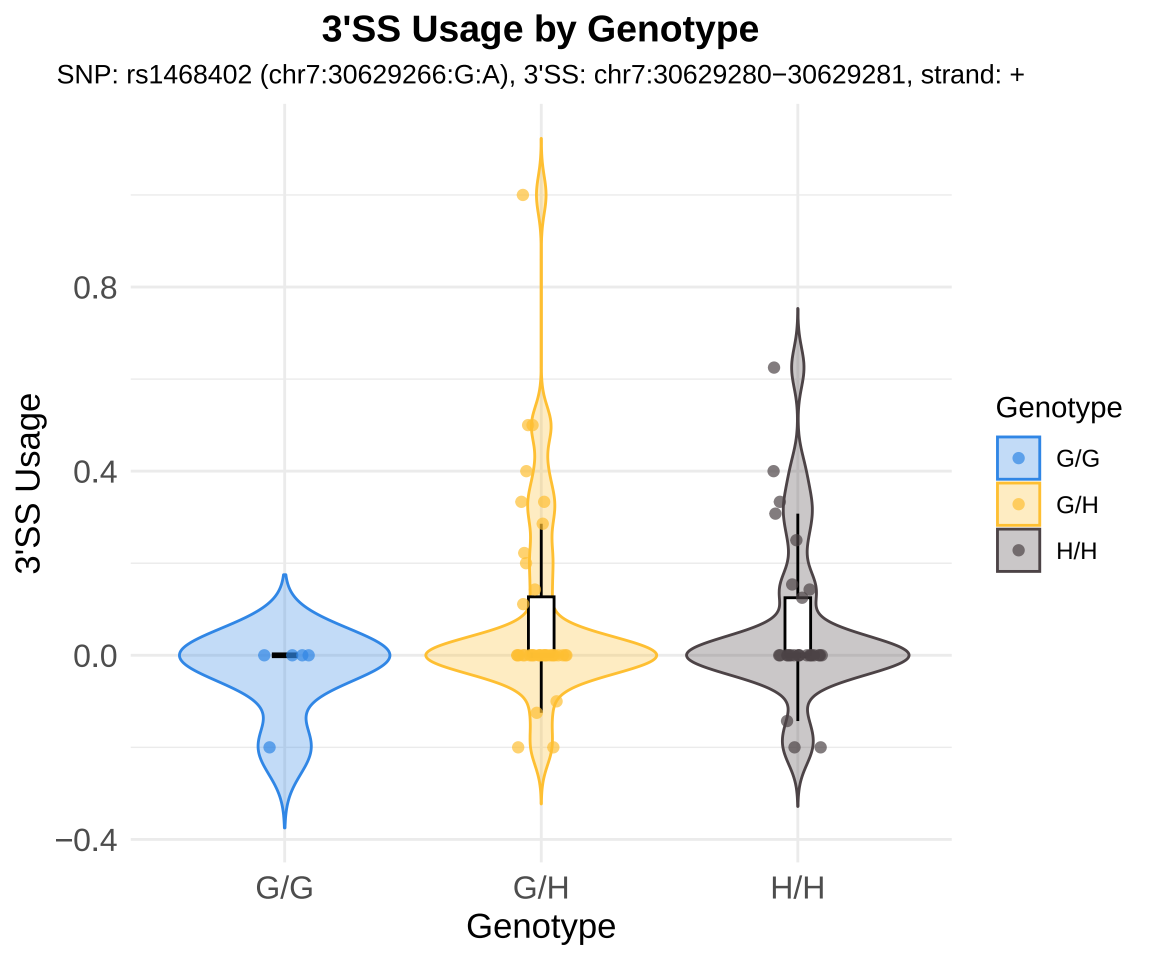
3'SS Usage by Genotype
SNP: rs1468402 (chr7:30629266:G:A), 3'SS: chr7:30629280−30629281, strand: +
3'SS Usage
0.8
0.4
0.0
−0.4
G/G
G/H
H/H
Genotype
Genotype
G/G
G/H
H/H

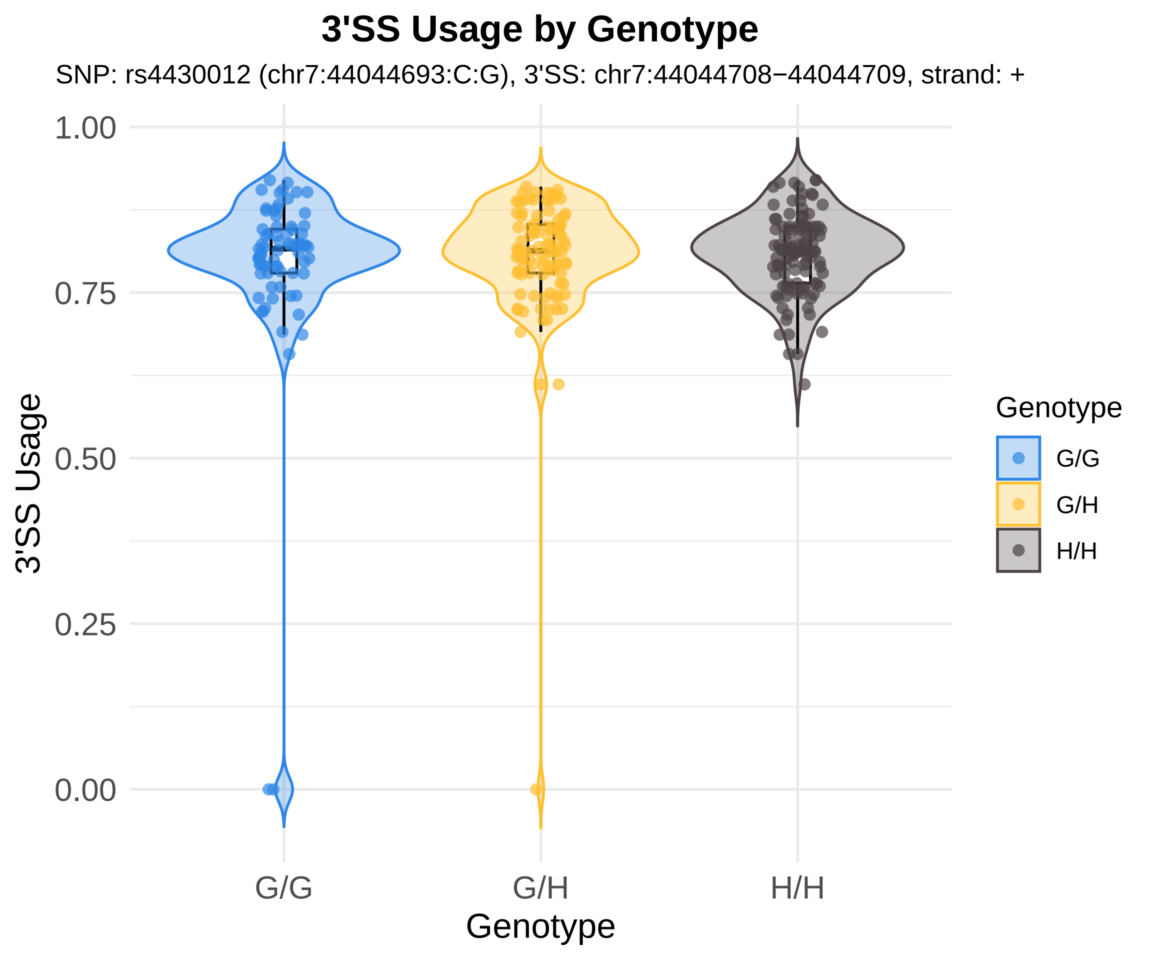
3'SS Usage by Genotype
SNP: rs4430012 (chr7:44044693:C:G), 3'SS: chr7:44044708−44044709, strand: +
3'SS Usage
1.00
0.75
0.50
0.25
0.00
G/G
G/H
H/H
Genotype
Genotype
G/G
G/H
H/H

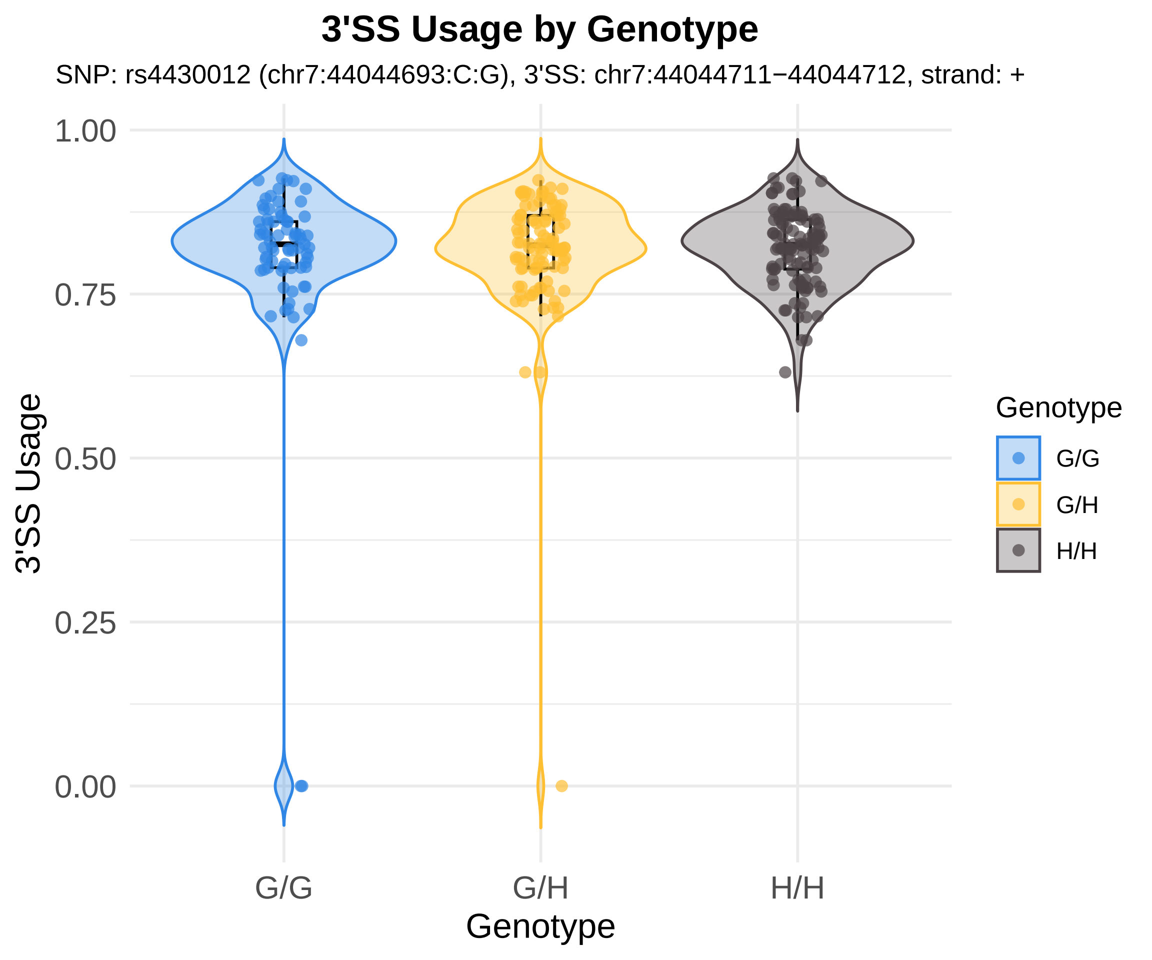
3'SS Usage by Genotype
SNP: rs4430012 (chr7:44044693:C:G), 3'SS: chr7:44044711−44044712, strand: +
1.00
0.75
0.50
0.25
0.00
3'SS Usage
G/G
G/H
H/H
Genotype
Genotype
G/G
G/H
H/H

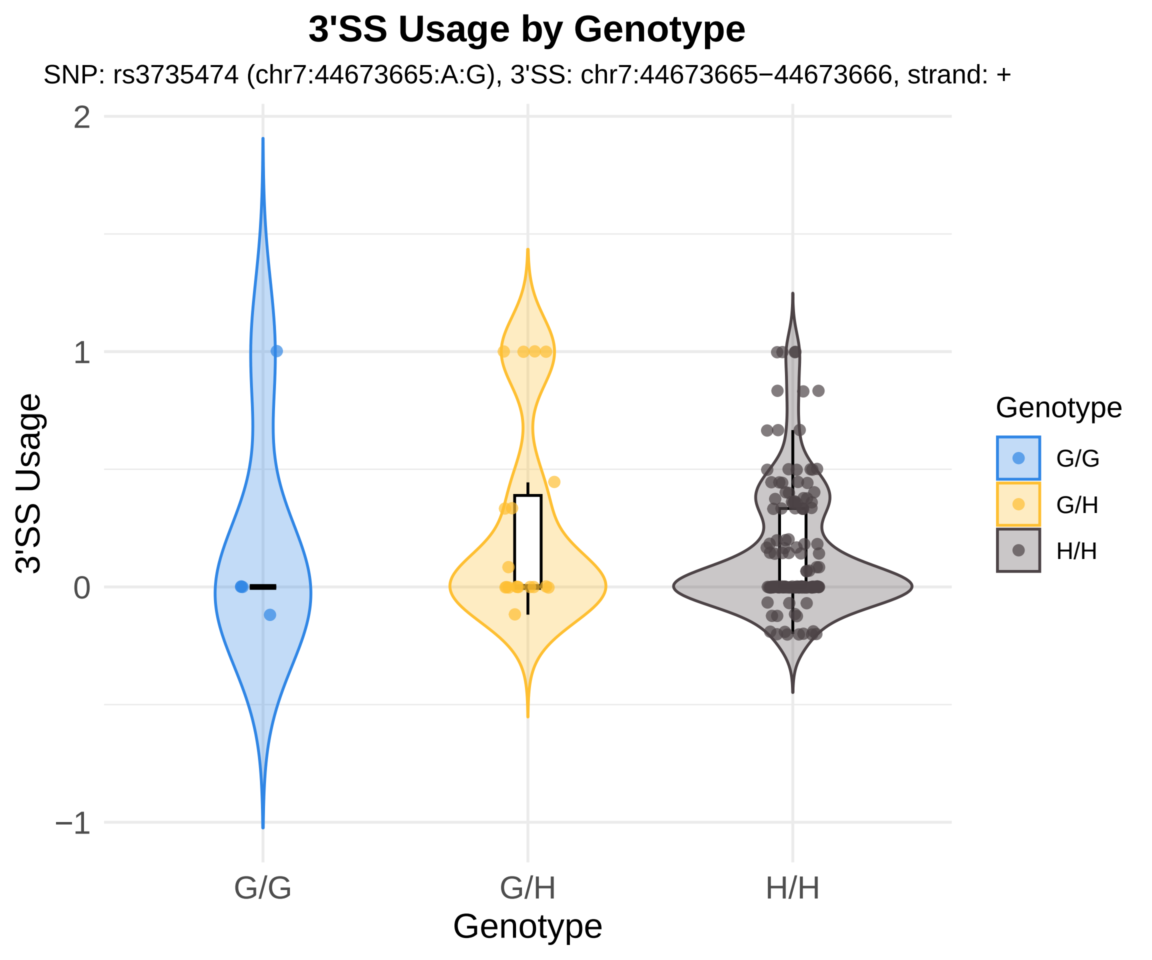

3'SS Usage by Genotype
SNP: rs3735474 (chr7:44673665:A:G), 3'SS: chr7:44673665−44673666, strand: +
3'SS Usage
2
1
0
−1
G/G
G/H
H/H
Genotype
Genotype
G/G
G/H
H/H

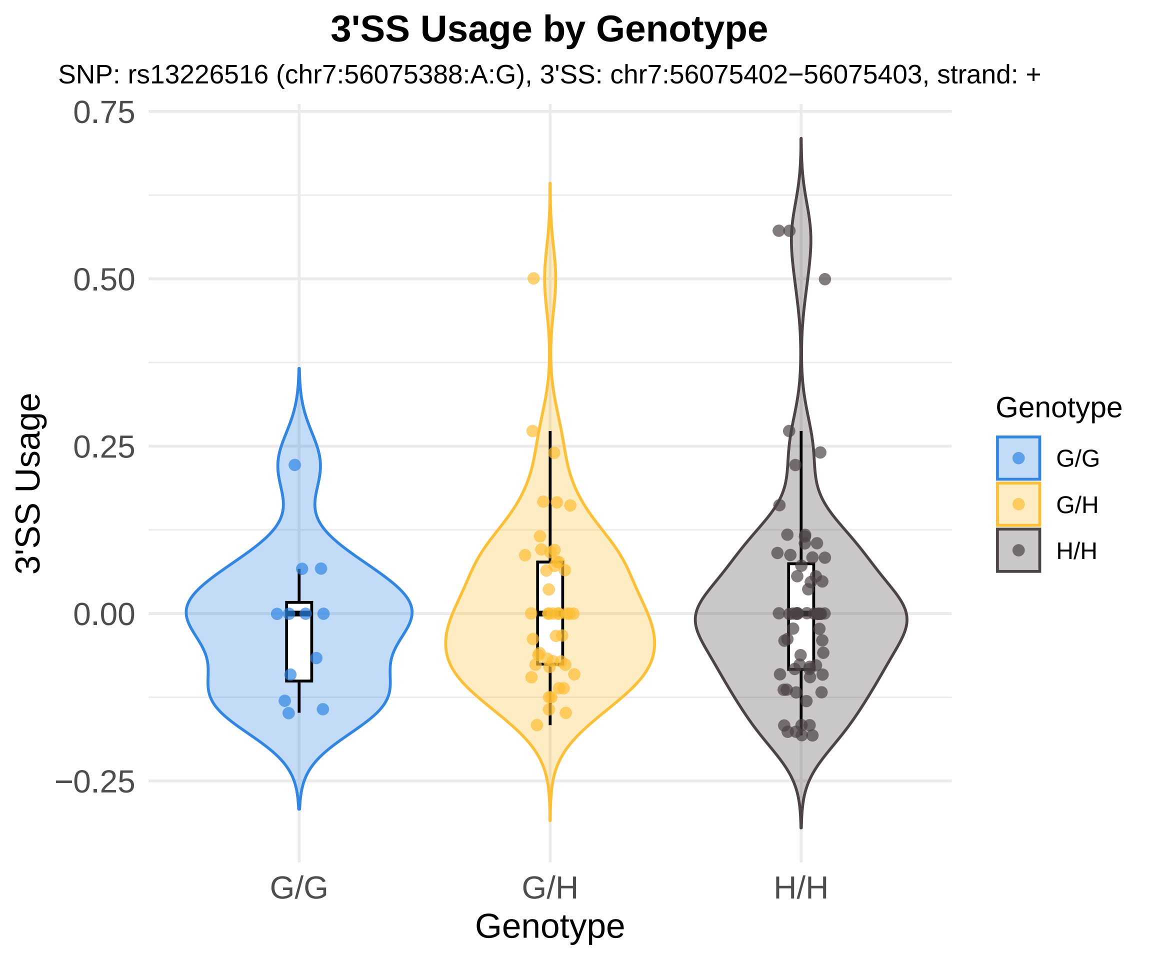
3'SS Usage by Genotype
SNP: rs13226516 (chr7:56075388:A:G), 3'SS: chr7:56075402−56075403, strand: +
3'SS Usage
0.75
0.50
0.25
0.00
−0.25
G/G
G/H
H/H
Genotype
Genotype
G/G
G/H
H/H

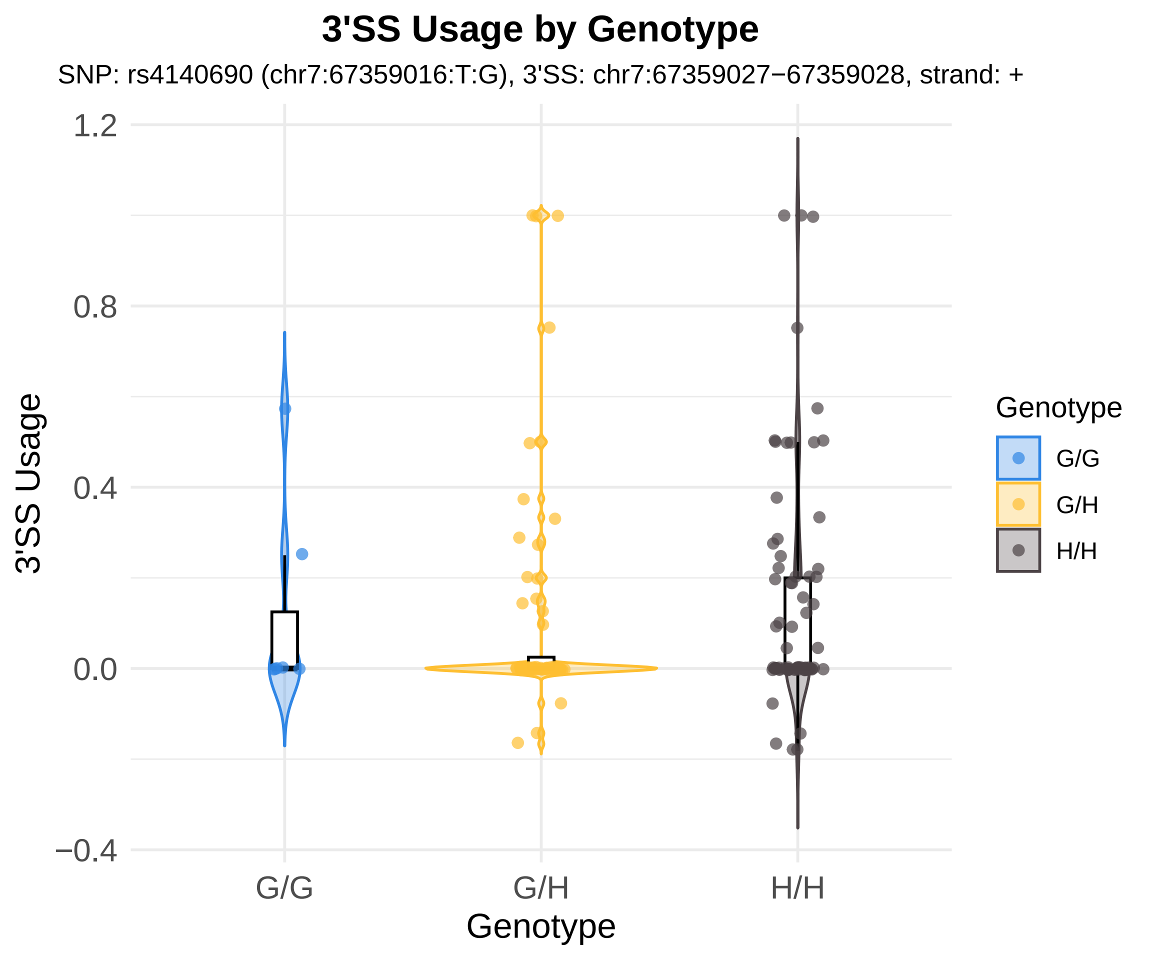

3'SS Usage by Genotype
SNP: rs4140690 (chr7:67359016:T:G), 3'SS: chr7:67359027−67359028, strand: +
3'SS Usage
1.2
0.8
0.4
0.0
−0.4
G/G
G/H
H/H
Genotype
Genotype
G/G
G/H
H/H

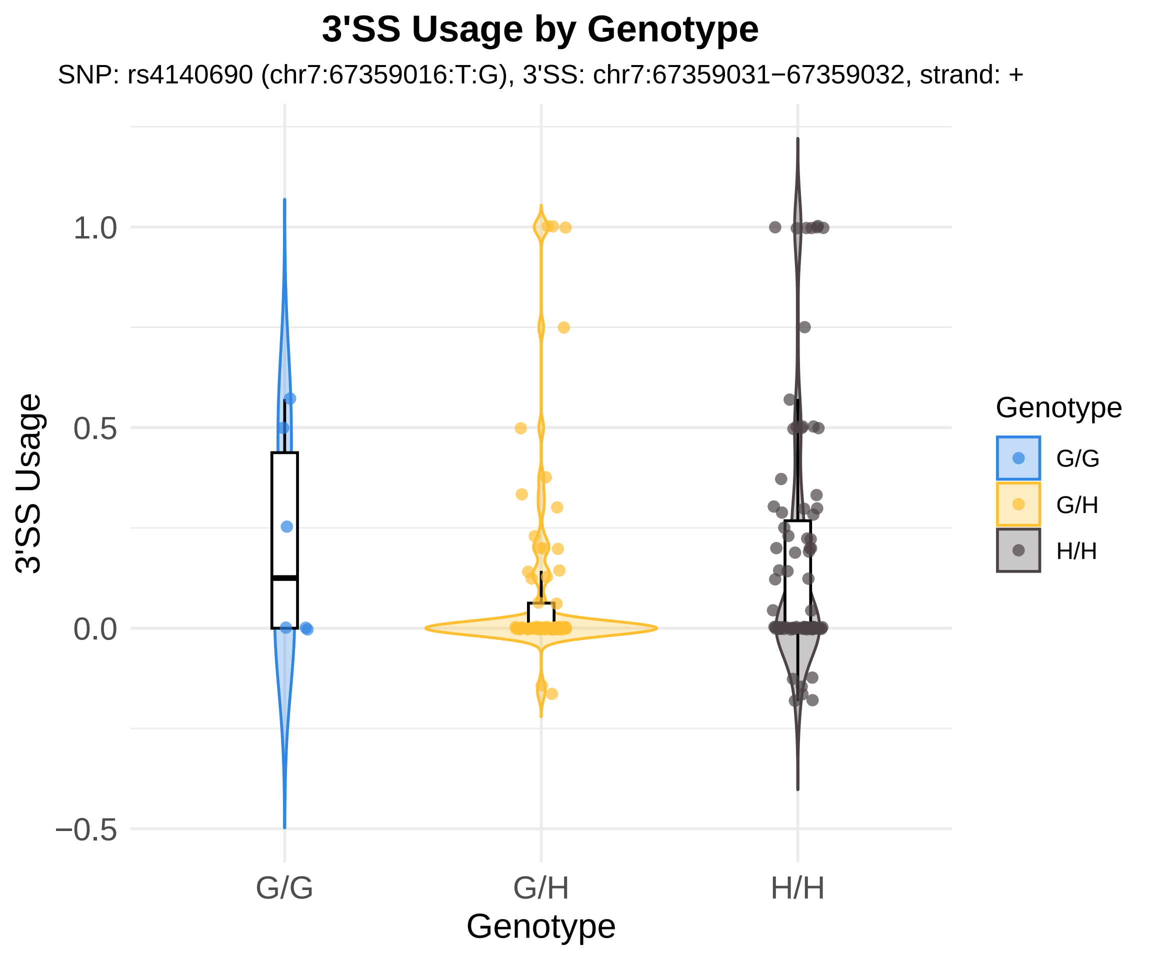
3'SS Usage by Genotype
SNP: rs4140690 (chr7:67359016:T:G), 3'SS: chr7:67359031−67359032, strand: +
3'SS Usage
1.0
0.5
0.0
−0.5
G/G
G/H
H/H
Genotype
Genotype
G/G
G/H
H/H

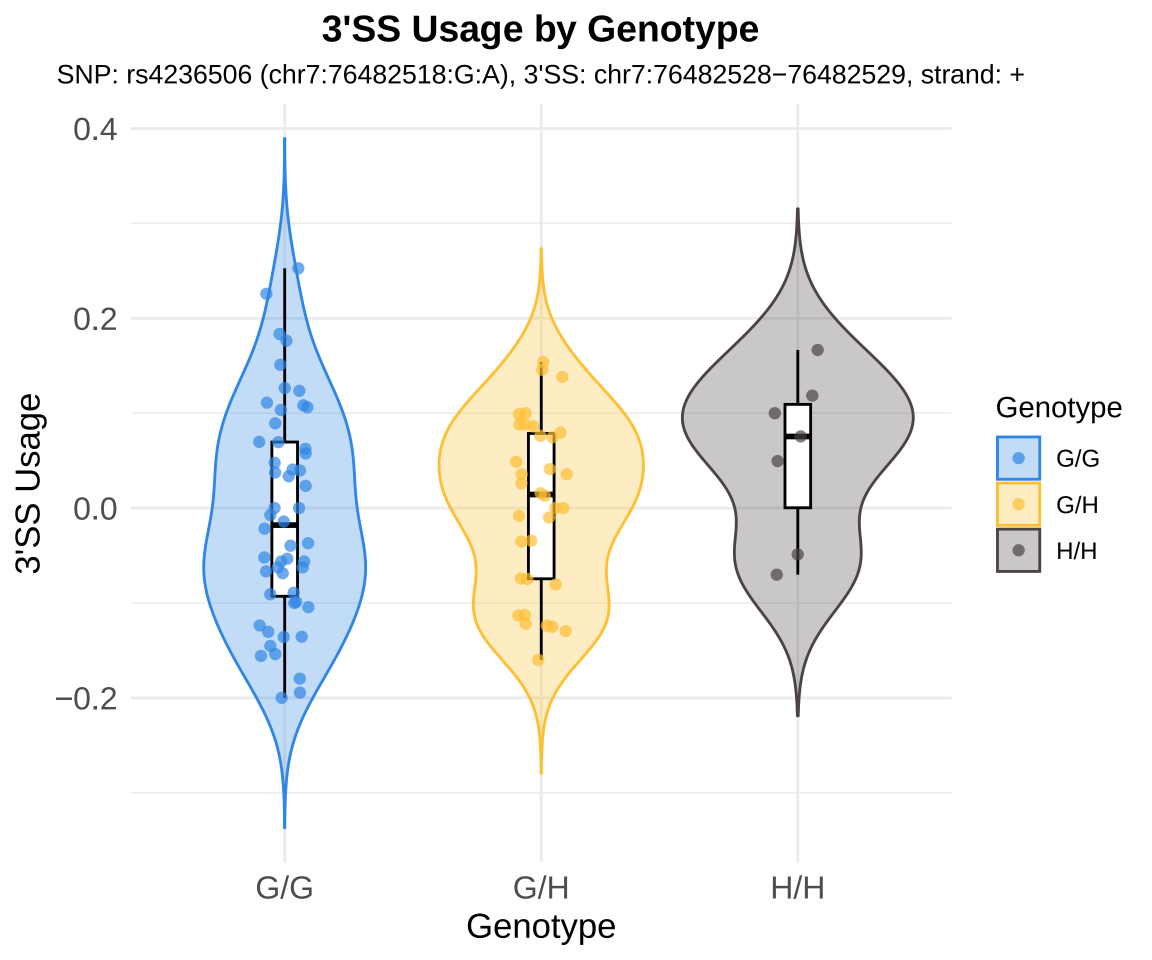
3'SS Usage by Genotype
SNP: rs4236506 (chr7:76482518:G:A), 3'SS: chr7:76482528−76482529, strand: +
3'SS Usage
0.4
0.2
0.0
−0.2
G/G
G/H
H/H
Genotype
Genotype
G/G
G/H
H/H

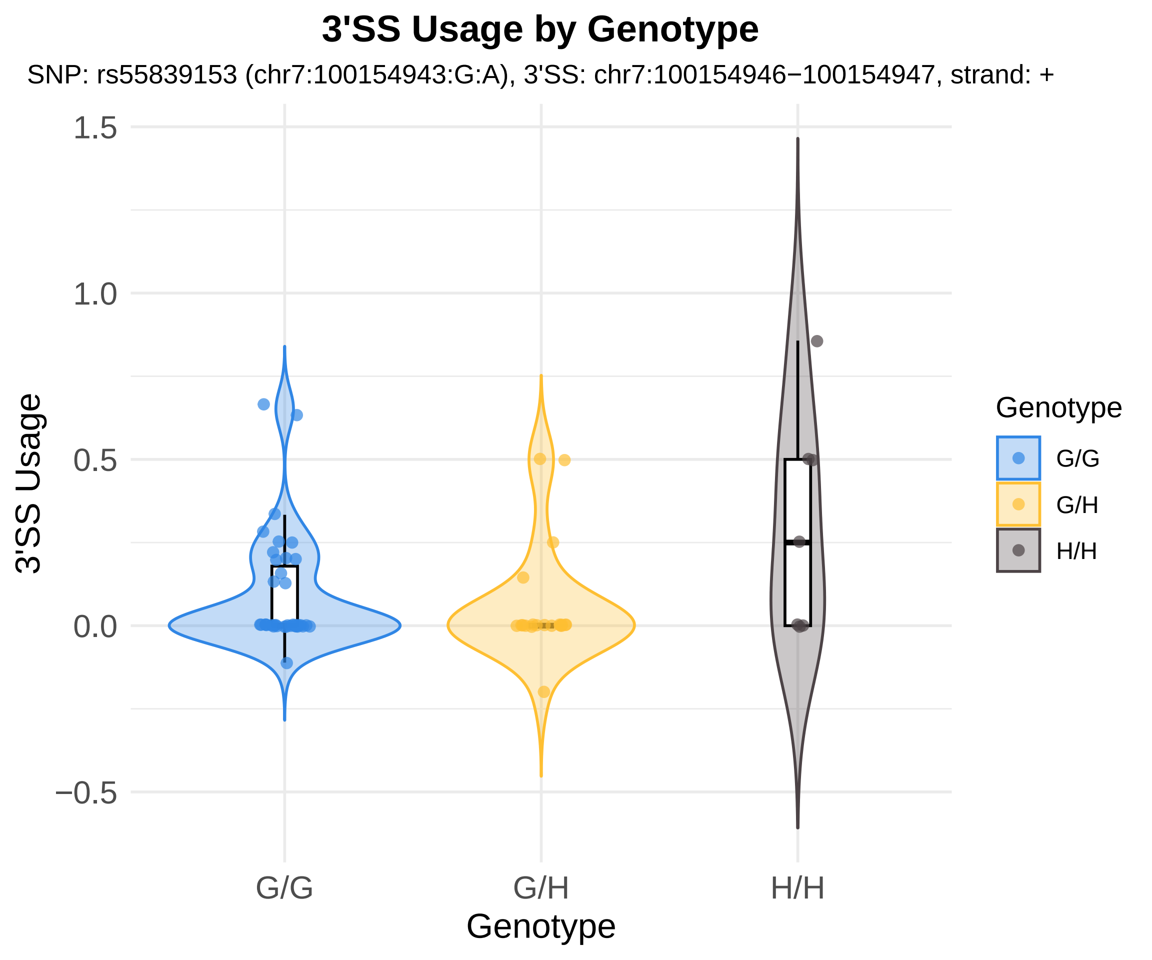

3'SS Usage by Genotype
SNP: rs55839153 (chr7:100154943:G:A), 3'SS: chr7:100154946−100154947, strand: +
1.5
1.0
0.5
0.0
−0.5
3'SS Usage
G/G
G/H
H/H
Genotype
Genotype
G/G
G/H
H/H

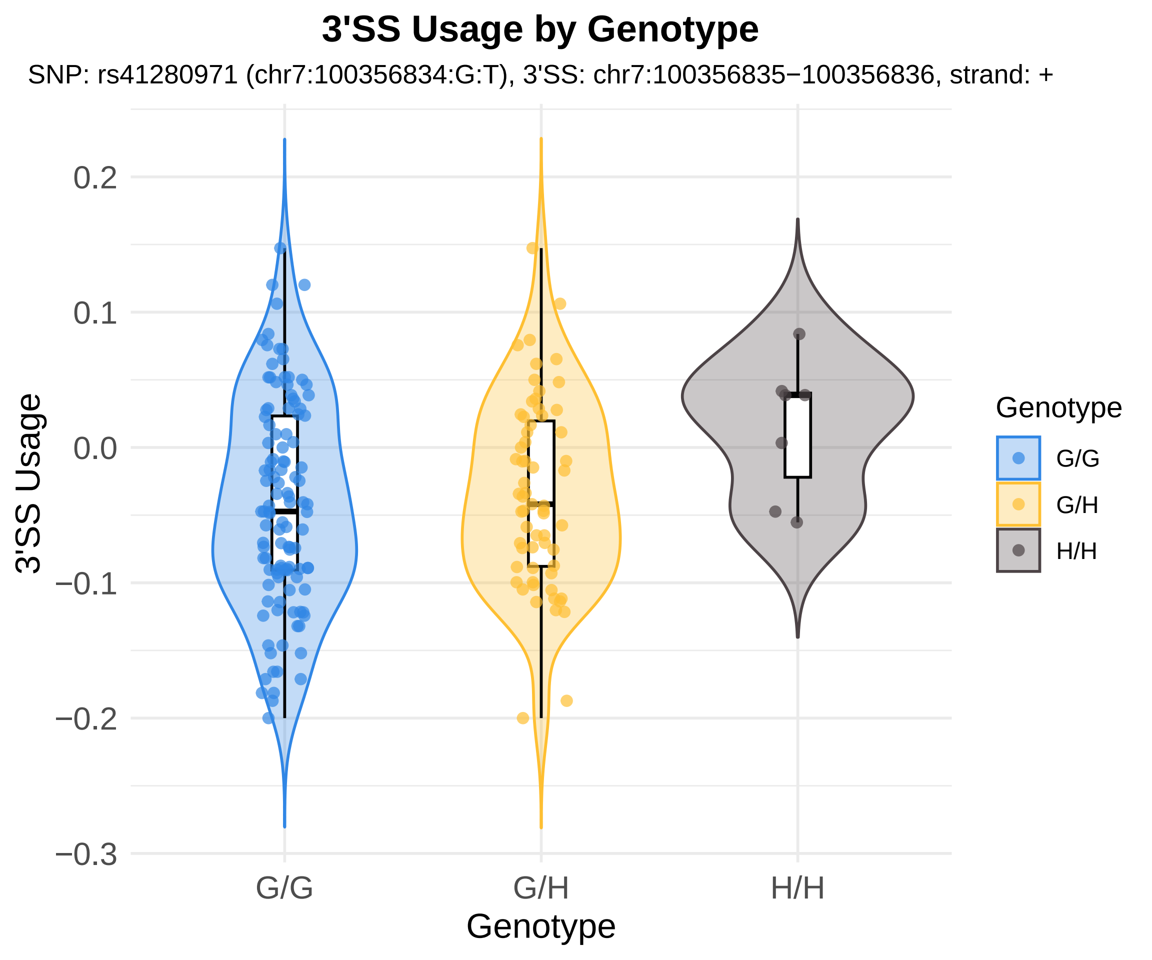

3'SS Usage by Genotype
SNP: rs41280971 (chr7:100356834:G:T), 3'SS: chr7:100356835−100356836, strand: +
3'SS Usage
0.2
0.1
0.0
−0.1
−0.2
−0.3
G/G
G/H
H/H
Genotype
Genotype
G/G
G/H
H/H

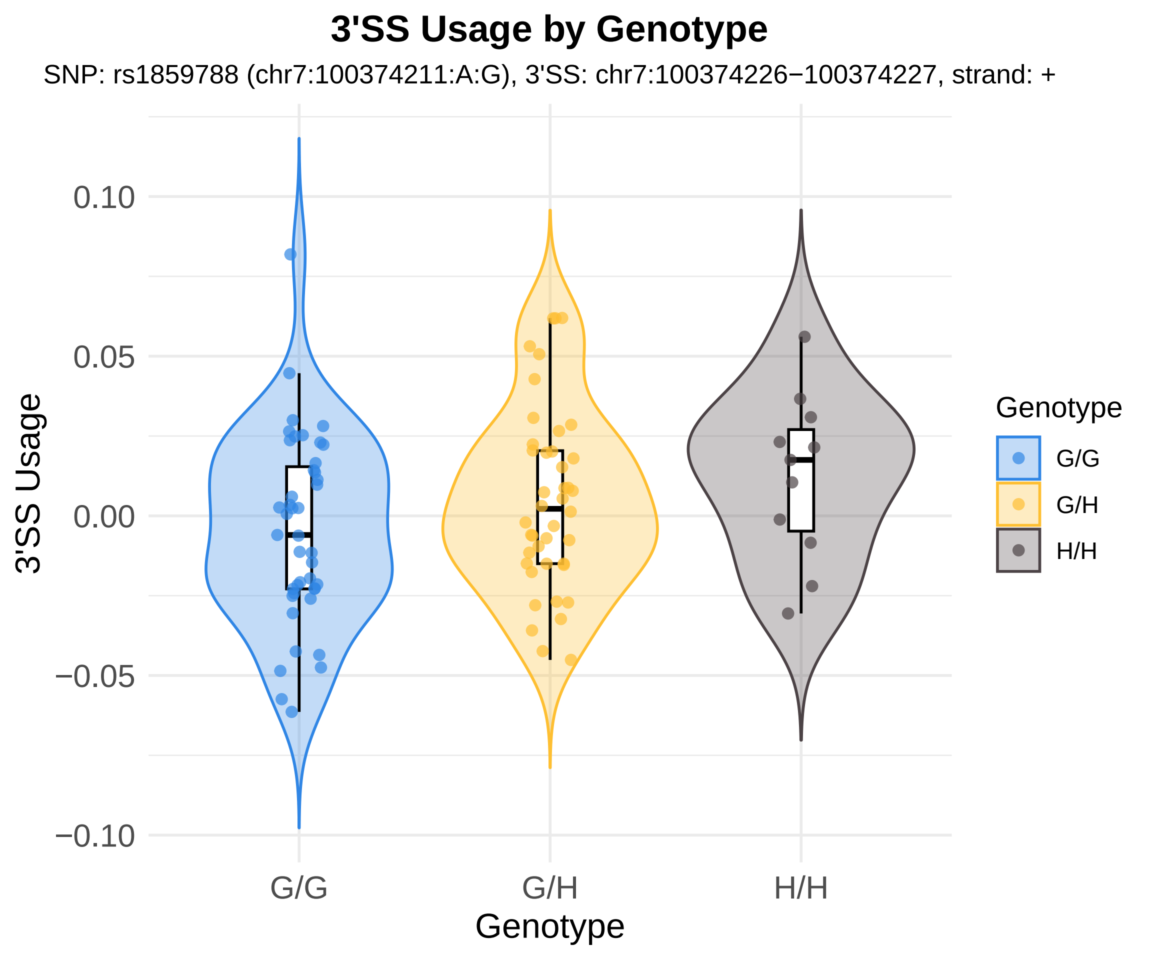
3'SS Usage by Genotype
SNP: rs1859788 (chr7:100374211:A:G), 3'SS: chr7:100374226−100374227, strand: +
3'SS Usage
0.10
0.05
0.00
−0.05
−0.10
G/G
G/H
H/H
Genotype
Genotype
G/G
G/H
H/H

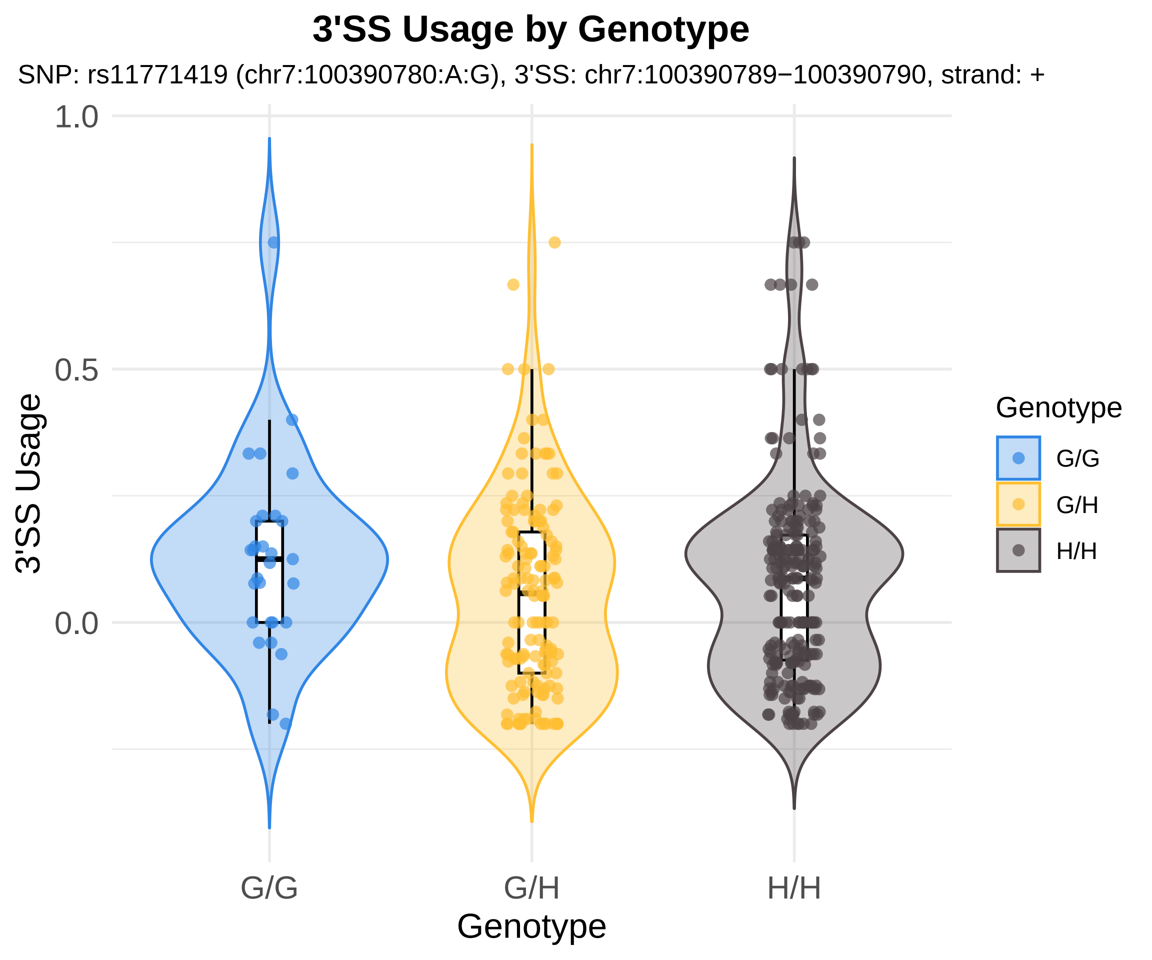
3'SS Usage by Genotype
SNP: rs11771419 (chr7:100390780:A:G), 3'SS: chr7:100390789−100390790, strand: +
1.0
0.5
0.0
3'SS Usage
G/G
G/H
H/H
Genotype
Genotype
G/G
G/H
H/H

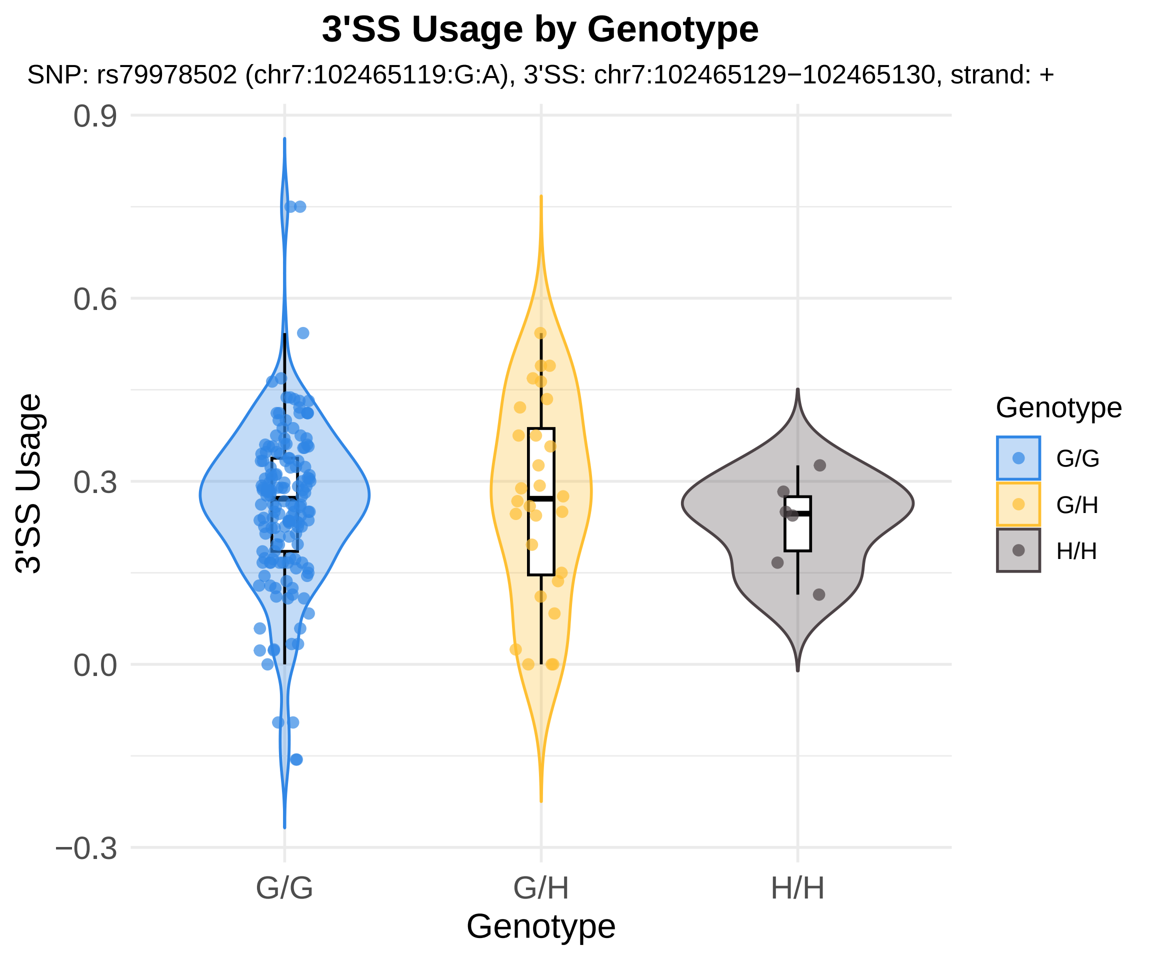
3'SS Usage by Genotype
SNP: rs79978502 (chr7:102465119:G:A), 3'SS: chr7:102465129−102465130, strand: +
3'SS Usage
0.9
0.6
0.3
0.0
−0.3
G/G
G/H
H/H
Genotype
Genotype
G/G
G/H
H/H

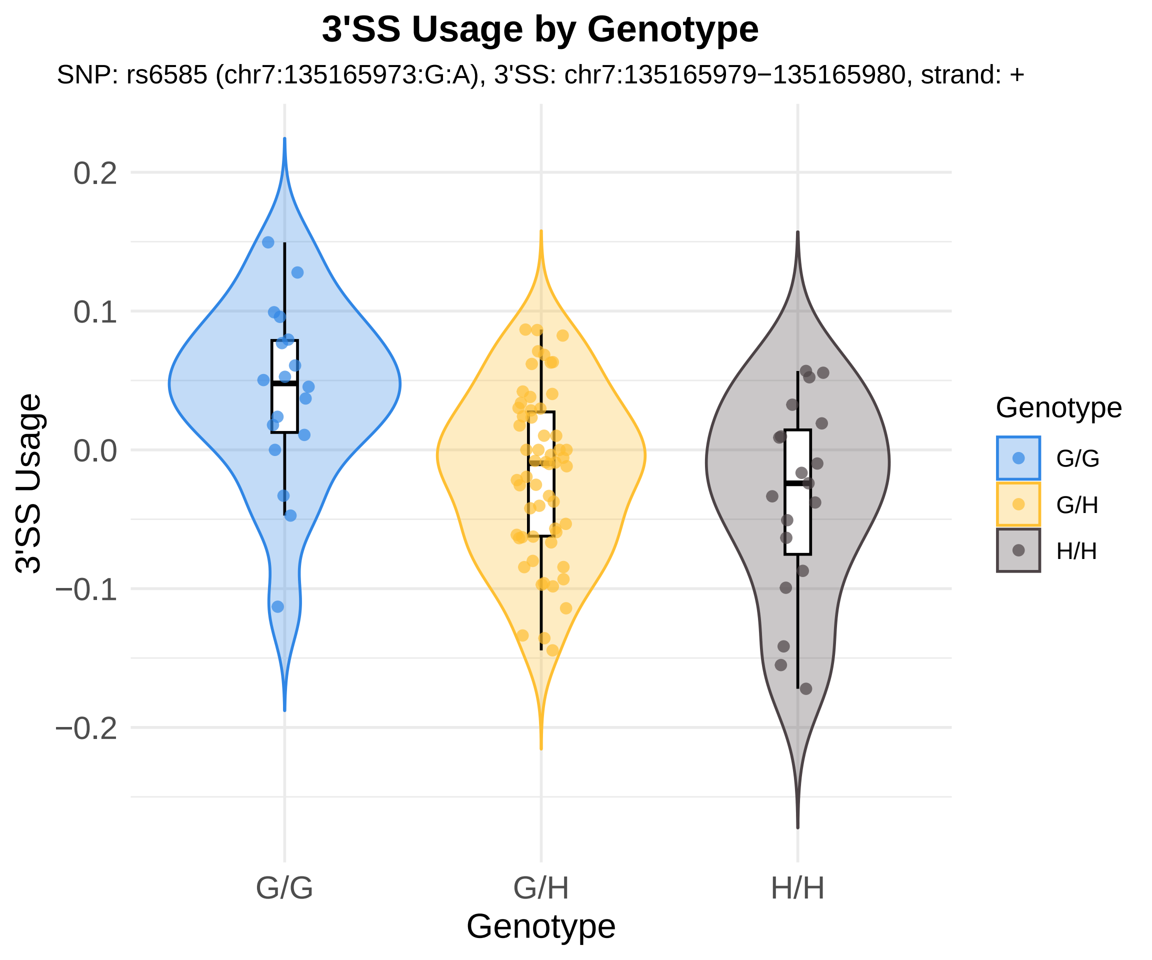

3'SS Usage by Genotype
SNP: rs6585 (chr7:135165973:G:A), 3'SS: chr7:135165979−135165980, strand: +
3'SS Usage
0.2
0.1
0.0
−0.1
−0.2
G/G
G/H
H/H
Genotype
Genotype
G/G
G/H
H/H

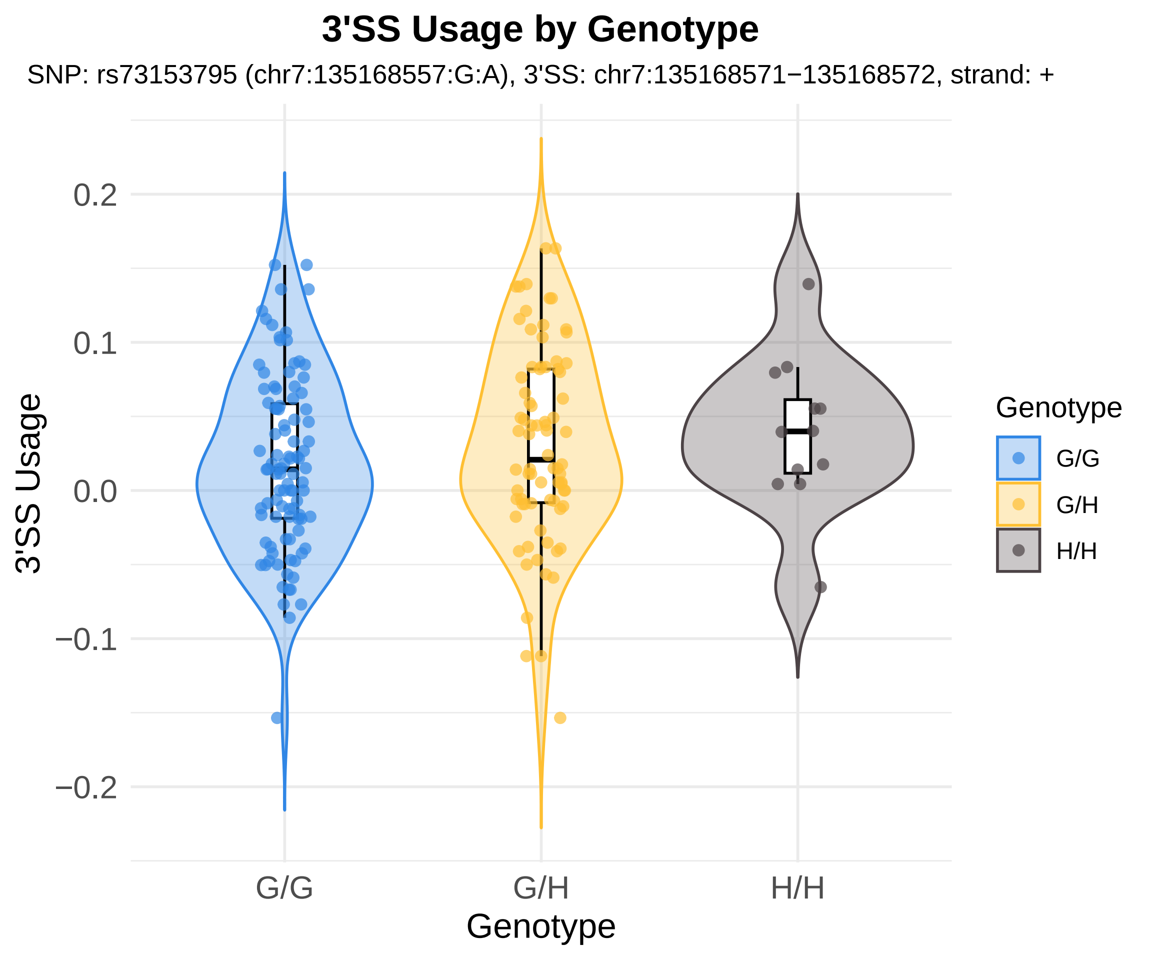
3'SS Usage by Genotype
SNP: rs73153795 (chr7:135168557:G:A), 3'SS: chr7:135168571−135168572, strand: +
3'SS Usage
0.2
0.1
0.0
−0.1
−0.2
G/G
G/H
H/H
Genotype
Genotype
G/G
G/H
H/H

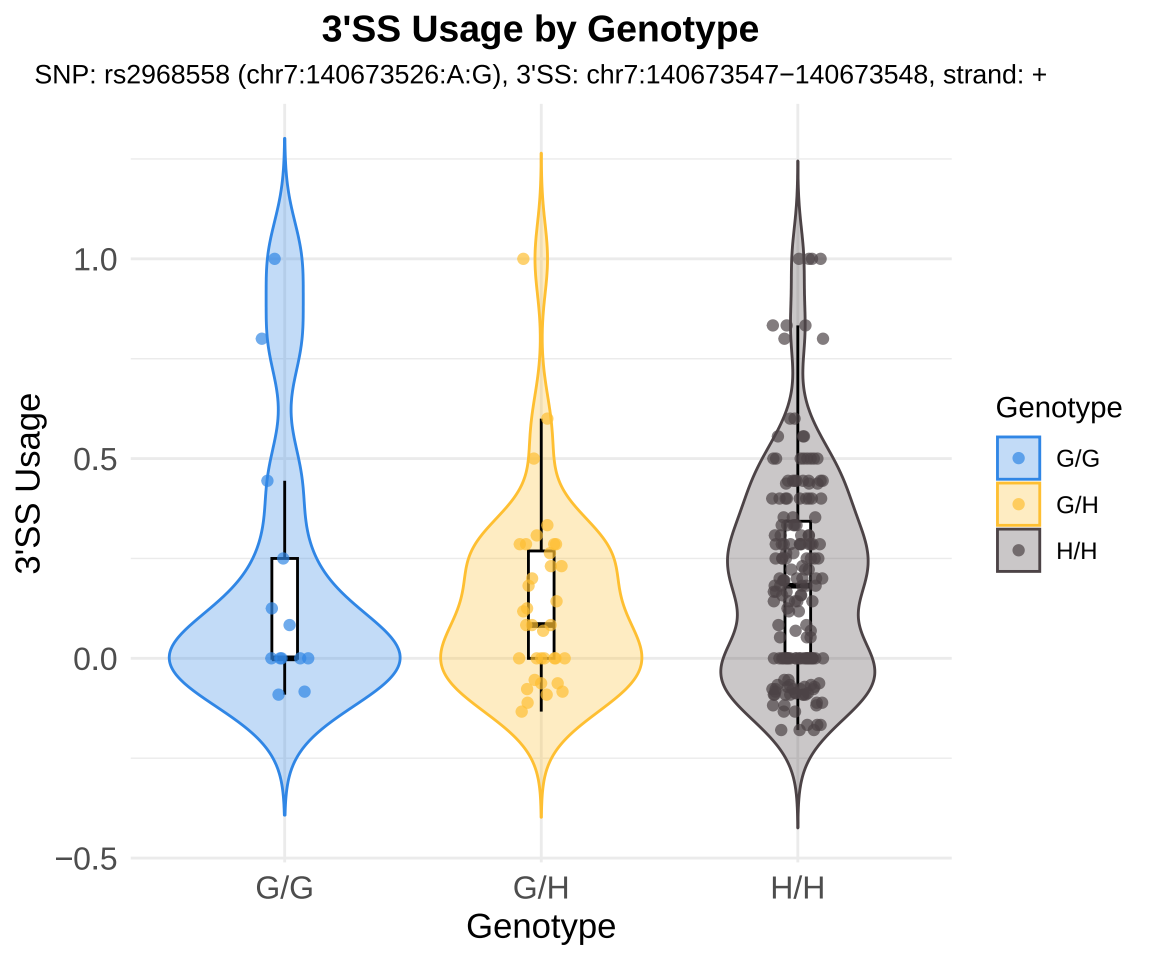

3'SS Usage by Genotype
SNP: rs2968558 (chr7:140673526:A:G), 3'SS: chr7:140673547−140673548, strand: +
3'SS Usage
1.0
0.5
0.0
−0.5
G/G
G/H
H/H
Genotype
Genotype
G/G
G/H
H/H

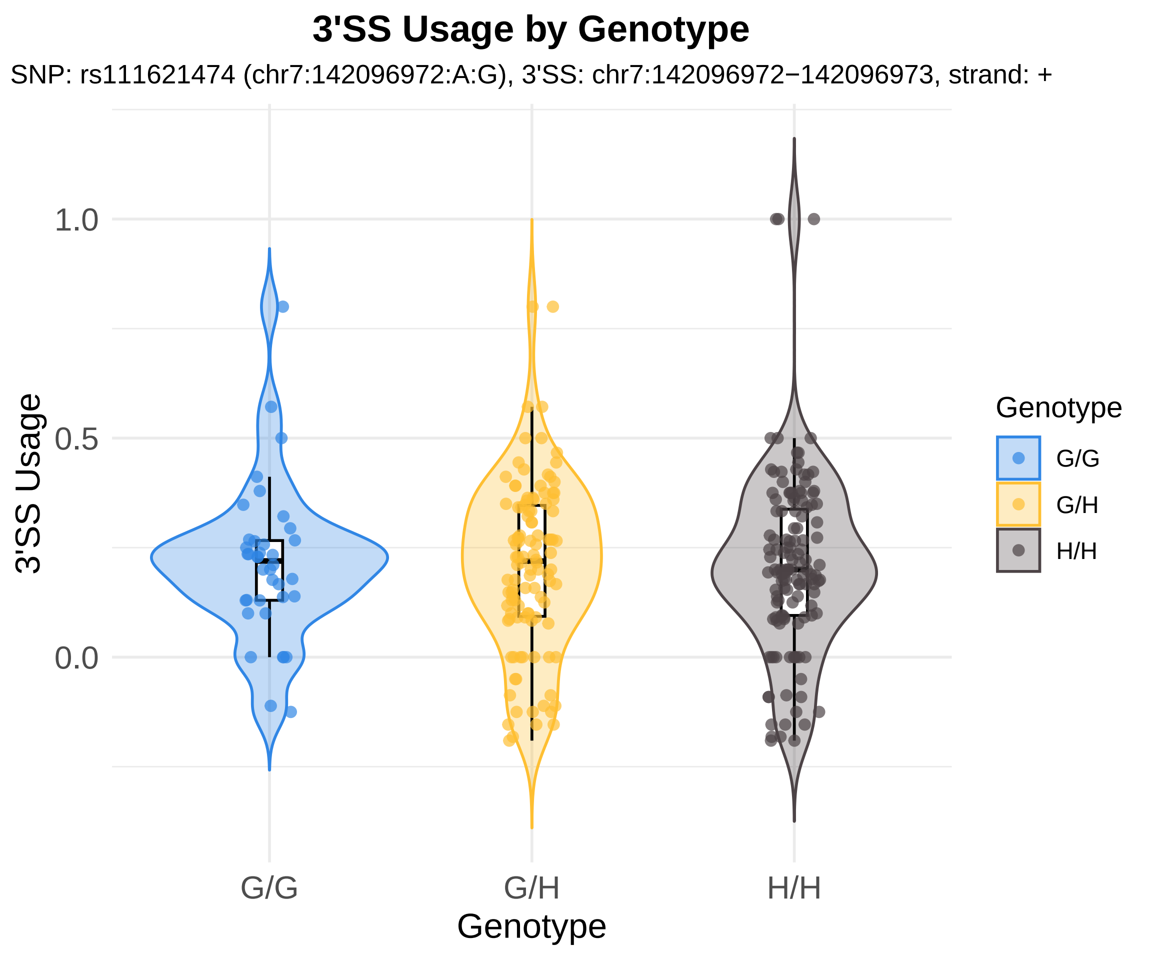
3'SS Usage by Genotype
SNP: rs111621474 (chr7:142096972:A:G), 3'SS: chr7:142096972−142096973, strand: +
3'SS Usage
1.0
0.5
0.0
G/G
G/H
H/H
Genotype
Genotype
G/G
G/H
H/H

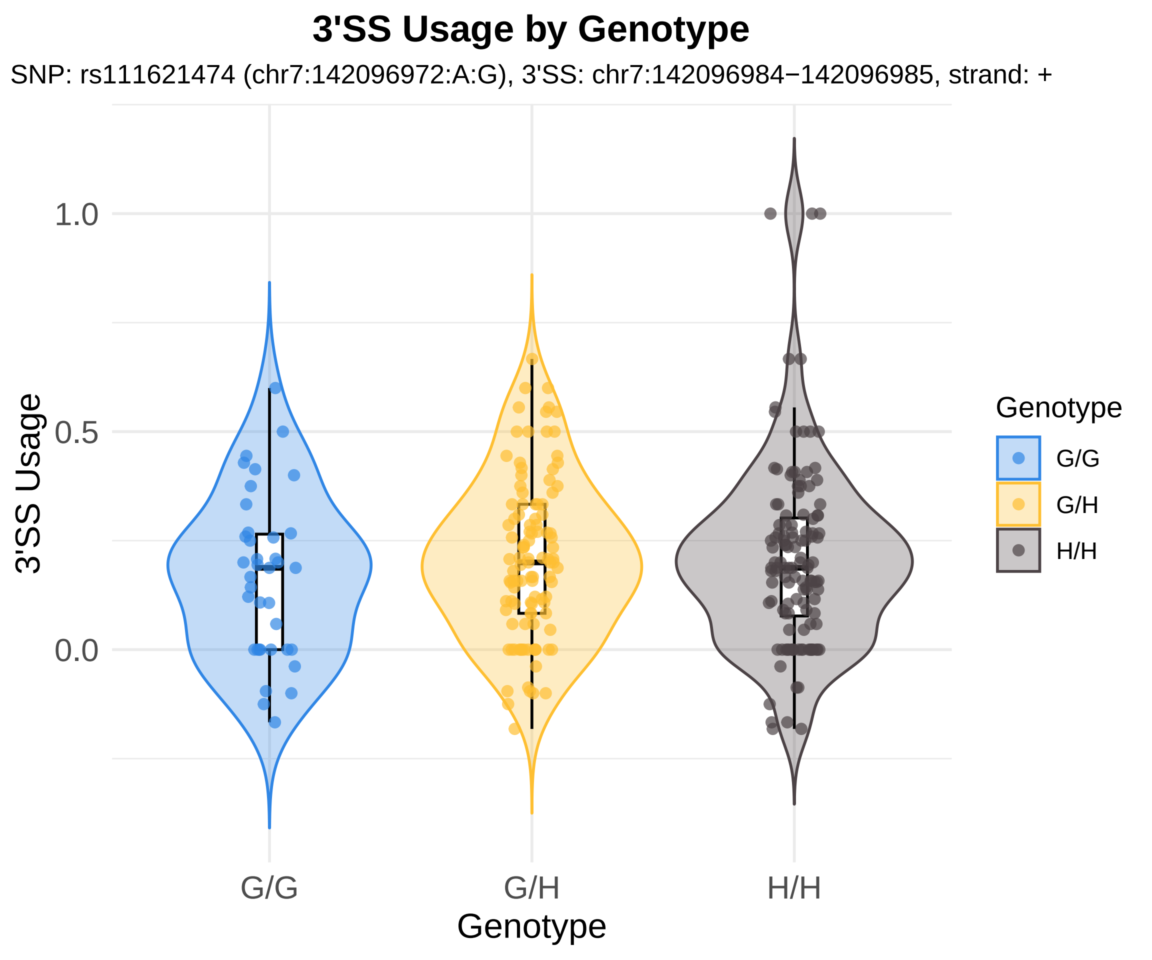
3'SS Usage by Genotype
SNP: rs111621474 (chr7:142096972:A:G), 3'SS: chr7:142096984−142096985, strand: +
3'SS Usage
1.0
0.5
0.0
G/G
G/H
H/H
Genotype
Genotype
G/G
G/H
H/H

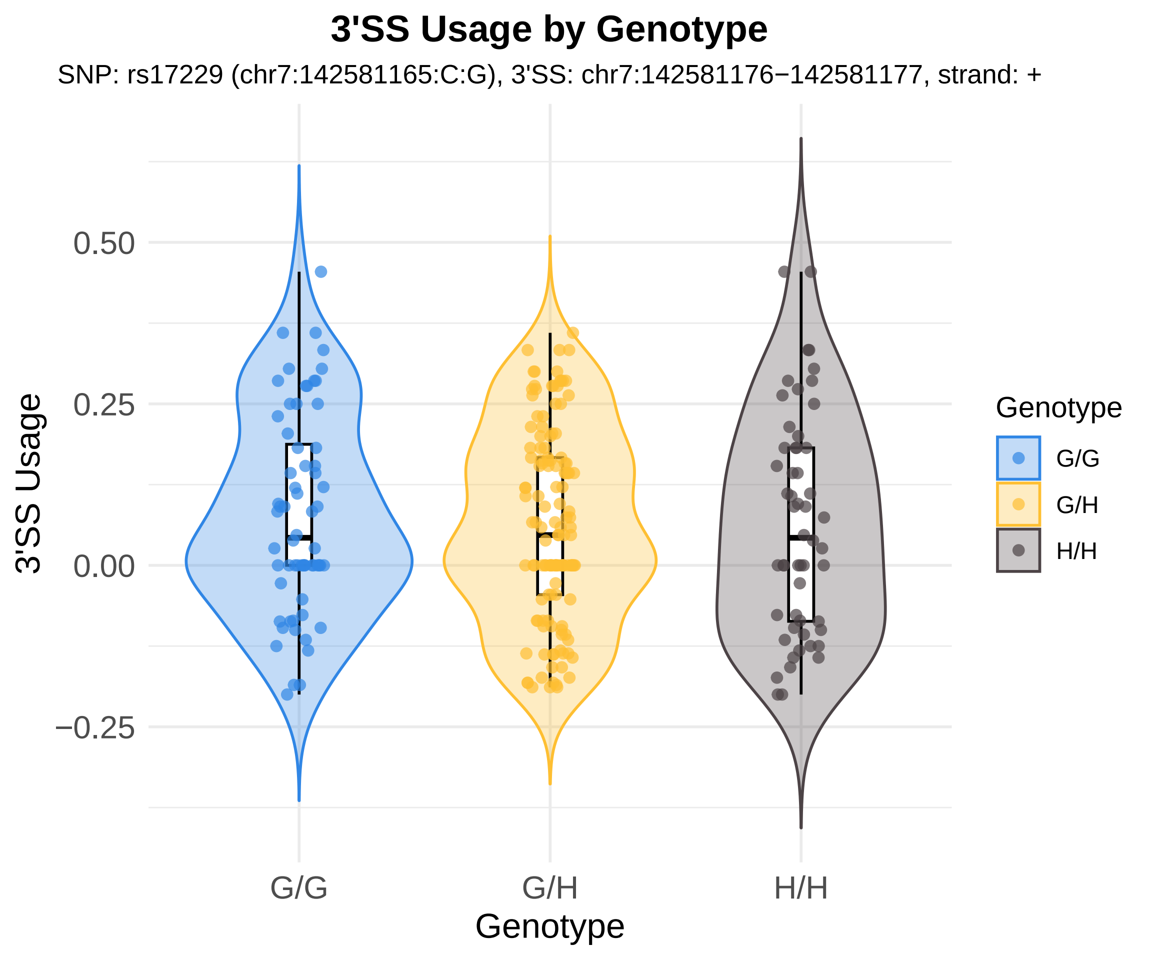

3'SS Usage by Genotype
SNP: rs17229 (chr7:142581165:C:G), 3'SS: chr7:142581176−142581177, strand: +
3'SS Usage
0.50
0.25
0.00
−0.25
G/G
G/H
H/H
Genotype
Genotype
G/G
G/H
H/H

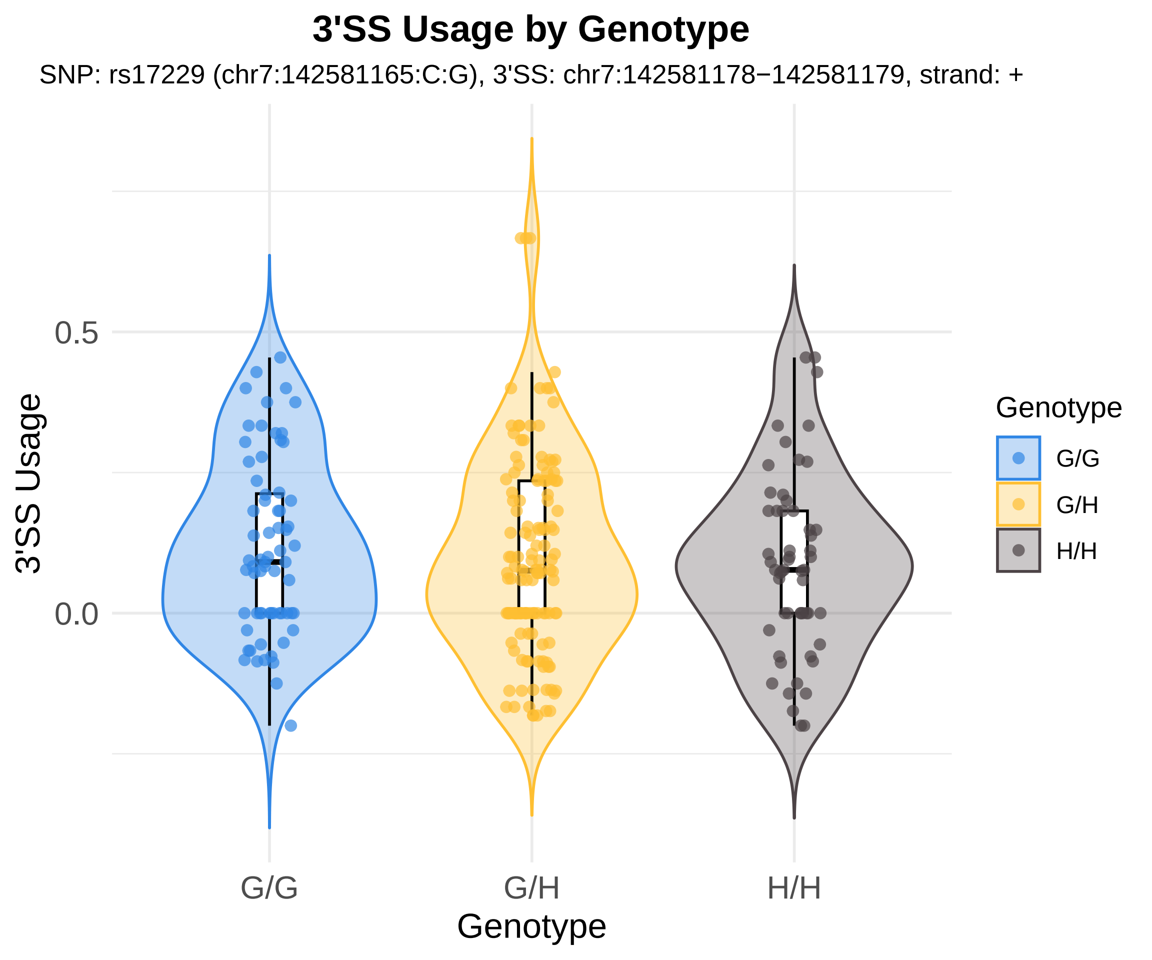
3'SS Usage by Genotype
SNP: rs17229 (chr7:142581165:C:G), 3'SS: chr7:142581178−142581179, strand: +
3'SS Usage
0.5
0.0
G/G
G/H
H/H
Genotype
Genotype
G/G
G/H
H/H

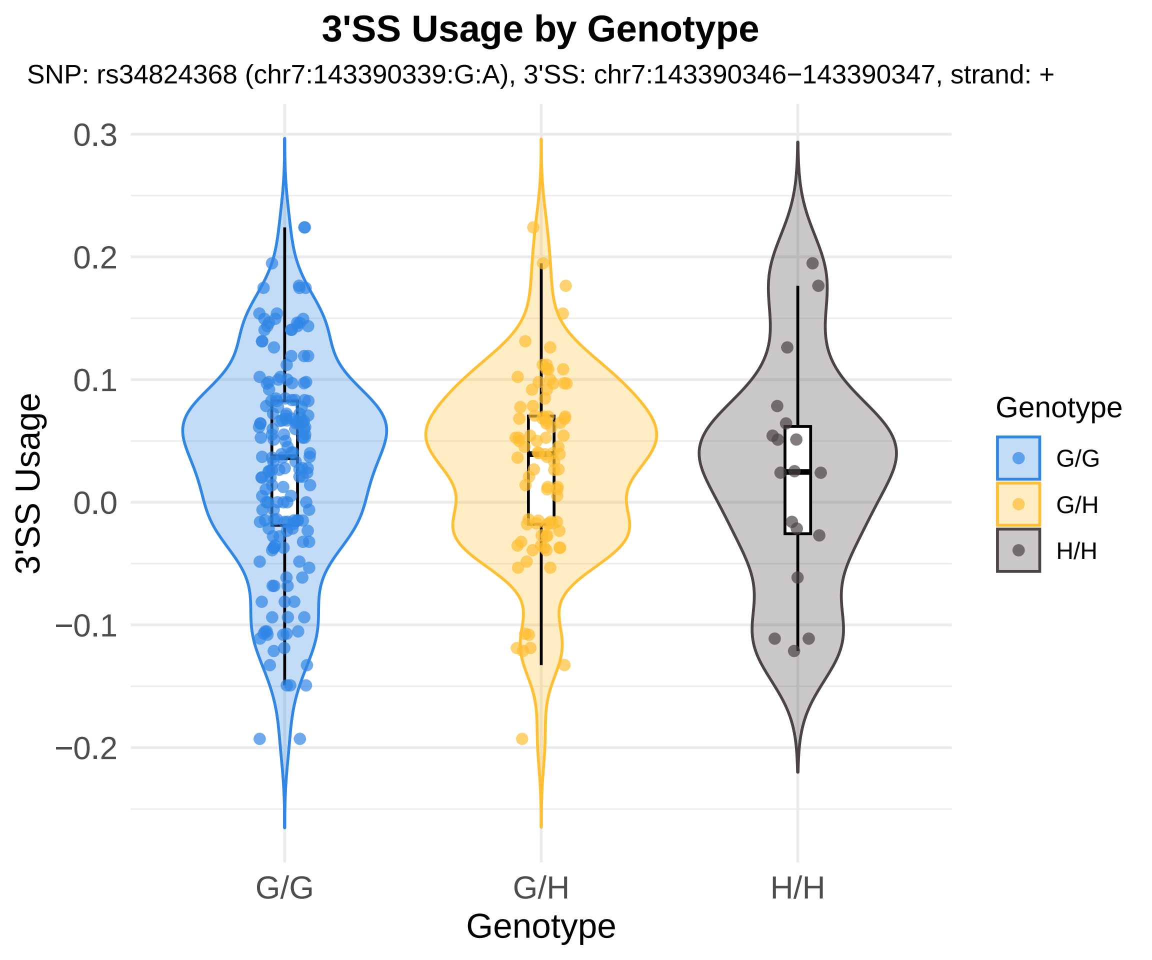

3'SS Usage by Genotype
SNP: rs34824368 (chr7:143390339:G:A), 3'SS: chr7:143390346−143390347, strand: +
3'SS Usage
0.3
0.2
0.1
0.0
−0.1
−0.2
G/G
G/H
H/H
Genotype
Genotype
G/G
G/H
H/H

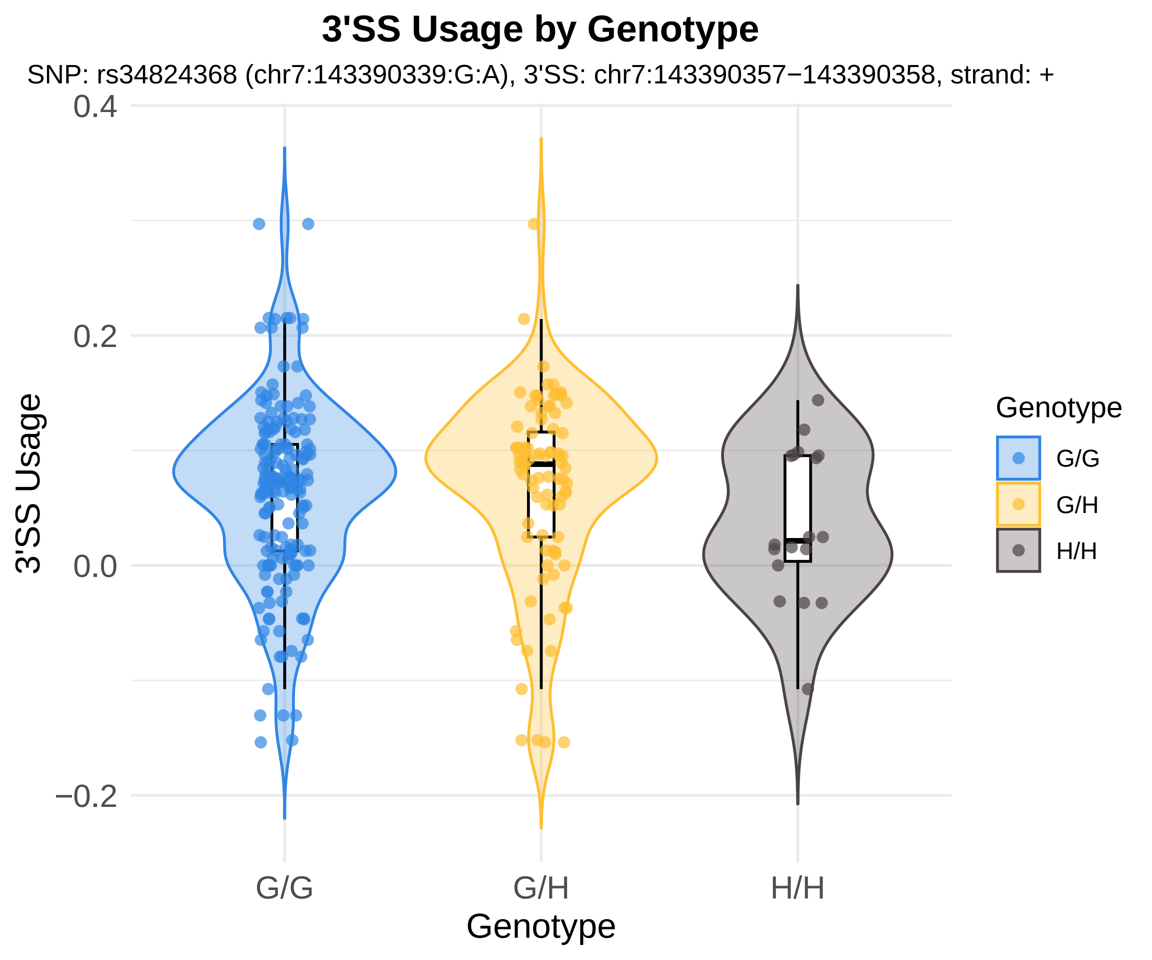
3'SS Usage by Genotype
SNP: rs34824368 (chr7:143390339:G:A), 3'SS: chr7:143390357−143390358, strand: +
3'SS Usage
0.4
0.2
0.0
−0.2
G/G
G/H
H/H
Genotype
Genotype
G/G
G/H
H/H

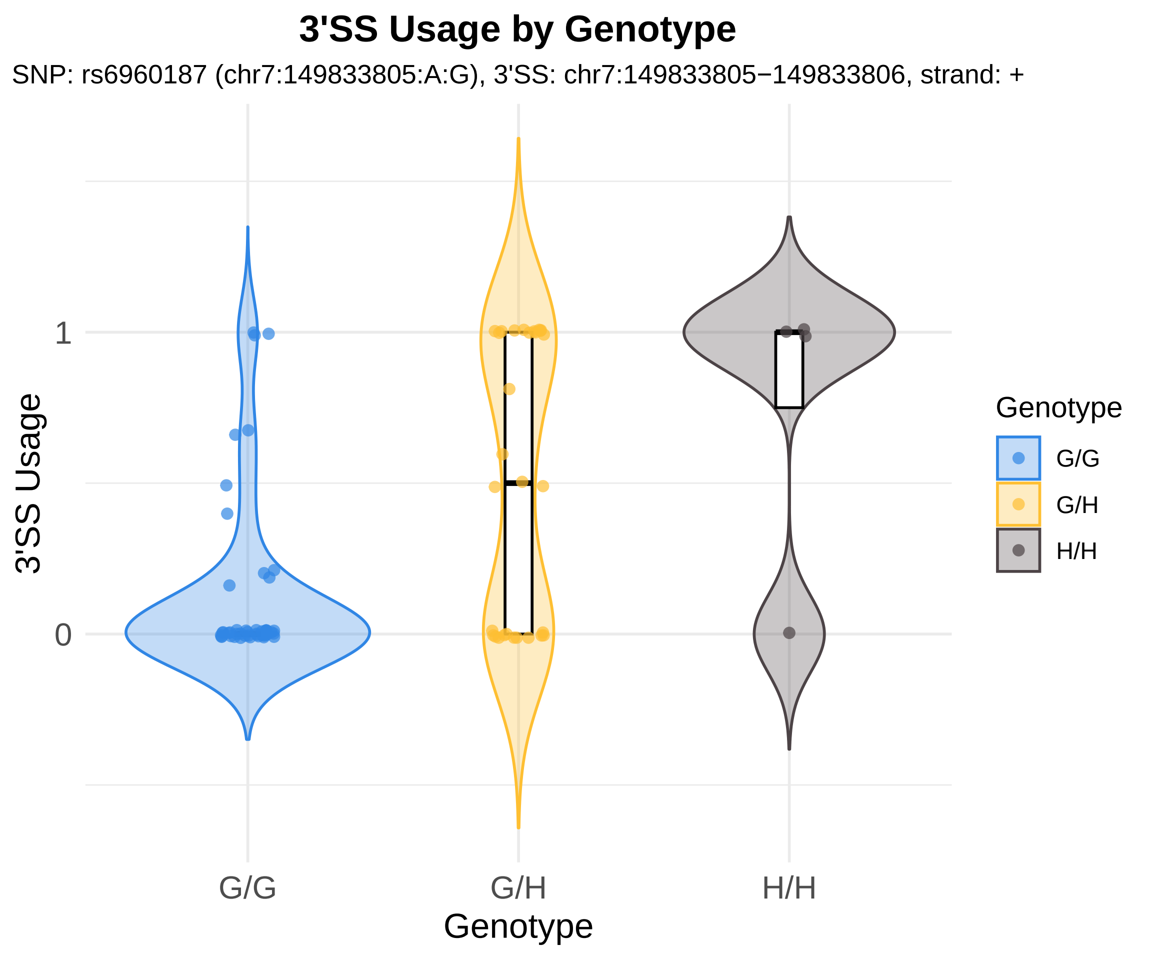
3'SS Usage by Genotype
SNP: rs6960187 (chr7:149833805:A:G), 3'SS: chr7:149833805−149833806, strand: +
3'SS Usage
1
0
G/G
G/H
H/H
Genotype
Genotype
G/G
G/H
H/H

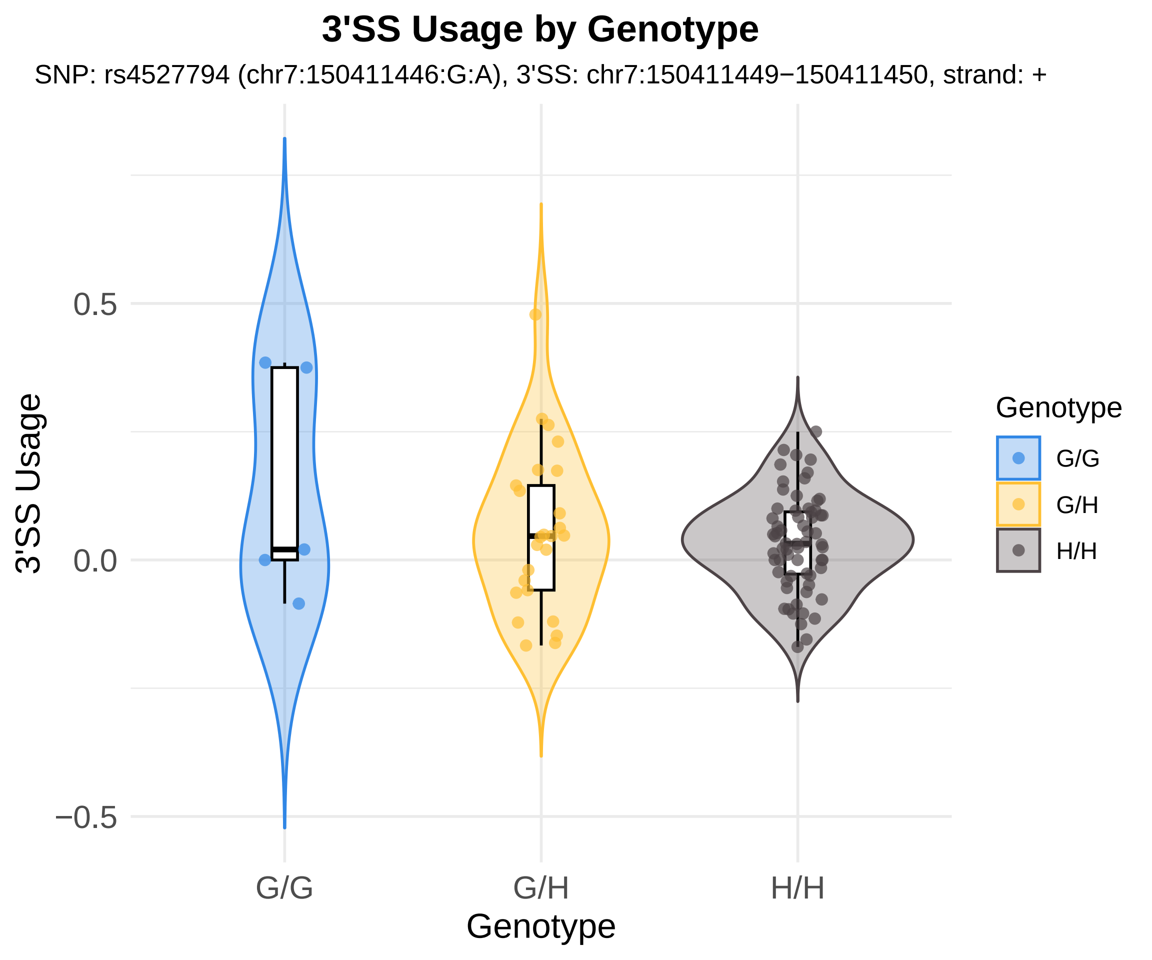
3'SS Usage by Genotype
SNP: rs4527794 (chr7:150411446:G:A), 3'SS: chr7:150411449−150411450, strand: +
3'SS Usage
0.5
0.0
−0.5
G/G
G/H
H/H
Genotype
Genotype
G/G
G/H
H/H

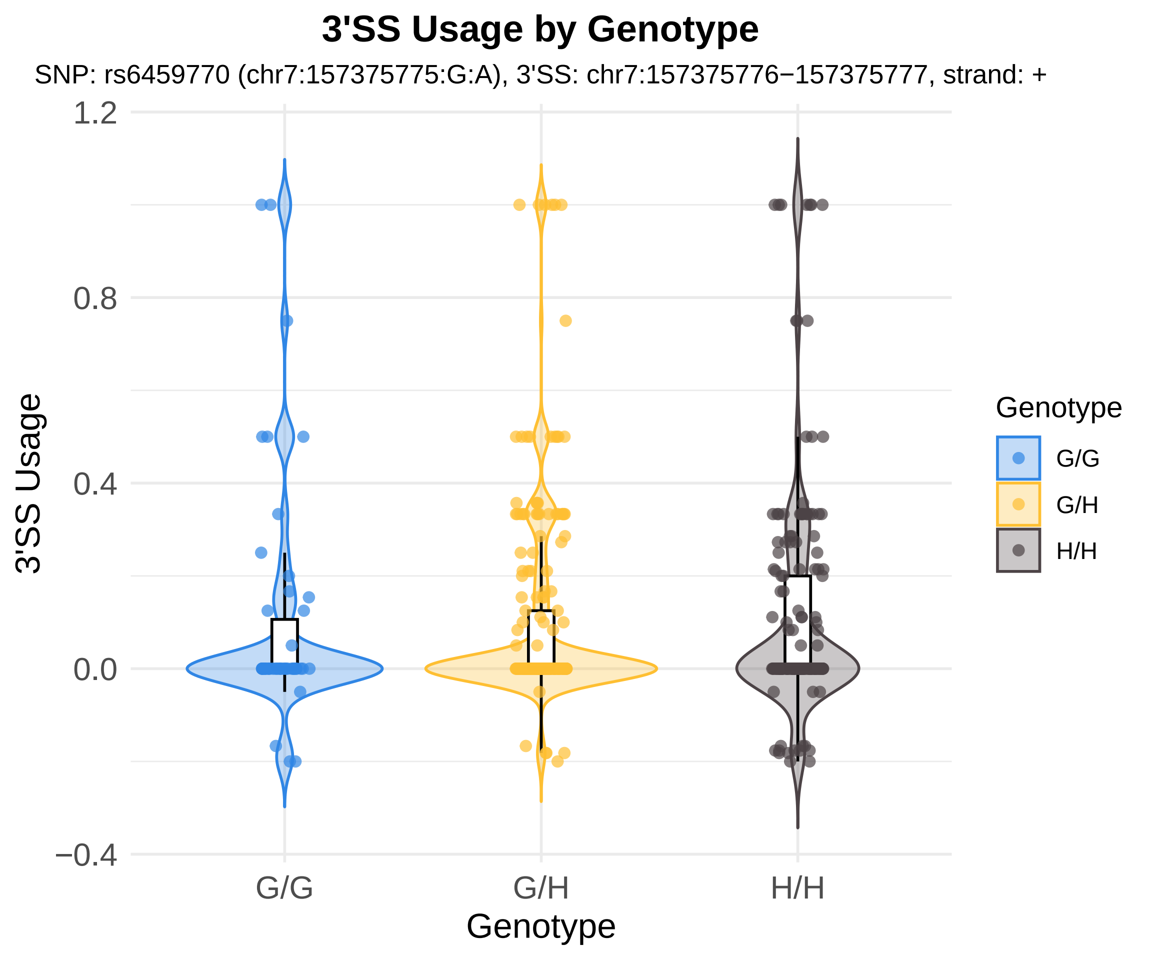

3'SS Usage by Genotype
SNP: rs6459770 (chr7:157375775:G:A), 3'SS: chr7:157375776−157375777, strand: +
1.2
0.8
0.4
0.0
−0.4
3'SS Usage
G/G
G/H
H/H
Genotype
Genotype
G/G
G/H
H/H

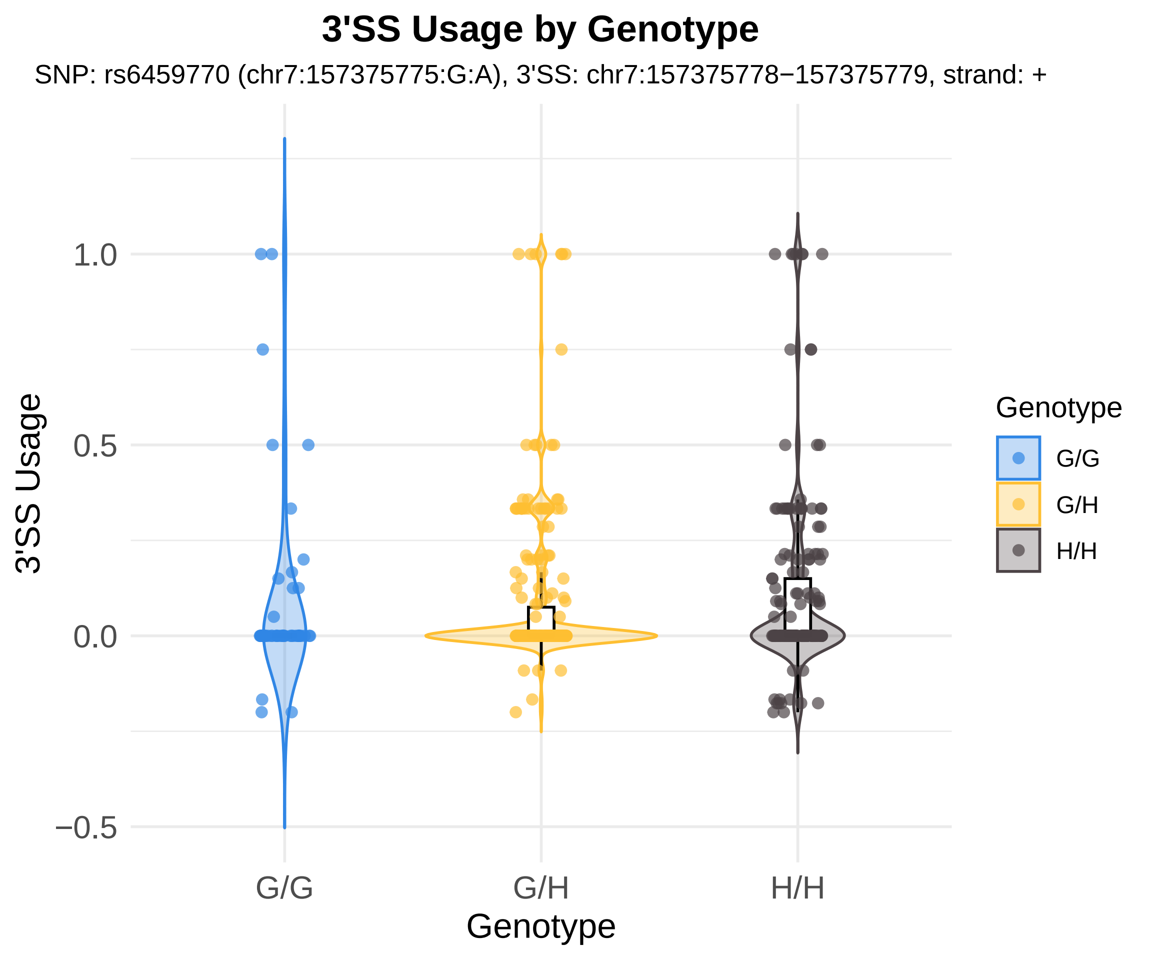

3'SS Usage by Genotype
SNP: rs6459770 (chr7:157375775:G:A), 3'SS: chr7:157375778−157375779, strand: +
3'SS Usage
1.0
0.5
0.0
−0.5
G/G
G/H
H/H
Genotype
Genotype
G/G
G/H
H/H

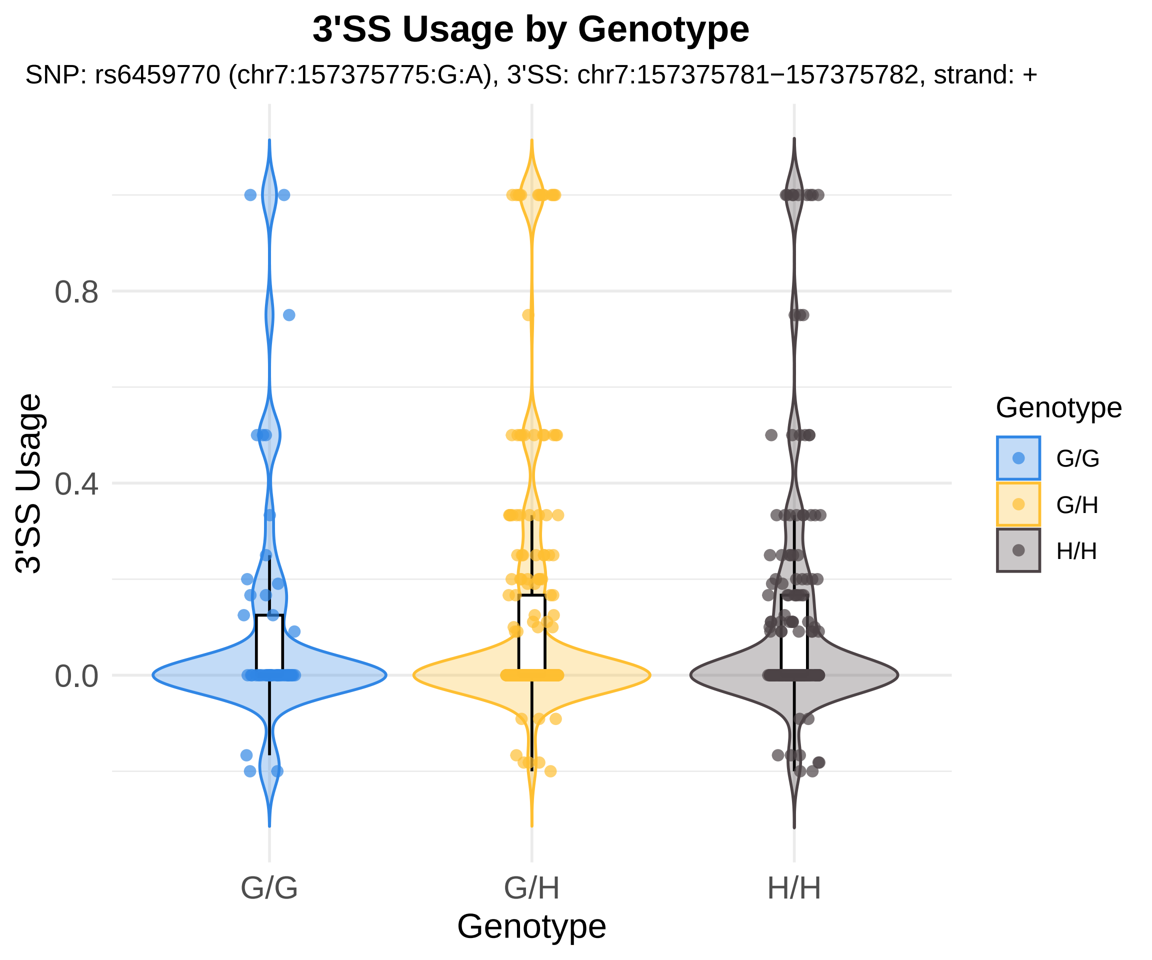
3'SS Usage by Genotype
SNP: rs6459770 (chr7:157375775:G:A), 3'SS: chr7:157375781−157375782, strand: +
3'SS Usage
0.8
0.4
0.0
G/G
G/H
H/H
Genotype
Genotype
G/G
G/H
H/H

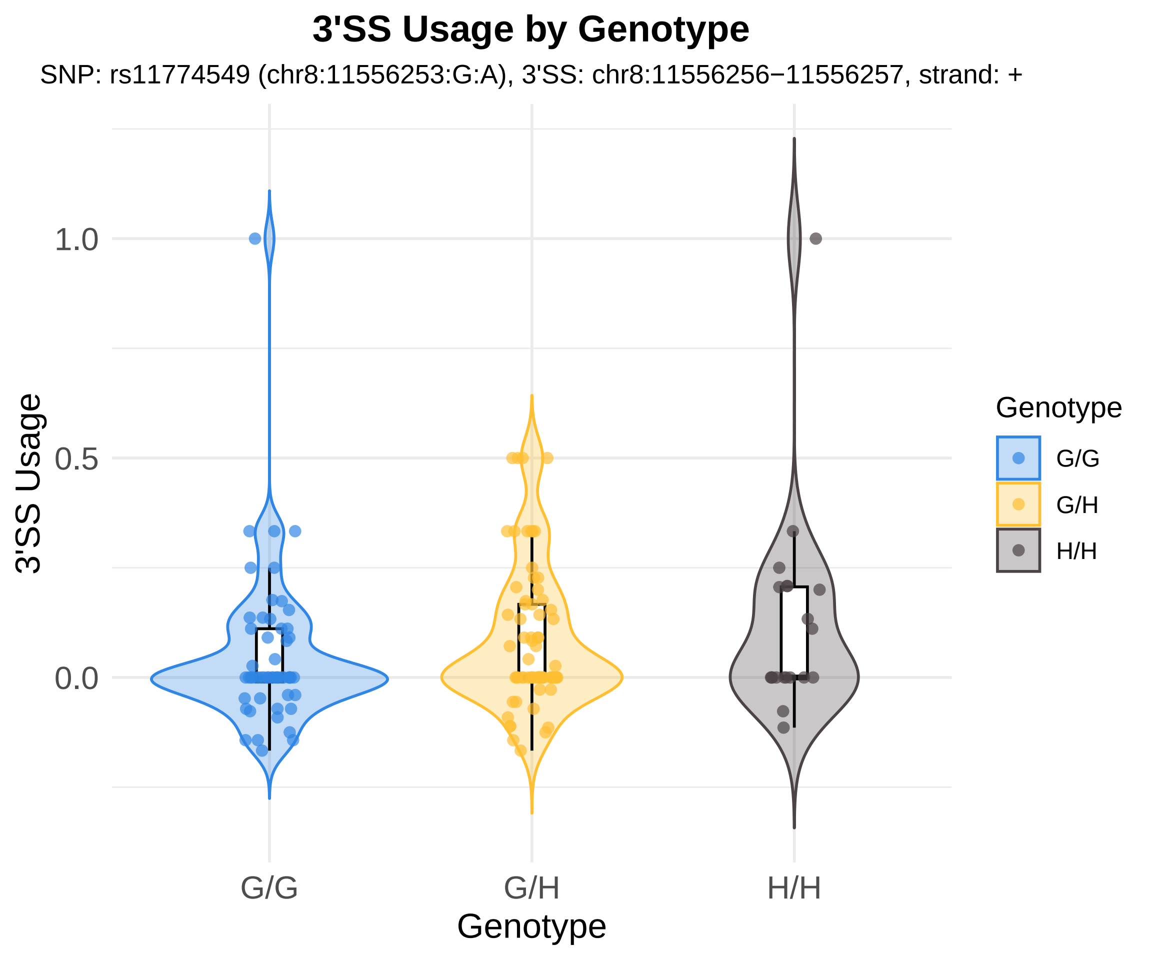
3'SS Usage by Genotype
SNP: rs11774549 (chr8:11556253:G:A), 3'SS: chr8:11556256−11556257, strand: +
3'SS Usage
1.0
0.5
0.0
G/G
G/H
H/H
Genotype
Genotype
G/G
G/H
H/H

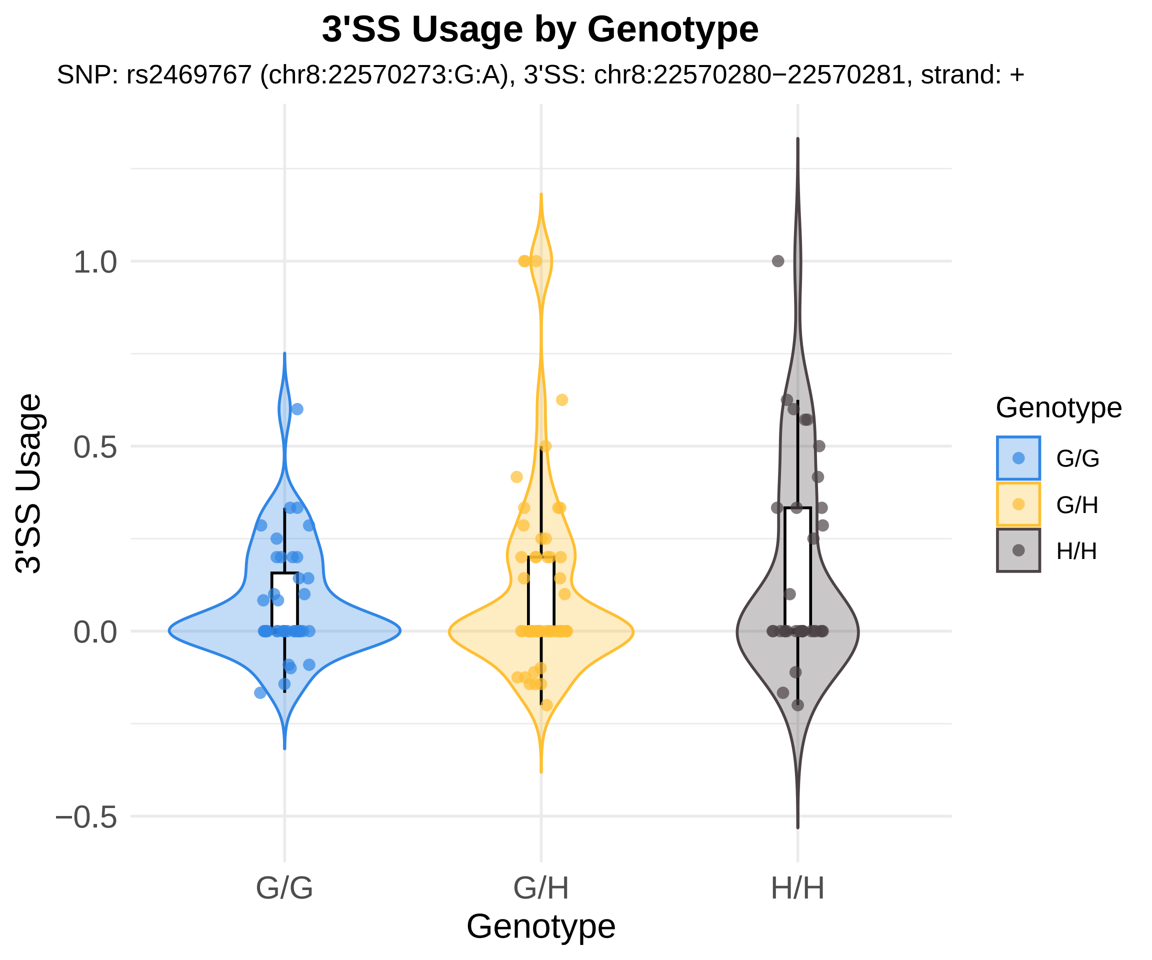
3'SS Usage by Genotype
SNP: rs2469767 (chr8:22570273:G:A), 3'SS: chr8:22570280−22570281, strand: +
3'SS Usage
1.0
0.5
0.0
−0.5
G/G
G/H
H/H
Genotype
Genotype
G/G
G/H
H/H

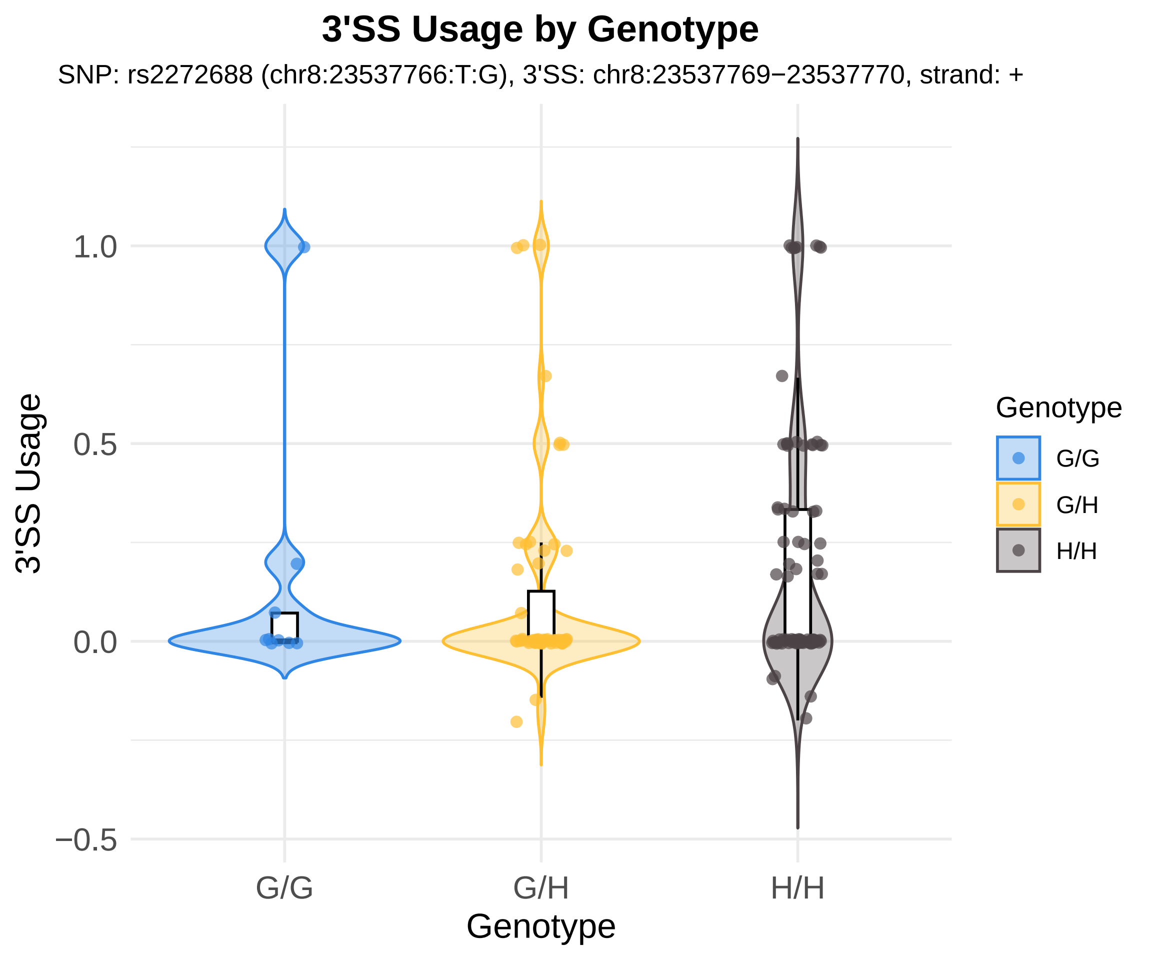
3'SS Usage by Genotype
SNP: rs2272688 (chr8:23537766:T:G), 3'SS: chr8:23537769−23537770, strand: +
3'SS Usage
1.0
0.5
0.0
−0.5
G/G
G/H
H/H
Genotype
Genotype
G/G
G/H
H/H

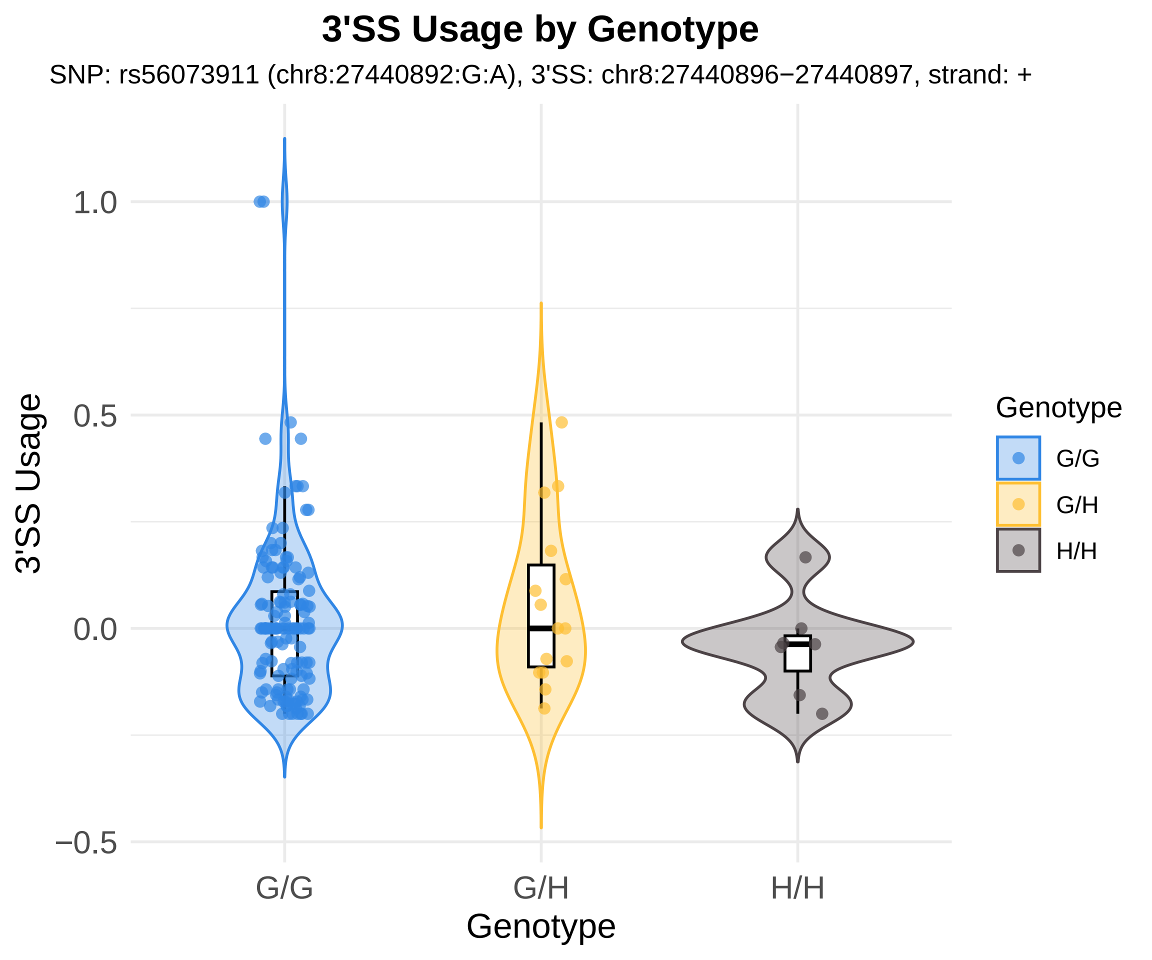
3'SS Usage by Genotype
SNP: rs56073911 (chr8:27440892:G:A), 3'SS: chr8:27440896−27440897, strand: +
1.0
0.5
0.0
−0.5
3'SS Usage
G/G
G/H
H/H
Genotype
Genotype
G/G
G/H
H/H

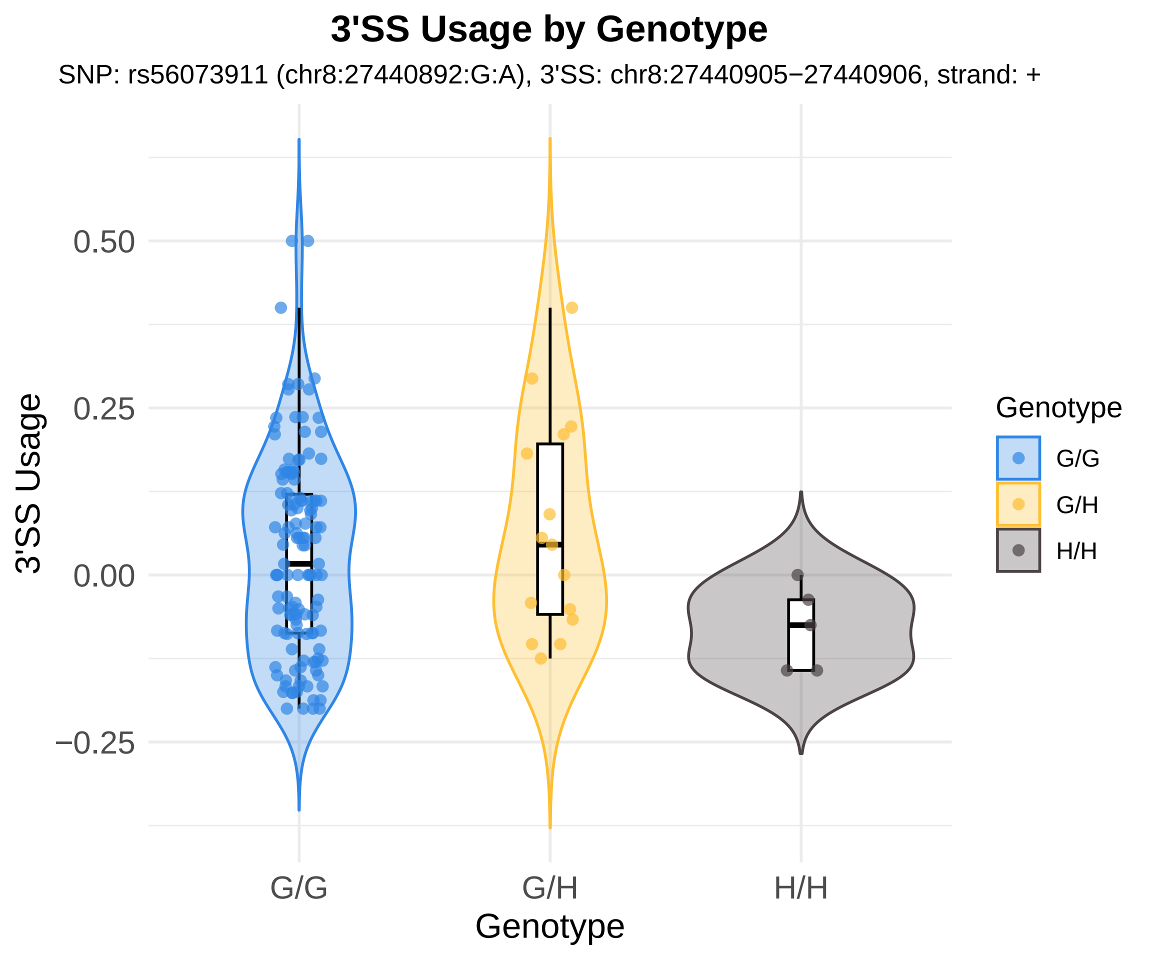
3'SS Usage by Genotype
SNP: rs56073911 (chr8:27440892:G:A), 3'SS: chr8:27440905−27440906, strand: +
3'SS Usage
0.50
0.25
0.00
−0.25
G/G
G/H
H/H
Genotype
Genotype
G/G
G/H
H/H

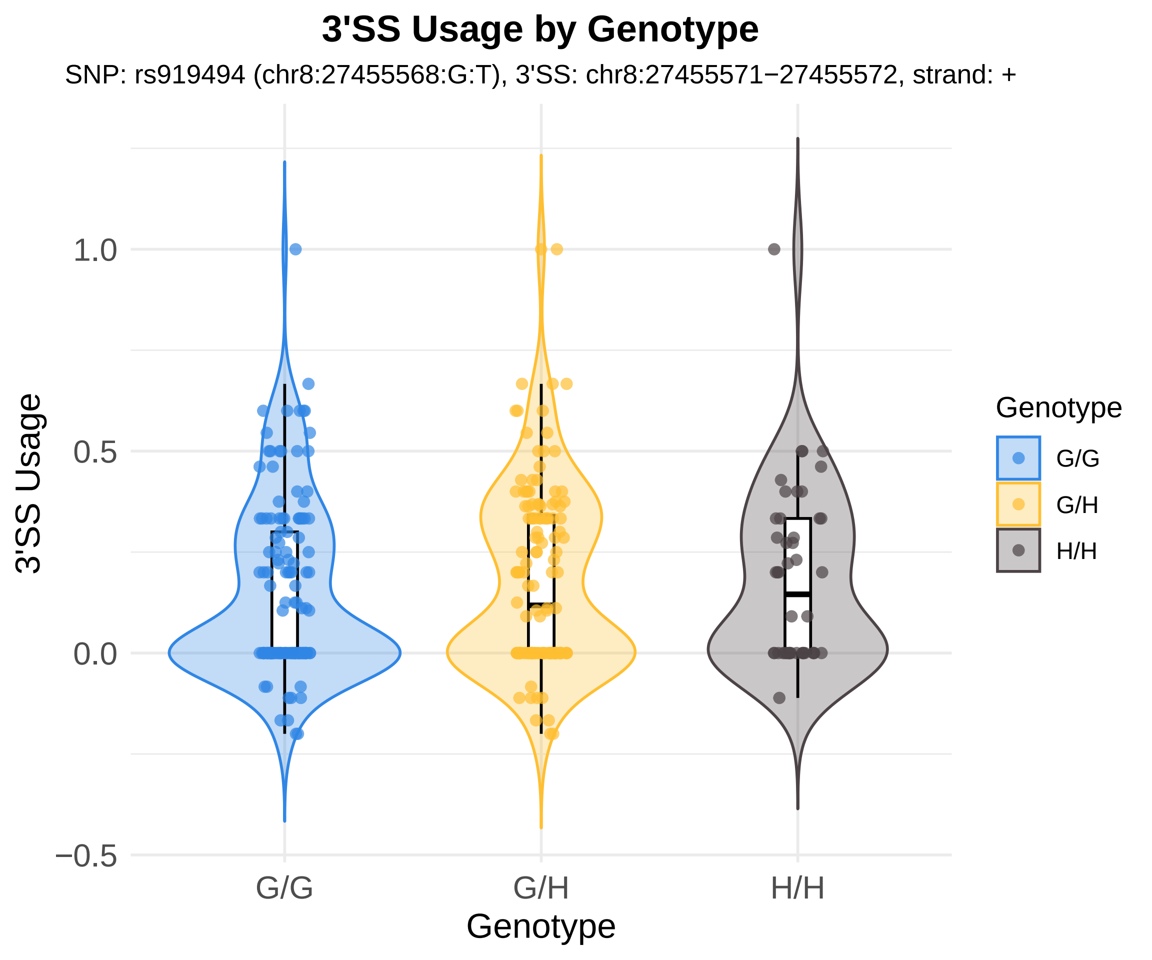

3'SS Usage by Genotype
SNP: rs919494 (chr8:27455568:G:T), 3'SS: chr8:27455571−27455572, strand: +
3'SS Usage
1.0
0.5
0.0
−0.5
G/G
G/H
H/H
Genotype
Genotype
G/G
G/H
H/H

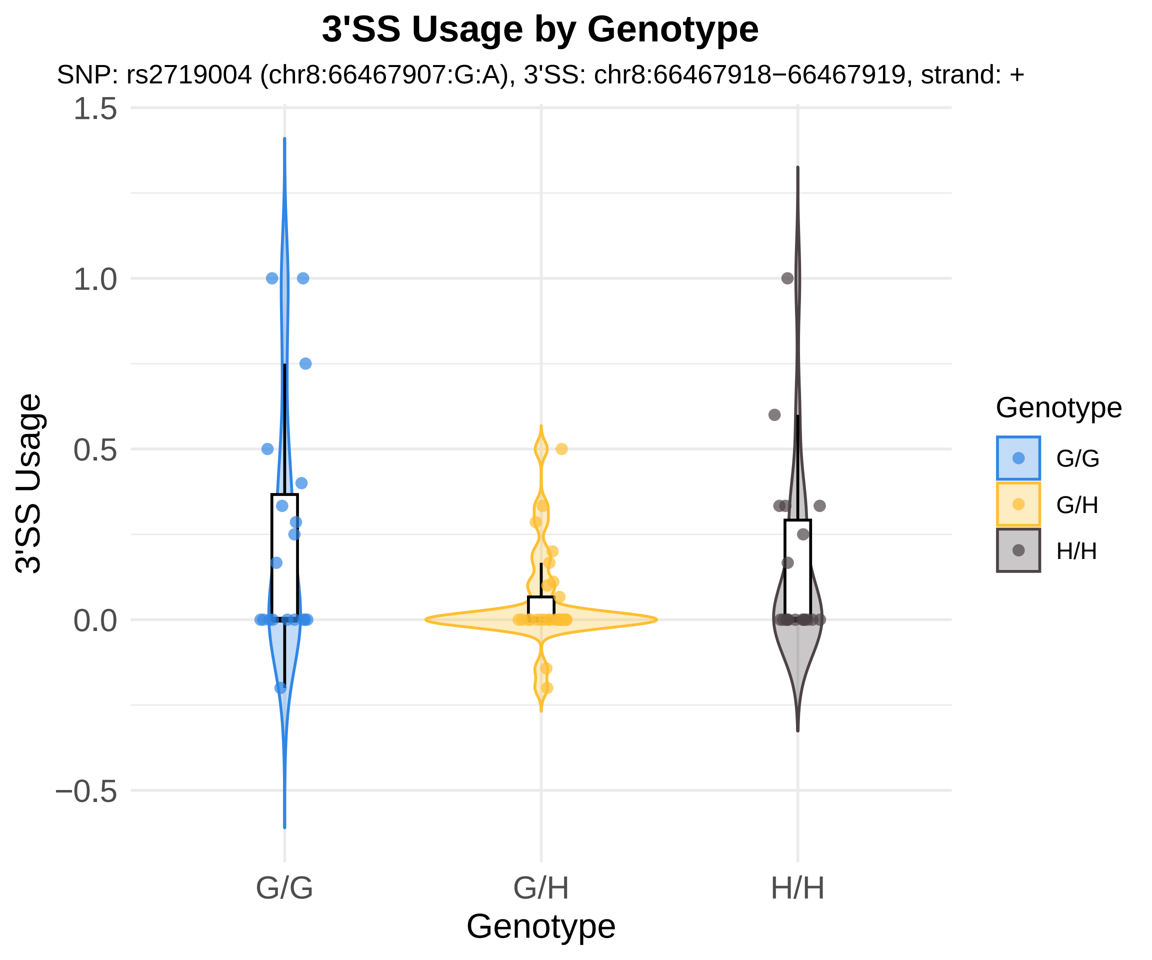
3'SS Usage by Genotype
SNP: rs2719004 (chr8:66467907:G:A), 3'SS: chr8:66467918−66467919, strand: +
1.5
1.0
0.5
0.0
−0.5
3'SS Usage
G/G
G/H
H/H
Genotype
Genotype
G/G
G/H
H/H

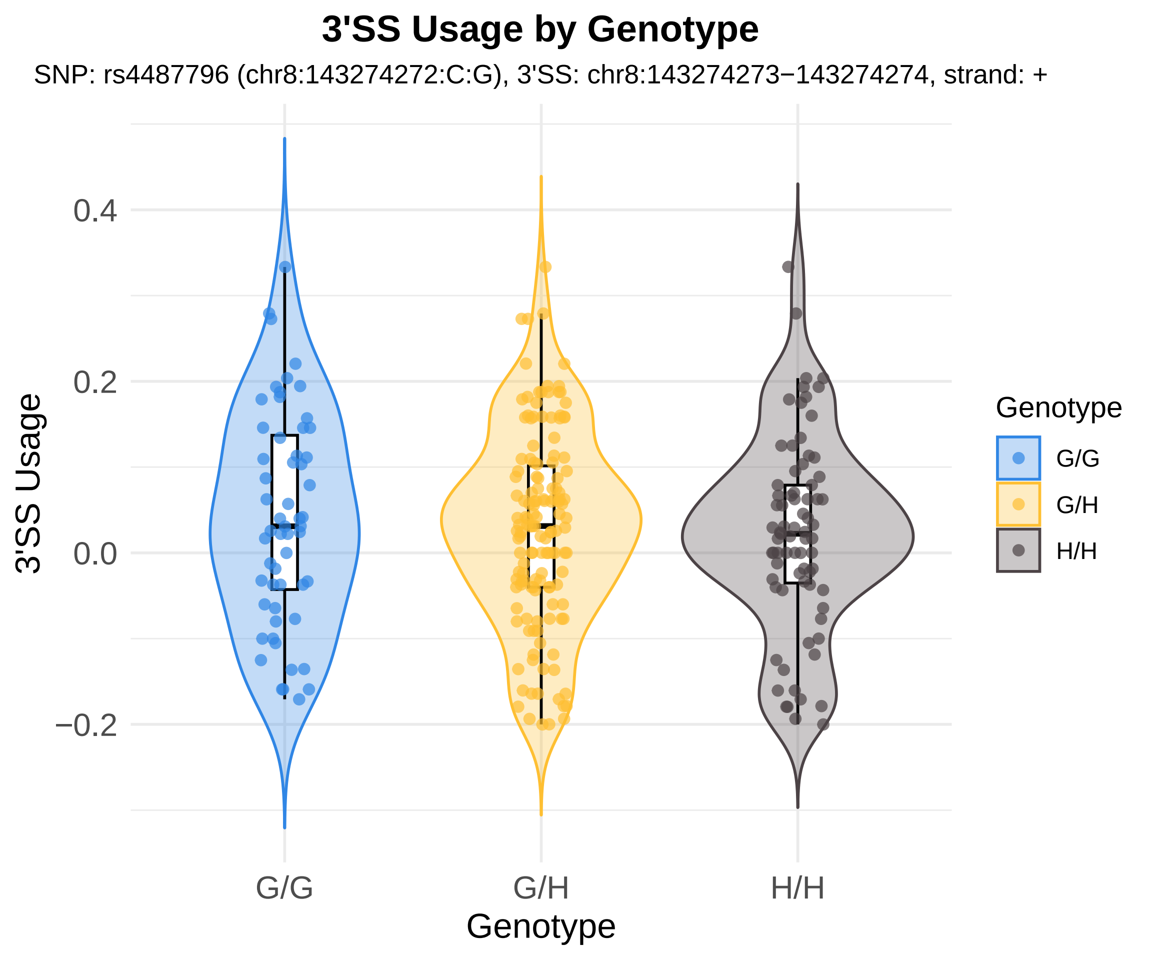
3'SS Usage by Genotype
SNP: rs4487796 (chr8:143274272:C:G), 3'SS: chr8:143274273−143274274, strand: +
3'SS Usage
0.4
0.2
0.0
−0.2
G/G
G/H
H/H
Genotype
Genotype
G/G
G/H
H/H

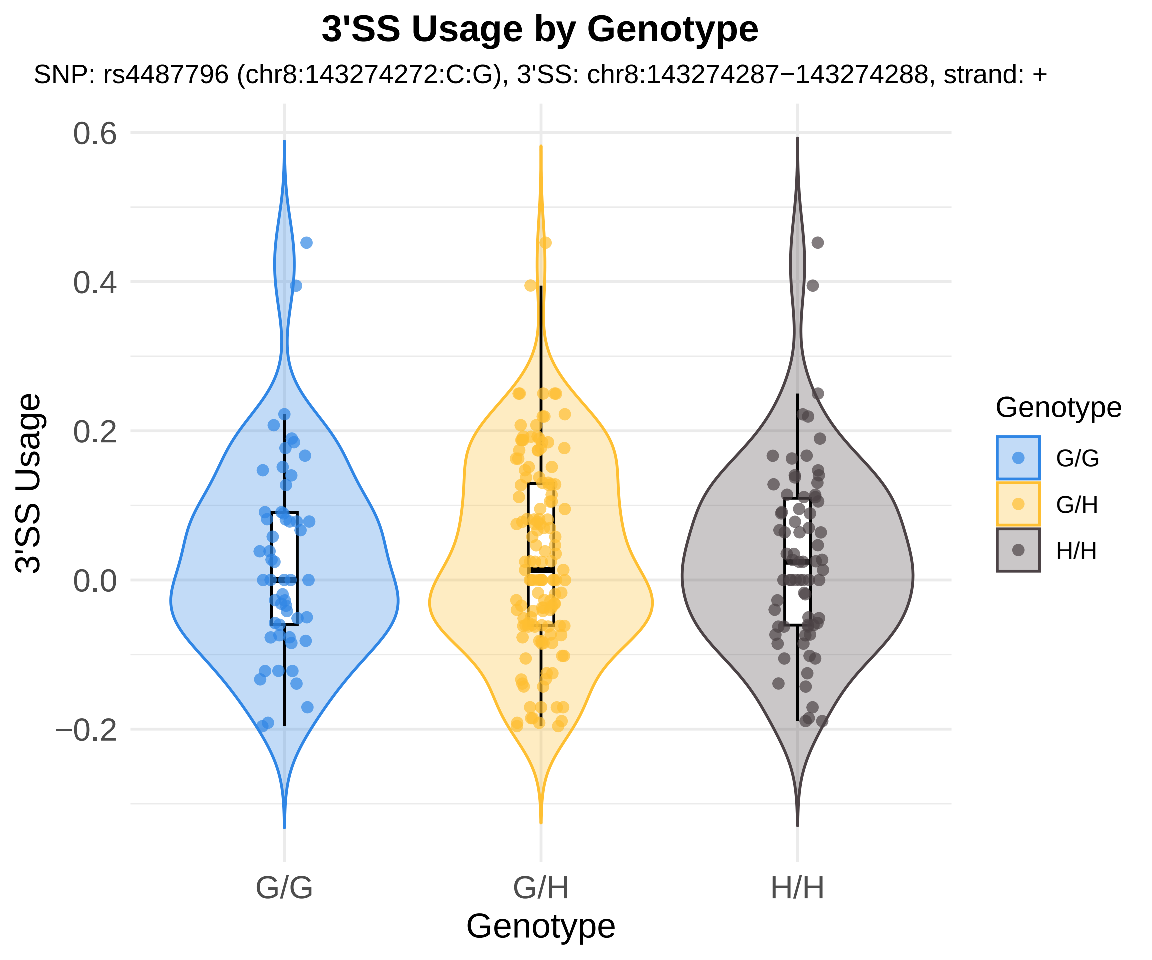
3'SS Usage by Genotype
SNP: rs4487796 (chr8:143274272:C:G), 3'SS: chr8:143274287−143274288, strand: +
3'SS Usage
0.6
0.4
0.2
0.0
−0.2
G/G
G/H
H/H
Genotype
Genotype
G/G
G/H
H/H

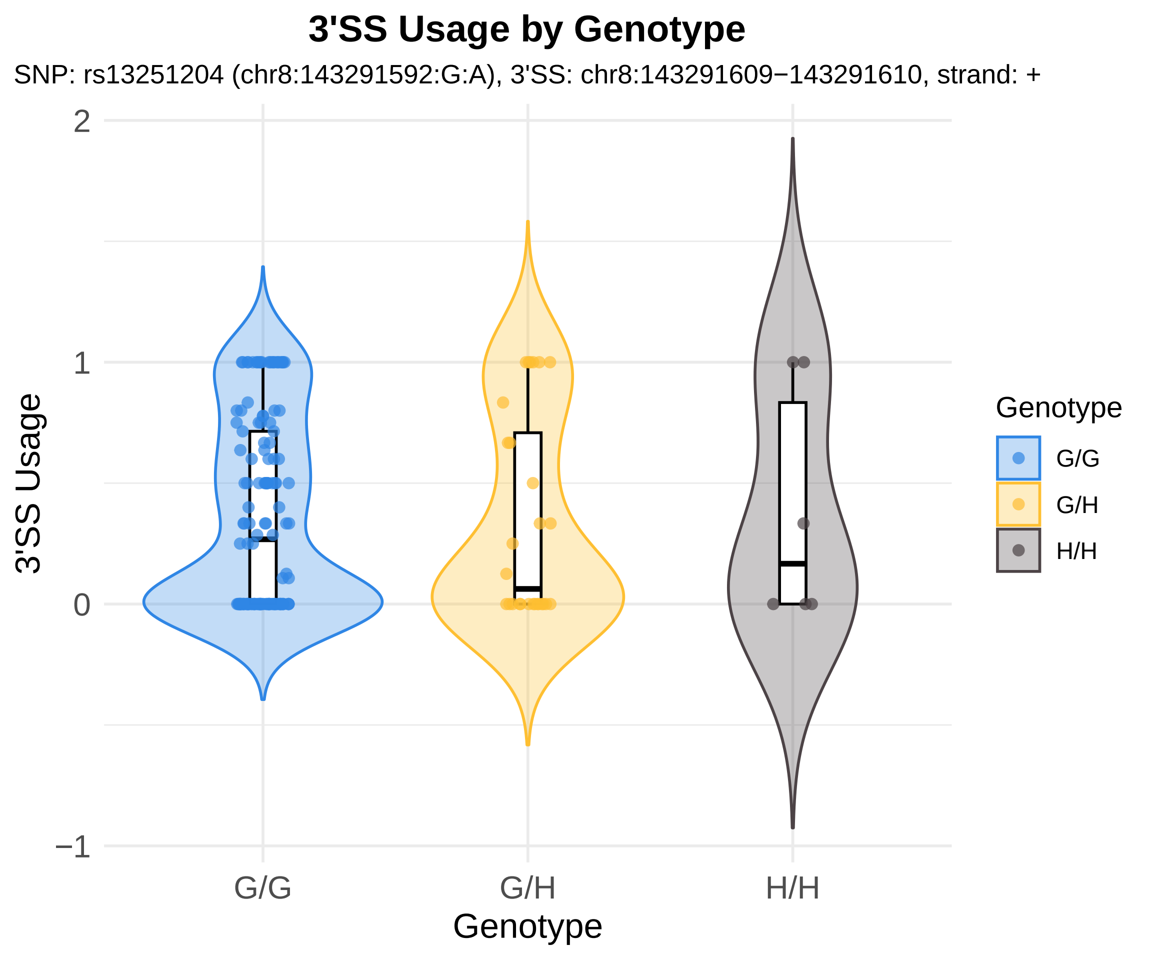
3'SS Usage by Genotype
SNP: rs13251204 (chr8:143291592:G:A), 3'SS: chr8:143291609−143291610, strand: +
3'SS Usage
2
1
0
−1
G/G
G/H
H/H
Genotype
Genotype
G/G
G/H
H/H

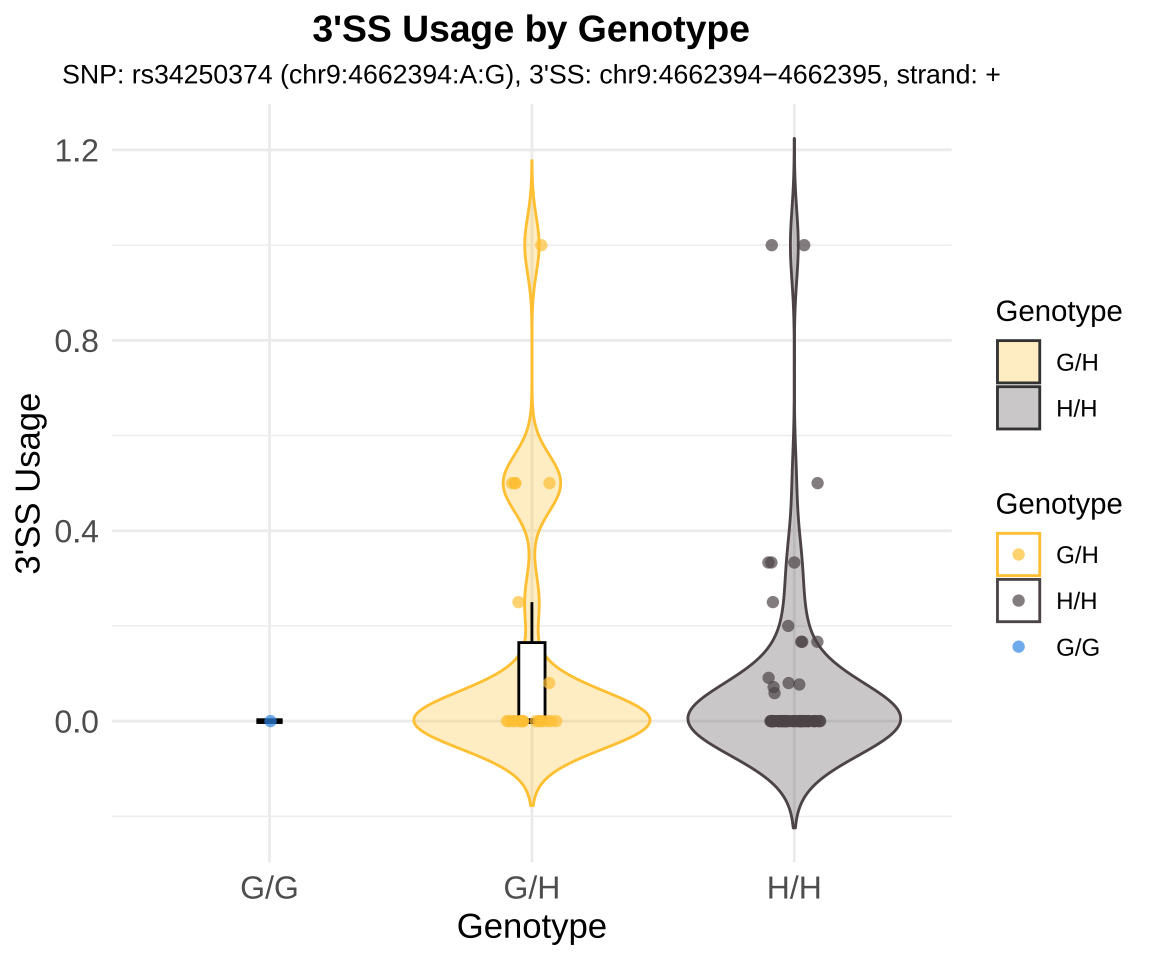
3'SS Usage by Genotype
SNP: rs34250374 (chr9:4662394:A:G), 3'SS: chr9:4662394−4662395, strand: +
3'SS Usage
1.2
0.8
0.4
0.0
G/G
G/H
H/H
Genotype
Genotype
G/H
H/H
Genotype
G/H
H/H
G/G

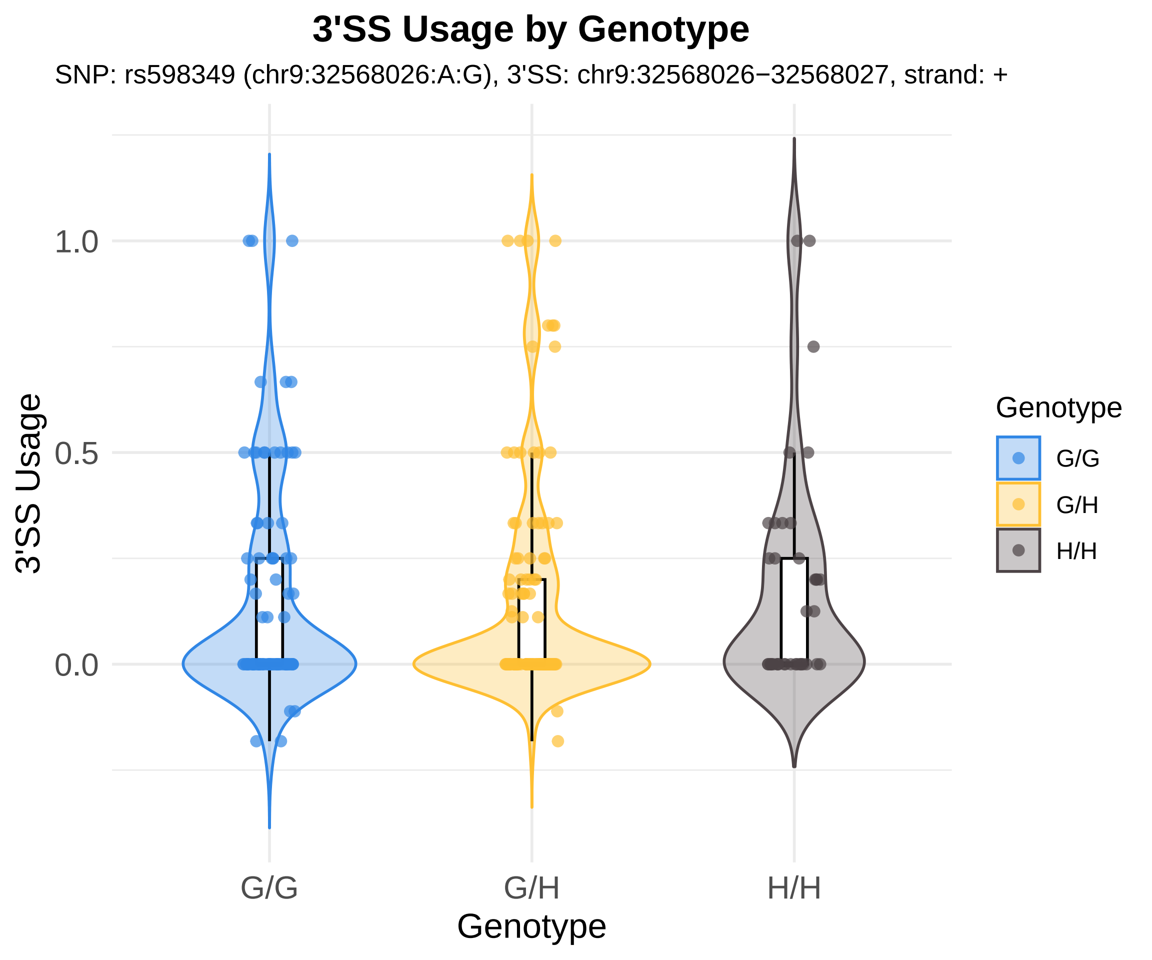
3'SS Usage by Genotype
SNP: rs598349 (chr9:32568026:A:G), 3'SS: chr9:32568026−32568027, strand: +
3'SS Usage
1.0
0.5
0.0
G/G
G/H
H/H
Genotype
Genotype
G/G
G/H
H/H

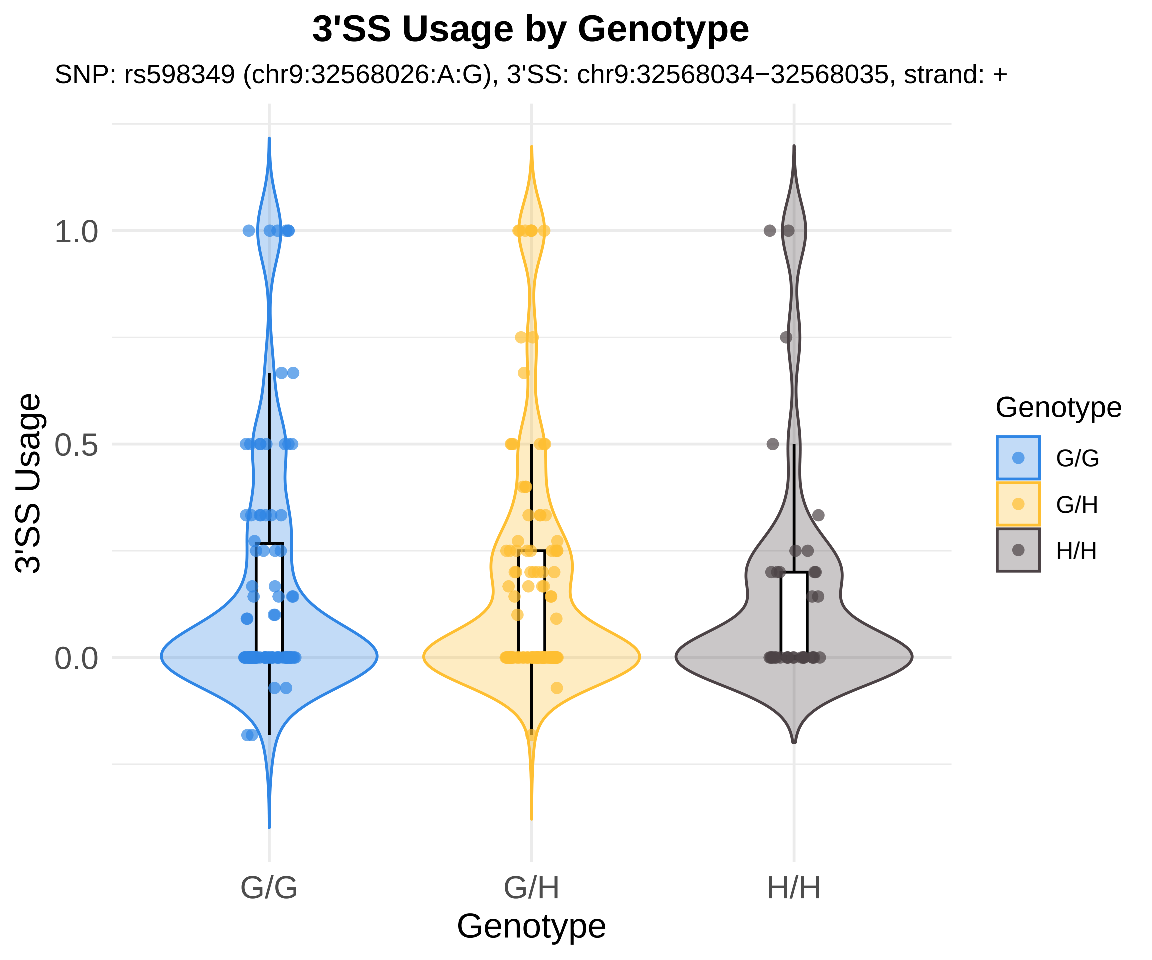
3'SS Usage by Genotype
SNP: rs598349 (chr9:32568026:A:G), 3'SS: chr9:32568034−32568035, strand: +
3'SS Usage
1.0
0.5
0.0
G/G
G/H
H/H
Genotype
Genotype
G/G
G/H
H/H

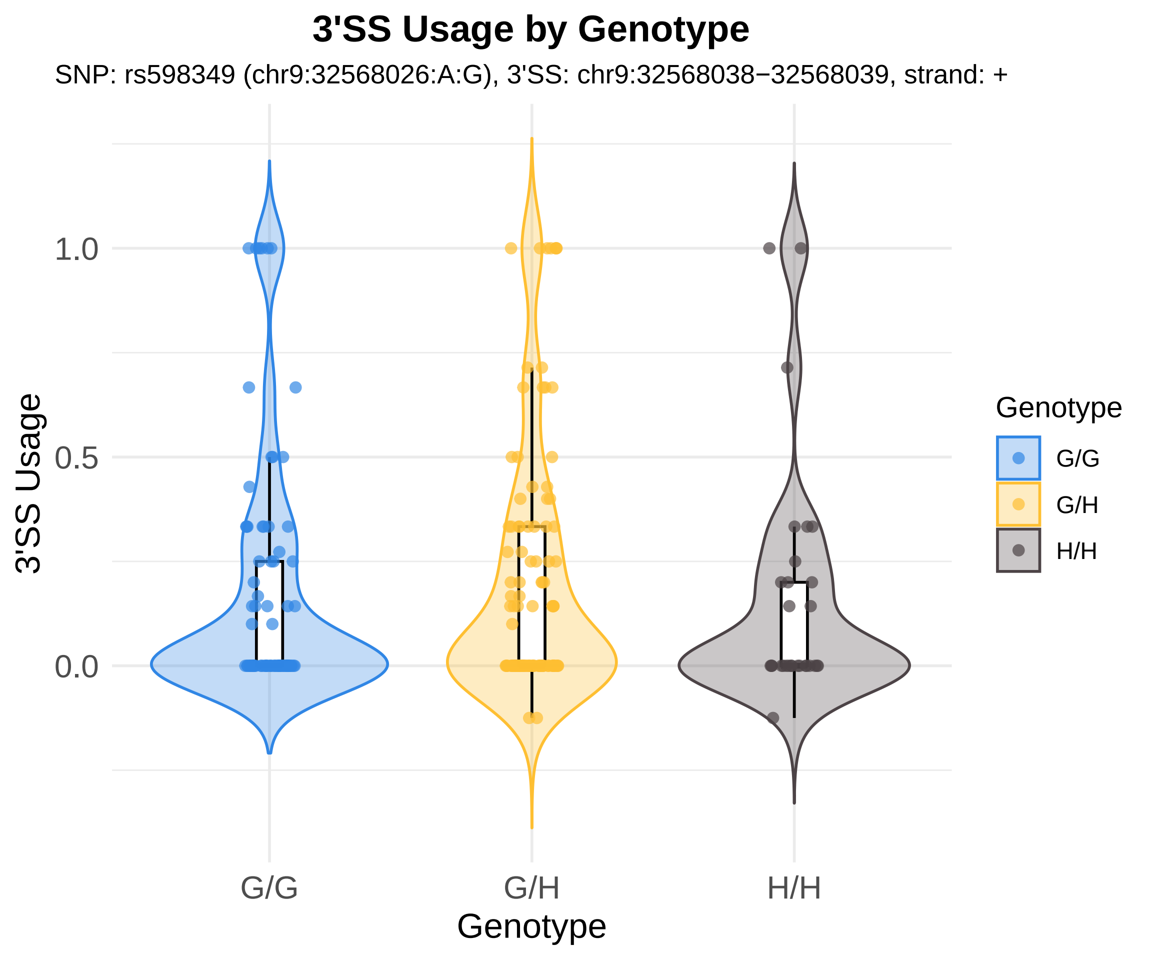

3'SS Usage by Genotype
SNP: rs598349 (chr9:32568026:A:G), 3'SS: chr9:32568038−32568039, strand: +
3'SS Usage
1.0
0.5
0.0
G/G
G/H
H/H
Genotype
Genotype
G/G
G/H
H/H

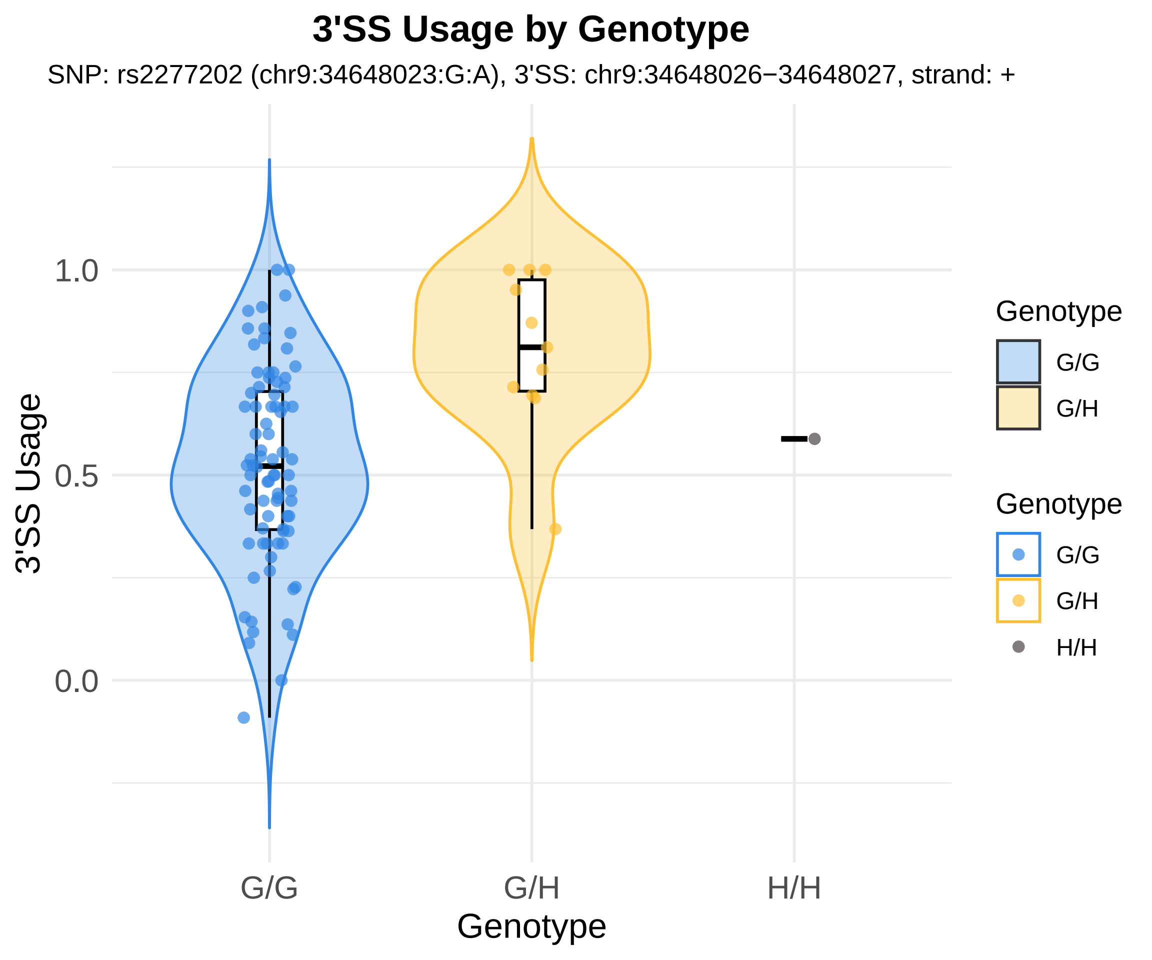
3'SS Usage by Genotype
SNP: rs2277202 (chr9:34648023:G:A), 3'SS: chr9:34648026−34648027, strand: +
3'SS Usage
1.0
0.5
0.0
G/G
G/H
H/H
Genotype
Genotype
G/G
G/H
Genotype
G/G
G/H
H/H

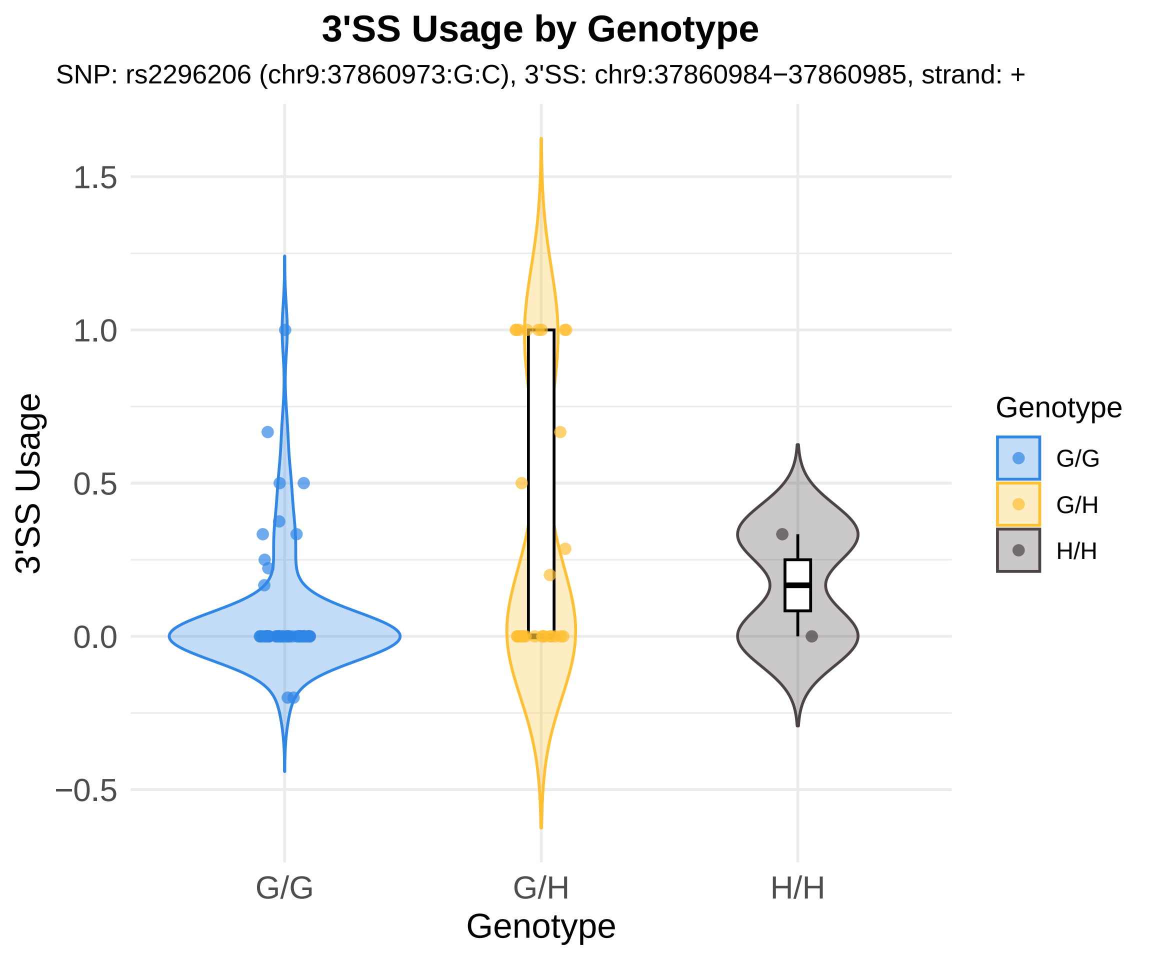
3'SS Usage by Genotype
SNP: rs2296206 (chr9:37860973:G:C), 3'SS: chr9:37860984−37860985, strand: +
3'SS Usage
1.5
1.0
0.5
0.0
−0.5
G/G
G/H
H/H
Genotype
Genotype
G/G
G/H
H/H

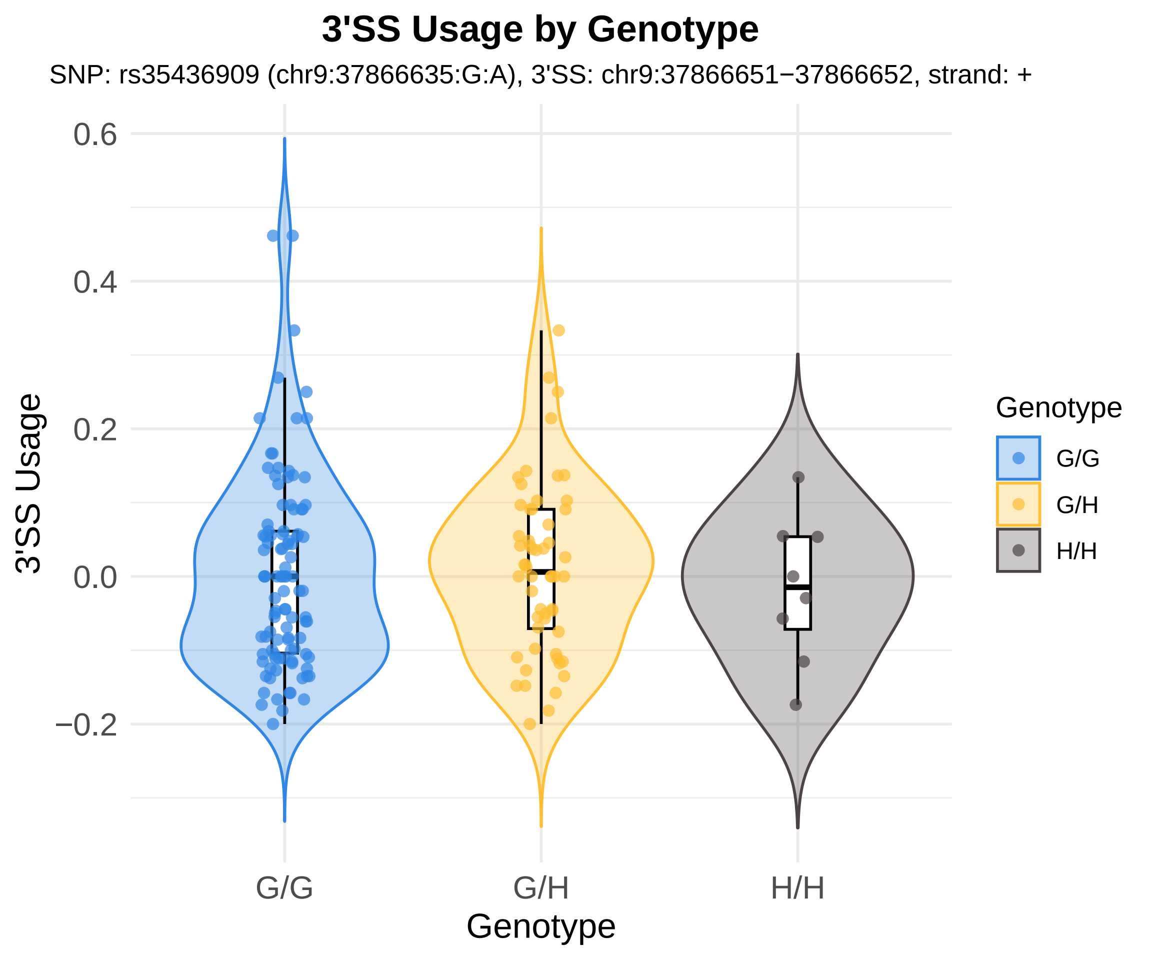
3'SS Usage by Genotype
SNP: rs35436909 (chr9:37866635:G:A), 3'SS: chr9:37866651−37866652, strand: +
0.6
0.4
0.2
0.0
−0.2
3'SS Usage
G/G
G/H
H/H
Genotype
Genotype
G/G
G/H
H/H

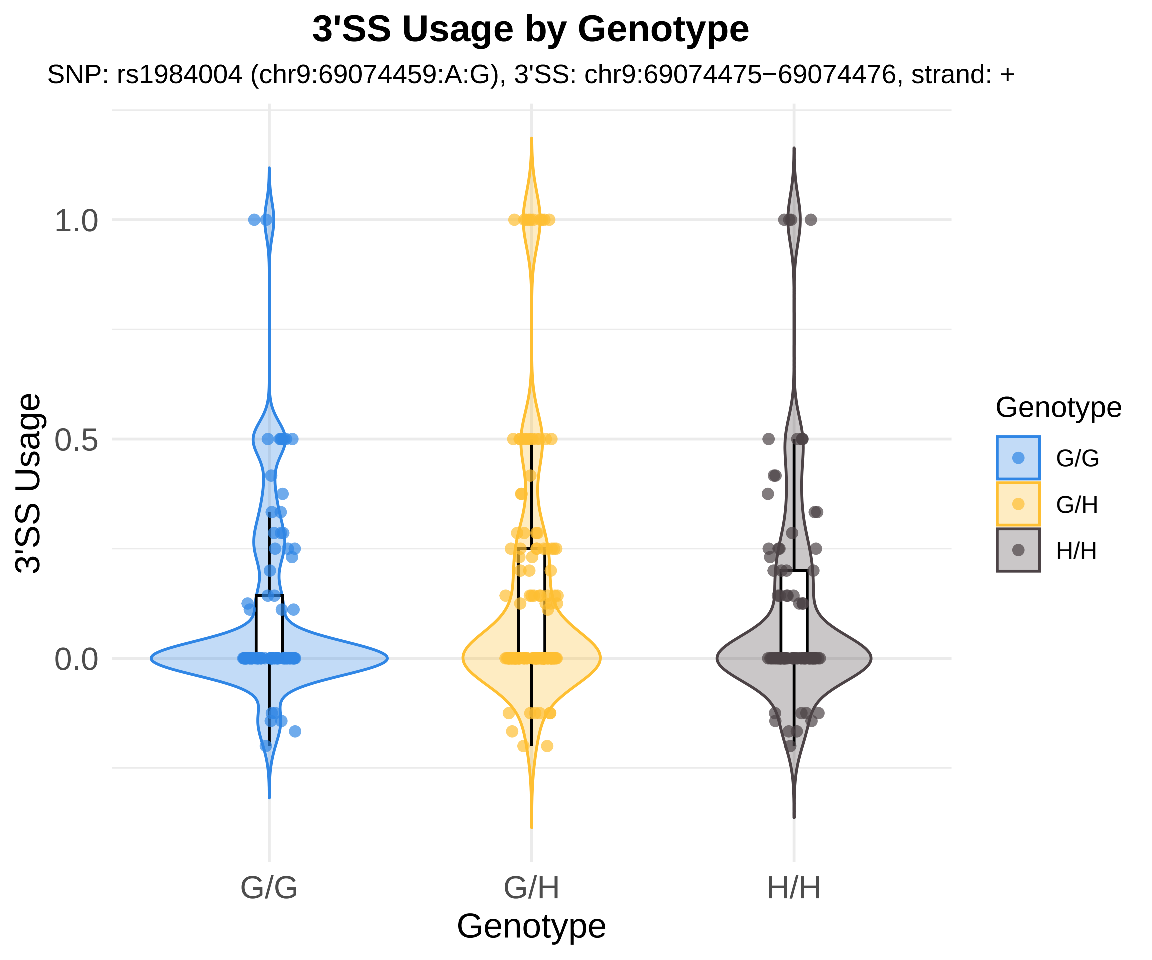

3'SS Usage by Genotype
SNP: rs1984004 (chr9:69074459:A:G), 3'SS: chr9:69074475−69074476, strand: +
1.0
0.5
0.0
3'SS Usage
G/G
G/H
H/H
Genotype
Genotype
G/G
G/H
H/H

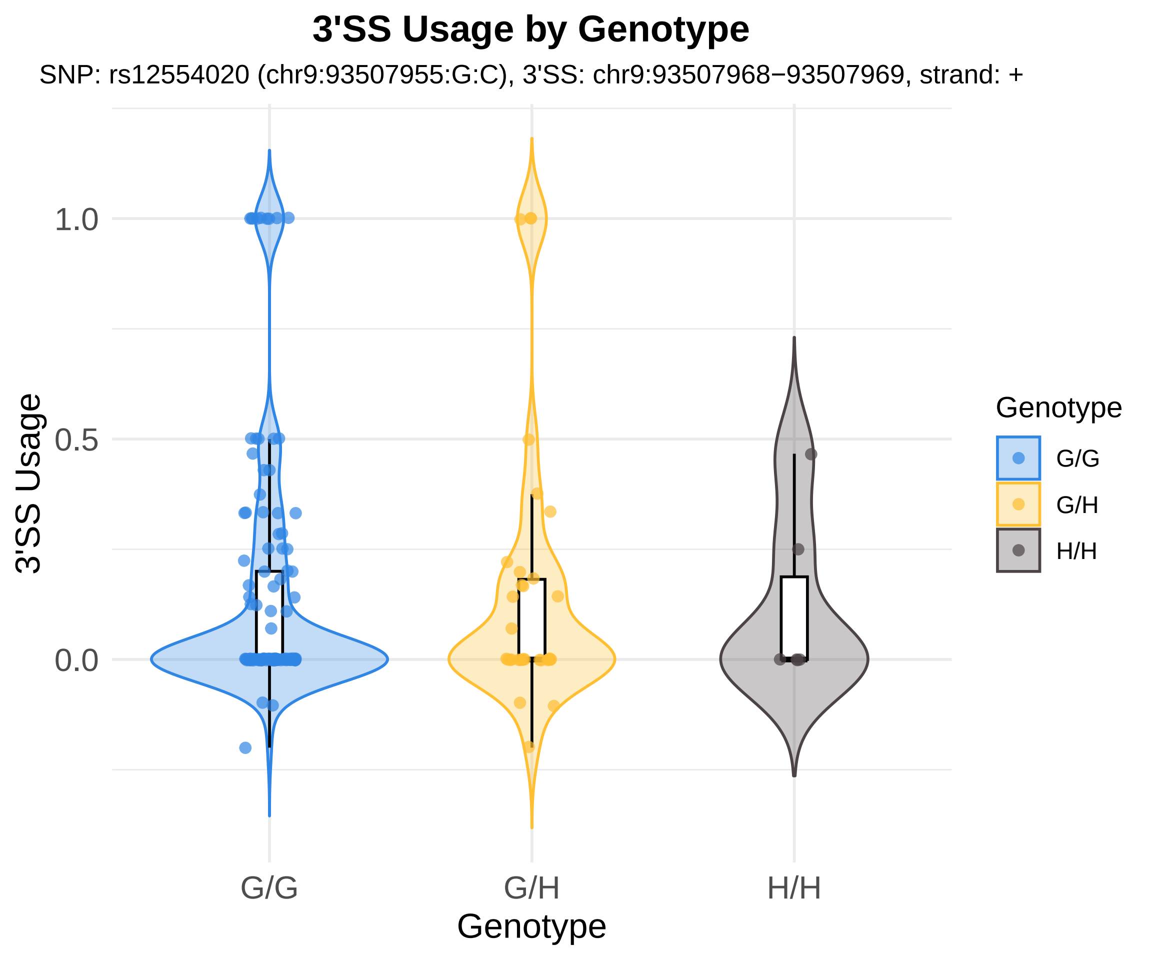

3'SS Usage by Genotype
SNP: rs12554020 (chr9:93507955:G:C), 3'SS: chr9:93507968−93507969, strand: +
3'SS Usage
1.0
0.5
0.0
G/G
G/H
H/H
Genotype
Genotype
G/G
G/H
H/H

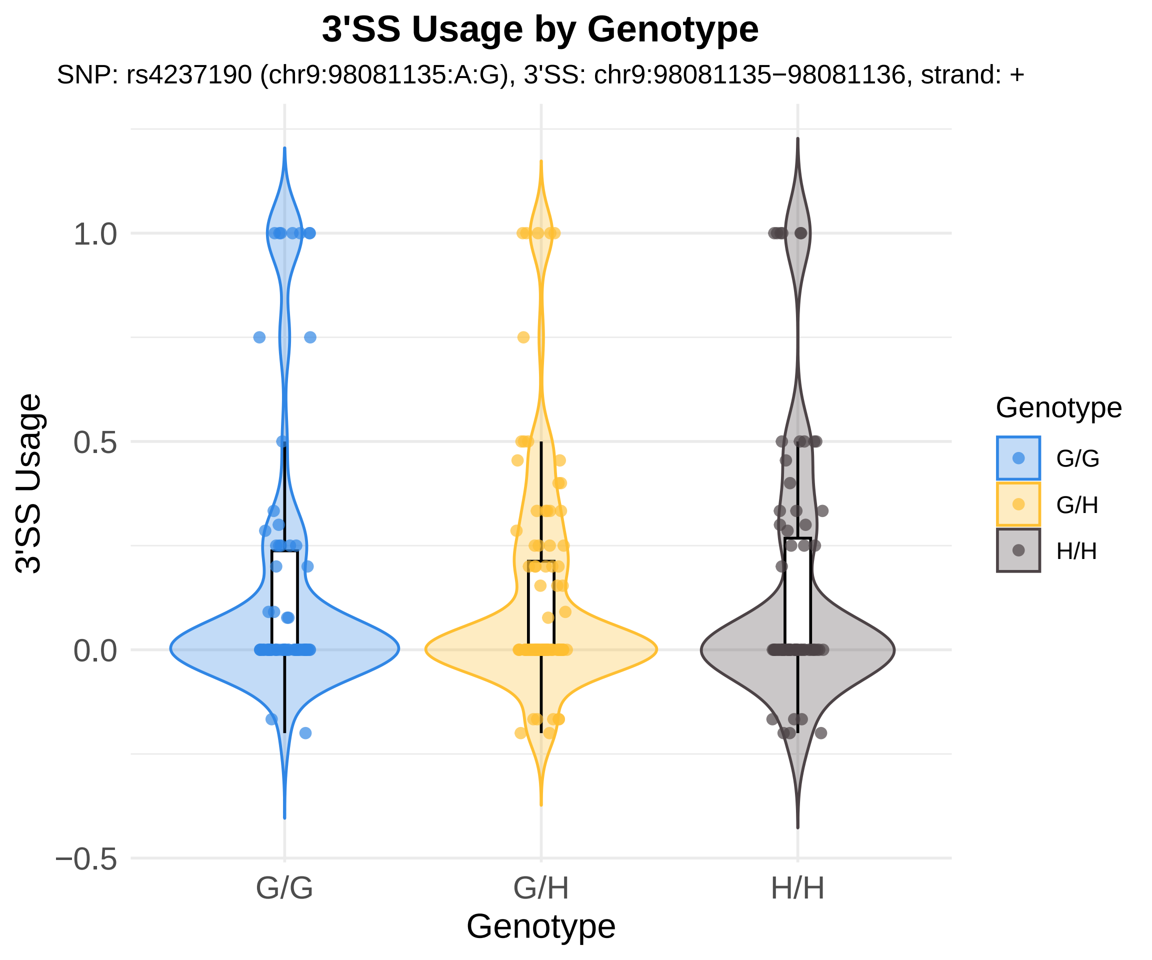
3'SS Usage by Genotype
SNP: rs4237190 (chr9:98081135:A:G), 3'SS: chr9:98081135−98081136, strand: +
3'SS Usage
1.0
0.5
0.0
−0.5
G/G
G/H
H/H
Genotype
Genotype
G/G
G/H
H/H

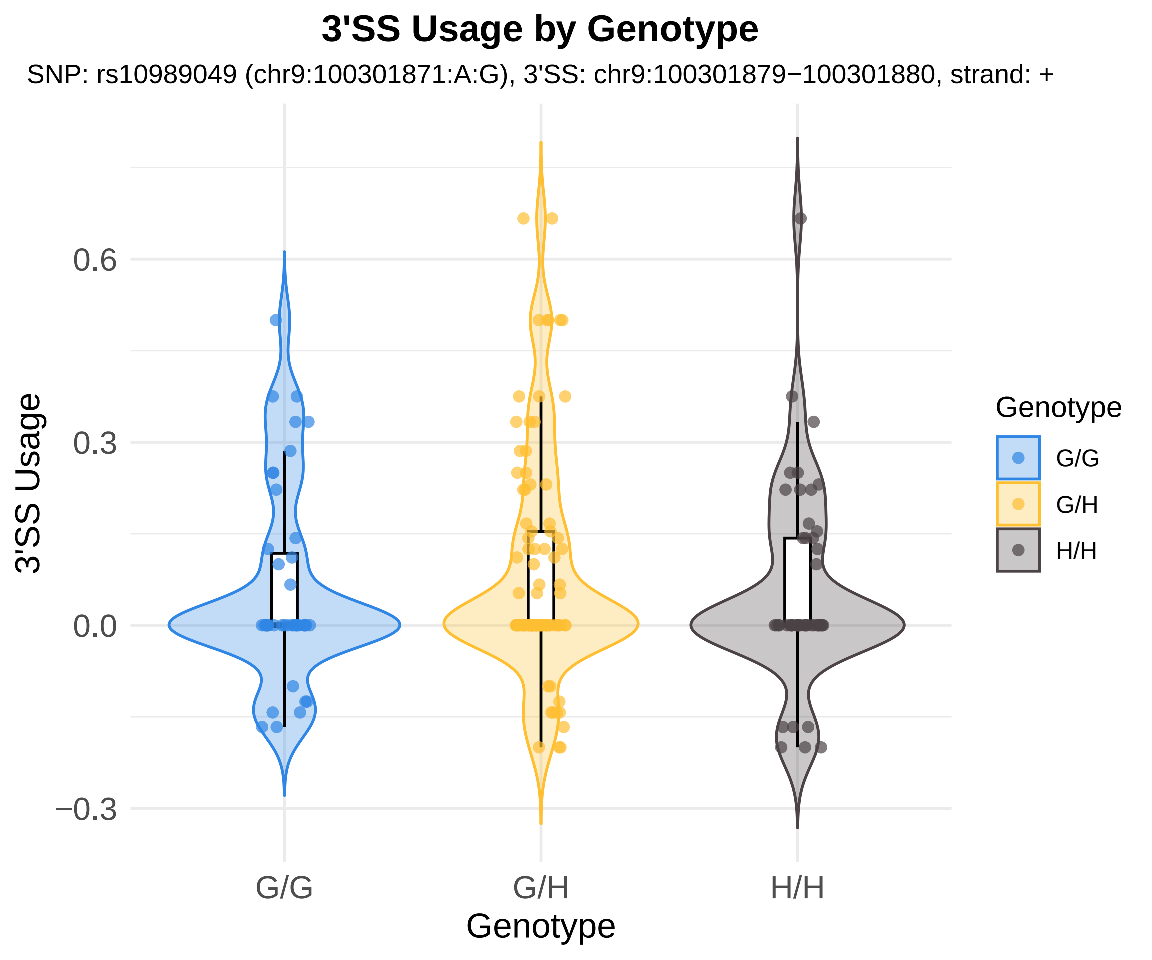
3'SS Usage by Genotype
SNP: rs10989049 (chr9:100301871:A:G), 3'SS: chr9:100301879−100301880, strand: +
3'SS Usage
0.6
0.3
0.0
−0.3
G/G
G/H
H/H
Genotype
Genotype
G/G
G/H
H/H

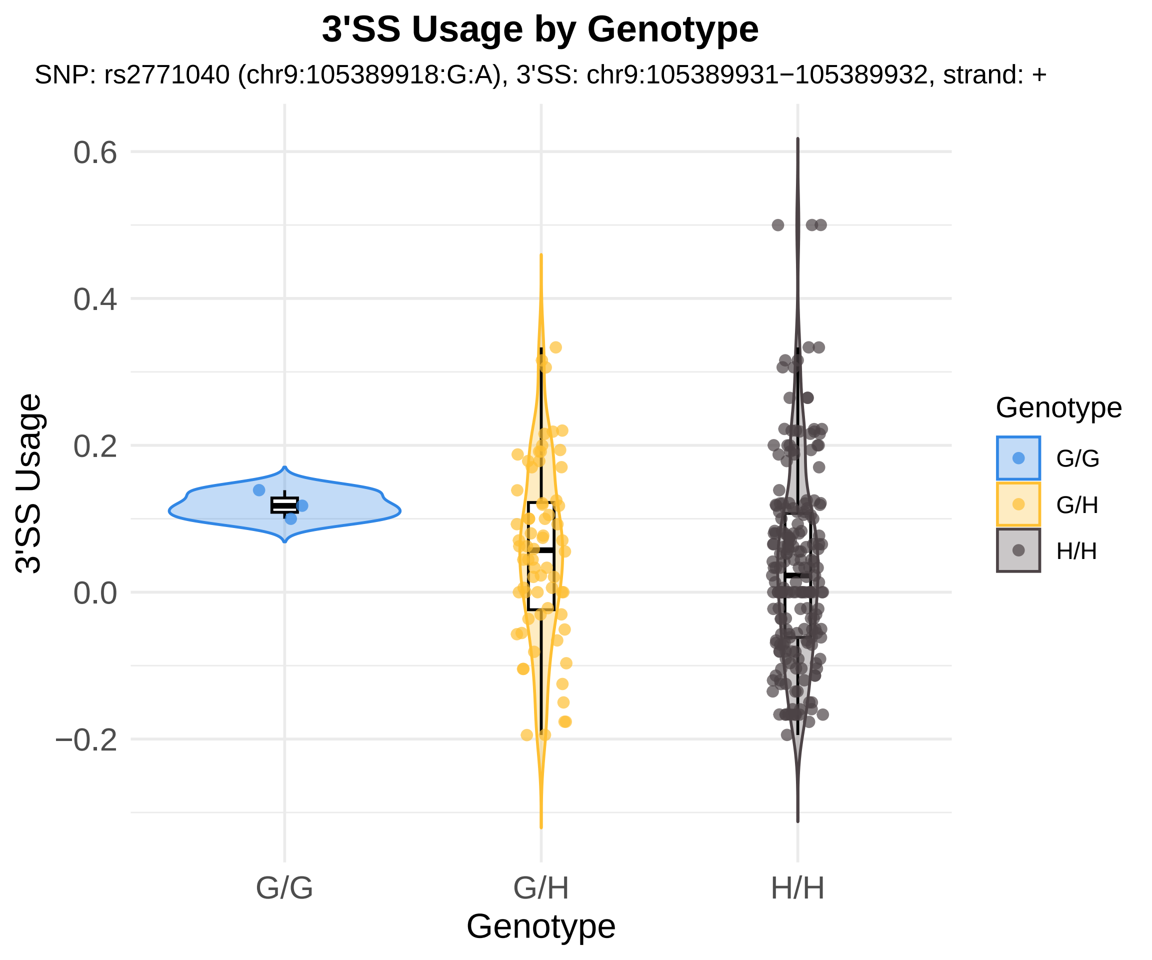
3'SS Usage by Genotype
SNP: rs2771040 (chr9:105389918:G:A), 3'SS: chr9:105389931−105389932, strand: +
3'SS Usage
0.6
0.4
0.2
0.0
−0.2
G/G
G/H
H/H
Genotype
Genotype
G/G
G/H
H/H

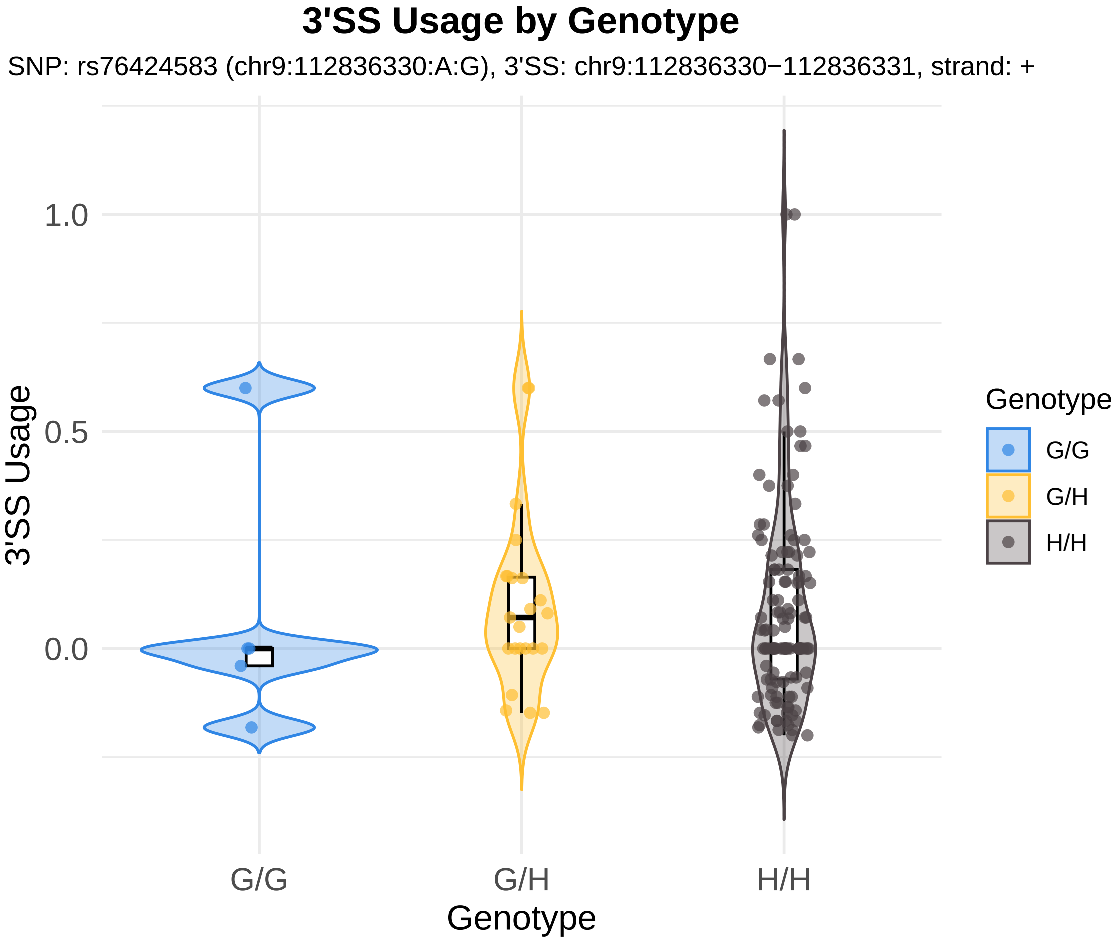
3'SS Usage by Genotype
SNP: rs76424583 (chr9:112836330:A:G), 3'SS: chr9:112836330−112836331, strand: +
1.0
0.5
0.0
3'SS Usage
G/G
G/H
H/H
Genotype
Genotype
G/G
G/H
H/H

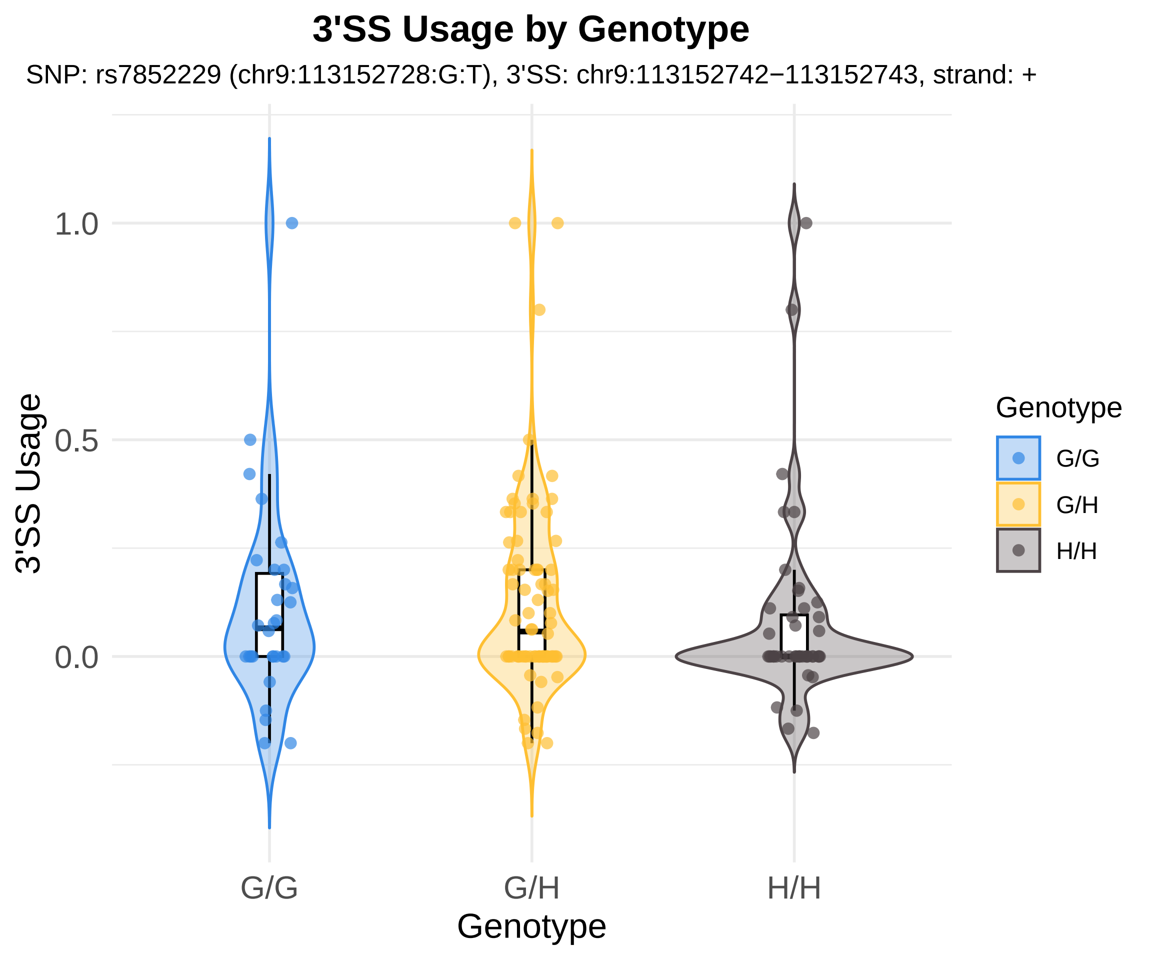
3'SS Usage by Genotype
SNP: rs7852229 (chr9:113152728:G:T), 3'SS: chr9:113152742−113152743, strand: +
3'SS Usage
1.0
0.5
0.0
G/G
G/H
H/H
Genotype
Genotype
G/G
G/H
H/H

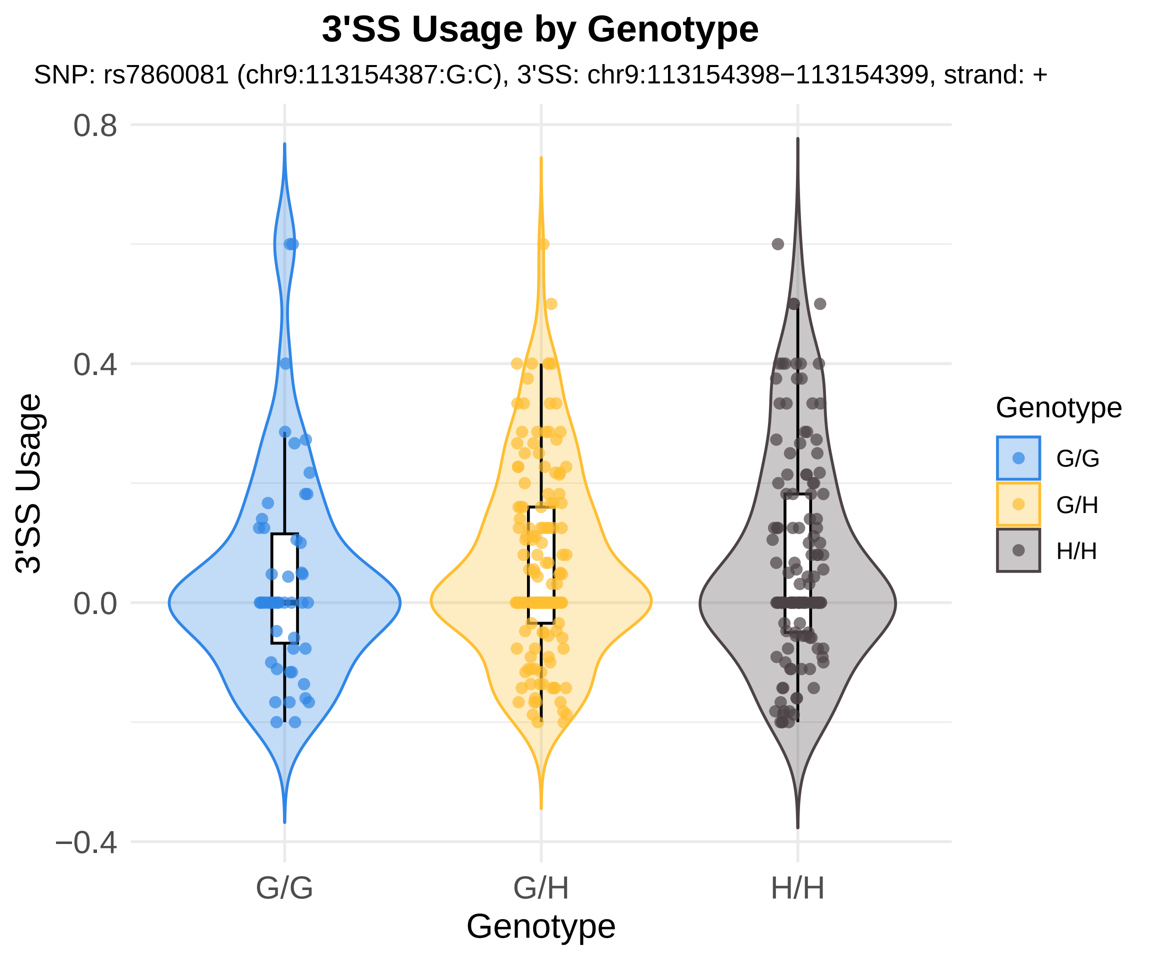
3'SS Usage by Genotype
SNP: rs7860081 (chr9:113154387:G:C), 3'SS: chr9:113154398−113154399, strand: +
0.8
0.4
0.0
−0.4
3'SS Usage
G/G
G/H
H/H
Genotype
Genotype
G/G
G/H
H/H

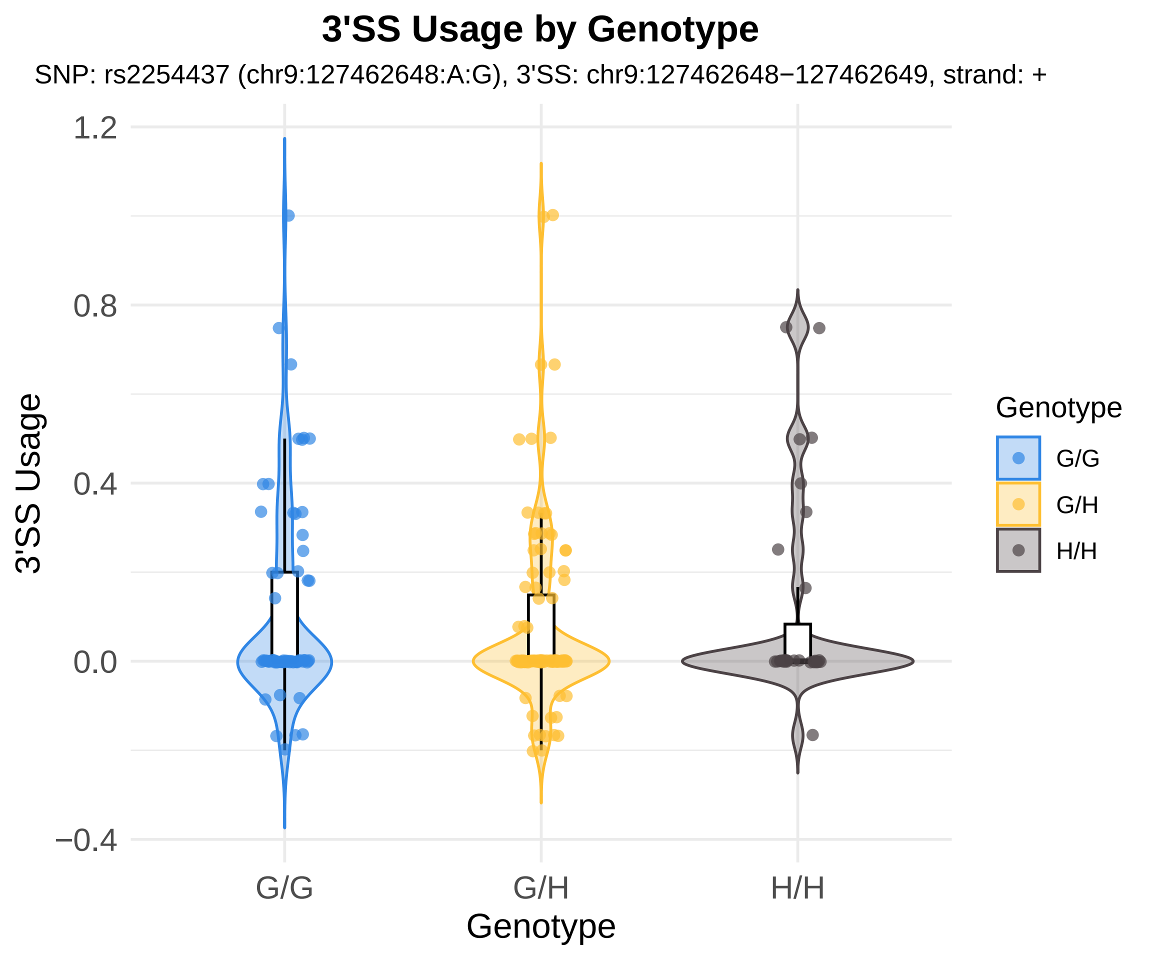

3'SS Usage by Genotype
SNP: rs2254437 (chr9:127462648:A:G), 3'SS: chr9:127462648−127462649, strand: +
3'SS Usage
1.2
0.8
0.4
0.0
−0.4
G/G
G/H
H/H
Genotype
Genotype
G/G
G/H
H/H

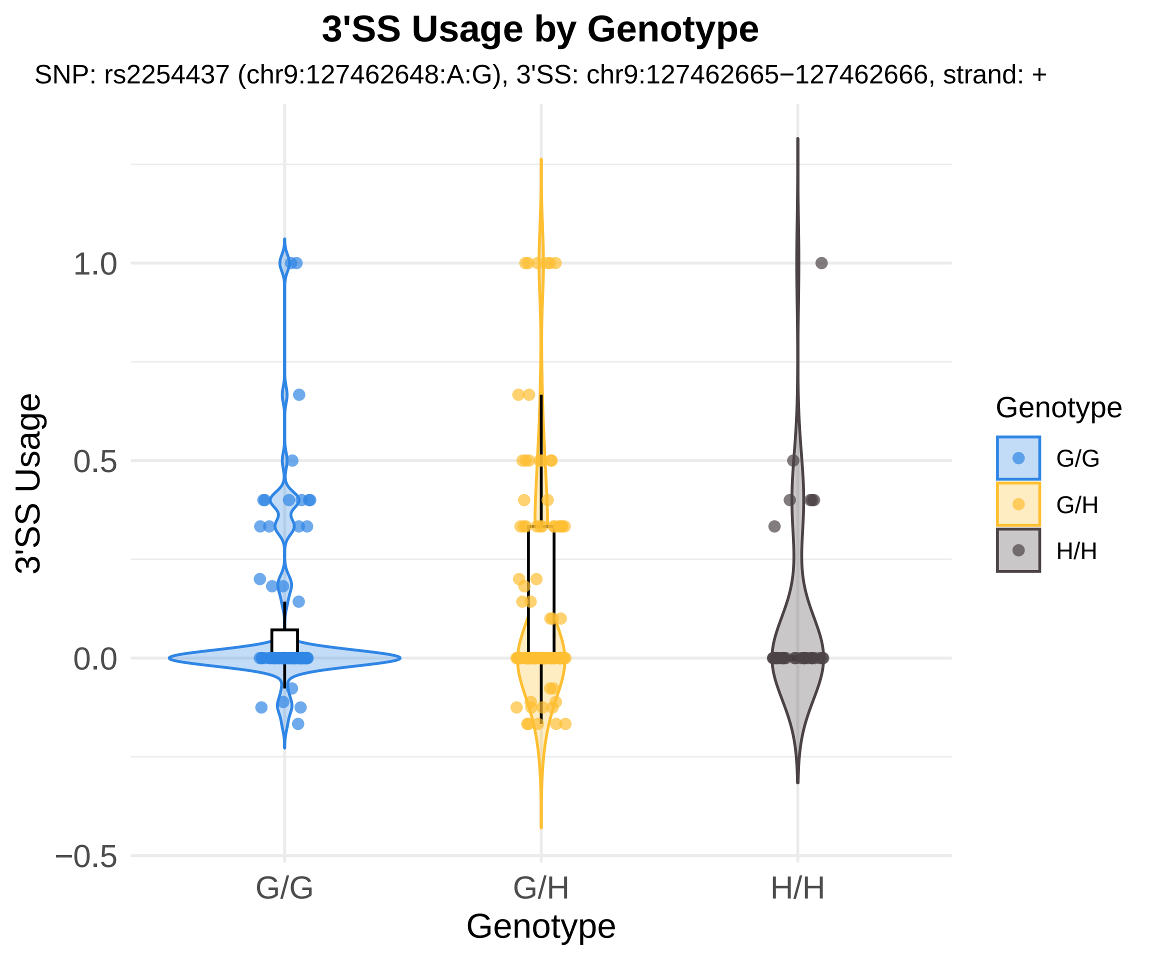
3'SS Usage by Genotype
SNP: rs2254437 (chr9:127462648:A:G), 3'SS: chr9:127462665−127462666, strand: +
3'SS Usage
1.0
0.5
0.0
−0.5
G/G
G/H
H/H
Genotype
Genotype
G/G
G/H
H/H

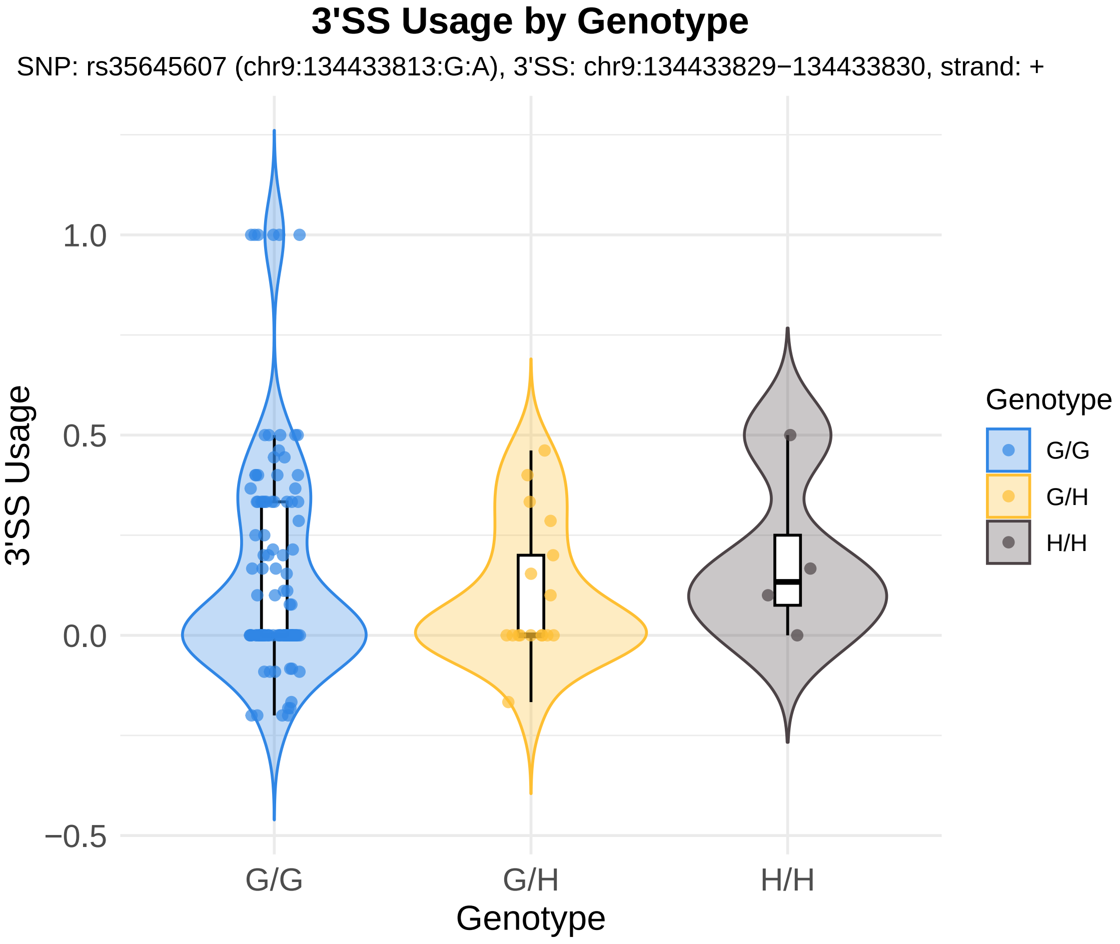
3'SS Usage by Genotype
SNP: rs35645607 (chr9:134433813:G:A), 3'SS: chr9:134433829−134433830, strand: +
3'SS Usage
1.0
0.5
0.0
−0.5
G/G
G/H
H/H
Genotype
Genotype
G/G
G/H
H/H

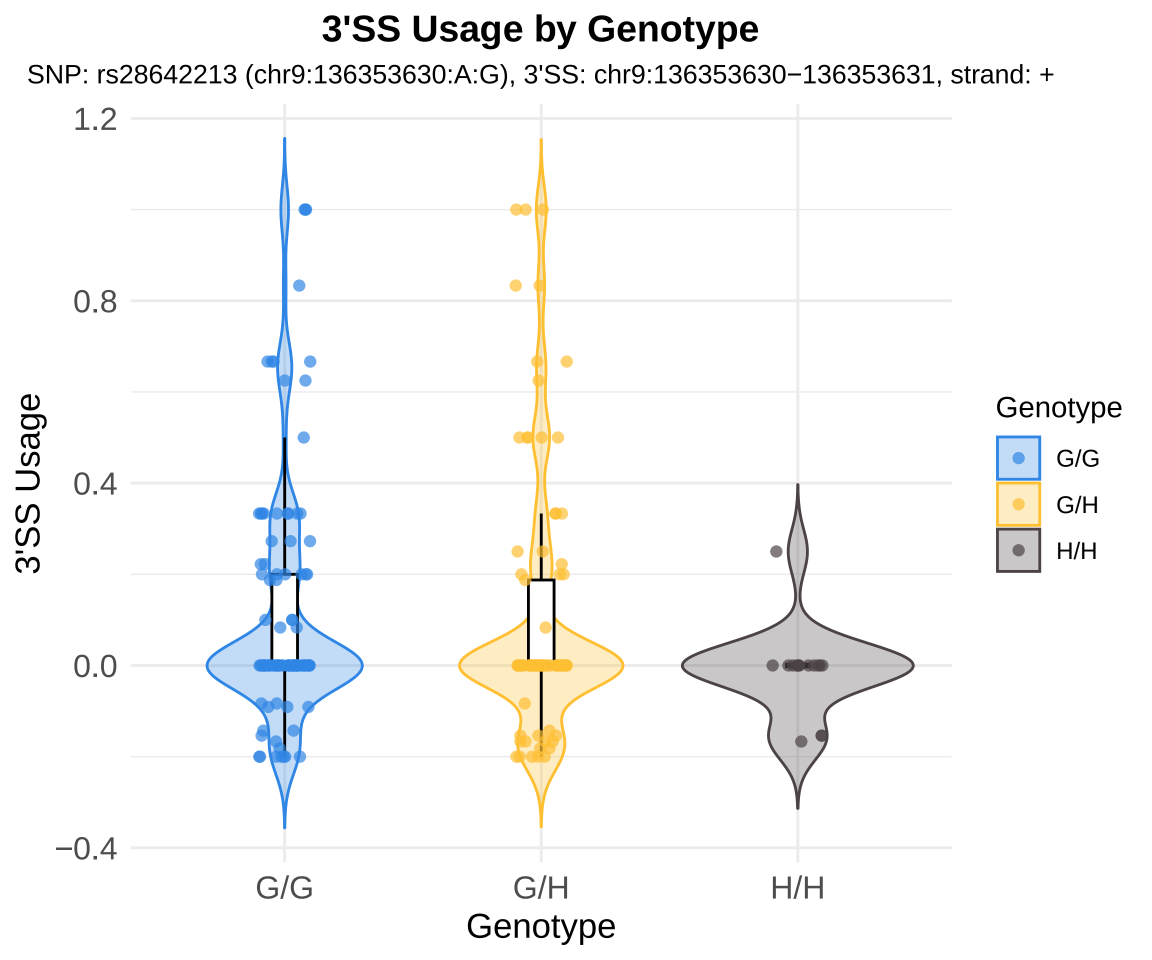
3'SS Usage by Genotype
SNP: rs28642213 (chr9:136353630:A:G), 3'SS: chr9:136353630−136353631, strand: +
3'SS Usage
1.2
0.8
0.4
0.0
−0.4
G/G
G/H
H/H
Genotype
Genotype
G/G
G/H
H/H

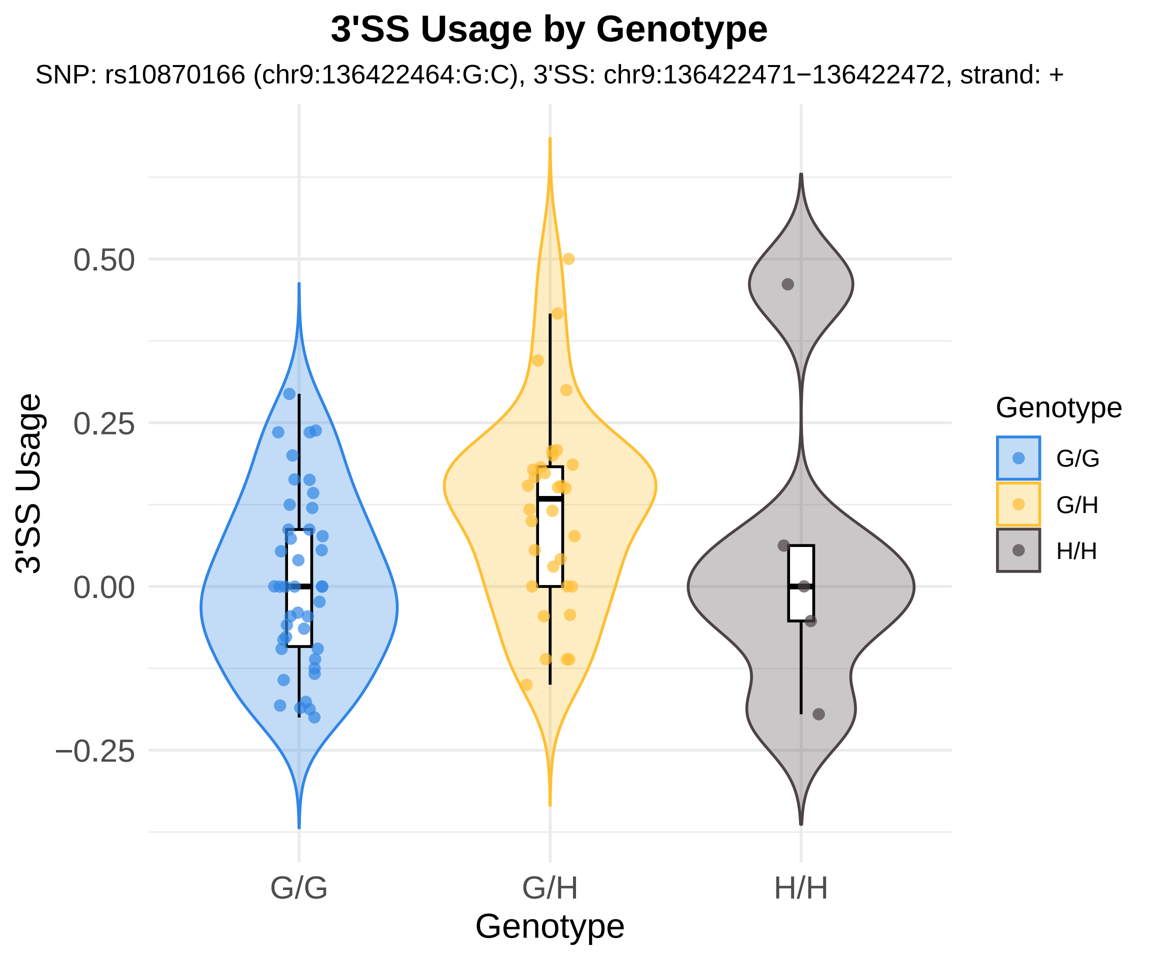

3'SS Usage by Genotype
SNP: rs10870166 (chr9:136422464:G:C), 3'SS: chr9:136422471−136422472, strand: +
3'SS Usage
0.50
0.25
0.00
−0.25
G/G
G/H
H/H
Genotype
Genotype
G/G
G/H
H/H

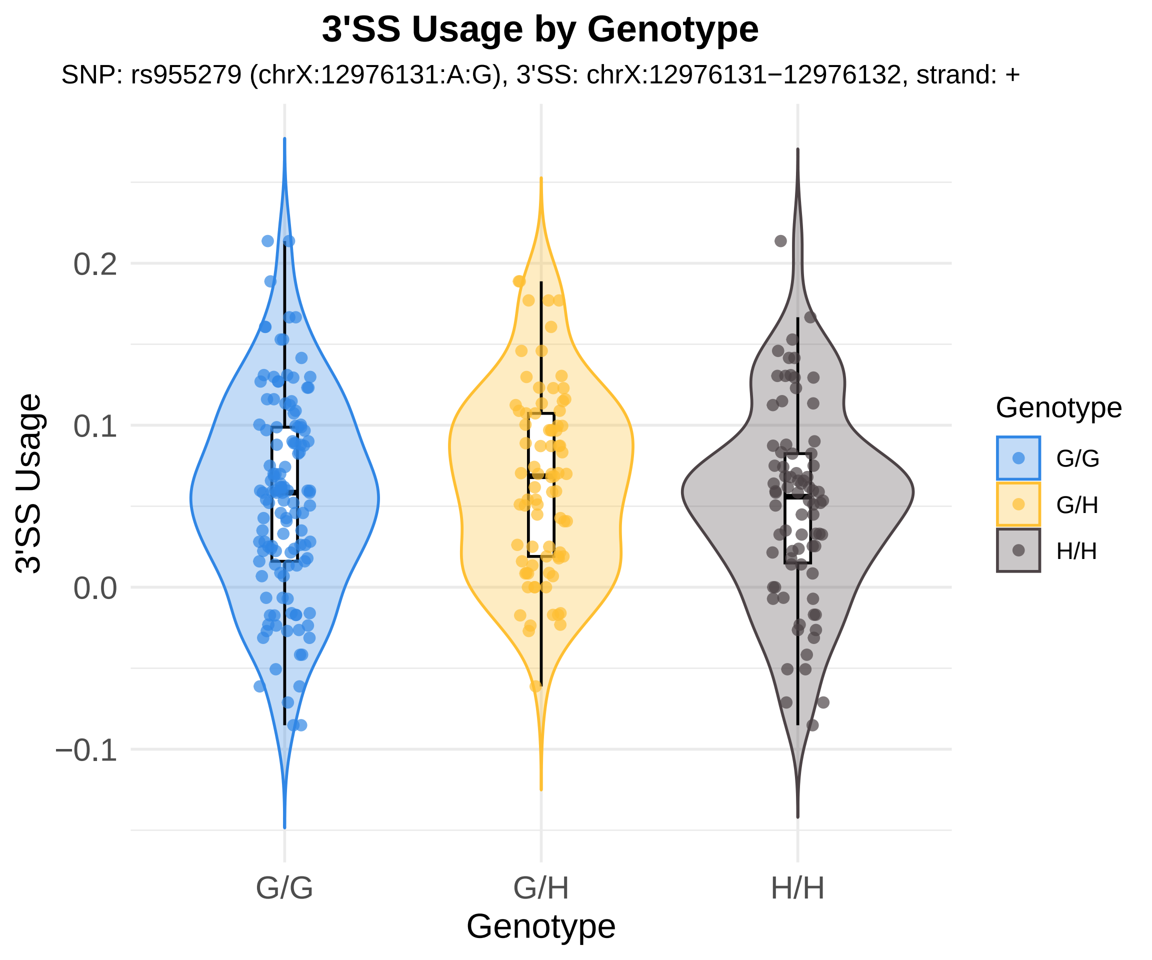

3'SS Usage by Genotype
SNP: rs955279 (chrX:12976131:A:G), 3'SS: chrX:12976131−12976132, strand: +
3'SS Usage
0.2
0.1
0.0
−0.1
G/G
G/H
H/H
Genotype
Genotype
G/G
G/H
H/H

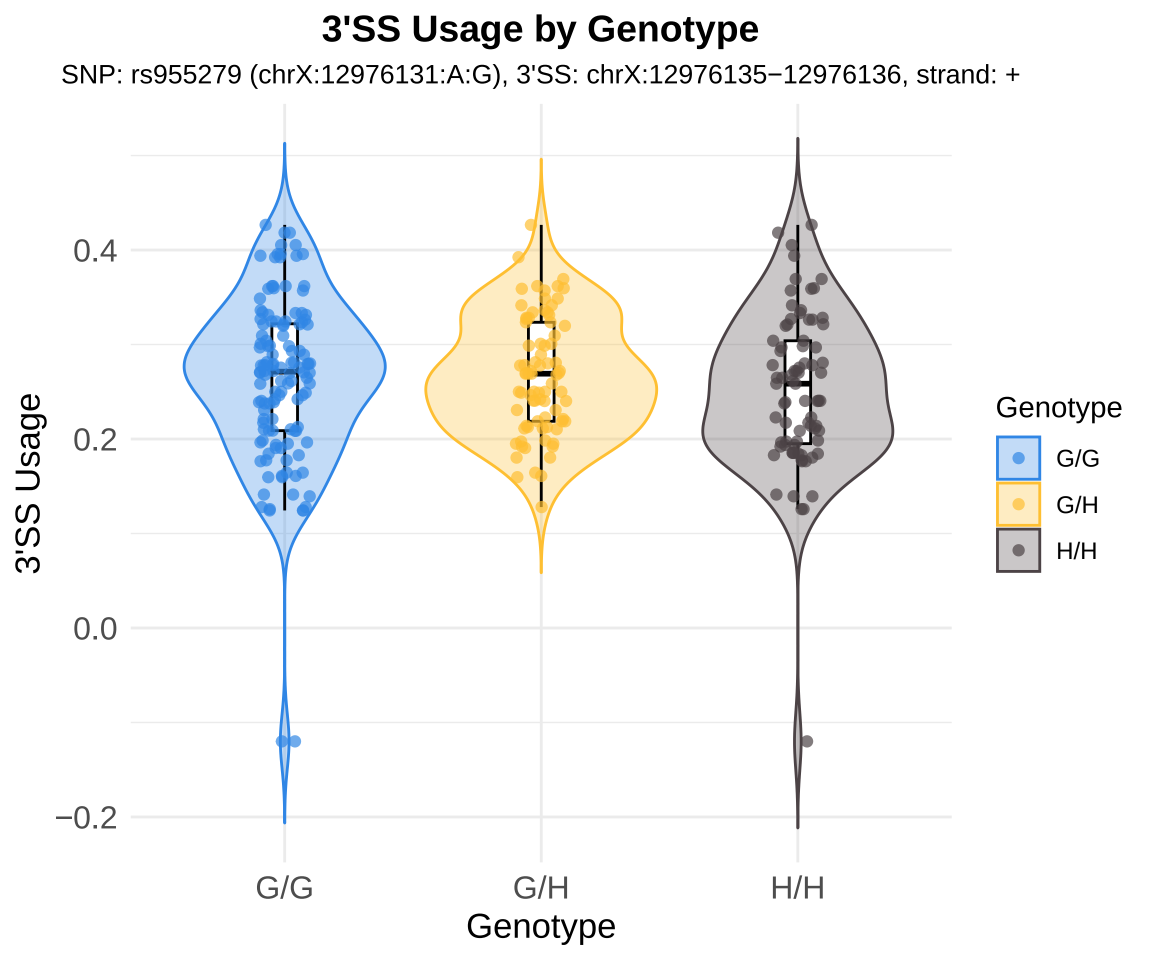

3'SS Usage by Genotype
SNP: rs955279 (chrX:12976131:A:G), 3'SS: chrX:12976135−12976136, strand: +
0.4
0.2
0.0
−0.2
3'SS Usage
G/G
G/H
H/H
Genotype
Genotype
G/G
G/H
H/H

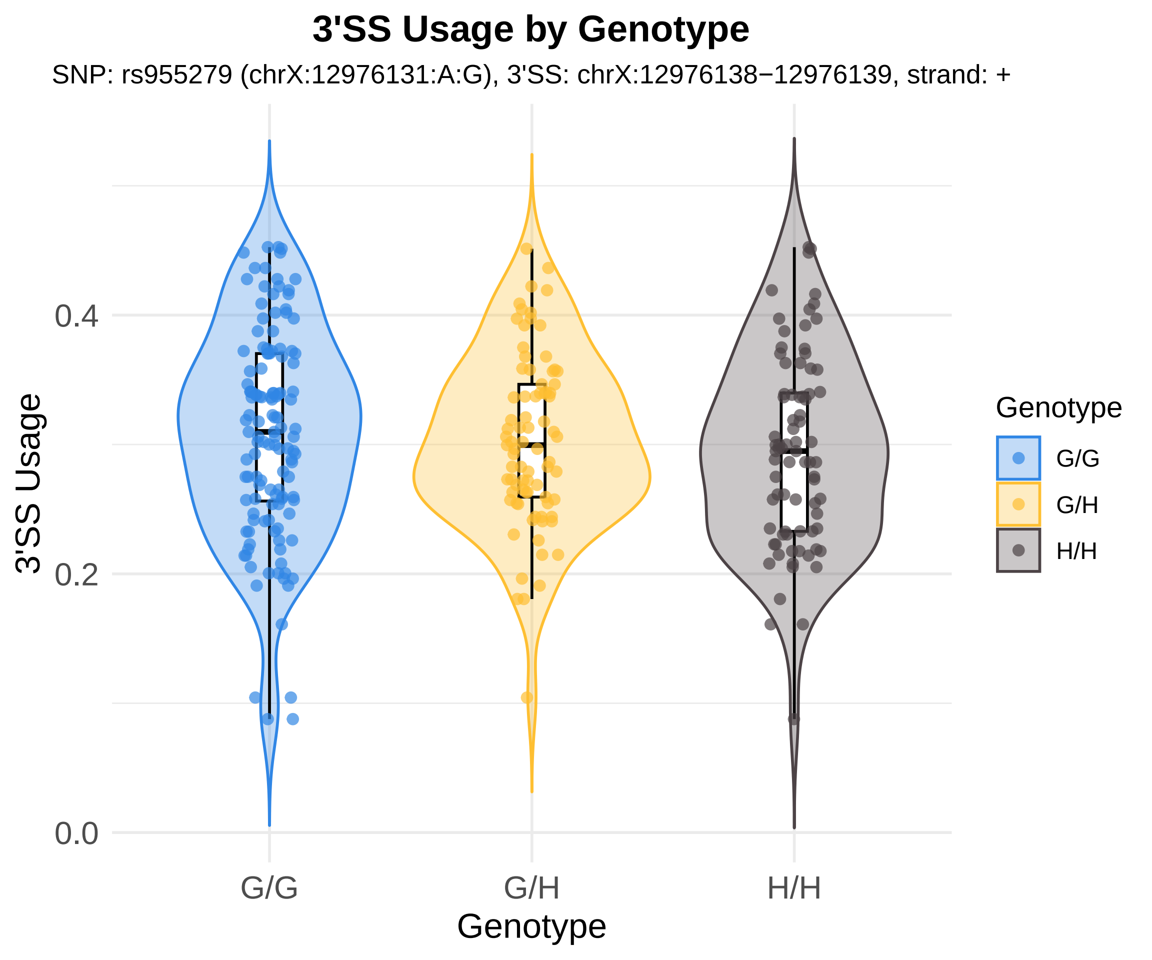
3'SS Usage by Genotype
SNP: rs955279 (chrX:12976131:A:G), 3'SS: chrX:12976138−12976139, strand: +
3'SS Usage
0.0
0.2
0.4
G/G
G/H
H/H
Genotype
Genotype
G/G
G/H
H/H

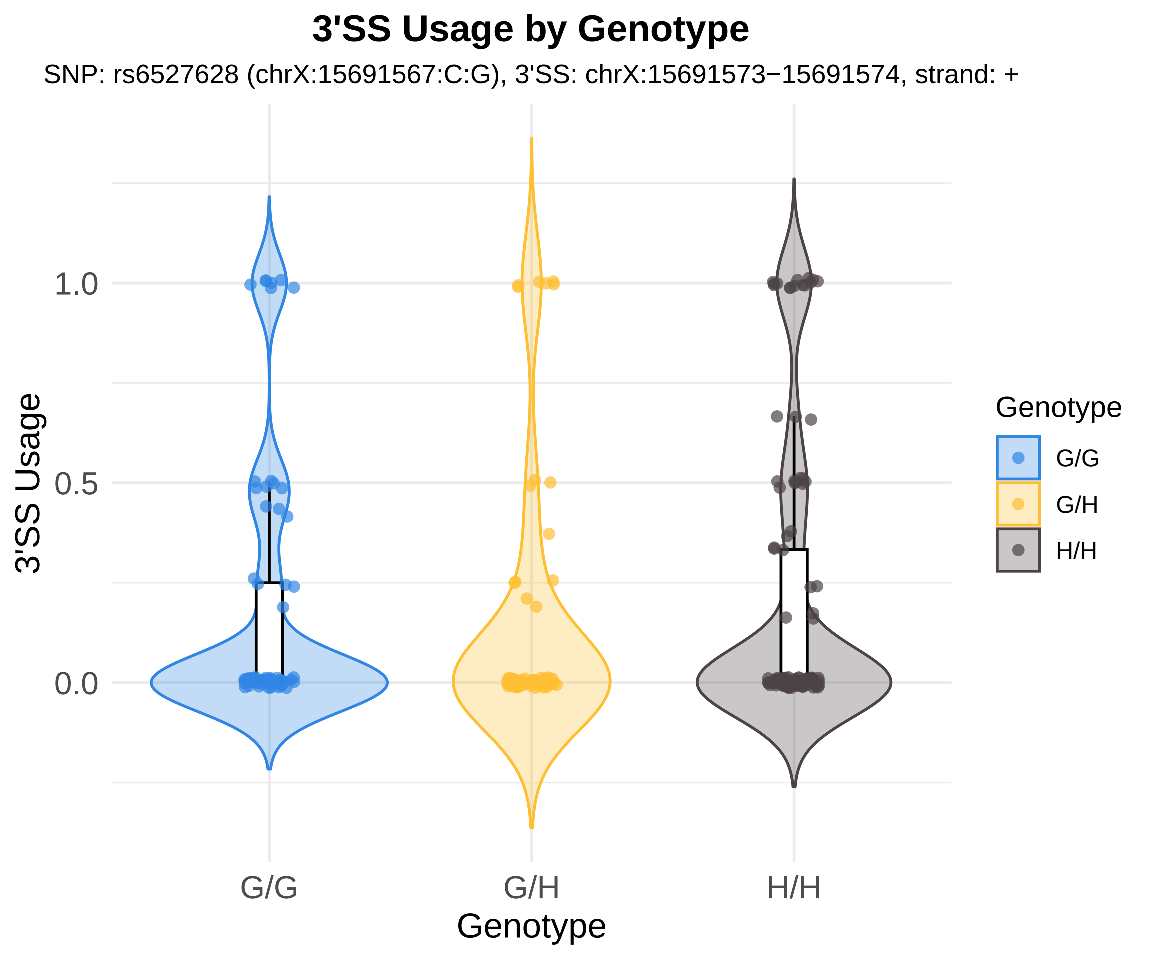

3'SS Usage by Genotype
SNP: rs6527628 (chrX:15691567:C:G), 3'SS: chrX:15691573−15691574, strand: +
3'SS Usage
1.0
0.5
0.0
G/G
G/H
H/H
Genotype
Genotype
G/G
G/H
H/H

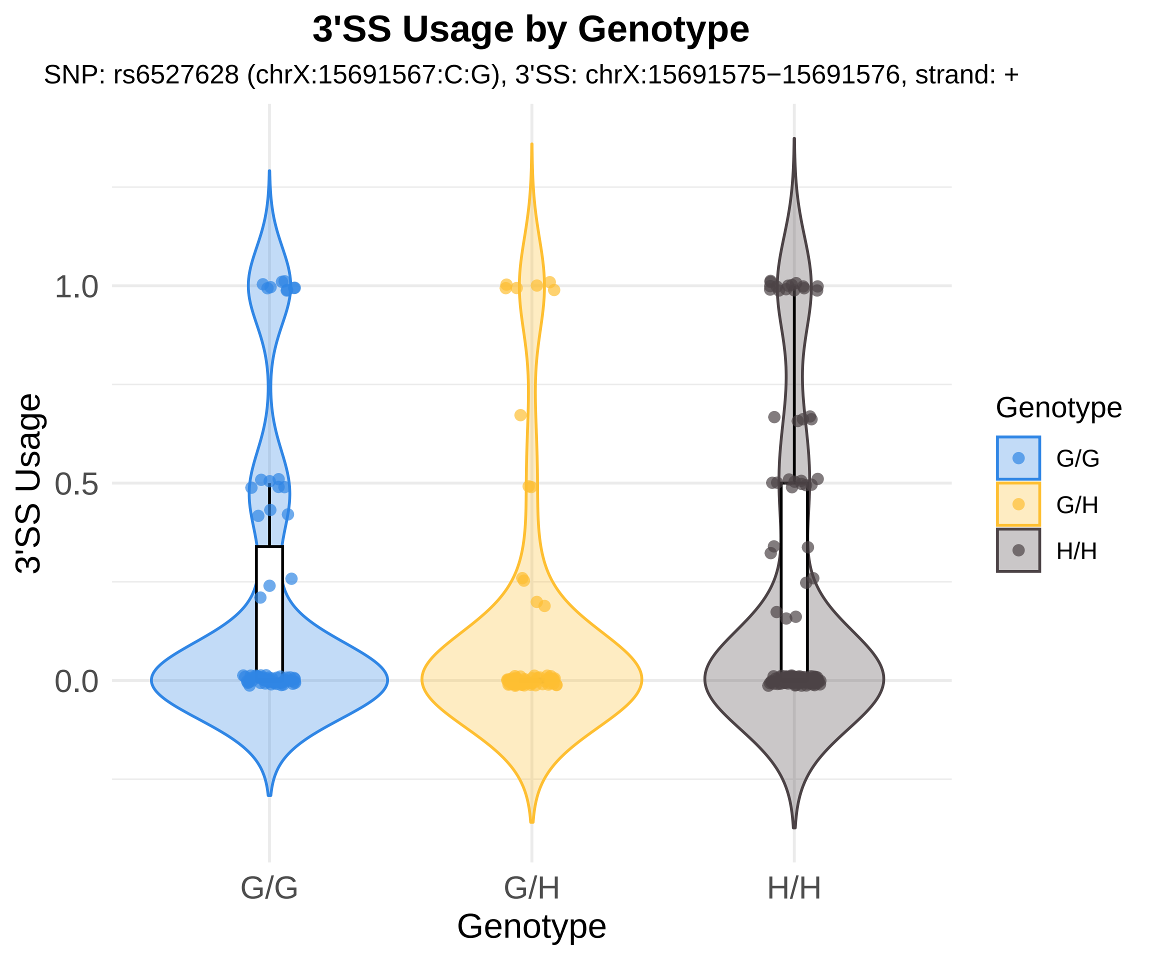
3'SS Usage by Genotype
SNP: rs6527628 (chrX:15691567:C:G), 3'SS: chrX:15691575−15691576, strand: +
3'SS Usage
1.0
0.5
0.0
G/G
G/H
H/H
Genotype
Genotype
G/G
G/H
H/H

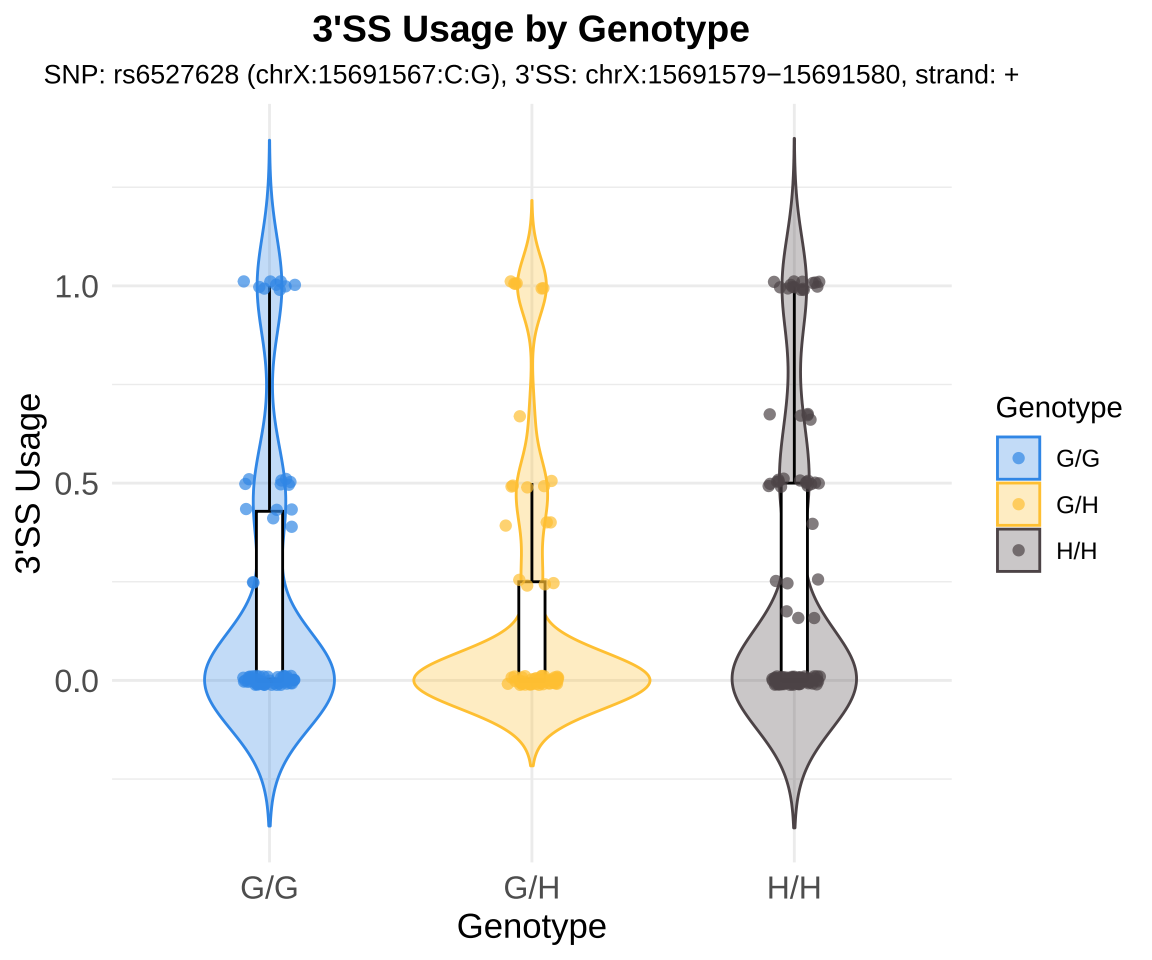

3'SS Usage by Genotype
SNP: rs6527628 (chrX:15691567:C:G), 3'SS: chrX:15691579−15691580, strand: +
3'SS Usage
1.0
0.5
0.0
G/G
G/H
H/H
Genotype
Genotype
G/G
G/H
H/H

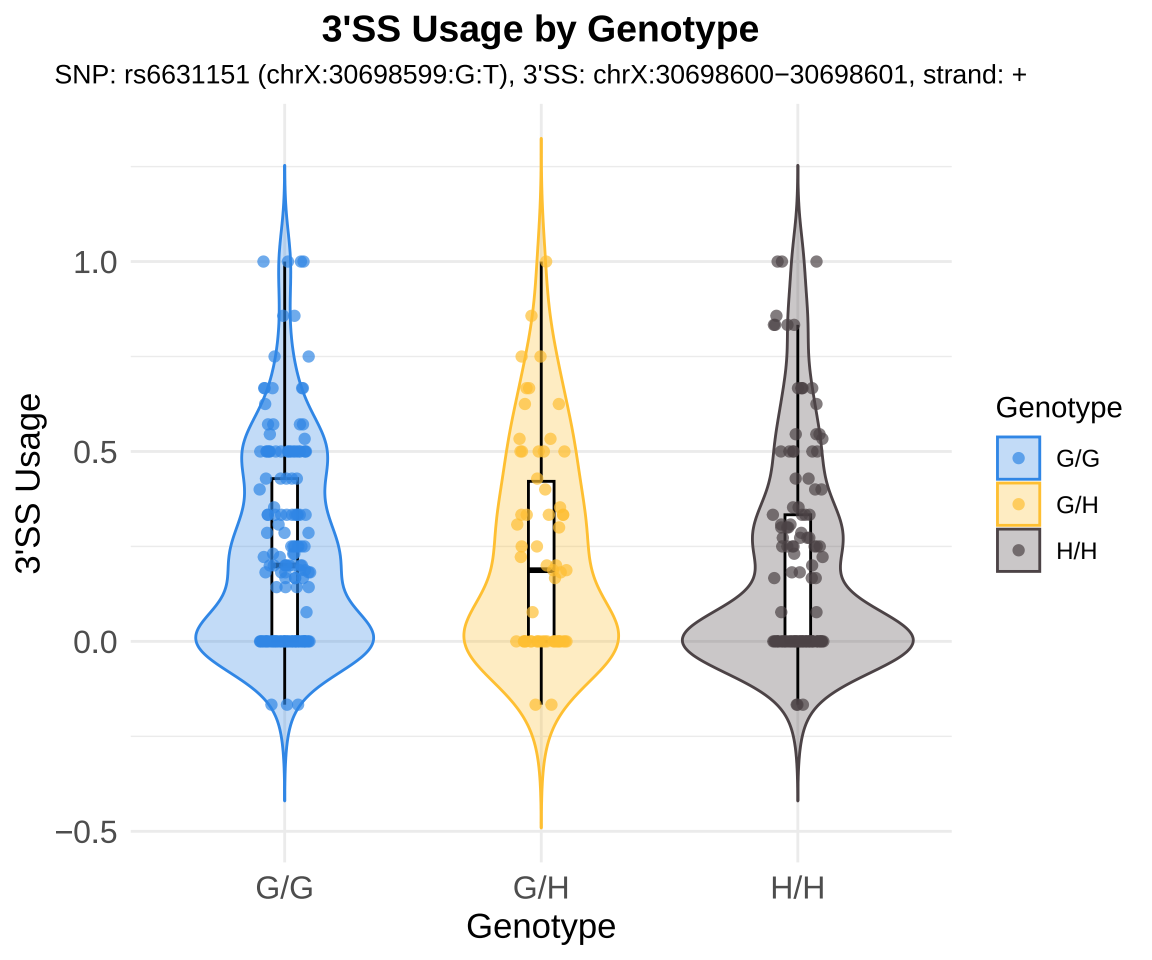
3'SS Usage by Genotype
SNP: rs6631151 (chrX:30698599:G:T), 3'SS: chrX:30698600−30698601, strand: +
3'SS Usage
1.0
0.5
0.0
−0.5
G/G
G/H
H/H
Genotype
Genotype
G/G
G/H
H/H

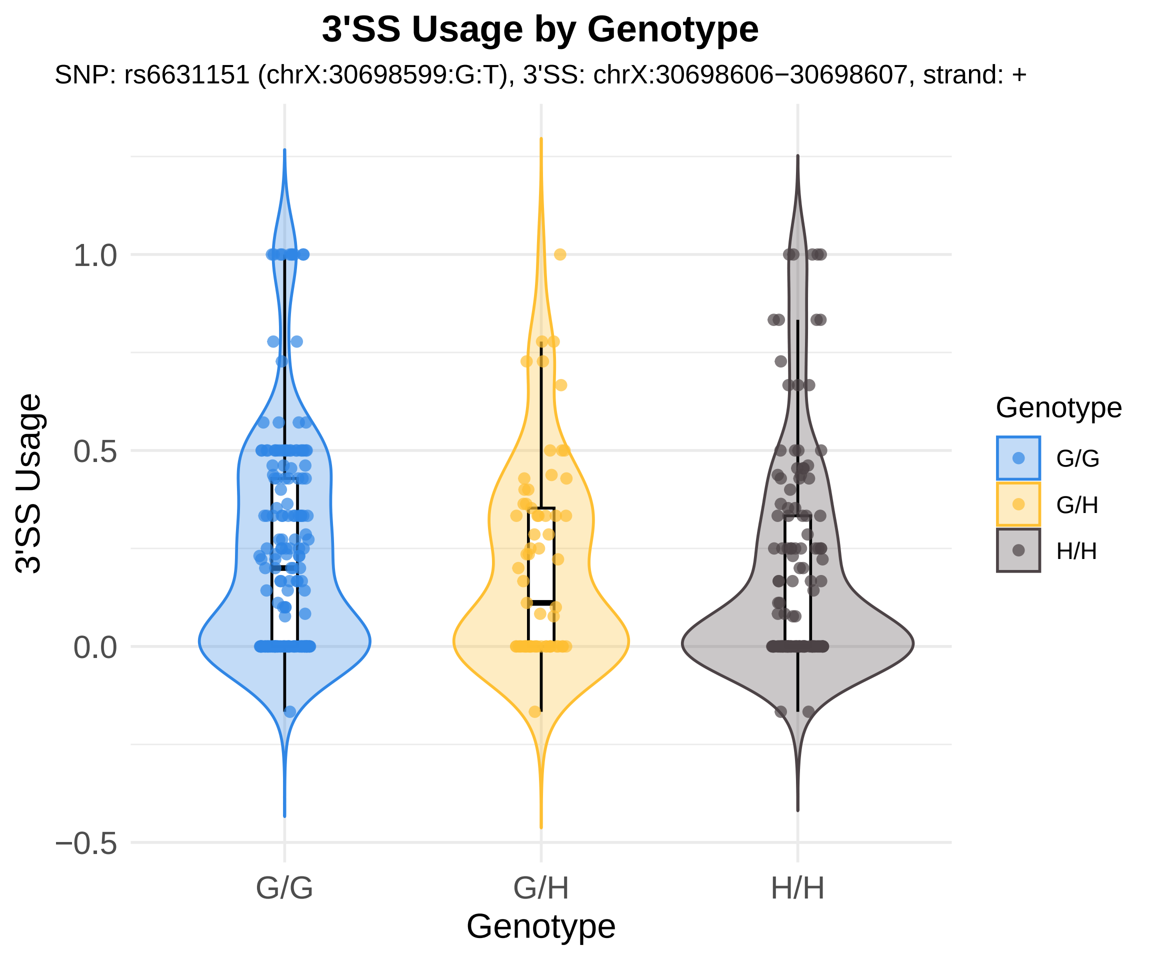
3'SS Usage by Genotype
SNP: rs6631151 (chrX:30698599:G:T), 3'SS: chrX:30698606−30698607, strand: +
3'SS Usage
1.0
0.5
0.0
−0.5
G/G
G/H
H/H
Genotype
Genotype
G/G
G/H
H/H

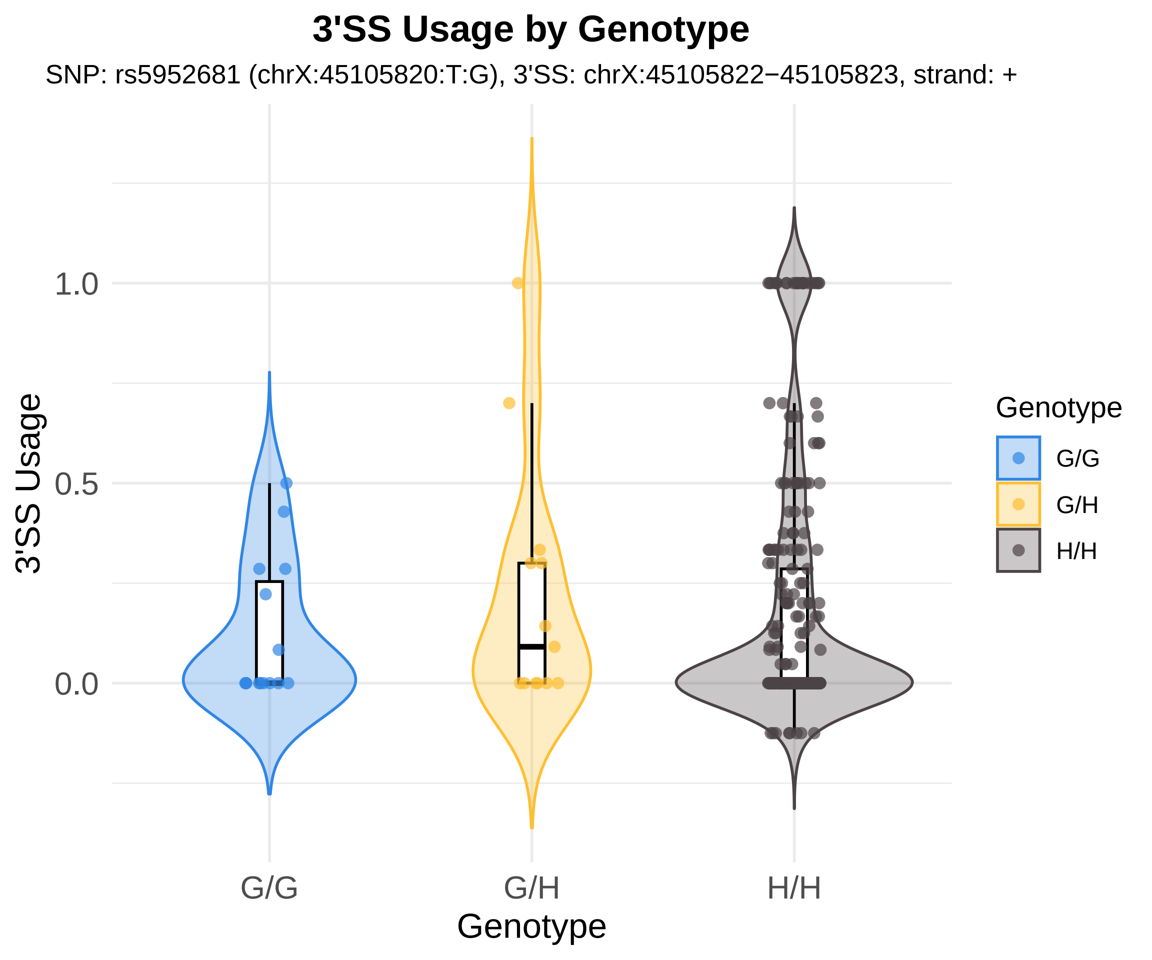

3'SS Usage by Genotype
SNP: rs5952681 (chrX:45105820:T:G), 3'SS: chrX:45105822−45105823, strand: +
1.0
0.5
0.0
3'SS Usage
G/G
G/H
H/H
Genotype
Genotype
G/G
G/H
H/H

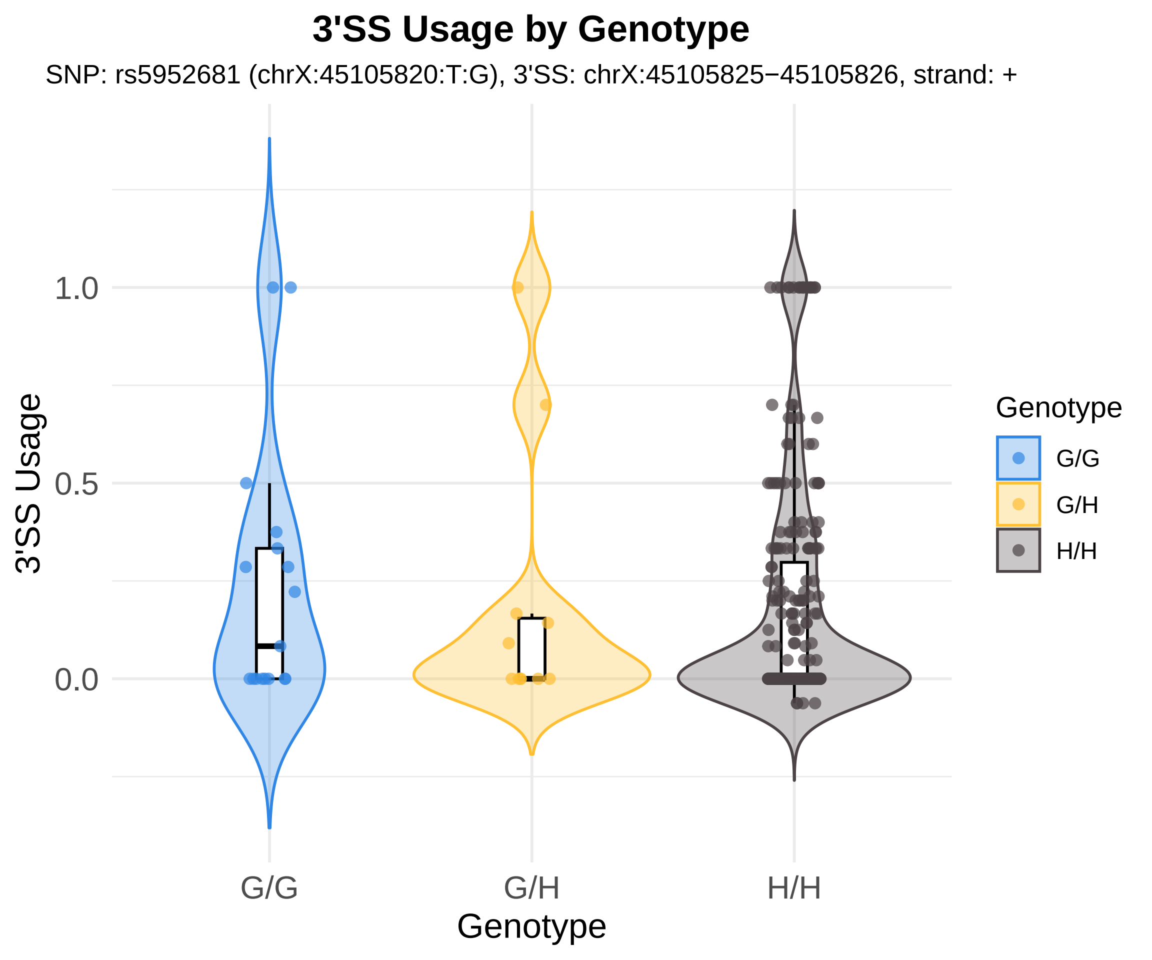
3'SS Usage by Genotype
SNP: rs5952681 (chrX:45105820:T:G), 3'SS: chrX:45105825−45105826, strand: +
3'SS Usage
1.0
0.5
0.0
G/G
G/H
H/H
Genotype
Genotype
G/G
G/H
H/H

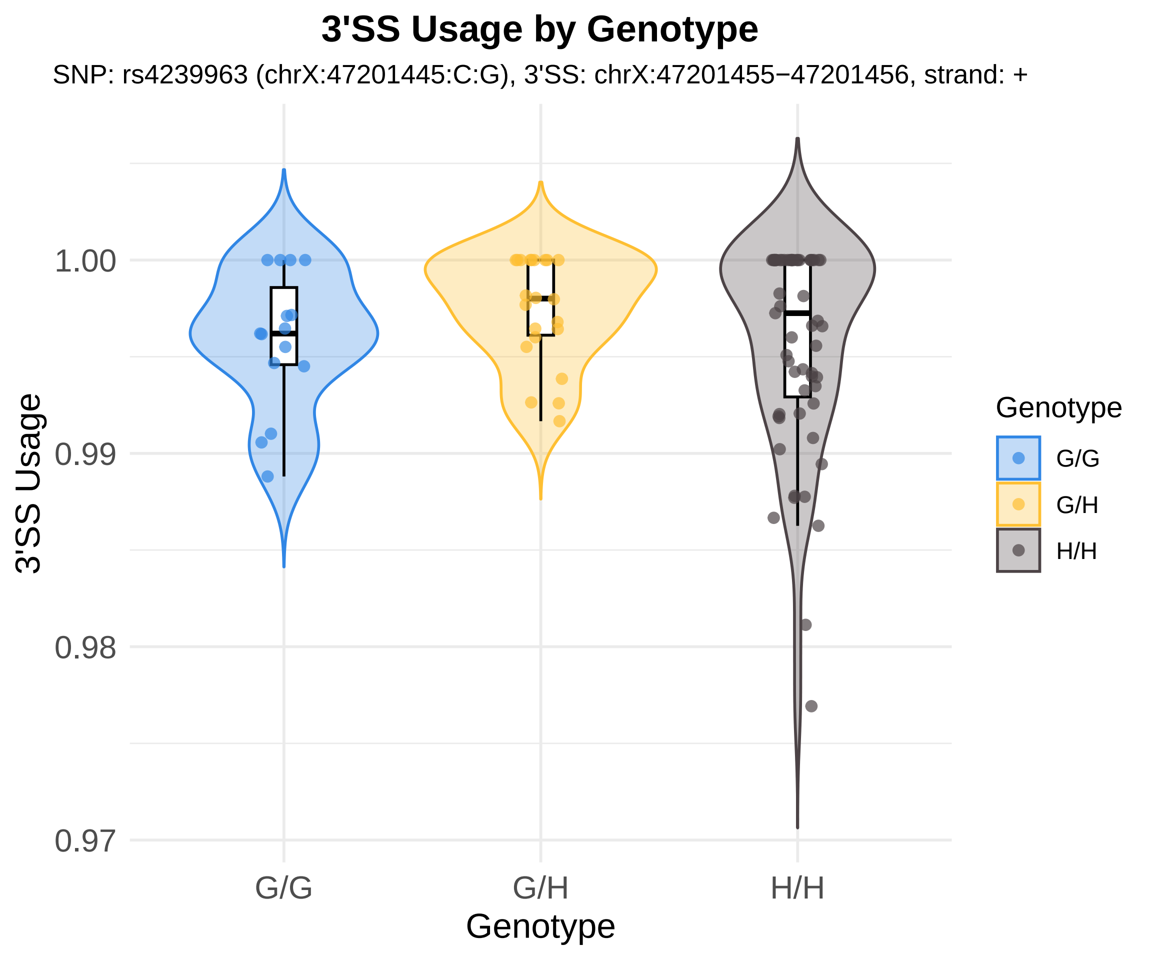
3'SS Usage by Genotype
SNP: rs4239963 (chrX:47201445:C:G), 3'SS: chrX:47201455−47201456, strand: +
3'SS Usage
1.00
0.99
0.98
0.97
G/G
G/H
H/H
Genotype
Genotype
G/G
G/H
H/H

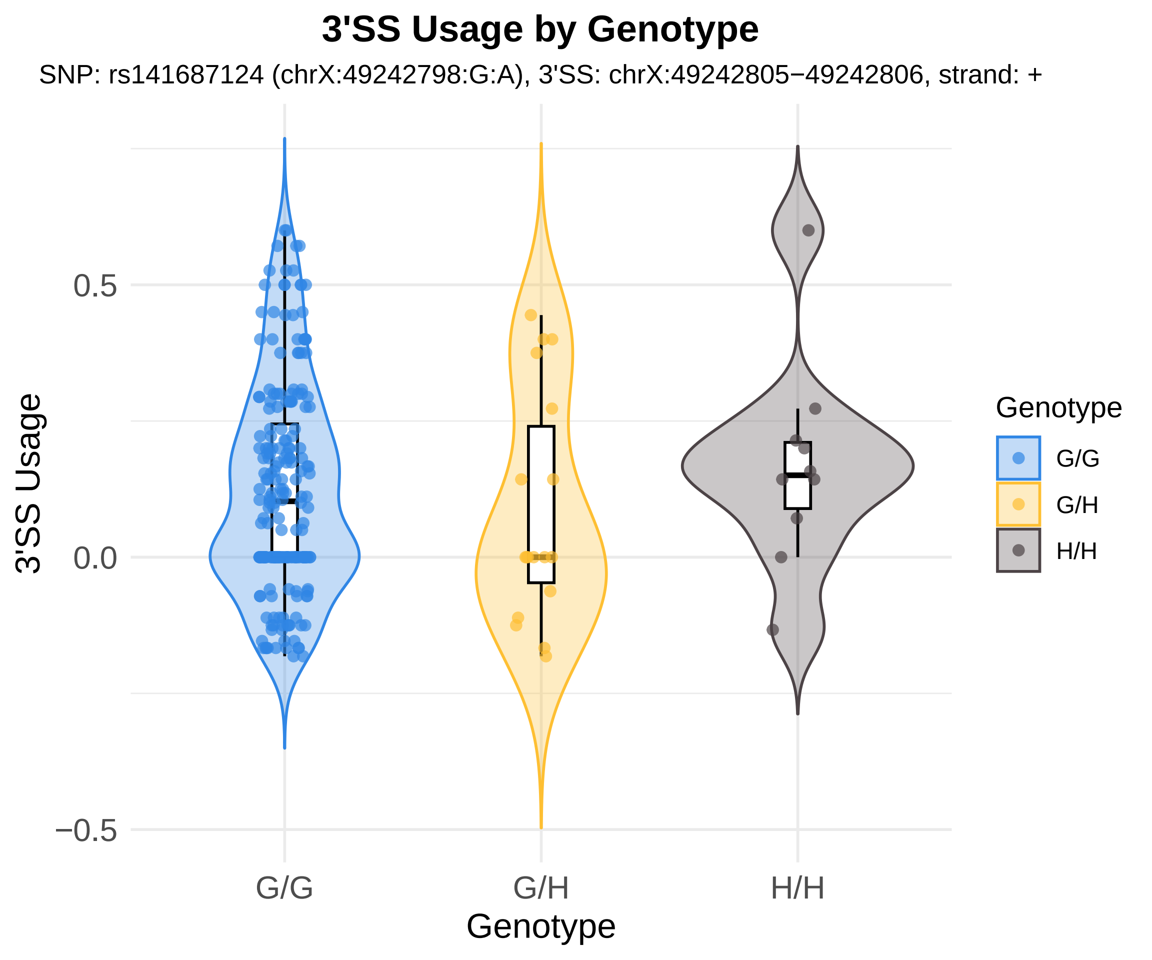
3'SS Usage by Genotype
SNP: rs141687124 (chrX:49242798:G:A), 3'SS: chrX:49242805−49242806, strand: +
3'SS Usage
0.5
0.0
−0.5
G/G
G/H
H/H
Genotype
Genotype
G/G
G/H
H/H

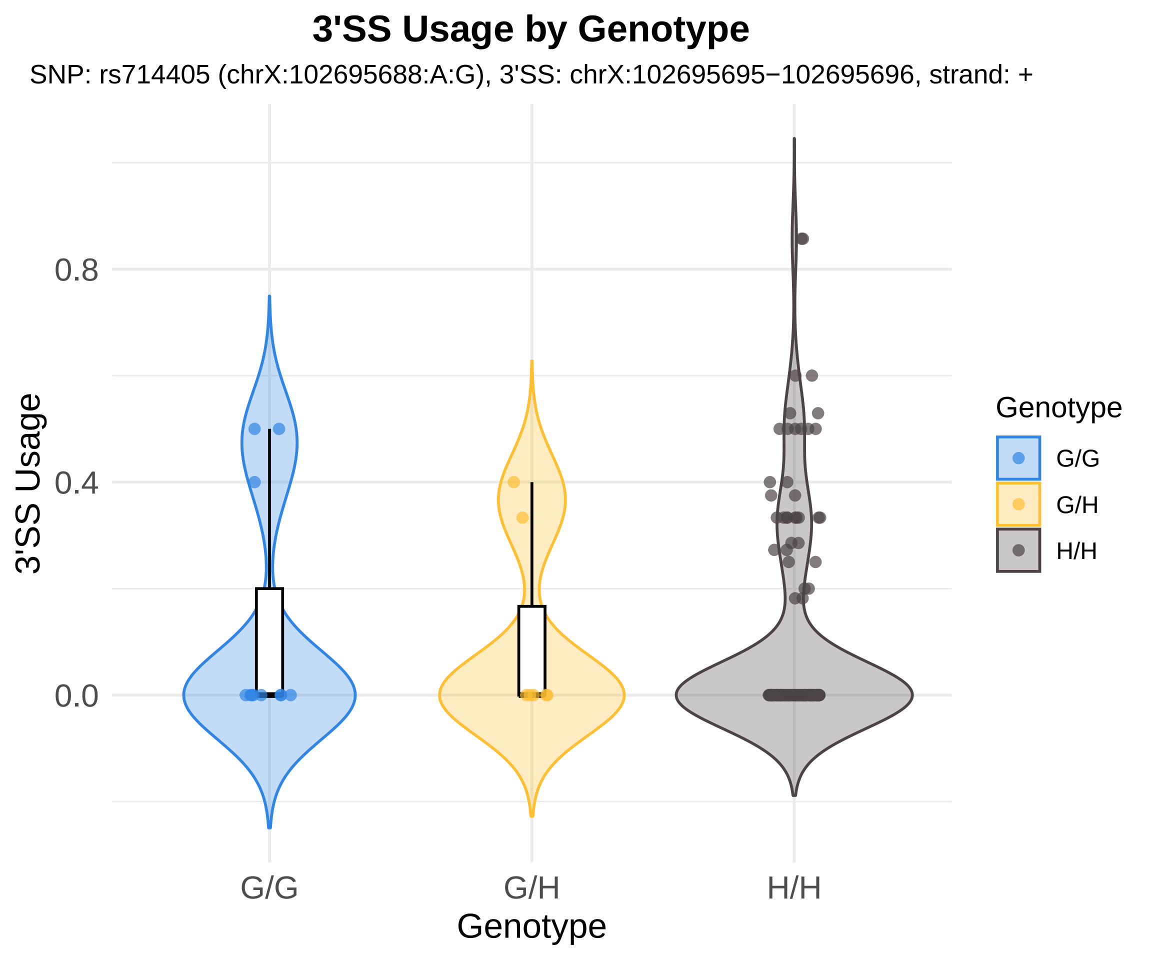
3'SS Usage by Genotype
SNP: rs714405 (chrX:102695688:A:G), 3'SS: chrX:102695695−102695696, strand: +
3'SS Usage
0.8
0.4
0.0
G/G
G/H
H/H
Genotype
Genotype
G/G
G/H
H/H
